## Supplementary material for "_Altered neutrophil extracellular traps in response to *Mycobacterium tuberculosis* in persons living with HIV with no previous TB and negative TST and IGRA_"

### Supporting information captions

#### S1 Table: Antibodies used for Flow Cytometry Analysis of Contaminating Cell Populations

| Staining step | Fluorophore | Specificity | Supplier | Clone | Catalogue Number |
|---|---|---|---|---|---|
| **Surface Staining** | BUV496 | CD16 | BD | 3G8 | 612945 |
| | V450 | CD66b | BD | G10F5 | 561649 |
| **Staining After Perm/Wash Incubation** | BV650 | CD15 | BioLegend | W6D3 | 323034 |
| | Spark Blue 550 | CD45 | BioLegend | 2D1 | 368549 |
| | PE-eFluor-610 | CD14 | Thermo Fisher (Invitrogen) | 61D3 | 61-0149-42 |
| | PE-Cy5 | CD3 | BD | UCHT1 | 561007 |

#### S2 Table: Cell population distribution of isolated PMN from HITTIN and HIT, as determined by flow cytometry

| Characteristic | HITTIN[a], N = 14 (3 missing values)[*] | HIT[b], N = 10 (1 missing value)[*] | p-value[**] |
|---|---|---|---|
| **CD15+ CD66b+** | 92.70% (87.05, 93.53) | 94.35% (93.65, 97.08) | 0.02** |
| **CD16+ (Neutrophils)** | 86.90 (80.28, 92.90) | 90.45 (86.10, 93.25) | 0.28 |
| **CD16- $CD14_{low}$ (Eosinophils)** | 4.33 (1.51, 7.23) | 3.63 (2.11, 6.60) | 0.93 |
| **CD15- CD66b-** | 7.31% (6.44, 12.95) | 5.67% (2.92, 6.32) | 0.03** |
| **CD3- CD14- (Other)** | 0.30 (0.17, 0.74) | 0.39 (0.18, 0.58) | 0.66 |
| **CD3- CD14+ (Monocytes)** | 0.07 (0.04, 0.12) | 0.11 (0.07, 0.19) | 0.22 |
| **CD3+ (T-cells)** | 6.70 (6.10, 12.43) | 4.55 (2.27, 6.00) | 0.02** |

[*]Median (IQR); [**]Wilcoxon rank sum test

[a] HITTIN (HIV-1-infected persistently TB, tuberculin and IGRA negative), [b] HIT (HIV-1-infected IGRA positive tuberculin positive)

#### S3 Table: Differential gene expression testing results

Included as a separate document due to size.

#### S4 Table: Sample characteristics

| **Characteristic** | **$PMN_{HITTIN}$** [a], N = 17[*] | **$PMN_{HIT}$** [b], N = 11[*] | **p-value**[**] |
|---|---|---|---|
| **Sequencer** | | | 0.4 |
| HiSeq | 3/ 17 (18%) | 4/ 11 (36%) | |
| NovaSeq | 14/ 17 (82%) | 7/ 11 (64%) | |

[*]n/ N (%); Mean (SD) [**]Fisher's exact test; Wilcoxon rank sum test

[a] $PMN_{HITTIN}$ (neutrophils from HIV-1-infected persistently TB, tuberculin and IGRA negative), [b] $PMN_{HIT}$ (neutrophils from HIV-1-infected IGRA positive tuberculin positive))

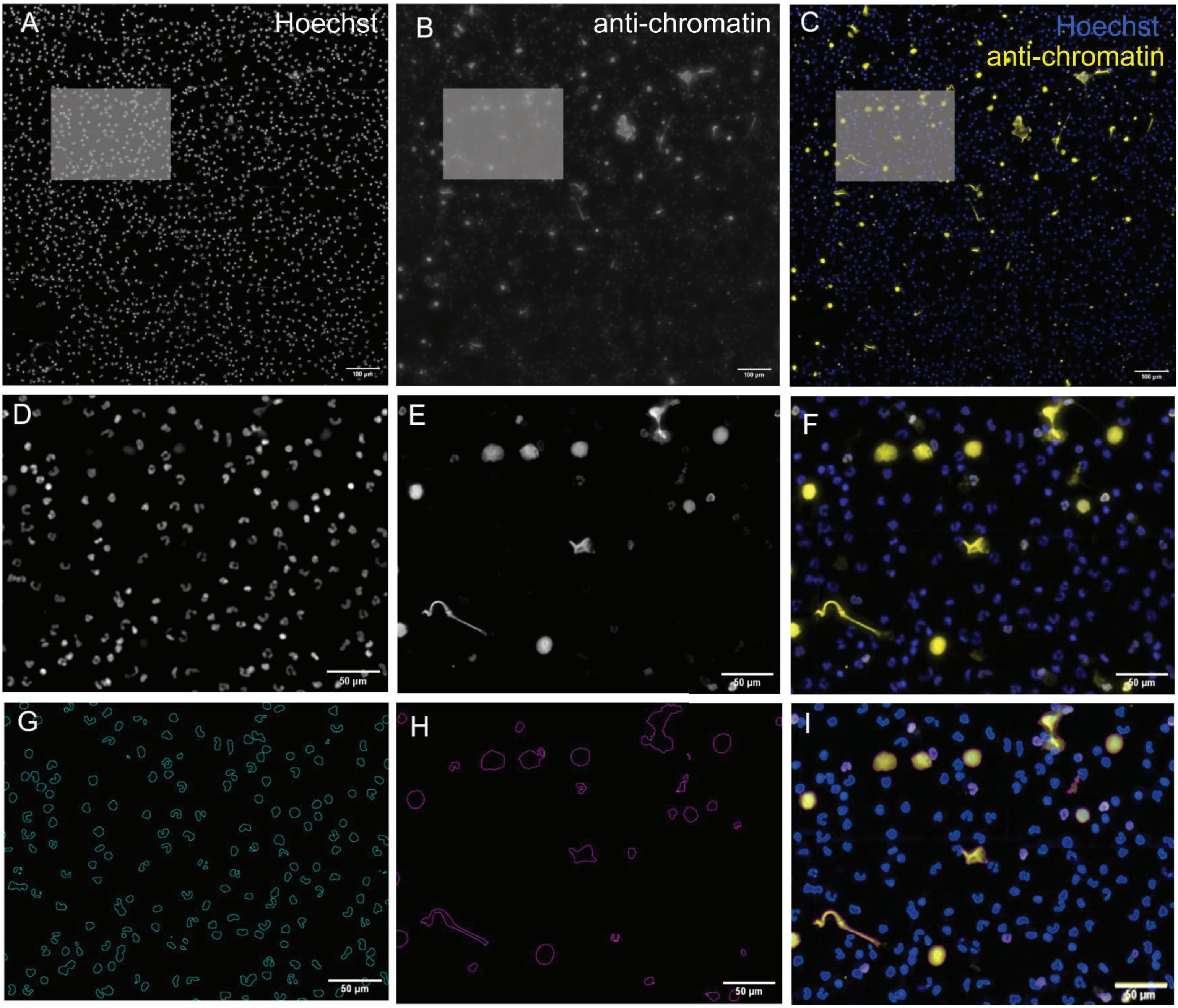


**S1 Fig: Fluorescence imaging process overview to calculate neutrophil nuclei and neutrophil extracellular trap (chromatin) area. Scale bars: 100 µm (A-C) and 50 µm (D-I).**

Isolated neutrophils were stimulated with *Mtb* and stained with Hoechst 33342 **(A and boxed area D)** and PL2-3 **(B and boxed area E)**. **C and the boxed area F** show the overlap (blue, DNA; yellow, chromatin). The segmentation of fluorescent signals used to calculate the neutrophil nuclei area **(G)** and neutrophil extracellular trap area **(H)** with the overlap shown in **(I)**. The scale bars represent 100 µm **(A-C)** and 50 µm **(D-I)**.

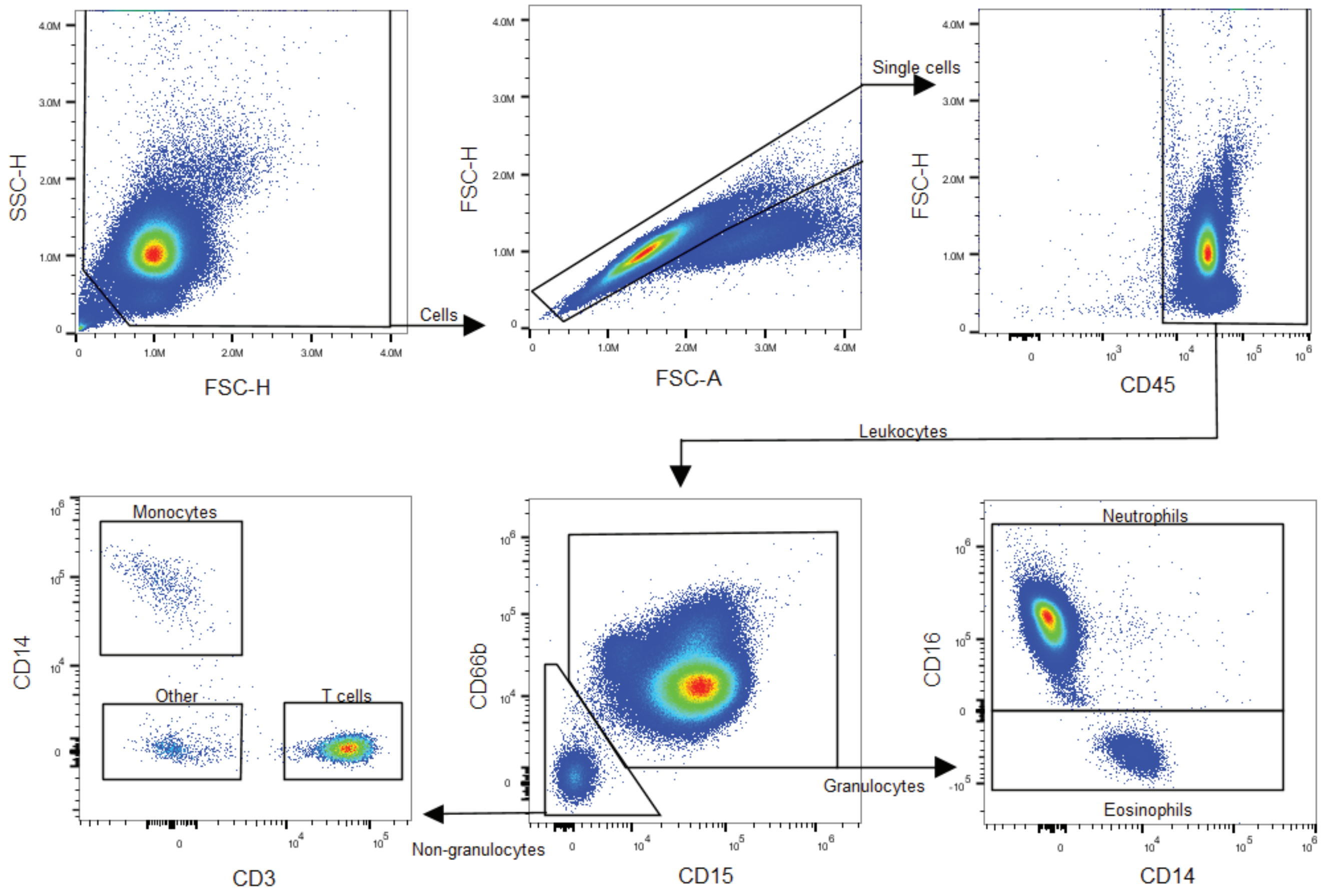


**S2 Fig: Flow Cytometry Analysis of Cell Populations**

The first gate was applied to exclude debris (the low SSC-H, FSC-H values in the bottom left corner). Single cells were separated and gated by a CD45+ marker for leukocytes. After single cell gating, CD45+ cells were then grouped into CD15+CD66b+ (granulocytes) and CD15-CD66b-(non-granulocytes) cells. The CD15+CD66b+ cells were further classified as CD14- CD16+ (Neutrophils) and CD14- CD16- (Eosinophils). CD15-CD66b- were stratified as CD3+ (T-cells), CD3- CD14+ (Monocytes) and CD14- CD3- (Other) cells.

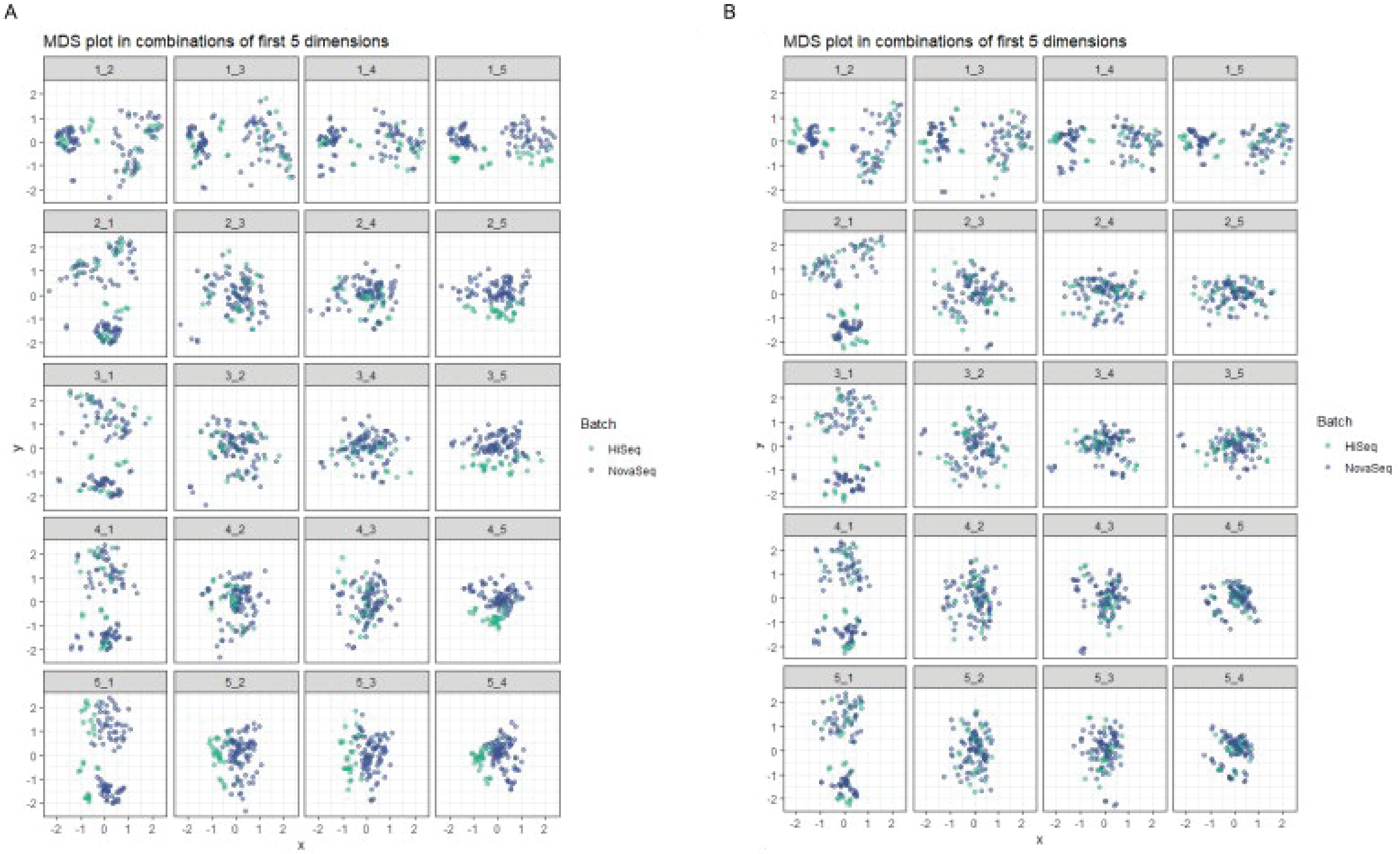


**S3 Fig: Multidimensional scaling (MDS) plot before (A) and after (B) ComBat-seq batch correction**

The multidimensional scaling (MDS) plots the Euclidian distances between samples with the x and y axis representing the sample distances between samples of read counts normalized by depth but not covariates. Each row of plots in **(A)** and **(B)** represent dimension 1 to 5 respectively (represented by the x-axis) and shown with the combination of the other dimensions on the y-axis. Samples are colored for batch effect with green showing samples sequenced on Illumina HiSeq2500 and blue for samples sequenced on Illumina NovaSeq6000. **(A)** shows the multiple combination of plots by dimension for the normalized counts prior to batch correction. Clear separation of samples by batch can be seen and is most notable in dimensions 4 and 5. **(B)** Appropriate batch correction after correction of raw counts with ComBat-seq. After batch correct outlier samples are most notably observed in dimension 3.

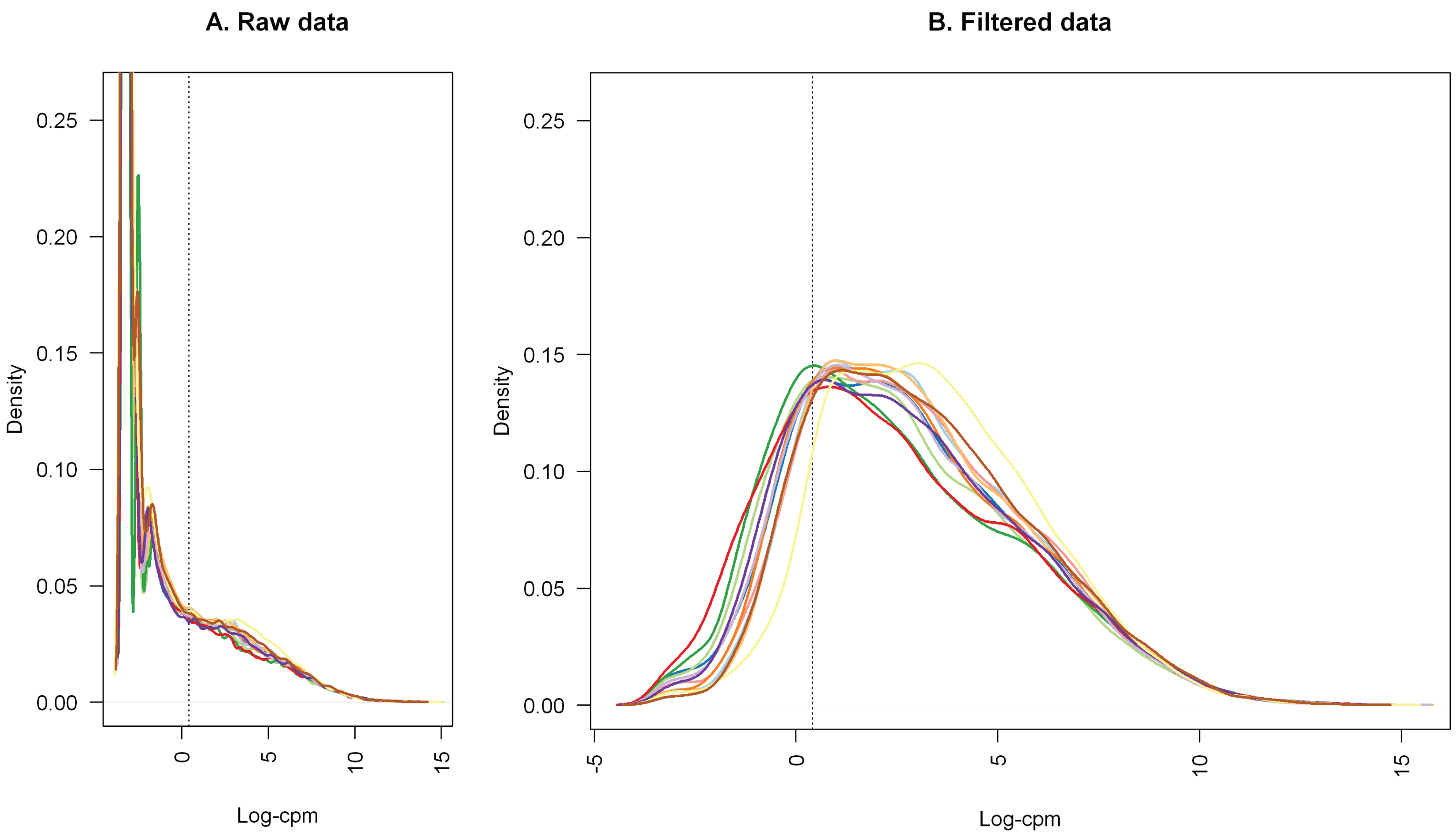


**S4 Fig: Density plot showing successful filtering applied to counts.**

Density plot with the density represented on the y-axis for the log-cpm values before **(A)** and after **(B)** filtering. The dotted vertical line is equivalent to the counts per million (CPM) threshold of 1.32 which was used in the filtering step.

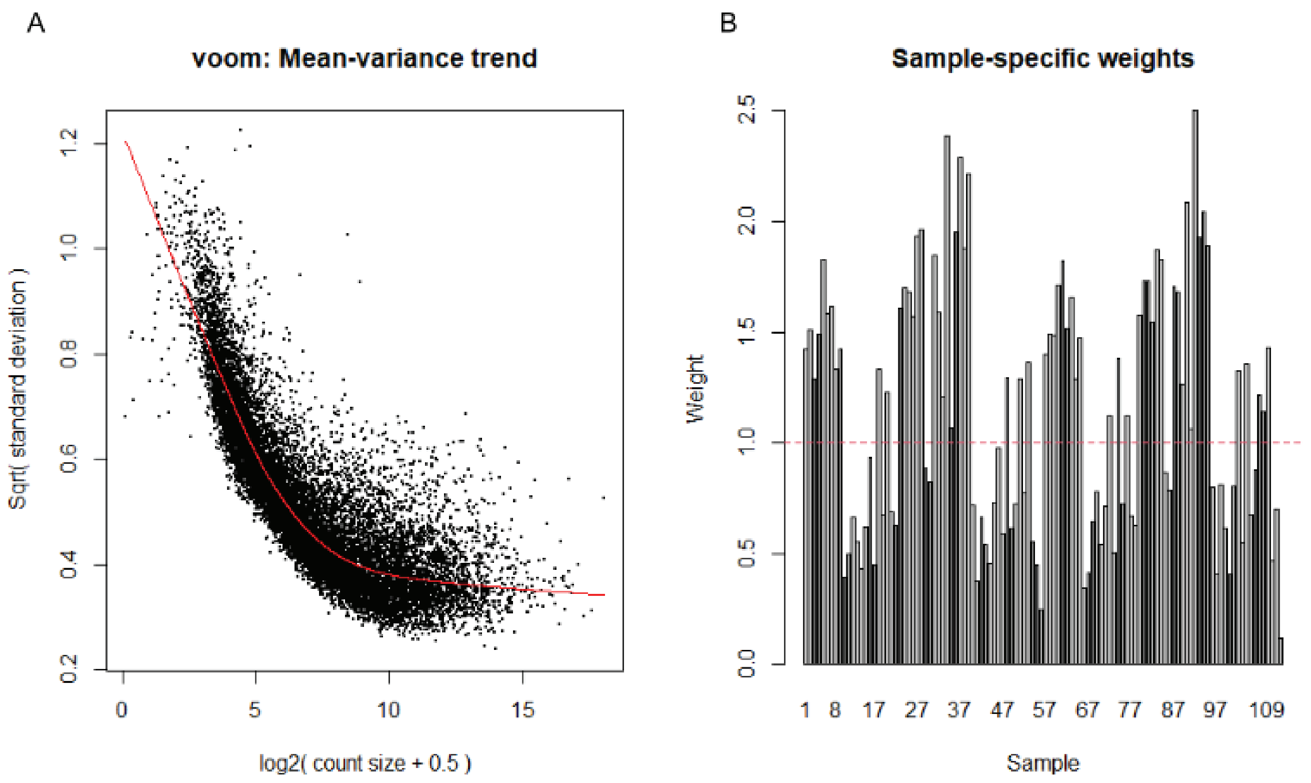


**S5 Fig: Voom observational mean-variance trend and sample-specific weights**

**(A)** shows rescaled (square-root of standard deviations) residual variances plotted against the mean expression (log2 transformed with an offset of 2) of each gene. A decreasing trend is seen between the mean gene expression and the variance with higher expressed genes showing less variation. The sample specific weight for each sample used in the analysis is shown in **(B)**. A total of 112 samples were analysed (28 participants and 4 conditions, uninfected and infected after 1 and 6h).

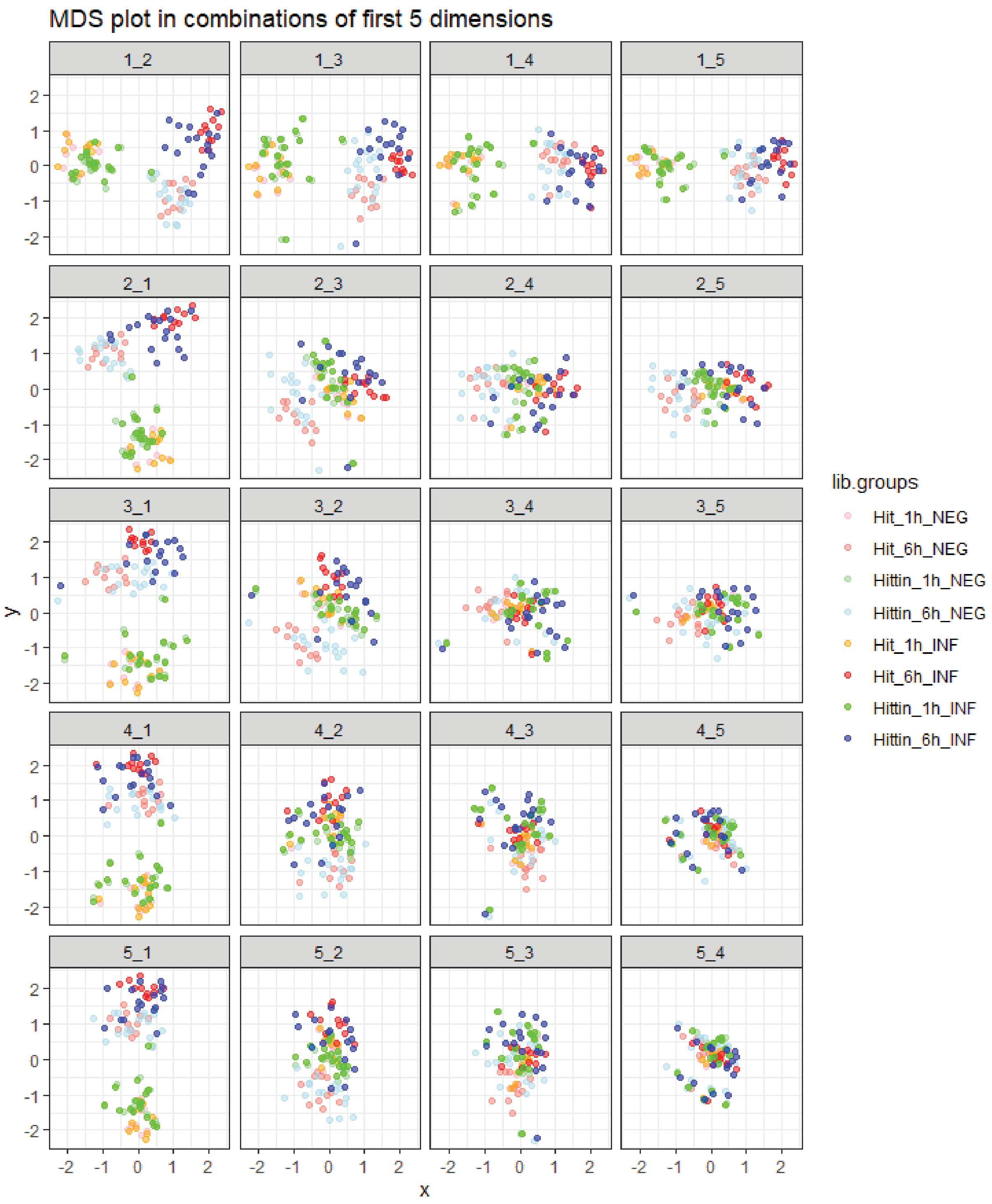


**S6 Fig: Multidimensional scaling (MDS) plot of group separation**

The multidimensional scaling (MDS) plots the Euclidian distances between samples with the x and y axis representing the sample distances between samples of read counts normalized by depth but not covariates. Each row of plots in represent dimension 1 to 5 respectively (represented by the x-axis) and shown with the combination of the other dimensions on the y-axis. Samples are colored for phenotype (HITTIN and HIT), timepoint (1h and 6h) and infection status (uninfected [NEG] and infected [INF]) and as depicted in the legend. Separation of the groups by time can be seen in dimension 1, by infection in dimension 2 and the phenotype in dimension 3. Additional separation is seen in dimension 4 and 5 due to sex and possibly smoking (see S8 and S9 Figs).

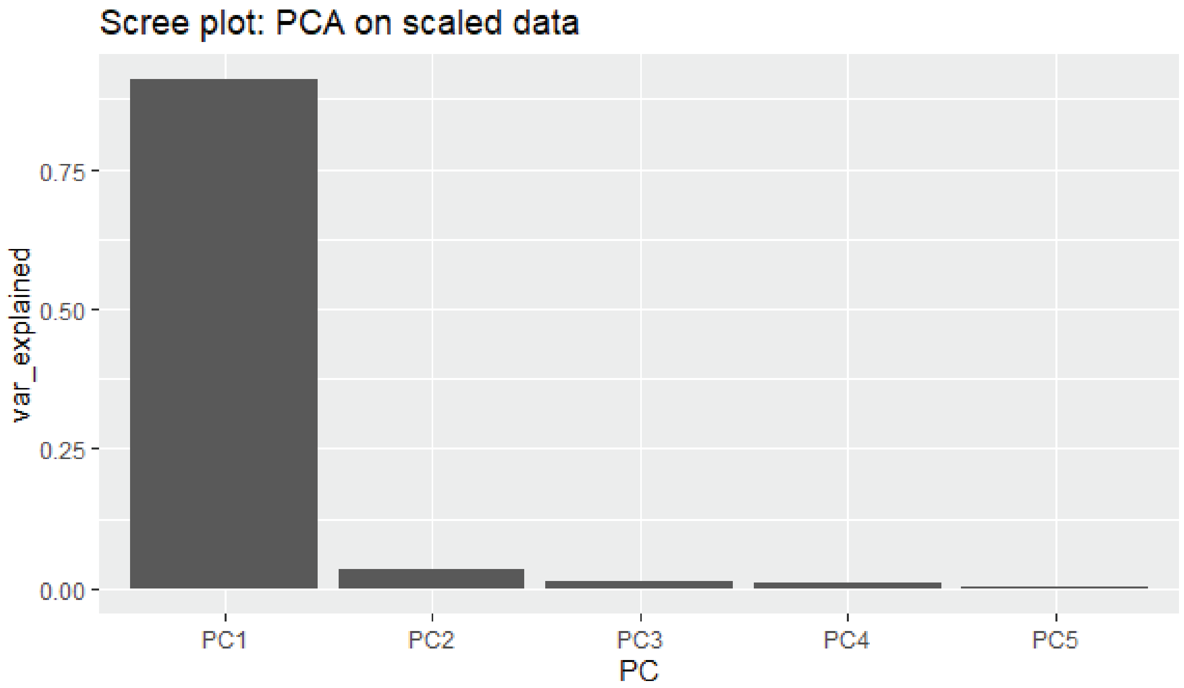


**S7 Fig: Scree plot**

The top 5 principal components are represented on the x-axis and the variance explained by each component on the y-axis. Principal component (PC) 1 contributes to most of the variance seen and represents time. PC2 represent infection effect and PC3 the phenotype groups.

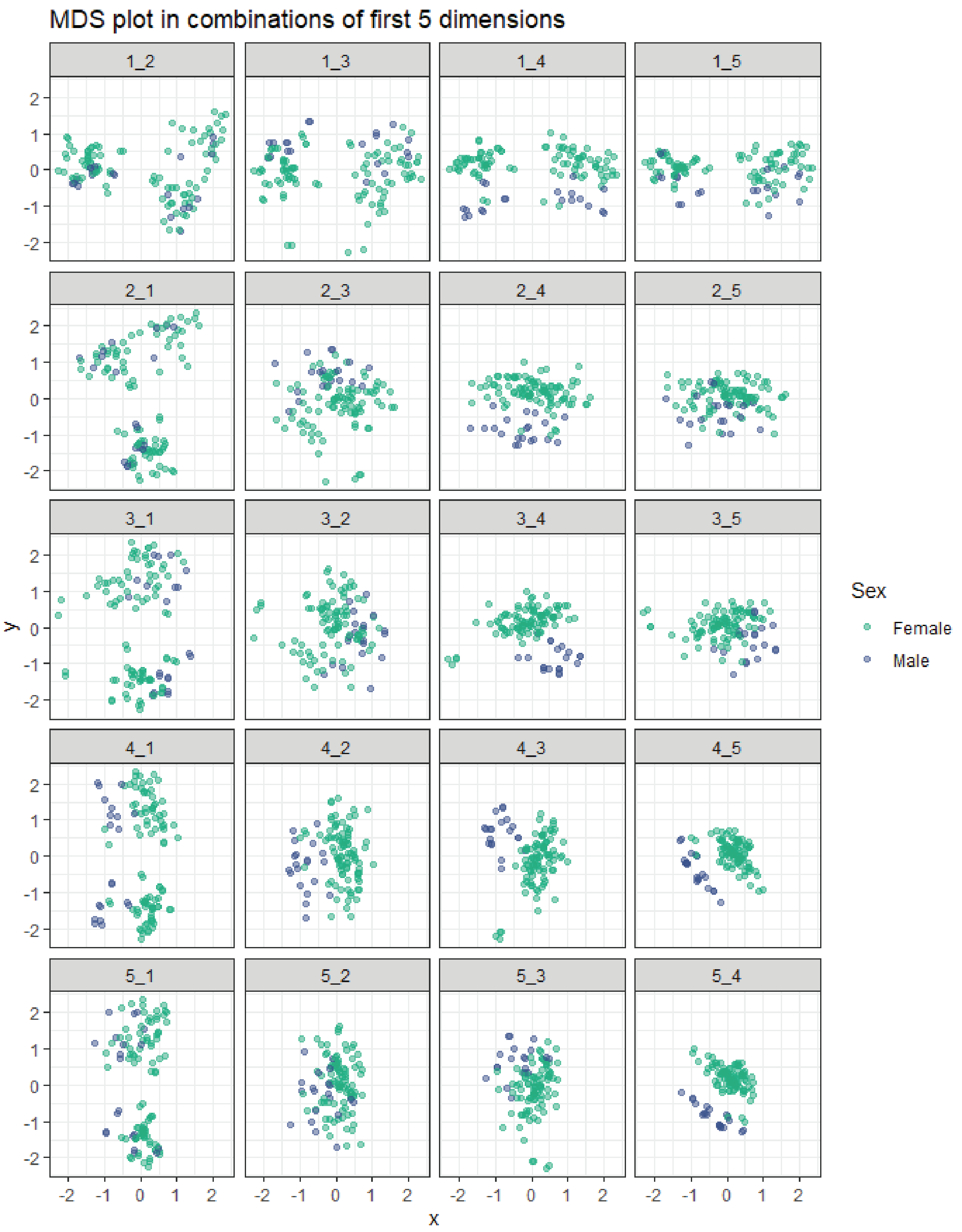


**S8 Fig: Multidimensional scaling (MDS) plot of participant sex**

The multidimensional scaling (MDS) plots the Euclidian distances between samples with the x and y axis representing the sample distances between samples of read counts normalized by depth but not covariates. Each row of plots in represent dimension 1 to 5 respectively (represented by the x-axis) and shown with the combination of the other dimensions on the y-axis. Samples are colored for sex (female and male) as depicted in the legend. Separation is seen in dimension 4 and 5.

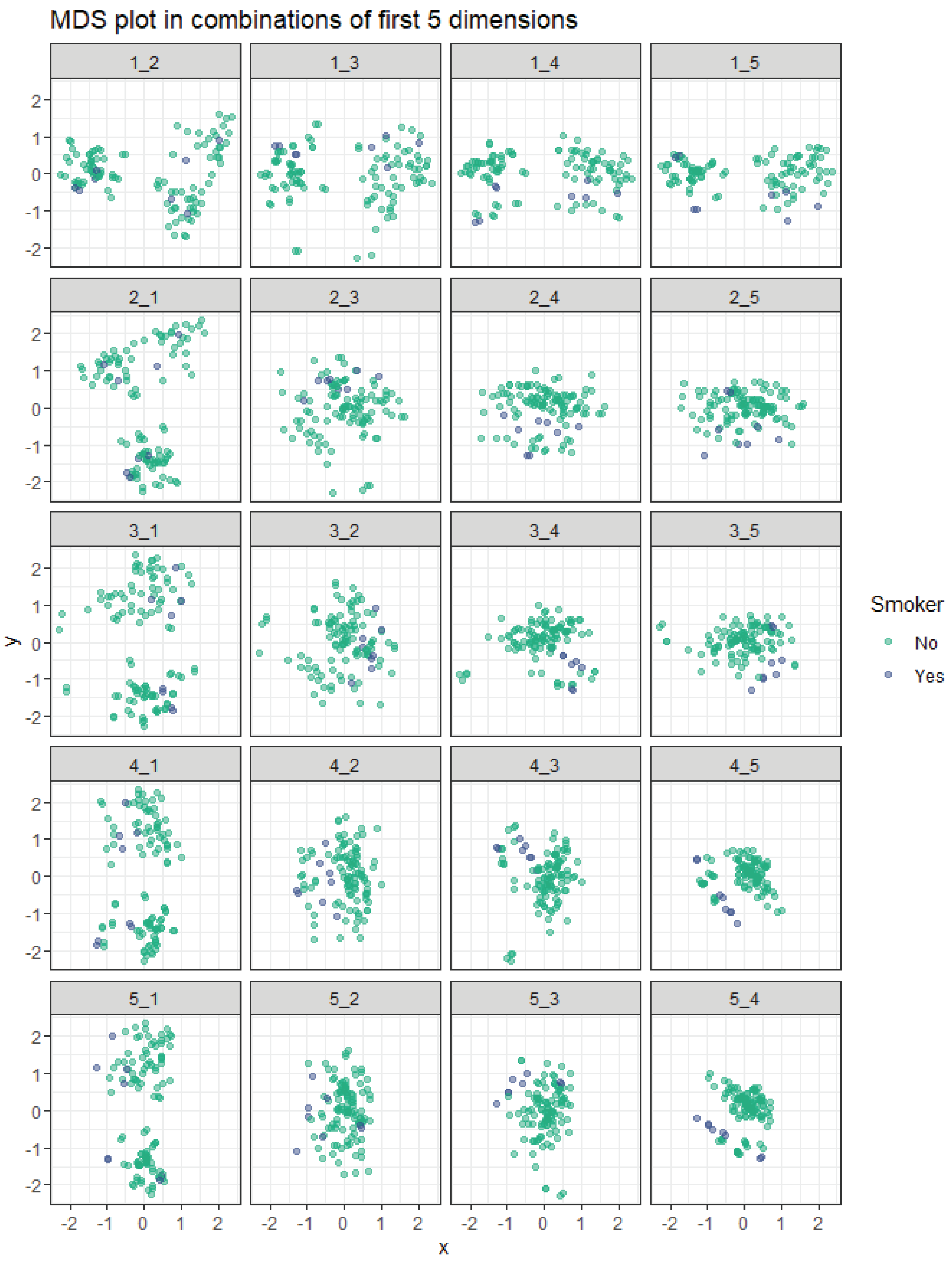


**S9 Fig: Multidimensional scaling (MDS) plot of participant smoking habit**

The multidimensional scaling (MDS) plots the Euclidian distances between samples with the x and y axis representing the sample distances between samples of read counts normalized by depth but not covariates. Each row of plots in represent dimension 1 to 5 respectively (represented by the x-axis) and shown with the combination of the other dimensions on the y-axis. Samples are colored for participants classified as smokers or not as depicted in the legend. Each participant is represented by 4 dots (1h uninfected and infected; 6h uninfected and infected). There are only two smokers making it difficult to comment on separation due to smoking.

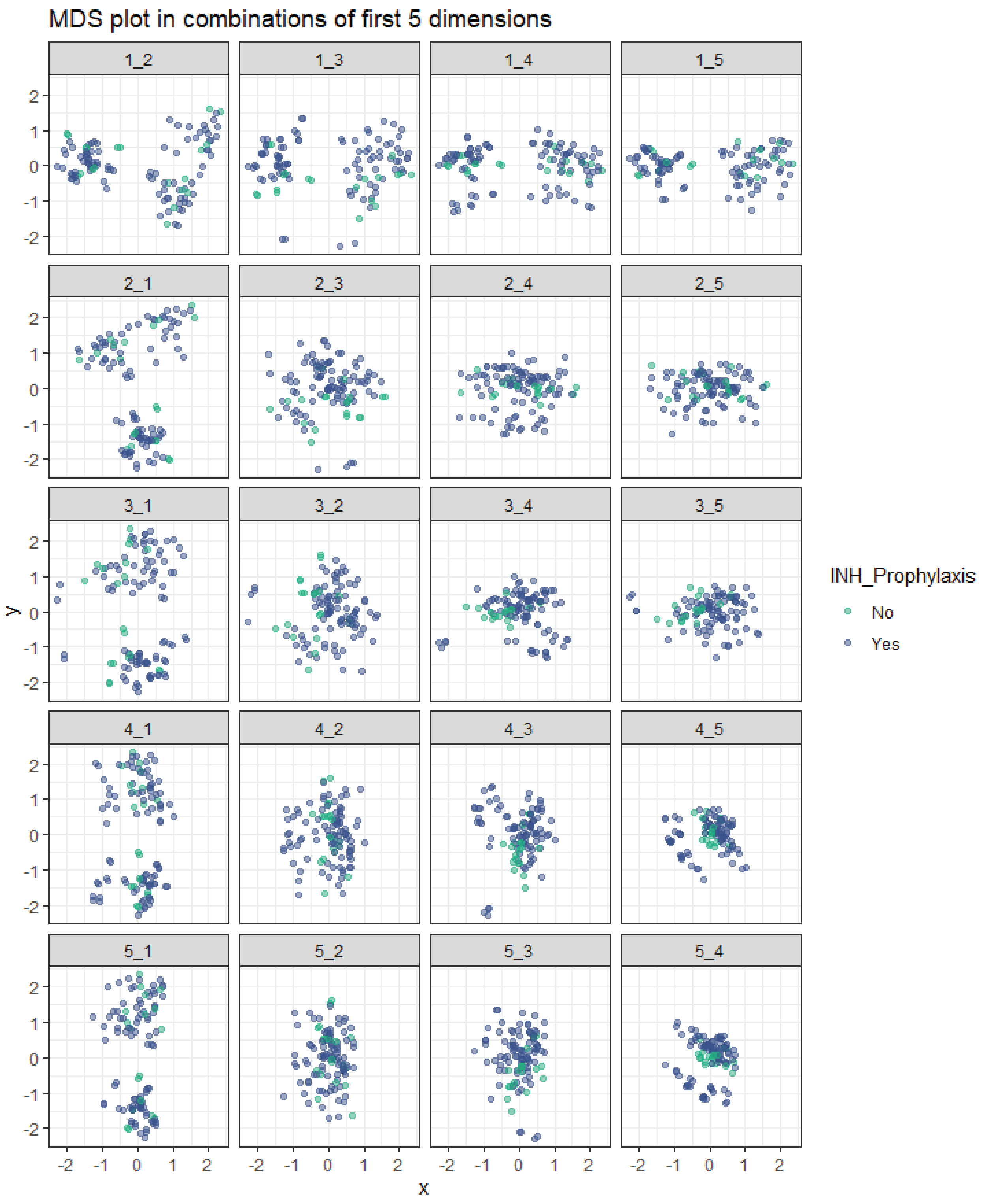


**S10 Fig: Multidimensional scaling (MDS) plot of participant isoniazid (INH) prophylaxis use**

The multidimensional scaling (MDS) plots the Euclidian distances between samples with the x and y axis representing the sample distances between samples of read counts normalized by depth but not covariates. Each row of plots in represent dimension 1 to 5 respectively (represented by the x-axis) and shown with the combination of the other dimensions on the y-axis. Samples are colored for participants classified as either having used INH prophylaxis previously/currently or not, as depicted in the legend. There is no clear separation based on INH prophylaxis use.

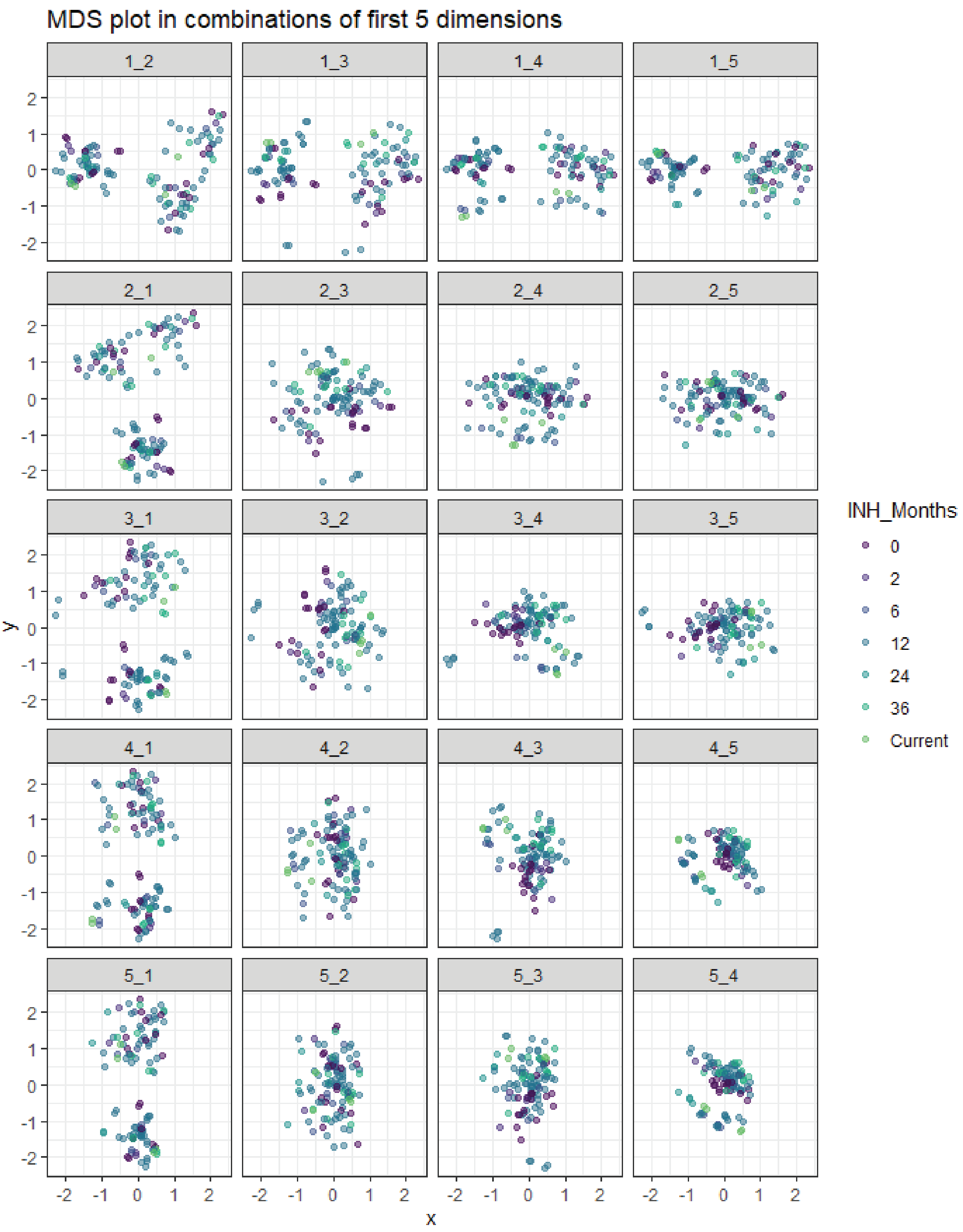


**S11 Fig: Multidimensional scaling (MDS) plot of duration of participant isoniazid (INH) prophylaxis use in months**

The multidimensional scaling (MDS) plots the Euclidian distances between samples with the x and y axis representing the sample distances between samples of read counts normalized by depth but not covariates. Each row of plots in represent dimension 1 to 5 respectively (represented by the x-axis) and shown with the combination of the other dimensions on the y-axis. Samples are colored for the numbers of months participants previously used INH (0 or none, 2, 6, 12, 24 or 36 months) or participants who are currently using INH, as depicted in the legend. There is no clear separation based on duration of INH prophylaxis use.

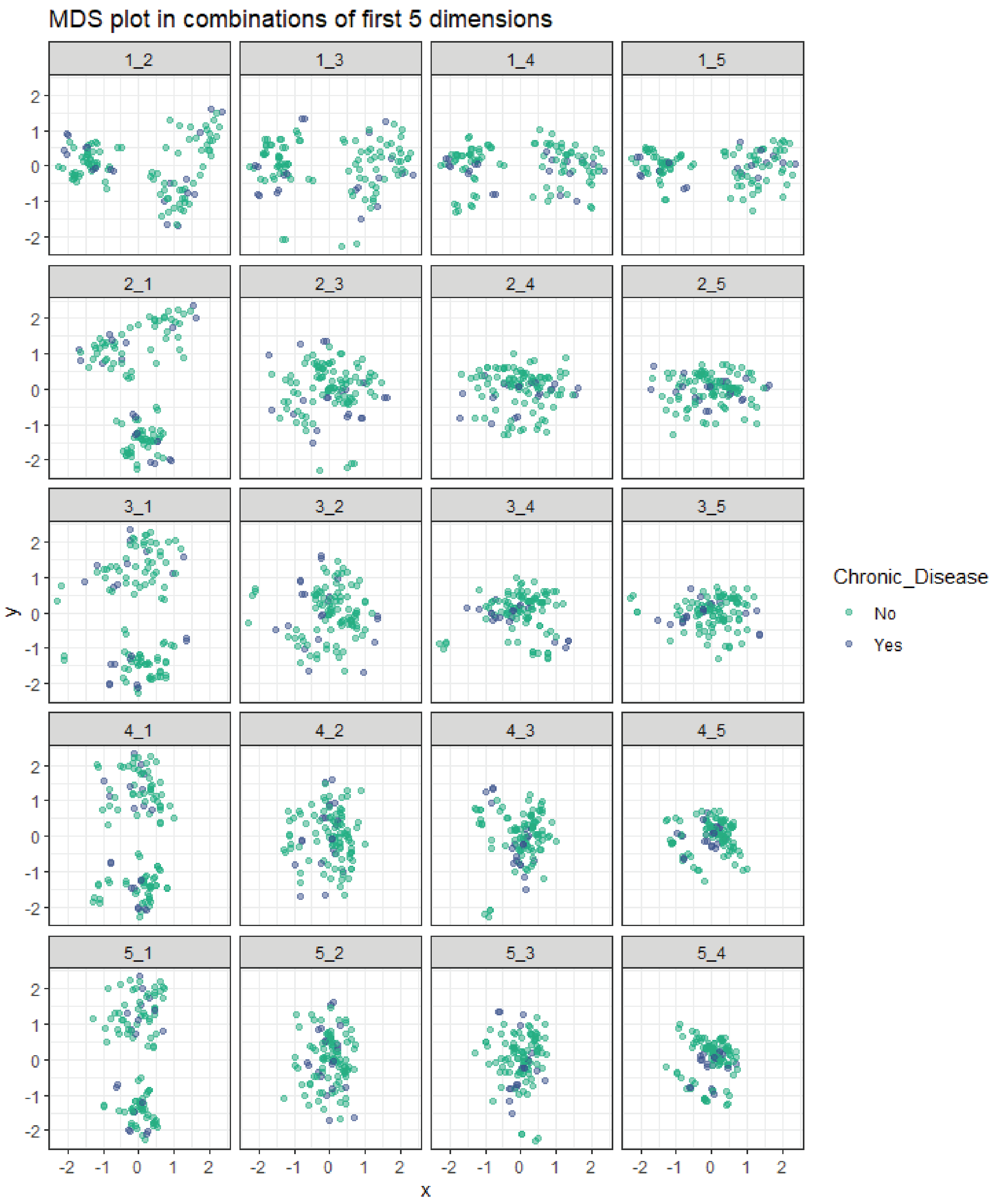


**S12 Fig: Multidimensional scaling (MDS) plot of participants with a known history of chronic disease**

The multidimensional scaling (MDS) plots the Euclidian distances between samples with the x and y axis representing the sample distances between samples of read counts normalized by depth but not covariates. Each row of plots in represent dimension 1 to 5 respectively (represented by the x-axis) and shown with the combination of the other dimensions on the y-axis. Samples are colored for participants classified as either having a chronic disease or not, as depicted in the legend. There is no clear separation based on participants with chronic disease.

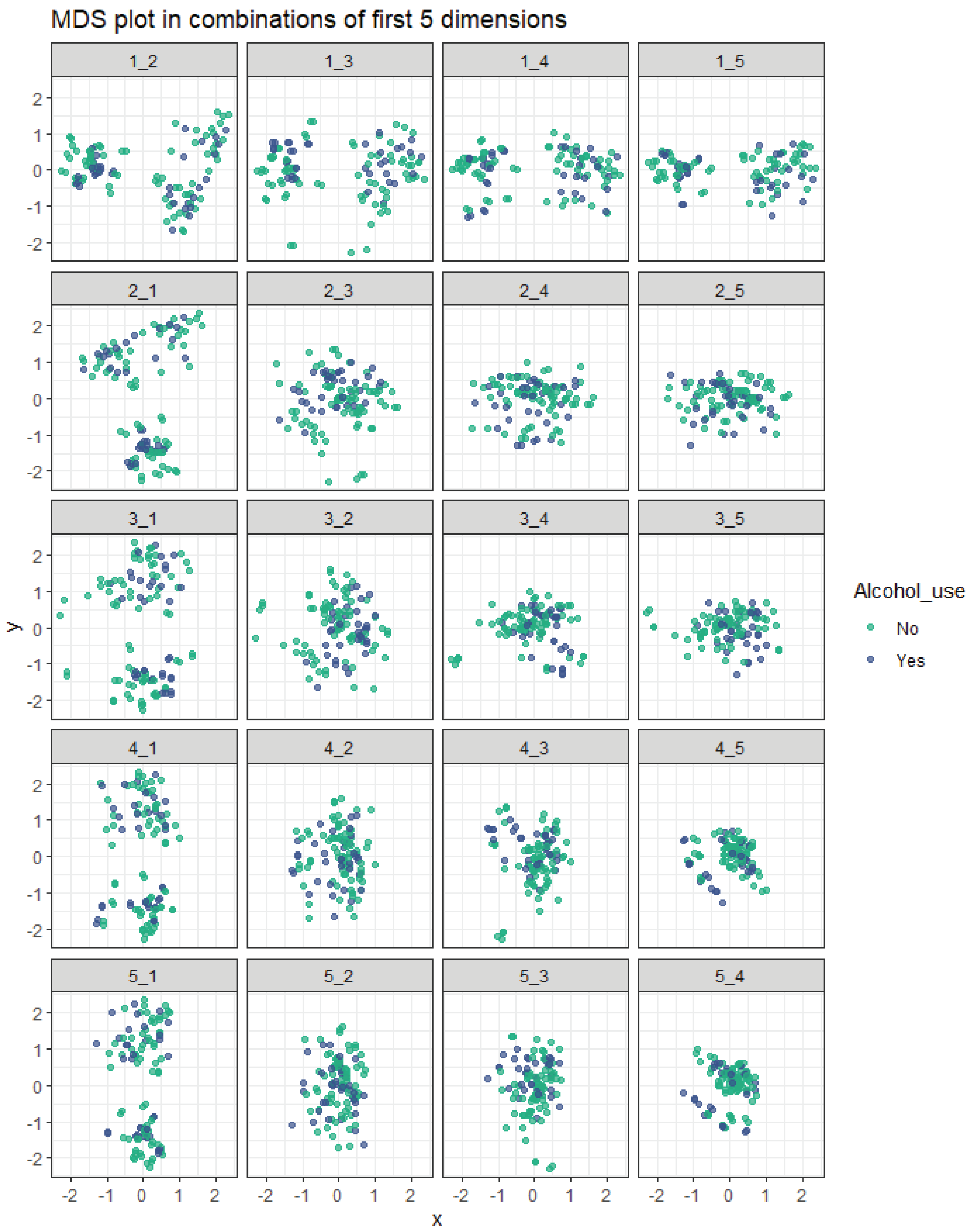


**S13 Fig: Multidimensional scaling (MDS) plot of participant social alcohol use**

The multidimensional scaling (MDS) plots the Euclidian distances between samples with the x and y axis representing the sample distances between samples of read counts normalized by depth but not covariates. Each row of plots in represent dimension 1 to 5 respectively (represented by the x-axis) and shown with the combination of the other dimensions on the y-axis. Samples are colored for participants classified as either using alcohol or not, as depicted in the legend. There is no clear separation based on participant alcohol use.

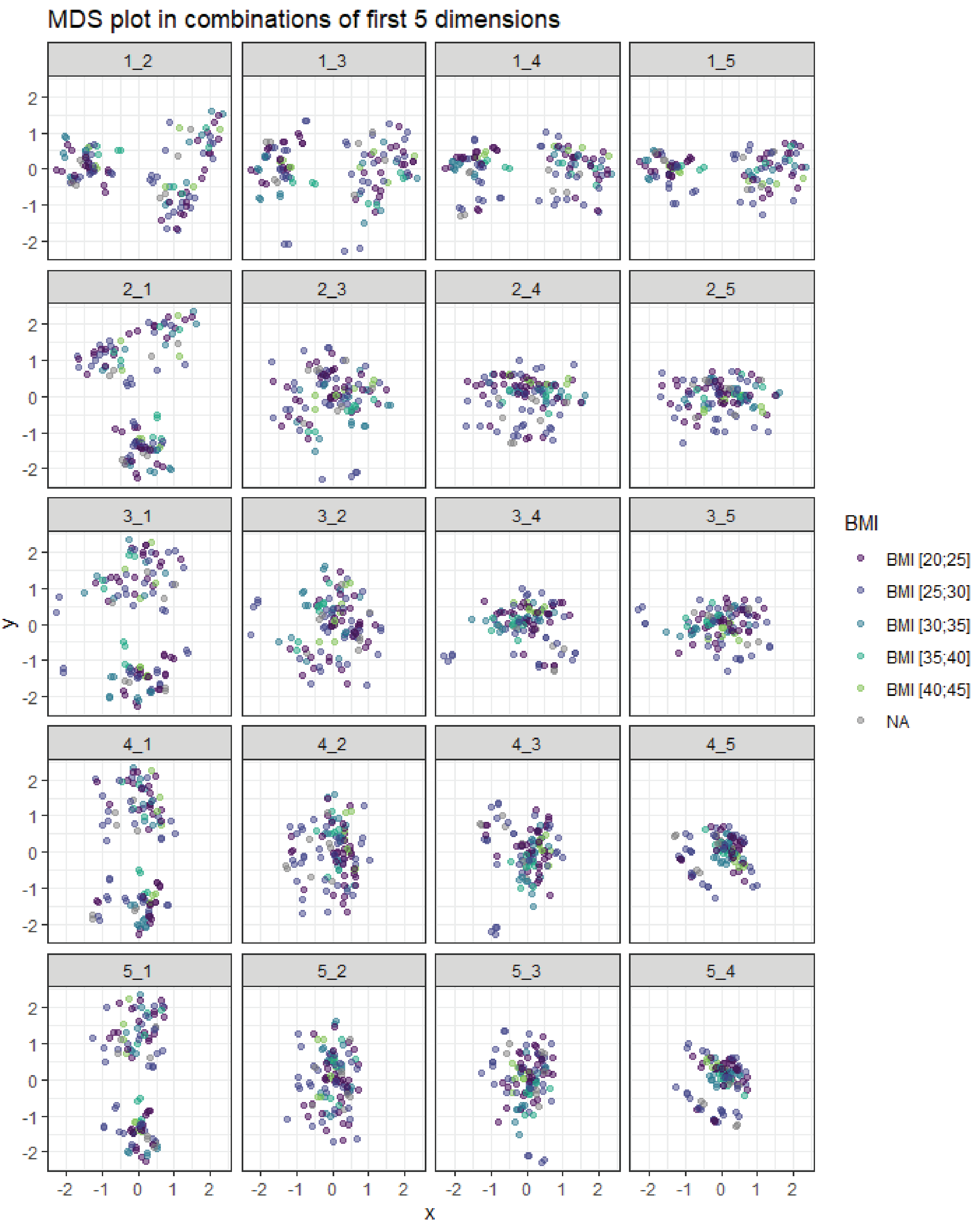


**S14 Fig: Multidimensional scaling (MDS) plot of participant body mass index (BMI)**

The multidimensional scaling (MDS) plots the Euclidian distances between samples with the x and y axis representing the sample distances between samples of read counts normalized by depth but not covariates. Each row of plots in represent dimension 1 to 5 respectively (represented by the x-axis) and shown with the combination of the other dimensions on the y-axis. Samples are colored for participants classified according to their BMI, as depicted in the legend. NA represents samples with missing values and is shown in grey. There is no clear separation based on participant BMI.

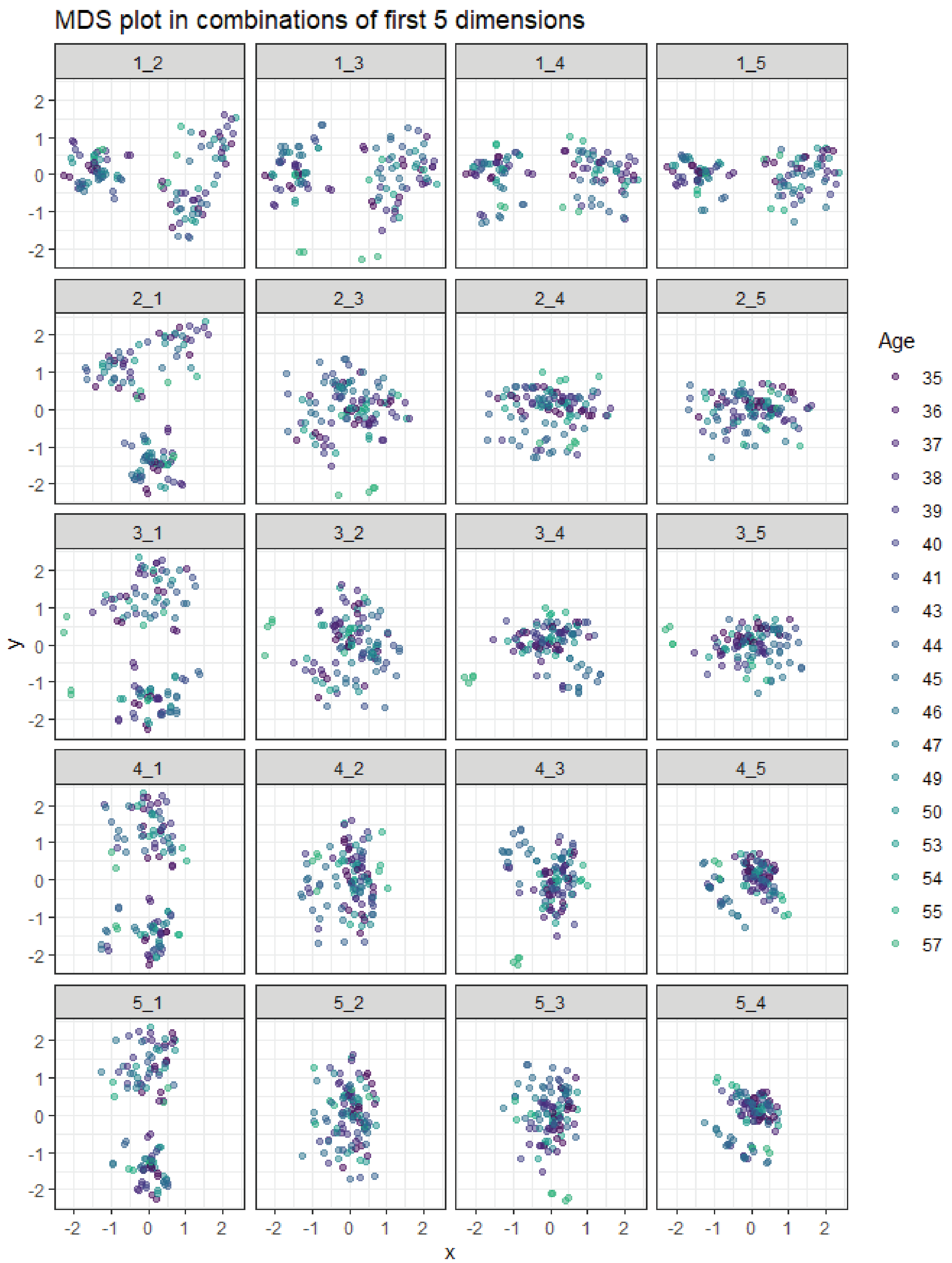


**S15 Fig: Multidimensional scaling (MDS) plot of participant age**

The multidimensional scaling (MDS) plots the Euclidian distances between samples with the x and y axis representing the sample distances between samples of read counts normalized by depth but not covariates. Each row of plots in represent dimension 1 to 5 respectively (represented by the x-axis) and shown with the combination of the other dimensions on the y-axis. Samples are colored for participants classified by age, as depicted in the legend. There is no clear separation based on participant age, although there is an older outlier seen in dimensions 3.
