## Supplementary table 3 for "_Altered neutrophil extracellular traps in response to *Mycobacterium tuberculosis* in persons living with HIV with no previous TB and negative TST and IGRA_"

### Supplementary table 3 : Differential gene expression testing results

| | | Average Expression | Log2FC Expression values | | | Adj.P.Value | Log2FC Expression values | | | Adj.P.Value |
|---|---|---|---|---|---|---|---|---|---|---|
| Ensembl_ID | Gene | All | HITTIN 1h inf | HIT 1h inf | HITTINxHIT 1h inf | HITTINxHIT 1h inf | HITTIN 6h inf | HIT 6h inf | HITTINxHIT 6h inf | HITTINxHIT 6h inf |
| ENSG00000013810 | *TACC3* | 6.290964045 | -0.002851691 | -0.012062479 | 0.009210788 | 0.999933887 | -0.920121901 | -2.064248081 | 1.14412618 | 4.0e-08 |
| ENSG00000198728 | *LDB1* | 6.566089821 | -0.018328008 | -0.022618037 | 0.004290029 | 0.999933887 | -0.537086423 | -1.179098795 | 0.642012372 | 4.1e-08 |
| ENSG00000163950 | *SLBP* | 4.391521096 | 0.007184897 | -0.067021321 | 0.074206218 | 0.999933887 | -1.092490973 | -2.271706574 | 1.179215602 | 4.1e-08 |
| ENSG00000114770 | *ABCC5* | 4.598229697 | -0.04577187 | -0.007035407 | -0.038736463 | 0.999933887 | -0.38288356 | -1.107346016 | 0.724462456 | 1.55e-07 |
| ENSG00000154889 | *MPPE1* | 4.986227645 | -0.060510226 | -0.04892043 | -0.011589796 | 0.999933887 | -0.956374604 | -2.120342821 | 1.163968216 | 1.55e-07 |
| ENSG00000148411 | *NACC2* | 4.429865479 | 0.028911995 | -0.028333317 | 0.057245312 | 0.999933887 | -0.686865258 | -1.720275735 | 1.033410477 | 1.55e-07 |
| ENSG00000115525 | *ST3GAL5* | 3.212199503 | 0.024936133 | 0.056258708 | -0.031322575 | 0.999933887 | -0.909640132 | -2.028942914 | 1.119302783 | 1.55e-07 |
| ENSG00000114650 | *SCAP* | 5.894223641 | 0.00407177 | 0.000219602 | 0.003852167 | 0.999933887 | -0.693582353 | -1.491069581 | 0.797487227 | 2.04e-07 |
| ENSG00000186469 | *GNG2* | 7.636126174 | -0.086645275 | -0.039984134 | -0.046661142 | 0.999933887 | 1.689878598 | 2.786825127 | -1.096946529 | 2.83e-07 |
| ENSG00000136490 | *LIMD2* | 8.224782826 | 0.046063834 | 0.020552663 | 0.02551117 | 0.999933887 | -0.334981953 | -1.088005044 | 0.753023091 | 3.77e-07 |
| ENSG00000175115 | *PACS1* | 7.156963363 | 0.017294307 | 0.023147182 | -0.005852875 | 0.999933887 | -0.389540031 | -1.115206526 | 0.725666495 | 4.56e-07 |
| ENSG00000149177 | *PTPRJ* | 8.105117631 | 0.003543127 | -0.02488696 | 0.028430087 | 0.999933887 | 0.993095255 | 1.778970509 | -0.785875254 | 5.85e-07 |
| ENSG00000176454 | *LPCAT4* | 3.751378867 | -0.076903478 | 0.084847201 | -0.161750679 | 0.999933887 | -0.215394507 | -0.926666514 | 0.711272007 | 7.08e-07 |
| ENSG00000097033 | *SH3GLB1* | 7.561685911 | -0.037566006 | -0.061047052 | 0.023481046 | 0.999933887 | 0.055818994 | 0.609377741 | -0.553558747 | 7.28e-07 |
| ENSG00000142599 | *RERE* | 7.111115304 | -0.001125043 | 0.025483521 | -0.026608564 | 0.999933887 | -0.879698007 | -1.831043409 | 0.951345402 | 7.28e-07 |
| ENSG00000126882 | *FAM78A* | 4.354335938 | 0.035313925 | 0.020032206 | 0.015281719 | 0.999933887 | -0.360049992 | -1.140014822 | 0.77996483 | 8.67e-07 |
| ENSG00000077420 | *APBB1IP* | 7.011032353 | 0.068260876 | 0.038463138 | 0.029797738 | 0.999933887 | -0.644671435 | -1.358051197 | 0.713379763 | 1.247e-06 |
| ENSG00000148660 | *CAMK2G* | 6.00175391 | -0.019914597 | 0.019206591 | -0.039121189 | 0.999933887 | -0.485044854 | -1.247596411 | 0.762551557 | 1.247e-06 |
| ENSG00000130475 | *FCHO1* | 5.369262812 | 0.067658605 | 0.054474871 | 0.013183734 | 0.999933887 | -0.62047067 | -1.578634166 | 0.958163497 | 1.403e-06 |
| ENSG00000168906 | *MAT2A* | 5.788810601 | -0.12898742 | -0.053301496 | -0.075685924 | 0.999933887 | -0.404587622 | -1.0040006 | 0.599412978 | 1.405e-06 |
| ENSG00000132359 | *RAP1GAP2* | 6.965216034 | -0.055247579 | 0.034752219 | -0.089999798 | 0.999933887 | -0.5328804 | -1.334797713 | 0.801917313 | 2.145e-06 |
| ENSG00000198909 | *MAP3K3* | 6.925593001 | 0.006309852 | 0.032385554 | -0.026075702 | 0.999933887 | -0.923243447 | -1.751441154 | 0.828197708 | 2.245e-06 |
| ENSG00000065809 | *FAM107B* | 7.626215891 | -0.049455645 | -0.058092221 | 0.008636576 | 0.999933887 | 0.950935916 | 1.58013648 | -0.629200565 | 2.469e-06 |
| ENSG00000173020 | *GRK2* | 8.540296719 | 0.045699309 | 0.06557417 | -0.019874861 | 0.999933887 | -0.358906437 | -0.959921588 | 0.601015151 | 2.469e-06 |
| ENSG00000169220 | *RGS14* | 6.20992342 | 0.041852281 | 0.115571347 | -0.073719066 | 0.999933887 | -0.548504028 | -1.459109152 | 0.910605124 | 2.469e-06 |
| ENSG00000172375 | *C2CD2L* | 3.480366741 | 0.013879221 | 0.174006408 | -0.160127187 | 0.999933887 | -0.334832564 | -1.034191303 | 0.699358739 | 2.469e-06 |
| ENSG00000082212 | *ME2* | 4.529195555 | 0.022371314 | 0.023416201 | -0.001044887 | 0.999933887 | -0.846288138 | -1.593121487 | 0.746833348 | 3.069e-06 |
| ENSG00000106608 | *URGCP* | 3.230348518 | -0.04418902 | 0.036972839 | -0.081161858 | 0.999933887 | 1.405187847 | 2.295210356 | -0.890022509 | 3.854e-06 |
| ENSG00000130699 | *TAF4* | 3.786223363 | 0.015236879 | 0.008683517 | 0.006553362 | 0.999933887 | -0.455188327 | -1.220182054 | 0.764993727 | 3.854e-06 |
| ENSG00000168918 | *INPP5D* | 7.718757251 | 0.005114308 | 0.009818274 | -0.004703966 | 0.999933887 | -0.323458024 | -0.807702399 | 0.484244374 | 4.0e-06 |
| ENSG00000178104 | *PDE4DIP* | 4.264688646 | 0.032511927 | -0.143211394 | 0.175723321 | 0.999933887 | 1.015127346 | 1.794306972 | -0.779179626 | 4.628e-06 |
| ENSG00000198231 | *DDX42* | 5.363265192 | -0.034808736 | -0.074098825 | 0.039290089 | 0.999933887 | -0.321010591 | -0.881219363 | 0.560208773 | 4.65e-06 |
| ENSG00000146083 | *RNF44* | 7.444290972 | 0.008016684 | 0.030679566 | -0.022662882 | 0.999933887 | -0.499370932 | -1.077023227 | 0.577652295 | 4.65e-06 |
| ENSG00000167895 | *TMC8* | 6.777299354 | -0.001634845 | 0.034679133 | -0.036313978 | 0.999933887 | -0.472444916 | -1.143139929 | 0.670695014 | 5.27e-06 |
| ENSG00000133943 | *DGLUCY* | 6.132360015 | -0.001855521 | 0.005003189 | -0.006858711 | 0.999933887 | -0.218912398 | -0.742483871 | 0.523571473 | 5.325e-06 |
| ENSG00000123329 | *ARHGAP9* | 8.165973367 | -0.007638712 | -0.00276802 | -0.004870692 | 0.999933887 | -0.66738706 | -1.185433618 | 0.518046558 | 5.845e-06 |
| ENSG00000119844 | *AFTPH* | 7.209865128 | -0.046276039 | -0.115350946 | 0.069074907 | 0.999933887 | 0.439159178 | 0.924488316 | -0.485329138 | 6.046e-06 |
| ENSG00000088833 | *NSFL1C* | 5.661026915 | 0.002455419 | -0.011287075 | 0.013742495 | 0.999933887 | -0.458833826 | -0.987479167 | 0.528645342 | 6.357e-06 |
| ENSG00000075420 | *FNDC3B* | 8.237662676 | -0.0133526 | 0.004402784 | -0.017755384 | 0.999933887 | 1.197128388 | 1.898007669 | -0.700879281 | 6.798e-06 |
| ENSG00000169231 | *THBS3* | 3.535581999 | -0.001929168 | 0.006431848 | -0.008361016 | 0.999933887 | -0.389862815 | -1.030149936 | 0.640287121 | 7.538e-06 |
| ENSG00000172932 | *ANKRD13D* | 7.146472498 | 0.057239192 | 0.097639191 | -0.040399999 | 0.999933887 | -0.490647134 | -1.121310456 | 0.630663322 | 7.652e-06 |
| ENSG00000172794 | *RAB37* | 5.656575342 | 0.017405499 | -0.017140746 | 0.034546245 | 0.999933887 | -0.393957541 | -1.127790842 | 0.733833302 | 7.652e-06 |
| ENSG00000197142 | *ACSL5* | 6.375869596 | 0.011310521 | -0.055101964 | 0.066412485 | 0.999933887 | 1.221588824 | 1.841295156 | -0.619706332 | 1.005e-05 |
| ENSG00000136280 | *CCM2* | 6.106257077 | -0.03415799 | 0.054801455 | -0.088959445 | 0.999933887 | -0.777418987 | -1.50719577 | 0.729776783 | 1.0254e-05 |
| ENSG00000084070 | *SMAP2* | 8.502520365 | 0.021460197 | 0.014435331 | 0.007024866 | 0.999933887 | -0.571759356 | -1.248351251 | 0.676591896 | 1.045e-05 |
| ENSG00000061938 | *TNK2* | 6.416439929 | 0.03391351 | 0.075237887 | -0.041324376 | 0.999933887 | -0.456161307 | -1.179131658 | 0.722970352 | 1.057e-05 |
| ENSG00000072736 | *NFATC3* | 4.746903342 | 0.005816827 | 0.016761045 | -0.010944217 | 0.999933887 | -0.708766503 | -1.584622945 | 0.875856442 | 1.0571e-05 |
| ENSG00000143418 | *CERS2* | 5.242249155 | 0.076751762 | 0.088002973 | -0.011251211 | 0.999933887 | -0.503960036 | -1.108604218 | 0.604644181 | 1.0619e-05 |
| ENSG00000122515 | *ZMIZ2* | 6.094058829 | 0.056423391 | 0.023522079 | 0.032901312 | 0.999933887 | 0.987495903 | 1.606907823 | -0.619411921 | 1.0619e-05 |
| ENSG00000127663 | *KDM4B* | 6.902984478 | -0.036326011 | 0.021069235 | -0.057395246 | 0.999933887 | -0.3991895 | -1.045417531 | 0.646228031 | 1.0619e-05 |
| ENSG00000149289 | *ZC3H12C* | 4.31250693 | -0.02985739 | -0.04336922 | 0.01351183 | 0.999933887 | 1.120532112 | 1.944860009 | -0.824327897 | 1.0619e-05 |
| ENSG00000170776 | *AKAP13* | 7.855959673 | -0.033706668 | -0.039203124 | 0.005496456 | 0.999933887 | 0.404817359 | 1.014002547 | -0.609185187 | 1.1278e-05 |
| ENSG00000124496 | *TRERF1* | 5.660039947 | -0.002085982 | 0.036342486 | -0.038428468 | 0.999933887 | -0.2960214 | -0.862413658 | 0.566392257 | 1.1278e-05 |
| ENSG00000164938 | *TP53INP1* | 5.951823499 | -0.03912612 | -0.012460146 | -0.026665974 | 0.999933887 | -0.821019442 | -1.388679632 | 0.56766019 | 1.1755e-05 |
| ENSG00000108479 | *GALK1* | 2.938215054 | 0.035637383 | 0.010113401 | 0.025523982 | 0.999933887 | -0.840456606 | -1.821683638 | 0.981227032 | 1.1755e-05 |
| ENSG00000114446 | *IFT57* | 3.614690425 | -0.067203241 | -0.15600716 | 0.088803919 | 0.999933887 | 1.22506003 | 2.118661648 | -0.893601618 | 1.3402e-05 |
| ENSG00000197024 | *ZNF398* | 3.661984467 | -0.060360522 | -0.013476937 | -0.046883585 | 0.999933887 | -1.060979036 | -2.001919911 | 0.940940875 | 1.4634e-05 |
| ENSG00000124067 | *SLC12A4* | 4.159691512 | 0.051831849 | 0.03428606 | 0.017545789 | 0.999933887 | -0.736896536 | -1.559236391 | 0.822339855 | 1.4772e-05 |
| ENSG00000187109 | *NAP1L1* | 6.509610016 | -0.077393259 | -0.114708373 | 0.037315114 | 0.999933887 | 0.227178643 | 0.685234027 | -0.458055384 | 1.7612e-05 |
| ENSG00000129911 | *KLF16* | 4.350480746 | 0.113846363 | 0.167121941 | -0.053275578 | 0.999933887 | -1.068564571 | -2.102400418 | 1.033853847 | 1.8637e-05 |
| ENSG00000072274 | *TFRC* | 6.715511047 | -0.052872077 | -0.072672126 | 0.019800048 | 0.999933887 | 1.066976556 | 1.83596925 | -0.768992693 | 1.8637e-05 |
| ENSG00000167522 | *ANKRD11* | 7.652069794 | -0.008438045 | -0.025374392 | 0.016936347 | 0.999933887 | -0.403440064 | -0.784640018 | 0.381199955 | 1.8637e-05 |
| ENSG00000178764 | *ZHX2* | 6.062754902 | -0.022794807 | -0.014171798 | -0.008623009 | 0.999933887 | 1.048461858 | 1.636783233 | -0.588321375 | 1.9029e-05 |
| ENSG00000261221 | *ZNF865* | 4.581461258 | 0.088814707 | -0.017832338 | 0.106647045 | 0.999933887 | -0.670236674 | -1.443582102 | 0.773345428 | 1.934e-05 |
| ENSG00000170266 | *GLB1* | 3.989794072 | -0.010154649 | -0.029577025 | 0.019422376 | 0.999933887 | -0.574528202 | -1.170385833 | 0.595857631 | 1.934e-05 |
| ENSG00000185963 | *BICD2* | 5.246868351 | 0.003997328 | 0.074577185 | -0.070579856 | 0.999933887 | -0.857017 | -1.604623298 | 0.747606298 | 1.934e-05 |
| ENSG00000161980 | *POLR3K* | 1.896842638 | 0.069435916 | -0.155579532 | 0.225015448 | 0.999933887 | -1.102369579 | -2.147970051 | 1.045600472 | 2.0533e-05 |
| ENSG00000158470 | *B4GALT5* | 7.967916792 | 0.027272438 | 0.03112881 | -0.003856373 | 0.999933887 | 1.582174739 | 2.163203298 | -0.581028559 | 2.1043e-05 |
| ENSG00000175463 | *TBC1D10C* | 6.640421337 | 0.046697737 | 0.091331436 | -0.044633699 | 0.999933887 | -0.37231127 | -1.134078509 | 0.761767239 | 2.1522e-05 |
| ENSG00000164088 | *PPM1M* | 5.089046667 | -0.035798282 | -0.056602142 | 0.020803861 | 0.999933887 | -0.692823402 | -1.333613478 | 0.640790076 | 2.4268e-05 |
| ENSG00000133874 | *RNF122* | 3.512510895 | -0.069419139 | 0.005922269 | -0.075341408 | 0.999933887 | -0.651886704 | -1.364651695 | 0.712764991 | 2.5203e-05 |
| ENSG00000100605 | *ITPK1* | 7.041676962 | -0.014329097 | -0.000833266 | -0.013495831 | 0.999933887 | -0.625830863 | -1.479689141 | 0.853858278 | 2.5203e-05 |
| ENSG00000171522 | *PTGER4* | 9.112463595 | 0.09305502 | 0.028950047 | 0.064104972 | 0.999933887 | 0.985578739 | 1.515878348 | -0.530299609 | 2.5572e-05 |
| ENSG00000136161 | *RCBTB2* | 4.270111034 | -0.1322119 | -0.03888944 | -0.09332246 | 0.999933887 | -1.075706597 | -1.952215626 | 0.876509029 | 2.6108e-05 |
| ENSG00000107099 | *DOCK8* | 8.241036158 | -0.039974702 | -0.043777231 | 0.003802529 | 0.999933887 | -0.254823142 | -0.724429524 | 0.469606382 | 2.6108e-05 |
| ENSG00000148158 | *SNX30* | 4.459368284 | -0.05897294 | -0.128370587 | 0.069397646 | 0.999933887 | -0.632186835 | -1.321057384 | 0.688870549 | 2.6108e-05 |
| ENSG00000170265 | *ZNF282* | 3.061189479 | -0.024343724 | -0.032856269 | 0.008512545 | 0.999933887 | -0.268789324 | -1.028135314 | 0.759345989 | 2.6108e-05 |
| ENSG00000262370 | *AC108134.3* | 4.170000055 | -0.020055202 | -0.016020937 | -0.004034265 | 0.999933887 | -0.117373798 | -0.71117137 | 0.593797572 | 2.689e-05 |
| ENSG00000104312 | *RIPK2* | 5.982029547 | 0.059481317 | 0.029719938 | 0.029761379 | 0.999933887 | 1.48382956 | 2.091243852 | -0.607414292 | 2.7693e-05 |
| ENSG00000111554 | *MDM1* | 2.958056966 | -0.090142581 | -0.177234868 | 0.087092287 | 0.999933887 | 0.674620612 | 1.68400263 | -1.009382018 | 2.8417e-05 |
| ENSG00000178927 | *CYBC1* | 6.512412249 | 0.009366682 | -0.002301262 | 0.011667944 | 0.999933887 | -0.454298837 | -1.019453827 | 0.56515499 | 3.1216e-05 |
| ENSG00000185033 | *SEMA4B* | 4.65768804 | 0.008379295 | 0.028968947 | -0.020589652 | 0.999933887 | -0.976389375 | -1.809334944 | 0.832945568 | 3.1467e-05 |
| ENSG00000142208 | *AKT1* | 6.856448615 | 0.038309326 | 0.050906596 | -0.01259727 | 0.999933887 | -0.428805041 | -0.999983904 | 0.571178863 | 3.1468e-05 |
| ENSG00000182095 | *TNRC18* | 7.292012628 | 0.029069332 | 0.087921854 | -0.058852522 | 0.999933887 | -0.715395605 | -1.540606797 | 0.825211193 | 3.3674e-05 |
| ENSG00000010322 | *NISCH* | 5.941426818 | -0.007555315 | 0.032740067 | -0.040295382 | 0.999933887 | -0.287112182 | -0.699489625 | 0.412377443 | 3.4586e-05 |
| ENSG00000196535 | *MYO18A* | 5.31585968 | 0.052711226 | -0.000782399 | 0.053493624 | 0.999933887 | -0.211186292 | -0.726004074 | 0.514817782 | 3.739e-05 |
| ENSG00000105063 | *PPP6R1* | 6.873124656 | 0.02717147 | 0.028572972 | -0.001401502 | 0.999933887 | -0.359550968 | -0.845333588 | 0.48578262 | 3.739e-05 |
| ENSG00000197852 | *INKA2* | 5.611720601 | -0.059848755 | -0.039763265 | -0.02008549 | 0.999933887 | -1.519672723 | -2.384656789 | 0.864984065 | 4.0313e-05 |
| ENSG00000165671 | *NSD1* | 6.08950729 | -0.02836576 | -0.077824133 | 0.049458372 | 0.999933887 | -0.612874938 | -1.15168642 | 0.538811483 | 4.2202e-05 |
| ENSG00000183323 | *CCDC125* | 3.439679479 | -0.129614031 | -0.040640955 | -0.088973076 | 0.999933887 | -0.853773109 | -1.891981439 | 1.038208329 | 4.3566e-05 |
| ENSG00000161203 | *AP2M1* | 5.99528036 | 0.022793665 | -0.024275211 | 0.047068876 | 0.999933887 | -0.303901376 | -0.726249466 | 0.422348091 | 4.676e-05 |
| ENSG00000110911 | *SLC11A2* | 5.095201037 | 0.000682727 | -0.126628398 | 0.127311125 | 0.999933887 | 2.077431823 | 2.955633337 | -0.878201515 | 4.7381e-05 |
| ENSG00000168936 | *TMEM129* | 2.87590367 | 0.116278274 | 0.086295423 | 0.029982852 | 0.999933887 | -0.220205142 | -0.94808347 | 0.727878328 | 4.8639e-05 |
| ENSG00000137642 | *SORL1* | 10.159646271 | -0.052035552 | -0.038401783 | -0.013633769 | 0.999933887 | -0.289240127 | -0.748679516 | 0.459439388 | 5.021e-05 |
| ENSG00000128791 | *TWSG1* | 1.764435395 | -0.097533723 | -0.095952561 | -0.001581162 | 0.999933887 | 0.666664111 | 1.526086403 | -0.859422292 | 5.021e-05 |
| ENSG00000068024 | *HDAC4* | 6.01381678 | -0.009347047 | 0.021609004 | -0.030956051 | 0.999933887 | -0.334793181 | -0.841529845 | 0.506736664 | 5.054e-05 |
| ENSG00000103569 | *AQP9* | 11.438079296 | -0.062899547 | -0.099385257 | 0.03648571 | 0.999933887 | 1.430266544 | 1.994875776 | -0.564609232 | 5.1462e-05 |
| ENSG00000206344 | *HCG27* | 3.853383578 | -0.253277799 | -0.151009364 | -0.102268436 | 0.999933887 | -1.647544731 | -2.68769071 | 1.040145979 | 5.1573e-05 |
| ENSG00000168887 | *C2orf68* | 4.376881428 | -0.115748588 | -0.122008639 | 0.006260051 | 0.999933887 | -0.512220704 | -1.039203996 | 0.526983292 | 5.1573e-05 |
| ENSG00000127314 | *RAP1B* | 6.617679922 | -0.060923537 | -0.09583743 | 0.034913893 | 0.999933887 | 0.388562847 | 0.877727719 | -0.489164872 | 5.1573e-05 |
| ENSG00000141524 | *TMC6* | 6.747463663 | 0.071311779 | 0.09011674 | -0.01880496 | 0.999933887 | -0.310375703 | -0.933315519 | 0.622939816 | 5.1573e-05 |

| | | | | | | | | | | |
|---|---|---|---|---|---|---|---|---|---|---|
| ENSG00000116685 | *KIAA2013* | 5.196244863 | 0.072549181 | 0.014480546 | 0.058068635 | 0.999933887 | -0.601427321 | -1.324402741 | 0.722975421 | 5.1573e-05 |
| ENSG00000106144 | *CASP2* | 4.334164006 | -0.011239216 | -0.071730569 | 0.060491353 | 0.999933887 | -0.218176748 | -0.77462884 | 0.556452093 | 5.1573e-05 |
| ENSG00000031081 | *ARHGAP31* | 2.971786361 | 0.017044277 | 0.028884149 | -0.011839871 | 0.999933887 | 1.852292041 | 2.916988934 | -1.064696893 | 5.1573e-05 |
| ENSG00000091542 | *ALKBH5* | 5.248493509 | 0.058733965 | 0.016972187 | 0.041761778 | 0.999933887 | -0.318851633 | -0.883247477 | 0.564395844 | 5.2874e-05 |
| ENSG00000181192 | *DHTKD1* | 3.810553508 | -0.024623433 | -0.032545574 | 0.007922141 | 0.999933887 | -0.86826476 | -1.660973322 | 0.792708562 | 5.3299e-05 |
| ENSG00000122359 | *ANXA11* | 8.184550833 | -0.017245939 | 0.001235212 | -0.018481151 | 0.999933887 | -0.32552756 | -0.774590883 | 0.449063322 | 5.4974e-05 |
| ENSG00000180992 | *MRPL14* | 2.299509152 | 0.016393501 | 0.096547399 | -0.080153898 | 0.999933887 | 1.293195975 | 2.057455967 | -0.764259992 | 5.4974e-05 |
| ENSG00000070413 | *DGCR2* | 5.979323245 | 0.009887424 | -0.007767047 | 0.017654471 | 0.999933887 | -0.453665715 | -1.14662462 | 0.692958905 | 5.4974e-05 |
| ENSG00000275131 | *AC241952.1* | 2.358888181 | -0.104409698 | -0.013737424 | -0.090672273 | 0.999933887 | 0.435835947 | 1.033259323 | -0.597423376 | 5.6213e-05 |
| ENSG00000023892 | *DEF6* | 6.218974363 | 0.029219153 | 0.069277476 | -0.040058323 | 0.999933887 | -0.238078285 | -0.77677757 | 0.538699284 | 5.6213e-05 |
| ENSG00000279369 | *AC046185.3* | 2.630421058 | 0.02849853 | 0.062661143 | -0.034162613 | 0.999933887 | -0.449390993 | -1.335433753 | 0.886042759 | 5.6213e-05 |
| ENSG00000104915 | *STX10* | 5.332392688 | 0.04363254 | 0.026144329 | 0.017488211 | 0.999933887 | -0.562711525 | -1.398504423 | 0.835792898 | 5.6302e-05 |
| ENSG00000152804 | *HHEX* | 3.003777197 | -0.119276793 | 0.011368368 | -0.130645161 | 0.999933887 | -1.346938269 | -2.434663502 | 1.087725233 | 5.6527e-05 |
| ENSG00000108306 | *FBXL20* | 4.689349271 | -0.029796973 | 0.109142475 | -0.138939448 | 0.999933887 | -0.663461024 | -1.302255225 | 0.638794201 | 5.959e-05 |
| ENSG00000089159 | *PXN* | 7.629933757 | 0.010099607 | -0.024240581 | 0.034340189 | 0.999933887 | -0.823778252 | -1.376827166 | 0.553048914 | 6.0851e-05 |
| ENSG00000189339 | *SLC35E2B* | 5.428709483 | 0.026469614 | -0.055181045 | 0.081650659 | 0.999933887 | -0.221047606 | -0.593639688 | 0.372592082 | 6.1934e-05 |
| ENSG00000263528 | *IKBKE* | 3.867450284 | -0.013214348 | -0.075093642 | 0.061879295 | 0.999933887 | 1.181051084 | 2.188857082 | -1.007805997 | 6.3123e-05 |
| ENSG00000242247 | *ARFGAP3* | 6.822814623 | 0.005662773 | -0.090122351 | 0.095785125 | 0.999933887 | 0.886759053 | 1.357889561 | -0.471130508 | 6.3676e-05 |
| ENSG00000141401 | *IMPA2* | 4.690593141 | -0.007905947 | -0.021418954 | 0.013513007 | 0.999933887 | -1.158802175 | -2.345322103 | 1.186519927 | 6.3676e-05 |
| ENSG00000144597 | *EAF1* | 6.603504729 | -0.007865281 | -0.019469745 | 0.011604464 | 0.999933887 | 1.198690172 | 1.736121725 | -0.537431553 | 6.5127e-05 |
| ENSG00000112576 | *CCND3* | 6.348766398 | 0.089999466 | 0.072026484 | 0.017972981 | 0.999933887 | -0.450282406 | -1.068861119 | 0.618578713 | 6.6879e-05 |
| ENSG00000153029 | *MR1* | 5.037120279 | -0.008860535 | -0.018056652 | 0.009196117 | 0.999933887 | 0.435331069 | 0.913862978 | -0.478531909 | 6.6908e-05 |
| ENSG00000123610 | *TNFAIP6* | 7.671591934 | -0.006760387 | 0.057575992 | -0.064336379 | 0.999933887 | 3.099319565 | 4.070842057 | -0.971522492 | 6.6908e-05 |
| ENSG00000110697 | *PITPNM1* | 6.636056624 | 0.112352551 | 0.053770388 | 0.058582163 | 0.999933887 | -0.351277263 | -0.893326514 | 0.542049251 | 6.8159e-05 |
| ENSG00000166523 | *CLEC4E* | 7.983767091 | -0.01256623 | 0.027780944 | -0.040347174 | 0.999933887 | 1.993090714 | 2.5572695 | -0.564178786 | 6.8159e-05 |
| ENSG00000271601 | *LIX1L* | 3.074129974 | -0.01591519 | 0.009927599 | -0.025842789 | 0.999933887 | -0.188866361 | -0.823331048 | 0.634464686 | 7.1027e-05 |
| ENSG00000125730 | *C3* | 4.290919655 | 0.059465496 | -0.049403284 | 0.10886878 | 0.999933887 | 2.166974178 | 3.034212042 | -0.867237864 | 7.2381e-05 |
| ENSG00000204713 | *TRIM27* | 4.667633302 | 0.036192169 | -0.086008445 | 0.122200614 | 0.999933887 | -0.388922443 | -0.956906405 | 0.567983961 | 7.6819e-05 |
| ENSG00000187257 | *RSBN1L* | 4.572085456 | 0.011871371 | 0.023859848 | -0.011988476 | 0.999933887 | -0.822891909 | -1.548249568 | 0.725357658 | 7.9437e-05 |
| ENSG00000121578 | *B4GALT4* | 3.361789483 | -0.03500484 | -0.083963073 | 0.048958233 | 0.999933887 | 0.925886906 | 1.481551297 | -0.555664391 | 7.9813e-05 |
| ENSG00000123240 | *OPTN* | 4.964283456 | 0.019374357 | -0.155232538 | 0.174606895 | 0.999933887 | 0.548645245 | 1.086644648 | -0.537999403 | 8.1278e-05 |
| ENSG00000167552 | *TUBA1A* | 7.94142701 | -0.042793738 | -0.088039737 | 0.045245998 | 0.999933887 | -0.715743104 | -1.398062696 | 0.682319592 | 8.14e-05 |
| ENSG00000139613 | *SMARCC2* | 5.539934014 | 0.019094683 | 0.074318512 | -0.055223829 | 0.999933887 | -0.406135569 | -0.837696565 | 0.431560997 | 8.14e-05 |
| ENSG00000160285 | *LSS* | 4.285526782 | -0.018440629 | 0.090762359 | -0.109202988 | 0.999933887 | 2.109705668 | 3.229635513 | -1.119929846 | 8.14e-05 |
| ENSG00000104974 | *LILRA1* | 4.290693708 | 0.001177589 | -0.038137169 | 0.039314758 | 0.999933887 | -1.28278991 | -2.261615323 | 0.978825413 | 8.14e-05 |
| ENSG00000196782 | *MAML3* | 4.001320772 | -0.073558127 | 0.031949396 | -0.105507522 | 0.999933887 | -0.393930238 | -0.98721309 | 0.593282852 | 8.1771e-05 |
| ENSG00000113368 | *LMNB1* | 6.680646682 | -0.046376509 | -0.065779898 | 0.01940339 | 0.999933887 | 0.651980356 | 1.147768305 | -0.495787949 | 8.2386e-05 |
| ENSG00000110934 | *BIN2* | 7.406867867 | 0.003393237 | -0.010796694 | 0.014189931 | 0.999933887 | -0.184319121 | -0.681826507 | 0.497507386 | 8.2386e-05 |
| ENSG00000163050 | *COQ8A* | 5.021133219 | -0.018896949 | 0.026221704 | -0.045118654 | 0.999933887 | -0.586293826 | -1.238320704 | 0.652026878 | 8.3487e-05 |
| ENSG00000197555 | *SIPA1L1* | 6.957835753 | 0.003095838 | -0.049461715 | 0.052557552 | 0.999933887 | 0.338868748 | 0.820855588 | -0.48198684 | 8.3487e-05 |
| ENSG00000091640 | *SPAG7* | 4.472679426 | 0.051224162 | 0.041107685 | 0.010116477 | 0.999933887 | 0.730351348 | 1.203792348 | -0.473441 | 8.444e-05 |
| ENSG00000099991 | *CABIN1* | 5.67584602 | 0.020933027 | 0.049385252 | -0.028452225 | 0.999933887 | -0.098379209 | -0.617258916 | 0.518879707 | 8.5579e-05 |
| ENSG00000105939 | *ZC3HAV1* | 6.109419084 | 0.012615292 | -0.09398765 | 0.106602941 | 0.999933887 | -0.385110172 | -0.838310077 | 0.453199905 | 8.5579e-05 |
| ENSG00000066294 | *CD84* | 5.440471952 | -0.018768682 | -0.120810176 | 0.102041494 | 0.999933887 | 0.294825836 | 0.940508819 | -0.645682983 | 8.6003e-05 |
| ENSG00000240849 | *PEDS1* | 2.338072948 | -0.009268574 | 0.041826672 | -0.051095246 | 0.999933887 | 0.399813848 | 1.120367051 | -0.720553203 | 8.7406e-05 |
| ENSG00000103502 | *CDIPT* | 5.141461283 | 0.051635765 | -0.026973659 | 0.078609424 | 0.999933887 | -0.4831706 | -1.110995899 | 0.627825299 | 8.9675e-05 |
| ENSG00000124570 | *SERPINB6* | 3.394423387 | -0.068544206 | -0.071909973 | 0.003365768 | 0.999933887 | 0.272745849 | 1.061919284 | -0.789173434 | 8.9843e-05 |
| ENSG00000186298 | *PPP1CC* | 5.197340673 | -0.039660488 | 0.033208108 | -0.072868596 | 0.999933887 | -0.376431953 | -0.832881125 | 0.456449171 | 9.0564e-05 |
| ENSG00000100342 | *APOL1* | 5.066475479 | 0.020492725 | -0.06172811 | 0.082220835 | 0.999933887 | -0.881600206 | -1.511286315 | 0.629686109 | 9.1108e-05 |
| ENSG00000088832 | *FKBP1A* | 5.905197263 | -0.068163957 | -0.041480064 | -0.026683893 | 0.999933887 | -0.383000097 | -0.965413562 | 0.582413465 | 9.2691e-05 |
| ENSG00000124256 | *ZBP1* | 5.318297747 | 0.015451319 | 0.028634268 | -0.013182949 | 0.999933887 | -0.367032979 | -0.786119739 | 0.41908676 | 9.2691e-05 |
| ENSG00000080371 | *RAB21* | 8.036346702 | 0.011944323 | 0.037905317 | -0.025960994 | 0.999933887 | 1.516672385 | 2.010171464 | -0.49349908 | 9.516e-05 |
| ENSG00000214753 | *HNRNPUL2* | 3.586903115 | -0.075287413 | -0.055959908 | -0.019327505 | 0.999933887 | -0.457184203 | -1.098218715 | 0.641034512 | 0.000100004 |
| ENSG00000100299 | *ARSA* | 5.837957626 | 0.032980544 | 0.010816811 | 0.022163733 | 0.999933887 | -0.288934212 | -0.747442864 | 0.458508652 | 0.000100004 |
| ENSG00000170542 | *SERPINB9* | 6.820558712 | 0.003676961 | 0.113572317 | -0.109895357 | 0.999933887 | 2.565527197 | 3.518728957 | -0.95320176 | 0.000100004 |
| ENSG00000178719 | *GRINA* | 7.662975626 | 0.038646411 | 0.033642533 | 0.005003877 | 0.999933887 | 1.500660243 | 2.082451341 | -0.581791098 | 0.000101059 |
| ENSG00000125731 | *SH2D3A* | 3.661076595 | 0.020906176 | 0.150335957 | -0.129429781 | 0.999933887 | 1.626789012 | 2.519512345 | -0.892723333 | 0.000101059 |
| ENSG00000197912 | *SPG7* | 5.831660434 | -0.030046924 | 0.030674696 | -0.06072162 | 0.999933887 | -0.220672906 | -0.565327021 | 0.344654114 | 0.000102073 |
| ENSG00000180891 | *CUEDC1* | 3.634562755 | 0.009476964 | 0.065754293 | -0.056277329 | 0.999933887 | -0.392863375 | -1.136433405 | 0.74357003 | 0.000102073 |
| ENSG00000096996 | *IL12RB1* | 4.505343621 | 0.093402075 | 0.010827342 | 0.082574734 | 0.999933887 | -0.38277599 | -0.975666368 | 0.592890378 | 0.000104397 |
| ENSG00000111912 | *NCOA7* | 3.700613356 | 0.089662344 | -0.082227976 | 0.171890321 | 0.999933887 | -0.707243749 | -1.593417553 | 0.886173805 | 0.000104397 |
| ENSG00000100307 | *CBX7* | 5.363080207 | 0.057896487 | -0.004923848 | 0.062820336 | 0.999933887 | -0.638016176 | -1.264594038 | 0.626577862 | 0.000109289 |
| ENSG00000023445 | *BIRC3* | 6.421204193 | -0.088746623 | -0.144759827 | 0.056013204 | 0.999933887 | 0.779340429 | 1.798279315 | -1.018938886 | 0.000114085 |
| ENSG00000163082 | *SGPP2* | 3.402664208 | 0.072169955 | 0.134228532 | -0.062058577 | 0.999933887 | 2.267611196 | 3.136170412 | -0.868559217 | 0.000114085 |
| ENSG00000116213 | *WRAP73* | 3.532723107 | 0.131608409 | 0.107729019 | 0.02387939 | 0.999933887 | -0.323011888 | -0.824377784 | 0.501365896 | 0.000118976 |
| ENSG00000197183 | *NOL4L* | 5.112875044 | 0.071452455 | 0.056459665 | 0.01499279 | 0.999933887 | -0.393198451 | -0.836542711 | 0.44334426 | 0.000118976 |
| ENSG00000104880 | *ARHGEF18* | 5.135943592 | 0.064915441 | 0.033448324 | 0.031467117 | 0.999933887 | -0.26551078 | -0.786976691 | 0.521465911 | 0.000121859 |
| ENSG00000140443 | *IGF1R* | 6.948582491 | -0.060027299 | -0.01588316 | -0.044144139 | 0.999933887 | -0.780791648 | -1.275370769 | 0.494579121 | 0.000123754 |
| ENSG00000232934 | *AL157786.1* | 3.026417215 | 0.041150464 | 0.133862996 | -0.092712532 | 0.999933887 | 1.576550003 | 2.268449334 | -0.691899331 | 0.000123754 |
| ENSG00000010295 | *IFFO1* | 3.970754318 | -0.011498515 | 0.019905397 | -0.031403912 | 0.999933887 | -0.221712888 | -0.763315457 | 0.541602569 | 0.000136424 |
| ENSG00000132819 | *RBM38* | 5.991726698 | 0.062907836 | 0.021624543 | 0.041283293 | 0.999933887 | -0.344132886 | -0.824140168 | 0.480007282 | 0.000139671 |
| ENSG00000235750 | *KIAA0040* | 5.364402475 | 0.100591763 | 0.032912034 | 0.067679729 | 0.999933887 | -0.753789444 | -1.375603873 | 0.62181443 | 0.000142944 |
| ENSG00000121281 | *ADCY7* | 6.004016262 | 0.05978705 | 0.076236836 | -0.016449786 | 0.999933887 | -0.508883676 | -0.92914088 | 0.420257204 | 0.000142944 |
| ENSG00000074319 | *TSG101* | 6.097345071 | 0.022472082 | -0.039837091 | 0.062309174 | 0.999933887 | 0.205614885 | 0.567317354 | -0.361702469 | 0.000143164 |
| ENSG00000115896 | *PLCL1* | 3.646926842 | 0.013248703 | -0.053434519 | 0.066683222 | 0.999933887 | 0.151010362 | 0.671340931 | -0.520330569 | 0.000146958 |
| ENSG00000048828 | *FAM120A* | 6.34425053 | 0.011503481 | -0.022970873 | 0.034474354 | 0.999933887 | -0.239483254 | -0.599813164 | 0.36032991 | 0.000149532 |
| ENSG00000198863 | *RUNDC1* | 3.648199713 | -0.062143697 | 0.016964836 | -0.079108532 | 0.999933887 | -0.638412895 | -1.267145479 | 0.628732584 | 0.000152314 |
| ENSG00000135439 | *AGAP2* | 5.161490322 | 0.060030877 | 0.056880968 | 0.00314991 | 0.999933887 | -0.20874256 | -0.750502793 | 0.541760233 | 0.000153201 |
| ENSG00000172725 | *CORO1B* | 4.762069943 | 0.08879845 | 0.058170796 | 0.030627654 | 0.999933887 | -0.747969281 | -1.433017059 | 0.685047778 | 0.000156565 |
| ENSG00000197965 | *MPZL1* | 4.888727371 | 0.011504512 | -0.076379593 | 0.087884104 | 0.999933887 | -0.933464548 | -1.992756526 | 1.059291978 | 0.000158508 |
| ENSG00000180354 | *MTURN* | 4.277437722 | -0.080573611 | -0.039359851 | -0.04121376 | 0.999933887 | -0.43242999 | -0.926530197 | 0.494100207 | 0.000172528 |
| ENSG00000165169 | *DYNLT3* | 4.023396564 | -0.114430981 | -0.06350791 | -0.050923071 | 0.999933887 | 0.901744338 | 1.602722374 | -0.700978036 | 0.000175985 |
| ENSG00000010818 | *HIVEP2* | 7.533875946 | 0.024267247 | -0.002534529 | 0.026801777 | 0.999933887 | 1.346339833 | 1.994733087 | -0.648393254 | 0.000180899 |
| ENSG00000167613 | *LAIR1* | 3.499649536 | -0.012112946 | 0.128921423 | -0.141034369 | 0.999933887 | 1.149612424 | 2.109860592 | -0.960248168 | 0.000184766 |
| ENSG00000111276 | *CDKN1B* | 6.332204871 | -0.045795577 | -0.200279819 | 0.154484241 | 0.999933887 | -0.773917318 | -1.415389256 | 0.641471939 | 0.000185381 |
| ENSG00000141905 | *NFIC* | 4.712526922 | 0.025646017 | -0.040679834 | 0.066325851 | 0.999933887 | -0.80936435 | -1.438005626 | 0.628641276 | 0.000186201 |
| ENSG00000103490 | *PYCARD* | 4.3769037 | 0.033902641 | -0.005391161 | 0.039293802 | 0.999933887 | -0.758729503 | -1.643990439 | 0.885260936 | 0.000186782 |
| ENSG00000100068 | *LRP5L* | 3.100461582 | 0.006849981 | -0.048149771 | 0.054999753 | 0.999933887 | 0.677048315 | 1.324148887 | -0.647100572 | 0.000186782 |
| ENSG00000124587 | *PEX6* | 3.578012649 | 0.003161306 | 0.086497605 | -0.0833363 | 0.999933887 | -0.248720005 | -0.712705445 | 0.46398544 | 0.000186782 |
| ENSG00000186687 | *LYRM7* | 2.29513875 | 0.02550131 | -0.067437314 | 0.092938624 | 0.999933887 | -0.079707415 | 0.687291178 | -0.766998593 | 0.000191707 |
| ENSG00000124702 | *KLHDC3* | 3.790460894 | 0.010600706 | -0.060145579 | 0.070746285 | 0.999933887 | -0.239328912 | -0.741042468 | 0.501713556 | 0.000192577 |
| ENSG00000143507 | *DUSP10* | 4.982359534 | -0.031765178 | -0.005886149 | -0.025879029 | 0.999933887 | 0.910562651 | 1.459647185 | -0.549084535 | 0.000197108 |
| ENSG00000119950 | *MXI1* | 4.822318728 | 0.09962565 | 0.020452526 | 0.079173124 | 0.999933887 | -1.05002184 | -1.832587214 | 0.782565374 | 0.00019954 |
| ENSG00000127666 | *TICAM1* | 6.008258797 | 0.078513534 | 0.042936764 | 0.03557677 | 0.999933887 | 2.014155525 | 2.632645442 | -0.618489917 | 0.00019954 |
| ENSG00000133805 | *AMPD3* | 6.947313633 | -0.083981354 | -0.009213496 | -0.074767858 | 0.999933887 | 1.872941879 | 2.502154351 | -0.629212472 | 0.000201389 |
| ENSG00000142197 | *DOP1B* | 4.418206273 | -0.035808556 | 0.048208942 | -0.084017497 | 0.999933887 | -0.719442141 | -1.264569401 | 0.545127261 | 0.000201389 |
| ENSG00000130340 | *SNX9* | 4.1904169 | 0.032847287 | -0.078637676 | 0.111484963 | 0.999933887 | 0.781112874 | 1.466774496 | -0.685661622 | 0.000201389 |
| ENSG00000076242 | *MLH1* | 3.731248342 | -0.058820782 | 0.079538146 | -0.138358928 | 0.999933887 | 1.097328401 | 1.790026518 | -0.692698116 | 0.000205097 |
| ENSG00000197779 | *ZNF81* | 2.839199618 | -0.063000261 | -0.058524756 | -0.004475504 | 0.999933887 | 0.105650043 | 0.810010081 | -0.704360038 | 0.000211925 |
| ENSG00000144401 | *METTL21A* | 3.543826341 | 0.042827908 | 0.117403515 | -0.074575607 | 0.999933887 | 0.59696454 | 1.118630297 | -0.521665757 | 0.000211925 |
| ENSG00000090020 | *SLC9A1* | 6.281828726 | 0.009281339 | 0.026961341 | -0.017680002 | 0.999933887 | -0.18598371 | -0.599601541 | 0.413617831 | 0.000211925 |
| ENSG00000263264 | *AC119396.1* | 3.21210731 | 0.077452603 | -0.002382725 | 0.079835327 | 0.999933887 | -0.137448377 | -0.788321593 | 0.650873216 | 0.000213046 |
| ENSG00000254087 | *LYN* | 10.502904072 | -0.011637782 | -0.039977884 | 0.028340103 | 0.999933887 | 0.852734655 | 1.302740101 | -0.450005446 | 0.000213046 |
| ENSG00000171443 | *ZNF524* | 2.22337691 | 0.152550734 | 0.016627653 | 0.135923081 | 0.999933887 | -0.376794998 | -1.315647808 | 0.93885281 | 0.000213281 |
| ENSG00000122733 | *PHF24* | 1.577472214 | -0.06604745 | 0.011915885 | -0.077963335 | 0.999933887 | 1.334187996 | 2.14514802 | -0.810960024 | 0.000214438 |

| | | | | | | | | | | |
|---|---|---|---|---|---|---|---|---|---|---|
| ENSG00000238227 | *TMEM250* | 3.747208982 | 0.065702932 | -0.023351946 | 0.089054878 | 0.999933887 | -0.504419353 | -1.249701776 | 0.745282423 | 0.000215256 |
| ENSG00000166477 | *LEO1* | 2.066744886 | -0.071753102 | -0.064423821 | -0.007329281 | 0.999933887 | 0.566903971 | 1.284676792 | -0.717772821 | 0.000223747 |
| ENSG00000262246 | *CORO7* | 5.21504005 | 0.015307925 | 0.038076806 | -0.02276888 | 0.999933887 | -0.173068285 | -0.497947414 | 0.324879129 | 0.000223747 |
| ENSG00000164032 | *H2AZ1* | 5.766563792 | 0.002520891 | -0.07981449 | 0.082335381 | 0.999933887 | -0.20648341 | -0.67029474 | 0.46381133 | 0.000223747 |
| ENSG00000204839 | *MROH6* | 3.579884365 | 0.116811623 | 0.060763397 | 0.056048226 | 0.999933887 | -0.78492525 | -1.56131998 | 0.77639473 | 0.000225643 |
| ENSG00000166398 | *GARRE1* | 3.657842062 | 0.027310631 | 0.11871207 | -0.091401439 | 0.999933887 | -0.139213279 | -0.593874348 | 0.454661069 | 0.000225643 |
| ENSG00000196209 | *SIRPB2* | 4.433157527 | -0.03982362 | -0.001358977 | -0.038464643 | 0.999933887 | -0.722619928 | -1.461170698 | 0.73855077 | 0.000227422 |
| ENSG00000100060 | *MFNG* | 6.055318048 | -0.020099496 | -0.028167576 | 0.008068079 | 0.999933887 | -0.902707106 | -1.534194412 | 0.631487306 | 0.000227699 |
| ENSG00000165474 | *GJB2* | 4.018471658 | 0.184063499 | 0.070394379 | 0.11366912 | 0.999933887 | 1.678213457 | 2.502046137 | -0.82383268 | 0.000228996 |
| ENSG00000175137 | *SH3BP5L* | 4.99787884 | -0.065102059 | -0.010215797 | -0.054886262 | 0.999933887 | -0.847646549 | -1.458054322 | 0.610407773 | 0.000228996 |
| ENSG00000123416 | *TUBA1B* | 3.855245703 | 0.054544998 | 0.022145335 | 0.032399663 | 0.999933887 | -0.354632706 | -1.025497907 | 0.670865201 | 0.000228996 |
| ENSG00000114353 | *GNAI2* | 9.158639486 | 0.04023449 | 0.01855277 | 0.02168172 | 0.999933887 | -0.476032843 | -1.002195513 | 0.52616267 | 0.000228996 |
| ENSG00000108406 | *DHX40* | 5.550744809 | -0.024207967 | -0.090557877 | 0.06634991 | 0.999933887 | 0.36465064 | 0.764964832 | -0.400314192 | 0.000228996 |
| ENSG00000122986 | *HVCN1* | 3.250315266 | -0.00690454 | 0.004096068 | -0.011000608 | 0.999933887 | -0.731850894 | -1.461740116 | 0.729889222 | 0.000230904 |
| ENSG00000124659 | *TBCC* | 4.457067302 | -0.058493622 | -0.175751266 | 0.117257644 | 0.999933887 | -0.427415617 | -0.890889827 | 0.46347421 | 0.000233644 |
| ENSG00000021355 | *SERPINB1* | 7.089210047 | 0.051527359 | -0.044536822 | 0.096064181 | 0.999933887 | 1.493446749 | 2.117840147 | -0.624393398 | 0.000239943 |
| ENSG00000048740 | *CELF2* | 7.782255196 | 0.055473523 | -0.027244395 | 0.082717919 | 0.999933887 | -0.726154195 | -1.298467065 | 0.572312869 | 0.000241325 |
| ENSG00000135842 | *NIBAN1* | 9.195632183 | 0.019526462 | -0.028262603 | 0.047789065 | 0.999933887 | 1.126985936 | 1.717986609 | -0.591000673 | 0.000241325 |
| ENSG00000275302 | *CCL4* | 9.130925942 | 0.379507849 | 0.38471106 | -0.005203211 | 0.999933887 | 3.717792632 | 4.862261055 | -1.144468424 | 0.000246643 |
| ENSG00000119335 | *SET* | 6.321347653 | -0.066031836 | -0.04504822 | -0.020983616 | 0.999933887 | -0.356648609 | -0.826923858 | 0.470275249 | 0.000246643 |
| ENSG00000164620 | *RELL2* | 2.201792452 | -0.095371238 | 0.099071222 | -0.19444246 | 0.999933887 | -0.768903198 | -1.521731526 | 0.752828328 | 0.000246643 |
| ENSG00000146828 | *SLC12A9* | 6.695554705 | 0.03973703 | 0.074704003 | -0.034966973 | 0.999933887 | -0.120766451 | -0.687442116 | 0.566675665 | 0.000246643 |
| ENSG00000138964 | *PARVG* | 5.878723171 | 0.018340927 | 0.041109021 | -0.022768095 | 0.999933887 | -0.183683898 | -0.561071913 | 0.377388016 | 0.000246643 |
| ENSG00000145779 | *TNFAIP8* | 5.4141109 | -0.032590625 | -0.008176978 | -0.024413647 | 0.999933887 | 1.0323358 | 1.664727325 | -0.632391525 | 0.000246901 |
| ENSG00000198740 | *ZNF652* | 4.731758649 | 0.001417028 | -0.037529531 | 0.038946559 | 0.999933887 | -1.01688726 | -1.774359002 | 0.757471741 | 0.000250265 |
| ENSG00000115956 | *PLEK* | 12.023675126 | 0.075079335 | 0.037220306 | 0.037859029 | 0.999933887 | 1.597892467 | 2.061758473 | -0.463866005 | 0.000252492 |
| ENSG00000196684 | *HSH2D* | 6.88649446 | -0.042401097 | -0.001750812 | -0.040650285 | 0.999933887 | -0.437522359 | -0.900319364 | 0.462797006 | 0.000252492 |
| ENSG00000180198 | *RCC1* | 2.912076758 | 0.040071405 | 0.173637518 | -0.133566114 | 0.999933887 | 0.577798494 | 1.307192435 | -0.729393941 | 0.000252492 |
| ENSG00000205189 | *ZBTB10* | 3.938061568 | 0.024748339 | -0.012201382 | 0.036949721 | 0.999933887 | 1.089135579 | 1.985584247 | -0.896448668 | 0.000252492 |
| ENSG00000100647 | *SUSD6* | 9.45735636 | 0.010436892 | 0.036888069 | -0.026451177 | 0.999933887 | 0.91140039 | 1.309033844 | -0.397633454 | 0.000252492 |
| ENSG00000136026 | *CKAP4* | 7.607937659 | 0.004202874 | 0.014878684 | -0.010675811 | 0.999933887 | 1.270295966 | 1.965786258 | -0.695490292 | 0.000252492 |
| ENSG00000198911 | *SREBF2* | 5.854413947 | 0.02169526 | -0.037149774 | 0.058845034 | 0.999933887 | 0.565988174 | 1.027213129 | -0.461224955 | 0.000257374 |
| ENSG00000169925 | *BRD3* | 4.164455573 | 0.010431938 | 0.020369462 | -0.009937525 | 0.999933887 | -0.651337506 | -1.327132039 | 0.675794534 | 0.000257374 |
| ENSG00000164251 | *F2RL1* | 2.759032672 | -0.037616033 | 0.117203937 | -0.15481997 | 0.999933887 | -2.413552025 | -4.098744666 | 1.685192641 | 0.000257464 |
| ENSG00000159314 | *ARHGAP27* | 7.668155691 | 0.047550806 | 0.088952478 | -0.041401672 | 0.999933887 | -0.595777113 | -1.089746916 | 0.493969803 | 0.000262944 |
| ENSG00000109654 | *TRIM2* | 0.00120883 | -0.078697902 | 0.451130405 | -0.529828307 | 0.999933887 | 0.532934931 | 1.580905084 | -1.047970153 | 0.000263444 |
| ENSG00000130560 | *UBAC1* | 3.695429311 | 0.032003507 | 0.043271158 | -0.011267651 | 0.999933887 | -0.216301756 | -0.784712087 | 0.568410332 | 0.000263767 |
| ENSG00000163110 | *PDLIM5* | 4.578542113 | 0.03823955 | 0.112531303 | -0.074291753 | 0.999933887 | 0.756530142 | 1.166392234 | -0.409862092 | 0.000272337 |
| ENSG00000102871 | *TRADD* | 4.735924686 | 0.071324524 | 0.013234666 | 0.058089857 | 0.999933887 | -0.290282686 | -0.894060355 | 0.603777668 | 0.000276518 |
| ENSG00000110844 | *PRPF40B* | -0.122007712 | -0.108806516 | 0.201079919 | -0.309886435 | 0.999933887 | 1.484003775 | 3.008363348 | -1.524359573 | 0.000290446 |
| ENSG00000165168 | *CYBB* | 6.738982697 | -0.070108547 | -0.031186093 | -0.038922455 | 0.999933887 | 0.602982377 | 1.080449674 | -0.477467297 | 0.00029364 |
| ENSG00000112339 | *HBS1L* | 4.4047058 | -0.061257071 | -0.064107555 | 0.002850484 | 0.999933887 | 0.574541671 | 1.0764278 | -0.50188613 | 0.00029364 |
| ENSG00000145569 | *OTULINL* | 3.706960469 | -0.094460687 | 0.005058795 | -0.099519482 | 0.999933887 | 0.263474523 | 0.833424946 | -0.569950423 | 0.000295879 |
| ENSG00000157514 | *TSC22D3* | 9.134040049 | -0.011916777 | -0.104292187 | 0.09237541 | 0.999933887 | -0.77628887 | -1.228715918 | 0.452427048 | 0.000296261 |
| ENSG00000168246 | *UBTD2* | 4.18125607 | -0.075763025 | -0.228083976 | 0.152320952 | 0.999933887 | 1.503379877 | 2.098187896 | -0.594808019 | 0.000297103 |
| ENSG00000118046 | *STK11* | 5.313184241 | 0.064348163 | 0.112190306 | -0.047842143 | 0.999933887 | -0.308829625 | -0.871492187 | 0.562662562 | 0.000310927 |
| ENSG00000178623 | *GPR35* | 2.1171776 | -0.008914785 | -0.167741273 | 0.158826488 | 0.999933887 | 0.347240485 | 1.175361781 | -0.828121297 | 0.00031148 |
| ENSG00000127481 | *UBR4* | 9.070915414 | 0.018468391 | -0.00327772 | 0.021746111 | 0.999933887 | 0.825319599 | 1.29239217 | -0.467072571 | 0.0003116 |
| ENSG00000226479 | *TMEM185B* | 6.284323945 | 0.045320704 | 0.028668138 | 0.016652566 | 0.999933887 | 1.311359819 | 1.763462082 | -0.452102263 | 0.000320855 |
| ENSG00000275183 | *LENG9* | -0.003696788 | 0.407066343 | 0.36159922 | 0.045467124 | 0.999933887 | 0.923062846 | 2.367513803 | -1.444450957 | 0.000322891 |
| ENSG00000155307 | *SAMSN1* | 7.361672702 | -0.035232391 | -0.093150808 | 0.057918417 | 0.999933887 | 1.375488844 | 2.23342943 | -0.857940586 | 0.000328361 |
| ENSG00000120913 | *PDLIM2* | 5.763933627 | 0.046706991 | 0.058095803 | -0.011388813 | 0.999933887 | -0.592193905 | -1.071276454 | 0.479082548 | 0.000328961 |
| ENSG00000177963 | *RIC8A* | 6.06040413 | 0.049903089 | 0.051216703 | -0.001313614 | 0.999933887 | -0.27344288 | -0.673720428 | 0.400277548 | 0.000336036 |
| ENSG00000127311 | *HELB* | 3.07806817 | -0.074746483 | -0.080984999 | 0.006238516 | 0.999933887 | 0.920101277 | 1.56364098 | -0.643539703 | 0.000340246 |
| ENSG00000108651 | *UTP6* | 3.753831768 | -0.101602368 | -0.011021808 | -0.090580559 | 0.999933887 | -0.338873643 | -0.905998575 | 0.567124931 | 0.000342831 |
| ENSG00000064687 | *ABCA7* | 7.140526705 | 0.053165691 | 0.070036064 | -0.016870373 | 0.999933887 | -0.21182583 | -0.662920736 | 0.451094905 | 0.000342831 |
| ENSG00000162894 | *FCMR* | 5.566905152 | 0.018916731 | -0.040110136 | 0.059026867 | 0.999933887 | -0.379283865 | -0.980961236 | 0.601677372 | 0.000342831 |
| ENSG00000063854 | *HAGH* | 3.37349388 | -0.015526495 | 0.121366734 | -0.136893229 | 0.999933887 | -0.235978833 | -0.815820986 | 0.579842153 | 0.000342831 |
| ENSG00000096968 | *JAK2* | 4.829666692 | -0.065669532 | -0.074804238 | 0.009134706 | 0.999933887 | 0.140770073 | 0.872101852 | -0.731331779 | 0.00034346 |
| ENSG00000171206 | *TRIM8* | 6.149454054 | 0.051304978 | 0.039997705 | 0.011307272 | 0.999933887 | -0.375453055 | -0.832709426 | 0.457256371 | 0.000347638 |
| ENSG00000185010 | *F8* | 2.899725049 | -0.049423081 | 0.015389995 | -0.064813076 | 0.999933887 | 1.183654891 | 1.908150233 | -0.724495342 | 0.000351263 |
| ENSG00000163545 | *NUAK2* | 4.734755249 | -0.116094494 | -0.094025629 | -0.022068865 | 0.999933887 | -1.312199132 | -2.114633122 | 0.80243399 | 0.000363508 |
| ENSG00000247596 | *TWF2* | 4.720285522 | -0.00883377 | 0.057236771 | -0.06607054 | 0.999933887 | -0.942269909 | -1.618419363 | 0.676149455 | 0.000363508 |
| ENSG00000163513 | *TGFBR2* | 6.746729634 | -0.037999402 | -0.006431059 | -0.031568343 | 0.999933887 | -0.320455523 | -0.787632824 | 0.467177301 | 0.000365152 |
| ENSG00000078804 | *TP53INP2* | 7.470845163 | 0.101793777 | 0.016483737 | 0.08531004 | 0.999933887 | 1.015564483 | 1.625605549 | -0.610041065 | 0.000380181 |
| ENSG00000130723 | *PRRC2B* | 6.421379404 | 0.039470274 | 0.024987753 | 0.01448252 | 0.999933887 | -0.153940435 | -0.480676553 | 0.326736118 | 0.000380941 |
| ENSG00000104518 | *GSDMD* | 5.285067435 | 0.0620982 | 0.110069298 | -0.047971099 | 0.999933887 | -0.260582224 | -0.955633255 | 0.695051032 | 0.000380941 |
| ENSG00000124466 | *LYPD3* | 0.952918453 | 0.142397702 | 0.154981695 | -0.012583993 | 0.999933887 | 1.587694586 | 2.685839716 | -1.09814513 | 0.000382875 |
| ENSG00000177666 | *PNPLA2* | 6.112612197 | 0.023720823 | 0.030934588 | -0.007213765 | 0.999933887 | -0.357530952 | -0.767364966 | 0.409834014 | 0.000382875 |
| ENSG00000188559 | *RALGAPA2* | 6.850085421 | 0.017017115 | 0.034169465 | -0.01715235 | 0.999933887 | 0.382574072 | 0.771778747 | -0.389204675 | 0.000382875 |
| ENSG00000163393 | *SLC22A15* | 3.934525979 | -0.019241434 | -0.028810187 | 0.009568752 | 0.999933887 | -0.92340487 | -1.49638136 | 0.57297649 | 0.000382875 |
| ENSG00000007376 | *RPUSD1* | 2.748897741 | 0.144232633 | 0.072775968 | 0.071456665 | 0.999933887 | -0.219763514 | -0.866349871 | 0.646586357 | 0.000383359 |
| ENSG00000198771 | *RCSD1* | 5.690393357 | -0.037845445 | 0.053356796 | -0.091202241 | 0.999933887 | -0.379265409 | -0.884299882 | 0.505034473 | 0.000383359 |
| ENSG00000166527 | *CLEC4D* | 4.227749523 | 0.030030717 | -0.128974488 | 0.159005205 | 0.999933887 | 1.594116308 | 2.412598667 | -0.818482359 | 0.000383359 |
| ENSG00000196954 | *CASP4* | 7.761161173 | 0.004543991 | -0.084567162 | 0.089111153 | 0.999933887 | 0.53897292 | 0.959991855 | -0.421018934 | 0.000383359 |
| ENSG00000171720 | *HDAC3* | 3.999238741 | 0.053194648 | -0.021468723 | 0.074663371 | 0.999933887 | -0.404073892 | -0.810007824 | 0.405933932 | 0.000384019 |
| ENSG00000186407 | *CD300E* | 2.944192076 | -0.069054116 | -0.051933632 | -0.017120484 | 0.999933887 | -0.659544443 | -1.434269886 | 0.774725444 | 0.000386294 |
| ENSG00000115594 | *IL1R1* | 7.790224633 | -0.023434844 | -0.015918881 | -0.007515963 | 0.999933887 | 0.379607241 | 0.77954311 | -0.399935869 | 0.000389323 |
| ENSG00000152518 | *ZFP36L2* | 7.932307048 | -0.022566771 | 0.005889456 | -0.028456227 | 0.999933887 | -0.439024626 | -0.861787134 | 0.422762508 | 0.000389323 |
| ENSG00000128245 | *YWHAH* | 3.545677759 | -0.010536197 | 0.077245447 | -0.087781645 | 0.999933887 | -0.352069516 | -0.856682704 | 0.504613188 | 0.000398451 |
| ENSG00000104522 | *GFUS* | 3.637358339 | 0.082537477 | -0.0002963 | 0.082833777 | 0.999933887 | -0.127333287 | -0.660884069 | 0.533550781 | 0.000419257 |
| ENSG00000116815 | *CD58* | 6.27287418 | -0.034811517 | -0.045229629 | 0.010418111 | 0.999933887 | 1.036876948 | 1.528359243 | -0.491482295 | 0.000424593 |
| ENSG00000169032 | *MAP2K1* | 4.963971073 | 0.022843882 | 0.001441474 | 0.021402408 | 0.999933887 | 0.535697411 | 0.957248545 | -0.421551134 | 0.000429549 |
| ENSG00000027075 | *PRKCH* | 5.93754361 | 0.002291963 | 0.002888777 | -0.000596814 | 0.999933887 | 0.748574278 | 1.351965659 | -0.603391382 | 0.00043842 |
| ENSG00000128815 | *WDFY4* | 5.418324845 | 0.004615865 | 0.024106198 | -0.019490333 | 0.999933887 | -0.273189742 | -0.696552277 | 0.423362535 | 0.000441885 |
| ENSG00000100030 | *MAPK1* | 6.920482394 | -0.046051535 | -0.028984524 | -0.017067011 | 0.999933887 | -0.656273187 | -1.077412142 | 0.421138955 | 0.000442001 |
| ENSG00000101350 | *KIF3B* | 4.550306827 | -0.047295356 | -0.032874933 | -0.014420423 | 0.999933887 | 0.516258302 | 1.032159192 | -0.515900889 | 0.000442001 |
| ENSG00000127948 | *POR* | 6.064997002 | 0.079536915 | 0.010435683 | 0.069101232 | 0.999933887 | 0.789430653 | 1.261225254 | -0.471794602 | 0.000443174 |
| ENSG00000102172 | *SMS* | 4.029263704 | 0.052803808 | 0.028452808 | 0.024351 | 0.999933887 | 0.304335439 | 0.810299267 | -0.505963828 | 0.000443174 |
| ENSG00000180233 | *ZNRF2* | 4.063599814 | 0.171745845 | -0.028537575 | 0.200283419 | 0.999933887 | -0.450071691 | -1.074345674 | 0.624273983 | 0.000449751 |
| ENSG00000128487 | *SPECC1* | 2.78044192 | 0.002909515 | -0.003160547 | 0.006070062 | 0.999933887 | 0.23153459 | 0.95709718 | -0.72556259 | 0.000449751 |
| ENSG00000173221 | *GLRX* | 5.513715977 | -0.104411118 | -0.168002616 | 0.063591498 | 0.999933887 | -0.43955077 | -0.921778461 | 0.482227691 | 0.000451956 |
| ENSG00000105323 | *HNRNPUL1* | 7.201004861 | 0.022898354 | -0.015782767 | 0.038681121 | 0.999933887 | -0.503150111 | -0.873394095 | 0.370243984 | 0.000452872 |
| ENSG00000068650 | *ATP11A* | 6.540336762 | 0.00547934 | 0.011911333 | -0.006431993 | 0.999933887 | -0.28324549 | -0.648754681 | 0.365509191 | 0.000453076 |
| ENSG00000235385 | *LINC02154* | 2.068111595 | -0.127646427 | -0.173474972 | 0.045828545 | 0.999933887 | 1.699931992 | 2.378956931 | -0.679024938 | 0.000453222 |
| ENSG00000166797 | *CIAO2A* | 3.629525476 | 0.038839121 | -0.068898878 | 0.107737999 | 0.999933887 | -0.939918514 | -1.69863732 | 0.758718807 | 0.000453571 |
| ENSG00000140931 | *CMTM3* | 5.116688532 | 0.013726457 | -0.002136896 | 0.015863354 | 0.999933887 | -0.504204894 | -1.038553627 | 0.534348732 | 0.000453593 |
| ENSG00000162616 | *DNAJB4* | 2.139147296 | -0.222427696 | -0.08489688 | -0.137530816 | 0.999933887 | 1.3540377 | 2.165484192 | -0.811446492 | 0.000455605 |
| ENSG00000105771 | *SMG9* | 4.473063719 | 0.081676359 | 0.083399173 | -0.001722813 | 0.999933887 | 2.235076587 | 2.966552158 | -0.73147557 | 0.000455605 |
| ENSG00000125384 | *PTGER2* | 5.015057251 | 0.061045575 | 0.035178339 | 0.025867236 | 0.999933887 | 1.680499444 | 2.564656976 | -0.884157532 | 0.000455605 |
| ENSG00000156675 | *RAB11FIP1* | 7.228766406 | -0.03905698 | -0.023485442 | -0.015571538 | 0.999933887 | -1.426057154 | -2.078969782 | 0.652912628 | 0.000456712 |
| ENSG00000129245 | *FXR2* | 3.982898192 | 0.066477037 | 0.16269059 | -0.096213553 | 0.999933887 | -0.349901564 | -0.79284361 | 0.442942046 | 0.000456921 |
| ENSG00000110400 | *NECTIN1* | 2.975497652 | -0.098099612 | -0.049537529 | -0.048562083 | 0.999933887 | 1.479229522 | 2.286187203 | -0.806957681 | 0.00046129 |
| ENSG00000059378 | *PARP12* | 5.767711384 | 0.016420195 | 0.025727682 | -0.009307487 | 0.999933887 | -0.151420814 | -0.515575294 | 0.36415448 | 0.000461673 |
| ENSG00000100644 | *HIF1A* | 9.375688509 | -0.084711337 | -0.146426113 | 0.061714776 | 0.999933887 | 1.147411103 | 1.90057234 | -0.753161237 | 0.000463771 |

| | | | | | | | | | | |
|---|---|---|---|---|---|---|---|---|---|---|
| ENSG00000136059 | *VILL* | 3.904656882 | 0.040150453 | 0.06683193 | -0.026681477 | 0.999933887 | 2.086237209 | 2.910918988 | -0.824681779 | 0.000463771 |
| ENSG00000173868 | *PHOSPHO1* | 7.338655545 | 0.008237831 | -0.00488136 | 0.013119191 | 0.999933887 | -1.088414248 | -1.700667664 | 0.612251416 | 0.000463771 |
| ENSG00000079459 | *FDFT1* | 4.697173366 | 0.035090254 | -0.024609319 | 0.059699572 | 0.999933887 | -0.236137191 | -0.651858652 | 0.415721461 | 0.000465902 |
| ENSG00000099860 | *GADD45B* | 10.247449495 | 0.171428636 | 0.155579217 | 0.015849419 | 0.999933887 | 1.363104249 | 1.928093589 | -0.56498934 | 0.000474137 |
| ENSG00000173011 | *TADA2B* | 5.235598144 | 0.03179577 | 0.003956989 | 0.027838781 | 0.999933887 | -0.624137195 | -1.105875984 | 0.481738788 | 0.000474137 |
| ENSG00000117298 | *ECE1* | 9.089812894 | 0.014221799 | 0.005735823 | 0.008485975 | 0.999933887 | 1.742953283 | 2.317089955 | -0.574136672 | 0.000474137 |
| ENSG00000162433 | *AK4* | 0.490796962 | -0.235465689 | -0.327402117 | 0.091936428 | 0.999933887 | 2.188280806 | 2.949740164 | -0.761459358 | 0.000478436 |
| ENSG00000182511 | *FES* | 5.072112503 | 0.018858534 | 0.024954152 | -0.006095617 | 0.999933887 | -0.570558208 | -1.254934828 | 0.684376619 | 0.000478436 |
| ENSG00000118503 | *TNFAIP3* | 10.401396196 | 0.131013551 | 0.131600428 | -0.000586877 | 0.999933887 | 2.325472362 | 2.980843456 | -0.655371093 | 0.000481081 |
| ENSG00000120217 | *CD274* | 4.631863574 | 0.049397087 | 0.064695207 | -0.015298121 | 0.999933887 | 1.997103329 | 2.703446219 | -0.70634289 | 0.000493276 |
| ENSG00000099910 | *KLHL22* | 2.8548099 | -0.057505516 | 0.005317983 | -0.062823499 | 0.999933887 | -0.243420278 | -0.708316686 | 0.464896409 | 0.00049429 |
| ENSG00000232442 | *MHENCR* | 2.703767444 | 0.01094313 | -0.051906132 | 0.062849262 | 0.999933887 | -0.1354225 | -0.729809852 | 0.594387352 | 0.000498713 |
| ENSG00000166501 | *PRKCB* | 7.655081114 | -0.068956297 | -0.041310712 | -0.027645584 | 0.999933887 | -0.423642147 | -0.757976277 | 0.33433413 | 0.00049883 |
| ENSG00000130313 | *PGLS* | 4.008544025 | 0.074734564 | -0.010460198 | 0.085194763 | 0.999933887 | -0.532247002 | -1.150565308 | 0.618318307 | 0.00049883 |
| ENSG00000198315 | *ZKSCAN8* | 4.353249745 | -0.012552264 | -0.106695115 | 0.094142851 | 0.999933887 | 0.168265699 | 0.822822848 | -0.654557149 | 0.00049918 |
| ENSG00000110080 | *ST3GAL4* | 4.883979989 | 0.120873664 | 0.139814247 | -0.018940583 | 0.999933887 | 0.923438431 | 1.416935402 | -0.493496971 | 0.000509158 |
| ENSG00000073849 | *ST6GAL1* | 5.137139756 | 0.008299139 | -0.032144187 | 0.040443326 | 0.999933887 | -0.315752399 | -0.703174092 | 0.387421693 | 0.000509158 |
| ENSG00000115649 | *CNPPD1* | 5.224900151 | -0.036621252 | 0.000187896 | -0.036809148 | 0.999933887 | -0.706268412 | -1.173552934 | 0.467284523 | 0.000512618 |
| ENSG00000055208 | *TAB2* | 8.063379728 | -0.023003317 | -0.074135522 | 0.051132205 | 0.999933887 | 0.699293902 | 1.101446521 | -0.402152619 | 0.00051355 |
| ENSG00000160191 | *PDE9A* | 0.723217692 | 0.049836408 | 0.064928413 | -0.015092004 | 0.999933887 | 0.076881875 | 1.118035375 | -1.041153499 | 0.000521709 |
| ENSG00000095015 | *MAP3K1* | 6.831575066 | -0.030440972 | -0.049034851 | 0.018593878 | 0.999933887 | -0.168520606 | 0.297427913 | -0.46594852 | 0.000525996 |
| ENSG00000116729 | *WLS* | 5.197853349 | 0.028390501 | -0.010487496 | 0.038877997 | 0.999933887 | -0.787430261 | -1.367410468 | 0.579980207 | 0.000525996 |
| ENSG00000102554 | *KLF5* | 4.238786433 | 0.020212749 | 0.013776478 | 0.006436271 | 0.999933887 | 0.814897465 | 1.343313048 | -0.528415583 | 0.000525996 |
| ENSG00000167778 | *SPRYD3* | 4.086048572 | -0.004189092 | 0.04094156 | -0.045130652 | 0.999933887 | -0.392752689 | -0.927242874 | 0.534490185 | 0.000525996 |
| ENSG00000005844 | *ITGAL* | 7.968619303 | 0.026285021 | 0.022364862 | 0.003920159 | 0.999933887 | -0.054442639 | -0.349665831 | 0.295223192 | 0.000528974 |
| ENSG00000142405 | *NLRP12* | 4.998051598 | -0.076259096 | -0.083419673 | 0.007160577 | 0.999933887 | -1.253327554 | -1.91626857 | 0.662941016 | 0.000531247 |
| ENSG00000277406 | *SEC22B4P* | 2.696943022 | -0.117219606 | 0.132610968 | -0.249830574 | 0.999933887 | 1.625679534 | 2.193104622 | -0.567425088 | 0.000531247 |
| ENSG00000125656 | *CLPP* | 3.563830608 | 0.053177859 | -0.09749213 | 0.150669989 | 0.999933887 | -0.166994519 | -0.830337775 | 0.663343257 | 0.00054406 |
| ENSG00000147394 | *ZNF185* | 4.281441854 | 0.031550595 | -0.018498934 | 0.05004953 | 0.999933887 | -0.82086849 | -1.364822976 | 0.543954487 | 0.000553891 |
| ENSG00000125910 | *S1PR4* | 6.306404756 | 0.016717606 | -0.043721018 | 0.060438625 | 0.999933887 | -0.697309509 | -1.412671686 | 0.715362178 | 0.000558687 |
| ENSG00000099817 | *POLR2E* | 5.730153765 | 0.0226226 | 0.020026596 | 0.002596004 | 0.999933887 | -0.28996169 | -0.677846517 | 0.387884827 | 0.000558834 |
| ENSG00000111087 | *GLI1* | 2.624007187 | 0.049798876 | -0.044227185 | 0.094026061 | 0.999933887 | -0.553569614 | -1.140575758 | 0.587006145 | 0.000564584 |
| ENSG00000119403 | *PHF19* | 3.831820289 | -0.018326151 | 0.030036577 | -0.048362727 | 0.999933887 | 0.603947943 | 1.258501589 | -0.654553646 | 0.000564584 |
| ENSG00000163931 | *TKT* | 8.272076849 | 0.010518431 | -0.002207398 | 0.012725829 | 0.999933887 | -0.336796986 | -0.847302542 | 0.510505556 | 0.00056899 |
| ENSG00000266173 | *STRADA* | 3.326938532 | -0.010214006 | -0.134750543 | 0.124536537 | 0.999933887 | -0.518326979 | -1.100155504 | 0.581828525 | 0.00056899 |
| ENSG00000138614 | *INTS14* | 2.460627094 | -0.006507736 | 0.02537633 | -0.031884066 | 0.999933887 | 0.685669629 | 1.341883935 | -0.656214306 | 0.000569404 |
| ENSG00000286162 | *AL162253.2* | 1.64577002 | 0.13322837 | 0.087289541 | 0.045938829 | 0.999933887 | 1.57063582 | 2.467061202 | -0.896425382 | 0.000581413 |
| ENSG00000151465 | *CDC123* | 4.252510867 | -0.053746532 | -0.030487713 | -0.023258819 | 0.999933887 | -0.306430671 | -0.771693294 | 0.465262623 | 0.000581413 |
| ENSG00000068028 | *RASSF1* | 4.891462629 | -0.037162803 | 0.02853351 | -0.065696313 | 0.999933887 | -0.13494986 | -0.562207847 | 0.427257987 | 0.000581947 |
| ENSG00000110324 | *IL10RA* | 7.422934672 | 0.072740418 | 0.102350273 | -0.029609855 | 0.999933887 | 1.131844978 | 1.822963674 | -0.691118696 | 0.000582956 |
| ENSG00000116574 | *RHOU* | 2.726233695 | -0.102730842 | -0.214213816 | 0.111482974 | 0.999933887 | 0.78114959 | 1.839404265 | -1.058254675 | 0.000582956 |
| ENSG00000169756 | *LIMS1* | 6.915249345 | -0.058639423 | 0.005851231 | -0.064490654 | 0.999933887 | 1.638769898 | 2.220986789 | -0.582216891 | 0.000582956 |
| ENSG00000100596 | *SPTLC2* | 4.29971387 | -0.041050623 | -0.036920042 | -0.004130581 | 0.999933887 | -0.599764306 | -1.122627777 | 0.522863471 | 0.000582956 |
| ENSG00000154957 | *ZNF18* | 3.24696529 | -0.051323353 | 0.011618395 | -0.062941748 | 0.999933887 | -0.523975766 | -1.111147023 | 0.587171258 | 0.00059238 |
| ENSG00000173276 | *ZBTB21* | 5.822255805 | 0.036906955 | -0.091802073 | 0.128709028 | 0.999933887 | 0.541520042 | 1.044856748 | -0.503336707 | 0.00059238 |
| ENSG00000162924 | *REL* | 6.703556129 | -0.03931007 | -0.006483538 | -0.032826532 | 0.999933887 | 1.469172903 | 2.266581537 | -0.797408634 | 0.00059238 |
| ENSG00000167077 | *MEI1* | 3.664762043 | 0.065662383 | 0.03245316 | 0.033209223 | 0.999933887 | 0.640884165 | 1.308964043 | -0.668079877 | 0.000592893 |
| ENSG00000150054 | *MPP7* | 4.126406659 | -0.033362275 | 0.015469826 | -0.048832101 | 0.999933887 | 0.002872586 | 0.497844836 | -0.494972249 | 0.000598825 |
| ENSG00000198920 | *KIAA0753* | 2.833926859 | -0.071690608 | -0.122054894 | 0.050364285 | 0.999933887 | -0.641371467 | -1.201962916 | 0.56059145 | 0.000599905 |
| ENSG00000143207 | *COP1* | 6.046359176 | -0.044050497 | 0.022748502 | -0.066798999 | 0.999933887 | -0.463585687 | -0.90060972 | 0.437024033 | 0.000599905 |
| ENSG00000023041 | *ZDHHC6* | 5.006196152 | -0.005896617 | 0.003052504 | -0.00894912 | 0.999933887 | 0.653613196 | 1.038760046 | -0.385146849 | 0.000599905 |
| ENSG00000187994 | *RINL* | 3.434543152 | 0.093907251 | 0.16280181 | -0.068894559 | 0.999933887 | -0.102796442 | -0.660473923 | 0.557677481 | 0.000600908 |
| ENSG00000163156 | *SCNM1* | 4.239916091 | -0.04965325 | -0.039081249 | -0.010572001 | 0.999933887 | -0.215464228 | -0.634315967 | 0.418851739 | 0.000600908 |
| ENSG00000168763 | *CNNM3* | 4.382311719 | 0.050485755 | 0.029042453 | 0.021443301 | 0.999933887 | -0.337315054 | -0.735048783 | 0.397733729 | 0.000602361 |
| ENSG00000197070 | *ARRDC1* | 4.802213759 | 0.045818734 | 0.096673527 | -0.050854792 | 0.999933887 | -0.12106694 | -0.608540687 | 0.487473747 | 0.000602361 |
| ENSG00000172830 | *SSH3* | 4.489237102 | 0.025402329 | 0.071246487 | -0.045844158 | 0.999933887 | -0.977209232 | -1.613043141 | 0.635833909 | 0.000602361 |
| ENSG00000134070 | *IRAK2* | 8.465396421 | 0.007435236 | -0.058204457 | 0.065639693 | 0.999933887 | 2.076817628 | 2.654400998 | -0.577583369 | 0.000602361 |
| ENSG00000167261 | *DPEP2* | 6.704467166 | -0.051407234 | -0.039027158 | -0.012380076 | 0.999933887 | -1.456076039 | -2.074524735 | 0.618448696 | 0.000607851 |
| ENSG00000112419 | *PHACTR2* | 3.486976267 | -0.028876467 | -0.085627665 | 0.056751197 | 0.999933887 | 0.029435882 | 0.904048215 | -0.874612333 | 0.000607851 |
| ENSG00000186074 | *CD300LF* | 5.549016988 | -0.015508064 | 0.051273333 | -0.066781397 | 0.999933887 | -0.803353928 | -1.3474114 | 0.544057472 | 0.000607851 |
| ENSG00000173120 | *KDM2A* | 7.885612119 | 0.011230504 | 0.031453871 | -0.020223368 | 0.999933887 | -0.255324258 | -0.587376362 | 0.332052105 | 0.000625152 |
| ENSG00000105514 | *RAB3D* | 4.855767503 | -0.086710129 | -0.093705679 | 0.006995551 | 0.999933887 | -1.256174382 | -2.184413015 | 0.928238633 | 0.000636876 |
| ENSG00000138821 | *SLC39A8* | 4.32104294 | -0.034530771 | -0.005733192 | -0.028797579 | 0.999933887 | 1.769492516 | 2.698565707 | -0.92907319 | 0.000638366 |
| ENSG00000205339 | *IPO7* | 4.123677741 | -0.065744837 | -0.17258003 | 0.106835193 | 0.999933887 | 0.322654136 | 0.908415517 | -0.585761381 | 0.000639185 |
| ENSG00000173281 | *PPP1R3B* | 5.408562413 | -0.096006164 | -0.062085398 | -0.033920766 | 0.999933887 | -0.891823591 | -1.415668102 | 0.52384451 | 0.000651613 |
| ENSG00000107771 | *CCSER2* | 5.053372282 | -0.039765717 | -0.062880182 | 0.023114465 | 0.999933887 | 0.083338967 | 0.673996645 | -0.590657678 | 0.000651613 |
| ENSG00000075131 | *TIPIN* | -1.022044372 | -0.19702859 | -0.060792726 | -0.136235864 | 0.999933887 | 1.07889118 | 2.704298128 | -1.625406948 | 0.000665217 |
| ENSG00000144566 | *RAB5A* | 6.8954384 | 0.026013049 | 0.060220051 | -0.034207002 | 0.999933887 | 0.419443151 | 0.726072029 | -0.306628878 | 0.000668512 |
| ENSG00000104660 | *LEPROTL1* | 5.133251522 | 0.023259172 | -0.038540737 | 0.061799909 | 0.999933887 | -0.248828957 | -0.706030205 | 0.457201247 | 0.000668512 |
| ENSG00000187147 | *RNF220* | 4.278335805 | -0.048115165 | -0.030773336 | -0.017341829 | 0.999933887 | -0.307283804 | -0.664629197 | 0.357345393 | 0.000670753 |
| ENSG00000176393 | *RNPEP* | 3.849141579 | 0.040557681 | 0.042538101 | -0.00198042 | 0.999933887 | -0.567800472 | -1.205495463 | 0.637694991 | 0.000670753 |
| ENSG00000138670 | *RASGEF1B* | 6.587929909 | 0.075328306 | 0.052262228 | 0.023066078 | 0.999933887 | 0.57346098 | 1.022968733 | -0.449507753 | 0.000673004 |
| ENSG00000151116 | *UEVLD* | 2.383167553 | -0.050634726 | -0.051910165 | 0.001275439 | 0.999933887 | 0.328326911 | 0.950051241 | -0.62172433 | 0.000676611 |
| ENSG00000026751 | *SLAMF7* | 9.659828825 | -0.007269608 | 0.053395258 | -0.060664866 | 0.999933887 | 1.051754266 | 1.549022172 | -0.497267905 | 0.000677698 |
| ENSG00000179119 | *SPTY2D1* | 6.269354632 | 0.058595202 | -0.045067015 | 0.103662217 | 0.999933887 | 0.047757819 | 0.386816999 | -0.33905918 | 0.000678012 |
| ENSG00000162104 | *ADCY9* | 3.335054775 | 0.113220092 | 0.113348982 | -0.00012889 | 0.999933887 | 2.573194043 | 3.638345053 | -1.06515101 | 0.000678012 |
| ENSG00000261222 | *AC064805.1* | 1.645408369 | 0.03630679 | 0.015269093 | 0.021037697 | 0.999933887 | -0.847062844 | -1.84769918 | 1.000636336 | 0.000685315 |
| ENSG00000143815 | *LBR* | 6.380973389 | 0.00315357 | -0.021314533 | 0.024468103 | 0.999933887 | -0.897072195 | -1.427534376 | 0.530462181 | 0.000691622 |
| ENSG00000196547 | *MAN2A2* | 7.556823292 | -0.024102457 | 0.018677027 | -0.042779484 | 0.999933887 | -0.89009938 | -1.310174435 | 0.420075056 | 0.000691744 |
| ENSG00000149547 | *EI24* | 2.555658236 | 0.017082948 | 0.154428479 | -0.137345531 | 0.999933887 | 0.529022279 | 1.148405792 | -0.619383512 | 0.000710166 |
| ENSG00000177663 | *IL17RA* | 7.262781875 | -0.023919452 | -0.011395111 | -0.012524341 | 0.999933887 | -0.570077093 | -1.012984885 | 0.442907792 | 0.000721705 |
| ENSG00000180251 | *SLC9A4* | 1.371732711 | 0.207078331 | -0.070057644 | 0.277135975 | 0.999933887 | 2.079929732 | 2.951919755 | -0.871990023 | 0.000724211 |
| ENSG00000197943 | *PLCG2* | 7.266836129 | -0.026802566 | 0.007147802 | -0.033950368 | 0.999933887 | -0.507252827 | -0.83656692 | 0.329314093 | 0.000728783 |
| ENSG00000171314 | *PGAM1* | 4.506436938 | 0.055101634 | -0.016881678 | 0.071983311 | 0.999933887 | -0.339366049 | -0.811000454 | 0.471634405 | 0.000731801 |
| ENSG00000143772 | *ITPKB* | 7.056114795 | 0.035405451 | 0.020196007 | 0.015209444 | 0.999933887 | 0.209327187 | 0.579802154 | -0.370474967 | 0.000735093 |
| ENSG00000159339 | *PADI4* | 4.882174685 | -0.025149354 | 0.006251592 | -0.031400946 | 0.999933887 | -0.205395259 | -0.595604274 | 0.390209015 | 0.000735093 |
| ENSG00000129566 | *TEP1* | 6.232780882 | 0.014995888 | 0.061454487 | -0.046458599 | 0.999933887 | -0.556279079 | -0.973157754 | 0.416878675 | 0.000736532 |
| ENSG00000170458 | *CD14* | 8.828296574 | 0.007846503 | -0.018929823 | 0.026776326 | 0.999933887 | -0.329386631 | 0.265528554 | -0.594915184 | 0.000736532 |
| ENSG00000143401 | *ANP32E* | 4.531938743 | -0.050462953 | -0.124863838 | 0.074400885 | 0.999933887 | 0.375289572 | 0.871837943 | -0.496548371 | 0.000749858 |
| ENSG00000135250 | *SRPK2* | 5.891751389 | -0.033463949 | -0.001353639 | -0.03211031 | 0.999933887 | -0.413705414 | -0.918399866 | 0.504694452 | 0.000749858 |
| ENSG00000265491 | *RNF115* | 3.803413278 | -0.04844467 | -0.003376867 | -0.045067804 | 0.999933887 | 0.638162188 | 1.315038408 | -0.67687622 | 0.000749858 |
| ENSG00000134108 | *ARL8B* | 7.026558315 | -0.092369359 | -0.055571024 | -0.036798335 | 0.999933887 | 0.44095787 | 0.785664713 | -0.344706844 | 0.000750407 |
| ENSG00000123908 | *AGO2* | 7.766068705 | -0.049449539 | -0.047085628 | -0.00236391 | 0.999933887 | 0.366467311 | 0.721729036 | -0.355261725 | 0.000776341 |
| ENSG00000112146 | *FBXO9* | 4.362978734 | 0.02911889 | 0.086214681 | -0.057095791 | 0.999933887 | -0.364287062 | -0.852573109 | 0.488286048 | 0.000779597 |
| ENSG00000150977 | *RILPL2* | 9.535149527 | 0.009226912 | 0.051580092 | -0.042353179 | 0.999933887 | 2.180197083 | 2.756574651 | -0.576377568 | 0.000779597 |
| ENSG00000011243 | *AKAP8L* | 4.658927034 | -0.031900512 | -0.020904691 | -0.010995821 | 0.999933887 | -0.314531712 | -0.635171815 | 0.320640103 | 0.000782741 |
| ENSG00000179630 | *LACC1* | 2.352570178 | 0.051173076 | -0.164210185 | 0.215383261 | 0.999933887 | 0.705301748 | 1.667036183 | -0.961734435 | 0.00078338 |
| ENSG00000141956 | *PRDM15* | 2.547260076 | 0.034325687 | 0.048150192 | -0.013824506 | 0.999933887 | 0.23418849 | 0.744319459 | -0.510130969 | 0.000791789 |
| ENSG00000168067 | *MAP4K2* | 5.304019328 | 0.053344371 | 0.046629137 | 0.006715233 | 0.999933887 | -0.062772511 | -0.393381702 | 0.330609191 | 0.000792826 |
| ENSG00000078967 | *UBE2D4* | 1.073428295 | -0.090216824 | 0.092820023 | -0.183036848 | 0.999933887 | 0.724312503 | 1.614624345 | -0.890311842 | 0.000792826 |
| ENSG00000104081 | *BMF* | 5.535541113 | 0.009683369 | -0.000739349 | 0.010422717 | 0.999933887 | 1.416592799 | 2.09209501 | -0.675502211 | 0.000802845 |
| ENSG00000047056 | *WDR37* | 5.040737169 | -0.006846276 | -0.027311885 | 0.020465609 | 0.999933887 | -0.454609042 | -0.84878803 | 0.394178988 | 0.000806064 |
| ENSG00000138834 | *MAPK8IP3* | 7.416784767 | 0.024973187 | 0.087226296 | -0.062253109 | 0.999933887 | -0.515945747 | -0.993374984 | 0.477429238 | 0.000815355 |
| ENSG00000085449 | *WDFY1* | 5.472178937 | -0.073871905 | -0.049301209 | -0.024570696 | 0.999933887 | 0.318503369 | 0.743729148 | -0.425225779 | 0.000816688 |
| ENSG00000157764 | *BRAF* | 6.945800454 | 0.025834561 | 0.015216423 | 0.010618139 | 0.999933887 | 0.433873071 | 0.829533951 | -0.395660879 | 0.000816688 |

| | | | | | | | | | | |
|---|---|---|---|---|---|---|---|---|---|---|
| ENSG00000213398 | *LCAT* | 1.799986222 | 0.080204867 | 0.068330226 | 0.011874641 | 0.999933887 | -0.654952177 | -1.504975605 | 0.850023428 | 0.000831212 |
| ENSG00000160326 | *SLC2A6* | 2.282232105 | 0.161355104 | 0.009933379 | 0.151421726 | 0.999933887 | 3.271499775 | 4.489742289 | -1.218242514 | 0.000834147 |
| ENSG00000115602 | *IL1RL1* | 2.905921682 | -0.068124385 | -0.009300448 | -0.058823937 | 0.999933887 | -0.018412791 | 0.525336114 | -0.543748904 | 0.000851884 |
| ENSG00000151702 | *FLI1* | 5.65072938 | 0.04251064 | 0.048062083 | -0.005551444 | 0.999933887 | -0.266649821 | -0.702765479 | 0.436115658 | 0.000851884 |
| ENSG00000213625 | *LEPROT* | 6.756932198 | -0.06183044 | -0.023409061 | -0.038421379 | 0.999933887 | -0.294947933 | 0.156533862 | -0.451481795 | 0.000855648 |
| ENSG00000102879 | *CORO1A* | 9.099271042 | 0.029189919 | 0.023617169 | 0.00557275 | 0.999933887 | -0.628412864 | -1.155227762 | 0.526814898 | 0.000855648 |
| ENSG00000080815 | *PSEN1* | 7.967791916 | -0.089330885 | -0.046127572 | -0.043203314 | 0.999933887 | 1.167122172 | 1.552611796 | -0.385489624 | 0.000859511 |
| ENSG00000095319 | *NUP188* | 4.088413094 | -0.034942913 | -0.010234276 | -0.024708637 | 0.999933887 | 0.578225307 | 1.132824986 | -0.554599679 | 0.000859531 |
| ENSG00000151694 | *ADAM17* | 6.846157601 | -0.081503171 | -0.043276138 | -0.038227033 | 0.999933887 | 0.945624397 | 1.271658452 | -0.326034055 | 0.000861223 |
| ENSG00000131242 | *RAB11FIP4* | 7.029733623 | -0.096982499 | -0.052327907 | -0.044654592 | 0.999933887 | -1.253543634 | -1.794640701 | 0.541097067 | 0.000861223 |
| ENSG00000108262 | *GIT1* | 4.285374505 | 0.03413015 | 0.020001222 | 0.014128928 | 0.999933887 | -0.165869188 | -0.568420673 | 0.402551486 | 0.000861223 |
| ENSG00000215861 | *AC245297.1* | 1.500980679 | -0.129264089 | 0.413986395 | -0.543250484 | 0.999933887 | 1.258602537 | 1.912291656 | -0.653689119 | 0.000861223 |
| ENSG00000175471 | *MCTP1* | 5.880030353 | -0.014736746 | -0.023190288 | 0.008453543 | 0.999933887 | 1.225932422 | 1.82747085 | -0.601538428 | 0.000861223 |
| ENSG00000104973 | *MED25* | 6.499680055 | -0.004589642 | 0.023811591 | -0.028401233 | 0.999933887 | -0.907996878 | -1.381358628 | 0.473361751 | 0.000861223 |
| ENSG00000196498 | *NCOR2* | 7.457743169 | 0.090605118 | 0.061155803 | 0.029449315 | 0.999933887 | 1.383891049 | 1.951659956 | -0.567768907 | 0.000865401 |
| ENSG00000198879 | *SFMBT2* | 6.935850546 | -0.024493193 | 0.006517866 | -0.031011058 | 0.999933887 | 1.280894524 | 1.802098047 | -0.521203523 | 0.00086938 |
| ENSG00000026508 | *CD44* | 10.012102911 | -0.008263367 | 0.011141126 | -0.019404493 | 0.999933887 | 1.173283413 | 1.669825102 | -0.496541689 | 0.00086938 |
| ENSG00000105967 | *TFEC* | 3.262959541 | 0.134275547 | -0.016004502 | 0.150280049 | 0.999933887 | 2.592349255 | 3.634482164 | -1.042132909 | 0.000869393 |
| ENSG00000170448 | *NFXL1* | 3.001643868 | -0.117824547 | -0.023345124 | -0.094479423 | 0.999933887 | -0.429349902 | -1.080396134 | 0.651046232 | 0.000873123 |
| ENSG00000172732 | *MUS81* | 4.0947931 | 0.082357227 | 0.14626158 | -0.063904353 | 0.999933887 | -0.144765866 | -0.543353734 | 0.398587868 | 0.000873123 |
| ENSG00000137757 | *CASP5* | 3.749760443 | -0.08609178 | -0.182235994 | 0.096144214 | 0.999933887 | 1.512608821 | 2.273230805 | -0.760621985 | 0.000873123 |
| ENSG00000100811 | *YY1* | 6.093847832 | -0.005007753 | 0.044118022 | -0.049125775 | 0.999933887 | -0.478857033 | -0.840571919 | 0.361714886 | 0.000873123 |
| ENSG00000160766 | *GBAP1* | 0.903382235 | 0.107460104 | -0.254622086 | 0.36208219 | 0.999933887 | -0.698924426 | -1.772565537 | 1.073641111 | 0.000886957 |
| ENSG00000156873 | *PHKG2* | 4.013398329 | 0.032914119 | 0.020865958 | 0.012048161 | 0.999933887 | -0.26334179 | -0.675186363 | 0.411844573 | 0.000886957 |
| ENSG00000189319 | *FAM53B* | 5.863330618 | -0.035389795 | 0.00251613 | -0.037905926 | 0.999933887 | -0.545307394 | -1.078350952 | 0.533043558 | 0.000886957 |
| ENSG00000182287 | *AP1S2* | 4.709134929 | 0.012312835 | -0.171067645 | 0.18338048 | 0.999933887 | 1.070466726 | 1.564622083 | -0.494155356 | 0.000886957 |
| ENSG00000261804 | *AC007342.4* | 1.84888396 | -0.016519493 | -0.075906134 | 0.059386641 | 0.999933887 | -0.846757813 | -1.973536691 | 1.126778878 | 0.000888641 |
| ENSG00000169621 | *APLF* | 0.236054238 | -0.175551456 | 0.099786736 | -0.275338192 | 0.999933887 | 1.203355123 | 2.106635919 | -0.903280796 | 0.000892752 |
| ENSG00000102882 | *MAPK3* | 6.17902598 | 0.029810834 | 0.017738284 | 0.012072551 | 0.999933887 | -0.780748739 | -1.254757156 | 0.474008417 | 0.000903596 |
| ENSG00000111348 | *ARHGDIB* | 9.144516412 | 0.014736594 | -0.05683368 | 0.071570274 | 0.999933887 | -0.206022716 | -0.658814279 | 0.452791563 | 0.000904363 |
| ENSG00000151012 | *SLC7A11* | 6.158364357 | -0.064944797 | -0.057411143 | -0.007533654 | 0.999933887 | -0.073271427 | 0.432089152 | -0.50536058 | 0.000909441 |
| ENSG00000138646 | *HERC5* | 5.026477572 | -0.083749164 | -0.058704985 | -0.025044179 | 0.999933887 | -0.721078821 | -1.275293766 | 0.554214945 | 0.00091068 |
| ENSG00000084733 | *RAB10* | 7.088847968 | 0.047778287 | 0.063341354 | -0.015563067 | 0.999933887 | 0.491744298 | 0.876478915 | -0.384734617 | 0.000915528 |
| ENSG00000100106 | *TRIOBP* | 6.021055611 | 0.054558684 | 0.063812048 | -0.009253365 | 0.999933887 | -0.618894151 | -1.088074406 | 0.469180255 | 0.000915649 |
| ENSG00000104960 | *PTOV1* | 5.037908963 | 0.049903056 | 0.10887393 | -0.058970874 | 0.999933887 | -0.278479871 | -0.770054262 | 0.491574391 | 0.000926398 |
| ENSG00000111732 | *AICDA* | -1.068427222 | 0.311359986 | -0.549775674 | 0.86113566 | 0.999933887 | 1.557603969 | 2.740640977 | -1.183037008 | 0.000949765 |
| ENSG00000182973 | *CNOT10* | 3.548674091 | -0.0387956 | -0.026976516 | -0.011819084 | 0.999933887 | -0.29947756 | -0.80101289 | 0.501535331 | 0.000949765 |
| ENSG00000146826 | *MAP11* | 4.819692171 | 0.107134509 | 0.116923024 | -0.009788516 | 0.999933887 | -0.386154864 | -0.863227827 | 0.477072963 | 0.000952845 |
| ENSG00000274265 | *AC245297.3* | 2.355286057 | 0.062817848 | 0.074202031 | -0.011384183 | 0.999933887 | 0.783963714 | 1.313909565 | -0.529945851 | 0.000960953 |
| ENSG00000056558 | *TRAF1* | 6.65872219 | 0.043222643 | 0.027058333 | 0.01616431 | 0.999933887 | 2.604150443 | 3.449710752 | -0.845560309 | 0.000960953 |
| ENSG00000154237 | *LRRK1* | 4.15203469 | -0.025959454 | 0.027358265 | -0.053317719 | 0.999933887 | -0.370903319 | -0.93291908 | 0.562015761 | 0.000967348 |
| ENSG00000108861 | *DUSP3* | 5.991821097 | -0.058160242 | -0.087420092 | 0.029259849 | 0.999933887 | 0.935414221 | 1.362509467 | -0.427095247 | 0.000969831 |
| ENSG00000038210 | *PI4K2B* | 2.971242188 | -0.17442804 | 0.014339162 | -0.188767203 | 0.999933887 | 0.884343533 | 1.736433205 | -0.852089673 | 0.00097215 |
| ENSG00000126216 | *TUBGCP3* | 4.009229963 | -0.013935024 | 0.050176444 | -0.064111468 | 0.999933887 | -0.284708781 | -0.723243092 | 0.43853431 | 0.00097215 |
| ENSG00000139651 | *ZNF740* | 3.556275446 | -0.03534286 | -0.001449762 | -0.033893097 | 0.999933887 | -0.167301476 | -0.551628541 | 0.384327066 | 0.00097297 |
| ENSG00000123728 | *RAP2C* | 6.740101708 | 0.019185788 | 0.010604213 | 0.008581576 | 0.999933887 | 0.757529754 | 1.114401871 | -0.356872117 | 0.000987054 |
| ENSG00000122958 | *VPS26A* | 6.245553559 | -0.065660389 | -0.152938486 | 0.087278097 | 0.999933887 | -0.075710106 | 0.29289067 | -0.368600776 | 0.001006552 |
| ENSG00000160310 | *PRMT2* | 6.084208728 | 0.022518318 | -0.038198299 | 0.060716617 | 0.999933887 | 0.804957681 | 1.311656198 | -0.506698517 | 0.001006552 |
| ENSG00000179743 | *FLJ37453* | 2.086985545 | -0.07488705 | 0.045696202 | -0.120583252 | 0.999933887 | -0.311897845 | -0.915954757 | 0.604056912 | 0.001045357 |
| ENSG00000085978 | *ATG16L1* | 4.460021644 | -0.05552266 | -0.017953022 | -0.037569637 | 0.999933887 | -0.107672693 | -0.432284006 | 0.324611313 | 0.001057199 |
| ENSG00000131979 | *GCH1* | 5.598911626 | -0.050802097 | 0.024590322 | -0.075392419 | 0.999933887 | 2.958338265 | 3.869760131 | -0.911421867 | 0.001063286 |
| ENSG00000272501 | *AL662844.4* | 4.527859513 | -0.087503388 | -0.089438793 | 0.001935404 | 0.999933887 | -0.883104971 | -1.433593599 | 0.550488628 | 0.001072907 |
| ENSG00000116478 | *HDAC1* | 5.126536128 | 0.025448329 | -0.042998062 | 0.068446392 | 0.999933887 | -0.126543867 | -0.50503495 | 0.378491083 | 0.001089166 |
| ENSG00000137161 | *CNPY3* | 7.11298908 | 0.001965625 | -0.010919968 | 0.012885593 | 0.999933887 | -0.360322446 | -0.729515254 | 0.369192808 | 0.001089166 |
| ENSG00000120690 | *ELF1* | 9.074939424 | -0.061614246 | -0.144267855 | 0.082653609 | 0.999933887 | -0.053605827 | 0.293197105 | -0.346802932 | 0.001092888 |
| ENSG00000108622 | *ICAM2* | 3.194515294 | 0.071919981 | 0.012510454 | 0.059409527 | 0.999933887 | 0.368944739 | 0.810031012 | -0.441086273 | 0.00110226 |
| ENSG00000166145 | *SPINT1* | 2.364680459 | 0.021940463 | 0.063219973 | -0.041279509 | 0.999933887 | -0.958334028 | -1.938115495 | 0.979781468 | 0.00110226 |
| ENSG00000049239 | *H6PD* | 4.55803922 | 0.010937577 | -0.039203117 | 0.050140694 | 0.999933887 | -0.348712024 | -0.738936312 | 0.390224288 | 0.001113003 |
| ENSG00000185338 | *SOCS1* | 3.75060889 | 0.164057176 | 0.11755812 | 0.046499056 | 0.999933887 | 1.072436563 | 1.656482596 | -0.584046033 | 0.001126632 |
| ENSG00000146070 | *PLA2G7* | 1.070755343 | 0.17249263 | 0.184580699 | -0.012088069 | 0.999933887 | 0.488889488 | 1.141170228 | -0.65228074 | 0.001132075 |
| ENSG00000171163 | *ZNF692* | 3.702623715 | 0.028704578 | 0.056240757 | -0.027536179 | 0.999933887 | -0.356602666 | -0.827265932 | 0.470663267 | 0.001132075 |
| ENSG00000103415 | *HMOX2* | 4.43750536 | -0.036498042 | 0.001993042 | -0.038491084 | 0.999933887 | -0.43704414 | -1.004070863 | 0.567026722 | 0.001132167 |
| ENSG00000105122 | *RASAL3* | 6.389077992 | 0.021825805 | 0.043155708 | -0.021329902 | 0.999933887 | -0.23921628 | -0.574251043 | 0.335034762 | 0.001132167 |
| ENSG00000160214 | *RRP1* | 4.524148065 | 0.063491947 | 0.082693821 | -0.019201874 | 0.999933887 | 0.911728836 | 1.332408532 | -0.420679696 | 0.001136208 |
| ENSG00000141506 | *PIK3R5* | 9.557875607 | -0.007780963 | -0.001920783 | -0.00586018 | 0.999933887 | 0.527022804 | 0.78487653 | -0.257853726 | 0.00114191 |
| ENSG00000167034 | *NKX3-1* | 1.007559537 | 0.019750232 | -0.026367036 | 0.046117268 | 0.999933887 | -1.049529014 | -2.125213217 | 1.075684203 | 0.00114191 |
| ENSG00000017260 | *ATP2C1* | 4.291924181 | -0.052775512 | -0.092500352 | 0.03972484 | 0.999933887 | 0.274393355 | 0.783468934 | -0.509075578 | 0.001143123 |
| ENSG00000040275 | *SPDL1* | 1.757446492 | -0.151275343 | -0.371452951 | 0.220177608 | 0.999933887 | 0.234800241 | 0.839891104 | -0.605090863 | 0.001146891 |
| ENSG00000162222 | *TTC9C* | 2.044275448 | -0.020349162 | -0.071342337 | 0.050993175 | 0.999933887 | -0.312251873 | -0.937431068 | 0.625179195 | 0.001148212 |
| ENSG00000180539 | *C9orf139* | 1.824751912 | 0.008731282 | 0.130369365 | -0.121638083 | 0.999933887 | -0.376075397 | -1.331684145 | 0.955608747 | 0.00115915 |
| ENSG00000147813 | *NAPRT* | 4.931008226 | 0.056295556 | 0.015218807 | 0.041076749 | 0.999933887 | -1.496180663 | -2.196773914 | 0.700593251 | 0.001163393 |
| ENSG00000196839 | *ADA* | 2.861694027 | 0.07900337 | 0.007658935 | 0.071344435 | 0.999933887 | 0.913933388 | 1.601717051 | -0.687783663 | 0.001173433 |
| ENSG00000163171 | *CDC42EP3* | 10.39180724 | 0.010058429 | 0.012642135 | -0.002583705 | 0.999933887 | 0.272568063 | 0.587857049 | -0.315288986 | 0.001173433 |
| ENSG00000100883 | *SRP54* | 6.023390761 | -0.067691779 | 0.025815955 | -0.093507734 | 0.999933887 | 1.328188011 | 1.77615185 | -0.447963839 | 0.001178705 |
| ENSG00000075426 | *FOSL2* | 10.768004632 | 0.075277701 | 0.026471606 | 0.048806096 | 0.999933887 | 0.630111172 | 1.056156876 | -0.426045704 | 0.001193821 |
| ENSG00000168961 | *LGALS9* | 4.896452645 | 0.116125076 | 0.05471894 | 0.061406136 | 0.999933887 | -0.694263001 | -1.283483768 | 0.589220767 | 0.0012016 |
| ENSG00000114738 | *MAPKAPK3* | 5.897413329 | -0.020361045 | -0.014909332 | -0.005451713 | 0.999933887 | -1.091721876 | -1.593351009 | 0.501629134 | 0.001202766 |
| ENSG00000163154 | *TNFAIP8L2* | 2.470249233 | -0.203317114 | -0.204203308 | 0.000886194 | 0.999933887 | -0.633611626 | -1.405861342 | 0.772249716 | 0.001203375 |
| ENSG00000197646 | *PDCD1LG2* | 0.479990533 | 0.005199718 | -0.365802241 | 0.371001958 | 0.999933887 | 0.247597036 | 1.132495571 | -0.884898535 | 0.001204161 |
| ENSG00000147421 | *HMBOX1* | 4.24244837 | -0.018652999 | 0.004600376 | -0.023253375 | 0.999933887 | 0.159688325 | 0.59708195 | -0.437393625 | 0.00121255 |
| ENSG00000128340 | *RAC2* | 9.934842254 | 0.028523023 | -0.028474883 | 0.056997906 | 0.999933887 | 0.379539484 | 0.681054536 | -0.301515052 | 0.001214153 |
| ENSG00000074416 | *MGLL* | 4.795307858 | 0.047369024 | 0.213577207 | -0.166208183 | 0.999933887 | 0.664667768 | 1.15282506 | -0.488157292 | 0.001219929 |
| ENSG00000137710 | *RDX* | 2.830110131 | -0.150893298 | -0.178397965 | 0.027504667 | 0.999933887 | -0.012227297 | 0.521926042 | -0.534153339 | 0.001257358 |
| ENSG00000288156 | *AC104530.1* | 3.758414408 | -0.175033884 | -0.097406403 | -0.077627481 | 0.999933887 | -0.723844839 | -1.453764954 | 0.729920115 | 0.00125987 |
| ENSG00000204842 | *ATXN2* | 4.007151881 | 0.029050896 | 0.027529968 | 0.001520928 | 0.999933887 | -0.545126031 | -0.944641456 | 0.399515425 | 0.001263396 |
| ENSG00000145911 | *N4BP3* | 2.847106788 | 0.120893809 | 0.002678365 | 0.118215444 | 0.999933887 | 1.204535875 | 1.844475502 | -0.639939627 | 0.001280367 |
| ENSG00000104976 | *SNAPC2* | 2.844501046 | 0.037631356 | -0.112204886 | 0.149836242 | 0.999933887 | -0.722880718 | -1.244402894 | 0.521522175 | 0.001280706 |
| ENSG00000138663 | *COPS4* | 3.675239641 | -0.037775963 | -0.124585966 | 0.086810003 | 0.999933887 | -0.170443096 | 0.312967338 | -0.483410434 | 0.001286398 |
| ENSG00000149476 | *TKFC* | 2.355303561 | 0.098123379 | -0.021821671 | 0.11994505 | 0.999933887 | 0.684391297 | 1.198409423 | -0.514018126 | 0.001288706 |
| ENSG00000213024 | *NUP62* | 4.876136998 | -0.043611442 | -0.001391667 | -0.042219774 | 0.999933887 | 0.539259448 | 0.93562044 | -0.396360992 | 0.001288706 |
| ENSG00000157570 | *TSPAN18* | 2.208190651 | 0.115394062 | 0.096638306 | 0.018755756 | 0.999933887 | 0.637038653 | 1.370704979 | -0.733666326 | 0.001299133 |
| ENSG00000116731 | *PRDM2* | 7.153867851 | -0.032073041 | -0.080275769 | 0.048202728 | 0.999933887 | 0.482056475 | 0.803322457 | -0.321265981 | 0.001299133 |
| ENSG00000035687 | *ADSS2* | 4.642206822 | -0.032814184 | -0.054405914 | 0.02159173 | 0.999933887 | -1.015124797 | -1.59823131 | 0.583106514 | 0.001299882 |
| ENSG00000082397 | *EPB41L3* | 1.667384647 | 0.073630332 | -0.050037975 | 0.123668307 | 0.999933887 | 0.570728719 | 1.248389218 | -0.677660499 | 0.001304861 |
| ENSG00000197119 | *SLC25A29* | 4.7285602 | 0.039892733 | -0.030781565 | 0.070674299 | 0.999933887 | -0.760956199 | -1.207032119 | 0.44607592 | 0.001306118 |
| ENSG00000105520 | *PLPPR2* | 6.544207108 | 0.006071057 | 0.041838214 | -0.035767157 | 0.999933887 | -0.544997432 | -1.044841099 | 0.499843667 | 0.001306118 |
| ENSG00000155115 | *GTF3C6* | 2.655219403 | -0.138237231 | 0.019047033 | -0.157284264 | 0.999933887 | 2.001974612 | 2.838323193 | -0.836348582 | 0.001321337 |
| ENSG00000171860 | *C3AR1* | 7.852069919 | -0.001817445 | -0.054265615 | 0.05244817 | 0.999933887 | 1.060265875 | 1.63146335 | -0.571197476 | 0.001331046 |
| ENSG00000152558 | *TMEM123* | 7.716180565 | -0.032923437 | -0.115619034 | 0.082695597 | 0.999933887 | 0.028713314 | 0.474168258 | -0.445454945 | 0.001332219 |
| ENSG00000037637 | *FBXO42* | 4.781856494 | -0.024074987 | -0.010881739 | -0.013193247 | 0.999933887 | 0.722491813 | 1.204036415 | -0.481544603 | 0.001347282 |
| ENSG00000165516 | *KLHDC2* | 3.849440126 | -0.101499024 | -0.032446414 | -0.069052611 | 0.999933887 | -0.241229743 | -0.660661795 | 0.419432052 | 0.001351096 |
| ENSG00000140030 | *GPR65* | 6.292342717 | 0.025927402 | -0.031517234 | 0.057444636 | 0.999933887 | 0.962411716 | 1.506886099 | -0.544474383 | 0.001351096 |
| ENSG00000136560 | *TANK* | 8.228200072 | -0.014876622 | 0.022539326 | -0.037415949 | 0.999933887 | 0.801659315 | 1.333963831 | -0.532304516 | 0.001351096 |
| ENSG00000008516 | *MMP25* | 8.770077893 | 0.005702505 | 0.009882424 | -0.004179919 | 0.999933887 | -0.387193003 | -0.807040034 | 0.419847031 | 0.001351096 |
| ENSG00000110955 | *ATP5F1B* | 6.351470874 | -0.013482022 | -0.020443148 | 0.006961126 | 0.999933887 | -0.11678379 | -0.387918829 | 0.271135039 | 0.001360905 |

| | | | | | | | | | | |
|---|---|---|---|---|---|---|---|---|---|---|
| ENSG00000140932 | *CMTM2* | 6.650287658 | -0.038273753 | -0.064978863 | 0.02670511 | 0.999933887 | -0.43487889 | -1.038086622 | 0.603207733 | 0.001370793 |
| ENSG00000158201 | *ABHD3* | 5.36684154 | -0.029278807 | -0.032717582 | 0.003438775 | 0.999933887 | -0.444824932 | -0.859422976 | 0.414598044 | 0.001373997 |
| ENSG00000127328 | *RAB3IP* | 2.200302749 | -0.049835669 | -0.143462422 | 0.093626753 | 0.999933887 | 0.451126445 | 1.097024438 | -0.645897993 | 0.001390956 |
| ENSG00000067955 | *CBFB* | 4.357709591 | -0.037838468 | 0.003283496 | -0.041121963 | 0.999933887 | -0.490513422 | -0.865207711 | 0.374694289 | 0.001397599 |
| ENSG00000146476 | *ARMT1* | 4.057822996 | -0.06616615 | -0.065403908 | -0.000762242 | 0.999933887 | 0.115188619 | 0.489564042 | -0.374375422 | 0.001398212 |
| ENSG00000166128 | *RAB8B* | 8.422073549 | -0.082839163 | -0.138378184 | 0.055539021 | 0.999933887 | 0.468493465 | 0.953413518 | -0.484920052 | 0.001410206 |
| ENSG00000125779 | *PANK2* | 4.416017883 | 0.021439671 | 0.046380965 | -0.024941294 | 0.999933887 | -0.404803605 | -0.808672361 | 0.403868756 | 0.001417129 |
| ENSG00000015285 | *WAS* | 7.942319352 | 3.4829e-05 | -0.004711243 | 0.004746072 | 0.999933887 | -0.466864243 | -0.813630026 | 0.346765783 | 0.001419612 |
| ENSG00000104320 | *NBN* | 8.290914724 | -0.108007119 | -0.175803556 | 0.067796437 | 0.999933887 | 1.745953824 | 2.511749664 | -0.765795841 | 0.001421977 |
| ENSG00000155096 | *AZIN1* | 7.989130461 | -0.092308378 | -0.049106277 | -0.043202101 | 0.999933887 | 1.601183406 | 2.073971957 | -0.47278855 | 0.00142668 |
| ENSG00000104866 | *PPP1R37* | 3.434734634 | 0.075795769 | 0.058308487 | 0.017487282 | 0.999933887 | -0.304170148 | -0.788976893 | 0.484806745 | 0.001427275 |
| ENSG00000197860 | *SGTB* | 7.708349632 | -0.043899454 | -0.114931342 | 0.071031888 | 0.999933887 | 0.372939021 | 0.818817598 | -0.445878577 | 0.001427275 |
| ENSG00000257027 | *AC010186.3* | 2.200414169 | -0.034075521 | 0.087204356 | -0.121279877 | 0.999933887 | 0.839351153 | 1.729461296 | -0.890110143 | 0.001427275 |
| ENSG00000148290 | *SURF1* | 3.860321104 | 0.003662941 | -0.05473195 | 0.05839489 | 0.999933887 | -0.177253533 | -0.560297813 | 0.383044281 | 0.001427275 |
| ENSG00000128564 | *VGF* | 0.157970524 | -0.145556878 | 0.299168243 | -0.444725121 | 0.999933887 | -0.575043985 | -1.638403287 | 1.063359303 | 0.001445882 |
| ENSG00000103495 | *MAZ* | 4.029472511 | 0.032788355 | 0.104761989 | -0.071973634 | 0.999933887 | -0.413368229 | -0.956025557 | 0.542657328 | 0.001445882 |
| ENSG00000113916 | *BCL6* | 9.921030222 | 0.009469631 | -0.011243693 | 0.020713324 | 0.999933887 | 0.510006039 | 0.751846485 | -0.241840446 | 0.001445882 |
| ENSG00000035664 | *DAPK2* | 4.990465506 | -0.012765326 | 0.055061394 | -0.067826721 | 0.999933887 | -0.289636685 | -0.719050706 | 0.429414021 | 0.001445882 |
| ENSG00000043462 | *LCP2* | 10.941819185 | -0.006795142 | -0.001225155 | -0.005569987 | 0.999933887 | 0.921980152 | 1.258023786 | -0.336043634 | 0.001445882 |
| ENSG00000170638 | *TRABD* | 6.035562848 | 0.051131951 | 0.041558172 | 0.009573778 | 0.999933887 | -0.082725185 | -0.460201401 | 0.377476216 | 0.001447743 |
| ENSG00000134802 | *SLC43A3* | 6.398534944 | 0.036693287 | -0.006944167 | 0.043637654 | 0.999933887 | 1.589710385 | 2.031190276 | -0.441479891 | 0.001447743 |
| ENSG00000187266 | *EPOR* | 2.460028448 | 0.019134689 | 0.073763276 | -0.054628588 | 0.999933887 | -0.657894064 | -1.255716873 | 0.597822809 | 0.001459735 |
| ENSG00000112096 | *SOD2* | 12.211699964 | -0.040862904 | 0.022933325 | -0.063796229 | 0.999933887 | 2.031100578 | 2.529030648 | -0.49793007 | 0.001465509 |
| ENSG00000128594 | *LRRC4* | 5.863742671 | 0.020438484 | 0.039057027 | -0.018618544 | 0.999933887 | -1.568306981 | -2.163316753 | 0.595009773 | 0.001470514 |
| ENSG00000079805 | *DNM2* | 7.996709507 | 0.002575392 | 0.032184455 | -0.029609062 | 0.999933887 | -0.286296091 | -0.562380583 | 0.276084492 | 0.001524737 |
| ENSG00000148737 | *TCF7L2* | 4.718575352 | -0.038280177 | 0.105987416 | -0.144267593 | 0.999933887 | 0.571546136 | 0.941747091 | -0.370200954 | 0.001530305 |
| ENSG00000205356 | *TECPR1* | 5.493905349 | 0.037977813 | 0.053550041 | -0.015572228 | 0.999933887 | -0.289714419 | -0.681834273 | 0.392119854 | 0.00154636 |
| ENSG00000164603 | *BMT2* | 3.933299743 | -0.00577163 | -0.106498648 | 0.100727018 | 0.999933887 | 1.248532369 | 1.79381925 | -0.54528688 | 0.00154636 |
| ENSG00000165030 | *NFIL3* | 7.816060722 | 0.055898231 | -0.03946434 | 0.095362571 | 0.999933887 | 0.794924028 | 1.233732985 | -0.438808957 | 0.001546463 |
| ENSG00000069956 | *MAPK6* | 7.628388574 | -0.062497878 | -0.127723171 | 0.065225294 | 0.999933887 | 1.132372757 | 1.65203032 | -0.519657563 | 0.001546463 |
| ENSG00000119638 | *NEK9* | 4.793638018 | -0.03039132 | -0.024971584 | -0.005419736 | 0.999933887 | -0.203087138 | -0.47305837 | 0.269971232 | 0.001546463 |
| ENSG00000140990 | *NDUFB10* | 3.527518084 | 0.049236136 | 0.01203903 | 0.037197106 | 0.999933887 | -0.131783698 | -0.684326066 | 0.552542367 | 0.001546463 |
| ENSG00000156587 | *UBE2L6* | 7.024718755 | 0.039792329 | -0.011170747 | 0.050963076 | 0.999933887 | -0.423122335 | -0.846973328 | 0.423850993 | 0.001546463 |
| ENSG00000120437 | *ACAT2* | 4.236828779 | -0.037995823 | -0.032032712 | -0.005963112 | 0.999933887 | 1.672567215 | 2.152756971 | -0.480189756 | 0.001546463 |
| ENSG00000196588 | *MRTFA* | 5.977646559 | -0.041794038 | 0.05183244 | -0.093626477 | 0.999933887 | -0.230960892 | -0.624934249 | 0.393973358 | 0.001570613 |
| ENSG00000040933 | *INPP4A* | 5.824525316 | 0.016065675 | 0.006324758 | 0.009740917 | 0.999933887 | -0.323715569 | -0.72077693 | 0.397061361 | 0.001570613 |
| ENSG00000104154 | *SLC30A4* | 2.419907804 | -0.180495187 | -0.159962524 | -0.020532663 | 0.999933887 | 2.055480905 | 2.960444347 | -0.904963443 | 0.00160586 |
| ENSG00000067334 | *DNTTIP2* | 7.334978894 | -0.06754697 | -0.188661127 | 0.121114158 | 0.999933887 | 0.428847143 | 0.965127258 | -0.536280115 | 0.001615311 |
| ENSG00000119979 | *DENND10* | 4.309155494 | 0.061249476 | -0.028646492 | 0.089895968 | 0.999933887 | -0.359094367 | -0.742223497 | 0.38312913 | 0.001635524 |
| ENSG00000155099 | *PIP4P2* | 3.862429569 | 0.005404049 | 0.128901532 | -0.123497483 | 0.999933887 | -0.991107923 | -1.560253105 | 0.569145182 | 0.001640256 |
| ENSG00000198839 | *ZNF277* | 4.004704648 | -0.013454063 | -0.1308227 | 0.117368637 | 0.999933887 | 0.171511473 | 0.612101459 | -0.440589986 | 0.001649672 |
| ENSG00000279024 | *AC112255.1* | 2.743371378 | -0.127262741 | -0.124491866 | -0.002770875 | 0.999933887 | 0.209670404 | 0.713410165 | -0.503739762 | 0.001651684 |
| ENSG00000078081 | *LAMP3* | 2.401064945 | -0.004454341 | 0.107302014 | -0.111756354 | 0.999933887 | 1.57349036 | 2.452852864 | -0.879362504 | 0.001652104 |
| ENSG00000167508 | *MVD* | 4.498744565 | 0.054299967 | 0.109066419 | -0.054766452 | 0.999933887 | 1.592507882 | 2.415950217 | -0.823442335 | 0.00167496 |
| ENSG00000140995 | *DEF8* | 6.36772318 | -0.002342184 | 0.054379808 | -0.056721992 | 0.999933887 | -0.173883061 | -0.694981503 | 0.521098442 | 0.001675025 |
| ENSG00000274536 | *MIR223HG* | 4.723661018 | -0.117188132 | -0.057324685 | -0.059863447 | 0.999933887 | -0.251007326 | -0.771001632 | 0.519994305 | 0.001676007 |
| ENSG00000169598 | *DFFB* | 1.503030199 | -0.042271011 | 0.012382708 | -0.054653719 | 0.999933887 | -0.052257893 | -0.68361393 | 0.631356037 | 0.001682235 |
| ENSG00000163516 | *ANKZF1* | 4.564474014 | 0.016028951 | 0.056994774 | -0.040965823 | 0.999933887 | -0.123431033 | -0.478604096 | 0.355173063 | 0.001682235 |
| ENSG00000172590 | *MRPL52* | 2.630810029 | -0.098945318 | -0.145437301 | 0.046491983 | 0.999933887 | 1.187636568 | 1.873629861 | -0.685993293 | 0.001682453 |
| ENSG00000161542 | *PRPSAP1* | 3.282556893 | 0.056994165 | -0.216489195 | 0.273483359 | 0.999933887 | 1.197885226 | 1.804721453 | -0.606836228 | 0.001682453 |
| ENSG00000236939 | *BAALC-AS2* | -1.645688777 | -0.032896537 | -0.411022322 | 0.378125785 | 0.999933887 | 1.576544307 | 2.401167077 | -0.82462277 | 0.001682453 |
| ENSG00000109787 | *KLF3* | 6.818771052 | -0.032585855 | -0.019248699 | -0.013337156 | 0.999933887 | -1.122306337 | -1.564926625 | 0.442620288 | 0.00168609 |
| ENSG00000172531 | *PPP1CA* | 6.283589868 | 0.001727259 | -0.018459099 | 0.020186358 | 0.999933887 | -0.619176009 | -1.059594449 | 0.44041844 | 0.00168609 |
| ENSG00000213445 | *SIPA1* | 8.006244098 | 0.058689015 | 0.078346631 | -0.019657616 | 0.999933887 | -0.344905388 | -0.721700001 | 0.376794613 | 0.001693511 |
| ENSG00000213889 | *PPM1N* | 1.437522558 | -0.074964866 | 0.096032198 | -0.170997064 | 0.999933887 | 0.525201017 | 1.232943334 | -0.707742318 | 0.001698453 |
| ENSG00000137413 | *TAF8* | 3.562354095 | -0.01151045 | -0.038874328 | 0.027363878 | 0.999933887 | -0.565437652 | -1.049828056 | 0.484390404 | 0.001734874 |
| ENSG00000068831 | *RASGRP2* | 7.032998288 | 0.053206782 | 0.051136045 | 0.002070737 | 0.999933887 | -0.031169255 | -0.428350816 | 0.397181561 | 0.001742917 |
| ENSG00000127922 | *SEM1* | 3.33066589 | 0.001532115 | -0.061587509 | 0.063119624 | 0.999933887 | -0.034290185 | 0.337048787 | -0.371338973 | 0.001742917 |
| ENSG00000137497 | *NUMA1* | 6.810838926 | 0.110860981 | 0.063729587 | 0.047131394 | 0.999933887 | 0.573070869 | 0.92102668 | -0.347955811 | 0.001744885 |
| ENSG00000155304 | *HSPA13* | 4.779867695 | -0.020014688 | -0.155868825 | 0.135854137 | 0.999933887 | 0.316748662 | 0.812179621 | -0.495430959 | 0.001744885 |
| ENSG00000129071 | *MBD4* | 4.974586653 | -0.095242732 | -0.140746988 | 0.045504256 | 0.999933887 | -0.842956781 | -1.234858448 | 0.391901667 | 0.001745406 |
| ENSG00000220008 | *LINGO3* | 3.976532118 | -0.030148705 | -0.072090778 | 0.041942073 | 0.999933887 | -0.778475064 | -1.582952969 | 0.804477905 | 0.001754444 |
| ENSG00000237522 | *NONOP2* | 1.334090777 | -0.125249046 | -0.045135825 | -0.080113221 | 0.999933887 | 0.635544109 | 1.648867575 | -1.013323466 | 0.001769364 |
| ENSG00000115275 | *MOGS* | 4.361345817 | 0.016988322 | 0.066874978 | -0.049886655 | 0.999933887 | -0.045448379 | -0.349613572 | 0.304165193 | 0.001770791 |
| ENSG00000183091 | *NEB* | 1.243723659 | -0.143458762 | -0.160537684 | 0.017078922 | 0.999933887 | 0.218144842 | 1.139046996 | -0.920902154 | 0.001815722 |
| ENSG00000115825 | *PRKD3* | 3.82092395 | -0.086586725 | -0.097296291 | 0.010709566 | 0.999933887 | 0.219408738 | 0.848403476 | -0.628994738 | 0.001815722 |
| ENSG00000114450 | *GNB4* | 3.349327964 | -0.094815655 | 0.050125658 | -0.144941313 | 0.999933887 | -1.364311479 | -2.068364665 | 0.704053187 | 0.00181681 |
| ENSG00000287626 | *AL021978.1* | 4.553455939 | -0.090241026 | -0.050632908 | -0.039608118 | 0.999933887 | 0.387408228 | 0.748345576 | -0.360937348 | 0.001818128 |
| ENSG00000130309 | *COLGALT1* | 4.986161427 | 0.040774316 | -0.020313227 | 0.061087542 | 0.999933887 | -0.103504597 | -0.426274886 | 0.322770289 | 0.001845295 |
| ENSG00000139178 | *C1RL* | 4.969060745 | -0.021411455 | -0.000540953 | -0.020870501 | 0.999933887 | -0.244497221 | -0.575430012 | 0.330932792 | 0.001897128 |
| ENSG00000213390 | *ARHGAP19* | 3.207387212 | 0.027416621 | -0.017191398 | 0.044608018 | 0.999933887 | -0.52487679 | -1.072028802 | 0.547152013 | 0.001924838 |
| ENSG00000170260 | *ZNF212* | 2.518703359 | -0.096102545 | 0.015412133 | -0.111514678 | 0.999933887 | -0.436481145 | -0.977777484 | 0.54129634 | 0.001928964 |
| ENSG00000235531 | *MSC-AS1* | 1.188589313 | -0.180192987 | -0.249021129 | 0.068828142 | 0.999933887 | 0.882304661 | 1.588051113 | -0.705746452 | 0.001954053 |
| ENSG00000122299 | *ZC3H7A* | 5.562581208 | -0.062432777 | -0.070877402 | 0.008444625 | 0.999933887 | 0.266084778 | 0.638643423 | -0.372558646 | 0.001967748 |
| ENSG00000136167 | *LCP1* | 11.817379217 | -0.010931158 | -0.073358519 | 0.062427361 | 0.999933887 | 0.583538933 | 0.889995011 | -0.306456078 | 0.001972101 |
| ENSG00000031823 | *RANBP3* | 4.894394209 | -0.000882473 | 0.048524833 | -0.049407305 | 0.999933887 | -0.236444128 | -0.591476413 | 0.355032285 | 0.001978477 |
| ENSG00000113615 | *SEC24A* | 5.567674198 | -0.015896149 | -0.074655685 | 0.058759536 | 0.999933887 | 1.355859991 | 1.929300502 | -0.573440511 | 0.001983796 |
| ENSG00000106012 | *IQCE* | 4.196000849 | 0.025557008 | 0.054303213 | -0.028746205 | 0.999933887 | -0.594492432 | -1.092800004 | 0.498307572 | 0.002009638 |
| ENSG00000131149 | *GSE1* | 5.676094554 | 0.019070902 | 0.035535445 | -0.016464543 | 0.999933887 | 0.155162625 | 0.453772912 | -0.298610287 | 0.00203286 |
| ENSG00000076604 | *TRAF4* | 3.898016263 | 0.023472077 | 0.118726649 | -0.095254573 | 0.999933887 | 0.97498407 | 1.706579944 | -0.731595874 | 0.002036845 |
| ENSG00000161981 | *SNRNP25* | 2.454530324 | -0.010734354 | -0.056464322 | 0.045729968 | 0.999933887 | -1.598158662 | -2.357919646 | 0.759760984 | 0.002039838 |
| ENSG00000186591 | *UBE2H* | 7.384523721 | 0.0198901 | 0.004662352 | 0.015227748 | 0.999933887 | 0.760144646 | 1.116039636 | -0.355894991 | 0.002040707 |
| ENSG00000060138 | *YBX3* | 6.474325988 | 0.042843029 | 0.011879399 | 0.03096363 | 0.999933887 | 0.175458859 | 0.464249087 | -0.288790228 | 0.002045257 |
| ENSG00000139926 | *FRMD6* | -0.173424563 | -0.160899015 | -0.063105811 | -0.097793205 | 0.999933887 | 0.955739611 | 2.344841447 | -1.389101836 | 0.002051251 |
| ENSG00000164663 | *USP49* | 4.170606807 | -0.116472479 | -0.053365587 | -0.063106892 | 0.999933887 | 0.854068353 | 1.286387371 | -0.432319018 | 0.002069859 |
| ENSG00000187260 | *WDR86* | 0.741192358 | 0.119564893 | 0.189899136 | -0.069434243 | 0.999933887 | -0.012949353 | -0.762074215 | 0.749124862 | 0.002069859 |
| ENSG00000119048 | *UBE2B* | 8.280011437 | -0.008320474 | 0.03861762 | -0.046938094 | 0.999933887 | 0.291688005 | 0.544771809 | -0.253083805 | 0.002069859 |
| ENSG00000198719 | *DLL1* | 1.955332716 | 0.051584403 | 0.41679618 | -0.365211777 | 0.999933887 | 0.972495429 | 1.47933969 | -0.506844261 | 0.002074453 |
| ENSG00000198018 | *ENTPD7* | 2.931317446 | -0.031513051 | -0.009228969 | -0.022284082 | 0.999933887 | -0.085248294 | 0.518062498 | -0.603310792 | 0.002086623 |
| ENSG00000123358 | *NR4A1* | 6.017573788 | 0.113020453 | 0.159754566 | -0.046734113 | 0.999933887 | 0.921034983 | 1.517281211 | -0.596246228 | 0.002093108 |
| ENSG00000204231 | *RXRB* | 5.032917834 | 0.011054953 | 0.035551703 | -0.024496749 | 0.999933887 | -0.395799134 | -0.744731811 | 0.348932677 | 0.002093108 |
| ENSG00000105642 | *KCNN1* | -0.838271118 | -0.555806841 | -1.045159858 | 0.489353017 | 0.999933887 | 2.491481303 | 3.57200263 | -1.080521326 | 0.002132646 |
| ENSG00000174485 | *DENND4A* | 6.184723224 | -0.178570857 | -0.135404243 | -0.043166614 | 0.999933887 | 2.455678968 | 3.509553571 | -1.053874603 | 0.002132646 |
| ENSG00000177674 | *AGTRAP* | 6.729311301 | 0.031345516 | -0.019898506 | 0.051244022 | 0.999933887 | 1.130373771 | 1.565700437 | -0.435326666 | 0.002142636 |
| ENSG00000054983 | *GALC* | 6.727970803 | -0.00468086 | 0.00064332 | -0.005324179 | 0.999933887 | 1.070646535 | 1.502392566 | -0.431746031 | 0.002142636 |
| ENSG00000158669 | *GPAT4* | 5.417057677 | -0.002828899 | 0.008255858 | -0.011084756 | 0.999933887 | 0.636646093 | 0.98759597 | -0.350949877 | 0.002142636 |
| ENSG00000265975 | *AC002091.1* | 4.313103649 | 0.059894745 | 0.036353533 | 0.023541212 | 0.999933887 | 1.165251613 | 1.567638308 | -0.402386695 | 0.002148805 |
| ENSG00000224660 | *SH3BP5-AS1* | 6.557635944 | 0.00262507 | 0.014991054 | -0.012365984 | 0.999933887 | 0.735195924 | 1.151534479 | -0.416338555 | 0.002175136 |
| ENSG00000175857 | *GAPT* | 5.657648423 | -0.062486687 | -0.187784735 | 0.125298048 | 0.999933887 | 0.650090363 | 1.223817364 | -0.573727001 | 0.00217758 |
| ENSG00000104412 | *EMC2* | 3.666936231 | -0.110638244 | -0.099091639 | -0.011546605 | 0.999933887 | 0.422805377 | 0.951856687 | -0.52905131 | 0.002197043 |
| ENSG00000147202 | *DIAPH2* | 2.483957723 | -0.105999874 | 0.093352573 | -0.199352447 | 0.999933887 | 0.359764581 | 0.93787448 | -0.578109899 | 0.002221417 |
| ENSG00000119471 | *HSDL2* | 4.054889922 | -0.031345789 | -0.08494699 | 0.053601202 | 0.999933887 | -0.476417327 | -1.060097106 | 0.58367978 | 0.002266275 |
| ENSG00000141971 | *MVB12A* | 2.455715945 | 0.058243705 | -0.153097263 | 0.211340968 | 0.999933887 | -0.166544809 | -0.769240219 | 0.60269541 | 0.002297268 |
| ENSG00000104695 | *PPP2CB* | 3.475097998 | 0.038940457 | -0.077957879 | 0.116898336 | 0.999933887 | 0.286037128 | 0.635422303 | -0.349385176 | 0.002302011 |
| ENSG00000132507 | *EIF5A* | 7.050762932 | 0.047168854 | 0.034667849 | 0.012501004 | 0.999933887 | 1.209421965 | 1.642158968 | -0.432737004 | 0.002310861 |

| | | | | | | | | | | |
|---|---|---|---|---|---|---|---|---|---|---|
| ENSG00000108100 | *CCNY* | 6.394786948 | -0.023028271 | -0.012109427 | -0.010918843 | 0.999933887 | -0.733654533 | -1.128856564 | 0.395202031 | 0.002313026 |
| ENSG00000166548 | *TK2* | 2.963674087 | 0.006032504 | 0.082158369 | -0.076125865 | 0.999933887 | -0.575855302 | -1.266070035 | 0.690214733 | 0.002317187 |
| ENSG00000279833 | *AL031846.2* | 2.427553033 | -0.057611873 | -0.143792497 | 0.086180624 | 0.999933887 | -0.715010818 | -1.379967683 | 0.664956865 | 0.002320205 |
| ENSG00000135926 | *TMBIM1* | 7.588919248 | -0.012112756 | -0.005020194 | -0.007092562 | 0.999933887 | 0.493458359 | 0.857228672 | -0.363770313 | 0.002329934 |
| ENSG00000066583 | *ISOC1* | 2.769033518 | -0.138152847 | 0.083158771 | -0.221311618 | 0.999933887 | -0.237037245 | -0.83263441 | 0.595597165 | 0.002357699 |
| ENSG00000123374 | *CDK2* | 2.639281403 | -0.097077101 | 0.069074622 | -0.166151723 | 0.999933887 | 0.50529125 | 1.098018678 | -0.592727429 | 0.002383735 |
| ENSG00000266094 | *RASSF5* | 9.961578567 | 0.022709938 | 0.006430665 | 0.016279274 | 0.999933887 | 0.442366128 | 0.71643687 | -0.274070742 | 0.002383735 |
| ENSG00000160211 | *G6PD* | 6.814452304 | 0.011346899 | 0.005126583 | 0.006220316 | 0.999933887 | -0.667288606 | -1.087587773 | 0.420299167 | 0.002392071 |
| ENSG00000188986 | *NELFB* | 4.148545652 | 0.068294079 | 0.100060453 | -0.031766374 | 0.999933887 | -0.276523665 | -0.73414223 | 0.457618565 | 0.002394411 |
| ENSG00000125733 | *TRIP10* | 2.551926679 | 0.071028259 | -0.082433836 | 0.153462095 | 0.999933887 | 2.354667286 | 3.163407895 | -0.808740609 | 0.002394411 |
| ENSG00000092421 | *SEMA6A* | 0.600104887 | -0.047206097 | 0.378426768 | -0.425632865 | 0.999933887 | 0.63777397 | 1.636308074 | -0.998534105 | 0.002394411 |
| ENSG00000062194 | *GPBP1* | 6.751214588 | -0.080300895 | -0.121577691 | 0.041276796 | 0.999933887 | 0.551872992 | 0.982767261 | -0.430894269 | 0.002397925 |
| ENSG00000008513 | *ST3GAL1* | 7.796866626 | 0.010942557 | 0.000621332 | 0.010321225 | 0.999933887 | 0.916748062 | 1.329552178 | -0.412804116 | 0.002403977 |
| ENSG00000261971 | *MMP25-AS1* | 5.511641175 | -0.028590294 | -0.032033542 | 0.003443248 | 0.999933887 | -0.92362772 | -1.560659627 | 0.637031907 | 0.002418034 |
| ENSG00000133048 | *CHI3L1* | 7.298293087 | 0.07447316 | 0.012835717 | 0.061637443 | 0.999933887 | 1.787883781 | 2.443479662 | -0.65559588 | 0.002444117 |
| ENSG00000115604 | *IL18R1* | 3.944306597 | -0.062059126 | -0.119712524 | 0.057653398 | 0.999933887 | 0.055128983 | 0.490487167 | -0.435358184 | 0.002447732 |
| ENSG00000115840 | *SLC25A12* | 1.890968647 | -0.097724548 | -0.068900312 | -0.028824236 | 0.999933887 | 0.817886687 | 1.544617615 | -0.726730928 | 0.002447732 |
| ENSG00000135476 | *ESPL1* | 2.381463282 | -0.036280769 | -0.054239259 | 0.01795849 | 0.999933887 | 2.333748068 | 3.165769852 | -0.832021784 | 0.002447732 |
| ENSG00000095794 | *CREM* | 5.925942737 | 0.029096978 | -0.067529032 | 0.09662601 | 0.999933887 | 0.438736382 | 0.906444263 | -0.467707881 | 0.002449348 |
| ENSG00000118260 | *CREB1* | 6.037625303 | -0.019437578 | -0.080532162 | 0.061094584 | 0.999933887 | 0.251395108 | 0.682392041 | -0.430996934 | 0.002457077 |
| ENSG00000232713 | *RPS12P3* | 0.299621248 | -0.137212722 | -0.180441551 | 0.043228828 | 0.999933887 | 0.612072476 | 1.671274223 | -1.059201747 | 0.002494288 |
| ENSG00000165097 | *KDM1B* | 5.730215038 | -0.045918313 | -0.01787136 | -0.028046952 | 0.999933887 | 0.505307154 | 0.874465178 | -0.369158024 | 0.002494288 |
| ENSG00000137364 | *TPMT* | 2.959678014 | 0.04057513 | -0.007217559 | 0.047792689 | 0.999933887 | 1.024746576 | 1.555289262 | -0.530542686 | 0.002494288 |
| ENSG00000166189 | *HPS6* | 2.755354464 | 0.043906145 | 0.077237947 | -0.033331801 | 0.999933887 | -0.464453677 | -1.023353598 | 0.558899922 | 0.002545761 |
| ENSG00000163964 | *PIGX* | 4.874886232 | 0.011288634 | 0.033529869 | -0.022241235 | 0.999933887 | -0.66474325 | -1.223278077 | 0.558534828 | 0.002546999 |
| ENSG00000178149 | *DALRD3* | 2.935961504 | 0.084829019 | -0.058414468 | 0.143243487 | 0.999933887 | -0.227892376 | -0.660299131 | 0.432406754 | 0.002551886 |
| ENSG00000092094 | *OSGEP* | 3.993841008 | -0.114124466 | 0.057205296 | -0.171329762 | 0.999933887 | -0.347597496 | -0.77540424 | 0.427806744 | 0.002564032 |
| ENSG00000160209 | *PDXK* | 7.056255017 | -0.033226711 | -0.034768376 | 0.001541665 | 0.999933887 | 1.253242723 | 1.678335101 | -0.425092378 | 0.002564032 |
| ENSG00000160602 | *NEK8* | 0.975743057 | 0.040079949 | -0.047853426 | 0.087933375 | 0.999933887 | 0.387880868 | 1.112954394 | -0.725073526 | 0.002564032 |
| ENSG00000206562 | *METTL6* | 2.106586494 | 0.029820063 | -0.280162645 | 0.309982708 | 0.999933887 | 0.970812388 | 1.418153252 | -0.447340864 | 0.002564032 |
| ENSG00000124508 | *BTN2A2* | 5.712504717 | 0.037384723 | 0.026732331 | 0.010652392 | 0.999933887 | 2.008161977 | 2.479059055 | -0.470897078 | 0.002580204 |
| ENSG00000159840 | *ZYX* | 8.790142027 | 0.017056816 | 0.018616718 | -0.001559902 | 0.999933887 | -0.590649411 | -0.940623749 | 0.349974338 | 0.002592451 |
| ENSG00000139192 | *TAPBPL* | 4.596463977 | 0.043448015 | -0.002827944 | 0.046275959 | 0.999933887 | -0.312975242 | -0.754779517 | 0.441804275 | 0.002606478 |
| ENSG00000133313 | *CNDP2* | 4.608996209 | 0.022401523 | -0.05800886 | 0.080410383 | 0.999933887 | -0.253418521 | -0.591840123 | 0.338421602 | 0.002617961 |
| ENSG00000213918 | *DNASE1* | 3.599285017 | 0.013305263 | 0.056268572 | -0.042963309 | 0.999933887 | -0.191752147 | -0.568086273 | 0.376334126 | 0.002632533 |
| ENSG00000118564 | *FBXL5* | 8.633766787 | -0.064202114 | -0.123585657 | 0.059383543 | 0.999933887 | -0.092244007 | 0.215494442 | -0.307738449 | 0.002632813 |
| ENSG00000146802 | *TMEM168* | 3.201490893 | -0.133095364 | -0.148639298 | 0.015543934 | 0.999933887 | 0.32454347 | 0.826422399 | -0.501878928 | 0.002648581 |
| ENSG00000056586 | *RC3H2* | 5.353354228 | -0.005702929 | -0.048992279 | 0.04328935 | 0.999933887 | -0.259572243 | 0.137624426 | -0.397196669 | 0.002672593 |
| ENSG00000264364 | *DYNLL2* | 5.602166541 | 0.026052106 | -0.00544764 | 0.031499746 | 0.999933887 | -0.157730073 | -0.497280085 | 0.339550012 | 0.00268074 |
| ENSG00000155629 | *PIK3AP1* | 9.084389879 | -0.028339643 | -0.009231798 | -0.019107845 | 0.999933887 | 2.063245188 | 2.617862778 | -0.55461759 | 0.002684578 |
| ENSG00000148331 | *ASB6* | 3.84151362 | -0.016636622 | -0.041763586 | 0.025126964 | 0.999933887 | -0.156243358 | -0.529227845 | 0.372984487 | 0.002684578 |
| ENSG00000065613 | *SLK* | 6.710355186 | -0.113524791 | -0.142769087 | 0.029244296 | 0.999933887 | 0.91109654 | 1.704582594 | -0.793486054 | 0.002698269 |
| ENSG00000163932 | *PRKCD* | 9.270942921 | -0.033901815 | -0.017170908 | -0.016730907 | 0.999933887 | 0.597591874 | 0.916654917 | -0.319063043 | 0.002701117 |
| ENSG00000166188 | *ZNF319* | 4.129823931 | 0.108138889 | 0.033630597 | 0.074508293 | 0.999933887 | -0.623621009 | -1.08359353 | 0.459972521 | 0.002712351 |
| ENSG00000203880 | *PCMTD2* | 4.362727894 | -0.056751708 | -0.076231235 | 0.019479527 | 0.999933887 | -0.283602314 | -0.647479749 | 0.363877436 | 0.002712351 |
| ENSG00000090376 | *IRAK3* | 7.08762894 | -0.109851147 | -0.088190348 | -0.021660799 | 0.999933887 | 1.462172614 | 2.085982742 | -0.623810128 | 0.002712351 |
| ENSG00000137221 | *TJAP1* | 5.307584713 | 0.034370377 | 0.022032165 | 0.012338212 | 0.999933887 | 0.880159243 | 1.182856853 | -0.302697609 | 0.002712351 |
| ENSG00000185989 | *RASA3* | 6.326004302 | 0.02276308 | -0.007824158 | 0.030587238 | 0.999933887 | -0.193152516 | -0.442968951 | 0.249816435 | 0.002712351 |
| ENSG00000115738 | *ID2* | 7.060543458 | 0.011864998 | 0.093869521 | -0.082004522 | 0.999933887 | 1.898915125 | 2.670421608 | -0.771506483 | 0.002729525 |
| ENSG00000112679 | *DUSP22* | 5.039557682 | 0.033037801 | 0.024506447 | 0.008531353 | 0.999933887 | -0.290657429 | -0.633342924 | 0.342785495 | 0.002746223 |
| ENSG00000105483 | *CARD8* | 7.231027526 | -0.032960024 | -0.05330916 | 0.020349136 | 0.999933887 | -1.027603823 | -1.460434015 | 0.432830192 | 0.002746223 |
| ENSG00000100591 | *AHSA1* | 3.724631672 | -0.026611221 | 0.008534942 | -0.035146163 | 0.999933887 | -0.170148821 | -0.520793799 | 0.350644978 | 0.002754402 |
| ENSG00000173825 | *TIGD3* | 1.957627318 | -0.050490603 | -0.004024942 | -0.046465661 | 0.999933887 | -1.512773431 | -2.567321881 | 1.05454845 | 0.002780513 |
| ENSG00000133612 | *AGAP3* | 7.041055357 | 0.002562884 | -0.010903673 | 0.013466557 | 0.999933887 | 0.657611621 | 1.000854662 | -0.343243041 | 0.002780513 |
| ENSG00000182446 | *NPLOC4* | 6.591044263 | 0.021428801 | -0.024278539 | 0.045707341 | 0.999933887 | 0.686240662 | 1.041123049 | -0.354882387 | 0.002785955 |
| ENSG00000204396 | *VWA7* | 0.454746738 | -0.413431864 | -0.136013284 | -0.27741858 | 0.999933887 | -0.364773797 | -1.273658951 | 0.908885154 | 0.002793785 |
| ENSG00000101187 | *SLCO4A1* | 4.093211452 | 0.113688839 | 0.153083988 | -0.039395149 | 0.999933887 | 0.978266326 | 1.489573685 | -0.511307359 | 0.002793785 |
| ENSG00000100024 | *UPB1* | 5.38810352 | -0.033903264 | -0.042398388 | 0.008495123 | 0.999933887 | 1.719014537 | 2.151491105 | -0.432476568 | 0.002793785 |
| ENSG00000215041 | *NEURL4* | 3.178217245 | 0.114275605 | -0.017132127 | 0.131407732 | 0.999933887 | 0.044001523 | -0.388557283 | 0.432558806 | 0.002806915 |
| ENSG00000160593 | *JAML* | 8.583024809 | -0.064255283 | -0.044343219 | -0.019912064 | 0.999933887 | -0.285417048 | -0.706663836 | 0.421246788 | 0.002806915 |
| ENSG00000078269 | *SYNJ2* | 4.466846953 | 0.036701493 | 0.048406484 | -0.011704991 | 0.999933887 | 1.715611824 | 2.370756344 | -0.65514452 | 0.002806915 |
| ENSG00000154822 | *PLCL2* | 5.178191039 | -0.024172011 | -0.039577334 | 0.015405323 | 0.999933887 | 0.204718951 | 0.697552665 | -0.492833714 | 0.002886956 |
| ENSG00000271856 | *LINC01215* | 4.957943114 | -0.017005053 | -0.024492749 | 0.007487695 | 0.999933887 | 1.040497298 | 1.495126879 | -0.454629581 | 0.002886956 |
| ENSG00000130402 | *ACTN4* | 7.554141173 | 0.054999819 | 0.009640658 | 0.045359161 | 0.999933887 | -0.331311565 | -0.677221251 | 0.345909686 | 0.002888658 |
| ENSG00000118217 | *ATF6* | 5.161307879 | 0.020329401 | -0.059750836 | 0.080080237 | 0.999933887 | 0.266660409 | 0.629129001 | -0.362468593 | 0.002898289 |
| ENSG00000072110 | *ACTN1* | 8.613777521 | 0.03723982 | 0.011204435 | 0.026035385 | 0.999933887 | -0.646361342 | -0.963597903 | 0.317236561 | 0.002906618 |
| ENSG00000103769 | *RAB11A* | 5.851917115 | -0.025978462 | 0.004799202 | -0.030777663 | 0.999933887 | 0.338164488 | 0.666725785 | -0.328561298 | 0.002906618 |
| ENSG00000070081 | *NUCB2* | 3.000649652 | -0.180351207 | -0.212883365 | 0.032532158 | 0.999933887 | -0.009387797 | 0.678264262 | -0.687652058 | 0.00290939 |
| ENSG00000154642 | *C21orf91* | 3.842250963 | -0.074562692 | -0.008271299 | -0.066291393 | 0.999933887 | 0.397970982 | 1.03188628 | -0.633915298 | 0.002940265 |
| ENSG00000188186 | *LAMTOR4* | 6.434991117 | -0.005298481 | 0.001762322 | -0.007060802 | 0.999933887 | -0.211772971 | -0.600557219 | 0.388784248 | 0.002968487 |
| ENSG00000135269 | *TES* | 5.089620949 | -0.00682557 | -0.038205 | 0.03137943 | 0.999933887 | 0.504052215 | 0.931774202 | -0.427721987 | 0.002977149 |
| ENSG00000102886 | *GDPD3* | 2.299829629 | 0.111139961 | 0.130704226 | -0.019564265 | 0.999933887 | -0.844810002 | -1.470506 | 0.625695997 | 0.002977589 |
| ENSG00000147408 | *CSGALNACT1* | 4.797858225 | 0.006213115 | -0.015581149 | 0.021794264 | 0.999933887 | 0.401673126 | 0.836522243 | -0.434849117 | 0.002987845 |
| ENSG00000197467 | *COL13A1* | 0.763487812 | -0.153074966 | -0.012745611 | -0.140329355 | 0.999933887 | 1.176948209 | 1.945655645 | -0.768707437 | 0.002995387 |
| ENSG00000121858 | *TNFSF10* | 4.619030921 | -0.033122707 | -0.113683054 | 0.080560347 | 0.999933887 | -0.582761117 | -1.243798343 | 0.661037227 | 0.003040071 |
| ENSG00000178127 | *NDUFV2* | 1.709878155 | -0.074690731 | 0.003440681 | -0.078131412 | 0.999933887 | 0.634286374 | 1.366922359 | -0.732635985 | 0.003059272 |
| ENSG00000100077 | *GRK3* | 3.876261178 | -0.037920486 | 0.090502284 | -0.12842277 | 0.999933887 | 0.276887705 | 0.707388804 | -0.430501099 | 0.003071059 |
| ENSG00000141096 | *DPEP3* | 2.671454211 | -0.007766466 | -0.037632852 | 0.029866386 | 0.999933887 | -1.353168404 | -2.048699102 | 0.695530698 | 0.003080084 |
| ENSG00000160948 | *VPS28* | 5.874456191 | 0.010584045 | 0.068292243 | -0.057708198 | 0.999933887 | -0.122094578 | -0.485779459 | 0.36368488 | 0.003097567 |
| ENSG00000275023 | *MLLT6* | 8.40973129 | 0.073291844 | 0.006825311 | 0.066466532 | 0.999933887 | 0.818478099 | 1.193393885 | -0.374915786 | 0.00311743 |
| ENSG00000138433 | *CIR1* | 8.27537217 | -0.065367651 | -0.165627261 | 0.10025961 | 0.999933887 | 0.397014631 | 0.774458044 | -0.377443413 | 0.003165265 |
| ENSG00000103642 | *LACTB* | 5.475542665 | -0.0531549 | 0.008622308 | -0.061777208 | 0.999933887 | 0.261726119 | 0.56072236 | -0.298996241 | 0.003176267 |
| ENSG00000096070 | *BRPF3* | 6.501035093 | 0.012494675 | -0.021748329 | 0.034243004 | 0.999933887 | 0.751801616 | 1.093676909 | -0.341875293 | 0.003192123 |
| ENSG00000114796 | *KLHL24* | 7.170897649 | -0.043724696 | -0.113506495 | 0.0697818 | 0.999933887 | -0.281756457 | 0.01450277 | -0.296259227 | 0.003221368 |
| ENSG00000197380 | *DACT3* | -1.193473833 | 0.072252194 | 0.399155029 | -0.326902835 | 0.999933887 | 1.3165085 | 2.905758237 | -1.589249737 | 0.003221368 |
| ENSG00000178252 | *WDR6* | 5.075886236 | 0.01572241 | 0.014607375 | 0.001115034 | 0.999933887 | -0.063414644 | -0.362846932 | 0.299432288 | 0.003232836 |
| ENSG00000113163 | *CERT1* | 6.343021297 | -0.017651804 | -0.094503903 | 0.076852098 | 0.999933887 | 0.385857939 | 0.798932882 | -0.413074943 | 0.0032334 |
| ENSG00000165732 | *DDX21* | 6.978354673 | -0.01303621 | -0.120825888 | 0.107789678 | 0.999933887 | 0.656034543 | 1.168429553 | -0.51239501 | 0.003248952 |
| ENSG00000155329 | *ZCCHC10* | 2.957910642 | -0.139469138 | -0.009393906 | -0.130075232 | 0.999933887 | 0.615212194 | 1.020348128 | -0.405135934 | 0.003254615 |
| ENSG00000143067 | *ZNF697* | 3.057995518 | 0.058195358 | -0.08059986 | 0.138795218 | 0.999933887 | 0.792876165 | 1.3328386 | -0.539962435 | 0.003254615 |
| ENSG00000104365 | *IKBKB* | 6.754770093 | -0.006032322 | -0.034892098 | 0.028859776 | 0.999933887 | 0.639318151 | 0.951773271 | -0.312455119 | 0.003254615 |
| ENSG00000173801 | *JUP* | 2.716297688 | -0.022078172 | 0.060050328 | -0.0821285 | 0.999933887 | 0.073372832 | 0.58065293 | -0.507280098 | 0.003260597 |
| ENSG00000111144 | *LTA4H* | 5.674792975 | -0.079206354 | -0.093469441 | 0.014263087 | 0.999933887 | -0.546439098 | -0.976553341 | 0.430114242 | 0.003305158 |
| ENSG00000080345 | *RIF1* | 4.845051105 | -0.102258899 | -0.093815265 | -0.008443635 | 0.999933887 | 0.419699241 | 1.300946769 | -0.881247528 | 0.003305158 |
| ENSG00000132388 | *UBE2G1* | 4.957725699 | -0.033190513 | 0.040701246 | -0.073891759 | 0.999933887 | -0.706390996 | -1.109323293 | 0.402932297 | 0.003305158 |
| ENSG00000139636 | *LMBR1L* | 5.604191905 | -0.005821937 | 0.004587629 | -0.010409566 | 0.999933887 | -0.132107108 | -0.391469493 | 0.259362385 | 0.003305158 |
| ENSG00000162927 | *PUS10* | 1.923272167 | -0.120393565 | -0.215949266 | 0.095555701 | 0.999933887 | 0.511981395 | 1.446347168 | -0.934365772 | 0.003324819 |
| ENSG00000253167 | *WASHC5-AS1* | 0.363966404 | 0.196889281 | 0.000755321 | 0.19613396 | 0.999933887 | -0.143662675 | 0.636176088 | -0.779838763 | 0.003333652 |
| ENSG00000068366 | *ACSL4* | 5.59638782 | -0.099485818 | -0.133516279 | 0.034030462 | 0.999933887 | 0.165783074 | 0.856668519 | -0.690885445 | 0.003333652 |
| ENSG00000114648 | *KLHL18* | 4.698582137 | 0.01182571 | -0.002845558 | 0.014671268 | 0.999933887 | -0.309832624 | -0.679101443 | 0.369268819 | 0.003333652 |
| ENSG00000168002 | *POLR2G* | 3.836999916 | 0.001551403 | 0.033751502 | -0.032200099 | 0.999933887 | -0.126766252 | -0.514486662 | 0.38772041 | 0.003333652 |
| ENSG00000114388 | *NPRL2* | 3.365776518 | 0.028387123 | -0.028130939 | 0.056518062 | 0.999933887 | -0.193973414 | -0.589351858 | 0.395378444 | 0.003341921 |
| ENSG00000135124 | *P2RX4* | 5.847521242 | 0.000287346 | -0.053285032 | 0.053572378 | 0.999933887 | 1.055208232 | 1.482963237 | -0.427755004 | 0.003341921 |
| ENSG00000152291 | *TGOLN2* | 7.950654793 | -0.034978369 | -0.027632514 | -0.007345854 | 0.999933887 | -0.448969617 | -0.700380856 | 0.251411239 | 0.003354118 |
| ENSG00000122042 | *UBL3* | 6.168465714 | -0.043256924 | -0.056891521 | 0.013634598 | 0.999933887 | 0.112907707 | 0.423689139 | -0.310781433 | 0.003354118 |

| | | | | | | | | | | |
|---|---|---|---|---|---|---|---|---|---|---|
| ENSG00000134352 | *IL6ST* | 5.132124922 | -0.041119672 | -0.212383357 | 0.171263685 | 0.999933887 | 0.191960766 | 0.959876934 | -0.767916168 | 0.003354118 |
| ENSG00000035403 | *VCL* | 5.590052861 | 0.008567832 | -0.052107086 | 0.060674918 | 0.999933887 | 0.217201353 | 0.513499747 | -0.296298394 | 0.003354118 |
| ENSG00000253256 | *AC134043.2* | -0.390743575 | 0.018032636 | 0.191306471 | -0.173273835 | 0.999933887 | 0.572276871 | 1.409781174 | -0.837504304 | 0.003354118 |
| ENSG00000138600 | *SPPL2A* | 6.255438196 | -0.041055486 | -0.052152663 | 0.011097177 | 0.999933887 | 0.741317454 | 1.10603271 | -0.364715256 | 0.00335678 |
| ENSG00000168438 | *CDC40* | 3.741498982 | -0.084566015 | -0.013359504 | -0.071206511 | 0.999933887 | -0.61641209 | -1.091365564 | 0.474953474 | 0.00339547 |
| ENSG00000164142 | *FAM160A1* | 0.212239124 | -0.01797775 | 0.084684194 | -0.102661944 | 0.999933887 | 1.335667954 | 2.489065458 | -1.153397504 | 0.003397039 |
| ENSG00000176619 | *LMNB2* | 3.384593781 | 0.138846783 | 0.021949729 | 0.116897054 | 0.999933887 | 1.008870369 | 1.658325873 | -0.649455504 | 0.003422121 |
| ENSG00000088888 | *MAVS* | 4.804922006 | -0.00439618 | 0.009784215 | -0.014180396 | 0.999933887 | -0.205763358 | -0.581770556 | 0.376007199 | 0.003422121 |
| ENSG00000138760 | *SCARB2* | 3.176147309 | -0.010322383 | -0.0063092 | -0.004013183 | 0.999933887 | -0.482817267 | -0.982895718 | 0.500078452 | 0.003477086 |
| ENSG00000144283 | *PKP4* | 3.580449543 | -0.031581971 | 0.089436059 | -0.12101803 | 0.999933887 | -0.887849375 | -1.371557804 | 0.483708429 | 0.003477624 |
| ENSG00000033800 | *PIAS1* | 6.778675487 | -0.029146011 | -0.039512698 | 0.010366687 | 0.999933887 | 0.078456966 | 0.465827764 | -0.387370798 | 0.00350511 |
| ENSG00000142751 | *GPN2* | 3.604075023 | -0.060285754 | 0.131968099 | -0.192253853 | 0.944094537 | -0.146047919 | -0.453790895 | 0.307742977 | 0.003507203 |
| ENSG00000115091 | *ACTR3* | 9.497681864 | -0.016939671 | -0.004535351 | -0.01240432 | 0.999933887 | 0.744096652 | 1.016079013 | -0.271982361 | 0.003507203 |
| ENSG00000110756 | *HPS5* | 5.660160961 | -0.015438068 | -0.024960759 | 0.009522691 | 0.999933887 | 0.535529523 | 0.94244861 | -0.406919087 | 0.003520319 |
| ENSG00000286281 | *AL353152.2* | 1.380117748 | -0.0526744 | 0.381520884 | -0.434195283 | 0.999933887 | 0.525299865 | 1.118843439 | -0.593543575 | 0.003527351 |
| ENSG00000154310 | *TNIK* | 4.472668381 | -0.02589935 | -0.023064169 | -0.002835181 | 0.999933887 | 0.01825107 | 0.515539388 | -0.497288318 | 0.003530991 |
| ENSG00000234616 | *JRK* | 3.054569016 | -0.088559343 | -0.001256303 | -0.08730304 | 0.999933887 | -0.188885187 | -0.545397507 | 0.356512319 | 0.003548404 |
| ENSG00000160446 | *ZDHHC12* | 3.830894813 | 0.095166956 | -0.03379384 | 0.128960797 | 0.999933887 | -0.467119713 | -1.017273593 | 0.55015388 | 0.003548404 |
| ENSG00000151726 | *ACSL1* | 10.068648973 | -0.061427846 | -0.051561798 | -0.009866048 | 0.999933887 | 1.739207772 | 2.242398127 | -0.503190356 | 0.003548404 |
| ENSG00000186088 | *GSAP* | 5.36394477 | -0.025236702 | -0.003094197 | -0.022142505 | 0.999933887 | 1.081895612 | 1.611017854 | -0.529122243 | 0.003548404 |
| ENSG00000126561 | *STAT5A* | 8.046770933 | -0.028086008 | 0.004867381 | -0.03295339 | 0.999933887 | 2.540926236 | 3.164184309 | -0.623258073 | 0.00355842 |
| ENSG00000184640 | *SEPTIN9* | 7.773937361 | 0.012953249 | -0.018900762 | 0.031854011 | 0.999933887 | -0.303133668 | -0.665599737 | 0.362466069 | 0.003589577 |
| ENSG00000252172 | *RNU6-720P* | 0.092766874 | -0.406520971 | 0.488079678 | -0.894600649 | 0.944094537 | -0.484273349 | 0.189759258 | -0.674032606 | 0.003601601 |
| ENSG00000173465 | *ZNRD2* | 2.968275222 | -0.021048251 | -0.063113503 | 0.042065251 | 0.999933887 | 0.524675721 | 1.001415574 | -0.476739853 | 0.003601601 |
| ENSG00000121552 | *CSTA* | 4.267479694 | 0.002654431 | 0.0322774 | -0.029622969 | 0.999933887 | 0.628914949 | 1.066158298 | -0.437243349 | 0.003652967 |
| ENSG00000258926 | *AL355916.2* | 2.935949204 | 0.030359452 | -0.001884067 | 0.032243519 | 0.999933887 | 0.957629468 | 1.687258534 | -0.729629066 | 0.003668712 |
| ENSG00000112773 | *TENT5A* | 3.868766283 | -0.065597842 | -0.07544464 | 0.009846798 | 0.999933887 | -0.620939851 | -1.082991046 | 0.462051196 | 0.003675416 |
| ENSG00000196663 | *TECPR2* | 8.165887867 | -0.017928469 | -0.018410393 | 0.000481923 | 0.999933887 | 0.779724339 | 1.081531458 | -0.301807119 | 0.003675416 |
| ENSG00000065413 | *ANKRD44* | 7.109570287 | -0.032583703 | -0.053911519 | 0.021327816 | 0.999933887 | -0.477124701 | -0.980648931 | 0.50352423 | 0.00369443 |
| ENSG00000158773 | *USF1* | 6.062661124 | -0.041874777 | -0.002393976 | -0.039480801 | 0.999933887 | -0.422877948 | -0.773367211 | 0.350489263 | 0.003745502 |
| ENSG00000176597 | *B3GNT5* | 7.391567201 | 0.036345042 | -0.031855029 | 0.068200071 | 0.999933887 | 0.261578409 | 0.72214835 | -0.460569941 | 0.003745502 |
| ENSG00000065526 | *SPEN* | 6.913763594 | -0.033829708 | -0.061404342 | 0.027574634 | 0.999933887 | -0.860188755 | -1.307759648 | 0.447570892 | 0.003748876 |
| ENSG00000104093 | *DMXL2* | 7.115953014 | -0.166292375 | -0.144452313 | -0.021840062 | 0.999933887 | 1.793565171 | 2.763992888 | -0.970427717 | 0.003762073 |
| ENSG00000163444 | *TMEM183A* | 4.730113702 | 0.000413074 | 0.007900736 | -0.007487662 | 0.999933887 | -0.248013912 | -0.556390926 | 0.308377014 | 0.003762073 |
| ENSG00000131899 | *LLGL1* | 3.125475589 | 0.030769752 | -0.003366149 | 0.0341359 | 0.999933887 | -0.019131111 | -0.387669327 | 0.368538217 | 0.003765578 |
| ENSG00000167264 | *DUS2* | 3.472864256 | -0.134978211 | -0.016552567 | -0.118425643 | 0.999933887 | -1.691043981 | -2.380853979 | 0.689809998 | 0.003818742 |
| ENSG00000142046 | *TMEM91* | 3.430651765 | 0.055121604 | 0.03969476 | 0.015426844 | 0.999933887 | 0.048527165 | -0.456729106 | 0.505256271 | 0.003818742 |
| ENSG00000154153 | *RETREG1* | 3.045126496 | -0.096098633 | -0.006397073 | -0.08970156 | 0.999933887 | 0.524573059 | 1.080640489 | -0.55606743 | 0.003839672 |
| ENSG00000111371 | *SLC38A1* | 6.985376936 | -0.066231091 | -0.099041068 | 0.032809977 | 0.999933887 | 0.076549323 | 0.583517323 | -0.506967999 | 0.003884438 |
| ENSG00000163539 | *CLASP2* | 3.570597366 | -0.066273293 | -0.001564242 | -0.064709051 | 0.999933887 | 0.146597484 | 0.680268622 | -0.533671138 | 0.003888643 |
| ENSG00000159556 | *ISL2* | 0.587985461 | -0.176985224 | 0.290239474 | -0.467224698 | 0.944094537 | -0.986376159 | -2.088896472 | 1.102520314 | 0.003906181 |
| ENSG00000179163 | *FUCA1* | 3.328650257 | -0.201737404 | 0.028779466 | -0.230516869 | 0.999933887 | -0.757927105 | -1.25424809 | 0.496320984 | 0.003910623 |
| ENSG00000136044 | *APPL2* | 5.0402989 | -0.075619822 | -0.039592297 | -0.036027525 | 0.999933887 | -0.568732858 | -0.906048438 | 0.337315581 | 0.003910623 |
| ENSG00000153048 | *CARHSP1* | 3.003723714 | 0.089445724 | 0.048796532 | 0.040649192 | 0.999933887 | -0.139235879 | -0.675678001 | 0.536442122 | 0.003910623 |
| ENSG00000197948 | *FCHSD1* | 5.986270261 | 0.028791371 | 0.107656866 | -0.078865495 | 0.999933887 | -0.317715367 | -0.66094634 | 0.343230973 | 0.003910623 |
| ENSG00000100612 | *DHRS7* | 6.150034897 | -0.030013777 | -0.094386588 | 0.064372811 | 0.999933887 | -1.024403615 | -1.458835851 | 0.434432237 | 0.003910623 |
| ENSG00000142327 | *RNPEPL1* | 6.492476826 | -0.015655563 | 0.009361198 | -0.025016761 | 0.999933887 | -0.475347543 | -0.826474284 | 0.351126741 | 0.003910623 |
| ENSG00000067992 | *PDK3* | 5.215305824 | -0.091912754 | -0.09192784 | 1.5085e-05 | 0.999933887 | -1.322713757 | -1.808945742 | 0.486231985 | 0.003917996 |
| ENSG00000171368 | *TPPP* | 2.237900123 | -0.101150927 | -0.022793786 | -0.078357141 | 0.999933887 | 1.036882267 | 2.148356758 | -1.111474491 | 0.003927935 |
| ENSG00000255823 | *MTRNR2L8* | 1.76121405 | -0.19908938 | -0.122806857 | -0.076282523 | 0.999933887 | 2.036025584 | 2.829976688 | -0.793951103 | 0.003940125 |
| ENSG00000132589 | *FLOT2* | 7.777917681 | 0.023389402 | -0.046249787 | 0.06963919 | 0.999933887 | 0.419295446 | 0.7762401 | -0.356944655 | 0.003966922 |
| ENSG00000104450 | *SPAG1* | 2.417431086 | -0.00974323 | 0.027593931 | -0.037337161 | 0.999933887 | 0.574549554 | 1.233261211 | -0.658711656 | 0.003966922 |
| ENSG00000268581 | *SIGLEC18P* | -1.192952733 | 0.135592842 | -0.082993914 | 0.218586756 | 0.999933887 | 1.225787282 | 2.091602133 | -0.86581485 | 0.003990503 |
| ENSG00000115325 | *DOK1* | 4.17051304 | 0.015960695 | 0.008617542 | 0.007343153 | 0.999933887 | -0.68774336 | -1.170507141 | 0.482763781 | 0.003999376 |
| ENSG00000166670 | *MMP10* | 0.244724504 | 0.244365067 | 0.025311803 | 0.219053264 | 0.999933887 | 1.086922871 | 2.180863596 | -1.093940725 | 0.004012324 |
| ENSG00000188483 | *IER5L* | 2.012554191 | -0.226874126 | -0.133483449 | -0.093390678 | 0.999933887 | -1.463953544 | -2.238153769 | 0.774200225 | 0.004030639 |
| ENSG00000159055 | *MIS18A* | 0.059401798 | 0.076541772 | 0.07703551 | -0.000493738 | 0.999933887 | 1.051614328 | 2.093900411 | -1.042286083 | 0.004030639 |
| ENSG00000163625 | *WDFY3* | 7.528504268 | -0.034786371 | -0.0603605 | 0.025574129 | 0.999933887 | 0.566372105 | 0.963652467 | -0.397280362 | 0.004054681 |
| ENSG00000188921 | *HACD4* | 4.641438929 | 0.079396831 | -0.048243659 | 0.12764049 | 0.999933887 | -0.264116932 | -0.760208373 | 0.49609144 | 0.004056109 |
| ENSG00000139146 | *SINHCAF* | 3.611462925 | -0.067886744 | 0.016128765 | -0.084015508 | 0.999933887 | 0.132495295 | 0.477273128 | -0.344777833 | 0.00406207 |
| ENSG00000137486 | *ARRB1* | 5.343553556 | -0.014590191 | -0.00429512 | -0.010295071 | 0.999933887 | -0.866264838 | -1.336572911 | 0.470308073 | 0.00406207 |
| ENSG00000114268 | *PFKFB4* | 4.440363198 | -0.042253388 | 0.038297665 | -0.080551053 | 0.999933887 | -0.328299458 | -0.961046495 | 0.632747037 | 0.004077578 |
| ENSG00000131100 | *ATP6V1E1* | 7.064348553 | -0.033403212 | -0.051873715 | 0.018470503 | 0.999933887 | 0.157149539 | 0.42973883 | -0.272589291 | 0.004088839 |
| ENSG00000198189 | *HSD17B11* | 6.348771611 | -0.06404765 | -0.079305914 | 0.015258263 | 0.999933887 | -0.481993316 | -0.935804481 | 0.453811166 | 0.004091068 |
| ENSG00000115234 | *SNX17* | 5.046035121 | 0.060397094 | 0.031490657 | 0.028906437 | 0.999933887 | -0.166106617 | -0.498674125 | 0.332567508 | 0.004091108 |
| ENSG00000142765 | *SYTL1* | 5.983392265 | 0.043349882 | 0.060517577 | -0.017167694 | 0.999933887 | -0.009747115 | -0.338940648 | 0.329193533 | 0.004102848 |
| ENSG00000167703 | *SLC43A2* | 9.386692836 | 0.030732194 | 0.007083172 | 0.023649022 | 0.999933887 | 0.392319238 | 0.668043361 | -0.275724123 | 0.00413189 |
| ENSG00000092964 | *DPYSL2* | 1.516399616 | -0.081106434 | -0.05257484 | -0.028531594 | 0.999933887 | -0.199799268 | -0.81111552 | 0.611316251 | 0.004137556 |
| ENSG00000197208 | *SLC22A4* | 4.683140869 | 0.062518543 | 0.0028196 | 0.059698942 | 0.999933887 | -0.517467547 | -0.891277953 | 0.373810406 | 0.004141236 |
| ENSG00000110660 | *SLC35F2* | 1.950890288 | 0.153761983 | -0.087485099 | 0.241247082 | 0.999933887 | 1.64940502 | 2.24336956 | -0.59396454 | 0.004141236 |
| ENSG00000160213 | *CSTB* | 7.738179036 | 0.114107384 | 0.051101105 | 0.063006278 | 0.999933887 | 0.955346839 | 1.325025608 | -0.369678769 | 0.004174658 |
| ENSG00000108510 | *MED13* | 7.115606893 | -0.03125877 | -0.078293178 | 0.047034408 | 0.999933887 | 0.076062433 | 0.524697804 | -0.448635371 | 0.004179127 |
| ENSG00000265808 | *SEC22B* | 7.664608061 | -0.041581753 | -0.096160434 | 0.05457868 | 0.999933887 | 0.579136989 | 0.886647462 | -0.307510473 | 0.004196175 |
| ENSG00000263956 | *NBPF11* | 2.608090504 | -0.028086504 | -0.109324031 | 0.081237527 | 0.999933887 | 0.174767289 | 0.546358298 | -0.371591009 | 0.004196175 |
| ENSG00000064012 | *CASP8* | 7.594893196 | -0.097866557 | -0.018276108 | -0.07959045 | 0.999933887 | -0.11313839 | -0.37227454 | 0.25913615 | 0.004248238 |
| ENSG00000166949 | *SMAD3* | 5.458869924 | -0.042584509 | -0.005281068 | -0.037303441 | 0.999933887 | 0.320188827 | 0.693702865 | -0.373514038 | 0.004250116 |
| ENSG00000169504 | *CLIC4* | 4.510078149 | -0.055897519 | -0.161506464 | 0.105608945 | 0.999933887 | 3.977491053 | 4.861878502 | -0.884387449 | 0.004250116 |
| ENSG00000124813 | *RUNX2* | 4.055972266 | 0.005427641 | 0.045927809 | -0.040500168 | 0.999933887 | -0.468720616 | -1.018654057 | 0.549933441 | 0.004250116 |
| ENSG00000126003 | *PLAGL2* | 6.139464567 | -0.093536277 | -0.063655651 | -0.029880626 | 0.999933887 | 1.192170031 | 1.618603005 | -0.426432974 | 0.00427278 |
| ENSG00000137496 | *IL18BP* | 6.635925811 | 0.085186088 | 0.107141144 | -0.021955056 | 0.999933887 | 0.96717305 | 1.288500838 | -0.321327788 | 0.004297179 |
| ENSG00000099785 | *MARCHF2* | 4.2917905 | 0.008016814 | -0.001471297 | 0.009488111 | 0.999933887 | -0.546377095 | -0.963721227 | 0.417344132 | 0.004297179 |
| ENSG00000164880 | *INTS1* | 5.872242565 | 0.05663493 | 0.080148389 | -0.023513459 | 0.999933887 | -0.159006709 | -0.483690572 | 0.324683863 | 0.004304393 |
| ENSG00000140044 | *JDP2* | 4.48773761 | 0.026874471 | 0.056237175 | -0.029362703 | 0.999933887 | -0.980221363 | -1.601571592 | 0.621350229 | 0.004304393 |
| ENSG00000116752 | *BCAS2* | 4.77376395 | -0.021440885 | -0.108810876 | 0.087369991 | 0.999933887 | -0.048579552 | 0.349243503 | -0.397823055 | 0.004310283 |
| ENSG00000144843 | *ADPRH* | 1.892505495 | -0.141071941 | 0.00040303 | -0.141474972 | 0.999933887 | 0.669709639 | 1.196083552 | -0.526373914 | 0.004353579 |
| ENSG00000163399 | *ATP1A1* | 7.030357997 | -0.009724812 | -0.00335873 | -0.006366081 | 0.999933887 | 0.379876411 | 0.679011034 | -0.299134623 | 0.004353579 |
| ENSG00000119508 | *NR4A3* | 7.634489989 | 0.107843938 | 0.051490008 | 0.056353931 | 0.999933887 | 1.347395886 | 2.0845948 | -0.737198914 | 0.004370285 |
| ENSG00000120889 | *TNFRSF10B* | 6.237771822 | -0.016734895 | 0.052416505 | -0.0691514 | 0.999933887 | 0.058645806 | -0.314190666 | 0.372836472 | 0.004374451 |
| ENSG00000216864 | *GLT8D1* | 2.388476932 | -0.093258683 | -0.156709066 | 0.063450383 | 0.999933887 | -0.152731674 | -0.525334853 | 0.372603179 | 0.004375551 |
| ENSG00000103150 | *MLYCD* | 2.481426595 | 0.033698574 | 0.08310556 | -0.049406986 | 0.999933887 | -0.150284989 | -0.62660356 | 0.476318571 | 0.004375551 |
| ENSG00000152684 | *PELO* | 4.171883559 | -0.213959925 | -0.242257023 | 0.028297098 | 0.999933887 | -1.052612467 | -1.585578293 | 0.532965826 | 0.004390737 |
| ENSG00000164062 | *APEH* | 4.137031787 | 0.053189145 | -0.103094772 | 0.156283917 | 0.999933887 | -0.089372063 | -0.416364729 | 0.326992666 | 0.004460469 |
| ENSG00000100296 | *THOC5* | 4.557633942 | 0.024873011 | -0.011795149 | 0.03666816 | 0.999933887 | -0.10737941 | -0.553705598 | 0.446326188 | 0.004460469 |
| ENSG00000120438 | *TCP1* | 5.868573604 | -0.05577555 | -0.076042602 | 0.020267052 | 0.999933887 | 0.559316754 | 0.805410749 | -0.246093995 | 0.004472924 |
| ENSG00000196151 | *WDSUB1* | 1.948942152 | 0.002467891 | -0.163075699 | 0.16554359 | 0.999933887 | -0.012151442 | 0.467285015 | -0.479436457 | 0.004486055 |
| ENSG00000261371 | *PECAM1* | 7.336203771 | -0.009643566 | -0.093243474 | 0.083599908 | 0.999933887 | -0.619472616 | -1.261653589 | 0.642180973 | 0.004488162 |
| ENSG00000138279 | *ANXA7* | 6.43172484 | -0.033341708 | -0.048094076 | 0.014752368 | 0.999933887 | 0.683394696 | 0.977500258 | -0.294105562 | 0.004490805 |
| ENSG00000164068 | *RNF123* | 4.020266248 | 0.040683789 | 0.094729911 | -0.054046122 | 0.999933887 | -0.190072408 | -0.52365785 | 0.333585442 | 0.004494147 |
| ENSG00000111110 | *PPM1H* | 0.897314112 | -0.070395096 | 0.130588205 | -0.200983301 | 0.999933887 | -1.117987119 | -1.932987576 | 0.815000457 | 0.00450868 |
| ENSG00000109685 | *NSD2* | 4.207080316 | -0.016022124 | -0.069629645 | 0.053607521 | 0.999933887 | -0.335927815 | -0.714886254 | 0.378958439 | 0.004546145 |
| ENSG00000183020 | *AP2A2* | 4.30327739 | 0.022641418 | 0.019349055 | 0.003292363 | 0.999933887 | -0.221987335 | -0.550747118 | 0.328759783 | 0.004574549 |
| ENSG00000183597 | *TANGO2* | 4.441327121 | 0.017579204 | -0.08774071 | 0.105319914 | 0.999933887 | -0.568725213 | -0.88807401 | 0.319348797 | 0.004634346 |
| ENSG00000110713 | *NUP98* | 8.422234241 | 0.005848158 | 0.002818203 | 0.003029955 | 0.999933887 | 0.527991686 | 0.799590386 | -0.2715987 | 0.004639536 |
| ENSG00000102178 | *UBL4A* | 2.731043992 | -0.017341608 | -0.035717436 | 0.018375827 | 0.999933887 | -0.486665219 | -1.043201482 | 0.556536263 | 0.004659789 |
| ENSG00000118495 | *PLAGL1* | 5.401622104 | -0.016320392 | -0.086380684 | 0.070060292 | 0.999933887 | 0.25294981 | 0.674063877 | -0.421114066 | 0.004673418 |

| | | | | | | | | | | |
|---|---|---|---|---|---|---|---|---|---|---|
| ENSG00000103489 | *XYLT1* | 5.062924553 | -0.026349581 | 0.003066153 | -0.029415734 | 0.999933887 | -0.344490256 | -0.752094056 | 0.4076038 | 0.004722737 |
| ENSG00000187609 | *EXD3* | 2.602745193 | -0.002493101 | 0.063450139 | -0.06594324 | 0.999933887 | 0.300914588 | 0.732263144 | -0.431348556 | 0.004733331 |
| ENSG00000101040 | *ZMYND8* | 3.945362033 | -0.022805364 | -0.047364055 | 0.024558692 | 0.999933887 | -0.333446741 | -0.739928545 | 0.406481804 | 0.004746279 |
| ENSG00000100297 | *MCM5* | 3.852564574 | -0.008402128 | 0.03451027 | -0.042912399 | 0.999933887 | -0.132403369 | -0.476554511 | 0.344151142 | 0.004746279 |
| ENSG00000135604 | *STX11* | 8.553671189 | 0.089947586 | 0.0614051 | 0.028542487 | 0.999933887 | 1.000738584 | 1.294368115 | -0.293629531 | 0.004758562 |
| ENSG00000014216 | *CAPN1* | 5.698581057 | 0.050832359 | -0.008112252 | 0.058944611 | 0.999933887 | -0.231761412 | -0.592428705 | 0.360667293 | 0.004775836 |
| ENSG00000196814 | *MVB12B* | 2.794815731 | 0.013962397 | 0.035165084 | -0.021202688 | 0.999933887 | 0.549326375 | 1.153857527 | -0.604531152 | 0.004775836 |
| ENSG00000141569 | *TRIM65* | 2.189479855 | 0.099983792 | 0.001262756 | 0.098721036 | 0.999933887 | 0.13433308 | -0.315475577 | 0.449808657 | 0.004791884 |
| ENSG00000124102 | *PI3* | 9.348772601 | -0.043228343 | 0.037985421 | -0.081213764 | 0.999933887 | 2.287084983 | 2.981097968 | -0.694012985 | 0.004824025 |
| ENSG00000185475 | *TMEM179B* | 2.298995691 | 0.064466574 | -0.018624372 | 0.083090946 | 0.999933887 | -0.364158307 | -1.040802671 | 0.676644363 | 0.004858186 |
| ENSG00000004660 | *CAMKK1* | 4.438083525 | -0.052362933 | -0.022991985 | -0.029370948 | 0.999933887 | -1.459370694 | -1.95858571 | 0.499215015 | 0.004910284 |
| ENSG00000175634 | *RPS6KB2* | 4.327590061 | 0.038621863 | 0.044990272 | -0.006368409 | 0.999933887 | -0.282808719 | -0.698659893 | 0.415851173 | 0.004910284 |
| ENSG00000196776 | *CD47* | 6.739019085 | -0.030342216 | -0.030120892 | -0.000221324 | 0.999933887 | -0.122540245 | 0.229744233 | -0.352284478 | 0.004945664 |
| ENSG00000074800 | *ENO1* | 7.584134227 | -0.020785647 | -0.052975652 | 0.032190005 | 0.999933887 | -0.300560977 | -0.608818529 | 0.308257553 | 0.004946575 |
| ENSG00000104903 | *LYL1* | 3.990935002 | 0.019354455 | -0.05267793 | 0.072032385 | 0.999933887 | -1.575269475 | -2.38737183 | 0.812102355 | 0.004957942 |
| ENSG00000113638 | *TTC33* | 3.042753618 | -0.039524406 | 0.030252124 | -0.06977653 | 0.999933887 | 0.376347768 | 0.902533259 | -0.526185491 | 0.00495894 |
| ENSG00000087087 | *SRRT* | 6.263304862 | 0.020761633 | 0.042152569 | -0.021390935 | 0.999933887 | -0.097689933 | -0.34503947 | 0.247349537 | 0.004988891 |
| ENSG00000213380 | *COG8* | 0.64598216 | -0.08124276 | -0.027090226 | -0.054152535 | 0.999933887 | 0.069711273 | -0.607446735 | 0.677158008 | 0.00498895 |
| ENSG00000089916 | *GPATCH2L* | 6.783258397 | -0.035667778 | -0.05151284 | 0.015845062 | 0.999933887 | 0.342370139 | 0.623886087 | -0.281515948 | 0.00501099 |
| ENSG00000205220 | *PSMB10* | 4.469337967 | 0.043623405 | 0.035179032 | 0.008444372 | 0.999933887 | -0.657672371 | -1.044847027 | 0.387174656 | 0.005021317 |
| ENSG00000250565 | *ATP6V1E2* | 0.846054354 | 0.204333262 | -0.13460923 | 0.338942492 | 0.999933887 | 1.108724505 | 1.881508856 | -0.77278435 | 0.00502718 |
| ENSG00000223949 | *ROR1-AS1* | -1.203378618 | 0.398380387 | 0.206633096 | 0.191747291 | 0.999933887 | 1.618015552 | 2.446288972 | -0.828273421 | 0.005034651 |
| ENSG00000213937 | *CLDN9* | 1.360828733 | -0.012605435 | 0.134656573 | -0.147262009 | 0.999933887 | -1.344044502 | -2.275761333 | 0.931716832 | 0.005055411 |
| ENSG00000101337 | *TM9SF4* | 6.256751411 | 0.00238706 | -0.047819395 | 0.050206455 | 0.999933887 | 0.630395769 | 0.908810459 | -0.27841469 | 0.00508912 |
| ENSG00000111596 | *CNOT2* | 5.858534382 | -0.051334684 | -0.044537199 | -0.006797485 | 0.999933887 | 0.177032163 | 0.539233136 | -0.362200973 | 0.005090606 |
| ENSG00000171853 | *TRAPPC12* | 4.462001277 | 0.051588132 | 0.075693626 | -0.024105494 | 0.999933887 | -0.285771865 | -0.670155245 | 0.38438338 | 0.005134543 |
| ENSG00000110057 | *UNC93B1* | 6.421020481 | 0.046436902 | 0.025301735 | 0.021135167 | 0.999933887 | -1.065129749 | -1.480489186 | 0.415359437 | 0.00514807 |
| ENSG00000166920 | *C15orf48* | 6.50602043 | 0.278275277 | 0.396552714 | -0.118277438 | 0.999933887 | 2.314660568 | 2.970865014 | -0.656204446 | 0.005152533 |
| ENSG00000139718 | *SETD1B* | 6.480721792 | -0.030280919 | 0.097577012 | -0.127857931 | 0.999933887 | -0.776909253 | -1.182227947 | 0.405318694 | 0.005200615 |
| ENSG00000111786 | *SRSF9* | 5.180880374 | 0.013910436 | 0.028053443 | -0.014143007 | 0.999933887 | -0.272762627 | -0.604899107 | 0.33213648 | 0.005200615 |
| ENSG00000242048 | *AC093583.1* | -0.008556875 | 0.25594421 | 0.509314154 | -0.253369944 | 0.999933887 | 3.493669328 | 4.881390955 | -1.387721627 | 0.005202514 |
| ENSG00000196976 | *LAGE3* | 2.174520282 | 0.167482042 | -0.032455655 | 0.199937697 | 0.999933887 | 0.949864622 | 1.508699022 | -0.5588344 | 0.005208077 |
| ENSG00000010404 | *IDS* | 6.336953002 | 0.039868501 | -0.047712751 | 0.087581253 | 0.999933887 | -0.475901297 | -0.764996092 | 0.289094795 | 0.005208077 |
| ENSG00000182141 | *ZNF708* | 4.139606838 | -0.045324515 | -0.083826364 | 0.038501849 | 0.999933887 | -0.351467609 | 0.18128536 | -0.532752969 | 0.005208077 |
| ENSG00000175895 | *PLEKHF2* | 5.891265971 | -0.024245039 | -0.051368796 | 0.027123757 | 0.999933887 | 1.191822121 | 1.616971808 | -0.425149686 | 0.005208077 |
| ENSG00000100284 | *TOM1* | 9.589696082 | -0.024398324 | -0.014290426 | -0.010107898 | 0.999933887 | 0.501629205 | 0.750741119 | -0.249111914 | 0.00522581 |
| ENSG00000174151 | *CYB561D1* | 3.960124052 | -0.12061952 | -0.078980802 | -0.041638718 | 0.999933887 | -0.597616031 | -0.973349434 | 0.375733403 | 0.005282123 |
| ENSG00000069702 | *TGFBR3* | 4.715804446 | -0.033891681 | -0.005112945 | -0.028778737 | 0.999933887 | 0.364925695 | 0.848786971 | -0.483861276 | 0.005282123 |
| ENSG00000259668 | *AC066613.1* | 2.437570041 | -0.040137432 | -0.274346903 | 0.23420947 | 0.999933887 | 2.009144258 | 3.040766815 | -1.031622557 | 0.005282898 |
| ENSG00000138069 | *RAB1A* | 8.65850465 | -0.023636826 | -0.07208387 | 0.048447044 | 0.999933887 | 0.285795616 | 0.535041202 | -0.249245586 | 0.005285536 |
| ENSG00000172493 | *AFF1* | 6.683264873 | -0.026585459 | -0.008480053 | -0.018105407 | 0.999933887 | -1.176661978 | -1.652167671 | 0.475505693 | 0.005288241 |
| ENSG00000226328 | *NUP50-DT* | 2.95954831 | -0.002343468 | -0.001804447 | -0.000539021 | 0.999933887 | -1.126971128 | -1.698032742 | 0.571061613 | 0.005302994 |
| ENSG00000156313 | *RPGR* | 5.407542787 | -0.116947971 | -0.099300454 | -0.017647517 | 0.999933887 | 0.672142225 | 1.348900846 | -0.676758621 | 0.005307062 |
| ENSG00000109062 | *SLC9A3R1* | 6.632480186 | 0.064486622 | 0.065570487 | -0.001083864 | 0.999933887 | -0.28309175 | -0.642926982 | 0.359835232 | 0.005308939 |
| ENSG00000136715 | *SAP130* | 4.761810787 | 0.040341284 | 0.010911476 | 0.029429808 | 0.999933887 | -0.437181741 | -0.891045045 | 0.453863304 | 0.005308939 |
| ENSG00000187051 | *RPS19BP1* | 4.884794567 | -0.038896066 | 0.011782855 | -0.050678921 | 0.999933887 | -0.498200594 | -0.876364414 | 0.378163821 | 0.005308939 |
| ENSG00000251194 | *AL133330.1* | 3.281279122 | 0.018666898 | 0.017352564 | 0.001314334 | 0.999933887 | 1.427727303 | 2.252329696 | -0.824602393 | 0.005308939 |
| ENSG00000014138 | *POLA2* | 0.774821026 | -0.018873262 | 0.222194233 | -0.241067494 | 0.999933887 | 0.839955881 | 1.649312695 | -0.809356814 | 0.005308939 |
| ENSG00000102921 | *N4BP1* | 8.735485799 | -0.028986074 | -0.041621432 | 0.012635357 | 0.999933887 | 1.332331149 | 1.646515449 | -0.3141843 | 0.005326083 |
| ENSG00000185379 | *RAD51D* | 1.624291782 | -0.015850999 | -0.071258067 | 0.055407068 | 0.999933887 | 0.408256077 | 0.954542745 | -0.546286668 | 0.005326083 |
| ENSG00000180316 | *PNPLA1* | 4.765212967 | -0.063467594 | 0.060722817 | -0.124190411 | 0.999933887 | 2.059766114 | 2.540005319 | -0.480239206 | 0.00532927 |
| ENSG00000214413 | *BBIP1* | 4.917252797 | -0.069823367 | 0.034097678 | -0.103921045 | 0.999933887 | 0.307657982 | 0.664464477 | -0.356806496 | 0.00533851 |
| ENSG00000127483 | *HP1BP3* | 6.388703735 | 0.007512906 | 0.04896427 | -0.041451363 | 0.999933887 | -0.318180067 | -0.569134251 | 0.250954185 | 0.005394487 |
| ENSG00000146457 | *WTAP* | 7.933392765 | -0.050631115 | -0.035136722 | -0.015494393 | 0.999933887 | 1.746649698 | 2.131544034 | -0.384894336 | 0.005445704 |
| ENSG00000113657 | *DPYSL3* | 0.842000877 | -0.265497182 | -0.228131427 | -0.037365755 | 0.999933887 | 0.297884286 | 0.865800418 | -0.567916131 | 0.005479273 |
| ENSG00000091592 | *NLRP1* | 7.560174275 | 0.024294482 | 0.02152258 | 0.002771902 | 0.999933887 | -0.081421315 | -0.291994279 | 0.210572964 | 0.005479273 |
| ENSG00000253315 | *LINC01932* | 1.000142392 | 0.015737775 | 0.136530126 | -0.120792351 | 0.999933887 | 0.702911326 | 1.283046568 | -0.580135242 | 0.005479273 |
| ENSG00000088930 | *XRN2* | 6.068564492 | -0.051425286 | -0.041878584 | -0.009546702 | 0.999933887 | -0.466919356 | -0.731876863 | 0.264957507 | 0.005497848 |
| ENSG00000102096 | *PIM2* | 8.078718708 | 0.001125791 | -0.005720787 | 0.006846577 | 0.999933887 | 2.265407375 | 2.751990091 | -0.486582716 | 0.005497848 |
| ENSG00000154640 | *BTG3* | 2.356511009 | -0.030187071 | -0.11559977 | 0.085412699 | 0.999933887 | 1.003240743 | 1.667529146 | -0.664288403 | 0.00551054 |
| ENSG00000166900 | *STX3* | 9.246851234 | -0.044557061 | -0.034825861 | -0.0097312 | 0.999933887 | 0.220139424 | 0.414591919 | -0.194452495 | 0.00553225 |
| ENSG00000157191 | *NECAP2* | 5.789217535 | -0.011894609 | -0.000312918 | -0.011581691 | 0.999933887 | 0.45836293 | 0.71924131 | -0.26087838 | 0.005533342 |
| ENSG00000156411 | *ATP5MPL* | 4.167915537 | -0.029391496 | -0.098006863 | 0.068615367 | 0.999933887 | -0.34273701 | -0.671971341 | 0.329234331 | 0.005567498 |
| ENSG00000163735 | *CXCL5* | 0.55621579 | 0.334881485 | 0.152685604 | 0.182195881 | 0.999933887 | 0.46985402 | 1.385194906 | -0.915340886 | 0.00557473 |
| ENSG00000058272 | *PPP1R12A* | 7.432293738 | -0.109493413 | -0.167538337 | 0.058044924 | 0.999933887 | 0.87583261 | 1.680997194 | -0.805164583 | 0.005632858 |
| ENSG00000125912 | *NCLN* | 5.482451299 | 0.046668222 | 0.02683189 | 0.019836332 | 0.999933887 | -0.097106581 | -0.406826483 | 0.309719901 | 0.005661506 |
| ENSG00000133872 | *SARAF* | 8.221332769 | 0.025206086 | -0.039590122 | 0.064796208 | 0.999933887 | -0.339655099 | -0.649245259 | 0.30959016 | 0.005670121 |
| ENSG00000154767 | *XPC* | 4.655192273 | -0.023746302 | -0.038454358 | 0.014708056 | 0.999933887 | -0.261022726 | -0.567704338 | 0.306681611 | 0.005676414 |
| ENSG00000130244 | *FAM98C* | 3.580396406 | 0.092606895 | 0.118000995 | -0.0253941 | 0.999933887 | 0.013983647 | -0.384718591 | 0.398702239 | 0.005709642 |
| ENSG00000176390 | *CRLF3* | 4.733318426 | 0.042248276 | 0.067630508 | -0.025382232 | 0.999933887 | -0.646921196 | -0.987037255 | 0.340116059 | 0.005796659 |
| ENSG00000140455 | *USP3* | 5.072247854 | -0.060419862 | -0.064646145 | 0.004226282 | 0.999933887 | -0.331253005 | -0.593502926 | 0.262249922 | 0.005805739 |
| ENSG00000169926 | *KLF13* | 7.123561532 | 0.037730676 | -0.046750743 | 0.084481419 | 0.999933887 | -0.384847731 | -0.730830253 | 0.345982522 | 0.005818621 |
| ENSG00000185973 | *TMLHE* | 1.735407472 | 0.008160366 | -0.051200615 | 0.059360981 | 0.999933887 | -0.47619982 | -1.114144073 | 0.637944252 | 0.005818621 |
| ENSG00000163006 | *CCDC138* | 0.448494754 | 0.00419948 | 0.142019684 | -0.137820204 | 0.999933887 | 0.84140655 | 1.65459536 | -0.81318881 | 0.005911906 |
| ENSG00000065883 | *CDK13* | 5.57941409 | 0.050054387 | 0.0559366 | -0.005882213 | 0.999933887 | -0.434771711 | -0.723161629 | 0.288389917 | 0.005932049 |
| ENSG00000144802 | *NFKBIZ* | 9.553633073 | 0.278243582 | 0.233821579 | 0.044422003 | 0.999933887 | 3.006350961 | 3.588112948 | -0.581761987 | 0.005949376 |
| ENSG00000131473 | *ACLY* | 5.233086493 | 0.070239256 | 0.000716176 | 0.06952308 | 0.999933887 | -0.226950404 | -0.585909942 | 0.358959538 | 0.005963792 |
| ENSG00000072135 | *PTPN18* | 4.717568983 | 0.030505659 | 0.015615377 | 0.014890282 | 0.999933887 | -0.068002823 | -0.343944218 | 0.275941395 | 0.005963792 |
| ENSG00000123562 | *MORF4L2* | 6.954519264 | -0.014519255 | -0.107629731 | 0.093110476 | 0.999933887 | 0.336008323 | 0.672907562 | -0.33689924 | 0.005963792 |
| ENSG00000170142 | *UBE2E1* | 4.711799277 | -0.0227354 | 0.019905899 | -0.042641298 | 0.999933887 | 1.620806958 | 2.169583568 | -0.54877661 | 0.006012132 |
| ENSG00000136869 | *TLR4* | 7.731158654 | -0.04920278 | -0.071148254 | 0.021945474 | 0.999933887 | 0.841575215 | 1.223873474 | -0.382298259 | 0.006031687 |
| ENSG00000127824 | *TUBA4A* | 6.307991246 | 0.049632619 | -0.028040523 | 0.077673142 | 0.999933887 | -0.209618964 | -0.54196969 | 0.332350725 | 0.006079579 |
| ENSG00000222009 | *BTBD19* | 6.602120534 | 0.269965037 | 0.231341641 | 0.038623396 | 0.999933887 | 1.696406856 | 2.21741472 | -0.521007863 | 0.006091781 |
| ENSG00000080802 | *CNOT4* | 3.981804883 | 0.022664666 | -0.099831643 | 0.122496309 | 0.999933887 | 0.185644758 | 0.573714632 | -0.388069874 | 0.006095358 |
| ENSG00000165410 | *CFL2* | 1.006135279 | -0.225101433 | -0.032819812 | -0.192281622 | 0.999933887 | 0.487421059 | 1.177480661 | -0.690059602 | 0.006105698 |
| ENSG00000107140 | *TESK1* | 3.963576036 | 0.108690531 | 0.031537769 | 0.077152762 | 0.999933887 | 0.439545119 | 0.884534888 | -0.444989769 | 0.00612675 |
| ENSG00000166788 | *SAAL1* | 1.241209936 | -0.235972238 | -0.106793151 | -0.129179087 | 0.999933887 | 1.037635674 | 1.732287964 | -0.694652289 | 0.006135294 |
| ENSG00000099337 | *KCNK6* | 3.289405662 | -0.033921569 | -0.031639834 | -0.002281735 | 0.999933887 | -0.435430844 | -0.941348246 | 0.505917401 | 0.006135294 |
| ENSG00000073169 | *SELENOO* | 4.631852765 | 0.06509843 | -0.004914372 | 0.070012802 | 0.999933887 | -0.23994688 | -0.632822473 | 0.392875593 | 0.006138384 |
| ENSG00000237499 | *WAKMAR2* | 4.565138736 | 0.24840286 | 0.239789802 | 0.008613059 | 0.999933887 | 1.707198569 | 2.292091171 | -0.584892602 | 0.006150254 |
| ENSG00000163975 | *MELTF* | 4.905739832 | 0.068323046 | 0.204646558 | -0.136323512 | 0.999933887 | 0.484281926 | 0.978786493 | -0.494504567 | 0.006150254 |
| ENSG00000183044 | *ABAT* | 3.505722259 | -0.043020226 | 0.070485399 | -0.113505625 | 0.999933887 | -0.156422096 | -0.656935391 | 0.500513294 | 0.006150254 |
| ENSG00000165775 | *FUNDC2* | 5.134838445 | 0.025429247 | -0.061685899 | 0.087115146 | 0.999933887 | 0.222025126 | 0.534908799 | -0.312883672 | 0.006169686 |
| ENSG00000133624 | *ZNF767P* | 3.70912865 | 0.039541142 | -0.004550521 | 0.044091662 | 0.999933887 | -0.206884816 | -0.502287792 | 0.295402977 | 0.00618531 |
| ENSG00000180871 | *CXCR2* | 8.82889996 | -0.007982065 | -0.002784613 | -0.005197452 | 0.999933887 | -1.472834917 | -2.069921889 | 0.597086971 | 0.00619039 |
| ENSG00000146067 | *FAM193B* | 6.63448988 | 0.04672546 | 0.089590485 | -0.042865025 | 0.999933887 | -0.218256862 | -0.472880713 | 0.254623851 | 0.006230361 |
| ENSG00000015475 | *BID* | 8.520241359 | 0.034486041 | -0.054724785 | 0.089210827 | 0.999933887 | 1.008709351 | 1.316515849 | -0.307806499 | 0.006239497 |
| ENSG00000135912 | *TTLL4* | 5.560865639 | -0.017194627 | -0.04049637 | 0.023301743 | 0.999933887 | 0.625576054 | 1.050312437 | -0.424736383 | 0.006242407 |
| ENSG00000151502 | *VPS26B* | 4.359905571 | 0.037588386 | 0.000491753 | 0.037096633 | 0.999933887 | -0.157401015 | -0.518040561 | 0.360639546 | 0.006272701 |
| ENSG00000162521 | *RBBP4* | 3.741384568 | -0.082105547 | -0.109806641 | 0.027701094 | 0.999933887 | -0.065362114 | -0.364362829 | 0.299000715 | 0.006282121 |
| ENSG00000286067 | *AC004263.2* | 4.168369937 | -0.018211212 | 0.027902863 | -0.046114075 | 0.999933887 | -0.681350146 | -1.135474236 | 0.45412409 | 0.006303087 |
| ENSG00000162645 | *GBP2* | 9.789131942 | -0.037781691 | -0.192598971 | 0.15481728 | 0.999933887 | 0.711320768 | 1.16483777 | -0.453517002 | 0.00631032 |
| ENSG00000095951 | *HIVEP1* | 7.540204778 | -0.029606514 | -0.018033641 | -0.011572873 | 0.999933887 | 0.840474149 | 1.407458116 | -0.566983967 | 0.006311685 |
| ENSG00000178567 | *EPM2AIP1* | 5.145081989 | -0.229107687 | -0.159851236 | -0.069256451 | 0.999933887 | 2.049703894 | 2.861380254 | -0.81167636 | 0.006344947 |
| ENSG00000164144 | *ARFIP1* | 5.723284681 | -0.096612354 | -0.093660386 | -0.002951967 | 0.999933887 | 0.541817387 | 0.944196409 | -0.402379022 | 0.006344947 |

| | | | | | | | | | | |
|---|---|---|---|---|---|---|---|---|---|---|
| ENSG00000074621 | *SLC24A1* | 2.884744766 | -0.128811803 | -0.039682358 | -0.089129445 | 0.999933887 | 2.658607755 | 3.659176392 | -1.000568637 | 0.006344947 |
| ENSG00000007202 | *KIAA0100* | 5.474601498 | 0.010026177 | 0.008110114 | 0.001916062 | 0.999933887 | -0.173589481 | -0.451498986 | 0.277909506 | 0.006361713 |
| ENSG00000069399 | *BCL3* | 7.881679314 | 0.125306381 | 0.134548139 | -0.009241758 | 0.999933887 | 1.925062756 | 2.37272177 | -0.447659014 | 0.006372831 |
| ENSG00000103042 | *SLC38A7* | 2.995340358 | 0.009747041 | 0.095443595 | -0.085696554 | 0.999933887 | 0.218942103 | 0.549492454 | -0.330550351 | 0.006375408 |
| ENSG00000272886 | *DCP1A* | 5.855619853 | -0.029705772 | -0.039819823 | 0.010114051 | 0.999933887 | 0.415712134 | 0.665406204 | -0.24969407 | 0.006446897 |
| ENSG00000253372 | *AC016405.1* | 3.021059484 | 0.092433168 | 0.151734304 | -0.059301136 | 0.999933887 | 1.713509559 | 2.326621798 | -0.613112239 | 0.006447994 |
| ENSG00000090060 | *PAPOLA* | 7.338860248 | -0.04728022 | -0.012213238 | -0.035066982 | 0.999933887 | -0.126597492 | 0.154879372 | -0.281476864 | 0.006447994 |
| ENSG00000107882 | *SUFU* | 2.275292503 | -0.011786178 | 0.086915048 | -0.098701227 | 0.999933887 | -0.016284632 | -0.577332596 | 0.561047964 | 0.006447994 |
| ENSG00000113068 | *PFDN1* | 3.782800012 | -0.058923265 | -0.02215063 | -0.036772635 | 0.999933887 | 0.184487694 | 0.487496413 | -0.303008719 | 0.006477778 |
| ENSG00000198830 | *HMGN2* | 6.590506442 | 0.010273405 | -0.029921467 | 0.040194872 | 0.999933887 | -0.195271736 | -0.466271173 | 0.270999437 | 0.006496696 |
| ENSG00000177946 | *CENPBD1* | 0.497518121 | -0.048792623 | -0.140549499 | 0.091756877 | 0.999933887 | -0.488398765 | -1.289487003 | 0.801088238 | 0.0065302 |
| ENSG00000188886 | *ASTL* | 2.171631879 | 0.120680216 | 0.181303217 | -0.060623001 | 0.999933887 | 0.857349925 | 1.533352123 | -0.676002198 | 0.00654793 |
| ENSG00000129465 | *RIPK3* | 3.687668293 | -0.020641767 | 0.066828483 | -0.08747025 | 0.999933887 | -0.424291035 | -0.794782866 | 0.370491831 | 0.006553929 |
| ENSG00000164086 | *DUSP7* | 2.242424788 | 0.079707057 | 0.040002275 | 0.039704781 | 0.999933887 | -0.442252279 | -0.973264984 | 0.531012704 | 0.006590214 |
| ENSG00000039319 | *ZFYVE16* | 8.201307998 | -0.036642853 | -0.071754875 | 0.035112022 | 0.999933887 | -0.05030401 | 0.647873602 | -0.698177612 | 0.006620652 |
| ENSG00000247982 | *LINC00926* | 1.768623798 | 0.140282598 | -0.119649286 | 0.259931884 | 0.999933887 | 0.280657055 | 0.802388519 | -0.521731464 | 0.006663577 |
| ENSG00000277443 | *MARCKS* | 10.588034292 | -0.050743024 | 0.006325812 | -0.057068836 | 0.999933887 | 0.91064083 | 1.275207291 | -0.364566461 | 0.006700829 |
| ENSG00000237803 | *LINC00211* | 5.468515434 | 0.027517138 | -0.041337656 | 0.068854794 | 0.999933887 | 0.428366891 | 0.754362868 | -0.325995977 | 0.00674196 |
| ENSG00000152700 | *SAR1B* | 3.963489258 | -0.059083323 | 0.064936384 | -0.124019707 | 0.999933887 | 0.634157938 | 1.028427128 | -0.39426919 | 0.006748846 |
| ENSG00000101544 | *ADNP2* | 6.552911507 | -0.049785368 | -0.038895238 | -0.01089013 | 0.999933887 | 0.610806431 | 0.881984617 | -0.271178186 | 0.006782173 |
| ENSG00000267080 | *ASB16-AS1* | 1.495934419 | 0.100179724 | -0.030623523 | 0.130803247 | 0.999933887 | -0.154324931 | -0.82595192 | 0.671626989 | 0.006782173 |
| ENSG00000198791 | *CNOT7* | 4.598208731 | -0.065191945 | -0.043110503 | -0.022081442 | 0.999933887 | -0.412769357 | -0.709590916 | 0.296821559 | 0.006819588 |
| ENSG00000214106 | *PAXIP1-AS2* | 1.74221589 | -0.118981792 | -0.121252789 | 0.002270997 | 0.999933887 | 0.192442822 | 0.864665147 | -0.672222326 | 0.006832012 |
| ENSG00000163412 | *EIF4E3* | 5.265294779 | -0.024502009 | -0.010713307 | -0.013788703 | 0.999933887 | -0.35539035 | -0.63432577 | 0.27893542 | 0.006848499 |
| ENSG00000257354 | *AC048341.1* | 1.699784257 | -0.237111503 | -0.155169062 | -0.081942441 | 0.999933887 | 0.873040515 | 1.586036886 | -0.712996371 | 0.006854837 |
| ENSG00000106123 | *EPHB6* | 3.530529706 | 0.043263208 | 0.088080226 | -0.044817019 | 0.999933887 | -0.09260776 | -0.540944759 | 0.448336999 | 0.006854837 |
| ENSG00000204560 | *DHX16* | 4.686322628 | 0.025036849 | 0.01708145 | 0.0079554 | 0.999933887 | -0.161776749 | -0.418173501 | 0.256396752 | 0.006854837 |
| ENSG00000144040 | *SFXN5* | 3.147121299 | -0.076149705 | -0.027947486 | -0.048202219 | 0.999933887 | -0.193229946 | -0.587800113 | 0.394570167 | 0.006909551 |
| ENSG00000160305 | *DIP2A* | 5.817257846 | 0.043035339 | 0.034397579 | 0.008637761 | 0.999933887 | 0.458866881 | 0.911538112 | -0.452671231 | 0.006909551 |
| ENSG00000158062 | *UBXN11* | 4.33920628 | 0.014163768 | 0.045851489 | -0.031687721 | 0.999933887 | -0.030003181 | -0.337840501 | 0.30783732 | 0.006909551 |
| ENSG00000160584 | *SIK3* | 6.680228321 | 0.011602072 | 0.006360971 | 0.005241101 | 0.999933887 | 0.467667737 | 0.748908931 | -0.281241194 | 0.006909551 |
| ENSG00000138449 | *SLC40A1* | 3.486121028 | -0.147445441 | -0.038140701 | -0.109304739 | 0.999933887 | -0.885323251 | -1.45188929 | 0.566566038 | 0.006926248 |
| ENSG00000120742 | *SERP1* | 7.314791898 | 0.050324643 | -0.003048867 | 0.05337351 | 0.999933887 | 0.464237213 | 0.720348895 | -0.256111682 | 0.007013498 |
| ENSG00000085514 | *PILRA* | 8.545638083 | 0.072087313 | 0.042200261 | 0.029887052 | 0.999933887 | 1.72852057 | 2.157631989 | -0.429111419 | 0.007032458 |
| ENSG00000110711 | *AIP* | 4.541860858 | 0.020898758 | 0.0083998 | 0.012498958 | 0.999933887 | -0.119983724 | -0.510772832 | 0.390789107 | 0.007043205 |
| ENSG00000145088 | *EAF2* | 0.982045682 | 0.033362525 | 0.00998713 | 0.023375395 | 0.999933887 | 0.19693704 | 0.882585902 | -0.685648862 | 0.007051166 |
| ENSG00000197818 | *SLC9A8* | 7.725213442 | -0.024100132 | 0.024466359 | -0.048566491 | 0.999933887 | 1.019430285 | 1.327790824 | -0.308360538 | 0.007057086 |
| ENSG00000258919 | *AL049836.1* | 1.878473129 | -0.23338285 | 0.061368113 | -0.294750964 | 0.999933887 | 0.966110402 | 1.429399711 | -0.463289309 | 0.007084522 |
| ENSG00000175582 | *RAB6A* | 4.658521134 | -0.017938953 | 0.012459433 | -0.030398386 | 0.999933887 | -0.442529278 | -0.791492406 | 0.348963127 | 0.007149497 |
| ENSG00000237732 | *CT75* | 1.067569736 | 0.326871203 | 0.370437968 | -0.043566765 | 0.999933887 | 3.098517158 | 3.760554862 | -0.662037704 | 0.007192328 |
| ENSG00000115307 | *AUP1* | 5.391924138 | -0.002282138 | 0.038072387 | -0.040354525 | 0.999933887 | 0.090390154 | -0.149489116 | 0.23987927 | 0.007225737 |
| ENSG00000163346 | *PBXIP1* | 8.071159004 | 0.018397679 | 0.03246097 | -0.014063291 | 0.999933887 | -0.150090464 | -0.512718386 | 0.362627922 | 0.007244265 |
| ENSG00000133731 | *IMPA1* | 3.390049008 | -0.051834431 | -0.137686795 | 0.085852364 | 0.999933887 | -0.169772319 | 0.22073057 | -0.390502889 | 0.007250413 |
| ENSG00000254999 | *BRK1* | 5.84704191 | 0.064009315 | -0.019108719 | 0.083118033 | 0.999933887 | 0.629275872 | 0.940793637 | -0.311517764 | 0.007252815 |
| ENSG00000116701 | *NCF2* | 11.05621892 | -0.01562602 | -0.018236708 | 0.002610688 | 0.999933887 | -0.104238514 | 0.158283462 | -0.262521976 | 0.007252815 |
| ENSG00000185483 | *ROR1* | -0.932285495 | -0.013569945 | 0.630553648 | -0.644123594 | 0.999933887 | 1.895819651 | 2.687452698 | -0.791633046 | 0.007252815 |
| ENSG00000179833 | *SERTAD2* | 6.295920189 | -0.001014281 | -0.079093034 | 0.078078754 | 0.999933887 | 0.82606723 | 1.339529626 | -0.513462396 | 0.007252815 |
| ENSG00000197837 | *H4-16* | 1.428911281 | 0.087275109 | 0.217087744 | -0.129812635 | 0.999933887 | -0.067138335 | -0.670405377 | 0.603267042 | 0.007280029 |
| ENSG00000250571 | *GLI4* | 1.974715723 | -0.014462424 | 0.044077972 | -0.058540396 | 0.999933887 | -0.005822672 | -0.467228325 | 0.461405653 | 0.007280029 |
| ENSG00000075975 | *MKRN2* | 4.028781084 | -0.001215932 | 0.036264121 | -0.037480053 | 0.999933887 | -0.554881418 | -0.870486006 | 0.315604589 | 0.007280029 |
| ENSG00000147416 | *ATP6V1B2* | 10.461574598 | -0.076140959 | -0.066405538 | -0.009735422 | 0.999933887 | 0.215486371 | 0.492345461 | -0.27685909 | 0.007281496 |
| ENSG00000115486 | *GGCX* | 3.407708285 | 0.031573035 | -0.061205202 | 0.092778237 | 0.999933887 | -0.195001762 | -0.539611429 | 0.344609668 | 0.007295388 |
| ENSG00000115306 | *SPTBN1* | 7.142172345 | -0.004804384 | -0.057770385 | 0.052966001 | 0.999933887 | -0.009753102 | 0.23734287 | -0.247095972 | 0.007295388 |
| ENSG00000112659 | *CUL9* | 4.849573357 | 0.054484356 | 0.058748227 | -0.004263871 | 0.999933887 | 0.207333594 | 0.483460414 | -0.276126821 | 0.007336609 |
| ENSG00000127947 | *PTPN12* | 7.617749838 | -0.013053284 | -0.047316211 | 0.034262927 | 0.999933887 | 0.370199349 | 0.71470769 | -0.344508341 | 0.00735351 |
| ENSG00000067048 | *DDX3Y* | -1.64642325 | 0.048116252 | 0.079064675 | -0.030948423 | 0.999933887 | 0.408752915 | 0.958617198 | -0.549864283 | 0.007357975 |
| ENSG00000026297 | *RNASET2* | 7.893252123 | 0.050599568 | 0.069568919 | -0.018969351 | 0.999933887 | -0.340269927 | -0.799778813 | 0.459508886 | 0.007382221 |
| ENSG00000170892 | *TSEN34* | 5.501151433 | -0.046696609 | -0.046828413 | 0.000131804 | 0.999933887 | -1.227307582 | -1.728692947 | 0.501385365 | 0.007416857 |
| ENSG00000111671 | *SPSB2* | 1.749860455 | 0.034793305 | 0.021895294 | 0.01289801 | 0.999933887 | -0.322212775 | -0.853584348 | 0.531371573 | 0.007500167 |
| ENSG00000112110 | *MRPL18* | 3.457235575 | -0.045668324 | 0.045991678 | -0.091660002 | 0.999933887 | 1.019323114 | 1.439728846 | -0.420405733 | 0.00754233 |
| ENSG00000116337 | *AMPD2* | 7.984183341 | 0.03185726 | 0.083039707 | -0.051182447 | 0.999933887 | -0.400728105 | -0.830252893 | 0.429524788 | 0.007563416 |
| ENSG00000120008 | *WDR11* | 3.280297343 | -0.078646644 | -0.074039744 | -0.0046069 | 0.999933887 | 0.362658897 | 0.818372611 | -0.455713714 | 0.007653018 |
| ENSG00000138303 | *ASCC1* | 2.683066665 | 0.015202489 | 0.045827804 | -0.030625315 | 0.999933887 | 0.474291858 | 0.877612843 | -0.403320986 | 0.007653018 |
| ENSG00000102580 | *DNAJC3* | 7.73492038 | -0.058551561 | -0.066996207 | 0.008444646 | 0.999933887 | 0.623167815 | 1.012883588 | -0.389715773 | 0.007663374 |
| ENSG00000129625 | *REEP5* | 4.68815165 | -0.015369739 | -0.040943088 | 0.02557335 | 0.999933887 | -0.068925719 | -0.464082739 | 0.39515702 | 0.007675356 |
| ENSG00000103197 | *TSC2* | 5.616914762 | 0.051372522 | 0.09892222 | -0.047549698 | 0.999933887 | -0.165759329 | -0.444242681 | 0.278483352 | 0.007684693 |
| ENSG00000069493 | *CLEC2D* | 5.275837277 | -0.08070252 | -0.021057853 | -0.059644667 | 0.999933887 | 0.109765046 | 0.584422387 | -0.474657342 | 0.007684693 |
| ENSG00000185722 | *ANKFY1* | 5.601852858 | -0.014399458 | 0.070770167 | -0.085169625 | 0.999933887 | -0.328760717 | -0.633044376 | 0.304283659 | 0.007684693 |
| ENSG00000169891 | *REPS2* | 5.38951638 | -0.02983419 | -0.00514755 | -0.02468664 | 0.999933887 | 0.537120177 | 0.817268918 | -0.280148741 | 0.007698726 |
| ENSG00000180767 | *CHST13* | 1.723275454 | -0.01917439 | 0.043256669 | -0.062431059 | 0.999933887 | -1.393519558 | -2.280809698 | 0.887290141 | 0.007698726 |
| ENSG00000114383 | *TUSC2* | 4.560981395 | -0.024819705 | 0.007623866 | -0.032443571 | 0.999933887 | -0.502928932 | -0.854307704 | 0.351378772 | 0.007701037 |
| ENSG00000113742 | *CPEB4* | 8.135905938 | 0.001536958 | -0.023837838 | 0.025374796 | 0.999933887 | 0.656624727 | 1.060495094 | -0.403870367 | 0.007708398 |
| ENSG00000197774 | *EME2* | 2.892750048 | 0.046284251 | 0.166379475 | -0.120095224 | 0.999933887 | -0.050534085 | -0.518325547 | 0.467791462 | 0.007708851 |
| ENSG00000135862 | *LAMC1* | 0.741234602 | -0.046436732 | -0.090859175 | 0.044422444 | 0.999933887 | 0.956361629 | 1.591466213 | -0.635104584 | 0.007711634 |
| ENSG00000188056 | *TREML4* | 5.850977593 | 0.083740387 | -0.015807792 | 0.099548179 | 0.999933887 | 0.787773689 | 1.177304059 | -0.389530371 | 0.007788685 |
| ENSG00000065357 | *DGKA* | 6.339863522 | -0.039048387 | 0.011776475 | -0.050824862 | 0.999933887 | -0.061200757 | -0.249721082 | 0.188520324 | 0.007788685 |
| ENSG00000142227 | *EMP3* | 7.379945103 | 0.021853365 | -0.029390585 | 0.051243951 | 0.999933887 | 0.207941289 | 0.465236593 | -0.257295303 | 0.007788685 |
| ENSG00000101745 | *ANKRD12* | 7.616499703 | -0.223477793 | -0.194490488 | -0.028987304 | 0.999933887 | -0.271871632 | 0.466464365 | -0.738335997 | 0.007791588 |
| ENSG00000246705 | *H2AJ* | 2.502137642 | 0.073882861 | -0.075985569 | 0.14986843 | 0.999933887 | 0.074491981 | -0.391767161 | 0.466259143 | 0.007791588 |
| ENSG00000115866 | *DARS1* | 4.717697135 | -0.110450237 | -0.09873641 | -0.011713827 | 0.999933887 | -0.14740984 | 0.204208875 | -0.351618714 | 0.007792764 |
| ENSG00000166002 | *SMCO4* | 1.649482553 | 0.086452561 | 0.300408062 | -0.213955501 | 0.999933887 | 0.665315003 | 1.393601334 | -0.728286331 | 0.007871046 |
| ENSG00000124224 | *PPP4R1L* | 5.413614204 | 0.00027608 | 0.033558623 | -0.033282543 | 0.999933887 | 0.019492476 | 0.36184914 | -0.342356665 | 0.00789778 |
| ENSG00000198604 | *BAZ1A* | 7.627105422 | -0.111590618 | -0.11888956 | 0.007298941 | 0.999933887 | 1.276620776 | 1.772810568 | -0.496189791 | 0.007901347 |
| ENSG00000006634 | *DBF4* | 4.231712367 | -0.12276275 | -0.081076047 | -0.041686703 | 0.999933887 | 0.623144204 | 1.170856112 | -0.547711908 | 0.007901347 |
| ENSG00000130363 | *RSPH3* | 1.89835293 | -0.053921012 | 0.066101241 | -0.120022253 | 0.999933887 | -0.701122201 | -1.312836778 | 0.611714576 | 0.007901347 |
| ENSG00000189308 | *LIN54* | 5.220006374 | -0.037683838 | -0.067130834 | 0.029446995 | 0.999933887 | -0.413354687 | -0.033074289 | -0.380280398 | 0.007901347 |
| ENSG00000108639 | *SYNGR2* | 5.986150791 | 0.044440783 | -0.020447443 | 0.064888226 | 0.999933887 | 0.851842974 | 1.153051224 | -0.30120825 | 0.007909253 |
| ENSG00000139832 | *RAB20* | 6.321999612 | 0.027702875 | -0.052422961 | 0.080125836 | 0.999933887 | 0.627383219 | 1.097166336 | -0.469783116 | 0.007919104 |
| ENSG00000100084 | *HIRA* | 2.950524813 | 0.09221321 | -0.06853663 | 0.16074984 | 0.999933887 | -0.843250188 | -1.52319878 | 0.679948592 | 0.007971083 |
| ENSG00000011422 | *PLAUR* | 10.36394921 | 0.263083404 | 0.149132799 | 0.113950606 | 0.999933887 | 1.465440783 | 1.931537037 | -0.466096254 | 0.007987076 |
| ENSG00000130764 | *LRRC47* | 3.446088508 | 0.019268954 | 0.073730564 | -0.05446161 | 0.999933887 | -0.119459384 | -0.467908804 | 0.348449421 | 0.007992253 |
| ENSG00000028528 | *SNX1* | 5.531543624 | -0.088315639 | 0.012386117 | -0.100701756 | 0.999933887 | -0.78452461 | -1.125269412 | 0.340744801 | 0.008007374 |
| ENSG00000083799 | *CYLD* | 8.195987782 | -0.058994762 | -0.055966368 | -0.003028393 | 0.999933887 | 0.750196834 | 1.06666821 | -0.316471376 | 0.008013885 |
| ENSG00000101384 | *JAG1* | 5.3647317 | 0.074031906 | -0.048235084 | 0.122266989 | 0.999933887 | 2.874130023 | 3.632928836 | -0.758798813 | 0.00802485 |
| ENSG00000139163 | *ETNK1* | 4.8069626 | -0.032137168 | -0.050177931 | 0.018040763 | 0.999933887 | 0.148142585 | 0.626403696 | -0.47826111 | 0.00802485 |
| ENSG00000158122 | *PRXL2C* | 2.748814205 | -0.031437195 | -0.023070609 | -0.008366586 | 0.999933887 | 0.383216743 | 0.841380847 | -0.458164104 | 0.008102961 |
| ENSG00000115310 | *RTN4* | 8.570093176 | 0.003816174 | -0.00017732 | 0.003993493 | 0.999933887 | 0.242921413 | 0.472488965 | -0.229567551 | 0.008103924 |
| ENSG00000108669 | *CYTH1* | 8.970131961 | -0.025739655 | -0.005774955 | -0.0199647 | 0.999933887 | 0.464294605 | 0.721399684 | -0.25710508 | 0.008134299 |
| ENSG00000110315 | *RNF141* | 5.72532245 | -0.058814435 | -0.027778406 | -0.031036029 | 0.999933887 | 0.887723933 | 1.423184991 | -0.535461059 | 0.008142285 |
| ENSG00000102858 | *MGRN1* | 7.223779947 | 0.006459862 | 0.000424374 | 0.006035489 | 0.999933887 | -0.247268279 | -0.483711001 | 0.236442721 | 0.008188397 |
| ENSG00000141968 | *VAV1* | 7.48727601 | 0.020721786 | 0.021766063 | -0.001044277 | 0.999933887 | 0.754881636 | 1.114934685 | -0.360053048 | 0.00823616 |
| ENSG00000137185 | *ZSCAN9* | 1.454835106 | 0.000427955 | -0.269245257 | 0.269673212 | 0.999933887 | 0.236410659 | 0.723619081 | -0.487208422 | 0.008260023 |
| ENSG00000188167 | *TMPPE* | 2.817871281 | -0.032266097 | 0.033500703 | -0.065766799 | 0.999933887 | -0.820148509 | -1.287297206 | 0.467148698 | 0.008336163 |
| ENSG00000172113 | *NME6* | 2.422622624 | 0.045015251 | 0.096766577 | -0.051751326 | 0.999933887 | -0.304773696 | -0.755948744 | 0.451175048 | 0.0083739 |
| ENSG00000198876 | *DCAF12* | 3.983717148 | -0.092292788 | -0.039720018 | -0.052572771 | 0.999933887 | -0.444370503 | -0.89343553 | 0.449065028 | 0.008384176 |

| | | | | | | | | | | |
|---|---|---|---|---|---|---|---|---|---|---|
| ENSG00000111913 | *RIPOR2* | 8.685730181 | -0.010690329 | -0.012049778 | 0.001359449 | 0.999933887 | 0.190258877 | 0.474323939 | -0.284065063 | 0.008384176 |
| ENSG00000118689 | *FOXO3* | 7.415723562 | -0.071213079 | -0.007058266 | -0.064154813 | 0.999933887 | -0.455215748 | -0.770087462 | 0.314871714 | 0.008436443 |
| ENSG00000267344 | *AC003070.1* | 0.819858953 | 0.125340914 | 0.12361689 | 0.001724024 | 0.999933887 | -0.424995069 | -1.172749924 | 0.747754855 | 0.008445986 |
| ENSG00000197724 | *PHF2* | 5.552758163 | 0.015954168 | 0.054487198 | -0.03853303 | 0.999933887 | -0.204575333 | -0.489490494 | 0.284915161 | 0.008475058 |
| ENSG00000137075 | *RNF38* | 5.334514625 | 0.069113905 | 0.083392858 | -0.014278952 | 0.999933887 | -0.740417306 | -1.13255531 | 0.392138004 | 0.008477772 |
| ENSG00000198690 | *FAN1* | 3.878633311 | 0.029439222 | 0.03671495 | -0.007275729 | 0.999933887 | -0.285490501 | -0.610817983 | 0.325327482 | 0.008477772 |
| ENSG00000163867 | *ZMYM6* | 2.278822682 | -0.048520732 | -0.078308422 | 0.029787689 | 0.999933887 | -0.3568653 | 0.290111348 | -0.646976648 | 0.008488436 |
| ENSG00000164576 | *SAP30L* | 3.844219984 | 0.022565829 | -0.047387652 | 0.069953482 | 0.999933887 | -0.396230067 | -0.732024485 | 0.335794418 | 0.008521696 |
| ENSG00000240057 | *AC078785.1* | -1.105478127 | -0.221132418 | 0.14248819 | -0.363620608 | 0.999933887 | 0.430922284 | 1.523295352 | -1.092373068 | 0.008526681 |
| ENSG00000164091 | *WDR82* | 6.69767817 | -0.054472185 | -0.03603354 | -0.018438645 | 0.999933887 | -0.527502749 | -0.805395269 | 0.277892519 | 0.008535052 |
| ENSG00000137752 | *CASP1* | 7.729227293 | -0.073118065 | -0.166153155 | 0.09303509 | 0.999933887 | 1.027983393 | 1.400422697 | -0.372439303 | 0.008535052 |
| ENSG00000105732 | *ZNF574* | 3.26246829 | 0.047699629 | -0.051022353 | 0.098721982 | 0.999933887 | -0.171753679 | -0.518705708 | 0.346952029 | 0.008543685 |
| ENSG00000129968 | *ABHD17A* | 7.111969279 | 0.097078644 | 0.085275464 | 0.01180318 | 0.999933887 | -0.402052509 | -0.809047039 | 0.40699453 | 0.008574397 |
| ENSG00000087589 | *CASS4* | 5.548015034 | -0.023578992 | 0.118452036 | -0.142031028 | 0.999933887 | -1.36212561 | -1.987303183 | 0.625177573 | 0.008581508 |
| ENSG00000105705 | *SUGP1* | 3.752765879 | 0.029609197 | 0.028195013 | 0.001414184 | 0.999933887 | -0.268066994 | -0.626477605 | 0.358410611 | 0.008582344 |
| ENSG00000099622 | *CIRBP* | 7.344196253 | 0.009293952 | -0.018678479 | 0.027972431 | 0.999933887 | -0.358698205 | -0.617325433 | 0.258627228 | 0.00861444 |
| ENSG00000186141 | *POLR3C* | 3.129255218 | -0.071712083 | -0.123959647 | 0.052247564 | 0.999933887 | 0.984257653 | 1.660109197 | -0.675851544 | 0.008626883 |
| ENSG00000137574 | *TGS1* | 3.470335178 | -0.086385404 | -0.153104433 | 0.066719029 | 0.999933887 | 0.446737108 | 0.792291122 | -0.345554014 | 0.008643361 |
| ENSG00000140379 | *BCL2A1* | 9.502010607 | 0.067646164 | 0.018391678 | 0.049254486 | 0.999933887 | 1.112657558 | 1.519824694 | -0.407167136 | 0.008663522 |
| ENSG00000103381 | *CPPED1* | 7.383857608 | -0.022206055 | -0.028700114 | 0.006494059 | 0.999933887 | -0.797530593 | -1.235235777 | 0.437705184 | 0.008682849 |
| ENSG00000197283 | *SYNGAP1* | 3.433721363 | 0.054618727 | 0.022598673 | 0.032020053 | 0.999933887 | 0.554118236 | 1.012174742 | -0.458056506 | 0.008758927 |
| ENSG00000112851 | *ERBIN* | 8.867037715 | -0.059411405 | -0.113003686 | 0.053592281 | 0.999933887 | 0.280364547 | 0.920821974 | -0.640457427 | 0.008758927 |
| ENSG00000153936 | *HS2ST1* | 3.51864175 | -0.073638513 | 0.024600588 | -0.098239101 | 0.999933887 | -0.361557913 | 0.0930506 | -0.454608513 | 0.008791191 |
| ENSG00000112149 | *CD83* | 10.188261769 | 0.065820514 | 0.008129367 | 0.057691147 | 0.999933887 | 1.28479596 | 1.645788295 | -0.360992335 | 0.008821656 |
| ENSG00000144895 | *EIF2A* | 3.932884115 | -0.036175946 | -0.09252491 | 0.056348964 | 0.999933887 | 0.143724542 | 0.465548796 | -0.321824255 | 0.008821656 |
| ENSG00000108061 | *SHOC2* | 6.82873088 | -0.044717839 | -0.039424798 | -0.005293041 | 0.999933887 | -0.232896546 | 0.169893842 | -0.402790388 | 0.008821656 |
| ENSG00000015133 | *CCDC88C* | 6.049497525 | 0.025536269 | 0.080039596 | -0.054503327 | 0.999933887 | -0.102276724 | -0.33244024 | 0.230163516 | 0.008821656 |
| ENSG00000196544 | *BORCS6* | 1.568606373 | 0.053862687 | 0.045777657 | 0.00808503 | 0.999933887 | -0.498062414 | -1.145372366 | 0.647309952 | 0.008821656 |
| ENSG00000123815 | *COQ8B* | 4.074781103 | 0.021585558 | 0.076854979 | -0.055269421 | 0.999933887 | -0.071429719 | -0.40464554 | 0.333215821 | 0.008821656 |
| ENSG00000180398 | *MCFD2* | 4.230836099 | 0.019559114 | 0.028920252 | -0.009361138 | 0.999933887 | 0.433165359 | 0.747074417 | -0.313909058 | 0.008821656 |
| ENSG00000145901 | *TNIP1* | 9.417293046 | -0.015926756 | -0.019990994 | 0.004064238 | 0.999933887 | 1.564320029 | 1.959873311 | -0.395553282 | 0.008821656 |
| ENSG00000172053 | *QARS1* | 4.834482721 | 0.004996353 | -0.018015692 | 0.023012045 | 0.999933887 | 0.045931339 | -0.227504647 | 0.273435986 | 0.008860491 |
| ENSG00000109320 | *NFKB1* | 8.123360281 | -0.052092572 | 0.000575656 | -0.052668228 | 0.999933887 | 2.135915491 | 2.602510778 | -0.466595287 | 0.008890492 |
| ENSG00000260279 | *AC137932.1* | 3.19618909 | -0.102602773 | -0.076685242 | -0.025917531 | 0.999933887 | -0.366181806 | -0.818728769 | 0.452546963 | 0.008895359 |
| ENSG00000172724 | *CCL19* | -2.234237265 | 0.034191327 | 0.203420777 | -0.169229451 | 0.999933887 | 1.591513298 | 3.180011971 | -1.588498672 | 0.008895359 |
| ENSG00000093134 | *VNN3* | 6.520946651 | -0.080894637 | -0.108275103 | 0.027380466 | 0.999933887 | 0.389035417 | 0.788922221 | -0.399886805 | 0.008907669 |
| ENSG00000063244 | *U2AF2* | 6.950693277 | 0.029493899 | 0.041455657 | -0.011961759 | 0.999933887 | -0.289845887 | -0.550767859 | 0.260921972 | 0.008928526 |
| ENSG00000110925 | *CSRNP2* | 3.550893026 | 0.015878059 | 0.08850649 | -0.072628431 | 0.999933887 | 0.546140224 | 0.972949395 | -0.42680917 | 0.008964674 |
| ENSG00000196411 | *EPHB4* | 2.842045315 | 0.023721696 | 0.019943046 | 0.003778651 | 0.999933887 | -0.110639994 | -0.699281402 | 0.588641408 | 0.008967352 |
| ENSG00000241258 | *CRCP* | 3.876769748 | -0.086555022 | -0.003539725 | -0.083015297 | 0.999933887 | -0.194022562 | -0.557695036 | 0.363672474 | 0.008970024 |
| ENSG00000138073 | *PREB* | 3.542669637 | 0.010939319 | 0.03501886 | -0.024079542 | 0.999933887 | -0.172041623 | -0.486477305 | 0.314435682 | 0.009005826 |
| ENSG00000066697 | *MSANTD3* | 1.582945023 | -0.157287979 | -0.056899737 | -0.100388242 | 0.999933887 | 1.603539138 | 2.266420058 | -0.66288092 | 0.009015869 |
| ENSG00000184602 | *SNN* | 8.934781355 | -0.023253671 | -0.083323711 | 0.06007004 | 0.999933887 | 0.83123069 | 1.175244419 | -0.344013729 | 0.009017292 |
| ENSG00000164691 | *TAGAP* | 9.80561271 | 0.143195625 | 0.114807415 | 0.02838821 | 0.999933887 | 1.761879135 | 2.236951365 | -0.475072229 | 0.009058621 |
| ENSG00000089057 | *SLC23A2* | 6.329707518 | -0.098653133 | -0.047446687 | -0.051206446 | 0.999933887 | -1.094861131 | -1.405946363 | 0.311085231 | 0.00906111 |
| ENSG00000083937 | *CHMP2B* | 7.402111048 | -0.018762642 | -0.145378767 | 0.126616124 | 0.999933887 | 0.222524808 | 0.540164844 | -0.317640036 | 0.009074981 |
| ENSG00000108312 | *UBTF* | 5.342946152 | 0.01437657 | 0.043977123 | -0.029600553 | 0.999933887 | -0.090624339 | -0.358642992 | 0.268018653 | 0.009074981 |
| ENSG00000090316 | *MAEA* | 7.771152537 | 0.003000138 | -0.049523546 | 0.052523683 | 0.999933887 | 0.463651358 | 0.709301727 | -0.245650369 | 0.009133618 |
| ENSG00000168175 | *MAPK1IP1L* | 6.489435749 | -0.042010296 | -0.019640576 | -0.022369721 | 0.999933887 | 0.170908961 | 0.394934502 | -0.22402554 | 0.009195328 |
| ENSG00000096060 | *FKBP5* | 5.063923616 | 0.009505567 | -0.024003858 | 0.033509425 | 0.999933887 | 0.42109077 | 0.857416008 | -0.436325238 | 0.009195328 |
| ENSG00000182676 | *PPP1R27* | 0.657344415 | 0.257614549 | 0.329162542 | -0.071547994 | 0.999933887 | 1.630097755 | 2.605651107 | -0.975553352 | 0.009202286 |
| ENSG00000121749 | *TBC1D15* | 6.885800741 | -0.074316788 | -0.081202211 | 0.006885423 | 0.999933887 | -0.07454639 | 0.387762523 | -0.462308913 | 0.009228434 |
| ENSG00000198818 | *SFT2D1* | 5.545389777 | -0.001707519 | 0.007279126 | -0.008986644 | 0.999933887 | 0.278317621 | 0.514375997 | -0.236058375 | 0.009253392 |
| ENSG00000049249 | *TNFRSF9* | 3.321266406 | 0.040620289 | -0.08480377 | 0.125424059 | 0.999933887 | 1.173616968 | 1.669775228 | -0.49615826 | 0.009264876 |
| ENSG00000232912 | *RERE-AS1* | 0.073829276 | 0.058165456 | -0.097011972 | 0.155177428 | 0.999933887 | -0.772210625 | -1.712912145 | 0.940701521 | 0.009275755 |
| ENSG00000078369 | *GNB1* | 8.965924571 | 0.015812432 | 0.007155988 | 0.008656445 | 0.999933887 | 0.34639291 | 0.568311968 | -0.221919058 | 0.009341579 |
| ENSG00000197548 | *ATG7* | 6.633400633 | 0.036268487 | 0.094143831 | -0.057875344 | 0.999933887 | 1.452725447 | 1.80803993 | -0.355314483 | 0.009398771 |
| ENSG00000176542 | *USF3* | 7.836563386 | -0.049044488 | -0.086995249 | 0.037950761 | 0.999933887 | 0.666176245 | 1.197499062 | -0.531322816 | 0.009402896 |
| ENSG00000103222 | *ABCC1* | 6.02604141 | 0.032594555 | 0.037043395 | -0.004448841 | 0.999933887 | 0.713192601 | 1.039850184 | -0.326657583 | 0.009406642 |
| ENSG00000166164 | *BRD7* | 4.193765859 | -0.01204484 | -0.078326298 | 0.066281458 | 0.999933887 | -0.153738624 | -0.463833156 | 0.310094531 | 0.009406642 |
| ENSG00000089818 | *NECAP1* | 7.677512757 | -0.000935435 | 0.022632537 | -0.023567972 | 0.999933887 | 0.905468036 | 1.209450868 | -0.303982831 | 0.009406642 |
| ENSG00000144824 | *PHLDB2* | 1.021912735 | -0.134666333 | -0.00339066 | -0.131275673 | 0.999933887 | -0.198426138 | 0.574878096 | -0.773304235 | 0.009414184 |
| ENSG00000226137 | *BAIAP2-DT* | 1.071123398 | 0.079335706 | 0.130551061 | -0.051215355 | 0.999933887 | -0.752480221 | -1.64244499 | 0.889964769 | 0.009482901 |
| ENSG00000143851 | *PTPN7* | 5.978877678 | 0.005731372 | -0.003760213 | 0.009491584 | 0.999933887 | -0.278638068 | -0.520342213 | 0.241704145 | 0.009482901 |
| ENSG00000213073 | *CHP1P2* | 5.514519925 | 0.006988579 | 0.027512921 | -0.020524342 | 0.999933887 | -0.043870424 | 0.229529526 | -0.27339995 | 0.009494612 |
| ENSG00000157020 | *SEC13* | 5.281488213 | 0.026031187 | -0.007305728 | 0.033336915 | 0.999933887 | 0.921007004 | 1.210820858 | -0.289813854 | 0.00949591 |
| ENSG00000140553 | *UNC45A* | 3.843271187 | -0.066109764 | 0.012404157 | -0.078513921 | 0.999933887 | -0.403809327 | -0.748808291 | 0.344998964 | 0.009540465 |
| ENSG00000133703 | *KRAS* | 5.461825027 | -0.032985494 | -0.023658962 | -0.009326532 | 0.999933887 | 0.238819641 | 0.578703356 | -0.339883715 | 0.009570827 |
| ENSG00000229314 | *ORM1* | 6.265369327 | -0.083518107 | -0.0129869 | -0.070531207 | 0.999933887 | 1.906484722 | 2.454878383 | -0.54839366 | 0.009583212 |
| ENSG00000107679 | *PLEKHA1* | 3.111521934 | -0.06132005 | -0.098659563 | 0.037339513 | 0.999933887 | -0.13251171 | 0.270249838 | -0.402761548 | 0.009654826 |
| ENSG00000124391 | *IL17C* | -0.923840129 | -0.263117486 | 0.091984826 | -0.355102313 | 0.999933887 | 2.19907867 | 3.682139989 | -1.483061318 | 0.009667701 |
| ENSG00000214078 | *CPNE1* | 5.931749481 | -0.013730516 | -0.009508467 | -0.004222049 | 0.999933887 | -0.226117658 | -0.448782654 | 0.222664996 | 0.009670933 |
| ENSG00000160190 | *SLC37A1* | 4.079318244 | 0.091788901 | 0.022445373 | 0.069343528 | 0.999933887 | 0.2144093 | 0.496587174 | -0.282177874 | 0.009770337 |
| ENSG00000177542 | *SLC25A22* | 2.636201158 | 0.078608303 | 0.051326041 | 0.027282263 | 0.999933887 | -0.221748663 | -0.588552586 | 0.366803924 | 0.009770337 |
| ENSG00000153560 | *UBP1* | 4.650206307 | -0.043071164 | -0.041290543 | -0.001780621 | 0.999933887 | -0.13505712 | -0.357305131 | 0.222248012 | 0.009770337 |
| ENSG00000028137 | *TNFRSF1B* | 11.174381683 | -0.021473395 | -0.067566405 | 0.04609301 | 0.999933887 | 0.367778706 | 0.626660999 | -0.258882293 | 0.009808343 |
| ENSG00000167323 | *STIM1* | 5.270823792 | 0.027328536 | -0.015312105 | 0.042640641 | 0.999933887 | -0.082401828 | -0.398636699 | 0.316234871 | 0.009809565 |
| ENSG00000110958 | *PTGES3* | 5.462545915 | -0.046095511 | -0.066941526 | 0.020846015 | 0.999933887 | -0.296022828 | -0.533195599 | 0.237172771 | 0.009842122 |
| ENSG00000130584 | *ZBTB46* | 1.858074077 | -0.037413913 | -0.087096973 | 0.049683059 | 0.999933887 | 0.905896053 | 1.521510084 | -0.615614031 | 0.009842122 |
| ENSG00000160404 | *TOR2A* | 3.543413824 | 0.08592025 | 0.033963457 | 0.051956793 | 0.999933887 | -0.488581821 | -0.923544837 | 0.434963015 | 0.009848846 |
| ENSG00000104904 | *OAZ1* | 8.742387652 | 0.030133638 | 0.007685578 | 0.02244806 | 0.999933887 | -0.120888419 | -0.36036131 | 0.239472891 | 0.00988455 |
| ENSG00000125863 | *MKKS* | 1.770474631 | 0.18033562 | 0.117344542 | 0.062991077 | 0.999933887 | 1.379000312 | 1.985456542 | -0.606456231 | 0.009893026 |
| ENSG00000125304 | *TM9SF2* | 6.717711858 | -0.076129994 | -0.068458879 | -0.007671115 | 0.999933887 | -0.599544822 | -0.903973644 | 0.304428822 | 0.009944651 |
| ENSG00000152409 | *JMY* | 5.745621033 | -0.046111212 | -0.119029985 | 0.072918773 | 0.999933887 | 0.127976507 | 0.518526867 | -0.39055036 | 0.009953104 |
| ENSG00000120063 | *GNA13* | 8.876704312 | -0.049292756 | -0.074489612 | 0.025196855 | 0.999933887 | -0.348288931 | -0.073262741 | -0.27502619 | 0.00997099 |
| ENSG00000129646 | *QRICH2* | 1.398124915 | 0.077687199 | 0.16529589 | -0.087608691 | 0.999933887 | 0.827455381 | 1.568663662 | -0.741208281 | 0.009975333 |
| ENSG00000122694 | *GLIPR2* | 7.010974532 | -0.011302969 | -0.027946205 | 0.016643236 | 0.999933887 | -0.340342749 | -0.675142287 | 0.334799538 | 0.010041767 |
| ENSG00000236790 | *LINC00299* | 1.799289496 | 0.041228566 | -0.010158516 | 0.051387082 | 0.999933887 | 0.918919667 | 1.64162158 | -0.722701913 | 0.010088935 |
| ENSG00000152270 | *PDE3B* | 5.647294421 | 0.01997071 | -0.009712254 | 0.029682964 | 0.999933887 | -0.58367611 | -1.029507394 | 0.445831284 | 0.010090507 |
| ENSG00000278195 | *SSTR3* | 4.186693269 | 0.190153606 | 0.125898613 | 0.064254993 | 0.999933887 | 0.54445333 | 0.982243936 | -0.437790607 | 0.010133444 |
| ENSG00000251615 | *AC104825.1* | -0.023591424 | -0.253317254 | -0.029516397 | -0.223800857 | 0.999933887 | 0.759468217 | 1.552275775 | -0.792807558 | 0.010133444 |
| ENSG00000176788 | *BASP1* | 10.901470166 | 0.058799905 | 0.0649927 | -0.006192795 | 0.999933887 | 0.42565455 | 0.759005232 | -0.333350682 | 0.010133444 |
| ENSG00000258867 | *LINC01146* | 0.154541979 | -0.129548017 | 0.092530116 | -0.222078133 | 0.999933887 | 0.342286967 | 1.274965253 | -0.932678286 | 0.010148721 |
| ENSG00000103479 | *RBL2* | 6.146165119 | -0.011579048 | -0.036353435 | 0.024774388 | 0.999933887 | -0.322176561 | -0.581388196 | 0.259211635 | 0.010148721 |
| ENSG00000251023 | *AC114980.1* | 0.254325548 | 0.097403908 | -0.316571353 | 0.413975261 | 0.999933887 | -0.647488076 | 0.225279144 | -0.87276722 | 0.010153306 |
| ENSG00000068383 | *INPP5A* | 5.847255056 | 0.016687967 | 0.017312662 | -0.000624695 | 0.999933887 | 0.616073765 | 0.861324896 | -0.245251132 | 0.010229382 |
| ENSG00000129667 | *RHBDF2* | 7.101719398 | 0.007965546 | 0.019476173 | -0.011510627 | 0.999933887 | 1.157807091 | 1.564998487 | -0.407191396 | 0.010239963 |
| ENSG00000136731 | *UGGT1* | 5.686446829 | -0.046944151 | -0.055159896 | 0.008215744 | 0.999933887 | -0.150149625 | -0.377716083 | 0.227566458 | 0.010363449 |
| ENSG00000277130 | *AC073569.3* | 0.175215792 | -0.010036518 | 0.030645543 | -0.040682061 | 0.999933887 | 1.168727557 | 2.119671905 | -0.950944349 | 0.010363449 |
| ENSG00000102547 | *CAB39L* | 3.289958063 | -0.043269032 | -0.061952555 | 0.018683523 | 0.999933887 | -0.390628454 | -0.103091743 | -0.287536711 | 0.010372544 |
| ENSG00000185947 | *ZNF267* | 8.982942632 | -0.103472242 | -0.221857872 | 0.11838563 | 0.999933887 | 0.226989291 | 0.756069777 | -0.529080486 | 0.010401281 |
| ENSG00000109065 | *NAT9* | 4.689996478 | 0.037694899 | 0.048041382 | -0.010346483 | 0.999933887 | -0.294480414 | -0.612633296 | 0.318152882 | 0.010494747 |
| ENSG00000185728 | *YTHDF3* | 7.315253888 | -0.048455314 | -0.078716134 | 0.03026082 | 0.999933887 | 0.007688322 | 0.357356537 | -0.349668215 | 0.010506339 |
| ENSG00000162961 | *DPY30* | 2.505445363 | 0.058762763 | -0.068587064 | 0.127349827 | 0.999933887 | 0.092053026 | -0.44978681 | 0.541839836 | 0.010506339 |
| ENSG00000091527 | *CDV3* | 7.072038252 | -0.007418973 | -0.000884811 | -0.006534162 | 0.999933887 | -0.572257877 | -0.832656247 | 0.26039837 | 0.010506339 |

| | | | | | | | | | | |
|---|---|---|---|---|---|---|---|---|---|---|
| ENSG00000251136 | *AF117829.1* | 3.511940203 | 0.10538667 | 0.185128174 | -0.079741504 | 0.999933887 | 1.834988462 | 2.363214407 | -0.528225946 | 0.01050666 |
| ENSG00000131504 | *DIAPH1* | 7.842246107 | -0.028113308 | -0.01402163 | -0.014091678 | 0.999933887 | -0.412681204 | -0.648918313 | 0.236237109 | 0.010509856 |
| ENSG00000145860 | *RNF145* | 7.240198162 | -0.02325847 | -0.00711028 | -0.01614819 | 0.999933887 | 0.00213308 | 0.212190768 | -0.210057689 | 0.010509856 |
| ENSG00000197442 | *MAP3K5* | 5.858460184 | -0.011324201 | -0.027934916 | 0.016610715 | 0.999933887 | 0.709733419 | 1.118590911 | -0.408857492 | 0.010509856 |
| ENSG00000158161 | *EYA3* | 5.182168039 | -0.032539437 | -0.11595749 | 0.083418053 | 0.999933887 | 1.047202629 | 1.505236559 | -0.45803393 | 0.010524484 |
| ENSG00000163348 | *PYGO2* | 3.671006864 | -0.051331624 | -0.059485585 | 0.008153961 | 0.999933887 | -0.056798566 | -0.350647826 | 0.29384926 | 0.010541794 |
| ENSG00000101160 | *CTSZ* | 6.369313139 | 0.064040072 | 0.080247127 | -0.016207055 | 0.999933887 | -0.571192573 | -1.058611308 | 0.487418736 | 0.010541794 |
| ENSG00000152503 | *TRIM36* | -0.328209948 | -0.077627758 | 0.287528113 | -0.365155871 | 0.999933887 | 1.381463354 | 2.314578902 | -0.933115549 | 0.010541794 |
| ENSG00000185650 | *ZFP36L1* | 10.227713044 | 0.158110291 | 0.27203099 | -0.113920699 | 0.999933887 | -0.051854803 | 0.213379747 | -0.26523455 | 0.010563592 |
| ENSG00000213281 | *NRAS* | 5.33815715 | -0.055862738 | -0.062162689 | 0.006299951 | 0.999933887 | -0.134181905 | 0.122195606 | -0.256377511 | 0.010563592 |
| ENSG00000181790 | *ADGRB1* | 2.663837468 | 0.206042661 | -0.012829977 | 0.218872638 | 0.999933887 | 1.110438731 | 1.545072711 | -0.43463398 | 0.010563592 |
| ENSG00000185049 | *NELFA* | 3.177133043 | 0.060862976 | 0.020056231 | 0.040806745 | 0.999933887 | -0.002577353 | -0.354905388 | 0.352328035 | 0.010563592 |
| ENSG00000177105 | *RHOG* | 8.561865451 | 0.020831145 | -0.016672371 | 0.037503516 | 0.999933887 | 0.624062492 | 0.943118165 | -0.319055672 | 0.010617206 |
| ENSG00000115762 | *PLEKHB2* | 8.825658248 | -0.077090917 | -0.02254532 | -0.054545597 | 0.999933887 | 0.661424032 | 0.90188612 | -0.240462087 | 0.010632742 |
| ENSG00000240065 | *PSMB9* | 6.759828514 | 0.07329268 | -0.002422189 | 0.075714869 | 0.999933887 | -0.275809712 | -0.573482424 | 0.297672712 | 0.010632742 |
| ENSG00000005889 | *ZFX* | 6.097715613 | -0.072119743 | -0.052820689 | -0.019299053 | 0.999933887 | -0.102311731 | 0.268264034 | -0.370575766 | 0.010632742 |
| ENSG00000125430 | *HS3ST3B1* | 4.056506548 | 0.041534981 | 0.071983571 | -0.03044859 | 0.999933887 | 1.557413477 | 2.442758688 | -0.885345211 | 0.010632742 |
| ENSG00000167642 | *SPINT2* | 3.587560123 | 0.066144615 | 0.070899075 | -0.00475446 | 0.999933887 | 0.727382751 | 1.145069414 | -0.417686663 | 0.010688145 |
| ENSG00000113575 | *PPP2CA* | 6.793789624 | -0.025398411 | -0.059779993 | 0.034381583 | 0.999933887 | 0.082997606 | 0.2831465 | -0.200148894 | 0.010760612 |
| ENSG00000237765 | *FAM200B* | 5.135756596 | 0.029675396 | 0.121716292 | -0.092040896 | 0.999933887 | 0.109943254 | 0.448373787 | -0.338430534 | 0.010799218 |
| ENSG00000171490 | *RSL1D1* | 4.470106222 | -0.036120295 | -0.144780274 | 0.108659979 | 0.999933887 | 0.268775929 | 0.563937984 | -0.295162055 | 0.010807398 |
| ENSG00000090924 | *PLEKHG2* | 5.728202768 | 0.137574351 | 0.072999346 | 0.064575004 | 0.999933887 | 0.573412475 | 0.897490486 | -0.324078011 | 0.010832781 |
| ENSG00000126883 | *NUP214* | 6.1816606 | -0.028549281 | 0.021537208 | -0.050086489 | 0.999933887 | -0.447257297 | -0.764967384 | 0.317710087 | 0.010861629 |
| ENSG00000160791 | *CCR5* | 3.190003322 | -0.007802735 | -0.030219511 | 0.022416776 | 0.999933887 | 0.383318144 | 0.721141981 | -0.337823837 | 0.010946562 |
| ENSG00000112242 | *E2F3* | 6.98068466 | -0.073683169 | -0.059241002 | -0.014442167 | 0.999933887 | 0.007584752 | 0.237992072 | -0.23040732 | 0.010954779 |
| ENSG00000143878 | *RHOB* | 6.048418309 | 0.117438298 | 0.087354419 | 0.030083879 | 0.999933887 | -0.475420707 | -0.991000708 | 0.515580001 | 0.010997122 |
| ENSG00000153291 | *SLC25A27* | 1.941049561 | -0.062693327 | -0.157886435 | 0.095193108 | 0.999933887 | 0.243143129 | 0.683686901 | -0.440543772 | 0.010997122 |
| ENSG00000185753 | *CXorf38* | 4.915340699 | -0.086399895 | -0.031077622 | -0.055322273 | 0.999933887 | -0.746023256 | -1.111652666 | 0.36562941 | 0.011009003 |
| ENSG00000139645 | *ANKRD52* | 4.376852408 | 0.029742661 | -0.038055304 | 0.067797965 | 0.999933887 | -0.121205068 | -0.394055309 | 0.272850241 | 0.011009003 |
| ENSG00000128335 | *APOL2* | 6.351305992 | -0.023231552 | -0.040004792 | 0.016773239 | 0.999933887 | -0.759563268 | -1.074906867 | 0.315343599 | 0.011071997 |
| ENSG00000130305 | *NSUN5* | 3.237321084 | 0.022182889 | -0.106924196 | 0.129107085 | 0.999933887 | -0.293165105 | -0.671021667 | 0.377856562 | 0.011074296 |
| ENSG00000143390 | *RFX5* | 4.640711344 | 0.00883039 | 0.012053425 | -0.003223036 | 0.999933887 | 0.605288565 | 0.942140271 | -0.336851706 | 0.011104238 |
| ENSG00000173171 | *MTX1* | 3.917368937 | -0.058354905 | -0.059166409 | 0.000811504 | 0.999933887 | -0.783360805 | -1.252815606 | 0.469454801 | 0.011149479 |
| ENSG00000170035 | *UBE2E3* | 3.908368844 | -0.05525784 | -0.020217478 | -0.035040363 | 0.999933887 | -0.295338707 | -0.636020747 | 0.34068204 | 0.011149825 |
| ENSG00000102125 | *TAZ* | 4.775907872 | 0.02224218 | 0.04563642 | -0.023394241 | 0.999933887 | -0.183394325 | -0.469043099 | 0.285648774 | 0.011149825 |
| ENSG00000148516 | *ZEB1* | 5.856201958 | -0.045590037 | -0.025684079 | -0.019905958 | 0.999933887 | -0.514545561 | 0.087885488 | -0.602431048 | 0.011151931 |
| ENSG00000172578 | *KLHL6* | 7.012129582 | -0.032206794 | -0.063599164 | 0.03139237 | 0.999933887 | -0.175123658 | 0.07533749 | -0.250461148 | 0.011153532 |
| ENSG00000076053 | *RBM7* | 4.310456205 | -0.00887466 | 0.032067019 | -0.040941679 | 0.999933887 | 0.055396993 | 0.341338199 | -0.285941207 | 0.011199192 |
| ENSG00000113595 | *TRIM23* | 3.685239511 | -0.027321072 | -0.178534929 | 0.151213857 | 0.999933887 | -0.338639944 | 0.11749439 | -0.456134334 | 0.01121816 |
| ENSG00000184831 | *APOO* | 0.156078099 | 0.008194999 | 0.120595694 | -0.112400695 | 0.999933887 | 0.422874392 | 1.400346424 | -0.977472032 | 0.011242017 |
| ENSG00000084110 | *HAL* | 5.695709964 | -0.030051023 | 0.019022067 | -0.04907309 | 0.999933887 | -0.50368555 | -0.77137753 | 0.26769198 | 0.011255493 |
| ENSG00000149187 | *CELF1* | 7.327662126 | -0.060125646 | 0.000134557 | -0.060260203 | 0.999933887 | 0.608466972 | 0.920380812 | -0.31191384 | 0.011258303 |
| ENSG00000156127 | *BATF* | 3.396478853 | 0.051996095 | 0.038844835 | 0.01315126 | 0.999933887 | 0.541433306 | 1.0393963 | -0.497962994 | 0.01128075 |
| ENSG00000115590 | *IL1R2* | 7.753039878 | 0.043217129 | 0.018033767 | 0.025183362 | 0.999933887 | 0.780421163 | 1.065272306 | -0.284851143 | 0.011351218 |
| ENSG00000150961 | *SEC24D* | 4.44030428 | -0.01869759 | -0.019608248 | 0.000910658 | 0.999933887 | -0.416295045 | -0.729255341 | 0.312960297 | 0.011367767 |
| ENSG00000110330 | *BIRC2* | 6.057995984 | -0.010533721 | -0.06241607 | 0.051882349 | 0.999933887 | 0.673331849 | 1.033310622 | -0.359978773 | 0.011376317 |
| ENSG00000152484 | *USP12* | 4.168754619 | -0.051422108 | -0.067100269 | 0.015678161 | 0.999933887 | 0.452887055 | 0.827403671 | -0.374516617 | 0.01140366 |
| ENSG00000234191 | *LINC01283* | -1.860567304 | -0.131239876 | -0.377429623 | 0.246189746 | 0.999933887 | 0.988421698 | 1.675927248 | -0.68750555 | 0.01140366 |
| ENSG00000263847 | *AP005899.1* | 0.166121991 | -0.103884757 | 0.294984807 | -0.398869564 | 0.999933887 | 0.479986905 | 1.286430532 | -0.806443627 | 0.011406925 |
| ENSG00000144711 | *IQSEC1* | 9.045221391 | -0.003316012 | -0.037473737 | 0.034157725 | 0.999933887 | 0.310579568 | 0.492637418 | -0.182057851 | 0.011531948 |
| ENSG00000105352 | *CEACAM4* | 3.518691199 | 0.035218863 | 0.003707399 | 0.031511465 | 0.999933887 | -0.218814357 | -0.851654138 | 0.632839781 | 0.011554888 |
| ENSG00000139629 | *GALNT6* | 3.066037494 | 0.013376745 | 0.108963584 | -0.095586839 | 0.999933887 | 0.202425564 | 0.513430819 | -0.311005255 | 0.011556622 |
| ENSG00000131323 | *TRAF3* | 7.46318198 | -0.052816301 | -0.045652897 | -0.007163403 | 0.999933887 | 0.634785433 | 0.893741738 | -0.258956305 | 0.011556807 |
| ENSG00000272079 | *AC004233.2* | 1.339885517 | 0.019292209 | 0.062170771 | -0.042878562 | 0.999933887 | -1.315140047 | -2.101058578 | 0.785918531 | 0.011576407 |
| ENSG00000197632 | *SERPINB2* | 5.538252544 | 0.143361014 | 0.198751284 | -0.05539027 | 0.999933887 | 0.877009034 | 1.407894885 | -0.530885851 | 0.011709949 |
| ENSG00000146205 | *ANO7* | -2.348720182 | -0.019785324 | 0.465266785 | -0.485052109 | 0.999933887 | -0.875806674 | 0.924822962 | -1.800629636 | 0.011709949 |
| ENSG00000172977 | *KAT5* | 4.726366166 | 0.000621441 | 0.03773906 | -0.037117619 | 0.999933887 | 0.159889125 | 0.407412697 | -0.247523572 | 0.011709949 |
| ENSG00000075651 | *PLD1* | 4.008056231 | -0.010642916 | -0.023742894 | 0.013099978 | 0.999933887 | 2.862938065 | 3.656083757 | -0.793145692 | 0.011722331 |
| ENSG00000101190 | *TCFL5* | 3.844685828 | 0.117050929 | 0.075564003 | 0.041486926 | 0.999933887 | 3.325746097 | 4.011249057 | -0.685502959 | 0.011722647 |
| ENSG00000173575 | *CHD2* | 9.155684252 | -0.087246768 | -0.061866589 | -0.025380178 | 0.999933887 | 0.58640516 | 0.909698626 | -0.323293467 | 0.011737192 |
| ENSG00000273812 | *BX640514.2* | 3.673831059 | 0.183881758 | 0.266708982 | -0.082827223 | 0.999933887 | 1.345592779 | 1.875113966 | -0.529521188 | 0.011750009 |
| ENSG00000256771 | *ZNF253* | 0.101054558 | 0.127940137 | -0.12304191 | 0.250982048 | 0.999933887 | -0.383632841 | 0.299534375 | -0.683167216 | 0.011792753 |
| ENSG00000138413 | *IDH1* | 1.690641087 | -0.060699693 | 0.031933731 | -0.092633424 | 0.999933887 | -0.376482319 | -0.841612536 | 0.465130217 | 0.011797255 |
| ENSG00000042445 | *RETSAT* | 3.093894669 | -0.028548546 | 0.088169596 | -0.116718142 | 0.999933887 | -0.065055047 | -0.430469624 | 0.365414577 | 0.011797255 |
| ENSG00000008226 | *DLEC1* | 3.238949677 | -0.089540199 | 0.021738152 | -0.111278351 | 0.999933887 | 0.836721362 | 1.575353356 | -0.738631994 | 0.011862587 |
| ENSG00000178385 | *PLEKHM3* | 5.823228419 | 0.075183411 | 0.089173644 | -0.013990233 | 0.999933887 | 1.496096554 | 1.924641339 | -0.428544785 | 0.011887427 |
| ENSG00000204852 | *TCTN1* | 2.13737168 | -0.107884891 | -0.019194182 | -0.08869071 | 0.999933887 | -0.692186726 | -1.250062052 | 0.557875327 | 0.011909977 |
| ENSG00000233578 | *EIF4EP1* | 3.168272062 | -0.020807394 | 0.098122156 | -0.11892955 | 0.999933887 | -0.139265927 | 0.455845357 | -0.595111284 | 0.011917758 |
| ENSG00000133059 | *DSTYK* | 3.602250813 | 0.028138586 | -0.006084825 | 0.034223411 | 0.999933887 | -0.494435267 | -0.959232194 | 0.464796927 | 0.011931489 |
| ENSG00000151715 | *TMEM45B* | 2.51879103 | 0.023122651 | 0.033327269 | -0.010204618 | 0.999933887 | -0.809321269 | -1.351506015 | 0.542184746 | 0.01195218 |
| ENSG00000147894 | *C9orf72* | 8.228645741 | -0.079204303 | -0.083149461 | 0.003945157 | 0.999933887 | 0.215626813 | 0.654578968 | -0.438952155 | 0.011961847 |
| ENSG00000144560 | *VGLL4* | 2.250880804 | 0.05659976 | 0.032246939 | 0.024352821 | 0.999933887 | 0.062687756 | 0.49084828 | -0.428160525 | 0.011961847 |
| ENSG00000158545 | *ZC3H18* | 6.232387196 | 0.046870524 | 0.058222232 | -0.011351708 | 0.999933887 | 0.677823455 | 1.057625599 | -0.379802144 | 0.011961847 |
| ENSG00000131503 | *ANKHD1* | 3.085394198 | -0.082840432 | -0.014066653 | -0.06877378 | 0.999933887 | -0.109060126 | 0.278109584 | -0.387169711 | 0.011997124 |
| ENSG00000130939 | *UBE4B* | 5.722794814 | -0.013660968 | 0.004921279 | -0.018582247 | 0.999933887 | -0.489003254 | -0.707440305 | 0.218437051 | 0.012004854 |
| ENSG00000103152 | *MPG* | 2.260185302 | 0.162801136 | 0.150530253 | 0.012270883 | 0.999933887 | -0.66827048 | -1.296693545 | 0.628423064 | 0.012135237 |
| ENSG00000169554 | *ZEB2* | 9.28070514 | 0.029194384 | 0.124775573 | -0.095581189 | 0.999933887 | 1.203070252 | 1.55313599 | -0.350065738 | 0.012266245 |
| ENSG00000103510 | *KAT8* | 5.308247131 | -0.039821947 | -0.020185569 | -0.019636379 | 0.999933887 | -0.230297527 | -0.481281858 | 0.250984331 | 0.012276005 |
| ENSG00000241886 | *AC112496.1* | 3.520233279 | -0.110503506 | -0.034189323 | -0.076314184 | 0.999933887 | 0.067392579 | 0.580081475 | -0.512688896 | 0.012282594 |
| ENSG00000105221 | *AKT2* | 6.423437025 | 0.059159866 | 0.024092124 | 0.035067742 | 0.999933887 | -0.199226542 | -0.411609669 | 0.212383127 | 0.012297505 |
| ENSG00000116017 | *ARID3A* | 5.930210051 | 0.091386017 | 0.096936953 | -0.005550936 | 0.999933887 | 0.558969554 | 0.854016172 | -0.295046618 | 0.012309628 |
| ENSG00000147533 | *GOLGA7* | 4.881614707 | 0.080935032 | -0.017521754 | 0.098456786 | 0.999933887 | 0.908377052 | 1.240924794 | -0.332547743 | 0.01231622 |
| ENSG00000107897 | *ACBD5* | 3.387071544 | 0.010718137 | 0.039105214 | -0.028387077 | 0.999933887 | -0.486648973 | -0.885015981 | 0.398367008 | 0.012355476 |
| ENSG00000166938 | *DIS3L* | 2.059533307 | -0.042643307 | 0.063471157 | -0.106114465 | 0.999933887 | 0.054856683 | 0.417181952 | -0.362325269 | 0.01243997 |
| ENSG00000172366 | *MCRIP2* | 3.532468191 | 0.071251005 | 0.130110436 | -0.058859431 | 0.999933887 | 1.578141102 | 2.113796327 | -0.535655225 | 0.012448108 |
| ENSG00000165219 | *GAPVD1* | 5.304796898 | 0.01115629 | -0.07973968 | 0.090895969 | 0.999933887 | -0.396204603 | -0.708157437 | 0.311952834 | 0.012453929 |
| ENSG00000141232 | *TOB1* | 4.91367584 | 0.088243049 | 0.166141401 | -0.077898352 | 0.999933887 | -0.186265208 | -0.478733031 | 0.292467824 | 0.012499695 |
| ENSG00000144635 | *DYNC1LI1* | 5.232149402 | -0.03399081 | 0.042004799 | -0.075995609 | 0.999933887 | -0.829334255 | -1.113533274 | 0.284199019 | 0.012583477 |
| ENSG00000139436 | *GIT2* | 5.338432749 | -0.060501593 | -0.058751343 | -0.00175025 | 0.999933887 | -0.335012682 | -0.562582187 | 0.227569505 | 0.012589684 |
| ENSG00000215644 | *GCGR* | -2.711519817 | 0.192334257 | -0.6118585 | 0.804192756 | 0.999933887 | 3.756058346 | 5.510160314 | -1.754101968 | 0.012589684 |
| ENSG00000125457 | *MIF4GD* | 2.418931233 | -0.121716603 | 0.084437411 | -0.206154014 | 0.999933887 | -0.171402944 | -0.617052548 | 0.445649604 | 0.012634613 |
| ENSG00000076928 | *ARHGEF1* | 8.675629157 | 0.045045061 | 0.066361138 | -0.021316077 | 0.999933887 | -0.116954625 | -0.372541575 | 0.25558695 | 0.012634613 |
| ENSG00000001084 | *GCLC* | 4.639552566 | -0.055968176 | -0.01418645 | -0.041781726 | 0.999933887 | 0.936289066 | 1.424820087 | -0.488531021 | 0.012634613 |
| ENSG00000247853 | *AC006064.2* | 0.349552401 | 0.131784787 | 0.006005442 | 0.125779345 | 0.999933887 | 0.042480658 | 0.909395985 | -0.866915327 | 0.012643189 |
| ENSG00000154359 | *LONRF1* | 5.709431304 | 0.091855171 | 0.102907852 | -0.01105268 | 0.999933887 | 0.129105435 | -0.210379564 | 0.339484999 | 0.012644379 |
| ENSG00000153201 | *RANBP2* | 8.240332071 | -0.197514877 | -0.149126969 | -0.048387908 | 0.999933887 | 0.358045071 | 1.132569633 | -0.774524562 | 0.012644379 |
| ENSG00000114302 | *PRKAR2A* | 4.164728642 | -0.057249133 | -0.000882371 | -0.056366762 | 0.999933887 | -0.189111892 | -0.516971604 | 0.327859711 | 0.012644379 |
| ENSG00000204397 | *CARD16* | 6.968261626 | 0.060163279 | -0.033339035 | 0.093502314 | 0.999933887 | 1.079579167 | 1.416385788 | -0.33680662 | 0.012644379 |
| ENSG00000182541 | *LIMK2* | 9.839064238 | 0.022723855 | -0.005119695 | 0.027843551 | 0.999933887 | 1.449915822 | 1.717432614 | -0.267516792 | 0.012644379 |
| ENSG00000204308 | *RNF5* | 3.082459261 | -0.01645846 | 0.046878031 | -0.063336491 | 0.999933887 | -0.042228247 | -0.453744212 | 0.411515965 | 0.012644379 |
| ENSG00000099968 | *BCL2L13* | 4.366125375 | 0.022047595 | 0.05758192 | -0.035534325 | 0.999933887 | 0.360947505 | 0.620755781 | -0.259808276 | 0.012654086 |
| ENSG00000059915 | *PSD* | 2.435739418 | 0.245408713 | 0.299338864 | -0.053930151 | 0.999933887 | 1.730772705 | 2.306519998 | -0.575747293 | 0.012663439 |
| ENSG00000188786 | *MTF1* | 7.053393435 | -0.055756272 | -0.032230066 | -0.023526206 | 0.999933887 | 2.209454543 | 2.681010067 | -0.471555524 | 0.012663439 |
| ENSG00000196396 | *PTPN1* | 6.106365671 | -0.007997898 | -0.017392393 | 0.009394495 | 0.999933887 | 0.377877589 | 0.714916587 | -0.337038998 | 0.012663439 |
| ENSG00000007168 | *PAFAH1B1* | 7.433795082 | -0.043818491 | -0.031120466 | -0.012698026 | 0.999933887 | 0.364209187 | 0.590393965 | -0.226184778 | 0.012750032 |

| | | | | | | | | | | |
|---|---|---|---|---|---|---|---|---|---|---|
| ENSG00000105825 | *TFPI2* | -1.826989997 | -0.198870711 | 0.081314654 | -0.280185365 | 0.999933887 | 0.815095839 | 1.743640737 | -0.928544898 | 0.012831371 |
| ENSG00000116525 | *TRIM62* | 3.631261264 | 0.141556324 | 0.080426244 | 0.06113008 | 0.999933887 | -0.316363662 | -0.664986702 | 0.34862304 | 0.012900138 |
| ENSG00000186431 | *FCAR* | 8.125259563 | 0.157449278 | 0.136168643 | 0.021280635 | 0.999933887 | 1.82801463 | 2.226175811 | -0.398161181 | 0.012900138 |
| ENSG00000185112 | *FAM43A* | 1.714431669 | -0.061971234 | -0.05237038 | -0.009600854 | 0.999933887 | -0.069812518 | -0.544380571 | 0.474568053 | 0.012919341 |
| ENSG00000110436 | *SLC1A2* | -0.610501627 | 0.025844316 | 0.469128589 | -0.443284273 | 0.999933887 | 0.498128198 | 1.6035353 | -1.105407102 | 0.013040327 |
| ENSG00000148356 | *LRSAM1* | 3.905584776 | -0.026697711 | 0.10499416 | -0.131691871 | 0.999933887 | -0.13452494 | -0.485645377 | 0.351120437 | 0.013050847 |
| ENSG00000225422 | *RBMS1P1* | 1.967570959 | -0.109990479 | -0.176211863 | 0.066221384 | 0.999933887 | 1.518252441 | 2.191621258 | -0.673368818 | 0.013073263 |
| ENSG00000151689 | *INPP1* | 2.618727644 | -0.053628484 | -0.063140363 | 0.009511879 | 0.999933887 | -0.06118912 | 0.29969403 | -0.36088315 | 0.013119632 |
| ENSG00000143149 | *ALDH9A1* | 3.858763117 | -0.129485151 | 0.072154846 | -0.201639996 | 0.969206975 | -0.390335497 | -0.688606519 | 0.298271021 | 0.013241847 |
| ENSG00000111802 | *TDP2* | 8.224673971 | -0.034839047 | -0.101299432 | 0.066460384 | 0.999933887 | 0.06272639 | 0.302388638 | -0.239662247 | 0.013248345 |
| ENSG00000163519 | *TRAT1* | 3.600766461 | -0.05711455 | -0.156843984 | 0.099729434 | 0.999933887 | -0.017975896 | 0.441237722 | -0.459213619 | 0.013248345 |
| ENSG00000139190 | *VAMP1* | 4.865581627 | 0.012092209 | -0.03882217 | 0.050914379 | 0.999933887 | -0.141649874 | -0.439095165 | 0.297445291 | 0.013254284 |
| ENSG00000010361 | *FUZ* | 3.109482473 | 0.059110494 | -0.053564727 | 0.112675221 | 0.999933887 | -0.290151694 | -0.717195444 | 0.427043749 | 0.013510771 |
| ENSG00000171467 | *ZNF318* | 5.448708698 | 0.019378325 | -0.104715805 | 0.12409413 | 0.999933887 | 0.358033471 | 0.646776074 | -0.288742603 | 0.013534104 |
| ENSG00000205208 | *C4orf46* | 2.301216159 | -0.12156041 | -0.103145916 | -0.018414494 | 0.999933887 | -0.051112871 | 0.376669152 | -0.427782023 | 0.013555936 |
| ENSG00000176101 | *SSNA1* | 4.178158686 | 0.111742571 | 0.080319349 | 0.031423223 | 0.999933887 | -0.233473815 | -0.607981136 | 0.374507321 | 0.013555936 |
| ENSG00000168528 | *SERINC2* | 1.185925047 | -0.015113147 | -0.123136798 | 0.108023651 | 0.999933887 | -0.377488068 | 0.017266953 | -0.394755021 | 0.013559372 |
| ENSG00000108798 | *ABI3* | 4.28440785 | 0.042817407 | 0.039928519 | 0.002888888 | 0.999933887 | -0.211565507 | -0.5397714 | 0.328205892 | 0.013563059 |
| ENSG00000165997 | *ARL5B* | 7.614909991 | 0.11578085 | -0.036761365 | 0.152542215 | 0.999933887 | 1.620074593 | 2.219936846 | -0.599862253 | 0.013574174 |
| ENSG00000203644 | *AC083799.1* | 4.886542407 | -0.088546309 | -0.110623317 | 0.022077008 | 0.999933887 | -0.043355902 | 0.238575944 | -0.281931845 | 0.013575728 |
| ENSG00000214194 | *SMIM30* | 0.008631567 | -0.047772845 | -0.278872805 | 0.23109996 | 0.999933887 | -0.003152548 | 0.855045822 | -0.85819837 | 0.013660005 |
| ENSG00000173846 | *PLK3* | 8.676020753 | 0.187103647 | 0.094618976 | 0.092484672 | 0.999933887 | 1.84232184 | 2.268848973 | -0.426527134 | 0.013666711 |
| ENSG00000183513 | *COA5* | 3.080959882 | -0.023887789 | -0.230872137 | 0.206984348 | 0.999933887 | -0.295419809 | -0.608113333 | 0.312693524 | 0.013716153 |
| ENSG00000101266 | *CSNK2A1* | 4.22192711 | -0.055050817 | 0.000551998 | -0.055602815 | 0.999933887 | -0.2914478 | -0.574945141 | 0.283497341 | 0.013770704 |
| ENSG00000138495 | *COX17* | 3.798060225 | -0.008113505 | -0.033528736 | 0.025415232 | 0.999933887 | 0.671133903 | 0.936164405 | -0.265030502 | 0.013770704 |
| ENSG00000166716 | *ZNF592* | 6.431577473 | -0.004139076 | -0.002162699 | -0.001976377 | 0.999933887 | -0.548080853 | -0.784711919 | 0.236631066 | 0.013770704 |
| ENSG00000149679 | *CABLES2* | 3.479721356 | 0.028012312 | 0.022101308 | 0.005911004 | 0.999933887 | 0.504382016 | 0.862741485 | -0.358359468 | 0.013851156 |
| ENSG00000142347 | *MYO1F* | 10.339761823 | -0.011919759 | 0.020869143 | -0.032788902 | 0.999933887 | -0.732455661 | -1.048125121 | 0.31566946 | 0.013891473 |
| ENSG00000091972 | *CD200* | -0.025862591 | 0.212285212 | -0.415687584 | 0.627972796 | 0.999933887 | 1.907100073 | 2.744887274 | -0.837787201 | 0.013933931 |
| ENSG00000114904 | *NEK4* | 1.560331549 | 0.030464031 | -0.07362812 | 0.104092151 | 0.999933887 | 0.279077899 | 0.763116694 | -0.484038795 | 0.013933931 |
| ENSG00000077150 | *NFKB2* | 9.661266513 | 0.0569005 | 0.119924853 | -0.063024353 | 0.999933887 | 2.142827091 | 2.524923529 | -0.382096438 | 0.013938749 |
| ENSG00000100439 | *ABHD4* | 6.294281111 | -0.005065012 | -0.018673714 | 0.013608702 | 0.999933887 | 0.305518615 | 0.557029806 | -0.251511191 | 0.014001786 |
| ENSG00000090863 | *GLG1* | 6.171215998 | -0.022471069 | -0.060152466 | 0.037681397 | 0.999933887 | -0.181495332 | -0.406223598 | 0.224728266 | 0.014039157 |
| ENSG00000105835 | *NAMPT* | 12.55791678 | -0.012940281 | -0.070751979 | 0.057811699 | 0.999933887 | 0.352708477 | 0.658821444 | -0.306112967 | 0.014085244 |
| ENSG00000099942 | *CRKL* | 5.746057521 | -0.118666986 | -0.032895113 | -0.085771872 | 0.999933887 | -0.512913172 | -0.790977943 | 0.278064771 | 0.014088054 |
| ENSG00000164609 | *SLU7* | 6.647631096 | -0.061002597 | -0.095386246 | 0.034383649 | 0.999933887 | 0.004973742 | 0.29442794 | -0.289454198 | 0.014088054 |
| ENSG00000155961 | *RAB39B* | 2.91797255 | -0.041521502 | -0.09986885 | 0.058347348 | 0.999933887 | 0.407896818 | 0.834561771 | -0.426664954 | 0.014156326 |
| ENSG00000135679 | *MDM2* | 6.095659033 | -0.024178395 | 0.046473336 | -0.070651731 | 0.999933887 | 0.269246703 | 0.617810098 | -0.348563395 | 0.014164296 |
| ENSG00000160013 | *PTGIR* | 0.184208693 | 0.056070289 | -0.236905926 | 0.292976215 | 0.999933887 | 2.112590223 | 3.206958746 | -1.094368523 | 0.014167611 |
| ENSG00000134452 | *FBH1* | 4.187715419 | 0.02505346 | 0.024787722 | 0.000265737 | 0.999933887 | -0.118455227 | -0.401722522 | 0.283267295 | 0.014219225 |
| ENSG00000178028 | *DMAP1* | 3.971919006 | -0.073080324 | -0.007018206 | -0.066062119 | 0.999933887 | 0.066433665 | -0.177568112 | 0.244001778 | 0.014262378 |
| ENSG00000136048 | *DRAM1* | 5.486728626 | -0.007692011 | -0.063379066 | 0.055687054 | 0.999933887 | 1.825290721 | 2.347123607 | -0.521832886 | 0.014262378 |
| ENSG00000161204 | *ABCF3* | 4.539952492 | -0.043404046 | 0.012957997 | -0.056362044 | 0.999933887 | -0.05849678 | -0.37330926 | 0.314812479 | 0.014306387 |
| ENSG00000106628 | *POLD2* | 2.309096681 | 0.055568851 | -0.070504216 | 0.126073067 | 0.999933887 | 0.354669295 | 0.814513982 | -0.459844687 | 0.014306387 |
| ENSG00000118496 | *FBXO30* | 4.373865764 | -0.123343248 | -0.081851281 | -0.041491967 | 0.999933887 | 0.131062452 | 0.468501807 | -0.337439355 | 0.014361194 |
| ENSG00000229337 | *AC079305.2* | 2.502359385 | 0.247419817 | 0.00685206 | 0.240567757 | 0.999933887 | 0.815790147 | 0.175588664 | 0.640201483 | 0.014365099 |
| ENSG00000185043 | *CIB1* | 5.70124605 | 0.009008074 | 0.017635034 | -0.00862696 | 0.999933887 | -0.082936328 | -0.297298598 | 0.21436227 | 0.014395085 |
| ENSG00000196502 | *SULT1A1* | 5.791296186 | -0.112840811 | -0.059684733 | -0.053156078 | 0.999933887 | -1.124476633 | -1.471605528 | 0.347128895 | 0.014402877 |
| ENSG00000111186 | *WNT5B* | -1.764529404 | 0.374535542 | 0.183505859 | 0.191029683 | 0.999933887 | 2.560481492 | 4.272027757 | -1.711546265 | 0.014402877 |
| ENSG00000218565 | *AL592429.1* | -1.178158512 | -0.091449694 | 0.474436729 | -0.565886424 | 0.999933887 | 1.465527278 | 2.556874894 | -1.091347616 | 0.014402877 |
| ENSG00000184384 | *MAML2* | 6.886173578 | 0.041396872 | 0.081048547 | -0.039651675 | 0.999933887 | 0.689266616 | 0.983666194 | -0.294399577 | 0.014413097 |
| ENSG00000109381 | *ELF2* | 6.122993817 | -0.021792107 | -0.053518585 | 0.031726478 | 0.999933887 | 0.474902619 | 0.771583961 | -0.296681342 | 0.014425974 |
| ENSG00000161267 | *BDH1* | 1.128901392 | 0.021378715 | 0.099330027 | -0.077951312 | 0.999933887 | 0.079632253 | 0.579335273 | -0.49970302 | 0.014459402 |
| ENSG00000118985 | *ELL2* | 6.098135726 | -0.022700552 | -0.030158119 | 0.007457567 | 0.999933887 | 0.537534687 | 0.848326162 | -0.310791475 | 0.014513967 |
| ENSG00000164332 | *UBLCP1* | 4.992347883 | -0.079690337 | -0.148260745 | 0.068570408 | 0.999933887 | -0.07444164 | 0.467770122 | -0.542211762 | 0.014548207 |
| ENSG00000116266 | *STXBP3* | 5.500003717 | -0.072745559 | -0.117450856 | 0.044705297 | 0.999933887 | -0.177978362 | 0.219159316 | -0.397137678 | 0.014563595 |
| ENSG00000169299 | *PGM2* | 3.35978018 | -0.052553489 | -0.08827763 | 0.035724141 | 0.999933887 | -0.117854231 | -0.556121739 | 0.438267507 | 0.014611988 |
| ENSG00000242732 | *RTL5* | 1.716497299 | -0.100216677 | 0.260746142 | -0.360962819 | 0.944094537 | -0.221982354 | -0.918634462 | 0.696652108 | 0.014636942 |
| ENSG00000141458 | *NPC1* | 6.851303176 | -0.001284045 | -0.024381072 | 0.023097027 | 0.999933887 | 0.329886351 | 0.560811495 | -0.230925143 | 0.014636942 |
| ENSG00000183495 | *EP400* | 5.300325502 | 0.042002401 | -0.001438515 | 0.043440916 | 0.999933887 | -0.078888422 | -0.299883096 | 0.220994674 | 0.014663879 |
| ENSG00000168286 | *THAP11* | 2.729223279 | 0.009633889 | 0.034010145 | -0.024376256 | 0.999933887 | -0.042043874 | -0.439847027 | 0.397803154 | 0.014665368 |
| ENSG00000180773 | *SLC36A4* | 4.082084686 | -0.019563208 | 0.013377082 | -0.03294029 | 0.999933887 | 0.431054067 | 0.862179146 | -0.431125079 | 0.014747037 |
| ENSG00000075413 | *MARK3* | 6.773547546 | -0.006157124 | 0.020864359 | -0.027021483 | 0.999933887 | 0.205003126 | 0.397912638 | -0.192909512 | 0.014865658 |
| ENSG00000134627 | *PIWIL4* | 2.549170873 | 0.00346155 | -0.096042674 | 0.099504224 | 0.999933887 | 1.13534203 | 1.566389885 | -0.431047855 | 0.014865658 |
| ENSG00000138092 | *CENPO* | 1.132166257 | -0.000129372 | 0.161600734 | -0.161730106 | 0.999933887 | 0.834334737 | 1.406881487 | -0.57254675 | 0.014865658 |
| ENSG00000173621 | *LRFN4* | 0.091436031 | 0.090888531 | 0.060937401 | 0.02995113 | 0.999933887 | -0.491221959 | -1.249819434 | 0.758597475 | 0.014909641 |
| ENSG00000148296 | *SURF6* | 3.672569983 | 0.021804498 | 0.100119749 | -0.078315251 | 0.999933887 | -0.085413446 | -0.48727741 | 0.401863964 | 0.014973278 |
| ENSG00000011007 | *ELOA* | 3.578408208 | -0.020624119 | -0.100218615 | 0.079594497 | 0.999933887 | 0.222571127 | 0.485562374 | -0.262991246 | 0.014990028 |
| ENSG00000185745 | *IFIT1* | 5.486816474 | -0.15952655 | -0.231432483 | 0.071905933 | 0.999933887 | -1.137975363 | -1.675818018 | 0.537842655 | 0.014996076 |
| ENSG00000133884 | *DPF2* | 5.089303583 | -0.079802132 | 0.011581994 | -0.091384126 | 0.999933887 | -0.428248725 | -0.724012848 | 0.295764123 | 0.015016873 |
| ENSG00000133063 | *CHIT1* | 2.89009541 | 0.028179297 | 0.07187853 | -0.043699233 | 0.999933887 | 0.831752175 | 1.241969428 | -0.410217252 | 0.015016873 |
| ENSG00000105855 | *ITGB8* | -0.208703392 | -0.031175201 | 0.316308357 | -0.347483558 | 0.999933887 | 2.034246776 | 2.850108641 | -0.815861866 | 0.015016873 |
| ENSG00000087302 | *RTRAF* | 4.355835822 | 0.021748919 | -0.092372072 | 0.114120991 | 0.999933887 | 1.005803963 | 1.45437842 | -0.448574457 | 0.01504698 |
| ENSG00000269836 | *AC020913.3* | -0.08573025 | -0.014705484 | 0.098827715 | -0.1135332 | 0.999933887 | -0.592143895 | 0.126811654 | -0.718955549 | 0.01504698 |
| ENSG00000090621 | *PABPC4* | 5.194573098 | 0.049420982 | 0.030170125 | 0.019250858 | 0.999933887 | 0.215479159 | 0.439545485 | -0.224066326 | 0.015049948 |
| ENSG00000108883 | *EFTUD2* | 4.609530947 | -0.031260013 | -0.093238315 | 0.061978302 | 0.999933887 | -0.088424579 | -0.320028779 | 0.2316042 | 0.015049948 |
| ENSG00000196812 | *ZSCAN16* | 2.557499248 | -0.179131877 | -0.215776902 | 0.036645024 | 0.999933887 | -0.134789146 | 0.357249398 | -0.492038544 | 0.01510437 |
| ENSG00000171608 | *PIK3CD* | 8.374602765 | 0.024684011 | 0.034115873 | -0.009431862 | 0.999933887 | -0.091795383 | -0.35545425 | 0.263658868 | 0.01510437 |
| ENSG00000126456 | *IRF3* | 4.801855358 | 0.054777255 | 0.130915265 | -0.07613801 | 0.999933887 | -0.050310525 | -0.421933064 | 0.37162254 | 0.01525338 |
| ENSG00000136250 | *AOAH* | 5.74365993 | -0.020439828 | -0.039907651 | 0.019467823 | 0.999933887 | 0.205889066 | 0.490670039 | -0.284780972 | 0.01525338 |
| ENSG00000186111 | *PIP5K1C* | 4.535792988 | 0.047539498 | 0.043480772 | 0.004058726 | 0.999933887 | -0.383720364 | -0.682972058 | 0.299251694 | 0.015273117 |
| ENSG00000110876 | *SELPLG* | 8.413014359 | 0.029160618 | -0.05473248 | 0.083893099 | 0.999933887 | -1.523520912 | -2.018286703 | 0.494765791 | 0.015273117 |
| ENSG00000145214 | *DGKQ* | 4.375252855 | 0.071434371 | 0.082666874 | -0.011232502 | 0.999933887 | 0.035427425 | -0.358423992 | 0.393851417 | 0.015274048 |
| ENSG00000129933 | *MAU2* | 6.79062712 | 0.023920879 | 0.009521405 | 0.014399475 | 0.999933887 | -0.488631542 | -0.726837025 | 0.238205483 | 0.015282567 |
| ENSG00000164929 | *BAALC* | -0.910292163 | -0.715765615 | -0.158072801 | -0.557692814 | 0.999933887 | 1.523038148 | 2.249406651 | -0.726368503 | 0.015292463 |
| ENSG00000225572 | *DOCK4-AS1* | 1.689672934 | 0.113224996 | 0.363311404 | -0.250086409 | 0.999933887 | 0.496251091 | 1.033179091 | -0.536928001 | 0.015292463 |
| ENSG00000101421 | *CHMP4B* | 8.709476274 | -0.030208547 | -0.018669829 | -0.011538718 | 0.999933887 | 0.69271367 | 0.939915688 | -0.247202018 | 0.015302451 |
| ENSG00000126945 | *HNRNPH2* | 7.65538199 | -0.08006137 | -0.08879204 | 0.00873067 | 0.999933887 | 0.299526404 | 0.525307544 | -0.22578114 | 0.015305919 |
| ENSG00000111261 | *MANSC1* | 3.731976424 | -0.021175088 | -0.047782463 | 0.026607375 | 0.999933887 | -0.591810814 | -1.067733108 | 0.475922294 | 0.015333609 |
| ENSG00000137462 | *TLR2* | 9.797260779 | -0.079425299 | -0.129966127 | 0.050540828 | 0.999933887 | 0.346705065 | 0.755677563 | -0.408972498 | 0.015383854 |
| ENSG00000271821 | *AL662844.3* | -0.986104636 | -0.418844981 | -0.422675195 | 0.003830214 | 0.999933887 | -1.317887596 | -2.968712084 | 1.650824488 | 0.015450215 |
| ENSG00000185813 | *PCYT2* | 3.807973297 | 0.115990165 | 0.160557858 | -0.044567692 | 0.999933887 | 0.615791382 | 0.992185422 | -0.376394039 | 0.015494584 |
| ENSG00000197457 | *STMN3* | 3.518820389 | 0.013836933 | 0.025893147 | -0.012056214 | 0.999933887 | 0.06250467 | -0.274299538 | 0.336804208 | 0.015494584 |
| ENSG00000052795 | *FNIP2* | 6.622923636 | -0.09038131 | 0.063127703 | -0.153509013 | 0.999933887 | -0.152057477 | 0.1421583 | -0.294215776 | 0.015571179 |
| ENSG00000105499 | *PLA2G4C* | 1.4309254 | -0.078510991 | -0.198338659 | 0.119827668 | 0.999933887 | 0.943603093 | 1.422081948 | -0.478478855 | 0.015571179 |
| ENSG00000211450 | *SELENOH* | 1.03792792 | -0.111210765 | -0.119017681 | 0.007806916 | 0.999933887 | -0.054445233 | -0.493167422 | 0.43872219 | 0.015634624 |
| ENSG00000078237 | *TIGAR* | 0.940634458 | -0.064135974 | 0.143519583 | -0.207655557 | 0.999933887 | -0.386742987 | -1.136095364 | 0.749352377 | 0.015637747 |
| ENSG00000105576 | *TNPO2* | 5.083962922 | 0.025122471 | 0.036794733 | -0.011672262 | 0.999933887 | 0.663565947 | 0.978370677 | -0.31480473 | 0.015637747 |
| ENSG00000174405 | *LIG4* | 4.738603718 | -0.132017511 | -0.226316597 | 0.094299087 | 0.999933887 | 0.073090033 | 0.684315988 | -0.611225955 | 0.015677493 |
| ENSG00000255398 | *HCAR3* | 11.072643251 | 0.114728099 | 0.049456042 | 0.065272057 | 0.999933887 | 0.706859952 | 1.032292916 | -0.325432964 | 0.015692258 |
| ENSG00000125637 | *PSD4* | 7.162385062 | 0.017198438 | -0.000113646 | 0.017312083 | 0.999933887 | 0.103068846 | -0.186290863 | 0.28935971 | 0.015705526 |
| ENSG00000167670 | *CHAF1A* | 2.961129301 | 0.045573172 | -0.046228168 | 0.09180134 | 0.999933887 | 0.728069926 | 1.128767217 | -0.400697291 | 0.015748188 |
| ENSG00000005302 | *MSL3* | 6.199370397 | -0.045302236 | -0.063926472 | 0.018624236 | 0.999933887 | -0.049824417 | -0.250027899 | 0.200203482 | 0.015851672 |
| ENSG00000151092 | *NGLY1* | 4.666161151 | 0.043310463 | 0.017374281 | 0.025936182 | 0.999933887 | -0.65570063 | -1.038644909 | 0.382944279 | 0.015851672 |
| ENSG00000182768 | *NGRN* | 2.024485534 | 0.03826814 | -0.164851399 | 0.20311954 | 0.999933887 | -0.031751526 | -0.439731685 | 0.407980159 | 0.015851672 |

| | | | | | | | | | | |
|---|---|---|---|---|---|---|---|---|---|---|
| ENSG00000115828 | *QPCT* | 6.840130742 | 0.014180721 | -0.043042696 | 0.057223417 | 0.999933887 | 0.121890955 | 0.513026845 | -0.391135891 | 0.015851672 |
| ENSG00000135519 | *KCNH3* | 3.203373382 | 0.080351999 | 0.009527872 | 0.070824126 | 0.999933887 | -0.151962922 | -0.622121734 | 0.470158812 | 0.015915181 |
| ENSG00000119922 | *IFIT2* | 8.328032815 | -0.196746265 | -0.197820648 | 0.001074383 | 0.999933887 | -1.368412893 | -1.863504737 | 0.495091844 | 0.015989053 |
| ENSG00000124107 | *SLPI* | 6.813368997 | -0.028954057 | -0.034994363 | 0.006040306 | 0.999933887 | 0.983239957 | 1.339382842 | -0.356142886 | 0.015989053 |
| ENSG00000126391 | *FRMD8* | 6.331794645 | -0.014128532 | -0.031278923 | 0.01715039 | 0.999933887 | -0.594312079 | -0.847964957 | 0.253652879 | 0.015989053 |
| ENSG00000155363 | *MOV10* | 4.915073189 | 0.061468894 | 0.052946739 | 0.008522154 | 0.999933887 | -0.001514948 | -0.281975899 | 0.280460951 | 0.016008536 |
| ENSG00000105612 | *DNASE2* | 4.662103394 | -0.112064597 | -0.033335094 | -0.078729503 | 0.999933887 | -1.416570992 | -1.837733544 | 0.421162552 | 0.016012638 |
| ENSG00000110696 | *C11orf58* | 5.749405505 | -0.107889139 | -0.084057877 | -0.023831262 | 0.999933887 | -0.018184571 | 0.179593712 | -0.197778283 | 0.016013602 |
| ENSG00000174165 | *ZDHHC24* | 2.648838928 | 0.075005597 | 0.071588671 | 0.003416926 | 0.999933887 | -0.051483155 | -0.441920734 | 0.390437579 | 0.016013602 |
| ENSG00000125753 | *VASP* | 10.04952972 | 0.044570694 | 0.050189151 | -0.005618457 | 0.999933887 | 0.459158402 | 0.725919663 | -0.26676126 | 0.016013602 |
| ENSG00000124151 | *NCOA3* | 6.082704409 | 0.033001457 | -0.011041119 | 0.044042575 | 0.999933887 | -0.509322951 | -0.760087628 | 0.250764677 | 0.016013602 |
| ENSG00000145868 | *FBXO38* | 5.894349417 | -0.03782571 | -0.050351197 | 0.012525487 | 0.999933887 | 0.235534032 | 0.533319114 | -0.297785083 | 0.016013602 |
| ENSG00000072518 | *MARK2* | 6.943436178 | 0.030518088 | 0.085191611 | -0.054673523 | 0.999933887 | -0.422675553 | -0.69194266 | 0.269267107 | 0.016066721 |
| ENSG00000151304 | *SRFBP1* | 0.003955739 | -0.049500459 | 0.108101361 | -0.15760182 | 0.999933887 | -0.076491916 | 0.748828255 | -0.825320171 | 0.016156499 |
| ENSG00000173638 | *SLC19A1* | 6.737395798 | 0.069371539 | 0.015028005 | 0.054343533 | 0.999933887 | -0.755512772 | -1.096506008 | 0.340993236 | 0.016156612 |
| ENSG00000259354 | *AC025580.2* | 2.960912065 | 0.124413873 | 0.099786014 | 0.024627859 | 0.999933887 | 2.395114657 | 3.022808836 | -0.627694179 | 0.016156612 |
| ENSG00000188419 | *CHM* | 1.829254738 | 0.072001805 | -0.095278522 | 0.167280327 | 0.999933887 | -0.014676482 | 0.560406618 | -0.5750831 | 0.016156612 |
| ENSG00000107104 | *KANK1* | 1.07576382 | -0.099505923 | -0.046004811 | -0.053501112 | 0.999933887 | 1.205516245 | 2.004090324 | -0.798574078 | 0.016156612 |
| ENSG00000163162 | *RNF149* | 8.929363348 | 0.019836431 | 0.040636421 | -0.020799991 | 0.999933887 | -0.554935669 | -0.782577434 | 0.227641765 | 0.016156612 |
| ENSG00000161791 | *FMNL3* | 5.105190266 | 0.044147563 | -0.057340276 | 0.101487839 | 0.999933887 | 2.35342265 | 2.965421585 | -0.611998934 | 0.016156612 |
| ENSG00000125734 | *GPR108* | 7.261163676 | -0.003924045 | 0.000686722 | -0.004610766 | 0.999933887 | 1.471987199 | 1.861429118 | -0.389441919 | 0.016156612 |
| ENSG00000257923 | *CUX1* | 5.905766741 | -0.001754593 | -0.012852926 | 0.011098333 | 0.999933887 | 0.193382697 | 0.424406137 | -0.231023439 | 0.016156612 |
| ENSG00000225828 | *FAM229A* | 2.4399994 | 0.026082457 | 0.055627329 | -0.029544872 | 0.999933887 | -0.015288881 | -0.417227233 | 0.401938352 | 0.016237211 |
| ENSG00000106009 | *BRAT1* | 5.861094439 | 0.004256916 | 0.070492218 | -0.066235302 | 0.999933887 | -0.468171609 | -0.809824665 | 0.341653056 | 0.016237211 |
| ENSG00000165879 | *FRAT1* | 5.450188905 | -0.140478018 | -0.254616366 | 0.114138348 | 0.999933887 | -1.371269236 | -1.89390903 | 0.522639794 | 0.01627742 |
| ENSG00000182054 | *IDH2* | 4.695546007 | -0.041028251 | 0.012529639 | -0.05355789 | 0.999933887 | -0.187284705 | -0.411879515 | 0.22459481 | 0.01627742 |
| ENSG00000113448 | *PDE4D* | 4.836060579 | 0.020903217 | -0.111704883 | 0.1326081 | 0.999933887 | -0.233968444 | 0.155372849 | -0.389341293 | 0.01627742 |
| ENSG00000059728 | *MXD1* | 10.981261541 | -0.000416339 | 0.016621724 | -0.017038063 | 0.999933887 | 0.508145395 | 0.7098321 | -0.201686705 | 0.016342314 |
| ENSG00000122482 | *ZNF644* | 4.248658072 | -0.048165372 | -0.124529102 | 0.07636373 | 0.999933887 | 0.197954536 | 0.822675753 | -0.624721217 | 0.016395869 |
| ENSG00000178537 | *SLC25A20* | 1.854661903 | 0.037109463 | -0.012096124 | 0.049205586 | 0.999933887 | -0.090795239 | -0.567765795 | 0.476970556 | 0.016423264 |
| ENSG00000184207 | *PGP* | 2.888588982 | 0.162993421 | 0.040087041 | 0.12290638 | 0.999933887 | -0.115288938 | -0.573950823 | 0.458661884 | 0.016445691 |
| ENSG00000145365 | *TIFA* | 3.621265918 | 0.079263216 | 0.026075106 | 0.05318811 | 0.999933887 | 1.970353794 | 2.546078695 | -0.575724902 | 0.016475589 |
| ENSG00000166619 | *BLCAP* | 4.422926105 | 0.002151323 | -0.00855692 | 0.010708243 | 0.999933887 | -0.266515493 | -0.5187875 | 0.252272007 | 0.016475589 |
| ENSG00000090487 | *SPG21* | 6.88275736 | -0.031637023 | -0.038834618 | 0.007197595 | 0.999933887 | 0.333704091 | 0.514903269 | -0.181199178 | 0.016570464 |
| ENSG00000001617 | *SEMA3F* | -0.173194469 | 0.201201913 | 0.003480034 | 0.19772188 | 0.999933887 | -0.108484604 | 0.659965718 | -0.768450322 | 0.016570464 |
| ENSG00000064201 | *TSPAN32* | 4.239712826 | 0.044326904 | 0.057071902 | -0.012744998 | 0.999933887 | -0.150884414 | -0.480377953 | 0.329493539 | 0.016570464 |
| ENSG00000125510 | *OPRL1* | 0.168755098 | 0.010442415 | 0.26458645 | -0.254144035 | 0.999933887 | -0.881322709 | -1.825348711 | 0.944026002 | 0.016570464 |
| ENSG00000237576 | *LINC01888* | 2.605887108 | 0.091985328 | 0.000230014 | 0.091755314 | 0.999933887 | 0.958816549 | 1.378621278 | -0.419804729 | 0.016575242 |
| ENSG00000133961 | *NUMB* | 8.53167444 | 0.00194292 | -0.029724439 | 0.031667359 | 0.999933887 | 0.470019789 | 0.677308987 | -0.207289198 | 0.01673452 |
| ENSG00000198554 | *WDHD1* | -0.661910829 | -0.139569474 | -0.225340589 | 0.085771114 | 0.999933887 | -0.324777012 | 0.385015325 | -0.709792337 | 0.016786919 |
| ENSG00000175643 | *RMI2* | 1.948856457 | -0.038255995 | -0.190172232 | 0.151916236 | 0.999933887 | 0.656975492 | 1.155920742 | -0.498945249 | 0.016822687 |
| ENSG00000130830 | *MPP1* | 8.812309549 | -0.078402632 | -0.083795004 | 0.005392372 | 0.999933887 | -0.118565628 | 0.117058417 | -0.235624045 | 0.016829965 |
| ENSG00000126804 | *ZBTB1* | 6.172508566 | -0.050980696 | -0.095836325 | 0.044855629 | 0.999933887 | 0.022874045 | 0.338783135 | -0.315909089 | 0.016851768 |
| ENSG00000197930 | *ERO1A* | 5.199945674 | -0.017416136 | -0.039981362 | 0.022565226 | 0.999933887 | -0.424933454 | -0.693356035 | 0.268422581 | 0.016904787 |
| ENSG00000149182 | *ARFGAP2* | 5.528844315 | -0.015513088 | -0.027389125 | 0.011876037 | 0.999933887 | -0.169587082 | -0.359192929 | 0.189605846 | 0.016918715 |
| ENSG00000063978 | *RNF4* | 5.574145689 | -0.016257166 | -0.034002243 | 0.017745077 | 0.999933887 | -0.36239631 | -0.594772221 | 0.232375911 | 0.016918715 |
| ENSG00000214253 | *FIS1* | 4.376267212 | 0.02662835 | 0.001440691 | 0.02518766 | 0.999933887 | -0.077098246 | -0.354709493 | 0.277611246 | 0.016931578 |
| ENSG00000103591 | *AAGAB* | 3.745645738 | -0.055794411 | -0.110008666 | 0.054214255 | 0.999933887 | 0.38656415 | 0.70485943 | -0.31829528 | 0.016933643 |
| ENSG00000117713 | *ARID1A* | 7.170482015 | 0.008019646 | 0.052705716 | -0.04468607 | 0.999933887 | -0.274785511 | -0.518151416 | 0.243365905 | 0.016933643 |
| ENSG00000165006 | *UBAP1* | 7.781948812 | -0.012110561 | -0.069685313 | 0.057574752 | 0.999933887 | 0.290591212 | 0.503935565 | -0.213344353 | 0.016964 |
| ENSG00000167967 | *E4F1* | 4.805199585 | 0.085181787 | 0.092829385 | -0.007647598 | 0.999933887 | -0.179818919 | -0.514802009 | 0.334983091 | 0.016972079 |
| ENSG00000224597 | *SVIL-AS1* | 1.70998374 | 0.003028045 | -0.118747502 | 0.121775547 | 0.999933887 | 0.105964687 | 0.622290856 | -0.516326169 | 0.016972079 |
| ENSG00000111490 | *TBC1D30* | 1.87205188 | -0.012105418 | -0.14981231 | 0.137706892 | 0.999933887 | 1.050055357 | 1.501257733 | -0.451202376 | 0.017072827 |
| ENSG00000157227 | *MMP14* | 1.862174206 | 0.097212361 | 0.079961439 | 0.017250922 | 0.999933887 | 2.2346455 | 3.017633171 | -0.782987671 | 0.017124722 |
| ENSG00000122882 | *ECD* | 4.671038551 | -0.001766268 | -0.112570655 | 0.110804387 | 0.999933887 | 0.253211017 | 0.492171546 | -0.23896053 | 0.017169896 |
| ENSG00000161921 | *CXCL16* | 9.6272892 | 0.042644582 | 0.048879361 | -0.006234779 | 0.999933887 | -0.228239394 | 0.064873768 | -0.293113162 | 0.017376857 |
| ENSG00000171503 | *ETFDH* | 2.364607084 | -0.047931326 | 0.016445283 | -0.064376609 | 0.999933887 | -0.150254984 | 0.238246953 | -0.388501937 | 0.017393786 |
| ENSG00000196189 | *SEMA4A* | 6.862010081 | 0.018567925 | -0.019950122 | 0.038518047 | 0.999933887 | 0.799364886 | 1.187636678 | -0.388271792 | 0.017393786 |
| ENSG00000269293 | *ZSCAN16-AS1* | 1.14159728 | -0.134611632 | -0.302356866 | 0.167745234 | 0.999933887 | 0.533580464 | 1.026241622 | -0.492661158 | 0.017401471 |
| ENSG00000255443 | *CD44-AS1* | 3.248959645 | -0.061606083 | 0.055920638 | -0.11752672 | 0.999933887 | 1.535030452 | 2.164282915 | -0.629252463 | 0.017401471 |
| ENSG00000126822 | *PLEKHG3* | 7.404829493 | 0.001353658 | 0.090749738 | -0.08939608 | 0.999933887 | -0.396495094 | -0.70787563 | 0.311380536 | 0.017401471 |
| ENSG00000172575 | *RASGRP1* | 4.515447625 | 0.001058812 | -0.054238212 | 0.055297024 | 0.999933887 | 0.098281327 | 0.441789741 | -0.343508415 | 0.017401471 |
| ENSG00000153551 | *CMTM7* | 3.876838142 | 0.069963296 | 0.116582498 | -0.046619202 | 0.999933887 | -0.271184947 | -0.679435879 | 0.408250931 | 0.017431346 |
| ENSG00000160877 | *NACC1* | 4.529563634 | 0.023301524 | -0.009608228 | 0.032909753 | 0.999933887 | -0.543444317 | -0.856592906 | 0.31314859 | 0.017433638 |
| ENSG00000172273 | *HINFP* | 2.717924283 | -0.068306521 | -0.035797478 | -0.032509043 | 0.999933887 | -0.2501128 | -0.645782191 | 0.395669391 | 0.017467316 |
| ENSG00000281420 | *AP001052.1* | -0.453497025 | 0.052120443 | 0.177935291 | -0.125814849 | 0.999933887 | 1.239290162 | 2.096161941 | -0.856871779 | 0.017500888 |
| ENSG00000122008 | *POLK* | 2.434520606 | -0.115156035 | -0.261966835 | 0.1468108 | 0.999933887 | -0.205110224 | 0.514134463 | -0.719244687 | 0.017577709 |
| ENSG00000154978 | *VOPP1* | 6.858034054 | -0.040094085 | -0.053341108 | 0.013247023 | 0.999933887 | 0.270587053 | 0.582389862 | -0.311802809 | 0.01765558 |
| ENSG00000175029 | *CTBP2* | 4.996736822 | -0.011806026 | -0.00616824 | -0.005637786 | 0.999933887 | -0.416636703 | -0.693509083 | 0.27687238 | 0.017685912 |
| ENSG00000172939 | *OXSR1* | 7.87765093 | -0.031916936 | -0.0516001 | 0.019683164 | 0.999933887 | 0.189895921 | 0.456424442 | -0.266528521 | 0.017696496 |
| ENSG00000155097 | *ATP6V1C1* | 7.205479725 | -0.081636388 | -0.014927913 | -0.066708475 | 0.999933887 | 0.372425954 | 0.660704255 | -0.288278301 | 0.017717231 |
| ENSG00000128563 | *PRKRIP1* | 4.097600251 | -0.014884517 | 0.017426728 | -0.032311245 | 0.999933887 | -0.113923448 | -0.403689236 | 0.289765788 | 0.017717231 |
| ENSG00000156110 | *ADK* | 2.582987346 | -0.005675743 | -0.124096954 | 0.11842121 | 0.999933887 | -0.385282968 | -0.675858379 | 0.290575411 | 0.017738962 |
| ENSG00000188820 | *CALHM6* | 2.124001631 | 0.16216525 | 0.110056527 | 0.052108723 | 0.999933887 | -1.151540648 | -2.251474981 | 1.099934333 | 0.017758889 |
| ENSG00000137171 | *KLC4* | 3.145504221 | -0.056033527 | 0.050499838 | -0.106533365 | 0.999933887 | -0.32509515 | -0.664873351 | 0.339778202 | 0.017772396 |
| ENSG00000136816 | *TOR1B* | 5.135532336 | -0.067266225 | -0.100347618 | 0.033081393 | 0.999933887 | -0.638563714 | -0.930785155 | 0.29222144 | 0.017810927 |
| ENSG00000162613 | *FUBP1* | 5.981012589 | -0.039026643 | -0.054864539 | 0.015837896 | 0.999933887 | -0.11744795 | 0.136777776 | -0.254225726 | 0.01782574 |
| ENSG00000143157 | *POGK* | 4.120039319 | -0.027601324 | -0.074709917 | 0.047108594 | 0.999933887 | 0.507917453 | 0.818211639 | -0.310294186 | 0.01782574 |
| ENSG00000078699 | *CBFA2T2* | 4.067294005 | -0.019212497 | -0.001148747 | -0.01806375 | 0.999933887 | -0.45719375 | -0.762912163 | 0.305718413 | 0.01782574 |
| ENSG00000285639 | *AL133259.1* | 1.824221931 | 0.010297942 | 0.269661259 | -0.259363317 | 0.999933887 | 0.69138229 | 1.328502344 | -0.637120054 | 0.01782574 |
| ENSG00000278090 | *LUNAR1* | 0.35169648 | -0.167373807 | -0.209354704 | 0.041980896 | 0.999933887 | -1.24497685 | -1.966716365 | 0.721739515 | 0.017857807 |
| ENSG00000068120 | *COASY* | 4.264885701 | -0.115130216 | -0.15098125 | 0.035851034 | 0.999933887 | -0.357051961 | -0.638278065 | 0.281226104 | 0.017879039 |
| ENSG00000107551 | *RASSF4* | 1.080527406 | -0.031720701 | 0.006091444 | -0.037812145 | 0.999933887 | 0.563711784 | 1.403698113 | -0.839986328 | 0.017924782 |
| ENSG00000117143 | *UAP1* | 3.542033472 | -0.05234677 | -0.114407276 | 0.062060507 | 0.999933887 | 0.047896916 | 0.416666262 | -0.368769346 | 0.01796644 |
| ENSG00000164889 | *SLC4A2* | 4.866644893 | 0.08783217 | 0.090051551 | -0.00221938 | 0.999933887 | 0.489399499 | 0.800085693 | -0.310686194 | 0.018012382 |
| ENSG00000253522 | *MIR3142HG* | 1.317249424 | 0.219109798 | 0.439303628 | -0.22019383 | 0.999933887 | 2.01661299 | 2.892408301 | -0.875795311 | 0.018095581 |
| ENSG00000133816 | *MICAL2* | 5.651343333 | -0.011764062 | -0.01127276 | -0.000491302 | 0.999933887 | -0.382967366 | -0.707790791 | 0.324823425 | 0.018095581 |
| ENSG00000169230 | *PRELID1* | 5.614936646 | 0.028672632 | -0.003922359 | 0.03259499 | 0.999933887 | -0.07308246 | -0.388124234 | 0.315041774 | 0.01810546 |
| ENSG00000155330 | *C16orf87* | 2.166986434 | -0.043446289 | 0.031054592 | -0.074500882 | 0.999933887 | 0.135204171 | 0.498497023 | -0.363292853 | 0.01810546 |
| ENSG00000169692 | *AGPAT2* | 3.890326544 | 0.055667388 | 0.031581208 | 0.024086181 | 0.999933887 | -0.668779122 | -1.082692815 | 0.413913693 | 0.018120765 |
| ENSG00000070961 | *ATP2B1* | 9.657755321 | -0.047125124 | -0.093966328 | 0.046841204 | 0.999933887 | 1.011433213 | 1.605007617 | -0.593574404 | 0.01813517 |
| ENSG00000123595 | *RAB9A* | 3.855979721 | -0.104640928 | -0.142568932 | 0.037928004 | 0.999933887 | 1.079195001 | 1.4330866 | -0.353891599 | 0.018230834 |
| ENSG00000145675 | *PIK3R1* | 6.481332601 | -0.077963167 | -0.144467835 | 0.066504668 | 0.999933887 | -0.112871612 | 0.253040668 | -0.36591228 | 0.018230834 |
| ENSG00000115919 | *KYNU* | 2.733567677 | 0.024913732 | 0.062581105 | -0.037667373 | 0.999933887 | 1.141604746 | 1.628621634 | -0.487016888 | 0.018230834 |
| ENSG00000169375 | *SIN3A* | 6.87773346 | -0.002487782 | -0.064342248 | 0.061854466 | 0.999933887 | 0.705715134 | 0.963865819 | -0.258150685 | 0.018232605 |
| ENSG00000181631 | *P2RY13* | 6.166215201 | -0.090281674 | -0.137658718 | 0.047377044 | 0.999933887 | -1.922379216 | -2.424072464 | 0.501693248 | 0.018268492 |
| ENSG00000228382 | *ITPKB-IT1* | 2.490660025 | 0.127012555 | -0.180811338 | 0.307823893 | 0.999933887 | 0.838489288 | 1.218322119 | -0.37983283 | 0.018279956 |
| ENSG00000175550 | *DRAP1* | 5.680569199 | 0.08709309 | 0.024292417 | 0.062800673 | 0.999933887 | -0.184460677 | -0.684249082 | 0.499788405 | 0.018291641 |
| ENSG00000174738 | *NR1D2* | 5.569859303 | -0.072836478 | -0.163098196 | 0.090261718 | 0.999933887 | 0.245009468 | 0.618813782 | -0.373804313 | 0.018366142 |
| ENSG00000135940 | *COX5B* | 5.103062838 | -0.009787786 | -0.056378118 | 0.046590332 | 0.999933887 | -0.106944863 | -0.408146322 | 0.301201459 | 0.018366142 |
| ENSG00000127125 | *PPCS* | 4.050701423 | -0.116660122 | -0.135955439 | 0.019295317 | 0.999933887 | -0.416858776 | -0.716052524 | 0.299193748 | 0.01837878 |
| ENSG00000168769 | *TET2* | 7.714501632 | -0.026975585 | -0.090125314 | 0.063149728 | 0.999933887 | -0.586737885 | -0.102202549 | -0.484535335 | 0.018441867 |
| ENSG00000150867 | *PIP4K2A* | 6.12008218 | -0.004094582 | -0.033256216 | 0.029161634 | 0.999933887 | -0.192870685 | -0.414441512 | 0.221570827 | 0.018441867 |
| ENSG00000135541 | *AHI1* | 2.089545794 | 0.007519495 | 0.168680333 | -0.161160838 | 0.999933887 | -0.129412531 | 0.363382732 | -0.492795263 | 0.018441867 |
| ENSG00000104497 | *SNX16* | 3.176357007 | -0.074732528 | 0.019291255 | -0.094023783 | 0.999933887 | -0.25611046 | 0.15432947 | -0.41043993 | 0.018457725 |

| | | | | | | | | | | |
|---|---|---|---|---|---|---|---|---|---|---|
| ENSG00000169660 | *HEXD* | 3.870144027 | 0.04377214 | 0.087020576 | -0.043248436 | 0.999933887 | -0.058997893 | -0.447669 | 0.388671107 | 0.018457725 |
| ENSG00000121716 | *PILRB* | 3.114439211 | -0.017252151 | 0.079942538 | -0.097194689 | 0.999933887 | 0.771906259 | 1.338368339 | -0.56646208 | 0.018495435 |
| ENSG00000183484 | *GPR132* | 5.279336332 | 0.145827322 | 0.167252862 | -0.021425539 | 0.999933887 | 1.351945106 | 1.910675124 | -0.558730018 | 0.018521393 |
| ENSG00000108788 | *MLX* | 5.117458866 | -0.085929284 | -0.018074333 | -0.067854951 | 0.999933887 | -0.933021953 | -1.27039623 | 0.337374277 | 0.018521393 |
| ENSG00000108604 | *SMARCD2* | 5.5600537 | 0.044645688 | 0.020965683 | 0.023680005 | 0.999933887 | -0.367725566 | -0.637940443 | 0.270214877 | 0.018545522 |
| ENSG00000140153 | *WDR20* | 4.604527072 | -0.032978282 | -0.044198009 | 0.011219727 | 0.999933887 | 0.169654961 | 0.411168957 | -0.241513997 | 0.018551277 |
| ENSG00000102572 | *STK24* | 5.898806274 | -0.034504682 | -0.031770308 | -0.002734373 | 0.999933887 | -0.816577347 | -1.136658678 | 0.320081331 | 0.018630133 |
| ENSG00000102401 | *ARMCX3* | 5.158048029 | 0.072420393 | -0.06754099 | 0.139961383 | 0.999933887 | 0.50467542 | 0.818798303 | -0.314122883 | 0.018648127 |
| ENSG00000149532 | *CPSF7* | 7.191010307 | -0.041304772 | -0.042953966 | 0.001649194 | 0.999933887 | 0.100761667 | 0.277655837 | -0.176894171 | 0.018648472 |
| ENSG00000149480 | *MTA2* | 5.014892398 | 0.051551782 | 0.050843344 | 0.000708438 | 0.999933887 | 0.100172448 | -0.138180798 | 0.238353246 | 0.018648472 |
| ENSG00000181649 | *PHLDA2* | 3.362493764 | 0.451824118 | 0.412295341 | 0.039528777 | 0.999933887 | 1.84595073 | 2.353553145 | -0.507602414 | 0.018695933 |
| ENSG00000139437 | *TCHP* | 3.812637021 | -0.001608103 | -0.126650989 | 0.125042885 | 0.999933887 | -1.346660438 | -1.864483208 | 0.517822771 | 0.018716206 |
| ENSG00000137801 | *THBS1* | 4.933753946 | 0.068990931 | 0.086414886 | -0.017423955 | 0.999933887 | 0.424581297 | 1.035330095 | -0.610748798 | 0.018726434 |
| ENSG00000131236 | *CAP1* | 8.234651079 | -0.022291912 | -0.06093648 | 0.038644568 | 0.999933887 | -0.578626669 | -0.883084927 | 0.304458258 | 0.018873697 |
| ENSG00000166295 | *ANAPC16* | 5.049145075 | 0.022437962 | 0.031697254 | -0.009259292 | 0.999933887 | 0.331555964 | 0.574081909 | -0.242525945 | 0.018942994 |
| ENSG00000070476 | *ZXDC* | 5.964922511 | -0.005664819 | -0.006277239 | 0.00061242 | 0.999933887 | -0.611377648 | -0.842060319 | 0.230682671 | 0.018955865 |
| ENSG00000072310 | *SREBF1* | 6.705017434 | 0.037467879 | 0.041821256 | -0.004353376 | 0.999933887 | 0.451288066 | 0.700960448 | -0.249672382 | 0.018959397 |
| ENSG00000071553 | *ATP6AP1* | 8.547507946 | -0.049416281 | -0.043532103 | -0.005884179 | 0.999933887 | 0.613830034 | 0.828749808 | -0.214919774 | 0.018972246 |
| ENSG00000165131 | *LLCFC1* | -0.403338406 | 0.164377422 | -0.267319254 | 0.431696676 | 0.999933887 | -0.766688144 | -1.547469807 | 0.780781664 | 0.019007582 |
| ENSG00000112511 | *PHF1* | 7.623057645 | 0.028664365 | 0.058208193 | -0.029543828 | 0.999933887 | 0.536606046 | 0.756219155 | -0.219613109 | 0.019064876 |
| ENSG00000140853 | *NLRC5* | 8.360759594 | 0.053785191 | 0.032723777 | 0.021061414 | 0.999933887 | -0.321709437 | -0.551475334 | 0.229765897 | 0.01909151 |
| ENSG00000149231 | *CCDC82* | 5.163056616 | -0.107776268 | -0.154585757 | 0.046809489 | 0.999933887 | 0.638824301 | 1.375021965 | -0.736197664 | 0.01909151 |
| ENSG00000270457 | *AC093424.1* | -0.47289177 | -0.112905812 | -0.037430392 | -0.07547542 | 0.999933887 | 1.017316664 | 1.867021903 | -0.849705239 | 0.01909151 |
| ENSG00000197147 | *LRRC8B* | 3.011017673 | -0.041704958 | 0.002070592 | -0.04377555 | 0.999933887 | 0.484503593 | 0.932848152 | -0.448344559 | 0.01909151 |
| ENSG00000120314 | *WDR55* | 4.628243982 | -0.01678596 | -0.00478506 | -0.012000899 | 0.999933887 | -0.230349892 | -0.468609535 | 0.238259644 | 0.01909151 |
| ENSG00000013275 | *PSMC4* | 5.338842624 | -0.011339558 | 0.011590991 | -0.022930549 | 0.999933887 | 0.392373193 | 0.652396881 | -0.260023688 | 0.01909151 |
| ENSG00000181135 | *ZNF707* | 2.467970492 | 0.059802304 | 0.133406838 | -0.073604535 | 0.999933887 | -0.034971636 | -0.418819493 | 0.383847857 | 0.019144384 |
| ENSG00000159388 | *BTG2* | 9.768035326 | 0.364412936 | 0.369364415 | -0.004951478 | 0.999933887 | 1.006843983 | 1.428214917 | -0.421370934 | 0.019165546 |
| ENSG00000046653 | *GPM6B* | 0.625083964 | -0.20935902 | -0.14486986 | -0.06448916 | 0.999933887 | 0.099428037 | 0.88067843 | -0.781250393 | 0.019165546 |
| ENSG00000161036 | *LRWD1* | 5.100663282 | 0.044671423 | 0.110745287 | -0.066073863 | 0.999933887 | -0.112165853 | -0.402792394 | 0.290626541 | 0.019165546 |
| ENSG00000196387 | *ZNF140* | 2.997554288 | -0.040229198 | -0.073584611 | 0.033355413 | 0.999933887 | 0.326560611 | 0.639966281 | -0.31340567 | 0.019165546 |
| ENSG00000036054 | *TBC1D23* | 5.500642358 | -0.042190038 | -0.092685936 | 0.050495898 | 0.999933887 | -0.023184439 | 0.335968075 | -0.359152515 | 0.019207797 |
| ENSG00000064703 | *DDX20* | 3.402223121 | -0.03289515 | -0.026514664 | -0.006380486 | 0.999933887 | -0.111538102 | -0.435629225 | 0.324091123 | 0.019207797 |
| ENSG00000164054 | *SHISA5* | 7.866543889 | 0.079431504 | 0.011970259 | 0.067461245 | 0.999933887 | -0.467220802 | -0.802588581 | 0.335367778 | 0.019248971 |
| ENSG00000165915 | *SLC39A13* | 4.070149649 | 0.075108475 | -0.021643629 | 0.096752104 | 0.999933887 | 0.280941034 | 0.594202237 | -0.313261203 | 0.019248971 |
| ENSG00000118308 | *IRAG2* | 6.522404716 | -0.061007314 | -0.118640624 | 0.05763331 | 0.999933887 | -0.574142323 | -0.886398668 | 0.312256345 | 0.019248971 |
| ENSG00000100461 | *RBM23* | 7.169319836 | -0.021093341 | 0.005499167 | -0.026592508 | 0.999933887 | 0.596925805 | 0.828713909 | -0.231788104 | 0.019248971 |
| ENSG00000188191 | *PRKAR1B* | 1.351835343 | 0.059136973 | 0.120212394 | -0.061075421 | 0.999933887 | 0.336267742 | 0.961478547 | -0.625210805 | 0.019248971 |
| ENSG00000175470 | *PPP2R2D* | 5.813431072 | -0.012599575 | -0.024096375 | 0.0114968 | 0.999933887 | 0.192814972 | 0.373324792 | -0.18050982 | 0.019248971 |
| ENSG00000172869 | *DMXL1* | 3.953068142 | -0.0036904 | -0.13805764 | 0.13436724 | 0.999933887 | -0.195860582 | 0.328307403 | -0.524167985 | 0.019248971 |
| ENSG00000159348 | *CYB5R1* | 5.022593551 | -0.015590881 | -0.02116686 | 0.00557598 | 0.999933887 | -0.636745273 | -0.920631769 | 0.283886496 | 0.019253943 |
| ENSG00000114030 | *KPNA1* | 5.57091542 | -0.053229444 | -0.084659394 | 0.031429949 | 0.999933887 | -0.202870508 | 0.005202331 | -0.208072839 | 0.0192958 |
| ENSG00000273604 | *EPOP* | 1.39809035 | 0.086009837 | 0.176167135 | -0.090157298 | 0.999933887 | 2.092359671 | 2.719842871 | -0.6274832 | 0.019296374 |
| ENSG00000111837 | *MAK* | 3.678491864 | -0.045635743 | 0.02153147 | -0.067167213 | 0.999933887 | 0.184171303 | 0.495434492 | -0.311263189 | 0.019300053 |
| ENSG00000141337 | *ARSG* | 4.065019449 | -0.084239974 | -0.074205444 | -0.01003453 | 0.999933887 | -0.149541873 | -0.466557528 | 0.317015655 | 0.019322219 |
| ENSG00000238018 | *AC093110.1* | 1.027340254 | -0.099935186 | -0.396057772 | 0.296122586 | 0.999933887 | 0.020164712 | 0.441353918 | -0.421189206 | 0.019402191 |
| ENSG00000047249 | *ATP6V1H* | 4.996827613 | -0.034360211 | 0.021861437 | -0.056221648 | 0.999933887 | 0.662150731 | 0.964425358 | -0.302274627 | 0.019402191 |
| ENSG00000116044 | *NFE2L2* | 8.495838126 | -0.016574664 | -0.03301314 | 0.016438476 | 0.999933887 | 0.736253006 | 1.043966152 | -0.307713145 | 0.019402191 |
| ENSG00000137767 | *SQOR* | 5.041867824 | 0.01163546 | -0.05801786 | 0.06965332 | 0.999933887 | 0.828033237 | 1.188550845 | -0.360517607 | 0.019402191 |
| ENSG00000136485 | *DCAF7* | 4.995975259 | -0.08825825 | 0.022044737 | -0.110302987 | 0.999933887 | -0.337501475 | -0.613610075 | 0.2761086 | 0.019440913 |
| ENSG00000073417 | *PDE8A* | 2.616537674 | 0.013637068 | 0.008546076 | 0.005090993 | 0.999933887 | 0.285013193 | 0.702565637 | -0.417552445 | 0.019552784 |
| ENSG00000138835 | *RGS3* | 4.11939002 | 0.093105624 | 0.000956858 | 0.092148766 | 0.999933887 | -0.250446524 | -0.654597281 | 0.404150757 | 0.019589301 |
| ENSG00000075618 | *FSCN1* | 2.919385715 | 0.064231206 | -0.089866135 | 0.154097342 | 0.999933887 | 2.475703757 | 3.221965262 | -0.746261504 | 0.019589301 |
| ENSG00000051523 | *CYBA* | 9.178312278 | 0.068204279 | 0.015301427 | 0.052902851 | 0.999933887 | 0.985319545 | 1.324408758 | -0.339089214 | 0.01961462 |
| ENSG00000226648 | *PLCG1-AS1* | 2.020068743 | 0.093300693 | 0.005745698 | 0.087554994 | 0.999933887 | -0.080930057 | 0.468117651 | -0.549047708 | 0.019636955 |
| ENSG00000215458 | *AATBC* | 1.060080035 | 0.036298143 | 0.040859884 | -0.004561742 | 0.999933887 | 0.82532022 | 1.495409542 | -0.670089322 | 0.019730598 |
| ENSG00000107738 | *VSIR* | 9.442960556 | -0.009362462 | -0.008176713 | -0.001185749 | 0.999933887 | -0.430183561 | -0.734444608 | 0.304261047 | 0.019730598 |
| ENSG00000164111 | *ANXA5* | 7.443725353 | 0.002587578 | -0.040585495 | 0.043173073 | 0.999933887 | 1.346094054 | 1.687759226 | -0.341665173 | 0.019730598 |
| ENSG00000112379 | *ARFGEF3* | -1.240230178 | 0.098290083 | 0.625327037 | -0.527036955 | 0.999933887 | 0.874449681 | 1.59444942 | -0.71999974 | 0.019746995 |
| ENSG00000171298 | *GAA* | 6.184341239 | 0.048233169 | 0.033168487 | 0.015064682 | 0.999933887 | -0.244606991 | -0.50848487 | 0.263877879 | 0.019752552 |
| ENSG00000213639 | *PPP1CB* | 7.529225162 | -0.087492396 | -0.063194267 | -0.024298129 | 0.999933887 | -0.084130884 | 0.220868428 | -0.304999311 | 0.01975449 |
| ENSG00000213347 | *MXD3* | 3.663905161 | 0.050537919 | 0.059588258 | -0.00905034 | 0.999933887 | -0.107534301 | -0.53860925 | 0.43107495 | 0.019770664 |
| ENSG00000105127 | *AKAP8* | 5.390287569 | -0.000665917 | 0.022641165 | -0.023307082 | 0.999933887 | -0.161694085 | -0.366675515 | 0.204981429 | 0.019770664 |
| ENSG00000255921 | *AC026310.1* | -2.384324374 | -0.409115763 | -0.083318626 | -0.325797137 | 0.999933887 | 2.102207969 | 3.430342166 | -1.328134197 | 0.019790423 |
| ENSG00000047315 | *POLR2B* | 5.469007736 | -0.110725692 | -0.109447225 | -0.001278466 | 0.999933887 | -0.296412061 | -0.536326266 | 0.239914205 | 0.019801029 |
| ENSG00000128294 | *TPST2* | 6.104632222 | 0.023371912 | 0.00956458 | 0.013807332 | 0.999933887 | -0.745557514 | -1.055479409 | 0.309921895 | 0.019801029 |
| ENSG00000146376 | *ARHGAP18* | 3.063857264 | -0.147628357 | -0.033345322 | -0.114283035 | 0.999933887 | -0.455504539 | -0.081703464 | -0.373801076 | 0.019808519 |
| ENSG00000224609 | *HSD52* | 3.092397928 | 0.252506261 | 0.129432837 | 0.123073424 | 0.999933887 | 1.271381906 | 1.79488294 | -0.523501034 | 0.019829039 |
| ENSG00000103876 | *FAH* | 1.600446303 | -0.187019501 | 0.315149629 | -0.502169129 | 0.944094537 | -0.432121033 | -0.814786413 | 0.38266538 | 0.019829039 |
| ENSG00000114861 | *FOXP1* | 4.799667441 | -0.063940731 | 0.028919249 | -0.09285998 | 0.999933887 | 0.383399453 | 0.785204656 | -0.401805204 | 0.019829039 |
| ENSG00000146063 | *TRIM41* | 4.494919472 | 0.076656442 | 0.052599808 | 0.024056634 | 0.999933887 | -0.216844162 | -0.50711072 | 0.290266558 | 0.019960797 |
| ENSG00000156508 | *EEF1A1* | 10.934526471 | -0.033064696 | -0.177567366 | 0.14450267 | 0.999933887 | 0.304104004 | 0.590076109 | -0.285972106 | 0.0199651 |
| ENSG00000205155 | *PSENEN* | 2.960210957 | 0.077489544 | 0.064292578 | 0.013196966 | 0.999933887 | -0.295833015 | -0.676107393 | 0.380274379 | 0.020010856 |
| ENSG00000153827 | *TRIP12* | 7.827206781 | -0.047762142 | -0.072044065 | 0.024281924 | 0.999933887 | 0.144853835 | 0.377125703 | -0.232271868 | 0.020010856 |
| ENSG00000165650 | *PDZD8* | 5.182799183 | 0.062851676 | -0.011950824 | 0.074802499 | 0.999933887 | -0.663255004 | -1.116759472 | 0.453504468 | 0.020023433 |
| ENSG00000087470 | *DNM1L* | 3.235622457 | -0.13247971 | -0.202723808 | 0.070244098 | 0.999933887 | 0.173111658 | 0.733913738 | -0.56080208 | 0.020075975 |
| ENSG00000224531 | *SMIM13* | 3.081667061 | -0.074616869 | 0.008938266 | -0.083555134 | 0.999933887 | -0.165802097 | 0.144762117 | -0.310564213 | 0.020075975 |
| ENSG00000188313 | *PLSCR1* | 5.755929162 | -0.019445299 | -0.041224873 | 0.021779575 | 0.999933887 | 0.560610483 | 0.838077857 | -0.277467373 | 0.020075975 |
| ENSG00000166401 | *SERPINB8* | 3.913431967 | -0.03297302 | -0.008258085 | -0.024714935 | 0.999933887 | 0.75610363 | 1.195009891 | -0.438906261 | 0.020075975 |
| ENSG00000008282 | *SYPL1* | 3.655534044 | -0.014008712 | -0.067564074 | 0.053555363 | 0.999933887 | 0.187919412 | 0.481829362 | -0.29390995 | 0.020075975 |
| ENSG00000136950 | *ARPC5L* | 3.997404512 | -0.009336957 | -0.018396651 | 0.009059693 | 0.999933887 | -0.04601525 | -0.31466831 | 0.268653059 | 0.020075975 |
| ENSG00000138031 | *ADCY3* | 4.51306348 | -0.003038673 | -0.016775283 | 0.01373661 | 0.999933887 | 0.935513868 | 1.277800921 | -0.342287053 | 0.020075975 |
| ENSG00000142512 | *SIGLEC10* | 6.485211668 | 0.035629744 | 0.034243942 | 0.001385802 | 0.999933887 | 0.503827034 | 0.825163316 | -0.321336282 | 0.020077272 |
| ENSG00000250510 | *GPR162* | 1.194064892 | 0.017838049 | -0.027193643 | 0.045031692 | 0.999933887 | -0.597128838 | -1.440928942 | 0.843800103 | 0.020091685 |
| ENSG00000079387 | *SENP1* | 3.544185974 | 0.002023954 | -0.011707187 | 0.013731141 | 0.999933887 | 0.385858661 | 0.671694014 | -0.285835353 | 0.020091685 |
| ENSG00000082014 | *SMARCD3* | 3.685998976 | 0.029157404 | 0.022443815 | 0.006713589 | 0.999933887 | 0.100598502 | -0.244528115 | 0.345126617 | 0.020118608 |
| ENSG00000100994 | *PYGB* | 4.003969774 | -0.019738212 | 0.01860471 | -0.038342922 | 0.999933887 | -0.061629173 | -0.322423956 | 0.260794783 | 0.020118608 |
| ENSG00000168216 | *LMBRD1* | 5.230369397 | -0.037352683 | -0.049353221 | 0.012000538 | 0.999933887 | -0.324918392 | -0.62040295 | 0.295484558 | 0.02012639 |
| ENSG00000181444 | *ZNF467* | 6.066515759 | 0.04910024 | 0.088357531 | -0.039257291 | 0.999933887 | -0.950207923 | -1.386350653 | 0.43614273 | 0.020164442 |
| ENSG00000075303 | *SLC25A40* | 3.662648893 | -0.029617852 | -0.119588279 | 0.089970427 | 0.999933887 | 0.159516654 | 0.559277187 | -0.399760533 | 0.020164442 |
| ENSG00000269713 | *NBPF9* | 2.322581916 | 0.003103494 | -0.097797007 | 0.100900501 | 0.999933887 | 0.425631082 | 0.846542927 | -0.420911844 | 0.020164442 |
| ENSG00000137575 | *SDCBP* | 10.809163669 | -0.091083008 | -0.125275727 | 0.034192719 | 0.999933887 | 0.044818186 | 0.295292549 | -0.250474363 | 0.020254093 |
| ENSG00000141503 | *MINK1* | 6.44440546 | 0.026865534 | 0.019415549 | 0.007449985 | 0.999933887 | -0.152751992 | -0.361293676 | 0.208541684 | 0.020297625 |
| ENSG00000137193 | *PIM1* | 8.492363704 | -0.100666548 | -0.103544323 | 0.002877775 | 0.999933887 | 0.134409866 | 0.45186177 | -0.317451904 | 0.02044054 |
| ENSG00000126070 | *AGO3* | 4.94584566 | -0.001417687 | -0.012006222 | 0.010588536 | 0.999933887 | 0.218024818 | 0.515477541 | -0.297452723 | 0.020453532 |
| ENSG00000095139 | *ARCN1* | 6.012466305 | -0.032118458 | -0.005210168 | -0.02690829 | 0.999933887 | 0.009782335 | 0.213826789 | -0.204044454 | 0.020456012 |
| ENSG00000178803 | *ADORA2A-AS* | 5.952960649 | 0.230950781 | 0.146156546 | 0.084794236 | 0.999933887 | 1.873465116 | 2.281641848 | -0.408176732 | 0.020471595 |
| ENSG00000131203 | *IDO1* | 1.019822559 | -0.093396073 | -0.008443149 | -0.084952924 | 0.999933887 | 0.381876303 | 1.034919864 | -0.653043561 | 0.02049399 |
| ENSG00000261324 | *AC010168.2* | 1.680549125 | -0.056586218 | 0.070793249 | -0.127379467 | 0.999933887 | -0.084035071 | -0.60231618 | 0.518281109 | 0.02049399 |
| ENSG00000115137 | *DNAJC27* | 2.617162893 | -0.030207911 | -0.032721976 | 0.002514065 | 0.999933887 | 0.114532724 | 0.542345169 | -0.427812446 | 0.02049399 |
| ENSG00000261546 | *AC135782.3* | -0.255661167 | -0.00080661 | -0.044059023 | 0.043252413 | 0.999933887 | -0.619699209 | -1.563644579 | 0.94394537 | 0.020581865 |
| ENSG00000104064 | *GABPB1* | 3.955777106 | -0.00275484 | 0.073498877 | -0.076253716 | 0.999933887 | 0.883797244 | 1.20828844 | -0.324491196 | 0.020589562 |
| ENSG00000139625 | *MAP3K12* | 3.526309563 | -0.004323837 | 0.060417548 | -0.064741384 | 0.999933887 | -0.519118669 | -0.84398728 | 0.324868611 | 0.02061159 |
| ENSG00000159202 | *UBE2Z* | 6.26912915 | -0.004969917 | -0.095335599 | 0.090365682 | 0.999933887 | 0.560531518 | 0.766413441 | -0.205881923 | 0.020651994 |
| ENSG00000135766 | *EGLN1* | 4.181372167 | 0.116029719 | 0.090419305 | 0.025610414 | 0.999933887 | -0.641712598 | -1.204682769 | 0.562970172 | 0.020769419 |

| | | | | | | | | | | |
|---|---|---|---|---|---|---|---|---|---|---|
| ENSG00000071054 | *MAP4K4* | 9.783929495 | -0.044631056 | -0.007489439 | -0.037141617 | 0.999933887 | 0.40181685 | 0.601935254 | -0.200118403 | 0.020769419 |
| ENSG00000185404 | *SP140L* | 4.27168288 | -0.045178488 | -0.066552537 | 0.021374049 | 0.999933887 | -0.290678511 | -0.526930075 | 0.236251565 | 0.020769419 |
| ENSG00000148985 | *PGAP2* | 3.920419495 | 0.045041952 | -0.028538742 | 0.073580694 | 0.999933887 | 0.742987668 | 1.128182164 | -0.385194496 | 0.020769419 |
| ENSG00000158850 | *B4GALT3* | 4.142032927 | 0.021300574 | 0.052098421 | -0.030797847 | 0.999933887 | -0.115279216 | -0.351928715 | 0.236649499 | 0.020769419 |
| ENSG00000163482 | *STK36* | 2.510255551 | 0.032198235 | -0.080244451 | 0.112442686 | 0.999933887 | 0.53284275 | 0.877792783 | -0.344950033 | 0.020769419 |
| ENSG00000246575 | *AC093162.1* | 2.611078876 | -0.017235689 | -0.097210843 | 0.079975154 | 0.999933887 | -0.317062831 | -0.691655654 | 0.374592822 | 0.020769419 |
| ENSG00000102531 | *FNDC3A* | 6.345639422 | -0.021782003 | -0.035760261 | 0.013978258 | 0.999933887 | -0.463675762 | -0.089479343 | -0.374196419 | 0.020769419 |
| ENSG00000212907 | *MT-ND4L* | 6.03153988 | 0.001316329 | -0.082926244 | 0.084242573 | 0.999933887 | -0.02058827 | 0.310825815 | -0.331414085 | 0.020769419 |
| ENSG00000250616 | *AC012645.1* | 1.644984498 | -0.009308995 | 0.008481018 | -0.017790013 | 0.999933887 | -1.157019153 | -2.020096229 | 0.863077077 | 0.020839212 |
| ENSG00000244405 | *ETV5* | -0.218188178 | -0.025154275 | -0.087542271 | 0.062387996 | 0.999933887 | 0.756971673 | 1.800482725 | -1.043511052 | 0.021072907 |
| ENSG00000070423 | *RNF126* | 4.699187934 | -0.002497975 | 0.02920998 | -0.031707956 | 0.999933887 | -0.078912277 | -0.338690204 | 0.259777928 | 0.02109467 |
| ENSG00000219891 | *ZSCAN12P1* | 0.065464981 | 0.147285497 | 0.187875615 | -0.040590118 | 0.999933887 | -0.548424548 | 0.209671003 | -0.758095552 | 0.021123327 |
| ENSG00000198873 | *GRK5* | 4.720698901 | 0.016751839 | 0.004927354 | 0.011824485 | 0.999933887 | 0.24316487 | 0.489236397 | -0.246071528 | 0.021139399 |
| ENSG00000180448 | *ARHGAP45* | 9.123306221 | 0.052783902 | 0.08581531 | -0.033031408 | 0.999933887 | -0.359422 | -0.661418438 | 0.301996438 | 0.02114403 |
| ENSG00000105928 | *GSDME* | -0.544871663 | -0.218137776 | 0.162879203 | -0.381016978 | 0.999933887 | 0.713177304 | 1.388482027 | -0.675304724 | 0.021255058 |
| ENSG00000204681 | *GABBR1* | 3.845949769 | 0.052095781 | 0.059581284 | -0.007485503 | 0.999933887 | 0.286308036 | 0.595234641 | -0.308926605 | 0.021255058 |
| ENSG00000077585 | *GPR137B* | 2.701959553 | 0.030320366 | 0.028306093 | 0.002014273 | 0.999933887 | 0.219838828 | 0.541872023 | -0.322033195 | 0.021255058 |
| ENSG00000241764 | *AC002467.1* | 0.050713245 | -0.029911955 | -0.323573787 | 0.293661833 | 0.999933887 | -0.262838269 | -1.097446111 | 0.834607842 | 0.021255058 |
| ENSG00000085511 | *MAP3K4* | 3.022467502 | -0.009244642 | -0.074602375 | 0.065357733 | 0.999933887 | 0.170160987 | 0.536403125 | -0.366242138 | 0.021308422 |
| ENSG00000127616 | *SMARCA4* | 4.824265169 | -0.007692931 | 0.020414668 | -0.028107599 | 0.999933887 | -0.027579585 | -0.232042432 | 0.204462847 | 0.021378552 |
| ENSG00000168229 | *PTGDR* | 0.791261722 | 0.1080188 | -0.04982437 | 0.157843169 | 0.999933887 | 0.677828123 | 1.343511889 | -0.665683766 | 0.021407437 |
| ENSG00000166507 | *NDST2* | 0.288285456 | 0.046565782 | -0.215610204 | 0.262175986 | 0.999933887 | 0.008933907 | -0.728433878 | 0.737367785 | 0.021407437 |
| ENSG00000173320 | *STOX2* | 2.779059777 | 0.003941399 | 0.134053974 | -0.130112575 | 0.999933887 | -0.139410994 | 0.159902871 | -0.299313865 | 0.021436896 |
| ENSG00000249115 | *HAUS5* | 2.170714725 | 0.035473096 | -0.068864405 | 0.104337501 | 0.999933887 | 0.351757243 | 0.844424059 | -0.492666816 | 0.021481812 |
| ENSG00000165102 | *HGSNAT* | 4.588575037 | -0.054612007 | -0.023279478 | -0.031332529 | 0.999933887 | -0.095670335 | -0.353234957 | 0.257564621 | 0.021484232 |
| ENSG00000170852 | *KBTBD2* | 6.855151098 | -0.039398921 | -0.00852532 | -0.030873601 | 0.999933887 | 0.130081246 | 0.304000662 | -0.173919416 | 0.021484232 |
| ENSG00000175970 | *UNC119B* | 2.807857932 | -0.014445896 | -0.110075215 | 0.095629319 | 0.999933887 | -0.335259134 | -0.697238201 | 0.361979067 | 0.021484232 |
| ENSG00000241058 | *NSUN6* | 2.345770631 | 0.01485619 | 0.099131033 | -0.084274843 | 0.999933887 | 0.679020859 | 1.2047099 | -0.525689041 | 0.021484232 |
| ENSG00000109270 | *LAMTOR3* | 6.517294642 | 0.004573746 | -0.029917696 | 0.034491442 | 0.999933887 | 0.299794765 | 0.514633092 | -0.214838327 | 0.021484232 |
| ENSG00000102265 | *TIMP1* | 6.445366441 | 0.016434206 | -0.016491347 | 0.032925553 | 0.999933887 | 0.271437901 | 0.517914342 | -0.246476442 | 0.021562148 |
| ENSG00000167797 | *CDK2AP2* | 3.506216841 | 0.050184372 | 0.06763557 | -0.017451198 | 0.999933887 | 0.023619068 | -0.283367207 | 0.306986274 | 0.021624344 |
| ENSG00000023330 | *ALAS1* | 3.604017252 | 0.057615818 | 0.030958125 | 0.026657693 | 0.999933887 | -0.106602742 | -0.432366432 | 0.325763689 | 0.021624344 |
| ENSG00000243927 | *MRPS6* | 3.281086532 | 0.019242015 | 0.047232809 | -0.027990794 | 0.999933887 | 0.591195143 | 1.04876631 | -0.457571167 | 0.021624344 |
| ENSG00000204131 | *NHSL2* | 5.999584909 | -0.025985246 | -0.020970814 | -0.005014432 | 0.999933887 | -1.004109881 | -1.470713647 | 0.466603766 | 0.021656055 |
| ENSG00000001561 | *ENPP4* | 2.023684473 | -0.088408923 | -0.104755889 | 0.016346966 | 0.999933887 | 0.001978517 | 0.569760979 | -0.567782462 | 0.0217522 |
| ENSG00000126653 | *NSRP1* | 4.364150075 | -0.093979464 | -0.124873884 | 0.03089442 | 0.999933887 | -0.125414723 | 0.349594937 | -0.47500966 | 0.02181991 |
| ENSG00000112531 | *QKI* | 8.194449323 | -0.030407263 | -0.027399598 | -0.003007665 | 0.999933887 | 0.776022792 | 1.139704357 | -0.363681565 | 0.021821932 |
| ENSG00000119681 | *LTBP2* | 2.956072599 | 0.059205244 | 0.013441881 | 0.045763363 | 0.999933887 | -0.812510906 | -1.206811421 | 0.394300515 | 0.021822793 |
| ENSG00000011405 | *PIK3C2A* | 4.437591582 | -0.05368187 | -0.176731135 | 0.123049265 | 0.999933887 | -0.315104799 | 0.475191649 | -0.790296448 | 0.021822793 |
| ENSG00000113302 | *IL12B* | 0.678584304 | -0.204672788 | 0.373749128 | -0.578421916 | 0.999933887 | 1.03756536 | 1.640018116 | -0.602452756 | 0.021879525 |
| ENSG00000124541 | *RRP36* | 2.988451499 | -0.081537929 | -0.021287675 | -0.060250255 | 0.999933887 | -0.193790047 | -0.497728269 | 0.303992221 | 0.021953682 |
| ENSG00000096384 | *HSP90AB1* | 7.255156533 | -0.050095198 | -0.106557232 | 0.056462034 | 0.999933887 | 0.45118936 | 0.695090194 | -0.243900835 | 0.021958818 |
| ENSG00000171606 | *ZNF274* | 6.261814262 | 0.014549656 | 0.028574265 | -0.014024609 | 0.999933887 | 0.146737937 | 0.351404573 | -0.204666637 | 0.022073026 |
| ENSG00000160813 | *PPP1R35* | 3.852824149 | 0.105103188 | 0.170213245 | -0.065110057 | 0.999933887 | -0.316687053 | -0.651396564 | 0.334709512 | 0.022078172 |
| ENSG00000196739 | *COL27A1* | -0.479095701 | 0.036736729 | 0.143804012 | -0.107067283 | 0.999933887 | 1.377179921 | 2.39786742 | -1.020687498 | 0.022078172 |
| ENSG00000066135 | *KDM4A* | 2.612522408 | 0.060567217 | 0.051651955 | 0.008915262 | 0.999933887 | -0.325120079 | -0.735039329 | 0.40991925 | 0.02215695 |
| ENSG00000183864 | *TOB2* | 4.493294148 | 0.069320089 | 0.036251837 | 0.033068253 | 0.999933887 | -0.316665908 | -0.603701672 | 0.287035764 | 0.022173102 |
| ENSG00000160679 | *CHTOP* | 5.661422984 | -0.026156336 | 0.013228891 | -0.039385227 | 0.999933887 | -0.294392337 | -0.514056642 | 0.219664306 | 0.022222936 |
| ENSG00000082996 | *RNF13* | 8.300169145 | -0.078166101 | -0.088397071 | 0.01023097 | 0.999933887 | -0.392877959 | -0.152635265 | -0.240242694 | 0.022354492 |
| ENSG00000167962 | *ZNF598* | 4.450833575 | 0.098908892 | 0.123692617 | -0.024783724 | 0.999933887 | 0.539089484 | 0.863503265 | -0.324413782 | 0.02236655 |
| ENSG00000009413 | *REV3L* | 5.588997551 | -0.134714366 | -0.095028015 | -0.039686351 | 0.999933887 | 0.358497964 | 1.00773918 | -0.649241216 | 0.02236655 |
| ENSG00000183621 | *ZNF438* | 6.039782024 | -0.036065171 | -0.019890823 | -0.016174348 | 0.999933887 | 0.054664148 | 0.279559375 | -0.224895227 | 0.02236655 |
| ENSG00000215790 | *SLC35E2A* | 2.416355402 | 0.095850559 | 0.119124067 | -0.023273508 | 0.999933887 | -0.392157106 | -0.787862036 | 0.395704929 | 0.022407411 |
| ENSG00000006007 | *GDE1* | 3.867410195 | 0.009006753 | -0.138521262 | 0.147528016 | 0.999933887 | -0.321898368 | -0.757698201 | 0.435799832 | 0.022407411 |
| ENSG00000169026 | *SLC49A3* | 2.620983215 | 0.081721998 | -0.050630519 | 0.132352516 | 0.999933887 | -0.995474448 | -1.536828682 | 0.541354233 | 0.022428371 |
| ENSG00000123268 | *ATF1* | 3.520753755 | -0.079287758 | -0.108913591 | 0.029625833 | 0.999933887 | -0.064298541 | 0.198735964 | -0.263034505 | 0.022446524 |
| ENSG00000242125 | *SNHG3* | 3.73929552 | -0.072843018 | 0.048216262 | -0.12105928 | 0.999933887 | 0.125647109 | -0.149683118 | 0.275330227 | 0.022447436 |
| ENSG00000115053 | *NCL* | 6.815759375 | -0.039705205 | -0.084097448 | 0.044392243 | 0.999933887 | 0.25689313 | 0.510360455 | -0.253467325 | 0.022618078 |
| ENSG00000163249 | *CCNYL1* | 3.706411568 | 0.037807874 | 0.06128459 | -0.023476715 | 0.999933887 | 0.147952109 | 0.451111073 | -0.303158964 | 0.022653606 |
| ENSG00000149716 | *LTO1* | 3.052593633 | -0.066488355 | -0.100238924 | 0.033750569 | 0.999933887 | -0.08628084 | -0.339105077 | 0.252824237 | 0.02266467 |
| ENSG00000076770 | *MBNL3* | 2.48803395 | -0.097642002 | -0.063386232 | -0.034255771 | 0.999933887 | -0.243690347 | 0.406415708 | -0.650106054 | 0.02266467 |
| ENSG00000178761 | *FAM219B* | 2.015898794 | 0.007895325 | -0.109823563 | 0.117718888 | 0.999933887 | 0.07226678 | -0.317547201 | 0.389813981 | 0.022676565 |
| ENSG00000159128 | *IFNGR2* | 7.826886407 | -0.002372084 | 0.010233647 | -0.012605731 | 0.999933887 | 1.449021766 | 1.770795556 | -0.32177379 | 0.022723894 |
| ENSG00000124762 | *CDKN1A* | 7.414633029 | 0.243007898 | 0.262830201 | -0.019822303 | 0.999933887 | 1.183007137 | 1.540861108 | -0.357853971 | 0.022730871 |
| ENSG00000101493 | *ZNF516* | 6.640030957 | 0.017393572 | 0.055325585 | -0.037932013 | 0.999933887 | -0.653520203 | -0.924181689 | 0.270661486 | 0.022761432 |
| ENSG00000137547 | *MRPL15* | 1.208977996 | -0.085466993 | 0.182904216 | -0.268371209 | 0.999933887 | -0.043229626 | 0.44719916 | -0.490428786 | 0.022799205 |
| ENSG00000141497 | *ZMYND15* | 5.375091627 | -0.001911084 | -0.010651286 | 0.008740202 | 0.999933887 | -0.426785109 | -0.123576616 | -0.303208494 | 0.022806892 |
| ENSG00000164823 | *OSGIN2* | 6.233409878 | -0.053112935 | 0.018068846 | -0.07118178 | 0.999933887 | 0.048070076 | 0.327222011 | -0.279151935 | 0.022819217 |
| ENSG00000159069 | *FBXW5* | 5.802320848 | 0.05153199 | 0.018852072 | 0.032679918 | 0.999933887 | 0.00502816 | -0.25907571 | 0.26410387 | 0.022827491 |
| ENSG00000135114 | *OASL* | 5.275420502 | 0.030987304 | -0.035576628 | 0.066563932 | 0.999933887 | -0.157731873 | -0.495024982 | 0.337293109 | 0.022878301 |
| ENSG00000134243 | *SORT1* | 3.661349691 | -0.024932795 | -0.042876248 | 0.017943454 | 0.999933887 | -0.801399596 | -1.199217137 | 0.397817542 | 0.022878301 |
| ENSG00000105290 | *APLP1* | 0.725012185 | 0.194861567 | 0.304887418 | -0.110025852 | 0.999933887 | 2.411641629 | 3.287586503 | -0.875944873 | 0.023066406 |
| ENSG00000242259 | *C22orf39* | 3.175329976 | 0.044108197 | -0.048755075 | 0.092863272 | 0.999933887 | -0.588707889 | -0.945364265 | 0.356656376 | 0.023067299 |
| ENSG00000113194 | *FAF2* | 4.822265855 | -0.015152997 | -0.02742058 | 0.012267583 | 0.999933887 | 0.002833555 | 0.31658866 | -0.313755105 | 0.023101173 |
| ENSG00000120539 | *MASTL* | 3.672999739 | -0.036897065 | 0.071812292 | -0.108709357 | 0.999933887 | 0.707753333 | 1.019064258 | -0.311310925 | 0.023193251 |
| ENSG00000048649 | *RSF1* | 5.567964676 | -0.083939181 | -0.119560051 | 0.03562087 | 0.999933887 | -0.45735586 | -0.008451585 | -0.448904275 | 0.023212559 |
| ENSG00000113648 | *MACROH2A1* | 7.579486308 | -0.015912061 | -0.025743892 | 0.009831831 | 0.999933887 | -1.246224936 | -1.561348324 | 0.315123388 | 0.023364746 |
| ENSG00000013374 | *NUB1* | 5.919378907 | 0.015787953 | -0.003834453 | 0.019622406 | 0.999933887 | -0.31121456 | -0.568308187 | 0.257093626 | 0.023383096 |
| ENSG00000138029 | *HADHB* | 5.104268526 | -0.074352791 | -0.020468811 | -0.05388398 | 0.999933887 | -0.325841628 | -0.525591621 | 0.199749994 | 0.023406732 |
| ENSG00000091009 | *RBM27* | 3.700437637 | -0.063595366 | -0.153291425 | 0.089696058 | 0.999933887 | -0.413628414 | -0.045636258 | -0.367992157 | 0.023452472 |
| ENSG00000198952 | *SMG5* | 6.06932519 | 0.029951477 | 0.013375473 | 0.016576004 | 0.999933887 | -0.236970062 | -0.428213389 | 0.191243327 | 0.02347914 |
| ENSG00000144815 | *NXPE3* | 3.53666304 | 0.027201637 | -0.012749593 | 0.039951229 | 0.999933887 | -0.128193676 | -0.499199306 | 0.37100563 | 0.023696902 |
| ENSG00000159023 | *EPB41* | 7.439547853 | -0.076569878 | -0.032642082 | -0.043927797 | 0.999933887 | -0.414858038 | -0.656030747 | 0.241172709 | 0.02370665 |
| ENSG00000166592 | *RRAD* | 0.618266061 | 0.415859511 | 0.792326547 | -0.376467035 | 0.999933887 | 2.013966698 | 2.797266411 | -0.783299713 | 0.023725344 |
| ENSG00000090520 | *DNAJB11* | 4.773925218 | -0.022375785 | 0.034118561 | -0.056494345 | 0.999933887 | 0.268103294 | 0.526954707 | -0.258851413 | 0.023845456 |
| ENSG00000070047 | *PHRF1* | 5.784357864 | 0.012228217 | 0.063743662 | -0.051515445 | 0.999933887 | -0.17026656 | -0.441692527 | 0.271425967 | 0.023892146 |
| ENSG00000214046 | *SMIM7* | 2.867963938 | -0.057099051 | -0.033082712 | -0.024016339 | 0.999933887 | -0.200258001 | -0.480363225 | 0.280105223 | 0.02397898 |
| ENSG00000136379 | *ABHD17C* | -0.240883853 | 0.007284352 | -0.254379116 | 0.261663468 | 0.999933887 | -0.383617892 | -1.21582658 | 0.832208688 | 0.023984236 |
| ENSG00000157954 | *WIPI2* | 6.274227127 | -0.045360442 | -0.036979276 | -0.008381166 | 0.999933887 | -0.53293127 | -0.758319664 | 0.225388394 | 0.024041665 |
| ENSG00000104714 | *ERICH1* | 5.257304794 | 0.044361065 | 0.045498978 | -0.001137913 | 0.999933887 | -0.244205625 | -0.54732976 | 0.303124135 | 0.024072729 |
| ENSG00000142552 | *RCN3* | 4.062651025 | 0.03880702 | 0.161420454 | -0.122613433 | 0.999933887 | -0.315592516 | -0.830184151 | 0.514591635 | 0.024108155 |
| ENSG00000074695 | *LMAN1* | 2.930005217 | -0.051396618 | -0.096119752 | 0.044723134 | 0.999933887 | -0.062863372 | 0.406444029 | -0.469307401 | 0.024182826 |
| ENSG00000103066 | *PLA2G15* | 1.304072589 | 0.054542052 | -0.141388685 | 0.195930737 | 0.999933887 | -0.174511275 | -0.790180397 | 0.615669122 | 0.024182826 |
| ENSG00000171448 | *ZBTB26* | 3.033684326 | -0.091835958 | -0.218526758 | 0.126690801 | 0.999933887 | 0.662779614 | 1.002445169 | -0.339665555 | 0.024187191 |
| ENSG00000197045 | *GMFB* | 4.667228332 | -0.075979189 | -0.223853785 | 0.147874596 | 0.999933887 | -0.331373736 | 0.066158649 | -0.397532385 | 0.024231784 |
| ENSG00000286555 | *AP003065.1* | 4.476968629 | -0.046724989 | -0.071317791 | 0.024592803 | 0.999933887 | -0.167168068 | 0.136720992 | -0.30388906 | 0.024231784 |
| ENSG00000100281 | *HMGXB4* | 4.420952603 | -0.015679245 | 0.067842731 | -0.083521976 | 0.999933887 | 0.345409503 | 0.605331974 | -0.259922471 | 0.024250928 |
| ENSG00000186827 | *TNFRSF4* | 2.459879159 | -0.02871214 | 0.023056595 | -0.051768735 | 0.999933887 | 1.079071174 | 1.67783039 | -0.598759216 | 0.024268317 |
| ENSG00000110768 | *GTF2H1* | 5.701562729 | -0.036467983 | 0.029123121 | -0.065591104 | 0.999933887 | 0.283908188 | 0.529987578 | -0.24607939 | 0.024326791 |
| ENSG00000110047 | *EHD1* | 8.626710668 | -0.054442244 | 0.009863708 | -0.064305951 | 0.999933887 | 2.400045175 | 2.979681881 | -0.579636705 | 0.024368204 |
| ENSG00000148187 | *MRRF* | 2.981826754 | -0.064903674 | -0.015596269 | -0.049307405 | 0.999933887 | 0.196286277 | 0.508070731 | -0.311784453 | 0.024412527 |
| ENSG00000054965 | *FAM168A* | 5.719181154 | 0.007809861 | 0.015528093 | -0.007718232 | 0.999933887 | -0.76251863 | -1.053271983 | 0.290753353 | 0.024412527 |
| ENSG00000256229 | *ZNF486* | 1.633290229 | -0.058559498 | 0.033707005 | -0.092266503 | 0.999933887 | -0.823452282 | -1.374648588 | 0.551196306 | 0.024421223 |
| ENSG00000148297 | *MED22* | 4.57165746 | -0.082613131 | -0.018959592 | -0.063653538 | 0.999933887 | -0.319671922 | -0.613415387 | 0.293743465 | 0.024437538 |
| ENSG00000204310 | *AGPAT1* | 3.329253403 | -0.021677647 | 0.001974505 | -0.023652152 | 0.999933887 | -0.286832738 | -0.639073079 | 0.35224034 | 0.024462764 |

| | | | | | | | | | | |
|---|---|---|---|---|---|---|---|---|---|---|
| ENSG00000156042 | *CFAP70* | 1.161647234 | -0.010034756 | -0.089791407 | 0.079756651 | 0.999933887 | 0.644948646 | 1.263096133 | -0.618147488 | 0.024556015 |
| ENSG00000166016 | *ABTB2* | 3.60282547 | -0.009854434 | -0.016361941 | 0.006507507 | 0.999933887 | 0.398362422 | 0.822377569 | -0.424015147 | 0.024557495 |
| ENSG00000011304 | *PTBP1* | 6.846958324 | 0.006967807 | 0.021464646 | -0.014496839 | 0.999933887 | -0.142386932 | -0.322286109 | 0.179899176 | 0.024619635 |
| ENSG00000178234 | *GALNT11* | 4.369141516 | -0.003557189 | 0.010278807 | -0.013835996 | 0.999933887 | 0.200912848 | 0.4191982 | -0.218285352 | 0.024630167 |
| ENSG00000067798 | *NAV3* | 0.293640316 | -0.059439534 | -0.117392842 | 0.057953308 | 0.999933887 | 2.101244205 | 2.906460765 | -0.80521656 | 0.024644695 |
| ENSG00000134780 | *DAGLA* | 1.161775735 | -0.052288096 | 0.169576294 | -0.22186439 | 0.999933887 | -0.129467061 | 0.391969574 | -0.521436635 | 0.024644695 |
| ENSG00000197136 | *PCNX3* | 6.586279521 | 0.005037677 | 0.048708832 | -0.043671156 | 0.999933887 | -0.203493045 | -0.405888726 | 0.202395681 | 0.024685697 |
| ENSG00000143933 | *CALM2* | 7.088392152 | -0.089079777 | -0.116892058 | 0.02781228 | 0.999933887 | -0.459278338 | -0.698757111 | 0.239478773 | 0.024687679 |
| ENSG00000068079 | *IFI35* | 4.696236008 | 0.017791075 | -0.071643147 | 0.089434221 | 0.999933887 | -0.439368513 | -0.776158311 | 0.336789798 | 0.024687679 |
| ENSG00000072062 | *PRKACA* | 5.547969984 | 0.006991971 | 0.030958846 | -0.023966875 | 0.999933887 | -0.104862117 | -0.377898683 | 0.273036566 | 0.024744546 |
| ENSG00000160741 | *CRTC2* | 7.621386629 | 0.018632686 | 0.051251198 | -0.032618512 | 0.999933887 | 0.902106845 | 1.120626418 | -0.218519573 | 0.024799411 |
| ENSG00000171867 | *PRNP* | 5.786261098 | -0.000415512 | 0.017275544 | -0.017691057 | 0.999933887 | 0.267889901 | 0.51280759 | -0.244917689 | 0.024799411 |
| ENSG00000066336 | *SPI1* | 10.297134615 | 0.061479693 | 0.03714822 | 0.024331473 | 0.999933887 | 0.340383037 | 0.539251966 | -0.19886893 | 0.02480964 |
| ENSG00000197302 | *ZNF720* | 2.722967887 | 0.04151557 | 0.056571857 | -0.015056287 | 0.999933887 | -0.031808697 | 0.382888186 | -0.414696883 | 0.02480964 |
| ENSG00000140391 | *TSPAN3* | 4.592059049 | 0.007433155 | -0.008509737 | 0.015942892 | 0.999933887 | 0.361016531 | 0.655999559 | -0.294983028 | 0.02480964 |
| ENSG00000273611 | *ZNHIT3* | 2.855074334 | -0.018159318 | -0.072632352 | 0.054473033 | 0.999933887 | -0.12761981 | -0.43579885 | 0.30817904 | 0.024997937 |
| ENSG00000143409 | *MINDY1* | 4.822238837 | -0.056047948 | 0.071428513 | -0.127476461 | 0.999933887 | -0.660291252 | -1.066521575 | 0.406230322 | 0.025037515 |
| ENSG00000196371 | *FUT4* | 4.040090106 | 0.08140056 | 0.025137139 | 0.056263421 | 0.999933887 | 0.985194358 | 1.322342881 | -0.337148523 | 0.025081883 |
| ENSG00000128463 | *EMC4* | 3.49932858 | 0.016813819 | -0.131740147 | 0.148553966 | 0.999933887 | -0.029529075 | -0.294678355 | 0.26514928 | 0.025105783 |
| ENSG00000154832 | *CXXC1* | 4.318801307 | 0.054639303 | 0.018798127 | 0.035841175 | 0.999933887 | 0.008369355 | -0.28380952 | 0.292178875 | 0.025153686 |
| ENSG00000092330 | *TINF2* | 6.683428219 | 0.039240232 | 0.034832234 | 0.004407998 | 0.999933887 | 0.476429883 | 0.682701315 | -0.206271432 | 0.025153686 |
| ENSG00000120899 | *PTK2B* | 9.265902746 | 0.031668734 | 0.027924544 | 0.003744191 | 0.999933887 | 0.887982091 | 1.198641481 | -0.310659389 | 0.025153686 |
| ENSG00000277224 | *H2BC7* | 0.430432395 | 0.084342343 | 0.20558706 | -0.121244717 | 0.999933887 | -0.601941891 | -1.606067436 | 1.004125546 | 0.025242413 |
| ENSG00000171421 | *MRPL36* | 1.002463798 | 0.001378873 | 0.257687748 | -0.256308875 | 0.999933887 | 0.053143699 | -0.598120633 | 0.651264332 | 0.025250244 |
| ENSG00000006062 | *MAP3K14* | 4.129330585 | -0.006898154 | 0.061534008 | -0.068432162 | 0.999933887 | -0.149399982 | -0.437084362 | 0.28768438 | 0.025259189 |
| ENSG00000104529 | *EEF1D* | 6.389684508 | 0.002939793 | 0.00534265 | -0.002402857 | 0.999933887 | -0.229188093 | -0.454157324 | 0.224969231 | 0.025259189 |
| ENSG00000198900 | *TOP1* | 7.290191045 | -0.039742497 | -0.063337699 | 0.023595201 | 0.999933887 | 0.20006838 | 0.446973934 | -0.246905554 | 0.025310409 |
| ENSG00000147383 | *NSDHL* | 0.633414954 | -0.053566345 | 0.01284078 | -0.066407125 | 0.999933887 | 0.034674094 | -0.553339635 | 0.588013729 | 0.025310409 |
| ENSG00000273284 | *AP001033.2* | 2.582458378 | -0.148558747 | -0.25165716 | 0.103098413 | 0.999933887 | -0.259445645 | 0.362898458 | -0.622344103 | 0.025339684 |
| ENSG00000070540 | *WIPI1* | 3.663910632 | -0.050732165 | 0.075745919 | -0.126478085 | 0.999933887 | -0.685082634 | -1.023906478 | 0.338823843 | 0.025381782 |
| ENSG00000023909 | *GCLM* | 5.028918478 | -0.070366451 | 0.007311233 | -0.077677683 | 0.999933887 | 0.418167465 | 0.726408467 | -0.308241002 | 0.025381782 |
| ENSG00000282851 | *BISPR* | 1.852966571 | 0.172313679 | 0.213505488 | -0.041191809 | 0.999933887 | -0.140055942 | -0.61258099 | 0.472525048 | 0.025498332 |
| ENSG00000242485 | *MRPL20* | 3.404949719 | -0.085871701 | -0.122411677 | 0.036539976 | 0.999933887 | -0.158064891 | -0.391634629 | 0.233569737 | 0.025534178 |
| ENSG00000106261 | *ZKSCAN1* | 5.125820913 | -0.061029748 | -0.078371533 | 0.017341785 | 0.999933887 | -0.003401401 | 0.230651109 | -0.23405251 | 0.025534178 |
| ENSG00000083814 | *ZNF671* | 1.783826485 | 0.065676555 | 0.1017377 | -0.036061144 | 0.999933887 | 0.007907934 | -0.390084826 | 0.39799276 | 0.025534178 |
| ENSG00000133657 | *ATP13A3* | 6.866518609 | -0.049299833 | -0.029377399 | -0.019922434 | 0.999933887 | 0.49193398 | 0.922473504 | -0.430539525 | 0.025534178 |
| ENSG00000078403 | *MLLT10* | 3.845262964 | -0.034187032 | -0.059592825 | 0.025405793 | 0.999933887 | -0.498429571 | -0.823051345 | 0.324621774 | 0.025534178 |
| ENSG00000169429 | *CXCL8* | 13.85870056 | 0.547314938 | 0.527723852 | 0.019591086 | 0.999933887 | 2.034739193 | 2.530100787 | -0.495361595 | 0.025544397 |
| ENSG00000164081 | *TEX264* | 2.808516155 | 0.036228811 | -0.074145854 | 0.110374665 | 0.999933887 | 0.028325383 | -0.350160232 | 0.378485616 | 0.025559094 |
| ENSG00000198743 | *SLC5A3* | 3.949093873 | -0.034692545 | -0.111916861 | 0.077224316 | 0.999933887 | 0.653963885 | 1.319041524 | -0.665077639 | 0.025644012 |
| ENSG00000105717 | *PBX4* | 2.156319287 | -0.034802808 | 0.013291502 | -0.048094311 | 0.999933887 | 0.35594664 | 0.74859507 | -0.39264843 | 0.025681919 |
| ENSG00000106460 | *TMEM106B* | 4.160864273 | 0.048769767 | -0.163182297 | 0.211952064 | 0.999933887 | -0.473407925 | 0.030549015 | -0.50395694 | 0.025681919 |
| ENSG00000116793 | *PHTF1* | 5.017663339 | -0.043605556 | 0.017584691 | -0.061190247 | 0.999933887 | -0.359403324 | -0.585409858 | 0.226006535 | 0.025726915 |
| ENSG00000110848 | *CD69* | 5.737848119 | -0.031999641 | -0.079268009 | 0.047268368 | 0.999933887 | 0.132295768 | 0.484214393 | -0.351918625 | 0.025726915 |
| ENSG00000140519 | *RHCG* | -1.645547746 | -0.001035673 | 0.246876825 | -0.247912497 | 0.999933887 | 3.830612409 | 5.173221195 | -1.342608786 | 0.025726915 |
| ENSG00000197405 | *C5AR1* | 11.056494624 | 0.038543226 | 0.018102939 | 0.020440286 | 0.999933887 | -0.221799028 | -0.005561784 | -0.216237244 | 0.025760935 |
| ENSG00000126698 | *DNAJC8* | 6.503319004 | -0.04719119 | -0.122167909 | 0.07497672 | 0.999933887 | 0.547045452 | 0.785856474 | -0.238811022 | 0.025850622 |
| ENSG00000171310 | *CHST11* | 7.557069469 | 0.078594715 | 0.105614376 | -0.027019661 | 0.999933887 | -0.197590086 | -0.47382233 | 0.276232245 | 0.025875988 |
| ENSG00000175505 | *CLCF1* | 1.986654196 | 0.155913208 | 0.105429302 | 0.050483906 | 0.999933887 | 1.505106886 | 2.136462712 | -0.631355827 | 0.025875988 |
| ENSG00000072958 | *AP1M1* | 5.254712139 | 0.042091884 | -0.013114231 | 0.055206116 | 0.999933887 | -0.298687981 | -0.523272583 | 0.224584602 | 0.025875988 |
| ENSG00000092036 | *HAUS4* | 2.236151178 | 0.057606191 | -0.02113264 | 0.07873883 | 0.999933887 | -0.232154156 | -0.825730575 | 0.593576419 | 0.025875988 |
| ENSG00000187713 | *TMEM203* | 2.546677114 | 0.047105592 | -0.090257756 | 0.137363348 | 0.999933887 | -0.033294763 | -0.425732028 | 0.392437265 | 0.025875988 |
| ENSG00000070759 | *TESK2* | 4.538527994 | 0.014112336 | -0.010094773 | 0.024207109 | 0.999933887 | 0.248167706 | 0.513597817 | -0.26543011 | 0.025894986 |
| ENSG00000176973 | *FAM89B* | 1.169459282 | 0.1319522 | -0.119335002 | 0.251287202 | 0.999933887 | 0.247245192 | -0.455877809 | 0.703123001 | 0.025907486 |
| ENSG00000227502 | *MROCKI* | 2.482655805 | -0.036971025 | -0.053111843 | 0.016140817 | 0.999933887 | 0.841189536 | 1.446872428 | -0.605682892 | 0.025907486 |
| ENSG00000132141 | *CCT6B* | -0.201177554 | 0.005317428 | 0.186358207 | -0.181040779 | 0.999933887 | 0.136415499 | 1.052950539 | -0.916535039 | 0.025907486 |
| ENSG00000224051 | *CPTP* | 3.024342933 | 0.00708862 | 0.005591613 | 0.001497006 | 0.999933887 | -0.53060376 | -0.930373513 | 0.399769753 | 0.025930573 |
| ENSG00000179044 | *EXOC3L1* | -0.214381433 | -0.097852028 | 0.091570051 | -0.189422079 | 0.999933887 | -1.080882254 | -2.640032112 | 1.559149857 | 0.026032014 |
| ENSG00000231074 | *HCG18* | 4.381739555 | 0.063537359 | 0.035003279 | 0.02853408 | 0.999933887 | 0.145869753 | 0.45530225 | -0.309432498 | 0.02607577 |
| ENSG00000072163 | *LIMS2* | -0.015467429 | 0.051930046 | 0.088574169 | -0.036644123 | 0.999933887 | -0.246007841 | -1.169751623 | 0.923743782 | 0.026155447 |
| ENSG00000118454 | *ANKRD13C* | 2.574310966 | -0.014943729 | 0.011404743 | -0.026348472 | 0.999933887 | -0.10913838 | 0.245136607 | -0.354274987 | 0.026253579 |
| ENSG00000139990 | *DCAF5* | 5.522377349 | 0.005721116 | 0.013808126 | -0.00808701 | 0.999933887 | -0.437852177 | -0.669358515 | 0.231506338 | 0.026253579 |
| ENSG00000272084 | *AL137127.1* | 1.888004171 | 0.080529295 | -0.012294855 | 0.09282415 | 0.999933887 | 0.833972639 | 1.307727784 | -0.473755145 | 0.026286115 |
| ENSG00000131669 | *NINJ1* | 10.103021777 | 0.041006428 | -0.002092013 | 0.043098442 | 0.999933887 | 1.087995885 | 1.315806062 | -0.227810177 | 0.026299314 |
| ENSG00000124209 | *RAB22A* | 5.745629625 | -0.053004702 | -0.102104103 | 0.049099401 | 0.999933887 | -0.180266063 | -0.016125804 | -0.164140259 | 0.026316651 |
| ENSG00000198093 | *ZNF649* | 0.33017136 | 0.010278495 | -0.241823414 | 0.252101909 | 0.999933887 | 0.039804853 | 0.639460624 | -0.599655771 | 0.026316651 |
| ENSG00000071243 | *ING3* | 5.490253882 | -0.023889175 | -0.054958687 | 0.031069512 | 0.999933887 | 0.413601964 | 0.690553763 | -0.276951799 | 0.026436753 |
| ENSG00000102974 | *CTCF* | 4.913413067 | -0.082976308 | -0.015175104 | -0.067801204 | 0.999933887 | -0.382114775 | -0.661204919 | 0.279090144 | 0.02644315 |
| ENSG00000262884 | *AC015921.1* | -1.048490788 | 0.043664211 | 0.052605055 | -0.008940844 | 0.999933887 | -0.537573753 | -1.952912812 | 1.41533906 | 0.02644315 |
| ENSG00000107021 | *TBC1D13* | 3.741818305 | -0.023407089 | -0.026295925 | 0.002888835 | 0.999933887 | -0.347592589 | -0.652061348 | 0.304468759 | 0.026472065 |
| ENSG00000247081 | *BAALC-AS1* | -0.60122028 | -0.500550673 | 0.232060799 | -0.732611472 | 0.999933887 | 1.90344695 | 2.750698431 | -0.847251481 | 0.026631524 |
| ENSG00000113621 | *TXNDC15* | 2.443594812 | 0.002463546 | -0.159146024 | 0.161609571 | 0.999933887 | -0.161398383 | -0.535753581 | 0.374355198 | 0.026638275 |
| ENSG00000153406 | *NMRAL1* | 3.856645127 | -0.026796152 | -0.091056091 | 0.064259939 | 0.999933887 | -0.518192576 | -0.855299937 | 0.337107361 | 0.026675481 |
| ENSG00000005100 | *DHX33* | 1.788411771 | -0.155842197 | 0.13116633 | -0.287008527 | 0.999933887 | 0.096387763 | -0.280797292 | 0.377185056 | 0.026791545 |
| ENSG00000151151 | *IPMK* | 6.728415477 | -0.049204988 | -0.088970953 | 0.039765965 | 0.999933887 | -0.512739097 | -0.131420161 | -0.381318936 | 0.026820293 |
| ENSG00000146278 | *PNRC1* | 10.459089763 | 0.109929652 | 0.07259832 | 0.037331331 | 0.999933887 | 1.371562797 | 1.658290075 | -0.286727278 | 0.026902793 |
| ENSG00000105856 | *HBP1* | 8.117233362 | -0.085388418 | -0.123681719 | 0.038293301 | 0.999933887 | 0.124165855 | 0.350220747 | -0.226054892 | 0.026902793 |
| ENSG00000154370 | *TRIM11* | 4.509733445 | 0.016130051 | 0.075294788 | -0.059164737 | 0.999933887 | -0.314261085 | -0.613455879 | 0.299194794 | 0.026902793 |
| ENSG00000138814 | *PPP3CA* | 5.680245235 | 0.008010606 | 0.048530679 | -0.040520072 | 0.999933887 | -0.323485953 | -0.624826418 | 0.301340464 | 0.026903095 |
| ENSG00000107968 | *MAP3K8* | 7.591966985 | 0.062296146 | 0.048671303 | 0.013624843 | 0.999933887 | 1.76873989 | 2.194677308 | -0.425937418 | 0.026942319 |
| ENSG00000169105 | *CHST14* | 1.091991654 | 0.012836983 | -0.047801299 | 0.060638283 | 0.999933887 | 0.262053607 | -0.296006858 | 0.558060465 | 0.026942319 |
| ENSG00000162444 | *RBP7* | 2.358970921 | -0.04277004 | -0.078648212 | 0.035878172 | 0.999933887 | -0.590756145 | -1.171690857 | 0.580934713 | 0.026971504 |
| ENSG00000160219 | *GAB3* | 5.503338752 | -0.021016642 | -0.073889777 | 0.052873134 | 0.999933887 | -0.016563766 | 0.222251048 | -0.238814815 | 0.026971504 |
| ENSG00000204520 | *MICA* | 2.551691045 | 0.019675998 | -0.080160418 | 0.099836415 | 0.999933887 | -0.294649024 | -0.691647521 | 0.396998497 | 0.026971504 |
| ENSG00000134851 | *TMEM165* | 5.128588089 | -0.013729915 | 0.104786961 | -0.118516876 | 0.999933887 | 1.029178462 | 1.343504284 | -0.314325822 | 0.026971504 |
| ENSG00000166747 | *AP1G1* | 7.860590566 | -0.069164987 | -0.066192801 | -0.002972186 | 0.999933887 | 0.098607458 | 0.284901199 | -0.186293741 | 0.026993043 |
| ENSG00000114902 | *SPCS1* | 4.278573917 | -0.004801375 | -0.03759499 | 0.032793615 | 0.999933887 | -0.031234419 | -0.294295768 | 0.263061349 | 0.026993043 |
| ENSG00000124731 | *TREM1* | 10.885557178 | 0.054139734 | 0.02914062 | 0.024999114 | 0.999933887 | 0.379856177 | 0.610148168 | -0.230291991 | 0.027052382 |
| ENSG00000118257 | *NRP2* | -0.610079471 | -0.248380656 | 0.214974495 | -0.46335515 | 0.999933887 | 0.562641771 | 1.348427311 | -0.78578554 | 0.027077296 |
| ENSG00000043093 | *DCUN1D1* | 4.438150416 | -0.042943384 | -0.117947206 | 0.075003822 | 0.999933887 | -0.03039731 | 0.308020942 | -0.338418252 | 0.027081253 |
| ENSG00000114354 | *TFG* | 5.135269487 | -0.050984538 | -0.063541182 | 0.012556645 | 0.999933887 | 0.387722278 | 0.610147048 | -0.222424771 | 0.027087473 |
| ENSG00000135316 | *SYNCRIP* | 5.503108504 | -0.018135577 | -0.059278408 | 0.041142831 | 0.999933887 | -0.052593017 | 0.178016238 | -0.230609255 | 0.027087473 |
| ENSG00000170921 | *TANC2* | 5.22522522 | -0.076949965 | -0.003914107 | -0.073035858 | 0.999933887 | -0.593731716 | -0.889864145 | 0.296132429 | 0.027106315 |
| ENSG00000277632 | *CCL3* | 6.970312984 | 0.541102414 | 0.593214974 | -0.05211256 | 0.999933887 | 3.172144794 | 3.837581046 | -0.665436251 | 0.027136165 |
| ENSG00000131370 | *SH3BP5* | 6.625175386 | 0.080974846 | 0.048867211 | 0.032107635 | 0.999933887 | 0.472542427 | 0.75698316 | -0.284440733 | 0.027136165 |
| ENSG00000110075 | *PPP6R3* | 6.395514127 | -0.038109171 | -0.015056086 | -0.023053085 | 0.999933887 | 0.040994802 | 0.290705017 | -0.249710215 | 0.027166212 |
| ENSG00000111679 | *PTPN6* | 9.479280143 | 0.000594374 | 3.2031e-05 | 0.000562342 | 0.999933887 | 0.20051351 | 0.430133755 | -0.229620246 | 0.027166212 |
| ENSG00000055147 | *FAM114A2* | 2.558892233 | -0.031340753 | -0.069781272 | 0.038440519 | 0.999933887 | -0.324657957 | -0.023287826 | -0.301370132 | 0.027177008 |
| ENSG00000163870 | *TPRA1* | 6.002320644 | -0.030557018 | 0.062552528 | -0.093109546 | 0.999933887 | 1.192727646 | 1.5079946 | -0.315266954 | 0.027177008 |
| ENSG00000126767 | *ELK1* | 4.325625707 | -0.008917352 | 0.016551807 | -0.025469159 | 0.999933887 | 0.929261502 | 1.28210434 | -0.352842838 | 0.027228949 |
| ENSG00000168090 | *COPS6* | 3.461055164 | 0.013009222 | -0.000578588 | 0.01358781 | 0.999933887 | 0.130268819 | -0.158207134 | 0.288475953 | 0.027254581 |
| ENSG00000120705 | *ETF1* | 8.804603343 | -0.010529974 | -0.080043392 | 0.069513418 | 0.999933887 | 0.387608409 | 0.631896222 | -0.244287813 | 0.027254581 |
| ENSG00000100425 | *BRD1* | 5.951272799 | -0.015032943 | 0.021334551 | -0.036367494 | 0.999933887 | -0.240114742 | -0.435912664 | 0.195797921 | 0.027258939 |
| ENSG00000101199 | *ARFGAP1* | 6.023487788 | -0.01145438 | 0.07517744 | -0.086631821 | 0.999933887 | -0.12324538 | -0.353292367 | 0.230046987 | 0.027258939 |
| ENSG00000149294 | *NCAM1* | -0.207077974 | -0.032731394 | 0.116311079 | -0.149042473 | 0.999933887 | -0.65253443 | 0.157747801 | -0.810282231 | 0.027283129 |

| | | | | | | | | | | |
|---|---|---|---|---|---|---|---|---|---|---|
| ENSG00000168421 | *RHOH* | 5.40316308 | 0.054389957 | 0.068127777 | -0.013737821 | 0.999933887 | 1.062607681 | 1.552281386 | -0.489673705 | 0.027340245 |
| ENSG00000136603 | *SKIL* | 7.349384158 | -0.030730424 | -0.100450071 | 0.069719647 | 0.999933887 | 0.707214195 | 1.1687041 | -0.461489905 | 0.027342257 |
| ENSG00000148834 | *GSTO1* | 6.091533262 | -0.075993072 | -0.063553838 | -0.012439233 | 0.999933887 | 0.311680183 | 0.570153739 | -0.258473556 | 0.027406255 |
| ENSG00000139641 | *ESYT1* | 5.564356861 | -0.032744769 | 0.023775868 | -0.056520637 | 0.999933887 | -0.120556724 | -0.318812088 | 0.198255364 | 0.027542216 |
| ENSG00000125827 | *TMX4* | 6.158531538 | 0.00388676 | -0.038646253 | 0.042533014 | 0.999933887 | -0.249618507 | -0.5186946 | 0.269076093 | 0.027611924 |
| ENSG00000137726 | *FXYD6* | -0.216868585 | 0.084992056 | -0.405310786 | 0.490302841 | 0.999933887 | -0.093404796 | 0.425676389 | -0.519081185 | 0.027723641 |
| ENSG00000160325 | *CACFD1* | 1.427055126 | 0.148589717 | -0.017495681 | 0.166085398 | 0.999933887 | 1.680826764 | 2.248976041 | -0.568149277 | 0.02772805 |
| ENSG00000141452 | *RMC1* | 4.939689969 | 0.022249148 | 0.018942518 | 0.003306631 | 0.999933887 | -0.291330898 | -0.492384792 | 0.201053894 | 0.02772805 |
| ENSG00000104140 | *RHOV* | 1.952570912 | -0.060530652 | 0.174096466 | -0.234627118 | 0.999933887 | 2.043778145 | 2.512495086 | -0.468716941 | 0.027763728 |
| ENSG00000167604 | *NFKBID* | 8.809550491 | 0.106863916 | 0.127620512 | -0.020756595 | 0.999933887 | 2.398350392 | 2.784424553 | -0.386074161 | 0.027770019 |
| ENSG00000049541 | *RFC2* | 2.745078108 | -0.04851762 | -0.097061657 | 0.048544038 | 0.999933887 | -0.014174973 | -0.347154229 | 0.332979257 | 0.02782417 |
| ENSG00000013288 | *MAN2B2* | 5.497002783 | 0.051292712 | 0.039515064 | 0.011777647 | 0.999933887 | -0.266264832 | -0.509802992 | 0.24353816 | 0.027824341 |
| ENSG00000003402 | *CFLAR* | 9.557156309 | -0.012662443 | 0.003943402 | -0.016605845 | 0.999933887 | 1.15427913 | 1.400873582 | -0.246594452 | 0.027825157 |
| ENSG00000135924 | *DNAJB2* | 5.640683026 | 0.054836178 | 0.077009433 | -0.022173256 | 0.999933887 | -0.162562632 | -0.476474085 | 0.313911453 | 0.027858324 |
| ENSG00000095787 | *WAC* | 8.223328264 | 0.00997741 | -0.027289734 | 0.037267144 | 0.999933887 | -0.012335491 | 0.169346438 | -0.181681929 | 0.027858324 |
| ENSG00000074370 | *ATP2A3* | 8.534718474 | 0.030157918 | 0.011710801 | 0.018447117 | 0.999933887 | -0.541342727 | -0.782863321 | 0.241520594 | 0.027936005 |
| ENSG00000166130 | *IKBIP* | 3.439451933 | 0.048479382 | -0.032290152 | 0.080769534 | 0.999933887 | -0.602752585 | -1.064880189 | 0.462127604 | 0.028020988 |
| ENSG00000010244 | *ZNF207* | 6.992243118 | -0.058766135 | -0.051635428 | -0.007130707 | 0.999933887 | 0.015039012 | 0.208463403 | -0.193424391 | 0.028031789 |
| ENSG00000184887 | *BTBD6* | 3.541812218 | 0.012339247 | 0.048293042 | -0.035953795 | 0.999933887 | 0.147792988 | -0.173905464 | 0.321698452 | 0.028040745 |
| ENSG00000115520 | *COQ10B* | 5.994261081 | 0.110059368 | 0.123296811 | -0.013237443 | 0.999933887 | 0.249369898 | 0.446253059 | -0.196883161 | 0.028060422 |
| ENSG00000250771 | *AC106865.1* | 4.448528007 | 0.044058202 | 0.134306684 | -0.090248482 | 0.999933887 | 0.884946589 | 1.327256973 | -0.442310384 | 0.028060422 |
| ENSG00000134222 | *PSRC1* | 0.343393473 | 0.066309649 | 0.376637615 | -0.310327965 | 0.999933887 | -0.510188468 | -1.722279833 | 1.212091365 | 0.028101224 |
| ENSG00000131238 | *PPT1* | 6.27248025 | 0.035783463 | -0.061847244 | 0.097630707 | 0.999933887 | -0.614297314 | -0.938035364 | 0.323738051 | 0.028112042 |
| ENSG00000089775 | *ZBTB25* | 4.522176567 | -0.063782218 | 0.104141037 | -0.167923255 | 0.999933887 | 0.232247385 | 0.479930202 | -0.247682817 | 0.028215892 |
| ENSG00000236213 | *AC006369.1* | 1.994725247 | 0.135233136 | 0.180665245 | -0.045432109 | 0.999933887 | 1.201677449 | 1.729393947 | -0.527716498 | 0.028215892 |
| ENSG00000198216 | *CACNA1E* | 5.032366565 | 0.021060631 | 0.020392424 | 0.000668207 | 0.999933887 | 0.803465737 | 1.042391003 | -0.238925266 | 0.028215892 |
| ENSG00000242299 | *AC073861.1* | 0.205001331 | -0.022864523 | -0.042374357 | 0.019509834 | 0.999933887 | -0.272102461 | 0.367696506 | -0.639798967 | 0.028340164 |
| ENSG00000133055 | *MYBPH* | 1.694343932 | 0.133173781 | -0.092445742 | 0.225619523 | 0.999933887 | 1.712189398 | 2.199746854 | -0.487557456 | 0.028452773 |
| ENSG00000114416 | *FXR1* | 5.306143983 | -0.116557524 | -0.155696737 | 0.039139214 | 0.999933887 | 0.031239161 | 0.292158111 | -0.26091895 | 0.028480497 |
| ENSG00000101367 | *MAPRE1* | 7.956613587 | -0.028235899 | -0.082003284 | 0.053767385 | 0.999933887 | 0.196021641 | 0.422548761 | -0.226527121 | 0.028480497 |
| ENSG00000123415 | *SMUG1* | 1.889630448 | -0.076763986 | 0.009895813 | -0.086659799 | 0.999933887 | -0.222850855 | -0.629354801 | 0.406503946 | 0.028520165 |
| ENSG00000101158 | *NELFCD* | 4.171445121 | -0.005479111 | -0.010453755 | 0.004974645 | 0.999933887 | -0.10846592 | -0.350165671 | 0.241699751 | 0.028521233 |
| ENSG00000198925 | *ATG9A* | 3.773298005 | 0.057750658 | -0.042903769 | 0.100654427 | 0.999933887 | -0.392024758 | -0.704474304 | 0.312449546 | 0.028671753 |
| ENSG00000250033 | *SLC7A11-AS1* | 2.610803083 | -0.030247365 | 0.185742071 | -0.215989437 | 0.999933887 | -0.019635046 | 0.426647312 | -0.446282358 | 0.028718519 |
| ENSG00000126246 | *IGFLR1* | 1.322443301 | -0.137328229 | -0.239556649 | 0.102228421 | 0.999933887 | -0.379627241 | -0.826703857 | 0.447076616 | 0.02875246 |
| ENSG00000135525 | *MAP7* | 0.807114741 | -0.099615267 | -0.093235051 | -0.006380217 | 0.999933887 | -0.500134181 | -1.243541429 | 0.743407247 | 0.02875246 |
| ENSG00000251022 | *THAP9-AS1* | 6.430508475 | 0.075661395 | 0.00098762 | 0.074673776 | 0.999933887 | 0.666483766 | 1.165797116 | -0.49931335 | 0.02875246 |
| ENSG00000148841 | *ITPRIP* | 8.966548808 | 0.096305839 | 0.071171126 | 0.025134714 | 0.999933887 | 0.762385575 | 0.983205361 | -0.220819786 | 0.028765847 |
| ENSG00000123545 | *NDUFAF4* | 0.407645214 | -0.114470336 | -0.153485062 | 0.039014726 | 0.999933887 | 0.122645475 | 0.778750452 | -0.656104977 | 0.028772679 |
| ENSG00000265982 | *AC103810.3* | 2.143310667 | -0.039830038 | -0.070380988 | 0.03055095 | 0.999933887 | -0.273220889 | -0.716563744 | 0.443342855 | 0.028772679 |
| ENSG00000247400 | *DNAJC3-DT* | 4.816031175 | 0.02726392 | -0.050817099 | 0.078081019 | 0.999933887 | 1.467904279 | 1.825182538 | -0.357278258 | 0.028776839 |
| ENSG00000151876 | *FBXO4* | 0.212189291 | -0.303611683 | -0.104915471 | -0.198696211 | 0.999933887 | -0.122787779 | 0.518851501 | -0.64163928 | 0.028840794 |
| ENSG00000105246 | *EBI3* | -1.74598186 | 0.301893463 | -0.068250629 | 0.370144092 | 0.999933887 | 4.701810297 | 6.261026685 | -1.559216388 | 0.028840794 |
| ENSG00000017797 | *RALBP1* | 6.356855476 | -0.000493404 | 0.012410464 | -0.012903867 | 0.999933887 | -0.812255612 | -1.172699427 | 0.360443815 | 0.028840794 |
| ENSG00000173905 | *GOLIM4* | 0.835105923 | 0.07346057 | -0.20003845 | 0.27349902 | 0.999933887 | -0.193257221 | 0.41198753 | -0.605244752 | 0.028923733 |
| ENSG00000276317 | *AL357033.3* | 2.221139989 | 0.062012133 | -0.037249821 | 0.099261954 | 0.999933887 | 0.435852168 | 0.811986432 | -0.376134264 | 0.028979131 |
| ENSG00000221926 | *TRIM16* | -0.448005131 | 0.137178699 | 0.020268362 | 0.116910337 | 0.999933887 | 0.590216499 | 1.249948749 | -0.659732249 | 0.029002709 |
| ENSG00000148154 | *UGCG* | 5.618767513 | -0.075993108 | 0.007474217 | -0.083467325 | 0.999933887 | -0.034195366 | 0.191698586 | -0.225893952 | 0.02904515 |
| ENSG00000151327 | *FAM177A1* | 5.803380246 | 0.090084916 | 0.056450104 | 0.033634813 | 0.999933887 | 0.583986037 | 0.830689317 | -0.24670328 | 0.029182524 |
| ENSG00000101773 | *RBBP8* | 2.209892558 | -0.05774038 | -0.158761993 | 0.101021614 | 0.999933887 | 0.308066307 | 0.768575795 | -0.460509488 | 0.029182524 |
| ENSG00000176170 | *SPHK1* | 6.75625957 | 0.037925572 | 0.11779101 | -0.079865437 | 0.999933887 | 1.800685557 | 2.182460614 | -0.381775057 | 0.029182524 |
| ENSG00000204574 | *ABCF1* | 6.851088902 | -0.002733976 | 0.004786921 | -0.007520897 | 0.999933887 | 0.589862089 | 0.807216923 | -0.217354835 | 0.029182524 |
| ENSG00000197249 | *SERPINA1* | 10.491007963 | 0.036663322 | -0.01320071 | 0.049864032 | 0.999933887 | 1.548245535 | 1.876318792 | -0.328073256 | 0.029258576 |
| ENSG00000168010 | *ATG16L2* | 9.118356401 | -0.010168396 | 0.06009889 | -0.070267285 | 0.999933887 | -0.667749244 | -0.942173032 | 0.274423788 | 0.02929866 |
| ENSG00000130038 | *CRACR2A* | 3.324695522 | 0.052579586 | 0.025619637 | 0.026959948 | 0.999933887 | 0.116585435 | -0.230685424 | 0.347270859 | 0.029375059 |
| ENSG00000197562 | *RAB40C* | 3.626183309 | 0.038360712 | 0.002539285 | 0.035821427 | 0.999933887 | -0.18926136 | -0.53945635 | 0.35019499 | 0.029392546 |
| ENSG00000106615 | *RHEB* | 6.581974685 | -0.003394579 | 0.030440336 | -0.033834914 | 0.999933887 | 0.111824145 | 0.291762063 | -0.179937919 | 0.029392546 |
| ENSG00000125772 | *GPCPD1* | 9.430215003 | -0.082600497 | -0.061042364 | -0.021558133 | 0.999933887 | -0.279509257 | 0.031249494 | -0.310758751 | 0.029397973 |
| ENSG00000092020 | *PPP2R3C* | 4.126277672 | -0.021405736 | -0.093431351 | 0.072025614 | 0.999933887 | -0.071162042 | 0.156418194 | -0.227580236 | 0.029397973 |
| ENSG00000091436 | *MAP3K20* | 2.469289441 | 0.037203941 | 0.030037119 | 0.007166822 | 0.999933887 | 0.898134845 | 1.43438978 | -0.536254936 | 0.029397973 |
| ENSG00000070159 | *PTPN3* | -3.252609989 | -0.226081784 | -0.060462637 | -0.165619147 | 0.999933887 | -0.874714948 | -0.0262616 | -0.848453348 | 0.029401057 |
| ENSG00000149308 | *NPAT* | 4.477799457 | -0.057919657 | -0.169817927 | 0.11189827 | 0.999933887 | -0.067039551 | 0.534800028 | -0.601839579 | 0.029401057 |
| ENSG00000182871 | *COL18A1* | 6.51769146 | 0.081875897 | 0.129682583 | -0.047806686 | 0.999933887 | -0.130906709 | -0.496900174 | 0.365993465 | 0.029551184 |
| ENSG00000134287 | *ARF3* | 5.811557704 | 0.010137517 | 0.00081584 | 0.009321677 | 0.999933887 | -0.243067545 | -0.466805499 | 0.223737954 | 0.029551184 |
| ENSG00000142186 | *SCYL1* | 6.407137758 | 0.011207648 | 0.048934971 | -0.037727323 | 0.999933887 | -0.430725319 | -0.665578534 | 0.234853216 | 0.029551184 |
| ENSG00000269246 | *AC011445.1* | -0.178264623 | 0.0071602 | -0.694551223 | 0.701711422 | 0.594252256 | -0.553123744 | -1.444404263 | 0.891280519 | 0.029551184 |
| ENSG00000266389 | *AC002091.2* | 4.476780084 | 0.080422154 | 0.002702195 | 0.07771996 | 0.999933887 | 1.158755984 | 1.472927459 | -0.314171474 | 0.029564592 |
| ENSG00000077157 | *PPP1R12B* | 5.824619487 | -0.066193398 | -0.04551162 | -0.020681778 | 0.999933887 | -1.381305562 | -1.814788695 | 0.433483133 | 0.029575298 |
| ENSG00000276953 | *TRBV12-4* | -0.149253739 | -0.017828485 | 0.059664381 | -0.077492865 | 0.999933887 | 0.130304524 | -0.4568163 | 0.587120824 | 0.029601018 |
| ENSG00000133104 | *SPART* | 6.195975559 | -0.080226997 | -0.145966327 | 0.06573933 | 0.999933887 | -0.057823996 | 0.190310638 | -0.248134633 | 0.029635691 |
| ENSG00000111670 | *GNPTAB* | 4.368380081 | -0.035893417 | -0.090560055 | 0.054666638 | 0.999933887 | 0.069703811 | 0.393384332 | -0.323680521 | 0.029635691 |
| ENSG00000154813 | *DPH3* | 4.449071074 | 0.012722233 | -0.052572351 | 0.065294584 | 0.999933887 | 0.048559684 | 0.276748098 | -0.228188414 | 0.029635691 |
| ENSG00000145287 | *PLAC8* | 3.630684071 | -0.022136938 | 0.043792665 | -0.065929603 | 0.999933887 | 0.528729085 | 1.136379666 | -0.607650581 | 0.029688437 |
| ENSG00000141527 | *CARD14* | 2.982068116 | 0.005844668 | 0.184450045 | -0.178605377 | 0.999933887 | 0.131466918 | 0.413491802 | -0.282024884 | 0.029724969 |
| ENSG00000175727 | *MLXIP* | 5.404973492 | -0.018095759 | -0.042370617 | 0.024274858 | 0.999933887 | -0.133270035 | -0.355548616 | 0.222278581 | 0.029798453 |
| ENSG00000213676 | *ATF6B* | 6.046270069 | 0.026934465 | 0.055763758 | -0.028829293 | 0.999933887 | -0.299661885 | -0.491454895 | 0.19179301 | 0.029831133 |
| ENSG00000226945 | *AC098935.1* | -0.811114854 | 0.04535464 | 0.364131553 | -0.318776913 | 0.999933887 | -0.307645279 | -1.21920951 | 0.91156423 | 0.029831133 |
| ENSG00000184588 | *PDE4B* | 7.627776399 | 0.088417281 | 0.035841002 | 0.052576279 | 0.999933887 | 1.377118239 | 1.687446562 | -0.310328323 | 0.029837734 |
| ENSG00000100241 | *SBF1* | 7.207655609 | 0.050587285 | 0.098301886 | -0.047714601 | 0.999933887 | -0.088471547 | -0.299379203 | 0.210907656 | 0.029837734 |
| ENSG00000253276 | *CCDC71L* | 6.93111786 | 0.139728299 | 0.115192033 | 0.024536266 | 0.999933887 | 0.423775035 | 0.738480908 | -0.314705873 | 0.029994791 |
| ENSG00000100749 | *VRK1* | 1.86790528 | -0.140481854 | -0.155012976 | 0.014531122 | 0.999933887 | 0.077558959 | 0.486441878 | -0.408882919 | 0.030131048 |
| ENSG00000136997 | *MYC* | 3.066514224 | 0.061635595 | -0.010331072 | 0.071966667 | 0.999933887 | 0.38478685 | 0.799750985 | -0.414964135 | 0.030131048 |
| ENSG00000143624 | *INTS3* | 5.138486273 | -0.03022232 | 0.002861462 | -0.033084782 | 0.999933887 | -0.22560212 | -0.468376462 | 0.242774343 | 0.030131048 |
| ENSG00000188404 | *SELL* | 9.535069683 | -0.026502114 | -0.046908192 | 0.020406078 | 0.999933887 | -0.237825055 | -0.551318987 | 0.313493932 | 0.030174511 |
| ENSG00000123609 | *NMI* | 6.347427027 | -0.068132668 | -0.095599105 | 0.027466437 | 0.999933887 | 0.440616639 | 0.67899953 | -0.238382891 | 0.030186564 |
| ENSG00000162881 | *OXER1* | 3.070812076 | -0.181535865 | -0.06548601 | -0.116050825 | 0.999933887 | -0.861488752 | -1.432416763 | 0.570928011 | 0.030189651 |
| ENSG00000127054 | *INTS11* | 5.398276242 | 0.051869323 | 0.067333203 | -0.01546388 | 0.999933887 | -0.061292297 | -0.28378137 | 0.222489073 | 0.03024247 |
| ENSG00000100902 | *PSMA6* | 2.695934081 | -0.045047747 | -0.14949503 | 0.104447283 | 0.999933887 | 1.099580844 | 1.567779088 | -0.468198244 | 0.03024247 |
| ENSG00000129993 | *CBFA2T3* | 4.638933919 | 0.00426477 | 0.071959525 | -0.067694754 | 0.999933887 | -0.601249183 | -0.976873605 | 0.375624422 | 0.03024247 |
| ENSG00000132912 | *DCTN4* | 5.644158704 | -0.04662814 | -0.083676408 | 0.037048268 | 0.999933887 | 0.236247978 | 0.452947497 | -0.216699519 | 0.030249847 |
| ENSG00000132003 | *ZSWIM4* | 3.652604341 | 0.045345148 | 0.050436892 | -0.005091744 | 0.999933887 | 1.347839812 | 1.693327065 | -0.345487253 | 0.030262795 |
| ENSG00000138386 | *NAB1* | 5.220743104 | -0.04882057 | -0.023211196 | -0.025609374 | 0.999933887 | 0.003470625 | 0.238141591 | -0.234670965 | 0.030273009 |
| ENSG00000090615 | *GOLGA3* | 5.03234074 | -0.008448793 | 0.037997909 | -0.046446702 | 0.999933887 | -0.13111075 | -0.330263048 | 0.199152298 | 0.030324857 |
| ENSG00000173928 | *SWSAP1* | 0.380528576 | -0.029835021 | 0.046122941 | -0.075957962 | 0.999933887 | -0.185752882 | -0.962765475 | 0.777012593 | 0.030367456 |
| ENSG00000197321 | *SVIL* | 8.20503359 | 0.01862287 | 0.007849457 | 0.010773413 | 0.999933887 | 0.287176911 | 0.450437436 | -0.163260525 | 0.030377514 |
| ENSG00000188315 | *C3orf62* | 5.002783116 | -0.045950064 | -0.028993483 | -0.016956582 | 0.999933887 | -0.354579222 | -0.539853473 | 0.185274251 | 0.030413302 |
| ENSG00000126934 | *MAP2K2* | 6.406683396 | 0.044918935 | 0.006352133 | 0.038566803 | 0.999933887 | -0.281771037 | -0.516884937 | 0.2351139 | 0.030413302 |
| ENSG00000131626 | *PPFIA1* | 5.784713007 | 0.000470384 | 0.009807997 | -0.009337613 | 0.999933887 | 0.138639082 | 0.347934825 | -0.209295742 | 0.030420229 |
| ENSG00000189164 | *ZNF527* | 0.027149899 | -0.195944944 | -0.212663552 | 0.016718608 | 0.999933887 | -0.218234077 | 0.503902691 | -0.722136767 | 0.03045706 |
| ENSG00000178860 | *MSC* | 3.355388297 | 0.028019461 | 0.009911545 | 0.018107916 | 0.999933887 | 1.312478375 | 1.71803912 | -0.405560745 | 0.030466237 |
| ENSG00000163823 | *CCR1* | 9.887855017 | 0.116845512 | 0.135949091 | -0.019103579 | 0.999933887 | 0.200112948 | 0.405687156 | -0.205574208 | 0.030561833 |
| ENSG00000117174 | *ZNHIT6* | 1.435039469 | -0.05986946 | -0.209476509 | 0.149607049 | 0.999933887 | -0.084431053 | 0.386950219 | -0.471381272 | 0.030561833 |
| ENSG00000165288 | *BRWD3* | 6.071382739 | -0.056201544 | -0.163365213 | 0.107163668 | 0.999933887 | -0.224603342 | 0.286555526 | -0.511158868 | 0.030572177 |
| ENSG00000272333 | *KMT2B* | 7.266641007 | 0.014651933 | 0.05511462 | -0.040462688 | 0.999933887 | -0.429195755 | -0.639089096 | 0.209893341 | 0.030572177 |
| ENSG00000068097 | *HEATR6* | 2.78811974 | -0.032165173 | -0.02765741 | -0.004507763 | 0.999933887 | -0.400587076 | -0.706329957 | 0.305742881 | 0.030640215 |

| | | | | | | | | | | |
|---|---|---|---|---|---|---|---|---|---|---|
| ENSG00000143486 | *EIF2D* | 5.125092421 | 0.10296422 | 0.073975873 | 0.028988347 | 0.999933887 | 0.522330802 | 0.768106375 | -0.245775573 | 0.030661016 |
| ENSG00000113597 | *TRAPPC13* | 3.158021068 | -0.216399067 | -0.171866865 | -0.044532203 | 0.999933887 | -0.078333601 | 0.396424622 | -0.474758223 | 0.030674304 |
| ENSG00000180113 | *TDRD6* | -0.951157947 | -0.062327903 | -0.328412333 | 0.26608443 | 0.999933887 | 0.269340226 | 1.077118459 | -0.807778234 | 0.030674304 |
| ENSG00000064490 | *RFXANK* | 3.648508049 | -0.002236937 | -0.058191314 | 0.055954377 | 0.999933887 | -0.04459746 | -0.346583776 | 0.301986316 | 0.030717464 |
| ENSG00000158104 | *HPD* | 1.880739506 | 0.057423179 | 0.032056591 | 0.025366588 | 0.999933887 | 0.776056846 | 1.223339651 | -0.447282804 | 0.030756729 |
| ENSG00000164463 | *CREBRF* | 8.284095001 | -0.100230442 | -0.165004254 | 0.064773812 | 0.999933887 | -0.378218187 | -0.044683075 | -0.333535112 | 0.030875324 |
| ENSG00000126353 | *CCR7* | 4.728193281 | 0.061610991 | -0.020233436 | 0.081844427 | 0.999933887 | 0.56544941 | 0.854919357 | -0.289469947 | 0.030875406 |
| ENSG00000265206 | *AC004687.1* | 4.43902005 | 0.120426656 | 0.175505343 | -0.055078687 | 0.999933887 | -0.539431129 | -1.157966436 | 0.618535307 | 0.030875406 |
| ENSG00000116161 | *CACYBP* | 3.591687865 | -0.032662175 | -0.079585947 | 0.046923771 | 0.999933887 | -0.161750035 | -0.449492074 | 0.287742039 | 0.030875406 |
| ENSG00000260404 | *AC110079.1* | 4.037165187 | 0.022747009 | 0.010467008 | 0.012280001 | 0.999933887 | 0.205612093 | 0.4931569 | -0.287544807 | 0.030875406 |
| ENSG00000100416 | *TRMU* | 3.525792501 | 0.013391232 | 0.051018317 | -0.037627086 | 0.999933887 | 0.029476253 | -0.261710638 | 0.291186891 | 0.030875406 |
| ENSG00000023287 | *RB1CC1* | 7.404155789 | -0.068859107 | -0.12460343 | 0.055744323 | 0.999933887 | -0.23895776 | 0.340688184 | -0.579645944 | 0.030970209 |
| ENSG00000221978 | *CCNL2* | 6.739280948 | -0.000862874 | 0.04715874 | -0.048021614 | 0.999933887 | -0.176847378 | -0.342706621 | 0.165859242 | 0.031038883 |
| ENSG00000068976 | *PYGM* | 2.971480952 | 0.087189577 | 0.071150922 | 0.016038655 | 0.999933887 | -0.254880934 | -0.605500097 | 0.350619163 | 0.031071888 |
| ENSG00000173812 | *EIF1* | 10.560953674 | 0.163682669 | 0.149047151 | 0.014635518 | 0.999933887 | 0.86208882 | 1.128233456 | -0.266144636 | 0.031091103 |
| ENSG00000110002 | *VWA5A* | 2.782872511 | 0.000537552 | -0.103613162 | 0.104150715 | 0.999933887 | 0.435053775 | 0.770811872 | -0.335758097 | 0.031091934 |
| ENSG00000137996 | *RTCA* | 3.524186051 | 0.010809353 | -0.073147185 | 0.083956539 | 0.999933887 | -0.391354878 | -0.691003084 | 0.299648206 | 0.031143373 |
| ENSG00000051620 | *HEBP2* | 4.738362949 | 0.036307674 | -0.015966665 | 0.052274339 | 0.999933887 | -0.222960836 | -0.492732819 | 0.269771983 | 0.031157431 |
| ENSG00000102699 | *PARP4* | 6.344026168 | -0.090863095 | -0.108051751 | 0.017188656 | 0.999933887 | -0.753251001 | -1.025124539 | 0.271873538 | 0.031199273 |
| ENSG00000197182 | *MIRLET7BHG* | 0.675226229 | 0.126010172 | -0.062958627 | 0.188968799 | 0.999933887 | -0.549343585 | -1.296481909 | 0.747138323 | 0.031204046 |
| ENSG00000111581 | *NUP107* | 2.670833011 | -0.04484749 | -0.172126003 | 0.127278513 | 0.999933887 | 0.058208546 | 0.444786009 | -0.386577463 | 0.031214639 |
| ENSG00000183049 | *CAMK1D* | 6.495419366 | -0.036428086 | 0.024764079 | -0.061192165 | 0.999933887 | -0.107108287 | -0.303018094 | 0.195909807 | 0.031253072 |
| ENSG00000158825 | *CDA* | 5.845519777 | 0.058583924 | -0.022348429 | 0.080932352 | 0.999933887 | -0.411154587 | -0.701087279 | 0.289932692 | 0.031440161 |
| ENSG00000137200 | *CMTR1* | 5.649555729 | -0.007925756 | 0.071506616 | -0.079432372 | 0.999933887 | -0.080518023 | -0.30512602 | 0.224607996 | 0.031586254 |
| ENSG00000182810 | *DDX28* | 3.15567132 | -0.327471802 | -0.412901535 | 0.085429733 | 0.999933887 | -1.592551503 | -2.31114578 | 0.718594277 | 0.031615872 |
| ENSG00000279741 | *AC007342.8* | -0.261931872 | -0.208995225 | -0.032124115 | -0.17687111 | 0.999933887 | -0.949662134 | -2.133972132 | 1.184309998 | 0.031651392 |
| ENSG00000122643 | *NT5C3A* | 4.043036051 | 0.027639687 | -0.018872736 | 0.046512422 | 0.999933887 | -0.315047657 | -0.622061424 | 0.307013766 | 0.031689765 |
| ENSG00000139572 | *GPR84* | 5.074439977 | 0.353099694 | 0.46972728 | -0.116627587 | 0.999933887 | 2.721309665 | 3.137253606 | -0.415943941 | 0.031706079 |
| ENSG00000142546 | *NOSIP* | 5.149252803 | 0.062804639 | 0.003325159 | 0.059479479 | 0.999933887 | -0.110214957 | -0.363184067 | 0.25296911 | 0.031753288 |
| ENSG00000270231 | *NBPF8* | 3.205712594 | -0.120043311 | -0.158822818 | 0.038779507 | 0.999933887 | 0.228881887 | 0.576100562 | -0.347218675 | 0.031774918 |
| ENSG00000136021 | *SCYL2* | 5.940746255 | -0.020927549 | -0.067878976 | 0.046951427 | 0.999933887 | -0.003997967 | 0.396921447 | -0.400919413 | 0.031774918 |
| ENSG00000196917 | *HCAR1* | 5.471291761 | 0.12171565 | 0.119124437 | 0.002591213 | 0.999933887 | 1.898639183 | 2.388828955 | -0.490189771 | 0.031814115 |
| ENSG00000052749 | *RRP12* | 7.667383879 | 0.011831932 | 0.039310446 | -0.027478514 | 0.999933887 | 0.114536402 | -0.096999275 | 0.211535677 | 0.031814115 |
| ENSG00000127527 | *EPS15L1* | 7.788438959 | -0.007028247 | 0.016542213 | -0.02357046 | 0.999933887 | 0.152030242 | 0.324624316 | -0.172594075 | 0.031814115 |
| ENSG00000196352 | *CD55* | 9.525884635 | 0.090040482 | 0.047575783 | 0.042464699 | 0.999933887 | 1.222835069 | 1.556775375 | -0.333940306 | 0.031842452 |
| ENSG00000117616 | *RSRP1* | 7.134819469 | -0.019460637 | 0.04944082 | -0.068901457 | 0.999933887 | 0.08340127 | -0.103622188 | 0.187023458 | 0.031868235 |
| ENSG00000172602 | *RND1* | -0.720960056 | -0.05473168 | -0.086013215 | 0.031281536 | 0.999933887 | 1.338412735 | 2.507995608 | -1.169582873 | 0.031870719 |
| ENSG00000166439 | *RNF169* | 5.298875956 | -0.042162694 | -0.038978111 | -0.003184582 | 0.999933887 | -0.921481219 | -1.216154283 | 0.294673064 | 0.031890046 |
| ENSG00000171703 | *TCEA2* | 2.483083681 | 0.100829968 | 0.007469584 | 0.093360384 | 0.999933887 | -0.237913719 | -0.620111147 | 0.382197428 | 0.031899961 |
| ENSG00000163510 | *CWC22* | 4.446005931 | -0.098252348 | -0.187918676 | 0.089666328 | 0.999933887 | -0.234931996 | 0.243265804 | -0.4781978 | 0.031922949 |
| ENSG00000187650 | *VMAC* | 1.636787854 | -0.070969114 | -0.037020537 | -0.033948576 | 0.999933887 | 0.049434905 | -0.435003347 | 0.484438251 | 0.031922949 |
| ENSG00000257093 | *DENND11* | 4.382207543 | -0.003812484 | -0.022270113 | 0.018457629 | 0.999933887 | 0.022601894 | -0.182899578 | 0.205501472 | 0.031958267 |
| ENSG00000105568 | *PPP2R1A* | 5.543344637 | -0.020066416 | 0.031055605 | -0.051122021 | 0.999933887 | -0.218771884 | -0.440261081 | 0.221489197 | 0.032132022 |
| ENSG00000129235 | *TXNDC17* | 2.610314683 | -0.031772634 | 0.043728313 | -0.075500948 | 0.999933887 | -0.579734628 | -0.954316034 | 0.374581406 | 0.032235007 |
| ENSG00000258860 | *AL133163.2* | -1.260133867 | 0.389372033 | 0.289717709 | 0.099654324 | 0.999933887 | 2.784121999 | 3.725865026 | -0.941743027 | 0.032292883 |
| ENSG00000141522 | *ARHGDIA* | 8.407799192 | 0.103066505 | 0.041205167 | 0.061861337 | 0.999933887 | 0.544344274 | 0.783712167 | -0.239367893 | 0.032294642 |
| ENSG00000124795 | *DEK* | 5.779506005 | -0.055497458 | -0.146996148 | 0.09149869 | 0.999933887 | -0.191547385 | 0.221420496 | -0.412967881 | 0.032294642 |
| ENSG00000169180 | *XPO6* | 10.244333267 | 0.005452308 | 0.020243984 | -0.014791677 | 0.999933887 | -0.770925672 | -0.99514492 | 0.224219247 | 0.032379945 |
| ENSG00000176715 | *ACSF3* | 3.936286557 | 0.058841824 | -0.017182052 | 0.076023876 | 0.999933887 | -0.194806724 | -0.479468604 | 0.28466188 | 0.032399414 |
| ENSG00000198886 | *MT-ND4* | 9.598860048 | 0.040284852 | -0.071142358 | 0.111427209 | 0.999933887 | -0.094365939 | 0.286135772 | -0.380501711 | 0.032424679 |
| ENSG00000115008 | *IL1A* | 4.112924074 | 0.537032283 | 0.432105003 | 0.10492728 | 0.999933887 | 3.002128554 | 4.011219782 | -1.009091228 | 0.03245272 |
| ENSG00000234484 | *AL032821.1* | 0.696247802 | -0.343246624 | -0.061391628 | -0.281854996 | 0.999933887 | 0.049330925 | 0.810385937 | -0.761055012 | 0.032521781 |
| ENSG00000062716 | *VMP1* | 8.492880863 | 0.014124036 | -0.016068283 | 0.03019232 | 0.999933887 | -0.167925934 | -0.467285809 | 0.299359875 | 0.032543379 |
| ENSG00000184634 | *MED12* | 7.353043176 | 0.026411981 | 0.047130675 | -0.020718695 | 0.999933887 | 0.476077344 | 0.670108672 | -0.194031327 | 0.032563459 |
| ENSG00000225830 | *ERCC6* | 3.715701941 | -0.084142813 | -0.107300281 | 0.023157468 | 0.999933887 | -0.438211507 | -0.690101586 | 0.251890079 | 0.032579325 |
| ENSG00000115556 | *PLCD4* | 0.892268742 | -0.049759841 | -0.096199646 | 0.046439805 | 0.999933887 | 0.396276966 | 0.945523415 | -0.54924645 | 0.032600032 |
| ENSG00000179456 | *ZBTB18* | 6.223004576 | -0.152154276 | -0.094001906 | -0.058152371 | 0.999933887 | -0.848993684 | -1.16332353 | 0.314329846 | 0.032650145 |
| ENSG00000099381 | *SETD1A* | 4.55505046 | -0.012512798 | 0.02853925 | -0.041052048 | 0.999933887 | -0.254744506 | -0.457453163 | 0.202708656 | 0.032668259 |
| ENSG00000158050 | *DUSP2* | 7.121739564 | 0.356820646 | 0.474767661 | -0.117947015 | 0.999933887 | 1.059084881 | 1.628936835 | -0.569851954 | 0.032705545 |
| ENSG00000119414 | *PPP6C* | 5.63716158 | -0.049590635 | -0.04284541 | -0.006745225 | 0.999933887 | -0.11901644 | 0.056684873 | -0.175701314 | 0.032711833 |
| ENSG00000104325 | *DECR1* | 4.705664546 | -0.05541657 | -0.074814458 | 0.019397888 | 0.999933887 | -0.218317898 | -0.43392291 | 0.215605012 | 0.03278014 |
| ENSG00000115548 | *KDM3A* | 7.353734537 | -0.029364699 | -0.047568495 | 0.018203796 | 0.999933887 | -0.824220284 | -1.08170249 | 0.257482206 | 0.03278014 |
| ENSG00000103126 | *AXIN1* | 5.682043402 | 0.021190607 | 0.105824695 | -0.084634087 | 0.999933887 | -0.411480538 | -0.688987814 | 0.277507275 | 0.032781629 |
| ENSG00000115839 | *RAB3GAP1* | 4.509472726 | -0.109668939 | -0.148926654 | 0.039257715 | 0.999933887 | -0.034963939 | 0.167377077 | -0.202341016 | 0.032791564 |
| ENSG00000126814 | *TRMT5* | 2.573919286 | -0.006153883 | -0.215041568 | 0.208887684 | 0.999933887 | 0.023098533 | 0.3332367 | -0.310138168 | 0.032791564 |
| ENSG00000279716 | *AC006128.1* | 0.55966482 | -0.172170397 | -0.187511661 | 0.015341264 | 0.999933887 | -0.131025913 | -0.618148239 | 0.487122326 | 0.032920806 |
| ENSG00000005483 | *KMT2E* | 7.784650824 | -0.016152491 | -0.080589709 | 0.064437218 | 0.999933887 | -0.107479115 | 0.313810347 | -0.421289462 | 0.032920806 |
| ENSG00000145907 | *G3BP1* | 5.206032487 | -0.084883817 | -0.049574079 | -0.035309738 | 0.999933887 | 0.211245804 | 0.449328012 | -0.238082208 | 0.032921696 |
| ENSG00000104859 | *CLASRP* | 6.272203908 | 0.056697649 | 0.069195389 | -0.01249774 | 0.999933887 | -0.017802431 | -0.266337189 | 0.248534758 | 0.033183932 |
| ENSG00000116852 | *KIF21B* | 6.983516068 | 0.040140616 | 0.024676104 | 0.015464512 | 0.999933887 | -0.075506137 | -0.280133955 | 0.204627818 | 0.033223384 |
| ENSG00000265298 | *AC132812.1* | 2.283757366 | 0.06709234 | 0.049026855 | 0.018065485 | 0.999933887 | -0.068573999 | -0.437998504 | 0.369424506 | 0.033337589 |
| ENSG00000170525 | *PFKFB3* | 10.320894231 | 0.079238496 | 0.069699578 | 0.009538918 | 0.999933887 | 0.280513497 | 0.580569397 | -0.3000559 | 0.033650337 |
| ENSG00000164164 | *OTUD4* | 4.699515268 | -0.01810288 | -0.13790587 | 0.11980299 | 0.999933887 | 0.14363249 | 0.563580262 | -0.419947771 | 0.033650337 |
| ENSG00000283050 | *GTF2IP12* | 2.120218574 | 0.122512026 | -0.040241754 | 0.162753779 | 0.999933887 | 0.337817375 | 0.852839335 | -0.515021959 | 0.03368597 |
| ENSG00000186496 | *ZNF396* | 0.718600898 | 0.003424146 | -0.211543047 | 0.214967193 | 0.999933887 | 0.978419485 | 1.397509507 | -0.419090022 | 0.03375094 |
| ENSG00000169379 | *ARL13B* | 1.535103426 | -0.080747459 | -0.060937594 | -0.019809864 | 0.999933887 | 0.460998631 | 0.940870161 | -0.47987153 | 0.03377842 |
| ENSG00000117000 | *RLF* | 6.773840677 | -0.092585283 | -0.171904625 | 0.079319342 | 0.999933887 | 0.015431915 | 0.470239387 | -0.454807472 | 0.033812514 |
| ENSG00000204843 | *DCTN1* | 5.304422027 | -0.005661576 | 0.044225917 | -0.049887492 | 0.999933887 | -0.294241772 | -0.571410843 | 0.27716907 | 0.033823386 |
| ENSG00000141380 | *SS18* | 3.659771253 | -0.061829748 | 0.034058362 | -0.095888109 | 0.999933887 | 0.024596371 | 0.264426669 | -0.239830297 | 0.033829313 |
| ENSG00000177951 | *BET1L* | 4.613604417 | -0.047387222 | 0.000344978 | -0.0477322 | 0.999933887 | -0.379055823 | -0.628836822 | 0.249780998 | 0.033864645 |
| ENSG00000231881 | *AL109615.2* | -0.718744561 | 0.356431339 | 0.479236488 | -0.122805149 | 0.999933887 | 1.457342726 | 2.405896081 | -0.948553355 | 0.033876274 |
| ENSG00000183401 | *CCDC159* | 3.963065832 | 0.04498864 | -0.034121733 | 0.079110374 | 0.999933887 | 0.446259191 | 0.692444992 | -0.246185801 | 0.033943928 |
| ENSG00000186222 | *BLOC1S4* | 2.563751902 | 0.046272157 | 0.042268113 | 0.004004044 | 0.999933887 | -0.27501136 | -0.702998873 | 0.427987514 | 0.033993912 |
| ENSG00000163785 | *RYK* | 3.757428399 | -0.092157685 | 0.052109101 | -0.144266786 | 0.999933887 | -0.431535419 | -0.697841662 | 0.266306243 | 0.03410294 |
| ENSG00000161533 | *ACOX1* | 5.536474023 | -0.093248206 | -0.028797616 | -0.064450591 | 0.999933887 | -0.925636175 | -1.269990123 | 0.344353948 | 0.03410294 |
| ENSG00000185825 | *BCAP31* | 6.805333697 | -0.025051967 | -0.078659242 | 0.053607275 | 0.999933887 | -0.540617823 | -0.777063939 | 0.236446116 | 0.034121331 |
| ENSG00000172831 | *CES2* | 3.410937211 | 0.026244797 | 0.094952837 | -0.06870804 | 0.999933887 | -0.216607689 | -0.480394977 | 0.263787288 | 0.034167854 |
| ENSG00000168152 | *THAP9* | 3.11556583 | 0.037303917 | -0.199346689 | 0.236650605 | 0.999933887 | 0.818472363 | 1.420647525 | -0.602175162 | 0.034232015 |
| ENSG00000276600 | *RAB7B* | -1.338277705 | 0.22714424 | -0.283729729 | 0.510873969 | 0.999933887 | -1.97347026 | -3.121916597 | 1.148446336 | 0.034285671 |
| ENSG00000139218 | *SCAF11* | 7.614144916 | -0.065241804 | -0.06917817 | 0.003936367 | 0.999933887 | -0.665888666 | -0.151371608 | -0.514517058 | 0.03433521 |
| ENSG00000122862 | *SRGN* | 12.531072875 | 0.016650551 | -0.035627056 | 0.052277608 | 0.999933887 | 0.619955581 | 0.845272171 | -0.22531659 | 0.034362135 |
| ENSG00000164134 | *NAA15* | 3.305556749 | -0.002808893 | -0.134892528 | 0.132083636 | 0.999933887 | -0.037690135 | 0.339037735 | -0.37672787 | 0.034378872 |
| ENSG00000089351 | *GRAMD1A* | 7.963173787 | 0.094810986 | 0.119646003 | -0.024835017 | 0.999933887 | 2.713272322 | 3.167904002 | -0.45463168 | 0.034390081 |
| ENSG00000154265 | *ABCA5* | 3.728682701 | -0.172871752 | -0.059093517 | -0.113778235 | 0.999933887 | 0.221031046 | 0.842818016 | -0.62178697 | 0.034402516 |
| ENSG00000091164 | *TXNL1* | 5.330234733 | -0.039259066 | -0.007628648 | -0.031630418 | 0.999933887 | 0.057231768 | 0.239781699 | -0.182549931 | 0.034402516 |
| ENSG00000134571 | *MYBPC3* | 5.196233955 | 0.025833465 | 0.038962581 | -0.013129115 | 0.999933887 | 0.748319979 | 1.026578085 | -0.278258106 | 0.034446167 |
| ENSG00000131471 | *AOC3* | 2.939038824 | 0.063774986 | 0.030605585 | 0.0331694 | 0.999933887 | 1.269166022 | 0.618975502 | 0.65019052 | 0.034503966 |
| ENSG00000101336 | *HCK* | 8.739991495 | 0.083492994 | 0.042541808 | 0.040951186 | 0.999933887 | 1.334102286 | 1.62566326 | -0.291560975 | 0.03451867 |
| ENSG00000120694 | *HSPH1* | 4.19923435 | -0.071188027 | -0.145410315 | 0.074222288 | 0.999933887 | 0.068296555 | 0.515727572 | -0.447431017 | 0.034540634 |
| ENSG00000160703 | *NLRX1* | 3.201496224 | -0.041157476 | -0.068072757 | 0.026915281 | 0.999933887 | -0.414472557 | -0.803304453 | 0.388831895 | 0.034540634 |
| ENSG00000077044 | *DGKD* | 6.053586367 | 0.015700919 | 0.035389359 | -0.01968844 | 0.999933887 | -0.092609564 | -0.27443322 | 0.181824756 | 0.034540634 |
| ENSG00000197121 | *PGAP1* | 1.864337581 | -0.060931634 | -0.058581943 | -0.002349691 | 0.999933887 | 0.60829257 | 1.276565648 | -0.668273077 | 0.034540634 |
| ENSG00000165029 | *ABCA1* | 8.964618122 | -0.028826346 | -0.086289809 | 0.057463463 | 0.999933887 | 0.058624941 | 0.402301722 | -0.343676781 | 0.034540634 |
| ENSG00000267317 | *AC027307.2* | 0.891423977 | -0.037728309 | 0.069902752 | -0.107631061 | 0.999933887 | -0.179872025 | 0.278388702 | -0.458260727 | 0.034540634 |
| ENSG00000279672 | *AP006621.5* | -0.084844248 | 0.156512865 | 0.086192183 | 0.070320682 | 0.999933887 | -0.375462024 | -1.036845194 | 0.66138317 | 0.034540983 |

| | | | | | | | | | | |
|---|---|---|---|---|---|---|---|---|---|---|
| ENSG00000185420 | *SMYD3* | 0.477878309 | 0.035779436 | -0.057722036 | 0.093501472 | 0.999933887 | 0.235599299 | -0.499172626 | 0.734771925 | 0.034540983 |
| ENSG00000175482 | *POLD4* | 3.013585577 | 0.155009581 | 0.000511213 | 0.154498368 | 0.999933887 | 0.133661211 | -0.197964158 | 0.331625369 | 0.034559396 |
| ENSG00000138767 | *CNOT6L* | 6.054222846 | 0.048142846 | -0.031513635 | 0.079656481 | 0.999933887 | -0.436054445 | -0.772943971 | 0.336889526 | 0.034559396 |
| ENSG00000068724 | *TTC7A* | 4.181492799 | 0.091762807 | 0.029968056 | 0.06179475 | 0.999933887 | 0.462171696 | 0.724922122 | -0.262750427 | 0.034637154 |
| ENSG00000123143 | *PKN1* | 6.435825321 | 0.044488761 | 0.052645064 | -0.008156303 | 0.999933887 | -0.12661696 | -0.417004351 | 0.290387391 | 0.034637154 |
| ENSG00000174837 | *ADGRE1* | 6.026691148 | -0.014774505 | -0.003589188 | -0.011185317 | 0.999933887 | 1.346116092 | 1.785783328 | -0.439667236 | 0.034637154 |
| ENSG00000122547 | *EEPD1* | 5.792467514 | 0.003449602 | -0.021936231 | 0.025385833 | 0.999933887 | -0.309655773 | -0.091284854 | -0.21837092 | 0.034836165 |
| ENSG00000174132 | *FAM174A* | 3.530542798 | 0.12366471 | 0.144910919 | -0.021246209 | 0.999933887 | -0.933442756 | -1.371725408 | 0.438282652 | 0.034872204 |
| ENSG00000214872 | *SMTNL1* | 2.270307061 | -0.02850355 | -0.004194898 | -0.024308652 | 0.999933887 | -0.276190826 | -0.768276116 | 0.49208529 | 0.034931127 |
| ENSG00000256683 | *ZNF350* | 4.437753193 | -0.106549857 | -0.146722158 | 0.040172301 | 0.999933887 | -0.191468285 | 0.056552041 | -0.248020326 | 0.034962796 |
| ENSG00000133773 | *CCDC59* | 4.497016273 | -0.043168173 | -0.024759242 | -0.018408932 | 0.999933887 | 0.113044172 | 0.351820066 | -0.238775894 | 0.034962796 |
| ENSG00000162695 | *SLC30A7* | 3.945889345 | -0.075361611 | 0.018844274 | -0.094205886 | 0.999933887 | 0.431873255 | 0.791464411 | -0.359591156 | 0.034962796 |
| ENSG00000269556 | *TMEM185A* | 1.352592215 | -0.034706914 | 0.090487902 | -0.125194816 | 0.999933887 | -0.292723892 | -0.726549092 | 0.4338252 | 0.034962796 |
| ENSG00000136286 | *MYO1G* | 6.6781501 | 0.029811128 | 0.048349565 | -0.018538437 | 0.999933887 | 0.010008911 | -0.208898372 | 0.218907283 | 0.03502988 |
| ENSG00000108671 | *PSMD11* | 4.709674895 | -0.09816317 | -0.128349121 | 0.030185951 | 0.999933887 | 0.027365935 | 0.237892553 | -0.210526618 | 0.035039773 |
| ENSG00000271614 | *ATP2B1-AS1* | 8.017007494 | 0.271278348 | 0.111808059 | 0.15947029 | 0.999933887 | 1.649722673 | 2.115827284 | -0.46610461 | 0.035040639 |
| ENSG00000134086 | *VHL* | 6.142697693 | -0.059179605 | 0.01132719 | -0.070506795 | 0.999933887 | 1.098531806 | 1.392351407 | -0.293819601 | 0.035040639 |
| ENSG00000111269 | *CREBL2* | 3.599416038 | 0.047184659 | -0.13863836 | 0.185823019 | 0.999933887 | 0.198584233 | 0.502718124 | -0.304133891 | 0.035040639 |
| ENSG00000132612 | *VPS4A* | 3.69762588 | -0.037034851 | 0.056003207 | -0.093038059 | 0.999933887 | -0.084275621 | -0.352253013 | 0.267977391 | 0.035040639 |
| ENSG00000118707 | *TGIF2* | 5.280637076 | -0.015865584 | 0.014623008 | -0.030488593 | 0.999933887 | 1.031059356 | 1.306356335 | -0.27529698 | 0.035040639 |
| ENSG00000203993 | *ARRDC1-AS1* | 2.092810245 | 0.022294256 | 0.136361612 | -0.114067356 | 0.999933887 | -0.13425199 | -0.488870865 | 0.354618875 | 0.035040639 |
| ENSG00000030419 | *IKZF2* | 2.520092419 | -0.111976588 | -0.192373227 | 0.080396639 | 0.999933887 | -0.220495237 | 0.268445855 | -0.488941091 | 0.035204207 |
| ENSG00000160932 | *LY6E* | 5.971800443 | 0.064501432 | 0.004006975 | 0.060494457 | 0.999933887 | -0.189086245 | -0.556943462 | 0.367857217 | 0.035204207 |
| ENSG00000104856 | *RELB* | 7.532186948 | 0.037629261 | 0.014075023 | 0.023554238 | 0.999933887 | 0.945722556 | 1.17267544 | -0.226952883 | 0.035266601 |
| ENSG00000001167 | *NFYA* | 5.913879378 | 0.065310351 | -0.078970987 | 0.144281338 | 0.944094537 | 0.196279932 | 0.384912608 | -0.188632676 | 0.03534404 |
| ENSG00000136381 | *IREB2* | 5.257687957 | -0.056852794 | -0.084254932 | 0.027402137 | 0.999933887 | -0.306885286 | -0.041820142 | -0.265065144 | 0.03534404 |
| ENSG00000135677 | *GNS* | 7.936315518 | -0.042442186 | -0.019985209 | -0.022456977 | 0.999933887 | 0.070503485 | 0.280657235 | -0.210153751 | 0.03534404 |
| ENSG00000284690 | *CD300H* | 1.944277336 | -0.047599805 | -0.017067398 | -0.030532407 | 0.999933887 | -0.98094749 | -1.808529744 | 0.827582254 | 0.03534404 |
| ENSG00000213976 | *AC010615.1* | 0.285649835 | -0.22032318 | -0.222168732 | 0.001845553 | 0.999933887 | -0.69834931 | 0.207028189 | -0.905377499 | 0.035426851 |
| ENSG00000003400 | *CASP10* | 3.297098696 | -0.057325526 | 0.03504834 | -0.092373866 | 0.999933887 | -0.330042601 | -0.663929466 | 0.333886864 | 0.035426851 |
| ENSG00000285865 | *AC010285.3* | 2.26729959 | -0.009801275 | 0.182595943 | -0.192397218 | 0.999933887 | -0.019101319 | 0.38428057 | -0.403381889 | 0.035426851 |
| ENSG00000119139 | *TJP2* | -0.019803427 | 0.097834984 | -0.120801504 | 0.218636488 | 0.999933887 | 1.296947239 | 2.043029297 | -0.746082058 | 0.035528714 |
| ENSG00000133243 | *BTBD2* | 4.478071572 | 0.020887573 | 0.069286165 | -0.048398592 | 0.999933887 | -0.040058721 | -0.308441956 | 0.268383235 | 0.035528714 |
| ENSG00000134265 | *NAPG* | 4.309746782 | -0.054853765 | -0.072512149 | 0.017658384 | 0.999933887 | -0.691309802 | -1.001970832 | 0.31066103 | 0.035627819 |
| ENSG00000071889 | *FAM3A* | 3.601735092 | 0.018272186 | 0.069484334 | -0.051212148 | 0.999933887 | 0.119743769 | -0.129668508 | 0.249412277 | 0.035714739 |
| ENSG00000177683 | *THAP5* | 3.753969918 | 0.081750803 | -0.089778389 | 0.171529192 | 0.999933887 | -0.346486915 | 0.096842308 | -0.443329224 | 0.035730428 |
| ENSG00000140612 | *SEC11A* | 4.864168437 | -0.016919666 | -0.034753261 | 0.017833595 | 0.999933887 | -0.305723815 | -0.54953751 | 0.243813696 | 0.035808948 |
| ENSG00000111859 | *NEDD9* | 8.238145784 | 0.000857952 | 0.011489001 | -0.010631049 | 0.999933887 | 0.100388419 | 0.286211538 | -0.185823119 | 0.035880303 |
| ENSG00000254979 | *AP000781.2* | -1.148804937 | 0.28139041 | 0.387950834 | -0.106560424 | 0.999933887 | 1.05840113 | 1.857192495 | -0.798791366 | 0.035889278 |
| ENSG00000183307 | *TMEM121B* | 1.674257012 | 0.02159567 | -0.142906208 | 0.164501878 | 0.999933887 | -0.941508449 | -1.738557024 | 0.797048575 | 0.035889278 |
| ENSG00000113387 | *SUB1* | 6.633090459 | -0.032801491 | -0.024302802 | -0.008498689 | 0.999933887 | 0.186601636 | 0.382796276 | -0.19619464 | 0.035922918 |
| ENSG00000250274 | *AC034199.1* | 5.529156836 | 0.204361681 | 0.236763877 | -0.032402196 | 0.999933887 | 1.64185734 | 2.021444266 | -0.379586926 | 0.035996623 |
| ENSG00000179361 | *ARID3B* | 5.929178922 | 0.054177983 | 0.019986111 | 0.034191872 | 0.999933887 | -1.172068958 | -1.483901204 | 0.311832246 | 0.036021068 |
| ENSG00000145016 | *RUBCN* | 4.005101322 | 0.084715372 | 0.082029669 | 0.002685703 | 0.999933887 | -0.316068437 | -0.619552957 | 0.30348452 | 0.036028327 |
| ENSG00000172663 | *TMEM134* | 2.916224148 | 0.056838364 | -0.016105916 | 0.07294428 | 0.999933887 | -0.03281855 | -0.288132898 | 0.255314348 | 0.036028327 |
| ENSG00000115652 | *UXS1* | 3.174509607 | -0.033771729 | -0.123583631 | 0.089811902 | 0.999933887 | 0.363190923 | 0.691846286 | -0.328655363 | 0.036028327 |
| ENSG00000276980 | *AC008760.2* | 0.50598261 | -0.144570551 | 0.213278408 | -0.357848959 | 0.999933887 | 3.059205157 | 3.915064563 | -0.855859405 | 0.036078161 |
| ENSG00000266053 | *NDUFV2-AS1* | 2.860557894 | -0.095096892 | 0.015888762 | -0.110985655 | 0.999933887 | 0.662034539 | 0.986039439 | -0.3240049 | 0.036153348 |
| ENSG00000102962 | *CCL22* | 0.521486115 | -0.230904106 | -0.123525459 | -0.107378647 | 0.999933887 | 0.726268078 | 1.766411885 | -1.040143807 | 0.036153348 |
| ENSG00000166197 | *NOLC1* | 4.705767196 | 0.024608924 | -0.056283014 | 0.080891938 | 0.999933887 | 0.529167617 | 0.842695843 | -0.313528226 | 0.036153348 |
| ENSG00000169155 | *ZBTB43* | 5.679438911 | -0.058623939 | -0.048168772 | -0.010455167 | 0.999933887 | 0.274996469 | 0.501018509 | -0.22602204 | 0.036164762 |
| ENSG00000103168 | *TAF1C* | 4.586824292 | 0.083043367 | 0.057875238 | 0.025168129 | 0.999933887 | 0.5624122 | 0.989613973 | -0.427201773 | 0.036164762 |
| ENSG00000185670 | *ZBTB3* | 0.082383335 | -0.046094893 | -0.044493556 | -0.001601338 | 0.999933887 | -0.34654289 | -1.011851223 | 0.665308333 | 0.036164762 |
| ENSG00000152689 | *RASGRP3* | 1.680447094 | -0.052669597 | -0.042734105 | -0.009935492 | 0.999933887 | 0.334135012 | 0.76617972 | -0.432044709 | 0.036222941 |
| ENSG00000126266 | *FFAR1* | 0.172151599 | 0.04515907 | 0.256191935 | -0.211032865 | 0.999933887 | 0.407099685 | 0.889673025 | -0.48257334 | 0.03633022 |
| ENSG00000135090 | *TAOK3* | 5.936358036 | -0.002199146 | -0.028337809 | 0.026138663 | 0.999933887 | -0.470718084 | -0.694907128 | 0.224189045 | 0.03633022 |
| ENSG00000125871 | *MGME1* | 3.763804335 | -0.089433184 | -0.098920113 | 0.009486929 | 0.999933887 | -1.124019113 | -1.452533352 | 0.328514239 | 0.036380737 |
| ENSG00000135953 | *MFSD9* | 2.587313217 | 0.068627474 | 0.103057294 | -0.03442982 | 0.999933887 | -0.22034721 | -0.594181169 | 0.373833959 | 0.036389234 |
| ENSG00000126759 | *CFP* | 4.488146278 | 0.074533103 | 0.048286599 | 0.026246504 | 0.999933887 | 1.446841454 | 1.866351101 | -0.419509647 | 0.036389234 |
| ENSG00000174197 | *MGA* | 6.421327088 | -0.054877192 | -0.027579936 | -0.027297255 | 0.999933887 | -0.229358407 | 0.084761406 | -0.314119813 | 0.036389234 |
| ENSG00000173511 | *VEGFB* | 3.619746456 | -0.006003471 | -0.069466866 | 0.063463394 | 0.999933887 | -0.540024674 | -0.835290049 | 0.295265375 | 0.036389234 |
| ENSG00000144840 | *RABL3* | 2.488460914 | -0.053857976 | -0.025689467 | -0.028168509 | 0.999933887 | 0.215836649 | 0.573196891 | -0.357360242 | 0.036410497 |
| ENSG00000231925 | *TAPBP* | 8.897363357 | 0.029527643 | 0.011752425 | 0.017775218 | 0.999933887 | 0.686997601 | 0.887085924 | -0.200088324 | 0.036410497 |
| ENSG00000128513 | *POT1* | 1.89655697 | -0.030783386 | -0.249492271 | 0.218708885 | 0.999933887 | -0.070708989 | 0.462083685 | -0.532792674 | 0.036477901 |
| ENSG00000160785 | *SLC25A44* | 5.641907676 | -0.061547147 | -0.009324 | -0.052223147 | 0.999933887 | -0.470715149 | -0.743813038 | 0.273097888 | 0.036628458 |
| ENSG00000127957 | *PMS2P3* | 0.977790349 | 0.004147857 | 0.165128816 | -0.160980959 | 0.999933887 | -0.061103414 | -0.494325185 | 0.433221771 | 0.036632372 |
| ENSG00000126705 | *AHDC1* | 3.264292264 | 0.02905989 | -0.060801364 | 0.089861255 | 0.999933887 | 0.28565415 | 0.589093628 | -0.303439479 | 0.036694605 |
| ENSG00000136630 | *HLX* | 3.406635524 | -0.07872767 | -0.025147541 | -0.053580129 | 0.999933887 | -0.409245426 | -0.840642713 | 0.431397286 | 0.036873835 |
| ENSG00000110048 | *OSBP* | 5.742851804 | -0.000743092 | -0.058363664 | 0.057620572 | 0.999933887 | 0.223695706 | 0.404522254 | -0.180826548 | 0.03690754 |
| ENSG00000198431 | *TXNRD1* | 8.74840389 | -0.045867875 | -0.040116862 | -0.005751013 | 0.999933887 | -0.032461986 | 0.210828298 | -0.243290284 | 0.036916268 |
| ENSG00000167645 | *YIF1B* | 2.81328713 | 0.085374992 | 0.063660788 | 0.021714203 | 0.999933887 | -0.007212787 | -0.338898987 | 0.331686201 | 0.036980533 |
| ENSG00000197150 | *ABCB8* | 3.118203869 | 0.019737541 | 0.064351855 | -0.044614314 | 0.999933887 | 0.219133144 | 0.494513059 | -0.275379915 | 0.036980533 |
| ENSG00000130254 | *SAFB2* | 6.619125599 | 0.01148201 | 0.062609902 | -0.051127893 | 0.999933887 | -0.298713028 | -0.518879644 | 0.220166616 | 0.036983222 |
| ENSG00000162302 | *RPS6KA4* | 4.117005598 | 0.110373939 | 0.079230294 | 0.031143645 | 0.999933887 | -0.059731316 | -0.381497837 | 0.321766522 | 0.037149672 |
| ENSG00000123684 | *LPGAT1* | 6.082704723 | -0.038910041 | -0.032265897 | -0.006644144 | 0.999933887 | -0.442041553 | -0.675698936 | 0.233657383 | 0.037164507 |
| ENSG00000078401 | *EDN1* | 6.663003544 | 0.382842799 | 0.335520828 | 0.047321972 | 0.999933887 | 1.167521557 | 1.735906169 | -0.568384612 | 0.03723299 |
| ENSG00000137817 | *PARP6* | 5.194070126 | 0.004952859 | -0.001800615 | 0.006753474 | 0.999933887 | 0.547521367 | 0.771977225 | -0.224455858 | 0.037246429 |
| ENSG00000137601 | *NEK1* | 2.070152974 | -0.020538074 | 0.069118967 | -0.089657041 | 0.999933887 | -0.136056341 | 0.337958091 | -0.474014432 | 0.037346794 |
| ENSG00000145740 | *SLC30A5* | 3.544827299 | -0.070862044 | -0.000474833 | -0.070387211 | 0.999933887 | 0.361722207 | 0.619078802 | -0.257356595 | 0.037351418 |
| ENSG00000081041 | *CXCL2* | 8.448447411 | 0.70231544 | 0.853543872 | -0.151228432 | 0.999933887 | 2.433314688 | 3.089181656 | -0.655866968 | 0.037392822 |
| ENSG00000287691 | *AL031602.2* | 2.413446423 | -0.102938852 | -0.066711133 | -0.036227719 | 0.999933887 | 1.180267817 | 0.708150622 | 0.472117195 | 0.037392822 |
| ENSG00000135932 | *CAB39* | 6.231970294 | 0.014117076 | -0.00782669 | 0.021943767 | 0.999933887 | -0.570453483 | -0.82634509 | 0.255891607 | 0.037392822 |
| ENSG00000105186 | *ANKRD27* | 3.876808217 | 0.007166047 | 0.053791371 | -0.046625323 | 0.999933887 | -0.323668628 | -0.578426908 | 0.25475828 | 0.037392822 |
| ENSG00000099804 | *CDC34* | 5.400532216 | 0.034355363 | 0.042491285 | -0.008135922 | 0.999933887 | -0.167408147 | -0.454955927 | 0.28754778 | 0.037435286 |
| ENSG00000169136 | *ATF5* | 2.622096221 | 0.013088227 | 0.027529441 | -0.014441215 | 0.999933887 | 0.455693876 | 0.802411059 | -0.346717183 | 0.037594335 |
| ENSG00000119866 | *BCL11A* | 2.624993466 | -0.011388697 | 0.016274719 | -0.027663417 | 0.999933887 | 1.906474438 | 2.53014438 | -0.623669942 | 0.037613669 |
| ENSG00000160991 | *ORAI2* | 7.204821415 | -0.004219047 | 0.022653503 | -0.02687255 | 0.999933887 | -0.526423444 | -0.789114828 | 0.262691384 | 0.037613669 |
| ENSG00000113593 | *PPWD1* | 4.23542264 | -0.122979855 | -0.158752668 | 0.035772813 | 0.999933887 | -0.325432457 | 0.11833213 | -0.443764587 | 0.037650585 |
| ENSG00000172292 | *CERS6* | 2.083345583 | 0.093267572 | -0.020820414 | 0.114087986 | 0.999933887 | 0.253664774 | 0.590191397 | -0.336526623 | 0.037728313 |
| ENSG00000099875 | *MKNK2* | 8.508315248 | 0.033212001 | 0.049671519 | -0.016459518 | 0.999933887 | 0.013142593 | -0.176586005 | 0.189728598 | 0.037728313 |
| ENSG00000259717 | *LINC00677* | 1.542234371 | 0.046171118 | -0.026493873 | 0.072664991 | 0.999933887 | 0.540492697 | 0.095519413 | 0.444973284 | 0.037728313 |
| ENSG00000232915 | *RNF14P1* | -1.075489474 | 0.20097768 | 0.237311018 | -0.036333338 | 0.999933887 | 0.97907046 | 1.772780049 | -0.793709589 | 0.037728313 |
| ENSG00000115165 | *CYTIP* | 9.00177402 | 0.017558581 | -0.024233434 | 0.041792016 | 0.999933887 | -0.539795758 | -0.304115887 | -0.235679871 | 0.037728313 |
| ENSG00000136930 | *PSMB7* | 4.881497601 | 0.01618617 | -0.013252777 | 0.029438947 | 0.999933887 | -0.340164902 | -0.565999213 | 0.225834311 | 0.037728313 |
| ENSG00000181704 | *YIPF6* | 4.680317866 | 0.060812235 | 0.068596366 | -0.007784131 | 0.999933887 | 0.406197415 | 0.632018642 | -0.225821227 | 0.03777165 |
| ENSG00000109689 | *STIM2* | 4.75232685 | 0.008854757 | -0.000827891 | 0.009682648 | 0.999933887 | -0.320580698 | -0.629934186 | 0.309353488 | 0.03777165 |
| ENSG00000197102 | *DYNC1H1* | 7.208123998 | -0.046294563 | -0.061795863 | 0.0155013 | 0.999933887 | -0.477248359 | -0.67232722 | 0.195078861 | 0.037804339 |
| ENSG00000285165 | *AC124014.1* | 0.414543395 | -0.102672942 | -0.022461855 | -0.080211087 | 0.999933887 | -0.06789036 | 0.613328824 | -0.681219183 | 0.037804339 |
| ENSG00000262152 | *LINC00514* | 0.13297036 | -0.093489591 | -0.08147739 | -0.012012201 | 0.999933887 | -1.63405975 | -2.517579286 | 0.883519536 | 0.037804339 |
| ENSG00000173517 | *PEAK1* | 3.091997903 | -0.123028776 | 0.016478223 | -0.139506999 | 0.999933887 | -0.287492525 | 0.048584113 | -0.336076638 | 0.037859091 |
| ENSG00000187605 | *TET3* | 8.197580847 | -0.019470546 | -0.009426605 | -0.010043941 | 0.999933887 | 0.409646747 | 0.60882552 | -0.199178773 | 0.037859091 |
| ENSG00000136052 | *SLC41A2* | -0.383998529 | -0.033658306 | 0.15133915 | -0.184997457 | 0.999933887 | 0.144511724 | 0.727497304 | -0.58298558 | 0.037859091 |
| ENSG00000030110 | *BAK1* | 3.274567674 | 0.000667453 | -0.070380232 | 0.071047685 | 0.999933887 | -0.210152146 | -0.525876034 | 0.315723888 | 0.037859091 |
| ENSG00000105404 | *RABAC1* | 2.449520563 | -0.044156136 | 0.046810855 | -0.090966991 | 0.999933887 | -0.192730226 | -0.620667288 | 0.427937062 | 0.037872775 |
| ENSG00000075785 | *RAB7A* | 9.459309857 | 0.012884341 | -0.036303167 | 0.049187508 | 0.999933887 | 0.205268462 | 0.398078566 | -0.192810104 | 0.038036248 |

| | | | | | | | | | | |
|---|---|---|---|---|---|---|---|---|---|---|
| ENSG00000005187 | *ACSM3* | 0.277885133 | -0.074689603 | -0.097587878 | 0.022898275 | 0.999933887 | 0.226338802 | -0.35005954 | 0.576398342 | 0.038049241 |
| ENSG00000109079 | *TNFAIP1* | 2.867449544 | -0.109706168 | -0.029920669 | -0.079785499 | 0.999933887 | -0.057215478 | -0.304894116 | 0.247678639 | 0.038097499 |
| ENSG00000111737 | *RAB35* | 7.255998033 | 0.028381388 | 0.017071335 | 0.011310052 | 0.999933887 | -0.083484494 | 0.110281069 | -0.193765564 | 0.038097499 |
| ENSG00000124074 | *ENKD1* | 1.655704725 | 0.024813061 | 0.20978052 | -0.184967459 | 0.999933887 | -0.277730142 | -0.809097991 | 0.531367849 | 0.038097499 |
| ENSG00000124784 | *RIOK1* | 2.874801743 | -0.06170514 | 0.081176438 | -0.142881578 | 0.999933887 | 0.377240329 | 0.726311089 | -0.34907076 | 0.038105685 |
| ENSG00000135999 | *EPC2* | 5.309796269 | -0.034101474 | -0.013550215 | -0.020551259 | 0.999933887 | -0.294927607 | -0.080822101 | -0.214105506 | 0.038171621 |
| ENSG00000092439 | *TRPM7* | 5.988256836 | -0.083190281 | -0.057643561 | -0.02554672 | 0.999933887 | 0.124626766 | 0.536724422 | -0.412097656 | 0.038227686 |
| ENSG00000108588 | *CCDC47* | 4.757043993 | -0.033140555 | -0.099564472 | 0.066423917 | 0.999933887 | -0.573334514 | -0.837558986 | 0.264224472 | 0.038262277 |
| ENSG00000112053 | *SLC26A8* | 3.53621163 | -0.029877158 | 0.032416122 | -0.062293281 | 0.999933887 | -0.138848953 | 0.19098233 | -0.329831283 | 0.038262277 |
| ENSG00000165684 | *SNAPC4* | 3.326377313 | 0.073049617 | 0.105210833 | -0.032161216 | 0.999933887 | 0.095078717 | -0.234356016 | 0.329434734 | 0.038292207 |
| ENSG00000234925 | *ATP5PDP4* | 0.030319068 | 0.11989642 | 0.193801045 | -0.073904625 | 0.999933887 | 0.246397199 | 1.136873845 | -0.890476646 | 0.038320123 |
| ENSG00000061337 | *LZTS1* | 0.969788753 | 0.06612284 | 0.284414972 | -0.218292133 | 0.999933887 | 0.690449165 | 1.427043334 | -0.73659417 | 0.038376024 |
| ENSG00000076003 | *MCM6* | 4.36610633 | -0.177817119 | -0.10643899 | -0.071378129 | 0.999933887 | -1.268824971 | -1.631702366 | 0.362877394 | 0.03841788 |
| ENSG00000110011 | *DNAJC4* | 3.193641641 | 0.018225868 | -0.021447406 | 0.039673275 | 0.999933887 | -0.132822533 | -0.417226511 | 0.284403978 | 0.038454421 |
| ENSG00000187808 | *SOWAHD* | -0.384144231 | -0.017584507 | 0.059172054 | -0.076756561 | 0.999933887 | -0.641646846 | -1.572986781 | 0.931339935 | 0.038476081 |
| ENSG00000101972 | *STAG2* | 7.445464208 | -0.097877795 | -0.15957699 | 0.061699195 | 0.999933887 | -0.315180883 | 0.185017066 | -0.500197949 | 0.038519063 |
| ENSG00000280620 | *SCAANT1* | -0.368163024 | -0.002841994 | 0.10922265 | -0.112064644 | 0.999933887 | 0.495862184 | -0.46919347 | 0.965055654 | 0.038519063 |
| ENSG00000117385 | *P3H1* | 2.940931826 | 0.083303322 | 0.134230498 | -0.050927176 | 0.999933887 | 0.059625707 | -0.293165534 | 0.352791241 | 0.038657756 |
| ENSG00000172548 | *NIPAL4* | 1.320426973 | 0.074932376 | -0.173419924 | 0.248352299 | 0.999933887 | 2.763948884 | 3.391276579 | -0.627327695 | 0.038657756 |
| ENSG00000117748 | *RPA2* | 5.148720862 | 0.000549882 | -0.055592321 | 0.056142203 | 0.999933887 | 0.235506869 | 0.451948847 | -0.216441978 | 0.03874718 |
| ENSG00000152056 | *AP1S3* | 1.516415363 | -0.090446299 | -0.44014185 | 0.349695552 | 0.999933887 | 2.966835908 | 3.466615723 | -0.499779815 | 0.03876282 |
| ENSG00000177606 | *JUN* | 8.300620413 | 0.263597669 | 0.269293096 | -0.005695427 | 0.999933887 | 0.260876615 | 0.585337295 | -0.32446068 | 0.038862418 |
| ENSG00000276070 | *CCL4L2* | 9.240247983 | 0.517400761 | 0.444964577 | 0.072436184 | 0.999933887 | 3.149059254 | 3.782706858 | -0.633647604 | 0.03888051 |
| ENSG00000168288 | *MMADHC* | 5.654450953 | -0.06632344 | -0.016317388 | -0.050006052 | 0.999933887 | 0.049563731 | 0.228385176 | -0.178821446 | 0.038936861 |
| ENSG00000054267 | *ARID4B* | 7.163592179 | -0.073857167 | -0.146021202 | 0.072164036 | 0.999933887 | -0.132874137 | 0.375958723 | -0.508832861 | 0.038961611 |
| ENSG00000169972 | *PUSL1* | 1.550363113 | 0.01057681 | 0.106495885 | -0.095919076 | 0.999933887 | -0.003280717 | -0.513436373 | 0.510155656 | 0.039009377 |
| ENSG00000175354 | *PTPN2* | 4.530904234 | -0.035098984 | -0.077724741 | 0.042625757 | 0.999933887 | 0.046125063 | 0.326145077 | -0.280020014 | 0.039078823 |
| ENSG00000135052 | *GOLM1* | 0.787334351 | -0.053172483 | -0.240511138 | 0.187338655 | 0.999933887 | -0.096344011 | 0.507783495 | -0.604127506 | 0.039112766 |
| ENSG00000197498 | *RPF2* | 2.251992715 | -0.033161832 | -0.113851404 | 0.080689572 | 0.999933887 | 0.775751418 | 1.322788314 | -0.547036896 | 0.039118603 |
| ENSG00000186891 | *TNFRSF18* | 1.083264534 | 0.265002688 | -0.030257316 | 0.295260003 | 0.999933887 | 1.010703254 | 1.650913468 | -0.640210214 | 0.039187757 |
| ENSG00000166200 | *COPS2* | 4.10972635 | -0.104013038 | -0.237506536 | 0.133493497 | 0.999933887 | -0.295140462 | 0.049870507 | -0.345010968 | 0.039187757 |
| ENSG00000121481 | *RNF2* | 2.550880347 | -0.014746844 | -0.05758981 | 0.042842966 | 0.999933887 | 0.050683897 | 0.316687059 | -0.266003162 | 0.039276597 |
| ENSG00000196187 | *TMEM63A* | 5.725801727 | 0.008805595 | 0.031972157 | -0.023166562 | 0.999933887 | -0.037845172 | -0.222946161 | 0.185100989 | 0.039299501 |
| ENSG00000112658 | *SRF* | 6.896233605 | 0.196875694 | 0.172283555 | 0.024592139 | 0.999933887 | 0.387309731 | 0.687307433 | -0.299997702 | 0.039460153 |
| ENSG00000033100 | *CHPF2* | 6.39921544 | -0.036790984 | -0.021051681 | -0.015739303 | 0.999933887 | -0.643125454 | -0.88271561 | 0.239590155 | 0.039460153 |
| ENSG00000104852 | *SNRNP70* | 7.50641917 | 0.043014542 | 0.124523575 | -0.081509033 | 0.999933887 | 0.000879929 | -0.241256966 | 0.242136894 | 0.039525813 |
| ENSG00000255274 | *SMIM35* | 1.026464795 | 0.104423562 | -0.095536376 | 0.199959937 | 0.999933887 | 0.780888859 | 1.544843022 | -0.763954163 | 0.039525813 |
| ENSG00000078114 | *NEBL* | 2.500892749 | 0.054074988 | -0.129794906 | 0.183869893 | 0.999933887 | -0.035889176 | 0.354707982 | -0.390597158 | 0.039525813 |
| ENSG00000177173 | *NAP1L4P1* | 3.911429808 | 0.00785226 | 0.008097347 | -0.000245087 | 0.999933887 | 1.205954837 | 1.738760705 | -0.532805868 | 0.039525813 |
| ENSG00000169508 | *GPR183* | 4.446049788 | 0.039562246 | -0.000854527 | 0.040416772 | 0.999933887 | 0.014480125 | 0.309135286 | -0.294655161 | 0.039553484 |
| ENSG00000160949 | *TONSL* | 1.390734075 | -0.014192159 | 0.209249135 | -0.223441294 | 0.999933887 | 0.455052638 | 1.008601101 | -0.553548463 | 0.039553484 |
| ENSG00000153395 | *LPCAT1* | 10.223361221 | -0.017967587 | -0.01711852 | -0.000849067 | 0.999933887 | 0.852770588 | 1.093707331 | -0.240936743 | 0.039596599 |
| ENSG00000140474 | *ULK3* | 3.777135799 | -0.04683857 | 0.037897435 | -0.084736005 | 0.999933887 | 0.079243137 | -0.147815891 | 0.227059027 | 0.03979001 |
| ENSG00000117475 | *BLZF1* | 3.659908892 | -0.134053568 | -0.240320817 | 0.106267249 | 0.999933887 | -0.519389397 | -0.176528346 | -0.342861051 | 0.039810051 |
| ENSG00000275560 | *AC008115.3* | 1.815142288 | -0.119468404 | -0.175743955 | 0.056275551 | 0.999933887 | -0.438434603 | -0.923041191 | 0.484606587 | 0.039810051 |
| ENSG00000142082 | *SIRT3* | 2.54198914 | 0.082936543 | 0.021188376 | 0.061748167 | 0.999933887 | 0.03813146 | -0.287196679 | 0.32532814 | 0.039810051 |
| ENSG00000178585 | *CTNNBIP1* | 1.804158121 | 0.099011135 | 0.126232964 | -0.027221829 | 0.999933887 | -0.491415817 | -1.099654782 | 0.608238965 | 0.039810051 |
| ENSG00000087301 | *TXNDC16* | 3.366498884 | -0.068294982 | 0.064814373 | -0.133109355 | 0.999933887 | 0.415022303 | 0.937670102 | -0.5226478 | 0.039810051 |
| ENSG00000225783 | *MIAT* | 5.870358628 | -0.024328177 | 0.035715832 | -0.060044009 | 0.999933887 | -0.114608284 | -0.352350962 | 0.237742678 | 0.039810051 |
| ENSG00000169403 | *PTAFR* | 9.741022084 | -0.087680156 | -0.09846753 | 0.010787373 | 0.999933887 | 1.351019414 | 1.686256763 | -0.335237349 | 0.040017099 |
| ENSG00000285103 | *AL451123.1* | 4.148794353 | -0.060817246 | -0.039467755 | -0.021349491 | 0.999933887 | -0.032888036 | 0.500748183 | -0.533636219 | 0.040106542 |
| ENSG00000065882 | *TBC1D1* | 5.883739899 | -0.012693322 | 0.003110894 | -0.015804216 | 0.999933887 | 0.051768636 | -0.135383789 | 0.187152425 | 0.040106542 |
| ENSG00000124422 | *USP22* | 6.019585763 | -0.037312767 | -0.017598012 | -0.019714755 | 0.999933887 | -0.423346415 | -0.626751949 | 0.203405534 | 0.040125712 |
| ENSG00000150995 | *ITPR1* | 5.652554442 | -0.036004531 | -0.104596996 | 0.068592464 | 0.999933887 | 0.222244421 | 0.425384611 | -0.20314019 | 0.040125712 |
| ENSG00000198915 | *RASGEF1A* | 3.360194456 | 0.03388617 | -0.13370993 | 0.1675961 | 0.999933887 | -1.467139227 | -1.869881802 | 0.402742575 | 0.040125712 |
| ENSG00000165609 | *NUDT5* | 3.774675915 | -0.011491062 | -0.07671821 | 0.065227149 | 0.999933887 | -0.150423325 | -0.423882323 | 0.273458998 | 0.040170964 |
| ENSG00000162888 | *C1orf147* | -0.473412741 | -0.034344238 | 0.289298661 | -0.323642899 | 0.999933887 | 1.052139932 | 2.082189607 | -1.030049674 | 0.040170964 |
| ENSG00000156860 | *FBRS* | 9.474317995 | 0.099483077 | 0.085439843 | 0.014043234 | 0.999933887 | 0.696879323 | 0.903928994 | -0.207049671 | 0.04018161 |
| ENSG00000089041 | *P2RX7* | 2.902054572 | -0.010810949 | -0.028747524 | 0.017936575 | 0.999933887 | 1.274496291 | 1.740922545 | -0.466426254 | 0.040233858 |
| ENSG00000128159 | *TUBGCP6* | 5.877894949 | 0.026893456 | 0.07149006 | -0.044596604 | 0.999933887 | 0.042204039 | -0.146773238 | 0.188977277 | 0.040364118 |
| ENSG00000059377 | *TBXAS1* | 7.293073321 | -0.048148296 | -0.042745835 | -0.005402461 | 0.999933887 | -1.096822545 | -1.451976683 | 0.355154138 | 0.040412442 |
| ENSG00000130150 | *MOSPD2* | 5.224793381 | -0.034408722 | -0.117237553 | 0.082828831 | 0.999933887 | -0.60468588 | -0.242697153 | -0.361988728 | 0.040511434 |
| ENSG00000137312 | *FLOT1* | 8.472147632 | 0.097559232 | 0.053794224 | 0.043765008 | 0.999933887 | 1.135471223 | 1.434217035 | -0.298745812 | 0.040607178 |
| ENSG00000136819 | *C9orf78* | 5.47815194 | -0.034609451 | -0.076253557 | 0.041644106 | 0.999933887 | -0.380176138 | -0.576043252 | 0.195867114 | 0.040607178 |
| ENSG00000164236 | *ANKRD33B* | 5.06283045 | -0.06874268 | -0.029948871 | -0.038793809 | 0.999933887 | 2.255693094 | 2.762951176 | -0.507258081 | 0.040607178 |
| ENSG00000109736 | *MFSD10* | 5.380968908 | 0.048373649 | 0.035107985 | 0.013265664 | 0.999933887 | 0.038475094 | -0.17764839 | 0.216123484 | 0.040647603 |
| ENSG00000110665 | *C11orf21* | 5.518938847 | 0.047746241 | 0.052266176 | -0.004519935 | 0.999933887 | -0.391188664 | -0.678628482 | 0.287439819 | 0.040647603 |
| ENSG00000144747 | *TMF1* | 5.419610713 | -0.033117495 | -0.169523961 | 0.136406467 | 0.999933887 | -0.377501143 | 0.15714931 | -0.534650453 | 0.040647603 |
| ENSG00000182782 | *HCAR2* | 11.4216375 | 0.065221714 | 0.06651399 | -0.001292277 | 0.999933887 | 0.119994925 | 0.400234183 | -0.280239257 | 0.040719783 |
| ENSG00000103496 | *STX4* | 7.97558332 | 0.109626848 | 0.092625179 | 0.017001669 | 0.999933887 | 0.590246755 | 0.820935123 | -0.230688368 | 0.040817686 |
| ENSG00000198646 | *NCOA6* | 5.659842401 | -0.038127725 | -0.013376733 | -0.024750992 | 0.999933887 | -0.639110015 | -0.888171946 | 0.249061931 | 0.040817686 |
| ENSG00000156026 | *MCU* | 3.28620708 | 0.018158975 | 0.085619482 | -0.067460507 | 0.999933887 | -0.099652294 | -0.339761305 | 0.240109011 | 0.040838871 |
| ENSG00000178971 | *CTC1* | 6.138302612 | -0.021386618 | 0.044121042 | -0.06550766 | 0.999933887 | 0.123461422 | 0.302170354 | -0.178708932 | 0.040876562 |
| ENSG00000175130 | *MARCKSL1* | 6.435479167 | -0.000290789 | 0.090286978 | -0.090577767 | 0.999933887 | -0.210781271 | -0.515645197 | 0.304863926 | 0.041059821 |
| ENSG00000284669 | *AC092053.3* | 5.7601866 | 0.388172045 | 0.422271128 | -0.034099083 | 0.999933887 | 1.040390791 | 1.477586486 | -0.437195695 | 0.041151811 |
| ENSG00000198961 | *PJA2* | 6.985176333 | -0.054287115 | -0.063099139 | 0.008812024 | 0.999933887 | -0.19583257 | 0.104610256 | -0.300442826 | 0.041151811 |
| ENSG00000197746 | *PSAP* | 11.381219877 | -0.030049026 | -0.045489602 | 0.015440577 | 0.999933887 | 0.402119662 | 0.584366104 | -0.182246442 | 0.041309234 |
| ENSG00000107864 | *CPEB3* | 1.80383323 | 0.027257167 | 0.086117832 | -0.058860665 | 0.999933887 | -0.199990608 | -0.752845905 | 0.552855297 | 0.041311718 |
| ENSG00000141627 | *DYM* | 3.958456809 | 0.027795238 | -0.035306407 | 0.063101645 | 0.999933887 | -0.089090164 | 0.139560551 | -0.228650715 | 0.041323539 |
| ENSG00000227155 | *AL161725.1* | 3.230775142 | -0.00319414 | 0.010787857 | -0.013981997 | 0.999933887 | -0.237488248 | -0.795480351 | 0.557992103 | 0.041327894 |
| ENSG00000171617 | *ENC1* | 4.515763518 | -0.065841858 | -0.083333078 | 0.017491219 | 0.999933887 | -0.075048599 | -0.314985868 | 0.239937269 | 0.041433282 |
| ENSG00000167258 | *CDK12* | 6.408223858 | -0.013358219 | -0.04043177 | 0.02707355 | 0.999933887 | 0.061125108 | 0.226328028 | -0.16520292 | 0.041433282 |
| ENSG00000044090 | *CUL7* | 2.445426605 | 0.054847625 | 0.119763833 | -0.064916207 | 0.999933887 | -0.013541614 | -0.366265876 | 0.352724262 | 0.041487595 |
| ENSG00000112378 | *PERP* | 0.990973951 | -0.038567534 | -0.018933281 | -0.019634254 | 0.999933887 | 0.440666202 | 1.060795386 | -0.620129184 | 0.041497961 |
| ENSG00000183576 | *SETD3* | 3.922019932 | 0.012988405 | -0.058791039 | 0.071779444 | 0.999933887 | -0.223572605 | -0.474581897 | 0.251009291 | 0.041497961 |
| ENSG00000234511 | *C5orf58* | 4.663176417 | 0.072624111 | 0.083489742 | -0.010865631 | 0.999933887 | 0.806155386 | 1.357717869 | -0.551562483 | 0.041511416 |
| ENSG00000203364 | *AL390760.1* | 0.269913999 | 0.211292672 | -0.43385886 | 0.645151532 | 0.999933887 | 2.994154164 | 3.586146014 | -0.59199185 | 0.041565071 |
| ENSG00000169567 | *HINT1* | 5.288763982 | -0.007480124 | -0.021592412 | 0.014112289 | 0.999933887 | 0.271582298 | 0.512981638 | -0.241399341 | 0.041613456 |
| ENSG00000287839 | *AL353807.5* | 1.976298596 | 0.006766694 | -0.175316196 | 0.18208289 | 0.999933887 | 0.036504191 | -0.500827686 | 0.537331877 | 0.041640203 |
| ENSG00000187796 | *CARD9* | 0.454261603 | -0.063389598 | 0.060138392 | -0.12352799 | 0.999933887 | 0.026343447 | -0.691553498 | 0.717896945 | 0.041696353 |
| ENSG00000007129 | *CEACAM21* | 2.076305347 | 0.019230896 | 0.010929591 | 0.008301305 | 0.999933887 | 0.017638232 | -0.425155102 | 0.442793334 | 0.041696353 |
| ENSG00000139233 | *LLPH* | 3.314720467 | -0.196260489 | -0.120420872 | -0.075839617 | 0.999933887 | 0.045885072 | 0.310680096 | -0.264795024 | 0.041749447 |
| ENSG00000133318 | *RTN3* | 7.918592188 | -0.016514861 | -0.028373333 | 0.011858472 | 0.999933887 | -0.462611103 | -0.731191964 | 0.26858086 | 0.041764887 |
| ENSG00000103994 | *ZNF106* | 5.417171986 | -0.070924131 | -0.061253971 | -0.00967016 | 0.999933887 | -0.3725747 | -0.699473117 | 0.326898417 | 0.041824148 |
| ENSG00000012211 | *PRICKLE3* | 1.241142614 | 0.111866935 | 0.069043274 | 0.042823661 | 0.999933887 | -0.07907766 | -0.627656603 | 0.548578942 | 0.042071161 |
| ENSG00000120616 | *EPC1* | 5.655848772 | -0.041155517 | -0.025305819 | -0.015849698 | 0.999933887 | 0.009949519 | 0.201371792 | -0.191422273 | 0.042284865 |
| ENSG00000204569 | *PPP1R10* | 8.718473652 | 0.040093391 | 0.056842573 | -0.016749182 | 0.999933887 | 0.44696004 | 0.6544902 | -0.207530159 | 0.042527292 |
| ENSG00000149948 | *HMGA2* | -2.193312488 | -0.423295874 | -0.156691933 | -0.266603941 | 0.999933887 | 1.347664301 | 2.169750613 | -0.822086312 | 0.042561331 |
| ENSG00000198580 | *AC073343.1* | 0.899769816 | -0.149546398 | -0.552858026 | 0.403311628 | 0.999933887 | 0.11634482 | 0.554120479 | -0.437775659 | 0.042566563 |
| ENSG00000173535 | *TNFRSF10C* | 6.938295405 | -0.005392919 | -0.034560313 | 0.029167395 | 0.999933887 | -2.068368088 | -2.669798513 | 0.601430425 | 0.042631783 |
| ENSG00000166377 | *ATP9B* | 3.493387421 | -0.041414379 | 0.081855682 | -0.123270061 | 0.999933887 | -0.023550944 | 0.217042545 | -0.240593489 | 0.042674702 |
| ENSG00000121297 | *TSHZ3* | 2.944628722 | 0.124548412 | 0.074319329 | 0.050229083 | 0.999933887 | -0.83828668 | -1.201354928 | 0.363068249 | 0.042750877 |
| ENSG00000116014 | *KISS1R* | 0.435498909 | 0.191605297 | 0.016634567 | 0.17497073 | 0.999933887 | -0.677187433 | -1.434049569 | 0.756862136 | 0.042750877 |
| ENSG00000048471 | *SNX29* | 3.373083399 | 0.023295265 | 0.052364651 | -0.029069386 | 0.999933887 | 0.023941678 | -0.238958716 | 0.262900395 | 0.042750877 |
| ENSG00000157259 | *GATAD1* | 3.10234984 | -0.025332726 | 0.05655556 | -0.081888286 | 0.999933887 | -0.382690419 | -0.631015526 | 0.248325107 | 0.042780108 |

| | | | | | | | | | | |
|---|---|---|---|---|---|---|---|---|---|---|
| ENSG00000116698 | *SMG7* | 6.780309579 | -0.014122558 | -0.032144402 | 0.018021845 | 0.999933887 | 0.61049898 | 0.945984195 | -0.335485215 | 0.042780108 |
| ENSG00000132823 | *OSER1* | 7.970656719 | 0.00393199 | -0.044270387 | 0.048202377 | 0.999933887 | -0.203902735 | 0.015535781 | -0.219438516 | 0.04278054 |
| ENSG00000078061 | *ARAF* | 5.95607212 | 0.016045076 | 0.023514593 | -0.007469517 | 0.999933887 | 0.282239561 | 0.472312813 | -0.190073252 | 0.04287799 |
| ENSG00000010671 | *BTK* | 4.041103786 | -0.024621428 | -0.015977413 | -0.008644015 | 0.999933887 | -0.47656056 | -0.764407097 | 0.287846537 | 0.042950139 |
| ENSG00000278097 | *AC243829.5* | -1.935916471 | 0.225928777 | -0.002664324 | 0.228593101 | 0.999933887 | 0.239785708 | 1.37622597 | -1.136440262 | 0.042950446 |
| ENSG00000175220 | *ARHGAP1* | 5.207279429 | -0.040977191 | 0.032875474 | -0.073852665 | 0.999933887 | -0.293623189 | -0.491915007 | 0.198291819 | 0.042950446 |
| ENSG00000100031 | *GGT1* | 2.883322375 | 0.070739862 | 0.233337945 | -0.162598083 | 0.999933887 | 0.493660883 | 0.895931686 | -0.402270802 | 0.042960923 |
| ENSG00000197301 | *HMGA2-AS1* | -1.367110607 | -0.229260357 | 0.496529772 | -0.725790128 | 0.999933887 | 2.04930821 | 2.893890908 | -0.844582698 | 0.042963042 |
| ENSG00000100364 | *KIAA0930* | 5.141469307 | 0.018700582 | 0.035960739 | -0.017260157 | 0.999933887 | -1.49714068 | -1.915285135 | 0.418144455 | 0.042963042 |
| ENSG00000101868 | *POLA1* | 1.441280555 | -0.002162327 | 0.067895217 | -0.070057544 | 0.999933887 | 0.025382781 | 0.544556946 | -0.519174165 | 0.042979407 |
| ENSG00000100523 | *DDHD1* | 6.236118623 | -0.021337935 | 0.014702172 | -0.036040107 | 0.999933887 | 0.199240486 | 0.542090777 | -0.342850291 | 0.042983266 |
| ENSG00000057704 | *TMCC3* | 7.215542075 | -0.045500985 | -0.037836118 | -0.007664866 | 0.999933887 | -0.411069536 | -0.156850575 | -0.254218961 | 0.043169705 |
| ENSG00000239998 | *LILRA2* | 7.609432247 | 0.027143989 | -0.032795729 | 0.059939719 | 0.999933887 | -1.167770882 | -1.477903421 | 0.310132539 | 0.043169705 |
| ENSG00000167083 | *GNGT2* | 1.491630986 | -0.164642653 | -0.11746818 | -0.047174473 | 0.999933887 | 0.171156972 | -0.329730679 | 0.500887651 | 0.04322391 |
| ENSG00000113761 | *ZNF346* | 2.472520811 | 0.065326534 | -0.039346659 | 0.104673193 | 0.999933887 | -0.445587603 | -0.842636184 | 0.397048582 | 0.04322391 |
| ENSG00000164330 | *EBF1* | -1.075755845 | 0.080478653 | 0.263860735 | -0.183382082 | 0.999933887 | 0.340206499 | 1.207275516 | -0.867069016 | 0.04322391 |
| ENSG00000237181 | *PRKAR1B-AS* | 2.053977532 | -0.004095619 | 0.023939762 | -0.028035381 | 0.999933887 | 1.384145957 | 1.739732637 | -0.35558668 | 0.04322391 |
| ENSG00000148291 | *SURF2* | 1.558657488 | -0.001694903 | 0.062934369 | -0.064629272 | 0.999933887 | -0.000492855 | -0.355780732 | 0.355287876 | 0.04322391 |
| ENSG00000167487 | *KLHL26* | 1.981485431 | 0.14092495 | 0.065570358 | 0.075354593 | 0.999933887 | 0.149792127 | -0.247382109 | 0.397174235 | 0.043265754 |
| ENSG00000116830 | *TTF2* | 2.293861749 | -0.052204493 | -0.021616599 | -0.030587894 | 0.999933887 | 0.337960999 | 0.676004733 | -0.338043735 | 0.043265754 |
| ENSG00000258302 | *AC025034.1* | 3.877425882 | -0.13273152 | 0.128494752 | -0.261226272 | 0.999933887 | 0.971725681 | 1.794604255 | -0.822878574 | 0.043353676 |
| ENSG00000271383 | *NBPF19* | 1.169091639 | 0.020472862 | 0.088513376 | -0.068040514 | 0.999933887 | 0.513238441 | 0.977705295 | -0.464466854 | 0.043353676 |
| ENSG00000281195 | *AC007878.1* | 2.030327627 | 0.176123952 | 0.030995166 | 0.145128786 | 0.999933887 | -0.324263194 | 0.241470436 | -0.56573363 | 0.04341981 |
| ENSG00000101966 | *XIAP* | 5.914549093 | -0.049429348 | 0.009526519 | -0.058955867 | 0.999933887 | 0.14882588 | 0.470647033 | -0.321821153 | 0.043452148 |
| ENSG00000066322 | *ELOVL1* | 4.974844228 | -0.017291843 | -0.020040746 | 0.002748902 | 0.999933887 | -0.48685857 | -0.728003704 | 0.241145134 | 0.043467769 |
| ENSG00000089505 | *CMTM1* | 1.079746636 | 0.013423225 | -0.204641967 | 0.218065192 | 0.999933887 | -1.593468287 | -2.354038098 | 0.760569812 | 0.043467769 |
| ENSG00000100288 | *CHKB* | 0.631385614 | -0.202596846 | -0.004651301 | -0.197945545 | 0.999933887 | -0.11497576 | -0.663521588 | 0.548545828 | 0.043492498 |
| ENSG00000085185 | *BCORL1* | 3.196569822 | -0.053831981 | 0.080568469 | -0.13440045 | 0.999933887 | -0.291551226 | -0.60745644 | 0.315905215 | 0.043533756 |
| ENSG00000183723 | *CMTM4* | 0.406033936 | -0.112866308 | -0.130062669 | 0.017196361 | 0.999933887 | -0.402854131 | -1.159651242 | 0.756797111 | 0.043533756 |
| ENSG00000105991 | *HOXA1* | -0.005153012 | -0.035050038 | -0.238484222 | 0.203434185 | 0.999933887 | -0.646180473 | -1.707354965 | 1.061174493 | 0.043583401 |
| ENSG00000130382 | *MLLT1* | 5.9353508 | 0.050020788 | -0.006276064 | 0.056296852 | 0.999933887 | -0.248452442 | -0.464348365 | 0.215895923 | 0.04360068 |
| ENSG00000139132 | *FGD4* | 4.928647815 | -0.044075148 | -0.076927391 | 0.032852243 | 0.999933887 | 0.264216124 | 0.681679462 | -0.417463338 | 0.043643491 |
| ENSG00000130741 | *EIF2S3* | 7.076446039 | -0.021503115 | -0.011130792 | -0.010372323 | 0.999933887 | 0.126750062 | 0.279626851 | -0.152876789 | 0.043689744 |
| ENSG00000132661 | *NXT1* | 3.533508754 | 0.128372431 | 0.048413551 | 0.07995888 | 0.999933887 | 0.832906137 | 1.127578774 | -0.294672638 | 0.04376953 |
| ENSG00000108405 | *P2RX1* | 5.805197829 | -0.018994933 | 0.032489071 | -0.051484004 | 0.999933887 | -0.610594125 | -0.82984244 | 0.219248315 | 0.043782057 |
| ENSG00000123353 | *ORMDL2* | 2.465185426 | -0.0632595 | 0.042595947 | -0.105855447 | 0.999933887 | -0.568808512 | -0.901307519 | 0.332499008 | 0.043860227 |
| ENSG00000165417 | *GTF2A1* | 6.865212048 | -0.066378187 | -0.114379214 | 0.048001027 | 0.999933887 | 0.195441258 | 0.491331739 | -0.295890481 | 0.043921649 |
| ENSG00000198742 | *SMURF1* | 6.248955236 | 0.002605556 | -0.011875343 | 0.014480899 | 0.999933887 | 0.395095374 | 0.556665501 | -0.161570127 | 0.043945351 |
| ENSG00000183648 | *NDUFB1* | 3.880279182 | -0.034166009 | -0.05370381 | 0.019537801 | 0.999933887 | -0.101959294 | -0.386359674 | 0.28440038 | 0.043999532 |
| ENSG00000198804 | *MT-CO1* | 9.841663293 | 0.013682868 | -0.040331145 | 0.054014013 | 0.999933887 | -0.01870394 | 0.267416739 | -0.286120679 | 0.043999532 |
| ENSG00000144504 | *ANKMY1* | 3.032870954 | -0.062728075 | 0.056366114 | -0.119094188 | 0.999933887 | -0.071237815 | -0.335481114 | 0.2642433 | 0.04405078 |
| ENSG00000171862 | *PTEN* | 8.361259958 | -0.020467225 | -0.003695929 | -0.016771295 | 0.999933887 | -0.535005858 | -0.78138172 | 0.246375862 | 0.04405078 |
| ENSG00000258730 | *ITPK1-AS1* | -0.258670459 | -0.041694112 | 0.0731622 | -0.114856312 | 0.999933887 | -0.874726852 | -2.086118417 | 1.211391565 | 0.044136921 |
| ENSG00000141404 | *GNAL* | 1.041975506 | -0.149843888 | -0.115118064 | -0.034725824 | 0.999933887 | -0.450530018 | -0.967333042 | 0.516803025 | 0.044137619 |
| ENSG00000111863 | *ADTRP* | 0.435522556 | 0.00217819 | -0.254536302 | 0.256714492 | 0.999933887 | 0.420273068 | 0.957453269 | -0.537180202 | 0.044171545 |
| ENSG00000112624 | *BICRAL* | 5.267308708 | -0.037638237 | 0.053980712 | -0.091618949 | 0.999933887 | -0.668835456 | -0.9531906 | 0.284355143 | 0.04421323 |
| ENSG00000143622 | *RIT1* | 8.047026347 | -0.050731513 | -0.062851897 | 0.012120383 | 0.999933887 | 0.222614286 | 0.410353187 | -0.187738901 | 0.044217906 |
| ENSG00000090989 | *EXOC1* | 4.694782506 | -0.018336331 | -0.120156912 | 0.101820581 | 0.999933887 | -0.375228062 | -0.001755258 | -0.373472804 | 0.04431757 |
| ENSG00000232810 | *TNF* | 5.930030268 | 0.687892381 | 1.050933967 | -0.363041587 | 0.999933887 | 2.553222565 | 3.198183223 | -0.644960658 | 0.044483039 |
| ENSG00000134698 | *AGO4* | 7.280138115 | -0.077307954 | -0.059283571 | -0.018024384 | 0.999933887 | -0.824228945 | -1.056013759 | 0.231784813 | 0.044483039 |
| ENSG00000115155 | *OTOF* | -0.009304876 | 0.05642628 | -0.106249811 | 0.162676091 | 0.999933887 | 0.255988738 | -0.363061033 | 0.61904977 | 0.044505018 |
| ENSG00000233791 | *LINC01136* | 3.777709075 | 0.172309065 | 0.155760525 | 0.016548539 | 0.999933887 | 1.497606107 | 1.864800244 | -0.367194137 | 0.044652564 |
| ENSG00000228109 | *MELTF-AS1* | -0.7551082 | 0.24720064 | 0.300757892 | -0.053557252 | 0.999933887 | 0.714677141 | 1.516765598 | -0.802088457 | 0.044652564 |
| ENSG00000232630 | *PRPS1P2* | 0.88203665 | 0.013813381 | 0.067782386 | -0.053969005 | 0.999933887 | -0.313016898 | 0.198442352 | -0.51145925 | 0.044652564 |
| ENSG00000178952 | *TUFM* | 4.835918286 | -0.005053668 | -0.047040929 | 0.041987261 | 0.999933887 | -7.1678e-05 | -0.222227563 | 0.222155886 | 0.044652564 |
| ENSG00000163874 | *ZC3H12A* | 8.688105141 | 0.047867219 | 0.070284444 | -0.022417225 | 0.999933887 | 1.896313348 | 2.168763163 | -0.272449815 | 0.044936541 |
| ENSG00000183087 | *GAS6* | 1.784914318 | 0.036808615 | 0.098090424 | -0.061281809 | 0.999933887 | 0.432303092 | 0.81777841 | -0.385475318 | 0.044936541 |
| ENSG00000128604 | *IRF5* | 4.108829803 | 0.031044539 | 0.023270531 | 0.007774008 | 0.999933887 | 1.507344905 | 1.915962913 | -0.408618009 | 0.044936541 |
| ENSG00000129673 | *AANAT* | 0.64440941 | -0.098876011 | 0.089573085 | -0.188449097 | 0.999933887 | 1.180587235 | 1.970686721 | -0.790099485 | 0.045018285 |
| ENSG00000075624 | *ACTB* | 12.44988677 | 0.026838974 | 0.038609065 | -0.011770091 | 0.999933887 | -0.328817951 | -0.548319841 | 0.21950189 | 0.045018285 |
| ENSG00000166946 | *CCNDBP1* | 6.460661125 | -0.049790836 | -0.057645206 | 0.007854371 | 0.999933887 | -0.569150033 | -0.759571828 | 0.190421795 | 0.045038269 |
| ENSG00000118515 | *SGK1* | 11.375549727 | 0.191871894 | 0.215341141 | -0.023469246 | 0.999933887 | 0.564336437 | 0.800392981 | -0.236056544 | 0.04523552 |
| ENSG00000109066 | *TMEM104* | 4.24186213 | -0.053816124 | 0.001142311 | -0.054958435 | 0.999933887 | -0.664368731 | -0.907865812 | 0.24349708 | 0.04523552 |
| ENSG00000133985 | *TTC9* | 3.531508557 | 0.012858941 | 0.05453478 | -0.041675839 | 0.999933887 | -0.536905941 | -0.850363693 | 0.313457751 | 0.04523552 |
| ENSG00000174796 | *THAP6* | 3.063657549 | -0.040913657 | 0.038711838 | -0.079625495 | 0.999933887 | -0.152312074 | 0.179915692 | -0.332227766 | 0.045409382 |
| ENSG00000138592 | *USP8* | 5.58333298 | -0.041939716 | -0.124472991 | 0.082533276 | 0.999933887 | -0.172056626 | 0.233511833 | -0.405568459 | 0.045409382 |
| ENSG00000141027 | *NCOR1* | 6.306823779 | 0.003800716 | -0.023901953 | 0.027702669 | 0.999933887 | -0.221060688 | -0.43319951 | 0.212138821 | 0.045409382 |
| ENSG00000139154 | *AEBP2* | 3.781429483 | 0.000984588 | -0.06767354 | 0.068658127 | 0.999933887 | 0.203920629 | 0.499897608 | -0.295976979 | 0.045409382 |
| ENSG00000168884 | *TNIP2* | 5.094220714 | 0.024980721 | -0.005825097 | 0.030805818 | 0.999933887 | 0.668452453 | 0.950538717 | -0.282086264 | 0.045416465 |
| ENSG00000084734 | *GCKR* | -2.911788345 | -0.146755024 | 0.23942716 | -0.386182184 | 0.999933887 | 2.759667504 | 4.397955088 | -1.638287584 | 0.045545305 |
| ENSG00000101940 | *WDR13* | 4.576929576 | 0.02969057 | 0.049295147 | -0.019604578 | 0.999933887 | -0.112924411 | -0.356417312 | 0.243492901 | 0.045654959 |
| ENSG00000243273 | *AC020636.1* | 0.164783832 | 0.22429215 | -0.030373179 | 0.254665328 | 0.999933887 | 0.983021256 | 1.761370522 | -0.778349267 | 0.045691083 |
| ENSG00000237513 | *AC007384.1* | 2.85878649 | -0.079167391 | 0.052961528 | -0.132128919 | 0.999933887 | 2.156506838 | 2.77645028 | -0.619943442 | 0.045691083 |
| ENSG00000114745 | *GORASP1* | 4.452039726 | 0.010462471 | 0.065999486 | -0.055537015 | 0.999933887 | -0.289175568 | -0.542424442 | 0.253248873 | 0.045691083 |
| ENSG00000172232 | *AZU1* | -0.079639871 | -0.006215748 | 0.184764804 | -0.190980552 | 0.999933887 | -0.233399971 | 0.461469018 | -0.69486899 | 0.045691083 |
| ENSG00000164941 | *INTS8* | 3.891068676 | -0.032607811 | -0.018059469 | -0.014548342 | 0.999933887 | -0.375624917 | -0.653090398 | 0.277465481 | 0.045722637 |
| ENSG00000141179 | *PCTP* | 2.525486715 | -0.018255938 | -0.034063271 | 0.015807333 | 0.999933887 | -0.037755691 | 0.272206449 | -0.30996214 | 0.04579357 |
| ENSG00000250929 | *LINC01181* | -1.567563909 | -0.473363029 | -0.539547284 | 0.066184255 | 0.999933887 | 1.620705892 | 2.33712723 | -0.716421337 | 0.046029611 |
| ENSG00000168209 | *DDIT4* | 4.651289441 | 0.276332987 | 0.387251737 | -0.11091875 | 0.999933887 | 1.499252784 | 1.905457821 | -0.406205037 | 0.046057631 |
| ENSG00000181896 | *ZNF101* | 3.415425365 | -0.048003477 | -0.042992133 | -0.005011344 | 0.999933887 | -0.084667301 | -0.386560687 | 0.301893387 | 0.046057631 |
| ENSG00000119242 | *CCDC92* | 3.359938415 | 0.014865038 | -0.10268675 | 0.117551788 | 0.999933887 | -0.130974617 | -0.387910527 | 0.25693591 | 0.046057631 |
| ENSG00000217128 | *FNIP1* | 7.978963376 | 0.000404566 | 0.0141069 | -0.013702334 | 0.999933887 | -0.117374474 | 0.145620573 | -0.262995047 | 0.046057631 |
| ENSG00000141574 | *SECTM1* | 7.663564564 | 0.136716431 | 0.039525343 | 0.097191088 | 0.999933887 | -0.691202933 | -1.08507282 | 0.393869887 | 0.046161368 |
| ENSG00000105726 | *ATP13A1* | 4.5918479 | 0.06036476 | 0.025646671 | 0.034718089 | 0.999933887 | 0.020292637 | -0.169817379 | 0.190110017 | 0.046161368 |
| ENSG00000117091 | *CD48* | 7.436993403 | -0.019378942 | -0.061265795 | 0.041886853 | 0.999933887 | 0.437884948 | 0.645204974 | -0.207320026 | 0.046161368 |
| ENSG00000121316 | *PLBD1* | 5.734694769 | -0.014429214 | -0.028607408 | 0.014178194 | 0.999933887 | -0.207493931 | -0.54143926 | 0.333945329 | 0.046161368 |
| ENSG00000265241 | *RBM8A* | 4.61866475 | 0.009770268 | -0.006122555 | 0.015892823 | 0.999933887 | -0.414117073 | -0.685696284 | 0.271579211 | 0.046161368 |
| ENSG00000050130 | *JKAMP* | 3.34039957 | -0.003203252 | 0.01077817 | -0.013981422 | 0.999933887 | -0.9136072 | -1.282318712 | 0.368711512 | 0.046219461 |
| ENSG00000076382 | *SPAG5* | 2.646110792 | 0.070356377 | -0.126139293 | 0.196495671 | 0.999933887 | 0.823931888 | 1.242880229 | -0.418948341 | 0.046328859 |
| ENSG00000106603 | *COA1* | 4.030076828 | -0.005560436 | -0.200339895 | 0.194779459 | 0.999933887 | -0.150620176 | -0.461159747 | 0.310539572 | 0.046328859 |
| ENSG00000164754 | *RAD21* | 6.928248211 | -0.093396345 | -0.060867152 | -0.032529193 | 0.999933887 | -0.099785057 | 0.165464313 | -0.26524937 | 0.046417923 |
| ENSG00000126107 | *HECTD3* | 4.168072975 | 0.007213387 | 0.037939757 | -0.03072637 | 0.999933887 | -0.28112436 | -0.49806896 | 0.2169446 | 0.046442352 |
| ENSG00000254995 | *STX16-NPEPL* | 1.85243559 | 0.014479812 | 0.073049586 | -0.058569774 | 0.999933887 | 0.183439908 | -0.174461389 | 0.357901297 | 0.046450375 |
| ENSG00000177427 | *MIEF2* | 1.679408453 | 0.126944608 | 0.03817168 | 0.088772928 | 0.999933887 | -0.605518021 | -1.125496586 | 0.519978565 | 0.046636977 |
| ENSG00000123106 | *CCDC91* | 2.052130892 | -0.137623635 | -0.21681568 | 0.079192045 | 0.999933887 | -0.196389811 | 0.228442709 | -0.42483252 | 0.04668796 |
| ENSG00000136888 | *ATP6V1G1* | 7.207268603 | 0.001241401 | -0.000874143 | 0.002115544 | 0.999933887 | 0.32706051 | 0.485293978 | -0.158233468 | 0.04668796 |
| ENSG00000100075 | *SLC25A1* | 3.027263624 | 0.043994409 | 0.016275867 | 0.027718543 | 0.999933887 | 0.083476689 | -0.281116536 | 0.364593225 | 0.046708851 |
| ENSG00000176018 | *LYSMD3* | 4.615081479 | -0.046420553 | -0.214218955 | 0.167798402 | 0.999933887 | -0.401177927 | 0.012471812 | -0.413649738 | 0.046708851 |
| ENSG00000131042 | *LILRB2* | 7.738276012 | 0.017401226 | 0.007478702 | 0.009922524 | 0.999933887 | -0.268551746 | -0.562834962 | 0.294283216 | 0.046881621 |
| ENSG00000101577 | *LPIN2* | 7.608265381 | -0.088332408 | 0.005167674 | -0.093500082 | 0.999933887 | -1.139101814 | -1.45386169 | 0.314759876 | 0.046922633 |
| ENSG00000185515 | *BRCC3* | 2.084378555 | 0.034021499 | -0.056362361 | 0.09038386 | 0.999933887 | -0.553100929 | -1.053278791 | 0.500177862 | 0.046922633 |
| ENSG00000125741 | *OPA3* | 4.215085021 | 0.0215921 | 0.179975956 | -0.158383857 | 0.999933887 | 0.689277312 | 0.962497215 | -0.273219903 | 0.046985169 |
| ENSG00000196911 | *KPNA5* | 2.095440248 | -0.067474505 | -0.193060316 | 0.125585811 | 0.999933887 | -0.326036639 | 0.176978449 | -0.503015089 | 0.046988989 |
| ENSG00000037749 | *MFAP3* | 3.127671731 | -0.124800619 | 0.011401095 | -0.136201713 | 0.999933887 | -0.259645764 | -0.010018542 | -0.249627222 | 0.047061096 |
| ENSG00000126226 | *PCID2* | 3.342197638 | -0.020654666 | -0.007443942 | -0.013210724 | 0.999933887 | 0.23877773 | 0.520322009 | -0.281544279 | 0.047061096 |

| | | | | | | | | | | |
|---|---|---|---|---|---|---|---|---|---|---|
| ENSG00000162144 | *CYB561A3* | 5.300086359 | 0.001255323 | 0.024489747 | -0.023234424 | 0.999933887 | 0.916961119 | 1.116526792 | -0.199565674 | 0.047061096 |
| ENSG00000174004 | *NRROS* | 5.067230672 | -0.1051133 | -0.093628305 | -0.011484994 | 0.999933887 | -0.972843666 | -1.362366133 | 0.389522468 | 0.047104943 |
| ENSG00000238260 | *AL513320.1* | -0.518662995 | -0.155600461 | -0.166374774 | 0.010774314 | 0.999933887 | -1.114105587 | -2.045305257 | 0.93119967 | 0.04716322 |
| ENSG00000141867 | *BRD4* | 8.377072076 | -0.008693065 | 0.005296715 | -0.01398978 | 0.999933887 | 0.133205577 | 0.273760347 | -0.14055477 | 0.04716322 |
| ENSG00000103966 | *EHD4* | 4.74215198 | 0.020607194 | -0.008227129 | 0.028834323 | 0.999933887 | 0.135043417 | 0.391523254 | -0.256479837 | 0.047177372 |
| ENSG00000164323 | *CFAP97* | 2.10949713 | -0.10004251 | -0.050057124 | -0.049985386 | 0.999933887 | -0.065950307 | 0.381681306 | -0.447631613 | 0.047254931 |
| ENSG00000132471 | *WBP2* | 9.519211088 | 0.016723075 | 0.021244353 | -0.004521279 | 0.999933887 | 0.473790041 | 0.632864451 | -0.15907441 | 0.04728428 |
| ENSG00000223959 | *AFG3L1P* | 2.795168548 | -0.007012911 | 0.008094993 | -0.015107904 | 0.999933887 | 0.012610819 | -0.26896092 | 0.281571739 | 0.04728428 |
| ENSG00000077327 | *SPAG6* | -0.367941606 | 0.148451158 | 0.07993435 | 0.068516809 | 0.999933887 | 0.301953033 | 1.054128788 | -0.752175755 | 0.047379211 |
| ENSG00000100426 | *ZBED4* | 3.679444776 | 0.007242534 | 0.053791885 | -0.046549351 | 0.999933887 | -0.062379338 | -0.329830589 | 0.267451251 | 0.04758416 |
| ENSG00000228340 | *MIR646HG* | 3.623064964 | 0.003654146 | 0.019739623 | -0.016085477 | 0.999933887 | 0.513434589 | 0.843111811 | -0.329677223 | 0.047706502 |
| ENSG00000198727 | *MT-CYB* | 9.151402594 | 0.080402483 | -0.035546663 | 0.115949146 | 0.999933887 | -0.009882913 | 0.344440332 | -0.354323245 | 0.047803569 |
| ENSG00000003056 | *M6PR* | 7.474442343 | 0.031374118 | 0.017348409 | 0.014025709 | 0.999933887 | 0.160747357 | 0.327821935 | -0.167074579 | 0.047803569 |
| ENSG00000086758 | *HUWE1* | 7.16821569 | -0.024681082 | 0.004142226 | -0.028823308 | 0.999933887 | 0.162985201 | 0.331707766 | -0.168722565 | 0.047803569 |
| ENSG00000111652 | *COPS7A* | 3.74238591 | -0.003943742 | -0.073728146 | 0.069784403 | 0.999933887 | -0.322723572 | -0.530340754 | 0.207617183 | 0.047803569 |
| ENSG00000169733 | *RFNG* | 3.139860979 | 0.00346737 | 0.05016977 | -0.0467024 | 0.999933887 | -0.128773767 | -0.425910248 | 0.297136481 | 0.048180143 |
| ENSG00000198846 | *TOX* | 2.041559218 | 0.012258472 | -0.128935747 | 0.141194219 | 0.999933887 | -0.102684074 | 0.270618712 | -0.373302786 | 0.048222927 |
| ENSG00000148175 | *STOM* | 5.059972923 | -0.013161915 | -0.005111025 | -0.00805089 | 0.999933887 | 0.119374824 | 0.32559125 | -0.206216425 | 0.048264664 |
| ENSG00000108771 | *DHX58* | 3.663925604 | 0.083633842 | 0.012780102 | 0.070853741 | 0.999933887 | 0.027460193 | -0.261918154 | 0.289378346 | 0.048334771 |
| ENSG00000173334 | *TRIB1* | 10.381047556 | 0.04976718 | 0.063750853 | -0.013983673 | 0.999933887 | -0.051617686 | 0.170612473 | -0.222230159 | 0.048342036 |
| ENSG00000140090 | *SLC24A4* | 2.943963967 | -0.027411589 | -0.085769686 | 0.058358096 | 0.999933887 | -0.354882377 | -0.754845 | 0.399962624 | 0.048342036 |
| ENSG00000168883 | *USP39* | 3.89166192 | 0.018388869 | -0.077723961 | 0.09611283 | 0.999933887 | -0.30556877 | -0.582218458 | 0.276649688 | 0.048342036 |
| ENSG00000164985 | *PSIP1* | 4.511276463 | 0.030709915 | -0.026855017 | 0.057564931 | 0.999933887 | -0.38616712 | -0.757169656 | 0.371002535 | 0.048361051 |
| ENSG00000153006 | *SREK1IP1* | 2.458937929 | 0.011679292 | -0.117089309 | 0.128768601 | 0.999933887 | -0.38805638 | 0.116791581 | -0.504847961 | 0.048361051 |
| ENSG00000111897 | *SERINC1* | 8.567560347 | -0.053689999 | -0.032183098 | -0.021506901 | 0.999933887 | -0.008597734 | 0.197705239 | -0.206302973 | 0.0484995 |
| ENSG00000108587 | *GOSR1* | 4.328880127 | 0.010204383 | 0.061448473 | -0.051244091 | 0.999933887 | 0.274218705 | 0.473088761 | -0.198870055 | 0.048534931 |
| ENSG00000181274 | *FRAT2* | 7.225521319 | -0.250853326 | -0.290088326 | 0.039235 | 0.999933887 | -1.874752034 | -2.327573639 | 0.452821605 | 0.048778683 |
| ENSG00000100908 | *EMC9* | 1.414606591 | -0.294786903 | 0.13568295 | -0.430469852 | 0.944094537 | 0.097049108 | -0.356419312 | 0.45346842 | 0.048892328 |
| ENSG00000065548 | *ZC3H15* | 5.121514541 | -0.104502725 | -0.129278751 | 0.024776026 | 0.999933887 | -0.172334128 | 0.10829455 | -0.280628677 | 0.048892328 |
| ENSG00000130517 | *PGPEP1* | 5.468642327 | -0.088236159 | -0.046572068 | -0.041664091 | 0.999933887 | -0.799122887 | -1.081587123 | 0.282464236 | 0.048892328 |
| ENSG00000117523 | *PRRC2C* | 8.51473349 | -0.0738817 | -0.11336149 | 0.03947979 | 0.999933887 | 0.32082656 | 0.673625121 | -0.352798561 | 0.048892328 |
| ENSG00000106268 | *NUDT1* | 1.729920579 | 0.105610125 | 0.047666714 | 0.057943412 | 0.999933887 | 0.573661963 | 0.98734053 | -0.413678568 | 0.048918155 |
| ENSG00000169957 | *ZNF768* | 1.735534689 | 0.02547368 | 0.08021655 | -0.05474287 | 0.999933887 | -0.944592976 | -1.421630438 | 0.477037463 | 0.048918155 |
| ENSG00000168310 | *IRF2* | 7.560315469 | 0.013650505 | -0.019022695 | 0.0326732 | 0.999933887 | 0.040226599 | 0.221542504 | -0.181315905 | 0.048919709 |
| ENSG00000113811 | *SELENOK* | 5.953729116 | 0.096384056 | 0.121898405 | -0.025514349 | 0.999933887 | 0.716452088 | 0.995644513 | -0.279192425 | 0.048927348 |
| ENSG00000196865 | *NHLRC2* | 2.352462155 | 0.045495273 | -0.059165458 | 0.104660732 | 0.999933887 | -0.215180992 | 0.182001237 | -0.397182229 | 0.048927348 |
| ENSG00000164266 | *SPINK1* | 2.573579337 | 0.097167712 | 0.192352218 | -0.095184505 | 0.999933887 | 0.505411904 | 0.862924989 | -0.357513085 | 0.049028853 |
| ENSG00000136040 | *PLXNC1* | 9.420029698 | -0.009451468 | -0.005930145 | -0.003521324 | 0.999933887 | 0.424671222 | 0.620311858 | -0.195640636 | 0.049083263 |
| ENSG00000140262 | *TCF12* | 3.0136615 | -0.006195564 | -0.159994643 | 0.153799078 | 0.999933887 | -0.138388999 | 0.238762716 | -0.377151716 | 0.049132456 |
| ENSG00000183486 | *MX2* | 8.055989464 | 0.015507967 | 0.023085139 | -0.007577172 | 0.999933887 | -0.123696007 | -0.297553039 | 0.173857032 | 0.049228998 |
| ENSG00000149657 | *LSM14B* | 5.158423979 | -0.007166976 | 0.014379013 | -0.02154599 | 0.999933887 | -0.876733338 | -1.139237737 | 0.262504399 | 0.049557793 |
| ENSG00000223901 | *AP001469.1* | -0.973255627 | 0.1723289 | -0.092751104 | 0.265080004 | 0.999933887 | 2.088912395 | 3.186518783 | -1.097606389 | 0.049581726 |
| ENSG00000196455 | *PIK3R4* | 3.90388956 | -0.049304072 | -0.016121777 | -0.033182296 | 0.999933887 | 0.109057573 | 0.340943536 | -0.231885963 | 0.049712897 |
| ENSG00000135185 | *TMEM243* | 2.761260828 | -0.108401287 | 0.028343802 | -0.136745089 | 0.999933887 | -0.027491296 | -0.280743954 | 0.253252658 | 0.049783801 |
| ENSG00000153922 | *CHD1* | 7.910178756 | -0.062483515 | -0.177849775 | 0.11536626 | 0.999933887 | -0.108444358 | 0.430309701 | -0.538754059 | 0.049820842 |
| ENSG00000128311 | *TST* | 1.377177948 | -0.067935233 | 0.087280535 | -0.155215768 | 0.999933887 | -0.813111716 | -1.54479975 | 0.731688035 | 0.049897614 |
| ENSG00000117036 | *ETV3* | 5.230587146 | 0.008069824 | -0.018645102 | 0.026714926 | 0.999933887 | 0.153144422 | 0.333471466 | -0.180327044 | 0.049897614 |
| ENSG00000128829 | *EIF2AK4* | 2.300442152 | -0.005157486 | -0.083522027 | 0.078364541 | 0.999933887 | -0.173496608 | -0.534922667 | 0.361426059 | 0.049897614 |
| ENSG00000174840 | *PDE12* | 4.345892594 | 0.041210417 | -0.056035462 | 0.097245879 | 0.999933887 | 0.264800424 | 0.482852959 | -0.218052535 | 0.049990821 |
| ENSG00000187608 | *ISG15* | 6.241678647 | 0.091983984 | 0.04396306 | 0.048020924 | 0.999933887 | -0.123894872 | -0.447884564 | 0.323989692 | 0.050039717 |
| ENSG00000109956 | *B3GAT1* | 1.774042872 | -0.013763835 | 0.065645116 | -0.07940895 | 0.999933887 | 0.22969448 | -0.137006594 | 0.366701074 | 0.05008292 |
| ENSG00000117450 | *PRDX1* | 3.801123387 | -0.07470135 | 0.095382215 | -0.170083565 | 0.999933887 | 0.381484651 | 0.643592853 | -0.262108202 | 0.050255923 |
| ENSG00000287630 | *AC011370.1* | -0.30648871 | 0.196848111 | 0.60011255 | -0.403264439 | 0.999933887 | -0.37449303 | 0.268196733 | -0.642689763 | 0.050255923 |
| ENSG00000184203 | *PPP1R2* | 6.427636436 | -0.036945867 | -0.011554925 | -0.025390942 | 0.999933887 | -0.04555947 | 0.101438401 | -0.146997871 | 0.050352977 |
| ENSG00000244945 | *RUFY1-AS1* | 0.724580004 | 0.012259057 | 0.080169622 | -0.067910565 | 0.999933887 | 0.105089102 | 0.627265703 | -0.5221766 | 0.050352977 |
| ENSG00000142875 | *PRKACB* | 3.645933806 | -0.121887728 | -0.215691563 | 0.093803835 | 0.999933887 | -0.187931513 | 0.256189763 | -0.444121276 | 0.050427846 |
| ENSG00000198951 | *NAGA* | 2.292099656 | -0.031050967 | -8.7561e-05 | -0.030963406 | 0.999933887 | -0.292593946 | -0.648706921 | 0.356112975 | 0.050427846 |
| ENSG00000143554 | *SLC27A3* | 3.441086698 | 0.101911894 | 0.013684606 | 0.088227288 | 0.999933887 | -0.196339363 | -0.529690366 | 0.333351004 | 0.050534741 |
| ENSG00000182957 | *SPATA13* | 6.424455962 | -0.054223613 | -0.021679194 | -0.032544418 | 0.999933887 | 0.148491697 | 0.347010274 | -0.198518577 | 0.050613572 |
| ENSG00000162976 | *SLC66A3* | 2.598453744 | -0.006578875 | -0.100823832 | 0.094244957 | 0.999933887 | -0.131632928 | -0.520417027 | 0.388784099 | 0.050654531 |
| ENSG00000176783 | *RUFY1* | 5.331423743 | 0.003799377 | 0.042507572 | -0.038708195 | 0.999933887 | 0.114839129 | 0.303305107 | -0.188465979 | 0.050732996 |
| ENSG00000241839 | *PLEKHO2* | 9.493031998 | -0.036100647 | -0.115169344 | 0.079068697 | 0.999933887 | 0.485994078 | 0.697272911 | -0.211278833 | 0.050853094 |
| ENSG00000139826 | *ABHD13* | 4.702317779 | -0.023257989 | -0.098394447 | 0.075136457 | 0.999933887 | 0.009060852 | 0.260019349 | -0.250958497 | 0.0508586 |
| ENSG00000115568 | *ZNF142* | 3.921087696 | 0.072585848 | 0.029855992 | 0.042729856 | 0.999933887 | 0.329975618 | 0.599845096 | -0.269869478 | 0.050948782 |
| ENSG00000145414 | *NAF1* | 2.015618337 | -0.01446949 | -0.075673363 | 0.061203873 | 0.999933887 | 0.831917113 | 1.357102756 | -0.525185644 | 0.051018243 |
| ENSG00000125970 | *RALY* | 7.232923295 | 0.044485067 | 0.02483719 | 0.019647878 | 0.999933887 | -0.199825469 | -0.400939252 | 0.201113784 | 0.05103601 |
| ENSG00000198160 | *MIER1* | 6.326949891 | -0.050829477 | -0.124412764 | 0.073583287 | 0.999933887 | -0.436129983 | -0.122336583 | -0.3137934 | 0.05103601 |
| ENSG00000173473 | *SMARCC1* | 5.302324921 | -0.012126765 | 0.010011522 | -0.022138287 | 0.999933887 | -0.086855972 | -0.255285421 | 0.168429449 | 0.051200849 |
| ENSG00000152359 | *POC5* | 1.157453994 | -0.036636595 | -0.325697365 | 0.28906077 | 0.999933887 | 0.560497348 | 1.064507062 | -0.504009714 | 0.051264542 |
| ENSG00000162702 | *ZNF281* | 6.661315861 | -0.101038568 | -0.123673386 | 0.022634818 | 0.999933887 | 0.336542022 | 0.552312887 | -0.215770866 | 0.051526891 |
| ENSG00000176371 | *ZSCAN2* | -0.267324889 | -0.056214277 | 0.172249915 | -0.228464192 | 0.999933887 | -0.004578705 | -0.59955887 | 0.594980166 | 0.051569377 |
| ENSG00000158869 | *FCER1G* | 8.704431339 | -0.013404366 | 0.0005016 | -0.013905966 | 0.999933887 | 0.63233732 | 0.835128183 | -0.202790863 | 0.051611659 |
| ENSG00000104419 | *NDRG1* | 7.085507121 | 0.00283816 | 0.045228203 | -0.042390043 | 0.999933887 | 0.077282016 | 0.293418743 | -0.216136727 | 0.051611827 |
| ENSG00000144655 | *CSRNP1* | 10.034656736 | 0.243753309 | 0.198314262 | 0.045439047 | 0.999933887 | 0.927867988 | 1.2112019 | -0.283333912 | 0.051664385 |
| ENSG00000071189 | *SNX13* | 5.762580947 | -0.009619321 | -0.071980059 | 0.062360738 | 0.999933887 | -0.63747624 | -0.240873017 | -0.396603222 | 0.051666486 |
| ENSG00000132294 | *EFR3A* | 5.843832749 | -0.073740485 | -0.102972208 | 0.029231724 | 0.999933887 | -0.303434278 | 0.052178702 | -0.35561298 | 0.051695458 |
| ENSG00000197329 | *PELI1* | 9.664967774 | -0.00908764 | -0.024756146 | 0.015668507 | 0.999933887 | -0.058714516 | 0.205546118 | -0.264260634 | 0.051695458 |
| ENSG00000287644 | *AC080112.4* | 2.390280989 | 0.11214846 | 0.097714841 | 0.014433619 | 0.999933887 | 0.407975423 | 0.860204904 | -0.452229481 | 0.051756348 |
| ENSG00000224596 | *ZMIZ1-AS1* | 0.792868125 | 0.121049838 | -0.095637207 | 0.216687045 | 0.999933887 | 0.507006502 | 0.97157474 | -0.464568238 | 0.051863609 |
| ENSG00000114933 | *INO80D* | 5.182191741 | 0.022002309 | -0.106668607 | 0.128670916 | 0.999933887 | -0.178007261 | 0.346827791 | -0.524835052 | 0.051863609 |
| ENSG00000116668 | *SWT1* | 4.394725726 | -0.033630138 | -0.086248147 | 0.052618009 | 0.999933887 | -0.084520246 | 0.343934865 | -0.428455111 | 0.051873362 |
| ENSG00000189430 | *NCR1* | 1.590654029 | -0.024755405 | 0.019862262 | -0.044617667 | 0.999933887 | 1.042405698 | 1.497153665 | -0.454747968 | 0.052068414 |
| ENSG00000119878 | *CRIPT* | 3.505607037 | -0.11961115 | -0.057645038 | -0.061966112 | 0.999933887 | -0.277826635 | -0.534431754 | 0.256605119 | 0.052095091 |
| ENSG00000008018 | *PSMB1* | 5.870794534 | -0.040730408 | -0.074131927 | 0.033401519 | 0.999933887 | 0.101978825 | 0.273815691 | -0.171836867 | 0.052176045 |
| ENSG00000198833 | *UBE2J1* | 6.552147375 | -0.005425091 | -0.022021572 | 0.016596482 | 0.999933887 | -0.305223538 | -0.523799321 | 0.218575783 | 0.052186915 |
| ENSG00000286276 | *AL390066.2* | 0.159607746 | 0.224919808 | 0.22360431 | 0.001315499 | 0.999933887 | 0.935287719 | 1.378691987 | -0.443404268 | 0.052233773 |
| ENSG00000151229 | *SLC2A13* | 1.720252382 | -0.029676227 | -0.083840155 | 0.054163928 | 0.999933887 | -1.065434428 | -1.456126555 | 0.390692127 | 0.052233773 |
| ENSG00000026103 | *FAS* | 6.317371794 | -0.07413799 | -0.086911565 | 0.012773575 | 0.999933887 | 0.90493047 | 1.21396481 | -0.30903434 | 0.052354556 |
| ENSG00000104164 | *BLOC1S6* | 6.384361551 | -0.021615165 | -0.007991319 | -0.013623845 | 0.999933887 | 0.033860504 | 0.332066161 | -0.298205657 | 0.052413982 |
| ENSG00000080189 | *SLC35C2* | 5.629643629 | 0.011139793 | 0.009264147 | 0.001875647 | 0.999933887 | 0.664663818 | 0.869144906 | -0.204481088 | 0.052413982 |
| ENSG00000155093 | *PTPRN2* | 3.59733287 | -0.066031813 | -0.036689721 | -0.029342092 | 0.999933887 | -0.447469504 | -0.765658105 | 0.318188601 | 0.052432139 |
| ENSG00000241163 | *LINC00877* | 6.377744484 | 0.116579694 | 0.083714081 | 0.032865613 | 0.999933887 | 1.184126084 | 1.483113201 | -0.298987117 | 0.052477597 |
| ENSG00000125735 | *TNFSF14* | 8.594179704 | 0.069622742 | 0.049294359 | 0.020328384 | 0.999933887 | 0.461058597 | 0.716204328 | -0.255145731 | 0.052477597 |
| ENSG00000144791 | *LIMD1* | 1.87943653 | -0.056292128 | -0.050423317 | -0.00586881 | 0.999933887 | 0.02458704 | -0.318898425 | 0.343485465 | 0.052477597 |
| ENSG00000214087 | *ARL16* | 2.344258265 | -0.024702063 | -0.058239965 | 0.033537902 | 0.999933887 | 0.178158494 | -0.097200387 | 0.275358881 | 0.052477597 |
| ENSG00000124120 | *TTPAL* | 7.664524649 | -0.075111077 | -0.04654091 | -0.028570167 | 0.999933887 | -0.100831888 | 0.114313275 | -0.215145163 | 0.052501405 |
| ENSG00000178691 | *SUZ12* | 5.020785406 | -0.003706997 | -0.132976519 | 0.129269522 | 0.999933887 | -0.243247108 | 0.175793796 | -0.419040903 | 0.052569827 |
| ENSG00000130726 | *TRIM28* | 6.491569653 | 0.007614903 | 0.064951619 | -0.057336717 | 0.999933887 | 0.0489872 | -0.12374605 | 0.172733251 | 0.052840694 |
| ENSG00000141504 | *SAT2* | 3.454582581 | 0.037800853 | -0.028893386 | 0.066694239 | 0.999933887 | -0.295207045 | -0.564976642 | 0.269769597 | 0.052854784 |
| ENSG00000095209 | *TMEM38B* | 3.094639042 | 0.013546794 | -0.048230262 | 0.061777055 | 0.999933887 | 0.759232393 | 1.063750535 | -0.304518141 | 0.052854784 |
| ENSG00000172062 | *SMN1* | 0.450256412 | -0.123857331 | 0.22704352 | -0.350900851 | 0.999933887 | -0.053545771 | 0.535605071 | -0.589150842 | 0.052900039 |
| ENSG00000254837 | *AP001372.2* | -1.219411234 | 0.203992354 | 0.006857003 | 0.197135351 | 0.999933887 | 1.645326416 | 2.41004182 | -0.764715405 | 0.05305383 |
| ENSG00000115109 | *EPB41L5* | 1.222015092 | 0.042964439 | -0.062065295 | 0.105029734 | 0.999933887 | 0.209845248 | 0.785834384 | -0.575989136 | 0.05305383 |
| ENSG00000196205 | *EEF1A1P5* | 2.598757535 | -0.036607184 | -0.305332335 | 0.26872515 | 0.999933887 | 0.264120373 | 0.641421469 | -0.377301096 | 0.05305383 |
| ENSG00000101574 | *METTL4* | 2.568911343 | 0.033929178 | -0.069676443 | 0.103605621 | 0.999933887 | -0.736535693 | -1.090459623 | 0.353923931 | 0.053105871 |

| | | | | | | | | | | |
|---|---|---|---|---|---|---|---|---|---|---|
| ENSG00000151849 | *CENPJ* | 0.652631864 | 0.00256264 | -0.101060109 | 0.103622749 | 0.999933887 | -0.276201233 | 0.299297092 | -0.575498325 | 0.05310712 |
| ENSG00000182087 | *TMEM259* | 7.577772022 | 0.076335158 | 0.105522884 | -0.029187726 | 0.999933887 | 0.042463853 | -0.16092943 | 0.203393283 | 0.053119877 |
| ENSG00000160959 | *LRRC14* | 2.771841464 | 0.061822032 | 0.060381635 | 0.001440397 | 0.999933887 | 0.068765002 | -0.256058421 | 0.324823423 | 0.053119877 |
| ENSG00000198689 | *SLC9A6* | 2.975293095 | -0.068659154 | 0.019017174 | -0.087676328 | 0.999933887 | -0.208356773 | -0.57808077 | 0.369723997 | 0.053194228 |
| ENSG00000075240 | *GRAMD4* | 3.054086348 | -0.046450065 | 0.107880258 | -0.154330323 | 0.999933887 | -0.152027892 | -0.456635304 | 0.304607411 | 0.053194228 |
| ENSG00000147457 | *CHMP7* | 4.403693108 | -0.024283274 | 0.048481323 | -0.072764597 | 0.999933887 | -0.115079056 | -0.315150384 | 0.200071328 | 0.053194228 |
| ENSG00000198964 | *SGMS1* | 4.401766027 | 0.021939439 | -0.030475329 | 0.052414768 | 0.999933887 | 0.303228557 | 0.535118787 | -0.23189023 | 0.053194228 |
| ENSG00000174373 | *RALGAPA1* | 3.650137724 | -0.019080613 | -0.094252732 | 0.075172119 | 0.999933887 | 0.143280552 | 0.511937293 | -0.368656741 | 0.053217547 |
| ENSG00000100092 | *SH3BP1* | 4.651959345 | 0.067552662 | 0.08645631 | -0.018903649 | 0.999933887 | -0.246452649 | -0.458441315 | 0.211988666 | 0.053305413 |
| ENSG00000140416 | *TPM1* | 1.006970665 | -0.069148709 | -0.048323848 | -0.020824861 | 0.999933887 | -0.574855332 | -1.019603793 | 0.444748462 | 0.053319898 |
| ENSG00000144554 | *FANCD2* | 3.484704765 | -0.029917277 | -0.139049911 | 0.109132634 | 0.999933887 | 0.244955462 | 0.498078461 | -0.253123 | 0.053783269 |
| ENSG00000171316 | *CHD7* | 6.423295397 | -0.054083891 | -0.06876365 | 0.01467976 | 0.999933887 | -0.881012875 | -1.142051007 | 0.261038132 | 0.053915951 |
| ENSG00000071626 | *DAZAP1* | 5.922739242 | 0.010182218 | 0.020189776 | -0.010007558 | 0.999933887 | -0.226677598 | -0.384206973 | 0.157529375 | 0.053948756 |
| ENSG00000255857 | *PXN-AS1* | 1.406685792 | 0.11945469 | 0.035868108 | 0.083586583 | 0.999933887 | -0.501795201 | -1.032623874 | 0.530828672 | 0.054067939 |
| ENSG00000168615 | *ADAM9* | 5.655770341 | -0.026468694 | -0.096707057 | 0.070238363 | 0.999933887 | -0.442232311 | -0.129525751 | -0.31270656 | 0.054277053 |
| ENSG00000164163 | *ABCE1* | 2.882033722 | -0.077009682 | -0.154205937 | 0.077196255 | 0.999933887 | 0.006153135 | 0.373691648 | -0.367538513 | 0.054330591 |
| ENSG00000178694 | *NSUN3* | 3.317965317 | -0.098904387 | 0.039728827 | -0.138633213 | 0.999933887 | -0.135302912 | 0.100352415 | -0.235655327 | 0.054513498 |
| ENSG00000175782 | *SLC35E3* | 4.936154474 | -0.065306985 | 0.0143419 | -0.079648885 | 0.999933887 | 0.330519829 | 0.528447984 | -0.197928156 | 0.054513498 |
| ENSG00000168944 | *CEP120* | 3.463259953 | 0.067512717 | 0.027160596 | 0.040352121 | 0.999933887 | -0.138418733 | 0.288404746 | -0.426823479 | 0.054513498 |
| ENSG00000105849 | *POLR1F* | 3.311850866 | -0.101532351 | -0.182794359 | 0.081262008 | 0.999933887 | 0.780337456 | 1.264345698 | -0.484008243 | 0.054516861 |
| ENSG00000064999 | *ANKS1A* | 4.483797205 | 0.003460537 | 0.04228548 | -0.038824943 | 0.999933887 | -0.200406437 | -0.42152649 | 0.221120052 | 0.054516861 |
| ENSG00000149091 | *DGKZ* | 6.854598258 | 0.002375279 | 0.050531351 | -0.048156072 | 0.999933887 | -0.172165501 | -0.363649387 | 0.191483886 | 0.054516861 |
| ENSG00000105559 | *PLEKHA4* | -1.04025098 | 0.269203329 | 0.101450252 | 0.167753077 | 0.999933887 | 1.081428147 | 1.859304355 | -0.777876208 | 0.054634561 |
| ENSG00000124772 | *CPNE5* | 1.342462746 | 0.020443506 | 0.026109939 | -0.005666433 | 0.999933887 | 0.825598889 | 1.223025838 | -0.397426949 | 0.054651279 |
| ENSG00000064601 | *CTSA* | 7.662847139 | 0.008250334 | -0.021847132 | 0.030097466 | 0.999933887 | -0.476560481 | -0.654516655 | 0.177956174 | 0.054713018 |
| ENSG00000136536 | *MARCHF7* | 6.573976308 | -0.070911901 | -0.149637756 | 0.078725855 | 0.999933887 | -0.503571124 | -0.183049613 | -0.32052151 | 0.054847274 |
| ENSG00000149418 | *ST14* | 1.498269261 | 0.06452191 | 0.027452508 | 0.037069402 | 0.999933887 | -0.603242911 | -1.104933896 | 0.501690985 | 0.054847274 |
| ENSG00000119673 | *ACOT2* | 2.373118212 | 0.031499187 | 0.207917673 | -0.176418486 | 0.999933887 | 0.644812129 | 0.993994958 | -0.349182829 | 0.054847274 |
| ENSG00000255050 | *AC067930.4* | -0.572477713 | -0.019864737 | 0.107472827 | -0.127337564 | 0.999933887 | -0.611350141 | -1.333107358 | 0.721757218 | 0.054847274 |
| ENSG00000083838 | *ZNF446* | 1.038575445 | 0.007440298 | -0.030390531 | 0.037830828 | 0.999933887 | 0.171161791 | -0.471410048 | 0.642571839 | 0.05492458 |
| ENSG00000130940 | *CASZ1* | 3.843583096 | 0.037857362 | 0.035479255 | 0.002378107 | 0.999933887 | 0.534650804 | 0.843602593 | -0.30895179 | 0.054961631 |
| ENSG00000087053 | *MTMR2* | 1.938353696 | -0.030188435 | 0.024737317 | -0.054925752 | 0.999933887 | 0.222832865 | 0.525326192 | -0.302493327 | 0.054978312 |
| ENSG00000117862 | *TXNDC12* | 2.489721768 | -0.080078895 | -0.033413421 | -0.046665474 | 0.999933887 | -0.066099848 | -0.447164663 | 0.381064815 | 0.055089803 |
| ENSG00000174574 | *AKIRIN1* | 7.269584277 | 0.053764322 | 0.024605071 | 0.029159251 | 0.999933887 | 0.202563667 | 0.383415615 | -0.180851948 | 0.055117093 |
| ENSG00000243968 | *RN7SL402P* | -1.352339622 | -0.259440709 | 0.083714176 | -0.343154885 | 0.999933887 | 2.253106898 | 3.274636765 | -1.021529867 | 0.055203062 |
| ENSG00000155463 | *OXA1L* | 5.506829012 | -0.009843274 | -0.046493573 | 0.0366503 | 0.999933887 | -0.087154481 | -0.265560044 | 0.178405563 | 0.055223615 |
| ENSG00000160111 | *CPAMD8* | -1.025569332 | -0.097868235 | 0.193199092 | -0.291067327 | 0.999933887 | 0.476986476 | 1.556943154 | -1.079956678 | 0.055358491 |
| ENSG00000170909 | *OSCAR* | 5.681153696 | 0.055746294 | -0.057739341 | 0.113485635 | 0.999933887 | 0.248105873 | 0.474160375 | -0.226054502 | 0.055365144 |
| ENSG00000169826 | *CSGALNACT2* | 8.16244016 | -0.036717952 | -0.075705159 | 0.038987206 | 0.999933887 | 0.14042351 | 0.372265503 | -0.231841994 | 0.055365144 |
| ENSG00000137073 | *UBAP2* | 4.233170427 | 0.028844993 | 0.029346735 | -0.000501742 | 0.999933887 | 0.063711072 | 0.251411737 | -0.187700665 | 0.055405514 |
| ENSG00000182185 | *RAD51B* | 1.4116626 | 0.026939146 | -0.009698736 | 0.036637882 | 0.999933887 | 0.054668392 | 0.473886462 | -0.41921807 | 0.055464425 |
| ENSG00000077097 | *TOP2B* | 5.679757567 | -0.049508867 | -0.165953373 | 0.116444506 | 0.999933887 | -0.104952086 | 0.378393376 | -0.483345462 | 0.055482981 |
| ENSG00000171475 | *WIPF2* | 7.313967506 | -0.044797432 | -0.001045281 | -0.043752151 | 0.999933887 | 0.00027506 | 0.149793442 | -0.149518382 | 0.055528601 |
| ENSG00000102103 | *PQBP1* | 4.09068335 | 0.077588681 | 0.033358549 | 0.044230133 | 0.999933887 | 0.643756548 | 0.924546618 | -0.28079007 | 0.055528601 |
| ENSG00000171262 | *FAM98B* | 1.650059192 | -0.051054548 | -0.040997693 | -0.010056855 | 0.999933887 | -0.141828266 | 0.305452723 | -0.447280989 | 0.055528601 |
| ENSG00000077235 | *GTF3C1* | 5.112989462 | 0.002027249 | 0.01699451 | -0.01496726 | 0.999933887 | -0.03680285 | -0.222098238 | 0.185295388 | 0.055528601 |
| ENSG00000128656 | *CHN1* | -0.020622396 | -0.08253555 | -0.084155765 | 0.001620215 | 0.999933887 | -0.148199261 | 0.591692294 | -0.739891555 | 0.055532034 |
| ENSG00000073803 | *MAP3K13* | 3.651710779 | 0.045790684 | 0.039570209 | 0.006220475 | 0.999933887 | 0.599419467 | 0.923730522 | -0.324311055 | 0.055532034 |
| ENSG00000139687 | *RB1* | 4.295975762 | 0.054843032 | -0.030432018 | 0.08527505 | 0.999933887 | -0.401471124 | -0.085963342 | -0.315507782 | 0.055584601 |
| ENSG00000287263 | *AC008875.3* | 4.28471554 | 0.022401968 | 0.014456598 | 0.00794537 | 0.999933887 | -0.111818245 | -0.378698814 | 0.266880569 | 0.055724824 |
| ENSG00000158828 | *PINK1* | 3.831985217 | -0.026826156 | -0.010440398 | -0.016385758 | 0.999933887 | -0.767382041 | -1.083027246 | 0.315645205 | 0.055773375 |
| ENSG00000169592 | *INO80E* | 4.077652121 | 0.050658412 | 0.037056969 | 0.013601443 | 0.999933887 | -0.055850784 | -0.34294899 | 0.287098206 | 0.055789596 |
| ENSG00000148248 | *SURF4* | 6.458532051 | 0.019490333 | 0.002222414 | 0.01726792 | 0.999933887 | -0.227600852 | -0.420369528 | 0.192768676 | 0.055808397 |
| ENSG00000090238 | *YPEL3* | 8.624407456 | 0.008251654 | 0.030405041 | -0.022153387 | 0.999933887 | -0.639444794 | -0.857167501 | 0.217722707 | 0.055808397 |
| ENSG00000162869 | *PPP1R21* | 4.340895246 | -0.13104745 | -0.013891713 | -0.117155737 | 0.999933887 | -0.540449306 | -0.745052675 | 0.204603369 | 0.055916462 |
| ENSG00000166750 | *SLFN5* | 5.07551299 | -0.037350959 | -0.185070607 | 0.147719647 | 0.999933887 | 0.066835766 | 0.464842962 | -0.398007196 | 0.055916462 |
| ENSG00000172594 | *SMPDL3A* | 1.034254371 | -0.142979176 | 0.004516152 | -0.147495327 | 0.999933887 | 0.704185133 | 1.177565086 | -0.473379953 | 0.056233726 |
| ENSG00000198918 | *RPL39* | 4.933699128 | 0.057427201 | 0.00735854 | 0.050068661 | 0.999933887 | 0.065603166 | 0.390708377 | -0.32510521 | 0.056365902 |
| ENSG00000148926 | *ADM* | 7.395216228 | 0.23413581 | 0.259541285 | -0.025405475 | 0.999933887 | 0.920498274 | 1.248923524 | -0.32842525 | 0.056413352 |
| ENSG00000163738 | *MTHFD2L* | 1.982457128 | -0.166630292 | -0.082683314 | -0.083946977 | 0.999933887 | 0.088824149 | 0.455935172 | -0.367111023 | 0.056413352 |
| ENSG00000085117 | *CD82* | 8.463865003 | 0.007617665 | 0.019258567 | -0.011640902 | 0.999933887 | 1.254014903 | 1.525047389 | -0.271032486 | 0.056431498 |
| ENSG00000275238 | *MIR4734* | -0.387867907 | 0.487061001 | 0.363342339 | 0.123718662 | 0.999933887 | 2.269259792 | 2.984181056 | -0.714921264 | 0.05644008 |
| ENSG00000137501 | *SYTL2* | 3.474862502 | -0.112470622 | -0.043797682 | -0.06867294 | 0.999933887 | -0.140998379 | 0.187885476 | -0.328883855 | 0.056640463 |
| ENSG00000154582 | *ELOC* | 4.567270518 | -0.029951847 | -0.000720524 | -0.029231324 | 0.999933887 | -0.10654878 | 0.098117421 | -0.204666201 | 0.056640463 |
| ENSG00000196204 | *RNF216P1* | 2.158779526 | 0.018531307 | 0.016257787 | 0.00227352 | 0.999933887 | 0.007298867 | -0.32757826 | 0.334877127 | 0.056672243 |
| ENSG00000138678 | *GPAT3* | 4.75795477 | -0.109398836 | -0.076300956 | -0.03309788 | 0.999933887 | -1.283046291 | -1.646056388 | 0.363010097 | 0.056754077 |
| ENSG00000133056 | *PIK3C2B* | 4.816532107 | -0.02152344 | -0.066698916 | 0.045175476 | 0.999933887 | -2.521256914 | -3.033679205 | 0.512422291 | 0.056754338 |
| ENSG00000102908 | *NFAT5* | 6.950908662 | -0.009639308 | -0.059577518 | 0.049938211 | 0.999933887 | -0.208126006 | 0.227266491 | -0.435392496 | 0.056943583 |
| ENSG00000203747 | *FCGR3A* | 7.701159835 | -0.063133178 | -0.121541177 | 0.058408 | 0.999933887 | -0.453908537 | -0.743933777 | 0.290025241 | 0.057003094 |
| ENSG00000164758 | *MED30* | 3.877650103 | 0.033073336 | 0.122470665 | -0.089397329 | 0.999933887 | 0.138072833 | 0.37083399 | -0.232761157 | 0.057003094 |
| ENSG00000107262 | *BAG1* | 5.747967864 | -0.002149587 | 0.095434099 | -0.097583686 | 0.999933887 | -0.169056252 | -0.472857407 | 0.303801155 | 0.057003094 |
| ENSG00000130312 | *MRPL34* | 2.222071245 | 0.099031646 | -0.063168039 | 0.162199686 | 0.999933887 | 0.003783639 | -0.351238524 | 0.355022163 | 0.057029923 |
| ENSG00000134308 | *YWHAQ* | 5.740743064 | -0.038411568 | -0.087381648 | 0.04897008 | 0.999933887 | 0.074266525 | 0.241346224 | -0.167079699 | 0.05710324 |
| ENSG00000176444 | *CLK2* | 4.5040483 | -0.014807597 | -0.074918764 | 0.060111167 | 0.999933887 | -0.2823464 | -0.466053865 | 0.183707465 | 0.057199923 |
| ENSG00000236453 | *AC003092.1* | -2.3908742 | -0.101619594 | -0.503311854 | 0.40169226 | 0.999933887 | -0.036750786 | 0.638906904 | -0.67565769 | 0.057261634 |
| ENSG00000184371 | *CSF1* | 9.030867871 | 0.045765964 | 0.042896212 | 0.002869752 | 0.999933887 | 0.786239235 | 1.048009321 | -0.261770085 | 0.057263221 |
| ENSG00000155545 | *MIER3* | 2.372875115 | -0.02901318 | 0.002629084 | -0.031642264 | 0.999933887 | -0.129548177 | 0.346909221 | -0.476457398 | 0.057282506 |
| ENSG00000184992 | *BRI3BP* | 2.270142437 | -0.088797034 | 0.085032463 | -0.173829497 | 0.999933887 | -0.362022503 | -0.774825686 | 0.412803183 | 0.057521037 |
| ENSG00000165264 | *NDUFB6* | 3.530533971 | 0.004147485 | 0.061454884 | -0.057307399 | 0.999933887 | -0.046518287 | -0.450429533 | 0.403911246 | 0.057641643 |
| ENSG00000164683 | *HEY1* | 0.750912366 | 0.053071212 | -0.151967071 | 0.205038283 | 0.999933887 | -0.072434924 | -0.677148107 | 0.604713183 | 0.057708403 |
| ENSG00000093000 | *NUP50* | 6.170466448 | -0.118567325 | -0.078960531 | -0.039606794 | 0.999933887 | -0.879394659 | -1.192893157 | 0.313498498 | 0.0577424 |
| ENSG00000123933 | *MXD4* | 5.381072371 | 0.027585334 | 0.08166725 | -0.054081916 | 0.999933887 | 0.033404653 | -0.186632894 | 0.220037547 | 0.057756447 |
| ENSG00000135968 | *GCC2* | 5.587919935 | -0.135034739 | -0.204013051 | 0.068978312 | 0.999933887 | -0.1569041 | 0.467361343 | -0.624265444 | 0.057769986 |
| ENSG00000196730 | *DAPK1* | 5.095533678 | -0.055058549 | 0.095049557 | -0.150108106 | 0.999933887 | -0.940050027 | -1.21476644 | 0.274716413 | 0.057769986 |
| ENSG00000260256 | *AP001063.1* | 0.70755853 | 0.254698388 | 0.112217777 | 0.142480611 | 0.999933887 | 1.743547929 | 2.285673705 | -0.542125775 | 0.057799751 |
| ENSG00000158864 | *NDUFS2* | 4.626938733 | -0.077165754 | -0.020133219 | -0.057032535 | 0.999933887 | -0.109893603 | -0.317255255 | 0.207361652 | 0.057799751 |
| ENSG00000287387 | *AC010883.3* | -0.136598131 | -0.087028069 | -0.180539401 | 0.093511332 | 0.999933887 | -0.433309472 | -1.085288702 | 0.651979229 | 0.057799751 |
| ENSG00000141446 | *ESCO1* | 5.754684402 | -0.084445752 | -0.112032629 | 0.027586877 | 0.999933887 | -0.386451983 | 0.070326031 | -0.456778014 | 0.057908471 |
| ENSG00000054967 | *RELT* | 8.051449405 | 0.042475726 | 0.037559181 | 0.004916545 | 0.999933887 | -0.28695498 | -0.541642757 | 0.254687777 | 0.057908471 |
| ENSG00000280739 | *EIF1B-AS1* | 0.219216351 | -0.065104165 | -0.132694894 | 0.067590729 | 0.999933887 | 0.037132445 | 0.739674586 | -0.702542141 | 0.057908471 |
| ENSG00000248527 | *MTATP6P1* | 6.138805734 | -0.000451118 | -0.149131914 | 0.148680796 | 0.999933887 | -0.017138576 | 0.231542556 | -0.248681132 | 0.057909756 |
| ENSG00000081087 | *OSTM1* | 6.277784545 | 0.060412217 | 0.046956253 | 0.013455965 | 0.999933887 | -0.043426756 | 0.182416642 | -0.225843397 | 0.057918481 |
| ENSG00000132716 | *DCAF8* | 5.477334221 | -0.002868816 | -0.028471555 | 0.025602739 | 0.999933887 | 0.142095925 | 0.35207725 | -0.209981326 | 0.057977055 |
| ENSG00000088205 | *DDX18* | 6.044616019 | -0.011347628 | -0.060308746 | 0.048961119 | 0.999933887 | 0.281397832 | 0.560079578 | -0.278681746 | 0.058033055 |
| ENSG00000239697 | *TNFSF12* | 2.605713151 | 0.070475803 | 0.075748304 | -0.005272481 | 0.999933887 | -0.038380836 | -0.3869828 | 0.348601963 | 0.058311889 |
| ENSG00000100221 | *JOSD1* | 7.530871127 | 0.031884976 | 0.025765319 | 0.006119658 | 0.999933887 | 0.116398954 | 0.306518821 | -0.190119867 | 0.058538736 |
| ENSG00000100242 | *SUN2* | 7.798020492 | 0.048428293 | 0.035651015 | 0.012777278 | 0.999933887 | -0.123319149 | -0.334540391 | 0.211221241 | 0.058604376 |
| ENSG00000153317 | *ASAP1* | 8.61782687 | 0.006297766 | -0.024828156 | 0.031125922 | 0.999933887 | 0.107658215 | 0.27782427 | -0.170166055 | 0.058617999 |
| ENSG00000120832 | *MTERF2* | 1.57760094 | -0.051166349 | -0.106020852 | 0.054854503 | 0.999933887 | -0.232177152 | -0.652752783 | 0.420575631 | 0.058673449 |
| ENSG00000277496 | *AL357033.4* | 4.127884059 | 0.021945686 | 0.017394364 | 0.004551323 | 0.999933887 | 0.413327944 | 0.695638389 | -0.282310445 | 0.058676144 |
| ENSG00000176563 | *CNTD1* | 0.849921622 | -0.024523219 | -0.063970762 | 0.039447543 | 0.999933887 | -0.165159997 | 0.375820703 | -0.5409807 | 0.058676144 |
| ENSG00000130827 | *PLXNA3* | 3.509834435 | 0.032325222 | 0.106290811 | -0.073965588 | 0.999933887 | 0.17022771 | 0.397839203 | -0.227611493 | 0.058688994 |
| ENSG00000101888 | *NXT2* | 1.771333879 | -0.027098827 | -0.159631627 | 0.1325328 | 0.999933887 | 0.498995051 | 0.901013812 | -0.402018761 | 0.058688994 |
| ENSG00000023608 | *SNAPC1* | 5.926061147 | 0.144434737 | -0.080947383 | 0.225382119 | 0.999933887 | 0.744101137 | 1.094634169 | -0.350533032 | 0.058793625 |
| ENSG00000081377 | *CDC14B* | 1.584956802 | -0.130861296 | 0.015745982 | -0.146607279 | 0.999933887 | 0.438471812 | 0.815797886 | -0.377326074 | 0.058807619 |

| | | | | | | | | | | |
|---|---|---|---|---|---|---|---|---|---|---|
| ENSG00000279149 | *AL356750.1* | -0.071284424 | -0.047617387 | 0.237706467 | -0.285323854 | 0.999933887 | -0.868424058 | -0.347563752 | -0.520860306 | 0.058807619 |
| ENSG00000179094 | *PER1* | 8.641527567 | 0.080245304 | 0.052149885 | 0.028095419 | 0.999933887 | 0.313926714 | 0.514776128 | -0.200849415 | 0.058877162 |
| ENSG00000158552 | *ZFAND2B* | 4.77322461 | -0.024760051 | 0.016034699 | -0.04079475 | 0.999933887 | -0.271473565 | -0.476774195 | 0.20530063 | 0.059033122 |
| ENSG00000176208 | *ATAD5* | 0.264543424 | -0.161064572 | -0.068061804 | -0.093002768 | 0.999933887 | -0.373565535 | 0.305764052 | -0.679329587 | 0.059073313 |
| ENSG00000182093 | *GET1* | 1.409492335 | -0.093981941 | -0.009135318 | -0.084846623 | 0.999933887 | -0.853551784 | -1.281076065 | 0.427524281 | 0.059073313 |
| ENSG00000165929 | *TC2N* | 4.354385192 | -0.066664959 | -0.19106189 | 0.124396931 | 0.999933887 | -0.162567801 | 0.182388014 | -0.344955815 | 0.059073313 |
| ENSG00000175984 | *DENND2C* | 0.209043659 | 0.071236502 | -0.102262672 | 0.173499175 | 0.999933887 | -0.455287315 | 0.215557297 | -0.670844612 | 0.059073313 |
| ENSG00000251661 | *AC136475.1* | 1.731188977 | 0.062797212 | -0.022953705 | 0.085750917 | 0.999933887 | -0.424980628 | -1.086388283 | 0.661407655 | 0.059073313 |
| ENSG00000047188 | *YTHDC2* | 4.427827909 | -0.061869945 | -0.048199342 | -0.013670602 | 0.999933887 | -0.160237745 | 0.222999637 | -0.383237382 | 0.059076565 |
| ENSG00000175573 | *C11orf68* | 3.857983646 | 0.087698865 | -0.106897307 | 0.194596172 | 0.999933887 | -0.592447242 | -0.912026554 | 0.319579312 | 0.059078515 |
| ENSG00000183255 | *PTTG1IP* | 6.834363011 | 0.022574501 | -0.006096969 | 0.028671469 | 0.999933887 | 0.973924311 | 1.337779868 | -0.363855557 | 0.059078515 |
| ENSG00000134852 | *CLOCK* | 1.600355806 | -0.040162725 | -0.170432162 | 0.130269437 | 0.999933887 | 0.116818648 | 0.675487317 | -0.558668669 | 0.059130661 |
| ENSG00000100354 | *TNRC6B* | 6.717597879 | 0.014115646 | 0.001631295 | 0.012484351 | 0.999933887 | -0.68225419 | -0.93803592 | 0.25578173 | 0.059208073 |
| ENSG00000269066 | *AC020908.3* | 0.96262286 | 0.126830351 | 0.150408749 | -0.023578398 | 0.999933887 | -0.241356266 | 0.272447491 | -0.513803757 | 0.05930969 |
| ENSG00000118900 | *UBN1* | 7.550583366 | -0.033784487 | 0.011173709 | -0.044958196 | 0.999933887 | -0.536780577 | -0.718964422 | 0.182183845 | 0.05930969 |
| ENSG00000231233 | *CFAP58-DT* | 5.903288991 | 0.03209865 | 0.025203626 | 0.006895024 | 0.999933887 | 1.963455767 | 2.285593584 | -0.322137817 | 0.05930969 |
| ENSG00000115902 | *SLC1A4* | 3.295663791 | -0.015346539 | -0.016131719 | 0.00078518 | 0.999933887 | -0.370539958 | -0.757236061 | 0.386696103 | 0.05930969 |
| ENSG00000100711 | *ZFYVE21* | 1.766824361 | -0.064056724 | 0.137703845 | -0.201760569 | 0.999933887 | -0.403852337 | -0.738175829 | 0.334323492 | 0.059325353 |
| ENSG00000173926 | *MARCHF3* | 1.925062733 | 0.068118591 | 0.035044296 | 0.033074295 | 0.999933887 | 0.902695372 | 1.401086018 | -0.498390646 | 0.059325353 |
| ENSG00000145354 | *CISD2* | 1.566738669 | -0.060688559 | -0.020650572 | -0.040037988 | 0.999933887 | -0.324232549 | -0.819050364 | 0.494817815 | 0.059325451 |
| ENSG00000175155 | *YPEL2* | 3.81132491 | -0.058360686 | 0.004940291 | -0.063300977 | 0.999933887 | -0.672330398 | -0.950266183 | 0.277935785 | 0.05934181 |
| ENSG00000188938 | *FAM120AOS* | 5.022446332 | 0.040705592 | 0.040306736 | 0.000398857 | 0.999933887 | -0.16688055 | -0.347450332 | 0.180569782 | 0.059368466 |
| ENSG00000071794 | *HLTF* | 0.119667686 | -0.210778101 | -0.110268155 | -0.100509946 | 0.999933887 | -0.327912543 | 0.38075324 | -0.708665783 | 0.059443278 |
| ENSG00000163812 | *ZDHHC3* | 5.217216036 | 0.022688487 | 0.030963834 | -0.008275347 | 0.999933887 | -0.169245755 | -0.394255391 | 0.225009636 | 0.059500683 |
| ENSG00000182578 | *CSF1R* | 2.40575839 | 0.024587521 | -0.06350981 | 0.088097331 | 0.999933887 | -1.205076514 | -1.794991626 | 0.589915112 | 0.059501464 |
| ENSG00000164305 | *CASP3* | 4.915918345 | -0.116647922 | -0.102034427 | -0.014613495 | 0.999933887 | -0.082145712 | 0.109940098 | -0.192085809 | 0.05953527 |
| ENSG00000175215 | *CTDSP2* | 7.336803709 | -0.052589477 | -0.002983176 | -0.0496063 | 0.999933887 | -0.538839596 | -0.758125628 | 0.219286032 | 0.059590792 |
| ENSG00000058668 | *ATP2B4* | 6.562464284 | -0.007567808 | -0.014802512 | 0.007234704 | 0.999933887 | -0.33572792 | -0.54821205 | 0.21248413 | 0.059590792 |
| ENSG00000225339 | *AL354740.1* | 1.257309833 | -0.010997442 | -0.197609543 | 0.186612101 | 0.999933887 | 0.160436594 | 0.526692052 | -0.366255458 | 0.059590792 |
| ENSG00000196591 | *HDAC2* | 4.061753944 | -0.070064912 | -0.019521204 | -0.050543708 | 0.999933887 | 0.220312752 | 0.478997598 | -0.258684845 | 0.059621239 |
| ENSG00000228329 | *LINC01890* | 2.665614179 | 0.038105368 | 0.103566294 | -0.065460927 | 0.999933887 | 0.327155756 | 0.660088035 | -0.33293228 | 0.059621239 |
| ENSG00000141424 | *SLC39A6* | 4.363878776 | -0.055141139 | -0.083246253 | 0.028105114 | 0.999933887 | -0.453100435 | -0.687528984 | 0.234428549 | 0.059662123 |
| ENSG00000146556 | *WASH2P* | 2.389745298 | -0.012313511 | 0.099925176 | -0.112238687 | 0.999933887 | -0.145159127 | -0.447861882 | 0.302702756 | 0.059677596 |
| ENSG00000277978 | *AC010542.5* | -0.377246622 | -0.468900118 | -0.548373881 | 0.079473762 | 0.999933887 | -1.014536205 | -2.016157428 | 1.001621224 | 0.059927367 |
| ENSG00000148719 | *DNAJB12* | 6.018189657 | -0.008033895 | -0.001736088 | -0.006297807 | 0.999933887 | 0.129844225 | 0.279090308 | -0.149246083 | 0.059927367 |
| ENSG00000172771 | *EFCAB12* | 1.340594791 | -0.108349377 | 0.048021651 | -0.156371028 | 0.999933887 | -0.317642809 | -0.825833032 | 0.508190223 | 0.059938379 |
| ENSG00000121742 | *GJB6* | 6.108709385 | -0.077026718 | -0.049863282 | -0.027163436 | 0.999933887 | -0.381388392 | -0.15304301 | -0.228345382 | 0.060141788 |
| ENSG00000105662 | *CRTC1* | 2.991979198 | 0.059246964 | 0.082563185 | -0.023316221 | 0.999933887 | -0.130375431 | -0.392034684 | 0.261659253 | 0.060141788 |
| ENSG00000119574 | *ZBTB45* | 2.334401111 | -0.00680056 | 0.010675979 | -0.017476539 | 0.999933887 | -0.252941612 | -0.57252562 | 0.319584007 | 0.060161669 |
| ENSG00000159335 | *PTMS* | 3.336328121 | 0.014928685 | 0.047920844 | -0.032992159 | 0.999933887 | -0.018212076 | -0.381293418 | 0.363081342 | 0.060295248 |
| ENSG00000188677 | *PARVB* | 1.108770068 | 0.02018897 | 0.148732501 | -0.128543531 | 0.999933887 | 0.652263689 | 1.133926251 | -0.481662562 | 0.060349298 |
| ENSG00000179588 | *ZFPM1* | 3.106251715 | 0.047874259 | 0.086336772 | -0.038462513 | 0.999933887 | 0.050449841 | -0.278723089 | 0.32917293 | 0.06038871 |
| ENSG00000266709 | *AC005224.3* | 1.884970272 | 0.032779881 | -0.102734786 | 0.135514667 | 0.999933887 | 1.37056786 | 2.01456329 | -0.64399543 | 0.060395897 |
| ENSG00000073050 | *XRCC1* | 3.332548986 | 0.049064257 | -0.004050494 | 0.053114751 | 0.999933887 | -0.222049534 | -0.506263683 | 0.284214148 | 0.060403903 |
| ENSG00000170903 | *MSANTD4* | 1.5203384 | -0.120254979 | -0.129417407 | 0.009162428 | 0.999933887 | -0.179694661 | 0.240286525 | -0.419981186 | 0.060477256 |
| ENSG00000124222 | *STX16* | 5.924803208 | 0.023943197 | 0.065028378 | -0.041085181 | 0.999933887 | -0.249247001 | -0.426046178 | 0.176799177 | 0.060477256 |
| ENSG00000127220 | *ABHD8* | 1.750607468 | 0.212177091 | 0.070909801 | 0.14126729 | 0.999933887 | -0.094233932 | -0.566682365 | 0.472448433 | 0.060519482 |
| ENSG00000150347 | *ARID5B* | 4.556726981 | -0.015117552 | -0.100523154 | 0.085405602 | 0.999933887 | 0.17693684 | 0.488694731 | -0.31175789 | 0.060519482 |
| ENSG00000125538 | *IL1B* | 8.683670028 | 0.66009871 | 0.850344427 | -0.190245718 | 0.999933887 | 2.548933538 | 3.265567783 | -0.716634245 | 0.060538246 |
| ENSG00000250138 | *AC139495.2* | 2.197596364 | 0.160502695 | 0.099587209 | 0.060915486 | 0.999933887 | -0.755420314 | -1.389545355 | 0.634125041 | 0.060538246 |
| ENSG00000100239 | *PPP6R2* | 5.810256303 | -0.011297873 | 0.070474753 | -0.081772626 | 0.999933887 | -0.181690343 | -0.357334309 | 0.175643966 | 0.060538246 |
| ENSG00000182179 | *UBA7* | 5.608369997 | 0.040243296 | 0.073311008 | -0.033067712 | 0.999933887 | 0.039652991 | -0.198074864 | 0.237727855 | 0.060551554 |
| ENSG00000146425 | *DYNLT1* | 4.352696149 | 0.017400013 | 0.02135904 | -0.003959028 | 0.999933887 | -0.468014974 | -0.720083731 | 0.252068757 | 0.060551554 |
| ENSG00000182749 | *PAQR7* | 0.572458915 | 0.025969926 | 0.00784789 | 0.018122036 | 0.999933887 | -0.366922071 | -1.004772425 | 0.637850354 | 0.060578053 |
| ENSG00000231507 | *LINC01353* | 3.610751096 | 0.18960387 | 0.171578043 | 0.018025827 | 0.999933887 | 1.402807182 | 1.778674741 | -0.375867559 | 0.060601738 |
| ENSG00000159792 | *PSKH1* | 2.8525903 | 0.018182074 | -0.076360673 | 0.094542747 | 0.999933887 | -0.286671522 | -0.561967789 | 0.275296267 | 0.060601738 |
| ENSG00000165995 | *CACNB2* | 0.165553225 | -0.241059025 | -0.308194942 | 0.067135917 | 0.999933887 | 0.09943097 | 1.067271244 | -0.967840274 | 0.060672815 |
| ENSG00000034713 | *GABARAPL2* | 8.43051702 | -0.017708119 | 0.007722552 | -0.02543067 | 0.999933887 | 1.026861504 | 1.243662058 | -0.216800553 | 0.060672815 |
| ENSG00000246526 | *LINC02481* | 1.437897349 | -0.119060651 | -0.009033255 | -0.110027396 | 0.999933887 | 0.220635007 | 0.616993155 | -0.396358148 | 0.060687333 |
| ENSG00000140749 | *IGSF6* | 8.869906985 | -0.012274821 | -0.078176395 | 0.065901574 | 0.999933887 | 0.820771416 | 1.02768063 | -0.206909213 | 0.060806593 |
| ENSG00000131848 | *ZSCAN5A* | -0.195292778 | -0.010956336 | 0.07503412 | -0.085990456 | 0.999933887 | 0.241956475 | 0.798766566 | -0.55681009 | 0.060853653 |
| ENSG00000100911 | *PSME2* | 5.345907174 | 0.019665832 | -0.013377404 | 0.033043236 | 0.999933887 | 0.042229809 | -0.162291051 | 0.20452086 | 0.060862777 |
| ENSG00000267321 | *SNHG30* | 2.718914315 | -0.010911206 | -0.041379873 | 0.030468667 | 0.999933887 | -0.148417387 | -0.477749699 | 0.329332313 | 0.060903003 |
| ENSG00000162298 | *SYVN1* | 5.081001471 | 0.093643021 | 0.025164175 | 0.068478846 | 0.999933887 | -0.271716833 | -0.45967847 | 0.187961637 | 0.060907198 |
| ENSG00000155090 | *KLF10* | 8.313400343 | 0.239500191 | 0.253011262 | -0.013511072 | 0.999933887 | 1.318958424 | 1.583954447 | -0.264996022 | 0.060932122 |
| ENSG00000234883 | *MIR155HG* | 1.846814388 | 0.052004178 | 0.325287796 | -0.273283618 | 0.999933887 | 1.971293789 | 2.57518998 | -0.603896192 | 0.061152639 |
| ENSG00000166347 | *CYB5A* | 0.826570296 | -0.076275552 | 0.127260684 | -0.203536237 | 0.999933887 | 0.423429165 | 0.86971814 | -0.446288975 | 0.061285755 |
| ENSG00000100083 | *GGA1* | 7.82245223 | 0.051177969 | 0.059422161 | -0.008244193 | 0.999933887 | 0.637421663 | 0.834469221 | -0.197047559 | 0.061288802 |
| ENSG00000080503 | *SMARCA2* | 5.978365671 | 0.006325543 | -0.034239412 | 0.040564954 | 0.999933887 | -0.115076107 | -0.333245019 | 0.218168912 | 0.061567397 |
| ENSG00000184489 | *PTP4A3* | 5.746185341 | 0.092380725 | 0.058994316 | 0.033386409 | 0.999933887 | 0.450161708 | 0.108965686 | 0.341196023 | 0.061809249 |
| ENSG00000188295 | *ZNF669* | 2.28710075 | 0.015459547 | -0.000536503 | 0.01599605 | 0.999933887 | -0.071026612 | -0.40721334 | 0.336186728 | 0.061917301 |
| ENSG00000008294 | *SPAG9* | 9.449316124 | -0.013737472 | -0.01223278 | -0.001504692 | 0.999933887 | 0.266964441 | 0.483989832 | -0.21702539 | 0.062054893 |
| ENSG00000114850 | *SSR3* | 3.888865677 | -0.057028044 | -0.036375637 | -0.020652407 | 0.999933887 | 0.032382713 | -0.187120307 | 0.21950302 | 0.06211776 |
| ENSG00000153094 | *BCL2L11* | 5.622478607 | -0.095559337 | -0.052279805 | -0.043279531 | 0.999933887 | -0.79861379 | -1.092414635 | 0.293800845 | 0.062167971 |
| ENSG00000230417 | *LINC00595* | 1.272081954 | 0.257115039 | 0.126436137 | 0.130678902 | 0.999933887 | 0.674703145 | 1.020756039 | -0.346052895 | 0.062167971 |
| ENSG00000230417 | *LINC00595* | 1.272081954 | 0.257115039 | 0.126436137 | 0.130678902 | 0.999933887 | 0.674703145 | 1.020756039 | -0.346052895 | 0.062167971 |
| ENSG00000115211 | *EIF2B4* | 3.313277565 | -0.051147733 | 0.055478429 | -0.106626162 | 0.999933887 | -0.374607667 | -0.678723584 | 0.304115916 | 0.062420794 |
| ENSG00000004864 | *SLC25A13* | 6.782723919 | 0.134327356 | 0.140721792 | -0.006394436 | 0.999933887 | 0.282036104 | 0.512504997 | -0.230468893 | 0.062498977 |
| ENSG00000103154 | *NECAB2* | 3.260732849 | -0.038761461 | -0.030218691 | -0.00854277 | 0.999933887 | -0.874634895 | -1.235108699 | 0.360473804 | 0.062498977 |
| ENSG00000087502 | *ERGIC2* | 4.389767642 | -0.022843129 | -0.128712245 | 0.105869116 | 0.999933887 | -0.571958298 | -0.289907729 | -0.282050569 | 0.06256791 |
| ENSG00000100906 | *NFKBIA* | 11.251387314 | 0.470374894 | 0.522283926 | -0.051909033 | 0.999933887 | 2.712002909 | 3.187404676 | -0.475401767 | 0.062675731 |
| ENSG00000175538 | *KCNE3* | 4.313408472 | -0.153082849 | -0.226663899 | 0.07358105 | 0.999933887 | -0.244331782 | -0.501584142 | 0.25725236 | 0.062831367 |
| ENSG00000254004 | *ZNF260* | 0.438637006 | -0.05563473 | -0.054402562 | -0.001232168 | 0.999933887 | -0.209234017 | 0.496086934 | -0.705320952 | 0.06294127 |
| ENSG00000228010 | *AC073343.2* | 0.988839419 | -0.160879602 | 0.16797998 | -0.328859582 | 0.999933887 | 0.216724954 | 0.602072265 | -0.385347311 | 0.0629444 |
| ENSG00000120306 | *CYSTM1* | 7.372798632 | 0.02405108 | 0.025963434 | -0.001912355 | 0.999933887 | 0.43995024 | 0.606805442 | -0.166855202 | 0.063171682 |
| ENSG00000140987 | *ZSCAN32* | 2.927204902 | -0.131448721 | -0.166725041 | 0.03527632 | 0.999933887 | -0.284092214 | -0.536028633 | 0.251936419 | 0.063180108 |
| ENSG00000130203 | *APOE* | 0.25643152 | -0.075793886 | 0.462671406 | -0.538465292 | 0.999933887 | 0.05832458 | 1.041053425 | -0.982728845 | 0.06318655 |
| ENSG00000182923 | *CEP63* | 6.652284312 | -0.049254209 | -0.116633292 | 0.067379083 | 0.999933887 | -0.827539733 | -0.547015573 | -0.28052416 | 0.063208645 |
| ENSG00000123338 | *NCKAP1L* | 6.590561902 | -0.008639435 | 0.008522736 | -0.017162171 | 0.999933887 | -0.174446953 | -0.349364726 | 0.174917773 | 0.063270918 |
| ENSG00000125351 | *UPF3B* | 2.945801199 | -0.133075387 | -0.051107149 | -0.081968237 | 0.999933887 | 0.804146823 | 1.15201042 | -0.347863597 | 0.063338109 |
| ENSG00000230869 | *AGAP10P* | -0.590573215 | 0.022676326 | 0.037862732 | -0.015186406 | 0.999933887 | -0.240078925 | 0.445508375 | -0.6855873 | 0.063864309 |
| ENSG00000100150 | *DEPDC5* | 3.024660622 | 0.038134981 | -0.039975921 | 0.078110902 | 0.999933887 | -0.285954258 | -0.593073517 | 0.307119259 | 0.063913323 |
| ENSG00000137714 | *FDX1* | 2.671646483 | 0.030075001 | 0.065394341 | -0.03531934 | 0.999933887 | -0.084826671 | 0.226281983 | -0.311108654 | 0.063913323 |
| ENSG00000125945 | *ZNF436* | 1.602633013 | -0.025169369 | 0.133683561 | -0.15885293 | 0.999933887 | -0.190812984 | -0.540983428 | 0.350170443 | 0.063916453 |
| ENSG00000133574 | *GIMAP4* | 5.138412501 | 0.000216319 | -0.059433956 | 0.059650275 | 0.999933887 | -0.121720985 | -0.40442461 | 0.282703625 | 0.063916453 |
| ENSG00000109861 | *CTSC* | 6.443100407 | 0.001412165 | -0.036070765 | 0.03748293 | 0.999933887 | 0.434904938 | 0.853773356 | -0.418868418 | 0.063928689 |
| ENSG00000157540 | *DYRK1A* | 7.624490451 | -0.048232857 | -0.059447629 | 0.011214772 | 0.999933887 | -0.027453171 | 0.113999822 | -0.141452993 | 0.064266208 |
| ENSG00000237036 | *ZEB1-AS1* | 4.010593004 | 0.015376482 | 0.202256437 | -0.186879955 | 0.999291887 | 0.193448049 | 0.420489099 | -0.227041049 | 0.06429652 |
| ENSG00000172530 | *BANP* | 6.442021183 | 0.033840827 | -0.010775577 | 0.044616404 | 0.999933887 | 0.633515275 | 0.820004672 | -0.186489397 | 0.064496971 |
| ENSG00000111642 | *CHD4* | 4.826496749 | -0.002753366 | 0.014723027 | -0.017476393 | 0.999933887 | 0.550538658 | 0.75350433 | -0.202965672 | 0.064501612 |
| ENSG00000106605 | *BLVRA* | 3.057824942 | -0.179882601 | -0.034989737 | -0.144892864 | 0.999933887 | -1.161597823 | -1.563889726 | 0.402291903 | 0.064542267 |
| ENSG00000245148 | *ARAP1-AS2* | 4.514584002 | 0.000969875 | 0.006520644 | -0.005550768 | 0.999933887 | -0.052672646 | 0.294766626 | -0.347439271 | 0.064542267 |
| ENSG00000042980 | *ADAM28* | 2.914264954 | -0.056227042 | -0.063752983 | 0.007525941 | 0.999933887 | -0.111212705 | 0.321796766 | -0.433009472 | 0.064697908 |
| ENSG00000273188 | *AL022328.3* | 1.251493974 | -0.040213659 | 0.022970995 | -0.063184654 | 0.999933887 | -0.070806227 | -0.58449514 | 0.513688913 | 0.064751551 |
| ENSG00000074071 | *MRPS34* | 1.996562375 | 0.056894089 | -0.086207985 | 0.143102074 | 0.999933887 | -0.037868847 | -0.410849023 | 0.372980176 | 0.064769938 |

| | | | | | | | | | | |
|---|---|---|---|---|---|---|---|---|---|---|
| ENSG00000196562 | *SULF2* | 7.051126062 | 0.01116693 | -0.000967723 | 0.012134653 | 0.999933887 | -1.201882384 | -1.462112946 | 0.260230562 | 0.064769938 |
| ENSG00000111328 | *CDK2AP1* | 2.645059649 | 0.078011274 | 0.004342571 | 0.073668703 | 0.999933887 | 0.079054215 | -0.205182868 | 0.284237082 | 0.064912997 |
| ENSG00000204283 | *LINC01973* | 1.901602316 | 0.039281032 | 0.103922899 | -0.064641866 | 0.999933887 | 1.307732354 | 1.895051043 | -0.587318689 | 0.065341411 |
| ENSG00000257702 | *LBX2-AS1* | 0.346913954 | 0.198657578 | 0.415056248 | -0.216398671 | 0.999933887 | -0.184387257 | 0.169420756 | -0.353808013 | 0.065532953 |
| ENSG00000161996 | *WDR90* | 1.639236179 | 0.081301832 | 0.193669342 | -0.11236751 | 0.999933887 | 0.29350306 | 0.654595828 | -0.361092768 | 0.065532953 |
| ENSG00000164190 | *NIPBL* | 7.179741251 | -0.084149157 | -0.15815503 | 0.074005874 | 0.999933887 | -0.695228691 | -0.179230471 | -0.51599822 | 0.065589326 |
| ENSG00000125462 | *MIR9-1HG* | 3.490534373 | 0.02238218 | 0.008609951 | 0.01377223 | 0.999933887 | 2.041030033 | 2.414326996 | -0.373296963 | 0.065589326 |
| ENSG00000079819 | *EPB41L2* | 0.405182601 | 0.030756294 | -0.123925913 | 0.154682207 | 0.999933887 | 0.166791381 | 0.711585116 | -0.544793735 | 0.06559643 |
| ENSG00000258404 | *LINC02320* | 1.9138707 | 0.119505023 | 0.122788796 | -0.003283773 | 0.999933887 | 1.268199254 | 1.616847782 | -0.348648527 | 0.065630025 |
| ENSG00000100266 | *PACSIN2* | 9.250750944 | -0.000326068 | 0.002279681 | -0.002605749 | 0.999933887 | 0.032936566 | 0.186703377 | -0.153766812 | 0.065728378 |
| ENSG00000183604 | *SMG1P5* | 2.255164223 | 0.070898558 | 0.059737128 | 0.01116143 | 0.999933887 | -0.265296806 | -0.603593059 | 0.338296253 | 0.065828176 |
| ENSG00000196968 | *FUT11* | 2.801028062 | -0.156019291 | 0.000587806 | -0.156607097 | 0.999933887 | -0.179621764 | -0.419172179 | 0.239550416 | 0.065868662 |
| ENSG00000261218 | *AC099524.1* | 0.051414871 | -0.150756764 | 0.163483982 | -0.314240746 | 0.999933887 | -0.606827765 | -1.435978367 | 0.829150602 | 0.065974568 |
| ENSG00000165678 | *GHITM* | 6.851173528 | -0.02351366 | -0.036781502 | 0.013267841 | 0.999933887 | 0.188395506 | 0.344406631 | -0.156011125 | 0.065990783 |
| ENSG00000234127 | *TRIM26* | 5.23955782 | 0.033917977 | 0.043485172 | -0.009567195 | 0.999933887 | -0.125735141 | -0.369031476 | 0.243296336 | 0.066085134 |
| ENSG00000104756 | *KCTD9* | 3.841392539 | -0.124289741 | -0.065028862 | -0.059260879 | 0.999933887 | -0.4488999 | -0.212354109 | -0.236545791 | 0.066097896 |
| ENSG00000167595 | *PROSER3* | 3.081079205 | -0.051181652 | -0.039813695 | -0.011367958 | 0.999933887 | -0.275730876 | -0.57293647 | 0.297205594 | 0.066097896 |
| ENSG00000176142 | *TMEM39A* | 3.423392654 | -0.042118087 | 0.022129419 | -0.064247506 | 0.999933887 | 0.22878986 | 0.471380159 | -0.242590298 | 0.066097896 |
| ENSG00000237943 | *PRKCQ-AS1* | 3.114383538 | -0.127614745 | -0.191511984 | 0.063897238 | 0.999933887 | 0.134137967 | 0.389307659 | -0.255169691 | 0.066131955 |
| ENSG00000168439 | *STIP1* | 5.416357058 | -0.053103375 | -0.054361791 | 0.001258417 | 0.999933887 | 0.257675668 | 0.452766506 | -0.195090838 | 0.066131955 |
| ENSG00000102738 | *MRPS31* | 1.071371923 | -0.088877199 | -0.147316381 | 0.058439181 | 0.999933887 | -0.032911299 | 0.34224374 | -0.375155039 | 0.066131955 |
| ENSG00000180530 | *NRIP1* | 2.935064038 | -0.087748355 | -0.154412389 | 0.066664034 | 0.999933887 | 0.339165213 | 0.918290314 | -0.5791251 | 0.066131955 |
| ENSG00000154529 | *CNTNAP3B* | 0.164338136 | -0.045204698 | 0.044197634 | -0.089402332 | 0.999933887 | -0.841090328 | -1.512343899 | 0.67125357 | 0.066131955 |
| ENSG00000141854 | *MISP3* | -0.218478687 | 0.036532576 | -0.069508272 | 0.106040848 | 0.999933887 | 0.240340463 | -0.391906096 | 0.632246559 | 0.066131955 |
| ENSG00000159640 | *ACE* | -0.097848197 | 0.021114042 | 0.00977857 | 0.011335472 | 0.999933887 | -0.120522284 | 0.490756946 | -0.611279231 | 0.066131955 |
| ENSG00000116991 | *SIPA1L2* | 7.059765456 | -0.002465766 | -0.003095441 | 0.000629675 | 0.999933887 | -0.722332292 | -0.948477267 | 0.226144975 | 0.066131955 |
| ENSG00000136897 | *MRPL50* | 1.808272665 | 0.070278626 | -0.096709158 | 0.166987784 | 0.999933887 | 0.183995357 | 0.539572285 | -0.355576928 | 0.066150787 |
| ENSG00000121774 | *KHDRBS1* | 6.541624255 | -0.041004416 | 0.003454652 | -0.044459067 | 0.999933887 | 0.113627072 | 0.261051673 | -0.147424601 | 0.066165184 |
| ENSG00000072364 | *AFF4* | 8.045205172 | -0.042744971 | -0.107255514 | 0.064510544 | 0.999933887 | -0.051182006 | 0.218082019 | -0.269264025 | 0.066353345 |
| ENSG00000105963 | *ADAP1* | 3.930915143 | 0.057806309 | 0.079872752 | -0.022066443 | 0.999933887 | -0.153165749 | -0.490374273 | 0.337208524 | 0.066395632 |
| ENSG00000234518 | *PTGES3P1* | 3.509969296 | -0.170689144 | -0.187617196 | 0.016928052 | 0.999933887 | 0.973024903 | 1.518535905 | -0.545511002 | 0.066449356 |
| ENSG00000131873 | *CHSY1* | 6.11151017 | 0.051567797 | 0.128321269 | -0.076753471 | 0.999933887 | -0.438070194 | -0.706946803 | 0.268876609 | 0.066449356 |
| ENSG00000160179 | *ABCG1* | 5.906283735 | 0.094765889 | 0.030084932 | 0.064680957 | 0.999933887 | -0.262173079 | 0.102352325 | -0.364525404 | 0.066513155 |
| ENSG00000149557 | *FEZ1* | -0.107759909 | 0.01380753 | 0.013545309 | 0.000262222 | 0.999933887 | 0.559559603 | 1.149006274 | -0.589446671 | 0.066541392 |
| ENSG00000101346 | *POFUT1* | 2.366953622 | 0.029307104 | 0.017982233 | 0.011324871 | 0.999933887 | 0.312721323 | 0.63023067 | -0.317509347 | 0.066708357 |
| ENSG00000104290 | *FZD3* | -0.289785452 | -0.04854393 | -0.052333323 | 0.003789393 | 0.999933887 | -0.552032707 | 0.18293912 | -0.734971828 | 0.066812067 |
| ENSG00000258289 | *CHURC1* | 5.236348311 | -0.060218 | -0.148289308 | 0.088071307 | 0.999933887 | -0.11637824 | 0.139920283 | -0.256298523 | 0.066864349 |
| ENSG00000264522 | *OTUD7B* | 1.398080655 | 0.060173302 | 0.070896856 | -0.010723555 | 0.999933887 | -0.190182929 | 0.198183883 | -0.388366813 | 0.066864349 |
| ENSG00000014164 | *ZC3H3* | 5.205053215 | 0.02215315 | 0.019218869 | 0.002934281 | 0.999933887 | -0.255357502 | -0.465219423 | 0.20986192 | 0.066864349 |
| ENSG00000156804 | *FBXO32* | 2.964133122 | -0.100449807 | -0.001483316 | -0.09896649 | 0.999933887 | -0.202630785 | -0.475027904 | 0.272397119 | 0.066903895 |
| ENSG00000133789 | *SWAP70* | 4.098286265 | 0.017338611 | 0.059155479 | -0.041816868 | 0.999933887 | -0.021769304 | 0.197579917 | -0.219349221 | 0.066940078 |
| ENSG00000198814 | *GK* | 8.2581606 | -0.027820714 | 0.000365881 | -0.028186594 | 0.999933887 | 0.248390072 | 0.501679908 | -0.253289836 | 0.067211581 |
| ENSG00000227268 | *KLLN* | 1.746037093 | -0.123505445 | -0.083330769 | -0.040174676 | 0.999933887 | -0.266519989 | -0.791532211 | 0.525012222 | 0.067228411 |
| ENSG00000076641 | *PAG1* | 8.053093852 | -0.040026465 | -0.040498131 | 0.000471666 | 0.999933887 | -0.274732765 | -0.071643118 | -0.203089646 | 0.06724967 |
| ENSG00000087008 | *ACOX3* | 2.826783796 | 0.138576311 | -0.094523924 | 0.233100235 | 0.999933887 | 1.111083131 | 1.472174008 | -0.361090877 | 0.067398475 |
| ENSG00000074660 | *SCARF1* | 9.461748187 | 0.064796023 | 0.09041919 | -0.025623167 | 0.999933887 | 0.962387772 | 1.183461021 | -0.221073248 | 0.067398475 |
| ENSG00000145632 | *PLK2* | 2.687443616 | 0.194982484 | 0.083302375 | 0.111680109 | 0.999933887 | 1.604479365 | 2.034844136 | -0.430364771 | 0.067404541 |
| ENSG00000137700 | *SLC37A4* | 1.184019635 | 0.155248845 | 0.124247728 | 0.031001117 | 0.999933887 | 0.204449469 | -0.133656298 | 0.338105767 | 0.067404541 |
| ENSG00000197885 | *NKIRAS1* | 1.717916596 | 0.1108849 | 0.119576378 | -0.008691478 | 0.999933887 | 1.428841164 | 1.856735448 | -0.427894284 | 0.067404541 |
| ENSG00000225648 | *SBDSP1* | 2.985684641 | -0.10642342 | -0.145239261 | 0.038815841 | 0.999933887 | 0.191795876 | 0.488435998 | -0.296640123 | 0.067493997 |
| ENSG00000182606 | *TRAK1* | 4.695660797 | -0.025041292 | 0.002158429 | -0.027199721 | 0.999933887 | 0.575320503 | 0.881403324 | -0.306082821 | 0.067493997 |
| ENSG00000070190 | *DAPP1* | 5.673982029 | -0.002708992 | 0.015632137 | -0.018341129 | 0.999933887 | 0.584962349 | 0.84256917 | -0.257606821 | 0.067493997 |
| ENSG00000117153 | *KLHL12* | 4.642522108 | -0.049396822 | 0.022798255 | -0.072195077 | 0.999933887 | 0.283019322 | 0.553768277 | -0.270748954 | 0.067585486 |
| ENSG00000198756 | *COLGALT2* | 0.298282343 | 0.070961274 | -0.021321014 | 0.092282288 | 0.999933887 | 0.018032324 | 0.600678962 | -0.582646638 | 0.067927742 |
| ENSG00000154188 | *ANGPT1* | 0.418991699 | 0.192276723 | -0.021370044 | 0.213646766 | 0.999933887 | -0.696817123 | -1.592553252 | 0.895736129 | 0.068006062 |
| ENSG00000168137 | *SETD5* | 7.135247208 | -0.037151764 | -0.03940406 | 0.002252296 | 0.999933887 | 0.187749084 | 0.351823874 | -0.16407479 | 0.068263371 |
| ENSG00000130487 | *KLHDC7B* | 1.289505553 | 0.09213815 | -0.003453631 | 0.095591781 | 0.999933887 | -0.290740631 | -0.73643318 | 0.445692549 | 0.068271276 |
| ENSG00000143373 | *ZNF687* | 6.110674874 | -0.0215566 | -0.017190585 | -0.004366015 | 0.999933887 | 0.006641405 | -0.162553357 | 0.169194762 | 0.068313406 |
| ENSG00000162747 | *FCGR3B* | 10.584646055 | -0.026751794 | -0.136641597 | 0.109889804 | 0.999933887 | -0.618652933 | -0.890860845 | 0.272207912 | 0.068313406 |
| ENSG00000121104 | *FAM117A* | 4.372921863 | 0.049366921 | -0.046185674 | 0.095552594 | 0.999933887 | -0.163061246 | -0.430290841 | 0.267229595 | 0.068376463 |
| ENSG00000159692 | *CTBP1* | 5.325010077 | 0.057153131 | -0.026092891 | 0.083246022 | 0.999933887 | -0.086725778 | -0.248701187 | 0.161975409 | 0.068383165 |
| ENSG00000100784 | *RPS6KA5* | 4.66171742 | -0.043597818 | -0.024316522 | -0.019281296 | 0.999933887 | -0.521283241 | -0.858617162 | 0.337333922 | 0.068383165 |
| ENSG00000140465 | *CYP1A1* | 0.319149994 | -0.14962225 | 0.090113133 | -0.239735384 | 0.999933887 | 0.59449895 | 0.967644837 | -0.373145888 | 0.068505494 |
| ENSG00000153234 | *NR4A2* | 7.492828573 | 0.120028068 | 0.116934472 | 0.003093596 | 0.999933887 | 0.376978055 | 0.658115654 | -0.2811376 | 0.068518743 |
| ENSG00000249786 | *EAF1-AS1* | -0.118307768 | -0.067962055 | -0.047624817 | -0.020337238 | 0.999933887 | 0.886008462 | 1.561386676 | -0.675378214 | 0.068518743 |
| ENSG00000071994 | *PDCD2* | 3.557510001 | -0.011248017 | -0.023765891 | 0.012517875 | 0.999933887 | -0.068387427 | -0.269703744 | 0.201316316 | 0.068537483 |
| ENSG00000215845 | *TSTD1* | 3.255420379 | -0.055131423 | -0.024627705 | -0.030503717 | 0.999933887 | -0.118328229 | -0.40064237 | 0.282314141 | 0.068546903 |
| ENSG00000138185 | *ENTPD1* | 5.474851589 | -0.045025579 | -0.086672159 | 0.04164658 | 0.999933887 | -0.633944499 | -0.92761732 | 0.29367282 | 0.068870989 |
| ENSG00000178093 | *TSSK6* | 0.683746446 | -0.182457443 | -0.077456298 | -0.105001145 | 0.999933887 | 0.093787619 | -0.44930928 | 0.543096899 | 0.068914124 |
| ENSG00000120278 | *PLEKHG1* | 1.320949263 | -0.053169694 | 0.106419311 | -0.159589005 | 0.999933887 | 0.319169905 | 0.721999651 | -0.402829746 | 0.069040763 |
| ENSG00000082641 | *NFE2L1* | 5.273086283 | -0.008822494 | -0.007575427 | -0.001247067 | 0.999933887 | -0.457477787 | -0.695842076 | 0.238364289 | 0.069074164 |
| ENSG00000160551 | *TAOK1* | 6.403524069 | 0.002193909 | -0.089381076 | 0.091574984 | 0.999933887 | -0.549027765 | -0.212553298 | -0.336474468 | 0.069234986 |
| ENSG00000182327 | *GLTPD2* | 0.835674144 | 0.134093381 | 0.071259209 | 0.062834173 | 0.999933887 | 0.175732775 | 0.780715073 | -0.604982299 | 0.069381298 |
| ENSG00000101986 | *ABCD1* | 4.26413475 | 0.059732319 | 0.043190455 | 0.016541865 | 0.999933887 | -0.865975827 | -1.138432502 | 0.272456674 | 0.069381298 |
| ENSG00000211694 | *TRGV10* | 0.564055262 | -0.005502933 | -0.044847427 | 0.039344494 | 0.999933887 | 0.032616409 | 0.497256998 | -0.464640589 | 0.069391441 |
| ENSG00000163788 | *SNRK* | 6.810766155 | -0.035188081 | -0.009358166 | -0.025829915 | 0.999933887 | -0.255890041 | -0.064346719 | -0.191543322 | 0.069398446 |
| ENSG00000125107 | *CNOT1* | 7.218253436 | -0.045717655 | -0.118661175 | 0.07294352 | 0.999933887 | 0.165524027 | 0.347409773 | -0.181885746 | 0.069461114 |
| ENSG00000243055 | *GK-AS1* | 3.721928708 | -0.084790452 | 0.087975006 | -0.172765458 | 0.999933887 | -0.027431929 | 0.378722097 | -0.406154026 | 0.069461114 |
| ENSG00000181404 | *WASHC1* | 1.984190246 | 0.10136423 | -0.001021592 | 0.102385822 | 0.999933887 | -0.318254315 | -0.709016109 | 0.390761794 | 0.069493162 |
| ENSG00000119927 | *GPAM* | 0.841642061 | -0.020775352 | -0.256741605 | 0.235966253 | 0.999933887 | 0.191877394 | 0.690123006 | -0.498245612 | 0.069502599 |
| ENSG00000071967 | *CYBRD1* | 2.167408865 | -0.061847844 | 0.130366962 | -0.192214806 | 0.999933887 | -0.756422003 | -1.272440575 | 0.516018571 | 0.069575649 |
| ENSG00000113555 | *PCDH12* | 1.253882536 | 0.21874856 | -0.128495584 | 0.347244144 | 0.981460244 | -0.01841572 | 0.424392484 | -0.442808205 | 0.069586432 |
| ENSG00000107872 | *FBXL15* | 2.613797311 | 0.164675494 | 0.087117814 | 0.07755768 | 0.999933887 | 0.72008978 | 1.073954364 | -0.353864585 | 0.069693775 |
| ENSG00000203710 | *CR1* | 7.451891882 | -0.038194792 | -0.072688619 | 0.034493827 | 0.999933887 | -0.182761159 | -0.328035014 | 0.145273855 | 0.069737276 |
| ENSG00000158769 | *F11R* | 5.482735667 | -0.046735145 | 0.006949321 | -0.053684467 | 0.999933887 | -0.469346427 | -0.777001258 | 0.307654832 | 0.069741859 |
| ENSG00000137857 | *DUOX1* | -0.086912018 | -0.298894389 | 0.180803186 | -0.479697575 | 0.999933887 | 0.475478579 | 1.013450121 | -0.537971542 | 0.069806796 |
| ENSG00000153774 | *CFDP1* | 3.142035822 | 0.005000587 | 0.002580002 | 0.002420585 | 0.999933887 | 0.019649114 | 0.235795579 | -0.216146466 | 0.069957981 |
| ENSG00000215305 | *VPS16* | 4.60060343 | 0.01649529 | 0.054276105 | -0.037780815 | 0.999933887 | -0.070171989 | -0.276835098 | 0.206663109 | 0.070051184 |
| ENSG00000038358 | *EDC4* | 5.057720152 | 0.024360808 | 0.093196651 | -0.068835844 | 0.999933887 | -0.092684995 | -0.272005111 | 0.179320115 | 0.070060142 |
| ENSG00000171603 | *CLSTN1* | 5.285193977 | 0.004854657 | 0.013274761 | -0.008420104 | 0.999933887 | -0.071435096 | -0.250303145 | 0.178868049 | 0.070217106 |
| ENSG00000132953 | *XPO4* | 2.977932849 | -0.099282193 | -0.197662795 | 0.098380602 | 0.999933887 | -0.108248552 | 0.249552998 | -0.35780155 | 0.070335538 |
| ENSG00000143319 | *ISG20L2* | 5.689154658 | 0.041365023 | 0.022853958 | 0.018511066 | 0.999933887 | 0.112164554 | -0.05127491 | 0.163439464 | 0.070335538 |
| ENSG00000131002 | *TXLNGY* | -3.058917679 | 0.060247978 | 0.286962011 | -0.226714033 | 0.999933887 | -0.601699178 | 0.101350652 | -0.70304983 | 0.070388792 |
| ENSG00000156471 | *PTDSS1* | 4.272446362 | -0.004076968 | -0.027652869 | 0.0235759 | 0.999933887 | -0.114090083 | -0.334290365 | 0.220200282 | 0.070532836 |
| ENSG00000276644 | *DACH1* | 1.701148365 | 0.019550347 | 0.049184455 | -0.029634108 | 0.999933887 | -0.331397749 | -0.760377237 | 0.428979488 | 0.070533343 |
| ENSG00000121797 | *CCRL2* | 9.27961563 | 0.177088224 | 0.150688287 | 0.026399937 | 0.999933887 | 0.968519999 | 1.220824993 | -0.252304994 | 0.070614513 |
| ENSG00000140948 | *ZCCHC14* | 4.406145007 | -0.002775292 | 0.060360712 | -0.063136003 | 0.999933887 | 0.164161574 | 0.359367897 | -0.195206323 | 0.070629268 |
| ENSG00000116670 | *MAD2L2* | 3.319725389 | 0.085765228 | 0.074380516 | 0.011384711 | 0.999933887 | 0.269117426 | 0.585012771 | -0.315895345 | 0.070665163 |
| ENSG00000279080 | *AL022322.2* | 0.663344599 | -0.026206628 | -0.025828465 | -0.000378162 | 0.999933887 | 0.574202654 | 1.100419569 | -0.526216914 | 0.070678741 |
| ENSG00000089693 | *MLF2* | 7.258258098 | 0.031915006 | 0.020386153 | 0.011528853 | 0.999933887 | -0.150060393 | -0.31182961 | 0.161769218 | 0.070776572 |
| ENSG00000100422 | *CERK* | 6.788342663 | 0.020748829 | 0.010059439 | 0.01068939 | 0.999933887 | 0.352702 | 0.540414495 | -0.187712494 | 0.070997003 |
| ENSG00000272645 | *GTF2IP20* | 2.214093417 | 0.045021252 | -0.019615347 | 0.064636599 | 0.999933887 | 0.468902894 | 0.857676745 | -0.38877385 | 0.071193228 |
| ENSG00000257122 | *RRN3P3* | 3.173837298 | -0.044007268 | -0.005300036 | -0.038707232 | 0.999933887 | -0.150561425 | 0.244477461 | -0.395038886 | 0.071193228 |
| ENSG00000196358 | *NTNG2* | 6.080018809 | -0.013336111 | 0.083847647 | -0.097183758 | 0.999933887 | 0.049422385 | -0.125319618 | 0.174742003 | 0.071239573 |
| ENSG00000183431 | *SF3A3* | 4.41264409 | -0.068105899 | -0.027742753 | -0.040363145 | 0.999933887 | -0.211604512 | -0.414504645 | 0.202900133 | 0.07137411 |

| | | | | | | | | | | |
|---|---|---|---|---|---|---|---|---|---|---|
| ENSG00000179134 | *SAMD4B* | 7.366603729 | 0.021938984 | -0.041086181 | 0.063025165 | 0.999933887 | 0.343874946 | 0.519650456 | -0.17577551 | 0.07154832 |
| ENSG00000261416 | *AC012645.3* | 1.106662166 | -0.073589172 | -0.063084925 | -0.010504246 | 0.999933887 | -0.605006662 | -1.197425773 | 0.59241911 | 0.071577488 |
| ENSG00000186130 | *ZBTB6* | 2.904495848 | -0.245662808 | -0.384441136 | 0.138778327 | 0.999933887 | -0.287314073 | 0.033118342 | -0.320432415 | 0.07184797 |
| ENSG00000111817 | *DSE* | 6.164804694 | -0.067888782 | -0.018787702 | -0.04910108 | 0.999933887 | 0.891811264 | 1.199093922 | -0.307282658 | 0.07184797 |
| ENSG00000085063 | *CD59* | 5.803927278 | -0.040911262 | -0.068781911 | 0.027870649 | 0.999933887 | 0.831704217 | 1.082961044 | -0.251256827 | 0.071929796 |
| ENSG00000115685 | *PPP1R7* | 3.698670207 | 0.043128307 | -0.055368816 | 0.098497123 | 0.999933887 | -0.050618514 | -0.322122406 | 0.271503893 | 0.072081434 |
| ENSG00000099821 | *POLRMT* | 4.063876085 | 0.05097835 | 0.046033752 | 0.004944598 | 0.999933887 | 0.160683922 | -0.085386887 | 0.246070808 | 0.072140107 |
| ENSG00000232618 | *AL355304.1* | 2.088166132 | -0.025589664 | 0.043146526 | -0.06873619 | 0.999933887 | 2.139970189 | 2.540792917 | -0.400822728 | 0.072149212 |
| ENSG00000198898 | *CAPZA2* | 7.305577892 | -0.079863647 | -0.117014271 | 0.037150624 | 0.999933887 | -0.222859873 | -0.021441723 | -0.20141815 | 0.072213057 |
| ENSG00000119321 | *FKBP15* | 7.252149514 | -0.046142154 | -0.068968468 | 0.022826314 | 0.999933887 | -0.241034771 | -0.058492706 | -0.182542064 | 0.072220829 |
| ENSG00000122033 | *MTIF3* | 2.262526335 | 0.04057186 | 0.003471688 | 0.037100172 | 0.999933887 | -0.2831393 | -0.655736333 | 0.372597032 | 0.072445183 |
| ENSG00000070495 | *JMJD6* | 7.979107113 | 0.175145053 | 0.128947379 | 0.046197674 | 0.999933887 | 0.620035392 | 0.902184003 | -0.282148611 | 0.072508371 |
| ENSG00000050327 | *ARHGEF5* | 0.22209739 | 0.007491473 | 0.221246224 | -0.21375475 | 0.999933887 | -0.265472352 | 0.341776403 | -0.607248755 | 0.072645451 |
| ENSG00000132330 | *SCLY* | 0.317544245 | 0.120925085 | -0.211760808 | 0.332685894 | 0.999933887 | 0.013799822 | -0.512138533 | 0.525938355 | 0.072665162 |
| ENSG00000198888 | *MT-ND1* | 8.980604584 | 0.061688084 | -0.010728153 | 0.072416237 | 0.999933887 | 0.107772864 | 0.384820907 | -0.277048043 | 0.072686694 |
| ENSG00000233429 | *HOTAIRM1* | 2.456812701 | -0.234744469 | -0.084648198 | -0.15009627 | 0.999933887 | -1.4480322 | -1.900113827 | 0.452081627 | 0.072696158 |
| ENSG00000196428 | *TSC22D2* | 7.372765204 | 0.096551867 | 0.168098114 | -0.071546247 | 0.999933887 | 0.575365664 | 0.77003258 | -0.194666916 | 0.072696158 |
| ENSG00000272837 | *OR10AE3P* | 2.291622168 | 0.247987111 | 0.214418516 | 0.033568595 | 0.999933887 | 0.96438017 | 1.327085401 | -0.362705232 | 0.07273399 |
| ENSG00000136754 | *ABI1* | 6.584156638 | -0.03510435 | -0.04447133 | 0.00936698 | 0.999933887 | -0.108298421 | 0.040619585 | -0.148918006 | 0.07273399 |
| ENSG00000060642 | *PIGV* | 2.855831322 | -0.047773484 | 0.059576829 | -0.107350313 | 0.999933887 | 1.339954065 | 1.694561156 | -0.354607091 | 0.07273399 |
| ENSG00000270170 | *NCBP2AS2* | 2.415080808 | -0.04205049 | -0.157743622 | 0.115693132 | 0.999933887 | -0.140623621 | -0.439569253 | 0.298945632 | 0.072838902 |
| ENSG00000134330 | *IAH1* | 3.265824252 | -0.039019056 | -0.115412425 | 0.07639337 | 0.999933887 | 0.618152737 | 0.85794467 | -0.239791933 | 0.072878109 |
| ENSG00000122218 | *COPA* | 6.949514049 | -0.022279094 | -0.018796175 | -0.003482919 | 0.999933887 | 0.04704086 | 0.184343895 | -0.137303035 | 0.072938647 |
| ENSG00000253218 | *KLF3P1* | -0.989694179 | 0.687726242 | -0.011133039 | 0.698859281 | 0.999933887 | 1.600797469 | 2.39031111 | -0.789513641 | 0.073060658 |
| ENSG00000129675 | *ARHGEF6* | 5.500274411 | -0.021174216 | -0.039543986 | 0.018369771 | 0.999933887 | -0.181040082 | -0.354928575 | 0.173888492 | 0.073153573 |
| ENSG00000173821 | *RNF213* | 9.941437057 | 0.039610917 | -0.015700628 | 0.055311545 | 0.999933887 | -0.307940691 | -0.521540085 | 0.213599394 | 0.073249396 |
| ENSG00000053254 | *FOXN3* | 6.752509 | 0.002875589 | -0.013346438 | 0.016222027 | 0.999933887 | -0.536452324 | -0.726608199 | 0.190155875 | 0.073273741 |
| ENSG00000134463 | *ECHDC3* | -0.444256528 | -0.004895119 | 0.112052951 | -0.11694807 | 0.999933887 | -0.353689731 | 0.562846825 | -0.916536557 | 0.073273741 |
| ENSG00000269837 | *IPO5P1* | 0.594013458 | -0.166938211 | 0.001104878 | -0.168043089 | 0.999933887 | -0.047137889 | -0.544971255 | 0.497833367 | 0.073431992 |
| ENSG00000151743 | *AMN1* | 3.357948148 | -0.090433068 | -0.108767199 | 0.018334131 | 0.999933887 | -0.03704372 | 0.174914099 | -0.211957819 | 0.073458979 |
| ENSG00000119596 | *YLPM1* | 4.899197273 | 0.001063039 | -0.028603281 | 0.02966632 | 0.999933887 | -0.228919562 | -0.412635856 | 0.183716294 | 0.073458979 |
| ENSG00000141568 | *FOXK2* | 3.735550946 | -0.004356859 | 0.078720285 | -0.083077143 | 0.999933887 | -0.169626842 | -0.390840354 | 0.221213512 | 0.073474449 |
| ENSG00000105701 | *FKBP8* | 8.357538712 | 0.061255898 | 0.039608028 | 0.02164787 | 0.999933887 | -0.213838173 | -0.492695008 | 0.278856835 | 0.073671867 |
| ENSG00000159082 | *SYNJ1* | 5.448584922 | -0.022065995 | -0.061512126 | 0.03944613 | 0.999933887 | -0.533135464 | -0.21222609 | -0.320909374 | 0.073767647 |
| ENSG00000163902 | *RPN1* | 6.83426712 | -0.037010222 | -0.038553427 | 0.001543205 | 0.999933887 | -0.501773186 | -0.695053381 | 0.193280195 | 0.073775017 |
| ENSG00000000938 | *FGR* | 10.396319838 | 0.040178539 | 0.040135454 | 4.3084e-05 | 0.999933887 | 0.511760109 | 0.681251555 | -0.169491446 | 0.073815165 |
| ENSG00000162601 | *MYSM1* | 5.69542748 | -0.060743138 | -0.182370784 | 0.121627646 | 0.999933887 | -0.315077013 | 0.30471781 | -0.619794824 | 0.073815165 |
| ENSG00000149483 | *TMEM138* | 3.860073476 | -0.019330892 | 0.033903958 | -0.05323485 | 0.999933887 | 0.543039504 | 0.762450798 | -0.219411294 | 0.073815165 |
| ENSG00000120071 | *KANSL1* | 6.399923103 | -0.026818833 | 0.040208966 | -0.067027798 | 0.999933887 | -0.491182444 | -0.668418272 | 0.177235829 | 0.074005086 |
| ENSG00000130755 | *GMFG* | 8.405344904 | -0.01303103 | -0.036528928 | 0.023497898 | 0.999933887 | -0.240672261 | -0.428161413 | 0.187489152 | 0.074066285 |
| ENSG00000072818 | *ACAP1* | 7.015680698 | 0.028126674 | 0.049471298 | -0.021344625 | 0.999933887 | 0.002294416 | -0.201398283 | 0.203692699 | 0.074182239 |
| ENSG00000167100 | *SAMD14* | -0.864445926 | 0.471203907 | -0.226028913 | 0.69723282 | 0.999933887 | 2.169822174 | 2.85126605 | -0.681443876 | 0.07430869 |
| ENSG00000166483 | *WEE1* | 2.2445774 | -0.072734927 | -0.082541169 | 0.009806242 | 0.999933887 | 0.232013659 | 0.531322509 | -0.29930885 | 0.07430869 |
| ENSG00000158195 | *WASF2* | 7.06743894 | -0.021695868 | -0.011115247 | -0.010580621 | 0.999933887 | -0.408441015 | -0.575557124 | 0.167116109 | 0.07430869 |
| ENSG00000135272 | *MDFIC* | 3.323635118 | -0.015200505 | -0.016133097 | 0.000932591 | 0.999933887 | 0.065329167 | 0.427024631 | -0.361695464 | 0.07430869 |
| ENSG00000130813 | *SHFL* | 4.308028998 | 0.068001986 | -0.016736792 | 0.084738778 | 0.999933887 | -0.351618557 | -0.617174196 | 0.265555639 | 0.074498924 |
| ENSG00000136169 | *SETDB2* | 5.339848715 | 0.067023573 | 0.078218672 | -0.011195099 | 0.999933887 | 0.238403055 | 0.484422764 | -0.246019708 | 0.074583932 |
| ENSG00000255112 | *CHMP1B* | 7.148321043 | -0.025998632 | -0.041697754 | 0.015699122 | 0.999933887 | 0.024480259 | -0.150731191 | 0.175211451 | 0.074583932 |
| ENSG00000099795 | *NDUFB7* | 2.732125879 | 0.102822402 | -0.02256411 | 0.125386512 | 0.999933887 | 0.120633971 | -0.234372566 | 0.355006537 | 0.074611564 |
| ENSG00000117408 | *IPO13* | 2.822253328 | 0.068887036 | 0.102941 | -0.034053964 | 0.999933887 | -0.082619999 | -0.369149355 | 0.286529357 | 0.074668978 |
| ENSG00000140983 | *RHOT2* | 6.55917094 | 0.052413787 | 0.101870332 | -0.049456545 | 0.999933887 | 0.030285365 | -0.16844624 | 0.198731604 | 0.074724517 |
| ENSG00000103657 | *HERC1* | 6.84917979 | -0.033184562 | -0.045035569 | 0.011851007 | 0.999933887 | -0.154248738 | -0.349203254 | 0.194954516 | 0.074724517 |
| ENSG00000138942 | *RNF185* | 5.844334453 | -0.022637705 | -0.043848405 | 0.0212107 | 0.999933887 | 0.074744146 | 0.261504334 | -0.186760188 | 0.074819991 |
| ENSG00000081237 | *PTPRC* | 10.370218142 | -0.061863045 | -0.2108921 | 0.149029054 | 0.999933887 | -0.138102866 | 0.294877193 | -0.432980059 | 0.074849221 |
| ENSG00000124882 | *EREG* | 2.92901325 | 0.504787474 | 0.614340346 | -0.109552872 | 0.999933887 | 1.810807195 | 2.571385346 | -0.760578151 | 0.075073322 |
| ENSG00000275052 | *PPP4R3B* | 5.96618128 | -0.025842373 | -0.071460146 | 0.045617772 | 0.999933887 | -0.370746443 | -0.127657736 | -0.243088707 | 0.07519984 |
| ENSG00000076984 | *MAP2K7* | 5.688874687 | 0.044865145 | 0.063293853 | -0.018428708 | 0.999933887 | -0.273008545 | -0.490920206 | 0.217911661 | 0.075231217 |
| ENSG00000010610 | *CD4* | 4.440126905 | 0.036688251 | -0.008103776 | 0.044792027 | 0.999933887 | 0.04887548 | -0.196424263 | 0.245299743 | 0.075231217 |
| ENSG00000161904 | *LEMD2* | 5.719522373 | -0.029906107 | 0.046504779 | -0.076410886 | 0.999933887 | -0.238395697 | -0.427898939 | 0.189503242 | 0.075989295 |
| ENSG00000237190 | *CDKN2AIPNL* | 1.769277323 | 0.087156143 | 0.097675652 | -0.010519509 | 0.999933887 | 0.084903311 | 0.471711899 | -0.386808588 | 0.07602411 |
| ENSG00000121964 | *GTDC1* | 4.478742104 | -0.015865023 | -0.064216941 | 0.048351917 | 0.999933887 | 0.066823864 | 0.242631743 | -0.17580788 | 0.07614709 |
| ENSG00000171988 | *JMJD1C* | 9.327119403 | -0.098373818 | -0.140726147 | 0.042352329 | 0.999933887 | -0.593018379 | -0.054917122 | -0.538101256 | 0.07616359 |
| ENSG00000174720 | *LARP7* | 4.421011863 | -0.06067613 | -0.28457339 | 0.22389726 | 0.999933887 | -0.096673663 | 0.315686815 | -0.412360478 | 0.076222275 |
| ENSG00000234678 | *ELF3-AS1* | 0.481730902 | 0.032210124 | -0.135786607 | 0.167996731 | 0.999933887 | -0.603076058 | -1.193070401 | 0.589994343 | 0.076222275 |
| ENSG00000172348 | *RCAN2* | -0.698716885 | -0.110778764 | -0.149540095 | 0.038761331 | 0.999933887 | -0.056061834 | 0.565994847 | -0.622056681 | 0.07641288 |
| ENSG00000130734 | *ATG4D* | 4.135295663 | 0.095296745 | 0.117231643 | -0.021934898 | 0.999933887 | -0.064768038 | -0.337784829 | 0.273016791 | 0.076568709 |
| ENSG00000144021 | *CIAO1* | 5.019215158 | -0.005791715 | -0.010122979 | 0.004331264 | 0.999933887 | -0.189326838 | -0.36593209 | 0.176605252 | 0.076624773 |
| ENSG00000103266 | *STUB1* | 3.198104171 | 0.059864966 | 0.072720801 | -0.012855835 | 0.999933887 | -0.114736702 | -0.479937084 | 0.365200382 | 0.076714828 |
| ENSG00000276809 | *AL138955.1* | 0.899839553 | -0.016205076 | 0.377377247 | -0.393582323 | 0.999933887 | 1.072956572 | 1.563495584 | -0.490539012 | 0.076714828 |
| ENSG00000090339 | *ICAM1* | 9.146844484 | 0.173766103 | 0.147916766 | 0.025849337 | 0.999933887 | 2.317859041 | 2.639269358 | -0.321410317 | 0.076906361 |
| ENSG00000243660 | *ZNF487* | 2.596871708 | -0.025615216 | -0.064460399 | 0.038845183 | 0.999933887 | -0.518445665 | -0.795867986 | 0.277422321 | 0.077295655 |
| ENSG00000173674 | *EIF1AX* | 4.046784186 | -0.078805595 | -0.118842905 | 0.04003731 | 0.999933887 | 0.027851706 | 0.271514083 | -0.243662376 | 0.077339698 |
| ENSG00000180573 | *H2AC6* | 6.886647227 | 0.012679842 | -0.022369214 | 0.035049056 | 0.999933887 | -0.630424632 | -0.915873476 | 0.285448843 | 0.077369867 |
| ENSG00000125744 | *RTN2* | 4.148278245 | 0.141908103 | 0.039477945 | 0.102430158 | 0.999933887 | 0.761752953 | 1.085140285 | -0.323387333 | 0.077399235 |
| ENSG00000174276 | *ZNHIT2* | 0.128115771 | 0.158668241 | -0.105402393 | 0.264070634 | 0.999933887 | 0.298557404 | -0.3412088 | 0.639766204 | 0.077399235 |
| ENSG00000166888 | *STAT6* | 9.146789959 | -0.015089137 | 6.4151e-05 | -0.015153287 | 0.999933887 | -0.572907873 | -0.733745303 | 0.16083743 | 0.077403716 |
| ENSG00000104679 | *R3HCC1* | 2.451684981 | -0.075927902 | -0.12291581 | 0.046987908 | 0.999933887 | 0.017833877 | -0.285776984 | 0.303610861 | 0.077426312 |
| ENSG00000175309 | *PHYKPL* | 5.21525258 | 0.009226611 | 0.036098707 | -0.026872096 | 0.999933887 | -0.263396565 | -0.438745362 | 0.175348798 | 0.077426312 |
| ENSG00000170779 | *CDCA4* | 2.315532622 | 0.044566231 | -0.152196314 | 0.196762545 | 0.999933887 | -0.273827152 | -0.616372241 | 0.342545089 | 0.077549176 |
| ENSG00000137841 | *PLCB2* | 7.382253873 | 0.028722786 | 0.088618105 | -0.059895319 | 0.999933887 | -0.033669505 | -0.216234395 | 0.18256489 | 0.077643225 |
| ENSG00000172840 | *PDP2* | 1.528740082 | 0.039720892 | 0.133606672 | -0.09388578 | 0.999933887 | -0.477360626 | -0.849174009 | 0.371813383 | 0.077643225 |
| ENSG00000187239 | *FNBP1* | 7.054889915 | 0.001043335 | -0.06917665 | 0.070219985 | 0.999933887 | 0.324839846 | 0.554379396 | -0.229539551 | 0.077762462 |
| ENSG00000110321 | *EIF4G2* | 8.590448758 | -0.037156532 | -0.073417835 | 0.036261304 | 0.999933887 | -0.29872794 | -0.433882908 | 0.135154968 | 0.077776622 |
| ENSG00000111845 | *PAK1IP1* | 1.241078461 | -0.10851592 | -0.093116066 | -0.015399854 | 0.999933887 | 0.18734766 | 0.546475515 | -0.359127856 | 0.077986305 |
| ENSG00000165406 | *MARCHF8* | 6.574376188 | -0.058806815 | -0.005671192 | -0.053135623 | 0.999933887 | -0.755098746 | -0.957387644 | 0.202288899 | 0.077986305 |
| ENSG00000070061 | *ELP1* | 3.751935374 | -0.032207858 | -0.124613799 | 0.09240594 | 0.999933887 | -0.050407611 | 0.183923255 | -0.234330866 | 0.078056865 |
| ENSG00000220305 | *HNRNPH1P1* | 0.317824866 | -0.126559879 | -0.082064847 | -0.044495032 | 0.999933887 | 1.357770108 | 1.927671583 | -0.569901475 | 0.078416354 |
| ENSG00000256660 | *CLEC12B* | 2.080038988 | -0.020939656 | -0.042797711 | 0.021858055 | 0.999933887 | 0.754332448 | 1.296692756 | -0.542360308 | 0.07848573 |
| ENSG00000111696 | *NT5DC3* | 4.202468875 | 0.059073263 | 0.06638582 | -0.007312557 | 0.999933887 | 0.567697706 | 0.792858478 | -0.225160772 | 0.078895267 |
| ENSG00000157600 | *TMEM164* | 6.61257457 | -0.050604252 | 0.024345122 | -0.074949374 | 0.999933887 | -0.70895676 | -1.024396739 | 0.31543998 | 0.078970192 |
| ENSG00000197362 | *ZNF786* | 0.62073953 | -0.079746469 | -0.013722224 | -0.066024245 | 0.999933887 | 0.084258502 | -0.351308222 | 0.435566724 | 0.079169705 |
| ENSG00000182685 | *BRICD5* | 0.868216506 | 0.090052601 | 0.357993216 | -0.267940615 | 0.999933887 | -0.060914622 | -0.541170355 | 0.480255733 | 0.079203204 |
| ENSG00000132024 | *CC2D1A* | 3.74215623 | 0.029993653 | 0.012677526 | 0.017316127 | 0.999933887 | -0.075364735 | -0.313406746 | 0.238042011 | 0.079324044 |
| ENSG00000140259 | *MFAP1* | 4.204191911 | -0.089559535 | -0.108332146 | 0.018772611 | 0.999933887 | 0.197870075 | 0.431626569 | -0.233756494 | 0.079460411 |
| ENSG00000144455 | *SUMF1* | 2.915190422 | 0.048645955 | -0.072057604 | 0.120703559 | 0.999933887 | -0.25617649 | -0.505139227 | 0.248962737 | 0.079514304 |
| ENSG00000134058 | *CDK7* | 4.168742207 | -0.061521808 | -0.098708245 | 0.037186437 | 0.999933887 | 1.102229399 | 1.371777237 | -0.269547838 | 0.079680012 |
| ENSG00000167106 | *FAM102A* | 5.941288506 | -0.000464001 | 0.049390922 | -0.049854923 | 0.999933887 | 0.032887467 | -0.105163308 | 0.138050775 | 0.079800417 |
| ENSG00000115816 | *CEBPZ* | 3.437103359 | -0.095766593 | -0.245708572 | 0.149941979 | 0.999933887 | -0.112163045 | 0.271251438 | -0.383414484 | 0.079816791 |
| ENSG00000197956 | *S100A6* | 8.381663824 | 0.001174624 | -0.064886265 | 0.066060889 | 0.999933887 | -0.182461194 | -0.370647602 | 0.188186409 | 0.079816791 |
| ENSG00000204923 | *FBXO48* | 1.404790874 | -0.010391505 | 0.286705644 | -0.297097149 | 0.999933887 | 0.746488372 | 1.120590098 | -0.374101727 | 0.079876883 |
| ENSG00000142634 | *EFHD2* | 9.996269323 | 0.069768899 | 0.031463843 | 0.038305056 | 0.999933887 | 0.367497661 | 0.57751811 | -0.210020449 | 0.079952386 |
| ENSG00000116497 | *S100PBP* | 3.52929275 | 0.016101024 | 0.000421156 | 0.015679868 | 0.999933887 | -0.20026535 | -0.462163319 | 0.26189797 | 0.079956638 |
| ENSG00000119929 | *CUTC* | 2.806771337 | 0.015545314 | -0.125004961 | 0.140550274 | 0.999933887 | -0.428359708 | -0.727989967 | 0.299630259 | 0.079956638 |
| ENSG00000145919 | *BOD1* | 3.194586688 | -0.021794819 | -0.057043807 | 0.035248988 | 0.999933887 | 0.521687458 | 0.791952684 | -0.270265226 | 0.080018524 |
| ENSG00000189050 | *RNFT1* | 3.012647519 | -0.103849497 | -0.036577048 | -0.067272449 | 0.999933887 | -0.236339486 | 0.002627496 | -0.238966983 | 0.080060434 |

| | | | | | | | | | | |
|---|---|---|---|---|---|---|---|---|---|---|
| ENSG00000198087 | *CD2AP* | 2.686988846 | -0.000932304 | -0.117649063 | 0.116716759 | 0.999933887 | -0.22751151 | 0.260131587 | -0.487643097 | 0.0801872 |
| ENSG00000061656 | *SPAG4* | 2.004534627 | 0.072406894 | 0.114275313 | -0.041868419 | 0.999933887 | 0.221997833 | 0.59905615 | -0.377058317 | 0.080306088 |
| ENSG00000177731 | *FLII* | 7.498768741 | 0.005267815 | 0.025317196 | -0.020049382 | 0.999933887 | -0.642334497 | -0.821461315 | 0.179126818 | 0.080383142 |
| ENSG00000077684 | *JADE1* | 4.571211898 | 0.053855725 | -0.018476889 | 0.072332614 | 0.999933887 | -0.288506498 | -0.545468496 | 0.256961998 | 0.080467073 |
| ENSG00000235106 | *BRD3OS* | 0.932210158 | 0.085334473 | 0.024219058 | 0.061115415 | 0.999933887 | -0.094634623 | -0.526002508 | 0.431367885 | 0.080467073 |
| ENSG00000136146 | *MED4* | 3.953036812 | -0.058017245 | -0.038861204 | -0.019156041 | 0.999933887 | -0.190486492 | -0.002985107 | -0.187501386 | 0.080478592 |
| ENSG00000144567 | *RETREG2* | 7.051682774 | 0.008661852 | -0.023822357 | 0.032484209 | 0.999933887 | -0.763320863 | -0.974511308 | 0.211190445 | 0.080515465 |
| ENSG00000119729 | *RHOQ* | 6.195673601 | -0.057281236 | -0.003845815 | -0.053435421 | 0.999933887 | 0.168863224 | 0.402521662 | -0.233658439 | 0.080529093 |
| ENSG00000103051 | *COG4* | 3.329783634 | 0.00708321 | -0.068885835 | 0.075969046 | 0.999933887 | -0.168392979 | -0.396572072 | 0.228179093 | 0.080529093 |
| ENSG00000092820 | *EZR* | 7.131316091 | -0.056622886 | -0.002219225 | -0.054403661 | 0.999933887 | 0.153936248 | 0.314446013 | -0.160509765 | 0.081117656 |
| ENSG00000186105 | *LRRC70* | 3.683260443 | -0.035197006 | 0.059887039 | -0.095084045 | 0.999933887 | 0.084967859 | 0.385981583 | -0.301013723 | 0.081152743 |
| ENSG00000187840 | *EIF4EBP1* | 1.667288106 | -0.015865632 | 0.083895867 | -0.099761499 | 0.999933887 | -0.654515459 | -1.147169897 | 0.492654438 | 0.081288345 |
| ENSG00000100368 | *CSF2RB* | 9.710640926 | 0.055330881 | 0.072076888 | -0.016746008 | 0.999933887 | 0.48371395 | 0.298007924 | 0.185706026 | 0.08156402 |
| ENSG00000287561 | *AL121899.4* | 3.272192498 | 0.26357207 | 0.309588619 | -0.046016549 | 0.999933887 | 0.349657261 | 0.721902816 | -0.372245555 | 0.08166956 |
| ENSG00000109332 | *UBE2D3* | 9.397169272 | 0.009847078 | -0.011103509 | 0.020950587 | 0.999933887 | -0.001379167 | 0.109683862 | -0.111063029 | 0.08166956 |
| ENSG00000143384 | *MCL1* | 10.51752024 | 0.108846884 | 0.150557153 | -0.041710269 | 0.999933887 | 0.079473914 | -0.115695362 | 0.195169276 | 0.081825545 |
| ENSG00000141098 | *GFOD2* | 2.900155792 | 0.00270307 | 0.005995311 | -0.003292241 | 0.999933887 | -0.753156152 | -1.168049753 | 0.414893601 | 0.081825545 |
| ENSG00000185361 | *TNFAIP8L1* | 2.939060593 | -0.079655452 | -0.065420604 | -0.014234848 | 0.999933887 | -0.198959234 | -0.508986163 | 0.310026929 | 0.081854666 |
| ENSG00000231856 | *AL162377.1* | 1.789461482 | 0.058255695 | 0.127719853 | -0.069464157 | 0.999933887 | 1.954160106 | 2.470049666 | -0.51588956 | 0.081870335 |
| ENSG00000157557 | *ETS2* | 8.946814827 | 0.214395643 | 0.213153731 | 0.001241911 | 0.999933887 | 1.67698242 | 1.990346957 | -0.313364537 | 0.08214628 |
| ENSG00000236200 | *KDM4A-AS1* | -0.197930636 | 0.22451804 | -0.081067386 | 0.305585426 | 0.999933887 | 0.082071219 | -0.764452852 | 0.846524071 | 0.08214628 |
| ENSG00000066117 | *SMARCD1* | 4.412487987 | -0.010667003 | -0.052085073 | 0.04141807 | 0.999933887 | -0.208363912 | -0.410127913 | 0.201764001 | 0.082181858 |
| ENSG00000186652 | *PRG2* | -0.507773972 | -0.088927638 | 0.254983194 | -0.343910832 | 0.999933887 | 1.282486089 | 1.880609956 | -0.598123867 | 0.082182978 |
| ENSG00000177889 | *UBE2N* | 4.960509159 | -0.01141612 | -0.026341868 | 0.014925747 | 0.999933887 | 0.187744273 | 0.358970817 | -0.171226544 | 0.082223984 |
| ENSG00000104870 | *FCGRT* | 7.909274817 | 0.024688019 | 0.013796872 | 0.010891148 | 0.999933887 | -0.312748289 | -0.531243604 | 0.218495315 | 0.082261214 |
| ENSG00000144674 | *GOLGA4* | 4.923088039 | -0.117445064 | -0.123472306 | 0.006027242 | 0.999933887 | -0.52232597 | 0.041152561 | -0.563478531 | 0.082277946 |
| ENSG00000123124 | *WWP1* | 3.569825502 | -0.033217861 | -0.147278919 | 0.114061058 | 0.999933887 | -0.15121154 | 0.084352417 | -0.235563957 | 0.082395481 |
| ENSG00000113303 | *BTNL8* | 4.548861581 | -0.009758908 | 0.025805147 | -0.035564056 | 0.999933887 | 0.209237907 | 0.412921575 | -0.203683668 | 0.082440874 |
| ENSG00000280080 | *TBC1D22A-AS* | 0.471353466 | 0.172912929 | -0.035900997 | 0.208813926 | 0.999933887 | 0.391274956 | 0.993539971 | -0.602265015 | 0.082556861 |
| ENSG00000198712 | *MT-CO2* | 9.16940856 | 0.029562096 | -0.06510153 | 0.094663626 | 0.999933887 | 0.03050244 | 0.334493705 | -0.303991265 | 0.082754203 |
| ENSG00000160117 | *ANKLE1* | -0.026168927 | 0.036440327 | 0.067806619 | -0.031366292 | 0.999933887 | 1.208506802 | 1.939734284 | -0.731227482 | 0.082754203 |
| ENSG00000134900 | *TPP2* | 5.572023732 | -0.004872675 | -0.12901651 | 0.124143835 | 0.999933887 | -1.02345795 | -1.380551728 | 0.357093778 | 0.082791095 |
| ENSG00000198218 | *QRICH1* | 5.816359665 | 0.001554505 | 0.027994966 | -0.026440461 | 0.999933887 | -0.192056939 | -0.351381765 | 0.159324826 | 0.082875323 |
| ENSG00000112357 | *PEX7* | -0.358042132 | -0.163638657 | 0.397443768 | -0.561082425 | 0.999933887 | 1.0627258 | 1.700354596 | -0.637628796 | 0.082907765 |
| ENSG00000102226 | *USP11* | 4.285975856 | -0.037045354 | -0.030711686 | -0.006333668 | 0.999933887 | 0.149261265 | 0.358081406 | -0.208820141 | 0.082907765 |
| ENSG00000178053 | *MLF1* | 0.437521251 | 0.05422811 | -0.256521854 | 0.310749964 | 0.999933887 | -0.008622347 | 0.566781099 | -0.575403446 | 0.082907765 |
| ENSG00000176595 | *KBTBD11* | 2.243324219 | -0.037603149 | -0.004630741 | -0.032972407 | 0.999933887 | -0.089532365 | -0.396455953 | 0.306923588 | 0.082907765 |
| ENSG00000135778 | *NTPCR* | 1.806208424 | -0.02734669 | 0.168911178 | -0.196257868 | 0.999933887 | -0.021889787 | -0.345189647 | 0.32329986 | 0.082907765 |
| ENSG00000089006 | *SNX5* | 4.269915971 | -0.002942856 | -0.044786741 | 0.041843886 | 0.999933887 | -0.08157075 | -0.255715614 | 0.174144864 | 0.082907765 |
| ENSG00000064763 | *FAR2* | 3.506073837 | -0.001255331 | -0.033530275 | 0.032274945 | 0.999933887 | -0.267257954 | -0.525444973 | 0.258187019 | 0.08291344 |
| ENSG00000226942 | *IL9RP3* | 1.298032608 | -0.278987558 | -0.184600442 | -0.094387115 | 0.999933887 | -1.427244212 | -1.850926553 | 0.423682342 | 0.083022978 |
| ENSG00000247287 | *AL359220.1* | 1.094922586 | 0.029104504 | -0.183806454 | 0.212910958 | 0.999933887 | -0.257263065 | 0.192675844 | -0.449938909 | 0.083022978 |
| ENSG00000152495 | *CAMK4* | 4.043641136 | 0.022203031 | -0.075820435 | 0.098023466 | 0.999933887 | -0.114408153 | 0.222616486 | -0.337024639 | 0.083057628 |
| ENSG00000140525 | *FANCI* | 2.030753733 | -0.043458828 | -0.036236897 | -0.007221932 | 0.999933887 | -0.009929975 | 0.363876519 | -0.373806494 | 0.083221995 |
| ENSG00000203879 | *GDI1* | 8.219053409 | 0.010629496 | 0.029870833 | -0.019241337 | 0.999933887 | 0.232696597 | 0.373503877 | -0.14080728 | 0.083221995 |
| ENSG00000101138 | *CSTF1* | 3.364221654 | -0.047493821 | -0.018007558 | -0.029486263 | 0.999933887 | -0.374038342 | -0.633127375 | 0.259089034 | 0.083246855 |
| ENSG00000261884 | *AC040162.1* | 3.140688962 | -0.076903795 | 0.017850477 | -0.094754272 | 0.999933887 | -0.529870579 | -0.785930816 | 0.256060237 | 0.083323161 |
| ENSG00000167207 | *NOD2* | 5.469518198 | 0.059432602 | -0.051807107 | 0.111239709 | 0.999933887 | 1.122940991 | 1.488205568 | -0.365264578 | 0.083338462 |
| ENSG00000105723 | *GSK3A* | 5.843319629 | 0.025656807 | 0.070250273 | -0.044593466 | 0.999933887 | -0.015700424 | 0.156631161 | -0.172331586 | 0.083338462 |
| ENSG00000009335 | *UBE3C* | 4.292297605 | -0.043915731 | -0.001126936 | -0.042788795 | 0.999933887 | -0.100668063 | -0.317114129 | 0.216446065 | 0.083353739 |
| ENSG00000039650 | *PNKP* | 5.17091932 | 0.03330873 | 0.060419503 | -0.027110773 | 0.999933887 | -0.204555824 | -0.428825733 | 0.224269909 | 0.083444817 |
| ENSG00000132017 | *DCAF15* | 5.864203973 | 0.039693071 | 0.059255422 | -0.019562351 | 0.999933887 | -0.302793659 | -0.523005471 | 0.220211812 | 0.083482224 |
| ENSG00000112249 | *ASCC3* | 2.894962664 | -0.012395598 | -0.110511834 | 0.098116236 | 0.999933887 | -0.100371057 | 0.261722297 | -0.362093354 | 0.083489954 |
| ENSG00000237886 | *NALT1* | 1.158609236 | -0.065784912 | -0.068556144 | 0.002771232 | 0.999933887 | -0.768516006 | -1.195598729 | 0.427082723 | 0.083559587 |
| ENSG00000105401 | *CDC37* | 6.819609154 | 0.05427355 | 0.013319353 | 0.040954196 | 0.999933887 | 0.376559703 | 0.564263571 | -0.187703868 | 0.083636891 |
| ENSG00000127445 | *PIN1* | 3.950836791 | 0.11381173 | 0.074537933 | 0.039273797 | 0.999933887 | -0.077319747 | -0.340091362 | 0.262771615 | 0.083648635 |
| ENSG00000037757 | *MRI1* | 1.895176818 | 0.082927915 | 0.134916776 | -0.051988862 | 0.999933887 | 0.060485656 | -0.269531977 | 0.330017633 | 0.083648635 |
| ENSG00000197114 | *ZGPAT* | 3.015301907 | 0.075508254 | 0.053815022 | 0.021693232 | 0.999933887 | -0.048509735 | -0.410876605 | 0.362366871 | 0.083893348 |
| ENSG00000183734 | *ASCL2* | 1.832329269 | 0.085935327 | 0.1461795 | -0.060244173 | 0.999933887 | 0.045615353 | 0.515644965 | -0.470029612 | 0.083893348 |
| ENSG00000130052 | *STARD8* | 4.148521086 | 0.112318429 | -0.032082129 | 0.144400558 | 0.999933887 | 1.09305787 | 1.3994452 | -0.306387329 | 0.084033495 |
| ENSG00000065135 | *GNAI3* | 7.187336532 | -0.039390804 | -0.047534362 | 0.008143558 | 0.999933887 | -0.311429103 | -0.137020662 | -0.174408441 | 0.084033495 |
| ENSG00000136999 | *CCN3* | 4.097613987 | -0.069675944 | -0.064893561 | -0.004782383 | 0.999933887 | -1.383094703 | -1.708306339 | 0.325211636 | 0.084033495 |
| ENSG00000077782 | *FGFR1* | 2.274791447 | 0.045288088 | 0.11391372 | -0.068625633 | 0.999933887 | -0.292823583 | -0.553281825 | 0.260458242 | 0.084039884 |
| ENSG00000168066 | *SF1* | 8.494133603 | 0.008912047 | 0.007744666 | 0.001167381 | 0.999933887 | -0.179347406 | -0.310901592 | 0.131554186 | 0.0840681 |
| ENSG00000105245 | *NUMBL* | 1.736837282 | 0.135673508 | -0.042421207 | 0.178094714 | 0.999933887 | -0.139479823 | -0.49779702 | 0.358317198 | 0.084115399 |
| ENSG00000127022 | *CANX* | 7.366698983 | -0.044612927 | -0.1185419 | 0.073928973 | 0.999933887 | 0.04050678 | 0.182780708 | -0.142273928 | 0.084115399 |
| ENSG00000186283 | *TOR3A* | 2.987429518 | 0.027149507 | 0.098076073 | -0.070926567 | 0.999933887 | 0.783039127 | 1.078374144 | -0.295335017 | 0.084126529 |
| ENSG00000130714 | *POMT1* | 3.432068489 | -0.031963791 | -0.033660285 | 0.001696494 | 0.999933887 | 0.180472598 | -0.037547367 | 0.218019965 | 0.084292877 |
| ENSG00000142959 | *BEST4* | 0.155447837 | -0.164572686 | 0.11947917 | -0.284051855 | 0.999933887 | 0.486964309 | 0.930265645 | -0.443301337 | 0.084318919 |
| ENSG00000189159 | *JPT1* | 5.397265201 | 0.056692193 | -0.036291698 | 0.092983891 | 0.999933887 | -0.04362089 | -0.338317526 | 0.294696637 | 0.084497382 |
| ENSG00000252391 | *RNU6-638P* | 0.389448081 | -0.033951239 | 0.054137164 | -0.088088403 | 0.999933887 | 0.639894168 | 1.096086051 | -0.456191883 | 0.084497382 |
| ENSG00000287763 | *AC068279.2* | -1.755103097 | 0.236873647 | 0.36027218 | -0.123398533 | 0.999933887 | 0.092675251 | -0.535017711 | 0.627692963 | 0.084635225 |
| ENSG00000077238 | *IL4R* | 8.288266041 | 0.015960666 | 0.010128904 | 0.005831762 | 0.999933887 | 0.54726258 | 0.737906114 | -0.190643534 | 0.08477547 |
| ENSG00000091483 | *FH* | 1.499630153 | -0.044373129 | -0.014140578 | -0.030232552 | 0.999933887 | 0.395189952 | 0.762293701 | -0.367103749 | 0.084783483 |
| ENSG00000101901 | *ALG13* | 3.617102817 | 0.007906474 | -0.008180272 | 0.016086747 | 0.999933887 | -0.022696678 | 0.24591389 | -0.268610568 | 0.084894001 |
| ENSG00000065559 | *MAP2K4* | 5.123879678 | -0.068807945 | -0.036008148 | -0.032799798 | 0.999933887 | -0.87979774 | -1.118506319 | 0.238708578 | 0.085233296 |
| ENSG00000171700 | *RGS19* | 7.027455144 | 0.07873626 | 0.070417591 | 0.008318669 | 0.999933887 | -0.917106208 | -1.21384323 | 0.296737022 | 0.085233296 |
| ENSG00000163251 | *FZD5* | 1.71807898 | -0.096780579 | -0.075956592 | -0.020823987 | 0.999933887 | 0.841261465 | 1.178185707 | -0.336924242 | 0.085233296 |
| ENSG00000165175 | *MID1IP1* | 7.636377249 | 0.192086709 | 0.200498357 | -0.008411648 | 0.999933887 | 0.342724381 | 0.599531066 | -0.256806685 | 0.085313828 |
| ENSG00000131759 | *RARA* | 7.83384247 | 0.06646499 | 0.044774969 | 0.021690021 | 0.999933887 | 0.363792601 | 0.563663251 | -0.19987065 | 0.085313828 |
| ENSG00000189350 | *TOGARAM2* | 2.242874555 | 0.239588827 | 0.268730512 | -0.029141685 | 0.999933887 | -0.455264305 | -0.819621617 | 0.364357312 | 0.085459525 |
| ENSG00000068697 | *LAPTM4A* | 6.517842602 | 0.033750133 | -0.003613446 | 0.03736358 | 0.999933887 | -0.332042909 | -0.54355991 | 0.211517001 | 0.085459525 |
| ENSG00000082805 | *ERC1* | 3.291203974 | 0.000524979 | 0.098625769 | -0.09810079 | 0.999933887 | 0.015613372 | 0.250970047 | -0.235356674 | 0.085459525 |
| ENSG00000188396 | *TCTEX1D4* | 3.561811665 | 0.30507307 | 0.328285347 | -0.023212277 | 0.999933887 | 1.644528092 | 2.074508364 | -0.429980273 | 0.08557155 |
| ENSG00000125740 | *FOSB* | 9.268129828 | 0.295840982 | 0.154454419 | 0.141386563 | 0.999933887 | 0.750744756 | 1.144415125 | -0.39367037 | 0.08557155 |
| ENSG00000186660 | *ZFP91* | 5.43012791 | -0.053392183 | -0.041623029 | -0.011769155 | 0.999933887 | -0.154369089 | -0.320278224 | 0.165909135 | 0.08557155 |
| ENSG00000101654 | *RNMT* | 5.814616322 | -0.077067281 | -0.13317749 | 0.056110209 | 0.999933887 | 0.056170775 | 0.326709491 | -0.270538717 | 0.08557155 |
| ENSG00000138629 | *UBL7* | 4.01029996 | 0.053560632 | 0.064819846 | -0.011259215 | 0.999933887 | -0.688130276 | -0.973765733 | 0.285635457 | 0.08557155 |
| ENSG00000060558 | *GNA15* | 4.451363642 | 0.029700536 | 0.047430033 | -0.017729496 | 0.999933887 | 0.921132569 | 1.168029504 | -0.246896935 | 0.08557155 |
| ENSG00000230943 | *LINC02541* | 0.626471896 | 0.045234287 | 0.212500784 | -0.167266497 | 0.999933887 | 3.207116675 | 3.87581984 | -0.668703164 | 0.08557155 |
| ENSG00000167977 | *KCTD5* | 4.332639431 | -0.044129092 | 0.019334159 | -0.063463251 | 0.999933887 | -0.638216064 | -0.852204164 | 0.2139881 | 0.085852636 |
| ENSG00000135047 | *CTSL* | 4.212821983 | -0.044001743 | -0.176011734 | 0.132009991 | 0.999933887 | 0.439264876 | 0.692594878 | -0.253330002 | 0.085852636 |
| ENSG00000071051 | *NCK2* | 4.971065671 | 0.095027908 | 0.087577957 | 0.00744995 | 0.999933887 | 0.313370711 | 0.502362915 | -0.188992204 | 0.085855344 |
| ENSG00000120860 | *WASHC3* | 2.228454668 | -0.068431388 | -0.085763673 | 0.017332285 | 0.999933887 | -0.005682976 | 0.301560028 | -0.307243004 | 0.085855344 |
| ENSG00000131480 | *AOC2* | 3.666327952 | 0.155779891 | -0.022740414 | 0.178520304 | 0.999933887 | 1.195670213 | 0.708357118 | 0.487313095 | 0.085940877 |
| ENSG00000203326 | *ZNF525* | 0.535340903 | -0.36049767 | -0.140027219 | -0.220470451 | 0.999933887 | -0.152669637 | 0.420344532 | -0.573014169 | 0.086094428 |
| ENSG00000159322 | *ADPGK* | 7.363904595 | -0.035886774 | 0.015655736 | -0.051542509 | 0.999933887 | 0.345737428 | 0.509539054 | -0.163801626 | 0.086096989 |
| ENSG00000166546 | *BEAN1* | -0.111386644 | -0.108198872 | -0.005248847 | -0.102950026 | 0.999933887 | -0.545399766 | -1.271531705 | 0.726131939 | 0.086290805 |
| ENSG00000173786 | *CNP* | 3.465769853 | 0.033453197 | -0.007807179 | 0.041260376 | 0.999933887 | -0.301166245 | -0.563316867 | 0.262150622 | 0.08630493 |
| ENSG00000110931 | *CAMKK2* | 9.036289411 | -0.009693254 | -0.006050991 | -0.003642262 | 0.999933887 | 0.490227931 | 0.651638725 | -0.161410794 | 0.08630493 |
| ENSG00000277895 | *AC135279.3* | -0.448223445 | 0.127511309 | 0.122679781 | 0.004831528 | 0.999933887 | 2.643360535 | 3.532218941 | -0.888858406 | 0.086325192 |
| ENSG00000181800 | *CELF2-AS1* | 3.613900728 | 0.062499023 | -0.04500544 | 0.107504463 | 0.999933887 | -0.885972036 | -1.475947266 | 0.589975231 | 0.086588726 |
| ENSG00000254685 | *FPGT* | 0.75724841 | -0.130674769 | -0.217610615 | 0.086935846 | 0.999933887 | -0.488809894 | -1.049570767 | 0.560760873 | 0.086691886 |
| ENSG00000228278 | *ORM2* | 2.99308938 | -0.438702892 | 0.233496986 | -0.672199878 | 0.999933887 | 2.031898618 | 2.542636247 | -0.510737629 | 0.086928715 |
| ENSG00000177054 | *ZDHHC13* | 3.314836524 | -0.07492564 | 0.081571246 | -0.156496886 | 0.999933887 | -0.308967283 | -0.074053329 | -0.234913954 | 0.086937213 |

| | | | | | | | | | | |
|---|---|---|---|---|---|---|---|---|---|---|
| ENSG00000100504 | *PYGL* | 6.318384433 | -0.031690825 | -0.02669161 | -0.004999215 | 0.999933887 | -0.151296991 | -0.339263069 | 0.187966078 | 0.086951674 |
| ENSG00000165804 | *ZNF219* | 2.957213228 | 0.152697593 | 0.184427605 | -0.031730012 | 0.999933887 | -0.640155872 | -0.975628714 | 0.335472842 | 0.087048498 |
| ENSG00000183291 | *SELENOF* | 5.979156874 | 0.023402258 | -0.053207708 | 0.076609966 | 0.999933887 | -0.047122014 | 0.137499997 | -0.184622011 | 0.087048498 |
| ENSG00000070950 | *RAD18* | 1.969332891 | -0.069279147 | -0.190498354 | 0.121219207 | 0.999933887 | 0.217481389 | 0.650061604 | -0.432580216 | 0.087199454 |
| ENSG00000163563 | *MNDA* | 8.468525323 | -0.101381442 | -0.221746493 | 0.120365051 | 0.999933887 | -0.876606644 | -1.32301839 | 0.446411747 | 0.087283987 |
| ENSG00000081320 | *STK17B* | 8.238268932 | 0.054049476 | -0.005581058 | 0.059630533 | 0.999933887 | -0.453984461 | -0.650332284 | 0.196347824 | 0.087317925 |
| ENSG00000111530 | *CAND1* | 4.267539911 | -0.065740702 | -0.043911165 | -0.021829537 | 0.999933887 | -0.049041378 | 0.178660593 | -0.227701971 | 0.087341644 |
| ENSG00000268119 | *AC010615.2* | 1.682647559 | 0.008241365 | 0.081528896 | -0.073287531 | 0.999933887 | -0.259694216 | 0.149392409 | -0.409086625 | 0.087405274 |
| ENSG00000154001 | *PPP2R5E* | 3.93301641 | 0.023185229 | -0.058055289 | 0.081240518 | 0.999933887 | -0.266133615 | -0.47379398 | 0.207660365 | 0.08746701 |
| ENSG00000272668 | *AL590560.3* | 1.325103375 | 0.008228972 | -0.00141398 | 0.009642953 | 0.999933887 | 0.009792 | -0.425868412 | 0.435660412 | 0.08746701 |
| ENSG00000182858 | *ALG12* | 2.358087855 | 0.065166911 | -0.047474628 | 0.112641539 | 0.999933887 | -0.191877237 | -0.497355262 | 0.305478026 | 0.087519354 |
| ENSG00000139618 | *BRCA2* | 1.449268476 | -0.083068891 | -0.265767105 | 0.182698214 | 0.999933887 | 0.08196565 | 0.848282725 | -0.766317075 | 0.087519354 |
| ENSG00000185022 | *MAFF* | 9.144944873 | 0.328448665 | 0.271996597 | 0.056452068 | 0.999933887 | 1.88116178 | 2.228695099 | -0.347533319 | 0.087603193 |
| ENSG00000138744 | *NAAA* | 3.255639582 | -0.024157249 | -0.01092079 | -0.013236459 | 0.999933887 | -0.197655301 | -0.472000222 | 0.274344921 | 0.087603193 |
| ENSG00000133112 | *TPT1* | 9.553028321 | 0.002512479 | 0.019844908 | -0.017332429 | 0.999933887 | 0.188972715 | 0.371375457 | -0.182402742 | 0.087603193 |
| ENSG00000063169 | *BICRA* | 4.646743574 | 0.056282975 | 0.044000192 | 0.012282783 | 0.999933887 | -0.413190663 | -0.671721568 | 0.258530905 | 0.087847671 |
| ENSG00000122873 | *CISD1* | 0.233628907 | 0.017157827 | 0.064739177 | -0.04758135 | 0.999933887 | -0.124218873 | 0.521058823 | -0.645277697 | 0.088022826 |
| ENSG00000143420 | *ENSA* | 6.762572882 | -0.005601015 | -0.009522418 | 0.003921404 | 0.999933887 | 0.252232321 | 0.423209241 | -0.17097692 | 0.088022826 |
| ENSG00000238241 | *CCR12P* | -0.240717369 | 0.222866372 | -0.146653267 | 0.369519639 | 0.999933887 | 0.789078288 | 1.453431284 | -0.664352996 | 0.088122923 |
| ENSG00000230982 | *DSTNP1* | 0.074032771 | -0.064314488 | -0.179471161 | 0.115156674 | 0.999933887 | 0.750577112 | 1.364783219 | -0.614206107 | 0.08819215 |
| ENSG00000179218 | *CALR* | 6.207791914 | 0.004102573 | -0.046264975 | 0.050367548 | 0.999933887 | -0.016193251 | -0.183517189 | 0.167323938 | 0.08819215 |
| ENSG00000123130 | *ACOT9* | 6.536666868 | -0.062354019 | -0.018819011 | -0.043535008 | 0.999933887 | -0.000695858 | 0.169927371 | -0.170623229 | 0.088237579 |
| ENSG00000088298 | *EDEM2* | 4.845591586 | -0.02311061 | -0.030554474 | 0.007443864 | 0.999933887 | 0.037544034 | -0.156118478 | 0.193662512 | 0.088321368 |
| ENSG00000108389 | *MTMR4* | 5.422287547 | -0.042267387 | -0.054745123 | 0.012477735 | 0.999933887 | 1.011105753 | 1.280183046 | -0.269077293 | 0.088534992 |
| ENSG00000197961 | *ZNF121* | 3.070451416 | -0.066260355 | -0.031224185 | -0.03503617 | 0.999933887 | -0.138748731 | 0.131233911 | -0.269982642 | 0.088539038 |
| ENSG00000135828 | *RNASEL* | 4.5408833 | -0.037752291 | -0.110557579 | 0.072805288 | 0.999933887 | -0.408446071 | -0.603183824 | 0.194737753 | 0.088701205 |
| ENSG00000140943 | *MBTPS1* | 5.050620205 | -0.01037912 | -0.030222983 | 0.019843863 | 0.999933887 | -0.011178352 | -0.174288595 | 0.163110243 | 0.088978636 |
| ENSG00000237753 | *AC079922.2* | 1.79009424 | -0.161920255 | 0.182895939 | -0.344816194 | 0.999933887 | -0.409228467 | -0.771835712 | 0.362607245 | 0.088990362 |
| ENSG00000196453 | *ZNF777* | 2.797200448 | 0.063508239 | 0.00481799 | 0.058690249 | 0.999933887 | -0.196972077 | -0.455910377 | 0.2589383 | 0.08906901 |
| ENSG00000279164 | *AL118508.2* | 0.551915841 | 0.108551741 | 0.108291826 | 0.000259915 | 0.999933887 | -0.315987554 | 0.484317213 | -0.800304767 | 0.089170294 |
| ENSG00000263731 | *AC145207.5* | -0.188896838 | 0.11989764 | 0.117930163 | 0.001967477 | 0.999933887 | -0.110450036 | 0.696064938 | -0.806514974 | 0.089270559 |
| ENSG00000053524 | *MCF2L2* | 2.946356251 | 0.053334642 | 0.131575344 | -0.078240702 | 0.999933887 | 0.670413675 | 0.980372833 | -0.309959159 | 0.089762208 |
| ENSG00000105711 | *SCN1B* | 3.623993305 | 0.058506056 | -0.004946064 | 0.063452121 | 0.999933887 | 3.492270411 | 4.04487305 | -0.552602639 | 0.089762208 |
| ENSG00000230149 | *AL021707.3* | 1.529365595 | 0.031526486 | -0.027096315 | 0.058622801 | 0.999933887 | 0.195573393 | -0.23500353 | 0.430576923 | 0.089762208 |
| ENSG00000234389 | *AC007278.1* | 2.914828906 | -0.030341417 | -0.066142068 | 0.035800651 | 0.999933887 | 0.049413305 | 0.547953046 | -0.498539741 | 0.089762208 |
| ENSG00000096401 | *CDC5L* | 4.917919814 | -0.010678049 | -0.090058944 | 0.079380895 | 0.999933887 | -0.100151041 | 0.120908751 | -0.221059792 | 0.089762208 |
| ENSG00000112137 | *PHACTR1* | 10.351358763 | 0.088815031 | 0.069130819 | 0.019684211 | 0.999933887 | 0.699661313 | 0.924008172 | -0.224346858 | 0.08991288 |
| ENSG00000170322 | *NFRKB* | 4.857745056 | -0.025415965 | -0.045559528 | 0.020143563 | 0.999933887 | -0.226148803 | -0.399845266 | 0.173696463 | 0.089937542 |
| ENSG00000253738 | *OTUD6B-AS1* | 2.473322769 | -0.001126183 | -0.24097798 | 0.239851797 | 0.999933887 | -0.351062026 | 0.069784877 | -0.420846903 | 0.089951539 |
| ENSG00000009307 | *CSDE1* | 8.674625145 | -0.032351173 | -0.119305898 | 0.086954725 | 0.999933887 | -0.087297807 | 0.047423302 | -0.134721109 | 0.089991863 |
| ENSG00000163840 | *DTX3L* | 5.966129666 | -0.066292395 | -0.033269648 | -0.033022748 | 0.999933887 | -0.364394459 | -0.55547802 | 0.19108356 | 0.090019787 |
| ENSG00000124164 | *VAPB* | 4.031419193 | -0.055228392 | -0.046301275 | -0.008927116 | 0.999933887 | -0.628809838 | -0.855291391 | 0.226481553 | 0.090039665 |
| ENSG00000120370 | *GORAB* | 0.778272495 | 0.005452714 | -0.290465393 | 0.295918107 | 0.999933887 | 0.023807292 | 0.53974989 | -0.515942597 | 0.090243096 |
| ENSG00000155508 | *CNOT8* | 5.493679624 | -0.094770652 | 0.009110961 | -0.103881613 | 0.999933887 | -0.323441318 | -0.487381218 | 0.1639399 | 0.09033252 |
| ENSG00000068001 | *HYAL2* | 1.933998517 | 0.109418553 | -0.09510989 | 0.204528443 | 0.999933887 | -0.674773836 | -1.066158794 | 0.391384958 | 0.09033252 |
| ENSG00000133246 | *PRAM1* | 6.155998462 | -0.045200235 | -0.06062861 | 0.015428374 | 0.999933887 | -1.475403663 | -1.725075466 | 0.249671803 | 0.09033252 |
| ENSG00000132424 | *PNISR* | 7.413861153 | -0.06309474 | -0.224391852 | 0.161297112 | 0.999933887 | -0.378608442 | 0.098712495 | -0.477320937 | 0.090358251 |
| ENSG00000163512 | *AZI2* | 2.531876806 | -0.027385576 | -0.043874154 | 0.016488578 | 0.999933887 | -0.2892801 | 0.011034041 | -0.30031414 | 0.090358251 |
| ENSG00000185220 | *PGBD2* | 0.832764551 | -0.030878969 | -0.045957174 | 0.015078205 | 0.999933887 | -0.167849525 | -0.545791671 | 0.377942146 | 0.090358251 |
| ENSG00000196268 | *ZNF493* | 4.852968065 | -0.015436322 | -0.039440921 | 0.024004598 | 0.999933887 | -0.258633958 | 0.126687454 | -0.385321412 | 0.090358251 |
| ENSG00000178188 | *SH2B1* | 4.559238735 | 0.010280034 | 0.069766109 | -0.059486075 | 0.999933887 | -0.166242052 | -0.377684311 | 0.211442259 | 0.090571907 |
| ENSG00000167536 | *DHRS13* | 2.622486563 | 0.141763218 | 0.019965794 | 0.121797424 | 0.999933887 | 1.457779025 | 2.110556467 | -0.652777441 | 0.090588411 |
| ENSG00000006530 | *AGK* | 2.665845581 | 0.049124657 | 0.042255627 | 0.00686903 | 0.999933887 | 0.240369937 | 0.480220536 | -0.239850599 | 0.090609722 |
| ENSG00000271795 | *AC011337.1* | -1.179465613 | -0.035244288 | -0.497647692 | 0.462403404 | 0.999933887 | -0.847470378 | -0.195824411 | -0.651645968 | 0.090627104 |
| ENSG00000089127 | *OAS1* | 3.70051178 | -0.043613849 | 0.041076881 | -0.08469073 | 0.999933887 | -0.504848416 | -0.834344847 | 0.329496431 | 0.091073872 |
| ENSG00000207736 | *MIR657* | 0.579412334 | 0.069200125 | 0.070572683 | -0.001372558 | 0.999933887 | 0.182042666 | 0.664737387 | -0.48269472 | 0.09110639 |
| ENSG00000181481 | *RNF135* | 2.279561254 | -0.019171589 | 0.047783648 | -0.066955237 | 0.999933887 | -0.523837164 | -0.951405968 | 0.427568804 | 0.091140191 |
| ENSG00000181634 | *TNFSF15* | 7.665598257 | -0.071498384 | 0.017729375 | -0.089227759 | 0.999933887 | -0.112559446 | 0.172191334 | -0.28475078 | 0.091469857 |
| ENSG00000174243 | *DDX23* | 5.083236712 | -0.026895481 | -0.060768342 | 0.033872861 | 0.999933887 | -0.083473571 | -0.227233091 | 0.143759521 | 0.091469857 |
| ENSG00000155313 | *USP25* | 6.062414917 | -0.008866381 | -0.099959985 | 0.091093604 | 0.999933887 | -0.252934055 | 0.054827576 | -0.307761631 | 0.09149167 |
| ENSG00000092199 | *HNRNPC* | 9.154834639 | -0.053195403 | -0.057639236 | 0.004443833 | 0.999933887 | 1.383206203 | 1.635651983 | -0.25244578 | 0.09159359 |
| ENSG00000141219 | *C17orf80* | 0.782073795 | -0.110170056 | -0.110674316 | 0.000504259 | 0.999933887 | -0.196759619 | -0.587162725 | 0.390403106 | 0.09159359 |
| ENSG00000134884 | *ARGLU1* | 7.404164894 | -0.023082305 | 0.037694179 | -0.060776483 | 0.999933887 | -0.173774629 | -0.329056799 | 0.15528217 | 0.09159359 |
| ENSG00000221821 | *C6orf226* | 1.270861471 | 0.025432119 | -0.02425519 | 0.049687309 | 0.999933887 | 0.000512318 | -0.448982681 | 0.449494999 | 0.09159359 |
| ENSG00000237172 | *B3GNT9* | 1.299667997 | -0.008292321 | 0.065132017 | -0.073424338 | 0.999933887 | -0.503006764 | -0.924238256 | 0.421231492 | 0.09159359 |
| ENSG00000100767 | *PAPLN* | 1.649276662 | 0.008559188 | 0.087111729 | -0.078552541 | 0.999933887 | 0.472387404 | 0.886878036 | -0.414490632 | 0.091947522 |
| ENSG00000006459 | *KDM7A* | 6.147640473 | 0.042077682 | -0.057446177 | 0.09952386 | 0.999933887 | -1.272448623 | -1.597863491 | 0.325414868 | 0.091948898 |
| ENSG00000213923 | *CSNK1E* | 4.102602611 | 0.01995183 | 0.099942035 | -0.079990205 | 0.999933887 | 0.232412304 | 0.442322411 | -0.209910107 | 0.092267053 |
| ENSG00000125818 | *PSMF1* | 5.31567053 | 0.016349185 | 0.017242055 | -0.00089287 | 0.999933887 | -0.203150527 | -0.407303285 | 0.204152759 | 0.092329192 |
| ENSG00000188549 | *CCDC9B* | 2.805171277 | 0.134099817 | 0.23459394 | -0.100494123 | 0.999933887 | 0.839651488 | 1.106801604 | -0.267150116 | 0.092348208 |
| ENSG00000270120 | *AC007728.3* | 0.455027542 | 0.089746593 | -0.038004546 | 0.127751139 | 0.999933887 | 1.061841014 | 1.730919304 | -0.66907829 | 0.092396489 |
| ENSG00000111676 | *ATN1* | 3.575550111 | -0.013532575 | 0.13121085 | -0.144743425 | 0.999933887 | 0.098098696 | 0.305292638 | -0.207193943 | 0.092396489 |
| ENSG00000176903 | *PNMA1* | 3.653377072 | 0.089150429 | -0.009384193 | 0.098534622 | 0.999933887 | -0.090704038 | -0.272378606 | 0.181674568 | 0.092440851 |
| ENSG00000135070 | *ISCA1* | 4.593878285 | 0.052463932 | 0.032917989 | 0.019545943 | 0.999933887 | -0.043049705 | 0.173075794 | -0.216125499 | 0.092440851 |
| ENSG00000083896 | *YTHDC1* | 7.933972162 | -0.056279957 | -0.096408769 | 0.040128812 | 0.999933887 | 0.048786995 | 0.266658109 | -0.217871114 | 0.092440851 |
| ENSG00000286691 | *AL590009.1* | -0.74970361 | -0.118771123 | -0.130283626 | 0.011512503 | 0.999933887 | -1.25239567 | -2.088908805 | 0.836513134 | 0.092440851 |
| ENSG00000229931 | *ATXN1-AS1* | 2.410712214 | 0.04624186 | -0.151758654 | 0.198000514 | 0.999933887 | 0.28304282 | 0.710103622 | -0.427060802 | 0.092440851 |
| ENSG00000228794 | *LINC01128* | 3.988855069 | -0.01512257 | -0.047115903 | 0.031993333 | 0.999933887 | -0.010896801 | 0.227857096 | -0.238753897 | 0.092440851 |
| ENSG00000197429 | *IPP* | 1.412838649 | 0.015213695 | -0.167847147 | 0.183060842 | 0.999933887 | 0.1012437 | 0.475464844 | -0.374221144 | 0.092440851 |
| ENSG00000088247 | *KHSRP* | 5.689901986 | 0.05151325 | 0.046244594 | 0.005268655 | 0.999933887 | -0.190920717 | -0.37207408 | 0.181153363 | 0.092462752 |
| ENSG00000156171 | *DRAM2* | 3.00712037 | -0.081056093 | -0.093415373 | 0.01235928 | 0.999933887 | -0.496549499 | -0.790489774 | 0.293940275 | 0.092462752 |
| ENSG00000173480 | *ZNF417* | 3.482883203 | -0.023403885 | -0.108426144 | 0.085022259 | 0.999933887 | -0.032422157 | 0.179563175 | -0.211985332 | 0.092462752 |
| ENSG00000132646 | *PCNA* | 3.332702106 | 0.084777543 | -0.039322014 | 0.124099556 | 0.999933887 | -0.493175906 | -0.786166316 | 0.292990409 | 0.092589075 |
| ENSG00000259342 | *AC025580.1* | 1.999852423 | 0.334767457 | 0.530590798 | -0.195823342 | 0.999933887 | 2.557732959 | 3.114255154 | -0.556522196 | 0.092661109 |
| ENSG00000179295 | *PTPN11* | 4.965621254 | -0.052919131 | -0.00411878 | -0.048800351 | 0.999933887 | -0.087523832 | 0.122191379 | -0.209715211 | 0.092855332 |
| ENSG00000138443 | *ABI2* | 1.871263499 | -0.052663516 | -0.11120747 | 0.058543954 | 0.999933887 | -0.21604967 | 0.094451494 | -0.310501164 | 0.092951852 |
| ENSG00000011454 | *RABGAP1* | 5.075294561 | -0.037679966 | -0.061091761 | 0.023411795 | 0.999933887 | -0.06404191 | 0.127146124 | -0.191188034 | 0.09301689 |
| ENSG00000005436 | *GCFC2* | 0.568024012 | -0.230892284 | -0.116575491 | -0.114316793 | 0.999933887 | -0.118744779 | 0.442258948 | -0.561003727 | 0.093231597 |
| ENSG00000165629 | *ATP5F1C* | 3.638367857 | -0.070056832 | -0.098372887 | 0.028316055 | 0.999933887 | -0.259681844 | -0.473583713 | 0.213901868 | 0.093231597 |
| ENSG00000128512 | *DOCK4* | 7.502603755 | -0.026394478 | -0.018279606 | -0.008114872 | 0.999933887 | 0.259394685 | 0.454950993 | -0.195556308 | 0.093312078 |
| ENSG00000026652 | *AGPAT4* | 3.977001153 | -0.051907217 | 0.024695373 | -0.07660259 | 0.999933887 | 0.196220984 | 0.411096956 | -0.214875972 | 0.093416232 |
| ENSG00000104897 | *SF3A2* | 4.719644423 | 0.023148368 | 0.017793175 | 0.005355193 | 0.999933887 | -0.020778193 | -0.215170231 | 0.194392038 | 0.093666764 |
| ENSG00000117868 | *ESYT2* | 6.427061375 | 0.003993112 | 0.033003564 | -0.029010451 | 0.999933887 | 0.287515837 | 0.445975091 | -0.158459254 | 0.093666764 |
| ENSG00000087111 | *PIGS* | 4.087104545 | 0.002410183 | 0.041707684 | -0.039297501 | 0.999933887 | -0.043865478 | -0.248450464 | 0.204584986 | 0.093672335 |
| ENSG00000197223 | *C1D* | 3.62669313 | -0.076070678 | -0.077381929 | 0.001311251 | 0.999933887 | -0.067901598 | 0.140403938 | -0.208305535 | 0.093685139 |
| ENSG00000272402 | *AL031775.2* | 0.305558403 | 0.130815322 | 0.038078722 | 0.0927366 | 0.999933887 | 0.355983565 | 0.810446437 | -0.454462871 | 0.093685139 |
| ENSG00000072042 | *RDH11* | 2.640970535 | -0.084899255 | -0.047647739 | -0.037251515 | 0.999933887 | -0.017299231 | 0.251172713 | -0.268471944 | 0.093685139 |
| ENSG00000127511 | *SIN3B* | 4.927775759 | 0.040747143 | 0.039223793 | 0.00152335 | 0.999933887 | 0.032548149 | -0.151707252 | 0.184255401 | 0.093685139 |
| ENSG00000113532 | *ST8SIA4* | 5.876836957 | -0.045251672 | -0.034280742 | -0.01097093 | 0.999933887 | -0.819411208 | -1.112860694 | 0.293449486 | 0.093685139 |
| ENSG00000164654 | *MIOS* | 3.293479152 | 0.050348725 | 0.004586948 | 0.045761777 | 0.999933887 | 0.165481171 | 0.372575243 | -0.207094072 | 0.093733497 |
| ENSG00000177425 | *PAWR* | -0.604456649 | 0.022364334 | -0.055284772 | 0.077649106 | 0.999933887 | 1.106508722 | 1.906763419 | -0.800254697 | 0.093806051 |
| ENSG00000111271 | *ACAD10* | 2.920032175 | 0.068794197 | 0.008128722 | 0.060665475 | 0.999933887 | -0.187167485 | -0.472109238 | 0.284941753 | 0.093809819 |
| ENSG00000086232 | *EIF2AK1* | 4.250688544 | -0.031148309 | -0.105536624 | 0.074388316 | 0.999933887 | -0.1310363 | -0.337250181 | 0.206213881 | 0.093984336 |
| ENSG00000127838 | *PNKD* | 3.984407768 | 0.058987163 | -0.131979723 | 0.190966887 | 0.999933887 | 0.48314524 | 0.89342585 | -0.41028061 | 0.093984336 |
| ENSG00000156273 | *BACH1* | 9.437785403 | -0.166775077 | -0.177109372 | 0.010334296 | 0.999933887 | -0.342811743 | -0.064260641 | -0.278551102 | 0.094107056 |

| | | | | | | | | | | |
|---|---|---|---|---|---|---|---|---|---|---|
| ENSG00000236778 | *INTS6-AS1* | 0.290461703 | -0.252003603 | 0.123774574 | -0.375778177 | 0.999933887 | -0.431153134 | 0.019023393 | -0.450176527 | 0.094450608 |
| ENSG00000197969 | *VPS13A* | 4.026424359 | -0.128316881 | -0.219097447 | 0.090780566 | 0.999933887 | -0.272381536 | 0.398667213 | -0.671048749 | 0.094492521 |
| ENSG00000122884 | *P4HA1* | 3.643410485 | -0.051649268 | -0.100905237 | 0.049255968 | 0.999933887 | 0.625710551 | 0.907672192 | -0.281961641 | 0.094492521 |
| ENSG00000164949 | *GEM* | 0.665028597 | -0.006432961 | 0.266874822 | -0.273307784 | 0.999933887 | 0.767816211 | 1.140260521 | -0.37244431 | 0.09458739 |
| ENSG00000134516 | *DOCK2* | 7.865379873 | -0.019831525 | -0.053524027 | 0.033692502 | 0.999933887 | -0.053244419 | -0.178727057 | 0.125482638 | 0.094651574 |
| ENSG00000108846 | *ABCC3* | 0.101634695 | 0.034093354 | 0.304372881 | -0.270279527 | 0.999933887 | 0.8597978 | 1.587614478 | -0.727816678 | 0.09474837 |
| ENSG00000110422 | *HIPK3* | 6.732459062 | -0.031152672 | -0.071810566 | 0.040657894 | 0.999933887 | -0.134842092 | 0.152190865 | -0.287032957 | 0.094942326 |
| ENSG00000166024 | *R3HCC1L* | 4.176826355 | -0.124675646 | -0.093360278 | -0.031315369 | 0.999933887 | -0.162627525 | 0.069619262 | -0.232246788 | 0.095002543 |
| ENSG00000173545 | *ZNF622* | 4.840045161 | -0.029491642 | -0.094735407 | 0.065243765 | 0.999933887 | -0.008285229 | 0.166781285 | -0.175066514 | 0.095075505 |
| ENSG00000126821 | *SGPP1* | 3.211976783 | -0.107408507 | 0.072541563 | -0.17995007 | 0.999933887 | -1.154872742 | -1.545313245 | 0.390440503 | 0.095076984 |
| ENSG00000138286 | *FAM149B1* | 2.240286873 | -0.01629435 | 0.02794232 | -0.04423667 | 0.999933887 | 0.025293765 | 0.314209741 | -0.288915977 | 0.095086117 |
| ENSG00000170017 | *ALCAM* | 3.903786421 | -0.079526946 | -0.061828073 | -0.017698873 | 0.999933887 | 0.072403744 | 0.467708137 | -0.395304392 | 0.095373679 |
| ENSG00000084092 | *NOA1* | 2.45579761 | 0.053602794 | 0.207500528 | -0.153897734 | 0.999933887 | -0.011780098 | -0.294998607 | 0.283218509 | 0.095419669 |
| ENSG00000169951 | *ZNF764* | 2.280349269 | 0.051769044 | 0.162133436 | -0.110364392 | 0.999933887 | -0.227652421 | -0.537574093 | 0.309921672 | 0.095419669 |
| ENSG00000114374 | *USP9Y* | -2.967694079 | 0.000108542 | 0.057759265 | -0.057650723 | 0.999933887 | -0.459474312 | 0.147072155 | -0.606546467 | 0.095419669 |
| ENSG00000198899 | *MT-ATP6* | 8.353753722 | 0.047786475 | -0.044887061 | 0.092673536 | 0.999933887 | -0.054255463 | 0.250990568 | -0.305246032 | 0.09556972 |
| ENSG00000156256 | *USP16* | 5.116454494 | -0.120639711 | -0.202623358 | 0.081983647 | 0.999933887 | -0.168825876 | 0.256233814 | -0.42505969 | 0.095605699 |
| ENSG00000111231 | *GPN3* | 1.10578999 | 0.024922914 | 0.087024933 | -0.062102019 | 0.999933887 | -0.230541238 | -0.616507199 | 0.385965961 | 0.095802876 |
| ENSG00000136295 | *TTYH3* | 5.98846648 | 0.008029791 | -0.020129042 | 0.028158833 | 0.999933887 | -0.292648339 | -0.076073929 | -0.216574409 | 0.095802876 |
| ENSG00000136051 | *WASHC4* | 6.718639487 | -0.038908703 | -0.163854431 | 0.124945728 | 0.999933887 | -0.563163707 | -0.08841809 | -0.474745617 | 0.095860651 |
| ENSG00000168159 | *RNF187* | 4.054361953 | 0.021920548 | 0.139216963 | -0.117296415 | 0.999933887 | 0.037364025 | -0.189931686 | 0.227295711 | 0.095903008 |
| ENSG00000285999 | *AC025442.2* | 1.448000068 | -0.195962555 | 0.101114154 | -0.297076709 | 0.999933887 | 0.730530561 | 1.078503457 | -0.347972896 | 0.095911327 |
| ENSG00000108515 | *ENO3* | 3.416585685 | 0.241749631 | 0.151084759 | 0.090664872 | 0.999933887 | 0.797960125 | 1.159582514 | -0.361622389 | 0.095970813 |
| ENSG00000164329 | *TENT2* | 5.533276934 | -0.010673067 | -0.047728865 | 0.037055799 | 0.999933887 | -0.131367093 | 0.116522742 | -0.247889835 | 0.095970813 |
| ENSG00000263072 | *ZNF213-AS1* | 1.450124884 | 0.015086916 | -0.073051813 | 0.088138729 | 0.999933887 | -0.096903566 | -0.474161774 | 0.377258207 | 0.095970813 |
| ENSG00000087303 | *NID2* | -0.753821211 | -0.024386926 | 0.302717604 | -0.327104529 | 0.999933887 | 1.0680676 | 1.998055583 | -0.929987984 | 0.095970813 |
| ENSG00000169439 | *SDC2* | 2.66486533 | 0.012676756 | -0.031800789 | 0.044477545 | 0.999933887 | -1.169928433 | -1.632591102 | 0.462662669 | 0.095970813 |
| ENSG00000138658 | *ZGRF1* | 1.042846616 | -0.089534897 | -0.04258977 | -0.046945127 | 0.999933887 | -0.225332778 | 0.376209756 | -0.601542535 | 0.096050224 |
| ENSG00000065675 | *PRKCQ* | 4.133615597 | -0.036795722 | 0.050107601 | -0.086903323 | 0.999933887 | 0.163616813 | 0.374521393 | -0.21090458 | 0.096050224 |
| ENSG00000103343 | *ZNF174* | 1.002791562 | 0.062502816 | -0.045576481 | 0.108079296 | 0.999933887 | -0.310759635 | -0.756339555 | 0.445579919 | 0.096050224 |
| ENSG00000229390 | *MICD* | 0.501214167 | -0.031986317 | -0.135578904 | 0.103592587 | 0.999933887 | -0.672317634 | -1.138090257 | 0.465772623 | 0.096050224 |
| ENSG00000260007 | *AC107871.1* | 2.910131436 | -0.000561538 | -0.104534053 | 0.103972515 | 0.999933887 | 0.064679161 | 0.461433882 | -0.396754721 | 0.096050224 |
| ENSG00000140740 | *UQCRC2* | 7.304196335 | -0.03395154 | -0.055719898 | 0.021768359 | 0.999933887 | -0.182144329 | -0.018217525 | -0.163926804 | 0.096361706 |
| ENSG00000116030 | *SUMO1* | 5.598185831 | -0.020055406 | -0.086726488 | 0.066671082 | 0.999933887 | -0.096264609 | 0.07340171 | -0.169666319 | 0.096361706 |
| ENSG00000204482 | *LST1* | 7.957469376 | 0.025562592 | 0.020968904 | 0.004593688 | 0.999933887 | 0.090348289 | -0.129850618 | 0.220198907 | 0.096593298 |
| ENSG00000172661 | *WASHC2C* | 7.209985843 | -0.065168454 | -0.006300457 | -0.058867996 | 0.999933887 | 0.251431814 | 0.397505973 | -0.14607416 | 0.096620053 |
| ENSG00000116791 | *CRYZ* | 0.477653871 | -0.051588077 | -0.138670636 | 0.087082559 | 0.999933887 | -0.094494497 | 0.399766286 | -0.494260783 | 0.096620053 |
| ENSG00000132357 | *CARD6* | 2.606612013 | -0.115064211 | 0.0007363 | -0.115800511 | 0.999933887 | -0.674220129 | -0.965477297 | 0.291257168 | 0.09684281 |
| ENSG00000256235 | *SMIM3* | 3.031289098 | -0.027609618 | 0.062421485 | -0.090031103 | 0.999933887 | 1.351340888 | 1.711743096 | -0.360402208 | 0.09684281 |
| ENSG00000272341 | *AL137003.1* | 2.564537445 | -0.2610983 | -0.375408659 | 0.11431036 | 0.999933887 | 0.157550369 | 0.710773065 | -0.553222696 | 0.096854897 |
| ENSG00000233058 | *LINC00884* | 1.739806234 | 0.153859547 | 0.084933913 | 0.068925634 | 0.999933887 | 1.351877746 | 1.714682054 | -0.362804309 | 0.096962178 |
| ENSG00000183475 | *ASB7* | 3.117235902 | -0.08384783 | -0.034954342 | -0.048893488 | 0.999933887 | -0.472477395 | -0.718692106 | 0.246214711 | 0.096962178 |
| ENSG00000228140 | *AL031283.1* | 1.862085304 | -0.042890492 | -0.015157932 | -0.027732561 | 0.999933887 | 1.543134004 | 1.962036022 | -0.418902017 | 0.096990606 |
| ENSG00000146094 | *DOK3* | 9.302720956 | 0.004112653 | -0.003551277 | 0.00766393 | 0.999933887 | -0.321653571 | -0.150806702 | -0.170846869 | 0.097048151 |
| ENSG00000101665 | *SMAD7* | 5.02683054 | 0.053878065 | 0.077868489 | -0.023990424 | 0.999933887 | 0.457084695 | 0.716280851 | -0.259196156 | 0.0971222 |
| ENSG00000158805 | *ZNF276* | 5.843887401 | 0.058369395 | 0.077920076 | -0.019550681 | 0.999933887 | -0.114452998 | -0.287907884 | 0.173454886 | 0.097165628 |
| ENSG00000108239 | *TBC1D12* | -0.85972803 | -0.291742491 | 0.082878915 | -0.374621406 | 0.999933887 | 0.721358331 | 1.475339272 | -0.753980941 | 0.097172089 |
| ENSG00000196967 | *ZNF585A* | 1.082473898 | -0.190367089 | -0.175925621 | -0.014441468 | 0.999933887 | -0.325077553 | 0.063586716 | -0.388664269 | 0.097425944 |
| ENSG00000188994 | *ZNF292* | 7.41644241 | -0.109838947 | -0.131164372 | 0.021325426 | 0.999933887 | -0.427166042 | 0.055607728 | -0.48277377 | 0.097506193 |
| ENSG00000152465 | *NMT2* | 2.412663356 | -0.032322691 | 0.08123548 | -0.113558172 | 0.999933887 | 0.045016072 | 0.31007812 | -0.265062048 | 0.097506193 |
| ENSG00000280434 | *AL031595.3* | -0.802601088 | -0.059346197 | 0.483179019 | -0.542525216 | 0.999933887 | 0.304844226 | 1.034323133 | -0.729478907 | 0.097614188 |
| ENSG00000270049 | *AC009061.2* | 0.999450363 | -0.0656029 | -0.212306766 | 0.146703866 | 0.999933887 | -0.272301174 | -0.791326116 | 0.519024942 | 0.097618946 |
| ENSG00000161179 | *YDJC* | 3.305755489 | 0.017163657 | 0.071775859 | -0.054612202 | 0.999933887 | -0.246719507 | -0.506227811 | 0.259508304 | 0.097809228 |
| ENSG00000163558 | *PRKCI* | 2.426375083 | -0.008097508 | -0.016684227 | 0.008586719 | 0.999933887 | -0.092731731 | 0.20639284 | -0.299124571 | 0.09781528 |
| ENSG00000255874 | *PRECSIT* | 0.66845516 | 0.455774392 | 0.117452972 | 0.33832142 | 0.999933887 | 2.546752047 | 3.20354061 | -0.656788563 | 0.097980503 |
| ENSG00000150779 | *TIMM8B* | 2.060495484 | -0.112285971 | 0.007011122 | -0.119297093 | 0.999933887 | -0.06325891 | -0.429977322 | 0.366718412 | 0.09798863 |
| ENSG00000186020 | *ZNF529* | 2.267035064 | -0.087656065 | -0.07542715 | -0.012228915 | 0.999933887 | -0.188643049 | 0.16479766 | -0.353440708 | 0.09798863 |
| ENSG00000127980 | *PEX1* | 2.301354782 | 0.048464701 | -0.117663214 | 0.166127914 | 0.999933887 | -0.052252211 | -0.380403704 | 0.328151493 | 0.09811011 |
| ENSG00000166508 | *MCM7* | 5.23007153 | -0.031726298 | 0.032618002 | -0.0643443 | 0.999933887 | 0.076642128 | -0.095132806 | 0.171774933 | 0.098118059 |
| ENSG00000055483 | *USP36* | 7.017357851 | -0.029954151 | 0.002942386 | -0.032896537 | 0.999933887 | 0.043636648 | 0.234912582 | -0.191275934 | 0.098118059 |
| ENSG00000171100 | *MTM1* | 3.07575815 | -0.024250767 | -0.02620929 | 0.001958523 | 0.999933887 | -0.587445786 | -0.870437444 | 0.282991658 | 0.098118059 |
| ENSG00000106105 | *GARS1* | 5.959167649 | 0.019690584 | -0.034713011 | 0.054403595 | 0.999933887 | 0.347782102 | 0.568572541 | -0.220790439 | 0.098118059 |
| ENSG00000157617 | *C2CD2* | 0.609823098 | -0.030098307 | -0.031590114 | 0.001491807 | 0.999933887 | 0.047940963 | 0.547756105 | -0.499815141 | 0.098266235 |
| ENSG00000266320 | *MIR3909* | 2.10736128 | -0.080034188 | -0.004263736 | -0.075770453 | 0.999933887 | 0.593860923 | 0.950182748 | -0.356321825 | 0.098284619 |
| ENSG00000161202 | *DVL3* | 5.83102012 | 0.015889805 | 0.033861923 | -0.017972118 | 0.999933887 | -0.139176202 | -0.288008764 | 0.148832563 | 0.098416817 |
| ENSG00000054219 | *LY75* | 2.640404814 | -0.033083203 | -0.091406302 | 0.058323099 | 0.999933887 | -0.4358761 | -0.864938627 | 0.429062526 | 0.098548689 |
| ENSG00000198848 | *CES1* | -0.573756927 | 0.018908957 | 0.313443773 | -0.294534817 | 0.999933887 | 0.887795053 | 1.487109245 | -0.599314192 | 0.098577044 |
| ENSG00000120137 | *PANK3* | 4.456594851 | 0.012253721 | -0.14668509 | 0.158938811 | 0.999933887 | -0.377006577 | 0.022293258 | -0.399299835 | 0.098706841 |
| ENSG00000038382 | *TRIO* | 3.385034663 | 0.080968156 | -0.007239462 | 0.088207618 | 0.999933887 | 0.189123233 | 0.421128908 | -0.232005675 | 0.098733673 |
| ENSG00000116906 | *GNPAT* | 3.267593809 | -0.054846484 | -0.052048592 | -0.002797892 | 0.999933887 | -0.281735765 | -0.526773352 | 0.245037587 | 0.098733673 |
| ENSG00000130724 | *CHMP2A* | 7.814680744 | 0.027564828 | -0.014879121 | 0.042443948 | 0.999933887 | 0.418708607 | 0.580047711 | -0.161339104 | 0.098733673 |
| ENSG00000133794 | *ARNTL* | 5.964463607 | 0.005194723 | -0.019856023 | 0.025050746 | 0.999933887 | -0.620276249 | -0.796574765 | 0.176298516 | 0.098733673 |
| ENSG00000188486 | *H2AX* | 2.728019269 | 0.066765858 | 0.12479021 | -0.058024352 | 0.999933887 | 0.137845829 | -0.125860676 | 0.263706506 | 0.098805333 |
| ENSG00000137747 | *TMPRSS13* | -0.292601772 | 0.012960954 | -0.274895978 | 0.287856932 | 0.999933887 | 0.472220167 | 1.052268819 | -0.580048652 | 0.098936256 |
| ENSG00000230551 | *AC021078.1* | 5.034773524 | -0.001951428 | -0.008835343 | 0.006883915 | 0.999933887 | -0.160388471 | 0.184610521 | -0.344998991 | 0.099008835 |
| ENSG00000124875 | *CXCL6* | 1.62289214 | 0.040419894 | 0.066427881 | -0.026007987 | 0.999933887 | -0.838372319 | -1.253426061 | 0.415053742 | 0.099239851 |
| ENSG00000143653 | *SCCPDH* | 0.855216616 | -0.010855157 | 0.075089242 | -0.085944399 | 0.999933887 | -0.095812607 | -0.460695804 | 0.364883198 | 0.099278641 |
| ENSG00000137563 | *GGH* | 1.990900711 | -0.03435326 | 0.086634092 | -0.120987352 | 0.999933887 | 0.59676925 | 0.945385065 | -0.348615815 | 0.099522068 |
| ENSG00000157045 | *NTAN1* | 2.848145999 | 0.192468998 | 0.012743409 | 0.179725589 | 0.999933887 | -0.070380931 | -0.370665439 | 0.300284508 | 0.099523943 |
| ENSG00000163607 | *GTPBP8* | 1.362157272 | 0.03139998 | 0.019246804 | 0.012153176 | 0.999933887 | -0.08583765 | 0.29940946 | -0.38524711 | 0.099543943 |
| ENSG00000186806 | *VSIG10L* | -0.055326846 | 0.087055212 | 0.344845168 | -0.257789956 | 0.999933887 | 0.721635082 | 1.294276366 | -0.572641284 | 0.099638048 |
| ENSG00000149743 | *TRPT1* | 2.42715873 | 0.084799201 | -0.074848621 | 0.159647822 | 0.999933887 | 0.051504661 | -0.237371521 | 0.288876183 | 0.099827201 |
| ENSG00000283839 | *AC096667.1* | 2.892391438 | -0.123780378 | -0.080765498 | -0.04301488 | 0.999933887 | 0.407961266 | 0.679353603 | -0.271392338 | 0.09993016 |
| ENSG00000227398 | *KIF9-AS1* | -0.224121762 | 0.191584208 | 0.230421506 | -0.038837299 | 0.999933887 | -0.192967037 | -0.953220141 | 0.760253105 | 0.09993016 |
| ENSG00000274180 | *NATD1* | 6.582120851 | -0.082973691 | -0.125056902 | 0.042083211 | 0.999933887 | -1.549317605 | -1.856553507 | 0.307235901 | 0.09993016 |
| ENSG00000070831 | *CDC42* | 8.690592154 | -0.017616243 | -0.013555697 | -0.004060547 | 0.999933887 | 0.098751205 | 0.241599755 | -0.14284855 | 0.099991931 |
| ENSG00000279838 | *AL356273.3* | 4.365345061 | -0.062979227 | 0.061166666 | -0.124145893 | 0.999933887 | -0.055108442 | 0.457254602 | -0.512363044 | 0.100122643 |
| ENSG00000164400 | *CSF2* | -0.57053444 | 1.028429173 | 1.07997508 | -0.051545908 | 0.999933887 | 3.973544366 | 4.933436777 | -0.959892411 | 0.100211801 |
| ENSG00000141759 | *TXNL4A* | 3.616474125 | -0.025882924 | -0.008305118 | -0.017577806 | 0.999933887 | -0.063365523 | -0.27595641 | 0.212590887 | 0.100244648 |
| ENSG00000005022 | *SLC25A5* | 4.906690288 | -0.043603027 | -0.052216869 | 0.008613842 | 0.999933887 | -0.120077535 | -0.309583274 | 0.189505739 | 0.100382707 |
| ENSG00000116679 | *IVNS1ABP* | 11.599785201 | -0.023883288 | -0.051322551 | 0.027439263 | 0.999933887 | 0.289374829 | 0.482591992 | -0.193217164 | 0.10047511 |
| ENSG00000149761 | *NUDT22* | 2.63002669 | 0.024646594 | 0.128416508 | -0.103769914 | 0.999933887 | -0.292183584 | -0.616516721 | 0.324333137 | 0.100751261 |
| ENSG00000121931 | *LRIF1* | 3.932228252 | -0.078068544 | -0.056109052 | -0.021959492 | 0.999933887 | -0.068124508 | 0.187266146 | -0.255390654 | 0.100789514 |
| ENSG00000111880 | *RNGTT* | 3.490096367 | -0.047474639 | -0.08623958 | 0.038764941 | 0.999933887 | 0.102370837 | 0.334801979 | -0.232431142 | 0.100789514 |
| ENSG00000163904 | *SENP2* | 4.853951957 | -0.017607972 | 0.045482821 | -0.063090794 | 0.999933887 | 0.119026977 | 0.284763318 | -0.165736341 | 0.100803808 |
| ENSG00000160570 | *DEDD2* | 8.489739363 | 0.014377287 | -0.002637832 | 0.01701512 | 0.999933887 | -0.119216533 | 0.072171605 | -0.191388137 | 0.100870811 |
| ENSG00000229017 | *LINC01277* | -0.41983696 | -0.179390532 | 0.095748975 | -0.275139507 | 0.999933887 | 1.291414622 | 1.974970831 | -0.68355621 | 0.100936722 |
| ENSG00000172890 | *NADSYN1* | 6.335727214 | -0.016150965 | 0.000625387 | -0.016776352 | 0.999933887 | -0.092161406 | -0.222700564 | 0.130539159 | 0.100983938 |
| ENSG00000100813 | *ACIN1* | 7.298529526 | -0.028921805 | 0.028077587 | -0.056999392 | 0.999933887 | -0.209628623 | -0.345375349 | 0.135746726 | 0.101075262 |
| ENSG00000286219 | *NOTCH2NLC* | 3.744162951 | -0.035590308 | -0.001514076 | -0.034076233 | 0.999933887 | 0.715273783 | 1.096388563 | -0.381114779 | 0.101126903 |
| ENSG00000168374 | *ARF4* | 6.544945982 | -0.015064447 | -0.013326754 | -0.001737692 | 0.999933887 | 0.260192102 | 0.420350668 | -0.160158565 | 0.101278962 |
| ENSG00000076662 | *ICAM3* | 6.277953409 | -0.006469568 | -0.029749551 | 0.023279982 | 0.999933887 | -0.198944332 | -0.413182232 | 0.2142379 | 0.101278962 |
| ENSG00000177556 | *ATOX1* | 3.260593556 | 0.005104356 | 0.039338732 | -0.034234377 | 0.999933887 | 0.600649527 | 0.840355852 | -0.239706325 | 0.101278962 |
| ENSG00000285888 | *Z85996.2* | -0.982587404 | 0.034390928 | -0.226075683 | 0.260466611 | 0.999933887 | 0.82531145 | 1.389033977 | -0.563722527 | 0.101347999 |
| ENSG00000266074 | *BAHCC1* | -0.274883097 | 0.012840642 | -0.020223763 | 0.033064404 | 0.999933887 | -0.018156006 | 0.629828247 | -0.647984253 | 0.101518518 |

| | | | | | | | | | | |
|---|---|---|---|---|---|---|---|---|---|---|
| ENSG00000065911 | *MTHFD2* | 3.761201318 | 0.012861256 | 0.121635325 | -0.108774068 | 0.999933887 | 0.234656109 | 0.495647936 | -0.260991827 | 0.101581224 |
| ENSG00000184307 | *ZDHHC23* | 1.968467142 | -0.26224443 | 0.064925674 | -0.327170104 | 0.999933887 | 0.011141919 | 0.256430442 | -0.245288524 | 0.1015866 |
| ENSG00000056972 | *TRAF3IP2* | 2.99670158 | 0.051588521 | 0.043912837 | 0.007675684 | 0.999933887 | 0.873772338 | 1.202212692 | -0.328440354 | 0.1015866 |
| ENSG00000028277 | *POU2F2* | 6.296189466 | 0.000193189 | 0.013530669 | -0.01333748 | 0.999933887 | 0.164196838 | 0.429723951 | -0.265527113 | 0.101748948 |
| ENSG00000242294 | *STAG3L5P* | 2.437003002 | 0.041325004 | 0.092551711 | -0.051226707 | 0.999933887 | 0.461665567 | 0.822210567 | -0.360545 | 0.101764703 |
| ENSG00000172403 | *SYNPO2* | -0.648960019 | 0.030646303 | 0.160645039 | -0.129998736 | 0.999933887 | -0.125910483 | 0.73824921 | -0.864159693 | 0.101940346 |
| ENSG00000164181 | *ELOVL7* | -0.361962519 | 0.201953563 | 0.34620934 | -0.144255777 | 0.999933887 | 2.305640725 | 3.091062431 | -0.785421705 | 0.102156738 |
| ENSG00000164366 | *CCDC127* | 1.347470768 | 0.006013577 | 0.070264369 | -0.064250792 | 0.999933887 | 0.100463974 | -0.266295005 | 0.366758979 | 0.102234553 |
| ENSG00000150045 | *KLRF1* | 0.804985155 | -0.089058073 | -0.23154668 | 0.142488607 | 0.999933887 | -0.211577308 | 0.173199653 | -0.384776961 | 0.102255258 |
| ENSG00000064419 | *TNPO3* | 4.906314681 | -0.049765502 | -0.010968837 | -0.038796666 | 0.999933887 | -0.193220194 | -0.39078936 | 0.197569166 | 0.102335121 |
| ENSG00000257285 | *AL132780.1* | 0.817126754 | -0.020533404 | 0.019624359 | -0.040157763 | 0.999933887 | -0.468386775 | -1.109736033 | 0.641349258 | 0.102335121 |
| ENSG00000135387 | *CAPRIN1* | 5.716031418 | -0.01855653 | -0.049316897 | 0.030760367 | 0.999933887 | -0.473422624 | -0.625368107 | 0.151945483 | 0.102628661 |
| ENSG00000286190 | *AC055839.2* | 3.104578688 | 0.047263218 | 0.007419954 | 0.039843263 | 0.999933887 | -0.121033489 | -0.454226102 | 0.333192613 | 0.102794244 |
| ENSG00000165813 | *CCDC186* | 5.178159628 | -0.040436457 | -0.15279249 | 0.112356033 | 0.999933887 | -0.813151138 | -0.253150418 | -0.56000072 | 0.102794244 |
| ENSG00000205890 | *AC108134.1* | -0.679375612 | 0.229886812 | 0.04796903 | 0.181917783 | 0.999933887 | -0.443874124 | -1.397670953 | 0.953796829 | 0.102853273 |
| ENSG00000135899 | *SP110* | 7.040199518 | -0.033905682 | -0.089189072 | 0.05528339 | 0.999933887 | -0.599694432 | -0.770668849 | 0.170974417 | 0.102853273 |
| ENSG00000112701 | *SENP6* | 5.463626279 | -0.007512322 | -0.096024513 | 0.088512191 | 0.999933887 | -0.57203874 | -0.989734849 | 0.417696109 | 0.102853273 |
| ENSG00000230438 | *SERPINB9P1* | 4.701156391 | -0.072610442 | -0.008410658 | -0.064199784 | 0.999933887 | 2.583924887 | 3.006617752 | -0.422692864 | 0.10303842 |
| ENSG00000198265 | *HELZ* | 5.620476042 | -0.002069772 | -0.114576619 | 0.112506847 | 0.999933887 | -0.309516641 | 0.002634715 | -0.312151355 | 0.103201787 |
| ENSG00000113749 | *HRH2* | 4.467795348 | 0.02604853 | 0.012049613 | 0.013998917 | 0.999933887 | 0.053876476 | -0.242830195 | 0.296706671 | 0.103225921 |
| ENSG00000151014 | *NOCT* | 1.941522227 | 0.028378979 | 0.056998608 | -0.028619629 | 0.999933887 | 1.034160754 | 1.419806069 | -0.385645315 | 0.103323198 |
| ENSG00000113504 | *SLC12A7* | 4.047554905 | 0.086707404 | -0.002655205 | 0.089362609 | 0.999933887 | 0.75895985 | 1.111851579 | -0.35289173 | 0.103329704 |
| ENSG00000174799 | *CEP135* | 2.881172232 | -0.177229616 | -0.164004532 | -0.013225084 | 0.999933887 | -0.14555432 | 0.415735082 | -0.561289402 | 0.103510089 |
| ENSG00000064393 | *HIPK2* | 5.830765813 | -0.058502689 | -0.035676767 | -0.022825922 | 0.999933887 | -0.961264916 | -1.19114111 | 0.229876194 | 0.103510089 |
| ENSG00000260135 | *MMP2-AS1* | 1.026550495 | 0.083909551 | -0.020963692 | 0.104873243 | 0.999933887 | 2.505556154 | 3.122178889 | -0.616622736 | 0.103510089 |
| ENSG00000114209 | *PDCD10* | 3.871921466 | -0.026384063 | -0.110285726 | 0.083901663 | 0.999933887 | -0.350848599 | -0.123823019 | -0.22702558 | 0.103630863 |
| ENSG00000009780 | *FAM76A* | 2.384779895 | -0.05765018 | 0.135456674 | -0.193106854 | 0.999933887 | -0.210226157 | -0.442606222 | 0.232380066 | 0.103694869 |
| ENSG00000157933 | *SKI* | 7.213083218 | 0.006571805 | -0.029313918 | 0.035885722 | 0.999933887 | 0.387002806 | 0.567869749 | -0.180866943 | 0.103694869 |
| ENSG00000186047 | *DLEU7* | 0.835749623 | -0.09578177 | -0.139147363 | 0.043365593 | 0.999933887 | 1.209089208 | 1.747620196 | -0.538530988 | 0.103724509 |
| ENSG00000228300 | *FAM174C* | 3.608013101 | 0.048422117 | 0.082576669 | -0.034154552 | 0.999933887 | -0.150920735 | -0.406366145 | 0.25544541 | 0.103724509 |
| ENSG00000196072 | *BLOC1S2* | 5.816562349 | -0.08919446 | -0.038127778 | -0.051066681 | 0.999933887 | -0.253713143 | -0.084905512 | -0.168807631 | 0.103805489 |
| ENSG00000184545 | *DUSP8* | 4.084314304 | 0.053659961 | 0.073640753 | -0.019980791 | 0.999933887 | 0.721510881 | 1.036577775 | -0.315066894 | 0.103805489 |
| ENSG00000170412 | *GPRC5C* | 0.600508248 | -0.086532575 | 0.023959952 | -0.110492527 | 0.999933887 | 0.555786095 | -0.079466731 | 0.635252826 | 0.103805489 |
| ENSG00000173039 | *RELA* | 7.449891307 | 0.032568881 | 0.016607561 | 0.015961321 | 0.999933887 | 0.809018934 | 1.004253149 | -0.195234215 | 0.103805489 |
| ENSG00000110799 | *VWF* | 2.348192963 | -0.106861318 | -0.079372792 | -0.027488526 | 0.999933887 | -2.347241049 | -2.795237036 | 0.447995987 | 0.103805489 |
| ENSG00000287562 | *AL109615.4* | -0.530871267 | -0.130226632 | 0.335758168 | -0.4659848 | 0.999933887 | 1.12461336 | 1.744571447 | -0.619958087 | 0.103906175 |
| ENSG00000144029 | *MRPS5* | 2.71926261 | -0.072607812 | -0.123906991 | 0.05129918 | 0.999933887 | 0.022741978 | -0.194335045 | 0.217077024 | 0.104048867 |
| ENSG00000143515 | *ATP8B2* | 5.060379196 | 0.036727807 | 0.05999308 | -0.023265274 | 0.999933887 | 0.14851783 | 0.351254835 | -0.202737005 | 0.104048867 |
| ENSG00000243147 | *MRPL33* | 3.432199424 | 0.062869835 | 0.08499511 | -0.022125275 | 0.999933887 | 0.121133232 | 0.335959247 | -0.214826015 | 0.104211557 |
| ENSG00000184900 | *SUMO3* | 4.489361116 | -0.012282905 | -0.088461279 | 0.076178375 | 0.999933887 | -0.049797013 | 0.132936721 | -0.182733735 | 0.104214947 |
| ENSG00000162378 | *ZYG11B* | 3.841452702 | 0.037532211 | 0.043824414 | -0.006292202 | 0.999933887 | -0.898126012 | -1.179063871 | 0.280937859 | 0.104224408 |
| ENSG00000232118 | *BACH1-AS1* | 1.434472517 | -0.054151366 | 0.094430521 | -0.148581887 | 0.999933887 | -0.253278925 | 0.139887034 | -0.393165959 | 0.104224408 |
| ENSG00000274286 | *ADRA2B* | 0.694589978 | 0.06796318 | 0.170919564 | -0.102956384 | 0.999933887 | 0.398822776 | 1.209197778 | -0.810375002 | 0.104252176 |
| ENSG00000135686 | *KLHL36* | 5.748572382 | -0.019795261 | -0.043907931 | 0.02411267 | 0.999933887 | -0.356910196 | -0.546805022 | 0.189894825 | 0.104440668 |
| ENSG00000218283 | *MORF4L1P1* | 2.845615247 | -0.093897222 | -0.024950317 | -0.068946906 | 0.999933887 | -0.212501082 | 0.060228815 | -0.272729896 | 0.104452239 |
| ENSG00000166275 | *BORCS7* | 2.816186233 | 0.042520407 | 0.032125506 | 0.010394901 | 0.999933887 | 0.012021222 | 0.282636827 | -0.270615605 | 0.104520941 |
| ENSG00000122861 | *PLAU* | 10.763219043 | 0.146545597 | 0.240247849 | -0.093702252 | 0.999933887 | 1.419441181 | 1.692566859 | -0.273125678 | 0.104532299 |
| ENSG00000267121 | *AC008105.3* | 4.83199317 | -0.086160657 | -0.025877074 | -0.060283582 | 0.999933887 | -0.04265716 | -0.217328985 | 0.174671824 | 0.104541654 |
| ENSG00000121671 | *CRY2* | 4.500061676 | -0.011924893 | 0.023727304 | -0.035652197 | 0.999933887 | 0.002229263 | -0.208735026 | 0.21096429 | 0.104541654 |
| ENSG00000237892 | *KLF7-IT1* | 1.453089024 | 0.011659634 | 0.034644265 | -0.022984631 | 0.999933887 | 0.581360522 | 1.124344946 | -0.542984424 | 0.104541654 |
| ENSG00000130119 | *GNL3L* | 4.996265598 | -0.051534607 | -0.048171014 | -0.003363593 | 0.999933887 | -0.277670367 | -0.486405708 | 0.208735341 | 0.104783203 |
| ENSG00000165801 | *ARHGEF40* | 6.624190349 | 0.071026825 | 0.093923248 | -0.022896422 | 0.999933887 | -0.408157897 | -0.618006107 | 0.20984821 | 0.104789883 |
| ENSG00000116863 | *ADPRS* | 3.1059031 | -0.054479832 | 0.082841915 | -0.137321747 | 0.999933887 | -0.148068876 | -0.434235727 | 0.28616685 | 0.104791612 |
| ENSG00000163714 | *U2SURP* | 5.091456213 | -0.075668727 | -0.149761282 | 0.074092555 | 0.999933887 | -0.435479152 | -0.04405112 | -0.391428032 | 0.104840181 |
| ENSG00000141429 | *GALNT1* | 4.927113348 | -0.046457732 | -0.157587831 | 0.1111301 | 0.999933887 | -0.053404041 | 0.153969 | -0.207373041 | 0.105105042 |
| ENSG00000082258 | *CCNT2* | 5.918120195 | -0.052078409 | -0.135292823 | 0.083214414 | 0.999933887 | -0.379651133 | -0.008465218 | -0.371185915 | 0.105105042 |
| ENSG00000287978 | *AC245407.2* | 2.565957116 | 0.018633028 | 0.051877185 | -0.033244158 | 0.999933887 | 1.067255598 | 1.413994939 | -0.346739342 | 0.105105042 |
| ENSG00000204103 | *MAFB* | 6.969120217 | 0.249003476 | 0.327514823 | -0.078511346 | 0.999933887 | 0.265223995 | 0.466692269 | -0.201468274 | 0.105200848 |
| ENSG00000198763 | *MT-ND2* | 8.951473459 | 0.114766957 | 0.001876953 | 0.112890004 | 0.999933887 | 0.048301938 | 0.381645467 | -0.333343529 | 0.105323806 |
| ENSG00000135637 | *CCDC142* | 1.390873516 | -0.048858867 | 0.210606003 | -0.25946487 | 0.999933887 | -0.109000139 | -0.475244565 | 0.366244426 | 0.105323806 |
| ENSG00000273729 | *AC007686.3* | 0.610574574 | 0.006499488 | 0.012534639 | -0.006035151 | 0.999933887 | -0.227412407 | -0.772932493 | 0.545520086 | 0.105369064 |
| ENSG00000270210 | *AC104695.2* | 0.841900565 | 0.15009608 | 0.209212239 | -0.059116159 | 0.999933887 | 0.90850828 | 1.390126755 | -0.481618475 | 0.105393171 |
| ENSG00000162407 | *PLPP3* | 6.264472817 | -0.072405286 | -0.026390436 | -0.04601485 | 0.999933887 | -0.048715458 | 0.132004158 | -0.180719616 | 0.105629691 |
| ENSG00000118873 | *RAB3GAP2* | 4.665852705 | 0.005568292 | -0.10529624 | 0.110864532 | 0.999933887 | -0.274794034 | -0.008203333 | -0.266590701 | 0.105629691 |
| ENSG00000101974 | *ATP11C* | 3.598742026 | 0.009402048 | -0.041861892 | 0.05126394 | 0.999933887 | 0.319044422 | 0.714862342 | -0.39581792 | 0.10570842 |
| ENSG00000217275 | *AL031777.1* | -0.959199157 | 0.099752619 | 0.229326131 | -0.129573512 | 0.999933887 | -0.638642206 | -1.392275299 | 0.753633093 | 0.105798883 |
| ENSG00000198382 | *UVRAG* | 5.72821318 | -0.02288402 | 0.034839331 | -0.057723351 | 0.999933887 | 0.340872898 | 0.48603329 | -0.145160392 | 0.105798883 |
| ENSG00000148341 | *SH3GLB2* | 4.503806908 | 0.056704073 | 0.06268359 | -0.005979517 | 0.999933887 | 0.110067948 | -0.086041864 | 0.196109812 | 0.105825566 |
| ENSG00000224950 | *AL390066.1* | 2.345103325 | -0.055371942 | 0.029707292 | -0.085079234 | 0.999933887 | 0.892442114 | 1.169178371 | -0.276736257 | 0.105993293 |
| ENSG00000136448 | *NMT1* | 4.503496155 | -0.033669259 | -0.093764609 | 0.060095349 | 0.999933887 | -0.17232479 | -0.362929711 | 0.19060492 | 0.106348994 |
| ENSG00000131378 | *RFTN1* | 3.731653675 | -0.040045266 | 0.100501283 | -0.140546548 | 0.999933887 | 0.296445161 | 0.540639711 | -0.24419455 | 0.10648322 |
| ENSG00000174953 | *DHX36* | 5.031566476 | -0.061858983 | -0.091699335 | 0.029840351 | 0.999933887 | -0.066988639 | 0.259075131 | -0.32606377 | 0.106519599 |
| ENSG00000032444 | *PNPLA6* | 6.400332511 | 0.00210606 | 0.033841738 | -0.031735679 | 0.999933887 | -0.098321177 | -0.249050548 | 0.150729371 | 0.106544659 |
| ENSG00000122390 | *NAA60* | 5.120171656 | -0.008574607 | 0.024259702 | -0.032834309 | 0.999933887 | -0.403372457 | -0.593471779 | 0.190099322 | 0.106670166 |
| ENSG00000169914 | *OTUD3* | 2.366331723 | -0.016738795 | 0.097340455 | -0.114079249 | 0.999933887 | 0.001013545 | -0.232218352 | 0.233231897 | 0.106672634 |
| ENSG00000144228 | *SPOPL* | 6.900455426 | 0.015441196 | 0.00061603 | 0.014825166 | 0.999933887 | 0.051910078 | 0.324789981 | -0.272879902 | 0.106825355 |
| ENSG00000225402 | *AC010878.1* | 1.300061228 | 0.061937853 | 0.096326598 | -0.034388745 | 0.999933887 | 0.564993039 | 1.01204296 | -0.447049922 | 0.106876444 |
| ENSG00000260233 | *ZNRD2-AS1* | 2.484646571 | 0.133472416 | -0.038796338 | 0.172268754 | 0.999933887 | 1.366402924 | 1.655567309 | -0.289164385 | 0.106910591 |
| ENSG00000162129 | *CLPB* | 1.258396911 | 0.111854107 | 0.072198537 | 0.03965557 | 0.999933887 | 0.12235993 | 0.503615291 | -0.381255362 | 0.106931995 |
| ENSG00000142867 | *BCL10* | 5.513804878 | 0.021956646 | 0.064836104 | -0.042879458 | 0.999933887 | 0.042485947 | 0.204243265 | -0.161757318 | 0.107137774 |
| ENSG00000163395 | *IGFN1* | -2.873480007 | -0.234616251 | -0.440740014 | 0.206123763 | 0.999933887 | -0.427970682 | -1.228157374 | 0.800186692 | 0.107137774 |
| ENSG00000270980 | *AC060764.1* | -2.261467299 | -0.171065064 | 0.308620799 | -0.479685863 | 0.999933887 | 0.619563332 | 1.22144941 | -0.601886078 | 0.107137774 |
| ENSG00000260361 | *AC106028.3* | 0.758694522 | 0.168022596 | 0.034635479 | 0.133387118 | 0.999933887 | 0.442014484 | 0.794706506 | -0.352692021 | 0.107560395 |
| ENSG00000169087 | *HSPBAP1* | 5.746218373 | -0.028630888 | -0.062788131 | 0.034157243 | 0.999933887 | -0.312853608 | -0.12055209 | -0.192301518 | 0.107679315 |
| ENSG00000108010 | *GLRX3* | 2.952102907 | -0.007638483 | -0.038164498 | 0.030526016 | 0.999933887 | 0.112915969 | 0.318839961 | -0.205923991 | 0.10786858 |
| ENSG00000015676 | *NUDCD3* | 3.920216863 | -0.063780487 | 0.082242364 | -0.146022851 | 0.999933887 | -0.06175419 | -0.245704434 | 0.183950244 | 0.10792566 |
| ENSG00000178988 | *MRFAP1L1* | 4.357368939 | -0.076104699 | -0.176454398 | 0.100349699 | 0.999933887 | -0.320338895 | -0.543871251 | 0.223532356 | 0.108130518 |
| ENSG00000244038 | *DDOST* | 4.751065769 | 0.026354538 | -0.00601123 | 0.032365768 | 0.999933887 | -0.048837953 | -0.227428991 | 0.178591038 | 0.108377548 |
| ENSG00000170043 | *TRAPPC1* | 4.488984369 | 0.034834241 | -0.017860513 | 0.052694754 | 0.999933887 | -0.133734678 | -0.35183625 | 0.218101572 | 0.108397447 |
| ENSG00000032219 | *ARID4A* | 6.007065076 | -0.03439945 | -0.159365421 | 0.12496597 | 0.999933887 | -0.789782819 | -0.287183765 | -0.502599054 | 0.108877923 |
| ENSG00000152229 | *PSTPIP2* | 5.315719079 | 0.010755345 | -0.027148026 | 0.037903371 | 0.999933887 | 1.19856329 | 1.61038969 | -0.4118264 | 0.108877923 |
| ENSG00000138101 | *DTNB* | 2.587516576 | 0.029743655 | 0.194588823 | -0.164845168 | 0.999933887 | 0.599254362 | 0.887490695 | -0.288236333 | 0.108965914 |
| ENSG00000025039 | *RRAGD* | 7.06243667 | -0.059583306 | 0.000908161 | -0.060491467 | 0.999933887 | -0.089214611 | 0.075360799 | -0.16457541 | 0.108993262 |
| ENSG00000183562 | *AC131971.1* | -0.563875964 | -0.077789623 | 0.556412769 | -0.634202392 | 0.999933887 | 0.26828017 | -0.390828077 | 0.659108247 | 0.109062076 |
| ENSG00000113013 | *HSPA9* | 7.4816381 | 0.010598182 | -0.009965593 | 0.020563774 | 0.999933887 | 0.373368828 | 0.574388961 | -0.201020133 | 0.109440359 |
| ENSG00000215196 | *BASP1-AS1* | 2.836485205 | 0.123129635 | 0.053913095 | 0.069216539 | 0.999933887 | 0.878664622 | 1.182360533 | -0.303695911 | 0.1098193 |
| ENSG00000226979 | *LTA* | 0.990898492 | 0.046153505 | -0.185768041 | 0.231921546 | 0.999933887 | 0.765195601 | 1.149214292 | -0.384018692 | 0.1098193 |
| ENSG00000167695 | *TLCD3A* | 0.465197457 | 0.069176136 | 0.009913332 | 0.059262804 | 0.999933887 | 1.416231658 | 2.05064836 | -0.634416702 | 0.1098193 |
| ENSG00000182919 | *C11orf54* | 2.586380121 | -0.050828674 | -0.02817495 | -0.022653724 | 0.999933887 | -0.476388361 | -0.7656831 | 0.289294738 | 0.11001911 |
| ENSG00000196646 | *ZNF136* | 3.185743919 | -0.106185062 | -0.12344821 | 0.017263148 | 0.999933887 | -0.09608644 | 0.23244791 | -0.32853435 | 0.110148302 |
| ENSG00000170385 | *SLC30A1* | 5.063648413 | -0.075301858 | -0.068810281 | -0.006491577 | 0.999933887 | -0.508799145 | -0.219926335 | -0.28887281 | 0.110148302 |
| ENSG00000121879 | *PIK3CA* | 4.928721561 | -0.031898959 | -0.124860107 | 0.092961148 | 0.999933887 | -0.50199291 | -0.157159895 | -0.344833015 | 0.110148302 |
| ENSG00000175206 | *NPPA* | 0.140840348 | 0.058465498 | -0.386782505 | 0.445248003 | 0.999933887 | -0.299156794 | 0.096489403 | -0.395646197 | 0.110148302 |
| ENSG00000172673 | *THEMIS* | 3.384420302 | -0.012705777 | -0.041379904 | 0.028674127 | 0.999933887 | -0.036409318 | 0.293510881 | -0.329920198 | 0.110148302 |
| ENSG00000164970 | *FAM219A* | 3.857188601 | 0.050759674 | -0.029630308 | 0.080389981 | 0.999933887 | -0.472944673 | -0.673305444 | 0.200360771 | 0.110156155 |

| | | | | | | | | | | |
|---|---|---|---|---|---|---|---|---|---|---|
| ENSG00000233223 | *AC016876.1* | 3.660824166 | 0.378420177 | 0.359423413 | 0.018996764 | 0.999933887 | 0.550242915 | 0.805419856 | -0.255176941 | 0.110181365 |
| ENSG00000261207 | *AL031717.1* | 0.836140432 | 0.00914956 | -0.051709311 | 0.060858871 | 0.999933887 | -0.787098426 | -1.25866879 | 0.471570365 | 0.110181365 |
| ENSG00000137478 | *FCHSD2* | 6.890564274 | -0.030571481 | -0.0318562 | 0.001284719 | 0.999933887 | -0.080849982 | 0.116388652 | -0.197238634 | 0.110211669 |
| ENSG00000014824 | *SLC30A9* | 3.271794986 | -0.082787392 | -0.050500601 | -0.032286791 | 0.999933887 | -0.164905944 | 0.095844972 | -0.260750916 | 0.110405925 |
| ENSG00000183386 | *FHL3* | 4.920374048 | 0.032778862 | 0.061411531 | -0.028632669 | 0.999933887 | -0.84978757 | -1.131941331 | 0.282153761 | 0.110405925 |
| ENSG00000153214 | *TMEM87B* | 2.918030614 | 0.028953596 | 0.049650786 | -0.02069719 | 0.999933887 | -0.28890184 | -0.612778139 | 0.323876298 | 0.110405925 |
| ENSG00000168385 | *SEPTIN2* | 6.233345626 | -0.042161024 | -0.031032651 | -0.011128373 | 0.999933887 | -0.071753721 | 0.062335973 | -0.134089693 | 0.11049517 |
| ENSG00000078140 | *UBE2K* | 4.363035835 | -0.049388245 | -0.130522636 | 0.08113439 | 0.999933887 | -0.427612325 | -0.625095909 | 0.197483584 | 0.110609072 |
| ENSG00000273033 | *LINC02035* | 4.810917527 | -0.105109355 | -0.075671636 | -0.029437719 | 0.999933887 | -0.378899021 | -0.144380543 | -0.234518479 | 0.110699653 |
| ENSG00000050730 | *TNIP3* | 0.169333978 | -0.012169778 | -0.108310603 | 0.096140825 | 0.999933887 | 0.047606857 | 0.599397094 | -0.551790236 | 0.110699653 |
| ENSG00000109790 | *KLHL5* | 4.384320875 | -0.030842313 | 0.013928157 | -0.044770471 | 0.999933887 | 0.137357056 | 0.377096715 | -0.239739659 | 0.110869099 |
| ENSG00000151422 | *FER* | 0.478874277 | 0.117194526 | -0.257041627 | 0.374236153 | 0.999933887 | -0.260374452 | 0.215852969 | -0.476227422 | 0.111137067 |
| ENSG00000103363 | *ELOB* | 5.685184878 | 0.027019645 | 0.006089802 | 0.020929843 | 0.999933887 | -0.082990002 | -0.266713712 | 0.18372371 | 0.111160707 |
| ENSG00000109756 | *RAPGEF2* | 7.829495897 | 0.009582703 | -0.027999562 | 0.037582266 | 0.999933887 | 0.196264153 | 0.388926247 | -0.192662094 | 0.111160707 |
| ENSG00000011132 | *APBA3* | 4.779723626 | 0.104257389 | 0.022951952 | 0.081305437 | 0.999933887 | 0.673329912 | 0.886179149 | -0.212849237 | 0.111620517 |
| ENSG00000217930 | *PAM16* | 1.09121647 | -0.062807923 | -0.095620657 | 0.032812734 | 0.999933887 | -0.14928443 | -0.590824758 | 0.441540328 | 0.111620517 |
| ENSG00000171681 | *ATF7IP* | 5.943906405 | -0.041501467 | -0.074737688 | 0.033236221 | 0.999933887 | -0.069319448 | 0.19389197 | -0.263211418 | 0.111708108 |
| ENSG00000091039 | *OSBPL8* | 7.531291714 | -0.067868098 | -0.107680213 | 0.039812115 | 0.999933887 | -0.712736498 | -0.331458939 | -0.381277559 | 0.111793028 |
| ENSG00000037474 | *NSUN2* | 4.469048293 | -0.026648009 | 0.052148985 | -0.078796994 | 0.999933887 | 0.133500942 | 0.342302395 | -0.208801453 | 0.11208977 |
| ENSG00000253227 | *AC090192.2* | -0.115192263 | -0.288310978 | 0.471425983 | -0.75973696 | 0.999933887 | 0.606824135 | 1.035049636 | -0.428225501 | 0.112148816 |
| ENSG00000008952 | *SEC62* | 6.535544668 | -0.089747461 | -0.19524102 | 0.105493559 | 0.999933887 | -0.550821175 | -0.338058716 | -0.212762458 | 0.11227655 |
| ENSG00000184164 | *CRELD2* | 3.434099262 | 0.007178729 | 0.002554225 | 0.004624504 | 0.999933887 | 0.29770171 | 0.553108671 | -0.255406962 | 0.112568189 |
| ENSG00000272391 | *POM121C* | 5.677768602 | 0.003920197 | 0.029620606 | -0.025700409 | 0.999933887 | -0.341751487 | -0.498757998 | 0.157006511 | 0.112568189 |
| ENSG00000135241 | *PNPLA8* | 7.006515735 | -0.003730426 | -0.194700366 | 0.19096994 | 0.999933887 | 0.744741367 | 1.112081238 | -0.36733987 | 0.112568189 |
| ENSG00000196937 | *FAM3C* | 1.115694249 | -0.029530422 | -0.057716159 | 0.028185737 | 0.999933887 | -0.112249877 | 0.240739654 | -0.35298953 | 0.112613591 |
| ENSG00000089820 | *ARHGAP4* | 7.81702536 | 0.076235726 | 0.147424682 | -0.071188956 | 0.999933887 | 0.077423984 | -0.137539022 | 0.214963006 | 0.112669217 |
| ENSG00000112592 | *TBP* | 2.862370974 | -0.028872875 | 0.046100621 | -0.074973496 | 0.999933887 | -0.142942658 | -0.360483042 | 0.217540384 | 0.112669217 |
| ENSG00000112584 | *FAM120B* | 4.650717992 | -0.048440581 | 0.069004658 | -0.117445239 | 0.999933887 | -0.003276535 | 0.158538267 | -0.161814803 | 0.11277243 |
| ENSG00000262580 | *AC087741.1* | 4.39139774 | -0.126334924 | -0.033012308 | -0.093322616 | 0.999933887 | 0.167349858 | 0.336358239 | -0.16900838 | 0.112785701 |
| ENSG00000263001 | *GTF2I* | 2.820368037 | -0.102430306 | 0.041583139 | -0.144013445 | 0.999933887 | 0.01141702 | 0.224124682 | -0.212707662 | 0.112785701 |
| ENSG00000175121 | *WFDC5* | 0.634233891 | 0.310748449 | 0.465579372 | -0.154830924 | 0.999933887 | 1.873114845 | 2.302515202 | -0.429400358 | 0.112785701 |
| ENSG00000210082 | *MT-RNR2* | 12.320455824 | 0.046262892 | 0.041245334 | 0.005017557 | 0.999933887 | 0.248459487 | 0.444125573 | -0.195666086 | 0.112785701 |
| ENSG00000204345 | *CD300LD* | -0.534601288 | -0.146760872 | 0.098473165 | -0.245234037 | 0.999933887 | -0.739123083 | -1.678522084 | 0.939399001 | 0.112893745 |
| ENSG00000170801 | *HTRA3* | 1.184604328 | -0.132653743 | -0.134937776 | 0.002284033 | 0.999933887 | 1.401777336 | 1.711636665 | -0.309859329 | 0.11289729 |
| ENSG00000072415 | *MPP5* | 5.39270516 | 0.052214272 | 0.039080396 | 0.013133876 | 0.999933887 | 0.06707224 | 0.467059586 | -0.399987346 | 0.112999135 |
| ENSG00000256806 | *C17orf100* | -0.327904654 | 0.25153681 | -0.045490045 | 0.297026856 | 0.999933887 | -0.152679831 | -0.858731489 | 0.706051657 | 0.112999436 |
| ENSG00000138674 | *SEC31A* | 5.227983544 | -0.027897622 | 0.019287704 | -0.047185326 | 0.999933887 | 0.099825357 | 0.275213036 | -0.175387679 | 0.112999436 |
| ENSG00000120686 | *UFM1* | 4.348353054 | -0.073731004 | -0.158967445 | 0.085236442 | 0.999933887 | -0.091414006 | 0.207551821 | -0.298965827 | 0.113079494 |
| ENSG00000137094 | *DNAJB5* | 2.332854321 | 0.065585205 | 0.036060407 | 0.029524798 | 0.999933887 | 0.21662051 | 0.647730182 | -0.431109672 | 0.113098302 |
| ENSG00000151725 | *CENPU* | 1.024069018 | 0.115056645 | 0.277765436 | -0.162708791 | 0.999933887 | 0.190492239 | 0.784510779 | -0.59401854 | 0.113119668 |
| ENSG00000285448 | *AL136961.1* | 0.600972381 | -0.073695427 | 0.164440131 | -0.238135558 | 0.999933887 | 2.318479039 | 2.789347938 | -0.470868899 | 0.113126736 |
| ENSG00000261654 | *AL360270.2* | 0.307071748 | 0.047135025 | -0.065224772 | 0.112359798 | 0.999933887 | -0.313279902 | 0.622178175 | -0.935458077 | 0.113126736 |
| ENSG00000174839 | *DENND6A* | 3.679382537 | 0.046952314 | 0.013256054 | 0.03369626 | 0.999933887 | -0.330827934 | -0.595045765 | 0.264217832 | 0.11346909 |
| ENSG00000007541 | *PIGQ* | 3.006422365 | 0.020254808 | 0.135703397 | -0.11544859 | 0.999933887 | 0.081342482 | -0.180521928 | 0.26186441 | 0.11346909 |
| ENSG00000115170 | *ACVR1* | 2.063273656 | -0.026914259 | 0.054476638 | -0.081390897 | 0.999933887 | -0.397265245 | -0.695731824 | 0.298466579 | 0.11346909 |
| ENSG00000112640 | *PPP2R5D* | 2.653597712 | 0.017670034 | -0.009477865 | 0.027147899 | 0.999933887 | 0.084243992 | -0.197261235 | 0.281505226 | 0.11346909 |
| ENSG00000111850 | *SMIM8* | 2.295305739 | -0.061591981 | -0.054107546 | -0.007484435 | 0.999933887 | -1.288402903 | -1.640221954 | 0.351819051 | 0.113481492 |
| ENSG00000163683 | *SMIM14* | 3.491566521 | -0.042094873 | 0.14982454 | -0.191919414 | 0.999933887 | -0.308265138 | -0.606603284 | 0.298338146 | 0.113543144 |
| ENSG00000155465 | *SLC7A7* | 5.762588532 | -0.107568458 | -0.01764452 | -0.089923938 | 0.999933887 | 1.627320799 | 1.97909545 | -0.351774651 | 0.11360349 |
| ENSG00000145782 | *ATG12* | 5.899528268 | -0.040070063 | -0.060460557 | 0.020390494 | 0.999933887 | -0.356088873 | -0.187970624 | -0.168118249 | 0.113628219 |
| ENSG00000078142 | *PIK3C3* | 4.769875582 | -0.0816068 | -0.095484226 | 0.013877426 | 0.999933887 | -0.138552579 | 0.069966763 | -0.208519342 | 0.113643433 |
| ENSG00000159461 | *AMFR* | 4.503398507 | -0.00862869 | -0.045993378 | 0.037364689 | 0.999933887 | -0.434199391 | -0.686771967 | 0.252572576 | 0.113711981 |
| ENSG00000163464 | *CXCR1* | 8.205008192 | -0.029982155 | -0.055179184 | 0.025197029 | 0.999933887 | -1.820009999 | -2.180362392 | 0.360352393 | 0.113731991 |
| ENSG00000171222 | *SCAND1* | 4.457766775 | 0.006970128 | -0.062495696 | 0.069465823 | 0.999933887 | -0.32863218 | -0.578680636 | 0.250048456 | 0.113731991 |
| ENSG00000226950 | *DANCR* | 1.234572119 | -0.0012077 | -0.086539454 | 0.085331754 | 0.999933887 | 0.215575104 | -0.183688711 | 0.399263815 | 0.113731991 |
| ENSG00000176155 | *CCDC57* | 4.868394551 | 0.026985239 | 0.023349157 | 0.003636083 | 0.999933887 | 0.320094136 | 0.503125434 | -0.183031298 | 0.113753072 |
| ENSG00000169964 | *TMEM42* | 1.31049505 | -0.098212761 | -0.01290935 | -0.085303411 | 0.999933887 | 0.193309447 | 0.493223762 | -0.299914315 | 0.113909042 |
| ENSG00000221823 | *PPP3R1* | 5.542557954 | -0.022569376 | 0.058836518 | -0.081405895 | 0.999933887 | -0.529246458 | -0.713534367 | 0.184287909 | 0.113909042 |
| ENSG00000159164 | *SV2A* | 0.335051631 | -0.101073211 | -0.183002542 | 0.081929332 | 0.999933887 | -0.016244035 | 0.535823977 | -0.552068012 | 0.113994411 |
| ENSG00000170456 | *DENND5B* | -0.506293463 | -0.179577025 | -0.292563223 | 0.112986198 | 0.999933887 | -0.819809819 | -0.200745832 | -0.619063986 | 0.114103687 |
| ENSG00000166855 | *CLPX* | 3.787588881 | -0.037843678 | -0.146301844 | 0.108458165 | 0.999933887 | -0.444349713 | -0.6313678 | 0.187018088 | 0.114187916 |
| ENSG00000134996 | *OSTF1* | 6.390401761 | 0.015098344 | 0.027163017 | -0.012064673 | 0.999933887 | -0.702025686 | -0.894614452 | 0.192588766 | 0.114232591 |
| ENSG00000100228 | *RAB36* | 0.627775682 | 0.067810306 | 0.086607616 | -0.01879731 | 0.999933887 | -0.434486731 | -1.12172548 | 0.687238748 | 0.114234562 |
| ENSG00000187231 | *SESTD1* | 3.707620512 | -0.042496221 | -0.060806297 | 0.018310075 | 0.999933887 | 0.495091361 | 0.803353269 | -0.308261907 | 0.114337998 |
| ENSG00000251474 | *RPL32P3* | 3.433069941 | -0.018940171 | -0.001815526 | -0.017124645 | 0.999933887 | -0.219359798 | -0.410366424 | 0.191006626 | 0.114337998 |
| ENSG00000086288 | *NME8* | 0.246417147 | -0.006479885 | -0.179843866 | 0.173363981 | 0.999933887 | -0.108543679 | 0.56097914 | -0.669522819 | 0.114337998 |
| ENSG00000227500 | *SCAMP4* | 5.218576285 | -0.035090245 | -0.034353245 | -0.000737 | 0.999933887 | -1.105269281 | -1.339040135 | 0.233770854 | 0.114383477 |
| ENSG00000112877 | *CEP72* | 0.17787788 | 0.189210489 | -0.172970673 | 0.362181162 | 0.999933887 | 0.083057313 | 0.664559568 | -0.581502256 | 0.114468581 |
| ENSG00000163389 | *POGLUT1* | 2.0906616 | 0.059210714 | -0.143612983 | 0.202823697 | 0.999933887 | 0.159935715 | 0.45419608 | -0.294260366 | 0.114580358 |
| ENSG00000169727 | *GPS1* | 4.056080178 | 0.016686048 | 0.065167943 | -0.048481895 | 0.999933887 | 0.05231531 | -0.15056261 | 0.20287792 | 0.114649318 |
| ENSG00000198840 | *MT-ND3* | 7.473521289 | 0.02495813 | -0.0113073 | 0.03626543 | 0.999933887 | 0.090838485 | 0.315985104 | -0.225146618 | 0.1148211 |
| ENSG00000256249 | *AC026333.3* | 3.984240447 | 0.222397882 | 0.236686052 | -0.01428817 | 0.999933887 | 0.826055666 | 1.197644468 | -0.371588802 | 0.114822464 |
| ENSG00000112234 | *FBXL4* | 2.34361323 | 0.051342282 | -0.005026056 | 0.056368338 | 0.999933887 | -0.539801211 | -0.846346493 | 0.306545282 | 0.115076184 |
| ENSG00000144445 | *KANSL1L* | 3.682882328 | -0.015430501 | -0.080078955 | 0.064648454 | 0.999933887 | -0.254323324 | -0.04364622 | -0.210677104 | 0.115141866 |
| ENSG00000183741 | *CBX6* | 6.631648737 | -0.060034926 | 0.0037768 | -0.063811726 | 0.999933887 | -0.260589429 | -0.423757345 | 0.163167915 | 0.115196061 |
| ENSG00000112972 | *HMGCS1* | 5.336666201 | 0.021437244 | -0.067867076 | 0.08930432 | 0.999933887 | 0.623750801 | 0.840382515 | -0.216631714 | 0.11529042 |
| ENSG00000131143 | *COX4I1* | 6.931286124 | -0.013636083 | -0.052796031 | 0.039159948 | 0.999933887 | 0.368488343 | 0.513974112 | -0.145485769 | 0.11529042 |
| ENSG00000248309 | *MEF2C-AS1* | 1.228413207 | -0.103944252 | 0.106316384 | -0.210260636 | 0.999933887 | 0.554268505 | 0.908987651 | -0.354719146 | 0.115355374 |
| ENSG00000005007 | *UPF1* | 7.46308019 | -0.020755513 | 0.024480154 | -0.045235667 | 0.999933887 | -0.288009775 | -0.438532546 | 0.150522771 | 0.115357968 |
| ENSG00000115267 | *IFIH1* | 7.090559157 | 0.018666317 | -0.085713328 | 0.104379645 | 0.999933887 | 0.69538453 | 0.933188942 | -0.237804412 | 0.115359428 |
| ENSG00000184708 | *EIF4ENIF1* | 4.004948955 | -0.067559084 | 0.026474152 | -0.094033236 | 0.999933887 | 0.289221305 | 0.525987176 | -0.236765872 | 0.115652477 |
| ENSG00000143127 | *ITGA10* | -0.91991926 | -0.071749916 | -0.348553862 | 0.276803945 | 0.999933887 | -0.002807369 | -0.68300258 | 0.680195211 | 0.115652477 |
| ENSG00000006555 | *TTC22* | 1.532210898 | 0.102245464 | 0.054021303 | 0.048224161 | 0.999933887 | 0.052457384 | -0.262366016 | 0.3148234 | 0.115753855 |
| ENSG00000123689 | *G0S2* | 10.673364821 | 0.732704413 | 0.667785604 | 0.064918809 | 0.999933887 | 2.008472164 | 2.480931412 | -0.472459249 | 0.115891391 |
| ENSG00000155111 | *CDK19* | 4.784567117 | -0.055078664 | -0.033510269 | -0.021568395 | 0.999933887 | -0.530944473 | -0.773541354 | 0.242596881 | 0.115922268 |
| ENSG00000126067 | *PSMB2* | 3.860238894 | -0.040851322 | -0.029494694 | -0.011356628 | 0.999933887 | -0.163694821 | -0.389294164 | 0.225599343 | 0.11604271 |
| ENSG00000136868 | *SLC31A1* | 3.444551085 | 0.000549608 | -0.0662549 | 0.066804508 | 0.999933887 | -0.325054659 | -0.050732758 | -0.274321901 | 0.116156376 |
| ENSG00000084073 | *ZMPSTE24* | 3.534886526 | -0.014036763 | -0.124564903 | 0.11052814 | 0.999933887 | -0.246919419 | -0.511964528 | 0.265045109 | 0.11626822 |
| ENSG00000134317 | *GRHL1* | -0.041770919 | -0.040339304 | -0.134773827 | 0.094434524 | 0.999933887 | -0.050412869 | 0.619511458 | -0.669924327 | 0.116281335 |
| ENSG00000197620 | *EOLA1* | 2.899850695 | 0.029224884 | -0.070772414 | 0.099997298 | 0.999933887 | -0.148370614 | -0.424374174 | 0.27600356 | 0.116315983 |
| ENSG00000182004 | *SNRPE* | 3.058413431 | -0.092414845 | -0.075618541 | -0.016796303 | 0.999933887 | 0.021696651 | -0.245979762 | 0.267676412 | 0.116319501 |
| ENSG00000115966 | *ATF2* | 4.473920342 | -0.044587258 | -0.099266777 | 0.054679519 | 0.999933887 | -0.465077912 | -0.215931037 | -0.249146875 | 0.116357077 |
| ENSG00000077454 | *LRCH4* | 5.657185688 | 0.054073513 | 0.043337023 | 0.01073649 | 0.999933887 | -0.071609998 | -0.232780096 | 0.161170098 | 0.116450708 |
| ENSG00000164327 | *RICTOR* | 8.003805841 | -0.08356935 | -0.165234651 | 0.081665301 | 0.999933887 | -0.961512853 | -0.453763222 | -0.507749631 | 0.116450708 |
| ENSG00000070614 | *NDST1* | 6.727488658 | -0.103779661 | -0.085586266 | -0.018193395 | 0.999933887 | -1.099946737 | -1.333008051 | 0.233061313 | 0.116462746 |
| ENSG00000130449 | *ZSWIM6* | 6.300179647 | 0.036714912 | 0.046279738 | -0.009564826 | 0.999933887 | 0.097881447 | 0.291093857 | -0.19321241 | 0.116462746 |
| ENSG00000100105 | *PATZ1* | 3.641496379 | -0.003084522 | 0.011377568 | -0.014462089 | 0.999933887 | 0.068847982 | -0.122470838 | 0.191318819 | 0.11654885 |
| ENSG00000171208 | *NETO2* | 0.362967757 | 0.016462312 | 0.024538449 | -0.008076137 | 0.999933887 | 0.175521863 | 0.625205276 | -0.449683413 | 0.116607313 |
| ENSG00000125810 | *CD93* | 6.676611179 | -0.045486309 | 0.00088145 | -0.046367758 | 0.999933887 | -0.968233226 | -1.322725953 | 0.354492727 | 0.116698242 |
| ENSG00000127507 | *ADGRE2* | 7.506315674 | -0.016564153 | 0.022274796 | -0.03883895 | 0.999933887 | -0.382893322 | -0.611438553 | 0.228545231 | 0.116698242 |
| ENSG00000166340 | *TPP1* | 8.294950498 | 0.016495132 | -0.020465266 | 0.036960398 | 0.999933887 | 0.357594312 | 0.52466894 | -0.167074628 | 0.116959882 |
| ENSG00000270195 | *AC016773.1* | -0.865655797 | -0.204895667 | -0.017988713 | -0.186906954 | 0.999933887 | -0.748293916 | -1.616329649 | 0.868035733 | 0.117103212 |
| ENSG00000136758 | *YME1L1* | 6.531551556 | -0.01816096 | -0.034643278 | 0.016482318 | 0.999933887 | 0.236689878 | 0.374758872 | -0.138068994 | 0.117146087 |
| ENSG00000271913 | *AL035530.2* | 5.509090239 | 0.46910624 | 0.605547938 | -0.136441697 | 0.999933887 | 1.666360957 | 2.083465256 | -0.417104299 | 0.117480881 |
| ENSG00000132824 | *SERINC3* | 8.935073608 | -0.043198946 | -0.007933643 | -0.035265302 | 0.999933887 | -0.103780578 | 0.028206377 | -0.131986956 | 0.117507751 |

| | | | | | | | | | | |
|---|---|---|---|---|---|---|---|---|---|---|
| ENSG00000167434 | *CA4* | 3.729393979 | -0.039517196 | -0.079013496 | 0.0394963 | 0.999933887 | -0.668355551 | -0.964359726 | 0.296004175 | 0.117985742 |
| ENSG00000100324 | *TAB1* | 2.553588187 | 0.020203002 | -0.006747209 | 0.026950211 | 0.999933887 | -0.007083853 | -0.296931654 | 0.289847801 | 0.117997583 |
| ENSG00000100603 | *SNW1* | 5.750011995 | -0.041104167 | -0.101880772 | 0.060776604 | 0.999933887 | 0.106905108 | 0.283276221 | -0.176371113 | 0.118098305 |
| ENSG00000100325 | *ASCC2* | 4.750249535 | 0.022630382 | -0.034394755 | 0.057025137 | 0.999933887 | 0.148261559 | 0.324684578 | -0.176423018 | 0.118205289 |
| ENSG00000157916 | *RER1* | 5.856336708 | -0.01039251 | -0.044901954 | 0.034509444 | 0.999933887 | -0.188262047 | -0.321097203 | 0.132835156 | 0.118212941 |
| ENSG00000158716 | *DUSP23* | 3.403485437 | -0.160628492 | -0.107711805 | -0.052916687 | 0.999933887 | -0.811743954 | -1.166659096 | 0.354915142 | 0.118325264 |
| ENSG00000134871 | *COL4A2* | 1.373641028 | 0.176641993 | 0.175227075 | 0.001414918 | 0.999933887 | 1.006639236 | 1.444798517 | -0.43815928 | 0.118709616 |
| ENSG00000138448 | *ITGAV* | 2.220588531 | -0.139942874 | -0.217858981 | 0.077916108 | 0.999933887 | -0.300649995 | 0.041099114 | -0.341749109 | 0.118709616 |
| ENSG00000198301 | *SDAD1* | 2.343323978 | -0.087351087 | 0.035682385 | -0.123033472 | 0.999933887 | -0.105631729 | 0.194172701 | -0.29980443 | 0.118774083 |
| ENSG00000132549 | *VPS13B* | 5.753288366 | 0.004423632 | -0.011932594 | 0.016356226 | 0.999933887 | -0.129000362 | 0.187870765 | -0.316871127 | 0.118837325 |
| ENSG00000148634 | *HERC4* | 4.974059769 | -0.024902361 | -0.107824556 | 0.082922195 | 0.999933887 | -0.450086912 | -0.635228583 | 0.185141671 | 0.119085351 |
| ENSG00000108829 | *LRRC59* | 5.163162916 | 0.000682046 | -0.069929412 | 0.070611458 | 0.999933887 | 0.168477442 | 0.32522437 | -0.156746928 | 0.119118959 |
| ENSG00000126858 | *RHOT1* | 3.862463662 | 0.009893814 | -0.065371054 | 0.075264869 | 0.999933887 | -0.558945563 | -0.811185024 | 0.252239461 | 0.119216505 |
| ENSG00000101608 | *MYL12A* | 6.949736387 | 0.012890738 | 0.019034616 | -0.006143878 | 0.999933887 | -0.095868989 | -0.270299474 | 0.174430486 | 0.119484671 |
| ENSG00000260807 | *CEROX1* | 0.362540587 | -0.24810013 | 0.024403391 | -0.272503521 | 0.999933887 | 0.194331068 | -0.270902117 | 0.465233185 | 0.119509992 |
| ENSG00000089060 | *SLC8B1* | 5.214992296 | 0.025445345 | 0.010544347 | 0.014900998 | 0.999933887 | -0.288182658 | -0.460476186 | 0.172293528 | 0.119509992 |
| ENSG00000127124 | *HIVEP3* | 3.531866482 | 0.023956258 | 0.057834926 | -0.033878668 | 0.999933887 | 0.225707144 | 0.46964876 | -0.243941616 | 0.119509992 |
| ENSG00000171552 | *BCL2L1* | 3.583933089 | 0.017241206 | 0.102048653 | -0.084807447 | 0.999933887 | 0.939756126 | 1.344793234 | -0.405037109 | 0.119509992 |
| ENSG00000137845 | *ADAM10* | 7.436953638 | -0.056059614 | -0.047090092 | -0.008969523 | 0.999933887 | -0.773832575 | -0.546579757 | -0.227252818 | 0.119667688 |
| ENSG00000159496 | *RGL4* | 5.396603887 | 0.12766711 | 0.086856237 | 0.040810873 | 0.999933887 | 0.133102615 | 0.320950745 | -0.18784813 | 0.119813409 |
| ENSG00000232803 | *SLCO4A1-AS1* | -0.167468148 | 0.331691172 | -0.146082882 | 0.477774054 | 0.999933887 | 1.041656168 | 1.617383157 | -0.57572699 | 0.119813409 |
| ENSG00000287737 | *AC132872.5* | 1.813576146 | 0.066912215 | -0.055995038 | 0.122907253 | 0.999933887 | -0.13183397 | -0.469593726 | 0.337759756 | 0.119813409 |
| ENSG00000137337 | *MDC1* | 3.194617918 | -0.027642408 | -0.025856721 | -0.001785687 | 0.999933887 | 0.149162449 | 0.338709951 | -0.189547502 | 0.119813409 |
| ENSG00000092067 | *CEBPE* | 1.053903268 | -0.21540638 | 0.256476136 | -0.471882516 | 0.999933887 | -0.324062568 | -0.717710702 | 0.393648135 | 0.119859893 |
| ENSG00000281530 | *AC004461.2* | 0.371182788 | -0.086128573 | 0.089830993 | -0.175959565 | 0.999933887 | -0.388319209 | -1.006418177 | 0.618098968 | 0.119887643 |
| ENSG00000159111 | *MRPL10* | 3.972241305 | -0.054673226 | -0.031245228 | -0.023427997 | 0.999933887 | 0.00758924 | -0.191612864 | 0.199202104 | 0.11989341 |
| ENSG00000173588 | *CEP83* | 1.820153925 | -0.082865221 | -0.130345415 | 0.047480194 | 0.999933887 | 0.062300851 | 0.545037504 | -0.482736653 | 0.119900285 |
| ENSG00000138593 | *SECISBP2L* | 4.881867393 | 0.042501183 | -0.02063151 | 0.063132693 | 0.999933887 | -0.538980712 | -0.188227555 | -0.350753157 | 0.120000158 |
| ENSG00000110344 | *UBE4A* | 5.58505566 | -0.028494776 | -0.064019994 | 0.035525219 | 0.999933887 | -0.13463349 | 0.043848541 | -0.178482031 | 0.120012602 |
| ENSG00000172197 | *MBOAT1* | 5.312908633 | -0.016256604 | 0.010091958 | -0.026348562 | 0.999933887 | -0.367742256 | -0.528010196 | 0.160267939 | 0.120012602 |
| ENSG00000125503 | *PPP1R12C* | 7.246324362 | 0.089012699 | 0.146203705 | -0.057191006 | 0.999933887 | 0.001775699 | -0.211147282 | 0.212922981 | 0.120039527 |
| ENSG00000159885 | *ZNF222* | 2.904360319 | 0.042684225 | -0.016992228 | 0.059676453 | 0.999933887 | 0.15585936 | 0.39697669 | -0.24111733 | 0.12004278 |
| ENSG00000162959 | *MEMO1* | 1.161129539 | 0.156436031 | 0.230458861 | -0.07402283 | 0.999933887 | -0.330201583 | -0.761820022 | 0.431618439 | 0.120088869 |
| ENSG00000153815 | *CMIP* | 6.817618086 | -0.021884901 | 0.055196879 | -0.07708178 | 0.999933887 | -0.12972986 | -0.305734533 | 0.176004673 | 0.120088869 |
| ENSG00000160796 | *NBEAL2* | 9.322782875 | 0.054918114 | 0.054145514 | 0.000772599 | 0.999933887 | 0.349401008 | 0.500083846 | -0.150682839 | 0.120134619 |
| ENSG00000177697 | *CD151* | 2.190454583 | 0.055352121 | -0.001647225 | 0.056999346 | 0.999933887 | -0.34512672 | -0.682436571 | 0.337309851 | 0.120227403 |
| ENSG00000152492 | *CCDC50* | 1.893495936 | 0.023366368 | 0.013288358 | 0.01007801 | 0.999933887 | 0.124134336 | 0.445944899 | -0.321810563 | 0.120289173 |
| ENSG00000186767 | *SPIN4* | -0.315830061 | 0.107209126 | -0.086222607 | 0.193431733 | 0.999933887 | -0.439231224 | 0.212168931 | -0.651400156 | 0.120497302 |
| ENSG00000057608 | *GDI2* | 5.913505141 | -0.070063299 | -0.067747369 | -0.002315931 | 0.999933887 | -0.451985293 | -0.637340194 | 0.185354901 | 0.12061387 |
| ENSG00000178467 | *P4HTM* | 3.183517709 | 0.077807975 | 0.128454426 | -0.050646451 | 0.999933887 | 0.134102909 | -0.092182367 | 0.226285276 | 0.12061387 |
| ENSG00000152219 | *ARL14EP* | 2.18345939 | -0.070989757 | -0.19099261 | 0.120002853 | 0.999933887 | 0.106574199 | 0.401941003 | -0.295366804 | 0.120720268 |
| ENSG00000261460 | *AC009690.2* | 2.188807273 | 0.037553073 | 0.069893111 | -0.032340038 | 0.999933887 | 0.027946678 | 0.347281763 | -0.319335085 | 0.121462023 |
| ENSG00000165682 | *CLEC1B* | 1.502572721 | -0.141190599 | 0.089992857 | -0.231183456 | 0.999933887 | 1.098146157 | 1.606350808 | -0.508204651 | 0.121721554 |
| ENSG00000225300 | *AL591623.1* | -0.350796777 | 0.063661399 | 0.175474941 | -0.111813542 | 0.999933887 | 0.615355179 | 1.168734239 | -0.553379059 | 0.121721554 |
| ENSG00000138381 | *ASNSD1* | 1.525422418 | -0.072749039 | -0.30529865 | 0.232549611 | 0.999933887 | 0.238072857 | 0.58702157 | -0.348948713 | 0.121799923 |
| ENSG00000074356 | *NCBP3* | 5.296766589 | -0.045916045 | 0.002018515 | -0.04793456 | 0.999933887 | -0.303627858 | -0.460135571 | 0.156507713 | 0.121976844 |
| ENSG00000197122 | *SRC* | 4.106528664 | 0.016412994 | 0.06364315 | -0.047230155 | 0.999933887 | 1.920281813 | 2.34901685 | -0.428735036 | 0.122040223 |
| ENSG00000165458 | *INPPL1* | 6.238909568 | 0.059545181 | 0.085250632 | -0.025705451 | 0.999933887 | -0.077652725 | -0.221604792 | 0.143952066 | 0.122116566 |
| ENSG00000163719 | *MTMR14* | 7.491575328 | 0.004624077 | -0.006129748 | 0.010753825 | 0.999933887 | 0.129633268 | 0.291107524 | -0.161474257 | 0.122116566 |
| ENSG00000184056 | *VPS33B* | -0.179381988 | -0.318174323 | 0.007439429 | -0.325613752 | 0.999933887 | -0.172129893 | 0.357864602 | -0.529994495 | 0.122137955 |
| ENSG00000113272 | *THG1L* | 0.747638845 | 0.060477137 | -0.150287888 | 0.210765025 | 0.999933887 | 0.598606589 | 1.044186045 | -0.445579456 | 0.122137955 |
| ENSG00000197483 | *ZNF628* | 3.184382574 | 0.092036266 | 0.096070683 | -0.004034417 | 0.999933887 | -0.251943855 | -0.562280624 | 0.310336768 | 0.122164108 |
| ENSG00000172780 | *RAB43* | 3.720159638 | 0.118620494 | 0.063611055 | 0.055009438 | 0.999933887 | -0.373197815 | -0.693190717 | 0.319992902 | 0.122233267 |
| ENSG00000170604 | *IRF2BP1* | 2.805846596 | 0.098572973 | 0.156455525 | -0.057882552 | 0.999933887 | -0.070612724 | -0.38498728 | 0.314374556 | 0.122233267 |
| ENSG00000137270 | *GCM1* | 0.384653523 | -0.235911847 | 0.046666676 | -0.282578523 | 0.999933887 | -0.969342252 | -1.527602801 | 0.558260548 | 0.122297954 |
| ENSG00000143258 | *USP21* | 3.198374122 | 0.044817942 | 0.08560087 | -0.040782929 | 0.999933887 | -0.02390551 | -0.262468374 | 0.238562864 | 0.122297954 |
| ENSG00000151612 | *ZNF827* | 2.050253717 | -0.110395013 | 0.037168489 | -0.147563501 | 0.999933887 | 0.020580433 | 0.324559509 | -0.303979076 | 0.122556499 |
| ENSG00000136738 | *STAM* | 3.608347036 | 0.050229211 | 0.119649867 | -0.069420656 | 0.999933887 | 0.057074745 | 0.253699679 | -0.196624934 | 0.122556679 |
| ENSG00000108244 | *KRT23* | 5.329220898 | 0.042241054 | -0.008598858 | 0.050839913 | 0.999933887 | 0.936603936 | 1.135458371 | -0.198854435 | 0.122577457 |
| ENSG00000114127 | *XRN1* | 6.410894427 | 0.000321724 | -0.094745882 | 0.095067606 | 0.999933887 | -0.6464633 | -0.206220427 | -0.440242873 | 0.122667394 |
| ENSG00000047644 | *WWC3* | 6.508034863 | 0.049888175 | 0.085609656 | -0.035721482 | 0.999933887 | -0.005489586 | -0.181627711 | 0.176138125 | 0.122695023 |
| ENSG00000278129 | *ZNF8* | 2.347624806 | 0.0160897 | -0.118805203 | 0.134894903 | 0.999933887 | -0.187246474 | -0.451666887 | 0.264420413 | 0.122816649 |
| ENSG00000189051 | *RNF222* | 0.037985613 | 0.142243053 | 0.358691137 | -0.216448084 | 0.999933887 | 0.293111765 | -0.31146535 | 0.604577115 | 0.122917817 |
| ENSG00000171488 | *LRRC8C* | 3.883878815 | -0.014079541 | -0.019313015 | 0.005233474 | 0.999933887 | -0.084700795 | 0.16924424 | -0.253945034 | 0.122972913 |
| ENSG00000226330 | *AL606489.1* | -1.440908946 | 0.348672111 | -0.54952387 | 0.898195981 | 0.999933887 | -1.784388165 | -2.552461228 | 0.768073063 | 0.1230916 |
| ENSG00000105519 | *CAPS* | 1.593572459 | 0.019020341 | -0.068231659 | 0.087252 | 0.999933887 | 0.062702943 | -0.341662071 | 0.404365015 | 0.123154166 |
| ENSG00000176749 | *CDK5R1* | 0.981814328 | -0.047371117 | 0.092191669 | -0.139562786 | 0.999933887 | -0.216385695 | -0.614402358 | 0.398016663 | 0.123194813 |
| ENSG00000126351 | *THRA* | 5.865209306 | -0.025212714 | -0.038021426 | 0.012808712 | 0.999933887 | -0.331571737 | -0.131635786 | -0.199935951 | 0.12325158 |
| ENSG00000073969 | *NSF* | 2.453768485 | -0.077613176 | -0.015031932 | -0.062581244 | 0.999933887 | -0.234341662 | -0.520341626 | 0.285999964 | 0.123255393 |
| ENSG00000155876 | *RRAGA* | 4.446580459 | 0.015567976 | 0.002394815 | 0.013173161 | 0.999933887 | -0.337177374 | -0.505065445 | 0.167888071 | 0.123255393 |
| ENSG00000170876 | *TMEM43* | 6.427766037 | -0.030722534 | -0.026105101 | -0.004617433 | 0.999933887 | -0.534354775 | -0.694457483 | 0.160102707 | 0.123392767 |
| ENSG00000173681 | *BCLAF3* | 3.678848619 | 0.054571443 | -0.033714368 | 0.088285811 | 0.999933887 | -0.308119315 | -0.017253638 | -0.290865676 | 0.123454877 |
| ENSG00000176171 | *BNIP3* | 2.893382246 | 0.036134796 | 0.059472339 | -0.023337543 | 0.999933887 | -0.098118069 | 0.134737048 | -0.232855117 | 0.123477181 |
| ENSG00000198040 | *ZNF84* | 2.553145993 | -0.079019401 | 0.007312201 | -0.086331602 | 0.999933887 | -0.258643615 | 0.044428377 | -0.303071992 | 0.123679931 |
| ENSG00000111911 | *HINT3* | 5.03302609 | -0.042687528 | 0.007624048 | -0.050311576 | 0.999933887 | -0.699799731 | -0.465820373 | -0.233979358 | 0.123679931 |
| ENSG00000103888 | *CEMIP* | 0.04687767 | 0.210884225 | -0.058875006 | 0.269759231 | 0.999933887 | -0.450945839 | -0.979575542 | 0.528629703 | 0.123855634 |
| ENSG00000205138 | *SDHAF1* | 1.199423486 | 0.1350885 | 0.176526033 | -0.041437532 | 0.999933887 | -0.26409815 | -0.57236338 | 0.30826523 | 0.123855634 |
| ENSG00000215301 | *DDX3X* | 10.432557352 | -0.006548367 | -0.040315765 | 0.033767397 | 0.999933887 | 0.203263693 | 0.394001603 | -0.190737911 | 0.123855634 |
| ENSG00000229644 | *NAMPTP1* | 5.791907432 | -0.037209598 | -0.073466027 | 0.036256429 | 0.999933887 | 0.398090742 | 0.660810495 | -0.262719753 | 0.12408535 |
| ENSG00000183891 | *TTC32* | 4.740078707 | 0.114175858 | 0.140186395 | -0.026010537 | 0.999933887 | 0.808379813 | 0.998755218 | -0.190375405 | 0.124209501 |
| ENSG00000135503 | *ACVR1B* | 5.381305577 | -0.058748512 | -0.0094097 | -0.049338812 | 0.999933887 | -0.355864465 | -0.148011607 | -0.207852858 | 0.124209501 |
| ENSG00000178177 | *LCORL* | 2.487515958 | -0.025514513 | 0.007242191 | -0.032756704 | 0.999933887 | -0.40130367 | -0.06831464 | -0.33298903 | 0.124252284 |
| ENSG00000229152 | *ANKRD10-IT1* | 2.576500469 | -0.095461784 | -0.213967241 | 0.118505457 | 0.999933887 | -0.083388719 | 0.347608841 | -0.43099756 | 0.12448161 |
| ENSG00000104951 | *IL4I1* | 0.776125445 | -0.037357313 | -0.0086906 | -0.028666713 | 0.999933887 | 0.851614352 | 1.367354283 | -0.515739931 | 0.12448161 |
| ENSG00000086300 | *SNX10* | 6.076169835 | 0.023417526 | -0.054550881 | 0.077968407 | 0.999933887 | 1.255849854 | 1.634383104 | -0.37853325 | 0.124531881 |
| ENSG00000076067 | *RBMS2* | 1.03192729 | -0.065966194 | -0.09506159 | 0.029095397 | 0.999933887 | -0.246714164 | 0.260176222 | -0.506890386 | 0.124603374 |
| ENSG00000153310 | *CYRIB* | 8.949346205 | -0.012526417 | -0.004039553 | -0.008486865 | 0.999933887 | -0.085189498 | 0.046999818 | -0.132189316 | 0.124635694 |
| ENSG00000092203 | *TOX4* | 6.911098384 | -0.009618842 | -0.030565012 | 0.02094617 | 0.999933887 | 0.111141639 | 0.255913178 | -0.144771538 | 0.124818959 |
| ENSG00000287733 | *AC083862.2* | 5.69974831 | -0.002033641 | -0.082895961 | 0.08086232 | 0.999933887 | -0.166379132 | 0.045099772 | -0.211478904 | 0.124867556 |
| ENSG00000279946 | *AC244034.3* | -1.994152684 | 0.187937642 | 1.230400671 | -1.042463029 | 0.999933887 | 2.186828089 | 3.463783671 | -1.276955582 | 0.12487743 |
| ENSG00000104549 | *SQLE* | 2.7344362 | -0.019778537 | 0.081450212 | -0.101228749 | 0.999933887 | 0.648918498 | 0.907465255 | -0.258546757 | 0.12487743 |
| ENSG00000122729 | *ACO1* | 2.870524653 | 0.049613809 | 0.085881414 | -0.036267605 | 0.999933887 | -0.35709593 | 0.00502292 | -0.36211885 | 0.125059525 |
| ENSG00000129128 | *SPCS3* | 5.522900165 | 0.030978521 | 0.016608281 | 0.014370241 | 0.999933887 | -0.569379291 | -0.813837933 | 0.244458641 | 0.125084836 |
| ENSG00000183726 | *TMEM50A* | 6.923694609 | 0.0070499 | -0.014582823 | 0.021632723 | 0.999933887 | -0.340124536 | -0.498718269 | 0.158593733 | 0.125317285 |
| ENSG00000185507 | *IRF7* | 6.798625497 | 0.113788095 | 0.104614472 | 0.009173623 | 0.999933887 | -0.025547709 | -0.245653608 | 0.220105899 | 0.125375505 |
| ENSG00000170340 | *B3GNT2* | 4.304435825 | 0.056731054 | 0.226863845 | -0.170132791 | 0.999933887 | -0.377840906 | -0.588644604 | 0.210803697 | 0.125375505 |
| ENSG00000142039 | *CCDC97* | 4.968793952 | -0.001492755 | 0.001422461 | -0.002915216 | 0.999933887 | -0.474606557 | -0.703541528 | 0.228934971 | 0.125449982 |
| ENSG00000010256 | *UQCRC1* | 4.196434723 | 0.00807474 | -0.038600444 | 0.046675184 | 0.999933887 | -0.092044193 | -0.293480055 | 0.201435862 | 0.125467634 |
| ENSG00000152193 | *OBI1* | 0.752511207 | 0.078990135 | -0.509127099 | 0.588117234 | 0.944094537 | -0.407149661 | -0.019453623 | -0.387696038 | 0.125481234 |
| ENSG00000075415 | *SLC25A3* | 6.681885632 | 0.00834399 | -0.027624494 | 0.035968484 | 0.999933887 | -0.122925924 | -0.248113319 | 0.125187395 | 0.125548004 |
| ENSG00000272221 | *AL645933.2* | 0.63350834 | -0.176138446 | 0.010443114 | -0.18658156 | 0.999933887 | -0.04086628 | -0.458140733 | 0.417274453 | 0.125643803 |
| ENSG00000067369 | *TP53BP1* | 5.85228135 | -0.019713109 | -0.048907372 | 0.029194263 | 0.999933887 | 0.263187721 | 0.434318295 | -0.171130574 | 0.125643803 |
| ENSG00000226200 | *SGMS1-AS1* | 2.176194754 | -0.139434116 | 0.100142662 | -0.239576779 | 0.999933887 | 0.604880918 | 0.847949607 | -0.243068688 | 0.125933618 |
| ENSG00000142002 | *DPP9* | 5.173207347 | -0.017992288 | -0.005436855 | -0.012555433 | 0.999933887 | -0.055031437 | -0.193302616 | 0.13827118 | 0.125933618 |
| ENSG00000112343 | *TRIM38* | 6.411593152 | -0.0460308 | -0.030106914 | -0.015923885 | 0.999933887 | -0.288325687 | -0.427889036 | 0.13956335 | 0.126191015 |

| | | | | | | | | | | |
|---|---|---|---|---|---|---|---|---|---|---|
| ENSG00000156639 | *ZFAND3* | 7.214211626 | 0.006610564 | 0.023516595 | -0.016906031 | 0.999933887 | 0.240333999 | 0.370423521 | -0.130089522 | 0.126266535 |
| ENSG00000078487 | *ZCWPW1* | 4.326463881 | -0.053189431 | 0.045043541 | -0.098232972 | 0.999933887 | 1.132330655 | 1.369946577 | -0.237615922 | 0.126302178 |
| ENSG00000080493 | *SLC4A4* | -0.160883269 | -0.095503326 | 0.071823765 | -0.16732709 | 0.999933887 | -0.155997287 | 0.473219339 | -0.629216626 | 0.126373718 |
| ENSG00000160131 | *VMA21* | 2.163593343 | -0.001678249 | 0.108642776 | -0.110321025 | 0.999933887 | -0.07584337 | -0.38612195 | 0.310278579 | 0.12648322 |
| ENSG00000244687 | *UBE2V1* | 1.549623715 | 0.035640765 | -0.136514602 | 0.172155367 | 0.999933887 | -0.07310535 | 0.239060664 | -0.312166014 | 0.126573736 |
| ENSG00000124134 | *KCNS1* | 4.528710425 | -0.028965899 | 0.044618162 | -0.073584061 | 0.999933887 | -0.348721883 | -0.139691435 | -0.209030448 | 0.12661267 |
| ENSG00000235217 | *TSPY26P* | 1.766328587 | -0.003606131 | 0.037109222 | -0.040715353 | 0.999933887 | 1.084031931 | 1.487351816 | -0.403319885 | 0.126723203 |
| ENSG00000139323 | *POC1B* | 3.602069194 | -0.001984684 | -0.02223919 | 0.020254506 | 0.999933887 | 0.349124294 | 0.595700994 | -0.246576699 | 0.12672697 |
| ENSG00000285976 | *AL135905.2* | 2.127978175 | 0.171775961 | 0.317866608 | -0.146090647 | 0.999933887 | 0.573605562 | 0.071557883 | 0.50204768 | 0.126733605 |
| ENSG00000167548 | *KMT2D* | 8.637677011 | 0.03651769 | 0.080580248 | -0.044062558 | 0.999933887 | -0.611332038 | -0.788263005 | 0.176930966 | 0.126756457 |
| ENSG00000128590 | *DNAJB9* | 6.224200412 | 0.10073216 | 0.055291017 | 0.045441143 | 0.999933887 | 0.067764614 | 0.298301587 | -0.230536973 | 0.126864727 |
| ENSG00000260565 | *ERVK13-1* | 4.714616374 | 0.037054177 | -0.027725234 | 0.064779411 | 0.999933887 | -0.626571018 | -0.85900771 | 0.232436693 | 0.127040523 |
| ENSG00000186654 | *PRR5* | 2.894947679 | 0.056291349 | 0.023972909 | 0.03231844 | 0.999933887 | 0.041023906 | 0.312987135 | -0.271963229 | 0.127171507 |
| ENSG00000063176 | *SPHK2* | 1.740831198 | 0.099272755 | 0.20452201 | -0.105249256 | 0.999933887 | -0.105469862 | -0.457551852 | 0.35208199 | 0.127260287 |
| ENSG00000172331 | *BPGM* | 2.962354501 | -0.130888383 | -0.089240504 | -0.041647879 | 0.999933887 | -0.433292148 | -0.676963091 | 0.243670942 | 0.127266471 |
| ENSG00000175352 | *NRIP3* | 6.401416981 | 0.020522648 | 0.062522086 | -0.041999438 | 0.999933887 | 0.056936364 | 0.317429145 | -0.26049278 | 0.127266471 |
| ENSG00000155636 | *RBM45* | 0.683339674 | -0.02717538 | -0.171207263 | 0.144031883 | 0.999933887 | -0.118073969 | -0.572696659 | 0.454622689 | 0.127266471 |
| ENSG00000157014 | *TATDN2* | 4.852805979 | 0.010411284 | 0.063885565 | -0.053474281 | 0.999933887 | 1.06415544 | 1.289574046 | -0.225418606 | 0.127266471 |
| ENSG00000196139 | *AKR1C3* | 1.081396691 | 0.023084231 | -0.132599633 | 0.155683864 | 0.999933887 | -0.727798151 | -1.097617308 | 0.369819157 | 0.127266471 |
| ENSG00000155957 | *TMBIM4* | 4.11418331 | -0.00229534 | -0.064020946 | 0.061725607 | 0.999933887 | 0.768891679 | 0.992704027 | -0.223812348 | 0.127268375 |
| ENSG00000187498 | *COL4A1* | 0.123499182 | -0.056384177 | 0.060338061 | -0.116722238 | 0.999933887 | -0.1415484 | 0.351441584 | -0.492989984 | 0.127335856 |
| ENSG00000260542 | *AL499627.1* | -1.59499495 | 0.160106973 | 0.019437953 | 0.140669021 | 0.999933887 | 0.411862166 | 1.058649204 | -0.646787039 | 0.127379246 |
| ENSG00000196154 | *S100A4* | 7.825153019 | -0.044917626 | -0.046970787 | 0.002053161 | 0.999933887 | -0.111006861 | -0.345437556 | 0.234430695 | 0.127552893 |
| ENSG00000125386 | *FAM193A* | 5.508951095 | -0.020636176 | 0.007459833 | -0.028096009 | 0.999933887 | -0.307267199 | -0.482361627 | 0.175094428 | 0.127945108 |
| ENSG00000198912 | *C1orf174* | 2.987633172 | -0.051081659 | -0.113004926 | 0.061923267 | 0.999933887 | -0.321812608 | -0.521849674 | 0.200037066 | 0.127981302 |
| ENSG00000112078 | *KCTD20* | 6.152374737 | -0.00974303 | -0.065419089 | 0.055676059 | 0.999933887 | 1.126465074 | 1.458337203 | -0.331872129 | 0.127981302 |
| ENSG00000170275 | *CRTAP* | 3.868689879 | 0.049477115 | -0.024640183 | 0.074117298 | 0.999933887 | -0.116137541 | -0.369160585 | 0.253023045 | 0.128068898 |
| ENSG00000276570 | *AC010327.6* | 1.442659056 | 0.067847396 | 0.279970729 | -0.212123334 | 0.999933887 | -0.501029183 | -0.894215068 | 0.393185885 | 0.128068898 |
| ENSG00000255284 | *AP006621.3* | -0.121487646 | 0.106587987 | -0.220393247 | 0.326981233 | 0.999933887 | -0.272908853 | -0.837503993 | 0.56459514 | 0.128068898 |
| ENSG00000132274 | *TRIM22* | 7.514967907 | 0.024560744 | -0.01671227 | 0.041273014 | 0.999933887 | -0.131559058 | -0.299569799 | 0.168010741 | 0.128068898 |
| ENSG00000157106 | *SMG1* | 7.837289213 | -0.022713456 | -0.071597924 | 0.048884468 | 0.999933887 | -0.167742153 | 0.162921092 | -0.330663245 | 0.128068898 |
| ENSG00000120318 | *ARAP3* | 7.222514552 | -0.020525293 | 0.071188324 | -0.091713617 | 0.999933887 | -0.591895063 | -0.792481674 | 0.200586611 | 0.128201787 |
| ENSG00000186529 | *CYP4F3* | 7.467335148 | -0.044079009 | 0.036661131 | -0.08074014 | 0.999933887 | -0.178479239 | -0.294710877 | 0.116231638 | 0.128330097 |
| ENSG00000140299 | *BNIP2* | 6.602111143 | -0.079895771 | -0.041637799 | -0.038257973 | 0.999933887 | -0.396052492 | -0.197797665 | -0.198254827 | 0.128426789 |
| ENSG00000005075 | *POLR2J* | 4.645984888 | 0.057589283 | 0.048694112 | 0.008895171 | 0.999933887 | -0.105656032 | -0.27287402 | 0.167217988 | 0.128426789 |
| ENSG00000281357 | *ARRDC3-AS1* | 1.625839308 | -0.141415272 | -0.156059867 | 0.014644595 | 0.999933887 | 0.38745389 | 0.670640445 | -0.283186555 | 0.128436425 |
| ENSG00000167904 | *TMEM68* | 1.335502744 | -0.054201772 | 0.003582249 | -0.057784022 | 0.999933887 | 0.464231722 | 0.840326592 | -0.37609487 | 0.128436425 |
| ENSG00000172466 | *ZNF24* | 6.955472574 | -0.108054665 | -0.164950182 | 0.056895517 | 0.999933887 | 0.195595694 | 0.388549736 | -0.192954042 | 0.128480911 |
| ENSG00000134049 | *IER3IP1* | 0.325758836 | -0.2228851 | 0.413371991 | -0.636257091 | 0.944094537 | -0.142969908 | 0.429828666 | -0.572798574 | 0.128480911 |
| ENSG00000160014 | *CALM3* | 7.205837862 | 0.058209273 | 0.005671651 | 0.052537621 | 0.999933887 | 0.071695922 | 0.212750451 | -0.141054529 | 0.128480911 |
| ENSG00000118965 | *WDR35* | 2.11696405 | -0.126302274 | -0.178937256 | 0.052634981 | 0.999933887 | 0.040962465 | 0.366148594 | -0.325186129 | 0.128480911 |
| ENSG00000196456 | *ZNF775* | 1.048743641 | 0.07027043 | 0.104832203 | -0.034561772 | 0.999933887 | 0.844170621 | 1.218512309 | -0.374341688 | 0.128517344 |
| ENSG00000196182 | *STK40* | 9.47251234 | 0.013898282 | 0.001221273 | 0.012677009 | 0.999933887 | -0.058162395 | 0.064450965 | -0.12261336 | 0.128568144 |
| ENSG00000169992 | *NLGN2* | -0.030924254 | -0.094417408 | 0.240788258 | -0.335205665 | 0.999933887 | -0.101034346 | 0.559695039 | -0.660729385 | 0.12872819 |
| ENSG00000126749 | *EMG1* | 2.315759619 | -0.034145299 | -0.008601703 | -0.025543596 | 0.999933887 | 0.1865438 | 0.455109489 | -0.268565689 | 0.12872819 |
| ENSG00000135336 | *ORC3* | 2.216645271 | -0.021781226 | -0.146310998 | 0.124529771 | 0.999933887 | 0.023759642 | 0.338523624 | -0.314763982 | 0.12872819 |
| ENSG00000070444 | *MNT* | 6.319272009 | 0.007264432 | -0.006902931 | 0.014167363 | 0.999933887 | -0.583014427 | -0.739634679 | 0.156620252 | 0.12872819 |
| ENSG00000101182 | *PSMA7* | 6.982103095 | 0.006989953 | -0.023724707 | 0.03071466 | 0.999933887 | -0.176552451 | -0.324997166 | 0.148444716 | 0.12872819 |
| ENSG00000124198 | *ARFGEF2* | 5.193353861 | -0.042238716 | -0.115850947 | 0.073612231 | 0.999933887 | -0.108574721 | 0.050602463 | -0.159177184 | 0.128743221 |
| ENSG00000177575 | *CD163* | 3.447776666 | 0.129118925 | 0.136399039 | -0.007280114 | 0.999933887 | -0.864948989 | -1.177190594 | 0.312241605 | 0.128796951 |
| ENSG00000138378 | *STAT4* | 4.011873062 | -0.063358556 | -0.156763356 | 0.0934048 | 0.999933887 | 0.201896809 | 0.435352974 | -0.233456165 | 0.128965966 |
| ENSG00000165898 | *ISCA2* | 1.786777652 | -0.114842049 | -0.01228848 | -0.102553569 | 0.999933887 | -0.324208396 | -0.672525975 | 0.348317579 | 0.129040936 |
| ENSG00000170242 | *USP47* | 4.737544032 | -0.078778362 | -0.080642656 | 0.001864294 | 0.999933887 | -0.125232609 | 0.278616663 | -0.403849272 | 0.129040936 |
| ENSG00000198959 | *TGM2* | 1.000979611 | 0.07462047 | 0.207658658 | -0.133038188 | 0.999933887 | -0.018518915 | 0.526096898 | -0.544615814 | 0.129040936 |
| ENSG00000139597 | *N4BP2L1* | 3.130795266 | -0.069874303 | -0.07444161 | 0.004567307 | 0.999933887 | -0.021194802 | 0.249085831 | -0.270280633 | 0.129082391 |
| ENSG00000085872 | *CHERP* | 5.863898734 | 0.033087862 | -0.002774038 | 0.0358619 | 0.999933887 | 0.267299987 | 0.453212586 | -0.185912599 | 0.129252705 |
| ENSG00000004059 | *ARF5* | 6.134953715 | -0.009490695 | 0.010321079 | -0.019811774 | 0.999933887 | -0.211317829 | -0.402081814 | 0.190763985 | 0.129318773 |
| ENSG00000169413 | *RNASE6* | 2.512410198 | -0.020035459 | -0.001971856 | -0.018063603 | 0.999933887 | -0.419305668 | -1.111567927 | 0.692262259 | 0.12944918 |
| ENSG00000092098 | *RNF31* | 1.292182792 | 0.002828231 | 0.076500739 | -0.073672508 | 0.999933887 | -0.675605442 | -1.115729066 | 0.440123624 | 0.129464244 |
| ENSG00000263766 | *AC025682.1* | 0.126890341 | 0.068979682 | 0.079464278 | -0.010484596 | 0.999933887 | 0.653170609 | 1.285239328 | -0.632068719 | 0.129482371 |
| ENSG00000186501 | *TMEM222* | 4.734278283 | -0.022023911 | -0.001530502 | -0.020493409 | 0.999933887 | -0.395459069 | -0.564842142 | 0.169383072 | 0.129650883 |
| ENSG00000175938 | *ORAI3* | 3.551547386 | -0.104640299 | -0.014881632 | -0.089758666 | 0.999933887 | -0.899234623 | -1.149974915 | 0.250740292 | 0.129668394 |
| ENSG00000130811 | *EIF3G* | 5.430054565 | 0.014017152 | -0.026430214 | 0.040447366 | 0.999933887 | -0.020612994 | -0.186449657 | 0.165836662 | 0.129745363 |
| ENSG00000254777 | *AC022182.1* | 1.19997669 | 0.102202167 | 0.202567937 | -0.10036577 | 0.999933887 | 0.176484111 | -0.335238586 | 0.511722698 | 0.129815573 |
| ENSG00000137393 | *RNF144B* | 6.344176609 | -0.231192418 | -0.080807504 | -0.150384914 | 0.999933887 | 2.547298785 | 2.935105338 | -0.387806552 | 0.129868743 |
| ENSG00000167637 | *ZNF283* | -0.1826437 | 0.251031366 | -0.156016155 | 0.407047522 | 0.999933887 | -0.372553059 | 0.091083127 | -0.463636186 | 0.129868743 |
| ENSG00000226711 | *FAM66C* | -0.401006565 | -0.005206181 | 0.137440296 | -0.142646477 | 0.999933887 | 0.29638198 | 0.97268069 | -0.676298711 | 0.129868743 |
| ENSG00000262879 | *AC005670.3* | 1.45184196 | -0.0090604 | 0.066230175 | -0.075290576 | 0.999933887 | -0.335893239 | -0.618466204 | 0.282572965 | 0.129879904 |
| ENSG00000106290 | *TAF6* | 2.51076887 | -0.061584097 | 0.047165391 | -0.108749488 | 0.999933887 | 0.075831968 | -0.159676219 | 0.235508186 | 0.129889885 |
| ENSG00000287494 | *AC069368.2* | 2.852478223 | 0.107962457 | -0.010893156 | 0.118855613 | 0.999933887 | 1.043433935 | 1.375075108 | -0.331641173 | 0.129917072 |
| ENSG00000217555 | *CKLF* | -0.140377018 | -0.259004946 | -0.255373284 | -0.003631662 | 0.999933887 | -0.969873655 | -1.80103026 | 0.831156605 | 0.129948683 |
| ENSG00000177225 | *GATD1* | 3.408721708 | -0.084639268 | 0.015407357 | -0.100046625 | 0.999933887 | -0.328638921 | -0.553911912 | 0.225272991 | 0.129948683 |
| ENSG00000188185 | *LINC00265* | 1.275627006 | -0.136774235 | 0.082228708 | -0.219002943 | 0.999933887 | 0.256357713 | 0.612075579 | -0.355717867 | 0.129998969 |
| ENSG00000118922 | *KLF12* | 4.009126158 | -0.079427935 | -0.185924413 | 0.106496478 | 0.999933887 | -0.285724475 | 0.105525235 | -0.39124971 | 0.130203859 |
| ENSG00000205084 | *TMEM231* | 2.217942222 | 0.059242963 | 0.006059831 | 0.053183132 | 0.999933887 | 0.860286337 | 1.122932844 | -0.262646507 | 0.130203859 |
| ENSG00000165689 | *ENTR1* | 3.052230359 | 0.031675917 | -0.060728955 | 0.092404873 | 0.999933887 | -0.077597332 | -0.321567401 | 0.243970069 | 0.130203859 |
| ENSG00000111860 | *CEP85L* | 3.862350065 | -0.015257309 | -0.099674258 | 0.084416949 | 0.999933887 | -0.650911752 | -1.112249971 | 0.461338218 | 0.130203859 |
| ENSG00000153391 | *INO80C* | 2.279743937 | -0.1182441 | 0.045381886 | -0.163625986 | 0.999933887 | 0.367753567 | 0.659861362 | -0.292107795 | 0.13023857 |
| ENSG00000095397 | *WHRN* | 4.327268714 | 0.095328438 | 0.039772355 | 0.055556083 | 0.999933887 | 0.321990874 | 0.543339328 | -0.221348454 | 0.13023857 |
| ENSG00000228223 | *HCG11* | 0.926832871 | 0.045455666 | 0.225114179 | -0.179658512 | 0.999933887 | -0.397516275 | -0.750413033 | 0.352896758 | 0.13023857 |
| ENSG00000079393 | *DUSP13* | 1.247732767 | -0.023959064 | 0.241034002 | -0.264993066 | 0.999933887 | -1.160476406 | -1.514536477 | 0.354060071 | 0.13023857 |
| ENSG00000114378 | *HYAL1* | -0.008695784 | -0.120102016 | -0.281284862 | 0.161182846 | 0.999933887 | -0.532431315 | -1.046365268 | 0.513933953 | 0.130261133 |
| ENSG00000143061 | *IGSF3* | -1.754068161 | 0.034270158 | -0.223967682 | 0.258237841 | 0.999933887 | 3.101752406 | 4.086322624 | -0.984570218 | 0.130261133 |
| ENSG00000163472 | *TMEM79* | 1.937069025 | -0.007880689 | -0.270868884 | 0.262988195 | 0.999933887 | -0.379116009 | -0.660534122 | 0.281418113 | 0.130341498 |
| ENSG00000169905 | *TOR1AIP2* | 5.79741532 | -0.04365056 | -0.028565437 | -0.015085123 | 0.999933887 | -0.465069014 | -0.2812242 | -0.183844813 | 0.130649991 |
| ENSG00000150527 | *MIA2* | 4.667996678 | -0.03857174 | -0.096897144 | 0.058325405 | 0.999933887 | -0.687501004 | -0.898133249 | 0.210632245 | 0.130686288 |
| ENSG00000177125 | *ZBTB34* | 5.388058924 | -0.124174401 | 0.025661432 | -0.149835833 | 0.999933887 | -1.023734534 | -1.310041349 | 0.286306815 | 0.130810051 |
| ENSG00000228526 | *MIR34AHG* | -1.620714339 | -0.202798565 | 0.270849776 | -0.473648341 | 0.999933887 | -0.307907815 | 0.257916829 | -0.565824644 | 0.131129847 |
| ENSG00000022976 | *ZNF839* | 2.808647423 | 0.087161946 | 0.059143852 | 0.028018094 | 0.999933887 | 0.096941208 | -0.128638379 | 0.225579587 | 0.131190484 |
| ENSG00000177239 | *MAN1B1* | 4.929367383 | 0.026382565 | -0.003870931 | 0.030253496 | 0.999933887 | -0.028784425 | -0.189920648 | 0.161136223 | 0.131190484 |
| ENSG00000151176 | *PLBD2* | 2.733617583 | 0.03609334 | -0.128847719 | 0.164941058 | 0.999933887 | -0.464077119 | -0.700728854 | 0.236651735 | 0.131190484 |
| ENSG00000249096 | *LINC02362* | 4.090340097 | 0.015130344 | 0.049510166 | -0.034379822 | 0.999933887 | 0.690147375 | 0.947720026 | -0.25757265 | 0.131190484 |
| ENSG00000138750 | *NUP54* | 2.855109564 | -0.00565533 | -0.188075765 | 0.182420434 | 0.999933887 | -0.113848406 | 0.145714965 | -0.259563371 | 0.131190484 |
| ENSG00000149930 | *TAOK2* | 5.206888725 | 0.029454027 | 0.057917798 | -0.028463771 | 0.999933887 | -0.220090486 | -0.400636585 | 0.180546099 | 0.131251106 |
| ENSG00000214595 | *EML6* | 1.563589044 | -0.000967813 | 0.220865923 | -0.221833736 | 0.999933887 | 0.041305825 | 0.561891586 | -0.520585761 | 0.131251106 |
| ENSG00000080822 | *CLDND1* | 5.283815571 | -0.05847197 | -0.089517573 | 0.031045602 | 0.999933887 | -0.031008577 | 0.159059362 | -0.190067939 | 0.131289285 |
| ENSG00000196668 | *LINC00173* | 4.494682693 | -0.155259386 | -0.072016712 | -0.083242674 | 0.999933887 | -0.488322951 | -0.699938851 | 0.2116159 | 0.13207783 |
| ENSG00000138398 | *PPIG* | 5.421675251 | -0.079879968 | -0.221649562 | 0.141769594 | 0.999933887 | -0.523584135 | -0.147047276 | -0.376536859 | 0.13207783 |
| ENSG00000244459 | *AC147067.1* | -0.460760805 | 0.076370873 | -0.143870462 | 0.220241335 | 0.999933887 | -0.318092708 | -1.114947731 | 0.796855022 | 0.132181639 |
| ENSG00000183784 | *DOCK8-AS1* | 0.521218348 | 0.182086111 | 0.037623889 | 0.144462222 | 0.999933887 | -0.813124734 | -1.620885336 | 0.807760602 | 0.13227524 |
| ENSG00000124831 | *LRRFIP1* | 8.219934114 | 0.007229309 | -0.073483002 | 0.08071231 | 0.999933887 | -0.574306603 | -0.826329226 | 0.252022622 | 0.13227524 |
| ENSG00000134285 | *FKBP11* | 3.253046772 | -0.038851852 | -0.059054592 | 0.02020274 | 0.999933887 | -0.064102248 | -0.242348348 | 0.1782461 | 0.132463306 |
| ENSG00000241489 | *AC244197.3* | 1.47965341 | -0.028900347 | -0.077159448 | 0.048259101 | 0.999933887 | -0.270904923 | -0.615515424 | 0.344610501 | 0.132463306 |
| ENSG00000099139 | *PCSK5* | 0.804771807 | 0.167716005 | -0.167701715 | 0.335417721 | 0.944094537 | -0.202030417 | 0.2582565 | -0.460286918 | 0.132528275 |
| ENSG00000232811 | *AL360270.1* | -0.118326532 | 0.134924699 | 0.199995477 | -0.065070777 | 0.999933887 | -0.301183044 | 0.217755331 | -0.518938375 | 0.132528275 |

| | | | | | | | | | | |
|---|---|---|---|---|---|---|---|---|---|---|
| ENSG00000160993 | *ALKBH4* | 2.759762912 | 0.026086163 | 0.019040244 | 0.007045919 | 0.999933887 | 0.03462345 | -0.258098171 | 0.292721621 | 0.132528275 |
| ENSG00000058673 | *ZC3H11A* | 0.581727556 | -0.016962795 | -0.122779754 | 0.105816958 | 0.999933887 | 0.362222023 | 0.734030447 | -0.371808423 | 0.132528275 |
| ENSG00000163703 | *CRELD1* | 0.077445357 | -0.178625614 | 0.083381793 | -0.262007407 | 0.999933887 | -0.390302127 | -0.77305136 | 0.382749233 | 0.132773397 |
| ENSG00000262089 | *AC040977.1* | 3.26622783 | -0.136735128 | 0.001998743 | -0.138733872 | 0.999933887 | 0.112095524 | 0.306352903 | -0.194257379 | 0.132801888 |
| ENSG00000260630 | *SNAI3-AS1* | 1.721312188 | 0.142419484 | 0.338820218 | -0.196400734 | 0.999933887 | 0.674076767 | 1.005096892 | -0.331020125 | 0.132801888 |
| ENSG00000271032 | *AC020907.4* | 1.244271949 | 0.05109105 | 0.274306228 | -0.223215177 | 0.999933887 | 2.840051532 | 3.331257289 | -0.491205757 | 0.132801888 |
| ENSG00000135636 | *DYSF* | 8.336059033 | -0.00099744 | 0.092194126 | -0.093191565 | 0.999933887 | -0.179602329 | -0.40289625 | 0.223293922 | 0.132801888 |
| ENSG00000242086 | *MUC20-OT1* | 3.177317491 | -0.035124684 | 0.096695623 | -0.131820307 | 0.999933887 | -0.153408965 | 0.068644259 | -0.222053224 | 0.13307667 |
| ENSG00000279884 | *AC010894.4* | 0.430662464 | -0.213933738 | -0.117219786 | -0.096713952 | 0.999933887 | -0.661346978 | -1.346496939 | 0.685149961 | 0.133161831 |
| ENSG00000237491 | *LINC01409* | 0.428551243 | -0.100966491 | -0.073270002 | -0.027696489 | 0.999933887 | -0.213553223 | 0.261372174 | -0.474925398 | 0.133394046 |
| ENSG00000244879 | *GABPB1-AS1* | 3.902909026 | -0.089790565 | -0.026376115 | -0.063414451 | 0.999933887 | 0.086333141 | 0.266261328 | -0.179928186 | 0.133568236 |
| ENSG00000198176 | *TFDP1* | 5.884116711 | -0.048715423 | 0.024717465 | -0.073432889 | 0.999933887 | 0.639754473 | 0.817893137 | -0.178138664 | 0.133568236 |
| ENSG00000198053 | *SIRPA* | 9.537137177 | -0.015404544 | -0.033628563 | 0.018224019 | 0.999933887 | 0.412521106 | 0.576382536 | -0.163861431 | 0.133568236 |
| ENSG00000100280 | *AP1B1* | 4.815556136 | 0.01552863 | 0.024626652 | -0.009098022 | 0.999933887 | 0.316862162 | 0.48682299 | -0.169960828 | 0.133621717 |
| ENSG00000154548 | *SRSF12* | 1.626549311 | -0.003664021 | 0.073657774 | -0.077321795 | 0.999933887 | 0.750611589 | 1.066516513 | -0.315904924 | 0.133672733 |
| ENSG00000121775 | *TMEM39B* | 1.719513883 | 0.090714883 | 0.028406413 | 0.06230847 | 0.999933887 | -0.007592861 | -0.369502795 | 0.361909933 | 0.133740731 |
| ENSG00000163155 | *LYSMD1* | 0.916708229 | -0.16957122 | -0.07838524 | -0.09118598 | 0.999933887 | -0.419881611 | -0.834178323 | 0.414296712 | 0.133891865 |
| ENSG00000120051 | *CFAP58* | 2.537757996 | -0.113549425 | -0.163201775 | 0.049652349 | 0.999933887 | 1.019468645 | 1.346311234 | -0.326842589 | 0.133891865 |
| ENSG00000014919 | *COX15* | 2.313961564 | -0.182206293 | -0.109104375 | -0.073101918 | 0.999933887 | -0.247105816 | -0.477306997 | 0.23020118 | 0.133938626 |
| ENSG00000185252 | *ZNF74* | 0.598594202 | 0.022712594 | 0.147804756 | -0.125092163 | 0.999933887 | 0.158724787 | 0.502844531 | -0.344119744 | 0.133938626 |
| ENSG00000186810 | *CXCR3* | 2.885020942 | -0.032441074 | -0.074117189 | 0.041676115 | 0.999933887 | 0.156949996 | 0.395798898 | -0.238848902 | 0.134000893 |
| ENSG00000163961 | *RNF168* | 5.254341029 | -0.132657266 | -0.025044729 | -0.107612537 | 0.999933887 | -0.194671517 | -0.348456343 | 0.153784826 | 0.134046448 |
| ENSG00000261094 | *AC007066.2* | 1.143656285 | -0.041258654 | -0.156592505 | 0.115333851 | 0.999933887 | 0.844841621 | 1.171544408 | -0.326702786 | 0.134046448 |
| ENSG00000236675 | *MTX1P1* | 3.118909719 | 0.011688455 | -0.074104512 | 0.085792967 | 0.999933887 | -0.43062122 | -0.678542084 | 0.247920864 | 0.134053369 |
| ENSG00000160255 | *ITGB2* | 9.443084137 | 0.044067352 | 0.000313822 | 0.043753531 | 0.999933887 | -0.188274821 | -0.359416227 | 0.171141406 | 0.134081898 |
| ENSG00000156052 | *GNAQ* | 6.245718581 | 0.02670285 | 0.015869573 | 0.010833277 | 0.999933887 | -0.615772998 | -0.856830784 | 0.241057786 | 0.13414887 |
| ENSG00000143498 | *TAF1A* | -0.368117839 | -0.285792875 | 0.430676588 | -0.716469462 | 0.944094537 | 0.03700242 | 0.58089257 | -0.54389015 | 0.134169863 |
| ENSG00000185621 | *LMLN* | 0.53395075 | -0.120001583 | -0.03594989 | -0.084051693 | 0.999933887 | -0.210158998 | 0.24051623 | -0.450675229 | 0.134169863 |
| ENSG00000280332 | *AC020917.4* | 3.153820298 | 0.052518208 | 0.045101756 | 0.007416451 | 0.999933887 | -0.021097579 | 0.183852427 | -0.204950005 | 0.134227808 |
| ENSG00000112715 | *VEGFA* | 10.559120951 | 0.180800346 | 0.244035953 | -0.063235607 | 0.999933887 | 0.523144879 | 0.720675023 | -0.197530144 | 0.13439979 |
| ENSG00000110274 | *CEP164* | 4.367150057 | 0.046245247 | 0.05550337 | -0.009258123 | 0.999933887 | -0.054729858 | -0.223290555 | 0.168560697 | 0.134556605 |
| ENSG00000029153 | *ARNTL2* | -1.167599415 | -0.489639976 | -0.427815317 | -0.06182466 | 0.999933887 | 0.215859399 | 0.861454478 | -0.645595079 | 0.134775773 |
| ENSG00000168291 | *PDHB* | 2.522311904 | 0.024566792 | 0.052609009 | -0.028042217 | 0.999933887 | -0.07900096 | -0.307192759 | 0.2281918 | 0.134845697 |
| ENSG00000111653 | *ING4* | 4.07438491 | -0.113018279 | -0.067582746 | -0.045435533 | 0.999933887 | -0.310369486 | -0.499044645 | 0.18867516 | 0.13493934 |
| ENSG00000221817 | *PPP3CB-AS1* | 0.978100848 | -0.026642791 | -0.004655369 | -0.021987421 | 0.999933887 | -0.111647192 | 0.220520539 | -0.332167731 | 0.13493934 |
| ENSG00000115827 | *DCAF17* | 1.595629323 | 0.010869541 | -0.133621004 | 0.144490545 | 0.999933887 | -0.176933624 | 0.129828355 | -0.306761979 | 0.135125876 |
| ENSG00000130222 | *GADD45G* | 3.985131317 | -0.086415047 | -0.149488632 | 0.063073586 | 0.999933887 | -0.122620507 | -0.489115461 | 0.366494954 | 0.135467069 |
| ENSG00000087074 | *PPP1R15A* | 11.72942688 | 0.369578552 | 0.351393575 | 0.018184977 | 0.999933887 | 1.579993507 | 1.866464287 | -0.28647078 | 0.13547322 |
| ENSG00000178026 | *LRRC75B* | 2.853921439 | 0.111005281 | 0.099493297 | 0.011511985 | 0.999933887 | -0.361319219 | -0.107926449 | -0.25339277 | 0.135578709 |
| ENSG00000156735 | *BAG4* | 2.592036828 | 0.028911164 | 0.073369356 | -0.044458192 | 0.999933887 | -0.475047909 | -0.799809296 | 0.324761387 | 0.135578709 |
| ENSG00000236977 | *ANKRD44-IT1* | 1.237834319 | 0.069805865 | 0.05438269 | 0.015423174 | 0.999933887 | -0.241670038 | -0.775766931 | 0.534096893 | 0.135578876 |
| ENSG00000159110 | *IFNAR2* | 2.229065544 | -0.154876879 | -0.074713303 | -0.080163576 | 0.999933887 | -0.175650049 | -0.532496883 | 0.356846835 | 0.135739602 |
| ENSG00000108528 | *SLC25A11* | 4.009575119 | 0.071485044 | 0.03035484 | 0.041130204 | 0.999933887 | 0.087189182 | -0.100246048 | 0.18743523 | 0.135739602 |
| ENSG00000130303 | *BST2* | 4.281188282 | 0.125738856 | 0.059083924 | 0.066654931 | 0.999933887 | -0.22738765 | -0.536007063 | 0.308619413 | 0.135861121 |
| ENSG00000253653 | *AC009185.1* | 0.652740873 | 0.040184357 | 0.043989104 | -0.003804747 | 0.999933887 | -0.412761856 | 0.080322248 | -0.493084104 | 0.13608266 |
| ENSG00000164978 | *NUDT2* | 0.934040835 | -0.065567815 | 0.053728723 | -0.119296538 | 0.999933887 | -0.036011281 | -0.417964934 | 0.381953653 | 0.136363487 |
| ENSG00000187824 | *TMEM220* | -0.430853442 | -0.119705049 | -0.10074403 | -0.018961019 | 0.999933887 | -0.36944169 | 0.306913974 | -0.676355664 | 0.136482813 |
| ENSG00000092871 | *RFFL* | 6.074820896 | -0.021982877 | 0.029310392 | -0.051293269 | 0.999933887 | 0.965651102 | 1.162230794 | -0.196579691 | 0.136731506 |
| ENSG00000156697 | *UTP14A* | 1.724496752 | -0.24934154 | -0.140156482 | -0.109185058 | 0.999933887 | 0.01952046 | 0.285520463 | -0.266000003 | 0.136970844 |
| ENSG00000095906 | *NUBP2* | 3.35565564 | 0.066795627 | 0.10735296 | -0.040557333 | 0.999933887 | 0.0158313 | -0.226631603 | 0.242462904 | 0.136970844 |
| ENSG00000136891 | *TEX10* | 4.342956417 | -0.057716566 | 0.059144951 | -0.116861517 | 0.999933887 | 0.100145809 | 0.280509766 | -0.180363958 | 0.136970844 |
| ENSG00000107566 | *ERLIN1* | 4.859993219 | -0.04306147 | -0.049077707 | 0.006016237 | 0.999933887 | 0.04741962 | 0.283191047 | -0.235771427 | 0.136970844 |
| ENSG00000100949 | *RABGGTA* | 4.878714127 | 0.042845769 | 0.076789911 | -0.033944142 | 0.999933887 | 0.736451782 | 0.929057931 | -0.19260615 | 0.136970844 |
| ENSG00000151650 | *VENTX* | 1.262636503 | 0.107422569 | -0.010792323 | 0.118214892 | 0.999933887 | 0.016788799 | 0.372694735 | -0.355905936 | 0.13726186 |
| ENSG00000119669 | *IRF2BPL* | 4.530304604 | 0.058959834 | 0.029292896 | 0.029666938 | 0.999933887 | -0.101066683 | -0.338145823 | 0.23707914 | 0.13726186 |
| ENSG00000105355 | *PLIN3* | 5.947348757 | 0.051226435 | 0.002892852 | 0.048333583 | 0.999933887 | -0.067646016 | 0.159865303 | -0.22751132 | 0.13726186 |
| ENSG00000184840 | *TMED9* | 5.044349752 | -0.01630641 | -0.008957138 | -0.007349272 | 0.999933887 | -0.307658168 | -0.525526332 | 0.217868163 | 0.13726186 |
| ENSG00000100726 | *TELO2* | 3.469145562 | 0.109455638 | 0.126656443 | -0.017200805 | 0.999933887 | -0.00530964 | -0.237955581 | 0.232645941 | 0.137408886 |
| ENSG00000149084 | *HSD17B12* | 2.763426408 | 0.15395625 | 0.01076256 | 0.14319369 | 0.999933887 | -0.223390694 | 0.018693066 | -0.24208376 | 0.137572723 |
| ENSG00000125633 | *CCDC93* | 7.468202066 | -0.064488947 | -0.126792732 | 0.062303785 | 0.999933887 | -0.018412677 | 0.147996141 | -0.166408818 | 0.137572723 |
| ENSG00000121644 | *DESI2* | 3.40390332 | -0.03247766 | -0.03994517 | 0.00746751 | 0.999933887 | -0.336551392 | -0.539016416 | 0.202465024 | 0.137572723 |
| ENSG00000144224 | *UBXN4* | 7.536896441 | -0.031166283 | -0.101720446 | 0.070554162 | 0.999933887 | -0.412422082 | -0.184675168 | -0.227746914 | 0.137572723 |
| ENSG00000146192 | *FGD2* | 1.28033119 | 0.014145002 | 0.142488132 | -0.12834313 | 0.999933887 | 0.081510578 | 0.425173841 | -0.343663263 | 0.137572723 |
| ENSG00000273133 | *AC116651.1* | 1.292931858 | -0.01739245 | 0.324931345 | -0.342323796 | 0.944094537 | 0.050485044 | 0.473584405 | -0.42309936 | 0.137625439 |
| ENSG00000138175 | *ARL3* | 2.12508656 | 0.023309243 | 0.130862622 | -0.107553379 | 0.999933887 | -0.604371398 | -0.954080423 | 0.349709025 | 0.137694325 |
| ENSG00000123091 | *RNF11* | 6.679097485 | 0.0338894 | 0.010617584 | 0.023271816 | 0.999933887 | 0.011124579 | 0.157222128 | -0.146097549 | 0.137855401 |
| ENSG00000165118 | *C9orf64* | 0.217458789 | -0.11995271 | -0.070592968 | -0.049359741 | 0.999933887 | -0.295135204 | -0.676894508 | 0.381759304 | 0.13787568 |
| ENSG00000170236 | *USP50* | 0.820689404 | -0.09287602 | -0.092911563 | 3.5543e-05 | 0.999933887 | -0.213492722 | 0.305512286 | -0.519005009 | 0.13787568 |
| ENSG00000198346 | *ZNF813* | -0.208523807 | -0.044946905 | -0.194728469 | 0.149781565 | 0.999933887 | -0.332980687 | 0.210135546 | -0.543116233 | 0.13787568 |
| ENSG00000187479 | *C11orf96* | 1.670980594 | 0.288223575 | 0.626835928 | -0.338612353 | 0.999933887 | 2.049082065 | 2.369273385 | -0.320191321 | 0.137974392 |
| ENSG00000072840 | *EVC* | -0.00882689 | 0.112292717 | -0.007704542 | 0.119997259 | 0.999933887 | -0.202814783 | 0.347566484 | -0.550381267 | 0.137974392 |
| ENSG00000237550 | *AC243919.1* | 3.029823346 | -0.042830438 | -0.068326861 | 0.025496423 | 0.999933887 | 0.148484845 | 0.398641822 | -0.250156978 | 0.137974392 |
| ENSG00000249859 | *PVT1* | 2.075569112 | -0.085472112 | 0.125409826 | -0.210881937 | 0.999933887 | 0.101008209 | 0.36358493 | -0.262576721 | 0.138120408 |
| ENSG00000169955 | *ZNF747* | 0.568794224 | -0.145202388 | 0.013244778 | -0.158447165 | 0.999933887 | -0.091964092 | -0.540504259 | 0.448540167 | 0.138207123 |
| ENSG00000149792 | *MRPL49* | 3.412825928 | -0.028966385 | 0.053115627 | -0.082082013 | 0.999933887 | -0.081335407 | -0.281234733 | 0.199899327 | 0.138210925 |
| ENSG00000213341 | *CHUK* | 5.338522925 | -0.022238511 | 0.034564484 | -0.056802995 | 0.999933887 | -0.152552167 | 0.04576285 | -0.198315017 | 0.13828656 |
| ENSG00000174885 | *NLRP6* | 6.051756986 | -0.002701213 | -0.017378865 | 0.014677652 | 0.999933887 | -2.145375254 | -2.550280354 | 0.404905099 | 0.138303754 |
| ENSG00000115271 | *GCA* | 7.274958247 | -0.024516002 | -0.017806509 | -0.006709493 | 0.999933887 | -0.183886009 | -0.380285335 | 0.196399326 | 0.138357057 |
| ENSG00000129355 | *CDKN2D* | 6.103489354 | 0.222234888 | 0.258276815 | -0.036041927 | 0.999933887 | -0.060177913 | -0.343737096 | 0.283559183 | 0.138360342 |
| ENSG00000120458 | *MSANTD2* | 0.778770198 | -0.180253456 | -0.036516582 | -0.143736874 | 0.999933887 | 0.270959552 | -0.131291986 | 0.402251538 | 0.13838791 |
| ENSG00000102796 | *DHRS12* | 2.84528177 | 0.059090423 | -0.069443828 | 0.128534252 | 0.999933887 | 0.553741362 | 0.816240232 | -0.26249887 | 0.138447819 |
| ENSG00000010292 | *NCAPD2* | 4.197328252 | -0.018446233 | -0.026692249 | 0.008246016 | 0.999933887 | -0.440799984 | -0.686178219 | 0.245378235 | 0.138544243 |
| ENSG00000248476 | *BACH1-IT1* | 2.68500061 | -0.049362959 | -0.010893208 | -0.038469751 | 0.999933887 | -0.043537239 | 0.189939241 | -0.23347648 | 0.138899699 |
| ENSG00000170144 | *HNRNPA3* | 8.183357398 | 0.002436671 | 0.016886056 | -0.014449385 | 0.999933887 | 0.307564585 | 0.479601218 | -0.172036633 | 0.138899699 |
| ENSG00000162910 | *MRPL55* | 2.668379198 | 0.062761571 | 0.062721485 | 4.0086e-05 | 0.999933887 | 0.104383784 | -0.164209669 | 0.268593453 | 0.139029983 |
| ENSG00000121486 | *TRMT1L* | 3.344901711 | -0.067208911 | 0.005010852 | -0.072219763 | 0.999933887 | -0.038802414 | 0.139674397 | -0.178476811 | 0.139029983 |
| ENSG00000182240 | *BACE2* | -0.089471435 | 0.008355684 | -0.152016197 | 0.160371881 | 0.999933887 | -0.270521127 | 0.302243467 | -0.572764593 | 0.139029983 |
| ENSG00000179523 | *EIF3J-DT* | 1.597223008 | 0.053589263 | -0.135621326 | 0.189210589 | 0.999933887 | -0.356272678 | -0.684002717 | 0.32773004 | 0.13909779 |
| ENSG00000070731 | *ST6GALNAC2* | 3.615279447 | 0.064685445 | 0.048900409 | 0.015785036 | 0.999933887 | -0.781470094 | -1.038267088 | 0.256796993 | 0.139108408 |
| ENSG00000120733 | *KDM3B* | 6.923641358 | -0.027510556 | 0.012336563 | -0.039847119 | 0.999933887 | -0.638390923 | -0.814748837 | 0.176357915 | 0.139108408 |
| ENSG00000015153 | *YAF2* | 4.055061239 | -0.018966933 | -0.007542441 | -0.011424492 | 0.999933887 | 0.089429877 | 0.255382619 | -0.165952742 | 0.139119391 |
| ENSG00000198938 | *MT-CO3* | 9.470977802 | 0.085656512 | 0.005170785 | 0.080485727 | 0.999933887 | 0.045109932 | 0.325128598 | -0.280018666 | 0.139285365 |
| ENSG00000215908 | *CROCCP2* | 4.246040413 | 0.05691688 | 0.097875882 | -0.040959002 | 0.999933887 | 0.050126002 | -0.135854151 | 0.185980153 | 0.139285365 |
| ENSG00000183718 | *TRIM52* | 3.19516821 | -0.042089318 | -0.021922909 | -0.020166408 | 0.999933887 | -0.215757944 | -0.419978094 | 0.20422015 | 0.139285365 |
| ENSG00000089048 | *ESF1* | 1.402579167 | -0.276099591 | -0.245610531 | -0.03048906 | 0.999933887 | -0.215096903 | 0.209883233 | -0.424980136 | 0.139450653 |
| ENSG00000100815 | *TRIP11* | 3.298431415 | -0.132631488 | -0.103139318 | -0.029492171 | 0.999933887 | -0.356319975 | 0.017833862 | -0.374153837 | 0.139525219 |
| ENSG00000268043 | *NBPF12* | 3.144927478 | -0.076392259 | -0.132762654 | 0.056370395 | 0.999933887 | -0.008342069 | 0.229181256 | -0.237523325 | 0.139525219 |
| ENSG00000111731 | *C2CD5* | 3.970800127 | -0.064585867 | -0.000732726 | -0.06385114 | 0.999933887 | 0.037422804 | 0.318353069 | -0.280930265 | 0.139525219 |
| ENSG00000134508 | *CABLES1* | 0.529093955 | -0.040736731 | 0.00526158 | -0.045998311 | 0.999933887 | 0.163122622 | 0.515561115 | -0.352438493 | 0.139525219 |
| ENSG00000002330 | *BAD* | 1.785394999 | 0.012887839 | 0.158137998 | -0.145250159 | 0.999933887 | 0.026734962 | -0.308241154 | 0.334976116 | 0.139525219 |
| ENSG00000273344 | *PAXIP1-AS1* | 1.242121707 | 0.015538826 | 0.217480854 | -0.201942028 | 0.999933887 | 0.509493186 | 0.88203889 | -0.372545704 | 0.139525219 |
| ENSG00000105829 | *BET1* | 0.985429567 | -0.00715525 | -0.019178129 | 0.012022879 | 0.999933887 | 0.035055451 | 0.390822612 | -0.355767162 | 0.139525219 |
| ENSG00000158636 | *EMSY* | 4.514410906 | 0.00285356 | -0.073427114 | 0.076280674 | 0.999933887 | -0.427615056 | -0.19776106 | -0.229853995 | 0.139525219 |
| ENSG00000241370 | *RPP21* | -0.111392783 | 0.147626299 | 0.217832003 | -0.070205704 | 0.999933887 | 0.320843355 | -0.221673794 | 0.542517149 | 0.139639796 |
| ENSG00000244036 | *AC073320.1* | -0.070612438 | 0.09416649 | 0.110311695 | -0.016145205 | 0.999933887 | 0.526425215 | 1.027649447 | -0.501224232 | 0.139639796 |

| | | | | | | | | | | |
|---|---|---|---|---|---|---|---|---|---|---|
| ENSG00000188342 | *GTF2F2* | 2.783457724 | 0.009636157 | -0.076887162 | 0.086523319 | 0.999933887 | -0.000302815 | 0.215642746 | -0.215945561 | 0.139808268 |
| ENSG00000092931 | *MFSD11* | 4.182747901 | 0.069309247 | 0.184215983 | -0.114906736 | 0.999933887 | 0.517515846 | 0.695877651 | -0.178361805 | 0.140112267 |
| ENSG00000163161 | *ERCC3* | 3.990156306 | 0.037421372 | -0.064009245 | 0.101430617 | 0.999933887 | 0.008907647 | -0.14781382 | 0.156721467 | 0.140112267 |
| ENSG00000184381 | *PLA2G6* | 4.521988687 | 0.13611294 | 0.1686793 | -0.03256636 | 0.999933887 | 0.757773159 | 0.975993013 | -0.218219853 | 0.140157435 |
| ENSG00000183844 | *FAM3B* | -1.163815916 | 0.165891311 | -0.131960912 | 0.297852223 | 0.999933887 | -0.056525824 | -0.778898335 | 0.722372511 | 0.140162169 |
| ENSG00000169223 | *LMAN2* | 6.053745775 | 0.024797787 | -0.02343405 | 0.048231837 | 0.999933887 | -0.194908622 | -0.349323414 | 0.154414792 | 0.140393419 |
| ENSG00000183309 | *ZNF623* | 2.278664015 | 0.040652844 | -0.035743079 | 0.076395923 | 0.999933887 | -0.298687389 | -0.556003283 | 0.257315894 | 0.140437846 |
| ENSG00000130856 | *ZNF236* | 3.255730603 | 0.052689342 | -0.028041636 | 0.080730978 | 0.999933887 | -0.310534777 | -0.491393226 | 0.180858449 | 0.140461428 |
| ENSG00000287878 | *AC022382.2* | 3.603950793 | -0.026291515 | 0.07769475 | -0.103986265 | 0.999933887 | 0.078205865 | 0.319185582 | -0.240979717 | 0.140496983 |
| ENSG00000033178 | *UBA6* | 4.595281828 | -0.059054887 | -0.067606604 | 0.008551717 | 0.999933887 | -0.286130633 | 0.124732986 | -0.410863619 | 0.14049984 |
| ENSG00000115718 | *PROC* | 0.652521018 | -0.187141105 | -0.187793303 | 0.000652198 | 0.999933887 | -1.919772235 | -2.525106684 | 0.605334448 | 0.140776085 |
| ENSG00000117020 | *AKT3* | 3.037839795 | -0.040493895 | 0.022623215 | -0.06311711 | 0.999933887 | 0.041009229 | 0.327079244 | -0.286070015 | 0.140803705 |
| ENSG00000203872 | *C6orf163* | 0.240677534 | 0.105640513 | -0.148564447 | 0.25420496 | 0.999933887 | -1.624021617 | -2.193559335 | 0.569537719 | 0.141025084 |
| ENSG00000127578 | *WFIKKN1* | -0.516718571 | 0.049033548 | 0.160003165 | -0.110969617 | 0.999933887 | 0.050636287 | -0.579428436 | 0.630064723 | 0.141209706 |
| ENSG00000266028 | *SRGAP2* | 4.078647114 | 0.00138314 | 0.071910609 | -0.070527469 | 0.999933887 | -0.082496031 | 0.149148191 | -0.231644222 | 0.141263664 |
| ENSG00000111424 | *VDR* | 6.248811632 | 0.003266135 | 0.063274755 | -0.06000862 | 0.999933887 | 0.070512989 | -0.106221261 | 0.176734249 | 0.141362506 |
| ENSG00000182010 | *RTKN2* | 0.374971205 | 0.164051911 | -0.419039941 | 0.583091852 | 0.999933887 | 0.263071928 | 0.698920686 | -0.435848758 | 0.141588505 |
| ENSG00000074696 | *HACD3* | 1.75368762 | 0.048560389 | -0.215486903 | 0.264047292 | 0.999933887 | 0.038085522 | 0.367996381 | -0.329910859 | 0.141648559 |
| ENSG00000287631 | *AC005071.1* | 2.169389223 | -0.246314285 | -0.043004352 | -0.203309933 | 0.999933887 | 2.34995168 | 2.875744035 | -0.525792355 | 0.141858036 |
| ENSG00000139514 | *SLC7A1* | 4.037865879 | 0.043440765 | -0.068128643 | 0.111569408 | 0.999933887 | 0.282929013 | 0.473433564 | -0.190504551 | 0.14202508 |
| ENSG00000139278 | *GLIPR1* | 5.573927046 | -0.009848429 | -0.123417721 | 0.113569292 | 0.999933887 | -0.386118521 | -0.657704453 | 0.271585931 | 0.142070682 |
| ENSG00000214765 | *SEPTIN7P2* | 2.961335468 | -0.007914022 | -0.00432555 | -0.003588472 | 0.999933887 | 0.115026343 | 0.380367207 | -0.265340863 | 0.142072279 |
| ENSG00000106952 | *TNFSF8* | 5.499256268 | 0.00657426 | -0.017911786 | 0.024486046 | 0.999933887 | 0.081168354 | 0.27676141 | -0.195593056 | 0.142072279 |
| ENSG00000232254 | *CSF2RBP1* | -0.727710449 | -0.080689636 | 0.315802856 | -0.396492492 | 0.999933887 | 0.159510485 | -0.849090091 | 1.008600576 | 0.142240927 |
| ENSG00000256594 | *AC010186.2* | 1.889345922 | 0.020229086 | 0.059739717 | -0.039510631 | 0.999933887 | 0.563297136 | 0.857732322 | -0.294435186 | 0.142240927 |
| ENSG00000127995 | *CASD1* | 2.00833274 | -0.078438879 | -0.226344942 | 0.147906063 | 0.999933887 | -0.271708117 | 0.056394073 | -0.32810219 | 0.142246568 |
| ENSG00000164010 | *ERMAP* | 1.710025654 | 0.069581676 | -0.054790031 | 0.124371706 | 0.999933887 | -0.153775595 | 0.151472161 | -0.305247756 | 0.142246568 |
| ENSG00000174945 | *AMZ1* | -0.750762502 | 0.124316045 | 0.257487528 | -0.133171483 | 0.999933887 | 0.22390381 | 0.823216977 | -0.599313167 | 0.142246568 |
| ENSG00000125686 | *MED1* | 4.799972147 | -0.006422021 | -0.149839645 | 0.143417623 | 0.999933887 | 0.064266688 | 0.226855404 | -0.162588716 | 0.142246568 |
| ENSG00000124523 | *SIRT5* | 2.008190978 | -0.006365252 | 0.080846382 | -0.087211634 | 0.999933887 | -0.418568546 | -0.653177671 | 0.234609125 | 0.142348562 |
| ENSG00000157212 | *PAXIP1* | 2.261001357 | -0.102291359 | 0.022900799 | -0.125192158 | 0.999933887 | 0.381897285 | 0.729372266 | -0.347474981 | 0.142623682 |
| ENSG00000079385 | *CEACAM1* | 6.326475404 | -0.019097549 | 0.020866054 | -0.039963603 | 0.999933887 | -0.087473895 | 0.122348469 | -0.209822363 | 0.142840806 |
| ENSG00000100567 | *PSMA3* | 5.142588473 | -0.089563977 | -0.126625221 | 0.037061245 | 0.999933887 | -0.122771505 | 0.023649311 | -0.146420816 | 0.142900798 |
| ENSG00000204084 | *INPP5B* | 3.570999346 | -0.007590236 | 0.006494841 | -0.014085076 | 0.999933887 | 0.091157657 | -0.077561866 | 0.168719523 | 0.142900798 |
| ENSG00000186480 | *INSIG1* | 7.975238772 | 0.088006264 | 0.087347059 | 0.000659205 | 0.999933887 | 1.422541426 | 1.763653474 | -0.341112048 | 0.142929833 |
| ENSG00000130640 | *TUBGCP2* | 5.601491917 | 0.007855603 | 0.024189866 | -0.016334262 | 0.999933887 | -0.046689733 | -0.170361318 | 0.123671585 | 0.142929833 |
| ENSG00000105248 | *YJU2* | 3.319818632 | -0.009832628 | 0.032594792 | -0.04242742 | 0.999933887 | 0.261802996 | 0.525212966 | -0.26340997 | 0.14332617 |
| ENSG00000177732 | *SOX12* | 1.738215156 | 0.068117506 | 0.027807441 | 0.040310065 | 0.999933887 | -0.188738824 | -0.545519755 | 0.356780932 | 0.143390962 |
| ENSG00000158882 | *TOMM40L* | 4.611589771 | 0.028147904 | 0.028062083 | 8.5821e-05 | 0.999933887 | 0.822277854 | 1.022693511 | -0.200415657 | 0.143461402 |
| ENSG00000065150 | *IPO5* | 3.575927972 | -0.000722936 | -0.123764308 | 0.123041373 | 0.999933887 | 0.354042396 | 0.564371644 | -0.210329248 | 0.144197887 |
| ENSG00000260051 | *AL031600.1* | 3.494174835 | -0.010015434 | -0.029159417 | 0.019143983 | 0.999933887 | 0.110086577 | 0.364303547 | -0.254216971 | 0.144255914 |
| ENSG00000118292 | *C1orf54* | -0.147427492 | 0.026046142 | 0.00064285 | 0.025403291 | 0.999933887 | -0.021760697 | 0.417806216 | -0.439566913 | 0.144859094 |
| ENSG00000178726 | *THBD* | 4.389051964 | 0.136350752 | 0.12240289 | 0.013947862 | 0.999933887 | -1.863862586 | -2.267850553 | 0.403987967 | 0.145069471 |
| ENSG00000136156 | *ITM2B* | 10.528344329 | -0.006818302 | -0.028151972 | 0.02133367 | 0.999933887 | -0.384752341 | -0.548812471 | 0.16406013 | 0.145069471 |
| ENSG00000163359 | *COL6A3* | -0.362695309 | -0.030113286 | 0.157382564 | -0.18749585 | 0.999933887 | -0.457138934 | 0.368675879 | -0.825814813 | 0.145794852 |
| ENSG00000272758 | *AC083798.2* | 1.106787324 | -0.168285745 | -0.053468978 | -0.114816768 | 0.999933887 | -0.139295351 | 0.203830901 | -0.343126252 | 0.145812278 |
| ENSG00000196655 | *TRAPPC4* | 3.666361242 | -0.073657909 | -0.033872414 | -0.039785495 | 0.999933887 | -0.077358215 | -0.242745512 | 0.165387297 | 0.145812278 |
| ENSG00000168488 | *ATXN2L* | 6.988827128 | 0.001896722 | 0.066737557 | -0.064840835 | 0.999933887 | 0.050858064 | 0.173206353 | -0.122348289 | 0.145812278 |
| ENSG00000273243 | *Z82243.1* | -0.388000008 | -0.219778063 | -0.042818919 | -0.176959144 | 0.999933887 | -0.954380154 | -1.600775784 | 0.64639563 | 0.146062252 |
| ENSG00000196705 | *ZNF431* | 3.10753668 | -0.042565277 | -0.199519841 | 0.156954564 | 0.999933887 | -0.395219567 | -0.057168074 | -0.338051494 | 0.146077263 |
| ENSG00000197037 | *ZSCAN25* | 3.358136008 | 0.061737111 | 0.051414846 | 0.010322264 | 0.999933887 | -0.111075331 | -0.345399747 | 0.234324416 | 0.146126903 |
| ENSG00000149541 | *B3GAT3* | 3.985501211 | 0.081157829 | -0.018922864 | 0.100080693 | 0.999933887 | 0.798830258 | 1.130114336 | -0.331284078 | 0.146157273 |
| ENSG00000135835 | *KIAA1614* | 0.220540814 | 0.031644886 | -0.30459832 | 0.336243207 | 0.999933887 | -0.525700533 | -1.014482599 | 0.488782066 | 0.146157273 |
| ENSG00000148840 | *PPRC1* | 3.821765316 | -0.046108856 | -0.026081642 | -0.020027214 | 0.999933887 | 0.548949393 | 0.752013309 | -0.203063916 | 0.146422222 |
| ENSG00000067646 | *ZFY* | -3.421724204 | -0.159379079 | 0.051879907 | -0.211258986 | 0.999933887 | -0.051130051 | 0.444249078 | -0.495379129 | 0.146438709 |
| ENSG00000253320 | *MAILR* | 6.694393316 | 0.095042724 | 0.204637545 | -0.109594821 | 0.999933887 | 3.71131949 | 4.21966943 | -0.508349941 | 0.146870927 |
| ENSG00000149346 | *SLX4IP* | 1.858564461 | 0.047212635 | -0.09686463 | 0.144077265 | 0.999933887 | 0.63569404 | 0.984268058 | -0.348574017 | 0.146870927 |
| ENSG00000215267 | *AKR1C7P* | 0.75359138 | 0.177093798 | -0.046449941 | 0.223543739 | 0.999933887 | 0.100644369 | 0.421256519 | -0.32061215 | 0.147214245 |
| ENSG00000173065 | *FAM222B* | 2.918427611 | -0.006509284 | -0.068266155 | 0.061756871 | 0.999933887 | 0.14005393 | 0.355245231 | -0.2151913 | 0.147315614 |
| ENSG00000261526 | *AC012615.1* | 1.277912123 | -0.106910105 | -0.032433863 | -0.074476242 | 0.999933887 | -0.855123582 | -1.305235273 | 0.450111691 | 0.147484265 |
| ENSG00000071242 | *RPS6KA2* | 3.116842561 | -0.010720909 | -0.017570746 | 0.006849837 | 0.999933887 | -1.077911473 | -1.384341998 | 0.306430525 | 0.147714616 |
| ENSG00000112983 | *BRD8* | 3.914468345 | -0.114571813 | -0.111517193 | -0.00305462 | 0.999933887 | -0.351786509 | -0.571751574 | 0.219965065 | 0.147724171 |
| ENSG00000113649 | *TCERG1* | 3.728845496 | -0.041451185 | -0.101912495 | 0.06046131 | 0.999933887 | -0.049108403 | 0.2026182 | -0.251726603 | 0.147724171 |
| ENSG00000180758 | *GPR157* | 3.014800567 | -0.090871641 | 0.102623371 | -0.193495012 | 0.999933887 | 0.895894371 | 1.286194902 | -0.390300531 | 0.147762332 |
| ENSG00000084774 | *CAD* | 3.287447343 | 0.027047107 | 0.062829344 | -0.035782237 | 0.999933887 | 0.108998084 | -0.087712185 | 0.196710269 | 0.147838936 |
| ENSG00000258921 | *AC009779.4* | -0.468997016 | 0.012021549 | -0.041235269 | 0.053256818 | 0.999933887 | 1.382002741 | 1.860580973 | -0.478578232 | 0.148081502 |
| ENSG00000146776 | *ATXN7L1* | 2.834423794 | -0.003192605 | -0.019505838 | 0.016313233 | 0.999933887 | 0.176331324 | 0.412933378 | -0.236602054 | 0.148081502 |
| ENSG00000204619 | *PPP1R11* | 6.701932121 | 0.036891942 | 0.002369565 | 0.034522377 | 0.999933887 | 0.392686255 | 0.545109069 | -0.152422814 | 0.148230089 |
| ENSG00000187741 | *FANCA* | 2.16837568 | 0.139861035 | 0.127600826 | 0.01226021 | 0.999933887 | -0.349793865 | -0.735488403 | 0.385694538 | 0.148467486 |
| ENSG00000142961 | *MOB3C* | 4.149834255 | -0.011959976 | 0.014032988 | -0.025992964 | 0.999933887 | 0.247278649 | 0.524634397 | -0.277355748 | 0.148477574 |
| ENSG00000224614 | *TNK2-AS1* | 1.204128271 | -0.039474785 | -0.096668373 | 0.057193588 | 0.999933887 | -1.019960189 | -1.455728118 | 0.435767929 | 0.148802784 |
| ENSG00000224307 | *AL161785.1* | 1.046517319 | -0.069731287 | -0.055466387 | -0.0142649 | 0.999933887 | -1.250705784 | -1.758363507 | 0.507657723 | 0.148811419 |
| ENSG00000151914 | *DST* | 0.218376341 | -0.066742865 | 0.118075888 | -0.184818753 | 0.999933887 | -0.351176333 | 0.269785729 | -0.620962062 | 0.148811419 |
| ENSG00000174886 | *NDUFA11* | 2.138171414 | 0.135647394 | 0.041149959 | 0.094497435 | 0.999933887 | 0.071841549 | -0.207796338 | 0.279637888 | 0.148907387 |
| ENSG00000133106 | *EPSTI1* | 5.078541582 | -0.051252457 | -0.053637751 | 0.002385294 | 0.999933887 | -0.092079263 | -0.251696518 | 0.159617256 | 0.148907387 |
| ENSG00000170759 | *KIF5B* | 6.027375994 | -0.021209917 | -0.119199359 | 0.097989442 | 0.999933887 | -0.606170831 | -0.273674155 | -0.332496675 | 0.149035621 |
| ENSG00000165185 | *KIAA1958* | 1.212158406 | 0.003484773 | -0.041903446 | 0.045388219 | 0.999933887 | -0.268935113 | -0.753131985 | 0.484196872 | 0.149147597 |
| ENSG00000165832 | *TRUB1* | 0.509468608 | -0.056417937 | -0.08226133 | 0.025843393 | 0.999933887 | -0.101111682 | 0.352376325 | -0.453488007 | 0.149172154 |
| ENSG00000149591 | *TAGLN* | 2.791973582 | 0.027395027 | 0.073067965 | -0.045672938 | 0.999933887 | 0.072069965 | -0.173300407 | 0.245370372 | 0.149220517 |
| ENSG00000146904 | *EPHA1* | 2.31095607 | 0.155011538 | 0.184463181 | -0.029451643 | 0.999933887 | -0.359921471 | -0.720556818 | 0.360635346 | 0.149241409 |
| ENSG00000258441 | *LINC00641* | 5.410605811 | -0.017380388 | -0.00412999 | -0.013250398 | 0.999933887 | 1.424545687 | 1.825250569 | -0.400704883 | 0.149345453 |
| ENSG00000100162 | *CENPM* | -0.30335497 | -0.052664362 | 0.1269947 | -0.179659062 | 0.999933887 | 0.243433425 | -0.29731733 | 0.540750755 | 0.149368536 |
| ENSG00000041802 | *LSG1* | 3.903682982 | -0.041803079 | 0.033647901 | -0.07545098 | 0.999933887 | 0.108573017 | 0.271980454 | -0.163407437 | 0.149415071 |
| ENSG00000012171 | *SEMA3B* | 1.924123934 | 0.031617733 | 0.126048154 | -0.094430421 | 0.999933887 | 0.035797365 | -0.373521239 | 0.409318605 | 0.149415071 |
| ENSG00000084090 | *STARD7* | 4.364613919 | -0.021897715 | -0.00799051 | -0.013907205 | 0.999933887 | 0.086117062 | 0.296648695 | -0.210531633 | 0.149415071 |
| ENSG00000115295 | *CLIP4* | 3.174598706 | -0.025105148 | -0.107899561 | 0.082794413 | 0.999933887 | -0.357711611 | -0.657787092 | 0.300075481 | 0.149415071 |
| ENSG00000206418 | *RAB12* | 2.252509941 | 0.02034683 | 0.059974552 | -0.039627722 | 0.999933887 | 0.688321589 | 0.972555913 | -0.284234325 | 0.149415071 |
| ENSG00000107929 | *LARP4B* | 6.25400346 | -0.02697087 | 0.039314438 | -0.066285307 | 0.999933887 | -0.387562036 | -0.513093928 | 0.125531892 | 0.149449412 |
| ENSG00000049759 | *NEDD4L* | 0.431386481 | -0.121021079 | 0.021935766 | -0.142956845 | 0.999933887 | 0.446794213 | 0.797255404 | -0.350461191 | 0.149504886 |
| ENSG00000269825 | *AC022150.4* | 0.00640683 | 0.115518317 | 0.163749634 | -0.048231317 | 0.999933887 | -0.297035945 | 0.187146734 | -0.484182679 | 0.149622169 |
| ENSG00000164442 | *CITED2* | 5.352821438 | 0.03123852 | 0.033277527 | -0.002039007 | 0.999933887 | 0.089501721 | 0.283157883 | -0.193656162 | 0.149622169 |
| ENSG00000130881 | *LRP3* | -0.11347176 | -0.052803439 | 0.005972903 | -0.058776342 | 0.999933887 | -0.192770608 | -0.836897309 | 0.644126701 | 0.149877134 |
| ENSG00000021762 | *OSBPL5* | 2.906424413 | 0.004837567 | 0.212897096 | -0.208059528 | 0.999933887 | -0.074409324 | -0.323604564 | 0.24919524 | 0.149877134 |
| ENSG00000253710 | *ALG11* | 1.84230728 | -0.004637557 | -0.089155394 | 0.084517837 | 0.999933887 | -0.003856283 | -0.311319286 | 0.307463003 | 0.149974297 |
| ENSG00000147403 | *RPL10* | 8.351173928 | 0.050950151 | -0.02223084 | 0.073180991 | 0.999933887 | 0.218747023 | 0.380282685 | -0.161535662 | 0.149992749 |
| ENSG00000153363 | *LINC00467* | 0.021148686 | -0.03065179 | 0.021115409 | -0.051767199 | 0.999933887 | -0.025961508 | 0.552265197 | -0.578226705 | 0.150094166 |
| ENSG00000101310 | *SEC23B* | 4.747441944 | 0.007053917 | -0.019303234 | 0.026357151 | 0.999933887 | 0.37551035 | 0.547531126 | -0.172020776 | 0.150421826 |
| ENSG00000221963 | *APOL6* | 6.592990245 | 0.007715664 | -0.021401894 | 0.029117558 | 0.999933887 | -0.470156965 | -0.655229342 | 0.185072376 | 0.150469436 |
| ENSG00000159842 | *ABR* | 7.716961895 | 0.017980145 | 0.015460894 | 0.002519251 | 0.999933887 | 0.088143649 | 0.215024756 | -0.126881107 | 0.150725338 |
| ENSG00000179921 | *GPBAR1* | -0.046382135 | 0.017932836 | 0.031812831 | -0.013879995 | 0.999933887 | 0.029406699 | -0.596165352 | 0.625572051 | 0.15083128 |
| ENSG00000285730 | *Z94721.3* | 1.232922958 | 0.097197639 | -0.067030901 | 0.16422854 | 0.999933887 | -0.419029569 | -0.85914107 | 0.440111502 | 0.150947639 |
| ENSG00000162819 | *BROX* | 5.46184249 | -0.159535924 | -0.066106363 | -0.093429561 | 0.999933887 | -0.439449321 | -0.226959757 | -0.212489565 | 0.151086304 |
| ENSG00000167674 | *HDGFL2* | 3.932342824 | 0.082412231 | 0.143606855 | -0.061194624 | 0.999933887 | -0.133258456 | -0.41079045 | 0.277531994 | 0.151396732 |
| ENSG00000230521 | *AL645929.1* | -0.100610568 | -0.048128548 | 0.393815375 | -0.441943922 | 0.999933887 | 0.901275595 | 0.374856367 | 0.526419228 | 0.151446913 |
| ENSG00000172667 | *ZMAT3* | 3.121094356 | 0.006858628 | 0.018169542 | -0.011310913 | 0.999933887 | -0.01267997 | 0.287998586 | -0.300678556 | 0.151446913 |

| | | | | | | | | | | |
|---|---|---|---|---|---|---|---|---|---|---|
| ENSG00000114013 | CD86 | 0.673401877 | 0.00913733 | -0.126100337 | 0.135237667 | 0.999933887 | -0.443048895 | 0.119798658 | -0.562847553 | 0.1514647 |
| ENSG00000138768 | USO1 | 4.003340524 | -0.035216224 | -0.141015422 | 0.105799198 | 0.999933887 | -0.314016846 | -0.012112226 | -0.301904619 | 0.151591045 |
| ENSG00000244754 | N4BP2L2 | 6.805177846 | -0.054101043 | -0.087926797 | 0.033825754 | 0.999933887 | -0.009535533 | 0.334410172 | -0.343945704 | 0.151595764 |
| ENSG00000168101 | NUDT16L1 | 3.594020695 | 0.026956641 | -0.050846072 | 0.077802714 | 0.999933887 | 0.032576912 | -0.224299411 | 0.256876323 | 0.151595764 |
| ENSG00000113296 | THBS4 | -0.527727599 | 0.066951114 | 0.175814023 | -0.10886291 | 0.999933887 | -0.300920432 | 0.384795981 | -0.685716413 | 0.151669291 |
| ENSG00000157353 | FCSK | 2.938484096 | 0.043634017 | 0.198958257 | -0.155324241 | 0.999933887 | 0.049225936 | -0.162527649 | 0.211753585 | 0.151707712 |
| ENSG00000124145 | SDC4 | 1.148010384 | -0.170208506 | -0.090136005 | -0.080072501 | 0.999933887 | 1.631012368 | 2.185558833 | -0.554546465 | 0.151707712 |
| ENSG00000099624 | ATP5F1D | 3.556715219 | 0.041838934 | -0.043865414 | 0.085704347 | 0.999933887 | -0.10002802 | -0.360475789 | 0.260447769 | 0.151707712 |
| ENSG00000198862 | LTN1 | 4.792231937 | -0.072562746 | -0.156003679 | 0.083440933 | 0.999933887 | -0.519230108 | -0.068053579 | -0.451176529 | 0.151707712 |
| ENSG00000204946 | ZNF783 | 2.589317188 | 0.037625243 | 0.085821404 | -0.048196161 | 0.999933887 | 0.065434593 | -0.167174654 | 0.232609247 | 0.151707712 |
| ENSG00000034152 | MAP2K3 | 8.701344868 | -0.027327068 | -0.008627197 | -0.01869987 | 0.999933887 | 1.187668351 | 1.379308672 | -0.191640321 | 0.151707712 |
| ENSG00000021776 | AQR | 3.869758218 | 0.00664505 | -0.064060038 | 0.070705088 | 0.999933887 | -0.188786981 | 0.052867161 | -0.241654141 | 0.151707712 |
| ENSG00000118246 | FASTKD2 | 2.180896683 | -0.139784545 | -0.054460519 | -0.085324026 | 0.999933887 | 0.010457991 | 0.321010773 | -0.310552782 | 0.151726246 |
| ENSG00000166889 | PATL1 | 8.121588279 | 0.00631347 | 0.016578376 | -0.010264906 | 0.999933887 | 0.499430378 | 0.671088004 | -0.171657627 | 0.151726246 |
| ENSG00000179144 | GIMAP7 | 3.890955126 | -0.094982377 | -0.184467733 | 0.089485356 | 0.999933887 | 0.105867724 | 0.298529501 | -0.192661777 | 0.151749122 |
| ENSG00000204960 | BLACE | -0.653758995 | 0.154014417 | 0.39985454 | -0.245840122 | 0.999933887 | 2.456220623 | 3.046439932 | -0.59021931 | 0.151749122 |
| ENSG00000114626 | ABTB1 | 9.436386599 | 0.007573989 | 0.008718563 | -0.001144574 | 0.999933887 | -0.17417909 | -0.327015395 | 0.152836304 | 0.151762201 |
| ENSG00000121988 | ZRANB3 | -0.277020161 | -0.02285781 | -0.133721572 | 0.110863762 | 0.999933887 | -0.476237899 | 0.178065063 | -0.654302962 | 0.151762201 |
| ENSG00000108785 | HSD17B1P1 | 0.009302307 | 0.132507825 | 0.107915098 | 0.024592727 | 0.999933887 | -0.171767182 | -0.725015488 | 0.553248306 | 0.151778607 |
| ENSG00000225630 | MTND2P28 | 4.056775559 | 0.128388373 | 0.059963539 | 0.068424834 | 0.999933887 | 0.051851245 | 0.319968375 | -0.26811713 | 0.151882137 |
| ENSG00000111052 | LIN7A | 4.387342231 | -0.061882457 | -0.021415076 | -0.040467381 | 0.999933887 | -0.894723967 | -1.218201664 | 0.323477697 | 0.151891014 |
| ENSG00000181019 | NQO1 | 0.354869481 | 0.137111385 | 0.220508367 | -0.083396982 | 0.999933887 | 0.189303015 | 0.549205263 | -0.359902248 | 0.15194206 |
| ENSG00000175826 | CTDNEP1 | 5.282275677 | -0.043685466 | 0.007161791 | -0.050847256 | 0.999933887 | -0.075330702 | -0.21792476 | 0.142594058 | 0.152046131 |
| ENSG00000157601 | MX1 | 7.284294973 | -0.021050288 | 0.008557263 | -0.029607551 | 0.999933887 | -0.397725436 | -0.571056469 | 0.173331033 | 0.152046131 |
| ENSG00000164073 | MFSD8 | 2.474373876 | -0.027836406 | -0.027578418 | -0.000257988 | 0.999933887 | -0.357786571 | -0.616794282 | 0.259007711 | 0.152101527 |
| ENSG00000223482 | NUTM2A-AS1 | 3.172183016 | -0.057249379 | -0.022921007 | -0.034328372 | 0.999933887 | -0.312196108 | -0.521828518 | 0.20963241 | 0.152108116 |
| ENSG00000204673 | AKT1S1 | 3.883269879 | 0.044609642 | 0.068746069 | -0.024136427 | 0.999933887 | -0.558933771 | -0.748614289 | 0.189680518 | 0.152108116 |
| ENSG00000055044 | NOP58 | 3.828811234 | -0.067039957 | -0.192678232 | 0.125638275 | 0.999933887 | 0.124353193 | 0.329846694 | -0.2054935 | 0.152278364 |
| ENSG00000205744 | DENND1C | 6.127017033 | 0.027342512 | 0.04894585 | -0.021603338 | 0.999933887 | -0.068754732 | -0.189915522 | 0.121160791 | 0.152334828 |
| ENSG00000129353 | SLC44A2 | 8.597467412 | -0.029043713 | -0.035990207 | 0.006946494 | 0.999933887 | -0.826639463 | -1.022499341 | 0.195859878 | 0.152334828 |
| ENSG00000011275 | RNF216 | 5.520553874 | -0.018176687 | 0.021570393 | -0.039747081 | 0.999933887 | -0.046778773 | -0.160562475 | 0.113783702 | 0.152368068 |
| ENSG00000124713 | GNMT | 0.234554024 | -0.023547174 | 0.014275398 | -0.037822572 | 0.999933887 | -0.172013251 | -0.713432246 | 0.541418996 | 0.152530602 |
| ENSG00000172046 | USP19 | 6.397972773 | -0.049047254 | 0.037970375 | -0.087017629 | 0.999933887 | -0.376954854 | -0.51686125 | 0.139906397 | 0.152675819 |
| ENSG00000276085 | CCL3L3 | 10.130244606 | 0.59802638 | 0.609265036 | -0.011238656 | 0.999933887 | 2.721293567 | 3.188856757 | -0.46756319 | 0.152728443 |
| ENSG00000276085 | CCL3L3 | 10.130244606 | 0.59802638 | 0.609265036 | -0.011238656 | 0.999933887 | 2.721293567 | 3.188856757 | -0.46756319 | 0.152728443 |
| ENSG00000120265 | PCMT1 | 3.954563164 | 0.032137706 | -0.038246744 | 0.07038445 | 0.999933887 | -0.38017006 | -0.600791512 | 0.220621452 | 0.153014005 |
| ENSG00000280020 | AC116407.3 | -0.766404744 | 0.012214854 | -0.091310154 | 0.103525008 | 0.999933887 | -0.164928069 | -1.115771801 | 0.950843732 | 0.153104927 |
| ENSG00000286584 | AC023494.1 | 0.244078491 | -0.223678348 | 0.070663066 | -0.294341415 | 0.999933887 | 0.354960722 | 0.781380646 | -0.426419924 | 0.153249624 |
| ENSG00000248323 | LUCAT1 | 9.124534462 | 0.057227236 | 0.090927037 | -0.033699801 | 0.999933887 | 0.511284116 | 0.691762481 | -0.180478365 | 0.153609554 |
| ENSG00000287269 | Z82215.1 | 0.546144657 | 0.057646813 | 0.171729683 | -0.11408287 | 0.999933887 | 0.431416961 | 0.951149132 | -0.519732171 | 0.153609554 |
| ENSG00000286004 | AC124944.3 | 1.177819361 | -0.055365738 | 0.058790567 | -0.114156305 | 0.999933887 | -0.465449283 | -0.902739659 | 0.437290376 | 0.153674043 |
| ENSG00000175467 | SART1 | 5.581016468 | 0.029547096 | 0.058382572 | -0.028835476 | 0.999933887 | -0.20981851 | -0.460757476 | 0.250938966 | 0.153674043 |
| ENSG00000167182 | SP2 | 5.646117354 | 0.006760707 | -0.041727227 | 0.048487934 | 0.999933887 | 0.068089155 | 0.202288257 | -0.134199102 | 0.153873003 |
| ENSG00000142669 | SH3BGRL3 | 8.131526663 | 0.028368968 | -0.009126834 | 0.037495802 | 0.999933887 | -0.258981221 | -0.418968699 | 0.159987479 | 0.15399776 |
| ENSG00000166685 | COG1 | 3.635495506 | 0.082450654 | 0.065197069 | 0.017253586 | 0.999933887 | -0.085424461 | -0.293218646 | 0.207794185 | 0.15404243 |
| ENSG00000151748 | SAV1 | 1.563265394 | 0.087167958 | 0.216625307 | -0.129457349 | 0.999933887 | 0.463921358 | 0.766937069 | -0.303015711 | 0.15404243 |
| ENSG00000251127 | AC091173.1 | 3.878099471 | 0.186001847 | 0.115971139 | 0.070030708 | 0.999933887 | 1.333414354 | 1.684441882 | -0.351027529 | 0.154227046 |
| ENSG00000041988 | THAP3 | 1.95074799 | -0.146466958 | -0.280729935 | 0.134262977 | 0.999933887 | -0.105169321 | -0.34025731 | 0.235087989 | 0.154290925 |
| ENSG00000173918 | C1QTNF1 | -1.95464785 | -0.245324469 | 1.227233421 | -1.47255789 | 0.999933887 | 1.389338773 | 2.048938096 | -0.659599323 | 0.154290925 |
| ENSG00000005339 | CREBBP | 9.07465888 | 0.01301701 | 0.055141958 | -0.042124948 | 0.999933887 | 0.142783755 | 0.253549032 | -0.110765276 | 0.154290925 |
| ENSG00000047634 | SCML1 | 3.685666347 | -0.012169954 | -0.143482909 | 0.131312955 | 0.999933887 | 0.263988876 | 0.64597006 | -0.381981184 | 0.154290925 |
| ENSG00000104408 | EIF3E | 5.177948861 | -0.091901411 | -0.168020589 | 0.076119178 | 0.999933887 | 0.023322058 | 0.245786228 | -0.22246417 | 0.154315413 |
| ENSG00000156515 | HK1 | 5.143489744 | 0.00934078 | -0.015477108 | 0.024817889 | 0.999933887 | -0.089716563 | -0.238144214 | 0.148427652 | 0.154315413 |
| ENSG00000247828 | TMEM161B-A | 1.880861043 | 0.046217251 | -0.286979613 | 0.333196865 | 0.999933887 | -0.319242289 | 0.117936384 | -0.437178674 | 0.154347596 |
| ENSG00000259807 | AC009093.1 | -0.44836692 | -0.041817265 | -0.184559782 | 0.142742516 | 0.999933887 | -0.194503638 | 0.461110267 | -0.655613905 | 0.154365792 |
| ENSG00000198836 | OPA1 | 4.224676961 | 0.016086383 | -0.056679371 | 0.072765754 | 0.999933887 | -0.229767758 | 0.10723014 | -0.336997897 | 0.15480416 |
| ENSG00000100429 | HDAC10 | 0.646843116 | 0.110474341 | -0.011525399 | 0.121999739 | 0.999933887 | 0.010971574 | -0.447321645 | 0.458293219 | 0.15482473 |
| ENSG00000100258 | LMF2 | 6.226127874 | 0.048660097 | 0.057130157 | -0.008470061 | 0.999933887 | -0.03578996 | -0.188155283 | 0.152365323 | 0.154970282 |
| ENSG00000255801 | AC092746.1 | 1.394257435 | 0.019160521 | -0.009191044 | 0.028351565 | 0.999933887 | 1.456565728 | 1.881262933 | -0.424697205 | 0.154970282 |
| ENSG00000092201 | SUPT16H | 4.685347861 | -0.087112192 | -0.226182846 | 0.139070654 | 0.999933887 | -0.203205232 | -0.365023944 | 0.161818712 | 0.155038369 |
| ENSG00000147168 | IL2RG | 6.762997652 | 0.042140148 | 0.067793001 | -0.025652853 | 0.999933887 | 0.522407263 | 0.670185422 | -0.147778159 | 0.155038369 |
| ENSG00000187595 | ZNF385C | -2.18342944 | -0.030292859 | -0.098525627 | 0.068232768 | 0.999933887 | 2.588301905 | 3.806247951 | -1.217946046 | 0.155038369 |
| ENSG00000018189 | RUFY3 | 2.040368243 | -0.034657498 | -0.036295122 | 0.001637624 | 0.999933887 | -0.118425193 | 0.164007441 | -0.282432634 | 0.155114557 |
| ENSG00000266338 | NBPF15 | 2.826045287 | -0.014248325 | -0.024786482 | 0.010538157 | 0.999933887 | 0.169203645 | 0.416690078 | -0.247486433 | 0.15513726 |
| ENSG00000271335 | AL117336.2 | -0.659708531 | -0.03010294 | 0.257710781 | -0.287813722 | 0.999933887 | -0.417632668 | -1.038948647 | 0.621315979 | 0.15513726 |
| ENSG00000077254 | USP33 | 4.517318309 | 0.005449516 | -0.01985044 | 0.025299956 | 0.999933887 | -0.337666302 | -0.563950478 | 0.226284176 | 0.15513726 |
| ENSG00000103202 | NME4 | 2.716243728 | 0.074165161 | 0.016036722 | 0.058128439 | 0.999933887 | 0.006598049 | -0.240380568 | 0.246978617 | 0.155144254 |
| ENSG00000133997 | MED6 | 3.294227428 | -0.034260593 | 0.00966443 | -0.043925023 | 0.999933887 | -0.117467911 | 0.05169929 | -0.169167201 | 0.155144254 |
| ENSG00000091181 | IL5RA | 2.501150156 | -0.047341981 | 0.113288668 | -0.160630649 | 0.999933887 | -0.084557564 | 0.137031143 | -0.221588707 | 0.155221971 |
| ENSG00000167291 | TBC1D16 | 0.974213991 | 0.023226386 | 0.084776977 | -0.061550591 | 0.999933887 | 0.920245079 | 1.35443853 | -0.434193451 | 0.155327453 |
| ENSG00000264304 | AC024267.3 | 0.349406715 | -0.007961114 | 0.001765034 | -0.009726148 | 0.999933887 | 0.605748611 | 1.254219186 | -0.648470575 | 0.155327453 |
| ENSG00000168476 | REEP4 | 3.005349507 | 0.013288897 | 0.012702384 | 0.000586514 | 0.999933887 | -0.075622408 | -0.338572021 | 0.262949613 | 0.155477574 |
| ENSG00000204632 | HLA-G | 4.029116289 | 0.084878431 | 0.002572141 | 0.082306291 | 0.999933887 | 0.369554809 | -0.058701957 | 0.428256766 | 0.155504213 |
| ENSG00000165891 | E2F7 | -0.261287748 | -0.046421309 | 0.088666731 | -0.13508804 | 0.999933887 | 0.438500981 | 1.033397873 | -0.594896892 | 0.155524118 |
| ENSG00000179981 | TSHZ1 | 4.860394747 | -0.054344968 | 0.006175215 | -0.060520183 | 0.999933887 | -0.972771009 | -1.215728194 | 0.242957185 | 0.155550206 |
| ENSG00000224397 | PELATON | 7.620070107 | 0.043806584 | 0.026847143 | 0.016959441 | 0.999933887 | 0.432356074 | 0.585752751 | -0.153396677 | 0.155569838 |
| ENSG00000131355 | ADGRE3 | 7.77703752 | -0.026165933 | -0.016402108 | -0.009763825 | 0.999933887 | -0.850618601 | -1.047633364 | 0.197014763 | 0.155569838 |
| ENSG00000136810 | TXN | 6.690016368 | -0.011809805 | -0.022832305 | 0.0110225 | 0.999933887 | 0.580951686 | 0.755161859 | -0.174210173 | 0.155621151 |
| ENSG00000164022 | AIMP1 | 3.14102872 | -0.094980914 | -0.106394342 | 0.011413427 | 0.999933887 | -0.109223696 | 0.132067323 | -0.241291019 | 0.155927184 |
| ENSG00000087206 | UIMC1 | 5.902255081 | -0.039474593 | -0.027952259 | -0.011522334 | 0.999933887 | -0.707375589 | -0.862135956 | 0.154760367 | 0.155927184 |
| ENSG00000189042 | ZNF567 | 0.370423957 | -0.136636615 | -0.311551981 | 0.174915366 | 0.999933887 | -0.053820223 | 0.470745589 | -0.524565812 | 0.156028452 |
| ENSG00000196460 | RFX8 | -0.715842515 | 0.278384773 | 0.141795824 | 0.136588949 | 0.999933887 | 2.674441585 | 3.366557712 | -0.692116127 | 0.156368636 |
| ENSG00000196628 | TCF4 | 0.673379441 | -0.078693869 | 0.054070535 | -0.132764404 | 0.999933887 | -0.450191378 | -0.016546556 | -0.433644821 | 0.156368636 |
| ENSG00000160208 | RRP1B | 3.2357012 | -0.070371261 | -0.045065779 | -0.025305482 | 0.999933887 | 0.221545741 | 0.41666163 | -0.195115888 | 0.156394541 |
| ENSG00000204186 | ZDBF2 | 2.05327961 | -0.110411882 | -0.196275524 | 0.085863642 | 0.999933887 | -0.223313442 | 0.345803119 | -0.569116561 | 0.156394541 |
| ENSG00000179362 | HMGN2P46 | -1.270099294 | -0.237811123 | -0.419797901 | 0.181986778 | 0.999933887 | 1.807241129 | 2.54231196 | -0.735070831 | 0.156414972 |
| ENSG00000116539 | ASH1L | 6.242810068 | -0.040374979 | -0.053699799 | 0.013324819 | 0.999933887 | -0.271494354 | -0.013428117 | -0.258066236 | 0.156420525 |
| ENSG00000168283 | BMI1 | 2.866783005 | 0.016142493 | -0.005820737 | 0.02196323 | 0.999933887 | -0.275885663 | -0.005038806 | -0.270846857 | 0.156429367 |
| ENSG00000178695 | KCTD12 | 4.969390161 | -0.198750392 | -0.198503916 | -0.000246476 | 0.999933887 | -1.015567247 | -1.311927711 | 0.296360464 | 0.156520031 |
| ENSG00000185561 | TLCD2 | 1.388787297 | -0.019390551 | 0.040480747 | -0.059871298 | 0.999933887 | -0.086787395 | 0.225867235 | -0.31265463 | 0.156827783 |
| ENSG00000087338 | GMCL1 | 3.864549791 | 0.054920533 | 7.6742e-05 | 0.054843791 | 0.999933887 | -0.422874248 | -0.647652181 | 0.224777933 | 0.156929443 |
| ENSG00000237596 | AL138828.1 | -1.366475113 | 0.611443938 | 0.782307903 | -0.170863964 | 0.999933887 | 1.03247625 | 1.526944803 | -0.494468553 | 0.157028935 |
| ENSG00000103035 | PSMD7 | 4.66242009 | -0.059121326 | -0.128217954 | 0.069096628 | 0.999933887 | -0.036856997 | 0.103317436 | -0.140174433 | 0.157028935 |
| ENSG00000177868 | SVBP | 2.145013755 | -0.098173815 | -0.100780532 | 0.002606717 | 0.999933887 | -0.29171672 | -0.005962299 | -0.285754421 | 0.157278212 |
| ENSG00000157873 | TNFRSF14 | 10.030765685 | 0.047919465 | 0.058877755 | -0.01095829 | 0.999933887 | 0.670235919 | 0.824220639 | -0.153984721 | 0.157278212 |
| ENSG00000135951 | TSGA10 | 0.670992292 | 0.027391532 | 0.073280815 | -0.045889283 | 0.999933887 | 0.46962932 | 0.845137924 | -0.375508604 | 0.15730306 |
| ENSG00000141367 | CLTC | 8.024607357 | -0.118822703 | -0.143035243 | 0.02421254 | 0.999933887 | -0.0783 | 0.111258642 | -0.189558643 | 0.157395135 |
| ENSG00000214706 | IFRD2 | 2.348946332 | 0.020211129 | -0.069383034 | 0.089594163 | 0.999933887 | 0.665540229 | 0.948013366 | -0.282473137 | 0.157404456 |
| ENSG00000286940 | MICB-DT | 3.041256122 | 0.02156592 | 0.029987668 | -0.008421748 | 0.999933887 | 0.993359037 | 1.274442146 | -0.28108311 | 0.157534564 |
| ENSG00000069667 | RORA | 5.224014197 | -0.111580684 | -0.117101547 | 0.005520863 | 0.999933887 | -0.293750872 | 0.124357058 | -0.41810793 | 0.158052651 |
| ENSG00000116095 | PLEKHA3 | 3.463683804 | 0.003164076 | 0.104807712 | -0.101643635 | 0.999933887 | -0.251817649 | -0.454894424 | 0.203076775 | 0.158052651 |
| ENSG00000171497 | PPID | 4.23761206 | -0.14178747 | -0.119071574 | -0.022715896 | 0.999933887 | 0.180273521 | 0.337768444 | -0.157494922 | 0.158099731 |
| ENSG00000186265 | BTLA | 0.678265216 | -0.11603346 | -0.489204884 | 0.373171424 | 0.999933887 | -0.014553071 | 0.351356447 | -0.365909518 | 0.158205299 |
| ENSG00000239948 | RN7SL368P | 3.55141622 | 0.286173169 | 0.160684242 | 0.125488927 | 0.999933887 | 1.322763319 | 1.807406303 | -0.484642984 | 0.158580596 |
| ENSG00000160712 | IL6R | 9.268635389 | -0.032447104 | -0.031001381 | -0.001445723 | 0.999933887 | -0.205172332 | -0.038959103 | -0.166213229 | 0.158580596 |

| | | | | | | | | | | |
|---|---|---|---|---|---|---|---|---|---|---|
| ENSG00000119720 | *NRDE2* | 3.836205892 | -0.081261059 | -0.051178732 | -0.030082327 | 0.999933887 | -0.231096543 | -0.404782801 | 0.173686259 | 0.158663911 |
| ENSG00000155287 | *SLC25A28* | 4.407152903 | 0.118563931 | 0.051941042 | 0.066622889 | 0.999933887 | 0.024085349 | -0.2291079 | 0.253193249 | 0.158800929 |
| ENSG00000147905 | *ZCCHC7* | 2.947000803 | -0.020170102 | -0.116937054 | 0.096766952 | 0.999933887 | -0.01045924 | -0.211389527 | 0.200930287 | 0.158800929 |
| ENSG00000167461 | *RAB8A* | 4.725745595 | -0.02760791 | -0.006220097 | -0.021387813 | 0.999933887 | -0.326066592 | -0.502813538 | 0.176746945 | 0.159221423 |
| ENSG00000027847 | *B4GALT7* | 2.702976481 | 0.007016999 | 0.230346861 | -0.223329863 | 0.999933887 | 0.51968674 | 0.76583257 | -0.24614583 | 0.159413832 |
| ENSG00000050405 | *LIMA1* | 0.758146415 | -0.012214196 | 0.068661015 | -0.08087521 | 0.999933887 | -0.13351771 | 0.249469871 | -0.382987581 | 0.159466978 |
| ENSG00000157551 | *KCNJ15* | 8.804238678 | -0.048968082 | -0.033371542 | -0.01559654 | 0.999933887 | -0.531881846 | -0.369396346 | -0.162485499 | 0.15967798 |
| ENSG00000111711 | *GOLT1B* | 2.974906168 | -0.126357353 | -0.114335797 | -0.012021557 | 0.999933887 | -0.225687628 | 0.0362114 | -0.261899028 | 0.159975042 |
| ENSG00000184106 | *TREML3P* | 5.075255847 | 0.014772457 | -0.043827023 | 0.05859948 | 0.999933887 | 0.558227194 | 0.75171268 | -0.193485486 | 0.160066378 |
| ENSG00000158480 | *SPATA2* | 4.85597875 | 0.066226163 | 0.00632606 | 0.059900103 | 0.999933887 | 0.370873223 | 0.544307026 | -0.173433803 | 0.160171466 |
| ENSG00000185477 | *GPRIN3* | 4.744285639 | -0.024839705 | -0.254675828 | 0.229836123 | 0.999933887 | -0.333077954 | 0.07126583 | -0.404343784 | 0.160171466 |
| ENSG00000233276 | *GPX1* | 6.170466486 | -0.054587908 | -0.06788634 | 0.013298431 | 0.999933887 | -0.122465676 | -0.339259677 | 0.216794001 | 0.160182584 |
| ENSG00000116205 | *TCEANC2* | 2.59467553 | 0.094413595 | -0.064747868 | 0.159161464 | 0.999933887 | -0.013410946 | -0.20902158 | 0.195610635 | 0.16021872 |
| ENSG00000128271 | *ADORA2A* | 4.754548031 | 0.31873947 | 0.182726043 | 0.136013427 | 0.999933887 | 1.989422872 | 2.381790948 | -0.392368075 | 0.160318555 |
| ENSG00000007520 | *TSR3* | 3.776125621 | 0.054410988 | 0.086634107 | -0.032223119 | 0.999933887 | -0.138099942 | -0.376940164 | 0.238840222 | 0.160318555 |
| ENSG00000108932 | *SLC16A6* | 5.137145824 | -0.137726838 | -0.040984748 | -0.09674209 | 0.999933887 | -0.148468827 | 0.054445931 | -0.202914758 | 0.160477575 |
| ENSG00000131263 | *RLIM* | 6.870254875 | -0.012323865 | -0.089125015 | 0.07680115 | 0.999933887 | -0.169070457 | 0.069472452 | -0.238542909 | 0.160523818 |
| ENSG00000163882 | *POLR2H* | 1.921291022 | 0.012773263 | -0.003293614 | 0.016066877 | 0.999933887 | 0.15472329 | -0.112242159 | 0.266965449 | 0.160523818 |
| ENSG00000223722 | *IFITM3P2* | 2.129389531 | 0.225343406 | 0.046343482 | 0.178999924 | 0.999933887 | 1.180059664 | 1.536758264 | -0.356698599 | 0.160556796 |
| ENSG00000253194 | *AL137009.1* | -0.125747972 | -0.063142635 | -0.05449761 | -0.008645024 | 0.999933887 | -0.218889069 | 0.549101347 | -0.767990416 | 0.160669596 |
| ENSG00000165525 | *NEMF* | 4.406900757 | 0.002042069 | -0.110488986 | 0.112531056 | 0.999933887 | -0.335949137 | 0.045686408 | -0.381635545 | 0.160678219 |
| ENSG00000102100 | *SLC35A2* | 4.81232209 | 0.037279955 | 0.066296703 | -0.029016748 | 0.999933887 | 1.519839992 | 1.826923554 | -0.307083562 | 0.160737302 |
| ENSG00000169499 | *PLEKHA2* | 7.738224032 | -0.083798043 | 0.001466979 | -0.085265023 | 0.999933887 | -0.321345534 | -0.158955767 | -0.162389767 | 0.160904192 |
| ENSG00000168685 | *IL7R* | 7.007448576 | -0.005164961 | -0.114242595 | 0.109077634 | 0.999933887 | 0.122487692 | 0.25519128 | -0.132703588 | 0.161332093 |
| ENSG00000072694 | *FCGR2B* | 3.02994491 | 0.022213818 | -0.0869438 | 0.109157617 | 0.999933887 | -0.159700258 | -0.423540802 | 0.263840544 | 0.16160197 |
| ENSG00000108465 | *CDK5RAP3* | 3.998276691 | -0.005234761 | 0.035896686 | -0.041131446 | 0.999933887 | 0.023887012 | -0.157091508 | 0.18097852 | 0.161804543 |
| ENSG00000162600 | *OMA1* | 2.002704132 | -0.046527756 | -0.042426549 | -0.004101206 | 0.999933887 | -0.062057924 | -0.316733417 | 0.254675493 | 0.161805694 |
| ENSG00000090273 | *NUDC* | 4.277730574 | -0.002312824 | -0.001061773 | -0.001251052 | 0.999933887 | -0.209061669 | -0.411733398 | 0.202671729 | 0.161805694 |
| ENSG00000211689 | *TRGC1* | 2.150069979 | -0.108510016 | -0.024596059 | -0.083913957 | 0.999933887 | 0.029068701 | 0.301197817 | -0.272129116 | 0.161823148 |
| ENSG00000031698 | *SARS1* | 4.903998316 | -0.017348021 | -0.009075557 | -0.008272463 | 0.999933887 | -0.040821219 | -0.189261121 | 0.148439902 | 0.162128337 |
| ENSG00000103264 | *FBXO31* | 2.924085817 | 0.030657978 | 0.043677905 | -0.013019928 | 0.999933887 | 0.071707593 | -0.141231891 | 0.212939484 | 0.16216114 |
| ENSG00000267680 | *ZNF224* | 2.633110261 | 0.027953805 | -0.114639644 | 0.142593449 | 0.999933887 | -0.239259784 | -0.546661816 | 0.307402032 | 0.16216114 |
| ENSG00000213337 | *ANKRD39* | 0.193415555 | -0.125899301 | -0.208969689 | 0.083070387 | 0.999933887 | -0.122949757 | -0.596021656 | 0.473071899 | 0.162592754 |
| ENSG00000136147 | *PHF11* | 5.76386695 | -0.029253179 | -0.03378273 | 0.004529551 | 0.999933887 | -0.175556312 | -0.051316926 | -0.124239386 | 0.162592754 |
| ENSG00000095321 | *CRAT* | 1.42944817 | -0.034750817 | 0.029965145 | -0.064715961 | 0.999933887 | -0.067193385 | -0.39064172 | 0.323448335 | 0.162657362 |
| ENSG00000225111 | *AC083900.1* | -0.010255455 | 0.077713752 | 0.350702673 | -0.272988921 | 0.999933887 | 1.704944152 | 2.122024712 | -0.417080561 | 0.162692383 |
| ENSG00000188735 | *TMEM120B* | 2.795989894 | 0.048348207 | 0.15185605 | -0.103507843 | 0.999933887 | 1.517554251 | 1.859325196 | -0.341770945 | 0.162829139 |
| ENSG00000225963 | *AC009950.1* | 2.11078961 | -0.052205041 | -0.047297813 | -0.004907228 | 0.999933887 | -0.788741936 | -1.153720805 | 0.364978869 | 0.163066226 |
| ENSG00000179041 | *RRS1* | 1.249755979 | -0.016569068 | 0.018191118 | -0.034760187 | 0.999933887 | 0.206796793 | 0.54475004 | -0.337953246 | 0.163080517 |
| ENSG00000163597 | *SNHG16* | 4.075314831 | -0.066184828 | -0.087198183 | 0.021013355 | 0.999933887 | -0.237785348 | -0.453216213 | 0.215430865 | 0.163225906 |
| ENSG00000105223 | *PLD3* | 4.513436463 | 0.118243013 | -0.060783396 | 0.17902641 | 0.944094537 | -0.058317632 | -0.227992105 | 0.169674474 | 0.163285189 |
| ENSG00000273117 | *AC144652.1* | 1.976465754 | 0.213623652 | 0.126133137 | 0.087490515 | 0.999933887 | 0.881895829 | 1.301334822 | -0.419438993 | 0.163285189 |
| ENSG00000248734 | *AC008906.1* | 1.126952167 | -0.102212457 | 0.035871778 | -0.138084235 | 0.999933887 | -0.618662844 | -0.104592072 | -0.514070772 | 0.163285189 |
| ENSG00000278017 | *AC064801.1* | 0.362519152 | -0.067490024 | 0.030129872 | -0.097619896 | 0.999933887 | 0.365873319 | 0.877452301 | -0.511578981 | 0.163474065 |
| ENSG00000254887 | *AC010247.1* | 0.428779697 | -0.013879049 | 0.436844457 | -0.450723506 | 0.999933887 | 0.385699636 | 0.719584332 | -0.333884696 | 0.163563378 |
| ENSG00000275074 | *NUDT18* | 2.058544928 | 0.027822096 | -0.010033389 | 0.037855485 | 0.999933887 | -0.159805067 | -0.433535047 | 0.273729979 | 0.163638522 |
| ENSG00000139496 | *NUP58* | 6.263129368 | -0.023220127 | -0.027773901 | 0.004553774 | 0.999933887 | 0.211090134 | 0.412721571 | -0.201631437 | 0.163696317 |
| ENSG00000235944 | *ZNF815P* | 1.062797218 | 0.152730718 | 0.290899657 | -0.13816894 | 0.999933887 | 0.117319008 | -0.275541913 | 0.392860921 | 0.163721183 |
| ENSG00000068308 | *OTUD5* | 7.280536145 | 0.037633207 | 0.020561421 | 0.017071787 | 0.999933887 | 0.418434224 | 0.56630929 | -0.147875066 | 0.163788969 |
| ENSG00000185344 | *ATP6V0A2* | 3.645483132 | -0.020376863 | -0.018674745 | -0.001702118 | 0.999933887 | -0.093023699 | -0.29740685 | 0.20438315 | 0.163788969 |
| ENSG00000183696 | *UPP1* | 6.83658304 | -0.038523464 | -0.004891202 | -0.033632262 | 0.999933887 | 0.739836551 | 0.910476463 | -0.170639912 | 0.163950339 |
| ENSG00000198668 | *CALM1* | 7.448914695 | 0.015479078 | -0.070034931 | 0.085514009 | 0.999933887 | -0.023892593 | -0.154812084 | 0.13091949 | 0.163950339 |
| ENSG00000100142 | *POLR2F* | 3.209193017 | 0.04670558 | -0.011998846 | 0.058704426 | 0.999933887 | -0.161836422 | -0.398565675 | 0.236729253 | 0.16410975 |
| ENSG00000106400 | *ZNHIT1* | 4.533475458 | 0.004056347 | 0.033424672 | -0.029368325 | 0.999933887 | -0.198774468 | -0.372450382 | 0.173675914 | 0.16410975 |
| ENSG00000143947 | *RPS27A* | 7.768833755 | -0.038940419 | -0.051562504 | 0.012622085 | 0.999933887 | 0.18207037 | 0.33799942 | -0.15592905 | 0.164141745 |
| ENSG00000215717 | *TMEM167B* | 6.812054554 | -0.056870274 | -0.044604698 | -0.012265575 | 0.999933887 | 0.012750839 | 0.143777486 | -0.131026647 | 0.164219168 |
| ENSG00000198492 | *YTHDF2* | 4.307808443 | -0.043852592 | -0.1011873 | 0.057334708 | 0.999933887 | -0.27784318 | -0.445866177 | 0.168022997 | 0.164348382 |
| ENSG00000067829 | *IDH3G* | 4.144783375 | 0.07618991 | -0.003585585 | 0.079775495 | 0.999933887 | -0.238005735 | -0.404196692 | 0.166190958 | 0.164366048 |
| ENSG00000204389 | *HSPA1A* | 5.551846764 | 0.038676375 | 0.032205551 | 0.006470824 | 0.999933887 | 0.69334642 | 1.030084026 | -0.336737606 | 0.164852794 |
| ENSG00000146834 | *MEPCE* | 6.745041824 | 0.096606901 | 0.117481429 | -0.020874527 | 0.999933887 | 0.225277576 | 0.384794971 | -0.159517395 | 0.165055845 |
| ENSG00000034510 | *TMSB10* | 7.847556443 | 0.011431077 | -0.064823381 | 0.076254458 | 0.999933887 | 0.032991443 | -0.118218616 | 0.151210058 | 0.165055845 |
| ENSG00000103507 | *BCKDK* | 5.157297172 | -0.022511054 | 0.019167232 | -0.041678287 | 0.999933887 | -0.658758007 | -0.849577846 | 0.190819839 | 0.165074964 |
| ENSG00000198786 | *MT-ND5* | 8.288225626 | 0.009004829 | -0.029462681 | 0.038467509 | 0.999933887 | 0.024484933 | 0.249308807 | -0.224823873 | 0.165074964 |
| ENSG00000108094 | *CUL2* | 3.817796161 | -0.098255811 | -0.127170332 | 0.028914522 | 0.999933887 | -0.351059382 | -0.126961511 | -0.224097872 | 0.165308352 |
| ENSG00000093010 | *COMT* | 2.026381423 | -0.029922311 | 0.178344312 | -0.208266623 | 0.999933887 | 0.02732074 | -0.213060854 | 0.240381594 | 0.165308352 |
| ENSG00000176953 | *NFATC2IP* | 5.187939059 | -0.048496882 | -0.002973182 | -0.045523701 | 0.999933887 | 0.079692606 | 0.222299853 | -0.142607247 | 0.165340798 |
| ENSG00000181450 | *ZNF678* | 0.419191058 | -0.056810807 | -0.261145344 | 0.204334537 | 0.999933887 | -0.290840793 | 0.250483627 | -0.54132442 | 0.165519511 |
| ENSG00000068903 | *SIRT2* | 6.316457954 | 0.025235606 | 0.00434272 | 0.020892886 | 0.999933887 | 0.266667438 | 0.394364459 | -0.127697021 | 0.165540782 |
| ENSG00000279133 | *AC018628.1* | 1.437137204 | -0.048769566 | 0.114379111 | -0.163148677 | 0.999933887 | 0.223689871 | 0.658319438 | -0.434629567 | 0.165540782 |
| ENSG00000004961 | *HCCS* | 2.58428543 | 0.021642651 | 0.058425461 | -0.036782809 | 0.999933887 | 0.046484222 | -0.170765942 | 0.217250164 | 0.165987102 |
| ENSG00000140941 | *MAP1LC3B* | 8.590791287 | 0.037890441 | 0.002209295 | 0.035681146 | 0.999933887 | 0.032676359 | 0.154264305 | -0.121587946 | 0.166040855 |
| ENSG00000103245 | *CIAO3* | 2.161629858 | 0.052034934 | 0.084300988 | -0.032266053 | 0.999933887 | 0.084435584 | -0.17657361 | 0.261009194 | 0.166067792 |
| ENSG00000170075 | *GPR37L1* | 1.033503484 | -0.104099597 | -0.00775678 | -0.096342817 | 0.999933887 | -0.437628156 | -0.074294351 | -0.363333806 | 0.166077014 |
| ENSG00000123836 | *PFKFB2* | 5.324714618 | -0.012049048 | 0.064652259 | -0.076701307 | 0.999933887 | 0.040088355 | -0.188818579 | 0.228906934 | 0.166132388 |
| ENSG00000132434 | *LANCL2* | 1.985340944 | -0.033471258 | -0.110953879 | 0.077482621 | 0.999933887 | 0.368575482 | 0.639705936 | -0.271130454 | 0.166374001 |
| ENSG00000274528 | *AC090970.2* | 2.2101638 | 0.200183731 | 0.152018189 | 0.048165542 | 0.999933887 | 2.297669292 | 2.675088304 | -0.377419012 | 0.166402414 |
| ENSG00000008517 | *IL32* | 6.062189682 | 0.021446175 | -0.011033686 | 0.032479862 | 0.999933887 | 0.260576276 | 0.413080643 | -0.152504367 | 0.166402414 |
| ENSG00000226312 | *CFLAR-AS1* | 4.0942492 | -0.101022558 | -0.083457962 | -0.017564596 | 0.999933887 | 1.244516426 | 1.555397472 | -0.310881046 | 0.166885271 |
| ENSG00000111875 | *ASF1A* | 1.729746154 | -0.095156135 | -0.021912072 | -0.073244063 | 0.999933887 | -0.098907767 | 0.195542844 | -0.294450611 | 0.166932868 |
| ENSG00000196642 | *RABL6* | 4.340029741 | 0.062117072 | 0.088685019 | -0.026567947 | 0.999933887 | -0.029064837 | -0.219044063 | 0.189979226 | 0.167195941 |
| ENSG00000120594 | *PLXDC2* | 6.222109369 | -0.036358907 | -0.033128987 | -0.00322992 | 0.999933887 | -0.86866252 | -1.100213449 | 0.231550929 | 0.167195941 |
| ENSG00000099377 | *HSD3B7* | 1.737541196 | 0.048723223 | 0.085726093 | -0.03700287 | 0.999933887 | -0.840546821 | -0.53739554 | -0.30315128 | 0.167195941 |
| ENSG00000150776 | *NKAPD1* | 2.469714552 | -0.012349214 | -0.038475023 | 0.026125809 | 0.999933887 | -0.067016823 | -0.286465476 | 0.219448653 | 0.167195941 |
| ENSG00000168765 | *GSTM4* | 2.154164385 | -0.004834312 | -0.039964383 | 0.035130071 | 0.999933887 | -0.813940895 | -1.20902836 | 0.395087465 | 0.167195941 |
| ENSG00000093167 | *LRRFIP2* | 5.327372322 | -0.056337819 | -0.047704098 | -0.008633721 | 0.999933887 | 0.093721164 | 0.234241278 | -0.140520114 | 0.167398057 |
| ENSG00000122779 | *TRIM24* | 3.674631016 | -0.005369875 | 0.005901876 | -0.01127175 | 0.999933887 | -0.774312419 | -1.007790609 | 0.23347819 | 0.167591722 |
| ENSG00000049769 | *PPP1R3F* | 0.357697655 | 0.170822157 | 0.081370888 | 0.089451269 | 0.999933887 | -0.063797986 | -0.466732223 | 0.402934237 | 0.167986553 |
| ENSG00000109323 | *MANBA* | 6.528274168 | -0.043159443 | -0.077960817 | 0.034801374 | 0.999933887 | -0.111028959 | 0.020815479 | -0.131844438 | 0.167986553 |
| ENSG00000105053 | *VRK3* | 5.283668129 | 0.01391266 | 0.005280814 | 0.008631845 | 0.999933887 | -0.171080358 | -0.028622065 | -0.142458293 | 0.167986553 |
| ENSG00000265972 | *TXNIP* | 10.512277695 | -0.17702616 | -0.197161798 | 0.020135638 | 0.999933887 | -1.03258688 | -1.354015515 | 0.321428635 | 0.168037676 |
| ENSG00000254236 | *AP003550.1* | 0.024215259 | 0.300718149 | 0.780841915 | -0.480123766 | 0.999933887 | 0.933927519 | 1.442062065 | -0.508134546 | 0.168152777 |
| ENSG00000170293 | *CMTM8* | 0.040564734 | 0.181903914 | 0.165601609 | 0.016302305 | 0.999933887 | -0.309279019 | -0.710702607 | 0.401423588 | 0.168184299 |
| ENSG00000141560 | *FN3KRP* | 2.016283148 | -0.03716632 | -0.111219911 | 0.074053591 | 0.999933887 | 0.050005094 | 0.293095296 | -0.243090201 | 0.168496568 |
| ENSG00000234338 | *AC073349.2* | -1.528467227 | 0.005325493 | 0.032345813 | -0.02702032 | 0.999933887 | -0.223166978 | 0.578394364 | -0.801561342 | 0.168496568 |
| ENSG00000125458 | *NT5C* | 2.025674963 | 0.103247971 | 0.074707996 | 0.028539975 | 0.999933887 | -0.243042021 | -0.536484573 | 0.293442552 | 0.168625766 |
| ENSG00000132670 | *PTPRA* | 4.744921874 | 0.037856943 | 0.012997131 | 0.024859812 | 0.999933887 | -0.113568224 | -0.287171249 | 0.173603026 | 0.168632907 |
| ENSG00000075223 | *SEMA3C* | 1.668887345 | -0.075022802 | -0.241659107 | 0.166636305 | 0.999933887 | -1.002752159 | -1.372392505 | 0.369640345 | 0.168632907 |
| ENSG00000278231 | *AL133342.1* | 3.452509851 | -0.033275498 | 0.004501708 | -0.037777205 | 0.999933887 | -0.235408509 | -0.426307502 | 0.190898993 | 0.168632907 |
| ENSG00000198586 | *TLK1* | 5.529394472 | -0.031533926 | -0.044791821 | 0.013257895 | 0.999933887 | -0.505003718 | -0.299012634 | -0.205991085 | 0.168632907 |
| ENSG00000139505 | *MTMR6* | 5.728584314 | -0.066083066 | -0.116541927 | 0.050458861 | 0.999933887 | -0.332328988 | -0.096863347 | -0.235465641 | 0.168719835 |
| ENSG00000185596 | *WASH3P* | 2.70463859 | 0.013962856 | 0.134912447 | -0.120949591 | 0.999933887 | -0.107250106 | -0.333206423 | 0.225956316 | 0.168719835 |
| ENSG00000203999 | *LINC01270* | 3.730721436 | 0.031994255 | -0.005764908 | 0.037759164 | 0.999933887 | 0.5881969 | 0.795556364 | -0.207359464 | 0.168853464 |
| ENSG00000254901 | *BORCS8* | 4.001338622 | -0.07176352 | 0.00453343 | -0.07629695 | 0.999933887 | -0.47241228 | -0.646850246 | 0.174437966 | 0.168854627 |
| ENSG00000153975 | *ZUP1* | 2.107948158 | -0.068455187 | -0.08978608 | 0.021330893 | 0.999933887 | 0.327366788 | 0.617116291 | -0.289749503 | 0.168854627 |
| ENSG00000132463 | *GRSF1* | 3.384127107 | 0.037791937 | -0.040226614 | 0.078018551 | 0.999933887 | 0.061448839 | -0.146799643 | 0.208248482 | 0.168854627 |

| | | | | | | | | | | |
|---|---|---|---|---|---|---|---|---|---|---|
| ENSG00000248483 | *POU5F2* | 0.356912665 | 0.230306749 | 0.54674455 | -0.316437801 | 0.999933887 | -0.078699941 | 0.641639865 | -0.720339806 | 0.168886807 |
| ENSG00000170348 | *TMED10* | 5.337252734 | -0.029133123 | 0.022284368 | -0.051417491 | 0.999933887 | -0.420517258 | -0.584433981 | 0.163916723 | 0.168958981 |
| ENSG00000166794 | *PPIB* | 3.830802316 | 0.030919949 | 0.00158061 | 0.02933934 | 0.999933887 | -0.108454279 | -0.307186972 | 0.198732693 | 0.169125077 |
| ENSG00000100023 | *PPIL2* | 4.033603191 | 0.065326025 | 0.08392385 | -0.018597825 | 0.999933887 | -0.095654637 | -0.248887183 | 0.153232547 | 0.169224134 |
| ENSG00000010017 | *RANBP9* | 5.268612133 | 0.014913242 | 0.002386254 | 0.012526988 | 0.999933887 | -0.0140471 | 0.106525316 | -0.120572416 | 0.169436978 |
| ENSG00000170088 | *TMEM192* | 2.796648395 | -0.02951041 | -0.042797372 | 0.013286962 | 0.999933887 | -0.344441077 | -0.152393217 | -0.19204786 | 0.169436978 |
| ENSG00000181163 | *NPM1* | 6.95700912 | -0.023035073 | -0.049137826 | 0.026102753 | 0.999933887 | 0.249595944 | 0.416822562 | -0.167226618 | 0.169622904 |
| ENSG00000260236 | *AC099778.1* | 1.527451659 | 0.154982741 | 0.233368401 | -0.07838566 | 0.999933887 | 0.456430201 | -0.011213855 | 0.467644056 | 0.169807669 |
| ENSG00000036828 | *CASR* | -1.342220193 | -0.215533354 | 0.406629513 | -0.622162867 | 0.999933887 | 1.028319228 | 1.618891059 | -0.59057183 | 0.170141057 |
| ENSG00000203896 | *LIME1* | 0.645662941 | 0.062528037 | -0.090966535 | 0.153494573 | 0.999933887 | -0.199098589 | -0.577995038 | 0.378896449 | 0.170141057 |
| ENSG00000247199 | *AC011346.1* | 0.060381407 | 0.025574829 | 0.140457196 | -0.114882367 | 0.999933887 | 0.714982754 | 1.332345427 | -0.617362673 | 0.170229485 |
| ENSG00000066926 | *FECH* | 1.716388638 | -0.155409073 | -0.106594245 | -0.048814828 | 0.999933887 | -0.349744626 | -0.643193802 | 0.293449175 | 0.170258221 |
| ENSG00000080986 | *NDC80* | 0.44428372 | 0.034606381 | 0.051448696 | -0.016842315 | 0.999933887 | -0.555508535 | -1.074347997 | 0.518839462 | 0.170439923 |
| ENSG00000113282 | *CLINT1* | 5.39119104 | -0.020143585 | -0.072537429 | 0.052393844 | 0.999933887 | 0.014451374 | 0.160941467 | -0.146490093 | 0.17052161 |
| ENSG00000198937 | *CCDC167* | 0.806206901 | 0.016065715 | 0.054657306 | -0.03859159 | 0.999933887 | 0.032871558 | 0.430301868 | -0.39743031 | 0.17052161 |
| ENSG00000143537 | *ADAM15* | 3.27448644 | 0.005463239 | -0.080241346 | 0.085704585 | 0.999933887 | -0.309190624 | -0.551397768 | 0.242207143 | 0.170616731 |
| ENSG00000173208 | *ABCD2* | 0.911082131 | -0.039312768 | -0.274967718 | 0.23565495 | 0.999933887 | -0.066297161 | 0.319940127 | -0.386237289 | 0.170638557 |
| ENSG00000153898 | *MCOLN2* | 2.914891338 | 0.010716467 | -0.014456463 | 0.02517293 | 0.999933887 | 0.094107711 | 0.296303442 | -0.202195731 | 0.170815074 |
| ENSG00000062582 | *MRPS24* | 0.049457478 | 0.064656444 | -0.071334418 | 0.135990862 | 0.999933887 | 2.340626823 | 2.961658755 | -0.621031932 | 0.170890798 |
| ENSG00000174915 | *PTDSS2* | 1.693374924 | 0.126475794 | 0.001449717 | 0.125026077 | 0.999933887 | 0.0459621 | -0.271951332 | 0.317913432 | 0.171143429 |
| ENSG00000168214 | *RBPJ* | 7.469689036 | -0.07110556 | -0.073711231 | 0.00260567 | 0.999933887 | -0.649864043 | -0.460056424 | -0.189807619 | 0.171143429 |
| ENSG00000119906 | *SLF2* | 3.196934159 | -0.049966188 | -0.062202703 | 0.012236515 | 0.999933887 | -0.582287406 | -0.920522646 | 0.33823524 | 0.171143429 |
| ENSG00000260249 | *AC007608.3* | 0.32120543 | -0.035538739 | 0.013123423 | -0.048662161 | 0.999933887 | 0.20126053 | 0.718793218 | -0.517532688 | 0.171143429 |
| ENSG00000279605 | *AC067930.5* | 0.976794767 | -0.00724727 | 0.240667007 | -0.247914277 | 0.999933887 | -0.802176632 | -1.214709658 | 0.412533026 | 0.171189159 |
| ENSG00000187953 | *PMS2CL* | 3.113592145 | 0.070645786 | -0.008811135 | 0.079456921 | 0.999933887 | -0.238582617 | -0.451489159 | 0.212906542 | 0.171295225 |
| ENSG00000284052 | *AC006460.2* | 2.051849237 | 0.233443224 | 0.014156732 | 0.219286491 | 0.999933887 | 0.656574402 | 0.906836537 | -0.250262135 | 0.171446857 |
| ENSG00000106348 | *IMPDH1* | 7.755210333 | 0.03649565 | -0.003103892 | 0.039599542 | 0.999933887 | 0.724230139 | 0.913069374 | -0.188839236 | 0.171467481 |
| ENSG00000151883 | *PARP8* | 7.451058323 | -0.044291597 | -0.09330855 | 0.049016953 | 0.999933887 | -0.521796953 | -0.726671517 | 0.204874564 | 0.171788771 |
| ENSG00000119616 | *FCF1* | 4.411126394 | -0.028639696 | 0.068917348 | -0.097557045 | 0.999933887 | 0.24635988 | 0.411596689 | -0.165236809 | 0.171900575 |
| ENSG00000100926 | *TM9SF1* | 1.292612121 | 0.082119156 | 0.159972264 | -0.077853108 | 0.999933887 | -0.023124866 | -0.340235103 | 0.317110236 | 0.171909809 |
| ENSG00000179454 | *KLHL28* | 4.112935731 | -0.036942749 | -0.119921707 | 0.082978958 | 0.999933887 | -0.411624755 | -0.11291862 | -0.298706134 | 0.172017033 |
| ENSG00000158019 | *BABAM2* | 6.001529588 | 0.167953621 | 0.205390773 | -0.037437152 | 0.999933887 | 0.78001031 | 1.038456503 | -0.258446193 | 0.172124277 |
| ENSG00000197272 | *IL27* | 2.478647467 | 0.106489161 | 0.089435457 | 0.017053704 | 0.999933887 | 0.295262992 | 0.661335411 | -0.366072419 | 0.172280103 |
| ENSG00000108509 | *CAMTA2* | 5.505328765 | 0.043249368 | 0.004301525 | 0.038947842 | 0.999933887 | 0.289484589 | 0.416227794 | -0.126743205 | 0.172280103 |
| ENSG00000136867 | *SLC31A2* | 3.024695837 | 0.045827467 | -0.061187925 | 0.107015392 | 0.999933887 | 0.420486322 | 0.658236247 | -0.237749925 | 0.172280103 |
| ENSG00000228804 | *AC072022.1* | 0.180303678 | -0.023298689 | 0.232459456 | -0.255758145 | 0.999933887 | 0.605994784 | 1.130258484 | -0.5242637 | 0.172280103 |
| ENSG00000116667 | *C1orf21* | 2.826400629 | -0.01195602 | -0.0077624 | -0.00419362 | 0.999933887 | 0.042499081 | 0.246272326 | -0.203773245 | 0.172420599 |
| ENSG00000117242 | *PINK1-AS* | 0.234941419 | 0.117810822 | -0.106219131 | 0.224029954 | 0.999933887 | -0.268780995 | -0.739538991 | 0.470757996 | 0.172639932 |
| ENSG00000134371 | *CDC73* | 4.996110391 | -0.006964085 | -0.019121158 | 0.012157073 | 0.999933887 | -0.29322606 | -0.083344306 | -0.209881754 | 0.172676238 |
| ENSG00000120333 | *MRPS14* | 1.567961948 | 0.063780009 | -0.171528171 | 0.235308179 | 0.999933887 | -0.166672323 | -0.450776648 | 0.284104325 | 0.172734468 |
| ENSG00000144589 | *STK11IP* | 4.304915855 | 0.083258522 | 0.039559997 | 0.043698525 | 0.999933887 | -0.001242584 | -0.165486625 | 0.164244041 | 0.172756613 |
| ENSG00000197021 | *EOLA2* | 3.404578414 | -0.047108409 | -0.021211366 | -0.025897043 | 0.999933887 | -0.021089453 | -0.230462085 | 0.209372632 | 0.172756613 |
| ENSG00000014123 | *UFL1* | 2.894594901 | -0.09075108 | -0.193359211 | 0.10260813 | 0.999933887 | -0.363076156 | 0.065812645 | -0.428888801 | 0.172756613 |
| ENSG00000163743 | *RCHY1* | 4.022364394 | -0.040498886 | -0.149492242 | 0.108993356 | 0.999933887 | 0.043059143 | 0.230925369 | -0.187866226 | 0.172756613 |
| ENSG00000161835 | *TAMALIN* | 2.9853007 | 0.109652561 | 0.131222016 | -0.021569455 | 0.999933887 | 0.383166713 | 0.659032441 | -0.275865728 | 0.172779036 |
| ENSG00000170325 | *PRDM10* | 3.790180812 | -0.032203394 | -0.114669045 | 0.08246565 | 0.999933887 | -0.326609062 | -0.513779238 | 0.187170176 | 0.172855447 |
| ENSG00000176463 | *SLCO3A1* | 6.723549235 | -0.000506741 | 0.012553812 | -0.013060553 | 0.999933887 | -0.026987406 | -0.15369573 | 0.126708324 | 0.172855447 |
| ENSG00000114544 | *SLC41A3* | 2.190289269 | -0.012380377 | 0.038451771 | -0.050832147 | 0.999933887 | 0.124080769 | -0.103486938 | 0.227567707 | 0.173271278 |
| ENSG00000241155 | *ARHGAP31-A* | -0.514177727 | 0.174332697 | -0.369114351 | 0.543447048 | 0.999933887 | 2.812653789 | 3.447450386 | -0.634796596 | 0.17327716 |
| ENSG00000198829 | *SUCNR1* | 5.154505779 | -0.150894327 | -0.165921656 | 0.015027329 | 0.999933887 | -1.29408358 | -0.977962588 | -0.316120992 | 0.173323939 |
| ENSG00000197020 | *ZNF100* | 4.29303886 | -0.060771795 | -0.071568933 | 0.010797138 | 0.999933887 | -0.326822847 | 0.041583357 | -0.368406204 | 0.173411624 |
| ENSG00000079263 | *SP140* | 5.968425512 | 0.023366888 | -0.027628544 | 0.050995431 | 0.999933887 | -0.16952754 | -0.015657986 | -0.153869554 | 0.173519136 |
| ENSG00000287011 | *AC003077.1* | 0.043917954 | -0.10483928 | 0.43430625 | -0.53914553 | 0.999933887 | 0.401170218 | 0.807658062 | -0.406487844 | 0.173519136 |
| ENSG00000197381 | *ADARB1* | 4.067770607 | 0.03143532 | 0.084894418 | -0.053459098 | 0.999933887 | -0.373976772 | -0.530888711 | 0.156911939 | 0.17377709 |
| ENSG00000164007 | *CLDN19* | -0.567447497 | 0.207135868 | 0.574129394 | -0.366993526 | 0.999933887 | 0.297923223 | 0.800340151 | -0.502416928 | 0.173839322 |
| ENSG00000028203 | *VEZT* | 2.441495877 | -0.069666685 | -0.411811341 | 0.342144656 | 0.944094537 | 0.049412403 | 0.328849526 | -0.279437123 | 0.173958452 |
| ENSG00000260367 | *AC109460.1* | 1.258986061 | 0.05133462 | 0.143138601 | -0.091803981 | 0.999933887 | 0.333501512 | 0.66174683 | -0.328245318 | 0.173958452 |
| ENSG00000214331 | *AC009053.1* | 3.348365461 | -0.016972667 | -0.107425117 | 0.09045245 | 0.999933887 | -0.062901453 | 0.131960907 | -0.19486236 | 0.173958452 |
| ENSG00000285921 | *AP000779.1* | -0.971631252 | 0.061969587 | 0.684713904 | -0.622744317 | 0.999933887 | 0.739292834 | 1.322243684 | -0.58295085 | 0.173958452 |
| ENSG00000152683 | *SLC30A6* | 2.760945817 | 0.009425923 | 0.002313608 | 0.007112315 | 0.999933887 | -0.097852164 | 0.085506491 | -0.183358655 | 0.173958452 |
| ENSG00000147162 | *OGT* | 7.990133434 | -0.044930006 | -0.035775475 | -0.009154531 | 0.999933887 | -0.084490675 | -0.210495376 | 0.126004701 | 0.173985245 |
| ENSG00000167863 | *ATP5PD* | 3.729991456 | -0.07639306 | 0.004449876 | -0.080842936 | 0.999933887 | -0.128435788 | -0.31492804 | 0.186492252 | 0.174052063 |
| ENSG00000139324 | *TMTC3* | 0.733129559 | -0.103428292 | -0.37430475 | 0.270876458 | 0.999933887 | -0.334162641 | 0.179799768 | -0.513962408 | 0.174302672 |
| ENSG00000028310 | *BRD9* | 4.983004942 | -0.021330102 | -0.018055744 | -0.003274358 | 0.999933887 | -0.0487657 | -0.190363242 | 0.141597543 | 0.174493198 |
| ENSG00000082153 | *BZW1* | 6.533477806 | -0.049794588 | -0.131612089 | 0.081817501 | 0.999933887 | 0.078981155 | 0.222618591 | -0.143637436 | 0.174525816 |
| ENSG00000008735 | *MAPK8IP2* | -0.578159324 | 0.056772892 | -0.022434843 | 0.079207735 | 0.999933887 | -0.999853997 | -1.535703091 | 0.535849094 | 0.174581823 |
| ENSG00000184988 | *TMEM106A* | 1.726811353 | -0.107851915 | 0.357540199 | -0.465392114 | 0.999933887 | 0.745843981 | 1.055738233 | -0.309894252 | 0.174954733 |
| ENSG00000197599 | *CCDC154* | 3.074701434 | 0.231553512 | 0.330611383 | -0.099057871 | 0.999933887 | 0.56553175 | 0.83578364 | -0.270251889 | 0.175348201 |
| ENSG00000260078 | *AC007342.3* | 4.866318222 | -0.107766711 | -0.113456358 | 0.005689648 | 0.999933887 | -0.600629756 | -0.838264492 | 0.237634735 | 0.175434428 |
| ENSG00000171174 | *RBKS* | 4.783055551 | 0.385312577 | 0.224536636 | 0.160775941 | 0.999933887 | 1.082647059 | 1.6014202 | -0.518773142 | 0.175478339 |
| ENSG00000183283 | *DAZAP2* | 9.956114324 | -0.005797227 | -0.015550024 | 0.009752797 | 0.999933887 | 0.127980123 | 0.249752436 | -0.121772312 | 0.175478339 |
| ENSG00000232774 | *AL355916.1* | -0.477952261 | -0.075459097 | -0.14447892 | 0.069019823 | 0.999933887 | 0.613340653 | 1.264756338 | -0.651415685 | 0.175783702 |
| ENSG00000260271 | *AL132996.1* | -0.28335596 | 0.46792952 | 0.482977756 | -0.015048236 | 0.999933887 | 0.575230906 | 1.002322075 | -0.427091169 | 0.175955849 |
| ENSG00000240230 | *COX19* | 3.582877234 | -0.061250618 | 0.031128451 | -0.092379069 | 0.999933887 | -0.233655969 | -0.437751973 | 0.204096004 | 0.175955849 |
| ENSG00000061987 | *MON2* | 5.651692099 | 0.027151842 | 0.035040409 | -0.007888566 | 0.999933887 | -0.199698772 | 0.023838576 | -0.223537349 | 0.175998577 |
| ENSG00000218739 | *CEBPZOS* | 2.030272082 | -0.002057965 | -0.064202898 | 0.062144933 | 0.999933887 | -0.187919579 | 0.109344509 | -0.297264089 | 0.176111578 |
| ENSG00000135801 | *TAF5L* | 3.16434771 | -0.068715205 | 0.028658634 | -0.097373839 | 0.999933887 | -0.019413885 | -0.202330472 | 0.182916587 | 0.176140951 |
| ENSG00000171101 | *SIGLEC17P* | 0.457197775 | 0.268359015 | -0.136217052 | 0.404576067 | 0.944094537 | -0.017558634 | 0.398210029 | -0.415768663 | 0.176225663 |
| ENSG00000260475 | *AL353719.1* | -0.665367825 | 0.159465204 | 0.227009439 | -0.067544235 | 0.999933887 | 0.016667696 | 0.915574683 | -0.898906987 | 0.176225663 |
| ENSG00000131016 | *AKAP12* | 0.882509179 | 0.171220403 | -0.358395017 | 0.529615421 | 0.999933887 | -1.186380897 | -1.563695999 | 0.377315102 | 0.176225663 |
| ENSG00000214174 | *AMZ2P1* | 4.392394319 | 0.11397349 | 0.036984458 | 0.076989032 | 0.999933887 | 0.503184188 | 0.706496026 | -0.203311838 | 0.176374486 |
| ENSG00000286302 | *AC084724.1* | -1.069427725 | -0.179634575 | -0.028613899 | -0.151020676 | 0.999933887 | -0.176950554 | 0.7365537 | -0.913504254 | 0.176584169 |
| ENSG00000125952 | *MAX* | 6.352412625 | -0.025014594 | 0.00492264 | -0.029937234 | 0.999933887 | -0.498430811 | -0.641033806 | 0.142602995 | 0.176811193 |
| ENSG00000274736 | *CCL23* | -0.29898285 | 0.033078283 | -0.169087896 | 0.202166179 | 0.999933887 | 0.600981679 | 1.317934389 | -0.71695271 | 0.177067816 |
| ENSG00000138756 | *BMP2K* | 4.453322627 | -0.013311387 | 0.038087316 | -0.051398703 | 0.999933887 | -0.925891884 | -1.135674927 | 0.209783043 | 0.177381562 |
| ENSG00000229236 | *TTTY10* | -3.83077962 | 0.128762019 | 0.312187821 | -0.183425803 | 0.999933887 | -0.501920466 | 0.184622592 | -0.686543058 | 0.177534315 |
| ENSG00000100294 | *MCAT* | 2.952646283 | 0.17410915 | 0.065690895 | 0.108418255 | 0.999933887 | 0.236531402 | 0.508024202 | -0.2714928 | 0.177633924 |
| ENSG00000131188 | *PRR7* | 5.972431432 | 0.197424939 | 0.139629399 | 0.057795541 | 0.999933887 | 1.002234727 | 1.220566619 | -0.218331893 | 0.177873913 |
| ENSG00000274070 | *CASTOR2* | 2.942883207 | -0.016736406 | -0.058884038 | 0.042147632 | 0.999933887 | -0.463985802 | -0.69153247 | 0.227546668 | 0.178056519 |
| ENSG00000133119 | *RFC3* | 0.785796998 | -0.305788025 | -0.068667551 | -0.237120475 | 0.999933887 | -0.059768836 | 0.491134271 | -0.550903107 | 0.178084379 |
| ENSG00000143924 | *EML4* | 7.520737537 | -0.06468975 | -0.03664517 | -0.02804458 | 0.999933887 | -0.038784197 | 0.194243489 | -0.233027686 | 0.178084379 |
| ENSG00000228817 | *BACH1-IT2* | 2.551703061 | 0.103561397 | 0.054267863 | 0.049293534 | 0.999933887 | 0.059168396 | 0.296682275 | -0.237513879 | 0.17810489 |
| ENSG00000072786 | *STK10* | 9.616683992 | 0.002004767 | -0.00939541 | 0.011400178 | 0.999933887 | 0.271797917 | 0.375703899 | -0.103905982 | 0.17810489 |
| ENSG00000156831 | *NSMCE2* | 3.978863249 | 0.053786466 | 0.06183083 | -0.008044364 | 0.999933887 | 0.310171443 | 0.502409292 | -0.192237848 | 0.178147436 |
| ENSG00000110880 | *CORO1C* | 7.994146465 | -0.02729961 | -0.04555386 | 0.018254249 | 0.999933887 | -0.832022994 | -0.640749074 | -0.19127392 | 0.178157581 |
| ENSG00000185787 | *MORF4L1* | 7.42413206 | -0.038498211 | -0.080127794 | 0.041629584 | 0.999933887 | -0.182306255 | -0.069412187 | -0.112894068 | 0.178313468 |
| ENSG00000147050 | *KDM6A* | 6.740721469 | 0.019379124 | 0.010098599 | 0.009280525 | 0.999933887 | 0.031854371 | 0.205320746 | -0.173466375 | 0.178313468 |
| ENSG00000010318 | *PHF7* | 1.338482601 | 0.034922316 | -0.087713419 | 0.122635735 | 0.999933887 | -0.270884263 | -0.597925835 | 0.327041572 | 0.178313468 |
| ENSG00000239732 | *TLR9* | -0.588572654 | -0.127387464 | 0.143259345 | -0.270646809 | 0.999933887 | -0.28651211 | -1.086737323 | 0.800225213 | 0.178389187 |
| ENSG00000166783 | *MARF1* | 7.696292775 | -0.04652181 | -0.066429402 | 0.019907592 | 0.999933887 | -0.129767773 | 0.020941915 | -0.150709688 | 0.178440911 |
| ENSG00000100319 | *ZMAT5* | 2.012507125 | -0.064765619 | 0.120246758 | -0.185012377 | 0.999933887 | -0.462361023 | -0.786407045 | 0.324046021 | 0.178609502 |
| ENSG00000177854 | *TMEM187* | 1.400391199 | -0.00888758 | -0.260344847 | 0.251457267 | 0.999933887 | 0.686164432 | 0.981769603 | -0.29560517 | 0.178739202 |
| ENSG00000275791 | *TRBV10-3* | -0.339504024 | 0.150381547 | 0.062801484 | 0.087580063 | 0.999933887 | 0.187296373 | 0.729165112 | -0.541868738 | 0.178749941 |
| ENSG00000204348 | *DXO* | 4.088379671 | 0.099298347 | 0.039995769 | 0.059302578 | 0.999933887 | 0.536879992 | 0.773482581 | -0.236602589 | 0.17875345 |
| ENSG00000109103 | *UNC119* | 7.046232332 | 0.079702333 | 0.024314978 | 0.055387355 | 0.999933887 | 1.479700126 | 1.687413959 | -0.207713833 | 0.17875345 |

| | | | | | | | | | | |
|---|---|---|---|---|---|---|---|---|---|---|
| ENSG00000177989 | *ODF3B* | 6.376780037 | 0.172970771 | 0.178212189 | -0.005241419 | 0.999933887 | -0.077253213 | -0.310512681 | 0.233259468 | 0.179062202 |
| ENSG00000065989 | *PDE4A* | 2.633718399 | -0.064717127 | 0.056881409 | -0.121598536 | 0.999933887 | 0.124631033 | 0.390920239 | -0.266289206 | 0.179100033 |
| ENSG00000184220 | *CMSS1* | 0.315331842 | -0.088557121 | -0.083397911 | -0.00515921 | 0.999933887 | 0.120868235 | 0.437883071 | -0.317014836 | 0.179121649 |
| ENSG00000134602 | *STK26* | 5.610555809 | -0.013804754 | -0.019278369 | 0.005473615 | 0.999933887 | -0.14769771 | 0.073812206 | -0.221509917 | 0.179277364 |
| ENSG00000108055 | *SMC3* | 4.0600093 | -0.10260886 | -0.127839803 | 0.025230943 | 0.999933887 | -0.843246817 | -0.493752413 | -0.349494404 | 0.179495832 |
| ENSG00000163406 | *SLC15A2* | 0.719994677 | -0.17470852 | -0.01486629 | -0.15984223 | 0.999933887 | 0.005618467 | 0.366931921 | -0.361313454 | 0.179605983 |
| ENSG00000251323 | *LINC02728* | 1.437673425 | 0.213977451 | 0.415740545 | -0.201763094 | 0.999933887 | 2.470800605 | 2.964190381 | -0.493389777 | 0.179859648 |
| ENSG00000147548 | *NSD3* | 6.089558109 | -0.018856483 | -0.065964576 | 0.047108093 | 0.999933887 | -0.470602279 | -0.640048033 | 0.169445754 | 0.180205846 |
| ENSG00000203865 | *ATP1A1-AS1* | 1.326045524 | 0.023314509 | 0.086193231 | -0.062878722 | 0.999933887 | 0.312445028 | 0.716715036 | -0.404270008 | 0.180285708 |
| ENSG00000106479 | *ZNF862* | 3.249981881 | -0.011677418 | -0.014835292 | 0.003157874 | 0.999933887 | -0.019694508 | -0.226581116 | 0.206886608 | 0.180399105 |
| ENSG00000107020 | *PLGRKT* | 1.662012881 | 0.051312019 | -0.05708135 | 0.108393369 | 0.999933887 | 0.341490819 | 0.711678171 | -0.370187353 | 0.180759907 |
| ENSG00000133028 | *SCO1* | 2.89554024 | -0.032504243 | 0.020841553 | -0.053345796 | 0.999933887 | -0.059900149 | -0.283514351 | 0.223614202 | 0.180759907 |
| ENSG00000172000 | *ZNF556* | -2.853330483 | -0.149840275 | -0.529393997 | 0.379553721 | 0.999933887 | 0.543756038 | 1.301532825 | -0.757776787 | 0.180759907 |
| ENSG00000181827 | *RFX7* | 3.189730406 | 0.076862112 | -0.190882673 | 0.267744785 | 0.999933887 | -0.262447714 | 0.104889573 | -0.367337287 | 0.180765703 |
| ENSG00000144426 | *NBEAL1* | 1.559591248 | 0.059478303 | 0.066589472 | -0.007111169 | 0.999933887 | 0.225582799 | 0.750997981 | -0.525415182 | 0.180847507 |
| ENSG00000103091 | *WDR59* | 3.66041074 | 0.012111577 | -0.018608998 | 0.030720574 | 0.999933887 | -0.050782698 | -0.216313711 | 0.165531013 | 0.181059169 |
| ENSG00000211753 | *TRBV28* | 0.6315717 | 0.054778486 | 0.232843492 | -0.178065006 | 0.999933887 | 0.143803878 | -0.201593825 | 0.345397704 | 0.181119605 |
| ENSG00000228784 | *LINC00954* | 3.576443095 | 0.078943383 | 0.019344797 | 0.059598586 | 0.999933887 | 1.183571075 | 1.465631157 | -0.282060082 | 0.181333979 |
| ENSG00000184014 | *DENND5A* | 9.892119275 | -0.037084846 | 0.009331009 | -0.046415856 | 0.999933887 | 1.102781914 | 1.246631291 | -0.143849376 | 0.181333979 |
| ENSG00000119685 | *TTLL5* | 2.210058384 | 0.028988714 | -0.096858809 | 0.125847523 | 0.999933887 | -0.194080093 | 0.131814221 | -0.325894314 | 0.181333979 |
| ENSG00000258907 | *PSMB3P1* | 2.157245527 | 0.22740673 | 0.471004721 | -0.243597991 | 0.999933887 | 1.196200287 | 1.483926363 | -0.287726076 | 0.181382578 |
| ENSG00000068323 | *TFE3* | 8.357767755 | 0.038739989 | 0.019775298 | 0.018964691 | 0.999933887 | 0.19955477 | 0.331040433 | -0.131485663 | 0.181382578 |
| ENSG00000172123 | *SLFN12* | 1.850002766 | -0.033650061 | 0.07258509 | -0.106235151 | 0.999933887 | -0.448456192 | -0.760520155 | 0.312063963 | 0.181447458 |
| ENSG00000261737 | *CLCA4-AS1* | 0.450152391 | 0.005723942 | -0.032793004 | 0.038516946 | 0.999933887 | 0.525675934 | 0.952077758 | -0.426401824 | 0.181502439 |
| ENSG00000108799 | *EZH1* | 6.195777161 | -0.042332222 | 0.00830109 | -0.050633312 | 0.999933887 | -0.323206335 | -0.442755608 | 0.119549273 | 0.181519669 |
| ENSG00000161011 | *SQSTM1* | 10.502997176 | 0.034663942 | 0.026990082 | 0.007673861 | 0.999933887 | 0.540812299 | 0.681131759 | -0.14031946 | 0.181519669 |
| ENSG00000163145 | *C1QTNF7* | -0.572732477 | 0.021730635 | 0.358173437 | -0.336442803 | 0.999933887 | 0.047389989 | 0.716544501 | -0.669154512 | 0.181519669 |
| ENSG00000273559 | *CWC25* | 7.866754096 | 0.010694945 | 0.012899751 | -0.002204806 | 0.999933887 | 0.598264228 | 0.739558054 | -0.141293826 | 0.181587107 |
| ENSG00000103145 | *HCFC1R1* | 1.688180614 | -0.020210277 | -0.043531855 | 0.023321578 | 0.999933887 | -0.38181192 | -0.665131793 | 0.283319873 | 0.181674881 |
| ENSG00000086061 | *DNAJA1* | 7.245092205 | -0.018454669 | -0.11081259 | 0.092357921 | 0.999933887 | 0.700202767 | 0.866635772 | -0.166433005 | 0.181764186 |
| ENSG00000166225 | *FRS2* | 3.471800598 | -0.062998065 | -0.032102296 | -0.030895769 | 0.999933887 | 0.01071239 | 0.233639053 | -0.222926663 | 0.181929992 |
| ENSG00000135870 | *RC3H1* | 6.511335337 | -0.054885762 | -0.017215 | -0.037670762 | 0.999933887 | -0.705683487 | -0.476994429 | -0.228689058 | 0.181929992 |
| ENSG00000119431 | *HDHD3* | 1.682961839 | -0.064772537 | -0.049229026 | -0.015543511 | 0.999933887 | 0.175687388 | -0.081624736 | 0.257312124 | 0.181929992 |
| ENSG00000243335 | *KCTD7* | 5.356766768 | 0.034468516 | -0.020012559 | 0.054481075 | 0.999933887 | 0.189622812 | 0.340070113 | -0.150447301 | 0.181929992 |
| ENSG00000073111 | *MCM2* | 3.062284711 | 0.013787219 | 0.080591129 | -0.066803911 | 0.999933887 | 0.704947055 | 0.935204194 | -0.230257139 | 0.181929992 |
| ENSG00000258704 | *SRP54-AS1* | 0.158428895 | -0.019797523 | 0.516893711 | -0.536691234 | 0.999933887 | 1.04864719 | 1.513869866 | -0.465222676 | 0.181929992 |
| ENSG00000198081 | *ZBTB14* | 2.229574643 | -0.006892281 | -0.145004316 | 0.138112034 | 0.999933887 | 0.031315933 | -0.202230199 | 0.233546132 | 0.181929992 |
| ENSG00000139697 | *SBNO1* | 6.285749271 | -0.078688735 | -0.132120018 | 0.053431283 | 0.999933887 | -0.226252932 | -0.00815448 | -0.218098452 | 0.181959553 |
| ENSG00000116120 | *FARSB* | 1.463143201 | -0.142338568 | -0.167944506 | 0.025605938 | 0.999933887 | 0.245353743 | 0.515775603 | -0.270421859 | 0.181974864 |
| ENSG00000101347 | *SAMHD1* | 5.47958537 | -0.026503228 | -0.07596972 | 0.049466492 | 0.999933887 | -0.008359597 | -0.157290263 | 0.148930666 | 0.182142553 |
| ENSG00000253795 | *MTND1P5* | -1.417281792 | 0.445227633 | 0.245394028 | 0.199833605 | 0.999933887 | 2.237472123 | 3.157872024 | -0.920399901 | 0.182209603 |
| ENSG00000105639 | *JAK3* | 6.86800901 | 0.038707133 | 0.038195905 | 0.000511228 | 0.999933887 | 0.24633491 | 0.106707047 | 0.139627863 | 0.182231713 |
| ENSG00000104805 | *NUCB1* | 6.328768563 | 0.056676656 | 0.027923852 | 0.028752805 | 0.999933887 | -0.176187656 | -0.333230152 | 0.157042497 | 0.182320843 |
| ENSG00000101236 | *RNF24* | 7.267335991 | -0.059756647 | -0.011366468 | -0.048390179 | 0.999933887 | -0.858286022 | -1.114309146 | 0.256023125 | 0.182370404 |
| ENSG00000135148 | *TRAFD1* | 5.651181026 | -0.011862147 | 0.013639009 | -0.025501156 | 0.999933887 | -0.309831896 | -0.504694507 | 0.19486261 | 0.182499692 |
| ENSG00000275070 | *Metazoa_SRP* | 0.59090211 | 0.081978883 | -0.046619567 | 0.12859845 | 0.999933887 | -0.063742098 | 0.419967378 | -0.483709476 | 0.182837464 |
| ENSG00000161955 | *TNFSF13* | 1.553114066 | -0.003977322 | 0.197711939 | -0.201689261 | 0.999933887 | 0.071749371 | -0.237158518 | 0.308907889 | 0.183399783 |
| ENSG00000197771 | *MCMBP* | 4.597409762 | -0.026879001 | 0.076874398 | -0.103753399 | 0.999933887 | 0.032058858 | -0.128015985 | 0.160074844 | 0.183523858 |
| ENSG00000023191 | *RNH1* | 5.506951155 | 0.069697268 | 0.049175514 | 0.020521754 | 0.999933887 | -0.132350582 | -0.301597 | 0.169246418 | 0.183707113 |
| ENSG00000091262 | *ABCC6* | 1.014959991 | -0.099501249 | -0.194903379 | 0.095402131 | 0.999933887 | -0.41883731 | 0.013916256 | -0.432753565 | 0.183760983 |
| ENSG00000257743 | *MGAM2* | 2.956233852 | 0.03162847 | -0.160740534 | 0.192369004 | 0.999933887 | -0.168389083 | 0.202548205 | -0.370937289 | 0.183760983 |
| ENSG00000156345 | *CDK20* | 3.228940981 | 0.129646417 | 0.172892173 | -0.043245755 | 0.999933887 | -0.015582363 | 0.177223408 | -0.192805771 | 0.183768477 |
| ENSG00000131748 | *STARD3* | 6.266426256 | 0.012015929 | 0.038684408 | -0.026668479 | 0.999933887 | -0.128833308 | -0.259352568 | 0.13051926 | 0.183768477 |
| ENSG00000116260 | *QSOX1* | 6.832929673 | 0.013755866 | 0.008936274 | 0.004819592 | 0.999933887 | 0.06917529 | 0.222868721 | -0.153693431 | 0.183768477 |
| ENSG00000158874 | *APOA2* | 2.23070346 | 0.108328676 | 0.079806799 | 0.028521877 | 0.999933887 | 0.753790705 | 1.139090991 | -0.385300286 | 0.18382937 |
| ENSG00000107249 | *GLIS3* | -0.04978007 | -0.274934666 | -0.00620992 | -0.268724746 | 0.999933887 | -0.227897518 | -0.67281915 | 0.444921632 | 0.183837741 |
| ENSG00000136244 | *IL6* | 0.016161984 | 0.760927628 | 1.458129661 | -0.697202034 | 0.999933887 | 3.561017055 | 4.486722749 | -0.925705694 | 0.183837741 |
| ENSG00000230989 | *HSBP1* | 4.85004988 | -0.02980246 | -0.094846511 | 0.065044051 | 0.999933887 | -0.433823913 | -0.614048696 | 0.180224783 | 0.183837741 |
| ENSG00000174175 | *SELP* | 0.51036522 | 0.038622809 | -0.083318797 | 0.121941606 | 0.999933887 | -0.385525445 | -0.830535817 | 0.445010371 | 0.183837741 |
| ENSG00000101849 | *TBL1X* | 6.808825556 | -0.053802895 | -0.026267423 | -0.027535472 | 0.999933887 | -0.220559954 | -0.343961559 | 0.123401605 | 0.18384731 |
| ENSG00000198131 | *ZNF544* | 1.118511069 | -0.075647845 | -0.045295677 | -0.030352168 | 0.999933887 | -0.060947652 | -0.391835483 | 0.330887831 | 0.18384731 |
| ENSG00000076555 | *ACACB* | 2.657004719 | 0.130113732 | -0.067748508 | 0.19786224 | 0.999933887 | -0.05927546 | 0.166342434 | -0.225617894 | 0.183856506 |
| ENSG00000138641 | *HERC3* | 1.114431943 | 0.107056633 | 0.137017608 | -0.029960974 | 0.999933887 | -0.279184413 | -0.794280807 | 0.515096393 | 0.183856506 |
| ENSG00000171219 | *CDC42BPG* | 2.705915289 | 0.076185742 | 0.027634174 | 0.048551568 | 0.999933887 | 0.5610029 | 0.888613609 | -0.327610709 | 0.183856506 |
| ENSG00000064545 | *TMEM161A* | 1.618741653 | -0.061799436 | -0.073951205 | 0.01215177 | 0.999933887 | 0.197904127 | 0.474138066 | -0.276233938 | 0.183856506 |
| ENSG00000125352 | *RNF113A* | 3.093779466 | -0.026266869 | -0.20761094 | 0.181344071 | 0.999933887 | -0.373647016 | -0.613771128 | 0.240124112 | 0.183856506 |
| ENSG00000137842 | *TMEM62* | 2.143357386 | -0.036083638 | 0.064151871 | -0.100235509 | 0.999933887 | 0.076286121 | -0.330536925 | 0.406823046 | 0.184455379 |
| ENSG00000183150 | *GPR19* | -0.303793775 | -0.188357167 | -0.246556474 | 0.058199307 | 0.999933887 | -0.73252546 | -1.275609811 | 0.543084351 | 0.184524022 |
| ENSG00000197093 | *GAL3ST4* | 1.234189757 | 0.022836377 | -0.048266327 | 0.071102704 | 0.999933887 | -0.022353672 | -0.454500853 | 0.432147182 | 0.184592676 |
| ENSG00000186350 | *RXRA* | 8.10906012 | -0.013262276 | 0.025937052 | -0.039199328 | 0.999933887 | -0.747633421 | -0.936955816 | 0.189322395 | 0.184856796 |
| ENSG00000156709 | *AIFM1* | 1.849471844 | -0.092827915 | -0.001646134 | -0.091181781 | 0.999933887 | -0.095197255 | -0.359571303 | 0.264374048 | 0.184869261 |
| ENSG00000113810 | *SMC4* | 3.904434 | -0.050213273 | -0.309237628 | 0.259024355 | 0.999933887 | -0.44155308 | 0.110534358 | -0.552087438 | 0.184869261 |
| ENSG00000173264 | *GPR137* | 3.949200184 | 0.058983481 | 0.014374509 | 0.044608972 | 0.999933887 | 0.456570473 | 0.625022866 | -0.168452394 | 0.185187362 |
| ENSG00000083812 | *ZNF324* | 3.425178011 | 0.104493294 | 0.001587371 | 0.102905923 | 0.999933887 | -0.042824057 | -0.249200709 | 0.206376652 | 0.185717593 |
| ENSG00000237938 | *AL450998.2* | 2.992869723 | -0.08609283 | -0.080403896 | -0.005688934 | 0.999933887 | 0.286671208 | 0.467370181 | -0.180698974 | 0.185771243 |
| ENSG00000175105 | *ZNF654* | 4.01722546 | -0.172873457 | -0.195695789 | 0.022822332 | 0.999933887 | -0.283222593 | 0.079789989 | -0.363012581 | 0.186131254 |
| ENSG00000243156 | *MICAL3* | 1.894903314 | 0.184980252 | 0.071267236 | 0.113713017 | 0.999933887 | 0.409564327 | 0.733487214 | -0.323922887 | 0.186205687 |
| ENSG00000111906 | *HDDC2* | 1.624229636 | 0.037773464 | -0.05848795 | 0.096261414 | 0.999933887 | -0.07602063 | 0.145569977 | -0.221590607 | 0.186205687 |
| ENSG00000196878 | *LAMB3* | 6.268445034 | 0.059899626 | 0.022208523 | 0.037691103 | 0.999933887 | 1.707740514 | 1.938203041 | -0.230462528 | 0.186491974 |
| ENSG00000163754 | *GYG1* | 4.059260335 | -0.048561699 | -0.092640957 | 0.044079259 | 0.999933887 | -0.28369017 | -0.459328847 | 0.175638678 | 0.186601883 |
| ENSG00000198673 | *TAFA2* | 3.39827038 | 0.052295018 | 0.103622136 | -0.051327118 | 0.999933887 | 0.265167859 | 0.508766141 | -0.243598281 | 0.186601883 |
| ENSG00000131381 | *RBSN* | 4.868285704 | -0.00279801 | -0.015774458 | 0.012976448 | 0.999933887 | -0.086219011 | 0.055613028 | -0.141832039 | 0.186601883 |
| ENSG00000183878 | *UTY* | -2.683781403 | 0.109669248 | 0.122254147 | -0.012584899 | 0.999933887 | -0.319837425 | -0.035853184 | -0.283984241 | 0.186604325 |
| ENSG00000171456 | *ASXL1* | 6.113512006 | -0.011798731 | -0.015231033 | 0.003432302 | 0.999933887 | 0.173806906 | 0.287898762 | -0.114091856 | 0.186606524 |
| ENSG00000166126 | *AMN* | -0.047544036 | 0.015052588 | 0.273533766 | -0.258481178 | 0.999933887 | 1.190881919 | 1.672872595 | -0.481990677 | 0.186606524 |
| ENSG00000133275 | *CSNK1G2* | 8.072552564 | 0.070227231 | 0.063049572 | 0.007177659 | 0.999933887 | -0.004053618 | -0.152710171 | 0.148656553 | 0.186681513 |
| ENSG00000163219 | *ARHGAP25* | 9.10721606 | -0.086194202 | -0.042192046 | -0.044002156 | 0.999933887 | -0.635029599 | -0.449932322 | -0.185097276 | 0.18672216 |
| ENSG00000235724 | *AC009299.2* | 1.630510751 | -0.17302697 | 0.083591172 | -0.256618141 | 0.999933887 | 0.687258717 | 1.211104629 | -0.523845912 | 0.18672216 |
| ENSG00000107960 | *STN1* | 2.973232091 | 0.068541762 | -0.029634923 | 0.098176685 | 0.999933887 | 0.133499262 | -0.158864822 | 0.292364083 | 0.18672216 |
| ENSG00000089053 | *ANAPC5* | 5.211005677 | -0.013104439 | 0.005515646 | -0.018620085 | 0.999933887 | 0.22036077 | 0.351764464 | -0.131403694 | 0.186901129 |
| ENSG00000101161 | *PRPF6* | 5.50687943 | 0.002453395 | -0.023304683 | 0.025758078 | 0.999933887 | -0.160361064 | -0.308501138 | 0.148140073 | 0.187038286 |
| ENSG00000081014 | *AP4E1* | 2.135478254 | -0.060028727 | -0.059775437 | -0.00025329 | 0.999933887 | 0.071421914 | 0.347291856 | -0.275869942 | 0.187082452 |
| ENSG00000280832 | *GSEC* | 2.574466017 | -0.045016529 | 0.201850723 | -0.246867252 | 0.999933887 | 0.413684968 | 0.660188868 | -0.2465039 | 0.187138536 |
| ENSG00000234797 | *RPS3AP6* | 0.339134836 | -0.032554825 | -0.494045976 | 0.461491151 | 0.999933887 | 0.142522601 | 0.571973193 | -0.429450591 | 0.187328357 |
| ENSG00000236859 | *NIFK-AS1* | 0.66874756 | -0.042607418 | 0.071223749 | -0.113831167 | 0.999933887 | -0.09771223 | -0.444968017 | 0.347255787 | 0.187445303 |
| ENSG00000138785 | *INTS12* | 2.846597745 | -0.011913276 | -0.13501573 | 0.123102454 | 0.999933887 | -0.037847521 | 0.172528159 | -0.210375681 | 0.187465582 |
| ENSG00000173597 | *SULT1B1* | 3.694438434 | 0.003007335 | 0.023868754 | -0.020861419 | 0.999933887 | -0.226286322 | -0.625228322 | 0.398942 | 0.187465582 |
| ENSG00000238160 | *LINC02863* | 2.002964814 | -0.024458266 | 0.025695178 | -0.050153444 | 0.999933887 | -0.126555955 | -0.537167098 | 0.410611142 | 0.187654811 |
| ENSG00000259600 | *AC066616.2* | 3.790440321 | 0.342918097 | 0.714833014 | -0.371914918 | 0.999933887 | 2.000964737 | 2.609509898 | -0.608545162 | 0.187825005 |
| ENSG00000182199 | *SHMT2* | 2.788134499 | 0.027919774 | 0.14546985 | -0.117550075 | 0.999933887 | 0.383853293 | 0.617659435 | -0.233806143 | 0.187916883 |
| ENSG00000039123 | *MTREX* | 3.296831769 | -0.13098811 | -0.076106268 | -0.054881842 | 0.999933887 | -0.17352027 | 0.088430194 | -0.261950464 | 0.187959522 |
| ENSG00000204856 | *FAM216A* | -0.272856561 | 0.076993618 | -0.317135764 | 0.394129382 | 0.999933887 | -0.060718833 | -0.510718526 | 0.449999692 | 0.187959522 |
| ENSG00000069188 | *SDK2* | -0.619057669 | -0.105790004 | 0.104089719 | -0.209879723 | 0.999933887 | -0.491654756 | 0.225924329 | -0.717579085 | 0.187971885 |
| ENSG00000135074 | *ADAM19* | 5.764349831 | 0.000394853 | 0.035253069 | -0.034858216 | 0.999933887 | 0.020390036 | -0.104771849 | 0.125161884 | 0.188341703 |
| ENSG00000240288 | *GHRLOS* | 1.424094059 | 0.115959691 | -0.074641965 | 0.190601656 | 0.999933887 | 1.674244089 | 2.112100446 | -0.437856357 | 0.188595294 |

| | | | | | | | | | | |
|---|---|---|---|---|---|---|---|---|---|---|
| ENSG00000261220 | *AC103706.1* | 2.175309903 | 0.021612376 | 0.03773597 | -0.016123594 | 0.999933887 | 0.948810338 | 1.236702235 | -0.287891897 | 0.188595294 |
| ENSG00000136541 | *ERMN* | 3.484827207 | -0.146350954 | -0.041210901 | -0.105140053 | 0.999933887 | -0.541499769 | -0.229627455 | -0.311872314 | 0.188671181 |
| ENSG00000152061 | *RABGAP1L* | 5.353477011 | 0.021144371 | -0.0754234 | 0.096567771 | 0.999933887 | -0.685023473 | -0.926773227 | 0.241749754 | 0.188774881 |
| ENSG00000127837 | *AAMP* | 4.044647571 | 0.074028624 | 0.102365564 | -0.028336941 | 0.999933887 | 0.101306895 | -0.065779031 | 0.167085926 | 0.189261209 |
| ENSG00000110237 | *ARHGEF17* | -0.366256938 | -0.034094834 | 0.05683234 | -0.090927174 | 0.999933887 | -0.314754215 | -0.946390525 | 0.631636309 | 0.18941553 |
| ENSG00000148288 | *GBGT1* | 1.951347101 | 0.097538154 | 0.144111452 | -0.046573298 | 0.999933887 | -0.229407874 | -0.507274806 | 0.277866933 | 0.189431182 |
| ENSG00000108468 | *CBX1* | 3.160556198 | 0.018420545 | -0.133008241 | 0.151428786 | 0.999933887 | 0.021531582 | -0.233302428 | 0.25483401 | 0.189431182 |
| ENSG00000110497 | *AMBRA1* | 4.143314763 | -0.005449251 | 0.081931024 | -0.087380275 | 0.999933887 | -0.07802557 | 0.064197238 | -0.142222809 | 0.189431182 |
| ENSG00000211456 | *SACM1L* | 4.523775604 | 0.01280508 | 0.020113495 | -0.007308415 | 0.999933887 | -0.468486992 | -0.703230887 | 0.234743896 | 0.189449711 |
| ENSG00000053918 | *KCNQ1* | 4.076907808 | 0.080361664 | 0.029269947 | 0.051091718 | 0.999933887 | -0.057708298 | -0.279039552 | 0.221331253 | 0.189458212 |
| ENSG00000125826 | *RBCK1* | 6.682452775 | 0.037828127 | 0.019101291 | 0.018726836 | 0.999933887 | -0.287875829 | -0.43171363 | 0.143837801 | 0.190033391 |
| ENSG00000119698 | *PPP4R4* | -0.898660693 | -0.124956213 | 0.111116571 | -0.236072784 | 0.999933887 | 2.359235975 | 2.997889973 | -0.638653999 | 0.190161273 |
| ENSG00000224187 | *LINC01991* | 0.853918925 | 0.226047171 | 0.378382195 | -0.152335023 | 0.999933887 | 1.93712091 | 2.37482496 | -0.43770405 | 0.190234321 |
| ENSG00000166478 | *ZNF143* | 5.106911191 | -0.022337575 | -0.035236861 | 0.012899286 | 0.999933887 | -0.234339057 | -0.096941496 | -0.137397562 | 0.190234321 |
| ENSG00000163848 | *ZNF148* | 5.157257564 | -0.070732334 | -0.043974031 | -0.026758304 | 0.999933887 | -0.823483019 | -0.557770957 | -0.265712062 | 0.190527174 |
| ENSG00000162734 | *PEA15* | 3.6410487 | 0.008910731 | -0.171200736 | 0.180111467 | 0.999933887 | 0.329046625 | 0.505607387 | -0.176560762 | 0.190527174 |
| ENSG00000159899 | *NPR2* | -1.028326909 | -0.418416607 | -0.211139006 | -0.207277601 | 0.999933887 | 0.341155881 | -0.190062925 | 0.531218806 | 0.190543255 |
| ENSG00000134460 | *IL2RA* | 1.266715303 | -0.079350256 | 0.191440499 | -0.270790756 | 0.999933887 | 0.99675974 | 1.395325381 | -0.398565642 | 0.190543255 |
| ENSG00000117682 | *DHDDS* | 3.363444088 | -0.025099119 | -0.062466294 | 0.037367175 | 0.999933887 | -0.178941654 | -0.342490388 | 0.163548734 | 0.190543255 |
| ENSG00000012223 | *LTF* | 3.300143326 | -0.016795668 | -0.076280847 | 0.059485179 | 0.999933887 | 0.580836344 | 0.780861145 | -0.2000248 | 0.190543255 |
| ENSG00000170027 | *YWHAG* | 5.551651615 | 0.002163772 | 0.027470149 | -0.025306376 | 0.999933887 | -0.480502151 | -0.629824152 | 0.149322001 | 0.190543255 |
| ENSG00000262001 | *DLGAP1-AS2* | 3.815569034 | -0.083652627 | 0.171896539 | -0.255549166 | 0.981460244 | 0.444911639 | 0.634146679 | -0.18923504 | 0.190822664 |
| ENSG00000005156 | *LIG3* | 3.942474639 | 0.012734445 | -0.105599875 | 0.11833432 | 0.999933887 | 0.43398652 | 0.615513907 | -0.181527387 | 0.190822664 |
| ENSG00000011566 | *MAP4K3* | 1.234893629 | -0.045955092 | -0.007510109 | -0.038444983 | 0.999933887 | -0.536811686 | -0.891870775 | 0.355059089 | 0.190843641 |
| ENSG00000162775 | *RBM15* | 4.427260877 | -0.003225333 | 0.060817775 | -0.064043108 | 0.999933887 | -0.123009001 | -0.267406119 | 0.144397118 | 0.191053822 |
| ENSG00000070882 | *OSBPL3* | 3.475278207 | -0.043527461 | -0.049756039 | 0.006228578 | 0.999933887 | -0.026488024 | 0.13532325 | -0.161811274 | 0.191177979 |
| ENSG00000102760 | *RGCC* | 5.299659925 | 0.022817036 | -0.019733055 | 0.042550091 | 0.999933887 | -0.164095504 | -0.323278382 | 0.159182878 | 0.191586116 |
| ENSG00000100376 | *FAM118A* | 4.250697881 | -0.051116807 | -0.025209937 | -0.02590687 | 0.999933887 | 0.109074707 | -0.036977962 | 0.146052669 | 0.191591718 |
| ENSG00000007968 | *E2F2* | 3.965358268 | -0.081538495 | -0.116235771 | 0.034697276 | 0.999933887 | -0.401816509 | -0.184860803 | -0.216955707 | 0.191591718 |
| ENSG00000153113 | *CAST* | 7.587309754 | -0.002370379 | 0.010103365 | -0.012473744 | 0.999933887 | -0.375894998 | -0.221295567 | -0.154599431 | 0.191669166 |
| ENSG00000185838 | *GNB1L* | 1.117320829 | -0.100949466 | 0.162158108 | -0.263107574 | 0.999933887 | -0.063252996 | -0.430132361 | 0.366879365 | 0.191675662 |
| ENSG00000245149 | *RNF139-AS1* | 0.556662892 | -0.049947918 | -0.136335404 | 0.086387486 | 0.999933887 | 0.165543935 | -0.303976501 | 0.469520436 | 0.191940765 |
| ENSG00000166147 | *FBN1* | 0.25619395 | -0.191256911 | 0.065625606 | -0.256882517 | 0.999933887 | 0.550839619 | 1.045397753 | -0.494558134 | 0.192033427 |
| ENSG00000272384 | *AC016405.3* | -0.957809547 | 0.259650856 | 0.532398624 | -0.272747768 | 0.999933887 | 2.441712018 | 3.327446891 | -0.885734873 | 0.192146618 |
| ENSG00000196510 | *ANAPC7* | 2.984354176 | -0.090904161 | -0.025057741 | -0.06584642 | 0.999933887 | -0.180442228 | -0.352577911 | 0.172135683 | 0.192219758 |
| ENSG00000213588 | *ZBTB9* | 0.862028466 | 0.052019348 | -0.018986191 | 0.071005539 | 0.999933887 | 0.10318981 | 0.416373788 | -0.313183978 | 0.192365686 |
| ENSG00000105429 | *MEGF8* | 2.128004162 | 0.077208586 | 0.090327659 | -0.013119073 | 0.999933887 | -0.020724978 | 0.235271919 | -0.255996896 | 0.192414872 |
| ENSG00000125746 | *EML2* | 4.214535944 | 0.013657983 | 0.111287683 | -0.097629699 | 0.999933887 | 0.258250112 | 0.46790859 | -0.209658478 | 0.192586484 |
| ENSG00000267385 | *AC011498.4* | -1.084610637 | -0.122921618 | 0.104539333 | -0.22746095 | 0.999933887 | -0.142928175 | 0.82130449 | -0.964232665 | 0.192920622 |
| ENSG00000247556 | *OIP5-AS1* | 4.050477191 | -0.087639361 | -0.070016086 | -0.017623274 | 0.999933887 | -0.126744876 | -0.297121986 | 0.170377109 | 0.193191833 |
| ENSG00000185453 | *ZSWIM9* | 1.261518116 | 0.150398448 | 0.1165768 | 0.033821648 | 0.999933887 | -0.025345089 | -0.403046385 | 0.377701296 | 0.193191833 |
| ENSG00000113240 | *CLK4* | 5.231118164 | -0.09244776 | -0.138386655 | 0.045938895 | 0.999933887 | 0.084288836 | 0.342820156 | -0.258531319 | 0.193191833 |
| ENSG00000002549 | *LAP3* | 5.473066924 | -0.00245799 | -0.062430155 | 0.059972165 | 0.999933887 | -0.324098692 | -0.518221474 | 0.194122781 | 0.19353446 |
| ENSG00000164398 | *ACSL6* | 0.325665777 | -0.039296087 | -0.046801894 | 0.007505808 | 0.999933887 | -0.283592449 | 0.133934545 | -0.417526995 | 0.193726321 |
| ENSG00000087274 | *ADD1* | 6.844693262 | -0.043723531 | -0.005072221 | -0.03865131 | 0.999933887 | -0.157270405 | -0.277290318 | 0.120019913 | 0.193834652 |
| ENSG00000258667 | *HIF1A-AS3* | 4.619712846 | 0.084588197 | 0.318560982 | -0.233972785 | 0.999933887 | 1.009466909 | 1.35464718 | -0.345180271 | 0.193842009 |
| ENSG00000135253 | *KCP* | 0.633048877 | 0.14848317 | 0.332604508 | -0.184121338 | 0.999933887 | 0.417728876 | 0.813013331 | -0.395284455 | 0.193977764 |
| ENSG00000197044 | *ZNF441* | 1.703447733 | -0.291757411 | -0.160433869 | -0.131323542 | 0.999933887 | -0.389009832 | 0.002180526 | -0.391190358 | 0.194049511 |
| ENSG00000196405 | *EVL* | 6.61292162 | 0.006332811 | 0.054608653 | -0.048275843 | 0.999933887 | 0.00541217 | -0.117671736 | 0.123083906 | 0.194118072 |
| ENSG00000130479 | *MAP1S* | 4.779337765 | -0.016429334 | -0.033352991 | 0.016923657 | 0.999933887 | -0.958231187 | -1.167664742 | 0.209433555 | 0.194123952 |
| ENSG00000196712 | *NF1* | 5.202762455 | -0.04755642 | -0.031973805 | -0.015582616 | 0.999933887 | -0.245134542 | -0.017990105 | -0.227144437 | 0.194266329 |
| ENSG00000237772 | *LINC02631* | 1.570826029 | -0.032032661 | 0.002736353 | -0.034769014 | 0.999933887 | 0.174858092 | 0.59344807 | -0.418589978 | 0.19428112 |
| ENSG00000121417 | *ZNF211* | 3.355298123 | -0.021694326 | -0.035003705 | 0.013309378 | 0.999933887 | -0.009735385 | -0.170062405 | 0.160327021 | 0.194307756 |
| ENSG00000197057 | *DTHD1* | 1.817580655 | 0.075977475 | -0.031261739 | 0.107239214 | 0.999933887 | -0.192900896 | 0.197235112 | -0.390136008 | 0.194334688 |
| ENSG00000160094 | *ZNF362* | 2.617646831 | 0.079214801 | 0.068826176 | 0.010388625 | 0.999933887 | -0.160520723 | -0.402229639 | 0.241708916 | 0.194382582 |
| ENSG00000148481 | *MINDY3* | 3.012348109 | -0.053416006 | -0.190354362 | 0.136938356 | 0.999933887 | 0.512619442 | 0.828501888 | -0.315882446 | 0.194382582 |
| ENSG00000106948 | *AKNA* | 9.130536172 | 0.04273028 | 0.029888317 | 0.012841963 | 0.999933887 | 0.290729555 | 0.41434944 | -0.123619885 | 0.194527043 |
| ENSG00000171943 | *SRGAP2C* | 1.644519703 | 0.052347014 | 0.007924177 | 0.044422837 | 0.999933887 | -0.023882282 | 0.339230427 | -0.363112708 | 0.194560218 |
| ENSG00000133142 | *TCEAL4* | 1.634519793 | 0.014748219 | -0.186489155 | 0.201237374 | 0.999933887 | 0.142552486 | 0.432457517 | -0.289905031 | 0.194560218 |
| ENSG00000186432 | *KPNA4* | 6.708511875 | -0.05863671 | -0.017137682 | -0.041499029 | 0.999933887 | -0.391189462 | -0.252762968 | -0.138426495 | 0.194866424 |
| ENSG00000130204 | *TOMM40* | 2.541539774 | 0.077354526 | 0.116132242 | -0.038777715 | 0.999933887 | 0.080025794 | 0.307691941 | -0.227666147 | 0.194866424 |
| ENSG00000151366 | *NDUFC2* | -0.256571517 | 0.057263968 | 0.289061898 | -0.23179793 | 0.999933887 | 0.122874292 | -0.31941586 | 0.442290153 | 0.194866424 |
| ENSG00000117597 | *UTP25* | 2.346642729 | -0.147359788 | -0.18142258 | 0.034062792 | 0.999933887 | -0.12385642 | 0.105259393 | -0.229115813 | 0.195237847 |
| ENSG00000223768 | *LINC00205* | 1.885491319 | 0.089831665 | 0.110472397 | -0.020640732 | 0.999933887 | -0.029454571 | -0.299659601 | 0.27020503 | 0.195237847 |
| ENSG00000197536 | *IRF1-AS1* | 4.609300455 | 0.125492589 | 0.109651049 | 0.01584154 | 0.999933887 | -0.180651203 | -0.363209111 | 0.182557908 | 0.195246429 |
| ENSG00000232573 | *RPL3P4* | 1.110125329 | 0.006524843 | -0.176833685 | 0.183358529 | 0.999933887 | 0.017955108 | 0.332677269 | -0.314722161 | 0.195246429 |
| ENSG00000273341 | *AC004921.1* | -0.541623447 | 0.1803312 | -0.011499198 | 0.191830397 | 0.999933887 | 1.687213486 | 2.0778867 | -0.390673214 | 0.195361482 |
| ENSG00000169764 | *UGP2* | 4.550540938 | -0.060101867 | -0.174809679 | 0.114707811 | 0.999933887 | -0.213355123 | -0.376658394 | 0.163303271 | 0.195369851 |
| ENSG00000177302 | *TOP3A* | 5.406005787 | -4.6527e-05 | 0.007852654 | -0.007899181 | 0.999933887 | -0.525634033 | -0.693980742 | 0.168346709 | 0.195573101 |
| ENSG00000185721 | *DRG1* | 2.769939668 | -0.040790252 | -0.083974003 | 0.043183751 | 0.999933887 | -0.029732677 | 0.170228986 | -0.199961663 | 0.195588445 |
| ENSG00000259626 | *MTND3P12* | 1.705217849 | 0.445543901 | 0.762015135 | -0.316471234 | 0.999933887 | 1.976246946 | 2.796163864 | -0.819916918 | 0.195731689 |
| ENSG00000229727 | *AC013460.1* | -0.749103291 | -0.280433634 | -0.152657806 | -0.127775828 | 0.999933887 | -0.344490358 | 0.138385475 | -0.482875833 | 0.195731689 |
| ENSG00000115539 | *PDCL3* | 1.488956049 | -0.025967974 | -0.151315226 | 0.125347252 | 0.999933887 | -0.05008093 | -0.400688946 | 0.350608016 | 0.195807139 |
| ENSG00000048707 | *VPS13D* | 6.0222617 | -0.03660012 | -0.055402942 | 0.018802822 | 0.999933887 | 0.200039527 | 0.350347941 | -0.150308414 | 0.19598738 |
| ENSG00000211653 | *IGLV1-40* | -1.667620815 | -0.197364011 | -0.020067388 | -0.177296623 | 0.999933887 | 0.213819735 | -0.89632234 | 1.110142076 | 0.19598738 |
| ENSG00000136319 | *TTC5* | 1.1479107 | 0.054156561 | -0.267014133 | 0.321170694 | 0.999933887 | 0.060363163 | -0.262394037 | 0.3227572 | 0.19598738 |
| ENSG00000077147 | *TM9SF3* | 6.562663315 | -0.038967465 | -0.072734641 | 0.033767176 | 0.999933887 | 0.015589555 | 0.188278521 | -0.172688966 | 0.196012299 |
| ENSG00000169359 | *SLC33A1* | 0.943706444 | -0.177112531 | -0.120251181 | -0.05686135 | 0.999933887 | -0.405325602 | -0.072478613 | -0.332846989 | 0.19606161 |
| ENSG00000188322 | *SBK1* | 2.776939717 | 0.052598175 | 0.026596282 | 0.026001893 | 0.999933887 | 0.042752602 | -0.170910314 | 0.213662916 | 0.19606161 |
| ENSG00000135931 | *ARMC9* | 0.09461823 | -0.422467336 | -0.518068188 | 0.095600853 | 0.999933887 | 0.50287414 | 0.836022259 | -0.333148119 | 0.196073865 |
| ENSG00000239306 | *RBM14* | 4.486638618 | -0.033106411 | -0.008327313 | -0.024779098 | 0.999933887 | 0.008454831 | -0.156459692 | 0.164914523 | 0.196073865 |
| ENSG00000135426 | *TESPA1* | 3.775325896 | -0.018113114 | -0.126766377 | 0.108653264 | 0.999933887 | 0.116834539 | 0.25163555 | -0.134801012 | 0.196073865 |
| ENSG00000106785 | *TRIM14* | 4.126977412 | 0.012022918 | -0.003247303 | 0.015270221 | 0.999933887 | -0.124766351 | -0.299431019 | 0.174664668 | 0.196123385 |
| ENSG00000075292 | *ZNF638* | 5.459637749 | -0.069003028 | -0.102419136 | 0.033416108 | 0.999933887 | -0.372395106 | -0.064982194 | -0.307412912 | 0.196180834 |
| ENSG00000268705 | *BNIP3P26* | -0.016659485 | 0.089236777 | 0.146766935 | -0.057530158 | 0.999933887 | -0.100067405 | 0.468031162 | -0.568098567 | 0.196180834 |
| ENSG00000105866 | *SP4* | 4.170566432 | 0.032658337 | 0.025126284 | 0.007532052 | 0.999933887 | -0.263506152 | 0.04553576 | -0.309041912 | 0.196320165 |
| ENSG00000122965 | *RBM19* | 3.545715682 | -3.7927e-05 | 0.054327618 | -0.054365545 | 0.999933887 | 0.238399942 | 0.41669044 | -0.178290498 | 0.196323964 |
| ENSG00000083312 | *TNPO1* | 6.203737491 | -0.041022639 | -0.102217565 | 0.061194926 | 0.999933887 | -0.585446824 | -0.336614621 | -0.248832203 | 0.196380423 |
| ENSG00000275339 | *Z99129.1* | 0.595158023 | 0.052207492 | -0.053636452 | 0.105843944 | 0.999933887 | -0.240702069 | 0.228372186 | -0.469074254 | 0.196539261 |
| ENSG00000233175 | *AC008105.1* | 1.556217538 | -0.070534545 | -0.044114697 | -0.026419847 | 0.999933887 | -0.518084468 | -0.190757148 | -0.327327321 | 0.196574605 |
| ENSG00000033627 | *ATP6V0A1* | 6.18618972 | -0.100894582 | -0.028980693 | -0.071913889 | 0.999933887 | -0.739423617 | -0.917836329 | 0.178412712 | 0.196621915 |
| ENSG00000167173 | *C15orf39* | 9.773225456 | -0.0460779 | -0.140970242 | 0.094892342 | 0.999933887 | 0.881874478 | 1.042122336 | -0.160247858 | 0.196793496 |
| ENSG00000164659 | *ELAPOR2* | 0.363163024 | -0.040814858 | -0.33849075 | 0.297675892 | 0.999933887 | -0.145615425 | 0.230456424 | -0.376071849 | 0.196793496 |
| ENSG00000198892 | *SHISA4* | -0.829418696 | 0.217601653 | 0.223562395 | -0.005960742 | 0.999933887 | -0.05902966 | -0.713373111 | 0.654343451 | 0.197048193 |
| ENSG00000129680 | *MAP7D3* | 0.731112094 | -0.070674236 | -0.223720891 | 0.153046654 | 0.999933887 | -0.192813491 | 0.123237361 | -0.316050852 | 0.197048193 |
| ENSG00000218018 | *RBM38-AS1* | 0.8251089 | -0.019044751 | 0.009599444 | -0.028644195 | 0.999933887 | -0.044967001 | -0.436037947 | 0.391070946 | 0.197183763 |
| ENSG00000169683 | *LRRC45* | 1.562848916 | -0.000126332 | 0.217850756 | -0.217977088 | 0.999933887 | 0.162189339 | -0.102169134 | 0.264358473 | 0.197472695 |
| ENSG00000065618 | *COL17A1* | 1.762367852 | -0.048064942 | 0.090483846 | -0.138548788 | 0.999933887 | 3.834598555 | 4.400063359 | -0.565464803 | 0.197577067 |
| ENSG00000118197 | *DDX59* | 4.313960224 | -0.058789195 | 0.110094996 | -0.168884191 | 0.999933887 | -0.307080388 | -0.155565911 | -0.151514477 | 0.197607008 |
| ENSG00000257621 | *PSMA3-AS1* | 6.300888875 | -0.017886405 | -0.107351892 | 0.089465487 | 0.999933887 | -0.218388253 | -0.008500697 | -0.209887556 | 0.197607008 |
| ENSG00000100393 | *EP300* | 9.093806431 | -0.009834847 | -0.016463801 | 0.006628955 | 0.999933887 | -0.150932101 | -0.024328685 | -0.126603416 | 0.197607008 |
| ENSG00000164209 | *SLC25A46* | 3.660844706 | -0.012825063 | -0.135959797 | 0.123134734 | 0.999933887 | -0.22058637 | 0.050579804 | -0.271166174 | 0.197956608 |
| ENSG00000079999 | *KEAP1* | 4.785558611 | -0.006959434 | -0.132467313 | 0.125507879 | 0.999933887 | 0.343562363 | 0.511403883 | -0.167841519 | 0.197984176 |
| ENSG00000198695 | *MT-ND6* | 6.690793266 | 0.005390806 | 0.027065013 | -0.021674208 | 0.999933887 | -0.010477852 | 0.208377045 | -0.218854897 | 0.198062285 |
| ENSG00000227032 | *RPS2P36* | 1.270450085 | 0.116559889 | 0.256060715 | -0.139500826 | 0.999933887 | 2.476949956 | 2.919852004 | -0.442902048 | 0.198133737 |

| | | | | | | | | | | |
|---|---|---|---|---|---|---|---|---|---|---|
| ENSG00000226233 | *Z82188.1* | -0.807766546 | 0.360578279 | -0.175614105 | 0.536192384 | 0.999933887 | 0.76006762 | 1.249062805 | -0.488995185 | 0.198341898 |
| ENSG00000102119 | *EMD* | 7.531002598 | 0.106829157 | 0.113959646 | -0.007130489 | 0.999933887 | 0.569594302 | 0.728938574 | -0.159344273 | 0.198589712 |
| ENSG00000141480 | *ARRB2* | 9.725086708 | -0.001775364 | 0.016768644 | -0.018544009 | 0.999933887 | -0.400598251 | -0.525489439 | 0.124891187 | 0.198660463 |
| ENSG00000276718 | *AC005840.4* | 0.282184149 | -0.155032269 | 0.062809704 | -0.217841972 | 0.999933887 | -0.125559455 | -0.603876575 | 0.47831712 | 0.198693911 |
| ENSG00000145734 | *BDP1* | 4.189067537 | -0.103884877 | -0.03939343 | -0.064491447 | 0.999933887 | -0.343500335 | 0.08298176 | -0.426482096 | 0.198693911 |
| ENSG00000102978 | *POLR2C* | 4.390580802 | -0.039684002 | 0.030490702 | -0.070174705 | 0.999933887 | -0.092011955 | -0.271254046 | 0.179242091 | 0.198693911 |
| ENSG00000132405 | *TBC1D14* | 6.573201191 | -0.023342315 | -0.057941123 | 0.034598809 | 0.999933887 | -0.592947046 | -0.742202941 | 0.149255895 | 0.198693911 |
| ENSG00000164180 | *TMEM161B* | 2.626080322 | -0.040622389 | -0.104829085 | 0.064206696 | 0.999933887 | -0.347832067 | -0.037982815 | -0.309849251 | 0.198944725 |
| ENSG00000251230 | *MIR3945HG* | 2.412585636 | 0.13672564 | 0.098264635 | 0.038461005 | 0.999933887 | 2.752940384 | 3.175819509 | -0.422879124 | 0.199009695 |
| ENSG00000196757 | *ZNF700* | 3.148859006 | -0.107093259 | -0.096980125 | -0.010113134 | 0.999933887 | -0.433877074 | -0.707841539 | 0.273964465 | 0.199009695 |
| ENSG00000173530 | *TNFRSF10D* | 1.116496837 | -0.066993073 | 0.071476142 | -0.138469215 | 0.999933887 | -0.312465691 | -0.718320416 | 0.405854725 | 0.199009695 |
| ENSG00000259820 | *AC083843.2* | 2.386712739 | -0.168164408 | -0.212282161 | 0.044117753 | 0.999933887 | 0.059140819 | 0.537342828 | -0.478202008 | 0.199266532 |
| ENSG00000244625 | *MIATNB* | 3.206703678 | 0.045198423 | -0.029994338 | 0.07519276 | 0.999933887 | -0.369262675 | -0.540518451 | 0.171255776 | 0.199266532 |
| ENSG00000145012 | *LPP* | 5.084687729 | 0.005459727 | -0.007768245 | 0.013227972 | 0.999933887 | -0.1953495 | 0.052906044 | -0.248255544 | 0.199266532 |
| ENSG00000181038 | *METTL23* | 3.908292021 | 0.049242829 | 0.084602961 | -0.035360132 | 0.999933887 | 0.612300411 | 0.79994858 | -0.187648169 | 0.199306018 |
| ENSG00000177150 | *FAM210A* | 3.516441912 | 0.017323055 | 0.028100849 | -0.010777794 | 0.999933887 | 0.199819191 | 0.469008427 | -0.269189236 | 0.199317422 |
| ENSG00000155827 | *RNF20* | 4.217015335 | -0.037477898 | -0.123332817 | 0.085854919 | 0.999933887 | -0.475985006 | -0.658788982 | 0.182803975 | 0.199606608 |
| ENSG00000091136 | *LAMB1* | 1.029658159 | -0.234004299 | -0.035599891 | -0.198404408 | 0.999933887 | 0.073180579 | 0.397285416 | -0.324104837 | 0.199628604 |
| ENSG00000162604 | *TM2D1* | 2.722576197 | 0.043747427 | 0.043422334 | 0.000325094 | 0.999933887 | -0.224610954 | -0.044346674 | -0.18026428 | 0.199628604 |
| ENSG00000272047 | *GTF2H5* | 1.09055742 | -0.00932243 | -0.170927579 | 0.161605149 | 0.999933887 | -0.193407541 | 0.145815724 | -0.339223265 | 0.199628604 |
| ENSG00000229473 | *RGS17P1* | 2.326719132 | 0.015999966 | 0.011538988 | 0.004460978 | 0.999933887 | 0.495219924 | 0.786402108 | -0.291182184 | 0.19963107 |
| ENSG00000163608 | *NEPRO* | 3.939331577 | -0.114140842 | -0.171605724 | 0.057464881 | 0.999933887 | -0.052057972 | 0.108775218 | -0.16083319 | 0.199657821 |
| ENSG00000248996 | *AC145098.1* | 4.68856727 | -0.02260588 | -0.010343894 | -0.012261986 | 0.999933887 | -0.306881884 | -0.120246008 | -0.186635876 | 0.199657821 |
| ENSG00000143612 | *C1orf43* | 7.658755053 | -0.014442514 | -0.053664418 | 0.039221903 | 0.999933887 | 0.05424652 | 0.198061844 | -0.143815325 | 0.199722321 |
| ENSG00000128606 | *LRRC17* | -0.364921159 | 0.230409757 | 0.141596008 | 0.088813749 | 0.999933887 | -0.303310868 | 0.197400323 | -0.500711191 | 0.199782955 |
| ENSG00000108784 | *NAGLU* | 3.749597044 | -0.088685196 | -0.202818764 | 0.114133568 | 0.999933887 | -0.430951706 | -0.623458461 | 0.192506755 | 0.199840367 |
| ENSG00000254317 | *AC022973.5* | 1.63219855 | 0.034469463 | 0.19559467 | -0.161125207 | 0.999933887 | -0.237250295 | 0.194893115 | -0.43214341 | 0.199840367 |
| ENSG00000157181 | *ODR4* | 3.188717631 | 0.001801913 | -0.123476739 | 0.125278652 | 0.999933887 | -0.102645261 | 0.226904478 | -0.329549739 | 0.199840367 |
| ENSG00000196821 | *ILRUN* | 7.10877493 | -0.065713416 | -0.032588163 | -0.033125254 | 0.999933887 | 0.014273187 | 0.140089949 | -0.125816761 | 0.199964441 |
| ENSG00000136689 | *IL1RN* | 9.211585529 | 0.063303932 | 0.054122856 | 0.009181076 | 0.999933887 | 2.197090949 | 2.646006395 | -0.448915446 | 0.199994474 |
| ENSG00000214922 | *HLA-F-AS1* | 2.151527292 | 0.100430986 | 0.013893842 | 0.086537144 | 0.999933887 | 0.730462896 | 1.036522546 | -0.306059651 | 0.20013866 |
| ENSG00000100365 | *NCF4* | 7.939531781 | 0.042184221 | 0.007992592 | 0.034191629 | 0.999933887 | 0.672031555 | 0.486279544 | 0.185752011 | 0.20013866 |
| ENSG00000167981 | *ZNF597* | 2.499887945 | -0.048754182 | -0.066079831 | 0.017325649 | 0.999933887 | 0.459861006 | 0.776856786 | -0.31699578 | 0.20013866 |
| ENSG00000168785 | *TSPAN5* | 0.990364079 | 0.015717593 | -0.016326627 | 0.03204422 | 0.999933887 | 0.065931856 | 0.40487113 | -0.338939274 | 0.200219493 |
| ENSG00000169062 | *UPF3A* | 3.510572986 | -0.075901183 | 0.00804436 | -0.083945543 | 0.999933887 | -0.177423386 | -0.368965285 | 0.191541898 | 0.200351112 |
| ENSG00000146729 | *NIPSNAP2* | 2.735961631 | 0.006722523 | 0.035240306 | -0.028517783 | 0.999933887 | 0.106088404 | 0.322173133 | -0.216084729 | 0.200725708 |
| ENSG00000141977 | *CIB3* | -1.048114633 | -0.046224654 | -0.085701288 | 0.039476634 | 0.999933887 | -0.124192741 | -1.103842398 | 0.979649657 | 0.200747001 |
| ENSG00000111011 | *RSRC2* | 6.563669984 | -0.037611675 | -0.054317997 | 0.016706322 | 0.999933887 | -0.076577537 | 0.13032767 | -0.206905207 | 0.201116779 |
| ENSG00000168016 | *TRANK1* | 7.727611046 | -0.027803953 | -0.077883088 | 0.050079134 | 0.999933887 | -0.105941056 | -0.259809702 | 0.153868646 | 0.201193581 |
| ENSG00000061794 | *MRPS35* | 1.407632782 | -0.042204248 | -0.107042319 | 0.064838071 | 0.999933887 | 0.109170097 | 0.40025914 | -0.291089043 | 0.201193581 |
| ENSG00000280213 | *UCKL1-AS1* | 2.117387605 | 0.031283425 | 0.139521061 | -0.108237635 | 0.999933887 | 0.388247226 | 0.684263386 | -0.296016161 | 0.201193581 |
| ENSG00000143669 | *LYST* | 8.383569106 | -0.030702861 | -0.082215066 | 0.051512205 | 0.999933887 | -0.126255382 | 0.118294154 | -0.244549537 | 0.201193581 |
| ENSG00000165312 | *OTUD1* | 5.45079299 | -0.010137281 | 0.020168576 | -0.030305858 | 0.999933887 | -0.467875306 | -0.637434167 | 0.169558861 | 0.201193581 |
| ENSG00000236577 | *SNRPGP14* | 0.40251416 | -0.093738928 | 0.449709806 | -0.543448734 | 0.999933887 | 0.324383784 | 0.604296928 | -0.279913144 | 0.201263205 |
| ENSG00000163466 | *ARPC2* | 9.236149927 | -0.020218877 | -0.006702324 | -0.013516553 | 0.999933887 | -0.101397558 | -0.19784244 | 0.096444882 | 0.201352469 |
| ENSG00000106397 | *PLOD3* | 3.103260916 | 0.055235086 | 0.016414653 | 0.038820432 | 0.999933887 | -0.175999675 | -0.385713687 | 0.209714012 | 0.201418898 |
| ENSG00000269352 | *PTOV1-AS2* | 1.966491718 | 0.033552975 | 0.099993956 | -0.066440981 | 0.999933887 | -0.212090778 | -0.528982328 | 0.31689155 | 0.201418898 |
| ENSG00000139793 | *MBNL2* | 2.324878572 | 0.035336488 | -0.14231105 | 0.177647538 | 0.999933887 | -0.157465169 | 0.275443703 | -0.432908872 | 0.201418898 |
| ENSG00000179958 | *DCTPP1* | 0.584673836 | 0.116707587 | -0.13952088 | 0.256228467 | 0.999933887 | 0.191468466 | 0.578654503 | -0.387186037 | 0.201419794 |
| ENSG00000237854 | *LINC00674* | 2.271509983 | 0.109021298 | -0.012672305 | 0.121693602 | 0.999933887 | 0.075342876 | -0.125952115 | 0.201294991 | 0.201645777 |
| ENSG00000109670 | *FBXW7* | 6.40244251 | 0.00478782 | 0.012113516 | -0.007325697 | 0.999933887 | 0.141446988 | 0.324448611 | -0.183001624 | 0.201702726 |
| ENSG00000230844 | *ZNF674-AS1* | 4.787185561 | 0.086546227 | 0.126491847 | -0.03994562 | 0.999933887 | 0.601114272 | 0.803205677 | -0.202091404 | 0.201854762 |
| ENSG00000166886 | *NAB2* | 4.512122108 | 0.026011743 | -0.191806702 | 0.217818445 | 0.999933887 | -1.287523721 | -1.543178596 | 0.255654875 | 0.201854762 |
| ENSG00000136504 | *KAT7* | 5.597170939 | -0.021952605 | -0.028679045 | 0.00672644 | 0.999933887 | -0.300243885 | -0.445148572 | 0.144904687 | 0.201862142 |
| ENSG00000204388 | *HSPA1B* | 3.272113394 | -0.048492296 | -0.010540374 | -0.037951922 | 0.999933887 | 0.513891948 | 0.870866323 | -0.356974374 | 0.201862142 |
| ENSG00000095574 | *IKZF5* | 4.930826966 | -0.024359614 | -0.033623883 | 0.009264269 | 0.999933887 | -0.296402101 | -0.12726751 | -0.169134591 | 0.202003879 |
| ENSG00000084623 | *EIF3I* | 5.30007952 | 0.050832502 | -0.012553759 | 0.06338626 | 0.999933887 | 0.18855323 | 0.336333329 | -0.147780098 | 0.202049585 |
| ENSG00000117090 | *SLAMF1* | 3.243848839 | -0.071535933 | -0.118891589 | 0.047355656 | 0.999933887 | 0.276520365 | 0.473542327 | -0.197021963 | 0.202063266 |
| ENSG00000129535 | *NRL* | -0.042732824 | 0.242788418 | -0.18320434 | 0.425992758 | 0.999933887 | -0.279541853 | -0.837102057 | 0.557560204 | 0.202136416 |
| ENSG00000081721 | *DUSP12* | 1.582771609 | -0.063391629 | -0.058504186 | -0.004887444 | 0.999933887 | 0.149889532 | -0.150819239 | 0.300708771 | 0.202136416 |
| ENSG00000069869 | *NEDD4* | 0.803384494 | 0.029363073 | -0.173705995 | 0.203069068 | 0.999933887 | 0.153520398 | 0.570956471 | -0.417436073 | 0.202136416 |
| ENSG00000054116 | *TRAPPC3* | 5.356175229 | 0.004036009 | -0.0561287 | 0.060164708 | 0.999933887 | -0.079508613 | 0.074172059 | -0.153680671 | 0.202136416 |
| ENSG00000015479 | *MATR3* | 2.492096148 | 0.033041235 | 0.052455374 | -0.019414138 | 0.999933887 | -0.152856033 | 0.239045541 | -0.391901574 | 0.20218443 |
| ENSG00000104894 | *CD37* | 8.285953673 | 0.026106531 | 0.012249807 | 0.013856724 | 0.999933887 | -0.104258131 | -0.238297366 | 0.134039235 | 0.202189559 |
| ENSG00000170382 | *LRRN2* | 3.453589991 | -0.118938856 | 0.101570418 | -0.220509273 | 0.999933887 | -0.522802242 | -0.735522968 | 0.212720726 | 0.202368017 |
| ENSG00000167995 | *BEST1* | 7.856505041 | 0.057066946 | 0.057535442 | -0.000468496 | 0.999933887 | 0.327675267 | 0.456636583 | -0.128961316 | 0.202368017 |
| ENSG00000106052 | *TAX1BP1* | 6.726240257 | -0.069927763 | -0.136784673 | 0.06685691 | 0.999933887 | -0.308348275 | -0.124192848 | -0.184155427 | 0.202368017 |
| ENSG00000196507 | *TCEAL3* | 0.91557481 | 0.182549539 | 0.020297647 | 0.162251892 | 0.999933887 | 0.318757785 | 0.649030193 | -0.330272408 | 0.202636302 |
| ENSG00000196961 | *AP2A1* | 5.983901361 | 0.011533926 | 0.001449195 | 0.010084731 | 0.999933887 | -0.409133012 | -0.537915989 | 0.128782977 | 0.202738688 |
| ENSG00000122644 | *ARL4A* | 3.726489523 | 0.077241871 | 0.039729404 | 0.037512467 | 0.999933887 | 0.600150178 | 0.817530383 | -0.217380204 | 0.202749692 |
| ENSG00000223705 | *NSUN5P1* | 2.7940126 | 0.032502671 | 0.071699098 | -0.039196428 | 0.999933887 | 0.04411699 | -0.157095007 | 0.201211997 | 0.202749692 |
| ENSG00000066557 | *LRRC40* | 1.352237073 | -0.151755507 | -0.163822154 | 0.012066648 | 0.999933887 | -0.082558811 | 0.337498548 | -0.420057359 | 0.202883127 |
| ENSG00000253556 | *MTCO1P4* | -0.75750417 | 0.385872235 | 0.868876603 | -0.483004368 | 0.999933887 | 2.081671533 | 2.749226772 | -0.667555239 | 0.202923389 |
| ENSG00000272316 | *AL021368.2* | -1.064827532 | -0.202180821 | -0.351951912 | 0.149771092 | 0.999933887 | -0.387763214 | 0.160222238 | -0.547985452 | 0.20301073 |
| ENSG00000129691 | *ASH2L* | 3.354853478 | 0.044375922 | 0.018145526 | 0.026230395 | 0.999933887 | -0.392161747 | -0.580768582 | 0.188606835 | 0.20301073 |
| ENSG00000073331 | *ALPK1* | 6.667106283 | -0.031224944 | -0.046983066 | 0.015758122 | 0.999933887 | 0.305921357 | 0.448961308 | -0.143039951 | 0.20301073 |
| ENSG00000186575 | *NF2* | 2.921943542 | 0.008952619 | 0.066028217 | -0.057075597 | 0.999933887 | -0.079253392 | -0.370065452 | 0.29081206 | 0.20301073 |
| ENSG00000265656 | *AC106037.3* | -0.858252135 | -0.008476665 | 0.000417807 | -0.008894472 | 0.999933887 | -0.7082289 | -1.280582098 | 0.572353198 | 0.20301073 |
| ENSG00000116675 | *DNAJC6* | -0.968898966 | -0.324627537 | -0.059544947 | -0.26508259 | 0.999933887 | 0.056342816 | 0.530903836 | -0.47456102 | 0.203103104 |
| ENSG00000117281 | *CD160* | 0.766610422 | -0.066612112 | -0.268519929 | 0.201907817 | 0.999933887 | 0.065173211 | 0.434575055 | -0.369401845 | 0.203337807 |
| ENSG00000007392 | *LUC7L* | 4.659595495 | -0.023394382 | -0.000618629 | -0.022775752 | 0.999933887 | 0.074543002 | -0.04146528 | 0.116008282 | 0.204086508 |
| ENSG00000143858 | *SYT2* | 0.417151308 | -0.090129118 | 0.083548138 | -0.173677255 | 0.999933887 | -1.159539867 | -1.649512506 | 0.489972639 | 0.204088963 |
| ENSG00000169689 | *CENPX* | 1.490196508 | 0.140810654 | -0.014421735 | 0.155232389 | 0.999933887 | 0.216359565 | -0.136674436 | 0.353034001 | 0.204156746 |
| ENSG00000168894 | *RNF181* | 4.180966032 | -0.025494163 | 0.024462001 | -0.049956164 | 0.999933887 | -0.265890864 | -0.43188433 | 0.165993465 | 0.204204694 |
| ENSG00000184557 | *SOCS3* | 7.322245178 | 0.495545422 | 0.518487132 | -0.02294171 | 0.999933887 | 0.563328158 | 0.873553519 | -0.310225361 | 0.20422717 |
| ENSG00000161714 | *PLCD3* | 1.391745208 | 0.024126822 | -0.038271148 | 0.062397971 | 0.999933887 | -0.544583231 | -0.86484689 | 0.320263659 | 0.20422717 |
| ENSG00000275385 | *CCL18* | -0.184975449 | 0.386090249 | 0.219971606 | 0.166118643 | 0.999933887 | 1.364881081 | 1.779914993 | -0.415033912 | 0.204461643 |
| ENSG00000115207 | *GTF3C2* | 3.625354181 | -0.048057783 | 0.008772985 | -0.056830768 | 0.999933887 | -0.506960311 | -0.675867842 | 0.168907531 | 0.204461643 |
| ENSG00000173598 | *NUDT4* | 4.397737858 | -0.034592103 | 0.03112653 | -0.065718632 | 0.999933887 | -0.048864204 | 0.120426688 | -0.169290892 | 0.204461643 |
| ENSG00000119383 | *PTPA* | 3.49040645 | 0.013056298 | 0.040261412 | -0.027205114 | 0.999933887 | -0.027244822 | -0.23797464 | 0.210729818 | 0.204461643 |
| ENSG00000124193 | *SRSF6* | 6.27521514 | -0.051711769 | 0.008733463 | -0.060445232 | 0.999933887 | -0.092857731 | -0.203274228 | 0.110416497 | 0.204599989 |
| ENSG00000198168 | *SVIP* | 3.142754333 | -0.050260428 | -0.128469556 | 0.078209128 | 0.999933887 | 0.008067223 | 0.242884535 | -0.234817313 | 0.204599989 |
| ENSG00000145391 | *SETD7* | 2.278996187 | 0.047393826 | 0.088261645 | -0.040867819 | 0.999933887 | -0.559984785 | -0.881953456 | 0.321968671 | 0.204599989 |
| ENSG00000123405 | *NFE2* | 4.850055256 | 0.019338608 | 0.050944177 | -0.031605569 | 0.999933887 | -0.041722227 | -0.25215937 | 0.210437143 | 0.204701456 |
| ENSG00000241399 | *CD302* | 2.284790487 | -0.067786218 | 0.025983424 | -0.093769642 | 0.999933887 | -0.803588115 | -1.189962526 | 0.386374411 | 0.204995443 |
| ENSG00000278845 | *MRPL45* | 1.174916901 | -0.039196795 | 0.00704524 | -0.046242035 | 0.999933887 | 0.056899271 | 0.36185084 | -0.304951569 | 0.205047192 |
| ENSG00000146859 | *TMEM140* | 8.069630565 | -0.014091124 | -0.055108662 | 0.041017538 | 0.999933887 | -0.300678593 | -0.131622651 | -0.169055942 | 0.205077737 |
| ENSG00000225663 | *MCRIP1* | 5.01952148 | 0.052247814 | -0.007988881 | 0.060236694 | 0.999933887 | 0.280878336 | 0.436202091 | -0.155323755 | 0.205164696 |
| ENSG00000128607 | *KLHDC10* | 3.366528759 | -0.000833254 | -0.055347602 | 0.054514347 | 0.999933887 | -0.493586843 | -0.688702553 | 0.195115709 | 0.205164696 |
| ENSG00000150401 | *DCUN1D2* | 3.474580894 | -0.06623733 | -0.0314665 | -0.03477083 | 0.999933887 | -0.661202587 | -0.855875783 | 0.194673196 | 0.20525403 |
| ENSG00000261505 | *AL031714.1* | 2.890887975 | -0.068300464 | -0.013916871 | -0.054383593 | 0.999933887 | 0.170134669 | 0.372783115 | -0.202648446 | 0.20525403 |
| ENSG00000122783 | *CYREN* | 7.294068143 | -0.018928228 | -0.002334754 | -0.016593474 | 0.999933887 | -0.229304676 | -0.087210445 | -0.142094231 | 0.20525403 |
| ENSG00000075275 | *CELSR1* | 0.267663794 | 0.125547599 | -0.030695447 | 0.156243047 | 0.999933887 | -0.363137817 | -0.007467006 | -0.35567081 | 0.205493125 |
| ENSG00000143614 | *GATAD2B* | 6.216365285 | -0.082594755 | -0.011011335 | -0.07158342 | 0.999933887 | -0.540780537 | -0.687596906 | 0.146816369 | 0.205555199 |
| ENSG00000103257 | *SLC7A5* | 6.350055892 | 0.076280809 | 0.041320739 | 0.034960069 | 0.999933887 | 1.595486851 | 1.985164352 | -0.389677501 | 0.205555199 |

| | | | | | | | | | | |
|---|---|---|---|---|---|---|---|---|---|---|
| ENSG00000269303 | *AC020907.3* | -0.442434602 | 0.121334496 | 0.439116347 | -0.317781851 | 0.999933887 | 2.671951782 | 3.384398589 | -0.712446808 | 0.205565536 |
| ENSG00000138606 | *SHF* | -0.086826206 | 0.010895211 | -0.219973404 | 0.230868614 | 0.999933887 | 0.593847666 | 0.949743956 | -0.355896291 | 0.205611947 |
| ENSG00000109680 | *TBC1D19* | -0.101250652 | 0.12233879 | -0.342488575 | 0.464827365 | 0.999933887 | -0.070777602 | 0.427773586 | -0.498551188 | 0.205825341 |
| ENSG00000176095 | *IP6K1* | 7.144338014 | -0.065493552 | -0.042016838 | -0.023476713 | 0.999933887 | -0.137985201 | -0.011271896 | -0.126713305 | 0.205884958 |
| ENSG00000167978 | *SRRM2* | 10.056826482 | 0.007319141 | -0.009240226 | 0.016559368 | 0.999933887 | -0.09202052 | 0.004622242 | -0.096642762 | 0.206476137 |
| ENSG00000205423 | *CNEP1R1* | 4.11609446 | 0.001367387 | 0.043707798 | -0.042340411 | 0.999933887 | -0.120104111 | 0.070464239 | -0.19056835 | 0.206569407 |
| ENSG00000238105 | *GOLGA2P5* | 2.43671913 | -0.004731366 | -0.001595097 | -0.003136268 | 0.999933887 | 0.060906066 | -0.127837968 | 0.188744034 | 0.206642147 |
| ENSG00000166971 | *AKTIP* | 2.700341696 | 0.0513076 | 0.104820075 | -0.053512476 | 0.999933887 | -0.220272459 | -0.481933511 | 0.261661053 | 0.206644168 |
| ENSG00000225855 | *RUSC1-AS1* | 0.506272167 | -0.028943487 | 0.106016794 | -0.134960282 | 0.999933887 | 0.091772509 | -0.295081698 | 0.386854208 | 0.206717658 |
| ENSG00000157796 | *WDR19* | 3.371354337 | -0.067725669 | -0.096307954 | 0.028582284 | 0.999933887 | -0.248224438 | -0.412700064 | 0.164475626 | 0.206864637 |
| ENSG00000073350 | *LLGL2* | 3.538834269 | 0.010811172 | 0.04640992 | -0.035598748 | 0.999933887 | 0.116375266 | -0.070481798 | 0.186857064 | 0.207060691 |
| ENSG00000204291 | *COL15A1* | 1.608881251 | 0.076596099 | 0.170978547 | -0.094382448 | 0.999933887 | 1.727781734 | 2.182884049 | -0.455102314 | 0.207132267 |
| ENSG00000285693 | *AP002381.2* | 3.115356322 | -0.095635711 | -0.102574397 | 0.006938686 | 0.999933887 | -0.81526546 | -1.075397294 | 0.260131834 | 0.207299667 |
| ENSG00000176014 | *TUBB6* | 1.101766193 | -0.059163005 | -0.247514014 | 0.188351009 | 0.999933887 | -0.162242859 | 0.123465224 | -0.285708083 | 0.208154186 |
| ENSG00000069998 | *HDHD5* | 1.8194702 | -0.020016365 | 0.215214076 | -0.235230442 | 0.999933887 | 0.13062737 | -0.114082501 | 0.244709871 | 0.208154186 |
| ENSG00000260086 | *AC007611.1* | 1.918678797 | 0.277575951 | 0.442380211 | -0.16480426 | 0.999933887 | 1.786853826 | 1.467816069 | 0.319037757 | 0.208292773 |
| ENSG00000064115 | *TM7SF3* | 2.290603047 | 0.079971804 | 0.090971518 | -0.010999714 | 0.999933887 | -0.221576281 | -0.494234199 | 0.272657918 | 0.208292773 |
| ENSG00000178573 | *MAF* | 4.439105388 | 0.021225464 | 0.01607909 | 0.005146374 | 0.999933887 | 0.030233142 | -0.124107863 | 0.154341004 | 0.208292773 |
| ENSG00000235859 | *AC006978.1* | 2.541567731 | 0.00558867 | -0.058101361 | 0.063690031 | 0.999933887 | -0.444488557 | -0.815249562 | 0.370761005 | 0.208715828 |
| ENSG00000204020 | *LIPN* | 5.285532734 | -0.005014143 | -0.056353636 | 0.051339493 | 0.999933887 | 0.793226129 | 1.106231766 | -0.313005637 | 0.208715828 |
| ENSG00000270055 | *AC127502.2* | 0.680883054 | -0.076298673 | 0.457444723 | -0.533743396 | 0.999933887 | -0.156858945 | 0.288345771 | -0.445204716 | 0.209006869 |
| ENSG00000183250 | *LINC01547* | 1.209043008 | -0.12455397 | 0.009000159 | -0.133554129 | 0.999933887 | 0.979373763 | 1.520919254 | -0.54154549 | 0.209144882 |
| ENSG00000286256 | *AC084871.3* | 2.627079471 | 0.035594952 | 0.053282833 | -0.017687881 | 0.999933887 | 2.740514292 | 3.125407972 | -0.38489368 | 0.209144882 |
| ENSG00000170113 | *NIPA1* | 1.941682273 | 0.039773212 | 0.048021646 | -0.008248435 | 0.999933887 | 0.069003487 | 0.303514019 | -0.234510532 | 0.209255892 |
| ENSG00000149781 | *FERMT3* | 8.984504227 | 0.060899751 | 0.04955454 | 0.01134521 | 0.999933887 | 0.197741755 | 0.350781472 | -0.153039717 | 0.209260934 |
| ENSG00000198574 | *SH2D1B* | 0.402245452 | -0.044293042 | -0.07840315 | 0.034110107 | 0.999933887 | 0.392322849 | 0.792800754 | -0.400477905 | 0.209368001 |
| ENSG00000174437 | *ATP2A2* | 5.819159113 | -0.000634895 | -0.010638821 | 0.010003926 | 0.999933887 | -0.0967436 | -0.202171797 | 0.105428197 | 0.209390997 |
| ENSG00000104324 | *CPQ* | 6.264197659 | -0.000358913 | -0.049997071 | 0.049638159 | 0.999933887 | -0.146540654 | -0.280625179 | 0.134084525 | 0.209400863 |
| ENSG00000114737 | *CISH* | 1.666091077 | -0.090744791 | -0.109440842 | 0.018696052 | 0.999933887 | 0.085516648 | -0.303672002 | 0.38918865 | 0.209419523 |
| ENSG00000140403 | *DNAJA4* | 2.596098751 | -0.010496249 | -0.026106309 | 0.01561006 | 0.999933887 | 0.127803172 | 0.342266126 | -0.214462954 | 0.209534681 |
| ENSG00000072121 | *ZFYVE26* | 6.089405533 | -0.026366112 | 0.017072031 | -0.043438143 | 0.999933887 | 0.006096137 | 0.162272033 | -0.156175895 | 0.209624983 |
| ENSG00000251442 | *LINC01094* | 1.799487076 | -0.085141014 | -0.138994408 | 0.053853395 | 0.999933887 | -0.725790835 | -1.10465184 | 0.378861005 | 0.209952292 |
| ENSG00000257218 | *GATC* | 1.688325016 | 0.002659262 | -0.043060416 | 0.045719677 | 0.999933887 | 0.011653568 | -0.282633503 | 0.294287071 | 0.209952292 |
| ENSG00000171786 | *NHLH1* | 0.012003794 | -0.104185355 | 0.292937998 | -0.397123353 | 0.944094537 | -0.306013111 | 0.256739803 | -0.562752914 | 0.210117899 |
| ENSG00000007341 | *ST7L* | 3.525712695 | 0.094908949 | 0.081869098 | 0.013039852 | 0.999933887 | 0.403255346 | 0.586033032 | -0.182777686 | 0.210140885 |
| ENSG00000170832 | *USP32* | 7.525524438 | -0.044594507 | -0.08039628 | 0.035801773 | 0.999933887 | -0.671339416 | -0.826806027 | 0.155466611 | 0.210140885 |
| ENSG00000225931 | *AL139246.2* | 4.188291812 | 0.337292302 | 0.451600417 | -0.114308115 | 0.999933887 | 1.440095157 | 1.762292622 | -0.322197465 | 0.210571279 |
| ENSG00000086730 | *LAT2* | 8.05287523 | 0.02933135 | 0.002241776 | 0.027089574 | 0.999933887 | -0.187268803 | -0.352572457 | 0.165303653 | 0.210586225 |
| ENSG00000008853 | *RHOBTB2* | 0.65904543 | 0.240464003 | 0.215967363 | 0.02449664 | 0.999933887 | -0.164727922 | -0.543469232 | 0.37874131 | 0.210626477 |
| ENSG00000257315 | *ZBED6* | 0.96228462 | 0.093687088 | 0.063616462 | 0.030070627 | 0.999933887 | -0.154892688 | 0.117697494 | -0.272590181 | 0.210919769 |
| ENSG00000105204 | *DYRK1B* | 3.826701816 | 0.034368679 | 0.045671983 | -0.011303304 | 0.999933887 | -0.422214081 | -0.585613141 | 0.163399059 | 0.210919769 |
| ENSG00000145428 | *RNF175* | 4.269331754 | -0.138154662 | 0.032614693 | -0.170769355 | 0.999933887 | 0.320838866 | 0.608071291 | -0.287232425 | 0.210996174 |
| ENSG00000140350 | *ANP32A* | 8.640934803 | -0.058681862 | -0.054468659 | -0.004213204 | 0.999933887 | 0.850815782 | 0.990014258 | -0.139198476 | 0.211136931 |
| ENSG00000148180 | *GSN* | 7.41764167 | -0.045468261 | 0.037858301 | -0.083326562 | 0.999933887 | -0.51153821 | -0.736024376 | 0.224486166 | 0.211144111 |
| ENSG00000257605 | *MYG1-AS1* | 0.671234136 | -0.023107208 | -0.118499072 | 0.095391864 | 0.999933887 | -0.138755786 | 0.296191804 | -0.434947589 | 0.211144111 |
| ENSG00000275066 | *SYNRG* | 5.866459677 | -0.023820579 | -0.032730174 | 0.008909595 | 0.999933887 | -0.239794442 | -0.121464316 | -0.118330127 | 0.211228175 |
| ENSG00000162711 | *NLRP3* | 3.867420876 | 0.185260778 | 0.173198398 | 0.01206238 | 0.999933887 | 1.350080047 | 1.817186357 | -0.46710631 | 0.211259098 |
| ENSG00000170837 | *GPR27* | 3.568413941 | 0.020667335 | 0.060063798 | -0.039396463 | 0.999933887 | -0.587373755 | -0.896701274 | 0.309327519 | 0.211259098 |
| ENSG00000114423 | *CBLB* | 4.551189251 | -0.050275496 | 0.106139407 | -0.156414903 | 0.999933887 | -0.037254859 | 0.16227091 | -0.199525769 | 0.211505812 |
| ENSG00000229638 | *RPL4P4* | -0.756844348 | 0.070124822 | 0.163426257 | -0.093301435 | 0.999933887 | 0.137799459 | -0.387829076 | 0.525628535 | 0.211541619 |
| ENSG00000175040 | *CHST2* | 3.376776869 | 0.057676396 | -0.000643088 | 0.058319484 | 0.999933887 | 0.201689204 | 0.50143171 | -0.299742506 | 0.211643255 |
| ENSG00000178996 | *SNX18* | 6.402132813 | -0.009469373 | 0.045844184 | -0.055313557 | 0.999933887 | -0.962579027 | -1.243580746 | 0.281001718 | 0.211777011 |
| ENSG00000235081 | *AC245052.3* | -0.972438835 | 0.179126637 | -0.028740258 | 0.207866895 | 0.999933887 | -0.639670332 | 0.201425261 | -0.841095592 | 0.211871914 |
| ENSG00000110619 | *CARS1* | 5.676116628 | -0.032781833 | 0.030580671 | -0.063362504 | 0.999933887 | 0.14750271 | 0.255024896 | -0.107522186 | 0.211871914 |
| ENSG00000254996 | *ANKHD1-EIF4* | 2.015838751 | 0.083418265 | 0.041629245 | 0.041789021 | 0.999933887 | 0.044809066 | 0.43059895 | -0.385789884 | 0.211871914 |
| ENSG00000168038 | *ULK4* | 2.295705627 | 0.005656046 | 0.028215813 | -0.022559767 | 0.999933887 | -0.051478562 | 0.220782923 | -0.272261485 | 0.211871914 |
| ENSG00000105617 | *LENG1* | 2.194909898 | 0.097055371 | 0.124880889 | -0.027825518 | 0.999933887 | -0.343165415 | -0.609022242 | 0.265856827 | 0.21195424 |
| ENSG00000260711 | *AL121839.2* | 0.080759043 | 0.007386226 | 0.200942079 | -0.193555853 | 0.999933887 | -0.29519921 | 0.238994032 | -0.534193242 | 0.212183806 |
| ENSG00000147669 | *POLR2K* | 2.815064837 | -0.065268549 | -0.060985661 | -0.004282888 | 0.999933887 | 0.354733445 | 0.611812597 | -0.257079152 | 0.212198323 |
| ENSG00000277595 | *AC007546.1* | 0.958502726 | -0.048652221 | -0.020552293 | -0.028099928 | 0.999933887 | 0.692766341 | 1.163189451 | -0.47042311 | 0.212198323 |
| ENSG00000167968 | *DNASE1L2* | -0.452044284 | 0.155399462 | 0.089300878 | 0.066098584 | 0.999933887 | -0.194278573 | -0.908783749 | 0.714505176 | 0.212441007 |
| ENSG00000102081 | *FMR1* | 6.228552573 | -0.048676588 | -0.091233471 | 0.042556884 | 0.999933887 | -0.517165932 | -0.270755527 | -0.246410405 | 0.21293742 |
| ENSG00000004838 | *ZMYND10* | -0.83884943 | 0.059107948 | 0.090988349 | -0.031880401 | 0.999933887 | -0.543178454 | -1.224766792 | 0.681588338 | 0.21293742 |
| ENSG00000213079 | *SCAF8* | 5.667227789 | 0.013325426 | -0.020116408 | 0.033441834 | 0.999933887 | -0.197124626 | -0.075850601 | -0.121274025 | 0.212946491 |
| ENSG00000088970 | *KIZ* | 3.116210404 | -0.021359196 | -0.05331719 | 0.031957994 | 0.999933887 | -0.199064511 | -0.4182598 | 0.219195289 | 0.212955218 |
| ENSG00000197323 | *TRIM33* | 6.942944657 | -0.006806565 | -0.083666555 | 0.07685999 | 0.999933887 | -0.943658186 | -0.731074055 | -0.212584131 | 0.213453737 |
| ENSG00000171425 | *ZNF581* | 2.610023595 | 0.005767643 | -0.182954479 | 0.188722123 | 0.999933887 | -0.235948262 | -0.437174161 | 0.201225899 | 0.213539619 |
| ENSG00000228918 | *LINC01344* | -0.284919139 | -0.009647847 | -0.410803663 | 0.401155815 | 0.999933887 | 0.031615171 | 0.568863355 | -0.537248184 | 0.213570014 |
| ENSG00000275139 | *AL133492.1* | -0.779206134 | -0.286490956 | -0.114161379 | -0.172329577 | 0.999933887 | -0.189400804 | 0.307294624 | -0.496695428 | 0.213620424 |
| ENSG00000164011 | *ZNF691* | 0.1213705 | 0.077304209 | -0.26017885 | 0.337483059 | 0.999933887 | -0.068710152 | -0.396747974 | 0.328037822 | 0.213910589 |
| ENSG00000228763 | *LIMS1-AS1* | 0.280335583 | -0.082338207 | 0.184675086 | -0.267013294 | 0.999933887 | 1.861182273 | 2.325862088 | -0.464679815 | 0.213951323 |
| ENSG00000126777 | *KTN1* | 4.567497489 | -0.156362929 | -0.223170808 | 0.06680788 | 0.999933887 | -0.498145381 | -0.148971653 | -0.349173728 | 0.214115557 |
| ENSG00000060339 | *CCAR1* | 4.394097191 | -0.051006285 | -0.139235423 | 0.088229138 | 0.999933887 | -0.179910466 | 0.044501699 | -0.224412166 | 0.214115557 |
| ENSG00000111266 | *DUSP16* | 5.780068989 | -0.03443027 | -0.005407206 | -0.029023064 | 0.999933887 | 1.103882965 | 1.409337949 | -0.305454985 | 0.214115557 |
| ENSG00000099725 | *PRKY* | -3.049772494 | -0.00748937 | -0.026134675 | 0.018645306 | 0.999933887 | -0.156217412 | -0.466384162 | 0.31016675 | 0.214115557 |
| ENSG00000130522 | *JUND* | 8.868936533 | 0.109273971 | 0.077879656 | 0.031394315 | 0.999933887 | -0.037976032 | -0.243779719 | 0.205803687 | 0.214183941 |
| ENSG00000131844 | *MCCC2* | 1.942208631 | -0.117405733 | -0.005754171 | -0.111651562 | 0.999933887 | 0.090095599 | 0.316181446 | -0.226085846 | 0.214183941 |
| ENSG00000198837 | *DENND4B* | 7.778338528 | 0.030372452 | 0.10840944 | -0.078036988 | 0.999933887 | 0.06386089 | -0.070919019 | 0.134779909 | 0.214183941 |
| ENSG00000133101 | *CCNA1* | -1.014024091 | -0.140729538 | -0.156247526 | 0.015517988 | 0.999933887 | 1.282509666 | 2.068293424 | -0.785783758 | 0.214183941 |
| ENSG00000074935 | *TUBE1* | 1.373084312 | 0.033264288 | -0.15436234 | 0.187626628 | 0.999933887 | 0.037946123 | 0.291756862 | -0.253810739 | 0.214183941 |
| ENSG00000254477 | *AP000640.1* | 3.939046447 | 0.01311293 | -0.06396819 | 0.077081121 | 0.999933887 | 0.521437428 | 0.666322725 | -0.144885296 | 0.214183941 |
| ENSG00000117676 | *RPS6KA1* | 7.737003776 | 0.004619033 | 0.002072498 | 0.002546535 | 0.999933887 | 0.044050523 | -0.061697451 | 0.105747974 | 0.214183941 |
| ENSG00000140471 | *LINS1* | 3.162477008 | -0.006748917 | -0.087616246 | 0.080867329 | 0.999933887 | -0.149387756 | 0.021730359 | -0.171118114 | 0.214183941 |
| ENSG00000172671 | *ZFAND4* | 2.315879574 | 0.044433965 | -0.104758984 | 0.149192949 | 0.999933887 | -0.051764728 | 0.201285036 | -0.253049763 | 0.214242746 |
| ENSG00000211455 | *STK38L* | 5.491241288 | -0.095480992 | -0.076026544 | -0.019454447 | 0.999933887 | -0.825020279 | -1.077573708 | 0.252553429 | 0.214306277 |
| ENSG00000166887 | *VPS39* | 6.218411033 | -0.042349128 | -0.020163894 | -0.022185234 | 0.999933887 | -0.156299381 | -0.261137596 | 0.104838214 | 0.214460145 |
| ENSG00000123200 | *ZC3H13* | 5.668153139 | -0.079028979 | -0.15287557 | 0.073846592 | 0.999933887 | -0.894975087 | -0.58627232 | -0.308702767 | 0.214460145 |
| ENSG00000120837 | *NFYB* | 2.242723529 | -0.015473238 | 0.051250566 | -0.066723804 | 0.999933887 | 0.30618189 | 0.505476681 | -0.199294791 | 0.215473197 |
| ENSG00000140406 | *TLNRD1* | 4.950741705 | 0.146528726 | 0.186228626 | -0.0396999 | 0.999933887 | 0.519311087 | 0.769690981 | -0.250379894 | 0.215818278 |
| ENSG00000064886 | *CHI3L2* | 2.023988906 | 0.134673263 | 0.338163612 | -0.203490349 | 0.999933887 | 0.511289687 | 0.274671999 | 0.236617688 | 0.215818278 |
| ENSG00000165806 | *CASP7* | 2.55063681 | -0.05968786 | -0.004001437 | -0.055686423 | 0.999933887 | 0.243320107 | 0.454054684 | -0.210734578 | 0.215952263 |
| ENSG00000186184 | *POLR1D* | 4.727206984 | 0.036937852 | 0.080623938 | -0.043686086 | 0.999933887 | -0.162125582 | -0.306240534 | 0.144114952 | 0.216011695 |
| ENSG00000168434 | *COG7* | 1.801507013 | -0.037606718 | 0.082663679 | -0.120270397 | 0.999933887 | -0.093464694 | -0.360191285 | 0.266726591 | 0.216011695 |
| ENSG00000163872 | *YEATS2* | 3.507959787 | 0.039799423 | 0.024722875 | 0.015076548 | 0.999933887 | -0.181012651 | -0.329573328 | 0.148560676 | 0.216304763 |
| ENSG00000179526 | *SHARPIN* | 3.871693207 | 0.055558335 | 0.048393435 | 0.007164899 | 0.999933887 | -0.004432705 | -0.168987222 | 0.164554516 | 0.216573195 |
| ENSG00000004455 | *AK2* | 4.621844254 | -0.042792803 | 0.019096842 | -0.061889645 | 0.999933887 | 0.516568956 | 0.69043917 | -0.173870214 | 0.216604859 |
| ENSG00000107651 | *SEC23IP* | 4.09591294 | -0.075093133 | 0.011623537 | -0.08671667 | 0.999933887 | -0.180220855 | -0.050321202 | -0.129899653 | 0.217131584 |
| ENSG00000109458 | *GAB1* | 3.682584599 | -0.086647122 | -0.023070551 | -0.063576572 | 0.999933887 | -1.242950848 | -1.560140486 | 0.317189638 | 0.217232212 |
| ENSG00000105810 | *CDK6* | 3.31171396 | -0.061513593 | -0.130043595 | 0.068530003 | 0.999933887 | -0.142169388 | 0.112913341 | -0.255082729 | 0.217393128 |
| ENSG00000106077 | *ABHD11* | 1.207962003 | 0.211360766 | -0.118810039 | 0.330170804 | 0.944094537 | -0.194606276 | -0.511600934 | 0.316994658 | 0.217404213 |
| ENSG00000214212 | *C19orf38* | 4.563138394 | 0.095754921 | 0.056839902 | 0.03891502 | 0.999933887 | -0.116609011 | -0.340792179 | 0.224183168 | 0.217710111 |
| ENSG00000143119 | *CD53* | 10.294225686 | -0.008410999 | -0.040890272 | 0.032479273 | 0.999933887 | 0.21786001 | 0.34193714 | -0.12407713 | 0.218020636 |
| ENSG00000185905 | *C16orf54* | 6.94920641 | -0.012930814 | -0.05721048 | 0.044279666 | 0.999933887 | -0.808589552 | -0.983587171 | 0.174997619 | 0.218059683 |
| ENSG00000141034 | *GID4* | 1.808393201 | -0.084619922 | 0.121688212 | -0.206308133 | 0.999933887 | -0.331638238 | -0.608875784 | 0.277237545 | 0.218075888 |
| ENSG00000116809 | *ZBTB17* | 5.898344568 | 0.05109796 | 0.041183177 | 0.009914783 | 0.999933887 | 0.447160681 | 0.621547143 | -0.174386462 | 0.218144408 |

| | | | | | | | | | | |
|---|---|---|---|---|---|---|---|---|---|---|
| ENSG00000157654 | *PALM2AKAP2* | 3.109099923 | -0.069137064 | 0.001035579 | -0.070172644 | 0.999933887 | 0.534258521 | 0.747914319 | -0.213655798 | 0.218733498 |
| ENSG00000125148 | *MT2A* | 4.005395807 | 0.055040709 | -0.050379884 | 0.105420593 | 0.999933887 | 0.577882953 | 0.848976627 | -0.271093675 | 0.218733498 |
| ENSG00000170873 | *MTSS1* | 3.463107085 | -0.038958459 | 0.083832326 | -0.122790785 | 0.999933887 | 0.02004673 | 0.196633022 | -0.176586292 | 0.218788434 |
| ENSG00000166226 | *CCT2* | 3.963734763 | 0.024624378 | -0.071639377 | 0.096263755 | 0.999933887 | 0.25919549 | 0.447898392 | -0.188702902 | 0.218819072 |
| ENSG00000139112 | *GABARAPL1* | 8.693798455 | 0.010195445 | -0.024680601 | 0.034876045 | 0.999933887 | -0.190391045 | -0.057955493 | -0.132435552 | 0.218843024 |
| ENSG00000188227 | *ZNF793* | -0.431901112 | -0.277631685 | -0.41043191 | 0.132800225 | 0.999933887 | -0.138740392 | 0.402294427 | -0.541034819 | 0.219033369 |
| ENSG00000118960 | *HS1BP3* | 6.10684051 | -0.012639452 | -0.03589455 | 0.023255098 | 0.999933887 | -0.334946951 | -0.186450964 | -0.148495987 | 0.219176841 |
| ENSG00000183160 | *TMEM119* | 2.528007058 | -0.050709921 | -0.019998965 | -0.030710956 | 0.999933887 | -0.386993153 | -0.69751398 | 0.310520827 | 0.219266668 |
| ENSG00000099364 | *FBXL19* | 2.860711554 | 0.055394576 | 0.171389328 | -0.115994751 | 0.999933887 | -0.242927836 | -0.496601947 | 0.253674111 | 0.219266668 |
| ENSG00000139974 | *SLC38A6* | 1.513385555 | -0.378497153 | 0.048585706 | -0.427082859 | 0.999933887 | -0.051599452 | 0.187181034 | -0.238780486 | 0.219292803 |
| ENSG00000241343 | *RPL36A* | 1.612171075 | 0.146063249 | 0.148279368 | -0.002216119 | 0.999933887 | 0.575431992 | 0.930330346 | -0.354898354 | 0.219301185 |
| ENSG00000198521 | *ZNF43* | 2.841322461 | -0.132768317 | -0.063333959 | -0.069434358 | 0.999933887 | -0.244375727 | 0.021543806 | -0.265919533 | 0.219301185 |
| ENSG00000173230 | *GOLGB1* | 6.452027756 | -0.139425424 | -0.124514692 | -0.014910732 | 0.999933887 | -0.642289706 | -0.211206941 | -0.431082765 | 0.219301185 |
| ENSG00000168487 | *BMP1* | 1.169656541 | 0.11672113 | 0.074669042 | 0.042052088 | 0.999933887 | -0.000380177 | 0.32440189 | -0.324782066 | 0.219301185 |
| ENSG00000107372 | *ZFAND5* | 8.6358189 | 0.027092702 | 0.060590872 | -0.03349817 | 0.999933887 | -0.216972869 | -0.092491751 | -0.124481118 | 0.219301185 |
| ENSG00000173818 | *ENDOV* | 2.427955687 | 0.025302371 | -0.002818619 | 0.028120991 | 0.999933887 | -0.007252071 | 0.217262986 | -0.224515058 | 0.219301185 |
| ENSG00000169122 | *FAM110B* | 1.3558417 | 0.129690747 | 0.062831477 | 0.06685927 | 0.999933887 | 0.238189685 | 0.558923793 | -0.320734109 | 0.219313346 |
| ENSG00000145494 | *NDUFS6* | 2.96657084 | 0.052255963 | -0.025205744 | 0.077461707 | 0.999933887 | -0.038445742 | -0.229693609 | 0.191247867 | 0.219424567 |
| ENSG00000171155 | *C1GALT1C1* | 2.649284705 | -0.055321511 | 0.063817238 | -0.119138749 | 0.999933887 | -0.558876449 | -0.825216712 | 0.266340263 | 0.21948097 |
| ENSG00000205476 | *CCDC85C* | 4.136672959 | 0.011853686 | 0.010667143 | 0.001186544 | 0.999933887 | -0.047799753 | 0.130867012 | -0.178666766 | 0.21948097 |
| ENSG00000171365 | *CLCN5* | 2.052612287 | -0.108392737 | 0.074469207 | -0.182861944 | 0.999933887 | -0.16161636 | 0.092120703 | -0.253737063 | 0.219557035 |
| ENSG00000099365 | *STX1B* | 1.607999604 | 0.029757482 | -0.034819828 | 0.064577309 | 0.999933887 | 0.050875678 | 0.295926718 | -0.24505104 | 0.219773345 |
| ENSG00000119280 | *C1orf198* | -0.184234353 | -0.11296507 | 0.355546933 | -0.468512002 | 0.999933887 | 0.024899543 | 0.458421453 | -0.43352191 | 0.220018331 |
| ENSG00000155100 | *OTUD6B* | 0.981910148 | -0.024742037 | -0.237860515 | 0.213118478 | 0.999933887 | -0.404552508 | 0.065457012 | -0.47000952 | 0.220018331 |
| ENSG00000008118 | *CAMK1G* | 2.373980682 | 0.145629008 | 0.285781208 | -0.1401522 | 0.999933887 | 2.765854393 | 3.188419514 | -0.422565121 | 0.220022477 |
| ENSG00000178977 | *LINC00324* | 1.611589587 | 0.01274244 | -0.113380597 | 0.126123037 | 0.999933887 | -0.324210623 | -0.659238618 | 0.335027996 | 0.220051409 |
| ENSG00000161265 | *U2AF1L4* | 2.545841849 | 0.015415359 | -0.012430111 | 0.02784547 | 0.999933887 | 0.058012473 | -0.165303307 | 0.22331578 | 0.220073447 |
| ENSG00000177082 | *WDR73* | 1.263793017 | -0.115629361 | -0.042157457 | -0.073471904 | 0.999933887 | -0.107243627 | -0.396071223 | 0.288827596 | 0.220288586 |
| ENSG00000197535 | *MYO5A* | 5.156083749 | -0.007214585 | -0.07157093 | 0.064356345 | 0.999933887 | -0.285524514 | -0.525096345 | 0.239571831 | 0.220288586 |
| ENSG00000161847 | *RAVER1* | 4.016369561 | -0.013003472 | -0.039521293 | 0.026517821 | 0.999933887 | -0.109152435 | -0.271336193 | 0.162183758 | 0.220441609 |
| ENSG00000007080 | *CCDC124* | 3.232727401 | 0.1485641 | 0.077889765 | 0.070674336 | 0.999933887 | 0.226027854 | 0.446821072 | -0.220793218 | 0.220482732 |
| ENSG00000090470 | *PDCD7* | 2.341002788 | 0.047051847 | 0.093158728 | -0.046106881 | 0.999933887 | 0.188310734 | -0.00649032 | 0.194801054 | 0.220482732 |
| ENSG00000180228 | *PRKRA* | 2.146003316 | -0.045743441 | 0.063515301 | -0.109258742 | 0.999933887 | -0.196189361 | -0.412827922 | 0.216638561 | 0.220482732 |
| ENSG00000130695 | *CEP85* | 3.182553838 | 0.040821479 | 0.023380398 | 0.017441081 | 0.999933887 | -0.418992758 | -0.60193466 | 0.182941902 | 0.220482732 |
| ENSG00000181852 | *RNF41* | 4.757459318 | 0.017006299 | 0.084222969 | -0.06721667 | 0.999933887 | 0.002319079 | 0.146668359 | -0.14434928 | 0.220482732 |
| ENSG00000185414 | *MRPL30* | 1.729449186 | 0.019338843 | -0.112618747 | 0.13195759 | 0.999933887 | 0.138649176 | -0.091082321 | 0.229731497 | 0.220482732 |
| ENSG00000101181 | *MTG2* | 4.262483288 | -0.004487157 | -0.026115748 | 0.021628591 | 0.999933887 | -0.155734209 | -0.301291236 | 0.145557027 | 0.220482732 |
| ENSG00000154678 | *PDE1C* | -1.054496793 | -0.005228171 | -0.260131481 | 0.25490331 | 0.999933887 | 0.00312977 | 0.693402008 | -0.690272238 | 0.220482732 |
| ENSG00000135535 | *CD164* | 7.848524822 | -0.068870441 | -0.079997399 | 0.011126958 | 0.999933887 | -0.578831193 | -0.444012973 | -0.13481822 | 0.220715512 |
| ENSG00000151136 | *BTBD11* | 2.400922165 | 0.00362963 | 0.002365555 | 0.001264075 | 0.999933887 | 0.257297012 | 0.003313701 | 0.253983311 | 0.22103979 |
| ENSG00000181090 | *EHMT1* | 4.771278693 | 0.032633608 | 0.065956364 | -0.033322755 | 0.999933887 | -0.044486173 | -0.191584553 | 0.14709838 | 0.221159679 |
| ENSG00000166323 | *C11orf65* | -0.898392946 | 0.229969765 | 0.283347478 | -0.053377713 | 0.999933887 | -0.439130946 | 0.275765411 | -0.714896358 | 0.221322853 |
| ENSG00000124733 | *MEA1* | 2.94904805 | 0.047074293 | -0.080504961 | 0.127579254 | 0.999933887 | -0.153740327 | -0.384380704 | 0.230640377 | 0.221322853 |
| ENSG00000186716 | *BCR* | 5.005331233 | 0.033083387 | -0.011338339 | 0.044421726 | 0.999933887 | 0.008421193 | -0.165095238 | 0.17351643 | 0.221322853 |
| ENSG00000177853 | *ZNF518A* | 3.649872098 | -0.076693307 | -0.152036904 | 0.075343597 | 0.999933887 | -0.683367956 | -0.247392479 | -0.435975476 | 0.221322853 |
| ENSG00000108448 | *TRIM16L* | -1.024620282 | -0.141664773 | 0.096147505 | -0.237812278 | 0.999933887 | 0.776067208 | 1.279388034 | -0.503320826 | 0.221322853 |
| ENSG00000178184 | *PARD6G* | 1.525128647 | -0.014922128 | 0.261665245 | -0.276587373 | 0.999933887 | 0.790762445 | 1.105410735 | -0.314648289 | 0.221322853 |
| ENSG00000134028 | *ADAMDEC1* | -1.099869093 | -0.409392558 | 0.05386199 | -0.463254548 | 0.999933887 | 1.288292153 | 1.913079915 | -0.624787763 | 0.221402888 |
| ENSG00000187742 | *SECISBP2* | 6.014768217 | -0.002497817 | -0.022218344 | 0.019720526 | 0.999933887 | 0.023856405 | 0.13380553 | -0.109949125 | 0.221402888 |
| ENSG00000114026 | *OGG1* | 2.173236138 | 0.033942803 | 0.068087487 | -0.034144683 | 0.999933887 | -0.168311932 | -0.369077789 | 0.200765857 | 0.221515353 |
| ENSG00000185122 | *HSF1* | 5.180798245 | 0.05804897 | 0.073355963 | -0.015306993 | 0.999933887 | -0.020035561 | -0.150460019 | 0.130424458 | 0.221743179 |
| ENSG00000188211 | *NCR3LG1* | 5.742352781 | -0.169230572 | 0.010260156 | -0.179490728 | 0.999933887 | 3.024943994 | 3.369204968 | -0.344260973 | 0.221743179 |
| ENSG00000065457 | *ADAT1* | 5.465431561 | -0.062347563 | -0.03552577 | -0.026821793 | 0.999933887 | 1.388782008 | 1.600947981 | -0.212165973 | 0.221743179 |
| ENSG00000105993 | *DNAJB6* | 7.346926465 | 0.012860838 | 0.026811587 | -0.01395075 | 0.999933887 | -0.106771024 | 0.024653756 | -0.13142478 | 0.221743179 |
| ENSG00000153250 | *RBMS1* | 6.627171754 | -0.004808187 | -0.017064906 | 0.012256719 | 0.999933887 | -0.4684893 | -0.321097795 | -0.147391505 | 0.22201348 |
| ENSG00000206337 | *HCP5* | 8.31806591 | 0.014256146 | 0.022352027 | -0.00809588 | 0.999933887 | 0.825096506 | 0.960675971 | -0.135579465 | 0.222049373 |
| ENSG00000099840 | *IZUMO4* | 0.917372129 | -0.010744935 | 0.080333817 | -0.091078752 | 0.999933887 | 0.216915623 | -0.11502772 | 0.331943343 | 0.222049373 |
| ENSG00000183735 | *TBK1* | 6.226871708 | -0.141926028 | -0.182193537 | 0.040267509 | 0.999933887 | 1.11432196 | 1.393394434 | -0.279072474 | 0.222239833 |
| ENSG00000104983 | *CCDC61* | 2.03721637 | 0.071214201 | 0.070818525 | 0.000395675 | 0.999933887 | -0.026838758 | -0.296557509 | 0.269718751 | 0.222239833 |
| ENSG00000215021 | *PHB2* | 4.664776247 | 0.027172958 | 0.036234553 | -0.009061596 | 0.999933887 | 0.020383914 | -0.123447119 | 0.143831033 | 0.222239833 |
| ENSG00000205181 | *LINC00654* | 2.09744427 | 0.045164168 | 0.137433267 | -0.0922691 | 0.999933887 | -0.144327703 | 0.050199259 | -0.194526962 | 0.222239833 |
| ENSG00000128309 | *MPST* | 2.644843599 | 0.039104559 | 0.045298918 | -0.006194359 | 0.999933887 | -0.007276206 | -0.265463675 | 0.258187469 | 0.222518788 |
| ENSG00000134955 | *SLC37A2* | 3.675921609 | -0.014598763 | 0.104088258 | -0.118687022 | 0.999933887 | -0.486539222 | -0.329936205 | -0.156603017 | 0.22268021 |
| ENSG00000159720 | *ATP6V0D1* | 9.151414177 | -0.048479645 | -0.040657568 | -0.007822077 | 0.999933887 | 0.307693712 | 0.432814828 | -0.125121116 | 0.222918861 |
| ENSG00000112425 | *EPM2A* | -0.165599321 | -0.030012239 | -0.184341319 | 0.154329081 | 0.999933887 | 0.346958496 | -0.151862747 | 0.498821244 | 0.222965489 |
| ENSG00000230587 | *LINC02580* | 1.018189129 | 0.081902807 | -0.104900223 | 0.18680303 | 0.999933887 | -0.393105076 | -0.744147624 | 0.351042548 | 0.223014107 |
| ENSG00000162591 | *MEGF6* | 5.339239523 | 0.113503923 | 0.126254806 | -0.012750883 | 0.999933887 | 0.026855508 | -0.157794966 | 0.184650474 | 0.223035611 |
| ENSG00000276231 | *PIK3R6* | 3.584416219 | 0.09191452 | 0.046736854 | 0.045177666 | 0.999933887 | 0.176438758 | 0.385687631 | -0.209248873 | 0.223075582 |
| ENSG00000173214 | *MFSD4B* | 2.79778972 | -0.056264334 | -0.045692516 | -0.010571818 | 0.999933887 | -0.026951807 | 0.201048941 | -0.228000748 | 0.223075582 |
| ENSG00000144485 | *HES6* | -0.057939177 | 0.159410764 | 0.060025335 | 0.099385428 | 0.999933887 | 0.304479697 | -0.160640918 | 0.465120615 | 0.22313598 |
| ENSG00000138182 | *KIF20B* | 1.212244148 | -0.244926658 | -0.19082073 | -0.054105928 | 0.999933887 | -0.313199091 | 0.11349715 | -0.42669624 | 0.223174775 |
| ENSG00000164961 | *WASHC5* | 5.462542 | 0.035616632 | -0.022533753 | 0.058150385 | 0.999933887 | -0.145591689 | 0.047089283 | -0.192680972 | 0.223181955 |
| ENSG00000174718 | *RESF1* | 9.946383699 | -0.152241814 | -0.168438653 | 0.016196839 | 0.999933887 | -0.563677955 | -0.228087909 | -0.335590046 | 0.223563887 |
| ENSG00000213064 | *SFT2D2* | 4.994756033 | -0.022136678 | 0.00672571 | -0.028862388 | 0.999933887 | -0.172161694 | -0.374412447 | 0.202250754 | 0.223593925 |
| ENSG00000229474 | *PATL2* | 5.356681531 | -0.00528931 | 0.052147978 | -0.057437288 | 0.999933887 | 0.33066486 | 0.46509762 | -0.13443276 | 0.223712385 |
| ENSG00000171953 | *ATPAF2* | 1.671884251 | -0.052442406 | 0.089006137 | -0.141448544 | 0.999933887 | -0.054076195 | -0.302704877 | 0.248628682 | 0.223865162 |
| ENSG00000115540 | *MOB4* | 2.48366919 | -0.057932389 | -0.003560732 | -0.054371657 | 0.999933887 | -0.396454492 | -0.115156867 | -0.281297625 | 0.224375836 |
| ENSG00000074054 | *CLASP1* | 5.300759236 | 0.032665885 | -0.022889313 | 0.055555198 | 0.999933887 | -0.20980091 | -0.356457222 | 0.146656312 | 0.224470944 |
| ENSG00000100490 | *CDKL1* | -0.147604135 | -0.038019386 | 0.013598476 | -0.051617862 | 0.999933887 | -0.360663501 | 0.170459231 | -0.531122733 | 0.224470944 |
| ENSG00000213551 | *DNAJC9* | 1.168748334 | 0.105279006 | -0.160078174 | 0.265357179 | 0.999933887 | 0.10096818 | -0.308767811 | 0.40973599 | 0.224607124 |
| ENSG00000158402 | *CDC25C* | 0.333483579 | 0.193152826 | -0.107060955 | 0.300213781 | 0.999933887 | 0.033594053 | -0.50472449 | 0.538318543 | 0.224711306 |
| ENSG00000137522 | *RNF121* | 2.904095567 | 0.07796524 | 0.046577727 | 0.031387513 | 0.999933887 | 0.299866152 | 0.459166291 | -0.159300139 | 0.224711306 |
| ENSG00000089737 | *DDX24* | 6.530452822 | -0.048339668 | -0.078301139 | 0.029961472 | 0.999933887 | 0.241995422 | 0.37586416 | -0.133868738 | 0.224711306 |
| ENSG00000128915 | *ICE2* | 2.348289092 | -0.011975456 | -0.219132608 | 0.207157152 | 0.999933887 | -0.112230128 | 0.198181883 | -0.310412012 | 0.224813007 |
| ENSG00000088448 | *ANKRD10* | 5.670751453 | -0.030538337 | -0.071523607 | 0.040985269 | 0.999933887 | 0.207997185 | 0.412194602 | -0.204197417 | 0.224952253 |
| ENSG00000165156 | *ZHX1* | 2.154667194 | -0.137658932 | -0.141358557 | 0.003699624 | 0.999933887 | -0.326715283 | 0.014302346 | -0.341017629 | 0.225109902 |
| ENSG00000136193 | *SCRN1* | 1.05427393 | 0.040524857 | -0.143894586 | 0.184419443 | 0.999933887 | -0.16898795 | 0.123469127 | -0.292457077 | 0.225109902 |
| ENSG00000152457 | *DCLRE1C* | 3.176222321 | 0.011928531 | -0.076677746 | 0.088606277 | 0.999933887 | -0.565344369 | -0.791189484 | 0.225845116 | 0.225109902 |
| ENSG00000231711 | *LINC00899* | -0.473782536 | -0.011490281 | -0.087096627 | 0.075606346 | 0.999933887 | -0.336727404 | -0.889073013 | 0.552345609 | 0.225109902 |
| ENSG00000105552 | *BCAT2* | 0.935793881 | 0.038183677 | -0.088492577 | 0.126676254 | 0.999933887 | 0.116279885 | 0.423450397 | -0.307170512 | 0.225272067 |
| ENSG00000212694 | *LINC01089* | 2.728631247 | 0.037950059 | -0.032577518 | 0.070527577 | 0.999933887 | -0.072784135 | -0.241262001 | 0.168477866 | 0.22530063 |
| ENSG00000247708 | *STX18-AS1* | -0.072302856 | 0.036795816 | 0.052621331 | -0.015825515 | 0.999933887 | 0.197809322 | 0.659962696 | -0.462153374 | 0.225439458 |
| ENSG00000227543 | *SPAG5-AS1* | 1.511758829 | 0.120388781 | 0.043229454 | 0.077159327 | 0.999933887 | 0.733062069 | 0.991304447 | -0.258242378 | 0.225454148 |
| ENSG00000086712 | *TXLNG* | 3.118594467 | 0.041978525 | -0.054547914 | 0.09652644 | 0.999933887 | 0.048491487 | 0.241541306 | -0.193049819 | 0.225561735 |
| ENSG00000107937 | *GTPBP4* | 3.7730535 | -0.088817492 | -0.065164387 | -0.023653105 | 0.999933887 | 0.389394799 | 0.538763007 | -0.149368209 | 0.225599785 |
| ENSG00000273759 | *AL117379.1* | -0.015752114 | -0.399227266 | -0.169997567 | -0.229229699 | 0.999933887 | -0.351828354 | -0.727147712 | 0.375319359 | 0.225600188 |
| ENSG00000273272 | *U62317.3* | 1.865572929 | 0.280263686 | 0.342820381 | -0.062556695 | 0.999933887 | 0.486537141 | 0.100274746 | 0.386262394 | 0.225659388 |
| ENSG00000109667 | *SLC2A9* | 0.560943725 | 0.078355827 | 0.076887487 | 0.00146834 | 0.999933887 | 0.258038829 | -0.109138046 | 0.367176875 | 0.225659388 |
| ENSG00000198169 | *ZNF251* | 1.70253195 | -0.114742564 | 0.037059421 | -0.151801985 | 0.999933887 | 0.155389361 | -0.090830707 | 0.246220068 | 0.225754466 |
| ENSG00000176092 | *CRYBG2* | 0.082246308 | 0.046311775 | 0.248643089 | -0.202331314 | 0.999933887 | -0.05683112 | -0.489633103 | 0.432801983 | 0.225853944 |
| ENSG00000152926 | *ZNF117* | 6.118179695 | -0.127649219 | -0.169317681 | 0.041668462 | 0.999933887 | -0.70301143 | -0.321026217 | -0.381985213 | 0.225854087 |
| ENSG00000103978 | *TMEM87A* | 3.79291642 | -0.03783468 | -0.033300524 | -0.004534156 | 0.999933887 | -0.174486039 | -0.346812567 | 0.172326528 | 0.22595505 |
| ENSG00000231006 | *RPL7P32* | -0.281558741 | 0.391624452 | 0.485923326 | -0.094298874 | 0.999933887 | 0.531128474 | 1.652164099 | -1.121035624 | 0.22596074 |
| ENSG00000104221 | *BRF2* | 1.018283542 | -0.040088994 | -0.10551778 | 0.065428785 | 0.999933887 | -0.057818473 | -0.458385922 | 0.400567449 | 0.22596074 |
| ENSG00000135678 | *CPM* | 0.096639732 | 0.020315292 | -0.026709821 | 0.047025113 | 0.999933887 | 0.326723277 | 0.762827087 | -0.43610381 | 0.226034797 |

| | | | | | | | | | | |
|---|---|---|---|---|---|---|---|---|---|---|
| ENSG00000182107 | *TMEM30B* | -0.164382455 | -0.2569466 | -0.142431619 | -0.114514981 | 0.999933887 | 0.338136674 | -0.18573591 | 0.523872584 | 0.226173351 |
| ENSG00000198399 | *ITSN2* | 7.293194482 | -0.005514488 | -0.145804558 | 0.140290069 | 0.999933887 | -0.327129046 | 0.002783969 | -0.329913015 | 0.226173351 |
| ENSG00000144320 | *LNPK* | 2.64495814 | 0.003511401 | -0.152374184 | 0.155885585 | 0.999933887 | -0.430672837 | -0.123375109 | -0.307297728 | 0.226233963 |
| ENSG00000227507 | *LTB* | 7.187123742 | 0.146480226 | 0.164340986 | -0.01786076 | 0.999933887 | 0.167601235 | -0.149984697 | 0.317585933 | 0.226408617 |
| ENSG00000185669 | *SNAI3* | 2.14294324 | 0.072416953 | -0.133176499 | 0.205593452 | 0.999933887 | 0.217892824 | 0.479508268 | -0.261615444 | 0.226409795 |
| ENSG00000279453 | *Z99129.4* | 2.738305493 | -0.036166205 | -0.003344918 | -0.032821287 | 0.999933887 | -0.077628109 | 0.187623676 | -0.265251784 | 0.226441642 |
| ENSG00000204287 | *HLA-DRA* | 6.652120263 | -0.022054515 | -0.018891451 | -0.003163065 | 0.999933887 | 1.083762593 | 1.424404829 | -0.340642236 | 0.226441642 |
| ENSG00000112473 | *SLC39A7* | 4.891833492 | 0.010608498 | 0.012384991 | -0.001776493 | 0.999933887 | -0.295345957 | -0.464167003 | 0.168821046 | 0.226441642 |
| ENSG00000170684 | *ZNF296* | 1.567223572 | 0.023018972 | -0.006630024 | 0.029648996 | 0.999933887 | -0.291560666 | -0.619711459 | 0.328150794 | 0.226447435 |
| ENSG00000222898 | *RN7SKP97* | 0.253630465 | 0.155818634 | 0.033533639 | 0.122284995 | 0.999933887 | -0.388976847 | 0.078059654 | -0.467036502 | 0.226583314 |
| ENSG00000269951 | *AC090181.2* | -0.059316191 | 0.246093137 | 0.061000384 | 0.185092752 | 0.999933887 | 0.880188454 | 1.221067293 | -0.34087884 | 0.226820581 |
| ENSG00000179021 | *C3orf38* | 3.11600442 | -0.018491549 | -0.076040574 | 0.057549025 | 0.999933887 | -0.132158777 | 0.040716835 | -0.172875613 | 0.226963976 |
| ENSG00000198015 | *MRPL42* | 2.263410535 | -0.059193001 | -0.10545339 | 0.046260389 | 0.999933887 | -0.152512145 | 0.071517228 | -0.224029372 | 0.227144762 |
| ENSG00000282988 | *AL031777.2* | -0.466456386 | 0.035213569 | 0.11637193 | -0.08115836 | 0.999933887 | -0.809325282 | -1.375589145 | 0.566263863 | 0.227144762 |
| ENSG00000172322 | *CLEC12A* | 5.75470992 | -0.006051057 | -0.035833983 | 0.029782926 | 0.999933887 | 1.141349311 | 1.467431411 | -0.3260821 | 0.227468597 |
| ENSG00000271936 | *AC012073.1* | -0.47320171 | 0.121768234 | 0.205706392 | -0.083938157 | 0.999933887 | 1.027124602 | 1.50582053 | -0.478695928 | 0.227618189 |
| ENSG00000240905 | *RN7SL798P* | 2.030099349 | 0.107363361 | 0.228183524 | -0.120820164 | 0.999933887 | 1.305626627 | 1.693381151 | -0.387754524 | 0.227687311 |
| ENSG00000167333 | *TRIM68* | 0.895260775 | 0.122055833 | -0.032139286 | 0.154195119 | 0.999933887 | 0.190493857 | -0.105566151 | 0.296060008 | 0.227687311 |
| ENSG00000105656 | *ELL* | 7.78406305 | 0.032707058 | 0.051955889 | -0.019248831 | 0.999933887 | 0.020251002 | 0.134217804 | -0.113966803 | 0.227687311 |
| ENSG00000079257 | *LXN* | 1.248850527 | 0.053696466 | 0.104823453 | -0.051126988 | 0.999933887 | -0.681160153 | -1.003462296 | 0.322302143 | 0.22780962 |
| ENSG00000067900 | *ROCK1* | 7.240742722 | -0.055689751 | -0.179554584 | 0.123864533 | 0.999933887 | -0.733300883 | -0.327721239 | -0.405579644 | 0.228044187 |
| ENSG00000249915 | *PDCD6* | 3.472842492 | 0.025185009 | -0.088266479 | 0.113451488 | 0.999933887 | -0.191288007 | -0.343766506 | 0.152478499 | 0.228066994 |
| ENSG00000157911 | *PEX10* | 0.71853649 | 0.036810086 | -0.018654772 | 0.055464858 | 0.999933887 | 0.004445041 | -0.301636023 | 0.306081064 | 0.228101209 |
| ENSG00000130066 | *SAT1* | 12.281217993 | 0.067255902 | 0.02685472 | 0.040401182 | 0.999933887 | 0.53409719 | 0.689945315 | -0.155848124 | 0.228130322 |
| ENSG00000187010 | *RHD* | 2.368661331 | -0.126036043 | -0.010107571 | -0.115928471 | 0.999933887 | -0.162280507 | -0.404512422 | 0.242231915 | 0.228272166 |
| ENSG00000202538 | *RNU4-2* | 0.222315398 | 0.163489024 | -0.815744824 | 0.979233848 | 0.999933887 | 2.079131716 | 0.05297284 | 2.026158876 | 0.22848546 |
| ENSG00000067533 | *RRP15* | 1.480423379 | -0.202943076 | -0.252723532 | 0.049780456 | 0.999933887 | -0.076986185 | 0.225558492 | -0.302544677 | 0.228539045 |
| ENSG00000229740 | *U91324.1* | -1.869401856 | 0.392894657 | 0.728329068 | -0.335434411 | 0.999933887 | 3.092115951 | 3.875106913 | -0.782990962 | 0.22856947 |
| ENSG00000239779 | *WBP1* | 2.015355923 | 0.058545306 | 0.086151354 | -0.027606048 | 0.999933887 | 0.123829724 | -0.127458263 | 0.251287987 | 0.22856947 |
| ENSG00000114742 | *WDR48* | 5.914491883 | -0.059180702 | -0.059390061 | 0.00020936 | 0.999933887 | 0.272563401 | 0.39021344 | -0.117650039 | 0.228674558 |
| ENSG00000135929 | *CYP27A1* | 4.880354203 | 0.004667435 | 0.053115582 | -0.048448147 | 0.999933887 | 0.119100784 | 0.340430629 | -0.221329845 | 0.228824361 |
| ENSG00000010810 | *FYN* | 6.258099936 | -0.049281969 | -0.038971053 | -0.010310915 | 0.999933887 | 0.099551778 | 0.208096392 | -0.108544614 | 0.228901802 |
| ENSG00000104643 | *MTMR9* | 2.584875563 | -0.141422539 | -0.065074526 | -0.076348013 | 0.999933887 | -0.194361928 | 0.04144326 | -0.235805188 | 0.229108597 |
| ENSG00000258826 | *LINC01147* | -0.850907659 | -0.148595187 | -0.026424421 | -0.122170767 | 0.999933887 | 0.87486864 | 1.65495952 | -0.78009088 | 0.229108597 |
| ENSG00000107863 | *ARHGAP21* | 2.186466606 | -0.032124006 | -0.257919315 | 0.225795309 | 0.999933887 | -0.063598875 | 0.232107115 | -0.29570599 | 0.229108597 |
| ENSG00000112561 | *TFEB* | 5.732300805 | 0.005871045 | 0.032717408 | -0.026846363 | 0.999933887 | -0.678875904 | -0.889282612 | 0.210406707 | 0.229203194 |
| ENSG00000167272 | *POP5* | 1.431549351 | 0.047108057 | -0.032766953 | 0.07987501 | 0.999933887 | 0.204992011 | -0.088703844 | 0.293695855 | 0.229240064 |
| ENSG00000077514 | *POLD3* | 2.866833089 | -0.010602233 | -0.071020339 | 0.060418107 | 0.999933887 | 0.019165691 | 0.229906293 | -0.210740602 | 0.229240064 |
| ENSG00000140263 | *SORD* | 2.589723362 | -0.105323724 | 0.086405487 | -0.191729211 | 0.999933887 | -0.055193884 | -0.246817578 | 0.191623694 | 0.229270799 |
| ENSG00000266993 | *AL050343.2* | 0.077393312 | 0.153586897 | 0.054666072 | 0.098920825 | 0.999933887 | -0.296693372 | 0.218062357 | -0.514755729 | 0.229270799 |
| ENSG00000175711 | *B3GNTL1* | 4.589244896 | -0.012302876 | 0.057403092 | -0.069705968 | 0.999933887 | -0.170401006 | -0.301262619 | 0.130861613 | 0.229270799 |
| ENSG00000187446 | *CHP1* | 6.352834598 | -0.032703851 | -0.027542574 | -0.005161277 | 0.999933887 | -0.021481588 | 0.109800901 | -0.131282489 | 0.229330215 |
| ENSG00000140526 | *ABHD2* | 7.87520672 | -0.102122032 | -0.01485443 | -0.087267603 | 0.999933887 | -0.838146637 | -1.024109654 | 0.185963017 | 0.229389357 |
| ENSG00000178038 | *ALS2CL* | 1.62731872 | -0.025517268 | -0.035374368 | 0.0098571 | 0.999933887 | 0.082047845 | 0.368101972 | -0.286054127 | 0.229389357 |
| ENSG00000164023 | *SGMS2* | 1.928334168 | -0.007503145 | -0.087869151 | 0.080366006 | 0.999933887 | -0.121443146 | 0.231363892 | -0.352807038 | 0.229389357 |
| ENSG00000131788 | *PIAS3* | 2.816649411 | 0.052193569 | 0.054914333 | -0.002720764 | 0.999933887 | 0.048079294 | -0.109527138 | 0.157606433 | 0.229404431 |
| ENSG00000114853 | *ZBTB47* | 1.023881459 | -0.069285813 | 0.073151426 | -0.142437239 | 0.999933887 | -0.612342076 | -1.145257178 | 0.532915102 | 0.229404431 |
| ENSG00000152464 | *RPP38* | 1.034488639 | 0.04625275 | -0.129293472 | 0.175546223 | 0.999933887 | 0.007588922 | -0.315146576 | 0.322735497 | 0.229465646 |
| ENSG00000244486 | *SCARF2* | 1.151170535 | 0.088799471 | 0.231402136 | -0.142602665 | 0.999933887 | -0.186919403 | -0.568638275 | 0.381718872 | 0.229478989 |
| ENSG00000244065 | *MARK2P17* | -0.911515038 | -0.026933071 | 0.173784024 | -0.200717095 | 0.999933887 | 0.490280393 | 1.085211175 | -0.594930782 | 0.229527074 |
| ENSG00000266947 | *AC022916.1* | 0.351716169 | -0.005571364 | 0.049193644 | -0.054765008 | 0.999933887 | 0.370166752 | 0.801020648 | -0.430853897 | 0.229533767 |
| ENSG00000140265 | *ZSCAN29* | 3.489179969 | -0.111082487 | 0.005730776 | -0.116813263 | 0.999933887 | -0.170883587 | -0.341062859 | 0.170179272 | 0.229684474 |
| ENSG00000287920 | *AL022098.1* | 5.095673133 | 0.064446594 | 0.278945792 | -0.214499198 | 0.999933887 | 0.595471613 | 0.859178775 | -0.263707162 | 0.229874403 |
| ENSG00000203950 | *RTL8A* | 1.405693327 | 0.116636941 | -0.171419507 | 0.288056448 | 0.999933887 | -0.118471273 | -0.467564597 | 0.349093324 | 0.230102336 |
| ENSG00000174365 | *SNHG11* | 0.558933031 | -0.036435688 | 0.316088751 | -0.35252444 | 0.999933887 | 0.347123592 | -0.029600412 | 0.376724004 | 0.230284359 |
| ENSG00000196227 | *FAM217B* | 3.064147454 | -0.026981742 | -0.281538072 | 0.254556331 | 0.999933887 | -0.291909943 | -0.581161586 | 0.289251643 | 0.230354353 |
| ENSG00000151806 | *GUF1* | 1.802883684 | -0.002436255 | -0.255503933 | 0.253067678 | 0.999933887 | -0.165468401 | 0.170272969 | -0.33574137 | 0.230354353 |
| ENSG00000070610 | *GBA2* | 5.854182406 | 0.002431808 | 0.067688128 | -0.06525632 | 0.999933887 | -0.02930113 | -0.140192171 | 0.110891042 | 0.230364837 |
| ENSG00000100029 | *PES1* | 3.256686307 | 0.008060405 | 0.061400813 | -0.053340409 | 0.999933887 | 0.175557733 | 0.340192861 | -0.164635128 | 0.230603078 |
| ENSG00000047365 | *ARAP2* | 5.077033113 | -0.058171744 | -0.099318937 | 0.041147193 | 0.999933887 | -0.321946922 | 0.063132269 | -0.385079191 | 0.230724162 |
| ENSG00000172775 | *PSME3IP1* | 6.169439179 | -0.040059851 | 0.001001916 | -0.041061767 | 0.999933887 | -0.078452877 | 0.014768762 | -0.093221639 | 0.230769715 |
| ENSG00000064666 | *CNN2* | 8.673683212 | 0.060937534 | 0.000570582 | 0.060366952 | 0.999933887 | -0.738477452 | -0.912839443 | 0.174361991 | 0.230769715 |
| ENSG00000067606 | *PRKCZ* | 3.239140753 | -0.015241137 | 0.002106964 | -0.017348101 | 0.999933887 | -0.136556656 | -0.325891139 | 0.189334483 | 0.230769715 |
| ENSG00000198356 | *GET3* | 4.966741491 | 0.027422913 | -0.029189468 | 0.056612381 | 0.999933887 | 0.03989823 | 0.174290504 | -0.134392273 | 0.230865264 |
| ENSG00000139117 | *CPNE8* | 0.356884507 | -0.205244405 | 0.007985884 | -0.213230288 | 0.999933887 | 0.243470809 | 0.651467357 | -0.407996548 | 0.231039069 |
| ENSG00000235453 | *SMIM27* | 1.841990716 | 0.077833563 | 0.057247675 | 0.020585887 | 0.999933887 | -0.150500204 | -0.430661545 | 0.280161341 | 0.231039069 |
| ENSG00000222020 | *HDAC4-AS1* | 1.491085034 | -0.018362044 | -0.184547682 | 0.166185638 | 0.999933887 | 0.405590329 | 0.157179364 | 0.248410965 | 0.231039069 |
| ENSG00000196924 | *FLNA* | 11.249699033 | 0.073511841 | 0.076940231 | -0.003428389 | 0.999933887 | 0.250546898 | 0.378246585 | -0.127699688 | 0.231172185 |
| ENSG00000197858 | *GPAA1* | 3.902035034 | 0.024728006 | 0.041122733 | -0.016394727 | 0.999933887 | 0.176113858 | 0.015577676 | 0.160536182 | 0.231172185 |
| ENSG00000102471 | *NDFIP2* | 0.711093038 | -0.138628463 | 0.02465166 | -0.163280123 | 0.999933887 | -0.093668881 | 0.153777379 | -0.24744626 | 0.231517491 |
| ENSG00000141564 | *RPTOR* | 3.679292239 | 0.0521026 | 0.043929212 | 0.008173388 | 0.999933887 | -0.120922606 | 0.04389581 | -0.164818415 | 0.231636175 |
| ENSG00000227392 | *HPN-AS1* | -1.705640796 | 0.114898045 | 0.164746443 | -0.049848398 | 0.999933887 | 3.252104291 | 4.382684418 | -1.130580127 | 0.231880594 |
| ENSG00000163704 | *PRRT3* | 1.898722445 | 0.108510968 | 0.121702951 | -0.013191983 | 0.999933887 | -0.58336795 | -0.806537255 | 0.223169305 | 0.232186648 |
| ENSG00000259005 | *AC005479.1* | 0.942356756 | -0.030003559 | -0.127310124 | 0.097306565 | 0.999933887 | -0.018293265 | 0.221121157 | -0.239414422 | 0.232307821 |
| ENSG00000170827 | *CELP* | -0.903828445 | 0.070795374 | 0.218920798 | -0.148125424 | 0.999933887 | 0.682940741 | 1.200886745 | -0.517946004 | 0.232435065 |
| ENSG00000126464 | *PRR12* | 4.118526324 | 0.060782998 | 0.031793818 | 0.028989179 | 0.999933887 | -0.111398173 | -0.279646481 | 0.168248308 | 0.232722802 |
| ENSG00000167515 | *TRAPPC2L* | 3.451785999 | 0.109624829 | -0.106467931 | 0.21609276 | 0.944094537 | -0.060515196 | -0.219071847 | 0.158556651 | 0.232810188 |
| ENSG00000163527 | *STT3B* | 6.136670694 | -0.046686698 | -0.049000183 | 0.002313484 | 0.999933887 | -0.426245589 | -0.559049064 | 0.132803475 | 0.232810188 |
| ENSG00000091140 | *DLD* | 4.571971143 | -0.043041772 | -0.072795219 | 0.029753447 | 0.999933887 | -0.086423341 | 0.065089336 | -0.151512677 | 0.232810188 |
| ENSG00000101624 | *CEP76* | 0.091226871 | -0.075147062 | 0.084495309 | -0.159642371 | 0.999933887 | -0.036667348 | -0.390917616 | 0.354250268 | 0.232810188 |
| ENSG00000188612 | *SUMO2* | 5.118527681 | -0.021546521 | -0.01999022 | -0.001556301 | 0.999933887 | -0.051532315 | -0.18432763 | 0.132795315 | 0.232810188 |
| ENSG00000156384 | *SFR1* | 2.770379725 | 0.084879461 | 0.062519129 | 0.022360332 | 0.999933887 | 2.801710804 | 3.106833513 | -0.305122709 | 0.232811354 |
| ENSG00000147677 | *EIF3H* | 5.746595155 | -0.044056737 | -0.097950604 | 0.053893867 | 0.999933887 | -0.013945632 | 0.108020514 | -0.121966147 | 0.232926554 |
| ENSG00000072849 | *DERL2* | 4.636364345 | -0.043470498 | -0.044194632 | 0.000724134 | 0.999933887 | -0.199692674 | -0.32013772 | 0.120445046 | 0.232926554 |
| ENSG00000167986 | *DDB1* | 4.955767126 | -0.035946253 | 0.060881515 | -0.096827768 | 0.999933887 | 0.232846777 | 0.368825032 | -0.135978254 | 0.232926554 |
| ENSG00000055332 | *EIF2AK2* | 5.838943421 | -0.038089871 | -0.032042041 | -0.00604783 | 0.999933887 | -0.423260949 | -0.576085789 | 0.15282484 | 0.232926554 |
| ENSG00000099849 | *RASSF7* | 2.504387369 | 0.039896144 | 0.040723512 | -0.000827368 | 0.999933887 | 0.067902669 | -0.174997806 | 0.242900475 | 0.232926554 |
| ENSG00000214783 | *POLR2J4* | 0.265265903 | -0.008100897 | 0.103463887 | -0.111564784 | 0.999933887 | 0.34319244 | 0.788061235 | -0.444868795 | 0.232926554 |
| ENSG00000196218 | *RYR1* | 0.029637081 | -0.24779572 | -0.068890044 | -0.178905676 | 0.999933887 | -0.356902831 | 0.169240075 | -0.526142906 | 0.23306673 |
| ENSG00000119953 | *SMNDC1* | 5.255781506 | -0.066588391 | -0.048654319 | -0.017934072 | 0.999933887 | -0.021077033 | 0.080934364 | -0.102011397 | 0.23310471 |
| ENSG00000177479 | *ARIH2* | 4.884075107 | -0.06635151 | 0.016287295 | -0.082638805 | 0.999933887 | -0.088234543 | -0.195991656 | 0.107757113 | 0.23310471 |
| ENSG00000184857 | *TMEM186* | 0.328272484 | 0.05182955 | 0.009243175 | 0.042586375 | 0.999933887 | -0.127471737 | -0.542560993 | 0.415089256 | 0.23310471 |
| ENSG00000075914 | *EXOSC7* | 1.636935552 | 0.029800074 | -0.143416427 | 0.173216502 | 0.999933887 | 0.134504728 | 0.412630012 | -0.278125284 | 0.23310471 |
| ENSG00000123349 | *PFDN5* | 7.150197751 | -0.0064528 | -0.027227592 | 0.020774793 | 0.999933887 | 0.03954209 | 0.167158283 | -0.127616193 | 0.23310471 |
| ENSG00000102897 | *LYRM1* | 4.383406991 | -0.010087368 | -0.03722521 | 0.027137842 | 0.999933887 | -0.25917862 | -0.421989509 | 0.162810889 | 0.233316123 |
| ENSG00000197448 | *GSTK1* | 6.256362161 | 0.005636178 | 0.001190253 | 0.004445925 | 0.999933887 | 0.077659453 | -0.041588112 | 0.119247565 | 0.233484896 |
| ENSG00000183762 | *KREMEN1* | 6.069600774 | -0.026037986 | 0.039083619 | -0.065121605 | 0.999933887 | -0.242997786 | -0.417754367 | 0.174756581 | 0.233661929 |
| ENSG00000125505 | *MBOAT7* | 9.292205823 | 0.012953136 | -0.001636437 | 0.014589573 | 0.999933887 | -0.849609705 | -1.032887916 | 0.183278211 | 0.233837867 |
| ENSG00000173614 | *NMNAT1* | 1.368784744 | 0.000319952 | -0.02626928 | 0.026589232 | 0.999933887 | -0.550033457 | -0.806821157 | 0.256787699 | 0.23440533 |
| ENSG00000012124 | *CD22* | 4.234154691 | -0.022990251 | 0.227345834 | -0.250336086 | 0.999933887 | 0.081590448 | 0.226602228 | -0.14501178 | 0.234685061 |
| ENSG00000099937 | *SERPIND1* | 0.795835561 | 0.083746937 | 0.037426922 | 0.046320014 | 0.999933887 | 0.077802925 | 0.458521434 | -0.380718509 | 0.234849077 |
| ENSG00000019144 | *PHLDB1* | 2.274697057 | 0.042529265 | -0.11828009 | 0.160809355 | 0.999933887 | 0.43332534 | 0.645113321 | -0.211787981 | 0.23549024 |
| ENSG00000260916 | *CCPG1* | 4.170548144 | -0.102362653 | -0.132485429 | 0.030122777 | 0.999933887 | -1.052026088 | -1.33878979 | 0.286763701 | 0.235866944 |
| ENSG00000166311 | *SMPD1* | 5.034368474 | 0.008425964 | -0.019699786 | 0.02812575 | 0.999933887 | 0.312412671 | 0.449323823 | -0.136911151 | 0.235866944 |
| ENSG00000143363 | *PRUNE1* | 3.42781289 | -0.049791594 | -0.044884714 | -0.004906881 | 0.999933887 | -0.320691311 | -0.533222607 | 0.212531296 | 0.235901274 |

| | | | | | | | | | | |
|---|---|---|---|---|---|---|---|---|---|---|
| ENSG00000253352 | *TUG1* | 7.408519758 | 0.000384848 | -0.00552474 | 0.005909588 | 0.999933887 | -0.093525588 | 0.110936173 | -0.204461761 | 0.236030409 |
| ENSG00000184677 | *ZBTB40* | 4.665025644 | -0.007362473 | -0.018480394 | 0.011117921 | 0.999933887 | -0.027790444 | 0.093788177 | -0.121578622 | 0.236228429 |
| ENSG00000253719 | *ATXN7L3B* | 4.43880817 | -0.060975257 | -0.050142027 | -0.010833231 | 0.999933887 | 0.007659624 | -0.123451738 | 0.131111363 | 0.236377475 |
| ENSG00000205765 | *C5orf51* | 4.481604503 | -0.113368046 | -0.100996498 | -0.012371547 | 0.999933887 | -0.237263839 | -0.021645451 | -0.215618388 | 0.236377475 |
| ENSG00000108344 | *PSMD3* | 4.489163519 | 0.029762289 | -0.060645677 | 0.090407966 | 0.999933887 | -0.154037334 | -0.284425364 | 0.13038803 | 0.236377475 |
| ENSG00000132570 | *PCBD2* | 0.763570968 | 0.029660983 | -0.195645353 | 0.225306336 | 0.999933887 | -0.138186639 | -0.45621249 | 0.31802585 | 0.236377475 |
| ENSG00000236320 | *SLFN14* | 0.403050343 | 0.010379396 | -0.162990177 | 0.173369574 | 0.999933887 | -0.054585572 | 0.368305038 | -0.42289061 | 0.236377475 |
| ENSG00000223496 | *EXOSC6* | 2.925002789 | -0.001309624 | 0.034233627 | -0.03554325 | 0.999933887 | 0.093375998 | -0.147151415 | 0.240527413 | 0.236577008 |
| ENSG00000204351 | *SKIV2L* | 6.383092959 | 0.018159399 | 0.045274156 | -0.027114756 | 0.999933887 | 0.063741079 | -0.062044001 | 0.125785081 | 0.236637658 |
| ENSG00000163811 | *WDR43* | 3.219495365 | -0.008796185 | -0.045767191 | 0.036971006 | 0.999933887 | 0.124409235 | 0.311340539 | -0.186931304 | 0.237059877 |
| ENSG00000141699 | *RETREG3* | 4.923129365 | -0.049817335 | -0.059689449 | 0.009872115 | 0.999933887 | -0.292448056 | -0.433348642 | 0.140900585 | 0.237060737 |
| ENSG00000158156 | *XKR8* | 4.870593798 | -0.014186797 | -0.066743208 | 0.052556411 | 0.999933887 | -0.242372223 | -0.50553625 | 0.263164026 | 0.237060737 |
| ENSG00000137404 | *NRM* | 4.344457259 | -0.018749505 | -0.002635221 | -0.016114284 | 0.999933887 | 0.29506072 | 0.4340982 | -0.13903748 | 0.237458285 |
| ENSG00000105953 | *OGDH* | 5.644058786 | -0.023121622 | 0.023131137 | -0.046252759 | 0.999933887 | -0.220928933 | -0.349446813 | 0.128517879 | 0.237553522 |
| ENSG00000088682 | *COQ9* | 1.780307752 | -0.04455333 | 0.058515855 | -0.103069185 | 0.999933887 | -0.012336896 | -0.244351195 | 0.232014299 | 0.237831898 |
| ENSG00000135521 | *LTV1* | 1.034580566 | -0.114349139 | -0.368536004 | 0.254186866 | 0.999933887 | 0.180128056 | 0.481429535 | -0.301301478 | 0.237937629 |
| ENSG00000105072 | *C19orf44* | 0.359366415 | 0.149668085 | 0.383473151 | -0.233805066 | 0.999933887 | 0.041547545 | 0.4245472 | -0.382999655 | 0.237937629 |
| ENSG00000177191 | *B3GNT8* | 5.71269006 | -0.048209634 | -0.151759329 | 0.103549695 | 0.999933887 | -0.689311651 | -0.47636424 | -0.212947411 | 0.237937629 |
| ENSG00000266472 | *MRPS21* | 3.058220914 | -0.015191694 | -0.080477715 | 0.06528602 | 0.999933887 | 0.086131493 | -0.095160567 | 0.181292059 | 0.237937629 |
| ENSG00000168397 | *ATG4B* | 5.549164024 | 0.000194897 | 0.050382052 | -0.050187155 | 0.999933887 | 0.109623753 | -0.014931798 | 0.12455555 | 0.237937629 |
| ENSG00000236255 | *AC009404.1* | 2.652970803 | -0.007858723 | 0.001077597 | -0.008936319 | 0.999933887 | 0.123932686 | 0.416773929 | -0.292841242 | 0.238245525 |
| ENSG00000175265 | *GOLGA8A* | 3.825676351 | -0.02417841 | 0.001167823 | -0.025346234 | 0.999933887 | -0.031948647 | 0.149006903 | -0.180955551 | 0.238309377 |
| ENSG00000134202 | *GSTM3* | 0.46330991 | -0.048874012 | 0.158530432 | -0.207404445 | 0.999933887 | 0.025331032 | 0.473107759 | -0.447776727 | 0.238383629 |
| ENSG00000105341 | *DMAC2* | 3.324473972 | -0.085586925 | 0.009145641 | -0.094732565 | 0.999933887 | -0.029841227 | 0.147670073 | -0.177511301 | 0.238388961 |
| ENSG00000234284 | *ZNF879* | -0.532860116 | -0.069863804 | 0.178299977 | -0.248163781 | 0.999933887 | -0.111321604 | 0.38620667 | -0.497528274 | 0.238518916 |
| ENSG00000134256 | *CD101* | 1.843885236 | 0.088482998 | 0.083994983 | 0.004488016 | 0.999933887 | -0.121420544 | -0.429953502 | 0.308532958 | 0.238736625 |
| ENSG00000176124 | *DLEU1* | 2.112728246 | 0.136114317 | 0.205030598 | -0.068916281 | 0.999933887 | 0.278873767 | 0.67724014 | -0.398366373 | 0.238870669 |
| ENSG00000135245 | *HILPDA* | -0.83696805 | -0.054936789 | 0.102626348 | -0.157563137 | 0.999933887 | 1.096974689 | 1.584376035 | -0.487401346 | 0.239064594 |
| ENSG00000153037 | *SRP19* | 2.734547379 | -0.096498063 | -0.087554538 | -0.008943524 | 0.999933887 | -0.028015927 | 0.133169204 | -0.161185131 | 0.2390863 |
| ENSG00000135823 | *STX6* | 4.638912074 | -0.097063378 | -0.053201995 | -0.043861383 | 0.999933887 | -0.706811353 | -0.890699033 | 0.18388768 | 0.239149012 |
| ENSG00000149932 | *TMEM219* | 4.044276268 | -0.028339737 | -0.043188084 | 0.014848348 | 0.999933887 | -0.085838307 | -0.266016534 | 0.180178226 | 0.239189277 |
| ENSG00000121350 | *PYROXD1* | 2.356191021 | -0.153022757 | -0.114619972 | -0.038402785 | 0.999933887 | -0.102545505 | 0.221777505 | -0.32432301 | 0.239249136 |
| ENSG00000181915 | *ADO* | 3.99604311 | -0.085227999 | -0.136027545 | 0.050799546 | 0.999933887 | -0.254775787 | -0.463856093 | 0.209080306 | 0.239423814 |
| ENSG00000101916 | *TLR8* | 5.998082979 | -0.085624102 | -0.098206527 | 0.012582425 | 0.999933887 | -1.03910086 | -1.298435906 | 0.259335046 | 0.239423814 |
| ENSG00000000457 | *SCYL3* | 3.004546847 | -0.050082408 | -0.090733992 | 0.040651584 | 0.999933887 | -0.187289762 | 0.00384429 | -0.191134053 | 0.239423814 |
| ENSG00000140368 | *PSTPIP1* | 6.89343696 | -0.016153737 | 0.046901102 | -0.063054838 | 0.999933887 | -0.106399293 | -0.254598496 | 0.148199203 | 0.239423814 |
| ENSG00000135698 | *MPHOSPH6* | 1.223636139 | 0.008265199 | -0.011790179 | 0.020055378 | 0.999933887 | 0.002251992 | 0.325169833 | -0.322917841 | 0.239423814 |
| ENSG00000133858 | *ZFC3H1* | 6.749574552 | 0.057106725 | 0.004434054 | 0.052672671 | 0.999933887 | -0.034992113 | 0.243485638 | -0.27847775 | 0.239475486 |
| ENSG00000148943 | *LIN7C* | 2.917222607 | -0.08140238 | -0.025295077 | -0.056107303 | 0.999933887 | -0.263270441 | 0.046025596 | -0.309296037 | 0.239844443 |
| ENSG00000256464 | *YWHABP2* | 0.079246289 | 0.045766631 | 0.099325028 | -0.053558398 | 0.999933887 | 0.139601966 | 0.601709024 | -0.462107058 | 0.239844443 |
| ENSG00000197566 | *ZNF624* | -0.225787531 | -0.177428209 | 0.072328397 | -0.249756605 | 0.999933887 | -0.26546686 | 0.271277145 | -0.536744005 | 0.240002719 |
| ENSG00000145416 | *MARCHF1* | 0.736813788 | -0.199806858 | -0.024123211 | -0.175683647 | 0.999933887 | -0.459895873 | -0.138540356 | -0.321355518 | 0.240146555 |
| ENSG00000166451 | *CENPN* | 0.232107243 | -0.012645955 | 0.045531468 | -0.058177424 | 0.999933887 | 0.076943659 | -0.39934136 | 0.476285019 | 0.240594825 |
| ENSG00000084463 | *WBP11* | 5.532383847 | -0.030691204 | -0.034519611 | 0.003828407 | 0.999933887 | 0.134407274 | 0.251284749 | -0.116877475 | 0.240634991 |
| ENSG00000130985 | *UBA1* | 7.822508439 | 0.008411879 | 0.028941499 | -0.02052962 | 0.999933887 | 0.203052382 | 0.299367573 | -0.096315191 | 0.240702678 |
| ENSG00000167113 | *COQ4* | 2.520618958 | -0.008307756 | 0.048755384 | -0.05706314 | 0.999933887 | 0.169239207 | -0.013266327 | 0.182505534 | 0.240702678 |
| ENSG00000011451 | *WIZ* | 3.729189366 | 0.0749915 | 0.008728537 | 0.066262963 | 0.999933887 | -0.029293426 | -0.205071595 | 0.17577817 | 0.240782274 |
| ENSG00000188158 | *NHS* | 1.792640401 | -0.073775627 | -0.192603549 | 0.118827922 | 0.999933887 | -0.566484371 | -1.153763759 | 0.587279389 | 0.240782274 |
| ENSG00000203965 | *EFCAB7* | 0.770467027 | -0.115847119 | -0.066981159 | -0.04886596 | 0.999933887 | 0.114900419 | 0.469108837 | -0.354208418 | 0.240828802 |
| ENSG00000076685 | *NT5C2* | 6.54435891 | -0.046495017 | -0.033808909 | -0.012686108 | 0.999933887 | -0.110259001 | 0.035943003 | -0.146202004 | 0.240903748 |
| ENSG00000116983 | *HPCAL4* | 2.130914257 | -0.041545696 | -0.080842019 | 0.039296323 | 0.999933887 | 0.123990785 | -0.099253703 | 0.223244488 | 0.240903748 |
| ENSG00000175387 | *SMAD2* | 5.475017775 | -0.064944435 | 0.004933687 | -0.069878121 | 0.999933887 | -0.654231506 | -0.788800533 | 0.134569027 | 0.241046966 |
| ENSG00000008311 | *AASS* | -0.830272901 | -0.183742278 | 0.028053124 | -0.211795402 | 0.999933887 | -0.38559059 | 0.134484822 | -0.520075412 | 0.241046966 |
| ENSG00000139266 | *MARCHF9* | 2.791051643 | 0.058618607 | 0.021547178 | 0.037071429 | 0.999933887 | 0.129658414 | -0.056263744 | 0.185922158 | 0.241046966 |
| ENSG00000211584 | *SLC48A1* | 4.077025245 | 0.030395603 | -0.028779596 | 0.059175198 | 0.999933887 | -0.578486553 | -0.807548486 | 0.229061933 | 0.241046966 |
| ENSG00000163606 | *CD200R1* | 0.750599769 | 0.042814689 | -0.057098499 | 0.099913188 | 0.999933887 | -0.092893249 | 0.250614032 | -0.343507281 | 0.241303506 |
| ENSG00000133704 | *IPO8* | 3.698178608 | 0.001359667 | -0.033072695 | 0.034432362 | 0.999933887 | -0.061034157 | -0.230922529 | 0.169888372 | 0.241514368 |
| ENSG00000204304 | *PBX2* | 7.792067758 | 0.043833417 | 0.0304153 | 0.013418117 | 0.999933887 | 0.013766934 | -0.097002741 | 0.110769675 | 0.241534568 |
| ENSG00000171530 | *TBCA* | 3.568119575 | 0.021498453 | -0.014610418 | 0.036108871 | 0.999933887 | 0.068294506 | -0.101301459 | 0.169595965 | 0.241534568 |
| ENSG00000136718 | *IMP4* | 3.33230214 | 0.017903807 | -0.027022457 | 0.044926264 | 0.999933887 | 0.202189549 | 0.044439484 | 0.157750065 | 0.241915853 |
| ENSG00000236936 | *AL031005.1* | 0.352891774 | 0.024016025 | -0.013599924 | 0.037615949 | 0.999933887 | 2.226029064 | 3.002955135 | -0.776926071 | 0.241915853 |
| ENSG00000230753 | *ZNF341-AS1* | -0.311424819 | -0.019269615 | 0.39180285 | -0.411072465 | 0.999933887 | 1.05831182 | 1.422180455 | -0.363868635 | 0.241928367 |
| ENSG00000169609 | *C15orf40* | 2.098415627 | -0.047631332 | -0.019161363 | -0.028469968 | 0.999933887 | 0.017481124 | -0.181892777 | 0.199373901 | 0.24221276 |
| ENSG00000104957 | *CCDC130* | 4.894888945 | 0.000747652 | -0.004668937 | 0.00541659 | 0.999933887 | 0.023687826 | -0.102806648 | 0.126494474 | 0.24252744 |
| ENSG00000255197 | *AC090559.1* | 6.816044574 | 0.012660709 | 0.034175601 | -0.021514892 | 0.999933887 | 0.226132901 | 0.355527314 | -0.129394412 | 0.242529234 |
| ENSG00000119787 | *ATL2* | 3.589899871 | 0.002645331 | -0.159516723 | 0.162162054 | 0.999933887 | -0.005890869 | 0.157745501 | -0.16363637 | 0.242529234 |
| ENSG00000153786 | *ZDHHC7* | 5.976649211 | -0.095270778 | -0.070549342 | -0.024721436 | 0.999933887 | -0.797850452 | -0.966336418 | 0.168485965 | 0.242708126 |
| ENSG00000254858 | *MPV17L2* | 1.753044268 | 0.159745462 | 0.002246553 | 0.157498909 | 0.999933887 | -0.112964845 | -0.409046047 | 0.296081202 | 0.242708126 |
| ENSG00000141858 | *SAMD1* | 2.517514438 | -0.007368056 | 0.15093937 | -0.158307425 | 0.999933887 | 0.175963117 | 0.401443949 | -0.225480832 | 0.242708126 |
| ENSG00000166595 | *CIAO2B* | 3.80360706 | 0.019229674 | 0.030586466 | -0.011356793 | 0.999933887 | -0.046280393 | -0.221673702 | 0.175393308 | 0.242845729 |
| ENSG00000136153 | *LMO7* | 1.185298851 | -0.016675448 | -0.168553405 | 0.151877957 | 0.999933887 | 0.019576114 | 0.332714365 | -0.313138251 | 0.242845729 |
| ENSG00000010072 | *SPRTN* | 3.385633671 | -0.060705467 | -0.047562388 | -0.013143079 | 0.999933887 | -0.164273364 | 0.01858525 | -0.182858613 | 0.242949038 |
| ENSG00000073008 | *PVR* | 0.509769701 | 0.078057942 | 0.102022119 | -0.023964178 | 0.999933887 | -0.104235066 | -0.541326179 | 0.437091113 | 0.242949038 |
| ENSG00000279862 | *AC092964.1* | 0.437437933 | 0.02529584 | 0.252106063 | -0.226810223 | 0.999933887 | 1.477528183 | 2.150149928 | -0.672621745 | 0.242949038 |
| ENSG00000278818 | *Metazoa_SRP* | -0.961860904 | 0.500092918 | 0.249979726 | 0.250113192 | 0.999933887 | -0.242361147 | 0.503289005 | -0.745650152 | 0.242969918 |
| ENSG00000230013 | *CT70* | 1.470814998 | 0.358601241 | 0.024958046 | 0.333643195 | 0.999933887 | 0.018602639 | -0.45932233 | 0.477924969 | 0.242969918 |
| ENSG00000184979 | *USP18* | 0.140076273 | 0.114564165 | 0.087692105 | 0.02687206 | 0.999933887 | -0.175950737 | -0.582547646 | 0.406596909 | 0.242969918 |
| ENSG00000058600 | *POLR3E* | 4.104433589 | 0.037387234 | 0.090179982 | -0.052792748 | 0.999933887 | 0.10035538 | 0.244843306 | -0.144487927 | 0.242969918 |
| ENSG00000225889 | *AC012368.1* | 6.729594239 | -0.025657068 | -0.005226363 | -0.020430705 | 0.999933887 | 0.107327508 | -0.015035974 | 0.122363483 | 0.243256434 |
| ENSG00000254470 | *AP5B1* | 4.84928763 | -0.059619824 | -0.039980373 | -0.01963945 | 0.999933887 | -0.621650509 | -0.756385171 | 0.134734662 | 0.243426178 |
| ENSG00000237094 | *AL732372.2* | 3.469099398 | 0.023430927 | -0.008379638 | 0.031810564 | 0.999933887 | 0.195198065 | 0.364761133 | -0.169563069 | 0.24348952 |
| ENSG00000132881 | *CPLANE2* | -0.563244951 | 0.495157856 | 0.17453552 | 0.320622335 | 0.999933887 | 0.227289833 | -0.34630807 | 0.573597903 | 0.243662825 |
| ENSG00000239569 | *KMT2E-AS1* | 4.247120025 | 0.155525762 | 0.17045568 | -0.014929917 | 0.999933887 | 0.687251593 | 0.923001093 | -0.2357495 | 0.24376341 |
| ENSG00000089902 | *RCOR1* | 6.811626348 | -0.076860812 | 0.03361649 | -0.110477302 | 0.999933887 | -0.79545424 | -0.999030167 | 0.203575927 | 0.244044175 |
| ENSG00000058262 | *SEC61A1* | 5.988535324 | 0.00159468 | -0.021540285 | 0.023134966 | 0.999933887 | 0.267308981 | 0.374465399 | -0.107156418 | 0.244148416 |
| ENSG00000259438 | *MAPK6-DT* | 1.119025083 | 0.272428704 | 0.05327954 | 0.219149164 | 0.999933887 | 2.227924926 | 2.603875344 | -0.375950419 | 0.244160852 |
| ENSG00000107731 | *UNC5B* | -1.682413883 | 0.264856407 | -0.185468933 | 0.45032534 | 0.999933887 | 1.300456331 | 1.785182053 | -0.484725722 | 0.24427361 |
| ENSG00000188511 | *C22orf34* | 1.671987355 | -0.011514428 | 0.089129361 | -0.100643789 | 0.999933887 | -0.112989186 | -0.378244519 | 0.265255333 | 0.24427361 |
| ENSG00000155850 | *SLC26A2* | 2.58236106 | -0.026393989 | -0.159507878 | 0.133113889 | 0.999933887 | -0.075714273 | 0.154833879 | -0.230548152 | 0.244520288 |
| ENSG00000166167 | *BTRC* | 1.939052816 | -0.061619624 | -0.122866476 | 0.061246852 | 0.999933887 | -0.032438156 | 0.200922171 | -0.233360326 | 0.244764798 |
| ENSG00000236449 | *AC010894.2* | 0.834365541 | 0.061477833 | 0.177518726 | -0.116040893 | 0.999933887 | -0.615474741 | -0.990359129 | 0.374884388 | 0.244900579 |
| ENSG00000172340 | *SUCLG2* | 2.235982191 | 0.038190219 | 0.028654733 | 0.009535486 | 0.999933887 | -0.014476713 | 0.226463707 | -0.24094042 | 0.244900579 |
| ENSG00000273018 | *FAM106A* | -0.012678446 | 0.050242387 | -0.457662078 | 0.507904465 | 0.999933887 | -0.403393438 | 0.029321353 | -0.432714791 | 0.244900579 |
| ENSG00000139880 | *CDH24* | -0.29961568 | -0.029222482 | 0.11085122 | -0.140073702 | 0.999933887 | 0.230077335 | 0.723551318 | -0.493473983 | 0.244900579 |
| ENSG00000274020 | *LINC01138* | 2.060683803 | 0.035234915 | 0.06241416 | -0.027179246 | 0.999933887 | -0.173001002 | -0.395473909 | 0.222472908 | 0.244921844 |
| ENSG00000230224 | *PHBP9* | 1.718325976 | -0.056069048 | 0.094018422 | -0.15008747 | 0.999933887 | -0.025140234 | 0.206903252 | -0.232043485 | 0.244964071 |
| ENSG00000223697 | *AF230666.1* | 0.053245605 | -0.26333539 | -0.067110974 | -0.196224416 | 0.999933887 | -0.676052261 | -0.035006946 | -0.641045314 | 0.245124187 |
| ENSG00000102393 | *GLA* | 7.880531575 | 0.075756663 | -0.01334327 | 0.089099932 | 0.999933887 | 1.145575496 | 1.37326686 | -0.227691364 | 0.245124187 |
| ENSG00000229833 | *PET100* | 1.978050118 | -0.150150424 | -0.142854252 | -0.007296172 | 0.999933887 | -0.131995013 | -0.363576266 | 0.231581253 | 0.245229535 |
| ENSG00000111669 | *TPI1* | 6.202601176 | 0.027186206 | -0.035711687 | 0.062897892 | 0.999933887 | 0.010756465 | -0.135845188 | 0.146601654 | 0.245229535 |
| ENSG00000139714 | *MORN3* | 1.69916546 | -0.006846427 | -0.024533298 | 0.017686871 | 0.999933887 | 0.046395654 | -0.187958128 | 0.234353782 | 0.245229535 |
| ENSG00000215421 | *ZNF407* | 5.441222401 | 0.086025822 | 0.215247971 | -0.129222149 | 0.999933887 | 0.06086461 | 0.220047524 | -0.159182914 | 0.24526117 |
| ENSG00000131779 | *PEX11B* | 2.087139105 | -0.176539468 | -0.097773584 | -0.078765885 | 0.999933887 | -0.514373766 | -0.806193045 | 0.291819279 | 0.24526117 |
| ENSG00000155755 | *TMEM237* | -0.855642361 | 0.097898846 | -0.129646618 | 0.227545464 | 0.999933887 | -0.367599449 | 0.23304272 | -0.600642169 | 0.24526117 |
| ENSG00000115993 | *TRAK2* | 3.705444288 | -0.039061074 | -0.082133945 | 0.043072871 | 0.999933887 | -0.116180135 | -0.262363446 | 0.146183311 | 0.24530323 |

| | | | | | | | | | | |
|---|---|---|---|---|---|---|---|---|---|---|
| ENSG00000225190 | PLEKHM1 | 7.722207243 | -0.017256147 | -0.005612173 | -0.011643974 | 0.999933887 | -0.06576827 | 0.037394221 | -0.103162491 | 0.245319491 |
| ENSG00000187556 | NANOS3 | 0.751029633 | 0.209206345 | -0.113555549 | 0.322761894 | 0.999933887 | 1.550770547 | 1.893689846 | -0.342919299 | 0.245343423 |
| ENSG00000269553 | U62631.1 | -1.011304884 | 0.161534324 | -0.06192836 | 0.223462684 | 0.999933887 | 0.069487756 | 0.41025231 | -0.340764554 | 0.246040207 |
| ENSG00000152767 | FARP1 | 2.016834661 | 0.132000625 | 0.199408041 | -0.067407416 | 0.999933887 | -0.724671588 | -0.995458079 | 0.270786491 | 0.246149094 |
| ENSG00000157077 | ZFYVE9 | -0.192620261 | -0.088910727 | 0.078941414 | -0.167852142 | 0.999933887 | -0.008767952 | -0.429242244 | 0.420474292 | 0.246149094 |
| ENSG00000134262 | AP4B1 | 3.086759807 | -0.03072525 | -0.060737675 | 0.030012424 | 0.999933887 | 0.112646294 | -0.081362063 | 0.194008357 | 0.2461969 |
| ENSG00000148358 | GPR107 | 5.873480588 | -0.050798024 | -0.008903379 | -0.041894645 | 0.999933887 | 0.149399277 | 0.328844474 | -0.179445197 | 0.246349525 |
| ENSG00000134686 | PHC2 | 7.319659006 | -0.035714047 | -0.016810153 | -0.018903894 | 0.999933887 | -1.33820463 | -1.553409951 | 0.215205321 | 0.246378869 |
| ENSG00000124233 | SEMG1 | 0.91179455 | 0.049706299 | -0.035560611 | 0.08526691 | 0.999933887 | -0.062955877 | 0.233398146 | -0.296354023 | 0.246378869 |
| ENSG00000114956 | DGUOK | 3.65983173 | -0.060884169 | 0.049181468 | -0.110065637 | 0.999933887 | 0.048764984 | 0.19527547 | -0.146510487 | 0.246405044 |
| ENSG00000138621 | PPCDC | 7.699834655 | 0.003416423 | -0.030145619 | 0.033562042 | 0.999933887 | 1.157354369 | 1.31202137 | -0.154667002 | 0.246405044 |
| ENSG00000175376 | EIF1AD | 3.738886522 | -0.097781088 | 0.011301945 | -0.109083033 | 0.999933887 | -0.323872171 | -0.467337525 | 0.143465355 | 0.24647985 |
| ENSG00000142409 | ZNF787 | 5.344783316 | 0.095763212 | 0.127413519 | -0.031650307 | 0.999933887 | -0.213919882 | -0.441308991 | 0.227389108 | 0.24647985 |
| ENSG00000177337 | DLGAP1-AS1 | 4.14766104 | -0.03613244 | -0.147085521 | 0.110953081 | 0.999933887 | 0.300624678 | 0.454311705 | -0.153687028 | 0.24647985 |
| ENSG00000175203 | DCTN2 | 5.101488593 | -0.016155044 | -0.000192719 | -0.015962325 | 0.999933887 | -0.157873924 | -0.045811982 | -0.112061942 | 0.24647985 |
| ENSG00000041353 | RAB27B | -0.097657329 | 0.043641055 | -0.065124188 | 0.108765243 | 0.999933887 | -0.236715141 | 0.209055014 | -0.445770155 | 0.24647985 |
| ENSG00000146263 | MMS22L | 0.648843606 | -0.019366206 | -0.341077648 | 0.321711442 | 0.999933887 | -0.094846655 | 0.380197541 | -0.475044196 | 0.246526851 |
| ENSG00000141084 | RANBP10 | 3.472110808 | -0.009837408 | -0.029153534 | 0.019316127 | 0.999933887 | 0.044137214 | -0.108372025 | 0.152509239 | 0.246674077 |
| ENSG00000240211 | PPP1R35-AS1 | -0.724308689 | 0.09764095 | 0.341689248 | -0.244048298 | 0.999933887 | 0.642283075 | 1.238561431 | -0.596278356 | 0.246727302 |
| ENSG00000118058 | KMT2A | 5.396438948 | -0.026807472 | -0.033192367 | 0.006384895 | 0.999933887 | -0.070270894 | 0.078558828 | -0.148829721 | 0.246792119 |
| ENSG00000165661 | QSOX2 | 3.690651122 | -0.038565711 | 0.080067523 | -0.118633233 | 0.999933887 | 0.019040241 | -0.128772796 | 0.147813037 | 0.24741361 |
| ENSG00000162236 | STX5 | 5.479107703 | 0.018303768 | -0.023777889 | 0.042081656 | 0.999933887 | 0.223242305 | 0.333979819 | -0.110737515 | 0.24741361 |
| ENSG00000170365 | SMAD1 | 0.060878494 | -0.033538832 | -0.124460071 | 0.090921239 | 0.999933887 | -0.017151479 | 0.385202834 | -0.402354313 | 0.24741361 |
| ENSG00000151292 | CSNK1G3 | 2.915317178 | -0.008103554 | -0.267721522 | 0.259617968 | 0.999933887 | -0.217069219 | 0.091553981 | -0.3086232 | 0.247492535 |
| ENSG00000172586 | CHCHD1 | 2.146912597 | 0.04788579 | -0.268823269 | 0.316709058 | 0.999933887 | -0.397198174 | -0.63612369 | 0.238925516 | 0.247554737 |
| ENSG00000197647 | ZNF433 | 1.399001056 | -0.329442605 | -0.089223928 | -0.240218677 | 0.999933887 | -0.747552931 | -0.522736139 | -0.224816792 | 0.247611707 |
| ENSG00000091073 | DTX2 | 5.50621606 | 0.009192467 | 0.011908107 | -0.002715641 | 0.999933887 | 0.405567366 | 0.535104536 | -0.12953717 | 0.247611707 |
| ENSG00000180667 | YOD1 | 4.140571976 | -0.08109496 | -0.212036673 | 0.130941713 | 0.999933887 | -0.170200732 | 0.090405283 | -0.260606015 | 0.247662633 |
| ENSG00000180509 | KCNE1 | 2.188964634 | 0.193130503 | 0.161120163 | 0.03201034 | 0.999933887 | 1.085912389 | 1.586353639 | -0.50044125 | 0.247796476 |
| ENSG00000177595 | PIDD1 | 2.767705837 | 0.051205324 | 0.077143964 | -0.025938639 | 0.999933887 | 0.099864429 | -0.116161357 | 0.216025786 | 0.247796476 |
| ENSG00000148842 | CNNM2 | 3.194420715 | -0.014939563 | -0.046553815 | 0.031614252 | 0.999933887 | -0.063543147 | 0.117591532 | -0.181134679 | 0.247796476 |
| ENSG00000214113 | LYRM4 | 1.519198763 | -0.02830405 | -0.073241063 | 0.044937013 | 0.999933887 | 0.331676976 | 0.604713898 | -0.273036922 | 0.247796476 |
| ENSG00000052841 | TTC17 | 4.82021902 | 0.000119973 | -0.081576364 | 0.081696337 | 0.999933887 | -0.210625477 | -0.089410922 | -0.121214555 | 0.247796476 |
| ENSG00000182208 | MOB2 | 4.490966879 | 0.022881306 | -0.026406125 | 0.04928743 | 0.999933887 | -0.080002726 | -0.24034967 | 0.160346945 | 0.247923924 |
| ENSG00000117500 | TMED5 | 6.391549143 | -0.034646373 | -0.14466039 | 0.110014018 | 0.999933887 | -0.354330081 | -0.120218098 | -0.234111983 | 0.248003555 |
| ENSG00000163661 | PTX3 | 3.84656883 | 0.25098728 | 0.254162448 | -0.003175168 | 0.999933887 | 1.949971787 | 2.307487107 | -0.35751532 | 0.248058229 |
| ENSG00000253977 | MTCO2P4 | -2.564837008 | 0.393512783 | -0.118511849 | 0.512024632 | 0.999933887 | 2.017615384 | 2.926771231 | -0.909155846 | 0.248146609 |
| ENSG00000012660 | ELOVL5 | 8.022349656 | -0.025189673 | 0.003667515 | -0.028857188 | 0.999933887 | -0.618920275 | -0.740188011 | 0.121267736 | 0.248146609 |
| ENSG00000286319 | AL135978.1 | 1.676238942 | -0.069710257 | 0.077345573 | -0.14705583 | 0.999933887 | 0.004858537 | 0.393135676 | -0.388277139 | 0.248147311 |
| ENSG00000144746 | ARL6IP5 | 6.016158462 | -0.025606097 | -0.092533171 | 0.066927074 | 0.999933887 | -0.070665518 | -0.239067165 | 0.168401648 | 0.248284267 |
| ENSG00000113558 | SKP1 | 5.993568678 | -0.042618465 | -0.024728223 | -0.017890242 | 0.999933887 | -0.002738077 | 0.098960401 | -0.101698478 | 0.24838636 |
| ENSG00000275665 | AC006449.3 | -0.435276806 | -0.154658018 | 0.128105014 | -0.282763032 | 0.999933887 | 0.965867141 | 1.319818076 | -0.353950936 | 0.24838636 |
| ENSG00000198715 | GLMP | 4.747025416 | -0.06235213 | -0.096888612 | 0.034536482 | 0.999933887 | -0.406093515 | -0.556830225 | 0.15073671 | 0.248454731 |
| ENSG00000152818 | UTRN | 6.266542981 | 0.061169689 | -0.01286474 | 0.074034428 | 0.999933887 | -0.23046754 | 0.101151672 | -0.331619212 | 0.248621208 |
| ENSG00000156966 | B3GNT7 | 1.282384716 | 0.158481402 | 0.128019553 | 0.030461849 | 0.999933887 | 2.791419378 | 3.181742118 | -0.39032274 | 0.248675125 |
| ENSG00000166986 | MARS1 | 6.496905109 | -0.019690911 | 0.001613629 | -0.02130454 | 0.999933887 | 0.016218851 | -0.08211038 | 0.098329231 | 0.248675125 |
| ENSG00000124641 | MED20 | 2.670817775 | -0.037727947 | -0.056886686 | 0.019158739 | 0.999933887 | 0.469583083 | 0.656300197 | -0.186717114 | 0.248675125 |
| ENSG00000144306 | SCRN3 | 3.379326491 | -0.020331626 | -0.164642354 | 0.144310728 | 0.999933887 | -0.1568738 | 0.028135671 | -0.185009471 | 0.248675125 |
| ENSG00000044446 | PHKA2 | 4.389737492 | -0.024981749 | 0.027156137 | -0.052137886 | 0.999933887 | -0.018807634 | -0.164559678 | 0.145752045 | 0.248739366 |
| ENSG00000102317 | RBM3 | 6.611778508 | -0.021482932 | 0.005222661 | -0.026705593 | 0.999933887 | 0.113220314 | 0.231222536 | -0.118002221 | 0.248770293 |
| ENSG00000177885 | GRB2 | 8.878790496 | 0.013781792 | 0.020350883 | -0.006569091 | 0.999933887 | 0.3555361 | 0.467855701 | -0.112319602 | 0.249042833 |
| ENSG00000125089 | SH3TC1 | 2.787310839 | 0.037391474 | -0.001040641 | 0.038432115 | 0.999933887 | 0.005268531 | -0.242652675 | 0.247921206 | 0.249079498 |
| ENSG00000165685 | TMEM52B | 0.134572055 | -0.041609167 | -0.041449055 | -0.000160112 | 0.999933887 | 1.099683559 | 1.710760906 | -0.611077347 | 0.249085453 |
| ENSG00000152234 | ATP5F1A | 5.101624655 | -0.025870212 | -0.127413009 | 0.101542797 | 0.999933887 | 0.052891598 | -0.095889405 | 0.148781003 | 0.249094089 |
| ENSG00000259834 | AL365361.1 | 2.500830346 | -0.122444223 | -0.305397261 | 0.182953038 | 0.999933887 | -0.323435345 | 0.130206819 | -0.453642164 | 0.249171824 |
| ENSG00000177426 | TGIF1 | 7.702795341 | 0.1103923 | 0.114590751 | -0.004198451 | 0.999933887 | 0.10006836 | 0.224530501 | -0.124462142 | 0.249498429 |
| ENSG00000183625 | CCR3 | 4.582986747 | -0.082975209 | -0.000258152 | -0.082717057 | 0.999933887 | -0.288566409 | -0.47139773 | 0.182831321 | 0.249645025 |
| ENSG00000108443 | RPS6KB1 | 3.614223383 | -0.055761041 | -0.123987025 | 0.068225984 | 0.999933887 | -0.295688385 | -0.049695703 | -0.245992682 | 0.249645025 |
| ENSG00000279141 | LINC01451 | 1.249196372 | 0.031784845 | -0.25110367 | 0.282888515 | 0.999933887 | -1.600412472 | -1.948848334 | 0.348435862 | 0.249645025 |
| ENSG00000120800 | UTP20 | 1.832178656 | 0.00110853 | -0.150518759 | 0.151627289 | 0.999933887 | -0.150002824 | 0.123869798 | -0.273872623 | 0.249825094 |
| ENSG00000227741 | AL121987.2 | -0.330361581 | 0.187715556 | -0.05003202 | 0.237747576 | 0.999933887 | 0.551913973 | 0.976151378 | -0.424237405 | 0.24987066 |
| ENSG00000008405 | CRY1 | 5.457394232 | -0.02697344 | -0.073320557 | 0.046347118 | 0.999933887 | 0.207796292 | 0.387284582 | -0.17948829 | 0.24987066 |
| ENSG00000130758 | MAP3K10 | 2.963980012 | 0.027428206 | 0.156414095 | -0.128985889 | 0.999933887 | 0.120283469 | -0.101840496 | 0.222123965 | 0.24987066 |
| ENSG00000138463 | SLC49A4 | 5.755589162 | -0.003206275 | 0.000311746 | -0.00351802 | 0.999933887 | 0.043539889 | 0.148426816 | -0.104886927 | 0.24990195 |
| ENSG00000286696 | AC090826.3 | 0.356419243 | -0.080625146 | 0.160887921 | -0.241513068 | 0.999933887 | 0.041856982 | -0.400930136 | 0.442787118 | 0.25023928 |
| ENSG00000135341 | MAP3K7 | 4.572953458 | -0.050411342 | -0.122332302 | 0.07192096 | 0.999933887 | -0.361185354 | -0.208296429 | -0.152888925 | 0.25023928 |
| ENSG00000177406 | NINJ2-AS1 | 4.33011703 | -0.031569484 | -0.023409732 | -0.008159752 | 0.999933887 | 0.48679017 | 0.65906793 | -0.172277761 | 0.25023928 |
| ENSG00000182952 | HMGN4 | 4.922276311 | -0.009528358 | -0.022734204 | 0.013205846 | 0.999933887 | -0.142346834 | -0.275190952 | 0.132844118 | 0.25023928 |
| ENSG00000198824 | CHAMP1 | 2.084685514 | -0.010462797 | -0.095715252 | 0.085252455 | 0.999933887 | -0.392174089 | -0.640620486 | 0.248446396 | 0.25023928 |
| ENSG00000166272 | WBP1L | 6.237150294 | -0.090115613 | -0.018114862 | -0.072000751 | 0.999933887 | -0.113557711 | 0.045390577 | -0.158948288 | 0.250250037 |
| ENSG00000215012 | RTL10 | 2.984830309 | -0.114544149 | -0.093212854 | -0.021331294 | 0.999933887 | -0.234359514 | -0.42913729 | 0.194777777 | 0.250494937 |
| ENSG00000122912 | SLC25A16 | 3.745460569 | -0.020062037 | 0.013561183 | -0.03362322 | 0.999933887 | -0.118206248 | 0.032864388 | -0.151070636 | 0.250602347 |
| ENSG00000061273 | HDAC7 | 6.808276139 | 0.035323836 | 0.085539982 | -0.050216147 | 0.999933887 | 0.062083095 | -0.056968702 | 0.119051797 | 0.250613965 |
| ENSG00000182944 | EWSR1 | 7.869517459 | -0.008735406 | 0.021602863 | -0.030338269 | 0.999933887 | -0.007979571 | 0.088312396 | -0.096291966 | 0.250617913 |
| ENSG00000156973 | PDE6D | 2.583119079 | 0.053131111 | 0.045293049 | 0.007838062 | 0.999933887 | 0.469212541 | 0.662193818 | -0.192981277 | 0.250651003 |
| ENSG00000124486 | USP9X | 8.097883148 | -0.036900774 | -0.067895308 | 0.030994534 | 0.999933887 | -0.323734986 | -0.119616701 | -0.204118285 | 0.250651003 |
| ENSG00000171827 | ZNF570 | 0.765149552 | 0.10128834 | -0.15459385 | 0.255882191 | 0.999933887 | -0.160540907 | 0.235834434 | -0.396375342 | 0.250870295 |
| ENSG00000169139 | UBE2V2 | 2.961566692 | -0.007439205 | -0.103318108 | 0.095878903 | 0.999933887 | -0.243900521 | -0.058480644 | -0.185419876 | 0.250870295 |
| ENSG00000100109 | TFIP11 | 5.828148683 | -0.037819891 | -0.044496252 | 0.006676361 | 0.999933887 | -0.011843719 | 0.092277851 | -0.10412157 | 0.250878664 |
| ENSG00000134077 | THUMPD3 | 4.451441801 | -0.075059691 | -0.067860968 | -0.007198724 | 0.999933887 | -0.24039677 | -0.080324957 | -0.160071813 | 0.250982021 |
| ENSG00000114023 | FAM162A | 1.689872131 | -0.078191994 | -0.075401416 | -0.002790577 | 0.999933887 | 0.198561058 | -0.03346191 | 0.232022968 | 0.251320452 |
| ENSG00000122068 | FYTTD1 | 4.204989577 | -0.000751382 | -0.059901281 | 0.059149899 | 0.999933887 | -0.272844782 | -0.101031119 | -0.171813663 | 0.251320452 |
| ENSG00000182150 | ERCC6L2 | 2.699957663 | -0.054925329 | -0.258399837 | 0.203474508 | 0.999933887 | -0.187870465 | 0.192019574 | -0.379890039 | 0.251384568 |
| ENSG00000183918 | SH2D1A | 3.591322342 | -0.085851786 | -0.204299322 | 0.118447536 | 0.999933887 | 0.098855046 | 0.274967115 | -0.176112069 | 0.251384664 |
| ENSG00000205808 | PLPP6 | 1.176091444 | 0.027537195 | -0.210366324 | 0.237903519 | 0.999933887 | 0.696639251 | 0.353814302 | 0.342824948 | 0.251384664 |
| ENSG00000230569 | AC114763.2 | -1.296628132 | -0.298683184 | 1.004887334 | -1.303570518 | 0.999933887 | 0.969471554 | 1.443846138 | -0.474374584 | 0.251400757 |
| ENSG00000184277 | TM2D3 | 6.078624464 | 0.028242606 | 0.029877685 | -0.001635079 | 0.999933887 | -0.051864169 | 0.067570433 | -0.119434602 | 0.251400757 |
| ENSG00000072506 | HSD17B10 | 2.467805091 | -0.027206296 | -0.017922651 | -0.009283644 | 0.999933887 | 0.137193799 | -0.072629489 | 0.209823288 | 0.25140547 |
| ENSG00000198105 | ZNF248 | 0.327593888 | -0.213865114 | 0.051571224 | -0.265436337 | 0.999933887 | -0.149688848 | 0.196373688 | -0.346062536 | 0.251677643 |
| ENSG00000171798 | KNDC1 | -0.543252342 | 0.134217114 | -0.038872586 | 0.173089701 | 0.999933887 | 0.919984687 | 1.38389285 | -0.463908164 | 0.251834907 |
| ENSG00000100554 | ATP6V1D | 7.004702264 | 0.000598166 | -0.025371805 | 0.025969971 | 0.999933887 | 0.102989255 | 0.263547742 | -0.160558486 | 0.251834907 |
| ENSG00000149499 | EML3 | 5.635907405 | 0.04227735 | 0.095498661 | -0.053221311 | 0.999933887 | 0.369070801 | 0.550906748 | -0.181835947 | 0.251850199 |
| ENSG00000285868 | AC008676.3 | 0.85855022 | 0.041296481 | -0.085550381 | 0.126846862 | 0.999933887 | -0.399157555 | -0.029822675 | -0.36933488 | 0.252085977 |
| ENSG00000198520 | ARMH1 | 3.001798214 | 0.123871714 | 0.029108765 | 0.094762948 | 0.999933887 | -0.003776788 | -0.203674163 | 0.199897375 | 0.252094401 |
| ENSG00000273356 | LINC02019 | 1.271768408 | -0.092185664 | -0.049952954 | -0.04223271 | 0.999933887 | -0.652115843 | -0.989877759 | 0.337761916 | 0.252094401 |
| ENSG00000110719 | TCIRG1 | 9.73619722 | 0.047133643 | 0.063762521 | -0.016628878 | 0.999933887 | -0.297326173 | -0.427568564 | 0.130242391 | 0.252231725 |
| ENSG00000012232 | EXTL3 | 5.331098713 | -0.061104431 | -0.042682794 | -0.018421637 | 0.999933887 | -0.894873475 | -1.096491802 | 0.201618327 | 0.252231725 |
| ENSG00000182173 | TSEN54 | 3.551921334 | 0.053559228 | 0.068962473 | -0.015403245 | 0.999933887 | 0.17411875 | 0.012742345 | 0.161376405 | 0.252385262 |
| ENSG00000228253 | MT-ATP8 | 4.599706188 | 0.053139016 | -0.068283522 | 0.121422538 | 0.999933887 | -0.037410668 | 0.21142213 | -0.248832798 | 0.252385262 |
| ENSG00000198826 | ARHGAP11A | -0.582986989 | -0.014890586 | 0.2356783 | -0.250568886 | 0.999933887 | -0.195661615 | 0.380190531 | -0.575852146 | 0.252385262 |
| ENSG00000142168 | SOD1 | 4.379956038 | 0.027789309 | 0.017669245 | 0.010120063 | 0.999933887 | 0.165477406 | 0.015865786 | 0.14961162 | 0.252541241 |
| ENSG00000265354 | TIMM23 | 2.306743655 | -0.050490553 | -0.076506774 | 0.02601622 | 0.999933887 | 0.071859447 | 0.255725687 | -0.183866241 | 0.252860121 |
| ENSG00000274173 | AL035661.1 | 2.269508669 | -0.051964107 | -0.006775145 | -0.045188962 | 0.999933887 | 0.351109685 | 0.651692871 | -0.300583186 | 0.252860121 |
| ENSG00000135378 | PRRG4 | 3.734579038 | 0.032789539 | 0.053674704 | -0.020885164 | 0.999933887 | 0.521501724 | 0.743845417 | -0.222343693 | 0.252860121 |
| ENSG00000099866 | MADCAM1 | 0.473971354 | -0.053575188 | -0.133189792 | 0.079614604 | 0.999933887 | -0.104712729 | -0.474385393 | 0.369672664 | 0.252893771 |

| | | | | | | | | | | |
|---|---|---|---|---|---|---|---|---|---|---|
| ENSG00000271933 | *AL603756.1* | 1.999367955 | 0.146092939 | 0.174573751 | -0.028480812 | 0.999933887 | 0.246400759 | 0.557231136 | -0.310830377 | 0.252911027 |
| ENSG00000173153 | *ESRRA* | 5.777376163 | 0.09962337 | 0.010341838 | 0.089281532 | 0.999933887 | -0.102360375 | 0.083500781 | -0.185861156 | 0.252911027 |
| ENSG00000237310 | *GS1-124K5.4* | -0.346742182 | 0.200355855 | 0.023368984 | 0.176986872 | 0.999933887 | 0.37374644 | 0.820410089 | -0.446663649 | 0.252911027 |
| ENSG00000224137 | *LINC01857* | 0.412392475 | -0.180068652 | -0.035316644 | -0.144752009 | 0.999933887 | 1.012007809 | 1.261613453 | -0.249605644 | 0.252911027 |
| ENSG00000104142 | *VPS18* | 9.269916501 | 0.03207161 | 0.014038016 | 0.018033594 | 0.999933887 | 0.373583966 | 0.523179659 | -0.149595693 | 0.252911027 |
| ENSG00000278867 | *AC090616.6* | 0.403470944 | 0.066050697 | -0.083189299 | 0.149239996 | 0.999933887 | -0.481310603 | -0.157512708 | -0.323797895 | 0.252911027 |
| ENSG00000115464 | *USP34* | 6.574384054 | -0.048509621 | -0.094718538 | 0.046208917 | 0.999933887 | -0.437912712 | -0.134291021 | -0.303621692 | 0.252911027 |
| ENSG00000107485 | *GATA3* | 3.561352223 | -0.003391547 | -0.06958019 | 0.066188643 | 0.999933887 | 0.132489575 | 0.290977629 | -0.158488054 | 0.252911027 |
| ENSG00000173402 | *DAG1* | 2.093903305 | 0.003322207 | -0.100756776 | 0.104078983 | 0.999933887 | -0.136569618 | -0.38903472 | 0.252465102 | 0.252911027 |
| ENSG00000214013 | *GANC* | 2.125418649 | -0.000563826 | -0.135758766 | 0.135194939 | 0.999933887 | -0.022340878 | -0.259185776 | 0.236844899 | 0.252911027 |
| ENSG00000165071 | *TMEM71* | 6.322009841 | -0.039291587 | -0.060653434 | 0.021361847 | 0.999933887 | -0.222872847 | -0.363356701 | 0.140483854 | 0.25297253 |
| ENSG00000224086 | *PPM1F-AS1* | 3.831650287 | -0.216878773 | -0.237286014 | 0.020407241 | 0.999933887 | 0.487056511 | 0.722969281 | -0.23591277 | 0.252974786 |
| ENSG00000271254 | *AC240274.1* | 1.400447512 | -0.090671685 | -0.013629908 | -0.077041777 | 0.999933887 | 0.048642436 | 0.270164803 | -0.221522367 | 0.252974786 |
| ENSG00000196172 | *ZNF681* | -0.795557699 | -0.248442396 | 0.373413288 | -0.621855684 | 0.999933887 | -0.716546025 | -0.221494932 | -0.495051093 | 0.252974786 |
| ENSG00000261008 | *LINC01572* | -1.116776423 | -0.376641589 | -0.302387774 | -0.074253815 | 0.999933887 | 1.040941667 | 1.554392934 | -0.513451267 | 0.252974786 |
| ENSG00000083099 | *LYRM2* | 4.102764474 | -0.045050429 | -0.000342397 | -0.044708032 | 0.999933887 | -0.170638839 | -0.030926632 | -0.139712208 | 0.252974786 |
| ENSG00000153914 | *SREK1* | 6.044705626 | -0.048083307 | -0.194990954 | 0.146907647 | 0.999933887 | -0.594882074 | -0.200787514 | -0.39409456 | 0.252974786 |
| ENSG00000167874 | *TMEM88* | 4.483753913 | 0.227741287 | 0.140006291 | 0.087734996 | 0.999933887 | 1.143710578 | 1.345346634 | -0.201636056 | 0.253118333 |
| ENSG00000282100 | *HSP90AB4P* | 0.673818975 | -0.010150828 | 0.08563397 | -0.095784798 | 0.999933887 | 0.051410952 | -0.38370093 | 0.435111882 | 0.253118333 |
| ENSG00000121413 | *ZSCAN18* | 3.293309107 | 0.007917211 | -0.046676507 | 0.054593718 | 0.999933887 | -0.079010777 | 0.109029549 | -0.188040326 | 0.253511706 |
| ENSG00000067208 | *EVI5* | 2.258004137 | -0.086471871 | -0.10920256 | 0.022730689 | 0.999933887 | -0.644835709 | -1.01365439 | 0.368818681 | 0.253514194 |
| ENSG00000225138 | *SLC9A3-AS1* | 1.832564671 | -0.04119131 | -0.009952778 | -0.031238532 | 0.999933887 | 0.277273619 | 0.482713564 | -0.205439945 | 0.253514194 |
| ENSG00000237927 | *AL078604.2* | 0.436640649 | -0.131581763 | -0.087148092 | -0.044433672 | 0.999933887 | 1.300691709 | 1.6215609 | -0.320869191 | 0.25364839 |
| ENSG00000113263 | *ITK* | 5.790654067 | -0.026894444 | -0.037737043 | 0.010842599 | 0.999933887 | 0.142850478 | 0.255337626 | -0.112487148 | 0.253707741 |
| ENSG00000277203 | *F8A1* | 3.024160019 | -0.012536247 | 0.009760281 | -0.022296528 | 0.999933887 | 1.047379327 | 1.34538165 | -0.298002323 | 0.253789373 |
| ENSG00000165688 | *PMPCA* | 4.347184567 | -0.038815521 | 0.013391891 | -0.052207412 | 0.999933887 | -0.030811359 | -0.164016349 | 0.13320499 | 0.253880076 |
| ENSG00000224195 | *AC022400.1* | -0.422073828 | 0.163455823 | 0.090148853 | 0.07330697 | 0.999933887 | -0.589077524 | -1.17538667 | 0.586309147 | 0.253897313 |
| ENSG00000228606 | *AL139011.1* | -0.14930991 | 0.142625218 | -0.098421579 | 0.241046798 | 0.999933887 | 0.508791485 | 0.861819758 | -0.353028273 | 0.253897313 |
| ENSG00000237551 | *AC096775.1* | -0.231139474 | 0.042910454 | 0.279365688 | -0.236455234 | 0.999933887 | 0.516985942 | 1.05797841 | -0.540992467 | 0.253897313 |
| ENSG00000175087 | *PDIK1L* | 0.646293497 | -0.14728428 | 0.106933848 | -0.254218127 | 0.999933887 | -0.278534923 | 0.088902304 | -0.367437227 | 0.253955163 |
| ENSG00000110092 | *CCND1* | -0.599651607 | 0.032146734 | 0.016984916 | 0.015161818 | 0.999933887 | 0.215297946 | 0.665399498 | -0.450101552 | 0.253955163 |
| ENSG00000104299 | *INTS9* | 2.506559377 | 0.076633299 | -0.020454017 | 0.097087316 | 0.999933887 | 0.434657708 | 0.680273167 | -0.245615459 | 0.253957182 |
| ENSG00000091127 | *PUS7* | 0.097733048 | -0.135409772 | 0.014394894 | -0.149804666 | 0.999933887 | 0.049325168 | -0.363826908 | 0.413152076 | 0.253957182 |
| ENSG00000188229 | *TUBB4B* | 6.589822627 | 0.045852824 | -0.012437375 | 0.058290199 | 0.999933887 | -0.45114869 | -0.623966424 | 0.172817734 | 0.253957182 |
| ENSG00000185009 | *AP3M1* | 3.664803598 | -0.050937506 | -0.013306218 | -0.037631287 | 0.999933887 | -0.643377174 | -0.807645658 | 0.164268484 | 0.253957182 |
| ENSG00000101290 | *CDS2* | 6.805055581 | -0.026425047 | 0.00044391 | -0.026868957 | 0.999933887 | -0.990332694 | -1.158893865 | 0.168561171 | 0.253957182 |
| ENSG00000179915 | *NRXN1* | -0.649661643 | -0.046208231 | 0.172858417 | -0.219066647 | 0.999933887 | -0.647497643 | 0.192671935 | -0.840169578 | 0.253957182 |
| ENSG00000163755 | *HPS3* | 3.040218573 | 0.015167749 | -0.081136875 | 0.096304624 | 0.999933887 | -0.584048132 | -0.761702429 | 0.177654297 | 0.253957182 |
| ENSG00000081386 | *ZNF510* | 1.178393201 | -0.027200327 | -0.301846088 | 0.274645761 | 0.999933887 | 0.059307738 | 0.404975511 | -0.345667773 | 0.253957182 |
| ENSG00000144535 | *DIS3L2* | 2.493116584 | 0.006343652 | -0.082078393 | 0.088422045 | 0.999933887 | 0.11990502 | -0.056168855 | 0.176073874 | 0.253957182 |
| ENSG00000164542 | *KIAA0895* | -0.270054086 | 0.012461689 | 0.023556911 | -0.011095221 | 0.999933887 | 0.289337623 | 0.759868262 | -0.470530639 | 0.253957182 |
| ENSG00000276649 | *AL117335.1* | 4.217305933 | 0.010517436 | 0.053791967 | -0.04327453 | 0.999933887 | 0.814422839 | 0.967605899 | -0.15318306 | 0.254050997 |
| ENSG00000141384 | *TAF4B* | 0.972549261 | 0.011288433 | -0.083006229 | 0.094294661 | 0.999933887 | 0.116430158 | 0.407650698 | -0.29122054 | 0.254050997 |
| ENSG00000081870 | *HSPB11* | 1.851977295 | -0.03065164 | 0.033496521 | -0.064148161 | 0.999933887 | -0.013452565 | -0.240596444 | 0.227143879 | 0.254119588 |
| ENSG00000237169 | *RPL12P27* | 0.29105623 | -0.108538703 | 0.020997143 | -0.129535845 | 0.999933887 | -1.20670675 | -1.653390387 | 0.446683637 | 0.254177244 |
| ENSG00000077458 | *FAM76B* | 4.940174824 | -0.009708542 | -0.026567469 | 0.016858927 | 0.999933887 | -0.342561852 | -0.105815706 | -0.236746147 | 0.254401794 |
| ENSG00000117983 | *MUC5B* | 1.639681115 | -0.04475443 | -0.033161302 | -0.011593128 | 0.999933887 | 0.9254053 | 1.28768235 | -0.36227705 | 0.254974199 |
| ENSG00000146109 | *ABT1* | 4.087536376 | -0.012982622 | 0.011710568 | -0.02469319 | 0.999933887 | -0.117038092 | -0.285836419 | 0.168798326 | 0.255045423 |
| ENSG00000286532 | *PARTICL* | 0.683301261 | -0.069187199 | -0.393829101 | 0.324641903 | 0.999933887 | -0.962355207 | -1.446483341 | 0.484128134 | 0.255059508 |
| ENSG00000179195 | *ZNF664* | 3.111654615 | -0.035199003 | 0.013574221 | -0.048773224 | 0.999933887 | -0.079651126 | -0.234020544 | 0.154369418 | 0.255123147 |
| ENSG00000169249 | *ZRSR2* | 3.448362971 | -0.069015755 | -0.091021181 | 0.022005426 | 0.999933887 | -0.008750735 | -0.179497335 | 0.1707466 | 0.25548312 |
| ENSG00000082269 | *FAM135A* | -0.029120792 | 0.108410134 | -0.316211476 | 0.42462161 | 0.999933887 | -0.732906432 | -0.31346554 | -0.419440892 | 0.25548312 |
| ENSG00000199405 | *SNORA1B* | -0.084590515 | 0.123108042 | 0.113365896 | 0.009742147 | 0.999933887 | 0.98305983 | 1.617940756 | -0.634880926 | 0.255551747 |
| ENSG00000159658 | *EFCAB14* | 5.490944822 | 0.000155085 | -0.03885358 | 0.039008665 | 0.999933887 | -0.147776535 | -0.264408061 | 0.116631526 | 0.255666453 |
| ENSG00000180815 | *MAP3K15* | -0.639747961 | -0.086062888 | -0.043739772 | -0.042323116 | 0.999933887 | -0.63272825 | 0.086039447 | -0.718767696 | 0.255911754 |
| ENSG00000132436 | *FIGNL1* | 0.438022119 | 0.034835838 | 0.187034954 | -0.152199116 | 0.999933887 | -0.091358376 | 0.300909357 | -0.392267733 | 0.256108687 |
| ENSG00000134759 | *ELP2* | 3.964120659 | -0.101362074 | -0.010865306 | -0.090496768 | 0.999933887 | 0.03513563 | -0.090509997 | 0.125645627 | 0.256179533 |
| ENSG00000204086 | *RPA4* | -1.069204337 | 0.432798387 | -0.009801182 | 0.44259957 | 0.999933887 | 1.136636744 | 1.605435028 | -0.468798284 | 0.256194337 |
| ENSG00000164284 | *GRPEL2* | 1.177533581 | -0.085722277 | -0.243685742 | 0.157963465 | 0.999933887 | -0.234702501 | -0.538324295 | 0.303621794 | 0.256194337 |
| ENSG00000137054 | *POLR1E* | 1.747497516 | -0.023080326 | -0.073245238 | 0.050164912 | 0.999933887 | 0.147446517 | 0.377271143 | -0.229824626 | 0.256389421 |
| ENSG00000157036 | *EXOG* | 1.533717356 | -0.015199736 | -0.092354378 | 0.077154643 | 0.999933887 | 0.025401446 | 0.248456411 | -0.223054966 | 0.256389421 |
| ENSG00000103342 | *GSPT1* | 4.882631858 | -0.025853426 | -0.013108897 | -0.012744528 | 0.999933887 | 0.091036183 | 0.205665602 | -0.114629419 | 0.256739456 |
| ENSG00000198178 | *CLEC4C* | -1.958163524 | 0.09534865 | -0.051802827 | 0.147151477 | 0.999933887 | -0.114510619 | -1.054563853 | 0.940053234 | 0.256874495 |
| ENSG00000132485 | *ZRANB2* | 4.848247159 | -0.118706974 | -0.145596074 | 0.0268891 | 0.999933887 | -0.228259948 | 0.089995673 | -0.318255621 | 0.256893353 |
| ENSG00000169221 | *TBC1D10B* | 5.930117421 | 0.079529952 | 0.061833176 | 0.017696776 | 0.999933887 | -0.750046066 | -0.949667602 | 0.199621536 | 0.257278905 |
| ENSG00000146576 | *C7orf26* | 5.321807067 | -0.037852917 | -0.068238919 | 0.030386002 | 0.999933887 | -0.422325334 | -0.557625084 | 0.13529975 | 0.257306876 |
| ENSG00000255139 | *AP000442.1* | 1.566324143 | -0.130708038 | 0.122096746 | -0.252804784 | 0.999933887 | 0.291178862 | 0.552369579 | -0.261190717 | 0.25740934 |
| ENSG00000130511 | *SSBP4* | 4.121803212 | 0.079086755 | 0.034057602 | 0.045029154 | 0.999933887 | 0.035289292 | -0.134509701 | 0.169798993 | 0.257594788 |
| ENSG00000279633 | *AL137918.1* | -1.239683501 | 0.053919256 | 0.39110096 | -0.337181703 | 0.999933887 | 0.44697574 | 0.994837014 | -0.547861274 | 0.257594788 |
| ENSG00000047578 | *KATNIP* | 5.213114173 | 0.003882951 | 0.015118793 | -0.011235841 | 0.999933887 | 0.009598765 | -0.102574533 | 0.112173298 | 0.257594788 |
| ENSG00000175591 | *P2RY2* | 2.815205078 | -0.023467127 | 0.006374251 | -0.029841378 | 0.999933887 | 0.161608472 | 0.431756378 | -0.270147905 | 0.257675536 |
| ENSG00000228043 | *AC114763.1* | 0.587933041 | 0.181002104 | 0.083503737 | 0.097498367 | 0.999933887 | 1.334648678 | 1.608565854 | -0.273917176 | 0.258045338 |
| ENSG00000161048 | *NAPEPLD* | 0.632011712 | -0.269763684 | -0.216678879 | -0.053084805 | 0.999933887 | -0.29802822 | 0.046605547 | -0.344633767 | 0.258110157 |
| ENSG00000153443 | *UBALD1* | 5.862272353 | 0.031437805 | 0.032504439 | -0.001066633 | 0.999933887 | -0.428068825 | -0.612728185 | 0.18465936 | 0.258310554 |
| ENSG00000182310 | *SPACA6* | 4.150661681 | 0.043893188 | 0.163398433 | -0.119505245 | 0.999933887 | 0.920980183 | 1.089979676 | -0.168999494 | 0.258333701 |
| ENSG00000086589 | *RBM22* | 6.071677097 | 0.00833963 | 0.008387402 | -4.7772e-05 | 0.999933887 | 0.279866651 | 0.387217039 | -0.107350388 | 0.258333701 |
| ENSG00000078177 | *N4BP2* | 2.606986703 | 0.02164348 | -0.163678793 | 0.185322273 | 0.999933887 | -0.274885799 | 0.138516751 | -0.41340255 | 0.258333701 |
| ENSG00000095261 | *PSMD5* | 2.790970235 | -0.013291001 | 0.044941675 | -0.058232676 | 0.999933887 | 0.275325203 | 0.513035202 | -0.237709998 | 0.258333701 |
| ENSG00000196810 | *CTBP1-DT* | 2.452281391 | 0.139693141 | 0.109703045 | 0.029990096 | 0.999933887 | 0.664164299 | 0.913645747 | -0.249481449 | 0.258451406 |
| ENSG00000120910 | *PPP3CC* | 3.399779411 | 0.003221701 | -0.004298496 | 0.007520197 | 0.999933887 | 0.292900657 | 0.447909852 | -0.155009196 | 0.258451406 |
| ENSG00000101473 | *ACOT8* | 2.645057399 | 0.075653723 | 0.021032039 | 0.054621684 | 0.999933887 | -0.269590714 | -0.491054003 | 0.221463289 | 0.258482332 |
| ENSG00000277449 | *CEBPB-AS1* | 1.641977555 | 0.00087961 | -0.047137307 | 0.048016917 | 0.999933887 | 0.179582265 | 0.470284038 | -0.290701773 | 0.258482332 |
| ENSG00000213983 | *AP1G2* | 4.598427449 | 0.054127073 | 0.093000442 | -0.038873369 | 0.999933887 | -0.024844031 | -0.182425633 | 0.157581602 | 0.258667233 |
| ENSG00000082701 | *GSK3B* | 6.122414554 | -0.022952933 | 0.011429407 | -0.034382339 | 0.999933887 | -0.023736719 | -0.142602207 | 0.118865488 | 0.258667233 |
| ENSG00000154122 | *ANKH* | 4.221764854 | -0.087561268 | 0.049015981 | -0.136577249 | 0.999933887 | -0.043187269 | -0.163362957 | 0.120175688 | 0.258947004 |
| ENSG00000180089 | *TMEM86B* | 1.119556436 | 0.117511545 | -0.018431326 | 0.135942871 | 0.999933887 | -0.336584067 | -0.66652736 | 0.329943294 | 0.258947004 |
| ENSG00000167555 | *ZNF528* | 0.760950837 | -0.033350712 | 0.023945984 | -0.057296697 | 0.999933887 | -0.257016563 | 0.051685885 | -0.308702449 | 0.258947004 |
| ENSG00000165630 | *PRPF18* | 1.868416104 | -0.061036111 | -0.127156804 | 0.066120693 | 0.999933887 | -0.336530238 | -0.106899671 | -0.229630567 | 0.259039694 |
| ENSG00000010030 | *ETV7* | 1.806353915 | 0.053497685 | 0.027532594 | 0.025965091 | 0.999933887 | -0.139114539 | -0.436912277 | 0.297797738 | 0.259199401 |
| ENSG00000114978 | *MOB1A* | 7.563499047 | -0.043933284 | 0.005645596 | -0.04957888 | 0.999933887 | -0.11157583 | 0.004151712 | -0.115727542 | 0.259310854 |
| ENSG00000100427 | *MLC1* | -0.490446854 | -0.036103351 | 0.026671707 | -0.062775058 | 0.999933887 | 0.20201217 | -0.218443465 | 0.420455634 | 0.259396899 |
| ENSG00000259956 | *RBM15B* | 4.722057338 | 0.031967815 | 0.004423438 | 0.027544377 | 0.999933887 | -0.317813718 | -0.467732311 | 0.149918593 | 0.259649996 |
| ENSG00000132376 | *INPP5K* | 6.77252597 | -0.012736646 | 0.013668376 | -0.026405022 | 0.999933887 | 0.015753719 | -0.094903154 | 0.110656873 | 0.259649996 |
| ENSG00000136205 | *TNS3* | 0.874410538 | 0.315063994 | 0.21387588 | 0.101188114 | 0.999933887 | -0.577441163 | -0.231402462 | -0.3460387 | 0.259887085 |
| ENSG00000140564 | *FURIN* | 7.395672235 | 0.038750186 | 0.036975363 | 0.001774823 | 0.999933887 | -0.811546362 | -0.98891419 | 0.177367828 | 0.259890869 |
| ENSG00000117620 | *SLC35A3* | 2.20490282 | 0.062618534 | -0.158851936 | 0.22147047 | 0.999933887 | -0.226508128 | 0.113006819 | -0.339514947 | 0.259890869 |
| ENSG00000007944 | *MYLIP* | 8.061033939 | 0.069556497 | 0.054874796 | 0.014681702 | 0.999933887 | 0.302546972 | 0.414450371 | -0.111903398 | 0.259959364 |
| ENSG00000139719 | *VPS33A* | 2.884196416 | -0.02007701 | -0.089880779 | 0.069803769 | 0.999933887 | -0.185088533 | -0.036825507 | -0.148263026 | 0.259985751 |
| ENSG00000091317 | *CMTM6* | 8.458886746 | -0.051005394 | -0.003439041 | -0.047566353 | 0.999933887 | 0.169761975 | 0.278195719 | -0.108433744 | 0.260025142 |
| ENSG00000259924 | *AC011939.1* | -0.063243474 | 0.201286469 | 0.489256975 | -0.287970506 | 0.999933887 | 2.787512001 | 3.390908443 | -0.603396442 | 0.260025142 |
| ENSG00000180346 | *TIGD2* | -0.590085163 | -0.147578156 | 0.006558266 | -0.154136422 | 0.999933887 | 0.122804949 | -0.346425255 | 0.469230204 | 0.260025142 |
| ENSG00000260152 | *AC113208.2* | 0.295514393 | -0.151161789 | 0.037177481 | -0.188339269 | 0.999933887 | 1.530539109 | 2.049714227 | -0.519175119 | 0.260025142 |
| ENSG00000162769 | *FLVCR1* | 3.184700582 | -0.045501269 | -0.145378358 | 0.099877088 | 0.999933887 | -0.041090492 | 0.118051411 | -0.159141902 | 0.260025142 |
| ENSG00000076201 | *PTPN23* | 5.649348561 | 0.025395964 | 0.00366964 | 0.021726324 | 0.999933887 | -0.605344628 | -0.738162262 | 0.132817634 | 0.260025142 |
| ENSG00000211598 | *IGKV4-1* | -0.630195326 | 0.025110263 | 0.144683777 | -0.119573514 | 0.999933887 | 0.405503139 | -0.197822134 | 0.603325273 | 0.260088802 |

| | | | | | | | | | | |
|---|---|---|---|---|---|---|---|---|---|---|
| ENSG00000169738 | *DCXR* | 3.283672476 | 0.008667507 | 0.120186792 | -0.111519286 | 0.999933887 | 0.173283639 | -0.047397972 | 0.220681611 | 0.260178983 |
| ENSG00000196912 | *ANKRD36B* | 1.119880309 | -0.089908348 | -0.248237332 | 0.158328985 | 0.999933887 | -0.159986441 | 0.233402432 | -0.393388873 | 0.260199691 |
| ENSG00000260101 | *AC008074.2* | 0.917302412 | -0.189246217 | -0.17590234 | -0.013343877 | 0.999933887 | 1.772645197 | 2.26760237 | -0.494957173 | 0.260219204 |
| ENSG00000287613 | *AC069549.2* | -0.412947575 | 0.182957884 | 0.164305454 | 0.01865243 | 0.999933887 | -0.765804779 | -1.686387457 | 0.920582677 | 0.260322229 |
| ENSG00000151657 | *KIN* | 3.457237686 | -0.10209046 | -0.24228726 | 0.1401968 | 0.999933887 | -0.091818138 | 0.065869861 | -0.157688 | 0.260530623 |
| ENSG00000166394 | *CYB5R2* | 0.003486621 | -0.062750777 | -0.079459692 | 0.016708915 | 0.999933887 | 2.462213098 | 3.029348384 | -0.567135286 | 0.260530623 |
| ENSG00000237672 | *KRR1P1* | 0.328626845 | 0.021964676 | 0.124063923 | -0.102099247 | 0.999933887 | -0.310710649 | 0.08802341 | -0.398734059 | 0.260530623 |
| ENSG00000185261 | *KIAA0825* | 3.475406632 | -0.001770767 | 0.024080113 | -0.02585088 | 0.999933887 | -0.374065181 | -0.065513406 | -0.308551775 | 0.260530623 |
| ENSG00000273142 | *LINC02604* | 2.632973269 | 0.044862524 | -0.065135691 | 0.109998215 | 0.999933887 | 0.410474986 | 0.62857956 | -0.218104573 | 0.26061061 |
| ENSG00000177272 | *KCNA3* | 3.203200357 | 0.03999609 | -0.045973338 | 0.085969428 | 0.999933887 | -0.052443159 | 0.201540478 | -0.253983636 | 0.26061061 |
| ENSG00000083444 | *PLOD1* | 3.553677997 | 0.012497438 | 0.02731341 | -0.014815972 | 0.999933887 | -0.243727486 | -0.473145396 | 0.22941791 | 0.26082082 |
| ENSG00000117525 | *F3* | 4.154552386 | 0.353855453 | 0.468323516 | -0.114468063 | 0.999933887 | 1.668017472 | 2.103821078 | -0.435803606 | 0.261247738 |
| ENSG00000240219 | *AL512306.2* | 2.286377369 | -0.010998622 | 0.104683931 | -0.115682553 | 0.999933887 | -0.545990345 | -0.752124161 | 0.206133816 | 0.261247738 |
| ENSG00000285016 | *AC017002.3* | -2.353046832 | 0.896208329 | 0.671734573 | 0.224473756 | 0.999933887 | 1.950972295 | 2.540011812 | -0.589039517 | 0.261691347 |
| ENSG00000273141 | *AP001269.4* | 1.086167633 | -0.086463481 | -0.162890088 | 0.076426607 | 0.999933887 | -0.594416293 | -1.008873536 | 0.414457243 | 0.261887428 |
| ENSG00000172500 | *FIBP* | 2.681735607 | 0.035757888 | -0.053836048 | 0.089593936 | 0.999933887 | -0.036214876 | -0.246752578 | 0.210537701 | 0.261887428 |
| ENSG00000136514 | *RTP4* | 0.506545224 | -0.189038863 | 0.008205802 | -0.197244665 | 0.999933887 | -0.360569252 | -0.732072586 | 0.371503333 | 0.261927904 |
| ENSG00000166704 | *ZNF606* | 1.11304888 | -0.188017091 | -0.150525577 | -0.037491513 | 0.999933887 | -0.180653769 | -0.439672396 | 0.259018628 | 0.262030463 |
| ENSG00000176105 | *YES1* | 1.735184792 | -0.091552049 | -0.124417084 | 0.032865034 | 0.999933887 | -0.226253128 | 0.149243968 | -0.375497096 | 0.262030463 |
| ENSG00000139675 | *HNRNPA1L2* | -0.005043076 | -0.134429847 | 0.060233836 | -0.194663682 | 0.999933887 | 0.099325489 | 0.455834826 | -0.356509337 | 0.262030463 |
| ENSG00000236194 | *AC099811.1* | 3.13895677 | 0.056761025 | 0.034073387 | 0.022687638 | 0.999933887 | 1.230328765 | 1.553127242 | -0.322798477 | 0.262030463 |
| ENSG00000168268 | *NT5DC2* | -0.726170414 | -0.085687819 | -0.256536506 | 0.170848687 | 0.999933887 | 0.022443872 | -0.472245199 | 0.494689071 | 0.262030463 |
| ENSG00000131013 | *PPIL4* | 3.949939115 | -0.036665479 | -0.014051643 | -0.022613836 | 0.999933887 | -0.343230067 | -0.116575307 | -0.22665476 | 0.262030463 |
| ENSG00000048405 | *ZNF800* | 4.8001644 | 0.034751316 | -0.106199605 | 0.140950921 | 0.999933887 | -0.328524428 | -0.027362819 | -0.30116161 | 0.262030463 |
| ENSG00000287742 | *AC068282.2* | 0.468227778 | -0.010468459 | -0.178859497 | 0.168391037 | 0.999933887 | -0.980156923 | -1.294998527 | 0.314841604 | 0.262030463 |
| ENSG00000197712 | *FAM114A1* | -0.123454588 | -0.145336767 | -0.217693086 | 0.07235632 | 0.999933887 | -1.02116743 | -1.533733306 | 0.512565876 | 0.262291183 |
| ENSG00000167987 | *VPS37C* | 5.936455082 | -0.032191484 | -0.005226756 | -0.026964728 | 0.999933887 | -0.110242777 | 0.023691458 | -0.133934235 | 0.262376517 |
| ENSG00000134242 | *PTPN22* | 4.433539476 | -0.054137706 | -0.178497708 | 0.124360002 | 0.999933887 | -0.209868021 | 0.079192389 | -0.28906041 | 0.262377218 |
| ENSG00000122126 | *OCRL* | 0.241464006 | -0.020494114 | -0.194621145 | 0.17412703 | 0.999933887 | -0.178577148 | 0.125528006 | -0.304105154 | 0.262377218 |
| ENSG00000160007 | *ARHGAP35* | 4.397634074 | 0.022107831 | 0.036637646 | -0.014529815 | 0.999933887 | -0.242302816 | -0.378086388 | 0.135783572 | 0.262400567 |
| ENSG00000178741 | *COX5A* | 3.029674148 | 0.088779709 | -0.091544854 | 0.180324563 | 0.999933887 | -0.144520344 | -0.355086115 | 0.210565771 | 0.262983286 |
| ENSG00000174238 | *PITPNA* | 7.7874424 | -0.020920233 | -0.050699562 | 0.029779329 | 0.999933887 | 0.053070497 | 0.151269697 | -0.0981992 | 0.263045661 |
| ENSG00000162522 | *KIAA1522* | -0.510385436 | -0.057417376 | -0.214045928 | 0.156628552 | 0.999933887 | 0.239343208 | -0.143465528 | 0.382808735 | 0.26326015 |
| ENSG00000167612 | *ANKRD33* | 2.403511691 | 0.049016121 | 0.145379164 | -0.096363043 | 0.999933887 | -1.159010675 | -1.371366144 | 0.212355469 | 0.26326015 |
| ENSG00000123159 | *GIPC1* | 2.1669082 | 0.148708349 | 0.009610241 | 0.139098108 | 0.999933887 | 0.116752103 | -0.107228732 | 0.223980834 | 0.263273159 |
| ENSG00000167202 | *TBC1D2B* | 5.802101253 | -0.002622409 | 0.060946328 | -0.063568737 | 0.999933887 | 0.300973614 | 0.483332797 | -0.182359183 | 0.263526722 |
| ENSG00000123975 | *CKS2* | 1.764067974 | 0.103191099 | 0.032863534 | 0.070327565 | 0.999933887 | 0.391381503 | 0.710966282 | -0.319584778 | 0.263956459 |
| ENSG00000285596 | *AC017116.2* | -0.16853513 | -0.067019435 | -0.592778067 | 0.525758632 | 0.999933887 | -0.082593897 | 0.307378715 | -0.389972612 | 0.264310507 |
| ENSG00000062650 | *WAPL* | 6.583474142 | -0.066690407 | -0.088105251 | 0.021414844 | 0.999933887 | -0.260974366 | -0.126432535 | -0.134541831 | 0.264401685 |
| ENSG00000181991 | *MRPS11* | 1.220247059 | -0.07975607 | 0.063440602 | -0.143196672 | 0.999933887 | -0.045825385 | -0.307554233 | 0.261728847 | 0.264401685 |
| ENSG00000125652 | *ALKBH7* | 3.292294305 | 0.058882898 | 0.048511783 | 0.010371115 | 0.999933887 | -0.147700477 | -0.376546094 | 0.228845617 | 0.264401685 |
| ENSG00000082293 | *COL19A1* | -1.259340851 | -0.149589426 | 0.136102238 | -0.285691665 | 0.999933887 | -0.361550152 | 0.126798768 | -0.48834892 | 0.26444703 |
| ENSG00000131697 | *NPHP4* | 3.377883411 | 0.007716379 | 0.037439466 | -0.029723087 | 0.999933887 | -0.172264105 | -0.338616397 | 0.166352292 | 0.26444703 |
| ENSG00000136688 | *IL36G* | -2.242390043 | 0.399621435 | -0.134800847 | 0.534422282 | 0.999933887 | 4.180979671 | 5.169417813 | -0.988438143 | 0.264541549 |
| ENSG00000143756 | *FBXO28* | 3.833173239 | -0.058051747 | 0.042368855 | -0.100420602 | 0.999933887 | -0.212122014 | -0.075639324 | -0.13648269 | 0.264603837 |
| ENSG00000143344 | *RGL1* | 2.180995805 | -0.07973155 | 0.033055117 | -0.112786666 | 0.999933887 | 0.28808934 | 0.509562827 | -0.221473487 | 0.264603837 |
| ENSG00000081059 | *TCF7* | 6.24035252 | -0.004632485 | 0.042311812 | -0.046944296 | 0.999933887 | 0.198983178 | 0.310384297 | -0.111401118 | 0.264603837 |
| ENSG00000150782 | *IL18* | 0.191901289 | -0.069877065 | -0.230869715 | 0.16099265 | 0.999933887 | 0.377276726 | -0.129678029 | 0.506954756 | 0.264793082 |
| ENSG00000255355 | *AP000640.2* | 1.500169793 | 0.285751188 | 0.327750599 | -0.041999411 | 0.999933887 | 1.800760302 | 2.077339295 | -0.276578993 | 0.264886827 |
| ENSG00000131375 | *CAPN7* | 5.038191227 | 0.048746931 | -0.008988565 | 0.057735496 | 0.999933887 | -0.003965773 | 0.197202326 | -0.201168099 | 0.264886827 |
| ENSG00000161277 | *THAP8* | 1.01217966 | -0.084703273 | 0.089547621 | -0.174250894 | 0.999933887 | -0.85202916 | -1.106675889 | 0.254646729 | 0.264886827 |
| ENSG00000079335 | *CDC14A* | 4.267827927 | 0.018489124 | -0.035469436 | 0.05395856 | 0.999933887 | -0.027416432 | 0.139106408 | -0.16652284 | 0.264886827 |
| ENSG00000248559 | *AC109454.2* | 0.105814626 | 0.157852543 | -0.015934938 | 0.173787481 | 0.999933887 | 0.334139513 | 0.960268361 | -0.626128848 | 0.264896198 |
| ENSG00000134748 | *PRPF38A* | 5.545002454 | -0.025257729 | -0.066285897 | 0.041028168 | 0.999933887 | 0.075307032 | 0.180353778 | -0.105046746 | 0.265367559 |
| ENSG00000070214 | *SLC44A1* | 2.611425698 | -0.042512784 | 0.064881482 | -0.107394265 | 0.999933887 | -0.526224802 | -0.725447858 | 0.199223056 | 0.265367559 |
| ENSG00000089248 | *ERP29* | 4.820199946 | 0.031873759 | -0.032860435 | 0.064734195 | 0.999933887 | -0.060378656 | -0.21898419 | 0.158605534 | 0.265367559 |
| ENSG00000260339 | *HEXA-AS1* | 1.093256482 | 0.141946221 | 0.018107749 | 0.123838472 | 0.999933887 | 0.559098198 | 0.869965305 | -0.310867107 | 0.265537451 |
| ENSG00000132475 | *H3-3B* | 11.92396565 | 0.070118697 | 0.059120028 | 0.010998669 | 0.999933887 | 0.081977881 | 0.205119105 | -0.123141223 | 0.265933868 |
| ENSG00000179409 | *GEMIN4* | 2.563314279 | -0.020042855 | 0.050855434 | -0.070898289 | 0.999933887 | 0.490787853 | 0.718228596 | -0.227440743 | 0.265965265 |
| ENSG00000126522 | *ASL* | 2.635529239 | 0.022998934 | 0.156798807 | -0.133799873 | 0.999933887 | 0.046539501 | -0.144506532 | 0.191046032 | 0.266011137 |
| ENSG00000198585 | *NUDT16* | 3.238781882 | 0.012536927 | -0.019476871 | 0.032013797 | 0.999933887 | -0.177105854 | -0.393803245 | 0.216697391 | 0.266049856 |
| ENSG00000111142 | *METAP2* | 3.054187841 | -0.081889837 | -0.085422372 | 0.003532535 | 0.999933887 | 0.104469766 | 0.285469196 | -0.18099943 | 0.266113226 |
| ENSG00000252908 | *RNU6-1003P* | 0.447916348 | 0.164252499 | 0.539631012 | -0.375378513 | 0.999933887 | 0.675428933 | 1.201290308 | -0.525861374 | 0.266113226 |
| ENSG00000101265 | *RASSF2* | 10.383930598 | -0.077215854 | -0.076571011 | -0.000644843 | 0.999933887 | 0.022454586 | 0.126243591 | -0.103789006 | 0.266122791 |
| ENSG00000225492 | *GBP1P1* | -0.850516783 | -0.004789218 | 0.246181448 | -0.250970666 | 0.999933887 | 0.780547924 | 1.575997159 | -0.795449234 | 0.266122791 |
| ENSG00000130177 | *CDC16* | 3.666213527 | -0.099626911 | -0.04217463 | -0.057452281 | 0.999933887 | -0.116491858 | -0.245728009 | 0.129236152 | 0.26612303 |
| ENSG00000215375 | *MYL5* | 2.176828348 | 0.022208269 | 0.144793134 | -0.122584865 | 0.999933887 | -0.163883802 | -0.378432116 | 0.214548314 | 0.266638202 |
| ENSG00000129351 | *ILF3* | 7.768365671 | 0.035138561 | 0.018375251 | 0.01676331 | 0.999933887 | 0.299585358 | 0.435700169 | -0.136114811 | 0.266646789 |
| ENSG00000152127 | *MGAT5* | 5.096230922 | -0.025262967 | 0.024733028 | -0.049995994 | 0.999933887 | 0.126432889 | 0.272758951 | -0.146326062 | 0.266700056 |
| ENSG00000230322 | *AL136320.1* | 0.16829944 | -0.05228489 | 0.032737012 | -0.085021901 | 0.999933887 | -0.374928322 | -1.256141405 | 0.881213083 | 0.266808489 |
| ENSG00000151470 | *C4orf33* | 2.49198216 | -0.037342219 | -0.107684976 | 0.070342757 | 0.999933887 | -0.25668978 | 0.057965341 | -0.314655121 | 0.266861922 |
| ENSG00000163626 | *COX18* | 1.678828098 | -0.018962592 | -0.066924731 | 0.047962138 | 0.999933887 | 0.151545371 | -0.064256669 | 0.215802039 | 0.266910643 |
| ENSG00000122641 | *INHBA* | 4.371739503 | 0.060288034 | -0.104598568 | 0.164886602 | 0.999933887 | 0.543999168 | 0.775587668 | -0.2315885 | 0.267009531 |
| ENSG00000170364 | *SETMAR* | 0.783383388 | 0.066617319 | -0.05071327 | 0.117330589 | 0.999933887 | 0.170365974 | -0.159595832 | 0.329961806 | 0.267009531 |
| ENSG00000162739 | *SLAMF6* | 3.514317629 | 0.010516399 | -0.093027113 | 0.103543512 | 0.999933887 | 0.076174704 | 0.236084756 | -0.159910053 | 0.267009531 |
| ENSG00000158417 | *EIF5B* | 3.866023764 | -0.178049404 | -0.212462587 | 0.034413183 | 0.999933887 | -0.139413884 | 0.135948881 | -0.275362765 | 0.267012521 |
| ENSG00000258044 | *AC073569.1* | 0.00063561 | 0.171428258 | 0.082255692 | 0.089172566 | 0.999933887 | 2.068062065 | 2.61513357 | -0.547071504 | 0.267012521 |
| ENSG00000143379 | *SETDB1* | 4.192328216 | -0.033884233 | -0.011752805 | -0.022131429 | 0.999933887 | -0.363851959 | -0.497759121 | 0.133907162 | 0.267201757 |
| ENSG00000133740 | *E2F5* | 0.012008668 | -0.238732006 | 0.094694254 | -0.333426261 | 0.999933887 | -0.142080152 | 0.243723206 | -0.385803358 | 0.267612074 |
| ENSG00000229780 | *UBE2Q1-AS1* | -0.396000412 | -0.042150782 | -0.230981613 | 0.188830831 | 0.999933887 | -0.077112422 | 0.273369988 | -0.35048241 | 0.267612074 |
| ENSG00000136709 | *WDR33* | 4.848596613 | -0.011921833 | 0.033959841 | -0.045881674 | 0.999933887 | -0.187258358 | -0.289675569 | 0.102417211 | 0.267628799 |
| ENSG00000260708 | *AL118516.1* | 4.200799781 | 0.212559423 | 0.178792217 | 0.033767206 | 0.999933887 | 1.643545716 | 1.85092108 | -0.207375365 | 0.267809233 |
| ENSG00000273654 | *AC020904.2* | 0.470619217 | -0.103141834 | -0.551704632 | 0.448562798 | 0.999933887 | 0.27801415 | 0.47878329 | -0.200769139 | 0.267819057 |
| ENSG00000254325 | *AC018607.1* | 2.270226053 | -0.046158044 | 0.129135531 | -0.175293575 | 0.999933887 | 1.134125007 | 1.540544256 | -0.406419249 | 0.267819057 |
| ENSG00000259225 | *LINC02345* | -1.098934113 | 0.093938079 | -0.23492956 | 0.32886764 | 0.999933887 | 0.593440873 | 1.134412906 | -0.540972033 | 0.267819057 |
| ENSG00000120156 | *TEK* | -0.191252688 | -0.175225714 | -0.268650291 | 0.093424577 | 0.999933887 | -0.496124553 | -1.086173819 | 0.590049266 | 0.26784826 |
| ENSG00000168903 | *BTNL3* | 0.643034476 | -0.052059588 | -0.02714119 | -0.024918398 | 0.999933887 | 0.121153114 | 0.391537794 | -0.27038468 | 0.268167537 |
| ENSG00000143499 | *SMYD2* | 2.482022431 | -0.053048334 | -0.081721821 | 0.028673487 | 0.999933887 | -0.028887351 | 0.164352803 | -0.193240154 | 0.268440149 |
| ENSG00000008441 | *NFIX* | 0.176900947 | 0.109543838 | -0.012749991 | 0.122293829 | 0.999933887 | -0.204469453 | -0.648154845 | 0.443685393 | 0.268440149 |
| ENSG00000204253 | *HNRNPCP2* | -0.030632061 | -0.133305889 | -0.102951604 | -0.030354286 | 0.999933887 | 1.216325794 | 1.695917665 | -0.479591871 | 0.268440149 |
| ENSG00000147439 | *BIN3* | 6.141072056 | -0.021202434 | -0.053318623 | 0.032116189 | 0.999933887 | 0.161629615 | 0.275702304 | -0.114072688 | 0.268450304 |
| ENSG00000156958 | *GALK2* | 1.354195946 | -0.077402011 | 0.026791997 | -0.104194008 | 0.999933887 | -0.187059564 | -0.452157401 | 0.265097837 | 0.26859582 |
| ENSG00000088876 | *ZNF343* | 0.442584944 | 0.01752839 | -0.281149078 | 0.298677468 | 0.999933887 | 0.305370145 | 0.00081848 | 0.304551664 | 0.268899096 |
| ENSG00000184428 | *TOP1MT* | 2.221722967 | 0.098297474 | -0.035708566 | 0.13400604 | 0.999933887 | 0.165159192 | -0.04428328 | 0.209442472 | 0.269212325 |
| ENSG00000136436 | *CALCOCO2* | 7.005696381 | -0.061182693 | -0.07999871 | 0.018816017 | 0.999933887 | 0.062299344 | 0.153276437 | -0.090977094 | 0.269363502 |
| ENSG00000069275 | *NUCKS1* | 4.40167743 | -0.042571352 | -0.037688462 | -0.00488289 | 0.999933887 | 0.053848465 | 0.192191149 | -0.138342685 | 0.269781635 |
| ENSG00000083520 | *DIS3* | 3.933419187 | -0.111691454 | -0.196387006 | 0.084695552 | 0.999933887 | -0.341189516 | -0.09274106 | -0.248448456 | 0.27005471 |
| ENSG00000148090 | *AUH* | 2.103426415 | 0.068576128 | -0.001408555 | 0.069984683 | 0.999933887 | -0.145107648 | -0.340807904 | 0.195700255 | 0.270123756 |
| ENSG00000100888 | *CHD8* | 6.277800517 | -0.04198681 | -0.127902507 | 0.085915697 | 0.999933887 | -0.34449646 | -0.466031199 | 0.121534739 | 0.270123756 |
| ENSG00000135083 | *CCNJL* | 3.3020445 | 0.037613609 | 0.013141895 | 0.024471713 | 0.999933887 | 0.282123689 | 0.005100131 | 0.277023558 | 0.270123756 |
| ENSG00000279748 | *AC008764.9* | -0.369872791 | 0.051280057 | 0.381456774 | -0.330176717 | 0.999933887 | 0.0662154 | 0.561450928 | -0.495235528 | 0.270123756 |
| ENSG00000100532 | *CGRRF1* | 2.098702934 | -0.019566337 | -0.103786608 | 0.08422027 | 0.999933887 | -0.094224945 | 0.126367038 | -0.220591983 | 0.270323065 |
| ENSG00000157500 | *APPL1* | 2.614839314 | 0.028556925 | -0.061304375 | 0.0898613 | 0.999933887 | -0.187435247 | 0.080195755 | -0.267631001 | 0.270458519 |
| ENSG00000198805 | *PNP* | 5.155111644 | 0.094812927 | 0.145300524 | -0.050487597 | 0.999933887 | -0.104320317 | 0.09233085 | -0.196651167 | 0.270725658 |
| ENSG00000167702 | *KIFC2* | 4.62170921 | 0.114160785 | 0.206307893 | -0.092147107 | 0.999933887 | 0.778125362 | 1.038790752 | -0.26066539 | 0.270725658 |

| | | | | | | | | | | |
|---|---|---|---|---|---|---|---|---|---|---|
| ENSG00000066455 | *GOLGA5* | 3.789396785 | -0.045253677 | -0.117032838 | 0.071779161 | 0.999933887 | -0.266661683 | -0.121675438 | -0.144986245 | 0.270725658 |
| ENSG00000196367 | *TRRAP* | 5.64019612 | 0.057914801 | 0.011690048 | 0.046224753 | 0.999933887 | -0.026625089 | -0.14584745 | 0.119222361 | 0.271145385 |
| ENSG00000152749 | *GPR180* | 0.016872896 | -0.085070818 | -0.194629465 | 0.109558646 | 0.999933887 | -0.205041037 | 0.190348059 | -0.395389096 | 0.271146852 |
| ENSG00000185262 | *UBALD2* | 8.699214568 | 0.035826867 | 0.025236953 | 0.010589914 | 0.999933887 | 0.014732661 | 0.155426119 | -0.140693459 | 0.271210195 |
| ENSG00000175866 | *BAIAP2* | 3.08536533 | 0.029116131 | 0.015175547 | 0.013940584 | 0.999933887 | 0.087755867 | 0.300374513 | -0.212618646 | 0.27158916 |
| ENSG00000259984 | *AL928711.1* | -0.910191433 | -0.271788034 | -0.160186514 | -0.11160152 | 0.999933887 | 0.320604028 | -0.185630514 | 0.506234542 | 0.271989589 |
| ENSG00000090776 | *EFNB1* | 0.727527035 | -0.070537608 | 0.164713258 | -0.235250865 | 0.999933887 | 0.158056249 | 0.510766181 | -0.352709932 | 0.2720204 |
| ENSG00000166432 | *ZMAT1* | 2.880367181 | -0.085239278 | -0.301149167 | 0.215909889 | 0.999933887 | -0.420734249 | 0.073065601 | -0.493799849 | 0.2720204 |
| ENSG00000134107 | *BHLHE40* | 8.590299863 | -0.006890664 | -0.060909942 | 0.054019278 | 0.999933887 | 0.507483543 | 0.632269187 | -0.124785644 | 0.2720204 |
| ENSG00000135837 | *CEP350* | 5.548484521 | -0.026149825 | -0.103576947 | 0.077427121 | 0.999933887 | -0.488940508 | -0.129196217 | -0.359744291 | 0.272043983 |
| ENSG00000183018 | *SPNS2* | 1.873273131 | -0.021788628 | -0.088051063 | 0.066262435 | 0.999933887 | -0.786076394 | -1.089627831 | 0.303551437 | 0.272245967 |
| ENSG00000267278 | *MAP3K14-AS1* | 1.832058079 | 0.02036937 | 0.102227125 | -0.081857755 | 0.999933887 | 0.219743255 | -0.095781164 | 0.31552442 | 0.272281334 |
| ENSG00000135655 | *USP15* | 8.691994775 | -0.079356461 | -0.183078807 | 0.103722346 | 0.999933887 | -0.538356505 | -0.257457589 | -0.280898916 | 0.272548391 |
| ENSG00000116747 | *RO60* | 4.344272725 | -0.046789909 | -0.039433108 | -0.007356801 | 0.999933887 | -0.551956184 | -0.265372863 | -0.286583321 | 0.272548391 |
| ENSG00000154222 | *CC2D1B* | 6.089022171 | -0.006266615 | 0.055263456 | -0.061530071 | 0.999933887 | -0.115856939 | -0.009418295 | -0.106438644 | 0.272548391 |
| ENSG00000168298 | *H1-4* | 0.148197138 | 0.000574479 | -0.201356393 | 0.201930873 | 0.999933887 | 0.320062618 | -0.182473621 | 0.502536239 | 0.272548391 |
| ENSG00000116984 | *MTR* | 3.92563646 | -0.027422821 | -0.030945743 | 0.003522922 | 0.999933887 | 0.040535778 | 0.190520576 | -0.149984799 | 0.272720982 |
| ENSG00000181523 | *SGSH* | 7.562273086 | -0.053122333 | -0.076130755 | 0.023008422 | 0.999933887 | 0.115257367 | 0.226965805 | -0.111708438 | 0.272787045 |
| ENSG00000148690 | *FRA10AC1* | 1.916475916 | 0.094054542 | -0.053808013 | 0.147862555 | 0.999933887 | -0.21934168 | 0.063870346 | -0.283212026 | 0.27338867 |
| ENSG00000075151 | *EIF4G3* | 5.573682417 | 0.001884505 | -0.064160019 | 0.066044524 | 0.999933887 | -0.083788799 | 0.028467335 | -0.112256134 | 0.273449153 |
| ENSG00000246982 | *Z84485.1* | 0.268584496 | 0.185257449 | 0.225120089 | -0.03986264 | 0.999933887 | 0.301845883 | 0.63096238 | -0.329116497 | 0.273494603 |
| ENSG00000162227 | *TAF6L* | 1.126526493 | 0.137366473 | -0.073984886 | 0.211351359 | 0.999933887 | 0.082697898 | -0.166500008 | 0.249197906 | 0.274390559 |
| ENSG00000070371 | *CLTCL1* | 0.866214969 | -0.026714941 | -0.066892298 | 0.040177357 | 0.999933887 | -0.465089275 | -0.905311322 | 0.440222047 | 0.274390559 |
| ENSG00000134470 | *IL15RA* | 1.440561588 | -0.183521103 | 0.050617966 | -0.23413907 | 0.999933887 | 0.865671962 | 1.203554711 | -0.337882749 | 0.274734375 |
| ENSG00000261442 | *AC023830.1* | 0.13140331 | 0.222889337 | -0.062243485 | 0.285132822 | 0.999933887 | -0.005129887 | -0.50752498 | 0.502395093 | 0.27516543 |
| ENSG00000233230 | *AC079807.1* | 0.035336511 | -0.043819436 | 0.014883549 | -0.058702985 | 0.999933887 | 0.344674688 | -0.000355216 | 0.345029904 | 0.275212397 |
| ENSG00000144290 | *SLC4A10* | -0.695977701 | -0.170928294 | -0.315127071 | 0.144198776 | 0.999933887 | -0.286215927 | 0.328251181 | -0.614467109 | 0.275213185 |
| ENSG00000162461 | *SLC25A34* | 2.754326534 | -0.01490416 | 0.124407456 | -0.139311616 | 0.999933887 | 0.615413471 | 0.845142408 | -0.229728937 | 0.275697734 |
| ENSG00000249898 | *MCPH1-AS1* | 0.104234291 | 0.108410545 | -0.174194317 | 0.282604862 | 0.999933887 | 1.581000226 | 2.106359877 | -0.525359651 | 0.27590668 |
| ENSG00000227591 | *HSD11B1-AS1* | 2.938427847 | 0.173304641 | 0.060155752 | 0.113148888 | 0.999933887 | 1.11643837 | 1.315289153 | -0.198850783 | 0.275995132 |
| ENSG00000189266 | *PNRC2* | 5.726782622 | -0.086564176 | -0.072039587 | -0.014524589 | 0.999933887 | -0.031256292 | 0.080127982 | -0.111384273 | 0.276132028 |
| ENSG00000225831 | *RPS18P1* | -0.639894608 | 0.104328317 | 0.351719415 | -0.247391098 | 0.999933887 | -0.448755518 | 0.069364582 | -0.5181201 | 0.276132028 |
| ENSG00000135486 | *HNRNPA1* | 7.598945592 | -0.068140163 | -0.138570109 | 0.070429945 | 0.999933887 | 0.076229263 | 0.179948687 | -0.103719424 | 0.27639565 |
| ENSG00000013441 | *CLK1* | 8.607286562 | -0.055440299 | -0.058946659 | 0.003506361 | 0.999933887 | -0.021349205 | 0.136076268 | -0.157425472 | 0.27639565 |
| ENSG00000148334 | *PTGES2* | 2.896927484 | 0.031954781 | 0.133658806 | -0.101704025 | 0.999933887 | 0.161153978 | 0.00410187 | 0.157052108 | 0.27639565 |
| ENSG00000254789 | *AC073172.1* | 1.342482479 | 0.011451712 | 0.046436452 | -0.03498474 | 0.999933887 | -0.920157315 | -1.294241532 | 0.374084217 | 0.27639565 |
| ENSG00000114098 | *ARMC8* | 4.125817928 | -0.030796984 | -0.061957497 | 0.031160513 | 0.999933887 | -0.154515829 | -0.306430997 | 0.151915168 | 0.276656979 |
| ENSG00000205622 | *AP001042.1* | 0.943034243 | 0.286378263 | 0.566788787 | -0.280410524 | 0.999933887 | 1.975932007 | 2.511175118 | -0.535243111 | 0.276722527 |
| ENSG00000140950 | *MEAK7* | 1.364490072 | -0.0442772 | -0.105964806 | 0.061687606 | 0.999933887 | 0.170131246 | 0.446350768 | -0.276219522 | 0.276789347 |
| ENSG00000126746 | *ZNF384* | 4.460491468 | -0.037034908 | -0.029848892 | -0.007186016 | 0.999933887 | -0.360440306 | -0.497399259 | 0.136958954 | 0.276824331 |
| ENSG00000228037 | *AL139246.3* | 2.038037642 | -0.093379546 | 0.015001069 | -0.108380615 | 0.999933887 | 1.584395528 | 1.81257836 | -0.228182833 | 0.276824331 |
| ENSG00000183617 | *MRPL54* | 3.367115144 | 0.010694756 | -0.011157104 | 0.02185186 | 0.999933887 | 0.201166368 | 0.413368736 | -0.212202368 | 0.277026512 |
| ENSG00000250069 | *AC011379.1* | 2.476347991 | 0.233957957 | 0.428158292 | -0.194200335 | 0.999933887 | 0.527626614 | 0.898812058 | -0.371185444 | 0.27720024 |
| ENSG00000278259 | *MYO19* | 2.912830039 | -0.12432286 | 0.104358342 | -0.228681202 | 0.944094537 | 0.067031264 | -0.081922339 | 0.148953602 | 0.27720024 |
| ENSG00000221949 | *LINC01465* | 1.3591687 | 0.264758064 | 0.209442147 | 0.055315918 | 0.999933887 | 2.12002447 | 2.524159275 | -0.404134805 | 0.27720024 |
| ENSG00000128272 | *ATF4* | 8.426616977 | 0.054700143 | 0.100064516 | -0.045364373 | 0.999933887 | -0.002466212 | -0.113192775 | 0.110726564 | 0.27720024 |
| ENSG00000198663 | *C6orf89* | 6.044223312 | -0.04626023 | -0.022292366 | -0.023967864 | 0.999933887 | -0.171466408 | -0.055348061 | -0.116118348 | 0.27720024 |
| ENSG00000227671 | *AL390728.4* | 4.645705531 | -0.051957885 | -0.121370062 | 0.069412177 | 0.999933887 | 0.127473867 | 0.42897958 | -0.301505714 | 0.27720024 |
| ENSG00000172748 | *ZNF596* | 1.14092899 | 0.027590543 | 0.047268036 | -0.019677493 | 0.999933887 | 0.176719184 | 0.438053992 | -0.261334808 | 0.27720024 |
| ENSG00000187775 | *DNAH17* | 5.559287431 | 0.013982777 | 0.034100562 | -0.020117785 | 0.999933887 | 0.653024369 | 0.904226113 | -0.251201744 | 0.27720024 |
| ENSG00000125445 | *MRPS7* | 2.018318304 | 0.000261868 | -0.008610974 | 0.008872843 | 0.999933887 | 0.100865131 | -0.090659789 | 0.191524921 | 0.27720024 |
| ENSG00000100290 | *BIK* | 1.078509623 | -0.086723958 | -0.148917383 | 0.062193425 | 0.999933887 | 0.2227059 | 0.542871858 | -0.320165958 | 0.27771342 |
| ENSG00000144043 | *TEX261* | 4.120118334 | -0.00021371 | 0.034466633 | -0.034680342 | 0.999933887 | -0.200684104 | -0.340423514 | 0.13973941 | 0.27771342 |
| ENSG00000099385 | *BCL7C* | 1.603636698 | 0.139731115 | 0.154665295 | -0.01493418 | 0.999933887 | 0.130071657 | -0.175878084 | 0.305949741 | 0.277788783 |
| ENSG00000172270 | *BSG* | 7.559088295 | 0.049190588 | 0.00649081 | 0.042699778 | 0.999933887 | -0.088898815 | -0.217026819 | 0.128128004 | 0.277887968 |
| ENSG00000143919 | *CAMKMT* | 0.229910135 | -0.049278195 | -0.242887427 | 0.193609231 | 0.999933887 | -0.169359333 | 0.269964673 | -0.439324006 | 0.277913063 |
| ENSG00000239264 | *TXNDC5* | 0.059866445 | 0.061817282 | 0.054399411 | 0.00741787 | 0.999933887 | 0.085861404 | -0.403864259 | 0.489725663 | 0.278200723 |
| ENSG00000179406 | *LINC00174* | 3.299993653 | 0.007192623 | -0.036292748 | 0.043485371 | 0.999933887 | 0.419267219 | 0.628465178 | -0.209197958 | 0.278394724 |
| ENSG00000171659 | *GPR34* | 0.428731112 | -0.08590618 | -0.332454825 | 0.246548645 | 0.999933887 | -0.275170683 | 0.040921807 | -0.316092491 | 0.278457362 |
| ENSG00000274184 | *AC011815.2* | -0.441257193 | 0.417561044 | 0.10414167 | 0.313419375 | 0.999933887 | -1.137043692 | -0.71645253 | -0.420591161 | 0.278522778 |
| ENSG00000182551 | *ADI1* | 3.058108024 | -0.023581492 | 0.002149583 | -0.025731075 | 0.999933887 | -0.063020631 | -0.270929808 | 0.207909177 | 0.278522778 |
| ENSG00000197343 | *ZNF655* | 6.079793294 | 0.016353273 | -0.026995355 | 0.043348628 | 0.999933887 | -0.178403777 | -0.070842291 | -0.107561486 | 0.278651684 |
| ENSG00000131408 | *NR1H2* | 7.519665253 | 0.073806263 | 0.038133428 | 0.035672835 | 0.999933887 | 0.312337947 | 0.428185545 | -0.115847598 | 0.278760383 |
| ENSG00000135723 | *FHOD1* | 5.48551065 | -0.050962938 | 0.023781368 | -0.074744306 | 0.999933887 | 0.050061608 | -0.104322379 | 0.154383986 | 0.278896867 |
| ENSG00000175356 | *SCUBE2* | -0.51355272 | -0.043597953 | 0.164466187 | -0.208064141 | 0.999933887 | 0.899164984 | 1.334101099 | -0.434936115 | 0.278901251 |
| ENSG00000129083 | *COPB1* | 5.469658968 | -0.085030792 | -0.09539937 | 0.010368578 | 0.999933887 | -0.328843271 | -0.183548822 | -0.14529445 | 0.278970443 |
| ENSG00000221838 | *AP4M1* | 1.394602786 | -0.030446055 | 0.139396294 | -0.169842349 | 0.999933887 | 0.236871918 | 0.45981808 | -0.222946162 | 0.27917681 |
| ENSG00000133302 | *SLF1* | 3.193802257 | 0.109160182 | -0.06239379 | 0.171553973 | 0.999933887 | -0.439113323 | -0.087319788 | -0.351793535 | 0.279345469 |
| ENSG00000240445 | *FOXO3B* | 0.057114933 | -0.018959543 | 0.449071651 | -0.468031194 | 0.999933887 | -0.184094307 | 0.257978188 | -0.442072495 | 0.280022919 |
| ENSG00000172336 | *POP7* | 1.274768664 | -0.036724121 | 0.092131721 | -0.128855842 | 0.999933887 | 0.112434111 | -0.193084166 | 0.305518277 | 0.280096223 |
| ENSG00000143633 | *C1orf131* | 1.741368877 | -0.018362033 | -0.256056138 | 0.237694104 | 0.999933887 | -0.177129682 | -0.378203466 | 0.201073784 | 0.280096223 |
| ENSG00000168004 | *PLAAT5* | 0.393607791 | -0.12473551 | 0.215643925 | -0.340379435 | 0.944094537 | -0.132622261 | 0.307862149 | -0.44048441 | 0.280207593 |
| ENSG00000120727 | *PAIP2* | 6.031989062 | -0.018777199 | -0.12422656 | 0.105449361 | 0.999933887 | -0.180295207 | -0.057896168 | -0.12239904 | 0.280222681 |
| ENSG00000075884 | *ARHGAP15* | 5.957455887 | -0.013598791 | -0.073469106 | 0.059870315 | 0.999933887 | 0.00625548 | 0.148528925 | -0.142273445 | 0.280288323 |
| ENSG00000113580 | *NR3C1* | 8.041959291 | -0.024632676 | -0.039424375 | 0.014791699 | 0.999933887 | 0.075700525 | 0.204056908 | -0.128356384 | 0.280650011 |
| ENSG00000278133 | *AC135050.6* | 2.65553361 | -0.01322511 | -0.012364711 | -0.000860399 | 0.999933887 | -0.139242433 | -0.35067596 | 0.211433527 | 0.280650011 |
| ENSG00000182985 | *CADM1* | -0.23796074 | 0.003485361 | 0.283178884 | -0.279693523 | 0.999933887 | -0.362140781 | 0.10752351 | -0.469664291 | 0.280834459 |
| ENSG00000130023 | *ERMARD* | 1.992167778 | -0.001947077 | 0.208938992 | -0.210886069 | 0.999933887 | 0.033380689 | -0.146219167 | 0.179599856 | 0.280981326 |
| ENSG00000113318 | *MSH3* | 1.791626264 | -0.02459746 | 0.013178702 | -0.037776162 | 0.999933887 | -0.043091548 | -0.290488884 | 0.247397337 | 0.281030444 |
| ENSG00000115339 | *GALNT3* | 3.788402556 | -0.104306341 | -0.066493984 | -0.037812358 | 0.999933887 | -0.474357114 | -0.679223368 | 0.204866254 | 0.28103047 |
| ENSG00000035681 | *NSMAF* | 8.591189648 | 0.021735162 | 0.013338094 | 0.008397068 | 0.999933887 | 0.711157443 | 0.861195741 | -0.150038298 | 0.281101419 |
| ENSG00000278969 | *AC026310.2* | -0.121546147 | -0.000638088 | 0.079424147 | -0.080062235 | 0.999933887 | 0.31723304 | 0.711217126 | -0.393984086 | 0.281181739 |
| ENSG00000106714 | *CNTNAP3* | 3.767438667 | 0.017113768 | 0.058708594 | -0.041594826 | 0.999933887 | -0.329434961 | -0.484650558 | 0.155215597 | 0.281243516 |
| ENSG00000163378 | *EOGT* | 1.890325282 | 0.041473844 | -0.126268595 | 0.167742439 | 0.999933887 | 0.054742211 | 0.360402769 | -0.305660558 | 0.281564565 |
| ENSG00000168280 | *KIF5C* | 1.300661339 | 0.021178623 | -0.249153457 | 0.27033208 | 0.999933887 | -0.160539899 | 0.106139536 | -0.266679435 | 0.281564565 |
| ENSG00000162599 | *NFIA* | 0.458125347 | -0.061469193 | -0.217846625 | 0.156377432 | 0.999933887 | -0.679747219 | -0.097499677 | -0.582247542 | 0.281916543 |
| ENSG00000255538 | *OR10V2P* | 1.013450476 | 0.027362843 | 0.097012784 | -0.069649941 | 0.999933887 | 0.340400843 | 0.601097515 | -0.260696672 | 0.282358822 |
| ENSG00000121807 | *CCR2* | 0.970973056 | -0.004282144 | -0.048714709 | 0.044432564 | 0.999933887 | -0.134177339 | 0.095140157 | -0.229317496 | 0.282358822 |
| ENSG00000184162 | *NR2C2AP* | 1.021146402 | 0.057639535 | 0.220467997 | -0.162828461 | 0.999933887 | 0.164351592 | -0.092767946 | 0.257119537 | 0.282676006 |
| ENSG00000110172 | *CHORDC1* | 4.264199685 | -0.039865545 | -0.051213071 | 0.011347526 | 0.999933887 | -0.134154061 | 0.192069828 | -0.326223889 | 0.282676006 |
| ENSG00000100263 | *RHBDD3* | 1.863303365 | -0.018044368 | 0.024531299 | -0.042575667 | 0.999933887 | -0.205472123 | -0.405981332 | 0.200509209 | 0.282676006 |
| ENSG00000273841 | *TAF9* | 4.655405699 | 0.013981518 | -0.085322028 | 0.099303545 | 0.999933887 | 0.060910198 | 0.205818088 | -0.144907889 | 0.282915232 |
| ENSG00000187912 | *CLEC17A* | 3.036858135 | -0.097720618 | -0.066314182 | -0.031406437 | 0.999933887 | -0.575109586 | -0.800035709 | 0.224926123 | 0.283055333 |
| ENSG00000186001 | *LRCH3* | 4.162500204 | -0.003070693 | -0.009291751 | 0.006221057 | 0.999933887 | 0.177008898 | 0.295887531 | -0.118878632 | 0.283055333 |
| ENSG00000064995 | *TAF11* | 3.342228569 | 0.010020859 | 0.074284774 | -0.064263915 | 0.999933887 | -0.080517206 | -0.231842046 | 0.15132484 | 0.283056818 |
| ENSG00000046651 | *OFD1* | 4.88694655 | -0.038934009 | -0.116763702 | 0.077829693 | 0.999933887 | 0.191079487 | 0.397912407 | -0.20683292 | 0.283209976 |
| ENSG00000283355 | *AC074194.2* | -2.258740839 | -0.035678623 | -0.159941971 | 0.124263348 | 0.999933887 | -0.353083034 | 0.09233581 | -0.445418844 | 0.283307596 |
| ENSG00000165280 | *VCP* | 7.701638793 | 0.012238808 | -0.021834002 | 0.03407281 | 0.999933887 | -0.150118653 | -0.032252298 | -0.117866355 | 0.283328145 |
| ENSG00000121274 | *TENT4B* | 3.749683542 | 0.065642284 | 0.067801155 | -0.002158871 | 0.999933887 | -0.281130956 | -0.494825538 | 0.213694581 | 0.28370033 |
| ENSG00000225313 | *AL513327.1* | 3.547219897 | -0.170252248 | -0.126183983 | -0.044068265 | 0.999933887 | -1.185318051 | -1.412152483 | 0.226834432 | 0.283960695 |
| ENSG00000055118 | *KCNH2* | 0.778055298 | 0.00441588 | 0.048252679 | -0.043836799 | 0.999933887 | -0.103529627 | 0.263982348 | -0.367511975 | 0.284022567 |
| ENSG00000262919 | *CCNQ* | 2.032299512 | 0.05054627 | -0.068184037 | 0.118730307 | 0.999933887 | 0.083433718 | -0.181511669 | 0.264945387 | 0.284495462 |
| ENSG00000260260 | *SNHG19* | 0.195732722 | 0.006071682 | 0.081639096 | -0.075567414 | 0.999933887 | 0.101301701 | -0.193732044 | 0.295033745 | 0.28488271 |
| ENSG00000119705 | *SLIRP* | 1.717603668 | -0.057894492 | -0.093933803 | 0.036039311 | 0.999933887 | -0.057046054 | 0.188272124 | -0.245318178 | 0.284914845 |
| ENSG00000258738 | *AL121603.2* | 3.887913445 | 0.111283099 | 0.194711243 | -0.083428145 | 0.999933887 | 1.747816878 | 1.930069065 | -0.182252187 | 0.284932823 |

| | | | | | | | | | | |
|---|---|---|---|---|---|---|---|---|---|---|
| ENSG00000174442 | *ZWILCH* | 0.363244912 | -0.093025416 | -0.089761235 | -0.00326418 | 0.999933887 | -0.020382651 | 0.417954169 | -0.43833682 | 0.284932823 |
| ENSG00000174705 | *SH3PXD2B* | -0.567571259 | -0.053450565 | -0.231440451 | 0.177989887 | 0.999933887 | 1.703897754 | 2.198589052 | -0.494691298 | 0.284932823 |
| ENSG00000256667 | *KLRA1P* | 1.083901385 | 0.138142872 | -0.209513468 | 0.347656339 | 0.999933887 | -0.064225821 | 0.267008478 | -0.331234298 | 0.285468443 |
| ENSG00000149489 | *ROM1* | 0.567511893 | 0.091708279 | 0.057376944 | 0.034331335 | 0.999933887 | 1.383089031 | 1.816578283 | -0.433489252 | 0.285468443 |
| ENSG00000003147 | *ICA1* | 0.212099652 | -0.052893325 | -0.117353748 | 0.064460423 | 0.999933887 | 0.113144476 | 0.495725685 | -0.382581208 | 0.285468443 |
| ENSG00000198498 | *TMA16* | 0.09839524 | 0.066210317 | 0.005155323 | 0.061054994 | 0.999933887 | 0.198359938 | 0.501365315 | -0.303005377 | 0.285614516 |
| ENSG00000136279 | *DBNL* | 7.61258067 | -0.01857991 | 0.00905171 | -0.027631621 | 0.999933887 | -0.191500001 | -0.296985684 | 0.105485683 | 0.285614516 |
| ENSG00000241772 | *AC092620.1* | 0.305231002 | -0.040820441 | 0.281571899 | -0.32239234 | 0.999933887 | -0.023677799 | 0.430917531 | -0.45459533 | 0.285614516 |
| ENSG00000187735 | *TCEA1* | 3.510029847 | -0.019726973 | -0.01143308 | -0.008293893 | 0.999933887 | -0.032912487 | 0.111254076 | -0.144166563 | 0.285628632 |
| ENSG00000166902 | *MRPL16* | 3.29524444 | -0.12043595 | 0.049625367 | -0.170061317 | 0.999933887 | -0.009898612 | 0.151173184 | -0.161071797 | 0.285799451 |
| ENSG00000226029 | *LINC01772* | 0.917017397 | -0.201702105 | 0.151697622 | -0.353399727 | 0.999933887 | 0.227370939 | 0.495733328 | -0.26836239 | 0.285799451 |
| ENSG00000104388 | *RAB2A* | 6.707991462 | -0.0289408 | -0.020776189 | -0.008164611 | 0.999933887 | -0.560869406 | -0.447639431 | -0.113229975 | 0.285799451 |
| ENSG00000130749 | *ZC3H4* | 5.711489037 | 0.013693819 | 0.04023933 | -0.026545511 | 0.999933887 | 0.227949906 | 0.339780324 | -0.111830418 | 0.285799451 |
| ENSG00000123342 | *MMP19* | 2.492035941 | -0.156082505 | -0.281403004 | 0.125320499 | 0.999933887 | -0.851402728 | -1.074649341 | 0.223246613 | 0.285830904 |
| ENSG00000215712 | *TMEM242* | 1.363236744 | -0.171313089 | -0.198617872 | 0.027304783 | 0.999933887 | 0.177955278 | 0.423088133 | -0.245132855 | 0.286391497 |
| ENSG00000177410 | *ZFAS1* | 4.907893026 | -0.072562302 | -0.060672571 | -0.011889731 | 0.999933887 | -0.060295072 | -0.179663258 | 0.119368186 | 0.286391497 |
| ENSG00000121966 | *CXCR4* | 12.034696704 | 0.071801421 | 0.110887076 | -0.039085655 | 0.999933887 | -0.394820122 | -0.254847859 | -0.139972263 | 0.286391497 |
| ENSG00000196756 | *SNHG17* | 3.181348503 | -0.053107209 | -0.002610999 | -0.05049621 | 0.999933887 | 0.077888133 | -0.08611698 | 0.164005113 | 0.286391497 |
| ENSG00000108691 | *CCL2* | 3.114443579 | 0.037710391 | 0.139692266 | -0.101981874 | 0.999933887 | -1.226499004 | -1.554262295 | 0.327763291 | 0.286391497 |
| ENSG00000124201 | *ZNFX1* | 8.578014237 | -0.036434054 | -0.05553424 | 0.019100187 | 0.999933887 | 0.47702346 | 0.589744832 | -0.112721373 | 0.286606187 |
| ENSG00000160271 | *RALGDS* | 8.679815513 | 0.095933701 | 0.119098 | -0.023164298 | 0.999933887 | 1.032931589 | 1.199806249 | -0.16687466 | 0.286701828 |
| ENSG00000112335 | *SNX3* | 5.621069583 | 0.041321037 | -0.06258451 | 0.103905547 | 0.999933887 | -0.595358571 | -0.736907923 | 0.141549352 | 0.286701828 |
| ENSG00000135390 | *ATP5MC2* | 4.657020735 | 0.026261903 | -0.013848223 | 0.040110126 | 0.999933887 | -0.011888465 | -0.151320983 | 0.139432517 | 0.286701828 |
| ENSG00000140396 | *NCOA2* | 6.670166789 | -0.022363531 | -0.071170295 | 0.048806764 | 0.999933887 | -0.713903122 | -0.917927349 | 0.204024226 | 0.286701828 |
| ENSG00000241743 | *XACT* | -1.495236294 | -1.219171742 | -0.206594579 | -1.012577163 | 0.999933887 | -1.489208745 | -0.635027316 | -0.85418143 | 0.28675871 |
| ENSG00000148218 | *ALAD* | 2.704261374 | -0.022091543 | 0.034417386 | -0.056508929 | 0.999933887 | 0.012522564 | -0.184580755 | 0.197103318 | 0.28675871 |
| ENSG00000144741 | *SLC25A26* | 2.14451982 | -0.076814389 | -0.148511197 | 0.071696809 | 0.999933887 | 0.168416951 | 0.369454293 | -0.201037341 | 0.286821154 |
| ENSG00000154114 | *TBCEL* | 0.768193556 | 0.122687335 | 0.120902389 | 0.001784947 | 0.999933887 | -0.183026264 | 0.152001786 | -0.33502805 | 0.286821154 |
| ENSG00000270000 | *AC005479.2* | 0.014793036 | -0.088787697 | 0.262806171 | -0.351593868 | 0.999933887 | -0.019107802 | 0.28801648 | -0.307124282 | 0.286821154 |
| ENSG00000150768 | *DLAT* | 1.36548977 | 0.003674588 | 0.058772536 | -0.055097948 | 0.999933887 | -0.008902176 | 0.232313739 | -0.241215914 | 0.286926577 |
| ENSG00000162585 | *FAAP20* | 3.839035338 | 0.078495375 | 0.000876057 | 0.077619319 | 0.999933887 | -0.112355446 | -0.274645282 | 0.162289835 | 0.286936361 |
| ENSG00000166321 | *NUDT13* | 0.524521751 | 0.210111815 | 0.116313304 | 0.093798511 | 0.999933887 | 0.143311688 | 0.693282987 | -0.549971299 | 0.286936361 |
| ENSG00000110917 | *MLEC* | 3.368206233 | -0.038764077 | -0.044398683 | 0.005634606 | 0.999933887 | -0.061463335 | -0.209213746 | 0.147750411 | 0.286936361 |
| ENSG00000138613 | *APH1B* | 3.842148338 | -0.049064662 | 0.043598146 | -0.092662808 | 0.999933887 | -0.227443673 | -0.364719785 | 0.137276113 | 0.286955831 |
| ENSG00000102218 | *RP2* | 6.272979765 | 0.029390544 | 0.044545116 | -0.015154572 | 0.999933887 | -0.225834837 | -0.069922977 | -0.155911861 | 0.286955831 |
| ENSG00000183955 | *KMT5A* | 4.426561532 | 0.067530442 | 0.058362026 | 0.009168415 | 0.999933887 | 0.511412363 | 0.659729376 | -0.148317012 | 0.2870818 |
| ENSG00000176490 | *DIRAS1* | -1.336713655 | 0.250160347 | -0.164013675 | 0.414174022 | 0.999933887 | -0.750897801 | -0.134538388 | -0.616359413 | 0.2870818 |
| ENSG00000205268 | *PDE7A* | 5.638469883 | 0.013626704 | 0.018079798 | -0.004453094 | 0.999933887 | 0.085456348 | 0.226944344 | -0.141487996 | 0.2870818 |
| ENSG00000129749 | *CHRNA10* | 2.79740796 | 0.038122286 | 0.093995742 | -0.055873457 | 0.999933887 | 0.59236709 | 0.823874162 | -0.231507072 | 0.287168449 |
| ENSG00000204439 | *C6orf47* | 3.516982796 | 0.083190708 | 0.06308054 | 0.020110168 | 0.999933887 | -0.504483658 | -0.696891087 | 0.192407428 | 0.287317707 |
| ENSG00000125245 | *GPR18* | 1.687502256 | -0.147079891 | -0.056525507 | -0.090554384 | 0.999933887 | -0.002447234 | 0.266211395 | -0.268658629 | 0.287507271 |
| ENSG00000128708 | *HAT1* | 3.697193641 | -0.088660734 | -0.163597075 | 0.074936342 | 0.999933887 | 0.194264073 | 0.437340401 | -0.243076328 | 0.287507271 |
| ENSG00000276337 | *AC105429.1* | -0.661304824 | 0.201543681 | 0.114581561 | 0.08696212 | 0.999933887 | 0.339659609 | 0.791544618 | -0.451885009 | 0.287507271 |
| ENSG00000075407 | *ZNF37A* | 1.610759869 | -0.064157431 | -0.024601356 | -0.039556075 | 0.999933887 | -0.268721456 | 0.118059805 | -0.386781261 | 0.287507271 |
| ENSG00000108953 | *YWHAE* | 6.090339472 | -0.054708259 | -0.031894439 | -0.02281382 | 0.999933887 | 0.335366816 | 0.458651076 | -0.12328426 | 0.287519859 |
| ENSG00000092200 | *RPGRIP1* | 1.294972907 | -0.084041221 | -0.015489822 | -0.068551399 | 0.999933887 | -0.864851802 | -1.347429697 | 0.482577896 | 0.287554317 |
| ENSG00000154803 | *FLCN* | 7.881159032 | -0.054300159 | -0.010950737 | -0.043349422 | 0.999933887 | 0.133940688 | 0.245229869 | -0.111289181 | 0.287726654 |
| ENSG00000281026 | *N4BP2L2-IT2* | 2.094553362 | -0.242168803 | 0.011944637 | -0.254113441 | 0.999933887 | -0.008732291 | 0.32456412 | -0.333296411 | 0.287731847 |
| ENSG00000273112 | *AL590385.2* | 1.066949222 | -0.1197242 | -0.019848866 | -0.099875315 | 0.999933887 | 0.915676962 | 1.265110193 | -0.349433231 | 0.287898439 |
| ENSG00000226067 | *LINC00623* | 2.500103182 | -0.041894184 | -0.093852685 | 0.051958501 | 0.999933887 | 0.003264587 | -0.230313974 | 0.233578561 | 0.28795106 |
| ENSG00000083223 | *TUT7* | 8.244547187 | -0.07170838 | -0.085719241 | 0.014010861 | 0.999933887 | -0.550957142 | -0.363665316 | -0.187291826 | 0.288173242 |
| ENSG00000111666 | *CHPT1* | 3.19915736 | 0.029692592 | 0.035113519 | -0.005420927 | 0.999933887 | -0.067768703 | -0.239417579 | 0.171648876 | 0.288249213 |
| ENSG00000151287 | *TEX30* | 0.602214968 | 0.031060316 | -0.095556249 | 0.126616564 | 0.999933887 | 0.021026714 | -0.320348403 | 0.341375117 | 0.288249213 |
| ENSG00000053438 | *NNAT* | 0.196577742 | 0.224243643 | -0.109467743 | 0.333711386 | 0.999933887 | -0.439935357 | -0.742185028 | 0.302249671 | 0.288372668 |
| ENSG00000197579 | *TOPORS* | 4.666928309 | -0.112473257 | -0.198303193 | 0.085829936 | 0.999933887 | -0.792218381 | -1.004505695 | 0.212287315 | 0.288599766 |
| ENSG00000215039 | *CD27-AS1* | 0.831826575 | -0.069634947 | -0.140048062 | 0.070413116 | 0.999933887 | -0.124600345 | -0.459233769 | 0.334633424 | 0.288599766 |
| ENSG00000211696 | *TRGV8* | -0.794009336 | 0.163498637 | -0.172786444 | 0.336285081 | 0.999933887 | 0.181406876 | 0.641334993 | -0.459928117 | 0.288665157 |
| ENSG00000101557 | *USP14* | 3.175478047 | -0.136637687 | -0.038035067 | -0.09860262 | 0.999933887 | 0.056465914 | 0.216455114 | -0.1599892 | 0.28873381 |
| ENSG00000168795 | *ZBTB5* | 3.097828943 | -0.050693006 | -0.082994461 | 0.032301455 | 0.999933887 | 0.378759968 | 0.568161292 | -0.189401324 | 0.28873381 |
| ENSG00000126259 | *KIRREL2* | -0.757110984 | 0.171116006 | 0.00228962 | 0.168826386 | 0.999933887 | 2.733374514 | 3.371110113 | -0.637735599 | 0.288948221 |
| ENSG00000114125 | *RNF7* | 2.869750962 | 0.033516976 | -0.045852529 | 0.079369505 | 0.999933887 | -0.190547012 | -0.391495613 | 0.200948602 | 0.289462553 |
| ENSG00000131653 | *TRAF7* | 4.839495041 | 0.058000597 | 0.046555213 | 0.011445384 | 0.999933887 | -0.211585728 | -0.342093754 | 0.130508026 | 0.289905984 |
| ENSG00000012983 | *MAP4K5* | 2.904763333 | -0.070070807 | -0.114341084 | 0.044270277 | 0.999933887 | -0.126589724 | 0.159271211 | -0.285860935 | 0.290165997 |
| ENSG00000102781 | *KATNAL1* | 0.52264102 | -0.076836412 | -0.257092591 | 0.180255778 | 0.999933887 | -0.142458306 | 0.251168008 | -0.393626314 | 0.290171288 |
| ENSG00000110851 | *PRDM4* | 3.891034862 | 0.015631648 | 0.007900282 | 0.007731367 | 0.999933887 | -0.267829767 | -0.41603025 | 0.148200483 | 0.290251101 |
| ENSG00000089558 | *KCNH4* | -0.790428772 | 0.062624738 | -0.287384593 | 0.350009331 | 0.999933887 | 0.150671309 | 0.708721727 | -0.558050418 | 0.290537006 |
| ENSG00000273311 | *DGCR11* | 1.326019582 | -0.005620673 | -0.072643356 | 0.067022683 | 0.999933887 | -0.185121157 | -0.596402622 | 0.411281465 | 0.290560694 |
| ENSG00000185432 | *METTL7A* | 1.77605804 | -0.144827932 | -0.016658044 | -0.128169888 | 0.999933887 | 0.195244948 | 0.475883985 | -0.280639037 | 0.290583249 |
| ENSG00000091490 | *SEL1L3* | 4.248062843 | -0.011354666 | 0.026950076 | -0.038304743 | 0.999933887 | 0.13944928 | 0.267896534 | -0.128447254 | 0.290600652 |
| ENSG00000056277 | *ZNF280C* | -0.153473481 | -0.255647435 | -0.000524517 | -0.255122918 | 0.999933887 | -0.021910694 | 0.4418342 | -0.463744894 | 0.290992846 |
| ENSG00000114062 | *UBE3A* | 3.763472716 | 0.033266204 | -0.039762586 | 0.07302879 | 0.999933887 | -0.120009901 | 0.023744057 | -0.143753959 | 0.290992846 |
| ENSG00000108924 | *HLF* | -0.224693161 | 0.04962882 | -0.341417543 | 0.391046363 | 0.999933887 | 0.138699779 | 0.549579372 | -0.410879593 | 0.290992846 |
| ENSG00000100335 | *MIEF1* | 3.576476048 | -0.068078723 | -0.174495492 | 0.106416769 | 0.999933887 | -0.259877706 | -0.416926968 | 0.157049263 | 0.291205353 |
| ENSG00000029363 | *BCLAF1* | 7.142459828 | -0.042293744 | -0.190051239 | 0.147757495 | 0.999933887 | -0.357748079 | -0.07940069 | -0.278347389 | 0.291317819 |
| ENSG00000108960 | *MMD* | 0.564241552 | 0.101714683 | 0.160188903 | -0.058474221 | 0.999933887 | -0.131279937 | 0.207432173 | -0.33871211 | 0.291323958 |
| ENSG00000236914 | *LINC01852* | -1.054550973 | 0.364275064 | 0.359449728 | 0.004825336 | 0.999933887 | 0.682072203 | 1.427988325 | -0.745916121 | 0.291330202 |
| ENSG00000156050 | *FAM161B* | -0.079755105 | -0.257388068 | -0.150073411 | -0.107314657 | 0.999933887 | 0.110961194 | 0.552580911 | -0.441619716 | 0.291330202 |
| ENSG00000185504 | *FAAP100* | 3.910613582 | 0.046640393 | 0.029269998 | 0.017370395 | 0.999933887 | 0.632074683 | 0.79390046 | -0.161825777 | 0.291393238 |
| ENSG00000196611 | *MMP1* | 0.997572015 | -0.147930837 | -0.550297153 | 0.402366316 | 0.999933887 | -0.260670015 | 0.016609214 | -0.277279228 | 0.291393238 |
| ENSG00000154146 | *NRGN* | 1.693138415 | 0.039005853 | -0.014109628 | 0.053115481 | 0.999933887 | 0.003971361 | -0.264567459 | 0.26853882 | 0.291393238 |
| ENSG00000163159 | *VPS72* | 2.939848858 | -0.011816771 | 0.062178414 | -0.073995185 | 0.999933887 | -0.227573239 | -0.403093537 | 0.175520298 | 0.291393238 |
| ENSG00000158615 | *PPP1R15B* | 9.09924218 | 0.075659622 | 0.075967828 | -0.000308206 | 0.999933887 | 0.325259246 | 0.419367871 | -0.094108625 | 0.291590646 |
| ENSG00000262155 | *LINC02175* | 0.174851936 | 0.094681973 | -0.180664451 | 0.275346423 | 0.999933887 | 0.446435138 | 0.824895854 | -0.378460716 | 0.291590646 |
| ENSG00000228363 | *AC015971.1* | 3.916274292 | -0.027761656 | 0.067267697 | -0.095029353 | 0.999933887 | 0.180163723 | 0.376970445 | -0.196806722 | 0.291590646 |
| ENSG00000174579 | *MSL2* | 6.696183222 | -0.060103682 | -0.031771376 | -0.028332306 | 0.999933887 | -0.247913856 | -0.350800552 | 0.102886696 | 0.291692367 |
| ENSG00000175398 | *OR10P1* | -0.092161883 | 0.268394403 | 0.321341281 | -0.052946878 | 0.999933887 | 1.205384299 | 1.58001717 | -0.374632871 | 0.291692367 |
| ENSG00000070269 | *TMEM260* | 2.969673418 | 0.034734651 | -0.034878297 | 0.069612948 | 0.999933887 | -0.165133007 | -0.374127802 | 0.208994795 | 0.291692367 |
| ENSG00000140386 | *SCAPER* | 2.543093435 | 0.021868411 | -0.151444649 | 0.173313061 | 0.999933887 | -0.116548324 | 0.109474481 | -0.226022805 | 0.291692367 |
| ENSG00000142937 | *RPS8* | 7.623218405 | 0.00247836 | -0.038071159 | 0.040549519 | 0.999933887 | 0.106590683 | 0.232167128 | -0.125576445 | 0.291692367 |
| ENSG00000130598 | *TNNI2* | 5.248733761 | -0.054385007 | -0.032832899 | -0.021552108 | 0.999933887 | -0.546070153 | -0.734968841 | 0.188898688 | 0.291891071 |
| ENSG00000171204 | *TMEM126B* | 3.100907118 | -0.095206364 | -0.201423985 | 0.106217621 | 0.999933887 | -0.416045216 | -0.57489722 | 0.158852005 | 0.291960637 |
| ENSG00000184584 | *STING1* | 3.601090489 | 0.076292926 | -0.000196021 | 0.076488948 | 0.999933887 | 0.129532186 | -0.019049648 | 0.148581835 | 0.291978354 |
| ENSG00000065183 | *WDR3* | 2.100249573 | -0.062573995 | -0.030551954 | -0.032022041 | 0.999933887 | 0.084788372 | 0.339639472 | -0.2548511 | 0.291978354 |
| ENSG00000213096 | *ZNF254* | 3.860671196 | -0.091693526 | -0.16810895 | 0.076415424 | 0.999933887 | -0.850135454 | -0.606141863 | -0.243993591 | 0.292203829 |
| ENSG00000157303 | *SUSD3* | 2.305586584 | 0.029981674 | -0.071228143 | 0.101209817 | 0.999933887 | -0.256513053 | -0.46717753 | 0.210664477 | 0.292290841 |
| ENSG00000048140 | *TSPAN17* | 3.024361955 | -0.000137055 | -0.060149938 | 0.060012884 | 0.999933887 | -0.067723327 | -0.242572906 | 0.174849579 | 0.292345791 |
| ENSG00000286242 | *AC018797.3* | 2.287423924 | 0.123978397 | 0.152570005 | -0.028591608 | 0.999933887 | -0.290358021 | 0.08053046 | -0.370888482 | 0.292453834 |
| ENSG00000183154 | *AC138356.1* | 0.967477779 | 0.130059958 | 0.314680409 | -0.184620451 | 0.999933887 | 0.494499092 | -0.098244728 | 0.59274382 | 0.292565604 |
| ENSG00000151623 | *NR3C2* | 1.26950213 | -0.116399115 | -0.054358229 | -0.062040887 | 0.999933887 | 0.032825084 | 0.309217761 | -0.276392677 | 0.29259419 |
| ENSG00000139410 | *SDSL* | -0.442444938 | 0.29558083 | 0.403235983 | -0.107655152 | 0.999933887 | -0.637148551 | -0.317839898 | -0.319308653 | 0.29259419 |
| ENSG00000141258 | *SGSM2* | 5.244204496 | 0.042796995 | 0.053793787 | -0.010996792 | 0.999933887 | 0.058097865 | -0.058310765 | 0.11640863 | 0.292600654 |
| ENSG00000081307 | *UBA5* | 2.084171598 | -0.027606984 | -0.139185247 | 0.111578263 | 0.999933887 | -0.161953067 | 0.104648391 | -0.266601458 | 0.292629844 |
| ENSG00000239305 | *RNF103* | 6.933818964 | -0.088716365 | -0.024692859 | -0.064023507 | 0.999933887 | -0.254248882 | -0.119246661 | -0.135002221 | 0.292721309 |
| ENSG00000147231 | *RADX* | 0.703487816 | -0.133206367 | -0.174313729 | 0.041107361 | 0.999933887 | -0.109914347 | 0.262442334 | -0.37235668 | 0.292965261 |
| ENSG00000146143 | *PRIM2* | 0.026832867 | 0.12312992 | 0.089862714 | 0.033267206 | 0.999933887 | -0.234198124 | 0.168528183 | -0.402726307 | 0.292989684 |

| | | | | | | | | | | |
|---|---|---|---|---|---|---|---|---|---|---|
| ENSG00000132792 | *CTNNBL1* | 4.139105815 | 0.025635455 | -0.035599537 | 0.061234992 | 0.999933887 | -0.173461751 | -0.310257148 | 0.136795397 | 0.293119072 |
| ENSG00000241852 | *C8orf58* | 2.23805294 | 0.047438042 | 0.041730881 | 0.005707161 | 0.999933887 | -0.000113522 | -0.272567446 | 0.272453923 | 0.293168643 |
| ENSG00000173114 | *LRRN3* | 0.981323411 | 0.055880948 | -0.023996891 | 0.07987784 | 0.999933887 | -0.215110318 | 0.036173267 | -0.251283585 | 0.293246365 |
| ENSG00000129636 | *ITFG1* | 2.823090703 | 0.019966801 | 0.045433547 | -0.025466746 | 0.999933887 | -0.229380466 | -0.414428174 | 0.185047708 | 0.293246365 |
| ENSG00000219545 | *UMAD1* | 2.505501195 | -0.010475053 | 0.022049882 | -0.032524935 | 0.999933887 | -0.177227004 | 0.028676855 | -0.205903859 | 0.293246365 |
| ENSG00000168876 | *ANKRD49* | 4.311356417 | -0.158045204 | -0.210528447 | 0.052483243 | 0.999933887 | -0.116744061 | 0.053266781 | -0.170010842 | 0.293425672 |
| ENSG00000169251 | *NMD3* | 3.308864504 | -0.056460667 | -0.07617867 | 0.019718003 | 0.999933887 | -0.091767006 | 0.132518844 | -0.22428585 | 0.294205937 |
| ENSG00000110446 | *SLC15A3* | 9.238425703 | 0.06182005 | 0.082405427 | -0.020585377 | 0.999933887 | 0.997437639 | 1.110494007 | -0.113056367 | 0.294276092 |
| ENSG00000175213 | *ZNF408* | 2.620030124 | -0.071134857 | 0.002478424 | -0.073613282 | 0.999933887 | -0.345399527 | -0.532163229 | 0.186763702 | 0.294423464 |
| ENSG00000111785 | *RIC8B* | 0.890560441 | -0.133365811 | -0.040418748 | -0.092947063 | 0.999933887 | -0.030563424 | 0.258736426 | -0.289299849 | 0.294573791 |
| ENSG00000172216 | *CEBPB* | 9.125505557 | 0.263699909 | 0.273323652 | -0.009623743 | 0.999933887 | 0.112493981 | 0.329989708 | -0.217495727 | 0.295061266 |
| ENSG00000228446 | *ANKRD49P1* | -0.078526098 | 0.222841789 | 0.241550657 | -0.018708868 | 0.999933887 | -0.299238542 | 0.222956293 | -0.522194835 | 0.295250122 |
| ENSG00000255145 | *STX17-AS1* | -0.845624211 | 0.116130462 | 0.266442378 | -0.150311916 | 0.999933887 | 0.232244644 | 1.007871669 | -0.775627025 | 0.295250122 |
| ENSG00000087152 | *ATXN7L3* | 7.112451034 | 0.031246822 | 0.005802008 | 0.025444813 | 0.999933887 | -0.238008708 | -0.359180535 | 0.121171827 | 0.295250122 |
| ENSG00000234912 | *SNHG20* | 4.142675418 | -0.090296072 | -0.036672353 | -0.05362372 | 0.999933887 | 0.270105473 | 0.127886452 | 0.142219021 | 0.29532307 |
| ENSG00000134697 | *GNL2* | 4.76519878 | -0.04035273 | -0.120391542 | 0.080038812 | 0.999933887 | 0.322513685 | 0.490658901 | -0.168145216 | 0.29545569 |
| ENSG00000158169 | *FANCC* | -0.165125259 | -0.001210672 | 0.070869933 | -0.072080605 | 0.999933887 | 0.060325161 | 0.470194823 | -0.409869662 | 0.295511449 |
| ENSG00000134152 | *KATNBL1* | 8.729749927 | -0.00562564 | 0.079305922 | -0.084931562 | 0.999933887 | 0.151255026 | 0.280868227 | -0.129613201 | 0.295623254 |
| ENSG00000259865 | *AL390728.6* | 2.19187607 | -0.130442392 | 0.050590328 | -0.18103272 | 0.999933887 | 0.055475 | 0.437527572 | -0.382052572 | 0.295716464 |
| ENSG00000234617 | *SNRK-AS1* | 0.744578604 | 0.056613205 | -0.142440429 | 0.199053634 | 0.999933887 | 0.026233283 | 0.460243589 | -0.434010306 | 0.295716464 |
| ENSG00000184068 | *SREBF2-AS1* | 2.182230176 | 0.105326223 | 0.027437159 | 0.077889064 | 0.999933887 | 0.808835608 | 1.048115649 | -0.239280041 | 0.295928053 |
| ENSG00000172716 | *SLFN11* | 3.725787442 | -0.020510638 | -0.082420693 | 0.061910055 | 0.999933887 | -0.108919111 | -0.255051104 | 0.146131994 | 0.295928053 |
| ENSG00000105750 | *ZNF85* | -0.376300655 | 0.059650487 | -0.094522106 | 0.154172593 | 0.999933887 | -0.631736226 | -0.051434888 | -0.580301338 | 0.295928053 |
| ENSG00000161960 | *EIF4A1* | 4.726107279 | 0.24296503 | 0.349228059 | -0.106263029 | 0.999933887 | 0.205727887 | 0.393082105 | -0.187354218 | 0.295984317 |
| ENSG00000066379 | *POLR1H* | 2.550313701 | 0.042915536 | 0.023125606 | 0.01978993 | 0.999933887 | 0.01014449 | -0.199783031 | 0.209927521 | 0.296073037 |
| ENSG00000241878 | *PISD* | 7.25198499 | -0.052347571 | -0.071627587 | 0.019280016 | 0.999933887 | -0.252117623 | -0.368005739 | 0.115888116 | 0.296519192 |
| ENSG00000135317 | *SNX14* | 4.010962043 | -0.02218791 | -0.040378816 | 0.018149906 | 0.999933887 | -0.325884098 | -0.523505465 | 0.197621368 | 0.296519192 |
| ENSG00000262468 | *LINC01569* | -0.863561475 | -0.212957011 | 0.46936833 | -0.68232534 | 0.999933887 | -0.243576292 | 0.283596417 | -0.527172709 | 0.296555117 |
| ENSG00000054654 | *SYNE2* | 7.286963854 | -0.0628977 | -0.004760319 | -0.058137381 | 0.999933887 | -0.403876362 | -0.156182484 | -0.247693878 | 0.296555117 |
| ENSG00000166707 | *ZCCHC18* | -0.634455483 | -0.228049702 | -0.114559634 | -0.113490068 | 0.999933887 | 0.129699211 | 0.545223682 | -0.415524471 | 0.296751747 |
| ENSG00000121454 | *LHX4* | -0.414884686 | 0.388732541 | -0.237318691 | 0.626051232 | 0.944094537 | -0.126943928 | 0.286650533 | -0.413594461 | 0.29731488 |
| ENSG00000121542 | *SEC22A* | 0.730875523 | 0.154333543 | 0.014401968 | 0.139931575 | 0.999933887 | 0.404405906 | 0.709032856 | -0.304626951 | 0.297399553 |
| ENSG00000075399 | *VPS9D1* | 5.713361497 | 0.043475967 | 0.071681496 | -0.028205529 | 0.999933887 | 0.111364787 | -0.015711861 | 0.127076647 | 0.297399553 |
| ENSG00000106477 | *CEP41* | -0.421471578 | -0.122262961 | -0.020324652 | -0.101938309 | 0.999933887 | -0.087677604 | 0.317205187 | -0.404882791 | 0.297399553 |
| ENSG00000100345 | *MYH9* | 10.672429126 | 0.014336789 | -0.016892051 | 0.03122884 | 0.999933887 | 0.381882232 | 0.510848429 | -0.128966198 | 0.297399553 |
| ENSG00000231160 | *KLF3-AS1* | 1.597034894 | 0.021063488 | -0.033970516 | 0.055034004 | 0.999933887 | -0.273522182 | -0.522124091 | 0.248601909 | 0.297399553 |
| ENSG00000080546 | *SESN1* | 3.039795166 | -0.018567284 | -0.099029464 | 0.08046218 | 0.999933887 | 0.087689868 | 0.278614654 | -0.190924786 | 0.29748176 |
| ENSG00000205578 | *POM121B* | -0.711603033 | 0.080745797 | -0.066569886 | 0.147315684 | 0.999933887 | -0.235451241 | 0.306285055 | -0.541736296 | 0.297749785 |
| ENSG00000251381 | *LINC00958* | 2.072548651 | 0.166764308 | 0.370017036 | -0.203252728 | 0.999933887 | 0.310821574 | 0.580932162 | -0.270110587 | 0.297795667 |
| ENSG00000279277 | *AC012020.1* | 2.296718958 | 0.035600603 | 0.091559217 | -0.055958614 | 0.999933887 | 0.195738345 | 0.406374163 | -0.210635818 | 0.297851446 |
| ENSG00000115159 | *GPD2* | 4.405591045 | -0.028288764 | -0.099674742 | 0.071385978 | 0.999933887 | -1.033741054 | -0.815593106 | -0.218147948 | 0.298226407 |
| ENSG00000103061 | *SLC7A6OS* | 1.870887929 | 0.073703197 | -0.057204708 | 0.130907905 | 0.999933887 | -0.26866557 | -0.482304829 | 0.213639259 | 0.298570377 |
| ENSG00000142065 | *ZFP14* | 0.442191209 | 0.012369495 | -0.080398432 | 0.092767927 | 0.999933887 | -0.231910887 | 0.075548539 | -0.307459426 | 0.298597094 |
| ENSG00000105402 | *NAPA* | 8.100610235 | 0.057433165 | 0.073716705 | -0.01628354 | 0.999933887 | 0.236702937 | 0.332184467 | -0.095481529 | 0.299120989 |
| ENSG00000228393 | *LINC01004* | 2.348519824 | 0.134469139 | 0.190484003 | -0.056014864 | 0.999933887 | 0.960473814 | 1.180605779 | -0.220131965 | 0.299120989 |
| ENSG00000140691 | *ARMC5* | 3.608018511 | 0.092201453 | 0.15041461 | -0.058213157 | 0.999933887 | 0.47513476 | 0.687759387 | -0.212624626 | 0.299120989 |
| ENSG00000198198 | *SZT2* | 5.691959201 | 0.036944639 | 0.049122965 | -0.012178326 | 0.999933887 | -0.068472026 | -0.174460366 | 0.105988339 | 0.299120989 |
| ENSG00000149212 | *SESN3* | 4.333366859 | -0.044054292 | -0.157117242 | 0.11306295 | 0.999933887 | -0.366930486 | -0.093996418 | -0.272934068 | 0.299120989 |
| ENSG00000104267 | *CA2* | 3.859937273 | 0.079300167 | 0.054361182 | 0.024938985 | 0.999933887 | 0.663338528 | 0.842110089 | -0.178771561 | 0.299434453 |
| ENSG00000132510 | *KDM6B* | 11.141986875 | 0.132918862 | 0.108933785 | 0.023985077 | 0.999933887 | 1.046044448 | 1.180106822 | -0.134062374 | 0.299446701 |
| ENSG00000112033 | *PPARD* | 4.91811744 | -0.037381366 | -0.061026301 | 0.023644935 | 0.999933887 | 0.657633698 | 0.808080194 | -0.150446496 | 0.299919919 |
| ENSG00000150907 | *FOXO1* | 4.311436619 | 0.032838413 | -0.013408995 | 0.046247408 | 0.999933887 | -0.055887989 | -0.205130646 | 0.149242658 | 0.299927919 |
| ENSG00000107625 | *DDX50* | 3.711575824 | 0.018735893 | -0.132840537 | 0.151576429 | 0.999933887 | 0.147379318 | 0.287566376 | -0.140187058 | 0.299962007 |
| ENSG00000138594 | *TMOD3* | 5.182834405 | 0.026257524 | 0.0595571 | -0.033299576 | 0.999933887 | 0.068870712 | 0.217663753 | -0.14879304 | 0.299995528 |
| ENSG00000055130 | *CUL1* | 3.830311108 | 0.02273919 | -0.139112488 | 0.161851678 | 0.999933887 | 0.030680607 | 0.16769195 | -0.137011343 | 0.300006892 |
| ENSG00000140332 | *TLE3* | 9.097672469 | 0.015379966 | 0.046507567 | -0.031127601 | 0.999933887 | 0.147189455 | 0.26755112 | -0.120361666 | 0.30003225 |
| ENSG00000023734 | *STRAP* | 5.262559601 | -0.054336182 | -0.076073134 | 0.021736952 | 0.999933887 | 0.058424922 | 0.160133636 | -0.101708714 | 0.30012988 |
| ENSG00000179632 | *MAF1* | 5.783214925 | 0.029040148 | 0.028996944 | 4.3204e-05 | 0.999933887 | -0.164792322 | -0.289057482 | 0.12426516 | 0.30012988 |
| ENSG00000165283 | *STOML2* | 2.979053949 | 0.008540299 | -0.066458456 | 0.074998754 | 0.999933887 | -0.04444009 | -0.191434583 | 0.146994493 | 0.30012988 |
| ENSG00000118690 | *ARMC2* | 1.866484617 | -0.055636171 | -0.146732867 | 0.091096696 | 0.999933887 | 0.039779109 | 0.284362122 | -0.244583014 | 0.300337604 |
| ENSG00000131368 | *MRPS25* | 2.667310285 | -0.007776974 | 0.065278113 | -0.073055088 | 0.999933887 | -0.048313983 | 0.106574919 | -0.154888901 | 0.300337604 |
| ENSG00000284948 | *AC107959.4* | 2.016307841 | 0.127581012 | -0.024971831 | 0.152552842 | 0.999933887 | -0.8825226 | -1.290849758 | 0.408327158 | 0.3007583 |
| ENSG00000153283 | *CD96* | 5.06080328 | -0.083018212 | -0.06543164 | -0.017586572 | 0.999933887 | -0.025762509 | 0.108603324 | -0.134365833 | 0.300833622 |
| ENSG00000181754 | *AMIGO1* | 1.343459702 | -0.002448984 | 0.064899762 | -0.067348746 | 0.999933887 | 0.176498002 | -0.054012584 | 0.230510586 | 0.300833622 |
| ENSG00000149658 | *YTHDF1* | 4.749880778 | 0.008601413 | -0.023382179 | 0.031983593 | 0.999933887 | -0.186941316 | -0.296651765 | 0.109710449 | 0.301078407 |
| ENSG00000114346 | *ECT2* | 0.058503358 | -0.055427287 | -0.240691217 | 0.18526393 | 0.999933887 | 0.334783519 | 0.920440506 | -0.585656986 | 0.301125479 |
| ENSG00000231721 | *LINC-PINT* | 5.69203179 | 0.020039829 | 0.153307422 | -0.133267593 | 0.999933887 | -0.320567287 | -0.476299901 | 0.155732615 | 0.301125479 |
| ENSG00000173611 | *SCAI* | 0.968833549 | -0.022519251 | -0.042433338 | 0.019914087 | 0.999933887 | -0.339819576 | 0.016947155 | -0.35676673 | 0.301125479 |
| ENSG00000261716 | *H2BC20P* | 3.465730539 | -0.130420067 | -0.046074839 | -0.084345229 | 0.999933887 | -0.626532998 | -0.823058986 | 0.196525988 | 0.301204438 |
| ENSG00000156983 | *BRPF1* | 4.62300381 | 0.125711765 | 0.044725114 | 0.080986651 | 0.999933887 | -0.449976468 | -0.589840028 | 0.13986356 | 0.301254379 |
| ENSG00000261888 | *AC144831.1* | 2.421604573 | 0.058996512 | -0.019743314 | 0.078739825 | 0.999933887 | -0.042306028 | -0.29918108 | 0.256875052 | 0.301254379 |
| ENSG00000146463 | *ZMYM4* | 2.280050578 | -0.062940906 | -0.14155207 | 0.078611164 | 0.999933887 | -0.131409437 | 0.086583378 | -0.217992815 | 0.301254379 |
| ENSG00000185104 | *FAF1* | 2.475020716 | -0.05734582 | -0.108178295 | 0.050832475 | 0.999933887 | 0.01091249 | 0.184793769 | -0.173881278 | 0.301403313 |
| ENSG00000134882 | *UBAC2* | 4.4724491 | 0.008469569 | -0.044871609 | 0.053341179 | 0.999933887 | -0.167816078 | -0.279788791 | 0.111972713 | 0.301421984 |
| ENSG00000124767 | *GLO1* | 2.636047469 | 0.011314875 | -0.179635155 | 0.19095003 | 0.999933887 | -0.163111814 | 0.033645595 | -0.196757409 | 0.301421984 |
| ENSG00000185624 | *P4HB* | 7.279233212 | 0.053625423 | 0.017084536 | 0.036540887 | 0.999933887 | 0.584336626 | 0.723183551 | -0.138846925 | 0.301823432 |
| ENSG00000070501 | *POLB* | 4.215158525 | 0.008303231 | 0.024989122 | -0.016685891 | 0.999933887 | -0.199754125 | -0.341059962 | 0.141305837 | 0.301987303 |
| ENSG00000114554 | *PLXNA1* | 2.817673411 | 0.014697828 | 0.136887551 | -0.122189723 | 0.999933887 | 0.38664588 | 0.567494789 | -0.180848909 | 0.302177819 |
| ENSG00000136824 | *SMC2* | 0.230267956 | -0.007963233 | 0.051307107 | -0.05927034 | 0.999933887 | -0.277897636 | 0.160698477 | -0.438596113 | 0.3026933 |
| ENSG00000248544 | *AC008676.1* | 1.488920181 | 0.169483817 | 0.03065897 | 0.138824847 | 0.999933887 | -0.489405011 | -0.19173256 | -0.297672451 | 0.302765744 |
| ENSG00000155719 | *OTOA* | 0.949656282 | -0.195970468 | -0.404536662 | 0.208566194 | 0.999933887 | -0.092271454 | 0.128364666 | -0.220636119 | 0.302765744 |
| ENSG00000135506 | *OS9* | 7.745248323 | -0.001060663 | -0.004570646 | 0.003509983 | 0.999933887 | -0.284338694 | -0.402250444 | 0.117911749 | 0.302765744 |
| ENSG00000163281 | *GNPDA2* | 0.862751859 | -0.235207006 | -0.381802055 | 0.146595049 | 0.999933887 | -0.141786485 | 0.175985099 | -0.317771585 | 0.30278964 |
| ENSG00000237719 | *Z95152.1* | 2.369565852 | 0.002186799 | -0.078689057 | 0.080875856 | 0.999933887 | -0.293091622 | 0.017924559 | -0.311016181 | 0.30278964 |
| ENSG00000154781 | *CCDC174* | 4.958155293 | -0.082808071 | -0.05603709 | -0.026770981 | 0.999933887 | 0.104553388 | 0.198773597 | -0.094220209 | 0.303157862 |
| ENSG00000155366 | *RHOC* | 2.664640401 | 0.077961697 | -0.141364879 | 0.219326576 | 0.999933887 | -0.005062135 | -0.200099545 | 0.195037411 | 0.303169253 |
| ENSG00000159267 | *HLCS* | 1.328601503 | -0.0680263 | -0.093499361 | 0.025473061 | 0.999933887 | -0.091991036 | 0.165307893 | -0.257298928 | 0.303169253 |
| ENSG00000147138 | *GPR174* | 3.656452366 | -0.067081076 | -0.094625296 | 0.02754422 | 0.999933887 | -0.03722257 | 0.203809676 | -0.241032247 | 0.303169253 |
| ENSG00000136827 | *TOR1A* | 5.199715362 | -0.046582221 | -0.058032873 | 0.011450652 | 0.999933887 | -0.972416651 | -1.149734535 | 0.177317884 | 0.303169253 |
| ENSG00000172175 | *MALT1* | 4.259424399 | -0.044982726 | -0.076583523 | 0.031600797 | 0.999933887 | -0.185739609 | 0.009767475 | -0.195507084 | 0.303169253 |
| ENSG00000133619 | *KRBA1* | 2.05108466 | -0.014086076 | 0.114343916 | -0.128429992 | 0.999933887 | 0.230873855 | 0.425147699 | -0.194273844 | 0.303169253 |
| ENSG00000248719 | *AC021127.1* | 1.639830074 | 0.009992719 | 0.180335377 | -0.170342658 | 0.999933887 | 0.261300921 | -0.024771649 | 0.28607257 | 0.303169253 |
| ENSG00000241553 | *ARPC4* | 6.962898247 | -0.007972868 | 0.028057703 | -0.036030571 | 0.999933887 | -0.084400679 | -0.190546208 | 0.106145529 | 0.303502598 |
| ENSG00000231105 | *ECE1-AS1* | 2.44046279 | 0.191832627 | 0.19818664 | -0.006354012 | 0.999933887 | 2.143996589 | 2.453247232 | -0.309250643 | 0.303623595 |
| ENSG00000145741 | *BTF3* | 6.42232234 | -0.022261136 | -0.110093746 | 0.08783261 | 0.999933887 | 0.105405819 | 0.212868952 | -0.107463133 | 0.303623595 |
| ENSG00000189195 | *BTBD8* | 0.418132501 | -0.12184242 | -0.201072769 | 0.07923035 | 0.999933887 | -0.745582149 | -0.355370009 | -0.390212141 | 0.303636099 |
| ENSG00000163577 | *EIF5A2* | 2.043030431 | 0.051441825 | -0.130509603 | 0.181951428 | 0.999933887 | -0.333031936 | -0.137870122 | -0.195161814 | 0.303693079 |
| ENSG00000151690 | *MFSD6* | 3.345641168 | -0.079141203 | -0.022174521 | -0.056966682 | 0.999933887 | 0.031640344 | -0.123660391 | 0.155300734 | 0.303730857 |
| ENSG00000134313 | *KIDINS220* | 6.460977747 | -0.017763628 | -0.053964371 | 0.036200743 | 0.999933887 | -0.353983278 | -0.21189641 | -0.142086867 | 0.303730857 |
| ENSG00000107951 | *MTPAP* | 1.835857406 | -0.108566369 | -0.083856124 | -0.024710245 | 0.999933887 | 0.076538973 | 0.27719555 | -0.200656577 | 0.303847995 |
| ENSG00000279570 | *AC099804.1* | 1.0171755 | -0.038021138 | 0.237305992 | -0.27532713 | 0.999933887 | 0.600041658 | 0.947199993 | -0.347158334 | 0.303993955 |
| ENSG00000164930 | *FZD6* | -0.923144898 | -0.023513161 | 0.138494031 | -0.162007192 | 0.999933887 | 0.81041168 | 1.338273525 | -0.527861846 | 0.30404276 |
| ENSG00000237604 | *AP001056.1* | -2.50570992 | -0.215322332 | 0.318097544 | -0.533419876 | 0.999933887 | 2.662073994 | 3.288861089 | -0.626787095 | 0.304062752 |
| ENSG00000134709 | *HOOK1* | 0.399916819 | -0.37113301 | -0.095941673 | -0.275191337 | 0.999933887 | -0.185140502 | 0.265366482 | -0.450506985 | 0.304104117 |
| ENSG00000104853 | *CLPTM1* | 5.62816866 | 0.027777817 | 0.007120467 | 0.020657349 | 0.999933887 | -0.072013458 | -0.177675042 | 0.105661584 | 0.304104117 |

| | | | | | | | | | | |
|---|---|---|---|---|---|---|---|---|---|---|
| ENSG00000204209 | *DAXX* | 4.697949522 | -0.002742045 | -0.042562139 | 0.039820094 | 0.999933887 | 0.017792327 | -0.099261626 | 0.117053953 | 0.304186757 |
| ENSG00000147475 | *ERLIN2* | 5.414451358 | 0.033892709 | -0.036687576 | 0.070580286 | 0.999933887 | -0.715038345 | -0.890564773 | 0.175526427 | 0.304281984 |
| ENSG00000164244 | *PRRC1* | 2.924306808 | 0.014088865 | -0.115493942 | 0.129582807 | 0.999933887 | -0.186865666 | -0.035841482 | -0.151024184 | 0.304281984 |
| ENSG00000155380 | *SLC16A1* | 1.544718511 | 0.00545963 | 0.043314007 | -0.037854378 | 0.999933887 | 0.16200753 | 0.440333703 | -0.278326173 | 0.304284326 |
| ENSG00000250995 | *TMEM30BP1* | -0.150542743 | 0.106910236 | -0.026372507 | 0.133282744 | 0.999933887 | 0.674344052 | 0.174507983 | 0.49983607 | 0.30469462 |
| ENSG00000242474 | *AC093627.5* | 0.255814313 | -0.16438442 | 0.027985415 | -0.192369835 | 0.999933887 | -0.562921454 | -0.878862943 | 0.315941489 | 0.304827282 |
| ENSG00000136141 | *LRCH1* | 2.898471307 | -0.066615127 | -0.023955498 | -0.042659629 | 0.999933887 | 0.08349576 | 0.257555777 | -0.174060017 | 0.304827282 |
| ENSG00000183665 | *TRMT12* | 2.569667405 | -0.068834688 | -0.142715213 | 0.073880525 | 0.999933887 | -0.287767849 | -0.490007978 | 0.202240129 | 0.304827282 |
| ENSG00000029639 | *TFB1M* | 0.388641905 | -0.051025959 | 0.194806616 | -0.245832575 | 0.999933887 | -0.067911158 | 0.212781986 | -0.280693145 | 0.304827282 |
| ENSG00000167553 | *TUBA1C* | 2.8826497 | 0.016218907 | -0.071802254 | 0.088021161 | 0.999933887 | -0.034056537 | -0.220375642 | 0.186319105 | 0.305414123 |
| ENSG00000174891 | *RSRC1* | 1.343213048 | 0.133681178 | -0.070972272 | 0.204653451 | 0.999933887 | -0.062104079 | 0.256307764 | -0.318411842 | 0.30560508 |
| ENSG00000166348 | *USP54* | 0.246657566 | -0.316541804 | 0.071165711 | -0.387707515 | 0.999933887 | 0.083870636 | -0.310359586 | 0.394230221 | 0.305642982 |
| ENSG00000244720 | *NT5C3AP2* | 0.277659503 | -0.291367652 | -0.111765385 | -0.179602267 | 0.999933887 | -0.299807849 | 0.148177145 | -0.447984994 | 0.30587961 |
| ENSG00000114395 | *CYB561D2* | 3.128862285 | 0.017013272 | -0.003361052 | 0.020374323 | 0.999933887 | -0.23875225 | -0.404293484 | 0.165541234 | 0.306084574 |
| ENSG00000272994 | *AC012360.3* | 0.383634503 | 0.143905003 | 0.172592141 | -0.028687138 | 0.999933887 | 0.619917479 | 0.239530326 | 0.380387153 | 0.306112733 |
| ENSG00000151611 | *MMAA* | 0.121301338 | -0.047238819 | -0.319333991 | 0.272095171 | 0.999933887 | -0.080931091 | 0.25272399 | -0.333655081 | 0.306138201 |
| ENSG00000251405 | *AC016571.1* | 0.629497176 | 0.035198189 | 0.056064115 | -0.020865925 | 0.999933887 | -0.566068374 | -0.223742047 | -0.342326328 | 0.306639006 |
| ENSG00000130175 | *PRKCSH* | 6.803855555 | 0.042696068 | 0.014601903 | 0.028094166 | 0.999933887 | -0.033159892 | -0.148374781 | 0.11521489 | 0.306707273 |
| ENSG00000163001 | *CFAP36* | 2.385927174 | -0.038012108 | -0.061566447 | 0.023554339 | 0.999933887 | -0.004036765 | 0.188537685 | -0.19257445 | 0.306782565 |
| ENSG00000084652 | *TXLNA* | 4.208744391 | 0.000342918 | 0.043574284 | -0.043231366 | 0.999933887 | 0.060089899 | -0.096509034 | 0.156598934 | 0.306782565 |
| ENSG00000143761 | *ARF1* | 9.120348572 | 0.018616286 | 0.005714634 | 0.012901652 | 0.999933887 | 0.15214363 | 0.242258091 | -0.090114462 | 0.30694257 |
| ENSG00000132522 | *GPS2* | 2.346244887 | 0.012038361 | 0.039760252 | -0.027721891 | 0.999933887 | 0.259196033 | 0.053539477 | 0.205656556 | 0.30694257 |
| ENSG00000177058 | *SLC38A9* | 1.162577422 | 0.116023814 | -0.002623963 | 0.118647778 | 0.999933887 | -0.207293676 | 0.0652812 | -0.272574877 | 0.307239947 |
| ENSG00000101193 | *GID8* | 5.099465444 | -0.072853542 | -0.108084115 | 0.035230573 | 0.999933887 | -0.43965668 | -0.567514534 | 0.127857854 | 0.307244598 |
| ENSG00000258890 | *CEP95* | 4.318976597 | -0.032165899 | -0.092141448 | 0.059975549 | 0.999933887 | -0.085571281 | 0.054164313 | -0.139735594 | 0.307291753 |
| ENSG00000123983 | *ACSL3* | 4.575405319 | -0.028375607 | -0.02828332 | -9.2287e-05 | 0.999933887 | -0.55178247 | -0.707489509 | 0.155707038 | 0.307291753 |
| ENSG00000121579 | *NAA50* | 6.66932333 | -0.094764234 | -0.11111627 | 0.016352036 | 0.999933887 | -0.276291925 | -0.147673206 | -0.128618719 | 0.307338154 |
| ENSG00000256223 | *ZNF10* | 1.107354247 | -0.173441253 | -0.141474898 | -0.031966355 | 0.999933887 | -0.300825058 | 0.137071514 | -0.437896572 | 0.307338154 |
| ENSG00000100442 | *FKBP3* | 1.436327078 | 0.056219846 | -0.026678973 | 0.082898819 | 0.999933887 | -0.077057405 | 0.199942365 | -0.276999769 | 0.307338154 |
| ENSG00000136828 | *RALGPS1* | 2.502081137 | 0.026924371 | 0.025005775 | 0.001918596 | 0.999933887 | -0.145005309 | -0.356511137 | 0.211505827 | 0.307520293 |
| ENSG00000237788 | *AL162615.1* | 1.525050745 | 0.135904608 | 0.051860579 | 0.084044028 | 0.999933887 | 1.243656773 | 1.55513896 | -0.311482187 | 0.307729237 |
| ENSG00000131018 | *SYNE1* | 6.849130562 | -0.062119919 | -0.020504698 | -0.041615221 | 0.999933887 | -0.076394577 | 0.087649485 | -0.164044062 | 0.307729237 |
| ENSG00000115524 | *SF3B1* | 9.367109048 | -0.070950952 | -0.132099917 | 0.061148965 | 0.999933887 | -0.184419587 | 0.00753021 | -0.191949797 | 0.307729237 |
| ENSG00000106991 | *ENG* | 1.887165334 | 0.100474724 | -0.050099732 | 0.150574456 | 0.999933887 | -0.679864449 | -0.913183607 | 0.233319158 | 0.307729237 |
| ENSG00000259330 | *INAFM2* | 7.005262551 | 0.039092015 | 0.06499629 | -0.025904275 | 0.999933887 | 0.404367209 | 0.565980864 | -0.161613655 | 0.307729237 |
| ENSG00000115425 | *PECR* | 0.690019222 | 0.008616621 | -0.102934355 | 0.111550977 | 0.999933887 | -0.063449823 | -0.424308218 | 0.360858395 | 0.307730705 |
| ENSG00000213066 | *CEP43* | 2.711726721 | -0.020236222 | -0.06860776 | 0.048371538 | 0.999933887 | -0.366891655 | -0.566617774 | 0.199726119 | 0.307731986 |
| ENSG00000231113 | *AL035587.1* | 1.860144449 | -0.119817724 | -0.077978041 | -0.041839683 | 0.999933887 | -0.482946624 | -0.789796033 | 0.306849409 | 0.307846515 |
| ENSG00000156787 | *TBC1D31* | 0.502694386 | -0.114872877 | -0.138481767 | 0.02360889 | 0.999933887 | -0.252351881 | 0.096627673 | -0.348979554 | 0.307846515 |
| ENSG00000125676 | *THOC2* | 4.05955616 | -0.080054958 | -0.145613685 | 0.065558728 | 0.999933887 | -0.377148774 | -0.03939318 | -0.337755594 | 0.308012884 |
| ENSG00000198258 | *UBL5* | 5.860487104 | 0.009213462 | -0.026272727 | 0.03548619 | 0.999933887 | 0.021844571 | -0.084118484 | 0.105963055 | 0.308077185 |
| ENSG00000077800 | *FKBP6* | 0.160763901 | 0.156149286 | -0.070613198 | 0.226762484 | 0.999933887 | 0.511137331 | 0.85651822 | -0.345380889 | 0.30827461 |
| ENSG00000147164 | *SNX12* | 4.431241421 | -0.019101139 | -0.057074825 | 0.037973686 | 0.999933887 | -0.545576949 | -0.40656078 | -0.139016169 | 0.308353459 |
| ENSG00000135913 | *USP37* | 2.141462302 | -0.007779718 | -0.017126645 | 0.009346927 | 0.999933887 | -0.202945655 | 0.037535917 | -0.240481572 | 0.308380557 |
| ENSG00000276916 | *AL442125.2* | -0.175994355 | -0.177177351 | -0.566133532 | 0.388956182 | 0.999933887 | -1.03877656 | -0.689466603 | -0.349309957 | 0.308392663 |
| ENSG00000081154 | *PCNP* | 5.147363878 | -0.026873288 | -0.140946679 | 0.114073391 | 0.999933887 | -0.142178554 | -0.00484876 | -0.137329793 | 0.308602856 |
| ENSG00000159140 | *SON* | 8.548814551 | -0.052370256 | -0.064666733 | 0.012296477 | 0.999933887 | -0.080596659 | 0.005068313 | -0.085664972 | 0.308641111 |
| ENSG00000159409 | *CELF3* | -0.169553046 | -0.042520246 | 0.226936213 | -0.26945646 | 0.999933887 | -0.364719656 | -0.040815252 | -0.323904404 | 0.308704764 |
| ENSG00000174697 | *LEP* | 0.549093674 | -0.039720429 | -0.022448804 | -0.017271625 | 0.999933887 | -0.0501095 | 0.172943114 | -0.223052615 | 0.308704764 |
| ENSG00000246596 | *AC139795.1* | -0.381623755 | -0.019556784 | 0.20280406 | -0.222360845 | 0.999933887 | -0.540122324 | -0.076533869 | -0.463588456 | 0.308774759 |
| ENSG00000116678 | *LEPR* | 0.82446939 | 0.031112634 | -0.158605481 | 0.189718114 | 0.999933887 | -0.353550137 | -0.094224773 | -0.259325364 | 0.309169204 |
| ENSG00000058799 | *YIPF1* | 3.561553404 | 0.015084475 | -0.109374344 | 0.124458819 | 0.999933887 | -0.086758659 | -0.295771208 | 0.209012549 | 0.309589548 |
| ENSG00000170946 | *DNAJC24* | 0.812221738 | -0.260890385 | 0.006866574 | -0.267756959 | 0.999933887 | -0.100106205 | 0.178668995 | -0.2787752 | 0.309958105 |
| ENSG00000163993 | *S100P* | 7.542199024 | 0.029882556 | -0.003934728 | 0.033817284 | 0.999933887 | 0.426726283 | 0.579443451 | -0.152717167 | 0.310245231 |
| ENSG00000251259 | *AC004069.1* | 1.291162419 | 0.029934679 | 0.017837356 | 0.012097324 | 0.999933887 | -0.067898194 | 0.278960125 | -0.346858319 | 0.310444697 |
| ENSG00000279159 | *AC003681.1* | 5.377343344 | 0.049923435 | 0.066182327 | -0.016258893 | 0.999933887 | 0.05110481 | 0.213108479 | -0.162003669 | 0.310511668 |
| ENSG00000279530 | *AC092881.1* | 1.50695491 | 0.090278841 | 0.01335659 | 0.076922251 | 0.999933887 | 0.136221403 | 0.624464355 | -0.488242952 | 0.310511668 |
| ENSG00000073737 | *DHRS9* | 4.080613686 | -0.085341624 | -0.07554033 | -0.009801295 | 0.999933887 | -1.908485059 | -2.173582649 | 0.265097591 | 0.310653568 |
| ENSG00000115241 | *PPM1G* | 4.960694502 | -0.030564513 | 0.022242068 | -0.052806581 | 0.999933887 | -0.259154495 | -0.383507332 | 0.124352836 | 0.311177273 |
| ENSG00000060069 | *CTDP1* | 6.083198514 | 0.008364537 | 0.019895572 | -0.011531035 | 0.999933887 | -0.354297758 | -0.490131665 | 0.135833907 | 0.311193637 |
| ENSG00000235173 | *HGH1* | 0.957240393 | 0.160817859 | -0.160080712 | 0.320898571 | 0.999933887 | 0.208370604 | -0.032167498 | 0.240538103 | 0.31154297 |
| ENSG00000254760 | *AC008750.1* | -0.455885203 | 0.017172301 | -0.183682266 | 0.200854567 | 0.999933887 | 0.964833286 | 1.358536334 | -0.393703049 | 0.311622642 |
| ENSG00000177888 | *ZBTB41* | 1.228203412 | -0.220399202 | -0.208120415 | -0.012278787 | 0.999933887 | -0.288581143 | 0.144003475 | -0.432584618 | 0.311777209 |
| ENSG00000128298 | *BAIAP2L2* | -0.66744515 | 0.218819177 | 0.111901112 | 0.106918065 | 0.999933887 | 0.32085029 | 0.747724815 | -0.426874525 | 0.311798425 |
| ENSG00000110107 | *PRPF19* | 3.672844477 | 0.027904033 | -0.017166866 | 0.045070899 | 0.999933887 | 0.156641802 | 0.0168636 | 0.139778202 | 0.311798425 |
| ENSG00000173273 | *TNKS* | 4.805486476 | -0.019164668 | -0.070383588 | 0.05121892 | 0.999933887 | -0.07397056 | 0.057050994 | -0.131021554 | 0.311798425 |
| ENSG00000162676 | *GFI1* | 3.129846514 | -0.012029652 | -0.084939908 | 0.072910256 | 0.999933887 | 0.19944382 | 0.06069719 | 0.13874663 | 0.311866112 |
| ENSG00000251301 | *LINC02384* | 1.91249618 | 0.012471212 | 0.013382112 | -0.0009109 | 0.999933887 | 0.140768413 | 0.335174 | -0.194405587 | 0.311866112 |
| ENSG00000129518 | *EAPP* | 4.758368845 | -0.058252944 | -0.112849951 | 0.054597007 | 0.999933887 | -0.38218241 | -0.233002333 | -0.149180077 | 0.311977901 |
| ENSG00000232284 | *GNG12-AS1* | 0.097287676 | 0.089314473 | -0.057029953 | 0.146344426 | 0.999933887 | -1.246172204 | -1.682232757 | 0.436060553 | 0.311980954 |
| ENSG00000257557 | *PPP1R12A-AS* | -1.044490286 | 0.212161123 | 0.038884932 | 0.173276191 | 0.999933887 | 1.850374317 | 2.292936287 | -0.442561969 | 0.311980954 |
| ENSG00000227372 | *TP73-AS1* | 2.09133804 | 0.051075118 | -0.054447854 | 0.105522972 | 0.999933887 | 0.01366042 | -0.180057806 | 0.193718227 | 0.311980954 |
| ENSG00000144130 | *NT5DC4* | 2.228686899 | -0.015983011 | 0.101809344 | -0.117792355 | 0.999933887 | -0.222798441 | -0.391918029 | 0.169119588 | 0.311980954 |
| ENSG00000100889 | *PCK2* | 2.439267763 | 0.003432607 | -0.019629498 | 0.023062105 | 0.999933887 | -0.349878408 | -0.530308152 | 0.180429744 | 0.311980954 |
| ENSG00000198945 | *L3MBTL3* | 2.814041948 | -0.026471529 | -0.136799382 | 0.110327854 | 0.999933887 | -0.241547255 | -0.453952667 | 0.212405412 | 0.311984172 |
| ENSG00000170653 | *ATF7* | 2.57823457 | 0.090755162 | 0.022151102 | 0.06860406 | 0.999933887 | -0.273931875 | -0.455671284 | 0.181739409 | 0.312116754 |
| ENSG00000196313 | *POM121* | 5.124712745 | 0.041426782 | 0.069549341 | -0.028122559 | 0.999933887 | 0.303290806 | 0.436554788 | -0.133263983 | 0.312239711 |
| ENSG00000147459 | *DOCK5* | 9.048850269 | -0.013255508 | -0.034304568 | 0.02104906 | 0.999933887 | -0.442351862 | -0.563222896 | 0.120871034 | 0.312239711 |
| ENSG00000160961 | *ZNF333* | 5.280986404 | -0.037820542 | -0.001182376 | -0.036638166 | 0.999933887 | -0.15867918 | -0.05891901 | -0.09976017 | 0.31224018 |
| ENSG00000087088 | *BAX* | 6.205475265 | 0.020330051 | 0.07730465 | -0.0569746 | 0.999933887 | -0.092978165 | -0.196106573 | 0.103128407 | 0.312484924 |
| ENSG00000134910 | *STT3A* | 4.239798083 | -0.019407734 | -0.011978656 | -0.007429078 | 0.999933887 | 0.085712913 | 0.205708312 | -0.119995398 | 0.312771408 |
| ENSG00000140521 | *POLG* | 5.829067756 | 0.030287131 | 0.068335887 | -0.038048756 | 0.999933887 | 0.164412873 | 0.263149739 | -0.098736866 | 0.312957702 |
| ENSG00000101278 | *RPS10P5* | 0.39715961 | -0.215706931 | -0.080208832 | -0.135498099 | 0.999933887 | 0.579938509 | 0.88491379 | -0.304975281 | 0.313030214 |
| ENSG00000186866 | *POFUT2* | 6.293069613 | 0.028584043 | 0.00488577 | 0.023698273 | 0.999933887 | 0.013771293 | -0.092615669 | 0.106386962 | 0.313030214 |
| ENSG00000222389 | *RNU2-28P* | -1.801022039 | 0.078990957 | 0.137055685 | -0.058064728 | 0.999933887 | 1.0849111 | 1.622689247 | -0.537778148 | 0.313030214 |
| ENSG00000126254 | *RBM42* | 5.493023705 | 0.050187707 | 0.023029346 | 0.027158361 | 0.999933887 | 0.231768717 | 0.357197841 | -0.125429125 | 0.313191136 |
| ENSG00000228486 | *C2orf92* | 0.169517226 | 0.018819047 | 0.296644002 | -0.277824955 | 0.999933887 | 0.911209575 | 1.239587423 | -0.328377848 | 0.313191136 |
| ENSG00000138081 | *FBXO11* | 5.545937931 | -0.068638721 | -0.049204466 | -0.019434255 | 0.999933887 | -0.180034353 | -0.06581612 | -0.114218233 | 0.313360992 |
| ENSG00000143324 | *XPR1* | 4.140348866 | -0.043071937 | -0.003968983 | -0.039102954 | 0.999933887 | -0.045547427 | 0.104029266 | -0.149576693 | 0.313360992 |
| ENSG00000068971 | *PPP2R5B* | 3.076161509 | 0.038178563 | 0.010866755 | 0.027311808 | 0.999933887 | 0.007298325 | -0.189819223 | 0.197117548 | 0.313360992 |
| ENSG00000204257 | *HLA-DMA* | 3.150404487 | 0.035190823 | 0.046506901 | -0.011316078 | 0.999933887 | -0.055606881 | -0.265026754 | 0.209419873 | 0.313360992 |
| ENSG00000106066 | *CPVL* | 0.349806768 | -0.052075838 | -0.081731165 | 0.029655327 | 0.999933887 | 0.051573954 | -0.46554337 | 0.517117324 | 0.313493662 |
| ENSG00000163312 | *HELQ* | 3.545291612 | -0.045438225 | -0.041783805 | -0.003654419 | 0.999933887 | 0.106183909 | 0.246313376 | -0.140129467 | 0.313707922 |
| ENSG00000139946 | *PELI2* | 6.173464434 | -0.093309815 | -0.012167438 | -0.081142377 | 0.999933887 | -0.819190161 | -0.974744104 | 0.155553943 | 0.314063353 |
| ENSG00000151131 | *C12orf45* | 0.35521721 | -0.185940782 | 0.073928512 | -0.259869294 | 0.999933887 | -0.294698681 | 0.031143752 | -0.325842433 | 0.314063353 |
| ENSG00000128203 | *ASPHD2* | 1.75775837 | 0.032288219 | -0.10500746 | 0.137295679 | 0.999933887 | -0.36111131 | -0.630618754 | 0.269507443 | 0.314063353 |
| ENSG00000135821 | *GLUL* | 12.329701529 | 0.058900992 | 0.053039173 | 0.005861819 | 0.999933887 | -0.519119779 | -0.390031047 | -0.129088732 | 0.314073325 |
| ENSG00000107789 | *MINPP1* | 0.021978669 | -0.086095657 | -0.063700525 | -0.022395132 | 0.999933887 | -0.01552326 | -0.36941065 | 0.35388739 | 0.314073325 |
| ENSG00000226419 | *SLC16A1-AS1* | 0.012850316 | -0.145332867 | -0.047264903 | -0.098067964 | 0.999933887 | -0.168445011 | 0.246679649 | -0.41512466 | 0.314791627 |
| ENSG00000140961 | *OSGIN1* | 5.232075232 | 0.098195029 | 0.00239486 | 0.095800169 | 0.999933887 | 0.372694657 | 0.546516899 | -0.173822241 | 0.314922028 |
| ENSG00000273899 | *NOL12* | 2.068090282 | -0.092068779 | -0.04660765 | -0.04546113 | 0.999933887 | -0.071855119 | -0.248289535 | 0.176434416 | 0.314922028 |
| ENSG00000100219 | *XBP1* | 5.202833331 | 0.014796002 | 0.073355211 | -0.05855921 | 0.999933887 | 1.387549779 | 1.659090153 | -0.271540373 | 0.315005272 |
| ENSG00000127530 | *OR7C1* | -0.533981898 | -0.301493618 | -0.035265211 | -0.266228406 | 0.999933887 | -0.85068449 | -0.299265318 | -0.551419172 | 0.315166309 |
| ENSG00000174130 | *TLR6* | 5.960527456 | -0.094863833 | -0.108931511 | 0.014067678 | 0.999933887 | -1.458260221 | -1.687946372 | 0.229686151 | 0.315440179 |
| ENSG00000246089 | *AC016065.1* | 1.18373491 | 0.010065256 | 0.470724452 | -0.460659196 | 0.944094537 | 0.606788533 | 0.334986262 | 0.271802271 | 0.315513034 |

| | | | | | | | | | | |
|---|---|---|---|---|---|---|---|---|---|---|
| ENSG00000197614 | *MFAP5* | -2.531273955 | 0.45167895 | 0.221284131 | 0.230394819 | 0.999933887 | 1.887870258 | 2.428091473 | -0.540221216 | 0.315550802 |
| ENSG00000256060 | *TRAPPC2B* | 0.893416259 | 0.006049363 | 0.223416138 | -0.217366776 | 0.999933887 | 0.320668367 | 0.666825583 | -0.346157216 | 0.315550802 |
| ENSG00000130202 | *NECTIN2* | 2.994084766 | -0.061957528 | -0.059986924 | -0.001970604 | 0.999933887 | 0.84209734 | 1.150666588 | -0.308569248 | 0.315572544 |
| ENSG00000102755 | *FLT1* | 3.84496603 | 0.035794004 | -0.005257617 | 0.04105162 | 0.999933887 | 1.00263634 | 1.240115818 | -0.237479478 | 0.315572544 |
| ENSG00000287586 | *Z98886.1* | -0.960807839 | 0.186893901 | 0.057483039 | 0.129410862 | 0.999933887 | -0.162838987 | -0.653093634 | 0.490254647 | 0.315779991 |
| ENSG00000276842 | *AC023510.2* | 2.219098589 | -0.071066552 | -0.11089133 | 0.039824778 | 0.999933887 | -0.685900778 | -1.110559419 | 0.424658641 | 0.315831381 |
| ENSG00000109321 | *AREG* | 2.480506121 | 0.140583578 | 0.097568925 | 0.043014653 | 0.999933887 | -0.077572977 | -0.398926085 | 0.321353108 | 0.316065705 |
| ENSG00000185716 | *MOSMO* | 1.821184226 | -0.13436881 | -0.048338011 | -0.086030799 | 0.999933887 | -0.589104231 | -0.395840068 | -0.193264163 | 0.316065705 |
| ENSG00000226091 | *LINC00937* | 1.776595121 | 0.071610952 | -0.104175794 | 0.175786746 | 0.999933887 | 1.150675511 | 1.452156422 | -0.301480912 | 0.316113324 |
| ENSG00000076554 | *TPD52* | 0.495997839 | -0.138438749 | 0.058109005 | -0.196547754 | 0.999933887 | -0.072588865 | 0.263135561 | -0.335724426 | 0.316156598 |
| ENSG00000142330 | *CAPN10* | 3.905131582 | 0.059719053 | 0.127216547 | -0.067497494 | 0.999933887 | -0.04630784 | -0.186460559 | 0.14015272 | 0.316156598 |
| ENSG00000183258 | *DDX41* | 5.32767766 | -0.022562558 | -0.021430116 | -0.001132442 | 0.999933887 | -0.27333029 | -0.385069947 | 0.111739657 | 0.316156598 |
| ENSG00000188878 | *FBF1* | 0.281006165 | -0.056588277 | 0.135719533 | -0.192307809 | 0.999933887 | 0.132930339 | -0.167547152 | 0.300477492 | 0.316156598 |
| ENSG00000177189 | *RPS6KA3* | 6.237008822 | -0.028529832 | -0.090209845 | 0.061680014 | 0.999933887 | -0.008353852 | 0.168384985 | -0.176738837 | 0.316156598 |
| ENSG00000233913 | *RPL10P9* | 4.003630998 | -0.020745083 | 0.060347072 | -0.081092155 | 0.999933887 | 0.203795382 | 0.351983515 | -0.148188133 | 0.316156598 |
| ENSG00000113658 | *SMAD5* | 2.049347664 | 0.055134932 | -0.012782724 | 0.067917656 | 0.999933887 | -0.124489246 | 0.069846399 | -0.194335645 | 0.316188355 |
| ENSG00000102898 | *NUTF2* | 3.018569163 | 0.034643372 | -0.009393905 | 0.044037277 | 0.999933887 | -0.021520316 | -0.186879406 | 0.165359091 | 0.316188355 |
| ENSG00000215493 | *AC007731.1* | -1.01702913 | -0.217680745 | 0.255399649 | -0.473080394 | 0.999933887 | -0.037802604 | -0.430021778 | 0.392219174 | 0.316205877 |
| ENSG00000136840 | *ST6GALNAC4* | 1.543747113 | 0.025843821 | -0.099638499 | 0.12548232 | 0.999933887 | -0.125605599 | -0.337455444 | 0.211849845 | 0.316205877 |
| ENSG00000264757 | *MIR3198-1* | 0.585288127 | -0.034837966 | 0.082991599 | -0.117829565 | 0.999933887 | 1.573335993 | 2.003661236 | -0.430325243 | 0.316205877 |
| ENSG00000100147 | *CCDC134* | 1.469595584 | 0.018342968 | 0.003491228 | 0.01485174 | 0.999933887 | 0.078062831 | 0.382021014 | -0.303958183 | 0.316205877 |
| ENSG00000106462 | *EZH2* | 2.806555543 | -0.013260537 | -0.004498836 | -0.008761701 | 0.999933887 | 0.6203832 | 0.861193953 | -0.240810753 | 0.316205877 |
| ENSG00000168282 | *MGAT2* | -0.375244806 | -0.07182599 | 0.205482843 | -0.277308834 | 0.999933887 | 0.011802332 | -0.478687347 | 0.490489679 | 0.316223992 |
| ENSG00000268403 | *AC132192.2* | 1.175899228 | 0.164531675 | 0.034789917 | 0.129741758 | 0.999933887 | 0.665740968 | 0.388620784 | 0.277120184 | 0.31629929 |
| ENSG00000090432 | *MUL1* | 4.846969661 | -0.030194846 | -0.021006194 | -0.009188653 | 0.999933887 | -0.129757213 | -0.007891659 | -0.121865553 | 0.316905311 |
| ENSG00000029725 | *RABEP1* | 5.01386457 | -0.101395431 | -0.068730601 | -0.03266483 | 0.999933887 | -0.20168234 | -0.073874651 | -0.127807689 | 0.317017701 |
| ENSG00000228686 | *AL590723.1* | 1.082165657 | -0.082233108 | 0.053671223 | -0.135904332 | 0.999933887 | -0.633653112 | -1.052117466 | 0.418464354 | 0.317017701 |
| ENSG00000067840 | *PDZD4* | 3.271059213 | 0.023611264 | -0.008278507 | 0.031889771 | 0.999933887 | 0.197919204 | 0.002026209 | 0.195892995 | 0.317017701 |
| ENSG00000013588 | *GPRC5A* | -1.199766287 | -0.095125877 | -0.100965553 | 0.005839676 | 0.999933887 | -0.247351036 | 0.196055326 | -0.443406362 | 0.317022265 |
| ENSG00000254986 | *DPP3* | 2.283184117 | 0.062779734 | 0.017881474 | 0.04489826 | 0.999933887 | -0.010172505 | 0.179040858 | -0.189213363 | 0.317167363 |
| ENSG00000223478 | *AL441992.1* | 0.061869542 | 0.106509323 | 0.081991927 | 0.024517395 | 0.999933887 | 0.048562306 | -0.328648887 | 0.377211193 | 0.317205763 |
| ENSG00000116786 | *PLEKHM2* | 8.438795292 | 0.024353874 | 0.030385795 | -0.006031921 | 0.999933887 | 0.303357459 | 0.411067143 | -0.107709684 | 0.317205763 |
| ENSG00000181652 | *ATG9B* | -0.436566434 | 0.020422538 | -0.157635745 | 0.178058283 | 0.999933887 | -0.449731487 | -0.053887683 | -0.395843804 | 0.317205763 |
| ENSG00000134453 | *RBM17* | 4.827094474 | -0.032012886 | 0.007990437 | -0.040003322 | 0.999933887 | 0.324312468 | 0.448422987 | -0.124110519 | 0.31743379 |
| ENSG00000143630 | *HCN3* | -0.568637064 | -0.034746067 | -0.189686482 | 0.154940415 | 0.999933887 | -0.059180968 | 0.3499049 | -0.409085868 | 0.317455784 |
| ENSG00000163104 | *SMARCAD1* | 1.792722543 | -0.008076391 | -0.193758977 | 0.185682585 | 0.999933887 | -0.267081228 | 0.031074552 | -0.29815578 | 0.318093067 |
| ENSG00000122678 | *POLM* | 4.750514404 | 0.030848354 | 0.021630205 | 0.009218149 | 0.999933887 | -0.012728893 | -0.125746662 | 0.113017769 | 0.318200868 |
| ENSG00000185697 | *MYBL1* | 3.46860944 | -0.053311245 | -0.11802412 | 0.064712875 | 0.999933887 | -0.204695077 | 0.072552349 | -0.277247426 | 0.318229926 |
| ENSG00000185897 | *FFAR3* | 0.341051862 | 0.1194629 | 0.057977947 | 0.061484953 | 0.999933887 | 0.691469715 | 1.124002958 | -0.432533243 | 0.318241158 |
| ENSG00000162377 | *COA7* | 1.621146103 | 0.016258048 | -0.009906044 | 0.026164093 | 0.999933887 | 0.01029768 | 0.233375356 | -0.223077676 | 0.318567601 |
| ENSG00000078053 | *AMPH* | 1.185701051 | -0.095465691 | 0.167230768 | -0.26269646 | 0.999933887 | 1.081078891 | 1.367330001 | -0.28625111 | 0.318759485 |
| ENSG00000181744 | *DIPK2A* | 3.134058802 | 0.053559571 | 0.066168187 | -0.012608616 | 0.999933887 | -0.465203825 | -0.646633402 | 0.181429577 | 0.318759485 |
| ENSG00000020922 | *MRE11* | 3.06868973 | 0.013620547 | -0.113834225 | 0.127454772 | 0.999933887 | -0.291562846 | -0.039414741 | -0.252148106 | 0.318759485 |
| ENSG00000132199 | *ENOSF1* | 3.030429243 | -0.051813347 | -0.032113288 | -0.019700059 | 0.999933887 | 0.03469413 | 0.17848536 | -0.14379123 | 0.319088173 |
| ENSG00000145833 | *DDX46* | 4.619552216 | -0.001304432 | -0.183732935 | 0.182428502 | 0.999933887 | -0.234369926 | -0.414732036 | 0.180362109 | 0.319242184 |
| ENSG00000167186 | *COQ7* | 2.942590032 | 0.097156195 | 0.032732589 | 0.064423606 | 0.999933887 | 0.332449473 | 0.140192465 | 0.192257008 | 0.319334151 |
| ENSG00000159147 | *DONSON* | 1.768806839 | -0.002530949 | 0.044229353 | -0.046760301 | 0.999933887 | -0.01497624 | 0.24205933 | -0.25703557 | 0.319419905 |
| ENSG00000229873 | *OGFR-AS1* | 0.840389528 | 0.321376234 | 0.257432666 | 0.063943568 | 0.999933887 | 0.455625398 | 0.716935766 | -0.261310368 | 0.320068416 |
| ENSG00000138639 | *ARHGAP24* | 1.393098365 | 0.035955263 | -0.051120128 | 0.087075391 | 0.999933887 | 0.425022775 | 0.75428356 | -0.329260784 | 0.320123854 |
| ENSG00000015532 | *XYLT2* | 2.590780884 | -0.00837841 | -0.018593849 | 0.010215439 | 0.999933887 | -0.016531027 | -0.199414629 | 0.182883602 | 0.320190447 |
| ENSG00000112667 | *DNPH1* | 1.721086764 | 0.129122016 | 0.067768123 | 0.061353894 | 0.999933887 | 0.386024019 | 0.589485062 | -0.203461043 | 0.320316312 |
| ENSG00000285280 | *AL390957.1* | 1.604308981 | -0.053101294 | 0.15090771 | -0.204009004 | 0.999933887 | -0.236776589 | -0.584737966 | 0.347961377 | 0.320316312 |
| ENSG00000130304 | *SLC27A1* | 0.4730001 | -0.025930818 | 0.318297865 | -0.344228684 | 0.999933887 | 0.016347701 | -0.297042553 | 0.313390254 | 0.320316312 |
| ENSG00000137309 | *HMGA1* | 7.262404805 | 0.073506697 | 0.009481766 | 0.06402493 | 0.999933887 | -0.027205561 | 0.086286892 | -0.113492454 | 0.320341169 |
| ENSG00000234684 | *SDCBP2-AS1* | 1.326569086 | -0.084046231 | 0.014400322 | -0.098446552 | 0.999933887 | -0.114356611 | -0.376091709 | 0.261735098 | 0.320341169 |
| ENSG00000246250 | *AC087521.2* | -0.129366152 | -0.113149787 | -0.28520043 | 0.172050643 | 0.999933887 | 0.896150817 | 1.50084013 | -0.604689313 | 0.320341169 |
| ENSG00000116691 | *MIIP* | 3.954969578 | 0.091326068 | 0.081044308 | 0.01028176 | 0.999933887 | 0.219905849 | 0.057769008 | 0.162136841 | 0.32057648 |
| ENSG00000156876 | *SASS6* | 0.839134312 | 0.140934286 | -0.13596087 | 0.276895157 | 0.999933887 | -0.565825265 | -0.152742349 | -0.413082917 | 0.32057648 |
| ENSG00000166046 | *TCP11L2* | 5.789160153 | -0.078955695 | -0.026303924 | -0.052651771 | 0.999933887 | -0.443475556 | -0.552773626 | 0.10929807 | 0.321017956 |
| ENSG00000114120 | *SLC25A36* | 4.322755349 | -0.084216479 | -0.14440987 | 0.060193391 | 0.999933887 | -0.25477767 | 0.01718194 | -0.27195961 | 0.321017956 |
| ENSG00000070785 | *EIF2B3* | 0.171390491 | -0.12198664 | -0.008307807 | -0.113678834 | 0.999933887 | -0.079262903 | 0.278100509 | -0.357363413 | 0.321017956 |
| ENSG00000182899 | *RPL35A* | 6.935212694 | -0.001225001 | -0.033557039 | 0.032332038 | 0.999933887 | 0.102489924 | 0.223025486 | -0.120535562 | 0.321017956 |
| ENSG00000242244 | *ATP5MC1P3* | 0.470163226 | 0.4555511 | 0.476527276 | -0.020976176 | 0.999933887 | 1.051778485 | 1.587393574 | -0.535615089 | 0.321077014 |
| ENSG00000166881 | *NEMP1* | 3.882219585 | -0.076696533 | -0.001131943 | -0.07556459 | 0.999933887 | -0.411944924 | -0.291170247 | -0.120774677 | 0.321130712 |
| ENSG00000112299 | *VNN1* | 5.380943966 | -0.07081812 | -0.048397517 | -0.022420602 | 0.999933887 | -0.315531099 | -0.139405645 | -0.176125454 | 0.321130712 |
| ENSG00000286122 | *AC016074.2* | -0.017459404 | -0.109587608 | 0.381740104 | -0.491327712 | 0.999933887 | -0.265336622 | 0.140380179 | -0.405716801 | 0.321130712 |
| ENSG00000176209 | *SMIM19* | 1.859241559 | -0.023429706 | -0.087900392 | 0.064470686 | 0.999933887 | 0.149770835 | 0.349056434 | -0.199285599 | 0.321130712 |
| ENSG00000159593 | *NAE1* | 2.368766339 | -0.017094925 | -0.079325701 | 0.062230775 | 0.999933887 | -0.068028308 | 0.113242672 | -0.181270981 | 0.32119649 |
| ENSG00000125149 | *C16orf70* | 3.405066141 | -0.06126666 | -0.076792207 | 0.015525547 | 0.999933887 | -0.798704699 | -0.983945101 | 0.185240402 | 0.321339327 |
| ENSG00000143819 | *EPHX1* | 2.279781481 | 0.066233584 | 0.092761759 | -0.026528175 | 0.999933887 | 0.035547487 | -0.149435568 | 0.184983054 | 0.321469242 |
| ENSG00000185278 | *ZBTB37* | 5.415973502 | -0.004626042 | -0.108804772 | 0.104178729 | 0.999933887 | -0.59034781 | -0.329929101 | -0.260418709 | 0.321469242 |
| ENSG00000168538 | *TRAPPC11* | 3.61984693 | -0.062688083 | -0.080385383 | 0.0176973 | 0.999933887 | -0.17201934 | -0.331714815 | 0.159695475 | 0.321528884 |
| ENSG00000109390 | *NDUFC1* | 2.213288745 | -0.023647757 | -0.101710953 | 0.078063196 | 0.999933887 | -0.052099703 | -0.244041045 | 0.191941342 | 0.321528884 |
| ENSG00000213316 | *LTC4S* | 1.419442019 | 0.02177887 | 0.078670034 | -0.056891164 | 0.999933887 | -0.080964803 | -0.312980385 | 0.232015582 | 0.321528884 |
| ENSG00000159247 | *TUBBP5* | 0.118858898 | 0.000132537 | 0.159597614 | -0.159465077 | 0.999933887 | -1.292194742 | -1.619664338 | 0.327469596 | 0.321528884 |
| ENSG00000273253 | *AL022328.4* | 2.40222602 | 0.104517987 | -0.017406863 | 0.12192485 | 0.999933887 | -0.296449906 | -0.486129607 | 0.189679701 | 0.32231521 |
| ENSG00000118242 | *MREG* | 0.289280248 | -0.136918151 | -0.075156315 | -0.061761836 | 0.999933887 | 0.455105436 | 0.7759927 | -0.320887264 | 0.322660399 |
| ENSG00000112062 | *MAPK14* | 6.818665062 | -0.015573465 | -0.024910652 | 0.009337186 | 0.999933887 | -0.33401568 | -0.453858035 | 0.119842355 | 0.323279582 |
| ENSG00000164715 | *LMTK2* | 6.044285205 | -0.059252457 | -0.038755345 | -0.020497112 | 0.999933887 | -0.339017286 | -0.445745814 | 0.106728528 | 0.323289144 |
| ENSG00000213373 | *LINC00671* | 0.278954297 | 0.024520681 | 0.089822328 | -0.065301648 | 0.999933887 | -0.463362217 | -0.948005734 | 0.484643518 | 0.323289144 |
| ENSG00000049618 | *ARID1B* | 5.260100077 | 0.058776733 | 0.015349255 | 0.043427478 | 0.999933887 | -0.011007536 | -0.115919144 | 0.104911608 | 0.323350149 |
| ENSG00000101361 | *NOP56* | 4.50838189 | -0.013379461 | 0.033550061 | -0.046929522 | 0.999933887 | 0.189024204 | 0.064644829 | 0.124379375 | 0.323378096 |
| ENSG00000119397 | *CNTRL* | 4.564812054 | -0.090919348 | -0.125395311 | 0.034475963 | 0.999933887 | -0.362324383 | -0.048541817 | -0.313782566 | 0.323399599 |
| ENSG00000125870 | *SNRPB2* | 3.60990147 | -0.065664488 | -0.074948158 | 0.00928367 | 0.999933887 | 0.132437708 | 0.26052358 | -0.128085872 | 0.323616652 |
| ENSG00000107130 | *NCS1* | -0.643266119 | 0.054520817 | -0.459026848 | 0.513547666 | 0.999933887 | -0.152879574 | -0.594729397 | 0.441849823 | 0.323616652 |
| ENSG00000212176 | *RNA5SP207* | 1.260514966 | 0.14373361 | 0.213218028 | -0.069484418 | 0.999933887 | 0.48121928 | 0.860112148 | -0.378892868 | 0.323849544 |
| ENSG00000103024 | *NME3* | 2.626580002 | 0.145168604 | 0.002862344 | 0.14230626 | 0.999933887 | 0.25507087 | 0.039427432 | 0.215643438 | 0.324028357 |
| ENSG00000286974 | *AL662884.5* | 1.327300235 | 0.086624008 | 0.091935841 | -0.005311833 | 0.999933887 | -0.103233991 | -0.441389868 | 0.338155877 | 0.324302843 |
| ENSG00000109906 | *ZBTB16* | 1.338424449 | 0.071266721 | 0.085791277 | -0.014524556 | 0.999933887 | -0.386909012 | -0.661900496 | 0.274991484 | 0.324302843 |
| ENSG00000064547 | *LPAR2* | 6.749247419 | 0.046358728 | 0.039403555 | 0.006955173 | 0.999933887 | -0.152834915 | -0.253683218 | 0.100848303 | 0.324345989 |
| ENSG00000073921 | *PICALM* | 8.605419223 | 0.014372038 | 0.00784498 | 0.006527059 | 0.999933887 | -0.30172911 | -0.171840975 | -0.129888135 | 0.324569808 |
| ENSG00000091732 | *ZC3HC1* | 1.787541095 | 0.00379838 | -0.287153927 | 0.290952307 | 0.999933887 | 0.145893254 | 0.31712001 | -0.171226756 | 0.324569808 |
| ENSG00000171823 | *FBXL14* | 2.947175781 | 0.184355371 | -0.005353793 | 0.189709164 | 0.999933887 | 0.174187415 | 0.417299073 | -0.243111658 | 0.324588255 |
| ENSG00000160299 | *PCNT* | 4.393851639 | -0.041843184 | 0.022493416 | -0.0643366 | 0.999933887 | 0.056292925 | 0.182813605 | -0.12652068 | 0.324683501 |
| ENSG00000160999 | *SH2B2* | 6.644102117 | 0.030503595 | -0.026548655 | 0.057052251 | 0.999933887 | 0.654474805 | 0.795853208 | -0.141378403 | 0.324683501 |
| ENSG00000288596 | *C8orf44* | 0.235659949 | -0.049960867 | 0.11437881 | -0.164339677 | 0.999933887 | -0.46907504 | -0.147548558 | -0.321526482 | 0.324683501 |
| ENSG00000131148 | *EMC8* | 2.669937797 | -0.0146378 | 0.059383269 | -0.074021069 | 0.999933887 | -0.136265575 | -0.299205231 | 0.162939656 | 0.324890582 |
| ENSG00000130818 | *ZNF426* | 2.540393902 | -0.005852177 | 0.032233356 | -0.038085534 | 0.999933887 | 0.053474779 | 0.237011328 | -0.183536549 | 0.325242621 |
| ENSG00000163492 | *CCDC141* | -0.384386241 | -0.008287615 | 0.453673123 | -0.461960738 | 0.999933887 | -0.388385177 | 0.174302922 | -0.562688099 | 0.325560066 |
| ENSG00000167657 | *DAPK3* | 5.172863057 | 0.060644981 | 0.070165712 | -0.009520731 | 0.999933887 | -0.292153939 | -0.461663693 | 0.169509754 | 0.325586722 |
| ENSG00000023902 | *PLEKHO1* | 6.470145705 | 0.000795257 | 0.023018015 | -0.022222758 | 0.999933887 | -0.079508916 | -0.262867184 | 0.183358268 | 0.325586722 |
| ENSG00000100592 | *DAAM1* | 1.342170834 | -0.126605285 | -0.007154391 | -0.119450894 | 0.999933887 | -0.241241215 | 0.027528383 | -0.268769598 | 0.325625014 |
| ENSG00000117602 | *RCAN3* | 3.667111219 | -0.012436933 | 0.082682704 | -0.095119638 | 0.999933887 | 0.086492359 | -0.030503195 | 0.116995554 | 0.325656682 |
| ENSG00000272086 | *AC025181.2* | -0.322528396 | -0.067771031 | -0.302816786 | 0.235045756 | 0.999933887 | -0.288685012 | -0.760684959 | 0.471999947 | 0.325810121 |
| ENSG00000220378 | *KRT8P42* | 3.375161882 | 0.167662648 | 0.092347637 | 0.075315011 | 0.999933887 | 1.164363045 | 1.431888635 | -0.267525591 | 0.326000169 |

| | | | | | | | | | | |
|---|---|---|---|---|---|---|---|---|---|---|
| ENSG00000086666 | *ZFAND6* | 6.414275374 | -0.068349342 | -0.076903733 | 0.008554391 | 0.999933887 | -0.207869145 | -0.109585552 | -0.098283593 | 0.326000169 |
| ENSG00000215252 | *GOLGA8B* | 3.944716283 | -0.05476358 | -0.047618727 | -0.007144853 | 0.999933887 | -0.045585388 | 0.112092763 | -0.157678151 | 0.326000169 |
| ENSG00000177051 | *FBXO46* | 4.531706561 | 0.03122424 | 0.047864455 | -0.016640215 | 0.999933887 | 0.357777288 | 0.474956609 | -0.117179321 | 0.326000169 |
| ENSG00000272768 | *AC004854.2* | 1.040485724 | 0.18533914 | 0.294285609 | -0.108946469 | 0.999933887 | 1.327933442 | 0.934200372 | 0.393733069 | 0.326008705 |
| ENSG00000233214 | *AC002511.2* | 0.743522741 | -0.122462289 | -0.00879024 | -0.113672049 | 0.999933887 | 2.10967082 | 2.508495557 | -0.398824737 | 0.326151457 |
| ENSG00000125037 | *EMC3* | 5.694896567 | -0.040184979 | -0.031651144 | -0.008533835 | 0.999933887 | -0.616049974 | -0.46610908 | -0.149940894 | 0.326151457 |
| ENSG00000213995 | *NAXD* | 3.096913304 | -0.001705899 | 0.038309586 | -0.040015485 | 0.999933887 | 0.318743723 | 0.475999267 | -0.157255544 | 0.326151457 |
| ENSG00000174749 | *FAM241A* | 1.969902787 | 0.00300896 | 0.1360456 | -0.13303664 | 0.999933887 | 0.005659441 | 0.209552273 | -0.203892832 | 0.326153621 |
| ENSG00000147955 | *SIGMAR1* | 2.58637279 | 0.097993904 | -0.038991691 | 0.136985595 | 0.999933887 | 0.010431197 | -0.182348453 | 0.19277965 | 0.326196313 |
| ENSG00000106635 | *BCL7B* | 5.534462122 | 0.046343345 | 0.028735691 | 0.017607654 | 0.999933887 | 0.356331373 | 0.475631016 | -0.119299643 | 0.326201209 |
| ENSG00000101464 | *PIGU* | 1.349514124 | -0.018914786 | 0.10268128 | -0.121596066 | 0.999933887 | 0.258425294 | 0.498058183 | -0.23963289 | 0.326330433 |
| ENSG00000159459 | *UBR1* | 4.134332112 | 0.031196755 | -0.109043461 | 0.140240217 | 0.999933887 | -0.023350584 | 0.174975525 | -0.198326109 | 0.32635824 |
| ENSG00000175054 | *ATR* | 3.331212195 | -0.035160875 | -0.07365392 | 0.038493045 | 0.999933887 | -0.215144755 | 0.091113523 | -0.306258279 | 0.32635971 |
| ENSG00000197937 | *ZNF347* | 0.150419608 | -0.197340626 | 0.052756447 | -0.250097073 | 0.999933887 | -0.231885199 | -0.532409124 | 0.300523925 | 0.326360802 |
| ENSG00000243716 | *NPIPB5* | 3.508951875 | -0.006643569 | 0.004011819 | -0.010655388 | 0.999933887 | 0.091697265 | 0.254358092 | -0.162660827 | 0.326360802 |
| ENSG00000205786 | *LINC01531* | -0.067205137 | -0.081604369 | -0.112978526 | 0.031374157 | 0.999933887 | 1.247896303 | 1.821657267 | -0.573760963 | 0.326595396 |
| ENSG00000086544 | *ITPKC* | 4.668808193 | 0.067281508 | 0.024142737 | 0.043138771 | 0.999933887 | 0.766145088 | 0.903458288 | -0.1373132 | 0.326662086 |
| ENSG00000165792 | *METTL17* | 2.590535626 | 0.039765608 | 0.059287159 | -0.019521552 | 0.999933887 | 0.119038116 | -0.062732516 | 0.181770633 | 0.326662086 |
| ENSG00000129255 | *MPDU1* | 3.529494834 | 0.107218533 | 0.124006863 | -0.01678833 | 0.999933887 | 0.495785235 | 0.678907598 | -0.183122363 | 0.327082789 |
| ENSG00000134874 | *DZIP1* | -0.332857878 | -0.1358559 | 0.088989453 | -0.224845353 | 0.999933887 | 0.964715794 | 1.196949433 | -0.232233638 | 0.327096352 |
| ENSG00000244134 | *RPS12P20* | 1.723726924 | -0.011608084 | -0.14692705 | 0.135318967 | 0.999933887 | 0.027290575 | 0.326376496 | -0.299085921 | 0.327112185 |
| ENSG00000267523 | *AC008735.2* | 1.112379431 | 0.162400266 | -0.018501371 | 0.180901638 | 0.999933887 | -0.4073164 | -0.654632514 | 0.247316114 | 0.327442707 |
| ENSG00000162496 | *DHRS3* | 1.848360274 | -0.098488813 | 0.034948181 | -0.133436994 | 0.999933887 | 0.264434256 | 0.455420465 | -0.190986209 | 0.327480333 |
| ENSG00000180035 | *ZNF48* | 0.40379677 | 0.123316454 | 0.046182901 | 0.077133553 | 0.999933887 | 0.172388935 | -0.101309083 | 0.273698018 | 0.327480333 |
| ENSG00000122877 | *EGR2* | 7.617477208 | 0.288349865 | 0.26118325 | 0.027166615 | 0.999933887 | 0.321429183 | 0.495920018 | -0.174490835 | 0.327527987 |
| ENSG00000250116 | *RHOQ-AS1* | 2.556392339 | 0.008662117 | 0.083045402 | -0.074383284 | 0.999933887 | 0.377409378 | 0.610924993 | -0.233515614 | 0.327527987 |
| ENSG00000125447 | *GGA3* | 6.051841058 | -0.03649329 | 0.05461188 | -0.09110517 | 0.999933887 | -0.197116958 | -0.277568917 | 0.080451959 | 0.327812421 |
| ENSG00000111647 | *UHRF1BP1L* | 5.737186018 | -0.040256812 | -0.113974512 | 0.073717699 | 0.999933887 | -0.595757007 | -0.402471853 | -0.193285154 | 0.327827494 |
| ENSG00000167792 | *NDUFV1* | 4.51733889 | 0.025414527 | 0.00407619 | 0.021338337 | 0.999933887 | 0.147722195 | 0.002642638 | 0.145079558 | 0.327964052 |
| ENSG00000166676 | *TVP23A* | -0.571712128 | 0.09806023 | -0.009164373 | 0.107224603 | 0.999933887 | -0.212667776 | 0.225859781 | -0.438527558 | 0.327969975 |
| ENSG00000120896 | *SORBS3* | 2.356590021 | 0.004976837 | 0.085385574 | -0.080408737 | 0.999933887 | 0.099622034 | 0.274296483 | -0.174674449 | 0.327969975 |
| ENSG00000142207 | *URB1* | 2.47969446 | 0.047717991 | -0.002524416 | 0.050242407 | 0.999933887 | 0.227987686 | 0.406536366 | -0.17854868 | 0.328052327 |
| ENSG00000160695 | *VPS11* | 5.329431364 | 0.004196278 | -0.043647619 | 0.047843897 | 0.999933887 | -0.009989634 | -0.127837668 | 0.117848034 | 0.328060635 |
| ENSG00000075391 | *RASAL2* | -0.73540456 | 0.035757013 | 0.219182891 | -0.183425878 | 0.999933887 | -0.919816443 | -0.492871254 | -0.426945189 | 0.32815189 |
| ENSG00000169519 | *METTL15* | 0.288703078 | -0.024683504 | 0.172786751 | -0.197470255 | 0.999933887 | -0.246248007 | 0.211856197 | -0.458104204 | 0.328226075 |
| ENSG00000170100 | *ZNF778* | 2.169938079 | -0.194124927 | 0.075404454 | -0.269529382 | 0.999933887 | -0.119036579 | -0.299523598 | 0.180487019 | 0.328356945 |
| ENSG00000272444 | *AL118558.4* | -0.6967068 | -0.042721108 | -0.119233267 | 0.076512159 | 0.999933887 | 0.229658779 | 0.607618898 | -0.377960119 | 0.328402506 |
| ENSG00000171791 | *BCL2* | 4.345427458 | 0.035730265 | -0.027656166 | 0.063386431 | 0.999933887 | 0.187540027 | 0.314686391 | -0.127146364 | 0.328491483 |
| ENSG00000179476 | *C14orf28* | 1.315935026 | -0.024826754 | -0.161930799 | 0.137104045 | 0.999933887 | -0.392447136 | -0.646701311 | 0.254254175 | 0.328566694 |
| ENSG00000160185 | *UBASH3A* | 3.066755244 | -0.06408002 | 0.101775642 | -0.165855662 | 0.999933887 | 0.351413885 | 0.517039385 | -0.165625501 | 0.328936628 |
| ENSG00000118518 | *RNF146* | 4.912574883 | -0.068571327 | -0.096843846 | 0.028272519 | 0.999933887 | -0.29573261 | -0.166422906 | -0.129309704 | 0.32928822 |
| ENSG00000179918 | *SEPHS2* | 5.481982601 | 0.031398626 | -0.102784306 | 0.134182932 | 0.999933887 | 1.12987129 | 0.881525022 | 0.248346269 | 0.329307816 |
| ENSG00000135625 | *EGR4* | -0.422727745 | 0.205971308 | 0.130865547 | 0.07510576 | 0.999933887 | 1.050657597 | 1.6491997 | -0.598542103 | 0.329358883 |
| ENSG00000283286 | *AC093277.1* | -0.622042305 | -0.137520277 | -0.34281803 | 0.205297753 | 0.999933887 | 1.669480342 | 2.079803985 | -0.410323643 | 0.329358883 |
| ENSG00000138430 | *OLA1* | 2.322737305 | -0.002692989 | -0.24095434 | 0.238261351 | 0.999933887 | -0.064431161 | 0.141331719 | -0.20576288 | 0.329358883 |
| ENSG00000136830 | *NIBAN2* | 2.08262265 | 0.005413675 | -0.073880086 | 0.079293761 | 0.999933887 | -0.770063284 | -0.991061394 | 0.22099811 | 0.329401125 |
| ENSG00000179840 | *PIK3CD-AS1* | 1.045120834 | -0.005686569 | 0.14286734 | -0.148553909 | 0.999933887 | 0.946774566 | 1.336497164 | -0.389722598 | 0.329498164 |
| ENSG00000265205 | *AC010761.3* | 1.143839332 | -0.012705358 | -0.095004283 | 0.082298925 | 0.999933887 | 0.044149481 | 0.441774299 | -0.397624818 | 0.329515158 |
| ENSG00000165416 | *SUGT1* | 2.377911991 | -0.110443093 | 0.068074105 | -0.178517198 | 0.999933887 | -0.045229392 | 0.163958127 | -0.209187519 | 0.329524782 |
| ENSG00000172534 | *HCFC1* | 5.516458494 | 0.031615106 | 0.152813759 | -0.121198652 | 0.999933887 | 0.225191098 | 0.331871736 | -0.106680637 | 0.32953575 |
| ENSG00000163798 | *SLC4A1AP* | 3.300643928 | 0.007666096 | 0.0404819 | -0.032815803 | 0.999933887 | -0.094729208 | 0.059152222 | -0.15388143 | 0.32953575 |
| ENSG00000163635 | *ATXN7* | 6.401520444 | 0.043113072 | 0.010861063 | 0.032252009 | 0.999933887 | -0.249041605 | -0.393464001 | 0.144422396 | 0.329689054 |
| ENSG00000165476 | *REEP3* | 2.558756503 | 0.025615913 | -0.005225485 | 0.030841398 | 0.999933887 | -0.587622434 | -0.823708556 | 0.236086123 | 0.329945955 |
| ENSG00000115145 | *STAM2* | 4.045496155 | -0.013983925 | 0.017813059 | -0.031796984 | 0.999933887 | -0.581931391 | -0.41945601 | -0.162475381 | 0.329965408 |
| ENSG00000163453 | *IGFBP7* | -0.065372688 | -0.00106157 | 0.182341326 | -0.183402896 | 0.999933887 | 0.096055705 | -0.343904907 | 0.439960612 | 0.329965408 |
| ENSG00000288093 | *AL592295.5* | -0.232262627 | -0.144421535 | 0.19935466 | -0.343776195 | 0.999933887 | 0.094806526 | 0.639207413 | -0.544400886 | 0.330059208 |
| ENSG00000139116 | *KIF21A* | 1.916517712 | -0.11802535 | -0.099310397 | -0.018714953 | 0.999933887 | -0.102221509 | 0.190446059 | -0.292667568 | 0.330378616 |
| ENSG00000256294 | *ZNF225* | -0.25576226 | -0.010681114 | 0.12277857 | -0.133459685 | 0.999933887 | 0.055215208 | -0.249887311 | 0.305102519 | 0.330558523 |
| ENSG00000164053 | *ATRIP* | 2.161632629 | 0.038801222 | 0.042014948 | -0.003213726 | 0.999933887 | 0.043982086 | -0.150854442 | 0.194836529 | 0.330770747 |
| ENSG00000171735 | *CAMTA1* | 2.740539043 | -0.045277409 | 0.131424861 | -0.17670227 | 0.999933887 | 0.168549155 | 0.334537377 | -0.165988222 | 0.331003492 |
| ENSG00000235919 | *ASH1L-AS1* | 1.232310373 | -0.022832763 | -0.009513327 | -0.013319436 | 0.999933887 | 0.834915472 | 1.056114039 | -0.221198567 | 0.331003492 |
| ENSG00000106682 | *EIF4H* | 7.406738012 | 0.008904878 | 0.017272234 | -0.008367356 | 0.999933887 | 0.077973868 | 0.170436606 | -0.092462737 | 0.331114758 |
| ENSG00000232671 | *ZNF687-AS1* | 0.540368078 | 0.063102256 | 0.01326526 | 0.049836996 | 0.999933887 | 0.942393321 | 0.598613233 | 0.343780088 | 0.331321536 |
| ENSG00000116586 | *LAMTOR2* | 2.657155404 | 0.050732987 | 0.034308173 | 0.016424815 | 0.999933887 | 0.006041911 | -0.230290648 | 0.236332559 | 0.331389868 |
| ENSG00000197013 | *ZNF429* | 5.639091939 | 0.044182741 | 0.008019518 | 0.036163223 | 0.999933887 | 0.410808898 | 0.635260253 | -0.224451355 | 0.331389868 |
| ENSG00000148110 | *MFSD14B* | 4.553773774 | -0.007295718 | -0.007411172 | 0.000115454 | 0.999933887 | -0.2696697 | -0.402803001 | 0.133133301 | 0.331742375 |
| ENSG00000232434 | *AJM1* | 1.665258457 | 0.176687805 | 0.142843111 | 0.033844694 | 0.999933887 | 0.130245807 | 0.385311885 | -0.255066078 | 0.33183483 |
| ENSG00000162063 | *CCNF* | 0.206870723 | -0.080432794 | -0.114249676 | 0.033816882 | 0.999933887 | 0.309800826 | 0.602187961 | -0.292387135 | 0.33183483 |
| ENSG00000119541 | *VPS4B* | 7.157127537 | -0.032459164 | -0.025341047 | -0.007118117 | 0.999933887 | -0.03392053 | 0.07263134 | -0.10655187 | 0.331938995 |
| ENSG00000254634 | *SMG1P6* | -0.701746982 | -0.217820057 | 0.022746364 | -0.240566422 | 0.999933887 | -0.122034599 | 0.266548581 | -0.388583181 | 0.331940938 |
| ENSG00000198453 | *ZNF568* | 0.250893502 | -0.02714334 | -0.072249838 | 0.045106498 | 0.999933887 | -0.03522239 | 0.297546683 | -0.332769072 | 0.331940938 |
| ENSG00000126264 | *HCST* | 2.842524596 | 0.130523928 | 0.153497007 | -0.022973079 | 0.999933887 | 0.69012643 | 0.887795673 | -0.197669242 | 0.331951559 |
| ENSG00000230492 | *AL049651.1* | -0.53722594 | -0.058694153 | -0.004823895 | -0.053870258 | 0.999933887 | -0.066261598 | 0.456847718 | -0.523109316 | 0.332111088 |
| ENSG00000131467 | *PSME3* | 6.62156325 | 0.015672853 | 0.004927203 | 0.01074565 | 0.999933887 | 0.512930756 | 0.34677779 | 0.166152966 | 0.332646496 |
| ENSG00000007516 | *BAIAP3* | 4.666910992 | 0.058838732 | 0.068744042 | -0.00990531 | 0.999933887 | 0.10608473 | -0.033097684 | 0.139182414 | 0.333357518 |
| ENSG00000182463 | *TSHZ2* | 1.152566889 | -0.098726289 | 0.028042104 | -0.126768392 | 0.999933887 | -0.1623195 | 0.108050729 | -0.270370229 | 0.333494241 |
| ENSG00000143179 | *UCK2* | -0.290540822 | -0.009386875 | 0.028674811 | -0.038061686 | 0.999933887 | -0.021340494 | 0.388826903 | -0.410167397 | 0.333494241 |
| ENSG00000181751 | *MACIR* | 0.695575895 | -0.193356008 | -0.068878698 | -0.12447731 | 0.999933887 | -0.48146526 | -0.762157141 | 0.28069188 | 0.333675355 |
| ENSG00000138399 | *FASTKD1* | 0.878042952 | -0.141322251 | -0.156598548 | 0.015276297 | 0.999933887 | -0.062696919 | 0.175043215 | -0.237740134 | 0.333823078 |
| ENSG00000163956 | *LRPAP1* | 6.701964154 | 0.025720597 | 0.020369936 | 0.005350661 | 0.999933887 | -0.209738741 | -0.341684429 | 0.131945688 | 0.333823442 |
| ENSG00000116212 | *LRRC42* | 2.225469 | -0.001619942 | 0.06186898 | -0.063488922 | 0.999933887 | -0.504422595 | -0.732509567 | 0.228086972 | 0.33399772 |
| ENSG00000163877 | *SNIP1* | 5.410829119 | 0.02013787 | -0.000294599 | 0.020432468 | 0.999933887 | 0.208367173 | 0.309531822 | -0.101164649 | 0.334069453 |
| ENSG00000110318 | *CEP126* | -0.834772158 | -0.228238464 | 0.04080991 | -0.269048374 | 0.999933887 | -0.414992292 | 0.129844869 | -0.544837161 | 0.334097617 |
| ENSG00000114480 | *GBE1* | 6.611067153 | 0.155799498 | 0.236605344 | -0.080805846 | 0.999933887 | 1.555062256 | 1.770500338 | -0.215438082 | 0.334107454 |
| ENSG00000260852 | *FBXL19-AS1* | 0.176039928 | 0.004049564 | -0.012765259 | 0.016814823 | 0.999933887 | -0.253176058 | 0.045368136 | -0.298544194 | 0.334144004 |
| ENSG00000164120 | *HPGD* | 1.26208231 | -0.144579932 | -0.090935145 | -0.053644787 | 0.999933887 | -0.279394879 | -0.044528464 | -0.234866415 | 0.334150831 |
| ENSG00000185129 | *PURA* | 3.06568979 | 0.065789138 | -0.039467421 | 0.105256559 | 0.999933887 | -0.270893473 | -0.450184656 | 0.179291183 | 0.334150831 |
| ENSG00000269843 | *AC008537.2* | 0.513423803 | -0.052675884 | -0.351930461 | 0.299254578 | 0.999933887 | -0.151017048 | 0.150671819 | -0.301688867 | 0.334456217 |
| ENSG00000197134 | *ZNF257* | -1.029678156 | 0.110970157 | 0.185732551 | -0.074762394 | 0.999933887 | -0.27245064 | 0.223322414 | -0.495773054 | 0.334480637 |
| ENSG00000239351 | *NPM1P29* | 1.146700382 | -0.152802979 | -0.129925662 | -0.022877317 | 0.999933887 | -0.14485623 | 0.20004902 | -0.34490525 | 0.33454561 |
| ENSG00000058063 | *ATP11B* | 7.164121083 | -0.042404632 | -0.094060944 | 0.051656312 | 0.999933887 | -0.557538707 | -0.364688151 | -0.192850557 | 0.33454561 |
| ENSG00000267293 | *AC012569.1* | 0.821624879 | -0.065139256 | 0.019131682 | -0.084270938 | 0.999933887 | 1.872836816 | 2.353302908 | -0.480466092 | 0.33454561 |
| ENSG00000265148 | *TSPOAP1-AS1* | 2.540770673 | 0.159361398 | 0.049247114 | 0.110114284 | 0.999933887 | 0.225070028 | 0.027137973 | 0.197932056 | 0.334616545 |
| ENSG00000167881 | *SRP68* | 3.829878648 | -0.017396799 | -0.109916217 | 0.092519418 | 0.999933887 | 0.077505346 | 0.201278921 | -0.123773575 | 0.334796045 |
| ENSG00000116688 | *MFN2* | 6.541925563 | -0.014600026 | 0.059632409 | -0.074232436 | 0.999933887 | -0.039242076 | 0.064915739 | -0.104157816 | 0.335108565 |
| ENSG00000231346 | *LINC01160* | -0.208747872 | -0.042476034 | -0.066066108 | 0.023590073 | 0.999933887 | 0.112600427 | -0.222902291 | 0.335502719 | 0.33517383 |
| ENSG00000148019 | *CEP78* | 3.093383579 | -0.053109807 | -0.070913775 | 0.017803967 | 0.999933887 | -0.207825899 | 0.029888808 | -0.237714707 | 0.335206062 |
| ENSG00000145495 | *MARCHF6* | 7.992892481 | -0.047627554 | -0.027007515 | -0.020620039 | 0.999933887 | 0.112445996 | 0.197621705 | -0.085175709 | 0.335323566 |
| ENSG00000172301 | *COPRS* | 1.622790807 | 0.232347266 | 0.130884115 | 0.101463152 | 0.999933887 | 0.008499927 | -0.203125499 | 0.211625426 | 0.335565302 |
| ENSG00000116824 | *CD2* | 4.766589055 | -0.021516244 | -0.106638964 | 0.08512272 | 0.999933887 | 0.16161441 | 0.288649054 | -0.127034644 | 0.335565302 |
| ENSG00000087460 | *GNAS* | 8.745986019 | 0.015346163 | -0.00490456 | 0.020250723 | 0.999933887 | -0.181803349 | -0.283847693 | 0.102044343 | 0.335565302 |
| ENSG00000145439 | *CBR4* | 1.850625501 | -0.075898981 | -0.125814849 | 0.049915869 | 0.999933887 | -0.571725738 | -0.821913703 | 0.250187965 | 0.335940346 |
| ENSG00000274270 | *AL137060.3* | 1.051810949 | 0.176014928 | 0.130837993 | 0.045176935 | 0.999933887 | 0.88388121 | 1.165171149 | -0.281289938 | 0.336111577 |
| ENSG00000261098 | *AP000766.1* | -0.439203465 | 0.221042635 | 0.33190195 | -0.110859315 | 0.999933887 | -0.311969001 | 0.267625003 | -0.579594004 | 0.336111577 |
| ENSG00000172172 | *MRPL13* | 0.517810536 | -0.147635917 | -0.518169642 | 0.370533725 | 0.999933887 | 0.072833434 | 0.347755632 | -0.274922198 | 0.336116428 |

| | | | | | | | | | | |
|---|---|---|---|---|---|---|---|---|---|---|
| ENSG00000137145 | *DENND4C* | 3.992415389 | -0.046243294 | -0.000902677 | -0.045340617 | 0.999933887 | -0.305167344 | -0.108073631 | -0.197093713 | 0.336218604 |
| ENSG00000124217 | *MOCS3* | 1.351067341 | -0.125265921 | -0.145870379 | 0.020604457 | 0.999933887 | -0.127016951 | -0.346073905 | 0.219056954 | 0.336306623 |
| ENSG00000070367 | *EXOC5* | 3.753649425 | 0.035428455 | -0.19777316 | 0.233201616 | 0.999933887 | -0.72056318 | -0.45616271 | -0.26440047 | 0.336332624 |
| ENSG00000154727 | *GABPA* | 3.610443625 | -0.083678824 | -0.123204146 | 0.039525322 | 0.999933887 | -0.607039453 | -0.356208632 | -0.250830822 | 0.336572944 |
| ENSG00000065600 | *PACC1* | 0.983835811 | -0.049067862 | -0.074286497 | 0.025218635 | 0.999933887 | -0.197668611 | 0.03304707 | -0.230715682 | 0.336605497 |
| ENSG00000113552 | *GNPDA1* | 7.859820952 | -0.056766586 | -0.014207251 | -0.042559336 | 0.999933887 | 0.177491571 | 0.304608209 | -0.127116638 | 0.336613287 |
| ENSG00000235092 | *ID2-AS1* | -0.334901709 | -0.130011504 | -0.286789116 | 0.156777611 | 0.999933887 | 0.279825047 | 0.656044094 | -0.376219046 | 0.336613287 |
| ENSG00000111674 | *ENO2* | 3.421895049 | -0.030051541 | -0.139063734 | 0.109012193 | 0.999933887 | 0.159157564 | 0.003136394 | 0.156021169 | 0.336613287 |
| ENSG00000253669 | *GASAL1* | 1.38173175 | 0.288892857 | 0.346664076 | -0.05777122 | 0.999933887 | 1.347924838 | 1.653480688 | -0.30555585 | 0.336677763 |
| ENSG00000172115 | *CYCS* | 4.294115147 | -0.028971137 | -0.047313716 | 0.018342579 | 0.999933887 | 0.352310048 | 0.506063299 | -0.153753251 | 0.336998633 |
| ENSG00000270108 | *AL049840.6* | 0.405379445 | -0.182205196 | 0.054762874 | -0.236968071 | 0.999933887 | -0.191741468 | -0.555457767 | 0.3637163 | 0.337448109 |
| ENSG00000127452 | *FBXL12* | 4.221584186 | -0.023233148 | -0.065033763 | 0.041800615 | 0.999933887 | -0.29584045 | -0.422126244 | 0.126285794 | 0.337599203 |
| ENSG00000189077 | *TMEM120A* | 5.597779666 | 0.065181358 | 0.020607748 | 0.04457361 | 0.999933887 | 0.505806895 | 0.683225275 | -0.17741838 | 0.337667766 |
| ENSG00000246790 | *AP000977.1* | 0.021576329 | -0.084597925 | -0.377501764 | 0.292903839 | 0.999933887 | -0.023526768 | -0.40727344 | 0.383746672 | 0.337667766 |
| ENSG00000092010 | *PSME1* | 7.443098474 | -0.025257916 | -0.094225464 | 0.068967549 | 0.999933887 | 0.045011491 | -0.055443008 | 0.100454499 | 0.337667766 |
| ENSG00000272888 | *CHASERR* | 6.909805833 | -0.039266925 | -0.018982219 | -0.020284706 | 0.999933887 | 0.311110978 | 0.412748716 | -0.101637738 | 0.337894538 |
| ENSG00000255587 | *RAB44* | 4.155131005 | -0.052030984 | 0.07986985 | -0.131900834 | 0.999933887 | 0.032986562 | 0.18282858 | -0.149842019 | 0.337894538 |
| ENSG00000106346 | *USP42* | 4.606552461 | 0.079856055 | 0.109336663 | -0.029480608 | 0.999933887 | -0.100755762 | -0.2398888 | 0.139133038 | 0.338054044 |
| ENSG00000152207 | *CYSLTR2* | 2.178767249 | -0.034688354 | -0.087998788 | 0.053310434 | 0.999933887 | -0.072427942 | 0.137069687 | -0.209497629 | 0.338070391 |
| ENSG00000170860 | *LSM3* | 3.385833706 | 0.033705232 | -0.060903133 | 0.094608365 | 0.999933887 | 0.045512497 | 0.221947261 | -0.176434765 | 0.338070391 |
| ENSG00000105677 | *TMEM147* | 2.081580558 | 0.012247792 | 0.114734591 | -0.102486799 | 0.999933887 | 0.099411489 | 0.281296859 | -0.18188537 | 0.338070391 |
| ENSG00000211776 | *TRAV2* | -0.060856339 | 0.096579334 | 0.175265582 | -0.078686247 | 0.999933887 | 0.004382928 | -0.404904928 | 0.409287857 | 0.338111537 |
| ENSG00000114573 | *ATP6V1A* | 7.182042024 | -0.112604952 | -0.011249441 | -0.10135551 | 0.999933887 | -0.228964142 | -0.112101824 | -0.116862318 | 0.338295593 |
| ENSG00000154719 | *MRPL39* | 0.785505827 | -0.062726287 | -0.095320401 | 0.032594114 | 0.999933887 | 0.174416238 | 0.421290421 | -0.246874183 | 0.338295593 |
| ENSG00000104047 | *DTWD1* | 0.33387022 | -0.187755709 | -0.021580481 | -0.166175228 | 0.999933887 | 0.006195369 | 0.327235885 | -0.321040516 | 0.338307457 |
| ENSG00000100227 | *POLDIP3* | 5.901445744 | -0.034669296 | -0.017544208 | -0.017125087 | 0.999933887 | -0.186079085 | -0.282378098 | 0.096299013 | 0.338307457 |
| ENSG00000267394 | *AC004596.1* | 0.16108696 | 0.058433784 | 0.281376007 | -0.222942223 | 0.999933887 | 0.146851072 | -0.24599524 | 0.392846312 | 0.338307457 |
| ENSG00000100505 | *TRIM9* | 0.813847796 | -0.034013112 | 0.009399311 | -0.043412422 | 0.999933887 | -0.069912485 | -0.458655597 | 0.388743113 | 0.338307457 |
| ENSG00000136527 | *TRA2B* | 6.640017976 | 0.012480947 | 0.015701372 | -0.003220425 | 0.999933887 | 0.11978608 | 0.207642042 | -0.087855962 | 0.338307457 |
| ENSG00000184983 | *NDUFA6* | 3.8355635 | -0.009613723 | -0.036807128 | 0.027193404 | 0.999933887 | -0.194620063 | -0.33679429 | 0.142174226 | 0.338307457 |
| ENSG00000167693 | *NXN* | -0.995940045 | 0.044605549 | -0.077619881 | 0.12222543 | 0.999933887 | 1.843565773 | 2.402838885 | -0.559273111 | 0.338398211 |
| ENSG00000100003 | *SEC14L2* | 0.974425595 | 0.062945333 | -0.123596606 | 0.186541938 | 0.999933887 | 0.188224354 | -0.099891056 | 0.28811541 | 0.338512359 |
| ENSG00000178105 | *DDX10* | 1.203369418 | -0.074061077 | -0.111941965 | 0.037880888 | 0.999933887 | 0.103371458 | 0.349512922 | -0.246141464 | 0.338512359 |
| ENSG00000211795 | *TRAV8-6* | -0.319719669 | 0.002181678 | -0.287413887 | 0.289595565 | 0.999933887 | 0.186516854 | -0.156596627 | 0.343113481 | 0.338512359 |
| ENSG00000123360 | *PDE1B* | 2.53300511 | -0.033653039 | 0.094673303 | -0.128326342 | 0.999933887 | -0.005620006 | -0.213382838 | 0.207762831 | 0.338542907 |
| ENSG00000020577 | *SAMD4A* | 0.007235702 | 0.175064088 | 0.502813341 | -0.327749254 | 0.999933887 | 0.166306546 | 0.52702844 | -0.360721895 | 0.338795787 |
| ENSG00000187554 | *TLR5* | 2.2157949 | 0.003542624 | -0.055867017 | 0.059409641 | 0.999933887 | 0.263373157 | 0.570388296 | -0.307015139 | 0.338795787 |
| ENSG00000125450 | *NUP85* | 3.935195736 | -0.044292447 | 0.036901368 | -0.081193814 | 0.999933887 | 0.042173346 | -0.073899561 | 0.116072907 | 0.338849688 |
| ENSG00000283597 | *FAM169B* | -0.092872869 | 0.2234248 | -0.024831379 | 0.248256179 | 0.999933887 | -0.20792278 | -0.604616597 | 0.396693817 | 0.339069522 |
| ENSG00000277117 | *FP565260.3* | -1.137130057 | -0.035507573 | 0.245948512 | -0.281456085 | 0.999933887 | 1.226995509 | 1.677375514 | -0.450380005 | 0.339190279 |
| ENSG00000112182 | *BACH2* | 4.11905326 | -0.0176433 | 0.082686965 | -0.100330265 | 0.999933887 | -0.319138018 | -0.458659403 | 0.139521385 | 0.339320608 |
| ENSG00000143570 | *SLC39A1* | -0.010207982 | 0.244509777 | 0.169379796 | 0.075129981 | 0.999933887 | 1.095975168 | 1.44020381 | -0.344228642 | 0.339331085 |
| ENSG00000126895 | *AVPR2* | 1.058746855 | 0.037253503 | 0.143787554 | -0.106534051 | 0.999933887 | 0.030748282 | 0.289442402 | -0.25869412 | 0.339829412 |
| ENSG00000122435 | *TRMT13* | 1.404726832 | -0.163714788 | -0.17226943 | 0.008554642 | 0.999933887 | -0.285583906 | 0.01774919 | -0.303333097 | 0.339863653 |
| ENSG00000101365 | *IDH3B* | 4.783918958 | -0.03089855 | 0.014611284 | -0.045509834 | 0.999933887 | -0.136570168 | -0.24057835 | 0.104008182 | 0.340222924 |
| ENSG00000136932 | *TRMO* | 4.99045778 | -0.044892826 | -0.035977143 | -0.008915683 | 0.999933887 | 0.17174632 | 0.277159563 | -0.105413243 | 0.340230285 |
| ENSG00000131462 | *TUBG1* | 0.021113652 | 0.039920324 | 0.086677879 | -0.046757556 | 0.999933887 | 0.132926448 | -0.201384066 | 0.334310514 | 0.340230285 |
| ENSG00000232043 | *AL133230.1* | 1.029198323 | -0.031839942 | -0.098163715 | 0.066323773 | 0.999933887 | 0.456769287 | 0.782671771 | -0.325902484 | 0.340280792 |
| ENSG00000159592 | *GPBP1L1* | 6.267673797 | -0.039352402 | -0.001555851 | -0.037796552 | 0.999933887 | -0.360319277 | -0.455870901 | 0.095551624 | 0.340584677 |
| ENSG00000065923 | *SLC9A7* | 4.338047056 | 0.009144098 | 0.010121753 | -0.000977655 | 0.999933887 | 0.173158712 | 0.331204317 | -0.158045605 | 0.340713137 |
| ENSG00000131127 | *ZNF141* | 2.089467524 | 0.078584234 | -0.122450484 | 0.201034717 | 0.999933887 | -0.508430365 | -0.191919217 | -0.316511148 | 0.340796842 |
| ENSG00000129315 | *CCNT1* | 6.020839651 | -0.057525117 | -0.044159745 | -0.013365372 | 0.999933887 | 0.099340128 | 0.208793322 | -0.109453194 | 0.340949609 |
| ENSG00000067082 | *KLF6* | 10.320545264 | 0.030150795 | 0.029254187 | 0.000896608 | 0.999933887 | 0.077157205 | 0.181288405 | -0.1041312 | 0.340949609 |
| ENSG00000135299 | *ANKRD6* | 1.815867557 | -0.060364203 | -0.017406719 | -0.042957484 | 0.999933887 | -0.546275403 | -0.73070691 | 0.184431507 | 0.340949609 |
| ENSG00000131389 | *SLC6A6* | 10.256554915 | -0.012716973 | -0.012087158 | -0.000629816 | 0.999933887 | 0.142605135 | 0.234625675 | -0.09202054 | 0.340949609 |
| ENSG00000181826 | *RELL1* | 4.671257364 | -0.106261425 | -0.011334593 | -0.094926831 | 0.999933887 | -0.312484568 | -0.45443083 | 0.141946262 | 0.341262589 |
| ENSG00000235706 | *DICER1-AS1* | 0.528653179 | 0.035047563 | -0.350708657 | 0.38575622 | 0.999933887 | -0.181319796 | -0.486402637 | 0.305082842 | 0.341262589 |
| ENSG00000112312 | *GMNN* | 1.528535017 | 0.015594277 | -0.012343577 | 0.027937854 | 0.999933887 | 0.076137655 | 0.275141275 | -0.19900362 | 0.341262589 |
| ENSG00000170581 | *STAT2* | 6.897219487 | 0.003386041 | 0.012423465 | -0.009037424 | 0.999933887 | -0.256231239 | -0.377783515 | 0.121552276 | 0.341262589 |
| ENSG00000244045 | *TMEM199* | 3.266625078 | 0.035713683 | 0.001313175 | 0.034400509 | 0.999933887 | -0.014440495 | 0.122981566 | -0.137422061 | 0.341714729 |
| ENSG00000103226 | *NOMO3* | 0.652521392 | -0.087404141 | -0.000279697 | -0.087124445 | 0.999933887 | -0.025229746 | -0.426268217 | 0.401038471 | 0.341963187 |
| ENSG00000176853 | *FAM91A1* | 6.748810771 | -0.017359551 | -0.099749199 | 0.082389648 | 0.999933887 | -0.247662667 | -0.071938737 | -0.17572393 | 0.341997012 |
| ENSG00000213722 | *DDAH2* | 5.292440434 | 0.098061807 | 0.120730513 | -0.022668706 | 0.999933887 | -0.766053829 | -0.957563298 | 0.191509469 | 0.342062328 |
| ENSG00000084764 | *MAPRE3* | 2.696213156 | 0.058167189 | 0.088162069 | -0.02999488 | 0.999933887 | -0.622934416 | -0.780282413 | 0.157338997 | 0.342062328 |
| ENSG00000127337 | *YEATS4* | 1.203956456 | -0.05395606 | -0.182763492 | 0.128807432 | 0.999933887 | 0.264776933 | 0.557961485 | -0.293184552 | 0.342098864 |
| ENSG00000251161 | *AC020661.1* | 3.518126309 | 0.155736862 | 0.057841769 | 0.097895093 | 0.999933887 | 0.90931874 | 1.118625381 | -0.209306641 | 0.342109117 |
| ENSG00000140374 | *ETFA* | 2.698510615 | -0.0360116 | -0.17355999 | 0.137548391 | 0.999933887 | -0.157652054 | -0.352029787 | 0.194377733 | 0.342162645 |
| ENSG00000112514 | *CUTA* | 4.303739466 | 0.01921681 | 0.044932547 | -0.025715737 | 0.999933887 | 0.01981866 | -0.112380517 | 0.132199178 | 0.342162645 |
| ENSG00000285799 | *AL645929.2* | -0.197604522 | 0.053140033 | 0.05411942 | -0.000979387 | 0.999933887 | 0.60042733 | 0.176195155 | 0.424232175 | 0.342162645 |
| ENSG00000101216 | *GMEB2* | 5.389642328 | -0.061396735 | 0.002138106 | -0.063534841 | 0.999933887 | -0.254089006 | -0.362215889 | 0.108126882 | 0.342240513 |
| ENSG00000116299 | *ELAPOR1* | 5.159066123 | 0.035171633 | -0.030120606 | 0.065292239 | 0.999933887 | -0.247803715 | -0.39304562 | 0.145241905 | 0.342347242 |
| ENSG00000167985 | *SDHAF2* | 2.987390091 | -0.016340384 | -0.08232698 | 0.065986596 | 0.999933887 | -0.02690015 | 0.131159115 | -0.158059265 | 0.342623362 |
| ENSG00000274051 | *AC005837.3* | 0.514605023 | 0.182055679 | 0.351958456 | -0.169902777 | 0.999933887 | 0.576897425 | 0.945866346 | -0.368968922 | 0.342853848 |
| ENSG00000164631 | *ZNF12* | 5.160034232 | -0.048788328 | -0.113486199 | 0.06469787 | 0.999933887 | -0.297811351 | -0.102610488 | -0.195200863 | 0.342853848 |
| ENSG00000180549 | *FUT7* | 2.15704491 | 0.031982102 | 0.088322985 | -0.056340883 | 0.999933887 | -0.481167782 | -0.851914407 | 0.370746625 | 0.342853848 |
| ENSG00000126262 | *FFAR2* | 8.82550716 | 0.026354543 | -0.001260145 | 0.027614687 | 0.999933887 | 2.082068701 | 2.303909946 | -0.221841245 | 0.342853848 |
| ENSG00000237637 | *FRY-AS1* | -0.091096072 | -0.016279703 | 0.083497151 | -0.099776855 | 0.999933887 | 0.222261804 | 0.678265361 | -0.456003557 | 0.342853848 |
| ENSG00000263335 | *AF001548.2* | 1.170760767 | 0.076041625 | -0.044367072 | 0.120408697 | 0.999933887 | 0.212422664 | 0.487900745 | -0.275478081 | 0.342964912 |
| ENSG00000183172 | *SMDT1* | 3.157052421 | 0.03951473 | 0.04539815 | -0.00588342 | 0.999933887 | -0.154611603 | -0.311321434 | 0.15670983 | 0.342964912 |
| ENSG00000285737 | *LINC02680* | 0.393253368 | 0.081601168 | 0.196961401 | -0.115360233 | 0.999933887 | 0.120379878 | -0.234370408 | 0.354750286 | 0.342964912 |
| ENSG00000148908 | *RGS10* | 3.336659768 | -0.022591168 | -0.120541889 | 0.097950721 | 0.999933887 | 0.034901364 | 0.177349619 | -0.142448255 | 0.342964912 |
| ENSG00000198246 | *SLC29A3* | 0.438144173 | 0.007629916 | 0.191278152 | -0.183648236 | 0.999933887 | -0.031997995 | -0.328784345 | 0.296786351 | 0.342964912 |
| ENSG00000067560 | *RHOA* | 9.266940396 | -0.022324988 | -0.055628891 | 0.033303903 | 0.999933887 | 0.054949554 | 0.153064642 | -0.098115088 | 0.343059112 |
| ENSG00000020129 | *NCDN* | 2.37443561 | 0.032496462 | 0.123563283 | -0.091066821 | 0.999933887 | 0.417141507 | 0.601629485 | -0.184487978 | 0.343059112 |
| ENSG00000110395 | *CBL* | 7.129070292 | -0.060662064 | 0.027163119 | -0.087825183 | 0.999933887 | -0.937214767 | -1.112631981 | 0.175417214 | 0.343088828 |
| ENSG00000115808 | *STRN* | 4.985957516 | -0.003678852 | -0.06331348 | 0.059634628 | 0.999933887 | -0.806140507 | -0.997674043 | 0.191533536 | 0.343088828 |
| ENSG00000235910 | *APOA1-AS* | -0.354774261 | -0.013064512 | -0.00937102 | -0.003693493 | 0.999933887 | 0.285522544 | 0.771272098 | -0.485749554 | 0.343301716 |
| ENSG00000197857 | *ZNF44* | 3.877039267 | -0.085321407 | -0.008719693 | -0.076601714 | 0.999933887 | 0.170613468 | 0.333167249 | -0.162553781 | 0.343360246 |
| ENSG00000229848 | *AC139149.1* | -0.648685394 | 0.237955212 | 0.206543097 | 0.031412115 | 0.999933887 | 1.04107486 | 0.474606216 | 0.566468643 | 0.343388302 |
| ENSG00000005194 | *CIAPIN1* | 2.363782848 | -0.054827404 | -0.135163615 | 0.080336211 | 0.999933887 | 0.060023208 | -0.104500094 | 0.164523302 | 0.343427592 |
| ENSG00000174501 | *ANKRD36C* | 2.105908151 | -0.049952603 | 0.059582951 | -0.109535554 | 0.999933887 | -0.174235118 | 0.133052678 | -0.307287796 | 0.343427592 |
| ENSG00000123066 | *MED13L* | 9.125799283 | -0.030732701 | -0.029845496 | -0.000887205 | 0.999933887 | -0.182576621 | -0.091379847 | -0.091196774 | 0.343559288 |
| ENSG00000196329 | *GIMAP5* | 2.250181759 | -0.138447039 | -0.009199392 | -0.129247648 | 0.999933887 | 0.271391261 | 0.082093966 | 0.189297295 | 0.343892865 |
| ENSG00000258769 | *RAP1AP* | 1.184157771 | 0.140362019 | -0.112817852 | 0.25317987 | 0.999933887 | -0.189980433 | 0.054269344 | -0.244249778 | 0.343939212 |
| ENSG00000100372 | *SLC25A17* | 0.93200595 | 0.086863499 | 0.00546912 | 0.081394379 | 0.999933887 | -0.086074531 | -0.297258473 | 0.211183941 | 0.344006796 |
| ENSG00000125124 | *BBS2* | 3.138906874 | -0.046696557 | 0.078193467 | -0.124890024 | 0.999933887 | 0.144299608 | 0.012244629 | 0.132054979 | 0.34409437 |
| ENSG00000160714 | *UBE2Q1* | 5.974629053 | 0.01618447 | 0.022801035 | -0.006616565 | 0.999933887 | -0.069391612 | 0.021917651 | -0.091309263 | 0.34409437 |
| ENSG00000171049 | *FPR2* | 7.843899173 | 0.035753563 | -0.001268938 | 0.037022501 | 0.999933887 | 0.692962334 | 0.817828462 | -0.124866128 | 0.344113797 |
| ENSG00000174842 | *GLMN* | 0.719702315 | 0.018460421 | -0.242015761 | 0.260476182 | 0.999933887 | -0.133840873 | 0.16963605 | -0.303476923 | 0.344290012 |
| ENSG00000184402 | *SS18L1* | 4.249788199 | 0.024752875 | -9.7332e-05 | 0.024850207 | 0.999933887 | -0.008021596 | -0.132714295 | 0.124692699 | 0.344394113 |
| ENSG00000198780 | *FAM169A* | 0.579030145 | -0.21668549 | -0.151341663 | -0.065343827 | 0.999933887 | -0.214164167 | 0.145148152 | -0.359312318 | 0.344483074 |
| ENSG00000083635 | *NUFIP1* | 2.052615656 | 0.010152165 | -0.117901928 | 0.128054093 | 0.999933887 | 0.547331589 | 0.834867778 | -0.287536189 | 0.344483074 |
| ENSG00000275022 | *MIR6753* | 1.509318352 | 0.045278996 | 0.039045387 | 0.006233609 | 0.999933887 | -0.40546831 | -0.153553763 | -0.251914547 | 0.344492098 |
| ENSG00000163686 | *ABHD6* | 0.672135939 | -0.013674361 | -0.049300938 | 0.035626576 | 0.999933887 | 0.114303951 | 0.342703054 | -0.228399102 | 0.344492098 |
| ENSG00000173578 | *XCR1* | -0.311360443 | -0.022115598 | -0.017026215 | -0.005089383 | 0.999933887 | -1.08787244 | -1.390447009 | 0.302574569 | 0.344492098 |

| | | | | | | | | | | |
|---|---|---|---|---|---|---|---|---|---|---|
| ENSG00000179818 | *PCBP1-AS1* | 6.979802649 | 0.017883693 | 0.107584674 | -0.089700981 | 0.999933887 | 0.489072048 | 0.602003142 | -0.112931094 | 0.34460163 |
| ENSG00000242113 | *RN7SL124P* | 0.629289823 | 0.263977351 | 0.418946373 | -0.154969023 | 0.999933887 | 3.492248646 | 4.054980505 | -0.562731858 | 0.344636012 |
| ENSG00000130227 | *XPO7* | 4.143394428 | -0.047765028 | 0.00424709 | -0.052012117 | 0.999933887 | -0.169875096 | -0.298661183 | 0.128786087 | 0.344636012 |
| ENSG00000139318 | *DUSP6* | 7.122415804 | -0.048689708 | -0.015388571 | -0.033301138 | 0.999933887 | 0.075470905 | 0.263522613 | -0.188051708 | 0.344636012 |
| ENSG00000286599 | *AC110611.1* | -0.186521137 | -0.040706726 | -0.00994367 | -0.030763056 | 0.999933887 | -0.122285012 | 0.353922216 | -0.476207228 | 0.344636012 |
| ENSG00000272374 | *Z97832.2* | -0.220182229 | 0.082210207 | -0.187519715 | 0.269729922 | 0.999933887 | 0.188172801 | 0.619913653 | -0.431740852 | 0.344777713 |
| ENSG00000156136 | *DCK* | 3.370513322 | -0.107672052 | 0.033822805 | -0.141494857 | 0.999933887 | -0.388722812 | -0.543427408 | 0.154704596 | 0.344882174 |
| ENSG00000184967 | *NOC4L* | 3.109908244 | 0.064654293 | 0.036389308 | 0.028264985 | 0.999933887 | 0.263357425 | 0.433777734 | -0.17042031 | 0.344884631 |
| ENSG00000179820 | *MYADM* | 8.417082576 | 0.045772684 | 0.068356639 | -0.022583955 | 0.999933887 | -0.033169935 | -0.162522147 | 0.129352212 | 0.345108809 |
| ENSG00000080824 | *HSP90AA1* | 8.093110987 | -0.147444109 | -0.20766965 | 0.060225541 | 0.999933887 | 0.348555321 | 0.559116428 | -0.210561107 | 0.345193997 |
| ENSG00000102984 | *ZNF821* | 1.520209715 | 0.080125481 | -0.003036118 | 0.083161599 | 0.999933887 | 0.063004475 | 0.284826053 | -0.221821578 | 0.34522745 |
| ENSG00000163541 | *SUCLG1* | 3.310226978 | 0.022724045 | -0.146500142 | 0.169224187 | 0.999933887 | 0.033109798 | 0.172478009 | -0.139368211 | 0.34525022 |
| ENSG00000276728 | *AC142472.1* | -0.195812751 | 0.187689674 | -0.023884237 | 0.211573911 | 0.999933887 | -0.179864834 | 0.141082524 | -0.320947358 | 0.345318873 |
| ENSG00000156802 | *ATAD2* | 2.433923508 | -0.101611118 | 0.009184827 | -0.110795945 | 0.999933887 | -0.24196467 | 0.111798632 | -0.353763302 | 0.345318873 |
| ENSG00000120992 | *LYPLA1* | 4.235766183 | -0.072829228 | -0.0765628 | 0.003733572 | 0.999933887 | -0.225355188 | -0.100737779 | -0.124617409 | 0.345661848 |
| ENSG00000160229 | *ZNF66* | 1.758874479 | -0.012717288 | -0.048495015 | 0.035777727 | 0.999933887 | -0.575063069 | -0.980470823 | 0.405407754 | 0.345661848 |
| ENSG00000261468 | *AC096921.2* | 1.238622465 | 0.047788863 | 0.001363283 | 0.046425581 | 0.999933887 | -0.118496848 | -0.482521695 | 0.364024847 | 0.345800449 |
| ENSG00000176225 | *RTTN* | 1.87093078 | 0.039104267 | -0.11163721 | 0.150741477 | 0.999933887 | -0.107337205 | 0.131458405 | -0.23879561 | 0.345813316 |
| ENSG00000125814 | *NAPB* | 2.750062861 | -0.09365231 | 0.013576055 | -0.107228365 | 0.999933887 | -0.209024122 | -0.394822623 | 0.185798501 | 0.34640681 |
| ENSG00000157152 | *SYN2* | -0.381303369 | 0.126828111 | -0.014272431 | 0.141100542 | 0.999933887 | -0.081645303 | -0.576387003 | 0.4947417 | 0.34640681 |
| ENSG00000235958 | *UBOX5-AS1* | -0.262089249 | -0.143420306 | -0.316313761 | 0.172893455 | 0.999933887 | 0.090010896 | 0.522909723 | -0.432898827 | 0.34640681 |
| ENSG00000160221 | *GATD3A* | -0.082545364 | -0.14809618 | -0.230256222 | 0.082160042 | 0.999933887 | 0.248629015 | -0.041149382 | 0.289778397 | 0.347091517 |
| ENSG00000141562 | *NARF* | 6.349602605 | -0.009726325 | 0.034301292 | -0.044027617 | 0.999933887 | -0.041296149 | -0.161730546 | 0.120434397 | 0.347241815 |
| ENSG00000125648 | *SLC25A23* | 0.979330555 | 0.128554744 | 0.05236438 | 0.076190363 | 0.999933887 | 0.082667149 | -0.216393897 | 0.299061045 | 0.347357337 |
| ENSG00000005700 | *IBTK* | 2.286625158 | -0.053381364 | -0.068147837 | 0.014766474 | 0.999933887 | -0.128375222 | 0.141089327 | -0.269464549 | 0.347357337 |
| ENSG00000172086 | *KRCC1* | 3.453247984 | 0.028156318 | -0.115771601 | 0.143927919 | 0.999933887 | -0.138137496 | 0.015509472 | -0.153646967 | 0.347357337 |
| ENSG00000260302 | *LINC01882* | -0.860276536 | 0.024222019 | -0.45384835 | 0.478070369 | 0.999933887 | 0.015923744 | 0.498475574 | -0.482551831 | 0.347357337 |
| ENSG00000172239 | *PAIP1* | 2.45648233 | -0.071584469 | -0.024307606 | -0.047276863 | 0.999933887 | 0.006254548 | -0.163025621 | 0.169280169 | 0.34740548 |
| ENSG00000165916 | *PSMC3* | 3.804741076 | -0.043173784 | 0.005157019 | -0.048330803 | 0.999933887 | 0.078580484 | 0.206055508 | -0.127475024 | 0.34740548 |
| ENSG00000145945 | *FAM50B* | -0.616955929 | 0.131641183 | 0.024392557 | 0.107248626 | 0.999933887 | 0.174090818 | -0.248762041 | 0.422852859 | 0.34740548 |
| ENSG00000226696 | *LENG8-AS1* | 0.823062216 | -0.045303368 | -0.036831456 | -0.008471911 | 0.999933887 | 0.004648604 | -0.209196671 | 0.213845275 | 0.34740548 |
| ENSG00000137504 | *CREBZF* | 5.276315989 | -0.012179737 | 0.049805142 | -0.061984879 | 0.999933887 | -0.193860945 | -0.30716143 | 0.113300485 | 0.347467454 |
| ENSG00000078668 | *VDAC3* | 4.363814656 | -0.069357172 | -0.162682519 | 0.093325347 | 0.999933887 | 0.098512295 | -0.023314012 | 0.121826307 | 0.347495649 |
| ENSG00000170522 | *ELOVL6* | -0.4540306 | -0.081610931 | -0.107978049 | 0.026367118 | 0.999933887 | -0.268356924 | 0.1712062 | -0.439563124 | 0.347495649 |
| ENSG00000181830 | *SLC35C1* | 2.224583919 | 0.075510383 | -0.144748441 | 0.220258824 | 0.999933887 | -0.206134537 | -0.437081948 | 0.230947411 | 0.347630955 |
| ENSG00000258682 | *AL132989.1* | 1.356083052 | 0.085619033 | -0.314856752 | 0.400475785 | 0.999933887 | -0.248155389 | -0.010803868 | -0.237351522 | 0.347630955 |
| ENSG00000267731 | *AC005332.2* | -0.639571254 | -0.260222432 | 0.238731446 | -0.498953877 | 0.999933887 | -0.180929051 | 0.276307035 | -0.457236086 | 0.347678901 |
| ENSG00000101146 | *RAE1* | 3.935619718 | -0.126854067 | -0.156549961 | 0.029695894 | 0.999933887 | -0.257657791 | -0.37179342 | 0.114135629 | 0.347776948 |
| ENSG00000057757 | *PITHD1* | 4.322216725 | 0.053275979 | -0.014336983 | 0.067612962 | 0.999933887 | -0.302081945 | -0.442281731 | 0.140199786 | 0.348765561 |
| ENSG00000125356 | *NDUFA1* | 4.686651911 | -0.001428745 | -0.020187553 | 0.018758808 | 0.999933887 | 0.09557118 | -0.032253741 | 0.127824921 | 0.348765561 |
| ENSG00000183638 | *RP1L1* | -2.659235282 | 0.146377212 | 0.351148653 | -0.204771441 | 0.999933887 | -0.158854446 | -1.076324898 | 0.917470453 | 0.349179103 |
| ENSG00000178458 | *H3P16* | 0.125337931 | -0.068737433 | 0.01094287 | -0.079680303 | 0.999933887 | -0.451217877 | -0.058542516 | -0.392675362 | 0.349307626 |
| ENSG00000177169 | *ULK1* | 7.961077405 | -0.012827991 | 0.019976819 | -0.03280481 | 0.999933887 | -0.728154058 | -0.858173506 | 0.130019448 | 0.349307626 |
| ENSG00000226210 | *WASH8P* | 3.527076066 | 0.010175101 | 0.056778261 | -0.046603161 | 0.999933887 | -0.207325486 | -0.337800138 | 0.130474652 | 0.349373608 |
| ENSG00000137871 | *ZNF280D* | 0.756016469 | -0.072761412 | -0.040314538 | -0.032446875 | 0.999933887 | -0.309261145 | 0.051440082 | -0.360701227 | 0.349655685 |
| ENSG00000186364 | *NUDT17* | -0.454660185 | 0.079556104 | -0.207063654 | 0.286619758 | 0.999933887 | 0.267979965 | 0.600416658 | -0.332436693 | 0.349808807 |
| ENSG00000122085 | *MTERF4* | 2.899105092 | -0.028470245 | -0.068642515 | 0.04017227 | 0.999933887 | 0.051424166 | -0.094304714 | 0.145728881 | 0.349831648 |
| ENSG00000143162 | *CREG1* | 6.94431823 | -0.076642703 | -0.115982394 | 0.03933969 | 0.999933887 | -0.322582368 | -0.192559833 | -0.130022536 | 0.349887003 |
| ENSG00000198000 | *NOL8* | 3.746456039 | -0.079163672 | -0.199083556 | 0.119919885 | 0.999933887 | -0.375776917 | -0.135436004 | -0.240340913 | 0.349887003 |
| ENSG00000151893 | *CACUL1* | 6.939697518 | -0.028234355 | 0.018394951 | -0.046629307 | 0.999933887 | -0.642884789 | -0.755531404 | 0.112646615 | 0.349887003 |
| ENSG00000141933 | *TPGS1* | 1.359644594 | 0.1892375 | -0.022398036 | 0.211635536 | 0.999933887 | -0.209424394 | 0.039879895 | -0.249304289 | 0.350042714 |
| ENSG00000135930 | *EIF4E2* | 4.618245173 | -0.025962256 | -0.099892489 | 0.073930233 | 0.999933887 | -0.221921749 | -0.329410664 | 0.107488915 | 0.350042714 |
| ENSG00000125775 | *SDCBP2* | -0.183952476 | 0.073631809 | 0.159568954 | -0.085937144 | 0.999933887 | -0.102263268 | -0.567234132 | 0.464970863 | 0.350147959 |
| ENSG00000117528 | *ABCD3* | 2.073559942 | -0.146976691 | 0.016199621 | -0.163176312 | 0.999933887 | -0.2769444 | -0.529872156 | 0.252927756 | 0.350343958 |
| ENSG00000287237 | *AC073655.3* | 1.717267772 | -0.141142613 | 0.179825231 | -0.320967844 | 0.999933887 | 0.970286859 | 1.223250633 | -0.252963774 | 0.350377122 |
| ENSG00000095303 | *PTGS1* | 2.978624938 | 0.035338059 | -0.026158939 | 0.061496998 | 0.999933887 | -0.103537713 | 0.065361845 | -0.168899558 | 0.350377122 |
| ENSG00000005206 | *SPPL2B* | 4.014336128 | 0.025578015 | 0.059539998 | -0.033961983 | 0.999933887 | 0.027833067 | -0.132575454 | 0.160408521 | 0.350377122 |
| ENSG00000106733 | *NMRK1* | 3.585486124 | -0.006898455 | 0.028756068 | -0.035654522 | 0.999933887 | -0.266654168 | -0.407671187 | 0.141017019 | 0.350377122 |
| ENSG00000243704 | *RN7SL105P* | -0.106888984 | 0.00873719 | -0.140248059 | 0.14898525 | 0.999933887 | 0.263774124 | 0.667268215 | -0.403494091 | 0.350377122 |
| ENSG00000261101 | *AC234775.2* | 0.263115654 | 0.058476462 | -0.066254827 | 0.124731289 | 0.999933887 | 0.555904374 | 0.953980806 | -0.398076432 | 0.350445309 |
| ENSG00000160352 | *ZNF714* | 1.689309793 | -0.031574057 | -0.017876203 | -0.013697854 | 0.999933887 | -0.126672785 | 0.128818798 | -0.255491583 | 0.35045011 |
| ENSG00000064651 | *SLC12A2* | 2.275467716 | 0.019137415 | -0.034859711 | 0.053997125 | 0.999933887 | -0.190430931 | 0.062884062 | -0.253314993 | 0.350639373 |
| ENSG00000168003 | *SLC3A2* | 9.393148447 | -0.008991904 | 0.005865828 | -0.014857732 | 0.999933887 | -0.023399128 | 0.088505146 | -0.111904274 | 0.350847136 |
| ENSG00000160408 | *ST6GALNAC6* | -0.786098414 | 0.148956922 | 0.536166367 | -0.387209445 | 0.999933887 | 0.041941792 | -0.408155161 | 0.450096953 | 0.350875564 |
| ENSG00000111707 | *SUDS3* | 5.63520046 | -0.030389697 | 0.050239654 | -0.080629351 | 0.999933887 | -0.403499263 | -0.509188284 | 0.105689021 | 0.350875564 |
| ENSG00000135315 | *CEP162* | 1.809359156 | -0.227113178 | -0.347464064 | 0.120350886 | 0.999933887 | -0.448611842 | -0.156379648 | -0.292232195 | 0.35124101 |
| ENSG00000099904 | *ZDHHC8* | 2.913592741 | 0.066427171 | 0.024401642 | 0.042025529 | 0.999933887 | 0.073590536 | -0.080896951 | 0.154487487 | 0.351242345 |
| ENSG00000071655 | *MBD3* | 2.670761763 | 0.045699563 | 0.057109798 | -0.011410235 | 0.999933887 | -0.032433243 | -0.193906087 | 0.161472844 | 0.351242345 |
| ENSG00000004975 | *DVL2* | 4.971026376 | 0.042550274 | 0.12617549 | -0.083625215 | 0.999933887 | -0.500664721 | -0.641593416 | 0.140928696 | 0.351242345 |
| ENSG00000122257 | *RBBP6* | 6.545793942 | 0.010495939 | -0.070558632 | 0.081054571 | 0.999933887 | -0.160627216 | -0.011363072 | -0.149264143 | 0.351242345 |
| ENSG00000164684 | *ZNF704* | -1.5133409 | 0.009050141 | -0.423629294 | 0.432679434 | 0.999933887 | -0.316252831 | 0.07513073 | -0.391383561 | 0.351242345 |
| ENSG00000108599 | *AKAP10* | 5.086703254 | -0.053214154 | -0.044435827 | -0.008778327 | 0.999933887 | -0.41118291 | -0.507733803 | 0.096550893 | 0.351256165 |
| ENSG00000148429 | *USP6NL* | 1.944864469 | -0.023325319 | 0.006713567 | -0.030038886 | 0.999933887 | -0.332221962 | -0.56917474 | 0.236952777 | 0.351378471 |
| ENSG00000162430 | *SELENON* | 2.550944076 | 0.11673335 | 0.103784468 | 0.012948881 | 0.999933887 | -0.099520564 | -0.300424373 | 0.200903809 | 0.351408501 |
| ENSG00000161091 | *MFSD12* | 6.911023415 | 0.063799752 | 0.068187767 | -0.004388015 | 0.999933887 | 0.318822555 | 0.444594779 | -0.125772223 | 0.351408501 |
| ENSG00000287091 | *AC009961.3* | 0.556507958 | 0.037167824 | 0.116305506 | -0.079137682 | 0.999933887 | -0.896046827 | -1.197591956 | 0.30154513 | 0.351444194 |
| ENSG00000228261 | *ITPRIP-AS1* | 3.279307051 | 0.213578328 | 0.246144124 | -0.032565796 | 0.999933887 | 1.287922097 | 1.589997711 | -0.302075614 | 0.35148982 |
| ENSG00000135049 | *AGTPBP1* | 6.708864306 | -0.046787549 | -0.107681515 | 0.060893966 | 0.999933887 | 0.054523822 | 0.249396062 | -0.19487224 | 0.351557808 |
| ENSG00000262903 | *AC027796.4* | 0.1780978 | 0.050860698 | -0.026352509 | 0.077213207 | 0.999933887 | 0.07181965 | -0.247946006 | 0.319765657 | 0.351557808 |
| ENSG00000148484 | *RSU1* | 3.990600694 | 0.042714604 | 0.007297013 | 0.035417591 | 0.999933887 | -0.025671257 | 0.115008344 | -0.140679601 | 0.351608515 |
| ENSG00000137802 | *MAPKBP1* | 3.743560151 | -0.018697367 | 0.093248075 | -0.111945442 | 0.999933887 | -0.082637113 | -0.18356716 | 0.100930047 | 0.351608515 |
| ENSG00000249379 | *AL033397.2* | -1.113143044 | 0.075993458 | -0.071816346 | 0.147809804 | 0.999933887 | 1.207746869 | 1.602089278 | -0.394342409 | 0.351608515 |
| ENSG00000254632 | *AP003119.1* | -0.569905368 | 0.379390363 | 0.777838473 | -0.39844811 | 0.999933887 | 1.029968865 | 1.325867191 | -0.295898325 | 0.351799078 |
| ENSG00000224078 | *SNHG14* | 3.234431898 | -0.057026138 | 0.027227155 | -0.084253294 | 0.999933887 | -0.233886852 | -0.03320948 | -0.200677371 | 0.351799078 |
| ENSG00000120254 | *MTHFD1L* | 1.239075089 | -0.033824557 | 0.008777977 | -0.042602534 | 0.999933887 | 0.109913197 | 0.317095879 | -0.207182682 | 0.352110272 |
| ENSG00000255097 | *AP006259.1* | 0.047991689 | -0.066831876 | 0.050322612 | -0.117154487 | 0.999933887 | 0.902021586 | 1.342963678 | -0.440942092 | 0.352161969 |
| ENSG00000166454 | *ATMIN* | 3.415805902 | 0.071020042 | -0.15756811 | 0.228588152 | 0.944094537 | -0.05423138 | -0.201254519 | 0.147023139 | 0.352521291 |
| ENSG00000142192 | *APP* | 4.131568789 | 0.032813067 | -0.018491544 | 0.051304611 | 0.999933887 | -0.121088453 | -0.249862764 | 0.12877431 | 0.352720904 |
| ENSG00000185864 | *NPIPB4* | 4.313147683 | 0.034776182 | -0.016644389 | 0.051420571 | 0.999933887 | 0.058381884 | 0.18115887 | -0.122776986 | 0.352772147 |
| ENSG00000152380 | *FAM151B* | 1.079325306 | -0.014868752 | -0.184768377 | 0.169899626 | 0.999933887 | 0.005557112 | 0.213545132 | -0.20798802 | 0.352772147 |
| ENSG00000078319 | *PMS2P1* | 0.764239792 | -0.12340407 | -0.105639572 | -0.017764498 | 0.999933887 | 0.095592231 | 0.378384183 | -0.282791952 | 0.35290127 |
| ENSG00000279078 | *SND1-IT1* | -0.409351536 | -0.016753824 | -0.062887261 | 0.046133437 | 0.999933887 | -0.751224253 | -1.272104518 | 0.520880264 | 0.352913778 |
| ENSG00000143889 | *HNRNPLL* | 4.192811192 | -0.008833143 | 0.01561078 | -0.024443923 | 0.999933887 | -0.027489692 | -0.149198741 | 0.121709049 | 0.352925843 |
| ENSG00000250696 | *AC111000.4* | -0.203645374 | -0.211005101 | -0.144125114 | -0.066879987 | 0.999933887 | -0.502791383 | -0.079245604 | -0.423545779 | 0.352936352 |
| ENSG00000140199 | *SLC12A6* | 9.438144699 | -0.008477561 | -0.001646773 | -0.006830788 | 0.999933887 | -0.151171368 | -0.063795286 | -0.087376082 | 0.353061853 |
| ENSG00000137713 | *PPP2R1B* | 2.976928779 | -0.015529356 | 0.011112822 | -0.026642178 | 0.999933887 | 0.102264042 | 0.274046706 | -0.171782664 | 0.353123988 |
| ENSG00000164040 | *PGRMC2* | 3.138218539 | 0.054345289 | 0.000671071 | 0.053674218 | 0.999933887 | -0.227795854 | -0.394330958 | 0.166535103 | 0.353939412 |
| ENSG00000121067 | *SPOP* | 5.269632138 | -0.096360545 | -0.092702938 | -0.003657607 | 0.999933887 | -0.445831988 | -0.565463109 | 0.119631121 | 0.353975889 |
| ENSG00000172757 | *CFL1* | 8.701399177 | 0.014597035 | -0.008805603 | 0.023402638 | 0.999933887 | -0.139559525 | -0.234750452 | 0.095190927 | 0.354142356 |
| ENSG00000152582 | *SPEF2* | 0.367782365 | -0.191341122 | -0.497923184 | 0.306582062 | 0.999933887 | -0.124293906 | 0.208560955 | -0.332854861 | 0.35421251 |
| ENSG00000132002 | *DNAJB1* | 7.066580199 | 0.060968987 | 0.005955945 | 0.055013042 | 0.999933887 | 0.153670655 | 0.340548631 | -0.186877975 | 0.354216864 |
| ENSG00000172057 | *ORMDL3* | 4.221535806 | 0.04159434 | 0.104647967 | -0.063053627 | 0.999933887 | 0.064777248 | -0.079590974 | 0.144368222 | 0.354216864 |
| ENSG00000114812 | *VIPR1* | 1.294519302 | 0.024252319 | 0.049598838 | -0.025346519 | 0.999933887 | -0.061106001 | 0.20481783 | -0.265923831 | 0.354216864 |
| ENSG00000022556 | *NLRP2* | 0.721392727 | -0.128956001 | 0.13798825 | -0.266944251 | 0.999933887 | -0.042906582 | 0.217242998 | -0.26014958 | 0.35451793 |
| ENSG00000198208 | *RPS6KL1* | -0.70218667 | 0.129872237 | -0.272542504 | 0.40241474 | 0.999933887 | -0.179580976 | 0.291210756 | -0.470791732 | 0.35451793 |

| | | | | | | | | | | |
|---|---|---|---|---|---|---|---|---|---|---|
| ENSG00000161618 | *ALDH16A1* | 2.812605969 | 0.022277659 | 0.1006259 | -0.078348241 | 0.999933887 | 0.005828021 | -0.204236424 | 0.210064445 | 0.35451793 |
| ENSG00000152348 | *ATG10* | 0.099883273 | -0.024490114 | 0.022737343 | -0.047227457 | 0.999933887 | 0.010635194 | -0.408065193 | 0.418700388 | 0.35451793 |
| ENSG00000109911 | *ELP4* | 0.681165284 | 0.233802019 | 0.010274759 | 0.223527261 | 0.999933887 | -0.001206608 | -0.338271795 | 0.337065187 | 0.354566477 |
| ENSG00000280254 | *AC233723.2* | 0.760334952 | -0.199707381 | -0.205908431 | 0.006201049 | 0.999933887 | 0.035804938 | 0.40290067 | -0.367095732 | 0.354970622 |
| ENSG00000285850 | *AC009152.3* | -0.184502454 | -0.107053254 | 0.097352373 | -0.204405627 | 0.999933887 | -0.090340091 | -0.585827757 | 0.495487665 | 0.354998195 |
| ENSG00000267898 | *AC026803.2* | 1.789180905 | 0.126597921 | -0.039399003 | 0.165996924 | 0.999933887 | 0.811521225 | 1.063667931 | -0.252146706 | 0.355079573 |
| ENSG00000171055 | *FEZ2* | 2.815488667 | -0.046845014 | -0.081922563 | 0.035077549 | 0.999933887 | -0.29374194 | -0.475995258 | 0.182253318 | 0.355079573 |
| ENSG00000127914 | *AKAP9* | 4.987613431 | -0.124039896 | -0.144475465 | 0.020435569 | 0.999933887 | -0.62351261 | -0.232415137 | -0.391097473 | 0.355317641 |
| ENSG00000090674 | *MCOLN1* | 8.106500044 | 0.007366072 | 0.010157061 | -0.002790989 | 0.999933887 | 0.201300657 | 0.307622492 | -0.106321835 | 0.355323826 |
| ENSG00000101782 | *RIOK3* | 6.543352794 | -0.078801406 | -0.082030565 | 0.003229159 | 0.999933887 | -0.275937254 | -0.18684425 | -0.089093005 | 0.355346346 |
| ENSG00000149531 | *FRG1BP* | 0.333922598 | -0.221310347 | 0.032213041 | -0.253523388 | 0.999933887 | -0.070386486 | 0.220856251 | -0.291242737 | 0.355346346 |
| ENSG00000283573 | *AL157371.2* | 2.02113488 | 0.122541947 | 0.172474207 | -0.049932261 | 0.999933887 | 0.400816371 | 0.613886998 | -0.213070627 | 0.355346346 |
| ENSG00000171813 | *PWWP2B* | 3.666612294 | 0.023350744 | -0.01202148 | 0.035372224 | 0.999933887 | -0.231895667 | -0.407841638 | 0.175945972 | 0.355346346 |
| ENSG00000232545 | *AC253536.3* | -0.698536236 | -0.001136879 | 0.059735856 | -0.060872735 | 0.999933887 | -0.023701704 | 0.416296277 | -0.43999798 | 0.355387896 |
| ENSG00000175787 | *ZNF169* | 1.368350021 | -0.054719128 | -0.083509637 | 0.028790509 | 0.999933887 | -0.055271926 | -0.263687431 | 0.208415505 | 0.355977015 |
| ENSG00000226742 | *HSBP1L1* | 1.548288812 | -0.028847669 | -0.114471769 | 0.0856241 | 0.999933887 | -0.605510601 | -0.820801394 | 0.215290792 | 0.355977015 |
| ENSG00000227218 | *AL157935.1* | -0.560209144 | 0.019605321 | 0.143325607 | -0.123720286 | 0.999933887 | -0.109391653 | -0.52027171 | 0.410880057 | 0.355977015 |
| ENSG00000103707 | *MTFMT* | 1.875024486 | 0.035969944 | 0.223492924 | -0.187522979 | 0.999933887 | -0.393715934 | -0.594404943 | 0.200689009 | 0.356041685 |
| ENSG00000068438 | *FTSJ1* | 3.195099048 | -0.015461418 | 0.05703249 | -0.072493908 | 0.999933887 | 0.14868579 | 0.294829944 | -0.146144153 | 0.356041685 |
| ENSG00000103005 | *USB1* | 6.165794939 | 0.045070264 | 0.0269466 | 0.018123663 | 0.999933887 | -0.070924893 | -0.187725662 | 0.116800769 | 0.356378959 |
| ENSG00000161671 | *EMC10* | 3.697993462 | 0.022698738 | 0.060372044 | -0.037673306 | 0.999933887 | 0.031315625 | -0.112421964 | 0.14373759 | 0.356378959 |
| ENSG00000072501 | *SMC1A* | 4.844964679 | -9.2528e-05 | -0.077804213 | 0.077711685 | 0.999933887 | -0.014743461 | -0.116655983 | 0.101912522 | 0.356378959 |
| ENSG00000063245 | *EPN1* | 6.241490577 | 0.037017346 | 0.005527575 | 0.031489771 | 0.999933887 | -0.430328304 | -0.540582635 | 0.110254331 | 0.35652937 |
| ENSG00000090054 | *SPTLC1* | 4.510396311 | -0.06780571 | -0.055823863 | -0.011981847 | 0.999933887 | -0.119922231 | -0.010489238 | -0.109432993 | 0.356880292 |
| ENSG00000184221 | *OLIG1* | 5.638544008 | -0.258780094 | -0.306452096 | 0.047672002 | 0.999933887 | -0.60374426 | -0.328075446 | -0.275668814 | 0.35694399 |
| ENSG00000076513 | *ANKRD13A* | 7.018866299 | -0.026798023 | -0.025715512 | -0.001082512 | 0.999933887 | 0.263277896 | 0.382618525 | -0.119340629 | 0.357025064 |
| ENSG00000131061 | *ZNF341* | 2.112106855 | 0.031154473 | -0.031282152 | 0.062436625 | 0.999933887 | -0.133438198 | -0.38574931 | 0.252311113 | 0.357025064 |
| ENSG00000228923 | *AP000355.1* | 1.765231273 | -0.096498151 | 0.103294003 | -0.199792154 | 0.999933887 | 1.968624991 | 2.249900755 | -0.281275764 | 0.357174819 |
| ENSG00000140848 | *CPNE2* | 4.023965172 | 0.024705316 | 0.00972188 | 0.014983437 | 0.999933887 | -0.690544489 | -0.838503031 | 0.147958542 | 0.357174819 |
| ENSG00000151150 | *ANK3* | 1.923752449 | 0.015036146 | -0.008849466 | 0.023885612 | 0.999933887 | -0.045169135 | 0.180316065 | -0.2254852 | 0.357296442 |
| ENSG00000163682 | *RPL9* | 7.4794904 | -0.005732044 | -0.036273427 | 0.030541383 | 0.999933887 | 0.1380829 | 0.249871946 | -0.111789046 | 0.357504145 |
| ENSG00000198734 | *F5* | 5.494285285 | -0.049137982 | -0.119938299 | 0.070800317 | 0.999933887 | -0.420356974 | -0.565716671 | 0.145359697 | 0.357509568 |
| ENSG00000145241 | *CENPC* | 3.529254886 | -0.017327101 | -0.073479436 | 0.056152335 | 0.999933887 | -0.20254082 | 0.157502246 | -0.360043066 | 0.35774515 |
| ENSG00000177494 | *ZBED2* | -0.020405615 | 0.116259734 | 0.163319303 | -0.047059569 | 0.999933887 | -0.101493386 | 0.320941056 | -0.422434442 | 0.357790074 |
| ENSG00000116128 | *BCL9* | 4.088692084 | -0.002239492 | 0.024705664 | -0.026945156 | 0.999933887 | -0.321257685 | -0.184073033 | -0.137184653 | 0.357806613 |
| ENSG00000144136 | *SLC20A1* | 8.433244065 | -0.047789401 | 0.032636562 | -0.080425963 | 0.999933887 | -0.453723701 | -0.585037194 | 0.131313492 | 0.35820432 |
| ENSG00000172071 | *EIF2AK3* | 5.033233361 | 0.009211509 | -0.131237738 | 0.140449247 | 0.999933887 | 0.163493722 | 0.38559389 | -0.222100168 | 0.358472074 |
| ENSG00000143153 | *ATP1B1* | -0.258498293 | 0.218374714 | -0.089295073 | 0.307669787 | 0.999933887 | -0.004198694 | 0.480391954 | -0.484590648 | 0.358484809 |
| ENSG00000103274 | *NUBP1* | 1.269119027 | 0.056236631 | -0.130287945 | 0.186524576 | 0.999933887 | -0.108001496 | 0.138663979 | -0.246665474 | 0.358484809 |
| ENSG00000120868 | *APAF1* | 5.255846987 | -0.064928051 | 0.028652483 | -0.093580534 | 0.999933887 | -0.423471673 | -0.569511886 | 0.146040213 | 0.358549544 |
| ENSG00000116455 | *WDR77* | 1.305458704 | -0.106922209 | 0.109383712 | -0.21630592 | 0.999933887 | 0.101801431 | -0.117352415 | 0.219153846 | 0.358549544 |
| ENSG00000153574 | *RPIA* | 2.581139018 | -0.009964451 | 0.038503585 | -0.048468036 | 0.999933887 | 0.157780371 | -0.012423241 | 0.170203612 | 0.358549544 |
| ENSG00000171148 | *TADA3* | 5.323937642 | 0.02251014 | 0.014110001 | 0.008400139 | 0.999933887 | -0.279752331 | -0.39390943 | 0.114157099 | 0.358549932 |
| ENSG00000105875 | *WDR91* | 7.082617896 | 0.01728914 | 0.077849118 | -0.060559977 | 0.999933887 | -0.239554751 | -0.144830353 | -0.094724398 | 0.358549932 |
| ENSG00000083454 | *P2RX5* | 0.706745193 | 0.053110915 | -0.141534896 | 0.194645811 | 0.999933887 | 0.129661833 | -0.146842368 | 0.276504201 | 0.358839724 |
| ENSG00000196781 | *TLE1* | 1.902813071 | 0.000907916 | 0.06679776 | -0.065889843 | 0.999933887 | 0.588658092 | 0.817375565 | -0.228717473 | 0.358839724 |
| ENSG00000267493 | *CIRBP-AS1* | 1.281881005 | -0.1052197 | -0.062682495 | -0.042537205 | 0.999933887 | -0.275979699 | -0.518016512 | 0.242036813 | 0.358843136 |
| ENSG00000049247 | *UTS2* | -2.806000659 | 0.149347421 | -0.565476276 | 0.714823697 | 0.999933887 | -0.144771686 | 0.392184635 | -0.536956322 | 0.358904523 |
| ENSG00000286786 | *AC116158.3* | 1.340154862 | -0.199388199 | 0.055667184 | -0.255055383 | 0.999933887 | -0.194449322 | 0.16084007 | -0.355289392 | 0.360594336 |
| ENSG00000186187 | *ZNRF1* | 1.321117236 | -0.125816528 | 0.218670863 | -0.344487391 | 0.999933887 | -0.156465008 | -0.43664758 | 0.280182573 | 0.360594336 |
| ENSG00000163617 | *CCDC191* | 1.141949104 | 0.014744216 | -0.17229402 | 0.187038236 | 0.999933887 | -0.192964846 | 0.12831462 | -0.321279466 | 0.360594336 |
| ENSG00000213585 | *VDAC1* | 3.509860399 | 0.036712655 | -0.089529155 | 0.126241809 | 0.999933887 | -0.008701901 | 0.128590862 | -0.137292763 | 0.360607712 |
| ENSG00000081791 | *DELE1* | 5.398371039 | -0.02172351 | -0.00616273 | -0.01556078 | 0.999933887 | -0.082535393 | -0.16659284 | 0.084057447 | 0.360607712 |
| ENSG00000163659 | *TIPARP* | 6.112379112 | -0.028103057 | -0.042090203 | 0.013987147 | 0.999933887 | 0.071715333 | 0.166491387 | -0.094776054 | 0.36064981 |
| ENSG00000128567 | *PODXL* | 0.054050208 | 0.125775057 | -0.037664373 | 0.16343943 | 0.999933887 | -0.212051343 | -0.615713617 | 0.403662274 | 0.360686344 |
| ENSG00000161912 | *ADCY10P1* | 3.051880221 | -0.019639035 | -0.099811001 | 0.080171966 | 0.999933887 | 0.090652065 | 0.289476314 | -0.198824249 | 0.360686344 |
| ENSG00000282826 | *FRG1CP* | 1.375804985 | -0.026195596 | -0.019268751 | -0.006926845 | 0.999933887 | 0.091364884 | -0.07759564 | 0.168960524 | 0.360687317 |
| ENSG00000094975 | *SUCO* | 5.585018969 | -0.004673814 | -0.120641431 | 0.115967617 | 0.999933887 | -0.068809011 | 0.189914488 | -0.258723499 | 0.360687317 |
| ENSG00000170004 | *CHD3* | 7.248221749 | 0.031350621 | 0.042607206 | -0.011256585 | 0.999933887 | 0.027104235 | -0.050198964 | 0.077303198 | 0.36072979 |
| ENSG00000177200 | *CHD9* | 3.696866315 | -0.082049359 | -0.154350548 | 0.072301189 | 0.999933887 | -0.432254558 | -0.162560459 | -0.269694099 | 0.36072979 |
| ENSG00000049245 | *VAMP3* | 5.78958427 | -0.032237109 | -0.000358051 | -0.031879058 | 0.999933887 | -0.709990168 | -0.84001985 | 0.130029682 | 0.36072979 |
| ENSG00000107890 | *ANKRD26* | 0.545235764 | -0.088757886 | -0.176348997 | 0.087591111 | 0.999933887 | -0.266654613 | 0.182068861 | -0.448723473 | 0.36072979 |
| ENSG00000281990 | *IGHV1-69-2* | -1.878267035 | 0.167143795 | 0.268361777 | -0.101217983 | 0.999933887 | 0.217371417 | -0.311811257 | 0.529182673 | 0.360797226 |
| ENSG00000258034 | *AC012157.2* | -0.351626018 | -0.063381374 | -0.02081815 | -0.042563224 | 0.999933887 | 0.161262507 | 0.670891176 | -0.509628669 | 0.360797226 |
| ENSG00000176386 | *CDC26* | 3.306479031 | 0.013597292 | -0.100143875 | 0.113741167 | 0.999933887 | -0.062845636 | 0.079908313 | -0.142753949 | 0.360797226 |
| ENSG00000102109 | *PCSK1N* | 0.23750683 | 0.046573191 | 0.092319779 | -0.045746588 | 0.999933887 | 0.23530347 | 0.574932345 | -0.339628875 | 0.36080581 |
| ENSG00000025293 | *PHF20* | 6.326554385 | -0.061903132 | -0.042342038 | -0.019561094 | 0.999933887 | -0.235867138 | -0.334220568 | 0.09835343 | 0.360878293 |
| ENSG00000233431 | *LINC02596* | 0.363072893 | -0.01036763 | 0.133133513 | -0.143501142 | 0.999933887 | 0.341349071 | -0.144145512 | 0.485494583 | 0.360878293 |
| ENSG00000106683 | *LIMK1* | 4.207537664 | 0.002143857 | 0.043842888 | -0.04169903 | 0.999933887 | -0.053186958 | -0.190130075 | 0.136916117 | 0.360878293 |
| ENSG00000155846 | *PPARGC1B* | 0.083817235 | 0.013335446 | -0.180845469 | 0.194180915 | 0.999933887 | -0.274624438 | 0.069454132 | -0.34407857 | 0.36092614 |
| ENSG00000159210 | *SNF8* | 3.953925884 | -0.030632071 | -0.025485385 | -0.005146686 | 0.999933887 | 0.322267876 | 0.431570044 | -0.109302168 | 0.36101882 |
| ENSG00000197299 | *BLM* | 1.066485812 | -0.233347064 | -0.072165999 | -0.161181065 | 0.999933887 | -0.180214539 | 0.040006558 | -0.220221098 | 0.361105283 |
| ENSG00000259962 | *AC007342.2* | -0.931597219 | -0.175998165 | -0.780959706 | 0.604961541 | 0.999933887 | -1.011621916 | -1.544172633 | 0.532550717 | 0.361199691 |
| ENSG00000259671 | *MTCYBP23* | 3.724417097 | 0.331451237 | 0.611899405 | -0.280448168 | 0.999933887 | 2.06420905 | 2.524689447 | -0.460480397 | 0.361352137 |
| ENSG00000204178 | *MACO1* | 3.710330426 | -0.013506816 | -0.115331309 | 0.101824493 | 0.999933887 | -0.102067934 | 0.011509361 | -0.113577295 | 0.361352137 |
| ENSG00000232888 | *RPS11P5* | -0.555938203 | -0.049398236 | 0.166949599 | -0.216347835 | 0.999933887 | -0.415627137 | -0.009054701 | -0.406572435 | 0.361352137 |
| ENSG00000268061 | *NAPA-AS1* | 2.522260848 | 0.131019405 | 0.267772895 | -0.13675349 | 0.999933887 | 0.309770388 | 0.562596305 | -0.252825917 | 0.361394836 |
| ENSG00000136367 | *ZFHX2* | 1.10371479 | -0.10452435 | 0.148625165 | -0.253149515 | 0.999933887 | 2.01505463 | 2.358230583 | -0.343175953 | 0.361472733 |
| ENSG00000073584 | *SMARCE1* | 1.448085843 | -0.165917921 | -0.034862923 | -0.131054998 | 0.999933887 | -0.384239118 | -0.120389413 | -0.263849705 | 0.36152472 |
| ENSG00000175455 | *CCDC14* | 2.477902514 | -0.12289013 | -0.306324955 | 0.183434825 | 0.999933887 | -0.309640865 | 0.040196457 | -0.349837322 | 0.36152472 |
| ENSG00000280205 | *AC009716.2* | 0.442062769 | 0.073263455 | 0.296857202 | -0.223593747 | 0.999933887 | -0.051700632 | -0.446110318 | 0.394409687 | 0.36152472 |
| ENSG00000186462 | *NAP1L2* | 0.03076592 | -0.068307155 | 0.244674532 | -0.312981688 | 0.999933887 | 0.075999885 | 0.45435371 | -0.378353826 | 0.36152472 |
| ENSG00000141577 | *CEP131* | 2.891563382 | 0.087747764 | 0.188092675 | -0.100344911 | 0.999933887 | 0.075702025 | 0.220146674 | -0.144444649 | 0.361781234 |
| ENSG00000103194 | *USP10* | 6.14833005 | -0.053305307 | -0.07203572 | 0.018730413 | 0.999933887 | -0.045026335 | 0.062604837 | -0.107631172 | 0.361781234 |
| ENSG00000010278 | *CD9* | 5.645110563 | -0.109933115 | 0.00920388 | -0.119136995 | 0.999933887 | -1.576421192 | -1.829326259 | 0.252905067 | 0.361892865 |
| ENSG00000169045 | *HNRNPH1* | 8.358918157 | -0.025019441 | -0.028016341 | 0.002996901 | 0.999933887 | 0.00357076 | 0.132026616 | -0.128455856 | 0.361986766 |
| ENSG00000081913 | *PHLPP1* | 3.940471875 | 0.037823268 | 0.028865847 | 0.008957421 | 0.999933887 | 0.498110556 | 0.652582183 | -0.154471628 | 0.362101545 |
| ENSG00000162777 | *DENND2D* | 5.917967184 | 0.033126259 | -0.007676245 | 0.040802503 | 0.999933887 | 0.304542099 | 0.421588157 | -0.117046058 | 0.362111968 |
| ENSG00000007264 | *MATK* | 4.193213542 | 0.025981775 | 0.076087899 | -0.050106125 | 0.999933887 | -0.034366065 | -0.162608294 | 0.128242229 | 0.362163054 |
| ENSG00000108839 | *ALOX12* | 4.902523687 | 0.110150747 | 0.118316098 | -0.008165351 | 0.999933887 | 0.227351133 | 0.336883461 | -0.109532328 | 0.362181082 |
| ENSG00000270164 | *LINC01480* | 0.427695897 | 0.13979113 | 0.090534609 | 0.049256521 | 0.999933887 | -0.497730146 | -0.016710764 | -0.481019382 | 0.362181082 |
| ENSG00000137414 | *FAM8A1* | 6.950513556 | -0.031488255 | -0.004570588 | -0.026917668 | 0.999933887 | -0.373747307 | -0.49503959 | 0.121292283 | 0.362181082 |
| ENSG00000184956 | *MUC6* | -0.742070872 | 0.028893865 | 0.072484536 | -0.043590671 | 0.999933887 | -0.111723646 | -0.715737002 | 0.604013356 | 0.362265116 |
| ENSG00000253616 | *AC107959.3* | 2.031289071 | 0.017447956 | -0.14949837 | 0.166946326 | 0.999933887 | -1.481906547 | -1.806813841 | 0.324907294 | 0.362265116 |
| ENSG00000268471 | *MIR4453HG* | 1.754664051 | 0.008148223 | 0.047392107 | -0.039243884 | 0.999933887 | 0.251615168 | 0.469386426 | -0.217771257 | 0.362265189 |
| ENSG00000213523 | *SRA1* | 6.742021569 | 0.067603516 | 0.04177951 | 0.025824006 | 0.999933887 | 0.099474753 | 0.205980213 | -0.10650546 | 0.362349628 |
| ENSG00000149577 | *SIDT2* | 4.441469327 | 0.057514336 | 0.007112512 | 0.050401824 | 0.999933887 | 0.042319958 | -0.049900539 | 0.092220497 | 0.362529395 |
| ENSG00000245156 | *AP001107.1* | 0.69864762 | -0.158672699 | 0.049826859 | -0.208499558 | 0.999933887 | -0.234658453 | 0.034422707 | -0.26908116 | 0.362867594 |
| ENSG00000101247 | *NDUFAF5* | 1.706433352 | -0.023735327 | 0.063145076 | -0.086880403 | 0.999933887 | -0.021586654 | -0.243762442 | 0.222175787 | 0.362867594 |
| ENSG00000155849 | *ELMO1* | 5.404428118 | 0.031447949 | 0.008523961 | 0.022923987 | 0.999933887 | -0.034247043 | -0.116973728 | 0.082726685 | 0.362919482 |
| ENSG00000112081 | *SRSF3* | 7.863614024 | 0.057426822 | 0.080143911 | -0.022717089 | 0.999933887 | 0.536205132 | 0.65259501 | -0.116389877 | 0.363074282 |
| ENSG00000253552 | *HOXA-AS2* | -1.088530908 | -0.060783543 | -0.066613386 | 0.005829843 | 0.999933887 | -0.803093361 | -1.429317111 | 0.626223751 | 0.363337606 |
| ENSG00000106636 | *YKT6* | 6.235661808 | -0.003957204 | -0.019258547 | 0.015301342 | 0.999933887 | -0.050241971 | 0.045769701 | -0.096011672 | 0.363860597 |
| ENSG00000143252 | *SDHC* | 3.469185499 | -0.046284174 | 0.015198875 | -0.061483049 | 0.999933887 | 0.074956127 | 0.21791288 | -0.142956753 | 0.363904868 |
| ENSG00000109111 | *SUPT6H* | 8.047464078 | -0.001374843 | -0.023363213 | 0.021988369 | 0.999933887 | 0.236642383 | 0.324009792 | -0.087367409 | 0.363904868 |

| | | | | | | | | | | |
|---|---|---|---|---|---|---|---|---|---|---|
| ENSG00000159256 | *MORC3* | 5.469482085 | -0.140518157 | -0.148689955 | 0.008171798 | 0.999933887 | -0.603085655 | -0.417152086 | -0.185933569 | 0.363906253 |
| ENSG00000231365 | *WARS2-AS1* | 0.213713671 | 0.019972425 | 0.143490882 | -0.123518457 | 0.999933887 | -0.109776491 | 0.200818077 | -0.310594568 | 0.363906253 |
| ENSG00000162139 | *NEU3* | 4.016449184 | -0.113104665 | -0.067964766 | -0.045139899 | 0.999933887 | -0.454023327 | -0.586036948 | 0.132013622 | 0.363948648 |
| ENSG00000166484 | *MAPK7* | 3.752722877 | 0.061723175 | 0.068301171 | -0.006577996 | 0.999933887 | -0.09518382 | -0.244286157 | 0.149102337 | 0.363948648 |
| ENSG00000100664 | *EIF5* | 8.535056685 | -0.014393258 | -0.035378461 | 0.020985203 | 0.999933887 | 0.382525519 | 0.486357975 | -0.103832456 | 0.364242949 |
| ENSG00000101343 | *CRNKL1* | 5.152064679 | -0.108995097 | -0.1140649 | 0.005069803 | 0.999933887 | -0.201440496 | -0.083225443 | -0.118215053 | 0.364459845 |
| ENSG00000179148 | *ALOXE3* | -0.978657269 | -0.129851249 | 0.376594288 | -0.506445537 | 0.999933887 | 0.442018914 | 0.778390463 | -0.336371548 | 0.364459845 |
| ENSG00000176476 | *SGF29* | 3.117667057 | -0.065579328 | -0.020119913 | -0.045459415 | 0.999933887 | -0.276550574 | -0.425380487 | 0.148829914 | 0.364527732 |
| ENSG00000103249 | *CLCN7* | 8.368751915 | 0.050797438 | 0.066755248 | -0.01595781 | 0.999933887 | 0.254225687 | 0.382386081 | -0.128160394 | 0.364710384 |
| ENSG00000116560 | *SFPQ* | 7.668325219 | -0.023515009 | -0.010376533 | -0.013138475 | 0.999933887 | 0.110047322 | 0.185558612 | -0.07551129 | 0.364710384 |
| ENSG00000127463 | *EMC1* | 3.106484377 | -0.010414883 | 0.025706233 | -0.036121116 | 0.999933887 | 0.153439934 | 0.31503612 | -0.161596187 | 0.364710384 |
| ENSG00000211460 | *TSN* | 3.739625819 | -0.063760145 | -0.048572056 | -0.015188089 | 0.999933887 | -0.109093596 | -0.232580928 | 0.123487331 | 0.364762608 |
| ENSG00000120093 | *HOXB3* | -0.559439483 | -0.169950739 | 0.032544071 | -0.20249481 | 0.999933887 | -0.415859473 | -0.014503392 | -0.401356081 | 0.364845737 |
| ENSG00000235192 | *AC009495.3* | 0.198212184 | 0.074141457 | -0.115098297 | 0.189239754 | 0.999933887 | 0.103259745 | -0.288857429 | 0.392117174 | 0.364845737 |
| ENSG00000253720 | *AC022973.3* | 3.639832241 | -0.060712192 | 0.101844457 | -0.162556648 | 0.999933887 | -0.085352347 | 0.198098505 | -0.283450853 | 0.364845737 |
| ENSG00000249884 | *RNF103-CHMI* | 2.211071345 | -0.014000303 | -0.065259124 | 0.05125882 | 0.999933887 | 0.085364715 | -0.096678012 | 0.182042727 | 0.364845737 |
| ENSG00000148399 | *DPH7* | 2.725876704 | 0.000287412 | 0.008703637 | -0.008416224 | 0.999933887 | 0.083681087 | -0.07679434 | 0.160475427 | 0.364845737 |
| ENSG00000068796 | *KIF2A* | 3.987042588 | -0.106116571 | -0.083486306 | -0.022630265 | 0.999933887 | -0.388965726 | -0.229509885 | -0.15945584 | 0.364940828 |
| ENSG00000245748 | *AC097382.2* | 0.173711045 | -0.083544646 | -0.208863289 | 0.125318643 | 0.999933887 | -0.583300721 | -1.02254509 | 0.439244369 | 0.364940828 |
| ENSG00000076356 | *PLXNA2* | 2.483909402 | 0.03779273 | 0.005074516 | 0.032718215 | 0.999933887 | 0.047305782 | -0.136554563 | 0.183860345 | 0.365004335 |
| ENSG00000138119 | *MYOF* | 1.365054692 | -0.021010531 | 0.068911495 | -0.089922026 | 0.999933887 | -0.260181515 | -0.026488028 | -0.233693487 | 0.365292603 |
| ENSG00000143751 | *SDE2* | 6.353893466 | 0.089687904 | 0.126668786 | -0.036980882 | 0.999933887 | -0.71506658 | -0.598336615 | -0.116729965 | 0.365540825 |
| ENSG00000162368 | *CMPK1* | 4.553337222 | -0.062609556 | 0.065267378 | -0.127876934 | 0.999933887 | -0.071521989 | 0.077092512 | -0.148614502 | 0.365694385 |
| ENSG00000166233 | *ARIH1* | 7.192679767 | -0.039530419 | -0.066059066 | 0.026528647 | 0.999933887 | 0.148224614 | 0.247596278 | -0.099371663 | 0.365694385 |
| ENSG00000139637 | *MYG1* | 2.7115491 | 0.035234375 | -0.064912087 | 0.100146462 | 0.999933887 | -0.328841253 | -0.496511606 | 0.167670353 | 0.365694385 |
| ENSG00000239300 | *AC080162.1* | 0.761365287 | -0.025599717 | 0.093012998 | -0.118612715 | 0.999933887 | 0.922383532 | 1.186393154 | -0.264009622 | 0.365694385 |
| ENSG00000198919 | *DZIP3* | 1.711805516 | -0.066459902 | -0.268267714 | 0.201807812 | 0.999933887 | 0.015397191 | 0.347619789 | -0.332222598 | 0.365718922 |
| ENSG00000147144 | *CCDC120* | 1.858412269 | -0.059768726 | 0.039640343 | -0.099409069 | 0.999933887 | 0.009542925 | -0.174637104 | 0.18418003 | 0.36581505 |
| ENSG00000237037 | *NDUFA6-DT* | 0.226532146 | 0.0927844 | -0.042885067 | 0.135669467 | 0.999933887 | -0.010335361 | 0.315928327 | -0.326263687 | 0.36581505 |
| ENSG00000148346 | *LCN2* | 0.940134419 | -0.111972622 | 0.070117283 | -0.182089904 | 0.999933887 | 0.826796766 | 1.088856904 | -0.262060139 | 0.366828627 |
| ENSG00000136710 | *CCDC115* | 3.512716487 | 0.05167643 | -0.020261004 | 0.071937435 | 0.999933887 | 0.073430284 | -0.067109691 | 0.140539975 | 0.366947919 |
| ENSG00000274561 | *AC005332.3* | 0.498350864 | 0.039722599 | -0.310926676 | 0.350649276 | 0.999933887 | -0.302365778 | -0.583853277 | 0.281487499 | 0.367068542 |
| ENSG00000260908 | *AC009093.3* | -0.267204477 | 0.092876251 | -0.068486257 | 0.161362508 | 0.999933887 | -0.098332008 | 0.252760043 | -0.351092051 | 0.367615392 |
| ENSG00000105501 | *SIGLEC5* | 3.903317228 | 0.064358789 | 0.024318833 | 0.040039956 | 0.999933887 | 0.726796412 | 0.579532615 | 0.147263797 | 0.367993007 |
| ENSG00000051825 | *MPHOSPH9* | 2.563549138 | 0.011476115 | -0.142735348 | 0.154211463 | 0.999933887 | -0.272802041 | -0.016373845 | -0.256428196 | 0.367993007 |
| ENSG00000101150 | *TPD52L2* | 9.48797349 | -0.034782345 | -0.050756454 | 0.015974109 | 0.999933887 | -0.105595922 | -0.016792402 | -0.08880352 | 0.368118952 |
| ENSG00000100201 | *DDX17* | 9.787003508 | -0.060647175 | -0.054393844 | -0.006253331 | 0.999933887 | -0.142450876 | -0.080764009 | -0.061686867 | 0.368381894 |
| ENSG00000277969 | *AC006449.6* | 4.368803151 | 0.157386804 | 0.110212489 | 0.047174314 | 0.999933887 | 2.153858215 | 2.340643443 | -0.186785228 | 0.368381894 |
| ENSG00000123552 | *USP45* | 0.683454735 | -0.210419605 | -0.345099133 | 0.134679528 | 0.999933887 | -0.210905656 | 0.193636305 | -0.404541961 | 0.368381894 |
| ENSG00000185798 | *WDR53* | 1.491238655 | 0.0452173 | -0.003384555 | 0.048601855 | 0.999933887 | -0.02214203 | -0.226057035 | 0.203915005 | 0.368381894 |
| ENSG00000174007 | *CEP19* | 3.910493899 | -0.025570049 | 0.045239313 | -0.070809362 | 0.999933887 | -0.714084537 | -0.899884345 | 0.185799807 | 0.368381894 |
| ENSG00000262777 | *AC032044.1* | 1.023150279 | -0.001175054 | 0.121028448 | -0.122203502 | 0.999933887 | -0.201966822 | 0.158766051 | -0.360732873 | 0.368381894 |
| ENSG00000064313 | *TAF2* | 4.213699883 | -4.4986e-05 | -0.069304894 | 0.069259907 | 0.999933887 | -0.05739552 | 0.122088246 | -0.179483766 | 0.368381894 |
| ENSG00000125449 | *ARMC7* | 4.170606945 | -0.005634587 | -0.131602068 | 0.125967481 | 0.999933887 | -0.437372958 | -0.570575973 | 0.133203015 | 0.368426058 |
| ENSG00000152223 | *EPG5* | 7.057133794 | -0.055738766 | -0.011752175 | -0.043986591 | 0.999933887 | 0.438320252 | 0.556378727 | -0.118058475 | 0.368450453 |
| ENSG00000283959 | *AP002851.1* | -0.55527548 | -0.21857555 | -0.846347385 | 0.627771835 | 0.999933887 | 2.11624527 | 1.762232205 | 0.354013065 | 0.368450453 |
| ENSG00000108840 | *HDAC5* | 7.823910508 | -0.001456307 | 0.021145381 | -0.022601688 | 0.999933887 | -0.340331396 | -0.436009098 | 0.095677702 | 0.368591693 |
| ENSG00000137055 | *PLAA* | 4.512938918 | -0.035769743 | -0.067457669 | 0.031687926 | 0.999933887 | -0.063226786 | 0.036252049 | -0.099478836 | 0.369028409 |
| ENSG00000197180 | *CH17-340M24* | 0.513830333 | -0.340069567 | -0.240384903 | -0.099684664 | 0.999933887 | 0.389753286 | 0.628812001 | -0.239058715 | 0.369099729 |
| ENSG00000182473 | *EXOC7* | 5.253924349 | 0.072005518 | -0.042313964 | 0.114319482 | 0.999933887 | -0.091875887 | -0.192249754 | 0.100373867 | 0.369795094 |
| ENSG00000101460 | *MAP1LC3A* | 6.36211131 | 0.074233739 | 0.095400031 | -0.021166292 | 0.999933887 | 1.127306007 | 1.268703512 | -0.141397505 | 0.369877318 |
| ENSG00000268154 | *Metazoa_SRP* | -0.239224878 | -0.063048655 | -0.124337609 | 0.061288954 | 0.999933887 | -0.51833434 | -0.129489609 | -0.388844732 | 0.369877318 |
| ENSG00000167549 | *CORO6* | 0.551345291 | -0.066817596 | 0.255092691 | -0.321910287 | 0.999933887 | 0.011170746 | -0.28035401 | 0.291524755 | 0.369877318 |
| ENSG00000148335 | *NTMT1* | 2.188645246 | -0.021595686 | 0.055407221 | -0.077002906 | 0.999933887 | 0.213414705 | -0.005990317 | 0.219405022 | 0.369877318 |
| ENSG00000100104 | *SRRD* | 2.176876129 | -0.012643655 | -0.054296939 | 0.041653284 | 0.999933887 | -0.017064563 | 0.187500031 | -0.204564593 | 0.369877318 |
| ENSG00000244165 | *P2RY11* | 0.593884999 | 0.022022967 | 0.010365695 | 0.011657271 | 0.999933887 | 0.125941369 | -0.200135017 | 0.326076386 | 0.370040162 |
| ENSG00000145545 | *SRD5A1* | 0.658931541 | -0.080663703 | 0.158552316 | -0.239216019 | 0.999933887 | -0.162411692 | 0.15100481 | -0.313416501 | 0.370143087 |
| ENSG00000287974 | *Z84486.1* | -0.539504172 | -0.19717829 | -0.336185462 | 0.139007171 | 0.999933887 | -0.018823516 | 0.306324406 | -0.325147921 | 0.37021375 |
| ENSG00000141469 | *SLC14A1* | 0.677726222 | -0.147398368 | 0.020686048 | -0.168084416 | 0.999933887 | -0.008353537 | 0.245181413 | -0.25353495 | 0.370595106 |
| ENSG00000272282 | *LINC02084* | 0.840063038 | -0.273219037 | -0.09486794 | -0.178351097 | 0.999933887 | 0.082981139 | 0.325189508 | -0.242208369 | 0.370734274 |
| ENSG00000124091 | *GCNT7* | 2.993760338 | 0.124773369 | 0.148893572 | -0.024120204 | 0.999933887 | -0.030070699 | 0.183648944 | -0.213719643 | 0.370734274 |
| ENSG00000137822 | *TUBGCP4* | 1.928927164 | 0.077248286 | 0.154569101 | -0.077320815 | 0.999933887 | -0.006788255 | 0.178892808 | -0.185681063 | 0.370734274 |
| ENSG00000285967 | *NIPBL-DT* | 3.46345297 | -0.072732672 | -0.061291506 | -0.011441166 | 0.999933887 | -0.405536175 | -0.556062197 | 0.150526022 | 0.370734274 |
| ENSG00000132963 | *POMP* | 4.900097725 | -0.018915761 | -0.058641937 | 0.039726176 | 0.999933887 | 0.12937169 | 0.227908959 | -0.098537269 | 0.370734274 |
| ENSG00000179832 | *MROH1* | 4.919609217 | 0.006657 | 0.013948916 | -0.007291916 | 0.999933887 | -0.154736145 | -0.261224788 | 0.106488643 | 0.370734274 |
| ENSG00000196338 | *NLGN3* | 4.140438326 | 0.094009698 | -0.091310927 | 0.185320625 | 0.981460244 | 0.499772189 | 0.630152827 | -0.130380638 | 0.370794933 |
| ENSG00000177646 | *ACAD9* | 2.859261656 | -0.042693046 | 0.136619445 | -0.179312492 | 0.999933887 | -0.097590179 | -0.226689548 | 0.12909937 | 0.371267928 |
| ENSG00000107554 | *DNMBP* | 4.198786706 | -0.107302706 | -0.012397307 | -0.094905399 | 0.999933887 | -0.824112892 | -0.971348856 | 0.147235964 | 0.37164658 |
| ENSG00000249138 | *SLED1* | 2.257430437 | 0.125731172 | 0.128731127 | -0.002999955 | 0.999933887 | 2.235949435 | 1.810525522 | 0.425423913 | 0.371850248 |
| ENSG00000172922 | *RNASEH2C* | 4.023713011 | 0.091356528 | 0.059511046 | 0.031845482 | 0.999933887 | 0.173252283 | 0.296157768 | -0.122905485 | 0.371956121 |
| ENSG00000095059 | *DHPS* | 1.131322036 | -0.061242104 | 0.16105992 | -0.222302024 | 0.999933887 | -0.088422655 | -0.351667664 | 0.263245009 | 0.371956121 |
| ENSG00000171476 | *HOPX* | 2.513463727 | 0.055587764 | 0.098111151 | -0.042523387 | 0.999933887 | 0.041300112 | 0.205162415 | -0.163862303 | 0.372215488 |
| ENSG00000067836 | *ROGDI* | 0.553130624 | 0.323886909 | 0.05244132 | 0.271445588 | 0.999933887 | 0.047845459 | -0.343599546 | 0.391445005 | 0.372407784 |
| ENSG00000173077 | *DELEC1* | 1.215529739 | 0.299355582 | 0.602792516 | -0.303436935 | 0.999933887 | 0.447940986 | 0.708464918 | -0.260523933 | 0.372407784 |
| ENSG00000065057 | *NTHL1* | -0.424926111 | 0.189827882 | -0.00982622 | 0.199654101 | 0.999933887 | 0.235045573 | -0.205709034 | 0.440754607 | 0.372407784 |
| ENSG00000107949 | *BCCIP* | 2.37646899 | -0.160315022 | -0.081095042 | -0.07921998 | 0.999933887 | 0.005301557 | 0.19992387 | -0.194622312 | 0.372798759 |
| ENSG00000273340 | *MICE* | -0.816529719 | 0.271688668 | -0.126233249 | 0.397921916 | 0.999933887 | -0.337013465 | 0.104942662 | -0.441956127 | 0.373125755 |
| ENSG00000106113 | *CRHR2* | -0.664702178 | 0.125723365 | 0.21793021 | -0.092206845 | 0.999933887 | 0.496334828 | 0.076646406 | 0.419688422 | 0.373125755 |
| ENSG00000123213 | *NLN* | 4.576596684 | -0.034205582 | 0.015164003 | -0.049369586 | 0.999933887 | -0.10303822 | 0.027005987 | -0.130044207 | 0.373379262 |
| ENSG00000160445 | *ZER1* | 5.588116733 | -0.029708047 | -0.025833001 | -0.003875046 | 0.999933887 | -0.53300639 | -0.643976473 | 0.110970083 | 0.373440892 |
| ENSG00000114942 | *EEF1B2* | 6.160099324 | -0.00578463 | -0.056269301 | 0.050484671 | 0.999933887 | 0.109229487 | 0.223757939 | -0.114528453 | 0.373700145 |
| ENSG00000166529 | *ZSCAN21* | 1.033910891 | 0.035977367 | -0.283677702 | 0.319655069 | 0.999933887 | 0.387408259 | 0.698929544 | -0.311521286 | 0.373982971 |
| ENSG00000116489 | *CAPZA1* | 8.246678056 | -0.090879542 | -0.062448548 | -0.028430994 | 0.999933887 | -0.194895944 | -0.088140422 | -0.106755522 | 0.374119562 |
| ENSG00000196605 | *ZNF846* | 0.059893101 | -0.172702522 | -0.200228491 | 0.027525969 | 0.999933887 | -0.144729239 | 0.143423449 | -0.288152688 | 0.374119562 |
| ENSG00000160218 | *TRAPPC10* | 6.174930517 | -0.027556088 | 0.02658477 | -0.054140858 | 0.999933887 | -0.08096723 | -0.004748724 | -0.076218505 | 0.374172342 |
| ENSG00000183340 | *JRKL* | 2.77374686 | -0.045845437 | 0.068541334 | -0.114386771 | 0.999933887 | 1.000544914 | 1.218124993 | -0.217580079 | 0.374172342 |
| ENSG00000223534 | *HLA-DQB1-AS* | -0.331917848 | -0.002466037 | 0.417829963 | -0.420296 | 0.999933887 | 1.497090189 | 1.963914676 | -0.466824487 | 0.374269278 |
| ENSG00000269343 | *ZNF587B* | 2.824033083 | 0.021292177 | 0.009806596 | 0.011485581 | 0.999933887 | 0.024718117 | -0.148884353 | 0.17360247 | 0.374428041 |
| ENSG00000204580 | *DDR1* | 0.361503295 | 0.056090212 | -0.274976378 | 0.33106659 | 0.999933887 | 0.280931732 | 0.541579172 | -0.26064744 | 0.374779213 |
| ENSG00000236349 | *SUCLG2P2* | -0.909910089 | 0.156993399 | 0.268678448 | -0.11168505 | 0.999933887 | -0.701486857 | 0.025989134 | -0.727475991 | 0.374889026 |
| ENSG00000140320 | *BAHD1* | 3.940762052 | 0.039642831 | -0.011727742 | 0.051370573 | 0.999933887 | -0.057894523 | -0.177705304 | 0.11981078 | 0.374889026 |
| ENSG00000143198 | *MGST3* | 2.188262771 | 0.065338861 | -0.009134744 | 0.074473605 | 0.999933887 | 0.097719882 | -0.112912066 | 0.210631948 | 0.375021046 |
| ENSG00000138735 | *PDE5A* | 0.572978855 | 0.006078248 | -0.020340539 | 0.026418787 | 0.999933887 | -0.44385956 | 0.040903341 | -0.4847629 | 0.375298107 |
| ENSG00000234975 | *FTH1P2* | 4.100635736 | -0.029650447 | 0.010966521 | -0.040616969 | 0.999933887 | 0.5418545 | 0.702328298 | -0.160473797 | 0.375485899 |
| ENSG00000014257 | *ACP3* | 3.466879421 | -0.082559805 | 0.049077123 | -0.131636928 | 0.999933887 | -0.36456193 | -0.646071522 | 0.281509592 | 0.375586193 |
| ENSG00000163534 | *FCRL1* | 1.076574914 | 0.011578103 | 0.067886773 | -0.056308669 | 0.999933887 | -0.12209811 | -0.374882941 | 0.252784831 | 0.375749598 |
| ENSG00000005884 | *ITGA3* | 0.610749712 | -0.014233097 | 0.040380178 | -0.054613275 | 0.999933887 | 0.054029087 | 0.369354742 | -0.315325655 | 0.376195433 |
| ENSG00000109113 | *RAB34* | 0.383627074 | 0.161946189 | -0.082716578 | 0.244662768 | 0.999933887 | 0.294809401 | -0.018154609 | 0.31296401 | 0.376276728 |
| ENSG00000073711 | *PPP2R3A* | -0.045484615 | -0.183941562 | -0.120758963 | -0.063182599 | 0.999933887 | -0.56574603 | -0.119582715 | -0.446163315 | 0.376899573 |
| ENSG00000152332 | *UHMK1* | 5.389817851 | 0.05161173 | -0.009231916 | 0.060843645 | 0.999933887 | -0.329155875 | -0.181668707 | -0.147487168 | 0.376899573 |
| ENSG00000150938 | *CRIM1* | 2.37914164 | 0.050996413 | 0.150095153 | -0.09909874 | 0.999933887 | 0.157358102 | 0.31913981 | -0.161781708 | 0.376899573 |
| ENSG00000147443 | *DOK2* | 4.713165953 | 0.060239557 | 0.000953122 | 0.059286436 | 0.999933887 | -0.065822539 | -0.189935341 | 0.124112802 | 0.376972341 |
| ENSG00000113732 | *ATP6V0E1* | 8.090158561 | -0.003658517 | -0.060471099 | 0.056812582 | 0.999933887 | 0.066179434 | 0.163979936 | -0.097800502 | 0.376974037 |
| ENSG00000166839 | *ANKDD1A* | 4.454321501 | -0.047218641 | -0.109410041 | 0.0621914 | 0.999933887 | 0.380969387 | 0.552529139 | -0.171559752 | 0.377198657 |
| ENSG00000179344 | *HLA-DQB1* | 5.351165141 | -0.0738818 | 0.046361135 | -0.120242936 | 0.999933887 | 1.182614356 | 1.466902111 | -0.284287755 | 0.377536172 |

| | | | | | | | | | | |
|---|---|---|---|---|---|---|---|---|---|---|
| ENSG00000114503 | *NCBP2* | 3.6993436 | -0.060041361 | 0.000490683 | -0.060532044 | 0.999933887 | 0.051908714 | 0.161987953 | -0.110079239 | 0.377795701 |
| ENSG00000171291 | *ZNF439* | 0.597094643 | -0.079471236 | 0.12796122 | -0.207432456 | 0.999933887 | -0.283591006 | -0.605233802 | 0.321642795 | 0.377795701 |
| ENSG00000124275 | *MTRR* | 3.158104122 | -0.129551424 | -0.165464664 | 0.035913241 | 0.999933887 | 0.015063765 | 0.147233642 | -0.132169877 | 0.378035441 |
| ENSG00000104812 | *GYS1* | 3.41949693 | 0.05499024 | -0.031636326 | 0.086626566 | 0.999933887 | 0.11109698 | -0.011481421 | 0.122578401 | 0.378035441 |
| ENSG00000272419 | *LINC01145* | 1.418442993 | -0.000690544 | -0.170704014 | 0.17001347 | 0.999933887 | -0.134930093 | -0.391558607 | 0.256628514 | 0.378035441 |
| ENSG00000267796 | *LIN37* | 1.039713059 | -0.116709479 | 0.126473452 | -0.243182931 | 0.999933887 | -0.506719416 | -0.759563961 | 0.252844544 | 0.378269623 |
| ENSG00000116584 | *ARHGEF2* | 7.724243173 | 0.002873438 | 0.018783891 | -0.015910454 | 0.999933887 | 0.302560114 | 0.395795056 | -0.093234942 | 0.378269623 |
| ENSG00000134294 | *SLC38A2* | 7.394747794 | -0.035171989 | -0.085637359 | 0.05046537 | 0.999933887 | -0.480473853 | -0.298212752 | -0.182261102 | 0.37854352 |
| ENSG00000006327 | *TNFRSF12A* | 3.225359528 | -0.102892072 | -0.161232949 | 0.058340877 | 0.999933887 | -1.643643547 | -1.954847409 | 0.311203862 | 0.378550461 |
| ENSG00000178980 | *SELENOW* | 3.292866181 | 0.056629476 | 0.014943391 | 0.041686085 | 0.999933887 | 0.216872977 | 0.052256419 | 0.164616558 | 0.378699312 |
| ENSG00000166908 | *PIP4K2C* | 5.295106153 | -0.085580772 | 0.022063155 | -0.107643927 | 0.999933887 | -0.516446857 | -0.616997797 | 0.100550941 | 0.378853753 |
| ENSG00000005893 | *LAMP2* | 8.665768927 | -0.004552788 | -0.053196975 | 0.048644187 | 0.999933887 | 0.011844566 | 0.096331836 | -0.08448727 | 0.378917794 |
| ENSG00000107816 | *LZTS2* | 1.705599241 | 0.144016989 | 0.040788503 | 0.103228487 | 0.999933887 | -0.007944771 | -0.215950776 | 0.208006005 | 0.379294838 |
| ENSG00000197081 | *IGF2R* | 11.01422367 | -0.028434258 | -0.003597299 | -0.024836959 | 0.999933887 | -0.197073138 | -0.11142838 | -0.085644757 | 0.379294838 |
| ENSG00000083828 | *ZNF586* | 4.437113287 | -0.079517113 | -0.068225191 | -0.011291922 | 0.999933887 | -0.280968216 | -0.392004919 | 0.111036702 | 0.379345946 |
| ENSG00000125454 | *SLC25A19* | 0.736690482 | 0.189057027 | -0.024225594 | 0.213282621 | 0.999933887 | 0.09104383 | -0.174620628 | 0.265664458 | 0.379345946 |
| ENSG00000124104 | *SNX21* | -0.270859692 | 0.168447846 | -0.043179416 | 0.211627262 | 0.999933887 | -0.215653991 | -0.575287331 | 0.359633341 | 0.37935159 |
| ENSG00000100528 | *CNIH1* | 2.53865695 | -0.056750867 | -0.065716808 | 0.008965941 | 0.999933887 | -0.098140826 | 0.062936626 | -0.161077451 | 0.37935159 |
| ENSG00000247925 | *AL139807.1* | 0.858850205 | -0.091848658 | 0.40265906 | -0.494507718 | 0.944094537 | -0.104917666 | 0.271795859 | -0.376713526 | 0.37935159 |
| ENSG00000104885 | *DOT1L* | 6.940421428 | 0.036643413 | -0.009324892 | 0.045968305 | 0.999933887 | 1.56724835 | 1.784352272 | -0.217103922 | 0.37935159 |
| ENSG00000103313 | *MEFV* | 7.446818147 | -0.038381309 | 0.017267778 | -0.055649087 | 0.999933887 | -0.029295517 | -0.180725125 | 0.151429608 | 0.379402196 |
| ENSG00000204386 | *NEU1* | 7.767864153 | 0.096946745 | 0.049268347 | 0.047678398 | 0.999933887 | 0.298189931 | 0.406268465 | -0.108078534 | 0.379531709 |
| ENSG00000078098 | *FAP* | -1.174151871 | -0.493735099 | -0.596786423 | 0.103051325 | 0.999933887 | 0.800126832 | 1.201035813 | -0.400908981 | 0.379590599 |
| ENSG00000177830 | *CHID1* | 1.720491345 | -0.001151132 | -0.147368363 | 0.14621723 | 0.999933887 | 0.056848607 | -0.160532186 | 0.217380793 | 0.379802765 |
| ENSG00000129292 | *PHF20L1* | 6.857645857 | -0.03787017 | -0.035121688 | -0.002748481 | 0.999933887 | -0.241184373 | -0.097141385 | -0.144042988 | 0.379894157 |
| ENSG00000262714 | *AC007342.5* | 1.316421687 | -0.038689849 | 0.028026603 | -0.066716453 | 0.999933887 | -1.272340439 | -1.73047583 | 0.458135391 | 0.379972053 |
| ENSG00000259398 | *AC090970.1* | -0.372334467 | -0.038359207 | 0.349365917 | -0.387725124 | 0.999933887 | 1.038287797 | 1.526746164 | -0.488458367 | 0.380188829 |
| ENSG00000143164 | *DCAF6* | 3.411384722 | 0.008196 | -0.213332063 | 0.221528063 | 0.999291887 | -0.446506252 | -0.615785179 | 0.169278928 | 0.380347662 |
| ENSG00000157890 | *MEGF11* | 0.719603768 | -0.155368909 | -0.032936656 | -0.122432254 | 0.999933887 | 0.680135512 | 0.908600627 | -0.228465115 | 0.38044301 |
| ENSG00000197386 | *HTT* | 4.879418027 | 0.029504681 | 0.052791643 | -0.023286962 | 0.999933887 | 0.057398561 | -0.041129222 | 0.098527784 | 0.38044301 |
| ENSG00000158717 | *RNF166* | 6.90614422 | 0.053435582 | 0.032354283 | 0.021081299 | 0.999933887 | -0.084959978 | -0.190769752 | 0.105809774 | 0.38060018 |
| ENSG00000135473 | *PAN2* | 3.38072249 | -0.03421153 | 0.030811634 | -0.065023164 | 0.999933887 | 0.26736312 | 0.42044701 | -0.15308389 | 0.380704108 |
| ENSG00000189420 | *ZFP92* | 2.417525829 | -0.050216324 | -0.201946491 | 0.151730167 | 0.999933887 | -0.971901644 | -1.168368905 | 0.196467261 | 0.380718642 |
| ENSG00000103540 | *CCP110* | 2.142730556 | -0.044484602 | -0.061049882 | 0.01656528 | 0.999933887 | -0.416757767 | -0.058710436 | -0.358047331 | 0.380718642 |
| ENSG00000151239 | *TWF1* | 3.464165263 | -0.034072646 | -0.186138454 | 0.152065808 | 0.999933887 | -0.536807069 | -0.324442723 | -0.212364347 | 0.380777679 |
| ENSG00000105607 | *GCDH* | 1.983243793 | 0.030672509 | -0.045373673 | 0.076046182 | 0.999933887 | 0.094715785 | -0.09568521 | 0.190400995 | 0.380777679 |
| ENSG00000122565 | *CBX3* | 5.787085993 | -0.046973836 | -0.114375841 | 0.067402005 | 0.999933887 | -0.479959138 | -0.584585931 | 0.104626793 | 0.38096682 |
| ENSG00000072778 | *ACADVL* | 6.799847763 | 0.071405485 | 0.053135373 | 0.018270113 | 0.999933887 | -0.171372161 | -0.280579495 | 0.109207334 | 0.381027968 |
| ENSG00000269743 | *SLC25A53* | 2.70028341 | -0.073792198 | 0.122008587 | -0.195800785 | 0.999933887 | -0.075023246 | 0.052541451 | -0.127564697 | 0.381071298 |
| ENSG00000242960 | *FTH1P23* | 0.262732854 | -0.059272354 | -0.341528149 | 0.282255795 | 0.999933887 | 0.396849768 | 0.670544485 | -0.273694716 | 0.381071298 |
| ENSG00000104763 | *ASAH1* | 10.305378173 | -0.076675857 | -0.079284477 | 0.00260862 | 0.999933887 | -0.24100979 | -0.155568831 | -0.085440959 | 0.38115011 |
| ENSG00000149016 | *TUT1* | 1.110456634 | -0.188367653 | 0.140112044 | -0.328479697 | 0.999933887 | 0.019790605 | -0.215492774 | 0.235283378 | 0.381207458 |
| ENSG00000092068 | *SLC7A8* | -0.698943028 | -0.094403739 | 0.168889709 | -0.263293448 | 0.999933887 | -0.604992088 | -0.111010749 | -0.493981339 | 0.381207458 |
| ENSG00000058091 | *CDK14* | 4.231312167 | -0.000722575 | 0.007499272 | -0.008221846 | 0.999933887 | 0.419949057 | 0.269052102 | 0.150896955 | 0.381207458 |
| ENSG00000211810 | *TRAV29DV5* | -0.706489094 | 0.063688157 | 0.179116063 | -0.115427906 | 0.999933887 | -0.124262899 | 0.29649444 | -0.420757339 | 0.381234535 |
| ENSG00000231794 | *AC009542.1* | 2.026779168 | 0.076929308 | 0.108675957 | -0.031746649 | 0.999933887 | -0.190898678 | 0.081445804 | -0.272344482 | 0.381868101 |
| ENSG00000173542 | *MOB1B* | 1.559615076 | -0.058834734 | -0.150967069 | 0.092132335 | 0.999933887 | -0.331049311 | -0.025499353 | -0.305549958 | 0.381983074 |
| ENSG00000273148 | *LINC00653* | 0.313817549 | -0.223799798 | -0.079027547 | -0.144772251 | 0.999933887 | 0.072902415 | 0.406435334 | -0.333532919 | 0.382091206 |
| ENSG00000127951 | *FGL2* | 6.29646366 | -0.035509321 | -0.156626057 | 0.121116736 | 0.999933887 | -0.494042089 | -0.717069181 | 0.223027093 | 0.382123807 |
| ENSG00000171865 | *RNASEH1* | 2.508943219 | -0.030233294 | -0.146159446 | 0.115926152 | 0.999933887 | -0.071789547 | 0.089620742 | -0.161410289 | 0.382131174 |
| ENSG00000102181 | *CD99L2* | 2.445666831 | 0.05451297 | 0.014951716 | 0.039561254 | 0.999933887 | -0.028331154 | -0.231388497 | 0.203057343 | 0.382171697 |
| ENSG00000184009 | *ACTG1* | 10.349632366 | 0.044226033 | 0.022810069 | 0.021415964 | 0.999933887 | -0.182765079 | -0.281125807 | 0.098360728 | 0.382220866 |
| ENSG00000146592 | *CREB5* | 5.816777582 | -0.029019984 | -0.025523401 | -0.003496582 | 0.999933887 | -0.935602654 | -1.13845047 | 0.202847816 | 0.382258854 |
| ENSG00000102245 | *CD40LG* | 1.983078327 | -0.072930201 | -0.208885991 | 0.13595579 | 0.999933887 | 0.208433518 | 0.412913192 | -0.204479674 | 0.382414632 |
| ENSG00000180357 | *ZNF609* | 4.431092604 | 0.021062097 | 0.035639478 | -0.01457738 | 0.999933887 | -0.029529789 | 0.080777304 | -0.110307093 | 0.382414632 |
| ENSG00000167766 | *ZNF83* | 3.345115819 | -0.090776945 | -0.049175308 | -0.041601638 | 0.999933887 | -0.142667302 | 0.066896525 | -0.209563827 | 0.382441297 |
| ENSG00000155506 | *LARP1* | 6.134940474 | 0.005278305 | 0.086688688 | -0.081410383 | 0.999933887 | 0.001813239 | 0.116595338 | -0.1147821 | 0.382441297 |
| ENSG00000122223 | *CD244* | 5.133626918 | -0.033470538 | -0.063812895 | 0.030342357 | 0.999933887 | -0.810754711 | -0.652614323 | -0.158140388 | 0.382862442 |
| ENSG00000103932 | *RPAP1* | 1.986674762 | 0.017144659 | 0.034813466 | -0.017668807 | 0.999933887 | 0.124258882 | 0.290649414 | -0.166390532 | 0.382992475 |
| ENSG00000243806 | *RPL7P18* | 0.030434381 | -0.257494505 | -0.035932123 | -0.221562382 | 0.999933887 | 1.500037705 | 1.875527737 | -0.375490031 | 0.383647413 |
| ENSG00000259635 | *AC100830.1* | 0.057180124 | -0.143930877 | -0.063550126 | -0.080380752 | 0.999933887 | 0.281295666 | 0.602183371 | -0.320887705 | 0.383789656 |
| ENSG00000139370 | *SLC15A4* | 8.931341969 | -0.073241718 | -0.035787602 | -0.037454116 | 0.999933887 | -0.265937564 | -0.169097999 | -0.096839565 | 0.384213048 |
| ENSG00000250318 | *AC003072.1* | 1.117551135 | -0.125125658 | 0.033147813 | -0.158273471 | 0.999933887 | 0.107523897 | -0.144579226 | 0.252103124 | 0.384213048 |
| ENSG00000108774 | *RAB5C* | 6.164366881 | 0.011036165 | -0.032953513 | 0.043989678 | 0.999933887 | 0.058621529 | 0.154876856 | -0.096255327 | 0.384213048 |
| ENSG00000123600 | *METTL8* | 0.849570795 | -0.016807612 | 0.068966362 | -0.085773974 | 0.999933887 | -0.013922002 | 0.235299458 | -0.249221461 | 0.384213048 |
| ENSG00000123179 | *EBPL* | 1.460654345 | 0.030780455 | -0.153200705 | 0.18398116 | 0.999933887 | -0.331557233 | -0.619568975 | 0.288011742 | 0.384282822 |
| ENSG00000175606 | *TMEM70* | 4.321381871 | -0.094288504 | -0.067461848 | -0.026826656 | 0.999933887 | -0.867357386 | -1.019890483 | 0.152533097 | 0.384393035 |
| ENSG00000181690 | *PLAG1* | -0.010813796 | -0.170554963 | -0.522813352 | 0.352258388 | 0.999933887 | -0.158015518 | 0.144684193 | -0.302699711 | 0.384393035 |
| ENSG00000196597 | *ZNF782* | 0.42847924 | 0.083537772 | 0.229350018 | -0.145812245 | 0.999933887 | -0.116294678 | 0.128565438 | -0.244860117 | 0.384438975 |
| ENSG00000040341 | *STAU2* | 2.637115849 | 0.0549952 | -0.172174649 | 0.22716985 | 0.999933887 | -0.397340349 | -0.603685985 | 0.206345636 | 0.384438975 |
| ENSG00000129467 | *ADCY4* | 3.625529153 | 0.09024456 | 0.017725855 | 0.072518706 | 0.999933887 | -0.14471757 | -0.290951509 | 0.146233939 | 0.384562469 |
| ENSG00000178449 | *COX14* | 2.509742067 | -0.007312025 | 0.028241081 | -0.035553106 | 0.999933887 | -0.086152284 | -0.267602304 | 0.18145002 | 0.384897548 |
| ENSG00000121073 | *SLC35B1* | 3.69738859 | 0.012154708 | -0.018710539 | 0.030865247 | 0.999933887 | -0.032994862 | -0.163339792 | 0.13034493 | 0.38502062 |
| ENSG00000162704 | *ARPC5* | 8.968147536 | -0.04299571 | -0.05773258 | 0.01473687 | 0.999933887 | -0.375744783 | -0.278778147 | -0.096966635 | 0.385170301 |
| ENSG00000187105 | *HEATR4* | 0.289127706 | -0.26573577 | 0.02318411 | -0.28891988 | 0.999933887 | 0.996959231 | 1.260218363 | -0.263259132 | 0.385270104 |
| ENSG00000266498 | *AC055811.3* | 1.428255905 | 0.079925292 | 0.145818998 | -0.065893706 | 0.999933887 | 0.538656381 | 0.810260979 | -0.271604598 | 0.385270104 |
| ENSG00000110074 | *FOXRED1* | 2.080912045 | 0.013958393 | -0.070302674 | 0.084261067 | 0.999933887 | 0.150768713 | 0.335575454 | -0.184806741 | 0.385270104 |
| ENSG00000119917 | *IFIT3* | 7.926532732 | -0.137513989 | -0.174656998 | 0.037143009 | 0.999933887 | -0.969163602 | -1.133014233 | 0.163850631 | 0.38536829 |
| ENSG00000235863 | *B3GALT4* | 2.839378644 | -0.104223627 | 0.107312014 | -0.211535641 | 0.999933887 | -0.504923404 | -0.660529051 | 0.155605648 | 0.38536829 |
| ENSG00000187860 | *CCDC157* | 1.348021218 | 0.131061632 | 0.176967592 | -0.04590596 | 0.999933887 | 0.238475609 | 0.505052684 | -0.266577075 | 0.385561603 |
| ENSG00000166228 | *PCBD1* | -0.175392578 | 0.015991759 | 0.116960238 | -0.100968479 | 0.999933887 | 0.219362718 | -0.149084787 | 0.368447505 | 0.385561603 |
| ENSG00000102931 | *ARL2BP* | 0.020919388 | 0.083987953 | -0.277083261 | 0.361071214 | 0.999933887 | 0.230535478 | -0.140503636 | 0.371039113 | 0.385899824 |
| ENSG00000125347 | *IRF1* | 8.546380233 | 0.053334309 | 0.033129351 | 0.020204957 | 0.999933887 | 0.244692815 | 0.128179624 | 0.116513191 | 0.386006309 |
| ENSG00000164414 | *SLC35A1* | 0.564880149 | 0.168326359 | 0.085050699 | 0.08327566 | 0.999933887 | -0.66064849 | -1.001141665 | 0.340493176 | 0.386069564 |
| ENSG00000229331 | *GK-IT1* | 3.344537548 | 0.041777205 | 0.06539262 | -0.023615416 | 0.999933887 | 0.284021784 | 0.437929307 | -0.153907523 | 0.386186424 |
| ENSG00000245532 | *NEAT1* | 11.836750613 | 0.127632586 | 0.114400182 | 0.013232404 | 0.999933887 | 0.345530746 | 0.459743591 | -0.114212846 | 0.386532364 |
| ENSG00000262312 | *AC004494.1* | 0.21082752 | -0.150338181 | 0.127015409 | -0.27735359 | 0.999933887 | 0.033029438 | 0.344159189 | -0.311129751 | 0.386778177 |
| ENSG00000213903 | *LTB4R* | 5.199285646 | 0.011132074 | -0.069097883 | 0.080229958 | 0.999933887 | -0.835372835 | -1.021004904 | 0.18563207 | 0.386848078 |
| ENSG00000109445 | *ZNF330* | 3.663224061 | -0.060125798 | -0.098613032 | 0.038487234 | 0.999933887 | 0.043059985 | 0.150894474 | -0.107834488 | 0.386952481 |
| ENSG00000142444 | *TIMM29* | 1.511037666 | 0.210066924 | 0.130959725 | 0.079107199 | 0.999933887 | -0.019280726 | -0.217375498 | 0.198094772 | 0.387049776 |
| ENSG00000179922 | *ZNF784* | 1.318016984 | 0.137894286 | 0.172441289 | -0.034547003 | 0.999933887 | -0.410622048 | -0.65820786 | 0.247585812 | 0.38715418 |
| ENSG00000154589 | *LY96* | 5.811579667 | 0.002361747 | 0.003926606 | -0.001564859 | 0.999933887 | -0.212284051 | -0.340673985 | 0.128389934 | 0.387366564 |
| ENSG00000080561 | *MID2* | -0.481097287 | 0.085616193 | -0.10349546 | 0.189111653 | 0.999933887 | -0.114376148 | 0.218759613 | -0.333135762 | 0.387459659 |
| ENSG00000204160 | *ZDHHC18* | 9.896033081 | 0.027706456 | 0.031117797 | -0.00341134 | 0.999933887 | 0.317288395 | 0.402784677 | -0.085496282 | 0.387486535 |
| ENSG00000117335 | *CD46* | 8.160643245 | -0.048233708 | -0.106058229 | 0.057824521 | 0.999933887 | -0.289415597 | -0.394186582 | 0.104770985 | 0.387619602 |
| ENSG00000105643 | *ARRDC2* | 7.729696806 | -0.067785512 | -0.055570671 | -0.012214841 | 0.999933887 | -0.201189279 | -0.064140119 | -0.13704916 | 0.387619602 |
| ENSG00000183605 | *SFXN4* | -0.177161177 | -0.129556404 | 0.073282302 | -0.202838706 | 0.999933887 | 0.285125359 | -0.098529648 | 0.383655007 | 0.387619602 |
| ENSG00000100056 | *ESS2* | 4.05576622 | 0.02637396 | 0.033694617 | -0.007320657 | 0.999933887 | -0.072886711 | -0.211706227 | 0.138819516 | 0.387619602 |
| ENSG00000140854 | *KATNB1* | 3.830718976 | 0.040714578 | 0.081858286 | -0.041143708 | 0.999933887 | -0.068826467 | -0.222646021 | 0.153819554 | 0.387648165 |
| ENSG00000263934 | *SNORD3A* | 0.357973119 | -0.314656544 | -0.769332566 | 0.454676022 | 0.999933887 | 0.686300218 | 0.110236114 | 0.576064104 | 0.387707388 |
| ENSG00000214135 | *AC132008.2* | 2.181430449 | -0.010984589 | -0.003631624 | -0.007352966 | 0.999933887 | -0.221263167 | -0.401445358 | 0.180182191 | 0.387789111 |
| ENSG00000272636 | *DOC2B* | -0.351155701 | 0.160026136 | 0.265851097 | -0.105824961 | 0.999933887 | 1.175065644 | 1.718324656 | -0.543259012 | 0.387934232 |
| ENSG00000109929 | *SC5D* | 2.54086778 | -0.11238772 | -0.067479462 | -0.044908259 | 0.999933887 | -0.187087627 | -0.004039483 | -0.183048145 | 0.388056535 |
| ENSG00000236383 | *CCDC200* | 0.336525355 | -0.050809337 | 0.078184593 | -0.128993929 | 0.999933887 | 0.433410686 | 0.779285135 | -0.345874449 | 0.388056535 |
| ENSG00000163605 | *PPP4R2* | 6.996170102 | -0.08816864 | -0.057767775 | -0.030400865 | 0.999933887 | 0.404226805 | 0.571391186 | -0.167164381 | 0.388211862 |

| | | | | | | | | | | |
|---|---|---|---|---|---|---|---|---|---|---|
| ENSG00000012048 | *BRCA1* | 2.110855657 | -0.045712183 | -0.110438776 | 0.064726592 | 0.999933887 | -0.301000293 | -0.04048232 | -0.260517973 | 0.388211862 |
| ENSG00000100379 | *KCTD17* | 1.498883114 | 0.006799007 | -0.041877879 | 0.048676886 | 0.999933887 | 0.28700348 | 0.090132607 | 0.196870873 | 0.388211862 |
| ENSG00000110435 | *PDHX* | 1.454710354 | -0.146791474 | -0.214700103 | 0.067908629 | 0.999933887 | 0.13581347 | 0.329082951 | -0.19326948 | 0.388818478 |
| ENSG00000104687 | *GSR* | 6.106962076 | -0.063798355 | 0.022360049 | -0.086158404 | 0.999933887 | -0.304930553 | -0.422028088 | 0.117097535 | 0.388931669 |
| ENSG00000133606 | *MKRN1* | 7.203557322 | -0.073188351 | -0.082871284 | 0.009682933 | 0.999933887 | -0.568590404 | -0.679148043 | 0.110557639 | 0.388944861 |
| ENSG00000246082 | *NUDT16P1* | -1.619843875 | 0.155818744 | 0.117833122 | 0.037985622 | 0.999933887 | -0.726014904 | -1.411398496 | 0.685383592 | 0.38898874 |
| ENSG00000267141 | *AC012615.4* | -1.336924417 | -0.113774703 | -0.168666608 | 0.054891905 | 0.999933887 | -0.485661358 | -1.17214686 | 0.686485502 | 0.38898874 |
| ENSG00000258439 | *AC007956.2* | 0.616789807 | -0.111149822 | 0.161960032 | -0.273109854 | 0.999933887 | 0.098281286 | 0.421732928 | -0.323451641 | 0.38907558 |
| ENSG00000139970 | *RTN1* | 1.005769503 | 0.101716472 | 0.191595159 | -0.089878687 | 0.999933887 | 0.071653285 | -0.285400587 | 0.357053872 | 0.38907558 |
| ENSG00000228594 | *FNDC10* | 0.41275377 | 0.156168373 | -0.213161058 | 0.369329432 | 0.999933887 | -0.345421309 | -0.628408976 | 0.282987666 | 0.389083283 |
| ENSG00000150477 | *KIAA1328* | 0.400992093 | -0.028898228 | -0.133967492 | 0.105069263 | 0.999933887 | -0.296941273 | 0.034943479 | -0.331884752 | 0.389445218 |
| ENSG00000173692 | *PSMD1* | 4.674412882 | -0.082197744 | -0.129264585 | 0.047066841 | 0.999933887 | 0.015970865 | 0.122462195 | -0.106491329 | 0.389534719 |
| ENSG00000153207 | *AHCTF1* | 6.574048305 | -0.100359474 | -0.209935049 | 0.109575575 | 0.999933887 | -1.582010737 | -1.869997823 | 0.287987086 | 0.389534719 |
| ENSG00000107614 | *TRDMT1* | 2.02686489 | 0.039161921 | -0.249113057 | 0.288274978 | 0.944094537 | -0.257236649 | -0.05521781 | -0.202018839 | 0.389534719 |
| ENSG00000122566 | *HNRNPA2B1* | 9.014550548 | -0.02276689 | -0.021833354 | -0.000933536 | 0.999933887 | 0.154671553 | 0.258558435 | -0.103886882 | 0.389534719 |
| ENSG00000036672 | *USP2* | 0.943335283 | 0.037678269 | 0.112105281 | -0.074427012 | 0.999933887 | -0.049176099 | 0.320480035 | -0.369656134 | 0.389534719 |
| ENSG00000092208 | *GEMIN2* | -0.417369299 | 0.059106751 | 0.267024757 | -0.207918007 | 0.999933887 | -0.058453925 | 0.318741362 | -0.377195287 | 0.389534719 |
| ENSG00000123700 | *KCNJ2* | 5.653375357 | 0.000933634 | 0.102163454 | -0.10122982 | 0.999933887 | 3.305401548 | 3.643217397 | -0.337815849 | 0.389534719 |
| ENSG00000069849 | *ATP1B3* | 4.421520609 | -0.042015723 | -0.08437172 | 0.042355997 | 0.999933887 | 0.096904678 | 0.21507568 | -0.118171003 | 0.389856611 |
| ENSG00000189046 | *ALKBH2* | 1.928903219 | 0.007232239 | 0.027569 | -0.020336761 | 0.999933887 | 0.442640279 | 0.159025518 | 0.283614761 | 0.390077135 |
| ENSG00000211893 | *IGHG2* | 0.750101257 | -0.102772025 | -0.109759124 | 0.006987099 | 0.999933887 | 0.248781616 | 0.714277099 | -0.465495483 | 0.390097339 |
| ENSG00000214193 | *SH3D21* | 7.00155958 | 0.021569494 | 0.02093194 | 0.000637553 | 0.999933887 | 0.265279173 | 0.376378357 | -0.111099184 | 0.390097339 |
| ENSG00000077348 | *EXOSC5* | 0.61377402 | -0.113577917 | 0.049054383 | -0.1626323 | 0.999933887 | 0.136468462 | -0.108633728 | 0.24510219 | 0.390327519 |
| ENSG00000198700 | *IPO9* | 3.836378056 | -0.059692258 | -0.009479567 | -0.050212691 | 0.999933887 | -0.233314985 | -0.340390401 | 0.107075416 | 0.390611284 |
| ENSG00000183520 | *UTP11* | 1.846018203 | -0.1126591 | -0.161316111 | 0.048657011 | 0.999933887 | -0.198024364 | -0.058166123 | -0.139858241 | 0.390629413 |
| ENSG00000119862 | *LGALSL* | 3.962119735 | -0.063794913 | 0.016379819 | -0.080174732 | 0.999933887 | -0.850068832 | -1.00025845 | 0.150189617 | 0.390629413 |
| ENSG00000135077 | *HAVCR2* | 2.659146536 | -0.162034292 | -0.098418376 | -0.063615916 | 0.999933887 | -0.499253063 | -0.670843064 | 0.171590001 | 0.390975964 |
| ENSG00000111727 | *HCFC2* | 1.847466941 | -0.070314875 | -0.272491675 | 0.2021768 | 0.999933887 | 0.452631822 | 0.702481 | -0.249849178 | 0.39103307 |
| ENSG00000110700 | *RPS13* | 6.587906868 | -0.015936597 | -0.032311214 | 0.016374617 | 0.999933887 | 0.143146404 | 0.249436602 | -0.106290199 | 0.39103307 |
| ENSG00000110203 | *FOLR3* | 4.947869815 | 0.059947874 | 0.017069637 | 0.042878237 | 0.999933887 | 0.0013743 | -0.161004474 | 0.162378774 | 0.391159837 |
| ENSG00000111684 | *LPCAT3* | 2.401571196 | 0.06306674 | -0.015549925 | 0.078616665 | 0.999933887 | -0.033804458 | 0.195471661 | -0.229276119 | 0.391159837 |
| ENSG00000165943 | *MOAP1* | 5.492832066 | 0.046880505 | 0.047080559 | -0.000200054 | 0.999933887 | 0.294695347 | 0.167545382 | 0.127149965 | 0.391159837 |
| ENSG00000279789 | *AC120114.3* | 1.23718092 | -0.103798986 | -0.337239899 | 0.233440913 | 0.999933887 | 0.019696467 | 0.27338858 | -0.253692113 | 0.391159837 |
| ENSG00000204577 | *LILRB3* | 8.720109075 | 0.022450544 | 0.020608645 | 0.001841899 | 0.999933887 | -0.398173778 | -0.31167063 | -0.086503148 | 0.391159837 |
| ENSG00000103275 | *UBE2I* | 5.539406869 | -0.018140513 | -0.030304038 | 0.012163525 | 0.999933887 | 0.144520726 | 0.221280315 | -0.076759588 | 0.391159837 |
| ENSG00000132635 | *PCED1A* | 3.495743674 | 0.025661447 | 0.036747806 | -0.011086359 | 0.999933887 | 0.100426171 | -0.013058557 | 0.113484728 | 0.391159837 |
| ENSG00000237356 | *AL365295.1* | -0.446897804 | -0.134093372 | 0.030133106 | -0.164226479 | 0.999933887 | 0.201750216 | 0.549873922 | -0.348123706 | 0.391159837 |
| ENSG00000118997 | *DNAH7* | -0.296413828 | -0.058857622 | -0.073616051 | 0.014758429 | 0.999933887 | -0.4766272 | 0.05627566 | -0.53290286 | 0.391159837 |
| ENSG00000177688 | *SUMO4* | 1.239798118 | 0.034337623 | 0.148804237 | -0.114466613 | 0.999933887 | 0.728157926 | 1.008989347 | -0.280831421 | 0.391159837 |
| ENSG00000096093 | *EFHC1* | 2.289571592 | 0.023288904 | -0.037246292 | 0.060535196 | 0.999933887 | 0.110030547 | 0.327504394 | -0.217473847 | 0.391159837 |
| ENSG00000133317 | *LGALS12* | 2.956018672 | 0.008784368 | 0.049287394 | -0.040503026 | 0.999933887 | 0.028017773 | 0.22528436 | -0.197266588 | 0.391159837 |
| ENSG00000141002 | *TCF25* | 7.172767071 | 0.025235507 | 0.012459796 | 0.01277571 | 0.999933887 | 0.026426246 | 0.105181545 | -0.078755299 | 0.391172484 |
| ENSG00000266378 | *AC005224.2* | -1.020695862 | 0.112786487 | 0.246337444 | -0.133550957 | 0.999933887 | 1.423820742 | 2.108928279 | -0.685107536 | 0.391252784 |
| ENSG00000236409 | *NRADDP* | -0.067692715 | -0.042511576 | 0.059091335 | -0.101602911 | 0.999933887 | 0.408305703 | 0.794929393 | -0.386623689 | 0.391252784 |
| ENSG00000142188 | *TMEM50B* | 3.317359979 | -0.064078722 | -0.059384196 | -0.004694526 | 0.999933887 | 0.123783912 | 0.250431739 | -0.126647827 | 0.391375525 |
| ENSG00000108733 | *PEX12* | 1.1468614 | -0.243115924 | -0.088336592 | -0.154779332 | 0.999933887 | -0.238433815 | -0.442328821 | 0.203895006 | 0.391496613 |
| ENSG00000214279 | *SCART1* | 0.768384304 | 0.092754988 | 0.268861387 | -0.176106399 | 0.999933887 | 0.015436483 | 0.262755344 | -0.247318861 | 0.391496613 |
| ENSG00000224080 | *UBE2FP1* | 0.065950991 | -0.075780104 | 0.10596344 | -0.181743544 | 0.999933887 | 0.172395929 | 0.632556249 | -0.460160319 | 0.391635957 |
| ENSG00000248712 | *CCDC153* | 0.726207416 | -0.114978144 | 0.05523489 | -0.170213034 | 0.999933887 | -0.229560079 | -0.649220857 | 0.419660779 | 0.391838052 |
| ENSG00000280721 | *LINC01943* | -0.653900346 | -0.350052952 | 0.056206481 | -0.406259433 | 0.999933887 | -0.081418987 | -0.48959587 | 0.408176883 | 0.39184136 |
| ENSG00000196693 | *ZNF33B* | 1.821640278 | -0.088149164 | -0.269205357 | 0.181056193 | 0.999933887 | -0.250928357 | 0.035678241 | -0.286606598 | 0.391883245 |
| ENSG00000103253 | *HAGHL* | 1.329284266 | 0.117667742 | 0.261362388 | -0.143694646 | 0.999933887 | 0.103468172 | -0.124180521 | 0.227648693 | 0.392333752 |
| ENSG00000088538 | *DOCK3* | -0.044260941 | 0.009281791 | -0.141218214 | 0.150500006 | 0.999933887 | 0.279290028 | -0.048460782 | 0.32775081 | 0.392358837 |
| ENSG00000117859 | *OSBPL9* | 5.622956163 | -0.050918442 | -0.112540385 | 0.061621943 | 0.999933887 | -0.124557435 | 0.024908569 | -0.149466004 | 0.392463399 |
| ENSG00000268027 | *AC243960.1* | 1.324412473 | -0.044741428 | 0.027147227 | -0.071888655 | 0.999933887 | 0.089554389 | -0.131540663 | 0.221095052 | 0.3926314 |
| ENSG00000126524 | *SBDS* | 4.384686062 | -0.001089319 | -0.003199505 | 0.002110187 | 0.999933887 | 0.123271146 | 0.241394521 | -0.118123376 | 0.393025405 |
| ENSG00000175567 | *UCP2* | 5.593533885 | -0.015925933 | -0.047902122 | 0.03197619 | 0.999933887 | 0.065457333 | -0.042959627 | 0.108416959 | 0.393149761 |
| ENSG00000099326 | *MZF1* | 4.028085084 | 0.059197442 | 0.031607176 | 0.027590266 | 0.999933887 | -0.294085563 | -0.403702063 | 0.1096165 | 0.393191988 |
| ENSG00000145649 | *GZMA* | 3.707372663 | 0.025229851 | -0.089389501 | 0.114619353 | 0.999933887 | 0.0602235 | 0.208178546 | -0.147955046 | 0.393210415 |
| ENSG00000119402 | *FBXW2* | 4.737839111 | -0.084764374 | -0.031422361 | -0.053342013 | 0.999933887 | 0.080744108 | 0.190924252 | -0.110180143 | 0.393387483 |
| ENSG00000167635 | *ZNF146* | 3.058040887 | -0.080398616 | -0.236178459 | 0.155779842 | 0.999933887 | -0.178607088 | 0.026869782 | -0.20547687 | 0.393461717 |
| ENSG00000244071 | *RPL9P33* | 1.859426316 | -0.069491814 | 0.035449066 | -0.10494088 | 0.999933887 | 0.750669916 | 0.526137251 | 0.224532665 | 0.393719049 |
| ENSG00000242622 | *AC092910.3* | 1.232977065 | -0.11456175 | -0.247674262 | 0.133112512 | 0.999933887 | 0.763513914 | 0.506011614 | 0.2575023 | 0.393719049 |
| ENSG00000160917 | *CPSF4* | 2.892726495 | 0.030444733 | 0.030661003 | -0.000216271 | 0.999933887 | 0.042393891 | -0.101091167 | 0.143485058 | 0.393844161 |
| ENSG00000110200 | *ANAPC15* | 1.460063691 | 0.127661011 | 0.047976385 | 0.079684626 | 0.999933887 | 0.175238034 | -0.105026418 | 0.280264451 | 0.394165866 |
| ENSG00000279469 | *AC109454.4* | -0.083020404 | -0.160953674 | -0.071195037 | -0.089758638 | 0.999933887 | 0.052965709 | 0.480085329 | -0.427119621 | 0.394304437 |
| ENSG00000143971 | *ETAA1* | 0.628386502 | -0.239033303 | -0.179859767 | -0.059173536 | 0.999933887 | -0.403826605 | -0.039093148 | -0.364733457 | 0.394561425 |
| ENSG00000256720 | *BTG1P1* | -0.504513304 | -0.187254556 | -0.137103779 | -0.050150777 | 0.999933887 | -0.295442692 | 0.120348991 | -0.415791683 | 0.394614093 |
| ENSG00000138867 | *GUCD1* | 7.172487083 | -0.053597676 | -0.041589524 | -0.012008152 | 0.999933887 | -0.110036457 | -0.013796556 | -0.0962399 | 0.394614093 |
| ENSG00000155660 | *PDIA4* | 3.526778358 | -0.007096106 | -0.036393149 | 0.029297043 | 0.999933887 | 0.013879424 | 0.133316508 | -0.119437084 | 0.394614093 |
| ENSG00000158987 | *RAPGEF6* | 3.537465745 | -0.075979594 | -0.104125698 | 0.028146104 | 0.999933887 | -0.221849631 | 0.049685934 | -0.271535565 | 0.395010806 |
| ENSG00000169607 | *CKAP2L* | 0.221398075 | -0.196901804 | 0.00905695 | -0.205958754 | 0.999933887 | -0.424402809 | -0.719881894 | 0.295479084 | 0.395393345 |
| ENSG00000131732 | *ZCCHC9* | 2.518331936 | -0.029471522 | -0.111607145 | 0.082135624 | 0.999933887 | -0.240487442 | -0.09100911 | -0.149478332 | 0.395393345 |
| ENSG00000055609 | *KMT2C* | 8.159105151 | -0.006269076 | -0.05280542 | 0.046536345 | 0.999933887 | -0.431777231 | -0.258610701 | -0.173166529 | 0.395393345 |
| ENSG00000243438 | *LARP1BP2* | -0.638401342 | 0.004354543 | 0.108805738 | -0.104451195 | 0.999933887 | 1.018943663 | 1.541412381 | -0.522468718 | 0.395412553 |
| ENSG00000141576 | *RNF157* | 1.469160622 | 0.221260569 | 0.098795442 | 0.122465127 | 0.999933887 | 0.093895008 | -0.122121984 | 0.216016992 | 0.395415748 |
| ENSG00000156475 | *PPP2R2B* | 1.599462206 | -0.09717396 | -0.020199887 | -0.076974073 | 0.999933887 | -0.047470336 | 0.136695938 | -0.184166275 | 0.395915564 |
| ENSG00000204256 | *BRD2* | 8.953686016 | 0.047686198 | 0.043994434 | 0.003691765 | 0.999933887 | 0.723936155 | 0.826909918 | -0.102973763 | 0.396224467 |
| ENSG00000032742 | *IFT88* | 2.360635531 | 0.038480307 | -0.001212273 | 0.03969258 | 0.999933887 | -0.462942584 | -0.233364313 | -0.229578271 | 0.396473386 |
| ENSG00000225093 | *RPL3P7* | -1.42783911 | -0.521270934 | 0.143845114 | -0.665116048 | 0.999933887 | 0.021087702 | 0.561826048 | -0.540738346 | 0.396555812 |
| ENSG00000163960 | *UBXN7* | 5.867010693 | -0.083587526 | -0.075557514 | -0.008030012 | 0.999933887 | -0.3967221 | -0.240554919 | -0.156167181 | 0.396555812 |
| ENSG00000100300 | *TSPO* | 6.584629101 | 0.057475807 | 0.026724835 | 0.030750972 | 0.999933887 | 0.019416782 | -0.109135274 | 0.128552056 | 0.396555812 |
| ENSG00000172845 | *SP3* | 7.719800701 | -0.031573799 | -0.044331811 | 0.012758012 | 0.999933887 | -0.895578189 | -0.75428141 | -0.141296779 | 0.396555812 |
| ENSG00000100629 | *CEP128* | 0.913078363 | 0.022879696 | -0.305000905 | 0.327880601 | 0.999933887 | 0.263355356 | 0.539428603 | -0.276073247 | 0.396555812 |
| ENSG00000087269 | *NOP14* | 2.920171252 | -0.005501932 | -0.037837727 | 0.032335795 | 0.999933887 | 0.203144783 | 0.354346569 | -0.151201786 | 0.396555812 |
| ENSG00000242553 | *AP001432.1* | -0.605803205 | 0.037293006 | 0.183199923 | -0.145906916 | 0.999933887 | -0.521625087 | -0.948529421 | 0.426904335 | 0.396587616 |
| ENSG00000130717 | *UCK1* | 3.774897772 | -0.034424042 | 0.015217512 | -0.049641553 | 0.999933887 | -0.276970198 | -0.391336556 | 0.114366357 | 0.396706932 |
| ENSG00000254910 | *AC136475.2* | 0.877679636 | -0.124452194 | -0.320298961 | 0.195846767 | 0.999933887 | -0.400226841 | -0.700080827 | 0.299853986 | 0.397036466 |
| ENSG00000278831 | *AL513190.1* | -0.787843835 | 0.06496948 | -0.132476421 | 0.197445901 | 0.999933887 | -0.196618001 | -0.794553753 | 0.597935753 | 0.397036466 |
| ENSG00000241468 | *ATP5MF* | 2.732361634 | 0.007031267 | 0.027568971 | -0.020537704 | 0.999933887 | -0.033072507 | -0.213951469 | 0.180878962 | 0.397036466 |
| ENSG00000196220 | *SRGAP3* | -0.193091804 | 0.059872803 | 0.114420096 | -0.054547293 | 0.999933887 | -0.378810506 | 0.16859726 | -0.547407767 | 0.397080112 |
| ENSG00000145332 | *KLHL8* | 2.766749282 | 0.029027653 | 0.002785399 | 0.026242254 | 0.999933887 | -0.347336709 | -0.537569888 | 0.190233179 | 0.397080112 |
| ENSG00000100298 | *APOBEC3H* | -0.308514383 | 0.137386953 | 0.031262994 | 0.106123959 | 0.999933887 | -0.013422131 | -0.302240461 | 0.28881833 | 0.397234128 |
| ENSG00000149571 | *KIRREL3* | -0.182646159 | 0.028295375 | 0.008581247 | 0.019714128 | 0.999933887 | 0.417869837 | 0.900189059 | -0.482319222 | 0.397287083 |
| ENSG00000239911 | *PRKAG2-AS1* | -0.453810935 | -0.144955085 | 0.01680864 | -0.161763725 | 0.999933887 | 1.003039427 | 1.338185607 | -0.33514618 | 0.397324266 |
| ENSG00000104825 | *NFKBIB* | 6.598130406 | 0.120450899 | 0.093361381 | 0.027089518 | 0.999933887 | 1.084692444 | 1.2361224 | -0.151429956 | 0.397396787 |
| ENSG00000116521 | *SCAMP3* | 3.482909549 | -0.042482031 | -0.027230839 | -0.015251192 | 0.999933887 | 0.049422563 | -0.063364204 | 0.112786767 | 0.397396787 |
| ENSG00000048544 | *MRPS10* | 2.893110927 | -0.043051471 | -0.101968425 | 0.058916955 | 0.999933887 | -0.151527953 | -0.019512155 | -0.132015798 | 0.397396787 |
| ENSG00000100181 | *TPTEP1* | 0.632734792 | 0.070883161 | 0.045627002 | 0.025256159 | 0.999933887 | -0.834181195 | -0.585249497 | -0.248931698 | 0.397396787 |
| ENSG00000135898 | *GPR55* | -0.097321374 | -0.061982196 | -0.010829024 | -0.051153172 | 0.999933887 | 0.387515112 | 0.721333668 | -0.333818556 | 0.397396787 |
| ENSG00000118508 | *RAB32* | 6.168489882 | 0.014657905 | -0.062085388 | 0.076743293 | 0.999933887 | 0.083437213 | 0.193022108 | -0.109584895 | 0.397396787 |
| ENSG00000128524 | *ATP6V1F* | 6.470606173 | 0.0125219 | -0.040293636 | 0.052815536 | 0.999933887 | 0.268995404 | 0.388636302 | -0.119640897 | 0.397396787 |
| ENSG00000003393 | *ALS2* | 3.610006962 | 0.013023046 | -0.033837451 | 0.046860496 | 0.999933887 | -0.236738146 | -0.108133593 | -0.128604553 | 0.397396787 |
| ENSG00000112406 | *HECA* | 7.948795342 | -0.006424647 | 0.031224813 | -0.03764946 | 0.999933887 | -0.278707147 | -0.175891942 | -0.102815205 | 0.397396787 |
| ENSG00000108592 | *FTSJ3* | 3.727299649 | -0.005497833 | 0.048575704 | -0.054073536 | 0.999933887 | 0.009144473 | -0.117268131 | 0.126412604 | 0.397396787 |

| | | | | | | | | | | |
|---|---|---|---|---|---|---|---|---|---|---|
| ENSG00000169918 | *OTUD7A* | -0.090235338 | 0.153445531 | 0.035151806 | 0.118293725 | 0.999933887 | 0.097307314 | 0.421016507 | -0.323709193 | 0.397440292 |
| ENSG00000127080 | *IPPK* | 1.990337457 | -0.046528104 | 0.036161318 | -0.082689423 | 0.999933887 | 0.628090176 | 0.852981523 | -0.224891347 | 0.397440292 |
| ENSG00000166710 | *B2M* | 13.388422422 | -0.002253769 | -0.043354406 | 0.041100637 | 0.999933887 | 0.299354463 | 0.384217641 | -0.084863178 | 0.39744833 |
| ENSG00000164919 | *COX6C* | 2.665202494 | -0.091752996 | -0.062260824 | -0.029492172 | 0.999933887 | 0.150609975 | 0.312432017 | -0.161822042 | 0.397548969 |
| ENSG00000078687 | *TNRC6C* | 5.170909987 | 0.025105311 | 0.044702883 | -0.019597572 | 0.999933887 | -0.209506471 | -0.099791895 | -0.109714575 | 0.397632222 |
| ENSG00000106610 | *STAG3L4* | 2.901376332 | 0.000272935 | 0.036124467 | -0.035851532 | 0.999933887 | -0.071375553 | 0.090213343 | -0.161588896 | 0.397632222 |
| ENSG00000243646 | *IL10RB* | 2.687894188 | -0.021501198 | -0.001003203 | -0.020497994 | 0.999933887 | 0.424486497 | 0.251805556 | 0.172680941 | 0.397857484 |
| ENSG00000148572 | *NRBF2* | 6.949409664 | 0.024480375 | -0.012317669 | 0.036798045 | 0.999933887 | -0.340488325 | -0.240212548 | -0.100275777 | 0.398258937 |
| ENSG00000231249 | *ITPR1-DT* | 0.14059268 | 0.108641578 | -0.068640559 | 0.177282137 | 0.999933887 | 0.332385071 | 0.562435578 | -0.230050507 | 0.398258937 |
| ENSG00000175662 | *TOM1L2* | 4.344456875 | 0.010103806 | 0.059812204 | -0.049708398 | 0.999933887 | -0.18128889 | -0.301799769 | 0.120510879 | 0.398258937 |
| ENSG00000092140 | *G2E3* | 2.79792797 | -0.000968522 | -0.052287305 | 0.051318783 | 0.999933887 | -0.406382423 | -0.121309712 | -0.285072712 | 0.398897839 |
| ENSG00000176731 | *RBIS* | 1.77093704 | -0.043747888 | 0.000173132 | -0.04392102 | 0.999933887 | 0.038247041 | -0.153956583 | 0.192203625 | 0.399276838 |
| ENSG00000257433 | *AC004241.1* | 1.084222433 | 0.044622779 | -0.188166658 | 0.232789437 | 0.999933887 | -0.06172666 | -0.339228832 | 0.277502173 | 0.399276838 |
| ENSG00000211799 | *TRAV19* | 0.17272202 | 0.086912959 | -0.302600238 | 0.389513196 | 0.999933887 | 0.215514955 | 0.545158107 | -0.329643152 | 0.399287442 |
| ENSG00000256039 | *LINC02446* | 1.354923085 | 0.065508738 | -0.029005337 | 0.094514075 | 0.999933887 | 0.005999436 | 0.212985404 | -0.206985968 | 0.399316381 |
| ENSG00000042429 | *MED17* | 1.823208548 | -0.094600179 | 0.004539506 | -0.099139685 | 0.999933887 | -0.273324576 | -0.462211988 | 0.188887412 | 0.399512499 |
| ENSG00000118620 | *ZNF430* | 3.243626897 | 0.08899169 | -0.055217307 | 0.144208997 | 0.999933887 | -0.385425931 | -0.165795812 | -0.219630119 | 0.399512499 |
| ENSG00000172164 | *SNTB1* | 3.402021299 | 0.060080802 | 0.16923701 | -0.109156209 | 0.999933887 | -0.304770033 | -0.485067797 | 0.180297764 | 0.399512499 |
| ENSG00000213366 | *GSTM2* | 1.936883833 | 0.042118097 | 0.001654656 | 0.040463441 | 0.999933887 | -0.333820597 | -0.572265112 | 0.238444515 | 0.399512499 |
| ENSG00000277791 | *PSMB3* | 6.052291836 | -0.008872347 | -0.034414437 | 0.02554209 | 0.999933887 | -0.21655787 | -0.329027684 | 0.112469814 | 0.399512499 |
| ENSG00000288398 | *AL109627.1* | -0.226950565 | 0.149243344 | -0.107410017 | 0.256653361 | 0.999933887 | 0.09943571 | 0.507360409 | -0.407924699 | 0.399678327 |
| ENSG00000103248 | *MTHFSD* | 2.703088704 | -0.151246429 | 0.014853076 | -0.166099506 | 0.999933887 | 0.196913668 | 0.334806796 | -0.137893128 | 0.399695599 |
| ENSG00000162385 | *MAGOH* | 4.389513208 | 0.0271235 | -0.017785212 | 0.044908712 | 0.999933887 | 0.190580979 | 0.310224769 | -0.11964379 | 0.399695599 |
| ENSG00000272277 | *AL031963.3* | 0.490012388 | 0.068228385 | 0.33195912 | -0.263730736 | 0.999933887 | 0.053113085 | -0.293136855 | 0.34624994 | 0.399710615 |
| ENSG00000174099 | *MSRB3* | 1.470858717 | -0.148628988 | 0.028385481 | -0.177014469 | 0.999933887 | -0.071335819 | 0.180347084 | -0.251682903 | 0.399766473 |
| ENSG00000113328 | *CCNG1* | 3.563885982 | -0.118694148 | -0.243686817 | 0.12499267 | 0.999933887 | -0.114757341 | 0.03554317 | -0.150300511 | 0.399766473 |
| ENSG00000122705 | *CLTA* | 5.643209226 | -0.077653152 | -0.152668822 | 0.07501567 | 0.999933887 | -0.357839039 | -0.243118383 | -0.114720655 | 0.399766473 |
| ENSG00000189269 | *DRICH1* | -0.410207254 | -0.175473276 | 0.067797456 | -0.243270732 | 0.999933887 | 0.357943208 | -0.071450916 | 0.429394125 | 0.399766473 |
| ENSG00000167302 | *TEPSIN* | 4.715338638 | -0.035228 | 0.027726351 | -0.06295435 | 0.999933887 | -0.113054948 | -0.209500053 | 0.096445104 | 0.399766473 |
| ENSG00000142235 | *LMTK3* | 0.972874861 | 0.063601315 | -0.051154336 | 0.11475565 | 0.999933887 | -0.006551099 | 0.281390283 | -0.287941382 | 0.399766473 |
| ENSG00000131023 | *LATS1* | 3.722821082 | -0.009979104 | -0.005306178 | -0.004672926 | 0.999933887 | -0.373659175 | -0.218515603 | -0.155143572 | 0.399766473 |
| ENSG00000246548 | *LINC02288* | -0.941253711 | -0.126259992 | 0.104138114 | -0.230398106 | 0.999933887 | -0.430871466 | -1.100685406 | 0.66981394 | 0.399953371 |
| ENSG00000240710 | *AL512306.3* | 1.043820177 | -0.12536514 | -0.107605114 | -0.017760026 | 0.999933887 | -0.378219646 | -0.610629127 | 0.232409482 | 0.399953371 |
| ENSG00000172893 | *DHCR7* | 2.858806845 | -0.005372731 | -0.043678385 | 0.038305655 | 0.999933887 | 0.103045828 | 0.298558981 | -0.195513153 | 0.399953371 |
| ENSG00000105767 | *CADM4* | 2.573649613 | 0.079173958 | -0.044158184 | 0.123332141 | 0.999933887 | 0.683201461 | 0.856971183 | -0.173769722 | 0.400174272 |
| ENSG00000101191 | *DIDO1* | 6.124748458 | -0.019913391 | -0.004269954 | -0.015643437 | 0.999933887 | -0.143441781 | -0.217299124 | 0.073857342 | 0.400174272 |
| ENSG00000152778 | *IFIT5* | 5.333502424 | -0.150263423 | -0.139000687 | -0.011262736 | 0.999933887 | -0.616616447 | -0.799281635 | 0.182665189 | 0.400427698 |
| ENSG00000168813 | *ZNF507* | 3.789374925 | -0.145707507 | -0.164111186 | 0.018403679 | 0.999933887 | -0.484018955 | -0.295754475 | -0.18826448 | 0.400560758 |
| ENSG00000103056 | *SMPD3* | 3.383682438 | 0.005166978 | 0.087951389 | -0.082784411 | 0.999933887 | -0.17948226 | -0.372082649 | 0.192600389 | 0.400560758 |
| ENSG00000261373 | *VPS9D1-AS1* | 1.907983872 | 0.125312897 | 0.236987241 | -0.111674344 | 0.999933887 | 0.117444821 | -0.18714622 | 0.304591041 | 0.40075414 |
| ENSG00000143106 | *PSMA5* | 4.617754995 | -0.018851234 | -0.066798694 | 0.04794746 | 0.999933887 | 0.264722068 | 0.363802162 | -0.099080094 | 0.40075414 |
| ENSG00000173145 | *NOC3L* | 1.006330962 | -0.155742638 | -0.131040485 | -0.024702153 | 0.999933887 | 0.106461173 | 0.319239439 | -0.212778265 | 0.401104949 |
| ENSG00000142230 | *SAE1* | 3.873891848 | -0.025796936 | 0.035961247 | -0.061758184 | 0.999933887 | -0.196094356 | -0.316364777 | 0.120270421 | 0.401104949 |
| ENSG00000267980 | *AC007292.1* | -1.043094908 | 0.277437042 | 0.144495997 | 0.132941045 | 0.999933887 | 0.04179712 | 0.67201139 | -0.63021427 | 0.401116965 |
| ENSG00000144749 | *LRIG1* | 3.532729755 | -0.025691991 | 0.106339323 | -0.132031314 | 0.999933887 | 0.119245498 | 0.234961967 | -0.115716469 | 0.401177861 |
| ENSG00000204138 | *PHACTR4* | 2.695670901 | -0.002875965 | -0.026244765 | 0.0233688 | 0.999933887 | -0.152253386 | 0.003143883 | -0.155397269 | 0.401177861 |
| ENSG00000147601 | *TERF1* | 2.583848905 | 0.030564784 | -0.091870589 | 0.122435373 | 0.999933887 | -0.219955753 | -0.043181652 | -0.176774101 | 0.401197865 |
| ENSG00000139289 | *PHLDA1* | 4.818905915 | 0.122235114 | 0.296860184 | -0.174625069 | 0.999933887 | 0.395630625 | 0.541806366 | -0.146175741 | 0.401302165 |
| ENSG00000128917 | *DLL4* | -0.003403981 | 0.125467698 | 0.292287892 | -0.166820194 | 0.999933887 | 0.485692093 | 0.866978158 | -0.381286065 | 0.401462955 |
| ENSG00000189298 | *ZKSCAN3* | 0.003052783 | -0.036289626 | -0.093947736 | 0.057658111 | 0.999933887 | 0.031448693 | -0.260105863 | 0.291554555 | 0.401462955 |
| ENSG00000221988 | *PPT2* | -0.773885672 | -0.037042998 | 0.442966707 | -0.480009704 | 0.999933887 | 0.045798306 | 0.526721672 | -0.480923366 | 0.401462955 |
| ENSG00000143226 | *FCGR2A* | 8.9676917 | 0.007650373 | -0.041953517 | 0.04960389 | 0.999933887 | 0.586130751 | 0.697865016 | -0.111734265 | 0.401462955 |
| ENSG00000278829 | *AC099811.5* | 2.621825764 | 0.200732083 | 0.12356717 | 0.077164913 | 0.999933887 | 1.398212828 | 1.663970529 | -0.265757701 | 0.401516707 |
| ENSG00000239857 | *GET4* | 2.613697729 | 0.163313222 | 0.145535196 | 0.017778026 | 0.999933887 | 0.125374439 | -0.064337382 | 0.189711821 | 0.401516707 |
| ENSG00000070087 | *PFN2* | 1.187062548 | -0.036632579 | 0.03520318 | -0.071835759 | 0.999933887 | -0.627816783 | -0.857629109 | 0.229812326 | 0.401728457 |
| ENSG00000116106 | *EPHA4* | 3.045250227 | -0.062317902 | -0.117463462 | 0.055145559 | 0.999933887 | -0.03045507 | 0.110309344 | -0.140764414 | 0.402000906 |
| ENSG00000172765 | *TMCC1* | 6.309514582 | -0.003885475 | 0.002429109 | -0.006314584 | 0.999933887 | -0.221526619 | -0.120673912 | -0.100852707 | 0.402755934 |
| ENSG00000198612 | *COPS8* | 2.620218192 | -0.010067376 | -0.087743297 | 0.077675921 | 0.999933887 | 0.013969247 | 0.181253344 | -0.167284098 | 0.402848813 |
| ENSG00000170291 | *ELP5* | 2.526642888 | -0.070723327 | 0.076550903 | -0.147274229 | 0.999933887 | 0.293530614 | 0.129765214 | 0.1637654 | 0.402939058 |
| ENSG00000103043 | *VAC14* | 4.51144786 | -0.012297516 | 0.185647178 | -0.197944694 | 0.944094537 | -0.069199996 | -0.156075581 | 0.086875585 | 0.402939058 |
| ENSG00000221944 | *TIGD1* | 1.396362281 | -0.032614927 | -0.120076554 | 0.087461627 | 0.999933887 | 0.134346968 | -0.074154539 | 0.208501507 | 0.40304495 |
| ENSG00000249936 | *RAC1P2* | -0.290281765 | 0.017877224 | 0.150153571 | -0.132276347 | 0.999933887 | 0.370032133 | 0.071023435 | 0.299008698 | 0.40306369 |
| ENSG00000005812 | *FBXL3* | 5.077230146 | -0.142293296 | -0.213010354 | 0.070717057 | 0.999933887 | -0.66125204 | -0.532550747 | -0.128701294 | 0.403081496 |
| ENSG00000165113 | *GKAP1* | 1.13994411 | 0.095798443 | -0.190659591 | 0.286458034 | 0.999933887 | -0.525088367 | -0.721674993 | 0.196586626 | 0.403081496 |
| ENSG00000134982 | *APC* | 5.131834449 | -0.044782621 | -0.111142474 | 0.066359852 | 0.999933887 | -0.912165798 | -0.627890917 | -0.284274882 | 0.403081496 |
| ENSG00000111728 | *ST8SIA1* | 1.088018715 | 0.24284265 | 0.272506655 | -0.029664005 | 0.999933887 | 0.041291757 | 0.227754324 | -0.186462567 | 0.403134551 |
| ENSG00000252904 | *AC010359.1* | -0.730965894 | -0.180873554 | 0.239314112 | -0.420187667 | 0.999933887 | 0.997102479 | 1.463976452 | -0.466873973 | 0.403134551 |
| ENSG00000038219 | *BOD1L1* | 7.834311808 | -0.111186731 | -0.186869777 | 0.075683046 | 0.999933887 | -0.905319985 | -0.651305063 | -0.254014922 | 0.403276497 |
| ENSG00000130826 | *DKC1* | 5.084719847 | -0.02608809 | -0.081437251 | 0.055349161 | 0.999933887 | -0.070276868 | 0.013229809 | -0.083506677 | 0.403276497 |
| ENSG00000144048 | *DUSP11* | 3.782190131 | -0.012308932 | 0.005801709 | -0.018110641 | 0.999933887 | -0.323563137 | -0.449041617 | 0.125478479 | 0.403276497 |
| ENSG00000132773 | *TOE1* | 2.574086065 | 0.040396531 | 0.020272375 | 0.020124155 | 0.999933887 | 0.190712764 | 0.041058037 | 0.149654727 | 0.403394952 |
| ENSG00000118513 | *MYB* | 1.152967588 | -0.140152769 | -0.267858216 | 0.127705447 | 0.999933887 | 0.086366936 | 0.243779896 | -0.15741296 | 0.403633689 |
| ENSG00000133687 | *TMTC1* | -0.191527272 | -0.076106793 | 0.082333173 | -0.158439965 | 0.999933887 | -0.623005186 | -1.024432444 | 0.401427258 | 0.403633689 |
| ENSG00000174946 | *GPR171* | 2.771047324 | -0.076285001 | 0.051186777 | -0.127471778 | 0.999933887 | 0.009738724 | 0.175541331 | -0.165802607 | 0.403722661 |
| ENSG00000006576 | *PHTF2* | 3.691842123 | -0.047810251 | -0.092403821 | 0.04459357 | 0.999933887 | -0.396548346 | -0.222508911 | -0.174039436 | 0.403722661 |
| ENSG00000001036 | *FUCA2* | 1.509733165 | 0.142941842 | -0.022110637 | 0.165052479 | 0.999933887 | -0.163883169 | -0.361586405 | 0.197703236 | 0.403794834 |
| ENSG00000163297 | *ANTXR2* | 6.703502957 | 0.024009137 | 0.048259469 | -0.024250332 | 0.999933887 | -0.467283313 | -0.570686532 | 0.103403219 | 0.403794834 |
| ENSG00000132906 | *CASP9* | 3.17417455 | -0.001135785 | 0.071747037 | -0.072882821 | 0.999933887 | -0.066873452 | -0.216953836 | 0.150080383 | 0.403803404 |
| ENSG00000141552 | *ANAPC11* | 3.507290795 | 0.081808333 | 0.093772407 | -0.011964074 | 0.999933887 | 0.131317972 | -0.03772662 | 0.169044592 | 0.403862683 |
| ENSG00000243302 | *AC018638.4* | 4.043281015 | 0.082707873 | 0.059794373 | 0.0229135 | 0.999933887 | -0.121496911 | 0.037479819 | -0.15897673 | 0.404555234 |
| ENSG00000113119 | *TMCO6* | 3.242960492 | -0.013630448 | 0.10058194 | -0.114212388 | 0.999933887 | -0.313480617 | -0.437828346 | 0.124347729 | 0.404857881 |
| ENSG00000148824 | *MTG1* | 0.309089123 | 0.108582515 | -0.136169033 | 0.244751548 | 0.999933887 | 0.135792517 | -0.119124529 | 0.254917045 | 0.404995395 |
| ENSG00000154710 | *RABGEF1* | 4.386371929 | 0.117662046 | 0.067585283 | 0.050076764 | 0.999933887 | 0.871281311 | 1.013492641 | -0.14221133 | 0.405002796 |
| ENSG00000167528 | *ZNF641* | 5.496147512 | -0.099704136 | 0.052902455 | -0.152606591 | 0.999933887 | -0.34220898 | -0.456853025 | 0.114644046 | 0.405002796 |
| ENSG00000152413 | *HOMER1* | 1.578780595 | 0.093760384 | 0.075618748 | 0.018141636 | 0.999933887 | 0.474006113 | 0.725992457 | -0.251986344 | 0.405002796 |
| ENSG00000009694 | *TENM1* | 2.282727761 | -0.112996544 | -0.149262289 | 0.036265746 | 0.999933887 | -0.192172619 | 0.145083632 | -0.337256251 | 0.405002796 |
| ENSG00000130021 | *PUDP* | 0.797878703 | 0.081116635 | -0.142869493 | 0.223986128 | 0.999933887 | -0.021214924 | -0.276379734 | 0.25516481 | 0.405002796 |
| ENSG00000108395 | *TRIM37* | 3.017075323 | -0.046338719 | -0.095876042 | 0.049537323 | 0.999933887 | -0.344468848 | -0.494518221 | 0.150049373 | 0.405002796 |
| ENSG00000151532 | *VTI1A* | 4.84901986 | 0.025962897 | -0.011167961 | 0.037130858 | 0.999933887 | 0.464540411 | 0.564689193 | -0.100148783 | 0.405002796 |
| ENSG00000116237 | *ICMT* | 1.716190552 | 0.048210289 | -0.184748174 | 0.232958463 | 0.999933887 | 0.094880972 | -0.10698198 | 0.201862952 | 0.405002796 |
| ENSG00000186448 | *ZNF197* | 0.848657842 | 0.037850269 | -0.132282722 | 0.170132991 | 0.999933887 | -0.270257615 | -0.042828397 | -0.227429218 | 0.405002796 |
| ENSG00000068305 | *MEF2A* | 5.222027478 | -0.006769427 | 0.016673348 | -0.023442775 | 0.999933887 | -0.284647946 | -0.405879122 | 0.121231177 | 0.405002796 |
| ENSG00000165527 | *ARF6* | 6.395168552 | 0.079179509 | 0.011170908 | 0.068008602 | 0.999933887 | -0.392253558 | -0.493631337 | 0.101377779 | 0.405165086 |
| ENSG00000107736 | *CDH23* | 4.983649706 | 0.048224089 | 0.054673901 | -0.006449812 | 0.999933887 | 0.471512241 | 0.579258977 | -0.107746736 | 0.405165086 |
| ENSG00000165572 | *KBTBD6* | 1.332619251 | 0.028370549 | -0.205933765 | 0.234304314 | 0.999933887 | 0.344173972 | 0.580843299 | -0.236669327 | 0.405165086 |
| ENSG00000197056 | *ZMYM1* | 0.450595275 | 0.006479466 | -0.173797696 | 0.180277162 | 0.999933887 | -0.244136071 | 0.043151873 | -0.287287944 | 0.405294489 |
| ENSG00000126088 | *UROD* | 3.076676974 | 0.017973466 | 0.164346587 | -0.146373121 | 0.999933887 | 0.184576811 | 0.295422151 | -0.11084534 | 0.405520726 |
| ENSG00000137507 | *LRRC32* | -0.997003014 | 0.081721862 | -0.18124692 | 0.262968782 | 0.999933887 | 1.665095594 | 2.231595654 | -0.56650006 | 0.405520726 |
| ENSG00000135108 | *FBXO21* | 2.778526475 | -0.114425561 | 0.105890173 | -0.220315734 | 0.944094537 | 0.018257807 | 0.138036051 | -0.119778245 | 0.4058403 |
| ENSG00000102786 | *INTS6* | 5.39869072 | 0.024045129 | 0.032118244 | -0.008073116 | 0.999933887 | 0.267274551 | 0.415926871 | -0.14865232 | 0.4058403 |
| ENSG00000106266 | *SNX8* | 5.164972794 | -0.006954264 | 0.055044997 | -0.061999261 | 0.999933887 | 0.241273259 | 0.395932529 | -0.154659271 | 0.4058403 |
| ENSG00000185803 | *SLC52A2* | 2.337949113 | 0.005228902 | -0.033183468 | 0.03841237 | 0.999933887 | 0.009573195 | -0.160385929 | 0.169959124 | 0.4058403 |
| ENSG00000168060 | *NAALADL1* | -0.251660168 | -0.003695955 | 0.060463973 | -0.064159928 | 0.999933887 | 0.258636356 | -0.077302813 | 0.33593917 | 0.405849431 |
| ENSG00000181031 | *RPH3AL* | -0.182124615 | -0.099285106 | 0.022557421 | -0.121842527 | 0.999933887 | -0.194666568 | 0.185899231 | -0.380565799 | 0.405923021 |
| ENSG00000286071 | *AL136171.2* | 0.609490487 | 0.035162385 | 0.28918546 | -0.254023074 | 0.999933887 | 0.336907922 | 0.92319135 | -0.586283428 | 0.405973076 |

| | | | | | | | | | | |
|---|---|---|---|---|---|---|---|---|---|---|
| ENSG00000224383 | *PRR29* | 1.451728328 | 0.076657012 | -0.011238787 | 0.087895799 | 0.999933887 | 0.416646148 | 0.670277126 | -0.253630978 | 0.406071602 |
| ENSG00000143442 | *POGZ* | 6.368361254 | 0.00249648 | -0.020423819 | 0.0229203 | 0.999933887 | 0.013726809 | 0.082403338 | -0.068676529 | 0.406166607 |
| ENSG00000168995 | *SIGLEC7* | 2.500241701 | -0.037640894 | -0.015128424 | -0.02251247 | 0.999933887 | -0.299483373 | -0.534135448 | 0.234652076 | 0.406529282 |
| ENSG00000180957 | *PITPNB* | 4.587425384 | -0.040057911 | -0.098002526 | 0.057944616 | 0.999933887 | 0.046384735 | 0.141965243 | -0.095580508 | 0.406697903 |
| ENSG00000143314 | *MRPL24* | 1.262324234 | 0.066257984 | 0.087863256 | -0.021605272 | 0.999933887 | 0.380723557 | 0.595969368 | -0.21524581 | 0.406700916 |
| ENSG00000089289 | *IGBP1* | 4.229233572 | -0.01824403 | -0.166494288 | 0.148250258 | 0.999933887 | -0.053907294 | 0.053448435 | -0.107355729 | 0.406700916 |
| ENSG00000251192 | *ZNF674* | 3.922627422 | -0.00761222 | 0.102082276 | -0.109694497 | 0.999933887 | -0.025637191 | 0.115915303 | -0.141552494 | 0.406700916 |
| ENSG00000160180 | *TFF3* | -1.482278117 | 0.12822988 | -0.11654847 | 0.24477835 | 0.999933887 | 0.216005785 | -0.514553981 | 0.730559766 | 0.406812231 |
| ENSG00000143643 | *TTC13* | 3.242153249 | -0.051758485 | 0.021567162 | -0.073325647 | 0.999933887 | -0.18680699 | -0.334526301 | 0.147719311 | 0.407089928 |
| ENSG00000092529 | *CAPN3* | 0.741318363 | 0.051451699 | -0.008505533 | 0.059957231 | 0.999933887 | 0.07103573 | -0.211642676 | 0.282678407 | 0.407089928 |
| ENSG00000189079 | *ARID2* | 4.589634707 | -0.072001961 | -0.079674385 | 0.007672425 | 0.999933887 | -0.053790946 | 0.112635273 | -0.166426219 | 0.407227055 |
| ENSG00000183647 | *ZNF530* | -0.155777656 | -0.162969146 | 0.124515812 | -0.287484958 | 0.999933887 | -0.036778429 | 0.262838903 | -0.299617333 | 0.407267848 |
| ENSG00000179091 | *CYC1* | 3.535948725 | -0.013394368 | -0.011471595 | -0.001922773 | 0.999933887 | 0.231526589 | 0.348937477 | -0.117410888 | 0.407267848 |
| ENSG00000258733 | *LINC02328* | -0.012553141 | -0.017187881 | 0.18361659 | -0.200804471 | 0.999933887 | -0.007199686 | 0.341969086 | -0.349168772 | 0.407286368 |
| ENSG00000230061 | *TRPM2-AS* | 0.460067722 | -0.076587192 | -0.158359611 | 0.081772418 | 0.999933887 | 0.976993768 | 1.207874519 | -0.230880751 | 0.407348886 |
| ENSG00000105298 | *CACTIN* | 3.514874457 | 0.045432772 | 0.16871533 | -0.123282558 | 0.999933887 | 0.005890698 | -0.157749681 | 0.16364038 | 0.407381375 |
| ENSG00000000419 | *DPM1* | 4.307588162 | -0.005036904 | -0.032891046 | 0.027854143 | 0.999933887 | 0.05269906 | 0.136969464 | -0.084270404 | 0.407403629 |
| ENSG00000162852 | *CNST* | 4.838249709 | -0.025383907 | -0.013069283 | -0.012314624 | 0.999933887 | -0.04417601 | 0.050662114 | -0.094838124 | 0.407826095 |
| ENSG00000130779 | *CLIP1* | 5.932587926 | 0.008925573 | -0.085231716 | 0.09415729 | 0.999933887 | -0.584026037 | -0.438664078 | -0.145361959 | 0.407826095 |
| ENSG00000054282 | *SDCCAG8* | 2.487326637 | 0.055826314 | 0.036875925 | 0.01895039 | 0.999933887 | -0.130825124 | -0.296167756 | 0.165342632 | 0.407877632 |
| ENSG00000086619 | *ERO1B* | 3.707606754 | 0.064656398 | 0.031958806 | 0.032697593 | 0.999933887 | 0.010916038 | -0.155329436 | 0.166245474 | 0.40792702 |
| ENSG00000202019 | *Y_RNA* | 2.795004321 | 0.309277177 | 0.444832195 | -0.135555019 | 0.999933887 | 1.730074188 | 2.049387531 | -0.319313343 | 0.408303917 |
| ENSG00000188157 | *AGRN* | 3.320918258 | 0.06711897 | 0.090892331 | -0.023773362 | 0.999933887 | 0.494634431 | 0.677436252 | -0.182801821 | 0.408493697 |
| ENSG00000135914 | *HTR2B* | -1.360124062 | -0.138856324 | -0.425562392 | 0.286706068 | 0.999933887 | 0.628176493 | 0.976173906 | -0.347997413 | 0.408493697 |
| ENSG00000111196 | *MAGOHB* | 0.646761501 | -0.004168294 | 0.064278274 | -0.068446567 | 0.999933887 | 0.024453416 | 0.291674838 | -0.267221422 | 0.408493697 |
| ENSG00000150457 | *LATS2* | 6.460778632 | 0.019391805 | 0.066529576 | -0.047137771 | 0.999933887 | -0.082491073 | 0.000318551 | -0.082809624 | 0.408983092 |
| ENSG00000281404 | *LINC01176* | 1.19392998 | -0.070837145 | -0.013109696 | -0.057727449 | 0.999933887 | -0.521680185 | -0.817196785 | 0.2955166 | 0.409025303 |
| ENSG00000271734 | *Z98200.1* | 0.469941728 | 0.029559067 | 0.020268155 | 0.009290912 | 0.999933887 | 0.231663275 | 0.438181321 | -0.206518045 | 0.409097912 |
| ENSG00000228477 | *AL663070.1* | -0.137149269 | -0.293636698 | 0.39237074 | -0.686007438 | 0.999933887 | 2.41987694 | 2.762587369 | -0.34271043 | 0.409210928 |
| ENSG00000102158 | *MAGT1* | 5.013329613 | -0.023095916 | -0.037227032 | 0.014131116 | 0.999933887 | 0.229692795 | 0.333844543 | -0.104151749 | 0.409210928 |
| ENSG00000238005 | *AL391832.2* | 2.905476904 | -0.143660044 | 0.041518139 | -0.185178183 | 0.999933887 | 1.426926015 | 1.636532423 | -0.209606407 | 0.409737508 |
| ENSG00000197372 | *ZNF675* | 1.977053414 | -0.000370279 | -0.09494412 | 0.09457384 | 0.999933887 | -0.67507888 | -0.420556895 | -0.254521985 | 0.409737508 |
| ENSG00000128609 | *NDUFA5* | 2.958492507 | -0.118983395 | 0.015602675 | -0.134586071 | 0.999933887 | -0.00888639 | 0.1819985 | -0.19088489 | 0.409903451 |
| ENSG00000078902 | *TOLLIP* | 6.492903424 | 0.026783487 | -0.034052015 | 0.060835503 | 0.999933887 | -0.063766984 | 0.02824578 | -0.092012765 | 0.409920144 |
| ENSG00000135018 | *UBQLN1* | 6.601045733 | -0.047535143 | -0.030348041 | -0.017187102 | 0.999933887 | -0.334055735 | -0.26439557 | -0.069660165 | 0.40992564 |
| ENSG00000198894 | *CIPC* | 2.702830118 | -0.080440744 | 0.183576079 | -0.264016823 | 0.981460244 | -0.61625711 | -0.770234687 | 0.153977577 | 0.40992564 |
| ENSG00000086015 | *MAST2* | 2.43237346 | 0.016189255 | 0.000760649 | 0.015428606 | 0.999933887 | -0.084337815 | 0.047706074 | -0.132043889 | 0.40992564 |
| ENSG00000109618 | *SEPSECS* | 2.482630723 | 0.00124292 | -0.262082369 | 0.263325289 | 0.999933887 | -0.006637217 | 0.191115558 | -0.197752776 | 0.4099505 |
| ENSG00000165115 | *KIF27* | 2.811714797 | -0.028786244 | -0.21479548 | 0.186009236 | 0.999933887 | -0.629549155 | -0.938865846 | 0.30931669 | 0.410231666 |
| ENSG00000169495 | *HTRA4* | 1.988433457 | -0.013791851 | 0.165043477 | -0.178835328 | 0.999933887 | 0.172244319 | 0.311738818 | -0.139494499 | 0.410231666 |
| ENSG00000102144 | *PGK1* | 8.118777071 | -0.005844116 | -0.013890239 | 0.008046123 | 0.999933887 | 0.053208431 | -0.056269611 | 0.109478042 | 0.410243942 |
| ENSG00000178163 | *ZNF518B* | 3.821275254 | -0.051019397 | -0.049298742 | -0.001720655 | 0.999933887 | -0.509918077 | -0.374179275 | -0.135738802 | 0.410672271 |
| ENSG00000135722 | *FBXL8* | 0.996340956 | 0.095735811 | -0.04577307 | 0.141508881 | 0.999933887 | -0.046992112 | -0.314347455 | 0.267355343 | 0.411062093 |
| ENSG00000079337 | *RAPGEF3* | 1.053581794 | -0.048031529 | -0.11806063 | 0.070029101 | 0.999933887 | -0.321030208 | -0.539776028 | 0.21874582 | 0.411605426 |
| ENSG00000186312 | *CA5BP1* | 2.221195622 | 0.026197858 | -0.056930231 | 0.083128089 | 0.999933887 | 0.112983944 | 0.308054594 | -0.19507065 | 0.411605426 |
| ENSG00000287104 | *AC097382.3* | 1.589727941 | -0.076251003 | -0.021394667 | -0.054856335 | 0.999933887 | 0.259196053 | 0.083909846 | 0.175286206 | 0.411874053 |
| ENSG00000133393 | *CEP20* | 2.421968347 | -0.031900961 | 0.111876738 | -0.143777698 | 0.999933887 | -0.053741194 | 0.079478878 | -0.133220072 | 0.411964078 |
| ENSG00000162923 | *WDR26* | 8.752050268 | -0.010469804 | -0.000886666 | -0.009583138 | 0.999933887 | -0.392642775 | -0.312644722 | -0.079998054 | 0.412015143 |
| ENSG00000173653 | *RCE1* | 3.306570569 | -0.028832004 | 0.008336833 | -0.037168837 | 0.999933887 | -0.065317954 | -0.190560864 | 0.125242909 | 0.412167504 |
| ENSG00000119812 | *FAM98A* | 1.570634132 | -0.033899527 | -0.089084735 | 0.055185208 | 0.999933887 | 0.123497804 | 0.338968487 | -0.215470683 | 0.412167504 |
| ENSG00000267074 | *AC015911.3* | 0.920914463 | -0.01716385 | 0.042419252 | -0.059583103 | 0.999933887 | -0.194406516 | 0.044243686 | -0.238650203 | 0.41216789 |
| ENSG00000133627 | *ACTR3B* | 0.606441812 | 0.075330086 | -0.21887985 | 0.294209936 | 0.999933887 | -0.048884503 | -0.39778451 | 0.348900007 | 0.412268408 |
| ENSG00000229180 | *AC006001.3* | 4.069638834 | 0.064571614 | 0.06436161 | 0.000210004 | 0.999933887 | -0.158525705 | -0.258189568 | 0.099663864 | 0.412306201 |
| ENSG00000176485 | *PLAAT3* | 0.343370809 | -0.182915082 | -0.204899212 | 0.021984129 | 0.999933887 | 0.034577394 | 0.331188308 | -0.296610914 | 0.412509598 |
| ENSG00000160818 | *GPATCH4* | 2.122919449 | -0.13155037 | -0.11424808 | -0.01730229 | 0.999933887 | 0.25870346 | 0.441353271 | -0.182649811 | 0.412509598 |
| ENSG00000166535 | *A2ML1* | 0.802490654 | -0.398720766 | -0.270623955 | -0.128096811 | 0.999933887 | -0.564783254 | -0.383792775 | -0.18099048 | 0.412509598 |
| ENSG00000130713 | *EXOSC2* | 2.31804448 | -0.075726576 | -0.076439314 | 0.000712738 | 0.999933887 | 0.227958894 | 0.078482178 | 0.149476716 | 0.412509598 |
| ENSG00000244436 | *AC023050.1* | 1.410287492 | 0.140255941 | 0.044458461 | 0.09579748 | 0.999933887 | 0.495885572 | 0.783249547 | -0.287363975 | 0.412509598 |
| ENSG00000284976 | *BX255925.3* | 3.155221776 | -0.02026483 | 0.22990538 | -0.25017021 | 0.594252256 | 0.006284006 | -0.128121998 | 0.134406004 | 0.412509598 |
| ENSG00000176531 | *PHLDB3* | 1.349439004 | 0.005947172 | 0.158243447 | -0.152296275 | 0.999933887 | -0.063691104 | 0.175416907 | -0.239108011 | 0.412509598 |
| ENSG00000160051 | *IQCC* | 0.680050692 | 0.025875266 | 0.038185482 | -0.012310216 | 0.999933887 | 0.135384392 | -0.143918277 | 0.279302669 | 0.412689421 |
| ENSG00000179335 | *CLK3* | 7.419658153 | 0.030312763 | 0.047835453 | -0.01752269 | 0.999933887 | 0.202035454 | 0.276643134 | -0.07460768 | 0.412933188 |
| ENSG00000266983 | *AC011444.1* | -0.905412683 | -0.022949819 | 0.294803307 | -0.317753125 | 0.999933887 | -0.413747749 | -1.039761038 | 0.626013289 | 0.412941082 |
| ENSG00000285909 | *AP002762.2* | 0.295228929 | 0.0166776 | 0.153475619 | -0.136798018 | 0.999933887 | -0.226246371 | 0.110052866 | -0.336299236 | 0.413116752 |
| ENSG00000140006 | *WDR89* | 0.828639565 | 0.109967756 | 0.026531638 | 0.083436118 | 0.999933887 | 0.097827706 | 0.355699804 | -0.257872099 | 0.413444721 |
| ENSG00000186908 | *ZDHHC17* | 5.228723557 | -0.053004019 | -0.118478455 | 0.065474436 | 0.999933887 | -0.516273527 | -0.300102949 | -0.216170579 | 0.413475831 |
| ENSG00000277715 | *AC079174.2* | 0.699623789 | -0.136961136 | -0.099932953 | -0.037028183 | 0.999933887 | 1.349020399 | 1.6795804 | -0.330560002 | 0.414034662 |
| ENSG00000273032 | *DGCR5* | -0.967128788 | -0.181531137 | 0.333307411 | -0.514838548 | 0.594252256 | 0.18947005 | -0.454480861 | 0.643950911 | 0.414569741 |
| ENSG00000182022 | *CHST15* | 8.44162772 | -0.020389029 | -0.008768116 | -0.011620912 | 0.999933887 | -0.414061915 | -0.329185378 | -0.084876537 | 0.414741309 |
| ENSG00000137216 | *TMEM63B* | 3.61737624 | 0.041265236 | 0.129268688 | -0.088003452 | 0.999933887 | 0.72460454 | 0.866986793 | -0.142382253 | 0.414752306 |
| ENSG00000267212 | *AC018761.3* | 0.685132907 | 0.205599436 | 0.258298998 | -0.052699562 | 0.999933887 | 0.585462798 | 0.836439317 | -0.250976519 | 0.415034046 |
| ENSG00000261114 | *AC012181.1* | 0.123975933 | 0.142486141 | 0.112490277 | 0.029995864 | 0.999933887 | 0.3438285 | -0.149735857 | 0.493564357 | 0.415034046 |
| ENSG00000105339 | *DENND3* | 9.548978313 | 0.0512974 | 0.030996987 | 0.020300413 | 0.999933887 | -1.145216073 | -1.282281984 | 0.137065911 | 0.415034046 |
| ENSG00000147649 | *MTDH* | 5.103871151 | -0.024553603 | -0.025636097 | 0.001082494 | 0.999933887 | -0.305403832 | -0.416228811 | 0.110824979 | 0.415034046 |
| ENSG00000049860 | *HEXB* | 6.874466615 | 0.000677249 | -0.00055155 | 0.001228798 | 0.999933887 | -0.593677043 | -0.689600256 | 0.095923213 | 0.415034046 |
| ENSG00000125651 | *GTF2F1* | 5.517300392 | 0.076413477 | 0.052374746 | 0.024038731 | 0.999933887 | -0.082103524 | 0.044649321 | -0.126752845 | 0.415186187 |
| ENSG00000249992 | *TMEM158* | 2.669153639 | 0.056724758 | 0.073357749 | -0.016632991 | 0.999933887 | -0.634858518 | -0.869543962 | 0.234685444 | 0.415186187 |
| ENSG00000121060 | *TRIM25* | 8.730081719 | -0.139767048 | -0.066140581 | -0.073626467 | 0.999933887 | -0.746475382 | -0.86826571 | 0.121790328 | 0.415413212 |
| ENSG00000246889 | *AP000487.1* | -0.144981414 | 0.212350405 | -0.053275187 | 0.265625593 | 0.999933887 | 0.117269806 | 0.432636271 | -0.315366465 | 0.415413212 |
| ENSG00000196743 | *GM2A* | 4.250096449 | -0.058144418 | -0.069100124 | 0.010955766 | 0.999933887 | 1.016231142 | 1.260028749 | -0.243797607 | 0.415413212 |
| ENSG00000274092 | *AC106739.1* | -1.150466273 | 0.23865879 | -0.096539725 | 0.335198515 | 0.999933887 | 0.599323284 | 1.203196687 | -0.603873403 | 0.415439214 |
| ENSG00000285852 | *AL353147.1* | -0.150880706 | 0.048853132 | 0.003359237 | 0.045493895 | 0.999933887 | -0.386744727 | -0.848148168 | 0.461403441 | 0.415439214 |
| ENSG00000038945 | *MSR1* | 0.183222322 | -0.270353759 | 0.025140706 | -0.295494465 | 0.999933887 | -1.502986528 | -1.896344837 | 0.393358309 | 0.415480262 |
| ENSG00000131845 | *ZNF304* | 1.291972821 | -0.105364402 | 0.013935627 | -0.119300028 | 0.999933887 | -0.033204117 | 0.210270118 | -0.243474235 | 0.415480262 |
| ENSG00000075711 | *DLG1* | 3.615064667 | 0.021213885 | -0.091956936 | 0.113170822 | 0.999933887 | -0.153582126 | 0.02421803 | -0.177800156 | 0.415492353 |
| ENSG00000110777 | *POU2AF1* | 0.288563124 | -0.127885196 | -0.063288628 | -0.064596568 | 0.999933887 | -0.14411546 | 0.085829802 | -0.229945263 | 0.415620968 |
| ENSG00000168405 | *CMAHP* | 2.706711876 | 0.056180129 | -0.158488888 | 0.214669017 | 0.999933887 | 0.016041029 | 0.205548693 | -0.189507664 | 0.415620968 |
| ENSG00000125354 | *SEPTIN6* | 5.72570065 | 0.011029652 | -0.02285082 | 0.033880472 | 0.999933887 | -0.010217528 | -0.105818579 | 0.095601051 | 0.415620968 |
| ENSG00000286681 | *AF235103.3* | 0.044516336 | -0.029604217 | 0.08431322 | -0.113917436 | 0.999933887 | 0.553097272 | 0.913544498 | -0.360447225 | 0.415620968 |
| ENSG00000282022 | *AC006581.2* | -0.780803793 | 0.142181145 | 0.04857165 | 0.093609495 | 0.999933887 | -0.000799944 | 0.21712187 | -0.217921814 | 0.415699209 |
| ENSG00000169976 | *SF3B5* | 5.149371463 | 0.045451397 | 0.010553491 | 0.034897906 | 0.999933887 | 0.100443338 | -0.009877984 | 0.110321323 | 0.41610153 |
| ENSG00000202502 | *RNA5SP151* | -0.005886161 | 0.467361403 | 0.393619523 | 0.07374188 | 0.999933887 | 0.542816722 | 1.252066359 | -0.709249637 | 0.416273928 |
| ENSG00000154814 | *OXNAD1* | 3.027337099 | 0.016844458 | -0.105280211 | 0.12212467 | 0.999933887 | -0.033974089 | 0.092279426 | -0.126253515 | 0.416380505 |
| ENSG00000116663 | *FBXO6* | 3.066402786 | 0.035188072 | 0.021263177 | 0.013924895 | 0.999933887 | 0.302716089 | 0.458468491 | -0.155752402 | 0.416474837 |
| ENSG00000278970 | *HEIH* | 3.634041092 | 0.028467112 | 0.046593716 | -0.018126603 | 0.999933887 | 0.560119577 | 0.696946931 | -0.136827354 | 0.416474837 |
| ENSG00000156875 | *MFSD14A* | 2.677517288 | 0.029229574 | -0.085219248 | 0.114448823 | 0.999933887 | 0.233706913 | 0.377263565 | -0.143556652 | 0.416698044 |
| ENSG00000129473 | *BCL2L2* | 0.557361681 | 0.060161806 | 0.371217466 | -0.31105566 | 0.999933887 | -0.213309734 | -0.428174684 | 0.214864951 | 0.417089965 |
| ENSG00000248049 | *UBA6-AS1* | -0.155197092 | -0.006943757 | 0.17390522 | -0.180848977 | 0.999933887 | 0.027074543 | -0.289032237 | 0.316106779 | 0.417607823 |
| ENSG00000066136 | *NFYC* | 6.076308337 | -0.00857586 | -0.022931517 | 0.014355656 | 0.999933887 | -0.21505254 | -0.134138695 | -0.080913844 | 0.417632855 |
| ENSG00000132879 | *FBXO44* | 2.691769682 | 0.013679588 | 0.09048903 | -0.076809442 | 0.999933887 | 0.152369049 | 0.018002025 | 0.134367024 | 0.417632855 |
| ENSG00000198799 | *LRIG2* | 2.607887233 | -0.015993223 | -0.042612474 | 0.026619252 | 0.999933887 | -0.046307075 | -0.183753405 | 0.13744633 | 0.417953139 |
| ENSG00000102934 | *PLLP* | -0.746990128 | -0.243233626 | 0.061795978 | -0.305029604 | 0.999933887 | -0.398457238 | 0.012697458 | -0.411154697 | 0.417957921 |
| ENSG00000169258 | *GPRIN1* | -0.193361395 | 0.152388843 | -0.233712563 | 0.386101407 | 0.999933887 | -0.253172265 | -0.688731831 | 0.435559566 | 0.418190779 |
| ENSG00000166140 | *ZFYVE19* | 2.545344709 | 0.019355435 | 0.109463692 | -0.090108257 | 0.999933887 | 0.054845367 | -0.104389934 | 0.159235301 | 0.418190779 |
| ENSG00000129534 | *MIS18BP1* | 2.968922918 | -0.038181972 | -0.178696427 | 0.140514455 | 0.999933887 | -0.818810034 | -1.081002002 | 0.262191968 | 0.418355377 |

| | | | | | | | | | | |
|---|---|---|---|---|---|---|---|---|---|---|
| ENSG00000129354 | *AP1M2* | 0.271288919 | -0.078522361 | 0.082568256 | -0.161090617 | 0.999933887 | -1.598241514 | -2.031655821 | 0.433414306 | 0.41854785 |
| ENSG00000233622 | *CYP2T1P* | 2.119380496 | 0.058063769 | 0.164661273 | -0.106597504 | 0.999933887 | 0.024867963 | 0.254814041 | -0.229946078 | 0.41854785 |
| ENSG00000133069 | *TMCC2* | 0.179546413 | -0.106487903 | -0.254017999 | 0.147530095 | 0.999933887 | 0.123739233 | -0.186459832 | 0.310199065 | 0.418814195 |
| ENSG00000143167 | *GPA33* | -0.052955639 | -0.085343692 | 0.05444446 | -0.139788151 | 0.999933887 | 0.076799645 | 0.41012975 | -0.333330105 | 0.418814195 |
| ENSG00000090971 | *NAT14* | 0.750286031 | 0.326205161 | 0.180145684 | 0.146059477 | 0.999933887 | -0.203456174 | -0.536281971 | 0.332825796 | 0.418954831 |
| ENSG00000169902 | *TPST1* | 2.914284028 | 0.110399706 | 0.128102487 | -0.017702782 | 0.999933887 | -0.159507979 | 0.000629443 | -0.160137422 | 0.418954831 |
| ENSG00000228863 | *AL121985.1* | 3.617478287 | 0.031464634 | 0.198234542 | -0.166769908 | 0.999933887 | 0.603845036 | 0.80586809 | -0.202023054 | 0.418954831 |
| ENSG00000177738 | *AC025171.1* | 1.582199328 | -0.009302552 | 0.151879872 | -0.161182424 | 0.999933887 | -0.036449779 | -0.249648564 | 0.213198785 | 0.418954831 |
| ENSG00000116604 | *MEF2D* | 8.385059644 | 0.051081732 | 0.097275799 | -0.046194067 | 0.999933887 | -0.059206294 | 0.020374622 | -0.079580915 | 0.418956694 |
| ENSG00000174851 | *YIF1A* | 2.321080261 | -0.023678411 | 0.049000767 | -0.072679178 | 0.999933887 | 0.139694339 | 0.008702453 | 0.130991886 | 0.418970147 |
| ENSG00000279942 | *AL353597.3* | -1.234055546 | 0.04701667 | -0.075800834 | 0.122817504 | 0.999933887 | 0.426243913 | 0.017317941 | 0.408925972 | 0.418970147 |
| ENSG00000253593 | *AC110741.1* | -0.83551364 | 0.334109778 | 0.483945349 | -0.149835572 | 0.999933887 | 1.617542099 | 2.056572174 | -0.439030075 | 0.419620288 |
| ENSG00000118816 | *CCNI* | 8.232649341 | -0.005443128 | -0.016543681 | 0.011100552 | 0.999933887 | 0.042120365 | 0.112076192 | -0.069955827 | 0.419726937 |
| ENSG00000128585 | *MKLN1* | 5.316187685 | -0.058772266 | -0.058042783 | -0.000729483 | 0.999933887 | -0.724512168 | -0.567171917 | -0.157340252 | 0.420185791 |
| ENSG00000282458 | *WASH5P* | 0.846667497 | 0.043936696 | 0.203946222 | -0.160009526 | 0.999933887 | -0.046190164 | -0.275448822 | 0.229258658 | 0.420185791 |
| ENSG00000160888 | *IER2* | 6.611099344 | 0.346861873 | 0.456306884 | -0.109445011 | 0.999933887 | 0.197634057 | -0.034348464 | 0.231982521 | 0.420478379 |
| ENSG00000131981 | *LGALS3* | 8.9632194 | -0.058003876 | -0.024361024 | -0.033642852 | 0.999933887 | 0.133034525 | 0.231568883 | -0.098534359 | 0.420669487 |
| ENSG00000136628 | *EPRS1* | 3.515730537 | -0.067776218 | -0.141097645 | 0.073321426 | 0.999933887 | 0.024759356 | 0.180257948 | -0.155498591 | 0.420669487 |
| ENSG00000008277 | *ADAM22* | 0.214776548 | -0.066311029 | -0.195940572 | 0.129629543 | 0.999933887 | -0.321684677 | 0.063779116 | -0.385463792 | 0.420868096 |
| ENSG00000228315 | *GUSBP11* | 4.189257718 | -0.00677704 | 0.015145982 | -0.021923022 | 0.999933887 | 0.123036225 | 0.236966618 | -0.113930393 | 0.420953378 |
| ENSG00000138738 | *PRDM5* | -0.010923751 | -0.031647026 | -0.055751219 | 0.024104194 | 0.999933887 | -0.378148996 | -0.764470679 | 0.386321683 | 0.420973309 |
| ENSG00000180008 | *SOCS4* | 3.872633265 | 0.019696192 | -0.180971107 | 0.200667299 | 0.999933887 | -0.429040756 | -0.21037787 | -0.218662885 | 0.421370576 |
| ENSG00000107862 | *GBF1* | 5.895502546 | -0.015807412 | 0.03185658 | -0.047663992 | 0.999933887 | 0.094258008 | 0.171121622 | -0.076863614 | 0.421591836 |
| ENSG00000280035 | *AC011676.5* | 4.829007373 | 0.027245958 | 0.021566329 | 0.005679629 | 0.999933887 | 0.146700017 | 0.305226641 | -0.158526623 | 0.421593064 |
| ENSG00000181873 | *IBA57* | 3.299007863 | 0.047851328 | 0.127723351 | -0.079872023 | 0.999933887 | 0.091209935 | 0.223910377 | -0.132700442 | 0.421805168 |
| ENSG00000135365 | *PHF21A* | 6.986451057 | 0.013635355 | -0.029650474 | 0.043285828 | 0.999933887 | 0.013423959 | -0.062157662 | 0.07558162 | 0.421805168 |
| ENSG00000160075 | *SSU72* | 7.210266976 | -0.010754156 | -0.030512568 | 0.019758412 | 0.999933887 | -0.088415569 | -0.015919419 | -0.072496151 | 0.421913089 |
| ENSG00000129932 | *DOHH* | 1.332178817 | 0.078651897 | 0.147228035 | -0.068576137 | 0.999933887 | 0.118611734 | 0.357529613 | -0.238917879 | 0.422246503 |
| ENSG00000167074 | *TEF* | 0.49015101 | 0.0524783 | -0.127397796 | 0.179876096 | 0.999933887 | 0.05558538 | -0.248796107 | 0.304381487 | 0.422601349 |
| ENSG00000160685 | *ZBTB7B* | 7.57600997 | 0.073884565 | 0.091360272 | -0.017475706 | 0.999933887 | -0.309764872 | -0.172693291 | -0.137071582 | 0.422697224 |
| ENSG00000163564 | *PYHIN1* | 3.224363586 | -0.080233605 | -0.124740019 | 0.044506414 | 0.999933887 | -0.125564897 | 0.068529417 | -0.194094313 | 0.422697224 |
| ENSG00000151923 | *TIAL1* | 5.231575354 | -0.015752838 | 0.047287246 | -0.063040084 | 0.999933887 | -0.272486575 | -0.37030392 | 0.097817345 | 0.422697224 |
| ENSG00000146963 | *LUC7L2* | 4.900580517 | -0.029524004 | -0.038859175 | 0.009335171 | 0.999933887 | -0.149941571 | -0.029424094 | -0.120517476 | 0.422846143 |
| ENSG00000101082 | *SLA2* | 3.359858945 | -0.002517885 | -0.017592704 | 0.015074819 | 0.999933887 | 0.053378942 | 0.186405378 | -0.133026436 | 0.423209564 |
| ENSG00000116209 | *TMEM59* | 6.748710914 | -0.023759276 | -0.067329515 | 0.043570239 | 0.999933887 | -0.167096221 | -0.261118145 | 0.094021924 | 0.423390636 |
| ENSG00000233896 | *PDYN-AS1* | 0.064685822 | 0.2007075 | 0.197169053 | 0.003538447 | 0.999933887 | 2.975996146 | 3.510385635 | -0.534389488 | 0.42416159 |
| ENSG00000144579 | *CTDSP1* | 6.926588914 | 0.030298605 | 0.029846699 | 0.000451906 | 0.999933887 | -0.201332205 | -0.295568927 | 0.094236722 | 0.42416159 |
| ENSG00000243679 | *AC018638.5* | 4.186724631 | 0.0623405 | -0.044632618 | 0.106973118 | 0.999933887 | -0.169927337 | -0.052537766 | -0.117389571 | 0.424269749 |
| ENSG00000220323 | *H2BC19P* | 1.968216404 | 0.056034568 | -0.086641659 | 0.142676227 | 0.999933887 | -0.692409566 | -0.874937387 | 0.18252782 | 0.424269749 |
| ENSG00000279170 | *TSTD3* | -0.287056488 | -0.095548836 | -0.052494026 | -0.04305481 | 0.999933887 | -0.128494654 | 0.288363671 | -0.416858325 | 0.424269749 |
| ENSG00000188070 | *C11orf95* | 0.270735584 | 0.068923826 | 0.294730991 | -0.225807165 | 0.999933887 | 0.215617485 | -0.076640998 | 0.292258483 | 0.424269749 |
| ENSG00000268081 | *AC123912.1* | 0.490695371 | 0.283311819 | 0.362521644 | -0.079209825 | 0.999933887 | 0.69242113 | 0.373675242 | 0.318745888 | 0.424352893 |
| ENSG00000173715 | *C11orf80* | 0.510039215 | -0.028119193 | -0.015214227 | -0.012904966 | 0.999933887 | -0.048602558 | -0.342114526 | 0.293511968 | 0.424352893 |
| ENSG00000157625 | *TAB3* | 4.954494624 | -0.003378347 | -0.079211218 | 0.075832871 | 0.999933887 | -0.371312478 | -0.199123448 | -0.17218903 | 0.42462857 |
| ENSG00000155368 | *DBI* | 4.54317514 | 0.017655599 | -0.059613908 | 0.077269508 | 0.999933887 | 0.436728861 | 0.306072931 | 0.13065593 | 0.424953729 |
| ENSG00000258623 | *AL121820.1* | 1.821148952 | 0.386309678 | 0.217304786 | 0.169004892 | 0.999933887 | 0.375139479 | 0.57238856 | -0.197249081 | 0.42495488 |
| ENSG00000223552 | *CCR5AS* | 3.365449631 | 0.074023746 | 0.244005478 | -0.169981732 | 0.999933887 | 0.462883168 | 0.596268795 | -0.133385627 | 0.42495488 |
| ENSG00000268583 | *AC011466.1* | 0.608531257 | 0.101077916 | 0.013025856 | 0.088052059 | 0.999933887 | -0.776665438 | -1.050496703 | 0.273831264 | 0.425197172 |
| ENSG00000144848 | *ATG3* | 6.773609491 | -0.022399463 | -0.014019876 | -0.008379587 | 0.999933887 | 0.193786243 | 0.26441469 | -0.070628447 | 0.425605006 |
| ENSG00000100353 | *EIF3D* | 5.95011214 | -0.044244959 | -0.073115102 | 0.028870143 | 0.999933887 | -0.068356999 | -0.141930787 | 0.073573788 | 0.425708339 |
| ENSG00000283525 | *AC005622.1* | 2.06892188 | 0.058823374 | 0.052876457 | 0.005946917 | 0.999933887 | 1.340126099 | 1.522675858 | -0.182549759 | 0.425708339 |
| ENSG00000111321 | *LTBR* | 5.733837862 | 0.057456294 | 0.0281742 | 0.029282094 | 0.999933887 | -0.913476688 | -1.02518364 | 0.111706952 | 0.4257142 |
| ENSG00000137814 | *HAUS2* | 1.75411252 | 0.090050554 | -0.151440581 | 0.241491135 | 0.999933887 | 0.103658563 | 0.277021084 | -0.173362521 | 0.425860856 |
| ENSG00000180953 | *ST20* | 1.414192712 | -0.022552742 | -0.08005169 | 0.057498948 | 0.999933887 | -0.277194746 | -0.014609559 | -0.262585187 | 0.425920399 |
| ENSG00000162688 | *AGL* | 2.542003888 | -0.179454606 | -0.413632585 | 0.234177978 | 0.999933887 | -0.336372026 | -0.098442961 | -0.237929065 | 0.426062347 |
| ENSG00000142687 | *KIAA0319L* | 5.629426713 | -0.076103462 | -0.047764431 | -0.028339031 | 0.999933887 | 1.145471753 | 1.367551934 | -0.222080181 | 0.426062347 |
| ENSG00000269220 | *LINC00528* | 6.643813863 | 0.158851478 | 0.120681005 | 0.038170473 | 0.999933887 | 1.704264381 | 1.824591788 | -0.120327407 | 0.426144503 |
| ENSG00000112695 | *COX7A2* | 5.06321995 | -0.06029965 | -0.0403601 | -0.01993955 | 0.999933887 | 0.18048489 | 0.27880925 | -0.098324361 | 0.426144503 |
| ENSG00000225973 | *PIGBOS1* | 1.507702554 | -0.046725883 | 0.066407215 | -0.113133097 | 0.999933887 | -0.209400033 | -0.398030155 | 0.188630122 | 0.426144503 |
| ENSG00000112144 | *CILK1* | 1.400294271 | 0.019099044 | -0.171199251 | 0.190298295 | 0.999933887 | -0.167669512 | 0.061834382 | -0.229503894 | 0.426144503 |
| ENSG00000133226 | *SRRM1* | 6.227440609 | -0.044718536 | -0.063711146 | 0.01899261 | 0.999933887 | -0.214840538 | -0.129855261 | -0.084985277 | 0.42663615 |
| ENSG00000204116 | *CHIC1* | 1.390942029 | -0.121754798 | -0.252991518 | 0.13123672 | 0.999933887 | -0.105075494 | 0.236045635 | -0.341121129 | 0.42663615 |
| ENSG00000161920 | *MED11* | 4.063326664 | 0.01461198 | -0.038482206 | 0.053094186 | 0.999933887 | -0.369038281 | -0.250767499 | -0.118270782 | 0.42663615 |
| ENSG00000163357 | *DCST1* | -0.154351611 | -0.027879166 | -0.011149567 | -0.016729598 | 0.999933887 | 0.049974114 | 0.423040877 | -0.373066763 | 0.42663615 |
| ENSG00000131374 | *TBC1D5* | 3.927259611 | -0.011150083 | -0.030518663 | 0.019368581 | 0.999933887 | -0.100774035 | -0.207313429 | 0.106539394 | 0.426685335 |
| ENSG00000204564 | *C6orf136* | 2.208911413 | 0.009693744 | -0.018807741 | 0.028501485 | 0.999933887 | -0.152667562 | -0.311998302 | 0.15933074 | 0.426703809 |
| ENSG00000007545 | *CRAMP1* | 3.162463371 | -0.106339986 | 0.029876778 | -0.136216764 | 0.999933887 | -0.184020037 | -0.298189358 | 0.114169321 | 0.426705257 |
| ENSG00000068745 | *IP6K2* | 5.591509663 | 0.040243877 | 0.00093994 | 0.039303937 | 0.999933887 | 0.046695492 | 0.136822631 | -0.090127139 | 0.426705257 |
| ENSG00000164440 | *TXLNB* | -0.140328828 | 0.02082027 | 0.017732743 | 0.003087528 | 0.999933887 | -0.084811325 | 0.1904113 | -0.275222625 | 0.426705257 |
| ENSG00000119718 | *EIF2B2* | 1.549177946 | -0.155819295 | -0.034497997 | -0.121321298 | 0.999933887 | 0.025136049 | 0.200195722 | -0.175059674 | 0.427169065 |
| ENSG00000125817 | *CENPB* | 3.581404182 | 0.06396701 | 0.064280325 | -0.000313315 | 0.999933887 | -0.018772143 | -0.156805707 | 0.138033564 | 0.427169065 |
| ENSG00000157778 | *PSMG3* | 1.7535503 | -0.055681614 | -0.060664654 | 0.004983041 | 0.999933887 | 0.176194704 | 0.010693459 | 0.165501245 | 0.427169065 |
| ENSG00000254827 | *SLC22A18AS* | 3.182130046 | 0.041618845 | -0.042961676 | 0.084580521 | 0.999933887 | 0.20207969 | 0.341935981 | -0.139856291 | 0.427169065 |
| ENSG00000269514 | *AC024257.3* | -0.449025688 | -0.059995969 | 0.032182637 | -0.092178606 | 0.999933887 | -0.376760229 | 0.098060921 | -0.47482115 | 0.427293039 |
| ENSG00000178789 | *CD300LB* | 5.600773302 | -0.024296698 | -0.004796358 | -0.019500341 | 0.999933887 | 0.05394884 | -0.108766749 | 0.16271559 | 0.42747245 |
| ENSG00000231177 | *LINC00852* | 1.273579983 | 0.061003199 | 0.039810509 | 0.02119269 | 0.999933887 | 1.170791659 | 1.433466897 | -0.262675238 | 0.427619642 |
| ENSG00000139668 | *WDFY2* | 4.211911343 | 0.002226946 | -0.030894676 | 0.033121623 | 0.999933887 | 0.113768487 | -0.00769553 | 0.121464017 | 0.428414644 |
| ENSG00000167447 | *SMG8* | 1.571427471 | -0.116254736 | -0.13196289 | 0.015708155 | 0.999933887 | -0.035107916 | -0.225818278 | 0.190710363 | 0.428822948 |
| ENSG00000117262 | *GPR89A* | -0.15153103 | -0.265394154 | -0.047250561 | -0.218143593 | 0.999933887 | 0.071794228 | -0.206543184 | 0.278337412 | 0.428933957 |
| ENSG00000168826 | *ZBTB49* | 2.987688522 | 0.09016085 | -0.085792511 | 0.175953361 | 0.999933887 | -0.150795672 | -0.264354556 | 0.113558884 | 0.428933957 |
| ENSG00000185324 | *CDK10* | 3.850130858 | 0.026443555 | 0.139091557 | -0.112648002 | 0.999933887 | 0.112246387 | 0.009848721 | 0.102397666 | 0.428933957 |
| ENSG00000166762 | *CATSPER2* | -0.988747999 | -0.310270014 | 0.253412425 | -0.563682438 | 0.999933887 | -0.16637568 | 0.242008694 | -0.408384374 | 0.428996813 |
| ENSG00000146909 | *NOM1* | 2.625678984 | -0.058280889 | 0.048168244 | -0.106449133 | 0.999933887 | 0.200850054 | 0.049649449 | 0.151200605 | 0.428996813 |
| ENSG00000161010 | *MRNIP* | 6.938306133 | 0.050560767 | 0.11103131 | -0.060470542 | 0.999933887 | 0.673772381 | 0.79807475 | -0.124302368 | 0.428996813 |
| ENSG00000138496 | *PARP9* | 7.250570676 | -0.017880747 | -0.058296606 | 0.04041586 | 0.999933887 | -0.302174136 | -0.393465491 | 0.091291355 | 0.42918102 |
| ENSG00000023516 | *AKAP11* | 3.443371836 | -0.053286058 | -0.151087278 | 0.09780122 | 0.999933887 | -0.360658966 | -0.088168729 | -0.272490237 | 0.42918102 |
| ENSG00000156642 | *NPTN* | 6.189859953 | -0.003125117 | -0.043254546 | 0.04012943 | 0.999933887 | -0.209741039 | -0.309188575 | 0.099447537 | 0.429236143 |
| ENSG00000183474 | *GTF2H2C* | 0.742508929 | -0.18971304 | -0.010151119 | -0.179561921 | 0.999933887 | -0.742741003 | -0.97847971 | 0.235738706 | 0.429445255 |
| ENSG00000232931 | *LINC00342* | 2.640958016 | -0.172293536 | -0.157399159 | -0.014894376 | 0.999933887 | -0.263344982 | 0.040565307 | -0.303910289 | 0.429624632 |
| ENSG00000111237 | *VPS29* | 4.462255849 | -0.017962493 | -0.014147863 | -0.00381463 | 0.999933887 | -0.244600324 | -0.339012948 | 0.094412624 | 0.429624632 |
| ENSG00000180769 | *WDFY3-AS2* | 2.352965681 | 0.121940374 | -0.000232431 | 0.122172805 | 0.999933887 | 0.766567124 | 0.988422107 | -0.221854983 | 0.42973878 |
| ENSG00000102870 | *ZNF629* | 0.626188144 | 0.00340058 | -0.088366949 | 0.091767529 | 0.999933887 | -0.00969915 | 0.213958888 | -0.223658038 | 0.429943373 |
| ENSG00000149823 | *VPS51* | 4.987548148 | 0.114615806 | 0.041489603 | 0.073126203 | 0.999933887 | 0.049701789 | -0.066066824 | 0.115768613 | 0.429948697 |
| ENSG00000159588 | *CCDC17* | 2.542881098 | 0.089934873 | 0.177504431 | -0.087569558 | 0.999933887 | -0.203269263 | -0.041314965 | -0.161954298 | 0.430064782 |
| ENSG00000232578 | *AC093311.1* | 0.871812761 | -0.024044864 | 0.000598616 | -0.024643481 | 0.999933887 | 0.06919742 | 0.394618745 | -0.325421325 | 0.430064782 |
| ENSG00000197582 | *GPX1P1* | -0.404353151 | 0.03823474 | -0.0057293 | 0.043964041 | 0.999933887 | -0.210755496 | -0.502000147 | 0.291244651 | 0.430246267 |
| ENSG00000182979 | *MTA1* | 3.672397577 | 0.071465667 | -0.033028465 | 0.104494131 | 0.999933887 | 0.071861823 | -0.053988186 | 0.125850009 | 0.430407231 |
| ENSG00000167136 | *ENDOG* | -0.674432157 | 0.077261675 | -0.199861091 | 0.277122766 | 0.999933887 | 0.307993587 | -0.066337723 | 0.374331309 | 0.430407231 |
| ENSG00000116288 | *PARK7* | 4.483632066 | 0.007340447 | -0.121836697 | 0.129177144 | 0.999933887 | 0.085880363 | -0.030258477 | 0.11613884 | 0.430407231 |
| ENSG00000280128 | *AL662795.2* | 0.985841949 | -0.038360996 | -0.059023698 | 0.020662702 | 0.999933887 | -0.102729587 | 0.330151255 | -0.432880842 | 0.430475 |
| ENSG00000152620 | *NADK2* | 1.707930852 | 0.039783537 | -0.046298278 | 0.086081815 | 0.999933887 | -0.249182755 | -0.438396013 | 0.189213258 | 0.430674832 |
| ENSG00000229325 | *ACAP2-IT1* | 0.981893615 | 0.115000141 | 0.059414739 | 0.055585403 | 0.999933887 | -0.481002563 | -0.154333328 | -0.326669235 | 0.430728967 |
| ENSG00000211788 | *TRAV13-1* | 0.398312715 | -0.021061019 | -0.05830531 | 0.037244291 | 0.999933887 | 0.024720857 | 0.386291538 | -0.361570681 | 0.430746482 |
| ENSG00000100578 | *KIAA0586* | 2.43998676 | 0.096833677 | -0.02326524 | 0.120098917 | 0.999933887 | -0.31516747 | -0.030998012 | -0.284169458 | 0.430783419 |
| ENSG00000110046 | *ATG2A* | 9.752182062 | 0.036809283 | 0.058095447 | -0.021286164 | 0.999933887 | 0.132757512 | 0.221322704 | -0.088565192 | 0.431044284 |

| | | | | | | | | | | |
|---|---|---|---|---|---|---|---|---|---|---|
| ENSG00000126461 | *SCAF1* | 6.024860416 | -0.005118846 | 0.105366028 | -0.110484874 | 0.999933887 | -0.160895246 | -0.278261071 | 0.117365826 | 0.431044284 |
| ENSG00000168389 | *MFSD2A* | 4.15236092 | -0.020759502 | 0.056733464 | -0.077492966 | 0.999933887 | 2.532932297 | 2.767768622 | -0.234836325 | 0.431083742 |
| ENSG00000087076 | *HSD17B14* | -0.298918243 | -0.897071737 | 0.062270514 | -0.959342251 | 0.999933887 | 0.369688376 | 0.563404678 | -0.193716302 | 0.43124129 |
| ENSG00000092295 | *TGM1* | 2.593388048 | 0.116981535 | 0.120738237 | -0.003756702 | 0.999933887 | 1.067649911 | 1.203067188 | -0.135417277 | 0.43124129 |
| ENSG00000083720 | *OXCT1* | 2.008405594 | 0.042066641 | -0.0287332 | 0.070799841 | 0.999933887 | 0.075911071 | 0.26301402 | -0.187102949 | 0.43124129 |
| ENSG00000167711 | *SERPINF2* | -0.099592343 | 0.070693144 | 0.064516185 | 0.006176958 | 0.999933887 | 0.135300289 | -0.145660142 | 0.280960431 | 0.43124129 |
| ENSG00000086570 | *FAT2* | -0.512992995 | -0.273079214 | 0.226158329 | -0.499237543 | 0.999933887 | -0.60982218 | -0.936742206 | 0.326920027 | 0.431302161 |
| ENSG00000133392 | *MYH11* | 3.453045942 | -0.007462881 | -0.025561977 | 0.018099097 | 0.999933887 | 0.366601504 | 0.530185778 | -0.163584273 | 0.431302161 |
| ENSG00000147889 | *CDKN2A* | -0.670272726 | 0.009395607 | -0.033568674 | 0.042964281 | 0.999933887 | 0.227177897 | -0.145524079 | 0.372701977 | 0.431302161 |
| ENSG00000133193 | *FAM104A* | 3.538605263 | -0.049216949 | -0.023554733 | -0.025662216 | 0.999933887 | -0.599519814 | -0.744345155 | 0.144825341 | 0.431439919 |
| ENSG00000279086 | *AC073130.3* | 0.396886098 | 0.032957087 | 0.214570182 | -0.181613094 | 0.999933887 | 0.196754823 | 0.545073393 | -0.34831857 | 0.431439919 |
| ENSG00000189180 | *ZNF33A* | 4.853511799 | -0.020301269 | -0.098033561 | 0.077732293 | 0.999933887 | -0.553006864 | -0.324019732 | -0.228987132 | 0.431439919 |
| ENSG00000104863 | *LIN7B* | -0.667760445 | 0.01727897 | -0.272938063 | 0.290217033 | 0.999933887 | -0.038216263 | -0.467878103 | 0.42966184 | 0.431586224 |
| ENSG00000261609 | *GAN* | 0.585549427 | 0.142774302 | 0.103314095 | 0.039460207 | 0.999933887 | -0.153506189 | -0.482166958 | 0.328660769 | 0.431594896 |
| ENSG00000198933 | *TBKBP1* | 5.058395722 | 0.061585883 | 0.048747951 | 0.012837933 | 0.999933887 | 0.098212048 | -0.0222291 | 0.120441148 | 0.431594896 |
| ENSG00000166224 | *SGPL1* | 4.222733729 | -0.053010905 | -0.048709837 | -0.004301068 | 0.999933887 | -0.318711323 | -0.433750075 | 0.115038753 | 0.431594896 |
| ENSG00000181029 | *TRAPPC5* | 0.619378992 | -0.04207379 | -0.124815049 | 0.082741259 | 0.999933887 | 0.124708707 | 0.374428951 | -0.249720243 | 0.431594896 |
| ENSG00000242372 | *EIF6* | 3.708092431 | -0.019669153 | -0.031142406 | 0.011473253 | 0.999933887 | -0.09941686 | -0.227420698 | 0.128003838 | 0.431594896 |
| ENSG00000143553 | *SNAPIN* | 2.087135563 | 0.050388227 | -0.16852966 | 0.218917887 | 0.999933887 | -0.017880016 | -0.188922529 | 0.171042513 | 0.431744863 |
| ENSG00000171658 | *NMRAL2P* | -0.058632408 | -0.237998758 | -0.119606824 | -0.118391934 | 0.999933887 | -0.02727659 | 0.169178553 | -0.196455143 | 0.431780438 |
| ENSG00000239521 | *CASTOR3* | 2.769677378 | 0.069258218 | -0.017629918 | 0.086888135 | 0.999933887 | -0.046372396 | -0.220017365 | 0.173644969 | 0.431785621 |
| ENSG00000214517 | *PPME1* | 2.395906709 | 0.034858404 | 0.018607563 | 0.016250841 | 0.999933887 | -0.09248481 | -0.254287779 | 0.161802969 | 0.431785621 |
| ENSG00000122490 | *SLC66A2* | 5.995223692 | 0.07761638 | 0.040137137 | 0.037479244 | 0.999933887 | 0.117066567 | 0.005680056 | 0.111386511 | 0.431825041 |
| ENSG00000148200 | *NR6A1* | 3.687621744 | -0.033308333 | -0.053685935 | 0.020377602 | 0.999933887 | 0.910766174 | 1.060540044 | -0.14977387 | 0.431825041 |
| ENSG00000006015 | *REX1BD* | 3.570798203 | 0.068119502 | -0.007228382 | 0.075347885 | 0.999933887 | 0.114223023 | -0.0542503 | 0.168473323 | 0.431835306 |
| ENSG00000269019 | *HOMER3-AS1* | -0.050529624 | 0.149197572 | 0.065011261 | 0.084186311 | 0.999933887 | 0.423772947 | 0.666669359 | -0.242896412 | 0.431976865 |
| ENSG00000131828 | *PDHA1* | 3.74148977 | 0.007047131 | -0.042975273 | 0.050022404 | 0.999933887 | 0.257744551 | 0.376497188 | -0.118752637 | 0.431976865 |
| ENSG00000143493 | *INTS7* | 1.009352564 | -0.095305316 | -0.325585821 | 0.230280505 | 0.999933887 | 0.010127055 | 0.235525421 | -0.225398366 | 0.432098567 |
| ENSG00000186416 | *NKRF* | 1.394334756 | -0.063537202 | -0.152535145 | 0.088997943 | 0.999933887 | 0.154080905 | -0.058426675 | 0.21250758 | 0.432098567 |
| ENSG00000177721 | *ANXA2R* | 2.277709618 | 0.045266438 | 0.091053242 | -0.045786804 | 0.999933887 | 0.173178459 | 0.005463549 | 0.167714911 | 0.432098567 |
| ENSG00000180964 | *TCEAL8* | 0.446541358 | 0.062161851 | 0.053876952 | 0.008284899 | 0.999933887 | 0.161265298 | 0.386966021 | -0.225700723 | 0.432521845 |
| ENSG00000173137 | *ADCK5* | 1.701906796 | -0.071289334 | -0.063842857 | -0.007446477 | 0.999933887 | 0.329634802 | 0.55304576 | -0.223410958 | 0.432521845 |
| ENSG00000151348 | *EXT2* | 3.073678434 | 0.01798516 | -0.067859388 | 0.085844548 | 0.999933887 | -0.002115688 | 0.135971166 | -0.138086854 | 0.432521845 |
| ENSG00000063322 | *MED29* | 5.398694973 | 0.060716837 | 0.002184282 | 0.058532555 | 0.999933887 | 0.206233162 | 0.098797551 | 0.10743561 | 0.432863249 |
| ENSG00000186132 | *C2orf76* | 1.072065393 | -0.045522749 | 0.143635466 | -0.189158215 | 0.999933887 | -0.104244088 | -0.302894436 | 0.198650348 | 0.432902934 |
| ENSG00000163463 | *KRTCAP2* | 0.223615614 | 0.032740779 | 0.131330773 | -0.098589994 | 0.999933887 | -0.082237074 | -0.368038309 | 0.285801235 | 0.432902934 |
| ENSG00000280069 | *AC127024.8* | -0.555486244 | -0.126676753 | 0.280948579 | -0.407625332 | 0.999933887 | -0.255830142 | 0.06254714 | -0.318377281 | 0.433065515 |
| ENSG00000156990 | *RPUSD3* | 1.947743751 | -0.029789756 | -0.020805687 | -0.008984069 | 0.999933887 | 0.022707637 | -0.130202068 | 0.152909705 | 0.433324174 |
| ENSG00000112305 | *SMAP1* | 2.145695966 | 0.09509895 | -0.028874593 | 0.123973543 | 0.999933887 | -0.096210631 | -0.282653545 | 0.186442914 | 0.433428515 |
| ENSG00000135372 | *NAT10* | 3.264762771 | -0.000898418 | 0.011804485 | -0.012702903 | 0.999933887 | 0.1984345 | 0.305162221 | -0.106727721 | 0.433537846 |
| ENSG00000230202 | *AL450405.1* | 2.56181285 | 0.115050791 | 0.070300963 | 0.044749828 | 0.999933887 | 0.018377671 | 0.200703113 | -0.182325442 | 0.433766885 |
| ENSG00000103811 | *CTSH* | 3.732284528 | 0.05607543 | -0.021935968 | 0.078011398 | 0.999933887 | 0.396921485 | 0.534422623 | -0.137501138 | 0.433766885 |
| ENSG00000131584 | *ACAP3* | 4.784696125 | 0.032005749 | 0.120583718 | -0.088577969 | 0.999933887 | 0.109024541 | -0.003056387 | 0.112080928 | 0.433771792 |
| ENSG00000121577 | *POPDC2* | 0.553731669 | 0.035906078 | 0.008312678 | 0.0275934 | 0.999933887 | 0.443046243 | 0.711758328 | -0.268712086 | 0.433781087 |
| ENSG00000138686 | *BBS7* | 0.458879486 | -0.179913522 | 0.065747691 | -0.245661213 | 0.999933887 | -0.028958016 | 0.250321983 | -0.279279999 | 0.433964539 |
| ENSG00000223891 | *OSER1-DT* | 4.285269325 | -0.023483179 | -0.132882142 | 0.109398963 | 0.999933887 | -0.594706797 | -0.456408931 | -0.138297866 | 0.433964539 |
| ENSG00000254554 | *AC080023.1* | 2.922314809 | 0.118762643 | 0.201541866 | -0.082779223 | 0.999933887 | 0.69138961 | 0.828785638 | -0.137396029 | 0.43402369 |
| ENSG00000012061 | *ERCC1* | 3.190817816 | 0.034489945 | 0.027718342 | 0.006771604 | 0.999933887 | 0.106302507 | 0.247531105 | -0.141228598 | 0.43402369 |
| ENSG00000185608 | *MRPL40* | 0.879777607 | -0.017652436 | 0.176537911 | -0.194190347 | 0.999933887 | -0.179294848 | -0.485594693 | 0.306299845 | 0.434236608 |
| ENSG00000158796 | *DEDD* | 4.724273216 | -0.040638418 | -0.039459345 | -0.001179072 | 0.999933887 | -0.194266778 | -0.105553192 | -0.088713587 | 0.43432134 |
| ENSG00000103254 | *ANTKMT* | 1.093415595 | 0.067010931 | -0.004807739 | 0.07181867 | 0.999933887 | 0.021181656 | -0.188363551 | 0.209545206 | 0.43432134 |
| ENSG00000162804 | *SNED1* | 1.599756314 | 0.025112743 | 0.151471266 | -0.126358523 | 0.999933887 | -0.009786263 | 0.159334345 | -0.169120607 | 0.434474819 |
| ENSG00000242588 | *AC108010.1* | 2.222219325 | 0.020319945 | 0.005655432 | 0.014664513 | 0.999933887 | -0.172865111 | -0.009837017 | -0.163028094 | 0.434571369 |
| ENSG00000149970 | *CNKSR2* | -0.045733064 | -0.254161059 | 0.10498724 | -0.359148299 | 0.999933887 | -0.267280885 | 0.066749394 | -0.334030279 | 0.434812779 |
| ENSG00000185246 | *PRPF39* | 3.580097755 | -0.088020499 | -0.171003559 | 0.08298306 | 0.999933887 | -0.030534481 | 0.158489349 | -0.189023829 | 0.434812779 |
| ENSG00000110514 | *MADD* | 5.430996417 | 0.001181764 | -0.035091648 | 0.036273412 | 0.999933887 | -0.037065724 | -0.114932853 | 0.077867129 | 0.434812779 |
| ENSG00000228716 | *DHFR* | 0.456068669 | 0.101563665 | -0.120762084 | 0.222325749 | 0.999933887 | -0.387158666 | -0.083909752 | -0.303248915 | 0.434829791 |
| ENSG00000215067 | *ALOX12-AS1* | 4.864984605 | 0.11028519 | 0.086713978 | 0.023571213 | 0.999933887 | 0.392774434 | 0.486174481 | -0.093400047 | 0.435059528 |
| ENSG00000251369 | *ZNF550* | 1.168804741 | -0.064620807 | -0.163509287 | 0.098888479 | 0.999933887 | 0.097904691 | -0.076494309 | 0.174399 | 0.435119114 |
| ENSG00000112699 | *GMDS* | 2.068591359 | 0.102342992 | 0.122799026 | -0.020456034 | 0.999933887 | -0.042150359 | 0.186578603 | -0.228728962 | 0.435419491 |
| ENSG00000102977 | *ACD* | 3.405384687 | 0.067515496 | 0.047860838 | 0.019654658 | 0.999933887 | 0.231873841 | 0.355583092 | -0.123709251 | 0.435432729 |
| ENSG00000065518 | *NDUFB4* | 3.74405201 | -0.037262911 | 0.009650247 | -0.046913158 | 0.999933887 | 0.043807526 | 0.155816803 | -0.112009276 | 0.435432729 |
| ENSG00000213462 | *ERV3-1* | 6.981770791 | -0.027264433 | -0.04377259 | 0.016508157 | 0.999933887 | -0.258687142 | -0.144166307 | -0.114520835 | 0.435432729 |
| ENSG00000205937 | *RNPS1* | 5.839797492 | -0.011674181 | 0.001948983 | -0.013623163 | 0.999933887 | -0.068065896 | -0.141431915 | 0.073366019 | 0.435432729 |
| ENSG00000139211 | *AMIGO2* | 2.496296722 | 0.010087121 | 6.234e-05 | 0.010024781 | 0.999933887 | 0.327937655 | 0.459715858 | -0.131778202 | 0.435432729 |
| ENSG00000104972 | *LILRB1* | 3.966999872 | 0.04092912 | -0.004767878 | 0.045696998 | 0.999933887 | -0.083693501 | -0.214135606 | 0.130442106 | 0.435549788 |
| ENSG00000112237 | *CCNC* | 2.072213805 | -0.008305332 | -0.280104046 | 0.271798714 | 0.999933887 | -0.001980273 | 0.197192837 | -0.19917311 | 0.435549788 |
| ENSG00000167550 | *RHEBL1* | -0.168095024 | 0.021319641 | -0.389544931 | 0.410864573 | 0.999933887 | 0.017410633 | -0.428136233 | 0.445546866 | 0.435691914 |
| ENSG00000073150 | *PANX2* | 5.729818791 | 0.001210012 | 0.069950211 | -0.068740199 | 0.999933887 | -0.344736376 | -0.45686494 | 0.112128564 | 0.435691914 |
| ENSG00000139908 | *TSSK4* | 2.26904951 | -0.090888997 | -0.057090881 | -0.033798116 | 0.999933887 | 0.09147737 | 0.237858116 | -0.146380745 | 0.435820145 |
| ENSG00000198373 | *WWP2* | 7.44136283 | -0.027706055 | -0.009280708 | -0.018425347 | 0.999933887 | -0.417081142 | -0.507742301 | 0.090661159 | 0.435820145 |
| ENSG00000198624 | *CCDC69* | 6.498340988 | 0.003196142 | -0.01927311 | 0.022469252 | 0.999933887 | -0.064529538 | -0.147528735 | 0.082999196 | 0.435950156 |
| ENSG00000086848 | *ALG9* | -0.240451794 | -0.366072647 | 0.031240157 | -0.397312804 | 0.999933887 | -0.105303223 | 0.231514306 | -0.336817529 | 0.436558769 |
| ENSG00000182325 | *FBXL6* | 2.619724797 | 0.124138171 | 0.087008538 | 0.037129633 | 0.999933887 | 0.276797037 | 0.097411876 | 0.179385161 | 0.436558769 |
| ENSG00000117009 | *KMO* | -0.230947534 | -0.101394682 | 0.290768444 | -0.392163127 | 0.999933887 | 1.128851879 | 1.450699863 | -0.321847984 | 0.436558769 |
| ENSG00000111145 | *ELK3* | 3.216217959 | 0.008389108 | 0.145369927 | -0.136980819 | 0.999933887 | -0.193276608 | -0.340607463 | 0.147330855 | 0.436558769 |
| ENSG00000233365 | *AL121956.1* | 1.863923288 | -0.033631623 | -0.131923451 | 0.098291829 | 0.999933887 | 0.507288153 | 0.706214842 | -0.198926688 | 0.436569498 |
| ENSG00000143368 | *SF3B4* | 5.596400786 | -0.003306414 | 0.007722881 | -0.011029295 | 0.999933887 | -0.163105527 | -0.071789756 | -0.091315771 | 0.436571338 |
| ENSG00000196267 | *ZNF836* | 0.846893153 | -0.114280709 | -0.080676569 | -0.03360414 | 0.999933887 | -0.285841288 | -0.518657099 | 0.232815811 | 0.436586748 |
| ENSG00000198677 | *TTC37* | 3.342249665 | -0.111132484 | -0.161231272 | 0.050098788 | 0.999933887 | -0.324467717 | -0.102637266 | -0.221830451 | 0.436801046 |
| ENSG00000011485 | *PPP5C* | 2.44089097 | -0.035605397 | 0.002378864 | -0.037984261 | 0.999933887 | 0.109402097 | 0.259826699 | -0.150424603 | 0.436801046 |
| ENSG00000160201 | *U2AF1* | 3.274935997 | 0.070297619 | 0.090209784 | -0.019912165 | 0.999933887 | -0.029785092 | 0.120574676 | -0.150359768 | 0.436834436 |
| ENSG00000119707 | *RBM25* | 7.563893181 | -0.068129443 | -0.179841558 | 0.111712114 | 0.999933887 | -0.504659153 | -0.238864594 | -0.265794559 | 0.436899904 |
| ENSG00000011009 | *LYPLA2* | 5.261349308 | 0.016054062 | -0.060345601 | 0.076399663 | 0.999933887 | -0.249565941 | -0.340488004 | 0.090922063 | 0.436899904 |
| ENSG00000182885 | *ADGRG3* | 8.39324188 | 0.008768714 | 0.015618478 | -0.006849764 | 0.999933887 | 0.02220641 | 0.105549116 | -0.083342706 | 0.436899904 |
| ENSG00000114491 | *UMPS* | 1.174000026 | -0.004914518 | 0.246548056 | -0.251462574 | 0.999933887 | 0.191633114 | 0.365108006 | -0.173474892 | 0.437101227 |
| ENSG00000112739 | *PRPF4B* | 5.450741692 | -0.083622335 | -0.154683846 | 0.071061511 | 0.999933887 | -0.435128086 | -0.224866253 | -0.210261833 | 0.437463803 |
| ENSG00000102901 | *CENPT* | 3.550958174 | -0.034156514 | 0.059928222 | -0.094084735 | 0.999933887 | 0.119713638 | -0.001636166 | 0.121349804 | 0.437466102 |
| ENSG00000156232 | *WHAMM* | 4.749004228 | -0.006946786 | -0.000366977 | -0.00657981 | 0.999933887 | -0.006464136 | -0.09932676 | 0.092862625 | 0.437562302 |
| ENSG00000140464 | *PML* | 6.044295949 | 0.031620963 | -0.007003956 | 0.038624918 | 0.999933887 | -0.037517538 | -0.128685502 | 0.091167964 | 0.437675341 |
| ENSG00000236008 | *LINC01814* | 0.36711483 | 0.078303595 | -0.014000572 | 0.092304167 | 0.999933887 | 0.41733737 | 0.746520091 | -0.329182721 | 0.437675341 |
| ENSG00000173212 | *MAB21L3* | -1.142792871 | -0.058342271 | -0.139706829 | 0.081364559 | 0.999933887 | -0.700969494 | -0.114654794 | -0.5863147 | 0.437675341 |
| ENSG00000099974 | *DDTL* | 0.546013157 | -0.069545003 | 0.108703573 | -0.178248575 | 0.999933887 | 0.078532342 | -0.159146292 | 0.237678634 | 0.437720445 |
| ENSG00000148362 | *PAXX* | 3.772858774 | 0.017840068 | 0.087249268 | -0.069409201 | 0.999933887 | 0.167563529 | 0.048401446 | 0.119162083 | 0.437720445 |
| ENSG00000268758 | *ADGRE4P* | 4.232281309 | -0.062462862 | 0.016967653 | -0.079430515 | 0.999933887 | -0.045548426 | 0.08083675 | -0.126385176 | 0.437724391 |
| ENSG00000160953 | *PWWP3A* | 2.213158326 | 0.034845974 | -0.009211201 | 0.044057176 | 0.999933887 | -0.081331195 | -0.258723634 | 0.177392439 | 0.437724391 |
| ENSG00000126756 | *UXT* | 3.581309839 | -0.006724188 | 0.043729119 | -0.050453307 | 0.999933887 | 0.256371984 | 0.373562988 | -0.117191004 | 0.437724391 |
| ENSG00000280295 | *AC099811.6* | -0.035378373 | 0.063368722 | 0.13722879 | -0.073860067 | 0.999933887 | 0.619223973 | 0.993006821 | -0.373782847 | 0.437789686 |
| ENSG00000165819 | *METTL3* | 3.995450214 | -0.031520137 | -0.033534363 | 0.002014227 | 0.999933887 | -0.094940169 | -0.190152145 | 0.095211977 | 0.437872014 |
| ENSG00000143443 | *C1orf56* | 1.420310101 | 0.009884985 | 0.187076727 | -0.177191742 | 0.999933887 | 0.473442859 | 0.275570714 | 0.197872144 | 0.437872014 |
| ENSG00000078304 | *PPP2R5C* | 8.028982969 | -0.027839104 | -0.053009239 | 0.025170135 | 0.999933887 | 0.217218716 | 0.290028737 | -0.072810021 | 0.437928367 |
| ENSG00000173889 | *PHC3* | 5.623971371 | -0.035714446 | -0.037164489 | 0.001450043 | 0.999933887 | -0.512493102 | -0.342512498 | -0.169980604 | 0.438153334 |
| ENSG00000120688 | *WBP4* | 2.797753998 | -0.031268603 | -0.119950023 | 0.088681419 | 0.999933887 | -0.251784251 | -0.114230171 | -0.13755408 | 0.438153334 |
| ENSG00000132334 | *PTPRE* | 10.027251754 | 0.067403965 | 0.056463368 | 0.010940597 | 0.999933887 | 0.086964951 | 0.161961363 | -0.074996412 | 0.4385266 |
| ENSG00000161813 | *LARP4* | 2.037500059 | -0.180725191 | -0.136584462 | -0.044140728 | 0.999933887 | -0.229607805 | -0.052856444 | -0.176751361 | 0.4385266 |

| | | | | | | | | | | |
|---|---|---|---|---|---|---|---|---|---|---|
| ENSG00000197903 | *H2BC12* | 4.321325703 | 0.066556603 | 0.014083527 | 0.052473076 | 0.999933887 | -0.128888175 | -0.26883151 | 0.139943334 | 0.438670247 |
| ENSG00000150637 | *CD226* | 3.421098679 | -0.053997606 | -0.074635011 | 0.020637404 | 0.999933887 | -0.163721456 | 0.0231733 | -0.186894756 | 0.438862344 |
| ENSG00000071537 | *SEL1L* | 6.496471917 | -0.100204531 | -0.107602806 | 0.007398275 | 0.999933887 | -0.505733071 | -0.375853574 | -0.129879497 | 0.438873136 |
| ENSG00000105821 | *DNAJC2* | 3.279690952 | -0.091854987 | -0.195104426 | 0.103249439 | 0.999933887 | -0.092633412 | 0.027836579 | -0.120469991 | 0.43899152 |
| ENSG00000125898 | *FAM110A* | 5.254600692 | -0.076128325 | -0.097167892 | 0.021039567 | 0.999933887 | 0.51829197 | 0.646057121 | -0.12776515 | 0.43899704 |
| ENSG00000124786 | *SLC35B3* | 2.485006796 | 0.030357023 | 0.170677669 | -0.140320646 | 0.999933887 | -0.097287519 | 0.038901475 | -0.136188994 | 0.43899704 |
| ENSG00000153071 | *DAB2* | -0.041959497 | -0.00903913 | -0.001992562 | -0.007046567 | 0.999933887 | -1.279884245 | -1.738966658 | 0.459082412 | 0.43899704 |
| ENSG00000152904 | *GGPS1* | 1.571603343 | 0.009951752 | -0.029828937 | 0.039780689 | 0.999933887 | -0.300800547 | -0.134080819 | -0.166719728 | 0.439377757 |
| ENSG00000024048 | *UBR2* | 7.585991983 | -0.056403891 | -0.07232285 | 0.015918959 | 0.999933887 | -0.967836603 | -1.111018327 | 0.143181724 | 0.43969078 |
| ENSG00000145388 | *METTL14* | 3.170035678 | -0.10923158 | -0.076637051 | -0.032594529 | 0.999933887 | -0.226829109 | -0.055380181 | -0.171448928 | 0.439701842 |
| ENSG00000111361 | *EIF2B1* | 3.407088655 | -0.056185891 | -0.156133837 | 0.099947945 | 0.999933887 | -0.056994993 | -0.180936777 | 0.123941784 | 0.439701842 |
| ENSG00000130311 | *DDA1* | 4.707631515 | 0.01215372 | 0.023842227 | -0.011688507 | 0.999933887 | 0.028099819 | 0.119956174 | -0.091856355 | 0.439701842 |
| ENSG00000139197 | *PEX5* | 2.921950741 | 0.043261548 | -0.081605928 | 0.124867476 | 0.999933887 | -0.085654014 | -0.220469555 | 0.134815541 | 0.439705448 |
| ENSG00000182183 | *SHISAL2A* | 2.104167001 | -0.05768774 | -0.010604293 | -0.047083447 | 0.999933887 | -0.158538714 | -0.305254501 | 0.146715787 | 0.439789632 |
| ENSG00000084234 | *APLP2* | 8.566714692 | -0.017130273 | -0.043603804 | 0.026473531 | 0.999933887 | 0.162093699 | 0.2535233 | -0.091429601 | 0.439789632 |
| ENSG00000278535 | *DHRS11* | -0.576507638 | -0.141311339 | -0.284617693 | 0.143306354 | 0.999933887 | 0.097253656 | -0.286119028 | 0.383372685 | 0.439941655 |
| ENSG00000164548 | *TRA2A* | 6.415977773 | -0.030324581 | -0.000143784 | -0.030180797 | 0.999933887 | 0.19041918 | 0.280061738 | -0.089642558 | 0.440084406 |
| ENSG00000107560 | *RAB11FIP2* | 3.213050337 | 0.025298661 | -0.15695177 | 0.182250431 | 0.999933887 | -0.839296403 | -0.602108958 | -0.237187445 | 0.440084406 |
| ENSG00000136478 | *TEX2* | 1.889779599 | 0.118794327 | 0.071312235 | 0.047482092 | 0.999933887 | -0.267086782 | -0.458201835 | 0.191115053 | 0.440138423 |
| ENSG00000102893 | *PHKB* | 4.048204041 | -0.052634873 | -0.086669966 | 0.034035093 | 0.999933887 | -0.342897817 | -0.472746441 | 0.129848624 | 0.440138423 |
| ENSG00000259657 | *PIGHP1* | 0.71230326 | 0.080741815 | 0.113500759 | -0.032758945 | 0.999933887 | -0.025814998 | 0.270032807 | -0.295847804 | 0.440138423 |
| ENSG00000225231 | *LINC02470* | 3.041073108 | 0.018609911 | -0.150005667 | 0.168615578 | 0.999933887 | 1.768593512 | 2.131737885 | -0.363144373 | 0.440138423 |
| ENSG00000279719 | *AC015871.5* | 1.268849667 | 0.359649133 | 0.238294432 | 0.121354702 | 0.999933887 | 1.647140741 | 1.282446425 | 0.364694316 | 0.440175391 |
| ENSG00000013016 | *EHD3* | 0.063343782 | -0.202736928 | 0.006330933 | -0.209067861 | 0.999933887 | -0.035827878 | -0.362951351 | 0.327123473 | 0.440175391 |
| ENSG00000125843 | *AP5S1* | 1.273977799 | 0.008694246 | -0.038416716 | 0.047110963 | 0.999933887 | -0.128060821 | 0.091254565 | -0.219315386 | 0.440227371 |
| ENSG00000088387 | *DOCK9* | 3.700928559 | -0.004277145 | -0.086743911 | 0.082466767 | 0.999933887 | -0.01069384 | 0.109490868 | -0.120184709 | 0.440227371 |
| ENSG00000134480 | *CCNH* | 4.920627687 | -0.002483947 | -0.015820939 | 0.013336991 | 0.999933887 | -0.021357976 | 0.156603407 | -0.177961384 | 0.440227371 |
| ENSG00000142166 | *IFNAR1* | 7.131455605 | -0.021095376 | -0.010061476 | -0.011033901 | 0.999933887 | 0.272211891 | 0.387135456 | -0.114923564 | 0.440575844 |
| ENSG00000184005 | *ST6GALNAC3* | 1.140718969 | -0.019210071 | 0.003440244 | -0.022650316 | 0.999933887 | -0.266581105 | -0.507766284 | 0.241185179 | 0.440680863 |
| ENSG00000146426 | *TIAM2* | 3.283728657 | -0.078692599 | -0.052481507 | -0.026211092 | 0.999933887 | -0.146190232 | -0.018446455 | -0.127743777 | 0.440710789 |
| ENSG00000197992 | *CLEC9A* | 1.916746514 | 0.117869953 | -0.012811698 | 0.130681652 | 0.999933887 | -0.93962745 | -1.216667785 | 0.277040335 | 0.440906742 |
| ENSG00000144218 | *AFF3* | -0.086264048 | -0.059531549 | 0.035328291 | -0.09485984 | 0.999933887 | -0.086396857 | 0.193237846 | -0.279634703 | 0.441063494 |
| ENSG00000196235 | *SUPT5H* | 7.313314958 | -0.013079047 | 0.022022513 | -0.03510156 | 0.999933887 | -0.00863792 | -0.08546549 | 0.07682757 | 0.441281543 |
| ENSG00000163684 | *RPP14* | 2.348122377 | -0.075516848 | 0.042921698 | -0.118438547 | 0.999933887 | 0.042438779 | -0.131260141 | 0.17369892 | 0.441557181 |
| ENSG00000183779 | *ZNF703* | -0.139892158 | 0.195716195 | 0.224177981 | -0.028461786 | 0.999933887 | -0.18942488 | -0.593699968 | 0.404275087 | 0.441855448 |
| ENSG00000177106 | *EPS8L2* | 0.111668528 | -0.226312471 | -0.251238771 | 0.024926299 | 0.999933887 | 0.040648096 | -0.301999415 | 0.34264751 | 0.441869754 |
| ENSG00000164117 | *FBXO8* | 2.269922727 | -0.038378122 | 0.038290698 | -0.07666882 | 0.999933887 | -0.039884371 | 0.095100182 | -0.134984553 | 0.441869754 |
| ENSG00000136986 | *DERL1* | 5.467730424 | 0.014732417 | 0.020963407 | -0.00623099 | 0.999933887 | -0.11129107 | -0.187490723 | 0.076199654 | 0.441869754 |
| ENSG00000181788 | *SIAH2* | 5.434428349 | 0.001281699 | 0.063116332 | -0.061834633 | 0.999933887 | 0.717475211 | 0.854430582 | -0.13695537 | 0.441869754 |
| ENSG00000182628 | *SKA2* | 0.793085433 | 0.061505105 | -0.077564361 | 0.139069466 | 0.999933887 | 0.006319872 | 0.252726216 | -0.246406344 | 0.442196293 |
| ENSG00000035928 | *RFC1* | 3.655478145 | 0.005787071 | -0.109433599 | 0.11522067 | 0.999933887 | -0.230429476 | -0.051564537 | -0.17886494 | 0.442238367 |
| ENSG00000197054 | *ZNF763* | 0.909294147 | -0.134940467 | 0.037929203 | -0.17286967 | 0.999933887 | -0.089662987 | -0.299675136 | 0.210012148 | 0.442351452 |
| ENSG00000143479 | *DYRK3* | 1.208346585 | 0.38466585 | 0.077215565 | 0.307450285 | 0.999933887 | 2.299599505 | 2.631547826 | -0.331948322 | 0.44241127 |
| ENSG00000168710 | *AHCYL1* | 5.791958324 | -0.040077521 | -0.055962465 | 0.015884944 | 0.999933887 | -0.249554002 | -0.171764414 | -0.077789588 | 0.44241127 |
| ENSG00000248905 | *FMN1* | 1.056137661 | -0.091018972 | -0.066402354 | -0.024616618 | 0.999933887 | -0.128480578 | 0.158018214 | -0.286498791 | 0.44241127 |
| ENSG00000240053 | *LY6G5B* | 2.409099014 | -0.041526079 | -7.5884e-05 | -0.041450195 | 0.999933887 | -0.476825459 | -0.317021607 | -0.159803852 | 0.44241127 |
| ENSG00000175602 | *CCDC85B* | 3.136530432 | 0.140649833 | 0.188189884 | -0.04754005 | 0.999933887 | 0.083323925 | -0.137727672 | 0.221051598 | 0.442411622 |
| ENSG00000235652 | *FBXO30-DT* | -0.797670267 | -0.242684652 | -0.034389768 | -0.208294885 | 0.999933887 | -0.054680096 | 0.379692236 | -0.434372331 | 0.44242094 |
| ENSG00000118418 | *HMGN3* | 1.810120894 | -0.178733207 | 0.093702863 | -0.27243607 | 0.999933887 | 0.134714708 | -0.064446409 | 0.199161117 | 0.442425842 |
| ENSG00000139180 | *NDUFA9* | 0.775407141 | 0.133109119 | -0.013965134 | 0.147074253 | 0.999933887 | 0.132393169 | -0.158036365 | 0.290429534 | 0.442537988 |
| ENSG00000151413 | *NUBPL* | -0.581571951 | 0.082521508 | -0.215338819 | 0.297860327 | 0.999933887 | -0.3910358 | -0.019573629 | -0.371462171 | 0.442566292 |
| ENSG00000168701 | *TMEM208* | 3.582315958 | -0.019022309 | -0.12767955 | 0.108657241 | 0.999933887 | 0.237259223 | 0.116935054 | 0.120324169 | 0.442566292 |
| ENSG00000272288 | *AL451165.2* | 2.782993941 | 0.032238519 | 0.256500102 | -0.224261583 | 0.999933887 | 1.005171095 | 0.862136864 | 0.143034232 | 0.442648341 |
| ENSG00000040633 | *PHF23* | 5.169385821 | -0.041154204 | -0.024108555 | -0.017045649 | 0.999933887 | -0.936054542 | -1.095682937 | 0.159628395 | 0.442727765 |
| ENSG00000175806 | *MSRA* | 3.202104911 | -0.014184922 | -0.024109655 | 0.009924733 | 0.999933887 | -0.112537885 | -0.25418542 | 0.141647535 | 0.442727765 |
| ENSG00000096746 | *HNRNPH3* | 7.784271865 | -0.090070152 | -0.096298913 | 0.006228761 | 0.999933887 | -0.27444416 | -0.166747308 | -0.107696852 | 0.442787425 |
| ENSG00000113712 | *CSNK1A1* | 6.708315653 | 0.014450558 | -0.025205135 | 0.039655693 | 0.999933887 | 0.070145097 | 0.132408439 | -0.062263342 | 0.44290307 |
| ENSG00000263753 | *LINC00667* | 2.357902607 | -0.080473448 | 0.075553287 | -0.156026736 | 0.999933887 | 0.022621482 | 0.151622173 | -0.129000691 | 0.443097506 |
| ENSG00000123384 | *LRP1* | 2.955063883 | 0.083590078 | 0.034740313 | 0.048849765 | 0.999933887 | -0.485467528 | -0.721958086 | 0.236490558 | 0.443108249 |
| ENSG00000125846 | *ZNF133* | 0.675771565 | -0.118376963 | -0.145781443 | 0.027404481 | 0.999933887 | 0.120574785 | -0.090338254 | 0.210913039 | 0.443474177 |
| ENSG00000101220 | *C20orf27* | 1.129446902 | -0.074091872 | 0.114151452 | -0.188243324 | 0.999933887 | 0.16116641 | -0.104631957 | 0.265798366 | 0.443474177 |
| ENSG00000215867 | *KRT18P57* | -0.163931597 | 0.085896601 | 0.221461583 | -0.135564982 | 0.999933887 | 0.014482508 | -0.417820619 | 0.432303127 | 0.443568213 |
| ENSG00000224870 | *MRPL20-AS1* | 2.916357001 | 0.001643708 | 0.079425774 | -0.077782066 | 0.999933887 | -0.236586291 | -0.374832627 | 0.138246336 | 0.443568213 |
| ENSG00000053501 | *USE1* | 2.802086088 | 0.044602166 | 0.100404655 | -0.055802489 | 0.999933887 | 0.271776644 | 0.421411758 | -0.149635114 | 0.443685319 |
| ENSG00000196586 | *MYO6* | 0.915469244 | -0.058195836 | -0.240760292 | 0.182564456 | 0.999933887 | -0.085270534 | 0.213644252 | -0.298914786 | 0.44390599 |
| ENSG00000274008 | *Metazoa_SRP* | 2.182488175 | 0.454227552 | 0.430739685 | 0.023487867 | 0.999933887 | 0.974457007 | 1.355338204 | -0.380881197 | 0.443911643 |
| ENSG00000078808 | *SDF4* | 6.719146091 | 0.047296016 | -0.023048541 | 0.070344557 | 0.999933887 | 0.176823089 | 0.264019656 | -0.087196567 | 0.443911643 |
| ENSG00000257913 | *DDN-AS1* | -0.319669527 | 0.106785734 | 0.131797762 | -0.025012028 | 0.999933887 | -0.058576815 | 0.373513671 | -0.432090486 | 0.443911643 |
| ENSG00000137720 | *C11orf1* | -0.344569991 | -0.112877112 | -0.049286567 | -0.063590545 | 0.999933887 | -0.078660422 | 0.297781341 | -0.376441763 | 0.443911643 |
| ENSG00000273749 | *CYFIP1* | 1.640941994 | -0.062952998 | -0.143996865 | 0.081043867 | 0.999933887 | 0.022729559 | 0.190016275 | -0.167286716 | 0.443911643 |
| ENSG00000140287 | *HDC* | 0.911800381 | -0.007061402 | 0.110689268 | -0.11775067 | 0.999933887 | 0.206152207 | 0.433266384 | -0.227114177 | 0.443911643 |
| ENSG00000163918 | *RFC4* | 0.702251066 | -0.001112711 | 0.197431851 | -0.198544562 | 0.999933887 | 0.033425324 | 0.274493731 | -0.241068406 | 0.444055938 |
| ENSG00000276386 | *CNTNAP3P2* | 0.025359603 | 0.156116047 | 0.032577677 | 0.12353837 | 0.999933887 | -1.026371322 | -1.32008656 | 0.293715238 | 0.444067477 |
| ENSG00000187091 | *PLCD1* | 3.98124933 | 0.092409003 | 0.087894076 | 0.004514928 | 0.999933887 | 0.396567662 | 0.545727825 | -0.149160163 | 0.444068658 |
| ENSG00000280138 | *AC027290.2* | 6.39685346 | 0.076060297 | 0.121491213 | -0.045430916 | 0.999933887 | 0.077039655 | 0.222848179 | -0.145808524 | 0.444584447 |
| ENSG00000171246 | *NPTX1* | -1.034834137 | 0.04725763 | -0.108937931 | 0.156195561 | 0.999933887 | -0.347871211 | -0.019974441 | -0.32789677 | 0.444830171 |
| ENSG00000257337 | *AC068888.1* | 1.311528151 | 0.005724096 | 0.05434279 | -0.048618695 | 0.999933887 | -0.042342612 | 0.156066635 | -0.198409247 | 0.445051338 |
| ENSG00000105136 | *ZNF419* | 0.713211923 | 0.003185941 | 0.239341259 | -0.236155318 | 0.999933887 | -0.117820387 | -0.362675677 | 0.244855289 | 0.445506896 |
| ENSG00000179583 | *CIITA* | 3.414458531 | 0.063678545 | 0.076546317 | -0.012867771 | 0.999933887 | -0.013517358 | -0.182448581 | 0.168931223 | 0.445560785 |
| ENSG00000134539 | *KLRD1* | 4.377350822 | -0.019266915 | -0.016895036 | -0.002371879 | 0.999933887 | -0.039987544 | 0.050350075 | -0.090337618 | 0.44561203 |
| ENSG00000010539 | *ZNF200* | 4.305425591 | -0.081456257 | -0.021781423 | -0.059674834 | 0.999933887 | -0.728711638 | -0.856568661 | 0.127857023 | 0.445952194 |
| ENSG00000225886 | *AL445490.1* | 0.323223597 | -0.152438368 | -0.18205862 | 0.029620252 | 0.999933887 | -0.357651573 | -0.693979762 | 0.33632819 | 0.445952194 |
| ENSG00000122696 | *SLC25A51* | 2.035821308 | 0.071935781 | 0.016359025 | 0.055576756 | 0.999933887 | -0.015844249 | -0.180170627 | 0.164326378 | 0.445952194 |
| ENSG00000091831 | *ESR1* | 0.579519491 | -0.081950647 | 0.135006459 | -0.216957106 | 0.999933887 | -0.204747562 | 0.102435045 | -0.307182606 | 0.446355635 |
| ENSG00000143437 | *ARNT* | 4.906049245 | -0.085055785 | 0.021573097 | -0.106628881 | 0.999933887 | -0.447759186 | -0.549529217 | 0.101770031 | 0.446623553 |
| ENSG00000118363 | *SPCS2* | 3.789917313 | -0.046598663 | -0.11511277 | 0.068514107 | 0.999933887 | 0.000398947 | 0.121433033 | -0.121034086 | 0.446623553 |
| ENSG00000088179 | *PTPN4* | 3.815375606 | -0.017330594 | -0.072917151 | 0.055586557 | 0.999933887 | -0.192792336 | 0.00742221 | -0.200214547 | 0.446623553 |
| ENSG00000283646 | *LINC02009* | 3.463274605 | 0.039785994 | 0.266991742 | -0.227205747 | 0.999933887 | 0.723118726 | 0.864839388 | -0.141720662 | 0.446879949 |
| ENSG00000230882 | *AC005077.4* | -1.721416287 | 0.259957031 | 0.122293093 | 0.137663938 | 0.999933887 | 1.017496176 | 1.407771499 | -0.390275323 | 0.447118021 |
| ENSG00000033050 | *ABCF2* | 1.179796829 | -0.019844323 | 0.105389783 | -0.125234106 | 0.999933887 | 0.038152138 | 0.209904938 | -0.1717528 | 0.447244235 |
| ENSG00000136111 | *TBC1D4* | 3.687116683 | -0.06360019 | -0.102340099 | 0.038739909 | 0.999933887 | -0.082408996 | 0.042288178 | -0.124697175 | 0.447343517 |
| ENSG00000117228 | *GBP1* | 7.878296784 | 0.042753646 | -0.048235638 | 0.090989284 | 0.999933887 | 0.727488079 | 0.863731728 | -0.136243648 | 0.447343517 |
| ENSG00000117480 | *FAAH* | 1.339697287 | 0.037990628 | 0.07180691 | -0.033816282 | 0.999933887 | 0.316981301 | 0.52100356 | -0.204022259 | 0.447352545 |
| ENSG00000235576 | *LINC01871* | 1.141566749 | -0.062316901 | -0.166070806 | 0.103753905 | 0.999933887 | 0.083221668 | 0.300737891 | -0.217516222 | 0.447471547 |
| ENSG00000102780 | *DGKH* | 2.008673048 | -0.047814531 | -0.058066502 | 0.01025197 | 0.999933887 | -0.30566228 | -0.017476602 | -0.288185678 | 0.447527534 |
| ENSG00000169994 | *MYO7B* | 1.909069486 | 0.074775077 | 0.045226364 | 0.029548713 | 0.999933887 | -0.337892309 | -0.666232689 | 0.32834038 | 0.44766618 |
| ENSG00000124920 | *MYRF* | -0.284612911 | -0.06057879 | 0.04628862 | -0.10686741 | 0.999933887 | 0.00136827 | -0.359476493 | 0.360844763 | 0.447698848 |
| ENSG00000124155 | *PIGT* | 4.788859712 | -0.055640342 | -0.055516893 | -0.00012345 | 0.999933887 | -0.128482781 | -0.214696668 | 0.086213886 | 0.447712705 |
| ENSG00000259712 | *AC023906.5* | 1.297760944 | 0.000261069 | 0.054798205 | -0.054537137 | 0.999933887 | 1.275646307 | 1.532117199 | -0.256470892 | 0.447712705 |
| ENSG00000242473 | *KIR2DP1* | -1.290520394 | -0.085128452 | 0.23170371 | -0.316832163 | 0.999933887 | 1.40318492 | 1.731460744 | -0.328275824 | 0.447763788 |
| ENSG00000270629 | *NBPF14* | 4.690473129 | -0.056233136 | -0.032547087 | -0.023686049 | 0.999933887 | 0.216249465 | 0.326077242 | -0.109827777 | 0.447806276 |
| ENSG00000105372 | *RPS19* | 7.6322358 | 0.027826255 | -0.027437818 | 0.055264073 | 0.999933887 | 0.197865623 | 0.298069517 | -0.100203894 | 0.447980062 |
| ENSG00000109854 | *HTATIP2* | 5.433476625 | -0.023375435 | -0.031985164 | 0.008609729 | 0.999933887 | -0.237920894 | -0.334002554 | 0.096081661 | 0.447980062 |
| ENSG00000111252 | *SH2B3* | 6.121529759 | -0.021325473 | 0.015734855 | -0.037060328 | 0.999933887 | 0.071125072 | -0.065282645 | 0.136407717 | 0.447980062 |
| ENSG00000140365 | *COMMD4* | 4.085938225 | 0.038367657 | 0.030618777 | 0.007748881 | 0.999933887 | 0.42526471 | 0.528807007 | -0.103542297 | 0.448372355 |

| | | | | | | | | | | |
|---|---|---|---|---|---|---|---|---|---|---|
| ENSG00000268279 | *AC090004.1* | 0.782960789 | -0.028137983 | -0.25440607 | 0.226268087 | 0.999933887 | -0.375392636 | -0.585454551 | 0.210061915 | 0.448372355 |
| ENSG00000122971 | *ACADS* | 1.178924913 | 0.091400146 | -0.152786619 | 0.244186765 | 0.999933887 | 0.050345966 | -0.177660762 | 0.228006728 | 0.448488656 |
| ENSG00000093144 | *ECHDC1* | 3.363186262 | 0.026514823 | -0.041957763 | 0.068472585 | 0.999933887 | -0.167523993 | -0.010459106 | -0.157064887 | 0.448488656 |
| ENSG00000175489 | *LRRC25* | 8.11325279 | -0.089675163 | -0.0670848 | -0.022590363 | 0.999933887 | -0.827450314 | -0.695647354 | -0.13180296 | 0.448867353 |
| ENSG00000229391 | *HLA-DRB6* | 0.557158333 | 0.130122123 | 0.171690562 | -0.041568438 | 0.999933887 | 0.993698212 | 1.322174851 | -0.328476639 | 0.448867353 |
| ENSG00000090061 | *CCNK* | 4.589688137 | 0.016054399 | 0.025218079 | -0.00916368 | 0.999933887 | 0.119456473 | 0.215088895 | -0.095632422 | 0.448931517 |
| ENSG00000003436 | *TFPI* | -1.597029895 | -0.124357056 | -0.475519784 | 0.351162728 | 0.999933887 | -0.422551505 | -0.773613104 | 0.351061599 | 0.449065114 |
| ENSG00000145191 | *EIF2B5* | 4.080412937 | -0.089661707 | 0.048126726 | -0.137788433 | 0.999933887 | 0.055233347 | -0.042638589 | 0.097871936 | 0.449125714 |
| ENSG00000130731 | *METTL26* | 2.600163822 | 0.103917331 | -0.041041813 | 0.144959144 | 0.999933887 | 0.148921235 | 0.00381321 | 0.145108025 | 0.449708799 |
| ENSG00000105926 | *MPP6* | 0.636394756 | 0.13367788 | 0.042239234 | 0.091438646 | 0.999933887 | -0.097176529 | 0.205388766 | -0.302565295 | 0.449708799 |
| ENSG00000158467 | *AHCYL2* | 2.061444824 | 0.032437518 | 0.019072802 | 0.013364716 | 0.999933887 | -0.331861921 | -0.509804062 | 0.177942141 | 0.449708799 |
| ENSG00000160753 | *RUSC1* | 1.910294837 | 0.01631173 | -0.022092215 | 0.038403945 | 0.999933887 | 0.178702367 | -0.022858811 | 0.201561178 | 0.449708799 |
| ENSG00000105321 | *CCDC9* | 6.56064689 | 0.089815633 | 0.098757345 | -0.008941712 | 0.999933887 | 0.285699703 | 0.425546457 | -0.139846754 | 0.449752785 |
| ENSG00000134905 | *CARS2* | 5.186100015 | 0.024515956 | 0.053098379 | -0.028582423 | 0.999933887 | -0.129268838 | -0.21940898 | 0.090140141 | 0.449982046 |
| ENSG00000085231 | *AK6* | 0.99473922 | -0.022546264 | -0.059649463 | 0.037103199 | 0.999933887 | -0.162884043 | 0.025961736 | -0.188845779 | 0.449982046 |
| ENSG00000185304 | *RGPD2* | 0.723880921 | -0.231312672 | -0.217912562 | -0.013400111 | 0.999933887 | 0.511012325 | 0.690567757 | -0.179555431 | 0.450238208 |
| ENSG00000149809 | *TM7SF2* | 0.683909688 | 0.116561948 | 0.171253835 | -0.054691887 | 0.999933887 | 0.031449053 | -0.197844937 | 0.229293989 | 0.450238208 |
| ENSG00000153147 | *SMARCA5* | 5.769495029 | -0.028347837 | -0.137623605 | 0.109275768 | 0.999933887 | -0.201723282 | -0.019459281 | -0.182264 | 0.450238208 |
| ENSG00000146416 | *AIG1* | 3.807493733 | -0.018208615 | 0.070785663 | -0.088994278 | 0.999933887 | 0.825293026 | 0.948636142 | -0.123343115 | 0.450238208 |
| ENSG00000179750 | *APOBEC3B* | 2.492388996 | 0.020597906 | 0.048596824 | -0.027998919 | 0.999933887 | 0.02654241 | 0.208133005 | -0.181590595 | 0.450238208 |
| ENSG00000074201 | *CLNS1A* | 2.533499582 | -0.031521316 | -0.062049989 | 0.030528673 | 0.999933887 | 0.052683943 | 0.200915935 | -0.148231993 | 0.450338827 |
| ENSG00000173327 | *MAP3K11* | 8.276404791 | 0.076549113 | 0.05376662 | 0.022782493 | 0.999933887 | 0.490434617 | 0.59524132 | -0.104806703 | 0.450695741 |
| ENSG00000180902 | *D2HGDH* | 2.48769711 | 0.073678873 | 0.084723981 | -0.011045108 | 0.999933887 | 0.210170984 | 0.371660863 | -0.161489878 | 0.450819875 |
| ENSG00000113140 | *SPARC* | 0.689761376 | 0.111405796 | 0.263892302 | -0.152486506 | 0.999933887 | 0.010736278 | 0.282887753 | -0.272151475 | 0.450819875 |
| ENSG00000189376 | *C8orf76* | -0.076352423 | -0.092826996 | 0.156499491 | -0.249326488 | 0.999933887 | -0.278483259 | -0.005205938 | -0.273277321 | 0.450819875 |
| ENSG00000108352 | *RAPGEFL1* | 3.387900882 | -0.005561729 | 0.088604174 | -0.094165903 | 0.999933887 | -0.215603898 | -0.337121509 | 0.121517612 | 0.450819875 |
| ENSG00000196126 | *HLA-DRB1* | 5.750035095 | 0.002821558 | 0.022727309 | -0.019905751 | 0.999933887 | 0.610118255 | 0.817355259 | -0.207237005 | 0.450819875 |
| ENSG00000186056 | *MATN1-AS1* | 0.950307025 | 0.135514348 | 0.100196421 | 0.035317927 | 0.999933887 | 0.202675842 | 0.490266256 | -0.287590415 | 0.451038059 |
| ENSG00000138134 | *STAMBPL1* | 1.871369996 | -0.020212585 | -0.026244457 | 0.006031871 | 0.999933887 | -0.009521007 | 0.146526336 | -0.156047343 | 0.451038059 |
| ENSG00000176473 | *WDR25* | 1.806532398 | 0.003187325 | -0.046171401 | 0.049358726 | 0.999933887 | -0.308208614 | -0.468774663 | 0.160566049 | 0.451038059 |
| ENSG00000116954 | *RRAGC* | 6.593941773 | -0.075778759 | -0.053584494 | -0.022194265 | 0.999933887 | 0.254981182 | 0.352964429 | -0.097983247 | 0.451109072 |
| ENSG00000116198 | *CEP104* | 3.249894459 | 0.065747317 | -0.006948767 | 0.072696084 | 0.999933887 | -0.135234746 | -0.269089389 | 0.133854644 | 0.451109072 |
| ENSG00000259379 | *MTND5P32* | 4.410233307 | 0.366135632 | 0.695346795 | -0.329211163 | 0.999933887 | 2.043678727 | 2.424605132 | -0.380926405 | 0.451591653 |
| ENSG00000144893 | *MED12L* | 0.080809264 | -0.232976854 | -0.140429734 | -0.092547119 | 0.999933887 | -0.421090303 | -0.080899714 | -0.340190589 | 0.451591653 |
| ENSG00000084112 | *SSH1* | 6.297950829 | -0.063407954 | 0.021289506 | -0.08469746 | 0.999933887 | -0.455973158 | -0.558783187 | 0.102810029 | 0.451591653 |
| ENSG00000160710 | *ADAR* | 9.51326002 | -0.034545001 | -0.057393329 | 0.022848328 | 0.999933887 | 0.211173044 | 0.28190391 | -0.070730866 | 0.451792869 |
| ENSG00000170160 | *CCDC144A* | -2.872688351 | -0.060693446 | 0.35520759 | -0.415901036 | 0.999933887 | -0.460881934 | -0.010600243 | -0.450281691 | 0.45194673 |
| ENSG00000119004 | *CYP20A1* | 3.054528252 | -0.055491527 | 0.041076275 | -0.096567802 | 0.999933887 | -0.103002922 | 0.015906275 | -0.118909197 | 0.452027914 |
| ENSG00000095370 | *SH2D3C* | 6.829291587 | 0.000822902 | 0.022875963 | -0.022053061 | 0.999933887 | -0.408993858 | -0.513911032 | 0.104917174 | 0.452027914 |
| ENSG00000140859 | *KIFC3* | 2.930068825 | 0.009740369 | 0.171212362 | -0.161471993 | 0.999933887 | 0.023069884 | 0.149831196 | -0.126761311 | 0.452426215 |
| ENSG00000173013 | *CCDC96* | 0.167547791 | 0.009134862 | 0.088685693 | -0.079550831 | 0.999933887 | -0.439619045 | -0.769579852 | 0.329960808 | 0.452426215 |
| ENSG00000122484 | *RPAP2* | 1.838091447 | 0.065802184 | -0.076685528 | 0.142487712 | 0.999933887 | -0.226581784 | 0.002987059 | -0.229568843 | 0.452453768 |
| ENSG00000101255 | *TRIB3* | 5.94181048 | -0.086598101 | -0.070192349 | -0.016405753 | 0.999933887 | 0.133113827 | 0.24199159 | -0.108877763 | 0.452530869 |
| ENSG00000144736 | *SHQ1* | 0.904197099 | 0.075944435 | -0.233426569 | 0.309371003 | 0.999933887 | -0.022898947 | 0.203112297 | -0.226011245 | 0.452530869 |
| ENSG00000184432 | *COPB2* | 5.53560194 | -0.079364795 | -0.227167765 | 0.14780297 | 0.999933887 | -0.017130532 | 0.078137114 | -0.095267646 | 0.452716301 |
| ENSG00000103187 | *COTL1* | 9.67635487 | -0.000888757 | -0.017694072 | 0.016805315 | 0.999933887 | -0.051991253 | -0.14978335 | 0.097792097 | 0.452908843 |
| ENSG00000273783 | *AL136040.1* | -0.201222567 | 0.240958355 | 0.173535286 | 0.067423069 | 0.999933887 | 1.195747079 | 0.834129546 | 0.361617533 | 0.452950105 |
| ENSG00000100802 | *C14orf93* | 1.734512719 | 0.084661334 | -0.099440755 | 0.184102089 | 0.999933887 | -0.434439049 | -0.650660931 | 0.216221882 | 0.452950105 |
| ENSG00000100243 | *CYB5R3* | 5.399114331 | 0.072912972 | 0.070165923 | 0.002747049 | 0.999933887 | 0.027663539 | 0.11121577 | -0.083552231 | 0.453198035 |
| ENSG00000282164 | *PEG13* | 1.034498208 | 0.099512022 | 0.195294323 | -0.0957823 | 0.999933887 | 0.121585271 | -0.160607228 | 0.282192499 | 0.453198035 |
| ENSG00000143185 | *XCL2* | 0.911381038 | -0.088447454 | 0.014874897 | -0.103322351 | 0.999933887 | 0.028238227 | 0.234379525 | -0.206141298 | 0.453198035 |
| ENSG00000074966 | *TXK* | 1.796382369 | -0.068433026 | -0.088645531 | 0.020212505 | 0.999933887 | -0.018207016 | 0.161133407 | -0.179340423 | 0.453198035 |
| ENSG00000196116 | *TDRD7* | 4.628306984 | 0.005590101 | -0.010155377 | 0.015745478 | 0.999933887 | -0.176958894 | -0.264938987 | 0.087980092 | 0.453198035 |
| ENSG00000196704 | *AMZ2* | 4.590969203 | -0.000950959 | -0.012720933 | 0.011769974 | 0.999933887 | -0.088092664 | -0.191205687 | 0.103113023 | 0.453198035 |
| ENSG00000252135 | *RNU1-155P* | -2.466719317 | -0.105209764 | 0.041669813 | -0.146879577 | 0.999933887 | -0.23294869 | 0.236173668 | -0.469122358 | 0.453219436 |
| ENSG00000175066 | *GK5* | 3.33522643 | -0.077910689 | 0.108292178 | -0.186202867 | 0.999933887 | -0.137312338 | 0.005250196 | -0.142562534 | 0.45322448 |
| ENSG00000267554 | *AC015911.7* | 0.983529684 | -0.110139625 | -0.414076216 | 0.303936591 | 0.999933887 | -0.388311422 | -0.11602093 | -0.272290492 | 0.45322448 |
| ENSG00000166669 | *ATF7IP2* | 1.460740894 | -0.069914407 | -0.185818227 | 0.11590382 | 0.999933887 | 0.241425941 | 0.462831943 | -0.221406003 | 0.45322448 |
| ENSG00000140319 | *SRP14* | 6.221441416 | 0.007237449 | -0.106355881 | 0.113593329 | 0.999933887 | 0.163808495 | 0.252736635 | -0.08892814 | 0.453604357 |
| ENSG00000288066 | *AC097448.1* | -0.375706354 | 0.1126811 | -0.07454065 | 0.187221749 | 0.999933887 | -0.24419793 | 0.15834724 | -0.40254517 | 0.453619896 |
| ENSG00000232528 | *AL109809.1* | 2.622147491 | 0.018462587 | 0.073974702 | -0.055512115 | 0.999933887 | 0.007428389 | 0.140356401 | -0.132928012 | 0.453619896 |
| ENSG00000137288 | *UQCC2* | 0.908761419 | 0.023452109 | -0.11346651 | 0.136918619 | 0.999933887 | 0.168097969 | -0.044810081 | 0.212908051 | 0.453654798 |
| ENSG00000095066 | *HOOK2* | 2.848804052 | -0.007307326 | -0.086175311 | 0.078867985 | 0.999933887 | 0.375366344 | 0.523710684 | -0.148344341 | 0.453795926 |
| ENSG00000114391 | *RPL24* | 6.932504385 | 0.018832904 | 0.002430861 | 0.016402043 | 0.999933887 | 0.158964821 | 0.257627651 | -0.098662831 | 0.453958048 |
| ENSG00000139055 | *ERP27* | 0.998181778 | 0.00982842 | -0.009070363 | 0.018898783 | 0.999933887 | -0.171248849 | -0.517258189 | 0.34600934 | 0.454270851 |
| ENSG00000132530 | *XAF1* | 4.157199497 | -0.018725993 | -0.12273813 | 0.104012137 | 0.999933887 | -0.306294194 | -0.457323482 | 0.151029288 | 0.454504022 |
| ENSG00000160867 | *FGFR4* | -0.384636376 | 0.161575545 | 0.057234206 | 0.104341339 | 0.999933887 | -0.145790395 | -0.560289059 | 0.414498664 | 0.454816926 |
| ENSG00000273319 | *AC058791.1* | 2.489014556 | 0.118558156 | 0.219872156 | -0.101313999 | 0.999933887 | 0.017587782 | -0.323269912 | 0.340857695 | 0.454894671 |
| ENSG00000111450 | *STX2* | 2.026306843 | -0.050195607 | 0.058809697 | -0.109005304 | 0.999933887 | 0.012726037 | -0.14194585 | 0.154671886 | 0.454894671 |
| ENSG00000107175 | *CREB3* | 3.748857259 | -0.013818943 | -0.02737638 | 0.013557437 | 0.999933887 | 0.666556048 | 0.799879377 | -0.133323329 | 0.454944507 |
| ENSG00000182117 | *NOP10* | 7.712924609 | 0.045884954 | -0.03640987 | 0.082294824 | 0.999933887 | 0.065353771 | 0.15647167 | -0.091117899 | 0.455095228 |
| ENSG00000285492 | *AL356417.3* | 3.720341545 | -0.089920033 | 0.208975587 | -0.29889562 | 0.999933887 | 3.861821037 | 4.163468757 | -0.30164772 | 0.45510873 |
| ENSG00000125375 | *DMAC2L* | 1.349309127 | -0.044497481 | 0.191497112 | -0.235994593 | 0.999933887 | -0.183633523 | 0.028568934 | -0.212202457 | 0.455172773 |
| ENSG00000259374 | *NDUFB4P11* | -0.521235615 | 0.265378601 | 0.452790101 | -0.1874115 | 0.999933887 | 0.973855778 | 1.461321224 | -0.487465446 | 0.45525749 |
| ENSG00000176978 | *DPP7* | 5.979710445 | 0.05919994 | 0.059131198 | 6.8741e-05 | 0.999933887 | -0.030874676 | -0.163291579 | 0.132416903 | 0.45525749 |
| ENSG00000074590 | *NUAK1* | -0.920192729 | -0.102501219 | -0.054020524 | -0.048480696 | 0.999933887 | 0.041830817 | -0.308556231 | 0.350387047 | 0.45525749 |
| ENSG00000168497 | *CAVIN2* | -0.601358929 | 0.054512437 | 0.110755237 | -0.0562428 | 0.999933887 | -0.038608348 | 0.386905435 | -0.425513783 | 0.45525749 |
| ENSG00000145687 | *SSBP2* | 2.427069438 | -0.004834485 | 0.044182782 | -0.049017267 | 0.999933887 | 0.05729132 | -0.093404639 | 0.150695959 | 0.45525749 |
| ENSG00000280228 | *AC079753.1* | -0.24812537 | 0.585328041 | 0.644199944 | -0.058871903 | 0.999933887 | 2.361276373 | 2.76688332 | -0.405606947 | 0.455363178 |
| ENSG00000182149 | *IST1* | 7.821176376 | -0.053866909 | -0.053559931 | -0.000306979 | 0.999933887 | 0.058000851 | 0.128724106 | -0.070723255 | 0.455363178 |
| ENSG00000136695 | *IL36RN* | -2.369888072 | -0.200623561 | -0.513784362 | 0.313160801 | 0.999933887 | 0.994262985 | 1.301260667 | -0.306997682 | 0.455363178 |
| ENSG00000022840 | *RNF10* | 7.5623103 | 0.007110274 | -0.004623455 | 0.011733729 | 0.999933887 | -0.155756767 | -0.071609799 | -0.084146969 | 0.455363178 |
| ENSG00000152520 | *PAN3* | 6.861187645 | -0.064493618 | -0.000944844 | -0.063548774 | 0.999933887 | -0.395414942 | -0.28050994 | -0.114905003 | 0.455678816 |
| ENSG00000224442 | *RPL4P7* | -0.558513495 | 0.356361529 | 0.240385337 | 0.115976192 | 0.999933887 | -0.274969293 | -0.7785015 | 0.503532207 | 0.455720288 |
| ENSG00000179914 | *ITLN1* | -0.900894213 | -0.412434416 | 0.014781232 | -0.427215647 | 0.999933887 | -0.033302806 | 0.216279316 | -0.249582121 | 0.455756536 |
| ENSG00000100938 | *GMPR2* | 6.398750329 | 0.049408927 | -0.023805531 | 0.073214458 | 0.999933887 | 0.227936245 | 0.152771167 | 0.075165079 | 0.455756536 |
| ENSG00000166681 | *BEX3* | -0.13179427 | -0.054478666 | 0.019549226 | -0.074027892 | 0.999933887 | 0.357262195 | 0.006780636 | 0.350481559 | 0.455756536 |
| ENSG00000240666 | *MME-AS1* | -0.620020761 | -0.023035884 | -0.134022106 | 0.110986222 | 0.999933887 | -1.591098759 | -0.92678675 | -0.664312009 | 0.455756536 |
| ENSG00000175691 | *ZNF77* | 0.62079604 | -0.012742929 | -0.126259802 | 0.113516873 | 0.999933887 | 0.285612011 | 0.026712284 | 0.258899727 | 0.455756536 |
| ENSG00000132383 | *RPA1* | 3.604876407 | -0.000145478 | -0.046080272 | 0.045934794 | 0.999933887 | -0.012094837 | -0.119517332 | 0.107422495 | 0.455756536 |
| ENSG00000204792 | *LINC01291* | -2.799989166 | 0.260775074 | 0.294098269 | -0.033323195 | 0.999933887 | -0.357048498 | 0.19072187 | -0.547770368 | 0.455932147 |
| ENSG00000125630 | *POLR1B* | 2.527953407 | -0.117946352 | 0.000695053 | -0.118641405 | 0.999933887 | 0.202198416 | 0.33823272 | -0.136034304 | 0.456131957 |
| ENSG00000228302 | *AL512770.1* | 0.783247697 | -0.086090235 | -0.046353781 | -0.039736454 | 0.999933887 | -0.144417084 | -0.473148226 | 0.328731142 | 0.456347758 |
| ENSG00000105287 | *PRKD2* | 7.318006007 | 0.039327669 | 0.019297094 | 0.020030575 | 0.999933887 | -0.31197417 | -0.402614318 | 0.090640148 | 0.456361743 |
| ENSG00000115507 | *OTX1* | -1.418469508 | -0.080891519 | 0.063160442 | -0.144051961 | 0.999933887 | -1.162854062 | -1.857488724 | 0.694634661 | 0.456372574 |
| ENSG00000117592 | *PRDX6* | 4.469038489 | -0.002699308 | -0.009008572 | 0.006309264 | 0.999933887 | 0.286264432 | 0.177919846 | 0.108344586 | 0.456559861 |
| ENSG00000149925 | *ALDOA* | 4.466751056 | 0.076877874 | 0.042521306 | 0.034356568 | 0.999933887 | -0.194239601 | -0.311907321 | 0.11766772 | 0.456603958 |
| ENSG00000154845 | *PPP4R1* | 7.545833013 | -0.053125992 | -0.058907735 | 0.005781743 | 0.999933887 | -1.109714589 | -1.229028025 | 0.119313435 | 0.456603958 |
| ENSG00000167081 | *PBX3* | 2.709381308 | -0.015357101 | -0.032146342 | 0.016789241 | 0.999933887 | -0.379859565 | -0.516519044 | 0.136659479 | 0.456603958 |
| ENSG00000238045 | *AC009133.1* | 1.220292678 | 0.020814568 | -0.021951007 | 0.042765576 | 0.999933887 | 0.165737595 | -0.045731839 | 0.211469434 | 0.456603958 |
| ENSG00000108342 | *CSF3* | -1.916670986 | -0.161945523 | 0.39691269 | -0.558858213 | 0.999933887 | 6.277553593 | 7.093363477 | -0.815809884 | 0.4566215 |
| ENSG00000147118 | *ZNF182* | 2.77350099 | -0.135626258 | -0.243112786 | 0.107486529 | 0.999933887 | 0.066315034 | 0.228305216 | -0.161990182 | 0.456764138 |
| ENSG00000197670 | *AL157838.1* | 1.876109995 | -0.233276713 | -0.381725796 | 0.148449083 | 0.999933887 | -0.062596612 | -0.230868419 | 0.168271807 | 0.456901055 |
| ENSG00000197345 | *MRPL21* | 2.181252821 | 0.017779413 | 0.108628716 | -0.090849303 | 0.999933887 | 0.411399883 | 0.604272888 | -0.192873005 | 0.456999593 |
| ENSG00000204356 | *NELFE* | 5.316793848 | 0.04667766 | 0.046960927 | -0.000283268 | 0.999933887 | 0.105291706 | 0.189338113 | -0.084046407 | 0.45702608 |

| | | | | | | | | | | |
|---|---|---|---|---|---|---|---|---|---|---|
| ENSG00000148450 | *MSRB2* | 2.139587578 | 0.064408819 | -0.053886697 | 0.118295516 | 0.999933887 | -0.126521779 | -0.358084931 | 0.231563153 | 0.457097987 |
| ENSG00000117010 | *ZNF684* | -0.209233911 | -0.080377744 | -0.111307551 | 0.030929807 | 0.999933887 | 0.200874452 | -0.186128784 | 0.387003236 | 0.457097987 |
| ENSG00000168890 | *TMEM150A* | 1.206889977 | 0.060443285 | 0.048494644 | 0.011948641 | 0.999933887 | 0.05880626 | -0.141987437 | 0.200793697 | 0.457140852 |
| ENSG00000086548 | *CEACAM6* | -0.276148934 | 0.130837782 | 0.190288887 | -0.059451105 | 0.999933887 | -0.357102876 | 0.015961645 | -0.37306452 | 0.457252813 |
| ENSG00000184281 | *TSSC4* | 4.864818007 | 0.01766396 | 0.063007558 | -0.045343598 | 0.999933887 | -0.134741014 | -0.266974979 | 0.132233965 | 0.457364638 |
| ENSG00000170802 | *FOXN2* | 4.6722822 | -0.057227533 | -0.160109054 | 0.102881521 | 0.999933887 | -0.686169471 | -0.473239005 | -0.212930467 | 0.457700483 |
| ENSG00000035141 | *FAM136A* | 2.557850445 | -0.111570682 | 0.074965218 | -0.1865359 | 0.999933887 | 0.089838364 | 0.222581141 | -0.132742777 | 0.457763804 |
| ENSG00000111364 | *DDX55* | 2.460875972 | -0.0533059 | -0.12136755 | 0.06806165 | 0.999933887 | 0.090701596 | -0.054977546 | 0.145679142 | 0.457763804 |
| ENSG00000168300 | *PCMTD1* | 5.089186466 | -0.025628944 | -0.005654811 | -0.019974134 | 0.999933887 | -0.648885512 | -0.517962725 | -0.130922786 | 0.457763804 |
| ENSG00000178381 | *ZFAND2A* | 4.76363391 | 0.046564306 | 0.026871265 | 0.019693042 | 0.999933887 | 0.334239102 | 0.427681278 | -0.093442177 | 0.458063003 |
| ENSG00000271870 | *AC024060.2* | -0.083424042 | -0.134606345 | -0.157615777 | 0.023009432 | 0.999933887 | 0.064982941 | -0.215783842 | 0.280766782 | 0.458063003 |
| ENSG00000280027 | *AC007342.9* | 0.358524997 | -0.140605191 | 0.020260957 | -0.160866148 | 0.999933887 | -1.374225722 | -1.876149659 | 0.501923937 | 0.458099675 |
| ENSG00000113845 | *TIMMDC1* | 2.715641318 | -0.015436853 | 0.131573462 | -0.147010315 | 0.999933887 | 0.072321724 | 0.197286207 | -0.124964483 | 0.458099675 |
| ENSG00000196369 | *SRGAP2B* | 0.945063625 | 0.011156088 | 0.187563704 | -0.176407616 | 0.999933887 | -0.212328569 | 0.059229737 | -0.271558305 | 0.458099675 |
| ENSG00000062485 | *CS* | 5.647696947 | -0.042637804 | -0.023865415 | -0.01877239 | 0.999933887 | 0.186337175 | 0.271204669 | -0.084867494 | 0.458143699 |
| ENSG00000103353 | *UBFD1* | 2.459409847 | -0.02071771 | 0.095172955 | -0.115890665 | 0.999933887 | 0.046849242 | -0.077361171 | 0.124210414 | 0.458197163 |
| ENSG00000026036 | *RTEL1-TNFRS* | 0.183212274 | -0.124050448 | 0.101458638 | -0.225509086 | 0.999933887 | 0.164313398 | -0.150236477 | 0.314549875 | 0.458366285 |
| ENSG00000173960 | *UBXN2A* | 1.768116836 | -0.047214514 | -0.057444286 | 0.010229772 | 0.999933887 | 0.146943188 | 0.351023576 | -0.204080388 | 0.458366285 |
| ENSG00000101004 | *NINL* | -1.253962586 | -0.25774707 | 0.337691799 | -0.595438869 | 0.999933887 | -0.044500519 | 0.326834262 | -0.371334781 | 0.458863068 |
| ENSG00000234494 | *SP2-AS1* | 0.364574442 | 0.193489232 | 0.26763518 | -0.074145949 | 0.999933887 | 0.166274087 | 0.444025804 | -0.277751717 | 0.45887709 |
| ENSG00000241983 | *RN7SL566P* | -0.280576526 | 0.056645112 | -0.173053022 | 0.229698134 | 0.999933887 | 0.005841931 | 0.441254068 | -0.435412137 | 0.459176637 |
| ENSG00000086200 | *IPO11* | 3.763767833 | -0.022556759 | -0.002082918 | -0.02047384 | 0.999933887 | -0.180709515 | -0.061287849 | -0.119421666 | 0.459176637 |
| ENSG00000146701 | *MDH2* | 4.17852923 | -0.028545504 | -0.071808451 | 0.043262947 | 0.999933887 | -0.112922606 | -0.220882763 | 0.107960156 | 0.459246486 |
| ENSG00000173213 | *TUBB8B* | -1.762197812 | 0.054332579 | -0.193409847 | 0.247742426 | 0.999933887 | -1.700996457 | -1.367573253 | -0.333423204 | 0.459304632 |
| ENSG00000129450 | *SIGLEC9* | 4.630073569 | -0.012873886 | -0.076118634 | 0.063244748 | 0.999933887 | 0.407992847 | 0.577630041 | -0.169637193 | 0.45934096 |
| ENSG00000055163 | *CYFIP2* | 8.363060834 | 0.021355631 | 0.019318027 | 0.002037604 | 0.999933887 | -0.300165547 | -0.219373989 | -0.080791558 | 0.459406799 |
| ENSG00000006194 | *ZNF263* | 5.365548633 | 0.000431395 | 0.03067381 | -0.030242414 | 0.999933887 | -0.060534878 | -0.169379355 | 0.108844478 | 0.459446352 |
| ENSG00000119820 | *YIPF4* | 5.102710299 | -0.062140496 | 0.043635346 | -0.105775842 | 0.999933887 | -0.320830577 | -0.238284376 | -0.082546201 | 0.459485106 |
| ENSG00000103522 | *IL21R* | 3.14566029 | -0.065617468 | 0.054535705 | -0.120153173 | 0.999933887 | 0.341580257 | 0.48674529 | -0.145165033 | 0.459485106 |
| ENSG00000169967 | *MAP3K2* | 8.747622876 | -0.055655111 | -0.104929056 | 0.049273944 | 0.999933887 | -0.993838437 | -0.806680827 | -0.18715761 | 0.459485106 |
| ENSG00000118655 | *DCLRE1B* | 0.478435483 | 0.068294425 | 0.101605236 | -0.03331081 | 0.999933887 | 0.113595498 | -0.231178492 | 0.34477399 | 0.459485106 |
| ENSG00000136878 | *USP20* | 6.312013666 | 0.016125402 | 0.030863288 | -0.014737886 | 0.999933887 | -0.210950678 | -0.310074018 | 0.09912334 | 0.459624159 |
| ENSG00000259475 | *AC036108.1* | -0.298813687 | 0.006221405 | 0.289387783 | -0.283166379 | 0.999933887 | -0.807817334 | -1.22798862 | 0.420171286 | 0.459761284 |
| ENSG00000279207 | *AC015813.6* | 1.554607278 | 0.026579926 | -0.06766512 | 0.094245047 | 0.999933887 | -0.144614982 | -0.391973621 | 0.247358639 | 0.459879271 |
| ENSG00000124608 | *AARS2* | 1.824036607 | -0.117264251 | 0.179853623 | -0.297117874 | 0.962755146 | 0.147955247 | -0.002829541 | 0.150784789 | 0.459896559 |
| ENSG00000106404 | *CLDN15* | 3.524631817 | 0.077937567 | -0.012853523 | 0.09079109 | 0.999933887 | -0.01762021 | -0.13017328 | 0.11255307 | 0.459896559 |
| ENSG00000267369 | *TAF5LP1* | 1.87493884 | -0.036488869 | 0.188627074 | -0.225115943 | 0.999933887 | -0.070488045 | -0.243491815 | 0.17300377 | 0.459896559 |
| ENSG00000160563 | *MED27* | 1.975244553 | 0.025465012 | 0.010383327 | 0.015081685 | 0.999933887 | 0.217684264 | 0.393578866 | -0.175894602 | 0.459896559 |
| ENSG00000065621 | *GSTO2* | 2.807777042 | 0.201662482 | 0.210581652 | -0.00891917 | 0.999933887 | 0.592881836 | 0.808952202 | -0.216070365 | 0.459968799 |
| ENSG00000149527 | *PLCH2* | 2.858316697 | 0.102082098 | 0.259411736 | -0.157329638 | 0.999933887 | 0.067545092 | 0.209394165 | -0.141849073 | 0.459968799 |
| ENSG00000101558 | *VAPA* | 7.011922619 | -0.028884256 | 0.014639969 | -0.043524224 | 0.999933887 | -0.109340197 | -0.037902883 | -0.071437315 | 0.459968799 |
| ENSG00000246130 | *AC107959.2* | 1.304157169 | -0.092846159 | 0.049651953 | -0.142498112 | 0.999933887 | -1.208616782 | -1.480333426 | 0.271716645 | 0.459968799 |
| ENSG00000025770 | *NCAPH2* | 5.035411403 | 0.060379654 | 0.095780171 | -0.035400517 | 0.999933887 | 0.280894341 | 0.380356566 | -0.099462225 | 0.459999388 |
| ENSG00000160223 | *ICOSLG* | -0.444391748 | 0.102366218 | 0.212657164 | -0.110290946 | 0.999933887 | 1.207683667 | 1.501067782 | -0.293384115 | 0.460184206 |
| ENSG00000140107 | *SLC25A47* | -1.398445758 | 0.054142527 | -0.078678457 | 0.132820985 | 0.999933887 | 1.149192067 | 0.413286177 | 0.73590589 | 0.460184206 |
| ENSG00000123104 | *ITPR2* | 5.000225622 | -0.004152388 | -0.044237359 | 0.040084971 | 0.999933887 | -0.084743117 | 0.103915889 | -0.188659005 | 0.460184206 |
| ENSG00000167085 | *PHB* | 1.879164641 | 0.046458063 | -0.020119115 | 0.066577178 | 0.999933887 | 0.284059567 | 0.439603155 | -0.155543589 | 0.460484496 |
| ENSG00000132964 | *CDK8* | 1.420214949 | -0.032295421 | -0.054797127 | 0.022501707 | 0.999933887 | -0.078772709 | -0.278595156 | 0.199822447 | 0.460484496 |
| ENSG00000133641 | *C12orf29* | 0.960382291 | -0.120610803 | -0.076451706 | -0.044159097 | 0.999933887 | -0.052725661 | 0.217986642 | -0.270712303 | 0.460731085 |
| ENSG00000149257 | *SERPINH1* | 0.010010658 | 0.196946924 | 0.043200512 | 0.153746412 | 0.999933887 | 0.125203247 | 0.352361371 | -0.227158124 | 0.461117044 |
| ENSG00000268087 | *AC008764.2* | -0.511119937 | -0.151645046 | -0.266204333 | 0.114559287 | 0.999933887 | -0.225938747 | -0.52886084 | 0.302922093 | 0.461117044 |
| ENSG00000124177 | *CHD6* | 4.205259807 | 0.047068829 | -0.032893545 | 0.079962374 | 0.999933887 | -0.000845048 | 0.13501273 | -0.135857778 | 0.461117044 |
| ENSG00000164399 | *IL3* | -2.830725274 | 0.115218025 | -0.003591772 | 0.118809797 | 0.999933887 | 0.594638792 | -0.247585094 | 0.842223885 | 0.461117044 |
| ENSG00000257335 | *MGAM* | 8.356101024 | -0.014752025 | -0.046738137 | 0.031986112 | 0.999933887 | 0.281101304 | 0.367905586 | -0.086804282 | 0.461117044 |
| ENSG00000275764 | *AC092747.4* | 1.524384534 | -0.027402629 | -0.075543224 | 0.048140595 | 0.999933887 | -0.411762423 | -0.701154684 | 0.28939226 | 0.461117044 |
| ENSG00000132182 | *NUP210* | 5.558907176 | 0.026287807 | 0.032489603 | -0.006201796 | 0.999933887 | 0.072466174 | -0.002652512 | 0.075118686 | 0.461409024 |
| ENSG00000148488 | *ST8SIA6* | -0.223669413 | 0.283755447 | 0.143043534 | 0.140711913 | 0.999933887 | -0.16811632 | 0.080753263 | -0.248869582 | 0.461530768 |
| ENSG00000259883 | *EHD4-AS1* | -0.850926224 | 0.149320659 | 0.296458876 | -0.147138216 | 0.999933887 | 0.206492704 | 0.764412163 | -0.55791946 | 0.461530768 |
| ENSG00000256628 | *ZBTB11-AS1* | -0.022823308 | 0.150574049 | 0.121651905 | 0.028922145 | 0.999933887 | 0.184658927 | 0.503257168 | -0.318598241 | 0.461530768 |
| ENSG00000135617 | *PRADC1* | -0.140150125 | 0.125802852 | -0.06648045 | 0.192283302 | 0.999933887 | 0.124076363 | -0.263445337 | 0.3875217 | 0.461530768 |
| ENSG00000109452 | *INPP4B* | 3.188704336 | -0.065432917 | -0.153755847 | 0.088322929 | 0.999933887 | -0.194298841 | 0.01316255 | -0.20746139 | 0.461530768 |
| ENSG00000286864 | *AC012076.1* | 0.829171587 | 0.073437506 | 0.033450001 | 0.039987505 | 0.999933887 | -0.648881985 | -1.138055442 | 0.489173456 | 0.461530768 |
| ENSG00000126838 | *PZP* | 1.034177133 | 0.094974222 | 0.337693754 | -0.242719532 | 0.999933887 | -0.108511987 | 0.218125361 | -0.326637348 | 0.461530768 |
| ENSG00000211667 | *IGLV3-12* | -2.252569817 | 0.124448791 | -0.181346057 | 0.305794848 | 0.999933887 | -0.206105545 | -0.557028196 | 0.35092265 | 0.461530768 |
| ENSG00000197265 | *GTF2E2* | 3.287779772 | 0.035670795 | 0.102661425 | -0.06699063 | 0.999933887 | -0.155771678 | -0.279793118 | 0.12402144 | 0.461530768 |
| ENSG00000102225 | *CDK16* | 5.92940624 | 0.019688255 | 0.084629277 | -0.064941022 | 0.999933887 | 0.217673934 | 0.304386732 | -0.086712798 | 0.461530768 |
| ENSG00000105707 | *HPN* | -1.040715543 | 0.074934606 | 0.107194185 | -0.032259579 | 0.999933887 | 3.597086467 | 4.158541478 | -0.561455012 | 0.461530768 |
| ENSG00000186577 | *SMIM29* | 6.452903048 | 0.00041874 | -0.070393209 | 0.070811949 | 0.999933887 | 0.282169226 | 0.175042883 | 0.107126342 | 0.461559664 |
| ENSG00000173915 | *ATP5MD* | 3.533595555 | 0.019827402 | 0.034091548 | -0.014264146 | 0.999933887 | -0.027071222 | 0.117813226 | -0.144884448 | 0.462447268 |
| ENSG00000115317 | *HTRA2* | 3.93742306 | 0.098671429 | 0.118827559 | -0.02015613 | 0.999933887 | 0.440099643 | 0.308241364 | 0.131858279 | 0.462572513 |
| ENSG00000259366 | *AC108449.2* | 0.989874234 | 0.023006749 | -0.206074607 | 0.229081356 | 0.999933887 | 0.231931492 | 0.452488996 | -0.220557504 | 0.462654333 |
| ENSG00000237232 | *ZNF295-AS1* | 0.035841628 | 0.245975868 | 0.192471381 | 0.053504487 | 0.999933887 | 0.636913759 | 0.904550262 | -0.267636503 | 0.46270627 |
| ENSG00000211710 | *TRBV4-1* | 0.174289754 | -0.124236464 | -0.116282196 | -0.007954268 | 0.999933887 | 0.235481408 | -0.011438313 | 0.246919721 | 0.46270627 |
| ENSG00000163599 | *CTLA4* | 1.919326338 | -0.089714251 | 0.236772546 | -0.326486797 | 0.944094537 | -0.000425823 | -0.163138763 | 0.162712941 | 0.462722449 |
| ENSG00000286196 | *AL162586.2* | 2.39579799 | -0.083084978 | 0.01272679 | -0.095811768 | 0.999933887 | -0.250103315 | -0.411266292 | 0.161162977 | 0.462722449 |
| ENSG00000239382 | *ALKBH6* | 0.448631629 | 0.180813593 | -0.229663447 | 0.41047704 | 0.999933887 | -0.012402106 | -0.288274737 | 0.275872631 | 0.462960334 |
| ENSG00000141499 | *WRAP53* | 2.113443503 | 0.101049674 | 0.027063556 | 0.073986118 | 0.999933887 | 0.271380136 | 0.096337469 | 0.175042668 | 0.462990667 |
| ENSG00000189068 | *VSTM1* | 2.095341738 | 0.02164532 | -0.033628343 | 0.055273663 | 0.999933887 | -0.114904857 | -0.297172198 | 0.182267341 | 0.462997864 |
| ENSG00000149115 | *TNKS1BP1* | 1.730348951 | 0.066417462 | 0.023411297 | 0.043006165 | 0.999933887 | 0.395527491 | 0.573693603 | -0.178166112 | 0.463274892 |
| ENSG00000183060 | *LYSMD4* | 1.461292441 | -0.042374078 | -0.171049617 | 0.128675539 | 0.999933887 | -0.001731351 | -0.169516014 | 0.167784662 | 0.463274892 |
| ENSG00000114166 | *KAT2B* | 5.891632166 | -0.000866086 | 0.017230816 | -0.018096902 | 0.999933887 | -0.541615853 | -0.63947291 | 0.097857057 | 0.46371367 |
| ENSG00000140575 | *IQGAP1* | 9.388004539 | -0.058742886 | -0.108559925 | 0.049817039 | 0.999933887 | -0.440561936 | -0.326639234 | -0.113922702 | 0.463963609 |
| ENSG00000261423 | *TMEM202-AS* | 1.370169746 | 0.114192359 | 0.194265502 | -0.080073143 | 0.999933887 | 0.537074216 | 0.291006124 | 0.246068092 | 0.464170965 |
| ENSG00000230076 | *RPL10P6* | 0.56956171 | 0.074077821 | 0.317222614 | -0.243144793 | 0.999933887 | 0.181334713 | -0.126342685 | 0.307677397 | 0.464264373 |
| ENSG00000119977 | *TCTN3* | 1.487564917 | -0.032991918 | -0.107686451 | 0.074694534 | 0.999933887 | 0.152279949 | -0.039198984 | 0.191478933 | 0.464264373 |
| ENSG00000246451 | *AL049840.2* | 2.928196324 | 0.101002404 | 0.109496458 | -0.008494054 | 0.999933887 | -0.283860769 | -0.167947885 | -0.115912884 | 0.464265661 |
| ENSG00000184719 | *RNLS* | 0.289763574 | 0.087119813 | 0.113568608 | -0.026448794 | 0.999933887 | 0.567989831 | 0.205927732 | 0.362062099 | 0.464265661 |
| ENSG00000168724 | *DNAJC21* | 3.699932361 | -0.036783683 | -0.103121881 | 0.066338199 | 0.999933887 | 0.011775314 | 0.135476729 | -0.123701415 | 0.464265661 |
| ENSG00000182568 | *SATB1* | 6.101339608 | -0.013109992 | 0.002227251 | -0.015337243 | 0.999933887 | -0.211449542 | -0.307970975 | 0.096521432 | 0.464265661 |
| ENSG00000143952 | *VPS54* | 2.771852296 | -0.004738225 | -0.136495415 | 0.13175719 | 0.999933887 | -0.417034255 | -0.231669993 | -0.185364262 | 0.464265661 |
| ENSG00000140688 | *RUSF1* | 2.561503756 | 0.037760006 | 0.144406768 | -0.106646762 | 0.999933887 | 0.106029233 | -0.044744714 | 0.150773947 | 0.46441848 |
| ENSG00000165672 | *PRDX3* | 3.140595607 | 0.035533404 | -0.087565324 | 0.123098728 | 0.999933887 | -0.188985493 | -0.329796719 | 0.140811226 | 0.464546956 |
| ENSG00000152382 | *TADA1* | 1.372297196 | -0.050191096 | 0.035180737 | -0.085371833 | 0.999933887 | 0.18700778 | 0.376514577 | -0.189506797 | 0.464546956 |
| ENSG00000169446 | *MMGT1* | 3.878444287 | -0.03351291 | -0.011597394 | -0.021915516 | 0.999933887 | -0.25813618 | -0.377887351 | 0.11975117 | 0.464549387 |
| ENSG00000187325 | *TAF9B* | 2.146648759 | 0.101613086 | 0.066022491 | 0.035590595 | 0.999933887 | -0.133476273 | 0.112018319 | -0.245494592 | 0.464576264 |
| ENSG00000211809 | *TRAV27* | -1.272395362 | -0.222179061 | 0.011457362 | -0.233636423 | 0.999933887 | 0.113387925 | 0.519250063 | -0.405862138 | 0.464981963 |
| ENSG00000259494 | *MRPL46* | -0.529526429 | -0.054019507 | 0.162357371 | -0.216376878 | 0.999933887 | -0.020199617 | -0.387360146 | 0.367160529 | 0.465140393 |
| ENSG00000143811 | *PYCR2* | 2.805808474 | -0.044548051 | 0.07297831 | -0.117526361 | 0.999933887 | -0.132790978 | -0.25441288 | 0.121621903 | 0.465174859 |
| ENSG00000158985 | *CDC42SE2* | 7.773267026 | -0.080614366 | -0.040352091 | -0.040262275 | 0.999933887 | -0.084047027 | 0.020956146 | -0.105003172 | 0.46524233 |
| ENSG00000125691 | *RPL23* | 7.330418645 | 0.028831076 | -0.016856999 | 0.045688075 | 0.999933887 | 0.107058167 | 0.202765207 | -0.09570704 | 0.465891704 |
| ENSG00000091157 | *WDR7* | 3.163336681 | 0.092960503 | -0.02635695 | 0.119317453 | 0.999933887 | -0.03692941 | -0.168266006 | 0.131336596 | 0.466303132 |
| ENSG00000122692 | *SMU1* | 5.280557586 | -0.085361739 | -0.039966251 | -0.045395488 | 0.999933887 | 0.130942701 | 0.206407668 | -0.075464968 | 0.466352576 |
| ENSG00000189127 | *ANKRD34B* | 0.469462176 | -0.063504775 | -0.144661156 | 0.081156381 | 0.999933887 | -0.778317974 | -0.474611511 | -0.303706463 | 0.466387528 |
| ENSG00000139626 | *ITGB7* | 4.739169149 | 0.024242698 | 0.02174127 | 0.002501428 | 0.999933887 | 0.087961763 | -0.014595341 | 0.102557103 | 0.466508069 |
| ENSG00000263393 | *AC011825.2* | -1.454821508 | 0.020233893 | 0.03131654 | -0.011082647 | 0.999933887 | -0.982489432 | -0.22025059 | -0.762238842 | 0.467094542 |

| | | | | | | | | | | |
|---|---|---|---|---|---|---|---|---|---|---|
| ENSG00000138095 | *LRPPRC* | 3.381043468 | -0.166607231 | -0.095207644 | -0.071399587 | 0.999933887 | -0.025506686 | 0.110945921 | -0.136452607 | 0.467328729 |
| ENSG00000268433 | *MTDHP3* | 3.64880396 | 0.121279004 | 0.0628522 | 0.058426804 | 0.999933887 | -0.102628752 | 0.088654651 | -0.191283403 | 0.467328729 |
| ENSG00000121766 | *ZCCHC17* | 1.851118845 | 0.059575771 | 0.063241253 | -0.003665481 | 0.999933887 | 0.201373737 | -0.007358661 | 0.208732398 | 0.467896568 |
| ENSG00000120159 | *CAAP1* | 1.456777178 | -0.128030014 | 0.179832483 | -0.307862497 | 0.999933887 | 0.175980691 | 0.368098911 | -0.19211822 | 0.468049394 |
| ENSG00000198060 | *MARCHF5* | 4.275703093 | 0.016125628 | -0.019592742 | 0.035718371 | 0.999933887 | -0.27326115 | -0.376112153 | 0.102851004 | 0.468090424 |
| ENSG00000229809 | *ZNF688* | 2.07964784 | 0.13246773 | 0.082736947 | 0.049730783 | 0.999933887 | -0.054965049 | -0.261398635 | 0.206433586 | 0.468443292 |
| ENSG00000117481 | *NSUN4* | 3.760142164 | -0.081076315 | 0.033042651 | -0.114118966 | 0.999933887 | 0.954854101 | 1.094958165 | -0.140104064 | 0.468563942 |
| ENSG00000280239 | *AC011498.7* | 0.319979454 | 0.101116437 | 0.096025647 | 0.00509079 | 0.999933887 | 0.170595743 | -0.071562262 | 0.242158004 | 0.468563942 |
| ENSG00000118298 | *CA14* | 0.11068671 | 0.048166744 | -0.082627396 | 0.130794141 | 0.999933887 | 0.090952793 | -0.288383288 | 0.379336081 | 0.468563942 |
| ENSG00000103174 | *NAGPA* | 2.101291041 | -0.056261975 | 0.010401577 | -0.066663553 | 0.999933887 | -0.077600947 | -0.226068632 | 0.148467685 | 0.468828551 |
| ENSG00000163376 | *KBTBD8* | 2.426154434 | -0.152284047 | -0.217823306 | 0.06553926 | 0.999933887 | -0.157192762 | 0.038035699 | -0.19522846 | 0.468928526 |
| ENSG00000202272 | *Y_RNA* | -0.774025372 | 0.331282114 | 0.482420904 | -0.15113879 | 0.999933887 | 1.271793548 | 0.563580603 | 0.708212945 | 0.469329816 |
| ENSG00000278745 | *AL391903.3* | 1.659887519 | 0.146925734 | 0.181744073 | -0.034818339 | 0.999933887 | 0.68808814 | 0.989215849 | -0.301127709 | 0.469361593 |
| ENSG00000258920 | *FOXN3-AS1* | 0.990765986 | 0.06872887 | 0.206169609 | -0.137440739 | 0.999933887 | -0.549262476 | -0.759427121 | 0.210164645 | 0.469361593 |
| ENSG00000112303 | *VNN2* | 8.436156051 | -0.002695991 | -0.090915869 | 0.088219878 | 0.999933887 | -0.413650564 | -0.281713272 | -0.131937292 | 0.469865531 |
| ENSG00000270959 | *LPP-AS2* | 2.719815477 | 0.047130406 | -0.051824594 | 0.098954999 | 0.999933887 | 0.671397699 | 0.814083473 | -0.142685775 | 0.470135436 |
| ENSG00000103196 | *CRISPLD2* | 5.006638679 | -0.056416031 | 0.016645247 | -0.073061278 | 0.999933887 | -0.494427627 | -0.633907162 | 0.139479535 | 0.470191192 |
| ENSG00000205702 | *CYP2D7* | 2.545348161 | 0.05674483 | 0.171475661 | -0.114730831 | 0.999933887 | -0.000633514 | -0.12820157 | 0.127568056 | 0.470191192 |
| ENSG00000163959 | *SLC51A* | 2.97583014 | 0.00970471 | -0.002926381 | 0.012631091 | 0.999933887 | -0.093199009 | 0.041877742 | -0.135076751 | 0.470191192 |
| ENSG00000183977 | *PP2D1* | 1.581762832 | -0.046601597 | -0.016518056 | -0.030083541 | 0.999933887 | 0.231475978 | 0.437296452 | -0.205820474 | 0.470321454 |
| ENSG00000072952 | *IRAG1* | 3.310928901 | -0.091084216 | -0.010041381 | -0.081042835 | 0.999933887 | -0.026942454 | -0.261313545 | 0.234371091 | 0.470515429 |
| ENSG00000196083 | *IL1RAP* | 7.339034021 | -0.023707998 | -0.016892154 | -0.006815844 | 0.999933887 | -0.242144854 | -0.123057944 | -0.11908691 | 0.470630454 |
| ENSG00000172009 | *THOP1* | 2.724440385 | 0.011949566 | 0.05439937 | -0.042449805 | 0.999933887 | 0.199762787 | 0.069411165 | 0.130351622 | 0.470630454 |
| ENSG00000272256 | *AC044849.1* | 1.103425911 | 0.162647601 | 0.192926788 | -0.030279187 | 0.999933887 | 0.25759176 | -0.042391803 | 0.299983563 | 0.470631792 |
| ENSG00000189144 | *ZNF573* | -0.692664508 | -0.188233164 | -0.32665053 | 0.138417366 | 0.999933887 | -0.246052189 | 0.133901818 | -0.379954007 | 0.470631792 |
| ENSG00000070366 | *SMG6* | 3.31720326 | -0.021663001 | -0.060768134 | 0.039105133 | 0.999933887 | 0.084528706 | -0.035400033 | 0.119928739 | 0.470631792 |
| ENSG00000145335 | *SNCA* | -1.630215482 | -0.454202074 | -0.015197755 | -0.439004319 | 0.999933887 | -0.191461889 | 0.353721395 | -0.545183284 | 0.470722072 |
| ENSG00000235328 | *AC006946.1* | -0.93156693 | -0.257621015 | -0.289300017 | 0.031679002 | 0.999933887 | -0.293156632 | -0.791563275 | 0.498406643 | 0.470722072 |
| ENSG00000166598 | *HSP90B1* | 5.766556572 | -0.074771157 | -0.208894468 | 0.134123311 | 0.999933887 | -0.008297629 | 0.11662528 | -0.124922909 | 0.470722072 |
| ENSG00000162636 | *FAM102B* | 3.113681437 | -0.093794681 | -0.159328881 | 0.0655342 | 0.999933887 | -0.32268717 | -0.128383444 | -0.194303727 | 0.470722072 |
| ENSG00000101997 | *CCDC22* | 2.588884707 | -0.021310027 | 0.117560825 | -0.138870851 | 0.999933887 | 0.170767769 | 0.030029524 | 0.140738245 | 0.470722072 |
| ENSG00000182934 | *SRPRA* | 6.83143201 | 0.007682616 | 0.006351664 | 0.001330952 | 0.999933887 | 0.116061867 | 0.188879877 | -0.07281801 | 0.470722072 |
| ENSG00000231964 | *AL731567.1* | 1.308840382 | 0.000887515 | -0.094883928 | 0.095771442 | 0.999933887 | 0.36819597 | 0.640404239 | -0.272208269 | 0.470722072 |
| ENSG00000104738 | *MCM4* | 1.240305118 | -0.063198129 | -0.021981199 | -0.04121693 | 0.999933887 | 0.058956693 | 0.27988019 | -0.220923497 | 0.470737258 |
| ENSG00000154451 | *GBP5* | 8.720262885 | -0.010787355 | -0.100984373 | 0.090197017 | 0.999933887 | 0.304721213 | 0.428078281 | -0.123357067 | 0.470737258 |
| ENSG00000286809 | *AC008964.1* | 0.524170022 | -0.006623538 | -0.073705205 | 0.067081668 | 0.999933887 | -0.76000402 | -0.518269265 | -0.241734755 | 0.470969519 |
| ENSG00000164080 | *RAD54L2* | 3.633713039 | 0.059081927 | -0.026731791 | 0.085813718 | 0.999933887 | 0.025305872 | -0.072823483 | 0.098129355 | 0.470998285 |
| ENSG00000223501 | *VPS52* | 3.304547717 | 0.002486849 | 0.165820643 | -0.163333794 | 0.999933887 | 0.032489904 | -0.088034331 | 0.120524234 | 0.470998285 |
| ENSG00000115504 | *EHBP1* | 0.919266448 | 0.120403158 | 0.037799402 | 0.082603756 | 0.999933887 | -0.073127115 | 0.172670895 | -0.245798011 | 0.471355465 |
| ENSG00000185811 | *IKZF1* | 6.782199886 | 0.030733564 | 0.034711613 | -0.003978049 | 0.999933887 | 0.032998847 | 0.119048582 | -0.086049735 | 0.471355465 |
| ENSG00000141753 | *IGFBP4* | 1.121804506 | -0.064573842 | 0.047835978 | -0.11240982 | 0.999933887 | 0.008429353 | 0.268956952 | -0.260527599 | 0.471498252 |
| ENSG00000135720 | *DYNC1LI2* | 5.170556928 | -0.057768446 | -0.057263642 | -0.000504804 | 0.999933887 | -0.402460453 | -0.495023656 | 0.092563203 | 0.471845901 |
| ENSG00000089050 | *RBBP9* | 0.742244505 | 0.108047045 | 0.095834682 | 0.012212363 | 0.999933887 | 0.483931496 | 0.737385424 | -0.253453929 | 0.471931274 |
| ENSG00000165240 | *ATP7A* | 4.526816848 | -0.006619617 | -0.02529466 | 0.018675043 | 0.999933887 | -0.721899597 | -0.867706168 | 0.145806571 | 0.472126748 |
| ENSG00000227782 | *AC002553.1* | -0.150592801 | 0.041822286 | 0.130229802 | -0.088407516 | 0.999933887 | -0.141800584 | 0.235565951 | -0.377366535 | 0.472283787 |
| ENSG00000099822 | *HCN2* | -0.638268358 | 0.105400449 | 0.182232541 | -0.076832093 | 0.999933887 | 0.293589979 | 0.704924312 | -0.411334332 | 0.472348063 |
| ENSG00000161677 | *JOSD2* | 2.041683987 | 0.135886775 | 0.203787663 | -0.067900888 | 0.999933887 | 0.111285788 | -0.062256677 | 0.173542465 | 0.47247283 |
| ENSG00000154874 | *CCDC144B* | 1.112433042 | 0.07594099 | -0.00453889 | 0.08047988 | 0.999933887 | -0.137375773 | 0.129951996 | -0.267327769 | 0.47247283 |
| ENSG00000161654 | *LSM12* | 2.089489356 | -0.016988939 | -0.049844379 | 0.03285544 | 0.999933887 | -0.044013819 | -0.200577004 | 0.156563185 | 0.47247283 |
| ENSG00000185946 | *RNPC3* | 4.058616713 | 0.017955916 | -0.100791535 | 0.118747451 | 0.999933887 | -0.387539517 | -0.200806654 | -0.186732863 | 0.47247283 |
| ENSG00000186815 | *TPCN1* | 2.837523623 | 0.009683284 | 0.16349306 | -0.153809776 | 0.999933887 | 0.0575854 | 0.184621303 | -0.127035903 | 0.47247283 |
| ENSG00000260661 | *AC116903.2* | 0.840569109 | 0.061137298 | 0.159372792 | -0.098235494 | 0.999933887 | 0.057473297 | -0.25410827 | 0.311581567 | 0.472609862 |
| ENSG00000102309 | *PIN4* | 1.022101426 | -0.063316964 | -0.007648348 | -0.055668616 | 0.999933887 | -0.202343314 | 0.058629207 | -0.260972521 | 0.472609862 |
| ENSG00000197766 | *CFD* | 6.871300449 | 0.10056605 | 0.040948846 | 0.059617204 | 0.999933887 | -0.088366917 | -0.236321047 | 0.14795413 | 0.472612194 |
| ENSG00000182504 | *CEP97* | 1.636472921 | -0.003539633 | -0.176646664 | 0.173107031 | 0.999933887 | -0.486598132 | -0.225992445 | -0.260605687 | 0.472665395 |
| ENSG00000164808 | *SPIDR* | 3.840121555 | 0.022237612 | 0.007567287 | 0.014670325 | 0.999933887 | -0.156418184 | -0.267817245 | 0.111399061 | 0.472741983 |
| ENSG00000074755 | *ZZEF1* | 7.418008757 | -0.035740283 | -0.027460297 | -0.008279986 | 0.999933887 | -0.113816363 | -0.183010342 | 0.069193979 | 0.4729293 |
| ENSG00000223956 | *LINC01767* | -1.655939125 | 0.201691409 | -0.001519888 | 0.203211297 | 0.999933887 | -0.297578893 | -0.485234279 | 0.187655387 | 0.4729293 |
| ENSG00000099995 | *SF3A1* | 7.565178962 | -0.00808707 | 0.022238488 | -0.030325558 | 0.999933887 | 0.056155311 | 0.11988605 | -0.063730739 | 0.472953951 |
| ENSG00000160803 | *UBQLN4* | 2.729181066 | 0.090722952 | 0.017559993 | 0.073162959 | 0.999933887 | 0.141048946 | 0.283711327 | -0.142662382 | 0.473118703 |
| ENSG00000214900 | *LINC01588* | 2.253878743 | -0.030760564 | 0.039982586 | -0.070743149 | 0.999933887 | 0.086408079 | 0.276454519 | -0.190046439 | 0.473118703 |
| ENSG00000152242 | *C18orf25* | 5.365688142 | 0.003909723 | 0.062835577 | -0.058925854 | 0.999933887 | -0.300027337 | -0.380590999 | 0.080563662 | 0.473118703 |
| ENSG00000224877 | *NDUFAF8* | 1.835262983 | -0.099736805 | 0.077907634 | -0.177644439 | 0.999933887 | 0.172259612 | 0.017388322 | 0.15487129 | 0.473586239 |
| ENSG00000150681 | *RGS18* | 4.648263834 | -0.076363166 | -0.10353557 | 0.027172404 | 0.999933887 | -0.784321384 | -0.97326602 | 0.188944636 | 0.473586239 |
| ENSG00000152439 | *ZNF773* | 0.104847979 | -0.044316038 | -0.419682394 | 0.375366356 | 0.999933887 | -0.109484628 | -0.361974715 | 0.252490087 | 0.473586239 |
| ENSG00000164308 | *ERAP2* | 4.41564466 | -0.070638846 | -0.105186094 | 0.034547248 | 0.999933887 | -0.156665608 | -0.278466849 | 0.121801241 | 0.473894945 |
| ENSG00000171103 | *TRMT61B* | 0.358017567 | -0.123471934 | -0.029013537 | -0.094458397 | 0.999933887 | 0.141597736 | 0.382435011 | -0.240837274 | 0.473952647 |
| ENSG00000225342 | *LRRK2-DT* | 3.752065033 | -0.210671311 | -0.29164122 | 0.080969909 | 0.999933887 | -0.394818193 | -0.556758648 | 0.161940455 | 0.474003006 |
| ENSG00000188234 | *AGAP4* | 1.256779868 | -0.076868186 | -0.047757576 | -0.02911061 | 0.999933887 | 0.406083855 | 0.18688547 | 0.219198385 | 0.474003006 |
| ENSG00000187566 | *NHLRC1* | 1.122232591 | 0.008138359 | 0.032105108 | -0.023966749 | 0.999933887 | 1.104148084 | 1.392749087 | -0.288601003 | 0.474003006 |
| ENSG00000280077 | *AL353763.2* | 0.328491531 | -0.208413538 | -0.379535758 | 0.171122219 | 0.999933887 | -0.570917065 | -0.306164026 | -0.264753039 | 0.474027257 |
| ENSG00000236829 | *Z97634.1* | 0.839371903 | -0.013464718 | -0.377382869 | 0.363918151 | 0.999933887 | 0.304382453 | 0.11797781 | 0.186404643 | 0.474051429 |
| ENSG00000198075 | *SULT1C4* | -0.969771105 | 0.347089658 | -0.135246395 | 0.482336053 | 0.999933887 | 1.637514486 | 1.945498999 | -0.307984513 | 0.474110391 |
| ENSG00000223522 | *AC093690.1* | 0.46601962 | 0.033484583 | -0.256317828 | 0.289802411 | 0.999933887 | 0.04141753 | 0.368995755 | -0.327578225 | 0.474110391 |
| ENSG00000240143 | *AL023653.1* | 1.887986915 | -0.027052475 | -0.008995701 | -0.018056774 | 0.999933887 | 0.191086021 | 0.377108836 | -0.186022816 | 0.474291987 |
| ENSG00000179950 | *PUF60* | 5.438681011 | 0.027182461 | -0.011711392 | 0.038893853 | 0.999933887 | -0.090975671 | -0.184240787 | 0.093265116 | 0.47459318 |
| ENSG00000162194 | *LBHD1* | 0.560380611 | 0.011165906 | 0.092116215 | -0.08095031 | 0.999933887 | 0.54339109 | 0.806175792 | -0.262784701 | 0.474861381 |
| ENSG00000182180 | *MRPS16* | 3.248751142 | -0.142173499 | -0.067565636 | -0.074607863 | 0.999933887 | -0.130838012 | -0.250807197 | 0.119969185 | 0.474925826 |
| ENSG00000214353 | *VAC14-AS1* | 1.89062497 | -0.072611959 | 0.04975995 | -0.122371909 | 0.999933887 | 0.137196233 | 0.309831394 | -0.172635162 | 0.475071905 |
| ENSG00000182400 | *TRAPPC6B* | 1.819280423 | -0.007955902 | -0.023692483 | 0.015736581 | 0.999933887 | -0.186406895 | -0.344878774 | 0.158471879 | 0.475076722 |
| ENSG00000154479 | *CCDC173* | -0.523515933 | -0.068924344 | 0.025218337 | -0.094142681 | 0.999933887 | 0.025227086 | -0.371912984 | 0.39714007 | 0.475254195 |
| ENSG00000134072 | *CAMK1* | -0.017200116 | 0.15012721 | -0.031847247 | 0.181974456 | 0.999933887 | -0.022360195 | -0.308231284 | 0.285871089 | 0.475573634 |
| ENSG00000169962 | *TAS1R3* | 0.650682678 | 0.105167989 | -0.016183106 | 0.121351095 | 0.999933887 | 0.132715243 | -0.162696414 | 0.295411657 | 0.475573634 |
| ENSG00000226688 | *ENTPD1-AS1* | 2.090729564 | -0.005164277 | 0.13184617 | -0.137010447 | 0.999933887 | -0.69084052 | -0.930049377 | 0.239208856 | 0.475573634 |
| ENSG00000167046 | *AL357033.1* | 1.488857872 | 0.059360354 | 0.057504075 | 0.001856279 | 0.999933887 | 0.517449443 | 0.664642043 | -0.147192601 | 0.475653655 |
| ENSG00000101605 | *MYOM1* | 0.586163713 | -0.000454894 | -0.13388115 | 0.133426256 | 0.999933887 | -0.249199568 | -9.7692e-05 | -0.249101876 | 0.475789851 |
| ENSG00000148343 | *MIGA2* | 4.007774003 | 0.07347642 | 0.097866479 | -0.024390059 | 0.999933887 | 0.021922839 | -0.087394853 | 0.109317693 | 0.475918936 |
| ENSG00000105649 | *RAB3A* | 0.127948572 | -0.032619928 | -0.065068809 | 0.032448881 | 0.999933887 | -0.44344752 | -0.778192083 | 0.334744563 | 0.475929487 |
| ENSG00000071859 | *FAM50A* | 6.847880209 | 0.041642472 | 0.107222799 | -0.065580327 | 0.999933887 | 0.587311587 | 0.69086091 | -0.103549323 | 0.476251077 |
| ENSG00000160293 | *VAV2* | 0.50907056 | 0.21328414 | 0.081272344 | 0.132011797 | 0.999933887 | 0.45733761 | 0.698063759 | -0.240726149 | 0.47632275 |
| ENSG00000187626 | *ZKSCAN4* | 0.973148465 | -0.143557621 | -0.035181 | -0.10837662 | 0.999933887 | -0.449984051 | -0.622206366 | 0.172222315 | 0.47632275 |
| ENSG00000225032 | *AL162586.1* | -0.227594982 | 0.136319471 | -0.001322337 | 0.137641808 | 0.999933887 | 0.084053263 | -0.157659837 | 0.2417131 | 0.47632275 |
| ENSG00000173198 | *CYSLTR1* | 3.187826717 | -0.052924378 | -0.204690942 | 0.151766564 | 0.999933887 | -0.240653005 | -0.437967608 | 0.197314603 | 0.47632275 |
| ENSG00000227057 | *WDR46* | 3.189689331 | -0.031021485 | -0.027820659 | -0.003200826 | 0.999933887 | 0.024614889 | -0.074716302 | 0.099331191 | 0.47632275 |
| ENSG00000118579 | *MED28* | 6.961513881 | -0.087767969 | -0.087555811 | -0.000212158 | 0.999933887 | -0.09258054 | -0.017415753 | -0.075164786 | 0.476341888 |
| ENSG00000198355 | *PIM3* | 9.245477955 | 0.20396058 | 0.216364814 | -0.012404234 | 0.999933887 | 1.117685132 | 1.236277269 | -0.118592137 | 0.476445143 |
| ENSG00000051382 | *PIK3CB* | 4.96674403 | -0.045231161 | -0.047468129 | 0.002236968 | 0.999933887 | -0.637202041 | -0.741126245 | 0.103924204 | 0.476445143 |
| ENSG00000287351 | *AC092120.3* | -0.599280848 | 0.124773431 | 0.288171847 | -0.163398416 | 0.999933887 | 0.048562314 | -0.478699221 | 0.527261535 | 0.476445143 |
| ENSG00000108556 | *CHRNE* | 2.184074368 | 0.023830333 | 0.058537841 | -0.034707508 | 0.999933887 | 0.297543229 | 0.477458502 | -0.179915272 | 0.476549218 |
| ENSG00000182057 | *OGFRP1* | -0.112183398 | 0.129085057 | 0.094733009 | 0.034352048 | 0.999933887 | -0.012901257 | -0.2719158 | 0.259014542 | 0.476701653 |
| ENSG00000279382 | *AC018665.1* | 1.285028108 | -0.100398784 | -0.099917292 | -0.000481493 | 0.999933887 | -0.050386943 | -0.238593402 | 0.188206459 | 0.476872579 |
| ENSG00000158006 | *PAFAH2* | 3.636137771 | -0.015470379 | -0.001538699 | -0.01393168 | 0.999933887 | -0.024955146 | -0.11274261 | 0.087787464 | 0.476872579 |
| ENSG00000155229 | *MMS19* | 3.925150531 | 0.010714895 | 0.019379903 | -0.008665008 | 0.999933887 | -0.024512364 | -0.106298511 | 0.081786147 | 0.476872579 |
| ENSG00000156521 | *TYSND1* | 2.624080327 | -0.014986626 | -0.019550828 | 0.004564202 | 0.999933887 | 0.124891852 | -0.00851517 | 0.133407022 | 0.476955649 |
| ENSG00000165959 | *CLMN* | 4.751743447 | -0.037588219 | -0.043678634 | 0.006090415 | 0.999933887 | 0.223065202 | 0.333283274 | -0.110218073 | 0.477189009 |

| | | | | | | | | | | |
|---|---|---|---|---|---|---|---|---|---|---|
| ENSG00000169228 | *RAB24* | 4.325064793 | 0.145844234 | 0.199720401 | -0.053876167 | 0.999933887 | 0.493028736 | 0.292720164 | 0.200308572 | 0.477274191 |
| ENSG00000285979 | *AC009090.6* | 3.43377209 | -0.0021321 | 0.008527244 | -0.010659344 | 0.999933887 | -0.1008222 | 0.053203279 | -0.15402548 | 0.477274191 |
| ENSG00000107077 | *KDM4C* | 4.281932147 | -0.042114866 | -0.080577923 | 0.038463058 | 0.999933887 | 0.003243422 | -0.077480369 | 0.08072379 | 0.477382978 |
| ENSG00000090612 | *ZNF268* | 1.418979826 | -0.108605603 | -0.078559938 | -0.030045665 | 0.999933887 | -0.216593894 | 0.009559708 | -0.226153602 | 0.477382978 |
| ENSG00000237135 | *DDX10P1* | 1.718171581 | -0.104547798 | 0.072558567 | -0.177106365 | 0.999933887 | -0.069042436 | 0.192895735 | -0.261938171 | 0.477382978 |
| ENSG00000229124 | *VIM-AS1* | 4.811074548 | 0.012240853 | -0.041002856 | 0.053243709 | 0.999933887 | 0.276838998 | 0.389590667 | -0.112751669 | 0.477382978 |
| ENSG00000145781 | *COMMD10* | 0.928202231 | 0.00141909 | -0.099663235 | 0.101082324 | 0.999933887 | 0.036337778 | 0.275547002 | -0.239209223 | 0.477382978 |
| ENSG00000137965 | *IFI44* | 4.707453048 | -0.14672915 | -0.162387577 | 0.015658427 | 0.999933887 | -0.420291157 | -0.608741463 | 0.188450306 | 0.477707339 |
| ENSG00000176681 | *LRRC37A* | -0.875405782 | -0.207542051 | -0.051755773 | -0.155786279 | 0.999933887 | -0.159758007 | 0.201648036 | -0.361406043 | 0.478107565 |
| ENSG00000143013 | *LMO4* | 2.806167422 | -0.069102824 | 0.111764198 | -0.180867022 | 0.999933887 | -0.195749995 | -0.349877052 | 0.154127057 | 0.478278704 |
| ENSG00000171202 | *TMEM126A* | 1.3918651 | -0.028595969 | 0.056493921 | -0.08508989 | 0.999933887 | 0.128156835 | 0.310870605 | -0.18271377 | 0.478330976 |
| ENSG00000118181 | *RPS25* | 7.415819355 | -0.007427569 | -0.072725785 | 0.065298216 | 0.999933887 | 0.09645249 | 0.182400712 | -0.085948222 | 0.478330976 |
| ENSG00000105996 | *HOXA2* | -0.805321252 | -0.17759959 | -0.453424625 | 0.275825035 | 0.999933887 | -0.81884345 | -1.224684274 | 0.405840824 | 0.479035825 |
| ENSG00000134954 | *ETS1* | 7.35818214 | -0.026871617 | -0.024261793 | -0.002609823 | 0.999933887 | 0.124792021 | 0.205826679 | -0.081034658 | 0.479035825 |
| ENSG00000187037 | *GPR141* | 1.446123349 | 0.116251068 | -0.008148383 | 0.124399451 | 0.999933887 | 0.980814006 | 1.208597036 | -0.22778303 | 0.4791631 |
| ENSG00000119899 | *SLC17A5* | 3.540676111 | -0.075266391 | 0.001429809 | -0.0766962 | 0.999933887 | -0.717846358 | -0.569044789 | -0.148801569 | 0.4791631 |
| ENSG00000166387 | *PPFIBP2* | 3.149858166 | 0.037167356 | -0.016202408 | 0.053369765 | 0.999933887 | -0.214849256 | -0.123899124 | -0.090950133 | 0.4791631 |
| ENSG00000161905 | *ALOX15* | 3.750265002 | -0.00493622 | -0.006113415 | 0.001177195 | 0.999933887 | -0.060675448 | 0.056430729 | -0.117106177 | 0.4791631 |
| ENSG00000174171 | *AC020659.1* | 2.130098231 | -0.055494431 | 0.041815119 | -0.097309549 | 0.999933887 | -0.026560727 | 0.124395316 | -0.150956042 | 0.479218613 |
| ENSG00000101246 | *ARFRP1* | 3.754968628 | 0.04106195 | 0.058233893 | -0.017171943 | 0.999933887 | 0.059384379 | -0.050911515 | 0.110295894 | 0.479258173 |
| ENSG00000019995 | *ZRANB1* | 4.46643475 | 0.009701905 | -0.082945756 | 0.092647661 | 0.999933887 | -0.169933801 | -0.07395113 | -0.095982671 | 0.479267642 |
| ENSG00000094841 | *UPRT* | 1.627542784 | -0.079216994 | 0.026899084 | -0.106116078 | 0.999933887 | 0.244656418 | 0.439799384 | -0.195142965 | 0.479447185 |
| ENSG00000187097 | *ENTPD5* | 1.10282196 | -0.050629113 | -0.179148654 | 0.128519541 | 0.999933887 | -0.102414713 | 0.098717624 | -0.201132337 | 0.479447207 |
| ENSG00000206077 | *ZDHHC11B* | -0.207057972 | -0.072606762 | 0.20538836 | -0.277995123 | 0.999933887 | 0.689983298 | 1.149099114 | -0.459115816 | 0.479447207 |
| ENSG00000162174 | *ASRGL1* | 0.555838537 | -0.063393584 | -0.157669136 | 0.094275553 | 0.999933887 | -0.656505535 | -0.883596124 | 0.227090589 | 0.479463312 |
| ENSG00000128699 | *ORMDL1* | 4.232348478 | 0.011912691 | -0.065285811 | 0.077198502 | 0.999933887 | -0.04525428 | -0.14659541 | 0.101341129 | 0.479463312 |
| ENSG00000142303 | *ADAMTS10* | 1.867277188 | 0.012886195 | 0.061655183 | -0.048768988 | 0.999933887 | 0.084561325 | -0.068447203 | 0.153008528 | 0.479463312 |
| ENSG00000244187 | *TMEM141* | 0.168008591 | 0.004010896 | 0.364378767 | -0.360367871 | 0.999933887 | 0.13458113 | -0.134207937 | 0.268789068 | 0.479463312 |
| ENSG00000232593 | *KANTR* | 0.47777646 | -0.121804069 | -0.374739548 | 0.252935478 | 0.999933887 | -0.276455019 | 0.026401947 | -0.302856966 | 0.479521668 |
| ENSG00000106459 | *NRF1* | 3.870453694 | -0.032413207 | 0.076316848 | -0.108730055 | 0.999933887 | -0.312368892 | -0.413443749 | 0.101074857 | 0.479521668 |
| ENSG00000284977 | *AL160272.1* | 1.008078106 | -0.008434509 | -0.171624823 | 0.163190314 | 0.999933887 | 0.274794342 | 0.614951523 | -0.340157181 | 0.479521668 |
| ENSG00000253736 | *AC022217.3* | 2.589277696 | 0.157427889 | 0.114583689 | 0.0428442 | 0.999933887 | 0.980872983 | 1.167167038 | -0.186294054 | 0.479739403 |
| ENSG00000107164 | *FUBP3* | 3.334255093 | 0.006165225 | 0.018908023 | -0.012742798 | 0.999933887 | -0.080664056 | -0.179753733 | 0.099089677 | 0.479739403 |
| ENSG00000185305 | *ARL15* | 1.789892201 | -0.005004898 | -0.022961868 | 0.01795697 | 0.999933887 | -0.198070579 | -0.018108909 | -0.17996167 | 0.479821459 |
| ENSG00000155256 | *ZFYVE27* | 5.659879187 | 0.001487711 | 0.019652677 | -0.018164967 | 0.999933887 | -0.009488735 | -0.078637482 | 0.069148748 | 0.479821459 |
| ENSG00000113070 | *HBEGF* | 2.372560197 | 0.245795869 | 0.408750182 | -0.162954313 | 0.999933887 | 0.241084335 | 0.48187588 | -0.240791546 | 0.480069154 |
| ENSG00000085365 | *SCAMP1* | 2.681230478 | 0.026974981 | -0.123662934 | 0.150637914 | 0.999933887 | -0.315769698 | -0.478911741 | 0.163142043 | 0.480213414 |
| ENSG00000158793 | *NIT1* | 3.697878195 | -0.015164301 | -0.018315218 | 0.003150917 | 0.999933887 | -0.019554938 | -0.139498428 | 0.11994349 | 0.480545805 |
| ENSG00000204264 | *PSMB8* | 7.051329716 | -0.021872116 | -0.035338661 | 0.013466545 | 0.999933887 | 0.067060276 | -0.009093392 | 0.076153668 | 0.480558054 |
| ENSG00000078747 | *ITCH* | 6.282909992 | -0.06174908 | -0.053425615 | -0.008323465 | 0.999933887 | -0.323386009 | -0.235421238 | -0.087964771 | 0.480626287 |
| ENSG00000125657 | *TNFSF9* | 1.615977089 | 0.023517385 | -0.075788285 | 0.09930567 | 0.999933887 | 0.196092694 | 0.365800956 | -0.169708262 | 0.481059292 |
| ENSG00000250303 | *LINC02762* | 1.661459635 | 0.15547294 | 0.087687748 | 0.067785192 | 0.999933887 | 0.602809067 | 0.831329786 | -0.228520719 | 0.481154214 |
| ENSG00000261471 | *AC092145.1* | 2.531419934 | -0.110180096 | -0.053035429 | -0.057144666 | 0.999933887 | -0.612867133 | -0.814665147 | 0.201798014 | 0.481482692 |
| ENSG00000167664 | *TMIGD2* | -0.50936403 | 0.212546329 | 0.085517223 | 0.127029106 | 0.999933887 | 0.038229463 | -0.306498766 | 0.344728229 | 0.481482692 |
| ENSG00000164611 | *PTTG1* | 0.160537794 | 0.027715293 | -0.378523461 | 0.406238753 | 0.999933887 | 0.294019747 | 0.600678294 | -0.306658547 | 0.481482692 |
| ENSG00000274290 | *H2BC6* | -0.872413457 | 0.021881778 | -0.085925938 | 0.107807716 | 0.999933887 | -0.871426129 | -1.297763105 | 0.426336977 | 0.481482692 |
| ENSG00000250539 | *KRT8P33* | -0.122204846 | -0.133183855 | -0.248531355 | 0.1153475 | 0.999933887 | 0.071942441 | 0.358916837 | -0.286974396 | 0.481790667 |
| ENSG00000272145 | *NFYC-AS1* | 0.972011344 | 0.068325228 | 0.090225643 | -0.021900415 | 0.999933887 | 0.076951796 | 0.288107263 | -0.211155468 | 0.481881812 |
| ENSG00000126247 | *CAPNS1* | 5.780929449 | 0.083884692 | -0.036047352 | 0.119932044 | 0.999933887 | -0.143337403 | -0.24273886 | 0.099401457 | 0.481884062 |
| ENSG00000172262 | *ZNF131* | 4.170017177 | -0.043501368 | -0.105998134 | 0.062496766 | 0.999933887 | 0.026756644 | 0.141142935 | -0.11438629 | 0.482004444 |
| ENSG00000231731 | *AC010976.1* | 0.968418578 | -0.12652104 | 0.114033388 | -0.240554428 | 0.999933887 | -0.487895684 | -0.269651279 | -0.218244405 | 0.482038537 |
| ENSG00000170248 | *PDCD6IP* | 5.898888627 | 0.015729298 | -0.04894054 | 0.064669838 | 0.999933887 | -0.125067511 | -0.026398872 | -0.098668639 | 0.482038537 |
| ENSG00000087995 | *METTL2A* | 0.452093854 | -0.27778181 | -0.389717265 | 0.111935454 | 0.999933887 | 0.072527802 | 0.314415556 | -0.241887754 | 0.482074203 |
| ENSG00000275180 | *AC048341.2* | 0.975094022 | 0.025601334 | -0.182272596 | 0.20787393 | 0.999933887 | 1.572954004 | 1.797337623 | -0.224383619 | 0.482074203 |
| ENSG00000148308 | *GTF3C5* | 4.34979168 | -0.004587494 | 0.06363482 | -0.068222314 | 0.999933887 | -0.093049671 | -0.18566342 | 0.092613749 | 0.482074203 |
| ENSG00000196689 | *TRPV1* | -0.210764798 | -0.07179477 | 0.135766848 | -0.207561618 | 0.999933887 | -0.343734963 | -0.673244185 | 0.329509222 | 0.482306408 |
| ENSG00000271643 | *AC112220.2* | 0.210415421 | -0.222748736 | 0.206484192 | -0.429232929 | 0.999933887 | 0.106106398 | 0.33274729 | -0.226640892 | 0.482488961 |
| ENSG00000124602 | *UNC5CL* | 0.68420246 | -0.109337013 | 0.043183268 | -0.152520281 | 0.999933887 | -0.057481105 | -0.325360493 | 0.267879388 | 0.482514203 |
| ENSG00000143158 | *MPC2* | 3.00700804 | -0.067588556 | 0.133049269 | -0.200637825 | 0.944094537 | -0.262849969 | -0.411440634 | 0.148590665 | 0.482572699 |
| ENSG00000163328 | *GPR155* | 4.415159079 | 0.041465203 | 0.120038632 | -0.078573429 | 0.999933887 | -0.286243963 | -0.453956744 | 0.167712781 | 0.482572699 |
| ENSG00000233785 | *AC131011.1* | 0.675699703 | 0.250943233 | 0.274356951 | -0.023413718 | 0.999933887 | 0.713251324 | 0.998835399 | -0.285584074 | 0.482584377 |
| ENSG00000090661 | *CERS4* | 3.203895319 | 0.145055059 | 0.070795425 | 0.074259635 | 0.999933887 | 0.709410422 | 0.853686015 | -0.144275593 | 0.482584377 |
| ENSG00000280003 | *AC097637.2* | -0.495395676 | -0.145916005 | -0.010158866 | -0.135757139 | 0.999933887 | -0.225199343 | -0.667572934 | 0.442373591 | 0.482584377 |
| ENSG00000287597 | *AC108105.1* | -1.575348332 | -0.387803118 | 0.268506804 | -0.656309922 | 0.999933887 | 1.11382882 | 1.474386309 | -0.360557488 | 0.482584377 |
| ENSG00000064607 | *SUGP2* | 5.393655109 | -0.021888753 | 0.018186299 | -0.040075052 | 0.999933887 | -0.02718565 | -0.10400605 | 0.0768204 | 0.482584377 |
| ENSG00000067113 | *PLPP1* | -0.519834658 | -0.054821636 | -0.347348472 | 0.292526836 | 0.999933887 | 0.315690554 | 0.596911485 | -0.281220931 | 0.482584377 |
| ENSG00000011260 | *UTP18* | 4.021356721 | -0.007224424 | -0.060264605 | 0.053040181 | 0.999933887 | -0.164080009 | -0.255974331 | 0.091894321 | 0.482584377 |
| ENSG00000167378 | *IRGQ* | 5.142831131 | 0.006509981 | 0.032869624 | -0.026359644 | 0.999933887 | 0.496514275 | 0.631132588 | -0.134618313 | 0.482584377 |
| ENSG00000002834 | *LASP1* | 8.612672732 | -0.00450021 | -0.009942513 | 0.005442304 | 0.999933887 | -0.624184812 | -0.749459998 | 0.125275186 | 0.482584377 |
| ENSG00000134278 | *SPIRE1* | 1.610019313 | 0.003694471 | -0.295050176 | 0.298744647 | 0.999933887 | -0.40947281 | -0.283584756 | -0.125888054 | 0.482584377 |
| ENSG00000272556 | *GTF2IP13* | 1.291165599 | 0.113386387 | 0.133365392 | -0.019979005 | 0.999933887 | 0.305105685 | 0.518462698 | -0.213357013 | 0.482631078 |
| ENSG00000204590 | *GNL1* | 5.870978267 | 0.027263962 | 0.048744187 | -0.021480224 | 0.999933887 | 0.128096293 | 0.216072256 | -0.087975962 | 0.482631078 |
| ENSG00000127870 | *RNF6* | 3.69786672 | -0.013332832 | -0.176611312 | 0.16327848 | 0.999933887 | -0.528684747 | -0.369770102 | -0.158914645 | 0.482631078 |
| ENSG00000107643 | *MAPK8* | 4.385857191 | -0.002086609 | 0.007531384 | -0.009617993 | 0.999933887 | -0.125378021 | -0.001285704 | -0.124092317 | 0.482631078 |
| ENSG00000132694 | *ARHGEF11* | 6.171229184 | -0.020642458 | 0.038012295 | -0.058654753 | 0.999933887 | -0.442521592 | -0.535509877 | 0.092988285 | 0.482671786 |
| ENSG00000104969 | *SGTA* | 6.409717385 | -0.026145079 | -0.041075886 | 0.014930807 | 0.999933887 | 0.153126087 | 0.053063433 | 0.100062654 | 0.482681123 |
| ENSG00000101882 | *NKAP* | 3.121065911 | -0.076909661 | -0.024847939 | -0.052061722 | 0.999933887 | -0.045256777 | 0.065980461 | -0.111237238 | 0.482820773 |
| ENSG00000230482 | *ATP5MC2P3* | -0.491805444 | 0.028332125 | 0.066099242 | -0.037767118 | 0.999933887 | -0.252911518 | -0.71717043 | 0.464258912 | 0.482820773 |
| ENSG00000213928 | *IRF9* | 5.482750824 | -0.01401815 | -0.062030032 | 0.048011882 | 0.999933887 | 0.091574646 | 0.012350682 | 0.079223964 | 0.482975631 |
| ENSG00000174748 | *RPL15* | 7.338046805 | 0.008441578 | -0.057399327 | 0.065840906 | 0.999933887 | 0.179275234 | 0.267642983 | -0.088367749 | 0.483031309 |
| ENSG00000111725 | *PRKAB1* | 3.86052872 | -0.076388492 | 0.029587164 | -0.105975656 | 0.999933887 | -0.253024349 | -0.355500679 | 0.10247633 | 0.483143522 |
| ENSG00000168495 | *POLR3D* | 2.340010294 | 0.049735312 | -0.107764356 | 0.157499668 | 0.999933887 | 0.160655526 | 0.290647778 | -0.129992252 | 0.483143522 |
| ENSG00000011198 | *ABHD5* | 7.256276311 | -0.014656754 | -0.017836225 | 0.003179471 | 0.999933887 | -1.411506436 | -1.256914961 | -0.154591476 | 0.483240025 |
| ENSG00000131943 | *C19orf12* | 2.399278456 | 0.124485518 | -0.079939925 | 0.204425443 | 0.999933887 | -0.169677988 | -0.337006964 | 0.167328975 | 0.483272017 |
| ENSG00000183010 | *PYCR1* | -1.650582791 | 0.118839251 | -0.305322862 | 0.424162113 | 0.999933887 | 0.42165395 | 0.746291572 | -0.324637623 | 0.483541501 |
| ENSG00000271806 | *AL590822.2* | 1.091941801 | 0.142786928 | 0.090019782 | 0.052767146 | 0.999933887 | -0.491714369 | -0.191116858 | -0.300597512 | 0.483593455 |
| ENSG00000215068 | *AC025171.2* | 1.717404912 | 0.075073242 | -0.075453976 | 0.150527218 | 0.999933887 | -0.175168829 | -0.370635557 | 0.195466728 | 0.483593455 |
| ENSG00000119121 | *TRPM6* | 3.964642921 | -0.012452077 | -0.054519003 | 0.042066926 | 0.999933887 | -0.442408576 | -0.305756678 | -0.136651899 | 0.484009755 |
| ENSG00000251139 | *AC084871.1* | 2.127712305 | 0.28628208 | 0.255659422 | 0.030622658 | 0.999933887 | 1.79596637 | 2.136227155 | -0.340260785 | 0.484068561 |
| ENSG00000235945 | *AC002543.1* | 0.444026856 | 0.274304519 | 0.409103922 | -0.134799403 | 0.999933887 | -0.023785422 | 0.482063794 | -0.505849216 | 0.484189846 |
| ENSG00000119421 | *NDUFA8* | 0.714966948 | 0.29431049 | 0.048424708 | 0.245885782 | 0.999933887 | 0.186511859 | -0.063357496 | 0.249869355 | 0.484260369 |
| ENSG00000105397 | *TYK2* | 8.04486526 | 0.047395706 | 0.077076272 | -0.029680566 | 0.999933887 | 0.16358773 | 0.078092952 | 0.085494778 | 0.485126773 |
| ENSG00000169018 | *FEM1B* | 3.911991037 | -0.071922905 | -0.136135305 | 0.064212399 | 0.999933887 | -0.461381467 | -0.575867995 | 0.114486528 | 0.485206628 |
| ENSG00000205784 | *ARRDC5* | -0.080553664 | -0.301848661 | 0.0365727 | -0.338421361 | 0.999933887 | 0.023479942 | -0.264395418 | 0.287875361 | 0.485383383 |
| ENSG00000069345 | *DNAJA2* | 5.470724255 | -0.055382564 | -0.144281718 | 0.088899155 | 0.999933887 | -0.339707589 | -0.432385091 | 0.092677502 | 0.485738849 |
| ENSG00000269335 | *IKBKG* | 4.021829883 | 0.069194886 | 0.024166796 | 0.04502809 | 0.999933887 | 0.137853618 | 0.231209754 | -0.093356136 | 0.485876009 |
| ENSG00000164211 | *STARD4* | 2.86443814 | -0.131754934 | -0.202780286 | 0.071025352 | 0.999933887 | -0.184781042 | 0.029968707 | -0.214749749 | 0.485876009 |
| ENSG00000140367 | *UBE2Q2* | 2.910236043 | -0.137375611 | -0.09551727 | -0.041858341 | 0.999933887 | -0.106257339 | 0.010093714 | -0.116351053 | 0.485900676 |
| ENSG00000099949 | *LZTR1* | 2.395155387 | 0.082124802 | -0.02548544 | 0.107610243 | 0.999933887 | 0.141772163 | -0.017084956 | 0.158857118 | 0.485928762 |
| ENSG00000160799 | *CCDC12* | 3.212216457 | -0.036232997 | 0.048285974 | -0.08451897 | 0.999933887 | -0.030833037 | -0.148456666 | 0.117623629 | 0.485928762 |
| ENSG00000283994 | *AC092652.2* | 2.183274842 | 0.00864839 | 0.120910667 | -0.112262277 | 0.999933887 | 0.222003726 | 0.475580898 | -0.253577172 | 0.485928762 |
| ENSG00000090686 | *USP48* | 4.509445263 | 0.013935094 | -0.106528928 | 0.120464022 | 0.999933887 | -0.126438626 | -0.260058226 | 0.133619599 | 0.486839489 |
| ENSG00000164118 | *CEP44* | 1.739493239 | -0.014778265 | -0.095188727 | 0.080410461 | 0.999933887 | -0.417169814 | -0.172428235 | -0.24474158 | 0.487085013 |
| ENSG00000239665 | *AL157392.3* | 2.823865843 | 0.015574936 | 0.0169541 | -0.001379164 | 0.999933887 | -0.217519212 | -0.339257756 | 0.121738544 | 0.487107145 |
| ENSG00000146830 | *GIGYF1* | 7.309671171 | 0.023890028 | 0.12530061 | -0.101410582 | 0.999933887 | -0.185422402 | -0.275154288 | 0.089731886 | 0.487181244 |

| | | | | | | | | | | |
|---|---|---|---|---|---|---|---|---|---|---|
| ENSG00000197006 | *METTL9* | 6.042213068 | 0.051498151 | 0.007569264 | 0.043928887 | 0.999933887 | -0.018334304 | -0.102863318 | 0.084529015 | 0.48719309 |
| ENSG00000127561 | *SYNGR3* | -0.266257498 | 0.215052476 | -0.012710847 | 0.227763324 | 0.999933887 | 0.582789516 | 0.875184328 | -0.292394812 | 0.487244196 |
| ENSG00000131437 | *KIF3A* | 1.539058633 | -0.125029746 | -0.121203367 | -0.003826379 | 0.999933887 | -0.1614501 | 0.065770008 | -0.227220109 | 0.487511981 |
| ENSG00000085662 | *AKR1B1* | 2.929073051 | -0.030914638 | 0.141376917 | -0.172291555 | 0.999933887 | 0.231949925 | 0.122129799 | 0.109820126 | 0.487511981 |
| ENSG00000126261 | *UBA2* | 3.696192674 | -0.09171401 | -0.18354404 | 0.09183003 | 0.999933887 | -0.043736675 | 0.053520871 | -0.097257546 | 0.487557777 |
| ENSG00000141556 | *TBCD* | 4.526703661 | 0.010005536 | 0.017048806 | -0.007043271 | 0.999933887 | 0.027267202 | -0.061961169 | 0.089228371 | 0.487557777 |
| ENSG00000177548 | *RABEP2* | 2.983871712 | 0.076259719 | 0.033015616 | 0.043244104 | 0.999933887 | 0.015143903 | -0.126833272 | 0.141977175 | 0.48789841 |
| ENSG00000260276 | *AC022167.2* | -0.518683507 | 0.16290974 | -0.130366644 | 0.293276384 | 0.999933887 | 0.085967053 | -0.283698619 | 0.369665672 | 0.48789841 |
| ENSG00000115233 | *PSMD14* | 1.928097664 | -0.089444345 | -0.082294288 | -0.007150057 | 0.999933887 | 0.293454706 | 0.436870001 | -0.143415295 | 0.487938807 |
| ENSG00000145022 | *TCTA* | 3.202527957 | 0.006643459 | 0.036243581 | -0.029600122 | 0.999933887 | -0.036911791 | -0.143135569 | 0.106223778 | 0.487938807 |
| ENSG00000268030 | *AC005253.1* | 1.48020081 | -0.045001216 | 0.023591516 | -0.068592733 | 0.999933887 | -0.098455147 | 0.07332316 | -0.171778306 | 0.488105217 |
| ENSG00000009844 | *VTA1* | 3.101274328 | -0.051837312 | -0.150079348 | 0.098242035 | 0.999933887 | -0.339584881 | -0.468767863 | 0.129182982 | 0.488227334 |
| ENSG00000159346 | *ADIPOR1* | 8.232347514 | 0.070462483 | 0.040622603 | 0.029839881 | 0.999933887 | -0.098875667 | -0.017923068 | -0.080952599 | 0.488539347 |
| ENSG00000225756 | *DBH-AS1* | -1.328593185 | -0.034905802 | -0.046911941 | 0.012006139 | 0.999933887 | 0.064142221 | -0.335730382 | 0.399872603 | 0.488539347 |
| ENSG00000188215 | *DCUN1D3* | 4.409046874 | -0.028728299 | -0.015572139 | -0.01315616 | 0.999933887 | 0.160211445 | 0.28636134 | -0.126149896 | 0.488666322 |
| ENSG00000231758 | *ARHGAP15-A* | 0.522444531 | 0.386812278 | 0.218335802 | 0.168476476 | 0.999933887 | -0.106492099 | 0.466069054 | -0.572561153 | 0.488737542 |
| ENSG00000100722 | *ZC3H14* | 2.377089804 | -0.118277647 | -0.128016478 | 0.009738831 | 0.999933887 | -0.072263593 | 0.066548687 | -0.13881228 | 0.488774829 |
| ENSG00000237702 | *TRBV3-1* | -0.049355263 | -0.018291216 | -0.238105303 | 0.219814087 | 0.999933887 | -0.104525745 | 0.151037171 | -0.255562917 | 0.488774829 |
| ENSG00000089486 | *CDIP1* | 4.675514441 | 0.006369059 | 0.020277457 | -0.013908398 | 0.999933887 | 0.248410673 | 0.343417459 | -0.095006785 | 0.488774829 |
| ENSG00000156467 | *UQCRB* | 4.930959677 | -0.046659172 | -0.084889105 | 0.038229933 | 0.999933887 | -0.008200207 | 0.080265958 | -0.088466165 | 0.488810248 |
| ENSG00000132196 | *HSD17B7* | 2.007790488 | 0.098549951 | -0.059775832 | 0.158325784 | 0.999933887 | -0.077217104 | -0.215420528 | 0.138203423 | 0.48910632 |
| ENSG00000121022 | *COPS5* | 4.085644608 | -0.118750417 | -0.113097259 | -0.005653158 | 0.999933887 | -0.2555307 | -0.175431832 | -0.080098868 | 0.489145362 |
| ENSG00000166822 | *TMEM170A* | 4.876613202 | 0.01239322 | 0.005118549 | 0.00727467 | 0.999933887 | -0.109085638 | -0.022230873 | -0.086854766 | 0.489386579 |
| ENSG00000272008 | *AL139274.2* | 0.773867483 | 0.065398221 | -0.315161219 | 0.38055944 | 0.999933887 | 0.766057059 | 1.014271104 | -0.248214045 | 0.489454857 |
| ENSG00000211445 | *GPX3* | 3.272076304 | 0.085043638 | 0.096207502 | -0.011163864 | 0.999933887 | 1.471413697 | 1.662939044 | -0.191525347 | 0.489627859 |
| ENSG00000111775 | *COX6A1* | 2.887997002 | -0.046069886 | 0.012213037 | -0.058282923 | 0.999933887 | 0.023310264 | -0.10345734 | 0.126767605 | 0.490176223 |
| ENSG00000173875 | *ZNF791* | 3.732478539 | 0.022379093 | -0.140500802 | 0.162879896 | 0.999933887 | -0.216176232 | -0.322976094 | 0.106799862 | 0.490176223 |
| ENSG00000235016 | *SEMA3F-AS1* | 2.797831771 | -0.031977816 | 0.139409399 | -0.171387214 | 0.962755146 | 0.028137832 | 0.148214282 | -0.12007645 | 0.490222102 |
| ENSG00000254263 | *AC022973.4* | 1.237041793 | 0.036764013 | 0.1939866 | -0.157222588 | 0.999933887 | 0.038669458 | 0.342108764 | -0.303439306 | 0.490291187 |
| ENSG00000166435 | *XRRA1* | 2.555580829 | 0.114607235 | 0.087349965 | 0.02725727 | 0.999933887 | -0.095504277 | -0.258871658 | 0.163367381 | 0.490315298 |
| ENSG00000152133 | *GPATCH11* | 1.530585284 | -0.067159316 | -0.139322935 | 0.072163619 | 0.999933887 | -0.136841773 | 0.059935746 | -0.196777519 | 0.490315298 |
| ENSG00000163734 | *CXCL3* | 4.834718089 | 0.716158847 | 0.966262516 | -0.250103669 | 0.999933887 | 1.961179756 | 2.237087677 | -0.275907921 | 0.490315896 |
| ENSG00000125834 | *STK35* | 5.676223077 | 0.009608126 | -0.030247342 | 0.039855468 | 0.999933887 | -0.040482867 | -0.146814768 | 0.106331901 | 0.490315896 |
| ENSG00000156671 | *SAMD8* | 5.85173168 | -0.003418768 | 0.02403165 | -0.027450418 | 0.999933887 | -0.55398796 | -0.429128317 | -0.124859643 | 0.490315896 |
| ENSG00000001461 | *NIPAL3* | 2.714879197 | -0.016042288 | 0.077225483 | -0.093267771 | 0.999933887 | 0.026056268 | -0.107333005 | 0.133389272 | 0.49056007 |
| ENSG00000172354 | *GNB2* | 8.592488114 | 0.111221355 | 0.093622314 | 0.017599041 | 0.999933887 | -0.269351143 | -0.3729087 | 0.103557557 | 0.490675617 |
| ENSG00000135956 | *TMEM127* | 8.184712541 | -0.052529943 | -0.029045902 | -0.023484041 | 0.999933887 | -0.241919064 | -0.159371088 | -0.082547975 | 0.490675617 |
| ENSG00000130559 | *CAMSAP1* | 3.328732189 | -0.033450242 | -0.002640318 | -0.030809924 | 0.999933887 | -0.062061561 | -0.17061127 | 0.108549709 | 0.491123168 |
| ENSG00000198331 | *HYLS1* | -0.652500195 | -0.158746702 | 0.411012186 | -0.569758889 | 0.999933887 | 0.123326644 | 0.44679334 | -0.323466696 | 0.49113718 |
| ENSG00000108590 | *MED31* | 1.480899419 | -0.076382862 | 0.173729695 | -0.250112557 | 0.999933887 | -0.364521638 | -0.523507127 | 0.158985489 | 0.49113718 |
| ENSG00000111181 | *SLC6A12* | 1.102400584 | -0.09566546 | -0.084034826 | -0.011630634 | 0.999933887 | 2.27358502 | 2.50392784 | -0.23034282 | 0.491252624 |
| ENSG00000116032 | *GRIN3B* | 0.511329102 | 0.079725567 | 0.103053423 | -0.023327856 | 0.999933887 | -0.158992792 | -0.444080631 | 0.285087839 | 0.491618442 |
| ENSG00000286408 | *AC113410.4* | 0.139566356 | 0.289875745 | 0.187649013 | 0.102226732 | 0.999933887 | 0.499481987 | 0.234734141 | 0.264747847 | 0.491827032 |
| ENSG00000233038 | *PTPRN2-AS1* | -0.98643194 | 0.01243753 | 0.098525934 | -0.086088403 | 0.999933887 | -0.229405161 | -0.718194716 | 0.488789555 | 0.491890247 |
| ENSG00000103942 | *HOMER2* | -0.343608626 | -0.2116465 | -0.297881692 | 0.086235192 | 0.999933887 | -0.32317873 | -0.645919833 | 0.322741103 | 0.492077219 |
| ENSG00000174695 | *TMEM167A* | 5.12159276 | -0.060993384 | -0.08867816 | 0.027684776 | 0.999933887 | -0.50207887 | -0.393547881 | -0.108530989 | 0.492077219 |
| ENSG00000213613 | *RPL11P3* | 1.58977303 | -0.100397463 | -0.004372364 | -0.096025099 | 0.999933887 | -0.463157209 | -0.809080591 | 0.345923381 | 0.492427991 |
| ENSG00000129696 | *TTI2* | 1.497238602 | -0.065354506 | -0.118515443 | 0.053160937 | 0.999933887 | -0.201921264 | -0.378585097 | 0.176663833 | 0.492427991 |
| ENSG00000138326 | *RPS24* | 6.786519567 | -0.025027271 | -0.100529436 | 0.075502165 | 0.999933887 | 0.05985224 | 0.148390964 | -0.088538724 | 0.492588954 |
| ENSG00000137038 | *DMAC1* | 1.55686242 | -0.009201237 | -0.175961083 | 0.166759845 | 0.999933887 | 0.082889 | -0.07214346 | 0.15503246 | 0.492588954 |
| ENSG00000167283 | *ATP5MG* | 2.925260623 | -0.116316734 | -0.042001753 | -0.074314981 | 0.999933887 | 0.019129376 | -0.127253192 | 0.146382567 | 0.492613105 |
| ENSG00000108423 | *TUBD1* | 0.973579851 | 0.100085567 | 0.091071468 | 0.009014099 | 0.999933887 | -0.07355337 | 0.153226215 | -0.226779585 | 0.492686586 |
| ENSG00000280927 | *CTBP1-AS* | 1.740942223 | 0.152589793 | 0.017888835 | 0.134700957 | 0.999933887 | -0.248984007 | -0.075354768 | -0.173629239 | 0.492978019 |
| ENSG00000165949 | *IFI27* | -1.390444154 | -0.102594871 | -0.200447397 | 0.097852526 | 0.999933887 | 0.223651927 | -0.248760297 | 0.472412224 | 0.493099381 |
| ENSG00000066855 | *MTFR1* | 1.588290172 | -0.021982988 | -0.089438626 | 0.067455637 | 0.999933887 | -0.012490414 | 0.18223125 | -0.194721665 | 0.493134419 |
| ENSG00000262420 | *AC007613.1* | 1.30278892 | 0.174944631 | 0.23263579 | -0.057691159 | 0.999933887 | 0.623103729 | 0.902257224 | -0.279153495 | 0.493520863 |
| ENSG00000121957 | *GPSM2* | 0.616373922 | -0.143550897 | 0.018154147 | -0.161705043 | 0.999933887 | -0.410923426 | -0.67700111 | 0.266077683 | 0.493521035 |
| ENSG00000250462 | *LRRC37BP1* | -0.268550867 | 0.280964939 | -0.269677044 | 0.550641983 | 0.999933887 | -0.145863353 | -0.482125947 | 0.336262594 | 0.493676933 |
| ENSG00000075142 | *SRI* | 3.654573091 | -0.098943724 | -0.098999567 | 5.5843e-05 | 0.999933887 | -0.149193419 | -0.052806474 | -0.096386945 | 0.494021875 |
| ENSG00000029993 | *HMGB3* | -0.505209289 | -0.078205216 | 0.252520737 | -0.330725952 | 0.999933887 | -0.044032443 | -0.441891021 | 0.397858577 | 0.494021875 |
| ENSG00000243477 | *NAA80* | 1.433725012 | 0.073617353 | 0.047258743 | 0.02635861 | 0.999933887 | 0.264214791 | 0.089784889 | 0.174429901 | 0.494048467 |
| ENSG00000132704 | *FCRL2* | -0.75856619 | 0.016962047 | -0.167113771 | 0.184075818 | 0.999933887 | -0.223596841 | 0.063154007 | -0.286750848 | 0.494108737 |
| ENSG00000233334 | *FAM53B-AS1* | 0.269067632 | 0.156592935 | 0.204383011 | -0.047790075 | 0.999933887 | -0.203657024 | -0.724776567 | 0.521119543 | 0.494145005 |
| ENSG00000096006 | *CRISP3* | -0.528841823 | -0.1276447 | -0.117418178 | -0.010226522 | 0.999933887 | -0.28814158 | 0.065224303 | -0.353365884 | 0.494145005 |
| ENSG00000186918 | *ZNF395* | 2.800098595 | -0.076957621 | -0.094959708 | 0.018002087 | 0.999933887 | -0.011761315 | -0.117058954 | 0.105297639 | 0.49418814 |
| ENSG00000136518 | *ACTL6A* | 1.523052618 | -0.154925892 | -0.159973756 | 0.005047864 | 0.999933887 | 0.037932543 | 0.186161561 | -0.148229018 | 0.494230283 |
| ENSG00000104218 | *CSPP1* | 1.891387306 | -0.092078201 | 0.000420354 | -0.092498556 | 0.999933887 | -0.085341497 | 0.091386589 | -0.176728086 | 0.494325254 |
| ENSG00000169857 | *AVEN* | 0.25704978 | 0.069208644 | 0.117660987 | -0.048452343 | 0.999933887 | 0.217940728 | -0.033247704 | 0.251188432 | 0.494325254 |
| ENSG00000276517 | *AL133243.3* | 1.15160527 | -0.039284901 | 0.136350862 | -0.175635763 | 0.999933887 | 0.01252396 | -0.255225434 | 0.267749395 | 0.494325254 |
| ENSG00000276293 | *PIP4K2B* | 3.853274934 | 0.025150745 | 0.011221717 | 0.013929028 | 0.999933887 | -0.077907549 | -0.171172857 | 0.093265308 | 0.494476605 |
| ENSG00000117118 | *SDHB* | 4.842825092 | 0.041539863 | -0.034621775 | 0.076161638 | 0.999933887 | -0.091912733 | -0.190152903 | 0.09824017 | 0.494507368 |
| ENSG00000179115 | *FARSA* | 2.491384637 | 0.083626744 | 0.142470642 | -0.058843898 | 0.999933887 | -0.002079268 | -0.132902825 | 0.130823557 | 0.494638563 |
| ENSG00000140398 | *NEIL1* | 2.295230662 | 0.091333146 | 0.133853552 | -0.042520406 | 0.999933887 | -0.094722383 | 0.037439252 | -0.132161635 | 0.494879998 |
| ENSG00000164074 | *ABHD18* | 2.713659784 | 0.060307766 | 0.027159163 | 0.033148604 | 0.999933887 | -0.394391333 | -0.563691938 | 0.169300606 | 0.494879998 |
| ENSG00000182195 | *LDOC1* | -0.673621911 | 0.116330584 | 0.106314809 | 0.010015775 | 0.999933887 | 0.630165622 | 0.270709648 | 0.359455974 | 0.494879998 |
| ENSG00000243279 | *PRAF2* | 2.460670722 | 0.075267665 | 0.119374376 | -0.044106711 | 0.999933887 | -0.206857723 | -0.357201994 | 0.150344272 | 0.495046916 |
| ENSG00000215769 | *ARHGAP27P1* | 1.117126807 | -0.067806321 | -0.116580705 | 0.048774384 | 0.999933887 | -0.093090623 | -0.333294654 | 0.240204032 | 0.495046916 |
| ENSG00000246477 | *AF131216.1* | -0.25286279 | 0.030293276 | 0.171799204 | -0.141505928 | 0.999933887 | -0.052105223 | 0.226357016 | -0.278462239 | 0.495046916 |
| ENSG00000170296 | *GABARAP* | 3.130871657 | -0.012790772 | -0.014695905 | 0.001905133 | 0.999933887 | 0.365780165 | 0.492056113 | -0.126275948 | 0.495046916 |
| ENSG00000162971 | *TYW5* | 1.887446826 | -0.01751339 | -0.173379653 | 0.155866263 | 0.999933887 | -0.06718116 | 0.137544941 | -0.204726101 | 0.495046916 |
| ENSG00000260618 | *AC025917.1* | -0.039393207 | -0.011517189 | 0.089631429 | -0.101148619 | 0.999933887 | -0.511753194 | -0.924140913 | 0.412387719 | 0.495046916 |
| ENSG00000163833 | *FBXO40* | 0.395831396 | -0.000641104 | 0.378439245 | -0.379080349 | 0.944094537 | -0.263195513 | 0.01601197 | -0.279207483 | 0.495046916 |
| ENSG00000186166 | *CENATAC* | 4.19682081 | -0.074507669 | 0.082367303 | -0.156874972 | 0.944094537 | -0.115491244 | -0.189983739 | 0.074492495 | 0.495047347 |
| ENSG00000132155 | *RAF1* | 9.14395931 | -0.038894822 | -0.008778919 | -0.030115903 | 0.999933887 | -0.413174129 | -0.474653021 | 0.061478891 | 0.495075506 |
| ENSG00000258512 | *LINC00239* | 3.891267104 | 0.155631931 | 0.247224314 | -0.091592383 | 0.999933887 | 2.577695291 | 2.786826832 | -0.209131541 | 0.495255332 |
| ENSG00000109572 | *CLCN3* | 3.861001803 | -0.067715724 | -0.064041862 | -0.003673862 | 0.999933887 | -0.088344218 | 0.010920786 | -0.099265004 | 0.495255332 |
| ENSG00000113356 | *POLR3G* | -0.475273694 | 0.173394867 | -0.199730344 | 0.373125211 | 0.999933887 | 0.209045454 | -0.242741738 | 0.451787192 | 0.495255332 |
| ENSG00000091106 | *NLRC4* | 2.562664026 | -0.072358931 | -0.1527251 | 0.08036617 | 0.999933887 | -0.373688721 | -0.165290249 | -0.208398472 | 0.495255332 |
| ENSG00000205022 | *PABPN1L* | -0.979480997 | -0.06431504 | 0.191389857 | -0.255704897 | 0.999933887 | 0.054827379 | -0.384302197 | 0.439129577 | 0.495255332 |
| ENSG00000267412 | *AC092068.2* | -0.731925296 | -0.00626347 | 0.616983827 | -0.623247297 | 0.999933887 | 0.397082856 | -0.022059089 | 0.419141945 | 0.495255332 |
| ENSG00000196236 | *XPNPEP3* | 2.114669819 | -0.066314146 | -0.071648991 | 0.005334845 | 0.999933887 | -0.224241661 | -0.385128584 | 0.160886923 | 0.495316935 |
| ENSG00000275714 | *H3C1* | -0.584902775 | 0.179704442 | -0.167138675 | 0.346843117 | 0.999933887 | 0.070414151 | 0.512161981 | -0.441747829 | 0.495387186 |
| ENSG00000100036 | *SLC35E4* | 1.283182174 | 0.076234707 | -0.075552331 | 0.151787037 | 0.999933887 | 0.748345644 | 1.000653692 | -0.252308048 | 0.495782843 |
| ENSG00000163875 | *MEAF6* | 3.547492338 | 0.042328804 | 0.006934702 | 0.035394102 | 0.999933887 | -0.100006709 | -0.197109116 | 0.097102408 | 0.495855098 |
| ENSG00000204611 | *ZNF616* | 0.51439801 | -0.065653937 | -0.246108045 | 0.180454108 | 0.999933887 | -0.568987365 | -0.343155376 | -0.225831989 | 0.495855098 |
| ENSG00000114784 | *EIF1B* | 5.39850803 | 0.01410243 | -0.011640832 | 0.025743262 | 0.999933887 | 0.392807224 | 0.488870623 | -0.096063399 | 0.495855098 |
| ENSG00000259768 | *AC004943.2* | 0.379053865 | -0.032409784 | 0.002391006 | -0.03480079 | 0.999933887 | 0.090103811 | 0.289610598 | -0.199506787 | 0.495855098 |
| ENSG00000104375 | *STK3* | 2.165146105 | 0.063859762 | 0.068534328 | -0.004674566 | 0.999933887 | 0.137224161 | 0.326824983 | -0.189600822 | 0.496008676 |
| ENSG00000104490 | *NCALD* | 1.363678786 | 0.080015462 | -0.150150238 | 0.230165701 | 0.999933887 | 0.029216205 | 0.24518852 | -0.215972315 | 0.496008676 |
| ENSG00000145936 | *KCNMB1* | -0.493313337 | 0.096025323 | 0.035861561 | 0.060163762 | 0.999933887 | 0.735993271 | 1.085796633 | -0.349803363 | 0.496008676 |
| ENSG00000163428 | *LRRC58* | 3.654352226 | 0.045208354 | -0.155237222 | 0.200445576 | 0.999933887 | -0.509990627 | -0.296073219 | -0.213917408 | 0.496172135 |
| ENSG00000126215 | *XRCC3* | 2.899974364 | 0.04299559 | -0.044559127 | 0.087554717 | 0.999933887 | -0.198390063 | -0.320809621 | 0.122419557 | 0.496185202 |
| ENSG00000080845 | *DLGAP4* | 5.233023584 | 0.028159992 | -0.009823872 | 0.037983864 | 0.999933887 | 0.16511521 | 0.258768344 | -0.093653134 | 0.496501038 |
| ENSG00000138688 | *KIAA1109* | 6.700355284 | -0.07092154 | -0.079807926 | 0.008886386 | 0.999933887 | -0.499640133 | -0.227682282 | -0.271957851 | 0.496501038 |
| ENSG00000088986 | *DYNLL1* | 3.469689724 | 0.003386643 | 0.020801361 | -0.017414718 | 0.999933887 | -0.014421419 | 0.103638709 | -0.118060128 | 0.496539336 |

| | | | | | | | | | | |
|---|---|---|---|---|---|---|---|---|---|---|
| ENSG00000167397 | *VKORC1* | 0.375127244 | -0.032781773 | -0.146267985 | 0.113486211 | 0.999933887 | -0.210391297 | 0.046988402 | -0.257379699 | 0.496611333 |
| ENSG00000184990 | *SIVA1* | 4.524948434 | 0.075304556 | 0.01707589 | 0.058228666 | 0.999933887 | 0.079383473 | -0.02652415 | 0.105907623 | 0.496760799 |
| ENSG00000154723 | *ATP5PF* | 2.617105616 | 0.037273836 | 0.080839791 | -0.043565955 | 0.999933887 | -0.163671204 | -0.315997786 | 0.152326582 | 0.496760799 |
| ENSG00000171044 | *XKR6* | 1.17168196 | 0.089270328 | 0.103361433 | -0.014091104 | 0.999933887 | 0.322848869 | 0.52067812 | -0.197829251 | 0.496773428 |
| ENSG00000171916 | *LGALS9C* | -0.421263874 | -0.030058369 | 0.078863456 | -0.108921825 | 0.999933887 | -1.24413375 | -1.667594176 | 0.423460427 | 0.496773428 |
| ENSG00000011201 | *ANOS1* | -1.289336818 | -0.094015344 | 0.532583345 | -0.626598689 | 0.999933887 | -0.878265414 | -1.302971291 | 0.424705877 | 0.496926622 |
| ENSG00000125877 | *ITPA* | 2.322785031 | 0.091247835 | 0.054959845 | 0.03628799 | 0.999933887 | 0.101691103 | -0.038364083 | 0.140055186 | 0.497018518 |
| ENSG00000185800 | *DMWD* | 2.987900865 | 0.085523967 | -0.079603935 | 0.165127903 | 0.999933887 | 0.390764148 | 0.585323009 | -0.194558861 | 0.497089245 |
| ENSG00000102409 | *BEX4* | 2.389766013 | 0.001121019 | 0.124712316 | -0.123591296 | 0.999933887 | 0.183116947 | 0.054337722 | 0.128779225 | 0.497107045 |
| ENSG00000158747 | *NBL1* | 0.486250309 | 0.133766614 | -0.123193628 | 0.256960242 | 0.999933887 | 0.338502044 | 0.021617574 | 0.31688447 | 0.497115576 |
| ENSG00000186174 | *BCL9L* | 5.659977137 | 0.06029591 | 0.096197071 | -0.03590116 | 0.999933887 | 0.755966629 | 0.899417259 | -0.143450631 | 0.497116738 |
| ENSG00000132005 | *RFX1* | 6.173403948 | 0.01878893 | 0.054468383 | -0.035679453 | 0.999933887 | -0.039034525 | 0.047381458 | -0.086415983 | 0.497116738 |
| ENSG00000198890 | *PRMT6* | 0.830754139 | -0.039846947 | 0.191889836 | -0.231736783 | 0.999933887 | 0.156855507 | 0.370306018 | -0.213450511 | 0.497116738 |
| ENSG00000067167 | *TRAM1* | 5.415816395 | 0.006927656 | 0.001278743 | 0.005648912 | 0.999933887 | -0.261568148 | -0.356370598 | 0.09480245 | 0.497116738 |
| ENSG00000260060 | *AC009088.1* | -0.228318228 | 0.188023921 | 0.198678724 | -0.010654803 | 0.999933887 | -0.185713994 | 0.219941931 | -0.405655925 | 0.497781991 |
| ENSG00000156030 | *MIDEAS* | 7.358916264 | 0.031687955 | 0.079575908 | -0.047887953 | 0.999933887 | -0.377538258 | -0.469590973 | 0.092052715 | 0.497781991 |
| ENSG00000159445 | *THEM4* | 2.214424704 | -0.013240848 | 0.052751224 | -0.065992072 | 0.999933887 | -0.016095436 | 0.113495869 | -0.129591306 | 0.497781991 |
| ENSG00000182389 | *CACNB4* | 0.883132462 | -0.051187623 | -0.172272743 | 0.12108512 | 0.999933887 | -0.622039038 | -0.247141426 | -0.374897612 | 0.497966359 |
| ENSG00000166266 | *CUL5* | 2.508524128 | -0.217926581 | -0.05016653 | -0.167760051 | 0.999933887 | -0.376097194 | -0.193472022 | -0.182625172 | 0.498212765 |
| ENSG00000116473 | *RAP1A* | 7.22166791 | -0.031081581 | -0.049605223 | 0.018523642 | 0.999933887 | -0.292811754 | -0.372801448 | 0.079989694 | 0.498212765 |
| ENSG00000078596 | *ITM2A* | 4.206092762 | -0.021577416 | -0.123787222 | 0.102209807 | 0.999933887 | 0.114277217 | 0.223063407 | -0.108786191 | 0.498212765 |
| ENSG00000070814 | *TCOF1* | 4.451965473 | 0.040417461 | 0.094559405 | -0.054141944 | 0.999933887 | 0.041048973 | -0.056449981 | 0.097498954 | 0.498892697 |
| ENSG00000099985 | *OSM* | 8.586899258 | 0.7293827 | 0.626963386 | 0.102419314 | 0.999933887 | 2.530232093 | 2.798648284 | -0.268416191 | 0.499148432 |
| ENSG00000167996 | *FTH1* | 12.864689448 | 0.150606785 | 0.102868728 | 0.047738058 | 0.999933887 | 0.596843273 | 0.705317432 | -0.108474159 | 0.499148432 |
| ENSG00000105982 | *RNF32* | -0.320509166 | -0.214509324 | 0.0243237 | -0.238833024 | 0.999933887 | -0.463831471 | -0.154579428 | -0.309252043 | 0.499148432 |
| ENSG00000152404 | *CWF19L2* | 2.784663145 | -0.12929782 | -0.195088465 | 0.065790645 | 0.999933887 | -0.096593283 | 0.080953247 | -0.17754653 | 0.499148432 |
| ENSG00000164114 | *MAP9* | 0.089083895 | -0.130071256 | -0.195243959 | 0.065172703 | 0.999933887 | -0.140067014 | 0.213510363 | -0.353577377 | 0.499148432 |
| ENSG00000153046 | *CDYL* | 3.702420419 | 0.05640554 | 0.034956668 | 0.021448872 | 0.999933887 | -0.284000844 | -0.189366082 | -0.094634762 | 0.499148432 |
| ENSG00000233483 | *AC008105.2* | 0.512721484 | 0.042548112 | -0.063838991 | 0.106387103 | 0.999933887 | -0.289584538 | -0.015943236 | -0.273641302 | 0.499148432 |
| ENSG00000167680 | *SEMA6B* | 5.1475801 | 0.041203893 | 0.071983518 | -0.030779625 | 0.999933887 | 0.421938657 | 0.569708605 | -0.147769948 | 0.499329262 |
| ENSG00000258539 | *AC068896.1* | -0.947833477 | 0.137614655 | 0.117452094 | 0.020162561 | 0.999933887 | 0.153291117 | -0.283996079 | 0.437287196 | 0.499366246 |
| ENSG00000196345 | *ZKSCAN7* | -0.475264771 | 0.110223912 | -0.264002036 | 0.374225948 | 0.999933887 | -0.576877699 | -0.897106617 | 0.320228918 | 0.499366246 |
| ENSG00000211895 | *IGHA1* | 2.63314422 | 0.025335584 | -0.023950346 | 0.04928593 | 0.999933887 | 0.333993823 | 0.08699137 | 0.247002454 | 0.499366246 |
| ENSG00000181904 | *C5orf24* | 2.347219045 | 0.046487848 | 0.051533873 | -0.005046026 | 0.999933887 | -0.279502048 | -0.428971481 | 0.149469433 | 0.499366954 |
| ENSG00000071575 | *TRIB2* | 4.393174739 | -0.035987414 | 0.057358845 | -0.093346259 | 0.999933887 | -0.005585449 | -0.0896287 | 0.084043252 | 0.499416677 |
| ENSG00000187122 | *SLIT1* | 0.75960102 | -0.076713981 | 0.049027152 | -0.125741133 | 0.999933887 | 0.069252518 | 0.387825103 | -0.318572585 | 0.499662876 |
| ENSG00000069943 | *PIGB* | 4.224673673 | -0.060508425 | -0.028459004 | -0.032049421 | 0.999933887 | -0.180378211 | -0.286778572 | 0.106400362 | 0.499885043 |
| ENSG00000268355 | *AC243960.3* | 2.186389891 | 0.031959107 | -0.004502676 | 0.036461783 | 0.999933887 | -0.01269733 | 0.18922745 | -0.20192478 | 0.499885043 |
| ENSG00000141428 | *C18orf21* | 1.681579965 | -0.089213667 | -0.073440353 | -0.015773314 | 0.999933887 | -0.446438627 | -0.613089151 | 0.166650524 | 0.500033797 |
| ENSG00000198435 | *NRARP* | 2.217924339 | 0.072767974 | 0.056970346 | 0.015797628 | 0.999933887 | -0.255634459 | -0.062079082 | -0.193555377 | 0.500033797 |
| ENSG00000155561 | *NUP205* | 3.557896121 | -0.041101546 | -0.055417112 | 0.014315566 | 0.999933887 | 0.069813671 | 0.175389907 | -0.105576236 | 0.500033797 |
| ENSG00000204576 | *PRR3* | 1.915734987 | 0.093342979 | 0.099233454 | -0.005890475 | 0.999933887 | -0.050901733 | -0.207316122 | 0.156414389 | 0.500074033 |
| ENSG00000013364 | *MVP* | 8.005463255 | 0.051745676 | 0.051942453 | -0.000196777 | 0.999933887 | 0.178937606 | 0.087324894 | 0.091612712 | 0.500074033 |
| ENSG00000165637 | *VDAC2* | 4.199860154 | 0.03356231 | -0.00795988 | 0.04152219 | 0.999933887 | -0.088399101 | -0.175621217 | 0.087222116 | 0.500074033 |
| ENSG00000270589 | *AL158163.1* | 1.253680729 | -0.175244681 | 0.019426371 | -0.194671052 | 0.999933887 | 0.334777141 | 0.608182015 | -0.273404873 | 0.500279595 |
| ENSG00000104936 | *DMPK* | 2.548901876 | 0.031237436 | 0.002857099 | 0.028380337 | 0.999933887 | 0.356248873 | 0.533253001 | -0.177004128 | 0.500286697 |
| ENSG00000041880 | *PARP3* | 0.890282165 | 0.197923553 | -0.023263229 | 0.221186781 | 0.999933887 | 0.128911841 | -0.077890356 | 0.206802198 | 0.500460324 |
| ENSG00000018280 | *SLC11A1* | 10.21375508 | 0.025723341 | 0.068278905 | -0.042555564 | 0.999933887 | -0.118529478 | -0.190766473 | 0.072236995 | 0.500460324 |
| ENSG00000268670 | *AC016586.1* | 0.40291897 | -0.061288381 | 0.097454953 | -0.158743334 | 0.999933887 | -0.136510469 | 0.111363025 | -0.247873493 | 0.500539945 |
| ENSG00000178935 | *ZNF552* | 2.784098071 | -0.147570449 | -0.117413715 | -0.030156734 | 0.999933887 | -0.469062551 | -0.611840815 | 0.142778264 | 0.500639794 |
| ENSG00000130589 | *HELZ2* | 8.297198147 | 0.058524535 | 0.084023088 | -0.025498554 | 0.999933887 | 0.128717816 | 0.031232979 | 0.097484837 | 0.500639794 |
| ENSG00000070770 | *CSNK2A2* | 2.760923538 | 0.050367372 | 0.078046643 | -0.027679271 | 0.999933887 | 0.002175338 | 0.122118973 | -0.119943634 | 0.500659053 |
| ENSG00000205413 | *SAMD9* | 6.228931383 | -0.135162393 | -0.191000468 | 0.055838075 | 0.999933887 | -0.812341467 | -0.574957589 | -0.237383877 | 0.500952583 |
| ENSG00000228696 | *ARL17B* | -0.622381786 | 0.168137057 | 0.01790659 | 0.150230466 | 0.999933887 | -0.341533183 | 0.035273736 | -0.376806919 | 0.501004495 |
| ENSG00000182247 | *UBE2E2* | -0.840533112 | 0.142782793 | -0.006380406 | 0.149163198 | 0.999933887 | 0.526588951 | 0.819387648 | -0.292798697 | 0.501004495 |
| ENSG00000111412 | *SPRING1* | 4.5520569 | -0.015187886 | -0.065668118 | 0.050480232 | 0.999933887 | -0.284297678 | -0.182475748 | -0.10182193 | 0.501004495 |
| ENSG00000138794 | *CASP6* | 0.130223364 | -0.105708593 | -0.092963441 | -0.012745152 | 0.999933887 | -0.127870715 | 0.154224712 | -0.282095427 | 0.50101113 |
| ENSG00000089022 | *MAPKAPK5* | 2.934497142 | -0.09433718 | -0.039192483 | -0.055144697 | 0.999933887 | 0.054831822 | -0.040548913 | 0.095380735 | 0.501136654 |
| ENSG00000245910 | *SNHG6* | 3.769339957 | 0.035483044 | -0.06541128 | 0.100894323 | 0.999933887 | 0.075222815 | -0.024640948 | 0.099863762 | 0.501136654 |
| ENSG00000122642 | *FKBP9* | -0.569512472 | -0.171478982 | 0.036935888 | -0.20841487 | 0.999933887 | -0.263803245 | -0.733541827 | 0.469738583 | 0.501164309 |
| ENSG00000055957 | *ITIH1* | -1.668595504 | -0.312694584 | -0.081337061 | -0.231357523 | 0.999933887 | 2.290973827 | 2.678428974 | -0.387455147 | 0.501164309 |
| ENSG00000232818 | *RPS2P32* | -0.097719879 | 0.129630665 | -0.042549585 | 0.17218025 | 0.999933887 | -0.032050989 | 0.338897821 | -0.37094881 | 0.501164309 |
| ENSG00000258476 | *LINC02207* | 2.288696801 | 0.012139222 | -0.08851907 | 0.100658292 | 0.999933887 | 0.466046116 | 0.307793784 | 0.158252332 | 0.501164309 |
| ENSG00000132256 | *TRIM5* | 3.441777789 | 0.027141906 | -0.012432701 | 0.039574607 | 0.999933887 | 0.103509874 | -0.000503179 | 0.104013053 | 0.501302794 |
| ENSG00000233231 | *HNRNPA1P49* | 1.289217877 | 0.105659883 | -0.026360337 | 0.13202022 | 0.999933887 | 0.248704047 | 0.467595556 | -0.218891508 | 0.501431076 |
| ENSG00000105948 | *TTC26* | -0.799396049 | 0.181199921 | -0.096234999 | 0.27743492 | 0.999933887 | 0.202099661 | -0.186007772 | 0.388107433 | 0.501520562 |
| ENSG00000233261 | *FAM238A* | -0.268747849 | 0.057260909 | -0.164759084 | 0.222019993 | 0.999933887 | 0.107578829 | -0.31019992 | 0.41777875 | 0.501597927 |
| ENSG00000273742 | *MIR6075* | 1.358474719 | 0.157949533 | -0.042500961 | 0.200450494 | 0.999933887 | 0.314932241 | 0.112924494 | 0.202007747 | 0.501701176 |
| ENSG00000105254 | *TBCB* | 4.129944754 | 0.075798953 | 0.022732476 | 0.053066477 | 0.999933887 | 0.022045339 | -0.087113781 | 0.10915912 | 0.501874175 |
| ENSG00000286966 | *AL137800.1* | -0.278513819 | -0.130084628 | -0.129739778 | -0.000344851 | 0.999933887 | -0.709941709 | -0.931730348 | 0.221788639 | 0.501874175 |
| ENSG00000111788 | *AC009533.1* | -0.456125656 | 0.11098933 | -0.106482577 | 0.217471907 | 0.999933887 | -0.120943637 | 0.185739363 | -0.306683 | 0.501874175 |
| ENSG00000189241 | *TSPYL1* | 5.465299011 | -0.021522668 | 0.021111253 | -0.042633922 | 0.999933887 | 0.105556648 | 0.026003013 | 0.079553635 | 0.501874175 |
| ENSG00000145743 | *FBXL17* | 2.486049781 | 0.036875853 | 0.046799157 | -0.009923304 | 0.999933887 | -0.138242474 | 0.051365901 | -0.189608375 | 0.501874175 |
| ENSG00000083097 | *DOP1A* | 2.670721932 | 0.014221744 | -0.208613496 | 0.22283524 | 0.999933887 | -0.222185138 | -0.000517869 | -0.22166727 | 0.501874175 |
| ENSG00000179933 | *C14orf119* | 3.494208778 | -0.034845432 | 0.015003986 | -0.049849418 | 0.999933887 | 0.006827304 | 0.105629195 | -0.098801891 | 0.501937492 |
| ENSG00000148672 | *GLUD1* | 4.230430496 | -0.043447566 | -0.081391555 | 0.037943989 | 0.999933887 | -0.065243506 | -0.156774131 | 0.091530625 | 0.501983422 |
| ENSG00000232539 | *LINC00945* | -0.228483636 | -0.441307239 | -0.679726165 | 0.238418926 | 0.999933887 | -0.172004144 | 0.071937286 | -0.243941431 | 0.502030367 |
| ENSG00000198648 | *STK39* | 2.454658503 | -0.05418664 | 0.113936371 | -0.168123011 | 0.999933887 | 0.065360496 | 0.211909921 | -0.146549424 | 0.502030367 |
| ENSG00000259881 | *AC092384.2* | -0.119615063 | -0.109357592 | -0.004929675 | -0.104427918 | 0.999933887 | -0.804153892 | -1.11713205 | 0.312978158 | 0.502030367 |
| ENSG00000267940 | *AC022762.2* | -0.253744399 | 0.257767504 | -0.165102534 | 0.422870038 | 0.999933887 | 1.659224872 | 1.231106823 | 0.428118048 | 0.502194492 |
| ENSG00000158042 | *MRPL17* | 0.667292656 | -0.137079535 | -0.132818673 | -0.004260863 | 0.999933887 | -0.088152535 | 0.121830677 | -0.209983212 | 0.502194492 |
| ENSG00000143322 | *ABL2* | 4.085360612 | 0.046991794 | 0.069907595 | -0.022915801 | 0.999933887 | 0.468889417 | 0.596419145 | -0.127529728 | 0.502194492 |
| ENSG00000197815 | *AC122129.1* | 0.612033319 | 0.039178922 | 0.094952982 | -0.055774059 | 0.999933887 | -0.22657646 | 0.046066952 | -0.272643412 | 0.502194492 |
| ENSG00000139899 | *CBLN3* | -0.453677513 | -0.024361382 | 0.219604198 | -0.243965581 | 0.999933887 | 0.167463691 | 0.457645555 | -0.290181863 | 0.502194492 |
| ENSG00000172888 | *ZNF621* | 2.783797037 | -0.074964636 | -0.033503915 | -0.041460721 | 0.999933887 | 0.022942732 | -0.104913415 | 0.127856147 | 0.50221915 |
| ENSG00000170190 | *SLC16A5* | 6.315908043 | 0.047103051 | -0.05768531 | 0.104788361 | 0.999933887 | 0.006111598 | 0.081168246 | -0.075056648 | 0.50221915 |
| ENSG00000130653 | *PNPLA7* | 0.660678052 | 0.046978757 | 0.022977352 | 0.024001404 | 0.999933887 | 0.02178335 | 0.247603715 | -0.225820364 | 0.50221915 |
| ENSG00000261351 | *AC116913.1* | 0.313834857 | -0.112833029 | 0.082347406 | -0.195180435 | 0.999933887 | 0.595378466 | 0.859235896 | -0.26385743 | 0.50238545 |
| ENSG00000196505 | *GDAP2* | 3.775532973 | 0.007891685 | -0.059065087 | 0.066956773 | 0.999933887 | -0.615134384 | -0.763362147 | 0.148227763 | 0.502614349 |
| ENSG00000134690 | *CDCA8* | 0.505421779 | -0.180198869 | -0.27557952 | 0.095380651 | 0.999933887 | -0.158001571 | -0.365435087 | 0.207433515 | 0.502645694 |
| ENSG00000167005 | *NUDT21* | 3.93159098 | 0.022956195 | -0.06846096 | 0.091417156 | 0.999933887 | -0.176941498 | -0.280950539 | 0.104009041 | 0.502646978 |
| ENSG00000136122 | *BORA* | -0.364091141 | 0.283896023 | -0.399479988 | 0.683376011 | 0.944094537 | -0.060024536 | 0.238716857 | -0.298741393 | 0.502674747 |
| ENSG00000143569 | *UBAP2L* | 8.153463981 | -0.01416755 | 0.006154177 | -0.020321726 | 0.999933887 | 0.206274386 | 0.294756082 | -0.088481696 | 0.502674747 |
| ENSG00000163660 | *CCNL1* | 10.086549896 | 0.050855549 | 0.001363187 | 0.049492362 | 0.999933887 | 0.411324539 | 0.538185326 | -0.126860787 | 0.502708338 |
| ENSG00000253314 | *LINC00293* | -0.417888911 | 0.004186705 | 0.41710968 | -0.412922975 | 0.999933887 | 0.109556869 | 0.401121203 | -0.291564334 | 0.502708338 |
| ENSG00000102057 | *KCND1* | 2.122656242 | 0.058474799 | -0.033013839 | 0.091488638 | 0.999933887 | 0.285034733 | 0.449814927 | -0.164780194 | 0.502809692 |
| ENSG00000196470 | *SIAH1* | 4.06645623 | -0.047927757 | 0.031855209 | -0.079782966 | 0.999933887 | 0.421632589 | 0.529713043 | -0.108080454 | 0.502879388 |
| ENSG00000180611 | *MB21D2* | -0.609371813 | 0.040451591 | 0.049200053 | -0.008748461 | 0.999933887 | 0.359289504 | 0.820144468 | -0.460854965 | 0.502954405 |
| ENSG00000277147 | *LINC00869* | 2.525183477 | 0.064082978 | 0.135149851 | -0.071066873 | 0.999933887 | 0.18462014 | 0.323601042 | -0.138980902 | 0.503202641 |
| ENSG00000136874 | *STX17* | 3.766230647 | -0.018564572 | -0.051211333 | 0.032646761 | 0.999933887 | -0.228942671 | -0.0869694 | -0.141973271 | 0.503202641 |
| ENSG00000198171 | *DDRGK1* | 2.971070445 | 0.089434137 | 0.122507193 | -0.033073056 | 0.999933887 | 0.093276285 | -0.070755103 | 0.164031388 | 0.503206377 |
| ENSG00000178921 | *PFAS* | 2.432847631 | 0.065072265 | 0.033365714 | 0.031706551 | 0.999933887 | 0.1734279 | 0.310917104 | -0.137489204 | 0.503206377 |
| ENSG00000140992 | *PDPK1* | 6.406993182 | -0.017696553 | -0.03600259 | 0.018306036 | 0.999933887 | -0.676384386 | -0.764611419 | 0.088227033 | 0.503206377 |
| ENSG00000165704 | *HPRT1* | 2.092083839 | -0.004469075 | 0.072200338 | -0.076669413 | 0.999933887 | 0.055156313 | 0.203272186 | -0.148115874 | 0.5035824 |
| ENSG00000255040 | *MORF4L1P3* | -0.863782885 | 0.258400673 | 0.084508928 | 0.173891745 | 0.999933887 | -0.479079165 | -0.960945875 | 0.48186671 | 0.50362464 |

| | | | | | | | | | | |
|---|---|---|---|---|---|---|---|---|---|---|
| ENSG00000286808 | *AC092338.3* | -0.332939123 | -0.106418166 | 0.170960801 | -0.277378967 | 0.999933887 | 0.098084983 | -0.211187302 | 0.309272286 | 0.50362464 |
| ENSG00000157017 | *GHRL* | 2.76140388 | -0.070915411 | 0.046824961 | -0.117740372 | 0.999933887 | 1.138796146 | 1.300394894 | -0.161598749 | 0.503930217 |
| ENSG00000286076 | *AC005050.3* | 3.173407327 | 0.370895078 | 0.283893771 | 0.087001307 | 0.999933887 | 1.492894687 | 1.69104823 | -0.198153542 | 0.50411223 |
| ENSG00000166166 | *TRMT61A* | 1.600141299 | 0.120753933 | 0.234331202 | -0.113577269 | 0.999933887 | 0.183289037 | 0.332958235 | -0.149669198 | 0.504129248 |
| ENSG00000165521 | *EML5* | -0.919933931 | 0.167789511 | -0.179191522 | 0.346981033 | 0.999933887 | -0.171217069 | 0.19236762 | -0.363584689 | 0.504129248 |
| ENSG00000279253 | *AL121753.2* | 1.574197413 | 0.02050693 | 0.091768089 | -0.071261159 | 0.999933887 | -0.24708439 | -0.466433566 | 0.219349176 | 0.504129248 |
| ENSG00000108557 | *RAI1* | 3.159884442 | 0.070746453 | 0.163766418 | -0.093019965 | 0.999933887 | -0.22341199 | -0.323250575 | 0.099838585 | 0.504268111 |
| ENSG00000260452 | *TPRKBP2* | 0.01973934 | 0.152115923 | 0.027919974 | 0.124195949 | 0.999933887 | -1.570981851 | -1.11894572 | -0.452036131 | 0.504314855 |
| ENSG00000055950 | *MRPL43* | 2.317172638 | 0.068827907 | 0.0254041 | 0.043423807 | 0.999933887 | -0.017884622 | -0.168256033 | 0.150371411 | 0.504398234 |
| ENSG00000274256 | *AL391903.2* | 1.691911647 | 0.132796437 | 0.044621088 | 0.088175349 | 0.999933887 | 0.485316716 | 0.718383216 | -0.2330665 | 0.504537766 |
| ENSG00000111058 | *ACSS3* | 1.256757361 | -0.062908298 | -0.028041386 | -0.034866912 | 0.999933887 | -0.923870578 | -1.152103155 | 0.228232577 | 0.504823413 |
| ENSG00000259651 | *MTCO3P23* | 2.208241005 | 0.50935204 | 0.704501292 | -0.195149251 | 0.999933887 | 2.052939578 | 2.427598508 | -0.37465893 | 0.504854029 |
| ENSG00000164306 | *PRIMPOL* | 0.501255242 | -0.129059505 | 0.114422303 | -0.243481809 | 0.999933887 | -0.171115563 | 0.025336189 | -0.196451752 | 0.504854029 |
| ENSG00000101335 | *MYL9* | -0.125018008 | -0.14781952 | 0.321891535 | -0.469711055 | 0.999933887 | 0.195666745 | 0.466985981 | -0.271319235 | 0.504854029 |
| ENSG00000165359 | *INTS6L* | 4.443441401 | 0.019876058 | -0.090998289 | 0.110874347 | 0.999933887 | -0.030875129 | 0.088182009 | -0.119057139 | 0.504902368 |
| ENSG00000145703 | *IQGAP2* | 5.786352283 | -0.007726209 | -0.110808094 | 0.103081885 | 0.999933887 | -0.330579365 | -0.150166609 | -0.180412756 | 0.504902368 |
| ENSG00000136875 | *PRPF4* | 3.996880943 | -0.038690504 | -0.032120231 | -0.006570273 | 0.999933887 | 0.075760328 | 0.174541761 | -0.098781433 | 0.505128172 |
| ENSG00000101224 | *CDC25B* | 5.33204401 | 0.014409805 | -0.000616039 | 0.015025844 | 0.999933887 | 0.114163057 | 0.037516209 | 0.076646848 | 0.505256273 |
| ENSG00000105497 | *ZNF175* | 0.019764992 | -0.201534279 | 0.230675399 | -0.432209678 | 0.999933887 | 0.05046633 | -0.184312591 | 0.234778922 | 0.505513518 |
| ENSG00000112893 | *MAN2A1* | 4.886434227 | -0.005692681 | -0.104954097 | 0.099261416 | 0.999933887 | -0.340278684 | -0.508422136 | 0.168143451 | 0.505513518 |
| ENSG00000251453 | *HAUS1P1* | -0.58469624 | -0.5089487 | -0.093859473 | -0.415089227 | 0.999933887 | -0.966047884 | -0.574703854 | -0.39134403 | 0.505540972 |
| ENSG00000147896 | *IFNK* | -1.148216675 | -0.455081338 | 0.17426143 | -0.629342767 | 0.999933887 | 0.263924461 | 0.654783611 | -0.39085915 | 0.506157036 |
| ENSG00000204305 | *AGER* | 4.93240099 | -0.012713476 | 0.011948547 | -0.024662023 | 0.999933887 | 0.002722968 | -0.094732977 | 0.097455945 | 0.506157036 |
| ENSG00000121892 | *PDS5A* | 4.978110916 | -0.0266152 | -0.052658129 | 0.026042929 | 0.999933887 | -0.369761614 | -0.254561174 | -0.115200439 | 0.5064395 |
| ENSG00000174744 | *BRMS1* | 5.250638831 | -0.017097744 | 0.028756104 | -0.045853848 | 0.999933887 | -0.145268154 | -0.226737151 | 0.081468997 | 0.506476341 |
| ENSG00000153179 | *RASSF3* | 7.965029976 | -0.054801888 | 0.007937497 | -0.062739385 | 0.999933887 | -0.244381767 | -0.163257236 | -0.081124531 | 0.506643944 |
| ENSG00000134684 | *YARS1* | 5.461487815 | -0.041494681 | -0.108524517 | 0.067029836 | 0.999933887 | 0.011436534 | 0.086608178 | -0.075171644 | 0.506643944 |
| ENSG00000106609 | *TMEM248* | 5.532386866 | -0.02661097 | -0.044707294 | 0.018096323 | 0.999933887 | -0.210583549 | -0.282955221 | 0.072371671 | 0.506643944 |
| ENSG00000159131 | *GART* | 2.9952691 | -0.033436938 | 0.087870564 | -0.121307502 | 0.999933887 | 0.062139331 | 0.167965827 | -0.105826495 | 0.506643944 |
| ENSG00000230124 | *ACBD6* | 2.175084836 | 0.044286272 | 0.155127787 | -0.110841515 | 0.999933887 | 0.003976548 | 0.135060564 | -0.131084015 | 0.506706448 |
| ENSG00000141449 | *GREB1L* | 0.713664307 | -0.057646514 | 0.401284973 | -0.458931487 | 0.944094537 | -0.232721526 | 0.14127535 | -0.373996876 | 0.506706448 |
| ENSG00000112167 | *SAYSD1* | 1.379364961 | -0.056410431 | -0.119379817 | 0.062969386 | 0.999933887 | -0.070030887 | 0.082342481 | -0.152373367 | 0.506710645 |
| ENSG00000185900 | *POMK* | 0.293829071 | 0.049070521 | -0.011927351 | 0.060997872 | 0.999933887 | -0.172182668 | -0.426028348 | 0.253845681 | 0.506710645 |
| ENSG00000164543 | *STK17A* | 5.410886767 | -0.024541758 | 0.078223119 | -0.102764877 | 0.999933887 | 0.012474644 | -0.070792061 | 0.083266705 | 0.506754336 |
| ENSG00000115368 | *WDR75* | 2.703598638 | -0.054369066 | -0.11828663 | 0.063917565 | 0.999933887 | 0.112118001 | 0.270283449 | -0.158165448 | 0.506787232 |
| ENSG00000232388 | *SMIM26* | 1.528394389 | -0.186619782 | -0.087187068 | -0.099432713 | 0.999933887 | -0.154739114 | -0.347710262 | 0.192971148 | 0.506983094 |
| ENSG00000239280 | *AC108693.1* | -0.465735188 | -0.275941744 | 0.313533173 | -0.589474916 | 0.944094537 | -0.179349083 | 0.153271473 | -0.332620556 | 0.506999166 |
| ENSG00000138660 | *AP1AR* | 4.251953876 | -0.139060134 | -0.14160859 | 0.002548456 | 0.999933887 | -0.729832232 | -0.566208209 | -0.163624022 | 0.507088707 |
| ENSG00000011376 | *LARS2* | 1.950179727 | 0.045391713 | 0.22987923 | -0.184487518 | 0.999933887 | 0.057607587 | -0.100782747 | 0.158390334 | 0.507088707 |
| ENSG00000270344 | *POC1B-AS1* | 1.745343045 | 0.151386442 | 0.028969767 | 0.122416676 | 0.999933887 | 0.975054614 | 1.195476594 | -0.22042198 | 0.507161352 |
| ENSG00000179627 | *ZBTB42* | 0.856627726 | -0.011440181 | 0.112944856 | -0.124385037 | 0.999933887 | -0.182101714 | -0.375182856 | 0.193081143 | 0.50720444 |
| ENSG00000277687 | *AL139407.1* | 2.536732391 | -0.037815344 | 0.00016985 | -0.037985194 | 0.999933887 | -0.698444829 | -0.527610287 | -0.170834542 | 0.507868255 |
| ENSG00000164405 | *UQCRQ* | 2.803058354 | 0.055266285 | -0.057490683 | 0.112756968 | 0.999933887 | 0.069184931 | -0.060906799 | 0.13009173 | 0.507878496 |
| ENSG00000244274 | *DBNDD2* | -0.238310246 | 0.007875736 | -0.017148016 | 0.025023752 | 0.999933887 | 1.209114231 | 0.824956327 | 0.384157904 | 0.507878496 |
| ENSG00000267519 | *AC020916.1* | 9.498186257 | 0.315787628 | 0.153963974 | 0.161823654 | 0.999933887 | 1.095978776 | 1.223774549 | -0.127795772 | 0.50794983 |
| ENSG00000167842 | *MIS12* | 2.772859467 | 0.004952636 | 0.063205044 | -0.058252408 | 0.999933887 | 0.029911345 | 0.125023982 | -0.095112637 | 0.50794983 |
| ENSG00000155393 | *HEATR3* | 1.197282256 | -0.018902423 | 0.011717126 | -0.03061955 | 0.999933887 | -0.199685719 | -0.429351813 | 0.229666094 | 0.508007543 |
| ENSG00000087266 | *SH3BP2* | 8.482673546 | -0.007285988 | 0.0196591 | -0.026945088 | 0.999933887 | 0.091842978 | 0.166035939 | -0.074192961 | 0.508007543 |
| ENSG00000159479 | *MED8* | 3.244115759 | -0.114656087 | -0.128400487 | 0.013744401 | 0.999933887 | -0.44285124 | -0.555660335 | 0.112809095 | 0.508146372 |
| ENSG00000168411 | *RFWD3* | 3.783730631 | -0.025575916 | 0.08023122 | -0.105807137 | 0.999933887 | 0.055521205 | -0.058113966 | 0.113635171 | 0.508146372 |
| ENSG00000261267 | *AC026470.2* | 2.19449542 | 0.055855552 | 0.023334239 | 0.032521314 | 0.999933887 | 1.400253809 | 1.594238324 | -0.193984515 | 0.508522275 |
| ENSG00000133561 | *GIMAP6* | 3.762843219 | 0.034137734 | -0.021212451 | 0.055350185 | 0.999933887 | 0.158308045 | 0.256692791 | -0.098384746 | 0.508522275 |
| ENSG00000134014 | *ELP3* | 2.192043767 | 0.009656899 | -0.08002563 | 0.08968253 | 0.999933887 | 0.081127266 | 0.223609946 | -0.14248268 | 0.508882264 |
| ENSG00000143771 | *CNIH4* | 2.734669377 | -0.017574228 | -0.021265624 | 0.003691395 | 0.999933887 | 0.075930294 | -0.071988722 | 0.147919016 | 0.508931788 |
| ENSG00000171960 | *PPIH* | 0.908307613 | -0.110137124 | -0.122243261 | 0.012106137 | 0.999933887 | -0.075129659 | 0.113180791 | -0.18831045 | 0.508943212 |
| ENSG00000117543 | *DPH5* | 1.501390089 | -0.06909131 | -0.161885364 | 0.092794053 | 0.999933887 | 0.102778872 | 0.273891889 | -0.171113017 | 0.508943212 |
| ENSG00000278000 | *AC139100.2* | -0.243282239 | 0.090470475 | -0.022761012 | 0.113231487 | 0.999933887 | 0.956776296 | 1.200255667 | -0.243479372 | 0.508998881 |
| ENSG00000167384 | *ZNF180* | 1.608809122 | -0.207813316 | 0.083520689 | -0.291334005 | 0.999933887 | -0.291732148 | -0.477617698 | 0.185885551 | 0.509001758 |
| ENSG00000225470 | *JPX* | 0.597668589 | 0.209143995 | 0.10970273 | 0.099441265 | 0.999933887 | -0.36067166 | -0.679099844 | 0.318428183 | 0.509010248 |
| ENSG00000145817 | *YIPF5* | 3.459042963 | -0.013201822 | -0.033982317 | 0.020780495 | 0.999933887 | -0.108457551 | -0.010271652 | -0.098185899 | 0.509247988 |
| ENSG00000240356 | *RPL23AP7* | 0.422359332 | -0.012048552 | -0.12289732 | 0.110848768 | 0.999933887 | -0.499748895 | -0.682787967 | 0.183039072 | 0.509272311 |
| ENSG00000275888 | *AC132872.3* | 4.195444893 | 0.086884807 | 0.115896905 | -0.029012098 | 0.999933887 | -0.303192809 | -0.198691457 | -0.104501352 | 0.509310365 |
| ENSG00000167565 | *SERTAD3* | 4.560026859 | 0.120506312 | 0.090527539 | 0.029978772 | 0.999933887 | -0.801736138 | -0.935197448 | 0.13346131 | 0.509310365 |
| ENSG00000180817 | *PPA1* | 2.716345997 | -0.069149896 | -0.120715099 | 0.051565203 | 0.999933887 | 0.114210162 | 0.234112519 | -0.119902357 | 0.509310365 |
| ENSG00000117640 | *MTFR1L* | 2.700835995 | 0.043660165 | 0.038618998 | 0.005041167 | 0.999933887 | -0.176979458 | -0.302294849 | 0.125315391 | 0.509310365 |
| ENSG00000162300 | *ZFPL1* | 1.790846319 | -0.059307891 | 0.039798038 | -0.099105929 | 0.999933887 | 0.027426025 | -0.16509972 | 0.192525745 | 0.509310365 |
| ENSG00000137441 | *FGFBP2* | 4.030411868 | 0.03289071 | -0.110459656 | 0.143350367 | 0.999933887 | 0.038381162 | 0.14567296 | -0.107291798 | 0.509310365 |
| ENSG00000188707 | *ZBED6CL* | 2.190635261 | -0.045513594 | -0.000469477 | -0.045044117 | 0.999933887 | 0.698163568 | 0.535532866 | 0.162630702 | 0.509555954 |
| ENSG00000222808 | *RNU4-47P* | 0.902028444 | -0.20367981 | 0.078878586 | -0.282558395 | 0.999933887 | -0.710180076 | -0.472748693 | -0.237431383 | 0.510179851 |
| ENSG00000143222 | *UFC1* | 3.665744337 | -0.040920455 | 0.023311519 | -0.064231974 | 0.999933887 | -0.11065261 | -0.213268891 | 0.102616281 | 0.510213989 |
| ENSG00000198843 | *SELENOT* | 6.149118056 | -0.054094949 | -0.028866643 | -0.025228306 | 0.999933887 | -0.044937275 | 0.032011328 | -0.076948604 | 0.510732453 |
| ENSG00000251002 | *AC244502.1* | -0.345887607 | 0.159436418 | -0.094986965 | 0.254423382 | 0.999933887 | -0.27654248 | 0.055899462 | -0.332441942 | 0.510732453 |
| ENSG00000072858 | *SIDT1* | 3.375499841 | -0.036140274 | -0.019500887 | -0.016639387 | 0.999933887 | 0.047435695 | 0.142374212 | -0.094938517 | 0.510732453 |
| ENSG00000261308 | *FIGNL2* | 0.162602871 | 0.191808832 | 0.26682414 | -0.075015308 | 0.999933887 | -1.013206369 | -0.815772343 | -0.197434025 | 0.510745419 |
| ENSG00000138642 | *HERC6* | 3.219500971 | 0.06129968 | 0.025138432 | 0.036161248 | 0.999933887 | -0.160871986 | -0.300454198 | 0.139582211 | 0.510745419 |
| ENSG00000235602 | *POU5F1P3* | 3.115381747 | -0.111854515 | -0.142622652 | 0.030768137 | 0.999933887 | 0.128198709 | 0.265826657 | -0.137627948 | 0.510790676 |
| ENSG00000186517 | *ARHGAP30* | 9.109167637 | -0.035313949 | -0.018642192 | -0.016671757 | 0.999933887 | -0.640151621 | -0.745879811 | 0.105728189 | 0.510790676 |
| ENSG00000137310 | *TCF19* | 0.718824093 | -0.022104689 | -0.052418702 | 0.030314013 | 0.999933887 | 0.024830768 | -0.214951187 | 0.239781956 | 0.510790676 |
| ENSG00000143753 | *DEGS1* | 4.686225975 | 0.051397343 | -0.045084663 | 0.096482006 | 0.999933887 | 0.255403395 | 0.373001596 | -0.117598201 | 0.510834858 |
| ENSG00000279394 | *AC015871.4* | -1.001971841 | 0.043485049 | -0.046402317 | 0.089887365 | 0.999933887 | -0.21954978 | 0.381020481 | -0.600570261 | 0.510834858 |
| ENSG00000106665 | *CLIP2* | 1.761076605 | 0.113843701 | -0.080070075 | 0.193913776 | 0.999933887 | -0.061526372 | -0.250160693 | 0.188634321 | 0.510846206 |
| ENSG00000055211 | *GINM1* | 4.023576516 | -0.056469423 | -0.003983203 | -0.05248622 | 0.999933887 | -0.306093593 | -0.403261551 | 0.097167958 | 0.51088005 |
| ENSG00000129187 | *DCTD* | 2.900394579 | 0.079768707 | 0.004034126 | 0.075734581 | 0.999933887 | 0.11918861 | 0.231888486 | -0.112699876 | 0.51130893 |
| ENSG00000165409 | *TSHR* | -1.288982725 | -0.193176624 | -0.438908344 | 0.24573172 | 0.999933887 | -0.332271976 | 0.007235577 | -0.339507552 | 0.51130893 |
| ENSG00000081026 | *MAGI3* | -0.00185426 | -0.180979303 | -0.548009787 | 0.367030484 | 0.999933887 | -0.146226836 | -0.512024235 | 0.365797399 | 0.511359009 |
| ENSG00000182986 | *ZNF320* | 1.245571151 | -0.077169131 | -0.042658877 | -0.034510254 | 0.999933887 | -0.075334147 | -0.243766325 | 0.168432178 | 0.511359009 |
| ENSG00000182670 | *TTC3* | 4.670689662 | -0.056856534 | -0.141610674 | 0.084754141 | 0.999933887 | -0.214683936 | -0.047120895 | -0.167563041 | 0.511359009 |
| ENSG00000164077 | *MON1A* | 0.513078854 | -0.104456407 | 0.049872445 | -0.154328852 | 0.999933887 | 0.145516197 | -0.060651589 | 0.206167786 | 0.51146359 |
| ENSG00000166471 | *TMEM41B* | 4.67055504 | 0.021333489 | 0.011292698 | 0.010040792 | 0.999933887 | 0.598500252 | 0.710509702 | -0.11200945 | 0.51152398 |
| ENSG00000248208 | *WDR45P1* | 0.314838554 | 0.151802423 | -0.09311396 | 0.244916383 | 0.999933887 | 0.813339969 | 1.103724607 | -0.290384638 | 0.511590746 |
| ENSG00000167658 | *EEF2* | 9.240823827 | 0.045158856 | 0.022453 | 0.022705856 | 0.999933887 | 0.118957241 | 0.197761537 | -0.078804296 | 0.512173901 |
| ENSG00000197785 | *ATAD3A* | 1.671645483 | 0.042784245 | 0.069829388 | -0.027045142 | 0.999933887 | 0.217949246 | 0.065813213 | 0.152136034 | 0.512173901 |
| ENSG00000213553 | *RPLP0P6* | -0.250583877 | 0.056269168 | 0.098762827 | -0.042493659 | 0.999933887 | 0.081850502 | 0.355376887 | -0.273526385 | 0.512279791 |
| ENSG00000100348 | *TXN2* | 3.64653746 | 0.01222829 | 0.129006833 | -0.116778543 | 0.999933887 | -0.030146026 | -0.145136551 | 0.114990525 | 0.512279791 |
| ENSG00000198440 | *ZNF583* | -0.850562707 | -0.423678455 | -0.006886913 | -0.416791542 | 0.999933887 | -0.153573881 | 0.206850135 | -0.360424016 | 0.512436179 |
| ENSG00000188554 | *NBR1* | 6.603740022 | -0.066261429 | 0.011297348 | -0.077558777 | 0.999933887 | 0.19816726 | 0.29875267 | -0.100585409 | 0.512436179 |
| ENSG00000092841 | *MYL6* | 8.068841889 | -0.010801483 | -0.01878176 | 0.007980276 | 0.999933887 | -0.075215985 | -0.151708868 | 0.076492882 | 0.512436179 |
| ENSG00000167524 | *RSKR* | 2.265014345 | -0.052759972 | 0.041794266 | -0.094554238 | 0.999933887 | 0.292366834 | 0.464492235 | -0.172125401 | 0.512493302 |
| ENSG00000155592 | *ZKSCAN2* | 0.913452348 | 0.109716215 | -0.036707164 | 0.146423378 | 0.999933887 | 0.08570035 | 0.310726206 | -0.225025856 | 0.512493458 |
| ENSG00000232956 | *SNHG15* | 3.561900119 | -0.041649702 | 0.075115192 | -0.116764894 | 0.999933887 | 0.326821793 | 0.199355671 | 0.127466122 | 0.512591627 |
| ENSG00000258413 | *AL158801.2* | 0.361149127 | 0.274488028 | 0.248704779 | 0.025783249 | 0.999933887 | 0.74389885 | 0.399267177 | 0.344631673 | 0.512937013 |
| ENSG00000180901 | *KCTD2* | 4.690731421 | -0.059820993 | -0.084351712 | 0.02453072 | 0.999933887 | -0.411106269 | -0.497615808 | 0.086509539 | 0.512937013 |
| ENSG00000180479 | *ZNF571* | 0.178385312 | -0.12101242 | -0.255121551 | 0.134109131 | 0.999933887 | 0.046719337 | 0.345382497 | -0.298663161 | 0.512937013 |
| ENSG00000204301 | *NOTCH4* | 1.038121046 | 0.078118402 | 0.056564992 | 0.02155341 | 0.999933887 | 0.387112001 | 0.166340783 | 0.220771218 | 0.512937013 |
| ENSG00000196670 | *ZFP62* | 2.407381446 | 0.023674346 | -0.130938762 | 0.154613107 | 0.999933887 | -0.31834692 | -0.435705199 | 0.117358279 | 0.512937013 |

| | | | | | | | | | | |
|---|---|---|---|---|---|---|---|---|---|---|
| ENSG00000068489 | *PRR11* | 0.55559687 | -0.019489327 | -0.114055484 | 0.094566156 | 0.999933887 | -0.3325105 | -0.5876702 | 0.2551597 | 0.512937013 |
| ENSG00000164877 | *MICALL2* | 0.81430297 | -0.001846567 | -0.127396364 | 0.125549796 | 0.999933887 | 0.321500696 | 0.555450595 | -0.233949898 | 0.512937013 |
| ENSG00000143321 | *HDGF* | 6.211327598 | 0.035512352 | -0.013468113 | 0.048980466 | 0.999933887 | 1.17756007 | 1.341110353 | -0.163550282 | 0.512980934 |
| ENSG00000147454 | *SLC25A37* | 9.694263637 | -0.003255197 | 0.012127884 | -0.015383081 | 0.999933887 | 0.162557328 | 0.103076404 | 0.059480924 | 0.512980934 |
| ENSG00000259781 | *HMGB1P6* | 1.377975969 | -0.129194048 | 0.103435516 | -0.232629563 | 0.999933887 | -0.105957213 | 0.076109644 | -0.182066857 | 0.513210581 |
| ENSG00000120656 | *TAF12* | 4.252983977 | 0.005552893 | 0.066502663 | -0.06094977 | 0.999933887 | -0.317887001 | -0.420394814 | 0.102507814 | 0.513443782 |
| ENSG00000228903 | *RASA4CP* | 0.633193971 | 0.064259314 | 0.099839678 | -0.035580364 | 0.999933887 | 0.077736454 | -0.210551062 | 0.288287516 | 0.513478038 |
| ENSG00000106443 | *PHF14* | 2.018816118 | 0.038396737 | -0.036317162 | 0.074713899 | 0.999933887 | -0.131534746 | 0.013935736 | -0.145470483 | 0.513478038 |
| ENSG00000156886 | *ITGAD* | 2.093256388 | 0.041706141 | -0.074313651 | 0.116019792 | 0.999933887 | -0.102054881 | 0.060235194 | -0.162290075 | 0.513478038 |
| ENSG00000143376 | *SNX27* | 6.825024233 | 0.013491037 | 0.00317455 | 0.010316487 | 0.999933887 | -0.472451668 | -0.382666684 | -0.089784984 | 0.513478038 |
| ENSG00000106546 | *AHR* | 3.290194931 | -0.043329799 | 0.08219843 | -0.125528228 | 0.999933887 | 0.127900219 | 0.259955512 | -0.132055293 | 0.513546133 |
| ENSG00000002726 | *AOC1* | -1.470738634 | 0.08725452 | 0.243031608 | -0.155777088 | 0.999933887 | 0.68063956 | 1.233063531 | -0.552423971 | 0.513546133 |
| ENSG00000155744 | *FAM126B* | 5.888127794 | -0.077906905 | -0.092200656 | 0.014293751 | 0.999933887 | -0.191323359 | -0.036504784 | -0.154818575 | 0.513598393 |
| ENSG00000188026 | *RILPL1* | 1.196253866 | 0.011032826 | 0.071358116 | -0.06032529 | 0.999933887 | 0.072675546 | 0.286330858 | -0.213655312 | 0.513746814 |
| ENSG00000186063 | *AIDA* | 2.393297691 | -0.105780458 | 0.022719494 | -0.128499952 | 0.999933887 | -0.25655031 | -0.390278031 | 0.133727721 | 0.513785977 |
| ENSG00000115661 | *STK16* | 3.524729775 | 0.068801081 | -0.01291434 | 0.081715421 | 0.999933887 | -0.247968077 | -0.356886593 | 0.108918516 | 0.513785977 |
| ENSG00000233144 | *AL512656.1* | -0.812287197 | -0.255164246 | 0.074828158 | -0.329992404 | 0.999933887 | 1.291992925 | 1.599134454 | -0.307141528 | 0.513788437 |
| ENSG00000166261 | *ZNF202* | 1.417234599 | -0.017864811 | -0.129876258 | 0.112011447 | 0.999933887 | 0.232680397 | 0.41113909 | -0.178458693 | 0.513788437 |
| ENSG00000148730 | *EIF4EBP2* | 7.046645585 | 0.004715846 | -0.044509011 | 0.049224857 | 0.999933887 | -0.268091311 | -0.343481695 | 0.075390384 | 0.513788437 |
| ENSG00000137106 | *GRHPR* | 2.891721563 | 0.082480758 | 0.014284971 | 0.068195786 | 0.999933887 | 0.047322598 | -0.074845349 | 0.122167947 | 0.514214182 |
| ENSG00000013503 | *POLR3B* | 0.736097835 | -0.253401733 | 0.00170655 | -0.255108283 | 0.999933887 | -0.082630134 | 0.100403905 | -0.18303404 | 0.514293914 |
| ENSG00000164924 | *YWHAZ* | 9.604324744 | -0.040227154 | -0.052977796 | 0.012750641 | 0.999933887 | -0.095825214 | -0.031740114 | -0.0640851 | 0.514293914 |
| ENSG00000266405 | *CBX3P2* | 1.167298018 | -0.084573054 | -0.156442418 | 0.071869364 | 0.999933887 | -0.632789141 | -0.869493585 | 0.236704445 | 0.514293914 |
| ENSG00000165046 | *LETM2* | 2.217406306 | 0.126975569 | -0.068865388 | 0.195840957 | 0.999933887 | -0.712671612 | -0.898490618 | 0.185819006 | 0.514406316 |
| ENSG00000140307 | *GTF2A2* | 2.89288741 | 0.116651746 | 0.120870261 | -0.004218516 | 0.999933887 | -0.126289852 | -0.244515768 | 0.118225916 | 0.514877489 |
| ENSG00000163938 | *GNL3* | 3.405346533 | -0.116764225 | -0.096596915 | -0.02016731 | 0.999933887 | 0.150876359 | 0.266318941 | -0.115442582 | 0.514877489 |
| ENSG00000273004 | *AL078644.1* | 2.925482074 | 0.115962397 | 0.318006362 | -0.202043965 | 0.999933887 | 0.414417988 | 0.571304049 | -0.156886062 | 0.514877489 |
| ENSG00000213714 | *FAM209B* | 1.417339728 | -0.15223989 | -0.031660042 | -0.120579848 | 0.999933887 | 0.17340376 | -0.033892701 | 0.207296461 | 0.514885165 |
| ENSG00000068354 | *TBC1D25* | 4.633904325 | -0.026938771 | -0.018015473 | -0.008923299 | 0.999933887 | -0.120374831 | -0.192832139 | 0.072457308 | 0.514885165 |
| ENSG00000272941 | *AC083862.1* | 3.860077754 | -0.023895263 | 0.023185004 | -0.047080267 | 0.999933887 | -0.255111115 | -0.139277375 | -0.11583374 | 0.514885165 |
| ENSG00000143178 | *TBX19* | 0.982533868 | 0.023735512 | 0.295110925 | -0.271375414 | 0.999933887 | -0.115818601 | -0.297367906 | 0.181549305 | 0.514885165 |
| ENSG00000137942 | *FNBP1L* | -0.196537636 | -0.029471601 | -0.185090872 | 0.155619271 | 0.999933887 | -0.090529301 | 0.130531022 | -0.221060323 | 0.514976828 |
| ENSG00000286634 | *AC146507.3* | -0.668637682 | -0.103092528 | 0.126063487 | -0.229156015 | 0.999933887 | -0.585386215 | -0.134084441 | -0.451301774 | 0.51498321 |
| ENSG00000227376 | *FTH1P16* | 1.030847865 | -0.063020824 | 0.102069033 | -0.165089857 | 0.999933887 | 0.382036952 | 0.573568935 | -0.191531983 | 0.515263131 |
| ENSG00000109920 | *FNBP4* | 5.740264125 | -0.024852214 | -0.053007597 | 0.028155384 | 0.999933887 | -0.180400137 | -0.261526302 | 0.081126165 | 0.515530462 |
| ENSG00000261338 | *AC021016.1* | 0.774583027 | -0.025303284 | -0.034987876 | 0.009684593 | 0.999933887 | -0.206855856 | -0.495415451 | 0.288559594 | 0.515530462 |
| ENSG00000157823 | *AP3S2* | 3.097977985 | -0.140309137 | -0.11628182 | -0.024027318 | 0.999933887 | 0.039427075 | -0.064737206 | 0.104164281 | 0.515762514 |
| ENSG00000274272 | *AC069281.2* | 2.947991946 | 0.066227825 | 0.046464818 | 0.019763008 | 0.999933887 | -0.157216839 | -0.290793018 | 0.133092179 | 0.515762514 |
| ENSG00000101452 | *DHX35* | 1.588771142 | -0.021628675 | -0.193657716 | 0.172029041 | 0.999933887 | -0.021784794 | -0.187885218 | 0.166100424 | 0.515762514 |
| ENSG00000143977 | *SNRPG* | 3.579379471 | 0.036155325 | -0.018102787 | 0.054258113 | 0.999933887 | 0.210896081 | 0.116671662 | 0.094224419 | 0.51589104 |
| ENSG00000100580 | *TMED8* | 3.369183181 | 0.018083776 | 0.106598719 | -0.088514943 | 0.999933887 | -0.076938586 | -0.196451106 | 0.11951252 | 0.516288959 |
| ENSG00000280091 | *AC020895.2* | 0.791522309 | -0.077304983 | 0.143487845 | -0.220792829 | 0.999933887 | 1.677365391 | 1.933058657 | -0.255693266 | 0.516639006 |
| ENSG00000196407 | *THEM5* | -0.594879431 | -0.02148429 | -0.063706673 | 0.042222382 | 0.999933887 | 0.045433926 | 0.36065908 | -0.315225154 | 0.516639006 |
| ENSG00000131475 | *VPS25* | 2.815940499 | -0.009500124 | 0.12451168 | -0.134011804 | 0.999933887 | 0.016568642 | -0.097004436 | 0.113573077 | 0.516639006 |
| ENSG00000112308 | *C6orf62* | 7.729954567 | -0.06367134 | -0.033996915 | -0.029674425 | 0.999933887 | 0.296167325 | 0.362188172 | -0.066020847 | 0.516702268 |
| ENSG00000013392 | *RWDD2A* | -0.658574075 | 0.193383065 | 0.043846931 | 0.149536134 | 0.999933887 | 0.143213015 | 0.451428098 | -0.308215082 | 0.516877933 |
| ENSG00000196092 | *PAX5* | 1.090047118 | 0.210612678 | 0.160667388 | 0.049945289 | 0.999933887 | 0.072342075 | 0.231426653 | -0.159084578 | 0.517090455 |
| ENSG00000214846 | *AC114744.1* | 1.625590695 | -0.156926085 | -0.040935736 | -0.11599035 | 0.999933887 | -0.684191109 | -0.490672494 | -0.193518614 | 0.517090455 |
| ENSG00000167395 | *ZNF646* | 7.049062162 | 0.053564729 | 0.11858511 | -0.065020382 | 0.999933887 | -0.071280498 | 0.003705885 | -0.074986383 | 0.517090455 |
| ENSG00000203446 | *SUGCT-AS1* | -0.508760758 | 0.025755425 | -0.377170649 | 0.402926073 | 0.999933887 | -0.030636247 | 0.158007886 | -0.188644133 | 0.517090455 |
| ENSG00000167862 | *MRPL58* | 0.641937738 | 0.0827475 | -0.141404147 | 0.224151648 | 0.999933887 | 0.094108606 | 0.286260102 | -0.192151497 | 0.517238052 |
| ENSG00000245281 | *AC124242.1* | 0.016264256 | -0.038058444 | 0.138212455 | -0.176270898 | 0.999933887 | 0.089761829 | 0.306464893 | -0.216703064 | 0.517238052 |
| ENSG00000141526 | *SLC16A3* | 9.20323401 | 0.077869059 | 0.079399227 | -0.001530168 | 0.999933887 | -0.06605692 | 0.041213309 | -0.107270229 | 0.517288093 |
| ENSG00000263482 | *ANTXRLP1* | -1.030583737 | 0.00286554 | -0.311862981 | 0.31472852 | 0.999933887 | 0.170649495 | -0.21960273 | 0.390252225 | 0.517316499 |
| ENSG00000232273 | *FTH1P1* | -0.693420105 | 0.149042698 | 0.419565264 | -0.270522566 | 0.999933887 | 0.188307904 | 0.681599139 | -0.493291235 | 0.517411572 |
| ENSG00000115459 | *ELMOD3* | 3.370380456 | 0.022636481 | 0.057183239 | -0.034546758 | 0.999933887 | 0.167869272 | 0.268501698 | -0.100632426 | 0.517435774 |
| ENSG00000134755 | *DSC2* | 2.620189431 | -0.04955581 | 0.017133487 | -0.066689296 | 0.999933887 | -0.350603137 | -0.569351599 | 0.218748461 | 0.517462606 |
| ENSG00000273329 | *AC078846.1* | 1.381678041 | -0.03950827 | 0.105766976 | -0.145275246 | 0.999933887 | -0.391358132 | -0.230244773 | -0.161113359 | 0.517462606 |
| ENSG00000131931 | *THAP1* | 1.926304329 | -0.035423568 | 0.037632525 | -0.073056094 | 0.999933887 | -0.610282902 | -0.766224259 | 0.155941356 | 0.517462606 |
| ENSG00000115446 | *UNC50* | 4.637629868 | -0.01102491 | -0.088042612 | 0.077017702 | 0.999933887 | -0.018630442 | 0.054479106 | -0.073109548 | 0.517462606 |
| ENSG00000005249 | *PRKAR2B* | -0.879616342 | -0.008277733 | 0.035451112 | -0.043728845 | 0.999933887 | -0.071636314 | -0.422793371 | 0.351157057 | 0.51756896 |
| ENSG00000128342 | *LIF* | 4.154893208 | 0.301777365 | 0.280389673 | 0.021387692 | 0.999933887 | 0.87267222 | 1.0539358 | -0.18126358 | 0.517608679 |
| ENSG00000162654 | *GBP4* | 5.285727403 | -0.001252464 | 0.027731918 | -0.028984382 | 0.999933887 | 0.090831341 | -0.00746637 | 0.09829771 | 0.517633077 |
| ENSG00000126768 | *TIMM17B* | 4.461292077 | 0.089452273 | 0.018730724 | 0.070721549 | 0.999933887 | 0.837956334 | 0.942806677 | -0.104850342 | 0.518235287 |
| ENSG00000090905 | *TNRC6A* | 4.497348158 | -0.06525951 | -0.031487284 | -0.033772226 | 0.999933887 | -0.250623093 | -0.336490585 | 0.085867492 | 0.518235287 |
| ENSG00000186666 | *BCDIN3D* | 0.082334301 | -0.133167569 | -0.403745053 | 0.270577484 | 0.999933887 | -0.136311453 | 0.104646607 | -0.240958061 | 0.518427236 |
| ENSG00000169570 | *DTWD2* | -0.069187584 | -0.11339753 | -0.132958359 | 0.019560829 | 0.999933887 | -0.467201241 | -0.710269958 | 0.243068716 | 0.518579834 |
| ENSG00000242314 | *RPL12P32* | 0.429641676 | -0.153148283 | 0.006170157 | -0.15931844 | 0.999933887 | -0.141856945 | 0.167705636 | -0.309562581 | 0.518724301 |
| ENSG00000276557 | *TRBV18* | -0.02456065 | -0.184413369 | -0.022350878 | -0.162062491 | 0.999933887 | 0.014282439 | 0.290375834 | -0.276093395 | 0.51883721 |
| ENSG00000120708 | *TGFBI* | 1.285252489 | -0.137201194 | 0.034419538 | -0.171620732 | 0.999933887 | -0.578872786 | -0.849655431 | 0.270782645 | 0.51883721 |
| ENSG00000095739 | *BAMBI* | 0.430314919 | 0.082270864 | 0.204545866 | -0.122275003 | 0.999933887 | 0.322482259 | 0.566235875 | -0.243753615 | 0.51883721 |
| ENSG00000119943 | *PYROXD2* | 2.039845428 | 0.115144754 | -0.112645532 | 0.227790286 | 0.999933887 | 0.139206568 | -0.007051277 | 0.146257845 | 0.519232177 |
| ENSG00000175548 | *ALG10B* | 0.59375082 | -0.159818361 | -0.326030498 | 0.166212137 | 0.999933887 | -0.321867931 | -0.02567019 | -0.296197741 | 0.519232177 |
| ENSG00000158373 | *H2BC5* | 2.576512681 | 0.062414259 | 0.070544338 | -0.008130079 | 0.999933887 | -0.30738416 | -0.49695046 | 0.189566299 | 0.519232177 |
| ENSG00000166734 | *GOLM2* | 5.553309955 | 0.011812471 | -0.059300644 | 0.071113115 | 0.999933887 | -0.307599116 | -0.39769651 | 0.090097394 | 0.519232177 |
| ENSG00000066777 | *ARFGEF1* | 6.056366135 | -0.01151225 | -0.009288648 | -0.002223601 | 0.999933887 | -0.302642151 | -0.164315348 | -0.138326803 | 0.519232177 |
| ENSG00000137869 | *CYP19A1* | -2.654635758 | 0.049625241 | -0.6934194 | 0.743044641 | 0.999933887 | 3.796384535 | 4.392510181 | -0.596125646 | 0.519232177 |
| ENSG00000202533 | *Y_RNA* | -0.552149792 | 0.159058365 | -0.134026124 | 0.293084489 | 0.999933887 | -0.026610515 | -0.412527707 | 0.385917192 | 0.519451568 |
| ENSG00000261596 | *AC005632.3* | 2.411404845 | 0.037061036 | -0.014758709 | 0.051819745 | 0.999933887 | 0.313466441 | 0.166672675 | 0.146793766 | 0.519451568 |
| ENSG00000271581 | *AL671883.2* | 0.081636829 | 0.077011981 | 0.160866372 | -0.083854391 | 0.999933887 | 1.18582985 | 0.916343326 | 0.269486524 | 0.519464916 |
| ENSG00000189337 | *KAZN* | 1.049398521 | 0.065184207 | 0.066555433 | -0.001371227 | 0.999933887 | -0.395291971 | -0.121177867 | -0.274114104 | 0.519465888 |
| ENSG00000122378 | *PRXL2A* | -0.698160463 | -0.144739626 | -0.225620197 | 0.080880571 | 0.999933887 | -0.141490558 | 0.275595091 | -0.417085649 | 0.519610575 |
| ENSG00000284858 | *AC091185.2* | -0.444120143 | -0.470452736 | 0.105359036 | -0.575811772 | 0.944094537 | -1.17063324 | -0.798222122 | -0.372411118 | 0.519636528 |
| ENSG00000171643 | *S100Z* | -0.67713666 | -0.022890783 | 0.029935774 | -0.052826557 | 0.999933887 | -0.742937469 | -1.213015788 | 0.47007832 | 0.519872299 |
| ENSG00000169715 | *MT1E* | -0.736913977 | 0.176410452 | -0.292896353 | 0.469306806 | 0.999933887 | 0.583095307 | 0.93082762 | -0.347732312 | 0.519958527 |
| ENSG00000261799 | *AC007406.4* | 1.404917026 | -0.025957506 | 0.029119091 | -0.055076597 | 0.999933887 | -0.261228932 | -0.014556213 | -0.246672719 | 0.520037971 |
| ENSG00000143452 | *HORMAD1* | -0.002487571 | -0.055283926 | -0.167038181 | 0.111754254 | 0.999933887 | -0.212454496 | 0.221799721 | -0.434254217 | 0.520129029 |
| ENSG00000196418 | *ZNF124* | 0.967355995 | 0.126431833 | 0.032169506 | 0.094262327 | 0.999933887 | 0.532797753 | 0.756771006 | -0.223973253 | 0.520276062 |
| ENSG00000068784 | *SRBD1* | 3.050167144 | -0.045412025 | -0.131747058 | 0.086335032 | 0.999933887 | -0.234083805 | -0.103924491 | -0.130159315 | 0.520276062 |
| ENSG00000070729 | *CNGB1* | -0.584960153 | -0.135531796 | -0.126362464 | -0.009169332 | 0.999933887 | 0.212774337 | 0.606762959 | -0.393988622 | 0.520330651 |
| ENSG00000136940 | *PDCL* | 2.317170979 | -0.128630552 | 0.007868858 | -0.13649941 | 0.999933887 | -0.132841917 | -0.000436491 | -0.132405425 | 0.520463281 |
| ENSG00000162630 | *B3GALT2* | -0.528237173 | 0.357485255 | 0.701071685 | -0.34358643 | 0.999933887 | -0.252447088 | 0.284021483 | -0.536468571 | 0.520463281 |
| ENSG00000155966 | *AFF2* | 0.019430727 | -0.078414406 | 0.078214131 | -0.156628537 | 0.999933887 | 0.094357599 | -0.251742714 | 0.346100312 | 0.520463281 |
| ENSG00000224975 | *INE1* | 2.002412116 | -0.028565096 | 0.158027223 | -0.186592319 | 0.999933887 | 0.17062321 | 0.349416412 | -0.178793202 | 0.520463281 |
| ENSG00000138085 | *ATRAID* | 2.936869153 | 0.01360891 | -0.036293632 | 0.049902542 | 0.999933887 | -0.046990548 | -0.163854795 | 0.116864248 | 0.520463281 |
| ENSG00000115289 | *PCGF1* | 1.960344023 | -0.139091882 | -0.042360731 | -0.096731151 | 0.999933887 | -0.128824856 | -0.242269971 | 0.113445115 | 0.520718835 |
| ENSG00000005882 | *PDK2* | 0.743316265 | 0.072187837 | 0.002159037 | 0.0700288 | 0.999933887 | 0.168199114 | -0.064009209 | 0.232208323 | 0.520828673 |
| ENSG00000156738 | *MS4A1* | 2.256918192 | 0.014609727 | -0.042447899 | 0.057057626 | 0.999933887 | 0.052199551 | 0.189530679 | -0.137331127 | 0.520828673 |
| ENSG00000226608 | *FTLP3* | 2.509552225 | -0.032326312 | -0.092680836 | 0.060354524 | 0.999933887 | 0.327560907 | 0.190302716 | 0.137258192 | 0.520837974 |
| ENSG00000235568 | *NFAM1* | 8.579367426 | -0.001326359 | 0.001046156 | -0.002372515 | 0.999933887 | -0.018466471 | -0.11803346 | 0.099566989 | 0.520837974 |
| ENSG00000182481 | *KPNA2* | 2.925843198 | -0.030645264 | -0.197337126 | 0.166691862 | 0.999933887 | 0.104158673 | 0.223478824 | -0.119320151 | 0.520935763 |
| ENSG00000286307 | *AC017028.1* | -0.051121492 | 0.220613126 | 0.477144842 | -0.256531716 | 0.999933887 | -0.545804115 | -0.206164687 | -0.339639428 | 0.521003563 |
| ENSG00000167112 | *TRUB2* | 0.920297618 | 0.136517132 | -0.126154659 | 0.262671791 | 0.999933887 | 0.038926815 | 0.22270836 | -0.183781544 | 0.521003563 |
| ENSG00000165591 | *FAAH2* | -0.525435924 | -0.184657912 | 0.127260712 | -0.311918623 | 0.999933887 | 0.291903863 | 0.594787925 | -0.302884063 | 0.521003563 |
| ENSG00000154518 | *ATP5MC3* | 3.324031578 | 0.051489987 | -0.049198002 | 0.100687989 | 0.999933887 | 0.065031319 | -0.055879089 | 0.120910408 | 0.521416244 |

| | | | | | | | | | | |
|---|---|---|---|---|---|---|---|---|---|---|
| ENSG00000226450 | *CYP2D8P* | 1.576434409 | 0.06455335 | 0.211299498 | -0.146746148 | 0.999933887 | 0.00344349 | -0.16233545 | 0.165778941 | 0.521419936 |
| ENSG00000116985 | *BMP8B* | -0.4210712 | 0.109781162 | -0.003851162 | 0.113632324 | 0.999933887 | -0.123428597 | 0.134396398 | -0.257824995 | 0.521419936 |
| ENSG00000267546 | *AC015802.4* | -0.426383622 | -0.229071231 | -0.288471378 | 0.059400147 | 0.999933887 | 0.543147367 | 0.338717161 | 0.204430206 | 0.521771203 |
| ENSG00000204936 | *CD177* | 2.219089988 | 0.010012207 | -0.108349122 | 0.118361329 | 0.999933887 | -0.282822945 | -0.443681038 | 0.160858094 | 0.52219631 |
| ENSG00000101188 | *NTSR1* | 2.032007362 | 0.091967566 | -0.010853468 | 0.102821033 | 0.999933887 | 0.150252622 | 0.288088303 | -0.13783568 | 0.522274367 |
| ENSG00000146676 | *PURB* | 7.301351489 | 0.109393199 | 0.126758876 | -0.017365677 | 0.999933887 | 0.150973686 | 0.247502553 | -0.096528867 | 0.522294676 |
| ENSG00000100865 | *CINP* | 3.552501697 | -0.027492474 | 0.037239104 | -0.064731578 | 0.999933887 | 0.036101978 | 0.156446661 | -0.120344682 | 0.522294676 |
| ENSG00000197077 | *KIAA1671* | 2.077985375 | -0.053822468 | 0.08404627 | -0.137868739 | 0.999933887 | 0.006328612 | -0.122325985 | 0.128654597 | 0.522328735 |
| ENSG00000257226 | *AC079584.2* | -0.856488787 | -0.076291296 | -0.045877576 | -0.03041372 | 0.999933887 | -0.398792055 | 0.042151439 | -0.440943493 | 0.522503453 |
| ENSG00000275210 | *AC245884.11* | 0.917741162 | 0.063973948 | 0.112926524 | -0.048952576 | 0.999933887 | 0.306255708 | -0.006390301 | 0.312646008 | 0.52264761 |
| ENSG00000131558 | *EXOC4* | 3.090338134 | -0.030524837 | 0.012135934 | -0.04266077 | 0.999933887 | -0.002430798 | 0.119664231 | -0.122095029 | 0.523186559 |
| ENSG00000007255 | *TRAPPC6A* | 2.113439406 | 0.116749473 | -0.059228458 | 0.175977931 | 0.999933887 | 0.001859759 | -0.136323617 | 0.138183376 | 0.52338398 |
| ENSG00000224470 | *ATXN1L* | 6.584887786 | -0.001325838 | -0.035568192 | 0.034242355 | 0.999933887 | -0.12826395 | -0.042334967 | -0.085928983 | 0.52338398 |
| ENSG00000003756 | *RBM5* | 7.206799642 | -0.054905066 | -0.022674585 | -0.032230481 | 0.999933887 | -0.027024458 | 0.037797918 | -0.064822376 | 0.523566513 |
| ENSG00000108179 | *PPIF* | 11.495385398 | 0.121742201 | 0.116142683 | 0.005599518 | 0.999933887 | 0.514381502 | 0.597109632 | -0.08272813 | 0.523709647 |
| ENSG00000132623 | *ANKEF1* | -0.784195126 | -0.032740003 | -0.016916427 | -0.015823577 | 0.999933887 | 0.130194636 | 0.422449602 | -0.292254965 | 0.524149653 |
| ENSG00000286629 | *AC005803.1* | 1.437364094 | -0.083548782 | -0.090420201 | 0.006871419 | 0.999933887 | -0.20799695 | 0.037140298 | -0.245137247 | 0.524420004 |
| ENSG00000238825 | *RNVU1-2* | -1.792845644 | 0.128970132 | 0.256220258 | -0.127250126 | 0.999933887 | 0.20927309 | -0.245636618 | 0.454909709 | 0.524420004 |
| ENSG00000164597 | *COG5* | 3.717621483 | 0.018618007 | -0.012290409 | 0.030908417 | 0.999933887 | 0.121188181 | 0.225089911 | -0.10390173 | 0.524504884 |
| ENSG00000130703 | *OSBPL2* | 7.507235436 | -0.023128541 | 0.024481331 | -0.047609872 | 0.999933887 | -0.582063653 | -0.651605787 | 0.069542134 | 0.524705969 |
| ENSG00000225092 | *AL355336.1* | -0.655554202 | -0.058218433 | 0.273269145 | -0.331487577 | 0.999933887 | -0.739324166 | -0.434313707 | -0.305010459 | 0.524705969 |
| ENSG00000128973 | *CLN6* | 2.885115118 | -0.010618903 | 0.052768782 | -0.063387685 | 0.999933887 | -0.420146947 | -0.572453917 | 0.152306969 | 0.524705969 |
| ENSG00000213438 | *YBX2P1* | -0.072125165 | 0.03388275 | -0.082579042 | 0.116461792 | 0.999933887 | -0.284159638 | -0.739128383 | 0.454968745 | 0.524710976 |
| ENSG00000238164 | *TNFRSF14-AS* | 5.705140576 | 0.165409869 | 0.209109628 | -0.043699759 | 0.999933887 | 0.831424253 | 0.949725995 | -0.118301742 | 0.524798409 |
| ENSG00000198205 | *ZXDA* | 0.690098567 | 0.124660747 | 0.147962391 | -0.023301644 | 0.999933887 | 0.150762782 | -0.102678118 | 0.253440899 | 0.52484931 |
| ENSG00000123352 | *SPATS2* | -0.267198491 | 0.104964921 | 0.144396839 | -0.039431918 | 0.999933887 | -0.1020803 | 0.214344683 | -0.316424984 | 0.52484931 |
| ENSG00000079616 | *KIF22* | 3.651077799 | -0.019475571 | -0.018807914 | -0.000667658 | 0.999933887 | 0.135630428 | 0.033309107 | 0.102321322 | 0.52484931 |
| ENSG00000072134 | *EPN2* | 1.544748954 | -0.058537703 | 0.151378866 | -0.209916569 | 0.999933887 | 0.017932594 | -0.149144258 | 0.167076853 | 0.525235031 |
| ENSG00000273129 | *PACERR* | 0.982637913 | 0.355383502 | 0.356030625 | -0.000647123 | 0.999933887 | 0.708379922 | 0.380639197 | 0.327740725 | 0.525405887 |
| ENSG00000253328 | *SUMO2P19* | 2.280296175 | 0.001457641 | -0.010890989 | 0.01234863 | 0.999933887 | -0.48161379 | -0.350838583 | -0.130775206 | 0.525493786 |
| ENSG00000068912 | *ERLEC1* | 2.714257187 | -0.056870086 | 0.095834959 | -0.152705045 | 0.999933887 | -0.107646672 | 0.014600614 | -0.122247286 | 0.525671507 |
| ENSG00000264829 | *AC087749.2* | -0.443560561 | 0.118394343 | 0.332700841 | -0.214306498 | 0.999933887 | 0.34680311 | 0.826668008 | -0.479864898 | 0.525671507 |
| ENSG00000260943 | *LINC02555* | 0.476038689 | 0.067037424 | -0.21955332 | 0.286590745 | 0.999933887 | 0.194111234 | 0.476847562 | -0.282736329 | 0.525671507 |
| ENSG00000165181 | *SHOC1* | 0.790006281 | 0.078382505 | 0.266172418 | -0.187789913 | 0.999933887 | 0.100991291 | 0.500232472 | -0.399241181 | 0.525671507 |
| ENSG00000167325 | *RRM1* | 2.137536976 | 0.035103504 | -0.091777422 | 0.126880926 | 0.999933887 | 0.052627153 | 0.185225795 | -0.132598642 | 0.525671507 |
| ENSG00000102054 | *RBBP7* | 4.48347253 | 0.019778299 | -0.028548347 | 0.048326645 | 0.999933887 | 0.048073005 | 0.123867493 | -0.075794488 | 0.525671507 |
| ENSG00000100796 | *PPP4R3A* | 6.660323345 | -0.020559032 | -0.017739362 | -0.00281967 | 0.999933887 | -0.499433101 | -0.577843791 | 0.07841069 | 0.525671507 |
| ENSG00000177352 | *CCDC71* | 4.020096934 | -0.132521146 | -0.174895937 | 0.042374792 | 0.999933887 | -0.368011887 | -0.260072366 | -0.107939521 | 0.525718495 |
| ENSG00000267904 | *AC024075.1* | -0.105656223 | -0.177630677 | -0.062202159 | -0.115428518 | 0.999933887 | -0.429817053 | -0.132460136 | -0.297356917 | 0.52588913 |
| ENSG00000119699 | *TGFB3* | -0.066775194 | 0.109020437 | -0.054412302 | 0.163432739 | 0.999933887 | -0.390767219 | -0.116929761 | -0.273837458 | 0.52588913 |
| ENSG00000103423 | *DNAJA3* | 2.785416833 | -0.035389599 | -0.033984698 | -0.001404901 | 0.999933887 | 0.137461782 | 0.016804768 | 0.120657013 | 0.52588913 |
| ENSG00000269086 | *AC008555.2* | -0.492370636 | 0.164395314 | 0.433196319 | -0.268801005 | 0.999933887 | -0.027089317 | 0.240787221 | -0.267876538 | 0.52595401 |
| ENSG00000085563 | *ABCB1* | 1.949157371 | -0.174684385 | -0.088726438 | -0.085957947 | 0.999933887 | -0.119820402 | 0.0122417 | -0.132062102 | 0.52641766 |
| ENSG00000196998 | *WDR45* | 5.469781368 | -0.073708535 | 0.021145832 | -0.094854366 | 0.999933887 | -0.237687065 | -0.313694428 | 0.076007363 | 0.52641766 |
| ENSG00000126773 | *PCNX4* | 5.579718342 | -0.091096486 | -0.06665374 | -0.024442745 | 0.999933887 | -0.798268674 | -0.903668187 | 0.105399514 | 0.52641766 |
| ENSG00000105058 | *FAM32A* | 5.488111565 | -0.059400488 | -0.044581405 | -0.014819083 | 0.999933887 | -0.312473883 | -0.397377139 | 0.084903255 | 0.52641766 |
| ENSG00000277589 | *AC244093.4* | -0.387391306 | 0.084179312 | 0.003358811 | 0.0808205 | 0.999933887 | 0.20718077 | 0.552415232 | -0.345234462 | 0.52641766 |
| ENSG00000166532 | *RIMKLB* | 4.151523451 | -0.049750849 | -0.080799787 | 0.031048938 | 0.999933887 | -0.261420632 | -0.153877838 | -0.107542794 | 0.52641766 |
| ENSG00000100897 | *DCAF11* | 6.883631996 | 0.010500099 | 0.005673959 | 0.00482614 | 0.999933887 | -0.12352022 | -0.190943132 | 0.067422911 | 0.52641766 |
| ENSG00000127081 | *ZNF484* | 1.7926128 | -0.026718232 | -0.318646677 | 0.291928445 | 0.999933887 | -0.148645962 | 0.061643759 | -0.210289721 | 0.52641766 |
| ENSG00000237441 | *RGL2* | 7.373571088 | 0.015012634 | 0.082906303 | -0.067893669 | 0.999933887 | 0.207932656 | 0.146476636 | 0.06145602 | 0.526642699 |
| ENSG00000084754 | *HADHA* | 5.434318676 | -0.007304635 | -0.05108829 | 0.043783655 | 0.999933887 | -0.244824214 | -0.315763608 | 0.070939394 | 0.526642699 |
| ENSG00000073060 | *SCARB1* | 2.055533477 | 0.062914441 | 0.064610124 | -0.001695684 | 0.999933887 | -0.183944315 | -0.035144583 | -0.148799732 | 0.526756223 |
| ENSG00000146918 | *NCAPG2* | 0.990760038 | 0.033319898 | -0.237657953 | 0.270977851 | 0.999933887 | -0.031745134 | 0.250456183 | -0.282201317 | 0.526756223 |
| ENSG00000033327 | *GAB2* | 8.92749996 | 0.025200145 | 0.054532895 | -0.029332749 | 0.999933887 | -0.140508202 | -0.069360009 | -0.071148193 | 0.526830821 |
| ENSG00000150593 | *PDCD4* | 6.931190992 | -0.092834169 | -0.064180401 | -0.028653767 | 0.999933887 | 0.245178012 | 0.378292012 | -0.133114 | 0.526867567 |
| ENSG00000119446 | *RBM18* | 3.841801211 | -0.026187639 | 0.010903351 | -0.037090989 | 0.999933887 | -0.245466581 | -0.151921177 | -0.093545404 | 0.526867567 |
| ENSG00000138802 | *SEC24B* | 5.63878536 | -0.030694225 | -0.0434848 | 0.012790576 | 0.999933887 | -0.393183785 | -0.283898389 | -0.109285396 | 0.526960907 |
| ENSG00000124181 | *PLCG1* | 4.999744451 | 0.038505685 | 0.046220906 | -0.007715221 | 0.999933887 | 0.149054303 | 0.071522783 | 0.07753152 | 0.527056107 |
| ENSG00000105393 | *BABAM1* | 1.858601874 | -0.015073598 | 0.194578352 | -0.20965195 | 0.999933887 | 0.153112499 | -0.004333178 | 0.157445677 | 0.527081647 |
| ENSG00000120903 | *CHRNA2* | -0.283870347 | 0.173827766 | -0.210069536 | 0.383897302 | 0.999933887 | 1.44080393 | 1.774342429 | -0.333538499 | 0.527296524 |
| ENSG00000166575 | *TMEM135* | 1.04832297 | 0.003489545 | -0.114656443 | 0.118145989 | 0.999933887 | -0.16013792 | 0.079973909 | -0.240111828 | 0.527390324 |
| ENSG00000167721 | *TSR1* | 2.338425007 | 0.02506831 | -0.020853517 | 0.045921827 | 0.999933887 | 0.203560126 | 0.329070236 | -0.125510111 | 0.527425057 |
| ENSG00000276058 | *STMN1P1* | -0.108710172 | 0.059379152 | -0.264903113 | 0.324282264 | 0.999933887 | 0.231102299 | 0.522131088 | -0.291028789 | 0.527425057 |
| ENSG00000100604 | *CHGA* | -0.031030736 | 0.120749917 | 0.297114748 | -0.176364831 | 0.999933887 | 0.14193138 | -0.286259784 | 0.428191164 | 0.527580499 |
| ENSG00000151704 | *KCNJ1* | -1.27882575 | 0.074896804 | -0.07225117 | 0.147147974 | 0.999933887 | 0.134802064 | -0.387714185 | 0.522516249 | 0.527627732 |
| ENSG00000273320 | *AC007032.1* | 0.43468853 | 0.12953755 | 0.03037223 | 0.09916532 | 0.999933887 | 0.794817271 | 1.07348525 | -0.278667979 | 0.527641868 |
| ENSG00000128165 | *ADM2* | 0.287758646 | -0.134319104 | 0.019415243 | -0.153734347 | 0.999933887 | -0.323665034 | -0.086817255 | -0.236847779 | 0.527641868 |
| ENSG00000121594 | *CD80* | -1.378958052 | 0.224733691 | -0.619501282 | 0.844234974 | 0.999933887 | 1.941214945 | 2.368640659 | -0.427425714 | 0.527641868 |
| ENSG00000167232 | *ZNF91* | 2.731646223 | -0.014974461 | 0.043135089 | -0.058109551 | 0.999933887 | -0.180502086 | -0.016321731 | -0.164180355 | 0.527641868 |
| ENSG00000224003 | *YES1P1* | -1.155384774 | 0.017511966 | 0.545587437 | -0.528075472 | 0.999933887 | 0.77866637 | 1.099423403 | -0.320757034 | 0.527641868 |
| ENSG00000205581 | *HMGN1* | 6.289628042 | -0.021495323 | -0.012343084 | -0.009152239 | 0.999933887 | -0.225454167 | -0.290840545 | 0.065386378 | 0.527645099 |
| ENSG00000115904 | *SOS1* | 4.237268658 | 0.022419293 | -0.009063099 | 0.031482392 | 0.999933887 | -0.404192027 | -0.263772189 | -0.140419838 | 0.527686327 |
| ENSG00000163820 | *FYCO1* | 4.328821542 | -0.037691964 | 0.013504534 | -0.051196498 | 0.999933887 | -0.058247771 | 0.022063936 | -0.080311707 | 0.52781068 |
| ENSG00000196295 | *GARS1-DT* | 5.870335092 | 0.006607495 | -0.012578803 | 0.019186298 | 0.999933887 | -0.13142925 | -0.016820479 | -0.114608771 | 0.527849588 |
| ENSG00000164338 | *UTP15* | 2.24756614 | -0.076820872 | -0.122716128 | 0.045895256 | 0.999933887 | 0.134868338 | 0.300804177 | -0.16593584 | 0.528220325 |
| ENSG00000114541 | *FRMD4B* | 3.762064714 | 0.030647312 | -0.042362447 | 0.073009759 | 0.999933887 | 0.162368568 | 0.349131768 | -0.1867632 | 0.528220325 |
| ENSG00000147121 | *KRBOX4* | 2.536135653 | 0.011807795 | 0.008609401 | 0.003198394 | 0.999933887 | -0.014741525 | 0.088932327 | -0.103673852 | 0.528220325 |
| ENSG00000275895 | *U2AF1L5* | 2.880162162 | 0.008227293 | -0.014885716 | 0.023113009 | 0.999933887 | 0.003088665 | 0.13716794 | -0.134079275 | 0.528220325 |
| ENSG00000142185 | *TRPM2* | 6.222760427 | 0.081667812 | 0.030700568 | 0.050967245 | 0.999933887 | 0.873522556 | 0.97922543 | -0.105702874 | 0.5282805 |
| ENSG00000241889 | *AC079944.2* | -0.56807151 | -0.050254887 | -0.315004614 | 0.264749727 | 0.999933887 | -1.221039972 | -1.63031528 | 0.409275307 | 0.528509807 |
| ENSG00000182621 | *PLCB1* | 0.928622904 | -0.080961988 | -0.158095404 | 0.077133416 | 0.999933887 | -0.213969232 | -0.003524455 | -0.210444777 | 0.528534852 |
| ENSG00000259687 | *LINC01220* | 0.937812386 | 0.075417286 | 0.280869093 | -0.205451807 | 0.999933887 | 0.183215565 | -0.088363832 | 0.271579397 | 0.528551628 |
| ENSG00000106537 | *TSPAN13* | 0.733358736 | 0.096343935 | -0.130781319 | 0.227125254 | 0.999933887 | -0.050371477 | -0.316612169 | 0.266240692 | 0.529313547 |
| ENSG00000143641 | *GALNT2* | 4.187675405 | -0.028318264 | -0.035687575 | 0.007369311 | 0.999933887 | -0.292989415 | -0.372887656 | 0.079898241 | 0.529313547 |
| ENSG00000167685 | *ZNF444* | 2.453731437 | -0.005939242 | 0.080003839 | -0.08594308 | 0.999933887 | 0.109715817 | -0.032927982 | 0.142643799 | 0.529542928 |
| ENSG00000282936 | *AC004706.3* | -0.74460185 | 0.283624842 | 0.362926719 | -0.079301877 | 0.999933887 | -0.515405343 | -0.997145663 | 0.481740321 | 0.529594073 |
| ENSG00000164082 | *GRM2* | -0.280262436 | 0.105166913 | 0.206969074 | -0.101802162 | 0.999933887 | -0.171678879 | 0.157608523 | -0.329287403 | 0.529594073 |
| ENSG00000131043 | *AAR2* | 2.519289956 | -0.118193409 | -0.164766858 | 0.046573449 | 0.999933887 | 0.086825552 | 0.210746797 | -0.123921245 | 0.529647362 |
| ENSG00000173226 | *IQCB1* | 2.56585651 | -0.099574527 | -0.083853847 | -0.015720679 | 0.999933887 | -0.021334574 | 0.12166698 | -0.143001554 | 0.529647362 |
| ENSG00000155542 | *SETD9* | 0.695592039 | 0.113938559 | 0.086272881 | 0.027665679 | 0.999933887 | 0.055415241 | 0.248197235 | -0.192781994 | 0.529647362 |
| ENSG00000143797 | *MBOAT2* | 5.127699501 | -0.035894687 | -0.045035921 | 0.009141234 | 0.999933887 | -0.293142746 | -0.40272667 | 0.109583925 | 0.529770572 |
| ENSG00000020426 | *MNAT1* | -0.217018712 | -0.083623956 | -0.019947935 | -0.063676021 | 0.999933887 | -0.056106395 | 0.239049547 | -0.295155942 | 0.529770572 |
| ENSG00000175390 | *EIF3F* | 5.278059362 | -0.015170129 | -0.015243325 | 7.3195e-05 | 0.999933887 | 0.030542851 | 0.104645864 | -0.074103013 | 0.529770572 |
| ENSG00000106100 | *NOD1* | 2.501778758 | 0.030170353 | -0.015517221 | 0.045687574 | 0.999933887 | -0.123333303 | -0.285987643 | 0.16265434 | 0.529815983 |
| ENSG00000170425 | *ADORA2B* | 1.949467319 | 0.020044846 | 0.048682709 | -0.028637862 | 0.999933887 | -0.012379809 | -0.166044802 | 0.153664993 | 0.529815983 |
| ENSG00000273669 | *AC015819.1* | 3.065547757 | 0.013156205 | 0.039840887 | -0.026684681 | 0.999933887 | -0.154352216 | -0.030951777 | -0.123400439 | 0.529847546 |
| ENSG00000004777 | *ARHGAP33* | 1.964520087 | 0.026724544 | 0.160019587 | -0.133295043 | 0.999933887 | 0.053748117 | 0.194122289 | -0.140374172 | 0.529891715 |
| ENSG00000177119 | *ANO6* | 3.136358684 | 0.082704809 | -0.077982965 | 0.160687774 | 0.999933887 | -0.056192013 | 0.063818577 | -0.12001059 | 0.530037343 |
| ENSG00000029534 | *ANK1* | 0.725101593 | 0.167022772 | 0.089958439 | 0.077064333 | 0.999933887 | -0.174563715 | 0.03975099 | -0.214314705 | 0.530304342 |
| ENSG00000226833 | *AC092164.1* | 0.483488014 | 0.077365084 | 0.203718179 | -0.126353095 | 0.999933887 | 0.00029668 | 0.226948219 | -0.226651539 | 0.530369919 |
| ENSG00000167088 | *SNRPD1* | 1.995275691 | -0.026254064 | 0.009674258 | -0.035928323 | 0.999933887 | 0.106652226 | 0.240522129 | -0.133869904 | 0.530369919 |
| ENSG00000132326 | *PER2* | 3.225517594 | 0.002504984 | 0.032520646 | -0.030015663 | 0.999933887 | 0.219922746 | 0.333766455 | -0.113843709 | 0.530369919 |
| ENSG00000265625 | *AC104564.5* | 1.816965525 | -0.051167519 | -0.058767664 | 0.007600145 | 0.999933887 | -0.288793559 | -0.072318954 | -0.216474606 | 0.530667155 |
| ENSG00000271503 | *CCL5* | 7.24993542 | 0.015555383 | 0.006189691 | 0.009365692 | 0.999933887 | 0.112321512 | 0.206324559 | -0.094003047 | 0.530851357 |

| | | | | | | | | | | |
|---|---|---|---|---|---|---|---|---|---|---|
| ENSG00000167130 | *DOLPP1* | 0.2653315 | 0.081398485 | -0.13692618 | 0.218324665 | 0.999933887 | 0.055635826 | -0.160292462 | 0.215928289 | 0.530893984 |
| ENSG00000267534 | *S1PR2* | 1.266126991 | -0.060040178 | -0.068358788 | 0.00831861 | 0.999933887 | 0.280558353 | 0.489241546 | -0.208683193 | 0.530893984 |
| ENSG00000168610 | *STAT3* | 8.605660758 | -0.017703215 | -0.007342673 | -0.010360542 | 0.999933887 | 0.130315869 | 0.196171776 | -0.065855906 | 0.530893984 |
| ENSG00000100360 | *IFT27* | 2.120285953 | 0.031509948 | 0.166197522 | -0.134687574 | 0.999933887 | 0.00652816 | -0.121390132 | 0.127918292 | 0.530893984 |
| ENSG00000240970 | *RPL23AP64* | 0.90431075 | -0.036565062 | -0.110663383 | 0.074098322 | 0.999933887 | 0.006432349 | -0.174527263 | 0.180959612 | 0.530893984 |
| ENSG00000206149 | *HERC2P9* | 0.516302662 | 0.048483977 | 0.087572032 | -0.039088054 | 0.999933887 | -0.017653825 | 0.207169442 | -0.224823266 | 0.530893984 |
| ENSG00000166825 | *ANPEP* | 8.844843386 | -0.010805009 | -0.031765328 | 0.020960319 | 0.999933887 | -0.28967422 | -0.381946004 | 0.092271784 | 0.530893984 |
| ENSG00000130638 | *ATXN10* | 2.260880034 | -0.01332005 | -0.024848279 | 0.011528229 | 0.999933887 | -0.013994129 | 0.128954455 | -0.142948584 | 0.530893984 |
| ENSG00000119314 | *PTBP3* | 7.286209082 | -0.008522968 | -0.017496055 | 0.008973087 | 0.999933887 | -0.699632984 | -0.591857487 | -0.107775498 | 0.530893984 |
| ENSG00000205643 | *CDPF1* | -0.259712139 | -0.288670671 | 0.164808386 | -0.453479057 | 0.999933887 | 0.010305744 | -0.206652567 | 0.216958311 | 0.531033933 |
| ENSG00000127419 | *TMEM175* | 4.54628663 | 0.063993646 | 0.033258945 | 0.030734701 | 0.999933887 | 0.092132334 | 0.008907343 | 0.083224991 | 0.531033933 |
| ENSG00000164051 | *CCDC51* | 0.164885861 | -0.129947063 | -0.069626272 | -0.06032079 | 0.999933887 | 0.218278164 | -0.037850861 | 0.256129025 | 0.531033933 |
| ENSG00000006704 | *GTF2IRD1* | -1.088717141 | -0.386505105 | 0.18948914 | -0.575994245 | 0.999933887 | -0.497344336 | -0.216901621 | -0.280442714 | 0.531033933 |
| ENSG00000125482 | *TTF1* | 3.462057508 | -0.06293226 | -0.055330167 | -0.007602093 | 0.999933887 | -0.308497895 | -0.416415199 | 0.107917304 | 0.531033933 |
| ENSG00000153140 | *CETN3* | -0.893021359 | 0.141003527 | -0.327794279 | 0.468797806 | 0.999933887 | -0.249443317 | 0.061053088 | -0.310496405 | 0.531033933 |
| ENSG00000166446 | *CDYL2* | 2.560803581 | -0.020014626 | 0.040138831 | -0.060153457 | 0.999933887 | -0.058328083 | 0.067460396 | -0.12578848 | 0.531033933 |
| ENSG00000170584 | *NUDCD2* | 1.968292479 | 0.010916413 | -0.118791638 | 0.129708051 | 0.999933887 | 0.074951701 | 0.23136963 | -0.156417929 | 0.531033933 |
| ENSG00000282939 | *TRBV7-2* | 0.732973618 | -0.076686627 | -0.073128208 | -0.003558419 | 0.999933887 | 0.261595592 | 0.450530562 | -0.18893497 | 0.531139215 |
| ENSG00000165434 | *PGM2L1* | 1.668398019 | 0.064740572 | 0.011457771 | 0.053282802 | 0.999933887 | -0.292022545 | -0.083487449 | -0.208535096 | 0.531139215 |
| ENSG00000048342 | *CC2D2A* | 3.934603125 | 0.063826187 | 0.316123731 | -0.252297544 | 0.999933887 | -0.088125757 | 0.129298134 | -0.217423891 | 0.531171111 |
| ENSG00000104946 | *TBC1D17* | 5.48158862 | -0.004474926 | 0.080040291 | -0.084515217 | 0.999933887 | 0.058575115 | 0.122757796 | -0.06418268 | 0.531347555 |
| ENSG00000249042 | *AC008771.1* | 3.599384004 | 0.165277373 | 0.345496753 | -0.18021938 | 0.999933887 | 0.569901743 | 0.722926117 | -0.153024375 | 0.531435371 |
| ENSG00000187210 | *GCNT1* | 1.512289763 | -0.094605465 | 0.026368207 | -0.120973672 | 0.999933887 | -0.231179242 | -0.097198118 | -0.133981125 | 0.531487711 |
| ENSG00000179104 | *TMTC2* | 0.564132512 | 0.055993532 | 0.127259659 | -0.071266127 | 0.999933887 | 0.256865198 | -0.0420699 | 0.298935098 | 0.531670575 |
| ENSG00000109519 | *GRPEL1* | 3.985915678 | 0.01439431 | 0.115181059 | -0.100786749 | 0.999933887 | 0.121589928 | 0.235882546 | -0.114292618 | 0.531670575 |
| ENSG00000187624 | *C17orf97* | -0.403503818 | -0.172765992 | -0.238919607 | 0.066153615 | 0.999933887 | -0.134992761 | -0.415893693 | 0.280900932 | 0.531754369 |
| ENSG00000238024 | *DDX39BP2* | -2.151899277 | -0.069208684 | 0.498892258 | -0.568100941 | 0.999933887 | -0.763378992 | -0.441557321 | -0.321821671 | 0.532188661 |
| ENSG00000244313 | *AC024293.1* | -0.405192716 | -0.106062505 | 0.074566616 | -0.180629121 | 0.999933887 | -0.034456217 | -0.361214038 | 0.326757822 | 0.532358981 |
| ENSG00000210196 | *MT-TP* | 1.822861493 | -0.057939546 | -0.076978664 | 0.019039118 | 0.999933887 | 0.112177001 | 0.314318347 | -0.202141346 | 0.532358981 |
| ENSG00000130024 | *PHF10* | 3.879728014 | -0.024609033 | -0.035188955 | 0.010579922 | 0.999933887 | -0.017647787 | 0.088185967 | -0.105833754 | 0.532358981 |
| ENSG00000008256 | *CYTH3* | 3.49812401 | -0.008057465 | -0.027269247 | 0.019211782 | 0.999933887 | 0.246569269 | 0.36682961 | -0.120260341 | 0.532358981 |
| ENSG00000119326 | *CTNNAL1* | 1.788908255 | -0.013432183 | -0.054622934 | 0.041190751 | 0.999933887 | -0.055365748 | -0.214776552 | 0.159410804 | 0.532396934 |
| ENSG00000174775 | *HRAS* | 1.791852163 | 0.012214967 | 0.072915046 | -0.06070008 | 0.999933887 | 0.047647702 | -0.125988867 | 0.173636569 | 0.532396934 |
| ENSG00000178338 | *ZNF354B* | 0.548461419 | 0.130238231 | -0.115770077 | 0.246008307 | 0.999933887 | -0.088196284 | 0.117997856 | -0.206194139 | 0.532422682 |
| ENSG00000229097 | *CALM2P2* | 1.845753031 | 0.120897868 | 0.084470829 | 0.036427038 | 0.999933887 | 0.076393138 | 0.261806118 | -0.185412979 | 0.532602686 |
| ENSG00000163354 | *DCST2* | 0.640340024 | -0.075618971 | 0.16180858 | -0.237427551 | 0.999933887 | 0.107604899 | -0.144698073 | 0.252302972 | 0.532624457 |
| ENSG00000138166 | *DUSP5* | 5.69178993 | 0.111809839 | 0.156212762 | -0.044402923 | 0.999933887 | 0.81414945 | 0.941772838 | -0.127623389 | 0.53265516 |
| ENSG00000278997 | *AL662907.1* | -0.345183681 | 0.150014476 | 0.071818294 | 0.078196182 | 0.999933887 | 0.032610367 | -0.280033497 | 0.312643864 | 0.533008566 |
| ENSG00000211734 | *TRBV5-1* | 1.294477314 | 0.045527798 | -0.021436873 | 0.066964671 | 0.999933887 | 0.269994925 | 0.101205688 | 0.168789238 | 0.533008566 |
| ENSG00000239713 | *APOBEC3G* | 4.339547593 | -0.03610546 | -0.060817772 | 0.024712312 | 0.999933887 | -0.019283738 | -0.116538795 | 0.097255058 | 0.533409358 |
| ENSG00000159086 | *PAXBP1* | 4.802967737 | -0.078017006 | -0.161397459 | 0.083380453 | 0.999933887 | -0.338053717 | -0.171990116 | -0.166063601 | 0.533733195 |
| ENSG00000213199 | *ASIC3* | 0.131949904 | -0.033821262 | 0.468832561 | -0.502653823 | 0.999933887 | 0.161581807 | -0.049436664 | 0.211018471 | 0.533733195 |
| ENSG00000138385 | *SSB* | 3.032476077 | -0.083470771 | -0.107271481 | 0.023800709 | 0.999933887 | -0.037724919 | 0.08725275 | -0.124977669 | 0.533757682 |
| ENSG00000272578 | *AP000347.1* | 2.456843523 | -0.067469739 | -0.089361807 | 0.021892068 | 0.999933887 | -0.055993528 | 0.088916004 | -0.144909533 | 0.533757682 |
| ENSG00000140104 | *CLBA1* | 0.548193022 | 0.11004085 | -0.031677396 | 0.141718245 | 0.999933887 | -0.250962185 | -0.47560561 | 0.224643426 | 0.533757682 |
| ENSG00000144580 | *CNOT9* | 3.992700483 | -0.023962588 | 0.042805668 | -0.066768256 | 0.999933887 | 0.031667396 | 0.105438153 | -0.073770757 | 0.533757682 |
| ENSG00000110717 | *NDUFS8* | 3.545439137 | -0.007691415 | 0.042814533 | -0.050505948 | 0.999933887 | 0.113054953 | 0.027745441 | 0.085309512 | 0.533757682 |
| ENSG00000132640 | *BTBD3* | 1.176331402 | -0.114134577 | -0.192714628 | 0.078580051 | 0.999933887 | -0.917005214 | -0.735968916 | -0.181036298 | 0.533776035 |
| ENSG00000128059 | *PPAT* | 0.530907054 | -0.136510414 | -0.499871315 | 0.3633609 | 0.999933887 | -0.088468964 | 0.136553931 | -0.225022894 | 0.533776035 |
| ENSG00000109466 | *KLHL2* | 5.559220556 | -0.034091939 | -0.038224958 | 0.004133019 | 0.999933887 | -0.427439069 | -0.340804538 | -0.086634531 | 0.533776035 |
| ENSG00000234663 | *LINC01934* | 0.427583759 | -0.07578827 | -0.17695232 | 0.101164049 | 0.999933887 | -0.280167776 | -0.059427235 | -0.220740541 | 0.533776035 |
| ENSG00000226396 | *AL031727.1* | -0.280887528 | 0.035925194 | -0.041391106 | 0.0773163 | 0.999933887 | 0.319077794 | 0.007846448 | 0.311231346 | 0.533776035 |
| ENSG00000107833 | *NPM3* | 0.963671625 | 0.010849404 | -0.016982233 | 0.027831637 | 0.999933887 | 0.041252923 | -0.155502809 | 0.196755732 | 0.533776035 |
| ENSG00000144134 | *RABL2A* | 1.125839343 | -0.020760852 | -0.241960699 | 0.221199847 | 0.999933887 | -0.102552351 | -0.244942568 | 0.142390216 | 0.534101185 |
| ENSG00000256433 | *AC005840.2* | -0.516444203 | 0.312015496 | -0.012331583 | 0.324347079 | 0.999933887 | -0.423892916 | -0.864876154 | 0.440983238 | 0.534188357 |
| ENSG00000125434 | *SLC25A35* | -0.342963537 | 0.165914013 | -0.020926159 | 0.186840172 | 0.999933887 | 0.065114447 | -0.237001292 | 0.302115739 | 0.534188357 |
| ENSG00000073614 | *KDM5A* | 7.205639989 | -0.071833111 | -0.131001544 | 0.059168433 | 0.999933887 | -0.180492117 | -0.103686265 | -0.076805852 | 0.534355158 |
| ENSG00000147044 | *CASK* | 2.564667449 | -0.05526224 | -0.084572546 | 0.029310306 | 0.999933887 | -0.113525612 | -0.002200199 | -0.111325413 | 0.534355158 |
| ENSG00000132591 | *ERAL1* | 1.301072641 | -0.064658945 | 0.094018549 | -0.158677494 | 0.999933887 | 0.502537412 | 0.704572371 | -0.202034959 | 0.534355158 |
| ENSG00000182903 | *ZNF721* | 2.344603138 | -0.188797089 | -0.172291464 | -0.016505625 | 0.999933887 | -0.564845073 | -0.348594128 | -0.216250945 | 0.534684219 |
| ENSG00000088256 | *GNA11* | 0.374167166 | -0.031113132 | 0.089253074 | -0.120366206 | 0.999933887 | -0.067939849 | 0.145766205 | -0.213706054 | 0.534684219 |
| ENSG00000165490 | *DDIAS* | 0.020042625 | -0.029013104 | 0.088950653 | -0.117963756 | 0.999933887 | -0.153186677 | -0.487315445 | 0.334128768 | 0.534856324 |
| ENSG00000173914 | *RBM4B* | 1.722696629 | 0.031633687 | -0.052960879 | 0.084594566 | 0.999933887 | -0.165542507 | -0.307399698 | 0.141857191 | 0.535307366 |
| ENSG00000085274 | *MYNN* | 3.679909094 | -0.020165227 | -0.076879192 | 0.056713966 | 0.999933887 | 0.011242136 | 0.140344219 | -0.129102083 | 0.53553461 |
| ENSG00000240403 | *KIR3DL2* | 0.220425383 | 0.013513557 | 0.084976325 | -0.071462768 | 0.999933887 | 0.036253212 | 0.272993193 | -0.236739981 | 0.535568668 |
| ENSG00000279491 | *AP003733.3* | 5.060934196 | 0.285879348 | 0.199069232 | 0.086810115 | 0.999933887 | 1.094407467 | 1.22816112 | -0.133753653 | 0.535704774 |
| ENSG00000072071 | *ADGRL1* | 2.876446002 | 0.076736846 | 0.077791534 | -0.001054687 | 0.999933887 | 0.131339136 | 0.24236253 | -0.111023395 | 0.535704774 |
| ENSG00000111224 | *PARP11* | 2.598133796 | 0.06403045 | -0.026265074 | 0.090295524 | 0.999933887 | -0.291817552 | -0.16540186 | -0.126415692 | 0.535704774 |
| ENSG00000137513 | *NARS2* | 0.423226908 | -0.099813818 | -0.070735082 | -0.029078736 | 0.999933887 | 0.014943136 | 0.230878972 | -0.215935837 | 0.535704774 |
| ENSG00000126005 | *MMP24OS* | 5.072216257 | 0.018625863 | -0.002829662 | 0.021455525 | 0.999933887 | -0.2118434 | -0.295519628 | 0.083676228 | 0.535704774 |
| ENSG00000163565 | *IFI16* | 7.389644226 | -0.118408244 | -0.160978551 | 0.042570307 | 0.999933887 | -0.817587748 | -0.929144316 | 0.111556567 | 0.535722959 |
| ENSG00000266913 | *LINC01841* | -0.052613722 | 0.045551803 | 0.220543902 | -0.1749921 | 0.999933887 | 1.060520024 | 1.367239445 | -0.306719421 | 0.535799493 |
| ENSG00000118507 | *AKAP7* | -0.527380874 | -0.067897627 | 0.130343514 | -0.198241141 | 0.999933887 | -0.145205426 | 0.276675181 | -0.421880608 | 0.535821496 |
| ENSG00000229598 | *PRDX3P1* | 2.391150233 | 0.105031531 | 0.071888328 | 0.033143202 | 0.999933887 | 0.114794836 | 0.266842575 | -0.152047739 | 0.535854823 |
| ENSG00000203667 | *COX20* | 0.137768051 | -0.067650597 | 0.158758317 | -0.226408914 | 0.999933887 | 0.15597497 | -0.109565472 | 0.265540442 | 0.535897078 |
| ENSG00000180846 | *CSNK1G2-AS* | 2.394691461 | -0.101666392 | -0.008342341 | -0.093324051 | 0.999933887 | 0.050749368 | -0.155639338 | 0.206388706 | 0.535915083 |
| ENSG00000164902 | *PHAX* | 2.048460012 | -0.063999076 | -0.111895371 | 0.047896296 | 0.999933887 | -0.10477416 | 0.031009097 | -0.135783257 | 0.535915083 |
| ENSG00000236552 | *RPL13AP5* | 1.330227331 | -0.049211092 | -0.137871194 | 0.088660101 | 0.999933887 | 0.221429186 | 0.012877389 | 0.208551797 | 0.535915083 |
| ENSG00000188542 | *DUSP28* | 2.816701907 | -0.134481769 | -0.059170809 | -0.07531096 | 0.999933887 | -0.4705816 | -0.604140233 | 0.133558633 | 0.536045778 |
| ENSG00000185085 | *INTS5* | 2.732180864 | 0.049739661 | 0.009357535 | 0.040382126 | 0.999933887 | 0.242022374 | 0.117763737 | 0.124258637 | 0.536045778 |
| ENSG00000123146 | *ADGRE5* | 9.546762568 | 0.017721455 | -0.010283431 | 0.028004886 | 0.999933887 | 0.213080969 | 0.283749068 | -0.0706681 | 0.536045778 |
| ENSG00000225808 | *DNAJC19P5* | 2.232123786 | -0.079568172 | 0.145068028 | -0.2246362 | 0.999933887 | 0.487505058 | 0.696488327 | -0.208983269 | 0.536085289 |
| ENSG00000130821 | *SLC6A8* | 3.148749709 | 0.121177204 | 0.107820926 | 0.013356278 | 0.999933887 | -0.05211642 | -0.156627832 | 0.104511412 | 0.536706761 |
| ENSG00000164169 | *PRMT9* | 2.674291257 | -0.038440851 | -0.107585604 | 0.069144753 | 0.999933887 | -0.037452097 | -0.193316448 | 0.155864351 | 0.537026953 |
| ENSG00000164258 | *NDUFS4* | 0.578814243 | 0.013349053 | -0.099503232 | 0.112852286 | 0.999933887 | 0.001074302 | 0.20322965 | -0.202155348 | 0.537155668 |
| ENSG00000149554 | *CHEK1* | -0.419888025 | 0.125877784 | 0.024268885 | 0.1016089 | 0.999933887 | 0.296048758 | 0.627157684 | -0.331108927 | 0.537185553 |
| ENSG00000196850 | *PPTC7* | 6.643551414 | -0.020200118 | 0.018860244 | -0.039060362 | 0.999933887 | -0.248156412 | -0.342139239 | 0.093982827 | 0.537588977 |
| ENSG00000116001 | *TIA1* | 4.081241626 | 0.013324614 | -0.027053658 | 0.040378272 | 0.999933887 | -0.29908621 | -0.158204079 | -0.140882131 | 0.537588977 |
| ENSG00000173141 | *MRPL57* | 1.889758573 | 0.124254176 | -0.100581997 | 0.224836173 | 0.999933887 | 0.150214198 | 0.00682574 | 0.143388457 | 0.537601961 |
| ENSG00000173451 | *THAP2* | 1.713551038 | 0.098738844 | 0.157869966 | -0.059131122 | 0.999933887 | 0.595771562 | 0.84228784 | -0.246516278 | 0.537601961 |
| ENSG00000158473 | *CD1D* | -1.493756734 | 0.037502853 | -0.198997694 | 0.236500547 | 0.999933887 | -0.886856215 | -0.276318276 | -0.61053794 | 0.53785075 |
| ENSG00000197563 | *PIGN* | 1.562338307 | -0.028511256 | -0.096641158 | 0.068129903 | 0.999933887 | -0.172944975 | -0.365525635 | 0.19258066 | 0.538622202 |
| ENSG00000179085 | *DPM3* | 1.879267666 | -0.009413838 | -0.024684177 | 0.015270339 | 0.999933887 | -0.033491768 | -0.217109054 | 0.183617286 | 0.538892891 |
| ENSG00000122203 | *KIAA1191* | 3.587650263 | 0.025391738 | 0.038738761 | -0.013347023 | 0.999933887 | -0.042422089 | -0.143013026 | 0.100590937 | 0.539048536 |
| ENSG00000214029 | *ZNF891* | 1.363988543 | 0.012333443 | 0.007617729 | 0.004715714 | 0.999933887 | -0.223875663 | 0.012628199 | -0.236503862 | 0.539095215 |
| ENSG00000162642 | *C1orf52* | 4.262633368 | 0.006648666 | -0.020178831 | 0.026827497 | 0.999933887 | -0.161415028 | -0.090226356 | -0.071188672 | 0.539476704 |
| ENSG00000113300 | *CNOT6* | 3.472592269 | -0.119882788 | -0.062167173 | -0.057715616 | 0.999933887 | -0.20152934 | -0.094132848 | -0.107396492 | 0.539492111 |
| ENSG00000171311 | *EXOSC1* | 2.273888469 | -0.032264688 | -0.001024631 | -0.031240056 | 0.999933887 | -0.032975542 | -0.176658621 | 0.143683079 | 0.539492111 |
| ENSG00000174255 | *ZNF80* | -0.04344884 | 0.021367428 | -0.557490133 | 0.578857561 | 0.999933887 | -0.077448837 | 0.204151837 | -0.281600674 | 0.539612046 |
| ENSG00000227124 | *ZNF717* | -0.635826939 | -0.002012854 | 0.106390296 | -0.10840315 | 0.999933887 | -0.041845743 | -0.330160761 | 0.288315017 | 0.539627511 |
| ENSG00000205485 | *AC004980.1* | 2.474163142 | 0.043682804 | 0.074346653 | -0.03066385 | 0.999933887 | -0.021528885 | -0.184069568 | 0.162540683 | 0.539657763 |
| ENSG00000066654 | *THUMPD1* | 4.603188849 | 0.018050039 | -0.085581826 | 0.103631865 | 0.999933887 | -0.444869629 | -0.580998838 | 0.136129208 | 0.539657763 |
| ENSG00000128011 | *LRFN1* | 4.856870022 | 0.024007847 | 0.056656473 | -0.032648626 | 0.999933887 | -0.787069216 | -0.898068001 | 0.110998784 | 0.539807923 |
| ENSG00000225690 | *TREML5P* | 2.669277858 | 0.116163737 | 0.247993895 | -0.131830157 | 0.999933887 | 0.652638328 | 0.839096406 | -0.186458078 | 0.540298336 |
| ENSG00000231799 | *PA2G4P6* | 0.109307642 | 0.135017138 | 0.251545954 | -0.116528816 | 0.999933887 | 0.108341783 | -0.221222262 | 0.329564045 | 0.540298336 |

| | | | | | | | | | | |
|---|---|---|---|---|---|---|---|---|---|---|
| ENSG00000141510 | *TP53* | 3.054390174 | -0.047137704 | 0.125072478 | -0.172210182 | 0.999933887 | 0.19323591 | 0.097831953 | 0.095403957 | 0.540323893 |
| ENSG00000162783 | *IER5* | 7.907271112 | 0.165141407 | 0.203442677 | -0.03830127 | 0.999933887 | 2.24803212 | 2.414453499 | -0.166421378 | 0.540471919 |
| ENSG00000286608 | *AC010201.3* | 0.106943209 | -0.089798326 | -0.006351234 | -0.083447092 | 0.999933887 | 0.186171989 | 0.515155772 | -0.328983783 | 0.540471919 |
| ENSG00000151414 | *NEK7* | 5.692236488 | -0.053701332 | -0.071039844 | 0.017338512 | 0.999933887 | -0.62702446 | -0.487722992 | -0.139301468 | 0.540471919 |
| ENSG00000101203 | *COL20A1* | -0.646116046 | 0.296084602 | 0.18346942 | 0.112615182 | 0.999933887 | 1.529033025 | 1.765254048 | -0.236221023 | 0.540699403 |
| ENSG00000121851 | *POLR3GL* | 2.064554999 | -0.016406729 | -0.07683524 | 0.060428512 | 0.999933887 | -0.09847978 | -0.278528854 | 0.180049074 | 0.540813991 |
| ENSG00000174013 | *FBXO45* | 1.272020183 | 0.057295409 | 0.067710678 | -0.010415269 | 0.999933887 | -0.1491836 | -0.310059125 | 0.160875524 | 0.541150586 |
| ENSG00000026025 | *VIM* | 8.12292753 | 0.034022577 | 0.003419783 | 0.030602794 | 0.999933887 | 0.033415255 | 0.103819402 | -0.070404147 | 0.541212124 |
| ENSG00000224116 | *INHBA-AS1* | -0.611759192 | -0.181845136 | -0.287324046 | 0.10547891 | 0.999933887 | 0.824076409 | 1.124652831 | -0.300576422 | 0.541269997 |
| ENSG00000186230 | *ZNF749* | -0.233590695 | -0.01074155 | 0.139973919 | -0.150715469 | 0.999933887 | 0.237797191 | -0.012418684 | 0.250215876 | 0.541539986 |
| ENSG00000069696 | *DRD4* | -0.684047891 | 0.157738534 | -0.275933364 | 0.433671898 | 0.999933887 | -0.098862153 | 0.308087167 | -0.40694932 | 0.541645101 |
| ENSG00000237118 | *CYP2F2P* | 0.812755458 | -0.02090563 | -0.092100411 | 0.071194781 | 0.999933887 | 0.044271914 | 0.275393084 | -0.23112117 | 0.541645101 |
| ENSG00000196437 | *ZNF569* | 0.047573064 | 0.098300992 | -0.046268485 | 0.144569476 | 0.999933887 | -0.104825587 | 0.208344473 | -0.31317006 | 0.541680694 |
| ENSG00000196526 | *AFAP1* | -0.57916891 | -0.234871461 | -0.176371961 | -0.0584995 | 0.999933887 | -0.075250185 | 0.265942748 | -0.341192933 | 0.541726018 |
| ENSG00000185495 | *AC138393.1* | 1.008449191 | 0.086061092 | 0.154479038 | -0.068417946 | 0.999933887 | 0.03189945 | 0.272534942 | -0.240635492 | 0.541726018 |
| ENSG00000165121 | *AL353743.1* | 0.547576277 | -0.216933441 | -0.015666137 | -0.201267305 | 0.999933887 | -0.20564462 | 0.038207752 | -0.243852372 | 0.541778929 |
| ENSG00000119408 | *NEK6* | 5.874339181 | -0.05467874 | 0.037423968 | -0.092102708 | 0.999933887 | -1.023806988 | -1.131766521 | 0.107959533 | 0.541778929 |
| ENSG00000259079 | *AC005476.1* | -0.809202671 | 0.315421316 | 0.364293959 | -0.048872643 | 0.999933887 | 0.922412758 | 0.403500635 | 0.518912122 | 0.541841106 |
| ENSG00000280828 | *AC090114.3* | 0.645753984 | 0.215129554 | -0.013588508 | 0.228718062 | 0.999933887 | -0.125484223 | -0.328654557 | 0.203170334 | 0.541880849 |
| ENSG00000005379 | *TSPOAP1* | 4.768286396 | 0.07234599 | 0.10371541 | -0.03136942 | 0.999933887 | 0.155073369 | 0.057245682 | 0.097827686 | 0.542053667 |
| ENSG00000013619 | *MAMLD1* | 1.040982749 | 0.248368754 | 0.265693595 | -0.017324841 | 0.999933887 | -0.005522222 | 0.157358344 | -0.162880567 | 0.542053667 |
| ENSG00000173442 | *EHBP1L1* | 8.268678397 | 0.092643641 | 0.10457272 | -0.011929079 | 0.999933887 | 0.218118489 | 0.346623295 | -0.128504806 | 0.542053667 |
| ENSG00000019582 | *CD74* | 8.665207372 | 0.042609252 | 0.00133684 | 0.041272411 | 0.999933887 | 0.064301521 | 0.137912233 | -0.073610713 | 0.542053667 |
| ENSG00000100225 | *FBXO7* | 5.146802319 | -0.033758471 | -0.071692405 | 0.037933934 | 0.999933887 | -0.102817459 | -0.167264999 | 0.06444754 | 0.542053667 |
| ENSG00000198142 | *SOWAHC* | -0.27222738 | -0.276285213 | 0.12008533 | -0.396370543 | 0.999933887 | -0.764618443 | -0.98492049 | 0.220302048 | 0.542053667 |
| ENSG00000163221 | *S100A12* | 6.417815678 | 0.026173551 | 0.036973263 | -0.010799712 | 0.999933887 | 0.132066102 | 0.243409006 | -0.111342904 | 0.542053667 |
| ENSG00000085433 | *WDR47* | 3.840836273 | -0.014925618 | 0.030007146 | -0.044932764 | 0.999933887 | -0.138514328 | -0.240653198 | 0.10213887 | 0.542053667 |
| ENSG00000116127 | *ALMS1* | 2.638923065 | -0.020417677 | 0.074660603 | -0.09507828 | 0.999933887 | -0.164530724 | 0.054131052 | -0.218661776 | 0.542053667 |
| ENSG00000075539 | *FRYL* | 5.452431705 | 0.000875102 | -0.116104577 | 0.116979678 | 0.999933887 | -0.462970315 | -0.249130404 | -0.213839911 | 0.542053667 |
| ENSG00000038427 | *VCAN* | 1.693787103 | -0.000240796 | -0.036872733 | 0.036631937 | 0.999933887 | -2.688808249 | -3.161240944 | 0.472432695 | 0.542053667 |
| ENSG00000134597 | *RBMX2* | 2.113052495 | -0.079114827 | -0.055531129 | -0.023583698 | 0.999933887 | -0.080000422 | -0.230498793 | 0.150498371 | 0.542096648 |
| ENSG00000151576 | *QTRT2* | 3.021999995 | -0.020910386 | 0.035927049 | -0.056837435 | 0.999933887 | 0.002757992 | 0.105541802 | -0.10278381 | 0.542186471 |
| ENSG00000268173 | *AC007192.1* | 0.081540838 | 0.328191365 | 0.465941024 | -0.137749659 | 0.999933887 | 1.386249772 | 1.721688393 | -0.335438621 | 0.542331724 |
| ENSG00000119778 | *ATAD2B* | 4.152854199 | 0.060964782 | -0.036747985 | 0.097712767 | 0.999933887 | -0.182757413 | -0.022063378 | -0.160694035 | 0.542331724 |
| ENSG00000177873 | *ZNF619* | 0.682368356 | 0.187025448 | 0.054859943 | 0.132165505 | 0.999933887 | 0.0070228 | -0.195230782 | 0.202253582 | 0.542339935 |
| ENSG00000279873 | *LINC01126* | 1.469878928 | -0.198778598 | 0.206803081 | -0.405581679 | 0.944094537 | 0.255232566 | 0.40932804 | -0.154095474 | 0.542491694 |
| ENSG00000132825 | *PPP1R3D* | 4.121401098 | -0.064397165 | -0.086323796 | 0.021926631 | 0.999933887 | -0.360150968 | -0.477601893 | 0.117450925 | 0.543100022 |
| ENSG00000100034 | *PPM1F* | 7.233340241 | -0.018237575 | -0.027816567 | 0.009578992 | 0.999933887 | 0.026215605 | 0.132162141 | -0.105946536 | 0.543115253 |
| ENSG00000205537 | *AC121338.1* | 0.065000413 | 0.362993252 | -0.014245926 | 0.377239178 | 0.999933887 | 0.373860102 | 0.087607239 | 0.286252863 | 0.543281795 |
| ENSG00000267270 | *PARD6G-AS1* | 0.988314147 | 0.076474205 | 0.189678311 | -0.113204106 | 0.999933887 | 0.628327397 | 0.803440581 | -0.175113184 | 0.543281795 |
| ENSG00000168569 | *TMEM223* | 0.833390843 | 0.137008188 | -0.187706404 | 0.324714592 | 0.999933887 | -0.058475763 | -0.232830132 | 0.174354369 | 0.543338737 |
| ENSG00000132341 | *RAN* | 5.047252237 | -0.045102616 | -0.073366416 | 0.0282638 | 0.999933887 | 0.368771805 | 0.458602925 | -0.08983112 | 0.543394766 |
| ENSG00000156162 | *DPY19L4* | 0.252363398 | -0.041010183 | -0.234064703 | 0.193054521 | 0.999933887 | -0.10671343 | 0.153731856 | -0.260445286 | 0.543404325 |
| ENSG00000024862 | *CCDC28A* | 4.266219485 | 0.00342688 | -0.036827066 | 0.040253946 | 0.999933887 | -0.554080909 | -0.649762743 | 0.095681833 | 0.543404325 |
| ENSG00000106344 | *RBM28* | 4.503215832 | -0.040364408 | -0.078900452 | 0.038536044 | 0.999933887 | -0.239923891 | -0.164171084 | -0.075752808 | 0.54373604 |
| ENSG00000184205 | *TSPYL2* | 6.26076315 | 0.018967059 | 0.032813232 | -0.013846173 | 0.999933887 | 0.163296802 | 0.239697214 | -0.076400411 | 0.54373604 |
| ENSG00000136521 | *NDUFB5* | 2.631213226 | -0.085143926 | -0.031492691 | -0.053651235 | 0.999933887 | -0.126233198 | -0.011260373 | -0.114972826 | 0.543755553 |
| ENSG00000187607 | *ZNF286A* | -1.044222905 | -0.446323718 | 0.229495723 | -0.675819441 | 0.999933887 | -0.288959168 | 0.029351113 | -0.318310281 | 0.543760404 |
| ENSG00000279602 | *AC109326.1* | 2.121592383 | 0.260112154 | 0.215477324 | 0.044634831 | 0.999933887 | 0.519202615 | 0.754463141 | -0.235260526 | 0.543915769 |
| ENSG00000271303 | *SRXN1* | 2.946935863 | 0.262288956 | 0.034525614 | 0.227763342 | 0.999933887 | 0.243918338 | 0.431979799 | -0.188061462 | 0.543915769 |
| ENSG00000102805 | *CLN5* | 2.941537676 | -0.081525197 | -0.123812127 | 0.042286931 | 0.999933887 | -0.504305366 | -0.367457494 | -0.136847872 | 0.543915769 |
| ENSG00000169599 | *NFU1* | 1.415734487 | -0.060837199 | -0.164172551 | 0.103335352 | 0.999933887 | 0.001570491 | 0.221176578 | -0.219606087 | 0.543915769 |
| ENSG00000076248 | *UNG* | 1.148934673 | 0.034898186 | -0.027859724 | 0.06275791 | 0.999933887 | -0.033067173 | 0.151394357 | -0.18446153 | 0.543915769 |
| ENSG00000147274 | *RBMX* | 5.714978205 | -0.064278587 | -0.094770325 | 0.030491738 | 0.999933887 | 0.026171896 | 0.085750039 | -0.059578143 | 0.544261187 |
| ENSG00000207042 | *RNU6-196P* | 0.190222964 | 0.209110812 | 0.098827146 | 0.110283666 | 0.999933887 | 0.753984756 | 1.05677798 | -0.302793224 | 0.544261187 |
| ENSG00000175792 | *RUVBL1* | 2.361398223 | -0.042010246 | -0.025504191 | -0.016506055 | 0.999933887 | 0.159472337 | 0.303185899 | -0.143713562 | 0.544261187 |
| ENSG00000171634 | *BPTF* | 5.931487063 | 0.009994805 | -0.03910588 | 0.049100685 | 0.999933887 | -0.468066151 | -0.33311412 | -0.134952031 | 0.544261187 |
| ENSG00000167705 | *RILP* | 4.729897247 | 0.074774219 | 0.019465593 | 0.055308626 | 0.999933887 | 0.249462088 | 0.156036086 | 0.093426001 | 0.544321946 |
| ENSG00000182220 | *ATP6AP2* | 6.293448911 | -0.021795087 | -0.052079974 | 0.030284886 | 0.999933887 | 0.011589831 | 0.074031602 | -0.062441771 | 0.544321946 |
| ENSG00000111877 | *MCM9* | 3.307635811 | -0.042625635 | -0.045721369 | 0.003095734 | 0.999933887 | -0.287657631 | -0.392654077 | 0.104996445 | 0.544443677 |
| ENSG00000105671 | *DDX49* | 4.32165123 | -0.008274322 | 0.055441946 | -0.063716268 | 0.999933887 | -0.08830623 | -0.167101306 | 0.078795076 | 0.544572183 |
| ENSG00000135838 | *NPL* | 8.289062843 | -0.073437738 | -0.050658277 | -0.022779461 | 0.999933887 | -0.462921504 | -0.392191683 | -0.070729821 | 0.544728465 |
| ENSG00000236345 | *SCAT8* | -0.067123585 | -0.113429319 | -0.164707298 | 0.051277979 | 0.999933887 | -0.323983222 | -0.63654517 | 0.312561948 | 0.544762498 |
| ENSG00000245848 | *CEBPA* | 1.692038345 | 0.097505983 | -0.060517928 | 0.158023912 | 0.999933887 | -0.204374872 | -0.042130305 | -0.162244567 | 0.544766015 |
| ENSG00000004487 | *KDM1A* | 3.204282428 | -0.054818028 | -0.030553528 | -0.024264499 | 0.999933887 | 0.116750414 | 0.206757343 | -0.090006929 | 0.544826184 |
| ENSG00000178209 | *PLEC* | 8.249604755 | 0.056554025 | 0.073095302 | -0.016541277 | 0.999933887 | 0.155994673 | 0.245209341 | -0.089214668 | 0.544977277 |
| ENSG00000136933 | *RABEPK* | 1.647820918 | 0.135024848 | 0.099093731 | 0.035931118 | 0.999933887 | 0.101024045 | -0.077217435 | 0.17824148 | 0.545295672 |
| ENSG00000286379 | *AL591845.1* | 0.285468804 | 0.192843213 | 0.076602363 | 0.116240851 | 0.999933887 | 1.227081538 | 1.000936221 | 0.226145317 | 0.545313782 |
| ENSG00000168071 | *CCDC88B* | 8.762006199 | 0.122964867 | 0.141860914 | -0.018896047 | 0.999933887 | 0.328668061 | 0.46146752 | -0.132799459 | 0.54577016 |
| ENSG00000245680 | *ZNF585B* | 0.712409395 | -0.109319611 | -0.092871542 | -0.016448069 | 0.999933887 | -0.081696854 | 0.084314265 | -0.16601112 | 0.54577016 |
| ENSG00000104883 | *PEX11G* | 1.800097129 | 0.016149766 | 0.00729864 | 0.008851126 | 0.999933887 | 0.097671879 | 0.274184509 | -0.17651263 | 0.54577016 |
| ENSG00000105516 | *DBP* | 1.651137158 | -0.021058633 | -0.094398877 | 0.073340244 | 0.999933887 | -0.307257447 | -0.482930313 | 0.175672867 | 0.54577016 |
| ENSG00000255092 | *AC010768.2* | 2.312204289 | -0.123410211 | -0.093610681 | -0.029799531 | 0.999933887 | 1.911108564 | 2.078023706 | -0.166915142 | 0.545828559 |
| ENSG00000145685 | *LHFPL2* | 3.206666047 | -0.110756054 | -0.01487772 | -0.095878334 | 0.999933887 | 0.321341371 | 0.493769579 | -0.172428207 | 0.545828559 |
| ENSG00000168653 | *NDUFS5* | 3.223419938 | 0.051062313 | -0.123885346 | 0.174947659 | 0.999933887 | 0.141259174 | 0.252690867 | -0.111431693 | 0.545828559 |
| ENSG00000135587 | *SMPD2* | 4.142869453 | -0.051261963 | 0.005828062 | -0.057090025 | 0.999933887 | 1.031536069 | 1.134953317 | -0.103417247 | 0.545828559 |
| ENSG00000188566 | *NDOR1* | 4.263235836 | 0.024798704 | 0.068430426 | -0.043631722 | 0.999933887 | -0.071450044 | -0.159038849 | 0.087588805 | 0.545828559 |
| ENSG00000146386 | *ABRACL* | 3.991811596 | -0.045856087 | -0.061742405 | 0.015886318 | 0.999933887 | -0.235852917 | -0.346174934 | 0.110322017 | 0.545882384 |
| ENSG00000144152 | *FBLN7* | 0.585594101 | 0.05340189 | 0.016619808 | 0.036782081 | 0.999933887 | 0.106234867 | -0.100302191 | 0.206537058 | 0.545882384 |
| ENSG00000286531 | *AC097015.1* | 0.745503338 | -0.017719882 | 0.109078755 | -0.126798637 | 0.999933887 | 0.850912788 | 1.117890723 | -0.266977935 | 0.545882384 |
| ENSG00000123575 | *FAM199X* | 4.33692854 | -0.009582425 | -0.026061367 | 0.016478942 | 0.999933887 | -0.679833279 | -0.79709143 | 0.117258151 | 0.545882384 |
| ENSG00000101017 | *CD40* | 0.539220589 | 0.217977839 | -0.147609909 | 0.365587749 | 0.999933887 | 1.79243038 | 2.052459997 | -0.260029617 | 0.546037982 |
| ENSG00000135407 | *AVIL* | 2.401550918 | -0.003290126 | -0.051091642 | 0.047801516 | 0.999933887 | -0.156523187 | -0.285646534 | 0.129123346 | 0.546320641 |
| ENSG00000115020 | *PIKFYVE* | 5.967390029 | -0.081372521 | -0.12364609 | 0.042273569 | 0.999933887 | -0.555377863 | -0.402306595 | -0.153071269 | 0.546406826 |
| ENSG00000177700 | *POLR2L* | 3.509947545 | -0.021582965 | 0.045647144 | -0.067230109 | 0.999933887 | -0.166856372 | -0.280833872 | 0.1139775 | 0.546968998 |
| ENSG00000214026 | *MRPL23* | 0.758426503 | 0.114150543 | 0.059064986 | 0.055085556 | 0.999933887 | 0.020346933 | -0.165903419 | 0.186250352 | 0.547019002 |
| ENSG00000256705 | *AL137779.1* | 0.26188245 | -0.09154982 | 0.112334402 | -0.203884223 | 0.999933887 | 0.078644601 | -0.182661205 | 0.261305806 | 0.547019002 |
| ENSG00000108819 | *PPP1R9B* | 7.976307595 | 0.037042341 | 0.083225475 | -0.046183134 | 0.999933887 | 0.06452098 | -0.056569199 | 0.121090179 | 0.547019002 |
| ENSG00000179912 | *R3HDM2* | 4.753151112 | 0.010513217 | 0.111365112 | -0.100851895 | 0.999933887 | 0.019559461 | -0.049627918 | 0.069187379 | 0.547019002 |
| ENSG00000147119 | *CHST7* | 7.356671599 | -0.005339856 | -0.049825299 | 0.044485443 | 0.999933887 | -0.013990726 | 0.088214863 | -0.102205588 | 0.547019002 |
| ENSG00000138757 | *G3BP2* | 6.600215397 | -0.056070549 | -0.082997683 | 0.026927134 | 0.999933887 | -0.270610041 | -0.334039285 | 0.063429244 | 0.547059143 |
| ENSG00000169896 | *ITGAM* | 7.850145009 | 0.015680726 | 0.009922941 | 0.005757785 | 0.999933887 | 0.167635645 | 0.089941716 | 0.077693929 | 0.547059143 |
| ENSG00000161040 | *FBXL13* | 3.502090469 | 0.002709912 | -0.081242442 | 0.083952354 | 0.999933887 | -0.482971652 | -0.375300969 | -0.107670682 | 0.547320026 |
| ENSG00000211791 | *TRAV13-2* | -0.49445346 | 0.004066667 | -0.241391 | 0.245457667 | 0.999933887 | 0.017998146 | 0.352029074 | -0.334030928 | 0.547320026 |
| ENSG00000253829 | *AC067817.2* | -0.460815977 | 0.268037792 | -0.086444592 | 0.354482384 | 0.999933887 | 0.249235386 | -0.091837541 | 0.341072927 | 0.547467681 |
| ENSG00000214851 | *LINC00612* | -0.507473646 | 0.170109724 | 0.290595559 | -0.120485834 | 0.999933887 | -0.078323077 | 0.30102396 | -0.379347037 | 0.547486437 |
| ENSG00000111445 | *RFC5* | 1.409294594 | -0.035963119 | -0.192147092 | 0.156183973 | 0.999933887 | -0.189576561 | -0.339402294 | 0.149825733 | 0.547486437 |
| ENSG00000259673 | *IQCH-AS1* | -0.71382007 | -0.044796055 | -0.583510267 | 0.538714212 | 0.999933887 | -0.32554702 | 0.007991899 | -0.333538919 | 0.547486437 |
| ENSG00000116771 | *AGMAT* | -0.530518472 | 0.056348854 | -0.339131246 | 0.3954801 | 0.999933887 | 0.016494284 | 0.299056534 | -0.28256225 | 0.547557778 |
| ENSG00000280063 | *AC012676.5* | 0.673866687 | -0.072856829 | -0.234022973 | 0.161166144 | 0.999933887 | -0.2807379 | -0.482369944 | 0.201632044 | 0.547728654 |
| ENSG00000126016 | *AMOT* | -0.690043789 | -0.011424923 | -0.129440904 | 0.118015981 | 0.999933887 | 0.074431286 | -0.255021832 | 0.329453118 | 0.547865268 |
| ENSG00000271964 | *AC090948.1* | -0.804183973 | -0.420451312 | 0.03313048 | -0.453581792 | 0.999933887 | -0.155566515 | 0.254583357 | -0.410149871 | 0.547946361 |
| ENSG00000171492 | *LRRC8D* | 2.943481617 | -0.055102446 | 0.057830598 | -0.112933044 | 0.999933887 | 0.342658832 | 0.505849844 | -0.163191013 | 0.547946361 |
| ENSG00000267364 | *AC022706.1* | -0.692004903 | -0.05270851 | 0.044386675 | -0.097095185 | 0.999933887 | -0.141149507 | 0.204669699 | -0.345819207 | 0.547946361 |
| ENSG00000279568 | *AC093525.9* | 1.046162873 | -0.020036649 | -0.07089867 | 0.050862021 | 0.999933887 | -0.620945974 | -0.792669874 | 0.1717239 | 0.547946361 |

| | | | | | | | | | | |
|---|---|---|---|---|---|---|---|---|---|---|
| ENSG00000130803 | *ZNF317* | 4.811995925 | -0.029744006 | -0.029916446 | 0.000172439 | 0.999933887 | -0.174655371 | -0.095264481 | -0.07939089 | 0.548088663 |
| ENSG00000198400 | *NTRK1* | -0.558118015 | 0.146077042 | -0.235728731 | 0.381805773 | 0.999933887 | -0.119003016 | -0.467313248 | 0.348310233 | 0.548185439 |
| ENSG00000145604 | *SKP2* | 1.458926358 | -0.040655231 | -0.001157921 | -0.03949731 | 0.999933887 | 0.011193575 | -0.134079948 | 0.145273523 | 0.54845709 |
| ENSG00000187118 | *CMC1* | 2.770326881 | -0.06429184 | -0.043208046 | -0.021083795 | 0.999933887 | 0.029252779 | 0.137341131 | -0.108088352 | 0.548550934 |
| ENSG00000182318 | *ZSCAN22* | 0.606362228 | 0.054392058 | -0.010434543 | 0.064826602 | 0.999933887 | -0.012840543 | -0.241171449 | 0.228330906 | 0.548550934 |
| ENSG00000175334 | *BANF1* | 2.62667887 | 0.002335044 | -0.035473796 | 0.03780884 | 0.999933887 | 0.067076911 | -0.055948607 | 0.123025518 | 0.548665769 |
| ENSG00000261512 | *AC092368.3* | 1.357784997 | 0.130726165 | 0.090997901 | 0.039728264 | 0.999933887 | -0.767942939 | -0.989362476 | 0.221419536 | 0.548762017 |
| ENSG00000116750 | *UCHL5* | 2.354959647 | 0.005616725 | -0.205144985 | 0.21076171 | 0.999933887 | -0.273784449 | -0.088455275 | -0.185329174 | 0.548762017 |
| ENSG00000182774 | *RPS17* | 1.693037138 | -0.024396525 | -0.051582341 | 0.027185816 | 0.999933887 | 0.255563894 | 0.080603033 | 0.174960861 | 0.548834499 |
| ENSG00000178057 | *NDUFAF3* | 2.796321203 | -0.093220929 | 0.032261342 | -0.125482272 | 0.999933887 | -0.22449811 | -0.327747894 | 0.103249784 | 0.54888921 |
| ENSG00000197696 | *NMB* | 0.621597397 | 0.171636942 | 0.141349041 | 0.030287901 | 0.999933887 | 1.260241822 | 1.463507885 | -0.203266063 | 0.549333055 |
| ENSG00000228079 | *AC012368.2* | 0.641945106 | 0.021898311 | 0.241605308 | -0.219706997 | 0.999933887 | 0.00922495 | 0.253814888 | -0.244589938 | 0.549576916 |
| ENSG00000163319 | *MRPS18C* | 1.980122383 | 0.162238997 | 0.158719659 | 0.003519338 | 0.999933887 | -0.22304661 | -0.102276036 | -0.120770573 | 0.549727967 |
| ENSG00000183742 | *MACC1* | -0.884604311 | -0.191128886 | 0.623952439 | -0.815081325 | 0.999933887 | 1.284746847 | 1.593157894 | -0.308411047 | 0.549727967 |
| ENSG00000072864 | *NDE1* | 6.288428402 | 0.019367551 | 0.024795072 | -0.005427521 | 0.999933887 | 0.536846968 | 0.647357171 | -0.110510203 | 0.549727967 |
| ENSG00000236337 | *FMR1-IT1* | 0.001067633 | -0.036138316 | 0.093078375 | -0.129216692 | 0.999933887 | -0.365674024 | -0.075300704 | -0.290373319 | 0.549738433 |
| ENSG00000279641 | *AC120057.3* | -0.003407353 | -0.019951625 | -0.216612667 | 0.196661042 | 0.999933887 | 0.406484989 | 0.185376796 | 0.221108193 | 0.549767305 |
| ENSG00000048052 | *HDAC9* | 1.594252982 | 0.055460381 | -0.04260755 | 0.098067931 | 0.999933887 | 0.093951584 | 0.246084535 | -0.152132951 | 0.549857332 |
| ENSG00000105383 | *CD33* | 0.37312468 | -0.107166661 | 0.071040471 | -0.178207132 | 0.999933887 | -0.702405346 | -1.075335791 | 0.372930445 | 0.549987849 |
| ENSG00000253251 | *SHLD3* | 0.737209784 | -0.213879422 | -0.019782585 | -0.194096836 | 0.999933887 | 0.214167328 | 0.380142146 | -0.165974818 | 0.550200364 |
| ENSG00000203799 | *CCDC162P* | 0.560766294 | -0.209329156 | 0.187614489 | -0.396943645 | 0.999933887 | -0.684826508 | -0.477995374 | -0.206831134 | 0.550200364 |
| ENSG00000103227 | *LMF1* | 1.817335061 | 0.051922094 | 0.076524015 | -0.024601921 | 0.999933887 | 0.087043429 | 0.231908222 | -0.144864792 | 0.550200364 |
| ENSG00000181026 | *AEN* | 3.332877827 | -0.010727109 | -0.006554314 | -0.004172795 | 0.999933887 | 0.581922272 | 0.697820399 | -0.115898127 | 0.550200364 |
| ENSG00000132718 | *SYT11* | 3.437430778 | -0.003436222 | 0.002070219 | -0.005506441 | 0.999933887 | 0.005398046 | -0.093990085 | 0.099388132 | 0.550200364 |
| ENSG00000090263 | *MRPS33* | 0.805658856 | -0.005126397 | 0.221923821 | -0.227050219 | 0.999933887 | 0.056219583 | 0.245892031 | -0.189672449 | 0.550200364 |
| ENSG00000100814 | *CCNB1IP1* | 1.516244477 | 0.074729534 | 0.001213044 | 0.07351649 | 0.999933887 | 0.014398821 | 0.153869197 | -0.139470376 | 0.550331637 |
| ENSG00000173258 | *ZNF483* | 0.696587916 | -0.110151542 | -0.048433173 | -0.061718369 | 0.999933887 | -0.16916075 | 0.082714552 | -0.251875302 | 0.550331637 |
| ENSG00000085871 | *MGST2* | 2.312690504 | -0.126410039 | 0.016760343 | -0.143170383 | 0.999933887 | -1.838554526 | -2.036661585 | 0.198107059 | 0.550374606 |
| ENSG00000125944 | *HNRNPR* | 4.866964669 | -0.033703714 | -0.005818519 | -0.027885195 | 0.999933887 | -0.03191595 | -0.10393354 | 0.07201759 | 0.550374606 |
| ENSG00000278376 | *AP004609.3* | 0.200206377 | 0.009435907 | -0.203569729 | 0.213005636 | 0.999933887 | 0.037566037 | -0.156001004 | 0.19356704 | 0.550374606 |
| ENSG00000279957 | *AC110769.3* | 0.576592036 | 0.464220868 | 0.699350664 | -0.235129796 | 0.999933887 | 1.322383807 | 0.855667952 | 0.466715855 | 0.550408644 |
| ENSG00000173992 | *CCS* | 2.869165631 | 0.017119068 | 0.135295723 | -0.118176656 | 0.999933887 | -0.048865849 | -0.20630713 | 0.157441281 | 0.550408644 |
| ENSG00000169919 | *GUSB* | 4.652915197 | -0.0027474 | 0.010602374 | -0.013349774 | 0.999933887 | -0.176556116 | -0.255128974 | 0.078572858 | 0.550408644 |
| ENSG00000066044 | *ELAVL1* | 3.481483633 | -0.028063274 | -0.112303144 | 0.084239871 | 0.999933887 | -0.03369736 | -0.135158418 | 0.101461058 | 0.550439571 |
| ENSG00000125962 | *ARMCX5* | 0.671938415 | 0.011453732 | 0.099762919 | -0.088309188 | 0.999933887 | -0.109751046 | 0.072505718 | -0.182256764 | 0.550439571 |
| ENSG00000122591 | *FAM126A* | 2.559225268 | 0.004380038 | -0.030372431 | 0.034752469 | 0.999933887 | -0.072726809 | 0.095902371 | -0.16862918 | 0.550439571 |
| ENSG00000168899 | *VAMP5* | 2.712897497 | 0.043320633 | 0.000280619 | 0.043040014 | 0.999933887 | 0.059757048 | -0.089961839 | 0.149718887 | 0.550444924 |
| ENSG00000159884 | *CCDC107* | -0.016548178 | -0.051605825 | -0.269090526 | 0.217484701 | 0.999933887 | 0.329434095 | 0.064469469 | 0.264964626 | 0.550444924 |
| ENSG00000168404 | *MLKL* | 5.4089848 | 0.001393006 | 0.012781036 | -0.01138803 | 0.999933887 | 0.05655747 | -0.029476772 | 0.086034241 | 0.550444924 |
| ENSG00000180336 | *MEIOC* | -0.073051033 | -0.172188244 | 0.056305236 | -0.22849348 | 0.999933887 | -0.434231699 | -0.15610045 | -0.278131249 | 0.550570499 |
| ENSG00000234383 | *CTBP2P8* | -0.880367407 | -0.00089941 | 0.00909455 | -0.00999396 | 0.999933887 | -0.336391963 | -0.724498957 | 0.388106995 | 0.550752691 |
| ENSG00000143412 | *ANXA9* | 2.12882527 | -0.094702307 | 0.045629101 | -0.140331408 | 0.999933887 | -0.635746188 | -0.815655531 | 0.179909343 | 0.550936709 |
| ENSG00000108688 | *CCL7* | 1.165890026 | -0.049843003 | -0.593314569 | 0.543471566 | 0.999933887 | -1.710813698 | -1.927837059 | 0.217023361 | 0.550936709 |
| ENSG00000127528 | *KLF2* | 8.610655954 | 0.206388959 | 0.147586671 | 0.058802288 | 0.999933887 | 0.045840884 | -0.055523998 | 0.101364882 | 0.551062964 |
| ENSG00000179010 | *MRFAP1* | 6.634400531 | -0.054964773 | -0.111608332 | 0.056643558 | 0.999933887 | -0.220797851 | -0.149334325 | -0.071463525 | 0.551062964 |
| ENSG00000179152 | *TCAIM* | 1.3272632 | 0.082832888 | 0.10732076 | -0.024487872 | 0.999933887 | -0.1477452 | 0.099369329 | -0.247114529 | 0.551062964 |
| ENSG00000211899 | *IGHM* | 3.931355084 | 0.047612318 | 0.096164343 | -0.048552025 | 0.999933887 | 0.098061629 | -0.022212379 | 0.120274008 | 0.551062964 |
| ENSG00000184110 | *EIF3C* | -0.258930118 | -0.086009596 | -0.159188339 | 0.073178743 | 0.999933887 | 0.339783355 | 0.577287441 | -0.237504086 | 0.551062964 |
| ENSG00000141385 | *AFG3L2* | 2.998511863 | -0.014966569 | -0.132693881 | 0.117727311 | 0.999933887 | -0.193090547 | -0.081486104 | -0.111604443 | 0.551062964 |
| ENSG00000197841 | *ZNF181* | 0.458737868 | 0.028500017 | -0.135937163 | 0.16443718 | 0.999933887 | -0.064552218 | 0.129534654 | -0.194086872 | 0.551062964 |
| ENSG00000224852 | *MTND5P24* | -0.848362444 | 0.14344575 | -0.02738297 | 0.170828721 | 0.999933887 | 0.582379655 | 0.091669537 | 0.490710119 | 0.551144067 |
| ENSG00000183397 | *C19orf71* | 3.009391955 | 0.208954328 | 0.179119579 | 0.029834749 | 0.999933887 | 0.466238173 | 0.642957035 | -0.176718862 | 0.551959014 |
| ENSG00000101109 | *STK4* | 9.043917462 | -0.095005542 | -0.013376251 | -0.081629291 | 0.999933887 | -0.358811598 | -0.430809391 | 0.071997793 | 0.552217735 |
| ENSG00000143374 | *TARS2* | 2.102360134 | -0.063174269 | -0.050098705 | -0.013075564 | 0.999933887 | 0.097228783 | -0.040544817 | 0.1377736 | 0.552421457 |
| ENSG00000258928 | *AL358333.2* | 1.027476779 | 0.085247458 | 0.275670823 | -0.190423366 | 0.999933887 | 2.398587058 | 2.748086012 | -0.349498954 | 0.552421457 |
| ENSG00000089876 | *DHX32* | -0.412694554 | -0.101541467 | -0.048728948 | -0.052812519 | 0.999933887 | -0.145469792 | 0.064627958 | -0.21009775 | 0.552421457 |
| ENSG00000100138 | *SNU13* | 4.798014613 | 0.011440943 | -0.048616429 | 0.060057371 | 0.999933887 | -0.030136019 | -0.111574212 | 0.081438193 | 0.552421457 |
| ENSG00000188811 | *NHLRC3* | 1.827701535 | -0.0080576 | -0.01054958 | 0.002491981 | 0.999933887 | -0.158142618 | 0.02059114 | -0.178733758 | 0.552421457 |
| ENSG00000170234 | *PWWP2A* | 3.821978021 | 0.003154575 | 0.043084456 | -0.03992988 | 0.999933887 | 0.646030174 | 0.761374849 | -0.115344675 | 0.552421457 |
| ENSG00000167105 | *TMEM92* | 1.073125459 | -0.163674499 | -0.154361621 | -0.009312877 | 0.999933887 | -0.403519424 | -0.678041088 | 0.274521664 | 0.552601863 |
| ENSG00000147437 | *GNRH1* | 1.192125807 | -0.132592585 | 0.137376236 | -0.269968822 | 0.999933887 | -0.620509379 | -0.460707699 | -0.159801681 | 0.552689624 |
| ENSG00000101945 | *SUV39H1* | 0.880607496 | -0.063882845 | -0.210143201 | 0.146260356 | 0.999933887 | 0.019669803 | -0.189201311 | 0.208871114 | 0.552689624 |
| ENSG00000115350 | *POLE4* | 2.20075286 | -0.007140991 | -0.041270819 | 0.034129827 | 0.999933887 | 0.014693687 | -0.128527279 | 0.143220966 | 0.552689624 |
| ENSG00000138411 | *HECW2* | 1.890957976 | 0.003711738 | -0.066848389 | 0.070560127 | 0.999933887 | -0.436143392 | -0.651180161 | 0.215036769 | 0.552689624 |
| ENSG00000136104 | *RNASEH2B* | 3.627838867 | -0.054267218 | -0.110251613 | 0.055984395 | 0.999933887 | -0.074980307 | -0.164953394 | 0.089973087 | 0.552890884 |
| ENSG00000027001 | *MIPEP* | -0.571017394 | 0.068981119 | -0.030070474 | 0.099051593 | 0.999933887 | -0.280011629 | -0.000114468 | -0.279897161 | 0.55350745 |
| ENSG00000196792 | *STRN3* | 3.651360401 | 0.03218137 | 0.008750238 | 0.023431131 | 0.999933887 | -0.216731301 | -0.102835657 | -0.113895644 | 0.553544691 |
| ENSG00000130255 | *RPL36* | 6.292475359 | 0.056428511 | -0.028263066 | 0.084691577 | 0.999933887 | 0.103227778 | 0.18772637 | -0.084498592 | 0.553873817 |
| ENSG00000047662 | *FAM184B* | 1.167635321 | 0.117330202 | 0.11438308 | 0.002947122 | 0.999933887 | 0.133182097 | 0.333680623 | -0.200498526 | 0.553918887 |
| ENSG00000141040 | *ZNF287* | -0.461815712 | 0.148207335 | -0.008149064 | 0.156356399 | 0.999933887 | 0.170614839 | -0.054672298 | 0.225287137 | 0.553938233 |
| ENSG00000125875 | *TBC1D20* | 5.203629047 | -0.019448613 | -0.021751766 | 0.002303153 | 0.999933887 | -0.199529398 | -0.275802855 | 0.076273457 | 0.553938233 |
| ENSG00000247679 | *AC139795.2* | 0.550771274 | -0.035207516 | 0.123465133 | -0.158672649 | 0.999933887 | 0.128109128 | 0.335810075 | -0.207700946 | 0.553938233 |
| ENSG00000204920 | *ZNF155* | 1.291156242 | -0.139602141 | 0.123290403 | -0.262892544 | 0.999933887 | -0.29229379 | -0.134564781 | -0.15772901 | 0.554038187 |
| ENSG00000110921 | *MVK* | 1.886712409 | 0.049517818 | -0.087217605 | 0.136735423 | 0.999933887 | 0.518523931 | 0.687301402 | -0.16877747 | 0.554038187 |
| ENSG00000132849 | *PATJ* | 2.673548804 | -0.048703705 | -0.036924816 | -0.011778889 | 0.999933887 | -0.003255177 | 0.12265642 | -0.125911596 | 0.554257068 |
| ENSG00000223799 | *IL10RB-DT* | 2.677024702 | 0.074232611 | -0.015146141 | 0.089378752 | 0.999933887 | 1.210192067 | 1.054840062 | 0.155352006 | 0.554311895 |
| ENSG00000173110 | *HSPA6* | 6.353156523 | 0.049090843 | 0.043808052 | 0.005282791 | 0.999933887 | 0.683010358 | 0.576686551 | 0.106323807 | 0.554311895 |
| ENSG00000034677 | *RNF19A* | 6.082764316 | -0.025479051 | -0.021044122 | -0.004434929 | 0.999933887 | -0.300478066 | -0.204615114 | -0.095862952 | 0.554311895 |
| ENSG00000105327 | *BBC3* | 5.485617014 | -0.018519678 | 0.062004671 | -0.08052435 | 0.999933887 | -0.970113763 | -1.122746119 | 0.152632357 | 0.554311895 |
| ENSG00000253269 | *AC008680.1* | -0.161419635 | 0.171229146 | 0.226115303 | -0.054886157 | 0.999933887 | 0.104637911 | 0.398972599 | -0.294334688 | 0.554413726 |
| ENSG00000100139 | *MICALL1* | 3.773265014 | 0.035646375 | -0.020569484 | 0.056215859 | 0.999933887 | 0.038864951 | -0.071774136 | 0.110639087 | 0.55474832 |
| ENSG00000227232 | *WASH7P* | 0.822239974 | 0.073769026 | 0.001177949 | 0.072591077 | 0.999933887 | -0.131976613 | -0.334243021 | 0.202266407 | 0.55474832 |
| ENSG00000161526 | *SAP30BP* | 7.00530837 | -0.02804413 | 0.002762328 | -0.030806457 | 0.999933887 | 0.358506007 | 0.451500462 | -0.092994455 | 0.55474832 |
| ENSG00000268521 | *VN1R83P* | -0.570663874 | -0.037041063 | -0.525227812 | 0.488186748 | 0.999933887 | -0.354971291 | 0.000402265 | -0.355373556 | 0.55474832 |
| ENSG00000125520 | *SLC2A4RG* | 4.257880684 | 0.109957598 | 0.050358144 | 0.059599455 | 0.999933887 | 0.136898727 | 0.03172016 | 0.105178567 | 0.554799118 |
| ENSG00000136522 | *MRPL47* | 2.361083506 | -0.110934447 | -0.067408218 | -0.043526229 | 0.999933887 | 0.127473034 | 0.255394342 | -0.127921309 | 0.554799118 |
| ENSG00000179889 | *PDXDC1* | 4.731513202 | -0.0477826 | -0.01953769 | -0.02824491 | 0.999933887 | 0.054066989 | 0.114280266 | -0.060213278 | 0.554799118 |
| ENSG00000144161 | *ZC3H8* | 1.704347069 | -0.04013328 | -0.102863881 | 0.062730602 | 0.999933887 | -0.101276499 | 0.048258778 | -0.149535276 | 0.554799118 |
| ENSG00000228013 | *IL6R-AS1* | 1.61147445 | 0.193800011 | 0.109636001 | 0.08416401 | 0.999933887 | 1.222897226 | 1.454068907 | -0.231171681 | 0.554836677 |
| ENSG00000110108 | *TMEM109* | 3.616118179 | -0.029578477 | -0.035989031 | 0.006410553 | 0.999933887 | 0.154173608 | 0.054140014 | 0.100033594 | 0.555247782 |
| ENSG00000138758 | *SEPTIN11* | 3.035697715 | -0.052697973 | 0.032372852 | -0.085070825 | 0.999933887 | 0.040718327 | 0.161293705 | -0.120575378 | 0.555276441 |
| ENSG00000174871 | *CNIH2* | 0.696646442 | 0.088573539 | -0.0536513 | 0.142224839 | 0.999933887 | -0.026608237 | 0.242354093 | -0.26896233 | 0.555276441 |
| ENSG00000030582 | *GRN* | 9.14805352 | -0.024575855 | -0.016003435 | -0.00857242 | 0.999933887 | -0.275899319 | -0.200959606 | -0.074939712 | 0.555276441 |
| ENSG00000014914 | *MTMR11* | -0.510298127 | 0.019124143 | -0.124473943 | 0.143598086 | 0.999933887 | 0.147172846 | 0.415741984 | -0.268569138 | 0.555276441 |
| ENSG00000240869 | *RN7SL128P* | -0.218518979 | 0.080077439 | 0.2679498 | -0.187872361 | 0.999933887 | 0.574223427 | 0.319388828 | 0.254834599 | 0.555357078 |
| ENSG00000240344 | *PPIL3* | 0.981956615 | -0.173115003 | 0.161267966 | -0.334382969 | 0.999933887 | 0.1125251 | -0.069267539 | 0.181792639 | 0.555693404 |
| ENSG00000035115 | *SH3YL1* | 2.264465072 | -0.094443148 | -0.008688899 | -0.085754249 | 0.999933887 | 0.020243116 | 0.143611084 | -0.123367967 | 0.555693404 |
| ENSG00000114739 | *ACVR2B* | 0.539244595 | -0.064022124 | -0.110346195 | 0.046324072 | 0.999933887 | -0.010934604 | 0.141893724 | -0.152828329 | 0.555693404 |
| ENSG00000112304 | *ACOT13* | 3.931502638 | -0.024667996 | -0.019929888 | -0.004738108 | 0.999933887 | -0.192734538 | -0.101857193 | -0.090877345 | 0.555693404 |
| ENSG00000065802 | *ASB1* | 2.845696126 | -0.016763603 | 0.02198041 | -0.038744013 | 0.999933887 | 0.025742642 | 0.1247424 | -0.098999759 | 0.555693404 |
| ENSG00000205710 | *C17orf107* | 2.983573055 | 0.224636694 | 0.177817802 | 0.046818892 | 0.999933887 | 0.864281913 | 1.073098585 | -0.208816672 | 0.55578732 |
| ENSG00000126775 | *ATG14* | 3.714010499 | -0.091158575 | 0.03219195 | -0.123350525 | 0.999933887 | -0.157077823 | -0.238933769 | 0.081855945 | 0.55578732 |
| ENSG00000196312 | *MFSD14C* | 3.23661098 | 0.005664013 | 0.070380026 | -0.064716012 | 0.999933887 | -0.019274062 | -0.140227293 | 0.120953231 | 0.55578732 |
| ENSG00000157741 | *UBN2* | 3.670815405 | -0.006671274 | -0.055658361 | 0.048987087 | 0.999933887 | -0.264755009 | -0.135035634 | -0.129719375 | 0.555856619 |
| ENSG00000166313 | *APBB1* | 3.051673425 | 0.030650228 | 0.074835009 | -0.044184781 | 0.999933887 | 0.158223579 | 0.250664538 | -0.092440959 | 0.555889505 |

| | | | | | | | | | | |
|---|---|---|---|---|---|---|---|---|---|---|
| ENSG00000245750 | *DRAIC* | 2.428844363 | -0.004787893 | 0.080590039 | -0.085377932 | 0.999933887 | 0.31943968 | 0.471187679 | -0.151747999 | 0.555889505 |
| ENSG00000176714 | *CCDC121* | -0.789481869 | -0.104701235 | 0.012006416 | -0.116707651 | 0.999933887 | -0.536004418 | -0.989023856 | 0.453019437 | 0.556045274 |
| ENSG00000065243 | *PKN2* | 5.862065642 | -0.041786233 | -0.157595634 | 0.1158094 | 0.999933887 | -0.82920777 | -1.040962995 | 0.211755225 | 0.556131493 |
| ENSG00000140968 | *IRF8* | 2.802145049 | -0.004474411 | 0.08990174 | -0.094376151 | 0.999933887 | -0.006266283 | -0.110674534 | 0.104408251 | 0.556131493 |
| ENSG00000116350 | *SRSF4* | 6.352661212 | -0.02079234 | -0.034998026 | 0.014205686 | 0.999933887 | -0.309711523 | -0.372929614 | 0.063218091 | 0.556215758 |
| ENSG00000278869 | *BX539320.1* | 2.309922343 | 0.179993278 | 0.156548124 | 0.023445153 | 0.999933887 | 1.432537111 | 1.569589688 | -0.137052576 | 0.55628064 |
| ENSG00000135218 | *CD36* | 1.294920756 | 0.171103475 | -0.096001602 | 0.267105077 | 0.999933887 | -1.690425609 | -2.044614762 | 0.354189153 | 0.556600012 |
| ENSG00000184445 | *KNTC1* | 2.509557452 | 0.05579458 | -0.023927219 | 0.079721799 | 0.999933887 | -0.214911858 | -0.042232984 | -0.172678874 | 0.556842885 |
| ENSG00000114982 | *KANSL3* | 5.02380933 | -0.050842437 | -0.022807951 | -0.028034486 | 0.999933887 | -0.069285159 | -0.133323546 | 0.064038387 | 0.556968756 |
| ENSG00000165795 | *NDRG2* | 0.490832754 | -0.099237873 | -0.084985738 | -0.014252135 | 0.999933887 | -0.049415471 | 0.176991681 | -0.226407152 | 0.557544056 |
| ENSG00000058804 | *NDC1* | 0.950553552 | -0.265480064 | 0.239179381 | -0.504659445 | 0.944094537 | 0.112262571 | 0.306166344 | -0.193903774 | 0.557667305 |
| ENSG00000188529 | *SRSF10* | 5.732669717 | -0.053164387 | -0.08766491 | 0.034500523 | 0.999933887 | -0.052423072 | 0.032890247 | -0.08531332 | 0.557667305 |
| ENSG00000198838 | *RYR3* | -0.747492495 | -0.162341056 | -0.083604729 | -0.078736327 | 0.999933887 | -0.526927308 | -0.182596685 | -0.344330623 | 0.557667305 |
| ENSG00000023572 | *GLRX2* | 1.850950629 | 0.057916882 | 0.143921808 | -0.086004927 | 0.999933887 | -0.284931028 | -0.123028681 | -0.161902348 | 0.557667305 |
| ENSG00000071073 | *MGAT4A* | 5.148248286 | 0.041046242 | -0.052621664 | 0.093667906 | 0.999933887 | -0.62502322 | -0.760119151 | 0.135095931 | 0.557667305 |
| ENSG00000068654 | *POLR1A* | 2.999355547 | 0.011400543 | 0.010059461 | 0.001341081 | 0.999933887 | 0.068242384 | -0.033104651 | 0.101347035 | 0.557667305 |
| ENSG00000198711 | *SSBP3-AS1* | -0.792182168 | -0.000833102 | 0.315730798 | -0.3165639 | 0.999933887 | 0.157351218 | 0.441720301 | -0.284369083 | 0.557667305 |
| ENSG00000092108 | *SCFD1* | 3.430419596 | 3.217e-05 | -0.044436069 | 0.04446824 | 0.999933887 | -0.317747973 | -0.158052361 | -0.159695612 | 0.557729014 |
| ENSG00000278879 | *AP000560.1* | -0.602549006 | 0.114869344 | -0.040358653 | 0.155227998 | 0.999933887 | -0.830393784 | -0.430931069 | -0.399462715 | 0.557778362 |
| ENSG00000167110 | *GOLGA2* | 6.02091432 | -0.00045669 | 0.079737614 | -0.080194303 | 0.999933887 | -0.322649496 | -0.398506798 | 0.075857302 | 0.55810151 |
| ENSG00000162511 | *LAPTM5* | 11.121983734 | 0.054675589 | 0.006405407 | 0.048270182 | 0.999933887 | -0.019334076 | 0.046201286 | -0.065535362 | 0.558505899 |
| ENSG00000213015 | *ZNF580* | 3.561976913 | 0.04439293 | 0.072040791 | -0.027647861 | 0.999933887 | -0.149180419 | -0.237997281 | 0.088816862 | 0.558505899 |
| ENSG00000113734 | *BNIP1* | 0.769057077 | -0.067854759 | -0.048199423 | -0.019655336 | 0.999933887 | 0.191497758 | 0.397467645 | -0.205969887 | 0.558505899 |
| ENSG00000214837 | *LINC01347* | 0.158398658 | 0.06669006 | -0.17177509 | 0.23846515 | 0.999933887 | 0.054193356 | 0.278617933 | -0.224424577 | 0.558505899 |
| ENSG00000162664 | *ZNF326* | 3.480182584 | -0.02665662 | -0.08314959 | 0.05649297 | 0.999933887 | -0.213912311 | -0.028645893 | -0.185266417 | 0.558505899 |
| ENSG00000116754 | *SRSF11* | 6.492198778 | -0.036678586 | -0.098537448 | 0.061858861 | 0.999933887 | -0.274345359 | -0.126812436 | -0.147532923 | 0.558801535 |
| ENSG00000124491 | *F13A1* | 1.003918204 | -0.151720319 | -0.085689096 | -0.066031222 | 0.999933887 | -0.243254804 | -0.023909163 | -0.219345641 | 0.558905573 |
| ENSG00000205726 | *ITSN1* | 0.34173651 | -0.011121833 | -0.067228856 | 0.056107023 | 0.999933887 | -0.216472177 | -0.011044784 | -0.205427393 | 0.559105855 |
| ENSG00000101152 | *DNAJC5* | 7.95568386 | -0.103382201 | -0.128940156 | 0.025557955 | 0.999933887 | -0.089207092 | -0.011932898 | -0.077274195 | 0.559179847 |
| ENSG00000230487 | *PSMG3-AS1* | 0.269669489 | -0.153161488 | -0.165419057 | 0.012257569 | 0.999933887 | -0.020851488 | -0.19276055 | 0.171909062 | 0.559179847 |
| ENSG00000287680 | *AC093330.3* | 3.760459598 | 0.10707608 | 0.113827525 | -0.006751445 | 0.999933887 | -0.381540719 | -0.202890093 | -0.178650627 | 0.559179847 |
| ENSG00000111832 | *RWDD1* | 2.916249191 | -0.033188031 | 0.002129626 | -0.035317656 | 0.999933887 | 0.046756232 | 0.152551779 | -0.105795547 | 0.559179847 |
| ENSG00000215158 | *AC138409.2* | 0.096483482 | -0.003185633 | -0.028749945 | 0.025564312 | 0.999933887 | -0.347019206 | -0.020794764 | -0.326224442 | 0.559198761 |
| ENSG00000143702 | *CEP170* | 4.889921927 | 0.042321134 | 0.01970286 | 0.022618274 | 0.999933887 | -1.156579617 | -0.97410611 | -0.182473508 | 0.559256992 |
| ENSG00000238025 | *ZDHHC4P1* | -0.869219631 | -0.061595103 | 0.241350714 | -0.302945816 | 0.999933887 | -0.215401473 | 0.221062304 | -0.436463778 | 0.559589295 |
| ENSG00000205531 | *NAP1L4* | 5.464116904 | -0.009531436 | -0.069943141 | 0.060411704 | 0.999933887 | 0.064918893 | 0.011601319 | 0.053317574 | 0.559591276 |
| ENSG00000142396 | *ERVK3-1* | 3.28824721 | 0.108774603 | 0.077125988 | 0.031648615 | 0.999933887 | 0.003473881 | -0.111576655 | 0.115050537 | 0.559895077 |
| ENSG00000159352 | *PSMD4* | 5.985784817 | -0.031057358 | -0.070371443 | 0.039314085 | 0.999933887 | 0.382992872 | 0.459806453 | -0.076813581 | 0.559895077 |
| ENSG00000108654 | *DDX5* | 10.241222829 | 0.006995686 | -0.015116012 | 0.022111698 | 0.999933887 | 0.265859254 | 0.336796282 | -0.070937028 | 0.559966255 |
| ENSG00000146842 | *TMEM209* | 1.06037889 | -0.078277112 | 0.078651368 | -0.15692848 | 0.999933887 | -0.084883498 | 0.086871726 | -0.171755224 | 0.56011233 |
| ENSG00000115946 | *PNO1* | 1.623566699 | -0.074794719 | 0.116376225 | -0.191170944 | 0.999933887 | 0.291516105 | 0.448527937 | -0.157011833 | 0.560149373 |
| ENSG00000204954 | *C12orf73* | -0.088082094 | -0.125230717 | 0.006973612 | -0.132204329 | 0.999933887 | 0.125661662 | -0.143483298 | 0.26914496 | 0.560149373 |
| ENSG00000134013 | *LOXL2* | 0.264916915 | 0.027544391 | -0.138987381 | 0.166531772 | 0.999933887 | 0.194334253 | -0.057974241 | 0.252308493 | 0.560149373 |
| ENSG00000245025 | *AC107959.1* | 0.100768728 | -0.142046375 | -0.065842165 | -0.07620421 | 0.999933887 | 0.106726033 | -0.147403353 | 0.254129387 | 0.560374615 |
| ENSG00000137331 | *IER3* | 7.493463611 | 0.773085391 | 0.913163131 | -0.14007774 | 0.999933887 | 2.391680628 | 2.613741327 | -0.222060698 | 0.560429408 |
| ENSG00000092096 | *SLC22A17* | -1.129891652 | 0.095135722 | -0.025547645 | 0.120683367 | 0.999933887 | -0.005341649 | 0.315232436 | -0.320574084 | 0.560429408 |
| ENSG00000245060 | *LINC00847* | 3.974991289 | -0.02681225 | 0.013666495 | -0.040478745 | 0.999933887 | -0.027365698 | 0.055381836 | -0.082747534 | 0.560429408 |
| ENSG00000151651 | *ADAM8* | 10.414785529 | 0.087925181 | 0.069652016 | 0.018273165 | 0.999933887 | 0.524343495 | 0.591971293 | -0.067627799 | 0.560537649 |
| ENSG00000106617 | *PRKAG2* | 4.806693018 | 0.035006693 | 0.011876542 | 0.023130151 | 0.999933887 | 0.018372081 | 0.101485277 | -0.083113196 | 0.560537649 |
| ENSG00000184786 | *TCTE3* | 0.427239996 | -0.02179815 | 0.263501672 | -0.285299822 | 0.999933887 | -0.106713795 | 0.08244792 | -0.189161715 | 0.560537649 |
| ENSG00000126581 | *BECN1* | 5.414187203 | -0.006599936 | 0.020971823 | -0.027571759 | 0.999933887 | 0.055481676 | 0.12139826 | -0.065916584 | 0.560537649 |
| ENSG00000013563 | *DNASE1L1* | 3.61324005 | 0.005799914 | 0.081829809 | -0.076029895 | 0.999933887 | -0.169776277 | -0.068975968 | -0.100800309 | 0.560537649 |
| ENSG00000115009 | *CCL20* | 2.497830016 | 0.49179783 | 0.744759038 | -0.252961208 | 0.999933887 | 4.006886721 | 4.548872928 | -0.541986207 | 0.560983125 |
| ENSG00000188493 | *C19orf54* | 2.676965487 | 0.069886843 | -0.088841743 | 0.158728586 | 0.999933887 | 0.317362635 | 0.425168343 | -0.107805708 | 0.560983125 |
| ENSG00000169385 | *RNASE2* | 2.478254897 | 0.069556699 | -0.006768405 | 0.076325104 | 0.999933887 | -0.028063881 | 0.145032632 | -0.173096513 | 0.560983125 |
| ENSG00000100211 | *CBY1* | -0.403803878 | 0.053383474 | 0.211727187 | -0.158343713 | 0.999933887 | -0.215700061 | 0.063346341 | -0.279046402 | 0.560983125 |
| ENSG00000236254 | *MTND4P14* | -1.454007878 | -0.10879382 | 0.029724192 | -0.138518011 | 0.999933887 | 0.136508975 | 0.784816787 | -0.648307812 | 0.561014404 |
| ENSG00000162892 | *IL24* | 1.011010808 | 0.149527529 | -0.074854717 | 0.224382245 | 0.999933887 | -0.249032475 | -0.102931932 | -0.146100543 | 0.56108307 |
| ENSG00000176928 | *GCNT4* | 1.24311599 | -0.086127512 | -0.105744924 | 0.019617411 | 0.999933887 | -0.120456439 | 0.064046209 | -0.184502648 | 0.56108307 |
| ENSG00000286656 | *AC008875.2* | 1.652014745 | 0.046398032 | 0.08356288 | -0.037164849 | 0.999933887 | -0.02563128 | -0.231642836 | 0.206011556 | 0.56108307 |
| ENSG00000204070 | *SYS1* | 5.120783734 | -0.047037644 | 0.000819954 | -0.047857598 | 0.999933887 | 0.404289566 | 0.47868 | -0.074390434 | 0.561430113 |
| ENSG00000072609 | *CHFR* | 6.890191721 | 0.010623558 | 0.004526763 | 0.006096795 | 0.999933887 | -0.169833855 | -0.115386711 | -0.054447145 | 0.561430113 |
| ENSG00000197046 | *SIGLEC15* | -0.09450986 | -0.06501445 | 0.116125117 | -0.181139567 | 0.999933887 | -0.414722674 | -0.688706596 | 0.273983923 | 0.561656425 |
| ENSG00000260898 | *ADPGK-AS1* | 1.744660548 | 0.137241875 | 0.036884588 | 0.100357287 | 0.999933887 | 0.817977779 | 1.091469275 | -0.273491497 | 0.561798572 |
| ENSG00000115844 | *DLX2* | -0.672825282 | 0.570187503 | -0.28019723 | 0.850384733 | 0.144167088 | 1.400216405 | 1.162608319 | 0.237608086 | 0.562196353 |
| ENSG00000099290 | *WASHC2A* | 6.371186319 | -0.026524176 | -0.058708276 | 0.0321841 | 0.999933887 | 0.05660835 | 0.126804816 | -0.070196466 | 0.562196353 |
| ENSG00000184436 | *THAP7* | 3.883466006 | 0.013951083 | -0.119915044 | 0.133866127 | 0.999933887 | 0.256437418 | 0.135334235 | 0.121103183 | 0.562196353 |
| ENSG00000169641 | *LUZP1* | 3.651598424 | -0.026333846 | -0.027179633 | 0.000845787 | 0.999933887 | 0.010775754 | 0.12087173 | -0.110095976 | 0.56232508 |
| ENSG00000177432 | *NAP1L5* | 1.368789095 | 0.078052435 | 0.150366888 | -0.072314453 | 0.999933887 | 0.097819098 | -0.094851543 | 0.192670642 | 0.562549552 |
| ENSG00000108039 | *XPNPEP1* | 3.593888584 | -0.044655082 | 0.084881085 | -0.129536167 | 0.999933887 | 0.196087621 | 0.293526649 | -0.097439028 | 0.562549552 |
| ENSG00000160058 | *BSDC1* | 5.918952349 | -0.025432691 | 0.020355391 | -0.045788082 | 0.999933887 | -0.122472463 | -0.066337863 | -0.0561346 | 0.562549552 |
| ENSG00000100600 | *LGMN* | 0.686015018 | -0.018780935 | -0.067534202 | 0.048753267 | 0.999933887 | -0.193774456 | 0.080770709 | -0.274545164 | 0.562549552 |
| ENSG00000146555 | *SDK1* | -1.230384432 | 0.231691227 | -0.01168859 | 0.243379817 | 0.999933887 | -0.338166849 | 0.019175245 | -0.357342094 | 0.562980648 |
| ENSG00000118898 | *PPL* | 1.465695925 | -0.037421156 | -0.03496464 | -0.002456516 | 0.999933887 | -0.484591591 | -0.287648248 | -0.196943344 | 0.562980648 |
| ENSG00000132603 | *NIP7* | 3.171340591 | -0.082144406 | 0.020815338 | -0.102959745 | 0.999933887 | -0.034541102 | -0.137437393 | 0.102896291 | 0.563050611 |
| ENSG00000161055 | *SCGB3A1* | -2.141746029 | 0.235333214 | -0.3863466 | 0.621679814 | 0.999933887 | 0.375854007 | 0.018680562 | 0.357173445 | 0.563077771 |
| ENSG00000165487 | *MICU2* | 4.099271165 | 0.025503684 | -0.006913475 | 0.032417158 | 0.999933887 | -0.442047259 | -0.319054871 | -0.122992388 | 0.563077771 |
| ENSG00000073578 | *SDHA* | 1.621497085 | 0.034713254 | -0.057278818 | 0.091992071 | 0.999933887 | 0.05839533 | -0.093660881 | 0.152056211 | 0.563204084 |
| ENSG00000167608 | *TMC4* | 1.142634172 | 0.194414579 | 0.049811164 | 0.144603414 | 0.999933887 | -0.054184892 | -0.274756946 | 0.220572054 | 0.563261988 |
| ENSG00000117360 | *PRPF3* | 5.349143092 | -0.027066206 | 0.017191282 | -0.044257489 | 0.999933887 | 0.347442826 | 0.265463431 | 0.081979395 | 0.563261988 |
| ENSG00000060140 | *STYK1* | -0.678735529 | 0.068786163 | -0.702839325 | 0.771625488 | 0.944094537 | -0.069133506 | 0.222859042 | -0.291992548 | 0.563261988 |
| ENSG00000136870 | *ZNF189* | 3.287560637 | -0.048218714 | -0.1245654 | 0.076346686 | 0.999933887 | 0.029250013 | 0.126978325 | -0.097728312 | 0.563352441 |
| ENSG00000011295 | *TTC19* | 3.547705584 | -0.069208134 | -0.040057075 | -0.029151059 | 0.999933887 | 0.032736675 | -0.04184269 | 0.074579365 | 0.563591477 |
| ENSG00000248593 | *DSTNP2* | -0.116594005 | -0.056770003 | 0.006240994 | -0.063010997 | 0.999933887 | 0.135510231 | -0.14860467 | 0.284114901 | 0.5636489 |
| ENSG00000167770 | *OTUB1* | 5.603109796 | 0.062922002 | -0.02884781 | 0.091769811 | 0.999933887 | -0.026376395 | -0.089507752 | 0.063131357 | 0.56373233 |
| ENSG00000287771 | *AC007271.1* | 2.184513023 | 0.042521808 | 0.104080067 | -0.061558259 | 0.999933887 | 0.689457369 | 0.856810116 | -0.167352747 | 0.563852117 |
| ENSG00000159784 | *FAM131B* | -0.543843757 | 0.243727378 | -0.244088364 | 0.487815742 | 0.999933887 | -0.335640437 | -0.731397041 | 0.395756604 | 0.564004956 |
| ENSG00000115084 | *SLC35F5* | 4.74312492 | -0.017386139 | -0.011262184 | -0.006123955 | 0.999933887 | -0.465902552 | -0.35879717 | -0.107105383 | 0.564005472 |
| ENSG00000167543 | *TP53I13* | 3.511157858 | 0.008221598 | 0.21959445 | -0.211372852 | 0.999933887 | 0.094642748 | -0.006215231 | 0.100857979 | 0.564354962 |
| ENSG00000169184 | *MN1* | -0.465442883 | 0.1655865 | 0.168639124 | -0.003052624 | 0.999933887 | 0.19260331 | 0.498230978 | -0.305627669 | 0.56442543 |
| ENSG00000011258 | *MBTD1* | 4.132854868 | -0.039222544 | -0.092290091 | 0.053067547 | 0.999933887 | -0.629480758 | -0.748202908 | 0.118722151 | 0.564455386 |
| ENSG00000154473 | *BUB3* | 4.504072939 | -0.009660734 | -0.072479614 | 0.06281888 | 0.999933887 | -0.027971061 | -0.111324608 | 0.083353546 | 0.56469376 |
| ENSG00000168092 | *PAFAH1B2* | 5.632676947 | -0.113612196 | -0.027737218 | -0.085874978 | 0.999933887 | -0.59410867 | -0.6863414 | 0.09223273 | 0.56469706 |
| ENSG00000085733 | *CTTN* | -0.680426681 | -0.394103261 | 0.065304689 | -0.45940795 | 0.999933887 | -0.256598635 | 0.040882918 | -0.297481553 | 0.56469706 |
| ENSG00000248124 | *RRN3P1* | 5.060273886 | -0.029248388 | -0.059616042 | 0.030367653 | 0.999933887 | -0.020527216 | 0.067914934 | -0.088442151 | 0.56469706 |
| ENSG00000164331 | *ANKRA2* | 3.634290374 | -0.001805273 | 0.034185382 | -0.035990655 | 0.999933887 | -0.047397474 | 0.034128365 | -0.08152584 | 0.56469706 |
| ENSG00000167566 | *NCKAP5L* | 3.777376706 | 0.069145544 | -0.019397518 | 0.088543062 | 0.999933887 | 0.020672165 | -0.099307592 | 0.119979757 | 0.564737629 |
| ENSG00000147592 | *LACTB2* | 1.396755509 | 0.179434457 | 0.008653489 | 0.170780968 | 0.999933887 | 0.056749784 | -0.069138032 | 0.125887816 | 0.564744323 |
| ENSG00000272899 | *ATP6V1FNB* | 3.243042497 | 0.084925921 | 0.355019606 | -0.270093684 | 0.944094537 | 0.798262131 | 0.901234706 | -0.102972575 | 0.564756864 |
| ENSG00000171806 | *METTL18* | -0.041691795 | -0.339884261 | 0.140600393 | -0.480484654 | 0.999933887 | -0.072290719 | 0.141998887 | -0.214289606 | 0.565095154 |
| ENSG00000222043 | *AC079305.1* | 3.130071861 | 0.162885249 | 0.153406596 | 0.009478653 | 0.999933887 | 1.761551987 | 1.961331431 | -0.199779445 | 0.565242069 |
| ENSG00000237914 | *SIRPG-AS1* | -1.320040626 | 0.058964257 | -0.50777597 | 0.566740227 | 0.999933887 | -0.182793119 | 0.160165868 | -0.342958987 | 0.565383058 |
| ENSG00000124126 | *PREX1* | 10.485574979 | 0.024324251 | 0.043091359 | -0.018767107 | 0.999933887 | 0.009914935 | -0.040438008 | 0.050352943 | 0.565929937 |
| ENSG00000128694 | *OSGEPL1* | -0.594480308 | 0.052520571 | -0.072120336 | 0.124640907 | 0.999933887 | -0.145660736 | 0.14282931 | -0.288490046 | 0.565971592 |
| ENSG00000179715 | *PCED1B* | 3.107635063 | -0.019225145 | 0.005837986 | -0.025063132 | 0.999933887 | 0.099033785 | 0.011273526 | 0.087760259 | 0.565974227 |

| | | | | | | | | | | |
|---|---|---|---|---|---|---|---|---|---|---|
| ENSG00000156170 | *NDUFAF6* | 1.049500426 | -0.015188397 | 0.185036313 | -0.20022471 | 0.999933887 | 0.19421301 | 0.396704575 | -0.202491564 | 0.566199774 |
| ENSG00000130544 | *ZNF557* | 3.162681662 | -0.118877583 | -0.114844425 | -0.004033158 | 0.999933887 | -0.245711982 | -0.153565107 | -0.092146875 | 0.566261905 |
| ENSG00000263982 | *AC009716.1* | -0.145245261 | 0.218404327 | 0.005943362 | 0.212460965 | 0.999933887 | -0.158172264 | -0.433651331 | 0.275479067 | 0.566261905 |
| ENSG00000026950 | *BTN3A1* | 7.167291459 | 0.012539148 | 0.014593349 | -0.002054201 | 0.999933887 | -0.117092316 | -0.180782805 | 0.063690489 | 0.566261905 |
| ENSG00000182759 | *MAFA* | -0.78951907 | 0.093340896 | -0.487277866 | 0.580618763 | 0.999933887 | -0.508960043 | -0.854248017 | 0.345287974 | 0.566442075 |
| ENSG00000175756 | *AURKAIP1* | 6.228130577 | 0.094475153 | 0.141885575 | -0.047410422 | 0.999933887 | 0.02277609 | -0.099457076 | 0.122233166 | 0.566500352 |
| ENSG00000281398 | *SNHG4* | 0.35926767 | -0.209164583 | -0.264475003 | 0.05531042 | 0.999933887 | 0.733194882 | 0.941474405 | -0.208279523 | 0.566738862 |
| ENSG00000121058 | *COIL* | 2.90296333 | -0.078288046 | -0.061276059 | -0.017011987 | 0.999933887 | -0.395828388 | -0.28697147 | -0.108856918 | 0.566738862 |
| ENSG00000198792 | *TMEM184B* | 7.250241482 | -0.003979249 | 0.034295078 | -0.038274327 | 0.999933887 | -0.010053993 | 0.06112982 | -0.071183812 | 0.566738862 |
| ENSG00000087299 | *L2HGDH* | -0.704774498 | -0.023097658 | -0.135398031 | 0.112300374 | 0.999933887 | -0.24783561 | 0.132403263 | -0.380238872 | 0.566809675 |
| ENSG00000188042 | *ARL4C* | 7.468484542 | 0.022025465 | 0.007168945 | 0.01485652 | 0.999933887 | 0.09782645 | 0.169239143 | -0.071412693 | 0.566921082 |
| ENSG00000072954 | *TMEM38A* | -0.480096911 | 0.098816845 | 0.124502999 | -0.025686154 | 0.999933887 | -0.585298441 | -0.216149772 | -0.36914867 | 0.566984193 |
| ENSG00000285533 | *RELA-DT* | 1.285348957 | -0.04090477 | -0.048245266 | 0.007340497 | 0.999933887 | -0.138075633 | -0.317838511 | 0.179762878 | 0.567244328 |
| ENSG00000151491 | *EPS8* | -0.181424463 | -0.22741609 | -0.067314591 | -0.160101499 | 0.999933887 | -0.374870014 | -0.174070098 | -0.200799916 | 0.567380433 |
| ENSG00000157379 | *DHRS1* | 4.319071365 | 0.127229541 | 0.050273499 | 0.076956042 | 0.999933887 | 0.901197471 | 1.008921359 | -0.107723889 | 0.567584099 |
| ENSG00000166436 | *TRIM66* | 2.874796416 | 0.050406775 | -0.068004991 | 0.118411766 | 0.999933887 | -0.16324769 | -0.272630014 | 0.109382325 | 0.567673512 |
| ENSG00000094916 | *CBX5* | 3.142790172 | -0.085180904 | -0.072719262 | -0.012461642 | 0.999933887 | 0.003383866 | 0.112642055 | -0.109258189 | 0.567871478 |
| ENSG00000115944 | *COX7A2L* | 4.320247985 | -0.010323597 | -0.116150578 | 0.105826981 | 0.999933887 | -0.237346384 | -0.326041198 | 0.088694814 | 0.567873151 |
| ENSG00000168461 | *RAB31* | 7.009263517 | -0.064364411 | -0.041108951 | -0.02325546 | 0.999933887 | -0.782864412 | -0.682857203 | -0.100007209 | 0.567876769 |
| ENSG00000248571 | *AC106882.1* | 0.513814192 | -0.242624496 | -0.080495518 | -0.162128978 | 0.999933887 | -0.006521133 | -0.311401466 | 0.304880334 | 0.56791542 |
| ENSG00000205085 | *FAM71F2* | 1.624470165 | -0.06824608 | -0.05495993 | -0.013286151 | 0.999933887 | 0.068039911 | 0.258473831 | -0.19043392 | 0.56791542 |
| ENSG00000085832 | *EPS15* | 5.203554586 | 0.0117669 | -0.012983621 | 0.024750521 | 0.999933887 | -0.415927741 | -0.526074616 | 0.110146875 | 0.56791542 |
| ENSG00000117016 | *RIMS3* | -0.675897421 | 0.118567001 | 0.525261487 | -0.406694485 | 0.999933887 | 0.016306264 | 0.344826161 | -0.328519897 | 0.567968947 |
| ENSG00000162825 | *NBPF20* | -0.514150601 | -0.033022357 | 0.025513001 | -0.058535357 | 0.999933887 | 0.114616599 | 0.426381114 | -0.311764515 | 0.567968947 |
| ENSG00000100485 | *SOS2* | 6.29809357 | 0.032232705 | -0.081751156 | 0.113983861 | 0.999933887 | -0.621754711 | -0.488774328 | -0.132980383 | 0.568125869 |
| ENSG00000157827 | *FMNL2* | 1.067129355 | -0.01131511 | -0.007530901 | -0.003784209 | 0.999933887 | -0.579745901 | -0.772595699 | 0.192849798 | 0.568125869 |
| ENSG00000260306 | *AC092375.2* | 0.393527898 | 0.364225668 | 0.18712618 | 0.177099488 | 0.999933887 | 0.363721963 | 0.602352456 | -0.238630493 | 0.568239223 |
| ENSG00000177294 | *FBXO39* | -1.595489311 | -0.209375686 | -0.056705231 | -0.152670455 | 0.999933887 | 0.059479545 | -0.477721856 | 0.537201401 | 0.568239223 |
| ENSG00000259436 | *AC010247.2* | 0.689003971 | -0.290997953 | 0.132170371 | -0.423168324 | 0.999933887 | 0.362150335 | 0.528883075 | -0.166732739 | 0.568523334 |
| ENSG00000237973 | *MTCO1P12* | 2.657220577 | -0.046707539 | -0.040517014 | -0.006190524 | 0.999933887 | 0.027623083 | 0.159631642 | -0.132008559 | 0.568523334 |
| ENSG00000143365 | *RORC* | 0.233725422 | 0.189227042 | 0.035077402 | 0.154149639 | 0.999933887 | 0.133237238 | -0.114687005 | 0.247924243 | 0.568913638 |
| ENSG00000198538 | *ZNF28* | 1.103745098 | -0.00662194 | -0.246118782 | 0.239496842 | 0.999933887 | -0.32249473 | -0.148000928 | -0.174493802 | 0.568913638 |
| ENSG00000144596 | *GRIP2* | 0.349321166 | -0.177019376 | -0.018969974 | -0.158049401 | 0.999933887 | 0.204105688 | 0.454131521 | -0.250025833 | 0.568961365 |
| ENSG00000211459 | *MT-RNR1* | 9.717257386 | 0.116654584 | 0.177056378 | -0.060401794 | 0.999933887 | 0.314043947 | 0.436076916 | -0.122032969 | 0.568976133 |
| ENSG00000127603 | *MACF1* | 7.33027726 | -0.059855088 | -0.024873389 | -0.034981699 | 0.999933887 | -0.343855365 | -0.448512556 | 0.104657191 | 0.568976133 |
| ENSG00000274280 | *MIR6069* | 1.611574008 | 0.031480805 | 0.052895359 | -0.021414554 | 0.999933887 | 0.702087855 | 0.889806888 | -0.187719032 | 0.568987666 |
| ENSG00000237298 | *TTN-AS1* | 2.075569751 | -0.019097172 | 0.006072755 | -0.025169927 | 0.999933887 | -0.231786441 | -0.390047993 | 0.158261553 | 0.569247989 |
| ENSG00000281005 | *LINC00921* | 4.827250191 | -0.016300833 | 0.06770834 | -0.084009173 | 0.999933887 | -0.023861469 | 0.069209316 | -0.093070785 | 0.569350807 |
| ENSG00000198887 | *SMC5* | 3.802907565 | -0.022202388 | -0.130556307 | 0.108353919 | 0.999933887 | -0.378416717 | -0.167018298 | -0.211398419 | 0.569350807 |
| ENSG00000141448 | *GATA6* | -0.833963785 | 0.174853724 | 0.622827282 | -0.447973558 | 0.999933887 | 0.059068317 | -0.239712762 | 0.298781079 | 0.569381478 |
| ENSG00000273345 | *AC104109.4* | -0.268471656 | 0.026874342 | -0.076623833 | 0.103498175 | 0.999933887 | 0.018028553 | 0.29488567 | -0.276857118 | 0.569381478 |
| ENSG00000112977 | *DAP* | 5.750891097 | -0.002937211 | -0.034066374 | 0.031129163 | 0.999933887 | -0.036856219 | -0.11624957 | 0.07939335 | 0.569415654 |
| ENSG00000063015 | *SEZ6* | 2.862132983 | 0.101573359 | 0.080276001 | 0.021297359 | 0.999933887 | 0.707590826 | 0.851986354 | -0.144395527 | 0.569527472 |
| ENSG00000137818 | *RPLP1* | 8.222601129 | 0.009874096 | -0.035113358 | 0.044987454 | 0.999933887 | 0.174441765 | 0.249719871 | -0.075278105 | 0.569527472 |
| ENSG00000123453 | *SARDH* | -0.868969998 | 0.025141513 | 0.129834543 | -0.10469303 | 0.999933887 | -0.187946059 | 0.147008496 | -0.334954555 | 0.569839305 |
| ENSG00000149292 | *TTC12* | 1.540736335 | 0.108249474 | 0.038962472 | 0.069287003 | 0.999933887 | 0.001875971 | -0.133588851 | 0.135464822 | 0.569956359 |
| ENSG00000135045 | *C9orf40* | 3.306364165 | -0.083408419 | -0.144054399 | 0.06064598 | 0.999933887 | -0.662247043 | -0.568647178 | -0.093599865 | 0.569956359 |
| ENSG00000153339 | *TRAPPC8* | 5.714015199 | -0.079886276 | -0.111995125 | 0.03210885 | 0.999933887 | -0.653926713 | -0.531682471 | -0.122244243 | 0.570240065 |
| ENSG00000260033 | *AC023813.2* | 0.621122527 | 0.366792758 | 0.38891981 | -0.022127052 | 0.999933887 | 1.530694765 | 1.899030381 | -0.368335617 | 0.570460983 |
| ENSG00000284526 | *AC015802.6* | 1.249286283 | -0.112649066 | -0.017341437 | -0.095307629 | 0.999933887 | -0.810809438 | -0.961391426 | 0.150581988 | 0.570460983 |
| ENSG00000109536 | *FRG1* | 3.73603053 | -0.045490619 | -0.042073456 | -0.003417163 | 0.999933887 | -0.199476051 | -0.100301137 | -0.099174914 | 0.570460983 |
| ENSG00000167118 | *URM1* | 3.642712556 | -0.022919832 | -0.00696293 | -0.015956902 | 0.999933887 | 0.026658888 | -0.057642575 | 0.084301463 | 0.570460983 |
| ENSG00000113319 | *RASGRF2* | 1.606123854 | 0.023752786 | 0.101034246 | -0.07728146 | 0.999933887 | -0.004029564 | 0.137703983 | -0.141733547 | 0.570460983 |
| ENSG00000174444 | *RPL4* | 8.111058384 | -0.032523424 | -0.13617366 | 0.103650236 | 0.999933887 | 0.107289747 | 0.187437491 | -0.080147743 | 0.570571834 |
| ENSG00000008988 | *RPS20* | 7.611378184 | -0.00356769 | -0.062873249 | 0.059305559 | 0.999933887 | 0.166057133 | 0.244329051 | -0.078271918 | 0.570713792 |
| ENSG00000179029 | *TMEM107* | 1.421260505 | 0.032040688 | 0.151962667 | -0.119921979 | 0.999933887 | 0.107113138 | -0.106861262 | 0.2139744 | 0.570921711 |
| ENSG00000242324 | *AL049634.1* | 0.096602709 | -0.081898838 | -0.135633197 | 0.053734359 | 0.999933887 | -0.223224515 | -0.451664277 | 0.228439762 | 0.571222139 |
| ENSG00000160767 | *FAM189B* | 1.933311028 | 0.115452943 | -0.00305761 | 0.118510552 | 0.999933887 | -0.073017042 | -0.216065176 | 0.143048134 | 0.571247958 |
| ENSG00000172613 | *RAD9A* | 4.9856506 | 0.07798223 | 0.068320369 | 0.009661861 | 0.999933887 | 0.103231077 | 0.011511778 | 0.091719299 | 0.571405204 |
| ENSG00000138668 | *HNRNPD* | 6.902225375 | 0.005578614 | 0.018292616 | -0.012714002 | 0.999933887 | -0.187682034 | -0.249232374 | 0.061550341 | 0.571448182 |
| ENSG00000241975 | *ELOCP19* | 0.407720553 | 0.028693233 | -0.094422603 | 0.123115836 | 0.999933887 | -0.724622906 | -1.214032822 | 0.489409916 | 0.571674884 |
| ENSG00000285793 | *ANAPC1P2* | 1.670963229 | 0.145414104 | 0.125779006 | 0.019635099 | 0.999933887 | -0.752863634 | -0.98240473 | 0.229541096 | 0.57172616 |
| ENSG00000255320 | *AP000759.1* | 1.002134254 | -0.052261049 | -0.096002449 | 0.043741401 | 0.999933887 | -0.338081837 | -0.576678213 | 0.238596376 | 0.57172616 |
| ENSG00000170906 | *NDUFA3* | 3.085555927 | 0.036216905 | -0.025661408 | 0.061878314 | 0.999933887 | 0.031206228 | -0.077918865 | 0.109125093 | 0.57172616 |
| ENSG00000114107 | *CEP70* | -0.550199204 | -0.219825528 | -0.264725453 | 0.044899926 | 0.999933887 | -0.06755548 | 0.264278086 | -0.331833566 | 0.571796712 |
| ENSG00000062725 | *APPBP2* | 4.094963494 | -0.090424612 | -0.049678042 | -0.04074657 | 0.999933887 | -0.312526505 | -0.22538412 | -0.087142385 | 0.571861734 |
| ENSG00000137265 | *IRF4* | 4.090161946 | -0.086991433 | 0.045481364 | -0.132472797 | 0.999933887 | 0.326328054 | 0.415450534 | -0.089122479 | 0.572037927 |
| ENSG00000158528 | *PPP1R9A* | -1.684315186 | -0.413424794 | -0.42415531 | 0.010730516 | 0.999933887 | -1.22254922 | -1.477672156 | 0.255122936 | 0.572037927 |
| ENSG00000284697 | *AC006511.5* | 0.38237926 | -0.187467385 | 0.001150065 | -0.18861745 | 0.999933887 | 0.808403545 | 1.035329976 | -0.226926431 | 0.572037927 |
| ENSG00000166965 | *RCCD1* | 0.300824272 | 0.078434699 | -0.004275569 | 0.082710268 | 0.999933887 | 0.229069902 | 0.024130882 | 0.20493902 | 0.572037927 |
| ENSG00000274210 | *RNVU1-27* | -0.851814116 | -0.021701088 | 0.271909587 | -0.293610675 | 0.999933887 | 0.377132021 | 0.055948215 | 0.321183807 | 0.572037927 |
| ENSG00000118971 | *CCND2* | 4.828980693 | -0.062519831 | 0.028729427 | -0.091249258 | 0.999933887 | -0.100293601 | -0.180186288 | 0.079892687 | 0.572437936 |
| ENSG00000105369 | *CD79A* | 2.035844489 | 0.132244675 | -0.04526385 | 0.177508525 | 0.999933887 | -0.009073698 | 0.122519699 | -0.131593398 | 0.572470886 |
| ENSG00000162419 | *GMEB1* | 4.415546727 | -0.020564404 | 0.006130674 | -0.026695078 | 0.999933887 | -0.105227385 | -0.034348071 | -0.070879314 | 0.572621611 |
| ENSG00000196214 | *ZNF766* | 3.725470031 | -0.176581528 | -0.187988593 | 0.011407064 | 0.999933887 | -0.52860221 | -0.639599569 | 0.110997359 | 0.572782162 |
| ENSG00000171135 | *JAGN1* | 2.465831264 | -0.139757808 | -0.10305142 | -0.036706388 | 0.999933887 | 0.030443119 | -0.075757369 | 0.106200489 | 0.572782162 |
| ENSG00000265399 | *AP005329.2* | -0.168606598 | 0.213797549 | 0.319683607 | -0.105886059 | 0.999933887 | 0.045808411 | 0.270819608 | -0.225011196 | 0.572782162 |
| ENSG00000135205 | *CCDC146* | 1.945493296 | -0.121167015 | -0.191521885 | 0.07035487 | 0.999933887 | -0.196940826 | 0.042190876 | -0.239131702 | 0.572782162 |
| ENSG00000229515 | *FLT1P1* | 0.589264672 | 0.161414331 | -0.08405722 | 0.245471552 | 0.999933887 | -0.886886556 | -0.550362738 | -0.336523817 | 0.572782162 |
| ENSG00000269959 | *SPACA6P-AS* | -0.696989505 | 0.19563065 | 0.112193814 | 0.083436836 | 0.999933887 | 1.43728859 | 1.7541221 | -0.31683351 | 0.572782162 |
| ENSG00000119632 | *IFI27L2* | 1.881202022 | 0.067443119 | 0.008682299 | 0.05876082 | 0.999933887 | 0.570757338 | 0.752978685 | -0.182221347 | 0.572782162 |
| ENSG00000171777 | *RASGRP4* | 7.157490517 | 0.024372648 | 0.074659153 | -0.050286505 | 0.999933887 | -0.103254122 | -0.181129788 | 0.077875666 | 0.572782162 |
| ENSG00000249437 | *NAIP* | 4.0090528 | -0.028012262 | -0.031673497 | 0.003661235 | 0.999933887 | -0.953479878 | -1.066484051 | 0.113004174 | 0.572782162 |
| ENSG00000119686 | *FLVCR2* | 0.729425534 | -0.048241847 | -0.109747446 | 0.061505598 | 0.999933887 | 0.194753804 | 0.43452954 | -0.239775736 | 0.572782162 |
| ENSG00000288056 | *AL023806.4* | -1.569212785 | -0.026753742 | -0.293186965 | 0.266433222 | 0.999933887 | 0.090228524 | 0.304845074 | -0.21461655 | 0.57295526 |
| ENSG00000108021 | *TASOR2* | 4.380541027 | -0.02915051 | -0.167522627 | 0.138372117 | 0.999933887 | -0.335818269 | -0.168588926 | -0.167229343 | 0.573151185 |
| ENSG00000134490 | *TMEM241* | -0.68124367 | -0.389663105 | 0.236918908 | -0.626582013 | 0.999933887 | -0.288043928 | -0.018524872 | -0.269519056 | 0.573171562 |
| ENSG00000139537 | *CCDC65* | 1.103340177 | -0.049364968 | -0.02157774 | -0.027787228 | 0.999933887 | -0.072999729 | 0.079171982 | -0.152171711 | 0.573398829 |
| ENSG00000123080 | *CDKN2C* | 0.660342819 | 0.17815334 | 0.009796097 | 0.168357243 | 0.999933887 | 0.781413561 | 0.989070049 | -0.207656489 | 0.573518953 |
| ENSG00000260005 | *AC027601.1* | 1.975271982 | -0.009551325 | 0.144248773 | -0.153800097 | 0.999933887 | -0.164472365 | -0.003366604 | -0.161105761 | 0.574074781 |
| ENSG00000236088 | *COX10-AS1* | 0.85943727 | 0.091220918 | -0.224887104 | 0.316108022 | 0.999933887 | -0.244932702 | -0.062474586 | -0.182458116 | 0.574146362 |
| ENSG00000222046 | *DCDC2B* | -0.014090983 | 0.08206942 | -0.181778805 | 0.263848225 | 0.999933887 | 0.177771972 | -0.098740574 | 0.276512546 | 0.574146452 |
| ENSG00000131773 | *KHDRBS3* | -0.544480653 | -0.060943747 | 0.29891831 | -0.359862057 | 0.999933887 | 1.396932238 | 1.619357035 | -0.222424798 | 0.574149064 |
| ENSG00000148773 | *MKI67* | 0.124703761 | 0.020824774 | 0.094576485 | -0.073751711 | 0.999933887 | 0.221261258 | 0.499067255 | -0.277805997 | 0.574468447 |
| ENSG00000069329 | *VPS35* | 5.720197424 | -0.120036099 | -0.146946194 | 0.026910095 | 0.999933887 | -0.571128177 | -0.632317459 | 0.061189282 | 0.575286473 |
| ENSG00000141570 | *CBX8* | 1.001800413 | 0.092678191 | 0.12948333 | -0.036805139 | 0.999933887 | -0.11884108 | -0.348801853 | 0.229960773 | 0.575286473 |
| ENSG00000213600 | *U73169.1* | -0.52802878 | -0.061768676 | -0.243208409 | 0.181439733 | 0.999933887 | -0.585814708 | -0.904121249 | 0.318306541 | 0.575286473 |
| ENSG00000144036 | *EXOC6B* | 0.693264892 | 0.032119215 | -0.163385923 | 0.195505138 | 0.999933887 | -0.267713929 | -0.089535111 | -0.178178818 | 0.575286473 |
| ENSG00000004866 | *ST7* | -0.109380883 | -0.212852054 | 0.038405978 | -0.251258032 | 0.999933887 | -0.159183086 | 0.086377084 | -0.24556017 | 0.575391422 |
| ENSG00000171612 | *SLC25A33* | 4.149162284 | 0.006688122 | -0.027103935 | 0.033792057 | 0.999933887 | 0.456512015 | 0.572415645 | -0.11590363 | 0.575391422 |
| ENSG00000074582 | *BCS1L* | 1.505007344 | -0.013574233 | -0.17122967 | 0.157655436 | 0.999933887 | 0.411256919 | 0.544831726 | -0.133574808 | 0.575558851 |
| ENSG00000146707 | *POMZP3* | 1.926839433 | -0.119104449 | -0.014760596 | -0.104343852 | 0.999933887 | 0.751879496 | 0.933945865 | -0.182066369 | 0.575588347 |
| ENSG00000085788 | *DDHD2* | 2.926994666 | -0.078253993 | -0.20026648 | 0.122012487 | 0.999933887 | -0.009948052 | 0.096556769 | -0.10650482 | 0.575730251 |
| ENSG00000157350 | *ST3GAL2* | 7.213195815 | 0.050098852 | 0.064258677 | -0.014159825 | 0.999933887 | 0.28445554 | 0.377811552 | -0.093356012 | 0.575999234 |
| ENSG00000213965 | *NUDT19* | 1.307102304 | -0.049246314 | 0.122326299 | -0.171572614 | 0.999933887 | -0.041253694 | 0.140134509 | -0.181388203 | 0.575999234 |

| | | | | | | | | | | |
|---|---|---|---|---|---|---|---|---|---|---|
| ENSG00000282133 | *TRBJ1-3* | -0.481853579 | 0.161739052 | 0.058981965 | 0.102757087 | 0.999933887 | -0.019033068 | 0.220868068 | -0.239901136 | 0.576072414 |
| ENSG00000180694 | *TMEM64* | 1.664996866 | -0.084662256 | 0.029156756 | -0.113819012 | 0.999933887 | 0.095996276 | 0.27031783 | -0.174321554 | 0.576072414 |
| ENSG00000124383 | *MPHOSPH10* | 2.719202410 | -0.161404682 | -0.237357648 | 0.075952966 | 0.999933887 | -0.096468241 | 0.037521762 | -0.133990004 | 0.576200875 |
| ENSG00000147679 | *UTP23* | 2.29151121 | -0.077486663 | -0.134127942 | 0.056641279 | 0.999933887 | -0.303977771 | -0.137959896 | -0.166017875 | 0.576386738 |
| ENSG00000254676 | *AP000873.4* | 0.45053472 | 0.184420088 | 0.237072852 | -0.052652764 | 0.999933887 | 0.17026258 | 0.390425055 | -0.220162475 | 0.576469021 |
| ENSG00000158158 | *CNNM4* | 3.312896499 | 0.092660922 | 0.078845629 | 0.013815293 | 0.999933887 | -0.134732383 | -0.029099999 | -0.105632384 | 0.576469021 |
| ENSG00000282980 | *AC055839.1* | -0.451993646 | -0.229831507 | 0.040760015 | -0.270591522 | 0.999933887 | -0.070679658 | -0.488922132 | 0.418242474 | 0.576469021 |
| ENSG00000259719 | *LINC02284* | -0.489747442 | 0.110204588 | 0.114197654 | -0.003993065 | 0.999933887 | -0.588591003 | -0.148413667 | -0.440177336 | 0.576469021 |
| ENSG00000134215 | *VAV3* | 4.708047913 | 0.029766667 | -0.065686977 | 0.095453644 | 0.999933887 | -0.212800754 | -0.300995367 | 0.088194613 | 0.576469021 |
| ENSG00000181036 | *FCRL6* | 3.134879942 | -0.016402187 | -0.098894352 | 0.082492165 | 0.999933887 | 0.092620727 | -0.008938367 | 0.101559094 | 0.576469021 |
| ENSG00000080007 | *DDX43* | -0.021204931 | -0.038851303 | 0.259374951 | -0.298226254 | 0.999933887 | 0.47249668 | 0.712060319 | -0.23956364 | 0.576469021 |
| ENSG00000152454 | *ZNF256* | 0.129039501 | 0.025471363 | 0.437225088 | -0.411753725 | 0.999933887 | 0.337173242 | 0.119907253 | 0.21726599 | 0.576469021 |
| ENSG00000108506 | *INTS2* | 1.593706942 | -0.007139328 | -0.203336448 | 0.196197121 | 0.999933887 | -0.076852831 | 0.075342359 | -0.15219519 | 0.576469021 |
| ENSG00000204392 | *LSM2* | 1.016253269 | 0.001666583 | -0.111926764 | 0.113593347 | 0.999933887 | -0.045119138 | 0.151704144 | -0.196823282 | 0.576469021 |
| ENSG00000185024 | *BRF1* | 3.410992425 | 0.028497249 | 0.111494163 | -0.082996913 | 0.999933887 | 0.068178797 | -0.028883716 | 0.097062513 | 0.576992395 |
| ENSG00000163026 | *WDCP* | 0.086642408 | 0.205770626 | 0.190919188 | 0.014851438 | 0.999933887 | 0.204751012 | -0.002487477 | 0.207238489 | 0.577159371 |
| ENSG00000052126 | *PLEKHA5* | 0.376338376 | -0.153594911 | -0.093633138 | -0.059961773 | 0.999933887 | -0.334685286 | -0.074852229 | -0.259833057 | 0.577349983 |
| ENSG00000076944 | *STXBP2* | 7.792165593 | 0.045435686 | 0.074497757 | -0.029062071 | 0.999933887 | 0.276699033 | 0.347614945 | -0.070915912 | 0.577786042 |
| ENSG00000157216 | *SSBP3* | 4.060045569 | 0.049669663 | 0.027585121 | 0.022084542 | 0.999933887 | 0.12872812 | 0.222899442 | -0.094171322 | 0.577786042 |
| ENSG00000181778 | *TMEM252* | 0.033309511 | 0.088704952 | 0.064449635 | 0.024255318 | 0.999933887 | 0.639669056 | 0.862996507 | -0.223327451 | 0.57779728 |
| ENSG00000137078 | *SIT1* | 1.899152849 | 0.012650036 | 0.082866233 | -0.070216197 | 0.999933887 | 0.24519568 | 0.092887493 | 0.152308187 | 0.577811162 |
| ENSG00000276698 | *AL136295.6* | 0.138622483 | 0.021198626 | -0.157485724 | 0.17868435 | 0.999933887 | 0.203321413 | 0.013121042 | 0.190200371 | 0.577811796 |
| ENSG00000131351 | *HAUS8* | 2.417225333 | 0.050448664 | 0.058186296 | -0.007737632 | 0.999933887 | -0.164409686 | -0.062522268 | -0.101887418 | 0.577905855 |
| ENSG00000107341 | *UBE2R2* | 8.307491259 | 0.017570836 | 0.002886393 | 0.014684442 | 0.999933887 | -0.661662777 | -0.584282974 | -0.077379804 | 0.577953129 |
| ENSG00000131196 | *NFATC1* | 5.031377923 | -0.007964305 | 0.056409768 | -0.064374073 | 0.999933887 | 0.187214515 | 0.258632036 | -0.071417521 | 0.577966497 |
| ENSG00000120675 | *DNAJC15* | 1.814093381 | 0.007766383 | -0.081321931 | 0.089088314 | 0.999933887 | -0.128350768 | 0.041917167 | -0.170267935 | 0.577966497 |
| ENSG00000100151 | *PICK1* | 2.812322116 | 0.113579231 | 0.062760555 | 0.050818676 | 0.999933887 | 0.16340588 | 0.263092949 | -0.099687068 | 0.577984339 |
| ENSG00000256087 | *ZNF432* | 2.127716384 | -0.103868577 | -0.140687994 | 0.036819417 | 0.999933887 | -0.044936929 | 0.114011012 | -0.158947941 | 0.577984339 |
| ENSG00000130669 | *PAK4* | 2.044980785 | 0.03515011 | 0.323248743 | -0.288098632 | 0.999933887 | 0.224100291 | 0.347592431 | -0.12349214 | 0.577984339 |
| ENSG00000149196 | *HIKESHI* | 1.410460199 | 0.050876895 | 0.064029978 | -0.013153082 | 0.999933887 | 0.115177096 | 0.240718705 | -0.125541609 | 0.578070644 |
| ENSG00000122707 | *RECK* | 0.747465722 | 0.08656855 | -0.276930267 | 0.363498817 | 0.999933887 | -0.150220456 | 0.04667756 | -0.196898017 | 0.578470193 |
| ENSG00000163818 | *LZTFL1* | 2.737564358 | -0.012699885 | 0.002882247 | -0.015582131 | 0.999933887 | -0.721264865 | -0.573567153 | -0.147697712 | 0.578470193 |
| ENSG00000102312 | *PORCN* | 1.891602943 | 0.041769883 | 0.099724973 | -0.05795509 | 0.999933887 | 0.096665649 | 0.23811286 | -0.141447211 | 0.578472056 |
| ENSG00000242574 | *HLA-DMB* | 2.287137769 | 0.003720641 | -0.110022925 | 0.113743566 | 0.999933887 | -0.362319633 | -0.504868343 | 0.142548711 | 0.578472056 |
| ENSG00000147124 | *ZNF41* | 1.155210941 | -0.102309832 | 0.064482107 | -0.166791939 | 0.999933887 | 0.052128244 | -0.119652142 | 0.171780387 | 0.578500016 |
| ENSG00000198858 | *R3HDM4* | 9.260326339 | 0.038209896 | 0.01843845 | 0.019771447 | 0.999933887 | -0.305738606 | -0.377960101 | 0.072221495 | 0.578500016 |
| ENSG00000135845 | *PIGC* | 2.771170838 | -0.008949543 | -0.131887303 | 0.122937761 | 0.999933887 | -0.136342239 | -0.250137498 | 0.113795259 | 0.578500016 |
| ENSG00000213994 | *AL157395.1* | -1.306686269 | -0.015343761 | 0.243573739 | -0.2589175 | 0.999933887 | -0.211552796 | 0.357746436 | -0.569299233 | 0.578500016 |
| ENSG00000123612 | *ACVR1C* | -1.0242719 | -0.103262039 | 0.090616956 | -0.193878995 | 0.999933887 | -0.423098532 | -0.056563645 | -0.366534887 | 0.578959057 |
| ENSG00000211797 | *TRAV17* | -0.382161841 | -0.027415506 | 0.202107158 | -0.229522665 | 0.999933887 | 0.133399723 | 0.360356066 | -0.226956343 | 0.578959057 |
| ENSG00000187764 | *SEMA4D* | 8.215544589 | -0.020195574 | -0.003840227 | -0.016355347 | 0.999933887 | 0.236049994 | 0.143231487 | 0.092818506 | 0.579198648 |
| ENSG00000283674 | *AC068587.4* | -0.157712047 | -0.066239792 | 0.187008323 | -0.253248114 | 0.999933887 | -0.215592199 | 0.11212591 | -0.327718108 | 0.579273169 |
| ENSG00000236426 | *AL583824.1* | -0.279236554 | 0.2068294 | 0.03714299 | 0.16968641 | 0.999933887 | 0.69761883 | 0.933211291 | -0.235592461 | 0.579395657 |
| ENSG00000184752 | *NDUFA12* | 2.496362883 | -0.071852537 | 0.019471603 | -0.091324141 | 0.999933887 | 0.025346225 | -0.086941271 | 0.112287495 | 0.579395657 |
| ENSG00000256581 | *NLRP9P1* | 1.525390939 | -0.070396717 | -0.115353096 | 0.044956379 | 0.999933887 | -0.381547327 | -0.17506141 | -0.206485917 | 0.579395657 |
| ENSG00000134744 | *TUT4* | 4.800696863 | -0.042218442 | -0.076888933 | 0.034670491 | 0.999933887 | -0.292503046 | -0.166990797 | -0.12551225 | 0.57974535 |
| ENSG00000176387 | *HSD11B2* | 1.881682979 | 0.132057167 | 0.429012373 | -0.296955206 | 0.999933887 | 1.063340327 | 0.826322892 | 0.237017435 | 0.579845605 |
| ENSG00000254635 | *WAC-AS1* | 3.798699484 | -0.016666839 | 0.076778265 | -0.093445105 | 0.999933887 | -0.29077642 | -0.195238965 | -0.095537455 | 0.579868853 |
| ENSG00000167525 | *PROCA1* | 2.084636866 | 0.159545387 | 0.172545776 | -0.013000389 | 0.999933887 | 0.059069687 | -0.100875779 | 0.159945465 | 0.579977509 |
| ENSG00000080854 | *IGSF9B* | 2.832370444 | -0.090331349 | 0.03022567 | -0.120557019 | 0.999933887 | 0.020071677 | 0.120730314 | -0.100658637 | 0.579977509 |
| ENSG00000165861 | *ZFYVE1* | 5.710547918 | -0.082234491 | 0.052512152 | -0.134746643 | 0.999933887 | -0.056001129 | 0.014530075 | -0.070531204 | 0.580066163 |
| ENSG00000114316 | *USP4* | 7.546363848 | -0.046588929 | -0.022873778 | -0.023715152 | 0.999933887 | -0.175989828 | -0.222147319 | 0.04615749 | 0.580066163 |
| ENSG00000260467 | *AC018552.2* | 1.916789557 | -0.045356567 | 0.061036622 | -0.106393189 | 0.999933887 | -0.239389514 | 0.002643161 | -0.242032674 | 0.580066163 |
| ENSG00000107242 | *PIP5K1B* | 1.397796968 | -0.045737861 | -0.032798895 | -0.012938967 | 0.999933887 | -0.120548344 | -0.287101658 | 0.166553314 | 0.580160851 |
| ENSG00000180385 | *EMC3-AS1* | 0.999522829 | -0.004320186 | -0.27450382 | 0.270183634 | 0.999933887 | -0.603822884 | -0.454120104 | -0.14970278 | 0.580160851 |
| ENSG00000131876 | *SNRPA1* | 3.987077611 | -0.08894675 | -0.065045343 | -0.023901407 | 0.999933887 | 0.068295863 | 0.135747767 | -0.067451905 | 0.580344372 |
| ENSG00000106789 | *CORO2A* | 3.56094914 | 0.104913867 | 0.150433284 | -0.045519417 | 0.999933887 | -0.031288644 | 0.071793137 | -0.103081781 | 0.580344372 |
| ENSG00000163655 | *GMPS* | 2.61657835 | -0.044856486 | -0.137159043 | 0.092302557 | 0.999933887 | 0.049419745 | -0.049703504 | 0.099123249 | 0.580344372 |
| ENSG00000160972 | *PPP1R16A* | 2.041228888 | 0.053865432 | 0.142503863 | -0.088638431 | 0.999933887 | 0.291483539 | 0.13624049 | 0.15524305 | 0.580344372 |
| ENSG00000183935 | *HTR7P1* | 1.354285353 | 0.012880411 | 0.174008467 | -0.161128057 | 0.999933887 | -0.063290111 | 0.089928738 | -0.153218849 | 0.580507367 |
| ENSG00000106003 | *LFNG* | 5.698483983 | 0.00257957 | -0.025676923 | 0.028256493 | 0.999933887 | 0.281804196 | 0.380683661 | -0.098879465 | 0.580579222 |
| ENSG00000172508 | *CARNS1* | 1.30735398 | 0.094924361 | 0.109322327 | -0.014397966 | 0.999933887 | 0.042237746 | -0.124451689 | 0.166689435 | 0.580766234 |
| ENSG00000050344 | *NFE2L3* | 2.952940707 | -0.016937178 | 0.134612252 | -0.151549431 | 0.999933887 | -0.27432252 | -0.14527847 | -0.129044051 | 0.580766234 |
| ENSG00000161509 | *GRIN2C* | 0.747529848 | -0.105449921 | -0.05287363 | -0.052576291 | 0.999933887 | -0.196044239 | 0.034682821 | -0.23072706 | 0.580830142 |
| ENSG00000172171 | *TEFM* | -0.331971938 | -0.087228573 | 0.195978919 | -0.283207492 | 0.999933887 | -0.1059559 | 0.147384581 | -0.253340481 | 0.580890445 |
| ENSG00000167535 | *CACNB3* | 0.350667633 | 0.013053738 | 0.223520052 | -0.210466314 | 0.999933887 | 0.199379951 | 0.015870559 | 0.183509392 | 0.580890445 |
| ENSG00000260911 | *AC135050.3* | 2.218941211 | 0.091972127 | 0.223854821 | -0.131882694 | 0.999933887 | 0.518036966 | 0.634968711 | -0.116931745 | 0.581226211 |
| ENSG00000163291 | *PAQR3* | 2.068594268 | 0.020008139 | -0.045544692 | 0.065552831 | 0.999933887 | -0.257024284 | -0.051581121 | -0.205443163 | 0.581226211 |
| ENSG00000042832 | *TG* | 2.735796233 | -0.005430906 | -0.039945089 | 0.034514183 | 0.999933887 | 0.096727715 | -0.043735206 | 0.140462921 | 0.581350246 |
| ENSG00000283538 | *AC005972.3* | 0.295131453 | 0.283281715 | 0.588669515 | -0.3053878 | 0.999933887 | -0.519549729 | -0.78001054 | 0.260460811 | 0.581451905 |
| ENSG00000181409 | *AATK* | 8.681984464 | 0.060003583 | 0.037500079 | 0.022503505 | 0.999933887 | 0.100715579 | 0.206615084 | -0.105899505 | 0.581451905 |
| ENSG00000228492 | *RAB11FIP1P1* | -0.511414621 | -0.34383886 | -0.403421199 | 0.059582339 | 0.999933887 | -0.42993137 | -0.142311673 | -0.287619697 | 0.581497419 |
| ENSG00000255221 | *CARD17* | 4.793231737 | 0.107855359 | -0.049465497 | 0.157320856 | 0.999933887 | 1.131596407 | 1.272570165 | -0.140973758 | 0.581723793 |
| ENSG00000228113 | *AC003991.1* | 0.699196985 | -0.072666482 | -0.125848392 | 0.05318191 | 0.999933887 | -0.880002109 | -0.538673373 | -0.341328736 | 0.581723793 |
| ENSG00000157404 | *KIT* | 0.952708911 | -0.020397354 | -0.08871247 | 0.068315116 | 0.999933887 | -0.221133492 | -0.08620873 | -0.134924762 | 0.581723793 |
| ENSG00000144233 | *AMMECR1L* | 4.145466458 | -0.052899175 | 0.000621928 | -0.053521103 | 0.999933887 | -0.299822738 | -0.371012566 | 0.071189828 | 0.581737478 |
| ENSG00000165424 | *ZCCHC24* | 2.692287159 | -0.040749985 | 0.097778839 | -0.138528824 | 0.999933887 | -0.498027937 | -0.379186459 | -0.118841478 | 0.581737478 |
| ENSG00000286886 | *AJ011931.2* | -0.769281762 | -0.179154372 | 0.020310426 | -0.199464798 | 0.999933887 | 0.432961427 | 0.21592783 | 0.217033597 | 0.582205778 |
| ENSG00000174500 | *GCSAM* | 0.830096548 | 0.073185812 | -0.121133045 | 0.194318857 | 0.999933887 | 0.258878445 | 0.459151563 | -0.200273117 | 0.582302311 |
| ENSG00000162735 | *PEX19* | 3.183544555 | -0.019846951 | 0.095190021 | -0.115036972 | 0.999933887 | 0.111558133 | 0.21823533 | -0.106677197 | 0.582302311 |
| ENSG00000203668 | *CHML* | 1.262163461 | -0.020508531 | -0.238906117 | 0.218397586 | 0.999933887 | 0.182461908 | 0.409440529 | -0.22697862 | 0.582302311 |
| ENSG00000198522 | *GPN1* | 2.309473412 | -0.002403668 | -0.006363652 | 0.003959984 | 0.999933887 | 0.117642731 | 0.003340824 | 0.114301908 | 0.582302311 |
| ENSG00000164252 | *AGGF1* | 1.712603369 | -0.089300975 | -0.017221062 | -0.072079913 | 0.999933887 | -0.449277669 | -0.584609217 | 0.135331548 | 0.582477354 |
| ENSG00000281106 | *TMEM272* | 1.046113844 | -0.029538063 | 0.020843346 | -0.050381409 | 0.999933887 | 0.401445231 | 0.657196164 | -0.255750933 | 0.582737047 |
| ENSG00000181472 | *ZBTB2* | 4.581132961 | -0.056450619 | -0.028724372 | -0.027726247 | 0.999933887 | -0.219315136 | -0.290875628 | 0.071560492 | 0.583129267 |
| ENSG00000189043 | *NDUFA4* | 3.657612803 | 0.008476453 | -0.098660807 | 0.10713726 | 0.999933887 | 0.021186143 | -0.073528666 | 0.094714809 | 0.58317721 |
| ENSG00000131591 | *C1orf159* | 2.14376013 | 0.056750316 | 0.086240615 | -0.0294903 | 0.999933887 | 0.157351262 | 0.035077644 | 0.122273618 | 0.583228584 |
| ENSG00000231439 | *WASIR2* | -1.257444036 | -0.322669344 | -0.334173768 | 0.011504424 | 0.999933887 | -1.092774771 | -1.340617095 | 0.247842323 | 0.583420478 |
| ENSG00000175003 | *SLC22A1* | 3.021901341 | -0.080480741 | -0.00505635 | -0.07542439 | 0.999933887 | 0.483918254 | 0.600263679 | -0.116345425 | 0.583608435 |
| ENSG00000148339 | *SLC25A25* | 3.542883019 | 0.050562273 | 0.126403737 | -0.075841464 | 0.999933887 | -0.356606453 | -0.467653182 | 0.111046729 | 0.583608435 |
| ENSG00000211706 | *TRBV6-1* | -0.375496737 | -0.0874024 | -0.052602721 | -0.034799679 | 0.999933887 | 0.240894666 | -0.020909398 | 0.261804065 | 0.583688625 |
| ENSG00000073712 | *FERMT2* | -1.694810052 | -0.457480077 | 0.433840676 | -0.891320754 | 0.999933887 | 0.424628841 | 0.642371742 | -0.217742901 | 0.583758949 |
| ENSG00000164707 | *SLC13A4* | -0.816564434 | -0.122927851 | 0.085055311 | -0.207983162 | 0.999933887 | -0.108612644 | -0.354272119 | 0.245659476 | 0.583758949 |
| ENSG00000268658 | *LINC00664* | 5.631655716 | 0.028169109 | 0.104718513 | -0.076549405 | 0.999933887 | 0.999573339 | 1.139545157 | -0.139971819 | 0.583941304 |
| ENSG00000105270 | *CLIP3* | -0.386028938 | -0.025768939 | 0.185093391 | -0.210862331 | 0.999933887 | -0.081936613 | -0.371751036 | 0.289814424 | 0.58471721 |
| ENSG00000277161 | *PIGW* | 0.028948096 | -0.243080904 | -0.133481236 | -0.109599668 | 0.999933887 | 0.145790892 | 0.386902963 | -0.241112072 | 0.584794259 |
| ENSG00000219481 | *NBPF1* | 1.163070071 | 0.09923961 | -0.064895858 | 0.164135467 | 0.999933887 | 0.031089906 | -0.138652954 | 0.16974286 | 0.584794259 |
| ENSG00000120162 | *MOB3B* | -0.282313615 | 0.230072496 | -0.296893429 | 0.526965924 | 0.944094537 | -0.0189062 | 0.180290419 | -0.199196618 | 0.584941044 |
| ENSG00000204252 | *HLA-DOA* | 1.296922041 | 0.067684901 | 0.023153677 | 0.044531224 | 0.999933887 | 0.109177636 | 0.271596327 | -0.162418691 | 0.584971936 |
| ENSG00000188846 | *RPL14* | 7.395381656 | -0.002659006 | -0.109130691 | 0.106471685 | 0.999933887 | 0.080127264 | 0.158984976 | -0.078857713 | 0.585026451 |
| ENSG00000256751 | *PLBD1-AS1* | 0.265265136 | 0.145866942 | -0.061491643 | 0.207358585 | 0.999933887 | -0.443520962 | -0.786175481 | 0.342654519 | 0.585028334 |
| ENSG00000062282 | *DGAT2* | 7.329642842 | -0.000989285 | 0.031390898 | -0.032380184 | 0.999933887 | -0.081018311 | -0.155491136 | 0.074472826 | 0.585433792 |
| ENSG00000152795 | *HNRNPDL* | 6.835844828 | -0.063799369 | -0.053094038 | -0.010705331 | 0.999933887 | -0.108952963 | -0.166592915 | 0.057639952 | 0.585523609 |
| ENSG00000167785 | *ZNF558* | 1.845618071 | -0.074462666 | -0.149585167 | 0.075122501 | 0.999933887 | -0.115471247 | 0.009891112 | -0.125362359 | 0.585523609 |
| ENSG00000139624 | *CERS5* | 4.839184203 | -0.028952093 | -0.004817467 | -0.024134626 | 0.999933887 | -0.226760785 | -0.160361874 | -0.066398911 | 0.585523609 |

| | | | | | | | | | | |
|---|---|---|---|---|---|---|---|---|---|---|
| ENSG00000168297 | *PXK* | 5.203125648 | -0.031693283 | -0.042517288 | 0.010824005 | 0.999933887 | -0.469030635 | -0.549398385 | 0.08036775 | 0.585523609 |
| ENSG00000241769 | *LINC00893* | 0.067736479 | -0.031279002 | 0.085434652 | -0.116713654 | 0.999933887 | -0.359809738 | -0.597507761 | 0.237698024 | 0.585651699 |
| ENSG00000168140 | *VASN* | 2.333277998 | 0.036424287 | 0.035074737 | 0.00134955 | 0.999933887 | -0.118943408 | -0.265769232 | 0.146825824 | 0.585923954 |
| ENSG00000105609 | *LILRB5* | 2.28336474 | 0.073882627 | 0.047284173 | 0.026598454 | 0.999933887 | -0.233738728 | -0.337872046 | 0.104133318 | 0.586124687 |
| ENSG00000176700 | *SCAND2P* | 0.051291707 | -0.094600837 | 0.104427022 | -0.199027859 | 0.999933887 | -0.191326009 | -0.393073907 | 0.201747897 | 0.586124687 |
| ENSG00000139835 | *GRTP1* | 2.071732103 | 0.143515411 | 0.297607947 | -0.154092536 | 0.999933887 | 0.504864234 | 0.647311712 | -0.142447478 | 0.586511758 |
| ENSG00000104804 | *TULP2* | 2.250652269 | 0.427318646 | 0.509005096 | -0.08168645 | 0.999933887 | 1.938339188 | 2.213893793 | -0.275554604 | 0.586782905 |
| ENSG00000120949 | *TNFRSF8* | 4.79601977 | 0.12147477 | 0.110656753 | 0.010818016 | 0.999933887 | 0.252100569 | 0.346219343 | -0.094118774 | 0.586782905 |
| ENSG00000131171 | *SH3BGRL* | 5.673229765 | -0.053193666 | -0.164076285 | 0.110882619 | 0.999933887 | -0.086251647 | 0.004477097 | -0.090728744 | 0.586782905 |
| ENSG00000218713 | *AL512378.1* | 0.768009052 | -0.041378701 | -0.082869977 | 0.041491276 | 0.999933887 | -1.00386846 | -0.808352959 | -0.1955155 | 0.586841369 |
| ENSG00000143457 | *GOLPH3L* | 1.693722613 | -0.080366315 | -0.136607395 | 0.05624108 | 0.999933887 | -0.042577614 | 0.149721494 | -0.192299108 | 0.586914344 |
| ENSG00000254681 | *PKD1P5* | -2.252094938 | 0.034287296 | -0.01871349 | 0.053000786 | 0.999933887 | 0.122819264 | -0.092088077 | 0.214907341 | 0.586914344 |
| ENSG00000257698 | *GIHCG* | 1.286227457 | -0.016980309 | 0.022733299 | -0.039713608 | 0.999933887 | -0.30529638 | -0.454111153 | 0.148814774 | 0.586914344 |
| ENSG00000245275 | *SAP30L-AS1* | 1.750347536 | -0.037049313 | -0.007322525 | -0.029726787 | 0.999933887 | -0.230130831 | -0.381007136 | 0.150876306 | 0.587173032 |
| ENSG00000286760 | *AL022069.3* | 0.785373048 | 0.264758856 | 0.238103147 | 0.026655709 | 0.999933887 | 0.937685493 | 1.130139786 | -0.192454293 | 0.587228803 |
| ENSG00000010165 | *EEF1AKNMT* | 2.480061682 | 0.055875221 | 0.11002263 | -0.054147409 | 0.999933887 | -0.149995148 | -0.243837612 | 0.093842464 | 0.587228803 |
| ENSG00000176087 | *SLC35A4* | 5.295151186 | -0.001514362 | -0.050441419 | 0.048927058 | 0.999933887 | -0.04287281 | 0.027029486 | -0.069902295 | 0.587228803 |
| ENSG00000130748 | *TMEM160* | 1.591836276 | 0.001861986 | 0.16794135 | -0.166079364 | 0.999933887 | 0.116835372 | -0.065493359 | 0.182328731 | 0.587228803 |
| ENSG00000131187 | *F12* | 2.925481772 | 0.168569482 | -0.03346113 | 0.202030612 | 0.999933887 | 0.691088124 | 0.817178143 | -0.126090019 | 0.587636607 |
| ENSG00000114331 | *ACAP2* | 7.110820453 | -0.074157619 | -0.108538461 | 0.034380842 | 0.999933887 | -0.603103606 | -0.49201592 | -0.111087685 | 0.587636607 |
| ENSG00000083093 | *PALB2* | 0.371526063 | -0.126636877 | -0.028196621 | -0.098440256 | 0.999933887 | 0.081549914 | 0.250869717 | -0.169319803 | 0.587636607 |
| ENSG00000163320 | *CGGBP1* | 5.870224576 | -0.034054914 | -0.028779182 | -0.005275732 | 0.999933887 | -0.249581088 | -0.16992987 | -0.079651218 | 0.587636607 |
| ENSG00000122417 | *ODF2L* | 3.298392178 | -0.009966907 | -0.156269743 | 0.146302836 | 0.999933887 | -0.271393287 | -0.045466573 | -0.225926714 | 0.587693529 |
| ENSG00000271216 | *LINC01050* | -0.569471066 | -0.11660224 | 0.609217811 | -0.72582005 | 0.999933887 | 1.04220912 | 1.216457886 | -0.174248766 | 0.587781173 |
| ENSG00000111879 | *FAM184A* | -1.14782415 | -0.123646796 | -0.278474707 | 0.154827912 | 0.999933887 | -0.348083034 | 0.01465724 | -0.362740274 | 0.587796196 |
| ENSG00000104517 | *UBR5* | 7.316523891 | -0.057725401 | -0.089875726 | 0.032150325 | 0.999933887 | -0.097679566 | -0.006201082 | -0.091478484 | 0.587962552 |
| ENSG00000169241 | *SLC50A1* | 2.811719373 | -0.028872446 | 0.013229062 | -0.042101508 | 0.999933887 | 0.728518027 | 0.852883194 | -0.124365168 | 0.587985787 |
| ENSG00000137218 | *FRS3* | -0.086234459 | 0.123798318 | -0.074247398 | 0.198045717 | 0.999933887 | 0.093923131 | -0.114041365 | 0.207964496 | 0.587999825 |
| ENSG00000204463 | *BAG6* | 8.100092869 | 0.057828936 | 0.039383275 | 0.01844566 | 0.999933887 | 0.260803933 | 0.316964708 | -0.056160775 | 0.588257129 |
| ENSG00000107902 | *LHPP* | 1.880383592 | -0.07198457 | 0.05081658 | -0.12280115 | 0.999933887 | 0.003951155 | -0.170325269 | 0.174276424 | 0.588257129 |
| ENSG00000108578 | *BLMH* | 2.535739642 | 0.007644752 | 0.090137897 | -0.082493146 | 0.999933887 | 0.088555329 | 0.180180314 | -0.091624986 | 0.588257129 |
| ENSG00000279500 | *AC108704.2* | 4.566801656 | -0.075415622 | -0.034504179 | -0.040911443 | 0.999933887 | -0.257304993 | -0.161612408 | -0.095692584 | 0.588270813 |
| ENSG00000161914 | *ZNF653* | 0.990237078 | -0.056527928 | 0.041853499 | -0.098381427 | 0.999933887 | -0.020400896 | -0.189263549 | 0.168862653 | 0.588304796 |
| ENSG00000177917 | *ARL6IP6* | 2.831455889 | 0.115367867 | 0.062044309 | 0.053323558 | 0.999933887 | -0.184904485 | -0.314121724 | 0.129217238 | 0.588415871 |
| ENSG00000164687 | *FABP5* | 0.810708396 | -0.033549759 | -0.122861324 | 0.089311565 | 0.999933887 | -0.250962374 | -0.453858737 | 0.202896363 | 0.588415871 |
| ENSG00000100412 | *ACO2* | 4.076218217 | -0.001026561 | -0.009304162 | 0.0082776 | 0.999933887 | 0.006751188 | -0.069586975 | 0.076338163 | 0.588785757 |
| ENSG00000272914 | *AL359532.1* | 1.409280284 | -0.058881979 | -0.068098193 | 0.009216214 | 0.999933887 | -0.154349681 | -0.025297472 | -0.129052209 | 0.588943882 |
| ENSG00000286018 | *AF129075.3* | -0.425452151 | 0.221040949 | 0.105436481 | 0.115604468 | 0.999933887 | -0.090430808 | 0.161946121 | -0.252376929 | 0.588996124 |
| ENSG00000225518 | *LINC01703* | -0.00757587 | 0.039251436 | 0.26415204 | -0.224900604 | 0.999933887 | 0.441738866 | 0.029029033 | 0.412709833 | 0.589340504 |
| ENSG00000173209 | *AHSA2P* | 5.334986223 | 0.000940589 | -0.033032202 | 0.03397279 | 0.999933887 | -0.065876336 | -0.132869482 | 0.066993145 | 0.589408603 |
| ENSG00000204054 | *LINC00963* | 7.156335847 | -0.008836231 | -0.002407821 | -0.006428411 | 0.999933887 | 0.114038269 | 0.196539094 | -0.082500825 | 0.589494495 |
| ENSG00000177733 | *HNRNPA0* | 6.854885657 | 0.085149871 | 0.061948172 | 0.0232017 | 0.999933887 | -0.019512575 | -0.112502639 | 0.092990064 | 0.589584797 |
| ENSG00000103037 | *SETD6* | 1.681384603 | 0.100872176 | 0.046163015 | 0.054709161 | 0.999933887 | 0.2606662 | 0.138716571 | 0.121949629 | 0.589584797 |
| ENSG00000117322 | *CR2* | -0.37873525 | -0.180329347 | 0.159711397 | -0.340040744 | 0.999933887 | 0.333235273 | 0.154460131 | 0.178775142 | 0.589584797 |
| ENSG00000109220 | *CHIC2* | 5.422643243 | 0.041563828 | 0.098192599 | -0.056628771 | 0.999933887 | -0.127485095 | -0.054545829 | -0.072939266 | 0.589584797 |
| ENSG00000180370 | *PAK2* | 7.744100962 | -0.045104731 | 0.019499266 | -0.064603997 | 0.999933887 | -0.692140588 | -0.778601602 | 0.086461014 | 0.589584797 |
| ENSG00000118096 | *IFT46* | 0.547786777 | 0.051873045 | 0.057410158 | -0.005537113 | 0.999933887 | 0.14057121 | 0.311709667 | -0.171138457 | 0.589584797 |
| ENSG00000114520 | *SNX4* | 1.844105378 | -0.059435322 | -0.119958139 | 0.060522817 | 0.999933887 | -0.142331137 | -0.291991641 | 0.149660504 | 0.589642963 |
| ENSG00000135749 | *PCNX2* | 3.312794618 | -0.019148771 | 0.004789492 | -0.023938264 | 0.999933887 | -0.068366427 | 0.02420987 | -0.092576297 | 0.590311091 |
| ENSG00000101695 | *RNF125* | 3.841141886 | -0.072149315 | -0.109595083 | 0.037445769 | 0.999933887 | -0.088085664 | -0.202839448 | 0.114753784 | 0.590441122 |
| ENSG00000270972 | *AC136475.9* | -1.293120329 | -0.060552879 | -0.182477596 | 0.121924718 | 0.999933887 | 0.040703095 | -0.392930752 | 0.433633847 | 0.590523733 |
| ENSG00000106089 | *STX1A* | 1.727659734 | -0.105547602 | 0.092821962 | -0.198369564 | 0.999933887 | 0.006187366 | -0.161152043 | 0.167339409 | 0.59056313 |
| ENSG00000138468 | *SENP7* | 4.473055003 | -0.022899996 | -0.133105182 | 0.110205186 | 0.999933887 | -0.565786394 | -0.313725253 | -0.252061141 | 0.59056313 |
| ENSG00000198039 | *ZNF273* | 0.759727734 | 0.016035755 | 0.039694641 | -0.023658886 | 0.999933887 | -0.564188491 | -0.352675955 | -0.211512536 | 0.59056313 |
| ENSG00000121310 | *ECHDC2* | 3.072683862 | -0.081576529 | 0.085947911 | -0.167524441 | 0.999933887 | -0.002947087 | 0.101553587 | -0.104500674 | 0.590620513 |
| ENSG00000119013 | *NDUFB3* | 3.595173947 | -0.053640245 | -0.028258111 | -0.025382134 | 0.999933887 | 0.056953501 | -0.042786639 | 0.099740141 | 0.590872516 |
| ENSG00000119725 | *ZNF410* | 0.512364498 | 0.045136307 | 0.116875433 | -0.071739126 | 0.999933887 | 0.320723368 | 0.520373938 | -0.19965057 | 0.591015107 |
| ENSG00000274213 | *AC015912.3* | 5.296030542 | 0.125879503 | 0.073716339 | 0.052163164 | 0.999933887 | 0.191392539 | 0.065379662 | 0.126012877 | 0.591056053 |
| ENSG00000211829 | *TRDC* | 0.314651199 | -0.090186051 | -0.053978429 | -0.036207622 | 0.999933887 | 0.009181652 | 0.179311007 | -0.170129355 | 0.591056053 |
| ENSG00000116016 | *EPAS1* | 2.806557642 | -0.057215913 | 0.045636785 | -0.102852698 | 0.999933887 | -0.442206836 | -0.345677186 | -0.09652965 | 0.591357946 |
| ENSG00000104356 | *POP1* | -0.062606557 | 0.093939994 | 0.292240909 | -0.198300914 | 0.999933887 | 0.346930289 | 0.137977522 | 0.208952767 | 0.591357946 |
| ENSG00000114857 | *NKTR* | 6.447268965 | -0.048310218 | -0.195455003 | 0.147144784 | 0.999933887 | -0.516747945 | -0.272162579 | -0.244585366 | 0.591357946 |
| ENSG00000136732 | *GYPC* | 4.490347692 | 0.035219347 | -0.033578962 | 0.068798309 | 0.999933887 | 0.085637702 | -0.001449 | 0.087086702 | 0.591399284 |
| ENSG00000187792 | *ZNF70* | 0.005227895 | -0.003551289 | -0.055435993 | 0.051884705 | 0.999933887 | -0.010534145 | 0.22372274 | -0.234256885 | 0.591453868 |
| ENSG00000044574 | *HSPA5* | 8.280148799 | 0.0406526 | -0.033691021 | 0.07434362 | 0.999933887 | 0.31174396 | 0.384099626 | -0.072355666 | 0.59184642 |
| ENSG00000214688 | *C10orf105* | 0.233847568 | 0.055598936 | 0.051587122 | 0.004011814 | 0.999933887 | -0.292118065 | -0.661184665 | 0.3690666 | 0.592016903 |
| ENSG00000162836 | *ACP6* | 6.307476007 | 0.035940388 | 0.120375049 | -0.084434661 | 0.999933887 | -0.401535682 | -0.494480869 | 0.092945186 | 0.592085039 |
| ENSG00000124785 | *NRN1* | -0.292426849 | -0.161720656 | 0.186110925 | -0.34783158 | 0.958987651 | -0.442136182 | -0.142800842 | -0.29933534 | 0.592431423 |
| ENSG00000173262 | *SLC2A14* | 0.096918458 | -0.018947859 | 0.096010148 | -0.114958008 | 0.999933887 | -0.619303335 | -0.387368752 | -0.231934583 | 0.592431423 |
| ENSG00000272033 | *AL136984.1* | 0.166785766 | -0.040110695 | 0.089540153 | -0.129650847 | 0.999933887 | -0.458049355 | -0.174708854 | -0.283340501 | 0.59243454 |
| ENSG00000153107 | *ANAPC1* | 2.201053379 | -0.088509333 | -0.110020686 | 0.021511353 | 0.999933887 | 0.134973762 | 0.242106849 | -0.107133087 | 0.592936235 |
| ENSG00000055732 | *MCOLN3* | -0.838114487 | 0.143154537 | -0.104941647 | 0.248096184 | 0.999933887 | -0.163938935 | 0.127863802 | -0.291802736 | 0.592949119 |
| ENSG00000163935 | *SFMBT1* | 2.16812168 | -0.052493864 | 0.075224458 | -0.127718321 | 0.999933887 | 0.090560247 | 0.203629259 | -0.113069012 | 0.592949119 |
| ENSG00000227097 | *RPS28P7* | -0.100819316 | 0.11499697 | 0.152160646 | -0.037163676 | 0.999933887 | -0.031701388 | 0.252102468 | -0.283803856 | 0.592949119 |
| ENSG00000251648 | *AC008958.1* | 0.138769411 | 0.125610813 | 0.287472086 | -0.161861273 | 0.999933887 | 1.783615489 | 1.554643589 | 0.2289719 | 0.592949119 |
| ENSG00000123636 | *BAZ2B* | 6.110643139 | -0.039578565 | -0.127994357 | 0.088415791 | 0.999933887 | -0.749256221 | -0.563002755 | -0.186253466 | 0.592949119 |
| ENSG00000177932 | *ZNF354C* | 0.194897448 | -0.04611046 | 0.152143299 | -0.198253759 | 0.999933887 | 0.02926761 | 0.238855042 | -0.209587432 | 0.592949119 |
| ENSG00000178878 | *APOLD1* | 0.686164632 | 0.021690232 | 0.023295564 | -0.001605332 | 0.999933887 | -0.418161026 | -0.620124135 | 0.201963109 | 0.592949119 |
| ENSG00000232587 | *EEF1A1P3* | 0.410023159 | 0.138281386 | 0.312232674 | -0.173951288 | 0.999933887 | -0.29507925 | -0.034624735 | -0.260454516 | 0.59311888 |
| ENSG00000104814 | *MAP4K1* | 4.13084555 | -0.012483753 | 0.024446023 | -0.036929776 | 0.999933887 | 0.112175465 | 0.035419038 | 0.076756427 | 0.59311888 |
| ENSG00000203709 | *MIR29B2CHG* | 7.947146887 | 0.047073287 | 0.001436171 | 0.045637116 | 0.999933887 | -0.139553552 | -0.063742218 | -0.075811334 | 0.593190538 |
| ENSG00000184347 | *SLIT3* | -1.069008382 | -0.224645969 | -0.17100729 | -0.053638678 | 0.999933887 | -0.012748648 | 0.186280297 | -0.199028945 | 0.593190538 |
| ENSG00000272368 | *AC074032.1* | -0.199040802 | 0.087102875 | 0.047632034 | 0.039470841 | 0.999933887 | -0.052987695 | -0.333246919 | 0.280259224 | 0.593577758 |
| ENSG00000015171 | *ZMYND11* | 3.402874397 | -0.014131743 | 0.045967872 | -0.060099615 | 0.999933887 | -0.016118354 | 0.074160898 | -0.090279252 | 0.593727964 |
| ENSG00000105185 | *PDCD5* | 2.09891753 | -0.016424347 | -0.103675096 | 0.087250748 | 0.999933887 | 0.165463276 | 0.281171401 | -0.115708125 | 0.593727964 |
| ENSG00000136158 | *SPRY2* | 2.706876355 | -0.168094838 | -0.04250072 | -0.125594118 | 0.999933887 | -0.545437306 | -0.42396307 | -0.121474236 | 0.594184951 |
| ENSG00000273820 | *USP27X* | 0.504854671 | -0.119839756 | -0.002156504 | -0.117683252 | 0.999933887 | 0.130736178 | -0.026221857 | 0.156958036 | 0.594317934 |
| ENSG00000275395 | *FCGBP* | 2.300204805 | 0.017312541 | 0.126003384 | -0.108690843 | 0.999933887 | 0.312901377 | 0.412217114 | -0.099315737 | 0.594647693 |
| ENSG00000057663 | *ATG5* | 3.464965841 | -0.148144896 | 0.012768063 | -0.160912959 | 0.999933887 | -0.020214274 | 0.059820897 | -0.080035171 | 0.59484237 |
| ENSG00000130167 | *TSPAN16* | -0.084171647 | -0.078820877 | 0.153199288 | -0.232020164 | 0.999933887 | -0.706098363 | -1.047181713 | 0.34108335 | 0.59484237 |
| ENSG00000101843 | *PSMD10* | 2.34696848 | 0.043383482 | -0.118717665 | 0.162101147 | 0.999933887 | -0.138430582 | -0.024141037 | -0.114289545 | 0.59484237 |
| ENSG00000109046 | *WSB1* | 8.364977874 | -0.028240345 | 0.025202232 | -0.053442577 | 0.999933887 | 0.293482869 | 0.374138775 | -0.080655906 | 0.594916201 |
| ENSG00000221866 | *PLXNA4* | -0.74653649 | -0.017771792 | 0.073329518 | -0.09110131 | 0.999933887 | -0.132462669 | 0.190707118 | -0.323169787 | 0.594916201 |
| ENSG00000218809 | *AL391903.1* | 4.181776126 | 0.172248842 | 0.154230695 | 0.018018147 | 0.999933887 | 0.608705002 | 0.752458078 | -0.143753076 | 0.594927061 |
| ENSG00000164967 | *RPP25L* | 2.319663676 | -0.000493772 | -0.146871397 | 0.146377626 | 0.999933887 | -0.131275062 | -0.228506604 | 0.097231542 | 0.595321328 |
| ENSG00000137154 | *RPS6* | 8.403367959 | -0.028402557 | -0.133598216 | 0.105195658 | 0.999933887 | 0.066141479 | 0.137777615 | -0.071636137 | 0.595436617 |
| ENSG00000186106 | *ANKRD46* | 0.656276162 | 0.009264209 | 0.079068121 | -0.069803912 | 0.999933887 | -0.12792576 | 0.037296279 | -0.165222039 | 0.59587901 |
| ENSG00000145217 | *SLC26A1* | -0.204584828 | -0.156188204 | 0.017158844 | -0.173347047 | 0.999933887 | 0.099120295 | -0.103351436 | 0.202471731 | 0.596214916 |
| ENSG00000126803 | *HSPA2* | 1.07272921 | -0.030409632 | 0.113485293 | -0.143894925 | 0.999933887 | 0.524054442 | 0.334300411 | 0.189754031 | 0.596214916 |
| ENSG00000167315 | *ACAA2* | 1.703365005 | -0.021122462 | -0.024819862 | 0.0036974 | 0.999933887 | -0.241424845 | -0.10312887 | -0.138295975 | 0.596967301 |
| ENSG00000130338 | *TULP4* | 3.142723608 | -0.049536167 | 0.116379523 | -0.16591569 | 0.999933887 | -0.245845611 | -0.328362798 | 0.082517186 | 0.597050859 |
| ENSG00000186635 | *ARAP1* | 10.204659464 | 0.014293234 | 0.037300342 | -0.023007108 | 0.999933887 | 0.025905817 | -0.034327447 | 0.060233264 | 0.597171018 |
| ENSG00000260805 | *AC092803.1* | 2.8315967 | 0.208728217 | 0.144377718 | 0.0643505 | 0.999933887 | 0.973949157 | 1.113255504 | -0.139306347 | 0.597588826 |
| ENSG00000264931 | *MIR3138* | -1.630930885 | 0.31626762 | 0.629127595 | -0.312859975 | 0.999933887 | -0.07050854 | 0.464903379 | -0.535411919 | 0.597662867 |
| ENSG00000279811 | *AC093330.2* | 1.642152321 | 0.144816457 | -0.041987395 | 0.186803852 | 0.999933887 | -0.193086795 | -0.038601804 | -0.154484991 | 0.597662867 |

| | | | | | | | | | | |
|---|---|---|---|---|---|---|---|---|---|---|
| ENSG00000229619 | *MBNL1-AS1* | 0.584555836 | -0.103872184 | 0.007375741 | -0.111247924 | 0.999933887 | 0.111504863 | -0.16220289 | 0.273707753 | 0.597662867 |
| ENSG00000132763 | *MMACHC* | -0.389583511 | -0.408938396 | -0.223641489 | -0.185296906 | 0.999933887 | 0.329281236 | 0.551637746 | -0.22235651 | 0.597806778 |
| ENSG00000206013 | *IFITM5* | 0.043673938 | 0.278066364 | 0.066646844 | 0.21141952 | 0.999933887 | 0.171967391 | -0.082208586 | 0.254175977 | 0.597827103 |
| ENSG00000113360 | *DROSHA* | 2.468498638 | 0.058020862 | -0.025639872 | 0.083660733 | 0.999933887 | -0.082108017 | 0.013950572 | -0.09605859 | 0.597827103 |
| ENSG00000162714 | *ZNF496* | 2.499821597 | -0.081649589 | 0.104575724 | -0.186225313 | 0.999933887 | 0.095767652 | -0.015781615 | 0.111549267 | 0.597989847 |
| ENSG00000114473 | *IQCG* | 1.078045364 | 0.090785263 | 0.007935709 | 0.082849554 | 0.999933887 | 0.391312986 | 0.540686341 | -0.149373355 | 0.597989847 |
| ENSG00000239213 | *NCK1-DT* | -0.175374478 | -0.056546771 | 0.041723213 | -0.098269984 | 0.999933887 | -0.209136733 | -0.436061734 | 0.226925001 | 0.597989847 |
| ENSG00000129682 | *FGF13* | 2.542207022 | 0.168967604 | 0.306507753 | -0.137540149 | 0.999933887 | 0.323628768 | 0.158444728 | 0.16518404 | 0.598123738 |
| ENSG00000122406 | *RPL5* | 7.118425146 | -0.032220479 | -0.147972158 | 0.115751679 | 0.999933887 | 0.091654549 | 0.162810881 | -0.071156332 | 0.598123738 |
| ENSG00000269858 | *EGLN2* | 5.671420359 | 0.01197041 | 0.012126129 | -0.000155719 | 0.999933887 | 0.011996593 | 0.083787962 | -0.071791368 | 0.598123738 |
| ENSG00000260942 | *CAPN10-DT* | 1.499479003 | -0.259081527 | -0.18990305 | -0.069178477 | 0.999933887 | -0.007918699 | -0.192521202 | 0.184602503 | 0.598163741 |
| ENSG00000182872 | *RBM10* | 5.336057943 | 0.067196224 | 0.041781505 | 0.025414719 | 0.999933887 | -0.001710483 | -0.060733438 | 0.059022955 | 0.598163741 |
| ENSG00000091947 | *TMEM101* | 1.717714665 | -0.110406169 | -0.18597119 | 0.075565021 | 0.999933887 | -0.300990718 | -0.444288046 | 0.143297328 | 0.598163741 |
| ENSG00000131634 | *TMEM204* | 1.941605226 | 0.028855758 | 0.011090311 | 0.017765447 | 0.999933887 | 0.07163282 | -0.048603564 | 0.120236384 | 0.598163741 |
| ENSG00000227920 | *AL353597.1* | -1.41249047 | 0.040217228 | 0.506697488 | -0.466480261 | 0.999933887 | 0.232006627 | -0.063658908 | 0.295665536 | 0.598163741 |
| ENSG00000137944 | *KYAT3* | 3.426163716 | -0.134234221 | -0.027957018 | -0.106277203 | 0.999933887 | -0.021432578 | 0.057504106 | -0.078936684 | 0.5983948 |
| ENSG00000267174 | *AC011472.2* | 0.869705872 | 0.126039404 | 0.092788218 | 0.033251186 | 0.999933887 | -0.220881804 | -0.46073367 | 0.239851866 | 0.598812188 |
| ENSG00000113391 | *FAM172A* | 3.511696195 | -0.015315673 | 0.034713051 | -0.050028724 | 0.999933887 | -0.092167228 | 0.000988526 | -0.093155754 | 0.598812188 |
| ENSG00000182544 | *MFSD5* | 4.138942983 | -0.01115666 | -0.080593636 | 0.069436976 | 0.999933887 | -0.142336027 | -0.052662423 | -0.089673604 | 0.598812188 |
| ENSG00000100614 | *PPM1A* | 6.764061706 | -0.044798165 | -0.043612591 | -0.001185574 | 0.999933887 | -0.518499842 | -0.446473477 | -0.072026366 | 0.598818577 |
| ENSG00000176624 | *MEX3C* | 4.360364877 | 0.03800033 | 0.028483593 | 0.009516738 | 0.999933887 | -0.12318459 | -0.201290749 | 0.078106159 | 0.598818577 |
| ENSG00000137103 | *TMEM8B* | 0.601017758 | -0.067589983 | -0.0896321 | 0.022042118 | 0.999933887 | -0.048920276 | 0.131866514 | -0.18078679 | 0.598818577 |
| ENSG00000123064 | *DDX54* | 3.507821471 | 0.041289521 | 0.13721957 | -0.09593005 | 0.999933887 | 0.128929847 | 0.219923273 | -0.090993426 | 0.599117516 |
| ENSG00000086062 | *B4GALT1* | 7.418624243 | -0.050656993 | -0.007320866 | -0.043336126 | 0.999933887 | 0.944170985 | 1.04321597 | -0.099044985 | 0.599293241 |
| ENSG00000269952 | *AL117336.1* | 1.096836808 | 0.046708264 | 0.2012059 | -0.154497636 | 0.999933887 | 0.326267526 | 0.570811745 | -0.244544219 | 0.599420618 |
| ENSG00000258571 | *PTTG4P* | -0.064779886 | -0.016297793 | -0.09000194 | 0.073704147 | 0.999933887 | -0.608796478 | -0.26131363 | -0.347482848 | 0.599420618 |
| ENSG00000157483 | *MYO1E* | 0.348635495 | -0.065066758 | 0.116188537 | -0.181255295 | 0.999933887 | -0.361203985 | -0.201598251 | -0.159605734 | 0.599434099 |
| ENSG00000106701 | *FSD1L* | -0.418197812 | -0.053889794 | 0.021212775 | -0.075102569 | 0.999933887 | -0.680178198 | -0.450814275 | -0.229363922 | 0.599434099 |
| ENSG00000178814 | *OPLAH* | 3.61464313 | 0.026513328 | 0.086469073 | -0.059955745 | 0.999933887 | 0.201783938 | 0.101056395 | 0.100727543 | 0.599436077 |
| ENSG00000171150 | *SOCS5* | 2.686904781 | 0.051066085 | -0.153466266 | 0.204532351 | 0.999933887 | -0.216526565 | -0.328598583 | 0.112072018 | 0.599675964 |
| ENSG00000225611 | *LINC02158* | 2.523892535 | 0.032251131 | 0.10678318 | -0.074532049 | 0.999933887 | 0.620526528 | 0.739917854 | -0.119391326 | 0.599869649 |
| ENSG00000157637 | *SLC38A10* | 5.685670794 | 0.020992544 | -0.022993743 | 0.043986288 | 0.999933887 | -0.079184617 | -0.158579689 | 0.079395072 | 0.600041958 |
| ENSG00000151779 | *NBAS* | 2.584871037 | -0.004018268 | -0.055818114 | 0.051799846 | 0.999933887 | 0.083284235 | 0.224975352 | -0.141691117 | 0.600236095 |
| ENSG00000241560 | *ZBTB20-AS1* | -0.978201944 | -0.023464632 | 0.373284259 | -0.396748891 | 0.999933887 | -0.010800168 | 0.444946877 | -0.455747044 | 0.600433487 |
| ENSG00000229781 | *RRN3P4* | -1.298832669 | 0.212641431 | 0.003669417 | 0.208972014 | 0.999933887 | 1.859449032 | 2.212688987 | -0.353239955 | 0.600525512 |
| ENSG00000153044 | *CENPH* | -0.732540254 | -0.076575974 | -0.007754934 | -0.06882104 | 0.999933887 | 0.084252154 | 0.336195302 | -0.251943148 | 0.600559409 |
| ENSG00000196735 | *HLA-DQA1* | 4.230833441 | 0.026846592 | 0.096598947 | -0.069752354 | 0.999933887 | 1.671943346 | 1.928487497 | -0.256544151 | 0.600588304 |
| ENSG00000177954 | *RPS27* | 8.428246257 | 0.027468648 | -0.024055591 | 0.051524239 | 0.999933887 | 0.112770562 | 0.192457517 | -0.079686954 | 0.600731387 |
| ENSG00000108582 | *CPD* | 10.062907268 | -0.083857243 | -0.079786191 | -0.004071052 | 0.999933887 | 0.082153425 | 0.167206586 | -0.085053161 | 0.600959556 |
| ENSG00000249502 | *AC006160.1* | 1.487007232 | 0.104280966 | -0.038024636 | 0.142305602 | 0.999933887 | 0.130816041 | 0.283165015 | -0.152348975 | 0.600959556 |
| ENSG00000231993 | *EP300-AS1* | 0.748929536 | -0.02294896 | 0.147544912 | -0.170493871 | 0.999933887 | -0.184299566 | 0.04093288 | -0.225232446 | 0.600959556 |
| ENSG00000254531 | *FLJ20021* | -0.110324925 | 0.070194984 | 0.241488331 | -0.171293346 | 0.999933887 | -0.143957381 | -0.378350542 | 0.234393162 | 0.601053096 |
| ENSG00000100099 | *HPS4* | 4.900740224 | -0.078067841 | 0.014922704 | -0.092990545 | 0.999933887 | -0.204294774 | -0.141470798 | -0.062823975 | 0.601199109 |
| ENSG00000242802 | *AP5Z1* | 8.233450047 | 0.070761228 | 0.096950088 | -0.02618886 | 0.999933887 | 0.86472608 | 0.954269869 | -0.089543789 | 0.601199109 |
| ENSG00000139354 | *GAS2L3* | 0.71033557 | -0.183935908 | 0.240158603 | -0.424094511 | 0.999933887 | -0.517797451 | -0.359388672 | -0.158408778 | 0.601199109 |
| ENSG00000246263 | *UBR5-AS1* | 4.260375897 | -0.005983799 | -0.086763119 | 0.08077932 | 0.999933887 | -0.471421311 | -0.379396012 | -0.092025299 | 0.601876191 |
| ENSG00000145425 | *RPS3A* | 6.664855219 | -0.070684182 | -0.162305057 | 0.091620875 | 0.999933887 | 0.076518976 | 0.14472578 | -0.068206804 | 0.601944598 |
| ENSG00000174306 | *ZHX3* | 0.746015313 | 0.047238421 | 0.096565742 | -0.049327322 | 0.999933887 | -0.108624658 | -0.292749217 | 0.18412456 | 0.601944598 |
| ENSG00000184182 | *UBE2F* | 1.898254286 | 0.14346096 | 0.072355616 | 0.071105344 | 0.999933887 | -0.022608382 | 0.107417169 | -0.130025552 | 0.6020543 |
| ENSG00000134531 | *EMP1* | 3.358989355 | -0.186530799 | -0.123947126 | -0.062583673 | 0.999933887 | -1.520181861 | -1.660124379 | 0.139942519 | 0.6020543 |
| ENSG00000090006 | *LTBP4* | 4.246233589 | 0.019625761 | 0.05545757 | -0.035831809 | 0.999933887 | 0.091965485 | 0.006361798 | 0.085603687 | 0.602070109 |
| ENSG00000280347 | *AC000123.3* | 1.62790738 | -0.021409027 | 0.235953456 | -0.257362484 | 0.999933887 | -0.35192353 | -0.07970033 | -0.272223199 | 0.602070109 |
| ENSG00000168393 | *DTYMK* | 0.91678955 | 0.242406828 | -0.187743747 | 0.430150575 | 0.944094537 | 0.123970241 | -0.056111057 | 0.180081298 | 0.602318429 |
| ENSG00000259556 | *AC090971.3* | 0.649061994 | -0.057796995 | -0.179404803 | 0.121607808 | 0.999933887 | 0.359365129 | 0.575423352 | -0.216058224 | 0.602318429 |
| ENSG00000142544 | *CTU1* | 0.952343729 | 0.01515744 | 0.227852771 | -0.21269533 | 0.999933887 | 0.005442911 | -0.143458035 | 0.148900946 | 0.602330299 |
| ENSG00000213434 | *VTI1BP2* | -2.983975257 | 0.277834989 | 0.34715157 | -0.069316581 | 0.999933887 | 2.053479364 | 2.32383876 | -0.270359396 | 0.602401185 |
| ENSG00000229106 | *BTBD6P1* | 0.856262109 | -0.246698984 | -0.016035535 | -0.230663449 | 0.999933887 | -0.113909945 | 0.039879678 | -0.153789622 | 0.60296518 |
| ENSG00000005059 | *MCUB* | 3.437798905 | 0.019319702 | -0.06547222 | 0.084791922 | 0.999933887 | 0.037216937 | 0.119382704 | -0.082165767 | 0.603202521 |
| ENSG00000153561 | *RMND5A* | 7.642754499 | -0.106432196 | -0.017045072 | -0.089387124 | 0.999933887 | -0.754985601 | -0.831618126 | 0.076632525 | 0.603374504 |
| ENSG00000187555 | *USP7* | 7.026176869 | -0.038148175 | -0.033664887 | -0.004483288 | 0.999933887 | 0.041221716 | 0.088590086 | -0.04736837 | 0.603374504 |
| ENSG00000243402 | *AC022973.2* | 0.016946114 | 0.068197409 | 0.271200941 | -0.203003531 | 0.999933887 | -0.138938583 | 0.194993646 | -0.333932229 | 0.60375267 |
| ENSG00000044115 | *CTNNA1* | 4.800197668 | -0.039433592 | -0.05682627 | 0.017392678 | 0.999933887 | -0.189305027 | -0.119065732 | -0.070239295 | 0.604037737 |
| ENSG00000204899 | *MZT1* | 1.084590691 | 0.027016108 | 0.000200979 | 0.026815129 | 0.999933887 | -0.140452255 | 0.018573434 | -0.159025689 | 0.604253838 |
| ENSG00000255026 | *AC136475.3* | 4.926351505 | 0.108362443 | 0.131211437 | -0.022848995 | 0.999933887 | -0.233299837 | -0.330196813 | 0.096896976 | 0.604265952 |
| ENSG00000167193 | *CRK* | 6.325955665 | -0.027636821 | -0.110737392 | 0.083100572 | 0.999933887 | -0.160585372 | -0.106167532 | -0.054417841 | 0.604388135 |
| ENSG00000005020 | *SKAP2* | 6.663552152 | 0.022635021 | -0.040658064 | 0.063293085 | 0.999933887 | -0.769771694 | -0.868438657 | 0.098666964 | 0.604407591 |
| ENSG00000111335 | *OAS2* | 4.574813922 | -0.018531954 | -0.032766881 | 0.014234928 | 0.999933887 | -0.265009373 | -0.348727652 | 0.083718279 | 0.604407591 |
| ENSG00000116285 | *ERRFI1* | 0.474112318 | 0.036037651 | 0.099726376 | -0.063688726 | 0.999933887 | -0.403910198 | -0.649258981 | 0.245348783 | 0.604407591 |
| ENSG00000139546 | *TARBP2* | 1.963220239 | 0.003029257 | 0.060650213 | -0.057620956 | 0.999933887 | -0.279072586 | -0.410779238 | 0.131706652 | 0.604407591 |
| ENSG00000134198 | *TSPAN2* | 3.968371707 | -0.036840103 | -0.102515161 | 0.065675058 | 0.999933887 | 0.010766843 | 0.119338305 | -0.108571462 | 0.604796361 |
| ENSG00000115365 | *LANCL1* | 2.256438385 | -0.113351013 | -0.070145313 | -0.0432057 | 0.999933887 | -0.016862819 | 0.100585888 | -0.117448707 | 0.604950325 |
| ENSG00000013573 | *DDX11* | 1.532513544 | -0.095212808 | 0.319853827 | -0.415066635 | 0.594252256 | 0.137496959 | 0.022902097 | 0.114594862 | 0.604950325 |
| ENSG00000166452 | *AKIP1* | 0.297615756 | 0.102014131 | -0.277408098 | 0.379422229 | 0.999933887 | 0.166103608 | 0.000383955 | 0.165719653 | 0.604950325 |
| ENSG00000162434 | *JAK1* | 8.679477478 | -0.023797925 | -0.058863455 | 0.03506553 | 0.999933887 | 0.179144587 | 0.230621337 | -0.05147675 | 0.604950325 |
| ENSG00000258311 | *AC009779.3* | 1.982043022 | 0.152916282 | 0.001973036 | 0.150943246 | 0.999933887 | 0.037973064 | 0.168644257 | -0.130671193 | 0.60501947 |
| ENSG00000184611 | *KCNH7* | -0.550932213 | -0.129822237 | 0.153858771 | -0.283681008 | 0.999933887 | -0.286219817 | -0.625934582 | 0.339714765 | 0.60501947 |
| ENSG00000108786 | *HSD17B1* | 0.098624168 | -0.147614716 | -0.091146086 | -0.05646863 | 0.999933887 | -0.640869047 | -0.918908338 | 0.27803929 | 0.605355737 |
| ENSG00000100522 | *GNPNAT1* | 0.043599813 | -0.100036841 | 0.133080704 | -0.233117545 | 0.999933887 | 0.102034547 | 0.285020781 | -0.182986234 | 0.605355737 |
| ENSG00000170049 | *KCNAB3* | 1.849915153 | 0.021527876 | -0.160656757 | 0.182184634 | 0.999933887 | 0.173711533 | 0.032734899 | 0.140976633 | 0.605355737 |
| ENSG00000258082 | *AL391832.3* | 2.883709566 | 0.172704157 | 0.117968347 | 0.054735811 | 0.999933887 | 1.88420727 | 2.046028223 | -0.161820953 | 0.605619432 |
| ENSG00000074842 | *MYDGF* | 2.954263621 | 0.12772504 | -0.08203047 | 0.20975551 | 0.999933887 | 0.12868113 | 0.215424184 | -0.086743054 | 0.605619432 |
| ENSG00000059769 | *DNAJC25* | 1.566975816 | -0.12193588 | -0.08807922 | -0.03385666 | 0.999933887 | 0.139386034 | 0.324023923 | -0.184637889 | 0.605683536 |
| ENSG00000255310 | *AF131215.5* | -0.264989721 | -0.052304698 | -0.060720587 | 0.008415888 | 0.999933887 | 0.090365462 | 0.338593915 | -0.248228453 | 0.605683536 |
| ENSG00000188321 | *ZNF559* | 1.249480593 | -0.023430132 | -0.157400417 | 0.133970284 | 0.999933887 | -0.31073965 | -0.162947932 | -0.147791718 | 0.605707001 |
| ENSG00000156931 | *VPS8* | 6.76981894 | -0.086361677 | -0.089626182 | 0.003264505 | 0.999933887 | -1.079927945 | -1.171939195 | 0.09201125 | 0.605742635 |
| ENSG00000146757 | *ZNF92* | 1.47437848 | -0.195175777 | -0.290407954 | 0.095232177 | 0.999933887 | -0.448474794 | -0.241729107 | -0.206745687 | 0.606158547 |
| ENSG00000180900 | *SCRIB* | 3.806559034 | 0.0717835 | 0.052885334 | 0.018898167 | 0.999933887 | 0.070081563 | -0.030961295 | 0.101042858 | 0.606158547 |
| ENSG00000130725 | *UBE2M* | 6.00083289 | 0.105014909 | 0.123227049 | -0.018212141 | 0.999933887 | -0.315417072 | -0.396662641 | 0.081245569 | 0.606747961 |
| ENSG00000146872 | *TLK2* | 5.059086804 | -0.064532947 | 0.025933752 | -0.090466699 | 0.999933887 | -0.247308627 | -0.186355328 | -0.060953298 | 0.606785865 |
| ENSG00000169710 | *FASN* | 4.38111666 | 0.072656535 | 0.103434436 | -0.0307779 | 0.999933887 | 0.202555933 | 0.286780115 | -0.084224183 | 0.606785865 |
| ENSG00000178425 | *NT5DC1* | 2.011562923 | 0.077142461 | -0.142582617 | 0.219725078 | 0.999933887 | -0.18765733 | -0.332059386 | 0.144402057 | 0.606785865 |
| ENSG00000211793 | *TRAV9-2* | -0.926180883 | 0.171345724 | 0.380716272 | -0.209370549 | 0.999933887 | -0.086223158 | 0.201309001 | -0.287532159 | 0.606864181 |
| ENSG00000115128 | *SF3B6* | 4.655733033 | -0.036701193 | -0.040633216 | 0.003932023 | 0.999933887 | -0.166257463 | -0.094249734 | -0.072007729 | 0.607005977 |
| ENSG00000170558 | *CDH2* | -0.653644725 | -0.065538633 | 0.046942691 | -0.112481324 | 0.999933887 | -0.054291584 | 0.351014491 | -0.405306074 | 0.607005977 |
| ENSG00000130816 | *DNMT1* | 4.362298237 | -0.011750234 | 0.03547312 | -0.047223354 | 0.999933887 | 0.120353845 | 0.186268225 | -0.06591438 | 0.607005977 |
| ENSG00000158186 | *MRAS* | -0.374302923 | 0.22471631 | 0.185465754 | 0.039250556 | 0.999933887 | -0.457829758 | -0.679129131 | 0.221299373 | 0.607040481 |
| ENSG00000147684 | *NDUFB9* | 3.585964808 | -0.031369574 | -0.025191784 | -0.00617779 | 0.999933887 | -0.255483491 | -0.355401058 | 0.099917567 | 0.607040481 |
| ENSG00000213699 | *SLC35F6* | 4.157157361 | -0.156081004 | -0.084957053 | -0.071123951 | 0.999933887 | -0.682048894 | -0.761544194 | 0.079495299 | 0.607360188 |
| ENSG00000275532 | *AC006449.2* | 0.89481715 | 0.307561414 | 0.004295668 | 0.303265746 | 0.999933887 | 1.025609776 | 1.212974664 | -0.187364888 | 0.607360188 |
| ENSG00000261158 | *AC109597.2* | -0.396638899 | -0.002981667 | 0.080644241 | -0.083625908 | 0.999933887 | -0.401426433 | -0.852022649 | 0.450596216 | 0.607360188 |
| ENSG00000168564 | *CDKN2AIP* | 4.034925532 | -0.127064701 | -0.247176986 | 0.120112285 | 0.999933887 | 0.632421714 | 0.752533819 | -0.120112105 | 0.60752227 |
| ENSG00000126432 | *PRDX5* | 6.175369308 | 0.042044376 | 0.036404394 | 0.005639982 | 0.999933887 | -0.033962368 | -0.113439953 | 0.079477585 | 0.60752227 |
| ENSG00000128928 | *IVD* | 2.802080549 | 0.026303312 | -0.0398485 | 0.066151812 | 0.999933887 | 0.156413922 | 0.051937701 | 0.104476222 | 0.60752227 |
| ENSG00000126243 | *LRFN3* | 0.815875758 | -0.031635299 | -0.167922196 | 0.136286896 | 0.999933887 | 0.024458692 | 0.193321707 | -0.168863015 | 0.60752227 |
| ENSG00000237254 | *TRBV30* | 0.141060663 | 0.035590252 | -0.041680952 | 0.077271205 | 0.999933887 | 0.22497263 | 0.42018869 | -0.19521606 | 0.60752227 |

| | | | | | | | | | | |
|---|---|---|---|---|---|---|---|---|---|---|
| ENSG00000171723 | *GPHN* | -0.187388933 | -0.101746312 | -0.08378055 | -0.017965761 | 0.999933887 | 0.124195441 | -0.097819563 | 0.222015004 | 0.607804071 |
| ENSG00000211745 | *TRBV4-2* | -0.494783668 | 0.082096335 | 0.148587049 | -0.066490713 | 0.999933887 | 0.040831401 | 0.266801709 | -0.225970309 | 0.607804071 |
| ENSG00000171999 | *OR52P2P* | -0.557381468 | 0.065184051 | -0.184272151 | 0.249456202 | 0.999933887 | -0.111385626 | 0.32222797 | -0.433613596 | 0.607804071 |
| ENSG00000177728 | *TMEM94* | 4.162160248 | -0.013938534 | 0.0838914 | -0.097829934 | 0.999933887 | 0.004294213 | -0.08598722 | 0.090281433 | 0.60800331 |
| ENSG00000184481 | *FOXO4* | 5.735714245 | 0.010386778 | -0.020530996 | 0.030917775 | 0.999933887 | -0.539686045 | -0.626640029 | 0.086953983 | 0.60800331 |
| ENSG00000156253 | *RWDD2B* | -0.384479165 | -0.024135334 | -0.234512301 | 0.210376967 | 0.999933887 | 0.322729743 | 0.10114005 | 0.221589694 | 0.60800331 |
| ENSG00000279759 | *AC118344.2* | 0.307373247 | -0.22905917 | 0.126950692 | -0.356009861 | 0.999933887 | -0.392135124 | -0.196616147 | -0.195518977 | 0.608330037 |
| ENSG00000181513 | *ACBD4* | 1.653636183 | 0.06151171 | 0.08903686 | -0.027525149 | 0.999933887 | 0.172473541 | 0.03949354 | 0.132980001 | 0.608330037 |
| ENSG00000137947 | *GTF2B* | 5.139543098 | -0.023313333 | -0.052980162 | 0.029666828 | 0.999933887 | -0.128058072 | -0.059200219 | -0.068857853 | 0.608330037 |
| ENSG00000264493 | *MIR4298* | 0.519541961 | 0.026567051 | -0.102542206 | 0.129109257 | 0.999933887 | -0.261547532 | -0.023175885 | -0.238371647 | 0.608330037 |
| ENSG00000119537 | *KDSR* | 3.306102453 | -0.01046596 | -0.033575742 | 0.023109782 | 0.999933887 | -0.624200212 | -0.726931593 | 0.10273138 | 0.608330037 |
| ENSG00000124203 | *ZNF831* | 4.014188005 | 0.100115043 | 0.1619988 | -0.061883757 | 0.999933887 | 0.048136202 | -0.026355184 | 0.074491386 | 0.608740171 |
| ENSG00000223725 | *AC009226.1* | 0.965700644 | -0.047861795 | 0.302834441 | -0.350696236 | 0.999933887 | 1.228549133 | 1.408565808 | -0.180016675 | 0.608740171 |
| ENSG00000111667 | *USP5* | 2.974926908 | 0.006617216 | 0.022532609 | -0.015915393 | 0.999933887 | 0.170127705 | 0.073177774 | 0.096949931 | 0.608740171 |
| ENSG00000278030 | *TRBV7-9* | 0.814340236 | -0.109661326 | -0.242626885 | 0.13296556 | 0.999933887 | 0.136054957 | 0.271246336 | -0.135191379 | 0.608768489 |
| ENSG00000169682 | *SPNS1* | 2.638238624 | 0.043270905 | 0.193549642 | -0.150278737 | 0.999933887 | 0.494318359 | 0.611338065 | -0.117019706 | 0.60923981 |
| ENSG00000140939 | *NOL3* | 0.209601642 | -0.032372888 | 0.266218234 | -0.298591122 | 0.999933887 | 0.071136693 | 0.278610401 | -0.207473708 | 0.609294777 |
| ENSG00000100731 | *PCNX1* | 7.665626849 | 0.001583808 | -0.034080517 | 0.035664325 | 0.999933887 | -0.527421898 | -0.602186053 | 0.074764155 | 0.609347928 |
| ENSG00000139133 | *ALG10* | -0.413413717 | -0.148839991 | -0.133568306 | -0.015271685 | 0.999933887 | -0.195087284 | 0.046599622 | -0.241686907 | 0.609794726 |
| ENSG00000185482 | *STAC3* | 0.236394616 | 0.066888733 | -0.048363464 | 0.115252198 | 0.999933887 | 0.058849129 | 0.28205579 | -0.223206661 | 0.610004093 |
| ENSG00000233016 | *SNHG7* | 2.696861615 | 0.096228745 | -0.076697505 | 0.172926251 | 0.999933887 | 0.140479044 | 0.038263059 | 0.102215985 | 0.610034575 |
| ENSG00000184508 | *HDDC3* | -0.200528132 | 0.156326599 | -0.031931048 | 0.188257647 | 0.999933887 | 0.082296394 | -0.167794155 | 0.250090549 | 0.610034575 |
| ENSG00000268635 | *AP003680.1* | 0.504694295 | 0.339110552 | 0.579680816 | -0.240570264 | 0.999933887 | 1.726009152 | 1.970959639 | -0.244950487 | 0.610071544 |
| ENSG00000065029 | *ZNF76* | 3.988365077 | 0.039507566 | 0.113666184 | -0.074158618 | 0.999933887 | 0.212391689 | 0.282353857 | -0.069962167 | 0.610143823 |
| ENSG00000276966 | *H4C5* | -0.653654868 | -0.012235758 | -0.253655654 | 0.241419895 | 0.999933887 | -0.024125613 | -0.310462271 | 0.286336658 | 0.610155379 |
| ENSG00000241015 | *TPM3P9* | 1.194591305 | 0.08087008 | 0.085752908 | -0.004882828 | 0.999933887 | 0.123053494 | -0.025752692 | 0.148806186 | 0.610517183 |
| ENSG00000154743 | *TSEN2* | 0.012784751 | -0.18062034 | -0.05206334 | -0.128557 | 0.999933887 | 0.157240883 | 0.330441461 | -0.173200578 | 0.610555212 |
| ENSG00000224186 | *C5orf66* | 1.027649463 | -0.172745383 | -0.074108368 | -0.098637015 | 0.999933887 | -1.731648202 | -1.921554339 | 0.189906137 | 0.610590829 |
| ENSG00000211778 | *TRAV4* | -0.369384614 | 0.052310759 | 0.084708498 | -0.032397739 | 0.999933887 | 0.204157633 | 0.449119728 | -0.244962094 | 0.610590829 |
| ENSG00000090857 | *PDPR* | 6.134004082 | -0.06218982 | -0.081406365 | 0.019216545 | 0.999933887 | 0.09415481 | 0.173596258 | -0.079441448 | 0.610609895 |
| ENSG00000047597 | *XK* | -0.448254931 | -0.08998762 | 0.191636461 | -0.281624081 | 0.999933887 | -0.0472657 | 0.161031127 | -0.208296826 | 0.610756391 |
| ENSG00000135482 | *ZC3H10* | 2.097971145 | -0.029353243 | -0.001195594 | -0.028157649 | 0.999933887 | -0.419857331 | -0.306468515 | -0.113388816 | 0.610756391 |
| ENSG00000072803 | *FBXW11* | 4.659934112 | 0.016755239 | -0.016178881 | 0.03293412 | 0.999933887 | 0.073383676 | 0.152229073 | -0.078845397 | 0.610756391 |
| ENSG00000032389 | *EIPR1* | 1.922558482 | 0.020757829 | -0.019989031 | 0.040746859 | 0.999933887 | 0.039838831 | -0.108608158 | 0.148446989 | 0.610756391 |
| ENSG00000205133 | *TRIQK* | 2.597850967 | -0.057328794 | 0.014052659 | -0.071381453 | 0.999933887 | -0.495917234 | -0.385227952 | -0.110689281 | 0.610823601 |
| ENSG00000239480 | *AC073517.1* | -0.028409889 | 0.089632254 | 0.150965053 | -0.0613328 | 0.999933887 | -0.15559756 | -0.376588506 | 0.220990946 | 0.611049963 |
| ENSG00000222047 | *C10orf55* | 5.17407086 | 0.2634841 | 0.373451263 | -0.109967163 | 0.999933887 | 1.42652851 | 1.553156836 | -0.126628326 | 0.611205775 |
| ENSG00000108064 | *TFAM* | 3.5913429 | -0.043383927 | -0.129817721 | 0.086433794 | 0.999933887 | -0.137578782 | -0.232542102 | 0.094963319 | 0.611213345 |
| ENSG00000143742 | *SRP9* | 4.15450263 | -0.067727115 | -0.09515734 | 0.027430225 | 0.999933887 | -0.100543174 | -0.175618003 | 0.07507483 | 0.611533509 |
| ENSG00000168970 | *JMJD7-PLA2G* | 2.000372841 | 0.122986609 | 0.191822721 | -0.068836112 | 0.999933887 | -0.044606067 | -0.18173503 | 0.137128964 | 0.611533509 |
| ENSG00000198855 | *FICD* | 2.419923046 | 0.077701691 | 0.017998342 | 0.059703349 | 0.999933887 | 0.429081558 | 0.325649605 | 0.103431953 | 0.611623385 |
| ENSG00000205352 | *PRR13* | 6.705289726 | -0.042818124 | -0.01863064 | -0.024187484 | 0.999933887 | -0.152221282 | -0.215467297 | 0.063246015 | 0.611849286 |
| ENSG00000204228 | *HSD17B8* | 0.243741802 | 0.074421159 | 0.007580619 | 0.06684054 | 0.999933887 | 0.174094461 | 0.368932865 | -0.194838405 | 0.611961945 |
| ENSG00000101298 | *SNPH* | 1.388242894 | 0.037039272 | 0.043180045 | -0.006140774 | 0.999933887 | 0.084709981 | -0.08079126 | 0.165501241 | 0.612051474 |
| ENSG00000141682 | *PMAIP1* | 7.779261758 | 0.117658721 | 0.126901318 | -0.009242596 | 0.999933887 | -0.143761473 | -0.051860193 | -0.09190128 | 0.612090876 |
| ENSG00000155275 | *TRMT44* | 1.976633426 | 0.041057989 | -0.050286389 | 0.091344378 | 0.999933887 | 0.240331841 | 0.371420249 | -0.131088408 | 0.612231005 |
| ENSG00000187642 | *PERM1* | -0.885635343 | 0.065871187 | -0.142795022 | 0.20866621 | 0.999933887 | 1.499648497 | 1.097637229 | 0.402011268 | 0.612231005 |
| ENSG00000136720 | *HS6ST1* | 2.584181588 | 0.035647328 | 0.080481129 | -0.044833801 | 0.999933887 | 0.279107558 | 0.398565043 | -0.119457485 | 0.612304091 |
| ENSG00000055917 | *PUM2* | 7.059317844 | -0.051079719 | -0.05086435 | -0.000215369 | 0.999933887 | -0.409497074 | -0.339250787 | -0.070246287 | 0.612380734 |
| ENSG00000125885 | *MCM8* | -0.446992621 | -0.149046643 | -0.184780643 | 0.035734 | 0.999933887 | -0.012040166 | -0.331737127 | 0.319696961 | 0.612380734 |
| ENSG00000165055 | *METTL2B* | 1.005674742 | 0.027771772 | 0.034493874 | -0.006722102 | 0.999933887 | 0.138070994 | 0.284246778 | -0.146175784 | 0.612380734 |
| ENSG00000240853 | *RN7SL328P* | -0.224850257 | -0.008305101 | 0.14340076 | -0.151705862 | 0.999933887 | 0.185346552 | 0.470882491 | -0.285535939 | 0.612380734 |
| ENSG00000166912 | *MTMR10* | 6.233384318 | -0.093102014 | -0.092989029 | -0.000112985 | 0.999933887 | -1.363340311 | -1.471475981 | 0.108135669 | 0.612398915 |
| ENSG00000262766 | *AC135050.5* | 1.385066809 | 0.086466361 | 0.104977085 | -0.018510723 | 0.999933887 | -0.284306174 | -0.462443472 | 0.178137298 | 0.612398915 |
| ENSG00000286982 | *AC016931.1* | 0.065607502 | 0.154131919 | 0.37922314 | -0.225091221 | 0.999933887 | -0.14340618 | 0.084837781 | -0.228243961 | 0.612398915 |
| ENSG00000166123 | *GPT2* | -0.995019079 | 0.172364552 | 0.585379067 | -0.413014514 | 0.999933887 | -0.066809509 | 0.22779987 | -0.29460938 | 0.612398915 |
| ENSG00000100055 | *CYTH4* | 9.545040849 | -0.023510673 | 0.002365943 | -0.025876616 | 0.999933887 | -0.086787539 | -0.153412693 | 0.066625153 | 0.612398915 |
| ENSG00000258018 | *LINC02457* | -0.568329752 | 0.051887998 | 0.513311363 | -0.461423365 | 0.999933887 | 1.381573364 | 1.101432799 | 0.280140565 | 0.612398915 |
| ENSG00000084093 | *REST* | 4.690063165 | 0.098973828 | -0.063206843 | 0.162180672 | 0.999933887 | -0.903251955 | -1.04786224 | 0.144610285 | 0.612656653 |
| ENSG00000109475 | *RPL34* | 7.258683233 | -0.027093005 | -0.042367049 | 0.015274044 | 0.999933887 | 0.071223998 | 0.142138168 | -0.070914169 | 0.612656653 |
| ENSG00000048028 | *USP28* | 2.650304349 | -0.037649079 | -0.088174116 | 0.050533137 | 0.999933887 | -0.042587363 | 0.05260165 | -0.095189013 | 0.612656653 |
| ENSG00000175893 | *ZDHHC21* | 0.76089191 | -0.044027241 | -0.335968107 | 0.291940866 | 0.999933887 | -0.163171583 | 0.061794 | -0.224965583 | 0.612656653 |
| ENSG00000211771 | *TRBJ2-7* | -0.041770424 | 0.181386933 | 0.214396429 | -0.033009496 | 0.999933887 | 0.118658367 | -0.115771538 | 0.234429905 | 0.612770486 |
| ENSG00000180739 | *S1PR5* | 3.981627603 | 0.018472743 | -0.021023404 | 0.039496147 | 0.999933887 | 0.14765774 | 0.067093426 | 0.080564314 | 0.612770486 |
| ENSG00000225920 | *RIMKLBP2* | -0.923002813 | -0.144083342 | -0.779211848 | 0.635128507 | 0.999933887 | -1.315897945 | -0.96867785 | -0.347220096 | 0.612799775 |
| ENSG00000253214 | *AC079209.1* | 4.441752603 | -0.043934378 | 0.053819994 | -0.097754372 | 0.999933887 | 0.388719597 | 0.534405651 | -0.145686054 | 0.612861895 |
| ENSG00000100652 | *SLC10A1* | 1.034373775 | 0.157848217 | 0.171611908 | -0.01376369 | 0.999933887 | 0.498004443 | 0.679977793 | -0.18197335 | 0.613055169 |
| ENSG00000279089 | *AC005839.1* | 1.52743931 | -0.11787884 | 0.048965092 | -0.166843932 | 0.999933887 | -0.319879693 | -0.492706895 | 0.172827202 | 0.613066909 |
| ENSG00000258102 | *MAP1LC3B2* | 3.998637172 | 0.026789738 | 0.1092063 | -0.082416562 | 0.999933887 | 0.101966771 | 0.019544159 | 0.082422612 | 0.613066909 |
| ENSG00000184441 | *AP001062.1* | 0.559176009 | 0.018408328 | 0.194388541 | -0.175980213 | 0.999933887 | 0.063423035 | 0.238856319 | -0.175433284 | 0.613066909 |
| ENSG00000127526 | *SLC35E1* | 4.986907643 | -0.038059297 | -0.020072763 | -0.017986535 | 0.999933887 | -0.128712791 | -0.066282584 | -0.062430207 | 0.613077409 |
| ENSG00000155975 | *VPS37A* | 3.657644345 | 0.025626069 | 0.038133719 | -0.01250765 | 0.999933887 | -0.352937266 | -0.433153729 | 0.080216463 | 0.613077409 |
| ENSG00000025708 | *TYMP* | 7.48402929 | 0.036382616 | 0.081979582 | -0.045596966 | 0.999933887 | -0.73627262 | -0.82784574 | 0.09157312 | 0.613086441 |
| ENSG00000172456 | *FGGY* | 2.441512264 | 0.083582578 | -0.037271764 | 0.120854342 | 0.999933887 | 0.642626855 | 0.772342212 | -0.129715357 | 0.613414867 |
| ENSG00000103111 | *MON1B* | 6.359896 | -0.015935211 | -0.03720744 | 0.02127223 | 0.999933887 | -0.220648857 | -0.283205171 | 0.062556314 | 0.613421961 |
| ENSG00000267254 | *ZNF790-AS1* | -0.93489944 | -0.269098801 | 0.085923926 | -0.355022727 | 0.999933887 | -0.186066869 | 0.09059163 | -0.276658499 | 0.614326908 |
| ENSG00000215580 | *BCORP1* | -4.063275687 | 0.06910035 | 0.065409141 | 0.003691209 | 0.999933887 | 0.134730954 | 0.425294478 | -0.290563523 | 0.61453752 |
| ENSG00000211890 | *IGHA2* | 1.041889351 | 0.029679274 | 0.251826798 | -0.222147525 | 0.999933887 | 0.275262291 | 0.56049764 | -0.285235349 | 0.61453752 |
| ENSG00000133114 | *GPALPP1* | 2.838045642 | 0.089486452 | 0.08501644 | 0.004470012 | 0.999933887 | 0.252118207 | 0.382909499 | -0.130791293 | 0.615059068 |
| ENSG00000115421 | *PAPOLG* | 3.78723529 | -0.071210982 | -0.015101509 | -0.056109473 | 0.999933887 | -0.309834351 | -0.228547931 | -0.08128642 | 0.615059068 |
| ENSG00000197603 | *CPLANE1* | 0.987534428 | -0.031810319 | -0.023739303 | -0.008071017 | 0.999933887 | -0.293818414 | -0.061525455 | -0.232292959 | 0.615059068 |
| ENSG00000272812 | *AC004908.2* | 1.017535109 | 0.235885454 | 0.195630397 | 0.040255057 | 0.999933887 | 0.396750019 | 0.229519866 | 0.167230153 | 0.615136953 |
| ENSG00000059804 | *SLC2A3* | 8.867080036 | 0.041388803 | 0.074532956 | -0.033144153 | 0.999933887 | -0.361956497 | -0.285816521 | -0.076139976 | 0.615178865 |
| ENSG00000253978 | *CTB-178M22.2* | -0.87737406 | 0.06280515 | 0.062507052 | 0.000298098 | 0.999933887 | 0.908947289 | 1.123411878 | -0.214464589 | 0.615178865 |
| ENSG00000115947 | *ORC4* | 3.004205885 | -0.035968927 | -0.116810874 | 0.080841947 | 0.999933887 | -0.483051981 | -0.618337325 | 0.135285344 | 0.61530949 |
| ENSG00000123472 | *ATPAF1* | 2.074239437 | 0.075719773 | -0.068777085 | 0.144496858 | 0.999933887 | -0.011749692 | -0.119444729 | 0.107695037 | 0.615391379 |
| ENSG00000213619 | *NDUFS3* | 2.761098441 | -0.04949747 | 0.00850451 | -0.05800198 | 0.999933887 | 0.090654815 | -0.005126667 | 0.095781482 | 0.615391379 |
| ENSG00000271553 | *AC018638.7* | -0.440169791 | -0.041958591 | 0.219746272 | -0.261704863 | 0.999933887 | -0.157823175 | 0.155865369 | -0.313688544 | 0.615391379 |
| ENSG00000120451 | *SNX19* | 5.079928504 | -0.025650229 | -0.007370412 | -0.018279817 | 0.999933887 | -0.066638104 | -0.118968923 | 0.052330818 | 0.615597223 |
| ENSG00000258035 | *AC123567.2* | 3.366850226 | -0.004015889 | 0.184873395 | -0.188889284 | 0.999933887 | 0.792101025 | 0.942092397 | -0.149991372 | 0.615597223 |
| ENSG00000140694 | *PARN* | 3.650241756 | -0.045598558 | -0.04332889 | -0.002269667 | 0.999933887 | -0.15937374 | -0.243077363 | 0.083703623 | 0.615672042 |
| ENSG00000277149 | *TYW1B* | 0.989737653 | 0.05041427 | -0.060969833 | 0.111384103 | 0.999933887 | -0.050719981 | 0.112420898 | -0.16314088 | 0.615871569 |
| ENSG00000112282 | *MED23* | 4.480056995 | 0.025979225 | -0.043506575 | 0.0694858 | 0.999933887 | -0.316386183 | -0.43050219 | 0.114116007 | 0.615871569 |
| ENSG00000234358 | *AC003080.1* | 0.623211742 | -0.008778204 | 0.233164982 | -0.241943185 | 0.999933887 | 0.782701493 | 1.020762464 | -0.238060971 | 0.615871569 |
| ENSG00000115204 | *MPV17* | 3.166100231 | 0.12514604 | 0.105352054 | 0.019793986 | 0.999933887 | 0.074261746 | -0.003046321 | 0.077308067 | 0.616120924 |
| ENSG00000158887 | *MPZ* | 3.201122574 | 0.071439007 | -0.074343443 | 0.14578245 | 0.999933887 | 0.503169693 | 0.594947672 | -0.091777979 | 0.616120924 |
| ENSG00000167460 | *TPM4* | 8.036961329 | 0.043562727 | 0.026375601 | 0.017187126 | 0.999933887 | 0.086112394 | -0.002130291 | 0.088242685 | 0.616401862 |
| ENSG00000042753 | *AP2S1* | 3.337085913 | 0.034993695 | -0.094268345 | 0.129262039 | 0.999933887 | -0.005055471 | -0.106198259 | 0.101142788 | 0.616401862 |
| ENSG00000104205 | *SGK3* | 1.641551007 | -0.029794795 | 0.264008888 | -0.293803683 | 0.999933887 | -0.466652497 | -0.329228326 | -0.137424172 | 0.616567417 |
| ENSG00000135414 | *GDF11* | 1.866446159 | 0.006439777 | -0.082023039 | 0.088462816 | 0.999933887 | -0.110175031 | -0.220597455 | 0.110422424 | 0.616585207 |
| ENSG00000204677 | *FAM153CP* | -1.469592175 | -0.392128092 | 0.145494462 | -0.537622554 | 0.999933887 | -0.544278875 | -0.121569105 | -0.42270977 | 0.616917648 |
| ENSG00000182552 | *RWDD4* | 1.410960467 | 0.000838803 | -0.034568771 | 0.035407574 | 0.999933887 | -0.113768052 | -0.26814536 | 0.154377308 | 0.616917648 |
| ENSG00000118596 | *SLC16A7* | 2.677305722 | -0.026244994 | 0.064586757 | -0.090831751 | 0.999933887 | -0.522102985 | -0.393089528 | -0.129013457 | 0.616921241 |
| ENSG00000197355 | *UAP1L1* | 3.649661196 | 0.132440043 | 0.056571111 | 0.075868932 | 0.999933887 | -0.349650281 | -0.453906604 | 0.104256323 | 0.61718784 |
| ENSG00000119523 | *ALG2* | 2.64072723 | 0.028093458 | -0.085169146 | 0.113262604 | 0.999933887 | 0.135429239 | 0.042477361 | 0.092951878 | 0.617579717 |

| | | | | | | | | | | |
|---|---|---|---|---|---|---|---|---|---|---|
| ENSG00000245958 | *AC093752.1* | 2.550178719 | 0.104125749 | 0.035077121 | 0.069048628 | 0.999933887 | -0.049323391 | 0.108444028 | -0.157767419 | 0.617774156 |
| ENSG00000160588 | *MPZL3* | 6.24094965 | -0.087334412 | -0.025146981 | -0.062187431 | 0.999933887 | 0.075018068 | 0.001533534 | 0.073484534 | 0.618040165 |
| ENSG00000196576 | *PLXNB2* | 2.00436746 | 0.058488329 | 0.058345379 | 0.00014295 | 0.999933887 | -1.043808487 | -1.190636788 | 0.146828301 | 0.618089666 |
| ENSG00000111846 | *GCNT2* | -0.192363559 | -0.044998014 | -0.143408592 | 0.098410578 | 0.999933887 | 0.103024268 | -0.128497632 | 0.231521901 | 0.618233751 |
| ENSG00000176871 | *WSB2* | 5.802538331 | -0.088781075 | 0.003211311 | -0.091992386 | 0.999933887 | -0.301143676 | -0.230684635 | -0.070459041 | 0.618631494 |
| ENSG00000131966 | *ACTR10* | 4.166738567 | -0.061264401 | -0.041554577 | -0.019709824 | 0.999933887 | -0.212111289 | -0.140250523 | -0.071860766 | 0.618631494 |
| ENSG00000178982 | *EIF3K* | 5.411214132 | -0.043338424 | -0.042047535 | -0.001290889 | 0.999933887 | 0.053023127 | -0.007484908 | 0.060508035 | 0.618633032 |
| ENSG00000243943 | *ZNF512* | 2.67829872 | 0.003103882 | -0.0557238 | 0.058827682 | 0.999933887 | -0.173364869 | -0.262098438 | 0.088733569 | 0.618931069 |
| ENSG00000244617 | *ASPRV1* | 4.173198539 | -0.062430386 | 0.000328655 | -0.062759042 | 0.999933887 | 0.301140145 | 0.400607486 | -0.099467341 | 0.619026847 |
| ENSG00000207248 | *RNU6-1005P* | 0.363152125 | 0.143627474 | 0.265867572 | -0.122240097 | 0.999933887 | 0.554495429 | 0.779604296 | -0.225108867 | 0.619134842 |
| ENSG00000070018 | *LRP6* | -0.120949814 | -0.111473554 | 0.229678386 | -0.341151939 | 0.999933887 | -0.223104942 | 0.040787819 | -0.263892761 | 0.619232546 |
| ENSG00000116649 | *SRM* | 2.296555251 | 0.034210199 | 0.114942323 | -0.080732123 | 0.999933887 | 0.211869403 | 0.313619379 | -0.101749976 | 0.619280996 |
| ENSG00000037280 | *FLT4* | 0.131621656 | 0.038659612 | 0.196722871 | -0.158063259 | 0.999933887 | 0.085024351 | -0.145217283 | 0.230241634 | 0.619280996 |
| ENSG00000198960 | *ARMCX6* | 2.166774812 | 0.051394545 | 0.107535967 | -0.056141422 | 0.999933887 | -0.040749247 | 0.069082641 | -0.109831888 | 0.619561375 |
| ENSG00000151461 | *UPF2* | 5.327315014 | -0.015065629 | -0.164637926 | 0.149572297 | 0.999933887 | -0.877933319 | -1.003782038 | 0.125848719 | 0.619561375 |
| ENSG00000170345 | *FOS* | 9.075142236 | 0.526978282 | 0.448872848 | 0.078105435 | 0.999933887 | 0.135610132 | -0.00937729 | 0.144987422 | 0.61961244 |
| ENSG00000174917 | *MICOS13* | 2.17678388 | 0.099115498 | 0.075855746 | 0.023259752 | 0.999933887 | 0.093518864 | -0.010957818 | 0.104476682 | 0.619722041 |
| ENSG00000274215 | *AC106028.4* | 1.2546104 | 0.260763301 | -0.013124428 | 0.273887729 | 0.999933887 | 2.403173024 | 2.622787928 | -0.219608904 | 0.620050351 |
| ENSG00000223745 | *CCDC18-AS1* | 3.052829166 | -0.106454822 | -0.065769086 | -0.040685736 | 0.999933887 | -0.064084578 | -0.168981418 | 0.104896841 | 0.620050351 |
| ENSG00000145220 | *LYAR* | 3.284540143 | -0.050184339 | -0.045849217 | -0.004335122 | 0.999933887 | 0.065850215 | 0.147147927 | -0.081297712 | 0.620050351 |
| ENSG00000121680 | *PEX16* | 4.565621366 | -0.031307216 | 0.074719359 | -0.106026575 | 0.999933887 | -0.004241089 | -0.073296457 | 0.069055369 | 0.620050351 |
| ENSG00000009830 | *POMT2* | 2.007540934 | 0.024562484 | -0.126195013 | 0.150757497 | 0.999933887 | -0.20907466 | -0.085570723 | -0.123503937 | 0.620050351 |
| ENSG00000261270 | *AC012181.2* | 0.408069912 | 0.191964301 | 0.068448257 | 0.123516044 | 0.999933887 | 0.244248076 | -0.142757245 | 0.387005321 | 0.620116672 |
| ENSG00000011590 | *ZBTB32* | 0.672943353 | -0.090044976 | 0.13831277 | -0.228357746 | 0.999933887 | 0.324110579 | 0.132179048 | 0.191931531 | 0.620116672 |
| ENSG00000138380 | *CARF* | 0.696887018 | 0.093292145 | -0.2328966 | 0.326188745 | 0.999933887 | -0.182769466 | 0.013655746 | -0.196425213 | 0.620116672 |
| ENSG00000147324 | *MFHAS1* | 3.117479282 | -0.025212131 | 0.063186529 | -0.08839866 | 0.999933887 | -0.002932305 | -0.098152995 | 0.09522069 | 0.620116672 |
| ENSG00000111796 | *KLRB1* | 3.226213189 | -0.021230386 | -0.082543033 | 0.061312646 | 0.999933887 | 0.018684773 | 0.129724299 | -0.111039526 | 0.620116672 |
| ENSG00000148803 | *FUOM* | 0.021237611 | 0.038746005 | 0.128714955 | -0.08996895 | 0.999933887 | -0.031340426 | -0.223489642 | 0.192149217 | 0.620116672 |
| ENSG00000172809 | *RPL38* | 6.590778915 | -0.002216695 | -0.010695018 | 0.008478323 | 0.999933887 | 0.081656881 | 0.157168861 | -0.07551198 | 0.620116672 |
| ENSG00000163728 | *TTC14* | 4.183632382 | -0.098965375 | -0.252519208 | 0.153553833 | 0.999933887 | -0.424399601 | -0.246145953 | -0.178253649 | 0.620125905 |
| ENSG00000269961 | *AC010359.2* | 1.721608286 | 0.12563656 | 0.309860691 | -0.184224131 | 0.999933887 | 1.188092012 | 1.315310381 | -0.127218369 | 0.620267574 |
| ENSG00000100311 | *PDGFB* | -0.041868049 | 0.086638744 | 0.518808837 | -0.432170092 | 0.999933887 | 0.120532966 | 0.33804114 | -0.217508173 | 0.620267574 |
| ENSG00000272449 | *AL139246.5* | 7.080264469 | 0.200679278 | 0.18908537 | 0.011593909 | 0.999933887 | 0.981329965 | 1.12860399 | -0.147274025 | 0.620400777 |
| ENSG00000287265 | *AL365272.1* | 0.980950492 | 0.152245077 | -0.027830372 | 0.180075449 | 0.999933887 | -0.113358151 | -0.247576966 | 0.134218815 | 0.620400777 |
| ENSG00000146540 | *C7orf50* | 3.056600479 | 0.041693535 | 0.07915836 | -0.037464824 | 0.999933887 | -0.089423239 | -0.188690143 | 0.099266904 | 0.620400777 |
| ENSG00000274349 | *ZNF658* | -0.531345702 | -0.043270578 | -0.082091022 | 0.038820444 | 0.999933887 | -0.267236157 | -0.571296821 | 0.304060664 | 0.620400777 |
| ENSG00000113384 | *GOLPH3* | 5.280509036 | 0.001545164 | 0.004144029 | -0.002598865 | 0.999933887 | -0.336236696 | -0.403308168 | 0.067071472 | 0.620400777 |
| ENSG00000278743 | *AC087239.1* | 1.688988001 | 0.181147181 | 0.345100355 | -0.163953174 | 0.999933887 | 0.831642931 | 1.03419548 | -0.202552549 | 0.620519994 |
| ENSG00000184517 | *ZFP1* | 0.577037826 | -0.178681696 | -0.075082497 | -0.1035992 | 0.999933887 | 0.082746561 | -0.095646524 | 0.178393086 | 0.620531894 |
| ENSG00000112787 | *FBRSL1* | 6.669479279 | 0.034504214 | 0.049069002 | -0.014564788 | 0.999933887 | -0.057224953 | -0.11993848 | 0.062713527 | 0.620531894 |
| ENSG00000177409 | *SAMD9L* | 6.080398911 | -0.078322288 | -0.216337633 | 0.138015344 | 0.999933887 | -1.01753882 | -0.826961226 | -0.190577594 | 0.620531894 |
| ENSG00000138246 | *DNAJC13* | 4.580308515 | -0.037793904 | -0.186974502 | 0.149180598 | 0.999933887 | -0.994358962 | -1.127070949 | 0.132711987 | 0.620531894 |
| ENSG00000160789 | *LMNA* | 5.873693035 | -0.001362397 | 0.0014016 | -0.002763997 | 0.999933887 | 0.03402209 | 0.117961787 | -0.083939696 | 0.620531894 |
| ENSG00000231830 | *AC245140.1* | 1.325192225 | 0.048934856 | 0.026165898 | 0.022768957 | 0.999933887 | 0.32797101 | 0.501358879 | -0.173387869 | 0.62053791 |
| ENSG00000132604 | *TERF2* | 3.496515195 | 0.016369318 | -0.010738744 | 0.027108062 | 0.999933887 | -0.221956558 | -0.156634364 | -0.065322194 | 0.62053791 |
| ENSG00000205045 | *SLFN12L* | 1.67463911 | -0.110371073 | -0.024482239 | -0.085888834 | 0.999933887 | -0.177948615 | 0.016848574 | -0.194797189 | 0.620719002 |
| ENSG00000145730 | *PAM* | 3.472280941 | -0.02696452 | -0.001026527 | -0.025937993 | 0.999933887 | -0.008774572 | 0.078498709 | -0.087273282 | 0.620749372 |
| ENSG00000197050 | *ZNF420* | -0.181845981 | -0.170952354 | -0.125099774 | -0.04585258 | 0.999933887 | -0.039367933 | 0.174859406 | -0.214227339 | 0.620832051 |
| ENSG00000214226 | *C17orf67* | 1.264101857 | -0.100534123 | -0.012151202 | -0.088382921 | 0.999933887 | -0.394610112 | -0.544025778 | 0.149415666 | 0.620832051 |
| ENSG00000128654 | *MTX2* | -0.081035981 | 0.079743024 | 0.07774466 | 0.001998364 | 0.999933887 | 0.287284556 | 0.476712448 | -0.189427892 | 0.620832051 |
| ENSG00000158711 | *ELK4* | 4.598026605 | -0.018139021 | -0.066858948 | 0.048719927 | 0.999933887 | -0.277893729 | -0.150338968 | -0.127554761 | 0.620866835 |
| ENSG00000267046 | *E2F3P1* | 0.034860694 | 0.074166313 | -0.336881927 | 0.41104824 | 0.999933887 | -0.326595763 | -0.089639298 | -0.236956465 | 0.621204567 |
| ENSG00000077380 | *DYNC1I2* | 2.352032928 | -0.036649375 | -0.16073757 | 0.124088195 | 0.999933887 | 0.082481796 | 0.184709848 | -0.102228051 | 0.621204567 |
| ENSG00000197694 | *SPTAN1* | 6.487622036 | 0.000803139 | -0.04967189 | 0.050475029 | 0.999933887 | 0.023452495 | -0.030113044 | 0.053565539 | 0.621204567 |
| ENSG00000089472 | *HEPH* | -0.887537564 | -0.168653981 | -0.199396152 | 0.03074217 | 0.999933887 | -0.109657771 | 0.201040045 | -0.310697817 | 0.621345625 |
| ENSG00000213593 | *TMX2* | 3.033713381 | -0.049430117 | -0.005098654 | -0.044331463 | 0.999933887 | 0.062051198 | -0.022961644 | 0.085012842 | 0.621345625 |
| ENSG00000241170 | *AP001992.1* | 1.862231272 | -0.036992159 | 0.001921876 | -0.038914035 | 0.999933887 | -0.020296545 | 0.104572372 | -0.124868917 | 0.621942417 |
| ENSG00000119535 | *CSF3R* | 12.203206831 | 0.015563037 | 0.0118306 | 0.003732437 | 0.999933887 | -0.012656304 | -0.06108995 | 0.048433646 | 0.621942417 |
| ENSG00000186115 | *CYP4F2* | -2.83953 | -0.03570801 | -0.079358983 | 0.043650973 | 0.999933887 | 0.129231434 | -0.209784009 | 0.339015444 | 0.621942417 |
| ENSG00000086827 | *ZW10* | 0.614121482 | -0.054738637 | -0.11837201 | 0.063633373 | 0.999933887 | 0.151507779 | 0.309113706 | -0.157605927 | 0.622118117 |
| ENSG00000258839 | *MC1R* | 1.90303992 | -0.032433484 | 0.056342609 | -0.088776093 | 0.999933887 | 0.359890199 | 0.491064345 | -0.131174146 | 0.622118117 |
| ENSG00000036257 | *CUL3* | 6.16789017 | -0.028326851 | -0.102257134 | 0.073930283 | 0.999933887 | -0.368006989 | -0.296065603 | -0.071941386 | 0.622133237 |
| ENSG00000169203 | *NPIPB12* | -0.035444401 | 0.075420002 | -0.139293589 | 0.214713591 | 0.999933887 | -0.177847858 | 0.079474691 | -0.257322549 | 0.622133237 |
| ENSG00000256967 | *AC018653.3* | 1.531866422 | 0.078620735 | 0.024956694 | 0.053664041 | 0.999933887 | 0.178345708 | 0.007065289 | 0.171280419 | 0.62282165 |
| ENSG00000260144 | *AC100827.2* | -0.471934839 | 0.152588383 | 0.236349574 | -0.083761191 | 0.999933887 | 0.434578882 | 0.049849576 | 0.384729306 | 0.62282165 |
| ENSG00000232815 | *DUX4L50* | 1.842790313 | 0.054791753 | 0.164816653 | -0.110024901 | 0.999933887 | -0.363742364 | -0.551110846 | 0.187368482 | 0.62282165 |
| ENSG00000225964 | *NRIR* | 0.492590935 | -0.055697268 | -0.148580156 | 0.092882888 | 0.999933887 | 0.069869292 | 0.277592713 | -0.207723421 | 0.62282165 |
| ENSG00000100564 | *PIGH* | 2.613458405 | -0.119890876 | 0.099698741 | -0.219589617 | 0.999933887 | 0.313646872 | 0.449245827 | -0.135598954 | 0.622865118 |
| ENSG00000118162 | *KPTN* | 2.024207868 | 0.077846838 | 0.025863199 | 0.051983639 | 0.999933887 | 0.162545266 | 0.031935169 | 0.130610096 | 0.622865118 |
| ENSG00000100519 | *PSMC6* | 4.456908935 | -0.042595994 | -0.054753562 | 0.012157568 | 0.999933887 | -0.139158642 | -0.059431863 | -0.079726779 | 0.622865118 |
| ENSG00000152784 | *PRDM8* | 5.728352347 | 0.029834012 | 0.022824062 | 0.007009951 | 0.999933887 | -0.243015426 | -0.319689161 | 0.076673736 | 0.622865118 |
| ENSG00000124588 | *NQO2* | 5.637092583 | -0.053547232 | -0.081210818 | 0.027663586 | 0.999933887 | -0.12525739 | -0.209428642 | 0.084171252 | 0.623127003 |
| ENSG00000263712 | *MIR4639* | -1.396928883 | 0.064292778 | 0.018630195 | 0.045662583 | 0.999933887 | 0.076392846 | 0.564257004 | -0.487864157 | 0.623127003 |
| ENSG00000133398 | *MED10* | 3.684137336 | -0.038521716 | -0.022398412 | -0.016123304 | 0.999933887 | -0.059269636 | -0.128310126 | 0.069040491 | 0.623200711 |
| ENSG00000279765 | *AC013394.1* | 2.456986183 | 0.089788761 | 0.193002237 | -0.103213476 | 0.999933887 | 0.571568988 | 0.718308437 | -0.14673945 | 0.623200711 |
| ENSG00000136807 | *CDK9* | 5.59000903 | 0.0166879 | 0.024541251 | -0.007853352 | 0.999933887 | -0.223558929 | -0.287500441 | 0.063941513 | 0.623200711 |
| ENSG00000271064 | *AC027644.2* | 0.388197515 | 0.262675532 | 0.449360488 | -0.186684955 | 0.999933887 | 1.307069885 | 0.950145308 | 0.356924577 | 0.623227113 |
| ENSG00000204622 | *HLA-J* | 0.827673424 | 0.193814721 | 0.132119442 | 0.061695279 | 0.999933887 | 0.738851843 | 0.572312591 | 0.166539252 | 0.623227113 |
| ENSG00000142686 | *C1orf216* | 1.526137853 | -0.14629039 | 0.139647567 | -0.285937958 | 0.999933887 | 0.094674747 | -0.018790068 | 0.113464815 | 0.623227113 |
| ENSG00000102445 | *RUBCNL* | 6.325069821 | 0.009600923 | 0.021678166 | -0.012077243 | 0.999933887 | 0.000831193 | 0.068614854 | -0.067783661 | 0.623227113 |
| ENSG00000088812 | *ATRN* | 2.254954189 | 0.02662404 | 0.006437731 | 0.020186308 | 0.999933887 | -0.101031665 | -0.198996948 | 0.097965283 | 0.623334108 |
| ENSG00000265908 | *AC024267.5* | -0.480383897 | 0.027127699 | 0.019924553 | 0.007203146 | 0.999933887 | -0.416682985 | -0.071876225 | -0.344806759 | 0.623334108 |
| ENSG00000132952 | *USPL1* | 3.782997909 | -0.015640701 | -0.079241278 | 0.063600577 | 0.999933887 | -0.193063733 | -0.333057575 | 0.139993842 | 0.623481954 |
| ENSG00000131944 | *FAAP24* | 0.248768395 | -0.146287035 | -0.202209296 | 0.05592226 | 0.999933887 | -0.519645723 | -0.749032122 | 0.229386399 | 0.623580882 |
| ENSG00000185347 | *TEDC1* | 3.01159499 | 0.174745207 | 0.113310623 | 0.061434584 | 0.999933887 | 0.144706009 | 0.003532237 | 0.141173772 | 0.623585781 |
| ENSG00000073910 | *FRY* | 6.87726768 | -0.025259781 | -0.029273127 | 0.004013346 | 0.999933887 | -0.159146944 | -0.090981366 | -0.068165577 | 0.623585781 |
| ENSG00000081019 | *RSBN1* | 4.135912506 | 0.027138116 | -0.007532426 | 0.034670541 | 0.999933887 | -0.370356224 | -0.464416335 | 0.094060111 | 0.623585781 |
| ENSG00000196305 | *IARS1* | 4.074470685 | -0.023507553 | 0.053088348 | -0.076595902 | 0.999933887 | 0.055878576 | 0.127084946 | -0.07120637 | 0.623585781 |
| ENSG00000139631 | *CSAD* | 5.556779 | -0.013532768 | 0.01285566 | -0.026388429 | 0.999933887 | 0.092894112 | 0.011565449 | 0.081328663 | 0.623585781 |
| ENSG00000254415 | *SIGLEC14* | 6.835167331 | -0.002790439 | -0.054143564 | 0.051353125 | 0.999933887 | 0.824028418 | 0.904115692 | -0.080087273 | 0.623585781 |
| ENSG00000040531 | *CTNS* | 4.398513507 | 0.003442677 | -0.03165801 | 0.035100687 | 0.999933887 | -0.283008775 | -0.206314068 | -0.076694708 | 0.623757916 |
| ENSG00000135048 | *CEMIP2* | 6.456552264 | -0.037127726 | -0.020287396 | -0.01684033 | 0.999933887 | 0.001386849 | 0.064628635 | -0.063241786 | 0.623778511 |
| ENSG00000145819 | *ARHGAP26* | 8.683140974 | 0.029448843 | 0.026116522 | 0.003332321 | 0.999933887 | 0.107420269 | 0.063327223 | 0.044093046 | 0.624103708 |
| ENSG00000111790 | *FGFR1OP2* | 4.57380776 | -0.148002203 | -0.119400887 | -0.028601316 | 0.999933887 | -0.646587549 | -0.552704446 | -0.093883103 | 0.624161407 |
| ENSG00000167965 | *MLST8* | 1.954458385 | -0.113971398 | -0.014122771 | -0.099848627 | 0.999933887 | 0.157326872 | 0.050102214 | 0.107224658 | 0.62419816 |
| ENSG00000165606 | *DRGX* | -1.197269249 | 0.044574941 | -0.084913105 | 0.129488046 | 0.999933887 | -1.557974187 | -1.30368554 | -0.254288648 | 0.62419816 |
| ENSG00000050438 | *SLC4A8* | 1.255952298 | -0.011346266 | 0.132955681 | -0.144301946 | 0.999933887 | -0.214952332 | -0.059071566 | -0.155880766 | 0.62419816 |
| ENSG00000085382 | *HACE1* | 0.999322743 | -0.010637921 | -0.155429684 | 0.144791763 | 0.999933887 | 0.022089679 | 0.191977194 | -0.169887515 | 0.62419816 |
| ENSG00000103365 | *GGA2* | 5.202690512 | -0.040280165 | 0.032357174 | -0.072637339 | 0.999933887 | 0.001791155 | -0.058672482 | 0.060463637 | 0.624451984 |
| ENSG00000257838 | *OTOAP1* | -0.523039596 | -0.023383689 | 0.115348831 | -0.13873252 | 0.999933887 | -0.320010884 | -0.501439831 | 0.181428947 | 0.624487182 |
| ENSG00000232680 | *AC002511.1* | 1.18351925 | 0.053397945 | 0.058899456 | -0.00550151 | 0.999933887 | 2.268135246 | 2.511992349 | -0.243857103 | 0.624553346 |
| ENSG00000256069 | *A2MP1* | -0.117368972 | 0.008539057 | 0.00816831 | 0.000370747 | 0.999933887 | -0.363548366 | -0.122694582 | -0.240853784 | 0.624601477 |
| ENSG00000134046 | *MBD2* | 5.954092484 | 0.016835903 | -0.006062448 | 0.022898351 | 0.999933887 | -0.055065333 | 0.014815922 | -0.069881255 | 0.624625491 |
| ENSG00000070761 | *CFAP20* | 3.599026468 | 0.080429556 | -0.005100501 | 0.085530057 | 0.999933887 | 0.309863391 | 0.411233232 | -0.101369841 | 0.624679278 |
| ENSG00000162415 | *ZSWIM5* | -0.523908526 | 0.185321392 | 0.212609138 | -0.027287747 | 0.999933887 | -0.117733086 | 0.118901416 | -0.236634502 | 0.624679278 |

| | | | | | | | | | | |
|---|---|---|---|---|---|---|---|---|---|---|
| ENSG00000254093 | *PINX1* | 0.244112806 | 0.086229467 | 0.116181564 | -0.029952097 | 0.999933887 | 0.278841051 | 0.500420358 | -0.221579308 | 0.624679278 |
| ENSG00000225921 | *NOL7* | 3.636033594 | 0.034164002 | 0.022286682 | 0.011877321 | 0.999933887 | 0.023297715 | 0.103455107 | -0.080157392 | 0.624679278 |
| ENSG00000164674 | *SYTL3* | 4.44502437 | 0.003932112 | 0.016876984 | -0.012944872 | 0.999933887 | 0.096485252 | 0.013185326 | 0.083299926 | 0.624679278 |
| ENSG00000251634 | *NAIPP4* | -0.769956065 | 0.22765578 | -0.215140662 | 0.442796442 | 0.999933887 | -0.744377015 | -1.111975634 | 0.367598619 | 0.624792121 |
| ENSG00000170471 | *RALGAPB* | 6.082829374 | -0.018117825 | -0.051889787 | 0.033771962 | 0.999933887 | -0.345518091 | -0.280283075 | -0.065235016 | 0.62487509 |
| ENSG00000135974 | *C2orf49* | 3.808798658 | -0.020165022 | -0.07894094 | 0.058775918 | 0.999933887 | 0.005331208 | 0.086686546 | -0.081355338 | 0.62487509 |
| ENSG00000285667 | *AC012291.3* | 0.307941624 | -0.201633012 | -0.390620301 | 0.188987289 | 0.999933887 | -0.085277425 | -0.288419932 | 0.203142506 | 0.625127847 |
| ENSG00000170074 | *FAM153A* | -0.84538944 | 0.225908293 | -0.137943462 | 0.363851756 | 0.999933887 | -0.004677159 | 0.254574666 | -0.259251824 | 0.625127847 |
| ENSG00000115523 | *GNLY* | 7.000830551 | -0.01096925 | -0.023236519 | 0.012267269 | 0.999933887 | 0.109664397 | 0.166106521 | -0.056442124 | 0.625127847 |
| ENSG00000228107 | *AP000692.1* | -0.301209231 | -0.156203156 | 0.219026812 | -0.375229968 | 0.999933887 | -0.509444022 | -0.321921177 | -0.187522845 | 0.625133831 |
| ENSG00000100271 | *TTLL1* | -0.71433276 | 0.264229744 | -0.108143202 | 0.372372946 | 0.999933887 | -0.062803925 | -0.351902035 | 0.289098109 | 0.625218744 |
| ENSG00000197061 | *H4C3* | -0.873871833 | -0.229947531 | -0.391118796 | 0.161171265 | 0.999933887 | -0.09152049 | -0.586155316 | 0.494634826 | 0.625218744 |
| ENSG00000250130 | *GAPDHP43* | -0.05480078 | 0.161304801 | -0.157671191 | 0.318975992 | 0.999933887 | -0.313656265 | -0.572272012 | 0.258615747 | 0.625218744 |
| ENSG00000261455 | *LINC01003* | 4.459576469 | 0.068395372 | 0.023195835 | 0.045199537 | 0.999933887 | 0.26888972 | 0.368022074 | -0.099132354 | 0.625218744 |
| ENSG00000159377 | *PSMB4* | 6.95718056 | -0.033295839 | -0.023356849 | -0.00993899 | 0.999933887 | 0.189199974 | 0.243341835 | -0.05414186 | 0.625218744 |
| ENSG00000258352 | *IFITM3P6* | 0.10539492 | 0.106456643 | -0.161428589 | 0.267885232 | 0.999933887 | -0.201408205 | -0.005111817 | -0.196296388 | 0.625218744 |
| ENSG00000115286 | *NDUFS7* | 1.517468544 | -0.065849358 | 0.023317964 | -0.089167322 | 0.999933887 | -0.166125441 | -0.293734002 | 0.127608561 | 0.625218744 |
| ENSG00000118762 | *PKD2* | 1.634321111 | 0.062806775 | -0.008697983 | 0.071504759 | 0.999933887 | -0.111468596 | -0.265182403 | 0.153713808 | 0.625218744 |
| ENSG00000100632 | *ERH* | 3.225298974 | -0.109441482 | 0.075363027 | -0.184804509 | 0.999933887 | -0.090978534 | -0.181672697 | 0.090694163 | 0.625461441 |
| ENSG00000237489 | *C10orf143* | -0.054237951 | -0.204960412 | -0.140346017 | -0.064614395 | 0.999933887 | -0.226307999 | -0.027910535 | -0.198397464 | 0.625461441 |
| ENSG00000132128 | *LRRC41* | 3.381920118 | 0.082016882 | 0.005763992 | 0.076252891 | 0.999933887 | 0.026757079 | 0.098642252 | -0.071885173 | 0.62555875 |
| ENSG00000141012 | *GALNS* | 5.670330832 | 0.058639513 | -0.014302133 | 0.072941646 | 0.999933887 | 0.146745841 | 0.22216377 | -0.075417929 | 0.62585404 |
| ENSG00000254649 | *AP003086.2* | 0.099894104 | 0.223384259 | 0.228472959 | -0.0050887 | 0.999933887 | -0.198341101 | 0.1130256 | -0.311366701 | 0.62585404 |
| ENSG00000207870 | *MIR221* | -0.372999897 | 0.12962986 | 0.029841894 | 0.099787966 | 0.999933887 | 1.275880502 | 1.56663957 | -0.290759069 | 0.62585404 |
| ENSG00000167747 | *C19orf48* | 2.000733277 | 0.055062651 | -0.105908187 | 0.160970838 | 0.999933887 | 0.367083623 | 0.245465537 | 0.121618085 | 0.62585404 |
| ENSG00000137628 | *DDX60* | 4.053993494 | -0.055175837 | -0.102974196 | 0.04779836 | 0.999933887 | -0.443104812 | -0.286753124 | -0.156351688 | 0.62585404 |
| ENSG00000187134 | *AKR1C1* | 1.336608669 | 0.009535477 | 0.027611277 | -0.0180758 | 0.999933887 | 0.042163894 | 0.173679988 | -0.131516094 | 0.62585404 |
| ENSG00000283384 | *AL138694.1* | -0.350915837 | 0.002700385 | -0.262895841 | 0.265596225 | 0.999933887 | 1.01905344 | 1.286269924 | -0.267216484 | 0.62585404 |
| ENSG00000088899 | *LZTS3* | 1.566234231 | 0.089823329 | 0.170342996 | -0.080519668 | 0.999933887 | 0.207855976 | 0.349783284 | -0.141927308 | 0.625889246 |
| ENSG00000121933 | *TMIGD3* | 0.19044422 | 0.042910511 | -0.017277137 | 0.060187648 | 0.999933887 | -0.038910748 | -0.22131412 | 0.182403372 | 0.625889246 |
| ENSG00000166428 | *PLD4* | 0.472981478 | -0.040980434 | 0.061268435 | -0.102248869 | 0.999933887 | -0.099125565 | -0.323354867 | 0.224229302 | 0.626128171 |
| ENSG00000240857 | *RDH14* | 0.511077512 | -0.024011085 | -0.006308393 | -0.017702692 | 0.999933887 | 0.185677355 | 0.359956661 | -0.174279307 | 0.626210684 |
| ENSG00000214293 | *APTR* | 2.160782675 | 0.079694995 | -0.018456425 | 0.098151421 | 0.999933887 | -0.318488902 | -0.425888004 | 0.107399102 | 0.62621578 |
| ENSG00000174943 | *KCTD13* | 4.321269609 | 0.041962491 | 0.066753434 | -0.024790943 | 0.999933887 | 0.150247663 | 0.211326099 | -0.061078436 | 0.626424129 |
| ENSG00000144677 | *CTDSPL* | 1.321946733 | 0.24044567 | 0.17059324 | 0.06985243 | 0.999933887 | 3.265901687 | 3.047311935 | 0.218589752 | 0.62673988 |
| ENSG00000247240 | *UBL7-AS1* | 0.480078454 | -0.125014123 | -0.208339958 | 0.083325835 | 0.999933887 | -0.856329998 | -1.03353725 | 0.177207253 | 0.626835095 |
| ENSG00000211897 | *IGHG3* | 0.585752979 | 0.136377739 | -0.009107976 | 0.145485715 | 0.999933887 | -0.108278485 | 0.200177044 | -0.308455529 | 0.627162927 |
| ENSG00000173548 | *SNX33* | 0.245327285 | 0.004076702 | -0.057098941 | 0.061175643 | 0.999933887 | -0.121269364 | 0.086600588 | -0.207869952 | 0.62783298 |
| ENSG00000186638 | *KIF24* | 0.35970916 | 0.203892019 | 0.299057457 | -0.095165439 | 0.999933887 | 0.206751098 | 0.430098052 | -0.223346954 | 0.628022499 |
| ENSG00000275202 | *AL161421.1* | 4.20262322 | 0.079639051 | 0.085860407 | -0.006221356 | 0.999933887 | 0.083781482 | 0.179364463 | -0.095582981 | 0.628022499 |
| ENSG00000185485 | *SDHAP1* | 2.740471897 | 0.064078938 | -0.008415888 | 0.072494826 | 0.999933887 | -0.068941601 | -0.17033869 | 0.101397089 | 0.628022499 |
| ENSG00000078124 | *ACER3* | 2.715109286 | -0.048833121 | -0.104166488 | 0.055333368 | 0.999933887 | -0.275775527 | -0.169444379 | -0.106331147 | 0.628022499 |
| ENSG00000145996 | *CDKAL1* | 1.45689748 | -0.042331015 | 0.157415689 | -0.199746704 | 0.999933887 | 0.024528283 | 0.159339499 | -0.134811216 | 0.628022499 |
| ENSG00000108370 | *RGS9* | 0.299790565 | 0.018324579 | -0.174486498 | 0.192811077 | 0.999933887 | 0.143179964 | 0.333286672 | -0.190106708 | 0.628022499 |
| ENSG00000173041 | *ZNF680* | -0.349070212 | -0.002565424 | -0.032974502 | 0.030409078 | 0.999933887 | -0.338978391 | -0.05766156 | -0.281316831 | 0.628022499 |
| ENSG00000169635 | *HIC2* | 3.313754783 | 0.071919632 | 0.031205495 | 0.040714137 | 0.999933887 | -0.128798381 | -0.221569541 | 0.09277116 | 0.628094334 |
| ENSG00000100804 | *PSMB5* | 1.8188127 | -0.030929294 | -0.233620307 | 0.202691013 | 0.999933887 | -0.204693835 | -0.30995892 | 0.105265085 | 0.628275522 |
| ENSG00000174791 | *RIN1* | -0.42152293 | 0.195085461 | 0.038218641 | 0.15686682 | 0.999933887 | -0.160718207 | -0.385649095 | 0.224930889 | 0.628305183 |
| ENSG00000254420 | *AP003086.1* | 0.280297118 | -0.070781361 | 0.267463375 | -0.338244736 | 0.999933887 | 0.172880289 | 0.335757314 | -0.162877025 | 0.628305183 |
| ENSG00000145723 | *GIN1* | 0.466842812 | -0.167062418 | 0.06962161 | -0.236684028 | 0.999933887 | -0.413110299 | -0.280354605 | -0.132755694 | 0.628846945 |
| ENSG00000280433 | *FP565260.6* | 0.735900757 | -0.076694976 | -0.056136264 | -0.020558713 | 0.999933887 | -0.271319366 | -0.461622331 | 0.190302964 | 0.628846945 |
| ENSG00000107185 | *RGP1* | 4.489295981 | 0.048755425 | 0.00383771 | 0.044917715 | 0.999933887 | -0.007802838 | -0.083522705 | 0.075719867 | 0.628960316 |
| ENSG00000155906 | *RMND1* | 1.307873593 | -0.205711025 | 0.168303742 | -0.374014767 | 0.999933887 | 0.05385387 | -0.073281383 | 0.127135253 | 0.628971195 |
| ENSG00000172116 | *CD8B* | 4.133070862 | 0.056049739 | -0.094809036 | 0.150858775 | 0.999933887 | 0.08204735 | 0.170612884 | -0.088565535 | 0.628971195 |
| ENSG00000204524 | *ZNF805* | 2.526047118 | 0.027213692 | 0.039622944 | -0.012409252 | 0.999933887 | -0.17924452 | -0.042823281 | -0.136421239 | 0.629018829 |
| ENSG00000257446 | *ZNF878* | -1.570383944 | -0.528228967 | -0.341504146 | -0.18672482 | 0.999933887 | -0.849525518 | -0.553909391 | -0.295616127 | 0.629316506 |
| ENSG00000101413 | *RPRD1B* | 4.808700639 | -0.053640959 | -0.026097832 | -0.027543127 | 0.999933887 | -0.031986015 | 0.020291157 | -0.052277172 | 0.629867844 |
| ENSG00000181789 | *COPG1* | 5.707317276 | -0.030729803 | 0.050113816 | -0.080843619 | 0.999933887 | -0.345962745 | -0.402929943 | 0.056967198 | 0.629946049 |
| ENSG00000181982 | *CCDC149* | -0.719225708 | -0.033058342 | 0.118912836 | -0.151971178 | 0.999933887 | -0.753688864 | -1.085603223 | 0.331914358 | 0.629972333 |
| ENSG00000174106 | *LEMD3* | 5.391888147 | 0.013084278 | 0.030158717 | -0.017074438 | 0.999933887 | 0.059219677 | 0.114300996 | -0.055081319 | 0.630027572 |
| ENSG00000041357 | *PSMA4* | 5.454941224 | -0.052872936 | -0.119696108 | 0.066823172 | 0.999933887 | -0.026808771 | 0.032262506 | -0.059071277 | 0.630084121 |
| ENSG00000047410 | *TPR* | 6.541844446 | -0.107495384 | -0.185526966 | 0.078031582 | 0.999933887 | -0.792676288 | -0.574577677 | -0.218098611 | 0.630084121 |
| ENSG00000101417 | *PXMP4* | 0.259769987 | 0.068590008 | -0.048926408 | 0.117516416 | 0.999933887 | -0.107463801 | 0.031344382 | -0.138808183 | 0.630084121 |
| ENSG00000181220 | *ZNF746* | 7.6497113 | 0.015844491 | 0.097011294 | -0.081166803 | 0.999933887 | -0.326082588 | -0.247512569 | -0.078570019 | 0.630084121 |
| ENSG00000259250 | *AC018904.1* | -0.091056412 | -0.06919652 | -0.332869275 | 0.263672754 | 0.999933887 | -0.778848959 | -0.985025624 | 0.206176665 | 0.630114397 |
| ENSG00000244265 | *SIAH2-AS1* | -0.717018133 | -0.058876176 | 0.287872777 | -0.346748952 | 0.999933887 | 0.118987261 | 0.471113476 | -0.352126215 | 0.630114397 |
| ENSG00000224914 | *LINC00863* | 1.143845912 | 0.006329709 | 0.10072902 | -0.094399311 | 0.999933887 | 0.019845887 | 0.165380673 | -0.145534785 | 0.630114397 |
| ENSG00000267575 | *AC006504.5* | -0.588703511 | 0.051739053 | 0.372799725 | -0.321060672 | 0.999933887 | -0.51005443 | -0.247478445 | -0.262575985 | 0.630353551 |
| ENSG00000135976 | *ANKRD36* | 2.413820801 | -0.048799652 | 0.086369096 | -0.135168748 | 0.999933887 | 0.006240559 | 0.165901352 | -0.159660793 | 0.630359951 |
| ENSG00000174428 | *GTF2IRD2B* | 1.425559009 | 0.070911441 | -0.028866564 | 0.099778005 | 0.999933887 | 0.128722275 | 0.29104045 | -0.162318175 | 0.630407165 |
| ENSG00000142178 | *SIK1* | 0.78520919 | -0.014695882 | 0.028503808 | -0.04319969 | 0.999933887 | 0.174415092 | 0.023031076 | 0.151384016 | 0.630407165 |
| ENSG00000279649 | *AC020978.9* | -0.164270327 | 0.007159455 | 0.05991023 | -0.052750776 | 0.999933887 | -0.151771413 | 0.104326819 | -0.256098232 | 0.630407165 |
| ENSG00000211896 | *IGHG1* | 2.31450646 | 0.156510056 | 0.043152597 | 0.113357459 | 0.999933887 | 0.079681686 | -0.124386109 | 0.204067795 | 0.630444741 |
| ENSG00000170445 | *HARS1* | 4.024985104 | -0.036063443 | 0.005465905 | -0.041529348 | 0.999933887 | -0.00417181 | 0.058842644 | -0.063014454 | 0.630444741 |
| ENSG00000157388 | *CACNA1D* | 0.79635405 | -0.070759551 | 0.035258839 | -0.10601839 | 0.999933887 | -0.488187812 | -0.693517262 | 0.20532945 | 0.630598734 |
| ENSG00000134758 | *RNF138* | 4.889192866 | 0.043303748 | 0.034706113 | 0.008597635 | 0.999933887 | -0.41237183 | -0.532311498 | 0.119939668 | 0.630598734 |
| ENSG00000198783 | *ZNF830* | 4.13908109 | -0.009245339 | -0.044875037 | 0.035629698 | 0.999933887 | 0.188617647 | 0.263501325 | -0.074883679 | 0.630598734 |
| ENSG00000243176 | *AC092944.1* | 1.467952891 | 0.257874677 | 0.129935071 | 0.127939606 | 0.999933887 | 0.574776669 | 0.395637702 | 0.179138966 | 0.630667506 |
| ENSG00000163933 | *RFT1* | 4.253018845 | 0.043766763 | -0.068072597 | 0.111839359 | 0.999933887 | -0.06447682 | 0.027014942 | -0.091491761 | 0.630667506 |
| ENSG00000132801 | *ZSWIM3* | 0.633568711 | -0.099001513 | -0.3141587 | 0.215157187 | 0.999933887 | -0.098381988 | -0.235458733 | 0.137076746 | 0.630932034 |
| ENSG00000178562 | *CD28* | 4.529557072 | -0.029652808 | -0.058531615 | 0.028878807 | 0.999933887 | 0.061377649 | 0.125537577 | -0.064159928 | 0.63100311 |
| ENSG00000025772 | *TOMM34* | 2.458344273 | -0.047186548 | -0.069415621 | 0.022229074 | 0.999933887 | -0.013353343 | -0.116179053 | 0.10282571 | 0.631166528 |
| ENSG00000033011 | *ALG1* | 0.588835539 | 0.052233132 | 0.191433143 | -0.139200011 | 0.999933887 | 0.149089437 | -0.024802473 | 0.17389191 | 0.631333844 |
| ENSG00000124406 | *ATP8A1* | 4.460038181 | -0.031864611 | -0.094296991 | 0.06243238 | 0.999933887 | -0.300636311 | -0.119989182 | -0.180647129 | 0.631333844 |
| ENSG00000099246 | *RAB18* | 5.612519879 | -0.029364118 | 0.004771848 | -0.034135966 | 0.999933887 | -0.406891293 | -0.339987588 | -0.066903706 | 0.631456537 |
| ENSG00000138050 | *THUMPD2* | 0.762615832 | 0.035908244 | -0.072420123 | 0.108328367 | 0.999933887 | 0.07003228 | 0.234147834 | -0.164115555 | 0.631469237 |
| ENSG00000155666 | *KDM8* | 1.112727253 | -0.020616132 | 0.144476066 | -0.165092198 | 0.999933887 | 0.003358497 | 0.136943825 | -0.133585328 | 0.63162154 |
| ENSG00000082516 | *GEMIN5* | 1.390136919 | -0.074062611 | 0.016176288 | -0.090238899 | 0.999933887 | 0.184321702 | 0.067531894 | 0.116789808 | 0.63191358 |
| ENSG00000090554 | *FLT3LG* | -0.20019809 | 0.006627671 | 0.185478421 | -0.178850749 | 0.999933887 | -0.114524089 | 0.144370869 | -0.258894958 | 0.632123954 |
| ENSG00000187688 | *TRPV2* | 4.203893423 | 0.06309331 | 0.050349179 | 0.012744131 | 0.999933887 | -0.066493832 | -0.140695148 | 0.074201317 | 0.632204945 |
| ENSG00000108349 | *CASC3* | 7.808054711 | -0.016781476 | -0.021493496 | 0.004712021 | 0.999933887 | -0.38066113 | -0.43232735 | 0.05166622 | 0.632477705 |
| ENSG00000196262 | *PPIA* | 6.199775447 | -0.027502134 | -0.000296019 | -0.027206115 | 0.999933887 | 0.123901662 | 0.187341041 | -0.063439379 | 0.63261917 |
| ENSG00000104228 | *TRIM35* | 2.30094727 | 0.020361255 | 0.0746593 | -0.054298045 | 0.999933887 | 0.008496058 | 0.114768131 | -0.106272073 | 0.63261917 |
| ENSG00000100316 | *RPL3* | 8.899835056 | 0.010551243 | -0.033198173 | 0.043749416 | 0.999933887 | 0.105004148 | 0.172722466 | -0.067718319 | 0.63261917 |
| ENSG00000102390 | *PBDC1* | 2.392942512 | -0.04370399 | 0.107196346 | -0.150900335 | 0.999933887 | -0.148414256 | -0.259651153 | 0.111236897 | 0.632799056 |
| ENSG00000196110 | *ZNF699* | 2.642602923 | -0.038828365 | -0.00057273 | -0.038255635 | 0.999933887 | -0.253090411 | -0.360904893 | 0.107814483 | 0.632799056 |
| ENSG00000139998 | *RAB15* | 2.171524887 | -0.01728614 | -0.023737408 | 0.006451268 | 0.999933887 | -0.143151357 | -0.269508962 | 0.126357605 | 0.633162104 |
| ENSG00000087191 | *PSMC5* | 4.159432794 | 0.030670102 | -0.016779791 | 0.047449893 | 0.999933887 | -0.010360081 | -0.073128706 | 0.062768624 | 0.633433902 |
| ENSG00000101752 | *MIB1* | 4.361434262 | -0.002061726 | -0.070073572 | 0.068011846 | 0.999933887 | -0.372725651 | -0.213374899 | -0.159350752 | 0.633433902 |
| ENSG00000113407 | *TARS1* | 3.611105242 | -0.104021671 | -0.187444358 | 0.083422687 | 0.999933887 | 0.089265892 | 0.168669453 | -0.079403561 | 0.634012437 |
| ENSG00000117724 | *CENPF* | -0.644916015 | -0.200494181 | -0.154426538 | -0.046067642 | 0.999933887 | -0.136817752 | 0.204183699 | -0.341001452 | 0.634012437 |
| ENSG00000135900 | *MRPL44* | 3.480013647 | -0.100304199 | -0.052148273 | -0.048155926 | 0.999933887 | 0.342326851 | 0.425056179 | -0.082729328 | 0.634012437 |
| ENSG00000274512 | *TBC1D3L* | -3.156599491 | -0.204774601 | 0.431000598 | -0.635775199 | 0.594252256 | 0.104773702 | -0.104141729 | 0.208915432 | 0.634012437 |
| ENSG00000167210 | *LOXHD1* | 3.340457676 | -0.033053635 | -0.045912234 | 0.012858599 | 0.999933887 | 0.691261891 | 0.550436258 | 0.140825633 | 0.634012437 |
| ENSG00000270339 | *PPIAL4H* | -2.003125698 | -0.09133892 | 0.059035767 | -0.150374687 | 0.999933887 | 2.014730748 | 2.236647307 | -0.221916559 | 0.634012437 |

| | | | | | | | | | | |
|---|---|---|---|---|---|---|---|---|---|---|
| ENSG00000186152 | *LILRP1* | -0.277266245 | 0.121736744 | 0.091932231 | 0.029804512 | 0.999933887 | -0.428361804 | -0.224113432 | -0.204248373 | 0.634225201 |
| ENSG00000228323 | *MYADM-AS1* | 0.693037145 | 0.212542914 | 0.276165218 | -0.063622303 | 0.999933887 | 0.23312078 | -0.141405824 | 0.374526605 | 0.634254489 |
| ENSG00000153767 | *GTF2E1* | 2.119413262 | -0.096962345 | -0.068473961 | -0.028488384 | 0.999933887 | 0.074002219 | -0.010484761 | 0.084486979 | 0.634254489 |
| ENSG00000072682 | *P4HA2* | 1.079403894 | -0.070477711 | 0.025518751 | -0.095996462 | 0.999933887 | -0.396824567 | -0.211199229 | -0.185625338 | 0.634254489 |
| ENSG00000134899 | *ERCC5* | 2.269235673 | 0.030603685 | 0.005738422 | 0.024865263 | 0.999933887 | -0.177207121 | -0.310776306 | 0.133569185 | 0.634308891 |
| ENSG00000198034 | *RPS4X* | 7.86380355 | -0.011910117 | -0.06253234 | 0.050622223 | 0.999933887 | 0.097800817 | 0.170666616 | -0.072865799 | 0.634521853 |
| ENSG00000208028 | *MIR616* | 2.61528005 | 0.222939107 | 0.428808803 | -0.205869697 | 0.999933887 | 0.440902078 | 0.290871919 | 0.150030159 | 0.634619576 |
| ENSG00000157110 | *RBPMS* | 1.185070264 | 0.058029088 | 0.148711312 | -0.090682224 | 0.999933887 | -0.231138766 | -0.407464628 | 0.176325862 | 0.634619576 |
| ENSG00000179604 | *CDC42EP4* | 1.934469637 | -0.023896768 | 0.011043868 | -0.034940636 | 0.999933887 | -0.528083105 | -0.676179531 | 0.148096426 | 0.634619576 |
| ENSG00000167380 | *ZNF226* | 2.062324331 | -0.158159715 | -0.173765993 | 0.015606278 | 0.999933887 | 0.035133731 | 0.130775339 | -0.095641608 | 0.634630933 |
| ENSG00000183813 | *CCR4* | 2.851041946 | -0.011820778 | -0.02878758 | 0.016966802 | 0.999933887 | 0.188987398 | 0.086786542 | 0.102200857 | 0.634630933 |
| ENSG00000114631 | *PODXL2* | 2.009156021 | 0.184333524 | 0.292261445 | -0.107927921 | 0.999933887 | 0.129916713 | 0.000254403 | 0.12966231 | 0.634850136 |
| ENSG00000140326 | *CDAN1* | 2.50549635 | 0.097682081 | -0.095913318 | 0.193595399 | 0.999933887 | 0.080265593 | -0.036490255 | 0.116755848 | 0.634850136 |
| ENSG00000154099 | *DNAAF1* | -0.228978354 | 0.27379264 | 0.127144607 | 0.146648033 | 0.999933887 | 3.920150791 | 4.346999265 | -0.426848474 | 0.635036997 |
| ENSG00000100246 | *DNAL4* | 2.024966358 | 0.042651069 | 0.009858257 | 0.032792812 | 0.999933887 | -0.14732913 | -0.011382225 | -0.135946905 | 0.635036997 |
| ENSG00000100380 | *ST13* | 5.314739347 | -0.040655348 | -0.059082599 | 0.018427251 | 0.999933887 | -0.216427728 | -0.279654688 | 0.063226961 | 0.635079479 |
| ENSG00000112759 | *SLC29A1* | 3.450260893 | 0.038295401 | 0.03385186 | 0.004443541 | 0.999933887 | 0.058338339 | 0.160446334 | -0.102107995 | 0.635079479 |
| ENSG00000100836 | *PABPN1* | 5.90266472 | 0.013573273 | 0.02038298 | -0.006809707 | 0.999933887 | -0.128247578 | -0.179393495 | 0.051145916 | 0.63513379 |
| ENSG00000172215 | *CXCR6* | 0.904669468 | 0.1304549 | 0.087252224 | 0.043202676 | 0.999933887 | 0.078724804 | -0.074868378 | 0.153593182 | 0.635480584 |
| ENSG00000285420 | *PDGFRL2P* | 0.848424857 | -0.011516956 | -0.001712106 | -0.00980485 | 0.999933887 | -0.110840384 | -0.272872512 | 0.162032128 | 0.63556356 |
| ENSG00000228506 | *AL513550.1* | -0.081255079 | -0.225687747 | 0.017379687 | -0.243067434 | 0.999933887 | 0.175034604 | 0.368279058 | -0.193244454 | 0.635774369 |
| ENSG00000133422 | *MORC2* | 3.028685498 | -0.068177301 | 0.018867117 | -0.087044418 | 0.999933887 | -0.190504243 | -0.270605539 | 0.080101297 | 0.635774369 |
| ENSG00000198242 | *RPL23A* | 5.282564223 | -0.033307307 | 8.8949e-05 | -0.033396256 | 0.999933887 | 0.127359795 | 0.201936569 | -0.074576774 | 0.635774369 |
| ENSG00000186522 | *SEPTIN10* | -1.151296513 | -0.124559408 | -0.429003253 | 0.304443844 | 0.999933887 | -0.406008035 | -0.214997472 | -0.191010563 | 0.635774369 |
| ENSG00000155868 | *MED7* | 2.003682263 | 0.008200846 | -0.263580088 | 0.271780934 | 0.944094537 | -0.167766342 | -0.07284293 | -0.094923412 | 0.635774369 |
| ENSG00000117226 | *GBP3* | 3.278096741 | -0.00813801 | -0.086186361 | 0.078048351 | 0.999933887 | 0.432936565 | 0.575107345 | -0.14217078 | 0.635774369 |
| ENSG00000185909 | *KLHDC8B* | 6.618514717 | -0.234654788 | -0.199484995 | -0.035169793 | 0.999933887 | -1.438028013 | -1.572921056 | 0.134893043 | 0.635905772 |
| ENSG00000235245 | *AL360181.2* | 2.318700586 | 0.024286479 | -0.016833092 | 0.041119572 | 0.999933887 | -0.127195603 | -0.027335528 | -0.099860075 | 0.635913613 |
| ENSG00000161057 | *PSMC2* | 3.643454569 | -0.061042633 | -0.115903793 | 0.054861159 | 0.999933887 | -0.050799533 | 0.020066273 | -0.070865806 | 0.63615745 |
| ENSG00000151093 | *OXSM* | 1.918229681 | -0.028198971 | -0.071546736 | 0.043347765 | 0.999933887 | 0.25550765 | 0.387277123 | -0.131769472 | 0.63615745 |
| ENSG00000120833 | *SOCS2* | 2.163311102 | -0.14842332 | -0.031181725 | -0.117241595 | 0.999933887 | 0.12905841 | 0.266615925 | -0.137557514 | 0.636466643 |
| ENSG00000166801 | *FAM111A* | 4.394020077 | 0.011653214 | 0.028526115 | -0.0168729 | 0.999933887 | -0.215731346 | -0.311468805 | 0.095737459 | 0.636466643 |
| ENSG00000270426 | *AC099343.2* | 2.585533511 | -0.165718939 | -0.194229131 | 0.028510192 | 0.999933887 | 0.466685598 | 0.380443651 | 0.086241947 | 0.636730254 |
| ENSG00000165626 | *BEND7* | 0.038770691 | 0.118545092 | 0.059940271 | 0.058604821 | 0.999933887 | -0.736574984 | -0.943149788 | 0.206574804 | 0.63676072 |
| ENSG00000162066 | *AMDHD2* | 7.849459188 | 0.078700409 | 0.109839735 | -0.031139327 | 0.999933887 | -0.011614203 | -0.108828486 | 0.097214283 | 0.63676072 |
| ENSG00000234292 | *AC123595.1* | -0.320466727 | 0.183768543 | 0.161817048 | 0.021951496 | 0.999933887 | 1.244113857 | 1.018175044 | 0.225938814 | 0.63676072 |
| ENSG00000177981 | *ASB8* | 4.403525432 | -0.04185578 | -0.043878287 | 0.002022506 | 0.999933887 | -0.313359598 | -0.238840501 | -0.074519097 | 0.63676072 |
| ENSG00000197062 | *ZSCAN26* | 1.224444439 | 0.076482087 | 0.035131189 | 0.041350899 | 0.999933887 | -0.025208428 | -0.171155145 | 0.145946717 | 0.63676072 |
| ENSG00000053372 | *MRTO4* | 1.504906608 | 0.058886792 | 0.036892633 | 0.021994159 | 0.999933887 | 0.302952464 | 0.418241472 | -0.115289008 | 0.63682425 |
| ENSG00000152601 | *MBNL1* | 7.481489026 | 0.01448256 | -0.093848896 | 0.108331456 | 0.999933887 | -0.345908838 | -0.201943858 | -0.143964979 | 0.63690542 |
| ENSG00000119661 | *DNAL1* | -0.106506986 | 0.020264938 | -0.025637499 | 0.045902437 | 0.999933887 | -0.154892958 | 0.058489047 | -0.213382005 | 0.63739471 |
| ENSG00000159216 | *RUNX1* | 4.988775727 | 0.01345882 | -0.00860735 | 0.02206617 | 0.999933887 | 0.019195293 | 0.083148616 | -0.063953323 | 0.637701591 |
| ENSG00000128789 | *PSMG2* | 2.680243572 | -0.083754206 | 0.055871653 | -0.139625859 | 0.999933887 | 0.038717054 | -0.057233212 | 0.095950266 | 0.637748687 |
| ENSG00000141664 | *ZCCHC2* | 6.440000034 | -0.001305693 | -0.003596921 | 0.002291228 | 0.999933887 | -0.110347148 | -0.031895391 | -0.078451757 | 0.637751768 |
| ENSG00000196365 | *LONP1* | 5.74921643 | 0.091546276 | 0.10333493 | -0.011788654 | 0.999933887 | 0.341592761 | 0.439197891 | -0.09760513 | 0.637760383 |
| ENSG00000102007 | *PLP2* | 5.47664893 | 0.016165868 | 0.01962639 | -0.003460522 | 0.999933887 | -0.173230787 | -0.255433304 | 0.082202517 | 0.63785067 |
| ENSG00000114735 | *HEMK1* | 2.777320738 | -0.017044728 | -0.028203419 | 0.011158691 | 0.999933887 | 0.116228437 | 0.030068278 | 0.086160158 | 0.637884944 |
| ENSG00000215022 | *AL008729.1* | 6.12004821 | 0.209708522 | 0.308880497 | -0.099171975 | 0.999933887 | 0.81043141 | 0.910320633 | -0.099889223 | 0.637904217 |
| ENSG00000152147 | *GEMIN6* | -0.563293557 | -0.161290653 | -0.090827598 | -0.070463054 | 0.999933887 | 0.01281147 | 0.199312139 | -0.186500669 | 0.637904217 |
| ENSG00000135945 | *REV1* | 3.883906388 | -0.032611745 | -0.004705173 | -0.027906571 | 0.999933887 | -0.210326669 | -0.290350864 | 0.080024195 | 0.637964565 |
| ENSG00000184389 | *A3GALT2* | -0.927696641 | -0.059060756 | 0.012411047 | -0.071471803 | 0.999933887 | 0.00557114 | -0.400544256 | 0.406115396 | 0.637964565 |
| ENSG00000272078 | *AL139423.1* | 0.999491 | 0.082854566 | 0.074780382 | 0.008074184 | 0.999933887 | 0.566770566 | 0.743230513 | -0.176459947 | 0.638147309 |
| ENSG00000167625 | *ZNF526* | 1.883074734 | -0.060470684 | -0.107880755 | 0.047410071 | 0.999933887 | -0.245327168 | -0.358971359 | 0.113644191 | 0.638147309 |
| ENSG00000141101 | *NOB1* | 2.066453521 | -0.080633866 | -0.252878022 | 0.172244156 | 0.999933887 | 0.247747657 | 0.358668927 | -0.11092127 | 0.638266167 |
| ENSG00000255529 | *POLR2M* | 1.656281041 | -0.052050621 | 0.081426066 | -0.133476687 | 0.999933887 | 0.041596344 | 0.170251114 | -0.12865477 | 0.638266167 |
| ENSG00000144118 | *RALB* | 7.41285209 | 0.004973507 | -0.013410197 | 0.018383704 | 0.999933887 | 0.205213935 | 0.27690089 | -0.071686955 | 0.638474123 |
| ENSG00000228451 | *SDAD1P1* | 2.091793707 | 0.070984424 | 0.029565518 | 0.041418906 | 0.999933887 | 0.151451157 | 0.301845035 | -0.150393878 | 0.638614136 |
| ENSG00000234263 | *MAP3K5-AS1* | -0.439633228 | -0.012352804 | -0.073241712 | 0.060888908 | 0.999933887 | 1.472300075 | 1.726864765 | -0.25456469 | 0.638875898 |
| ENSG00000094804 | *CDC6* | 0.091333367 | 0.071321773 | 0.05336166 | 0.017960113 | 0.999933887 | 0.507052298 | 0.685742718 | -0.178690419 | 0.63911707 |
| ENSG00000010327 | *STAB1* | 1.42302964 | 0.095100013 | 0.009539342 | 0.085560671 | 0.999933887 | -0.275520823 | -0.412390991 | 0.136870168 | 0.639168477 |
| ENSG00000125812 | *GZF1* | 4.946290429 | 0.077288805 | -0.003695698 | 0.080984504 | 0.999933887 | -0.521511339 | -0.593981432 | 0.072470094 | 0.639245285 |
| ENSG00000099330 | *OCEL1* | 1.489148136 | 0.054149133 | -0.013559988 | 0.06770912 | 0.999933887 | -0.013245889 | -0.178736549 | 0.16549066 | 0.639245285 |
| ENSG00000122335 | *SERAC1* | 1.82539221 | -0.039449171 | -0.102294994 | 0.062845823 | 0.999933887 | -0.259011608 | -0.153794882 | -0.105216727 | 0.639245285 |
| ENSG00000108256 | *NUFIP2* | 6.536001146 | -0.015245046 | -0.073853211 | 0.058608165 | 0.999933887 | -0.84627122 | -0.7316591 | -0.114612119 | 0.63963097 |
| ENSG00000214894 | *LINC00243* | 1.686315311 | 0.135129474 | 0.020838201 | 0.114291273 | 0.999933887 | 1.162480572 | 1.304697579 | -0.142217008 | 0.639796626 |
| ENSG00000238113 | *LINC01410* | 2.398521183 | -0.051075862 | 0.039776444 | -0.090852306 | 0.999933887 | -0.597443049 | -0.736603958 | 0.139160909 | 0.639796626 |
| ENSG00000124160 | *NCOA5* | 3.541256371 | -0.038967258 | -0.021352128 | -0.01761513 | 0.999933887 | -0.099828127 | -0.035658576 | -0.064169551 | 0.639796626 |
| ENSG00000037042 | *TUBG2* | 0.506859286 | 0.066698737 | -0.00959436 | 0.076293098 | 0.999933887 | 0.154298285 | 0.012835523 | 0.141462762 | 0.639796626 |
| ENSG00000101844 | *ATG4A* | 1.274309047 | -0.029335336 | 0.011869257 | -0.041204593 | 0.999933887 | 0.122831861 | 0.300441833 | -0.177609972 | 0.639796626 |
| ENSG00000243797 | *AC004917.1* | 1.335303667 | -0.037776256 | 0.034895793 | -0.072672049 | 0.999933887 | 1.237875243 | 1.440315753 | -0.20244051 | 0.639796626 |
| ENSG00000130299 | *GTPBP3* | 1.526992752 | 0.066801069 | -0.111094591 | 0.17789566 | 0.999933887 | 0.114654971 | 0.003921048 | 0.110733923 | 0.639815833 |
| ENSG00000235852 | *AC005540.1* | -0.688030737 | 0.091324796 | 0.240323126 | -0.14899833 | 0.999933887 | 0.353957359 | 0.049011012 | 0.304946347 | 0.639815833 |
| ENSG00000188305 | *PEAK3* | 4.409474148 | -0.030460507 | -0.093614545 | 0.063154038 | 0.999933887 | -1.183524138 | -1.275446067 | 0.091921929 | 0.639815833 |
| ENSG00000214655 | *ZSWIM8* | 7.685773501 | 0.01254444 | 0.058660134 | -0.046115693 | 0.999933887 | -0.066714564 | -0.012743281 | -0.053971283 | 0.639815833 |
| ENSG00000196074 | *SYCP2* | 1.1659993 | -0.204246394 | -0.281188709 | 0.076942316 | 0.999933887 | -0.363973761 | -0.066949643 | -0.297024118 | 0.639939242 |
| ENSG00000236104 | *ZBTB22* | 3.820188856 | 0.090693423 | 0.072311402 | 0.018382021 | 0.999933887 | 0.248695279 | 0.326586611 | -0.077891333 | 0.639997775 |
| ENSG00000206199 | *ANKUB1* | 1.75316084 | -0.08681527 | 0.063567208 | -0.150382478 | 0.999933887 | 0.517449258 | 0.673382756 | -0.155933498 | 0.639997775 |
| ENSG00000287883 | *AC093627.22* | -0.209119214 | -0.034489109 | 0.026622497 | -0.061111606 | 0.999933887 | -0.091640197 | 0.088104807 | -0.179745005 | 0.640119526 |
| ENSG00000162408 | *NOL9* | 2.987061025 | -0.075784535 | 0.045028054 | -0.12081259 | 0.999933887 | 0.093538319 | 0.015075934 | 0.078462385 | 0.6401624 |
| ENSG00000184613 | *NELL2* | 3.881228642 | -0.033261028 | -0.145077607 | 0.111816579 | 0.999933887 | 0.121603325 | 0.194306643 | -0.072703317 | 0.64022778 |
| ENSG00000084731 | *KIF3C* | 2.845670173 | 0.035272181 | 0.115973517 | -0.080701336 | 0.999933887 | -0.053703214 | 0.037158354 | -0.090861568 | 0.640361976 |
| ENSG00000176022 | *B3GALT6* | 2.267279561 | 0.020480935 | 0.228304446 | -0.207823511 | 0.999933887 | 0.236582649 | 0.126011001 | 0.110571648 | 0.640427284 |
| ENSG00000067064 | *IDI1* | 5.513441042 | 0.029706103 | -0.044126633 | 0.073832736 | 0.999933887 | -0.014535427 | -0.084830314 | 0.070294887 | 0.640489119 |
| ENSG00000154127 | *UBASH3B* | 6.117329669 | -0.081293979 | -0.045153024 | -0.036140955 | 0.999933887 | -0.59797782 | -0.534027584 | -0.063950236 | 0.640639316 |
| ENSG00000129055 | *ANAPC13* | 5.728017647 | -0.083884796 | -0.07892048 | -0.004964315 | 0.999933887 | -0.648638374 | -0.576255084 | -0.07238329 | 0.640668077 |
| ENSG00000188092 | *GPR89B* | 0.470459208 | -0.15698238 | -0.033194108 | -0.123788273 | 0.999933887 | -0.094674535 | 0.064450314 | -0.159124849 | 0.640668077 |
| ENSG00000278540 | *ACACA* | 1.922864167 | -0.080113895 | -0.057876488 | -0.022237408 | 0.999933887 | 0.084615112 | 0.193948894 | -0.109333782 | 0.640668077 |
| ENSG00000163521 | *GLB1L* | 1.433865798 | -0.093782378 | -0.118622642 | 0.024840265 | 0.999933887 | -0.005075865 | -0.128798652 | 0.123722787 | 0.640668077 |
| ENSG00000168672 | *LRATD2* | 2.123480823 | -0.031946617 | -0.027850765 | -0.004095853 | 0.999933887 | 0.109224162 | 0.009352297 | 0.099871866 | 0.640668077 |
| ENSG00000197363 | *ZNF517* | 1.216397421 | 0.036815666 | 0.13332953 | -0.096513864 | 0.999933887 | 0.165786404 | 0.356512388 | -0.190725984 | 0.640668077 |
| ENSG00000147364 | *FBXO25* | 1.829930352 | -0.075652235 | -0.00998176 | -0.065670475 | 0.999933887 | -0.042519353 | 0.07417417 | -0.116693523 | 0.640898651 |
| ENSG00000122550 | *KLHL7* | 2.37214933 | -0.035981306 | -0.103968857 | 0.067987551 | 0.999933887 | -0.217695287 | -0.11203692 | -0.105658367 | 0.640898651 |
| ENSG00000179698 | *WDR97* | 0.471474098 | 0.060923808 | 0.193535043 | -0.132611235 | 0.999933887 | 0.058456766 | -0.12718414 | 0.185640906 | 0.640898651 |
| ENSG00000150712 | *MTMR12* | 3.735156685 | -0.021641578 | -0.054778716 | 0.033137137 | 0.999933887 | -0.173844826 | -0.256757606 | 0.08291278 | 0.640898651 |
| ENSG00000075239 | *ACAT1* | 1.239786612 | -0.028042228 | -0.018386949 | -0.009655279 | 0.999933887 | 0.041943882 | 0.185023521 | -0.14307964 | 0.640898651 |
| ENSG00000206560 | *ANKRD28* | 5.540791629 | -0.072721247 | -0.060161017 | -0.01256023 | 0.999933887 | -0.47227165 | -0.367242066 | -0.105029583 | 0.641106997 |
| ENSG00000134183 | *GNAT2* | 0.600415698 | 0.003713713 | 0.055883708 | -0.052169995 | 0.999933887 | -0.362496017 | -0.683997782 | 0.321501766 | 0.641200635 |
| ENSG00000113312 | *TTC1* | 4.623439175 | -0.020978266 | -0.044817381 | 0.023839116 | 0.999933887 | -0.047419204 | 0.014329803 | -0.061749007 | 0.641283886 |
| ENSG00000262049 | *AC139530.1* | 0.637198406 | 0.048144453 | 0.02531444 | 0.022830013 | 0.999933887 | -0.145989933 | 0.045693828 | -0.191683761 | 0.641283886 |
| ENSG00000089335 | *ZNF302* | 1.826925628 | -0.134562464 | -0.176622407 | 0.042059943 | 0.999933887 | -0.082196723 | 0.033612108 | -0.115808831 | 0.641338879 |
| ENSG00000131116 | *ZNF428* | 1.062866763 | 0.094376164 | 0.044625916 | 0.049750248 | 0.999933887 | 0.222606029 | 0.0533121 | 0.16929393 | 0.641347779 |
| ENSG00000172687 | *ZNF738* | 2.596729386 | 0.027691594 | 0.046722342 | -0.019030748 | 0.999933887 | 0.043033127 | 0.156430444 | -0.113397317 | 0.641385238 |
| ENSG00000173193 | *PARP14* | 7.833282839 | -0.001265818 | -0.046315582 | 0.045049764 | 0.999933887 | -0.160781655 | -0.060810975 | -0.099970681 | 0.641385238 |
| ENSG00000197629 | *MPEG1* | 7.136190188 | -0.103345275 | -0.072575388 | -0.030769888 | 0.999933887 | -0.650006841 | -0.75337486 | 0.103368018 | 0.641416035 |
| ENSG00000172965 | *MIR4435-2HG* | 4.572156638 | -0.063339328 | 0.00020068 | -0.063540008 | 0.999933887 | -0.286625967 | -0.206249084 | -0.080376883 | 0.641416035 |

| | | | | | | | | | | |
|---|---|---|---|---|---|---|---|---|---|---|
| ENSG00000271869 | *AC026979.2* | 2.19179764 | 0.137488695 | -0.090395544 | 0.227884239 | 0.999933887 | 0.04114172 | -0.08410873 | 0.125250451 | 0.641498543 |
| ENSG00000246273 | *SBF2-AS1* | 0.183196957 | -0.067073942 | 0.212180437 | -0.279254378 | 0.999933887 | 0.249607436 | 0.406779495 | -0.157172059 | 0.641498543 |
| ENSG00000228839 | *PIK3IP1-DT* | 0.241932784 | -0.044684737 | -0.337164589 | 0.292479852 | 0.999933887 | 0.889875046 | 1.040948925 | -0.151073879 | 0.641498543 |
| ENSG00000234420 | *ZNF37BP* | 3.079862478 | 0.018244071 | -0.058391285 | 0.076635355 | 0.999933887 | -0.157576336 | -0.032827118 | -0.124749218 | 0.641498543 |
| ENSG00000183688 | *RFLNB* | 6.85790862 | -0.104963713 | -0.039120344 | -0.06584337 | 0.999933887 | -0.960544705 | -1.060234875 | 0.09969017 | 0.641557918 |
| ENSG00000053371 | *AKR7A2* | 1.520701353 | 0.092012965 | -0.096116514 | 0.188129479 | 0.999933887 | 0.100422395 | -0.034656046 | 0.13507844 | 0.641863064 |
| ENSG00000157107 | *FCHO2* | 4.65907476 | -0.006483469 | -0.193866071 | 0.187382601 | 0.999933887 | -1.095910555 | -0.923469292 | -0.172441264 | 0.641932599 |
| ENSG00000237883 | *DGUOK-AS1* | 0.529512097 | -0.010730506 | -0.185318433 | 0.174587926 | 0.999933887 | 0.054638552 | 0.256629858 | -0.201991305 | 0.641974983 |
| ENSG00000152256 | *PDK1* | 4.368038377 | -0.055484735 | -0.010816721 | -0.044668015 | 0.999933887 | -0.419844882 | -0.495374199 | 0.075529317 | 0.642239382 |
| ENSG00000122512 | *PMS2* | 1.895453607 | -0.026720621 | -0.048176545 | 0.021455924 | 0.999933887 | -0.286932694 | -0.174800752 | -0.112131942 | 0.642261589 |
| ENSG00000268218 | *AC137932.3* | 2.647911143 | -0.02074224 | -0.017155031 | -0.003587209 | 0.999933887 | -0.438023356 | -0.562813667 | 0.12479031 | 0.642261589 |
| ENSG00000283199 | *C13orf46* | 1.622070231 | 0.119253788 | -0.021404888 | 0.140658676 | 0.999933887 | -0.13246514 | 0.027027873 | -0.159493013 | 0.642486108 |
| ENSG00000109436 | *TBC1D9* | 0.186243165 | 0.140335092 | 0.035858372 | 0.10447672 | 0.999933887 | 0.899616496 | 0.721403264 | 0.178213232 | 0.642543 |
| ENSG00000104524 | *PYCR3* | 0.013633326 | 0.025893624 | 0.009308209 | 0.016585415 | 0.999933887 | 0.170854557 | 0.342253787 | -0.17139923 | 0.642578561 |
| ENSG00000229692 | *SOS1-IT1* | -1.340275865 | -0.097846482 | -0.824417334 | 0.726570852 | 0.999933887 | -0.303304937 | -0.05017457 | -0.253130367 | 0.642684226 |
| ENSG00000178035 | *IMPDH2* | 4.117580311 | -0.049211502 | 0.008874625 | -0.058086127 | 0.999933887 | 0.119258574 | 0.050277951 | 0.068980623 | 0.642776056 |
| ENSG00000187950 | *OVCH1* | -0.863158692 | -0.232681962 | 0.351828151 | -0.584510113 | 0.999933887 | 0.609810539 | 0.380534284 | 0.229276255 | 0.642776056 |
| ENSG00000065615 | *CYB5R4* | 6.769342319 | -0.049916335 | -0.062672824 | 0.012756489 | 0.999933887 | -0.431292768 | -0.370908196 | -0.060384571 | 0.642824816 |
| ENSG00000161381 | *PLXDC1* | 1.330674242 | 0.023744839 | 0.087280781 | -0.063535942 | 0.999933887 | -0.085424144 | 0.050709521 | -0.136133665 | 0.642932801 |
| ENSG00000166136 | *NDUFB8* | 0.701468146 | -0.009633144 | 0.19932531 | -0.208958454 | 0.999933887 | 0.124308568 | 0.284764364 | -0.160455796 | 0.643054879 |
| ENSG00000120699 | *EXOSC8* | 1.765642418 | -0.062901792 | -0.110754574 | 0.047852782 | 0.999933887 | 0.136218183 | 0.245656492 | -0.109438309 | 0.64314144 |
| ENSG00000260917 | *AL158212.3* | 1.374580351 | -0.040240085 | -0.071278394 | 0.031038309 | 0.999933887 | -0.101339388 | 0.019403366 | -0.120742754 | 0.64344373 |
| ENSG00000151687 | *ANKAR* | -0.160719439 | -0.026600374 | 0.100887433 | -0.127487807 | 0.999933887 | -0.26239117 | -0.026904698 | -0.235486472 | 0.643803552 |
| ENSG00000140807 | *NKD1* | 1.596047882 | -0.177527772 | 0.11730267 | -0.294830442 | 0.999291887 | -0.036616044 | 0.10873682 | -0.145352864 | 0.643868867 |
| ENSG00000171566 | *PLRG1* | 3.228580543 | -0.016316388 | -0.106699652 | 0.090383264 | 0.999933887 | -0.099136797 | -0.0216727 | -0.077464098 | 0.643868867 |
| ENSG00000131652 | *THOC6* | 2.002085795 | 0.044287368 | -0.021714802 | 0.06600217 | 0.999933887 | -0.184870832 | -0.312936515 | 0.128065683 | 0.644132422 |
| ENSG00000155016 | *CYP2U1* | -0.727952936 | 0.01505719 | -0.09651493 | 0.111572121 | 0.999933887 | 0.046856212 | -0.177809349 | 0.224665561 | 0.644688125 |
| ENSG00000160226 | *CFAP410* | 2.146329569 | 0.124762617 | 0.224803204 | -0.100040587 | 0.999933887 | 0.13208448 | 0.236374916 | -0.104290436 | 0.644854267 |
| ENSG00000130520 | *LSM4* | 3.197215349 | 0.021883019 | -0.139447272 | 0.16133029 | 0.999933887 | 0.141587546 | 0.053521688 | 0.088065858 | 0.644939107 |
| ENSG00000167004 | *PDIA3* | 6.014132797 | -0.056957308 | -0.089456989 | 0.032499681 | 0.999933887 | 0.087624107 | 0.148653408 | -0.061029301 | 0.645123559 |
| ENSG00000064933 | *PMS1* | 1.546249882 | 0.10392523 | -0.214503558 | 0.318428788 | 0.999933887 | -0.268607816 | -0.130364155 | -0.138243662 | 0.645793615 |
| ENSG00000234534 | *CSNK1G2P1* | -1.767996286 | 0.033119323 | -0.022359025 | 0.055478348 | 0.999933887 | 0.932986126 | 0.353917832 | 0.579068294 | 0.645793615 |
| ENSG00000060491 | *OGFR* | 7.308877859 | 0.102929894 | 0.066079881 | 0.036850013 | 0.999933887 | 0.24828583 | 0.335063871 | -0.086778041 | 0.645796055 |
| ENSG00000134824 | *FADS2* | -0.096963161 | 0.127254763 | 0.137965499 | -0.010710736 | 0.999933887 | -0.245584369 | -0.026767133 | -0.218817236 | 0.645796055 |
| ENSG00000166133 | *RPUSD2* | 0.640895818 | 0.061628507 | -0.029907232 | 0.091535739 | 0.999933887 | -0.055718945 | 0.11259643 | -0.168315375 | 0.645796055 |
| ENSG00000100410 | *PHF5A* | 3.895323751 | -0.014187116 | 0.093339821 | -0.107526937 | 0.999933887 | -0.038647624 | -0.112414808 | 0.073767183 | 0.645796055 |
| ENSG00000187630 | *DHRS4L2* | -0.490091885 | -0.022024255 | -0.153040807 | 0.131016552 | 0.999933887 | -0.116507836 | 0.141128864 | -0.257636699 | 0.645796055 |
| ENSG00000213672 | *NCKIPSD* | 1.054387179 | 0.009992409 | -0.043148353 | 0.053140762 | 0.999933887 | 0.027791473 | -0.123874123 | 0.151665596 | 0.645796055 |
| ENSG00000178917 | *ZNF852* | 0.816521811 | -0.018781584 | -0.021015871 | 0.002234288 | 0.999933887 | -0.082229496 | 0.082055196 | -0.164284692 | 0.645810199 |
| ENSG00000168040 | *FADD* | 2.24002188 | -0.081864665 | -0.129933085 | 0.048068419 | 0.999933887 | -0.643752268 | -0.798753065 | 0.155000797 | 0.645854873 |
| ENSG00000264772 | *AC016876.2* | -0.902787829 | 0.022491095 | -0.066002514 | 0.088493609 | 0.999933887 | 0.721844293 | 1.027455379 | -0.305611086 | 0.645968214 |
| ENSG00000163041 | *H3-3A* | 9.350808237 | 0.036078867 | 0.047348846 | -0.011269979 | 0.999933887 | -0.265480622 | -0.209137885 | -0.056342737 | 0.646009477 |
| ENSG00000215256 | *DHRS4-AS1* | -0.080159513 | -0.297378231 | -0.030129758 | -0.267248472 | 0.999933887 | 0.134729917 | -0.07402815 | 0.208758067 | 0.646107154 |
| ENSG00000206053 | *JPT2* | 1.852070129 | -0.133288865 | 0.079307199 | -0.212596064 | 0.999933887 | 0.010310671 | -0.106512734 | 0.116823405 | 0.646107154 |
| ENSG00000231500 | *RPS18* | 8.082746848 | -0.000166521 | -0.033026128 | 0.032859607 | 0.999933887 | 0.118675677 | 0.186538796 | -0.067863119 | 0.646107154 |
| ENSG00000175221 | *MED16* | 5.204430224 | 0.036276207 | -0.00683254 | 0.043108747 | 0.999933887 | -0.023894831 | -0.092224463 | 0.068329633 | 0.646118957 |
| ENSG00000012822 | *CALCOCO1* | 6.887258003 | 0.005372388 | 0.011623152 | -0.006250764 | 0.999933887 | -0.533483461 | -0.59931343 | 0.065829968 | 0.646118957 |
| ENSG00000164944 | *VIRMA* | 4.333099862 | -0.037248902 | -0.113251157 | 0.076002255 | 0.999933887 | -0.394305031 | -0.318977594 | -0.075327437 | 0.646124752 |
| ENSG00000256053 | *COA8* | 1.053779895 | -0.02360034 | -0.052545282 | 0.028944942 | 0.999933887 | -0.274616494 | -0.418157252 | 0.143540758 | 0.646124752 |
| ENSG00000230099 | *TRBV5-4* | 0.010116364 | -0.0190569 | 0.126298802 | -0.145355702 | 0.999933887 | 0.284987303 | 0.112466461 | 0.172520842 | 0.646124752 |
| ENSG00000171159 | *C9orf16* | 4.069686893 | 0.069480695 | 0.030199751 | 0.039280944 | 0.999933887 | 0.096705193 | 0.013024243 | 0.08368095 | 0.646143318 |
| ENSG00000232859 | *LYRM9* | -0.075386915 | -0.136272135 | 0.197552864 | -0.333824999 | 0.999933887 | 0.039223648 | -0.163339371 | 0.202563019 | 0.646208399 |
| ENSG00000160447 | *PKN3* | 0.200449675 | 0.012886359 | 0.156454929 | -0.14356857 | 0.999933887 | -0.077616092 | 0.156466579 | -0.234082671 | 0.646270957 |
| ENSG00000212283 | *SNORD89* | 0.411317059 | 0.146147513 | -0.024063301 | 0.170210814 | 0.999933887 | -0.674619056 | -0.420657933 | -0.253961123 | 0.646383214 |
| ENSG00000280388 | *AC006330.1* | -0.038228697 | 0.13673933 | -0.366833826 | 0.503573156 | 0.999291887 | 0.168049806 | -0.05672313 | 0.224772935 | 0.64651468 |
| ENSG00000124942 | *AHNAK* | 9.156601098 | -0.04660613 | -0.086563257 | 0.039957127 | 0.999933887 | 0.039842237 | -0.035237211 | 0.075079448 | 0.646759335 |
| ENSG00000148700 | *ADD3* | 6.017231846 | 0.004442725 | -0.087040367 | 0.091483093 | 0.999933887 | -0.351864948 | -0.457596347 | 0.105731399 | 0.646759335 |
| ENSG00000234290 | *AC116366.1* | 4.23726952 | 0.070667817 | 0.055723751 | 0.014944066 | 0.999933887 | 0.025884896 | -0.062767423 | 0.088652319 | 0.647039105 |
| ENSG00000123395 | *ATG101* | 2.182019925 | 0.133813765 | 0.063825989 | 0.069987776 | 0.999933887 | 0.220778606 | 0.336567356 | -0.115788749 | 0.647039105 |
| ENSG00000236287 | *ZBED5* | 4.261297962 | -0.074937178 | -0.055491956 | -0.019445222 | 0.999933887 | -0.402446363 | -0.500933956 | 0.098487592 | 0.647039105 |
| ENSG00000103064 | *SLC7A6* | 3.093992482 | -0.024514612 | -0.102549271 | 0.078034659 | 0.999933887 | 0.029652708 | -0.043989182 | 0.07364189 | 0.647039105 |
| ENSG00000140743 | *CDR2* | 3.348767923 | 0.006119369 | 0.114417178 | -0.108297809 | 0.999933887 | -0.042709025 | 0.05784704 | -0.100556065 | 0.647039105 |
| ENSG00000204099 | *NEU4* | 0.699709845 | -0.008231994 | 0.072805906 | -0.081037901 | 0.999933887 | 1.641755456 | 1.828885035 | -0.187129579 | 0.647039105 |
| ENSG00000129493 | *HEATR5A* | 2.195400802 | -0.039983859 | -0.030772149 | -0.00921171 | 0.999933887 | -0.408806599 | -0.532361489 | 0.12355489 | 0.647053388 |
| ENSG00000080839 | *RBL1* | 1.301147427 | -0.142666299 | 0.055205283 | -0.197871582 | 0.999933887 | -0.138300498 | -0.003558652 | -0.134741846 | 0.647151965 |
| ENSG00000133030 | *MPRIP* | 5.724169324 | 0.041593177 | 0.056626807 | -0.01503363 | 0.999933887 | 0.034134519 | 0.082715278 | -0.048580759 | 0.647151965 |
| ENSG00000166411 | *IDH3A* | 1.236011137 | -0.089944694 | 0.241599517 | -0.331544211 | 0.981460244 | 0.092830358 | 0.210109923 | -0.117279565 | 0.647261534 |
| ENSG00000273828 | *AL133227.1* | 0.86000109 | 0.115415033 | 0.131225979 | -0.015810946 | 0.999933887 | -0.064147672 | -0.253441846 | 0.189294174 | 0.647261534 |
| ENSG00000170854 | *RIOX2* | 0.794751932 | -0.078643675 | 0.134431541 | -0.213075216 | 0.999933887 | 0.082519588 | 0.21978599 | -0.137266402 | 0.647261534 |
| ENSG00000115415 | *STAT1* | 8.758415737 | -0.057733438 | -0.076050108 | 0.01831667 | 0.999933887 | -0.387544323 | -0.447045067 | 0.059500744 | 0.647307443 |
| ENSG00000255306 | *AC004923.4* | -0.134110768 | -0.218190584 | -0.297922152 | 0.079731568 | 0.999933887 | -0.458761808 | -0.246744399 | -0.212017409 | 0.647307443 |
| ENSG00000257497 | *AC121761.1* | 0.475260987 | 0.128269759 | 0.409016321 | -0.280746562 | 0.999933887 | -0.054555972 | -0.278620726 | 0.224064754 | 0.647307443 |
| ENSG00000230280 | *HNRNPA1P59* | -0.211403659 | -0.095811565 | 0.077908988 | -0.173720553 | 0.999933887 | 0.57361169 | 0.729985072 | -0.156373381 | 0.647333552 |
| ENSG00000141456 | *PELP1* | 3.517728239 | -0.031175671 | 0.041082982 | -0.072258653 | 0.999933887 | 0.150846837 | 0.085115029 | 0.065731807 | 0.647426045 |
| ENSG00000166927 | *MS4A7* | 0.631089694 | 0.024640942 | 0.137167409 | -0.112526468 | 0.999933887 | -0.557751764 | -0.365877745 | -0.191874019 | 0.647426045 |
| ENSG00000122140 | *MRPS2* | 1.20416515 | -0.00328837 | -0.017058584 | 0.013770214 | 0.999933887 | 0.122481895 | 0.270129593 | -0.147647698 | 0.647426045 |
| ENSG00000229807 | *XIST* | 4.459609562 | -0.073098005 | -0.055804666 | -0.017293339 | 0.999933887 | -0.078730412 | 0.084704984 | -0.163435396 | 0.647645376 |
| ENSG00000169432 | *SCN9A* | -0.568197481 | -0.023965551 | 0.204587233 | -0.228552784 | 0.999933887 | -0.174980742 | 0.173958757 | -0.348939499 | 0.647645376 |
| ENSG00000188312 | *CENPP* | 0.848789203 | -0.019063011 | 0.102986628 | -0.122049639 | 0.999933887 | 0.376783814 | 0.540525704 | -0.163741891 | 0.647655261 |
| ENSG00000136937 | *NCBP1* | 3.603804345 | -0.027371845 | -0.003291331 | -0.024080514 | 0.999933887 | 0.001213303 | 0.076136849 | -0.074923545 | 0.647707737 |
| ENSG00000184232 | *OAF* | -0.457832383 | 0.207691698 | -0.005071183 | 0.212762881 | 0.999933887 | -0.383955993 | -0.172611415 | -0.211344577 | 0.64772267 |
| ENSG00000170502 | *NUDT9* | 0.732348744 | 0.061604755 | 0.146199661 | -0.084594906 | 0.999933887 | 0.013085822 | 0.162638047 | -0.149552225 | 0.64772267 |
| ENSG00000126709 | *IFI6* | 6.777794604 | 0.07443717 | 0.01550315 | 0.058934021 | 0.999933887 | -0.135770937 | -0.225033032 | 0.089262095 | 0.647775734 |
| ENSG00000178146 | *GK4P* | -0.053158507 | -0.176947233 | -0.076090751 | -0.100856482 | 0.999933887 | 0.242214241 | 0.440940124 | -0.198725883 | 0.648097132 |
| ENSG00000267002 | *AC060780.1* | 3.005925312 | -0.040191362 | -0.052357969 | 0.012166607 | 0.999933887 | -0.104450423 | -0.194548251 | 0.090097828 | 0.648160502 |
| ENSG00000075702 | *WDR62* | 0.24539964 | 0.043360406 | 0.018049515 | 0.025310891 | 0.999933887 | 0.156877628 | 0.358823301 | -0.201945673 | 0.648160502 |
| ENSG00000139726 | *DENR* | 4.773168801 | -0.066508867 | -0.050264469 | -0.016244398 | 0.999933887 | -0.362457723 | -0.304186717 | -0.058271006 | 0.648235457 |
| ENSG00000112309 | *B3GAT2* | -0.543570228 | 0.134029212 | 0.029964006 | 0.104065206 | 0.999933887 | -0.60461713 | -0.300344387 | -0.304272743 | 0.648235457 |
| ENSG00000163958 | *ZDHHC19* | 0.929678793 | 0.146687599 | -0.20346018 | 0.350147779 | 0.999933887 | -0.267603385 | -0.047333513 | -0.220269872 | 0.648265166 |
| ENSG00000160307 | *S100B* | -1.395269519 | 0.159020745 | -0.190639472 | 0.349660217 | 0.999933887 | 0.283536378 | 0.618006453 | -0.334470076 | 0.64830231 |
| ENSG00000117643 | *MAN1C1* | 1.810160947 | -0.077604856 | 0.129137459 | -0.206742315 | 0.999933887 | -0.131856184 | -0.224341559 | 0.092485375 | 0.64830231 |
| ENSG00000235313 | *HM13-IT1* | 1.187640258 | -0.067594034 | -0.131130485 | 0.063536452 | 0.999933887 | -0.101016675 | 0.049994104 | -0.151010779 | 0.64830231 |
| ENSG00000184787 | *UBE2G2* | 4.813445265 | -0.016383159 | 0.000165012 | -0.016548171 | 0.999933887 | 0.004027477 | -0.043620474 | 0.047647951 | 0.64830231 |
| ENSG00000082458 | *DLG3* | 1.529772908 | 0.054501941 | -0.056429328 | 0.110931269 | 0.999933887 | -0.124995955 | 0.009127577 | -0.134123532 | 0.648352867 |
| ENSG00000176102 | *CSTF3* | 1.84889248 | -0.08227305 | -0.057129984 | -0.025143067 | 0.999933887 | 0.118911619 | 0.228805989 | -0.10989437 | 0.648543107 |
| ENSG00000250831 | *AC074131.1* | -0.801550866 | 0.048365245 | 0.139911973 | -0.091546728 | 0.999933887 | 0.545623966 | 0.74013641 | -0.194512444 | 0.648543107 |
| ENSG00000084676 | *NCOA1* | 7.828479638 | -0.009485665 | -0.076101966 | 0.066616301 | 0.999933887 | -0.097677012 | -0.012887035 | -0.084789977 | 0.648543107 |
| ENSG00000137760 | *ALKBH8* | 0.705566047 | -0.06865131 | -0.124661181 | 0.056009871 | 0.999933887 | 0.030476756 | 0.17794651 | -0.147469754 | 0.648616516 |
| ENSG00000185651 | *UBE2L3* | 4.713173646 | -0.01725692 | 0.008545259 | -0.025802179 | 0.999933887 | -0.260320651 | -0.330471767 | 0.070151116 | 0.648800687 |
| ENSG00000226334 | *AL359182.1* | 1.291171791 | 0.122804112 | 0.394114608 | -0.271310496 | 0.999933887 | 0.400160424 | 0.238962502 | 0.161197922 | 0.648800716 |
| ENSG00000160957 | *RECQL4* | 0.416383843 | 0.096034315 | 0.04653352 | 0.049500795 | 0.999933887 | 0.356101601 | 0.1458679 | 0.210233701 | 0.648800716 |
| ENSG00000239557 | *PPP2R5CP* | 0.526011878 | 0.016510624 | -0.024730855 | 0.041241479 | 0.999933887 | 0.159482658 | -0.087386865 | 0.246869524 | 0.648928032 |
| ENSG00000203730 | *TEDDM1* | -0.127311717 | 0.160857722 | 0.221421843 | -0.060564121 | 0.999933887 | 0.806456565 | 0.552646786 | 0.25380978 | 0.648984204 |
| ENSG00000111886 | *GABRR2* | 2.650123102 | -0.055842828 | 0.003259294 | -0.059102122 | 0.999933887 | 0.107936247 | 0.204004413 | -0.096068166 | 0.648984204 |
| ENSG00000159685 | *CHCHD6* | -0.458509237 | 0.119988992 | 0.255776816 | -0.135787824 | 0.999933887 | 0.16345895 | -0.029431429 | 0.192890379 | 0.648984204 |

| | | | | | | | | | | |
|---|---|---|---|---|---|---|---|---|---|---|
| ENSG00000236278 | *PEBP1P3* | 1.503037657 | 0.100777513 | 0.318614034 | -0.21783652 | 0.999933887 | 0.467815004 | 0.70408337 | -0.236268366 | 0.649147084 |
| ENSG00000151929 | *BAG3* | 2.156885687 | 0.038885826 | -0.090378664 | 0.12926449 | 0.999933887 | 0.111671813 | 0.223249234 | -0.111577421 | 0.649147084 |
| ENSG00000198546 | *ZNF511* | 1.293786453 | -0.011216972 | -0.079854648 | 0.068637676 | 0.999933887 | -0.059038085 | -0.202651004 | 0.143612919 | 0.649147084 |
| ENSG00000164818 | *DNAAF5* | 1.722178931 | -0.083892634 | 0.020951978 | -0.104844612 | 0.999933887 | 0.06334861 | 0.165494642 | -0.102146032 | 0.649273826 |
| ENSG00000225720 | *AL031846.1* | -0.732206701 | 0.004431885 | 0.201915659 | -0.197483774 | 0.999933887 | -0.056238642 | 0.213105863 | -0.269344506 | 0.649273826 |
| ENSG00000085552 | *IGSF9* | 0.359985969 | 0.266688305 | 0.277264658 | -0.010576353 | 0.999933887 | 0.586788362 | 0.920186009 | -0.333397647 | 0.649307821 |
| ENSG00000242689 | *CNTF* | -0.419321377 | -0.238978626 | -0.139500009 | -0.099478617 | 0.999933887 | -0.03344826 | -0.322556624 | 0.289108363 | 0.649380513 |
| ENSG00000101391 | *CDK5RAP1* | 2.510174956 | -0.040172127 | 0.009146557 | -0.049318684 | 0.999933887 | 0.055104458 | -0.038779889 | 0.093884347 | 0.649610992 |
| ENSG00000113719 | *ERGIC1* | 8.090084261 | 0.012135442 | -0.004522107 | 0.016657549 | 0.999933887 | 0.216759479 | 0.271525221 | -0.054765742 | 0.649610992 |
| ENSG00000147123 | *NDUFB11* | 3.473198931 | 0.06753027 | -0.030943934 | 0.098474204 | 0.999933887 | 0.156512623 | 0.057575682 | 0.098936941 | 0.649957049 |
| ENSG00000203705 | *TATDN3* | 2.183750706 | -0.087165686 | -0.134809342 | 0.047643656 | 0.999933887 | -0.122616488 | -0.021558912 | -0.101057575 | 0.650077841 |
| ENSG00000259959 | *AC107068.1* | 0.129016227 | -0.136357733 | -0.067573564 | -0.068784169 | 0.999933887 | -0.362179706 | -0.63615969 | 0.273979984 | 0.650077841 |
| ENSG00000196504 | *PRPF40A* | 5.431111407 | -0.075037782 | -0.145624864 | 0.070587083 | 0.999933887 | -0.49691437 | -0.367040729 | -0.129873641 | 0.650077841 |
| ENSG00000146085 | *MMUT* | 2.390267251 | -0.019393545 | 0.032423297 | -0.051816842 | 0.999933887 | -0.130660898 | -0.033360437 | -0.09730046 | 0.650077841 |
| ENSG00000100292 | *HMOX1* | 8.57530175 | -0.024614562 | 0.007008037 | -0.031622599 | 0.999933887 | -0.800574063 | -0.654410925 | -0.146163138 | 0.650077841 |
| ENSG00000174282 | *ZBTB4* | 4.81884118 | 0.08557869 | 0.082922696 | 0.002655994 | 0.999933887 | -0.001018428 | -0.073601189 | 0.072582761 | 0.65014835 |
| ENSG00000109184 | *DCUN1D4* | 2.779177967 | -0.069359963 | 0.004934464 | -0.074294426 | 0.999933887 | -0.24752587 | -0.13425006 | -0.11327581 | 0.65014835 |
| ENSG00000103544 | *VPS35L* | 2.845035596 | -0.056456975 | -0.0035691 | -0.052887875 | 0.999933887 | 0.002930965 | 0.088595953 | -0.085664988 | 0.650455983 |
| ENSG00000229894 | *GK3P* | -0.276270321 | 0.019353838 | 0.286885138 | -0.2675313 | 0.999933887 | 0.072489003 | -0.196262243 | 0.268751246 | 0.650455983 |
| ENSG00000100079 | *LGALS2* | -1.458949269 | -0.007184328 | 0.049179081 | -0.056363409 | 0.999933887 | -0.150481769 | -0.488383596 | 0.337901827 | 0.650455983 |
| ENSG00000128872 | *TMOD2* | 3.384211376 | -0.002143742 | 0.072661008 | -0.07480475 | 0.999933887 | -0.199178977 | -0.102118889 | -0.097060088 | 0.650558538 |
| ENSG00000147133 | *TAF1* | 5.153655437 | -0.056875103 | -0.049875337 | -0.006999766 | 0.999933887 | -0.236162339 | -0.168736928 | -0.067425412 | 0.650642339 |
| ENSG00000257365 | *FNTB* | 0.672027717 | -0.004074994 | 0.142641062 | -0.146716056 | 0.999933887 | -0.756100759 | -0.922654754 | 0.166553995 | 0.650642339 |
| ENSG00000261067 | *AC109460.3* | 1.002561248 | 5.3472e-05 | 0.189304727 | -0.189251255 | 0.999933887 | 0.391672173 | 0.512921048 | -0.121248875 | 0.650642339 |
| ENSG00000196118 | *CCDC189* | 0.083363608 | -0.192586472 | 0.051034231 | -0.243620703 | 0.999933887 | -0.381786998 | -0.639874362 | 0.258087364 | 0.650999113 |
| ENSG00000168612 | *ZSWIM1* | 2.608808814 | -0.026723172 | -0.214485945 | 0.187762773 | 0.999933887 | -0.020063275 | 0.077211291 | -0.097274566 | 0.651092178 |
| ENSG00000008300 | *CELSR3* | 0.755765162 | 0.013172872 | -0.051458798 | 0.064631669 | 0.999933887 | 0.110623167 | 0.270498035 | -0.159874868 | 0.651369014 |
| ENSG00000164512 | *ANKRD55* | -0.035356232 | 0.115183741 | 0.153775633 | -0.038591891 | 0.999933887 | -0.330144429 | -0.093318511 | -0.236825918 | 0.651394821 |
| ENSG00000223935 | *LGALSL-DT* | 0.291276799 | 0.118081798 | -0.076735575 | 0.194817373 | 0.999933887 | -0.487287081 | -0.692452551 | 0.20516547 | 0.651470634 |
| ENSG00000004897 | *CDC27* | 3.522976191 | -0.037374316 | -0.093076468 | 0.055702151 | 0.999933887 | -0.285268149 | -0.197030081 | -0.088238068 | 0.651853454 |
| ENSG00000168026 | *TTC21A* | 3.245700948 | 0.175063227 | 0.260779392 | -0.085716166 | 0.999933887 | 0.027263821 | -0.100697537 | 0.127961358 | 0.651944312 |
| ENSG00000038274 | *MAT2B* | 5.869903533 | -0.087024908 | -0.015344955 | -0.071679953 | 0.999933887 | -0.287844097 | -0.345003528 | 0.057159431 | 0.651944312 |
| ENSG00000266891 | *AP000902.1* | 0.887095683 | -0.069545726 | 0.031497076 | -0.101042802 | 0.999933887 | -0.149704465 | 0.087969569 | -0.237674034 | 0.651944312 |
| ENSG00000164109 | *MAD2L1* | -0.220557907 | -0.14063813 | -0.125039593 | -0.015598537 | 0.999933887 | 0.125703456 | -0.119480081 | 0.245183537 | 0.651944312 |
| ENSG00000100218 | *RSPH14* | -1.13779651 | -0.157514909 | 0.118415467 | -0.275930376 | 0.999933887 | -0.283290371 | 0.165597624 | -0.448887995 | 0.651944312 |
| ENSG00000128050 | *PAICS* | 2.317614706 | -0.057628009 | -0.190700991 | 0.133072983 | 0.999933887 | 0.077453445 | 0.169676617 | -0.092223172 | 0.651944312 |
| ENSG00000115484 | *CCT4* | 3.803975425 | 0.013151349 | -0.153406486 | 0.166557835 | 0.999933887 | 0.159085798 | 0.232589628 | -0.07350383 | 0.651944312 |
| ENSG00000116151 | *MORN1* | 0.812269303 | -0.001180145 | -0.05531102 | 0.054130875 | 0.999933887 | 0.037561765 | -0.117124806 | 0.154686571 | 0.651944312 |
| ENSG00000226332 | *AL354836.1* | 0.632632672 | -0.052207554 | 0.164865788 | -0.217073341 | 0.999933887 | -0.096990255 | 0.035952446 | -0.132942701 | 0.651950052 |
| ENSG00000280161 | *AC022413.1* | -0.633392686 | 0.255721608 | -0.171845585 | 0.427567192 | 0.999933887 | 0.703941017 | 0.937648609 | -0.233707593 | 0.651987844 |
| ENSG00000260672 | *AC100827.4* | -1.302534324 | -0.052731981 | 0.243375019 | -0.296107 | 0.999933887 | 0.088253026 | 0.51742252 | -0.429169493 | 0.651987844 |
| ENSG00000159176 | *CSRP1* | 3.631517535 | -0.036791754 | 0.012405305 | -0.049197059 | 0.999933887 | -0.096479408 | -0.180813122 | 0.084333714 | 0.652295545 |
| ENSG00000186409 | *CCDC30* | 0.720236681 | -0.033102237 | 0.111238436 | -0.144340673 | 0.999933887 | -0.81324197 | -0.6201169 | -0.193125071 | 0.652295545 |
| ENSG00000124380 | *SNRNP27* | 2.168743457 | -0.010470897 | 0.06519906 | -0.075669958 | 0.999933887 | 0.003316176 | 0.099713355 | -0.096397179 | 0.652316527 |
| ENSG00000100453 | *GZMB* | 3.970346005 | -0.061730152 | -0.031202497 | -0.030527656 | 0.999933887 | 0.103564863 | 0.02661107 | 0.076953793 | 0.652510724 |
| ENSG00000177943 | *MAMDC4* | 1.20051602 | 0.072540587 | 0.102541301 | -0.030000713 | 0.999933887 | 0.102613524 | 0.248402643 | -0.145789118 | 0.652510724 |
| ENSG00000169118 | *CSNK1G1* | 3.60382041 | -0.099604591 | -0.129744702 | 0.030140111 | 0.999933887 | -0.355015127 | -0.444883114 | 0.089867987 | 0.652525561 |
| ENSG00000175063 | *UBE2C* | 0.238407075 | -0.133253279 | 0.150009827 | -0.283263106 | 0.999933887 | -0.135239724 | 0.081509956 | -0.21674968 | 0.652560575 |
| ENSG00000147536 | *GINS4* | 0.534137089 | 0.058777275 | 0.102093356 | -0.04331608 | 0.999933887 | 0.635181668 | 0.851670602 | -0.216488934 | 0.652739936 |
| ENSG00000260803 | *Z84723.1* | -1.717019172 | 0.016793625 | -0.411342845 | 0.42813647 | 0.999933887 | -1.399466128 | -1.693854097 | 0.294387969 | 0.65277847 |
| ENSG00000223750 | *SIRPB3P* | -0.390626716 | 0.093057696 | 0.048744894 | 0.044312802 | 0.999933887 | 0.03953706 | 0.311400647 | -0.271863587 | 0.652815725 |
| ENSG00000166402 | *TUB* | -0.503210484 | -0.039540959 | 0.003857718 | -0.043398677 | 0.999933887 | -0.108324561 | 0.141316168 | -0.249640729 | 0.652869219 |
| ENSG00000168234 | *TTC39C* | 3.536130985 | -0.069146157 | 0.010428361 | -0.079574518 | 0.999933887 | 0.140904444 | 0.207028486 | -0.066124042 | 0.653016674 |
| ENSG00000198363 | *ASPH* | 7.518241954 | -0.067297166 | 0.042099543 | -0.109396709 | 0.999933887 | -0.340767311 | -0.26074133 | -0.080025981 | 0.653016674 |
| ENSG00000119333 | *DYNC2I2* | 0.301857918 | -0.052864246 | -0.187635137 | 0.134770891 | 0.999933887 | 0.023165176 | -0.139195537 | 0.162360713 | 0.653016674 |
| ENSG00000125249 | *RAP2A* | 2.546534619 | -0.001079827 | -0.160808068 | 0.15972824 | 0.999933887 | -0.189383555 | -0.084658124 | -0.104725431 | 0.653016674 |
| ENSG00000174780 | *SRP72* | 4.186330489 | -0.047171989 | -0.138521221 | 0.091349232 | 0.999933887 | -0.047500352 | 0.031732295 | -0.079232647 | 0.653017894 |
| ENSG00000279155 | *AC233300.1* | 2.072104473 | -0.009907094 | 0.010201942 | -0.020109036 | 0.999933887 | 0.078448555 | 0.211985728 | -0.133537173 | 0.653145858 |
| ENSG00000205707 | *ETFRF1* | 2.013652288 | -0.053121901 | -0.094006161 | 0.04088426 | 0.999933887 | -0.026981523 | 0.109328994 | -0.136310517 | 0.653202405 |
| ENSG00000066422 | *ZBTB11* | 4.690019953 | -0.039450564 | -0.150656444 | 0.11120588 | 0.999933887 | -0.225827495 | -0.083547664 | -0.142279832 | 0.653202405 |
| ENSG00000226380 | *AC016831.1* | 5.281532819 | 0.112768884 | 0.093709238 | 0.019059646 | 0.999933887 | 1.097285216 | 1.22163647 | -0.124351254 | 0.653244526 |
| ENSG00000119801 | *YPEL5* | 9.556938214 | -0.007930418 | -0.068525059 | 0.060594641 | 0.999933887 | -0.194295902 | -0.143952595 | -0.050343307 | 0.653244526 |
| ENSG00000164168 | *TMEM184C* | 3.496877105 | 0.016443832 | -0.152637423 | 0.169081255 | 0.999933887 | -0.29887393 | -0.389716074 | 0.090842144 | 0.653477175 |
| ENSG00000115641 | *FHL2* | -0.130955886 | 0.026007129 | -0.100009694 | 0.126018623 | 0.999933887 | -0.109180039 | 0.093652977 | -0.202833016 | 0.653696064 |
| ENSG00000085465 | *OVGP1* | -0.18370849 | 0.066755033 | 0.257271092 | -0.190516059 | 0.999933887 | -0.020239055 | -0.231265954 | 0.211026899 | 0.653931093 |
| ENSG00000166579 | *NDEL1* | 7.121622275 | 0.064259259 | 0.040312546 | 0.023946713 | 0.999933887 | -0.519894691 | -0.57769061 | 0.057795919 | 0.654164192 |
| ENSG00000231128 | *AL137856.1* | -0.621881791 | 0.238804122 | 0.116761264 | 0.122042857 | 0.999933887 | -0.119056153 | 0.22230881 | -0.341364963 | 0.654164192 |
| ENSG00000075188 | *NUP37* | 0.200048938 | -0.140091507 | -0.180438186 | 0.040346679 | 0.999933887 | -0.218442946 | -0.043774266 | -0.17466868 | 0.654519161 |
| ENSG00000035862 | *TIMP2* | 7.264975449 | 0.013201194 | 0.011141937 | 0.002059257 | 0.999933887 | -1.334552095 | -1.433465222 | 0.098913128 | 0.654552881 |
| ENSG00000129422 | *MTUS1* | -0.905956295 | -0.18630146 | -0.104146874 | -0.082154587 | 0.999933887 | -0.208039319 | 0.042925477 | -0.250964796 | 0.65464143 |
| ENSG00000261490 | *AC005674.1* | 4.165060299 | -0.109684498 | -0.076768992 | -0.032915506 | 0.999933887 | -0.121955522 | -0.209293719 | 0.087338197 | 0.654731381 |
| ENSG00000146066 | *HIGD2A* | 3.045190203 | -0.007899641 | 0.002884028 | -0.010783668 | 0.999933887 | -0.050458056 | -0.149659171 | 0.099201115 | 0.655007297 |
| ENSG00000101166 | *PRELID3B* | 4.150040879 | -0.175040611 | -0.08608072 | -0.08895989 | 0.999933887 | -0.27809608 | -0.209117817 | -0.068978263 | 0.655199663 |
| ENSG00000115318 | *LOXL3* | 1.416920622 | 0.05416436 | -0.048289584 | 0.102453944 | 0.999933887 | 0.484926554 | 0.327176021 | 0.157750533 | 0.655199663 |
| ENSG00000237940 | *LINC01238* | 0.352679795 | -0.055864812 | -0.204355506 | 0.148490694 | 0.999933887 | 0.056985897 | 0.226227078 | -0.169241181 | 0.655199663 |
| ENSG00000156853 | *ZNF689* | 1.820027752 | -0.097018711 | -0.106644505 | 0.009625795 | 0.999933887 | 0.121991568 | 0.006463371 | 0.115528197 | 0.655201416 |
| ENSG00000278784 | *AL136295.7* | 2.024938968 | -0.013872758 | -0.016825049 | 0.002952291 | 0.999933887 | -0.020159465 | 0.085014391 | -0.105173856 | 0.655497244 |
| ENSG00000124357 | *NAGK* | 8.657108335 | -0.059745772 | -0.053235048 | -0.006510723 | 0.999933887 | -0.35428423 | -0.295422958 | -0.058861272 | 0.655781868 |
| ENSG00000129559 | *NEDD8* | 3.42742781 | -0.074263182 | 0.00579397 | -0.080057152 | 0.999933887 | -0.031610659 | -0.122498364 | 0.090887705 | 0.655781868 |
| ENSG00000179941 | *BBS10* | 0.914819718 | 0.146415328 | -0.122266325 | 0.268681653 | 0.999933887 | -0.267423379 | -0.397370486 | 0.129947107 | 0.655781868 |
| ENSG00000164885 | *CDK5* | 0.172212322 | -0.121302809 | 0.287483665 | -0.408786473 | 0.999933887 | 0.093410362 | -0.093459203 | 0.186869565 | 0.655781868 |
| ENSG00000135624 | *CCT7* | 4.469420927 | -0.037667358 | -0.024280087 | -0.013387271 | 0.999933887 | 0.128900406 | 0.068788354 | 0.060112052 | 0.655781868 |
| ENSG00000246922 | *UBAP1L* | 1.456930951 | 0.013579336 | 0.247896155 | -0.234316819 | 0.999933887 | -0.017191328 | 0.085481444 | -0.102672772 | 0.655781868 |
| ENSG00000108298 | *RPL19* | 8.128588081 | 0.001126851 | -0.061256333 | 0.062383184 | 0.999933887 | 0.050569505 | 0.111018018 | -0.060448513 | 0.655781868 |
| ENSG00000177685 | *CRACR2B* | 0.295315238 | 0.118543821 | 0.07742518 | 0.041118641 | 0.999933887 | -0.059110965 | -0.256531088 | 0.197420123 | 0.655786268 |
| ENSG00000117280 | *RAB29* | 4.394165636 | -0.02793047 | 0.082127912 | -0.110058382 | 0.999933887 | 0.032156582 | -0.024193598 | 0.05635018 | 0.65581725 |
| ENSG00000187676 | *B3GLCT* | 1.022392985 | 0.024953126 | -0.159277138 | 0.184230264 | 0.999933887 | -0.037067726 | -0.199545953 | 0.162478227 | 0.655811725 |
| ENSG00000183682 | *BMP8A* | 0.09546316 | 0.127557815 | 0.176397309 | -0.048839494 | 0.999933887 | -0.071191071 | 0.112917556 | -0.184108627 | 0.656252247 |
| ENSG00000204272 | *NBDY* | 1.090425542 | -0.032971298 | -0.155998201 | 0.123026903 | 0.999933887 | 0.119019371 | 0.257569501 | -0.13855013 | 0.656252247 |
| ENSG00000143190 | *POU2F1* | 5.345889546 | -0.09330034 | 0.001975818 | -0.095276158 | 0.999933887 | -0.444279359 | -0.34714872 | -0.097130638 | 0.656365763 |
| ENSG00000095002 | *MSH2* | 1.165827863 | 0.000479184 | -0.013935774 | 0.014414958 | 0.999933887 | -0.036824798 | 0.120715544 | -0.157540342 | 0.656636771 |
| ENSG00000182557 | *SPNS3* | 2.654005015 | -0.022074133 | 0.003899839 | -0.025973972 | 0.999933887 | -0.068652542 | -0.178322262 | 0.10966972 | 0.656642886 |
| ENSG00000258365 | *AC073655.2* | 1.618699283 | 0.207615083 | -0.021738902 | 0.229353985 | 0.999933887 | 0.737965838 | 0.895693899 | -0.157728062 | 0.65683246 |
| ENSG00000113812 | *ACTR8* | 2.481819265 | 0.087063908 | -0.024865075 | 0.111928983 | 0.999933887 | -0.06954646 | -0.157817873 | 0.088271413 | 0.65683246 |
| ENSG00000169131 | *ZNF354A* | 1.717858984 | -0.114876232 | -0.125822323 | 0.010946092 | 0.999933887 | -0.302705359 | -0.173519858 | -0.129185501 | 0.65683246 |
| ENSG00000205918 | *PDPK2P* | 0.552346092 | 0.103079899 | 0.083846311 | 0.019233588 | 0.999933887 | -0.213216952 | -0.378776559 | 0.165559607 | 0.65683246 |
| ENSG00000197713 | *RPE* | 1.334751328 | -0.108552029 | 0.061568626 | -0.170120654 | 0.999933887 | -0.075693203 | -0.214781677 | 0.139088474 | 0.657025986 |
| ENSG00000171604 | *CXXC5* | 0.695517473 | 0.067363354 | 0.061265941 | 0.006097412 | 0.999933887 | -0.056896985 | -0.22567177 | 0.168774785 | 0.657046329 |
| ENSG00000100483 | *VCPKMT* | 3.674423188 | -0.066621715 | -0.062072458 | -0.004549257 | 0.999933887 | -0.336136844 | -0.456397933 | 0.120261088 | 0.657263189 |
| ENSG00000271605 | *MILR1* | 1.130178573 | -0.053925697 | 0.002484122 | -0.056409819 | 0.999933887 | 0.054242778 | 0.264640129 | -0.210397351 | 0.657356768 |
| ENSG00000171469 | *ZNF561* | 2.312707967 | 0.121874707 | 0.027916155 | 0.093958552 | 0.999933887 | -0.168802973 | -0.278032839 | 0.109229867 | 0.657589896 |
| ENSG00000162543 | *UBXN10* | -0.984186913 | -0.167181187 | 0.068983672 | -0.236164858 | 0.999933887 | -0.330543185 | -0.044041106 | -0.28650208 | 0.657589896 |
| ENSG00000186376 | *ZNF75D* | 2.768941091 | 0.073504002 | 0.006966825 | 0.066537177 | 0.999933887 | -0.144831645 | -0.241826479 | 0.096994835 | 0.657589896 |
| ENSG00000188021 | *UBQLN2* | 5.445039427 | 0.029996121 | 0.015117535 | 0.014878587 | 0.999933887 | -0.285905611 | -0.346571692 | 0.060666081 | 0.657589896 |
| ENSG00000014641 | *MDH1* | 2.97380758 | -0.006190445 | -0.216867413 | 0.210676968 | 0.999933887 | 0.039106763 | 0.123702481 | -0.084595719 | 0.657589896 |

| | | | | | | | | | | |
|---|---|---|---|---|---|---|---|---|---|---|
| ENSG00000143458 | *GABPB2* | 2.722653036 | -0.003768357 | 0.121966731 | -0.125735088 | 0.999933887 | -0.014658903 | -0.108838156 | 0.094179253 | 0.657589896 |
| ENSG00000164008 | *C1orf50* | 0.367269793 | -0.005680819 | 0.327621692 | -0.333302511 | 0.999933887 | -0.01906601 | 0.159679468 | -0.178745478 | 0.657589896 |
| ENSG00000170571 | *EMB* | 5.623312011 | 0.040283127 | -0.124010174 | 0.164293301 | 0.999933887 | -0.179797848 | -0.087363272 | -0.092434576 | 0.657887078 |
| ENSG00000271888 | *AL136162.1* | 0.240067882 | -0.024053759 | 0.017750027 | -0.041803786 | 0.999933887 | 0.358501299 | 0.122886782 | 0.235614518 | 0.658248169 |
| ENSG00000235903 | *CPB2-AS1* | 3.192798625 | 0.24343985 | 0.412998364 | -0.169558513 | 0.999933887 | 0.830534727 | 1.029072611 | -0.198537883 | 0.658767384 |
| ENSG00000230539 | *AOAH-IT1* | 0.347287595 | 0.216643129 | 0.000427832 | 0.216215297 | 0.999933887 | 0.750691091 | 1.037791741 | -0.28710065 | 0.65878179 |
| ENSG00000008869 | *HEATR5B* | 4.478705033 | 0.042656114 | -0.140646259 | 0.183302373 | 0.999933887 | -0.10612691 | -0.184205811 | 0.078078901 | 0.65878179 |
| ENSG00000205250 | *E2F4* | 5.813035564 | 0.01922799 | 0.001793093 | 0.017434897 | 0.999933887 | -0.024629394 | -0.071439295 | 0.046809901 | 0.65878179 |
| ENSG00000241945 | *PWP2* | 1.443343374 | -0.040191997 | 0.152112379 | -0.192304376 | 0.999933887 | -0.034489492 | -0.193291632 | 0.15880214 | 0.65878179 |
| ENSG00000149428 | *HYOU1* | 4.357973185 | -0.005461423 | 0.032820299 | -0.038281722 | 0.999933887 | 0.164608035 | 0.227579307 | -0.062971272 | 0.65878179 |
| ENSG00000161929 | *SCIMP* | -0.97168561 | -0.179870878 | -0.01845988 | -0.161410998 | 0.999933887 | -0.461128769 | -0.217298232 | -0.243830537 | 0.658878298 |
| ENSG00000232295 | *AL589935.1* | 0.321687428 | 0.045270571 | -0.147694891 | 0.192965462 | 0.999933887 | 0.190421052 | -0.004502578 | 0.19492363 | 0.658878298 |
| ENSG00000107758 | *PPP3CB* | 4.029557981 | -0.032568577 | 0.109004284 | -0.14157286 | 0.999933887 | -0.138159465 | -0.203156199 | 0.064996734 | 0.658920515 |
| ENSG00000139182 | *CLSTN3* | 4.873323779 | -0.006448059 | 0.061916137 | -0.068364196 | 0.999933887 | 0.112580125 | 0.053893201 | 0.058686924 | 0.658920515 |
| ENSG00000282173 | *TRBJ1-5* | 0.494570503 | -0.258885772 | 0.146417307 | -0.405303079 | 0.999933887 | 0.245507473 | 0.084126096 | 0.161381377 | 0.659114583 |
| ENSG00000147155 | *EBP* | 1.670088875 | -0.112147573 | -0.109511825 | -0.002635748 | 0.999933887 | 0.031985792 | 0.213495721 | -0.181509929 | 0.659114583 |
| ENSG00000279108 | *AC008537.3* | 0.717614055 | -0.097579128 | 0.102821084 | -0.200400212 | 0.999933887 | -0.103993232 | 0.046087209 | -0.150080441 | 0.659114583 |
| ENSG00000033867 | *SLC4A7* | 3.804425761 | -0.003654142 | -0.106586416 | 0.102932274 | 0.999933887 | -0.413897627 | -0.269419814 | -0.144477813 | 0.659146694 |
| ENSG00000153487 | *ING1* | 3.981123551 | 0.143597161 | 0.099398178 | 0.044198983 | 0.999933887 | -0.186620513 | -0.25344729 | 0.066826777 | 0.659465202 |
| ENSG00000205903 | *ZNF316* | 9.737443674 | 0.086072965 | 0.049610363 | 0.036462601 | 0.999933887 | 0.51914286 | 0.608244254 | -0.089101393 | 0.659483874 |
| ENSG00000080819 | *CPOX* | 1.757777016 | 0.105936096 | -0.041066003 | 0.1470021 | 0.999933887 | 0.086753331 | 0.180642985 | -0.093889654 | 0.659508442 |
| ENSG00000120662 | *MTRF1* | 1.801769166 | -0.060952541 | -0.104412063 | 0.043459522 | 0.999933887 | -0.19706326 | -0.096440181 | -0.100623079 | 0.659508442 |
| ENSG00000158290 | *CUL4B* | 5.696572547 | -0.034187089 | -0.106166433 | 0.071979344 | 0.999933887 | -0.700936511 | -0.622389104 | -0.078547408 | 0.659508442 |
| ENSG00000123739 | *PLA2G12A* | 0.208490972 | 0.07778966 | -0.261592153 | 0.339381813 | 0.999933887 | 0.155883011 | -0.059910898 | 0.215793909 | 0.659508442 |
| ENSG00000186470 | *BTN3A2* | 6.569840275 | -0.004864589 | -0.051458316 | 0.046593727 | 0.999933887 | 0.015091541 | 0.072115036 | -0.057023495 | 0.659508442 |
| ENSG00000145349 | *CAMK2D* | 2.536232156 | -0.005438317 | -0.108221849 | 0.102783532 | 0.999933887 | -0.100846968 | -0.00354408 | -0.097302888 | 0.659531961 |
| ENSG00000242190 | *AC017015.2* | -0.771405418 | 0.43639207 | 0.292984007 | 0.143408063 | 0.999933887 | 2.082116402 | 1.778910788 | 0.303205614 | 0.659546756 |
| ENSG00000198121 | *LPAR1* | -0.904468749 | 0.064791719 | 0.303458528 | -0.238666809 | 0.999933887 | -0.575948979 | -0.268793132 | -0.307155848 | 0.659546756 |
| ENSG00000117691 | *NENF* | 1.362670267 | 0.052256146 | -0.088227805 | 0.140483951 | 0.999933887 | 0.277483573 | 0.158112234 | 0.119371339 | 0.659596906 |
| ENSG00000198468 | *FLVCR1-DT* | 0.594407727 | -0.018824665 | -0.043175935 | 0.02435127 | 0.999933887 | 0.030063115 | 0.19267141 | -0.162608296 | 0.659596906 |
| ENSG00000249353 | *NPM1P27* | 0.402773162 | -0.002645939 | 0.338304946 | -0.340950885 | 0.999933887 | 0.400596865 | 0.609420677 | -0.208823812 | 0.659724928 |
| ENSG00000244480 | *AC005154.2* | -0.080989955 | -0.105842511 | -0.134441219 | 0.028598707 | 0.999933887 | -0.114400948 | -0.3119317 | 0.197530752 | 0.65989362 |
| ENSG00000181016 | *LSMEM1* | 4.342935557 | -0.058118334 | 0.005288101 | -0.063406436 | 0.999933887 | 0.082049985 | 0.15381226 | -0.071762275 | 0.659966331 |
| ENSG00000090372 | *STRN4* | 6.62401897 | 0.043576078 | 0.075067017 | -0.031490939 | 0.999933887 | -0.277219699 | -0.344604129 | 0.067384429 | 0.660013785 |
| ENSG00000089012 | *SIRPG* | 2.333021905 | -0.001213402 | 0.087872767 | -0.089086169 | 0.999933887 | 0.085578762 | 0.182827736 | -0.097248974 | 0.660275889 |
| ENSG00000140264 | *SERF2* | 6.564945684 | 0.036182638 | 0.033701746 | 0.002480892 | 0.999933887 | 0.340744625 | 0.287654746 | 0.053089879 | 0.660307865 |
| ENSG00000267365 | *KCNJ2-AS1* | 2.928509551 | -0.131757745 | -0.110716557 | -0.021041187 | 0.999933887 | 3.647757798 | 3.835259138 | -0.18750134 | 0.660333969 |
| ENSG00000269243 | *AC008894.2* | -0.094902256 | 0.058098345 | -0.152605876 | 0.210704221 | 0.999933887 | -0.358015768 | -0.559345724 | 0.201329956 | 0.660333969 |
| ENSG00000276819 | *TRBV15* | -0.993696509 | -0.232675236 | -0.372979838 | 0.140304602 | 0.999933887 | 0.250523233 | 0.502495534 | -0.251972301 | 0.660417326 |
| ENSG00000157881 | *PANK4* | 5.73087863 | 0.006039437 | -0.057018097 | 0.063057533 | 0.999933887 | 0.084679069 | 0.143545139 | -0.058866069 | 0.660417326 |
| ENSG00000248971 | *KRT8P46* | 2.095175469 | -0.093346735 | -0.020528496 | -0.072818239 | 0.999933887 | 0.209014265 | 0.324915038 | -0.115900774 | 0.660518281 |
| ENSG00000196843 | *ARID5A* | 6.984454138 | 0.024258106 | 0.056025103 | -0.031766998 | 0.999933887 | 0.0949032 | 0.025532204 | 0.069370997 | 0.660518281 |
| ENSG00000177483 | *RBM44* | 0.765051577 | 0.056016452 | -0.124808742 | 0.180825194 | 0.999933887 | -0.431327414 | -0.138183028 | -0.293144386 | 0.660569379 |
| ENSG00000278730 | *AC005332.6* | 3.171393207 | -0.11194567 | -0.023583999 | -0.088361671 | 0.999933887 | -0.292150173 | -0.226393586 | -0.065756587 | 0.660849849 |
| ENSG00000250687 | *NAIPP1* | -0.121810139 | 0.129950828 | 0.020472209 | 0.109478619 | 0.999933887 | -0.391997242 | -0.661594761 | 0.269597519 | 0.660849849 |
| ENSG00000275709 | *AC090527.3* | -0.86580323 | 0.323818802 | 0.295370124 | 0.028448679 | 0.999933887 | 0.846755934 | 0.518190355 | 0.328565579 | 0.661056314 |
| ENSG00000273445 | *AC133644.1* | -0.842020042 | 0.174428519 | 0.196059708 | -0.021631189 | 0.999933887 | 0.149942303 | -0.137669733 | 0.287612036 | 0.661056314 |
| ENSG00000223642 | *AC008277.1* | -0.604277237 | 0.15693183 | 0.064335938 | 0.092595892 | 0.999933887 | -0.763743019 | -0.467939872 | -0.295803147 | 0.661056314 |
| ENSG00000049449 | *RCN1* | -0.466330998 | -0.08800758 | 0.180591148 | -0.268598727 | 0.999933887 | 0.223410146 | 0.004785298 | 0.218624848 | 0.661056314 |
| ENSG00000178913 | *TAF7* | 8.198807485 | 0.018816901 | -0.086633553 | 0.105450454 | 0.999933887 | 0.163659822 | 0.216685818 | -0.053025996 | 0.661056314 |
| ENSG00000280067 | *AC023818.1* | 1.04904606 | -0.049151892 | -0.040052968 | -0.009098924 | 0.999933887 | -0.020312324 | -0.166765676 | 0.146453352 | 0.661056314 |
| ENSG00000060688 | *SNRNP40* | 2.481057681 | 0.016663743 | -0.212673945 | 0.229337688 | 0.999933887 | 0.090078899 | 0.005551063 | 0.084527837 | 0.661056314 |
| ENSG00000279259 | *AC087741.3* | 0.587393569 | -0.018305151 | -0.086599909 | 0.068294757 | 0.999933887 | -0.164239821 | -0.348680537 | 0.184440715 | 0.661056314 |
| ENSG00000106211 | *HSPB1* | 2.05364166 | -0.000704061 | -0.040948539 | 0.040244478 | 0.999933887 | 0.176974779 | 0.301029225 | -0.124054446 | 0.661056314 |
| ENSG00000130770 | *ATP5IF1* | 4.095983805 | 0.056191632 | -0.071438411 | 0.127630042 | 0.999933887 | 0.108236511 | 0.17997785 | -0.071741338 | 0.661164008 |
| ENSG00000124782 | *RREB1* | 6.182428254 | -0.021219786 | 0.1013031 | -0.122522886 | 0.999933887 | -0.873225099 | -0.792089956 | -0.081135143 | 0.661164008 |
| ENSG00000189283 | *FHIT* | 0.965051689 | -0.01023491 | -0.005180475 | -0.005054436 | 0.999933887 | -0.040990962 | 0.100089465 | -0.141080427 | 0.661164008 |
| ENSG00000147251 | *DOCK11* | 7.012625751 | -0.009331713 | -0.126823749 | 0.117492036 | 0.999933887 | -0.806585956 | -0.683757397 | -0.122828559 | 0.661328112 |
| ENSG00000287510 | *AL137857.1* | -0.682733818 | 0.155072905 | 0.153800488 | 0.001272417 | 0.999933887 | -0.135924268 | 0.191746474 | -0.327670742 | 0.661500843 |
| ENSG00000163736 | *PPBP* | -0.107000213 | -0.044334633 | -0.061224571 | 0.016889938 | 0.999933887 | 0.019686162 | 0.241950562 | -0.2222644 | 0.661752189 |
| ENSG00000152642 | *GPD1L* | 1.523112729 | 0.013576653 | 0.11899546 | -0.105418807 | 0.999933887 | 0.0716936 | -0.049256715 | 0.120950316 | 0.661781861 |
| ENSG00000100014 | *SPECC1L* | 2.639736305 | 0.002678908 | -0.071844573 | 0.074523481 | 0.999933887 | 0.278509382 | 0.390277832 | -0.11176845 | 0.661781861 |
| ENSG00000181804 | *SLC9A9* | 0.99625866 | 0.048082778 | -0.037375713 | 0.085458491 | 0.999933887 | 0.012538029 | 0.156778106 | -0.144240077 | 0.661822893 |
| ENSG00000160439 | *RDH13* | 2.059884879 | 0.07848749 | 0.06589463 | 0.012592861 | 0.999933887 | 0.332905209 | 0.433357109 | -0.1004519 | 0.662010314 |
| ENSG00000220205 | *VAMP2* | 7.034849066 | 0.047606532 | 0.102733667 | -0.055127135 | 0.999933887 | 0.150251235 | 0.098222514 | 0.052028721 | 0.662300017 |
| ENSG00000260804 | *LINC01963* | 0.328144947 | -0.227534078 | 0.037722678 | -0.265256756 | 0.999933887 | -0.101984146 | 0.088750222 | -0.190734368 | 0.662371654 |
| ENSG00000111801 | *BTN3A3* | 4.953248049 | 0.022717243 | -0.053017487 | 0.075734729 | 0.999933887 | -0.013464764 | -0.092584767 | 0.079120003 | 0.662489176 |
| ENSG00000119655 | *NPC2* | 7.283350073 | -0.060512036 | -0.08462872 | 0.024116684 | 0.999933887 | -0.128061392 | -0.069205089 | -0.058856303 | 0.662529401 |
| ENSG00000120334 | *CENPL* | -0.089434479 | -0.072491441 | 0.098566901 | -0.171058342 | 0.999933887 | -0.014391857 | 0.16985118 | -0.184243036 | 0.662529401 |
| ENSG00000119689 | *DLST* | 5.544167659 | -0.005773855 | 0.006179211 | -0.011953066 | 0.999933887 | -0.070171143 | -0.113679405 | 0.043508262 | 0.662529401 |
| ENSG00000123737 | *EXOSC9* | 3.00534474 | -0.047805117 | -0.031480294 | -0.016324823 | 0.999933887 | 0.100024038 | 0.17573273 | -0.075708691 | 0.662735971 |
| ENSG00000120509 | *PDZD11* | 0.65273707 | -0.055802762 | 0.024656399 | -0.08045916 | 0.999933887 | 0.194897304 | 0.022821851 | 0.172075453 | 0.662894619 |
| ENSG00000087258 | *GNAO1* | 1.363559294 | 0.072575158 | -0.049029423 | 0.121604581 | 0.999933887 | -0.05479788 | -0.172159967 | 0.117362087 | 0.66294882 |
| ENSG00000262339 | *AC130371.1* | -1.096181115 | 0.121251427 | -0.195167469 | 0.316418896 | 0.999933887 | -0.244351536 | -0.627819666 | 0.383468131 | 0.66294882 |
| ENSG00000234585 | *CCT6P3* | 2.841212977 | -0.150730086 | -0.184893867 | 0.034163781 | 0.999933887 | -0.829448239 | -0.911726152 | 0.082277912 | 0.663113282 |
| ENSG00000141293 | *SKAP1* | 3.350004817 | -0.078450839 | 0.001992692 | -0.080443531 | 0.999933887 | 0.064725665 | 0.143495715 | -0.07877005 | 0.663113282 |
| ENSG00000050748 | *MAPK9* | 3.470645672 | -0.020350164 | 0.090504108 | -0.110854271 | 0.999933887 | 0.083923519 | 0.018382934 | 0.065540586 | 0.663113282 |
| ENSG00000017483 | *SLC38A5* | 1.459375771 | -0.027796234 | 0.050832874 | -0.078629108 | 0.999933887 | 0.347284897 | 0.231623161 | 0.115661736 | 0.663113282 |
| ENSG00000162004 | *CCDC78* | 0.341257453 | -0.039974905 | 0.171723724 | -0.211698629 | 0.999933887 | 0.035999384 | 0.207733522 | -0.171734138 | 0.663113282 |
| ENSG00000008283 | *CYB561* | 3.548782082 | 0.001405515 | 0.003589204 | -0.002183689 | 0.999933887 | 0.072711502 | -0.012147994 | 0.084859496 | 0.663113282 |
| ENSG00000173757 | *STAT5B* | 9.079457406 | -0.002366705 | 0.006140023 | -0.008506729 | 0.999933887 | 0.174247084 | 0.22272178 | -0.048474696 | 0.663318747 |
| ENSG00000221946 | *FXYD7* | -0.212339296 | 0.065302213 | 0.207903907 | -0.142601693 | 0.999933887 | 0.16057088 | -0.032944939 | 0.193515819 | 0.663329206 |
| ENSG00000248213 | *CICP16* | -0.004835715 | -0.026012044 | 0.03830719 | -0.064319235 | 0.999933887 | 0.072393152 | 0.311204184 | -0.238811032 | 0.663597828 |
| ENSG00000253439 | *CDC42P5* | 0.974916931 | 0.082749642 | 0.238918512 | -0.156168871 | 0.999933887 | 0.519078854 | 0.692048641 | -0.172969787 | 0.663642706 |
| ENSG00000187187 | *ZNF546* | -0.502020686 | -0.295658311 | -0.11928006 | -0.176378251 | 0.999933887 | -0.022137712 | 0.198321192 | -0.220458904 | 0.664882905 |
| ENSG00000177707 | *NECTIN3* | -0.269201337 | -0.195407173 | -0.383982083 | 0.18857491 | 0.999933887 | -0.026764504 | 0.195459702 | -0.222224206 | 0.664882905 |
| ENSG00000149474 | *KAT14* | 0.682046784 | -0.140517411 | 0.056614877 | -0.197132289 | 0.999933887 | 0.062173489 | -0.08454393 | 0.146717418 | 0.664882905 |
| ENSG00000173950 | *XXYLT1* | 0.425519564 | 0.012499611 | -0.102329265 | 0.114828876 | 0.999933887 | -0.017486289 | 0.193793664 | -0.211279954 | 0.664882905 |
| ENSG00000107317 | *PTGDS* | 0.107528242 | 0.252494503 | 0.187283634 | 0.065210869 | 0.999933887 | 0.07091658 | -0.097196192 | 0.168112772 | 0.66497446 |
| ENSG00000143514 | *TP53BP2* | 5.964912739 | 0.010121422 | -0.008548026 | 0.018669448 | 0.999933887 | 0.45844276 | 0.511248677 | -0.052805918 | 0.66497446 |
| ENSG00000112200 | *ZNF451* | 3.761212934 | -0.001618385 | 0.023487977 | -0.025106362 | 0.999933887 | -0.199434606 | -0.073552951 | -0.125881655 | 0.66497446 |
| ENSG00000128268 | *MGAT3* | 2.119261636 | -0.052050521 | -0.001490971 | -0.050559549 | 0.999933887 | -0.266898171 | -0.381216304 | 0.114318133 | 0.665283328 |
| ENSG00000268903 | *AL627309.6* | 5.211222976 | 0.080645481 | -0.010261772 | 0.090907252 | 0.999933887 | 0.279485592 | 0.190930076 | 0.088555516 | 0.665321775 |
| ENSG00000140830 | *TXNL4B* | 2.852751834 | -0.061275247 | -0.058296126 | -0.00297912 | 0.999933887 | 0.039629718 | -0.039797978 | 0.079427696 | 0.665321775 |
| ENSG00000185829 | *ARL17A* | -0.481672676 | 0.112197381 | -0.204623289 | 0.31682067 | 0.999933887 | -0.022258611 | 0.17271864 | -0.194977251 | 0.665321775 |
| ENSG00000213626 | *LBH* | 5.030197523 | -0.02738022 | -0.010809745 | -0.016570475 | 0.999933887 | 0.117781894 | 0.053988825 | 0.06379307 | 0.665321775 |
| ENSG00000254019 | *AC011676.2* | 2.128165168 | 0.054471975 | 0.137156382 | -0.082684408 | 0.999933887 | 0.05023683 | 0.22977925 | -0.17954242 | 0.665321775 |
| ENSG00000236056 | *GAPDHP14* | -0.700018413 | -0.04598248 | 0.141358577 | -0.187341057 | 0.999933887 | 0.126507106 | -0.117644973 | 0.244152078 | 0.665321775 |
| ENSG00000057252 | *SOAT1* | 4.130586994 | 0.012542197 | -0.005017661 | 0.017559858 | 0.999933887 | -0.082550093 | -0.011236156 | -0.071313938 | 0.665321775 |
| ENSG00000004766 | *VPS50* | 1.421030959 | -0.01066644 | -0.299316667 | 0.288650227 | 0.999933887 | -0.279616729 | -0.134678196 | -0.144938533 | 0.665321775 |
| ENSG00000229622 | *MTND5P2* | -0.345269604 | 0.01530131 | -0.138734984 | 0.154036294 | 0.999933887 | 0.82094432 | 1.070969917 | -0.250025597 | 0.665321775 |
| ENSG00000175198 | *PCCA* | 0.059101798 | -0.027876086 | -0.293977001 | 0.266100915 | 0.999933887 | 0.071066021 | 0.275042776 | -0.203976755 | 0.665415477 |
| ENSG00000102900 | *NUP93* | 3.400152047 | 0.026599256 | 0.016314278 | 0.010284977 | 0.999933887 | 0.040887963 | 0.128812392 | -0.087924429 | 0.665471716 |
| ENSG00000177463 | *NR2C2* | 6.262344892 | -0.06069997 | -0.041571167 | -0.019128803 | 0.999933887 | -0.260112208 | -0.311472076 | 0.051359868 | 0.665541467 |
| ENSG00000197016 | *ZNF470* | -0.83633238 | -0.033844961 | -0.296258857 | 0.262413896 | 0.999933887 | -0.056096214 | 0.18185307 | -0.237949285 | 0.666208268 |

| | | | | | | | | | | |
|---|---|---|---|---|---|---|---|---|---|---|
| ENSG00000008128 | *CDK11A* | 2.723676315 | 0.070937622 | 0.032939777 | 0.037997844 | 0.999933887 | -0.001767759 | -0.096185701 | 0.094417942 | 0.666721038 |
| ENSG00000132170 | *PPARG* | 0.309527971 | -0.137569152 | -0.040550257 | -0.097018895 | 0.999933887 | -1.280823673 | -1.426539076 | 0.145715403 | 0.666721038 |
| ENSG00000248508 | *SRP14-AS1* | -0.632128978 | 0.095630392 | -0.357277654 | 0.452908046 | 0.999933887 | 0.092286605 | 0.317603161 | -0.225316557 | 0.666721038 |
| ENSG00000123144 | *TRIR* | 6.754359055 | 0.077593608 | 0.105067614 | -0.027474006 | 0.999933887 | 0.135590303 | 0.193379916 | -0.057789613 | 0.666832243 |
| ENSG00000100385 | *IL2RB* | 5.00901077 | 0.035652124 | -0.097232744 | 0.132884868 | 0.999933887 | 0.098499216 | 0.03136807 | 0.067131146 | 0.666832243 |
| ENSG00000166289 | *PLEKHF1* | 2.446510035 | 0.055448332 | 0.137073491 | -0.081625159 | 0.999933887 | 0.039977801 | -0.073924323 | 0.113902124 | 0.666832243 |
| ENSG00000235381 | *AL596202.1* | -0.436793771 | 0.047002888 | -0.218840683 | 0.265843571 | 0.999933887 | -0.108736168 | -0.363652365 | 0.254916197 | 0.666832243 |
| ENSG00000066827 | *ZFAT* | 3.550221496 | 0.08315634 | 0.106196706 | -0.023040366 | 0.999933887 | 0.021346498 | 0.094017151 | -0.072670653 | 0.666955471 |
| ENSG00000127884 | *ECHS1* | 2.098745177 | -0.048339029 | -0.032306762 | -0.016032267 | 0.999933887 | 0.101135679 | -0.016376669 | 0.117512348 | 0.667068916 |
| ENSG00000267491 | *AC100788.1* | 0.770509457 | 0.062903548 | -0.290621502 | 0.35352505 | 0.999933887 | -0.682898185 | -0.878415358 | 0.195517173 | 0.667068916 |
| ENSG00000160284 | *SPATC1L* | -0.872689116 | -0.015207749 | 0.129256903 | -0.144464652 | 0.999933887 | 0.218789513 | -0.066346033 | 0.285135546 | 0.667068916 |
| ENSG00000089639 | *GMIP* | 8.700862555 | 0.012826907 | 0.042118489 | -0.029291581 | 0.999933887 | -0.298322122 | -0.351416251 | 0.053094129 | 0.6672222 |
| ENSG00000197989 | *SNHG12* | 2.954079167 | -0.053921514 | -0.104680291 | 0.050758778 | 0.999933887 | 0.133503706 | 0.041963746 | 0.09153996 | 0.667318469 |
| ENSG00000139168 | *ZCRB1* | 2.385741818 | -0.128863027 | -0.022650423 | -0.106212605 | 0.999933887 | -0.084612912 | 0.020360152 | -0.104973064 | 0.667351474 |
| ENSG00000104731 | *KLHDC4* | 3.803558585 | -0.014447706 | -0.004221785 | -0.010225921 | 0.999933887 | -0.020517255 | 0.045825953 | -0.066343208 | 0.667351474 |
| ENSG00000266714 | *MYO15B* | 6.017895373 | 0.045317398 | 0.080453267 | -0.035135869 | 0.999933887 | 0.068576285 | 0.010194078 | 0.058382207 | 0.667479294 |
| ENSG00000083844 | *ZNF264* | 3.793159393 | -0.024692725 | 0.039841389 | -0.064534114 | 0.999933887 | -0.343039721 | -0.412272 | 0.069232279 | 0.667479294 |
| ENSG00000272529 | *AC090948.3* | -1.065903574 | 0.0497813 | 0.434365258 | -0.384583959 | 0.999933887 | 0.00431693 | 0.279815923 | -0.275498993 | 0.667508814 |
| ENSG00000124251 | *TP53TG5* | 0.854231171 | 0.143919963 | 0.0375304 | 0.106389564 | 0.999933887 | 0.530749727 | 0.727258145 | -0.196508417 | 0.667605762 |
| ENSG00000197279 | *ZNF165* | 4.367102363 | 0.091153946 | 0.097188769 | -0.006034823 | 0.999933887 | -0.204820392 | -0.111279399 | -0.093540993 | 0.667605762 |
| ENSG00000100575 | *TIMM9* | 0.581090072 | -0.06554369 | 0.349890276 | -0.415433966 | 0.999933887 | 0.111316291 | 0.257445146 | -0.146128855 | 0.667605762 |
| ENSG00000254126 | *CD8B2* | -1.108096457 | 0.250935936 | -0.261455211 | 0.512391147 | 0.999933887 | -0.072633748 | 0.171484032 | -0.24411778 | 0.667624066 |
| ENSG00000086102 | *NFX1* | 5.158373514 | -0.055154869 | -0.074657349 | 0.01950248 | 0.999933887 | -0.047983447 | 0.002116391 | -0.050099837 | 0.667624066 |
| ENSG00000184319 | *RPL23AP82* | 0.733731687 | 0.105805151 | 0.225466234 | -0.119661083 | 0.999933887 | -0.278506424 | -0.392020481 | 0.113514057 | 0.667624066 |
| ENSG00000110013 | *SIAE* | -0.375805121 | -0.06662092 | -0.07058527 | 0.00396435 | 0.999933887 | 0.016868654 | -0.191222002 | 0.208090656 | 0.667624066 |
| ENSG00000137656 | *BUD13* | 3.03424419 | -0.003183597 | 0.037263254 | -0.040446851 | 0.999933887 | -0.188266646 | -0.112855336 | -0.075411311 | 0.667652347 |
| ENSG00000003987 | *MTMR7* | 0.279713889 | -0.011343983 | 0.039673731 | -0.051017713 | 0.999933887 | -0.208354799 | -0.437190259 | 0.228835461 | 0.667763485 |
| ENSG00000149136 | *SSRP1* | 3.680539728 | -0.056675181 | 0.041007007 | -0.097682188 | 0.999933887 | 0.157867278 | 0.230317988 | -0.07245071 | 0.667919245 |
| ENSG00000107854 | *TNKS2* | 6.301449213 | -0.046970949 | -0.026554303 | -0.020416646 | 0.999933887 | -0.413983434 | -0.330640273 | -0.083343161 | 0.667919245 |
| ENSG00000065427 | *KARS1* | 4.905300158 | -0.018557093 | -0.104373743 | 0.085816651 | 0.999933887 | 0.16013797 | 0.215483978 | -0.055346009 | 0.667919245 |
| ENSG00000128944 | *KNSTRN* | -0.292130348 | -0.030877154 | -0.152250557 | 0.121373403 | 0.999933887 | 0.176981118 | 0.365593343 | -0.188612225 | 0.667919245 |
| ENSG00000178074 | *C2orf69* | 2.464672944 | 0.201353215 | 0.05296044 | 0.148392775 | 0.999933887 | -0.421313002 | -0.239292261 | -0.182020741 | 0.668047926 |
| ENSG00000130590 | *SAMD10* | 1.351930715 | 0.165182531 | 0.151765064 | 0.013417467 | 0.999933887 | 0.207447152 | 0.335734201 | -0.128287049 | 0.668047926 |
| ENSG00000287950 | *AC009432.2* | -1.078130984 | 0.130511463 | 0.011702363 | 0.118809101 | 0.999933887 | -0.256452432 | 0.153362637 | -0.409815069 | 0.668047926 |
| ENSG00000279166 | *AC009951.4* | 4.021460777 | 0.079116409 | 0.466236796 | -0.387120387 | 0.999933887 | 1.380152861 | 1.581848024 | -0.201695162 | 0.668047926 |
| ENSG00000162517 | *PEF1* | 4.990244646 | 0.017204334 | -0.058558083 | 0.075762417 | 0.999933887 | 0.2646715 | 0.325696085 | -0.061024585 | 0.668047926 |
| ENSG00000115541 | *HSPE1* | 1.641875553 | -0.028625927 | 0.125600561 | -0.154226489 | 0.999933887 | 0.175995055 | 0.275233147 | -0.099238093 | 0.668047926 |
| ENSG00000251034 | *AC087854.1* | 0.179346798 | 0.01966878 | 0.011565226 | 0.008103555 | 0.999933887 | -0.137356696 | 0.095284645 | -0.232641341 | 0.668047926 |
| ENSG00000163795 | *ZNF513* | 4.567078514 | 0.119899588 | 0.140546964 | -0.020647375 | 0.999933887 | -0.152224342 | -0.231733349 | 0.079509006 | 0.668222915 |
| ENSG00000285938 | *AC072022.2* | 5.473181781 | 0.112048769 | 0.052765127 | 0.059283641 | 0.999933887 | 0.735978503 | 0.818734256 | -0.082755752 | 0.668222915 |
| ENSG00000198853 | *RUSC2* | 1.161394868 | 0.259859158 | -0.124244913 | 0.384104071 | 0.999933887 | -0.236696937 | -0.134039357 | -0.102657579 | 0.668222915 |
| ENSG00000011021 | *CLCN6* | 5.472810351 | 0.002394211 | -0.00726803 | 0.00966224 | 0.999933887 | -0.22717552 | -0.161086274 | -0.066089247 | 0.668222915 |
| ENSG00000164574 | *GALNT10* | 4.473697826 | 0.033218154 | 0.012131893 | 0.02108626 | 0.999933887 | -0.033094303 | -0.090001393 | 0.05690709 | 0.668350767 |
| ENSG00000145725 | *PPIP5K2* | 4.267774493 | -0.066625711 | -0.116722492 | 0.050096782 | 0.999933887 | -0.626477788 | -0.787663542 | 0.161185754 | 0.668350767 |
| ENSG00000183978 | *COA3* | 0.819993255 | 0.052827064 | -0.153242364 | 0.206069428 | 0.999933887 | 0.022308609 | 0.181111516 | -0.158802907 | 0.668350767 |
| ENSG00000163946 | *TASOR* | 4.394960914 | -0.016943201 | -0.050620289 | 0.033677088 | 0.999933887 | -0.368543335 | -0.260991534 | -0.107551802 | 0.668350767 |
| ENSG00000216937 | *CCDC7* | -0.007116963 | -0.102266926 | -0.467349276 | 0.36508235 | 0.999933887 | -0.174696399 | 0.012885294 | -0.187581694 | 0.668440194 |
| ENSG00000162402 | *USP24* | 5.705129532 | 0.010265758 | -0.08583393 | 0.096099688 | 0.999933887 | -0.232906532 | -0.319579639 | 0.086673107 | 0.668638977 |
| ENSG00000273096 | *AL021707.8* | 1.157328518 | 0.144557677 | 0.013120624 | 0.131437053 | 0.999933887 | -0.053942751 | 0.148748149 | -0.2026909 | 0.668718863 |
| ENSG00000197813 | *AC011450.1* | 1.336997772 | 0.208989792 | -0.004573333 | 0.213563124 | 0.999933887 | 0.334133692 | 0.227186647 | 0.106947045 | 0.668718863 |
| ENSG00000168228 | *ZCCHC4* | 0.712724728 | 0.050823702 | 0.29015374 | -0.239330038 | 0.999933887 | 0.218281519 | 0.06325073 | 0.155030789 | 0.668718863 |
| ENSG00000127540 | *UQCR11* | 2.506637846 | 0.027116866 | 0.026922093 | 0.000194773 | 0.999933887 | -0.294415186 | -0.397932496 | 0.10351731 | 0.668718863 |
| ENSG00000279457 | *WASH9P* | 1.177870967 | -0.030497792 | 0.000311466 | -0.030809258 | 0.999933887 | -0.118066582 | -0.258347554 | 0.140280972 | 0.668718863 |
| ENSG00000243811 | *APOBEC3D* | -0.779059498 | 0.038364084 | -0.06527614 | 0.103640224 | 0.999933887 | -0.117973267 | -0.402986221 | 0.285012955 | 0.668718863 |
| ENSG00000120942 | *UBIAD1* | 2.334753781 | 0.043878369 | -0.163938369 | 0.207816739 | 0.999933887 | 0.096129721 | -0.001351475 | 0.097481196 | 0.668739999 |
| ENSG00000103356 | *EARS2* | 0.826626618 | 0.001925292 | -0.037485687 | 0.039410979 | 0.999933887 | 0.128068776 | 0.001988384 | 0.126080392 | 0.668739999 |
| ENSG00000227827 | *PKD1P2* | -1.696455889 | -0.005165136 | 0.213022925 | -0.218188061 | 0.999933887 | -0.097740429 | 0.122618652 | -0.220359081 | 0.668781179 |
| ENSG00000204044 | *SLC12A5-AS1* | 0.609878303 | 0.136190898 | 0.118173253 | 0.018017646 | 0.999933887 | 0.369161305 | 0.567528089 | -0.198366784 | 0.668865529 |
| ENSG00000172967 | *XKR3* | -0.158680643 | 0.042310533 | 0.09851103 | -0.056200497 | 0.999933887 | -0.888847431 | -1.036013417 | 0.147165986 | 0.668891171 |
| ENSG00000185127 | *C6orf120* | 3.473330988 | -0.067797828 | -0.121041162 | 0.053243334 | 0.999933887 | -0.270688241 | -0.192521339 | -0.078166902 | 0.66907171 |
| ENSG00000204136 | *GGTA1* | -0.25216484 | -0.192668166 | -0.179205803 | -0.013462363 | 0.999933887 | 0.602996448 | 0.756615442 | -0.153618994 | 0.66907171 |
| ENSG00000185627 | *PSMD13* | 5.86112101 | -0.026582205 | -0.027984274 | 0.001402069 | 0.999933887 | -0.204598043 | -0.255814884 | 0.05121684 | 0.66907171 |
| ENSG00000132478 | *UNK* | 4.489421983 | -0.009412025 | 0.112887103 | -0.122299127 | 0.999933887 | 0.065101909 | 0.01222403 | 0.052877879 | 0.66907171 |
| ENSG00000197747 | *S100A10* | 5.059192863 | 0.004143222 | -0.156700257 | 0.160843479 | 0.999933887 | -0.046845034 | -0.111783157 | 0.064938123 | 0.66907171 |
| ENSG00000185522 | *LMNTD2* | 0.331181916 | 0.261707221 | 0.345824369 | -0.084117148 | 0.999933887 | 0.157252093 | 0.302640191 | -0.145388097 | 0.669112222 |
| ENSG00000273373 | *AL355488.1* | -0.380893617 | 0.162025713 | 0.080540307 | 0.081485406 | 0.999933887 | -0.192954672 | -0.43931069 | 0.246356018 | 0.669112222 |
| ENSG00000248503 | *AL356235.1* | 1.092221385 | -0.038326654 | 0.072419036 | -0.110745689 | 0.999933887 | -0.290608355 | -0.424825849 | 0.134217495 | 0.669112222 |
| ENSG00000235978 | *AC018816.1* | -0.158757394 | -0.234403826 | 0.22104786 | -0.455451686 | 0.999933887 | 1.025736782 | 1.218271114 | -0.192534331 | 0.669318589 |
| ENSG00000211698 | *TRGV4* | -0.142102223 | 0.118121091 | -0.210235259 | 0.32835635 | 0.999933887 | -0.119071971 | 0.03056843 | -0.149640401 | 0.669318589 |
| ENSG00000204370 | *SDHD* | 3.757897545 | -0.079736241 | -0.031203945 | -0.048532296 | 0.999933887 | -0.151335281 | -0.083480734 | -0.067854547 | 0.669487821 |
| ENSG00000169696 | *ASPSCR1* | 3.31784797 | 0.067288955 | 0.074268524 | -0.00697957 | 0.999933887 | 0.012766723 | 0.08243366 | -0.069666937 | 0.669786652 |
| ENSG00000129317 | *PUS7L* | 2.032503299 | -0.108484649 | -0.133820067 | 0.025335417 | 0.999933887 | -0.295886892 | -0.098953437 | -0.196933455 | 0.669981909 |
| ENSG00000138346 | *DNA2* | 0.95081428 | -0.15274167 | -0.261876506 | 0.109134836 | 0.999933887 | -0.352930902 | -0.199116987 | -0.153813914 | 0.670097666 |
| ENSG00000154174 | *TOMM70* | 2.887510794 | -0.143955645 | -0.022742499 | -0.121213146 | 0.999933887 | 0.071379301 | 0.145110536 | -0.073731236 | 0.670108316 |
| ENSG00000275432 | *AC022149.1* | 0.19049706 | -0.228589933 | 0.062403672 | -0.290993606 | 0.999933887 | -0.962021672 | -0.758751471 | -0.203270201 | 0.670108316 |
| ENSG00000162413 | *KLHL21* | 9.434332019 | -8.1347e-05 | -0.000219811 | 0.000138465 | 0.999933887 | 0.459158834 | 0.396038748 | 0.063120087 | 0.670108316 |
| ENSG00000230107 | *AL022316.1* | 0.841318426 | -0.124105977 | -0.507023812 | 0.382917835 | 0.999933887 | -0.056778567 | -0.179821403 | 0.123042835 | 0.67014134 |
| ENSG00000107443 | *CCNJ* | 1.386456535 | -0.077781464 | -0.178800187 | 0.101018723 | 0.999933887 | -0.096642368 | -0.23767922 | 0.141036852 | 0.67014134 |
| ENSG00000085721 | *RRN3* | 3.266365293 | 0.001694534 | -0.055310412 | 0.057004946 | 0.999933887 | 0.179299687 | 0.258707289 | -0.079407602 | 0.67014134 |
| ENSG00000255031 | *AP002807.1* | 3.473636897 | 0.181396025 | 0.235759919 | -0.054363894 | 0.999933887 | 0.035152577 | -0.083901889 | 0.119054466 | 0.670226823 |
| ENSG00000232022 | *FAAHP1* | -0.053984514 | -0.037950038 | 0.119484352 | -0.15743439 | 0.999933887 | -0.107405328 | 0.089218208 | -0.196623537 | 0.670273791 |
| ENSG00000108821 | *COL1A1* | 0.010950407 | 0.037403044 | -0.211637379 | 0.249040423 | 0.999933887 | 0.354282515 | 0.619194513 | -0.264911999 | 0.670393263 |
| ENSG00000188277 | *C15orf62* | 0.86197269 | -0.010593208 | 0.063201557 | -0.073794764 | 0.999933887 | -0.930698318 | -1.120981101 | 0.190282782 | 0.670393263 |
| ENSG00000198556 | *ZNF789* | 1.874661677 | -0.031559446 | 0.139076433 | -0.170635879 | 0.999933887 | 0.069934674 | -0.061237246 | 0.13117192 | 0.670509697 |
| ENSG00000247271 | *ZBED5-AS1* | 0.325264198 | -0.004688212 | 0.039433239 | -0.044121452 | 0.999933887 | -0.016386372 | 0.143983995 | -0.160370367 | 0.670509697 |
| ENSG00000198825 | *INPP5F* | 1.894200082 | -0.025678448 | 0.009285757 | -0.034964205 | 0.999933887 | -0.0226416 | 0.081468803 | -0.104110403 | 0.670524981 |
| ENSG00000126062 | *TMEM115* | 4.240356562 | -0.099353188 | 0.011264954 | -0.110618143 | 0.999933887 | -0.20128386 | -0.130478731 | -0.070805129 | 0.670528597 |
| ENSG00000196422 | *PPP1R26* | -0.046200794 | -0.108660327 | 0.201753316 | -0.310413643 | 0.999933887 | -0.270680122 | -0.089526428 | -0.181153693 | 0.670528597 |
| ENSG00000182796 | *TMEM198B* | 1.404006961 | -0.051625827 | 0.279897317 | -0.331523144 | 0.944094537 | -0.109905279 | -0.001521744 | -0.108383536 | 0.670528597 |
| ENSG00000266598 | *AC037487.2* | -0.115837122 | 0.056456688 | 0.194159129 | -0.137702441 | 0.999933887 | 0.313777879 | 0.557688868 | -0.243910989 | 0.670528597 |
| ENSG00000133460 | *SLC2A11* | 0.274530363 | -0.005008499 | 0.174846296 | -0.179854795 | 0.999933887 | 0.058642381 | 0.234852887 | -0.176210506 | 0.670528597 |
| ENSG00000187522 | *HSPA14* | 1.953243899 | -0.060727796 | -0.124091725 | 0.063363929 | 0.999933887 | 0.041977624 | -0.072261089 | 0.114238713 | 0.670797675 |
| ENSG00000116903 | *EXOC8* | 5.236791014 | -0.083397333 | -0.072612041 | -0.010785292 | 0.999933887 | -0.168635703 | -0.104450152 | -0.064185551 | 0.670844631 |
| ENSG00000149600 | *COMMD7* | 1.67877977 | -0.237320804 | 0.023722318 | -0.261043122 | 0.999933887 | 0.087716396 | -0.024104941 | 0.111821338 | 0.670865676 |
| ENSG00000258944 | *AC004846.2* | -0.052512258 | -0.275125849 | 0.006005652 | -0.281131501 | 0.999933887 | 0.64078632 | 0.905301228 | -0.264514908 | 0.670865676 |
| ENSG00000131969 | *ABHD12B* | -0.012768413 | 0.089309932 | 0.146523667 | -0.057213735 | 0.999933887 | -0.448235662 | -0.19562166 | -0.252614002 | 0.670865676 |
| ENSG00000172943 | *PHF8* | 5.457816079 | -0.01214032 | -0.040087903 | 0.027947583 | 0.999933887 | -0.08385832 | -0.129782172 | 0.045923851 | 0.670865676 |
| ENSG00000160294 | *MCM3AP* | 4.870348877 | -0.012394884 | 0.003839773 | -0.016234657 | 0.999933887 | 0.047725684 | -0.009954242 | 0.057679926 | 0.670865676 |
| ENSG00000251364 | *AC107884.1* | -0.180277581 | -0.351665466 | -0.051551099 | -0.300114367 | 0.999933887 | -0.005495096 | 0.190099994 | -0.19559509 | 0.671068626 |
| ENSG00000166848 | *TERF2IP* | 6.487792592 | 0.008467743 | 0.040959878 | -0.032492135 | 0.999933887 | 0.649016651 | 0.716481539 | -0.067464887 | 0.671108401 |
| ENSG00000004939 | *SLC4A1* | -1.955560348 | 7.6348e-05 | -0.312159781 | 0.312236129 | 0.999933887 | -0.302173584 | 0.112726151 | -0.414899735 | 0.671108401 |
| ENSG00000134369 | *NAV1* | 1.426042803 | -0.144786518 | -0.268461356 | 0.123674838 | 0.999933887 | -0.386601773 | -0.494162271 | 0.107560498 | 0.671190602 |
| ENSG00000132749 | *TESMIN* | 0.554058564 | -0.062972623 | -0.028106584 | -0.034866039 | 0.999933887 | -0.500377893 | -0.648937885 | 0.148559993 | 0.671236011 |
| ENSG00000062598 | *ELMO2* | 4.829217399 | -0.030070482 | 0.00019851 | -0.030268992 | 0.999933887 | -0.052438299 | 0.008485445 | -0.060923744 | 0.671247242 |
| ENSG00000204338 | *CYP21A1P* | -0.81804969 | -0.093292696 | -0.121347019 | 0.028054323 | 0.999933887 | 0.532170591 | 0.793407806 | -0.261237215 | 0.671247242 |

| | | | | | | | | | | |
|---|---|---|---|---|---|---|---|---|---|---|
| ENSG00000117151 | *CTBS* | 4.364131344 | -0.020848729 | -0.069193069 | 0.04834434 | 0.999933887 | -0.199197046 | -0.124077801 | -0.075119245 | 0.671247242 |
| ENSG00000145491 | *ROPN1L* | 2.647269482 | 0.014516488 | 0.025351429 | -0.01083494 | 0.999933887 | 0.062307898 | -0.073020089 | 0.135327987 | 0.671247242 |
| ENSG00000103160 | *HSDL1* | 4.362702616 | -0.01302621 | -0.041463948 | 0.028437738 | 0.999933887 | -0.064206054 | -0.137235462 | 0.073029408 | 0.671736188 |
| ENSG00000104231 | *ZFAND1* | 1.62475608 | 0.018344831 | -0.051274503 | 0.069619334 | 0.999933887 | 0.131287526 | 0.270033203 | -0.138745676 | 0.671736188 |
| ENSG00000079134 | *THOC1* | 2.939327173 | -0.118546201 | -0.104077191 | -0.01446901 | 0.999933887 | -0.123690582 | 0.008444399 | -0.132134981 | 0.671985995 |
| ENSG00000144559 | *TAMM41* | 0.652153171 | -0.13392147 | 0.024407405 | -0.158328876 | 0.999933887 | 0.126662994 | 0.260429576 | -0.133766582 | 0.671985995 |
| ENSG00000114126 | *TFDP2* | 3.084549317 | 0.048087631 | 0.053883384 | -0.005795753 | 0.999933887 | 0.016550671 | -0.0677332 | 0.08428387 | 0.671985995 |
| ENSG00000255717 | *SNHG1* | 4.328437422 | -0.038668275 | 0.072470243 | -0.111138519 | 0.999933887 | 0.292782478 | 0.222717941 | 0.070064537 | 0.671985995 |
| ENSG00000198625 | *MDM4* | 7.489797552 | -0.029886018 | 0.015323035 | -0.045209052 | 0.999933887 | -0.027067715 | 0.031773454 | -0.058841169 | 0.671985995 |
| ENSG00000087086 | *FTL* | 12.612230121 | 0.02643692 | -0.028418916 | 0.054855836 | 0.999933887 | 0.326550223 | 0.269611424 | 0.0569388 | 0.671985995 |
| ENSG00000189238 | *LINC00943* | -0.183107547 | -0.07503171 | 0.06543889 | -0.1404706 | 0.999933887 | 0.049508081 | 0.261016451 | -0.21150837 | 0.671985995 |
| ENSG00000140395 | *WDR61* | 1.772069182 | -0.032131913 | -0.048553428 | 0.016421515 | 0.999933887 | 0.061142657 | -0.05527234 | 0.116414997 | 0.671985995 |
| ENSG00000100351 | *GRAP2* | 3.982261113 | -0.014685582 | 0.055399061 | -0.070084644 | 0.999933887 | 0.117157006 | 0.053938376 | 0.06321863 | 0.671985995 |
| ENSG00000127586 | *CHTF18* | 2.841255225 | 0.093912187 | 0.06800762 | 0.025904567 | 0.999933887 | 0.144724834 | 0.043135813 | 0.10158902 | 0.672028791 |
| ENSG00000158246 | *TENT5B* | -0.891476472 | -0.130270838 | -0.164831011 | 0.034560173 | 0.999933887 | -0.278499946 | -0.546107938 | 0.267607992 | 0.672028791 |
| ENSG00000211789 | *TRAV12-2* | 0.247815526 | 0.062055395 | -0.070112988 | 0.132168384 | 0.999933887 | -0.014828548 | 0.14894585 | -0.163774398 | 0.672028791 |
| ENSG00000085377 | *PREP* | 2.287368545 | 0.022046829 | 0.033080006 | -0.011033178 | 0.999933887 | 0.039946311 | 0.139374002 | -0.09942769 | 0.672248319 |
| ENSG00000181381 | *DDX60L* | 8.08183235 | -0.042094159 | -0.090490014 | 0.048395855 | 0.999933887 | 0.430052745 | 0.594778616 | -0.164725871 | 0.672253052 |
| ENSG00000286288 | *AL109809.4* | 3.914865338 | 0.023037438 | 0.050223966 | -0.027186528 | 0.999933887 | 0.014604235 | 0.090114503 | -0.075510267 | 0.672397217 |
| ENSG00000027697 | *IFNGR1* | 9.229828696 | -0.080333209 | -0.094234997 | 0.013901788 | 0.999933887 | -0.200137292 | -0.135709752 | -0.06442754 | 0.672459188 |
| ENSG00000130595 | *TNNT3* | 4.449020312 | 0.070447482 | 0.059275034 | 0.011172448 | 0.999933887 | 0.285485708 | 0.218869335 | 0.066616373 | 0.672459188 |
| ENSG00000100097 | *LGALS1* | 4.170834359 | 0.06317832 | 0.02176436 | 0.04141396 | 0.999933887 | 0.17459559 | 0.259725888 | -0.085130298 | 0.672459188 |
| ENSG00000204381 | *LAYN* | -2.069562678 | 0.352316412 | -0.453283758 | 0.80560017 | 0.999933887 | 0.250313771 | -0.079375105 | 0.329688876 | 0.672565626 |
| ENSG00000142089 | *IFITM3* | 9.021279981 | 0.057995914 | 0.000523297 | 0.057472617 | 0.999933887 | 0.037429394 | -0.023553208 | 0.060982601 | 0.672793712 |
| ENSG00000124596 | *OARD1* | 2.869455825 | -0.047301784 | -0.052598786 | 0.005297001 | 0.999933887 | -0.047653871 | 0.044606818 | -0.092260689 | 0.673239288 |
| ENSG00000149311 | *ATM* | 6.234715954 | 0.003275996 | -0.058863147 | 0.062139143 | 0.999933887 | -0.579600546 | -0.428248073 | -0.151352473 | 0.67328765 |
| ENSG00000171596 | *NMUR1* | 0.613257427 | 0.043834511 | 0.196190969 | -0.152356458 | 0.999933887 | 0.125302067 | -0.022206729 | 0.147508796 | 0.673698737 |
| ENSG00000061936 | *SFSWAP* | 5.512350329 | -0.03566034 | 0.046911001 | -0.08257134 | 0.999933887 | 0.034352846 | 0.073552648 | -0.039199802 | 0.673706665 |
| ENSG00000163481 | *RNF25* | 2.684368011 | -0.059416673 | -0.041406816 | -0.018009857 | 0.999933887 | 0.134662051 | 0.218690959 | -0.084028908 | 0.673805553 |
| ENSG00000275801 | *AL121985.3* | -1.580440243 | 0.200530843 | 1.056747707 | -0.856216864 | 0.999933887 | 0.45551564 | 0.645251165 | -0.189735525 | 0.673805553 |
| ENSG00000239377 | *ZNF775-AS1* | -1.870139901 | 0.450426007 | -0.758533783 | 1.20895979 | 0.999933887 | 2.327628917 | 2.081515409 | 0.246113508 | 0.67396893 |
| ENSG00000165752 | *STK32C* | 1.099284096 | 0.13929517 | 0.137106501 | 0.00218867 | 0.999933887 | -0.381073749 | -0.529442927 | 0.148369177 | 0.67396893 |
| ENSG00000185418 | *TARS3* | 1.951503267 | -0.016061738 | 0.036455729 | -0.052517467 | 0.999933887 | -0.009029732 | 0.083545117 | -0.092574849 | 0.67396893 |
| ENSG00000251179 | *TMEM92-AS1* | 1.319058371 | -0.067850767 | -0.106146523 | 0.038295757 | 0.999933887 | -0.609351377 | -0.773165419 | 0.163814042 | 0.674396067 |
| ENSG00000213339 | *QTRT1* | 2.164680971 | 0.011827695 | -0.021731233 | 0.033558929 | 0.999933887 | 0.165551382 | 0.255649433 | -0.090098051 | 0.674396067 |
| ENSG00000181218 | *H2AW* | 0.631917305 | 0.076875775 | 0.253849593 | -0.176973818 | 0.999933887 | 0.092570111 | -0.06725346 | 0.159823571 | 0.674595269 |
| ENSG00000102145 | *GATA1* | 1.110895202 | 0.071846472 | 0.145463425 | -0.073616953 | 0.999933887 | 0.108331857 | 0.000741333 | 0.107590524 | 0.674595269 |
| ENSG00000184675 | *AMER1* | 0.209530803 | 0.041839609 | -0.27062842 | 0.312468029 | 0.999933887 | 0.104480052 | -0.062157122 | 0.166637174 | 0.674595269 |
| ENSG00000279434 | *AL049776.1* | 0.711790735 | -0.028439586 | 0.011087494 | -0.03952708 | 0.999933887 | 0.045610231 | 0.187577644 | -0.141967413 | 0.674595269 |
| ENSG00000128191 | *DGCR8* | 3.535189767 | 0.012637493 | 0.030281556 | -0.017644063 | 0.999933887 | -0.043369298 | -0.1215438 | 0.078174502 | 0.674595269 |
| ENSG00000255438 | *AL354813.1* | 0.131830335 | -0.026927543 | -0.054498088 | 0.027570545 | 0.999933887 | -1.555222052 | -1.865528768 | 0.310306716 | 0.674595269 |
| ENSG00000253200 | *AC037459.2* | 2.294131581 | -0.004124434 | 0.13882374 | -0.142948174 | 0.999933887 | 0.195757415 | 0.323245966 | -0.127488551 | 0.674595269 |
| ENSG00000256582 | *LINC02390* | -0.302661225 | -0.053011969 | 0.091247164 | -0.144259133 | 0.999933887 | -0.043328254 | 0.184971276 | -0.22829953 | 0.674637544 |
| ENSG00000254419 | *AL139349.1* | -0.030543122 | 0.067655059 | -0.067422706 | 0.135077765 | 0.999933887 | -0.052534234 | -0.273318753 | 0.22078452 | 0.674646554 |
| ENSG00000263941 | *RN7SL32P* | -0.88429156 | 0.345463027 | -0.163183393 | 0.50864642 | 0.999933887 | 0.326148092 | -0.052830678 | 0.378978769 | 0.674679182 |
| ENSG00000124191 | *TOX2* | 0.330961426 | 0.130528574 | 0.142034218 | -0.011505644 | 0.999933887 | -0.058878578 | -0.262074955 | 0.203196377 | 0.674679182 |
| ENSG00000198182 | *ZNF607* | -0.702427141 | 0.020914774 | 0.009144414 | 0.01177036 | 0.999933887 | 0.081423396 | -0.096864937 | 0.178288332 | 0.674679182 |
| ENSG00000244682 | *FCGR2C* | 4.405372092 | 0.004441068 | -0.186111834 | 0.190552902 | 0.999933887 | -0.001637278 | 0.143629209 | -0.145266487 | 0.674679182 |
| ENSG00000119777 | *TMEM214* | 3.591783632 | 0.114325625 | 0.059892 | 0.054433625 | 0.999933887 | -0.101406769 | -0.181391371 | 0.079984603 | 0.675385419 |
| ENSG00000125868 | *DSTN* | 3.533208005 | -0.016781804 | -0.098221547 | 0.081439742 | 0.999933887 | -0.045347112 | 0.035844895 | -0.081192007 | 0.675385419 |
| ENSG00000115514 | *TXNDC9* | 1.600815348 | -0.013785756 | 0.045790349 | -0.059576105 | 0.999933887 | -0.482665585 | -0.595389156 | 0.112723571 | 0.675385419 |
| ENSG00000182154 | *MRPL41* | 2.334944301 | 0.165404034 | 0.078535151 | 0.086868883 | 0.999933887 | 0.050999341 | -0.053615282 | 0.104614623 | 0.675414204 |
| ENSG00000139531 | *SUOX* | 1.40275835 | -0.10022091 | 0.028775044 | -0.128995954 | 0.999933887 | -0.056720765 | -0.192831386 | 0.13611062 | 0.675414204 |
| ENSG00000046647 | *GEMIN8* | 1.326886994 | -0.025210582 | 0.048488309 | -0.073698891 | 0.999933887 | -0.100625971 | 0.01310393 | -0.113729901 | 0.675414204 |
| ENSG00000235677 | *NPM1P26* | 1.00179452 | 0.088609036 | 0.027517757 | 0.061091279 | 0.999933887 | 0.442881504 | 0.275980657 | 0.166900847 | 0.675624471 |
| ENSG00000264659 | *AC064805.2* | -0.038945765 | -0.0093429 | -0.27999332 | 0.27065042 | 0.999933887 | -0.403773285 | -0.667582262 | 0.263808977 | 0.675624471 |
| ENSG00000231826 | *LINC01819* | -0.827918471 | 0.086881039 | -0.050489452 | 0.137370491 | 0.999933887 | -0.49291026 | -0.212405892 | -0.280504368 | 0.675816724 |
| ENSG00000211803 | *TRAV23DV6* | -0.934357943 | -0.158231109 | -0.157389822 | -0.000841287 | 0.999933887 | -0.173388691 | 0.09739505 | -0.270783741 | 0.675909668 |
| ENSG00000204428 | *LY6G5C* | 0.348699347 | -0.068808908 | 0.201081949 | -0.269890857 | 0.999933887 | -0.377290479 | -0.175208837 | -0.202081642 | 0.675979601 |
| ENSG00000213442 | *RPL18AP3* | -0.232185307 | 0.30215299 | -0.253684596 | 0.555837586 | 0.999933887 | 0.2762881 | 0.051345927 | 0.224942174 | 0.676107316 |
| ENSG00000108384 | *RAD51C* | 0.072629447 | -0.116010943 | -0.098848096 | -0.017162847 | 0.999933887 | 0.124639008 | 0.297488329 | -0.172849321 | 0.676107316 |
| ENSG00000099992 | *TBC1D10A* | 3.271980695 | 0.15922126 | 0.029051889 | 0.130169371 | 0.999933887 | 0.639997096 | 0.727714241 | -0.087717145 | 0.676130112 |
| ENSG00000101363 | *MANBAL* | 2.748431924 | -0.093377637 | -0.07651201 | -0.016865627 | 0.999933887 | 0.055808336 | 0.141003471 | -0.085195135 | 0.676130112 |
| ENSG00000106624 | *AEBP1* | -0.060203262 | 0.132549641 | 0.389616682 | -0.257067041 | 0.999933887 | 0.166328647 | 0.332930017 | -0.16660137 | 0.676130112 |
| ENSG00000108175 | *ZMIZ1* | 9.328852869 | 0.030755714 | 0.072793648 | -0.042037935 | 0.999933887 | 0.005774131 | 0.059440905 | -0.053666774 | 0.676130112 |
| ENSG00000178202 | *POGLUT3* | 0.660937405 | 0.04126809 | -0.124393201 | 0.16566129 | 0.999933887 | -0.076116083 | 0.070709723 | -0.146825805 | 0.676130112 |
| ENSG00000180096 | *SEPTIN1* | 4.316765123 | 0.014659837 | 0.043222632 | -0.028562795 | 0.999933887 | 0.149781985 | 0.073099169 | 0.076682816 | 0.676130112 |
| ENSG00000165178 | *NCF1C* | 7.182877581 | 0.010324008 | -0.008649264 | 0.018973273 | 0.999933887 | 0.907832334 | 0.825956524 | 0.08187581 | 0.676130112 |
| ENSG00000172409 | *CLP1* | 3.677788262 | -0.164451475 | -0.067389039 | -0.097062436 | 0.999933887 | -0.436222872 | -0.51168554 | 0.075462669 | 0.676187625 |
| ENSG00000079332 | *SAR1A* | 6.362694495 | -0.047283448 | -0.080488368 | 0.03320492 | 0.999933887 | 0.059362688 | 0.100430204 | -0.041067515 | 0.676296849 |
| ENSG00000082213 | *C5orf22* | 1.615461606 | -0.204259769 | 0.205779308 | -0.410039077 | 0.944094537 | 0.017448788 | -0.097618793 | 0.115067581 | 0.677133433 |
| ENSG00000176182 | *MYPOP* | 1.509281139 | 0.039353149 | 0.039740647 | -0.000387497 | 0.999933887 | 0.032361849 | 0.147656517 | -0.115294668 | 0.677133433 |
| ENSG00000218891 | *ZNF579* | 0.585907218 | -0.019193526 | 0.262181913 | -0.281375439 | 0.999933887 | -0.136286419 | -0.310840208 | 0.174553789 | 0.677133433 |
| ENSG00000116138 | *DNAJC16* | 2.122429039 | 0.002858175 | -0.115625798 | 0.118483973 | 0.999933887 | -0.009604981 | 0.094975487 | -0.104580468 | 0.677133433 |
| ENSG00000234426 | *AL139042.1* | -2.687131414 | 0.076596148 | -0.044172878 | 0.120769027 | 0.999933887 | 0.135495791 | -0.083467569 | 0.21896336 | 0.677163557 |
| ENSG00000134987 | *WDR36* | 2.499721811 | -0.15026911 | -0.327908314 | 0.177639204 | 0.999933887 | 0.050477063 | 0.169008578 | -0.118531515 | 0.677350633 |
| ENSG00000133706 | *LARS1* | 3.49606069 | -0.098892706 | -0.033706441 | -0.065186265 | 0.999933887 | 0.071355876 | 0.139958091 | -0.068602215 | 0.677350633 |
| ENSG00000286327 | *AL035427.2* | 0.361649902 | 0.349534649 | 0.101113083 | 0.248421566 | 0.999933887 | 0.05401994 | 0.206813126 | -0.152793186 | 0.677350633 |
| ENSG00000135905 | *DOCK10* | 5.114686868 | -0.076881466 | -0.081065638 | 0.004184171 | 0.999933887 | -0.035677599 | 0.070616983 | -0.106294582 | 0.677350633 |
| ENSG00000103121 | *CMC2* | 1.703650133 | -0.0538652 | 0.083371295 | -0.137236495 | 0.999933887 | -0.154619312 | -0.033374535 | -0.121244777 | 0.677350633 |
| ENSG00000280734 | *LINC01232* | 1.464351377 | 0.055034492 | -0.126788777 | 0.181823269 | 0.999933887 | -0.204652101 | -0.074801034 | -0.129851068 | 0.677350633 |
| ENSG00000100304 | *TTLL12* | 3.457512326 | 0.007136531 | 0.023693254 | -0.016556723 | 0.999933887 | 0.083265355 | 0.150799488 | -0.067534133 | 0.677350633 |
| ENSG00000160460 | *SPTBN4* | 0.845025803 | 0.147074122 | 0.201282895 | -0.054208773 | 0.999933887 | 0.084605441 | 0.202917639 | -0.118312198 | 0.677546374 |
| ENSG00000131470 | *PSMC3IP* | 1.105875667 | -0.254576374 | -0.13374942 | -0.120826954 | 0.999933887 | -0.630102023 | -0.799270515 | 0.169168492 | 0.677630807 |
| ENSG00000197714 | *ZNF460* | 2.400783937 | 0.168209766 | 0.050548827 | 0.117660938 | 0.999933887 | -0.120004446 | 0.01984138 | -0.139845827 | 0.677727287 |
| ENSG00000230795 | *HLA-K* | 0.302222598 | 0.069115687 | -0.412994099 | 0.482109787 | 0.999933887 | 0.674309343 | 0.482972059 | 0.191337284 | 0.677727287 |
| ENSG00000140750 | *ARHGAP17* | 4.15064343 | -0.003769354 | -0.005359572 | 0.001590217 | 0.999933887 | -0.05492244 | -0.002642981 | -0.052279458 | 0.677727287 |
| ENSG00000116918 | *TSNAX* | 2.975927851 | -0.05706163 | 0.01595252 | -0.07301415 | 0.999933887 | -0.377412563 | -0.498280576 | 0.120868013 | 0.677812555 |
| ENSG00000105135 | *ILVBL* | 2.155760934 | 0.034412541 | 0.089497712 | -0.055085171 | 0.999933887 | 0.139026168 | 0.248904491 | -0.109878322 | 0.677812555 |
| ENSG00000162946 | *DISC1* | 2.548202411 | -0.014854091 | 0.00184674 | -0.016700831 | 0.999933887 | 0.191354565 | 0.312864897 | -0.121510332 | 0.677812555 |
| ENSG00000006453 | *BAIAP2L1* | 3.527818649 | 0.07320749 | 0.176661731 | -0.103454241 | 0.999933887 | 0.206993595 | 0.297286611 | -0.090293015 | 0.678371616 |
| ENSG00000170037 | *CNTROB* | 3.528089369 | 0.026829795 | -0.008455546 | 0.035285341 | 0.999933887 | 0.234780036 | 0.163268933 | 0.071511103 | 0.678371616 |
| ENSG00000232533 | *AC093673.1* | 3.575811979 | 0.033451925 | 0.063492177 | -0.030040252 | 0.999933887 | -0.331733138 | -0.432013049 | 0.100279911 | 0.678371616 |
| ENSG00000136699 | *SMPD4* | 4.058514846 | 0.001410795 | 0.028997183 | -0.027586388 | 0.999933887 | 0.135676755 | 0.071029947 | 0.064646808 | 0.678486244 |
| ENSG00000118007 | *STAG1* | 4.374043054 | 0.027017093 | -0.102464803 | 0.129481896 | 0.999933887 | -0.310305075 | -0.405718025 | 0.09541295 | 0.678646799 |
| ENSG00000150967 | *ABCB9* | 0.364062672 | 0.056894143 | 0.062115397 | -0.005221254 | 0.999933887 | -0.373420142 | -0.513303254 | 0.139883112 | 0.678646799 |
| ENSG00000175348 | *TMEM9B* | 4.438839497 | 0.00750074 | 0.02662199 | -0.01912125 | 0.999933887 | -0.117624447 | -0.057110168 | -0.060514279 | 0.678646799 |
| ENSG00000164815 | *ORC5* | 0.526841096 | -0.002860885 | -0.062512012 | 0.059651127 | 0.999933887 | 0.040738302 | 0.186840132 | -0.14610183 | 0.678686162 |
| ENSG00000164241 | *C5orf63* | -0.706615272 | 0.057370294 | -0.133590699 | 0.190960993 | 0.999933887 | -0.133031227 | 0.077824848 | -0.210856076 | 0.678748637 |
| ENSG00000278607 | *AC015819.2* | 4.645783137 | 0.315585135 | 0.37622564 | -0.060640504 | 0.999933887 | 0.19606878 | 0.294152947 | -0.098084167 | 0.679058539 |
| ENSG00000177565 | *TBL1XR1* | 6.119895621 | -0.068902945 | -0.032483299 | -0.036419646 | 0.999933887 | -0.451426624 | -0.526629076 | 0.075202452 | 0.679058539 |
| ENSG00000272447 | *AL135925.1* | 0.549237917 | 0.135070634 | -0.05557244 | 0.190643074 | 0.999933887 | -0.08485285 | -0.260245938 | 0.175393088 | 0.67908631 |
| ENSG00000273017 | *AP000240.1* | 0.897040017 | 0.058450418 | -0.00360835 | 0.062058768 | 0.999933887 | -0.159239624 | -0.025427519 | -0.133812105 | 0.67934924 |
| ENSG00000139438 | *FAM222A* | 0.02103104 | 0.083090782 | 0.034526045 | 0.048564737 | 0.999933887 | 0.065236552 | -0.176497531 | 0.241734082 | 0.679352723 |
| ENSG00000254288 | *AC087672.2* | -0.46712427 | 0.082711732 | -0.282391795 | 0.365103527 | 0.999933887 | -0.253309582 | -0.549663502 | 0.29635392 | 0.679352723 |

| | | | | | | | | | | |
|---|---|---|---|---|---|---|---|---|---|---|
| ENSG00000203721 | *LINC00862* | 1.421394173 | -0.053712947 | -0.189802647 | 0.1360897 | 0.999933887 | 0.165413576 | 0.282614253 | -0.117200677 | 0.679352723 |
| ENSG00000108187 | *PBLD* | 3.245646819 | 0.011510466 | -0.119828819 | 0.131339285 | 0.999933887 | -0.339759354 | -0.232261162 | -0.107498192 | 0.679352723 |
| ENSG00000090013 | *BLVRB* | 3.558244158 | 0.120988502 | 0.039632292 | 0.081356209 | 0.999933887 | -0.243571065 | -0.331073109 | 0.087502044 | 0.67977545 |
| ENSG00000197782 | *ZNF780A* | 1.871134375 | -0.116484197 | 0.066003396 | -0.182487594 | 0.999933887 | -0.250272448 | -0.138046904 | -0.112225544 | 0.67977545 |
| ENSG00000162241 | *SLC25A45* | 3.73131849 | -0.05252297 | 0.011925903 | -0.064448873 | 0.999933887 | -0.074536085 | -0.140448663 | 0.065912578 | 0.67977545 |
| ENSG00000235314 | *LINC00957* | 0.283519638 | 0.123768437 | -0.061749185 | 0.185517622 | 0.999933887 | -0.376048051 | -0.571370841 | 0.195322791 | 0.67977545 |
| ENSG00000166503 | *HDGFL3* | 3.467512697 | -0.079914223 | -0.128912773 | 0.04899855 | 0.999933887 | -0.611182688 | -0.733037098 | 0.12185441 | 0.67977545 |
| ENSG00000103260 | *METRN* | 0.502938002 | 0.079446313 | 0.055025361 | 0.024420952 | 0.999933887 | -0.050047375 | 0.103893294 | -0.153940669 | 0.67977545 |
| ENSG00000141127 | *PRPSAP2* | 2.202237872 | 0.049164927 | 0.004780248 | 0.044384679 | 0.999933887 | 0.047906033 | -0.062392254 | 0.110298287 | 0.67977545 |
| ENSG00000126214 | *KLC1* | 4.958708813 | 0.017472701 | 0.051347563 | -0.033874862 | 0.999933887 | -0.20164352 | -0.252316629 | 0.050673109 | 0.67977545 |
| ENSG00000086598 | *TMED2* | 5.47752161 | -0.017564244 | -0.032586983 | 0.015022739 | 0.999933887 | -0.233485807 | -0.300159257 | 0.06667345 | 0.67977545 |
| ENSG00000269506 | *AC110792.2* | -0.045020713 | 0.05037274 | 0.11375126 | -0.06337852 | 0.999933887 | -0.068804324 | 0.216250352 | -0.285054676 | 0.67977545 |
| ENSG00000157870 | *PRXL2B* | 5.215221037 | 0.030479245 | 0.010240325 | 0.02023892 | 0.999933887 | -0.291096281 | -0.190931647 | -0.100164634 | 0.67977545 |
| ENSG00000136861 | *CDK5RAP2* | 5.393006898 | -0.006498205 | -0.08730993 | 0.080811725 | 0.999933887 | 0.204717933 | 0.270736123 | -0.066018191 | 0.67977545 |
| ENSG00000102753 | *KPNA3* | 4.077217279 | 0.01717037 | -0.001087046 | 0.018257416 | 0.999933887 | -0.460644579 | -0.377608259 | -0.08303632 | 0.679916907 |
| ENSG00000140092 | *FBLN5* | -0.492996743 | 0.021642544 | 0.325828014 | -0.30418547 | 0.999933887 | 0.078567954 | -0.120131031 | 0.198698985 | 0.679916907 |
| ENSG00000164253 | *WDR41* | 2.437395899 | -0.038780895 | -0.014550267 | -0.024230628 | 0.999933887 | -0.182053404 | -0.270255276 | 0.088201872 | 0.679977744 |
| ENSG00000198851 | *CD3E* | 6.26700514 | -0.019996389 | 0.001949391 | -0.02194578 | 0.999933887 | 0.094422276 | 0.147670862 | -0.053248586 | 0.680027391 |
| ENSG00000247315 | *ZCCHC3* | 3.722974124 | 0.104007463 | 0.01558997 | 0.088417492 | 0.999933887 | -0.3034831 | -0.376919237 | 0.073436137 | 0.680150505 |
| ENSG00000065060 | *UHRF1BP1* | 3.062824012 | -0.0364672 | -0.031546845 | -0.004920355 | 0.999933887 | -0.030988186 | 0.05795785 | -0.088946036 | 0.680150505 |
| ENSG00000162650 | *ATXN7L2* | 2.41946232 | 0.013901891 | 0.098676148 | -0.084774256 | 0.999933887 | 0.063393833 | -0.021657136 | 0.085050968 | 0.680193581 |
| ENSG00000076321 | *KLHL20* | 2.074448266 | -0.042461683 | 0.028507352 | -0.070969035 | 0.999933887 | -0.059275307 | 0.032893108 | -0.092168416 | 0.680513084 |
| ENSG00000173275 | *ZNF449* | 0.691989547 | 0.000465037 | -0.042038719 | 0.042503756 | 0.999933887 | -0.160920717 | -0.041956123 | -0.118964594 | 0.680513084 |
| ENSG00000201988 | *Y_RNA* | -0.755292338 | 0.22233291 | 0.154860569 | 0.067472341 | 0.999933887 | 1.199084521 | 1.416498488 | -0.217413967 | 0.680579633 |
| ENSG00000166260 | *COX11* | 1.841347819 | -0.032618869 | -0.062400956 | 0.029782087 | 0.999933887 | -0.034665709 | 0.071132207 | -0.105797916 | 0.680579633 |
| ENSG00000101104 | *PABPC1L* | 4.326343934 | -0.033999318 | 0.005575598 | -0.039574916 | 0.999933887 | 0.027917533 | 0.082331161 | -0.054413628 | 0.680643223 |
| ENSG00000203485 | *INF2* | 2.60814538 | 0.041478698 | 0.231288755 | -0.189810057 | 0.999933887 | 0.166777994 | 0.255984816 | -0.089206822 | 0.680643223 |
| ENSG00000141736 | *ERBB2* | 0.773274919 | 0.217877349 | 0.202322556 | 0.015554793 | 0.999933887 | 0.09601023 | 0.233477278 | -0.137467048 | 0.680897978 |
| ENSG00000173295 | *FAM86B3P* | 4.189872013 | 0.063963934 | 0.105174808 | -0.041210874 | 0.999933887 | -0.09222237 | -0.164396104 | 0.072173734 | 0.680945984 |
| ENSG00000272533 | *SNORA28* | -0.327209902 | 0.147705714 | 0.50015056 | -0.352444846 | 0.999933887 | 0.469885943 | 0.687126264 | -0.217240321 | 0.680945984 |
| ENSG00000225873 | *C3orf86* | 2.06118523 | 0.045029331 | 0.040195106 | 0.004834224 | 0.999933887 | 0.073346028 | -0.109308499 | 0.182654527 | 0.680945984 |
| ENSG00000137449 | *CPEB2* | 5.557477881 | 0.069363331 | 0.036893723 | 0.032469608 | 0.999933887 | -0.750901619 | -0.828047987 | 0.077146368 | 0.681209137 |
| ENSG00000263272 | *AC004148.1* | 0.490930795 | 0.182606919 | -0.12225018 | 0.304857099 | 0.999933887 | 0.076855298 | -0.074344802 | 0.1512001 | 0.681432495 |
| ENSG00000160256 | *FAM207A* | 2.701863908 | 0.073061811 | 0.151025789 | -0.077963979 | 0.999933887 | -0.080142404 | -0.200149327 | 0.120006924 | 0.681499305 |
| ENSG00000260528 | *FAM157C* | 4.109456675 | 0.081769902 | -0.006129666 | 0.087899568 | 0.999933887 | 0.414350923 | 0.498404553 | -0.08405363 | 0.681616608 |
| ENSG00000064787 | *BCAS1* | 0.223549227 | -0.044168884 | 0.143983592 | -0.188152476 | 0.999933887 | -0.13962004 | 0.050519492 | -0.190139531 | 0.681616608 |
| ENSG00000158092 | *NCK1* | 3.307350122 | -0.119307047 | -0.101525029 | -0.017782017 | 0.999933887 | -0.234210507 | -0.311025967 | 0.07681546 | 0.681990499 |
| ENSG00000270820 | *AC016727.1* | 1.055003317 | 0.024413396 | 0.084818521 | -0.060405125 | 0.999933887 | -0.048570707 | 0.077942356 | -0.126513064 | 0.681990499 |
| ENSG00000199695 | *RNU6-1128P* | 0.060031795 | 0.007432194 | 0.007874807 | -0.000442614 | 0.999933887 | 0.74430531 | 1.036473734 | -0.292168424 | 0.682067745 |
| ENSG00000279430 | *AL590560.4* | 1.656383699 | 0.079587429 | -0.005124745 | 0.084712174 | 0.999933887 | 0.212555304 | 0.401776213 | -0.189220909 | 0.682107742 |
| ENSG00000124214 | *STAU1* | 6.130429679 | -0.056312876 | -0.034518153 | -0.021794723 | 0.999933887 | -0.395520237 | -0.448049297 | 0.05252906 | 0.682381691 |
| ENSG00000151247 | *EIF4E* | 3.453802574 | 0.048773969 | 0.070607123 | -0.021833155 | 0.999933887 | 0.063113442 | 0.145764304 | -0.082650863 | 0.682381691 |
| ENSG00000120725 | *SIL1* | 1.098790399 | -0.123624409 | -0.050398995 | -0.073225414 | 0.999933887 | 0.139177521 | -0.035751025 | 0.174928546 | 0.682433034 |
| ENSG00000241112 | *RPL29P14* | -0.283266934 | 0.116539217 | -0.117883879 | 0.234423096 | 0.999933887 | 0.515702328 | 0.320927233 | 0.194775095 | 0.682511211 |
| ENSG00000171444 | *MCC* | 1.8220693 | -0.050748285 | -0.191890358 | 0.141142073 | 0.999933887 | -0.216486107 | -0.103088554 | -0.113397553 | 0.682511211 |
| ENSG00000249662 | *LINC02218* | 1.171524362 | -0.022783434 | -0.018794235 | -0.003989199 | 0.999933887 | 0.095290971 | -0.035579397 | 0.130870368 | 0.682511211 |
| ENSG00000100207 | *TCF20* | 6.851922539 | -0.039723358 | 0.069315364 | -0.109038722 | 0.999933887 | -0.297946182 | -0.35445671 | 0.056510528 | 0.682694126 |
| ENSG00000143294 | *PRCC* | 5.168932857 | -0.014010246 | -0.041012513 | 0.027002267 | 0.999933887 | 0.119075332 | 0.167431989 | -0.048356656 | 0.68281505 |
| ENSG00000111605 | *CPSF6* | 4.635659424 | -0.002639449 | -0.052551462 | 0.049912013 | 0.999933887 | -0.26701735 | -0.333041554 | 0.066024205 | 0.68281505 |
| ENSG00000237840 | *FAM21FP* | 1.365805944 | 0.260307294 | 0.136687133 | 0.123620161 | 0.999933887 | 0.623022684 | 0.490143985 | 0.132878699 | 0.682939866 |
| ENSG00000169814 | *BTD* | 3.588960103 | -0.053334424 | -0.034329616 | -0.019004808 | 0.999933887 | -0.349512037 | -0.2693321 | -0.080179937 | 0.683158785 |
| ENSG00000283849 | *AC092053.2* | 3.116261645 | 0.447643884 | 0.361078458 | 0.086565426 | 0.999933887 | 0.986963432 | 1.157578576 | -0.170615144 | 0.683214994 |
| ENSG00000227191 | *TRGC2* | 3.164865977 | -0.118707729 | -0.114152194 | -0.004555535 | 0.999933887 | 0.087168493 | 0.169102144 | -0.081933651 | 0.683214994 |
| ENSG00000287032 | *AC018647.3* | -0.370707902 | 0.151981654 | 0.179275274 | -0.02729362 | 0.999933887 | -0.282256069 | 0.040734477 | -0.322990546 | 0.683309705 |
| ENSG00000176896 | *TCEANC* | 1.65323815 | -0.054297839 | -0.099078916 | 0.044781076 | 0.999933887 | 0.146052106 | 0.268067491 | -0.122015385 | 0.683309705 |
| ENSG00000118412 | *CASP8AP2* | 2.321629835 | -0.057798552 | 0.006977934 | -0.064776487 | 0.999933887 | -0.515831281 | -0.334466576 | -0.181364705 | 0.683309705 |
| ENSG00000172269 | *DPAGT1* | 1.396958765 | 0.024446309 | -0.361112232 | 0.385558541 | 0.944094537 | 0.054457092 | -0.053855037 | 0.108312129 | 0.683309705 |
| ENSG00000148057 | *IDNK* | 1.308758387 | -0.009588581 | -0.311366203 | 0.301777623 | 0.999933887 | 0.181029665 | 0.293470065 | -0.1124404 | 0.683309705 |
| ENSG00000178342 | *KCNG2* | -0.082447352 | 0.046300214 | 0.203683746 | -0.157383532 | 0.999933887 | 0.099996923 | 0.384671152 | -0.284674229 | 0.683407282 |
| ENSG00000186352 | *ANKRD37* | 1.58632922 | 0.092847263 | 0.246909871 | -0.154062607 | 0.999933887 | 0.112240451 | 0.261184971 | -0.14894452 | 0.683892654 |
| ENSG00000265871 | *MIR3174* | -0.367666068 | -0.137775615 | 0.139688821 | -0.277464436 | 0.999933887 | 0.139433977 | 0.312122786 | -0.172688809 | 0.683892654 |
| ENSG00000175274 | *TP53I11* | 6.379124849 | -0.022447415 | 0.034624006 | -0.057071421 | 0.999933887 | -0.712776163 | -0.793735759 | 0.080959595 | 0.683952946 |
| ENSG00000189134 | *NKAPL* | -0.033017174 | 0.147147129 | -0.14278394 | 0.289931069 | 0.999933887 | -0.367447833 | -0.207503487 | -0.159944346 | 0.684943293 |
| ENSG00000161642 | *ZNF385A* | 4.46128849 | -0.017275016 | -0.027314794 | 0.010039778 | 0.999933887 | -1.178904049 | -1.271088303 | 0.092184254 | 0.684943293 |
| ENSG00000131069 | *ACSS2* | 3.831384366 | 0.011333483 | -0.00058633 | 0.011919813 | 0.999933887 | -0.562045356 | -0.65458726 | 0.092541904 | 0.685227502 |
| ENSG00000264548 | *AC132872.2* | 0.936793032 | 0.209250381 | 0.040207592 | 0.169042789 | 0.999933887 | 1.124506358 | 1.336603754 | -0.212097395 | 0.68528968 |
| ENSG00000186532 | *SMYD4* | 1.626845094 | -0.059198514 | 0.101671965 | -0.160870479 | 0.999933887 | 0.080756614 | -0.019995668 | 0.100752281 | 0.68528968 |
| ENSG00000089063 | *TMEM230* | 4.614088016 | 0.019167178 | -0.031394006 | 0.050561184 | 0.999933887 | -0.275400884 | -0.332280381 | 0.056879497 | 0.685592485 |
| ENSG00000184922 | *FMNL1* | 9.529647839 | 0.006149193 | 0.031226493 | -0.0250773 | 0.999933887 | -0.277023902 | -0.323712624 | 0.046688722 | 0.685679556 |
| ENSG00000119509 | *INVS* | 1.899375898 | 0.048399005 | -0.079737657 | 0.128136663 | 0.999933887 | -0.123196375 | -0.021330106 | -0.101866269 | 0.685817175 |
| ENSG00000178966 | *RMI1* | 1.155775244 | 0.057012769 | -0.045631507 | 0.102644277 | 0.999933887 | -0.819488148 | -0.953240533 | 0.133752384 | 0.685862631 |
| ENSG00000104219 | *ZDHHC2* | 3.594458821 | 0.090575335 | -0.164717206 | 0.255292541 | 0.594252256 | -0.321076888 | -0.395051399 | 0.07397451 | 0.686009881 |
| ENSG00000138190 | *EXOC6* | 2.962799698 | -0.080909804 | -0.043217812 | -0.037691992 | 0.999933887 | -0.470970927 | -0.373907166 | -0.097063761 | 0.686009881 |
| ENSG00000233954 | *UQCRHL* | 0.192418932 | 0.069368333 | -0.063829917 | 0.133198249 | 0.999933887 | -0.049043633 | -0.220118725 | 0.171075092 | 0.686150482 |
| ENSG00000278053 | *DDX52* | 2.587926835 | -0.067341526 | -0.065039215 | -0.002302311 | 0.999933887 | 0.060359123 | -0.023812989 | 0.084172112 | 0.686178796 |
| ENSG00000117410 | *ATP6V0B* | 9.212110946 | 0.018720565 | -0.008419018 | 0.027139583 | 0.999933887 | -0.073655273 | -0.122870626 | 0.049215353 | 0.686183138 |
| ENSG00000188243 | *COMMD6* | 3.927147935 | -0.013295741 | -0.005110651 | -0.00818509 | 0.999933887 | 0.030310113 | -0.037903626 | 0.068213739 | 0.686505991 |
| ENSG00000225972 | *MTND1P23* | 4.243129426 | 0.080581002 | 0.212749145 | -0.132168143 | 0.999933887 | 0.016133897 | 0.162951126 | -0.146817228 | 0.686802703 |
| ENSG00000092847 | *AGO1* | 5.925348281 | -0.043921875 | 0.014435327 | -0.058357203 | 0.999933887 | -0.239745927 | -0.319547487 | 0.07980156 | 0.686802703 |
| ENSG00000270127 | *AC027020.2* | -0.595531985 | 0.043010105 | 0.003244476 | 0.039765629 | 0.999933887 | -0.101294891 | 0.121878166 | -0.223173058 | 0.686802703 |
| ENSG00000083544 | *TDRD3* | 1.255452484 | 0.016863506 | -0.043343869 | 0.060207375 | 0.999933887 | -0.010854244 | 0.12932322 | -0.140177464 | 0.686802703 |
| ENSG00000135363 | *LMO2* | 6.563250644 | -0.093576186 | -0.14508738 | 0.051511194 | 0.999933887 | -0.468298086 | -0.395643752 | -0.072654334 | 0.686819895 |
| ENSG00000164916 | *FOXK1* | 4.630952914 | 0.004022826 | 0.082513327 | -0.078490501 | 0.999933887 | 0.177035507 | 0.238940388 | -0.061904881 | 0.687016877 |
| ENSG00000129521 | *EGLN3* | -0.729742969 | 0.007701096 | 0.285572698 | -0.277871601 | 0.999933887 | 0.100936958 | -0.113668373 | 0.214605331 | 0.68708419 |
| ENSG00000146247 | *PHIP* | 6.153176602 | -0.094003999 | -0.198436577 | 0.104432578 | 0.999933887 | -0.746447888 | -0.54518566 | -0.201262228 | 0.687193209 |
| ENSG00000170315 | *UBB* | 6.358999558 | -0.031496784 | -0.082078744 | 0.05058196 | 0.999933887 | 0.348859083 | 0.408939937 | -0.060080854 | 0.687339205 |
| ENSG00000185158 | *LRRC37B* | 4.246636988 | -0.024709308 | 0.056894702 | -0.081604011 | 0.999933887 | -0.377222306 | -0.42209593 | 0.044873624 | 0.687389849 |
| ENSG00000100744 | *GSKIP* | 2.994731192 | -0.032345646 | -0.122759308 | 0.090413662 | 0.999933887 | -0.490150641 | -0.600157996 | 0.110007355 | 0.687389849 |
| ENSG00000186834 | *HEXIM1* | 5.679226905 | 0.088779113 | 0.104378815 | -0.015599702 | 0.999933887 | -0.123963172 | -0.187896516 | 0.063933343 | 0.687513419 |
| ENSG00000184216 | *IRAK1* | 8.321991587 | 0.079326632 | 0.103376648 | -0.024050016 | 0.999933887 | 0.456252229 | 0.527063215 | -0.070810985 | 0.687513419 |
| ENSG00000221968 | *FADS3* | 2.324732462 | 0.054725348 | 0.081159073 | -0.026433726 | 0.999933887 | 0.032064221 | -0.040887229 | 0.07295145 | 0.687513419 |
| ENSG00000183569 | *SERHL2* | 0.06620416 | 0.107389377 | 0.103315168 | 0.004074208 | 0.999933887 | 0.209542619 | 0.026092934 | 0.183449685 | 0.687513419 |
| ENSG00000173559 | *NABP1* | 9.91505249 | 0.050426046 | 0.052298204 | -0.001872158 | 0.999933887 | 0.034416852 | 0.119919265 | -0.085502414 | 0.687513419 |
| ENSG00000089692 | *LAG3* | 2.648865734 | 0.032611805 | 0.107168218 | -0.074556413 | 0.999933887 | 0.040705505 | -0.042869805 | 0.08357531 | 0.687513419 |
| ENSG00000259096 | *SHLD2P2* | -0.472845666 | 0.00742423 | 0.232256446 | -0.224832216 | 0.999933887 | 1.43613375 | 1.791252716 | -0.355118966 | 0.687513419 |
| ENSG00000269925 | *Z98884.2* | -0.493661461 | 0.241024336 | 0.19478978 | 0.046234556 | 0.999933887 | -0.392249163 | -0.635913722 | 0.243664559 | 0.687553138 |
| ENSG00000189190 | *ZNF600* | 1.779508754 | -0.092599571 | -0.145740401 | 0.053140829 | 0.999933887 | -0.159126853 | -0.043529053 | -0.1155978 | 0.687705809 |
| ENSG00000277734 | *TRAC* | 6.169816769 | 0.011058907 | -0.035126703 | 0.04618561 | 0.999933887 | 0.139920682 | 0.195825709 | -0.055905027 | 0.687777738 |
| ENSG00000185591 | *SP1* | 7.3166999 | -0.068361343 | -0.004767118 | -0.063594225 | 0.999933887 | -0.763733818 | -0.830812356 | 0.067078538 | 0.687939058 |
| ENSG00000021300 | *PLEKHB1* | 1.402412897 | 0.047933438 | -0.121990031 | 0.169923469 | 0.999933887 | 0.028593314 | 0.169616754 | -0.14102344 | 0.688381207 |
| ENSG00000173482 | *PTPRM* | 1.284370685 | -0.040307915 | -0.276008135 | 0.235702022 | 0.999933887 | -0.063522279 | 0.057719565 | -0.121241844 | 0.688536872 |
| ENSG00000181315 | *ZNF322* | 1.032088199 | -0.038235011 | -0.051259576 | 0.013024565 | 0.999933887 | -0.41250084 | -0.517783008 | 0.105282168 | 0.688536872 |
| ENSG00000185000 | *DGAT1* | 5.896170672 | 0.059741257 | 0.052978779 | 0.006762478 | 0.999933887 | 0.220018961 | 0.147898339 | 0.072120621 | 0.688607397 |
| ENSG00000119688 | *ABCD4* | 3.418085916 | -0.04470968 | 0.001325744 | -0.046035424 | 0.999933887 | 0.084501895 | 0.003609158 | 0.080892737 | 0.688607397 |
| ENSG00000182487 | *NCF1B* | 6.149492079 | 0.034607466 | -0.011461792 | 0.046069258 | 0.999933887 | 0.990501522 | 0.908685955 | 0.081815567 | 0.688607397 |

| | | | | | | | | | | |
|---|---|---|---|---|---|---|---|---|---|---|
| ENSG00000142632 | *ARHGEF19* | 0.485828419 | 0.059233715 | -0.113062813 | 0.172296528 | 0.999933887 | 0.079467965 | -0.092440386 | 0.171908351 | 0.688607397 |
| ENSG00000007237 | *GAS7* | 7.674061614 | -0.048149135 | -0.003035812 | -0.045113324 | 0.999933887 | -0.447335189 | -0.390108742 | -0.057226447 | 0.688749815 |
| ENSG00000260278 | *AC098818.2* | 0.697855032 | 0.365184884 | 0.214535661 | 0.150649224 | 0.999933887 | 0.721292312 | 0.488800393 | 0.232491919 | 0.688824508 |
| ENSG00000206172 | *HBA1* | 3.88306826 | -0.102045946 | -0.116342481 | 0.014296535 | 0.999933887 | -0.038877463 | 0.064346333 | -0.103223796 | 0.688824508 |
| ENSG00000237711 | *MTCO3P11* | -1.515198471 | -0.167256152 | -0.197280002 | 0.03002385 | 0.999933887 | 0.419031658 | 0.908950733 | -0.489919075 | 0.688824508 |
| ENSG00000233527 | *ZNF529-AS1* | 0.093375148 | 0.068833924 | -0.016262009 | 0.085095933 | 0.999933887 | -0.265168664 | -0.114147191 | -0.151021472 | 0.688824508 |
| ENSG00000099203 | *TMED1* | 1.442499162 | 0.053502897 | 0.151208609 | -0.097705712 | 0.999933887 | -0.564438656 | -0.706136823 | 0.141698168 | 0.688824508 |
| ENSG00000104979 | *C19orf53* | 4.548506111 | 0.02054726 | 0.052570637 | -0.032023376 | 0.999933887 | 0.043152434 | -0.011314994 | 0.054467429 | 0.688824508 |
| ENSG00000287356 | *AL590822.3* | -1.052489368 | 0.126166639 | 0.221099738 | -0.094933099 | 0.999933887 | 0.203286039 | -0.113227898 | 0.316513936 | 0.689308684 |
| ENSG00000108797 | *CNTNAP1* | 1.403015712 | -0.021965881 | 0.124274288 | -0.146240168 | 0.999933887 | -0.131093877 | -0.253366578 | 0.122272701 | 0.689308684 |
| ENSG00000080298 | *RFX3* | 3.279087945 | -0.052035772 | -0.093083916 | 0.041048145 | 0.999933887 | -0.373670741 | -0.28077698 | -0.092893762 | 0.689630372 |
| ENSG00000196378 | *ZNF34* | 0.219039869 | -0.186848401 | 0.20332765 | -0.390176051 | 0.999933887 | 0.175710816 | 0.000306947 | 0.175403869 | 0.689764298 |
| ENSG00000163235 | *TGFA* | 2.728045295 | 0.115604817 | 0.10859878 | 0.007006037 | 0.999933887 | -0.071052255 | 0.040923853 | -0.111976109 | 0.689847258 |
| ENSG00000264769 | *AC145207.8* | 2.550084281 | 0.234231447 | 0.227157549 | 0.007073898 | 0.999933887 | 0.737680267 | 0.884344322 | -0.146664054 | 0.689847258 |
| ENSG00000178896 | *EXOSC4* | 1.36107855 | 0.104804488 | 0.090425949 | 0.014378538 | 0.999933887 | 0.462032408 | 0.304861248 | 0.15717116 | 0.689847258 |
| ENSG00000070182 | *SPTB* | 0.671418838 | -0.06155458 | -0.186321059 | 0.12476648 | 0.999933887 | -0.172676999 | -0.367126651 | 0.194449652 | 0.689847258 |
| ENSG00000114744 | *COMMD2* | 3.133039717 | -0.010648031 | -0.048680513 | 0.038032481 | 0.999933887 | -0.137325759 | -0.070762403 | -0.066563356 | 0.689942301 |
| ENSG00000147400 | *CETN2* | 2.251437601 | -0.112846328 | -0.18272611 | 0.069879782 | 0.999933887 | -0.059576597 | 0.06018878 | -0.119765377 | 0.6899623 |
| ENSG00000040199 | *PHLPP2* | 1.812151509 | -0.182576656 | -0.09009325 | -0.092483406 | 0.999933887 | 0.03900574 | 0.148700569 | -0.109694829 | 0.690308776 |
| ENSG00000158089 | *GALNT14* | 2.833156486 | 0.042547275 | -0.021569443 | 0.064116718 | 0.999933887 | 0.910584892 | 1.001040485 | -0.090455593 | 0.690308776 |
| ENSG00000188263 | *IL17REL* | 0.374066966 | 0.179650605 | 0.049386388 | 0.130264218 | 0.999933887 | 1.475443557 | 1.259468911 | 0.215974646 | 0.690419027 |
| ENSG00000155729 | *KCTD18* | 2.819897217 | -0.002930204 | -0.106546224 | 0.10361602 | 0.999933887 | -0.291862184 | -0.374800725 | 0.082938541 | 0.690449116 |
| ENSG00000138829 | *FBN2* | 0.532563793 | -0.068543538 | -0.278016405 | 0.209472867 | 0.999933887 | -0.450820091 | -0.282337221 | -0.168482869 | 0.690871994 |
| ENSG00000211746 | *TRBV19* | 0.976306277 | -0.150993111 | 0.178864187 | -0.329857298 | 0.999933887 | 0.10135733 | 0.255998177 | -0.154640846 | 0.690964141 |
| ENSG00000088827 | *SIGLEC1* | -1.107608039 | 0.101894114 | -0.151137094 | 0.253031208 | 0.999933887 | -0.136411412 | 0.221010825 | -0.357422237 | 0.690964141 |
| ENSG00000128000 | *ZNF780B* | 1.139930928 | -0.079384782 | -0.118145793 | 0.038761011 | 0.999933887 | -0.084212628 | 0.061761507 | -0.145974134 | 0.690964141 |
| ENSG00000137135 | *ARHGEF39* | -0.600821537 | -0.050268158 | -0.232619637 | 0.18235148 | 0.999933887 | 0.243466733 | 0.446112504 | -0.202645771 | 0.690964141 |
| ENSG00000110888 | *CAPRIN2* | 2.913396616 | -0.010060614 | -0.11918233 | 0.109121717 | 0.999933887 | -0.084554733 | -0.002060315 | -0.082494418 | 0.690964141 |
| ENSG00000226752 | *CUTALP* | 4.772948111 | -0.056902202 | -0.018834305 | -0.038067897 | 0.999933887 | 0.06822778 | 0.116532081 | -0.048304301 | 0.691052878 |
| ENSG00000152766 | *ANKRD22* | 1.034612892 | -0.188422699 | 0.115106096 | -0.303528794 | 0.944094537 | 0.302200899 | 0.53985127 | -0.237650371 | 0.691106759 |
| ENSG00000285728 | *AC098484.4* | 0.20975356 | -0.144048932 | 0.324982877 | -0.46903181 | 0.999933887 | 0.019704314 | 0.195059652 | -0.175355339 | 0.691116569 |
| ENSG00000165802 | *NSMF* | 3.900523958 | 0.063348627 | 0.075730733 | -0.012382106 | 0.999933887 | 0.213852008 | 0.288302758 | -0.074450751 | 0.691334481 |
| ENSG00000083290 | *ULK2* | 1.000950864 | 0.138421368 | 0.098048296 | 0.040373072 | 0.999933887 | 0.192939097 | 0.33497289 | -0.142033793 | 0.691634344 |
| ENSG00000146856 | *AGBL3* | 0.381277898 | -0.171784336 | 0.113924768 | -0.285709104 | 0.999933887 | -0.127533378 | -0.275701921 | 0.148168543 | 0.691634344 |
| ENSG00000150093 | *ITGB1* | 5.657278119 | -0.016242678 | -0.125648044 | 0.109405365 | 0.999933887 | -0.196705234 | -0.139842376 | -0.056862858 | 0.691634344 |
| ENSG00000272512 | *AL645608.7* | 4.115178893 | 0.299157204 | 0.297251952 | 0.001905252 | 0.999933887 | 1.329098203 | 1.180537756 | 0.148560447 | 0.69173477 |
| ENSG00000067182 | *TNFRSF1A* | 7.617693086 | -0.065662451 | -0.032078323 | -0.033584128 | 0.999933887 | -0.600992147 | -0.676166159 | 0.075174012 | 0.69173477 |
| ENSG00000124783 | *SSR1* | 5.351768087 | 0.032119431 | -0.081765707 | 0.113885138 | 0.999933887 | -0.274754233 | -0.33536224 | 0.060608007 | 0.69173477 |
| ENSG00000079277 | *MKNK1* | 6.008281351 | 0.024050726 | -0.014392147 | 0.038442873 | 0.999933887 | -0.301326056 | -0.233834608 | -0.067491447 | 0.691754538 |
| ENSG00000280202 | *AC005831.1* | 1.110931801 | -0.009970834 | -0.008007056 | -0.001963779 | 0.999933887 | -0.045383445 | -0.191463044 | 0.1460796 | 0.691791423 |
| ENSG00000023318 | *ERP44* | 6.0477495 | -0.046520134 | -0.038699829 | -0.007820305 | 0.999933887 | -0.083711749 | -0.034342744 | -0.049369005 | 0.692220631 |
| ENSG00000136643 | *RPS6KC1* | 2.600300625 | -0.067896342 | -0.095289059 | 0.027392718 | 0.999933887 | -0.179799666 | -0.092897821 | -0.086901846 | 0.692296529 |
| ENSG00000100941 | *PNN* | 5.614635858 | -0.059541867 | -0.14816821 | 0.088626343 | 0.999933887 | -0.269600772 | -0.362016207 | 0.092415435 | 0.692296529 |
| ENSG00000123095 | *BHLHE41* | 1.745439197 | -0.072276269 | -0.280479315 | 0.208203045 | 0.999933887 | -1.015916704 | -0.904818624 | -0.11109808 | 0.692296529 |
| ENSG00000164104 | *HMGB2* | 6.525645118 | -0.017357148 | -0.045839958 | 0.02848281 | 0.999933887 | 0.336944065 | 0.385391617 | -0.048447552 | 0.692296529 |
| ENSG00000012174 | *MBTPS2* | -0.112421818 | 0.028911666 | -0.043021193 | 0.07193286 | 0.999933887 | 0.033781868 | 0.19363081 | -0.159848942 | 0.692296529 |
| ENSG00000245293 | *CYP2U1-AS1* | -1.630288447 | 0.158903544 | 0.361134271 | -0.202230727 | 0.999933887 | 0.045282631 | 0.478579707 | -0.433297076 | 0.692496561 |
| ENSG00000100599 | *RIN3* | 7.183769543 | 0.026444778 | 0.026465709 | -2.0931e-05 | 0.999933887 | -0.089281588 | -0.15063607 | 0.061354481 | 0.692950142 |
| ENSG00000136936 | *XPA* | 1.674125829 | -0.062317586 | -0.169463932 | 0.107146346 | 0.999933887 | 0.108636701 | 0.218747045 | -0.110110344 | 0.692969073 |
| ENSG00000113971 | *NPHP3* | 1.009911176 | -0.189395031 | -0.111438746 | -0.077956285 | 0.999933887 | -0.277554648 | -0.412543516 | 0.134988868 | 0.692988431 |
| ENSG00000262655 | *SPON1* | 0.072206724 | 0.197833082 | 0.023617362 | 0.17421572 | 0.999933887 | 0.072177434 | 0.218215179 | -0.146037745 | 0.693049579 |
| ENSG00000240731 | *AL139287.1* | 0.324764069 | 0.131955019 | -0.202490197 | 0.334445217 | 0.999933887 | -0.052814333 | -0.208080201 | 0.155265868 | 0.69330527 |
| ENSG00000120129 | *DUSP1* | 10.107318972 | 0.515455262 | 0.583749325 | -0.068294063 | 0.999933887 | 1.122876046 | 1.247931578 | -0.125055532 | 0.69338341 |
| ENSG00000176834 | *VSIG10* | 0.788373457 | 0.113794349 | 0.143215506 | -0.029421157 | 0.999933887 | -0.166620916 | -0.374676772 | 0.208055857 | 0.69338341 |
| ENSG00000286025 | *FAM246C* | -0.637703783 | 0.155633399 | -0.015860483 | 0.171493882 | 0.999933887 | -0.331906602 | -0.015702494 | -0.316204108 | 0.693426661 |
| ENSG00000145244 | *CORIN* | -0.763333089 | -0.338094019 | -0.068477937 | -0.269616082 | 0.999933887 | -0.352921702 | -0.027922974 | -0.324998728 | 0.693550514 |
| ENSG00000108551 | *RASD1* | 0.818789007 | 0.272096365 | 0.542288694 | -0.270192329 | 0.999933887 | 1.546605898 | 1.382466194 | 0.164139704 | 0.693550514 |
| ENSG00000175575 | *PAAF1* | 0.434092168 | -0.175209909 | -0.105635499 | -0.06957441 | 0.999933887 | 0.031261961 | 0.190942399 | -0.159680438 | 0.693550514 |
| ENSG00000067704 | *IARS2* | 3.100694942 | -0.060910866 | -0.047274206 | -0.013636659 | 0.999933887 | -0.081535404 | -0.163369395 | 0.081833991 | 0.693550514 |
| ENSG00000196177 | *ACADSB* | 1.88690936 | 0.073083947 | -0.090672494 | 0.163756442 | 0.999933887 | -0.152949106 | -0.036209643 | -0.116739463 | 0.693550514 |
| ENSG00000239636 | *TFAP2E-AS1* | -0.197440042 | 0.093376848 | -0.041380325 | 0.134757173 | 0.999933887 | -0.417921871 | -0.62114103 | 0.203219159 | 0.693550514 |
| ENSG00000171763 | *SPATA5L1* | 3.416456733 | 0.025820795 | 0.064116901 | -0.038296106 | 0.999933887 | 0.122995039 | 0.044492927 | 0.078502112 | 0.693550514 |
| ENSG00000120805 | *ARL1* | 1.73723532 | -0.261995868 | -0.069132113 | -0.192863755 | 0.999933887 | 0.08555521 | 0.183101324 | -0.097546113 | 0.693620812 |
| ENSG00000228655 | *AC079793.1* | 2.296485839 | 0.409278311 | 0.464236682 | -0.054958371 | 0.999933887 | 0.124601118 | -0.180042686 | 0.304643804 | 0.693620812 |
| ENSG00000260229 | *PPIAP51* | -0.164452106 | -0.207246463 | -0.084217304 | -0.12302916 | 0.999933887 | 0.251777823 | 0.465643724 | -0.213865901 | 0.693620812 |
| ENSG00000156265 | *MAP3K7CL* | 0.401514377 | -0.093009982 | -0.027574327 | -0.065435654 | 0.999933887 | -0.588370953 | -0.755804888 | 0.167433935 | 0.693620812 |
| ENSG00000110442 | *COMMD9* | 2.914745303 | -0.025023523 | -0.041250518 | 0.016226995 | 0.999933887 | -0.298774745 | -0.373494305 | 0.07471956 | 0.693620812 |
| ENSG00000160908 | *ZNF394* | 6.866811346 | 0.010763412 | -0.024269336 | 0.035032748 | 0.999933887 | -0.020783439 | 0.026260467 | -0.047043905 | 0.693620812 |
| ENSG00000103326 | *CAPN15* | 4.169027883 | 0.058523794 | 0.047440518 | 0.011083276 | 0.999933887 | 0.037107131 | -0.032253601 | 0.069360732 | 0.693675188 |
| ENSG00000110721 | *CHKA* | 5.060508166 | 0.034164665 | 0.047844897 | -0.013680233 | 0.999933887 | -0.533988221 | -0.478084572 | -0.055903649 | 0.693675188 |
| ENSG00000100346 | *CACNA1I* | 2.428003676 | -0.028984808 | 0.168072043 | -0.197056851 | 0.999933887 | 0.054958046 | 0.136958912 | -0.082000867 | 0.693675188 |
| ENSG00000038002 | *AGA* | 1.824503011 | 0.009322509 | -0.000217854 | 0.009540363 | 0.999933887 | -0.08070471 | -0.181779133 | 0.101074423 | 0.694314083 |
| ENSG00000118113 | *MMP8* | -0.740406166 | 0.262393683 | 0.180332664 | 0.082061019 | 0.999933887 | 0.147889326 | 0.353178437 | -0.205289111 | 0.694448072 |
| ENSG00000170500 | *LONRF2* | -1.88379473 | -0.106260567 | 0.484567837 | -0.590828403 | 0.999933887 | -0.329039377 | -0.147006562 | -0.182032815 | 0.694712812 |
| ENSG00000181817 | *LSM10* | 5.251867697 | 0.052947833 | -0.045318001 | 0.098265834 | 0.999933887 | -0.357332904 | -0.421521527 | 0.064188623 | 0.694738216 |
| ENSG00000226121 | *AHCTF1P1* | -0.667051486 | 0.146553048 | 0.087707932 | 0.058845116 | 0.999933887 | -1.145974861 | -1.428734095 | 0.282759234 | 0.694738216 |
| ENSG00000068885 | *IFT80* | 0.812960005 | -0.126127059 | -0.0286889 | -0.097438159 | 0.999933887 | -0.046947874 | 0.161663075 | -0.208610949 | 0.694738216 |
| ENSG00000068400 | *GRIPAP1* | 6.172494843 | 0.02976954 | 0.043902123 | -0.014132582 | 0.999933887 | 0.131304418 | 0.181902304 | -0.050597886 | 0.694738216 |
| ENSG00000167850 | *CD300C* | 1.010009826 | 0.070440699 | -0.016491224 | 0.086931923 | 0.999933887 | -0.69738125 | -0.539987669 | -0.157393581 | 0.694738216 |
| ENSG00000119760 | *SUPT7L* | 3.915543369 | -0.0373747 | -0.061241173 | 0.023866474 | 0.999933887 | 0.154702888 | 0.220625448 | -0.065922559 | 0.694738216 |
| ENSG00000230069 | *LRRC37A15P* | -0.278225462 | -0.123285081 | 0.184996938 | -0.30828202 | 0.999933887 | 0.283185871 | 0.058484319 | 0.224701552 | 0.694738216 |
| ENSG00000273145 | *BX537318.1* | 0.004871363 | -0.136811889 | -0.175872212 | 0.039060323 | 0.999933887 | 0.38009095 | 0.251749009 | 0.128341941 | 0.694738216 |
| ENSG00000148688 | *RPP30* | 1.544646798 | 0.046003986 | 0.052610579 | -0.006606593 | 0.999933887 | 0.025279051 | 0.125385866 | -0.100106814 | 0.694738216 |
| ENSG00000137764 | *MAP2K5* | 1.992445706 | 0.033489872 | 0.042243553 | -0.00875368 | 0.999933887 | -0.048730906 | 0.056082015 | -0.10481292 | 0.694738216 |
| ENSG00000213071 | *LPAL2* | -0.808793969 | 0.067445625 | -0.032977688 | 0.100423313 | 0.999933887 | -0.395423673 | -0.163253442 | -0.232170231 | 0.694738216 |
| ENSG00000141741 | *MIEN1* | 4.006670755 | 0.013327523 | 0.048990271 | -0.035662748 | 0.999933887 | 0.191238924 | 0.258755653 | -0.067516729 | 0.694738216 |
| ENSG00000166004 | *CEP295* | 2.654222308 | -0.02720371 | -0.094081665 | 0.066877955 | 0.999933887 | -0.813078514 | -0.660596983 | -0.152481531 | 0.694738216 |
| ENSG00000082074 | *FYB1* | 9.093732544 | -0.01589984 | -0.10376285 | 0.08786301 | 0.999933887 | -0.901292122 | -0.810817024 | -0.090475098 | 0.694738216 |
| ENSG00000177034 | *MTX3* | 2.169778625 | -0.079138207 | 0.036709246 | -0.115847453 | 0.999933887 | -0.181063312 | -0.046841892 | -0.134221421 | 0.695161219 |
| ENSG00000273599 | *AL731571.1* | 2.720722404 | 0.060247184 | 0.18665035 | -0.126403167 | 0.999933887 | -0.400746554 | -0.483985696 | 0.083239142 | 0.695282216 |
| ENSG00000239672 | *NME1* | -0.714907141 | 0.027042253 | -0.379422977 | 0.40646523 | 0.999933887 | 0.030872748 | 0.239965553 | -0.209092806 | 0.695282216 |
| ENSG00000070669 | *ASNS* | -0.399698222 | -0.131393866 | 0.411888636 | -0.543282502 | 0.944094537 | 0.257070102 | 0.090194502 | 0.1668756 | 0.695654624 |
| ENSG00000204977 | *TRIM13* | 3.589039198 | -0.055737936 | 0.033287517 | -0.089025453 | 0.999933887 | 0.002503276 | 0.063062986 | -0.060559711 | 0.695753396 |
| ENSG00000198856 | *OSTC* | 2.404370551 | 0.041681492 | -0.025962703 | 0.067644195 | 0.999933887 | -0.015891432 | 0.080245494 | -0.096136926 | 0.695753396 |
| ENSG00000165244 | *ZNF367* | 0.139138299 | 0.082810624 | 0.189203298 | -0.106392674 | 0.999933887 | -0.016723118 | 0.18673234 | -0.203455458 | 0.695769076 |
| ENSG00000182158 | *CREB3L2* | 3.315002158 | -0.043996093 | -0.024330421 | -0.019665673 | 0.999933887 | -0.443312495 | -0.533771397 | 0.090458902 | 0.695901677 |
| ENSG00000117593 | *DARS2* | 1.034128086 | -0.028222883 | 0.075730815 | -0.103953698 | 0.999933887 | -0.00360712 | 0.111378443 | -0.114985563 | 0.695901677 |
| ENSG00000177000 | *MTHFR* | 5.933167442 | -0.007492108 | -0.002674014 | -0.004818095 | 0.999933887 | -0.393827104 | -0.460076278 | 0.066249174 | 0.695901677 |
| ENSG00000176148 | *TCP11L1* | 2.179633969 | 0.058834548 | 0.033476703 | 0.025357844 | 0.999933887 | 0.175111398 | 0.253492115 | -0.078380717 | 0.696047436 |
| ENSG00000180353 | *HCLS1* | 10.043929813 | -0.041155162 | -0.011916223 | -0.029238939 | 0.999933887 | -0.265619888 | -0.310769851 | 0.045149963 | 0.696072702 |
| ENSG00000144524 | *COPS7B* | 2.756467838 | 0.002174466 | -0.054713076 | 0.056887542 | 0.999933887 | -0.04163349 | -0.119887716 | 0.078254226 | 0.69620461 |
| ENSG00000130844 | *ZNF331* | 4.339649294 | 0.01307441 | -0.008192562 | 0.021266972 | 0.999933887 | 0.045146145 | 0.105260313 | -0.060114169 | 0.696298887 |
| ENSG00000088356 | *PDRG1* | 2.319086372 | -0.069516077 | -0.147404041 | 0.077887964 | 0.999933887 | 0.106995283 | 0.01663764 | 0.090357643 | 0.696364375 |
| ENSG00000066027 | *PPP2R5A* | 6.136905705 | -0.008271163 | 0.030572867 | -0.03884403 | 0.999933887 | -0.629784703 | -0.679906833 | 0.05012213 | 0.696364375 |
| ENSG00000100196 | *KDELR3* | 2.061047413 | 0.008689124 | 0.134911303 | -0.12622218 | 0.999933887 | 0.530375625 | 0.62978073 | -0.099405106 | 0.696364375 |

| | | | | | | | | | | |
|---|---|---|---|---|---|---|---|---|---|---|
| ENSG00000107796 | *ACTA2* | -0.117344576 | -0.104074566 | 0.168252223 | -0.272326789 | 0.999933887 | 0.295794054 | 0.501889601 | -0.206095547 | 0.696582156 |
| ENSG00000279413 | *AC112497.2* | -0.44099141 | -0.039687026 | 0.218994595 | -0.258681621 | 0.999933887 | -0.239179514 | 0.044180525 | -0.283360039 | 0.696783691 |
| ENSG00000172007 | *RAB33B* | 4.470890646 | -0.108684252 | -0.048617219 | -0.060067033 | 0.999933887 | -1.258414444 | -1.350962733 | 0.092548288 | 0.696943586 |
| ENSG00000179222 | *MAGED1* | 2.036912188 | -0.045243057 | -0.089502103 | 0.044259046 | 0.999933887 | 0.087341871 | 0.183012144 | -0.095670272 | 0.696943586 |
| ENSG00000272734 | *ADIRF-AS1* | -0.088087251 | -0.080742224 | 0.254258049 | -0.335000273 | 0.999933887 | 0.05760432 | 0.233375062 | -0.175770743 | 0.696943586 |
| ENSG00000105676 | *ARMC6* | 1.82699951 | -0.029621997 | 0.063836782 | -0.093458779 | 0.999933887 | 0.015295562 | -0.095775857 | 0.111071419 | 0.696943586 |
| ENSG00000179082 | *C9orf106* | 0.79863968 | -0.13343616 | 0.041454201 | -0.174890362 | 0.999933887 | -0.183964933 | -0.042142745 | -0.141822188 | 0.698004666 |
| ENSG00000109762 | *SNX25* | 1.086877817 | -0.067988863 | -0.190059903 | 0.12207104 | 0.999933887 | -0.018999938 | -0.132083079 | 0.11308314 | 0.698004666 |
| ENSG00000279476 | *AC092139.2* | 0.602704664 | -0.008662582 | -0.073772845 | 0.065110263 | 0.999933887 | -0.070250258 | 0.138512625 | -0.208762883 | 0.698004666 |
| ENSG00000177301 | *KCNA2* | -0.847337923 | -0.099224473 | 0.029315691 | -0.128540164 | 0.999933887 | -0.023452726 | 0.214847445 | -0.238300171 | 0.6981342 |
| ENSG00000071246 | *VASH1* | 0.736993874 | 0.042295511 | 0.056577513 | -0.014282002 | 0.999933887 | 0.096663395 | -0.056603261 | 0.153266656 | 0.6981342 |
| ENSG00000176809 | *LRRC37A3* | 0.584531094 | -0.019498574 | 0.209100046 | -0.228598619 | 0.999933887 | -0.405309308 | -0.226634195 | -0.178675113 | 0.698212223 |
| ENSG00000137815 | *RTF1* | 6.11990783 | -0.075235176 | -0.057648352 | -0.017586824 | 0.999933887 | -0.909527705 | -0.976342836 | 0.066815131 | 0.698221285 |
| ENSG00000181467 | *RAP2B* | 5.495051977 | -0.075112049 | -0.094568985 | 0.019456936 | 0.999933887 | -0.442650139 | -0.516880752 | 0.074230614 | 0.698437315 |
| ENSG00000188906 | *LRRK2* | 8.324888565 | -0.094537942 | -0.153603665 | 0.059065724 | 0.999933887 | -1.181733396 | -1.311431638 | 0.129698242 | 0.698437315 |
| ENSG00000243056 | *EIF4EBP3* | 1.041749917 | 0.046881819 | 0.097564114 | -0.050682295 | 0.999933887 | 0.066304309 | 0.229299982 | -0.162995673 | 0.698437315 |
| ENSG00000272072 | *AC004492.1* | 1.863679924 | -0.019054071 | 0.000266868 | -0.019320939 | 0.999933887 | 0.116033464 | 0.299684175 | -0.183650711 | 0.698437315 |
| ENSG00000084693 | *AGBL5* | 1.896489432 | 0.006565859 | 0.168590991 | -0.162025132 | 0.999933887 | -0.068991501 | -0.179308578 | 0.110317077 | 0.698437315 |
| ENSG00000109083 | *IFT20* | 2.780792901 | 0.019350412 | -0.030204296 | 0.049554708 | 0.999933887 | -0.176325314 | -0.256425972 | 0.080100658 | 0.698507828 |
| ENSG00000109180 | *OCIAD1* | 4.59268772 | -0.018246109 | -0.056342441 | 0.038096332 | 0.999933887 | -0.142934488 | -0.189834304 | 0.046899816 | 0.698600678 |
| ENSG00000163947 | *ARHGEF3* | 4.505540807 | -0.048291942 | -0.045174769 | -0.003117173 | 0.999933887 | 0.059175368 | 0.005131367 | 0.054044002 | 0.69861252 |
| ENSG00000204371 | *EHMT2* | 3.002138475 | 0.062814369 | 0.010218989 | 0.052595379 | 0.999933887 | 0.060736888 | -0.017210382 | 0.07794727 | 0.69861252 |
| ENSG00000110448 | *CD5* | 5.791137786 | -0.007507942 | -0.001505815 | -0.006002127 | 0.999933887 | 0.134947431 | 0.185551912 | -0.05060448 | 0.699044989 |
| ENSG00000252061 | *RNU6-415P* | -0.857556795 | -0.087390117 | 0.129753472 | -0.217143589 | 0.999933887 | -0.081996634 | 0.186653971 | -0.268650606 | 0.699066671 |
| ENSG00000158863 | *FAM160B2* | 4.08975605 | -0.010980547 | 0.148547376 | -0.159527923 | 0.999933887 | 0.045813529 | -0.01793847 | 0.063751999 | 0.699066671 |
| ENSG00000197226 | *TBC1D9B* | 5.429299752 | -0.007149801 | 0.016158305 | -0.023308106 | 0.999933887 | -0.32856288 | -0.385084702 | 0.056521822 | 0.699066671 |
| ENSG00000226167 | *AP4B1-AS1* | -0.273908476 | -0.149808993 | 0.016276189 | -0.166085182 | 0.999933887 | -0.154011208 | 0.068912569 | -0.222923777 | 0.699248394 |
| ENSG00000171657 | *GPR82* | -0.174040738 | -0.025585437 | -0.274716172 | 0.249130734 | 0.999933887 | -0.251929212 | -0.075000905 | -0.176928307 | 0.699248394 |
| ENSG00000175556 | *LONRF3* | 5.110881304 | 0.002815206 | -0.039878021 | 0.042693227 | 0.999933887 | -0.23759516 | -0.169378129 | -0.068217031 | 0.699248394 |
| ENSG00000256092 | *SBNO1-AS1* | 0.689176624 | -0.0300451 | 0.255072137 | -0.285117237 | 0.999933887 | 0.298608829 | 0.167152805 | 0.131456024 | 0.699314154 |
| ENSG00000221914 | *PPP2R2A* | 6.102186003 | -0.036913536 | -0.047416655 | 0.010503119 | 0.999933887 | -0.056573892 | -0.095376118 | 0.038802226 | 0.699620142 |
| ENSG00000142892 | *PIGK* | 1.662191951 | 0.032496436 | 0.01822511 | 0.014271326 | 0.999933887 | -0.017592791 | 0.120681639 | -0.13827443 | 0.699620142 |
| ENSG00000180229 | *HERC2P3* | 0.065233566 | -0.010594995 | 0.043437321 | -0.054032316 | 0.999933887 | -0.025513228 | 0.119891274 | -0.145404503 | 0.699620142 |
| ENSG00000138018 | *SELENOI* | 2.606460863 | -0.036682434 | -0.112347119 | 0.075664684 | 0.999933887 | -0.099677062 | -0.011850998 | -0.087826065 | 0.699627404 |
| ENSG00000133818 | *RRAS2* | 1.492727333 | -0.084640586 | -0.184255666 | 0.09961508 | 0.999933887 | 0.069372316 | 0.164967079 | -0.095594763 | 0.699628575 |
| ENSG00000066084 | *DIP2B* | 7.155654088 | 0.005438091 | -0.02075029 | 0.026188381 | 0.999933887 | -0.377893284 | -0.429362992 | 0.051469707 | 0.699628575 |
| ENSG00000165699 | *TSC1* | 5.828999197 | -0.10380458 | -0.14524271 | 0.04143813 | 0.999933887 | -0.420780114 | -0.368289777 | -0.052490337 | 0.699647601 |
| ENSG00000232160 | *RAP2C-AS1* | 0.337922936 | 0.21225864 | -0.035405396 | 0.247664037 | 0.999933887 | 1.019077641 | 1.166825767 | -0.147748125 | 0.699647601 |
| ENSG00000213190 | *MLLT11* | 1.19513841 | 0.090783526 | 0.225470075 | -0.134686548 | 0.999933887 | 1.417956868 | 1.595974553 | -0.178017684 | 0.699647601 |
| ENSG00000180138 | *CSNK1A1L* | 0.913011538 | -0.061408063 | -0.013284345 | -0.048123718 | 0.999933887 | 0.417837769 | 0.228776188 | 0.189061581 | 0.699647601 |
| ENSG00000164056 | *SPRY1* | 0.588881134 | 0.08080183 | -0.043170685 | 0.123972515 | 0.999933887 | -0.257343617 | -0.043510834 | -0.213832783 | 0.699647601 |
| ENSG00000164124 | *TMEM144* | -0.701690392 | 0.131320392 | 0.091160321 | 0.04016007 | 0.999933887 | -0.975883691 | -0.74245217 | -0.233431521 | 0.699647601 |
| ENSG00000175416 | *CLTB* | 5.351732029 | 0.028783752 | 0.027988858 | 0.000794894 | 0.999933887 | 0.098181402 | 0.034924177 | 0.063257224 | 0.699647601 |
| ENSG00000109084 | *TMEM97* | -0.773178739 | -0.08559302 | 0.582859813 | -0.668452833 | 0.969206975 | 0.054794383 | 0.277575057 | -0.222780674 | 0.699647601 |
| ENSG00000154945 | *ANKRD40* | 2.499125988 | -0.029430972 | -0.074579582 | 0.045148611 | 0.999933887 | -0.009887399 | 0.066360804 | -0.076248203 | 0.699647601 |
| ENSG00000145623 | *OSMR* | 0.488299996 | -0.073470407 | 0.35623094 | -0.429701347 | 0.999933887 | -0.57151398 | -0.406895642 | -0.164618339 | 0.699649141 |
| ENSG00000244482 | *LILRA6* | 7.127842414 | 0.014287834 | 0.040531144 | -0.02624331 | 0.999933887 | -0.497694258 | -0.451142656 | -0.046551601 | 0.699649141 |
| ENSG00000266777 | *SH3GL1P1* | 1.800172742 | -0.027700913 | 0.065421853 | -0.093122766 | 0.999933887 | -0.448276819 | -0.35918331 | -0.089093509 | 0.699649141 |
| ENSG00000142676 | *RPL11* | 7.71857764 | -0.004113287 | -0.063766012 | 0.059652725 | 0.999933887 | 0.086760232 | 0.145287467 | -0.058527235 | 0.699649141 |
| ENSG00000188897 | *AC099489.1* | 7.908405687 | 0.064712442 | 0.064684001 | 2.8441e-05 | 0.999933887 | 0.412107044 | 0.463360761 | -0.051253716 | 0.699810976 |
| ENSG00000179299 | *NSUN7* | 2.419921245 | 0.097033471 | -0.103651518 | 0.200684989 | 0.999933887 | -0.41377916 | -0.297472684 | -0.116306476 | 0.699810976 |
| ENSG00000185163 | *DDX51* | 2.963045373 | 0.063797025 | 0.165269452 | -0.101472428 | 0.999933887 | 0.132009117 | 0.210004754 | -0.077995638 | 0.699810976 |
| ENSG00000048162 | *NOP16* | 1.128776568 | 0.02407742 | 0.110114084 | -0.086036663 | 0.999933887 | 0.217490375 | 0.346976934 | -0.129486558 | 0.699810976 |
| ENSG00000187583 | *PLEKHN1* | -0.252300608 | 0.026007622 | -0.065012564 | 0.091020186 | 0.999933887 | 0.248079444 | 0.444975695 | -0.196896252 | 0.699810976 |
| ENSG00000234771 | *SLC25A25-AS1* | 2.254866235 | 0.010658915 | -0.053401121 | 0.064060036 | 0.999933887 | 0.022319428 | -0.066013758 | 0.088333186 | 0.699810976 |
| ENSG00000067221 | *STOML1* | 0.951101677 | 0.076194903 | -0.023060369 | 0.099255272 | 0.999933887 | 1.250666933 | 1.403126075 | -0.152459142 | 0.70001643 |
| ENSG00000273176 | *AL008635.1* | -0.072520924 | 0.271253765 | 0.394219747 | -0.122965981 | 0.999933887 | 1.947278664 | 2.12721384 | -0.179935177 | 0.700056161 |
| ENSG00000142733 | *MAP3K6* | 1.09693977 | 0.096785115 | 0.02957496 | 0.067210154 | 0.999933887 | -0.223285293 | -0.382369727 | 0.159084434 | 0.700056161 |
| ENSG00000280211 | *AC106886.4* | 0.112786327 | 0.023433544 | 0.030795695 | -0.007362151 | 0.999933887 | 0.138828009 | -0.08856216 | 0.227390168 | 0.700056161 |
| ENSG00000168824 | *NSG1* | 0.868160577 | -0.062632032 | -0.126269867 | 0.063637835 | 0.999933887 | -0.076120191 | -0.207287214 | 0.131167023 | 0.700068994 |
| ENSG00000116874 | *WARS2* | 0.414577409 | 0.163954924 | -0.111381099 | 0.275336022 | 0.999933887 | 0.306990065 | 0.45520882 | -0.148218755 | 0.700101361 |
| ENSG00000102189 | *EEA1* | 2.106812136 | -0.167417349 | -0.081793909 | -0.085623441 | 0.999933887 | -0.458271879 | -0.285477915 | -0.172793964 | 0.700101361 |
| ENSG00000277494 | *GPIHBP1* | -2.419676222 | 0.397250413 | 0.13982502 | 0.257425393 | 0.999933887 | 1.311003475 | 1.57631508 | -0.265311606 | 0.700101361 |
| ENSG00000101474 | *APMAP* | 6.885565823 | -0.025217173 | -0.039084996 | 0.013867823 | 0.999933887 | 0.118860387 | 0.166321105 | -0.047460718 | 0.700101361 |
| ENSG00000151500 | *THYN1* | 2.377915071 | -0.029829739 | -0.176819076 | 0.146989338 | 0.999933887 | 0.016457818 | -0.071612771 | 0.08807059 | 0.700483506 |
| ENSG00000230424 | *EMC1-AS1* | -0.294940233 | 0.026536893 | -0.213238443 | 0.239775336 | 0.999933887 | 0.79660333 | 1.015574771 | -0.218971441 | 0.700505597 |
| ENSG00000279447 | *AL118508.3* | -0.89867552 | -0.149181387 | 0.037133837 | -0.186315224 | 0.999933887 | -0.834068657 | -1.155009837 | 0.32094118 | 0.700641867 |
| ENSG00000106049 | *HIBADH* | 0.766286953 | -0.028237758 | -0.008098902 | -0.020138856 | 0.999933887 | -0.106889675 | 0.022884694 | -0.129774369 | 0.701096703 |
| ENSG00000159674 | *SPON2* | 3.473622232 | 0.002367055 | 0.065504905 | -0.06313785 | 0.999933887 | 0.103579121 | 0.031387833 | 0.072191289 | 0.701096703 |
| ENSG00000114767 | *RRP9* | 1.389873879 | -0.009292207 | 0.164683845 | -0.173976052 | 0.999933887 | 0.298654613 | 0.424001998 | -0.125347384 | 0.701132326 |
| ENSG00000270419 | *CAHM* | 2.139977689 | 0.108355227 | 0.137862844 | -0.029507617 | 0.999933887 | 0.606078619 | 0.499478822 | 0.106599797 | 0.701295049 |
| ENSG00000245164 | *LINC00861* | 3.419057897 | -0.050901424 | 0.039870602 | -0.090772026 | 0.999933887 | -0.040744944 | 0.045205013 | -0.085949957 | 0.701295049 |
| ENSG00000205269 | *TMEM170B* | 5.06733645 | -0.071450708 | -0.095411929 | 0.023961221 | 0.999933887 | -1.125270702 | -1.022455238 | -0.102815464 | 0.701563912 |
| ENSG00000226891 | *LINC01359* | 0.84536877 | -0.071627829 | -0.182892428 | 0.1112646 | 0.999933887 | -0.053627437 | 0.048478424 | -0.102105861 | 0.701563912 |
| ENSG00000120875 | *DUSP4* | 3.512267072 | -0.012569557 | -0.017950308 | 0.005380751 | 0.999933887 | 0.055968256 | 0.131429937 | -0.075461681 | 0.702045013 |
| ENSG00000264653 | *MIR5194* | 0.678874794 | 0.16789274 | 0.090712028 | 0.077180711 | 0.999933887 | 0.630427643 | 0.436688786 | 0.193738857 | 0.702213768 |
| ENSG00000087365 | *SF3B2* | 7.202161306 | -0.064850064 | -0.045185165 | -0.019664899 | 0.999933887 | -0.187351131 | -0.230080471 | 0.04272934 | 0.702297849 |
| ENSG00000172954 | *LCLAT1* | 1.812289195 | -0.14951142 | -0.058880812 | -0.090630608 | 0.999933887 | -0.385229081 | -0.495762356 | 0.110533275 | 0.702297849 |
| ENSG00000006715 | *VPS41* | 4.944700035 | -0.103366146 | -0.00497086 | -0.098395286 | 0.999933887 | -0.41345774 | -0.345355432 | -0.068102308 | 0.702297849 |
| ENSG00000081665 | *ZNF506* | 3.560195244 | -6.2026e-05 | 0.026168112 | -0.026230138 | 0.999933887 | -0.200339908 | -0.11128932 | -0.089050589 | 0.702297849 |
| ENSG00000081177 | *EXD2* | 0.020883324 | 0.163807147 | -0.007792118 | 0.171599265 | 0.999933887 | 0.050469879 | -0.098291832 | 0.148761711 | 0.702623613 |
| ENSG00000129197 | *RPAIN* | 2.924223247 | -0.071136878 | -0.161924506 | 0.090787628 | 0.999933887 | 0.100358886 | 0.165240692 | -0.064881806 | 0.702791892 |
| ENSG00000225975 | *LINC01534* | -0.175451408 | -0.156186512 | 0.006275701 | -0.162462214 | 0.999933887 | -0.050957497 | -0.212382496 | 0.161425 | 0.702791892 |
| ENSG00000108387 | *SEPTIN4* | -0.830324544 | 0.103955368 | -0.15832616 | 0.262281528 | 0.999933887 | 0.139983345 | -0.169753785 | 0.309737129 | 0.702791892 |
| ENSG00000211677 | *IGLC2* | 0.940151179 | -0.06878485 | 0.014879332 | -0.083664181 | 0.999933887 | 0.211198731 | 0.384771146 | -0.173572415 | 0.702791892 |
| ENSG00000142556 | *ZNF614* | 0.831989678 | -0.037965115 | -0.00928838 | -0.028676735 | 0.999933887 | -0.156884508 | -0.015973093 | -0.140911415 | 0.702791892 |
| ENSG00000087903 | *RFX2* | 5.240818004 | 0.009939504 | 0.053686572 | -0.043747067 | 0.999933887 | -0.412040747 | -0.485980676 | 0.073939929 | 0.702791892 |
| ENSG00000101280 | *ANGPT4* | -0.654802809 | 0.000763825 | -0.07901721 | 0.079781035 | 0.999933887 | 0.290952666 | 0.468317473 | -0.177364807 | 0.702791892 |
| ENSG00000285646 | *AL021155.4* | 0.335087651 | 0.178541388 | 0.352974689 | -0.174433301 | 0.999933887 | 0.496898893 | 0.321563108 | 0.175335785 | 0.702853727 |
| ENSG00000111962 | *UST* | 0.401425021 | 0.097929208 | -0.297819239 | 0.395748447 | 0.999933887 | 0.003065003 | -0.149248573 | 0.152313576 | 0.702853727 |
| ENSG00000105443 | *CYTH2* | 4.761327284 | 0.099248376 | 0.125984924 | -0.026736548 | 0.999933887 | -0.252966192 | -0.313749125 | 0.060782933 | 0.703215102 |
| ENSG00000224043 | *CCNT2-AS1* | -0.53998441 | 0.024430379 | 0.10934675 | -0.084916371 | 0.999933887 | -0.364050635 | -0.195106065 | -0.16894457 | 0.703215102 |
| ENSG00000109133 | *TMEM33* | 5.313649083 | 0.018603945 | 0.052469482 | -0.033865537 | 0.999933887 | -0.528610883 | -0.460588858 | -0.068022025 | 0.70323376 |
| ENSG00000269937 | *AC093525.7* | 0.569160418 | 0.17708298 | -0.132128182 | 0.309211162 | 0.999933887 | -0.614652685 | -0.428073345 | -0.18657934 | 0.703286791 |
| ENSG00000070010 | *UFD1* | 5.587950266 | -0.03412627 | -0.079519092 | 0.045392822 | 0.999933887 | -0.176517201 | -0.219721626 | 0.043204425 | 0.703286791 |
| ENSG00000212719 | *LINC02693* | 0.249304622 | 0.044325493 | -0.224140517 | 0.26846601 | 0.999933887 | -0.128557519 | -0.258424628 | 0.129867109 | 0.703286791 |
| ENSG00000224707 | *E2F3-IT1* | -0.153775781 | -0.028090355 | -0.113395654 | 0.085305298 | 0.999933887 | 0.682059994 | 0.867657831 | -0.185597837 | 0.703286791 |
| ENSG00000116977 | *LGALS8* | 5.093220164 | -0.047700593 | -0.064009508 | 0.016308915 | 0.999933887 | -0.472666148 | -0.528570001 | 0.055903853 | 0.703298385 |
| ENSG00000124226 | *RNF114* | 5.539944802 | -0.020714467 | -0.017175981 | -0.003538487 | 0.999933887 | 0.190398513 | 0.142077253 | 0.04832126 | 0.70334096 |
| ENSG00000186104 | *CYP2R1* | 2.485801439 | -0.04367588 | -0.202184648 | 0.158508768 | 0.999933887 | -0.125361945 | -0.218597687 | 0.093235742 | 0.703395006 |
| ENSG00000277763 | *AL138995.1* | -0.08128298 | -0.092271397 | -0.031148592 | -0.061122805 | 0.999933887 | -0.620556442 | -0.797474315 | 0.176917873 | 0.703418589 |
| ENSG00000285230 | *RALY-AS1* | 0.861166268 | -0.094734052 | -0.151049559 | 0.056315507 | 0.999933887 | -0.072082051 | -0.186504768 | 0.114422716 | 0.703492087 |
| ENSG00000127412 | *TRPV5* | 1.329562721 | -0.098260375 | -0.073009082 | -0.025251293 | 0.999933887 | -0.667028594 | -0.758899799 | 0.091871205 | 0.703523063 |
| ENSG00000163349 | *HIPK1* | 7.493365046 | -0.022168827 | 0.021954124 | -0.044122952 | 0.999933887 | -0.23546691 | -0.280952515 | 0.045485604 | 0.703785226 |
| ENSG00000005961 | *ITGA2B* | 0.776948581 | 0.110645952 | 0.123618517 | -0.012972565 | 0.999933887 | -0.043602847 | 0.129614224 | -0.173217071 | 0.704199972 |

| | | | | | | | | | | |
|---|---|---|---|---|---|---|---|---|---|---|
| ENSG00000097007 | *ABL1* | 3.577225283 | 0.047646454 | 0.049114528 | -0.001468074 | 0.999933887 | 0.011213607 | -0.054411689 | 0.065625296 | 0.704310298 |
| ENSG00000161395 | *PGAP3* | 2.198347015 | 0.005415715 | -0.029417664 | 0.034833379 | 0.999933887 | 0.140254672 | 0.049713967 | 0.090540705 | 0.704312182 |
| ENSG00000264773 | *MIR4420* | 1.327115852 | 0.157485876 | 0.340028981 | -0.182543106 | 0.999933887 | 0.068266584 | 0.282248519 | -0.213981934 | 0.70431238 |
| ENSG00000196247 | *ZNF107* | 2.975775595 | -0.093385678 | -0.070373499 | -0.023012179 | 0.999933887 | -0.493776146 | -0.336359313 | -0.157416834 | 0.704514451 |
| ENSG00000188706 | *ZDHHC9* | 1.407366013 | 0.084681711 | 0.036800394 | 0.047881317 | 0.999933887 | 0.243495218 | 0.379979454 | -0.136484236 | 0.704514451 |
| ENSG00000180376 | *CCDC66* | 2.019608598 | 0.072025489 | -0.029602588 | 0.101628077 | 0.999933887 | -0.028387243 | 0.081201341 | -0.109588584 | 0.704514451 |
| ENSG00000136935 | *GOLGA1* | 4.167435949 | 0.015042491 | -0.021100897 | 0.036143388 | 0.999933887 | -0.212770068 | -0.26648801 | 0.053717942 | 0.704514451 |
| ENSG00000213047 | *DENND1B* | 2.897626446 | 0.027346403 | -0.220286908 | 0.247633311 | 0.999933887 | -0.444372199 | -0.294356221 | -0.150015978 | 0.704514451 |
| ENSG00000225205 | *AC078883.1* | 0.216473288 | -0.092186352 | -0.07712393 | -0.015062422 | 0.999933887 | -0.287939643 | -0.476332736 | 0.188393093 | 0.704573185 |
| ENSG00000245648 | *KLRK1-AS1* | 1.867892699 | -0.080644199 | 0.031228439 | -0.111872638 | 0.999933887 | -0.05636241 | 0.061508253 | -0.117870664 | 0.704718003 |
| ENSG00000127804 | *METTL16* | 3.220582665 | -0.04393915 | -0.075356579 | 0.031417429 | 0.999933887 | 0.020839209 | 0.10298606 | -0.082146851 | 0.70482229 |
| ENSG00000182831 | *C16orf72* | 7.095011372 | -0.01123937 | 0.002374086 | -0.013613456 | 0.999933887 | -0.313409647 | -0.361876739 | 0.048467092 | 0.70482229 |
| ENSG00000130270 | *ATP8B3* | 0.918106637 | 0.040537778 | 0.115429444 | -0.074891665 | 0.999933887 | -0.354790141 | -0.51458112 | 0.159790979 | 0.704854919 |
| ENSG00000222011 | *FAM185A* | 0.845699373 | 0.09410902 | -0.203685459 | 0.297794479 | 0.999933887 | -0.188277441 | -0.323562108 | 0.135284667 | 0.705650154 |
| ENSG00000204516 | *MICB* | 3.671973046 | 0.052248596 | -0.066276506 | 0.118525101 | 0.999933887 | 0.326623637 | 0.400151441 | -0.073527804 | 0.705707991 |
| ENSG00000142453 | *CARM1* | 2.235483893 | -0.02671668 | 0.020057897 | -0.046774577 | 0.999933887 | 0.051442842 | -0.03396423 | 0.085407073 | 0.705707991 |
| ENSG00000140479 | *PCSK6* | -0.47135876 | -0.07371216 | -0.529363318 | 0.455651158 | 0.999933887 | -0.908483008 | -1.066699015 | 0.158216007 | 0.705707991 |
| ENSG00000128908 | *INO80* | 4.689052575 | -0.036690153 | -0.050889303 | 0.01419915 | 0.999933887 | -0.016924829 | -0.064772856 | 0.047848027 | 0.706398965 |
| ENSG00000203497 | *PDCD4-AS1* | 4.9032263 | 0.053865865 | 0.041107399 | 0.012758466 | 0.999933887 | 0.452373197 | 0.513772379 | -0.061399181 | 0.706430199 |
| ENSG00000224579 | *AC245052.2* | -0.181580618 | 0.304005886 | 0.269977792 | 0.034028094 | 0.999933887 | 0.461546048 | 0.129806266 | 0.331739782 | 0.706479445 |
| ENSG00000188807 | *TMEM201* | 1.362914848 | 0.078193572 | 0.026857806 | 0.051335766 | 0.999933887 | 0.158159738 | 0.262314318 | -0.10415458 | 0.706479445 |
| ENSG00000003509 | *NDUFAF7* | 2.757072085 | 0.061291516 | 0.063007095 | -0.001715579 | 0.999933887 | 0.007623736 | 0.081629033 | -0.074005297 | 0.706479445 |
| ENSG00000184924 | *PTRHD1* | 0.189660758 | 0.070638089 | -0.235561142 | 0.306199232 | 0.999933887 | -0.037858592 | 0.150111543 | -0.187970135 | 0.706479445 |
| ENSG00000248334 | *WHAMMP2* | 0.140053002 | -0.204016368 | 0.45712892 | -0.661145288 | 0.944094537 | 0.120303848 | 0.274738122 | -0.154434275 | 0.706503649 |
| ENSG00000054523 | *KIF1B* | 6.341885927 | -0.037679422 | -0.032752604 | -0.004926818 | 0.999933887 | 0.202255905 | 0.26280731 | -0.060551406 | 0.706606723 |
| ENSG00000181027 | *FKRP* | 3.31547442 | 0.060835977 | 0.116367606 | -0.055531628 | 0.999933887 | -0.136601574 | -0.07079506 | -0.065806514 | 0.706727281 |
| ENSG00000196440 | *ARMCX4* | 0.971502825 | 0.013089508 | -0.235459969 | 0.248549477 | 0.999933887 | 0.043907241 | 0.188819058 | -0.144911817 | 0.706727281 |
| ENSG00000168036 | *CTNNB1* | 6.882105153 | -0.066108593 | -0.059152712 | -0.006955881 | 0.999933887 | -0.050478497 | -0.016506762 | -0.033971735 | 0.706959635 |
| ENSG00000109107 | *ALDOC* | 2.506148369 | -0.118011567 | -0.057858218 | -0.060153349 | 0.999933887 | 0.538931925 | 0.63632392 | -0.097391995 | 0.706959635 |
| ENSG00000283378 | *CNTNAP3C* | -0.434576135 | 0.187418899 | 0.085967394 | 0.101451504 | 0.999933887 | 0.157604826 | -0.079616561 | 0.237221387 | 0.706959635 |
| ENSG00000050165 | *DKK3* | -0.21557177 | 0.192457666 | -0.130365548 | 0.322823214 | 0.999933887 | 0.145273513 | 0.296998988 | -0.151725475 | 0.706959635 |
| ENSG00000108424 | *KPNB1* | 7.28309505 | -0.037576135 | -0.068243942 | 0.030667808 | 0.999933887 | 0.090108779 | 0.136919246 | -0.046810468 | 0.706959635 |
| ENSG00000124172 | *ATP5F1E* | 6.417780424 | -0.015310774 | -0.033890588 | 0.018579815 | 0.999933887 | -0.068729587 | -0.12802837 | 0.059298783 | 0.706959635 |
| ENSG00000215533 | *LINC00189* | -0.741554854 | -0.023051634 | 0.124188857 | -0.147240491 | 0.999933887 | -0.097643746 | -0.352439764 | 0.254796018 | 0.706959635 |
| ENSG00000118946 | *PCDH17* | -1.804060607 | 0.207860688 | 0.2702585 | -0.062397813 | 0.999933887 | -0.18718564 | 0.016499548 | -0.203685187 | 0.707003148 |
| ENSG00000271119 | *AC026412.3* | -0.773719498 | -0.123276507 | 0.072243826 | -0.195520333 | 0.999933887 | -0.274623672 | -0.044436809 | -0.230186864 | 0.707003148 |
| ENSG00000132254 | *ARFIP2* | 3.531385395 | 0.017962574 | 0.032014444 | -0.01405187 | 0.999933887 | 0.160047715 | 0.101431244 | 0.058616471 | 0.707003148 |
| ENSG00000145916 | *RMND5B* | 2.773180701 | -0.005203018 | 0.140620589 | -0.145823607 | 0.999933887 | -0.038167291 | -0.105392271 | 0.06722498 | 0.707003148 |
| ENSG00000141030 | *COPS3* | 3.68312434 | -0.002775721 | -0.090753807 | 0.087978086 | 0.999933887 | 0.035655878 | -0.029041367 | 0.064697245 | 0.707003148 |
| ENSG00000115419 | *GLS* | 5.941133883 | -0.001793492 | -0.020340314 | 0.018546822 | 0.999933887 | -0.4366821 | -0.307839732 | -0.128842368 | 0.707003148 |
| ENSG00000198551 | *ZNF627* | 1.465103198 | -0.013298513 | -0.123256818 | 0.109958305 | 0.999933887 | -0.276411638 | -0.383532898 | 0.10712126 | 0.707015634 |
| ENSG00000151882 | *CCL28* | 0.061035541 | -0.047786363 | -0.256359536 | 0.208573173 | 0.999933887 | 0.138548255 | -0.000341602 | 0.138889857 | 0.70715721 |
| ENSG00000171045 | *TSNARE1* | 4.534791615 | 0.021214869 | 0.078987619 | -0.05777275 | 0.999933887 | 0.53859922 | 0.598554229 | -0.059955009 | 0.70715721 |
| ENSG00000158292 | *GPR153* | 0.7042171 | 0.153809094 | 0.150880586 | 0.002928507 | 0.999933887 | 0.158808348 | 0.009700907 | 0.149107441 | 0.707355059 |
| ENSG00000106591 | *MRPL32* | 2.827441254 | -0.065218978 | -0.19806638 | 0.132847402 | 0.999933887 | 0.139512597 | 0.058034018 | 0.081478579 | 0.707355059 |
| ENSG00000105738 | *SIPA1L3* | 2.993005571 | 0.0053299 | 0.052334659 | -0.047004759 | 0.999933887 | -0.091372117 | -0.174419171 | 0.083047054 | 0.707355059 |
| ENSG00000127084 | *FGD3* | 8.642832282 | -0.00791354 | -0.024870132 | 0.016956592 | 0.999933887 | -0.658275594 | -0.713594815 | 0.05531922 | 0.707513631 |
| ENSG00000280798 | *LINC00294* | 0.785490023 | -0.12585552 | 0.03116837 | -0.157023889 | 0.999933887 | -0.001821801 | 0.122249765 | -0.124071566 | 0.707711822 |
| ENSG00000176055 | *MBLAC2* | -0.224244983 | -0.131298074 | 0.211188236 | -0.34248631 | 0.999933887 | -0.050800586 | 0.129458373 | -0.180258959 | 0.707711822 |
| ENSG00000175073 | *VCPIP1* | 6.283958482 | -0.170224933 | -0.118053061 | -0.052171872 | 0.999933887 | -0.876638731 | -0.949231519 | 0.072592788 | 0.707816219 |
| ENSG00000287817 | *AL096701.4* | 0.276818705 | -0.137780234 | -0.030281698 | -0.107498536 | 0.999933887 | 0.941501305 | 0.75684615 | 0.184655155 | 0.707990782 |
| ENSG00000185187 | *SIGIRR* | 4.892617325 | 0.030880853 | 0.046915304 | -0.016034451 | 0.999933887 | 0.078350145 | 0.019176903 | 0.059173241 | 0.707990782 |
| ENSG00000169246 | *NPIPB3* | 1.761768481 | 0.04431311 | 0.101985269 | -0.05767216 | 0.999933887 | 0.017125201 | 0.126939057 | -0.109813857 | 0.707990782 |
| ENSG00000261026 | *AC105046.1* | 5.703527611 | 0.202358057 | 0.064738025 | 0.137620032 | 0.999933887 | 0.955203709 | 1.047283082 | -0.092079373 | 0.708061694 |
| ENSG00000257878 | *AC007298.2* | 1.451405172 | -0.119932569 | -0.148509453 | 0.028576884 | 0.999933887 | -0.474398039 | -0.605763651 | 0.131365612 | 0.708189091 |
| ENSG00000272054 | *AC007390.1* | -0.404348461 | -0.053101829 | 0.034896667 | -0.087998496 | 0.999933887 | -0.272416901 | -0.052972142 | -0.219444759 | 0.708189091 |
| ENSG00000113161 | *HMGCR* | 5.591817522 | -0.083982787 | -0.139612524 | 0.055629737 | 0.999933887 | 0.080863028 | 0.029964505 | 0.050898523 | 0.708301254 |
| ENSG00000250461 | *AC122718.1* | 0.194537023 | 0.255031614 | 0.14084766 | 0.114183954 | 0.999933887 | 0.293477532 | 0.518902333 | -0.225424801 | 0.708301254 |
| ENSG00000281103 | *TRG-AS1* | 2.406494585 | -0.071325148 | -0.15720819 | 0.085883042 | 0.999933887 | 0.06270309 | 0.14647122 | -0.08376813 | 0.708301254 |
| ENSG00000168566 | *SNRNP48* | 3.20213475 | 0.003829138 | -0.063187494 | 0.067016632 | 0.999933887 | -0.328693151 | -0.447233336 | 0.118540185 | 0.708301254 |
| ENSG00000214063 | *TSPAN4* | 1.202439821 | 0.005547307 | -0.04530744 | 0.050854747 | 0.999933887 | -0.936548858 | -1.089087013 | 0.152538155 | 0.708323483 |
| ENSG00000049089 | *COL9A2* | 2.573023145 | 0.079114342 | 0.251779237 | -0.172664895 | 0.999933887 | 0.136513884 | 0.029625153 | 0.106888731 | 0.708679475 |
| ENSG00000186205 | *MTARC1* | 1.431800531 | -0.105652832 | 0.073597304 | -0.179250135 | 0.999933887 | -0.648772405 | -0.832889315 | 0.18411691 | 0.708909311 |
| ENSG00000198736 | *MSRB1* | 7.865855305 | -0.073656927 | -0.056012608 | -0.017644318 | 0.999933887 | -1.007307556 | -1.096462884 | 0.089155328 | 0.708909311 |
| ENSG00000160633 | *SAFB* | 7.240821712 | -0.035061404 | 0.060378135 | -0.095439539 | 0.999933887 | -0.219547359 | -0.263805603 | 0.044258244 | 0.708909311 |
| ENSG00000242609 | *CCDC137P* | 0.06263823 | 0.106558636 | 0.253909175 | -0.147350538 | 0.999933887 | 1.989566892 | 2.277940987 | -0.288374095 | 0.708909311 |
| ENSG00000106366 | *SERPINE1* | 0.855675773 | 0.152859128 | 0.112576821 | 0.040282307 | 0.999933887 | 0.22071948 | 0.350026998 | -0.129307518 | 0.708909311 |
| ENSG00000173085 | *COQ2* | 3.428939077 | 0.04182087 | 0.070392912 | -0.028572041 | 0.999933887 | -0.657616023 | -0.729547879 | 0.071931856 | 0.708909311 |
| ENSG00000166974 | *MAPRE2* | 5.53653362 | -0.003911426 | -0.05168506 | 0.047773634 | 0.999933887 | -0.202288423 | -0.145367031 | -0.056921392 | 0.708909311 |
| ENSG00000029364 | *SLC39A9* | 4.210828284 | -0.013369626 | -0.057120545 | 0.043750919 | 0.999933887 | -0.143977795 | -0.09250343 | -0.051474365 | 0.708945503 |
| ENSG00000281162 | *LINC01127* | 2.162752563 | -0.021576733 | 0.017215468 | -0.038792201 | 0.999933887 | -0.336730013 | -0.200290636 | -0.136439377 | 0.709000628 |
| ENSG00000004139 | *SARM1* | 1.440624232 | 0.013845745 | 0.043824539 | -0.029978795 | 0.999933887 | 0.030412733 | 0.131231666 | -0.100818933 | 0.709000628 |
| ENSG00000116544 | *DLGAP3* | -1.152017648 | 0.227537144 | 0.345459324 | -0.11792218 | 0.999933887 | 0.505864397 | 0.241059664 | 0.264804733 | 0.709056264 |
| ENSG00000159374 | *M1AP* | -0.543979046 | -0.138709778 | -0.120284668 | -0.01842511 | 0.999933887 | -0.806044974 | -1.031050543 | 0.225005569 | 0.709056264 |
| ENSG00000279738 | *AL022311.1* | 0.861823208 | 0.024785899 | -0.070585702 | 0.095371601 | 0.999933887 | -0.306298999 | -0.150483991 | -0.155815008 | 0.709056264 |
| ENSG00000101407 | *TTI1* | 3.024686051 | -0.036109048 | -0.042178965 | 0.006069916 | 0.999933887 | -0.029603723 | 0.035364521 | -0.064968244 | 0.709261655 |
| ENSG00000170871 | *KIAA0232* | 6.82156945 | -0.070778966 | -0.10086469 | 0.030085724 | 0.999933887 | -0.758065857 | -0.688186263 | -0.069879594 | 0.709331256 |
| ENSG00000214022 | *REPIN1* | 7.345674947 | -0.0728686 | -0.065631624 | -0.007236976 | 0.999933887 | 1.59485886 | 1.688250823 | -0.093391963 | 0.709331256 |
| ENSG00000148400 | *NOTCH1* | 9.847176335 | 0.047759619 | 0.065353084 | -0.017593466 | 0.999933887 | -0.723916462 | -0.653193464 | -0.070722998 | 0.709331256 |
| ENSG00000242154 | *AC004884.2* | 0.103128282 | -0.038753704 | 0.149882514 | -0.188636218 | 0.999933887 | 0.400221248 | 0.587516369 | -0.18729512 | 0.709331256 |
| ENSG00000235655 | *H3P6* | 5.681533236 | 0.016439894 | -0.016839413 | 0.033279307 | 0.999933887 | -0.248003636 | -0.182714079 | -0.065289557 | 0.709331256 |
| ENSG00000143740 | *SNAP47* | 1.798766536 | -0.008011308 | 0.038079764 | -0.046091072 | 0.999933887 | 0.035049275 | -0.063215568 | 0.098264843 | 0.709331256 |
| ENSG00000119138 | *KLF9* | 3.385867348 | -0.025731143 | -0.0695051 | 0.043773957 | 0.999933887 | -0.104353389 | -0.174010617 | 0.069657229 | 0.709611495 |
| ENSG00000135318 | *NT5E* | 0.507817288 | -0.150864132 | -0.341516444 | 0.190652312 | 0.999933887 | -0.515719043 | -0.630394484 | 0.114675441 | 0.709657571 |
| ENSG00000147099 | *HDAC8* | 0.49052478 | -0.05210708 | -0.000192838 | -0.051914242 | 0.999933887 | -0.104048149 | 0.059317198 | -0.163365346 | 0.709657571 |
| ENSG00000287926 | *AC113208.5* | 1.654538695 | 0.170966102 | 0.203357803 | -0.032391701 | 0.999933887 | 1.231918029 | 1.40504541 | -0.173127381 | 0.709705873 |
| ENSG00000110060 | *PUS3* | 0.307189019 | -0.156823444 | 0.131032625 | -0.287856069 | 0.999933887 | -0.072727984 | 0.090787494 | -0.163515478 | 0.709780507 |
| ENSG00000116459 | *ATP5PB* | 4.826247705 | -0.069381474 | -0.101348356 | 0.031966882 | 0.999933887 | -0.183823997 | -0.129950558 | -0.053873439 | 0.70979478 |
| ENSG00000103148 | *NPRL3* | 2.52565128 | -0.083755692 | 0.093176867 | -0.176932559 | 0.999933887 | -0.07461544 | -0.155394839 | 0.080779399 | 0.709898512 |
| ENSG00000182308 | *DCAF4L1* | -0.534683484 | 0.009209316 | -0.259412632 | 0.268621948 | 0.999933887 | -0.039648582 | -0.238271558 | 0.198622976 | 0.709921595 |
| ENSG00000214367 | *HAUS3* | 2.920276721 | 0.007981332 | -0.056312493 | 0.064293825 | 0.999933887 | -0.195437542 | -0.291171584 | 0.095720042 | 0.710091958 |
| ENSG00000132275 | *RRP8* | 3.063102942 | -0.038047951 | -0.023603229 | -0.014444722 | 0.999933887 | 0.0230266 | -0.041557301 | 0.0645839 | 0.710150025 |
| ENSG00000153896 | *ZNF599* | -0.64550272 | 0.022238638 | 0.036946604 | -0.014707966 | 0.999933887 | 0.243096493 | 0.052589672 | 0.190506821 | 0.710150025 |
| ENSG00000171848 | *RRM2* | 0.659284054 | 0.148653785 | 0.234206088 | -0.085552303 | 0.999933887 | 0.10453829 | -0.110940989 | 0.215479279 | 0.710292241 |
| ENSG00000174292 | *TNK1* | -0.153182826 | -0.048701537 | 0.202478794 | -0.251180332 | 0.999933887 | -0.016351955 | 0.160158444 | -0.176510398 | 0.71056236 |
| ENSG00000197043 | *ANXA6* | 5.510069467 | -0.016329555 | -0.014694325 | -0.001635229 | 0.999933887 | 0.132258322 | 0.184756185 | -0.052497863 | 0.710611985 |
| ENSG00000211767 | *TRBJ2-3* | -0.511319474 | 0.139764849 | 0.089880337 | 0.049884512 | 0.999933887 | 0.143783912 | -0.058123348 | 0.201907259 | 0.710658471 |
| ENSG00000175592 | *FOSL1* | 4.346665732 | 0.309443158 | 0.156027519 | 0.153415639 | 0.999933887 | 1.349421233 | 1.457421996 | -0.108000763 | 0.710703329 |
| ENSG00000260196 | *AC124798.1* | 3.414109259 | -0.121086019 | -0.009392118 | -0.111693901 | 0.999933887 | 3.165477385 | 3.282444503 | -0.116967118 | 0.710703329 |
| ENSG00000100983 | *GSS* | 2.352749157 | -0.050951332 | 0.177852215 | -0.228803547 | 0.999933887 | 0.005760579 | 0.081018924 | -0.075258345 | 0.710703329 |
| ENSG00000100503 | *NIN* | 7.351647122 | -0.034224709 | -0.079553531 | 0.045328823 | 0.999933887 | -0.52446441 | -0.615450058 | 0.090985648 | 0.710703329 |
| ENSG00000283063 | *TRBV6-2* | -0.591751435 | 0.059705719 | 0.264748657 | -0.205042939 | 0.999933887 | 0.078563916 | 0.242712364 | -0.164148448 | 0.710918659 |
| ENSG00000006712 | *PAF1* | 6.214147497 | 0.048376897 | 0.040106675 | 0.008270221 | 0.999933887 | 0.196444563 | 0.247682631 | -0.051238068 | 0.710966342 |
| ENSG00000198954 | *KIFBP* | 1.907607726 | 0.063217614 | -0.057534691 | 0.120752305 | 0.999933887 | 0.187875627 | 0.295724833 | -0.107849205 | 0.710966342 |
| ENSG00000281649 | *EBLN3P* | 5.632499423 | 0.024948587 | -0.000742697 | 0.025691283 | 0.999933887 | -0.411567835 | -0.464281622 | 0.052713787 | 0.711137209 |

| | | | | | | | | | | |
|---|---|---|---|---|---|---|---|---|---|---|
| ENSG00000050426 | *LETMD1* | 3.265427427 | -0.026879237 | -0.033449699 | 0.006570462 | 0.999933887 | 0.23240103 | 0.290130273 | -0.057729243 | 0.711159094 |
| ENSG00000288082 | *AC023787.3* | -0.00729563 | 0.033944795 | -0.124525081 | 0.158469877 | 0.999933887 | 0.030632345 | 0.169054412 | -0.138422068 | 0.711198505 |
| ENSG00000142102 | *PGGHG* | 8.719245924 | 0.058554992 | 0.101719458 | -0.043164466 | 0.999933887 | -0.133855036 | -0.184985387 | 0.051130351 | 0.711226767 |
| ENSG00000104691 | *UBXN8* | 0.144962378 | -0.148210548 | -0.042414467 | -0.105796081 | 0.999933887 | -0.356382699 | -0.507336631 | 0.150953932 | 0.711280613 |
| ENSG00000186814 | *ZSCAN30* | 2.514548216 | -0.096523381 | -0.159451713 | 0.062928332 | 0.999933887 | -0.355888204 | -0.437806269 | 0.081918065 | 0.711364786 |
| ENSG00000181045 | *SLC26A11* | 7.253574779 | 0.036680708 | -0.024984555 | 0.061665264 | 0.999933887 | 0.252102657 | 0.312500033 | -0.060397376 | 0.711364786 |
| ENSG00000067057 | *PFKP* | 3.964391772 | -0.025729945 | 0.167317465 | -0.19304741 | 0.999933887 | -0.05308024 | -0.112585445 | 0.059505205 | 0.711364786 |
| ENSG00000165669 | *FAM204A* | 2.954663128 | -0.08284056 | -0.037966491 | -0.044874069 | 0.999933887 | -0.090555547 | -0.165747781 | 0.075192234 | 0.711436654 |
| ENSG00000165702 | *GFI1B* | 1.437852787 | 0.114395197 | 0.150193878 | -0.035798681 | 0.999933887 | -0.025805434 | 0.061530731 | -0.087336166 | 0.711508653 |
| ENSG00000281571 | *AC241585.2* | 0.81122814 | 0.091867915 | -0.224610596 | 0.316478511 | 0.999933887 | 0.319958779 | 0.184956489 | 0.13500229 | 0.711635182 |
| ENSG00000184083 | *FAM120C* | 0.430320268 | 0.037989051 | 0.18398672 | -0.145997669 | 0.999933887 | 0.027174931 | -0.129787359 | 0.15696229 | 0.711762869 |
| ENSG00000196873 | *CBWD3* | 0.950957393 | 0.1310137 | -0.173193561 | 0.304207261 | 0.944094537 | -0.019549559 | -0.195122545 | 0.175572986 | 0.71182171 |
| ENSG00000231329 | *AL031772.1* | 2.473582387 | 0.090650085 | 0.149663799 | -0.059013714 | 0.999933887 | -0.061537001 | 0.078793469 | -0.14033047 | 0.71182171 |
| ENSG00000143367 | *TUFT1* | 3.586024667 | -0.029177893 | 0.063075638 | -0.09225353 | 0.999933887 | -0.173898233 | -0.248960289 | 0.075062056 | 0.71182171 |
| ENSG00000152443 | *ZNF776* | 3.959943161 | -0.09130276 | -0.142407029 | 0.051104269 | 0.999933887 | -0.304587644 | -0.244094025 | -0.060493619 | 0.71184253 |
| ENSG00000076716 | *GPC4* | -0.045224031 | -0.149116884 | -0.069626354 | -0.07949053 | 0.999933887 | -0.913478716 | -1.165457003 | 0.251978287 | 0.71196344 |
| ENSG00000266401 | *AP002478.1* | -0.835640852 | -0.324569908 | -0.274466301 | -0.050103607 | 0.999933887 | -0.549334151 | -0.389007094 | -0.160327056 | 0.712155083 |
| ENSG00000196754 | *S100A2* | 0.289699311 | 0.126071017 | 0.089003821 | 0.037067196 | 0.999933887 | -0.280027816 | -0.118551415 | -0.161476401 | 0.712155083 |
| ENSG00000239483 | *RPS15AP16* | 0.650872372 | -0.055304692 | -0.12311838 | 0.067813688 | 0.999933887 | 0.33329235 | 0.147428954 | 0.185863396 | 0.712158566 |
| ENSG00000101846 | *STS* | 1.048226579 | -0.018577798 | -0.094255278 | 0.07567748 | 0.999933887 | -0.235948747 | -0.406175862 | 0.170227115 | 0.712386439 |
| ENSG00000268573 | *AC011815.1* | 0.020696802 | -0.092847853 | -0.032329566 | -0.060518287 | 0.999933887 | -0.541794432 | -0.6915984 | 0.149803968 | 0.712484477 |
| ENSG00000259699 | *HMGB1P8* | -0.797536823 | 0.073673617 | 0.137087768 | -0.063414151 | 0.999933887 | -0.710671903 | -0.379566735 | -0.331105168 | 0.712735674 |
| ENSG00000260563 | *AC132872.1* | 4.831397424 | 0.081613692 | 0.031320693 | 0.050292999 | 0.999933887 | 0.382558082 | 0.43883925 | -0.056281168 | 0.7130115 |
| ENSG00000104765 | *BNIP3L* | 6.868409788 | -0.017589756 | -0.073406117 | 0.055816361 | 0.999933887 | -0.424690555 | -0.379118831 | -0.045571724 | 0.713105643 |
| ENSG00000163138 | *PACRGL* | 0.239863802 | -0.02429623 | -0.045041559 | 0.020745328 | 0.999933887 | -0.038114087 | 0.123743486 | -0.161857573 | 0.713295252 |
| ENSG00000187790 | *FANCM* | 0.324245189 | -0.07322238 | -0.062129314 | -0.011093066 | 0.999933887 | -0.311165483 | -0.11656282 | -0.194602663 | 0.713383291 |
| ENSG00000132323 | *ILKAP* | 3.909470336 | -0.038749909 | 0.054664171 | -0.09341408 | 0.999933887 | 0.113328715 | 0.163639063 | -0.050310348 | 0.713639931 |
| ENSG00000226012 | *AP001434.1* | 3.054618408 | -0.018636612 | 0.181375244 | -0.200011856 | 0.999933887 | -0.459902384 | -0.290241749 | -0.169660635 | 0.713791131 |
| ENSG00000164307 | *ERAP1* | 5.09749908 | 0.042758603 | -0.03008689 | 0.072845494 | 0.999933887 | -0.457179791 | -0.382833535 | -0.074346256 | 0.714003342 |
| ENSG00000164187 | *LMBRD2* | 2.128877447 | 0.03021606 | -0.134250183 | 0.164466243 | 0.999933887 | -0.385174002 | -0.225700163 | -0.159473838 | 0.714127373 |
| ENSG00000188522 | *FAM83G* | 2.749714734 | 0.125571469 | -0.032131743 | 0.157703212 | 0.999933887 | -0.059339485 | 0.017314413 | -0.076653898 | 0.714206623 |
| ENSG00000276248 | *AL442125.1* | 1.20490478 | -0.242600065 | -0.184572338 | -0.058027727 | 0.999933887 | -0.870430456 | -0.73448782 | -0.135942636 | 0.714774142 |
| ENSG00000117595 | *IRF6* | -0.169345995 | -0.097156578 | -0.003954317 | -0.09320226 | 0.999933887 | -0.441341703 | -0.638423505 | 0.197081802 | 0.714785395 |
| ENSG00000164610 | *RP9* | 0.247005269 | -0.033737242 | -0.238171394 | 0.204434152 | 0.999933887 | 0.048844526 | 0.204168834 | -0.155324308 | 0.714881642 |
| ENSG00000170606 | *HSPA4* | 5.238308429 | -0.118549739 | -0.17318177 | 0.054632031 | 0.999933887 | -0.241003515 | -0.191742819 | -0.049260695 | 0.714929328 |
| ENSG00000110871 | *COQ5* | 0.630247844 | -0.06756658 | -0.061468419 | -0.006088161 | 0.999933887 | 0.061273274 | 0.180083763 | -0.11881049 | 0.714929328 |
| ENSG00000180182 | *MED14* | 5.043353414 | -0.029496613 | -0.058009938 | 0.028513325 | 0.999933887 | -0.165041591 | -0.219974798 | 0.054933206 | 0.714929328 |
| ENSG00000258315 | *C17orf49* | 1.704010611 | 0.2379052 | 0.251862492 | -0.013957292 | 0.999933887 | 0.626116714 | 0.744367709 | -0.118250995 | 0.715137131 |
| ENSG00000264270 | *AC087749.1* | 0.284006533 | 0.138199831 | 0.10535759 | 0.032842241 | 0.999933887 | 0.714079996 | 0.508064362 | 0.206015634 | 0.715137131 |
| ENSG00000215302 | *AC127502.1* | -0.199530621 | 0.088355735 | 0.214885914 | -0.12653018 | 0.999933887 | 0.028507287 | 0.223639203 | -0.195131916 | 0.715137131 |
| ENSG00000198785 | *GRIN3A* | -0.177919295 | -0.03094406 | -0.005374064 | -0.025569996 | 0.999933887 | -0.57516481 | -0.377879811 | -0.197284999 | 0.715137131 |
| ENSG00000180626 | *ZNF594* | -0.764690342 | -0.122948879 | 0.145770786 | -0.268719665 | 0.999933887 | -0.155969354 | 0.016899757 | -0.172869111 | 0.715282433 |
| ENSG00000237798 | *AC010894.3* | 0.239798183 | -0.031345894 | 0.01160419 | -0.042950084 | 0.999933887 | -0.168931747 | 0.02855788 | -0.197489627 | 0.715282433 |
| ENSG00000103319 | *EEF2K* | 2.473996927 | -0.012487914 | 0.127489754 | -0.139977668 | 0.999933887 | 0.050930908 | 0.136651348 | -0.08572044 | 0.715282433 |
| ENSG00000225101 | *OR52K3P* | 4.130524214 | 0.058605277 | -0.030876054 | 0.08948133 | 0.999933887 | -0.236551515 | -0.294243651 | 0.057692136 | 0.715309836 |
| ENSG00000249086 | *AC051649.1* | -1.169033137 | 0.241091781 | 0.118375205 | 0.122716576 | 0.999933887 | -0.493216709 | -0.036143779 | -0.45707293 | 0.715598951 |
| ENSG00000183527 | *PSMG1* | 0.996293301 | 0.07379192 | -0.000974188 | 0.074766108 | 0.999933887 | 0.177070973 | 0.312091294 | -0.135020321 | 0.715853122 |
| ENSG00000115806 | *GORASP2* | 3.790888884 | -0.022855083 | 0.017689376 | -0.040544459 | 0.999933887 | 0.131963945 | 0.183136907 | -0.051172962 | 0.715853122 |
| ENSG00000188352 | *FOCAD* | 2.51401443 | -0.01657882 | -0.100702857 | 0.084124036 | 0.999933887 | -0.013344841 | -0.100357316 | 0.087012475 | 0.715853122 |
| ENSG00000088451 | *TGDS* | 0.571696496 | -0.021651798 | -0.170061408 | 0.14840961 | 0.999933887 | -0.028521862 | 0.092343422 | -0.120865285 | 0.715878481 |
| ENSG00000155893 | *PXYLP1* | 1.933188804 | 0.000559507 | 0.124023697 | -0.12346419 | 0.999933887 | -0.044015968 | 0.056764772 | -0.100780741 | 0.715878481 |
| ENSG00000165282 | *PIGO* | 2.720508613 | 0.068773121 | 0.000647895 | 0.068125226 | 0.999933887 | 0.105340976 | 0.037689711 | 0.067651265 | 0.715883211 |
| ENSG00000156299 | *TIAM1* | 2.812480059 | -0.05123736 | -0.075645185 | 0.024407825 | 0.999933887 | 0.099803408 | 0.035681759 | 0.06412165 | 0.716194876 |
| ENSG00000094880 | *CDC23* | 1.774421216 | -0.042962011 | -0.116345217 | 0.073383206 | 0.999933887 | 0.177889913 | 0.091226434 | 0.086663479 | 0.716194876 |
| ENSG00000106772 | *PRUNE2* | -0.108149358 | -0.033151042 | 0.190217657 | -0.223368699 | 0.999933887 | -0.916324873 | -0.735435217 | -0.180889655 | 0.716194876 |
| ENSG00000153885 | *KCTD15* | 0.618540073 | 0.002781033 | 0.08204754 | -0.079266508 | 0.999933887 | -0.186891169 | -0.321917804 | 0.135026635 | 0.716194876 |
| ENSG00000148814 | *LRRC27* | -0.728889868 | 0.004105796 | 0.241766733 | -0.237660937 | 0.999933887 | -0.121371562 | -0.291698066 | 0.170326504 | 0.716194876 |
| ENSG00000171747 | *LGALS4* | -0.849003354 | -0.181963482 | -0.09784677 | -0.084116713 | 0.999933887 | -0.110875671 | 0.118157405 | -0.229033076 | 0.716319206 |
| ENSG00000185875 | *THNSL1* | -1.175917046 | 0.143298667 | -0.299097568 | 0.442396235 | 0.999933887 | 0.197692911 | 0.406088741 | -0.20839583 | 0.716611625 |
| ENSG00000105366 | *SIGLEC8* | 1.718356557 | 0.062890276 | 0.014975982 | 0.047914294 | 0.999933887 | 0.020708844 | 0.107331537 | -0.086622693 | 0.71664492 |
| ENSG00000284735 | *AL139424.3* | -0.331062308 | -0.201586737 | -0.318237209 | 0.116650472 | 0.999933887 | -0.322141148 | -0.582376161 | 0.260235014 | 0.716799959 |
| ENSG00000276291 | *FRG1HP* | 1.530329179 | 0.129404395 | -0.137645754 | 0.267050149 | 0.999933887 | -0.202754714 | -0.315281297 | 0.112526583 | 0.716799959 |
| ENSG00000142655 | *PEX14* | 1.753983111 | 0.025994122 | 0.008606982 | 0.017387141 | 0.999933887 | 0.092912656 | 0.181500609 | -0.088587953 | 0.716799959 |
| ENSG00000143793 | *C1orf35* | 2.403532402 | 0.090603543 | 0.058307996 | 0.032295547 | 0.999933887 | 0.021696759 | -0.076035639 | 0.097732398 | 0.716975034 |
| ENSG00000173409 | *ARV1* | -0.240898319 | -0.082666611 | -0.235446305 | 0.152779694 | 0.999933887 | -0.002170423 | 0.149113586 | -0.15124801 | 0.716975034 |
| ENSG00000273576 | *AC009283.1* | 0.258423405 | -0.031412202 | -0.195184142 | 0.16377194 | 0.999933887 | 0.246452492 | 0.117818047 | 0.128634446 | 0.716975034 |
| ENSG00000173706 | *HEG1* | 2.96284901 | -0.129942336 | 0.015494082 | -0.145436418 | 0.999933887 | 0.086639338 | 0.017016205 | 0.069623132 | 0.717011264 |
| ENSG00000263370 | *AC104564.1* | -0.427850296 | -0.118834098 | -0.009331156 | -0.109502942 | 0.999933887 | 0.088050107 | -0.13018896 | 0.218239067 | 0.717178857 |
| ENSG00000175283 | *DOLK* | 0.748431727 | 0.030469292 | 0.009115019 | 0.021354273 | 0.999933887 | 0.417004639 | 0.549345611 | -0.132340972 | 0.717178857 |
| ENSG00000244731 | *C4A* | -0.091176797 | 0.068391309 | 0.00796649 | 0.060424819 | 0.999933887 | 0.326295271 | 0.502242728 | -0.175947457 | 0.717198526 |
| ENSG00000034533 | *ASTE1* | 1.692819798 | -0.008850553 | -0.120043412 | 0.111192859 | 0.999933887 | 0.027936208 | 0.135151755 | -0.107215547 | 0.717271954 |
| ENSG00000188818 | *ZDHHC11* | -0.275617124 | -0.05294694 | -0.219742162 | 0.166795222 | 0.999933887 | 0.162957527 | 0.345560263 | -0.182602736 | 0.717399994 |
| ENSG00000270157 | *AC004918.3* | 0.438285612 | -0.015095199 | 0.159122582 | -0.174217781 | 0.999933887 | 0.015757081 | -0.111153498 | 0.12691058 | 0.717540724 |
| ENSG00000106415 | *GLCCI1* | 3.788025843 | 0.037281571 | -0.058675795 | 0.095957366 | 0.999933887 | -0.143150998 | -0.212119983 | 0.068968984 | 0.717686568 |
| ENSG00000132300 | *PTCD3* | 3.819797603 | -0.073618727 | -0.145643496 | 0.072024769 | 0.999933887 | -0.000169144 | -0.055517292 | 0.055348148 | 0.717758594 |
| ENSG00000260059 | *AC092620.2* | 0.045654247 | -0.226578726 | 0.042855243 | -0.269433969 | 0.999933887 | 0.392209087 | 0.585766381 | -0.193557295 | 0.717758594 |
| ENSG00000187862 | *TTC24* | 0.599613005 | -0.01904222 | 0.070442358 | -0.089484578 | 0.999933887 | 0.043182795 | -0.075501628 | 0.118684423 | 0.717758594 |
| ENSG00000137500 | *CCDC90B* | 2.567994004 | 0.045841031 | -0.026382443 | 0.072223474 | 0.999933887 | -0.131851547 | -0.061576212 | -0.070275335 | 0.717838345 |
| ENSG00000143333 | *RGS16* | 0.680692543 | -0.149988708 | -0.051422477 | -0.098566231 | 0.999933887 | 0.428075696 | 0.55448648 | -0.126410784 | 0.718035779 |
| ENSG00000174669 | *SLC29A2* | 0.506616492 | 0.055628239 | -0.128726343 | 0.184354582 | 0.999933887 | 0.089429697 | -0.03685091 | 0.126280608 | 0.718132503 |
| ENSG00000142864 | *SERBP1* | 5.537526612 | -0.065267304 | -0.04236712 | -0.022900184 | 0.999933887 | -0.154785109 | -0.192076338 | 0.037291229 | 0.71823322 |
| ENSG00000259953 | *AL138756.1* | 1.011861979 | 0.04306834 | -0.008898362 | 0.051966702 | 0.999933887 | 0.018797961 | 0.122268683 | -0.103470722 | 0.718342662 |
| ENSG00000107338 | *SHB* | -0.494363612 | 0.205222614 | 0.225348309 | -0.020125696 | 0.999933887 | -0.623071086 | -0.366059296 | -0.257011789 | 0.718491445 |
| ENSG00000145736 | *GTF2H2* | -0.147854792 | 0.220502689 | -0.152306671 | 0.37280936 | 0.999933887 | -0.723668086 | -0.927760906 | 0.204092821 | 0.718491445 |
| ENSG00000107798 | *LIPA* | 3.794693599 | 0.12410995 | 0.112914662 | 0.011195288 | 0.999933887 | -0.000781826 | -0.126997232 | 0.126215406 | 0.718491445 |
| ENSG00000267216 | *ZNF8-ERVK3-* | 0.138576573 | -0.07195357 | 0.024812507 | -0.096766077 | 0.999933887 | -0.211971601 | -0.365901822 | 0.153930221 | 0.718491445 |
| ENSG00000179761 | *PIPOX* | -0.231932295 | -0.146494369 | -0.200253785 | 0.053759417 | 0.999933887 | -0.00220046 | 0.173007509 | -0.175207969 | 0.718574235 |
| ENSG00000135116 | *HRK* | 0.532258698 | 0.015111825 | 0.020235051 | -0.005123226 | 0.999933887 | -0.194894125 | -0.377405907 | 0.182511782 | 0.71871303 |
| ENSG00000136682 | *CBWD2* | 2.605509413 | -0.004695796 | -0.082429631 | 0.077733835 | 0.999933887 | -0.335611206 | -0.43675027 | 0.101139064 | 0.718883132 |
| ENSG00000231551 | *AC245100.4* | 1.57088492 | 0.055500073 | -0.019862505 | 0.075362578 | 0.999933887 | -0.144525983 | -0.038082645 | -0.106443338 | 0.718897111 |
| ENSG00000250312 | *ZNF718* | 0.121951347 | -0.139908234 | -0.290088571 | 0.150180338 | 0.999933887 | -0.007407774 | -0.168608987 | 0.161201212 | 0.718930633 |
| ENSG00000260651 | *AF213884.3* | 0.487681069 | 0.296246303 | 0.18277506 | 0.113471242 | 0.999933887 | 2.172392073 | 1.991384889 | 0.181007184 | 0.719034408 |
| ENSG00000242616 | *GNG10* | 0.135084271 | -0.011889471 | 0.14795034 | -0.159839811 | 0.999933887 | 0.451800944 | 0.693903329 | -0.242102386 | 0.719060268 |
| ENSG00000166333 | *ILK* | 0.812401472 | -0.078045838 | 0.185248106 | -0.263293944 | 0.999933887 | 0.160422632 | 0.252285196 | -0.091862564 | 0.719138809 |
| ENSG00000155085 | *AK9* | 1.27297519 | -0.002830631 | 0.014781588 | -0.017612219 | 0.999933887 | -0.579196324 | -0.474155459 | -0.105040864 | 0.719138809 |
| ENSG00000107201 | *DDX58* | 5.354510849 | -0.120077139 | -0.124813436 | 0.004736297 | 0.999933887 | -0.444163057 | -0.372847268 | -0.071315789 | 0.719276327 |
| ENSG00000204618 | *RNF39* | 2.538005309 | 0.042698274 | 0.037290237 | 0.005408037 | 0.999933887 | 0.422726211 | 0.32761968 | 0.095106531 | 0.719276327 |
| ENSG00000272849 | *AC084018.1* | 0.897677659 | -0.137317747 | 0.000582464 | -0.137900211 | 0.999933887 | -0.192626067 | -0.094223737 | -0.09840233 | 0.719339337 |
| ENSG00000284194 | *SCO2* | 1.917511511 | 0.088855496 | 0.038537942 | 0.050317554 | 0.999933887 | -0.246175939 | -0.405431703 | 0.159255764 | 0.719471144 |
| ENSG00000286536 | *AL591506.1* | -0.279481316 | -0.230093625 | -0.089546372 | -0.140547254 | 0.999933887 | -0.059735287 | -0.182336079 | 0.122600792 | 0.719471144 |
| ENSG00000109534 | *GAR1* | 1.911125448 | 0.060967691 | 0.020310711 | 0.040656979 | 0.999933887 | -0.015655162 | -0.123129393 | 0.10747423 | 0.719471144 |
| ENSG00000226824 | *AC006001.2* | 0.737955401 | 0.056737491 | 0.049978383 | 0.006759108 | 0.999933887 | -0.108340998 | 0.018812833 | -0.127153831 | 0.719471144 |
| ENSG00000119650 | *IFT43* | 0.930958903 | 0.185657565 | -0.169847665 | 0.355505231 | 0.999933887 | 0.221019177 | 0.083421991 | 0.137597186 | 0.719471625 |
| ENSG00000215190 | *LINC00680* | 0.04926727 | -0.11933162 | 0.130825513 | -0.250157133 | 0.999933887 | -0.120489643 | 0.038658392 | -0.159148035 | 0.719471625 |
| ENSG00000171858 | *RPS21* | 6.513293907 | 0.029966611 | -0.016738448 | 0.04670506 | 0.999933887 | 0.113766693 | 0.171282166 | -0.057515473 | 0.719471625 |

| | | | | | | | | | | |
|---|---|---|---|---|---|---|---|---|---|---|
| ENSG00000144659 | *SLC25A38* | 3.29167427 | -0.028617754 | -0.114556872 | 0.085939117 | 0.999933887 | 0.153340347 | 0.217019833 | -0.063679487 | 0.719471625 |
| ENSG00000170145 | *SIK2* | 3.377472042 | -0.010482251 | 0.006234602 | -0.016716853 | 0.999933887 | -0.026320331 | 0.035846811 | -0.062167142 | 0.719471625 |
| ENSG00000174547 | *MRPL11* | 1.393919376 | 0.055573643 | 0.029265487 | 0.026308156 | 0.999933887 | 0.157428702 | 0.03900243 | 0.118426272 | 0.720039791 |
| ENSG00000256987 | *AC023050.5* | 1.259813475 | -0.081266331 | 0.030323267 | -0.111589597 | 0.999933887 | -0.011498175 | 0.159228084 | -0.170726259 | 0.720150793 |
| ENSG00000136206 | *SPDYE1* | 0.014937352 | -0.086206502 | -0.133815574 | 0.047609073 | 0.999933887 | 0.023325009 | 0.19385583 | -0.170530821 | 0.720487228 |
| ENSG00000115758 | *ODC1* | 5.463890749 | -0.00400753 | -0.023135835 | 0.019128305 | 0.999933887 | 0.141087986 | 0.194448392 | -0.053360406 | 0.720495923 |
| ENSG00000273702 | *AC091271.1* | 2.409291301 | 0.22777614 | 0.11534828 | 0.11242786 | 0.999933887 | 0.736459346 | 0.833175843 | -0.096716497 | 0.720998031 |
| ENSG00000183624 | *HMCES* | 4.892282401 | 0.061466195 | -0.039141948 | 0.100608143 | 0.999933887 | -0.111210853 | -0.05532634 | -0.055884513 | 0.720998031 |
| ENSG00000228264 | *PSMD8P1* | 1.429408515 | 0.142609899 | 0.099963037 | 0.042646862 | 0.999933887 | 0.858608431 | 1.03851479 | -0.179906359 | 0.720998031 |
| ENSG00000186854 | *TRABD2A* | 2.631552551 | 0.047099742 | 0.054746981 | -0.007647238 | 0.999933887 | 0.051276708 | -0.009215548 | 0.060492257 | 0.720998031 |
| ENSG00000138131 | *LOXL4* | -0.780620559 | 0.115095557 | 0.067047341 | 0.048048215 | 0.999933887 | 0.121721331 | -0.107026971 | 0.228748302 | 0.720998031 |
| ENSG00000277879 | *AL391988.1* | 0.7218305 | -0.068899634 | -0.103519124 | 0.03461949 | 0.999933887 | -0.316622078 | -0.524835798 | 0.20821372 | 0.720998031 |
| ENSG00000008083 | *JARID2* | 8.29506645 | -0.028197648 | 0.031036346 | -0.059233994 | 0.999933887 | -0.360823328 | -0.319219877 | -0.041603451 | 0.720998031 |
| ENSG00000249790 | *AC092490.1* | -1.290004301 | 0.061337107 | -0.249037452 | 0.310374559 | 0.999933887 | -0.731935849 | -0.518428958 | -0.213506891 | 0.720998031 |
| ENSG00000185436 | *IFNLR1* | 0.625844193 | -0.023134025 | -0.11774715 | 0.094613125 | 0.999933887 | 0.238288353 | 0.12448176 | 0.113806593 | 0.720998031 |
| ENSG00000259207 | *ITGB3* | -1.038186922 | -0.042546423 | -0.145581507 | 0.103035083 | 0.999933887 | -0.508922039 | -0.245668018 | -0.263254022 | 0.720998031 |
| ENSG00000253948 | *VPS13B-DT* | 0.217786323 | 0.014569645 | 0.054819819 | -0.040250174 | 0.999933887 | -0.122502759 | 0.081719768 | -0.204222527 | 0.720998031 |
| ENSG00000173744 | *AGFG1* | 5.773051851 | -0.000357624 | -0.027028228 | 0.026670604 | 0.999933887 | -0.175685872 | -0.130683226 | -0.045002646 | 0.720998031 |
| ENSG00000227039 | *ITGB2-AS1* | 5.170472667 | 0.020660822 | 0.0181417 | 0.002519123 | 0.999933887 | 0.215741035 | 0.267668593 | -0.051927559 | 0.721243048 |
| ENSG00000141447 | *OSBPL1A* | 2.843187851 | -0.002094579 | 0.045476098 | -0.047570677 | 0.999933887 | 0.062234579 | -0.02486626 | 0.087100839 | 0.721379435 |
| ENSG00000120885 | *CLU* | 1.118689784 | -0.04643479 | -0.075741062 | 0.029306272 | 0.999933887 | -0.089810997 | 0.0591688 | -0.148979797 | 0.72143778 |
| ENSG00000107874 | *CUEDC2* | 2.806493183 | 0.025862498 | -0.022447388 | 0.048309886 | 0.999933887 | 0.189298258 | 0.281225916 | -0.091927658 | 0.721686147 |
| ENSG00000197892 | *KIF13B* | 3.482458475 | -0.070841056 | 0.000852897 | -0.071693953 | 0.999933887 | 0.049069212 | -0.004908773 | 0.053977985 | 0.721798134 |
| ENSG00000163214 | *DHX57* | 1.065748398 | -0.069094058 | 0.116182586 | -0.185276645 | 0.999933887 | -0.129317975 | -0.000964964 | -0.128353011 | 0.721798134 |
| ENSG00000174611 | *KY* | 7.856388323 | 0.104851768 | 0.141574421 | -0.036722653 | 0.999933887 | 0.163483037 | 0.228574318 | -0.065091281 | 0.721921655 |
| ENSG00000177380 | *PPFIA3* | -0.635255808 | -0.10479743 | -0.04791257 | -0.05688486 | 0.999933887 | 0.025221049 | -0.17210958 | 0.197330628 | 0.721921655 |
| ENSG00000226660 | *TRBV2* | 0.650520428 | -0.084020564 | -0.118996973 | 0.034976408 | 0.999933887 | 0.020494625 | 0.135223055 | -0.11472843 | 0.722053337 |
| ENSG00000154165 | *GPR15* | 1.442777815 | -0.078093313 | -0.010741035 | -0.067352278 | 0.999933887 | 0.204631015 | 0.089289871 | 0.115341144 | 0.722053337 |
| ENSG00000180189 | *HMGB1P14* | -0.587194416 | 0.063819603 | 0.058419979 | 0.005399624 | 0.999933887 | 0.222815476 | 0.395407065 | -0.17259159 | 0.722166577 |
| ENSG00000079785 | *DDX1* | 2.547291043 | -0.057204131 | -0.09887709 | 0.041672959 | 0.999933887 | 0.093370279 | 0.00880047 | 0.084569808 | 0.722314013 |
| ENSG00000093183 | *SEC22C* | 4.131481887 | -0.056251256 | 0.024443992 | -0.080695248 | 0.999933887 | -0.045038844 | -0.098618845 | 0.053580001 | 0.722677185 |
| ENSG00000107959 | *PITRM1* | 3.897068586 | -0.02566402 | 0.121504607 | -0.147168627 | 0.999933887 | 0.003476118 | 0.066801753 | -0.063325635 | 0.722677185 |
| ENSG00000119318 | *RAD23B* | 6.679208385 | 0.008677147 | 0.004753811 | 0.003923337 | 0.999933887 | -0.336423586 | -0.295856652 | -0.040566934 | 0.722677185 |
| ENSG00000105329 | *TGFB1* | 8.590452757 | 0.026302139 | 0.016528756 | 0.009773383 | 0.999933887 | -0.2447255 | -0.304722604 | 0.059997104 | 0.722832028 |
| ENSG00000089157 | *RPLP0* | 7.48249216 | -0.002317327 | -0.040357203 | 0.038039876 | 0.999933887 | 0.131137736 | 0.182929741 | -0.051792005 | 0.723093989 |
| ENSG00000001460 | *STPG1* | -0.490226796 | -0.199532581 | -0.314379387 | 0.114846807 | 0.999933887 | -0.146837905 | 0.044260182 | -0.191098087 | 0.723275948 |
| ENSG00000204642 | *HLA-F* | 8.800286993 | 0.075294614 | 0.037820063 | 0.037474551 | 0.999933887 | 0.818450137 | 0.871035919 | -0.052585782 | 0.723546784 |
| ENSG00000116251 | *RPL22* | 3.733343003 | -0.017570969 | -0.110953682 | 0.093382713 | 0.999933887 | 0.084454728 | 0.155139693 | -0.070684966 | 0.723546784 |
| ENSG00000161298 | *ZNF382* | -0.101014963 | -0.004433554 | -0.0936155 | 0.089181946 | 0.999933887 | -0.026333559 | -0.20552411 | 0.179190551 | 0.723546784 |
| ENSG00000266970 | *AC061992.2* | 1.454127587 | 0.188130109 | 0.260030688 | -0.071900579 | 0.999933887 | 0.360897532 | 0.498561421 | -0.137663889 | 0.723582544 |
| ENSG00000167037 | *SGSM1* | -0.508701097 | 0.106084017 | -0.071182232 | 0.177266249 | 0.999933887 | -0.065733474 | 0.132753972 | -0.198487446 | 0.723582544 |
| ENSG00000177628 | *GBA* | 6.259669333 | 0.025159348 | -0.075214211 | 0.10037356 | 0.999933887 | -0.515790658 | -0.574841915 | 0.059051257 | 0.723582544 |
| ENSG00000235618 | *FAM21EP* | 0.237589523 | -0.055478847 | -0.036333822 | -0.019145025 | 0.999933887 | 0.376494913 | 0.511394585 | -0.134899671 | 0.723582544 |
| ENSG00000237541 | *HLA-DQA2* | 0.429479265 | -0.033805861 | 0.197461343 | -0.231267204 | 0.999933887 | 0.730602817 | 0.894141502 | -0.163538685 | 0.723582544 |
| ENSG00000136243 | *NUP42* | 1.441974814 | -0.019297524 | -0.000829919 | -0.018467605 | 0.999933887 | -0.267623421 | -0.379948547 | 0.112325126 | 0.72358557 |
| ENSG00000115364 | *MRPL19* | 1.033618745 | -0.110808689 | -0.129687302 | 0.018878613 | 0.999933887 | 0.164704696 | 0.279040977 | -0.114336281 | 0.72366304 |
| ENSG00000154760 | *SLFN13* | 3.645254113 | -0.004930104 | -0.042854089 | 0.037923985 | 0.999933887 | -0.141993078 | -0.082403274 | -0.059589804 | 0.724121613 |
| ENSG00000106244 | *PDAP1* | 5.368198945 | 1.1729e-05 | -0.016253433 | 0.016265162 | 0.999933887 | 0.035783849 | -0.01084914 | 0.046632989 | 0.724255343 |
| ENSG00000120262 | *CCDC170* | 0.81399594 | 0.134168278 | -0.216762878 | 0.350931157 | 0.944094537 | -0.059968595 | 0.099269902 | -0.159238497 | 0.724511441 |
| ENSG00000215193 | *PEX26* | 3.46780992 | -0.01402814 | -0.005200216 | -0.008827923 | 0.999933887 | -0.128592504 | -0.185635838 | 0.057043334 | 0.724587559 |
| ENSG00000143353 | *LYPLAL1* | 2.004802403 | -0.187351041 | -0.013684215 | -0.173666826 | 0.999933887 | -0.923710065 | -1.019513358 | 0.095803293 | 0.724645386 |
| ENSG00000117305 | *HMGCL* | 3.447739872 | 0.071698847 | -0.009256969 | 0.080955816 | 0.999933887 | -0.356272068 | -0.286981605 | -0.069290464 | 0.724645386 |
| ENSG00000056736 | *IL17RB* | -0.927103605 | 0.016645956 | 0.493546377 | -0.476900422 | 0.999933887 | 0.384917488 | 0.205953307 | 0.178964181 | 0.724645386 |
| ENSG00000081692 | *JMJD4* | -0.738820978 | 0.190036956 | 0.063262645 | 0.12677431 | 0.999933887 | 0.228491947 | 0.427745787 | -0.199253839 | 0.724791677 |
| ENSG00000119392 | *GLE1* | 4.056491576 | -0.014837251 | 0.044428769 | -0.059266019 | 0.999933887 | -0.379855132 | -0.437606594 | 0.057751462 | 0.724791677 |
| ENSG00000107371 | *EXOSC3* | 1.797221323 | -0.179876249 | -0.00195033 | -0.177925919 | 0.999933887 | -0.289998875 | -0.193711918 | -0.096286958 | 0.72495461 |
| ENSG00000236609 | *ZNF853* | 0.72596596 | 0.091876903 | -0.055583243 | 0.147460147 | 0.999933887 | -0.021237734 | -0.128203693 | 0.106965958 | 0.724966782 |
| ENSG00000259476 | *AC018904.2* | 1.772617384 | 0.281959695 | 0.421209596 | -0.139249901 | 0.999933887 | 0.802657277 | 0.943774401 | -0.141117124 | 0.72502786 |
| ENSG00000165548 | *TMEM63C* | 0.375333353 | 0.085157672 | -0.067868234 | 0.153025906 | 0.999933887 | -0.276438838 | -0.445196874 | 0.168758035 | 0.72502786 |
| ENSG00000069248 | *NUP133* | 3.545795198 | -0.020912786 | -0.016816141 | -0.004096646 | 0.999933887 | -0.102698733 | -0.033873497 | -0.068825237 | 0.72502786 |
| ENSG00000228474 | *OST4* | 4.419979283 | 0.016315927 | -0.049938844 | 0.06625477 | 0.999933887 | 0.094763724 | 0.041987041 | 0.052776683 | 0.72502786 |
| ENSG00000204438 | *GPANK1* | 3.814709772 | -0.003004692 | 0.093269377 | -0.096274068 | 0.999933887 | -0.371970197 | -0.445613538 | 0.073643341 | 0.72502786 |
| ENSG00000137207 | *YIPF3* | 6.745971581 | -0.051734993 | -0.094879839 | 0.043144846 | 0.999933887 | -0.190100622 | -0.146039312 | -0.04406131 | 0.725265521 |
| ENSG00000148337 | *CIZ1* | 4.257051601 | 0.034357455 | -0.020421008 | 0.054778462 | 0.999933887 | 0.097821302 | 0.151974769 | -0.054153467 | 0.725421435 |
| ENSG00000143870 | *PDIA6* | 3.773972494 | -0.040044127 | -0.093375107 | 0.05333098 | 0.999933887 | 0.018164152 | -0.041580446 | 0.059744598 | 0.725464872 |
| ENSG00000117751 | *PPP1R8* | 3.043806197 | -0.050948045 | -0.076293715 | 0.02534567 | 0.999933887 | 0.012859782 | -0.056291268 | 0.06915105 | 0.725464872 |
| ENSG00000063601 | *MTMR1* | 4.70816589 | 0.015574163 | 0.013732086 | 0.001842077 | 0.999933887 | -0.118735452 | -0.174439086 | 0.055703634 | 0.725464872 |
| ENSG00000160087 | *UBE2J2* | 5.501623059 | -0.000999235 | -0.068480239 | 0.067481004 | 0.999933887 | -0.121994491 | -0.079084299 | -0.042910192 | 0.725464872 |
| ENSG00000280326 | *AC067931.2* | -0.115262023 | -0.002111833 | -0.001358687 | -0.000753146 | 0.999933887 | 0.836879466 | 1.026727828 | -0.189848362 | 0.725464872 |
| ENSG00000262823 | *AC127521.1* | -0.351608402 | 0.095217776 | -0.131582064 | 0.22679984 | 0.999933887 | -0.081661203 | -0.245810538 | 0.164149336 | 0.725584758 |
| ENSG00000108559 | *NUP88* | 3.079617464 | -0.017426463 | -0.173150459 | 0.155723997 | 0.999933887 | 0.151966969 | 0.213142277 | -0.061175309 | 0.725584758 |
| ENSG00000067601 | *PMS2P4* | -0.026912845 | -0.010889487 | -0.180954723 | 0.170065236 | 0.999933887 | 0.073094789 | 0.237995953 | -0.164901164 | 0.725584758 |
| ENSG00000172803 | *SNX32* | 1.3018888 | 0.147069663 | 0.185556369 | -0.038486706 | 0.999933887 | 0.254822922 | 0.361802179 | -0.106979258 | 0.725627872 |
| ENSG00000197162 | *ZNF785* | -0.582551363 | 0.287263511 | 0.509096882 | -0.22183337 | 0.999933887 | -0.071481461 | 0.10783108 | -0.17931254 | 0.725627872 |
| ENSG00000099899 | *TRMT2A* | 5.557007557 | 0.029331263 | 0.008665067 | 0.020666196 | 0.999933887 | 0.194734823 | 0.258777036 | -0.064042213 | 0.725627872 |
| ENSG00000106351 | *AGFG2* | 2.640750896 | -0.003135844 | 0.089755125 | -0.092890969 | 0.999933887 | 0.037691834 | -0.03570461 | 0.073396444 | 0.725627872 |
| ENSG00000259456 | *ADNP-AS1* | 0.259342346 | -0.153044286 | -0.160543883 | 0.007499597 | 0.999933887 | 0.035006446 | -0.104232552 | 0.139238998 | 0.725672966 |
| ENSG00000155189 | *AGPAT5* | 2.119929028 | 0.043958708 | -0.006034654 | 0.049993363 | 0.999933887 | 0.313937246 | 0.227358792 | 0.086578454 | 0.725672966 |
| ENSG00000171928 | *TVP23B* | 3.148332108 | -0.071417471 | -0.220851256 | 0.149433785 | 0.999933887 | -0.23153464 | -0.15569285 | -0.075841789 | 0.725873022 |
| ENSG00000079739 | *PGM1* | 3.048839818 | -0.052693429 | 0.01602728 | -0.068720709 | 0.999933887 | -0.120704604 | -0.215744385 | 0.095039781 | 0.725964017 |
| ENSG00000183570 | *PCBP3* | 2.652778051 | -0.022001569 | 0.037565725 | -0.059567294 | 0.999933887 | 0.2850424 | 0.35178947 | -0.06674707 | 0.726022074 |
| ENSG00000228956 | *SATB1-AS1* | 1.07035655 | 0.178923796 | 0.179134509 | -0.000210713 | 0.999933887 | -0.069160342 | 0.066340226 | -0.135500569 | 0.72604531 |
| ENSG00000103044 | *HAS3* | -0.029445483 | 0.254541004 | -0.167664185 | 0.422205189 | 0.999933887 | -0.154824119 | 0.017939594 | -0.172763712 | 0.72604531 |
| ENSG00000161558 | *TMEM143* | 0.73299561 | 0.089155758 | 0.004828267 | 0.084327492 | 0.999933887 | -0.06518862 | 0.070382556 | -0.135571177 | 0.72604531 |
| ENSG00000286330 | *AL353660.1* | -0.735411079 | -0.077799756 | -0.512792274 | 0.434992518 | 0.999933887 | -0.219660152 | -0.006668799 | -0.212991352 | 0.72604531 |
| ENSG00000164038 | *SLC9B2* | 1.416534918 | -0.026506979 | -0.119744529 | 0.093237549 | 0.999933887 | 0.291977435 | 0.395312557 | -0.103335122 | 0.72604531 |
| ENSG00000154144 | *TBRG1* | 4.982545246 | -0.019905031 | -0.003163669 | -0.016741362 | 0.999933887 | -0.029949516 | 0.015398866 | -0.045348382 | 0.726425623 |
| ENSG00000240891 | *PLCXD2* | 3.236293149 | -0.094623003 | -0.009898246 | -0.084724757 | 0.999933887 | -0.064558986 | -0.132293983 | 0.067734997 | 0.726525992 |
| ENSG00000116191 | *RALGPS2* | 3.063682915 | -0.160432844 | -0.071537211 | -0.088895633 | 0.999933887 | -0.505944166 | -0.615976286 | 0.11003212 | 0.726535264 |
| ENSG00000211807 | *TRAV26-1* | -1.034573697 | 0.058282272 | 0.330331955 | -0.272049683 | 0.999933887 | 0.243015192 | 0.448213536 | -0.205198344 | 0.726535264 |
| ENSG00000169397 | *RNASE3* | -0.568852878 | 0.04037245 | 0.25479312 | -0.214420671 | 0.999933887 | 0.194750241 | 0.478921886 | -0.284171646 | 0.726535264 |
| ENSG00000134909 | *ARHGAP32* | -0.219852675 | -0.289523861 | 0.228751492 | -0.518275353 | 0.999933887 | 0.026968179 | 0.194667092 | -0.167698913 | 0.72663729 |
| ENSG00000088826 | *SMOX* | -0.749495471 | -0.023418752 | -0.127968358 | 0.104549606 | 0.999933887 | -0.307681118 | -0.500785492 | 0.193104373 | 0.72663729 |
| ENSG00000234750 | *AC012618.2* | 0.270005386 | -0.010886184 | 0.201399792 | -0.212285976 | 0.999933887 | 0.197667519 | 0.425658914 | -0.227991395 | 0.72663729 |
| ENSG00000183023 | *SLC8A1* | 4.013204786 | 0.020870351 | -0.015990319 | 0.03686067 | 0.999933887 | -0.039842314 | -0.126454959 | 0.086612645 | 0.726776388 |
| ENSG00000160233 | *LRRC3* | 1.074450977 | -0.029499478 | -0.060306689 | 0.030807211 | 0.999933887 | 0.566194764 | 0.683941818 | -0.117747054 | 0.726776388 |
| ENSG00000243696 | *AC006254.1* | -0.268767268 | -0.113542814 | 0.281556081 | -0.395098895 | 0.999933887 | 0.234277108 | 0.421944567 | -0.18766746 | 0.726806515 |
| ENSG00000025434 | *NR1H3* | 0.087679997 | 0.120424194 | 0.080967233 | 0.039456961 | 0.999933887 | 0.087051547 | -0.092509126 | 0.179560673 | 0.726976067 |
| ENSG00000123643 | *SLC36A1* | 6.176686765 | 0.015367152 | 0.025616474 | -0.010249321 | 0.999933887 | -0.768005847 | -0.710524395 | -0.057481452 | 0.727251847 |
| ENSG00000239350 | *AC105265.1* | -0.945483643 | -0.034026586 | 0.114773155 | -0.148799741 | 0.999933887 | 2.064864432 | 1.767407508 | 0.297456924 | 0.727251847 |
| ENSG00000189067 | *LITAF* | 12.030693643 | -0.056925377 | -0.045304321 | -0.011621056 | 0.999933887 | -0.085030948 | -0.041656338 | -0.04337461 | 0.727506568 |
| ENSG00000167523 | *SPATA33* | 0.511267446 | 0.123987416 | 0.194168327 | -0.07018091 | 0.999933887 | -0.114509658 | -0.24223236 | 0.127722702 | 0.727506568 |
| ENSG00000008710 | *PKD1* | 4.035547443 | 0.080069224 | 0.132348785 | -0.052279562 | 0.999933887 | 0.076364006 | 0.002968519 | 0.073395486 | 0.727668678 |
| ENSG00000249252 | *C1QTNF7-AS1* | -0.540522694 | 0.179996928 | -0.222814317 | 0.402811245 | 0.999933887 | -0.158723416 | -0.367510745 | 0.208787329 | 0.728100856 |
| ENSG00000213753 | *CENPBD1P1* | 4.646874084 | 0.051240752 | 0.037630594 | 0.013610158 | 0.999933887 | 0.003531012 | -0.051770795 | 0.055301807 | 0.728229216 |

| | | | | | | | | | | |
|---|---|---|---|---|---|---|---|---|---|---|
| ENSG00000282508 | *LINC01002* | 5.163174135 | 0.05147684 | -0.042347216 | 0.093824056 | 0.999933887 | 0.433667269 | 0.487405044 | -0.053737775 | 0.728437401 |
| ENSG00000220848 | *RPS18P9* | -0.13154467 | 0.043155 | -0.034620663 | 0.077775663 | 0.999933887 | -0.549658628 | -0.741107408 | 0.191448781 | 0.728437401 |
| ENSG00000173391 | *OLR1* | 3.838502921 | -0.020512684 | 0.047906253 | -0.068418937 | 0.999933887 | 1.0449918 | 1.213019305 | -0.168027505 | 0.728686407 |
| ENSG00000147419 | *CCDC25* | 1.869747344 | -0.060146596 | -0.161144689 | 0.100998092 | 0.999933887 | 0.099247711 | 0.183019517 | -0.083771807 | 0.728692189 |
| ENSG00000104907 | *TRMT1* | 3.445416871 | 0.052747625 | 0.137565969 | -0.084818344 | 0.999933887 | 0.124938719 | 0.052554315 | 0.072384403 | 0.728791276 |
| ENSG00000155962 | *CLIC2* | -0.269809204 | -0.030306584 | 0.134182109 | -0.164488693 | 0.999933887 | 0.476262331 | 0.651895486 | -0.175633155 | 0.728802441 |
| ENSG00000283633 | *AP000547.3* | 0.767016739 | 0.031722738 | 0.08663509 | -0.054912352 | 0.999933887 | -0.167042691 | -0.3106647 | 0.143622009 | 0.728902651 |
| ENSG00000287913 | *AL132986.1* | 1.922341386 | 0.193907547 | 0.137628112 | 0.056279436 | 0.999933887 | 0.636892932 | 0.766416656 | -0.129523724 | 0.729422636 |
| ENSG00000128923 | *MINDY2* | 4.086208947 | 0.064332154 | -0.065613263 | 0.129945417 | 0.999933887 | -1.128933519 | -1.016833089 | -0.11210043 | 0.729422636 |
| ENSG00000284642 | *AL139424.2* | -0.424522766 | -0.011190096 | 0.349035108 | -0.360225205 | 0.999933887 | -0.197185964 | -0.411467302 | 0.214281338 | 0.729422636 |
| ENSG00000164983 | *TMEM65* | 4.13412656 | 0.106075001 | 0.15931642 | -0.053241419 | 0.999933887 | -0.603349484 | -0.667549954 | 0.06420047 | 0.729569564 |
| ENSG00000071082 | *RPL31* | 7.305313223 | 0.020014424 | -0.06087105 | 0.080885474 | 0.999933887 | 0.097861275 | 0.15119379 | -0.053332515 | 0.729614564 |
| ENSG00000199753 | *SNORD104* | -0.34502939 | -0.089065459 | 0.298229393 | -0.387294851 | 0.999933887 | 0.076070661 | 0.303511332 | -0.227440671 | 0.729767348 |
| ENSG00000077721 | *UBE2A* | 5.726089759 | -0.023844191 | -0.032718623 | 0.008874433 | 0.999933887 | -0.168017551 | -0.128931313 | -0.039086238 | 0.729974584 |
| ENSG00000113569 | *NUP155* | 2.950314782 | 0.032359499 | 9.4272e-05 | 0.032265227 | 0.999933887 | 0.060451543 | -0.011981705 | 0.072433248 | 0.730287952 |
| ENSG00000177485 | *ZBTB33* | 3.797178995 | -0.026304234 | -0.157253847 | 0.130949613 | 0.999933887 | -0.890701056 | -0.787298184 | -0.103402872 | 0.730376311 |
| ENSG00000260257 | *AL035071.1* | 3.073154065 | -0.101585878 | 0.06464116 | -0.166227038 | 0.999933887 | -0.070900179 | 0.018419859 | -0.089320038 | 0.730615253 |
| ENSG00000206567 | *AC022007.1* | 0.894479786 | -0.045511603 | 0.174930293 | -0.220441896 | 0.999933887 | -0.938767506 | -1.05240742 | 0.113639914 | 0.730615253 |
| ENSG00000164105 | *SAP30* | 0.39362078 | 0.209506333 | 0.343180394 | -0.133674062 | 0.999933887 | 0.095288516 | -0.060189738 | 0.155478254 | 0.730801973 |
| ENSG00000036549 | *ZZZ3* | 2.874305253 | -0.127047534 | -0.177991067 | 0.050943533 | 0.999933887 | -0.317206848 | -0.237189103 | -0.080017745 | 0.730905641 |
| ENSG00000274986 | *MIR6750* | -0.143497586 | 0.165894926 | 0.164628973 | 0.001265953 | 0.999933887 | 0.26721774 | 0.059573072 | 0.207644668 | 0.730994868 |
| ENSG00000272918 | *AC005070.3* | 1.738614448 | -0.025523553 | 0.018441301 | -0.043964854 | 0.999933887 | -0.143085101 | -0.006350187 | -0.136734915 | 0.731020195 |
| ENSG00000130396 | *AFDN* | 3.338178059 | -0.004454164 | 0.003886759 | -0.008340923 | 0.999933887 | -0.097538003 | -0.157967106 | 0.060429103 | 0.731020195 |
| ENSG00000083642 | *PDS5B* | 4.365729559 | -0.080150909 | -0.095761474 | 0.015610565 | 0.999933887 | -0.485531802 | -0.35879889 | -0.126732912 | 0.731160214 |
| ENSG00000163344 | *PMVK* | 1.172059523 | 0.030074046 | 0.010891447 | 0.019182599 | 0.999933887 | 0.073081333 | 0.207903106 | -0.134821773 | 0.731160214 |
| ENSG00000117614 | *SYF2* | 6.175771244 | -0.095551051 | -0.178695023 | 0.083143972 | 0.999933887 | -0.236767223 | -0.18733293 | -0.049434293 | 0.731185865 |
| ENSG00000211749 | *TRBV23-1* | -0.55601509 | 0.110864796 | 0.358341139 | -0.247476343 | 0.999933887 | 0.141852719 | -0.020138582 | 0.161991302 | 0.731185865 |
| ENSG00000217644 | *AL355864.1* | -0.167896667 | -0.134072115 | 0.197324204 | -0.331396319 | 0.999933887 | 1.710445101 | 1.51326178 | 0.197183321 | 0.731213715 |
| ENSG00000069509 | *FUNDC1* | 0.326986045 | -0.083503908 | -0.102801593 | 0.019297685 | 0.999933887 | -0.225174334 | -0.351645353 | 0.126471018 | 0.731352183 |
| ENSG00000264269 | *AC016866.1* | -1.693017855 | 0.075551593 | 0.075538168 | 1.3425e-05 | 0.999978571 | 1.780164515 | 2.00419642 | -0.224031905 | 0.731560933 |
| ENSG00000233927 | *RPS28* | 7.064592306 | 0.057344785 | 0.008008005 | 0.049336779 | 0.999933887 | 0.221947143 | 0.268332653 | -0.046385509 | 0.731703475 |
| ENSG00000183530 | *PRR14L* | 5.378771164 | -0.080116743 | -0.042299378 | -0.037817364 | 0.999933887 | -0.892097534 | -0.973107968 | 0.081010433 | 0.731703475 |
| ENSG00000205542 | *TMSB4X* | 11.28846123 | 0.035224681 | -0.003059338 | 0.038284019 | 0.999933887 | 0.372441233 | 0.421119451 | -0.048678219 | 0.731703475 |
| ENSG00000221025 | *MIR1250* | 1.19307633 | 0.073528876 | -0.016941206 | 0.090470082 | 0.999933887 | 0.194289022 | 0.313433308 | -0.119144286 | 0.73197257 |
| ENSG00000101811 | *CSTF2* | 1.107239815 | -0.165828714 | 0.046481829 | -0.212310543 | 0.999933887 | 0.15391196 | 0.268066988 | -0.114155027 | 0.732051921 |
| ENSG00000213859 | *KCTD11* | 0.676635902 | 0.210395082 | 0.105947399 | 0.104447682 | 0.999933887 | 0.031441152 | 0.147929305 | -0.116488153 | 0.732188445 |
| ENSG00000089234 | *BRAP* | 4.52032067 | -0.068613236 | -0.000346525 | -0.068266711 | 0.999933887 | -0.310449428 | -0.256037632 | -0.054411797 | 0.732188445 |
| ENSG00000250548 | *LINC01303* | 1.691628028 | -0.005721135 | -0.136767167 | 0.131046032 | 0.999933887 | 1.100849051 | 1.22199295 | -0.121143899 | 0.732283679 |
| ENSG00000286314 | *AC010359.3* | 2.759828499 | 0.053490368 | 0.320712928 | -0.26722256 | 0.999933887 | 1.075646818 | 1.194726258 | -0.11907944 | 0.732326723 |
| ENSG00000135040 | *NAA35* | 2.742436751 | -0.021639913 | -0.065363737 | 0.043723825 | 0.999933887 | -0.017036492 | 0.061639459 | -0.078675951 | 0.732326723 |
| ENSG00000129219 | *PLD2* | 3.36590836 | -0.006429454 | 0.023352774 | -0.029782228 | 0.999933887 | -0.124136589 | -0.064622293 | -0.059514296 | 0.732368069 |
| ENSG00000218186 | *KRT8P43* | 0.369775413 | 0.115518514 | 0.056184446 | 0.059334069 | 0.999933887 | -0.534190647 | -0.418829408 | -0.11536124 | 0.732410641 |
| ENSG00000196323 | *ZBTB44* | 5.045737993 | -0.038195259 | -0.056277429 | 0.01808217 | 0.999933887 | -0.629369585 | -0.70305818 | 0.073688595 | 0.732664837 |
| ENSG00000205930 | *C21orf62-AS1* | -0.660210275 | 0.296805763 | -0.018804597 | 0.315610361 | 0.999933887 | -0.217895484 | -0.392714831 | 0.174819347 | 0.732884013 |
| ENSG00000179988 | *PSTK* | 1.542932171 | 0.001849843 | -0.014995025 | 0.016844868 | 0.999933887 | -0.081906634 | -0.187282606 | 0.105375972 | 0.732884013 |
| ENSG00000188825 | *LINC00910* | 2.224712381 | 0.00123033 | 0.030551649 | -0.029321319 | 0.999933887 | 0.189291571 | 0.11784697 | 0.071444602 | 0.733161627 |
| ENSG00000110944 | *IL23A* | 1.483275053 | -0.067701516 | 0.065363955 | -0.133065472 | 0.999933887 | 1.49298087 | 1.659812918 | -0.166832048 | 0.733165303 |
| ENSG00000125246 | *CLYBL* | -0.778738097 | -0.041420259 | -0.208448313 | 0.167028054 | 0.999933887 | 0.113211349 | -0.112248572 | 0.225459921 | 0.733165303 |
| ENSG00000115756 | *HPCAL1* | 7.726908009 | 0.075315889 | 0.043232787 | 0.032083103 | 0.999933887 | -1.013824323 | -0.948229022 | -0.065595301 | 0.733546998 |
| ENSG00000096092 | *TMEM14A* | -0.579158289 | 0.167610718 | -0.050782695 | 0.218393413 | 0.999933887 | 0.056933115 | 0.286747353 | -0.229814238 | 0.733546998 |
| ENSG00000228022 | *HCG20* | -0.068618987 | 0.036139907 | 0.233536524 | -0.197396617 | 0.999933887 | 1.057110579 | 1.192344836 | -0.135234257 | 0.733546998 |
| ENSG00000275993 | *SIK1B* | 1.573468581 | -0.005219329 | 0.035523206 | -0.040742536 | 0.999933887 | 0.216970957 | 0.0961677 | 0.120803257 | 0.733546998 |
| ENSG00000001629 | *ANKIB1* | 3.567337698 | 0.02671047 | -0.113144993 | 0.139855462 | 0.999933887 | -0.212687835 | -0.308603657 | 0.095915822 | 0.73373155 |
| ENSG00000059758 | *CDK17* | 5.727222996 | 0.01950247 | -0.016784753 | 0.036287224 | 0.999933887 | -0.439382517 | -0.504967509 | 0.065584991 | 0.733794106 |
| ENSG00000173163 | *COMMD1* | 0.674071517 | 0.012619036 | -0.21137494 | 0.223993977 | 0.999933887 | 0.151793831 | 0.278478783 | -0.126684952 | 0.733979837 |
| ENSG00000287405 | *AC017028.2* | -0.199919156 | 0.383008235 | 0.215109521 | 0.167898714 | 0.999933887 | -0.429287578 | -0.648389806 | 0.219102227 | 0.734013876 |
| ENSG00000275835 | *TUBGCP5* | 1.339110761 | 0.020309572 | 0.006709274 | 0.013600298 | 0.999933887 | 0.12085391 | 0.2467455 | -0.12589159 | 0.734089648 |
| ENSG00000115694 | *STK25* | 3.94658971 | 0.022908307 | 0.054150235 | -0.031241928 | 0.999933887 | 0.101486727 | 0.042126444 | 0.059360283 | 0.734173815 |
| ENSG00000185164 | *NOMO2* | 2.195837923 | 0.102071811 | -0.032799363 | 0.134871174 | 0.999933887 | -0.031904978 | -0.125678903 | 0.093773926 | 0.73454346 |
| ENSG00000173418 | *NAA20* | 2.296177075 | -0.086878234 | -0.020339297 | -0.066538937 | 0.999933887 | -0.09396078 | -0.011018073 | -0.082942707 | 0.73454346 |
| ENSG00000121989 | *ACVR2A* | 0.406455676 | -0.284640224 | 0.046339997 | -0.330980221 | 0.999933887 | 0.126790135 | 0.250575168 | -0.123785033 | 0.73482732 |
| ENSG00000117139 | *KDM5B* | 6.391901791 | -0.027833884 | -0.094291274 | 0.06645739 | 0.999933887 | -0.411878144 | -0.3698309 | -0.042047244 | 0.73482732 |
| ENSG00000172315 | *TP53RK* | 2.051695891 | -0.115749094 | -0.087473218 | -0.028275875 | 0.999933887 | 0.167640073 | 0.086768399 | 0.080871674 | 0.73523807 |
| ENSG00000170430 | *MGMT* | 0.676910726 | 0.031757515 | -0.085729997 | 0.117487512 | 0.999933887 | 0.127411853 | 0.006084047 | 0.121327806 | 0.735253902 |
| ENSG00000286887 | *AL355881.1* | 3.828587303 | 0.13651323 | 0.067433096 | 0.069080135 | 0.999933887 | 1.115799717 | 1.212192265 | -0.096392548 | 0.735448247 |
| ENSG00000138434 | *ITPRID2* | 6.344433352 | -0.008125989 | -0.005295234 | -0.002830755 | 0.999933887 | -0.476996388 | -0.537362478 | 0.06036609 | 0.735452481 |
| ENSG00000169239 | *CA5B* | 2.737005241 | -0.02142058 | -0.018765216 | -0.002655364 | 0.999933887 | 0.124714169 | 0.19213968 | -0.067425511 | 0.736218929 |
| ENSG00000232334 | *AL683842.1* | 0.970851789 | -0.106788067 | -0.061972902 | -0.044815164 | 0.999933887 | -0.467893691 | -0.62148712 | 0.153593429 | 0.736293823 |
| ENSG00000140548 | *ZNF710* | 6.601415067 | 0.027212275 | 0.049387058 | -0.022174784 | 0.999933887 | 0.033659696 | -0.00996598 | 0.043625676 | 0.736293823 |
| ENSG00000223865 | *HLA-DPB1* | 4.805908025 | 0.026177154 | 0.001911424 | 0.02426573 | 0.999933887 | 0.080047691 | 0.015183668 | 0.064864023 | 0.736293823 |
| ENSG00000158526 | *TSR2* | 3.396680049 | 0.03502936 | 0.088175905 | -0.053146545 | 0.999933887 | -0.088965753 | -0.016119123 | -0.072846631 | 0.736538506 |
| ENSG00000132467 | *UTP3* | 2.182795741 | -0.090877649 | -0.206939197 | 0.116061548 | 0.999933887 | 0.092348285 | 0.171596053 | -0.079247768 | 0.736682452 |
| ENSG00000095485 | *CWF19L1* | 4.530939352 | -0.019322512 | 0.016660422 | -0.035982933 | 0.999933887 | -0.035792283 | 0.022920905 | -0.058713187 | 0.736682452 |
| ENSG00000007038 | *PRSS21* | -0.632737503 | 0.238601859 | -0.066816465 | 0.305418324 | 0.999933887 | -0.154182243 | 0.01590888 | -0.170091123 | 0.736764606 |
| ENSG00000114021 | *NIT2* | 2.031379651 | -0.138832172 | -0.096294844 | -0.042537328 | 0.999933887 | 0.125569272 | 0.050297802 | 0.07527147 | 0.737146677 |
| ENSG00000167208 | *SNX20* | 6.106782438 | -0.005830942 | 0.004345521 | -0.010176464 | 0.999933887 | 0.292360963 | 0.340917219 | -0.048556255 | 0.737146677 |
| ENSG00000117560 | *FASLG* | 1.5304672 | -0.049319709 | -0.024713842 | -0.024605868 | 0.999933887 | 0.086164168 | 0.174708281 | -0.088544114 | 0.737256181 |
| ENSG00000008382 | *MPND* | 1.039722295 | 0.088694968 | -0.041548682 | 0.13024365 | 0.999933887 | 0.210359517 | 0.056790025 | 0.153569491 | 0.737308639 |
| ENSG00000164346 | *NSA2* | 4.068089694 | -0.017663647 | -0.087349376 | 0.069685728 | 0.999933887 | 0.070583891 | 0.129491872 | -0.05890798 | 0.737308639 |
| ENSG00000043143 | *JADE2* | 5.372925861 | 0.013991249 | -0.022439949 | 0.036431199 | 0.999933887 | 0.131821812 | 0.177317591 | -0.045495779 | 0.737308639 |
| ENSG00000171466 | *ZNF562* | 2.707759359 | -0.089420079 | -0.033910259 | -0.05550982 | 0.999933887 | -0.110267281 | -0.166040241 | 0.05577296 | 0.737311374 |
| ENSG00000135605 | *TEC* | 2.144944072 | -0.07666579 | -0.031033781 | -0.04563201 | 0.999933887 | -0.288265986 | -0.372002196 | 0.08373621 | 0.737311374 |
| ENSG00000215114 | *UBXN2B* | 5.799142309 | -0.051415253 | -0.069547308 | 0.018132054 | 0.999933887 | -0.811843493 | -0.87313742 | 0.061293927 | 0.737311374 |
| ENSG00000142794 | *NBPF3* | 1.29660567 | -0.091411298 | -0.051522234 | -0.039889064 | 0.999933887 | 0.239058385 | 0.353886143 | -0.114827758 | 0.737311374 |
| ENSG00000136238 | *RAC1* | 6.961292704 | -0.039730509 | -0.030215865 | -0.009514644 | 0.999933887 | 0.125310006 | 0.070670679 | 0.054639327 | 0.737311374 |
| ENSG00000174516 | *PELI3* | 1.003970599 | 0.070589548 | -0.044999645 | 0.115589193 | 0.999933887 | -0.164719155 | -0.26770835 | 0.102989195 | 0.737311374 |
| ENSG00000198496 | *NBR2* | 1.226085896 | -0.051649142 | -0.098533466 | 0.046884324 | 0.999933887 | -0.122608886 | -0.013076281 | -0.109532605 | 0.737311374 |
| ENSG00000196284 | *SUPT3H* | 0.330970279 | 0.048938161 | -0.111887512 | 0.160825672 | 0.999933887 | 0.065175925 | -0.070961761 | 0.136137686 | 0.737311374 |
| ENSG00000268947 | *AC002128.1* | -0.365782142 | 0.006117333 | 0.153242749 | -0.147125416 | 0.999933887 | -0.061975536 | 0.080653836 | -0.142629371 | 0.737311374 |
| ENSG00000155926 | *SLA* | 7.683759331 | -0.063910772 | -0.084479712 | 0.02056894 | 0.999933887 | 0.028190689 | -0.023342393 | 0.051533082 | 0.737334793 |
| ENSG00000134809 | *TIMM10* | 2.109201991 | -0.059453351 | 0.150291982 | -0.209745334 | 0.999933887 | 0.133791937 | 0.0540346 | 0.079757337 | 0.737334793 |
| ENSG00000162231 | *NXF1* | 7.12725089 | 0.014732878 | 0.031393667 | -0.016660789 | 0.999933887 | 0.148357202 | 0.10266662 | 0.045690582 | 0.737334793 |
| ENSG00000171817 | *ZNF540* | -0.238609994 | 0.056941134 | -0.256524191 | 0.313465325 | 0.999933887 | -0.182465178 | -0.017838018 | -0.16462716 | 0.737334793 |
| ENSG00000198755 | *RPL10A* | 7.107225029 | -0.004804377 | -0.040159716 | 0.035355339 | 0.999933887 | 0.126328981 | 0.177105208 | -0.050776227 | 0.737334793 |
| ENSG00000134830 | *C5AR2* | 7.265078062 | -0.078390451 | -0.104300512 | 0.02591006 | 0.999933887 | -0.746456216 | -0.799973606 | 0.05351739 | 0.737431291 |
| ENSG00000223884 | *AC068481.1* | 0.11615288 | -0.189968458 | 0.100022196 | -0.289990654 | 0.999933887 | -0.052848332 | -0.204681212 | 0.15183288 | 0.737431291 |
| ENSG00000173585 | *CCR9* | -1.001412628 | -0.013406949 | 0.132080285 | -0.145487235 | 0.999933887 | 0.407597327 | 0.218967057 | 0.18863027 | 0.737431291 |
| ENSG00000139083 | *ETV6* | 6.497421762 | -0.022613676 | -0.037039957 | 0.014426281 | 0.999933887 | 0.058899826 | 0.106448646 | -0.04754882 | 0.737694737 |
| ENSG00000085265 | *FCN1* | 5.773229922 | 0.05674082 | 0.03948768 | 0.01725314 | 0.999933887 | -0.111111811 | -0.188873286 | 0.077761474 | 0.73769625 |
| ENSG00000204584 | *AC027801.1* | 0.087329117 | 0.104111961 | 0.181688602 | -0.077576641 | 0.999933887 | -0.221953944 | -0.407494645 | 0.185540701 | 0.73769625 |
| ENSG00000175294 | *CATSPER1* | 0.065408809 | -0.044779673 | -0.182992639 | 0.138212966 | 0.999933887 | -0.814310398 | -0.973947061 | 0.159636663 | 0.73769625 |
| ENSG00000089091 | *DZANK1* | -0.160111456 | -0.029846635 | 0.094465746 | -0.124312381 | 0.999933887 | 0.022909474 | 0.20213124 | -0.179221766 | 0.737703346 |
| ENSG00000285851 | *AL359762.3* | 1.203269488 | -0.155312254 | 0.015784001 | -0.171096255 | 0.999933887 | -0.373951255 | -0.264645573 | -0.109305682 | 0.737933313 |
| ENSG00000162994 | *CLHC1* | -0.278011758 | -0.226490688 | -0.43102107 | 0.204530382 | 0.999933887 | -0.285290543 | -0.111959614 | -0.173330928 | 0.738308588 |
| ENSG00000274963 | *RN7SL600P* | 1.735790167 | 0.109976745 | 0.043835703 | 0.066141041 | 0.999933887 | 0.199614505 | 0.338282845 | -0.13866834 | 0.738427528 |

| | | | | | | | | | | |
|---|---|---|---|---|---|---|---|---|---|---|
| ENSG00000178295 | *GEN1* | 1.423689809 | -0.072139888 | -0.061762088 | -0.0103778 | 0.999933887 | -0.053494118 | 0.106238331 | -0.15973245 | 0.738671011 |
| ENSG00000129158 | *SERGEF* | 0.520640174 | 0.048237931 | -0.008659217 | 0.056897148 | 0.999933887 | 0.07948096 | 0.195685728 | -0.116204769 | 0.738701296 |
| ENSG00000119772 | *DNMT3A* | 6.006380137 | 0.023126879 | 0.090571313 | -0.067444434 | 0.999933887 | -0.440615161 | -0.490941998 | 0.050326837 | 0.738711268 |
| ENSG00000243836 | *WDR86-AS1* | -0.297112308 | 0.005380427 | 0.072893541 | -0.067513114 | 0.999933887 | 0.157109069 | -0.01299617 | 0.170105239 | 0.738831461 |
| ENSG00000166896 | *ATP23* | 0.878327472 | 0.109784434 | -0.176199404 | 0.285983838 | 0.999933887 | 0.219944879 | 0.353863459 | -0.13391858 | 0.739185815 |
| ENSG00000258634 | *AL160006.1* | 2.294099273 | -0.087829948 | 0.089107505 | -0.176937453 | 0.999933887 | -0.141105412 | -0.046718294 | -0.094387118 | 0.739542075 |
| ENSG00000254017 | *IGHEP2* | -0.193945386 | 0.046158716 | -0.132514926 | 0.178673642 | 0.999933887 | 0.237643464 | 0.511766916 | -0.274123452 | 0.739577722 |
| ENSG00000229926 | *IL9RP1* | -2.098262352 | -0.416839765 | -0.282242851 | -0.134596914 | 0.999933887 | -1.394493035 | -1.622096286 | 0.227603251 | 0.73978914 |
| ENSG00000047230 | *CTPS2* | -0.475824338 | -0.113989893 | -0.358974785 | 0.244984892 | 0.999933887 | 0.11467189 | -0.074351792 | 0.189023682 | 0.73978914 |
| ENSG00000125650 | *PSPN* | -0.2239958 | -0.15842639 | 0.350066579 | -0.508492969 | 0.999933887 | -0.072495544 | 0.086276166 | -0.158771709 | 0.739797955 |
| ENSG00000085224 | *ATRX* | 5.400542951 | -0.047194683 | -0.135534504 | 0.088339821 | 0.999933887 | -0.606982352 | -0.453197905 | -0.153784447 | 0.739934813 |
| ENSG00000099998 | *GGT5* | 0.440038645 | 0.033464785 | 0.187354773 | -0.153889988 | 0.999933887 | 0.7932002 | 0.961449228 | -0.168249028 | 0.740189191 |
| ENSG00000259877 | *AC009113.1* | 1.189723757 | -0.193993772 | -0.09847433 | -0.095519442 | 0.999933887 | -0.27970981 | -0.376016841 | 0.096307031 | 0.740335636 |
| ENSG00000205927 | *OLIG2* | 4.892813323 | -0.157010777 | -0.062528451 | -0.094482327 | 0.999933887 | -1.532332468 | -1.654757597 | 0.12242513 | 0.740430545 |
| ENSG00000159921 | *GNE* | 3.123382113 | 0.048099276 | 0.099332054 | -0.051232779 | 0.999933887 | -0.128846473 | -0.192583289 | 0.063736816 | 0.740621754 |
| ENSG00000165322 | *ARHGAP12* | 3.033758347 | -0.108730456 | -0.207842683 | 0.099112227 | 0.999933887 | -0.547435517 | -0.449394083 | -0.098041433 | 0.740710264 |
| ENSG00000253408 | *AC083973.1* | -0.743437195 | 0.313822783 | 0.478023855 | -0.164201072 | 0.999933887 | 1.954209787 | 2.1756278 | -0.221418012 | 0.740783821 |
| ENSG00000130368 | *MAS1* | -2.236674706 | 0.065301007 | -0.383747337 | 0.449048344 | 0.999933887 | 2.899413361 | 3.14508443 | -0.245671069 | 0.740783821 |
| ENSG00000127334 | *DYRK2* | 4.16935723 | 0.052766202 | -0.031139918 | 0.08390612 | 0.999933887 | -0.082299584 | -0.028076006 | -0.054223578 | 0.740822482 |
| ENSG00000224034 | *LINC02561* | -0.381701949 | 0.138468292 | -0.088467109 | 0.226935401 | 0.999933887 | -0.097675175 | 0.035238548 | -0.132913723 | 0.740843929 |
| ENSG00000105968 | *H2AZ2* | 5.618767568 | 0.013103195 | 0.005932409 | 0.007170786 | 0.999933887 | -0.120673375 | -0.084168366 | -0.03650501 | 0.740843929 |
| ENSG00000076650 | *GPATCH1* | 1.558910917 | 0.018996624 | 0.004153817 | 0.014842807 | 0.999933887 | -0.071776161 | -0.182988032 | 0.111211871 | 0.740880461 |
| ENSG00000159399 | *HK2* | 6.224262966 | 0.004357705 | 0.005004177 | -0.000646472 | 0.999933887 | -0.187073857 | -0.136946098 | -0.050127759 | 0.741414029 |
| ENSG00000146677 | *AC004453.1* | -0.26246464 | -0.28508034 | -0.191308846 | -0.093771494 | 0.999933887 | -0.037254892 | 0.190273202 | -0.227528094 | 0.741566297 |
| ENSG00000264112 | *AC015813.1* | 2.298772882 | -0.226639772 | -0.157935101 | -0.068704671 | 0.999933887 | -0.357518598 | -0.449944128 | 0.09242553 | 0.741821772 |
| ENSG00000137807 | *KIF23* | -0.889962658 | -0.377434065 | -0.137964079 | -0.239469987 | 0.999933887 | -0.351052136 | -0.590041793 | 0.238989657 | 0.741821772 |
| ENSG00000104043 | *ATP8B4* | 2.88925883 | -0.113867552 | 0.029045257 | -0.142912809 | 0.999933887 | -0.359840663 | -0.437130019 | 0.077289356 | 0.741821772 |
| ENSG00000131724 | *IL13RA1* | 6.576441799 | -0.054059699 | -0.020119355 | -0.033940345 | 0.999933887 | -0.143716817 | -0.215613841 | 0.071897024 | 0.741821772 |
| ENSG00000168994 | *PXDC1* | 1.489221915 | 0.089371891 | 0.192033449 | -0.102661558 | 0.999933887 | 0.702587119 | 0.766121909 | -0.06353479 | 0.741821772 |
| ENSG00000106554 | *CHCHD3* | 2.30595024 | -0.033116844 | -0.093621978 | 0.060505134 | 0.999933887 | 0.100596834 | 0.175332582 | -0.074735748 | 0.741821772 |
| ENSG00000125966 | *MMP24* | 2.027710322 | -0.036800878 | 0.087952689 | -0.124753567 | 0.999933887 | -0.254855239 | -0.162010079 | -0.09284516 | 0.741821772 |
| ENSG00000170473 | *PYM1* | 1.869011555 | -0.037116136 | 0.082724425 | -0.119840561 | 0.999933887 | -0.064310563 | -0.156756002 | 0.092445439 | 0.741821772 |
| ENSG00000144713 | *RPL32* | 7.621067456 | 0.017850862 | -0.052069651 | 0.069920513 | 0.999933887 | 0.089334909 | 0.141846764 | -0.052511855 | 0.741821772 |
| ENSG00000165392 | *WRN* | 1.9400575 | 0.009795891 | -0.270414271 | 0.280210162 | 0.999933887 | -0.189364272 | -0.051595274 | -0.137768999 | 0.741821772 |
| ENSG00000137267 | *TUBB2A* | 0.941253888 | -0.087757756 | -0.05722987 | -0.030527886 | 0.999933887 | 0.210692923 | 0.339067249 | -0.128374326 | 0.742002149 |
| ENSG00000242612 | *DECR2* | 0.034999708 | -0.114962016 | -0.251346775 | 0.136384759 | 0.999933887 | 0.065245653 | -0.072090875 | 0.137336528 | 0.742002149 |
| ENSG00000112208 | *BAG2* | -0.225893123 | -0.108146031 | -0.047678337 | -0.060467695 | 0.999933887 | 0.046034385 | 0.252588525 | -0.20655414 | 0.742002149 |
| ENSG00000170915 | *PAQR8* | 1.489483871 | 0.035278913 | 0.106762371 | -0.071483458 | 0.999933887 | 0.124229513 | 0.204471928 | -0.080242414 | 0.742002149 |
| ENSG00000151503 | *NCAPD3* | 2.719857623 | 0.021205905 | -0.053846034 | 0.075051939 | 0.999933887 | 0.084307312 | 0.016525114 | 0.067782198 | 0.742058807 |
| ENSG00000075336 | *TIMM21* | 0.540595554 | -0.139489254 | -0.147252277 | 0.007763023 | 0.999933887 | 0.091341172 | -0.029723393 | 0.121064564 | 0.742189262 |
| ENSG00000277548 | *AC018926.3* | -0.626710839 | -0.362782335 | 0.058258392 | -0.421040727 | 0.999933887 | -0.47953083 | -0.691857055 | 0.212326225 | 0.742427603 |
| ENSG00000245937 | *LINC01184* | 0.82433201 | -0.031424836 | -0.065265188 | 0.033840352 | 0.999933887 | 0.05916173 | -0.060504609 | 0.11966634 | 0.742427603 |
| ENSG00000213020 | *ZNF611* | 2.934671497 | -0.050294265 | 0.035006799 | -0.085301065 | 0.999933887 | -0.340907912 | -0.260589313 | -0.080318598 | 0.742463499 |
| ENSG00000099901 | *RANBP1* | 2.547560948 | 0.037450089 | -0.112042858 | 0.149492947 | 0.999933887 | 0.21689521 | 0.289069114 | -0.072173904 | 0.742463499 |
| ENSG00000128016 | *ZFP36* | 11.093066135 | 0.57665919 | 0.639322799 | -0.062663609 | 0.999933887 | 1.327856465 | 1.430957302 | -0.103100837 | 0.742570344 |
| ENSG00000238829 | *RNU7-45P* | -0.643928455 | -0.429123634 | 0.123927195 | -0.553050829 | 0.999933887 | 1.264812049 | 1.440183253 | -0.175371203 | 0.742570344 |
| ENSG00000241527 | *CA15P1* | -0.503871483 | -0.080227751 | -0.178869976 | 0.098642225 | 0.999933887 | -0.461735186 | -0.116488107 | -0.345247078 | 0.742570344 |
| ENSG00000163157 | *TMOD4* | -0.617774763 | 0.117869546 | -0.342270296 | 0.460139841 | 0.999933887 | -0.225424786 | -0.407784098 | 0.182359312 | 0.742570344 |
| ENSG00000175224 | *ATG13* | 5.976259194 | -0.062579172 | 0.013737396 | -0.076316568 | 0.999933887 | -0.212180862 | -0.169027028 | -0.043153835 | 0.742676743 |
| ENSG00000144115 | *THNSL2* | -0.261351795 | 0.173289135 | -0.165122023 | 0.338411158 | 0.999933887 | -0.649771218 | -0.828060279 | 0.178289061 | 0.742676743 |
| ENSG00000130766 | *SESN2* | 5.298748203 | 0.037897952 | 0.002547211 | 0.035350741 | 0.999933887 | 0.293079613 | 0.243829264 | 0.049250349 | 0.742676743 |
| ENSG00000107537 | *PHYH* | 0.40871803 | -0.071716285 | -0.028216087 | -0.043500198 | 0.999933887 | 0.151912213 | 0.284301109 | -0.132388896 | 0.742676743 |
| ENSG00000247572 | *CKMT2-AS1* | -0.549776226 | -0.097514464 | 0.155577179 | -0.253091643 | 0.999933887 | -0.04673115 | -0.235723592 | 0.188992442 | 0.742676743 |
| ENSG00000127399 | *LRRC61* | 0.134348636 | -0.049477023 | -0.005487088 | -0.043989934 | 0.999933887 | 0.761445638 | 0.923791154 | -0.162345517 | 0.742676743 |
| ENSG00000232517 | *CSPG4BP* | -0.408070152 | -0.094727905 | 0.023177054 | -0.11790496 | 0.999933887 | 5.253065415 | 5.005113873 | 0.247951542 | 0.742676743 |
| ENSG00000217078 | *AL021407.1* | -0.297772842 | -0.162168978 | 0.353384723 | -0.515553701 | 0.944094537 | 0.172117824 | 0.371720176 | -0.199602351 | 0.742745048 |
| ENSG00000205629 | *LCMT1* | 2.081406864 | -0.105712892 | -0.050494775 | -0.055218117 | 0.999933887 | -0.077741633 | -0.154821725 | 0.077080091 | 0.742794446 |
| ENSG00000164050 | *PLXNB1* | 0.142528895 | -0.037329015 | 0.254307482 | -0.291636497 | 0.999933887 | 0.011142997 | 0.175077252 | -0.163934254 | 0.742860869 |
| ENSG00000154447 | *SH3RF1* | 3.423904783 | 0.126094181 | 0.256211147 | -0.130116966 | 0.999933887 | -0.747027666 | -0.692480162 | -0.054547504 | 0.742897669 |
| ENSG00000259116 | *AL049869.2* | 0.552968101 | -0.172837074 | 0.110961089 | -0.283798163 | 0.999933887 | 0.114497654 | 0.272333157 | -0.157835503 | 0.74328827 |
| ENSG00000065970 | *FOXJ2* | 5.396418654 | -0.013127324 | 0.047015257 | -0.060142581 | 0.999933887 | 0.069905665 | 0.014682633 | 0.055223032 | 0.743419962 |
| ENSG00000109775 | *UFSP2* | 1.442693156 | -0.053153417 | -0.191274531 | 0.138121113 | 0.999933887 | 0.205435097 | 0.312158228 | -0.106723131 | 0.743476618 |
| ENSG00000119231 | *SENP5* | 5.048940064 | -0.055591499 | -0.001092962 | -0.054498537 | 0.999933887 | -0.02585311 | 0.016242832 | -0.042095942 | 0.743531951 |
| ENSG00000110583 | *NAA40* | 2.540187066 | 0.034448862 | 0.133752348 | -0.099303487 | 0.999933887 | 0.057030287 | -0.018191296 | 0.075221583 | 0.743531951 |
| ENSG00000156414 | *TDRD9* | 2.707471273 | 0.013597313 | -0.056082583 | 0.069679896 | 0.999933887 | -0.377454716 | -0.310226773 | -0.067227943 | 0.743531951 |
| ENSG00000110066 | *KMT5B* | 5.196933997 | -0.081426269 | -0.067044053 | -0.014382215 | 0.999933887 | -0.644289755 | -0.691545699 | 0.047255944 | 0.74354432 |
| ENSG00000175793 | *SFN* | 1.80526915 | -0.091384023 | 0.093961196 | -0.18534522 | 0.999933887 | 0.42683532 | 0.543531781 | -0.116696461 | 0.74354432 |
| ENSG00000198380 | *GFPT1* | 3.466315925 | -0.09305174 | -0.128926216 | 0.035874476 | 0.999933887 | -0.508576429 | -0.440550344 | -0.068026084 | 0.74354432 |
| ENSG00000128536 | *CDHR3* | -0.356744765 | 0.227447287 | -0.104397533 | 0.33184482 | 0.999933887 | -0.202152709 | -0.386684152 | 0.184531443 | 0.74354432 |
| ENSG00000103269 | *RHBDL1* | 0.151265889 | 0.109915786 | -0.207367638 | 0.317283425 | 0.999933887 | 0.074166824 | -0.10353147 | 0.177698295 | 0.74354432 |
| ENSG00000243317 | *STMP1* | 3.930504027 | -0.033570597 | 0.021006568 | -0.054577165 | 0.999933887 | -0.091697069 | -0.148926581 | 0.057229513 | 0.74354432 |
| ENSG00000097021 | *ACOT7* | 0.513296763 | 0.038165114 | 0.002746277 | 0.035418837 | 0.999933887 | 0.096652991 | -0.049306786 | 0.145959776 | 0.74354432 |
| ENSG00000162909 | *CAPN2* | 5.735446102 | 0.010880295 | -0.018194344 | 0.029074638 | 0.999933887 | 0.11993555 | 0.155257711 | -0.035322161 | 0.74354432 |
| ENSG00000265907 | *AP000919.1* | 1.301092168 | 0.025477691 | 0.198324166 | -0.172846475 | 0.999933887 | -0.883130214 | -1.019078267 | 0.135948053 | 0.74354432 |
| ENSG00000234571 | *H2BP2* | -1.534917906 | 0.011558776 | 0.102167985 | -0.090609209 | 0.999933887 | -0.542240875 | -0.180713055 | -0.36152782 | 0.74354432 |
| ENSG00000114270 | *COL7A1* | 1.408614288 | 0.008048065 | 0.27174284 | -0.263694775 | 0.999933887 | 0.231336572 | 0.10148376 | 0.129852812 | 0.74354432 |
| ENSG00000171792 | *RHNO1* | 1.62466459 | -0.003139481 | 0.06709825 | -0.070237731 | 0.999933887 | 0.131003856 | 0.036362638 | 0.094641218 | 0.74354432 |
| ENSG00000115355 | *CCDC88A* | 1.986357974 | 0.004099664 | -0.158153107 | 0.162252771 | 0.999933887 | -0.450593622 | -0.33932623 | -0.111267392 | 0.74354432 |
| ENSG00000049768 | *FOXP3* | -0.158494607 | 0.004358046 | 0.107845143 | -0.103487097 | 0.999933887 | 0.334895726 | 0.131639756 | 0.20325597 | 0.74354432 |
| ENSG00000213186 | *TRIM59* | 1.017625464 | 0.00341474 | -0.132292602 | 0.135707342 | 0.999933887 | -0.060646334 | 0.073738705 | -0.134385039 | 0.74354432 |
| ENSG00000171843 | *MLLT3* | 2.458462262 | 0.046473712 | -0.231607283 | 0.278080996 | 0.999933887 | -0.058770281 | 0.024142534 | -0.082912815 | 0.743648207 |
| ENSG00000108666 | *C17orf75* | 0.576853124 | -0.045035239 | 0.005013616 | -0.050048854 | 0.999933887 | 0.188695829 | 0.075939998 | 0.112755831 | 0.74372312 |
| ENSG00000123219 | *CENPK* | 0.781308141 | -0.151065991 | -0.057354301 | -0.093711691 | 0.999933887 | -0.020221791 | 0.124053298 | -0.14427509 | 0.743873609 |
| ENSG00000205981 | *DNAJC19* | 1.337042563 | 0.092057345 | -0.169067189 | 0.261124534 | 0.999933887 | 0.182177313 | 0.085247932 | 0.096929381 | 0.743873609 |
| ENSG00000173540 | *GMPPB* | 2.640050929 | 0.064467175 | 0.091423426 | -0.026956251 | 0.999933887 | 0.135795767 | 0.210077303 | -0.074281536 | 0.743873609 |
| ENSG00000108469 | *RECQL5* | 3.352533422 | 0.011360364 | 0.097121286 | -0.085760922 | 0.999933887 | 0.210556735 | 0.274955703 | -0.064398968 | 0.743873609 |
| ENSG00000133812 | *SBF2* | 5.555234654 | -0.007202106 | -0.025493227 | 0.018291121 | 0.999933887 | 0.05434452 | 0.099364025 | -0.045019506 | 0.743873609 |
| ENSG00000278881 | *FP325331.1* | -0.188431281 | 0.115765324 | 0.056448728 | 0.059316595 | 0.999933887 | 0.152234781 | 0.310107693 | -0.157872913 | 0.743991405 |
| ENSG00000129657 | *SEC14L1* | 10.225921101 | -0.035869291 | 0.009056643 | -0.044925933 | 0.999933887 | -0.818914179 | -0.761157335 | -0.057756845 | 0.743991405 |
| ENSG00000106367 | *AP1S1* | 1.484073417 | 0.168042971 | -0.152447307 | 0.320490278 | 0.999933887 | -0.298446495 | -0.397795929 | 0.099349434 | 0.744018409 |
| ENSG00000179240 | *GVQW3* | 0.185215003 | -0.185347291 | 0.16734099 | -0.352688281 | 0.999933887 | -0.131192164 | 0.005736232 | -0.136928396 | 0.744124372 |
| ENSG00000175899 | *A2M* | 2.152309469 | 0.324933354 | 0.498872519 | -0.173939165 | 0.999933887 | 0.020274705 | 0.207940876 | -0.187666171 | 0.744291981 |
| ENSG00000176410 | *DNAJC30* | 0.16777027 | -0.204809513 | 0.094191189 | -0.299000702 | 0.999933887 | 0.246476431 | 0.104426677 | 0.142049754 | 0.744291981 |
| ENSG00000173727 | *AP000769.1* | 1.543246615 | 0.167294005 | 0.170895265 | -0.00360126 | 0.999933887 | -0.374677599 | -0.264730334 | -0.109947264 | 0.744291981 |
| ENSG00000156304 | *SCAF4* | 4.767037284 | -0.055415194 | -0.093793014 | 0.038377819 | 0.999933887 | -0.227638969 | -0.184714502 | -0.042924467 | 0.744291981 |
| ENSG00000136273 | *HUS1* | 2.926019749 | -0.071486912 | 0.023157872 | -0.094644784 | 0.999933887 | 0.083638078 | 0.024822958 | 0.05881512 | 0.744291981 |
| ENSG00000111602 | *TIMELESS* | 1.071365719 | 0.114343448 | 0.132136201 | -0.017792753 | 0.999933887 | 0.055339729 | -0.05342614 | 0.108765869 | 0.744291981 |
| ENSG00000187815 | *ZFP69* | -0.216242417 | -0.109537749 | -0.068341278 | -0.04119647 | 0.999933887 | 0.008660474 | 0.160524296 | -0.151863822 | 0.744291981 |
| ENSG00000171302 | *CANT1* | 7.09321679 | -0.026417229 | -0.035714871 | 0.009297642 | 0.999933887 | -0.269286533 | -0.226478501 | -0.042808032 | 0.744291981 |
| ENSG00000188931 | *CFAP126* | -0.151477819 | 0.088966316 | 0.010810037 | 0.078156278 | 0.999933887 | -0.16375901 | -0.027742611 | -0.136016398 | 0.744291981 |
| ENSG00000111640 | *GAPDH* | 8.572733602 | 0.022779705 | -0.033769206 | 0.056548911 | 0.999933887 | -0.14353617 | -0.189763184 | 0.046227014 | 0.744291981 |
| ENSG00000121940 | *CLCC1* | 0.852171628 | -0.057395009 | 0.056730789 | -0.114125798 | 0.999933887 | -0.026818172 | 0.086102325 | -0.112920497 | 0.744291981 |
| ENSG00000134545 | *KLRC1* | -0.535367498 | -0.094927054 | 0.032785329 | -0.127712383 | 0.999933887 | 0.177401272 | 0.356120971 | -0.178719699 | 0.744291981 |
| ENSG00000250644 | *AC068580.4* | 5.559826822 | 0.03930389 | 0.024566122 | 0.014737768 | 0.999933887 | 0.560970527 | 0.638388044 | -0.077417517 | 0.744291981 |
| ENSG00000245904 | *AC025164.1* | 0.373979265 | -0.027967201 | 0.188814056 | -0.216781257 | 0.999933887 | -0.051992174 | 0.098781772 | -0.150773946 | 0.744291981 |

| | | | | | | | | | | |
|---|---|---|---|---|---|---|---|---|---|---|
| ENSG00000130921 | *C12orf65* | 2.087865831 | 0.008296265 | 0.040159644 | -0.03186338 | 0.999933887 | 0.215029982 | 0.290835823 | -0.075805841 | 0.744291981 |
| ENSG00000205758 | *CRYZL1* | 1.978367545 | -0.004448552 | 0.148234947 | -0.152683499 | 0.999933887 | -0.106726454 | -0.193851336 | 0.087124883 | 0.74506753 |
| ENSG00000278330 | *AC018529.2* | 6.147322527 | 0.115277246 | 0.205122253 | -0.089845007 | 0.999933887 | 0.2821945 | 0.353335055 | -0.071140555 | 0.745248299 |
| ENSG00000115282 | *TTC31* | 3.105134388 | -0.012274315 | -0.006001103 | -0.006273212 | 0.999933887 | 0.001394066 | -0.049761394 | 0.05115546 | 0.745248299 |
| ENSG00000277452 | *RN7SL473P* | 1.538756126 | 0.2114225 | 0.150288199 | 0.061134301 | 0.999933887 | 0.141017443 | 0.294031023 | -0.153013579 | 0.745350083 |
| ENSG00000260219 | *CD2BP2-DT* | 0.235185779 | 0.184745935 | -0.079251734 | 0.263997669 | 0.999933887 | 0.085384635 | -0.06194516 | 0.147329795 | 0.745627002 |
| ENSG00000151445 | *VIPAS39* | 3.041088203 | -0.042691016 | -0.128440103 | 0.085749086 | 0.999933887 | -0.27197253 | -0.327366752 | 0.055394221 | 0.745666853 |
| ENSG00000018699 | *TTC27* | 1.027193887 | -0.038851475 | 0.008339977 | -0.047191452 | 0.999933887 | 0.159283709 | 0.251522198 | -0.092238489 | 0.745883035 |
| ENSG00000277290 | *AC136475.10* | -0.209701029 | -0.040649317 | -0.26177275 | 0.221123433 | 0.999933887 | -0.395585273 | -0.203349907 | -0.192235366 | 0.74606879 |
| ENSG00000125755 | *SYMPK* | 4.845108011 | 0.022534897 | 0.013224935 | 0.009309962 | 0.999933887 | 0.009768102 | -0.028742824 | 0.038510926 | 0.746109065 |
| ENSG00000130433 | *CACNG6* | 0.363971944 | -0.062688347 | 0.287138504 | -0.349826851 | 0.999933887 | -0.04204991 | 0.080384195 | -0.122434105 | 0.746109065 |
| ENSG00000116514 | *RNF19B* | 9.00848978 | 0.005233704 | 0.038437007 | -0.033203303 | 0.999933887 | 1.007436333 | 0.944138477 | 0.063297856 | 0.746109065 |
| ENSG00000101445 | *PPP1R16B* | 5.722513972 | -0.00227784 | -0.042813047 | 0.040535208 | 0.999933887 | 0.109208497 | 0.163586992 | -0.054378495 | 0.746109065 |
| ENSG00000184178 | *SCFD2* | 0.462983508 | 0.06033219 | 0.058938348 | 0.001393843 | 0.999933887 | 0.122228916 | 0.004605573 | 0.117623343 | 0.746236609 |
| ENSG00000006282 | *SPATA20* | 1.294396801 | -0.020166295 | -0.11468113 | 0.094514834 | 0.999933887 | 0.045721821 | 0.1621412 | -0.116419379 | 0.746325312 |
| ENSG00000285366 | *AC008533.1* | 2.09839666 | -0.099855012 | 0.252488108 | -0.35234312 | 0.999933887 | 0.068877543 | 0.232521632 | -0.16364409 | 0.746637785 |
| ENSG00000287167 | *AL133313.1* | -0.75441628 | -0.209649408 | 0.034176904 | -0.243826312 | 0.999933887 | 0.455791918 | 0.278009129 | 0.177782788 | 0.746637785 |
| ENSG00000205730 | *ITPRIPL2* | 0.461748153 | 0.070025473 | -0.084711438 | 0.154736911 | 0.999933887 | -0.333076562 | -0.482326651 | 0.149250089 | 0.746954713 |
| ENSG00000049246 | *PER3* | 3.713891511 | 0.024508295 | 0.106837105 | -0.08232881 | 0.999933887 | -0.261962442 | -0.181218587 | -0.080743855 | 0.747286082 |
| ENSG00000197063 | *MAFG* | 8.442514275 | 0.087678806 | 0.02785949 | 0.059819316 | 0.999933887 | 0.562520218 | 0.611894149 | -0.049373931 | 0.747394125 |
| ENSG00000254750 | *CASP1P2* | 0.433207425 | 0.085781998 | -0.285829224 | 0.371611222 | 0.999933887 | 0.462388428 | 0.573271143 | -0.110882715 | 0.747394125 |
| ENSG00000260425 | *AL031709.1* | 0.618761717 | -0.012306488 | 0.029343915 | -0.041650403 | 0.999933887 | 0.428223738 | 0.521560869 | -0.093337132 | 0.747394125 |
| ENSG00000117758 | *STX12* | 4.488817198 | 0.003386422 | 0.036022105 | -0.032635683 | 0.999933887 | -0.534197155 | -0.478121126 | -0.056076029 | 0.747394125 |
| ENSG00000156504 | *FAM122B* | 3.567569643 | -0.035197343 | -0.035786439 | 0.000589096 | 0.999933887 | -0.144398312 | -0.199348023 | 0.054949711 | 0.748041494 |
| ENSG00000243650 | *RN7SL834P* | 1.265699365 | -0.029153357 | -0.106884518 | 0.077731161 | 0.999933887 | -0.224871625 | -0.390877118 | 0.166005493 | 0.748041494 |
| ENSG00000134333 | *LDHA* | 6.104695381 | -0.016295463 | -0.061614444 | 0.045318982 | 0.999933887 | 0.086619975 | 0.135326911 | -0.048706935 | 0.748377214 |
| ENSG00000163814 | *CDCP1* | 1.113823529 | 0.065648584 | 0.006953227 | 0.058695358 | 0.999933887 | -0.872180152 | -0.972087775 | 0.099907624 | 0.748447071 |
| ENSG00000183684 | *ALYREF* | 5.212341625 | 0.072749287 | -0.024048614 | 0.096797902 | 0.999933887 | 0.405388348 | 0.465834994 | -0.060446646 | 0.748513037 |
| ENSG00000141013 | *GAS8* | 0.173265453 | -0.086901963 | 0.361928464 | -0.448830427 | 0.981460244 | -0.127309223 | 0.010912842 | -0.138222065 | 0.748513037 |
| ENSG00000187726 | *DNAJB13* | -0.398895037 | 0.029959083 | -0.448891399 | 0.478850482 | 0.999933887 | 7.596e-05 | -0.239988401 | 0.24006436 | 0.748513037 |
| ENSG00000172936 | *MYD88* | 8.685984933 | -0.03076721 | -0.064564322 | 0.033797112 | 0.999933887 | -0.546595996 | -0.488062031 | -0.058533965 | 0.748625517 |
| ENSG00000180425 | *C11orf71* | 1.738890829 | 0.016364006 | -0.01890155 | 0.035265556 | 0.999933887 | 0.429822451 | 0.33463806 | 0.095184392 | 0.748625517 |
| ENSG00000256525 | *POLG2* | 2.752914603 | -0.097614344 | 0.060829228 | -0.158443572 | 0.999933887 | 0.067076745 | 0.006562729 | 0.060514016 | 0.748658397 |
| ENSG00000009724 | *MASP2* | 0.881851892 | -0.209978709 | -0.045834201 | -0.164144508 | 0.999933887 | -0.315352986 | -0.20102035 | -0.114332636 | 0.748658397 |
| ENSG00000231412 | *AC005392.2* | 0.339170485 | -0.105073509 | -0.016793618 | -0.088279892 | 0.999933887 | -0.746632007 | -0.885471551 | 0.138839544 | 0.748741297 |
| ENSG00000162438 | *CTRC* | 0.87860752 | 0.071196098 | 0.207919014 | -0.136722916 | 0.999933887 | 0.675255914 | 0.797468029 | -0.122212116 | 0.748741297 |
| ENSG00000087077 | *TRIP6* | 1.581025075 | 0.041321726 | 0.059930961 | -0.018609235 | 0.999933887 | -0.1509075 | -0.258623273 | 0.107715772 | 0.748741297 |
| ENSG00000006695 | *COX10* | 0.851101752 | 0.023097627 | 0.35714287 | -0.334045243 | 0.999933887 | 0.061835032 | -0.05910348 | 0.120938512 | 0.748741297 |
| ENSG00000245522 | *LINC02709* | -0.611106655 | 0.297107796 | 0.17465663 | 0.122451166 | 0.999933887 | 1.075960289 | 1.255245708 | -0.179285419 | 0.748778502 |
| ENSG00000115129 | *TP53I3* | 0.69130485 | 0.146526347 | 0.111255295 | 0.035271052 | 0.999933887 | -0.322243472 | -0.177415575 | -0.144827897 | 0.748851587 |
| ENSG00000227615 | *AP001324.1* | 1.158716422 | 0.073954213 | -0.228732746 | 0.30268696 | 0.999933887 | 0.066751112 | -0.028700777 | 0.095451889 | 0.748851587 |
| ENSG00000187961 | *KLHL17* | 2.287600912 | 0.020669265 | 0.139975612 | -0.119306347 | 0.999933887 | 0.004597797 | -0.073738258 | 0.078336055 | 0.748851587 |
| ENSG00000163141 | *BNIPL* | 0.000312969 | 0.071937967 | -0.05737214 | 0.129310107 | 0.999933887 | -0.067232727 | -0.208892582 | 0.141659855 | 0.7488971 |
| ENSG00000048392 | *RRM2B* | 5.214423719 | -0.114268812 | -0.104387294 | -0.009881518 | 0.999933887 | -0.972778318 | -0.888269503 | -0.084508815 | 0.748995991 |
| ENSG00000197451 | *HNRNPAB* | 4.139882821 | 0.064109841 | 0.06682618 | -0.002716339 | 0.999933887 | 0.128900125 | 0.193453604 | -0.064553479 | 0.748995991 |
| ENSG00000257221 | *AC007569.1* | 0.815328656 | 0.051705981 | -0.186159449 | 0.237865429 | 0.999933887 | -0.623668518 | -0.78083486 | 0.157166343 | 0.748995991 |
| ENSG00000104131 | *EIF3J* | 4.637565772 | -0.033975835 | -0.04013246 | 0.006156625 | 0.999933887 | -0.118082641 | -0.161880793 | 0.043798152 | 0.749103189 |
| ENSG00000160193 | *WDR4* | 1.087838689 | 0.058949214 | 0.296077963 | -0.237128749 | 0.999933887 | 0.366654144 | 0.276097322 | 0.090556822 | 0.749103189 |
| ENSG00000176340 | *COX8A* | 3.847228544 | 0.047290216 | -0.043461994 | 0.090752209 | 0.999933887 | 0.131984742 | 0.189546102 | -0.05756136 | 0.749149498 |
| ENSG00000122026 | *RPL21* | 7.461704583 | -0.036283104 | -0.109685904 | 0.0734028 | 0.999933887 | 0.078423609 | 0.124483964 | -0.046060355 | 0.749365019 |
| ENSG00000213463 | *SYNJ2BP* | 1.061674473 | -0.18106068 | -0.302412653 | 0.121351973 | 0.999933887 | 0.002773812 | -0.097970581 | 0.100744392 | 0.750058533 |
| ENSG00000090565 | *RAB11FIP3* | 2.990524604 | 0.112362365 | -0.015652317 | 0.128014682 | 0.999933887 | 0.067593639 | 0.003692928 | 0.063900711 | 0.750058533 |
| ENSG00000066230 | *SLC9A3* | -0.788803394 | 0.486260989 | 0.36772909 | 0.118531899 | 0.999933887 | 0.123680224 | 0.295909866 | -0.172229642 | 0.750058533 |
| ENSG00000196466 | *ZNF799* | -0.589650558 | 0.174061672 | -0.215673942 | 0.389735614 | 0.999933887 | -0.116347724 | -0.342682812 | 0.226335089 | 0.750058533 |
| ENSG00000213563 | *C8orf82* | 1.134590565 | 0.035831822 | 0.076966723 | -0.0411349 | 0.999933887 | 0.005393087 | -0.114523656 | 0.119916743 | 0.750058533 |
| ENSG00000168275 | *COA6* | 3.146726029 | -0.019516533 | -0.100762821 | 0.081246288 | 0.999933887 | -0.158344316 | -0.099949321 | -0.058394996 | 0.750058533 |
| ENSG00000181555 | *SETD2* | 6.05969792 | -0.002999736 | -0.127604979 | 0.124605244 | 0.999933887 | -0.737097349 | -0.820038882 | 0.082941534 | 0.750058533 |
| ENSG00000214941 | *ZSWIM7* | 1.79996716 | 0.198878368 | -0.042697983 | 0.241576351 | 0.999933887 | -0.061575274 | -0.143983701 | 0.082408426 | 0.750155195 |
| ENSG00000128185 | *DGCR6L* | 2.627994681 | 0.075969802 | -0.08362966 | 0.159599462 | 0.999933887 | 0.033575317 | -0.050991398 | 0.084566714 | 0.751016482 |
| ENSG00000128422 | *KRT17* | 0.658842524 | 0.08381369 | 0.278804779 | -0.194991089 | 0.999933887 | 1.105675561 | 1.286946494 | -0.181270933 | 0.751038496 |
| ENSG00000177030 | *DEAF1* | 3.686811285 | -0.012967585 | -0.062840951 | 0.049873366 | 0.999933887 | 0.075002739 | 0.12952151 | -0.054518771 | 0.751173533 |
| ENSG00000267481 | *AC011477.1* | -0.550525206 | 0.047359018 | 0.078643282 | -0.031284264 | 0.999933887 | -0.12893628 | -0.306551625 | 0.177615346 | 0.751522147 |
| ENSG00000087586 | *AURKA* | 0.323817738 | 0.030559938 | -0.007000402 | 0.03756034 | 0.999933887 | 0.177375745 | 0.334466754 | -0.157091009 | 0.751870444 |
| ENSG00000223547 | *ZNF844* | 2.484630836 | -0.125819277 | -0.098978473 | -0.026840804 | 0.999933887 | -0.141777642 | -0.069109187 | -0.072668455 | 0.751915127 |
| ENSG00000018510 | *AGPS* | 2.796746019 | -0.058212267 | -0.164949676 | 0.106737409 | 0.999933887 | -0.155137621 | -0.089112955 | -0.066024665 | 0.751915127 |
| ENSG00000132680 | *KHDC4* | 4.624239623 | -0.051505053 | 0.014461476 | -0.065966528 | 0.999933887 | 0.015552129 | -0.030534163 | 0.046086293 | 0.75198023 |
| ENSG00000259238 | *AC092755.2* | 0.632786164 | 0.220910562 | 0.354609354 | -0.133698792 | 0.999933887 | 0.579803323 | 0.752700833 | -0.173617509 | 0.752051015 |
| ENSG00000137692 | *DCUN1D5* | 1.997854246 | 0.077366106 | -0.134856893 | 0.212222999 | 0.999933887 | -0.000848398 | 0.075143138 | -0.075991536 | 0.752051015 |
| ENSG00000110031 | *LPXN* | 4.929260296 | -0.043727263 | -0.082972394 | 0.039245131 | 0.999933887 | 0.344959178 | 0.298840753 | 0.046118425 | 0.752051015 |
| ENSG00000165025 | *SYK* | 7.330498164 | -0.027804136 | -0.03288273 | 0.005078594 | 0.999933887 | 0.471155694 | 0.541326474 | -0.07017078 | 0.752051015 |
| ENSG00000059122 | *FLYWCH1* | 3.941513455 | 0.103133813 | 0.027859174 | 0.075274639 | 0.999933887 | 0.041458289 | 0.102290756 | -0.060832467 | 0.752398239 |
| ENSG00000185019 | *UBOX5* | 3.146580535 | -0.084046965 | -0.009738313 | -0.074308653 | 0.999933887 | -0.138658971 | -0.077978437 | -0.060680534 | 0.752398239 |
| ENSG00000146535 | *GNA12* | 3.295267324 | -0.064671252 | 0.041313684 | -0.105984936 | 0.999933887 | 0.000385274 | 0.061770659 | -0.061385385 | 0.752398239 |
| ENSG00000097046 | *CDC7* | 1.07704031 | 0.098190616 | -0.227790748 | 0.325981364 | 0.999933887 | 0.140300686 | 0.241842142 | -0.101541456 | 0.752398239 |
| ENSG00000224376 | *AC017104.1* | -1.34789945 | 0.298945962 | 0.316827593 | -0.017881632 | 0.999933887 | 1.953212097 | 2.202688329 | -0.249476232 | 0.752398239 |
| ENSG00000169718 | *DUS1L* | 4.514198929 | 0.043635203 | 0.103886261 | -0.060251057 | 0.999933887 | 0.151479097 | 0.097877984 | 0.053601113 | 0.752398239 |
| ENSG00000283602 | *AC116353.6* | 1.669123143 | -0.067652096 | -0.209996695 | 0.142344598 | 0.999933887 | -0.74798318 | -0.647565178 | -0.100418002 | 0.752398239 |
| ENSG00000255423 | *EBLN2* | 3.346533619 | 0.030850193 | 0.083532215 | -0.052682022 | 0.999933887 | 0.315525317 | 0.421764386 | -0.106239069 | 0.752398239 |
| ENSG00000170634 | *ACYP2* | 1.559467922 | -0.028400369 | 0.116153292 | -0.144553661 | 0.999933887 | 0.252239697 | 0.153356762 | 0.098882935 | 0.752398239 |
| ENSG00000259070 | *LINC00639* | 0.173463629 | -0.04751856 | 0.100561828 | -0.148080387 | 0.999933887 | -0.491756593 | -0.265313641 | -0.226442952 | 0.752398239 |
| ENSG00000239883 | *PARGP1* | 0.81932471 | 0.175283751 | -0.106966811 | 0.282250562 | 0.999933887 | 0.020612028 | 0.155853782 | -0.135241754 | 0.752457115 |
| ENSG00000106868 | *SUSD1* | 3.219550343 | -0.005567416 | -0.076511474 | 0.070944058 | 0.999933887 | -0.004926458 | 0.054128345 | -0.059054803 | 0.752457115 |
| ENSG00000176396 | *EID2* | 1.355957949 | 0.173546267 | 0.019993865 | 0.153552403 | 0.999933887 | -0.086030006 | 0.024921934 | -0.11095194 | 0.752605804 |
| ENSG00000153162 | *BMP6* | 4.889595916 | 0.073226001 | 0.026565532 | 0.046660469 | 0.999933887 | -0.85078288 | -0.782336395 | -0.068446485 | 0.752605804 |
| ENSG00000185869 | *ZNF829* | -0.31410273 | -0.118140768 | -0.093376685 | -0.024764083 | 0.999933887 | -0.1310059 | 0.045326591 | -0.17633249 | 0.752605804 |
| ENSG00000196639 | *HRH1* | -0.454877821 | -0.036346111 | -0.116079561 | 0.07973345 | 0.999933887 | 0.895342648 | 1.060225985 | -0.164883337 | 0.752605804 |
| ENSG00000120029 | *ARMH3* | 2.84607615 | -0.009651714 | 0.0255945 | -0.035246215 | 0.999933887 | -0.026095909 | -0.094485203 | 0.068389294 | 0.752605804 |
| ENSG00000047579 | *DTNBP1* | 5.626031365 | 0.002194708 | 0.055872503 | -0.053677795 | 0.999933887 | -0.109628786 | -0.064381056 | -0.045247729 | 0.752605804 |
| ENSG00000198815 | *FOXJ3* | 5.488593638 | -0.055079734 | 0.009612373 | -0.064692107 | 0.999933887 | -0.198722657 | -0.236964658 | 0.038242 | 0.752966808 |
| ENSG00000155158 | *TTC39B* | 2.818907661 | 0.006992188 | -0.045169745 | 0.052161933 | 0.999933887 | -7.3628e-05 | 0.071436574 | -0.071510203 | 0.752966808 |
| ENSG00000182134 | *TDRKH* | 0.238564808 | 0.044476055 | -0.06633838 | 0.110814436 | 0.999933887 | 0.286145305 | 0.436382912 | -0.150237607 | 0.75307934 |
| ENSG00000157837 | *SPPL3* | 5.649552909 | -0.049298686 | 0.06043896 | -0.109737645 | 0.999933887 | -0.302231604 | -0.263431848 | -0.038799755 | 0.753204307 |
| ENSG00000254177 | *AC079209.2* | 0.310309446 | 0.072624202 | 0.2593402 | -0.186715997 | 0.999933887 | 0.375153818 | 0.588268963 | -0.213115145 | 0.753555378 |
| ENSG00000159214 | *CCDC24* | 1.00479313 | 0.085651465 | 0.08845729 | -0.002805826 | 0.999933887 | 0.311074339 | 0.413765003 | -0.102690664 | 0.753838181 |
| ENSG00000104872 | *PIH1D1* | 3.485948467 | -0.034214282 | 0.002167926 | -0.036382207 | 0.999933887 | 0.106611972 | 0.049970928 | 0.056641043 | 0.753838181 |
| ENSG00000166037 | *CEP57* | 3.995854557 | -0.05262108 | -0.104602632 | 0.051981552 | 0.999933887 | -0.142239585 | -0.048654003 | -0.093585582 | 0.753838181 |
| ENSG00000139193 | *CD27* | 2.211916832 | 0.006382555 | -0.046998865 | 0.05338142 | 0.999933887 | 0.070079416 | 0.140346843 | -0.070267427 | 0.753838181 |
| ENSG00000165195 | *PIGA* | 4.232247172 | -0.001248275 | -0.040138502 | 0.038890227 | 0.999933887 | -0.01717343 | -0.090643258 | 0.073469827 | 0.753838181 |
| ENSG00000166479 | *TMX3* | 3.407703873 | -0.060856381 | -0.091981182 | 0.031124801 | 0.999933887 | -0.35223371 | -0.23314011 | -0.1190936 | 0.753934576 |
| ENSG00000167578 | *RAB4B* | 0.902940887 | -0.006100328 | -0.083398492 | 0.077298164 | 0.999933887 | -0.045442377 | 0.056435348 | -0.101877725 | 0.753958311 |
| ENSG00000176845 | *METRNL* | 7.980851901 | 0.109438907 | 0.14333205 | -0.033893142 | 0.999933887 | 0.009633982 | 0.071830883 | -0.062196901 | 0.754035004 |
| ENSG00000141646 | *SMAD4* | 4.920542241 | -0.06813297 | -0.010266083 | -0.057866887 | 0.999933887 | -0.567858944 | -0.625140084 | 0.05728114 | 0.754401243 |
| ENSG00000151490 | *PTPRO* | -1.033446106 | 0.131292955 | -0.017401605 | 0.14869456 | 0.999933887 | 0.017319173 | -0.164218731 | 0.181537904 | 0.754829757 |
| ENSG00000127993 | *RBM48* | 3.345079033 | -0.052366334 | -0.000543551 | -0.051822783 | 0.999933887 | -0.125217033 | -0.073163902 | -0.05205313 | 0.754944022 |
| ENSG00000129103 | *SUMF2* | 3.555246863 | -0.018578438 | -0.02151204 | 0.002933602 | 0.999933887 | 0.148129448 | 0.09056191 | 0.057567538 | 0.75498522 |

| | | | | | | | | | | |
|---|---|---|---|---|---|---|---|---|---|---|
| ENSG00000245146 | *MALINC1* | -0.200051172 | 0.240041829 | -0.020412488 | 0.260454317 | 0.999933887 | -0.289107627 | -0.145399784 | -0.143707844 | 0.755028372 |
| ENSG00000099994 | *SUSD2* | 0.157579036 | -0.02231509 | 0.249567381 | -0.27188247 | 0.999933887 | -0.055530899 | 0.116927712 | -0.172458611 | 0.755458163 |
| ENSG00000164221 | *CCDC112* | 0.912865974 | -0.196296258 | -0.094494524 | -0.101801734 | 0.999933887 | -0.300510709 | -0.411087204 | 0.110576494 | 0.755480626 |
| ENSG00000261693 | *AC134682.1* | 0.059628105 | 0.154263047 | 0.029094873 | 0.125168174 | 0.999933887 | 1.234773319 | 1.393662658 | -0.158889339 | 0.755480626 |
| ENSG00000137076 | *TLN1* | 9.684438696 | 0.004133129 | 0.01862498 | -0.014491851 | 0.999933887 | -0.234654995 | -0.276049418 | 0.041394423 | 0.755480626 |
| ENSG00000074266 | *EED* | 2.590040875 | -0.07175584 | -0.150361955 | 0.078606115 | 0.999933887 | 0.182670096 | 0.246073258 | -0.063403163 | 0.755545306 |
| ENSG00000181929 | *PRKAG1* | 4.520159554 | -0.029977585 | -0.073526244 | 0.04354866 | 0.999933887 | -0.163597247 | -0.120918279 | -0.042678968 | 0.755576227 |
| ENSG00000249476 | *AC008467.1* | 0.399932716 | 0.053205149 | 0.041637504 | 0.011567645 | 0.999933887 | -0.046329743 | 0.077492724 | -0.123822467 | 0.75567841 |
| ENSG00000153485 | *TMEM251* | 3.99788103 | 0.045029323 | -0.140231561 | 0.185260884 | 0.999933887 | 0.6800472 | 0.751562139 | -0.07151494 | 0.75567841 |
| ENSG00000254441 | *EIF2S2P6* | -0.828768735 | 0.065841266 | 0.104836543 | -0.038995277 | 0.999933887 | -0.125500842 | 0.212821239 | -0.338322081 | 0.75567841 |
| ENSG00000123178 | *SPRYD7* | -0.238284684 | -0.145556226 | 0.089922689 | -0.235478916 | 0.999933887 | -0.025965612 | 0.126961824 | -0.152927436 | 0.755946325 |
| ENSG00000185955 | *C7orf61* | 0.70003742 | 0.101177669 | -0.139684356 | 0.240862025 | 0.999933887 | 0.218774789 | 0.121701939 | 0.097072849 | 0.756076379 |
| ENSG00000277349 | *CICP25* | -1.993275889 | -0.040189292 | -0.007392056 | -0.032797236 | 0.999933887 | 0.179502183 | -0.062993391 | 0.242495573 | 0.756076379 |
| ENSG00000245105 | *A2M-AS1* | -0.270685861 | 0.302368351 | 0.569411878 | -0.267043528 | 0.999933887 | 0.07594695 | 0.302172432 | -0.226225481 | 0.756331325 |
| ENSG00000062822 | *POLD1* | 2.265180134 | 0.080350565 | 0.08008531 | 0.000265256 | 0.999933887 | 0.135134164 | 0.052521476 | 0.082612688 | 0.756331325 |
| ENSG00000214021 | *TTLL3* | 4.953876639 | -0.023620284 | 0.026452047 | -0.050072331 | 0.999933887 | -0.030067572 | 0.009501734 | -0.039569305 | 0.756331325 |
| ENSG00000276853 | *AC026124.2* | 0.400317502 | 0.071400842 | -0.178783602 | 0.250184444 | 0.999933887 | 0.426474753 | 0.286832468 | 0.139642285 | 0.756331325 |
| ENSG00000197798 | *FAM118B* | 1.32788154 | 0.034496181 | -0.226785845 | 0.261282025 | 0.999933887 | -0.096687954 | -0.008974196 | -0.087713757 | 0.756331325 |
| ENSG00000167851 | *CD300A* | 9.324930607 | 0.013037062 | -0.003612511 | 0.016649573 | 0.999933887 | -0.639668874 | -0.697279396 | 0.057610521 | 0.756331325 |
| ENSG00000101079 | *NDRG3* | 3.047040658 | -0.003508938 | -0.153150731 | 0.149641793 | 0.999933887 | -0.135889415 | -0.051466074 | -0.084423341 | 0.756331325 |
| ENSG00000236567 | *TCF3P1* | -0.033502568 | 0.212825485 | 0.18053635 | 0.032289135 | 0.999933887 | 0.53024009 | 0.842480666 | -0.312240576 | 0.756474969 |
| ENSG00000115526 | *CHST10* | -0.461059564 | 0.22583587 | 0.04028189 | 0.18555398 | 0.999933887 | 0.315663948 | 0.12926879 | 0.186395158 | 0.756657532 |
| ENSG00000147996 | *CBWD5* | 1.135803731 | -0.113934961 | 0.114316384 | -0.228251346 | 0.999933887 | -0.092846644 | -0.238592204 | 0.14574556 | 0.756657532 |
| ENSG00000138111 | *MFSD13A* | 2.586359486 | -0.054122026 | 0.012723218 | -0.066845244 | 0.999933887 | -0.613254111 | -0.718777818 | 0.105523707 | 0.756657532 |
| ENSG00000126217 | *MCF2L* | 0.383620388 | 0.007053757 | -0.067549783 | 0.07460354 | 0.999933887 | 0.209657318 | 0.33989888 | -0.130241561 | 0.756657532 |
| ENSG00000063046 | *EIF4B* | 7.419440196 | -0.060561936 | -0.065830782 | 0.005268846 | 0.999933887 | 0.026685456 | 0.062898841 | -0.036213385 | 0.75704849 |
| ENSG00000166913 | *YWHAB* | 7.370567926 | -0.064126368 | -0.100511289 | 0.036384921 | 0.999933887 | -0.142400953 | -0.179067658 | 0.036666705 | 0.757104842 |
| ENSG00000126970 | *ZC4H2* | -0.752581462 | 0.214162112 | 0.109962579 | 0.104199533 | 0.999933887 | -0.080924262 | 0.135308272 | -0.216232534 | 0.757129639 |
| ENSG00000272669 | *AL021707.6* | 3.971495916 | 0.031680042 | 0.07588904 | -0.044208997 | 0.999933887 | 0.14730705 | 0.07963904 | 0.06766801 | 0.757410555 |
| ENSG00000188917 | *TRMT2B* | 2.96554986 | 0.033655884 | -0.062922408 | 0.096578293 | 0.999933887 | -0.12805897 | -0.190539964 | 0.062480994 | 0.757470306 |
| ENSG00000204519 | *ZNF551* | 1.178127601 | -0.057514288 | 0.069085042 | -0.12659933 | 0.999933887 | 0.204254089 | 0.102238872 | 0.102015217 | 0.757810654 |
| ENSG00000206561 | *COLQ* | -0.022425909 | -0.069856879 | -0.091863937 | 0.022007059 | 0.999933887 | -0.06956951 | 0.096352665 | -0.165922175 | 0.758401889 |
| ENSG00000140678 | *ITGAX* | 11.557722995 | 0.072982687 | 0.076032655 | -0.003049969 | 0.999933887 | -0.059095247 | -0.020636652 | -0.038458595 | 0.758464034 |
| ENSG00000236779 | *AL512306.1* | -0.005599181 | 0.084715445 | 0.051507001 | 0.033208443 | 0.999933887 | 0.155079131 | -0.006421339 | 0.16150047 | 0.758503008 |
| ENSG00000178974 | *FBXO34* | 5.064467516 | -0.103959953 | -0.077772178 | -0.026187774 | 0.999933887 | -0.360986716 | -0.408776947 | 0.047790231 | 0.758864437 |
| ENSG00000258820 | *AF111167.1* | -0.199048644 | 0.248368142 | 0.079473798 | 0.168894344 | 0.999933887 | 0.578974978 | 0.352110786 | 0.226864192 | 0.758864437 |
| ENSG00000230730 | *AC074011.1* | -0.885756324 | 0.150032523 | 0.147069269 | 0.002963255 | 0.999933887 | 0.72047375 | 0.458416178 | 0.262057572 | 0.758864437 |
| ENSG00000286488 | *AC103858.3* | 1.283290537 | -0.0838919 | -0.075701583 | -0.008190317 | 0.999933887 | 0.602324918 | 0.718387537 | -0.116062619 | 0.758864437 |
| ENSG00000235290 | *HLA-W* | 0.980291828 | -0.052370172 | 0.201518059 | -0.253888231 | 0.999933887 | 0.252569658 | 0.116733245 | 0.135836412 | 0.758864437 |
| ENSG00000278022 | *AC118658.1* | 0.092938332 | 0.000799378 | 0.025177215 | -0.024377837 | 0.999933887 | 0.01274316 | 0.143769299 | -0.131026138 | 0.758864437 |
| ENSG00000092531 | *SNAP23* | 4.524439737 | 0.000536917 | -0.087153967 | 0.087690883 | 0.999933887 | 0.209550993 | 0.1467491 | 0.062801893 | 0.759143666 |
| ENSG00000115687 | *PASK* | 3.690306413 | 0.032616489 | 0.074396328 | -0.041779839 | 0.999933887 | 0.20296876 | 0.248766352 | -0.045797592 | 0.75919366 |
| ENSG00000116990 | *MYCL* | 0.422842061 | -0.471911965 | -0.26490693 | -0.207005035 | 0.999933887 | -0.834793828 | -0.951294058 | 0.116500229 | 0.759530027 |
| ENSG00000068878 | *PSME4* | 6.075219255 | -0.066776263 | -0.058190909 | -0.008585354 | 0.999933887 | -0.146049339 | -0.103190555 | -0.042858784 | 0.759530027 |
| ENSG00000179598 | *PLD6* | 3.958381104 | 0.08376443 | 0.159703387 | -0.075938957 | 0.999933887 | -0.270056052 | -0.34405401 | 0.073997958 | 0.759530027 |
| ENSG00000155008 | *APOOL* | 0.082416781 | -0.065029561 | -0.08708992 | 0.022060359 | 0.999933887 | -0.048259552 | 0.111384725 | -0.159644277 | 0.759530027 |
| ENSG00000188428 | *BLOC1S5* | 1.051245103 | -0.195664967 | -0.042426594 | -0.153238373 | 0.999933887 | -0.001786779 | -0.091145088 | 0.089358309 | 0.759772778 |
| ENSG00000184271 | *POU6F1* | 2.281413058 | -0.03689385 | -0.088866659 | 0.051972809 | 0.999933887 | 0.146176238 | 0.215400146 | -0.069223908 | 0.759772778 |
| ENSG00000232470 | *AL136368.1* | 0.850277976 | 0.059949803 | 0.054879688 | 0.005070115 | 0.999933887 | 0.258099956 | 0.376771984 | -0.118672028 | 0.759811222 |
| ENSG00000175166 | *PSMD2* | 6.095602356 | -0.01543009 | 0.003990601 | -0.019420691 | 0.999933887 | -0.079315116 | -0.113551319 | 0.034236203 | 0.759811222 |
| ENSG00000128383 | *APOBEC3A* | 7.635531097 | -0.004843198 | 0.018003617 | -0.022846815 | 0.999933887 | -0.116279758 | -0.159844205 | 0.043564446 | 0.760449504 |
| ENSG00000053770 | *AP5M1* | 3.559357585 | -0.122459555 | -0.099246822 | -0.023192732 | 0.999933887 | -0.360391775 | -0.302811309 | -0.057580467 | 0.760595002 |
| ENSG00000177613 | *CSTF2T* | 3.467253773 | -0.045385128 | 0.026741413 | -0.072126541 | 0.999933887 | 0.028583165 | -0.02983155 | 0.058414715 | 0.760595002 |
| ENSG00000139239 | *RPL14P1* | -0.138517695 | 0.082707536 | 0.119629559 | -0.036922022 | 0.999933887 | 0.052949835 | 0.199287938 | -0.146338103 | 0.760595002 |
| ENSG00000287697 | *Z99127.3* | 0.319738987 | 0.086428315 | 0.051256572 | 0.035171743 | 0.999933887 | 0.771488964 | 0.620921425 | 0.15056754 | 0.760595002 |
| ENSG00000040487 | *SLC66A1* | 4.088546081 | -0.031459358 | 0.015547037 | -0.047006395 | 0.999933887 | -0.512420124 | -0.57001841 | 0.057598286 | 0.760595002 |
| ENSG00000170464 | *DNAJC18* | 0.520529068 | -0.044166483 | -0.023670135 | -0.020496348 | 0.999933887 | 0.136615425 | -0.015317426 | 0.15193285 | 0.760595002 |
| ENSG00000130165 | *ELOF1* | 3.638282168 | -0.063172126 | 0.030858402 | -0.094030528 | 0.999933887 | -0.235003197 | -0.29376506 | 0.058761863 | 0.760718806 |
| ENSG00000104177 | *MYEF2* | 0.78966472 | -0.065776189 | -0.297755182 | 0.231978993 | 0.999933887 | -0.099191306 | 0.049572048 | -0.148763354 | 0.760729757 |
| ENSG00000150403 | *TMCO3* | 6.654091783 | -0.10364191 | -0.109613195 | 0.005971285 | 0.999933887 | -0.91851216 | -0.98160571 | 0.06309355 | 0.760842095 |
| ENSG00000107938 | *EDRF1* | 3.399260816 | 0.000364752 | -0.040202974 | 0.040567727 | 0.999933887 | -0.289033852 | -0.356127648 | 0.067093797 | 0.760867608 |
| ENSG00000280206 | *AC026401.3* | 3.328052283 | 0.097712896 | 0.096359481 | 0.001353415 | 0.999933887 | 0.680210988 | 0.753996538 | -0.073785551 | 0.761040268 |
| ENSG00000204120 | *GIGYF2* | 5.404852749 | -0.021348133 | -0.059875506 | 0.038527373 | 0.999933887 | -0.327232385 | -0.287193618 | -0.040038767 | 0.761067819 |
| ENSG00000259972 | *AC009120.2* | 1.548538784 | -0.138175634 | 0.093425405 | -0.231601039 | 0.999933887 | -0.071533441 | -0.162584625 | 0.091051183 | 0.761234933 |
| ENSG00000119900 | *OGFRL1* | 6.6489097 | -0.011106764 | -0.042267985 | 0.031161221 | 0.999933887 | -0.004384052 | -0.093422964 | 0.089038913 | 0.761234933 |
| ENSG00000218313 | *AL139274.1* | -0.012174295 | -0.038963063 | 0.386232067 | -0.42519513 | 0.999933887 | 0.303585118 | 0.546283598 | -0.24269848 | 0.761243888 |
| ENSG00000141295 | *SCRN2* | 2.381905086 | 0.042211433 | 0.08007544 | -0.037864008 | 0.999933887 | -0.00392573 | -0.091187718 | 0.087261988 | 0.761417303 |
| ENSG00000232615 | *AC026412.2* | -1.361627934 | -0.032063571 | 0.096912886 | -0.128976457 | 0.999933887 | 0.260476036 | 0.526347255 | -0.26587122 | 0.761558439 |
| ENSG00000226266 | *AC009961.1* | -0.006314481 | 0.035596793 | 0.131588605 | -0.095991812 | 0.999933887 | -0.610859316 | -0.426126385 | -0.184732931 | 0.761643874 |
| ENSG00000007402 | *CACNA2D2* | 1.778916996 | 0.063678923 | -0.019811928 | 0.083490851 | 0.999933887 | 0.077603267 | -0.009488253 | 0.087091521 | 0.761931891 |
| ENSG00000161692 | *DBF4B* | 0.739529361 | -0.061750791 | 0.022604082 | -0.084354873 | 0.999933887 | -0.114793913 | 0.004796259 | -0.119590172 | 0.761931891 |
| ENSG00000178950 | *GAK* | 7.953311609 | 0.040582595 | 0.006164947 | 0.034417648 | 0.999933887 | 0.215155511 | 0.174356135 | 0.040799376 | 0.761982151 |
| ENSG00000171132 | *PRKCE* | 2.868718248 | 0.003861731 | 0.152672025 | -0.148810294 | 0.999933887 | 0.14469016 | 0.077528346 | 0.067161813 | 0.761982151 |
| ENSG00000105486 | *LIG1* | 2.950047825 | 0.014699094 | 0.074843667 | -0.060144574 | 0.999933887 | 0.096141323 | 0.16077337 | -0.064632047 | 0.762007218 |
| ENSG00000285106 | *AC016831.6* | 4.716345387 | 0.0078415 | 0.107281954 | -0.099440454 | 0.999933887 | 0.690083688 | 0.757366021 | -0.067282333 | 0.762007218 |
| ENSG00000167900 | *TK1* | 0.255780713 | 0.204720594 | 0.130314006 | 0.074406589 | 0.999933887 | 0.739090135 | 0.878085931 | -0.138995795 | 0.762094352 |
| ENSG00000272602 | *ZNF595* | -0.640047185 | -0.217218369 | -0.134618505 | -0.082599864 | 0.999933887 | -0.344849652 | -0.161688598 | -0.183161054 | 0.762101038 |
| ENSG00000101425 | *BPI* | 0.245250266 | -0.161323862 | 0.241752542 | -0.403076403 | 0.999933887 | -0.015053469 | 0.123781902 | -0.138835371 | 0.762127896 |
| ENSG00000101751 | *POLI* | 2.33183107 | 0.014301486 | -0.042030299 | 0.056331785 | 0.999933887 | -0.319481846 | -0.213587709 | -0.105894137 | 0.762127896 |
| ENSG00000197324 | *LRP10* | 9.363062647 | 0.026585917 | -0.000227468 | 0.026813386 | 0.999933887 | -0.104830817 | -0.149620184 | 0.044789367 | 0.762309362 |
| ENSG00000160688 | *FLAD1* | 2.802300282 | 0.02810211 | -0.079034097 | 0.107136207 | 0.999933887 | -0.012409135 | -0.066891407 | 0.054482271 | 0.762433639 |
| ENSG00000178096 | *BOLA1* | 0.282660428 | 0.168720672 | 0.248635401 | -0.079914729 | 0.999933887 | 0.173244748 | 0.063118282 | 0.110126467 | 0.762692044 |
| ENSG00000225079 | *FTH1P22* | -0.633165398 | -0.085862889 | 0.556828132 | -0.642691021 | 0.999933887 | -0.089599289 | 0.113938043 | -0.203537332 | 0.762907579 |
| ENSG00000117569 | *PTBP2* | 3.130038025 | -0.015687031 | -0.055698065 | 0.040011034 | 0.999933887 | -0.11767048 | -0.030553469 | -0.08711701 | 0.763253123 |
| ENSG00000149313 | *AASDHPPT* | 2.135205679 | -0.162998698 | -0.027082704 | -0.135915993 | 0.999933887 | -0.038641533 | -0.123448068 | 0.084806536 | 0.763356663 |
| ENSG00000184208 | *C22orf46* | 2.283460372 | 0.035114655 | 0.055510835 | -0.02039618 | 0.999933887 | 0.115586475 | 0.195973733 | -0.080387258 | 0.763356663 |
| ENSG00000139620 | *KANSL2* | 3.70470429 | -0.034801307 | -0.079973596 | 0.045172289 | 0.999933887 | 0.071924233 | 0.017274525 | 0.054649708 | 0.763399227 |
| ENSG00000143337 | *TOR1AIP1* | 6.653236407 | -0.051053354 | -0.047563354 | -0.00349 | 0.999933887 | -0.17961278 | -0.224143021 | 0.044530241 | 0.763558653 |
| ENSG00000213397 | *HAUS7* | -0.639063774 | -0.252659112 | 0.00848253 | -0.261141643 | 0.999933887 | -0.061487568 | -0.270771448 | 0.209283881 | 0.763589223 |
| ENSG00000141294 | *LRRC46* | -0.264141541 | -0.094944388 | -0.207095988 | 0.1121516 | 0.999933887 | 0.035802647 | -0.158532642 | 0.194335289 | 0.763589223 |
| ENSG00000221957 | *KIR2DS4* | -0.796632257 | 0.105700147 | 0.223025034 | -0.117324887 | 0.999933887 | 0.965505564 | 1.145680735 | -0.180175171 | 0.763589223 |
| ENSG00000273015 | *AC008124.1* | 2.670351762 | 0.014082723 | 0.059750471 | -0.045667748 | 0.999933887 | 0.072950827 | 0.132651468 | -0.059700641 | 0.763589223 |
| ENSG00000185201 | *IFITM2* | 11.142784142 | 0.056212202 | 0.008782528 | 0.047429674 | 0.999933887 | -0.019516534 | -0.066092257 | 0.046575723 | 0.763826609 |
| ENSG00000178947 | *SMIM10L2A* | -0.952612666 | -0.157580049 | -0.00208167 | -0.155498379 | 0.999933887 | 0.006148046 | 0.171182829 | -0.165034783 | 0.763826711 |
| ENSG00000158445 | *KCNB1* | -0.324953821 | -0.050535359 | 0.242537787 | -0.293073145 | 0.999933887 | -0.249684487 | -0.422037091 | 0.172352605 | 0.763826711 |
| ENSG00000151729 | *SLC25A4* | 0.281619309 | -0.050973622 | 0.026607624 | -0.077581246 | 0.999933887 | 0.065890396 | 0.190710284 | -0.124819888 | 0.76395606 |
| ENSG00000233922 | *LINC01694* | -0.825996268 | 0.019205186 | 0.173700506 | -0.154495319 | 0.999933887 | 1.112088556 | 1.292085259 | -0.179996703 | 0.76395606 |
| ENSG00000164402 | *SEPTIN8* | 1.121516129 | 0.006476807 | 0.144333114 | -0.137856307 | 0.999933887 | 0.201982774 | 0.115648928 | 0.086333846 | 0.76395606 |
| ENSG00000211747 | *TRBV20-1* | 1.989914248 | -0.002423202 | -0.198423661 | 0.196000458 | 0.999933887 | 0.239803568 | 0.164181316 | 0.075622251 | 0.76395606 |
| ENSG00000086475 | *SEPHS1* | 1.964669473 | -0.02647897 | 0.028853904 | -0.055332873 | 0.999933887 | 0.015051435 | 0.099131941 | -0.084080506 | 0.763981502 |
| ENSG00000112079 | *STK38* | 6.076524604 | -0.0342539 | -0.061495114 | 0.027241214 | 0.999933887 | -0.027759278 | -0.066284643 | 0.038525366 | 0.764177051 |
| ENSG00000118785 | *SPP1* | 2.701361744 | 0.027533046 | -0.132020727 | 0.159553774 | 0.999933887 | -0.659053292 | -0.773540567 | 0.114487275 | 0.764209244 |
| ENSG00000183458 | *PKD1P3* | 1.03649158 | 0.128049705 | 0.275326816 | -0.147277111 | 0.999933887 | -0.108018717 | -0.011463451 | -0.096555266 | 0.764246591 |
| ENSG00000223704 | *LINC01422* | -0.722247368 | -0.117498334 | -0.134563091 | 0.017064756 | 0.999933887 | 0.133172699 | 0.315455113 | -0.182282415 | 0.764246591 |
| ENSG00000196388 | *INCA1* | -0.717243654 | -0.325961229 | -0.083722987 | -0.242238241 | 0.999933887 | 0.22444255 | 0.416031664 | -0.191589114 | 0.764724314 |

| | | | | | | | | | | |
|---|---|---|---|---|---|---|---|---|---|---|
| ENSG00000005238 | *FAM214B* | 6.469031167 | 0.035565265 | 0.081853187 | -0.046287922 | 0.999933887 | -0.502231632 | -0.451438282 | -0.05079335 | 0.76490757 |
| ENSG00000122557 | *HERPUD2* | 5.964479799 | 0.057903892 | 0.014068183 | 0.043835709 | 0.999933887 | -0.320425194 | -0.360258315 | 0.039833121 | 0.764940704 |
| ENSG00000159423 | *ALDH4A1* | 1.84859125 | 0.010067901 | 0.046678757 | -0.036610857 | 0.999933887 | 0.64804363 | 0.566563832 | 0.081479798 | 0.764940704 |
| ENSG00000174721 | *FGFBP3* | -0.85410843 | 0.27366428 | 0.198651632 | 0.075012648 | 0.999933887 | 0.017231017 | -0.181165603 | 0.19839662 | 0.765060293 |
| ENSG00000196275 | *GTF2IRD2* | 0.209478379 | -0.069543524 | 0.000715342 | -0.070258866 | 0.999933887 | 0.073029223 | 0.214701742 | -0.141672519 | 0.765060293 |
| ENSG00000101084 | *RAB5IF* | 4.515220607 | 0.136802719 | 0.038193738 | 0.09860898 | 0.999933887 | -0.099874248 | -0.170071618 | 0.07019737 | 0.765119766 |
| ENSG00000243466 | *IGKV1-5* | -0.926305665 | 0.084098758 | -0.516571561 | 0.600670319 | 0.999933887 | 0.217309121 | 0.495221232 | -0.277912111 | 0.765119766 |
| ENSG00000266852 | *MIR4482* | -0.047739305 | 0.006845991 | 0.152840491 | -0.1459945 | 0.999933887 | 0.774140465 | 0.634216111 | 0.139924355 | 0.765119766 |
| ENSG00000079950 | *STX7* | 6.426695413 | -0.081875123 | -0.032927354 | -0.048947769 | 0.999933887 | -0.983522443 | -1.039230886 | 0.055708443 | 0.765533917 |
| ENSG00000233029 | *AC244453.2* | 1.114360147 | 0.092291499 | -0.007285866 | 0.099577364 | 0.999933887 | 0.119462914 | 0.271948855 | -0.15248594 | 0.765686682 |
| ENSG00000197872 | *CYRIA* | 5.889865031 | -0.023512457 | -0.033687676 | 0.010175219 | 0.999933887 | 0.194491181 | 0.12827493 | 0.066216251 | 0.765686682 |
| ENSG00000104918 | *RETN* | -0.951445285 | 0.160409222 | 0.112203027 | 0.048206195 | 0.999933887 | -0.2015754 | 0.012849889 | -0.214425289 | 0.7658691 |
| ENSG00000123444 | *KBTBD4* | 1.824313129 | -0.164608622 | -0.194092648 | 0.029484026 | 0.999933887 | -0.106381994 | -0.024678137 | -0.081703858 | 0.766089155 |
| ENSG00000244267 | *RPL34P22* | -0.909130022 | -0.038261208 | 0.280698405 | -0.318959612 | 0.999933887 | 1.453508826 | 1.783873232 | -0.330364406 | 0.766294292 |
| ENSG00000145041 | *DCAF1* | 3.90288842 | -0.009817311 | -0.010130192 | 0.000312882 | 0.999933887 | 0.055164785 | 0.105641327 | -0.050476542 | 0.766567572 |
| ENSG00000285437 | *POLR2J3* | 1.160127516 | -0.034808243 | 0.07273643 | -0.107544673 | 0.999933887 | 0.049246186 | -0.047063827 | 0.096310013 | 0.766595554 |
| ENSG00000019485 | *PRDM11* | 1.501008308 | 0.059985648 | 0.003252371 | 0.056733277 | 0.999933887 | 0.114017154 | 0.010677452 | 0.103339702 | 0.767028515 |
| ENSG00000074657 | *ZNF532* | 0.27202046 | -0.096713243 | -0.034393553 | -0.062319689 | 0.999933887 | 0.093012224 | 0.218198657 | -0.125186433 | 0.767038139 |
| ENSG00000145216 | *FIP1L1* | 3.107581869 | -0.027734507 | -0.029382883 | 0.001648375 | 0.999933887 | -0.467569709 | -0.415871913 | -0.051697796 | 0.767038139 |
| ENSG00000284707 | *AC079781.5* | 0.007507056 | -0.03638379 | 0.088843807 | -0.125227598 | 0.999933887 | 0.071654562 | 0.20947019 | -0.137815628 | 0.767038139 |
| ENSG00000151353 | *TMEM18* | 2.55094211 | -0.008353014 | -0.044267614 | 0.0359146 | 0.999933887 | -0.038096932 | -0.094480476 | 0.056383543 | 0.767038139 |
| ENSG00000139725 | *RHOF* | 2.958752054 | 0.014212386 | -0.153870624 | 0.16808301 | 0.999933887 | 1.412314728 | 1.500711422 | -0.088396694 | 0.767038139 |
| ENSG00000181963 | *OR52K2* | 0.368704375 | 0.115104994 | -0.030569861 | 0.145674855 | 0.999933887 | 0.017940255 | 0.150647285 | -0.132707029 | 0.767646431 |
| ENSG00000138623 | *SEMA7A* | 4.839279073 | -0.041344302 | 0.047211715 | -0.088556017 | 0.999933887 | 0.195578819 | 0.257242816 | -0.061663997 | 0.767646431 |
| ENSG00000117054 | *ACADM* | 1.310677229 | -0.081067989 | 0.01392926 | -0.094997249 | 0.999933887 | -0.162271247 | -0.068275609 | -0.093995638 | 0.767646431 |
| ENSG00000267265 | *AC011476.3* | 0.131402875 | 0.101809175 | 0.000743431 | 0.101065744 | 0.999933887 | 0.520454754 | 0.643120202 | -0.122665447 | 0.767646431 |
| ENSG00000065268 | *WDR18* | 1.886554021 | 0.046849134 | 0.192160644 | -0.14531151 | 0.999933887 | 0.27011373 | 0.353896454 | -0.083782724 | 0.767646431 |
| ENSG00000223804 | *AC244669.1* | 0.665468129 | -0.030766666 | -0.310455668 | 0.279689003 | 0.999933887 | 0.32203844 | 0.444196632 | -0.122158192 | 0.767646431 |
| ENSG00000261556 | *SMG1P7* | 0.293470314 | -0.123199514 | 0.104446487 | -0.227646 | 0.999933887 | 0.127655927 | 0.01323889 | 0.114417036 | 0.767750384 |
| ENSG00000135457 | *TFCP2* | 3.436246067 | -0.022876164 | 0.057336817 | -0.080212981 | 0.999933887 | -0.10159815 | -0.149434803 | 0.047836653 | 0.767750384 |
| ENSG00000188011 | *RTP5* | 0.372157308 | -0.077888227 | -0.128914387 | 0.05102616 | 0.999933887 | 0.034242476 | 0.16768174 | -0.133439264 | 0.76792862 |
| ENSG00000179151 | *EDC3* | 2.497563599 | 0.027987442 | -0.003716699 | 0.031704141 | 0.999933887 | 0.049332092 | 0.10979834 | -0.060466249 | 0.76792862 |
| ENSG00000262962 | *KARS1P3* | -0.527112949 | -0.017666695 | -0.335338234 | 0.317671539 | 0.999933887 | -1.540826536 | -1.746666216 | 0.20583968 | 0.76792862 |
| ENSG00000251393 | *AC005280.1* | 1.935429942 | 0.110240152 | 0.099274874 | 0.010965278 | 0.999933887 | 1.60684361 | 1.703861189 | -0.097017579 | 0.768131736 |
| ENSG00000007923 | *DNAJC11* | 1.911143401 | -0.098705523 | -0.048884622 | -0.049820901 | 0.999933887 | 0.041984563 | 0.120909664 | -0.078925102 | 0.768399916 |
| ENSG00000273802 | *H2BC8* | 0.55865804 | -0.050234756 | 0.106250471 | -0.156485227 | 0.999933887 | -0.339010177 | -0.477228052 | 0.138217875 | 0.768399916 |
| ENSG00000196230 | *TUBB* | 5.523953875 | -0.013074101 | -0.050559698 | 0.037485597 | 0.999933887 | 0.295443373 | 0.341253842 | -0.045810469 | 0.768399916 |
| ENSG00000135362 | *PRR5L* | 5.434234991 | -0.051467863 | 0.034549074 | -0.086016937 | 0.999933887 | 0.038704506 | -0.009137069 | 0.047841575 | 0.768432203 |
| ENSG00000271270 | *TMCC1-AS1* | -0.397558926 | -0.119126812 | 0.085672162 | -0.204798974 | 0.999933887 | -0.639841239 | -0.503686009 | -0.13615523 | 0.768561749 |
| ENSG00000163629 | *PTPN13* | 0.086656459 | -0.242264558 | -0.112948981 | -0.129315578 | 0.999933887 | -0.039954941 | 0.159699692 | -0.199654633 | 0.768581498 |
| ENSG00000100601 | *ALKBH1* | 1.618342411 | -0.114769381 | -0.011112969 | -0.103656412 | 0.999933887 | -0.075940356 | 0.002130635 | -0.078070991 | 0.768727503 |
| ENSG00000093072 | *ADA2* | 3.952111421 | -0.041086793 | -0.012851689 | -0.028235104 | 0.999933887 | -0.037533762 | -0.084274626 | 0.046740864 | 0.768921362 |
| ENSG00000260105 | *AOC4P* | -0.777612631 | 0.120705346 | 0.108769763 | 0.011935584 | 0.999933887 | 1.07443053 | 0.832857588 | 0.241572943 | 0.769075238 |
| ENSG00000185880 | *TRIM69* | 5.848830845 | 0.042865348 | 0.036027077 | 0.006838271 | 0.999933887 | 0.238359513 | 0.281446294 | -0.04308678 | 0.769446972 |
| ENSG00000180011 | *ZADH2* | 5.172631584 | -0.121205466 | -0.048526667 | -0.072678799 | 0.999933887 | -0.706052874 | -0.761267978 | 0.055215103 | 0.769702827 |
| ENSG00000258086 | *AC079313.1* | 0.787433129 | -0.153853948 | -0.080085071 | -0.073768878 | 0.999933887 | -0.151692482 | -0.262216371 | 0.110523889 | 0.769702827 |
| ENSG00000160883 | *HK3* | 5.993327473 | 0.058024601 | 0.084501148 | -0.026476547 | 0.999933887 | 0.099773188 | 0.045757239 | 0.054015948 | 0.769702827 |
| ENSG00000080573 | *COL5A3* | 1.111301755 | 0.081332415 | 0.084366324 | -0.003033909 | 0.999933887 | -0.095375228 | -0.224591145 | 0.129215918 | 0.769702827 |
| ENSG00000103199 | *ZNF500* | 1.07955977 | -0.041123448 | 0.077856259 | -0.118979707 | 0.999933887 | 0.113558045 | 0.205295847 | -0.091737803 | 0.769702827 |
| ENSG00000008086 | *CDKL5* | 3.319089535 | 0.016924991 | -0.074492352 | 0.091417343 | 0.999933887 | -0.332350252 | -0.401668435 | 0.069318183 | 0.769702827 |
| ENSG00000280194 | *AD000864.1* | 2.987513141 | 0.316205196 | 0.20499619 | 0.111209006 | 0.999933887 | 1.750922654 | 1.855130675 | -0.104208021 | 0.769941192 |
| ENSG00000236753 | *MKLN1-AS* | 0.822068242 | -0.097657734 | 0.109582692 | -0.207240426 | 0.999933887 | -0.517533969 | -0.647794536 | 0.130260567 | 0.769941192 |
| ENSG00000259642 | *ST20-AS1* | 2.879427295 | 0.0269727 | 0.138002347 | -0.111029647 | 0.999933887 | 0.233103407 | 0.145565887 | 0.08753752 | 0.769941192 |
| ENSG00000170185 | *USP38* | 5.087261986 | 0.051452122 | 0.048320173 | 0.003131949 | 0.999933887 | -0.408643065 | -0.361184387 | -0.047458678 | 0.770036363 |
| ENSG00000132965 | *ALOX5AP* | 7.934049617 | -0.004726872 | -0.014856552 | 0.01012968 | 0.999933887 | 0.180254288 | 0.234602448 | -0.054348161 | 0.770036363 |
| ENSG00000105556 | *MIER2* | 1.721366935 | 0.130167235 | 0.157034349 | -0.026867114 | 0.999933887 | -0.104548277 | -0.186711806 | 0.082163529 | 0.770496597 |
| ENSG00000148925 | *BTBD10* | 3.548560199 | -0.076561906 | 0.015602505 | -0.092164411 | 0.999933887 | -0.462309742 | -0.409727059 | -0.052582683 | 0.770496597 |
| ENSG00000169224 | *GCSAML* | -0.172006333 | -0.230554837 | -0.086347114 | -0.144207723 | 0.999933887 | -0.168900762 | 0.019825873 | -0.188726636 | 0.770517278 |
| ENSG00000253385 | *AP003696.1* | -0.590588575 | 0.235888761 | 0.357755434 | -0.121866673 | 0.999933887 | 1.391154534 | 1.638436638 | -0.247282104 | 0.770689411 |
| ENSG00000259943 | *AL050341.2* | 1.080154599 | -0.271875191 | -0.167402368 | -0.104472824 | 0.999933887 | 0.016147575 | 0.096284674 | -0.080137099 | 0.771024678 |
| ENSG00000129226 | *CD68* | 0.013238014 | 0.287668054 | 0.17148117 | 0.116186884 | 0.999933887 | 0.689823466 | 0.863761842 | -0.173938376 | 0.771084313 |
| ENSG00000185950 | *IRS2* | 9.341709573 | -0.011227972 | 0.05038872 | -0.061616693 | 0.999933887 | -0.320935697 | -0.280053137 | -0.04088256 | 0.771289196 |
| ENSG00000167562 | *ZNF701* | 3.112089391 | -0.046465353 | -0.170676425 | 0.124211072 | 0.999933887 | -0.545481682 | -0.485615201 | -0.059866481 | 0.771550797 |
| ENSG00000167984 | *NLRC3* | 4.450060059 | -0.016854238 | 0.0822097 | -0.099063938 | 0.999933887 | 0.038442169 | -0.001020939 | 0.039463109 | 0.771665451 |
| ENSG00000260910 | *LINC00565* | -0.35247242 | -0.120362623 | -0.310626738 | 0.190264115 | 0.999933887 | -0.034687697 | 0.124693739 | -0.159381435 | 0.771688263 |
| ENSG00000106263 | *EIF3B* | 4.830965014 | 0.007346329 | 0.08631983 | -0.078973502 | 0.999933887 | 0.114034867 | 0.07529537 | 0.038739497 | 0.771688263 |
| ENSG00000082512 | *TRAF5* | 3.15499042 | -0.031252627 | -0.184276222 | 0.153023595 | 0.999933887 | -0.051157478 | 0.011377265 | -0.062534743 | 0.771884068 |
| ENSG00000119714 | *GPR68* | 3.33169672 | 0.119892986 | 0.007308675 | 0.112584312 | 0.999933887 | 0.133677544 | 0.195951727 | -0.062274183 | 0.771943241 |
| ENSG00000170629 | *DPY19L2P2* | -0.03494467 | -0.088661812 | 0.08819114 | -0.176852951 | 0.999933887 | 0.147525652 | 0.294873511 | -0.147347859 | 0.771943241 |
| ENSG00000005448 | *WDR54* | 1.555435951 | -0.027705836 | -0.021020077 | -0.006685758 | 0.999933887 | 0.12345809 | 0.207664582 | -0.084206492 | 0.772314853 |
| ENSG00000168477 | *TNXB* | 0.450762745 | 0.244492017 | 0.010294744 | 0.234197272 | 0.999933887 | -0.087563674 | -0.230882719 | 0.143319045 | 0.772512864 |
| ENSG00000279057 | *AC141586.4* | -0.292079675 | -0.019916566 | 0.079336138 | -0.099252704 | 0.999933887 | -0.652180833 | -0.486755805 | -0.165425028 | 0.772512864 |
| ENSG00000135829 | *DHX9* | 6.498520849 | -0.084317027 | -0.113534181 | 0.029217154 | 0.999933887 | -0.165465884 | -0.200057748 | 0.034591864 | 0.772513638 |
| ENSG00000134321 | *RSAD2* | 7.056415957 | -0.01433639 | -0.030742624 | 0.016406234 | 0.999933887 | 0.164991841 | 0.220858015 | -0.055866173 | 0.772513638 |
| ENSG00000277590 | *MIR7845* | -1.288625938 | 0.444717134 | 0.166399646 | 0.278317489 | 0.999933887 | 0.679458835 | 0.981050317 | -0.301591482 | 0.772532367 |
| ENSG00000258922 | *AC106028.2* | 0.120162997 | -0.048886728 | 0.035055906 | -0.083942633 | 0.999933887 | 1.050123217 | 1.178309236 | -0.12818602 | 0.772636878 |
| ENSG00000175414 | *ARL10* | 1.509921822 | -0.07231914 | 0.099600607 | -0.171919747 | 0.999933887 | 0.05241303 | -0.025547409 | 0.077960439 | 0.772891715 |
| ENSG00000204623 | *ZNRD1ASP* | 1.027597666 | -0.044136772 | 0.157530419 | -0.201667191 | 0.999933887 | -0.019994746 | 0.062788682 | -0.082783428 | 0.772891715 |
| ENSG00000004478 | *FKBP4* | 2.59775691 | 0.035419748 | 0.043599392 | -0.008179644 | 0.999933887 | 0.330886571 | 0.260701251 | 0.070185319 | 0.77309045 |
| ENSG00000174123 | *TLR10* | 2.925864448 | -0.030182076 | -0.156198054 | 0.126015978 | 0.999933887 | 0.130316428 | 0.012123981 | 0.118192447 | 0.77309045 |
| ENSG00000276023 | *DUSP14* | 0.451303018 | -0.023339155 | -0.067667343 | 0.044328188 | 0.999933887 | 0.112154129 | -0.018847873 | 0.131002001 | 0.77309045 |
| ENSG00000038532 | *CLEC16A* | 3.95793806 | -0.004893293 | 0.005210014 | -0.010103307 | 0.999933887 | -0.085353803 | -0.131200144 | 0.045846341 | 0.77309045 |
| ENSG00000174996 | *KLC2* | 2.296394975 | 0.10001265 | 0.233454575 | -0.133441926 | 0.999933887 | 0.303087284 | 0.380405823 | -0.07731854 | 0.773155223 |
| ENSG00000180329 | *CCDC43* | 2.706554909 | -0.128129217 | -0.22421071 | 0.096081493 | 0.999933887 | -0.126152399 | -0.062372866 | -0.063779533 | 0.77324581 |
| ENSG00000158411 | *MITD1* | 4.185293502 | -0.063737787 | -0.125815711 | 0.062077924 | 0.999933887 | 0.10925192 | 0.069789227 | 0.039462693 | 0.77324581 |
| ENSG00000146574 | *CCZ1B* | 1.622548199 | -0.093270723 | -0.043170676 | -0.050100048 | 0.999933887 | -0.29723935 | -0.368743007 | 0.071503657 | 0.77324581 |
| ENSG00000183828 | *NUDT14* | 0.318199575 | 0.166289401 | 0.147403306 | 0.018886095 | 0.999933887 | -0.176066922 | -0.288553719 | 0.112486797 | 0.77324581 |
| ENSG00000228663 | *PSMD10P1* | -0.14397151 | -0.15880242 | 0.348549876 | -0.507352295 | 0.999933887 | 0.070129533 | -0.128297619 | 0.198427152 | 0.77324581 |
| ENSG00000120158 | *RCL1* | 1.467058684 | -0.04976964 | 0.142995029 | -0.192764669 | 0.999933887 | 0.192574632 | 0.300642728 | -0.108068097 | 0.77324581 |
| ENSG00000145715 | *RASA1* | 4.439224139 | 0.052412422 | -0.022242854 | 0.074655276 | 0.999933887 | -0.391170974 | -0.502543526 | 0.111372552 | 0.77324581 |
| ENSG00000215417 | *MIR17HG* | 0.386201896 | -0.10561634 | -0.092272034 | -0.013344306 | 0.999933887 | 0.279270033 | 0.41509827 | -0.135828237 | 0.773454795 |
| ENSG00000006652 | *IFRD1* | 8.623401587 | 0.015321451 | 0.03362465 | -0.0183032 | 0.999933887 | 0.06923832 | 0.112643161 | -0.043404841 | 0.773454795 |
| ENSG00000180628 | *PCGF5* | 5.08433644 | 0.010327215 | -0.067452312 | 0.077779527 | 0.999933887 | -0.384713776 | -0.324469936 | -0.06024384 | 0.773454795 |
| ENSG00000168556 | *ING2* | 2.72317397 | 0.112994966 | 0.019520522 | 0.093474444 | 0.999933887 | 0.180443415 | 0.10405894 | 0.076384475 | 0.773534781 |
| ENSG00000170633 | *RNF34* | 4.843818432 | -0.043518767 | -0.038151278 | -0.005367489 | 0.999933887 | -0.063278804 | -0.026803564 | -0.03647524 | 0.773635224 |
| ENSG00000105755 | *ETHE1* | 0.571754916 | 0.070193038 | 0.35611248 | -0.285919442 | 0.999933887 | -0.024399017 | 0.079883514 | -0.104282531 | 0.773953894 |
| ENSG00000095564 | *BTAF1* | 6.309600492 | 0.003921223 | -0.103687811 | 0.107609034 | 0.999933887 | -0.136166625 | -0.062282585 | -0.07388404 | 0.773953894 |
| ENSG00000167699 | *GLOD4* | 3.64874249 | -0.053023158 | -0.118988538 | 0.06596538 | 0.999933887 | 0.070403795 | 0.109157396 | -0.038753601 | 0.774024271 |
| ENSG00000149218 | *ENDOD1* | 1.435813165 | 0.101059099 | -0.006659155 | 0.107718254 | 0.999933887 | 0.073658841 | -0.021514708 | 0.095173549 | 0.774024271 |
| ENSG00000110492 | *MDK* | 0.137011226 | 0.184678868 | 0.029129013 | 0.155549854 | 0.999933887 | 0.065784096 | 0.211192156 | -0.14540806 | 0.77412394 |
| ENSG00000059691 | *GATB* | -0.038784151 | -0.063537736 | -0.37149962 | 0.307961884 | 0.999933887 | 0.223517715 | 0.356958033 | -0.133440318 | 0.774125697 |
| ENSG00000249412 | *AC010285.1* | -0.252582327 | -0.040409123 | 0.183989521 | -0.224398643 | 0.999933887 | -0.32815018 | -0.225502265 | -0.102647915 | 0.774125697 |
| ENSG00000106399 | *RPA3* | 1.926950091 | -0.138404339 | 0.133451823 | -0.271856162 | 0.944094537 | 0.015306228 | 0.09249268 | -0.077186451 | 0.774293985 |
| ENSG00000197601 | *FAR1* | 6.158503491 | -0.085496111 | -0.218077937 | 0.132581827 | 0.999933887 | -0.902232992 | -0.771120481 | -0.131112511 | 0.774293985 |
| ENSG00000099956 | *SMARCB1* | 3.812624965 | 0.013377513 | -0.061422144 | 0.074799657 | 0.999933887 | 0.107282417 | 0.151842718 | -0.044560301 | 0.774293985 |

| | | | | | | | | | | |
|---|---|---|---|---|---|---|---|---|---|---|
| ENSG00000254363 | *AC011379.2* | -0.627973432 | -0.008444546 | 0.082280322 | -0.090724868 | 0.999933887 | -0.027116344 | 0.130232448 | -0.157348791 | 0.774293985 |
| ENSG00000114779 | *ABHD14B* | 2.900347838 | 0.07817903 | -0.053253692 | 0.131432722 | 0.999933887 | 0.109603174 | 0.173961233 | -0.064358059 | 0.77442889 |
| ENSG00000248643 | *RBM14-RBM4* | 0.187996427 | 0.191175351 | 0.253173064 | -0.061997713 | 0.999933887 | 0.167460791 | 0.284797138 | -0.117336347 | 0.774500371 |
| ENSG00000241157 | *AC104763.1* | 0.325791819 | 0.015297234 | -0.150468937 | 0.165766171 | 0.999933887 | -0.36404835 | -0.501762427 | 0.137714077 | 0.774551411 |
| ENSG00000120709 | *FAM53C* | 7.953520592 | 0.056739457 | 0.015830792 | 0.040908665 | 0.999933887 | -0.537603741 | -0.592196626 | 0.054592885 | 0.774563694 |
| ENSG00000139722 | *VPS37B* | 6.700100258 | -0.039969726 | -0.021689064 | -0.018280662 | 0.999933887 | -0.217921376 | -0.258437414 | 0.040516038 | 0.774563694 |
| ENSG00000264017 | *RN7SL336P* | 0.289457937 | 0.053949808 | 0.118422101 | -0.064472293 | 0.999933887 | 1.02286337 | 0.878005861 | 0.144857509 | 0.774563694 |
| ENSG00000213906 | *LTB4R2* | 0.504661386 | 0.043099821 | -0.05846649 | 0.101566311 | 0.999933887 | -0.124846584 | 0.022823801 | -0.147670385 | 0.774563694 |
| ENSG00000120925 | *RNF170* | 1.083227096 | 0.015786693 | 0.028886093 | -0.013099401 | 0.999933887 | -0.179762687 | -0.285618631 | 0.105855944 | 0.774563694 |
| ENSG00000227598 | *Z94721.1* | 0.457784996 | -0.010680009 | -0.08581137 | 0.075131361 | 0.999933887 | -0.709427497 | -0.884893929 | 0.175466433 | 0.774563694 |
| ENSG00000011478 | *QPCTL* | 2.945020435 | 0.078570541 | 0.12907348 | -0.050502939 | 0.999933887 | 0.257568583 | 0.324586119 | -0.067017536 | 0.77459624 |
| ENSG00000240038 | *AMY2B* | 2.897916141 | -0.054103984 | -0.173843413 | 0.119739429 | 0.999933887 | -0.160760804 | -0.061743549 | -0.099017255 | 0.77459624 |
| ENSG00000237296 | *SMG1P1* | -0.036464529 | -0.054375993 | 0.026367567 | -0.08074356 | 0.999933887 | 0.413987539 | 0.546293096 | -0.132305556 | 0.77459624 |
| ENSG00000168758 | *SEMA4C* | 3.004028737 | 0.042228368 | -0.014669682 | 0.05689805 | 0.999933887 | 0.089983032 | 0.150098913 | -0.060115881 | 0.774656165 |
| ENSG00000102763 | *VWA8* | 2.046948951 | 0.008798594 | -0.062052468 | 0.070851062 | 0.999933887 | -0.036171644 | 0.061510594 | -0.097682239 | 0.774656165 |
| ENSG00000279433 | *AC018529.3* | 3.859595812 | 0.170224597 | 0.314482585 | -0.144257987 | 0.999933887 | 0.131121502 | 0.223281649 | -0.092160147 | 0.774795196 |
| ENSG00000154330 | *PGM5* | 1.443492929 | 0.033321754 | -0.070242585 | 0.10356434 | 0.999933887 | -0.147663699 | -0.245858975 | 0.098195276 | 0.774795196 |
| ENSG00000188739 | *RBM34* | -0.454920189 | -0.198282278 | 0.008240542 | -0.20652282 | 0.999933887 | 0.080368452 | -0.088909652 | 0.169278104 | 0.775172981 |
| ENSG00000196372 | *ASB13* | -0.833616163 | -0.071097346 | -0.138796619 | 0.067699273 | 0.999933887 | 0.179713446 | -0.014032712 | 0.193746158 | 0.775172981 |
| ENSG00000157426 | *AASDH* | 0.872691004 | 0.014483075 | -0.207871383 | 0.222354458 | 0.999933887 | 0.089969467 | 0.198428312 | -0.108458845 | 0.775172981 |
| ENSG00000105819 | *PMPCB* | 4.411708509 | -0.098139292 | -0.10742328 | 0.009283988 | 0.999933887 | -0.039962024 | -0.002833544 | -0.037128479 | 0.775191063 |
| ENSG00000279520 | *AC093525.8* | 6.224872579 | 0.143917788 | 0.116640492 | 0.027277296 | 0.999933887 | 0.595391061 | 0.536702472 | 0.058688588 | 0.775191063 |
| ENSG00000279330 | *AJ003147.3* | 1.70145014 | -0.231535346 | 0.054160208 | -0.285695555 | 0.999933887 | -0.048260438 | -0.150240393 | 0.101979955 | 0.775191063 |
| ENSG00000156261 | *CCT8* | 4.100492715 | -0.130760118 | -0.085174642 | -0.045585476 | 0.999933887 | -0.097549592 | -0.046359178 | -0.051190414 | 0.775191063 |
| ENSG00000119203 | *CPSF3* | 2.007298418 | -0.166216459 | -0.109164643 | -0.057051816 | 0.999933887 | 0.060853132 | 0.125811455 | -0.064958324 | 0.775191063 |
| ENSG00000149262 | *INTS4* | 3.642695593 | -0.092135931 | 0.000854863 | -0.092990794 | 0.999933887 | 0.050803373 | 0.005713916 | 0.045089457 | 0.775191063 |
| ENSG00000134056 | *MRPS36* | 2.365360442 | 0.148635174 | 0.058770282 | 0.089864893 | 0.999933887 | 0.109110848 | 0.02829617 | 0.080814677 | 0.775191063 |
| ENSG00000132305 | *IMMT* | 3.408119524 | -0.085139242 | -0.107161232 | 0.022021991 | 0.999933887 | 0.018112324 | -0.031105323 | 0.049217647 | 0.775191063 |
| ENSG00000241106 | *HLA-DOB* | 0.479040589 | -0.185135606 | 0.008199788 | -0.193335394 | 0.999933887 | -0.0134026 | 0.134020848 | -0.147423448 | 0.775191063 |
| ENSG00000139517 | *LNX2* | 1.938622682 | 0.111475392 | 0.048277934 | 0.063197458 | 0.999933887 | -0.397074573 | -0.471654658 | 0.074580085 | 0.775191063 |
| ENSG00000141959 | *PFKL* | 5.626948431 | 0.079240965 | 0.038216307 | 0.041024658 | 0.999933887 | 0.266518598 | 0.321668337 | -0.055149739 | 0.775191063 |
| ENSG00000154328 | *NEIL2* | 0.730830697 | -0.128995406 | -0.208971793 | 0.079976387 | 0.999933887 | 0.046642787 | 0.14593299 | -0.099290203 | 0.775191063 |
| ENSG00000221909 | *FAM200A* | 0.572158987 | -0.179133758 | -0.142886786 | -0.036246972 | 0.999933887 | 0.025240105 | 0.146380774 | -0.121140669 | 0.775191063 |
| ENSG00000117505 | *DR1* | 5.655048028 | -0.05168978 | -0.008063278 | -0.043626502 | 0.999933887 | -0.350142529 | -0.392338814 | 0.042196285 | 0.775191063 |
| ENSG00000134644 | *PUM1* | 5.540135159 | -0.041506654 | -0.032654559 | -0.008852095 | 0.999933887 | -0.079183673 | -0.044408757 | -0.034774916 | 0.775191063 |
| ENSG00000197980 | *LEKR1* | -0.528176974 | 0.151658566 | 0.330441952 | -0.178783386 | 0.999933887 | 0.744111398 | 0.539337141 | 0.204774256 | 0.775191063 |
| ENSG00000188242 | *PP7080* | 0.385160562 | 0.135763858 | 0.159282131 | -0.023518273 | 0.999933887 | 0.066755857 | 0.18572566 | -0.118969802 | 0.775191063 |
| ENSG00000145555 | *MYO10* | 3.187458275 | -0.074084408 | 0.041523147 | -0.115607555 | 0.999933887 | -0.300617423 | -0.362307526 | 0.061690103 | 0.775191063 |
| ENSG00000247626 | *MARS2* | 0.95409941 | 0.091042823 | -0.070605064 | 0.161647887 | 0.999933887 | 0.158888752 | 0.277235093 | -0.118346341 | 0.775191063 |
| ENSG00000261456 | *TUBB8* | -2.158368029 | -0.199813953 | -0.570411072 | 0.370597119 | 0.999933887 | -1.629477801 | -1.428276344 | -0.201201456 | 0.775191063 |
| ENSG00000163806 | *SPDYA* | 0.834258067 | 0.065017151 | 0.052343966 | 0.012673185 | 0.999933887 | 0.055152831 | 0.210343178 | -0.155190348 | 0.775191063 |
| ENSG00000269119 | *HNRNPA1P52* | 0.380709151 | -0.065175028 | -0.24544751 | 0.180272483 | 0.999933887 | -0.345649183 | -0.506516653 | 0.16086747 | 0.775191063 |
| ENSG00000281691 | *RBM5-AS1* | -0.156117496 | -0.110392098 | 0.026432372 | -0.13682447 | 0.999933887 | -0.027449097 | 0.12297485 | -0.150423947 | 0.775191063 |
| ENSG00000170515 | *PA2G4* | 2.969779087 | -0.03499681 | 0.027412496 | -0.062409307 | 0.999933887 | 0.134755289 | 0.063773701 | 0.070981588 | 0.775191063 |
| ENSG00000229463 | *LYST-AS1* | -0.454141792 | 0.075709001 | -0.008243898 | 0.0839529 | 0.999933887 | -0.118846726 | 0.104680683 | -0.22352741 | 0.775191063 |
| ENSG00000257103 | *LSM14A* | 6.350519116 | -0.013569312 | -0.002710674 | -0.010858638 | 0.999933887 | -0.184737639 | -0.152260472 | -0.032477167 | 0.775191063 |
| ENSG00000198467 | *TPM2* | 1.958024173 | -0.030271425 | -0.01797036 | -0.012301065 | 0.999933887 | 0.049105356 | 0.131296888 | -0.082191532 | 0.775191063 |
| ENSG00000066468 | *FGFR2* | -0.011944207 | 0.03156454 | 0.025857969 | 0.005706571 | 0.999933887 | -0.142398476 | -0.014104179 | -0.128294296 | 0.775191063 |
| ENSG00000163584 | *RPL22L1* | 2.724345822 | 0.013373785 | -0.009812761 | 0.023186547 | 0.999933887 | 0.288969108 | 0.354093642 | -0.065124535 | 0.775191063 |
| ENSG00000196557 | *CACNA1H* | -0.057080513 | -0.029958676 | 0.392954829 | -0.422913506 | 0.999933887 | 0.005744016 | 0.159666048 | -0.153922032 | 0.775191063 |
| ENSG00000144867 | *SRPRB* | 1.631921791 | 0.015577943 | -0.086930631 | 0.102508574 | 0.999933887 | 0.268828952 | 0.3456113 | -0.076782348 | 0.775191063 |
| ENSG00000279700 | *AC131212.2* | 1.427430661 | -0.016120114 | -0.149056778 | 0.132936664 | 0.999933887 | -0.254687364 | -0.368356183 | 0.113668818 | 0.775191063 |
| ENSG00000132386 | *SERPINF1* | -0.879520551 | 0.030069016 | -0.395239007 | 0.425308023 | 0.999933887 | 0.110239155 | -0.083740228 | 0.193979383 | 0.775191063 |
| ENSG00000149573 | *MPZL2* | 1.419195597 | 0.011832097 | -0.025033708 | 0.036865805 | 0.999933887 | -0.323039479 | -0.414686522 | 0.091647043 | 0.775191063 |
| ENSG00000166979 | *EVA1C* | 0.565234698 | 0.013784141 | 0.017692415 | -0.003908274 | 0.999933887 | 0.21124829 | 0.334031848 | -0.122783558 | 0.775191063 |
| ENSG00000104884 | *ERCC2* | 0.933012312 | -0.012516895 | -0.027402525 | 0.01488563 | 0.999933887 | 0.020364456 | 0.120806991 | -0.100442535 | 0.775191063 |
| ENSG00000272906 | *AL353708.3* | 0.554718194 | 0.015837073 | 0.079150363 | -0.06331329 | 0.999933887 | 0.240970818 | 0.127318759 | 0.113652059 | 0.775191063 |
| ENSG00000049883 | *PTCD2* | 0.16677124 | -0.012875365 | -0.056108959 | 0.043233594 | 0.999933887 | -0.059906298 | 0.078155424 | -0.138061722 | 0.775191063 |
| ENSG00000151665 | *PIGF* | 0.74487587 | -0.005694645 | 0.249405899 | -0.255100544 | 0.999933887 | 0.029090996 | 0.130593908 | -0.101502912 | 0.775191063 |
| ENSG00000129194 | *SOX15* | -0.01070459 | 0.321997104 | 0.36069852 | -0.038701417 | 0.999933887 | 0.679539339 | 0.528309579 | 0.15122976 | 0.775233918 |
| ENSG00000164934 | *DCAF13* | 3.302710286 | -0.101125868 | 0.007850404 | -0.108976272 | 0.999933887 | -0.239004038 | -0.185914223 | -0.053089815 | 0.775233918 |
| ENSG00000106028 | *SSBP1* | 3.644128816 | -0.072099138 | -0.040453328 | -0.031645809 | 0.999933887 | -0.069524104 | -0.024688045 | -0.04483606 | 0.775233918 |
| ENSG00000251247 | *ZNF345* | 0.46520598 | 0.135341503 | -0.011579908 | 0.146921411 | 0.999933887 | -0.148454519 | -0.027674224 | -0.120780295 | 0.775233918 |
| ENSG00000247077 | *PGAM5* | 1.940144427 | -0.068303771 | 0.009159938 | -0.07746371 | 0.999933887 | 0.23700363 | 0.306359191 | -0.069355561 | 0.775233918 |
| ENSG00000281958 | *TRBJ1-4* | 0.222722455 | -0.02362256 | 0.213698525 | -0.237321085 | 0.999933887 | 0.165901934 | 0.056588431 | 0.109313502 | 0.775233918 |
| ENSG00000226272 | *ARHGAP26-A* | 0.802436661 | 0.291908134 | 0.597106851 | -0.305198717 | 0.999933887 | 0.364006564 | 0.084313039 | 0.279693526 | 0.775573053 |
| ENSG00000176248 | *ANAPC2* | 5.244398862 | 0.024847393 | 0.110859289 | -0.086011895 | 0.999933887 | 0.145953228 | 0.191228168 | -0.04527494 | 0.775573053 |
| ENSG00000060762 | *MPC1* | 3.72984044 | -0.077555691 | -0.019406426 | -0.058149265 | 0.999933887 | -0.13945789 | -0.088371245 | -0.051086645 | 0.775696593 |
| ENSG00000259912 | *AC023813.1* | 0.198583647 | 0.020776146 | 0.148790439 | -0.128014293 | 0.999933887 | 3.231196873 | 3.396033242 | -0.164836369 | 0.775860868 |
| ENSG00000122545 | *SEPTIN7* | 6.47279796 | -0.078082867 | -0.136583551 | 0.058500684 | 0.999933887 | -0.486287312 | -0.415903452 | -0.07038386 | 0.776183724 |
| ENSG00000136783 | *NIPSNAP3A* | 0.815323701 | -0.110882799 | 0.040053927 | -0.150936726 | 0.999933887 | -0.135430338 | -0.027894052 | -0.107536286 | 0.776183724 |
| ENSG00000071205 | *ARHGAP10* | 1.770584109 | -0.076453687 | 0.129172153 | -0.205625839 | 0.999933887 | -0.172840264 | -0.25591188 | 0.083071616 | 0.776183724 |
| ENSG00000242498 | *ARPIN* | -0.371127848 | 0.016296906 | 0.30069356 | -0.284396654 | 0.999933887 | -1.21776466 | -1.370933712 | 0.153169052 | 0.776183724 |
| ENSG00000146833 | *TRIM4* | 3.118498515 | -0.052137931 | 0.005803062 | -0.057940993 | 0.999933887 | -0.177762433 | -0.126860683 | -0.05090175 | 0.776364097 |
| ENSG00000125484 | *GTF3C4* | 2.618244121 | -0.014826135 | 0.022568553 | -0.037394688 | 0.999933887 | 0.074663162 | 0.011332145 | 0.063331017 | 0.776364097 |
| ENSG00000100058 | *CRYBB2P1* | 1.148447908 | 0.06638885 | -0.065940513 | 0.132329363 | 0.999933887 | 0.151401519 | 0.241810345 | -0.090408826 | 0.776459199 |
| ENSG00000089220 | *PEBP1* | 3.872132152 | 0.002845778 | 0.020458859 | -0.01761308 | 0.999933887 | 0.203538132 | 0.143272515 | 0.060265617 | 0.776624297 |
| ENSG00000149503 | *INCENP* | 2.206664573 | 0.047849158 | -0.067924818 | 0.115773977 | 0.999933887 | -0.154968226 | -0.08787061 | -0.067097615 | 0.776643178 |
| ENSG00000207171 | *AC079926.1* | -0.684349619 | 0.338756845 | 0.618268807 | -0.279511962 | 0.999933887 | 1.189289718 | 0.965949625 | 0.223340093 | 0.776674857 |
| ENSG00000109971 | *HSPA8* | 7.360212634 | -0.002713088 | -0.069497643 | 0.066784555 | 0.999933887 | 0.095577215 | 0.051593904 | 0.04398331 | 0.776731891 |
| ENSG00000167994 | *RAB3IL1* | 2.371156415 | -0.038882738 | -0.101221045 | 0.062338307 | 0.999933887 | -0.680903658 | -0.620570946 | -0.060332712 | 0.776738136 |
| ENSG00000237440 | *ZNF737* | 1.745556974 | -0.060620883 | -0.154927658 | 0.094306775 | 0.999933887 | -0.318185269 | -0.414871675 | 0.096686405 | 0.776946947 |
| ENSG00000186010 | *NDUFA13* | 0.174354192 | 0.125974093 | 0.062779016 | 0.063195078 | 0.999933887 | -0.114759537 | -0.224268561 | 0.109509024 | 0.77766106 |
| ENSG00000102981 | *PARD6A* | 0.843916288 | 0.166821509 | -0.170075085 | 0.336896594 | 0.999933887 | -0.442970791 | -0.600480349 | 0.157509558 | 0.777756211 |
| ENSG00000267416 | *AC025048.4* | -0.848604241 | -0.209013039 | -0.116790027 | -0.092223013 | 0.999933887 | -0.143156331 | -0.281604108 | 0.138477777 | 0.777816269 |
| ENSG00000188827 | *SLX4* | 2.502326621 | 0.066631572 | 0.049653699 | 0.016977873 | 0.999933887 | -0.103564568 | -0.181310936 | 0.077746368 | 0.777816269 |
| ENSG00000246223 | *LINC01550* | 0.335206363 | 0.097176262 | -0.250447196 | 0.347623458 | 0.999933887 | -0.004763861 | 0.110864622 | -0.115628484 | 0.777816269 |
| ENSG00000136854 | *STXBP1* | -0.780650185 | 0.078492182 | -0.054531563 | 0.133023744 | 0.999933887 | -0.098641534 | 0.068518338 | -0.167159872 | 0.777816269 |
| ENSG00000241860 | *AL627309.5* | 2.201360767 | -0.025156175 | 0.044985155 | -0.07014133 | 0.999933887 | 0.320743003 | 0.411919532 | -0.091176529 | 0.77783534 |
| ENSG00000183666 | *GUSBP1* | -0.238352722 | -0.045175343 | 0.122138874 | -0.167314217 | 0.999933887 | -0.352603951 | -0.200796761 | -0.15180719 | 0.778044921 |
| ENSG00000222057 | *RNU4-62P* | 1.104857787 | 0.086606568 | 0.053014095 | 0.033592473 | 0.999933887 | -0.334681036 | -0.493218497 | 0.158537461 | 0.778239456 |
| ENSG00000004468 | *CD38* | 1.254821433 | 0.028912613 | -0.00771558 | 0.036628193 | 0.999933887 | -0.216919672 | -0.120895365 | -0.096024306 | 0.778239456 |
| ENSG00000127774 | *EMC6* | -0.447415566 | -0.061942129 | -0.196345884 | 0.134403755 | 0.999933887 | -0.157197887 | -0.029307727 | -0.127890161 | 0.778255254 |
| ENSG00000157538 | *VPS26C* | 5.717549955 | -0.005658088 | -0.060371805 | 0.054713718 | 0.999933887 | -0.733310566 | -0.693020654 | -0.040289912 | 0.778374326 |
| ENSG00000197753 | *LHFPL5* | 4.668362487 | -0.073355253 | -0.00974712 | -0.063608132 | 0.999933887 | -0.762729216 | -0.698791693 | -0.063937523 | 0.778473457 |
| ENSG00000051128 | *HOMER3* | 3.297554039 | 0.058627918 | 0.02647662 | 0.032151298 | 0.999933887 | 0.131719639 | 0.193499452 | -0.061779813 | 0.778481033 |
| ENSG00000164047 | *CAMP* | 1.861817241 | -0.007505878 | 0.035950092 | -0.04345597 | 0.999933887 | -0.054770766 | -0.150964052 | 0.096193286 | 0.778481033 |
| ENSG00000211794 | *TRAV12-3* | -0.499265577 | -0.008228058 | 0.066857986 | -0.075086043 | 0.999933887 | 0.18330569 | 0.322417795 | -0.139112104 | 0.778481033 |
| ENSG00000181191 | *PJA1* | 1.344532297 | 0.004237208 | -0.233070441 | 0.237307649 | 0.999933887 | 0.091948874 | 0.175231797 | -0.083282923 | 0.778481033 |
| ENSG00000228649 | *SNHG26* | 0.138513306 | 0.155930222 | -0.198467397 | 0.354397619 | 0.999933887 | 0.18269202 | 0.313001313 | -0.130309294 | 0.778608028 |
| ENSG00000178199 | *ZC3H12D* | 4.448057637 | 0.037149553 | -0.00349275 | 0.040642303 | 0.999933887 | 0.27735648 | 0.325027259 | -0.047670779 | 0.778652553 |
| ENSG00000198286 | *CARD11* | 4.907195554 | -0.006623458 | 0.027406901 | -0.034030359 | 0.999933887 | 0.077814043 | 0.12259514 | -0.044781096 | 0.778652553 |
| ENSG00000274712 | *AC005332.4* | 1.753331219 | -0.010490802 | -0.038147135 | 0.027656333 | 0.999933887 | 0.062846631 | -0.020694894 | 0.083541525 | 0.778652553 |
| ENSG00000188389 | *PDCD1* | 1.994286616 | 0.108886962 | 0.063045933 | 0.045841029 | 0.999933887 | 0.148863798 | 0.071808181 | 0.077055617 | 0.778803207 |
| ENSG00000198146 | *ZNF770* | 2.543255963 | -0.122767519 | -0.042798658 | -0.079968861 | 0.999933887 | -0.302566792 | -0.196069616 | -0.106497176 | 0.779021065 |

| | | | | | | | | | | |
|---|---|---|---|---|---|---|---|---|---|---|
| ENSG00000162928 | *PEX13* | 1.689467404 | -0.087127598 | 0.007451888 | -0.094579485 | 0.999933887 | -0.129616581 | -0.203937204 | 0.074320623 | 0.779021065 |
| ENSG00000215788 | *TNFRSF25* | 4.101876276 | 0.050232661 | 0.065383921 | -0.01515126 | 0.999933887 | 0.260976171 | 0.201893398 | 0.059082773 | 0.779126514 |
| ENSG00000161328 | *LRRC56* | 0.127470843 | 0.038025631 | -0.057101421 | 0.095127052 | 0.999933887 | 0.058878994 | 0.185970485 | -0.127091491 | 0.779126514 |
| ENSG00000229413 | *AC018638.1* | 0.744872578 | 0.057809279 | -0.139748522 | 0.197557801 | 0.999933887 | -0.081097461 | -0.261360257 | 0.180262796 | 0.779132585 |
| ENSG00000276234 | *TADA2A* | 0.71127642 | -0.001626208 | -0.06609727 | 0.064471062 | 0.999933887 | 0.075334297 | -0.024561395 | 0.099895692 | 0.779132585 |
| ENSG00000170836 | *PPM1D* | 4.329540299 | 0.026380447 | 0.01814463 | 0.008235817 | 0.999933887 | -0.388733292 | -0.339163502 | -0.04956979 | 0.779294013 |
| ENSG00000253111 | *AC091114.1* | 1.074309684 | -0.10619804 | 0.266239621 | -0.37243766 | 0.999933887 | 1.512996168 | 1.636685686 | -0.123689518 | 0.779295515 |
| ENSG00000167107 | *ACSF2* | 1.762565163 | -0.023990888 | -0.020834013 | -0.003156875 | 0.999933887 | 0.024715508 | -0.060302125 | 0.085017633 | 0.779370715 |
| ENSG00000126870 | *DYNC2I1* | 2.200663342 | -0.012924841 | 0.037032704 | -0.049957544 | 0.999933887 | 0.145887209 | 0.208062755 | -0.062175546 | 0.779370715 |
| ENSG00000089094 | *KDM2B* | 4.616008508 | 0.025598464 | 0.067948146 | -0.042349681 | 0.999933887 | 0.06863167 | 0.113213273 | -0.044581604 | 0.779448897 |
| ENSG00000279801 | *AC111170.3* | 0.42159372 | -0.287640602 | -0.080796659 | -0.206843943 | 0.999933887 | -0.410045443 | -0.516358477 | 0.106313034 | 0.779611694 |
| ENSG00000288538 | *AP002985.1* | 0.351535402 | -0.053484902 | 0.029356223 | -0.082841126 | 0.999933887 | -0.248203648 | -0.120356674 | -0.127846974 | 0.779757629 |
| ENSG00000102030 | *NAA10* | 4.563915869 | 0.04994418 | 0.109257071 | -0.059312891 | 0.999933887 | 0.15702296 | 0.095629722 | 0.061393238 | 0.780004004 |
| ENSG00000115085 | *ZAP70* | 6.376097932 | 0.037205815 | 0.111372348 | -0.074166532 | 0.999933887 | 0.131656366 | 0.084714595 | 0.04694177 | 0.78003374 |
| ENSG00000225636 | *HNRNPA3P15* | -1.034015715 | 0.071877876 | 0.017368297 | 0.054509579 | 0.999933887 | -0.22580882 | -0.492266572 | 0.266457751 | 0.78003374 |
| ENSG00000140718 | *FTO* | 1.880285687 | -0.095095701 | -0.124075785 | 0.028980084 | 0.999933887 | 0.00388052 | 0.09210045 | -0.08821993 | 0.780068449 |
| ENSG00000144120 | *TMEM177* | -1.366761093 | -0.222436482 | -0.469034433 | 0.246597951 | 0.999933887 | -0.022908563 | -0.238000144 | 0.21509158 | 0.780155261 |
| ENSG00000073536 | *NLE1* | 0.909274894 | 0.113015662 | 0.069845772 | 0.04316989 | 0.999933887 | 0.357364624 | 0.263124168 | 0.094240456 | 0.780257668 |
| ENSG00000197019 | *SERTAD1* | 5.090920052 | 0.132637133 | 0.1547984 | -0.022161267 | 0.999933887 | -0.018794302 | 0.031784635 | -0.050578937 | 0.780350935 |
| ENSG00000168827 | *GFM1* | 2.606452936 | -0.104130866 | -0.100289385 | -0.003841481 | 0.999933887 | 0.00763586 | 0.066886417 | -0.059250557 | 0.780350935 |
| ENSG00000267430 | *AC036176.2* | 0.265380239 | 0.156119357 | 0.467091017 | -0.31097166 | 0.999933887 | 0.102567582 | 0.28895448 | -0.186386898 | 0.780350935 |
| ENSG00000177570 | *SAMD12* | -1.100735438 | 0.123284153 | -0.35011687 | 0.473401024 | 0.999933887 | -0.019371463 | -0.209471947 | 0.190100484 | 0.780350935 |
| ENSG00000157593 | *SLC35B2* | 5.324087738 | 0.065772781 | 0.100028751 | -0.03425597 | 0.999933887 | 1.711681986 | 1.792819826 | -0.08113784 | 0.780350935 |
| ENSG00000248098 | *BCKDHA* | -0.360280808 | -0.120284962 | -0.166997841 | 0.046712878 | 0.999933887 | -0.486540112 | -0.28095296 | -0.205587152 | 0.780350935 |
| ENSG00000204104 | *TRAF3IP1* | 0.482569319 | -0.076143235 | 0.033713484 | -0.109856719 | 0.999933887 | 0.106857157 | 0.221427218 | -0.11457006 | 0.780350935 |
| ENSG00000174206 | *C12orf66* | 0.389633233 | -0.296142261 | 0.245345848 | -0.541488109 | 0.940218856 | 0.327992714 | 0.224222861 | 0.103769853 | 0.780471716 |
| ENSG00000180488 | *MIGA1* | 1.809628528 | -0.069847081 | -0.005153514 | -0.064693568 | 0.999933887 | -0.172272992 | -0.070239765 | -0.102033227 | 0.780471716 |
| ENSG00000112763 | *BTN2A1* | 6.842851475 | 0.01971567 | 0.056034683 | -0.036319013 | 0.999933887 | 0.041870209 | 0.080136905 | -0.038266696 | 0.780725165 |
| ENSG00000159579 | *RSPRY1* | 3.798994786 | -0.02006945 | -0.111469168 | 0.091399718 | 0.999933887 | -0.403604166 | -0.356444138 | -0.047160028 | 0.78086707 |
| ENSG00000149639 | *SOGA1* | 0.750269728 | -0.032465162 | 0.033925931 | -0.066391093 | 0.999933887 | -0.907266554 | -1.038665766 | 0.131399212 | 0.781110236 |
| ENSG00000222267 | *RNU6-892P* | -0.09894163 | -0.283898593 | -0.019112133 | -0.26478646 | 0.999933887 | 0.30997085 | 0.46266797 | -0.15269712 | 0.781457311 |
| ENSG00000171161 | *ZNF672* | 5.355076078 | 0.050154344 | -0.029675847 | 0.079830191 | 0.999933887 | -0.197844622 | -0.240379462 | 0.04253484 | 0.781609694 |
| ENSG00000224032 | *EPB41L4A-AS1* | 2.305165095 | -0.039785051 | 0.039552414 | -0.079337465 | 0.999933887 | 0.048226995 | 0.121246165 | -0.073019169 | 0.781609694 |
| ENSG00000188895 | *MSL1* | 8.562490012 | -0.004784107 | 0.03567692 | -0.040461027 | 0.999933887 | -0.584456592 | -0.63172779 | 0.047271198 | 0.78171069 |
| ENSG00000164855 | *TMEM184A* | 1.488763337 | 0.001181584 | 0.105137617 | -0.103956033 | 0.999933887 | 0.053985985 | 0.147578367 | -0.093592382 | 0.781726898 |
| ENSG00000259248 | *USP3-AS1* | 1.769918608 | -0.033994694 | 0.026563435 | -0.060558129 | 0.999933887 | -0.359498225 | -0.207332768 | -0.152165457 | 0.781789279 |
| ENSG00000106070 | *GRB10* | 2.521102493 | -0.007038823 | 0.062914819 | -0.069953642 | 0.999933887 | -0.050537133 | -0.113042593 | 0.06250546 | 0.781805811 |
| ENSG00000107404 | *DVL1* | 4.18130137 | 0.056382252 | 0.053011637 | 0.003370615 | 0.999933887 | -0.011654948 | 0.046813283 | -0.058468231 | 0.781859779 |
| ENSG00000215014 | *AL645728.1* | 0.820993263 | -0.015747924 | 0.064416938 | -0.080164862 | 0.999933887 | 0.27755586 | 0.404932297 | -0.127376437 | 0.781966303 |
| ENSG00000083457 | *ITGAE* | 2.278803532 | -0.041393332 | -0.20613116 | 0.164737828 | 0.999933887 | -0.098914038 | -0.161118438 | 0.0622044 | 0.782228496 |
| ENSG00000154027 | *AK5* | -0.568359236 | 0.075170344 | -0.200326442 | 0.275496785 | 0.999933887 | -0.107821782 | 0.035414156 | -0.143235938 | 0.782233975 |
| ENSG00000002919 | *SNX11* | 4.318163308 | -0.020915632 | -0.036155025 | 0.015239394 | 0.999933887 | -0.020205926 | 0.037319893 | -0.057525819 | 0.782336232 |
| ENSG00000139746 | *RBM26* | 4.843301557 | -0.097455793 | -0.180099591 | 0.082643798 | 0.999933887 | -0.514869372 | -0.43183138 | -0.083037992 | 0.782410502 |
| ENSG00000151835 | *SACS* | 2.983441505 | -0.15701049 | -0.166221843 | 0.009211353 | 0.999933887 | -0.281884329 | -0.155081315 | -0.126803014 | 0.782784339 |
| ENSG00000258987 | *AL132642.1* | -0.919314645 | -0.052540395 | -0.004785012 | -0.047755383 | 0.999933887 | -0.330739857 | -0.109409503 | -0.221330354 | 0.782849731 |
| ENSG00000171649 | *ZIK1* | -0.707152098 | -0.435438455 | 0.236396482 | -0.671834937 | 0.969206975 | -0.180654705 | -0.039562844 | -0.141091861 | 0.782868513 |
| ENSG00000118894 | *EEF2KMT* | -0.991041026 | 0.120632824 | 0.332697968 | -0.212065145 | 0.999933887 | 0.305697815 | 0.480640046 | -0.174942231 | 0.782868513 |
| ENSG00000217165 | *ANKRD18EP* | 0.147012147 | -0.077166285 | -0.043035394 | -0.034130892 | 0.999933887 | -0.162945486 | -0.028195209 | -0.134750277 | 0.782879338 |
| ENSG00000170989 | *S1PR1* | 5.132166853 | -0.012046368 | -0.071975534 | 0.059929166 | 0.999933887 | 0.108048562 | 0.06236093 | 0.045687632 | 0.782879338 |
| ENSG00000136159 | *NUDT15* | 1.168222449 | 0.013189964 | 0.038003762 | -0.024813798 | 0.999933887 | 0.036073527 | -0.077027765 | 0.113101292 | 0.782879338 |
| ENSG00000155749 | *FLACC1* | -0.216328478 | 0.085093613 | -0.276315824 | 0.361409438 | 0.999933887 | -0.352929352 | -0.538643485 | 0.185714133 | 0.78301999 |
| ENSG00000143256 | *PFDN2* | 2.903984595 | 0.032690404 | 0.064434495 | -0.031744091 | 0.999933887 | -0.016311993 | 0.03832819 | -0.054640183 | 0.78301999 |
| ENSG00000126012 | *KDM5C* | 7.395873093 | 0.024272475 | 0.060076681 | -0.035804207 | 0.999933887 | -0.082640372 | -0.111904861 | 0.029264488 | 0.783122487 |
| ENSG00000133639 | *BTG1* | 8.625523225 | 0.018630428 | 0.023390713 | -0.004760286 | 0.999933887 | -0.032252697 | -0.066969766 | 0.034717069 | 0.783166408 |
| ENSG00000168675 | *LDLRAD4* | 2.574809703 | 0.029746199 | 0.133682669 | -0.103936471 | 0.999933887 | 0.004709732 | -0.052661321 | 0.057371053 | 0.783355618 |
| ENSG00000149150 | *SLC43A1* | 0.787077593 | -0.054612909 | -0.325893148 | 0.271280239 | 0.999933887 | 0.013466971 | 0.111595429 | -0.098128458 | 0.783826883 |
| ENSG00000230834 | *AC068580.2* | -1.070670085 | 0.255699152 | -0.000212106 | 0.255911259 | 0.999933887 | 0.291374295 | 0.475268865 | -0.183894569 | 0.783856072 |
| ENSG00000214300 | *SPDYE3* | 0.123867609 | 0.193760573 | -0.008736816 | 0.202497389 | 0.999933887 | 0.370548817 | 0.262316426 | 0.108232391 | 0.784074094 |
| ENSG00000108474 | *PIGL* | 1.55805361 | -0.019053247 | 0.04204194 | -0.061095187 | 0.999933887 | 0.066409857 | -0.014149647 | 0.080559505 | 0.784243831 |
| ENSG00000259479 | *SORD2P* | -0.043637674 | 0.035561303 | 0.025940337 | 0.009620966 | 0.999933887 | -0.20060137 | -0.319743816 | 0.119142446 | 0.784414078 |
| ENSG00000170540 | *ARL6IP1* | 5.181208092 | -0.084205303 | -0.077857912 | -0.006347391 | 0.999933887 | -0.291883487 | -0.341259671 | 0.049376184 | 0.784478366 |
| ENSG00000125247 | *TMTC4* | 1.120341141 | -0.0338748 | 0.061238962 | -0.095113763 | 0.999933887 | 0.04018869 | -0.04742294 | 0.08761163 | 0.784478366 |
| ENSG00000142528 | *ZNF473* | 2.424916891 | -0.041758162 | -0.059342404 | 0.017584243 | 0.999933887 | -0.02945375 | 0.049073699 | -0.078527449 | 0.784537545 |
| ENSG00000214456 | *PLIN5* | 4.214749954 | 0.056738954 | 0.068892636 | -0.012153682 | 0.999933887 | 0.077736375 | 0.016468554 | 0.061267821 | 0.784861199 |
| ENSG00000279908 | *AC026748.3* | 0.389493046 | -0.015554445 | 0.018191166 | -0.033745611 | 0.999933887 | 0.169977945 | 0.044159328 | 0.125818617 | 0.784861199 |
| ENSG00000176083 | *ZNF683* | 1.711508135 | 0.110918656 | -0.042115933 | 0.153034589 | 0.999933887 | -0.034705038 | 0.036867335 | -0.071572373 | 0.784908463 |
| ENSG00000233901 | *LINC01503* | 1.868512441 | 0.015208666 | -0.095004552 | 0.110213218 | 0.999933887 | -0.536439396 | -0.461341374 | -0.075098022 | 0.784908463 |
| ENSG00000241657 | *TRBV11-2* | -0.394277273 | 0.117344422 | 0.11322249 | 0.004121933 | 0.999933887 | 0.205022315 | 0.360226105 | -0.15520379 | 0.785001879 |
| ENSG00000267123 | *SCAT1* | -0.721488441 | -0.109149953 | -0.329617276 | 0.220467323 | 0.999933887 | 0.075249477 | -0.098372159 | 0.173621636 | 0.785001879 |
| ENSG00000245849 | *RAD51-AS1* | 2.769723059 | -0.000308812 | -0.044932318 | 0.044623506 | 0.999933887 | -0.077839074 | -0.02150193 | -0.056337144 | 0.785001879 |
| ENSG00000278434 | *AC023830.3* | 0.417934507 | 0.179953323 | 0.176495357 | 0.003457965 | 0.999933887 | 0.44972705 | 0.607721161 | -0.15799411 | 0.785083646 |
| ENSG00000138138 | *ATAD1* | 3.154310668 | -0.112391122 | -0.04314634 | -0.069244781 | 0.999933887 | -0.157342879 | -0.220154362 | 0.062811483 | 0.78509922 |
| ENSG00000204536 | *CCHCR1* | 2.174466101 | 0.084543811 | -0.030255515 | 0.114799326 | 0.999933887 | 0.066586705 | -0.010366868 | 0.076953573 | 0.78509922 |
| ENSG00000185710 | *SMG1P4* | -1.073880925 | 0.135704282 | 0.014869611 | 0.120834671 | 0.999933887 | -0.147503757 | 0.040176007 | -0.187679764 | 0.7851192 |
| ENSG00000189403 | *HMGB1* | 6.401744522 | -0.024312499 | -0.042506393 | 0.018193894 | 0.999933887 | -0.095932017 | -0.132708293 | 0.036776277 | 0.785227048 |
| ENSG00000113369 | *ARRDC3* | 7.476025709 | -0.316177649 | -0.415908295 | 0.099730646 | 0.999933887 | -0.269143942 | -0.203761844 | -0.065382098 | 0.785246073 |
| ENSG00000119041 | *GTF3C3* | 2.131015961 | -0.110106506 | -0.2468698 | 0.136763294 | 0.999933887 | -0.000429656 | 0.084803436 | -0.085233092 | 0.785246073 |
| ENSG00000196296 | *ATP2A1* | 0.322595022 | 0.050261806 | 0.171190392 | -0.120928587 | 0.999933887 | -0.280702395 | -0.434862733 | 0.154160338 | 0.785246073 |
| ENSG00000182272 | *B4GALNT4* | -0.813048531 | 0.153035369 | 0.176344292 | -0.023308923 | 0.999933887 | 0.128478369 | -0.087261548 | 0.215739918 | 0.785266702 |
| ENSG00000273797 | *AL133445.2* | 1.559147024 | 0.126235906 | 0.286455521 | -0.160219615 | 0.999933887 | 1.405804189 | 1.512311248 | -0.106507059 | 0.785460792 |
| ENSG00000213420 | *GPC2* | 0.947149043 | 0.116550515 | 0.035557788 | 0.080992727 | 0.999933887 | -0.226707292 | -0.348617859 | 0.121910567 | 0.785489795 |
| ENSG00000184350 | *MRGPRE* | -0.076494836 | 0.013072874 | -0.069433593 | 0.082506466 | 0.999933887 | -0.296971664 | -0.387412226 | 0.090440562 | 0.785489795 |
| ENSG00000203791 | *EEF1AKMT2* | -0.362685391 | 0.002476639 | -0.095459671 | 0.09793631 | 0.999933887 | -0.3833953 | -0.223489248 | -0.159906053 | 0.785489795 |
| ENSG00000184730 | *APOBR* | 8.787551313 | 0.074107279 | 0.16866735 | -0.094560071 | 0.999933887 | -0.42437688 | -0.504571373 | 0.080194493 | 0.78560797 |
| ENSG00000233750 | *CICP27* | 0.446502011 | -0.026966315 | -0.086418955 | 0.05945264 | 0.999933887 | 0.094071417 | -0.074728113 | 0.16879953 | 0.78560797 |
| ENSG00000264247 | *LINC00909* | 2.592057174 | -0.110585901 | -0.350615128 | 0.240029227 | 0.999933887 | -0.203420645 | -0.273560462 | 0.070139817 | 0.785635975 |
| ENSG00000160691 | *SHC1* | 5.345628421 | 0.034956911 | 0.025379923 | 0.009576988 | 0.999933887 | -0.157199358 | -0.195356767 | 0.038157409 | 0.785803675 |
| ENSG00000158234 | *FAIM* | -0.285085004 | -0.010348785 | -0.041512504 | 0.031163719 | 0.999933887 | -0.07599254 | 0.092006632 | -0.167999172 | 0.785803675 |
| ENSG00000171067 | *C11orf24* | 1.31234565 | -0.03301718 | 0.104787263 | -0.137804443 | 0.999933887 | 0.093825237 | 0.022439365 | 0.071385872 | 0.786138048 |
| ENSG00000182512 | *GLRX5* | 2.469105098 | 0.069110663 | 0.048536646 | 0.020574017 | 0.999933887 | -0.037512696 | -0.110821541 | 0.073308845 | 0.786295704 |
| ENSG00000253210 | *AC040970.1* | 3.899552005 | 0.056037442 | 0.033179714 | 0.022857728 | 0.999933887 | -0.467142184 | -0.405959733 | -0.061182451 | 0.786295704 |
| ENSG00000023228 | *NDUFS1* | 4.033232916 | -0.120460703 | -0.029026508 | -0.091434195 | 0.999933887 | -0.165014173 | -0.126029984 | -0.03898419 | 0.786576764 |
| ENSG00000100441 | *KHNYN* | 6.644599708 | 0.01307873 | -0.003791479 | 0.016870209 | 0.999933887 | -0.183206296 | -0.217471805 | 0.034265509 | 0.786660923 |
| ENSG00000265692 | *LINC01970* | -0.804894112 | 0.158967688 | 0.038291883 | 0.120675805 | 0.999933887 | -0.269444111 | -0.518538139 | 0.249094028 | 0.786673465 |
| ENSG00000068394 | *GPKOW* | 3.120301305 | 0.024520411 | -0.083256742 | 0.107777153 | 0.999933887 | -0.049075442 | 0.007205842 | -0.056281284 | 0.786673465 |
| ENSG00000230724 | *LINC01001* | 1.727821844 | 0.030431651 | -0.005992767 | 0.036424419 | 0.999933887 | 0.267492121 | 0.400099584 | -0.132607463 | 0.786673465 |
| ENSG00000146530 | *VWDE* | -0.485650503 | 0.039141044 | 0.171006781 | -0.131865738 | 0.999933887 | -0.442688874 | -0.565655938 | 0.122967064 | 0.786673465 |
| ENSG00000105983 | *LMBR1* | 4.134941946 | -0.008328303 | -0.02185836 | 0.013530058 | 0.999933887 | -0.130126491 | -0.170901156 | 0.040774665 | 0.786673465 |
| ENSG00000164483 | *SAMD3* | 2.891042415 | 0.010231236 | -0.141673464 | 0.1519047 | 0.999933887 | 0.081570379 | 0.13100475 | -0.049434371 | 0.786673465 |
| ENSG00000281831 | *HCP5B* | -1.029831128 | 0.015784154 | 0.143880825 | -0.128096671 | 0.999933887 | 0.549763257 | 0.405852004 | 0.143911253 | 0.786673465 |
| ENSG00000234238 | *AC022498.1* | -0.451042537 | 0.364877963 | 0.647447962 | -0.282569999 | 0.999933887 | 1.440851627 | 1.660316779 | -0.219465152 | 0.786790507 |
| ENSG00000105220 | *GPI* | 4.57647201 | 0.001950455 | -0.01285257 | 0.014803025 | 0.999933887 | 0.005003559 | -0.038726941 | 0.0437305 | 0.786790507 |
| ENSG00000258831 | *AC103996.2* | -0.601179666 | -0.205785093 | -0.209988758 | 0.004203665 | 0.999933887 | 0.480807393 | 0.314485938 | 0.166321455 | 0.786966529 |
| ENSG00000255571 | *MIR9-3HG* | -0.786653033 | -0.046039458 | -0.250736678 | 0.20469722 | 0.999933887 | 0.363411488 | 0.201597816 | 0.161813672 | 0.787087911 |
| ENSG00000171840 | *NINJ2* | 3.740983399 | 0.024285785 | 0.042597645 | -0.01831186 | 0.999933887 | 0.062411994 | 0.125035813 | -0.062623819 | 0.787392875 |

| | | | | | | | | | | |
|---|---|---|---|---|---|---|---|---|---|---|
| ENSG00000162065 | *TBC1D24* | 1.035425485 | -0.021927824 | -0.135576779 | 0.113648955 | 0.999933887 | 0.004012015 | -0.07738639 | 0.081398405 | 0.787392875 |
| ENSG00000131401 | *NAPSB* | -0.555521928 | 0.115173806 | -0.152399025 | 0.267572831 | 0.999933887 | -0.216366992 | -0.075399876 | -0.140967116 | 0.78748315 |
| ENSG00000102316 | *MAGED2* | 3.54431536 | 0.01869578 | 0.009374834 | 0.009320946 | 0.999933887 | 0.127441872 | 0.076053948 | 0.051387924 | 0.78748315 |
| ENSG00000198931 | *APRT* | 3.705450256 | 0.084507494 | 0.011619432 | 0.072888062 | 0.999933887 | 0.107974025 | 0.057886598 | 0.050087428 | 0.787570852 |
| ENSG00000126368 | *NR1D1* | 6.088687419 | 0.024399258 | 0.114854422 | -0.090455164 | 0.999933887 | 1.163116304 | 1.227472807 | -0.064356503 | 0.787766503 |
| ENSG00000132535 | *DLG4* | 0.824699999 | -0.064707277 | 0.130294232 | -0.195001509 | 0.999933887 | -0.061361343 | 0.034731841 | -0.096093184 | 0.788016404 |
| ENSG00000117394 | *SLC2A1* | 4.878690898 | 0.001174931 | 0.047478847 | -0.046303917 | 0.999933887 | -0.396469526 | -0.339583806 | -0.05688572 | 0.788636929 |
| ENSG00000236682 | *MAP3K2-DT* | 3.156991515 | -0.065809989 | 0.043934462 | -0.109744451 | 0.999933887 | -0.744737094 | -0.683768239 | -0.060968855 | 0.788881431 |
| ENSG00000169442 | *CD52* | 4.798565273 | -0.011321003 | -0.042752505 | 0.031431502 | 0.999933887 | 0.059557932 | 0.00737995 | 0.052177982 | 0.788881431 |
| ENSG00000025800 | *KPNA6* | 5.07820238 | -0.017506794 | -0.020387809 | 0.002881016 | 0.999933887 | -0.266317552 | -0.307261807 | 0.040944255 | 0.78890607 |
| ENSG00000104823 | *ECH1* | 2.638673146 | -0.033167739 | 0.075893271 | -0.10906101 | 0.999933887 | 0.068099555 | 0.123330224 | -0.055230669 | 0.789026724 |
| ENSG00000248092 | *NNT-AS1* | -0.325803716 | -0.324725759 | -0.570698724 | 0.245972965 | 0.999933887 | -0.198055537 | -0.081456291 | -0.116599245 | 0.789080675 |
| ENSG00000125901 | *MRPS26* | 1.364744243 | 0.144673637 | 0.146752707 | -0.00207907 | 0.999933887 | 0.043360329 | -0.046394552 | 0.089754881 | 0.789080675 |
| ENSG00000012779 | *ALOX5* | 7.565188602 | 0.045358578 | 0.025715879 | 0.019642699 | 0.999933887 | 0.122260278 | 0.160447976 | -0.038187698 | 0.789080675 |
| ENSG00000136938 | *ANP32B* | 5.192782372 | -0.047706402 | -0.039168794 | -0.008537608 | 0.999933887 | -0.038693643 | 0.000961184 | -0.039654828 | 0.789080675 |
| ENSG00000214176 | *PLEKHM1P1* | 6.56890235 | 0.034684024 | 0.014964832 | 0.019719192 | 0.999933887 | -0.194947661 | -0.225682364 | 0.030734703 | 0.789080675 |
| ENSG00000168781 | *PPIP5K1* | 0.65949413 | 0.101624576 | -0.0963621 | 0.197986676 | 0.999933887 | 0.145945892 | 0.235702209 | -0.089756317 | 0.789080675 |
| ENSG00000166793 | *YPEL4* | 0.961336558 | 0.109204843 | -0.079626949 | 0.188831792 | 0.999933887 | -0.871231213 | -0.976263114 | 0.105031901 | 0.789080675 |
| ENSG00000215424 | *MCM3AP-AS1* | 0.738562067 | -0.071564541 | 0.170829691 | -0.242394232 | 0.999933887 | 0.133100956 | 0.226435734 | -0.093334778 | 0.789080675 |
| ENSG00000086065 | *CHMP5* | 5.505707414 | 0.013654222 | 0.009846365 | 0.003807857 | 0.999933887 | 0.063787047 | 0.099734554 | -0.035947507 | 0.789080675 |
| ENSG00000104331 | *BPNT2* | 3.481339359 | 6.2674e-05 | 0.04053017 | -0.040467496 | 0.999933887 | -0.256376125 | -0.3095479 | 0.053171774 | 0.789080675 |
| ENSG00000136826 | *KLF4* | 5.171002304 | 0.208727129 | 0.242373532 | -0.033646403 | 0.999933887 | -0.48069985 | -0.405728516 | -0.074971335 | 0.789189009 |
| ENSG00000286813 | *AC015871.6* | 2.012046662 | 0.109705053 | 0.278992701 | -0.169287648 | 0.999933887 | 0.534805281 | 0.633372792 | -0.098567511 | 0.789189009 |
| ENSG00000241127 | *YAE1* | -1.226930791 | -0.083549903 | -0.370926059 | 0.287376156 | 0.999933887 | -0.123064114 | 0.045559129 | -0.168623243 | 0.789309565 |
| ENSG00000158023 | *CFAP251* | -0.249726796 | 0.237168451 | -0.194253352 | 0.431421803 | 0.999933887 | -0.401620159 | -0.241727266 | -0.159892893 | 0.789385287 |
| ENSG00000114656 | *CFAP92* | 3.323893374 | 0.05741192 | 0.096331298 | -0.038919379 | 0.999933887 | 0.127498632 | 0.073191613 | 0.054307019 | 0.789385287 |
| ENSG00000245213 | *AC105285.1* | 0.062114374 | 0.040447363 | 0.263051534 | -0.222604171 | 0.999933887 | 0.203017611 | 0.07202486 | 0.130992751 | 0.789385287 |
| ENSG00000269821 | *KCNQ1OT1* | 3.567711747 | 0.123533804 | 0.182449507 | -0.058915703 | 0.999933887 | 0.112607644 | 0.015506584 | 0.097101061 | 0.789601773 |
| ENSG00000170482 | *SLC23A1* | 0.499761503 | -0.236392259 | -0.016823317 | -0.219568942 | 0.999933887 | -0.137657898 | -0.252879909 | 0.11522201 | 0.789655594 |
| ENSG00000146007 | *ZMAT2* | 5.97603116 | -0.08763341 | -0.115236292 | 0.027602882 | 0.999933887 | -0.053385573 | -0.009099417 | -0.044286156 | 0.789655594 |
| ENSG00000148344 | *PTGES* | -0.848622849 | -0.314257002 | -0.116814386 | -0.197442616 | 0.999933887 | 3.145966697 | 2.912726654 | 0.233240043 | 0.789821856 |
| ENSG00000159363 | *ATP13A2* | 3.120604923 | -0.027975476 | 0.014555152 | -0.042530628 | 0.999933887 | 0.132032161 | 0.190463554 | -0.058431393 | 0.789821856 |
| ENSG00000232860 | *SMG7-AS1* | 0.545416954 | 0.233398673 | 0.458930741 | -0.225532068 | 0.999933887 | 1.347153248 | 1.461435798 | -0.11428255 | 0.789920255 |
| ENSG00000177879 | *AP3S1* | 5.338434725 | -0.016171532 | -0.040416652 | 0.02424512 | 0.999933887 | -0.412248771 | -0.45320431 | 0.040955539 | 0.789920255 |
| ENSG00000261592 | *AC010531.3* | 1.913960597 | 0.222391316 | 0.301850817 | -0.079459501 | 0.999933887 | 0.340930001 | 0.243178395 | 0.097751606 | 0.790184597 |
| ENSG00000257267 | *ZNF271P* | 3.079383411 | -0.183545897 | -0.114888109 | -0.068657787 | 0.999933887 | -0.239308485 | -0.298985634 | 0.059677149 | 0.79046132 |
| ENSG00000030066 | *NUP160* | 3.968446988 | -0.089102075 | 0.074982411 | -0.164084486 | 0.970123867 | -0.016056246 | 0.028231507 | -0.044287754 | 0.79057628 |
| ENSG00000092470 | *WDR76* | 0.222476142 | -0.114359065 | -0.106952465 | -0.007406601 | 0.999933887 | 0.101649152 | 0.000392135 | 0.101257017 | 0.79057628 |
| ENSG00000189306 | *RRP7A* | 3.507612976 | 0.046999359 | 0.0334191 | 0.01358026 | 0.999933887 | -0.028227952 | -0.073900234 | 0.045672282 | 0.79057628 |
| ENSG00000170962 | *PDGFD* | 0.842636541 | -0.02488813 | -0.121454595 | 0.096566465 | 0.999933887 | -0.183264758 | -0.090578591 | -0.092686167 | 0.79057628 |
| ENSG00000229153 | *EPHA1-AS1* | -0.100428879 | 0.157739334 | -0.042325436 | 0.20006477 | 0.999933887 | 0.015501488 | 0.193023244 | -0.177521756 | 0.790627673 |
| ENSG00000239653 | *PSMD6-AS2* | 1.758354553 | -0.131469498 | -0.204603677 | 0.07313418 | 0.999933887 | -0.615691994 | -0.764491624 | 0.14879963 | 0.790780856 |
| ENSG00000197375 | *SLC22A5* | 1.5329161 | -0.088539658 | 0.044754613 | -0.133294272 | 0.999933887 | 0.068253696 | 0.149005919 | -0.080752223 | 0.790780856 |
| ENSG00000168096 | *ANKS3* | 1.646062272 | 0.078645103 | 0.165091591 | -0.086446488 | 0.999933887 | 0.045793241 | 0.118584325 | -0.072791084 | 0.790780856 |
| ENSG00000173660 | *UQCRH* | 4.003294804 | -0.03654163 | 0.029810477 | -0.066352107 | 0.999933887 | 0.085408831 | 0.129319947 | -0.043911117 | 0.790780856 |
| ENSG00000170791 | *CHCHD7* | 2.801434879 | -0.038432386 | -0.065038746 | 0.02660636 | 0.999933887 | 0.163707263 | 0.093676058 | 0.070031205 | 0.790780856 |
| ENSG00000234444 | *ZNF736* | 2.343082558 | -0.012470026 | 0.117644709 | -0.130114735 | 0.999933887 | -0.119115419 | -0.052718186 | -0.066397234 | 0.790780856 |
| ENSG00000174456 | *C12orf76* | 2.071158867 | 0.012555681 | 0.154447864 | -0.141892183 | 0.999933887 | 0.049089162 | 0.125221348 | -0.076132186 | 0.790780856 |
| ENSG00000168924 | *LETM1* | 3.101735497 | 0.008460513 | -0.03786374 | 0.046324253 | 0.999933887 | 0.102832547 | 0.047862512 | 0.054970035 | 0.790780856 |
| ENSG00000240429 | *LRRFIP1P1* | -1.261209313 | 0.012541199 | 0.544480127 | -0.531938928 | 0.999933887 | -0.128683953 | -0.373243149 | 0.244559196 | 0.790780856 |
| ENSG00000090266 | *NDUFB2* | 2.085537104 | -0.048211974 | 0.07050187 | -0.118713844 | 0.999933887 | 0.049157297 | 0.109546163 | -0.060388866 | 0.790908938 |
| ENSG00000136463 | *TACO1* | 1.450528196 | -0.102096805 | 0.043060471 | -0.145157276 | 0.999933887 | -0.044102439 | -0.138523725 | 0.094421285 | 0.790959179 |
| ENSG00000080947 | *CROCCP3* | 1.777324062 | -0.066802174 | 0.006396048 | -0.073198223 | 0.999933887 | 0.069784696 | 0.137891481 | -0.068106786 | 0.790959179 |
| ENSG00000240023 | *AL133163.1* | -1.638451977 | -0.169151778 | 0.169089578 | -0.338241356 | 0.999933887 | 2.290188572 | 2.520414753 | -0.230226181 | 0.790959179 |
| ENSG00000266232 | *MIR3178* | -0.185955935 | 0.348415118 | 0.192972503 | 0.155442616 | 0.999933887 | 0.394494673 | 0.227371048 | 0.167123625 | 0.791046379 |
| ENSG00000279861 | *AC073548.1* | 5.052129707 | 0.080317353 | 0.132160591 | -0.051843239 | 0.999933887 | 0.194278587 | 0.247462598 | -0.053184011 | 0.791199266 |
| ENSG00000172183 | *ISG20* | 8.073990997 | 0.02870254 | 0.015389659 | 0.013312881 | 0.999933887 | 0.154632301 | 0.122913834 | 0.031718467 | 0.791199266 |
| ENSG00000183690 | *EFHC2* | 0.879274888 | 0.002012652 | -0.224932916 | 0.226945568 | 0.999933887 | 0.010450324 | 0.14156808 | -0.131117756 | 0.791199266 |
| ENSG00000213614 | *HEXA* | 6.076859075 | 0.008441533 | 0.07991777 | -0.071476237 | 0.999933887 | -0.02179618 | 0.034904162 | -0.056700342 | 0.79127598 |
| ENSG00000177971 | *IMP3* | 1.72416057 | -0.169144749 | 0.033469907 | -0.202614656 | 0.999933887 | 0.080767735 | 0.148863284 | -0.06809555 | 0.791972687 |
| ENSG00000263826 | *AC112907.3* | 0.877991006 | 0.116598583 | 0.359369494 | -0.242770911 | 0.999933887 | 0.363903554 | 0.470548844 | -0.10664529 | 0.791972687 |
| ENSG00000121064 | *SCPEP1* | 4.601468056 | -0.05103277 | -0.069175661 | 0.018142892 | 0.999933887 | -0.525986401 | -0.568310286 | 0.042323885 | 0.791972687 |
| ENSG00000254428 | *AP003392.1* | 1.080472439 | -0.085039395 | -0.007538267 | -0.077501128 | 0.999933887 | -0.150979562 | -0.060902969 | -0.090076594 | 0.791972687 |
| ENSG00000120647 | *CCDC77* | 1.04912214 | -0.087032135 | 0.072031944 | -0.159064079 | 0.999933887 | -0.333129604 | -0.426033119 | 0.092903515 | 0.791972687 |
| ENSG00000133169 | *BEX1* | 0.473831239 | -0.240644636 | 0.073645276 | -0.314289912 | 0.999933887 | 0.527163833 | 0.426004171 | 0.101159662 | 0.792049449 |
| ENSG00000176438 | *SYNE3* | 4.814305158 | -0.003807213 | 0.067840525 | -0.071647738 | 0.999933887 | -0.05843373 | -0.094079299 | 0.035645569 | 0.792049449 |
| ENSG00000132466 | *ANKRD17* | 5.930723809 | -0.006586468 | 0.015285296 | -0.021871764 | 0.999933887 | -0.191596473 | -0.151685699 | -0.039910774 | 0.792260602 |
| ENSG00000234745 | *HLA-B* | 13.079654663 | 0.024812836 | 0.001453488 | 0.023359348 | 0.999933887 | 0.315819552 | 0.346492092 | -0.03067254 | 0.792497333 |
| ENSG00000141644 | *MBD1* | 5.883326724 | 0.000113108 | -0.024490723 | 0.024603831 | 0.999933887 | -0.061010925 | -0.091725715 | 0.03071479 | 0.792497333 |
| ENSG00000130956 | *HABP4* | 1.604300222 | -0.07250175 | 0.100521658 | -0.173023408 | 0.999933887 | 0.090313906 | 0.168526225 | -0.07821232 | 0.792672776 |
| ENSG00000157349 | *DDX19B* | 1.80087464 | 0.019398668 | -0.0039651 | 0.023363768 | 0.999933887 | -0.053655972 | -0.133852619 | 0.080196647 | 0.792672776 |
| ENSG00000139372 | *TDG* | 3.560562204 | -0.006622776 | 0.019427705 | -0.026050481 | 0.999933887 | -0.332476118 | -0.29285598 | -0.039620139 | 0.792672776 |
| ENSG00000181722 | *ZBTB20* | 2.372787595 | 0.051507937 | 0.09499548 | -0.043487543 | 0.999933887 | -0.201777343 | -0.064128795 | -0.137648548 | 0.792754982 |
| ENSG00000169021 | *UQCRFS1* | 4.140479561 | -0.016815012 | -0.08284358 | 0.066028568 | 0.999933887 | 0.165428503 | 0.122394256 | 0.043034247 | 0.793307038 |
| ENSG00000129757 | *CDKN1C* | 2.388413596 | 0.145018348 | 0.037443628 | 0.107574721 | 0.999933887 | -0.469143207 | -0.564366494 | 0.095223287 | 0.793422138 |
| ENSG00000187535 | *IFT140* | 0.185455434 | -0.147108215 | 0.128923012 | -0.276031228 | 0.999933887 | 0.096819683 | -0.01226502 | 0.109084703 | 0.793422138 |
| ENSG00000089280 | *FUS* | 8.18681964 | 0.022673186 | 0.068862155 | -0.046188969 | 0.999933887 | 0.055929072 | 0.091330055 | -0.035400984 | 0.793462686 |
| ENSG00000285410 | *GABPB1-IT1* | 4.792557115 | 0.07379354 | 0.10618417 | -0.03239063 | 0.999933887 | 0.341982908 | 0.387914953 | -0.045932045 | 0.79349869 |
| ENSG00000258581 | *AL157871.3* | 1.201404011 | -0.083469407 | 0.132506761 | -0.215976168 | 0.999933887 | 0.134575744 | 0.227138198 | -0.092562454 | 0.79349869 |
| ENSG00000287216 | *AC106865.2* | -0.315770496 | 0.090582902 | 0.202059658 | -0.111476755 | 0.999933887 | 0.966369227 | 1.121418118 | -0.15504889 | 0.79349869 |
| ENSG00000196778 | *OR52K1* | -0.195031179 | 0.075674304 | 0.070004903 | 0.005669401 | 0.999933887 | -0.460218956 | -0.653789772 | 0.193570816 | 0.79349869 |
| ENSG00000171621 | *SPSB1* | 0.919970051 | -0.053692595 | -0.082295893 | 0.028603298 | 0.999933887 | 0.125031329 | 0.236525244 | -0.111493916 | 0.79349869 |
| ENSG00000198162 | *MAN1A2* | 4.012739202 | 0.124387416 | -0.126798911 | 0.251186327 | 0.999933887 | -0.28267037 | -0.358937544 | 0.076267174 | 0.793612478 |
| ENSG00000096433 | *ITPR3* | 4.544745131 | 0.052753417 | 0.049415916 | 0.003337502 | 0.999933887 | 0.157078885 | 0.119013732 | 0.038065153 | 0.793765619 |
| ENSG00000106246 | *PTCD1* | 1.540119347 | 0.041405588 | -0.007424316 | 0.048829903 | 0.999933887 | 0.076158351 | 0.150032396 | -0.073874046 | 0.793765619 |
| ENSG00000223749 | *MIR503HG* | -0.709620187 | 0.050892415 | 0.202213975 | -0.15132156 | 0.999933887 | -0.396743761 | -0.286861666 | -0.109882095 | 0.793979118 |
| ENSG00000073861 | *TBX21* | 3.830849392 | 0.003461677 | 0.023162322 | -0.019700645 | 0.999933887 | 0.095312213 | 0.140227307 | -0.044915094 | 0.794249837 |
| ENSG00000135144 | *DTX1* | 0.073858203 | 0.050219971 | 0.351509875 | -0.301289905 | 0.999933887 | 0.158648902 | 0.28494433 | -0.126295428 | 0.794302002 |
| ENSG00000170956 | *CEACAM3* | 6.912031174 | 0.008925886 | 0.043595061 | -0.034669175 | 0.999933887 | -0.215278895 | -0.174007111 | -0.041271785 | 0.794309705 |
| ENSG00000235865 | *GSN-AS1* | 1.624438556 | -0.180365728 | -0.002745309 | -0.17762042 | 0.999933887 | -0.197681242 | -0.122090397 | -0.075590845 | 0.794383629 |
| ENSG00000162882 | *HAAO* | 0.244376016 | 0.115493051 | 0.317702597 | -0.202209546 | 0.999933887 | -0.38171788 | -0.250632825 | -0.131085056 | 0.794383629 |
| ENSG00000166341 | *DCHS1* | 0.355649723 | -0.11168452 | -0.208049019 | 0.096364499 | 0.999933887 | -0.001587336 | 0.115114524 | -0.11670186 | 0.794383629 |
| ENSG00000181638 | *ZFP41* | -0.055361188 | 0.112577688 | -0.14535952 | 0.257937208 | 0.999933887 | 0.15372756 | 0.035995616 | 0.117731944 | 0.794383629 |
| ENSG00000280303 | *ERICD* | -0.216712798 | 0.084972331 | 0.228159861 | -0.14318753 | 0.999933887 | 1.454740859 | 1.606165824 | -0.151424965 | 0.794383629 |
| ENSG00000165275 | *TRMT10B* | 1.671397239 | -0.0518412 | -0.072711181 | 0.020869982 | 0.999933887 | -0.166783046 | -0.233973529 | 0.067190483 | 0.794383629 |
| ENSG00000102543 | *CDADC1* | 3.161789163 | -0.024530025 | -0.063696287 | 0.039166262 | 0.999933887 | 0.086410823 | 0.148571168 | -0.062160345 | 0.794383629 |
| ENSG00000164733 | *CTSB* | 6.940877819 | -0.014812258 | -0.019232629 | 0.004420371 | 0.999933887 | -0.118048985 | -0.082148276 | -0.035900709 | 0.794383629 |
| ENSG00000124766 | *SOX4* | 0.046136204 | -0.032667987 | 0.254258044 | -0.286926031 | 0.999933887 | -0.135186192 | -0.28629414 | 0.151107948 | 0.794383629 |
| ENSG00000122376 | *SHLD2* | 1.957202214 | -0.211025104 | -0.171750058 | -0.039275046 | 0.999933887 | -0.069440216 | 0.01532353 | -0.084763746 | 0.79465696 |
| ENSG00000260660 | *AC113208.3* | 2.681106859 | 0.145966815 | 0.265468171 | -0.119501356 | 0.999933887 | 1.072585766 | 1.175919517 | -0.103333751 | 0.79465696 |
| ENSG00000102804 | *TSC22D1* | 3.520991129 | 0.065050461 | -0.012739991 | 0.077790452 | 0.999933887 | 0.397759024 | 0.347992857 | 0.049766168 | 0.79465696 |
| ENSG00000205659 | *LIN52* | 1.090374398 | -0.114551455 | 0.080663704 | -0.195215159 | 0.999933887 | -0.113751325 | -0.010181521 | -0.103569804 | 0.79465696 |
| ENSG00000173040 | *EVC2* | -0.669232116 | 0.138167523 | 0.258235816 | -0.120068293 | 0.999933887 | 0.031860446 | 0.188063419 | -0.156202973 | 0.79465696 |
| ENSG00000108950 | *FAM20A* | 1.162816711 | -0.093279802 | -0.031047392 | -0.062232411 | 0.999933887 | -0.164547801 | -0.269970985 | 0.105423184 | 0.79465696 |
| ENSG00000250903 | *GMDS-DT* | 0.987603715 | 0.079261375 | -0.03356713 | 0.112828504 | 0.999933887 | -0.059849055 | -0.16952455 | 0.109675495 | 0.79465696 |

| | | | | | | | | | | |
|---|---|---|---|---|---|---|---|---|---|---|
| ENSG00000228242 | *XPC-AS1* | 3.542342324 | -0.042101265 | -0.056835221 | 0.014733956 | 0.999933887 | -0.839494643 | -0.901989205 | 0.062494562 | 0.79465696 |
| ENSG00000123989 | *CHPF* | 0.835937614 | 0.043137872 | -0.093918372 | 0.137056244 | 0.999933887 | 0.08230355 | 0.190608216 | -0.108304666 | 0.79465696 |
| ENSG00000286885 | *AL138824.1* | 0.096469295 | -0.038149274 | 0.148602575 | -0.186751849 | 0.999933887 | -0.070482954 | -0.203643778 | 0.133160824 | 0.79465696 |
| ENSG00000233355 | *CHRM3-AS2* | 2.76761138 | -0.002576928 | -0.184781441 | 0.182204513 | 0.999933887 | -0.124321704 | -0.058020165 | -0.066301539 | 0.79465696 |
| ENSG00000180917 | *CMTR2* | 1.834508336 | -0.102908139 | -0.157195868 | 0.054287729 | 0.999933887 | 0.014446603 | 0.095747177 | -0.081300574 | 0.794730918 |
| ENSG00000268041 | *ERFL* | 0.663441861 | 0.16296102 | 0.01769428 | 0.14526674 | 0.999933887 | -0.069921932 | 0.050440224 | -0.120362156 | 0.795010951 |
| ENSG00000198841 | *KTI12* | 0.109341202 | -0.127496402 | 0.287255104 | -0.414751506 | 0.999933887 | -0.205188945 | -0.324444648 | 0.119255704 | 0.795010951 |
| ENSG00000160072 | *ATAD3B* | 3.665347701 | 0.015850134 | 0.068109541 | -0.052259407 | 0.999933887 | 0.13581199 | 0.078752743 | 0.057059246 | 0.795010951 |
| ENSG00000138617 | *PARP16* | 2.711566917 | -0.007207226 | -0.006814574 | -0.000392652 | 0.999933887 | 0.202002171 | 0.259933134 | -0.057930963 | 0.795010951 |
| ENSG00000176383 | *B3GNT4* | -0.803241467 | -0.008974068 | 0.228561101 | -0.237535168 | 0.999933887 | 0.412513555 | 0.221567923 | 0.190945632 | 0.795010951 |
| ENSG00000229162 | *RUNX3-AS1* | -0.745550116 | 0.333588555 | 0.092370667 | 0.241217888 | 0.999933887 | 0.019355993 | 0.175043062 | -0.15568707 | 0.795051595 |
| ENSG00000176349 | *AC104129.1* | -0.740358017 | 0.040556286 | 0.122316487 | -0.081760202 | 0.999933887 | -0.226689071 | -0.048699888 | -0.177989183 | 0.795090774 |
| ENSG00000025796 | *SEC63* | 4.3746354 | -0.059382257 | -0.161069455 | 0.101687198 | 0.999933887 | -0.399129323 | -0.335708938 | -0.063420385 | 0.795226327 |
| ENSG00000179348 | *GATA2* | 1.523447875 | -0.005080919 | 0.179225145 | -0.184306064 | 0.999933887 | -0.025231162 | 0.050588552 | -0.075819714 | 0.795226327 |
| ENSG00000166170 | *BAG5* | 4.649644651 | -0.066970761 | -0.025650237 | -0.041320524 | 0.999933887 | -0.395549863 | -0.356096409 | -0.039453454 | 0.795632284 |
| ENSG00000140632 | *GLYR1* | 5.965152209 | -0.035115331 | 0.014962108 | -0.050077439 | 0.999933887 | -0.205580723 | -0.235733237 | 0.030152514 | 0.795632284 |
| ENSG00000124313 | *IQSEC2* | 1.652584646 | 0.061014257 | -0.032171536 | 0.093185793 | 0.999933887 | -0.226894509 | -0.143944376 | -0.082950133 | 0.795632284 |
| ENSG00000101400 | *SNTA1* | 0.416268282 | 0.038909109 | 0.197790961 | -0.158881852 | 0.999933887 | 0.128590274 | -0.00573618 | 0.134326454 | 0.795632284 |
| ENSG00000277972 | *CISD3* | 2.212904373 | 0.117060075 | 0.019848569 | 0.097211506 | 0.999933887 | 0.216356414 | 0.160597 | 0.055759414 | 0.795799772 |
| ENSG00000144647 | *POMGNT2* | 0.850301238 | -0.067758519 | 0.054927111 | -0.122685629 | 0.999933887 | 0.060224551 | 0.187997917 | -0.127773366 | 0.795799772 |
| ENSG00000214797 | *AP002358.1* | -0.853540307 | 0.064888347 | -0.125558147 | 0.190446494 | 0.999933887 | 0.324007156 | 0.574998267 | -0.250991111 | 0.795799772 |
| ENSG00000165905 | *LARGE2* | -0.333908232 | 0.133231341 | 0.128376108 | 0.004855233 | 0.999933887 | -0.075080612 | -0.197389842 | 0.12230923 | 0.796017408 |
| ENSG00000124549 | *BTN2A3P* | 0.974628889 | 0.068227911 | 0.116502829 | -0.048274918 | 0.999933887 | 0.166454971 | 0.077649388 | 0.088805583 | 0.796017408 |
| ENSG00000107745 | *MICU1* | 5.256612376 | 0.003057183 | 0.011260046 | -0.008202863 | 0.999933887 | 0.111871633 | 0.150877359 | -0.039005726 | 0.796017408 |
| ENSG00000181610 | *MRPS23* | 1.377260271 | 0.018958221 | -0.163134168 | 0.18209239 | 0.999933887 | 0.005289899 | 0.085211972 | -0.079922073 | 0.796261006 |
| ENSG00000152102 | *FAM168B* | 6.737737009 | -0.07943005 | -0.039942095 | -0.039487955 | 0.999933887 | 0.007145663 | 0.036000051 | -0.028854388 | 0.796264074 |
| ENSG00000259950 | *AC034105.3* | 1.993742778 | 0.190344853 | 0.450634439 | -0.260289585 | 0.999933887 | 0.893203004 | 1.00811459 | -0.114911586 | 0.796264074 |
| ENSG00000185236 | *RAB11B* | 5.967949109 | 0.044681501 | 0.032471709 | 0.012209792 | 0.999933887 | -0.164820975 | -0.200030757 | 0.035209782 | 0.796264074 |
| ENSG00000285053 | *AL357556.3* | -0.284204345 | -0.146040846 | -0.19835374 | 0.052312893 | 0.999933887 | -0.128863343 | -0.25436676 | 0.125503417 | 0.796264074 |
| ENSG00000119969 | *HELLS* | -0.091193518 | 0.102226582 | 0.034471334 | 0.067755248 | 0.999933887 | -0.140202384 | -0.007837053 | -0.132365331 | 0.796264074 |
| ENSG00000235499 | *AC073046.1* | 3.463376716 | 0.052239276 | 0.103063442 | -0.050824166 | 0.999933887 | 0.641508314 | 0.694860221 | -0.053351907 | 0.796264074 |
| ENSG00000187866 | *FAM122A* | 2.65862696 | 0.019853873 | -0.055328442 | 0.075182315 | 0.999933887 | -0.164856528 | -0.23108898 | 0.066232452 | 0.796264074 |
| ENSG00000197165 | *SULT1A2* | 0.261253065 | -0.270344743 | -0.382693144 | 0.112348401 | 0.999933887 | -0.943102514 | -1.048938916 | 0.105836402 | 0.796534478 |
| ENSG00000132356 | *PRKAA1* | 5.163624479 | -0.017325676 | -0.060081576 | 0.042755899 | 0.999933887 | -0.403748473 | -0.350177009 | -0.053571464 | 0.796534478 |
| ENSG00000162889 | *MAPKAPK2* | 9.225812742 | 0.007329771 | -0.004481636 | 0.011811407 | 0.999933887 | 0.405158117 | 0.436326436 | -0.031168319 | 0.796534478 |
| ENSG00000106524 | *ANKMY2* | 1.711134212 | -0.099935657 | 0.047686785 | -0.147622443 | 0.999933887 | 0.06930829 | -0.013798824 | 0.083107113 | 0.796760477 |
| ENSG00000121152 | *NCAPH* | 0.256104232 | -0.142315085 | -0.162371538 | 0.020056452 | 0.999933887 | 0.212068341 | 0.317988649 | -0.105920307 | 0.796898547 |
| ENSG00000132470 | *ITGB4* | 2.913020037 | 0.07115559 | 0.097598767 | -0.026443177 | 0.999933887 | -0.085080753 | -0.161023108 | 0.075942355 | 0.796898547 |
| ENSG00000273136 | *NBPF26* | 4.927266504 | -0.102688589 | -0.050441798 | -0.052246792 | 0.999933887 | 0.391367094 | 0.31320632 | 0.078160775 | 0.796898547 |
| ENSG00000115112 | *TFCP2L1* | -0.145338822 | -0.086139682 | 0.236759297 | -0.322898978 | 0.999933887 | -0.173358808 | -0.03699706 | -0.136361748 | 0.796898547 |
| ENSG00000163029 | *SMC6* | 2.391483025 | -0.058214203 | -0.151047055 | 0.092832852 | 0.999933887 | -0.201387756 | -0.089016884 | -0.112370871 | 0.796898547 |
| ENSG00000148843 | *PDCD11* | 3.11600284 | 0.028230935 | -0.016551043 | 0.044781978 | 0.999933887 | 0.160633414 | 0.112084228 | 0.048549186 | 0.796898547 |
| ENSG00000153187 | *HNRNPU* | 8.6420907 | -0.014248621 | -0.008711234 | -0.005537388 | 0.999933887 | -0.003852999 | 0.024773835 | -0.028626834 | 0.796898547 |
| ENSG00000053702 | *NRIP2* | 0.662515419 | -0.018637378 | 0.16606453 | -0.184701908 | 0.999933887 | 0.021445979 | -0.080326188 | 0.101772167 | 0.796898547 |
| ENSG00000137168 | *PPIL1* | 0.357970626 | -0.019305844 | 0.029554955 | -0.048860798 | 0.999933887 | 0.205281394 | 0.097736531 | 0.107544863 | 0.796898547 |
| ENSG00000180423 | *HARBI1* | 1.08720686 | -0.010019785 | -0.121437721 | 0.111417936 | 0.999933887 | 0.310069248 | 0.226955422 | 0.083113826 | 0.796898547 |
| ENSG00000283041 | *AC008038.1* | 0.263631194 | -0.117499715 | -0.009939627 | -0.107560088 | 0.999933887 | 0.17202982 | 0.279697563 | -0.107667743 | 0.797042916 |
| ENSG00000100445 | *SDR39U1* | 1.445475313 | 0.051466016 | 0.074919292 | -0.023453275 | 0.999933887 | 0.154779916 | 0.072297513 | 0.082482402 | 0.797042916 |
| ENSG00000107263 | *RAPGEF1* | 8.874505048 | 0.004047904 | 0.019934043 | -0.015886138 | 0.999933887 | 0.0356215 | 0.066774565 | -0.031153065 | 0.797042916 |
| ENSG00000235194 | *PPP1R3E* | 4.933549508 | 0.033032699 | 0.10590644 | -0.072873741 | 0.999933887 | 0.190617045 | 0.237414331 | -0.046797287 | 0.797148008 |
| ENSG00000140463 | *BBS4* | 2.584944905 | -0.070232245 | 0.031128455 | -0.101360701 | 0.999933887 | -0.000753742 | -0.051237533 | 0.050483791 | 0.797325512 |
| ENSG00000285518 | *AC004900.1* | -1.203626724 | -0.172986156 | -0.082054375 | -0.090931781 | 0.999933887 | 0.879712215 | 1.101384109 | -0.221671893 | 0.797619749 |
| ENSG00000136717 | *BIN1* | 4.382659086 | 0.049637412 | 0.020530534 | 0.029106879 | 0.999933887 | 0.050462173 | 0.004056071 | 0.046406102 | 0.797715008 |
| ENSG00000107223 | *EDF1* | 5.317514901 | 0.041599816 | 0.047383381 | -0.005783565 | 0.999933887 | 0.095628852 | 0.051394099 | 0.044234752 | 0.797715008 |
| ENSG00000198589 | *LRBA* | 3.706998059 | -0.041871558 | -0.013042071 | -0.028829487 | 0.999933887 | -0.051536289 | 0.013381346 | -0.064917636 | 0.797715008 |
| ENSG00000163508 | *EOMES* | 3.297644333 | -0.026223698 | -0.063143553 | 0.036919855 | 0.999933887 | 0.185573934 | 0.135817395 | 0.049756538 | 0.797715008 |
| ENSG00000164031 | *DNAJB14* | 4.33078675 | -0.03994773 | -0.203627142 | 0.163679412 | 0.999933887 | -0.799247182 | -0.694480688 | -0.104766494 | 0.797715008 |
| ENSG00000129824 | *RPS4Y1* | -2.454854904 | -0.020214717 | -0.165283911 | 0.145069194 | 0.999933887 | -0.084762244 | -0.15313365 | 0.068371406 | 0.797779716 |
| ENSG00000139914 | *FITM1* | 0.168451381 | 0.359851121 | -0.289251245 | 0.649102366 | 0.144167088 | -0.106044461 | -0.209757578 | 0.103713117 | 0.79779194 |
| ENSG00000132768 | *DPH2* | 1.621557607 | 0.10459765 | -0.156980453 | 0.261578104 | 0.999933887 | 0.278430265 | 0.20962986 | 0.068800405 | 0.797823174 |
| ENSG00000272555 | *AC009974.1* | -0.875105249 | -0.19528304 | -0.476641723 | 0.281358683 | 0.999933887 | 0.619991936 | 0.756320201 | -0.136328265 | 0.797823174 |
| ENSG00000255808 | *PDE2A-AS1* | -0.762401867 | 0.128147872 | 0.043282517 | 0.084865355 | 0.999933887 | 0.575817122 | 0.830365162 | -0.25454804 | 0.797823174 |
| ENSG00000165494 | *PCF11* | 7.975816777 | 0.042152328 | -0.066906441 | 0.109058769 | 0.999933887 | 0.264247502 | 0.309394582 | -0.04514708 | 0.798045325 |
| ENSG00000169057 | *MECP2* | 7.623209663 | 0.023656779 | 0.027798814 | -0.004142035 | 0.999933887 | 0.129968758 | 0.163175954 | -0.033207196 | 0.798045325 |
| ENSG00000139679 | *LPAR6* | 3.347543067 | -0.107119668 | -0.137244627 | 0.03012496 | 0.999933887 | -0.323720595 | -0.394313728 | 0.070593133 | 0.798644963 |
| ENSG00000231889 | *TRAF3IP2-AS1* | 2.049636921 | 0.066432618 | -0.035670647 | 0.102103265 | 0.999933887 | 0.80829399 | 0.88003202 | -0.071738031 | 0.798644963 |
| ENSG00000143502 | *SUSD4* | -0.876690097 | 0.077763052 | -0.149179955 | 0.226943007 | 0.999933887 | 0.091578183 | 0.239952837 | -0.148374654 | 0.798644963 |
| ENSG00000167700 | *MFSD3* | 0.650101757 | 0.042673707 | -0.034464516 | 0.077138223 | 0.999933887 | 0.344284165 | 0.239680208 | 0.104603957 | 0.798644963 |
| ENSG00000280135 | *AL096816.1* | 2.103075187 | -0.062071668 | -0.083957168 | 0.0218855 | 0.999933887 | 0.000240689 | -0.068060448 | 0.068301137 | 0.798964436 |
| ENSG00000204685 | *STARD7-AS1* | 3.371594517 | -0.078481539 | 0.014136386 | -0.092617925 | 0.999933887 | 0.123963927 | 0.060684096 | 0.063279831 | 0.799190091 |
| ENSG00000126903 | *SLC10A3* | 4.726698181 | 0.051898132 | -0.025509774 | 0.077407906 | 0.999933887 | -0.160531965 | -0.206081103 | 0.045549138 | 0.799197405 |
| ENSG00000186197 | *EDARADD* | -0.310783908 | -0.018380766 | 0.046341547 | -0.064722314 | 0.999933887 | 0.144032967 | 0.287458833 | -0.143425866 | 0.799197405 |
| ENSG00000112195 | *TREML2* | 6.815750088 | 0.005263083 | 0.010910233 | -0.00564715 | 0.999933887 | -0.263848867 | -0.210800343 | -0.053048524 | 0.799197405 |
| ENSG00000196497 | *IPO4* | 0.73136121 | 0.291164384 | 0.195359097 | 0.095805286 | 0.999933887 | 0.336141227 | 0.208960676 | 0.127180551 | 0.799352075 |
| ENSG00000128228 | *SDF2L1* | 2.478198821 | 0.168863442 | 0.154132752 | 0.01473069 | 0.999933887 | 0.187446425 | 0.117101496 | 0.070344928 | 0.799352075 |
| ENSG00000186594 | *MIR22HG* | 7.396713444 | 0.111451515 | 0.073359857 | 0.038091658 | 0.999933887 | 0.416313711 | 0.458533989 | -0.042220278 | 0.799352075 |
| ENSG00000183207 | *RUVBL2* | 2.312842876 | 0.106870083 | 0.020874214 | 0.085995869 | 0.999933887 | 0.117006643 | 0.054018692 | 0.062987951 | 0.799352075 |
| ENSG00000111203 | *ITFG2* | 2.739221673 | -0.039386405 | 0.126229592 | -0.165615997 | 0.999933887 | -0.052213117 | 0.003237636 | -0.055450752 | 0.799352075 |
| ENSG00000089009 | *RPL6* | 6.928605291 | -0.031314704 | -0.151013162 | 0.119698458 | 0.999933887 | 0.044028611 | 0.085618473 | -0.041589863 | 0.799352075 |
| ENSG00000186162 | *CIDECP1* | 3.420947627 | 0.035933386 | 0.083810506 | -0.04787712 | 0.999933887 | -0.128551368 | -0.070483502 | -0.058067866 | 0.799352075 |
| ENSG00000114686 | *MRPL3* | 1.939092697 | 0.02305923 | -0.133505933 | 0.156565163 | 0.999933887 | 0.170076833 | 0.235716469 | -0.065639636 | 0.799352075 |
| ENSG00000166326 | *TRIM44* | 4.03437459 | -0.012504719 | -0.122552981 | 0.110048262 | 0.999933887 | 0.024169715 | 0.066211066 | -0.042041351 | 0.799352075 |
| ENSG00000276045 | *ORAI1* | 4.648460775 | 0.070788705 | -0.023857477 | 0.094646182 | 0.999933887 | 0.008268477 | -0.038392591 | 0.046661068 | 0.799476086 |
| ENSG00000213799 | *ZNF845* | 1.942762236 | -0.155168019 | -0.275538129 | 0.12037011 | 0.999933887 | -0.580183926 | -0.484795755 | -0.095388171 | 0.799476086 |
| ENSG00000132676 | *DAP3* | 4.682122692 | -0.063682689 | 0.022448467 | -0.086131156 | 0.999933887 | 0.018228453 | 0.050846354 | -0.032617902 | 0.799490775 |
| ENSG00000280780 | *JAKMIP2-AS1* | -2.324042754 | -0.628539764 | -0.457838018 | -0.170701747 | 0.999933887 | -0.764288943 | -0.62011899 | -0.144169953 | 0.799490775 |
| ENSG00000214756 | *CSKMT* | 0.178344712 | 0.156224331 | 0.504261563 | -0.348037232 | 0.999933887 | 0.3637579 | 0.47469915 | -0.11094125 | 0.799490775 |
| ENSG00000141698 | *NT5C3B* | 1.149186335 | 0.084239712 | 0.124806979 | -0.040567267 | 0.999933887 | 0.168059483 | 0.245737884 | -0.077678401 | 0.799490775 |
| ENSG00000142534 | *RPS11* | 8.636852969 | 0.025609042 | -0.027754103 | 0.053363145 | 0.999933887 | 0.074690927 | 0.111165961 | -0.036475034 | 0.799490775 |
| ENSG00000279529 | *AC008764.8* | 1.739199116 | -0.020165694 | -0.060363229 | 0.040197536 | 0.999933887 | 0.031153808 | 0.112301288 | -0.08114748 | 0.799490775 |
| ENSG00000182866 | *LCK* | 5.414612611 | 0.070731556 | -0.003568474 | 0.07430003 | 0.999933887 | 0.096866932 | 0.058547331 | 0.038319601 | 0.799557281 |
| ENSG00000183354 | *KIAA2026* | 5.052505068 | 0.175315176 | 0.00560386 | 0.169711317 | 0.999933887 | -0.619366036 | -0.530401191 | -0.088964845 | 0.799658864 |
| ENSG00000197780 | *TAF13* | 4.048336812 | 0.026962337 | 0.206956968 | -0.179994631 | 0.944094537 | 0.538254307 | 0.587462801 | -0.049208494 | 0.799833667 |
| ENSG00000171962 | *DRC3* | 0.017493355 | -0.132289593 | -0.183239468 | 0.050949875 | 0.999933887 | -0.077171317 | 0.062897379 | -0.140068697 | 0.799890127 |
| ENSG00000072195 | *SPEG* | 0.33768812 | 0.073624851 | 0.239024563 | -0.165399712 | 0.999933887 | 0.025900138 | -0.088153003 | 0.114053141 | 0.799890127 |
| ENSG00000168056 | *LTBP3* | 4.364003824 | -0.001820862 | 0.061338344 | -0.063159207 | 0.999933887 | 0.11934603 | 0.072698641 | 0.046647389 | 0.80022529 |
| ENSG00000176986 | *SEC24C* | 5.442945645 | 0.00044428 | 0.032601723 | -0.032157443 | 0.999933887 | -0.063290283 | -0.031707678 | -0.031582606 | 0.8002938 |
| ENSG00000196531 | *NACA* | 6.743507847 | -0.108202808 | -0.180801906 | 0.072599098 | 0.999933887 | -0.163962581 | -0.1985381 | 0.034575519 | 0.800304159 |
| ENSG00000261253 | *AC137932.2* | -0.247733348 | 0.370312993 | 0.143067728 | 0.227245264 | 0.999933887 | -0.138325136 | -0.314588119 | 0.176262983 | 0.800304159 |
| ENSG00000163069 | *SGCB* | -0.41044152 | 0.196599681 | -0.159566782 | 0.356166463 | 0.999933887 | -0.182941677 | -0.036797939 | -0.146143738 | 0.800304159 |
| ENSG00000236901 | *MIR600HG* | 0.682626667 | -0.178394015 | -0.044759124 | -0.133634891 | 0.999933887 | -0.397664673 | -0.498205228 | 0.100540555 | 0.800304159 |
| ENSG00000115750 | *TAF1B* | 0.817798069 | -0.114718363 | -0.31926768 | 0.204549317 | 0.999933887 | -0.084623671 | 0.018161201 | -0.102784872 | 0.800304159 |
| ENSG00000135763 | *URB2* | 1.50465418 | -0.070483465 | 0.107778277 | -0.178261742 | 0.999933887 | 0.23108886 | 0.165393806 | 0.065695054 | 0.800304159 |
| ENSG00000137959 | *IFI44L* | 4.524428345 | -0.086330216 | -0.214030568 | 0.127700352 | 0.999933887 | -0.387536269 | -0.477485712 | 0.089949442 | 0.800304159 |
| ENSG00000276136 | *AC016957.2* | 3.000436792 | -0.059006043 | 0.027991942 | -0.086997985 | 0.999933887 | -0.221149294 | -0.291832164 | 0.07068287 | 0.800304159 |

| | | | | | | | | | | |
|---|---|---|---|---|---|---|---|---|---|---|
| ENSG00000133026 | *MYH10* | 3.749588668 | 0.047105701 | -0.019830117 | 0.066935818 | 0.999933887 | -0.029853295 | -0.104128837 | 0.074275542 | 0.800304159 |
| ENSG00000113758 | *DBN1* | 4.576108347 | 0.037470821 | 0.08373509 | -0.046264269 | 0.999933887 | 0.41569542 | 0.458585831 | -0.042890411 | 0.800304159 |
| ENSG00000127952 | *STYXL1* | 2.849619748 | 0.043236075 | -0.038462934 | 0.081699009 | 0.999933887 | 0.025207575 | 0.074164277 | -0.048956702 | 0.800304159 |
| ENSG00000179262 | *RAD23A* | 4.634916244 | -0.029034838 | 0.028940498 | -0.057975336 | 0.999933887 | -0.111254214 | -0.146567097 | 0.035312883 | 0.800304159 |
| ENSG00000286159 | *AL035106.1* | 3.232844788 | -0.030003561 | -0.075036529 | 0.045032968 | 0.999933887 | -0.097553182 | -0.028393135 | -0.069160047 | 0.800304159 |
| ENSG00000260231 | *KDM7A-DT* | 1.966715316 | 0.030946426 | 0.155625418 | -0.124678992 | 0.999933887 | 0.366399494 | 0.262814072 | 0.103585421 | 0.800304159 |
| ENSG00000167286 | *CD3D* | 4.945029344 | 0.009388365 | -0.027545152 | 0.036933517 | 0.999933887 | 0.12778374 | 0.168576312 | -0.040792572 | 0.800304159 |
| ENSG00000108433 | *GOSR2* | 2.964396376 | 0.010622982 | -0.017042424 | 0.027665407 | 0.999933887 | 0.058536677 | 0.107326171 | -0.048789494 | 0.800304159 |
| ENSG00000080200 | *CRYBG3* | -0.506010553 | 0.00693584 | 0.034061153 | -0.027125314 | 0.999933887 | -0.196377609 | 0.003046294 | -0.199423903 | 0.800304159 |
| ENSG00000279425 | *AC092279.2* | 1.232387281 | -0.002987763 | 0.13980243 | -0.142790193 | 0.999933887 | -0.481528507 | -0.571528085 | 0.090269578 | 0.800304159 |
| ENSG00000258521 | *AL157871.2* | 0.972078824 | -0.0014196 | -0.040365121 | 0.03894552 | 0.999933887 | 0.168584934 | 0.282470129 | -0.113885195 | 0.800304159 |
| ENSG00000126458 | *RRAS* | 0.751082696 | 0.062550825 | 0.025526202 | 0.037024623 | 0.999933887 | -0.133980472 | -0.023037963 | -0.110942509 | 0.800326581 |
| ENSG00000138801 | *PAPSS1* | 3.619233073 | 0.002834554 | -0.09362674 | 0.096461293 | 0.999933887 | -0.173173485 | -0.124571717 | -0.048601768 | 0.800329721 |
| ENSG00000272150 | *NBPF25P* | -0.831798722 | -0.069442246 | -0.459299053 | 0.389856807 | 0.999933887 | 0.341292743 | 0.191108038 | 0.150184705 | 0.800797278 |
| ENSG00000105197 | *TIMM50* | 2.701888098 | 0.063293047 | 0.099237565 | -0.035944518 | 0.999933887 | 0.099646089 | 0.160742563 | -0.061096474 | 0.800843298 |
| ENSG00000134186 | *PRPF38B* | 6.503823642 | 0.004382404 | -0.067749692 | 0.072132095 | 0.999933887 | -0.047780315 | 0.011690826 | -0.05947114 | 0.800843298 |
| ENSG00000095383 | *TBC1D2* | 7.923640771 | -0.076342665 | -0.068158894 | -0.008183771 | 0.999933887 | -0.422179856 | -0.371495202 | -0.050684653 | 0.801168876 |
| ENSG00000088766 | *CRLS1* | -0.609049975 | 0.1540938 | 0.161480315 | -0.007386516 | 0.999933887 | -0.090355537 | 0.088268695 | -0.178624232 | 0.801168876 |
| ENSG00000188536 | *HBA2* | 5.679225274 | -0.067999437 | -0.140515965 | 0.072516529 | 0.999933887 | 0.025335023 | 0.095693776 | -0.070358753 | 0.801168876 |
| ENSG00000204315 | *FKBPL* | 0.335969345 | 0.077758452 | -0.086442904 | 0.164201357 | 0.999933887 | -0.334164826 | -0.451583579 | 0.117418753 | 0.801186071 |
| ENSG00000241721 | *SUMO1P1* | 2.925022084 | -0.018067981 | 0.038107032 | -0.056175013 | 0.999933887 | 0.015512982 | -0.046729105 | 0.062242087 | 0.801300381 |
| ENSG00000205060 | *SLC35B4* | 1.250462733 | -0.132146341 | 0.048816886 | -0.180963227 | 0.999933887 | -0.156736826 | -0.07242811 | -0.084308716 | 0.801737273 |
| ENSG00000188647 | *PTAR1* | 5.09391902 | -0.011220974 | -0.060086035 | 0.048865061 | 0.999933887 | -0.543234107 | -0.46798611 | -0.075247997 | 0.801737273 |
| ENSG00000166165 | *CKB* | 1.478152602 | 0.119415579 | -0.03794057 | 0.157356149 | 0.999933887 | 0.911443762 | 1.025038316 | -0.113594554 | 0.801777601 |
| ENSG00000107819 | *SFXN3* | 4.753829175 | 0.037205116 | 0.011073019 | 0.026132097 | 0.999933887 | -0.141940645 | -0.180545804 | 0.03860516 | 0.801777601 |
| ENSG00000151135 | *TMEM263* | 2.060826642 | -0.058500523 | -0.171204943 | 0.11270442 | 0.999933887 | -0.073708395 | -0.000699533 | -0.073008861 | 0.801777601 |
| ENSG00000131495 | *NDUFA2* | 3.456929826 | -0.004652208 | 0.001480913 | -0.006133121 | 0.999933887 | -0.135666063 | -0.184681385 | 0.049015322 | 0.801777601 |
| ENSG00000138231 | *DBR1* | 1.998013577 | -0.00294137 | -0.17672076 | 0.17377939 | 0.999933887 | -0.139728356 | -0.21210646 | 0.072378104 | 0.801777601 |
| ENSG00000168818 | *STX18* | 2.576667382 | -0.088059668 | 0.157505407 | -0.245565075 | 0.944094537 | 0.059373679 | 0.103026572 | -0.043652893 | 0.801835092 |
| ENSG00000274767 | *AC243829.1* | 4.06386388 | -0.037413043 | -0.095327829 | 0.057914786 | 0.999933887 | 0.027983902 | -0.022578175 | 0.050562077 | 0.801835092 |
| ENSG00000153879 | *CEBPG* | 3.118519332 | -0.028830293 | -0.001255585 | -0.027574708 | 0.999933887 | 0.003843561 | 0.050261402 | -0.04641784 | 0.801835092 |
| ENSG00000084072 | *PPIE* | 2.784355735 | -0.031920226 | 0.00334088 | -0.035261106 | 0.999933887 | 0.097096638 | 0.041432757 | 0.055663882 | 0.801835092 |
| ENSG00000181396 | *OGFOD3* | 1.538768594 | 0.013938184 | -0.00170415 | 0.015642334 | 0.999933887 | 0.032048318 | 0.118747821 | -0.086699503 | 0.801835092 |
| ENSG00000131165 | *CHMP1A* | 7.103806118 | -0.00088213 | 0.021579402 | -0.022461532 | 0.999933887 | 0.053952034 | 0.088566449 | -0.034614415 | 0.801835092 |
| ENSG00000153944 | *MSI2* | 4.260219879 | 0.030490111 | 0.085616847 | -0.055126735 | 0.999933887 | -0.069024097 | -0.106860323 | 0.037836226 | 0.801839106 |
| ENSG00000071462 | *BUD23* | 4.119081051 | 0.002173171 | -0.033024681 | 0.035197851 | 0.999933887 | 0.157460902 | 0.122590277 | 0.034870624 | 0.801839106 |
| ENSG00000138002 | *IFT172* | 1.482954191 | 0.015810601 | 0.009883064 | 0.005927537 | 0.999933887 | -0.150533622 | -0.080253988 | -0.070279634 | 0.802371996 |
| ENSG00000170486 | *KRT72* | -1.181968495 | 0.025846118 | -0.0659516 | 0.091797718 | 0.999933887 | -0.104545894 | 0.045414759 | -0.149960653 | 0.802485978 |
| ENSG00000176593 | *AC008969.1* | 1.161620698 | -0.163383248 | -0.146901608 | -0.016481639 | 0.999933887 | -0.124810131 | -0.049076789 | -0.075733342 | 0.802495662 |
| ENSG00000117899 | *MESD* | 2.641688644 | -0.064839482 | -0.158053499 | 0.093214017 | 0.999933887 | 0.015581938 | -0.05274727 | 0.068329208 | 0.802495662 |
| ENSG00000218073 | *AL021407.2* | 0.650703239 | 0.030714835 | -0.01696063 | 0.047675465 | 0.999933887 | 0.244034773 | 0.391629121 | -0.147594347 | 0.802576406 |
| ENSG00000198835 | *GJC2* | -0.191633519 | 0.10974168 | 0.25347431 | -0.143732629 | 0.999933887 | 0.247662721 | 0.429737055 | -0.182074334 | 0.802650045 |
| ENSG00000204130 | *RUFY2* | 2.265225527 | -0.104031399 | -0.083427203 | -0.020604196 | 0.999933887 | -0.535647456 | -0.453449045 | -0.082198411 | 0.802736683 |
| ENSG00000113522 | *RAD50* | -0.828532928 | 0.156392034 | 0.094953016 | 0.061439018 | 0.999933887 | -0.002391096 | -0.15485121 | 0.152460115 | 0.802736683 |
| ENSG00000141378 | *PTRH2* | 2.557558412 | -0.070562729 | -0.02539005 | -0.04517268 | 0.999933887 | -0.123677485 | -0.188537096 | 0.064859611 | 0.802875477 |
| ENSG00000185808 | *PIGP* | -0.020403719 | 0.069826747 | 0.00743846 | 0.062388287 | 0.999933887 | 0.093492922 | -0.025745255 | 0.119238177 | 0.802875477 |
| ENSG00000276523 | *AC025287.3* | 3.613872239 | -0.00699357 | 0.111129427 | -0.118122997 | 0.999933887 | 0.830828148 | 0.890771028 | -0.05994288 | 0.802932166 |
| ENSG00000036448 | *MYOM2* | 1.214913836 | -0.095264783 | -0.032180395 | -0.063084388 | 0.999933887 | 0.256077299 | 0.151297538 | 0.104779761 | 0.803029896 |
| ENSG00000134970 | *TMED7* | 3.914811187 | -0.085813675 | 0.028202688 | -0.114016363 | 0.999933887 | -0.672191308 | -0.729350348 | 0.05715904 | 0.803529217 |
| ENSG00000099940 | *SNAP29* | 4.075034076 | -0.011704866 | 0.002003723 | -0.013708589 | 0.999933887 | -0.450254697 | -0.493813862 | 0.043559165 | 0.803529217 |
| ENSG00000279865 | *AC006511.3* | 0.540769327 | 0.110726341 | 0.212955737 | -0.102229396 | 0.999933887 | 1.41545194 | 1.308444136 | 0.107007804 | 0.803637186 |
| ENSG00000196549 | *MME* | 7.310883757 | -0.031985534 | -0.144317651 | 0.112332116 | 0.999933887 | -0.920323549 | -0.978873952 | 0.058550403 | 0.803860469 |
| ENSG00000260097 | *SPDYE6* | 0.411560175 | -0.186931053 | -0.037839668 | -0.149091385 | 0.999933887 | 0.126495016 | 0.22877071 | -0.102275693 | 0.804000145 |
| ENSG00000143257 | *NR1I3* | 2.474798192 | 0.09057824 | 0.095384624 | -0.004806384 | 0.999933887 | 1.32328409 | 1.407215591 | -0.083931501 | 0.804000145 |
| ENSG00000094631 | *HDAC6* | 4.258603442 | 0.021591416 | 0.114713522 | -0.093122106 | 0.999933887 | 0.179780378 | 0.224760605 | -0.044980227 | 0.804000145 |
| ENSG00000274114 | *ALOX15P1* | -0.74415804 | 0.026609981 | -0.164646312 | 0.191256294 | 0.999933887 | -0.129703114 | -0.296688734 | 0.16698562 | 0.804000145 |
| ENSG00000274750 | *H3C6* | -1.203829502 | 0.334851885 | -0.418836048 | 0.753687932 | 0.999933887 | -0.129432317 | -0.273699095 | 0.144266778 | 0.804052803 |
| ENSG00000144591 | *GMPPA* | 2.664262769 | 0.028785048 | -0.005969031 | 0.034754079 | 0.999933887 | -0.007070065 | -0.058187299 | 0.051117234 | 0.804070072 |
| ENSG00000002016 | *RAD52* | 1.612995809 | 0.008757011 | 0.004953025 | 0.003803986 | 0.999933887 | -0.041924068 | -0.123768835 | 0.081844767 | 0.804070072 |
| ENSG00000112782 | *CLIC5* | 0.193593642 | -0.048242373 | 0.065139054 | -0.113381426 | 0.999933887 | -0.029149483 | -0.127995438 | 0.098845955 | 0.804092021 |
| ENSG00000177706 | *FAM20C* | 1.340903292 | 0.000491198 | 0.034800439 | -0.034309241 | 0.999933887 | -0.578023756 | -0.476787134 | -0.101236621 | 0.804092021 |
| ENSG00000171105 | *INSR* | 0.755927956 | -0.117192557 | 0.133148724 | -0.250341281 | 0.999933887 | -0.731672584 | -0.610635334 | -0.12103725 | 0.804139901 |
| ENSG00000174032 | *SLC25A30* | 2.677625215 | -0.030249921 | -0.019417983 | -0.010831938 | 0.999933887 | -0.167726432 | -0.109345582 | -0.05838085 | 0.804139901 |
| ENSG00000067365 | *METTL22* | 4.686384042 | 0.065745428 | 0.051096289 | 0.014649139 | 0.999933887 | -0.145753098 | -0.184846794 | 0.039093695 | 0.804548534 |
| ENSG00000170092 | *SPDYE5* | -0.328313697 | -0.134566329 | 0.167933005 | -0.302499334 | 0.999933887 | -0.287366083 | -0.418740585 | 0.131374502 | 0.804548534 |
| ENSG00000089123 | *TASP1* | 0.751899011 | 0.050918045 | 0.162616808 | -0.111698763 | 0.999933887 | 0.01386824 | 0.092206206 | -0.078337966 | 0.804548534 |
| ENSG00000128482 | *RNF112* | -0.327336291 | 0.019942836 | 0.20565028 | -0.185707444 | 0.999933887 | 0.173667192 | 0.322390597 | -0.148723405 | 0.804548534 |
| ENSG00000104921 | *FCER2* | 0.89785308 | 0.054757632 | 0.083531701 | -0.028774069 | 0.999933887 | 0.536847772 | 0.449615962 | 0.08723181 | 0.804620515 |
| ENSG00000107611 | *CUBN* | 1.098358086 | 0.037797088 | 0.130704535 | -0.092907447 | 0.999933887 | -0.220003507 | -0.323064602 | 0.103061095 | 0.804620515 |
| ENSG00000274828 | *AC068473.5* | 0.764323158 | 0.079723303 | 0.439039111 | -0.359315807 | 0.999933887 | -0.153693942 | -0.224755584 | 0.071061642 | 0.805055589 |
| ENSG00000204420 | *MPIG6B* | 0.597119478 | 0.242223871 | 0.140508068 | 0.101715803 | 0.999933887 | -0.323578823 | -0.17741219 | -0.146166633 | 0.805091454 |
| ENSG00000225178 | *RPSAP58* | -0.951166883 | 0.288230515 | -0.113731939 | 0.401962455 | 0.999933887 | 0.096193589 | 0.234580454 | -0.138386865 | 0.805207607 |
| ENSG00000231047 | *GCNT1P3* | 1.024498557 | 0.050149682 | 0.187013301 | -0.136863619 | 0.999933887 | 0.212101946 | 0.462088731 | -0.249986785 | 0.805207607 |
| ENSG00000106799 | *TGFBR1* | 7.861987281 | -0.196638936 | -0.12348725 | -0.073151686 | 0.999933887 | -1.265991805 | -1.201060875 | -0.06493093 | 0.805387762 |
| ENSG00000196150 | *ZNF250* | 2.921595699 | 0.093085311 | 0.131040069 | -0.037954758 | 0.999933887 | 0.388481619 | 0.470298322 | -0.081816703 | 0.805400084 |
| ENSG00000124574 | *ABCC10* | 3.454917914 | -0.022030138 | 0.052812626 | -0.074842764 | 0.999933887 | 0.061375229 | 0.014535584 | 0.046839645 | 0.805400084 |
| ENSG00000174989 | *FBXW8* | 1.6613679 | 0.094477331 | 0.099066541 | -0.004589211 | 0.999933887 | 0.013931306 | -0.060927394 | 0.074858699 | 0.805774356 |
| ENSG00000113851 | *CRBN* | 3.895846634 | -0.041549244 | -0.110614842 | 0.069065598 | 0.999933887 | -0.274869971 | -0.224298811 | -0.050571159 | 0.805774356 |
| ENSG00000189227 | *C15orf61* | -0.592993104 | 0.025499658 | 0.018256984 | 0.007242674 | 0.999933887 | 0.042610815 | 0.200283735 | -0.15767292 | 0.805774356 |
| ENSG00000143799 | *PARP1* | 4.230665035 | 0.005696386 | -0.02826874 | 0.033965126 | 0.999933887 | 0.105453114 | 0.144518763 | -0.039065648 | 0.805774356 |
| ENSG00000154305 | *MIA3* | 5.061085967 | 0.022796422 | -0.065676334 | 0.088472757 | 0.999933887 | -0.419032534 | -0.474447793 | 0.055415258 | 0.805808121 |
| ENSG00000268575 | *AL031282.2* | 0.157775776 | 0.107046834 | 0.179539392 | -0.072492558 | 0.999933887 | -0.247685522 | -0.355569667 | 0.107884145 | 0.805811362 |
| ENSG00000256128 | *LINC00944* | 0.001757547 | -0.077687371 | 0.359819125 | -0.437506496 | 0.999933887 | 0.199568026 | 0.09552954 | 0.104038486 | 0.805811362 |
| ENSG00000149273 | *RPS3* | 8.530397117 | -0.025433286 | -0.050875585 | 0.025442299 | 0.999933887 | 0.080845044 | 0.043031506 | 0.037813538 | 0.805811362 |
| ENSG00000156928 | *MALSU1* | 1.338246117 | 0.045762773 | 0.133215868 | -0.087453095 | 0.999933887 | 0.134152467 | 0.040934697 | 0.093217769 | 0.805811362 |
| ENSG00000147224 | *PRPS1* | 3.243039013 | 0.00941077 | -0.017048884 | 0.026459653 | 0.999933887 | 0.093978033 | 0.1499883 | -0.056010267 | 0.805811362 |
| ENSG00000116221 | *MRPL37* | 2.102875385 | 0.002068579 | 0.027933139 | -0.02586456 | 0.999933887 | 0.225932505 | 0.159936791 | 0.065995714 | 0.806223066 |
| ENSG00000103355 | *PRSS33* | 3.50578363 | 0.061650143 | 0.07547267 | -0.013822527 | 0.999933887 | -0.028890194 | 0.026112043 | -0.055002237 | 0.8063262 |
| ENSG00000159228 | *CBR1* | -0.12536392 | 0.087204172 | 0.156730227 | -0.069526055 | 0.999933887 | -0.203660952 | -0.318163289 | 0.114502337 | 0.8063262 |
| ENSG00000158517 | *NCF1* | 7.59775237 | 0.026832459 | 0.001584652 | 0.025247807 | 0.999933887 | 0.888588638 | 0.838589155 | 0.049999483 | 0.8063262 |
| ENSG00000224546 | *EIF4BP3* | -1.24394067 | -0.092309285 | 0.085355464 | -0.177664749 | 0.999933887 | 0.143803587 | 0.301115967 | -0.157312381 | 0.80642519 |
| ENSG00000261997 | *AC007336.1* | 1.774662835 | 0.002243009 | -0.017937884 | 0.020180893 | 0.999933887 | -0.406412818 | -0.268161511 | -0.138251308 | 0.806511713 |
| ENSG00000196458 | *ZNF605* | 1.124755273 | 0.051435007 | -0.284499891 | 0.335934897 | 0.999933887 | -0.24346634 | -0.331121138 | 0.087654797 | 0.806910148 |
| ENSG00000242970 | *AC068522.1* | 1.905776495 | 0.309895494 | 0.429831828 | -0.119936334 | 0.999933887 | 0.975170273 | 1.084483995 | -0.109313721 | 0.807003277 |
| ENSG00000213793 | *ZNF888* | 0.983067572 | -0.160930401 | -0.04275292 | -0.118177481 | 0.999933887 | -0.419150066 | -0.320434334 | -0.098715732 | 0.807003277 |
| ENSG00000125845 | *BMP2* | 1.009468645 | 0.03983912 | 0.088153535 | -0.048314415 | 0.999933887 | 0.855215346 | 0.773829055 | 0.081386291 | 0.807299739 |
| ENSG00000172785 | *CBWD1* | 1.268767024 | -0.155155846 | -0.099135754 | -0.056020093 | 0.999933887 | -0.200655877 | -0.302706196 | 0.102050319 | 0.807312114 |
| ENSG00000188997 | *KCTD21* | 2.158754649 | -0.107369749 | -0.078340085 | -0.029029664 | 0.999933887 | -0.834538525 | -0.909427729 | 0.074889204 | 0.807312114 |
| ENSG00000189129 | *PLAC9* | 0.033308883 | -0.106263052 | 0.028720628 | -0.13498368 | 0.999933887 | -0.449478205 | -0.584982782 | 0.135504577 | 0.807312114 |
| ENSG00000167618 | *LAIR2* | -1.039736778 | -0.102485966 | -0.07994095 | -0.022545016 | 0.999933887 | 0.16540583 | 0.332968247 | -0.167562417 | 0.807312114 |
| ENSG00000139921 | *TMX1* | 2.75695553 | 0.038311797 | -0.040302905 | 0.078614702 | 0.999933887 | -0.148419113 | -0.07432714 | -0.074091974 | 0.807312114 |
| ENSG00000137177 | *KIF13A* | 6.57523732 | -0.029992274 | -0.034975697 | 0.004983423 | 0.999933887 | 0.290807584 | 0.356140065 | -0.06533248 | 0.807312114 |
| ENSG00000286646 | *AL121933.2* | 1.869413209 | -0.025386877 | 0.009415226 | -0.034802103 | 0.999933887 | 0.472426691 | 0.403608529 | 0.068818161 | 0.807312114 |
| ENSG00000223960 | *CHROMR* | 1.165948062 | 0.001612777 | -0.187438478 | 0.189051255 | 0.999933887 | -0.194595627 | -0.270044515 | 0.075448887 | 0.807312114 |
| ENSG00000150455 | *TIRAP* | 1.687777367 | 0.040945646 | -0.138821384 | 0.17976703 | 0.999933887 | 0.121632618 | 0.203052686 | -0.081420069 | 0.807379118 |

| | | | | | | | | | | |
|---|---|---|---|---|---|---|---|---|---|---|
| ENSG00000274370 | *AC130371.2* | 1.903376903 | -0.027639424 | 0.183620128 | -0.211259552 | 0.999933887 | -0.158866324 | -0.240554172 | 0.081687848 | 0.807379118 |
| ENSG00000204267 | *TAP2* | 5.998480388 | 0.035022368 | 0.026029334 | 0.008993034 | 0.999933887 | -0.034612075 | -0.066186603 | 0.031574528 | 0.80744505 |
| ENSG00000082515 | *MRPL22* | 1.362516954 | 0.013284713 | -0.108092913 | 0.121377626 | 0.999933887 | 0.109309052 | 0.030907067 | 0.078401985 | 0.80750109 |
| ENSG00000264853 | *AC011933.2* | 0.763512965 | -0.185221489 | 0.027972096 | -0.213193586 | 0.999933887 | 0.705961286 | 0.537396247 | 0.168565039 | 0.807862422 |
| ENSG00000260285 | *AL133367.1* | 0.771022374 | -0.106041226 | -0.031432357 | -0.074608869 | 0.999933887 | -0.615793657 | -0.530496324 | -0.085297332 | 0.80787768 |
| ENSG00000022267 | *FHL1* | 0.864788164 | -0.02081366 | 0.077641544 | -0.098455204 | 0.999933887 | 0.054852436 | -0.047351699 | 0.102204135 | 0.807947581 |
| ENSG00000108963 | *DPH1* | 2.148060367 | 0.003779077 | 0.052160143 | -0.048381066 | 0.999933887 | 0.146822564 | 0.215767163 | -0.068944599 | 0.80800488 |
| ENSG00000105655 | *ISYNA1* | 2.935514982 | 0.119109402 | 0.127351167 | -0.008241765 | 0.999933887 | 0.131024278 | 0.184261987 | -0.053237708 | 0.808121806 |
| ENSG00000124370 | *MCEE* | 0.07057171 | -0.282059687 | -0.208018817 | -0.07404087 | 0.999933887 | 0.125268477 | 0.031900407 | 0.09336807 | 0.808121806 |
| ENSG00000125257 | *ABCC4* | 1.232761067 | -0.058185228 | -0.250319826 | 0.192134598 | 0.999933887 | -0.036701842 | 0.030608897 | -0.067310739 | 0.808121806 |
| ENSG00000178409 | *BEND3* | 0.081948752 | 0.030295063 | 0.085342817 | -0.055047754 | 0.999933887 | 0.375672255 | 0.254409407 | 0.121262848 | 0.808121806 |
| ENSG00000186300 | *ZNF555* | 2.401148685 | -0.095126167 | -0.011097562 | -0.084028605 | 0.999933887 | 0.345041281 | 0.432584206 | -0.087542925 | 0.808232714 |
| ENSG00000145979 | *TBC1D7* | 6.584603205 | 0.017037108 | 0.053339122 | -0.036302014 | 0.999933887 | 0.599338347 | 0.646875434 | -0.047537087 | 0.808232714 |
| ENSG00000102010 | *BMX* | 1.302539121 | -0.019429867 | -0.098711542 | 0.079281674 | 0.999933887 | -0.877500679 | -0.765529791 | -0.111970888 | 0.808232714 |
| ENSG00000133103 | *COG6* | 1.146061356 | 0.01211883 | -0.007822632 | 0.019941462 | 0.999933887 | -0.013166012 | -0.11123909 | 0.098073077 | 0.808232714 |
| ENSG00000103671 | *TRIP4* | 3.384753885 | -0.001054867 | -0.104784188 | 0.103729321 | 0.999933887 | -0.141262136 | -0.180731537 | 0.0394694 | 0.808232714 |
| ENSG00000099797 | *TECR* | 3.810072559 | 0.075555439 | 0.033628821 | 0.041926619 | 0.999933887 | 0.165713241 | 0.209067397 | -0.043354156 | 0.808241121 |
| ENSG00000279584 | *AC005593.1* | 0.697344237 | 0.277484484 | 0.149800957 | 0.127683527 | 0.999933887 | 0.44242029 | 0.55611058 | -0.11369029 | 0.808293697 |
| ENSG00000147130 | *ZMYM3* | 2.234023401 | -0.053623893 | 0.060121271 | -0.113745165 | 0.999933887 | 0.039242219 | -0.014467209 | 0.053709427 | 0.808293697 |
| ENSG00000276168 | *RN7SL1* | 4.704535152 | -0.179411901 | -0.63527159 | 0.45585969 | 0.999933887 | 0.649505627 | 0.389223017 | 0.260282609 | 0.808351057 |
| ENSG00000171115 | *GIMAP8* | 2.522939599 | -0.084572122 | -0.062474323 | -0.022097799 | 0.999933887 | 0.143018583 | 0.19983849 | -0.056819907 | 0.808602118 |
| ENSG00000167394 | *ZNF668* | 2.92806386 | 0.064928209 | 0.065988429 | -0.00106022 | 0.999933887 | 0.006645606 | 0.067676483 | -0.061030878 | 0.808721468 |
| ENSG00000196290 | *NIF3L1* | 0.739163821 | -0.066798345 | -0.054719618 | -0.012078727 | 0.999933887 | 0.011296299 | -0.07944962 | 0.090745919 | 0.808788303 |
| ENSG00000279744 | *AC132938.5* | 0.480345579 | 0.060790002 | -0.088319085 | 0.149109087 | 0.999933887 | -0.47710351 | -0.595319197 | 0.118215687 | 0.808788303 |
| ENSG00000110104 | *CCDC86* | 0.855971862 | -0.070662422 | 0.064758412 | -0.135420834 | 0.999933887 | 0.115832006 | 0.03062739 | 0.085204615 | 0.80939566 |
| ENSG00000274922 | *AL139384.1* | 0.843040502 | 0.006786029 | -0.219557718 | 0.226343747 | 0.999933887 | -0.005539594 | -0.085890275 | 0.080350681 | 0.809568273 |
| ENSG00000156531 | *PHF6* | 1.941406712 | -0.146957527 | -0.128395415 | -0.018562112 | 0.999933887 | -0.048884911 | 0.027442803 | -0.076327715 | 0.809824182 |
| ENSG00000135469 | *COQ10A* | 1.457010944 | -0.118310883 | 0.223442005 | -0.341752887 | 0.999933887 | 0.234794423 | 0.157303431 | 0.077490992 | 0.809824182 |
| ENSG00000156500 | *FAM122C* | 2.539460529 | 0.038737406 | -0.004624498 | 0.043361904 | 0.999933887 | -0.086197179 | -0.143479177 | 0.057281999 | 0.809840761 |
| ENSG00000167815 | *PRDX2* | 2.351291607 | -0.039926315 | 0.024389218 | -0.064315534 | 0.999933887 | 0.392349227 | 0.464050124 | -0.071700897 | 0.809840761 |
| ENSG00000134324 | *LPIN1* | 4.643974785 | -0.019724293 | 0.012339095 | -0.032063388 | 0.999933887 | -0.003132982 | 0.036124664 | -0.039257646 | 0.809840761 |
| ENSG00000188732 | *FAM221A* | -0.546500925 | -0.084496102 | 0.018785868 | -0.10328197 | 0.999933887 | -0.231836131 | -0.083208016 | -0.148628116 | 0.809886795 |
| ENSG00000284024 | *AC069544.2* | 3.176139743 | -0.013037858 | 0.016999517 | -0.030037375 | 0.999933887 | -0.013266148 | 0.045585759 | -0.058851907 | 0.809886795 |
| ENSG00000204427 | *ABHD16A* | 1.950411445 | -0.059476114 | 0.106778626 | -0.166254741 | 0.999933887 | -0.456755204 | -0.380797183 | -0.075958021 | 0.809978815 |
| ENSG00000129038 | *LOXL1* | 2.139853006 | 0.078838958 | -0.006535863 | 0.085374821 | 0.999933887 | 0.034021956 | 0.137790324 | -0.103768367 | 0.810245984 |
| ENSG00000240418 | *AC020917.1* | 0.484513012 | -0.070474887 | -0.262384918 | 0.19191003 | 0.999933887 | 0.384963068 | 0.275996533 | 0.108966535 | 0.810277914 |
| ENSG00000158966 | *CACHD1* | -0.794631116 | 0.13367276 | 0.16519942 | -0.03152666 | 0.999933887 | -0.788175532 | -0.622186334 | -0.165989198 | 0.810615271 |
| ENSG00000115042 | *FAHD2A* | 0.916503028 | 0.011850365 | 0.067683162 | -0.055832797 | 0.999933887 | 0.151537157 | 0.069317497 | 0.08221966 | 0.810615271 |
| ENSG00000088053 | *GP6* | 0.23242254 | 0.248133802 | 0.223088558 | 0.025045244 | 0.999933887 | 0.660990972 | 0.505951183 | 0.155039789 | 0.810696005 |
| ENSG00000163002 | *NUP35* | -0.087730466 | -0.032724522 | -0.067556956 | 0.034832434 | 0.999933887 | 0.065950255 | 0.191920075 | -0.12596982 | 0.810696005 |
| ENSG00000047621 | *C12orf4* | 1.401145487 | 0.007365354 | -0.191758294 | 0.199123647 | 0.999933887 | -0.024956023 | 0.082760053 | -0.107716076 | 0.810867265 |
| ENSG00000162244 | *RPL29* | 6.866358702 | 0.074148116 | -0.013933252 | 0.088081367 | 0.999933887 | 0.06771682 | 0.111155332 | -0.043438512 | 0.811387251 |
| ENSG00000152475 | *ZNF837* | -1.230797086 | 0.264054438 | 0.18413499 | 0.079919448 | 0.999933887 | 0.321245017 | 0.16489724 | 0.156347778 | 0.811387251 |
| ENSG00000189091 | *SF3B3* | 4.710231523 | -0.086862544 | -0.011874631 | -0.074987913 | 0.999933887 | 0.086036519 | 0.119844664 | -0.033808145 | 0.811405788 |
| ENSG00000100330 | *MTMR3* | 5.005784822 | -0.042178119 | -0.001144691 | -0.041033428 | 0.999933887 | 0.019042142 | 0.053412959 | -0.034370817 | 0.811405788 |
| ENSG00000138796 | *HADH* | 1.62190349 | -0.074966554 | -0.224989283 | 0.150022729 | 0.999933887 | 0.024600285 | 0.087322421 | -0.062722136 | 0.811405788 |
| ENSG00000152284 | *TCF7L1* | -0.209408492 | -0.130352823 | -0.157597233 | 0.027244409 | 0.999933887 | -0.594273115 | -0.445397759 | -0.148875356 | 0.811405788 |
| ENSG00000124215 | *CDH26* | -0.426719905 | 0.095009383 | -0.255233523 | 0.350242906 | 0.999933887 | -0.20101819 | -0.391262803 | 0.190244613 | 0.811405788 |
| ENSG00000204628 | *RACK1* | 7.520050607 | -0.011166805 | -0.027190968 | 0.016024163 | 0.999933887 | 0.076328259 | 0.111112765 | -0.034784506 | 0.811405788 |
| ENSG00000134318 | *ROCK2* | 3.312149276 | -0.023664644 | 0.018971494 | -0.042636138 | 0.999933887 | -0.79293711 | -0.897629574 | 0.104692464 | 0.811405788 |
| ENSG00000158691 | *ZSCAN12* | 0.305336313 | 0.063985821 | -0.222746142 | 0.286731963 | 0.999933887 | -0.004867154 | 0.106507271 | -0.111374424 | 0.811723016 |
| ENSG00000125611 | *CHCHD5* | 2.613405957 | -0.077252286 | 0.061525825 | -0.138778111 | 0.999933887 | -0.262057083 | -0.316738072 | 0.054680989 | 0.811786829 |
| ENSG00000188290 | *HES4* | 7.305695269 | 0.104356167 | 0.091934446 | 0.012421721 | 0.999933887 | 0.794921342 | 0.726905191 | 0.068016151 | 0.81184016 |
| ENSG00000132361 | *CLUH* | 3.486450031 | 0.087312376 | 0.137764404 | -0.050452029 | 0.999933887 | 0.260586706 | 0.212214486 | 0.04837222 | 0.811851753 |
| ENSG00000172638 | *EFEMP2* | 1.8881005 | -0.010323124 | 0.257732475 | -0.268055599 | 0.944094537 | -0.211021865 | -0.274850557 | 0.063828692 | 0.811851753 |
| ENSG00000153563 | *CD8A* | 6.508228586 | -0.001187776 | -0.048715236 | 0.04752746 | 0.999933887 | 0.128442906 | 0.093004592 | 0.035438314 | 0.811851753 |
| ENSG00000183011 | *NAA38* | 3.779924598 | 0.125530518 | 0.101838026 | 0.023692492 | 0.999933887 | 0.496651619 | 0.549413204 | -0.052761584 | 0.811963085 |
| ENSG00000077984 | *CST7* | 6.989523432 | 0.014756481 | -0.029527218 | 0.044283699 | 0.999933887 | 0.046855343 | 0.084233894 | -0.03737855 | 0.811963085 |
| ENSG00000163507 | *CIP2A* | -0.571606617 | -0.018232189 | -0.088218905 | 0.069986716 | 0.999933887 | 0.102292297 | 0.268557767 | -0.16626547 | 0.811963085 |
| ENSG00000168916 | *ZNF608* | 1.288374149 | 0.125054957 | -0.056590987 | 0.181645945 | 0.999933887 | -1.133582216 | -1.238151227 | 0.104569011 | 0.812243027 |
| ENSG00000211787 | *TRAV8-3* | -0.263019328 | 0.082244431 | 0.149091005 | -0.066846574 | 0.999933887 | 0.208458553 | 0.302787734 | -0.094329181 | 0.812258339 |
| ENSG00000279693 | *AC099521.2* | -0.021766643 | -0.046664464 | 0.09032245 | -0.136986914 | 0.999933887 | -0.285011297 | -0.429344609 | 0.144333312 | 0.812258339 |
| ENSG00000233006 | *MIR3936HG* | -0.27525398 | 0.018581576 | -0.043081422 | 0.061662999 | 0.999933887 | -0.11702465 | 0.029050951 | -0.146075601 | 0.812258339 |
| ENSG00000100721 | *TCL1A* | -1.414133643 | 0.096539536 | -0.117430811 | 0.213970347 | 0.999933887 | -0.354411736 | -0.476229913 | 0.121818176 | 0.812315545 |
| ENSG00000141068 | *KSR1* | 5.093939182 | 0.033313043 | 0.05517583 | -0.021862787 | 0.999933887 | 0.078361722 | 0.112356221 | -0.033994499 | 0.812329238 |
| ENSG00000172426 | *RSPH9* | 0.495020128 | -0.080994018 | -0.126554289 | 0.045560271 | 0.999933887 | -0.412434772 | -0.522541216 | 0.110106444 | 0.812349573 |
| ENSG00000073756 | *PTGS2* | 8.752734756 | 0.531502347 | 0.510523294 | 0.020979053 | 0.999933887 | 1.072217417 | 1.169077004 | -0.096859587 | 0.812382943 |
| ENSG00000150753 | *CCT5* | 5.018697174 | -0.081241192 | -0.02993463 | -0.051306562 | 0.999933887 | -0.083241137 | -0.115906402 | 0.032665265 | 0.812382943 |
| ENSG00000207741 | *MIR590* | 0.041076788 | -0.238537119 | -0.256161963 | 0.017624844 | 0.999933887 | -0.042406643 | 0.086967146 | -0.129373789 | 0.812382943 |
| ENSG00000117308 | *GALE* | 2.197864034 | 0.160381864 | -0.178347663 | 0.338729527 | 0.944094537 | -0.56788722 | -0.629982632 | 0.062095412 | 0.812382943 |
| ENSG00000116273 | *PHF13* | 4.453096984 | -0.129349454 | -0.156939192 | 0.027589737 | 0.999933887 | -0.615442981 | -0.564801321 | -0.05064166 | 0.812382943 |
| ENSG00000087842 | *PIR* | -1.424665069 | 0.393814275 | -0.088162827 | 0.481977103 | 0.999933887 | 0.551975517 | 0.621728485 | -0.069752968 | 0.812382943 |
| ENSG00000280543 | *ASAP1-IT2* | 1.340972728 | 0.053126057 | 0.319678617 | -0.26655256 | 0.999933887 | 0.372642178 | 0.290757689 | 0.081884489 | 0.812382943 |
| ENSG00000188171 | *ZNF626* | -0.329213015 | 0.09712705 | 0.338234128 | -0.241107077 | 0.999933887 | -0.510339916 | -0.380452131 | -0.129887786 | 0.812382943 |
| ENSG00000111885 | *MAN1A1* | 5.960642011 | 0.015703157 | 0.012065557 | 0.0036376 | 0.999933887 | -0.008769609 | 0.030058231 | -0.03882784 | 0.812382943 |
| ENSG00000184860 | *SDR42E1* | -0.151388892 | -0.096508149 | 0.320138341 | -0.41664649 | 0.999933887 | -0.122215718 | -0.012671245 | -0.109544472 | 0.812383698 |
| ENSG00000241693 | *RN7SL704P* | -0.171591913 | 0.130743002 | 0.108073968 | 0.022669034 | 0.999933887 | 0.490008854 | 0.372777574 | 0.11723128 | 0.812383698 |
| ENSG00000102606 | *ARHGEF7* | 4.757650025 | -0.00877421 | 0.054197255 | -0.062971465 | 0.999933887 | -0.016652523 | -0.046623646 | 0.029971123 | 0.812383698 |
| ENSG00000158555 | *GDPD5* | 3.403515335 | 0.005708863 | 0.0389427 | -0.033233837 | 0.999933887 | 0.070788869 | 0.110335371 | -0.039546502 | 0.812383698 |
| ENSG00000163568 | *AIM2* | 2.569301987 | 0.000696003 | -0.07938277 | 0.080078773 | 0.999933887 | -0.054533102 | -0.113994601 | 0.059461498 | 0.812447255 |
| ENSG00000214530 | *STARD10* | 5.835868676 | 0.040948321 | 0.062773252 | -0.021824931 | 0.999933887 | 0.262358731 | 0.297391982 | -0.035033251 | 0.812498261 |
| ENSG00000135597 | *REPS1* | 3.485039186 | 0.015761267 | -0.053462419 | 0.069223686 | 0.999933887 | -0.079439801 | -0.120548244 | 0.041108442 | 0.812498261 |
| ENSG00000164896 | *FASTK* | 4.462578518 | -0.012311736 | 0.024161875 | -0.036473611 | 0.999933887 | 0.319460825 | 0.362563467 | -0.043102642 | 0.812498261 |
| ENSG00000186204 | *CYP4F12* | 1.473171049 | 0.011607896 | -0.039034177 | 0.050642073 | 0.999933887 | -0.168573713 | -0.264579087 | 0.096005373 | 0.812790453 |
| ENSG00000143748 | *NVL* | 1.542418961 | 0.053957615 | -0.035744148 | 0.089701763 | 0.999933887 | -0.143420323 | -0.0671757 | -0.076244623 | 0.812982746 |
| ENSG00000111678 | *C12orf57* | 2.22971254 | 0.086244694 | -0.012297391 | 0.098542084 | 0.999933887 | 0.377215118 | 0.304565591 | 0.072649527 | 0.813016997 |
| ENSG00000108518 | *PFN1* | 8.645742392 | 0.021091801 | -0.00238429 | 0.023476092 | 0.999933887 | 0.239111392 | 0.20117764 | 0.037933752 | 0.813376742 |
| ENSG00000275342 | *PRAG1* | 1.578199733 | -0.08922679 | 0.065676738 | -0.154903528 | 0.999933887 | 0.16126424 | 0.221809411 | -0.06054517 | 0.813388341 |
| ENSG00000139131 | *YARS2* | 0.585213495 | -0.010578154 | -0.079062501 | 0.068484347 | 0.999933887 | 0.075204076 | 0.174689317 | -0.099485241 | 0.813399962 |
| ENSG00000011638 | *TMEM159* | 2.976774794 | -0.110724133 | -0.135930723 | 0.02520659 | 0.999933887 | -0.437696289 | -0.388681739 | -0.04901455 | 0.813542077 |
| ENSG00000225434 | *LINC01504* | 1.493240005 | 0.008485608 | 0.227700749 | -0.219215141 | 0.999933887 | -0.150444115 | -0.078421166 | -0.072022949 | 0.813564246 |
| ENSG00000143774 | *GUK1* | 7.239818795 | 0.080844085 | 0.050849617 | 0.029994468 | 0.999933887 | 0.051101112 | 0.017459052 | 0.03364206 | 0.813586449 |
| ENSG00000225733 | *FGD5-AS1* | 6.130759977 | -0.055003294 | -0.049913402 | -0.005089892 | 0.999933887 | -0.40702003 | -0.438171063 | 0.031151033 | 0.813586449 |
| ENSG00000277511 | *AC116407.2* | 0.437169235 | 0.065264563 | -0.145430835 | 0.210695397 | 0.999933887 | -0.1177644 | -0.238853418 | 0.121089018 | 0.813586449 |
| ENSG00000233621 | *LINC01137* | 0.920508845 | -0.111834386 | 0.17902522 | -0.290859606 | 0.999933887 | 0.620051757 | 0.523488415 | 0.096563342 | 0.813586449 |
| ENSG00000213839 | *TMX2P1* | 0.451543564 | -0.026486521 | 0.215651945 | -0.242138466 | 0.999933887 | -0.193707306 | -0.054544958 | -0.139162348 | 0.813586449 |
| ENSG00000059145 | *UNKL* | 5.855708353 | -0.017531674 | -0.059998479 | 0.042466804 | 0.999933887 | -0.229845838 | -0.266705364 | 0.036859526 | 0.813600018 |
| ENSG00000171109 | *MFN1* | 4.95844509 | -0.065899756 | 0.015639986 | -0.081539742 | 0.999933887 | -0.310988367 | -0.252387456 | -0.05860091 | 0.813635055 |
| ENSG00000101439 | *CST3* | 5.525324795 | 0.03855085 | 0.024940664 | 0.013610185 | 0.999933887 | -0.282765164 | -0.340034727 | 0.057269563 | 0.813668325 |
| ENSG00000139291 | *TMEM19* | 0.826119358 | 0.013774059 | 0.098734535 | -0.084960476 | 0.999933887 | 0.032101493 | -0.052343679 | 0.084445172 | 0.813668325 |
| ENSG00000227345 | *PARG* | 1.981568636 | -0.009154157 | 0.000903494 | -0.01005765 | 0.999933887 | -0.129144396 | -0.19396337 | 0.064818974 | 0.813668325 |
| ENSG00000157985 | *AGAP1* | 0.916337717 | 0.044055131 | 0.070555883 | -0.026500753 | 0.999933887 | -0.078108878 | 0.011710348 | -0.089819226 | 0.814056545 |
| ENSG00000104442 | *ARMC1* | 1.696609349 | -0.077605857 | 0.023232757 | -0.100838614 | 0.999933887 | 0.08498313 | 0.155261378 | -0.070278248 | 0.814298015 |
| ENSG00000196419 | *XRCC6* | 5.993535247 | -0.033427802 | -0.02829194 | -0.005135862 | 0.999933887 | 0.013188132 | -0.019184473 | 0.032372605 | 0.814403834 |
| ENSG00000187801 | *ZFP69B* | 0.155135639 | 0.098349529 | -0.164882829 | 0.263232359 | 0.999933887 | 0.103704229 | -0.022973574 | 0.126677803 | 0.814403834 |

| | | | | | | | | | | |
|---|---|---|---|---|---|---|---|---|---|---|
| ENSG00000125084 | *WNT1* | -0.178257605 | 0.007783637 | -0.161286533 | 0.16907017 | 0.999933887 | -0.040982098 | 0.093179821 | -0.134161919 | 0.814403834 |
| ENSG00000165914 | *TTC7B* | 1.340458204 | 0.134431124 | 0.20143015 | -0.066999026 | 0.999933887 | 0.001360639 | 0.079263959 | -0.077903319 | 0.814404314 |
| ENSG00000087157 | *PGS1* | 7.237747866 | 0.046661025 | 0.039570609 | 0.007090416 | 0.999933887 | -0.195624898 | -0.160914783 | -0.034710115 | 0.814404314 |
| ENSG00000168028 | *RPSA* | 7.189608324 | -0.011320192 | -0.034384834 | 0.023064643 | 0.999933887 | 0.128517209 | 0.167731821 | -0.039214612 | 0.814404314 |
| ENSG00000148384 | *INPP5E* | 2.707709541 | -0.012763334 | 0.110138104 | -0.122901438 | 0.999933887 | 0.147426725 | 0.200858872 | -0.053432147 | 0.814404314 |
| ENSG00000247774 | *PCED1B-AS1* | 4.392808572 | 0.002159303 | -0.029529685 | 0.031688987 | 0.999933887 | 0.076845205 | 0.044558356 | 0.032286849 | 0.814404314 |
| ENSG00000067141 | *NEO1* | 1.478589732 | -0.104377499 | 0.084953462 | -0.189330961 | 0.999933887 | -0.046286254 | 0.040037199 | -0.086323453 | 0.814466423 |
| ENSG00000006625 | *GGCT* | -0.5148304 | -0.064451227 | -0.000758088 | -0.063693139 | 0.999933887 | -0.163514974 | -0.032178897 | -0.131336076 | 0.814466423 |
| ENSG00000138074 | *SLC5A6* | 3.785871069 | 0.010492039 | 0.031811217 | -0.021319178 | 0.999933887 | 0.156557893 | 0.198353301 | -0.041795408 | 0.814466423 |
| ENSG00000223573 | *TINCR* | -0.277293144 | 0.018238906 | 0.223119345 | -0.204880438 | 0.999933887 | -0.157226092 | -0.031433204 | -0.125792888 | 0.814466423 |
| ENSG00000135596 | *MICAL1* | 8.050807306 | 0.052178429 | 0.058352141 | -0.006173711 | 0.999933887 | -0.155165755 | -0.127326576 | -0.02783918 | 0.814478823 |
| ENSG00000273691 | *AC087284.1* | 0.87990639 | -0.102372098 | 0.002638062 | -0.10501016 | 0.999933887 | 0.116893816 | 0.027030116 | 0.0898637 | 0.814478823 |
| ENSG00000074603 | *DPP8* | 4.708262875 | -0.014120652 | -0.044265009 | 0.030144357 | 0.999933887 | -0.384475288 | -0.426357083 | 0.041881795 | 0.814478823 |
| ENSG00000279088 | *AC022400.6* | 2.202857685 | -0.140277471 | -0.180952935 | 0.040675464 | 0.999933887 | 0.093303223 | 0.039975665 | 0.053327558 | 0.814620149 |
| ENSG00000099338 | *CATSPERG* | 0.656379858 | 0.021264041 | -0.017406263 | 0.038670304 | 0.999933887 | -0.225741929 | -0.352951374 | 0.127209445 | 0.814806702 |
| ENSG00000226777 | *FAM30A* | -0.190945561 | -0.101758846 | 0.218669638 | -0.320428484 | 0.999933887 | -0.457121143 | -0.344472248 | -0.112648895 | 0.815287813 |
| ENSG00000145337 | *PYURF* | 2.927722115 | 0.087720212 | 0.083158236 | 0.004561977 | 0.999933887 | 0.049032408 | -0.012426994 | 0.061459402 | 0.815326412 |
| ENSG00000169813 | *HNRNPF* | 8.842777912 | -0.00679912 | 0.020495284 | -0.027294404 | 0.999933887 | -0.10358364 | -0.137145593 | 0.033561954 | 0.815522072 |
| ENSG00000135437 | *RDH5* | 1.736630345 | 0.051988114 | -0.12810212 | 0.180090233 | 0.999933887 | 0.026435778 | -0.029141538 | 0.055577316 | 0.815644869 |
| ENSG00000227512 | *AL592301.1* | 2.812315832 | -0.195779528 | -0.159981223 | -0.035798305 | 0.999933887 | -0.476523413 | -0.412047365 | -0.064476047 | 0.815662772 |
| ENSG00000117906 | *RCN2* | 2.668375406 | -0.044952536 | -0.040397152 | -0.004555384 | 0.999933887 | -0.170743108 | -0.111858704 | -0.058884404 | 0.815662772 |
| ENSG00000185219 | *ZNF445* | 3.670728674 | -0.013854541 | 0.004131705 | -0.017986246 | 0.999933887 | -0.078967366 | -0.034724457 | -0.044242909 | 0.815662772 |
| ENSG00000169714 | *CNBP* | 8.639849432 | -0.066242345 | -0.111701689 | 0.045459344 | 0.999933887 | -0.659340182 | -0.695810262 | 0.036470081 | 0.815670308 |
| ENSG00000165506 | *DNAAF2* | 1.898077055 | 0.160105793 | 0.017910634 | 0.142195158 | 0.999933887 | 0.186122041 | 0.115326491 | 0.070795551 | 0.81585173 |
| ENSG00000112039 | *FANCE* | -0.016790502 | -0.116863145 | -0.023276613 | -0.093586532 | 0.999933887 | 0.084736816 | -0.024229435 | 0.10896625 | 0.81585173 |
| ENSG00000184792 | *OSBP2* | -0.139065535 | -0.039244726 | 0.026759542 | -0.066004268 | 0.999933887 | 0.091182712 | -0.019367742 | 0.110550454 | 0.81585173 |
| ENSG00000197951 | *ZNF71* | 0.660948781 | 0.018894462 | 0.038953612 | -0.020059149 | 0.999933887 | 0.067833043 | -0.028537167 | 0.09637021 | 0.81585173 |
| ENSG00000166263 | *STXBP4* | -0.363191599 | -0.154281013 | -0.409687397 | 0.255406384 | 0.999933887 | -0.250018854 | -0.10852557 | -0.141493284 | 0.815883448 |
| ENSG00000214659 | *KRT8P26* | 0.501777457 | -0.057422068 | 0.008596186 | -0.066018255 | 0.999933887 | -0.446957342 | -0.571288401 | 0.124331059 | 0.815910874 |
| ENSG00000076924 | *XAB2* | 5.874799139 | -0.003198089 | 0.050917084 | -0.054115173 | 0.999933887 | 0.018135124 | -0.013934481 | 0.032069605 | 0.816132444 |
| ENSG00000128918 | *ALDH1A2* | 4.077718931 | 0.275342829 | 0.366489268 | -0.091146439 | 0.999933887 | 0.642998006 | 0.56714503 | 0.075852975 | 0.816253279 |
| ENSG00000165502 | *RPL36AL* | 6.285584732 | -0.034779462 | -0.075679349 | 0.040899887 | 0.999933887 | -0.077241212 | -0.046847493 | -0.030393719 | 0.816341491 |
| ENSG00000106330 | *MOSPD3* | 2.320609715 | -0.073619716 | 0.107567393 | -0.181187109 | 0.999933887 | -0.21216248 | -0.138960178 | -0.073202302 | 0.81637165 |
| ENSG00000203804 | *ADAMTSL4-AS* | 4.14919227 | 0.093710438 | 0.083959794 | 0.009750644 | 0.999933887 | 0.347640329 | 0.296870853 | 0.050769476 | 0.816747124 |
| ENSG00000281183 | *NPTN-IT1* | 3.002010063 | -0.046343105 | -0.090196893 | 0.043853788 | 0.999933887 | -0.427312263 | -0.376160272 | -0.051151992 | 0.816747124 |
| ENSG00000233806 | *LINC01237* | -0.641050807 | 0.046699949 | -0.058907863 | 0.105607811 | 0.999933887 | -0.029052426 | 0.12327973 | -0.152332156 | 0.816747124 |
| ENSG00000217527 | *RPS16P5* | 0.537697337 | 0.170903964 | 0.310888112 | -0.139984148 | 0.999933887 | 0.162491895 | -0.026634321 | 0.189126216 | 0.816759945 |
| ENSG00000148606 | *POLR3A* | 2.497957019 | -0.019432221 | -0.051010089 | 0.031577868 | 0.999933887 | -0.034565732 | 0.014461501 | -0.049027233 | 0.816759945 |
| ENSG00000105088 | *OLFM2* | -0.120538541 | -0.059984195 | 0.485183352 | -0.545167547 | 0.999933887 | 0.023031094 | 0.125470849 | -0.102439755 | 0.816759945 |
| ENSG00000104361 | *NIPAL2* | -1.607339119 | 0.247860677 | 0.233387427 | 0.01447325 | 0.999933887 | 0.2160853 | 0.401783465 | -0.185698165 | 0.816849032 |
| ENSG00000284428 | *AC092329.4* | 0.530547205 | -0.100986003 | 0.058940687 | -0.15992669 | 0.999933887 | -0.014902576 | -0.104708826 | 0.08980625 | 0.816930482 |
| ENSG00000239961 | *LILRA4* | -1.34236507 | -0.407034794 | -0.605152402 | 0.198117608 | 0.999933887 | -1.554975379 | -1.708832751 | 0.153857372 | 0.81733427 |
| ENSG00000166925 | *TSC22D4* | 8.301568497 | 0.052051105 | 0.05203001 | 2.1094e-05 | 0.999933887 | -0.134084478 | -0.170475162 | 0.036390683 | 0.81733427 |
| ENSG00000104447 | *TRPS1* | 2.836050487 | -0.094223249 | 0.087700978 | -0.181924226 | 0.999933887 | -0.178699084 | -0.113865376 | -0.064833708 | 0.81736372 |
| ENSG00000126001 | *CEP250* | 3.69072812 | -0.05057695 | 0.012156589 | -0.06273354 | 0.999933887 | -0.025898384 | -0.062416445 | 0.036518061 | 0.81736372 |
| ENSG00000138795 | *LEF1* | 4.28658501 | -0.042367782 | -0.041529615 | -0.000838166 | 0.999933887 | 0.114313496 | 0.077733772 | 0.036579725 | 0.81736372 |
| ENSG00000162148 | *PPP1R32* | 0.35507298 | 0.082638347 | 0.168352634 | -0.085714287 | 0.999933887 | -0.13704631 | -0.231430343 | 0.094384033 | 0.81736372 |
| ENSG00000101842 | *VSIG1* | 0.215156564 | 0.074322748 | 0.220853574 | -0.146530826 | 0.999933887 | 0.113859087 | 0.233391939 | -0.119532852 | 0.81736372 |
| ENSG00000235579 | *RPL38P5* | -0.681067449 | 0.12874662 | -0.178680002 | 0.307426622 | 0.999933887 | 1.102980547 | 1.255810813 | -0.152830266 | 0.81736372 |
| ENSG00000197958 | *RPL12* | 7.251054754 | 0.018046468 | -0.022352857 | 0.040399325 | 0.999933887 | 0.118251809 | 0.154308132 | -0.036056323 | 0.81736372 |
| ENSG00000254703 | *SENCR* | -0.339391428 | -0.051159402 | -0.041701794 | -0.009457608 | 0.999933887 | -0.178570268 | -0.347337653 | 0.168767386 | 0.81736372 |
| ENSG00000142507 | *PSMB6* | 3.850863358 | -0.019676527 | 0.049231328 | -0.068907856 | 0.999933887 | -0.168566899 | -0.124569745 | -0.043997154 | 0.81736372 |
| ENSG00000196872 | *CRACDL* | 0.671907899 | 0.03343312 | 0.168231907 | -0.134798787 | 0.999933887 | -0.550267553 | -0.438881019 | -0.111386534 | 0.81736372 |
| ENSG00000273218 | *AC005776.2* | 0.673984304 | 0.021333754 | -0.189185425 | 0.210519179 | 0.999933887 | 0.338927518 | 0.430188272 | -0.091260754 | 0.81736372 |
| ENSG00000260793 | *AC003102.1* | 0.791124232 | -0.018763281 | 0.262000313 | -0.280763594 | 0.999933887 | -0.072267068 | 0.021035367 | -0.093302436 | 0.81736372 |
| ENSG00000127184 | *COX7C* | 4.549938527 | 0.002464941 | -0.080821382 | 0.083286323 | 0.999933887 | 0.147967449 | 0.190885714 | -0.042918265 | 0.81736372 |
| ENSG00000050393 | *MCUR1* | 1.415835193 | -0.000276899 | 0.150926167 | -0.151203066 | 0.999933887 | -0.054359348 | 0.01405103 | -0.068410378 | 0.81736372 |
| ENSG00000279583 | *AC009086.2* | -0.159487366 | 0.385135208 | -0.211458867 | 0.596594075 | 0.594252256 | 0.101153908 | -0.002437378 | 0.103591286 | 0.817470877 |
| ENSG00000196923 | *PDLIM7* | 8.402562308 | 0.053228352 | 0.016633499 | 0.036594853 | 0.999933887 | -0.008693463 | 0.020809692 | -0.029503154 | 0.817555881 |
| ENSG00000182405 | *PGBD4* | 2.066661104 | -0.039509407 | -0.020394678 | -0.019114729 | 0.999933887 | 0.216695145 | 0.158612736 | 0.058082409 | 0.817555881 |
| ENSG00000215244 | *LINC02649* | 3.810884766 | 0.054998692 | 0.058026508 | -0.003027815 | 0.999933887 | 0.006426775 | -0.042334956 | 0.048761731 | 0.818045085 |
| ENSG00000150459 | *SAP18* | 5.741356273 | -0.041119167 | 0.03420962 | -0.075328787 | 0.999933887 | -0.125009785 | -0.093814825 | -0.03119496 | 0.818341876 |
| ENSG00000105373 | *NOP53* | 7.214112423 | 0.102094673 | 0.092213863 | 0.009880809 | 0.999933887 | 0.093416981 | 0.141822859 | -0.048405877 | 0.818859124 |
| ENSG00000285589 | *AC010422.8* | -0.484284868 | -0.187080149 | -0.063747743 | -0.123332406 | 0.999933887 | 0.098470679 | -0.039998217 | 0.138468896 | 0.818859124 |
| ENSG00000105447 | *GRWD1* | 2.759403075 | -0.063628146 | -0.104275177 | 0.040647031 | 0.999933887 | 0.299748462 | 0.35530023 | -0.055551768 | 0.818859124 |
| ENSG00000023839 | *ABCC2* | 1.70168364 | -0.064556651 | -0.074279951 | 0.0097233 | 0.999933887 | 0.152659206 | 0.072573639 | 0.080085567 | 0.818859124 |
| ENSG00000113441 | *LNPEP* | 5.444446114 | 0.048702958 | -0.009301095 | 0.058004052 | 0.999933887 | -0.42816986 | -0.361221874 | -0.066947986 | 0.818873089 |
| ENSG00000177084 | *POLE* | 3.74918338 | 0.002332957 | 0.068356196 | -0.066023239 | 0.999933887 | 0.17391735 | 0.222283793 | -0.048366443 | 0.819083064 |
| ENSG00000228705 | *LINC00659* | 1.621986645 | 0.1412158 | 0.388029244 | -0.246813444 | 0.999933887 | 0.595001797 | 0.666595938 | -0.071594141 | 0.819123372 |
| ENSG00000127418 | *FGFRL1* | 2.664112423 | 0.066235233 | 0.011642438 | 0.054592795 | 0.999933887 | 0.144183995 | 0.196608556 | -0.052424561 | 0.819269771 |
| ENSG00000166963 | *MAP1A* | 0.762744632 | 0.020660981 | 0.264438878 | -0.243777897 | 0.999933887 | -0.081565016 | -0.207906179 | 0.126341163 | 0.819269771 |
| ENSG00000180884 | *ZNF792* | 0.912025882 | 0.18678856 | 0.019545168 | 0.167243392 | 0.999933887 | 0.041651227 | 0.131428003 | -0.089776775 | 0.819303199 |
| ENSG00000162980 | *ARL5A* | 1.915182728 | -0.024268496 | -0.117160566 | 0.09289207 | 0.999933887 | -0.192677329 | -0.109850535 | -0.082826793 | 0.819303199 |
| ENSG00000151376 | *ME3* | -0.677130125 | 0.030677493 | -0.125712229 | 0.156389722 | 0.999933887 | 0.08422766 | -0.056731594 | 0.140959254 | 0.819303199 |
| ENSG00000261326 | *LINC01355* | -1.423880708 | -0.063582491 | -0.353383253 | 0.289800762 | 0.999933887 | -0.254851798 | -0.107975829 | -0.146875969 | 0.819473353 |
| ENSG00000117448 | *AKR1A1* | 4.160036749 | -0.05276096 | -0.147280628 | 0.094519668 | 0.999933887 | 0.124634593 | 0.084064527 | 0.040570066 | 0.819481479 |
| ENSG00000112992 | *NNT* | 3.015118384 | -0.023530651 | -0.053045015 | 0.029514364 | 0.999933887 | -0.03882732 | 0.000413753 | -0.039241073 | 0.819481479 |
| ENSG00000102794 | *ACOD1* | -2.381409195 | 0.703934944 | 0.66877877 | 0.035156173 | 0.999933887 | 3.877383152 | 4.209238458 | -0.331855305 | 0.819815869 |
| ENSG00000123838 | *C4BPA* | 1.40520972 | -0.101833199 | -0.17715285 | 0.075319651 | 0.999933887 | 0.041070132 | 0.121545052 | -0.080474919 | 0.819815869 |
| ENSG00000185245 | *GP1BA* | 1.397928327 | 0.040619193 | -0.106087293 | 0.146706486 | 0.999933887 | 0.077224542 | -0.026520674 | 0.103745216 | 0.819815869 |
| ENSG00000059588 | *TARBP1* | 2.601848046 | -0.003478821 | 0.133647079 | -0.1371259 | 0.999933887 | 0.043259337 | 0.095003709 | -0.051744372 | 0.819815869 |
| ENSG00000243179 | *AC110769.1* | -0.112677894 | -0.280109447 | -0.038783606 | -0.241325841 | 0.999933887 | 1.301395925 | 1.445492812 | -0.144096887 | 0.820422623 |
| ENSG00000267655 | *AC125437.1* | -0.256305451 | -0.170508084 | -0.173037883 | 0.002529799 | 0.999933887 | 0.110257245 | -0.069345864 | 0.179603109 | 0.820422623 |
| ENSG00000090861 | *AARS1* | 3.319282867 | -0.06013731 | -0.041307132 | -0.018830177 | 0.999933887 | 0.023969314 | -0.018521616 | 0.04249093 | 0.820422623 |
| ENSG00000121769 | *FABP3* | -0.902979819 | -0.170986812 | 0.377703337 | -0.548690149 | 0.999933887 | 0.064190307 | -0.059803146 | 0.123993453 | 0.820422623 |
| ENSG00000173848 | *NET1* | 1.192835449 | -0.058117155 | -0.026595465 | -0.03152169 | 0.999933887 | -0.222436625 | -0.132670853 | -0.089765772 | 0.820422623 |
| ENSG00000205078 | *SYCE1L* | -0.222848938 | 0.050897859 | -0.143288424 | 0.194186283 | 0.999933887 | 0.161814344 | 0.028543235 | 0.133271109 | 0.820422623 |
| ENSG00000143183 | *TMCO1* | 4.172354354 | -0.0706579 | -0.020307852 | -0.050350049 | 0.999933887 | -0.136849753 | -0.101330509 | -0.035519244 | 0.820501664 |
| ENSG00000156650 | *KAT6B* | 4.486045687 | 0.054505002 | -0.052297572 | 0.106802574 | 0.999933887 | -0.343422743 | -0.393517034 | 0.050094291 | 0.820501664 |
| ENSG00000123131 | *PRDX4* | 0.474963855 | -0.108967995 | 0.264033121 | -0.373001116 | 0.999933887 | 0.163206731 | 0.26501332 | -0.101806588 | 0.820501664 |
| ENSG00000147576 | *ADHFE1* | -0.467402195 | -0.123411689 | 0.070789853 | -0.194201542 | 0.999933887 | 0.021035578 | -0.10314712 | 0.124182698 | 0.820501664 |
| ENSG00000235272 | *RAMACL* | -1.458160583 | -0.416646531 | -0.332666477 | -0.083980055 | 0.999933887 | -0.479875331 | -0.637956636 | 0.158081304 | 0.821227343 |
| ENSG00000076108 | *BAZ2A* | 8.086654949 | 0.011008993 | 0.039264641 | -0.028255648 | 0.999933887 | -0.235152679 | -0.264403529 | 0.02925085 | 0.821310504 |
| ENSG00000121864 | *ZNF639* | 2.55022669 | 0.009556675 | 0.036458325 | -0.02690165 | 0.999933887 | 0.07697252 | 0.013089906 | 0.063882614 | 0.82134123 |
| ENSG00000279699 | *AC004264.2* | 1.222375128 | 0.227596298 | 0.019355936 | 0.208240363 | 0.999933887 | 2.239824951 | 2.339711647 | -0.099886696 | 0.821355445 |
| ENSG00000173762 | *CD7* | 5.324792736 | 0.030409665 | 0.051058692 | -0.020649027 | 0.999933887 | 0.193277341 | 0.229195268 | -0.035917926 | 0.821355445 |
| ENSG00000178015 | *GPR150* | -0.812506716 | 0.101399266 | -0.310275581 | 0.411674847 | 0.999933887 | -1.5039821 | -1.368861905 | -0.135120195 | 0.821355445 |
| ENSG00000117461 | *PIK3R3* | -0.20666038 | -0.218217 | -0.323088958 | 0.104871957 | 0.999933887 | -0.131135022 | -0.024243692 | -0.106891331 | 0.82137817 |
| ENSG00000146021 | *KLHL3* | 1.676706279 | -0.026864243 | -0.032099312 | 0.005235069 | 0.999933887 | -0.107133763 | -0.043728952 | -0.063404811 | 0.82137817 |
| ENSG00000152942 | *RAD17* | 2.192398437 | -0.049224838 | -0.163853317 | 0.114628479 | 0.999933887 | -0.274814041 | -0.221887406 | -0.052926635 | 0.821424876 |
| ENSG00000186193 | *SAPCD2* | -1.433882696 | 0.199030709 | -0.130031031 | 0.32906174 | 0.999933887 | -0.382530688 | -0.112939571 | -0.269591117 | 0.8214429 |
| ENSG00000063761 | *ADCK1* | -0.03908873 | -0.044375976 | 0.007300622 | -0.051676598 | 0.999933887 | -0.050572059 | -0.150642261 | 0.100070202 | 0.8214429 |
| ENSG00000223117 | *RN7SKP296* | 0.350870266 | -0.129366379 | -0.083319575 | -0.046046804 | 0.999933887 | -0.394030552 | -0.670225008 | 0.276194457 | 0.821702014 |
| ENSG00000254060 | *AC022778.1* | 0.45741044 | 0.184534024 | 0.287348293 | -0.10281427 | 0.999933887 | 0.476754176 | 0.616658546 | -0.139904369 | 0.82181086 |
| ENSG00000172543 | *CTSW* | 4.421059812 | 0.017412076 | 0.002265361 | 0.015146716 | 0.999933887 | 0.130169792 | 0.175669552 | -0.04549976 | 0.821842831 |

| | | | | | | | | | | |
|---|---|---|---|---|---|---|---|---|---|---|
| ENSG00000254328 | *AC008429.4* | -0.068496912 | 0.242894852 | 0.117186763 | 0.125708089 | 0.999933887 | 0.810887961 | 0.675710758 | 0.135177203 | 0.821898992 |
| ENSG00000211785 | *TRAV12-1* | 0.048840817 | -0.048354951 | 0.045293218 | -0.093648169 | 0.999933887 | 0.042867461 | 0.130666458 | -0.087798996 | 0.821925232 |
| ENSG00000123607 | *TTC21B* | 2.236430946 | -0.024930192 | -0.012642704 | -0.012287488 | 0.999933887 | -0.19248089 | -0.096359751 | -0.096121139 | 0.821925232 |
| ENSG00000287115 | *AC009951.6* | 0.678846642 | 0.192338439 | 0.016464747 | 0.175873692 | 0.999933887 | 1.541577303 | 1.463268004 | 0.078309299 | 0.821936918 |
| ENSG00000245888 | *NSMCE1-DT* | 0.985270413 | -0.142332481 | -0.172995402 | 0.030662921 | 0.999933887 | -0.365897605 | -0.463047668 | 0.097150063 | 0.822056577 |
| ENSG00000243609 | *RPS2P44* | 1.444377633 | 0.154668008 | 0.382216607 | -0.227548598 | 0.999933887 | 1.803248215 | 1.668162747 | 0.135085468 | 0.822099666 |
| ENSG00000099917 | *MED15* | 7.306884566 | 0.031197426 | 0.040158212 | -0.008960786 | 0.999933887 | 0.130519239 | 0.153601662 | -0.023082423 | 0.822108744 |
| ENSG00000187049 | *TMEM216* | 2.966119023 | 0.046222872 | -0.086345296 | 0.132568168 | 0.999933887 | -0.127267063 | -0.175981937 | 0.048714874 | 0.822108744 |
| ENSG00000084444 | *FAM234B* | 0.401081088 | -0.134910107 | 0.089093374 | -0.224003481 | 0.999933887 | -0.617773082 | -0.722785728 | 0.105012646 | 0.822387585 |
| ENSG00000180712 | *LINC02363* | 2.544561844 | 0.05289005 | 0.143404216 | -0.090514166 | 0.999933887 | 0.586919099 | 0.530742089 | 0.056177011 | 0.822425808 |
| ENSG00000180921 | *FAM83H* | -0.171209315 | 0.316324248 | -0.048407834 | 0.364732082 | 0.999933887 | 0.079130255 | -0.043266616 | 0.12239687 | 0.822453896 |
| ENSG00000164430 | *CGAS* | 1.791646147 | 0.038813963 | 0.185354402 | -0.146540439 | 0.999933887 | 0.195217858 | 0.2564277 | -0.061209842 | 0.822456841 |
| ENSG00000083750 | *RRAGB* | 2.117120305 | -0.040344891 | -0.074090538 | 0.033745647 | 0.999933887 | -0.062011056 | -0.010437482 | -0.051573574 | 0.822708256 |
| ENSG00000159733 | *ZFYVE28* | 3.022208455 | 0.142204551 | -0.009525149 | 0.1517297 | 0.999933887 | 0.160764232 | 0.208370214 | -0.047605981 | 0.822838547 |
| ENSG00000249249 | *AC010226.1* | -0.598530131 | 0.062119264 | -0.210187835 | 0.272307099 | 0.999933887 | 0.066268548 | 0.223027553 | -0.156759005 | 0.823029468 |
| ENSG00000250893 | *AC098869.2* | -0.082881986 | -0.078578733 | 0.129488301 | -0.208067034 | 0.999933887 | -0.324961625 | -0.198280762 | -0.126680863 | 0.823105949 |
| ENSG00000145780 | *FEM1C* | 5.31927889 | -0.015259253 | -0.032983949 | 0.017724696 | 0.999933887 | -0.452799473 | -0.498064362 | 0.045264889 | 0.823105949 |
| ENSG00000188372 | *ZP3* | -0.403945773 | -0.00241455 | 0.478471628 | -0.480886178 | 0.999933887 | 2.100240053 | 1.980937635 | 0.119302418 | 0.823105949 |
| ENSG00000280402 | *AC093525.10* | 0.065906214 | 0.19824217 | -0.003983559 | 0.202225729 | 0.999933887 | -0.776044586 | -0.89498559 | 0.118941004 | 0.823197465 |
| ENSG00000234545 | *FAM133B* | 2.519330277 | 0.078521106 | -0.021660338 | 0.100181444 | 0.999933887 | 0.006603204 | -0.06879805 | 0.075401254 | 0.823197465 |
| ENSG00000223662 | *SAMSN1-AS1* | -0.99809824 | -0.053711566 | -0.846740765 | 0.793029199 | 0.999933887 | 2.188371937 | 2.378624745 | -0.190252808 | 0.823197465 |
| ENSG00000196421 | *C20orf204* | 0.128622731 | 0.01403995 | 0.143839377 | -0.129799427 | 0.999933887 | -0.019509789 | 0.106276393 | -0.125786183 | 0.823197465 |
| ENSG00000080608 | *PUM3* | 1.616482847 | -0.199597368 | 0.045623416 | -0.245220784 | 0.999933887 | 0.037592693 | 0.10374352 | -0.066150827 | 0.823421468 |
| ENSG00000184361 | *SPATA32* | -0.219696742 | 0.237331395 | 0.043500734 | 0.193830661 | 0.999933887 | -0.095505097 | 0.05360068 | -0.149105777 | 0.823439541 |
| ENSG00000261118 | *AC092123.1* | -0.032484094 | -0.1237334 | -0.102667296 | -0.021066104 | 0.999933887 | -0.354014235 | -0.451603485 | 0.09758925 | 0.823439541 |
| ENSG00000286342 | *AC073210.3* | -0.982741265 | -0.19446671 | 0.471773172 | -0.666239882 | 0.999933887 | -0.720302736 | -0.88247203 | 0.162169294 | 0.823439541 |
| ENSG00000149269 | *PAK1* | 7.775408524 | -0.026299308 | -0.043857933 | 0.017558625 | 0.999933887 | 0.103280088 | 0.076639366 | 0.026640723 | 0.823439541 |
| ENSG00000146223 | *RPL7L1* | 3.454764483 | 0.025854016 | -0.049846548 | 0.075700564 | 0.999933887 | 0.084431535 | 0.047447738 | 0.036983798 | 0.823439541 |
| ENSG00000261349 | *AL031432.2* | 1.398359773 | 0.053536784 | 0.114907426 | -0.061370642 | 0.999933887 | 0.146793297 | 0.081975014 | 0.064818283 | 0.823439541 |
| ENSG00000257242 | *LINC01619* | 1.077490515 | 0.046810144 | -0.022439169 | 0.069249312 | 0.999933887 | 0.085220015 | 0.177908698 | -0.092688683 | 0.823439541 |
| ENSG00000164615 | *CAMLG* | 4.201687004 | -0.011950516 | -0.007239556 | -0.00471096 | 0.999933887 | -0.125922219 | -0.090408354 | -0.035513866 | 0.823476749 |
| ENSG00000099783 | *HNRNPM* | 7.319269227 | -0.022602742 | 0.006346687 | -0.028949429 | 0.999933887 | -0.048352535 | -0.023303248 | -0.025049287 | 0.823532635 |
| ENSG00000270062 | *AL606834.2* | 0.635930139 | 0.057845831 | -0.039862139 | 0.09770797 | 0.999933887 | -0.356378312 | -0.485523868 | 0.129145556 | 0.823532635 |
| ENSG00000170175 | *CHRNB1* | 0.863816303 | 0.064179337 | 0.184748841 | -0.120569504 | 0.999933887 | -0.254458706 | -0.357801757 | 0.103343051 | 0.823532635 |
| ENSG00000143126 | *CELSR2* | 0.62721894 | 0.079282208 | 0.150632725 | -0.071350517 | 0.999933887 | 0.027446135 | -0.0608697 | 0.088315834 | 0.823935864 |
| ENSG00000236911 | *AL137789.1* | -0.963010511 | 0.027667317 | 0.099321652 | -0.071654335 | 0.999933887 | -0.178400659 | 0.022981045 | -0.201381704 | 0.823935864 |
| ENSG00000174125 | *TLR1* | 6.742988896 | -0.162057805 | -0.173108569 | 0.011050764 | 0.999933887 | -1.324255928 | -1.396831397 | 0.072575469 | 0.823951105 |
| ENSG00000213891 | *RPL3P6* | 0.27134132 | 0.267824061 | 0.468612097 | -0.200788037 | 0.999933887 | 0.21464294 | 0.422150584 | -0.207507644 | 0.823951105 |
| ENSG00000131469 | *RPL27* | 7.241869032 | 0.032270539 | -0.060851323 | 0.093121862 | 0.999933887 | 0.080325008 | 0.11935161 | -0.039026603 | 0.823951105 |
| ENSG00000082068 | *WDR70* | 1.911620612 | 0.022292704 | -0.117722583 | 0.140015287 | 0.999933887 | 0.093379929 | 0.039850172 | 0.053529757 | 0.823951105 |
| ENSG00000006747 | *SCIN* | -1.625280405 | -0.789432947 | 0.115947637 | -0.905380583 | 0.999933887 | -0.966449253 | -0.83788967 | -0.128559583 | 0.82397279 |
| ENSG00000224578 | *HNRNPA1P48* | -0.073495687 | -0.249868462 | -0.337509218 | 0.087640756 | 0.999933887 | 0.205215053 | 0.113638623 | 0.091576431 | 0.82397279 |
| ENSG00000163374 | *YY1AP1* | 5.869799836 | -0.057045273 | -0.034178704 | -0.022866569 | 0.999933887 | 0.293054009 | 0.321808384 | -0.028754375 | 0.82397279 |
| ENSG00000255760 | *LINC02422* | 3.194054377 | -0.067443581 | -0.007631346 | -0.059812235 | 0.999933887 | 0.280945433 | 0.21793172 | 0.063013713 | 0.82397279 |
| ENSG00000258337 | *AC130895.1* | -0.474289991 | 0.142633984 | 0.230504872 | -0.087870888 | 0.999933887 | -0.008119613 | 0.163864196 | -0.17198381 | 0.82397279 |
| ENSG00000105700 | *KXD1* | 4.693426265 | 0.025517407 | 0.024371778 | 0.001145629 | 0.999933887 | -0.101038777 | -0.070529981 | -0.030508796 | 0.82397279 |
| ENSG00000139405 | *RITA1* | 0.407232524 | 0.089659989 | -0.1070721 | 0.196732089 | 0.999933887 | 0.054951649 | 0.175452378 | -0.12050073 | 0.82397279 |
| ENSG00000130513 | *GDF15* | 3.262806485 | 0.049633757 | -0.087431484 | 0.137065241 | 0.999933887 | 0.317810231 | 0.259913029 | 0.057897202 | 0.82397279 |
| ENSG00000143297 | *FCRL5* | -0.214939702 | -0.077593498 | -0.02640418 | -0.051189318 | 0.999933887 | 0.273571767 | 0.343569159 | -0.069997392 | 0.82397279 |
| ENSG00000254595 | *AC084337.1* | -0.10364534 | -0.045290246 | 0.521971508 | -0.567261754 | 0.594252256 | -0.155043456 | -0.270257921 | 0.115214465 | 0.82397279 |
| ENSG00000169629 | *RGPD8* | 0.269506066 | -0.046075266 | -0.101400683 | 0.055325417 | 0.999933887 | -0.388219282 | -0.275021263 | -0.113198018 | 0.82397279 |
| ENSG00000234171 | *RNASEH1-AS* | -0.193527218 | -0.042150617 | 0.084237845 | -0.126388461 | 0.999933887 | 0.101129073 | -0.013523099 | 0.114652171 | 0.82397279 |
| ENSG00000198553 | *KCNRG* | -0.700187162 | -0.036853604 | 0.420296814 | -0.457150418 | 0.999933887 | 0.174604071 | 0.044845965 | 0.129758105 | 0.82397279 |
| ENSG00000204394 | *VARS1* | 3.150539506 | 0.074830642 | 0.164405469 | -0.089574827 | 0.999933887 | 0.256006064 | 0.218136082 | 0.037869983 | 0.82407504 |
| ENSG00000183426 | *NPIPA1* | -0.398219043 | -0.105225348 | -0.169089895 | 0.063864546 | 0.999933887 | 0.05603138 | 0.169371479 | -0.113340099 | 0.824113925 |
| ENSG00000099194 | *SCD* | 2.43821644 | 0.056179633 | -0.07510803 | 0.131287664 | 0.999933887 | -0.396904973 | -0.465584844 | 0.06867987 | 0.824279751 |
| ENSG00000211786 | *TRAV8-2* | -0.183458824 | -0.281955701 | 0.116515757 | -0.398471458 | 0.999933887 | 0.1762534 | 0.306299511 | -0.130046111 | 0.8243847 |
| ENSG00000104824 | *HNRNPL* | 7.364123204 | 0.02189601 | 0.057386543 | -0.035490533 | 0.999933887 | -0.09291413 | -0.117946714 | 0.025032584 | 0.8243847 |
| ENSG00000224288 | *MTCYBP11* | -1.453213837 | -0.064951832 | 0.353027657 | -0.417979489 | 0.999933887 | 0.475902472 | 0.187398531 | 0.288503941 | 0.8243847 |
| ENSG00000119599 | *DCAF4* | 1.08610372 | 0.017503606 | 0.018784044 | -0.001280438 | 0.999933887 | 0.074412832 | 0.153124061 | -0.078711229 | 0.824743077 |
| ENSG00000244513 | *AC109587.1* | -0.207565377 | 0.097844785 | -0.033817229 | 0.131662014 | 0.999933887 | -0.056045231 | 0.072956772 | -0.129002003 | 0.824812614 |
| ENSG00000198720 | *ANKRD13B* | -0.505375241 | -0.283024974 | -0.294879742 | 0.011854769 | 0.999933887 | 0.023846876 | -0.097501641 | 0.121348517 | 0.824946203 |
| ENSG00000272655 | *POLR2J4* | 1.022147117 | 0.133939325 | 0.006884938 | 0.127054387 | 0.999933887 | 0.278169594 | 0.203704738 | 0.074464855 | 0.824946203 |
| ENSG00000135249 | *RINT1* | 2.362268022 | -0.046017128 | 0.155550212 | -0.20156734 | 0.999933887 | -0.007710099 | -0.061931445 | 0.054221346 | 0.824947968 |
| ENSG00000133466 | *C1QTNF6* | 3.083613473 | 0.037588052 | -0.005361443 | 0.042949495 | 0.999933887 | 0.245140491 | 0.289110172 | -0.043969682 | 0.824947968 |
| ENSG00000185156 | *MFSD6L* | 0.883980881 | 0.026010596 | 0.106239671 | -0.080229075 | 0.999933887 | 0.385786356 | 0.481037173 | -0.095250817 | 0.824947968 |
| ENSG00000000460 | *C1orf112* | 0.449761291 | -0.020255713 | -0.168482347 | 0.148226634 | 0.999933887 | -0.050256208 | 0.046653709 | -0.096909917 | 0.824947968 |
| ENSG00000261644 | *AC007728.2* | 2.890437953 | -0.033267435 | 0.089868805 | -0.12313624 | 0.999933887 | 0.553469766 | 0.602572489 | -0.049102723 | 0.825035313 |
| ENSG00000204220 | *PFDN6* | 2.831528467 | 0.028450256 | 0.061126268 | -0.032676012 | 0.999933887 | 0.164267249 | 0.123550157 | 0.040717092 | 0.825035313 |
| ENSG00000130193 | *THEM6* | 0.397198417 | 0.080717671 | 0.263570888 | -0.182853217 | 0.999933887 | 0.156384567 | 0.043285655 | 0.113098912 | 0.825314872 |
| ENSG00000127954 | *STEAP4* | 6.318469798 | -0.11997651 | -0.097030289 | -0.022946221 | 0.999933887 | -1.335863245 | -1.266004026 | -0.069859219 | 0.825349722 |
| ENSG00000276180 | *H4C9* | 2.331145338 | -0.007068915 | -0.058505661 | 0.051436747 | 0.999933887 | -0.104925491 | -0.165260072 | 0.060334581 | 0.825349722 |
| ENSG00000175611 | *LINC00476* | 0.605640832 | 0.249150107 | -0.148268053 | 0.39741816 | 0.999933887 | 0.031697924 | 0.125990011 | -0.094292087 | 0.825516364 |
| ENSG00000022567 | *SLC45A4* | 9.174283743 | -0.00938368 | -0.023308493 | 0.013924813 | 0.999933887 | -0.539505286 | -0.576086089 | 0.036580803 | 0.825629112 |
| ENSG00000143549 | *TPM3* | 9.356530333 | 0.002889277 | -0.024600613 | 0.027489889 | 0.999933887 | -0.171552831 | -0.143952157 | -0.027600674 | 0.825660887 |
| ENSG00000088992 | *TESC* | 2.08246636 | 0.001908965 | -0.103540526 | 0.105449492 | 0.999933887 | -0.05618776 | 0.008943069 | -0.06513083 | 0.825747705 |
| ENSG00000238366 | *Y_RNA* | -0.030840326 | 0.129276375 | 0.192290681 | -0.063014307 | 0.999933887 | 0.373930379 | 0.551821633 | -0.177891254 | 0.825750825 |
| ENSG00000042493 | *CAPG* | 5.222607407 | -0.042594615 | 0.014703459 | -0.057298074 | 0.999933887 | 0.362414023 | 0.325776224 | 0.0366378 | 0.826212955 |
| ENSG00000133315 | *MACROD1* | -0.27900163 | 0.082714386 | 0.08531593 | -0.002601544 | 0.999933887 | -0.020226663 | -0.145291624 | 0.125064961 | 0.826212955 |
| ENSG00000260273 | *AL359711.2* | 0.506348693 | 0.053146104 | -0.030280163 | 0.083426267 | 0.999933887 | 0.279379106 | 0.379669136 | -0.10029003 | 0.826212955 |
| ENSG00000224616 | *RTCA-AS1* | 0.365599258 | -0.362989378 | -0.324252508 | -0.038736871 | 0.999933887 | -1.365014668 | -1.2494502 | -0.115564467 | 0.826705983 |
| ENSG00000101307 | *SIRPB1* | 8.463274159 | -0.002770037 | -0.016747424 | 0.013977387 | 0.999933887 | -0.365400122 | -0.333618465 | -0.031781657 | 0.826705983 |
| ENSG00000198466 | *ZNF587* | 3.65394616 | -0.016281018 | 0.025805884 | -0.042086902 | 0.999933887 | -0.250137874 | -0.297649027 | 0.047511153 | 0.826798532 |
| ENSG00000013306 | *SLC25A39* | 3.733676118 | 0.078755935 | -0.001861692 | 0.080617627 | 0.999933887 | 0.1916536 | 0.153681763 | 0.037971837 | 0.826987617 |
| ENSG00000135776 | *ABCB10* | 2.145310383 | 0.023002724 | 0.000267002 | 0.022735722 | 0.999933887 | -0.086026422 | -0.021745047 | -0.064281376 | 0.827543702 |
| ENSG00000156482 | *RPL30* | 6.760218598 | 0.022666035 | -0.033926976 | 0.056593011 | 0.999933887 | 0.12306347 | 0.155341417 | -0.032277947 | 0.827557577 |
| ENSG00000082898 | *XPO1* | 5.259836689 | -0.057906642 | -0.044189987 | -0.013716655 | 0.999933887 | -0.332138034 | -0.286064235 | -0.046073798 | 0.827609897 |
| ENSG00000116741 | *RGS2* | 9.66834472 | 0.032761469 | -0.053292032 | 0.086053501 | 0.999933887 | -0.424528676 | -0.47410949 | 0.049580815 | 0.827616964 |
| ENSG00000213759 | *UGT2B11* | -0.640141336 | -0.001942726 | 0.339665077 | -0.341607803 | 0.999933887 | -0.200722426 | -0.087068329 | -0.113654098 | 0.827616964 |
| ENSG00000229891 | *LINC01315* | -0.361242493 | -0.197648456 | -0.298945209 | 0.101296753 | 0.999933887 | -0.281844853 | -0.392567992 | 0.110723139 | 0.82771522 |
| ENSG00000122122 | *SASH3* | 8.178633328 | -0.002435924 | -0.014539798 | 0.012103874 | 0.999933887 | -0.212907021 | -0.172811188 | -0.040095833 | 0.82771522 |
| ENSG00000087237 | *CETP* | 2.559081086 | -0.072672914 | -0.127013553 | 0.054340638 | 0.999933887 | -0.532094891 | -0.485902876 | -0.046192015 | 0.827800208 |
| ENSG00000124788 | *ATXN1* | 7.342609685 | -0.028043339 | -0.016469617 | -0.011573721 | 0.999933887 | -0.209586465 | -0.245466456 | 0.035879991 | 0.827800208 |
| ENSG00000104774 | *MAN2B1* | 6.438797093 | 0.008868388 | -0.058643428 | 0.067511817 | 0.999933887 | 0.046360058 | 0.016242013 | 0.030118045 | 0.827800208 |
| ENSG00000169193 | *CCDC126* | 2.149704401 | 0.102538557 | -0.004074178 | 0.106612735 | 0.999933887 | -0.696723615 | -0.625816274 | -0.070907341 | 0.827911691 |
| ENSG00000198502 | *HLA-DRB5* | 2.983195071 | 0.071109461 | 0.041188931 | 0.02992053 | 0.999933887 | 0.613672141 | 0.689582149 | -0.075910007 | 0.827911691 |
| ENSG00000181924 | *COA4* | 1.374246751 | 0.046194453 | 0.155857397 | -0.109662943 | 0.999933887 | 0.124514817 | 0.196597454 | -0.072082637 | 0.827911691 |
| ENSG00000107672 | *NSMCE4A* | 2.424880403 | 0.082674739 | -0.053030966 | 0.135705705 | 0.999933887 | 0.168588987 | 0.119069608 | 0.049519379 | 0.82792607 |
| ENSG00000180221 | *TPT1P10* | 0.089028526 | -0.033264039 | 0.192506165 | -0.225770204 | 0.999933887 | -0.167226689 | 0.01934835 | -0.186575039 | 0.82792607 |
| ENSG00000260920 | *AL031985.3* | -0.315743719 | -0.102574148 | -0.31079249 | 0.208218342 | 0.999933887 | -0.138322328 | 0.014168506 | -0.152490835 | 0.828180877 |
| ENSG00000064102 | *INTS13* | 1.672873649 | 0.061918908 | -0.029670964 | 0.091589872 | 0.999933887 | -0.010002149 | 0.059364076 | -0.069366225 | 0.828290659 |
| ENSG00000225450 | *AL021707.1* | -0.348508143 | 0.18346383 | 0.180709669 | 0.002754161 | 0.999933887 | -0.103366736 | -0.225796131 | 0.122429395 | 0.828573864 |
| ENSG00000043514 | *TRIT1* | 0.920430565 | -0.143334704 | 0.051214121 | -0.194548825 | 0.999933887 | 0.057117681 | 0.129099572 | -0.071981891 | 0.828573864 |
| ENSG00000264553 | *MIR4257* | -0.038359581 | 0.018475982 | 0.020127326 | -0.001651343 | 0.999933887 | 0.534858156 | 0.391664711 | 0.143193445 | 0.828642898 |
| ENSG00000103549 | *RNF40* | 6.418789856 | 0.028690778 | 0.026774533 | 0.001916245 | 0.999933887 | -0.22900832 | -0.264463982 | 0.035455661 | 0.828679379 |

| | | | | | | | | | | |
|---|---|---|---|---|---|---|---|---|---|---|
| ENSG00000145990 | *GFOD1* | 4.521265332 | -0.033229102 | 0.013438958 | -0.04666806 | 0.999933887 | -0.815975899 | -0.771841705 | -0.044134195 | 0.82873991 |
| ENSG00000168273 | *SMIM4* | 0.064501181 | 0.051133676 | -0.088568122 | 0.139701798 | 0.999933887 | -0.002435499 | -0.09853463 | 0.096099131 | 0.82873991 |
| ENSG00000267214 | *AC092068.1* | -0.510494598 | 0.052294389 | 0.140949799 | -0.08865541 | 0.999933887 | 0.216022842 | 0.09801874 | 0.118004102 | 0.82873991 |
| ENSG00000100979 | *PLTP* | -0.136797228 | -0.018532381 | -0.376677522 | 0.358145141 | 0.999933887 | -0.274604857 | -0.144224513 | -0.130380344 | 0.82873991 |
| ENSG00000061676 | *NCKAP1* | 0.408642355 | 0.093894616 | -0.052439932 | 0.146334548 | 0.999933887 | -0.306696763 | -0.398771089 | 0.092074327 | 0.829083148 |
| ENSG00000141425 | *RPRD1A* | 4.481935038 | -0.050843817 | -0.002662221 | -0.048181596 | 0.999933887 | -0.717053154 | -0.760728676 | 0.043675522 | 0.829163952 |
| ENSG00000185883 | *ATP6V0C* | 3.844390179 | 0.311177959 | 0.139869195 | 0.171308763 | 0.999933887 | 0.592219919 | 0.529775055 | 0.062444864 | 0.829365385 |
| ENSG00000100934 | *SEC23A* | 3.688216665 | 0.04815361 | 0.004041245 | 0.044112365 | 0.999933887 | -0.393179942 | -0.437112962 | 0.04393302 | 0.829365385 |
| ENSG00000163807 | *KIAA1143* | 3.197998766 | -0.104981478 | -0.09872162 | -0.006259858 | 0.999933887 | -0.064214395 | -0.021743931 | -0.042470464 | 0.829438414 |
| ENSG00000250959 | *GLUD1P3* | 0.31322239 | -0.14439128 | -0.093937032 | -0.050454249 | 0.999933887 | -0.049915634 | 0.03759491 | -0.087510544 | 0.829438414 |
| ENSG00000143951 | *WDPCP* | 0.463881996 | -0.112247967 | -0.540852544 | 0.428604578 | 0.944094537 | -0.071006814 | -0.166537333 | 0.095530519 | 0.829438414 |
| ENSG00000079435 | *LIPE* | 0.58260875 | 0.114581459 | 0.403319865 | -0.288738405 | 0.999933887 | -0.063221356 | -0.143401957 | 0.0801806 | 0.829438414 |
| ENSG00000163528 | *CHCHD4* | 0.294739047 | 0.099930163 | -0.058766179 | 0.158696342 | 0.999933887 | 0.114794622 | 0.022585383 | 0.09220924 | 0.829438414 |
| ENSG00000100226 | *GTPBP1* | 8.161634988 | 0.01056808 | 0.019487047 | -0.008918967 | 0.999933887 | 0.031310074 | 0.057658495 | -0.026348421 | 0.829438414 |
| ENSG00000163681 | *SLMAP* | 5.092597514 | 0.007855607 | -0.03091854 | 0.038774147 | 0.999933887 | -0.56165515 | -0.617301939 | 0.055646788 | 0.829438414 |
| ENSG00000287195 | *AC131568.1* | 3.491026071 | -0.121900076 | -0.00153028 | -0.120369796 | 0.999933887 | -0.02982291 | 0.064184297 | -0.094007207 | 0.829479248 |
| ENSG00000249850 | *KRT18P31* | -0.164064174 | 0.102752563 | 0.07486497 | 0.027887593 | 0.999933887 | 0.222622847 | 0.08820388 | 0.134418968 | 0.829728118 |
| ENSG00000085644 | *ZNF213* | 3.603238709 | 0.037935925 | -0.046758294 | 0.084694219 | 0.999933887 | -0.359150717 | -0.401053652 | 0.041902935 | 0.830041552 |
| ENSG00000113583 | *C5orf15* | 3.258177661 | -0.006354335 | -0.064344298 | 0.057989963 | 0.999933887 | -0.286501455 | -0.250147013 | -0.036354442 | 0.830123712 |
| ENSG00000180776 | *ZDHHC20* | 5.468962223 | 0.091208646 | 0.023986039 | 0.067222607 | 0.999933887 | -0.805026652 | -0.864691372 | 0.059664721 | 0.830123905 |
| ENSG00000075856 | *SART3* | 4.63596649 | -0.047744177 | -0.009663871 | -0.038080306 | 0.999933887 | 0.082569024 | 0.051074737 | 0.031494287 | 0.830212326 |
| ENSG00000167930 | *FAM234A* | 2.819477303 | 0.04199694 | 0.012188918 | 0.029808022 | 0.999933887 | -0.057094236 | -0.116781201 | 0.059686965 | 0.83042248 |
| ENSG00000135736 | *CCDC102A* | 0.378235776 | 0.157103694 | 0.123753138 | 0.033350556 | 0.999933887 | 0.183831796 | 0.280813074 | -0.096981279 | 0.830484581 |
| ENSG00000168904 | *LRRC28* | 1.838472849 | 0.001560372 | 0.105900929 | -0.104340557 | 0.999933887 | -0.03850933 | 0.023838168 | -0.062347498 | 0.83049125 |
| ENSG00000204227 | *RING1* | 3.676971141 | 0.130428966 | 0.055273542 | 0.075155424 | 0.999933887 | 0.079986037 | 0.033062884 | 0.046923152 | 0.830624989 |
| ENSG00000227201 | *CNN2P1* | -0.136449506 | 0.007012981 | 0.001853497 | 0.005159485 | 0.999933887 | 0.63057242 | 0.805273281 | -0.174700861 | 0.830624989 |
| ENSG00000124535 | *WRNIP1* | 3.460269655 | 0.084575617 | 0.21448517 | -0.129909553 | 0.999933887 | 0.121350237 | 0.081297184 | 0.040053053 | 0.830679665 |
| ENSG00000071894 | *CPSF1* | 4.746067779 | -0.031221192 | 0.033474359 | -0.06469555 | 0.999933887 | 0.14769269 | 0.187376569 | -0.039683878 | 0.830679665 |
| ENSG00000278864 | *AC055811.4* | 0.504602985 | -0.136476119 | -0.031603053 | -0.104873067 | 0.999933887 | 0.128317253 | 0.043295229 | 0.085022023 | 0.830945607 |
| ENSG00000198934 | *MAGEE1* | 0.071111257 | -0.043636155 | -0.044441469 | 0.000805314 | 0.999933887 | 0.057992344 | -0.043140046 | 0.101132389 | 0.830945607 |
| ENSG00000135069 | *PSAT1* | 0.679128961 | -0.024068997 | -0.120628252 | 0.096559255 | 0.999933887 | -0.038770642 | 0.032560914 | -0.071331557 | 0.831025279 |
| ENSG00000064961 | *HMG20B* | 5.320171945 | 0.054873695 | 0.046613796 | 0.0082599 | 0.999933887 | 0.007036855 | 0.037539779 | -0.030502924 | 0.831321386 |
| ENSG00000168778 | *TCTN2* | 0.000782838 | 0.219521902 | 0.134187644 | 0.085334258 | 0.999933887 | -0.11932577 | -0.206919654 | 0.087593885 | 0.831474723 |
| ENSG00000108523 | *RNF167* | 6.696968367 | 0.007207391 | 0.004394417 | 0.002812975 | 0.999933887 | 0.087330552 | 0.061149932 | 0.02618062 | 0.831569891 |
| ENSG00000261824 | *LINC00662* | 0.584771398 | -0.092986819 | 0.156392383 | -0.249379203 | 0.999933887 | -0.266993247 | -0.166402138 | -0.100591109 | 0.832191566 |
| ENSG00000058866 | *DGKG* | 1.471769858 | 0.077945734 | -0.018017407 | 0.095963141 | 0.999933887 | 0.128047645 | 0.047711132 | 0.080336513 | 0.832524486 |
| ENSG00000149582 | *TMEM25* | -0.155590396 | 0.126730261 | -0.132848488 | 0.25957875 | 0.999933887 | 0.159225537 | 0.04721554 | 0.112009997 | 0.832524486 |
| ENSG00000231327 | *LINC01816* | 0.900634671 | 0.272749787 | 0.52959782 | -0.256848033 | 0.999933887 | 0.957132684 | 1.055356049 | -0.098223365 | 0.832683956 |
| ENSG00000286511 | *Z95118.2* | 0.194138429 | -0.277999109 | -0.398781747 | 0.120782638 | 0.999933887 | -1.872434216 | -1.754458035 | -0.117976181 | 0.832683956 |
| ENSG00000161638 | *ITGA5* | 7.915851272 | -0.057952699 | -0.090203632 | 0.032250933 | 0.999933887 | -0.464577173 | -0.504357203 | 0.03978003 | 0.832683956 |
| ENSG00000160318 | *CLDND2* | 0.707643822 | -0.018880272 | 0.093894696 | -0.112774968 | 0.999933887 | 0.018737465 | 0.101631246 | -0.08289378 | 0.832683956 |
| ENSG00000103472 | *RRN3P2* | -0.117417379 | -0.184229333 | 0.025825693 | -0.210055027 | 0.999933887 | 0.100457862 | 0.188479276 | -0.088021414 | 0.832890474 |
| ENSG00000112306 | *RPS12* | 7.798646061 | -0.015336646 | -0.034740971 | 0.019404326 | 0.999933887 | 0.114252715 | 0.149507687 | -0.035254972 | 0.832890474 |
| ENSG00000138032 | *PPM1B* | 3.883566002 | -0.087511557 | -0.051873982 | -0.035637575 | 0.999933887 | -0.58567783 | -0.633128736 | 0.047450906 | 0.832914665 |
| ENSG00000259118 | *AL139022.1* | 1.244974522 | 0.120556698 | 0.017638545 | 0.102918153 | 0.999933887 | -0.380393167 | -0.471930038 | 0.09153687 | 0.833426347 |
| ENSG00000105205 | *CLC* | 5.35779413 | -0.060971016 | -0.007925503 | -0.053045513 | 0.999933887 | -0.097553793 | -0.147236602 | 0.049682809 | 0.833426347 |
| ENSG00000231181 | *AL954705.1* | -0.00688185 | -0.044751416 | -0.094083124 | 0.049331708 | 0.999933887 | 0.544828461 | 0.679279001 | -0.13445054 | 0.833426347 |
| ENSG00000251562 | *MALAT1* | 12.524352158 | 0.055451762 | 0.254226954 | -0.198775192 | 0.999933887 | 0.308817285 | 0.387869453 | -0.079052168 | 0.83349566 |
| ENSG00000188033 | *ZNF490* | 0.345484144 | 0.062519182 | -0.129267799 | 0.19178698 | 0.999933887 | -0.238896067 | -0.133765162 | -0.105130904 | 0.83349566 |
| ENSG00000189007 | *ADAT2* | 1.62259409 | -0.109142683 | -0.048446721 | -0.060695962 | 0.999933887 | 0.039517888 | 0.097133303 | -0.057615415 | 0.833629631 |
| ENSG00000157184 | *CPT2* | 1.26716504 | 0.083970729 | 0.080672318 | 0.003298411 | 0.999933887 | -0.236059014 | -0.307209513 | 0.0711505 | 0.833629631 |
| ENSG00000111275 | *ALDH2* | 1.485658808 | 0.028452255 | -0.125646066 | 0.154098322 | 0.999933887 | -0.101618368 | -0.211199145 | 0.109580777 | 0.833629631 |
| ENSG00000136144 | *RCBTB1* | 4.479185145 | -0.086141917 | -0.083635155 | -0.002506762 | 0.999933887 | -0.669989391 | -0.700724806 | 0.030735414 | 0.833789613 |
| ENSG00000211820 | *TRAV41* | -0.991136381 | -0.101940137 | -0.281647983 | 0.179707847 | 0.999933887 | 0.118252438 | 0.238178717 | -0.119926278 | 0.833789613 |
| ENSG00000162736 | *NCSTN* | 6.972441209 | -0.018731679 | 0.00243083 | -0.02116251 | 0.999933887 | -0.013999396 | 0.01221063 | -0.026210027 | 0.833789613 |
| ENSG00000006756 | *ARSD* | 1.697297042 | 0.147274781 | 0.018353572 | 0.128921209 | 0.999933887 | -0.031332355 | -0.104321105 | 0.072988751 | 0.833844742 |
| ENSG00000162552 | *WNT4* | 0.721934676 | 0.200687302 | 0.266189555 | -0.065502253 | 0.999933887 | 0.289913825 | 0.415510741 | -0.125596916 | 0.834208012 |
| ENSG00000106638 | *TBL2* | 1.963547182 | -0.05304608 | 0.036427299 | -0.089473379 | 0.999933887 | 0.104770854 | 0.052585802 | 0.052185052 | 0.834208012 |
| ENSG00000154358 | *OBSCN* | 3.171492949 | 0.092570267 | 0.07777037 | 0.014799898 | 0.999933887 | 0.075399622 | 0.130292441 | -0.054892819 | 0.834400208 |
| ENSG00000198590 | *C3orf35* | 0.45063743 | 0.128382875 | -0.116074054 | 0.24445693 | 0.999933887 | 0.039096337 | -0.04799763 | 0.087093967 | 0.834476616 |
| ENSG00000185215 | *TNFAIP2* | 10.995204246 | 0.078718042 | 0.093799324 | -0.015081282 | 0.999933887 | 1.096646219 | 1.13293185 | -0.036285632 | 0.834617168 |
| ENSG00000100216 | *TOMM22* | 3.067387094 | -0.015710279 | -0.012281205 | -0.003429074 | 0.999933887 | 0.103836945 | 0.149504484 | -0.045667539 | 0.834667298 |
| ENSG00000228327 | *AL669831.1* | 2.124858119 | 0.07669023 | 0.03928527 | 0.037404959 | 0.999933887 | 0.388810786 | 0.309827477 | 0.078983309 | 0.834726473 |
| ENSG00000232187 | *FTH1P7* | 1.786263196 | 0.031132872 | 0.025141329 | 0.005991543 | 0.999933887 | 0.607565 | 0.664080116 | -0.056515116 | 0.834873341 |
| ENSG00000182700 | *IGIP* | -0.099774147 | 0.025058743 | 0.124573538 | -0.099514795 | 0.999933887 | -0.487518559 | -0.588581607 | 0.101063048 | 0.834938915 |
| ENSG00000107669 | *ATE1* | 2.327436101 | -0.038954319 | -0.063586512 | 0.024632193 | 0.999933887 | -0.127768031 | -0.076193596 | -0.051574435 | 0.834997958 |
| ENSG00000286825 | *AL096870.10* | -0.196108402 | 0.454953515 | 0.54654439 | -0.091500876 | 0.999933887 | 1.35072105 | 1.482944907 | -0.132223856 | 0.835020223 |
| ENSG00000188375 | *H3-5* | 0.221775709 | 0.078528453 | 0.096225512 | -0.017697059 | 0.999933887 | 0.272718289 | 0.381195772 | -0.108477483 | 0.835020223 |
| ENSG00000134897 | *BIVM* | -0.277969787 | 0.025634107 | -0.057458144 | 0.083092251 | 0.999933887 | 0.156077775 | 0.020861319 | 0.135216455 | 0.835020223 |
| ENSG00000163737 | *PF4* | -0.837679687 | -0.018495914 | 0.025620252 | -0.044116166 | 0.999933887 | -0.09578744 | 0.046098508 | -0.141885949 | 0.835020223 |
| ENSG00000188177 | *ZC3H6* | 3.772931119 | 0.09567443 | -0.108114586 | 0.203789016 | 0.999933887 | -0.242587633 | -0.183518046 | -0.059069587 | 0.835034304 |
| ENSG00000136003 | *ISCU* | 6.111991189 | -0.008734519 | -0.058699731 | 0.049965212 | 0.999933887 | 0.047976841 | 0.019104499 | 0.028872342 | 0.835034304 |
| ENSG00000067191 | *CACNB1* | 1.165045212 | 0.025821229 | -0.134557036 | 0.160378265 | 0.999933887 | -0.052166382 | 0.015174905 | -0.067341287 | 0.835306869 |
| ENSG00000165983 | *PTER* | 1.076350886 | 0.079625365 | -0.066165206 | 0.145790571 | 0.999933887 | -0.252621789 | -0.17746024 | -0.075161549 | 0.835426731 |
| ENSG00000263002 | *ZNF234* | 0.349997395 | -0.213882551 | -0.052125159 | -0.161757392 | 0.999933887 | -0.034632563 | 0.040716741 | -0.075349304 | 0.835457339 |
| ENSG00000157927 | *RADIL* | -0.702643241 | 0.031987075 | 0.387079708 | -0.355092632 | 0.999933887 | 2.05119794 | 2.190030326 | -0.138832386 | 0.835457339 |
| ENSG00000110042 | *DTX4* | 4.585632706 | -0.141266085 | -0.127558354 | -0.013707731 | 0.999933887 | 0.257703993 | 0.189007443 | 0.06869655 | 0.83563846 |
| ENSG00000279432 | *AC015799.1* | 2.313073297 | -0.006225919 | 0.010543783 | -0.016769702 | 0.999933887 | 0.554417695 | 0.630576872 | -0.076159177 | 0.835945387 |
| ENSG00000054392 | *HHAT* | -0.501924616 | -0.16522742 | 0.090705716 | -0.255933135 | 0.999933887 | 0.040044731 | 0.193675008 | -0.153630277 | 0.836072385 |
| ENSG00000173531 | *MST1* | 0.342055362 | 0.097373123 | 0.097952052 | -0.000578928 | 0.999933887 | 0.061331781 | -0.040925754 | 0.102257535 | 0.836072385 |
| ENSG00000287812 | *AC105046.2* | 0.596433626 | 0.041545751 | -0.002627131 | 0.044172883 | 0.999933887 | 0.987466387 | 0.823863272 | 0.163603115 | 0.836072385 |
| ENSG00000272911 | *AC005696.3* | -0.01914648 | 0.324661187 | 0.365263894 | -0.040602707 | 0.999933887 | 1.201795265 | 1.310055467 | -0.108260203 | 0.836097857 |
| ENSG00000204387 | *SNHG32* | 2.479518049 | 0.173486737 | -0.017969365 | 0.191456102 | 0.999933887 | 0.226259418 | 0.169020784 | 0.057238634 | 0.836097857 |
| ENSG00000125459 | *MSTO1* | 1.457358863 | -0.160436204 | -0.10332734 | -0.057108864 | 0.999933887 | 0.101510511 | 0.164515115 | -0.063004604 | 0.836097857 |
| ENSG00000096063 | *SRPK1* | 7.219637542 | -0.056828131 | -0.007660752 | -0.04916738 | 0.999933887 | -0.48570465 | -0.521540636 | 0.035835986 | 0.836097857 |
| ENSG00000104472 | *CHRAC1* | 4.294822429 | 0.022791878 | 0.007822582 | 0.014969297 | 0.999933887 | -0.2575232 | -0.294348577 | 0.036825377 | 0.836097857 |
| ENSG00000189114 | *BLOC1S3* | 5.905358467 | 0.017465916 | 0.069120293 | -0.051654377 | 0.999933887 | -0.197256495 | -0.226749101 | 0.029492606 | 0.836097857 |
| ENSG00000285816 | *AP000944.5* | 0.37153934 | 0.005461672 | -0.024808227 | 0.030269899 | 0.999933887 | 0.90870185 | 1.034961972 | -0.126260122 | 0.836097857 |
| ENSG00000127152 | *BCL11B* | 5.025856738 | -0.001629583 | 0.049359829 | -0.050989412 | 0.999933887 | 0.042900122 | 0.006798978 | 0.036101144 | 0.836132812 |
| ENSG00000104626 | *ERI1* | 2.982722351 | -0.046090493 | -0.038698022 | -0.007392471 | 0.999933887 | -0.164604468 | -0.226035586 | 0.061431118 | 0.836205348 |
| ENSG00000251215 | *GOLGA5P1* | 0.665142344 | -0.404601814 | -0.321290434 | -0.083311381 | 0.999933887 | -0.676310953 | -0.590784071 | -0.085526882 | 0.836212673 |
| ENSG00000096080 | *MRPS18A* | 1.449385683 | 0.075937629 | 0.099371042 | -0.023433413 | 0.999933887 | 0.190801037 | 0.134407906 | 0.056393131 | 0.836212673 |
| ENSG00000127946 | *HIP1* | 6.64121787 | -0.069553571 | -0.038151274 | -0.031402297 | 0.999933887 | 0.901200814 | 0.965847124 | -0.064646309 | 0.836377812 |
| ENSG00000272908 | *AC006033.2* | -0.642793908 | -0.176023106 | 0.057389924 | -0.23341303 | 0.999933887 | 0.098041277 | -0.048064082 | 0.146105359 | 0.836377812 |
| ENSG00000124243 | *BCAS4* | 1.643293453 | 0.053896375 | 0.093774961 | -0.039878586 | 0.999933887 | 0.028101263 | 0.096999805 | -0.068898542 | 0.836377812 |
| ENSG00000282420 | *TRBJ1-2* | -1.069708116 | -0.121033435 | -0.096110166 | -0.024923269 | 0.999933887 | 0.126713595 | 0.257113901 | -0.130400306 | 0.836377812 |
| ENSG00000168256 | *NKIRAS2* | 3.934467497 | 0.034117977 | 0.136705346 | -0.102587369 | 0.999933887 | -0.085609804 | -0.126509722 | 0.040899918 | 0.836377812 |
| ENSG00000260032 | *NORAD* | 7.006846731 | -0.081804052 | -0.09355649 | 0.011752438 | 0.999933887 | -0.365373762 | -0.39657584 | 0.031202078 | 0.836511055 |
| ENSG00000089177 | *KIF16B* | 0.900105969 | 0.163406863 | -0.211584557 | 0.37499142 | 0.999933887 | 0.009918121 | 0.098617739 | -0.088699618 | 0.836511055 |
| ENSG00000244462 | *RBM12* | 5.603479609 | -0.046089839 | 0.012510465 | -0.058600304 | 0.999933887 | -0.15533952 | -0.129279506 | -0.026060015 | 0.836511055 |
| ENSG00000178700 | *DHFR2* | 0.985001665 | -0.119221018 | -0.113465461 | -0.005755557 | 0.999933887 | -0.206137181 | -0.146461341 | -0.05967584 | 0.836511055 |
| ENSG00000042286 | *AIFM2* | 0.667614734 | 0.095191564 | 0.256636909 | -0.161445345 | 0.999933887 | 0.080318097 | 0.148238453 | -0.067920357 | 0.836511055 |
| ENSG00000186818 | *LILRB4* | 2.566757383 | -0.058946805 | -0.029153112 | -0.029793693 | 0.999933887 | -0.495984093 | -0.548808481 | 0.052824387 | 0.836511055 |
| ENSG00000132205 | *EMILIN2* | 5.100761019 | 0.03696214 | 0.029308646 | 0.007653495 | 0.999933887 | -0.086476688 | -0.129738389 | 0.0432617 | 0.836511055 |
| ENSG00000167740 | *CYB5D2* | 1.626218367 | -0.057296147 | 0.103115677 | -0.160411824 | 0.999933887 | 0.063393661 | 0.00232146 | 0.061072201 | 0.836511055 |

| | | | | | | | | | | |
|---|---|---|---|---|---|---|---|---|---|---|
| ENSG00000176472 | *ZNF575* | -0.382337798 | -0.089161417 | -0.097008366 | 0.00784695 | 0.999933887 | 0.037243627 | 0.157036576 | -0.119792948 | 0.836511055 |
| ENSG00000159200 | *RCAN1* | 1.506689428 | 0.042978144 | 0.080663302 | -0.037685159 | 0.999933887 | -0.313109307 | -0.392515159 | 0.079405851 | 0.836511055 |
| ENSG00000132664 | *POLR3F* | 1.343092898 | -0.024066172 | 0.019768499 | -0.043834671 | 0.999933887 | 0.031137826 | -0.03355608 | 0.064693906 | 0.836511055 |
| ENSG00000197008 | *ZNF138* | 0.044851292 | -0.040666316 | -0.130140222 | 0.089473906 | 0.999933887 | -0.288000199 | -0.417331677 | 0.129331478 | 0.836548003 |
| ENSG00000247137 | *AP000873.2* | 1.656847248 | 0.161017383 | 0.025199138 | 0.135818245 | 0.999933887 | 0.100956603 | 0.032879251 | 0.068077352 | 0.836564134 |
| ENSG00000162772 | *ATF3* | 4.665826984 | 0.093765894 | 0.120422955 | -0.026657062 | 0.999933887 | -0.016331462 | -0.06760454 | 0.051273078 | 0.836564134 |
| ENSG00000185684 | *EP400P1* | 1.393056215 | 0.096347252 | 0.062058478 | 0.034288774 | 0.999933887 | -0.045102333 | 0.030767357 | -0.07586969 | 0.836564134 |
| ENSG00000120697 | *ALG5* | 0.976810475 | 0.044782493 | -0.21292672 | 0.257709213 | 0.999933887 | 0.193380059 | 0.119714006 | 0.073666054 | 0.836564134 |
| ENSG00000133256 | *PDE6B* | 0.532674174 | -0.054191114 | 0.055271104 | -0.109462218 | 0.999933887 | 0.193147328 | 0.106971153 | 0.086176175 | 0.836564134 |
| ENSG00000163634 | *THOC7* | 2.773763286 | -0.010073226 | -0.087491787 | 0.077418561 | 0.999933887 | -0.012329321 | 0.042490336 | -0.054819657 | 0.836564134 |
| ENSG00000133835 | *HSD17B4* | 4.983777026 | -0.202221516 | -0.049 | -0.153221515 | 0.999933887 | -1.272620637 | -1.324941301 | 0.052320664 | 0.836765116 |
| ENSG00000126860 | *EVI2A* | 6.480985362 | -0.102749578 | -0.19600967 | 0.093260092 | 0.999933887 | -1.103373123 | -1.050809513 | -0.05256361 | 0.836944773 |
| ENSG00000163531 | *NFASC* | 0.001961486 | 0.009824841 | 0.122542861 | -0.11271802 | 0.999933887 | 0.301315152 | 0.170984649 | 0.130330502 | 0.836967234 |
| ENSG00000110906 | *KCTD10* | 4.282947187 | -0.076935778 | -0.008544104 | -0.068391674 | 0.999933887 | -0.109276802 | -0.14030942 | 0.031032618 | 0.837019615 |
| ENSG00000271742 | *AL121992.2* | -0.240771185 | 0.301792324 | 0.188135626 | 0.113656698 | 0.999933887 | 0.939676018 | 1.050989145 | -0.111313128 | 0.837152973 |
| ENSG00000149328 | *GLB1L2* | 1.064724972 | 0.085695416 | -0.093106959 | 0.178802375 | 0.999933887 | -0.043734888 | 0.03621605 | -0.079950938 | 0.837476921 |
| ENSG00000143842 | *SOX13* | 1.808312398 | -0.056374925 | 0.033529727 | -0.089904651 | 0.999933887 | -0.192297571 | -0.24100145 | 0.048703878 | 0.837476921 |
| ENSG00000177311 | *ZBTB38* | 3.994982931 | -0.028328652 | -0.145717544 | 0.117388893 | 0.999933887 | -0.224604667 | -0.164844372 | -0.059760295 | 0.837476921 |
| ENSG00000234709 | *UPF3AP3* | -0.069051062 | 0.106591855 | 0.261529272 | -0.154937417 | 0.999933887 | -0.001435997 | 0.161101265 | -0.162537262 | 0.837611723 |
| ENSG00000130772 | *MED18* | 3.113845905 | -0.031627448 | -0.171399208 | 0.13977176 | 0.999933887 | 0.204291748 | 0.155578689 | 0.048713059 | 0.837611723 |
| ENSG00000105379 | *ETFB* | 1.835089444 | -0.017355051 | 0.069820506 | -0.087175556 | 0.999933887 | 0.192192655 | 0.2531599 | -0.060967245 | 0.837611723 |
| ENSG00000231535 | *LINC00278* | -3.761791396 | 0.148077183 | -0.287697425 | 0.435774608 | 0.999933887 | -0.458665263 | -0.556740845 | 0.098075582 | 0.837679461 |
| ENSG00000197461 | *PDGFA* | 0.236406574 | -0.145782512 | -0.235360082 | 0.08957757 | 0.999933887 | 0.313506183 | 0.419476903 | -0.105970719 | 0.837801469 |
| ENSG00000062524 | *LTK* | 1.236139678 | -0.048975306 | 0.128752319 | -0.177727625 | 0.999933887 | 0.112287734 | 0.192677055 | -0.080389321 | 0.837801469 |
| ENSG00000215883 | *CYB5RL* | -0.397204683 | -0.245483057 | 0.340321529 | -0.585804586 | 0.999933887 | -0.047243087 | -0.15626448 | 0.109021393 | 0.838062629 |
| ENSG00000175193 | *PARL* | 0.531531854 | -0.129303094 | -0.050099932 | -0.079203162 | 0.999933887 | 0.087451069 | -0.015848209 | 0.103299278 | 0.8381075 |
| ENSG00000100890 | *PRORP* | 0.745395159 | 0.147513988 | 0.003508068 | 0.14400592 | 0.999933887 | 0.107563332 | 0.032267409 | 0.075295923 | 0.838154449 |
| ENSG00000181061 | *HIGD1A* | 2.48021113 | -0.057114381 | 0.0609866 | -0.118100981 | 0.999933887 | 0.062978777 | 0.009825573 | 0.053153204 | 0.838154449 |
| ENSG00000094914 | *AAAS* | 3.312725466 | -0.051235724 | 0.033549631 | -0.084785355 | 0.999933887 | 0.007012245 | 0.048095602 | -0.041083358 | 0.838380616 |
| ENSG00000162086 | *ZNF75A* | 2.506622089 | -0.104543008 | -0.124511084 | 0.019968076 | 0.999933887 | -0.174782115 | -0.21999581 | 0.045213694 | 0.838628978 |
| ENSG00000170222 | *ADPRM* | 1.797844632 | 0.10971294 | -0.046653696 | 0.156366637 | 0.999933887 | -0.029584973 | 0.029989818 | -0.05957479 | 0.838657809 |
| ENSG00000225792 | *AC004540.2* | 0.48590996 | -0.165589816 | 0.07917022 | -0.244760036 | 0.999933887 | 2.278384546 | 2.416267884 | -0.137883338 | 0.838657809 |
| ENSG00000103995 | *CEP152* | 1.384211334 | -0.060553385 | -0.118324393 | 0.057771008 | 0.999933887 | -0.145249397 | -0.046975216 | -0.098274181 | 0.838815532 |
| ENSG00000226287 | *TMEM191A* | 0.545427565 | 0.015428267 | 0.023883722 | -0.008455455 | 0.999933887 | 0.078204147 | -0.010335365 | 0.088539513 | 0.839048114 |
| ENSG00000233461 | *AL445524.1* | 1.537223083 | 0.049253281 | -0.008747386 | 0.058000668 | 0.999933887 | 0.038803781 | -0.055289196 | 0.094092977 | 0.839058439 |
| ENSG00000104381 | *GDAP1* | 0.002728631 | -0.140637139 | 0.000843578 | -0.141480717 | 0.999933887 | 0.134507463 | 0.024590921 | 0.109916542 | 0.839305998 |
| ENSG00000114988 | *LMAN2L* | 1.159460384 | 0.060627833 | -0.034236362 | 0.094864195 | 0.999933887 | -0.157764245 | -0.103531104 | -0.054233142 | 0.839305998 |
| ENSG00000250739 | *LINC01262* | 1.199776628 | 0.221267808 | 0.298360402 | -0.077092594 | 0.999933887 | 1.853329963 | 1.960170499 | -0.106840536 | 0.839476536 |
| ENSG00000280407 | *AC132872.4* | 4.01591697 | 0.046393241 | 0.119509945 | -0.073116704 | 0.999933887 | 0.012926492 | -0.036062058 | 0.048988551 | 0.839476536 |
| ENSG00000136404 | *TM6SF1* | 3.280000294 | 0.072912681 | -0.060059138 | 0.132971819 | 0.999933887 | -0.576715598 | -0.628717545 | 0.052001947 | 0.839531134 |
| ENSG00000065491 | *TBC1D22B* | 2.26277401 | -0.0444452 | 0.006951795 | -0.051396995 | 0.999933887 | -0.005865522 | 0.047189444 | -0.053054966 | 0.839531134 |
| ENSG00000163382 | *NAXE* | 1.950168005 | -0.006711575 | -0.017440702 | 0.010729127 | 0.999933887 | 0.13153219 | 0.064850299 | 0.066681892 | 0.839531134 |
| ENSG00000260417 | *AC092127.1* | -0.237392886 | 0.040496825 | 0.157611989 | -0.117115164 | 0.999933887 | 0.483907272 | 0.63636976 | -0.152462488 | 0.839901277 |
| ENSG00000173890 | *GPR160* | 5.277136815 | 0.057750858 | 0.052300293 | 0.005450565 | 0.999933887 | -0.952768999 | -0.904669382 | -0.048099618 | 0.839997455 |
| ENSG00000042062 | *RIPOR3* | 0.765552032 | 0.06460444 | 0.068064153 | -0.003459713 | 0.999933887 | 0.369613419 | 0.518445857 | -0.148832437 | 0.840004167 |
| ENSG00000006025 | *OSBPL7* | 3.564398768 | 0.018931206 | -0.019368693 | 0.038299898 | 0.999933887 | 0.096771628 | 0.059715807 | 0.03705582 | 0.840004167 |
| ENSG00000277283 | *AC004812.2* | 0.444517406 | -0.052440437 | 0.125349504 | -0.177789941 | 0.999933887 | 0.478070624 | 0.374492049 | 0.103578575 | 0.840004167 |
| ENSG00000226791 | *LINC02611* | -0.613830809 | 0.045146068 | -0.209087077 | 0.254233145 | 0.999933887 | 0.161533212 | 0.281682307 | -0.120149096 | 0.840004167 |
| ENSG00000196169 | *KIF19* | 0.5328252 | 0.081881645 | -0.054603917 | 0.136485562 | 0.999933887 | 0.064129832 | -0.010427647 | 0.074557479 | 0.8400169 |
| ENSG00000230068 | *CDC42-IT1* | -0.307006488 | 0.079154831 | -0.010205105 | 0.089359936 | 0.999933887 | 0.181078643 | 0.311516672 | -0.130438028 | 0.8400169 |
| ENSG00000137460 | *FHDC1* | 3.456233788 | 0.016495026 | -0.001210246 | 0.017705272 | 0.999933887 | 0.403216426 | 0.439414372 | -0.036197946 | 0.8400169 |
| ENSG00000254211 | *LINC01485* | 0.078168792 | 0.020050522 | 0.205397019 | -0.185346497 | 0.999933887 | 0.464428031 | 0.385288004 | 0.079140027 | 0.8400169 |
| ENSG00000051009 | *FAM160A2* | 6.240408737 | 0.057464792 | 0.016244769 | 0.041220023 | 0.999933887 | 1.144311614 | 1.197644312 | -0.053332698 | 0.840070552 |
| ENSG00000285554 | *AC242988.2* | 0.468452868 | -0.003114808 | -0.00492244 | 0.001807632 | 0.999933887 | 0.287711402 | 0.388255651 | -0.100544249 | 0.840070552 |
| ENSG00000262870 | *CYCSP40* | 0.407415704 | -0.167656086 | -0.038734694 | -0.128921392 | 0.999933887 | -0.516826342 | -0.415406735 | -0.101419608 | 0.840078191 |
| ENSG00000183161 | *FANCF* | 0.694962108 | 0.027233465 | 0.054189623 | -0.026956158 | 0.999933887 | 0.069749955 | -0.002716903 | 0.072466858 | 0.840092193 |
| ENSG00000168014 | *C2CD3* | 4.846131185 | -0.014363096 | -0.040422407 | 0.02605931 | 0.999933887 | -0.097936171 | -0.068862259 | -0.029073912 | 0.840180281 |
| ENSG00000235172 | *LINC01366* | 4.352811525 | 0.177584521 | 0.160279212 | 0.017305309 | 0.999933887 | 1.479384472 | 1.533986817 | -0.054602345 | 0.840505524 |
| ENSG00000012963 | *UBR7* | 2.699787558 | -0.08193517 | 0.032315001 | -0.114250171 | 0.999933887 | 0.033926631 | 0.084353263 | -0.050426632 | 0.840585889 |
| ENSG00000077312 | *SNRPA* | 3.259690121 | -0.020367616 | -0.017074258 | -0.003293358 | 0.999933887 | 0.012894959 | -0.026920326 | 0.039815285 | 0.840585889 |
| ENSG00000109586 | *GALNT7* | 3.501120413 | 0.007214305 | -0.148560522 | 0.155774827 | 0.999933887 | -0.090857858 | -0.040745875 | -0.050111983 | 0.840585889 |
| ENSG00000269981 | *AL627309.7* | 4.16487672 | 0.073946681 | -0.03143523 | 0.10538191 | 0.999933887 | 0.312642026 | 0.261400732 | 0.051241294 | 0.840887473 |
| ENSG00000156858 | *PRR14* | 6.338256423 | 0.045033793 | 0.038223652 | 0.006810141 | 0.999933887 | 0.473594133 | 0.503584851 | -0.029990717 | 0.840887473 |
| ENSG00000211695 | *TRGV9* | -0.027409412 | -0.030964007 | -0.057019912 | 0.026055905 | 0.999933887 | 0.174892098 | 0.090626432 | 0.084265666 | 0.840887473 |
| ENSG00000139737 | *SLAIN1* | 1.480023109 | 0.115974585 | -0.007610961 | 0.123585546 | 0.999933887 | 0.05422502 | -0.008665859 | 0.062890878 | 0.840897401 |
| ENSG00000063177 | *RPL18* | 7.328951388 | 0.012261725 | -0.028659796 | 0.040921521 | 0.999933887 | 0.089924703 | 0.121693521 | -0.031768818 | 0.841043405 |
| ENSG00000268858 | *AL118506.1* | 4.384688318 | -0.010080192 | 0.109895205 | -0.119975397 | 0.999933887 | -0.342486787 | -0.301162773 | -0.041324014 | 0.8412405 |
| ENSG00000198113 | *TOR4A* | 4.806768508 | 0.064549704 | 0.089954698 | -0.025404993 | 0.999933887 | -0.462402955 | -0.506583589 | 0.044180633 | 0.841463086 |
| ENSG00000163913 | *IFT122* | 2.719709638 | -0.037633956 | 0.015231081 | -0.052865037 | 0.999933887 | 0.010178414 | -0.041250434 | 0.051428848 | 0.841624137 |
| ENSG00000226259 | *GTF2H2B* | -0.723716482 | 0.066728645 | -0.183737668 | 0.250466313 | 0.999933887 | -0.581732071 | -0.722746681 | 0.141014611 | 0.841624137 |
| ENSG00000235954 | *TTC28-AS1* | 1.61676261 | 0.023760323 | 0.203895812 | -0.180135489 | 0.999933887 | -0.053748061 | -0.119957395 | 0.066209334 | 0.841624137 |
| ENSG00000177374 | *HIC1* | 3.628198081 | -0.009149857 | -0.009694323 | 0.000544466 | 0.999933887 | 0.181656381 | 0.237408005 | -0.055751624 | 0.841624137 |
| ENSG00000099810 | *MTAP* | 0.934872239 | -0.006503686 | 0.289251953 | -0.295755639 | 0.999933887 | -0.051285832 | 0.020272401 | -0.071558233 | 0.841958546 |
| ENSG00000185189 | *NRBP2* | 1.733601024 | 0.173612322 | 0.056816377 | 0.116795945 | 0.999933887 | 0.013699809 | 0.067552354 | -0.053852546 | 0.841964782 |
| ENSG00000280587 | *LINC01348* | -0.717916492 | 0.108449182 | -0.201799667 | 0.310248849 | 0.999933887 | 2.378504417 | 2.21141497 | 0.167089447 | 0.842018816 |
| ENSG00000276961 | *MIR7848* | -0.118261659 | -0.028630286 | -0.095468545 | 0.06683826 | 0.999933887 | 0.155147069 | -0.007898999 | 0.163046069 | 0.84214854 |
| ENSG00000169189 | *NSMCE1* | 3.216562569 | -0.061070032 | -0.095496776 | 0.034426743 | 0.999933887 | -0.032236069 | 0.00875569 | -0.04099176 | 0.842151091 |
| ENSG00000005486 | *RHBDD2* | 5.670805785 | 0.034032512 | -0.000273581 | 0.034306093 | 0.999933887 | -0.013219418 | -0.042283238 | 0.029063821 | 0.842158553 |
| ENSG00000129472 | *RAB2B* | 2.803698822 | -0.091165024 | -0.075317921 | -0.015847103 | 0.999933887 | 0.12187064 | 0.073582735 | 0.048287906 | 0.842251015 |
| ENSG00000167840 | *ZNF232* | -0.6973944 | -0.158967666 | 0.191280861 | -0.350248527 | 0.999933887 | 0.300194728 | 0.417890179 | -0.117695452 | 0.842251015 |
| ENSG00000181894 | *ZNF329* | 1.29730142 | -0.099438106 | -0.193499961 | 0.094061855 | 0.999933887 | -0.074237239 | -0.005413723 | -0.068823516 | 0.842408964 |
| ENSG00000091536 | *MYO15A* | -0.382639443 | 0.098957322 | -0.019695939 | 0.118653261 | 0.999933887 | -0.206205747 | -0.075654919 | -0.130550828 | 0.842605936 |
| ENSG00000250317 | *SMIM20* | 1.389609619 | 0.037675965 | 0.000951533 | 0.036724432 | 0.999933887 | 0.059538823 | 0.131309038 | -0.071770215 | 0.842605936 |
| ENSG00000251586 | *TET2-AS1* | -0.071896071 | 0.173229255 | 0.490857017 | -0.317627763 | 0.999933887 | 0.021063467 | 0.176064246 | -0.155000779 | 0.84274191 |
| ENSG00000228719 | *AL022313.2* | -1.322920868 | 0.443040563 | 0.242094405 | 0.200946157 | 0.999933887 | 0.624753716 | 0.383606242 | 0.241147474 | 0.842866013 |
| ENSG00000175928 | *LRRN1* | 0.002871492 | 0.04092968 | -0.125060346 | 0.165990027 | 0.999933887 | -0.742779538 | -0.843223045 | 0.100443507 | 0.842955874 |
| ENSG00000170949 | *ZNF160* | 3.778970924 | -0.022802509 | -0.071987986 | 0.049185477 | 0.999933887 | -0.482622221 | -0.433304414 | -0.049317808 | 0.843007095 |
| ENSG00000175445 | *LPL* | 0.577297202 | -0.001127594 | -0.205433087 | 0.204305493 | 0.999933887 | -0.963415511 | -1.041053585 | 0.077638074 | 0.843101662 |
| ENSG00000109743 | *BST1* | 6.303015747 | -0.021455515 | 0.008597027 | -0.030052543 | 0.999933887 | -0.352699888 | -0.319521437 | -0.033178451 | 0.843140182 |
| ENSG00000198576 | *ARC* | -0.176409814 | -0.183637518 | -0.129984786 | -0.053652732 | 0.999933887 | -0.125159599 | -0.027443733 | -0.097715866 | 0.843153598 |
| ENSG00000225697 | *SLC26A6* | 3.522568911 | -0.052597647 | 0.015978737 | -0.068576384 | 0.999933887 | -0.063732162 | -0.110141632 | 0.04640947 | 0.843575364 |
| ENSG00000249673 | *NOP14-AS1* | 2.920011874 | 0.061714013 | -0.016840667 | 0.078554679 | 0.999933887 | 0.053842521 | 0.003799205 | 0.050043316 | 0.843676263 |
| ENSG00000173933 | *RBM4* | 3.541897527 | -0.013903466 | -0.039232883 | 0.025329417 | 0.999933887 | 0.063991214 | 0.028592048 | 0.035399165 | 0.843676263 |
| ENSG00000117362 | *APH1A* | 6.395187056 | -0.010424211 | -0.00657459 | -0.00384962 | 0.999933887 | -0.115137794 | -0.141009382 | 0.025871588 | 0.843676263 |
| ENSG00000244734 | *HBB* | 8.098226389 | -0.103908279 | -0.108651465 | 0.004743187 | 0.999933887 | 0.082009803 | 0.140273612 | -0.058263809 | 0.843830609 |
| ENSG00000139350 | *NEDD1* | 2.275750374 | 0.101229695 | -0.03579753 | 0.137027224 | 0.999933887 | -0.235871213 | -0.131827605 | -0.104043608 | 0.843830609 |
| ENSG00000141376 | *BCAS3* | 3.254805611 | 0.007953874 | 0.044906774 | -0.0369529 | 0.999933887 | 0.066123817 | 0.023748842 | 0.042374975 | 0.844336319 |
| ENSG00000161970 | *RPL26* | 3.695693902 | -0.101576097 | -0.115655701 | 0.014079604 | 0.999933887 | 0.100001295 | 0.14256634 | -0.042565045 | 0.84435925 |
| ENSG00000137266 | *SLC22A23* | 0.143508222 | 0.056017858 | 0.041315584 | 0.014702274 | 0.999933887 | 0.114722522 | 0.010838265 | 0.103884257 | 0.84435925 |
| ENSG00000135046 | *ANXA1* | 7.468454852 | -0.079879897 | -0.187629068 | 0.107749171 | 0.999933887 | -0.09551932 | -0.060213797 | -0.035305523 | 0.84482782 |
| ENSG00000149100 | *EIF3M* | 4.186076086 | -0.048307465 | -0.145317877 | 0.097010412 | 0.999933887 | 0.024384247 | 0.060229814 | -0.035845567 | 0.844939126 |
| ENSG00000268362 | *AC092279.1* | 5.071329704 | -0.011836177 | 0.058143209 | -0.069979386 | 0.999933887 | -0.294385134 | -0.356459365 | 0.062074231 | 0.844939126 |
| ENSG00000085982 | *USP40* | 1.168694827 | -0.051952289 | 0.002976248 | -0.054928537 | 0.999933887 | 0.059396654 | 0.127290437 | -0.067893782 | 0.8451025 |
| ENSG00000274523 | *RCC1L* | 2.264657448 | 0.023554249 | 0.12854822 | -0.104993971 | 0.999933887 | 0.206326079 | 0.16373662 | 0.042589459 | 0.8451025 |
| ENSG00000185896 | *LAMP1* | 8.814551765 | 0.020237368 | -0.003742779 | 0.023980147 | 0.999933887 | 0.148971807 | 0.172940396 | -0.023968589 | 0.845247275 |

| | | | | | | | | | | |
|---|---|---|---|---|---|---|---|---|---|---|
| ENSG00000213277 | *MARCKSL1P1* | 0.264459812 | -0.094593041 | -0.037247561 | -0.05734548 | 0.999933887 | 0.314445609 | 0.391789942 | -0.077344332 | 0.845280339 |
| ENSG00000251682 | *AC122718.2* | 0.407609251 | 0.260148187 | 0.492271371 | -0.232123184 | 0.999933887 | 0.450795966 | 0.279486585 | 0.171309381 | 0.845295155 |
| ENSG00000150687 | *PRSS23* | 1.720830146 | -0.047723581 | -0.194281573 | 0.146557992 | 0.999933887 | 0.017843778 | 0.073347953 | -0.055504175 | 0.845342578 |
| ENSG00000184007 | *PTP4A2* | 7.39635215 | -0.04205284 | -0.014661975 | -0.027390865 | 0.999933887 | -0.174679019 | -0.201211152 | 0.026532133 | 0.845490491 |
| ENSG00000172428 | *COPS9* | 2.502503573 | 0.068493284 | -0.055863403 | 0.124356687 | 0.999933887 | -0.01395911 | -0.068650679 | 0.054691569 | 0.845490491 |
| ENSG00000099250 | *NRP1* | 0.317975675 | -0.137302269 | 0.270445733 | -0.407748002 | 0.999933887 | -0.706548516 | -0.648087366 | -0.05846115 | 0.845706183 |
| ENSG00000196943 | *NOP9* | 2.966356706 | -0.000787378 | -0.064005042 | 0.063217664 | 0.999933887 | 0.16503876 | 0.124185968 | 0.040852792 | 0.845706183 |
| ENSG00000151240 | *DIP2C* | 0.004908462 | -0.401087107 | -0.166427266 | -0.234659841 | 0.999933887 | -0.190259517 | -0.08381347 | -0.106446047 | 0.845742665 |
| ENSG00000286724 | *AL732509.1* | 0.06052086 | 0.215091744 | 0.032973614 | 0.18211813 | 0.999933887 | -0.413138094 | -0.522550015 | 0.109411921 | 0.845742665 |
| ENSG00000116406 | *EDEM3* | 4.568783275 | 0.004274685 | -0.12673195 | 0.131006635 | 0.999933887 | -0.460682013 | -0.395921682 | -0.064760331 | 0.845742665 |
| ENSG00000141873 | *SLC39A3* | 2.596880936 | -0.033975517 | -0.074335745 | 0.040360228 | 0.999933887 | -0.359770841 | -0.409189978 | 0.049419136 | 0.845852696 |
| ENSG00000008130 | *NADK* | 9.749815879 | 0.000351641 | 0.016587051 | -0.016235409 | 0.999933887 | 0.141884239 | 0.121664667 | 0.020219572 | 0.845852696 |
| ENSG00000179862 | *CITED4* | 1.973016732 | 0.273573621 | 0.100017339 | 0.173556282 | 0.999933887 | 2.264423967 | 2.155712288 | 0.108711679 | 0.845978095 |
| ENSG00000081181 | *ARG2* | -0.724913244 | -0.042842176 | 0.079646052 | -0.122488228 | 0.999933887 | 0.264584327 | 0.413490388 | -0.148906062 | 0.846040153 |
| ENSG00000256043 | *CTSO* | 1.328694994 | -0.025407382 | -0.187353293 | 0.161945911 | 0.999933887 | -0.117574888 | -0.058198079 | -0.05937681 | 0.846173439 |
| ENSG00000260368 | *AC027373.1* | 0.142371194 | 0.190168278 | 0.393749865 | -0.203581586 | 0.999933887 | 0.034048103 | -0.079665463 | 0.113713565 | 0.846185861 |
| ENSG00000179364 | *PACS2* | 3.249900223 | 0.03429556 | 0.063995106 | -0.029699546 | 0.999933887 | 0.01395747 | -0.022037503 | 0.035994973 | 0.846185861 |
| ENSG00000204599 | *TRIM39* | 4.675339612 | -0.026941626 | 0.016557014 | -0.04349864 | 0.999933887 | 0.190068772 | 0.159330233 | 0.030738539 | 0.846185861 |
| ENSG00000159596 | *TMEM69* | 1.384577869 | 0.008725335 | 0.011927686 | -0.003202351 | 0.999933887 | 0.096028553 | 0.045549868 | 0.050478686 | 0.846185861 |
| ENSG00000213930 | *GALT* | 2.560787305 | -0.035881948 | 0.152113314 | -0.187995263 | 0.999933887 | 0.026193071 | 0.075170565 | -0.048977494 | 0.846442621 |
| ENSG00000261245 | *AC093520.2* | 3.111455904 | 0.011825796 | 0.097166616 | -0.085340821 | 0.999933887 | 0.017088867 | 0.07041329 | -0.053324423 | 0.846442621 |
| ENSG00000083817 | *ZNF416* | -0.309293956 | -0.020639313 | 0.04138708 | -0.062026392 | 0.999933887 | 0.302441165 | 0.401886516 | -0.099445351 | 0.846511676 |
| ENSG00000231527 | *FAM27C* | -1.276530719 | 0.078045268 | 0.059068437 | 0.018976831 | 0.999933887 | -0.328783734 | -0.435987957 | 0.107204223 | 0.846765492 |
| ENSG00000248489 | *LINC02062* | 1.231053335 | 0.122487169 | 0.202887241 | -0.080400072 | 0.999933887 | 0.413220083 | 0.488143356 | -0.074923273 | 0.846875389 |
| ENSG00000151881 | *TMEM267* | 0.353111003 | 0.137076088 | 0.33756449 | -0.200488402 | 0.999933887 | 0.021584987 | 0.117444475 | -0.095859488 | 0.846918163 |
| ENSG00000140451 | *PIF1* | -0.403824079 | 0.055577155 | 0.181211291 | -0.125634136 | 0.999933887 | 0.092310601 | 0.230064829 | -0.137754228 | 0.846964686 |
| ENSG00000140988 | *RPS2* | 8.213998115 | 0.070103502 | 0.007706852 | 0.06239665 | 0.999933887 | 0.092321108 | 0.057383611 | 0.034937496 | 0.847032 |
| ENSG00000090104 | *RGS1* | 4.928223245 | -0.059576252 | -0.094137725 | 0.034561472 | 0.999933887 | -0.022713488 | 0.025256361 | -0.047969849 | 0.847032 |
| ENSG00000250479 | *CHCHD10* | 1.698331848 | 0.090454646 | 0.049984241 | 0.040470404 | 0.999933887 | 0.066116946 | 0.135633464 | -0.069516518 | 0.847032 |
| ENSG00000132394 | *EEFSEC* | 1.129400735 | 0.089104254 | 0.006329566 | 0.082774688 | 0.999933887 | 0.18834191 | 0.099862761 | 0.088479149 | 0.847032 |
| ENSG00000285534 | *AL163541.1* | -0.07847177 | 0.048739995 | -0.000825278 | 0.049565273 | 0.999933887 | 0.04375125 | 0.140788453 | -0.097037203 | 0.847032 |
| ENSG00000211792 | *TRAV14DV4* | -0.286831552 | 0.045883006 | 0.132994832 | -0.087111826 | 0.999933887 | 0.284026071 | 0.18774912 | 0.096276951 | 0.847032 |
| ENSG00000120519 | *SLC10A7* | 0.760539306 | -0.021446853 | 0.054054026 | -0.075500879 | 0.999933887 | -0.143747627 | -0.06070482 | -0.083042807 | 0.847032 |
| ENSG00000099834 | *CDHR5* | -0.5437752 | 0.021684807 | -0.059942409 | 0.081627216 | 0.999933887 | -0.118060978 | 0.0072309 | -0.125291878 | 0.847032 |
| ENSG00000135404 | *CD63* | 10.284401809 | 0.001842246 | -0.011200275 | 0.013042522 | 0.999933887 | 0.089567612 | 0.061779924 | 0.027787688 | 0.847032 |
| ENSG00000178404 | *CEP295NL* | 3.247866322 | -0.002448577 | 0.001487516 | -0.003936093 | 0.999933887 | -0.844906648 | -0.787650411 | -0.057256236 | 0.847032 |
| ENSG00000130332 | *LSM7* | 2.968424268 | 0.001482482 | -0.058649791 | 0.060132273 | 0.999933887 | 0.05992489 | 0.019119474 | 0.040805415 | 0.847032 |
| ENSG00000122188 | *LAX1* | 3.220070042 | 0.028170697 | 0.02409508 | 0.004075617 | 0.999933887 | 0.081026014 | 0.118750587 | -0.037724573 | 0.847169072 |
| ENSG00000149179 | *C11orf49* | 1.44999256 | 0.089679582 | 0.157475558 | -0.067795977 | 0.999933887 | -0.164820569 | -0.107770181 | -0.057050387 | 0.847229582 |
| ENSG00000185386 | *MAPK11* | 0.356046353 | 0.006858858 | 0.033702729 | -0.026843871 | 0.999933887 | 0.310590206 | 0.228494324 | 0.082095882 | 0.847229582 |
| ENSG00000143382 | *ADAMTSL4* | 6.209249924 | 0.083538095 | 0.086167128 | -0.002629034 | 0.999933887 | 0.190226374 | 0.154183967 | 0.036042407 | 0.84732579 |
| ENSG00000006125 | *AP2B1* | 6.425233825 | -0.046823596 | -0.056778631 | 0.009955036 | 0.999933887 | -0.139558577 | -0.111802559 | -0.027756019 | 0.84732579 |
| ENSG00000120533 | *ENY2* | 4.787325874 | -0.038355019 | -0.06760592 | 0.029250901 | 0.999933887 | -0.112115534 | -0.0856834 | -0.026432134 | 0.84732579 |
| ENSG00000179841 | *AKAP5* | 0.000521616 | 0.099506594 | 0.359279493 | -0.259772899 | 0.999933887 | -0.119873885 | -0.019282109 | -0.100591776 | 0.84732579 |
| ENSG00000037241 | *RPL26L1* | 1.480423717 | -0.035418129 | -0.010005118 | -0.025413011 | 0.999933887 | 0.117037049 | 0.06376571 | 0.05327134 | 0.84732579 |
| ENSG00000196700 | *ZNF512B* | 2.698910201 | 0.029674262 | 0.136787362 | -0.1071131 | 0.999933887 | 0.104819509 | 0.146348067 | -0.041528558 | 0.847355117 |
| ENSG00000125631 | *HTR5BP* | 0.445172578 | 0.096377762 | 0.059194735 | 0.037183027 | 0.999933887 | 0.12128278 | 0.042537987 | 0.078744794 | 0.847375251 |
| ENSG00000028116 | *VRK2* | 1.910267306 | -0.113287653 | -0.011391252 | -0.1018964 | 0.999933887 | -0.296204439 | -0.376766674 | 0.080562235 | 0.847810274 |
| ENSG00000109606 | *DHX15* | 5.703712606 | -0.08359892 | -0.115308905 | 0.031709986 | 0.999933887 | -0.212807401 | -0.174219296 | -0.038588105 | 0.847834408 |
| ENSG00000243753 | *HLA-L* | 4.609554127 | 0.109670108 | 0.076865753 | 0.032804355 | 0.999933887 | 1.178129784 | 1.261657602 | -0.083527819 | 0.847834408 |
| ENSG00000237649 | *KIFC1* | -0.660236911 | -0.099209559 | -0.150304749 | 0.051095191 | 0.999933887 | 0.209399848 | 0.055487349 | 0.153912499 | 0.847834408 |
| ENSG00000287095 | *AF228727.1* | 1.179865904 | -0.072113078 | 0.129680978 | -0.201794056 | 0.999933887 | -0.709840937 | -0.606219962 | -0.103620976 | 0.847950961 |
| ENSG00000162069 | *BICDL2* | -0.356477856 | 0.058690253 | 0.097611625 | -0.038921372 | 0.999933887 | -0.355500111 | -0.495556836 | 0.140056725 | 0.847950961 |
| ENSG00000287185 | *AC041005.1* | 1.339513047 | -0.064274528 | -0.029077246 | -0.035197282 | 0.999933887 | -0.553719566 | -0.484765207 | -0.068954359 | 0.848064383 |
| ENSG00000228436 | *AL139260.1* | 1.582439681 | 0.022956197 | -0.333305531 | 0.356261728 | 0.999933887 | 0.270512245 | 0.219201482 | 0.051310763 | 0.848192543 |
| ENSG00000136877 | *FPGS* | 3.569710818 | 0.046728172 | 0.115089146 | -0.068360973 | 0.999933887 | 0.105305812 | 0.139377113 | -0.034071302 | 0.848370808 |
| ENSG00000153066 | *TXNDC11* | 5.376778937 | -0.016279017 | -0.009677045 | -0.006601971 | 0.999933887 | -0.028050689 | 0.003094438 | -0.031145127 | 0.848370808 |
| ENSG00000159199 | *ATP5MC1* | 1.64300991 | -0.003083151 | 0.143085879 | -0.146169031 | 0.999933887 | 0.2354695 | 0.29397743 | -0.058507929 | 0.848370808 |
| ENSG00000255521 | *AL356215.1* | 2.521464606 | 0.143810616 | 0.230382348 | -0.086571731 | 0.999933887 | 2.561831301 | 2.636675671 | -0.07484437 | 0.848378316 |
| ENSG00000104613 | *INTS10* | 3.511338586 | 0.000578643 | -0.002241979 | 0.002820622 | 0.999933887 | 0.089971818 | 0.05699126 | 0.032980558 | 0.848378316 |
| ENSG00000160613 | *PCSK7* | 5.212731068 | 0.038920903 | 0.038828144 | 9.2759e-05 | 0.999933887 | -0.005199021 | 0.020687968 | -0.025886989 | 0.848939375 |
| ENSG00000235897 | *TM4SF19-AS1* | -0.155756439 | -0.077504828 | -0.236365622 | 0.158860793 | 0.999933887 | 0.037564729 | -0.037440316 | 0.075005045 | 0.848939375 |
| ENSG00000109099 | *PMP22* | -0.520608422 | -0.187318901 | 0.068487788 | -0.255806689 | 0.999933887 | -0.009470639 | 0.149979649 | -0.159450288 | 0.848972774 |
| ENSG00000151948 | *GLT1D1* | 7.96626045 | -0.017298542 | -0.012965057 | -0.004333485 | 0.999933887 | -0.371291617 | -0.398461818 | 0.027170201 | 0.849246891 |
| ENSG00000102034 | *ELF4* | 7.40808578 | -0.005609895 | -0.010963992 | 0.005354097 | 0.999933887 | -0.507366774 | -0.480025656 | -0.027341118 | 0.849246891 |
| ENSG00000100852 | *ARHGAP5* | 1.563560971 | -0.040891068 | -0.203752055 | 0.162860988 | 0.999933887 | -0.304011903 | -0.194879321 | -0.109132582 | 0.849336503 |
| ENSG00000122674 | *CCZ1* | 1.560864059 | 0.032200591 | 0.030078921 | 0.00212167 | 0.999933887 | -0.347622025 | -0.412465731 | 0.064843706 | 0.849992043 |
| ENSG00000054598 | *FOXC1* | -0.198459672 | 0.070339193 | 0.109021981 | -0.038682788 | 0.999933887 | 0.113035631 | 0.233392002 | -0.12035637 | 0.850525955 |
| ENSG00000141551 | *CSNK1D* | 8.972499483 | -0.009512952 | 0.019385724 | -0.028898676 | 0.999933887 | -0.1570026 | -0.18104053 | 0.02403793 | 0.850525955 |
| ENSG00000124532 | *MRS2* | 1.254529905 | -0.100197601 | -0.108183092 | 0.007985492 | 0.999933887 | -0.057707589 | 0.00940428 | -0.067111868 | 0.850564355 |
| ENSG00000136856 | *SLC2A8* | 0.64646739 | 0.021271717 | -0.090628393 | 0.11190011 | 0.999933887 | -0.295883252 | -0.214444755 | -0.081438497 | 0.850564355 |
| ENSG00000259826 | *AC072061.1* | -0.240566971 | 0.215434008 | -0.061533966 | 0.276967974 | 0.999933887 | -0.208787009 | -0.328162497 | 0.119375488 | 0.850861535 |
| ENSG00000115761 | *NOL10* | 1.67885585 | -0.083240003 | -0.019760791 | -0.063479213 | 0.999933887 | 0.012111575 | -0.040881221 | 0.052992796 | 0.85086913 |
| ENSG00000186642 | *PDE2A* | 2.98619478 | 0.137306166 | 0.114364859 | 0.022941306 | 0.999933887 | 0.626197697 | 0.575394422 | 0.050803275 | 0.851188763 |
| ENSG00000170006 | *TMEM154* | 8.221837308 | -0.029423325 | -0.035666988 | 0.006243663 | 0.999933887 | -0.261707092 | -0.301089881 | 0.039382789 | 0.851188763 |
| ENSG00000182809 | *CRIP2* | 1.143384909 | -0.036165421 | 0.191394864 | -0.227560284 | 0.999933887 | 0.170724313 | 0.246922971 | -0.076198658 | 0.851188763 |
| ENSG00000111639 | *MRPL51* | 2.575288396 | -0.013931562 | -0.058609846 | 0.044678284 | 0.999933887 | -0.086437865 | -0.126646149 | 0.040208284 | 0.851272393 |
| ENSG00000129116 | *PALLD* | 0.542079238 | 0.347308649 | 0.133352334 | 0.213956315 | 0.999933887 | -0.010952878 | -0.081800197 | 0.070847319 | 0.851351137 |
| ENSG00000166105 | *GLB1L3* | 0.18770365 | 0.162985986 | 0.028686716 | 0.13429927 | 0.999933887 | 0.018044474 | -0.077708934 | 0.095753408 | 0.851351137 |
| ENSG00000234072 | *AC074117.1* | 0.322805249 | -0.183368299 | 0.330466741 | -0.513835039 | 0.944094537 | -0.423259858 | -0.354680056 | -0.068579803 | 0.851351137 |
| ENSG00000177370 | *TIMM22* | 1.782294367 | -0.119824014 | 0.043089049 | -0.162913063 | 0.999933887 | -0.054107838 | -0.002769269 | -0.051338569 | 0.851351137 |
| ENSG00000156869 | *FRRS1* | 1.694115449 | -0.104380792 | -0.003809764 | -0.100571028 | 0.999933887 | -0.431859391 | -0.499744826 | 0.067885436 | 0.851351137 |
| ENSG00000042317 | *SPATA7* | -1.016553854 | -0.166810025 | 0.399211316 | -0.566021341 | 0.999933887 | -1.005242161 | -0.896533136 | -0.108709025 | 0.851351137 |
| ENSG00000106993 | *CDC37L1* | 2.483005178 | -0.059296209 | -0.030355758 | -0.028940452 | 0.999933887 | -0.079456843 | -0.12406895 | 0.044612107 | 0.851351137 |
| ENSG00000105374 | *NKG7* | 6.666300906 | 0.035679859 | -0.047884451 | 0.08356431 | 0.999933887 | 0.055712431 | 0.086256907 | -0.030544476 | 0.851351137 |
| ENSG00000100418 | *DESI1* | 4.531447091 | -0.033246355 | -0.03688805 | 0.003641695 | 0.999933887 | -0.099485463 | -0.068574347 | -0.030911116 | 0.851351137 |
| ENSG00000232021 | *LEF1-AS1* | -0.979118786 | 0.120270063 | -0.189693151 | 0.309963213 | 0.999933887 | 0.119246776 | 0.024611031 | 0.094635745 | 0.851351137 |
| ENSG00000008438 | *PGLYRP1* | 4.112391816 | 0.008782544 | -0.005343665 | 0.014126209 | 0.999933887 | 0.033569252 | -0.010754651 | 0.044323902 | 0.851351137 |
| ENSG00000279412 | *AC020763.3* | 1.001078123 | 0.061866372 | -0.072990366 | 0.134856737 | 0.999933887 | -0.157279525 | -0.216359865 | 0.05908034 | 0.851450536 |
| ENSG00000006432 | *MAP3K9* | 1.244698225 | -0.115411131 | -0.02496993 | -0.0904412 | 0.999933887 | -0.064464496 | -0.132721232 | 0.068256735 | 0.851479147 |
| ENSG00000170855 | *TRIAP1* | 1.304018882 | -0.072342446 | -0.131229248 | 0.058886802 | 0.999933887 | 0.165883351 | 0.103754359 | 0.062128991 | 0.851566615 |
| ENSG00000169288 | *MRPL1* | 0.803521404 | 0.016270617 | -0.277537873 | 0.29380849 | 0.999933887 | 0.091706793 | 0.003342295 | 0.088364498 | 0.851566615 |
| ENSG00000163257 | *DCAF16* | 3.999925035 | -0.081411721 | -0.049373976 | -0.032037745 | 0.999933887 | -0.018271909 | -0.050730273 | 0.032458364 | 0.85164596 |
| ENSG00000198369 | *SPRED2* | 2.577318954 | -0.068663524 | -0.017924588 | -0.050738935 | 0.999933887 | 0.090163539 | 0.151308327 | -0.061144788 | 0.85164596 |
| ENSG00000285653 | *AC006238.2* | -0.51886164 | 0.023575206 | 0.148147199 | -0.124571993 | 0.999933887 | -0.497596011 | -0.361364418 | -0.136231593 | 0.85164596 |
| ENSG00000165389 | *SPTSSA* | 2.427006143 | -0.037611757 | 0.02020011 | -0.057811867 | 0.999933887 | -0.297074596 | -0.257167737 | -0.03990686 | 0.851729819 |
| ENSG00000238057 | *ZEB2-AS1* | 1.909066149 | 0.29424061 | 0.252723478 | 0.041517132 | 0.999933887 | 1.773465273 | 1.845454413 | -0.07198914 | 0.851849833 |
| ENSG00000270673 | *YTHDF3-AS1* | 0.074276235 | 0.214967792 | -0.056070016 | 0.271037808 | 0.999933887 | 1.012998924 | 1.154827382 | -0.141828457 | 0.851849833 |
| ENSG00000188001 | *TPRG1* | 1.008472342 | 0.087387485 | -0.33428007 | 0.421667555 | 0.999933887 | 0.064112269 | -0.000891851 | 0.065004121 | 0.851849833 |
| ENSG00000177455 | *CD19* | -0.641643649 | -0.10437393 | -0.045179528 | -0.059194401 | 0.999933887 | -0.124132154 | -0.00630779 | -0.117824364 | 0.852137011 |
| ENSG00000231123 | *SPATA20P1* | 3.215615601 | -0.018031526 | 0.217760829 | -0.235792355 | 0.999933887 | -0.447616114 | -0.352504058 | -0.095112055 | 0.852137011 |
| ENSG00000262898 | *AC139099.2* | 0.159490353 | 0.281313153 | 0.135464254 | 0.145848899 | 0.999933887 | 0.274273771 | 0.372305384 | -0.098031613 | 0.85257964 |
| ENSG00000107815 | *TWNK* | 0.802161678 | -0.087520759 | -0.090430048 | 0.002909289 | 0.999933887 | 0.242989649 | 0.311535819 | -0.06854617 | 0.85257964 |
| ENSG00000140511 | *HAPLN3* | 2.997170246 | -0.028154439 | 0.009381569 | -0.037536008 | 0.999933887 | 0.215737635 | 0.263897922 | -0.048160287 | 0.85257964 |
| ENSG00000120738 | *EGR1* | 6.622652301 | 0.639689151 | 0.674933952 | -0.035244801 | 0.999933887 | 0.693767569 | 0.791317444 | -0.097549875 | 0.852709354 |

| | | | | | | | | | | |
|---|---|---|---|---|---|---|---|---|---|---|
| ENSG00000160716 | *CHRNB2* | 0.551710113 | 0.199666837 | 0.158492832 | 0.041174005 | 0.999933887 | 0.333111282 | 0.431891924 | -0.098780642 | 0.852766171 |
| ENSG00000198133 | *TMEM229B* | 1.70303279 | 0.10803008 | 0.084897715 | 0.023132365 | 0.999933887 | -0.061868562 | -0.112450135 | 0.050581573 | 0.852774822 |
| ENSG00000117533 | *VAMP4* | 2.689986526 | -0.064636163 | -0.143143056 | 0.078506894 | 0.999933887 | -0.316875209 | -0.374806308 | 0.057931099 | 0.852864094 |
| ENSG00000149131 | *SERPING1* | 3.99193876 | 0.013048346 | -0.00745153 | 0.020499875 | 0.999933887 | -0.224153383 | -0.277791686 | 0.053638303 | 0.852864094 |
| ENSG00000236935 | *AP003774.2* | -0.487356114 | 0.017581356 | 0.168835559 | -0.151254203 | 0.999933887 | -0.183315083 | -0.064320652 | -0.118994431 | 0.852864094 |
| ENSG00000256843 | *AC023157.1* | 0.90134298 | 0.371078121 | 0.380076791 | -0.00899867 | 0.999933887 | 0.771387021 | 0.84403857 | -0.072651549 | 0.852885066 |
| ENSG00000013523 | *ANGEL1* | 1.717653098 | -0.026572155 | -0.037624754 | 0.011052599 | 0.999933887 | 0.083494044 | 0.02657479 | 0.056919254 | 0.85310696 |
| ENSG00000259293 | *LIPC-AS1* | 2.41281997 | 0.103918383 | 0.066385825 | 0.037532558 | 0.999933887 | 0.235305503 | 0.281675819 | -0.046370316 | 0.853133046 |
| ENSG00000137261 | *KIAA0319* | 4.181560159 | -0.121964186 | 0.006851579 | -0.128815765 | 0.999933887 | -0.401864028 | -0.436437457 | 0.034573429 | 0.853220036 |
| ENSG00000182600 | *SNORC* | -0.0991895 | 0.152849641 | 0.413626693 | -0.260777053 | 0.999933887 | -0.329176095 | -0.225772653 | -0.103403442 | 0.853220036 |
| ENSG00000152944 | *MED21* | 3.391155896 | 0.049949463 | -0.069904849 | 0.119854311 | 0.999933887 | -0.445630512 | -0.393246566 | -0.052383946 | 0.853220036 |
| ENSG00000274211 | *SOCS7* | 2.697480051 | -0.041406854 | 0.156091732 | -0.197498586 | 0.999933887 | -0.098724195 | -0.147710763 | 0.048986569 | 0.853220036 |
| ENSG00000139508 | *SLC46A3* | 2.16765405 | -0.024962943 | -0.108484035 | 0.083521092 | 0.999933887 | 0.082855188 | 0.138743441 | -0.055888253 | 0.853220036 |
| ENSG00000233668 | *AL353662.1* | 0.739399214 | 0.01250546 | 0.035940709 | -0.023435249 | 0.999933887 | 0.352168569 | 0.478141545 | -0.125972975 | 0.853220036 |
| ENSG00000197879 | *MYO1C* | 2.735476688 | 0.010591694 | 0.12870224 | -0.118110546 | 0.999933887 | 0.399359612 | 0.449815654 | -0.050456042 | 0.853220036 |
| ENSG00000164953 | *TMEM67* | -0.11535956 | 0.210760169 | 0.219072533 | -0.008312364 | 0.999933887 | -0.331240963 | -0.213709894 | -0.117531069 | 0.853640353 |
| ENSG00000276758 | *AC245884.12* | 0.506435193 | -0.033155885 | -0.071678994 | 0.038523108 | 0.999933887 | -0.117902759 | -0.23151614 | 0.113613381 | 0.853726522 |
| ENSG00000165209 | *STRBP* | 2.009453889 | -0.123315469 | -0.029264566 | -0.094050903 | 0.999933887 | -0.25017168 | -0.295657542 | 0.045485862 | 0.854372124 |
| ENSG00000280385 | *AP000648.4* | 0.67780374 | -0.092988386 | -0.135348146 | 0.04235976 | 0.999933887 | -0.258166513 | -0.120290029 | -0.137876484 | 0.854483321 |
| ENSG00000162076 | *FLYWCH2* | 0.5950672 | 0.043330387 | 0.027749256 | 0.015581132 | 0.999933887 | 0.040770169 | 0.132089452 | -0.091319283 | 0.854611949 |
| ENSG00000273824 | *AC008033.3* | -0.996923859 | 0.042286023 | -0.070451814 | 0.112737837 | 0.999933887 | 0.191200327 | 0.298457918 | -0.107257592 | 0.854716739 |
| ENSG00000164291 | *ARSK* | -0.336317676 | -0.217196582 | -0.36069972 | 0.143503139 | 0.999933887 | -0.342187982 | -0.431388593 | 0.089200612 | 0.854724219 |
| ENSG00000246334 | *PRR7-AS1* | 2.913756199 | 0.067966696 | 0.021191254 | 0.046775442 | 0.999933887 | -0.209728937 | -0.261938066 | 0.052209129 | 0.854724219 |
| ENSG00000204219 | *TCEA3* | -0.844037481 | -0.063731988 | 0.17698758 | -0.240719568 | 0.999933887 | 0.204909871 | 0.31072523 | -0.105815358 | 0.854724219 |
| ENSG00000160055 | *TMEM234* | 3.340344545 | -0.011982698 | -0.000380566 | -0.011602133 | 0.999933887 | -0.098231159 | -0.132535151 | 0.034303992 | 0.854724219 |
| ENSG00000259874 | *AC034105.2* | -0.814875483 | 0.225824294 | 0.514423336 | -0.288599042 | 0.999933887 | 1.957323737 | 1.81046858 | 0.146855157 | 0.854792353 |
| ENSG00000285781 | *AL117340.1* | 0.9915231 | 0.229694479 | 0.509439113 | -0.279744635 | 0.999933887 | 0.382227436 | 0.513894428 | -0.131666992 | 0.854792353 |
| ENSG00000267632 | *AC067852.3* | 1.510736565 | -0.12450321 | -0.039991351 | -0.084511859 | 0.999933887 | -0.483036894 | -0.421858723 | -0.061178171 | 0.854792353 |
| ENSG00000196436 | *NPIPB15* | 1.017332371 | 0.086241966 | 0.153640994 | -0.067399028 | 0.999933887 | 0.019578019 | 0.074986842 | -0.055408823 | 0.854792353 |
| ENSG00000113048 | *MRPS27* | 2.319639686 | 0.041206613 | -0.153961005 | 0.195167618 | 0.999933887 | 0.14819332 | 0.100270378 | 0.047922942 | 0.854792353 |
| ENSG00000182841 | *RRP7BP* | 1.919107847 | -0.046654297 | 0.239938174 | -0.286592471 | 0.999933887 | 0.156696588 | 0.104506797 | 0.052189792 | 0.854792353 |
| ENSG00000171130 | *ATP6V0E2* | 1.975128634 | 0.025061784 | 0.041727691 | -0.016665907 | 0.999933887 | 0.05774445 | -0.004981574 | 0.062726023 | 0.854792353 |
| ENSG00000124356 | *STAMBP* | 3.999118464 | -0.006265074 | -0.018317896 | 0.012052822 | 0.999933887 | -0.021880727 | -0.051895056 | 0.030014329 | 0.854792353 |
| ENSG00000234814 | *SVIL2P* | 1.609567696 | 0.00732395 | 0.051946902 | -0.044622952 | 0.999933887 | 0.293879087 | 0.348668012 | -0.054788925 | 0.854792353 |
| ENSG00000121289 | *CEP89* | 1.178151729 | 0.089525501 | -0.095783794 | 0.185309295 | 0.999933887 | -0.088394933 | -0.140158135 | 0.051763203 | 0.855025263 |
| ENSG00000162545 | *CAMK2N1* | 0.311477644 | -0.059213865 | 0.024657562 | -0.083871428 | 0.999933887 | 0.168318771 | 0.074827549 | 0.093491222 | 0.855025263 |
| ENSG00000128596 | *CCDC136* | 0.186206453 | 0.024465655 | 0.029018109 | -0.004552454 | 0.999933887 | 0.120148715 | 0.217087328 | -0.096938613 | 0.855025263 |
| ENSG00000005801 | *ZNF195* | 2.165999325 | -0.007994219 | -0.184193058 | 0.176198839 | 0.999933887 | -0.079234734 | -0.016598375 | -0.062636359 | 0.855025263 |
| ENSG00000103404 | *USP31* | 1.473454222 | -0.014706454 | -0.192346475 | 0.177640021 | 0.999933887 | -0.215359084 | -0.276908708 | 0.061549624 | 0.85524586 |
| ENSG00000171223 | *JUNB* | 9.742533054 | 0.439614977 | 0.632812176 | -0.193197199 | 0.999933887 | 1.000304674 | 1.060921551 | -0.060616877 | 0.855377583 |
| ENSG00000153130 | *SCOC* | 1.366693975 | -0.018505661 | -0.153728052 | 0.135222391 | 0.999933887 | -0.356611459 | -0.423933413 | 0.067321954 | 0.855377583 |
| ENSG00000010270 | *STARD3NL* | 4.828031222 | 0.038910267 | -0.028229083 | 0.067139349 | 0.999933887 | 0.640175065 | 0.683208577 | -0.043033512 | 0.855499083 |
| ENSG00000150756 | *ATPSCKMT* | -0.03411328 | 0.205445486 | 0.046479191 | 0.158966295 | 0.999933887 | -0.085544193 | 0.010416616 | -0.095960809 | 0.855718898 |
| ENSG00000116898 | *MRPS15* | 3.192007487 | 0.057737362 | 0.0532855 | 0.004451862 | 0.999933887 | -0.050959623 | -0.088937556 | 0.037977932 | 0.855718898 |
| ENSG00000105698 | *USF2* | 7.216664813 | 0.037980259 | 0.04693486 | -0.008954601 | 0.999933887 | 0.209129256 | 0.18382733 | 0.025301926 | 0.855718898 |
| ENSG00000101447 | *FAM83D* | -0.306148603 | -0.037777987 | -0.214331167 | 0.17655318 | 0.999933887 | -0.018238092 | 0.092659379 | -0.110897471 | 0.855718898 |
| ENSG00000287937 | *AC079753.2* | 0.429859566 | 0.230706839 | 0.208361302 | 0.022345536 | 0.999933887 | 1.549902145 | 1.653805923 | -0.103903778 | 0.85573835 |
| ENSG00000275552 | *AC243965.2* | -0.208836254 | 0.171518874 | 0.249618731 | -0.078099857 | 0.999933887 | 1.072634135 | 0.993777345 | 0.078856789 | 0.85573835 |
| ENSG00000106992 | *AK1* | 0.457266456 | -0.039593624 | -0.020296641 | -0.019296983 | 0.999933887 | 0.153824429 | 0.073939438 | 0.079884991 | 0.85573835 |
| ENSG00000110852 | *CLEC2B* | 7.811631229 | -0.144154793 | -0.124079711 | -0.020075082 | 0.999933887 | -0.412451152 | -0.449376263 | 0.036925111 | 0.85579564 |
| ENSG00000131871 | *SELENOS* | 3.328878672 | 0.026605715 | 0.019551275 | 0.007054439 | 0.999933887 | -0.089885884 | -0.051383112 | -0.038502772 | 0.85579564 |
| ENSG00000120685 | *PROSER1* | 3.295969109 | -0.011949974 | -0.041756694 | 0.02980672 | 0.999933887 | -0.105821535 | -0.064477111 | -0.041344424 | 0.85579564 |
| ENSG00000140577 | *CRTC3* | 4.221388697 | 0.018263844 | 0.079160461 | -0.060896617 | 0.999933887 | -0.011349046 | 0.015712097 | -0.027061143 | 0.855805085 |
| ENSG00000102890 | *ELMO3* | 2.565611513 | -0.043184768 | -0.056260191 | 0.013075424 | 0.999933887 | -0.017414983 | -0.058357786 | 0.040942803 | 0.855851966 |
| ENSG00000109189 | *USP46* | 1.173252642 | -0.0527232 | -0.287593566 | 0.234870367 | 0.999933887 | 0.011297402 | -0.045710782 | 0.057008184 | 0.855868132 |
| ENSG00000134109 | *EDEM1* | 7.519487316 | -0.063681758 | -0.039191517 | -0.02449024 | 0.999933887 | -0.070569558 | -0.047978792 | -0.022590766 | 0.855868786 |
| ENSG00000137770 | *CTDSPL2* | 3.72399562 | -0.02229026 | -0.098475849 | 0.076185589 | 0.999933887 | -0.308556139 | -0.250345022 | -0.058211117 | 0.855868786 |
| ENSG00000279554 | *AC130448.2* | 0.155021817 | 0.020865033 | 0.172966816 | -0.152101782 | 0.999933887 | -0.738556229 | -0.610986038 | -0.127570191 | 0.855868786 |
| ENSG00000132406 | *TMEM128* | 1.388658222 | -0.066863209 | -0.026002187 | -0.040861022 | 0.999933887 | -0.168217974 | -0.226292425 | 0.058074451 | 0.855913955 |
| ENSG00000250031 | *AC009927.1* | 1.516552741 | -0.009984378 | 0.06325696 | -0.073241338 | 0.999933887 | -0.684835539 | -0.783874312 | 0.099038773 | 0.855913955 |
| ENSG00000285863 | *AC009554.2* | -0.440217777 | 0.133241957 | -0.191993489 | 0.325235446 | 0.999933887 | -0.487599093 | -0.62142359 | 0.133824497 | 0.856021056 |
| ENSG00000129347 | *KRI1* | 3.319942821 | -0.007445 | -0.072187756 | 0.064742755 | 0.999933887 | 0.168608231 | 0.206557292 | -0.037949061 | 0.856021056 |
| ENSG00000155324 | *GRAMD2B* | -0.515033913 | -0.228992689 | 0.063036873 | -0.292029562 | 0.999933887 | 0.264574984 | 0.36494422 | -0.100369235 | 0.856033324 |
| ENSG00000119684 | *MLH3* | 1.30996668 | -0.131826686 | -0.024026555 | -0.107800131 | 0.999933887 | -0.141992382 | -0.085268673 | -0.056723708 | 0.856033324 |
| ENSG00000104689 | *TNFRSF10A* | 1.866626159 | -0.093355781 | 0.165110475 | -0.258466256 | 0.999933887 | -0.04700754 | 0.006496312 | -0.053503852 | 0.856033324 |
| ENSG00000261449 | *AC103724.3* | 0.701902603 | 0.050165206 | 0.310208151 | -0.260042944 | 0.999933887 | 0.262462084 | 0.125931103 | 0.136530981 | 0.856033324 |
| ENSG00000155438 | *NIFK* | 1.612069223 | 0.008026094 | -0.188412903 | 0.196438996 | 0.999933887 | 0.144308327 | 0.200970411 | -0.056662084 | 0.856033324 |
| ENSG00000132153 | *DHX30* | 4.088988141 | 0.003418626 | 0.029902295 | -0.026483669 | 0.999933887 | 0.149750864 | 0.18164137 | -0.031890506 | 0.856033324 |
| ENSG00000107438 | *PDLIM1* | 1.984705299 | -0.001540319 | -0.066134413 | 0.064594094 | 0.999933887 | 0.137419407 | 0.192458285 | -0.055038878 | 0.856033324 |
| ENSG00000131269 | *ABCB7* | 1.765346087 | 0.038409276 | -0.025731871 | 0.064141147 | 0.999933887 | -0.018995927 | 0.03556748 | -0.054563406 | 0.856092287 |
| ENSG00000105202 | *FBL* | 4.374376553 | 0.055277868 | 0.018706041 | 0.036571827 | 0.999933887 | 0.21566192 | 0.253212307 | -0.037550387 | 0.8561034 |
| ENSG00000230225 | *MTND5P14* | -1.532206012 | -0.046821401 | 0.132205179 | -0.179026581 | 0.999933887 | 0.397150093 | 0.130253981 | 0.266896112 | 0.856201435 |
| ENSG00000270060 | *AC090589.3* | 0.058119099 | -0.35129403 | -0.210953036 | -0.140340994 | 0.999933887 | -0.07974954 | -0.169839893 | 0.090090353 | 0.856257963 |
| ENSG00000106771 | *TMEM245* | 3.599306343 | -0.111882425 | 0.0599856 | -0.171868025 | 0.999933887 | -0.044924758 | -0.010033735 | -0.034891023 | 0.856257963 |
| ENSG00000249173 | *LINC01093* | 0.960718789 | 0.121112663 | 0.008124429 | 0.112988234 | 0.999933887 | 3.037246259 | 3.170022755 | -0.132776496 | 0.856283934 |
| ENSG00000216490 | *IFI30* | 6.746826295 | 0.193450909 | 0.327467706 | -0.134016796 | 0.999933887 | -0.066437643 | -0.108956988 | 0.042519344 | 0.85644199 |
| ENSG00000257284 | *AC013437.1* | -0.924247679 | 0.066726816 | 0.168314632 | -0.101587817 | 0.999933887 | -0.497172397 | -0.673330633 | 0.176158235 | 0.856490211 |
| ENSG00000110367 | *DDX6* | 7.449792405 | -0.071474572 | -0.045301037 | -0.026173536 | 0.999933887 | -0.309467985 | -0.286118233 | -0.023349752 | 0.85705485 |
| ENSG00000069974 | *RAB27A* | 6.286709962 | -0.077812108 | -0.05039501 | -0.027417099 | 0.999933887 | -0.320096491 | -0.291961882 | -0.028134609 | 0.85705485 |
| ENSG00000104341 | *LAPTM4B* | 0.718404877 | -0.116261078 | -0.104780585 | -0.011480493 | 0.999933887 | -0.171891569 | -0.10967046 | -0.062221109 | 0.85705485 |
| ENSG00000175182 | *FAM131A* | 4.784228978 | 0.04862946 | 0.022187812 | 0.026441648 | 0.999933887 | -0.30159826 | -0.334706741 | 0.033108482 | 0.85705485 |
| ENSG00000203875 | *SNHG5* | 4.516505653 | 0.048947701 | 0.107218901 | -0.058271201 | 0.999933887 | 0.207489929 | 0.244953902 | -0.037463974 | 0.85705485 |
| ENSG00000172819 | *RARG* | 2.699717697 | 0.060820861 | 0.058145637 | 0.002675225 | 0.999933887 | -0.034881412 | -0.078487637 | 0.043606226 | 0.85705485 |
| ENSG00000204444 | *APOM* | 0.563380483 | 0.036876851 | 0.091071196 | -0.054194345 | 0.999933887 | 0.147267703 | 0.067075098 | 0.080192605 | 0.85705485 |
| ENSG00000168734 | *PKIG* | 1.285724574 | 0.017663784 | 0.006795131 | 0.010868653 | 0.999933887 | -0.625424139 | -0.677377492 | 0.051953354 | 0.85705485 |
| ENSG00000169991 | *IFFO2* | 3.930637986 | 0.020342645 | 0.027662312 | -0.007319666 | 0.999933887 | 0.138950677 | 0.106385158 | 0.032565518 | 0.85705693 |
| ENSG00000198001 | *IRAK4* | 5.081620341 | -0.0056747 | -0.053922112 | 0.048247412 | 0.999933887 | -0.231466234 | -0.268843819 | 0.037377585 | 0.857149466 |
| ENSG00000137210 | *TMEM14B* | 2.844785622 | -0.063738833 | 0.044279444 | -0.108018278 | 0.999933887 | -0.012181525 | -0.05182768 | 0.039646154 | 0.85725693 |
| ENSG00000172432 | *GTPBP2* | 6.228921633 | 0.047456003 | 0.037802634 | 0.009653369 | 0.999933887 | 0.063636604 | 0.088109422 | -0.024472818 | 0.85762825 |
| ENSG00000140545 | *MFGE8* | 1.595015367 | -0.044982497 | 0.021423716 | -0.066406213 | 0.999933887 | 0.016538348 | 0.077507708 | -0.06096936 | 0.85762825 |
| ENSG00000164530 | *PI16* | -0.811147484 | 0.015593226 | 0.010886508 | 0.004706718 | 0.999933887 | 0.165592158 | 0.010368995 | 0.155223164 | 0.85762825 |
| ENSG00000196357 | *ZNF565* | 0.032078623 | -0.315219703 | -0.320579453 | 0.005359751 | 0.999933887 | 0.177952769 | 0.070976559 | 0.10697621 | 0.85765114 |
| ENSG00000085415 | *SEH1L* | 2.283581172 | -0.138641625 | -0.021935515 | -0.11670611 | 0.999933887 | 0.055425601 | 0.009207169 | 0.046218433 | 0.85765114 |
| ENSG00000168394 | *TAP1* | 8.778485752 | 0.055736866 | 0.039761764 | 0.015975103 | 0.999933887 | 0.303500884 | 0.327416737 | -0.023915853 | 0.85765114 |
| ENSG00000146453 | *PNLDC1* | -1.602799943 | -0.432806942 | -0.443810619 | 0.011003676 | 0.999933887 | 0.787643705 | 0.892323412 | -0.104679707 | 0.85765114 |
| ENSG00000271780 | *AL118558.3* | 3.203199984 | 0.121666394 | 0.146203154 | -0.024536759 | 0.999933887 | 0.686930301 | 0.633852778 | 0.053077523 | 0.85765114 |
| ENSG00000115073 | *ACTR1B* | 4.269735478 | 0.065793481 | 0.087467808 | -0.021674327 | 0.999933887 | 0.318584288 | 0.355230384 | -0.036646095 | 0.85765114 |
| ENSG00000100764 | *PSMC1* | 3.788377109 | -0.067192754 | -0.172472808 | 0.105280054 | 0.999933887 | -0.048889057 | -0.01373765 | -0.035151407 | 0.85765114 |
| ENSG00000175536 | *LIPT2* | -1.099144555 | -0.256638314 | -0.495255945 | 0.238617631 | 0.999933887 | 0.801942788 | 0.914118698 | -0.11217591 | 0.85765114 |
| ENSG00000128951 | *DUT* | 2.872041024 | -0.041884041 | 0.03109718 | -0.07298122 | 0.999933887 | 0.055635672 | 0.017333425 | 0.038302246 | 0.85765114 |
| ENSG00000254909 | *AP003392.2* | -0.033320211 | 0.100342992 | 0.052986651 | 0.047356341 | 0.999933887 | -0.216761611 | -0.125165834 | -0.091595777 | 0.85765114 |
| ENSG00000259182 | *AC019254.1* | -0.667984772 | 0.081289035 | -0.215537001 | 0.296826036 | 0.999933887 | -0.179884961 | -0.345891627 | 0.166006666 | 0.85765114 |
| ENSG00000152117 | *SMPD4BP* | 4.139312021 | 0.015055586 | 0.025515349 | -0.010459762 | 0.999933887 | 0.117262297 | 0.085311816 | 0.03195048 | 0.85765114 |
| ENSG00000181215 | *C4orf50* | 0.488611014 | 0.116017868 | -0.082006842 | 0.19802471 | 0.999933887 | 0.000582304 | 0.092612253 | -0.092029948 | 0.857656328 |
| ENSG00000233559 | *LINC00513* | 3.376966176 | 0.150613638 | 0.190964544 | -0.040350906 | 0.999933887 | 0.629856635 | 0.694604284 | -0.064747649 | 0.858871847 |

| | | | | | | | | | | |
|---|---|---|---|---|---|---|---|---|---|---|
| ENSG00000138709 | *LARP1B* | 1.52897632 | -0.088598607 | 0.062192327 | -0.150790933 | 0.999933887 | 0.125771426 | 0.073591341 | 0.052180086 | 0.858871847 |
| ENSG00000130158 | *DOCK6* | 0.843817127 | -0.097691709 | 0.012293869 | -0.109985579 | 0.999933887 | -0.038395533 | 0.034377904 | -0.072773438 | 0.858871847 |
| ENSG00000260145 | *AC023825.2* | 0.439332156 | -0.025023035 | 0.122087587 | -0.147110622 | 0.999933887 | -0.535102857 | -0.450631677 | -0.08447118 | 0.858871847 |
| ENSG00000115239 | *ASB3* | 2.31469109 | 0.082641188 | -0.074161376 | 0.156802564 | 0.999933887 | -0.189736297 | -0.139492116 | -0.050244181 | 0.858889646 |
| ENSG00000243449 | *C4orf48* | 2.342495664 | 0.023768271 | 0.043498671 | -0.0197304 | 0.999933887 | -0.16265755 | -0.230992136 | 0.068334586 | 0.859034749 |
| ENSG00000205885 | *C1RL-AS1* | 2.652771824 | 0.068371973 | 0.106763383 | -0.038391411 | 0.999933887 | 0.019077315 | 0.066605535 | -0.04752822 | 0.859118881 |
| ENSG00000286733 | *LINC02348* | -0.485457216 | 0.049262949 | 0.012952328 | 0.036310621 | 0.999933887 | 0.048164129 | 0.14610201 | -0.09793788 | 0.859118881 |
| ENSG00000170545 | *SMAGP* | 1.012614872 | 0.006601117 | 0.040048956 | -0.033447839 | 0.999933887 | -0.056630536 | 0.010938764 | -0.0675693 | 0.859345841 |
| ENSG00000184743 | *ATL3* | 2.864424747 | -0.074036072 | -0.020230316 | -0.053805757 | 0.999933887 | -0.202805829 | -0.162229912 | -0.040575917 | 0.859567078 |
| ENSG00000141279 | *NPEPPS* | 6.138158719 | -0.043950288 | -0.005395763 | -0.038554525 | 0.999933887 | -0.367279098 | -0.391292081 | 0.024012982 | 0.859626219 |
| ENSG00000155657 | *TTN* | 4.624008388 | 0.125547071 | 0.075751316 | 0.049795756 | 0.999933887 | -0.198199482 | -0.134194797 | -0.064004686 | 0.859912545 |
| ENSG00000176422 | *SPRYD4* | 0.830081989 | 0.096147325 | -0.001807294 | 0.097954618 | 0.999933887 | -0.031952247 | 0.026928571 | -0.058880818 | 0.859912545 |
| ENSG00000258056 | *AC009779.2* | 2.856681318 | 0.106070328 | 0.18742727 | -0.081356942 | 0.999933887 | 0.138532481 | 0.094881335 | 0.043651146 | 0.859912545 |
| ENSG00000170271 | *FAXDC2* | 0.579121563 | -0.12291046 | -0.0447781 | -0.07813236 | 0.999933887 | -0.05467027 | 0.02669617 | -0.08136644 | 0.859912545 |
| ENSG00000162441 | *LZIC* | 2.882369156 | -0.064762192 | -0.110802867 | 0.046040676 | 0.999933887 | -0.154571037 | -0.11102202 | -0.043549016 | 0.859912545 |
| ENSG00000279048 | *AC080080.1* | 0.401906539 | -0.150432876 | 0.181459351 | -0.331892227 | 0.999933887 | 0.509533321 | 0.425752732 | 0.083780589 | 0.859912545 |
| ENSG00000063587 | *ZNF275* | 3.02072614 | -0.044415618 | 0.061251408 | -0.105667026 | 0.999933887 | -0.255273967 | -0.288379149 | 0.033105182 | 0.859912545 |
| ENSG00000168005 | *SPINDOC* | 1.306076932 | -0.071753026 | -0.191239635 | 0.119486609 | 0.999933887 | 0.004490965 | -0.054305193 | 0.058796158 | 0.859912545 |
| ENSG00000105364 | *MRPL4* | 1.531820497 | 0.053526303 | 0.140639053 | -0.08711275 | 0.999933887 | 0.224839035 | 0.171757529 | 0.053081506 | 0.859912545 |
| ENSG00000172005 | *MAL* | 2.519312705 | 0.03275203 | -0.028112885 | 0.060864915 | 0.999933887 | 0.20752651 | 0.252968487 | -0.045441977 | 0.859912545 |
| ENSG00000269867 | *AC010326.3* | -0.003276947 | -0.058492616 | 0.384695898 | -0.443188514 | 0.981460244 | -0.182767837 | -0.271133555 | 0.088365718 | 0.859912545 |
| ENSG00000066427 | *ATXN3* | 3.402094045 | 0.019961408 | -0.028765783 | 0.048727192 | 0.999933887 | -0.486602022 | -0.52127522 | 0.034673198 | 0.859912545 |
| ENSG00000269313 | *MAGIX* | -0.278739951 | -0.020406663 | 0.102890322 | -0.123296985 | 0.999933887 | -0.06722001 | -0.189978277 | 0.122758267 | 0.859912545 |
| ENSG00000280387 | *AL109806.1* | 1.866290057 | -0.011959046 | 0.025889638 | -0.037848684 | 0.999933887 | -0.110269919 | -0.048762394 | -0.061507525 | 0.859912545 |
| ENSG00000163125 | *RPRD2* | 4.14097509 | 0.007437032 | 0.02690778 | -0.019470748 | 0.999933887 | -0.071349109 | -0.039677946 | -0.031671163 | 0.859912545 |
| ENSG00000156603 | *MED19* | 1.985496372 | -0.006945588 | -0.006624812 | -0.000320775 | 0.999933887 | 0.033716034 | -0.018821607 | 0.052537642 | 0.859938144 |
| ENSG00000189362 | *NEMP2* | 1.552923277 | -0.030163529 | -0.143187538 | 0.113024009 | 0.999933887 | -0.074177726 | -0.029341537 | -0.044836189 | 0.859968988 |
| ENSG00000287917 | *AP000944.7* | 0.383294546 | 0.066636779 | -0.002613203 | 0.069249981 | 0.999933887 | 0.963998834 | 0.887060813 | 0.076938021 | 0.85997749 |
| ENSG00000119723 | *COQ6* | 0.405482022 | -0.173381258 | -0.070680876 | -0.102700381 | 0.999933887 | 0.063690154 | -0.011189739 | 0.074879893 | 0.860019493 |
| ENSG00000186951 | *PPARA* | 2.67155566 | 0.041845607 | 0.034491976 | 0.007353631 | 0.999933887 | -0.173942065 | -0.214732841 | 0.040790776 | 0.860019493 |
| ENSG00000152240 | *HAUS1* | 0.406278016 | -0.063835286 | -0.159600505 | 0.095765219 | 0.999933887 | 0.082266477 | 0.166734784 | -0.084468307 | 0.860019493 |
| ENSG00000108826 | *MRPL27* | 2.039564927 | -0.014202508 | -0.056812683 | 0.042610175 | 0.999933887 | 0.072567134 | 0.121177922 | -0.048610788 | 0.860019493 |
| ENSG00000088038 | *CNOT3* | 5.735681908 | -0.004835706 | 0.000587552 | -0.005423259 | 0.999933887 | -0.337401162 | -0.362159892 | 0.02475873 | 0.860019493 |
| ENSG00000112697 | *TMEM30A* | 6.389621852 | -0.00595945 | -0.067707914 | 0.061748464 | 0.999933887 | -0.647070126 | -0.68836636 | 0.041296234 | 0.860019493 |
| ENSG00000145358 | *DDIT4L* | 2.130741671 | -0.012607103 | -0.122802231 | 0.110195128 | 0.999933887 | -0.290822171 | -0.236370681 | -0.05445149 | 0.86015966 |
| ENSG00000170379 | *TCAF2* | 2.565083571 | -0.123398439 | -0.050344909 | -0.07305353 | 0.999933887 | 0.215588843 | 0.170634777 | 0.044954066 | 0.860161002 |
| ENSG00000120053 | *GOT1* | 1.367881754 | -0.149088165 | 0.297386788 | -0.446474953 | 0.944094537 | 0.032545482 | -0.016071246 | 0.048616728 | 0.860161002 |
| ENSG00000124103 | *FAM209A* | 1.207242412 | -0.126744949 | 0.039703818 | -0.166448767 | 0.999933887 | -0.055961172 | -0.126350008 | 0.070388835 | 0.860161002 |
| ENSG00000130733 | *YIPF2* | 1.182885712 | 0.117810676 | 0.100239126 | 0.01757155 | 0.999933887 | 0.201216191 | 0.140136895 | 0.061079297 | 0.860161002 |
| ENSG00000234636 | *MED14OS* | 0.012075826 | -0.163948986 | -0.005852696 | -0.15809609 | 0.999933887 | 0.353028151 | 0.456863128 | -0.103834977 | 0.860161002 |
| ENSG00000025156 | *HSF2* | 2.030833296 | -0.043196953 | -0.082313941 | 0.039116988 | 0.999933887 | -0.093759255 | -0.14465332 | 0.050894065 | 0.860161002 |
| ENSG00000080603 | *SRCAP* | 2.498993048 | -0.012744412 | -0.035275704 | 0.022531292 | 0.999933887 | -0.216511781 | -0.174360672 | -0.04215111 | 0.860161002 |
| ENSG00000133321 | *PLAAT4* | 4.071923582 | 0.010045585 | -0.045791528 | 0.055837114 | 0.999933887 | 0.038327299 | 0.08103647 | -0.042709172 | 0.860161002 |
| ENSG00000063241 | *ISOC2* | -0.011107482 | 0.138429932 | 0.202339598 | -0.063909666 | 0.999933887 | 0.214286387 | 0.286713236 | -0.072426849 | 0.860174366 |
| ENSG00000125703 | *ATG4C* | -0.294529538 | -0.068292027 | -0.147451752 | 0.079159725 | 0.999933887 | -0.297542617 | -0.194397739 | -0.103144878 | 0.860214671 |
| ENSG00000168259 | *DNAJC7* | 4.535905202 | -0.081743382 | -0.029949316 | -0.051794066 | 0.999933887 | -0.158692304 | -0.182533404 | 0.0238411 | 0.860320981 |
| ENSG00000276568 | *Metazoa_SRP* | -0.697847896 | 0.117926727 | 0.280216896 | -0.16229017 | 0.999933887 | 1.319745048 | 1.428237418 | -0.10849237 | 0.861049176 |
| ENSG00000077549 | *CAPZB* | 8.251925724 | -0.007969048 | 0.00818041 | -0.016149458 | 0.999933887 | 0.008740319 | -0.011436213 | 0.020176533 | 0.861828427 |
| ENSG00000266980 | *AC087289.1* | -0.062717416 | 0.033311878 | 0.196153457 | -0.162841579 | 0.999933887 | 0.841309532 | 0.73802092 | 0.103288612 | 0.862256742 |
| ENSG00000185298 | *CCDC137* | 3.035726102 | 0.042630504 | 0.044727623 | -0.002097119 | 0.999933887 | 0.069876222 | 0.033232155 | 0.036644067 | 0.862349602 |
| ENSG00000164897 | *TMUB1* | 5.218243457 | 0.049799555 | 0.155404742 | -0.105605187 | 0.999933887 | 0.409658112 | 0.442785532 | -0.033127419 | 0.862403875 |
| ENSG00000164136 | *IL15* | 0.656415612 | -0.225418434 | -0.019703481 | -0.205714953 | 0.999933887 | 0.012280936 | -0.084327895 | 0.096608831 | 0.862415006 |
| ENSG00000154134 | *ROBO3* | 0.206318406 | 0.219493008 | 0.050727581 | 0.168765426 | 0.999933887 | 0.206301177 | 0.110691576 | 0.095609601 | 0.862415006 |
| ENSG00000274925 | *ZKSCAN2-DT* | 0.801534358 | -0.121841574 | 0.149144943 | -0.270986517 | 0.999933887 | -0.172419888 | -0.110508656 | -0.061911232 | 0.862415006 |
| ENSG00000163900 | *TMEM41A* | 1.408381356 | -0.077779593 | -0.072224586 | -0.005555007 | 0.999933887 | 0.15793185 | 0.100908839 | 0.057023011 | 0.862415006 |
| ENSG00000272595 | *OR10AH1P* | -1.16658988 | -0.095717502 | 0.188092372 | -0.283809874 | 0.999933887 | 0.0326661 | 0.170187577 | -0.137521477 | 0.862415006 |
| ENSG00000159433 | *STARD9* | 2.048554372 | 0.00934299 | -0.157252837 | 0.166595827 | 0.999933887 | -0.060140305 | -0.013844938 | -0.046295368 | 0.862415006 |
| ENSG00000173930 | *SLCO4C1* | 2.613017115 | -0.013223258 | 0.022668082 | -0.035891339 | 0.999933887 | -0.646242513 | -0.701866063 | 0.055623549 | 0.862418012 |
| ENSG00000230454 | *U73166.1* | -0.071955598 | 0.014752641 | -0.211509686 | 0.226262326 | 0.999933887 | -0.632342355 | -0.536952805 | -0.095389551 | 0.862418012 |
| ENSG00000146731 | *CCT6A* | 3.736739832 | -0.020722006 | -0.031826978 | 0.011104972 | 0.999933887 | 0.072539688 | 0.103282433 | -0.030742745 | 0.862739093 |
| ENSG00000174080 | *CTSF* | 0.864883371 | 0.105260592 | -0.023191128 | 0.12845172 | 0.999933887 | 0.185121225 | 0.121676885 | 0.06344434 | 0.86305464 |
| ENSG00000092758 | *COL9A3* | 1.312976408 | 0.086511485 | 0.147291758 | -0.060780274 | 0.999933887 | 2.920623904 | 3.017797953 | -0.097174049 | 0.863124721 |
| ENSG00000060971 | *ACAA1* | 5.948934605 | 0.010979448 | -0.003941198 | 0.014920647 | 0.999933887 | -0.186222554 | -0.224257298 | 0.038034744 | 0.863124721 |
| ENSG00000131725 | *WDR44* | 3.197354481 | -0.076215277 | -0.077423656 | 0.001208379 | 0.999933887 | -0.389576287 | -0.426673957 | 0.03709767 | 0.863147084 |
| ENSG00000169583 | *CLIC3* | 0.644464003 | -0.007263328 | 0.034657577 | -0.041920906 | 0.999933887 | 0.243904141 | 0.166844495 | 0.077059646 | 0.863147084 |
| ENSG00000198026 | *ZNF335* | 5.90164647 | 0.009716227 | 0.060296308 | -0.050580082 | 0.999933887 | 0.043048735 | 0.065794234 | -0.022745499 | 0.863217447 |
| ENSG00000258172 | *AC073655.1* | 0.964295954 | 0.220184736 | -0.003250606 | 0.223435342 | 0.999933887 | 0.854971467 | 0.948501869 | -0.093530402 | 0.863272791 |
| ENSG00000135972 | *MRPS9* | 0.303325598 | -0.108834808 | 0.078778406 | -0.187613214 | 0.999933887 | 0.14898247 | 0.072916876 | 0.076065594 | 0.863474728 |
| ENSG00000168517 | *HEXIM2* | 1.839391201 | 0.135962936 | -0.112829354 | 0.24879229 | 0.999933887 | 0.049614752 | 0.010989368 | 0.038625384 | 0.863494798 |
| ENSG00000118420 | *UBE3D* | -1.033121732 | -0.282805731 | 0.153628039 | -0.436433769 | 0.999933887 | -0.062232001 | -0.178015895 | 0.115783895 | 0.863494798 |
| ENSG00000102710 | *SUPT20H* | 4.761091601 | 0.057181608 | -0.050596887 | 0.107778495 | 0.999933887 | -0.27703815 | -0.249919544 | -0.027118606 | 0.863494798 |
| ENSG00000250848 | *AC021087.2* | -0.207371403 | 0.087721103 | -0.039209199 | 0.126930302 | 0.999933887 | -0.031326008 | -0.130209098 | 0.09888309 | 0.863494798 |
| ENSG00000168255 | *POLR2J3* | 0.787545425 | 0.034615486 | 0.056257479 | -0.021641994 | 0.999933887 | -0.112895096 | -0.186407492 | 0.073512395 | 0.863494798 |
| ENSG00000198393 | *ZNF26* | 2.75270236 | -0.094132448 | 0.013344037 | -0.107476485 | 0.999933887 | -0.028789176 | -0.068241857 | 0.039452682 | 0.863530987 |
| ENSG00000106333 | *PCOLCE* | 0.452951853 | -0.030196571 | 0.051618598 | -0.081815169 | 0.999933887 | -0.027815049 | -0.116572022 | 0.088756972 | 0.863614279 |
| ENSG00000175564 | *UCP3* | 0.527509291 | 0.073580288 | 0.051060339 | 0.022519949 | 0.999933887 | -0.345686768 | -0.42927089 | 0.083584121 | 0.863748462 |
| ENSG00000137274 | *BPHL* | -0.775299986 | 0.21147342 | 0.052486978 | 0.158986442 | 0.999933887 | 0.317195739 | 0.434331383 | -0.117135643 | 0.863955263 |
| ENSG00000102524 | *TNFSF13B* | 6.063869307 | -0.055357563 | -0.030335775 | -0.025021788 | 0.999933887 | -0.390030656 | -0.435147199 | 0.045116543 | 0.863955263 |
| ENSG00000197635 | *DPP4* | 1.860582055 | -0.023474898 | -0.219746761 | 0.196271864 | 0.999933887 | -0.068025223 | -0.010314915 | -0.057710308 | 0.863955263 |
| ENSG00000172349 | *IL16* | 6.686985249 | -0.01675598 | -0.088580588 | 0.071824608 | 0.999933887 | 0.64131541 | 0.679371489 | -0.038056079 | 0.863955263 |
| ENSG00000050628 | *PTGER3* | -1.0845152 | -0.012574984 | -0.12556831 | 0.112993326 | 0.999933887 | -0.2533146 | -0.082728357 | -0.170586243 | 0.863955263 |
| ENSG00000213638 | *ADAT3* | 1.670927305 | 0.017944066 | 0.049458588 | -0.031514522 | 0.999933887 | -0.838774956 | -0.900492071 | 0.061717115 | 0.863989143 |
| ENSG00000136100 | *VPS36* | 3.269967768 | 0.052316285 | -0.037569218 | 0.089885503 | 0.999933887 | -0.193894675 | -0.225835549 | 0.031940874 | 0.8640392 |
| ENSG00000187837 | *H1-2* | 5.596188098 | -0.000456312 | -0.063281174 | 0.062824862 | 0.999933887 | 0.028057719 | -0.007503597 | 0.035561315 | 0.864100148 |
| ENSG00000143155 | *TIPRL* | 3.657796949 | 0.047515374 | 0.085720806 | -0.038205432 | 0.999933887 | 0.182064158 | 0.154229298 | 0.027834861 | 0.864291204 |
| ENSG00000198019 | *FCGR1B* | 3.601447688 | 0.040561163 | -0.056843631 | 0.097404794 | 0.999933887 | -0.023165506 | 0.017184364 | -0.04034987 | 0.864376228 |
| ENSG00000128534 | *LSM8* | 3.919354653 | 0.027688276 | 0.037139088 | -0.009450812 | 0.999933887 | -0.174343378 | -0.13606545 | -0.038277929 | 0.86442154 |
| ENSG00000214114 | *MYCBP* | 4.300939723 | -0.067372385 | -0.127750719 | 0.060378334 | 0.999933887 | 0.121346516 | 0.090015433 | 0.031331083 | 0.864847674 |
| ENSG00000131459 | *GFPT2* | -0.118521913 | 0.212867127 | -0.024752528 | 0.237619654 | 0.999933887 | -0.080142237 | 0.018952009 | -0.099094246 | 0.86486804 |
| ENSG00000231890 | *DARS-AS1* | 4.862740777 | 0.239488988 | 0.285869371 | -0.046380383 | 0.999933887 | 0.488887847 | 0.433181875 | 0.055705972 | 0.865053507 |
| ENSG00000085719 | *CPNE3* | 4.14147453 | -0.036906643 | -0.006603813 | -0.03030283 | 0.999933887 | -0.245370258 | -0.279544392 | 0.034174134 | 0.865053507 |
| ENSG00000148396 | *SEC16A* | 7.303505239 | -0.018843983 | -0.01081824 | -0.008025744 | 0.999933887 | -0.064711146 | -0.04505325 | -0.019657896 | 0.865053507 |
| ENSG00000100413 | *POLR3H* | 2.174092058 | 0.160670453 | 0.156571644 | 0.004098809 | 0.999933887 | 0.052840829 | 0.100756781 | -0.047915952 | 0.865098849 |
| ENSG00000247121 | *AC009126.1* | 0.4068422 | 0.340174116 | 0.313488359 | 0.026685757 | 0.999933887 | -0.441454968 | -0.502547814 | 0.061092846 | 0.865205799 |
| ENSG00000101194 | *SLC17A9* | 1.319857303 | 0.205324503 | -0.012867069 | 0.218191573 | 0.999933887 | 0.177307695 | 0.240665032 | -0.063357337 | 0.865205799 |
| ENSG00000269480 | *AC020913.2* | 0.402019437 | 0.139027029 | 0.076951359 | 0.06207567 | 0.999933887 | -0.199993906 | -0.130668817 | -0.069325089 | 0.865205799 |
| ENSG00000151332 | *MBIP* | 1.336835839 | 0.03838162 | 0.039460556 | -0.001078936 | 0.999933887 | -0.103240263 | -0.170238848 | 0.066998585 | 0.865205799 |
| ENSG00000204745 | *AC083899.1* | -1.232262146 | -0.084618606 | 0.355635157 | -0.440253763 | 0.999933887 | -0.57813276 | -0.739429918 | 0.161297159 | 0.865205799 |
| ENSG00000006831 | *ADIPOR2* | 5.321560239 | -0.080056786 | -0.040994357 | -0.039062429 | 0.999933887 | -0.188263946 | -0.164158949 | -0.024104997 | 0.865279085 |
| ENSG00000278949 | *AC127070.4* | 1.348106368 | -0.145987338 | -0.231760862 | 0.085773524 | 0.999933887 | -0.172919972 | -0.114579283 | -0.058340689 | 0.865279085 |
| ENSG00000161547 | *SRSF2* | 7.151805137 | 0.058114252 | 0.101485735 | -0.043371483 | 0.999933887 | 0.426695996 | 0.460602948 | -0.033906952 | 0.865279085 |
| ENSG00000169020 | *ATP5ME* | 1.527756054 | -0.002954787 | 0.083723586 | -0.086678374 | 0.999933887 | -0.008878048 | -0.076182923 | 0.067304874 | 0.865279085 |
| ENSG00000134326 | *CMPK2* | 4.601863175 | 0.002096576 | -0.023841275 | 0.02593785 | 0.999933887 | 0.211646392 | 0.165854245 | 0.045792147 | 0.865486623 |
| ENSG00000137343 | *ATAT1* | 1.138531572 | 0.029024165 | -0.044530567 | 0.073554733 | 0.999933887 | 0.074016109 | 0.016189776 | 0.057826333 | 0.865609287 |
| ENSG00000110987 | *BCL7A* | -0.148387939 | 0.228548272 | -0.059728588 | 0.288276861 | 0.999933887 | -0.281644788 | -0.379001471 | 0.097356683 | 0.865832018 |

| | | | | | | | | | | |
|---|---|---|---|---|---|---|---|---|---|---|
| ENSG00000109917 | *ZPR1* | 4.479446061 | -0.037632731 | 0.003328789 | -0.04096152 | 0.999933887 | 0.188364478 | 0.215128598 | -0.02676412 | 0.865846334 |
| ENSG00000181104 | *F2R* | 3.129588889 | -0.037214203 | -0.113618855 | 0.076404652 | 0.999933887 | -0.032430645 | 0.004786284 | -0.03721693 | 0.865846334 |
| ENSG00000169188 | *APEX2* | 1.910338923 | -0.008253426 | -0.039429967 | 0.03117654 | 0.999933887 | 0.0143005 | 0.06105558 | -0.04675508 | 0.86610903 |
| ENSG00000137876 | *RSL24D1* | 4.843261804 | -0.075701721 | -0.058091126 | -0.017610595 | 0.999933887 | -0.036705592 | -0.006703871 | -0.030001722 | 0.866165118 |
| ENSG00000232112 | *TMA7* | 2.541295955 | -0.067396895 | 0.024639313 | -0.092036208 | 0.999933887 | 0.12571591 | 0.087968199 | 0.037747711 | 0.866165118 |
| ENSG00000134825 | *TMEM258* | 3.347958968 | 0.020667591 | -0.026458775 | 0.047126366 | 0.999933887 | 0.151822821 | 0.120138669 | 0.031684152 | 0.866165118 |
| ENSG00000184293 | *CLECL1* | -0.079171824 | 0.033147417 | -0.079295018 | 0.112442435 | 0.999933887 | 0.062555127 | 0.158705377 | -0.09615025 | 0.866165118 |
| ENSG00000244560 | *AC004890.2* | -0.332056026 | 0.008410067 | 0.049819611 | -0.041409544 | 0.999933887 | 0.133052543 | 0.042920747 | 0.090131796 | 0.86629096 |
| ENSG00000179869 | *ABCA13* | 1.130238169 | -0.212141836 | -0.043331718 | -0.168810118 | 0.999933887 | 0.224421213 | 0.27208064 | -0.047659427 | 0.866316984 |
| ENSG00000134250 | *NOTCH2* | 9.523944468 | -0.057348797 | -0.022107666 | -0.035241131 | 0.999933887 | -0.103734021 | -0.081913283 | -0.021820738 | 0.866500806 |
| ENSG00000102078 | *SLC25A14* | 1.81056972 | -0.022128244 | 0.025767213 | -0.047895458 | 0.999933887 | -0.108582153 | -0.165955716 | 0.057373563 | 0.866500806 |
| ENSG00000158435 | *CNOT11* | 4.479706305 | 0.000950783 | 0.111180396 | -0.110229613 | 0.999933887 | -0.191303133 | -0.164396555 | -0.026906578 | 0.866500806 |
| ENSG00000171863 | *RPS7* | 6.92699308 | 0.00825372 | -0.051870337 | 0.060124057 | 0.999933887 | 0.032963163 | 0.060214176 | -0.027251013 | 0.86657169 |
| ENSG00000106588 | *PSMA2* | 0.425907421 | -0.394659103 | -0.131478121 | -0.263180982 | 0.999933887 | -0.200923344 | -0.272580378 | 0.071657033 | 0.86665229 |
| ENSG00000162722 | *TRIM58* | -1.061447372 | 0.067248688 | 0.140316501 | -0.073067813 | 0.999933887 | 0.0115673 | 0.13217717 | -0.12060987 | 0.86665229 |
| ENSG00000135960 | *EDAR* | -1.166889064 | -0.316751939 | 0.429415984 | -0.746167923 | 0.969206975 | 0.281540093 | 0.394451454 | -0.112911361 | 0.866913574 |
| ENSG00000284930 | *AC005280.2* | 2.902129358 | 0.030948251 | -0.046444551 | 0.077392802 | 0.999933887 | 1.319557424 | 1.373809226 | -0.054251802 | 0.866913574 |
| ENSG00000106392 | *C1GALT1* | 4.107035848 | 0.028196853 | -0.05060675 | 0.078803603 | 0.999933887 | -0.436060039 | -0.37789213 | -0.058167909 | 0.866913574 |
| ENSG00000240449 | *REPIN1-AS1* | 0.558677366 | 0.036264511 | -0.31544542 | 0.351709931 | 0.999933887 | 1.499747253 | 1.440621705 | 0.059125548 | 0.866913574 |
| ENSG00000249471 | *ZNF324B* | -0.348596868 | -0.044341242 | -0.016832771 | -0.027508471 | 0.999933887 | 0.360163825 | 0.260167853 | 0.099995972 | 0.867082249 |
| ENSG00000286116 | *AL157394.3* | 1.782565784 | 0.16233875 | 0.190653274 | -0.028314524 | 0.999933887 | 1.6402878 | 1.563623976 | 0.076663824 | 0.867086377 |
| ENSG00000172985 | *SH3RF3* | 0.898480318 | 0.15506988 | 0.03946132 | 0.11560856 | 0.999933887 | -0.405300512 | -0.476830276 | 0.071529763 | 0.867372944 |
| ENSG00000184117 | *NIPSNAP1* | 2.325449963 | 0.102083482 | -0.029739437 | 0.131822919 | 0.999933887 | 0.179919085 | 0.124693368 | 0.055225717 | 0.8674099 |
| ENSG00000136925 | *TSTD2* | 1.964102099 | -0.098531104 | -0.013982054 | -0.08454905 | 0.999933887 | 0.028784602 | 0.076579586 | -0.047794983 | 0.8674099 |
| ENSG00000134419 | *RPS15A* | 5.233280142 | -0.046421956 | -0.00909354 | -0.037328417 | 0.999933887 | 0.067543313 | 0.098006641 | -0.030463327 | 0.8674099 |
| ENSG00000187098 | *MITF* | -1.088057202 | 0.12450624 | 0.140767435 | -0.016261196 | 0.999933887 | -2.080500018 | -2.188362892 | 0.107862874 | 0.8674099 |
| ENSG00000215154 | *AC141586.1* | 1.632900114 | 0.039608134 | 0.014076719 | 0.025531415 | 0.999933887 | 0.104982054 | 0.049647014 | 0.05533504 | 0.8674099 |
| ENSG00000158623 | *COPG2* | 0.659947456 | 0.043479396 | -0.080598755 | 0.124078151 | 0.999933887 | -0.122247544 | -0.032902328 | -0.089345216 | 0.8674099 |
| ENSG00000137100 | *DCTN3* | 4.030696216 | 0.013446079 | -0.018497788 | 0.031943867 | 0.999933887 | 0.04250865 | 0.0770142 | -0.03450555 | 0.8674099 |
| ENSG00000286616 | *AL008730.1* | -0.342037743 | -0.005956018 | -0.00440921 | -0.001546808 | 0.999933887 | 0.679792564 | 0.785653489 | -0.105860925 | 0.8674099 |
| ENSG00000189233 | *NUGGC* | 0.807857451 | -0.00185661 | 0.035444042 | -0.037300652 | 0.999933887 | 0.190102053 | 0.123123905 | 0.066978148 | 0.8674099 |
| ENSG00000138442 | *WDR12* | 0.790200087 | -0.02092553 | 0.056879186 | -0.077804716 | 0.999933887 | 0.075865587 | 0.131024608 | -0.05515902 | 0.867924644 |
| ENSG00000198464 | *ZNF480* | 1.404825384 | -0.046903773 | -0.124609122 | 0.077705349 | 0.999933887 | -0.155039738 | -0.215558344 | 0.060518605 | 0.867932867 |
| ENSG00000140839 | *CLEC18B* | -0.455555488 | 0.055678131 | 0.124663345 | -0.068985213 | 0.999933887 | -0.30165931 | -0.184178559 | -0.117480751 | 0.867968552 |
| ENSG00000116157 | *GPX7* | 0.843921271 | -0.090905557 | -0.216209309 | 0.125303752 | 0.999933887 | 0.148642556 | 0.085045031 | 0.063597524 | 0.867991655 |
| ENSG00000129595 | *EPB41L4A* | -1.185005393 | -0.19418994 | -0.337960856 | 0.143770916 | 0.999933887 | -0.064094068 | 0.049413478 | -0.113507545 | 0.868176764 |
| ENSG00000156599 | *ZDHHC5* | 6.381065778 | -0.039580934 | 0.027209609 | -0.066790542 | 0.999933887 | -0.244764299 | -0.218619957 | -0.026144342 | 0.868176764 |
| ENSG00000146373 | *RNF217* | -0.871722788 | -0.145056602 | -0.265457 | 0.120400398 | 0.999933887 | -0.770136294 | -0.920690957 | 0.150554662 | 0.868176764 |
| ENSG00000136213 | *CHST12* | 3.544038014 | -0.037218681 | -0.025267171 | -0.011951511 | 0.999933887 | -0.024133797 | -0.05878549 | 0.034651693 | 0.868176764 |
| ENSG00000211724 | *TRBV6-6* | -0.383724366 | -0.061518393 | -0.049045711 | -0.012472682 | 0.999933887 | 0.259310786 | 0.148793755 | 0.110517031 | 0.868176764 |
| ENSG00000131373 | *HACL1* | 2.094238441 | -0.017445069 | -0.089546417 | 0.072101348 | 0.999933887 | 0.052282857 | 0.09353124 | -0.041248383 | 0.868176764 |
| ENSG00000162383 | *SLC1A7* | -1.130791794 | 0.021832621 | 0.03356615 | -0.011733529 | 0.999933887 | 0.036565446 | 0.1239069 | -0.087341454 | 0.868176764 |
| ENSG00000011143 | *MKS1* | -0.130037812 | -0.196548153 | -0.032540082 | -0.164008071 | 0.999933887 | 0.163309737 | 0.241123977 | -0.07781424 | 0.868266185 |
| ENSG00000196155 | *PLEKHG4* | 1.191628198 | 0.00993096 | 0.255532943 | -0.245601983 | 0.999933887 | 0.167796994 | 0.11328755 | 0.054509444 | 0.868266185 |
| ENSG00000128626 | *MRPS12* | 1.44010144 | -0.005702136 | -0.012511771 | 0.006809635 | 0.999933887 | 0.132462276 | 0.185378898 | -0.052916622 | 0.868411217 |
| ENSG00000151062 | *CACNA2D4* | 1.758237285 | 0.0440131 | 0.049359057 | -0.005345957 | 0.999933887 | 0.161876647 | 0.208563451 | -0.046686804 | 0.868414455 |
| ENSG00000152455 | *SUV39H2* | -0.648273037 | 0.341747171 | -0.135178235 | 0.476925406 | 0.999933887 | -0.453820743 | -0.564812429 | 0.110991687 | 0.868471221 |
| ENSG00000066933 | *MYO9A* | 2.878735619 | 0.063229019 | -0.132862349 | 0.196091367 | 0.999933887 | -0.288288329 | -0.230825219 | -0.05746311 | 0.868471221 |
| ENSG00000135002 | *RFK* | 1.757298693 | 0.035392414 | -0.014632405 | 0.050024819 | 0.999933887 | -0.17700142 | -0.121567334 | -0.055434086 | 0.868471221 |
| ENSG00000077232 | *DNAJC10* | 3.69512517 | -0.013106299 | -0.179324501 | 0.166218202 | 0.999933887 | -0.131848127 | -0.078679199 | -0.053168929 | 0.868471221 |
| ENSG00000165632 | *TAF3* | 2.96690152 | -0.001304369 | -0.068827241 | 0.067522872 | 0.999933887 | 0.076850003 | 0.04134345 | 0.035506553 | 0.868471221 |
| ENSG00000224083 | *MTCO1P11* | -0.791665378 | 0.004033747 | 0.159783732 | -0.155749985 | 0.999933887 | 0.189659991 | 0.408468622 | -0.218808631 | 0.868471221 |
| ENSG00000272092 | *AC087623.2* | 0.17842775 | 0.28483382 | 0.062375853 | 0.222457967 | 0.999933887 | -0.222843053 | -0.296393882 | 0.073550829 | 0.868532214 |
| ENSG00000079462 | *PAFAH1B3* | 0.246346955 | -0.050568635 | 0.004435749 | -0.055004384 | 0.999933887 | 0.130391745 | 0.204687111 | -0.074295366 | 0.868532214 |
| ENSG00000276405 | *TRBV13* | -1.451871609 | -0.036995891 | -0.414108552 | 0.377112661 | 0.999933887 | 0.034892891 | 0.189433243 | -0.154540352 | 0.868532214 |
| ENSG00000162576 | *MXRA8* | -0.361415164 | 0.02784016 | 0.18774442 | -0.15990426 | 0.999933887 | 0.320472118 | 0.211632246 | 0.108839872 | 0.868532214 |
| ENSG00000286872 | *AC024270.4* | 0.258092218 | 0.04460573 | -0.155392954 | 0.199998684 | 0.999933887 | -0.3891084 | -0.495801495 | 0.106693095 | 0.868561919 |
| ENSG00000112941 | *TENT4A* | 4.786317017 | -0.019102246 | 0.015912965 | -0.035015211 | 0.999933887 | -0.045577289 | -0.070321153 | 0.024743864 | 0.86915984 |
| ENSG00000264577 | *AC010761.1* | 0.070633172 | -0.024835052 | -0.047434148 | 0.022599096 | 0.999933887 | 0.1022055 | 0.011135849 | 0.091069651 | 0.86915984 |
| ENSG00000123411 | *IKZF4* | 1.04438932 | 0.044279392 | 0.032871547 | 0.011407846 | 0.999933887 | 0.354787319 | 0.290427981 | 0.064359338 | 0.869217266 |
| ENSG00000265185 | *SNORD3B-1* | -0.496219848 | 0.00076065 | 0.086072034 | -0.085311384 | 0.999933887 | 0.72404525 | 0.838107531 | -0.114062281 | 0.869217266 |
| ENSG00000120280 | *CXorf21* | 4.437488314 | -0.116373607 | -0.128671564 | 0.012297957 | 0.999933887 | -0.050343418 | -0.087975242 | 0.037631824 | 0.869245515 |
| ENSG00000246067 | *RAB30-DT* | 0.808830351 | 0.055233281 | 0.015896307 | 0.039336974 | 0.999933887 | 0.208628227 | 0.152108107 | 0.05652012 | 0.869245515 |
| ENSG00000204390 | *HSPA1L* | 0.934111446 | 0.02279431 | 0.219605571 | -0.196811261 | 0.999933887 | 0.460169266 | 0.397688047 | 0.06248122 | 0.869245515 |
| ENSG00000173421 | *IHO1* | 1.540087539 | -0.147870589 | 0.013542339 | -0.161412928 | 0.999933887 | -0.468700521 | -0.408348842 | -0.060351679 | 0.869440195 |
| ENSG00000119682 | *AREL1* | 7.346100093 | -0.046651319 | -0.007060618 | -0.039590701 | 0.999933887 | 0.088312876 | 0.107463688 | -0.019150813 | 0.869440195 |
| ENSG00000066739 | *ATG2B* | 5.239226636 | -0.004735193 | -0.104672942 | 0.099937749 | 0.999933887 | -0.431153948 | -0.371273402 | -0.059880546 | 0.869472072 |
| ENSG00000235488 | *JARID2-AS1* | 3.067692473 | -0.033591036 | 0.115576026 | -0.149167062 | 0.999933887 | 0.375318164 | 0.421363468 | -0.046045304 | 0.869534476 |
| ENSG00000019991 | *HGF* | 0.311829016 | -0.082565127 | -0.040022814 | -0.042542314 | 0.999933887 | -1.216080479 | -1.314941761 | 0.098861282 | 0.869619484 |
| ENSG00000135164 | *DMTF1* | 5.014241217 | -0.047639192 | -0.010049387 | -0.037589805 | 0.999933887 | -0.403839168 | -0.45667655 | 0.052837383 | 0.869630296 |
| ENSG00000179639 | *FCER1A* | 0.337786363 | 0.194840373 | -0.304561762 | 0.499402135 | 0.944094537 | -0.188464824 | -0.123266204 | -0.06519862 | 0.869661709 |
| ENSG00000135709 | *KIAA0513* | 6.741315294 | -0.041748979 | -0.043221824 | 0.001472845 | 0.999933887 | -0.905049407 | -0.86420997 | -0.040839436 | 0.869925767 |
| ENSG00000256553 | *TRAV1-2* | -0.772768909 | -0.050779282 | -0.058813316 | 0.008034034 | 0.999933887 | 0.394051783 | 0.287802078 | 0.106249705 | 0.869971828 |
| ENSG00000198157 | *HMGN5* | -0.283008497 | -0.14474302 | -0.1122406 | -0.03250242 | 0.999933887 | -0.970933277 | -1.081814351 | 0.110881074 | 0.870212834 |
| ENSG00000088808 | *PPP1R13B* | 1.98609554 | -0.034957216 | 0.032755227 | -0.067712443 | 0.999933887 | 0.005150562 | -0.044512561 | 0.049663124 | 0.870212834 |
| ENSG00000124635 | *H2BC11* | 0.155637458 | -0.046615916 | -0.061522334 | 0.014906418 | 0.999933887 | 0.017590777 | -0.085678596 | 0.103269372 | 0.870415729 |
| ENSG00000197540 | *GZMM* | 4.267391712 | 0.100629676 | -0.028192555 | 0.128822231 | 0.999933887 | 0.160637133 | 0.123623409 | 0.037013724 | 0.870505375 |
| ENSG00000162929 | *KIAA1841* | 0.22536293 | 0.29056495 | -0.384384573 | 0.674949523 | 0.144167088 | -0.447338289 | -0.531590533 | 0.084252244 | 0.870526104 |
| ENSG00000108561 | *C1QBP* | 2.781063398 | 0.084742225 | 0.05031349 | 0.034428735 | 0.999933887 | 0.188282089 | 0.232336822 | -0.044054733 | 0.87053105 |
| ENSG00000178802 | *MPI* | 2.146525671 | 0.0090419 | 0.006249 | 0.002792899 | 0.999933887 | 0.17812365 | 0.225903678 | -0.047780028 | 0.87053105 |
| ENSG00000168395 | *ING5* | 3.442464574 | 0.004107394 | 0.081056884 | -0.076949489 | 0.999933887 | 0.176206898 | 0.13871841 | 0.037488488 | 0.87053105 |
| ENSG00000134030 | *CTIF* | 0.899019037 | 0.010847248 | -0.008759393 | 0.01960664 | 0.999933887 | -0.065322934 | 0.002829055 | -0.068151989 | 0.870670963 |
| ENSG00000132465 | *JCHAIN* | 0.44134625 | 0.02288801 | -0.059410294 | 0.082298304 | 0.999933887 | -0.043854297 | 0.053354054 | -0.097208351 | 0.870747641 |
| ENSG00000071564 | *TCF3* | 3.576486062 | 0.100079366 | 0.143109582 | -0.043030216 | 0.999933887 | 0.084860995 | 0.052969619 | 0.031891376 | 0.870899874 |
| ENSG00000136813 | *ECPAS* | 5.1902988 | -0.036623518 | 0.007984468 | -0.044607986 | 0.999933887 | -0.022654908 | -0.046550656 | 0.023895748 | 0.870899874 |
| ENSG00000232098 | *AC012313.1* | 0.805300618 | 0.260166429 | -0.162607845 | 0.422774274 | 0.999933887 | 0.062665732 | 0.011219926 | 0.051445806 | 0.870988098 |
| ENSG00000124207 | *CSE1L* | 2.924325002 | -0.087984217 | -0.002620527 | -0.08536369 | 0.999933887 | 0.006705525 | -0.032413751 | 0.039119276 | 0.870988098 |
| ENSG00000122034 | *GTF3A* | 4.786063546 | -0.024279413 | -0.097001292 | 0.072721879 | 0.999933887 | 0.044207634 | 0.017231307 | 0.026976327 | 0.870988098 |
| ENSG00000267758 | *AC099811.4* | 1.560347019 | -0.042447346 | 0.119792036 | -0.162239382 | 0.999933887 | 1.32621715 | 1.422045102 | -0.095827952 | 0.870988098 |
| ENSG00000116062 | *MSH6* | 3.336464522 | 0.015353682 | 0.01229887 | 0.003054813 | 0.999933887 | -0.112018078 | -0.076463918 | -0.035554161 | 0.870988098 |
| ENSG00000137509 | *PRCP* | 6.294895802 | -0.003403881 | -0.026931833 | 0.023527952 | 0.999933887 | -0.271776923 | -0.232151215 | -0.039625708 | 0.870988098 |
| ENSG00000140450 | *ARRDC4* | 4.735276148 | -0.145948087 | -0.112627719 | -0.033320369 | 0.999933887 | -0.233516868 | -0.181903433 | -0.051613435 | 0.871003879 |
| ENSG00000180574 | *EIF2S3B* | -1.770112482 | 0.274647295 | 0.006934395 | 0.2677129 | 0.999933887 | -0.004678668 | 0.103923161 | -0.108601829 | 0.871003879 |
| ENSG00000010219 | *DYRK4* | -0.454052378 | -0.112002859 | -0.018363825 | -0.093639034 | 0.999933887 | 0.20312507 | 0.108453155 | 0.094671916 | 0.871003879 |
| ENSG00000233170 | *AC138356.2* | 0.120778108 | -0.062483112 | -0.014248618 | -0.048234494 | 0.999933887 | -0.439476142 | -0.560970681 | 0.12149454 | 0.871003879 |
| ENSG00000269984 | *AC078795.1* | -0.443888 | 0.109509852 | -0.0057153 | 0.115225152 | 0.999933887 | -0.033726257 | 0.08465137 | -0.118377627 | 0.871124176 |
| ENSG00000261573 | *AL157402.2* | 1.219140718 | 0.047228054 | 0.452683807 | -0.405455753 | 0.999933887 | 0.871686428 | 0.725937259 | 0.145749169 | 0.871124176 |
| ENSG00000169330 | *MINAR1* | -0.682890308 | 0.025598317 | 0.08245597 | -0.056857652 | 0.999933887 | 0.018421596 | -0.064841801 | 0.083263397 | 0.871124176 |
| ENSG00000166405 | *RIC3* | 1.269731295 | 0.010305854 | 0.148831198 | -0.138525343 | 0.999933887 | -0.02298566 | 0.031583776 | -0.054569436 | 0.871124176 |
| ENSG00000143434 | *SEMA6C* | 0.715680753 | 0.008174611 | 0.133248332 | -0.125073721 | 0.999933887 | -0.118335406 | -0.189988605 | 0.0716532 | 0.871124176 |
| ENSG00000125355 | *TMEM255A* | 0.118950551 | 0.161628576 | -0.316641892 | 0.478270468 | 0.999933887 | -0.372512983 | -0.310874834 | -0.061638149 | 0.871177781 |
| ENSG00000102241 | *HTATSF1* | 3.397068239 | -0.035645415 | -0.178630498 | 0.142985083 | 0.999933887 | 0.043867473 | 0.004652467 | 0.039215006 | 0.871177781 |
| ENSG00000204514 | *ZNF814* | 2.337900813 | -0.064054019 | -0.09521414 | 0.031160121 | 0.999933887 | -0.196418336 | -0.157818173 | -0.038600162 | 0.871224564 |
| ENSG00000100321 | *SYNGR1* | -0.189471933 | 0.067366138 | -0.021470452 | 0.08883659 | 0.999933887 | 0.40790596 | 0.326281292 | 0.081624668 | 0.871224564 |
| ENSG00000137494 | *ANKRD42* | 3.964670463 | 0.077830604 | 0.043660674 | 0.03416993 | 0.999933887 | 0.437341829 | 0.473733969 | -0.03639214 | 0.871231854 |

| | | | | | | | | | | |
|---|---|---|---|---|---|---|---|---|---|---|
| ENSG00000171680 | *PLEKHG5* | -0.565976959 | 0.03570503 | 0.027008608 | 0.008696422 | 0.999933887 | -0.079012703 | -0.191213661 | 0.112200958 | 0.871231854 |
| ENSG00000244242 | *IFITM10* | 4.655531423 | 0.1377672 | 0.243334685 | -0.105567485 | 0.999933887 | 0.798328914 | 0.869718758 | -0.071389844 | 0.871288673 |
| ENSG00000237264 | *FTH1P11* | 0.818960404 | 0.059435975 | -0.009436569 | 0.068872543 | 0.999933887 | 0.529211797 | 0.585669325 | -0.056457528 | 0.871288673 |
| ENSG00000230701 | *FBXW4P1* | 0.281047965 | -0.044221066 | -0.060050286 | 0.01582922 | 0.999933887 | 0.246205399 | 0.144474431 | 0.101730968 | 0.871288673 |
| ENSG00000158710 | *TAGLN2* | 10.79183884 | 0.015803203 | -0.015823433 | 0.031626636 | 0.999933887 | 0.026101689 | 0.04943311 | -0.023331421 | 0.871288673 |
| ENSG00000145014 | *TMEM44* | 0.26192638 | -0.122341191 | -0.113233728 | -0.009107464 | 0.999933887 | 0.15013076 | 0.080577305 | 0.069553454 | 0.871347823 |
| ENSG00000130783 | *CCDC62* | -0.075407067 | -0.215742155 | -0.063994128 | -0.151748027 | 0.999933887 | -0.38383811 | -0.46966865 | 0.08583054 | 0.87146824 |
| ENSG00000174606 | *ANGEL2* | 3.654949531 | -0.048027277 | 0.004154176 | -0.052181453 | 0.999933887 | -0.207584797 | -0.168827271 | -0.038757526 | 0.87146824 |
| ENSG00000146281 | *PM20D2* | 3.448879949 | -0.058590507 | 0.022628594 | -0.081219101 | 0.999933887 | 0.263808602 | 0.223202788 | 0.040605813 | 0.87146824 |
| ENSG00000258777 | *HIF1A-AS1* | -2.141432959 | 0.006067397 | 0.266307313 | -0.260239916 | 0.999933887 | 2.377314463 | 2.516008621 | -0.138694157 | 0.87146824 |
| ENSG00000160199 | *PKNOX1* | 4.524079712 | -0.068318199 | -0.072127341 | 0.003809142 | 0.999933887 | -0.369858764 | -0.395685065 | 0.025826301 | 0.871572807 |
| ENSG00000121390 | *PSPC1* | 4.908999388 | -0.015610294 | -0.008409468 | -0.007200826 | 0.999933887 | -0.026099452 | -0.048298248 | 0.022198796 | 0.871572807 |
| ENSG00000110171 | *TRIM3* | 2.552018725 | 0.144847477 | 0.022861707 | 0.12198577 | 0.999933887 | 0.171945836 | 0.134075154 | 0.037870682 | 0.871647603 |
| ENSG00000111716 | *LDHB* | 0.657698427 | -0.070809986 | -0.185929343 | 0.115119357 | 0.999933887 | -0.001488071 | 0.074285766 | -0.075773838 | 0.871647603 |
| ENSG00000103047 | *TANGO6* | 1.619011112 | -0.038129872 | 0.095211623 | -0.133341494 | 0.999933887 | 0.088681172 | 0.143623078 | -0.054941905 | 0.871647603 |
| ENSG00000134153 | *EMC7* | 4.575860675 | 0.009042696 | 0.011803247 | -0.002760551 | 0.999933887 | -0.254115166 | -0.283293681 | 0.029178515 | 0.871647603 |
| ENSG00000125743 | *SNRPD2* | 4.370729279 | 0.021272204 | 0.019922914 | 0.00134929 | 0.999933887 | 0.14325566 | 0.174690486 | -0.031434826 | 0.871686358 |
| ENSG00000212743 | *LINC02656* | 2.24226644 | -0.004082414 | 0.197841901 | -0.201924315 | 0.999933887 | -0.159942073 | -0.105032561 | -0.054909513 | 0.871728514 |
| ENSG00000135297 | *MTO1* | 2.564314601 | 0.011602705 | 0.030280513 | -0.018677808 | 0.999933887 | 0.085217573 | 0.121055228 | -0.035837655 | 0.871991797 |
| ENSG00000153814 | *JAZF1* | 3.015574528 | -0.092074825 | -0.004162825 | -0.087912 | 0.999933887 | -0.144975492 | -0.179010652 | 0.03403516 | 0.872017737 |
| ENSG00000166135 | *HIF1AN* | 5.546204341 | -0.059206313 | -0.063359277 | 0.004152964 | 0.999933887 | -0.081269564 | -0.056661746 | -0.024607818 | 0.872017737 |
| ENSG00000228049 | *POLR2J2* | -0.157873125 | 0.086954263 | 0.017481413 | 0.069472849 | 0.999933887 | -0.06059929 | 0.020620693 | -0.081219983 | 0.872017737 |
| ENSG00000255328 | *AC136475.5* | -0.695501038 | -0.06033412 | 0.036699828 | -0.097033947 | 0.999933887 | 0.112146614 | 0.233974689 | -0.121828075 | 0.872017737 |
| ENSG00000199349 | *Y_RNA* | -1.474817383 | -0.069435517 | 0.018094224 | -0.087529741 | 0.999933887 | 0.481184106 | 0.245302283 | 0.235881823 | 0.872017737 |
| ENSG00000149636 | *DSN1* | 1.308055495 | -0.019623264 | 0.088872054 | -0.108495318 | 0.999933887 | -0.009168003 | 0.053491361 | -0.062659364 | 0.872017737 |
| ENSG00000163803 | *PLB1* | 5.192095237 | -0.009106128 | 0.025106062 | -0.03421219 | 0.999933887 | 0.109645741 | 0.134315709 | -0.024669968 | 0.872017737 |
| ENSG00000169217 | *CD2BP2* | 6.081635973 | 0.0104824 | 0.044855757 | -0.034373357 | 0.999933887 | -0.183230467 | -0.212179235 | 0.028948768 | 0.872017737 |
| ENSG00000078618 | *NRDC* | 7.081661471 | -0.000602654 | -0.083452983 | 0.082850329 | 0.999933887 | -0.160953561 | -0.139018551 | -0.02193501 | 0.872017737 |
| ENSG00000251441 | *RTEL1P1* | 1.694170596 | 0.066972423 | 0.057429579 | 0.009542844 | 0.999933887 | 0.634032094 | 0.539339847 | 0.094692247 | 0.872078866 |
| ENSG00000120215 | *MLANA* | -0.16233898 | 0.035289508 | -0.126048365 | 0.161337873 | 0.999933887 | -0.387736242 | -0.49811266 | 0.110376418 | 0.872109379 |
| ENSG00000150316 | *CWC15* | 3.912794529 | -0.143482954 | -0.165405299 | 0.021922345 | 0.999933887 | -0.112125551 | -0.083091719 | -0.029033832 | 0.872212032 |
| ENSG00000272462 | *U91328.1* | 0.915577271 | -0.037418125 | -0.161495989 | 0.124077864 | 0.999933887 | -0.016422534 | 0.061064182 | -0.077486715 | 0.872278976 |
| ENSG00000211750 | *TRBV24-1* | -0.427691773 | -0.12846519 | -0.437670025 | 0.309204835 | 0.999933887 | 0.22154672 | 0.31227262 | -0.090725899 | 0.872449293 |
| ENSG00000111537 | *IFNG* | 2.090062162 | -0.057039984 | -0.083547484 | 0.0265075 | 0.999933887 | 0.028860672 | -0.020395122 | 0.049255795 | 0.872449293 |
| ENSG00000279348 | *AC012513.3* | -0.369988246 | 0.03117557 | -0.335189548 | 0.366365118 | 0.999933887 | 0.094244909 | 0.173161412 | -0.078916503 | 0.872449293 |
| ENSG00000163694 | *RBM47* | 7.66009476 | 0.016404411 | 0.03455022 | -0.018145809 | 0.999933887 | -0.395164619 | -0.368385513 | -0.026779106 | 0.872490196 |
| ENSG00000242522 | *KLHL6-AS1* | 0.038112784 | -0.099524669 | -0.008018394 | -0.091506276 | 0.999933887 | 0.369829709 | 0.292365504 | 0.077464206 | 0.872493431 |
| ENSG00000171236 | *LRG1* | 6.107296672 | 0.087964846 | 0.017335104 | 0.070629742 | 0.999933887 | 0.568367169 | 0.538238377 | 0.030128792 | 0.872579034 |
| ENSG00000161999 | *JMJD8* | 1.374894351 | 0.140747026 | 0.064950632 | 0.075796393 | 0.999933887 | 0.036410737 | -0.022014932 | 0.058425669 | 0.872579034 |
| ENSG00000260971 | *AC119674.1* | -1.060615546 | 0.439744683 | -0.123059404 | 0.562804087 | 0.999933887 | 0.091058473 | 0.152010207 | -0.060951734 | 0.872579034 |
| ENSG00000237945 | *LINC00649* | 2.942873793 | 0.052822356 | -0.036944573 | 0.089766929 | 0.999933887 | 0.096866787 | 0.132787165 | -0.035920377 | 0.872579034 |
| ENSG00000224001 | *AL360081.1* | 0.311117561 | 0.096113862 | -0.024001242 | 0.120115104 | 0.999933887 | -0.115586815 | -0.034314073 | -0.081272742 | 0.872579034 |
| ENSG00000253982 | *AC100810.1* | 2.169021921 | -0.072175487 | -0.120995386 | 0.048819899 | 0.999933887 | -0.229479018 | -0.267460283 | 0.037981265 | 0.872579034 |
| ENSG00000250251 | *PKD1P6* | 1.525491694 | -0.058719233 | 0.107791764 | -0.166510997 | 0.999933887 | 0.060379909 | 0.119688325 | -0.059308416 | 0.872579034 |
| ENSG00000107281 | *NPDC1* | 2.579156932 | 0.041430617 | 0.165035763 | -0.123605146 | 0.999933887 | 0.061050091 | 0.011725948 | 0.049324142 | 0.872579034 |
| ENSG00000161800 | *RACGAP1* | 0.904885899 | -0.044215088 | -0.128525215 | 0.084310127 | 0.999933887 | -0.083366285 | -0.138878505 | 0.05551222 | 0.872579034 |
| ENSG00000134061 | *CD180* | -0.647765667 | 0.092795219 | 0.134679799 | -0.04188458 | 0.999933887 | -0.947884715 | -0.85060452 | -0.097280195 | 0.872579034 |
| ENSG00000004142 | *POLDIP2* | 4.145225149 | -0.016205514 | 0.015274031 | -0.031479546 | 0.999933887 | -0.088331164 | -0.116212636 | 0.027881472 | 0.872579034 |
| ENSG00000279407 | *AC007191.1* | 1.175190071 | 0.024443851 | 0.315418158 | -0.290974306 | 0.944094537 | 0.007685511 | -0.051014844 | 0.058700356 | 0.872579034 |
| ENSG00000256448 | *AP000763.3* | -0.32354979 | 0.018487913 | 0.0799902 | -0.061502287 | 0.999933887 | -0.963435122 | -1.10573113 | 0.142296009 | 0.872579034 |
| ENSG00000125971 | *DYNLRB1* | 1.425525 | 0.009385903 | 0.16452656 | -0.155140658 | 0.999933887 | -0.009294549 | 0.049711917 | -0.059006465 | 0.872579034 |
| ENSG00000226564 | *FTH1P20* | 3.02412795 | 0.008614396 | 0.009483451 | -0.000869054 | 0.999933887 | 0.660708177 | 0.617094889 | 0.043613289 | 0.872579034 |
| ENSG00000280195 | *AC245140.2* | 0.74003558 | 0.155062839 | 0.460460105 | -0.305397266 | 0.999933887 | 0.767447128 | 0.692428671 | 0.075018457 | 0.87287534 |
| ENSG00000267858 | *MZF1-AS1* | 1.705345011 | 0.206143903 | 0.303172997 | -0.097029094 | 0.999933887 | -0.050371368 | -0.130169654 | 0.079798286 | 0.872910206 |
| ENSG00000163421 | *PROK2* | 8.910788501 | 0.039681398 | 0.057231704 | -0.017550306 | 0.999933887 | -0.031560296 | -0.070684287 | 0.039123991 | 0.872975794 |
| ENSG00000137992 | *DBT* | 1.333683032 | -0.025079883 | 0.050251088 | -0.075330971 | 0.999933887 | -0.226106012 | -0.291118865 | 0.065012853 | 0.872992028 |
| ENSG00000175197 | *DDIT3* | 7.802334442 | 0.111690051 | 0.146027068 | -0.034337017 | 0.999933887 | 0.08779052 | 0.060236666 | 0.027553854 | 0.873177034 |
| ENSG00000259151 | *CAP2P1* | -0.168103733 | 0.186387893 | 0.059079654 | 0.127308239 | 0.999933887 | 0.036400414 | 0.126064497 | -0.089664083 | 0.873355089 |
| ENSG00000163602 | *RYBP* | 7.521928472 | -0.053937102 | -0.029799216 | -0.024137886 | 0.999933887 | 0.082611485 | 0.108689321 | -0.026077835 | 0.873355089 |
| ENSG00000284686 | *AC119674.2* | -1.536810601 | -0.276887362 | -0.513814672 | 0.236927309 | 0.999933887 | -0.081958995 | -0.015976251 | -0.065982744 | 0.873355089 |
| ENSG00000117479 | *SLC19A2* | 2.996285416 | -0.024452297 | -0.046929249 | 0.022476952 | 0.999933887 | -0.075979871 | -0.024425114 | -0.051554757 | 0.873355089 |
| ENSG00000027869 | *SH2D2A* | 4.330846318 | -0.035852123 | -0.006009255 | -0.029842868 | 0.999933887 | 0.165229334 | 0.193000293 | -0.027770958 | 0.873370287 |
| ENSG00000138640 | *FAM13A* | 2.093724923 | -0.03290455 | -0.115813797 | 0.082909247 | 0.999933887 | -0.298496014 | -0.236297251 | -0.062198763 | 0.873370287 |
| ENSG00000261172 | *AC133919.2* | 1.119424237 | 0.030961955 | -0.035895872 | 0.066857827 | 0.999933887 | 0.337438959 | 0.260273861 | 0.077165098 | 0.873370287 |
| ENSG00000253676 | *TAGLN2P1* | -0.195032815 | -0.002551204 | -0.126167276 | 0.123616072 | 0.999933887 | 0.202000917 | 0.103530555 | 0.098470362 | 0.873370287 |
| ENSG00000132600 | *PRMT7* | 2.748808541 | -0.000583171 | -0.041293478 | 0.040710306 | 0.999933887 | 0.026348823 | 0.060956182 | -0.034607359 | 0.873370287 |
| ENSG00000054611 | *TBC1D22A* | 5.134608127 | 0.038216071 | -0.0176652 | 0.055881271 | 0.999933887 | 0.273233192 | 0.296673846 | -0.023440655 | 0.873375936 |
| ENSG00000172824 | *CES4A* | -0.147964072 | -0.055936799 | 0.399384371 | -0.45532117 | 0.999933887 | -0.061417704 | -0.137850709 | 0.076433005 | 0.8733966 |
| ENSG00000204472 | *AIF1* | 6.819632149 | -0.032937377 | -0.039939936 | 0.007002559 | 0.999933887 | 0.098422785 | 0.0676827 | 0.030740085 | 0.873437224 |
| ENSG00000177383 | *MAGEF1* | -0.011070319 | 0.224205497 | 0.030990861 | 0.193214636 | 0.999933887 | -0.012414451 | 0.061106703 | -0.073521155 | 0.87375036 |
| ENSG00000138363 | *ATIC* | 2.456725618 | -0.047014359 | -0.192124265 | 0.145109906 | 0.999933887 | 0.17186843 | 0.205848812 | -0.033980382 | 0.87375036 |
| ENSG00000100918 | *REC8* | 6.85845196 | 0.020394527 | -0.018216587 | 0.038611114 | 0.999933887 | 0.193867484 | 0.217452645 | -0.023585161 | 0.87375036 |
| ENSG00000280047 | *AC091825.3* | 0.505484138 | -0.078519843 | -0.231127135 | 0.152607291 | 0.999933887 | -0.45556516 | -0.508268145 | 0.052702985 | 0.87375036 |
| ENSG00000101132 | *PFDN4* | 0.596501123 | -0.017118877 | -0.346707052 | 0.329588174 | 0.999933887 | -0.093468242 | -0.03421728 | -0.059250963 | 0.87375036 |
| ENSG00000012817 | *KDM5D* | -2.468918478 | -0.15448107 | 0.06372029 | -0.21820136 | 0.999933887 | -0.094441908 | -0.04928639 | -0.045155518 | 0.874199859 |
| ENSG00000136240 | *KDELR2* | 3.846800832 | -0.030160857 | 0.04555357 | -0.075714427 | 0.999933887 | -0.074668973 | -0.044503636 | -0.030165338 | 0.874199859 |
| ENSG00000279727 | *LINC02033* | -0.144511405 | 0.060088565 | 0.15272033 | -0.092631765 | 0.999933887 | 0.219663177 | 0.111186793 | 0.108476385 | 0.874199859 |
| ENSG00000004534 | *RBM6* | 5.041751726 | -0.011725774 | -0.042575182 | 0.030849407 | 0.999933887 | 0.046041757 | 0.024534861 | 0.021506896 | 0.874199859 |
| ENSG00000114054 | *PCCB* | 1.193231824 | 0.024611216 | 0.149068768 | -0.124457551 | 0.999933887 | -0.038897277 | 0.041315109 | -0.080212386 | 0.874199859 |
| ENSG00000224631 | *RPS27AP16* | -0.407682211 | 0.008847598 | 0.23672195 | -0.227874352 | 0.999933887 | 0.255079627 | 0.348349974 | -0.093270347 | 0.874199859 |
| ENSG00000231305 | *AC112484.1* | 1.226247386 | -0.005890948 | -0.019918195 | 0.014027247 | 0.999933887 | 0.580321334 | 0.633800301 | -0.053478966 | 0.874199859 |
| ENSG00000115998 | *C2orf42* | 2.464674995 | 0.062153426 | -0.126788412 | 0.188941838 | 0.999933887 | -0.004324454 | -0.039444638 | 0.035120185 | 0.874212347 |
| ENSG00000145293 | *ENOPH1* | 2.093984181 | -0.138588456 | -0.085837049 | -0.052751407 | 0.999933887 | -0.113780831 | -0.149061548 | 0.035280718 | 0.874267086 |
| ENSG00000280046 | *AC104581.4* | 1.871628506 | 0.256724261 | 0.335983437 | -0.079259176 | 0.999933887 | 1.078463759 | 1.143025462 | -0.064561703 | 0.874774864 |
| ENSG00000213145 | *CRIP1* | 2.624448114 | 0.069940816 | 0.020053958 | 0.049886858 | 0.999933887 | 0.157752116 | 0.108585643 | 0.049166472 | 0.874774864 |
| ENSG00000171984 | *SHLD1* | 1.106182123 | -0.095643618 | -0.029322612 | -0.066321006 | 0.999933887 | -0.247633996 | -0.299402544 | 0.051768548 | 0.874774864 |
| ENSG00000141543 | *EIF4A3* | 6.861728749 | 0.027509521 | -0.012278775 | 0.039788296 | 0.999933887 | 0.051220291 | 0.075298074 | -0.024077783 | 0.874774864 |
| ENSG00000123096 | *SSPN* | -0.200925082 | -0.158034834 | 0.063622438 | -0.221657272 | 0.999933887 | -1.783115428 | -1.864271772 | 0.081156344 | 0.874774864 |
| ENSG00000232629 | *HLA-DQB2* | -0.628673582 | 0.083890356 | -0.1486971 | 0.232587457 | 0.999933887 | 0.323422379 | 0.229328698 | 0.094093681 | 0.874774864 |
| ENSG00000049167 | *ERCC8* | 0.318857304 | -0.013557534 | -0.09795765 | 0.084400116 | 0.999933887 | 0.004633999 | 0.095809013 | -0.091175014 | 0.874774864 |
| ENSG00000280011 | *AL031595.2* | -0.252378924 | -0.016089655 | -0.069472647 | 0.053382992 | 0.999933887 | -0.179573814 | -0.28351265 | 0.103938836 | 0.874774864 |
| ENSG00000214425 | *LRRC37A4P* | 2.403204157 | -0.007754404 | 0.091716425 | -0.099470829 | 0.999933887 | -0.207031438 | -0.150377219 | -0.05665422 | 0.874774864 |
| ENSG00000236144 | *TMEM147-AS1* | -0.297793979 | -0.301759678 | 0.061804716 | -0.363564393 | 0.999933887 | 0.179743223 | 0.104548044 | 0.075195179 | 0.874914592 |
| ENSG00000273382 | *AL356488.3* | 1.526660617 | -0.132645352 | -0.10533569 | -0.027309663 | 0.999933887 | -0.218284505 | -0.158827563 | -0.059456942 | 0.874914592 |
| ENSG00000138439 | *FAM117B* | 2.710134811 | -0.054270398 | 0.059865416 | -0.114135815 | 0.999933887 | -0.16296785 | -0.211469318 | 0.048501467 | 0.874914592 |
| ENSG00000028839 | *TBPL1* | 2.609277194 | 0.04993924 | 0.027138576 | 0.022800664 | 0.999933887 | -0.189128688 | -0.222172008 | 0.033043319 | 0.874914592 |
| ENSG00000196233 | *LCOR* | 6.125425463 | -0.043789292 | -0.075922692 | 0.032133399 | 0.999933887 | -0.408202831 | -0.368595753 | -0.039607078 | 0.874914592 |
| ENSG00000114867 | *EIF4G1* | 7.163199786 | 0.005531337 | 0.027352049 | -0.021820712 | 0.999933887 | -0.049237737 | -0.066763648 | 0.017525911 | 0.874914592 |
| ENSG00000200814 | *RNU6-595P* | 0.459389177 | 0.001352632 | 0.103790305 | -0.102437674 | 0.999933887 | -0.026730493 | -0.117661243 | 0.09093075 | 0.874914592 |
| ENSG00000176623 | *RMDN1* | 2.342821154 | -0.080764659 | 0.055917917 | -0.136682576 | 0.999933887 | -0.089697074 | -0.050195953 | -0.039501121 | 0.875189859 |
| ENSG00000253893 | *FAM85B* | -1.049250251 | 0.169706037 | 0.007585108 | 0.162120929 | 0.999933887 | 0.187936522 | 0.06097301 | 0.126963512 | 0.875189859 |
| ENSG00000177156 | *TALDO1* | 9.555420377 | -0.020817099 | -0.034401838 | 0.013584739 | 0.999933887 | -0.345092588 | -0.314954106 | -0.030138481 | 0.875189859 |
| ENSG00000124098 | *FAM210B* | 2.566342298 | 0.018706425 | -0.126612416 | 0.145318841 | 0.999933887 | -0.171367611 | -0.214890361 | 0.04352275 | 0.875189859 |
| ENSG00000067596 | *DHX8* | 6.025597608 | -0.010442546 | -0.019843216 | 0.009400671 | 0.999933887 | -0.309366976 | -0.283006644 | -0.026360332 | 0.875189859 |
| ENSG00000117411 | *B4GALT2* | 0.273106971 | 0.028596621 | 0.224769545 | -0.196172925 | 0.999933887 | 0.113488047 | 0.188390877 | -0.07490283 | 0.875189859 |

| | | | | | | | | | | |
|---|---|---|---|---|---|---|---|---|---|---|
| ENSG00000152213 | *ARL11* | 5.369348871 | -0.011733519 | -0.114625447 | 0.102891927 | 0.999933887 | -0.769466939 | -0.73557841 | -0.033888529 | 0.875189859 |
| ENSG00000110063 | *DCPS* | 2.217391479 | 0.007309283 | 0.120805969 | -0.113496686 | 0.999933887 | 0.425754335 | 0.476803656 | -0.051049321 | 0.875189859 |
| ENSG00000144744 | *UBA3* | 4.39887586 | -0.039556467 | -0.00067253 | -0.038883938 | 0.999933887 | -0.083613598 | -0.10856566 | 0.024952062 | 0.87528306 |
| ENSG00000162885 | *B3GALNT2* | 0.757280602 | 0.091760929 | -0.179616091 | 0.27137702 | 0.999933887 | -0.267049963 | -0.200638296 | -0.066411667 | 0.875420763 |
| ENSG00000184260 | *H2AC20* | 1.545001731 | 0.08982348 | 0.186714971 | -0.096891491 | 0.999933887 | 0.28245254 | 0.212315216 | 0.070137324 | 0.875470424 |
| ENSG00000116857 | *TMEM9* | 1.424171383 | 0.087055223 | -0.059497199 | 0.146552422 | 0.999933887 | -0.017004239 | 0.047947588 | -0.064951827 | 0.875470424 |
| ENSG00000287576 | *AC067930.9* | -0.230839528 | 0.083048154 | 0.188203931 | -0.105155778 | 0.999933887 | 0.361630684 | 0.458912528 | -0.097281844 | 0.875470424 |
| ENSG00000263466 | *AC006441.1* | -0.023673076 | 0.056328779 | -0.178184983 | 0.234513762 | 0.999933887 | -0.257264212 | -0.335446974 | 0.078182762 | 0.875470424 |
| ENSG00000225217 | *HSPA7* | 3.986263416 | 0.017965808 | -0.047754857 | 0.065720665 | 0.999933887 | 0.032725721 | -0.018701089 | 0.051426809 | 0.875470424 |
| ENSG00000278918 | *AC080112.3* | 3.574536641 | 0.146616781 | 0.098224205 | 0.048392576 | 0.999933887 | 0.369518956 | 0.40852805 | -0.039009094 | 0.875529166 |
| ENSG00000165724 | *ZMYND19* | 1.289663903 | -0.0916441 | 0.114101615 | -0.205745715 | 0.999933887 | 0.221052948 | 0.171654715 | 0.049398233 | 0.875583315 |
| ENSG00000158428 | *CATIP* | 1.549193347 | 0.189109904 | 0.264273997 | -0.075164092 | 0.999933887 | 0.262391265 | 0.336736589 | -0.074345324 | 0.875696431 |
| ENSG00000108439 | *PNPO* | -0.548914975 | -0.115498993 | -0.548428525 | 0.432929532 | 0.999933887 | 0.291538624 | 0.204940927 | 0.086597698 | 0.875696431 |
| ENSG00000163576 | *EFHB* | -0.672490781 | -0.190120516 | 0.081136215 | -0.271256732 | 0.999933887 | 0.64986409 | 0.739634735 | -0.089770645 | 0.875814662 |
| ENSG00000235481 | *UBE2R2-AS1* | 2.515576204 | -0.052682293 | -0.121177528 | 0.068495235 | 0.999933887 | -0.05984177 | -0.015786358 | -0.044055412 | 0.87592751 |
| ENSG00000130787 | *HIP1R* | 4.755317623 | 0.057355842 | 0.141955086 | -0.084599244 | 0.999933887 | 0.110044288 | 0.137732126 | -0.027687838 | 0.876025873 |
| ENSG00000254838 | *GVINP1* | 4.510254878 | 0.104659581 | -0.092412015 | 0.197071596 | 0.999933887 | -0.347404532 | -0.405105571 | 0.057701039 | 0.876025873 |
| ENSG00000147854 | *UHRF2* | 4.820571231 | -0.048748378 | -0.100390463 | 0.051642085 | 0.999933887 | -0.164247164 | -0.201252335 | 0.037005171 | 0.876025873 |
| ENSG00000198746 | *GPATCH3* | 4.173427624 | -0.045645533 | -0.045934396 | 0.000288863 | 0.999933887 | 0.039086066 | 0.067118733 | -0.028032668 | 0.876025873 |
| ENSG00000143891 | *GALM* | 2.852900847 | -0.027858917 | 0.068546284 | -0.096405201 | 0.999933887 | 0.014077195 | -0.037019199 | 0.051096395 | 0.876025873 |
| ENSG00000258819 | *LINC02289* | 0.574521167 | -0.01751311 | -0.025267873 | 0.007754763 | 0.999933887 | -0.938266188 | -0.849238608 | -0.089027579 | 0.876025873 |
| ENSG00000213741 | *RPS29* | 7.278744652 | 0.010100077 | -0.016014709 | 0.026114786 | 0.999933887 | 0.130851013 | 0.159222185 | -0.028371172 | 0.876025873 |
| ENSG00000198276 | *UCKL1* | 3.414291795 | 0.026117601 | 0.036469141 | -0.010351539 | 0.999933887 | 0.072669254 | 0.105998302 | -0.033329048 | 0.876207006 |
| ENSG00000100478 | *AP4S1* | 1.771702151 | 0.013728392 | 0.083148046 | -0.069419654 | 0.999933887 | -0.134688121 | -0.088529298 | -0.046158823 | 0.876214382 |
| ENSG00000282608 | *ADORA3* | 1.359959232 | -0.142440747 | -0.074672514 | -0.067768233 | 0.999933887 | -0.017602403 | 0.028288015 | -0.045890417 | 0.876286569 |
| ENSG00000181007 | *ZFP82* | -0.008023285 | 0.113446141 | -0.365083831 | 0.478529972 | 0.999933887 | -0.004648827 | 0.080932616 | -0.085581443 | 0.876286569 |
| ENSG00000005108 | *THSD7A* | -1.452159367 | 0.147051264 | 0.049157916 | 0.097893348 | 0.999933887 | -0.722336823 | -0.628303104 | -0.094033719 | 0.8764804 |
| ENSG00000115232 | *ITGA4* | 6.122764319 | 0.011085042 | -0.15974494 | 0.170829982 | 0.999933887 | -0.296038408 | -0.346798415 | 0.050760007 | 0.8764804 |
| ENSG00000159882 | *ZNF230* | 3.983240978 | -0.032215925 | -0.08626576 | 0.054049835 | 0.999933887 | -0.510262782 | -0.539891384 | 0.029628602 | 0.876862464 |
| ENSG00000119401 | *TRIM32* | 0.556290096 | -0.108620663 | 0.018221064 | -0.126841727 | 0.999933887 | 0.233271835 | 0.291903343 | -0.058631508 | 0.876872998 |
| ENSG00000196417 | *ZNF765* | 1.383176708 | -0.089963749 | -0.082118739 | -0.007845009 | 0.999933887 | -0.621131757 | -0.557203081 | -0.063928676 | 0.876872998 |
| ENSG00000075945 | *KIFAP3* | 1.341388512 | -0.022081114 | -0.174007847 | 0.151926732 | 0.999933887 | 0.174744925 | 0.232440936 | -0.057696011 | 0.876872998 |
| ENSG00000236700 | *LINC01010* | 1.374134193 | -0.396073391 | -0.100674789 | -0.295398602 | 0.999933887 | -1.290310703 | -1.242024589 | -0.048286115 | 0.876941556 |
| ENSG00000124688 | *MAD2L1BP* | 4.097185709 | -0.069273458 | -0.109576085 | 0.040302627 | 0.999933887 | -0.142462755 | -0.115237731 | -0.027225025 | 0.87704154 |
| ENSG00000130812 | *ANGPTL6* | -0.787647703 | -0.025785654 | -0.021964002 | -0.003821652 | 0.999933887 | -0.160346962 | -0.260610893 | 0.100263931 | 0.877427697 |
| ENSG00000172059 | *KLF11* | 4.020923079 | 0.004496715 | 0.067496619 | -0.062999904 | 0.999933887 | -0.524737667 | -0.489725563 | -0.035012104 | 0.877653249 |
| ENSG00000080823 | *MOK* | 1.62814637 | 0.14873257 | 0.163058758 | -0.014326187 | 0.999933887 | 0.056997652 | 0.114612391 | -0.057614739 | 0.877736607 |
| ENSG00000153823 | *PID1* | 0.30512064 | -0.167084126 | 0.034923562 | -0.202007687 | 0.999933887 | -0.020986758 | 0.021411041 | -0.042397799 | 0.877875145 |
| ENSG00000124562 | *SNRPC* | 2.866403826 | 0.029401893 | -0.045043427 | 0.07444532 | 0.999933887 | 0.127317739 | 0.093192246 | 0.034125493 | 0.877875145 |
| ENSG00000185986 | *SDHAP3* | 2.193048215 | -0.042545705 | 0.094024151 | -0.136569856 | 0.999933887 | -0.095677694 | -0.138576687 | 0.042898993 | 0.877875145 |
| ENSG00000140563 | *MCTP2* | 6.508432242 | -0.007359414 | -0.039578117 | 0.032218703 | 0.999933887 | -0.390218404 | -0.360562407 | -0.029655998 | 0.877875145 |
| ENSG00000188976 | *NOC2L* | 3.668410643 | 0.043947527 | 0.033128279 | 0.010819248 | 0.999933887 | 0.117361045 | 0.086510562 | 0.030850483 | 0.877961825 |
| ENSG00000010803 | *SCMH1* | 1.663943757 | 0.076234806 | -0.007437028 | 0.083671834 | 0.999933887 | 0.02634503 | -0.019645079 | 0.04599011 | 0.8779643 |
| ENSG00000184575 | *XPOT* | 4.137872049 | -0.04190736 | -0.027217884 | -0.014689477 | 0.999933887 | -0.051871733 | -0.025810474 | -0.026061259 | 0.8779643 |
| ENSG00000253729 | *PRKDC* | 6.4780967 | -0.003621162 | -0.017942035 | 0.014320872 | 0.999933887 | -0.074155066 | -0.04410527 | -0.030049797 | 0.8779643 |
| ENSG00000119514 | *GALNT12* | 0.115707768 | 0.049978305 | 0.060039635 | -0.010061331 | 0.999933887 | 0.134682164 | 0.058307751 | 0.076374413 | 0.878002278 |
| ENSG00000175854 | *SWI5* | 0.765151398 | -0.041756 | -0.09568532 | 0.05392932 | 0.999933887 | -0.17930578 | -0.249652243 | 0.070346463 | 0.878002278 |
| ENSG00000119986 | *AVPI1* | 0.114846996 | -0.071134112 | -0.045478889 | -0.025655223 | 0.999933887 | -0.319752231 | -0.409097865 | 0.089345634 | 0.878118476 |
| ENSG00000149089 | *APIP* | 0.908075374 | -0.032296482 | -0.022309773 | -0.009986709 | 0.999933887 | -0.084941607 | -0.034964282 | -0.049977324 | 0.878280398 |
| ENSG00000165233 | *CARD19* | 7.077661649 | -0.015760176 | 0.019401399 | -0.035161576 | 0.999933887 | -0.434989714 | -0.408345418 | -0.026644295 | 0.878280398 |
| ENSG00000232373 | *MTCYBP3* | -0.211874779 | 0.045907976 | 0.175688719 | -0.129780743 | 0.999933887 | 1.282168352 | 1.395862507 | -0.113694155 | 0.878280398 |
| ENSG00000269845 | *CCNYL6* | 1.099545545 | 0.216594329 | 0.267792613 | -0.051198284 | 0.999933887 | 0.107422592 | 0.17856584 | -0.071143248 | 0.878383974 |
| ENSG00000196459 | *TRAPPC2* | 3.173783447 | -0.100964213 | -0.073073736 | -0.027890476 | 0.999933887 | 0.051837018 | 0.022362381 | 0.029474636 | 0.878383974 |
| ENSG00000078589 | *P2RY10* | 4.963380054 | -0.022874836 | -0.114288398 | 0.091413561 | 0.999933887 | -0.130705208 | -0.161742356 | 0.031037149 | 0.878383974 |
| ENSG00000176658 | *MYO1D* | 0.326719981 | 0.010130891 | 0.092309563 | -0.082178672 | 0.999933887 | 0.043416915 | 0.11867639 | -0.075259475 | 0.879001341 |
| ENSG00000153531 | *ADPRHL1* | 0.671389132 | -0.087717827 | 0.168872367 | -0.256590193 | 0.999933887 | 0.827667432 | 0.756251458 | 0.071415974 | 0.879174793 |
| ENSG00000175104 | *TRAF6* | 4.83671591 | -0.062050974 | -0.022138006 | -0.039912968 | 0.999933887 | -0.197590342 | -0.174683383 | -0.022906959 | 0.879379165 |
| ENSG00000149782 | *PLCB3* | 3.082797452 | 0.035633857 | 0.03251204 | 0.003121817 | 0.999933887 | 0.281640717 | 0.320898895 | -0.039258179 | 0.879379165 |
| ENSG00000105048 | *TNNT1* | -0.486037783 | -0.069150195 | 0.151196845 | -0.22034704 | 0.999933887 | -0.373978341 | -0.485109406 | 0.111131065 | 0.879453338 |
| ENSG00000129515 | *SNX6* | 5.674562051 | -0.049917435 | -0.119496015 | 0.069578579 | 0.999933887 | -0.488729739 | -0.463003181 | -0.025726558 | 0.879494188 |
| ENSG00000171169 | *NAIF1* | 3.770326936 | -0.020976273 | 0.023193979 | -0.044170252 | 0.999933887 | -0.181848755 | -0.144225344 | -0.03762341 | 0.879535626 |
| ENSG00000009954 | *BAZ1B* | 4.297611216 | -0.029531043 | -0.13568214 | 0.106151097 | 0.999933887 | 0.00494245 | 0.031918551 | -0.026976101 | 0.879719912 |
| ENSG00000138172 | *CALHM2* | 1.768501356 | -0.03577955 | -0.00713661 | -0.02864294 | 0.999933887 | -0.020889477 | 0.015659742 | -0.036549219 | 0.879719912 |
| ENSG00000102910 | *LONP2* | 4.094809493 | -0.014997415 | -0.047796095 | 0.03279868 | 0.999933887 | -0.022125135 | 0.006800214 | -0.028925349 | 0.879719912 |
| ENSG00000272498 | *AC090948.2* | -1.039151882 | 0.060132992 | 0.064353562 | -0.004220571 | 0.999933887 | -0.065947609 | 0.067628049 | -0.133575658 | 0.879719912 |
| ENSG00000279349 | *AC092692.1* | -0.164716607 | -0.053867337 | 0.277604313 | -0.33147165 | 0.999933887 | 0.232505786 | 0.169102287 | 0.063403499 | 0.879719912 |
| ENSG00000196696 | *PDXDC2P-NPIPB14P* | 1.416907143 | 0.009641928 | 0.07022593 | -0.060584002 | 0.999933887 | -0.055345611 | -0.103595313 | 0.048249702 | 0.879719912 |
| ENSG00000258424 | *AL512791.1* | 0.4059889 | -0.120991236 | -0.086018344 | -0.034972892 | 0.999933887 | -0.046443871 | -0.121004961 | 0.07456109 | 0.879732831 |
| ENSG00000169252 | *ADRB2* | 3.015150938 | 0.05247916 | 0.073286516 | -0.020807356 | 0.999933887 | 0.260673091 | 0.301604211 | -0.040931121 | 0.879792787 |
| ENSG00000128965 | *CHAC1* | -0.533442368 | -0.041369614 | -0.075177743 | 0.033808129 | 0.999933887 | 0.284572648 | 0.377273902 | -0.092701254 | 0.879864151 |
| ENSG00000224513 | *AC109309.1* | -1.792367188 | -0.262458507 | 0.126598568 | -0.389057074 | 0.999933887 | -0.168907592 | -0.272488035 | 0.103580443 | 0.879883218 |
| ENSG00000188747 | *NOXA1* | 0.109346857 | 0.168160389 | 0.284897353 | -0.116736964 | 0.999933887 | -0.042360389 | -0.122905553 | 0.080545164 | 0.879883218 |
| ENSG00000174231 | *PRPF8* | 7.741840091 | -0.016042175 | 0.004958758 | -0.021000932 | 0.999933887 | -0.010013977 | 0.007534691 | -0.017548668 | 0.879883218 |
| ENSG00000214439 | *FAM185BP* | -0.16908774 | -0.057541631 | -0.189882292 | 0.132340662 | 0.999933887 | -0.069165477 | -0.164176021 | 0.095010543 | 0.879883218 |
| ENSG00000090382 | *LYZ* | 7.247352636 | 0.015503258 | -0.024955579 | 0.040458837 | 0.999933887 | 0.018096271 | -0.008796384 | 0.026892655 | 0.879883218 |
| ENSG00000253307 | *AC011676.1* | 4.147661957 | -0.004029642 | -0.005756272 | 0.00172663 | 0.999933887 | -0.123530296 | -0.085028956 | -0.038501339 | 0.880247472 |
| ENSG00000174652 | *ZNF266* | 5.155370587 | -0.024611692 | -0.070571403 | 0.045959711 | 0.999933887 | 0.009775016 | -0.011155918 | 0.020930934 | 0.880552569 |
| ENSG00000249207 | *AC079921.1* | -0.275092456 | 0.023101192 | 0.171694644 | -0.148593452 | 0.999933887 | 0.179359815 | 0.26208994 | -0.082730125 | 0.880570465 |
| ENSG00000229512 | *AC068580.1* | 5.050922949 | 0.157073883 | 0.150846363 | 0.00622752 | 0.999933887 | 0.229923908 | 0.189773361 | 0.040150547 | 0.880707408 |
| ENSG00000244932 | *AL449212.1* | 0.034789404 | -0.057385535 | -0.248333687 | 0.190948152 | 0.999933887 | -0.313902365 | -0.253009198 | -0.060893167 | 0.880707408 |
| ENSG00000145375 | *SPATA5* | 0.57892197 | -0.036100853 | -0.007079897 | -0.029020956 | 0.999933887 | -0.064951413 | 0.00016325 | -0.065114663 | 0.880707408 |
| ENSG00000286177 | *AC011462.5* | 1.182415397 | -0.01645457 | 0.005140649 | -0.021595219 | 0.999933887 | -0.213994816 | -0.262886043 | 0.048891227 | 0.880707408 |
| ENSG00000106258 | *CYP3A5* | -0.17799084 | -0.020795108 | -0.646050567 | 0.625255459 | 0.999933887 | -0.283356011 | -0.192064246 | -0.091291765 | 0.880707408 |
| ENSG00000107521 | *HPS1* | 6.566220092 | -0.000479124 | 0.014433849 | -0.014912973 | 0.999933887 | 0.037267049 | 0.01963334 | 0.017633709 | 0.880707408 |
| ENSG00000156017 | *CARNMT1* | 1.47579602 | -0.058173395 | -0.048322976 | -0.009850419 | 0.999933887 | -0.153686154 | -0.100327541 | -0.053358612 | 0.881602629 |
| ENSG00000047932 | *GOPC* | 2.618822689 | -0.005870973 | -0.078292389 | 0.072421416 | 0.999933887 | -0.12341824 | -0.085853463 | -0.037564777 | 0.881776236 |
| ENSG00000140105 | *WARS1* | 7.68236929 | -0.029719821 | -0.021961 | -0.007758821 | 0.999933887 | -0.105742464 | -0.085644207 | -0.020098257 | 0.882009998 |
| ENSG00000169490 | *TM2D2* | 3.042640567 | -0.091074404 | -0.029194725 | -0.061879679 | 0.999933887 | -0.326687943 | -0.291342684 | -0.03534526 | 0.882014155 |
| ENSG00000173726 | *TOMM20* | 4.807193336 | -0.048607534 | -0.079264562 | 0.030657028 | 0.999933887 | 0.076156982 | 0.101474422 | -0.025317441 | 0.882014155 |
| ENSG00000072756 | *TRNT1* | 3.250121617 | 0.075939595 | 0.010110653 | 0.065828942 | 0.999933887 | 0.214897083 | 0.251355497 | -0.036458414 | 0.882014155 |
| ENSG00000232973 | *CYP1B1-AS1* | 0.522331138 | 0.054446205 | 0.10643327 | -0.051987065 | 0.999933887 | 0.330103192 | 0.274335177 | 0.055768015 | 0.882451287 |
| ENSG00000168591 | *TMUB2* | 6.84493152 | 0.046664314 | 0.013916578 | 0.032747736 | 0.999933887 | -0.127551996 | -0.107056753 | -0.020495243 | 0.882496451 |
| ENSG00000115598 | *IL1RL2* | 1.700663389 | -0.065938121 | 0.216898952 | -0.282837073 | 0.999933887 | 0.591662447 | 0.64370503 | -0.052042582 | 0.882496451 |
| ENSG00000225399 | *AC121247.1* | -0.275032161 | 0.052883151 | -0.005033658 | 0.057916809 | 0.999933887 | -0.430368976 | -0.325000807 | -0.105368169 | 0.882496451 |
| ENSG00000002822 | *MAD1L1* | 1.718922575 | -0.029588013 | 0.108338051 | -0.137926064 | 0.999933887 | 0.283218497 | 0.334021989 | -0.050803493 | 0.882496451 |
| ENSG00000174353 | *STAG3L3* | 0.572892537 | 0.014842115 | -0.065875126 | 0.080717241 | 0.999933887 | -0.050142126 | 0.013953277 | -0.064095402 | 0.882496451 |
| ENSG00000087095 | *NLK* | 2.135514419 | -0.009402308 | 0.180705292 | -0.1901076 | 0.999933887 | -0.09712918 | -0.146726237 | 0.049597057 | 0.882496451 |
| ENSG00000166780 | *BMERB1* | 0.9298983 | 0.013958294 | 0.140433768 | -0.126475474 | 0.999933887 | -0.327117707 | -0.257858271 | -0.069259436 | 0.882496451 |
| ENSG00000139194 | *RBP5* | 1.48796362 | 0.089649341 | -0.029169861 | 0.118819202 | 0.999933887 | 0.117399156 | 0.058258289 | 0.059140867 | 0.882886751 |
| ENSG00000165526 | *RPUSD4* | 2.373235253 | -0.06602896 | 0.087215965 | -0.153244925 | 0.999933887 | 0.193983016 | 0.160702756 | 0.03328026 | 0.882886751 |
| ENSG00000074706 | *IPCEF1* | 4.21088452 | -0.024372709 | -0.015992369 | -0.008380341 | 0.999933887 | -0.215338472 | -0.18855734 | -0.026781131 | 0.882886751 |
| ENSG00000287643 | *AC004825.3* | 0.498630052 | 0.13849966 | 0.079672931 | 0.058826729 | 0.999933887 | -0.092715173 | 0.001769022 | -0.094484195 | 0.882970447 |
| ENSG00000270554 | *AC069431.2* | -0.327396397 | 0.059548609 | -0.049949023 | 0.109497631 | 0.999933887 | -0.45459297 | -0.316866782 | -0.137726188 | 0.882970447 |
| ENSG00000164830 | *OXR1* | 3.088262798 | 0.019724962 | -0.009033068 | 0.02875803 | 0.999933887 | -0.533833554 | -0.49103179 | -0.042801763 | 0.882970447 |
| ENSG00000287261 | *AL139243.1* | 0.802098401 | 0.192305311 | 0.218242894 | -0.025937583 | 0.999933887 | 0.441633741 | 0.393450358 | 0.048183383 | 0.883207378 |

| | | | | | | | | | | |
|---|---|---|---|---|---|---|---|---|---|---|
| ENSG00000188603 | *CLN3* | 4.580791284 | 0.037770607 | -0.019367635 | 0.057138242 | 0.999933887 | -0.00391984 | 0.018567537 | -0.022487378 | 0.883207378 |
| ENSG00000204498 | *NFKBIL1* | 2.411690374 | 0.041769855 | -0.141397924 | 0.183167779 | 0.999933887 | -0.028547593 | 0.012617223 | -0.041164816 | 0.883207378 |
| ENSG00000101294 | *HM13* | 7.83017233 | 0.018428819 | -0.042977974 | 0.061406793 | 0.999933887 | -0.238590207 | -0.258655363 | 0.020065156 | 0.883207378 |
| ENSG00000162813 | *BPNT1* | 0.467675966 | -0.045651267 | 0.179504396 | -0.225155663 | 0.999933887 | 0.168751248 | 0.23033444 | -0.061583192 | 0.883207378 |
| ENSG00000155858 | *LSM11* | 0.618729916 | -0.398438738 | 0.006518527 | -0.404957265 | 0.999933887 | 0.050546171 | -0.012090145 | 0.062636316 | 0.883461887 |
| ENSG00000250608 | *AC010210.1* | -0.061632124 | -0.071175363 | -0.203739602 | 0.132564239 | 0.999933887 | -0.032302325 | 0.064760576 | -0.097062901 | 0.883670908 |
| ENSG00000211790 | *TRAV8-4* | -0.528059329 | -0.242852624 | -0.082225641 | -0.160626983 | 0.999933887 | 0.121407663 | 0.039085236 | 0.082322427 | 0.883706407 |
| ENSG00000233695 | *GAS6-AS1* | 2.281954486 | 0.125818813 | 0.244145849 | -0.118327036 | 0.999933887 | 0.37293989 | 0.417849702 | -0.044909812 | 0.883713361 |
| ENSG00000143575 | *HAX1* | 5.011963099 | 0.063520029 | 0.034242712 | 0.029277316 | 0.999933887 | 0.069941942 | 0.096055665 | -0.026113723 | 0.883775988 |
| ENSG00000082438 | *COBLL1* | -1.209302775 | -0.111786664 | 0.334675576 | -0.446462239 | 0.999933887 | 0.138887982 | 0.04006938 | 0.098818602 | 0.883790976 |
| ENSG00000135632 | *SMYD5* | 1.975060838 | -0.034730604 | 0.060411973 | -0.095142577 | 0.999933887 | 0.165255219 | 0.119003188 | 0.046252031 | 0.883790976 |
| ENSG00000283118 | *AC009951.5* | 1.593346185 | 0.052849579 | 0.289453974 | -0.236604395 | 0.999933887 | 1.638578758 | 1.726509391 | -0.087930633 | 0.883790976 |
| ENSG00000288393 | *AP003086.3* | 1.825913793 | 0.147106205 | 0.279035529 | -0.131929324 | 0.999933887 | 0.023808759 | 0.121112458 | -0.097303699 | 0.883794427 |
| ENSG00000089195 | *TRMT6* | 2.58416085 | -0.047429179 | -0.249277075 | 0.201847896 | 0.999933887 | 0.048327115 | 0.07988973 | -0.031562615 | 0.883794427 |
| ENSG00000160752 | *FDPS* | 2.749006301 | 0.095292478 | 0.040944601 | 0.054347877 | 0.999933887 | -0.062197047 | -0.099922909 | 0.037725862 | 0.883849595 |
| ENSG00000271204 | *AC016831.4* | 1.632071458 | 0.220431355 | 0.566875684 | -0.346444329 | 0.999933887 | -0.026921796 | -0.124901592 | 0.097979797 | 0.883849595 |
| ENSG00000145050 | *MANF* | 3.035752574 | 0.07796289 | -0.034260065 | 0.112222955 | 0.999933887 | 0.136984699 | 0.174415398 | -0.037430699 | 0.883849595 |
| ENSG00000179532 | *DNHD1* | 3.708361987 | 0.030077833 | 0.116909375 | -0.086831541 | 0.999933887 | -0.059424093 | -0.082014971 | 0.022590877 | 0.883849595 |
| ENSG00000145247 | *OCIAD2* | 2.184742273 | 0.047869261 | 0.015693979 | 0.032175282 | 0.999933887 | 0.116050471 | 0.078348378 | 0.037702093 | 0.883849595 |
| ENSG00000108946 | *PRKAR1A* | 7.014529232 | -0.022952847 | 0.055111318 | -0.078064165 | 0.999933887 | -0.26434346 | -0.286044008 | 0.021700548 | 0.883849595 |
| ENSG00000021574 | *SPAST* | 4.096306458 | -0.034473923 | -0.108525499 | 0.074051576 | 0.999933887 | -1.196778757 | -1.240621061 | 0.043842304 | 0.883849595 |
| ENSG00000140598 | *EFL1* | 1.713814459 | -0.037326576 | -0.09028659 | 0.052960014 | 0.999933887 | -0.118653788 | -0.072099537 | -0.046554251 | 0.883849595 |
| ENSG00000161013 | *MGAT4B* | 6.302191712 | -0.006294417 | 0.017043841 | -0.023338258 | 0.999933887 | 0.046200135 | 0.067348514 | -0.021148378 | 0.883849595 |
| ENSG00000160654 | *CD3G* | 3.885825634 | -0.06153608 | -0.102434701 | 0.040898621 | 0.999933887 | 0.067361253 | 0.093157534 | -0.025796281 | 0.883933767 |
| ENSG00000106723 | *SPIN1* | 2.85922424 | -0.063079707 | -0.014339542 | -0.048740165 | 0.999933887 | -0.004561908 | -0.0356607 | 0.031098791 | 0.884090546 |
| ENSG00000168958 | *MFF* | 3.847210354 | -0.06009108 | -0.003704264 | -0.056386816 | 0.999933887 | -0.190987543 | -0.215371256 | 0.024383714 | 0.884167055 |
| ENSG00000125977 | *EIF2S2* | 4.353389846 | -0.071962079 | -0.022769792 | -0.049192286 | 0.999933887 | -0.112028218 | -0.091100245 | -0.020927973 | 0.884215624 |
| ENSG00000214812 | *AL137792.1* | 0.856702767 | 0.070837708 | 0.376779836 | -0.305942128 | 0.999933887 | 1.946964623 | 1.999427533 | -0.052462909 | 0.884373185 |
| ENSG00000165140 | *FBP1* | 1.607448292 | 0.107139555 | 0.062333252 | 0.044806302 | 0.999933887 | 0.039690477 | 0.093221671 | -0.053531194 | 0.884393606 |
| ENSG00000287808 | *AL355999.1* | 0.119441553 | -0.12550161 | 0.109395058 | -0.234896667 | 0.999933887 | -0.534419385 | -0.468773078 | -0.065646307 | 0.884505789 |
| ENSG00000126457 | *PRMT1* | 3.559471076 | -0.012422187 | 0.062203267 | -0.074625453 | 0.999933887 | 0.286916033 | 0.317608498 | -0.030692465 | 0.884505789 |
| ENSG00000172940 | *SLC22A13* | 0.483724248 | 0.007140362 | 0.191959928 | -0.184819566 | 0.999933887 | 0.130105791 | 0.030489108 | 0.099616683 | 0.884645264 |
| ENSG00000006534 | *ALDH3B1* | 4.939994682 | -0.027772107 | -0.104537731 | 0.076765625 | 0.999933887 | -1.209986931 | -1.167303946 | -0.042682985 | 0.884761558 |
| ENSG00000198642 | *KLHL9* | 3.792676192 | -0.116694035 | -0.003135375 | -0.113558661 | 0.999933887 | -0.139701731 | -0.113360839 | -0.026340893 | 0.885282842 |
| ENSG00000211721 | *TRBV6-5* | 0.035489852 | -0.068909486 | -0.21624279 | 0.147333305 | 0.999933887 | 0.134774797 | 0.070403043 | 0.064371754 | 0.885282842 |
| ENSG00000238121 | *LINC00426* | 3.211608986 | 0.047523346 | 0.10318927 | -0.055665923 | 0.999933887 | -0.463029071 | -0.430818907 | -0.032210164 | 0.885341705 |
| ENSG00000283103 | *AC010642.2* | 2.312107798 | 0.089407666 | 0.070495003 | 0.018912663 | 0.999933887 | -0.082228126 | -0.03951161 | -0.042716517 | 0.885355023 |
| ENSG00000286145 | *AC068282.1* | 1.754985766 | -0.129838624 | 0.262265112 | -0.392103736 | 0.999933887 | -0.709315054 | -0.606136775 | -0.103178279 | 0.885355023 |
| ENSG00000274602 | *PI4KAP1* | 2.037047918 | -0.115455304 | 0.019176164 | -0.134631468 | 0.999933887 | 0.123631592 | 0.083108152 | 0.040523439 | 0.885395607 |
| ENSG00000183421 | *RIPK4* | -1.303712102 | -0.083691422 | 0.272288439 | -0.355979861 | 0.999933887 | -0.123426648 | 0.024950749 | -0.148377397 | 0.885395607 |
| ENSG00000227063 | *RPL41P1* | -0.221511774 | 0.050007498 | -0.095783783 | 0.145791281 | 0.999933887 | 0.180652682 | 0.112402755 | 0.068249927 | 0.885395607 |
| ENSG00000160336 | *ZNF761* | 1.014499873 | -0.032492861 | -0.124373243 | 0.091880382 | 0.999933887 | -0.225627727 | -0.289176705 | 0.063548978 | 0.885395607 |
| ENSG00000213762 | *ZNF134* | 2.813732496 | -0.110049594 | -0.116585641 | 0.006536047 | 0.999933887 | -0.173631739 | -0.203536058 | 0.029904319 | 0.88565001 |
| ENSG00000064932 | *SBNO2* | 8.892277852 | 0.106206146 | 0.105499133 | 0.000707013 | 0.999933887 | 0.254769652 | 0.279046313 | -0.024276661 | 0.885675932 |
| ENSG00000008394 | *MGST1* | 0.193635094 | 0.096838855 | 0.159655297 | -0.062816442 | 0.999933887 | -0.862513994 | -0.808815511 | -0.053698482 | 0.886007087 |
| ENSG00000160781 | *PAQR6* | 2.587003358 | 0.021545353 | 0.047202253 | -0.0256569 | 0.999933887 | -0.069803268 | -0.026885067 | -0.042918202 | 0.886007087 |
| ENSG00000134779 | *TPGS2* | 2.864323612 | -0.004257413 | -0.034917636 | 0.030660223 | 0.999933887 | 0.00938234 | -0.022439288 | 0.031821628 | 0.886007087 |
| ENSG00000148153 | *INIP* | 3.788713318 | -0.110576542 | -0.166958511 | 0.056381969 | 0.999933887 | 0.01273939 | -0.011852741 | 0.024592131 | 0.886138592 |
| ENSG00000140280 | *LYSMD2* | 4.973566356 | -0.009053791 | 0.061061258 | -0.070115049 | 0.999933887 | -0.000447826 | 0.026700847 | -0.027148673 | 0.886233194 |
| ENSG00000151952 | *TMEM132D* | 0.621093112 | 0.038437819 | -0.029575777 | 0.068013596 | 0.999933887 | -0.230511319 | -0.306831631 | 0.076320313 | 0.88630023 |
| ENSG00000278962 | *AC092645.1* | -0.957203801 | 0.152702587 | 0.04947496 | 0.103227627 | 0.999933887 | -0.029090949 | 0.071870291 | -0.10096124 | 0.886310209 |
| ENSG00000143303 | *RRNAD1* | 3.581107937 | 0.031923042 | 0.004801161 | 0.027121881 | 0.999933887 | 0.689538496 | 0.718877257 | -0.029338761 | 0.886712125 |
| ENSG00000181035 | *SLC25A42* | 2.223060188 | 0.054587884 | 0.09035247 | -0.035764585 | 0.999933887 | 0.026218711 | -0.009272837 | 0.035491548 | 0.88674699 |
| ENSG00000121406 | *ZNF549* | 1.381602121 | 0.1088405 | 0.070180861 | 0.038659639 | 0.999933887 | 0.006087029 | 0.055868383 | -0.049781354 | 0.886797254 |
| ENSG00000055070 | *SZRD1* | 6.297502817 | -0.013770312 | -0.005128406 | -0.008641906 | 0.999933887 | 0.075463774 | 0.09560386 | -0.020140086 | 0.886804057 |
| ENSG00000236438 | *FAM157A* | 5.752820735 | 0.069161704 | 0.025519955 | 0.043641748 | 0.999933887 | 0.490781456 | 0.464452156 | 0.0263293 | 0.886827513 |
| ENSG00000128989 | *ARPP19* | 5.60342643 | -0.019829352 | -0.025390224 | 0.005560871 | 0.999933887 | -0.352265313 | -0.328769994 | -0.023495319 | 0.886879696 |
| ENSG00000037897 | *METTL1* | -0.454123602 | 0.095420019 | -0.167240833 | 0.262660852 | 0.999933887 | 0.774105116 | 0.681799521 | 0.092305595 | 0.887593991 |
| ENSG00000167912 | *AC090152.1* | -0.368135855 | 0.045376982 | 0.025018629 | 0.020358354 | 0.999933887 | 0.138203857 | 0.037020001 | 0.101183856 | 0.887724918 |
| ENSG00000163322 | *ABRAXAS1* | 1.244034043 | 0.020650031 | 0.074912888 | -0.054262857 | 0.999933887 | -0.202203489 | -0.145688196 | -0.056515292 | 0.887774633 |
| ENSG00000139977 | *NAA30* | 2.934087898 | -0.047565068 | 0.082725448 | -0.130290517 | 0.999933887 | -0.229353953 | -0.270971549 | 0.041617596 | 0.887823457 |
| ENSG00000175216 | *CKAP5* | 2.78729048 | -0.053158587 | -0.173648994 | 0.120490407 | 0.999933887 | 0.018803663 | 0.058660082 | -0.03985642 | 0.88793933 |
| ENSG00000186260 | *MRTFB* | 1.13605566 | -0.079252041 | -0.282421222 | 0.203169181 | 0.999933887 | -0.198023139 | -0.146442361 | -0.051580778 | 0.888149586 |
| ENSG00000268170 | *AC073342.1* | -0.384076054 | 0.000134525 | 0.080731715 | -0.08059719 | 0.999933887 | 0.257017655 | 0.151955879 | 0.105061776 | 0.88820805 |
| ENSG00000130429 | *ARPC1B* | 2.837879109 | -0.064424419 | -0.029854054 | -0.034570365 | 0.999933887 | -0.037120738 | -0.074017324 | 0.036896586 | 0.888408384 |
| ENSG00000083168 | *KAT6A* | 7.732188224 | -0.02059161 | -0.024122635 | 0.003531025 | 0.999933887 | -0.647251979 | -0.671717599 | 0.02446562 | 0.888415318 |
| ENSG00000165060 | *FXN* | -0.331159847 | -0.122989022 | 0.067796917 | -0.190785939 | 0.999933887 | 0.127953273 | 0.214812032 | -0.086858758 | 0.888490677 |
| ENSG00000033030 | *ZCCHC8* | 4.084238049 | -0.008668441 | 0.017150136 | -0.025818577 | 0.999933887 | -0.062776686 | -0.088799989 | 0.026023303 | 0.888490677 |
| ENSG00000196363 | *WDR5* | 2.546757039 | -0.10522254 | 0.097548914 | -0.202771454 | 0.999933887 | 0.130844105 | 0.09708912 | 0.033754984 | 0.888492414 |
| ENSG00000166012 | *TAF1D* | 5.041654108 | -0.118721544 | -0.147835093 | 0.029113548 | 0.999933887 | -0.26579615 | -0.225142908 | -0.040653242 | 0.888492414 |
| ENSG00000157992 | *KRTCAP3* | 0.250787888 | -0.128781875 | 0.21142983 | -0.340211705 | 0.999933887 | 0.021213849 | -0.039956854 | 0.061170703 | 0.888492414 |
| ENSG00000213465 | *ARL2* | 0.460568011 | 0.027699936 | -0.012842813 | 0.040542749 | 0.999933887 | -0.018876031 | -0.074196336 | 0.055320306 | 0.888669289 |
| ENSG00000253043 | *RNU7-181P* | 0.20134461 | 0.166167178 | 0.391698373 | -0.225531195 | 0.999933887 | 0.238453478 | 0.303543536 | -0.065090058 | 0.889550681 |
| ENSG00000177192 | *PUS1* | 1.837015228 | 0.016685905 | 0.021177373 | -0.004491467 | 0.999933887 | 0.038799811 | 0.084913145 | -0.046113334 | 0.889550681 |
| ENSG00000152422 | *XRCC4* | 1.419362873 | -0.074829411 | -0.16802852 | 0.093199109 | 0.999933887 | -0.296040615 | -0.231218401 | -0.064822214 | 0.890278089 |
| ENSG00000198924 | *DCLRE1A* | 0.207497438 | 0.034495918 | -0.213404336 | 0.247900255 | 0.999933887 | 0.086674918 | 0.154711695 | -0.068036777 | 0.890278089 |
| ENSG00000175061 | *SNHG29* | 4.47887705 | -0.006476138 | -0.077021548 | 0.070545411 | 0.999933887 | 0.130859288 | 0.154187358 | -0.023328071 | 0.890278089 |
| ENSG00000151498 | *ACAD8* | 3.50517438 | -0.056145743 | -0.052792042 | -0.003353701 | 0.999933887 | -0.041108705 | -0.010685422 | -0.030423283 | 0.890289275 |
| ENSG00000234329 | *AL604028.2* | -0.085885877 | -0.209710801 | -0.237066627 | 0.027355826 | 0.999933887 | -0.30900626 | -0.245635817 | -0.063370443 | 0.890358202 |
| ENSG00000004700 | *RECQL* | 2.839969639 | -0.070003944 | -0.191594953 | 0.121591009 | 0.999933887 | -0.248710973 | -0.206217507 | -0.042493466 | 0.890358202 |
| ENSG00000154764 | *WNT7A* | -0.603139142 | 0.270269222 | 0.176760665 | 0.093508557 | 0.999933887 | 0.108938212 | 0.194383385 | -0.085445173 | 0.89050495 |
| ENSG00000056097 | *ZFR* | 4.980264983 | -0.047874288 | -0.066941828 | 0.01906754 | 0.999933887 | -0.328737388 | -0.352102938 | 0.02336555 | 0.890521975 |
| ENSG00000100220 | *RTCB* | 2.677057392 | -0.055064968 | -0.096562089 | 0.041497121 | 0.999933887 | 0.242121368 | 0.274022272 | -0.031900903 | 0.890521975 |
| ENSG00000167491 | *GATAD2A* | 8.002594158 | -0.017882789 | -0.001864772 | -0.016018017 | 0.999933887 | -0.16341424 | -0.184498191 | 0.021083951 | 0.890718527 |
| ENSG00000136371 | *MTHFS* | 2.803908553 | 0.16567924 | 0.16933973 | -0.003660491 | 0.999933887 | 0.27096713 | 0.312421151 | -0.041454021 | 0.890754756 |
| ENSG00000203761 | *MSTO2P* | 1.317820622 | 0.226868382 | -0.05527471 | 0.282143093 | 0.999933887 | 0.156050607 | 0.10386094 | 0.052189667 | 0.890754756 |
| ENSG00000223086 | *RNA5SP155* | 0.013824554 | 0.256530958 | 0.392647662 | -0.136116703 | 0.999933887 | 0.733666788 | 0.651071043 | 0.082595745 | 0.890754756 |
| ENSG00000280184 | *AL023806.3* | -0.794191314 | -0.697214709 | 0.385314274 | -1.082528983 | 0.999933887 | 0.193334469 | 0.26401158 | -0.07067711 | 0.890754756 |
| ENSG00000161217 | *PCYT1A* | 6.649349543 | -0.054315263 | -0.016855565 | -0.037459699 | 0.999933887 | -0.139925382 | -0.121455895 | -0.018469486 | 0.890754756 |
| ENSG00000166689 | *PLEKHA7* | -0.095521157 | 0.145960717 | -0.128670479 | 0.274631196 | 0.999933887 | -0.067117342 | 0.001728681 | -0.068846023 | 0.890754756 |
| ENSG00000187189 | *TSPYL4* | 3.091931 | -0.055889023 | -0.082996913 | 0.02710789 | 0.999933887 | 0.032894437 | 0.005859914 | 0.027034523 | 0.890754756 |
| ENSG00000226281 | *AL031123.1* | -0.557329604 | -0.210650285 | -0.352460152 | 0.141809867 | 0.999933887 | -0.068512849 | -0.164231308 | 0.095718458 | 0.890754756 |
| ENSG00000165028 | *NIPSNAP3B* | 0.039466808 | 0.167586892 | 0.180117047 | -0.012530155 | 0.999933887 | 0.049740021 | 0.153101523 | -0.103361502 | 0.890754756 |
| ENSG00000163939 | *PBRM1* | 4.609342044 | -0.052869701 | -0.020563904 | -0.032305797 | 0.999933887 | -0.352543451 | -0.325752138 | -0.026791313 | 0.890754756 |
| ENSG00000122483 | *CCDC18* | 2.289454264 | 0.0770412 | -0.022081359 | 0.099122559 | 0.999933887 | -0.281572284 | -0.340872366 | 0.059300082 | 0.890754756 |
| ENSG00000084207 | *GSTP1* | 3.81928046 | 0.045052324 | -0.046314293 | 0.091366618 | 0.999933887 | 0.132221044 | 0.101816493 | 0.030404551 | 0.890754756 |
| ENSG00000078246 | *TULP3* | 0.793565042 | 0.05852259 | -0.074459886 | 0.132982476 | 0.999933887 | 0.025819668 | 0.085557972 | -0.059738304 | 0.890754756 |
| ENSG00000143320 | *CRABP2* | 1.077607048 | 0.037141692 | -0.088299609 | 0.125441301 | 0.999933887 | 0.367759594 | 0.308414491 | 0.059345102 | 0.890754756 |
| ENSG00000117155 | *SSX2IP* | 1.400438569 | -0.045588567 | -0.085334325 | 0.039745758 | 0.999933887 | -0.264066515 | -0.201728047 | -0.062338468 | 0.890754756 |
| ENSG00000091986 | *CCDC80* | 0.265793497 | -0.033646807 | -0.069447488 | 0.035800681 | 0.999933887 | 0.06130132 | 0.14877904 | -0.08747772 | 0.890754756 |
| ENSG00000004799 | *PDK4* | -0.256593476 | -0.011780613 | -0.051027171 | 0.039246559 | 0.999933887 | -0.633695362 | -0.764039612 | 0.13034425 | 0.890754756 |
| ENSG00000226705 | *SDCBPP1* | -0.611309059 | -0.004492211 | 0.301194321 | -0.305686532 | 0.999933887 | 0.244180475 | 0.119487989 | 0.124692486 | 0.890754756 |
| ENSG00000149357 | *LAMTOR1* | 6.272042289 | 0.007993611 | 0.028245557 | -0.020251947 | 0.999933887 | -0.27654518 | -0.256488446 | -0.020056734 | 0.890756587 |
| ENSG00000101546 | *RBFA* | 0.195164608 | 0.142156149 | -0.091609536 | 0.233765686 | 0.999933887 | 0.168583815 | 0.225633815 | -0.05705 | 0.890813511 |
| ENSG00000185442 | *FAM174B* | 1.726117896 | -0.067617439 | 0.017849616 | -0.085467055 | 0.999933887 | 0.946548606 | 0.900778513 | 0.045770094 | 0.890832423 |

| | | | | | | | | | | |
|---|---|---|---|---|---|---|---|---|---|---|
| ENSG00000232133 | *IMPDH1P10* | 0.067324922 | 0.111758414 | 0.023108329 | 0.088650084 | 0.999933887 | 1.565478321 | 1.492284845 | 0.073193475 | 0.89096576 |
| ENSG00000250067 | *YJEFN3* | 0.209280063 | -0.045245376 | 0.232168521 | -0.277413897 | 0.999933887 | 0.128644129 | 0.056161289 | 0.072482839 | 0.89096576 |
| ENSG00000265666 | *RARA-AS1* | 3.420601721 | 0.14945513 | 0.013117498 | 0.136337632 | 0.999933887 | 0.038850652 | 0.078386281 | -0.039535629 | 0.891096774 |
| ENSG00000247950 | *SEC24B-AS1* | 0.080171349 | -0.014625406 | 0.276371319 | -0.290996725 | 0.999933887 | 0.088341743 | 0.018226159 | 0.070115585 | 0.891237097 |
| ENSG00000157450 | *RNF111* | 5.555939358 | 0.008629294 | 0.037108118 | -0.028478824 | 0.999933887 | -0.795793993 | -0.821997131 | 0.026203138 | 0.89141362 |
| ENSG00000124228 | *DDX27* | 4.766638691 | -0.005588277 | -0.006243469 | 0.000655191 | 0.999933887 | 0.015048484 | 0.035244146 | -0.020195662 | 0.891433575 |
| ENSG00000145284 | *SCD5* | -0.642151906 | -0.03858902 | -0.137762092 | 0.099173072 | 0.999933887 | -0.024671201 | -0.122272009 | 0.097600808 | 0.891657073 |
| ENSG00000166526 | *ZNF3* | 3.790777487 | -0.027615705 | -0.067026997 | 0.039411292 | 0.999933887 | 0.095211779 | 0.121289349 | -0.026077571 | 0.89172971 4 |
| ENSG00000198932 | *GPRASP1* | 2.63517214 | -0.020301094 | -0.092182142 | 0.071881048 | 0.999933887 | 0.095879983 | 0.12811254 | -0.032232558 | 0.891874481 |
| ENSG00000155621 | *C9orf85* | 0.245975278 | 0.19537291 | 0.191644742 | 0.003728168 | 0.999933887 | -0.223945465 | -0.152359343 | -0.071586122 | 0.89215572 |
| ENSG00000182108 | *DEXI* | -0.153789681 | 0.152538854 | 0.166254529 | -0.013715675 | 0.999933887 | -0.044384387 | -0.113121891 | 0.068737504 | 0.89215572 |
| ENSG00000264278 | *ZNF236-DT* | -0.319759966 | 0.130532192 | 0.102019749 | 0.028512444 | 0.999933887 | -0.336398258 | -0.423104978 | 0.08670672 | 0.89215572 |
| ENSG00000132432 | *SEC61G* | 2.568299962 | 0.039244796 | -0.05018379 | 0.089428586 | 0.999933887 | 0.131846557 | 0.098348545 | 0.033498013 | 0.892234999 |
| ENSG00000254206 | *NPIPB11* | 0.258523997 | 0.196742783 | -0.21175384 | 0.408496623 | 0.999933887 | -0.051845525 | 0.015226133 | -0.067071658 | 0.892254099 |
| ENSG00000141298 | *SSH2* | 8.602275907 | -0.055939015 | -0.023120579 | -0.032818436 | 0.999933887 | -0.293622485 | -0.318054036 | 0.02443155 | 0.892254099 |
| ENSG00000067225 | *PKM* | 8.130026074 | -0.032456536 | -0.003955904 | -0.028500632 | 0.999933887 | 0.228199446 | 0.210087105 | 0.018112341 | 0.892254099 |
| ENSG00000145592 | *RPL37* | 7.559296331 | 0.0182684 | -0.023843897 | 0.042112297 | 0.999933887 | 0.069669819 | 0.091467621 | -0.021797802 | 0.892254099 |
| ENSG00000112159 | *MDN1* | 4.666539837 | 0.015429901 | -0.05217208 | 0.067601981 | 0.999933887 | 0.021281022 | 0.049366525 | -0.028085502 | 0.892254099 |
| ENSG00000189060 | *H1-0* | 2.961278515 | 0.041231883 | -0.072581997 | 0.11381388 | 0.999933887 | -0.154188151 | -0.192203941 | 0.03801579 | 0.892369504 |
| ENSG00000197982 | *C1orf122* | 4.412590054 | 0.070925319 | 0.027843484 | 0.043081835 | 0.999933887 | 0.969824664 | 0.940786897 | 0.029037767 | 0.892424094 |
| ENSG00000106829 | *TLE4* | 6.957814068 | 0.006781398 | 0.048681054 | -0.041899656 | 0.999933887 | 0.198512264 | 0.22423491 | -0.025722647 | 0.892424094 |
| ENSG00000167196 | *FBXO22* | 1.695063343 | -0.057881756 | 0.083647607 | -0.141529363 | 0.999933887 | 0.032659327 | 0.07841536 | -0.045756033 | 0.892546002 |
| ENSG00000104980 | *TIMM44* | 3.973978939 | -0.021288435 | -0.005172233 | -0.016116202 | 0.999933887 | -0.031393717 | -0.007851676 | -0.02354204 | 0.892577324 |
| ENSG00000238083 | *LRRC37A2* | 0.916713897 | 0.033956787 | 0.03999604 | -0.006039253 | 0.999933887 | -0.247546354 | -0.186375077 | -0.061171277 | 0.892762904 |
| ENSG00000271843 | *AC012557.1* | -0.608731168 | -0.117839902 | -0.195991994 | 0.078152092 | 0.999933887 | -0.151059913 | -0.276439584 | 0.125379671 | 0.892836573 |
| ENSG00000230590 | *FTX* | 3.826310366 | 0.025344347 | 0.08192629 | -0.056581943 | 0.999933887 | -0.395014812 | -0.443588886 | 0.048574074 | 0.892947158 |
| ENSG00000236540 | *AC006547.1* | 1.583990738 | 0.003441523 | -0.220563002 | 0.224004525 | 0.999933887 | -0.660676392 | -0.706906887 | 0.046230495 | 0.892947158 |
| ENSG00000118705 | *RPN2* | 4.764503468 | 0.015288058 | -0.072629478 | 0.087917536 | 0.999933887 | 0.022787594 | 0.003046272 | 0.019741323 | 0.893053422 |
| ENSG00000083845 | *RPS5* | 6.653728287 | 0.027256705 | -0.003637415 | 0.03089412 | 0.999933887 | 0.131453213 | 0.154453479 | -0.023000266 | 0.893112199 |
| ENSG00000136960 | *ENPP2* | -0.279046758 | -0.067419564 | -0.12043995 | 0.053020386 | 0.999933887 | 0.04565957 | 0.135917068 | -0.090257498 | 0.893112199 |
| ENSG00000139684 | *ESD* | 2.284426924 | 0.020978343 | -0.129010314 | 0.149988657 | 0.999933887 | -0.058004418 | -0.024783927 | -0.033220491 | 0.893112199 |
| ENSG00000066651 | *TRMT11* | 1.485148603 | -0.025935433 | -0.013311207 | -0.012624226 | 0.999933887 | -0.006098925 | -0.048947152 | 0.042848227 | 0.893112199 |
| ENSG00000102349 | *KLF8* | 0.480965425 | -0.018716832 | -0.112053012 | 0.093336179 | 0.999933887 | -0.163878616 | -0.223629355 | 0.05975074 | 0.893112199 |
| ENSG00000114315 | *HES1* | 3.847863901 | 0.218872459 | 0.233127463 | -0.014255004 | 0.999933887 | 0.674747612 | 0.626097142 | 0.04865047 | 0.893267606 |
| ENSG00000280248 | *AC124319.3* | -0.271524199 | 0.158117353 | -0.028824878 | 0.186942231 | 0.999933887 | -0.456038795 | -0.549098536 | 0.093059741 | 0.893267606 |
| ENSG00000131067 | *GGT7* | 2.154582966 | 0.122452471 | 0.138709662 | -0.016257191 | 0.999933887 | 0.066019014 | 0.028108821 | 0.037910193 | 0.893359666 |
| ENSG00000271938 | *AC103724.4* | -0.278189079 | -0.013134493 | 0.186771511 | -0.199906004 | 0.999933887 | -0.767965813 | -0.662704664 | -0.105261149 | 0.893359666 |
| ENSG00000184898 | *RBM43* | 1.776785795 | 0.011565393 | -0.18040963 | 0.191975023 | 0.999933887 | 0.004101243 | 0.052811339 | -0.048710096 | 0.893359666 |
| ENSG00000101444 | *AHCY* | 1.929167646 | 0.007868832 | 0.014345585 | -0.006476753 | 0.999933887 | 0.119585759 | 0.076815216 | 0.042770543 | 0.893359666 |
| ENSG00000198860 | *TSEN15* | 2.108226242 | -0.064154256 | -0.109872198 | 0.045717942 | 0.999933887 | 0.100477817 | 0.135943453 | -0.035465636 | 0.89340106 |
| ENSG00000268812 | *AC004264.1* | -1.208907456 | 0.246700746 | 0.076721028 | 0.169979719 | 0.999933887 | 0.058162848 | -0.088397516 | 0.146560364 | 0.893501945 |
| ENSG00000133247 | *KMT5C* | 3.502383144 | 0.027600887 | 0.10417005 | -0.076569163 | 0.999933887 | 0.096120556 | 0.067097699 | 0.029022857 | 0.893558802 |
| ENSG00000215440 | *NPEPL1* | 3.467917952 | 0.094591328 | 0.044742945 | 0.049848384 | 0.999933887 | -0.117865052 | -0.153293838 | 0.035428786 | 0.893784223 |
| ENSG00000259363 | *AC090825.1* | -0.40596766 | 0.188464406 | -0.144594518 | 0.333058924 | 0.999933887 | 0.080461612 | -0.005503881 | 0.085965493 | 0.893784223 |
| ENSG00000133800 | *LYVE1* | -1.078443637 | 0.168938882 | -0.237646574 | 0.406585456 | 0.999933887 | 1.391328747 | 1.238223411 | 0.153105336 | 0.893950775 |
| ENSG00000269958 | *AL049840.5* | 2.778155759 | -0.076076409 | -0.029203143 | -0.046873265 | 0.999933887 | -0.290446497 | -0.258091855 | -0.032354641 | 0.894019694 |
| ENSG00000240342 | *RPS2P5* | 5.439726625 | 0.057352031 | 0.03208627 | 0.025265761 | 0.999933887 | 0.049928932 | 0.08152114 | -0.031592207 | 0.894019694 |
| ENSG00000164096 | *C4orf3* | 6.110221798 | -0.017281384 | 0.012990131 | -0.030271515 | 0.999933887 | 0.115745178 | 0.138829056 | -0.023083878 | 0.894019694 |
| ENSG00000269399 | *AC008764.6* | 0.35639315 | -0.343179 | -0.016605551 | -0.326573449 | 0.999933887 | 0.152411001 | 0.085660102 | 0.066750899 | 0.894032524 |
| ENSG00000138160 | *KIF11* | -0.855654082 | 0.10045119 | 0.11054041 | -0.010089219 | 0.999933887 | -0.330904684 | -0.219384672 | -0.111520012 | 0.894040085 |
| ENSG00000279891 | *FLJ42393* | -0.212132725 | 0.232165172 | 0.413027695 | -0.180862523 | 0.999933887 | 0.213760937 | 0.314018676 | -0.100257738 | 0.894044996 |
| ENSG00000162551 | *ALPL* | 8.411367893 | 0.028385914 | -0.014303309 | 0.042689224 | 0.999933887 | -0.102122186 | -0.126397946 | 0.02427576 | 0.894318867 |
| ENSG00000197157 | *SND1* | 4.423431417 | -0.017982197 | -0.020060535 | 0.002078338 | 0.999933887 | 0.085741807 | 0.106003662 | -0.020261855 | 0.894577903 |
| ENSG00000170619 | *COMMD5* | 6.282962255 | 0.038421119 | 0.046445457 | -0.008024338 | 0.999933887 | 0.130041177 | 0.156431016 | -0.026389839 | 0.894930689 |
| ENSG00000110077 | *MS4A6A* | 2.961792359 | 0.010497647 | -0.003134755 | 0.013632402 | 0.999933887 | 0.050376614 | 0.091586095 | -0.041209481 | 0.894963192 |
| ENSG00000272666 | *KLHDC7B-DT* | 1.227646643 | 0.256864854 | 0.298150282 | -0.041285428 | 0.999933887 | 0.090978066 | 0.153831571 | -0.062853505 | 0.894981308 |
| ENSG00000135916 | *ITM2C* | 3.060473014 | 0.129545286 | 0.026995319 | 0.102549968 | 0.999933887 | 0.206029047 | 0.171755964 | 0.034273082 | 0.894981308 |
| ENSG00000232926 | *AC000078.1* | 0.472282414 | 0.187338698 | 0.117274548 | 0.07006415 | 0.999933887 | 0.112383389 | 0.023097084 | 0.089286306 | 0.894981308 |
| ENSG00000167483 | *NIBAN3* | 1.098247389 | 0.077974989 | 0.099341666 | -0.021366677 | 0.999933887 | -0.156279157 | -0.213416503 | 0.057137346 | 0.894981308 |
| ENSG00000205560 | *CPT1B* | -0.338006934 | 0.135563087 | 0.324664805 | -0.189101717 | 0.999933887 | -0.242921981 | -0.32299556 | 0.080073579 | 0.894981308 |
| ENSG00000220267 | *ACTBP8* | -2.811239442 | 0.066398319 | 0.066816473 | -0.000418153 | 0.999933887 | 0.001438476 | 0.100996393 | -0.099557917 | 0.894981308 |
| ENSG00000164466 | *SFXN1* | 3.688874057 | -0.038906812 | -0.005893503 | -0.033013309 | 0.999933887 | 0.050898817 | 0.079286557 | -0.02838774 | 0.894981308 |
| ENSG00000136573 | *BLK* | 0.515447626 | 0.063996289 | -0.001368557 | 0.065364847 | 0.999933887 | 0.211731608 | 0.160087187 | 0.051644421 | 0.894981308 |
| ENSG00000112294 | *ALDH5A1* | 0.596463289 | 0.006770494 | 0.290807874 | -0.28403738 | 0.999933887 | 0.004034805 | 0.052588139 | -0.048553334 | 0.894981308 |
| ENSG00000243244 | *STON1* | 0.461715272 | 0.074078464 | 0.007052225 | 0.067026239 | 0.999933887 | -0.320764047 | -0.393066713 | 0.072302667 | 0.895096829 |
| ENSG00000260401 | *AP002761.4* | 1.375501961 | -0.140046606 | 0.14571512 | -0.285761727 | 0.999933887 | -0.074958498 | -0.015635152 | -0.059323346 | 0.895247372 |
| ENSG00000255198 | *SNHG9* | 1.063468421 | -0.105950129 | -0.010448207 | -0.095501922 | 0.999933887 | 0.106038766 | 0.053924277 | 0.05211449 | 0.895353254 |
| ENSG00000152217 | *SETBP1* | 1.689857754 | 0.13464014 | -0.058941887 | 0.193582027 | 0.999933887 | -0.112434464 | -0.166005286 | 0.053570822 | 0.895360322 |
| ENSG00000054148 | *PHPT1* | 1.949200044 | 0.075815094 | 0.121019865 | -0.045204771 | 0.999933887 | 0.055628379 | 0.096685063 | -0.041056684 | 0.895360322 |
| ENSG00000211772 | *TRBC2* | 6.399621623 | 0.007356772 | -0.052474293 | 0.059831065 | 0.999933887 | 0.164880288 | 0.144767717 | 0.020112571 | 0.895360322 |
| ENSG00000118640 | *VAMP8* | 2.979031326 | -0.055221472 | -0.05751487 | 0.002293398 | 0.999933887 | 0.046290488 | 0.079544743 | -0.033254254 | 0.895456128 |
| ENSG00000260669 | *AL096870.2* | 0.59104875 | 0.096122577 | 0.042906711 | 0.053215866 | 0.999933887 | 0.333346313 | 0.400712344 | -0.067366032 | 0.895456128 |
| ENSG00000163832 | *ELP6* | 1.008559975 | 0.046502041 | -0.136108129 | 0.18261017 | 0.999933887 | 0.145602737 | 0.089966958 | 0.05563578 | 0.895456128 |
| ENSG00000214160 | *ALG3* | 1.559780286 | -0.000145032 | -0.114295186 | 0.114150155 | 0.999933887 | 0.054167012 | 0.090121604 | -0.035954592 | 0.895456128 |
| ENSG00000173662 | *TAS1R1* | -0.292171254 | 0.244138102 | 0.141313832 | 0.102824269 | 0.999933887 | 0.675778632 | 0.583766588 | 0.092012044 | 0.895466022 |
| ENSG00000231971 | *CT69* | 0.114826984 | -0.285824906 | -0.062674211 | -0.223150695 | 0.999933887 | 0.096706657 | 0.149247276 | -0.052540619 | 0.895515454 |
| ENSG00000167671 | *UBXN6* | 5.890508222 | 0.055147087 | 0.053901451 | 0.001245636 | 0.999933887 | -0.118400496 | -0.14167177 | 0.023271275 | 0.895515454 |
| ENSG00000039523 | *RIPOR1* | 6.749588477 | 0.065426349 | 0.000255557 | 0.065170793 | 0.999933887 | 0.579533891 | 0.558197634 | 0.021336257 | 0.895855899 |
| ENSG00000145348 | *TBCK* | 1.355984651 | -0.104920023 | -0.032464179 | -0.072455844 | 0.999933887 | -0.04639099 | 0.005419784 | -0.051810774 | 0.895855899 |
| ENSG00000197971 | *MBP* | 9.436979123 | 0.025697501 | -0.014031611 | 0.039729112 | 0.999933887 | -0.577502052 | -0.556579904 | -0.020922149 | 0.895855899 |
| ENSG00000287214 | *AC021321.2* | 0.120185199 | 0.007391393 | 0.200848216 | -0.193456823 | 0.999933887 | -0.055595616 | 0.023021051 | -0.078616667 | 0.895855899 |
| ENSG00000148848 | *ADAM12* | -1.25619068 | -0.24386158 | -0.445833447 | 0.201971867 | 0.999933887 | -0.417706756 | -0.52641157 | 0.108704815 | 0.895860386 |
| ENSG00000251288 | *AC018797.2* | 0.784853529 | -0.177241164 | -0.0096611 | -0.167580064 | 0.999933887 | -0.06031458 | 0.010274623 | -0.070589203 | 0.895860386 |
| ENSG00000188785 | *ZNF548* | 2.59931892 | -0.081650294 | -0.010413274 | -0.07123702 | 0.999933887 | 0.126295588 | 0.160135964 | -0.033840376 | 0.895860386 |
| ENSG00000198730 | *CTR9* | 4.216505374 | -0.064530675 | -0.085848005 | 0.02131733 | 0.999933887 | -0.287061659 | -0.259398229 | -0.027663429 | 0.895860386 |
| ENSG00000150337 | *FCGR1A* | 4.036951441 | -0.035042983 | -0.09069453 | 0.055651547 | 0.999933887 | -0.359096484 | -0.319061885 | -0.040034598 | 0.895860386 |
| ENSG00000278600 | *AC015871.3* | 5.224064493 | -0.025496052 | 0.04151809 | -0.067014142 | 0.999933887 | 0.03687652 | 0.063348807 | -0.026472287 | 0.895860386 |
| ENSG00000142541 | *RPL13A* | 9.119562041 | -0.011261266 | -0.044476558 | 0.033215292 | 0.999933887 | 0.09515052 | 0.073294027 | 0.021856493 | 0.895860386 |
| ENSG00000145331 | *TRMT10A* | 0.203881346 | 0.019324643 | 0.027410514 | -0.008085871 | 0.999933887 | -0.238939538 | -0.329414864 | 0.090475326 | 0.895860386 |
| ENSG00000100450 | *GZMH* | 4.780829386 | -0.002366888 | -0.024541052 | 0.022174165 | 0.999933887 | 0.093104768 | 0.121702251 | -0.028597483 | 0.895860386 |
| ENSG00000245970 | *AP003352.1* | 0.188897222 | -0.126909347 | -0.190500372 | 0.063591025 | 0.999933887 | 0.04835163 | -0.030416609 | 0.078768239 | 0.895865495 |
| ENSG00000261226 | *AC092384.3* | -0.050075358 | -0.105334 | 0.018300848 | -0.123634848 | 0.999933887 | -0.289939628 | -0.362650252 | 0.072710624 | 0.895865495 |
| ENSG00000219200 | *RNASEK* | 5.036589406 | 0.131645684 | 0.17490301 | -0.043257325 | 0.999933887 | 0.639988161 | 0.664436295 | -0.024448135 | 0.896016221 |
| ENSG00000102678 | *FGF9* | -0.400223284 | -0.190846922 | -0.253290913 | 0.062443991 | 0.999933887 | 0.05654127 | 0.129480694 | -0.072939424 | 0.896016221 |
| ENSG00000126602 | *TRAP1* | 2.204578763 | -0.066596345 | -0.027762475 | -0.03883387 | 0.999933887 | 0.153152223 | 0.181359081 | -0.028206858 | 0.896016221 |
| ENSG00000248455 | *LINC02217* | 0.280654245 | 0.08588412 | 0.160042524 | -0.074158405 | 0.999933887 | 0.456003195 | 0.372123512 | 0.083879683 | 0.896016221 |
| ENSG00000146122 | *DAAM2* | -2.048806374 | -0.104077711 | 0.250835529 | -0.35491324 | 0.999933887 | -0.757798903 | -0.949973143 | 0.192174241 | 0.896016221 |
| ENSG00000164100 | *NDST3* | -0.661820341 | 0.115181744 | 0.040447856 | 0.074733888 | 0.999933887 | 0.674772145 | 0.603867594 | 0.070904551 | 0.896016221 |
| ENSG00000119711 | *ALDH6A1* | 0.42897829 | 0.050733205 | -0.632727386 | 0.683460591 | 0.944094537 | 0.0162887 | -0.046202707 | 0.062491407 | 0.896016221 |
| ENSG00000099889 | *ARVCF* | -0.464681869 | 0.037012359 | 0.239251148 | -0.202238789 | 0.999933887 | 0.067881063 | 0.002300336 | 0.065580727 | 0.896016221 |
| ENSG00000177600 | *RPLP2* | 7.707191358 | 0.008443559 | -0.021386001 | 0.029829561 | 0.999933887 | 0.098780987 | 0.121373454 | -0.022592467 | 0.896381949 |
| ENSG00000109171 | *SLAIN2* | 4.904297996 | 0.093834587 | 0.075056437 | 0.018778151 | 0.999933887 | -0.525571146 | -0.503015555 | -0.022555591 | 0.896646316 |
| ENSG00000141580 | *WDR45B* | 5.581076191 | 0.00587063 | -0.02913001 | 0.03500064 | 0.999933887 | -0.264394161 | -0.283456509 | 0.019062348 | 0.896646316 |
| ENSG00000156502 | *SUPV3L1* | 3.183066879 | 0.032744041 | 0.016593118 | 0.016150922 | 0.999933887 | 0.128550267 | 0.154622042 | -0.026071775 | 0.896651098 |
| ENSG00000111300 | *NAA25* | 2.48009869 | -0.074020994 | -0.042206249 | -0.031814744 | 0.999933887 | -0.016323224 | 0.019044012 | -0.035367236 | 0.89670095 |

| | | | | | | | | | | |
|---|---|---|---|---|---|---|---|---|---|---|
| ENSG00000175106 | *TVP23C* | -0.088094968 | -0.214386238 | 0.211196596 | -0.425582834 | 0.999933887 | -0.00477178 | 0.077353006 | -0.082124787 | 0.896809051 |
| ENSG00000105011 | *ASF1B* | 1.717231796 | 0.069107467 | -0.034088433 | 0.103195901 | 0.999933887 | 0.479066319 | 0.539065336 | -0.059999017 | 0.896809051 |
| ENSG00000214826 | *DDX12P* | -0.009909743 | -0.081857922 | 0.012821811 | -0.094679733 | 0.999933887 | -0.0550419 | 0.010303074 | -0.065344974 | 0.896809051 |
| ENSG00000063180 | *CA11* | 0.033535297 | 0.042993153 | 0.2043184 | -0.161325247 | 0.999933887 | -0.033608979 | 0.030878989 | -0.064487968 | 0.896809051 |
| ENSG00000164182 | *NDUFAF2* | 0.049931358 | -0.020253181 | -0.224212693 | 0.203959512 | 0.999933887 | 0.20381109 | 0.136829205 | 0.066981885 | 0.896809051 |
| ENSG00000102743 | *SLC25A15* | -0.083542169 | 0.002313714 | -0.284838436 | 0.28715215 | 0.999933887 | 0.091485377 | 0.024337249 | 0.067148128 | 0.896809051 |
| ENSG00000270871 | *AC015849.3* | -0.661602996 | 0.073229231 | 0.227274107 | -0.154044875 | 0.999933887 | -0.094901292 | -0.011509003 | -0.083392288 | 0.897291093 |
| ENSG00000235316 | *DUSP8P5* | 0.562454484 | 0.218707426 | 0.012115454 | 0.206591972 | 0.999933887 | -0.171468143 | -0.118831368 | -0.052636775 | 0.897473885 |
| ENSG00000165501 | *LRR1* | 0.39296087 | -0.097992467 | 0.138928797 | -0.236921264 | 0.999933887 | -0.097697117 | -0.15774966 | 0.060052543 | 0.897473885 |
| ENSG00000117834 | *SLC5A9* | -0.222042208 | -0.050123674 | -0.152297148 | 0.102173474 | 0.999933887 | -0.086220177 | -0.005370225 | -0.080849952 | 0.897473885 |
| ENSG00000053900 | *ANAPC4* | 3.598229183 | -0.015618183 | -0.018497244 | 0.002879061 | 0.999933887 | -0.204770681 | -0.239194529 | 0.034423847 | 0.897473885 |
| ENSG00000259473 | *LINC02205* | -0.526803414 | 0.169826558 | 0.079488824 | 0.090337734 | 0.999933887 | 0.16940997 | 0.268758398 | -0.099348428 | 0.897579247 |
| ENSG00000108854 | *SMURF2* | 4.954609139 | 0.031460974 | 0.007402049 | 0.024058925 | 0.999933887 | -0.470619028 | -0.448736965 | -0.021882063 | 0.897579247 |
| ENSG00000116711 | *PLA2G4A* | 0.190789942 | 0.045744199 | -0.134427859 | 0.180172058 | 0.999933887 | -0.621860783 | -0.702120039 | 0.080259256 | 0.897579247 |
| ENSG00000120915 | *EPHX2* | 1.209228904 | -0.070931506 | -0.055564156 | -0.01536735 | 0.999933887 | 0.232364428 | 0.275793204 | -0.043428776 | 0.897767801 |
| ENSG00000112029 | *FBXO5* | -0.319621986 | 0.120422294 | -0.158485345 | 0.278907639 | 0.999933887 | -0.12441538 | -0.039092483 | -0.085322896 | 0.897864062 |
| ENSG00000132967 | *HMGB1P5* | 0.410842809 | -0.060572937 | -0.320020862 | 0.259447925 | 0.999933887 | -0.081542732 | -0.154091535 | 0.072548803 | 0.897864062 |
| ENSG00000232869 | *TRBV29-1* | 0.850655527 | 0.047988052 | 0.00600804 | 0.041980012 | 0.999933887 | 0.083863674 | 0.036623166 | 0.047240508 | 0.898089501 |
| ENSG00000100714 | *MTHFD1* | 1.701195865 | 0.126547804 | 0.0951803 | 0.031367504 | 0.999933887 | 0.094292165 | 0.129568436 | -0.035276271 | 0.89819598 |
| ENSG00000278599 | *TBC1D3E* | -2.339179656 | 0.175928316 | -0.033022354 | 0.20895067 | 0.999933887 | 0.193180725 | 0.122657345 | 0.070523381 | 0.89833434 |
| ENSG00000173456 | *RNF26* | 1.779236101 | 0.101814721 | -0.122068981 | 0.223883702 | 0.999933887 | 0.158179192 | 0.198599347 | -0.040420154 | 0.89833434 |
| ENSG00000181126 | *HLA-V* | 2.121491028 | -0.060978953 | 0.055739365 | -0.116718318 | 0.999933887 | 0.729710807 | 0.758770395 | -0.029059589 | 0.89833434 |
| ENSG00000245571 | *FAM111A-DT* | -0.602898811 | -0.008936897 | 0.056669694 | -0.065606591 | 0.999933887 | 0.20275383 | 0.125624447 | 0.077129383 | 0.89833434 |
| ENSG00000264538 | *SUZ12P1* | 2.546277741 | 0.06125468 | 0.131829792 | -0.070575112 | 0.999933887 | 0.192450998 | 0.162947978 | 0.02950302 | 0.898342572 |
| ENSG00000273748 | *AL592183.1* | -0.602256533 | -0.043795296 | -0.36260202 | 0.318806724 | 0.999933887 | 0.075410225 | 0.141072432 | -0.065662207 | 0.898342572 |
| ENSG00000113643 | *RARS1* | 2.91297146 | -0.01347488 | -0.148919452 | 0.135444572 | 0.999933887 | -0.106997591 | -0.078889092 | -0.028108499 | 0.898342572 |
| ENSG00000286248 | *AC069209.2* | -2.177219221 | 0.044459725 | 0.773079156 | -0.728619431 | 0.999933887 | 1.156638156 | 1.248338447 | -0.091700291 | 0.898342572 |
| ENSG00000231084 | *RPL22P24* | -0.441021168 | -0.192214826 | 0.047245615 | -0.239460441 | 0.999933887 | 0.206161112 | 0.256858864 | -0.050697752 | 0.898386197 |
| ENSG00000158457 | *TSPAN33* | 0.776850303 | 0.223659349 | 0.235657331 | -0.011997982 | 0.999933887 | 2.808431909 | 2.9059751 | -0.097543191 | 0.898386197 |
| ENSG00000109814 | *UGDH* | 0.225576074 | -0.060506557 | 0.33463455 | -0.395141106 | 0.999933887 | -0.01397347 | -0.064469791 | 0.050496321 | 0.898386197 |
| ENSG00000105976 | *MET* | -0.573054386 | -0.581338452 | -0.429193496 | -0.152144956 | 0.999933887 | -0.088648709 | -0.032231575 | -0.056417134 | 0.898612671 |
| ENSG00000269688 | *AC008982.2* | 1.999164694 | 0.046534904 | 0.16127216 | -0.114737257 | 0.999933887 | -0.245813924 | -0.286297711 | 0.040483788 | 0.898612671 |
| ENSG00000127720 | *METTL25* | 0.8615275 | -0.067323395 | -0.195547948 | 0.128224554 | 0.999933887 | -0.218954998 | -0.271966076 | 0.053011078 | 0.898612671 |
| ENSG00000260517 | *AC009093.2* | -0.743409458 | 0.114865173 | 0.099520026 | 0.015345147 | 0.999933887 | 0.229937829 | 0.13298376 | 0.096954069 | 0.898612671 |
| ENSG00000100028 | *SNRPD3* | 2.511202768 | -0.067810694 | 0.044791124 | -0.112601817 | 0.999933887 | -0.13825249 | -0.106701041 | -0.031551449 | 0.898826214 |
| ENSG00000111319 | *SCNN1A* | -0.317549582 | -0.066868847 | 0.192461622 | -0.259330469 | 0.999933887 | -0.437414325 | -0.523795845 | 0.086381521 | 0.898917025 |
| ENSG00000143578 | *CREB3L4* | -0.575354012 | 0.066227686 | -0.063446859 | 0.129674546 | 0.999933887 | 0.358933602 | 0.280205722 | 0.07872788 | 0.898957863 |
| ENSG00000187240 | *DYNC2H1* | -0.130116224 | 0.133896765 | -0.357578514 | 0.491475279 | 0.999933887 | -0.310959102 | -0.207060452 | -0.10389865 | 0.898961089 |
| ENSG00000258227 | *CLEC5A* | 3.448098842 | -0.079319657 | -0.13086485 | 0.051545193 | 0.999933887 | -0.228225916 | -0.18052066 | -0.047705256 | 0.898961089 |
| ENSG00000136270 | *TBRG4* | 3.831025173 | 0.001297435 | 0.014986213 | -0.013688778 | 0.999933887 | 0.164016413 | 0.139882551 | 0.024133862 | 0.898961089 |
| ENSG00000233077 | *LINC01271* | -0.59000029 | 0.250953157 | 0.028360444 | 0.222592712 | 0.999933887 | 0.935875563 | 1.03805062 | -0.102175057 | 0.89896981 |
| ENSG00000136802 | *LRRC8A* | 4.691962372 | 0.076780728 | -0.041642716 | 0.118423443 | 0.999933887 | -0.38814979 | -0.361823843 | -0.026325947 | 0.89896981 |
| ENSG00000143398 | *PIP5K1A* | 4.975531877 | 0.063116288 | 0.131495261 | -0.068378973 | 0.999933887 | -0.065952783 | -0.045920297 | -0.020032486 | 0.89896981 |
| ENSG00000133131 | *MORC4* | 0.410138294 | -0.15092984 | 0.139716695 | -0.290646535 | 0.999933887 | 0.11113971 | 0.166213966 | -0.055074257 | 0.89896981 |
| ENSG00000187778 | *MCRS1* | 4.693584153 | -0.055657766 | 0.007902568 | -0.063560334 | 0.999933887 | -0.077677975 | -0.098462277 | 0.020784302 | 0.89896981 |
| ENSG00000204475 | *NCR3* | 1.391585582 | 0.030798375 | -0.047897276 | 0.078695651 | 0.999933887 | 0.021482726 | -0.020706924 | 0.04218965 | 0.89896981 |
| ENSG00000149485 | *FADS1* | 0.050360265 | -0.025792995 | 0.174790033 | -0.200583028 | 0.999933887 | -0.296866425 | -0.229983166 | -0.066883259 | 0.89896981 |
| ENSG00000213658 | *LAT* | 1.787458264 | 0.021669471 | 0.36538794 | -0.343718468 | 0.999933887 | 0.379780987 | 0.34410769 | 0.035673297 | 0.899407155 |
| ENSG00000270019 | *AC110769.2* | 1.311307798 | 0.388512897 | 0.277401043 | 0.111111854 | 0.999933887 | 1.799896696 | 1.71980079 | 0.080095906 | 0.899720745 |
| ENSG00000130518 | *IQCN* | 3.85883009 | 0.089942356 | 0.064495656 | 0.0254467 | 0.999933887 | -0.1390206 | -0.110388457 | -0.028632143 | 0.899720745 |
| ENSG00000184292 | *TACSTD2* | -0.17402679 | -0.280792303 | -0.212898894 | -0.067893409 | 0.999933887 | -0.328823029 | -0.394243829 | 0.0654208 | 0.899765298 |
| ENSG00000099331 | *MYO9B* | 9.648712392 | 0.012354519 | 0.037132504 | -0.024777984 | 0.999933887 | -0.164951069 | -0.179904264 | 0.014953195 | 0.899765298 |
| ENSG00000167615 | *LENG8* | 8.003677274 | 0.011611019 | 0.022353295 | -0.010742276 | 0.999933887 | -0.118782344 | -0.135467847 | 0.016685503 | 0.899765298 |
| ENSG00000165512 | *ZNF22* | 0.717677914 | 0.038614814 | -0.164673943 | 0.203288757 | 0.999933887 | 0.101652145 | 0.054528633 | 0.047123512 | 0.899894877 |
| ENSG00000204991 | *SPIRE2* | 0.489507713 | 0.007675133 | 0.143013339 | -0.135338206 | 0.999933887 | 0.284812533 | 0.343282439 | -0.058469906 | 0.899894877 |
| ENSG00000233834 | *AC005083.1* | -2.570292656 | 0.288493484 | 1.237258327 | -0.948764843 | 0.999933887 | 3.141061892 | 3.287655881 | -0.146593989 | 0.900088074 |
| ENSG00000170310 | *STX8* | 1.726158862 | -0.124821302 | 0.027645608 | -0.15246691 | 0.999933887 | 0.044794968 | 0.097088771 | -0.052293802 | 0.900204682 |
| ENSG00000198718 | *TOGARAM1* | 2.271211991 | 0.003962167 | -0.072401966 | 0.076364133 | 0.999933887 | -0.423904275 | -0.389951596 | -0.033952679 | 0.900204682 |
| ENSG00000181004 | *BBS12* | -0.662053974 | -0.120953848 | -0.183839694 | 0.062885846 | 0.999933887 | 0.120477282 | 0.042681694 | 0.077795588 | 0.900633113 |
| ENSG00000222041 | *CYTOR* | 4.070310063 | 0.089240778 | 0.093827635 | -0.004586857 | 0.999933887 | -0.605073067 | -0.56487608 | -0.040196987 | 0.900726129 |
| ENSG00000175544 | *CABP4* | -0.721396136 | 0.074887805 | -0.097410218 | 0.172298023 | 0.999933887 | -0.157313806 | -0.078488431 | -0.078825375 | 0.900726129 |
| ENSG00000177112 | *IRAG1-AS1* | -0.037923456 | -0.038061253 | -0.108501802 | 0.070440549 | 0.999933887 | 0.487214235 | 0.58524261 | -0.098028375 | 0.900726129 |
| ENSG00000198752 | *CDC42BPB* | 0.553653422 | 0.016845766 | 0.125668911 | -0.108823145 | 0.999933887 | -0.176043509 | -0.239987844 | 0.063944334 | 0.900726129 |
| ENSG00000261582 | *AL121753.1* | -0.521353472 | 0.003467772 | -0.02555285 | 0.029020622 | 0.999933887 | -0.293568899 | -0.372180717 | 0.078611818 | 0.900726129 |
| ENSG00000105171 | *POP4* | 3.275398386 | -0.006198992 | -0.084985563 | 0.078786571 | 0.999933887 | -0.027448047 | -0.003833893 | -0.023614154 | 0.900846877 |
| ENSG00000130414 | *NDUFA10* | 4.558115467 | -0.026518172 | -0.017108764 | -0.009409408 | 0.999933887 | 0.019989148 | 0.038199344 | -0.018210196 | 0.901187075 |
| ENSG00000064270 | *ATP2C2* | 1.482986899 | -0.076908868 | 0.0232416 | -0.100150468 | 0.999933887 | 0.401481922 | 0.436868512 | -0.03538659 | 0.901310177 |
| ENSG00000151233 | *GXYLT1* | 2.918075774 | -0.056642046 | -0.127268623 | 0.070626577 | 0.999933887 | -0.646421518 | -0.598817619 | -0.047603899 | 0.901318958 |
| ENSG00000170632 | *ARMC10* | 3.036819246 | -0.041134478 | 0.037362511 | -0.078496988 | 0.999933887 | -0.422202152 | -0.454887231 | 0.032685079 | 0.901318958 |
| ENSG00000275898 | *AC125603.4* | 0.164477929 | -0.047750956 | 0.195721415 | -0.24347237 | 0.999933887 | -0.336510732 | -0.408087476 | 0.071576744 | 0.901393981 |
| ENSG00000116704 | *SLC35D1* | 2.82465618 | -0.017940878 | -0.02956551 | 0.011624632 | 0.999933887 | -0.060721149 | -0.030757783 | -0.029963366 | 0.90205251 |
| ENSG00000113141 | *IK* | 6.543365905 | -0.05594384 | -0.13925898 | 0.083315141 | 0.999933887 | -0.275310753 | -0.293019037 | 0.017708284 | 0.902168771 |
| ENSG00000134864 | *GGACT* | -0.775534722 | -0.225148099 | -0.101771945 | -0.123376154 | 0.999933887 | 0.056019817 | -0.023696813 | 0.079716629 | 0.902292398 |
| ENSG00000274487 | *NPEPPSP1* | 1.883916488 | 0.088845536 | 0.053018217 | 0.035827319 | 0.999933887 | 0.09519442 | 0.056131951 | 0.039062469 | 0.902292398 |
| ENSG00000275557 | *AC242842.1* | 1.241057206 | 0.077034958 | 0.092685952 | -0.015650994 | 0.999933887 | 0.491769422 | 0.539059001 | -0.047289579 | 0.902292398 |
| ENSG00000166562 | *SEC11C* | 2.764974561 | -0.010762422 | -0.071062855 | 0.060300432 | 0.999933887 | 0.142109591 | 0.171399524 | -0.029289933 | 0.902292398 |
| ENSG00000164951 | *PDP1* | 3.288521148 | -0.007024233 | -0.037487636 | 0.030463403 | 0.999933887 | -0.216799839 | -0.243861567 | 0.027061728 | 0.902292398 |
| ENSG00000137070 | *IL11RA* | 3.085912005 | -0.005588222 | 0.126166588 | -0.13175481 | 0.999933887 | 0.059164169 | 0.031116267 | 0.028047903 | 0.902292398 |
| ENSG00000184209 | *SNRNP35* | 3.651909798 | -0.000840818 | 0.100426972 | -0.10126779 | 0.999933887 | 0.37209547 | 0.344907272 | 0.027188197 | 0.902292398 |
| ENSG00000066923 | *STAG3* | 1.916367166 | 0.085766478 | 0.065689425 | 0.020077053 | 0.999933887 | -0.200622896 | -0.160011285 | -0.040611611 | 0.902442421 |
| ENSG00000255987 | *TOMM20P2* | 0.06215781 | -0.068799434 | 0.219148484 | -0.287947918 | 0.999933887 | 0.023844752 | 0.086429153 | -0.062584402 | 0.902446951 |
| ENSG00000211949 | *IGHV3-23* | -1.404922698 | 0.062959293 | 0.128828731 | -0.065869439 | 0.999933887 | 0.135078086 | 0.274736568 | -0.139658482 | 0.902536583 |
| ENSG00000275700 | *AATF* | 5.408359698 | 0.008618275 | 0.011368819 | -0.002750544 | 0.999933887 | -0.193170211 | -0.21099956 | 0.017829349 | 0.902536583 |
| ENSG00000164587 | *RPS14* | 7.697605983 | 0.003589462 | -0.007172817 | 0.010762279 | 0.999933887 | 0.100057381 | 0.079261772 | 0.020795609 | 0.902536583 |
| ENSG00000083535 | *PIBF1* | 1.545825208 | 0.00073663 | -0.343664746 | 0.344401376 | 0.999933887 | -0.127437255 | -0.066735647 | -0.060701608 | 0.902536583 |
| ENSG00000281100 | *AC105749.1* | 1.060915832 | 0.072992622 | -0.185559375 | 0.258551997 | 0.999933887 | 0.157026681 | 0.098847106 | 0.058179576 | 0.902666332 |
| ENSG00000121691 | *CAT* | 6.203726269 | -0.065947004 | -0.008828642 | -0.057118362 | 0.999933887 | -0.04627796 | -0.022607658 | -0.023670302 | 0.902696306 |
| ENSG00000115677 | *HDLBP* | 6.603333444 | -0.035355663 | 0.006928812 | -0.042284474 | 0.999933887 | -0.272151987 | -0.287665178 | 0.015513191 | 0.902696306 |
| ENSG00000198089 | *SFI1* | 5.840986165 | 0.02530521 | 0.021498313 | 0.003806897 | 0.999933887 | 0.064244888 | 0.04481908 | 0.019425808 | 0.902696306 |
| ENSG00000275198 | *AL512791.2* | -0.553529806 | -0.060351829 | -0.081203279 | 0.020851451 | 0.999933887 | 0.404934353 | 0.320223782 | 0.084710571 | 0.902696306 |
| ENSG00000100197 | *CYP2D6* | 1.312573751 | -0.041780442 | 0.174497 | -0.216277442 | 0.999933887 | 0.053667578 | 0.00933796 | 0.044329617 | 0.902696306 |
| ENSG00000234741 | *GAS5* | 5.163587909 | 0.010854154 | -0.055794365 | 0.066648519 | 0.999933887 | 0.116811431 | 0.096539041 | 0.020272391 | 0.902696306 |
| ENSG00000184076 | *UQCR10* | 2.080399184 | 0.011425025 | -0.125109827 | 0.136534852 | 0.999933887 | -0.073711454 | -0.034306108 | -0.039405346 | 0.902696306 |
| ENSG00000288009 | *AL353135.2* | -0.157822517 | -0.283812184 | 0.096035003 | -0.379847187 | 0.999933887 | 0.347072344 | 0.399846 | -0.052773656 | 0.903105992 |
| ENSG00000287809 | *AC005632.6* | 1.975180995 | -0.026234846 | -0.021321143 | -0.004913702 | 0.999933887 | -0.01934831 | 0.028186574 | -0.047534884 | 0.903143807 |
| ENSG00000234118 | *RPL13AP6* | -0.90175447 | 0.01116546 | 0.139921674 | -0.128756214 | 0.999933887 | 0.325773347 | 0.154110284 | 0.171663063 | 0.903464397 |
| ENSG00000133111 | *RFXAP* | 0.155185639 | 0.159640251 | -0.033741794 | 0.193382046 | 0.999933887 | -0.155744387 | -0.103729877 | -0.05201451 | 0.903530887 |
| ENSG00000211770 | *TRBJ2-6* | -0.85701557 | 0.191067219 | 0.020717413 | 0.170349806 | 0.999933887 | 0.005023848 | 0.09615202 | -0.091128172 | 0.903530887 |
| ENSG00000244509 | *APOBEC3C* | 3.954894205 | -0.049665188 | -0.11158931 | 0.061924123 | 0.999933887 | -0.075714337 | -0.10356988 | 0.027855543 | 0.903555789 |
| ENSG00000232748 | *AC135050.1* | 0.453723754 | 0.044844858 | 0.247490291 | -0.202645433 | 0.999933887 | 0.171209935 | 0.245505546 | -0.074295611 | 0.903555789 |
| ENSG00000279913 | *AP001962.1* | 0.045718611 | -0.023438777 | 0.096058572 | -0.119497349 | 0.999933887 | 0.903368775 | 0.99803305 | -0.094664275 | 0.903555789 |
| ENSG00000153790 | *C7orf31* | 0.488841484 | -0.101696171 | -0.075418975 | -0.026277196 | 0.999933887 | 0.290877297 | 0.344137387 | -0.05326009 | 0.903671608 |
| ENSG00000120963 | *ZNF706* | 3.917526547 | 0.033246477 | -0.055148366 | 0.088394843 | 0.999933887 | 0.237026611 | 0.216876739 | 0.020149872 | 0.903671608 |
| ENSG00000140157 | *NIPA2* | 4.6757638 | 0.031249807 | -0.027063769 | 0.058313575 | 0.999933887 | -0.326273949 | -0.303935782 | -0.022338167 | 0.903671608 |
| ENSG00000174684 | *B4GAT1* | -0.84755259 | 0.11931717 | -0.086101921 | 0.205419091 | 0.999933887 | 0.040782298 | -0.034556812 | 0.075339111 | 0.90376476 |

| | | | | | | | | | | |
|---|---|---|---|---|---|---|---|---|---|---|
| ENSG00000170846 | *AC093323.1* | 2.41527224 | -0.019590371 | -0.051399686 | 0.031809315 | 0.999933887 | 0.08544493 | 0.051898509 | 0.033546421 | 0.90376476 |
| ENSG00000157800 | *SLC37A3* | 3.917694973 | -0.027021345 | -0.030971591 | 0.003950246 | 0.999933887 | -0.163829104 | -0.192915781 | 0.029086677 | 0.904044825 |
| ENSG00000174021 | *GNG5* | 4.501931665 | 0.104922388 | 0.025697572 | 0.079224816 | 0.999933887 | -0.103801016 | -0.128099328 | 0.024298311 | 0.904057846 |
| ENSG00000172795 | *DCP2* | 5.954933247 | -0.080394089 | -0.09647492 | 0.016080832 | 0.999933887 | -0.258951656 | -0.231275332 | -0.027676324 | 0.904057846 |
| ENSG00000183963 | *SMTN* | -0.0043137 | -0.149656797 | -0.591228558 | 0.441571761 | 0.999933887 | 0.322342374 | 0.259647825 | 0.062694548 | 0.904057846 |
| ENSG00000280649 | *AC245100.7* | 2.005715014 | 0.023424657 | -0.018273956 | 0.041698613 | 0.999933887 | 0.070869319 | 0.105300322 | -0.034431003 | 0.904403087 |
| ENSG00000241973 | *PI4KA* | 6.729763855 | 0.025045937 | -0.027592101 | 0.052638038 | 0.999933887 | -0.042732599 | -0.027016279 | -0.01571632 | 0.904448409 |
| ENSG00000174038 | *C9orf131* | -0.349708639 | -0.188202067 | -0.018700038 | -0.16950203 | 0.999933887 | 0.345827764 | 0.420497689 | -0.074669925 | 0.904476965 |
| ENSG00000258976 | *AC013451.2* | -0.036682238 | 0.152395643 | -0.294416848 | 0.446812492 | 0.999933887 | -0.9779298 | -0.91351464 | -0.06441516 | 0.90468999 |
| ENSG00000079432 | *CIC* | 7.992492294 | 0.038740465 | 0.056991967 | -0.018251502 | 0.999933887 | -0.00943669 | -0.028668825 | 0.019232135 | 0.90468999 |
| ENSG00000130810 | *PPAN* | 0.589172477 | -0.021205863 | 0.455185796 | -0.476391659 | 0.999291887 | 0.183333707 | 0.123560787 | 0.059772919 | 0.90468999 |
| ENSG00000141076 | *UTP4* | 2.381957649 | -0.000146507 | 0.007425711 | -0.007572218 | 0.999933887 | 0.253402848 | 0.291696097 | -0.03829325 | 0.90468999 |
| ENSG00000265681 | *RPL17* | 1.26924831 | 0.036845198 | 0.061284292 | -0.024439094 | 0.999933887 | -0.078749102 | -0.031240341 | -0.047508761 | 0.904864347 |
| ENSG00000155959 | *VBP1* | 3.356041756 | 0.062142389 | -0.014854921 | 0.076997309 | 0.999933887 | -0.049689616 | -0.076219461 | 0.026529845 | 0.904961346 |
| ENSG00000130348 | *QRSL1* | 1.069924224 | -0.031338622 | -0.126803586 | 0.095464964 | 0.999933887 | 0.064427366 | 0.104557204 | -0.040129838 | 0.90504771 |
| ENSG00000256427 | *AC010175.1* | -1.405373741 | 0.051004015 | 0.370341819 | -0.319337805 | 0.999933887 | 0.044384971 | 0.127105388 | -0.082720417 | 0.90504771 |
| ENSG00000143315 | *PIGM* | 1.057925678 | -0.100623001 | -0.044350424 | -0.056272576 | 0.999933887 | 0.094683768 | 0.137440154 | -0.042756386 | 0.905142499 |
| ENSG00000246422 | *DIAPH1-AS1* | -0.488796231 | 0.285434319 | 0.308525785 | -0.023091465 | 0.999933887 | -0.468793388 | -0.588909068 | 0.120115679 | 0.905275815 |
| ENSG00000118680 | *MYL12B* | 7.557896824 | -0.013012219 | -0.025466301 | 0.012454082 | 0.999933887 | -0.18456554 | -0.202363766 | 0.017798227 | 0.905275815 |
| ENSG00000145982 | *FARS2* | 0.702220153 | 0.086192944 | 0.254603426 | -0.168410482 | 0.999933887 | 0.174897557 | 0.122606237 | 0.05229132 | 0.905285146 |
| ENSG00000125166 | *GOT2* | 2.205939399 | -0.009630785 | -0.107679163 | 0.098048378 | 0.999933887 | 0.090689123 | 0.122788366 | -0.032099243 | 0.905285146 |
| ENSG00000147789 | *ZNF7* | 4.797475106 | 0.038602789 | 0.153807456 | -0.115204667 | 0.999933887 | 0.126266628 | 0.101972499 | 0.024294129 | 0.905399735 |
| ENSG00000243970 | *PPIEL* | 1.196783751 | 0.019002483 | -0.01822056 | 0.037223042 | 0.999933887 | 0.120630314 | 0.075402725 | 0.045227589 | 0.905429175 |
| ENSG00000233306 | *TRGV2* | 0.12893572 | -0.12920629 | 0.040250709 | -0.169456999 | 0.999933887 | 0.041511179 | 0.099267532 | -0.057756353 | 0.905707377 |
| ENSG00000164219 | *PGGT1B* | 3.421269422 | -0.089593596 | -0.033773422 | -0.055820174 | 0.999933887 | -0.563875006 | -0.532055353 | -0.031819654 | 0.905756917 |
| ENSG00000173917 | *HOXB2* | -0.158509842 | -0.013466171 | -0.15970121 | 0.146235039 | 0.999933887 | 0.100092427 | 0.162281046 | -0.062188619 | 0.905756917 |
| ENSG00000113716 | *HMGXB3* | 3.858377801 | -0.0128348 | 0.08143506 | -0.09426986 | 0.999933887 | -0.026931416 | -0.006690535 | -0.020240881 | 0.905757215 |
| ENSG00000148468 | *FAM171A1* | -0.996280488 | -0.045965214 | -0.090035946 | 0.044070732 | 0.999933887 | 0.261328606 | 0.178287682 | 0.083040924 | 0.90576535 |
| ENSG00000164024 | *METAP1* | 2.935858157 | -0.088193401 | -0.036269878 | -0.051923523 | 0.999933887 | -0.030409037 | -0.058537249 | 0.028128212 | 0.905768548 |
| ENSG00000104343 | *UBE2W* | 4.407683245 | -0.059799225 | 0.108730369 | -0.168529594 | 0.999933887 | -0.803282366 | -0.770618416 | -0.03266395 | 0.905791373 |
| ENSG00000182253 | *SYNM* | -0.436798616 | -0.052673649 | -0.085612505 | 0.032938856 | 0.999933887 | 0.106568599 | 0.043695792 | 0.062872807 | 0.905908077 |
| ENSG00000198055 | *GRK6* | 8.020670311 | 0.004298009 | 0.036373561 | -0.032075552 | 0.999933887 | -0.434517383 | -0.453509638 | 0.018992255 | 0.905908077 |
| ENSG00000229117 | *RPL41* | 6.923004689 | -0.030310232 | -0.063575086 | 0.033264854 | 0.999933887 | 0.062539485 | 0.083414336 | -0.020874851 | 0.905942323 |
| ENSG00000107957 | *SH3PXD2A* | 2.319500827 | 0.017650407 | -0.06683968 | 0.084490087 | 0.999933887 | 0.081236806 | 0.111178325 | -0.029941519 | 0.905942323 |
| ENSG00000106299 | *WASL* | 1.820179266 | 0.013230616 | -0.013261934 | 0.02649255 | 0.999933887 | -0.212373921 | -0.249516717 | 0.037142797 | 0.90635413 |
| ENSG00000132109 | *TRIM21* | 5.721124821 | -0.056412899 | -0.087606199 | 0.031193301 | 0.999933887 | -0.257589308 | -0.276727087 | 0.019137779 | 0.906454 |
| ENSG00000131174 | *COX7B* | 4.430702 | -0.021705308 | -0.012267892 | -0.009437416 | 0.999933887 | 0.10450371 | 0.123656669 | -0.019152959 | 0.906462732 |
| ENSG00000189223 | *PAX8-AS1* | 2.106657202 | -0.049670859 | 0.005404532 | -0.055075391 | 0.999933887 | 0.250676368 | 0.225156395 | 0.025519973 | 0.906477193 |
| ENSG00000117318 | *ID3* | 1.158673107 | -0.155387239 | 0.028702979 | -0.184090217 | 0.999933887 | 0.064516522 | 0.10070577 | -0.036189248 | 0.906613362 |
| ENSG00000201217 | *Y_RNA* | -0.891240762 | 0.038052723 | 0.192176767 | -0.154124044 | 0.999933887 | -0.109065836 | -0.202799689 | 0.093733852 | 0.906753343 |
| ENSG00000166086 | *JAM3* | -0.400077815 | -0.203885702 | 0.085177211 | -0.289062913 | 0.999933887 | -0.014413865 | -0.078459474 | 0.064045609 | 0.906843109 |
| ENSG00000160392 | *C19orf47* | 2.310549581 | 0.106954685 | -0.206638957 | 0.313593642 | 0.981460244 | -0.12144154 | -0.157198775 | 0.035757235 | 0.90713745 |
| ENSG00000174903 | *RAB1B* | 5.946715436 | 0.045237512 | 0.038460174 | 0.006777338 | 0.999933887 | -0.108976388 | -0.13039315 | 0.021416761 | 0.90713745 |
| ENSG00000258101 | *AC010173.1* | -0.245814643 | 0.081634069 | 0.234868815 | -0.153234746 | 0.999933887 | -0.014143634 | 0.093737243 | -0.107880877 | 0.90713745 |
| ENSG00000105193 | *RPS16* | 8.374629083 | 0.027818555 | -0.008861206 | 0.036679762 | 0.999933887 | 0.112584804 | 0.130087529 | -0.017502726 | 0.90713745 |
| ENSG00000143033 | *MTF2* | 3.503255805 | -0.03909617 | -0.101146747 | 0.062050577 | 0.999933887 | -0.323519064 | -0.296785209 | -0.026733855 | 0.90713745 |
| ENSG00000228775 | *WEE2-AS1* | 1.020970955 | -0.076904662 | -0.08938652 | 0.012481858 | 0.999933887 | -0.473008481 | -0.523379255 | 0.050370774 | 0.90713745 |
| ENSG00000166169 | *POLL* | 2.108581097 | -0.001946974 | 0.069801607 | -0.071748582 | 0.999933887 | 0.038298841 | 0.005577407 | 0.032721434 | 0.90713745 |
| ENSG00000184163 | *C1QTNF12* | 0.788489645 | 0.057386777 | -0.138875565 | 0.196262342 | 0.999933887 | -0.041188778 | -0.095620679 | 0.054431901 | 0.907252377 |
| ENSG00000113269 | *RNF130* | 8.084455413 | -0.003728623 | -0.017336827 | 0.013608204 | 0.999933887 | -0.343665385 | -0.324814355 | -0.01885103 | 0.907252377 |
| ENSG00000276141 | *WHAMMP3* | -0.419707475 | 0.011218364 | -0.410417416 | 0.42163578 | 0.999933887 | 0.110686086 | 0.182036566 | -0.07135048 | 0.907271916 |
| ENSG00000213694 | *S1PR3* | 0.877511558 | 0.006725841 | 0.051522853 | -0.044797012 | 0.999933887 | -0.828233133 | -0.769747159 | -0.058485974 | 0.907460178 |
| ENSG00000057019 | *DCBLD2* | 1.320577617 | -0.205352437 | -0.248117681 | 0.042765244 | 0.999933887 | -0.769584546 | -0.838073264 | 0.068488718 | 0.907741257 |
| ENSG00000105618 | *PRPF31* | 2.898465466 | 0.080755271 | 0.002513432 | 0.078241839 | 0.999933887 | 0.103147022 | 0.128007946 | -0.024860924 | 0.907741257 |
| ENSG00000279283 | *AC131009.4* | 2.401343156 | -0.04310715 | -0.074011661 | 0.030904511 | 0.999933887 | -0.893364785 | -0.93215463 | 0.038789845 | 0.907741257 |
| ENSG00000116661 | *FBXO2* | -0.928866773 | -0.038167286 | 0.077112895 | -0.115280182 | 0.999933887 | 0.348953731 | 0.447950851 | -0.09899712 | 0.907741257 |
| ENSG00000170881 | *RNF139* | 5.337843528 | 0.000293852 | -0.054493864 | 0.054787716 | 0.999933887 | -0.316368071 | -0.299371209 | -0.016996862 | 0.907741257 |
| ENSG00000150048 | *CLEC1A* | 2.780822934 | 0.024967579 | 0.092954842 | -0.067987263 | 0.999933887 | -0.338960167 | -0.373008585 | 0.034048418 | 0.907884191 |
| ENSG00000279549 | *AP000437.1* | 0.923856927 | -0.122225025 | 0.074740803 | -0.196965828 | 0.999933887 | 0.128242513 | 0.06563295 | 0.062609563 | 0.907944225 |
| ENSG00000173338 | *KCNK7* | 3.261311117 | 0.011792462 | 0.045950867 | -0.034158405 | 0.999933887 | 0.622839719 | 0.655508805 | -0.032669086 | 0.907944225 |
| ENSG00000124279 | *FASTKD3* | 0.330554308 | 0.032467021 | -0.027895684 | 0.060362706 | 0.999933887 | -0.04028765 | 0.021083731 | -0.06137138 | 0.908066694 |
| ENSG00000164506 | *STXBP5* | 6.048986004 | -0.065112337 | -0.03812803 | -0.026984307 | 0.999933887 | -0.266658161 | -0.292889424 | 0.026231263 | 0.908081015 |
| ENSG00000145882 | *PCYOX1L* | 2.24754883 | -0.004755458 | -0.050749899 | 0.045994441 | 0.999933887 | 0.020284784 | -0.012037275 | 0.032322059 | 0.908081015 |
| ENSG00000130349 | *MTRES1* | 0.073944318 | 0.071115474 | 0.105026238 | -0.033910764 | 0.999933887 | 0.022319053 | -0.027845482 | 0.050164534 | 0.908194397 |
| ENSG00000103351 | *CLUAP1* | 0.813671595 | -0.134619797 | 0.162996697 | -0.297616494 | 0.999933887 | -0.097240357 | -0.141030864 | 0.043790507 | 0.908304993 |
| ENSG00000133895 | *MEN1* | 2.999182966 | 0.087571987 | 0.18790948 | -0.100337493 | 0.999933887 | 0.170704758 | 0.197863585 | -0.027158827 | 0.908304993 |
| ENSG00000258355 | *AC079174.1* | 0.566312185 | 0.305465287 | 0.210865592 | 0.094599695 | 0.999933887 | 1.310826693 | 1.398242827 | -0.087416134 | 0.908318582 |
| ENSG00000255163 | *HSPE1P18* | 0.623542862 | -0.391759175 | -0.457168577 | 0.065409403 | 0.999933887 | -0.617103962 | -0.671982884 | 0.054878922 | 0.908318582 |
| ENSG00000268240 | *AC123912.2* | 3.527118081 | 0.231558103 | 0.194436944 | 0.037121159 | 0.999933887 | 0.517315775 | 0.474285579 | 0.043030196 | 0.908318582 |
| ENSG00000279841 | *AC092135.3* | 0.7281298 | 0.187472933 | 0.023108236 | 0.164364697 | 0.999933887 | 0.218643021 | 0.279233635 | -0.060590614 | 0.908318582 |
| ENSG00000211777 | *TRAV3* | -0.228191684 | -0.24954253 | -0.09697493 | -0.1525676 | 0.999933887 | 0.138338363 | 0.210573085 | -0.072234722 | 0.908318582 |
| ENSG00000280173 | *AC104447.1* | 0.020297176 | -0.134586251 | 0.012002219 | -0.14658847 | 0.999933887 | -0.725662636 | -0.80880942 | 0.083146784 | 0.908318582 |
| ENSG00000269427 | *AC024075.3* | -0.275266564 | 0.181291046 | -0.108685836 | 0.289976882 | 0.999933887 | -0.201668359 | -0.272052332 | 0.070383973 | 0.908318582 |
| ENSG00000164494 | *PDSS2* | 0.583091616 | -0.127806521 | 0.041693316 | -0.169499837 | 0.999933887 | 0.042247375 | -0.014259854 | 0.056507229 | 0.908318582 |
| ENSG00000115392 | *FANCL* | -0.255616403 | -0.183178902 | -0.34681868 | 0.163639779 | 0.999933887 | 0.048154273 | 0.133467532 | -0.085313259 | 0.908318582 |
| ENSG00000235027 | *AC068580.3* | 5.757904216 | 0.079851791 | 0.075911679 | 0.003940112 | 0.999933887 | 0.098146245 | 0.129015926 | -0.030869682 | 0.908318582 |
| ENSG00000268205 | *AC005261.1* | 2.134429927 | -0.06288663 | -0.012091518 | -0.050795112 | 0.999933887 | -0.074434966 | -0.113071195 | 0.038636229 | 0.908318582 |
| ENSG00000219507 | *FTH1P8* | 3.350170183 | 0.038929337 | -0.012501465 | 0.051430801 | 0.999933887 | 0.618289562 | 0.591131159 | 0.027158403 | 0.908318582 |
| ENSG00000135211 | *TMEM60* | 3.364478702 | -0.03724025 | -0.118206058 | 0.080965808 | 0.999933887 | -0.171511772 | -0.146856573 | -0.0246552 | 0.908318582 |
| ENSG00000113088 | *GZMK* | 1.879708584 | 0.027428533 | -0.053624366 | 0.081052899 | 0.999933887 | 0.158579967 | 0.197327583 | -0.038747617 | 0.908318582 |
| ENSG00000237436 | *CAMTA1-DT* | 0.19262456 | 0.072730005 | 0.288237713 | -0.215507708 | 0.999933887 | 1.100923829 | 1.032458572 | 0.068465258 | 0.908318582 |
| ENSG00000144645 | *OSBPL10* | -0.953733667 | -0.042307687 | -0.244843828 | 0.20253614 | 0.999933887 | -0.466889555 | -0.573862438 | 0.106972883 | 0.908318582 |
| ENSG00000139890 | *REM2* | 4.456454197 | -0.005695512 | -0.001962517 | -0.003732996 | 0.999933887 | 0.060393328 | 0.034148845 | 0.026244482 | 0.908318582 |
| ENSG00000249274 | *PDLIM1P4* | 1.302811131 | 0.004837855 | -0.206169539 | 0.211007394 | 0.999933887 | -0.513699845 | -0.552200938 | 0.038501093 | 0.908318582 |
| ENSG00000229023 | *RAB1AP1* | 0.375390459 | -0.002349178 | 0.229463516 | -0.231812694 | 0.999933887 | -0.210126489 | -0.297179241 | 0.087052752 | 0.908318582 |
| ENSG00000270069 | *MIR222HG* | 2.956612544 | 0.16282009 | 0.241945086 | -0.079124995 | 0.999933887 | 1.192578824 | 1.236263264 | -0.04368444 | 0.908505489 |
| ENSG00000287934 | *AP001885.3* | -0.640600337 | -0.106537628 | -0.063445059 | -0.043092569 | 0.999933887 | -0.224776536 | -0.313491971 | 0.088715435 | 0.908505489 |
| ENSG00000115216 | *NRBP1* | 7.126432726 | -0.002116371 | -0.029779416 | 0.027663045 | 0.999933887 | -0.355692582 | -0.339445927 | -0.016246656 | 0.908619469 |
| ENSG00000235908 | *RHOA-IT1* | 0.18931702 | -0.112491781 | -0.188847576 | 0.076355795 | 0.999933887 | 0.150839523 | 0.09114206 | 0.059697464 | 0.908896281 |
| ENSG00000179388 | *EGR3* | 8.729082053 | 0.224698447 | 0.159553013 | 0.065145434 | 0.999933887 | 1.045901066 | 1.015991944 | 0.029909123 | 0.909126471 |
| ENSG00000196123 | *KIAA0895L* | 1.28699786 | 0.042055572 | 0.026390891 | 0.015664681 | 0.999933887 | 0.139993267 | 0.092722266 | 0.047271001 | 0.909126471 |
| ENSG00000010626 | *LRRC23* | -0.011970265 | -0.041189165 | -0.042443543 | 0.001254378 | 0.999933887 | 0.060934822 | 0.000430833 | 0.060503989 | 0.909126471 |
| ENSG00000104635 | *SLC39A14* | 0.531706863 | -0.00070522 | 0.24878579 | -0.24949101 | 0.999933887 | 0.108666757 | 0.162060774 | -0.053394017 | 0.909126471 |
| ENSG00000204248 | *COL11A2* | 0.502908913 | 0.011090413 | 0.086106359 | -0.075015945 | 0.999933887 | -0.06667228 | -0.127970113 | 0.061297832 | 0.909212314 |
| ENSG00000198707 | *CEP290* | 1.566179056 | -0.160632889 | -0.096958348 | -0.063674541 | 0.999933887 | -0.267072325 | -0.200333684 | -0.066738641 | 0.909473345 |
| ENSG00000198455 | *ZXDB* | 2.028732646 | -0.016630344 | -0.101863211 | 0.085232867 | 0.999933887 | 0.163394978 | 0.197418997 | -0.034024019 | 0.909663724 |
| ENSG00000104998 | *IL27RA* | 3.97183599 | -0.000819981 | -0.021485252 | 0.02066527 | 0.999933887 | 0.146547573 | 0.12609057 | 0.020457002 | 0.909663724 |
| ENSG00000130768 | *SMPDL3B* | 0.382438222 | -0.085186192 | 0.026442985 | -0.111629177 | 0.999933887 | 0.319435398 | 0.377722412 | -0.058287014 | 0.910229989 |
| ENSG00000158109 | *TPRG1L* | 6.51472767 | -0.007870665 | -0.029584358 | 0.021713693 | 0.999933887 | -0.885678646 | -0.910401385 | 0.024722739 | 0.910229989 |
| ENSG00000100991 | *TRPC4AP* | 7.085196269 | -0.031861768 | 0.006670259 | -0.038532027 | 0.999933887 | -0.26347782 | -0.249338899 | -0.014138921 | 0.910559015 |
| ENSG00000198270 | *TMEM116* | 1.633171203 | -0.099489387 | 0.077669204 | -0.177158592 | 0.999933887 | 0.149053581 | 0.114180621 | 0.03487296 | 0.91057913 |
| ENSG00000155034 | *FBXL18* | 3.299366252 | 0.019805935 | -0.01677982 | 0.036585755 | 0.999933887 | -0.076921298 | -0.105865446 | 0.028944148 | 0.91057913 |
| ENSG00000132122 | *SPATA6* | 0.622051027 | 0.090796082 | 0.151926887 | -0.061130805 | 0.999933887 | -0.31874412 | -0.253241096 | -0.065503025 | 0.910642699 |
| ENSG00000181350 | *LRRC75A* | 0.642129775 | 0.010976005 | -0.015063114 | 0.026039119 | 0.999933887 | -0.177312963 | -0.111730356 | -0.065582607 | 0.910642699 |
| ENSG00000283787 | *PRR33* | 7.4150653 | 0.102084311 | 0.104133456 | -0.002049145 | 0.999933887 | -0.27888574 | -0.312900715 | 0.034014975 | 0.91065143 |

| | | | | | | | | | | |
|---|---|---|---|---|---|---|---|---|---|---|
| ENSG00000119737 | *GPR75* | 0.619348828 | -0.078213587 | 0.347258157 | -0.425471744 | 0.399697802 | -0.157966122 | -0.226366563 | 0.068400441 | 0.91065143 |
| ENSG00000089597 | *GANAB* | 6.214641896 | 0.003953915 | -0.073112978 | 0.077066893 | 0.999933887 | 0.105521487 | 0.091577667 | 0.01394382 | 0.91065143 |
| ENSG00000144028 | *SNRNP200* | 7.17294139 | -0.003019912 | -0.001194528 | -0.001825384 | 0.999933887 | -0.019351039 | -0.03323538 | 0.013884341 | 0.91065143 |
| ENSG00000147654 | *EBAG9* | 1.381204279 | -0.005658788 | -0.350734984 | 0.345076195 | 0.981460244 | -0.063283358 | -0.022089376 | -0.041193982 | 0.91065143 |
| ENSG00000158792 | *SPATA2L* | 5.052172191 | 0.100854255 | 0.035806959 | 0.065047297 | 0.999933887 | -0.126942829 | -0.156178369 | 0.02923554 | 0.910942505 |
| ENSG00000057935 | *MTA3* | 1.232608544 | -0.080983353 | 0.146523521 | -0.227506873 | 0.999933887 | -0.00792223 | -0.056420177 | 0.048497946 | 0.911121631 |
| ENSG00000287218 | *AL359962.3* | -0.689072655 | -0.017545419 | 0.149981477 | -0.167526896 | 0.999933887 | 0.023205772 | -0.069864321 | 0.093070093 | 0.911121631 |
| ENSG00000133401 | *PDZD2* | -0.043068436 | 0.003748785 | -0.218020872 | 0.221769657 | 0.999933887 | -0.176927658 | -0.250468664 | 0.073541006 | 0.911121631 |
| ENSG00000140993 | *TIGD7* | 0.810709646 | 0.002825183 | -0.10537237 | 0.108197553 | 0.999933887 | -0.196158192 | -0.240120851 | 0.04396266 | 0.911121631 |
| ENSG00000105426 | *PTPRS* | -0.07146387 | 0.09703471 | 0.107014457 | -0.009979747 | 0.999933887 | -0.248242926 | -0.319521662 | 0.071278736 | 0.911153039 |
| ENSG00000255150 | *EID3* | 3.206203552 | 0.039574207 | 0.177541158 | -0.137966952 | 0.999933887 | 0.388696864 | 0.420271863 | -0.031574999 | 0.911153039 |
| ENSG00000182704 | *TSKU* | -1.521196941 | -0.131160634 | 0.216321427 | -0.347482062 | 0.999933887 | -0.205792435 | -0.147818077 | -0.057974357 | 0.911153039 |
| ENSG00000175595 | *ERCC4* | 0.288646187 | 0.032964239 | -0.12827478 | 0.161239019 | 0.999933887 | 0.039497681 | 0.089460157 | -0.049962476 | 0.911635923 |
| ENSG00000197619 | *ZNF615* | 1.63953256 | -0.051028768 | -0.092085989 | 0.041057221 | 0.999933887 | -0.279279395 | -0.229433077 | -0.049846318 | 0.911687985 |
| ENSG00000147687 | *TATDN1* | 0.892743396 | -0.050811916 | 0.137471549 | -0.188283466 | 0.999933887 | -0.224327402 | -0.173666441 | -0.05066096 | 0.912014326 |
| ENSG00000276900 | *AC023157.2* | 8.32187284 | 0.188121373 | 0.171988071 | 0.016133301 | 0.999933887 | 0.573772722 | 0.600952807 | -0.027180085 | 0.912409693 |
| ENSG00000144909 | *OSBPL11* | 6.524233906 | -0.089173442 | -0.058821275 | -0.030352167 | 0.999933887 | -0.770188542 | -0.791850549 | 0.021662008 | 0.912409693 |
| ENSG00000204525 | *HLA-C* | 12.676787016 | 0.065977995 | 0.024255448 | 0.041722547 | 0.999933887 | 0.161284501 | 0.141912765 | 0.019371736 | 0.912409693 |
| ENSG00000148459 | *PDSS1* | -0.172269929 | -0.138526879 | 0.253153012 | -0.391679891 | 0.999933887 | 0.430716384 | 0.36170868 | 0.069007704 | 0.912409693 |
| ENSG00000211679 | *IGLC3* | -0.222999902 | -0.159948072 | -0.13201696 | -0.027931112 | 0.999933887 | 0.440433891 | 0.364552542 | 0.07588135 | 0.912409693 |
| ENSG00000067066 | *SP100* | 7.310305423 | -0.02862648 | -0.145259701 | 0.11663322 | 0.999933887 | -0.529808806 | -0.55140366 | 0.021594854 | 0.912409693 |
| ENSG00000073605 | *GSDMB* | 2.946322075 | 0.016397614 | 0.017004549 | -0.000606935 | 0.999933887 | 0.031881399 | 0.002488637 | 0.029392762 | 0.912409693 |
| ENSG00000131747 | *TOP2A* | -0.122616061 | -0.037408356 | -0.146355705 | 0.108947349 | 0.999933887 | 0.04907361 | 0.13129036 | -0.082216751 | 0.912409693 |
| ENSG00000109118 | *PHF12* | 7.128168555 | 0.010001046 | -0.013801648 | 0.023802694 | 0.999933887 | -0.314578609 | -0.331978043 | 0.017399434 | 0.912409693 |
| ENSG00000225614 | *ZNF469* | 0.039363904 | 0.357258571 | 0.261585888 | 0.095672683 | 0.999933887 | 0.124336395 | 0.068373929 | 0.055962467 | 0.912576716 |
| ENSG00000171861 | *MRM3* | 1.028509676 | 0.139563016 | 0.11314287 | 0.026420146 | 0.999933887 | 0.283145229 | 0.334143544 | -0.050998315 | 0.912576716 |
| ENSG00000279345 | *Z98885.3* | -0.061628102 | 0.170111786 | 0.042131186 | 0.1279806 | 0.999933887 | -0.278663758 | -0.34340758 | 0.064743822 | 0.912576716 |
| ENSG00000133134 | *BEX2* | 0.965898104 | 0.112236504 | 0.008149048 | 0.104087456 | 0.999933887 | 0.278410364 | 0.235136627 | 0.043273737 | 0.912576716 |
| ENSG00000111344 | *RASAL1* | -1.027517043 | 0.288343535 | 0.497552474 | -0.209208939 | 0.999933887 | -0.882239287 | -0.821394829 | -0.060844458 | 0.912576716 |
| ENSG00000133250 | *ZNF414* | 3.250433115 | 0.053764423 | 0.015838378 | 0.037926044 | 0.999933887 | -0.090636286 | -0.059354665 | -0.031281621 | 0.912576716 |
| ENSG00000109944 | *JHY* | -0.374027482 | 0.148576664 | -0.226176214 | 0.374752878 | 0.999933887 | -0.167813942 | -0.097476509 | -0.070337433 | 0.912576716 |
| ENSG00000203395 | *AC015969.1* | 1.353522657 | 0.594142003 | 0.869754158 | -0.275612156 | 0.999933887 | 1.768779934 | 1.661596892 | 0.107183042 | 0.912773846 |
| ENSG00000283208 | *AC001226.2* | -0.48434168 | 0.274012462 | 0.314435104 | -0.040422642 | 0.999933887 | -0.262353792 | -0.165995436 | -0.096358356 | 0.912773846 |
| ENSG00000152894 | *PTPRK* | -0.99847268 | -0.279438045 | -0.44152103 | 0.162082985 | 0.999933887 | 0.027717363 | 0.106035417 | -0.078318053 | 0.912773846 |
| ENSG00000146410 | *MTFR2* | 0.241957119 | -0.113616445 | -0.197066494 | 0.083450049 | 0.999933887 | 0.160965426 | 0.104379131 | 0.056586296 | 0.912773846 |
| ENSG00000126861 | *OMG* | 1.444669052 | -0.131992164 | -0.149343051 | 0.017350887 | 0.999933887 | -0.501854047 | -0.570210158 | 0.068356111 | 0.912773846 |
| ENSG00000268257 | *AIRN* | 2.469355494 | 0.142758668 | 0.312655209 | -0.169896541 | 0.999933887 | 0.368696424 | 0.299157077 | 0.069539347 | 0.912773846 |
| ENSG00000183655 | *KLHL25* | 1.054013834 | 0.072216161 | 0.029196725 | 0.043019435 | 0.999933887 | 0.214883724 | 0.255091704 | -0.040207981 | 0.912773846 |
| ENSG00000196636 | *SDHAF3* | 1.165278433 | -0.061817847 | 0.113113542 | -0.174931389 | 0.999933887 | 0.030162179 | 0.086262602 | -0.056100423 | 0.912773846 |
| ENSG00000137824 | *RMDN3* | 2.344461348 | 0.039235028 | -0.031174705 | 0.070409733 | 0.999933887 | 0.049830143 | 0.075561226 | -0.025731082 | 0.912773846 |
| ENSG00000261485 | *PAN3-AS1* | 1.158999157 | 0.063161781 | 0.021243478 | 0.041918303 | 0.999933887 | -0.354649162 | -0.317940897 | -0.036708265 | 0.912773846 |
| ENSG00000183337 | *BCOR* | 4.93778934 | -0.027043162 | -0.016541416 | -0.010501746 | 0.999933887 | -0.01324932 | -0.032280874 | 0.019031554 | 0.912773846 |
| ENSG00000262172 | *AC116025.2* | 0.098129004 | 0.071493159 | 0.464940815 | -0.393447656 | 0.999933887 | 1.77829244 | 1.83804932 | -0.059756879 | 0.912773846 |
| ENSG00000063660 | *GPC1* | -0.732394455 | -0.06846279 | 0.129505107 | -0.197967898 | 0.999933887 | 0.134061352 | 0.199757255 | -0.065695904 | 0.912773846 |
| ENSG00000163166 | *IWS1* | 4.358795985 | -0.020833811 | -0.119062012 | 0.098228201 | 0.999933887 | -0.218413415 | -0.196502487 | -0.021910928 | 0.912773846 |
| ENSG00000163596 | *ICA1L* | -0.383475842 | -0.058287036 | -0.077223514 | 0.018936478 | 0.999933887 | -0.140841028 | -0.226583458 | 0.085742431 | 0.912773846 |
| ENSG00000267551 | *AC005264.1* | 0.177984482 | -0.102329782 | -0.03727038 | -0.065059402 | 0.999933887 | 0.511736339 | 0.566350976 | -0.054614637 | 0.912854812 |
| ENSG00000233493 | *TMEM238* | 0.757151988 | 0.255606838 | -0.048825462 | 0.3044323 | 0.999933887 | 0.07903745 | 0.020429968 | 0.058607482 | 0.912872255 |
| ENSG00000096654 | *ZNF184* | 1.664836236 | 0.103894953 | -0.255576431 | 0.359471384 | 0.944094537 | -0.13217175 | -0.173491497 | 0.041319747 | 0.912872255 |
| ENSG00000011114 | *BTBD7* | 4.55347445 | -0.046672999 | -0.074992514 | 0.028319516 | 0.999933887 | -0.334580301 | -0.318445041 | -0.01613526 | 0.912872255 |
| ENSG00000131051 | *RBM39* | 8.835543868 | -0.041937733 | -0.095389489 | 0.053451756 | 0.999933887 | -0.058676531 | -0.075106571 | 0.01643004 | 0.912872255 |
| ENSG00000119965 | *C10orf88* | 0.285356511 | -0.129363467 | 0.20090734 | -0.330270807 | 0.999933887 | 0.049604498 | 0.098251294 | -0.048646796 | 0.912872255 |
| ENSG00000165238 | *WNK2* | -0.20506902 | 0.092098168 | 0.309878042 | -0.217779874 | 0.999933887 | 0.248775949 | 0.303465235 | -0.054689286 | 0.912872255 |
| ENSG00000277476 | *AC005332.5* | 1.85976818 | 0.044783812 | 0.079123145 | -0.034339333 | 0.999933887 | -0.308081613 | -0.271129339 | -0.036952274 | 0.912872255 |
| ENSG00000151474 | *FRMD4A* | 0.440156597 | -0.058869556 | 0.122819886 | -0.181689442 | 0.999933887 | -0.634219963 | -0.58632841 | -0.047891553 | 0.912872255 |
| ENSG00000174173 | *TRMT10C* | 1.855656269 | -0.031246807 | -0.120735153 | 0.089488347 | 0.999933887 | 0.012515829 | 0.043632977 | -0.031117149 | 0.912872255 |
| ENSG00000123154 | *WDR83* | 0.650990853 | -0.043895388 | -0.013892392 | -0.030002996 | 0.999933887 | 0.011306851 | 0.054203023 | -0.042896172 | 0.912872255 |
| ENSG00000178502 | *KLHL11* | 1.422283969 | 0.023527588 | 0.073960566 | -0.050432978 | 0.999933887 | -0.116923299 | -0.154758401 | 0.037835102 | 0.912872255 |
| ENSG00000158578 | *ALAS2* | -0.342021719 | -0.032890235 | -0.076502433 | 0.043612198 | 0.999933887 | 0.004665423 | 0.069136731 | -0.064471307 | 0.912872255 |
| ENSG00000247627 | *MTND4P12* | -0.344044188 | -0.025344894 | 0.160308941 | -0.185653836 | 0.999933887 | -0.126174974 | -0.075016684 | -0.05115829 | 0.912872255 |
| ENSG00000198324 | *PHETA1* | -0.115099515 | 0.120980319 | 0.068340188 | 0.052640131 | 0.999933887 | -0.133105754 | -0.18898463 | 0.055878876 | 0.912994675 |
| ENSG00000007312 | *CD79B* | -0.014986138 | -0.02581682 | 0.227030526 | -0.252847346 | 0.999933887 | -0.218162158 | -0.159855728 | -0.05830643 | 0.912994675 |
| ENSG00000058453 | *CROCC* | 3.65212397 | 0.090596914 | 0.157479008 | -0.066882095 | 0.999933887 | 0.013702069 | 0.046102446 | -0.032400378 | 0.913041877 |
| ENSG00000284691 | *AC073111.3* | 1.968881587 | 0.099575042 | 0.200036481 | -0.100461438 | 0.999933887 | 0.277622663 | 0.312878617 | -0.035255954 | 0.913041877 |
| ENSG00000170089 | *AC106795.1* | -0.13626501 | -0.034756789 | 0.087949812 | -0.122706601 | 0.999933887 | 0.070525371 | 0.124897987 | -0.054372616 | 0.913041877 |
| ENSG00000132581 | *SDF2* | 5.245659882 | 0.060287068 | -0.016399949 | 0.076687017 | 0.999933887 | 0.090964302 | 0.10916197 | -0.018197668 | 0.91311384 |
| ENSG00000274695 | *AC108704.1* | 0.57384916 | -0.150463124 | 0.013825273 | -0.164288397 | 0.999933887 | -0.449839149 | -0.401985635 | -0.047853514 | 0.91313934 |
| ENSG00000143436 | *MRPL9* | 2.025390613 | 0.019052665 | -0.03303662 | 0.052089285 | 0.999933887 | 0.174242678 | 0.145085017 | 0.029157661 | 0.91313934 |
| ENSG00000254826 | *AP001922.3* | 1.461363279 | 0.114057817 | 0.098683432 | 0.015374385 | 0.999933887 | -0.133608415 | -0.190914175 | 0.057305761 | 0.913190903 |
| ENSG00000178226 | *PRSS36* | -0.493267203 | -0.113651261 | -0.100351125 | -0.013300135 | 0.999933887 | -0.816079559 | -0.901974218 | 0.085894659 | 0.913190903 |
| ENSG00000274818 | *AC004825.2* | -0.313296361 | -0.055155324 | -0.005399281 | -0.049756043 | 0.999933887 | 0.072588444 | 0.160501504 | -0.08791306 | 0.913190903 |
| ENSG00000238197 | *PAXBP1-AS1* | 2.330195473 | -0.078915991 | -0.107129122 | 0.028213132 | 0.999933887 | -0.369460925 | -0.326297962 | -0.043162964 | 0.913290083 |
| ENSG00000135925 | *WNT10A* | -0.818089554 | -0.140645691 | 0.31519591 | -0.455841602 | 0.999933887 | 0.428796853 | 0.501362415 | -0.072565562 | 0.913290083 |
| ENSG00000269235 | *ZNF350-AS1* | 1.457586559 | -0.04751141 | 0.095388796 | -0.142900206 | 0.999933887 | 0.091739225 | 0.13789374 | -0.046154516 | 0.913290083 |
| ENSG00000272106 | *AL691432.2* | 2.444087984 | 0.033671784 | 0.036387705 | -0.002715921 | 0.999933887 | 0.013972256 | -0.018637232 | 0.032609488 | 0.913290083 |
| ENSG00000225439 | *BOLA3-AS1* | -0.607871464 | -0.004853396 | -0.476009096 | 0.4711557 | 0.999933887 | -0.198813632 | -0.133371429 | -0.065442203 | 0.913290083 |
| ENSG00000134717 | *BTF3L4* | 3.667278688 | -0.034104479 | 0.01580325 | -0.049907729 | 0.999933887 | -0.228851739 | -0.208472634 | -0.020379105 | 0.913496212 |
| ENSG00000267023 | *LRRC37A16P* | 1.977012188 | -0.068173565 | -0.14537795 | 0.077204386 | 0.999933887 | -0.288623971 | -0.3139386 | 0.025314629 | 0.913899444 |
| ENSG00000283215 | *LINC02537* | -0.851150141 | -0.100857434 | -0.115394509 | 0.014537075 | 0.999933887 | 0.061902502 | 0.114984801 | -0.053082299 | 0.913985627 |
| ENSG00000164649 | *CDCA7L* | 1.220125696 | -0.243719532 | -0.12481632 | -0.118903212 | 0.999933887 | -0.371074102 | -0.41610376 | 0.045029657 | 0.914149209 |
| ENSG00000170579 | *DLGAP1* | -0.075009938 | 0.259281052 | 0.011024106 | 0.248256946 | 0.999933887 | -1.255284612 | -1.308640578 | 0.053355965 | 0.914149209 |
| ENSG00000221869 | *CEBPD* | 7.33601521 | 0.051162942 | 0.049096967 | 0.002065975 | 0.999933887 | -0.24907567 | -0.221700105 | -0.027375565 | 0.914149209 |
| ENSG00000279996 | *AC004491.1* | 0.301649319 | 0.299347705 | 0.049292727 | 0.250054978 | 0.999933887 | 1.171560027 | 1.237617483 | -0.066057456 | 0.914618927 |
| ENSG00000255507 | *UVRAG-DT* | 1.039371802 | 0.026684883 | 0.106996054 | -0.080311171 | 0.999933887 | 1.377913545 | 1.32591379 | 0.051999755 | 0.914690208 |
| ENSG00000165119 | *HNRNPK* | 8.773102444 | -0.047868639 | -0.066311546 | 0.018442907 | 0.999933887 | -0.27786931 | -0.289732081 | 0.011862771 | 0.914786364 |
| ENSG00000269091 | *AC010624.3* | -1.199838126 | 0.068400064 | -0.056039418 | 0.124439482 | 0.999933887 | -0.043273159 | -0.148277566 | 0.105004407 | 0.914786364 |
| ENSG00000164828 | *SUN1* | 4.240113441 | -0.065062359 | -0.036342776 | -0.028719583 | 0.999933887 | 0.078875862 | 0.096493875 | -0.017618013 | 0.914834503 |
| ENSG00000205238 | *SPDYE2* | -0.442722984 | -0.159911738 | 0.235727557 | -0.395639295 | 0.999933887 | -0.037742911 | -0.12065382 | 0.08291091 | 0.914834503 |
| ENSG00000182718 | *ANXA2* | 4.827052669 | -0.030807125 | -0.138106994 | 0.107299869 | 0.999933887 | -0.01296962 | 0.00609496 | -0.01906458 | 0.914834503 |
| ENSG00000156976 | *EIF4A2* | 7.546125443 | -0.023321512 | -0.014316393 | -0.009005119 | 0.999933887 | 0.270671794 | 0.286337903 | -0.015666109 | 0.914834503 |
| ENSG00000198793 | *MTOR* | 4.819524671 | 0.020180009 | 0.009436362 | 0.010743647 | 0.999933887 | -0.04721221 | -0.029739312 | -0.017472898 | 0.914834503 |
| ENSG00000183751 | *TBL3* | 3.391081013 | -0.019944388 | 0.117699973 | -0.137644362 | 0.999933887 | 0.165769343 | 0.143863988 | 0.021905356 | 0.914834503 |
| ENSG00000129003 | *VPS13C* | 5.728527214 | -0.03996463 | -0.02347998 | -0.016484651 | 0.999933887 | -0.669280856 | -0.723573523 | 0.054292667 | 0.914834503 |
| ENSG00000170085 | *SIMC1* | 1.105852771 | -0.0137421 | 0.079483066 | -0.093225165 | 0.999933887 | -0.094558593 | -0.137390183 | 0.04283159 | 0.914834503 |
| ENSG00000184635 | *ZNF93* | 0.586347273 | -0.192334498 | -0.017270412 | -0.175064086 | 0.999933887 | -0.681597701 | -0.730563527 | 0.048965826 | 0.914931617 |
| ENSG00000163083 | *INHBB* | 1.157830602 | -0.129069245 | -0.047410831 | -0.081658413 | 0.999933887 | -0.017244324 | -0.061158506 | 0.043914182 | 0.914931617 |
| ENSG00000112578 | *BYSL* | 0.604209782 | 0.003235135 | -0.066558204 | 0.069793339 | 0.999933887 | 0.457600179 | 0.500448203 | -0.042848024 | 0.914931617 |
| ENSG00000186648 | *CARMIL3* | 0.310749936 | -0.017776136 | 0.147174911 | -0.164951047 | 0.999933887 | 0.030977534 | 0.080684327 | -0.049706793 | 0.915420378 |
| ENSG00000227262 | *HCG4B* | -1.21738882 | -0.041478249 | -0.43669003 | 0.395211781 | 0.999933887 | 1.182787353 | 1.113130196 | 0.069657157 | 0.915506694 |
| ENSG00000233030 | *AC243772.2* | 0.645411476 | 0.01593187 | 0.083867067 | -0.067935197 | 0.999933887 | -0.218141695 | -0.156427969 | -0.061713726 | 0.91558936 |
| ENSG00000164975 | *SNAPC3* | 1.443737593 | -0.10581974 | 0.16153714 | -0.26735688 | 0.999933887 | -0.004367549 | -0.039594736 | 0.035227188 | 0.915648403 |
| ENSG00000159403 | *C1R* | 1.130922107 | -0.035701 | 0.147061235 | -0.182762236 | 0.999933887 | 0.168644511 | 0.09225213 | 0.076392381 | 0.915760556 |
| ENSG00000173207 | *CKS1B* | -0.654425702 | -0.342383606 | 0.070225947 | -0.412609554 | 0.999933887 | 0.158378257 | 0.225772672 | -0.067394414 | 0.915813821 |
| ENSG00000244005 | *NFS1* | 1.122727405 | -0.075308326 | 0.03604926 | -0.111357586 | 0.999933887 | -0.314800124 | -0.350866823 | 0.036066699 | 0.915813821 |
| ENSG00000242992 | *FTH1P4* | 0.564746536 | -0.031093223 | 0.225915892 | -0.257009115 | 0.999933887 | 0.601094847 | 0.667500513 | -0.066405666 | 0.915813821 |

| | | | | | | | | | | |
|---|---|---|---|---|---|---|---|---|---|---|
| ENSG00000138078 | *PREPL* | 2.309511233 | -0.089612633 | -0.044859753 | -0.044752879 | 0.999933887 | -0.227388585 | -0.261395407 | 0.034006822 | 0.91591598 |
| ENSG00000171824 | *EXOSC10* | 3.311631732 | -0.01055493 | 0.037062711 | -0.047617641 | 0.999933887 | 0.113218134 | 0.134724305 | -0.021506171 | 0.91591598 |
| ENSG00000105699 | *LSR* | 2.966991426 | 0.074080813 | 0.028788459 | 0.045292354 | 0.999933887 | 0.099069937 | 0.123621048 | -0.024551111 | 0.916238262 |
| ENSG00000120802 | *TMPO* | 3.314307915 | 0.044795486 | -0.002721292 | 0.047516778 | 0.999933887 | 0.083462471 | 0.062024828 | 0.021437643 | 0.916238262 |
| ENSG00000134248 | *LAMTOR5* | 4.315797511 | 0.003519267 | -0.028144905 | 0.031664172 | 0.999933887 | -0.159751384 | -0.178498465 | 0.018747081 | 0.916238262 |
| ENSG00000176533 | *GNG7* | 1.904007431 | 0.101614253 | 0.072618126 | 0.028996127 | 0.999933887 | -0.419935024 | -0.380780802 | -0.039154222 | 0.91645235 |
| ENSG00000263465 | *SRSF8* | 3.126807354 | -0.042124274 | 0.07312706 | -0.115251334 | 0.999933887 | 0.147783748 | 0.124626847 | 0.023156901 | 0.91645235 |
| ENSG00000196449 | *YRDC* | 3.165953831 | 0.204948731 | 0.240541843 | -0.035593112 | 0.999933887 | 0.348563603 | 0.324241779 | 0.024321824 | 0.916922397 |
| ENSG00000072210 | *ALDH3A2* | 1.969666031 | -0.09509147 | 0.060280415 | -0.155371885 | 0.999933887 | -0.069351133 | -0.103632708 | 0.034281575 | 0.916922397 |
| ENSG00000213949 | *ITGA1* | 2.407718463 | 0.062639221 | -0.112512394 | 0.175151615 | 0.999933887 | -0.320863348 | -0.279437666 | -0.041425682 | 0.916922397 |
| ENSG00000136908 | *DPM2* | 2.268817898 | 0.013392719 | -0.013553415 | 0.026946134 | 0.999933887 | 0.024026279 | -0.001021249 | 0.025047528 | 0.916922397 |
| ENSG00000144959 | *NCEH1* | 0.427583619 | 0.166958652 | 0.106493709 | 0.060464943 | 0.999933887 | -0.675266499 | -0.742106546 | 0.066840048 | 0.917001088 |
| ENSG00000167257 | *RNF214* | 1.510366265 | -0.107055879 | -0.019711679 | -0.0873442 | 0.999933887 | -0.126587389 | -0.161450412 | 0.034863024 | 0.91702323 |
| ENSG00000245552 | *AP000787.1* | 0.053126717 | 0.176733787 | -0.078849068 | 0.255582855 | 0.999933887 | 0.061877618 | 0.01223937 | 0.049638248 | 0.917039277 |
| ENSG00000279511 | *AL356274.2* | 0.06542492 | 0.184752322 | 0.3237118 | -0.138959478 | 0.999933887 | 0.368131019 | 0.471190419 | -0.1030594 | 0.917039277 |
| ENSG00000085491 | *SLC25A24* | 1.969343111 | -0.07873445 | -0.063834176 | -0.014900273 | 0.999933887 | -0.289372944 | -0.254652634 | -0.03472031 | 0.917039277 |
| ENSG00000155903 | *RASA2* | 5.165234671 | -0.021867091 | -0.099715519 | 0.077848429 | 0.999933887 | -0.544471504 | -0.591887377 | 0.047415873 | 0.917039277 |
| ENSG00000263345 | *AC006435.2* | 0.099272154 | 0.00416406 | -0.221785646 | 0.225949706 | 0.999933887 | -0.172032258 | -0.103335836 | -0.068696422 | 0.917039277 |
| ENSG00000235111 | *Z97192.3* | -0.15698688 | -0.020114544 | 0.213497701 | -0.233612245 | 0.999933887 | -0.227830552 | -0.290550383 | 0.062719831 | 0.917396862 |
| ENSG00000158406 | *H4C8* | 2.492907254 | -0.003770755 | -0.014361147 | 0.010590393 | 0.999933887 | -0.249723521 | -0.27572134 | 0.025997819 | 0.917627039 |
| ENSG00000142856 | *ITGB3BP* | 0.478371661 | -0.093198952 | 0.049299898 | -0.142498851 | 0.999933887 | -0.167104405 | -0.123953538 | -0.043150866 | 0.917672501 |
| ENSG00000219163 | *HMGB1P20* | 0.963713208 | -0.047133513 | 0.034771113 | -0.081904626 | 0.999933887 | -0.333175551 | -0.283209825 | -0.049965726 | 0.917672501 |
| ENSG00000124159 | *MATN4* | 0.077571524 | 0.071138773 | 0.149511639 | -0.078372865 | 0.999933887 | 0.516814058 | 0.460263028 | 0.05655103 | 0.917674268 |
| ENSG00000132781 | *MUTYH* | -0.154764213 | -0.04255262 | 0.149053495 | -0.191606115 | 0.999933887 | 0.108698293 | 0.158659323 | -0.049961031 | 0.917674268 |
| ENSG00000196581 | *AJAP1* | -1.659274259 | 0.031997843 | 0.221448349 | -0.189450506 | 0.999933887 | -0.474907875 | -0.539892472 | 0.064984597 | 0.917674268 |
| ENSG00000196950 | *SLC39A10* | 2.199691126 | -0.139291182 | -0.112765761 | -0.026525421 | 0.999933887 | -0.349931268 | -0.312252909 | -0.037678359 | 0.917713916 |
| ENSG00000182518 | *FAM104B* | -0.272679699 | 0.045183385 | 0.062955711 | -0.017772325 | 0.999933887 | -0.13628681 | -0.077233663 | -0.059053146 | 0.917713916 |
| ENSG00000172766 | *NAA16* | 2.804596251 | -0.020585987 | -0.180981201 | 0.160395214 | 0.999933887 | -0.010046404 | 0.026745451 | -0.036791855 | 0.917713916 |
| ENSG00000254860 | *TMEM9B-AS1* | 0.804754684 | 0.243957168 | 0.14155092 | 0.102406247 | 0.999933887 | 0.132230809 | 0.079166883 | 0.053063925 | 0.917756779 |
| ENSG00000215251 | *FASTKD5* | 4.027103116 | -0.06926209 | -0.077966073 | 0.008703983 | 0.999933887 | -0.044963121 | -0.026082794 | -0.018880326 | 0.917826206 |
| ENSG00000165644 | *COMTD1* | 1.530764881 | 0.129445198 | 0.016087672 | 0.113357526 | 0.999933887 | -0.236351233 | -0.200173146 | -0.036178086 | 0.917826206 |
| ENSG00000227766 | *AL671277.1* | -1.897781676 | 0.634714191 | 0.471836254 | 0.162877936 | 0.999933887 | 1.072678622 | 0.964548689 | 0.108129933 | 0.917951579 |
| ENSG00000178307 | *TMEM11* | 3.38602472 | -0.116107057 | -0.211279821 | 0.095172765 | 0.999933887 | -0.868769176 | -0.894669979 | 0.025900802 | 0.917951579 |
| ENSG00000178607 | *ERN1* | 9.055140735 | -0.037886809 | 0.016539353 | -0.054426162 | 0.999933887 | -0.110125488 | -0.094698454 | -0.015427035 | 0.917951579 |
| ENSG00000223612 | *AC241585.1* | -0.528043865 | 0.264756428 | -0.414891628 | 0.679648056 | 0.999933887 | 0.39339662 | 0.446401362 | -0.053004742 | 0.918028667 |
| ENSG00000134255 | *CEPT1* | 3.920831543 | -0.045217295 | -0.063171112 | 0.017953817 | 0.999933887 | -0.21407943 | -0.185512844 | -0.028566586 | 0.91824849 |
| ENSG00000198723 | *TEX45* | 0.09642647 | 0.075756937 | 0.035285796 | 0.04047114 | 0.999933887 | 0.277773705 | 0.341109596 | -0.063335891 | 0.918336657 |
| ENSG00000116353 | *MECR* | -0.229964395 | 0.058801258 | 0.549620366 | -0.490819108 | 0.999933887 | 0.144284138 | 0.191689082 | -0.047404944 | 0.918336657 |
| ENSG00000215529 | *EFCAB8* | 0.953518797 | 0.029513118 | -0.032248073 | 0.061761192 | 0.999933887 | 0.121968972 | 0.083385096 | 0.038583876 | 0.918336657 |
| ENSG00000171453 | *POLR1C* | 1.847668231 | -0.081152338 | -0.145011804 | 0.063859467 | 0.999933887 | 0.197696797 | 0.167183931 | 0.030512867 | 0.918584111 |
| ENSG00000226644 | *AL121899.1* | 1.802612154 | 0.09691235 | 0.384073688 | -0.287161338 | 0.999933887 | 0.390958271 | 0.437556284 | -0.046598014 | 0.91881954 |
| ENSG00000169398 | *PTK2* | -0.07442925 | -0.036102193 | 0.040950369 | -0.077052563 | 0.999933887 | -0.116851915 | -0.063608965 | -0.05324295 | 0.918975767 |
| ENSG00000259146 | *AC005476.2* | -0.418089358 | 0.298247371 | 0.118627706 | 0.179619664 | 0.999933887 | 0.800124612 | 0.722787341 | 0.077337271 | 0.919022336 |
| ENSG00000149923 | *PPP4C* | 7.350203813 | 0.03731011 | 0.006625197 | 0.030684913 | 0.999933887 | -0.046738097 | -0.033135019 | -0.013603078 | 0.919022336 |
| ENSG00000237781 | *ADAMTSL4-A* | 3.040376991 | 0.050818247 | -0.008904473 | 0.059722719 | 0.999933887 | 0.281923628 | 0.252119426 | 0.029804202 | 0.919022336 |
| ENSG00000092621 | *PHGDH* | -0.030591696 | -0.13570578 | -0.080505215 | -0.055200565 | 0.999933887 | -0.3694572 | -0.409299294 | 0.039842094 | 0.919022336 |
| ENSG00000266412 | *NCOA4* | 7.563144733 | -0.045600994 | -0.040204085 | -0.005396909 | 0.999933887 | 0.763296708 | 0.800422832 | -0.037126124 | 0.919022336 |
| ENSG00000124299 | *PEPD* | 2.517020634 | 0.016151849 | 0.004051057 | 0.012100791 | 0.999933887 | 0.076885152 | 0.049275979 | 0.027609172 | 0.919022336 |
| ENSG00000206190 | *ATP10A* | 3.168034118 | -0.029999671 | 0.040522747 | -0.070522418 | 0.999933887 | -0.10059598 | -0.122753877 | 0.022157898 | 0.919047695 |
| ENSG00000115935 | *WIPF1* | 9.598163939 | -0.003861982 | -0.006212274 | 0.002350292 | 0.999933887 | -0.483762041 | -0.49613152 | 0.012369478 | 0.919167659 |
| ENSG00000204152 | *TIMM23B* | 0.018357097 | 0.074592793 | 0.354505909 | -0.279913117 | 0.999933887 | 0.090045996 | 0.037961884 | 0.052084112 | 0.919255306 |
| ENSG00000229951 | *FLJ31356* | 3.55562715 | 0.405401849 | 0.317080796 | 0.088321053 | 0.999933887 | 1.16287464 | 1.204344635 | -0.041469995 | 0.919376942 |
| ENSG00000174109 | *C16orf91* | 1.930173265 | 0.107436428 | -0.032553037 | 0.139989465 | 0.999933887 | 0.190257012 | 0.227676457 | -0.037419445 | 0.919376942 |
| ENSG00000179038 | *AP001885.1* | -0.671647765 | 0.281826372 | 0.243620416 | 0.038205956 | 0.999933887 | -0.44031492 | -0.542250738 | 0.101935818 | 0.919447249 |
| ENSG00000198517 | *MAFK* | 6.970709075 | 0.123538876 | 0.127975286 | -0.004436411 | 0.999933887 | -0.041094127 | -0.016422831 | -0.024671296 | 0.919617164 |
| ENSG00000162341 | *TPCN2* | 3.442769383 | -0.035238028 | 0.017101972 | -0.05234 | 0.999933887 | 0.05730674 | 0.033656483 | 0.023650256 | 0.919644757 |
| ENSG00000119559 | *C19orf25* | 4.496739116 | 0.007193013 | -0.020131388 | 0.027324402 | 0.999933887 | -0.065968222 | -0.086240408 | 0.020272186 | 0.919644757 |
| ENSG00000260267 | *AC026471.1* | 1.506864976 | 0.15859293 | 0.215162004 | -0.056569074 | 0.999933887 | 0.403704754 | 0.444402451 | -0.040697697 | 0.920433237 |
| ENSG00000116819 | *TFAP2E* | 0.153967821 | 0.211681336 | 0.264851525 | -0.053170189 | 0.999933887 | -0.354316436 | -0.397746251 | 0.043429815 | 0.920433237 |
| ENSG00000231683 | *AL033397.1* | -2.908152657 | -0.284254111 | 0.484012977 | -0.768267087 | 0.999933887 | 2.191296296 | 2.293126238 | -0.101829942 | 0.920521718 |
| ENSG00000147853 | *AK3* | 2.068176592 | -0.06652133 | 0.026557549 | -0.093078878 | 0.999933887 | -0.049770822 | -0.014358104 | -0.035412718 | 0.920578695 |
| ENSG00000266127 | *ZNF415P1* | -1.152126707 | 0.174683626 | 0.272713858 | -0.098030232 | 0.999933887 | -0.16391595 | -0.053058379 | -0.110857571 | 0.920610699 |
| ENSG00000196981 | *WDR5B* | 1.863894078 | -0.286040741 | -0.106776893 | -0.179263848 | 0.999933887 | -0.164178798 | -0.135759794 | -0.028419004 | 0.920726727 |
| ENSG00000163220 | *S100A9* | 11.619111856 | 0.060824056 | 0.030061588 | 0.030762467 | 0.999933887 | -0.001951413 | 0.020578264 | -0.022529678 | 0.920726727 |
| ENSG00000176024 | *ZNF613* | 0.395279073 | -0.148343662 | 0.013087608 | -0.161431269 | 0.999933887 | -0.102781925 | -0.141719437 | 0.038937511 | 0.920726727 |
| ENSG00000175701 | *MTLN* | -0.137516826 | -0.161915077 | -0.312238893 | 0.150323817 | 0.999933887 | 0.17043359 | 0.096561803 | 0.073871788 | 0.920726727 |
| ENSG00000088854 | *C20orf194* | 1.249725371 | -0.059638003 | 0.086283116 | -0.145921119 | 0.999933887 | -0.238242017 | -0.273295039 | 0.035053022 | 0.920726727 |
| ENSG00000115760 | *BIRC6* | 6.59217857 | 0.011626292 | -0.006993918 | 0.01862021 | 0.999933887 | -0.456882757 | -0.486047737 | 0.02916498 | 0.920726727 |
| ENSG00000197296 | *FITM2* | -0.094622223 | -0.153642435 | -0.021883932 | -0.131758503 | 0.999933887 | -0.106565558 | -0.051787163 | -0.054778395 | 0.92080875 |
| ENSG00000110693 | *SOX6* | 0.520401083 | -0.028672131 | 0.036733458 | -0.065405589 | 0.999933887 | -0.093338314 | -0.149078115 | 0.0557398 | 0.92080875 |
| ENSG00000103550 | *KNOP1* | 0.140404692 | -0.011424937 | 0.108913057 | -0.120337994 | 0.999933887 | 0.159343972 | 0.207285975 | -0.047942003 | 0.92080875 |
| ENSG00000006744 | *ELAC2* | 3.297285445 | -0.000775484 | 0.004123505 | -0.004898989 | 0.999933887 | 0.151327107 | 0.131209246 | 0.020117861 | 0.92080875 |
| ENSG00000244026 | *FAM86DP* | -0.327632285 | -0.053146921 | 0.276187228 | -0.329334149 | 0.999933887 | 0.140239337 | 0.187895788 | -0.047656452 | 0.920863753 |
| ENSG00000162542 | *TMCO4* | 1.748234991 | 0.018800543 | 0.187262583 | -0.16846204 | 0.999933887 | -0.004278488 | -0.036760144 | 0.032481656 | 0.920863753 |
| ENSG00000103512 | *NOMO1* | 2.470910761 | -0.006765414 | 0.010059501 | -0.016824915 | 0.999933887 | -0.026884338 | -0.05583057 | 0.028946233 | 0.920950314 |
| ENSG00000113108 | *APBB3* | 6.049360409 | 0.045029469 | 0.098193515 | -0.053164046 | 0.999933887 | 0.00673522 | 0.026508514 | -0.019773294 | 0.921192344 |
| ENSG00000144550 | *CPNE9* | 0.074410382 | 0.250930095 | 0.209234758 | 0.041695337 | 0.999933887 | 0.740134681 | 0.8002075 | -0.060072819 | 0.921250776 |
| ENSG00000013375 | *PGM3* | 1.289819267 | -0.078360539 | -0.179471342 | 0.101110803 | 0.999933887 | 0.098681727 | 0.05668921 | 0.041992517 | 0.921250776 |
| ENSG00000285704 | *AC004765.1* | -0.348935359 | 0.085940562 | 0.221375448 | -0.135434886 | 0.999933887 | 0.213982407 | 0.265731373 | -0.051748966 | 0.921250776 |
| ENSG00000164292 | *RHOBTB3* | 1.568310171 | -0.176561664 | 0.137619811 | -0.314181475 | 0.999933887 | 0.168645052 | 0.143102294 | 0.025542758 | 0.921251361 |
| ENSG00000136451 | *VEZF1* | 5.368850487 | -0.079546404 | -0.115620426 | 0.036074022 | 0.999933887 | -0.279656848 | -0.296056134 | 0.016399286 | 0.921251361 |
| ENSG00000161956 | *SENP3* | 2.373946793 | 0.074607737 | -0.015680022 | 0.090287759 | 0.999933887 | 0.006843288 | 0.035539784 | -0.028696496 | 0.921251361 |
| ENSG00000236776 | *RPL21P23* | -0.163418451 | 0.036378546 | 0.401020632 | -0.364642086 | 0.999933887 | 0.199882899 | 0.2431206 | -0.043237701 | 0.921251361 |
| ENSG00000273008 | *AC010864.1* | -0.00562473 | 0.350156864 | -0.047883214 | 0.398040077 | 0.999933887 | 0.424774395 | 0.358341334 | 0.06643306 | 0.92188656 |
| ENSG00000279162 | *AC141586.5* | 1.949624795 | 0.088371407 | -0.08577102 | 0.174142427 | 0.999933887 | -0.582569374 | -0.539996049 | -0.042573326 | 0.92188656 |
| ENSG00000167065 | *DUSP18* | 2.944490797 | -0.054425745 | 0.027317786 | -0.081743531 | 0.999933887 | 0.269317923 | 0.23876693 | 0.030550993 | 0.92188656 |
| ENSG00000106086 | *PLEKHA8* | 1.115385441 | -0.013175737 | -0.150970732 | 0.137794995 | 0.999933887 | -0.089749032 | -0.051222584 | -0.038526448 | 0.92188656 |
| ENSG00000176153 | *GPX2* | 0.342532872 | -0.121160133 | 0.112264844 | -0.233424978 | 0.999933887 | -0.025448293 | 0.025298082 | -0.050746375 | 0.921903985 |
| ENSG00000285872 | *AC007240.3* | 2.445596565 | -0.041289115 | 0.060358855 | -0.10164797 | 0.999933887 | -0.603552556 | -0.572124506 | -0.03142805 | 0.921903985 |
| ENSG00000106560 | *GIMAP2* | 3.345092159 | -0.023038689 | -0.115708698 | 0.092670009 | 0.999933887 | 0.033529858 | 0.012378441 | 0.021151417 | 0.921903985 |
| ENSG00000090447 | *TFAP4* | 0.670666051 | -0.004152769 | -0.285371245 | 0.281218475 | 0.999933887 | -0.038173894 | -0.005450841 | -0.032723052 | 0.921903985 |
| ENSG00000270605 | *AL353622.1* | 0.02253562 | -0.107570194 | -0.156514874 | 0.04894468 | 0.999933887 | 0.305264486 | 0.252802109 | 0.052462378 | 0.921905255 |
| ENSG00000143393 | *PI4KB* | 5.830063867 | -0.012041634 | 0.023202617 | -0.035244251 | 0.999933887 | 0.123646278 | 0.110965102 | 0.012681176 | 0.921905255 |
| ENSG00000138764 | *CCNG2* | 6.866618098 | -0.091251252 | 0.024433125 | -0.115684377 | 0.999933887 | -0.620113937 | -0.639007477 | 0.01889354 | 0.921970582 |
| ENSG00000278002 | *AL627171.1* | -0.738230726 | 0.461977619 | -0.163628014 | 0.625605633 | 0.999933887 | -0.214214996 | -0.315172771 | 0.100957775 | 0.922143217 |
| ENSG00000233452 | *STXBP5-AS1* | 1.264842234 | 0.201990996 | 0.056168568 | 0.145822428 | 0.999933887 | -0.340213851 | -0.298245633 | -0.041968217 | 0.922143217 |
| ENSG00000183019 | *MCEMP1* | 2.733999514 | -0.033304979 | 0.049859513 | -0.083164492 | 0.999933887 | 0.161194405 | 0.205855169 | -0.044660764 | 0.922143217 |
| ENSG00000129484 | *PARP2* | 0.3951301 | 0.030113604 | -0.061890507 | 0.092004111 | 0.999933887 | -0.094553497 | -0.04688896 | -0.047664537 | 0.922143217 |
| ENSG00000157734 | *SNX22* | 0.308572836 | 0.017132156 | 0.035836839 | -0.018704682 | 0.999933887 | -0.220552845 | -0.258505655 | 0.03795281 | 0.922143217 |
| ENSG00000115970 | *THADA* | 3.728769725 | 0.003429995 | -0.069946762 | 0.073376757 | 0.999933887 | -0.041277302 | -0.059263387 | 0.017986085 | 0.922143217 |
| ENSG00000185359 | *HGS* | 6.849453997 | 0.045926979 | 0.057716322 | -0.011789343 | 0.999933887 | 0.064323904 | 0.077332508 | -0.013008604 | 0.922213788 |
| ENSG00000140382 | *HMG20A* | 1.807105564 | 0.000182333 | -0.10905275 | 0.109235083 | 0.999933887 | 0.006097084 | -0.024439073 | 0.030536158 | 0.922247821 |
| ENSG00000226278 | *PSPHP1* | 0.527022104 | -0.037741001 | -0.132182006 | 0.094441004 | 0.999933887 | 0.185249858 | 0.237690194 | -0.052440337 | 0.922293122 |
| ENSG00000119328 | *ABITRAM* | 2.458036696 | -0.105180706 | -0.030395811 | -0.074784895 | 0.999933887 | -0.315196945 | -0.287022172 | -0.028174772 | 0.922336109 |
| ENSG00000076864 | *RAP1GAP* | 1.201522666 | 0.104749445 | 0.131644749 | -0.026895305 | 0.999933887 | 0.249998958 | 0.189086886 | 0.060912072 | 0.922336109 |

| | | | | | | | | | | |
|---|---|---|---|---|---|---|---|---|---|---|
| ENSG00000206503 | *HLA-A* | 11.947317057 | 0.055121286 | 0.033232258 | 0.021889028 | 0.999933887 | 0.519130753 | 0.536121849 | -0.016991096 | 0.922336109 |
| ENSG00000167861 | *HID1* | -0.701688916 | 0.155120309 | -0.475567989 | 0.630688299 | 0.944094537 | 0.013340501 | 0.072038419 | -0.058697918 | 0.922336109 |
| ENSG00000179886 | *TIGD5* | 0.808015509 | 0.115551653 | -0.127321121 | 0.242872773 | 0.999933887 | -0.150782683 | -0.186190491 | 0.035407808 | 0.922336109 |
| ENSG00000104886 | *PLEKHJ1* | 4.358904207 | -0.071406544 | 0.019860631 | -0.091267175 | 0.999933887 | 0.336429742 | 0.360902367 | -0.024472625 | 0.922336109 |
| ENSG00000148303 | *RPL7A* | 7.773206061 | -0.026433339 | -0.064756171 | 0.038322832 | 0.999933887 | 0.072981272 | 0.056677389 | 0.016303883 | 0.922336109 |
| ENSG00000285531 | *Z83840.1* | -0.067517996 | -0.057260955 | -0.102570495 | 0.04530954 | 0.999933887 | -0.029665222 | -0.093718213 | 0.064052992 | 0.922336109 |
| ENSG00000116962 | *NID1* | -0.411466632 | 0.070027678 | -0.099270642 | 0.169298321 | 0.999933887 | 0.588465437 | 0.528886371 | 0.059579066 | 0.922336109 |
| ENSG00000156097 | *GPR61* | -0.633843436 | 0.045863284 | -0.086118894 | 0.131982178 | 0.999933887 | 0.22024768 | 0.137656557 | 0.082591123 | 0.922336109 |
| ENSG00000273151 | *AC073957.3* | 3.003054819 | 0.019865276 | 0.010250574 | 0.009614702 | 0.999933887 | 0.023981193 | 5.5232e-05 | 0.02392596 | 0.922336109 |
| ENSG00000164167 | *LSM6* | 2.039215178 | 0.005777284 | -0.019234641 | 0.025011924 | 0.999933887 | 0.078371891 | 0.108459375 | -0.030087484 | 0.922336109 |
| ENSG00000160746 | *ANO10* | 3.616672541 | 0.002656228 | -0.029701037 | 0.032357264 | 0.999933887 | -0.350791384 | -0.375494404 | 0.024703019 | 0.922336109 |
| ENSG00000117395 | *EBNA1BP2* | 1.599411077 | -0.070814302 | 0.046630218 | -0.11744452 | 0.999933887 | 0.148255842 | 0.116946608 | 0.031309233 | 0.922422889 |
| ENSG00000119285 | *HEATR1* | 3.066287801 | -0.089358264 | -0.157808444 | 0.06845018 | 0.999933887 | 0.015642017 | 0.041834734 | -0.026192716 | 0.922426487 |
| ENSG00000146587 | *RBAK* | 2.129707485 | -0.12161426 | -0.269461166 | 0.147846906 | 0.999933887 | -0.360177755 | -0.305945845 | -0.05423191 | 0.922426487 |
| ENSG00000231663 | *COA6-AS1* | 0.454598011 | -0.242826228 | -0.285622636 | 0.042796408 | 0.999933887 | 0.064445041 | 0.021426523 | 0.043018518 | 0.922552172 |
| ENSG00000111961 | *SASH1* | -0.399352396 | 0.05249629 | -0.252917711 | 0.305414001 | 0.999933887 | -0.667695152 | -0.719672355 | 0.051977203 | 0.922712622 |
| ENSG00000274460 | *AC092119.2* | 1.399682283 | -0.238911832 | -0.302020379 | 0.063108547 | 0.999933887 | -0.302173808 | -0.343657871 | 0.041484062 | 0.923069264 |
| ENSG00000273270 | *AC090114.2* | 1.929784125 | 0.103032665 | -0.160243309 | 0.263275975 | 0.999933887 | 0.141829323 | 0.110360938 | 0.031468385 | 0.923069264 |
| ENSG00000134827 | *TCN1* | 1.943844408 | 0.015242003 | -0.138990566 | 0.154232569 | 0.999933887 | -0.154159737 | -0.11473692 | -0.039422818 | 0.923069264 |
| ENSG00000281344 | *HELLPAR* | -1.351940827 | -0.605687852 | -0.147039106 | -0.458648746 | 0.999933887 | -1.001743683 | -0.876500712 | -0.125242971 | 0.923100062 |
| ENSG00000137166 | *FOXP4* | 2.142869782 | 0.084566421 | 0.120174474 | -0.035608052 | 0.999933887 | 0.084465339 | 0.110008555 | -0.025543217 | 0.923100062 |
| ENSG00000104881 | *PPP1R13L* | -0.373651532 | 0.283814033 | 0.230497382 | 0.053316651 | 0.999933887 | -0.064743805 | -0.116473522 | 0.051729717 | 0.923155539 |
| ENSG00000154274 | *C4orf19* | 0.348186795 | 0.160764778 | 0.033607873 | 0.127156905 | 0.999933887 | -0.124655373 | -0.068616429 | -0.056038945 | 0.923155539 |
| ENSG00000136271 | *DDX56* | 4.227392567 | 0.006996872 | -0.036678692 | 0.043675564 | 0.999933887 | 0.231885322 | 0.249752352 | -0.01786703 | 0.923155539 |
| ENSG00000175581 | *MRPL48* | 0.623296939 | -0.098700482 | -0.148172329 | 0.049471847 | 0.999933887 | -0.101257526 | -0.161210591 | 0.059953065 | 0.923293589 |
| ENSG00000268996 | *MAN1B1-DT* | -0.459920411 | 0.103769875 | 0.053124944 | 0.050644932 | 0.999933887 | 0.014186286 | 0.076497794 | -0.062311508 | 0.923293589 |
| ENSG00000108984 | *MAP2K6* | 2.245027385 | 0.037304041 | -0.086138822 | 0.123442863 | 0.999933887 | -0.08872149 | -0.051930535 | -0.036790955 | 0.923293589 |
| ENSG00000147174 | *GCNA* | 1.162286947 | -0.04166387 | -0.066258279 | 0.024594409 | 0.999933887 | 0.117151545 | 0.158599764 | -0.041448219 | 0.923293589 |
| ENSG00000035720 | *STAP1* | -0.170331173 | -0.125869371 | -0.172963444 | 0.047094074 | 0.999933887 | -0.151662319 | -0.102108392 | -0.049553927 | 0.923364469 |
| ENSG00000187954 | *CYHR1* | 7.699645112 | 0.01722022 | 0.005062775 | 0.012157445 | 0.999933887 | 0.543524567 | 0.517065491 | 0.026459076 | 0.92379128 |
| ENSG00000130159 | *ECSIT* | 2.114774904 | 0.1400905 | -0.018778544 | 0.158869044 | 0.999933887 | 0.043814809 | 0.069698074 | -0.025883265 | 0.923887797 |
| ENSG00000147604 | *RPL7* | 7.218076545 | -0.057322989 | -0.145721558 | 0.088398569 | 0.999933887 | 0.045086491 | 0.06193087 | -0.016844379 | 0.923887797 |
| ENSG00000173436 | *MICOS10* | 1.302289397 | 0.010045766 | -0.228746201 | 0.238791967 | 0.999933887 | -0.205616735 | -0.171038704 | -0.034578031 | 0.923887797 |
| ENSG00000227383 | *RPL35AP2* | -0.631386435 | 0.071048661 | 0.090600427 | -0.019551766 | 0.999933887 | 0.348356036 | 0.268251546 | 0.080104489 | 0.923999627 |
| ENSG00000108848 | *LUC7L3* | 6.300371629 | -0.103968506 | -0.142208054 | 0.038239549 | 0.999933887 | -0.271604555 | -0.24639564 | -0.025208915 | 0.924052166 |
| ENSG00000230789 | *ARHGAP26-IT* | 0.346830381 | -0.052447133 | 0.067084159 | -0.119531291 | 0.999933887 | 0.02229258 | -0.05147607 | 0.07376865 | 0.924140561 |
| ENSG00000065154 | *OAT* | 4.573043953 | -0.082329614 | -0.047962346 | -0.034367268 | 0.999933887 | 0.046652429 | 0.061858813 | -0.015206384 | 0.924190483 |
| ENSG00000211699 | *TRGV3* | -0.488911841 | 0.293453754 | 0.215814856 | 0.077638897 | 0.999933887 | 0.095639989 | 0.152149661 | -0.056509672 | 0.924190483 |
| ENSG00000148300 | *REXO4* | 2.17614868 | 0.057656987 | 0.004052255 | 0.053604732 | 0.999933887 | 0.120953227 | 0.095764145 | 0.025189082 | 0.924190483 |
| ENSG00000168237 | *GLYCTK* | 2.910115286 | 0.045134334 | -0.020365025 | 0.06549936 | 0.999933887 | -0.119015187 | -0.095135195 | -0.023879992 | 0.924190483 |
| ENSG00000165138 | *ANKS6* | 0.944772494 | 0.030355552 | 0.251124097 | -0.220768545 | 0.999933887 | 0.05713273 | 0.100023672 | -0.042890942 | 0.924190483 |
| ENSG00000218226 | *TATDN2P2* | 2.241642208 | 0.003148982 | 0.051741452 | -0.04859247 | 0.999933887 | -0.02852607 | -0.057548589 | 0.02902252 | 0.924190483 |
| ENSG00000147535 | *PLPP5* | 3.069376754 | 0.00056595 | -0.028869706 | 0.029435656 | 0.999933887 | 0.090708977 | 0.069669721 | 0.021039256 | 0.924190483 |
| ENSG00000095380 | *NANS* | 3.185945057 | 0.046297268 | 0.106988988 | -0.06069172 | 0.999933887 | -0.31860559 | -0.344722803 | 0.026117213 | 0.924314482 |
| ENSG00000254505 | *CHMP4A* | 1.511642495 | 0.13742891 | 0.046924278 | 0.090504632 | 0.999933887 | 0.407317799 | 0.436256497 | -0.028938698 | 0.924462784 |
| ENSG00000009790 | *TRAF3IP3* | 6.513428065 | -0.028700534 | -0.079929154 | 0.05122862 | 0.999933887 | -0.293539431 | -0.307458774 | 0.013919342 | 0.924462784 |
| ENSG00000188343 | *CIBAR1* | 0.140554493 | -0.066937243 | -0.355164943 | 0.2882277 | 0.999933887 | -0.583810744 | -0.521881185 | -0.061929559 | 0.924462784 |
| ENSG00000211752 | *TRBV27* | 0.231963842 | 0.085629362 | -0.066804607 | 0.15243397 | 0.999933887 | 0.048714929 | 0.089926956 | -0.041212027 | 0.924637921 |
| ENSG00000132286 | *TIMM10B* | 3.894381294 | 0.015704828 | -0.076912793 | 0.092617621 | 0.999933887 | -0.020428696 | -0.038941756 | 0.018513061 | 0.924637921 |
| ENSG00000142025 | *DMRTC2* | 0.182981779 | -0.03567165 | 0.017255898 | -0.052927548 | 0.999933887 | -0.305213983 | -0.231902558 | -0.073311426 | 0.924761274 |
| ENSG00000115607 | *IL18RAP* | 5.916941867 | -0.025603268 | -0.050457601 | 0.024854333 | 0.999933887 | 0.166950294 | 0.180696109 | -0.013745814 | 0.924866199 |
| ENSG00000280208 | *GGT4P* | -0.173494606 | 0.126280636 | -0.365563419 | 0.491844054 | 0.999933887 | 0.685956423 | 0.7308276 | -0.044871178 | 0.924866199 |
| ENSG00000074181 | *NOTCH3* | -0.76767577 | -0.00131475 | -0.140420603 | 0.139105854 | 0.999933887 | -0.243880252 | -0.188684322 | -0.055195931 | 0.924866199 |
| ENSG00000179943 | *FIZ1* | 1.707740196 | 0.100564943 | 0.062869521 | 0.037695422 | 0.999933887 | -0.144045048 | -0.179434373 | 0.035389325 | 0.925254588 |
| ENSG00000188641 | *DPYD* | 4.749506234 | -0.019151085 | -0.001889474 | -0.01726161 | 0.999933887 | -0.147499713 | -0.16489892 | 0.017399208 | 0.925254588 |
| ENSG00000189343 | *RPS2P46* | -0.627064604 | -0.080579196 | -0.110371264 | 0.029792068 | 0.999933887 | 0.157750623 | 0.088237934 | 0.069512689 | 0.925254588 |
| ENSG00000172243 | *CLEC7A* | 9.750081753 | -0.053470549 | -0.005755926 | -0.047714623 | 0.999933887 | -0.337894802 | -0.323720619 | -0.014174183 | 0.925300722 |
| ENSG00000287789 | *AL158136.1* | -0.516987538 | 0.036222628 | 0.35209665 | -0.315874023 | 0.999933887 | -0.215259396 | -0.26443021 | 0.049170815 | 0.925395417 |
| ENSG00000124374 | *PAIP2B* | -0.4035554 | -0.072621434 | -0.612820447 | 0.540199013 | 0.999933887 | 0.099774898 | 0.14883646 | -0.049061562 | 0.925406065 |
| ENSG00000112365 | *ZBTB24* | 3.428174861 | -0.123123951 | -0.001986385 | -0.121137566 | 0.999933887 | -0.105985994 | -0.123243739 | 0.017257745 | 0.925441744 |
| ENSG00000267244 | *AC012615.6* | 2.568550582 | 0.103570253 | -0.043195124 | 0.146765377 | 0.999933887 | -0.063337739 | -0.037815381 | -0.025522358 | 0.925556412 |
| ENSG00000100568 | *VTI1B* | 3.025884972 | 0.064919128 | 0.092638921 | -0.027719792 | 0.999933887 | -0.264419946 | -0.282791627 | 0.018371681 | 0.925556412 |
| ENSG00000089154 | *GCN1* | 5.190275313 | -0.027514345 | -0.008750435 | -0.018763911 | 0.999933887 | 0.086403632 | 0.072451533 | 0.013952099 | 0.925556412 |
| ENSG00000204604 | *ZNF468* | 2.141406662 | -0.046452633 | -0.12327836 | 0.076825726 | 0.999933887 | -0.327045601 | -0.357829348 | 0.030783747 | 0.925556412 |
| ENSG00000271550 | *BNIP3P11* | 0.132742832 | -0.019569213 | 0.171357425 | -0.190926638 | 0.999933887 | -0.188264774 | -0.243153542 | 0.054888768 | 0.925556412 |
| ENSG00000185238 | *PRMT3* | 2.307457136 | -0.093647794 | -0.157391646 | 0.063743852 | 0.999933887 | -0.394702094 | -0.372981206 | -0.021720889 | 0.925929327 |
| ENSG00000006118 | *TMEM132A* | 2.748096702 | 0.250777643 | 0.269082623 | -0.01830498 | 0.999933887 | 1.18306958 | 1.148162491 | 0.034907089 | 0.925950535 |
| ENSG00000224114 | *AL591846.1* | 1.373812025 | 0.224662197 | 0.292233553 | -0.067571356 | 0.999933887 | 0.554036248 | 0.617904315 | -0.063868067 | 0.925950535 |
| ENSG00000251000 | *GGCTP1* | 0.296469169 | -0.272561015 | -0.695632862 | 0.423071847 | 0.999933887 | -0.354969629 | -0.304409855 | -0.050559775 | 0.925950535 |
| ENSG00000248275 | *TRIM52-AS1* | 1.756415477 | 0.043474398 | 0.081446039 | -0.037971641 | 0.999933887 | 0.01960209 | -0.008366239 | 0.027968328 | 0.925950535 |
| ENSG00000229320 | *KRT8P12* | 2.284050261 | -0.127492297 | -0.075318647 | -0.05217365 | 0.999933887 | -0.609731029 | -0.635886046 | 0.026155017 | 0.92599505 |
| ENSG00000269929 | *MIRLET7A1H(* | 2.68772232 | 0.062836717 | -0.012729672 | 0.07556639 | 0.999933887 | -0.02461306 | -0.046589003 | 0.021975943 | 0.92599505 |
| ENSG00000008324 | *SS18L2* | 3.090520992 | 0.022056581 | 0.045855356 | -0.023798775 | 0.999933887 | -0.133084481 | -0.156088515 | 0.023004035 | 0.926059484 |
| ENSG00000286186 | *AC018445.5* | 0.875874897 | -0.040270732 | -0.10392865 | 0.063657917 | 0.999933887 | 0.20089705 | 0.245003652 | -0.044106602 | 0.926059484 |
| ENSG00000164151 | *ICE1* | 4.245871204 | 0.018618769 | -0.076015486 | 0.094634256 | 0.999933887 | -0.311634888 | -0.281019617 | -0.030615271 | 0.926059484 |
| ENSG00000257017 | *HP* | -0.164270684 | -0.01918867 | -0.171791115 | 0.152602445 | 0.999933887 | 0.064223343 | 0.002813347 | 0.061409996 | 0.926084266 |
| ENSG00000167094 | *TTC16* | 1.809215661 | 0.04279566 | 0.049286666 | -0.006491006 | 0.999933887 | 0.000709104 | -0.029582533 | 0.030291637 | 0.926145092 |
| ENSG00000274376 | *ADAMTS7P1* | 0.062374629 | -0.121139862 | 0.177976249 | -0.299116111 | 0.999933887 | 0.045608803 | -0.01712698 | 0.062735783 | 0.926260565 |
| ENSG00000126091 | *ST3GAL3* | -0.41276352 | -0.187022143 | 0.124652551 | -0.311674694 | 0.999933887 | -0.139320854 | -0.209423227 | 0.070102373 | 0.926260565 |
| ENSG00000233730 | *LINC01765* | -0.1629722 | 0.069882466 | 0.019147881 | 0.050734585 | 0.999933887 | 0.403169005 | 0.335429533 | 0.067739472 | 0.926416674 |
| ENSG00000256537 | *SMIM10L1* | 1.520223572 | -0.15962997 | 0.017063672 | -0.176693641 | 0.999933887 | -0.044243364 | -0.01737955 | -0.026863814 | 0.926430176 |
| ENSG00000144357 | *UBR3* | 4.177087891 | 0.066433615 | -0.010102704 | 0.076536319 | 0.999933887 | -0.466342718 | -0.501485044 | 0.035142326 | 0.926430176 |
| ENSG00000278828 | *H3C10* | -0.300630724 | -0.036714731 | 0.1666293 | -0.203344031 | 0.999933887 | -0.336072176 | -0.400584483 | 0.064512307 | 0.926430176 |
| ENSG00000163739 | *CXCL1* | 8.939171249 | 0.440437877 | 0.596474159 | -0.156036282 | 0.999933887 | 1.162593501 | 1.189369428 | -0.026775928 | 0.926717503 |
| ENSG00000196652 | *ZKSCAN5* | 3.257184427 | 0.07014548 | -0.063864955 | 0.134010435 | 0.999933887 | 0.017911694 | 0.040578794 | -0.0226671 | 0.926717503 |
| ENSG00000196683 | *TOMM7* | 4.691865398 | -0.007109047 | -0.000794452 | -0.006314596 | 0.999933887 | 0.109567079 | 0.130348372 | -0.020781293 | 0.926736226 |
| ENSG00000128284 | *APOL3* | 3.735919874 | -0.04209705 | -0.019520093 | -0.022576957 | 0.999933887 | 0.185648142 | 0.167144514 | 0.018503627 | 0.926866973 |
| ENSG00000185862 | *EVI2B* | 7.935203458 | -0.1116516 | -0.210363431 | 0.098711831 | 0.999933887 | -1.390451486 | -1.424360647 | 0.033909161 | 0.926946836 |
| ENSG00000115268 | *RPS15* | 7.223560872 | 0.064323798 | -0.013552757 | 0.077876554 | 0.999933887 | 0.052376242 | 0.035161562 | 0.017214681 | 0.927026258 |
| ENSG00000279631 | *AL158211.5* | 0.155605758 | 0.146658381 | 0.296956242 | -0.150297862 | 0.999933887 | 0.08571986 | 0.128388949 | -0.042669089 | 0.927073902 |
| ENSG00000224745 | *AC063965.2* | 1.813834275 | 0.094963576 | 0.135545882 | -0.040582306 | 0.999933887 | -0.327116886 | -0.387491526 | 0.06037464 | 0.927267229 |
| ENSG00000117697 | *NSL1* | 2.596253418 | -0.000924863 | -0.010874163 | 0.0099493 | 0.999933887 | -0.213381373 | -0.241379409 | 0.027998036 | 0.927267229 |
| ENSG00000112245 | *PTP4A1* | -0.270457122 | 0.521212459 | 0.340309008 | 0.180903451 | 0.999933887 | 0.382586321 | 0.298267355 | 0.084318966 | 0.927399892 |
| ENSG00000233360 | *PDXP-DT* | -0.34690152 | 0.063762751 | -0.028322384 | 0.092085136 | 0.999933887 | -0.44068812 | -0.385114466 | -0.055573654 | 0.927399892 |
| ENSG00000187116 | *LILRA5* | 5.123742286 | 0.014620701 | 0.096229649 | -0.081608948 | 0.999933887 | -0.085696326 | -0.118286575 | 0.032590249 | 0.927399892 |
| ENSG00000143850 | *PLEKHA6* | -0.266394371 | -0.047914597 | 0.3472567 | -0.395171297 | 0.999933887 | 0.0005508 | 0.052833976 | -0.052283177 | 0.927430143 |
| ENSG00000182307 | *C8orf33* | 2.279388927 | 0.091458797 | -0.008710055 | 0.100168853 | 0.999933887 | 0.198837914 | 0.173223051 | 0.025614862 | 0.927477231 |
| ENSG00000175048 | *ZDHHC14* | 0.368834955 | -0.075984413 | 0.193503242 | -0.269487655 | 0.999933887 | -0.084172042 | -0.040333157 | -0.043838885 | 0.927477231 |
| ENSG00000279766 | *AC067931.1* | 0.216259309 | 0.062971072 | -0.125842536 | 0.188813608 | 0.999933887 | 0.761143841 | 0.810380439 | -0.049236598 | 0.927477231 |
| ENSG00000099341 | *PSMD8* | 5.685867968 | 0.029195514 | 0.034491716 | -0.005296201 | 0.999933887 | -0.056777988 | -0.043885426 | -0.012892561 | 0.927626979 |
| ENSG00000134489 | *HRH4* | 1.571651014 | -0.084990541 | 0.061515138 | -0.14650568 | 0.999933887 | 0.111839029 | 0.143950604 | -0.032111576 | 0.927660188 |
| ENSG00000123992 | *DNPEP* | 3.284805296 | 0.048623223 | -0.050225422 | 0.098848645 | 0.999933887 | 0.08761136 | 0.068858666 | 0.018752694 | 0.92771619 |
| ENSG00000223724 | *RAD17P2* | 0.053243243 | -0.084204652 | 0.060459491 | -0.144664143 | 0.999933887 | -0.175221615 | -0.103173904 | -0.072047711 | 0.92771619 |
| ENSG00000179859 | *RNF227* | 0.212733136 | -0.032631308 | 0.170190157 | -0.202821466 | 0.999933887 | 0.159717997 | 0.204298614 | -0.044580616 | 0.92771619 |
| ENSG00000197217 | *ENTPD4* | 6.431421825 | -0.043944304 | -0.075572747 | 0.031628444 | 0.999933887 | -0.156052311 | -0.167351995 | 0.011299683 | 0.92795009 |

| | | | | | | | | | | |
|---|---|---|---|---|---|---|---|---|---|---|
| ENSG00000265590 | *CFAP298-TCP* | -1.160754171 | -0.255306794 | -0.34794119 | 0.092634396 | 0.999933887 | -0.112776982 | -0.033158525 | -0.079618456 | 0.92795009 |
| ENSG00000167468 | *GPX4* | 6.209886209 | 0.052511145 | -0.041432962 | 0.093944107 | 0.999933887 | 0.431250883 | 0.416520037 | 0.014730847 | 0.92795009 |
| ENSG00000105229 | *PIAS4* | 5.331691572 | -0.046733281 | 0.04598802 | -0.092721301 | 0.999933887 | -0.274451461 | -0.260466097 | -0.013985364 | 0.92795009 |
| ENSG00000100985 | *MMP9* | 7.421891859 | 0.030805047 | -0.018054832 | 0.048859879 | 0.999933887 | -0.134198213 | -0.113754694 | -0.020443519 | 0.92795009 |
| ENSG00000101230 | *ISM1* | -1.25284795 | 0.058571306 | 0.015556491 | 0.043014815 | 0.999933887 | 0.001335474 | -0.096106976 | 0.097442451 | 0.92795009 |
| ENSG00000162692 | *VCAM1* | 0.007418984 | 0.039198498 | -0.008302904 | 0.047501402 | 0.999933887 | 0.051286414 | 0.107528104 | -0.05624169 | 0.92795009 |
| ENSG00000284882 | *AL359762.1* | 2.320526039 | -0.012790433 | -0.051146026 | 0.038355593 | 0.999933887 | -0.048252643 | -0.020717965 | -0.027534677 | 0.92795009 |
| ENSG00000100387 | *RBX1* | 3.557557547 | -0.007376369 | -0.061714738 | 0.054338369 | 0.999933887 | 0.00668965 | 0.025231374 | -0.018541724 | 0.92795009 |
| ENSG00000286997 | *AC005856.1* | -0.546078827 | -0.009775814 | 0.157640869 | -0.167416683 | 0.999933887 | 0.037010457 | 0.131646188 | -0.09463573 | 0.92795009 |
| ENSG00000269028 | *MTRNR2L12* | 0.963962935 | 0.015935598 | 0.133832863 | -0.117897265 | 0.999933887 | 0.351781173 | 0.316355222 | 0.035425951 | 0.928030738 |
| ENSG00000272247 | *AC080013.4* | -0.478709285 | 0.28305423 | 0.485115819 | -0.202061589 | 0.999933887 | 0.657287826 | 0.579368363 | 0.077919463 | 0.928106706 |
| ENSG00000285897 | *AC004687.2* | 0.073635527 | 0.138858479 | 0.171757714 | -0.032899235 | 0.999933887 | 0.120639421 | 0.064464431 | 0.05617499 | 0.928337918 |
| ENSG00000215397 | *SCRT2* | 0.697129452 | 0.114421485 | 0.225542864 | -0.111121379 | 0.999933887 | -0.273008256 | -0.307945051 | 0.034936794 | 0.928432104 |
| ENSG00000269564 | *AC008753.2* | -1.336873904 | 0.155100263 | 0.384933099 | -0.229832836 | 0.999933887 | -0.412061159 | -0.536148253 | 0.124087094 | 0.928488959 |
| ENSG00000223551 | *TMSB4XP4* | 1.149857781 | -0.086389157 | 0.065936536 | -0.152325693 | 0.999933887 | 0.431878824 | 0.392238056 | 0.039640768 | 0.928488959 |
| ENSG00000148204 | *CRB2* | -1.430313067 | -0.00543221 | 0.035679689 | -0.041111899 | 0.999933887 | -0.02822611 | 0.08070363 | -0.10892974 | 0.928488959 |
| ENSG00000168874 | *ATOH8* | -0.230713959 | -0.177864182 | 0.05621409 | -0.234078271 | 0.999933887 | 3.523120825 | 3.58010452 | -0.056983695 | 0.928572421 |
| ENSG00000117222 | *RBBP5* | 4.332959243 | -0.135958703 | 0.027361048 | -0.16331975 | 0.999933887 | -0.265959678 | -0.281937731 | 0.015978053 | 0.928578084 |
| ENSG00000129562 | *DAD1* | 5.306326351 | 0.050640332 | -0.015044901 | 0.065685233 | 0.999933887 | -0.080370346 | -0.097477231 | 0.017106884 | 0.928578084 |
| ENSG00000261055 | *AL450468.2* | 2.27902893 | 0.230511911 | 0.061263394 | 0.169248517 | 0.999933887 | 0.593259611 | 0.624512989 | -0.031253377 | 0.928608193 |
| ENSG00000110429 | *FBXO3* | 4.59177982 | -0.059897237 | -0.001281988 | -0.05861525 | 0.999933887 | -0.220356183 | -0.236081466 | 0.015725283 | 0.928608193 |
| ENSG00000105227 | *PRX* | 0.198620393 | 0.144578622 | 0.25052289 | -0.105944268 | 0.999933887 | 0.253772603 | 0.208222503 | 0.0455501 | 0.928608193 |
| ENSG00000233912 | *AC026202.2* | -0.762967059 | -0.00749631 | 0.291632984 | -0.299129294 | 0.999933887 | 0.691946797 | 0.63254675 | 0.059400048 | 0.92863075 |
| ENSG00000168496 | *FEN1* | 1.573852276 | -0.125896025 | -0.023948918 | -0.101947108 | 0.999933887 | 0.115384262 | 0.14757425 | -0.032189988 | 0.928960262 |
| ENSG00000107779 | *BMPR1A* | 0.196857583 | -0.080671205 | 0.159675797 | -0.240347003 | 0.999933887 | -0.162770753 | -0.208941192 | 0.046170439 | 0.928960262 |
| ENSG00000115977 | *AAK1* | 5.348570796 | 0.027405156 | 0.013870939 | 0.013534217 | 0.999933887 | -0.205929471 | -0.223336194 | 0.017406722 | 0.928960262 |
| ENSG00000151553 | *FAM160B1* | 5.269732764 | -0.031970962 | -0.081299142 | 0.04932818 | 0.999933887 | -0.70253291 | -0.724800521 | 0.02226761 | 0.928960262 |
| ENSG00000125122 | *LRRC29* | 1.665192035 | -0.116087499 | -0.050100353 | -0.065987147 | 0.999933887 | 0.152027405 | 0.120658535 | 0.03136887 | 0.929139827 |
| ENSG00000269737 | *AL691432.1* | 1.34104688 | -0.064945654 | 0.123285591 | -0.188231245 | 0.999933887 | -0.235978219 | -0.273228202 | 0.037249983 | 0.929139827 |
| ENSG00000101189 | *MRGBP* | 4.864691046 | 0.038128571 | 0.039853293 | -0.001724722 | 0.999933887 | 0.708550531 | 0.729080979 | -0.020530448 | 0.929139827 |
| ENSG00000104129 | *DNAJC17* | 1.431532685 | -0.002352599 | -0.091217279 | 0.08886468 | 0.999933887 | -0.09626109 | -0.126711698 | 0.030450608 | 0.929604043 |
| ENSG00000223969 | *AC002456.1* | 2.88744967 | 0.324203098 | 0.005941761 | 0.318261337 | 0.999933887 | 1.810543634 | 1.846170594 | -0.035626961 | 0.929738368 |
| ENSG00000225528 | *Z82206.1* | -0.555042766 | -0.212371417 | -0.256949431 | 0.044578014 | 0.999933887 | -0.065192139 | -0.126388771 | 0.061196632 | 0.929738368 |
| ENSG00000176049 | *JAKMIP2* | 2.449355812 | -0.201875634 | -0.016738732 | -0.185136903 | 0.999933887 | -0.778854479 | -0.802026095 | 0.023171616 | 0.929738368 |
| ENSG00000187601 | *MAGEH1* | 0.628617494 | 0.068333224 | 0.062735465 | 0.005597758 | 0.999933887 | 0.08335679 | 0.038758826 | 0.044597964 | 0.929738368 |
| ENSG00000110628 | *SLC22A18* | 6.095696321 | -0.003041566 | 0.017527081 | -0.020568647 | 0.999933887 | 0.309404274 | 0.293502829 | 0.015901444 | 0.929738368 |
| ENSG00000263606 | *CHORDC1P4* | 0.79949526 | -0.146528832 | 0.042118954 | -0.188647786 | 0.999933887 | -1.164972689 | -1.113459889 | -0.051512799 | 0.929741226 |
| ENSG00000153064 | *BANK1* | 0.953430212 | 0.052483356 | -0.220951916 | 0.273435272 | 0.999933887 | -0.208344971 | -0.24130899 | 0.032964019 | 0.929741226 |
| ENSG00000220804 | *LINC01881* | 1.260122367 | -0.019100389 | 0.152889734 | -0.171990124 | 0.999933887 | -0.213588721 | -0.186031602 | -0.027557119 | 0.929741226 |
| ENSG00000270504 | *AL391422.4* | -0.403889494 | -0.123563913 | 0.003073247 | -0.12663716 | 0.999933887 | 1.103600536 | 1.064594443 | 0.039006092 | 0.929750854 |
| ENSG00000274364 | *AL110115.1* | -0.359311441 | -0.145415855 | 0.169500656 | -0.314916511 | 0.999933887 | 0.1239292 | 0.058662255 | 0.065266945 | 0.929878335 |
| ENSG00000047346 | *FAM214A* | 4.96740301 | 0.052206006 | -0.014034881 | 0.066240887 | 0.999933887 | -0.628705709 | -0.596658744 | -0.032046965 | 0.929878335 |
| ENSG00000108773 | *KAT2A* | 3.090588194 | 0.062054972 | 0.009829938 | 0.052225033 | 0.999933887 | 0.126157173 | 0.14613863 | -0.019981457 | 0.929943985 |
| ENSG00000221890 | *NPTXR* | 0.129425416 | 0.117362744 | 0.0721165 | 0.045246243 | 0.999933887 | 0.088375477 | 0.039279457 | 0.04909602 | 0.929943985 |
| ENSG00000112118 | *MCM3* | 2.497312702 | -0.078178069 | 0.091940581 | -0.17011865 | 0.999933887 | 0.184866597 | 0.15931177 | 0.025554828 | 0.929943985 |
| ENSG00000143674 | *MAP3K21* | 0.430764442 | -0.10488133 | 0.138789304 | -0.243670634 | 0.999933887 | -0.263765254 | -0.31702465 | 0.053259396 | 0.929943985 |
| ENSG00000204650 | *LINC02210* | 0.558554104 | -0.016094798 | 0.029201252 | -0.04529605 | 0.999933887 | 0.166261285 | 0.127115079 | 0.039146206 | 0.929943985 |
| ENSG00000156239 | *N6AMT1* | -0.315048675 | -0.210254609 | 0.350724963 | -0.560979572 | 0.999933887 | -0.055237825 | -0.009291338 | -0.045946487 | 0.930060235 |
| ENSG00000089682 | *RBM41* | 0.390527503 | -0.135777638 | -0.258773492 | 0.122995854 | 0.999933887 | -0.160222575 | -0.2123465 | 0.052123925 | 0.930060235 |
| ENSG00000279821 | *AC145098.2* | 3.260451964 | 0.050254809 | 0.113364957 | -0.063110148 | 0.999933887 | 0.650261489 | 0.626847484 | 0.023414005 | 0.930060235 |
| ENSG00000138061 | *CYP1B1* | 4.025879229 | -0.051117836 | 0.062271374 | -0.11338921 | 0.999933887 | -0.367947846 | -0.396400116 | 0.02845227 | 0.930060235 |
| ENSG00000213654 | *GPSM3* | 9.225698403 | 0.012396424 | -0.021013326 | 0.03340975 | 0.999933887 | 0.07481784 | 0.062574267 | 0.012243572 | 0.930060235 |
| ENSG00000187017 | *ESPN* | -0.539654186 | -0.03833996 | 0.203397611 | -0.241737572 | 0.999933887 | 0.068286986 | 0.126060062 | -0.057773076 | 0.930292864 |
| ENSG00000001630 | *CYP51A1* | -0.266017845 | 0.239713304 | 0.054864944 | 0.184848359 | 0.999933887 | 0.028010905 | -0.04269461 | 0.070705515 | 0.930310413 |
| ENSG00000282572 | *AC215522.2* | 4.209905505 | 0.136961921 | 0.017135339 | 0.119826582 | 0.999933887 | 0.405831768 | 0.386182119 | 0.019649648 | 0.930333472 |
| ENSG00000269982 | *AC018809.2* | 1.922289176 | -0.126183753 | -0.118317537 | -0.007866217 | 0.999933887 | -0.5435613 | -0.514267546 | -0.029293754 | 0.930333472 |
| ENSG00000234769 | *WASH4P* | -0.740343583 | 0.188030898 | -0.286665065 | 0.474695963 | 0.999933887 | -0.741022218 | -0.790929911 | 0.049907693 | 0.930333472 |
| ENSG00000122507 | *BBS9* | 1.379310038 | 0.101306543 | -0.022788048 | 0.124094591 | 0.999933887 | -0.025803372 | 0.009013362 | -0.034816733 | 0.930521546 |
| ENSG00000162607 | *USP1* | 3.863329345 | -0.05143902 | -0.135397534 | 0.083958514 | 0.999933887 | -0.420772092 | -0.39469953 | -0.026072562 | 0.930521546 |
| ENSG00000172159 | *FRMD3* | 1.856569558 | -0.084036986 | 0.096656088 | -0.180693074 | 0.999933887 | -1.315180601 | -1.354145337 | 0.038964736 | 0.930521546 |
| ENSG00000155265 | *GOLGA7B* | 1.448692283 | 0.042693916 | 0.039904982 | 0.002788934 | 0.999933887 | 0.014856155 | -0.016511753 | 0.031367908 | 0.930521546 |
| ENSG00000116005 | *PCYOX1* | 1.684764683 | -0.033440956 | -0.170262808 | 0.136821852 | 0.999933887 | -0.025411552 | 0.004103681 | -0.029515233 | 0.930521546 |
| ENSG00000239900 | *ADSL* | -0.278235644 | -0.050630184 | -0.317908114 | 0.26727793 | 0.999933887 | -0.027693689 | -0.075703339 | 0.04800965 | 0.930521546 |
| ENSG00000264937 | *AC100830.2* | 1.269124626 | -0.031775456 | -0.134847422 | 0.103071966 | 0.999933887 | 0.329504904 | 0.295086291 | 0.034418613 | 0.930521546 |
| ENSG00000108828 | *VAT1* | 4.669727773 | 0.005618235 | -0.010313541 | 0.015931775 | 0.999933887 | -0.872689944 | -0.8958688 | 0.023178856 | 0.930521546 |
| ENSG00000271428 | *MPHOSPH6P1* | -0.553380291 | -0.006506938 | 0.227016712 | -0.23352365 | 0.999933887 | 0.251934181 | 0.338463052 | -0.086528871 | 0.930521546 |
| ENSG00000175322 | *ZNF519* | -0.817472536 | -0.348351147 | -0.180279019 | -0.168072127 | 0.999933887 | -0.055027075 | 0.012834135 | -0.067861209 | 0.930659767 |
| ENSG00000130592 | *LSP1* | 9.984119505 | 0.037566964 | 0.038493197 | -0.000926233 | 0.999933887 | -0.292086951 | -0.309443571 | 0.017356619 | 0.930697519 |
| ENSG00000157693 | *TMEM268* | 3.031188195 | -0.054456768 | -0.071805299 | 0.017348531 | 0.999933887 | -0.081645614 | -0.101858917 | 0.020213303 | 0.930763809 |
| ENSG00000077463 | *SIRT6* | 4.79202749 | 0.030083183 | 0.031926996 | -0.001843813 | 0.999933887 | 0.988168616 | 1.011054699 | -0.022886082 | 0.930763809 |
| ENSG00000148444 | *COMMD3* | 1.712937186 | -0.005399188 | -0.190282022 | 0.184882833 | 0.999933887 | -0.159693272 | -0.184926588 | 0.025233315 | 0.931096767 |
| ENSG00000225850 | *AL355490.1* | -0.319162746 | -0.264945821 | -0.11756373 | -0.147382091 | 0.999933887 | -1.469158552 | -1.541586634 | 0.072428083 | 0.93130303 |
| ENSG00000148229 | *POLE3* | 3.580522103 | -0.046019292 | -0.002417989 | -0.043601303 | 0.999933887 | 0.179115735 | 0.16258388 | 0.016531855 | 0.93130303 |
| ENSG00000153933 | *DGKE* | 3.26726716 | -0.055733814 | -0.003916519 | -0.051817296 | 0.999933887 | -0.617129002 | -0.590246483 | -0.026882519 | 0.931355051 |
| ENSG00000144034 | *TPRKB* | 1.712602414 | -0.115650684 | 0.077996865 | -0.193647549 | 0.999933887 | -0.116417637 | -0.088392011 | -0.028025626 | 0.931422379 |
| ENSG00000104964 | *TLE5* | 6.157058448 | 0.068211224 | 0.02896178 | 0.039249444 | 0.999933887 | 0.065111578 | 0.080137161 | -0.015025583 | 0.932038864 |
| ENSG00000204316 | *MRPL38* | 0.040370001 | -0.190942172 | 0.030226261 | -0.221168433 | 0.999933887 | 0.083655932 | 0.046141951 | 0.037513981 | 0.932181793 |
| ENSG00000111752 | *PHC1* | 5.064052509 | -0.051045447 | -0.069483371 | 0.018437924 | 0.999933887 | -0.257846333 | -0.242812126 | -0.015034207 | 0.932181793 |
| ENSG00000196141 | *SPATS2L* | 0.619902474 | -0.073678533 | 0.01806368 | -0.091742213 | 0.999933887 | -0.262786397 | -0.308034132 | 0.045247735 | 0.932181793 |
| ENSG00000011028 | *MRC2* | -0.885938998 | -0.103445568 | 0.116522868 | -0.219968436 | 0.999933887 | 0.034763749 | 0.101584699 | -0.06682095 | 0.932181793 |
| ENSG00000247373 | *TMED2-DT* | -0.018470752 | -0.035448466 | 0.049020533 | -0.084468999 | 0.999933887 | -0.219759381 | -0.176904428 | -0.042854953 | 0.932347917 |
| ENSG00000260034 | *LCMT1-AS2* | -0.8108901 | 0.195840937 | 0.061702856 | 0.134138081 | 0.999933887 | 0.10615132 | 0.174448964 | -0.068297643 | 0.932399574 |
| ENSG00000154217 | *PITPNC1* | 5.170568802 | 0.020360795 | 0.024143787 | -0.003782992 | 0.999933887 | -0.005071506 | -0.018666909 | 0.013595403 | 0.932697997 |
| ENSG00000206527 | *HACD2* | 2.530378319 | -0.015189629 | -0.079019021 | 0.063829392 | 0.999933887 | 0.021560221 | 0.045572662 | -0.024012441 | 0.932979558 |
| ENSG00000169871 | *TRIM56* | 6.712487542 | -0.014996406 | -0.0257554 | 0.010758994 | 0.999933887 | -0.039466927 | -0.054231795 | 0.014764868 | 0.933123104 |
| ENSG00000286545 | *AC092645.2* | -0.685649019 | -0.007331525 | 0.107129101 | -0.114460626 | 0.999933887 | -0.210135349 | -0.266434415 | 0.056299066 | 0.933123104 |
| ENSG00000263050 | *AC090617.5* | 0.558958046 | 0.052231938 | -0.03371112 | 0.085943058 | 0.999933887 | 0.911304206 | 0.967163023 | -0.055858817 | 0.933442544 |
| ENSG00000107742 | *SPOCK2* | 6.784501674 | 0.010555212 | 0.038343858 | -0.027788647 | 0.999933887 | 0.056118965 | 0.070109267 | -0.013990302 | 0.933442544 |
| ENSG00000140400 | *MAN2C1* | 4.991072366 | 0.070582958 | 0.107148798 | -0.03656584 | 0.999933887 | 0.154179446 | 0.140307555 | 0.013871891 | 0.933821092 |
| ENSG00000204592 | *HLA-E* | 12.368537423 | 0.04647761 | 0.012309039 | 0.03416857 | 0.999933887 | 0.12655634 | 0.138361188 | -0.011804848 | 0.933821092 |
| ENSG00000186566 | *GPATCH8* | 5.146422125 | -0.015825501 | -0.02905636 | 0.013230859 | 0.999933887 | -0.282741525 | -0.269789998 | -0.012951527 | 0.933821092 |
| ENSG00000135775 | *COG2* | 1.992778947 | -0.005978774 | -0.060526303 | 0.054547529 | 0.999933887 | 0.095521562 | 0.120073567 | -0.024552004 | 0.933821092 |
| ENSG00000105518 | *TMEM205* | 3.038383506 | -0.056116001 | -0.026339553 | -0.029776448 | 0.999933887 | 1.169883621 | 1.19213183 | -0.022248209 | 0.933842372 |
| ENSG00000231752 | *EMBP1* | 0.337341242 | -0.019217552 | 0.04980539 | -0.069022942 | 0.999933887 | 0.090176402 | 0.033746932 | 0.05642947 | 0.933922095 |
| ENSG00000164713 | *BRI3* | 9.390286805 | 0.02866621 | 0.003507111 | 0.025159099 | 0.999933887 | -0.082674618 | -0.095422201 | 0.012747583 | 0.933970893 |
| ENSG00000147140 | *NONO* | 6.810813067 | -0.033273363 | -0.02874872 | -0.004524643 | 0.999933887 | 0.036060336 | 0.045545553 | -0.009485217 | 0.934081655 |
| ENSG00000242028 | *HYPK* | 0.822880019 | -0.101251489 | 0.168940761 | -0.27019225 | 0.999933887 | 0.221963021 | 0.183318129 | 0.038644892 | 0.934157221 |
| ENSG00000198563 | *DDX39B* | 7.046381859 | -0.012276914 | 0.003553857 | -0.015830771 | 0.999933887 | -0.012537334 | -0.023685265 | 0.011147931 | 0.934423772 |
| ENSG00000189369 | *GSPT2* | -0.094663009 | -0.015605439 | -0.173914754 | 0.158309314 | 0.999933887 | 0.122842269 | 0.072348327 | 0.050493942 | 0.934548069 |
| ENSG00000260868 | *LINC01960* | -0.625494862 | -0.177262934 | 0.232122596 | -0.40938553 | 0.999933887 | -1.149499086 | -1.063308852 | -0.086190234 | 0.934588556 |
| ENSG00000236525 | *AC007278.2* | 0.95950898 | 0.006185595 | -0.116937486 | 0.123123081 | 0.999933887 | 0.024865607 | 0.11219026 | -0.087324654 | 0.934588556 |
| ENSG00000167969 | *ECI1* | 1.772752953 | 0.070637024 | 0.150485711 | -0.079848687 | 0.999933887 | -0.024083237 | -0.051290528 | 0.027207292 | 0.934608851 |
| ENSG00000228589 | *SPCS2P4* | -0.674997017 | -0.132942083 | 0.101322414 | -0.234264497 | 0.999933887 | 1.459064134 | 1.503629031 | -0.044564897 | 0.934749817 |
| ENSG00000266088 | *AC004585.1* | -0.927875291 | 0.025297544 | 0.061938067 | -0.036640523 | 0.999933887 | 0.007404731 | -0.049553077 | 0.056957808 | 0.934749817 |
| ENSG00000114439 | *BBX* | 4.865620148 | -0.012050544 | -0.051038891 | 0.038988346 | 0.999933887 | -0.30113011 | -0.330649723 | 0.029519613 | 0.935163653 |

| | | | | | | | | | | |
|---|---|---|---|---|---|---|---|---|---|---|
| ENSG00000070526 | *ST6GALNAC1* | -1.139696361 | -0.029021088 | -0.049082158 | 0.02006107 | 0.999933887 | 0.047612368 | 0.114954793 | -0.067342425 | 0.935224292 |
| ENSG00000133065 | *SLC41A1* | 2.700589778 | -0.040958788 | 0.048037856 | -0.088996645 | 0.999933887 | 0.115755374 | 0.095253513 | 0.020501861 | 0.935287826 |
| ENSG00000230084 | *AC006059.1* | 1.176669039 | 0.072811226 | -0.062171115 | 0.134982341 | 0.999933887 | -0.642777238 | -0.590521424 | -0.052255814 | 0.935351363 |
| ENSG00000273295 | *AP000350.6* | -1.547114734 | 0.039071477 | 0.241772461 | -0.202700984 | 0.999933887 | -0.084098866 | -0.162369678 | 0.078270812 | 0.935351363 |
| ENSG00000103653 | *CSK* | 7.926297137 | 0.014186303 | 0.026964265 | -0.012777962 | 0.999933887 | 0.030087981 | 0.042360962 | -0.012272981 | 0.935387513 |
| ENSG00000156711 | *MAPK13* | 8.106888896 | 0.013422331 | 0.054289626 | -0.040867294 | 0.999933887 | -0.053016012 | -0.040235788 | -0.012780224 | 0.935426616 |
| ENSG00000204217 | *BMPR2* | 2.838131184 | 0.102617825 | -0.056241118 | 0.158858943 | 0.999933887 | -0.385033629 | -0.41499805 | 0.029964421 | 0.935472479 |
| ENSG00000170734 | *POLH* | 2.605980338 | 0.005892646 | 0.090075688 | -0.084183042 | 0.999933887 | -0.057781972 | -0.076669079 | 0.018887108 | 0.935580854 |
| ENSG00000271109 | *AC008555.4* | 0.243184886 | 0.056987817 | -0.035192343 | 0.09218016 | 0.999933887 | -0.040545341 | -0.000639227 | -0.039906114 | 0.935789227 |
| ENSG00000228318 | *AP001610.2* | 0.703068645 | -0.036858388 | 0.148887393 | -0.185745782 | 0.999933887 | -0.206888657 | -0.166845807 | -0.040042849 | 0.935899542 |
| ENSG00000137275 | *RIPK1* | 5.414511293 | -0.053207311 | -0.051497273 | -0.001710038 | 0.999933887 | -0.251410882 | -0.264966843 | 0.013555961 | 0.936195323 |
| ENSG00000257277 | *AC092652.1* | 0.76039083 | 0.158432301 | 0.261947131 | -0.10351483 | 0.999933887 | 0.026076367 | 0.112163849 | -0.086087482 | 0.936195323 |
| ENSG00000185101 | *ANO9* | 3.804458436 | 0.11645864 | 0.115539762 | 0.000918878 | 0.999933887 | 0.185051404 | 0.167519179 | 0.017532225 | 0.936196588 |
| ENSG00000234028 | *EIF2AK3-DT* | 1.866692955 | 0.247338012 | 0.148325682 | 0.09901233 | 0.999933887 | 0.832941706 | 0.875533543 | -0.042591837 | 0.936196588 |
| ENSG00000287255 | *AC007877.1* | 0.790073865 | -0.103316031 | -0.014719035 | -0.088596995 | 0.999933887 | 0.170578226 | 0.131311946 | 0.039266279 | 0.936196588 |
| ENSG00000267169 | *AC022098.1* | 0.731765 | -0.087729368 | 0.157928631 | -0.245657999 | 0.999933887 | 0.430081404 | 0.463666382 | -0.033584978 | 0.936196588 |
| ENSG00000140470 | *ADAMTS17* | -0.627067271 | -0.056428769 | -0.232288581 | 0.175859812 | 0.999933887 | 0.012326583 | 0.064522983 | -0.052196401 | 0.936196588 |
| ENSG00000184343 | *SRPK3* | -0.761747287 | 0.067497771 | -0.033634287 | 0.101132057 | 0.999933887 | 0.082939949 | 0.028917888 | 0.054022061 | 0.936196588 |
| ENSG00000163930 | *BAP1* | 6.691503922 | 0.010092466 | 0.008276419 | 0.001816047 | 0.999933887 | 0.013136131 | 0.024560322 | -0.011424191 | 0.936196588 |
| ENSG00000249870 | *AC005920.2* | 0.907086664 | 0.142502839 | 0.364402172 | -0.221899333 | 0.999933887 | 0.909929828 | 0.949799875 | -0.039870047 | 0.936222942 |
| ENSG00000165714 | *BORCS5* | 3.255588798 | -0.030115402 | -0.031736665 | 0.001621263 | 0.999933887 | 0.217747873 | 0.241859963 | -0.024112091 | 0.936283283 |
| ENSG00000248936 | *AC027607.1* | -0.239962278 | -0.275558421 | 0.331967227 | -0.607525648 | 0.594252256 | 0.268708621 | 0.338568449 | -0.069859827 | 0.936430487 |
| ENSG00000261430 | *AL031600.2* | 1.296349643 | 0.131423623 | 0.140256572 | -0.008832949 | 0.999933887 | 0.889981395 | 0.926889758 | -0.036908363 | 0.936430487 |
| ENSG00000135424 | *ITGA7* | 3.746195027 | 0.030834571 | 0.022931711 | 0.007902859 | 0.999933887 | -0.13901602 | -0.155888657 | 0.016872637 | 0.936430487 |
| ENSG00000250734 | *AL391335.1* | 0.364874947 | 0.025677921 | -0.096440392 | 0.122118313 | 0.999933887 | 0.338883181 | 0.391572159 | -0.052688979 | 0.936430487 |
| ENSG00000180098 | *TRNAU1AP* | 2.31853665 | -0.080358953 | -0.042977977 | -0.037380977 | 0.999933887 | 0.049655888 | 0.028297079 | 0.021358808 | 0.93648854 |
| ENSG00000247092 | *SNHG10* | 0.457014977 | -0.081936333 | -0.239895319 | 0.157958986 | 0.999933887 | 0.248211506 | 0.284390211 | -0.036178706 | 0.936703649 |
| ENSG00000125995 | *ROMO1* | 2.832130702 | 0.042237304 | 0.00895051 | 0.033286793 | 0.999933887 | 0.101857546 | 0.121826044 | -0.019968497 | 0.93687145 |
| ENSG00000159259 | *CHAF1B* | 0.297462561 | 0.004406754 | -0.047096529 | 0.051503283 | 0.999933887 | -0.29017005 | -0.338488017 | 0.048317966 | 0.93687145 |
| ENSG00000011600 | *TYROBP* | 9.281110636 | 0.087804682 | 0.003488429 | 0.084316253 | 0.999933887 | 0.235880825 | 0.222661661 | 0.013219164 | 0.936902934 |
| ENSG00000236234 | *AC091132.2* | 0.952504663 | 0.08958136 | -0.132203862 | 0.221785222 | 0.999933887 | 0.053304583 | 0.087482591 | -0.034178007 | 0.936902934 |
| ENSG00000107036 | *RIC1* | 4.503620748 | -0.018454728 | -0.029442699 | 0.010987971 | 0.999933887 | -0.091317195 | -0.106552175 | 0.01523498 | 0.936902934 |
| ENSG00000254396 | *AL355432.1* | -0.774525434 | 0.00584029 | 0.185951936 | -0.180111646 | 0.999933887 | -0.309709954 | -0.359504093 | 0.04979414 | 0.936902934 |
| ENSG00000186468 | *RPS23* | 5.074802305 | 0.035317973 | -0.034589737 | 0.06990771 | 0.999933887 | 0.139740538 | 0.123109628 | 0.01663091 | 0.937050664 |
| ENSG00000197894 | *ADH5* | 2.734122499 | -0.011198838 | -0.114381513 | 0.103182675 | 0.999933887 | 0.056843996 | 0.075940645 | -0.01909665 | 0.937075962 |
| ENSG00000279930 | *AL032819.2* | 0.87390537 | 0.294164906 | 0.111111361 | 0.183053545 | 0.999933887 | 0.750087051 | 0.778961126 | -0.028874075 | 0.937093591 |
| ENSG00000254682 | *AP002387.1* | 0.552387426 | 0.138654744 | 0.036820377 | 0.101834367 | 0.999933887 | 0.023610497 | 0.065591578 | -0.041981081 | 0.93716166 |
| ENSG00000234436 | *AC245884.2* | -0.217676426 | -0.057506712 | -0.115641669 | 0.058134957 | 0.999933887 | 0.204250298 | 0.283565719 | -0.079315421 | 0.93716166 |
| ENSG00000185088 | *RPS27L* | 3.839207536 | -0.074981787 | -0.043367313 | -0.031614474 | 0.999933887 | -0.357921274 | -0.340569327 | -0.017351947 | 0.937205787 |
| ENSG00000196876 | *SCN8A* | -0.074916588 | -0.227993096 | -0.067478045 | -0.16051505 | 0.999933887 | -0.343542828 | -0.379525692 | 0.035982864 | 0.93736876 |
| ENSG00000135452 | *TSPAN31* | 2.524732069 | 0.033200586 | -0.025518174 | 0.05871876 | 0.999933887 | 0.206435001 | 0.185103229 | 0.021331772 | 0.937412599 |
| ENSG00000163781 | *TOPBP1* | 4.057192888 | -0.064586855 | -0.038217201 | -0.026369654 | 0.999933887 | -0.485811408 | -0.513405828 | 0.02759442 | 0.937545626 |
| ENSG00000186625 | *KATNA1* | 2.384344906 | -0.076338669 | 0.026232242 | -0.102570911 | 0.999933887 | -0.191074791 | -0.210872436 | 0.019797646 | 0.937733382 |
| ENSG00000151006 | *PRSS53* | 0.926076464 | 0.091734374 | 0.220771292 | -0.129036918 | 0.999933887 | 0.068483832 | 0.102353259 | -0.033869428 | 0.937733382 |
| ENSG00000278619 | *MRM1* | -0.720183399 | 0.127069997 | 0.321887245 | -0.194817247 | 0.999933887 | 0.284773387 | 0.243056143 | 0.041717244 | 0.937733382 |
| ENSG00000146674 | *IGFBP3* | 0.673093131 | -0.016477935 | 0.028033158 | -0.044511093 | 0.999933887 | 0.054195927 | 0.021354267 | 0.03284166 | 0.937733382 |
| ENSG00000228409 | *CCT6P1* | 1.133299361 | -0.170387321 | -0.248627876 | 0.078240555 | 0.999933887 | -0.137502153 | -0.108179072 | -0.029323081 | 0.938090596 |
| ENSG00000116871 | *MAP7D1* | 8.448976345 | 0.018824554 | 0.085072951 | -0.066248397 | 0.999933887 | -0.508739274 | -0.527009662 | 0.018270388 | 0.938090596 |
| ENSG00000187531 | *SIRT7* | 6.728514932 | 0.046696985 | 0.040312675 | 0.00638431 | 0.999933887 | 0.418468681 | 0.405060693 | 0.013407987 | 0.938332677 |
| ENSG00000245534 | *RORA-AS1* | -0.486642684 | 0.133889677 | 0.22183858 | -0.087948904 | 0.999933887 | 0.358216037 | 0.297264072 | 0.060951964 | 0.938421239 |
| ENSG00000286236 | *AC018755.5* | 2.149946002 | 0.244584083 | 0.238082878 | 0.006501205 | 0.999933887 | 1.563923124 | 1.526447214 | 0.03747591 | 0.938480591 |
| ENSG00000070718 | *AP3M2* | 3.702397873 | -0.078675528 | 0.023208068 | -0.101883596 | 0.999933887 | -0.110764124 | -0.128095564 | 0.01733144 | 0.938480591 |
| ENSG00000100417 | *PMM1* | 2.401474652 | 0.090107929 | -0.007638297 | 0.097746226 | 0.999933887 | -0.13558717 | -0.158291441 | 0.02270427 | 0.938480591 |
| ENSG00000287555 | *AC023424.2* | 1.387912955 | 0.240171672 | 0.505851389 | -0.265679717 | 0.999933887 | 0.383300607 | 0.461545414 | -0.078244807 | 0.938480591 |
| ENSG00000101412 | *E2F1* | -0.104715348 | 0.154653156 | 0.031304616 | 0.12334854 | 0.999933887 | 0.319403767 | 0.281381353 | 0.038022413 | 0.938480591 |
| ENSG00000198692 | *EIF1AY* | -2.85754512 | 0.082225844 | 0.227363313 | -0.145137469 | 0.999933887 | -0.49697925 | -0.528920669 | 0.031941419 | 0.938480591 |
| ENSG00000118263 | *KLF7* | 6.093299698 | -0.049166079 | -0.001917358 | -0.047248721 | 0.999933887 | 0.330039024 | 0.346849942 | -0.016810918 | 0.938480591 |
| ENSG00000279176 | *AC079316.2* | -0.015322151 | 0.1422099 | 0.213371244 | -0.071161344 | 0.999933887 | -0.059084667 | -0.122238359 | 0.063153692 | 0.938480591 |
| ENSG00000286388 | *AC026748.7* | 0.415495875 | 0.084463766 | 0.023707985 | 0.060755781 | 0.999933887 | 0.279346437 | 0.235865596 | 0.043480841 | 0.938480591 |
| ENSG00000143970 | *ASXL2* | 5.535579148 | -0.039723359 | -0.091458367 | 0.051735008 | 0.999933887 | -0.564561463 | -0.548478387 | -0.016083076 | 0.938480591 |
| ENSG00000227028 | *SLC8A1-AS1* | 0.485420454 | 0.096923023 | 0.489732162 | -0.392809139 | 0.999933887 | 0.07627503 | 0.16574508 | -0.08947005 | 0.938480591 |
| ENSG00000211728 | *TRBV5-6* | -0.334871835 | -0.097073784 | -0.311987587 | 0.214913803 | 0.999933887 | 0.184775925 | 0.131487907 | 0.053288018 | 0.938480591 |
| ENSG00000221983 | *UBA52* | 8.107933042 | 0.023359953 | -0.03817939 | 0.061539343 | 0.999933887 | 0.085324742 | 0.098210768 | -0.012886026 | 0.938480591 |
| ENSG00000173218 | *VANGL1* | -0.638066806 | -0.064571504 | -0.197747322 | 0.133175818 | 0.999933887 | -0.115328584 | -0.176103264 | 0.060774679 | 0.938480591 |
| ENSG00000105281 | *SLC1A5* | 3.645142083 | 0.022107635 | 0.066810585 | -0.044702951 | 0.999933887 | 0.297468459 | 0.277073821 | 0.020394638 | 0.938480591 |
| ENSG00000213809 | *KLRK1* | 1.01990428 | 0.026237832 | 0.174797321 | -0.148559488 | 0.999933887 | 0.023965331 | -0.005752938 | 0.029718268 | 0.938480591 |
| ENSG00000121897 | *LIAS* | 0.801501506 | -0.021132007 | -0.165406174 | 0.144274167 | 0.999933887 | -0.09354118 | -0.123805259 | 0.030264078 | 0.938480591 |
| ENSG00000157322 | *CLEC18A* | 0.356439373 | -0.008706545 | -0.088744431 | 0.080037886 | 0.999933887 | -0.129981176 | -0.085378138 | -0.044603038 | 0.938480591 |
| ENSG00000159079 | *CFAP298* | 0.099100491 | -0.002755115 | 0.024301555 | -0.02705667 | 0.999933887 | 0.034341799 | -0.005818732 | 0.040160531 | 0.938480591 |
| ENSG00000134291 | *TMEM106C* | 1.660653558 | 0.089929638 | -0.042698779 | 0.132628417 | 0.999933887 | 0.135687696 | 0.110978191 | 0.024709505 | 0.9384834 |
| ENSG00000176974 | *SHMT1* | 1.78728284 | 0.163934241 | 0.079469415 | 0.084464826 | 0.999933887 | -0.187911961 | -0.220400239 | 0.032488277 | 0.938816709 |
| ENSG00000112599 | *GUCA1B* | 0.258486925 | -0.086212245 | -0.109545705 | 0.023333459 | 0.999933887 | -0.193942437 | -0.15350916 | -0.040433276 | 0.938968029 |
| ENSG00000109063 | *MYH3* | 0.889443255 | -0.235465739 | 0.133372723 | -0.368838461 | 0.999933887 | -0.161857398 | -0.130215136 | -0.031642262 | 0.939006922 |
| ENSG00000181847 | *TIGIT* | 3.775440088 | -0.040935496 | -0.044876047 | 0.003940551 | 0.999933887 | 0.206944599 | 0.223470499 | -0.0165259 | 0.939006922 |
| ENSG00000159917 | *ZNF235* | -0.072139755 | -0.010089059 | -0.162075503 | 0.151986444 | 0.999933887 | 0.002202089 | -0.037219796 | 0.039421885 | 0.939006922 |
| ENSG00000111641 | *NOP2* | 1.403711264 | 0.183948254 | 0.102565666 | 0.081382588 | 0.999933887 | 0.617866262 | 0.661574038 | -0.043707776 | 0.939047917 |
| ENSG00000130775 | *THEMIS2* | 9.402828202 | -0.049431886 | -0.072690855 | 0.023258969 | 0.999933887 | -0.207226274 | -0.221692536 | 0.014466262 | 0.939047917 |
| ENSG00000269044 | *AC024075.2* | 0.813791408 | -0.117757841 | -0.214319328 | 0.096561487 | 0.999933887 | -0.320452043 | -0.283249519 | -0.037202524 | 0.939047917 |
| ENSG00000280181 | *AC025262.1* | 0.693559875 | -0.100210959 | 0.19679373 | -0.297004689 | 0.999933887 | 0.640099205 | 0.69109483 | -0.050995625 | 0.939047917 |
| ENSG00000131115 | *ZNF227* | 1.251541519 | -0.142930764 | -0.129322918 | -0.013607846 | 0.999933887 | 0.026521024 | 0.001447415 | 0.025073608 | 0.939053092 |
| ENSG00000109919 | *MTCH2* | 3.061911347 | -0.084314834 | -0.27403788 | 0.189723046 | 0.999933887 | -0.090007414 | -0.107308815 | 0.017301401 | 0.939053092 |
| ENSG00000226816 | *AC005082.1* | -1.42666851 | -0.137516932 | 0.451665662 | -0.589182594 | 0.999933887 | -1.425851438 | -1.366790352 | -0.059061086 | 0.939053092 |
| ENSG00000280383 | *Z95331.1* | -0.950082209 | -0.109377489 | 0.259314211 | -0.3686917 | 0.999933887 | -0.20503335 | -0.276687639 | 0.071654289 | 0.939160406 |
| ENSG00000143862 | *ARL8A* | 6.737461611 | 0.099892962 | 0.077335535 | 0.022557427 | 0.999933887 | -0.345767533 | -0.357306868 | 0.011539335 | 0.939194561 |
| ENSG00000122254 | *HS3ST2* | -1.780914873 | -0.596908367 | 0.650101903 | -1.247010271 | 0.999933887 | -0.03997541 | -0.088752848 | 0.048777438 | 0.939194561 |
| ENSG00000132780 | *NASP* | 4.085289447 | -0.055501424 | -0.104361685 | 0.048860262 | 0.999933887 | 0.12154264 | 0.10895297 | 0.01258967 | 0.939194561 |
| ENSG00000215148 | *PRSS41* | -1.313575645 | 0.229052656 | 0.308872469 | -0.079819813 | 0.999933887 | -0.076032274 | -0.030276601 | -0.045755673 | 0.939194561 |
| ENSG00000140829 | *DHX38* | 6.01575549 | 0.012229588 | 0.051818923 | -0.039589335 | 0.999933887 | 0.053333026 | 0.064582415 | -0.011249388 | 0.939194561 |
| ENSG00000244232 | *RN7SL698P* | -0.15083479 | 0.059863331 | 0.149901032 | -0.090037701 | 0.999933887 | 0.794413271 | 0.858336303 | -0.063923032 | 0.939194561 |
| ENSG00000118520 | *ARG1* | 2.589771103 | 0.026403466 | -0.033672713 | 0.060076179 | 0.999933887 | -0.009962319 | -0.039866753 | 0.029904434 | 0.939365243 |
| ENSG00000104783 | *KCNN4* | 2.243498148 | -0.048318564 | -0.117945821 | 0.069627256 | 0.999933887 | 0.101548639 | 0.12115066 | -0.019602021 | 0.93939498 |
| ENSG00000182197 | *EXT1* | 2.738529654 | 0.119213707 | -0.025075153 | 0.14428886 | 0.999933887 | 0.240832968 | 0.21897909 | 0.021853879 | 0.939414318 |
| ENSG00000116133 | *DHCR24* | 0.547149862 | -0.207155469 | -0.392803586 | 0.185648116 | 0.999933887 | 0.003134919 | 0.036974818 | -0.033839899 | 0.939414318 |
| ENSG00000168264 | *IRF2BP2* | 8.078004812 | 0.037609078 | 0.088562281 | -0.050953202 | 0.999933887 | 0.218103579 | 0.232153543 | -0.014049964 | 0.939414318 |
| ENSG00000181284 | *TMEM102* | 0.225178429 | -0.084044725 | 0.11381307 | -0.197857795 | 0.999933887 | 0.255946241 | 0.29132443 | -0.035378189 | 0.939414318 |
| ENSG00000166068 | *SPRED1* | 0.239561798 | -0.066236819 | 0.168564315 | -0.234801134 | 0.999933887 | -0.491282151 | -0.455840624 | -0.035441527 | 0.939414318 |
| ENSG00000165646 | *SLC18A2* | -0.300443938 | -0.044430685 | -0.107456973 | 0.063026287 | 0.999933887 | -0.419391897 | -0.4813239 | 0.061932003 | 0.939414318 |
| ENSG00000231389 | *HLA-DPA1* | 4.811763403 | 0.021222129 | -0.02458303 | 0.045805159 | 0.999933887 | 0.053679865 | 0.071447372 | -0.017767507 | 0.939414318 |
| ENSG00000078674 | *PCM1* | 5.492231275 | -0.015729611 | -0.110349899 | 0.094620288 | 0.999933887 | -0.85061838 | -0.818593078 | -0.032025302 | 0.939414318 |
| ENSG00000188690 | *UROS* | 1.99825757 | -0.008483167 | 0.054456058 | -0.062939225 | 0.999933887 | 0.116987582 | 0.13922326 | -0.022235679 | 0.939414318 |
| ENSG00000167716 | *WDR81* | 6.449740559 | -0.003572412 | 0.032132074 | -0.035704486 | 0.999933887 | -0.5057592 | -0.524586497 | 0.018827297 | 0.939414318 |
| ENSG00000136068 | *FLNB* | 3.412565813 | 0.050491746 | 0.009771163 | 0.040720583 | 0.999933887 | 0.075204445 | 0.056955168 | 0.018249277 | 0.939433512 |
| ENSG00000134815 | *DHX34* | 8.738311882 | 0.031910831 | 0.021044105 | 0.010866726 | 0.999933887 | -0.343521297 | -0.33010944 | -0.013411857 | 0.939433512 |
| ENSG00000274756 | *AC243732.1* | 0.606985256 | -0.134082875 | 0.000646365 | -0.13472924 | 0.999933887 | -0.013718288 | 0.026540572 | -0.04025886 | 0.93962447 |
| ENSG00000125821 | *DTD1* | 0.932680083 | -0.273868145 | 0.09645912 | -0.370327265 | 0.981460244 | -0.097385192 | -0.06088231 | -0.036502882 | 0.939766755 |

| | | | | | | | | | | |
|---|---|---|---|---|---|---|---|---|---|---|
| ENSG00000249456 | *AL731577.2* | 0.444421505 | 0.367820166 | 0.21467563 | 0.153144536 | 0.999933887 | -0.136931883 | -0.185419107 | 0.048487224 | 0.939766755 |
| ENSG00000198885 | *ITPRIPL1* | 1.264205707 | -0.047433164 | 0.06348197 | -0.110915133 | 0.999933887 | 0.026025496 | -0.001315395 | 0.027340891 | 0.939766755 |
| ENSG00000120253 | *NUP43* | 1.09419335 | 0.055702293 | -0.026885651 | 0.082587943 | 0.999933887 | -0.06191404 | -0.032202164 | -0.029711876 | 0.93979726 |
| ENSG00000136450 | *SRSF1* | 4.282335568 | -0.076466446 | -0.13307388 | 0.056607435 | 0.999933887 | -0.192202114 | -0.179850017 | -0.012352096 | 0.940033943 |
| ENSG00000287131 | *AC096970.1* | 1.68790154 | 0.094404305 | 0.118459122 | -0.024054816 | 0.999933887 | 0.715885098 | 0.681244829 | 0.034640268 | 0.940033943 |
| ENSG00000112394 | *SLC16A10* | -0.82185881 | 0.181805621 | -0.414140891 | 0.595946513 | 0.999933887 | -0.77985592 | -0.732279608 | -0.047576312 | 0.940033943 |
| ENSG00000179909 | *ZNF154* | 1.256281415 | -0.075324252 | -0.210198153 | 0.134873901 | 0.999933887 | 0.214412847 | 0.1887869 | 0.025625947 | 0.940033943 |
| ENSG00000112031 | *MTRF1L* | 1.712200627 | -0.003793555 | 0.011813178 | -0.015606733 | 0.999933887 | -0.115258442 | -0.091228941 | -0.024029501 | 0.940033943 |
| ENSG00000117115 | *PADI2* | 6.396476304 | 0.00178531 | 0.002126603 | -0.000341293 | 0.999933887 | 0.344760765 | 0.329742767 | 0.015017998 | 0.940033943 |
| ENSG00000275056 | *AC020663.3* | 0.812789418 | -0.08607884 | 0.145624737 | -0.231703577 | 0.999933887 | 0.344629954 | 0.377295865 | -0.032665911 | 0.940158725 |
| ENSG00000147872 | *PLIN2* | 6.880188541 | 0.041680886 | 0.120922014 | -0.079241127 | 0.999933887 | -0.39600798 | -0.412964423 | 0.016956443 | 0.940158725 |
| ENSG00000051596 | *THOC3* | 0.38894752 | 0.072973108 | 0.141164159 | -0.068191051 | 0.999933887 | 0.115729141 | 0.074674617 | 0.041054525 | 0.940158725 |
| ENSG00000211779 | *TRAV5* | -1.228823102 | 0.030052952 | 0.014855448 | 0.015197503 | 0.999933887 | 0.055278725 | 0.109236119 | -0.053957394 | 0.940158725 |
| ENSG00000011523 | *CEP68* | 4.78590009 | -0.007456665 | 0.073036391 | -0.080493056 | 0.999933887 | 0.262853171 | 0.280115811 | -0.01726264 | 0.940169938 |
| ENSG00000169372 | *CRADD* | 1.999844807 | -0.066893539 | 0.131507021 | -0.19840056 | 0.999933887 | -0.148358503 | -0.172060822 | 0.023702319 | 0.940363991 |
| ENSG00000165355 | *FBXO33* | 5.636322242 | -0.020120209 | -0.026895423 | 0.006775214 | 0.999933887 | -0.314343796 | -0.302227216 | -0.01211658 | 0.940363991 |
| ENSG00000264204 | *AGAP7P* | 0.861117952 | 0.059275216 | 0.069508884 | -0.010233668 | 0.999933887 | 0.104405848 | 0.156728526 | -0.052322678 | 0.94041877 |
| ENSG00000135392 | *DNAJC14* | 2.777527513 | -0.044449252 | 0.071982726 | -0.116431978 | 0.999933887 | -0.258721526 | -0.238968934 | -0.019752592 | 0.940524722 |
| ENSG00000273076 | *AL021707.7* | 1.115581857 | -0.094422374 | -0.000383659 | -0.094038715 | 0.999933887 | -0.042410239 | -0.007946289 | -0.03446395 | 0.940740069 |
| ENSG00000126953 | *TIMM8A* | 0.431598461 | -0.099926449 | -0.011854619 | -0.08807183 | 0.999933887 | -0.081935156 | -0.115385118 | 0.033449962 | 0.940740069 |
| ENSG00000153015 | *CWC27* | 1.299529461 | -0.046239849 | -0.15378378 | 0.107543931 | 0.999933887 | 0.012761304 | 0.045328596 | -0.032567292 | 0.940740069 |
| ENSG00000258875 | *AL135818.1* | 1.701039067 | 0.003943245 | 0.173787338 | -0.169844093 | 0.999933887 | 0.151841218 | 0.181834919 | -0.029993701 | 0.940740069 |
| ENSG00000065000 | *AP3D1* | 6.834956992 | -0.00705245 | -0.016750502 | 0.009698052 | 0.999933887 | -0.135056294 | -0.124491009 | -0.010565285 | 0.940768419 |
| ENSG00000287385 | *AL121820.3* | -1.259784383 | -0.219266154 | 0.249036646 | -0.4683028 | 0.999933887 | -0.425649141 | -0.475153478 | 0.049504337 | 0.940896827 |
| ENSG00000166482 | *MFAP4* | -0.248743398 | 0.061961536 | -0.13436328 | 0.196324816 | 0.999933887 | 0.58634949 | 0.520292194 | 0.066057296 | 0.940896827 |
| ENSG00000248727 | *LINC01948* | 1.524525348 | -0.041364979 | -0.095735151 | 0.054370172 | 0.999933887 | 0.007672328 | 0.038072563 | -0.030400235 | 0.940896827 |
| ENSG00000197587 | *DMBX1* | -1.878157041 | 0.177577695 | 0.420719847 | -0.243142152 | 0.999933887 | -0.58962286 | -0.548130287 | -0.041492574 | 0.940896827 |
| ENSG00000240132 | *ETF1P2* | 0.756060133 | 0.031338388 | 0.138029674 | -0.106691286 | 0.999933887 | 0.527411578 | 0.475265346 | 0.052146231 | 0.940896827 |
| ENSG00000184470 | *TXNRD2* | 3.176055283 | 0.039986186 | 0.173766338 | -0.133780152 | 0.999933887 | 0.078258886 | 0.061087779 | 0.017171107 | 0.940954604 |
| ENSG00000132313 | *MRPL35* | 0.860655488 | 0.17275688 | -0.016031122 | 0.188788002 | 0.999933887 | 0.114303394 | 0.139174938 | -0.024871544 | 0.94100158 |
| ENSG00000159713 | *TPPP3* | -0.799598283 | 0.170187678 | 0.179568073 | -0.009380395 | 0.999933887 | 0.791070335 | 0.713826881 | 0.077243454 | 0.94100158 |
| ENSG00000113273 | *ARSB* | 3.213415313 | 0.054413881 | 0.06793322 | -0.013519338 | 0.999933887 | 0.330744587 | 0.353126953 | -0.022382366 | 0.94100158 |
| ENSG00000255224 | *AC109322.1* | -0.306557717 | 0.079192912 | 0.393646242 | -0.314453329 | 0.999933887 | 0.29378811 | 0.360111499 | -0.066323388 | 0.94100158 |
| ENSG00000165675 | *ENOX2* | 0.904548673 | -0.026155887 | 0.204211468 | -0.230367355 | 0.999933887 | 0.023047423 | -0.005620377 | 0.0286678 | 0.94100158 |
| ENSG00000272356 | *AL080317.2* | 0.025661752 | 0.034606476 | -0.33381615 | 0.368422627 | 0.999933887 | 0.828997301 | 0.788699748 | 0.040297553 | 0.94100158 |
| ENSG00000151746 | *BICD1* | 1.133826463 | 0.002106599 | 0.08202829 | -0.079921692 | 0.999933887 | 0.021077295 | 0.046898693 | -0.025821398 | 0.94100158 |
| ENSG00000177369 | *FLJ40194* | -0.110128101 | -0.298086866 | -0.422125249 | 0.124038383 | 0.999933887 | -1.374472685 | -1.330692295 | -0.043780389 | 0.941031291 |
| ENSG00000225195 | *AC078962.1* | -0.478407712 | -0.208344439 | -0.092316157 | -0.116028282 | 0.999933887 | -0.137610047 | -0.093817491 | -0.043792556 | 0.941031291 |
| ENSG00000213085 | *CFAP45* | 3.939429694 | -0.046181079 | -0.018425665 | -0.027755413 | 0.999933887 | -0.015320245 | -0.033287543 | 0.017967298 | 0.941031291 |
| ENSG00000287202 | *AC008119.1* | -0.451987274 | -0.112524467 | -0.09572555 | -0.016798917 | 0.999933887 | -0.156951706 | -0.112259058 | -0.044692647 | 0.941031291 |
| ENSG00000271855 | *AC073195.1* | 0.034614068 | 0.057447924 | -0.031089857 | 0.088537781 | 0.999933887 | 1.690554564 | 1.651060657 | 0.039493908 | 0.941031291 |
| ENSG00000273247 | *AC097376.3* | 3.336347002 | -0.031021213 | -0.009543331 | -0.021477882 | 0.999933887 | -0.695298374 | -0.674964751 | -0.020333623 | 0.941197348 |
| ENSG00000197622 | *CDC42SE1* | 9.302859401 | 0.029378493 | 0.013762078 | 0.015616415 | 0.999933887 | 0.724476195 | 0.711882068 | 0.012594127 | 0.941350182 |
| ENSG00000075234 | *TTC38* | 3.091964022 | 0.056185112 | 0.120971267 | -0.064786155 | 0.999933887 | -0.135838428 | -0.115431811 | -0.020406617 | 0.941617057 |
| ENSG00000170468 | *RIOX1* | 4.272886248 | 0.125184425 | -0.025800646 | 0.150985071 | 0.999933887 | 0.97891014 | 0.9965325 | -0.01762236 | 0.942223382 |
| ENSG00000141252 | *VPS53* | 5.039576349 | -0.030288873 | -0.042495096 | 0.012206223 | 0.999933887 | -0.247523698 | -0.258185356 | 0.010661658 | 0.942223382 |
| ENSG00000269609 | *RPARP-AS1* | 1.860524399 | -0.22901373 | -0.487381388 | 0.258367658 | 0.999933887 | -0.38996997 | -0.36750966 | -0.022460309 | 0.942399463 |
| ENSG00000197756 | *RPL37A* | 7.226150764 | -0.015371603 | -0.027891554 | 0.012519951 | 0.999933887 | 0.099268923 | 0.112480759 | -0.013211836 | 0.942399463 |
| ENSG00000261236 | *BOP1* | 2.505181181 | 0.079716259 | 0.063091716 | 0.016624543 | 0.999933887 | 0.280531649 | 0.2589881 | 0.021543549 | 0.942442374 |
| ENSG00000141994 | *DUS3L* | 3.208340161 | 0.024311791 | -0.043086073 | 0.067397865 | 0.999933887 | 0.202193474 | 0.183448976 | 0.018744498 | 0.94245086 |
| ENSG00000246339 | *EXTL3-AS1* | -0.115872032 | -0.177120991 | 0.091673691 | -0.268794682 | 0.999933887 | -0.452412303 | -0.401889733 | -0.050522571 | 0.942612323 |
| ENSG00000005175 | *RPAP3* | 3.735663376 | -0.053505073 | -0.16576994 | 0.112264866 | 0.999933887 | -0.392035969 | -0.412635254 | 0.020599284 | 0.942681316 |
| ENSG00000285427 | *SOD2-OT1* | -1.280227786 | 0.521627556 | 0.156093108 | 0.365534449 | 0.999933887 | 2.295429041 | 2.364032079 | -0.068603038 | 0.942893103 |
| ENSG00000117133 | *RPF1* | 3.249132085 | -0.072384969 | -0.07022877 | -0.002156199 | 0.999933887 | -0.077529565 | -0.094436039 | 0.016906475 | 0.942893103 |
| ENSG00000176946 | *THAP4* | 3.175768341 | -0.048839597 | -0.006354724 | -0.042484873 | 0.999933887 | 0.299898515 | 0.318399749 | -0.018501233 | 0.942893103 |
| ENSG00000079246 | *XRCC5* | 6.817369986 | -0.104922204 | -0.1232959 | 0.018373696 | 0.999933887 | -0.189743046 | -0.178412223 | -0.011330824 | 0.943008692 |
| ENSG00000106355 | *LSM5* | 1.23584163 | -0.219017773 | -0.010689276 | -0.208328497 | 0.999933887 | 0.047947997 | 0.022609053 | 0.025338943 | 0.943229876 |
| ENSG00000221937 | *TAS2R40* | -1.083840803 | 0.155675232 | 0.194765139 | -0.039089908 | 0.999933887 | 0.707251698 | 0.63398723 | 0.073264468 | 0.943229876 |
| ENSG00000101162 | *TUBB1* | 0.792259266 | -0.120456418 | -0.098094952 | -0.022361466 | 0.999933887 | 0.110298041 | 0.079268039 | 0.031030002 | 0.943229876 |
| ENSG00000099999 | *RNF215* | 1.347679582 | 0.08849444 | 0.063504476 | 0.024989964 | 0.999933887 | 0.119146609 | 0.089895971 | 0.029250639 | 0.943229876 |
| ENSG00000167775 | *CD320* | 1.659791015 | -0.051232293 | 0.007977712 | -0.059210005 | 0.999933887 | 0.067887026 | 0.092287331 | -0.024400305 | 0.943229876 |
| ENSG00000076043 | *REXO2* | 1.555826584 | 0.037760324 | -0.268182234 | 0.305942558 | 0.999933887 | 0.093691437 | 0.070130785 | 0.023560652 | 0.943229876 |
| ENSG00000099251 | *HSD17B7P2* | -0.072534932 | -0.083521534 | -0.06286082 | -0.020660714 | 0.999933887 | -0.181121442 | -0.222787682 | 0.04166624 | 0.943229876 |
| ENSG00000176058 | *TPRN* | 1.088144905 | 0.038748792 | 0.0512259 | -0.012477108 | 0.999933887 | -0.04493183 | -0.016800692 | -0.028131137 | 0.943229876 |
| ENSG00000147804 | *SLC39A4* | 0.665204144 | 0.024060959 | 0.203485606 | -0.179424647 | 0.999933887 | 0.560697844 | 0.592899339 | -0.032201495 | 0.943229876 |
| ENSG00000280279 | *LINC02887* | -0.362407087 | -0.017529919 | -0.252146945 | 0.234617027 | 0.999933887 | 0.185248762 | 0.09796642 | 0.087282342 | 0.943284046 |
| ENSG00000261812 | *TUBB8P7* | -0.812922696 | 0.127992328 | -0.183750963 | 0.31174329 | 0.999933887 | -1.381931609 | -1.416765 | 0.034833391 | 0.943298504 |
| ENSG00000145476 | *CYP4V2* | 1.353167863 | 0.153641374 | -0.079395532 | 0.233036906 | 0.999933887 | 0.009744146 | -0.018449145 | 0.028193291 | 0.943429247 |
| ENSG00000056050 | *HPF1* | 0.037759992 | -0.246563057 | 0.054749182 | -0.301312239 | 0.999933887 | 0.177750427 | 0.138495992 | 0.039254435 | 0.943429247 |
| ENSG00000135919 | *SERPINE2* | 0.546053054 | -0.192821289 | -0.114423597 | -0.078397692 | 0.999933887 | 0.183724053 | 0.21162193 | -0.027897876 | 0.943429247 |
| ENSG00000166997 | *CNPY4* | 1.712428459 | 0.045234001 | 0.056073157 | -0.010839156 | 0.999933887 | 0.120221159 | 0.139858453 | -0.019637294 | 0.943429247 |
| ENSG00000136108 | *CKAP2* | 1.731066806 | -0.04057292 | -0.097834952 | 0.057262031 | 0.999933887 | -0.089085745 | -0.054624341 | -0.034461404 | 0.943429247 |
| ENSG00000185658 | *BRWD1* | 5.388378261 | -0.024659528 | -0.080312859 | 0.055653331 | 0.999933887 | -0.593504414 | -0.618728892 | 0.025224478 | 0.943429247 |
| ENSG00000178904 | *DPY19L3* | 3.900951487 | 0.020860127 | -0.071267819 | 0.092127946 | 0.999933887 | -0.567522537 | -0.594394659 | 0.026872121 | 0.943429247 |
| ENSG00000092148 | *HECTD1* | 5.361040693 | -0.011543818 | -0.092652679 | 0.081108861 | 0.999933887 | -0.499308549 | -0.479289297 | -0.020019252 | 0.943429247 |
| ENSG00000260077 | *AC104794.3* | 0.794209569 | 0.009504716 | 0.040919264 | -0.031414549 | 0.999933887 | 0.309621843 | 0.336312676 | -0.026690832 | 0.943429247 |
| ENSG00000134574 | *DDB2* | 3.208722833 | 0.003779951 | 0.150971103 | -0.147191152 | 0.999933887 | 0.134598154 | 0.117738957 | 0.016859196 | 0.943429247 |
| ENSG00000133265 | *HSPBP1* | 1.849479607 | 0.040326942 | -0.003082693 | 0.043409635 | 0.999933887 | 0.17975725 | 0.201890997 | -0.022133747 | 0.943703997 |
| ENSG00000147883 | *CDKN2B* | 1.223700885 | -0.118148098 | -0.195888678 | 0.07774058 | 0.999933887 | -0.19702996 | -0.227166044 | 0.030136085 | 0.943703997 |
| ENSG00000118482 | *PHF3* | 5.895260957 | -0.048809556 | -0.122386852 | 0.073577296 | 0.999933887 | -0.856726289 | -0.830242446 | -0.026483843 | 0.943703997 |
| ENSG00000204271 | *SPIN3* | 0.412670157 | 0.030176117 | 0.28784565 | -0.257669534 | 0.999933887 | -0.031984876 | -0.004764747 | -0.02722013 | 0.943703997 |
| ENSG00000004399 | *PLXND1* | 3.967280224 | 0.075751602 | 0.105550318 | -0.029798716 | 0.999933887 | 0.081521849 | 0.09655442 | -0.01503257 | 0.94375311 |
| ENSG00000211800 | *TRAV20* | -1.321858643 | -0.021579017 | -0.509895648 | 0.488316631 | 0.999933887 | 0.104237142 | 0.038825977 | 0.065411164 | 0.94375311 |
| ENSG00000261542 | *AC011978.2* | -0.162439957 | -0.240939734 | -0.210801465 | -0.030138269 | 0.999933887 | 0.093081211 | 0.044835111 | 0.0482461 | 0.943779371 |
| ENSG00000255838 | *AC069262.1* | -0.041750502 | 0.22483545 | 0.126035519 | 0.098799931 | 0.999933887 | 0.012289349 | -0.034578824 | 0.046868173 | 0.943779371 |
| ENSG00000275111 | *ZNF2* | -0.626920138 | -0.205185508 | -0.338893553 | 0.133708045 | 0.999933887 | -0.159297451 | -0.201342953 | 0.042045502 | 0.943779371 |
| ENSG00000165475 | *CRYL1* | 1.060856887 | 0.115619944 | -0.035556136 | 0.15117608 | 0.999933887 | -0.169707648 | -0.202288087 | 0.032580439 | 0.943779371 |
| ENSG00000176261 | *ZBTB8OS* | 2.941957672 | -0.049365413 | -0.120843989 | 0.071478576 | 0.999933887 | -0.107502796 | -0.123695864 | 0.016193068 | 0.943779371 |
| ENSG00000220785 | *MTMR9LP* | -0.414842859 | 0.112075718 | -0.098917522 | 0.210993241 | 0.999933887 | 0.02293704 | -0.038928561 | 0.0618656 | 0.943779371 |
| ENSG00000179051 | *RCC2* | 4.264586524 | -0.0134882 | -0.053431509 | 0.039943309 | 0.999933887 | 0.105904708 | 0.091414806 | 0.014489903 | 0.943779371 |
| ENSG00000279019 | *AC009090.4* | -0.209176755 | -0.037354967 | 0.094357087 | -0.131712053 | 0.999933887 | -0.509846833 | -0.460655537 | -0.049191296 | 0.943779371 |
| ENSG00000164048 | *ZNF589* | 2.36175295 | 0.009457951 | -0.045622901 | 0.055080852 | 0.999933887 | -0.008404397 | 0.011003053 | -0.01940745 | 0.943779371 |
| ENSG00000136444 | *RSAD1* | 2.886628017 | 0.005653543 | 0.094638577 | -0.088985034 | 0.999933887 | 0.097059778 | 0.114627877 | -0.017568098 | 0.943779371 |
| ENSG00000232656 | *IDI2-AS1* | 1.480762369 | 0.082288161 | -0.033585806 | 0.115873967 | 0.999933887 | -0.126473119 | -0.154009119 | 0.027535999 | 0.943961143 |
| ENSG00000275807 | *AC145285.6* | 0.214423653 | 0.158760193 | 0.435033259 | -0.276273066 | 0.999933887 | -0.003987436 | -0.033806245 | 0.029818809 | 0.943961143 |
| ENSG00000141985 | *SH3GL1* | 6.227334755 | 0.040714257 | 0.023521852 | 0.017192405 | 0.999933887 | -0.049796294 | -0.061705215 | 0.011908921 | 0.943961143 |
| ENSG00000063127 | *SLC6A16* | 2.933433273 | -0.062799067 | -0.044526876 | -0.018272191 | 0.999933887 | 0.310181997 | 0.295135283 | 0.015046714 | 0.943961143 |
| ENSG00000174446 | *SNAPC5* | 2.420302407 | 0.041177072 | 0.069100348 | -0.027923276 | 0.999933887 | -0.066119249 | -0.043925714 | -0.022193534 | 0.943961143 |
| ENSG00000164935 | *DCSTAMP* | 0.823005947 | -0.072183004 | 0.052975927 | -0.125158931 | 0.999933887 | -0.092633631 | -0.120077742 | 0.027444111 | 0.943961143 |
| ENSG00000125629 | *INSIG2* | 2.446947337 | 0.009840873 | -0.020470632 | 0.030311505 | 0.999933887 | -0.517535692 | -0.536633102 | 0.01909741 | 0.943961143 |
| ENSG00000111644 | *ACRBP* | 1.017289604 | -0.01171444 | -0.11798629 | 0.106271851 | 0.999933887 | -0.114271105 | -0.140058409 | 0.025787304 | 0.943961143 |
| ENSG00000041515 | *MYO16* | -1.330206508 | -0.012240072 | -0.687223917 | 0.674983845 | 0.999933887 | -0.752609881 | -0.709189662 | -0.043420219 | 0.943961143 |
| ENSG00000232119 | *MCTS1* | 3.228087395 | 0.006124803 | -0.232032609 | 0.238157413 | 0.944094537 | -0.303273363 | -0.32014595 | 0.016872587 | 0.944042 |
| ENSG00000105640 | *RPL18A* | 7.39892372 | 0.035270527 | 0.024701589 | 0.010568939 | 0.999933887 | 0.123780781 | 0.136843665 | -0.013062884 | 0.944116242 |
| ENSG00000120948 | *TARDBP* | 4.858357444 | -0.14663865 | -0.02789245 | -0.1187462 | 0.999933887 | -0.223516349 | -0.234153188 | 0.010636839 | 0.944175954 |

| | | | | | | | | | | |
|---|---|---|---|---|---|---|---|---|---|---|
| ENSG00000176659 | *C20orf197* | 0.788316264 | -0.126144484 | -0.069831802 | -0.056312682 | 0.999933887 | -0.165408635 | -0.208122357 | 0.042713722 | 0.944175954 |
| ENSG00000221923 | *ZNF880* | -0.02521221 | -0.196076689 | 0.20275435 | -0.398831039 | 0.999933887 | 0.052385835 | 0.092246584 | -0.039860749 | 0.944175954 |
| ENSG00000184897 | *H1-10* | 5.052914007 | 0.039673468 | 0.005213769 | 0.034459699 | 0.999933887 | 0.157502483 | 0.173095876 | -0.015593393 | 0.944175954 |
| ENSG00000133597 | *ADCK2* | 1.439165121 | 0.04538021 | 0.175915213 | -0.130535003 | 0.999933887 | 0.003897654 | 0.034412693 | -0.030515039 | 0.944175954 |
| ENSG00000078814 | *MYH7B* | 2.25191161 | 0.129057924 | 0.185150731 | -0.056092808 | 0.999933887 | -0.258147367 | -0.235470078 | -0.022677289 | 0.944234767 |
| ENSG00000286108 | *AL359916.1* | -0.544311741 | 0.15019553 | 0.191609105 | -0.041413575 | 0.999933887 | 0.607536534 | 0.668386615 | -0.060850081 | 0.944234767 |
| ENSG00000214401 | *KANSL1-AS1* | -0.150965761 | 0.101175275 | 0.21781526 | -0.116639985 | 0.999933887 | 0.102002102 | 0.053254869 | 0.048747233 | 0.944234767 |
| ENSG00000049656 | *CLPTM1L* | 6.004498723 | 0.037414498 | -0.005563893 | 0.042978391 | 0.999933887 | 0.083633184 | 0.093411809 | -0.009778625 | 0.944290609 |
| ENSG00000115942 | *ORC2* | 2.681898649 | -0.033500338 | -0.136948749 | 0.10344841 | 0.999933887 | -0.023283854 | -0.005563995 | -0.017719859 | 0.944290609 |
| ENSG00000276043 | *UHRF1* | 0.920868241 | -0.046430047 | -0.132817907 | 0.08638786 | 0.999933887 | -0.119436508 | -0.150207384 | 0.030770876 | 0.944290609 |
| ENSG00000205534 | *SMG1P2* | 1.757968994 | 0.011265658 | -0.01111332 | 0.022378978 | 0.999933887 | -0.047091783 | -0.065087913 | 0.017996129 | 0.944290609 |
| ENSG00000184154 | *LRTOMT* | 1.700814832 | -0.138584331 | 0.128045853 | -0.266630184 | 0.999933887 | -0.213865046 | -0.194595506 | -0.019269541 | 0.944671184 |
| ENSG00000173991 | *TCAP* | 0.191455962 | 0.195993592 | 0.227752906 | -0.031759314 | 0.999933887 | 0.128840018 | 0.087664394 | 0.041175625 | 0.944671184 |
| ENSG00000105325 | *FZR1* | 6.720697331 | 0.051460929 | 0.048923456 | 0.002537472 | 0.999933887 | 0.147993695 | 0.157581362 | -0.009587668 | 0.944671184 |
| ENSG00000018610 | *CXorf56* | 3.033013725 | -0.079615721 | 0.00210909 | -0.081724811 | 0.999933887 | -0.094998625 | -0.113117504 | 0.018118879 | 0.944671184 |
| ENSG00000241685 | *ARPC1A* | 2.526179746 | -0.044557133 | -0.077322418 | 0.032765284 | 0.999933887 | -0.182078561 | -0.164345386 | -0.017733175 | 0.944671184 |
| ENSG00000170270 | *GON7* | 1.302274958 | 0.053360222 | 0.204997632 | -0.15163741 | 0.999933887 | 0.197383841 | 0.223262608 | -0.025878767 | 0.944671184 |
| ENSG00000285756 | *BX890604.1* | 0.64106667 | 0.022081093 | -0.149728382 | 0.171809475 | 0.999933887 | 0.129352418 | 0.097856838 | 0.031495579 | 0.944671184 |
| ENSG00000124225 | *PMEPA1* | 1.141689614 | -0.000831844 | -0.230054621 | 0.229222777 | 0.999933887 | 0.060087116 | 0.026848549 | 0.033238567 | 0.944671184 |
| ENSG00000233154 | *LINC01762* | -0.985409358 | -0.324187491 | -0.093851387 | -0.230336105 | 0.999933887 | 0.961234809 | 0.915877058 | 0.045357751 | 0.944781302 |
| ENSG00000168301 | *KCTD6* | -0.282163313 | -0.068139593 | -0.083161947 | 0.015022354 | 0.999933887 | -0.105345701 | -0.061926805 | -0.043418896 | 0.944781302 |
| ENSG00000060749 | *QSER1* | 1.413915615 | -0.028423295 | -0.235915934 | 0.207492639 | 0.999933887 | -0.003022929 | -0.033652802 | 0.030629874 | 0.944781302 |
| ENSG00000118600 | *RXYLT1* | -0.154282557 | 0.026388968 | 0.157941121 | -0.131552152 | 0.999933887 | 0.028918099 | -0.014206278 | 0.043124377 | 0.944781302 |
| ENSG00000276791 | *AC092117.1* | 0.95886026 | 0.004180223 | 0.003706167 | 0.000474056 | 0.999933887 | 0.143071844 | 0.117681136 | 0.025390707 | 0.944781302 |
| ENSG00000257176 | *AC009318.1* | 1.966047406 | -0.021453513 | 0.080330686 | -0.101784198 | 0.999933887 | -0.457849515 | -0.487139143 | 0.029289628 | 0.944805485 |
| ENSG00000105509 | *HAS1* | 0.863646889 | 0.180712082 | -0.039847607 | 0.220559689 | 0.999933887 | 0.960867819 | 1.000906245 | -0.040038425 | 0.944830031 |
| ENSG00000143184 | *XCL1* | -0.30370997 | 0.066314715 | -0.236724501 | 0.303039216 | 0.999933887 | 0.043887338 | 0.000213506 | 0.043673832 | 0.944830031 |
| ENSG00000258469 | *CHMP4BP1* | 1.613506152 | 0.193075434 | 0.438479327 | -0.245403894 | 0.999933887 | 0.609394479 | 0.58156696 | 0.027827519 | 0.944831633 |
| ENSG00000169371 | *SNUPN* | 0.791817897 | -0.037500398 | -0.06075107 | 0.023250672 | 0.999933887 | -0.084982315 | -0.060493376 | -0.024488939 | 0.944831633 |
| ENSG00000249592 | *AC139887.2* | 1.682842086 | 0.005776209 | 0.004827392 | 0.000948817 | 0.999933887 | -0.008569858 | 0.01169471 | -0.020264568 | 0.944831633 |
| ENSG00000167280 | *ENGASE* | 3.358085856 | 0.048815995 | 0.126948728 | -0.078132733 | 0.999933887 | 0.13020572 | 0.109268346 | 0.020937374 | 0.944924598 |
| ENSG00000079156 | *OSBPL6* | -0.936598919 | -0.213832604 | 0.203506339 | -0.417338943 | 0.999933887 | -2.076541773 | -2.027710729 | -0.048831045 | 0.944939367 |
| ENSG00000268555 | *AC123912.4* | 4.030822607 | 0.109909003 | 0.034493295 | 0.075415708 | 0.999933887 | -0.004371657 | -0.020580877 | 0.01620922 | 0.945163278 |
| ENSG00000101096 | *NFATC2* | 4.66194251 | -0.031699418 | -0.038286062 | 0.006586644 | 0.999933887 | 0.216253382 | 0.232760044 | -0.016506663 | 0.945363164 |
| ENSG00000130935 | *NOL11* | 2.857846328 | -0.154914503 | -0.103504787 | -0.051409716 | 0.999933887 | 0.102781292 | 0.118272273 | -0.015490981 | 0.945396965 |
| ENSG00000272505 | *AC104964.3* | -0.533455798 | -0.018126161 | 0.094238604 | -0.112364765 | 0.999933887 | 0.019679811 | -0.033455288 | 0.053135099 | 0.945784408 |
| ENSG00000166337 | *TAF10* | 3.969729808 | 0.105298826 | 0.046545218 | 0.058753608 | 0.999933887 | 0.022587774 | 0.036429481 | -0.013841707 | 0.945864288 |
| ENSG00000146197 | *SCUBE3* | 1.170468456 | -0.002960652 | -0.044466122 | 0.04150547 | 0.999933887 | 0.440977113 | 0.406081791 | 0.034895322 | 0.945896879 |
| ENSG00000211892 | *IGHG4* | -1.998371498 | -0.43677688 | -0.501573956 | 0.064797075 | 0.999933887 | 0.492845781 | 0.577131794 | -0.084286013 | 0.946049285 |
| ENSG00000135441 | *BLOC1S1* | 3.941866583 | -0.073521146 | -0.039492403 | -0.034028743 | 0.999933887 | -0.011976448 | -0.024170518 | 0.01219407 | 0.946049285 |
| ENSG00000163810 | *TGM4* | -1.584371958 | 0.007588759 | -0.016669448 | 0.024258207 | 0.999933887 | -0.391597825 | -0.297017515 | -0.09458031 | 0.946049285 |
| ENSG00000198408 | *OGA* | 8.480074118 | -0.068780095 | -0.063307935 | -0.005472161 | 0.999933887 | -0.402250376 | -0.411105425 | 0.008855048 | 0.946059138 |
| ENSG00000114841 | *DNAH1* | 5.47295369 | 0.031399681 | 0.025849862 | 0.00554982 | 0.999933887 | 0.074587932 | 0.064451742 | 0.01013619 | 0.946242981 |
| ENSG00000048991 | *R3HDM1* | 2.510995636 | 0.012483904 | -0.095988047 | 0.108471951 | 0.999933887 | -0.03320262 | -0.052595826 | 0.019393206 | 0.946242981 |
| ENSG00000125498 | *KIR2DL1* | -0.923235509 | 0.011185022 | -0.174965077 | 0.186150099 | 0.999933887 | 0.764782842 | 0.807681812 | -0.042898971 | 0.946242981 |
| ENSG00000260727 | *SLC7A5P1* | 0.18633107 | 0.181553748 | 0.229238335 | -0.047684587 | 0.999933887 | 0.148882647 | 0.187496501 | -0.038613854 | 0.946305721 |
| ENSG00000183826 | *BTBD9* | 4.407303692 | 0.008440637 | 0.074305336 | -0.065864698 | 0.999933887 | -0.013160077 | -0.002040716 | -0.011119361 | 0.946576818 |
| ENSG00000164535 | *DAGLB* | 4.079908541 | -0.069060827 | 0.048734644 | -0.117795471 | 0.999933887 | 0.292427819 | 0.306542645 | -0.014114826 | 0.946683695 |
| ENSG00000151466 | *SCLT1* | 4.896089721 | -0.043499308 | -0.035015824 | -0.008483483 | 0.999933887 | -0.341156826 | -0.323868634 | -0.017288192 | 0.946683695 |
| ENSG00000231770 | *TMEM44-AS1* | 1.535065613 | 0.200819842 | 0.045075518 | 0.155744324 | 0.999933887 | 0.380104843 | 0.355988869 | 0.024115974 | 0.946769178 |
| ENSG00000203814 | *H2BC18* | 4.699492848 | 0.066242209 | -0.023402981 | 0.08964519 | 0.999933887 | 0.300853144 | 0.275086931 | 0.025766213 | 0.946769178 |
| ENSG00000147526 | *TACC1* | 7.377180634 | -0.048644599 | -0.031756135 | -0.016888464 | 0.999933887 | -0.159424353 | -0.148095635 | -0.011328718 | 0.946851234 |
| ENSG00000198721 | *ECI2* | 0.733125004 | -0.088798133 | -0.029262646 | -0.059535487 | 0.999933887 | 0.022992026 | 0.050390134 | -0.027398109 | 0.946861249 |
| ENSG00000092978 | *GPATCH2* | 1.40104605 | -0.165809656 | -0.220986899 | 0.055177243 | 0.999933887 | -0.060351549 | -0.030100321 | -0.030251228 | 0.946921801 |
| ENSG00000088727 | *KIF9* | 0.927232652 | 0.033360272 | 0.228969306 | -0.195609034 | 0.999933887 | -0.102155203 | -0.072347241 | -0.029807962 | 0.946921801 |
| ENSG00000180185 | *FAHD1* | 1.729183388 | -0.098045685 | -0.168048383 | 0.070002697 | 0.999933887 | -0.276284163 | -0.296011037 | 0.019726874 | 0.94700788 |
| ENSG00000272398 | *CD24* | 1.749601548 | 0.054281343 | 0.141137673 | -0.08685633 | 0.999933887 | 0.318634264 | 0.339591471 | -0.020957208 | 0.94703913 |
| ENSG00000151065 | *DCP1B* | 1.030770082 | -0.139224304 | -0.160152643 | 0.020928339 | 0.999933887 | 0.088731383 | 0.112661138 | -0.023929755 | 0.947114324 |
| ENSG00000142156 | *COL6A1* | 0.441729407 | 0.082080831 | -0.007985574 | 0.090066405 | 0.999933887 | 0.180202243 | 0.207347826 | -0.027145584 | 0.947198187 |
| ENSG00000177042 | *TMEM80* | 3.186837946 | -0.017633389 | 0.063433903 | -0.081067292 | 0.999933887 | 0.094202366 | 0.08028043 | 0.013921935 | 0.947352943 |
| ENSG00000287070 | *AL499604.1* | 1.44340013 | 0.262368853 | 0.165797326 | 0.096571527 | 0.999933887 | 0.612374551 | 0.570327114 | 0.042047437 | 0.94739741 |
| ENSG00000134001 | *EIF2S1* | 3.119275086 | -0.101896367 | -0.079675896 | -0.02222047 | 0.999933887 | 0.036357067 | 0.051385949 | -0.015028882 | 0.94739741 |
| ENSG00000260536 | *AL035071.2* | -0.273522393 | 0.237130337 | 0.377415689 | -0.140285352 | 0.999933887 | 0.166489222 | 0.103785134 | 0.062704088 | 0.94739741 |
| ENSG00000117625 | *RCOR3* | 4.729459202 | -0.030825548 | -0.042544 | 0.011718453 | 0.999933887 | -0.456335814 | -0.443289148 | -0.013046666 | 0.94739741 |
| ENSG00000120526 | *NUDCD1* | -0.005595785 | -0.096739547 | -0.198665766 | 0.10192622 | 0.999933887 | -0.005593976 | 0.032783851 | -0.038377828 | 0.94739741 |
| ENSG00000136942 | *RPL35* | 6.463912351 | 0.026233007 | -0.012984116 | 0.039217124 | 0.999933887 | 0.111493915 | 0.098142901 | 0.013351014 | 0.94739741 |
| ENSG00000132740 | *IGHMBP2* | 4.495457981 | 0.038821448 | 0.065433108 | -0.02661166 | 0.999933887 | 1.170399188 | 1.190523561 | -0.020124373 | 0.94739741 |
| ENSG00000169410 | *PTPN9* | 2.498921594 | 0.029631517 | 0.158222682 | -0.128591166 | 0.999933887 | -0.161070803 | -0.179883763 | 0.01881296 | 0.94739741 |
| ENSG00000177098 | *SCN4B* | -0.345257618 | -0.241958941 | 0.248739178 | -0.490698119 | 0.969206975 | -1.361211104 | -1.393900919 | 0.032689815 | 0.947453112 |
| ENSG00000177576 | *C18orf32* | 3.150272308 | 0.052081694 | 0.028079503 | 0.024002192 | 0.999933887 | -0.207847426 | -0.192685817 | -0.015161609 | 0.947453112 |
| ENSG00000278311 | *GGNBP2* | 6.377746681 | -0.00974028 | -0.064373708 | 0.054633429 | 0.999933887 | -0.122223495 | -0.131655823 | 0.009432328 | 0.947535018 |
| ENSG00000005513 | *SOX8* | -0.977342476 | 0.216981221 | 0.300359644 | -0.083378423 | 0.999933887 | 0.056184593 | 0.100847219 | -0.044662625 | 0.947598092 |
| ENSG00000088543 | *C3orf18* | 0.246574487 | 0.267553345 | 0.038960283 | 0.228593062 | 0.999933887 | 0.175655343 | 0.214693376 | -0.039038033 | 0.947688677 |
| ENSG00000087253 | *LPCAT2* | 6.10374854 | -0.071888738 | -0.057628914 | -0.014259823 | 0.999933887 | -0.340095062 | -0.353700536 | 0.013605474 | 0.9477066 |
| ENSG00000054118 | *THRAP3* | 7.506440862 | -0.042630418 | -0.003440402 | -0.039190016 | 0.999933887 | -0.045974532 | -0.038398693 | -0.007575839 | 0.9477066 |
| ENSG00000169895 | *SYAP1* | 6.084690292 | -0.039422491 | -0.021143674 | -0.018278817 | 0.999933887 | 0.194565922 | 0.20454922 | -0.009983298 | 0.9477066 |
| ENSG00000099800 | *TIMM13* | 1.979400959 | 0.072404974 | -0.035604368 | 0.108009342 | 0.999933887 | 0.085287925 | 0.065497589 | 0.019790336 | 0.9477066 |
| ENSG00000168807 | *SNTB2* | 3.106678098 | 0.039522772 | 0.038842379 | 0.000680393 | 0.999933887 | -0.266313785 | -0.283760205 | 0.017446419 | 0.9477066 |
| ENSG00000164751 | *PEX2* | 3.118171891 | -0.041018587 | -0.147055785 | 0.106037198 | 0.999933887 | -0.099056649 | -0.113277636 | 0.014220986 | 0.9477066 |
| ENSG00000153093 | *ACOXL* | 0.370833734 | 0.119595726 | -0.033700657 | 0.153296383 | 0.999933887 | 0.473982153 | 0.507800517 | -0.033818364 | 0.9477066 |
| ENSG00000215912 | *TTC34* | 1.22354987 | -0.086354607 | -0.00570139 | -0.080653217 | 0.999933887 | 0.093976154 | 0.112618845 | -0.018642691 | 0.9477066 |
| ENSG00000150991 | *UBC* | 10.993200286 | 0.018970867 | -0.103561864 | 0.122532731 | 0.999933887 | -0.178949911 | -0.193338386 | 0.014388475 | 0.9477066 |
| ENSG00000286341 | *AP002762.3* | 1.484701902 | 0.024357974 | 0.134846708 | -0.110488734 | 0.999933887 | -0.081086531 | -0.113056673 | 0.031970142 | 0.9477066 |
| ENSG00000159958 | *TNFRSF13C* | 2.048113216 | 0.017332148 | 0.072637826 | -0.055305678 | 0.999933887 | -0.169792722 | -0.14975572 | -0.020037002 | 0.9477066 |
| ENSG00000280789 | *PAGR1* | 0.981201104 | 0.017391963 | -0.016192656 | 0.033584619 | 0.999933887 | 0.147896317 | 0.122305349 | 0.025590968 | 0.9477066 |
| ENSG00000182873 | *PRKCZ-AS1* | 1.540286243 | -0.011347513 | 0.131247009 | -0.142594521 | 0.999933887 | -0.200803458 | -0.226746124 | 0.025942666 | 0.9477066 |
| ENSG00000154229 | *PRKCA* | 2.722575898 | -0.002648625 | 0.066180644 | -0.068829269 | 0.999933887 | -0.081631936 | -0.097740126 | 0.01610819 | 0.9477066 |
| ENSG00000211898 | *IGHD* | 1.323632492 | -0.024230207 | 0.000477049 | -0.024707256 | 0.999933887 | 0.04484999 | 0.065888849 | -0.021038859 | 0.947752935 |
| ENSG00000224940 | *PRRT4* | 0.996207934 | 0.076393901 | 0.278557817 | -0.202163915 | 0.999933887 | -0.732680442 | -0.71113019 | -0.021550252 | 0.947884554 |
| ENSG00000204344 | *STK19* | 3.799450063 | -0.006788959 | 0.062031026 | -0.068819985 | 0.999933887 | 0.337650038 | 0.350237365 | -0.012587327 | 0.947921041 |
| ENSG00000127415 | *IDUA* | 2.219233859 | 0.142103106 | 0.253221429 | -0.111118322 | 0.999933887 | 0.081105775 | 0.060120043 | 0.020985732 | 0.947959684 |
| ENSG00000135127 | *BICDL1* | 4.726334426 | 0.051691536 | 0.110669033 | -0.058977497 | 0.999933887 | 0.237045231 | 0.226625316 | 0.010419915 | 0.947959684 |
| ENSG00000079313 | *REXO1* | 5.958676235 | 0.035301631 | 0.132588696 | -0.097287065 | 0.999933887 | -0.13823049 | -0.1269524 | -0.01127809 | 0.947959684 |
| ENSG00000180644 | *PRF1* | 5.687772585 | -0.024681984 | -0.034958516 | 0.010276532 | 0.999933887 | 0.085797025 | 0.073948798 | 0.011848227 | 0.947959684 |
| ENSG00000255585 | *AL590627.1* | 0.451182335 | -0.010911043 | 0.029713311 | -0.040624354 | 0.999933887 | -0.310294886 | -0.345367754 | 0.035072868 | 0.947959684 |
| ENSG00000143224 | *PPOX* | 2.023715852 | -0.015214311 | 0.080713679 | -0.09592799 | 0.999933887 | 0.092117759 | 0.112921325 | -0.020803566 | 0.947962835 |
| ENSG00000139187 | *KLRG1* | 2.720464792 | -0.056224843 | -0.204885218 | 0.148660375 | 0.999933887 | 0.111271123 | 0.128827397 | -0.017556274 | 0.94809746 |
| ENSG00000156398 | *SFXN2* | 0.617342682 | -0.063110109 | 0.132743356 | -0.195853464 | 0.999933887 | 0.123110968 | 0.098919179 | 0.024191789 | 0.94809746 |
| ENSG00000160360 | *GPSM1* | 0.623488751 | 0.258484852 | -0.031643938 | 0.290128791 | 0.999933887 | 0.043069245 | 0.066202082 | -0.023132838 | 0.948205846 |
| ENSG00000267100 | *ILF3-DT* | 4.19255422 | 0.04109016 | -0.032884116 | 0.073974276 | 0.999933887 | 0.28518481 | 0.270424832 | 0.014759978 | 0.948205846 |
| ENSG00000100823 | *APEX1* | 3.857003778 | 0.03590774 | 0.069539619 | -0.033631878 | 0.999933887 | 0.183528129 | 0.197026035 | -0.013497906 | 0.948205846 |
| ENSG00000101856 | *PGRMC1* | 2.031815232 | 0.007050056 | -0.053058745 | 0.060108801 | 0.999933887 | -0.095415631 | -0.113043768 | 0.017628137 | 0.948205846 |
| ENSG00000258597 | *SERPINA2* | -0.824739428 | -0.270838105 | 0.185021936 | -0.45586004 | 0.999933887 | 2.974490868 | 3.041975695 | -0.067484827 | 0.948275964 |
| ENSG00000164776 | *PHKG1* | 0.696518509 | 0.11390333 | 0.20120643 | -0.0873031 | 0.999933887 | 0.182227582 | 0.152045996 | 0.030181586 | 0.948275964 |
| ENSG00000205517 | *RGL3* | -0.331002899 | -0.045099126 | -0.215786605 | 0.170687479 | 0.999933887 | -0.287023817 | -0.237107868 | -0.049915949 | 0.948275964 |

| | | | | | | | | | | |
|---|---|---|---|---|---|---|---|---|---|---|
| ENSG00000101126 | *ADNP* | 6.075507841 | -0.092286809 | -0.076192885 | -0.016093925 | 0.999933887 | -0.225075443 | -0.234236054 | 0.00916061 | 0.948369196 |
| ENSG00000125835 | *SNRPB* | 5.401672773 | -0.007028135 | 0.060897819 | -0.067925955 | 0.999933887 | 0.127522878 | 0.117971012 | 0.009551866 | 0.948380406 |
| ENSG00000184863 | *RBM33* | 6.828358958 | -0.003839796 | -0.043427599 | 0.039587803 | 0.999933887 | -0.330438057 | -0.339639387 | 0.00920133 | 0.948380406 |
| ENSG00000051108 | *HERPUD1* | 7.147189329 | 0.002125837 | 0.011979746 | -0.00985391 | 0.999933887 | 0.121431201 | 0.10870374 | 0.012727461 | 0.948642698 |
| ENSG00000163479 | *SSR2* | 6.151545493 | -0.008515095 | -0.047707261 | 0.039192167 | 0.999933887 | -0.041219005 | -0.050721384 | 0.009502379 | 0.948871892 |
| ENSG00000139428 | *MMAB* | 0.447297467 | 0.035210457 | 0.170094088 | -0.134883631 | 0.999933887 | 0.146655975 | 0.175648839 | -0.028992864 | 0.948964044 |
| ENSG00000259431 | *THTPA* | -0.386034592 | 0.001095235 | -0.065565608 | 0.066660842 | 0.999933887 | 0.191128961 | 0.156725784 | 0.034403177 | 0.948968457 |
| ENSG00000125844 | *RRBP1* | 5.241828003 | 0.052928542 | 0.054608397 | -0.001679855 | 0.999933887 | 0.083142361 | 0.094475392 | -0.01133303 | 0.949691277 |
| ENSG00000065665 | *SEC61A2* | 4.08124583 | -0.088816596 | 0.055861923 | -0.144678519 | 0.999933887 | -0.200614851 | -0.21192162 | 0.011306769 | 0.949930629 |
| ENSG00000267064 | *UXT-AS1* | -0.552757647 | 0.305935294 | 0.198622885 | 0.10731241 | 0.999933887 | 1.164161189 | 1.125817118 | 0.038344071 | 0.949930629 |
| ENSG00000197530 | *MIB2* | 5.408116767 | 0.12454725 | 0.152748539 | -0.028201289 | 0.999933887 | 0.145787684 | 0.128281851 | 0.017505834 | 0.949930629 |
| ENSG00000160410 | *SHKBP1* | 9.067354672 | 0.086881299 | 0.030024955 | 0.056856344 | 0.999933887 | -0.634646276 | -0.646924863 | 0.012278588 | 0.949930629 |
| ENSG00000287317 | *AC016769.6* | -0.994876693 | -0.148232751 | 0.087517568 | -0.235750319 | 0.999933887 | 0.353507866 | 0.401916767 | -0.0484089 | 0.949930629 |
| ENSG00000137040 | *RANBP6* | 3.117394967 | -0.080558815 | -0.214029231 | 0.133470415 | 0.999933887 | -0.158962278 | -0.179263566 | 0.020301288 | 0.949930629 |
| ENSG00000211714 | *TRBV7-3* | -0.857189854 | 0.107262513 | -0.166307366 | 0.273569878 | 0.999933887 | 0.168718581 | 0.207552307 | -0.038833725 | 0.949930629 |
| ENSG00000137776 | *SLTM* | 5.459733824 | -0.035156938 | -0.111722706 | 0.076565768 | 0.999933887 | -0.63740783 | -0.658768537 | 0.021360707 | 0.949930629 |
| ENSG00000163131 | *CTSS* | 9.984898271 | -0.014181002 | -0.072750568 | 0.058569566 | 0.999933887 | 0.134902739 | 0.124512243 | 0.010390496 | 0.949930629 |
| ENSG00000175931 | *UBE2O* | 4.030062305 | 0.008386051 | 0.023194634 | -0.014808583 | 0.999933887 | 0.207142734 | 0.221647051 | -0.014504316 | 0.949930629 |
| ENSG00000172992 | *DCAKD* | 1.658979528 | -0.027393888 | -0.046312631 | 0.018918743 | 0.999933887 | -0.079854241 | -0.09710068 | 0.017246439 | 0.950114404 |
| ENSG00000073792 | *IGF2BP2* | -0.530078221 | -0.291513305 | -0.224403778 | -0.067109527 | 0.999933887 | -0.245629142 | -0.211488938 | -0.034140204 | 0.950177045 |
| ENSG00000186395 | *KRT10* | 2.828936264 | 0.117621652 | 0.061900537 | 0.055721115 | 0.999933887 | 0.008378786 | -0.009440976 | 0.017819761 | 0.950177045 |
| ENSG00000144182 | *LIPT1* | 0.918490194 | -0.210569856 | -0.170185041 | -0.040384815 | 0.999933887 | 0.23279575 | 0.21023097 | 0.022564779 | 0.950208158 |
| ENSG00000114859 | *CLCN2* | -0.017241615 | 0.121206476 | -0.108736695 | 0.22994317 | 0.999933887 | 0.193227614 | 0.225744785 | -0.032517171 | 0.95029206 |
| ENSG00000213719 | *CLIC1* | 8.966249035 | 0.039184412 | -0.022369794 | 0.061554205 | 0.999933887 | -0.301693576 | -0.291159972 | -0.010533605 | 0.95029206 |
| ENSG00000223356 | *AL590666.1* | -0.792105123 | -0.031183933 | 0.217488688 | -0.248672621 | 0.999933887 | -0.113034298 | -0.170099338 | 0.057065041 | 0.95029206 |
| ENSG00000160856 | *FCRL3* | 3.691406517 | 0.01547312 | -0.085529867 | 0.101002986 | 0.999933887 | 0.13753529 | 0.147205794 | -0.009670503 | 0.950293365 |
| ENSG00000197471 | *SPN* | 4.601521363 | 0.010538547 | -0.024175315 | 0.034713861 | 0.999933887 | 0.087177491 | 0.098464125 | -0.011286633 | 0.950293365 |
| ENSG00000257599 | *OVCH1-AS1* | -0.227565496 | -0.215374968 | 0.195820427 | -0.411195394 | 0.999933887 | -0.308671113 | -0.33971686 | 0.031045747 | 0.950563732 |
| ENSG00000114999 | *TTL* | 4.154781306 | -0.091474647 | 0.048354231 | -0.139828878 | 0.999933887 | 1.156515084 | 1.176230308 | -0.019715224 | 0.950563732 |
| ENSG00000180596 | *H2BC4* | 5.528492838 | 0.070266897 | -0.012263602 | 0.082530499 | 0.999933887 | 0.06176994 | 0.074903914 | -0.013133975 | 0.950582282 |
| ENSG00000144026 | *ZNF514* | 1.189394548 | 0.049547825 | -0.16277504 | 0.212322865 | 0.999933887 | 0.14013024 | 0.119001059 | 0.021129181 | 0.950582282 |
| ENSG00000187514 | *PTMA* | 7.536795987 | -0.012329136 | -0.058781069 | 0.046451933 | 0.999933887 | 0.064928496 | 0.05544324 | 0.009485256 | 0.950582282 |
| ENSG00000125991 | *ERGIC3* | 4.632905135 | 0.003482339 | -0.014589821 | 0.018072161 | 0.999933887 | 0.003382021 | -0.007524342 | 0.010906364 | 0.95071707 |
| ENSG00000181666 | *ZNF875* | 2.423780712 | 0.003855138 | 0.149451532 | -0.145596394 | 0.999933887 | 0.104837749 | 0.12084936 | -0.016011611 | 0.95071707 |
| ENSG00000146282 | *RARS2* | 3.496712097 | -0.060549956 | -0.01162903 | -0.048920926 | 0.999933887 | -0.240662617 | -0.251827773 | 0.011165156 | 0.950782583 |
| ENSG00000144199 | *FAHD2B* | -0.155053472 | 0.178798788 | -0.211265445 | 0.390064233 | 0.999933887 | 0.190780483 | 0.228014735 | -0.037234253 | 0.950782583 |
| ENSG00000197111 | *PCBP2* | 6.674409144 | -0.033592627 | -0.007699551 | -0.025893076 | 0.999933887 | -0.093154215 | -0.100724723 | 0.007570508 | 0.950782583 |
| ENSG00000178229 | *ZNF543* | -0.001956869 | -0.157414356 | -0.101594008 | -0.055820348 | 0.999933887 | 0.019584359 | 0.053203661 | -0.033619302 | 0.950934019 |
| ENSG00000221962 | *TMEM14EP* | 0.235562102 | 0.160992648 | 0.469874434 | -0.308881786 | 0.999933887 | 0.248632895 | 0.304345953 | -0.055713058 | 0.950934019 |
| ENSG00000172878 | *METAP1D* | -0.635517621 | -0.079984285 | 0.164858287 | -0.244842571 | 0.999933887 | -0.120866991 | -0.164343894 | 0.043476903 | 0.950934019 |
| ENSG00000138459 | *SLC35A5* | 2.839249428 | 0.019643632 | -0.033886622 | 0.053530254 | 0.999933887 | 0.067960417 | 0.083631787 | -0.01567137 | 0.950934019 |
| ENSG00000173083 | *HPSE* | 6.190153325 | -0.019613051 | 0.013779458 | -0.033392508 | 0.999933887 | -1.061155103 | -1.045915171 | -0.015239932 | 0.950934019 |
| ENSG00000171051 | *FPR1* | 10.763814614 | 0.011735376 | -0.04111882 | 0.052854196 | 0.999933887 | 0.313509338 | 0.323808387 | -0.01029905 | 0.950934019 |
| ENSG00000101911 | *PRPS2* | 1.790133028 | 0.004225424 | -0.029978364 | 0.034203789 | 0.999933887 | 0.060438682 | 0.036218428 | 0.024220254 | 0.950934019 |
| ENSG00000112855 | *HARS2* | 4.485481863 | -0.103517905 | -0.038590686 | -0.064927218 | 0.999933887 | -0.077781186 | -0.067548194 | -0.010232991 | 0.950964728 |
| ENSG00000204469 | *PRRC2A* | 8.063523218 | 0.037177148 | 0.071526562 | -0.034349414 | 0.999933887 | 0.034046451 | 0.043404581 | -0.009358131 | 0.950964728 |
| ENSG00000099814 | *CEP170B* | 0.587479378 | 0.117069159 | 0.142171618 | -0.025102459 | 0.999933887 | 0.052883088 | 0.078277309 | -0.02539422 | 0.951070774 |
| ENSG00000157978 | *LDLRAP1* | 3.530036549 | 0.038863138 | 0.134560398 | -0.095697261 | 0.999933887 | 0.002649796 | -0.009246905 | 0.011896701 | 0.951070774 |
| ENSG00000186812 | *ZNF397* | 4.716835918 | -0.031832326 | -0.033251542 | 0.001419216 | 0.999933887 | -0.100180076 | -0.111705714 | 0.011525638 | 0.951070774 |
| ENSG00000183508 | *TENT5C* | 5.257571096 | -0.054788614 | -0.064493135 | 0.009704521 | 0.999933887 | 0.076096953 | 0.066551968 | 0.009544984 | 0.951157019 |
| ENSG00000147316 | *MCPH1* | 3.130193499 | 0.028784372 | 0.07565032 | -0.046865949 | 0.999933887 | 0.77235484 | 0.797414124 | -0.025059284 | 0.951157019 |
| ENSG00000213203 | *GIMAP1* | 2.846240191 | 0.106703834 | 0.04922279 | 0.057481044 | 0.999933887 | 0.140281715 | 0.12520044 | 0.015081276 | 0.951162738 |
| ENSG00000122035 | *RASL11A* | -0.303597833 | 0.134295957 | 0.325368756 | -0.191072798 | 0.999933887 | 0.353938701 | 0.311526526 | 0.042412174 | 0.951554843 |
| ENSG00000274717 | *AL049757.1* | 0.790239698 | 0.187557621 | 0.2579874 | -0.07042978 | 0.999933887 | 1.396951873 | 1.373468656 | 0.023483217 | 0.951610098 |
| ENSG00000177875 | *CCDC184* | -0.24748119 | 0.033960414 | 0.070383856 | -0.036423443 | 0.999933887 | 0.237034805 | 0.196183739 | 0.040851066 | 0.951610098 |
| ENSG00000100100 | *PIK3IP1* | 7.480890107 | 0.009720587 | 0.002276451 | 0.007444136 | 0.999933887 | 0.497603119 | 0.508638825 | -0.011035706 | 0.951610098 |
| ENSG00000198420 | *TCAF1* | -0.019385217 | 0.028534001 | -0.299552345 | 0.328086346 | 0.999933887 | -0.05014099 | -0.079906581 | 0.029765591 | 0.951610098 |
| ENSG00000241351 | *IGKV3-11* | -1.309942081 | -0.035238113 | -0.085120508 | 0.049882395 | 0.999933887 | 0.441680674 | 0.386954278 | 0.054726396 | 0.951610098 |
| ENSG00000120784 | *ZFP30* | 0.076974974 | -0.14095253 | 0.101420665 | -0.242373196 | 0.999933887 | -0.094292352 | -0.132159405 | 0.037867053 | 0.951691008 |
| ENSG00000259448 | *LINC02352* | 1.341017948 | 0.020682125 | 0.103174205 | -0.08249208 | 0.999933887 | -0.147331126 | -0.1189486 | -0.028382526 | 0.951691008 |
| ENSG00000155659 | *VSIG4* | -0.859669974 | 0.063789615 | 0.462885614 | -0.399095998 | 0.999933887 | 0.116961192 | 0.059552921 | 0.057408272 | 0.951736392 |
| ENSG00000134440 | *NARS1* | 6.06765241 | -0.070519751 | -0.091052375 | 0.020532624 | 0.999933887 | -0.220017419 | -0.231444373 | 0.011426953 | 0.951737716 |
| ENSG00000079308 | *TNS1* | -0.379798344 | -0.003265726 | 0.396882454 | -0.400148181 | 0.999933887 | -0.388431411 | -0.354294673 | -0.034136739 | 0.951737716 |
| ENSG00000074803 | *SLC12A1* | -0.076391963 | -0.211042373 | -0.093843754 | -0.117198619 | 0.999933887 | -0.16999544 | -0.11347845 | -0.05651699 | 0.951770366 |
| ENSG00000185885 | *IFITM1* | 6.947365428 | 0.123888042 | 0.02236676 | 0.101521282 | 0.999933887 | 0.06690853 | 0.049291552 | 0.017616978 | 0.951940223 |
| ENSG00000169019 | *COMMD8* | 2.484526587 | -0.132439271 | -0.230306417 | 0.097867147 | 0.999933887 | -0.516008129 | -0.535239641 | 0.019231511 | 0.951940223 |
| ENSG00000107331 | *ABCA2* | 6.582464586 | 0.059700842 | 0.058098254 | 0.001602588 | 0.999933887 | 0.151193341 | 0.160377038 | -0.009183697 | 0.951940223 |
| ENSG00000127191 | *TRAF2* | 2.926025359 | 0.068234902 | 0.071948796 | -0.003713894 | 0.999933887 | 0.242440244 | 0.229863825 | 0.012576419 | 0.951940223 |
| ENSG00000211592 | *IGKC* | 3.475080115 | 0.059808647 | -0.131960861 | 0.191769508 | 0.999933887 | 0.31414972 | 0.298828265 | 0.015321456 | 0.951940223 |
| ENSG00000155252 | *PI4K2A* | 4.309150847 | -0.040099387 | -0.023249119 | -0.016850267 | 0.999933887 | -0.321267846 | -0.30885032 | -0.012417526 | 0.951940223 |
| ENSG00000112297 | *CRYBG1* | 6.1811749 | -0.049004945 | -0.082546716 | 0.03354177 | 0.999933887 | 0.189952035 | 0.204497672 | -0.014545638 | 0.951940223 |
| ENSG00000141965 | *FEM1A* | 1.190871358 | -0.066946559 | 0.020162482 | -0.087109041 | 0.999933887 | -0.090320366 | -0.114253752 | 0.023933386 | 0.951940223 |
| ENSG00000160783 | *PMF1* | 2.857866571 | 0.001942608 | 0.034537371 | -0.032594763 | 0.999933887 | -0.178855938 | -0.198387441 | 0.019531503 | 0.951940223 |
| ENSG00000121741 | *ZMYM2* | 5.47388259 | 0.002535382 | -0.058508256 | 0.061043638 | 0.999933887 | -0.598566778 | -0.576315771 | -0.022251007 | 0.951940223 |
| ENSG00000165660 | *ABRAXAS2* | 2.465332989 | -0.087317023 | -0.047396322 | -0.039920701 | 0.999933887 | -0.17289862 | -0.188916297 | 0.016017676 | 0.952288266 |
| ENSG00000160050 | *CCDC28B* | 1.289527424 | 0.076130823 | 0.094429838 | -0.018299015 | 0.999933887 | 0.371993103 | 0.348796204 | 0.023196899 | 0.952288266 |
| ENSG00000106692 | *FKTN* | 0.546897474 | -0.32545684 | -0.303821904 | -0.021634936 | 0.999933887 | 0.029541937 | -0.000838325 | 0.030380262 | 0.952330865 |
| ENSG00000007350 | *TKTL1* | 3.290005338 | -0.135486844 | 0.068011262 | -0.203498106 | 0.999933887 | -0.957038399 | -0.940826505 | -0.016211894 | 0.952330865 |
| ENSG00000140650 | *PMM2* | 1.204516775 | 0.107000926 | 0.01530556 | 0.091695365 | 0.999933887 | 0.011875778 | 0.032093936 | -0.020218158 | 0.952330865 |
| ENSG00000235066 | *AC009303.2* | 2.329135394 | 0.115989544 | 0.340929961 | -0.224940417 | 0.999933887 | 0.418991863 | 0.442236698 | -0.023244834 | 0.952330865 |
| ENSG00000238035 | *AC138035.1* | 0.889225314 | -0.023630583 | 0.110571354 | -0.134201937 | 0.999933887 | 0.313065788 | 0.283679603 | 0.029386185 | 0.952330865 |
| ENSG00000279187 | *AC027601.4* | -0.443025768 | -0.056461568 | 0.060081092 | -0.11654266 | 0.999933887 | 0.019778369 | 0.061273821 | -0.041495452 | 0.952330865 |
| ENSG00000113456 | *RAD1* | 2.346422047 | 0.009415924 | -0.043922932 | 0.053338857 | 0.999933887 | -0.002071555 | 0.016018836 | -0.018090391 | 0.952330865 |
| ENSG00000060237 | *WNK1* | 7.914749112 | 0.004732266 | 0.027997377 | -0.02326511 | 0.999933887 | 0.00850347 | -0.004622559 | 0.013126029 | 0.952330865 |
| ENSG00000263069 | *RNF213-AS1* | 3.954982395 | 0.053980585 | 0.089964093 | -0.035983508 | 0.999933887 | -0.342214038 | -0.35744221 | 0.015228172 | 0.952413271 |
| ENSG00000116761 | *CTH* | -0.093127906 | -0.022258377 | -0.006884443 | -0.015373934 | 0.999933887 | 0.129425041 | 0.167274766 | -0.037849724 | 0.952413271 |
| ENSG00000234369 | *TATDN1P1* | -0.110651169 | 0.185273714 | 0.667328138 | -0.482054424 | 0.999933887 | 0.249529715 | 0.180195782 | 0.069333933 | 0.952489117 |
| ENSG00000151692 | *RNF144A* | 4.205832119 | -0.049992583 | 0.107666312 | -0.157658895 | 0.999933887 | -0.143290165 | -0.154530163 | 0.011239997 | 0.952489117 |
| ENSG00000182534 | *MXRA7* | 4.263378869 | 0.044418267 | -0.011289679 | 0.055707945 | 0.999933887 | -0.155880716 | -0.167955497 | 0.012074781 | 0.952489117 |
| ENSG00000261801 | *LOXL1-AS1* | 1.03286122 | 0.041964121 | 0.077582357 | -0.035618236 | 0.999933887 | -0.152443984 | -0.125776406 | -0.026667579 | 0.952489117 |
| ENSG00000171574 | *ZNF584* | -0.302298306 | -0.069346868 | 0.1164076 | -0.185754468 | 0.999933887 | 0.141055691 | 0.109579059 | 0.031476632 | 0.952489117 |
| ENSG00000091409 | *ITGA6* | 3.209437802 | -0.045763342 | 0.05037852 | -0.096141862 | 0.999933887 | -0.10852088 | -0.094197883 | -0.014322998 | 0.952641634 |
| ENSG00000263874 | *LINC00672* | 1.605543058 | -0.064837525 | 0.044826873 | -0.109664398 | 0.999933887 | -0.367822824 | -0.346607654 | -0.02121517 | 0.952641634 |
| ENSG00000273002 | *AL355388.1* | 0.693114703 | -0.102136205 | -0.081802321 | -0.020333883 | 0.999933887 | 0.664035759 | 0.637522003 | 0.026513756 | 0.952641634 |
| ENSG00000280604 | *AJ239328.1* | 0.323083058 | -0.082776443 | 0.065110871 | -0.147887315 | 0.999933887 | -0.151593209 | -0.128889394 | -0.022703815 | 0.952641634 |
| ENSG00000147234 | *FRMPD3* | 0.676815558 | 0.033102128 | -0.011435384 | 0.044537513 | 0.999933887 | -0.044101672 | -0.067134718 | 0.023033046 | 0.952641634 |
| ENSG00000249309 | *AC020703.1* | 0.03186179 | 0.021944827 | 0.160582803 | -0.138637976 | 0.999933887 | 0.677645334 | 0.711810384 | -0.03416505 | 0.952641634 |
| ENSG00000245112 | *SMARCA5-AS* | -0.431865674 | -0.015457544 | 0.144215469 | -0.159673013 | 0.999933887 | 0.367537415 | 0.325579684 | 0.041957731 | 0.952641634 |
| ENSG00000185115 | *NSMCE3* | 3.497920886 | 0.102186406 | -0.010802765 | 0.11298917 | 0.999933887 | 0.079473349 | 0.063782755 | 0.015690594 | 0.952662941 |
| ENSG00000124507 | *PACSIN1* | -0.561976512 | 0.109046968 | -0.13974855 | 0.248795518 | 0.999933887 | -0.074592191 | -0.030205528 | -0.044386663 | 0.952662941 |
| ENSG00000106327 | *TFR2* | 0.130611213 | 0.102429609 | 0.354668851 | -0.252239242 | 0.999933887 | -0.199382741 | -0.235025312 | 0.035642571 | 0.952662941 |
| ENSG00000187653 | *TMSB4XP8* | 2.452161559 | -0.020071797 | 0.013016228 | -0.033088025 | 0.999933887 | 0.361874407 | 0.381383208 | -0.019508801 | 0.952662941 |
| ENSG00000131831 | *RAI2* | 0.83231581 | -0.113801123 | -0.298306206 | 0.184505083 | 0.999933887 | -0.574971688 | -0.592355517 | 0.017383829 | 0.952737194 |
| ENSG00000163866 | *SMIM12* | 4.422733519 | -0.03665098 | -0.015673472 | -0.020977508 | 0.999933887 | -0.309497888 | -0.299349838 | -0.01014805 | 0.952832992 |
| ENSG00000287507 | *AP001053.1* | -0.74974774 | 0.038249058 | -0.129274234 | 0.167523292 | 0.999933887 | 2.050639408 | 2.088259623 | -0.037620215 | 0.952832992 |

| | | | | | | | | | | |
|---|---|---|---|---|---|---|---|---|---|---|
| ENSG00000273111 | *LYPD4* | -1.2861978 | 0.034485933 | -0.218309797 | 0.252795729 | 0.999933887 | -0.946746124 | -1.029888194 | 0.08314207 | 0.952832992 |
| ENSG00000235505 | *CASP4LP* | 0.99182953 | -0.037377878 | -0.168025869 | 0.130647992 | 0.999933887 | 0.034194228 | 0.013750802 | 0.020443426 | 0.953012139 |
| ENSG00000162390 | *ACOT11* | -0.573281027 | 0.02330739 | 0.222160415 | -0.198853025 | 0.999933887 | -0.239403189 | -0.197182183 | -0.042221007 | 0.953012139 |
| ENSG00000179899 | *PHC1P1* | 0.217631268 | 0.176002139 | 0.013784913 | 0.162217226 | 0.999933887 | -0.209084458 | -0.190131359 | -0.018953098 | 0.953171152 |
| ENSG00000132669 | *RIN2* | 0.766146471 | 0.05501205 | -0.006961332 | 0.061973382 | 0.999933887 | 1.06630712 | 1.042238098 | 0.024069022 | 0.95325376 |
| ENSG00000146232 | *NFKBIE* | 7.075156974 | -0.044376713 | -0.017139586 | -0.027237128 | 0.999933887 | 1.96054015 | 1.977207425 | -0.016667275 | 0.953456245 |
| ENSG00000134594 | *RAB33A* | 0.580379733 | 0.033263954 | -0.149999518 | 0.183263472 | 0.999933887 | -0.165592848 | -0.192374365 | 0.026781517 | 0.953627819 |
| ENSG00000079974 | *RABL2B* | 2.320610207 | 0.016152104 | -0.093722534 | 0.109874638 | 0.999933887 | -0.103672985 | -0.08821606 | -0.015456925 | 0.953627819 |
| ENSG00000165782 | *PIP4P1* | 8.069305057 | 0.104164523 | 0.129304952 | -0.025140429 | 0.999933887 | 0.291122675 | 0.303927599 | -0.012804924 | 0.953653605 |
| ENSG00000111729 | *CLEC4A* | 1.761790711 | -0.066956539 | -0.07940453 | 0.012447991 | 0.999933887 | 0.124930191 | 0.102709627 | 0.022220563 | 0.953653605 |
| ENSG00000164300 | *SERINC5* | 5.195212339 | -0.019340779 | 0.014613557 | -0.033954336 | 0.999933887 | -0.424636544 | -0.414754509 | -0.009882035 | 0.953653605 |
| ENSG00000259326 | *AC116158.1* | -0.471181288 | 0.038219195 | -0.092238484 | 0.13045768 | 0.999933887 | 0.176766772 | 0.136138807 | 0.040627965 | 0.953653605 |
| ENSG00000105851 | *PIK3CG* | 5.832044453 | -0.028694103 | 0.005642677 | -0.034336781 | 0.999933887 | -0.20931796 | -0.199754349 | -0.009563612 | 0.953854675 |
| ENSG00000185052 | *SLC24A3* | 0.648466014 | -0.065735605 | 0.100522847 | -0.166258452 | 0.999933887 | -0.200378873 | -0.174935857 | -0.025443015 | 0.953952022 |
| ENSG00000231793 | *DOC2GP* | -1.191617383 | 0.314134772 | 0.05408993 | 0.260044842 | 0.999933887 | 0.192356885 | 0.147824022 | 0.044532863 | 0.954069513 |
| ENSG00000286020 | *AC011997.2* | -1.021951772 | 0.30245517 | -0.206362219 | 0.508817389 | 0.999933887 | 1.066090401 | 1.020360245 | 0.045730156 | 0.954069513 |
| ENSG00000256937 | *KRT17P8* | -0.297520342 | 0.126936419 | 0.34555903 | -0.218622611 | 0.999933887 | 0.064179351 | 0.02998942 | 0.034189931 | 0.954069513 |
| ENSG00000106245 | *BUD31* | 4.270342774 | 0.025709348 | -0.028859695 | 0.054569042 | 0.999933887 | 0.058843334 | 0.068975346 | -0.010132012 | 0.954069513 |
| ENSG00000151458 | *ANKRD50* | 2.064925661 | 0.034887801 | -0.053802799 | 0.0886906 | 0.999933887 | -0.371512415 | -0.389736089 | 0.018223674 | 0.954069513 |
| ENSG00000132950 | *ZMYM5* | 2.737363725 | -0.017764201 | -0.005325409 | -0.012438791 | 0.999933887 | -0.35367076 | -0.368770335 | 0.015099574 | 0.954069513 |
| ENSG00000085831 | *TTC39A* | 1.476823831 | 0.04854969 | 0.111018877 | -0.062469187 | 0.999933887 | 0.305751697 | 0.33032113 | -0.024569433 | 0.954934965 |
| ENSG00000256576 | *LINC02361* | 0.361449575 | 0.12227563 | -0.336319027 | 0.458594657 | 0.944094537 | -0.089756795 | -0.069169659 | -0.020587136 | 0.954989585 |
| ENSG00000279227 | *AC009303.4* | 5.358437322 | -0.113702686 | -0.089016766 | -0.02468592 | 0.999933887 | -0.348220352 | -0.361005002 | 0.01278465 | 0.955195438 |
| ENSG00000176108 | *CHMP6* | 2.275278155 | 0.038966331 | -0.134201989 | 0.17316832 | 0.999933887 | -0.176422439 | -0.19123898 | 0.01481654 | 0.955195438 |
| ENSG00000135446 | *CDK4* | 2.4917992 | 0.027722013 | 0.031222269 | -0.003500257 | 0.999933887 | 0.073409225 | 0.059238643 | 0.014170583 | 0.955195438 |
| ENSG00000134590 | *RTL8C* | 2.6365341 | 0.005443247 | -0.003867975 | 0.009311221 | 0.999933887 | -0.0345938 | -0.018838682 | -0.015755118 | 0.955195438 |
| ENSG00000034693 | *PEX3* | 0.051701522 | -0.182294337 | -0.389449836 | 0.207155499 | 0.999933887 | 0.132607714 | 0.159048177 | -0.026440463 | 0.955197879 |
| ENSG00000112685 | *EXOC2* | 3.332522901 | 0.011413675 | -0.118915171 | 0.130328846 | 0.999933887 | -0.017845235 | -0.027971988 | 0.010126753 | 0.955197879 |
| ENSG00000102878 | *HSF4* | 1.027792914 | 0.028219273 | 0.076180215 | -0.047960942 | 0.999933887 | 0.044659744 | 0.071774678 | -0.027114935 | 0.955256161 |
| ENSG00000107581 | *EIF3A* | 6.784708662 | -0.047844777 | -0.104049384 | 0.056204607 | 0.999933887 | -0.21689112 | -0.229241936 | 0.012350816 | 0.95526546 |
| ENSG00000113763 | *UNC5A* | 1.107843374 | -0.031839587 | 0.070769013 | -0.1026086 | 0.999933887 | 0.356347881 | 0.398884956 | -0.042537075 | 0.95526546 |
| ENSG00000005469 | *CROT* | 0.867112551 | -0.080360513 | 0.011007673 | -0.091368186 | 0.999933887 | -0.172338407 | -0.196870864 | 0.024532456 | 0.955513525 |
| ENSG00000256269 | *HMBS* | 0.007134343 | 0.077906601 | 0.0116264 | 0.066280201 | 0.999933887 | 0.091439355 | 0.125064393 | -0.033625038 | 0.955628484 |
| ENSG00000007047 | *MARK4* | 4.774657848 | 0.004776495 | 0.05056112 | -0.045784625 | 0.999933887 | 0.00657481 | 0.016669034 | -0.010094224 | 0.955652226 |
| ENSG00000186153 | *WWOX* | 0.163718045 | -0.166750747 | 0.043127555 | -0.209878302 | 0.999933887 | 0.065652948 | 0.098530965 | -0.032878017 | 0.955664934 |
| ENSG00000133678 | *TMEM254* | -0.301230958 | -0.115464226 | -0.188023425 | 0.072559199 | 0.999933887 | 0.205345832 | 0.244037412 | -0.03869158 | 0.955666426 |
| ENSG00000223599 | *AL513523.1* | 0.252280885 | 0.169010986 | -0.048200113 | 0.217211098 | 0.999933887 | 0.062730385 | 0.094269279 | -0.031538894 | 0.955988514 |
| ENSG00000233013 | *FAM157B* | 2.514344862 | 0.076424602 | -0.119966996 | 0.196391598 | 0.999933887 | 0.604601623 | 0.585130596 | 0.019471027 | 0.956205902 |
| ENSG00000139579 | *NABP2* | 1.152150043 | -0.089687718 | 0.068790131 | -0.15847785 | 0.999933887 | 0.083593874 | 0.06278377 | 0.020810104 | 0.956205902 |
| ENSG00000203739 | *PRDX6-AS1* | 1.985028663 | -0.044875501 | -0.114556939 | 0.069681438 | 0.999933887 | 0.408791368 | 0.389979194 | 0.018812174 | 0.956205902 |
| ENSG00000125170 | *DOK4* | 0.585395277 | -0.045662437 | 0.006930275 | -0.052592712 | 0.999933887 | -0.039287696 | -0.079076899 | 0.039789203 | 0.956292034 |
| ENSG00000161405 | *IKZF3* | 5.114291519 | -0.051666579 | 0.033706589 | -0.085373168 | 0.999933887 | 0.135200972 | 0.143479045 | -0.008278073 | 0.956366357 |
| ENSG00000263167 | *AC115099.1* | 0.72838292 | -0.073870532 | 0.207295155 | -0.281165688 | 0.999933887 | 0.5670577 | 0.537403204 | 0.029654497 | 0.956366357 |
| ENSG00000136261 | *BZW2* | 1.686352472 | -0.042812515 | -0.090187667 | 0.047375152 | 0.999933887 | 0.287553746 | 0.269088725 | 0.018465021 | 0.956390782 |
| ENSG00000287736 | *PTCSC1* | 3.437260763 | -0.035042068 | 0.048527191 | -0.08356926 | 0.999933887 | 0.022898299 | 0.042111562 | -0.019213262 | 0.956430829 |
| ENSG00000235820 | *AL109935.1* | -0.101004789 | -0.178921907 | -0.16164187 | -0.017280037 | 0.999933887 | 0.07614578 | 0.109785481 | -0.033639701 | 0.956479179 |
| ENSG00000238086 | *PPP1R26P1* | -0.486808977 | -0.13037134 | 0.013830389 | -0.14420173 | 0.999933887 | -0.11998095 | -0.158046195 | 0.038065245 | 0.956479179 |
| ENSG00000089327 | *FXYD5* | 5.929658042 | -0.02370608 | -0.021888651 | -0.001817429 | 0.999933887 | 0.040727864 | 0.049922288 | -0.009194424 | 0.956479179 |
| ENSG00000257704 | *INAFM1* | 4.251261442 | 0.16872793 | 0.124536587 | 0.044191343 | 0.999933887 | -0.011375233 | -0.026625489 | 0.015250256 | 0.956710356 |
| ENSG00000171428 | *NAT1* | 2.075735854 | -0.140468717 | -0.135488685 | -0.004980032 | 0.999933887 | -0.219051066 | -0.202398419 | -0.016652647 | 0.956710356 |
| ENSG00000137571 | *SLCO5A1* | -0.191982202 | -0.108801423 | 0.062552761 | -0.171354184 | 0.999933887 | 2.015892726 | 2.053959764 | -0.038067038 | 0.956710356 |
| ENSG00000119640 | *ACYP1* | -0.317879113 | -0.06408025 | -0.256640042 | 0.192559791 | 0.999933887 | 0.096448164 | 0.130049024 | -0.03360086 | 0.956710356 |
| ENSG00000260378 | *AC109597.1* | -0.61572208 | -0.244598085 | 0.04791073 | -0.292508814 | 0.999933887 | -0.277351783 | -0.218344138 | -0.059007645 | 0.956749439 |
| ENSG00000058729 | *RIOK2* | 1.182762935 | -0.130313658 | -0.35426998 | 0.223956322 | 0.999933887 | 0.012901455 | 0.03638304 | -0.023481584 | 0.956749439 |
| ENSG00000168522 | *FNTA* | 4.5657756 | -0.060918535 | 0.001600807 | -0.062519342 | 0.999933887 | 0.076875934 | 0.066536764 | 0.01033917 | 0.956749439 |
| ENSG00000141837 | *CACNA1A* | 1.51844025 | -0.094540283 | -0.023815152 | -0.070725131 | 0.999933887 | -0.781495368 | -0.811466806 | 0.029971438 | 0.956749439 |
| ENSG00000277425 | *AL121890.4* | 0.316412474 | 0.174213065 | 0.0137075 | 0.160505564 | 0.999933887 | 0.441999832 | 0.469225098 | -0.027225266 | 0.956749439 |
| ENSG00000165272 | *AQP3* | 2.878578135 | 0.039525227 | 0.086024512 | -0.046499284 | 0.999933887 | 0.081466032 | 0.067244307 | 0.014221725 | 0.956749439 |
| ENSG00000127580 | *WDR24* | 1.929842512 | 0.052043908 | 0.012202145 | 0.039841763 | 0.999933887 | -0.081566848 | -0.098694735 | 0.017127887 | 0.956749439 |
| ENSG00000207034 | *Y_RNA* | 1.054091437 | 0.059274004 | 0.20297745 | -0.143703446 | 0.999933887 | -1.061390651 | -1.01667232 | -0.04471833 | 0.956749439 |
| ENSG00000136631 | *VPS45* | 1.297112564 | 0.034691161 | -0.319090476 | 0.353781637 | 0.999933887 | -0.049048817 | -0.067263643 | 0.018214826 | 0.956749439 |
| ENSG00000165271 | *NOL6* | 3.506215694 | 0.003380584 | 0.067560511 | -0.064179927 | 0.999933887 | 0.246103698 | 0.257116226 | -0.011012528 | 0.956749439 |
| ENSG00000136045 | *PWP1* | 2.76203156 | -0.076056609 | -0.186106655 | 0.110050046 | 0.999933887 | 0.099991662 | 0.08474347 | 0.015248192 | 0.956778693 |
| ENSG00000243022 | *MARK3P3* | -0.653778673 | 0.035961747 | 0.596900344 | -0.560938597 | 0.999933887 | 0.587106027 | 0.620209713 | -0.033103686 | 0.956914367 |
| ENSG00000146112 | *PPP1R18* | 9.294429288 | 0.040765557 | 0.047427591 | -0.006662034 | 0.999933887 | 0.179495786 | 0.170316931 | 0.009178855 | 0.95708171 |
| ENSG00000233406 | *AL162430.1* | 0.065864476 | -0.042894156 | 0.112309031 | -0.155203187 | 0.999933887 | 0.09517443 | 0.130187711 | -0.035013281 | 0.95708171 |
| ENSG00000164347 | *GFM2* | 2.376921961 | -0.136636666 | -0.03118232 | -0.105454346 | 0.999933887 | -0.334899175 | -0.349643301 | 0.014744126 | 0.957107861 |
| ENSG00000260997 | *AC004847.1* | 1.843053088 | -0.008135866 | -0.159400797 | 0.151264931 | 0.999933887 | 0.18431365 | 0.207900329 | -0.023586679 | 0.957130746 |
| ENSG00000116922 | *C1orf109* | 1.119016562 | 0.107382353 | 0.011397306 | 0.095985048 | 0.999933887 | 0.124334752 | 0.142581083 | -0.018246331 | 0.957182044 |
| ENSG00000130005 | *GAMT* | 0.246428895 | -0.061611145 | 0.021844431 | -0.083455576 | 0.999933887 | -0.03516921 | -0.064646322 | 0.029477112 | 0.957182044 |
| ENSG00000073464 | *CLCN4* | 1.254985061 | -0.006644863 | -0.000753207 | -0.005891656 | 0.999933887 | -0.31787596 | -0.337352136 | 0.019476175 | 0.957501387 |
| ENSG00000105669 | *COPE* | 5.99885924 | 0.024858252 | 0.026073485 | -0.001215233 | 0.999933887 | 0.077871161 | 0.08511555 | -0.007244389 | 0.957545384 |
| ENSG00000184939 | *ZFP90* | 2.372761165 | -0.032725529 | -0.09256729 | 0.05984176 | 0.999933887 | -0.010242932 | 0.002654143 | -0.012897075 | 0.957545384 |
| ENSG00000149806 | *FAU* | 7.699299715 | 0.021518928 | -0.008276464 | 0.029795392 | 0.999933887 | 0.097152929 | 0.105338472 | -0.008185543 | 0.957604262 |
| ENSG00000064726 | *BTBD1* | 4.199088427 | -0.043655678 | -0.043301781 | -0.000353896 | 0.999933887 | -0.064900459 | -0.078900218 | 0.013999759 | 0.957630271 |
| ENSG00000185917 | *SETD4* | 2.19013998 | -0.039258413 | 0.029042665 | -0.068301078 | 0.999933887 | -0.247344783 | -0.233139206 | -0.014205576 | 0.957630271 |
| ENSG00000136235 | *GPNMB* | 0.614899229 | 0.07601091 | 0.772412949 | -0.696402039 | 0.999933887 | -0.088476328 | -0.031345234 | -0.057131094 | 0.957802594 |
| ENSG00000260719 | *AC009133.3* | 1.949309107 | 0.038508741 | 0.119828024 | -0.081319284 | 0.999933887 | -0.138700912 | -0.11916188 | -0.019539033 | 0.957830032 |
| ENSG00000100401 | *RANGAP1* | 4.536010408 | -0.017063316 | 0.007316716 | -0.024380032 | 0.999933887 | 0.171082926 | 0.181420899 | -0.010337973 | 0.957830032 |
| ENSG00000105176 | *URI1* | 3.439349292 | -0.002983513 | -0.198957047 | 0.195973535 | 0.999933887 | -0.024656092 | -0.036228089 | 0.011571996 | 0.957982678 |
| ENSG00000196975 | *ANXA4* | 2.751467648 | -0.132265269 | 0.045417686 | -0.177682955 | 0.999933887 | -0.218843459 | -0.206317447 | -0.012526013 | 0.957983192 |
| ENSG00000213300 | *HNRNPA3P6* | 0.447093459 | -0.221462476 | -0.019004014 | -0.202458461 | 0.999933887 | 0.182405654 | 0.158308311 | 0.024097344 | 0.957983192 |
| ENSG00000127554 | *GFER* | 0.918042927 | 0.106104082 | 0.007595645 | 0.098508437 | 0.999933887 | 0.129390013 | 0.148789719 | -0.019399706 | 0.957983192 |
| ENSG00000182827 | *ACBD3* | 4.05651027 | -0.048108426 | -0.027528103 | -0.020580323 | 0.999933887 | -0.219468953 | -0.229234619 | 0.009765666 | 0.957983192 |
| ENSG00000286676 | *ACTBP4* | 0.533637072 | -0.023261776 | 0.182305853 | -0.20556763 | 0.999933887 | 0.175258549 | 0.148496393 | 0.026762157 | 0.957983192 |
| ENSG00000185664 | *PMEL* | -0.565641872 | -0.02697122 | -0.109887066 | 0.082915846 | 0.999933887 | -0.119515042 | -0.089139113 | -0.030375929 | 0.957983192 |
| ENSG00000178999 | *AURKB* | -0.317825479 | -0.021352616 | -0.26050779 | 0.239155174 | 0.999933887 | -0.307425419 | -0.265706606 | -0.041718813 | 0.957983192 |
| ENSG00000219626 | *FAM228B* | 0.819367852 | 0.011158244 | -0.106935166 | 0.11809341 | 0.999933887 | -0.284828289 | -0.25019589 | -0.034632399 | 0.957983192 |
| ENSG00000239951 | *IGKV3-20* | -0.255412205 | 0.007085917 | -0.05872457 | 0.065810487 | 0.999933887 | 0.168779118 | 0.199379495 | -0.030600377 | 0.957983192 |
| ENSG00000215271 | *HOMEZ* | 1.374530978 | -0.169051513 | -0.14162123 | -0.027430283 | 0.999933887 | -0.307639642 | -0.321178858 | 0.013539216 | 0.958114306 |
| ENSG00000108091 | *CCDC6* | 2.096884166 | -0.003299896 | -0.020617393 | 0.017317496 | 0.999933887 | 0.038518172 | 0.025097427 | 0.013420744 | 0.958114306 |
| ENSG00000013583 | *HEBP1* | 3.013493868 | -0.030858192 | -0.148983963 | 0.118125771 | 0.999933887 | -0.554259527 | -0.56710409 | 0.012844563 | 0.958189823 |
| ENSG00000182372 | *CLN8* | 3.85954628 | -0.061725527 | -0.102763165 | 0.041037638 | 0.999933887 | -0.464348654 | -0.452610166 | -0.011738487 | 0.958392959 |
| ENSG00000111358 | *GTF2H3* | 1.116059139 | -0.06615326 | -0.145759598 | 0.079606338 | 0.999933887 | 0.077530732 | 0.060526929 | 0.017003804 | 0.958392959 |
| ENSG00000106305 | *AIMP2* | 0.070770654 | 0.036743518 | 0.122127218 | -0.085383701 | 0.999933887 | 0.226989072 | 0.2481473 | -0.021158228 | 0.958392959 |
| ENSG00000169884 | *WNT10B* | 1.12952075 | 0.030110949 | 0.179083194 | -0.148972244 | 0.999933887 | 0.107452496 | 0.086688049 | 0.020764447 | 0.958392959 |
| ENSG00000148120 | *AOPEP* | 1.470890512 | 0.009375076 | 0.188545734 | -0.179170658 | 0.999933887 | 0.46066016 | 0.480555232 | -0.019895072 | 0.958392959 |
| ENSG00000231621 | *ANKRD44-AS1* | -0.603996312 | -0.104904113 | -0.104748418 | -0.000155695 | 0.999933887 | -0.026936179 | -0.064628246 | 0.037692067 | 0.958453904 |
| ENSG00000116199 | *FAM20B* | 2.699258299 | -0.014931774 | -0.141922536 | 0.126990762 | 0.999933887 | -0.005251189 | 0.007984842 | -0.013236032 | 0.958453904 |
| ENSG00000178075 | *GRAMD1C* | 2.629892994 | -0.166955743 | -0.115215575 | -0.051740168 | 0.999933887 | -1.042575859 | -1.022881026 | -0.019694833 | 0.958472055 |
| ENSG00000175164 | *ABO* | -1.246731105 | 0.14949995 | 0.180499369 | -0.030999419 | 0.999933887 | -0.099443819 | -0.134535959 | 0.03509214 | 0.958472055 |
| ENSG00000285994 | *AL731559.1* | 0.923482111 | 0.047062834 | 0.058614227 | -0.011551393 | 0.999933887 | -0.294576832 | -0.335892915 | 0.041316083 | 0.958472055 |
| ENSG00000120798 | *NR2C1* | 2.14235078 | -0.026960546 | -0.114588784 | 0.087628238 | 0.999933887 | 0.032309728 | 0.016190127 | 0.016119601 | 0.958472055 |
| ENSG00000065308 | *TRAM2* | 2.713774439 | 0.01516967 | -0.042907222 | 0.058076893 | 0.999933887 | -0.143122573 | -0.156942724 | 0.013820151 | 0.958472055 |
| ENSG00000224699 | *LAMTOR5-AS* | -0.218338371 | 0.00962248 | 0.309417969 | -0.299795489 | 0.999933887 | -0.230958458 | -0.20046304 | -0.030495417 | 0.958472055 |

| | | | | | | | | | | |
|---|---|---|---|---|---|---|---|---|---|---|
| ENSG00000179387 | *ELMOD2* | 2.652980753 | -0.074059098 | -0.053641159 | -0.020417939 | 0.999933887 | -0.629950478 | -0.613953997 | -0.01599648 | 0.958487536 |
| ENSG00000047849 | *MAP4* | 4.593931575 | -0.009850777 | -0.018464301 | 0.008613523 | 0.999933887 | 0.055728517 | 0.047293146 | 0.008435371 | 0.958487536 |
| ENSG00000143110 | *C1orf162* | 3.536888457 | 0.009431861 | -0.016943706 | 0.026375567 | 0.999933887 | 0.009992008 | -0.001592194 | 0.011584202 | 0.958487536 |
| ENSG00000100916 | *BRMS1L* | -0.215064073 | -0.304144912 | -0.115589205 | -0.188555707 | 0.999933887 | -0.224083373 | -0.18081921 | -0.043264163 | 0.958618737 |
| ENSG00000281490 | *CICP14* | 3.50104191 | 0.055249443 | 0.062466505 | -0.007217062 | 0.999933887 | -0.098417233 | -0.087120235 | -0.011296998 | 0.958618737 |
| ENSG00000088035 | *ALG6* | 1.983562901 | -0.075876902 | -0.079773201 | 0.003896299 | 0.999933887 | -0.276695841 | -0.259789494 | -0.016906348 | 0.958618737 |
| ENSG00000241837 | *ATP5PO* | 1.729640678 | 0.029614753 | 0.160655501 | -0.131040748 | 0.999933887 | 0.079318295 | 0.094989444 | -0.015671149 | 0.958618737 |
| ENSG00000162972 | *MAIP1* | 0.725024601 | -0.274406471 | -0.187126586 | -0.087279885 | 0.999933887 | 0.051040948 | 0.072259071 | -0.021218123 | 0.959258777 |
| ENSG00000186272 | *ZNF17* | -0.042816641 | -0.304693347 | -0.152633276 | -0.15206007 | 0.999933887 | -0.040763124 | -0.013019159 | -0.027743965 | 0.959258777 |
| ENSG00000229719 | *MIR194-2HG* | 2.037473988 | 0.156009483 | 0.101329418 | 0.054680065 | 0.999933887 | 1.311698947 | 1.288788325 | 0.022910622 | 0.959258777 |
| ENSG00000121210 | *TMEM131L* | 5.709284529 | -0.079031605 | -0.026270009 | -0.052761596 | 0.999933887 | -0.219170657 | -0.211622375 | -0.007548282 | 0.959258777 |
| ENSG00000229769 | *TRBV10-2* | -1.479499369 | -0.199325819 | 0.01611142 | -0.21543724 | 0.999933887 | 0.273349035 | 0.239137284 | 0.034211751 | 0.959258777 |
| ENSG00000142319 | *SLC6A3* | -0.363975511 | 0.26262269 | 0.926745165 | -0.664122475 | 0.999933887 | 1.562352058 | 1.59433647 | -0.031984412 | 0.959258777 |
| ENSG00000130164 | *LDLR* | 3.813174872 | 0.051044542 | 0.181721121 | -0.130676578 | 0.999933887 | 0.132289991 | 0.144722238 | -0.012432246 | 0.959258777 |
| ENSG00000171033 | *PKIA* | 0.612795967 | -0.089897735 | 0.176180986 | -0.266078722 | 0.999933887 | 0.04806788 | 0.067567974 | -0.019500093 | 0.959258777 |
| ENSG00000152556 | *PFKM* | 1.099795197 | -0.061036026 | 0.135141432 | -0.196177459 | 0.999933887 | 0.372019756 | 0.391052817 | -0.01903306 | 0.959258777 |
| ENSG00000197808 | *ZNF461* | 0.098243015 | -0.07368976 | 0.117001464 | -0.190691223 | 0.999933887 | 0.038190151 | 0.013979451 | 0.024210699 | 0.959258777 |
| ENSG00000211817 | *TRAV38-2DV8* | -1.089339241 | 0.098432564 | -0.177166843 | 0.275599407 | 0.999933887 | 0.194975776 | 0.155161316 | 0.03981446 | 0.959258777 |
| ENSG00000157895 | *C12orf43* | 1.578137883 | -0.048166257 | -0.075562211 | 0.027395955 | 0.999933887 | 0.080990555 | 0.0618074 | 0.019183154 | 0.959258777 |
| ENSG00000079215 | *SLC1A3* | 0.428916103 | 0.098486025 | 0.003679214 | 0.094806811 | 0.999933887 | 1.14777838 | 1.124298308 | 0.023480073 | 0.959258777 |
| ENSG00000122741 | *DCAF10* | 3.243807464 | -0.029337203 | -0.044969705 | 0.015632503 | 0.999933887 | -0.482713711 | -0.495338672 | 0.012624961 | 0.959258777 |
| ENSG00000176407 | *KCMF1* | 6.312329554 | -0.01637994 | 0.010344209 | -0.026724149 | 0.999933887 | -0.225874232 | -0.232522468 | 0.006648237 | 0.959258777 |
| ENSG00000135966 | *TGFBRAP1* | 3.639367467 | -0.049314685 | 0.056167616 | -0.105482301 | 0.999933887 | 0.598039583 | 0.617936988 | -0.019897405 | 0.959258777 |
| ENSG00000164172 | *MOCS2* | 1.349327641 | -0.038633425 | -0.001756924 | -0.036876501 | 0.999933887 | -0.000989354 | -0.019927421 | 0.018938067 | 0.959258777 |
| ENSG00000139433 | *GLTP* | 5.432490904 | -0.025197925 | -0.05558168 | 0.030383754 | 0.999933887 | -0.912605899 | -0.925656377 | 0.013050477 | 0.959258777 |
| ENSG00000100347 | *SAMM50* | 2.024987924 | 0.03283318 | 0.046797377 | -0.013964198 | 0.999933887 | 0.077821488 | 0.096044931 | -0.018223442 | 0.959258777 |
| ENSG00000034053 | *APBA2* | 3.093218551 | -0.013350571 | -0.091310466 | 0.077959895 | 0.999933887 | 0.045006665 | 0.033206936 | 0.011799728 | 0.959258777 |
| ENSG00000169047 | *IRS1* | 0.074036421 | -0.022010744 | 0.057448751 | -0.079459495 | 0.999933887 | 0.087889405 | 0.109983621 | -0.022094216 | 0.959258777 |
| ENSG00000196465 | *MYL6B* | -0.63562318 | -0.019820164 | -0.093682181 | 0.073862017 | 0.999933887 | -0.082205986 | -0.049672887 | -0.032533099 | 0.959258777 |
| ENSG00000185920 | *PTCH1* | 2.335502008 | 0.005459795 | -0.029891107 | 0.035350902 | 0.999933887 | -0.044345449 | -0.03062572 | -0.01371973 | 0.959258777 |
| ENSG00000124444 | *ZNF576* | 0.754824101 | -0.045890289 | 0.305458703 | -0.351348992 | 0.999933887 | -0.178574094 | -0.199702228 | 0.021128134 | 0.959313113 |
| ENSG00000162572 | *SCNN1D* | 0.313906038 | -0.001364957 | -0.189096422 | 0.187731465 | 0.999933887 | 0.155199178 | 0.178977018 | -0.02377784 | 0.959341423 |
| ENSG00000136383 | *ALPK3* | 0.633549766 | 0.141060299 | 0.059455944 | 0.081604355 | 0.999933887 | -0.218708705 | -0.246798748 | 0.028090043 | 0.959423082 |
| ENSG00000105722 | *ERF* | 6.842198799 | 0.079366723 | 0.066618717 | 0.012748007 | 0.999933887 | -0.453019989 | -0.443660577 | -0.009359412 | 0.959423082 |
| ENSG00000134575 | *ACP2* | 1.772753864 | -0.074491468 | -0.033480001 | -0.041011468 | 0.999933887 | 0.21701455 | 0.231225794 | -0.014211244 | 0.959423082 |
| ENSG00000008838 | *MED24* | 3.248970069 | -0.024849158 | -0.005158842 | -0.019690316 | 0.999933887 | 0.186818025 | 0.195421669 | -0.008603645 | 0.959423082 |
| ENSG00000010310 | *GIPR* | 1.817142819 | -0.042722705 | 0.013587443 | -0.056310148 | 0.999933887 | 0.012582784 | 0.02671904 | -0.014136256 | 0.959628071 |
| ENSG00000159788 | *RGS12* | 2.051052712 | -0.085753016 | 0.239371684 | -0.3251247 | 0.999933887 | -0.362905096 | -0.349493155 | -0.013411941 | 0.95981224 |
| ENSG00000278558 | *TMEM191B* | -1.783320353 | 0.015423783 | -0.078385836 | 0.093809619 | 0.999933887 | -0.271734108 | -0.226136436 | -0.045597672 | 0.95981224 |
| ENSG00000006451 | *RALA* | 2.672356815 | -0.003737759 | -0.133706298 | 0.129968539 | 0.999933887 | -0.092570365 | -0.080710484 | -0.011859881 | 0.95981224 |
| ENSG00000219665 | *ZNF433-AS1* | 3.2036995 | -0.199925525 | 0.067616938 | -0.267542463 | 0.999933887 | -0.42739826 | -0.439705075 | 0.012306815 | 0.960131867 |
| ENSG00000096717 | *SIRT1* | 5.63951753 | 0.034726428 | 0.037457182 | -0.002730754 | 0.999933887 | -0.203458995 | -0.213209774 | 0.00975078 | 0.960131867 |
| ENSG00000106803 | *SEC61B* | 4.450254093 | 0.057217811 | -0.001606843 | 0.058824654 | 0.999933887 | 0.039325315 | 0.047850433 | -0.008525117 | 0.960267872 |
| ENSG00000269001 | *AC092070.2* | 3.216567959 | -0.058412437 | -0.014142623 | -0.044269815 | 0.999933887 | -0.379819608 | -0.390545428 | 0.01072582 | 0.960267872 |
| ENSG00000124571 | *XPO5* | 2.742825691 | -0.051846706 | 0.153758958 | -0.205605664 | 0.999933887 | 0.044884976 | 0.032411857 | 0.012473118 | 0.960267872 |
| ENSG00000138376 | *BARD1* | 0.338797411 | 0.052886327 | 0.133463766 | -0.080577438 | 0.999933887 | -0.15558374 | -0.128223162 | -0.027360578 | 0.960267872 |
| ENSG00000096872 | *IFT74* | -0.126209336 | -0.240965067 | 0.128615518 | -0.369580585 | 0.999933887 | -0.111540393 | -0.080243451 | -0.031296942 | 0.960333497 |
| ENSG00000258484 | *SPESP1* | -0.495068348 | 0.232264258 | 0.319642419 | -0.087378161 | 0.999933887 | 0.956862621 | 0.92738536 | 0.029477261 | 0.960333497 |
| ENSG00000168792 | *ABHD15* | 1.760108207 | 0.079008369 | -0.111830341 | 0.190838711 | 0.999933887 | 0.042192836 | 0.056453768 | -0.014260932 | 0.960333497 |
| ENSG00000168303 | *MPLKIP* | 2.632626559 | 0.002101436 | -0.089680372 | 0.091781808 | 0.999933887 | -0.273398939 | -0.284622339 | 0.0112234 | 0.960333497 |
| ENSG00000115561 | *CHMP3* | 5.484139388 | -0.000649176 | -0.03383816 | 0.033188984 | 0.999933887 | -0.313674801 | -0.322006053 | 0.008331253 | 0.96035525 |
| ENSG00000137806 | *NDUFAF1* | 1.952048566 | -0.127434994 | -0.332711326 | 0.205276332 | 0.999933887 | -0.514367719 | -0.527192675 | 0.012824955 | 0.960525977 |
| ENSG00000160298 | *C21orf58* | -0.661721188 | -0.165627194 | -0.027429468 | -0.138197726 | 0.999933887 | -0.191419968 | -0.160802882 | -0.030617086 | 0.960525977 |
| ENSG00000164002 | *EXO5* | 0.153120094 | -0.126821835 | -0.084269063 | -0.042552772 | 0.999933887 | 0.1441967 | 0.12040013 | 0.02379657 | 0.960525977 |
| ENSG00000218510 | *LINC00339* | -0.546328119 | 0.071259502 | 0.019327791 | 0.051931711 | 0.999933887 | -0.26769609 | -0.299834343 | 0.032138253 | 0.960525977 |
| ENSG00000001497 | *LAS1L* | 2.763654337 | -0.110483132 | -0.021738037 | -0.088745095 | 0.999933887 | 0.17338929 | 0.184211782 | -0.010822492 | 0.960843497 |
| ENSG00000142408 | *CACNG8* | 1.261550404 | -0.092098508 | 0.092704513 | -0.184803021 | 0.999933887 | -0.117101339 | -0.1322545 | 0.015153161 | 0.960843497 |
| ENSG00000137936 | *BCAR3* | -0.588051439 | 0.163634722 | -0.331009218 | 0.49464394 | 0.999933887 | -0.371712583 | -0.393152667 | 0.021440084 | 0.960843497 |
| ENSG00000185513 | *L3MBTL1* | 0.615905768 | -0.040380747 | 0.185317999 | -0.225698746 | 0.999933887 | -0.163402464 | -0.137534403 | -0.025868062 | 0.960843497 |
| ENSG00000234493 | *RHOXF1P1* | -1.120934734 | 0.07559817 | -0.578804553 | 0.654402723 | 0.999933887 | -0.102648956 | -0.06280719 | -0.039841766 | 0.960890678 |
| ENSG00000126267 | *COX6B1* | 5.375820194 | 0.05483387 | 0.00113521 | 0.05369866 | 0.999933887 | 0.034024064 | 0.025412525 | 0.008611538 | 0.961031371 |
| ENSG00000128891 | *CCDC32* | 2.786830075 | 0.057488289 | -0.021882267 | 0.079370556 | 0.999933887 | 0.030441455 | 0.041397573 | -0.010956118 | 0.961035681 |
| ENSG00000033170 | *FUT8* | 1.259329063 | 0.06478196 | -0.037400964 | 0.102182924 | 0.999933887 | -0.059532779 | -0.040038392 | -0.019494387 | 0.961178159 |
| ENSG00000272037 | *AP002907.1* | 1.323196954 | 0.130546706 | 0.145366591 | -0.014819885 | 0.999933887 | -0.255991207 | -0.235866958 | -0.020124249 | 0.961366765 |
| ENSG00000130958 | *SLC35D2* | 0.196631353 | 0.232322223 | -0.099131006 | 0.331453229 | 0.999933887 | -0.044894485 | -0.073102813 | 0.028208329 | 0.961624303 |
| ENSG00000163121 | *NEURL3* | -0.652588066 | 0.225148885 | 0.481847462 | -0.256698578 | 0.999933887 | 3.405135506 | 3.366400697 | 0.03873481 | 0.961624303 |
| ENSG00000142784 | *WDTC1* | 6.770369892 | 0.040434581 | 0.002345281 | 0.0380893 | 0.999933887 | 0.352977498 | 0.34529554 | 0.007681958 | 0.96163238 |
| ENSG00000157833 | *GAREM2* | -0.503051278 | 0.267356797 | 0.065540227 | 0.20181657 | 0.999933887 | 0.146224211 | 0.178333138 | -0.032108928 | 0.961692047 |
| ENSG00000213782 | *DDX47* | 0.567603413 | -0.12336099 | -0.038312786 | -0.085048204 | 0.999933887 | 0.232521711 | 0.250489722 | -0.01796801 | 0.961954468 |
| ENSG00000276533 | *AC018926.2* | 1.964471041 | -0.035591835 | 0.275416761 | -0.311008596 | 0.944094537 | 0.292282608 | 0.270578776 | 0.021703832 | 0.961997232 |
| ENSG00000162191 | *UBXN1* | 6.771141406 | -0.009647783 | -0.017624958 | 0.007977175 | 0.999933887 | 0.362109417 | 0.370941059 | -0.008831642 | 0.96203962 |
| ENSG00000245556 | *SCAMP1-AS1* | 0.987777063 | -0.161472586 | -0.111257198 | -0.050215388 | 0.999933887 | 0.06491392 | 0.046524519 | 0.0183894 | 0.96218802 |
| ENSG00000243544 | *RN7SL172P* | 1.714884809 | 0.078182712 | 0.183489922 | -0.10530721 | 0.999933887 | -0.506908776 | -0.490143741 | -0.016765035 | 0.962290535 |
| ENSG00000167526 | *RPL13* | 8.600697663 | 0.039518493 | -0.009804544 | 0.049323038 | 0.999933887 | 0.096385223 | 0.105392252 | -0.00900703 | 0.962290535 |
| ENSG00000107829 | *FBXW4* | 3.257376283 | 0.037494509 | 0.131048779 | -0.09355427 | 0.999933887 | 0.155028092 | 0.164514021 | -0.009485929 | 0.962290535 |
| ENSG00000172728 | *FUT10* | 0.026752913 | -0.21411386 | 0.116605446 | -0.330719305 | 0.999933887 | -0.335078739 | -0.306643612 | -0.028435127 | 0.962434064 |
| ENSG00000139160 | *ETFBKMT* | 1.539434599 | -0.119683886 | -0.01133836 | -0.108345526 | 0.999933887 | 0.065003067 | 0.082414329 | -0.017411262 | 0.962434064 |
| ENSG00000116396 | *KCNC4* | 2.185088927 | 0.07346542 | 0.07585964 | -0.002394221 | 0.999933887 | -0.275120367 | -0.291163577 | 0.016043211 | 0.962434064 |
| ENSG00000100697 | *DICER1* | 6.584630454 | -0.057942506 | -0.106060823 | 0.048118318 | 0.999933887 | -0.412770049 | -0.39781393 | -0.014956119 | 0.962434064 |
| ENSG00000164039 | *BDH2* | 0.036139184 | 0.092498021 | -0.180198719 | 0.27269674 | 0.999933887 | 0.077250922 | 0.10271856 | -0.025467638 | 0.962434064 |
| ENSG00000133067 | *LGR6* | 1.831336633 | 0.032799043 | 0.013835181 | 0.018963861 | 0.999933887 | 0.012270136 | 0.024304284 | -0.012034148 | 0.962434064 |
| ENSG00000148835 | *TAF5* | 1.224820428 | 0.032796302 | -0.048207415 | 0.081003717 | 0.999933887 | -0.077411131 | -0.093390033 | 0.015978902 | 0.962434064 |
| ENSG00000234199 | *LINC01191* | 1.44653575 | -0.042320591 | 0.028864242 | -0.071184833 | 0.999933887 | 1.133793611 | 1.155047936 | -0.021254325 | 0.962434064 |
| ENSG00000279021 | *AC092139.1* | 0.455325991 | 0.047182691 | 0.286816845 | -0.239634155 | 0.999933887 | 0.40887239 | 0.380628614 | 0.028243776 | 0.962434064 |
| ENSG00000117632 | *STMN1* | 2.283864663 | -0.02016104 | -0.106972934 | 0.086811894 | 0.999933887 | -0.053349599 | -0.039772553 | -0.013577046 | 0.962434064 |
| ENSG00000161021 | *MAML1* | 6.628570023 | -0.007255775 | 0.036212832 | -0.043468608 | 0.999933887 | 0.117031644 | 0.110620538 | 0.006411106 | 0.962434064 |
| ENSG00000167925 | *GHDC* | 3.104498213 | -0.000465804 | -0.020549437 | 0.020083634 | 0.999933887 | 0.003591963 | -0.007909901 | 0.011501864 | 0.962434064 |
| ENSG00000111229 | *ARPC3* | 7.942057538 | -0.038822421 | -0.028062236 | -0.010760185 | 0.999933887 | -0.153265666 | -0.146761643 | -0.006504023 | 0.962731056 |
| ENSG00000256338 | *RPL41P2* | 0.052257629 | 0.008027854 | -0.118862429 | 0.126890283 | 0.999933887 | -0.119354666 | -0.085047283 | -0.034307383 | 0.962866942 |
| ENSG00000218426 | *AL590867.2* | 0.070944125 | -0.085604867 | -0.088103403 | 0.002498536 | 0.999933887 | 0.086762124 | 0.062624234 | 0.02413789 | 0.962949472 |
| ENSG00000280033 | *AC116407.4* | -0.529607519 | 0.119659324 | 0.176649889 | -0.056990565 | 0.999933887 | -0.160613785 | -0.192867193 | 0.032253408 | 0.96342123 |
| ENSG00000144827 | *ABHD10* | 1.665623912 | -0.048265493 | -0.273666516 | 0.225401024 | 0.999933887 | 0.019324731 | 0.004684252 | 0.014640479 | 0.963526612 |
| ENSG00000102575 | *ACP5* | 3.137219549 | 0.005977923 | 0.058900485 | -0.052922562 | 0.999933887 | -0.159392077 | -0.171436951 | 0.012044875 | 0.963526612 |
| ENSG00000101457 | *DNTTIP1* | 5.794495131 | 0.030211886 | 0.058488808 | -0.028276922 | 0.999933887 | 0.023620521 | 0.030815831 | -0.00719531 | 0.963671638 |
| ENSG00000125485 | *DDX31* | 1.155546549 | -0.105506762 | 0.087414447 | -0.192921209 | 0.999933887 | 0.302736749 | 0.318752953 | -0.016016204 | 0.963762114 |
| ENSG00000167470 | *MIDN* | 8.202493273 | 0.059130495 | 0.050184601 | 0.008945894 | 0.999933887 | -0.605941845 | -0.615491912 | 0.009550067 | 0.963894283 |
| ENSG00000141582 | *CBX4* | 6.134757168 | 0.184654431 | 0.192851833 | -0.008197402 | 0.999933887 | 0.201164979 | 0.19148366 | 0.009681319 | 0.963938222 |
| ENSG00000074276 | *CDHR2* | -0.025271427 | 0.071103794 | 0.065148372 | 0.005955422 | 0.999933887 | -0.088546023 | -0.060622078 | -0.027923946 | 0.964001598 |
| ENSG00000255302 | *EID1* | 4.07563405 | 0.029674412 | -0.058481772 | 0.088156185 | 0.999933887 | 0.01333016 | 0.004523905 | 0.008806255 | 0.964261593 |
| ENSG00000197312 | *DDI2* | 3.579427479 | -0.038455782 | -0.071391996 | 0.032936213 | 0.999933887 | -0.403181614 | -0.392859592 | -0.010322022 | 0.964261593 |
| ENSG00000145912 | *NHP2* | 1.403409044 | 0.031028734 | 0.193100787 | -0.162072053 | 0.999933887 | 0.241933939 | 0.256587013 | -0.014653074 | 0.964261593 |
| ENSG00000101019 | *UQCC1* | 1.740009673 | -0.066428926 | -0.18918204 | 0.122753114 | 0.999933887 | 0.062399052 | 0.049401335 | 0.012997717 | 0.964371076 |
| ENSG00000269956 | *MKNK1-AS1* | 0.438315104 | -0.073872998 | 0.002618796 | -0.076491793 | 0.999933887 | -0.243450028 | -0.217237609 | -0.026212419 | 0.964482793 |
| ENSG00000169016 | *E2F6* | 2.465350961 | -0.038343612 | -0.019284327 | -0.019059285 | 0.999933887 | -0.315745437 | -0.326533641 | 0.010788204 | 0.964663837 |
| ENSG00000205746 | *PKD1P4* | 0.430835073 | 0.279036304 | 0.246505763 | 0.032530541 | 0.999933887 | 0.09457743 | 0.114727149 | -0.020149719 | 0.964747361 |

| | | | | | | | | | | |
|---|---|---|---|---|---|---|---|---|---|---|
| ENSG00000272562 | *AL512343.2* | 0.773000584 | -0.117024487 | -0.215723854 | 0.098699367 | 0.999933887 | -0.653537114 | -0.63350419 | -0.020032923 | 0.964770221 |
| ENSG00000183808 | *RBM12B* | 3.067484195 | 0.040692936 | -0.068444655 | 0.109137591 | 0.999933887 | -0.500950477 | -0.512184541 | 0.011234064 | 0.964770221 |
| ENSG00000134283 | *PPHLN1* | 4.217742243 | -0.119873104 | 0.036402781 | -0.156275885 | 0.944094537 | -0.15157709 | -0.144363329 | -0.007213761 | 0.964847853 |
| ENSG00000279636 | *LINC00216* | -0.075796821 | -0.15353676 | 0.04666175 | -0.20019851 | 0.999933887 | 0.121020481 | 0.145581664 | -0.024561183 | 0.964847853 |
| ENSG00000258752 | *AL357093.2* | -0.304060205 | -0.160537598 | 0.087454145 | -0.247991743 | 0.999933887 | -0.226242943 | -0.263494794 | 0.03725185 | 0.964847853 |
| ENSG00000013561 | *RNF14* | 3.31359276 | -0.154294906 | -0.062588009 | -0.091706897 | 0.999933887 | -0.358474024 | -0.349612006 | -0.008862018 | 0.965134194 |
| ENSG00000107593 | *PKD2L1* | -1.257776759 | -0.079431884 | 0.033894409 | -0.113326293 | 0.999933887 | -1.229586508 | -1.254487244 | 0.024900736 | 0.965351086 |
| ENSG00000233184 | *AC093157.1* | 0.682554962 | 0.05627417 | 0.09464927 | -0.0383751 | 0.999933887 | 0.057474308 | 0.074285848 | -0.01681154 | 0.965516149 |
| ENSG00000105879 | *CBLL1* | 4.357441199 | -0.073152297 | -0.150584552 | 0.077432255 | 0.999933887 | -0.22373782 | -0.231815541 | 0.008077721 | 0.965697648 |
| ENSG00000183943 | *PRKX* | 4.365678167 | -0.068510671 | -0.043228097 | -0.025282574 | 0.999933887 | -0.02050428 | -0.027967045 | 0.007462765 | 0.965743117 |
| ENSG00000101935 | *AMMECR1* | 2.12079829 | 0.100693498 | -0.051343114 | 0.152036612 | 0.999933887 | -0.123272249 | -0.10866536 | -0.01460689 | 0.965743117 |
| ENSG00000160216 | *AGPAT3* | 4.200164585 | -0.049598465 | 0.037369244 | -0.086967708 | 0.999933887 | 0.09210441 | 0.084936253 | 0.007168157 | 0.965743117 |
| ENSG00000134184 | *GSTM1* | -0.225835836 | 0.176007237 | 0.260255739 | -0.084248502 | 0.999933887 | -0.205438088 | -0.238932739 | 0.033494651 | 0.965743117 |
| ENSG00000163517 | *HDAC11* | -0.012433032 | 0.098735323 | -0.211466603 | 0.310201926 | 0.999933887 | 0.02481427 | 0.001402483 | 0.023411787 | 0.965743117 |
| ENSG00000112996 | *MRPS30* | 2.273284462 | -0.034037925 | 0.127717251 | -0.161755176 | 0.999933887 | 0.097084943 | 0.084304124 | 0.012780819 | 0.965743117 |
| ENSG00000204261 | *PSMB8-AS1* | 4.696211644 | -0.005428354 | 0.054993335 | -0.06042169 | 0.999933887 | 0.054008086 | 0.046946304 | 0.007061782 | 0.965743117 |
| ENSG00000160124 | *CCDC58* | -0.23385011 | 0.343256269 | -0.132765721 | 0.47602199 | 0.999933887 | 0.232025575 | 0.254496102 | -0.022470527 | 0.965942255 |
| ENSG00000172006 | *ZNF554* | 2.843619427 | -0.099133487 | -0.18189322 | 0.082759733 | 0.999933887 | 0.262111631 | 0.272669768 | -0.010558137 | 0.966047241 |
| ENSG00000144381 | *HSPD1* | 4.60758584 | -0.048393592 | -0.084070135 | 0.035676543 | 0.999933887 | 0.109055988 | 0.11715162 | -0.008095632 | 0.966078841 |
| ENSG00000204406 | *MBD5* | 3.756113553 | 0.051125716 | 0.005303155 | 0.045822561 | 0.999933887 | -0.12224238 | -0.110546967 | -0.011695413 | 0.966078841 |
| ENSG00000283526 | *PRRT1B* | 0.139148984 | 0.04171354 | -0.051181439 | 0.092894979 | 0.999933887 | 0.243641085 | 0.223024462 | 0.020616623 | 0.966078841 |
| ENSG00000078043 | *PIAS2* | 1.957003662 | -0.025516972 | 0.008421154 | -0.033938126 | 0.999933887 | -0.244362837 | -0.263723772 | 0.019360935 | 0.966078841 |
| ENSG00000115266 | *APC2* | 2.523206734 | 0.015613046 | 0.040048341 | -0.024435295 | 0.999933887 | -0.05578476 | -0.067641998 | 0.011857238 | 0.966078841 |
| ENSG00000242861 | *AL591895.1* | 0.997888069 | -0.000119292 | -0.07284929 | 0.072729998 | 0.999933887 | -0.132442105 | -0.149913504 | 0.0174714 | 0.966078841 |
| ENSG00000100997 | *ABHD12* | 2.697159955 | -0.075323168 | 0.043328831 | -0.118651999 | 0.999933887 | -0.037000577 | -0.02775096 | -0.009249618 | 0.966321161 |
| ENSG00000257815 | *PRANCR* | 1.720496384 | -0.077577548 | 0.079861472 | -0.15743902 | 0.999933887 | 0.407262636 | 0.420540357 | -0.013277721 | 0.966322082 |
| ENSG00000270379 | *HEATR9* | -0.240939914 | 0.269049763 | 0.155039013 | 0.114010749 | 0.999933887 | -0.205762929 | -0.233035066 | 0.027272137 | 0.966335016 |
| ENSG00000196715 | *VKORC1L1* | 1.607729241 | 0.154375657 | 0.142087625 | 0.012288033 | 0.999933887 | -0.224015046 | -0.238750842 | 0.014735796 | 0.966541944 |
| ENSG00000198417 | *MT1F* | -0.120689577 | -0.221218759 | -0.260438926 | 0.039220167 | 0.999933887 | 0.514366706 | 0.487232972 | 0.027133734 | 0.966541944 |
| ENSG00000205302 | *SNX2* | 5.044252195 | -0.090449342 | -0.145763979 | 0.055314637 | 0.999933887 | -0.755466924 | -0.764792603 | 0.009325679 | 0.966541944 |
| ENSG00000249700 | *SRD5A3-AS1* | -0.412068242 | 0.048891267 | 0.056984264 | -0.008092996 | 0.999933887 | 1.077978013 | 1.107229149 | -0.029251136 | 0.966541944 |
| ENSG00000122224 | *LY9* | 4.067556943 | -0.008721114 | 0.042146912 | -0.050868026 | 0.999933887 | -0.007682681 | -0.015569671 | 0.007886991 | 0.966541944 |
| ENSG00000225484 | *NUTM2B-AS1* | 3.447566873 | -0.006027468 | 0.025772121 | -0.031799589 | 0.999933887 | -0.343445025 | -0.352891142 | 0.009446117 | 0.966541944 |
| ENSG00000077713 | *SLC25A43* | -0.292318357 | 0.000159284 | 0.33357823 | -0.333418946 | 0.999933887 | 0.168101839 | 0.143599831 | 0.024502008 | 0.966541944 |
| ENSG00000257135 | *ODC1-DT* | -0.888073659 | -0.057369135 | 0.265167474 | -0.322536609 | 0.999933887 | 0.096101478 | 0.125526627 | -0.029425148 | 0.966553564 |
| ENSG00000146285 | *SCML4* | 3.997588514 | 0.005941491 | -0.055544588 | 0.061486079 | 0.999933887 | -0.041216045 | -0.03386979 | -0.007346255 | 0.96672663 |
| ENSG00000205037 | *AC134312.1* | -0.769919116 | -0.187151265 | 0.046061037 | -0.233212302 | 0.999933887 | 0.722013314 | 0.753253997 | -0.031240684 | 0.966927224 |
| ENSG00000145246 | *ATP10D* | 2.013313806 | 0.092556081 | -0.134596738 | 0.227152819 | 0.999933887 | -0.389027891 | -0.376717679 | -0.012310213 | 0.967269743 |
| ENSG00000227417 | *AC114402.1* | -0.366375784 | 0.086881378 | -0.37444453 | 0.461325908 | 0.999933887 | 0.429304912 | 0.457465739 | -0.028160827 | 0.967514937 |
| ENSG00000136152 | *COG3* | 4.1677192 | -0.056749059 | -0.087827504 | 0.031078445 | 0.999933887 | -0.143730382 | -0.150481922 | 0.00675154 | 0.96793434 |
| ENSG00000154928 | *EPHB1* | 6.612362902 | 0.033809524 | 0.043178379 | -0.009368855 | 0.999933887 | 0.142209914 | 0.150735026 | -0.008525112 | 0.96793434 |
| ENSG00000117419 | *ERI3* | 1.502312022 | 0.20904169 | 0.042932433 | 0.166109257 | 0.999933887 | 0.075306518 | 0.091527789 | -0.016221271 | 0.968062896 |
| ENSG00000162639 | *HENMT1* | 1.897464263 | -0.082698997 | -0.186612907 | 0.10391391 | 0.999933887 | -0.214271028 | -0.200845443 | -0.013425585 | 0.968075892 |
| ENSG00000235162 | *C12orf75* | 2.463553672 | 0.161725448 | -0.02938422 | 0.191109668 | 0.999933887 | -0.003910882 | 0.006643176 | -0.010554058 | 0.968092273 |
| ENSG00000188283 | *ZNF383* | 1.143966745 | -0.15202493 | 0.170423241 | -0.322448171 | 0.999933887 | 0.147694935 | 0.132226917 | 0.015468018 | 0.968092273 |
| ENSG00000110090 | *CPT1A* | 3.592004432 | -0.035787735 | -0.036243756 | 0.000456021 | 0.999933887 | -0.101934675 | -0.111884044 | 0.009949369 | 0.968092273 |
| ENSG00000257802 | *MRS2P2* | 1.780121415 | -0.041658489 | 0.197576938 | -0.239235428 | 0.999933887 | 0.554591881 | 0.575282266 | -0.020690385 | 0.968092273 |
| ENSG00000233093 | *LINC00892* | -0.734819547 | -0.028829296 | -0.158446761 | 0.129617465 | 0.999933887 | 0.050437023 | 0.022062774 | 0.028374249 | 0.968092273 |
| ENSG00000176994 | *SMCR8* | 8.335497397 | -0.052608159 | -0.037704518 | -0.014903641 | 0.999933887 | 0.034525831 | 0.028033498 | 0.006492333 | 0.968209863 |
| ENSG00000279059 | *AC007485.2* | -1.319506115 | -0.386120216 | 0.757196634 | -1.143316849 | 0.969206975 | 0.096751388 | 0.064225495 | 0.032525893 | 0.968210213 |
| ENSG00000100129 | *EIF3L* | 4.829554108 | -0.013897182 | -0.119307227 | 0.105410045 | 0.999933887 | -0.07773455 | -0.085471546 | 0.007736996 | 0.968210213 |
| ENSG00000138698 | *RAP1GDS1* | 2.101963295 | -0.097374281 | -0.011381283 | -0.085992998 | 0.999933887 | -0.097463883 | -0.08674827 | -0.010715613 | 0.968221968 |
| ENSG00000228203 | *GRASLND* | 0.436408486 | -0.047965923 | 0.339713163 | -0.387679086 | 0.999291887 | -0.049332841 | -0.071944639 | 0.022611798 | 0.968221968 |
| ENSG00000138375 | *SMARCAL1* | 1.739508512 | 0.032863788 | -0.02904273 | 0.061906518 | 0.999933887 | 0.057292595 | 0.043750442 | 0.013542152 | 0.968221968 |
| ENSG00000147065 | *MSN* | 9.969724263 | -0.004549061 | -0.032822228 | 0.028273167 | 0.999933887 | 0.083656631 | 0.077143163 | 0.006513467 | 0.968221968 |
| ENSG00000147874 | *HAUS6* | 2.333057906 | 0.004409468 | -0.063668453 | 0.068077921 | 0.999933887 | -0.309100161 | -0.291773268 | -0.017326894 | 0.968221968 |
| ENSG00000271151 | *AC016737.1* | 0.856077855 | -0.233691502 | -0.303152495 | 0.069460994 | 0.999933887 | -1.084971791 | -1.068791409 | -0.016180382 | 0.968222678 |
| ENSG00000158941 | *CCAR2* | 5.735935474 | 0.049211651 | 0.08432698 | -0.035115329 | 0.999933887 | -0.053743881 | -0.04788859 | -0.00585529 | 0.968313463 |
| ENSG00000173113 | *TRMT112* | 6.301792908 | 0.06899047 | -0.006561065 | 0.075551535 | 0.999933887 | 0.209970403 | 0.202106611 | 0.007863793 | 0.968313463 |
| ENSG00000143543 | *JTB* | 2.026688359 | -0.106773987 | -0.05291628 | -0.053857708 | 0.999933887 | -0.143457676 | -0.156035345 | 0.012577669 | 0.968313463 |
| ENSG00000113460 | *BRIX1* | 1.451747474 | -0.087428979 | -0.031581588 | -0.055847391 | 0.999933887 | 0.070923533 | 0.056559669 | 0.014363863 | 0.968313463 |
| ENSG00000249738 | *AC008691.1* | 0.266247594 | -0.232988927 | 0.080331181 | -0.313320108 | 0.999933887 | 0.149629688 | 0.134518763 | 0.015110925 | 0.968313463 |
| ENSG00000111843 | *TMEM14C* | 2.528940121 | 0.075273442 | -0.007957824 | 0.083231266 | 0.999933887 | 0.075315489 | 0.090080785 | -0.014765296 | 0.968313463 |
| ENSG00000099219 | *ERMP1* | 2.037941871 | 0.069945235 | -0.088954541 | 0.158899776 | 0.999933887 | 0.001559422 | 0.014067721 | -0.012508299 | 0.968313463 |
| ENSG00000259291 | *ZNF710-AS1* | 2.950013769 | -0.066708211 | -0.097704829 | 0.030996618 | 0.999933887 | 0.136054573 | 0.146131997 | -0.010077423 | 0.968313463 |
| ENSG00000113721 | *PDGFRB* | -0.529934226 | -0.095726628 | 0.213866366 | -0.309592994 | 0.999933887 | 0.153074613 | 0.177225493 | -0.02415088 | 0.968313463 |
| ENSG00000146433 | *TMEM181* | 4.118662581 | 0.006647529 | -0.0080783 | 0.014725829 | 0.999933887 | -0.250018284 | -0.24138412 | -0.008634165 | 0.968313463 |
| ENSG00000129925 | *PGAP6* | 7.058172677 | 0.084887333 | 0.06883023 | 0.016057104 | 0.999933887 | 0.430259848 | 0.421405132 | 0.008854716 | 0.968317659 |
| ENSG00000183506 | *PI4KAP2* | 2.74557814 | 0.068630983 | 0.098384864 | -0.029753881 | 0.999933887 | 0.139814194 | 0.150112993 | -0.010298799 | 0.968387234 |
| ENSG00000204767 | *INSYN2B* | 2.847681378 | 0.05231942 | 0.402595888 | -0.350276468 | 0.999933887 | -0.062195118 | -0.07695989 | 0.014764772 | 0.968387234 |
| ENSG00000253438 | *PCAT1* | -0.272656264 | -0.084164274 | -0.001215515 | -0.082948759 | 0.999933887 | -0.360282363 | -0.390783838 | 0.030501475 | 0.968387234 |
| ENSG00000086504 | *MRPL28* | 4.926163985 | -0.011017162 | 0.001194891 | -0.012212053 | 0.999933887 | -0.102694244 | -0.109996594 | 0.00730235 | 0.968387234 |
| ENSG00000057657 | *PRDM1* | 6.99739195 | 0.013395694 | 0.08265707 | -0.069261376 | 0.999933887 | 0.717037645 | 0.726284197 | -0.009246551 | 0.968387234 |
| ENSG00000110651 | *CD81* | 4.987295414 | -0.00011832 | 0.003941517 | -0.004059837 | 0.999933887 | 0.097455133 | 0.090342724 | 0.007112409 | 0.968387234 |
| ENSG00000133935 | *ERG28* | 1.607359637 | -0.103987026 | -0.024353979 | -0.079633047 | 0.999933887 | 0.164470921 | 0.150652804 | 0.013818117 | 0.968562269 |
| ENSG00000178951 | *ZBTB7A* | 6.423114467 | 0.036299074 | 0.027898088 | 0.008400986 | 0.999933887 | -0.380883003 | -0.387635101 | 0.006752098 | 0.969211415 |
| ENSG00000196867 | *ZFP28* | -0.027748662 | -0.15798121 | 0.141625924 | -0.299607134 | 0.999933887 | 0.081595221 | 0.063759981 | 0.01783524 | 0.969373002 |
| ENSG00000180104 | *EXOC3* | 6.700074071 | -0.022980637 | 0.009885083 | -0.032865719 | 0.999933887 | -0.116217308 | -0.120760108 | 0.0045428 | 0.969373002 |
| ENSG00000100982 | *PCIF1* | 5.954052339 | -0.04357377 | 0.005219949 | -0.048793719 | 0.999933887 | -0.158608001 | -0.153525243 | -0.005082758 | 0.969417478 |
| ENSG00000136770 | *DNAJC1* | 3.143013643 | -0.019640109 | -0.019653823 | 1.3714e-05 | 0.999933887 | 0.092701646 | 0.082212095 | 0.010489551 | 0.969417478 |
| ENSG00000116717 | *GADD45A* | 3.591766094 | 0.119701755 | 0.076565189 | 0.043136566 | 0.999933887 | 0.640080732 | 0.629481433 | 0.010599299 | 0.96956042 |
| ENSG00000156381 | *ANKRD9* | 3.502180485 | 0.07938821 | 0.090782352 | -0.011394142 | 0.999933887 | 0.125971333 | 0.135958818 | -0.009987485 | 0.96956042 |
| ENSG00000174227 | *PIGG* | 5.320049638 | -0.028527873 | 0.00134578 | -0.029873653 | 0.999933887 | -0.037248267 | -0.042323359 | 0.005075091 | 0.96956042 |
| ENSG00000101052 | *IFT52* | 1.832946705 | -0.051619436 | 0.132259815 | -0.183879251 | 0.999933887 | 0.147486086 | 0.134667413 | 0.012818673 | 0.96956042 |
| ENSG00000105438 | *KDELR1* | 3.543765253 | -0.048029731 | -0.103649301 | 0.055619571 | 0.999933887 | -0.193077965 | -0.200317729 | 0.007239764 | 0.969670023 |
| ENSG00000153989 | *NUS1* | 2.587576575 | -0.095030923 | -0.139103506 | 0.044072583 | 0.999933887 | 0.033600226 | 0.023153848 | 0.010446378 | 0.969763916 |
| ENSG00000255455 | *AP003486.1* | 1.517427982 | -0.228575434 | -0.172523203 | -0.056052231 | 0.999933887 | -0.198302355 | -0.210554268 | 0.012251914 | 0.969933632 |
| ENSG00000124459 | *ZNF45* | 1.645073992 | -0.09882649 | 0.01226033 | -0.11108682 | 0.999933887 | 0.065586288 | 0.052466051 | 0.013120237 | 0.969933632 |
| ENSG00000197558 | *SSPOP* | 1.403406851 | 0.076308734 | 0.234897406 | -0.158588672 | 0.999933887 | -0.199548192 | -0.185803295 | -0.013744898 | 0.969933632 |
| ENSG00000272625 | *AP000919.3* | -0.056410374 | 0.072693545 | 0.251402264 | -0.178708719 | 0.999933887 | -0.383549436 | -0.408260012 | 0.024710575 | 0.969933632 |
| ENSG00000100628 | *ASB2* | 2.181838002 | 0.013821253 | 0.051921444 | -0.038100191 | 0.999933887 | 0.129339118 | 0.13969206 | -0.010352942 | 0.969933632 |
| ENSG00000196199 | *MPHOSPH8* | 5.390687298 | -0.007048324 | -0.072273618 | 0.065225294 | 0.999933887 | -0.392586265 | -0.384824001 | -0.007762263 | 0.969933632 |
| ENSG00000181222 | *POLR2A* | 7.995247034 | -0.002151077 | -0.007779393 | 0.005628316 | 0.999933887 | 0.012430989 | 0.006677298 | 0.00575369 | 0.969933632 |
| ENSG00000271425 | *NBPF10* | 4.203472408 | -0.11101449 | -0.025542662 | -0.085471828 | 0.999933887 | 0.197141579 | 0.189030589 | 0.00811099 | 0.970289523 |
| ENSG00000052802 | *MSMO1* | 1.453593729 | 0.051050959 | 0.090119289 | -0.03906833 | 0.999933887 | 0.07721828 | 0.063332853 | 0.013885427 | 0.970289523 |
| ENSG00000149634 | *SPATA25* | 0.305535257 | 0.196227289 | 0.094337345 | 0.101889944 | 0.999933887 | 0.180047108 | 0.19528151 | -0.015234402 | 0.970359144 |
| ENSG00000171940 | *ZNF217* | 6.830076014 | -0.245356756 | -0.188723173 | -0.056633583 | 0.999933887 | -0.290540584 | -0.299402678 | 0.008862094 | 0.970383583 |
| ENSG00000100395 | *L3MBTL2* | 2.441399853 | -0.133568563 | -0.053461386 | -0.080107177 | 0.999933887 | 0.138048431 | 0.127952914 | 0.010095516 | 0.970383583 |
| ENSG00000137877 | *SPTBN5* | 1.921574501 | 0.14078731 | 0.21211096 | -0.07132365 | 0.999933887 | 0.266245246 | 0.254751215 | 0.011494031 | 0.970383583 |
| ENSG00000125534 | *PPDPF* | 4.738001368 | 0.060757763 | 0.016869989 | 0.043887774 | 0.999933887 | 0.219872144 | 0.21221325 | 0.007658894 | 0.970383583 |
| ENSG00000267275 | *AC020911.2* | -0.353707547 | 0.133899331 | 0.194166 | -0.060266669 | 0.999933887 | 0.487790455 | 0.458062006 | 0.029728449 | 0.970383583 |
| ENSG00000123505 | *AMD1* | 6.040299625 | 0.04325732 | 0.001594468 | 0.041662852 | 0.999933887 | -0.127886025 | -0.120379664 | -0.007506361 | 0.970383583 |
| ENSG00000130706 | *ADRM1* | 5.008383694 | 0.041960558 | 0.056355637 | -0.014395079 | 0.999933887 | -0.00722045 | -0.013757651 | 0.006537201 | 0.970383583 |
| ENSG00000267041 | *ZNF850* | -0.626928543 | -0.138734907 | -0.229221175 | 0.090486268 | 0.999933887 | -0.295979242 | -0.319114338 | 0.023135096 | 0.970383583 |
| ENSG00000235437 | *LINC01278* | 1.374981831 | 0.072248834 | -0.0717694 | 0.144018234 | 0.999933887 | -0.037080329 | -0.051156573 | 0.014076243 | 0.970383583 |

| | | | | | | | | | | |
|---|---|---|---|---|---|---|---|---|---|---|
| ENSG00000138604 | *GLCE* | -0.578793283 | -0.072768134 | 0.16333779 | -0.236105924 | 0.999933887 | -0.056176422 | -0.083441938 | 0.027265516 | 0.970383583 |
| ENSG00000042088 | *TDP1* | 2.532643231 | 0.016716811 | -0.063529217 | 0.080246029 | 0.999933887 | 0.106917391 | 0.097542932 | 0.009374459 | 0.970383583 |
| ENSG00000166199 | *ALKBH3* | 0.898682666 | 0.014852469 | -0.054435168 | 0.069287637 | 0.999933887 | -0.019723225 | -0.032377506 | 0.012654281 | 0.970383583 |
| ENSG00000213025 | *COX20P1* | -1.201898549 | 0.10224449 | 0.124806565 | -0.022562075 | 0.999933887 | -0.270867417 | -0.245582844 | -0.025284573 | 0.970508184 |
| ENSG00000178685 | *PARP10* | 5.593745939 | 0.081724285 | 0.117080396 | -0.035356111 | 0.999933887 | 0.125427381 | 0.136129614 | -0.010702234 | 0.97092375 |
| ENSG00000180263 | *FGD6* | -0.305288581 | -0.149231348 | -0.199967486 | 0.050736138 | 0.999933887 | 0.018665111 | -0.007729684 | 0.026394795 | 0.97092375 |
| ENSG00000132842 | *AP3B1* | 4.770102052 | -0.006410162 | -0.082798095 | 0.076387933 | 0.999933887 | -0.205256133 | -0.196679703 | -0.00857643 | 0.971109544 |
| ENSG00000225880 | *LINC00115* | 2.290680225 | 0.028146791 | 0.024188581 | 0.00395821 | 0.999933887 | 0.424434459 | 0.413037432 | 0.011397027 | 0.971120024 |
| ENSG00000065717 | *TLE2* | 0.511090472 | -0.073867818 | 0.059489497 | -0.133357315 | 0.999933887 | 0.038843836 | 0.055741664 | -0.016897828 | 0.97112903 |
| ENSG00000183134 | *PTGDR2* | 3.151636041 | -0.003842117 | 0.005356671 | -0.009198788 | 0.999933887 | -0.09075333 | -0.081596008 | -0.009157321 | 0.971136789 |
| ENSG00000163013 | *FBXO41* | 2.607017506 | 0.021479167 | 0.148737714 | -0.127258547 | 0.999933887 | -0.000675852 | 0.008758982 | -0.009434834 | 0.97122061 |
| ENSG00000085760 | *MTIF2* | 1.791500679 | -0.12267893 | -0.064096002 | -0.058582928 | 0.999933887 | -0.107113019 | -0.119380368 | 0.012267349 | 0.971460831 |
| ENSG00000204922 | *UQCC3* | 0.392677593 | 0.097990806 | 0.266000797 | -0.168009991 | 0.999933887 | 0.670757285 | 0.688494493 | -0.017737208 | 0.971460831 |
| ENSG00000156795 | *NTAQ1* | -0.423984258 | 0.110328277 | -0.280917993 | 0.391246269 | 0.999933887 | -0.182608742 | -0.159542999 | -0.023065744 | 0.971460831 |
| ENSG00000175324 | *LSM1* | 1.832718391 | -0.007845236 | -0.05054668 | 0.042701444 | 0.999933887 | -0.524901117 | -0.511800305 | -0.013100813 | 0.971460831 |
| ENSG00000136247 | *ZDHHC4* | 1.225591436 | -0.026474005 | 0.1163305 | -0.142804505 | 0.999933887 | -0.02188491 | -0.010710269 | -0.011174641 | 0.971498955 |
| ENSG00000145107 | *TM4SF19* | 0.150385887 | -0.101846666 | 0.133226198 | -0.235072864 | 0.999933887 | 0.121323753 | 0.132892325 | -0.011568572 | 0.971525839 |
| ENSG00000197586 | *ENTPD6* | 3.113521476 | -0.069523201 | 0.109075403 | -0.178598604 | 0.999933887 | 0.077867936 | 0.069009137 | 0.008858799 | 0.971850725 |
| ENSG00000169740 | *ZNF32* | 0.397474616 | -0.071958919 | -0.026240269 | -0.04571865 | 0.999933887 | 0.198334787 | 0.17769311 | 0.020641677 | 0.971977962 |
| ENSG00000198252 | *STYX* | 3.141343567 | 0.094711619 | -0.058944813 | 0.153656432 | 0.999933887 | -0.406798507 | -0.395888773 | -0.010909734 | 0.971978561 |
| ENSG00000188725 | *SMIM15* | 2.191183082 | -0.029990417 | -0.03031807 | 0.000327653 | 0.999933887 | -0.386022773 | -0.397219412 | 0.011196639 | 0.971978561 |
| ENSG00000105793 | *GTPBP10* | 1.820876562 | -0.085998781 | -0.005038672 | -0.080960109 | 0.999933887 | -0.21224128 | -0.197571085 | -0.014670196 | 0.972144767 |
| ENSG00000276529 | *AP001505.1* | -1.060038098 | 0.112937095 | 0.167897199 | -0.054960103 | 0.999933887 | -0.092745097 | -0.055336451 | -0.037408646 | 0.972144767 |
| ENSG00000158604 | *TMED4* | 5.445527338 | 0.01077224 | 0.07635774 | -0.0655855 | 0.999933887 | 0.343693738 | 0.338394694 | 0.005299044 | 0.972260539 |
| ENSG00000104970 | *KIR3DX1* | -0.462329571 | 0.005586069 | -0.129056158 | 0.134642227 | 0.999933887 | -0.135504764 | -0.113195099 | -0.022309665 | 0.972260539 |
| ENSG00000173272 | *MZT2A* | 2.665925345 | 0.113856861 | -0.053635883 | 0.167492744 | 0.999933887 | -0.006737555 | -0.018602288 | 0.011864733 | 0.972846482 |
| ENSG00000100027 | *YPEL1* | 1.236229904 | 0.089770844 | -0.243330315 | 0.333101158 | 0.999933887 | -0.139166169 | -0.152902701 | 0.013736532 | 0.972846482 |
| ENSG00000185730 | *ZNF696* | 0.577369995 | 0.078170371 | 0.190599996 | -0.112429625 | 0.999933887 | 0.124602158 | 0.137528135 | -0.012925977 | 0.972935738 |
| ENSG00000221995 | *TIAF1* | 0.584368255 | 0.007751401 | 0.049691203 | -0.041939802 | 0.999933887 | -0.42608563 | -0.407657016 | -0.018428613 | 0.97301829 |
| ENSG00000184465 | *WDR27* | 1.398828014 | 0.014038538 | 0.025025359 | -0.010986821 | 0.999933887 | 0.061950671 | 0.049844088 | 0.012106583 | 0.973378996 |
| ENSG00000198130 | *HIBCH* | 0.666780015 | 0.258705323 | 0.071007948 | 0.187697376 | 0.999933887 | -0.183257999 | -0.160096725 | -0.023161274 | 0.973608957 |
| ENSG00000215030 | *RPL13P12* | 3.190302897 | 0.08741248 | -0.005546509 | 0.092958989 | 0.999933887 | 0.038467335 | 0.047603025 | -0.00913569 | 0.973608957 |
| ENSG00000270647 | *TAF15* | 5.116749725 | 0.058244619 | 0.091195581 | -0.032950962 | 0.999933887 | -0.030435312 | -0.038372137 | 0.007936825 | 0.973608957 |
| ENSG00000179271 | *GADD45GIP1* | 1.979237775 | 0.070369486 | 0.00214104 | 0.068228446 | 0.999933887 | 0.201606363 | 0.211936507 | -0.010330144 | 0.973608957 |
| ENSG00000163636 | *PSMD6* | 5.118361858 | -0.031495636 | -0.074277824 | 0.042782188 | 0.999933887 | -0.202007318 | -0.196957527 | -0.005049791 | 0.973608957 |
| ENSG00000112139 | *MDGA1* | 0.184509993 | -0.078599968 | 0.02967351 | -0.108273477 | 0.999933887 | -0.201948376 | -0.189332602 | -0.012615774 | 0.973608957 |
| ENSG00000255882 | *AC091814.1* | 0.15252843 | 0.074238629 | 0.527370348 | -0.453131719 | 0.999933887 | -0.164113732 | -0.193866164 | 0.029752432 | 0.973608957 |
| ENSG00000165886 | *UBTD1* | 3.638252527 | 0.03033206 | 0.011663445 | 0.018668615 | 0.999933887 | -0.807872051 | -0.797007866 | -0.010864185 | 0.973608957 |
| ENSG00000143621 | *ILF2* | 4.381943237 | -0.046365088 | -0.038111798 | -0.008253291 | 0.999933887 | 0.224354753 | 0.230427912 | -0.006073159 | 0.973680175 |
| ENSG00000163600 | *ICOS* | 3.773844719 | -0.031440419 | -0.058742019 | 0.0273016 | 0.999933887 | 0.058267838 | 0.064882397 | -0.006614559 | 0.973986977 |
| ENSG00000167632 | *TRAPPC9* | 3.898255885 | 0.0387579 | -0.115744688 | 0.154502589 | 0.999933887 | 0.077779637 | 0.070213107 | 0.00756653 | 0.974318908 |
| ENSG00000262999 | *AC099489.3* | 1.638530691 | 0.252841249 | 0.010164204 | 0.242677046 | 0.999933887 | 0.636365595 | 0.650586845 | -0.01422125 | 0.97432441 |
| ENSG00000100403 | *ZC3H7B* | 3.970354729 | 0.012767903 | 0.083154924 | -0.070387021 | 0.999933887 | 0.150180875 | 0.156636767 | -0.006455892 | 0.97432441 |
| ENSG00000261024 | *GS1-279B7.1* | 1.81868876 | 0.071621283 | 0.0289917 | 0.042629582 | 0.999933887 | 0.574461093 | 0.559843447 | 0.014617646 | 0.974535152 |
| ENSG00000165733 | *BMS1* | 2.721968895 | -0.067803818 | -0.049035495 | -0.018768323 | 0.999933887 | 0.177666645 | 0.185298646 | -0.007632001 | 0.97472717 |
| ENSG00000240767 | *RN7SL288P* | 1.113392229 | 0.358400446 | 0.541476835 | -0.183076389 | 0.999933887 | 0.894058777 | 0.866634189 | 0.027424588 | 0.974914546 |
| ENSG00000123136 | *DDX39A* | 7.46203267 | 0.082723652 | 0.042025253 | 0.040698398 | 0.999933887 | 0.19204011 | 0.197384898 | -0.005344788 | 0.974914546 |
| ENSG00000128595 | *CALU* | 3.262373862 | -0.052454211 | -0.044074874 | -0.008379337 | 0.999933887 | -0.178545755 | -0.184269142 | 0.005723387 | 0.974914546 |
| ENSG00000119912 | *IDE* | 1.915874387 | -0.06157227 | -0.24150296 | 0.179930689 | 0.999933887 | -0.147164443 | -0.157843597 | 0.010679154 | 0.974914546 |
| ENSG00000139644 | *TMBIM6* | 8.994553053 | -0.023914027 | -0.022047253 | -0.001866774 | 0.999933887 | -0.041170016 | -0.045887756 | 0.00471774 | 0.974914546 |
| ENSG00000168116 | *KIAA1586* | 0.004655592 | -0.006705708 | -0.195328987 | 0.188623279 | 0.999933887 | 0.005243621 | -0.012922211 | 0.018165832 | 0.974914546 |
| ENSG00000131797 | *CLUHP3* | 3.367513619 | 0.015000991 | 0.024262874 | -0.009261883 | 0.999933887 | 0.262792283 | 0.270871728 | -0.008079445 | 0.975034632 |
| ENSG00000105708 | *ZNF14* | 1.857900185 | 0.054620878 | -0.1984676 | 0.253088478 | 0.999933887 | -0.225882815 | -0.239055329 | 0.013172514 | 0.975178233 |
| ENSG00000058056 | *USP13* | 0.414930958 | 0.032278324 | -0.002131072 | 0.034409395 | 0.999933887 | -0.040609165 | -0.055280214 | 0.014671049 | 0.975263763 |
| ENSG00000184574 | *LPAR5* | 0.238668569 | 0.030523546 | -0.145343272 | 0.175866818 | 0.999933887 | 0.018906208 | 0.031095522 | -0.012189314 | 0.975263763 |
| ENSG00000164465 | *DCBLD1* | 0.195919022 | -0.022454941 | 0.309170415 | -0.331625356 | 0.999933887 | 0.127785399 | 0.113561169 | 0.014224231 | 0.975279865 |
| ENSG00000171295 | *ZNF440* | 1.599537807 | -0.123522947 | -0.100829011 | -0.022693936 | 0.999933887 | -0.295894113 | -0.282119897 | -0.013774216 | 0.975302949 |
| ENSG00000118200 | *CAMSAP2* | -0.322600782 | 0.092293793 | -0.063392839 | 0.155686632 | 0.999933887 | -0.437735675 | -0.455477686 | 0.017742011 | 0.975302949 |
| ENSG00000142173 | *COL6A2* | 3.507680121 | 0.034674663 | 0.111414581 | -0.076739917 | 0.999933887 | 0.095077985 | 0.10328572 | -0.008207735 | 0.975364214 |
| ENSG00000135093 | *USP30* | 2.873800375 | 0.003454677 | -0.07063776 | 0.074092437 | 0.999933887 | -0.154374243 | -0.161296313 | 0.00692207 | 0.975656167 |
| ENSG00000141026 | *MED9* | 1.91431254 | -0.011291275 | -0.049972733 | 0.038681459 | 0.999933887 | -0.075752321 | -0.06807317 | -0.007679151 | 0.975856612 |
| ENSG00000169762 | *TAPT1* | 3.756030853 | -0.071440766 | -0.0030669 | -0.068373866 | 0.999933887 | -0.129391901 | -0.122569217 | -0.006822684 | 0.97586365 |
| ENSG00000133477 | *FAM83F* | -0.791174026 | 0.118902901 | 0.057281528 | 0.061621372 | 0.999933887 | 0.169332205 | 0.149645672 | 0.019686533 | 0.975905973 |
| ENSG00000100650 | *SRSF5* | 9.207348785 | 0.020096918 | -0.006521728 | 0.026618646 | 0.999933887 | 0.281693619 | 0.286190874 | -0.004497255 | 0.975905973 |
| ENSG00000234608 | *MAPKAPK5-A* | 2.173769809 | -0.093968406 | 0.036532743 | -0.130501148 | 0.999933887 | -0.027830221 | -0.019029103 | -0.008801118 | 0.976106094 |
| ENSG00000168872 | *DDX19A* | 2.389920198 | 0.05525182 | -0.003082688 | 0.058334507 | 0.999933887 | 0.246441099 | 0.238007778 | 0.008433322 | 0.976106094 |
| ENSG00000204435 | *CSNK2B* | 3.190239571 | -0.048548186 | 0.021520097 | -0.070068283 | 0.999933887 | -0.176928612 | -0.170039336 | -0.006889276 | 0.976106094 |
| ENSG00000167972 | *ABCA3* | 1.059140267 | 0.08089944 | -0.013713377 | 0.094612817 | 0.999933887 | -0.001741536 | 0.010696312 | -0.012437848 | 0.976106094 |
| ENSG00000231999 | *LRRC8C-DT* | 1.09710363 | -0.14103499 | -0.086626673 | -0.054408317 | 0.999933887 | 0.119967557 | 0.131719077 | -0.011751521 | 0.976188612 |
| ENSG00000180787 | *ZFP3* | 0.142855756 | -0.017934102 | 0.045210854 | -0.063144956 | 0.999933887 | 0.0233554 | 0.013196576 | 0.010158824 | 0.976245386 |
| ENSG00000130684 | *ZNF337* | -0.485353367 | 0.088718445 | -0.017219956 | 0.105938401 | 0.999933887 | 0.174013002 | 0.155729583 | 0.018283419 | 0.976321662 |
| ENSG00000139842 | *CUL4A* | 4.775042547 | -0.021329092 | -0.064215416 | 0.042886323 | 0.999933887 | -0.161038707 | -0.165591585 | 0.004552878 | 0.976328015 |
| ENSG00000122679 | *RAMP3* | 2.655093775 | 0.197800674 | 0.261621568 | -0.063820894 | 0.999933887 | 1.226865545 | 1.240264129 | -0.013398584 | 0.976372799 |
| ENSG00000242114 | *MTFP1* | -0.475146335 | 0.264138653 | -0.526776957 | 0.790915609 | 0.944094537 | -0.052722927 | -0.031039471 | -0.021683456 | 0.976372799 |
| ENSG00000239282 | *CASTOR1* | -0.499928919 | 0.221360105 | -0.050505866 | 0.271865972 | 0.999933887 | 1.611052937 | 1.590045992 | 0.021006945 | 0.976372799 |
| ENSG00000081985 | *IL12RB2* | 0.622869261 | 0.092694734 | 0.130300825 | -0.037606091 | 0.999933887 | 0.112830329 | 0.098934843 | 0.013895486 | 0.976372799 |
| ENSG00000090975 | *PITPNM2* | 2.099370586 | -0.056964232 | -0.013723444 | -0.043240788 | 0.999933887 | 0.044279096 | 0.035776253 | 0.008502843 | 0.976372799 |
| ENSG00000275178 | *AC018529.1* | 2.679734605 | 0.069523598 | 0.227954721 | -0.158431123 | 0.999933887 | -0.019992393 | -0.009545406 | -0.010446987 | 0.976372799 |
| ENSG00000271646 | *IRF2-DT* | 4.358278676 | 0.056160086 | 0.065079369 | -0.008919283 | 0.999933887 | 1.054953422 | 1.04740027 | 0.007553152 | 0.976372799 |
| ENSG00000081189 | *MEF2C* | 4.476905628 | -0.066970725 | 0.038231029 | -0.105201754 | 0.999933887 | -1.304592085 | -1.313236676 | 0.00864459 | 0.976372799 |
| ENSG00000100124 | *ANKRD54* | 2.173177074 | -0.042284377 | 0.112463381 | -0.154747758 | 0.999933887 | -0.004159606 | 0.004445895 | -0.008605501 | 0.976372799 |
| ENSG00000162851 | *TFB2M* | 1.330353389 | 0.04761274 | -0.087536609 | 0.135149349 | 0.999933887 | 0.049894237 | 0.037983535 | 0.011910702 | 0.976372799 |
| ENSG00000211716 | *TRBV9* | -0.010225384 | 0.063455088 | 0.055725289 | 0.007729798 | 0.999933887 | 0.058135938 | 0.075144938 | -0.017009 | 0.976372799 |
| ENSG00000103429 | *BFAR* | 5.018056873 | -0.013571428 | -0.034016132 | 0.020444704 | 0.999933887 | -0.326660937 | -0.320706886 | -0.005954051 | 0.976372799 |
| ENSG00000221883 | *ARIH2OS* | -0.004326413 | 0.021375676 | 0.051551681 | -0.030176005 | 0.999933887 | -0.047699981 | -0.065167391 | 0.01746741 | 0.976372799 |
| ENSG00000284738 | *AL358472.5* | 0.090825987 | 0.190528545 | 0.105658528 | 0.084870017 | 0.999933887 | -0.154449418 | -0.128934452 | -0.025514966 | 0.976474949 |
| ENSG00000227081 | *AC005912.1* | 2.043146844 | -0.013342194 | -0.017174273 | 0.003832079 | 0.999933887 | 0.118660971 | 0.104273407 | 0.014387564 | 0.976474949 |
| ENSG00000168329 | *CX3CR1* | 3.271637017 | 0.086698442 | 0.037500912 | 0.04919753 | 0.999933887 | 0.122594291 | 0.114583252 | 0.008011039 | 0.976501528 |
| ENSG00000212978 | *AC016747.1* | -0.831233167 | -0.151095669 | 0.001828688 | -0.152924357 | 0.999933887 | 0.039251833 | 0.060286496 | -0.021034663 | 0.976501528 |
| ENSG00000133195 | *SLC39A11* | 1.428348848 | 0.039573767 | -0.02265456 | 0.062228327 | 0.999933887 | -0.101805067 | -0.115759253 | 0.013954186 | 0.976501528 |
| ENSG00000108106 | *UBE2S* | 4.69955983 | 0.176716223 | 0.253979445 | -0.077263222 | 0.999933887 | 0.437775553 | 0.428139365 | 0.009636188 | 0.976560315 |
| ENSG00000172367 | *PDZD3* | -0.404954066 | 0.202759203 | 0.146994601 | 0.055764602 | 0.999933887 | -0.589640908 | -0.611846151 | 0.022205243 | 0.976606957 |
| ENSG00000129460 | *NGDN* | 3.19787046 | -0.096135888 | -0.118629168 | 0.02249328 | 0.999933887 | 0.096852135 | 0.09081316 | 0.006038975 | 0.976626331 |
| ENSG00000160049 | *DFFA* | 1.832078376 | 0.10521173 | -0.066590651 | 0.171802381 | 0.999933887 | 0.058062259 | 0.049412282 | 0.008649977 | 0.976688256 |
| ENSG00000119487 | *MAPKAP1* | 4.247128699 | 0.050041655 | -0.002244801 | 0.052286455 | 0.999933887 | -0.176389367 | -0.181197295 | 0.004807928 | 0.976688256 |
| ENSG00000121895 | *TMEM156* | 0.995519908 | -0.051432967 | -0.184726087 | 0.133293121 | 0.999933887 | -0.029131662 | -0.017499374 | -0.011632289 | 0.976688256 |
| ENSG00000146858 | *ZC3HAV1L* | -1.661766416 | -0.336768922 | 0.643441828 | -0.98021075 | 0.999933887 | -0.199459364 | -0.2227476 | 0.023288236 | 0.976723091 |
| ENSG00000179562 | *GCC1* | 5.957959588 | -0.058658762 | -0.023506127 | -0.035152635 | 0.999933887 | -0.015517303 | -0.010736524 | -0.00478078 | 0.976755596 |
| ENSG00000162729 | *IGSF8* | 2.530022833 | 0.12522147 | 0.082306383 | 0.042915086 | 0.999933887 | 0.078652967 | 0.070545919 | 0.008107047 | 0.976845095 |
| ENSG00000273174 | *AC108673.2* | 1.640401988 | -0.039498824 | 0.114143032 | -0.153641856 | 0.999933887 | 0.022717001 | 0.036786311 | -0.01406931 | 0.977060799 |
| ENSG00000138035 | *PNPT1* | 2.983380874 | -0.034345668 | -0.112332633 | 0.077986965 | 0.999933887 | -0.226989398 | -0.219219013 | -0.007770385 | 0.977060799 |
| ENSG00000279198 | *AC008894.3* | 1.825982349 | 0.243925978 | 0.141974408 | 0.10195157 | 0.999933887 | 0.434517594 | 0.414450516 | 0.020067078 | 0.977061742 |
| ENSG00000279339 | *AC100788.2* | 2.374315041 | 0.144761789 | 0.020247432 | 0.124514357 | 0.999933887 | -0.897362208 | -0.909099881 | 0.011737673 | 0.977061742 |
| ENSG00000102032 | *RENBP* | 6.772144855 | -0.041381244 | -0.059431955 | 0.018050711 | 0.999933887 | -0.087733056 | -0.08118004 | -0.006553016 | 0.977061742 |
| ENSG00000244627 | *TPTEP2* | 1.381260305 | 0.062522961 | 0.25455637 | -0.192033409 | 0.999933887 | -0.594106955 | -0.584490961 | -0.009615994 | 0.977061742 |

| | | | | | | | | | | |
|---|---|---|---|---|---|---|---|---|---|---|
| ENSG00000189136 | *UBE2Q2P1* | -0.175829484 | -0.309805796 | -0.191379139 | -0.118426657 | 0.999933887 | -0.045539907 | -0.064320999 | 0.018781093 | 0.977176563 |
| ENSG00000271533 | *Z83843.1* | 1.517016247 | 0.124563342 | 0.174898704 | -0.050335362 | 0.999933887 | -0.324995755 | -0.311357072 | -0.013638683 | 0.977176563 |
| ENSG00000156469 | *MTERF3* | 0.504208864 | -0.075999405 | -0.096768014 | 0.020768609 | 0.999933887 | 0.111271111 | 0.097508428 | 0.013762683 | 0.977176563 |
| ENSG00000101639 | *CEP192* | 3.75550698 | 0.024339156 | -0.079177453 | 0.103516609 | 0.999933887 | -0.176617455 | -0.187707738 | 0.011090282 | 0.977176563 |
| ENSG00000128394 | *APOBEC3F* | 2.117438079 | -0.013453548 | 0.084882417 | -0.098335965 | 0.999933887 | -0.208476309 | -0.199286187 | -0.009190122 | 0.978037701 |
| ENSG00000279179 | *AL662907.2* | 0.779434137 | 0.146925917 | -0.115300222 | 0.26222614 | 0.999933887 | -0.323878687 | -0.339192752 | 0.015314065 | 0.978139158 |
| ENSG00000120696 | *KBTBD7* | 3.450779575 | 0.084460227 | -0.008298007 | 0.092758234 | 0.999933887 | 0.689902289 | 0.67854069 | 0.011361599 | 0.978267821 |
| ENSG00000271991 | *AC013400.1* | -1.144891714 | 0.502919112 | 0.062981223 | 0.43993789 | 0.999933887 | 2.669967646 | 2.642736954 | 0.027230692 | 0.978360383 |
| ENSG00000141642 | *ELAC1* | -0.652282525 | -0.018336191 | 0.047720206 | -0.066056396 | 0.999933887 | 0.075573366 | 0.093677912 | -0.018104546 | 0.978360383 |
| ENSG00000147010 | *SH3KBP1* | 7.314829835 | 8.876e-06 | 0.003878357 | -0.003869481 | 0.999933887 | 0.078912056 | 0.075330496 | 0.00358156 | 0.978360383 |
| ENSG00000206530 | *CFAP44* | 0.425463622 | -0.189642306 | -0.234537219 | 0.044894913 | 0.999933887 | -0.066734726 | -0.051360019 | -0.015374708 | 0.978877031 |
| ENSG00000152082 | *MZT2B* | 1.898035625 | 0.122880675 | 0.001377237 | 0.121503438 | 0.999933887 | -0.026073563 | -0.015590044 | -0.01048352 | 0.978877031 |
| ENSG00000163191 | *S100A11* | 9.598736224 | 0.055695687 | -0.018464646 | 0.074160333 | 0.999933887 | -0.578228059 | -0.573404963 | -0.004823096 | 0.978877031 |
| ENSG00000108679 | *LGALS3BP* | 3.313292432 | 0.056801737 | 0.029002731 | 0.027799006 | 0.999933887 | -0.059901489 | -0.052680656 | -0.007220833 | 0.978877031 |
| ENSG00000175110 | *MRPS22* | 2.191867716 | 0.049913093 | -0.112538314 | 0.162451408 | 0.999933887 | -0.006113079 | 0.001920429 | -0.008033508 | 0.978877031 |
| ENSG00000128881 | *TTBK2* | 2.911412244 | 0.047444778 | -0.015111521 | 0.062556299 | 0.999933887 | -0.054584829 | -0.043692913 | -0.010891916 | 0.978877031 |
| ENSG00000127989 | *MTERF1* | 1.419433977 | 0.052570926 | -0.254316979 | 0.306887905 | 0.999933887 | -0.276572985 | -0.287113709 | 0.010540724 | 0.978877031 |
| ENSG00000150787 | *PTS* | 2.568547733 | -0.022852167 | 0.076404518 | -0.099256685 | 0.999933887 | 0.090491578 | 0.083006672 | 0.007484906 | 0.978877031 |
| ENSG00000196152 | *ZNF79* | 1.815386125 | 0.033127547 | 0.068472559 | -0.035345013 | 0.999933887 | -0.104587546 | -0.112431975 | 0.007844429 | 0.978877031 |
| ENSG00000221829 | *FANCG* | 1.330334755 | 0.027385604 | 0.104991786 | -0.077606182 | 0.999933887 | -0.010757218 | -0.000988873 | -0.009768345 | 0.978877031 |
| ENSG00000100359 | *SGSM3* | -0.492826905 | -0.073422325 | -0.121211202 | 0.047788877 | 0.999933887 | -0.11200622 | -0.091834144 | -0.020172076 | 0.979120568 |
| ENSG00000213062 | *Z99572.1* | -0.798723404 | 0.156557896 | 0.114465902 | 0.042091994 | 0.999933887 | 0.095561085 | 0.125977549 | -0.030416465 | 0.979149204 |
| ENSG00000225774 | *SIRPAP1* | 0.843793216 | -0.06609707 | -0.0185298 | -0.047567269 | 0.999933887 | 0.6407808 | 0.62874713 | 0.01203367 | 0.979225723 |
| ENSG00000173457 | *PPP1R14B* | 2.35712467 | 0.037572235 | -0.017203048 | 0.054775283 | 0.999933887 | -0.007627568 | -0.01434563 | 0.006718062 | 0.979251889 |
| ENSG00000141540 | *TTYH2* | 2.111109312 | -0.025485847 | -0.045220068 | 0.019734221 | 0.999933887 | 0.125016823 | 0.134474768 | -0.009457945 | 0.979251889 |
| ENSG00000243566 | *UPK3B* | 0.809646145 | -0.125976193 | -0.127449451 | 0.001473258 | 0.999933887 | 0.198665502 | 0.211858642 | -0.01319314 | 0.979276155 |
| ENSG00000074855 | *ANO8* | 2.058148278 | 0.083221584 | 0.188694492 | -0.105472908 | 0.999933887 | 0.065995475 | 0.056538081 | 0.009457395 | 0.979301974 |
| ENSG00000109016 | *DHRS7B* | 2.567492346 | 0.063093898 | 0.075888412 | -0.012794514 | 0.999933887 | -0.053452237 | -0.059736555 | 0.006284319 | 0.97933366 |
| ENSG00000131446 | *MGAT1* | 8.765014923 | -0.047309796 | -0.045740017 | -0.001569779 | 0.999933887 | -0.779423387 | -0.784691227 | 0.00526784 | 0.97933366 |
| ENSG00000129295 | *LRRC6* | 0.631780544 | -0.055003765 | -0.092559326 | 0.037555561 | 0.999933887 | -0.237775857 | -0.223218183 | -0.014557674 | 0.97933366 |
| ENSG00000163923 | *RPL39L* | -0.567338558 | 0.127980172 | 0.034639651 | 0.093340521 | 0.999933887 | -0.278680233 | -0.291575072 | 0.012894839 | 0.97933366 |
| ENSG00000239839 | *DEFA3* | 0.806215167 | 0.087425866 | 0.049324943 | 0.038100923 | 0.999933887 | 0.073640397 | 0.063499393 | 0.010141004 | 0.979422251 |
| ENSG00000167114 | *SLC27A4* | 1.357335962 | 0.01688626 | 0.144852804 | -0.127966544 | 0.999933887 | 0.07037931 | 0.060963287 | 0.009416023 | 0.979517302 |
| ENSG00000265531 | *FCGR1CP* | 0.990494212 | 0.028350779 | -0.093606425 | 0.121957205 | 0.999933887 | 0.046352183 | 0.030074097 | 0.016278085 | 0.979757515 |
| ENSG00000237424 | *FOXD2-AS1* | -0.144366635 | 0.02572715 | 0.042730888 | -0.017003737 | 0.999933887 | 0.118192552 | 0.104298431 | 0.013894121 | 0.979785694 |
| ENSG00000178498 | *DTX3* | 2.078667656 | -0.141555304 | 0.137061641 | -0.278616945 | 0.944094537 | -0.027888714 | -0.035313382 | 0.007424668 | 0.979822784 |
| ENSG00000135334 | *AKIRIN2* | 8.956184283 | -0.012345094 | 0.008923029 | -0.021268122 | 0.999933887 | -0.659883877 | -0.664649601 | 0.004765725 | 0.979822784 |
| ENSG00000182902 | *SLC25A18* | 0.489261269 | 0.183853216 | 0.136803569 | 0.047049646 | 0.999933887 | 0.667700129 | 0.67654185 | -0.008841721 | 0.97995012 |
| ENSG00000228634 | *AL136115.1* | -0.162429463 | -0.157679532 | -0.000235251 | -0.157444282 | 0.999933887 | 0.164931738 | 0.149204787 | 0.015726951 | 0.97995012 |
| ENSG00000166181 | *API5* | 3.762335566 | -0.096684283 | -0.066963233 | -0.02972105 | 0.999933887 | -0.084706105 | -0.089821238 | 0.005115133 | 0.980173136 |
| ENSG00000060982 | *BCAT1* | 2.433962466 | -0.074826147 | 0.110318058 | -0.185144205 | 0.999933887 | -0.374746722 | -0.381754127 | 0.007007405 | 0.980200792 |
| ENSG00000204237 | *OXLD1* | 2.851212124 | -0.030274787 | -0.043498161 | 0.013223374 | 0.999933887 | 0.285791358 | 0.291726837 | -0.005935479 | 0.980200792 |
| ENSG00000275437 | *AL121832.3* | 1.770772733 | 0.036700468 | -0.048924186 | 0.085624654 | 0.999933887 | 0.760255731 | 0.771732378 | -0.011476648 | 0.980200792 |
| ENSG00000023171 | *GRAMD1B* | 3.123303525 | -0.021227948 | 0.000815119 | -0.022043067 | 0.999933887 | 0.053032091 | 0.05909562 | -0.00606353 | 0.980252906 |
| ENSG00000283142 | *AL049767.1* | -0.215962254 | -0.329832963 | -0.557241411 | 0.227408448 | 0.999933887 | -0.014373666 | -0.004829001 | -0.009544665 | 0.980328933 |
| ENSG00000287201 | *AC009682.1* | 2.141018467 | 0.079184219 | 0.072103798 | 0.007080421 | 0.999933887 | 0.155142785 | 0.169275104 | -0.014132319 | 0.980698601 |
| ENSG00000105778 | *AVL9* | 5.783808429 | -0.018211084 | -0.005513313 | -0.01269777 | 0.999933887 | -0.362718271 | -0.359049979 | -0.003668292 | 0.980698601 |
| ENSG00000154059 | *IMPACT* | 1.38141634 | -0.002663137 | 0.169469991 | -0.172133128 | 0.999933887 | -0.280181705 | -0.270555046 | -0.009626659 | 0.980848513 |
| ENSG00000196922 | *ZNF252P* | 1.385737065 | 0.050352506 | -0.149256216 | 0.199608722 | 0.999933887 | -0.129979971 | -0.115158938 | -0.014821033 | 0.980931071 |
| ENSG00000110218 | *PANX1* | 1.187068567 | 0.012500563 | 0.011913575 | 0.000586988 | 0.999933887 | 0.087177542 | 0.076206035 | 0.010971507 | 0.981201389 |
| ENSG00000196511 | *TPK1* | 1.685252078 | 0.041811018 | -0.083418527 | 0.125229545 | 0.999933887 | -0.011387297 | -0.021862436 | 0.010475139 | 0.981254414 |
| ENSG00000151208 | *DLG5* | 1.741839224 | 0.005665957 | 0.076182024 | -0.070516067 | 0.999933887 | -1.9461e-05 | -0.00823319 | 0.008213728 | 0.981504445 |
| ENSG00000161016 | *RPL8* | 8.323174273 | 0.026103889 | -0.011356818 | 0.037460706 | 0.999933887 | 0.083037384 | 0.087055907 | -0.004018523 | 0.981710334 |
| ENSG00000166582 | *CENPV* | 0.548309793 | 0.083665418 | -0.000524189 | 0.084189608 | 0.999933887 | 0.106032305 | 0.095963494 | 0.010068811 | 0.981775554 |
| ENSG00000276471 | *AC105265.3* | -0.122892845 | 0.070779632 | -0.073196044 | 0.143975677 | 0.999933887 | 1.807437625 | 1.828642199 | -0.021204573 | 0.981775554 |
| ENSG00000223509 | *AC135983.2* | 1.021228096 | 0.051263876 | 0.063071166 | -0.01180729 | 0.999933887 | 0.083756274 | 0.073128347 | 0.010627927 | 0.981775554 |
| ENSG00000020633 | *RUNX3* | 7.275072138 | 0.002793857 | 0.011749011 | -0.008955154 | 0.999933887 | 0.103375391 | 0.099958405 | 0.003416987 | 0.981775554 |
| ENSG00000168575 | *SLC20A2* | 2.654175075 | 0.004529771 | 0.067189406 | -0.062659635 | 0.999933887 | 0.092770134 | 0.098836043 | -0.006065909 | 0.981775554 |
| ENSG00000166557 | *TMED3* | 2.194175035 | 0.056191711 | -0.071745248 | 0.127936959 | 0.999933887 | 0.080012075 | 0.073479264 | 0.006532811 | 0.981836629 |
| ENSG00000151148 | *UBE3B* | 4.764494063 | -0.016873598 | 0.032203343 | -0.049076941 | 0.999933887 | -0.011917748 | -0.00833155 | -0.003586198 | 0.982040754 |
| ENSG00000269893 | *SNHG8* | 3.366552609 | 0.102538231 | 0.001522074 | 0.101016157 | 0.999933887 | 0.229102717 | 0.223639237 | 0.00546348 | 0.98204154 |
| ENSG00000100577 | *GSTZ1* | 0.171799877 | -0.221392616 | 0.002548779 | -0.223941395 | 0.999933887 | -0.120349402 | -0.109860517 | -0.010488886 | 0.982156505 |
| ENSG00000188822 | *CNR2* | 3.789836545 | -0.119682801 | -0.014367403 | -0.105315398 | 0.999933887 | -0.107911465 | -0.112815174 | 0.004903709 | 0.982156505 |
| ENSG00000090097 | *PCBP4* | 1.829469979 | 0.106407239 | 0.130343275 | -0.023936036 | 0.999933887 | 0.114928497 | 0.122663652 | -0.007735155 | 0.982156505 |
| ENSG00000124614 | *RPS10* | 0.256943918 | -0.106981623 | 0.061778057 | -0.16875968 | 0.999933887 | 0.108534991 | 0.120587471 | -0.01205248 | 0.982156505 |
| ENSG00000260853 | *AC109460.2* | 2.689963375 | 0.035986914 | 0.050498615 | -0.014511701 | 0.999933887 | 0.055456227 | 0.061205464 | -0.005749237 | 0.982156505 |
| ENSG00000233369 | *GTF2IP4* | 2.811052311 | -0.028206368 | 0.009824713 | -0.03803108 | 0.999933887 | -0.007116586 | -0.014736315 | 0.007619729 | 0.982156505 |
| ENSG00000006607 | *FARP2* | 2.625901859 | 0.015493561 | -0.009732958 | 0.025226519 | 0.999933887 | -0.119450498 | -0.112944469 | -0.006506029 | 0.982156505 |
| ENSG00000111077 | *TNS2* | 0.108286923 | -0.017641565 | 0.313790639 | -0.331432204 | 0.999933887 | -0.012576677 | -0.027077266 | 0.014500589 | 0.982156505 |
| ENSG00000256361 | *AC027544.1* | -0.373114561 | -0.068632019 | -0.108900004 | 0.040267986 | 0.999933887 | 0.217802748 | 0.194252461 | 0.023550287 | 0.982236846 |
| ENSG00000020256 | *ZFP64* | 0.737566032 | 0.1583546 | 0.089390965 | 0.068963635 | 0.999933887 | -0.0213587 | -0.030210848 | 0.008852148 | 0.982259617 |
| ENSG00000206651 | *Y_RNA* | -0.611643901 | -0.217236966 | -0.236106431 | 0.018869465 | 0.999933887 | 0.339003029 | 0.358730627 | -0.019727598 | 0.982259617 |
| ENSG00000280088 | *AC126474.2* | 1.527375649 | -0.031832104 | -0.130741681 | 0.098909577 | 0.999933887 | -0.633333978 | -0.623493669 | -0.009840309 | 0.982259617 |
| ENSG00000277075 | *H2AC8* | 0.120725159 | 0.055121763 | -0.266704121 | 0.321825884 | 0.999933887 | 0.147616468 | 0.132478373 | 0.015138096 | 0.982317442 |
| ENSG00000272325 | *NUDT3* | 2.655302086 | 0.144061416 | -0.050969986 | 0.195031402 | 0.999933887 | 0.35816429 | 0.34965934 | 0.00850495 | 0.98250006 |
| ENSG00000054793 | *ATP9A* | 1.452987679 | -0.022018716 | -0.047479886 | 0.02546117 | 0.999933887 | -0.065130862 | -0.058813337 | -0.006317525 | 0.982604465 |
| ENSG00000143387 | *CTSK* | 1.521761926 | -0.120926282 | 0.008354389 | -0.129280671 | 0.999933887 | -0.309452926 | -0.300906461 | -0.008546466 | 0.98262159 |
| ENSG00000254693 | *AC010768.1* | -0.538818796 | 0.134755818 | -0.163823068 | 0.298578886 | 0.999933887 | 2.044341223 | 2.028448225 | 0.015892998 | 0.98262159 |
| ENSG00000158321 | *AUTS2* | 3.712972631 | 0.019848501 | -0.021118628 | 0.040967129 | 0.999933887 | -0.026556168 | -0.031393407 | 0.004837239 | 0.98262159 |
| ENSG00000112130 | *RNF8* | 0.554247711 | 0.045313188 | 0.073326557 | -0.028013368 | 0.999933887 | 0.073125478 | 0.08329283 | -0.010167352 | 0.98262159 |
| ENSG00000271122 | *AC018647.2* | 0.095113798 | 0.043901376 | -0.266952132 | 0.310853508 | 0.999933887 | 0.14466339 | 0.133901512 | 0.010761878 | 0.98262159 |
| ENSG00000123191 | *ATP7B* | 0.48676454 | 0.034725505 | 0.055150339 | -0.020424833 | 0.999933887 | -0.04016383 | -0.056953474 | 0.016789644 | 0.98262159 |
| ENSG00000160200 | *CBS* | -1.120209613 | 0.161174108 | 0.363631538 | -0.20245743 | 0.999933887 | -1.025703901 | -1.012426105 | -0.013277796 | 0.982697523 |
| ENSG00000180257 | *ZNF816* | -0.504095652 | 0.227900643 | -0.205774541 | 0.433675184 | 0.999933887 | -0.177649149 | -0.164146866 | -0.013502282 | 0.982709495 |
| ENSG00000180747 | *SMG1P3* | 4.130467911 | 0.134230592 | 0.073733634 | 0.060496958 | 0.999933887 | 0.178003228 | 0.182399092 | -0.004395863 | 0.983041231 |
| ENSG00000213516 | *RBMXL1* | 4.008313105 | -0.096690068 | 0.012355839 | -0.109045906 | 0.999933887 | 0.169932734 | 0.166001459 | 0.003931275 | 0.983041231 |
| ENSG00000204859 | *ZBTB48* | 5.729384107 | 0.070131109 | 0.093694965 | -0.023563857 | 0.999933887 | 0.290173272 | 0.286662126 | 0.003511147 | 0.983041231 |
| ENSG00000171793 | *CTPS1* | 2.665623606 | -0.074702889 | 0.064093091 | -0.13879598 | 0.999933887 | 0.193045684 | 0.188062844 | 0.00498284 | 0.983041231 |
| ENSG00000124789 | *NUP153* | 7.468833789 | -0.064980221 | -0.042651573 | -0.022328648 | 0.999933887 | -0.689824178 | -0.685684381 | -0.004139797 | 0.983041231 |
| ENSG00000222112 | *RN7SKP16* | 0.28117641 | -0.119947297 | -0.097805878 | -0.02214142 | 0.999933887 | -1.420191213 | -1.43546404 | 0.015272827 | 0.983041231 |
| ENSG00000225676 | *AC002378.1* | -0.396776048 | 0.251094773 | 0.146944807 | 0.104149966 | 0.999933887 | 0.765331683 | 0.753096774 | 0.012234909 | 0.983041231 |
| ENSG00000174177 | *CTU2* | 1.862197902 | 0.073531709 | 0.06853767 | 0.004994039 | 0.999933887 | 0.330121197 | 0.323943564 | 0.006177632 | 0.983041231 |
| ENSG00000184678 | *H2BC21* | 6.854607816 | 0.054945472 | 0.020982838 | 0.033962634 | 0.999933887 | 0.099953928 | 0.104331634 | -0.004377707 | 0.983041231 |
| ENSG00000154429 | *CCSAP* | 3.848090072 | 0.065104234 | 0.111000976 | -0.045896741 | 0.999933887 | -0.764607185 | -0.770913548 | 0.006306362 | 0.983041231 |
| ENSG00000272211 | *AC114760.2* | 2.89014729 | 0.065059417 | 0.188193414 | -0.123133997 | 0.999933887 | -0.410085131 | -0.398756046 | -0.011329084 | 0.983041231 |
| ENSG00000108107 | *RPL28* | 8.465483274 | 0.028957583 | -0.01815966 | 0.047117243 | 0.999933887 | 0.055826603 | 0.052100908 | 0.003725694 | 0.983041231 |
| ENSG00000235703 | *LINC00894* | 1.604425028 | 0.076203078 | -0.058447825 | 0.134650902 | 0.999933887 | -0.207750703 | -0.198554167 | -0.009196536 | 0.983041231 |
| ENSG00000127585 | *FBXL16* | 1.327244308 | -0.051421359 | 0.148975885 | -0.200397244 | 0.999933887 | 0.095391814 | 0.086104729 | 0.009287085 | 0.983041231 |
| ENSG00000224429 | *LINC00539* | -0.576310943 | -0.079678114 | 0.098132382 | -0.177810496 | 0.999933887 | -0.131141647 | -0.115824779 | -0.015316868 | 0.983041231 |
| ENSG00000276550 | *HERC2P2* | 1.276350977 | -0.051493878 | -0.084139964 | 0.032646086 | 0.999933887 | 0.075021354 | 0.083659315 | -0.00863796 | 0.983041231 |
| ENSG00000211801 | *TRAV21* | -0.484556402 | -0.076566474 | 0.317161249 | -0.393727724 | 0.999933887 | 0.194262725 | 0.180226378 | 0.014036347 | 0.983041231 |
| ENSG00000067248 | *DHX29* | 2.539061314 | -0.039012071 | -0.184812074 | 0.145800003 | 0.999933887 | -0.234576191 | -0.227838962 | -0.006737229 | 0.983041231 |
| ENSG00000175395 | *ZNF25* | 1.128699867 | -0.05470727 | -0.071907442 | 0.017200172 | 0.999933887 | -0.067870707 | -0.077712316 | 0.009841609 | 0.983041231 |
| ENSG00000198682 | *PAPSS2* | 2.20718091 | 0.033120532 | -0.016001791 | 0.049122322 | 0.999933887 | -0.378899771 | -0.385030551 | 0.00613078 | 0.983041231 |
| ENSG00000022277 | *RTF2* | 7.379705275 | -0.016051037 | 0.001775473 | -0.01782651 | 0.999933887 | -0.057068487 | -0.060323445 | 0.003254958 | 0.983041231 |

| | | | | | | | | | | |
|---|---|---|---|---|---|---|---|---|---|---|
| ENSG00000180071 | *ANKRD18A* | -1.235076976 | 0.064901544 | 0.055915371 | 0.008986173 | 0.999933887 | 0.260865365 | 0.283878199 | -0.023012835 | 0.983041231 |
| ENSG00000253106 | *AC090198.1* | 0.233109924 | -0.033577526 | -0.114345966 | 0.08076844 | 0.999933887 | -0.459263282 | -0.446041823 | -0.013221459 | 0.983041231 |
| ENSG00000119771 | *KLHL29* | -0.041246538 | 0.029077699 | 0.134209952 | -0.105132253 | 0.999933887 | -0.011764514 | -0.023170011 | 0.011405497 | 0.983041231 |
| ENSG00000169612 | *RAMAC* | 2.140382198 | -0.020295608 | 0.033389043 | -0.053684652 | 0.999933887 | 0.051417771 | 0.05725757 | -0.005839799 | 0.983041231 |
| ENSG00000254197 | *AC011676.3* | 1.947143472 | 0.030025832 | 0.036783978 | -0.006758146 | 0.999933887 | -0.090080028 | -0.075267956 | -0.014812072 | 0.983041231 |
| ENSG00000113838 | *TBCCD1* | 0.958724844 | 0.024660978 | 0.249696368 | -0.22503539 | 0.999933887 | 0.0973228 | 0.106793331 | -0.009470531 | 0.983041231 |
| ENSG00000228549 | *BX284668.2* | -2.47887011 | -0.081940347 | -0.106393634 | 0.024453287 | 0.999933887 | -0.004438779 | -0.021161907 | 0.016723128 | 0.983041231 |
| ENSG00000196914 | *ARHGEF12* | 2.191385346 | -0.006615346 | -0.09133057 | 0.084715224 | 0.999933887 | -0.089112932 | -0.095498885 | 0.006385953 | 0.983041231 |
| ENSG00000106153 | *CHCHD2* | 6.003665742 | 0.046903226 | 0.080407888 | -0.033504662 | 0.999933887 | 0.16477104 | 0.160778292 | 0.003992748 | 0.983347066 |
| ENSG00000075568 | *TMEM131* | 5.223192585 | 0.01908745 | 0.002953218 | 0.016134233 | 0.999933887 | -0.236970936 | -0.241203059 | 0.004232123 | 0.983538626 |
| ENSG00000260314 | *MRC1* | -1.066980994 | 0.262300442 | 0.248980527 | 0.013319916 | 0.999933887 | 0.312088756 | 0.286078649 | 0.026010107 | 0.983927875 |
| ENSG00000138777 | *PPA2* | 3.253461911 | 0.031370963 | 0.075961626 | -0.044590663 | 0.999933887 | -0.090787248 | -0.086738797 | -0.004048452 | 0.983927875 |
| ENSG00000168172 | *HOOK3* | 4.67218973 | -0.004157638 | -0.031809613 | 0.027651975 | 0.999933887 | -0.586467605 | -0.580293703 | -0.006173902 | 0.983927875 |
| ENSG00000150990 | *DHX37* | 2.991729831 | 0.001108453 | 0.021285952 | -0.020177499 | 0.999933887 | 0.091732509 | 0.09624228 | -0.004509771 | 0.983927875 |
| ENSG00000198832 | *SELENOM* | 1.486126454 | -0.104397088 | 0.046008346 | -0.150405434 | 0.999933887 | 0.335377639 | 0.343922162 | -0.008544524 | 0.984056082 |
| ENSG00000139343 | *SNRPF* | 2.027144404 | -0.092329317 | -0.01698639 | -0.075342927 | 0.999933887 | 0.141105645 | 0.134171221 | 0.006934424 | 0.984167441 |
| ENSG00000122432 | *SPATA1* | 0.653977088 | -0.025905157 | -0.094772807 | 0.06886765 | 0.999933887 | -0.334612239 | -0.344473222 | 0.009860982 | 0.984167441 |
| ENSG00000138107 | *ACTR1A* | 5.748438069 | -0.007406848 | 0.028267564 | -0.035674413 | 0.999933887 | 0.032059847 | 0.035374096 | -0.003314249 | 0.984167441 |
| ENSG00000162437 | *RAVER2* | 1.435872256 | 0.005285309 | -0.008957687 | 0.014242996 | 0.999933887 | 0.129997032 | 0.118899538 | 0.011097494 | 0.984167441 |
| ENSG00000004779 | *NDUFAB1* | 1.862865572 | 0.004956496 | -0.154638594 | 0.15959509 | 0.999933887 | -0.001171181 | 0.005888694 | -0.007059875 | 0.984389942 |
| ENSG00000137074 | *APTX* | 2.967074585 | 0.023412066 | -0.023252075 | 0.046664141 | 0.999933887 | 0.096195631 | 0.101522284 | -0.005326654 | 0.984405899 |
| ENSG00000117425 | *PTCH2* | 0.127294248 | 0.165675274 | 0.060425746 | 0.105249528 | 0.999933887 | 0.485485532 | 0.496472648 | -0.010987116 | 0.98443403 |
| ENSG00000141349 | *G6PC3* | 1.103264447 | -0.014760309 | -0.000793267 | -0.013967042 | 0.999933887 | -0.13134697 | -0.138536027 | 0.007189056 | 0.98443403 |
| ENSG00000075089 | *ACTR6* | 1.786457792 | -0.06503258 | -0.131257358 | 0.066224778 | 0.999933887 | -0.037607507 | -0.043832137 | 0.006224631 | 0.984498268 |
| ENSG00000131791 | *PRKAB2* | 2.489953122 | -0.064426181 | -0.11347181 | 0.049045629 | 0.999933887 | -0.012614877 | -0.007180084 | -0.005434793 | 0.984498616 |
| ENSG00000181773 | *GPR3* | 0.565179719 | 0.160834836 | 0.129364337 | 0.031470499 | 0.999933887 | 1.875687324 | 1.896362838 | -0.020675515 | 0.984679316 |
| ENSG00000244723 | *ASLP1* | 0.155173684 | -0.056406535 | -0.010568112 | -0.045838422 | 0.999933887 | -0.12337742 | -0.135224948 | 0.011847528 | 0.984822951 |
| ENSG00000118961 | *LDAH* | 0.522562638 | -0.099055422 | -0.001845942 | -0.097209479 | 0.999933887 | 0.079909176 | 0.071717206 | 0.00819197 | 0.984829526 |
| ENSG00000135622 | *SEMA4F* | 0.913949883 | -0.009782006 | -0.00269689 | -0.007085116 | 0.999933887 | 0.14686138 | 0.139785124 | 0.007076256 | 0.985428308 |
| ENSG00000184047 | *DIABLO* | 2.089602559 | 0.115329109 | 0.162049293 | -0.046720184 | 0.999933887 | 0.319622592 | 0.31201732 | 0.007605273 | 0.985487863 |
| ENSG00000132382 | *MYBBP1A* | 3.419829389 | 0.04942476 | 0.137285282 | -0.087860523 | 0.999933887 | 0.232140538 | 0.235752117 | -0.003611579 | 0.985487863 |
| ENSG00000156374 | *PCGF6* | -0.147047848 | 0.14408165 | 0.148196274 | -0.004114624 | 0.999933887 | 0.027326739 | 0.038617695 | -0.011290956 | 0.985487863 |
| ENSG00000100462 | *PRMT5* | 1.685022996 | -0.048509342 | 0.006706972 | -0.055216314 | 0.999933887 | 0.120000322 | 0.1121804 | 0.007819922 | 0.985487863 |
| ENSG00000138382 | *METTL5* | 0.865261504 | 0.051418368 | -0.307328985 | 0.358747353 | 0.999933887 | 0.116111895 | 0.12368127 | -0.007569375 | 0.985487863 |
| ENSG00000197170 | *PSMD12* | 3.975985229 | -0.079723561 | -0.108944415 | 0.029220854 | 0.999933887 | -0.253274598 | -0.249211928 | -0.004062669 | 0.98553944 |
| ENSG00000158106 | *RHPN1* | 2.657242884 | 0.093450718 | 0.208384397 | -0.11493368 | 0.999933887 | 0.08884639 | 0.094413838 | -0.005567448 | 0.98553944 |
| ENSG00000118855 | *MFSD1* | 6.859063974 | 0.050824875 | 0.046198565 | 0.00462631 | 0.999933887 | -0.202614314 | -0.199040564 | -0.00357375 | 0.98553944 |
| ENSG00000129084 | *PSMA1* | 3.18068917 | -0.031900851 | -0.043400538 | 0.011499687 | 0.999933887 | 0.000180758 | 0.003958496 | -0.003777738 | 0.98553944 |
| ENSG00000180304 | *OAZ2* | 8.112090499 | -0.018082863 | -0.039579528 | 0.021496665 | 0.999933887 | 0.297656225 | 0.300716784 | -0.003060559 | 0.98553944 |
| ENSG00000146350 | *TBC1D32* | -0.073935957 | 0.034023265 | 0.183359699 | -0.149336435 | 0.999933887 | 0.142977611 | 0.1311797 | 0.011797911 | 0.98553944 |
| ENSG00000196656 | *AC004057.1* | -0.921266914 | 0.026808655 | -0.281186378 | 0.307995033 | 0.999933887 | 0.119000807 | 0.103805303 | 0.015195504 | 0.98553944 |
| ENSG00000246084 | *LINC02325* | -0.197754149 | -0.119658907 | -0.34170305 | 0.222044142 | 0.999933887 | -0.537597972 | -0.548508784 | 0.010910812 | 0.985569813 |
| ENSG00000116641 | *DOCK7* | 0.336884947 | 0.042399322 | -0.025591633 | 0.067990955 | 0.999933887 | -0.171979156 | -0.182163394 | 0.010184237 | 0.985569813 |
| ENSG00000005810 | *MYCBP2* | 5.994501639 | 0.022231998 | -0.055388517 | 0.077620515 | 0.999933887 | -0.422111576 | -0.416285221 | -0.005826355 | 0.985606763 |
| ENSG00000204161 | *TMEM273* | 6.520293358 | 0.107012461 | 0.145079714 | -0.038067253 | 0.999933887 | -0.665718713 | -0.661162548 | -0.004556165 | 0.985699124 |
| ENSG00000119396 | *RAB14* | 5.715953109 | -0.135179584 | -0.043165523 | -0.092014061 | 0.999933887 | -0.350000129 | -0.352562822 | 0.002562693 | 0.985810514 |
| ENSG00000273837 | *AC018755.4* | 2.599814504 | -0.069009378 | -0.093004667 | 0.023995288 | 0.999933887 | 0.617200848 | 0.622756642 | -0.005555794 | 0.985810514 |
| ENSG00000159063 | *ALG8* | 0.591939153 | -0.102440215 | -0.003982764 | -0.098457451 | 0.999933887 | 0.14733486 | 0.138139834 | 0.009195026 | 0.985810514 |
| ENSG00000085998 | *POMGNT1* | 2.203703368 | -0.011072813 | -0.031764062 | 0.020691249 | 0.999933887 | 0.150047581 | 0.154876788 | -0.004829206 | 0.985989211 |
| ENSG00000185761 | *ADAMTSL5* | -0.741865501 | -0.115436754 | -0.115821939 | 0.000385184 | 0.999933887 | 0.34888086 | 0.334582215 | 0.014298645 | 0.986009643 |
| ENSG00000183691 | *NOG* | -0.826344647 | 0.032797374 | -0.349243768 | 0.382041142 | 0.999933887 | -0.089455059 | -0.080668525 | -0.008786533 | 0.986009643 |
| ENSG00000144231 | *POLR2D* | 2.374272958 | -0.000266525 | 0.046687893 | -0.046954418 | 0.999933887 | -0.092132972 | -0.087281493 | -0.004851479 | 0.986009643 |
| ENSG00000052723 | *SIKE1* | 3.138934245 | 0.101773554 | -0.073382329 | 0.175155883 | 0.999933887 | -0.353753899 | -0.347699688 | -0.00605421 | 0.986152956 |
| ENSG00000103018 | *CYB5B* | 3.136432815 | -0.073445382 | 0.077002735 | -0.150448117 | 0.999933887 | 0.018918034 | 0.022918531 | -0.004000497 | 0.98657563 |
| ENSG00000166987 | *MBD6* | 7.892231969 | 0.067898664 | 0.087687743 | -0.019789079 | 0.999933887 | -0.100210886 | -0.102333007 | 0.002122122 | 0.986630673 |
| ENSG00000166441 | *RPL27A* | 7.832719384 | -0.01528838 | -0.052588146 | 0.037299765 | 0.999933887 | 0.093984844 | 0.096907282 | -0.002922438 | 0.986695656 |
| ENSG00000109943 | *CRTAM* | 2.60796581 | -0.091621174 | -0.143968438 | 0.052347264 | 0.999933887 | 0.020362052 | 0.015326051 | 0.005036002 | 0.986800084 |
| ENSG00000140009 | *ESR2* | 0.981304632 | 0.011587995 | 0.013068093 | -0.001480097 | 0.999933887 | -0.335139347 | -0.323589384 | -0.011549963 | 0.986800084 |
| ENSG00000163462 | *TRIM46* | 0.446841017 | 0.108027193 | 0.250263723 | -0.14223653 | 0.999933887 | 0.018921449 | 0.027371091 | -0.008449642 | 0.987005854 |
| ENSG00000069966 | *GNB5* | 2.240290908 | -0.010688237 | 0.073077633 | -0.08376587 | 0.999933887 | -0.022842578 | -0.018772268 | -0.00407031 | 0.987005854 |
| ENSG00000105289 | *TJP3* | 0.397220905 | 0.176171609 | 0.016307708 | 0.159863901 | 0.999933887 | 0.203465318 | 0.195123898 | 0.008341419 | 0.987094645 |
| ENSG00000131089 | *ARHGEF9* | 1.722969207 | -0.095684888 | -0.063603722 | -0.032081166 | 0.999933887 | 0.041065997 | 0.035867899 | 0.005198097 | 0.987204521 |
| ENSG00000229808 | *AL391825.1* | 0.219017776 | -0.123986669 | 0.003912874 | -0.127899543 | 0.999933887 | 0.732901211 | 0.719658965 | 0.013242245 | 0.987204521 |
| ENSG00000260052 | *AC023813.3* | -0.400209157 | -0.093259347 | -0.195873049 | 0.102613702 | 0.999933887 | 1.824429167 | 1.815832359 | 0.008596808 | 0.987271885 |
| ENSG00000069020 | *MAST4* | 2.856959251 | -0.05390486 | -0.068623034 | 0.014718174 | 0.999933887 | 0.148613504 | 0.144637751 | 0.003975754 | 0.987274323 |
| ENSG00000174600 | *CMKLR1* | -0.07115812 | 0.005261459 | 0.108462515 | -0.103201056 | 0.999933887 | -0.157795867 | -0.148556826 | -0.009239041 | 0.98730564 |
| ENSG00000128739 | *SNRPN* | 0.144536106 | -0.121920493 | 0.432916973 | -0.554837467 | 0.999291887 | 0.075191399 | 0.066322823 | 0.008868576 | 0.98733875 |
| ENSG00000277767 | *AL442128.2* | -0.291421341 | 0.05756166 | -0.089673202 | 0.147234862 | 0.999933887 | 0.868859889 | 0.879525032 | -0.010665144 | 0.98733875 |
| ENSG00000101928 | *MOSPD1* | 2.409988413 | -0.113206661 | -0.115947095 | 0.002740434 | 0.999933887 | -1.404585163 | -1.409886069 | 0.005300906 | 0.987342065 |
| ENSG00000173200 | *PARP15* | 3.705163201 | -0.100293028 | -0.071189565 | -0.029103463 | 0.999933887 | -0.047658153 | -0.043803079 | -0.003855074 | 0.987695001 |
| ENSG00000001631 | *KRIT1* | 0.673605476 | -0.107197866 | -0.157091426 | 0.04989356 | 0.999933887 | -0.116951179 | -0.125286774 | 0.008335596 | 0.987730046 |
| ENSG00000182362 | *YBEY* | 0.644492503 | 0.004304767 | -0.007720266 | 0.012025033 | 0.999933887 | 0.089128482 | 0.095791899 | -0.006663417 | 0.987730046 |
| ENSG00000131507 | *NDFIP1* | 6.131594555 | -0.050744626 | -0.031840475 | -0.01890415 | 0.999933887 | -0.352493721 | -0.35486751 | 0.002373789 | 0.988214559 |
| ENSG00000230257 | *NFE4* | 3.3849238 | -0.070593939 | -0.093705446 | 0.023111507 | 0.999933887 | -0.456155616 | -0.459658768 | 0.003503152 | 0.988214559 |
| ENSG00000075826 | *SEC31B* | 3.225714385 | -0.003379025 | 0.044395527 | -0.047774552 | 0.999933887 | 0.123527751 | 0.127487218 | -0.003959467 | 0.988214559 |
| ENSG00000156535 | *CD109* | 2.143430531 | -0.244853102 | -0.03335987 | -0.211493232 | 0.999933887 | -0.702356446 | -0.697919092 | -0.004437354 | 0.988229692 |
| ENSG00000184497 | *TMEM255B* | 1.8765399 | 0.226002848 | 0.121249051 | 0.104753797 | 0.999933887 | 0.632822557 | 0.626900935 | 0.005921622 | 0.988229692 |
| ENSG00000092929 | *UNC13D* | 8.04873051 | 0.035965018 | 0.038998729 | -0.003033711 | 0.999933887 | 0.203103762 | 0.205260818 | -0.002157056 | 0.988229692 |
| ENSG00000266236 | *NARF-IT1* | 0.566632177 | 0.07475436 | 0.042537911 | 0.032216449 | 0.999933887 | -0.373921972 | -0.361714033 | -0.012207939 | 0.988229692 |
| ENSG00000240682 | *ISY1* | 4.113838603 | -0.028371219 | -0.001511322 | -0.026859896 | 0.999933887 | -0.339523584 | -0.336110568 | -0.003413016 | 0.988229692 |
| ENSG00000162757 | *C1orf74* | 1.316516333 | -0.033756504 | -0.079262263 | 0.045505759 | 0.999933887 | -0.191736193 | -0.184024131 | -0.007712062 | 0.988229692 |
| ENSG00000160194 | *NDUFV3* | 2.349913292 | 0.018683555 | -0.064193748 | 0.082877303 | 0.999933887 | -0.136597188 | -0.132769626 | -0.003827562 | 0.988229692 |
| ENSG00000140497 | *SCAMP2* | 6.071556535 | -0.001109473 | -0.00472866 | 0.003619186 | 0.999933887 | -0.233343106 | -0.23546219 | 0.002119084 | 0.988229692 |
| ENSG00000157326 | *DHRS4* | -0.356694864 | 0.145688731 | -0.006201821 | 0.151890552 | 0.999933887 | 0.186189406 | 0.195423449 | -0.009234043 | 0.988456628 |
| ENSG00000108219 | *TSPAN14* | 7.433120996 | -0.031822796 | 0.003172575 | -0.034995371 | 0.999933887 | -0.255545876 | -0.257969947 | 0.002424071 | 0.988536525 |
| ENSG00000237272 | *OR51R1P* | 0.397577343 | 0.071263281 | 0.058357271 | 0.01290601 | 0.999933887 | 0.143423441 | 0.134555334 | 0.008868107 | 0.988536525 |
| ENSG00000121653 | *MAPK8IP1* | 0.468242583 | 0.01424465 | 0.277951174 | -0.263706525 | 0.999933887 | 0.363436701 | 0.355024297 | 0.008412403 | 0.988536525 |
| ENSG00000143727 | *ACP1* | 3.390330962 | 0.008122934 | -0.063897596 | 0.07202053 | 0.999933887 | 0.095785733 | 0.092615015 | 0.003170718 | 0.988536525 |
| ENSG00000205336 | *ADGRG1* | 5.099831291 | 0.045359991 | 0.082233128 | -0.036873137 | 0.999933887 | 0.046543506 | 0.043435884 | 0.003107622 | 0.988585524 |
| ENSG00000146409 | *SLC18B1* | 1.028503267 | -0.064074066 | 0.095193186 | -0.159267252 | 0.999933887 | 0.014014176 | 0.008109793 | 0.005904383 | 0.988585784 |
| ENSG00000185619 | *PCGF3* | 5.323256611 | 0.102081448 | 0.027430139 | 0.074651309 | 0.999933887 | -0.185855775 | -0.188646432 | 0.002790657 | 0.988660421 |
| ENSG00000231607 | *DLEU2* | 3.577577023 | 0.06356319 | 0.011349818 | 0.052213372 | 0.999933887 | 0.016115303 | 0.02000021 | -0.003884907 | 0.988660421 |
| ENSG00000162384 | *CZIB* | 2.444433418 | 0.040565279 | 0.036573144 | 0.003992135 | 0.999933887 | 0.015679734 | 0.019690154 | -0.00401042 | 0.988660421 |
| ENSG00000183347 | *GBP6* | -0.70255142 | -0.060609212 | -0.168048944 | 0.107439733 | 0.999933887 | -0.116442259 | -0.130356837 | 0.013914578 | 0.988660421 |
| ENSG00000253981 | *ALG1L13P* | 2.77903571 | 0.020775053 | 0.037824325 | -0.017049271 | 0.999933887 | -0.228605044 | -0.224749071 | -0.003855973 | 0.988660421 |
| ENSG00000159873 | *CCDC117* | 3.220307498 | -0.025426649 | -0.051736831 | 0.026310181 | 0.999933887 | 0.169032002 | 0.166006747 | 0.003025255 | 0.988660421 |
| ENSG00000164626 | *KCNK5* | 2.576729502 | -0.046764592 | -0.030798178 | -0.015966414 | 0.999933887 | 2.566533693 | 2.57296511 | -0.006431418 | 0.988660421 |
| ENSG00000111726 | *CMAS* | 1.350463556 | -0.004623152 | 0.071790009 | -0.076413161 | 0.999933887 | -0.038449238 | -0.03413914 | -0.004310098 | 0.988660421 |
| ENSG00000235961 | *PNMA6A* | -1.44974951 | -0.008321286 | -0.051233029 | 0.042911744 | 0.999933887 | -1.286476143 | -1.298411946 | 0.011935803 | 0.988660421 |
| ENSG00000197608 | *ZNF841* | 2.926653109 | -0.056741152 | -0.072522957 | 0.015781805 | 0.999933887 | -0.096838874 | -0.093142812 | -0.003696062 | 0.98870157 |
| ENSG00000088881 | *EBF4* | -1.166325437 | -0.115897119 | 0.262368585 | -0.378265704 | 0.999933887 | -0.176678007 | -0.164499913 | -0.012178094 | 0.98870157 |
| ENSG00000213058 | *AL365357.1* | -3.894285282 | 0.043459933 | -0.15492762 | 0.198387553 | 0.999933887 | 0.073006621 | 0.083085603 | -0.010078982 | 0.98870157 |
| ENSG00000197728 | *RPS26* | 6.789885669 | 0.006261789 | 0.036712648 | -0.030450859 | 0.999933887 | 0.047922694 | 0.045011328 | 0.002911366 | 0.98877087 |
| ENSG00000137409 | *MTCH1* | 7.768495369 | -0.002169852 | 0.049123103 | -0.051292955 | 0.999933887 | 0.209580898 | 0.211684798 | -0.002103901 | 0.98877087 |
| ENSG00000023697 | *DERA* | 1.415084878 | -0.009613947 | -0.046627972 | 0.037014025 | 0.999933887 | -0.165700279 | -0.172641241 | 0.006940962 | 0.988785361 |

| | | | | | | | | | | |
|---|---|---|---|---|---|---|---|---|---|---|
| ENSG00000258741 | *H2AZ2P1* | -0.630654618 | -0.242833831 | -0.064608666 | -0.178225165 | 0.999933887 | 0.56409488 | 0.555485466 | 0.008609415 | 0.988887326 |
| ENSG00000105173 | *CCNE1* | -0.561577834 | 0.150494777 | 0.147190538 | 0.003304239 | 0.999933887 | 0.246385839 | 0.256621124 | -0.010235284 | 0.988902054 |
| ENSG00000168062 | *BATF2* | 0.981767264 | 0.072793521 | 0.07825817 | -0.005464649 | 0.999933887 | -0.334709852 | -0.343591022 | 0.008881169 | 0.989061967 |
| ENSG00000128731 | *HERC2* | 4.085534498 | 0.032850316 | -0.025512608 | 0.058362924 | 0.999933887 | 0.001666202 | 0.005059939 | -0.003393737 | 0.989061967 |
| ENSG00000163468 | *CCT3* | 3.942871416 | -0.017823598 | -0.060982729 | 0.043159131 | 0.999933887 | 0.140498652 | 0.143250302 | -0.00275165 | 0.989061967 |
| ENSG00000163644 | *PPM1K* | 3.274429651 | -0.021110907 | -0.160500013 | 0.139389106 | 0.999933887 | -0.020797702 | -0.017463718 | -0.003333984 | 0.989061967 |
| ENSG00000279608 | *AL353795.3* | 0.273185021 | -0.042820752 | -0.248268848 | 0.205448096 | 0.999933887 | -0.102075523 | -0.111232559 | 0.009157036 | 0.989061967 |
| ENSG00000151552 | *QDPR* | 0.625426799 | 0.090448806 | 0.161384186 | -0.07093538 | 0.999933887 | 0.183204414 | 0.177114268 | 0.006090146 | 0.989084636 |
| ENSG00000179242 | *CDH4* | -0.167714623 | -0.105187725 | -0.016216009 | -0.088971716 | 0.999933887 | -0.296549445 | -0.287488585 | -0.00906086 | 0.989084636 |
| ENSG00000245954 | *LINC02273* | 0.013305548 | 0.040953485 | 0.08704058 | -0.046087095 | 0.999933887 | 0.014231098 | 0.021700171 | -0.007469073 | 0.989084636 |
| ENSG00000158856 | *DMTN* | 1.553682895 | -0.033159383 | 0.032663097 | -0.065822479 | 0.999933887 | 0.019928217 | 0.025925633 | -0.005997416 | 0.989101757 |
| ENSG00000188002 | *AC026412.1* | 2.106341764 | 0.116037784 | 0.053837704 | 0.062200079 | 0.999933887 | 0.294493608 | 0.299135167 | -0.004641559 | 0.989605168 |
| ENSG00000203666 | *EFCAB2* | 0.982328679 | -0.099886925 | -0.010138039 | -0.089748886 | 0.999933887 | -0.055170914 | -0.061099706 | 0.005928792 | 0.989767273 |
| ENSG00000110801 | *PSMD9* | 1.583718142 | 0.1591494 | 0.014011339 | 0.145138061 | 0.999933887 | -0.194216164 | -0.199921508 | 0.005705345 | 0.989932748 |
| ENSG00000078070 | *MCCC1* | 1.498602562 | -0.090501095 | -0.114872958 | 0.024371863 | 0.999933887 | 0.107878731 | 0.111776546 | -0.003897815 | 0.990010677 |
| ENSG00000256591 | *AP003108.2* | 1.074806714 | 0.128678292 | -0.046106966 | 0.174785258 | 0.999933887 | -0.007775672 | -0.001091435 | -0.006684237 | 0.990034278 |
| ENSG00000145390 | *USP53* | 2.156227451 | -0.0605892 | -0.256771987 | 0.196182787 | 0.999933887 | -0.385071288 | -0.379435995 | -0.005635293 | 0.990034278 |
| ENSG00000160282 | *FTCD* | -1.421021437 | 0.110469489 | 0.471662315 | -0.361192826 | 0.999933887 | 1.012956131 | 1.022200563 | -0.009244432 | 0.990034278 |
| ENSG00000243364 | *EFNA4* | 0.490608696 | -0.006554546 | 0.073426135 | -0.079980682 | 0.999933887 | -0.357335593 | -0.351258816 | -0.006076777 | 0.990034278 |
| ENSG00000142657 | *PGD* | 7.785338216 | -0.023781664 | -0.013398327 | -0.010383337 | 0.999933887 | -0.002395316 | 0.000110771 | -0.002506087 | 0.990455386 |
| ENSG00000206341 | *HLA-H* | 8.040472328 | 0.072737454 | 0.025423444 | 0.047314009 | 0.999933887 | 0.326041874 | 0.323678536 | 0.002363337 | 0.990486293 |
| ENSG00000154124 | *OTULIN* | 5.044299543 | -0.06933736 | 0.017808459 | -0.087145819 | 0.999933887 | -0.168459509 | -0.170709315 | 0.002249806 | 0.990491989 |
| ENSG00000149929 | *HIRIP3* | 0.959466407 | 0.094190131 | 0.310723849 | -0.216533718 | 0.999933887 | 0.174038449 | 0.180438653 | -0.006400204 | 0.990491989 |
| ENSG00000105085 | *MED26* | 3.103717491 | 0.022992149 | 0.086567528 | -0.063575379 | 0.999933887 | 0.189310286 | 0.186354454 | 0.002955832 | 0.990491989 |
| ENSG00000166343 | *MSS51* | 0.497361553 | -0.130035369 | 0.139819802 | -0.269855171 | 0.999933887 | -0.076971758 | -0.071555244 | -0.005416515 | 0.99057037 |
| ENSG00000072401 | *UBE2D1* | 5.734016277 | 0.02208349 | 0.029533558 | -0.007450068 | 0.999933887 | -0.188719364 | -0.18656095 | -0.002158414 | 0.99057037 |
| ENSG00000237483 | *RPS27AP8* | -0.448230464 | -0.162169439 | 0.353604438 | -0.515773877 | 0.999933887 | 1.8227468 | 1.831801833 | -0.009055034 | 0.990641715 |
| ENSG00000110455 | *ACCS* | 1.622592841 | 0.106990332 | -0.124651298 | 0.23164163 | 0.999933887 | -0.009802094 | -0.013871772 | 0.004069678 | 0.990655949 |
| ENSG00000159753 | *CARMIL2* | 4.069161452 | 0.06786344 | 0.159361947 | -0.091498507 | 0.999933887 | 0.119646328 | 0.121964283 | -0.002317955 | 0.990655949 |
| ENSG00000087448 | *KLHL42* | 1.411105682 | -0.063422721 | -0.163636097 | 0.100213376 | 0.999933887 | -0.100071644 | -0.096141147 | -0.003930497 | 0.990655949 |
| ENSG00000111615 | *KRR1* | 3.419921376 | -0.029854264 | -0.058930987 | 0.029076723 | 0.999933887 | -0.186340475 | -0.180391827 | -0.005948648 | 0.990772119 |
| ENSG00000169564 | *PCBP1* | 6.336777308 | 0.075637291 | 0.010963778 | 0.064673513 | 0.999933887 | -0.081077358 | -0.07820395 | -0.002873407 | 0.990779249 |
| ENSG00000275834 | *AC084368.1* | -0.660787536 | 0.373993177 | 0.331414928 | 0.042578248 | 0.999933887 | 0.793943129 | 0.782539738 | 0.011403391 | 0.990860063 |
| ENSG00000239503 | *MARK2P8* | 3.839013444 | 0.223255004 | 0.196082789 | 0.027172215 | 0.999933887 | 0.853338095 | 0.848461033 | 0.004877062 | 0.990967569 |
| ENSG00000101442 | *ACTR5* | 2.10182889 | -0.018598201 | 0.034614564 | -0.053212765 | 0.999933887 | 0.01640593 | 0.013030728 | 0.003375202 | 0.991140312 |
| ENSG00000260923 | *LINC02193* | -1.210672172 | 0.051655441 | 0.084002287 | -0.032346846 | 0.999933887 | 0.415224781 | 0.405672033 | 0.009552748 | 0.991187827 |
| ENSG00000154511 | *DIPK1A* | 0.588011034 | -0.039817898 | -0.188360881 | 0.148542983 | 0.999933887 | 0.118577907 | 0.124404647 | -0.005826741 | 0.991277682 |
| ENSG00000138071 | *ACTR2* | 8.507673792 | -0.093000474 | -0.054263761 | -0.038736713 | 0.999933887 | -0.329788889 | -0.331732061 | 0.001943172 | 0.99165002 |
| ENSG00000115138 | *POMC* | 0.006542579 | 0.236029689 | 0.215396272 | 0.020633417 | 0.999933887 | 0.035125387 | 0.029225311 | 0.005900077 | 0.99165002 |
| ENSG00000142459 | *EVI5L* | 2.079274018 | 0.123152143 | 0.196199183 | -0.07304704 | 0.999933887 | 0.18405154 | 0.179899115 | 0.004152425 | 0.99165002 |
| ENSG00000010704 | *HFE* | 0.489287046 | 0.149799963 | -0.040602362 | 0.190402324 | 0.999933887 | -0.243853875 | -0.250676791 | 0.006822916 | 0.99165002 |
| ENSG00000173064 | *HECTD4* | 5.128382196 | 0.045046095 | 0.086735719 | -0.041689624 | 0.999933887 | -0.103149342 | -0.105001853 | 0.001852511 | 0.99165002 |
| ENSG00000013725 | *CD6* | 6.370701314 | 0.038110812 | 0.022001551 | 0.016109261 | 0.999933887 | 0.150801397 | 0.152715957 | -0.00191456 | 0.99165002 |
| ENSG00000161513 | *FDXR* | 0.61938118 | -0.097437536 | 0.113633303 | -0.211070839 | 0.999933887 | 0.100764656 | 0.105798794 | -0.005034138 | 0.99165002 |
| ENSG00000170919 | *TPT1-AS1* | 3.485863975 | -0.039785193 | 0.03566131 | -0.075446503 | 0.999933887 | -0.090549075 | -0.087981273 | -0.002567802 | 0.99165002 |
| ENSG00000272667 | *AC012306.2* | 0.385206435 | 0.094701934 | -0.038301898 | 0.133003832 | 0.999933887 | -0.096895268 | -0.091525639 | -0.005369629 | 0.99165002 |
| ENSG00000277072 | *STAG3L2* | 1.690617175 | -0.035585279 | -0.060905563 | 0.025320284 | 0.999933887 | 0.098074779 | 0.101561838 | -0.003487059 | 0.99165002 |
| ENSG00000230923 | *LINC00309* | -0.809210948 | 0.068533644 | 0.359335928 | -0.290802284 | 0.999933887 | 1.45090477 | 1.440007396 | 0.010897374 | 0.99165002 |
| ENSG00000183837 | *PNMA3* | -0.259568258 | -0.044315121 | 0.071561537 | -0.115876657 | 0.999933887 | -0.045856029 | -0.039747083 | -0.006108946 | 0.99165002 |
| ENSG00000174944 | *P2RY14* | 3.571363339 | 0.01894501 | -0.145905984 | 0.164850995 | 0.999933887 | -0.266341959 | -0.269624875 | 0.003282916 | 0.99165002 |
| ENSG00000287040 | *AL590560.5* | -0.999503789 | 0.027841216 | 0.040595677 | -0.012754462 | 0.999933887 | -0.120814445 | -0.110842676 | -0.009971768 | 0.99165002 |
| ENSG00000250966 | *H3P14* | -0.024659672 | -0.033684208 | 0.21035707 | -0.244041278 | 0.999933887 | -0.185435061 | -0.192490083 | 0.007055022 | 0.99165002 |
| ENSG00000173852 | *DPY19L1* | 1.801238476 | 0.020442095 | -0.055220746 | 0.075662842 | 0.999933887 | -0.128226152 | -0.124089854 | -0.004136297 | 0.99165002 |
| ENSG00000123810 | *B9D2* | 3.197618624 | -0.012992082 | 0.001314323 | -0.014306405 | 0.999933887 | -0.010841865 | -0.007561609 | -0.003280256 | 0.99165002 |
| ENSG00000150433 | *TMEM218* | 2.212024989 | 0.001525242 | 0.008199304 | -0.006674062 | 0.999933887 | -0.10194275 | -0.104528718 | 0.002585968 | 0.99165002 |
| ENSG00000065978 | *YBX1* | 7.899446576 | 0.037641073 | -0.001231713 | 0.038872786 | 0.999933887 | -0.03543685 | -0.033881512 | -0.001555338 | 0.991684703 |
| ENSG00000282780 | *TRBJ1-6* | 0.490085644 | -0.113369813 | -0.007080948 | -0.106288865 | 0.999933887 | 0.136985801 | 0.132330532 | 0.004655269 | 0.992043528 |
| ENSG00000124216 | *SNAI1* | -0.106426959 | 0.162846715 | 0.378678036 | -0.215831321 | 0.999933887 | -0.357320924 | -0.366136844 | 0.00881592 | 0.992106195 |
| ENSG00000069424 | *KCNAB2* | 9.250966004 | 0.081695149 | 0.063390904 | 0.018304245 | 0.999933887 | -0.439374366 | -0.441610487 | 0.00223612 | 0.992165145 |
| ENSG00000265996 | *MIR3671* | 0.529288132 | -0.082510397 | -0.060618648 | -0.021891749 | 0.999933887 | 0.180765565 | 0.18438618 | -0.003620615 | 0.992282432 |
| ENSG00000136143 | *SUCLA2* | 1.033467907 | -0.185650987 | -0.20677183 | 0.021120844 | 0.999933887 | -0.274576888 | -0.27906708 | 0.004490192 | 0.992412021 |
| ENSG00000070756 | *PABPC1* | 9.515610719 | -0.019573332 | -0.048105607 | 0.028532275 | 0.999933887 | -0.002898013 | -0.001703246 | -0.001194766 | 0.992412021 |
| ENSG00000105619 | *TFPT* | 1.747094599 | -0.052064122 | 0.026998322 | -0.079062444 | 0.999933887 | -0.176850502 | -0.173651342 | -0.00319916 | 0.992412021 |
| ENSG00000117984 | *CTSD* | 9.130266227 | -0.031028276 | -0.033813352 | 0.002785076 | 0.999933887 | -0.388407067 | -0.386477783 | -0.001929284 | 0.992412021 |
| ENSG00000179978 | *NAIPP2* | 0.982267943 | -0.016548694 | 0.030193087 | -0.046741781 | 0.999933887 | -0.676305035 | -0.680698694 | 0.00439366 | 0.992412021 |
| ENSG00000179967 | *PPP1R14BP3* | 1.57684555 | 0.170637283 | 0.032095669 | 0.138541614 | 0.999933887 | 0.644130172 | 0.647644582 | -0.00351441 | 0.992417691 |
| ENSG00000264573 | *RN7SL15P* | 9.9155e-05 | 0.296514333 | 0.131091906 | 0.165422428 | 0.999933887 | 0.726135308 | 0.719809801 | 0.006325507 | 0.992540742 |
| ENSG00000131508 | *UBE2D2* | 6.217045486 | 0.03031948 | 0.035364547 | -0.005045067 | 0.999933887 | -0.065748949 | -0.064283371 | -0.001465577 | 0.992561217 |
| ENSG00000086189 | *DIMT1* | 1.651040394 | 0.006808299 | -0.005859962 | 0.012668262 | 0.999933887 | 0.09698774 | 0.094372364 | 0.002615376 | 0.992644653 |
| ENSG00000107290 | *SETX* | 7.435668633 | -0.093703778 | -0.155323831 | 0.061620052 | 0.999933887 | -0.908365071 | -0.905542329 | -0.002822741 | 0.992754417 |
| ENSG00000143093 | *STRIP1* | 5.403916028 | -0.011535178 | -0.016300996 | 0.004765818 | 0.999933887 | -0.146000032 | -0.147562684 | 0.001562652 | 0.992823012 |
| ENSG00000118369 | *USP35* | 3.053280002 | 0.027448534 | 0.212509059 | -0.185060526 | 0.999933887 | -0.253315044 | -0.256008117 | 0.002693073 | 0.99284556 |
| ENSG00000059573 | *ALDH18A1* | 2.034883419 | -0.049258891 | -0.141118447 | 0.091859556 | 0.999933887 | 0.241634471 | 0.238642114 | 0.002992357 | 0.992945634 |
| ENSG00000137101 | *CD72* | 0.142519762 | -0.07873075 | 0.288435036 | -0.367165786 | 0.999933887 | 0.038542124 | 0.034054555 | 0.004487569 | 0.992945634 |
| ENSG00000204172 | *AGAP9* | -0.573636762 | 0.200959503 | 0.063964192 | 0.13699531 | 0.999933887 | 0.139998266 | 0.145545663 | -0.005547397 | 0.993078728 |
| ENSG00000168118 | *RAB4A* | 2.061632289 | 0.031537776 | 0.1593922 | -0.127854424 | 0.999933887 | 0.042750956 | 0.045054989 | -0.002304033 | 0.993078728 |
| ENSG00000111481 | *COPZ1* | 4.602477934 | 0.067427135 | -0.008781138 | 0.076208273 | 0.999933887 | 0.163536023 | 0.165220447 | -0.001684424 | 0.993462138 |
| ENSG00000181588 | *MEX3D* | 0.531308471 | 0.110975973 | -0.099292995 | 0.210268968 | 0.999933887 | -0.148792048 | -0.1437366 | -0.005055448 | 0.993462138 |
| ENSG00000252759 | *Y_RNA* | 0.547151025 | 0.049698711 | 0.14581552 | -0.096116809 | 0.999933887 | 0.264488642 | 0.270324058 | -0.005835416 | 0.993504958 |
| ENSG00000168646 | *AXIN2* | 0.573146948 | 0.067502261 | 0.066601311 | 0.00090095 | 0.999933887 | 0.199277566 | 0.194794121 | 0.004483445 | 0.993801363 |
| ENSG00000103335 | *PIEZO1* | 5.534367273 | 0.074559934 | 0.113085939 | -0.038526005 | 0.999933887 | 0.073490877 | 0.072109311 | 0.001381565 | 0.993811903 |
| ENSG00000102967 | *DHODH* | 1.023986107 | 0.123858733 | -0.029198952 | 0.153057685 | 0.999933887 | 0.075246926 | 0.078915119 | -0.003668193 | 0.993811903 |
| ENSG00000253476 | *AC091185.1* | -0.537668741 | -0.027038848 | -0.248604664 | 0.221565816 | 0.999933887 | -0.585485489 | -0.578445197 | -0.007040292 | 0.993811903 |
| ENSG00000134375 | *TIMM17A* | 3.192322157 | -0.10816558 | -0.088066892 | -0.020098688 | 0.999933887 | -0.097806258 | -0.095984628 | -0.001821629 | 0.993899226 |
| ENSG00000108591 | *DRG2* | 2.997169165 | -0.061085088 | 0.155035854 | -0.216120943 | 0.999933887 | 0.073259378 | 0.071310213 | 0.001949165 | 0.993899226 |
| ENSG00000235253 | *AC010240.2* | 0.044259559 | 0.123413794 | 0.402615188 | -0.279201393 | 0.999933887 | 0.288808772 | 0.295490961 | -0.006682189 | 0.993899226 |
| ENSG00000175894 | *TSPEAR* | -1.622318273 | 0.116681896 | 0.032176061 | 0.084505835 | 0.999933887 | 0.345210082 | 0.336926382 | 0.008283699 | 0.993899226 |
| ENSG00000183155 | *RABIF* | 3.714581372 | 0.044070256 | -0.000268794 | 0.04433905 | 0.999933887 | -0.46261552 | -0.464402338 | 0.001786818 | 0.993899226 |
| ENSG00000142920 | *AZIN2* | -0.248020177 | 0.082506354 | -0.285613084 | 0.368119438 | 0.999933887 | 0.074123364 | 0.079231309 | -0.005107945 | 0.993899226 |
| ENSG00000106780 | *MEGF9* | 6.169170447 | 0.018522575 | -0.00082049 | 0.019343065 | 0.999933887 | -0.175338553 | -0.173632023 | -0.00170653 | 0.993899226 |
| ENSG00000139044 | *B4GALNT3* | -0.026937994 | -0.016414835 | 0.097188398 | -0.113603233 | 0.999933887 | -0.022550979 | -0.016975636 | -0.005575342 | 0.993899226 |
| ENSG00000280287 | *AC131212.3* | 3.45146523 | 0.063698213 | -0.055873293 | 0.119571506 | 0.999933887 | -0.1429248 | -0.140679793 | -0.002245008 | 0.994175957 |
| ENSG00000274021 | *AC024909.1* | 1.126474235 | 0.154290886 | 0.120648668 | 0.033642218 | 0.999933887 | 0.106362254 | 0.110135873 | -0.003773619 | 0.994272012 |
| ENSG00000147650 | *LRP12* | 0.155610161 | -0.193868574 | -0.22124348 | 0.027374906 | 0.999933887 | 0.136178618 | 0.132400774 | 0.003777845 | 0.994272012 |
| ENSG00000147471 | *PLPBP* | 4.495448461 | 0.048880315 | -0.057303542 | 0.106183856 | 0.999933887 | -0.520793222 | -0.519220561 | -0.00157266 | 0.994272012 |
| ENSG00000211751 | *TRBC1* | 5.583413329 | -0.035885614 | -0.060215064 | 0.02432945 | 0.999933887 | 0.17237036 | 0.171071173 | 0.001299187 | 0.994272012 |
| ENSG00000099308 | *MAST3* | 8.207218268 | -0.042764075 | 0.035934587 | -0.078698661 | 0.999933887 | -0.951364275 | -0.949506302 | -0.001857972 | 0.994272012 |
| ENSG00000133027 | *PEMT* | -0.48740443 | 0.105948438 | -0.04606697 | 0.152015408 | 0.999933887 | -0.105035228 | -0.09981804 | -0.005217188 | 0.994272012 |
| ENSG00000082146 | *STRADB* | 0.594165746 | -0.053593208 | -0.090393606 | 0.036800398 | 0.999933887 | -0.003928425 | -0.000178696 | -0.003749729 | 0.994272012 |
| ENSG00000198482 | *ZNF808* | 2.114602578 | -0.026639263 | -0.037212998 | 0.010573736 | 0.999933887 | -0.471893568 | -0.469195878 | -0.00269769 | 0.994272012 |
| ENSG00000144354 | *CDCA7* | 0.871971765 | -0.011825403 | 0.000623189 | -0.012448592 | 0.999933887 | 0.078372562 | 0.075658232 | 0.00271433 | 0.994272012 |
| ENSG00000087263 | *OGFOD1* | 1.797930676 | -0.135753637 | -0.082202404 | -0.053551233 | 0.999933887 | 0.091144429 | 0.089030408 | 0.002114021 | 0.994292069 |
| ENSG00000135631 | *RAB11FIP5* | 2.060583434 | 0.030970748 | 0.04938257 | -0.018411823 | 0.999933887 | 0.086990969 | 0.089800031 | -0.002809062 | 0.994298463 |
| ENSG00000186017 | *ZNF566* | -0.189840508 | -0.065926368 | -0.04841779 | -0.017508578 | 0.999933887 | -0.069584486 | -0.074246734 | 0.004662248 | 0.994298463 |
| ENSG00000285796 | *AL162458.1* | 2.243139294 | -0.021122513 | -0.011577145 | -0.009545368 | 0.999933887 | -0.067638743 | -0.065555833 | -0.00208291 | 0.994441024 |

| | | | | | | | | | | |
|---|---|---|---|---|---|---|---|---|---|---|
| ENSG00000104133 | *SPG11* | 6.400511008 | 0.016605064 | -0.036474777 | 0.053079841 | 0.999933887 | -0.340144586 | -0.341684127 | 0.00153954 | 0.994467381 |
| ENSG00000103485 | *QPRT* | -0.991854732 | 0.231264703 | 0.259769226 | -0.028504523 | 0.999933887 | 0.050254945 | 0.055136021 | -0.004881076 | 0.994801436 |
| ENSG00000198821 | *CD247* | 5.146101256 | -0.00563968 | -0.010315613 | 0.004675933 | 0.999933887 | 0.130234771 | 0.131459161 | -0.00122439 | 0.994801436 |
| ENSG00000144468 | *RHBDD1* | 2.12028446 | 0.050595005 | 0.028800915 | 0.02179409 | 0.999933887 | -0.029519796 | -0.027472407 | -0.002047389 | 0.994822633 |
| ENSG00000090581 | *GNPTG* | 6.332046848 | 0.032411432 | -0.04353406 | 0.075945492 | 0.999933887 | 0.301635081 | 0.300568202 | 0.001066879 | 0.995055185 |
| ENSG00000116580 | *GON4L* | 5.582094638 | -0.037173622 | -0.063473244 | 0.026299622 | 0.999933887 | -0.10657156 | -0.107787546 | 0.001215986 | 0.99506074 |
| ENSG00000134444 | *RELCH* | 5.04780747 | 0.011899899 | -0.031361534 | 0.043261434 | 0.999933887 | -0.412858449 | -0.410760657 | -0.002097792 | 0.99506074 |
| ENSG00000170631 | *ZNF16* | -0.056964164 | -0.123148839 | -0.022087309 | -0.10106153 | 0.999933887 | -0.060150016 | -0.064490486 | 0.00434047 | 0.99518552 |
| ENSG00000164379 | *FOXQ1* | -2.09385645 | 0.139052265 | 0.257303922 | -0.118251657 | 0.999933887 | -0.592423466 | -0.585449191 | -0.006974276 | 0.995253198 |
| ENSG00000248019 | *FAM13A-AS1* | 4.421380427 | 0.052449016 | -0.029463828 | 0.081912844 | 0.999933887 | -0.551416096 | -0.549329982 | -0.002086114 | 0.995253198 |
| ENSG00000206052 | *DOK6* | -0.338126937 | 0.097177448 | 0.091028675 | 0.006148773 | 0.999933887 | -0.046094896 | -0.051098251 | 0.005003355 | 0.995253198 |
| ENSG00000180879 | *SSR4* | 4.615981361 | -0.013685755 | -0.081232762 | 0.067547007 | 0.999933887 | 0.079257504 | 0.080361519 | -0.001104015 | 0.995253198 |
| ENSG00000126453 | *BCL2L12* | 1.892967114 | 0.008392647 | 0.117190882 | -0.108798235 | 0.999933887 | 0.05065545 | 0.053388705 | -0.002733255 | 0.995253198 |
| ENSG00000174327 | *SLC16A13* | 0.365901075 | -0.268058212 | 0.102228852 | -0.370287064 | 0.999933887 | 0.096664336 | 0.094393377 | 0.002270959 | 0.995289108 |
| ENSG00000054179 | *ENTPD2* | 0.888806182 | 0.032218642 | 0.072060908 | -0.039842265 | 0.999933887 | -0.099980182 | -0.096293085 | -0.003687097 | 0.995289108 |
| ENSG00000064225 | *ST3GAL6* | 5.339634629 | -0.117788141 | -0.129339835 | 0.011551695 | 0.999933887 | -0.479286417 | -0.477739713 | -0.001546704 | 0.995429348 |
| ENSG00000282431 | *TRBD1* | -0.971476015 | 0.36233452 | -0.356278756 | 0.718613276 | 0.999933887 | 0.116800375 | 0.112180747 | 0.004619628 | 0.995591112 |
| ENSG00000146670 | *CDCA5* | -0.810472874 | 0.184199436 | 0.242651701 | -0.058452265 | 0.999933887 | 0.447001424 | 0.452702744 | -0.00570132 | 0.995604373 |
| ENSG00000188636 | *RTL6* | 3.13028774 | -0.00609535 | -0.129005684 | 0.122910334 | 0.999933887 | 0.06791972 | 0.069467326 | -0.001547606 | 0.995690868 |
| ENSG00000158715 | *SLC45A3* | -0.297552294 | 0.268725087 | 0.185729205 | 0.082995882 | 0.999933887 | -0.07464263 | -0.077870083 | 0.003227452 | 0.995743989 |
| ENSG00000137955 | *RABGGTB* | 4.005330011 | -0.08886355 | -0.06726991 | -0.021593639 | 0.999933887 | -0.014049904 | -0.012982622 | -0.001067281 | 0.995826055 |
| ENSG00000081760 | *AACS* | 2.014181783 | -0.106384383 | -0.144186322 | 0.037801938 | 0.999933887 | 0.015286503 | 0.017145964 | -0.001859461 | 0.995826055 |
| ENSG00000143546 | *S100A8* | 10.180449895 | -0.000171396 | 0.014275348 | -0.014446744 | 0.999933887 | 0.121225332 | 0.122704243 | -0.001478911 | 0.995914868 |
| ENSG00000133316 | *WDR74* | 3.417100489 | 0.055007181 | 0.05779185 | -0.002784669 | 0.999933887 | 0.331479394 | 0.330066548 | 0.001412847 | 0.995915732 |
| ENSG00000137502 | *RAB30* | 1.900069337 | -0.048393129 | 0.01205486 | -0.060447989 | 0.999933887 | -0.276080869 | -0.274276397 | -0.001804472 | 0.995915732 |
| ENSG00000136811 | *ODF2* | 3.506465475 | -0.066296541 | -0.039270843 | -0.027025698 | 0.999933887 | -0.000847071 | 0.000254468 | -0.001101539 | 0.996087514 |
| ENSG00000224152 | *AC009506.1* | 2.90986291 | 0.064552042 | 0.083990387 | -0.019438345 | 0.999933887 | -0.431422321 | -0.433103061 | 0.00168074 | 0.996144571 |
| ENSG00000286699 | *AC084198.2* | -0.175847096 | 0.090710069 | -0.246963301 | 0.33767337 | 0.999933887 | 0.306905576 | 0.310164343 | -0.003258768 | 0.996144571 |
| ENSG00000163945 | *UVSSA* | 5.622285228 | 0.023517449 | 0.039685995 | -0.016168547 | 0.999933887 | 0.085664751 | 0.084696584 | 0.000968167 | 0.996144571 |
| ENSG00000154930 | *ACSS1* | 4.047635977 | 0.014212196 | 0.062287621 | -0.048075425 | 0.999933887 | 0.120068706 | 0.118961269 | 0.001107436 | 0.996144571 |
| ENSG00000233762 | *RPS15P4* | -0.560911129 | -0.040965124 | -0.145543836 | 0.104578712 | 0.999933887 | -0.057462071 | -0.054064604 | -0.003397467 | 0.996144571 |
| ENSG00000276805 | *AL133216.2* | 0.401814852 | -0.022576159 | -0.017326338 | -0.005249821 | 0.999933887 | 0.063391454 | 0.066750443 | -0.003358989 | 0.996144571 |
| ENSG00000270640 | *AC104695.3* | 4.361466022 | 0.352544651 | 0.439761569 | -0.087216918 | 0.999933887 | 1.172472764 | 1.170419008 | 0.002053756 | 0.99615725 |
| ENSG00000166847 | *DCTN5* | 4.012153487 | -0.059317804 | -0.046611976 | -0.012705828 | 0.999933887 | -0.071313174 | -0.072268407 | 0.000955233 | 0.99615725 |
| ENSG00000264947 | *MIR3181* | -0.15115846 | 0.312259175 | 0.403270004 | -0.09101083 | 0.999933887 | 1.155503467 | 1.151699733 | 0.003803734 | 0.996190922 |
| ENSG00000125967 | *NECAB3* | 1.418663844 | 0.112525637 | 0.05458705 | 0.057938587 | 0.999933887 | 0.130536627 | 0.132176292 | -0.001639664 | 0.996190922 |
| ENSG00000168806 | *LCMT2* | 1.056215746 | 0.050518551 | 0.054788932 | -0.004270381 | 0.999933887 | -0.058957288 | -0.061032932 | 0.002075644 | 0.996190922 |
| ENSG00000174529 | *TMEM81* | 0.166153169 | -0.096245039 | -0.125084728 | 0.028839689 | 0.999933887 | -0.368255831 | -0.365475602 | -0.002780228 | 0.996242619 |
| ENSG00000171307 | *ZDHHC16* | 1.619484629 | -0.035833761 | 0.038546449 | -0.074380211 | 0.999933887 | 0.055067888 | 0.056534359 | -0.001466471 | 0.996366116 |
| ENSG00000204568 | *MRPS18B* | 1.891687173 | -0.030568495 | -0.003502589 | -0.027065907 | 0.999933887 | 0.023950799 | 0.025491768 | -0.001540969 | 0.996416677 |
| ENSG00000230732 | *AC016949.1* | 0.559211841 | 0.158248684 | 0.082287307 | 0.075961377 | 0.999933887 | -0.33701122 | -0.333124248 | -0.003886972 | 0.996499459 |
| ENSG00000111540 | *RAB5B* | 6.551783253 | -0.045314315 | -0.01363117 | -0.031683146 | 0.999933887 | -0.244548083 | -0.243653667 | -0.000894415 | 0.996499459 |
| ENSG00000161551 | *ZNF577* | 0.594569317 | -0.057655619 | -0.113267773 | 0.055612155 | 0.999933887 | -0.090814103 | -0.093007341 | 0.002193238 | 0.996499459 |
| ENSG00000287575 | *AL390755.3* | -0.346473675 | -0.0514047 | 0.152646456 | -0.204051156 | 0.999933887 | 2.721452963 | 2.724950063 | -0.0034971 | 0.996499459 |
| ENSG00000265458 | *AC132938.3* | -0.057272628 | -0.030647124 | 0.409089064 | -0.439736188 | 0.999933887 | 0.233968652 | 0.23726033 | -0.003291678 | 0.996499459 |
| ENSG00000123870 | *ZNF137P* | 0.300612053 | 0.077814492 | 0.141891034 | -0.064076543 | 0.999933887 | -0.437459499 | -0.435079205 | -0.002380294 | 0.996507488 |
| ENSG00000115414 | *FN1* | -1.205555653 | -0.388539793 | 0.343078977 | -0.731618769 | 0.999933887 | -0.700952605 | -0.696835119 | -0.004117486 | 0.99656832 |
| ENSG00000112367 | *FIG4* | 4.040792622 | -0.061413152 | 0.083950966 | -0.145364118 | 0.999933887 | -0.530185797 | -0.529353195 | -0.000832602 | 0.99656832 |
| ENSG00000174010 | *KLHL15* | 4.081979278 | 0.045784576 | -0.006247217 | 0.052031793 | 0.999933887 | -0.113083661 | -0.111868787 | -0.001214874 | 0.996628165 |
| ENSG00000180822 | *PSMG4* | 1.460328472 | -0.019149878 | 0.163065175 | -0.182215053 | 0.999933887 | -0.082495729 | -0.080960438 | -0.001535291 | 0.996628165 |
| ENSG00000184060 | *ADAP2* | -0.850441704 | -0.190885474 | 0.270747524 | -0.461632998 | 0.999933887 | -0.971636899 | -0.966836068 | -0.004800831 | 0.996750723 |
| ENSG00000248333 | *CDK11B* | 6.544803192 | 0.011438253 | 0.048371572 | -0.03693332 | 0.999933887 | 0.083002432 | 0.083655824 | -0.000653392 | 0.996750723 |
| ENSG00000143801 | *PSEN2* | 0.297783879 | -0.251559811 | 0.343152441 | -0.594712252 | 0.944094537 | 0.099427148 | 0.10138492 | -0.001957772 | 0.996786307 |
| ENSG00000170889 | *RPS9* | 9.317002177 | 0.048975323 | 0.004520598 | 0.044454725 | 0.999933887 | 0.052162909 | 0.051433461 | 0.000729448 | 0.996786307 |
| ENSG00000111331 | *OAS3* | 6.28575697 | -0.009939705 | -0.001310853 | -0.008628852 | 0.999933887 | -0.061262645 | -0.060325859 | -0.000936786 | 0.996786307 |
| ENSG00000165934 | *CPSF2* | 4.039023948 | -0.110921357 | -0.065330486 | -0.045590871 | 0.999933887 | -0.361628516 | -0.360796235 | -0.000832281 | 0.996830554 |
| ENSG00000263676 | *MIR4632* | -0.165385582 | 0.316746464 | 0.180273008 | 0.136473456 | 0.999933887 | 0.680557092 | 0.677079511 | 0.003477582 | 0.996861883 |
| ENSG00000119185 | *ITGB1BP1* | 3.080699553 | 0.066175029 | -0.044390359 | 0.110565388 | 0.999933887 | -0.043164512 | -0.043991698 | 0.000827186 | 0.996861883 |
| ENSG00000275464 | *FP565260.1* | -0.603873777 | 0.102660878 | -0.197581125 | 0.300242003 | 0.999933887 | 0.092168722 | 0.095019753 | -0.002851031 | 0.996861883 |
| ENSG00000071127 | *WDR1* | 9.414666589 | 0.013360884 | 0.01147105 | 0.001889834 | 0.999933887 | 0.097421971 | 0.097882128 | -0.000460157 | 0.996861883 |
| ENSG00000233264 | *AC006042.2* | -0.478993085 | -0.062084815 | -0.437314134 | 0.375229319 | 0.999933887 | -0.135991246 | -0.132135889 | -0.003855357 | 0.996861883 |
| ENSG00000105887 | *MTPN* | 7.172973921 | -0.015151877 | 0.059027438 | -0.074179315 | 0.999933887 | -0.358635683 | -0.357920392 | -0.000715291 | 0.996861883 |
| ENSG00000215126 | *CBWD6* | 1.191486065 | -0.003230829 | -0.200721235 | 0.197490406 | 0.999933887 | -0.16822381 | -0.170460397 | 0.002236587 | 0.996861883 |
| ENSG00000116171 | *SCP2* | 3.121831183 | 0.000340129 | -0.137512632 | 0.137852761 | 0.999933887 | -0.006965513 | -0.007980758 | 0.001015245 | 0.996861883 |
| ENSG00000230715 | *AC018638.2* | 3.728078395 | 0.081190744 | 0.064772834 | 0.016417911 | 0.999933887 | -0.131971489 | -0.132842356 | 0.000870868 | 0.997012499 |
| ENSG00000255557 | *AP001266.2* | 1.332774274 | -0.168844254 | -0.065216584 | -0.10362767 | 0.999933887 | -0.469669653 | -0.467864464 | -0.001805189 | 0.997070945 |
| ENSG00000286638 | *AC123595.2* | 0.229071632 | -0.212983954 | 0.489445355 | -0.702429309 | 0.944094537 | 1.075188502 | 1.077075883 | -0.001887381 | 0.997070945 |
| ENSG00000157869 | *RAB28* | 2.593349706 | -0.067417332 | -0.004809843 | -0.06260749 | 0.999933887 | 0.00148846 | 0.002429355 | -0.000940895 | 0.997070945 |
| ENSG00000198874 | *TYW1* | 2.105451075 | 0.058036904 | -0.056737519 | 0.114774423 | 0.999933887 | 0.111871362 | 0.11091013 | 0.000961232 | 0.997070945 |
| ENSG00000217801 | *AL390719.1* | 1.667819276 | 0.0774602 | 0.170620762 | -0.093160562 | 0.999933887 | 0.516086751 | 0.514922301 | 0.00116445 | 0.997070945 |
| ENSG00000129250 | *KIF1C* | 3.887543358 | 0.01645168 | 0.055399942 | -0.038948262 | 0.999933887 | -0.083302934 | -0.082412061 | -0.000890873 | 0.997070945 |
| ENSG00000270441 | *LAMB2P1* | 0.170660662 | -0.00513447 | -0.240431729 | 0.235297259 | 0.999933887 | -0.60199096 | -0.603715988 | 0.001725027 | 0.997070945 |
| ENSG00000140836 | *ZFHX3* | 2.539517757 | 0.002411896 | -0.009838218 | 0.012250114 | 0.999933887 | 0.206977316 | 0.208218007 | -0.001240691 | 0.997070945 |
| ENSG00000137492 | *THAP12* | 5.186601484 | -0.084919694 | -0.131444105 | 0.046524412 | 0.999933887 | -0.367466706 | -0.366955609 | -0.000511097 | 0.997110398 |
| ENSG00000268001 | *CARD8-AS1* | 3.871119173 | -0.177567092 | -0.240984482 | 0.063417389 | 0.999933887 | -1.197141828 | -1.198230422 | 0.001088594 | 0.997110398 |
| ENSG00000104671 | *DCTN6* | 2.087488701 | -0.076772021 | -0.094746839 | 0.017974818 | 0.999933887 | 0.113295157 | 0.112171353 | 0.001123804 | 0.997110398 |
| ENSG00000101596 | *SMCHD1* | 9.568619302 | -0.044607186 | -0.149833759 | 0.105226573 | 0.999933887 | -1.132406208 | -1.131058514 | -0.001347694 | 0.997110398 |
| ENSG00000121057 | *AKAP1* | 2.417570634 | 0.017418553 | 0.017245396 | 0.000173157 | 0.999933887 | 0.074890413 | 0.075694812 | -0.000804399 | 0.997110398 |
| ENSG00000162623 | *TYW3* | 0.648226451 | -0.046889614 | -0.24713763 | 0.200248016 | 0.999933887 | 0.082562808 | 0.081128618 | 0.00143419 | 0.997180111 |
| ENSG00000126790 | *L3HYPDH* | 0.820337685 | 0.172700465 | 0.123240126 | 0.049460339 | 0.999933887 | 0.088714316 | 0.089849502 | -0.001135186 | 0.997426142 |
| ENSG00000270012 | *AC232271.1* | 0.548463294 | -0.09505425 | 0.319321463 | -0.414375713 | 0.999933887 | -0.213789403 | -0.21508984 | 0.001300437 | 0.997426142 |
| ENSG00000235532 | *LINC00402* | 0.818343591 | 0.045773203 | 0.051828217 | -0.006055014 | 0.999933887 | 0.05429643 | 0.055443576 | -0.001147146 | 0.997426142 |
| ENSG00000138772 | *ANXA3* | 4.525306037 | -0.008484833 | -0.103150304 | 0.094665471 | 0.999933887 | 0.272064734 | 0.271426457 | 0.000638276 | 0.997426142 |
| ENSG00000180979 | *LRRC57* | 2.423675208 | -0.093548038 | 0.049664714 | -0.143212751 | 0.999933887 | -0.004143218 | -0.00484137 | 0.000698153 | 0.997485807 |
| ENSG00000121039 | *RDH10* | 1.465279809 | 0.049551031 | 0.079900678 | -0.030349647 | 0.999933887 | -0.043640873 | -0.044591882 | 0.000951009 | 0.997573821 |
| ENSG00000259891 | *AC107375.1* | 1.248862151 | -0.082819429 | 0.128244578 | -0.211064007 | 0.999933887 | -0.209781635 | -0.210782237 | 0.001000603 | 0.99765007 |
| ENSG00000099204 | *ABLIM1* | 5.49893757 | -0.03384339 | -0.00906432 | -0.024779071 | 0.999933887 | 0.046076815 | 0.045682741 | 0.000394074 | 0.997679131 |
| ENSG00000157193 | *LRP8* | 2.021183106 | -0.014534174 | 0.148275196 | -0.16280937 | 0.999933887 | 0.06338544 | 0.062727053 | 0.000658387 | 0.997679131 |
| ENSG00000273137 | *AL022328.2* | 0.280791098 | 0.018790874 | 0.107724416 | -0.088933542 | 0.999933887 | -0.297765076 | -0.299118009 | 0.001352933 | 0.997790484 |
| ENSG00000249978 | *TRGV7* | -0.402632704 | -0.274039352 | 0.10733819 | -0.381377542 | 0.999933887 | 0.031391195 | 0.030336782 | 0.001054412 | 0.997974116 |
| ENSG00000261079 | *AC009053.2* | 0.000769589 | 0.1214744 | -0.017171912 | 0.138646312 | 0.999933887 | 0.682543349 | 0.681360374 | 0.001182975 | 0.997974116 |
| ENSG00000119522 | *DENND1A* | 4.226800522 | 0.031493344 | -0.012009716 | 0.04350306 | 0.999933887 | 0.111104706 | 0.110706412 | 0.000398294 | 0.997974116 |
| ENSG00000258546 | *CENPUP2* | -0.402545628 | -0.093895431 | -0.093092953 | -0.000802477 | 0.999933887 | -0.907890485 | -0.910314234 | 0.002423749 | 0.997974116 |
| ENSG00000159618 | *ADGRG5* | 3.46085037 | -0.053423077 | -0.018941238 | -0.034481839 | 0.999933887 | 0.171836986 | 0.171506768 | 0.000330219 | 0.998260028 |
| ENSG00000083123 | *BCKDHB* | 0.012644567 | 0.102244128 | -0.156018852 | 0.25826298 | 0.999933887 | -0.480746872 | -0.479594344 | -0.001152527 | 0.998260028 |
| ENSG00000185972 | *CCIN* | -0.932453691 | -0.073742314 | 0.031317211 | -0.105059525 | 0.999933887 | 0.249759907 | 0.252083194 | -0.002323287 | 0.998260028 |
| ENSG00000211666 | *IGLV2-14* | 0.099442435 | -0.124574094 | 0.181269066 | -0.30584316 | 0.999933887 | 0.237033225 | 0.238084234 | -0.001051009 | 0.998316198 |
| ENSG00000248099 | *INSL3* | 0.673336632 | 0.120390152 | -0.077739046 | 0.198129197 | 0.999933887 | 0.364802548 | 0.365937089 | -0.001134541 | 0.998379423 |
| ENSG00000159208 | *CIART* | 3.462147259 | 0.099575578 | 0.116750835 | -0.017175257 | 0.999933887 | 0.139125544 | 0.138705656 | 0.000419889 | 0.998379423 |
| ENSG00000008323 | *PLEKHG6* | 1.731560874 | 0.037985914 | 0.058402163 | -0.020416249 | 0.999933887 | -0.412401736 | -0.41156702 | -0.000834716 | 0.998379423 |
| ENSG00000211796 | *TRAV16* | -1.240919178 | 0.346667537 | -0.052392414 | 0.39905995 | 0.999933887 | 0.086180457 | 0.085152751 | 0.001027707 | 0.998713288 |
| ENSG00000146414 | *SHPRH* | 1.972210041 | 0.05864432 | 0.175707318 | -0.117062998 | 0.999933887 | -0.111007721 | -0.11036458 | -0.000643141 | 0.998713288 |
| ENSG00000206573 | *THUMPD3-AS* | 5.138015064 | -0.017318821 | 0.010163024 | -0.027481845 | 0.999933887 | 0.059036783 | 0.059223409 | -0.000186626 | 0.998713288 |
| ENSG00000234506 | *LINC01506* | 4.485228692 | -0.010023724 | 0.031397623 | -0.041421347 | 0.999933887 | 0.455013523 | 0.455262 | -0.000248478 | 0.998713288 |
| ENSG00000107362 | *ABHD17B* | 2.930801381 | -0.003788949 | 0.0164942 | -0.020283148 | 0.999933887 | -0.092414502 | -0.092674036 | 0.000259534 | 0.998713288 |

| | | | | | | | | | | |
|---|---|---|---|---|---|---|---|---|---|---|
| ENSG00000143156 | *NME7* | 0.132732804 | -0.003097129 | -0.068415622 | 0.065318493 | 0.999933887 | -0.070552433 | -0.071468207 | 0.000915774 | 0.998713288 |
| ENSG00000167676 | *PLIN4* | 5.132123838 | 0.070833053 | 0.061167771 | 0.009665282 | 0.999933887 | 0.242808403 | 0.242601209 | 0.000207194 | 0.999189691 |
| ENSG00000172469 | *MANEA* | -0.189229342 | 0.177192932 | -0.172586593 | 0.349779524 | 0.999933887 | -0.160833765 | -0.160346107 | -0.000487658 | 0.999189691 |
| ENSG00000198042 | *MAK16* | 1.143506695 | -0.133175329 | -0.245586344 | 0.112411016 | 0.999933887 | -0.041644247 | -0.041869868 | 0.000225622 | 0.999212988 |
| ENSG00000172081 | *MOB3A* | 8.544392904 | -0.083218534 | -0.082135677 | -0.001082857 | 0.999933887 | -0.249471067 | -0.24960789 | 0.000136823 | 0.999212988 |
| ENSG00000031003 | *FAM13B* | 5.307426426 | -0.050091322 | -0.039643422 | -0.0104479 | 0.999933887 | -0.440858457 | -0.440745349 | -0.000113108 | 0.999212988 |
| ENSG00000166860 | *ZBTB39* | 1.604299869 | 0.047997544 | 0.09498957 | -0.046992026 | 0.999933887 | 0.143790793 | 0.144007818 | -0.000217026 | 0.999212988 |
| ENSG00000115875 | *SRSF7* | 6.368229192 | -0.024615473 | 0.000305528 | -0.024921001 | 0.999933887 | 0.213481188 | 0.213578811 | -9.7623e-05 | 0.999212988 |
| ENSG00000089169 | *RPH3A* | 1.237995935 | -0.03066896 | -0.007834622 | -0.022834338 | 0.999933887 | -0.341116339 | -0.341412324 | 0.000295985 | 0.999212988 |
| ENSG00000176915 | *ANKLE2* | 6.243385322 | 0.003266403 | 0.031517499 | -0.028251096 | 0.999933887 | 0.804094635 | 0.804244774 | -0.000150138 | 0.999212988 |
| ENSG00000163590 | *PPM1L* | 1.362260688 | -0.037937884 | -0.014934189 | -0.023003694 | 0.999933887 | -0.043916893 | -0.043794189 | -0.000122704 | 0.999701 |
| ENSG00000204634 | *TBC1D8* | 3.304883736 | 0.028249195 | 0.028036464 | 0.00021273 | 0.999933887 | 0.275724986 | 0.275691925 | 3.306e-05 | 0.999878408 |
| ENSG00000224467 | *TANK-AS1* | 0.645760415 | 0.252398053 | 0.697448702 | -0.445050649 | 0.999933887 | 1.290038005 | 1.290078534 | -4.0528e-05 | 0.999904689 |

The overall average expression is shown for all of the groups.
**“HITTIN 1h inf”** and **“HITTIN 6h inf”** show the average log2FC *Mtb* infection compared to the uninfected response for neutrophils from HITTIN after 1 and 6 hours respectively.
**“HIT 1h inf”** and **“HIT 6h inf**” show the average log2FC *Mtb* infection compared to the uninfected response for neutrophils from HIT after 1 and 6 hours respectively.
**“HITTINxHIT 1h inf”** and **“HITTINxHIT 6h inf”** is the average log2FC response difference between neutrophils from HITTIN and HIT in response to *Mtb* infection at 1 hour and 6 hours respectively (interaction test).
**"Adj.P.Value"** is the adjusted p-value after the Benjamini Hochberg correction for multiple testing shown for both the 1 and 6 hour interaction tests
Significant genes were defined as genes with an absolute log2FC ≥ 0.2 and adjusted p value ≤ 0.05
